## Supplementary material for "Prenatal Arsenic Exposure and Gene Expression in Fetal Liver, Heart, Lung, and Placenta": Rychlik_Arsenic_microarray_Supplemental_Data

Author names & affiliations (with ORCIDs):

K.A. Rychlik[1,2], C. Kashiwagi[1], J. Liao[1], A. Mathur[1], E.J. Illingworth[1], S. Sanchez[1] (ORCID: 0000-0002-3649-778X), A. Kleensang[1], A. Maertens[1], F.C.M. Sillé[1] (ORCID: 0000-0003-4305-2049)

1 Department of Environmental Health and Engineering, Bloomberg School of Public Health, Johns Hopkins University, Baltimore, MD, USA

2 Public Health Program, School of Health Professions, Mayborn College of Health Sciences, University of Mary Hardin-Baylor, Belton, TX, USA

**Keywords**: arsenic, prenatal, mRNA, microarray, immune

**Running Head:** Prenatal Arsenic: Gene Expression in Multiple Fetal Tissues

**Supplemental Data**

**Table S1. Complete List of Differential Gene Expression Analysis for Heart Tissue.** Heart samples were obtained from gestational day (GD) 18 C57Bl/6J fetal mice exposed for 2 weeks prior to conception and during gestation to either 0 ppb (control) or 100 ppb (exposed) sodium (meta) arsenite. After microarray analysis, the expression of all genes from heart tissue were compared for arsenic exposed to unexposed controls. Data shown represents an N=3.

**Table S2. Complete List of Differential Gene Expression Analysis for Liver Tissue.** Liver samples were obtained from gestational day (GD) 18 C57Bl/6J fetal mice exposed for 2 weeks prior to conception and during gestation to either 0 ppb (control) or 100 ppb (exposed) sodium (meta) arsenite. After microarray analysis, the expression of all genes from liver tissue were compared for arsenic exposed to unexposed controls. Data shown represents an N=3.

**Table S3. Complete List of Differential Gene Expression Analysis for Lung Tissue.** Lung samples were obtained from gestational day (GD) 18 C57Bl/6J fetal mice exposed for 2 weeks prior to conception and during gestation to either 0 ppb (control) or 100 ppb (exposed) sodium (meta) arsenite. After microarray analysis, the expression of all genes from lung tissue were compared for arsenic exposed to unexposed controls. Data shown represents an N=3.

**Table S4. Complete List of Differential Gene Expression Analysis for Placenta Tissue.** Placenta samples were obtained from gestational day (GD) 18 C57Bl/6J fetal mice exposed for 2 weeks prior to conception and during gestation to either 0 ppb (control) or 100 ppb (exposed) sodium (meta) arsenite. After microarray analysis, the expression of all genes from placenta tissue were compared for arsenic exposed to unexposed controls. Data shown represents an N=3.

**Table S5. Complete List of Enriched Biological Processes in arsenic-exposed GD18 Liver Tissue.** Enriched biological processes were identified in STRING based on the 240 differentially expressed (comparing exposed to controls exhibited an unadjusted *p*-value <0.01) genes from the liver using gene ontology (GO). The specific GO number is reported along with its term description. Each protein from the differentially expressed input that are associated with that biological process are listed in the right column of the table. Data shown represents an N=3.

Rychlik_Arsenic_microarray_TableS1_Heart_100_vs_0_all_p_values

| | ID | logFC | CI.L | CI.R | AveExpr | t | P.Value | adj.P.Val | B |
|---|---|---|---|---|---|---|---|---|---|
| **A_51_P272553** | Bhlhe40 | -0.786096113 | -0.903681253 | -0.668510974 | 9.70348306 | -15.97836519 | 1.49E-06 | 0.028836071 | -0.453986073 |
| **A_52_P319438** | Ankrd37 | -0.621433468 | -0.754997891 | -0.487869045 | 10.09754107 | -11.12020973 | 1.54E-05 | 0.148494581 | -0.698637243 |
| **A_51_P151126** | Cd52 | -0.749209396 | -0.938996271 | -0.559422521 | 9.674489692 | -9.435092729 | 4.32E-05 | 0.278667429 | -0.865670567 |
| **A_55_P1996973** | Gvin1 | -0.503785296 | -0.649526792 | -0.358043799 | 8.779631163 | -8.261737563 | 9.83E-05 | 0.423359531 | -1.032426439 |
| **A_51_P226269** | Rgcc | -0.559678216 | -0.724459094 | -0.394897339 | 11.07352661 | -8.117844457 | 0.00010948 | 0.423359531 | -1.056772575 |
| **A_51_P102789** | C1qc | -0.455570436 | -0.609742798 | -0.301398075 | 9.653564141 | -7.062494786 | 0.000253756 | 0.705977075 | -1.346980217 |
| **A_55_P2027392** | Gpr146 | -0.349084455 | -0.469383302 | -0.228785608 | 11.21091296 | -6.93550641 | 0.00028266 | 0.705977075 | -1.299626482 |
| **A_51_P355753** | Hic1 | -0.436426327 | -0.593961122 | -0.278891533 | 11.88620151 | -6.621304847 | 0.000371653 | 0.705977075 | -1.379698798 |
| **A_52_P463977** | Tmem140 | -0.429410493 | -0.584844676 | -0.273976311 | 9.317362177 | -6.602908091 | 0.000377774 | 0.705977075 | -1.384626923 |
| **A_55_P2024888** | Ctss | -0.866619392 | -1.185136429 | -0.548102355 | 10.53935162 | -6.502866942 | 0.000413123 | 0.705977075 | -1.41191433 |
| **A_51_P359636** | Lgals3bp | -0.550083876 | -0.752669006 | -0.347498746 | 10.50385075 | -6.489787594 | 0.000418015 | 0.705977075 | -1.415543804 |
| **A_55_P1971889** | F3 | -0.309395867 | -0.424613756 | -0.194177979 | 9.53787301 | -6.418059348 | 0.000446036 | 0.705977075 | -1.43570831 |
| **A_55_P2137611** | Irgm2 | -0.324439551 | -0.447603764 | -0.201275338 | 9.245264488 | -6.295908218 | 0.00049878 | 0.705977075 | -1.47108558 |
| **A_55_P1966833** | Xaf1 | -0.324040735 | -0.447806616 | -0.200274854 | 8.519956602 | -6.257600053 | 0.000516746 | 0.705977075 | -1.482457064 |
| **A_51_P304170** | Rtp4 | -0.811952504 | -1.125694961 | -0.498210048 | 9.368251786 | -6.185381139 | 0.000552638 | 0.705977075 | -1.504263867 |
| **A_55_P2036567** | Vav1 | -0.332612425 | -0.462379043 | -0.202845806 | 8.695604838 | -6.126107479 | 0.000584207 | 0.705977075 | -1.522529448 |
| **A_55_P2163353** | Sox3 | 0.338007364 | 0.198933391 | 0.477081337 | 8.670119113 | 5.808840256 | 0.000791921 | 0.877509219 | -1.626234606 |
| **A_55_P2128032** | Hsf2 | 0.239975396 | 0.139768594 | 0.340182198 | 10.23399504 | 5.72372353 | 0.000861004 | 0.877509219 | -1.655851464 |
| **A_51_P438967** | Gpnmb | -0.593790231 | -0.841806134 | -0.345774328 | 8.428843677 | -5.722194967 | 0.000862305 | 0.877509219 | -1.656390601 |
| **A_51_P261517** | Tyrobp | -0.44429928 | -0.635076089 | -0.253522471 | 9.486204024 | -5.566204531 | 0.001007492 | 0.973992669 | -1.712794782 |
| **A_51_P129012** | B2m | -0.287324865 | -0.412291909 | -0.162357821 | 12.68094407 | -5.495242307 | 0.001082482 | 0.978744335 | -1.739385248 |
| **A_51_P503192** | Rbm15b | 0.302090528 | 0.169535264 | 0.434645792 | 9.307469863 | 5.44689824 | 0.001137156 | 0.978744335 | -1.757844306 |
| **A_55_P2151601** | Samd9l | -0.482061078 | -0.696512309 | -0.267609847 | 9.559630994 | -5.372577091 | 0.00122736 | 0.978744335 | -1.786777411 |
| **A_52_P351816** | Itgb1 | 0.417492636 | 0.231592452 | 0.603392819 | 10.26745064 | 5.367575133 | 0.001233714 | 0.978744335 | -1.788749173 |
| **A_55_P1962937** | Trem2 | -0.348583237 | -0.504514396 | -0.192652078 | 8.731733616 | -5.342969846 | 0.001265509 | 0.978744335 | -1.798494036 |
| **A_55_P1989772** | Sqor | -0.284265237 | -0.414566074 | -0.1539644 | 9.784206497 | -5.214175931 | 0.001447671 | 0.991835066 | -1.850758123 |
| **A_51_P345367** | Psmb8 | -0.284369054 | -0.416706613 | -0.152031495 | 8.322075451 | -5.135802861 | 0.001572814 | 0.991835066 | -1.88361942 |
| **A_66_P121787** | Samd9l | -0.357891118 | -0.524609894 | -0.191172342 | 9.190650782 | -5.130687219 | 0.001581394 | 0.991835066 | -1.885792831 |
| **A_52_P164136** | Arrdc3 | -0.326087611 | -0.479098555 | -0.173076666 | 9.243721924 | -5.093554237 | 0.001645267 | 0.991835066 | -1.901674856 |
| **A_55_P2002963** | Coro1a | -0.412371093 | -0.606101774 | -0.218640412 | 9.239199231 | -5.087435785 | 0.001656066 | 0.991835066 | -1.904309705 |
| **A_55_P1975560** | Ifi204 | -0.358089674 | -0.531297373 | -0.184881976 | 8.811527372 | -4.941214862 | 0.001939272 | 0.991835066 | -1.968818811 |
| **A_51_P514712** | Parp14 | -0.273631238 | -0.407057143 | -0.140205333 | 8.748646534 | -4.901563707 | 0.002025122 | 0.991835066 | -1.986831727 |
| **A_55_P2097279** | Pirb | -0.290553904 | -0.432323963 | -0.148783845 | 8.233008127 | -4.89836708 | 0.002032226 | 0.991835066 | -1.988293751 |
| **A_55_P2175245** | Tusc3 | -0.241613767 | -0.360551621 | -0.122675913 | 11.07404196 | -4.855240081 | 0.002130835 | 0.991835066 | -2.00816335 |
| **A_55_P2172619** | NA | -0.21392263 | -0.319904786 | -0.107940474 | 16.43484604 | -4.824286725 | 0.0022049 | 0.991835066 | -2.022591781 |
| **A_55_P2052834** | Lst1 | -0.405051307 | -0.605849184 | -0.204253429 | 9.855275538 | -4.821254435 | 0.002212309 | 0.991835066 | -2.024012821 |
| **A_55_P2102181** | NA | -0.239768402 | -0.358978663 | -0.12055814 | 10.47070187 | -4.807147416 | 0.002247143 | 0.991835066 | -2.03064176 |
| **A_51_P483280** | Prnp | -0.216637128 | -0.32494237 | -0.108331887 | 12.43146215 | -4.780711624 | 0.00231408 | 0.991835066 | -2.043143627 |
| **A_55_P2179413** | Lgals3bp | -0.415973582 | -0.624376788 | -0.207570375 | 9.753450103 | -4.770572334 | 0.00234034 | 0.991835066 | -2.047966319 |
| **A_55_P1961943** | Cd68 | -0.356638964 | -0.536577777 | -0.176700151 | 8.968518733 | -4.737105655 | 0.002429402 | 0.991835066 | -2.063994182 |
| **A_55_P1965000** | Ddx58 | -0.219690872 | -0.330983762 | -0.108397981 | 9.693802133 | -4.717954225 | 0.002482063 | 0.991835066 | -2.073242349 |
| **A_51_P189361** | Osgin1 | -0.214243635 | -0.322830534 | -0.105656735 | 8.10174124 | -4.715628946 | 0.002488543 | 0.991835066 | -2.074369013 |
| **A_55_P2114938** | Irf9 | -0.423757947 | -0.639882962 | -0.207632933 | 9.704730006 | -4.686212224 | 0.002572168 | 0.991835066 | -2.088693537 |
| **A_51_P316935** | Depp1 | -0.486742667 | -0.737363773 | -0.236121562 | 11.2234851 | -4.641847928 | 0.002704264 | 0.991835066 | -2.110548463 |
| **A_51_P390538** | Mpeg1 | -0.419739277 | -0.636082505 | -0.203396048 | 9.063528679 | -4.637089048 | 0.002718876 | 0.991835066 | -2.112910907 |

| | | | | | | | | | |
|---|---|---|---|---|---|---|---|---|---|
| **A_55_P2047800** | NA | -0.250722519 | -0.380666445 | -0.120778592 | 12.43578341 | -4.611545411 | 0.002798825 | 0.991835066 | -2.125651851 |
| **A_55_P2087240** | Mipep | -0.210873985 | -0.320375987 | -0.101371983 | 10.02858754 | -4.60267256 | 0.002827205 | 0.991835066 | -2.130101448 |
| **A_52_P56751** | Lcp1 | -0.303028564 | -0.461352721 | -0.144704407 | 9.399670633 | -4.574519834 | 0.002919393 | 0.991835066 | -2.144301635 |
| **A_55_P2156905** | NA | 0.254696204 | 0.121346298 | 0.38804611 | 13.38051835 | 4.564980128 | 0.002951385 | 0.991835066 | -2.149141861 |
| **A_51_P259296** | Lpl | -0.425873389 | -0.649117193 | -0.202629586 | 13.15632719 | -4.559426554 | 0.002970189 | 0.991835066 | -2.151966267 |
| **A_55_P2114953** | Usp18 | -0.249044196 | -0.380590285 | -0.117498108 | 7.986689806 | -4.524885751 | 0.003090172 | 0.991835066 | -2.16964318 |
| **A_55_P1961270** | Cd72 | -0.265128043 | -0.405591105 | -0.124664981 | 8.042621457 | -4.511309951 | 0.003138796 | 0.991835066 | -2.176643148 |
| **A_51_P405476** | Fcer1g | -0.473356297 | -0.72621442 | -0.220498174 | 9.986812079 | -4.47425139 | 0.003275906 | 0.991835066 | -2.195902661 |
| **A_55_P1954161** | Plagl2 | 0.182546527 | 0.084701747 | 0.280391307 | 8.161500067 | 4.459077742 | 0.003333954 | 0.991835066 | -2.203852762 |
| **A_55_P1958165** | Ms4a7 | -0.529120837 | -0.81462568 | -0.243615995 | 9.007042746 | -4.429457289 | 0.003450586 | 0.991835066 | -2.219480508 |
| **A_51_P246066** | Slamf9 | -0.364134775 | -0.560629316 | -0.167640234 | 8.439925153 | -4.429154188 | 0.003451803 | 0.991835066 | -2.219641167 |
| **A_51_P265495** | Ly6a | -0.449918623 | -0.693073435 | -0.20676381 | 8.783109449 | -4.422422517 | 0.003478946 | 0.991835066 | -2.223213205 |
| **A_55_P2158990** | Jun | 0.428853556 | 0.19214321 | 0.665563902 | 12.63102245 | 4.330129732 | 0.003875971 | 0.991835066 | -2.272944395 |
| **A_55_P1978216** | Cd300c2 | -0.31277276 | -0.485458545 | -0.140086976 | 8.355411038 | -4.328939013 | 0.003881411 | 0.991835066 | -2.273595307 |
| **A_52_P482203** | Tsen2 | 0.180573941 | 0.080270167 | 0.280877715 | 10.70981136 | 4.302758131 | 0.004003181 | 0.991835066 | -2.287967689 |
| **A_55_P2227355** | Ptpro | -0.208568102 | -0.325434526 | -0.091701677 | 8.366108533 | -4.26547366 | 0.004183958 | 0.991835066 | -2.308636215 |
| **A_55_P2123529** | NA | -0.198807993 | -0.310298662 | -0.087317324 | 11.08294074 | -4.26191174 | 0.004201695 | 0.991835066 | -2.310623155 |
| **A_55_P2011116** | L3mbtl3 | 0.212695856 | 0.093396004 | 0.331995708 | 9.475928938 | 4.261164053 | 0.004205429 | 0.991835066 | -2.311040511 |
| **A_52_P243054** | Nufip2 | 0.19188788 | 0.083908289 | 0.299867471 | 8.918947108 | 4.247320159 | 0.004275231 | 0.991835066 | -2.318785426 |
| **A_55_P2056729** | Igfbp2 | 0.274444358 | 0.119668268 | 0.429220449 | 8.791601568 | 4.237986484 | 0.004323014 | 0.991835066 | -2.324025672 |
| **A_55_P2048855** | Sprr2a2 | -0.273530201 | -0.427802659 | -0.119257743 | 8.73197744 | -4.237659114 | 0.004324701 | 0.991835066 | -2.32420974 |
| **A_52_P30451** | Ppp1r3c | -0.409818142 | -0.641993823 | -0.177642461 | 12.77969363 | -4.218748 | 0.00442338 | 0.991835066 | -2.334874113 |
| **A_52_P30989** | Cdkn3 | -0.263067897 | -0.412255587 | -0.113880207 | 9.617028983 | -4.21447998 | 0.004445993 | 0.991835066 | -2.337289472 |
| **A_66_P128593** | Lims1 | -0.215132236 | -0.338149836 | -0.092114635 | 10.92234575 | -4.179721744 | 0.004634982 | 0.991835066 | -2.357077352 |
| **A_52_P326354** | Erf | 0.274746153 | 0.11758194 | 0.431910366 | 12.25795391 | 4.178179466 | 0.004643571 | 0.991835066 | -2.357960235 |
| **A_55_P1998943** | Oas1a | -0.417881206 | -0.657193031 | -0.17856938 | 8.703784259 | -4.173474956 | 0.004669879 | 0.991835066 | -2.36065591 |
| **A_55_P1999838** | Cox7c | -0.238202026 | -0.374986638 | -0.101417414 | 15.14382918 | -4.162146294 | 0.004733908 | 0.991835066 | -2.367163053 |
| **A_55_P2017759** | Evi2a | -0.225188379 | -0.354572095 | -0.095804663 | 8.191162148 | -4.159829081 | 0.004747124 | 0.991835066 | -2.368496814 |
| **A_52_P281033** | Socs5 | -0.173495442 | -0.273829423 | -0.07316146 | 9.016181954 | -4.132845365 | 0.004904058 | 0.991835066 | -2.384097571 |
| **A_52_P266132** | Fgl2 | -0.398023169 | -0.62856089 | -0.167485448 | 10.01164561 | -4.126439567 | 0.004942149 | 0.991835066 | -2.38781989 |
| **A_55_P2472435** | Gbp3 | -0.327306591 | -0.5180784 | -0.136534783 | 8.442966081 | -4.100621519 | 0.005099013 | 0.991835066 | -2.40289573 |
| **A_55_P1978424** | Bcl2a1d | -0.331562651 | -0.525601696 | -0.137523605 | 8.432605524 | -4.083998796 | 0.005202911 | 0.991835066 | -2.412664563 |
| **A_55_P2143790** | Gm14410 | 0.216451725 | 0.089459019 | 0.343444431 | 9.709569784 | 4.073722169 | 0.005268311 | 0.991835066 | -2.418728455 |
| **A_52_P322181** | Adrb1 | -0.263562572 | -0.418296407 | -0.108828737 | 9.826095519 | -4.07106072 | 0.005285395 | 0.991835066 | -2.420301948 |
| **A_51_P280446** | Sdf2l1 | 0.246663261 | 0.101773856 | 0.391552667 | 9.600390798 | 4.068899576 | 0.005299313 | 0.991835066 | -2.42158058 |
| **A_55_P2066707** | Trim27 | 0.244795352 | 0.100911422 | 0.388679281 | 11.73525955 | 4.066305549 | 0.005316072 | 0.991835066 | -2.423116423 |
| **A_55_P2104798** | NA | 0.190582572 | 0.078277676 | 0.302887468 | 9.595883987 | 4.05595965 | 0.005383496 | 0.991835066 | -2.429253842 |
| **A_51_P154842** | Oas1f | -0.261172303 | -0.41530879 | -0.107035816 | 8.184135289 | -4.049774058 | 0.005424257 | 0.991835066 | -2.432932396 |
| **A_51_P128876** | Ifitm3 | -0.525074645 | -0.83508711 | -0.21506218 | 11.06969144 | -4.048098056 | 0.00543536 | 0.991835066 | -2.433930287 |
| **A_55_P2085142** | Spp1 | -0.251725691 | -0.400441336 | -0.103010045 | 8.041221322 | -4.045572732 | 0.005452137 | 0.991835066 | -2.435434812 |
| **A_51_P404084** | Psmb2 | -0.270555731 | -0.430412639 | -0.110698823 | 10.69116716 | -4.045148415 | 0.005454961 | 0.991835066 | -2.435687721 |
| **A_51_P181451** | C1qa | -0.335405548 | -0.53458722 | -0.136223876 | 8.739036829 | -4.02466744 | 0.005593229 | 0.991835066 | -2.447933513 |
| **A_55_P2064328** | Lama2 | -0.289120299 | -0.460921445 | -0.117319154 | 11.45822574 | -4.022181487 | 0.005610272 | 0.991835066 | -2.44942501 |
| **A_52_P522379** | Map3k20 | 0.251824718 | 0.101984214 | 0.401665221 | 8.598844403 | 4.016782026 | 0.005647489 | 0.991835066 | -2.452668349 |
| **A_55_P2131520** | Vash1 | 0.218388501 | 0.08832318 | 0.348453823 | 13.76040885 | 4.013075965 | 0.00567319 | 0.991835066 | -2.454897531 |
| **A_52_P638459** | Ccl5 | -0.197526984 | -0.315313616 | -0.079740352 | 7.954168466 | -4.008109354 | 0.005707836 | 0.991835066 | -2.457888805 |
| **A_51_P108108** | Lrrc25 | -0.173336088 | -0.27677372 | -0.069898456 | 8.095737211 | -4.005157061 | 0.00572854 | 0.991835066 | -2.459669007 |
| **A_55_P2006476** | NA | -0.293250296 | -0.468305072 | -0.11819552 | 12.05196027 | -4.003811538 | 0.005738004 | 0.991835066 | -2.460480864 |

| | | | | | | | | | |
|---|---|---|---|---|---|---|---|---|---|
| **A_55_P2094060** | Gzma | -0.581501495 | -0.929747761 | -0.23325523 | 8.336500191 | -3.990924394 | 0.005829522 | 0.991835066 | -2.468273202 |
| **A_51_P262171** | Irgm1 | -0.188065403 | -0.301102278 | -0.075028527 | 8.822440233 | -3.976471695 | 0.005934073 | 0.991835066 | -2.477047852 |
| **A_55_P2016105** | Errfi1 | -0.269404561 | -0.432044188 | -0.106764933 | 10.83576887 | -3.959020087 | 0.006063077 | 0.991835066 | -2.487693636 |
| **A_55_P1988202** | Ifi203 | -0.291851973 | -0.468209171 | -0.115494774 | 8.889599339 | -3.955292203 | 0.006091032 | 0.991835066 | -2.489974873 |
| **A_51_P259415** | Tmem70 | -0.181435247 | -0.291153921 | -0.071716573 | 12.16047293 | -3.952303081 | 0.006113549 | 0.991835066 | -2.491805857 |
| **A_55_P1992676** | Ubtf | 0.195769812 | 0.076552972 | 0.314986652 | 9.499467512 | 3.924797469 | 0.006325102 | 0.991835066 | -2.508730729 |
| **A_52_P202029** | Trak2 | -0.206175992 | -0.332192285 | -0.080159699 | 10.70509326 | -3.910394062 | 0.006439081 | 0.991835066 | -2.517648566 |
| **A_52_P393306** | Snap23 | 0.253635065 | 0.098428088 | 0.408842042 | 9.533150189 | 3.90577469 | 0.006476113 | 0.991835066 | -2.520516677 |
| **A_55_P2005813** | Abhd18 | -0.173031927 | -0.27895429 | -0.067109565 | 8.763367886 | -3.904340807 | 0.006487655 | 0.991835066 | -2.521407751 |
| **A_52_P682382** | Scd1 | -0.167278108 | -0.269814272 | -0.064741944 | 8.1341238 | -3.899161061 | 0.006529539 | 0.991835066 | -2.524629799 |
| **A_55_P1955039** | Myo1b | -0.214511315 | -0.346128528 | -0.082894102 | 9.061728297 | -3.895351458 | 0.006560533 | 0.991835066 | -2.527002691 |
| **A_51_P159352** | Sub1 | 0.240035472 | 0.092509542 | 0.387561402 | 8.939430851 | 3.888805264 | 0.00661417 | 0.991835066 | -2.531086346 |
| **A_55_P2071683** | NA | -0.16711935 | -0.26990424 | -0.064334461 | 14.53856647 | -3.88603401 | 0.006637021 | 0.991835066 | -2.532817483 |
| **A_51_P189746** | Pim3 | -0.282655032 | -0.457035454 | -0.108274609 | 9.772716872 | -3.874075926 | 0.006736625 | 0.991835066 | -2.540303599 |
| **A_55_P1961499** | Ly6c1 | -0.494905736 | -0.800607143 | -0.189204328 | 8.915395957 | -3.869315841 | 0.00677673 | 0.991835066 | -2.543290872 |
| **A_55_P2181356** | Cdh11 | 0.220797858 | 0.084004842 | 0.357590874 | 8.540630076 | 3.857803147 | 0.006874813 | 0.991835066 | -2.550533115 |
| **A_52_P674414** | Mkln1 | -0.240118962 | -0.388949856 | -0.091288068 | 9.65661092 | -3.856048595 | 0.006889898 | 0.991835066 | -2.55163899 |
| **A_55_P2103706** | NA | -0.194722907 | -0.315593487 | -0.073852326 | 10.85941293 | -3.850397515 | 0.006938729 | 0.991835066 | -2.555204662 |
| **A_55_P2086148** | NA | 0.196819569 | 0.073819913 | 0.319819224 | 11.12679791 | 3.82448986 | 0.007167513 | 0.991835066 | -2.571627209 |
| **A_51_P444447** | Cebpd | -0.216330174 | -0.351672814 | -0.080987534 | 11.77821919 | -3.82024828 | 0.007205751 | 0.991835066 | -2.57432773 |
| **A_55_P2114697** | Gcm1 | 0.440392397 | 0.205995073 | 0.674789721 | 12.10376396 | 5.419555748 | 0.007275757 | 0.991835066 | -3.430971153 |
| **A_55_P2051864** | Golgb1 | -0.331779407 | -0.54016336 | -0.123395454 | 13.52819518 | -3.805347186 | 0.007341869 | 0.991835066 | -2.583841379 |
| **A_51_P151433** | Ncbp2 | 0.190928504 | 0.070628224 | 0.311228784 | 9.278772885 | 3.793265706 | 0.0074543 | 0.991835066 | -2.591585106 |
| **A_51_P459350** | Dstn | -0.246567555 | -0.402180094 | -0.090955015 | 13.36151974 | -3.787045042 | 0.007512925 | 0.991835066 | -2.595582879 |
| **A_51_P150802** | Mpv17 | -0.305504318 | -0.498951078 | -0.112057557 | 11.17854912 | -3.774548623 | 0.007632226 | 0.991835066 | -2.603635607 |
| **A_55_P2182483** | Cd47 | -0.225818406 | -0.368897516 | -0.082739296 | 11.2589129 | -3.772178547 | 0.007655086 | 0.991835066 | -2.605166173 |
| **A_65_P05252** | Cdc25a | 0.195848119 | 0.071446601 | 0.320249637 | 9.981883563 | 3.762728284 | 0.007746984 | 0.991835066 | -2.611279458 |
| **A_52_P89064** | Dusp18 | -0.196730994 | -0.321782256 | -0.071679732 | 8.673383883 | -3.760051901 | 0.00777323 | 0.991835066 | -2.613013814 |
| **A_55_P2011341** | NA | -0.45681482 | -0.74802834 | -0.1656013 | 8.230625442 | -3.749192877 | 0.007880719 | 0.991835066 | -2.620064429 |
| **A_51_P189985** | Ppan | 0.321856119 | 0.116052742 | 0.527659495 | 10.55374153 | 3.737820194 | 0.00799504 | 0.991835066 | -2.627472194 |
| **A_55_P2151006** | C1qtnf7 | -0.220173229 | -0.360981084 | -0.079365373 | 9.103052078 | -3.737203604 | 0.00800129 | 0.991835066 | -2.62787451 |
| **A_65_P08227** | Snx5 | 0.16649221 | 0.059754301 | 0.273230118 | 8.615908277 | 3.728072594 | 0.00809447 | 0.991835066 | -2.633840705 |
| **A_51_P279552** | Cav2 | -0.216621492 | -0.355522864 | -0.07772012 | 9.954974346 | -3.727384016 | 0.008101545 | 0.991835066 | -2.634291255 |
| **A_55_P2144386** | Adgre1 | -0.234657748 | -0.385132051 | -0.084183445 | 8.372152704 | -3.727191349 | 0.008103526 | 0.991835066 | -2.634417336 |
| **A_51_P351860** | C1qb | -0.3595014 | -0.590077973 | -0.128924827 | 8.762033509 | -3.726443482 | 0.008111219 | 0.991835066 | -2.634906807 |
| **A_51_P139678** | Sprr1a | 0.478833698 | 0.170991971 | 0.786675425 | 8.258076752 | 3.717631611 | 0.008202476 | 0.991835066 | -2.640681983 |
| **A_55_P2023703** | Zfp386 | -0.160806627 | -0.264289627 | -0.057323628 | 8.628267593 | -3.714018549 | 0.008240217 | 0.991835066 | -2.643054142 |
| **A_51_P483473** | St3gal5 | 0.237680995 | 0.0845572 | 0.390804791 | 11.7212107 | 3.709888901 | 0.008283587 | 0.991835066 | -2.645768467 |
| **A_51_P160664** | Cox7b | -0.231568442 | -0.380909929 | -0.082226955 | 16.35911573 | -3.706022349 | 0.008324421 | 0.991835066 | -2.648312768 |
| **A_52_P539161** | Rdh11 | -0.203322238 | -0.335089664 | -0.071554813 | 9.690445565 | -3.687957884 | 0.008518132 | 0.991835066 | -2.660236913 |
| **A_51_P406583** | Myo1e | -0.265360432 | -0.437587895 | -0.09313297 | 9.265076894 | -3.68250118 | 0.008577611 | 0.991835066 | -2.663850889 |
| **A_55_P1964658** | NA | 0.211762457 | 0.074114319 | 0.349410595 | 12.45372071 | 3.676950289 | 0.008638583 | 0.991835066 | -2.667532996 |
| **A_66_P128537** | Isg15 | -0.242055329 | -0.399467904 | -0.084642753 | 8.022635159 | -3.675228581 | 0.00865759 | 0.991835066 | -2.668676245 |
| **A_55_P2005783** | Ifih1 | -0.203973949 | -0.336721409 | -0.071226488 | 8.415044821 | -3.672464557 | 0.0086882 | 0.991835066 | -2.670512783 |
| **A_55_P2031436** | Ly6e | -0.33607276 | -0.555702977 | -0.116442543 | 11.02160097 | -3.657213843 | 0.008859234 | 0.991835066 | -2.680671904 |
| **A_55_P2074414** | Gtf2h1 | 0.183461678 | 0.063466917 | 0.303456439 | 10.49443032 | 3.654199107 | 0.008893478 | 0.991835066 | -2.682685334 |
| **A_55_P2066578** | Ifi204 | -0.195585602 | -0.323877382 | -0.067293822 | 8.091551296 | -3.643738338 | 0.009013422 | 0.991835066 | -2.689684993 |
| **A_66_P112305** | Myo1f | -0.28696815 | -0.475344822 | -0.098591477 | 8.480890108 | -3.640958326 | 0.009045595 | 0.991835066 | -2.69154867 |

| | | | | | | | | | |
|---|---|---|---|---|---|---|---|---|---|
| **A_55_P2071596** | Ppara | -0.178868202 | -0.296328772 | -0.061407631 | 10.09158871 | -3.639570751 | 0.0090617 | 0.991835066 | -2.692479424 |
| **A_52_P489295** | Adamts1 | 0.293658154 | 0.100590401 | 0.486725907 | 11.10134002 | 3.635310026 | 0.009111348 | 0.991835066 | -2.695339696 |
| **A_55_P2057984** | Tbc1d19 | -0.157848289 | -0.261685842 | -0.054010735 | 9.408281494 | -3.633243986 | 0.009135529 | 0.991835066 | -2.696727887 |
| **A_55_P2053491** | Pdia6 | 0.208939038 | 0.071216238 | 0.346661838 | 14.39885487 | 3.625958927 | 0.009221355 | 0.991835066 | -2.70162922 |
| **A_55_P2086433** | Oasl1 | -0.274891308 | -0.456150832 | -0.093631784 | 8.248070665 | -3.624676934 | 0.009236549 | 0.991835066 | -2.702492773 |
| **A_55_P1990196** | NA | -0.233027948 | -0.386763748 | -0.079292149 | 12.45048767 | -3.622781425 | 0.009259065 | 0.991835066 | -2.703770162 |
| **A_55_P2192662** | Lepr | -0.18504919 | -0.307169187 | -0.062929194 | 7.928397556 | -3.621675524 | 0.009272229 | 0.991835066 | -2.704515745 |
| **A_55_P2317665** | E2f1 | 0.238354525 | 0.080895276 | 0.395813773 | 10.16398133 | 3.617964969 | 0.009316547 | 0.991835066 | -2.707019042 |
| **A_51_P215077** | Mgst3 | -0.444881504 | -0.73878525 | -0.150977757 | 12.21009651 | -3.617831704 | 0.009318143 | 0.991835066 | -2.707108996 |
| **A_55_P2145227** | Slmap | -0.183831303 | -0.305345136 | -0.06231747 | 13.18481654 | -3.615787309 | 0.009342663 | 0.991835066 | -2.708489396 |
| **A_52_P663413** | P4ha2 | 0.192962368 | 0.065383724 | 0.320541013 | 11.44473366 | 3.61496227 | 0.009352579 | 0.991835066 | -2.709046696 |
| **A_55_P2072228** | Cdk9 | 0.228527112 | 0.077294811 | 0.379759412 | 11.75675017 | 3.61162212 | 0.00939284 | 0.991835066 | -2.711304227 |
| **A_55_P1984103** | Gfra3 | -0.167248463 | -0.278164958 | -0.056331969 | 8.133167359 | -3.60391992 | 0.0094864 | 0.991835066 | -2.716518016 |
| **A_52_P225898** | Kcnj8 | -0.176781575 | -0.294022949 | -0.059540201 | 8.997014443 | -3.603837757 | 0.009487404 | 0.991835066 | -2.716573694 |
| **A_51_P235687** | Alox5ap | -0.215550273 | -0.35879786 | -0.072302685 | 8.706486284 | -3.596419905 | 0.009578483 | 0.991835066 | -2.721605736 |
| **A_55_P2076057** | Hmga1 | 0.226090618 | 0.075829021 | 0.376352215 | 8.480868808 | 3.596198646 | 0.009581214 | 0.991835066 | -2.721755992 |
| **A_55_P2070079** | Lgals9 | -0.282268052 | -0.470561629 | -0.093974476 | 9.074733063 | -3.582905504 | 0.009746877 | 0.991835066 | -2.730800285 |
| **A_55_P2002757** | Blnk | -0.24437716 | -0.407454234 | -0.081300086 | 8.065507804 | -3.581597975 | 0.00976334 | 0.991835066 | -2.731691702 |
| **A_51_P424221** | Irf2bp2 | -0.20267582 | -0.337946966 | -0.067404673 | 10.80309489 | -3.581013015 | 0.009770715 | 0.991835066 | -2.732090607 |
| **A_55_P2018934** | Adgre1 | -0.204977801 | -0.341919464 | -0.068036138 | 8.088249067 | -3.577505953 | 0.009815057 | 0.991835066 | -2.73448356 |
| **A_55_P1974636** | Gm16430 | 0.168914983 | 0.056026123 | 0.281803843 | 8.184695743 | 3.576236385 | 0.009831164 | 0.991835066 | -2.735350392 |
| **A_52_P420712** | Pja2 | -0.184625957 | -0.308263132 | -0.060988781 | 9.732729949 | -3.569051507 | 0.009922858 | 0.991835066 | -2.74026182 |
| **A_55_P2060487** | Zfp593 | 0.26671115 | 0.087736726 | 0.445685573 | 9.398758838 | 3.561716365 | 0.010017432 | 0.991835066 | -2.74528607 |
| **A_52_P392674** | Acot9 | -0.17477761 | -0.292167811 | -0.057387408 | 9.952814647 | -3.558468129 | 0.010059626 | 0.991835066 | -2.74751423 |
| **A_55_P2134581** | Papola | 0.215963258 | 0.070762949 | 0.361163567 | 9.804172532 | 3.554851491 | 0.010106833 | 0.991835066 | -2.749997455 |
| **A_55_P2051334** | H2-K2 | -0.225238533 | -0.376716979 | -0.073760087 | 9.964932615 | -3.55386546 | 0.010119745 | 0.991835066 | -2.750674907 |
| **A_51_P387845** | Fam120a | -0.196822276 | -0.329257831 | -0.064386721 | 10.34725219 | -3.552047675 | 0.010143596 | 0.991835066 | -2.751924297 |
| **A_52_P484081** | Ube2v1 | 0.207671567 | 0.067741737 | 0.347601397 | 9.543296832 | 3.547119887 | 0.010208563 | 0.991835066 | -2.755314393 |
| **A_51_P202633** | Ebi3 | -0.199198037 | -0.333446072 | -0.064950001 | 8.308233933 | -3.54638799 | 0.01021825 | 0.991835066 | -2.755818299 |
| **A_55_P2065113** | Gm11559 | -0.161755012 | -0.270945939 | -0.052564084 | 8.84758614 | -3.540628179 | 0.01029484 | 0.991835066 | -2.759787436 |
| **A_55_P2132549** | Cd48 | -0.160231279 | -0.268416953 | -0.052045604 | 7.867281703 | -3.539864785 | 0.010305037 | 0.991835066 | -2.76031397 |
| **A_51_P181517** | Fcgr4 | -0.28702378 | -0.481290383 | -0.092757177 | 8.288657624 | -3.531253257 | 0.010420839 | 0.991835066 | -2.766261253 |
| **A_51_P342652** | Cd79b | -0.159580829 | -0.267902506 | -0.051259151 | 8.106596913 | -3.521068473 | 0.010559628 | 0.991835066 | -2.773313244 |
| **A_55_P2022049** | Klf15 | -0.23055258 | -0.387420675 | -0.073684485 | 9.517207445 | -3.512727786 | 0.010674786 | 0.991835066 | -2.779103059 |
| **A_52_P255224** | NA | 0.180644628 | 0.057604465 | 0.303684792 | 8.401216991 | 3.509031648 | 0.010726254 | 0.991835066 | -2.781673018 |
| **A_55_P2064771** | Ly6c1 | -0.621148325 | -1.04446469 | -0.19783196 | 11.10466756 | -3.507029286 | 0.010754249 | 0.991835066 | -2.783066364 |
| **A_51_P172502** | Cxcl12 | -0.262051742 | -0.440859617 | -0.083243868 | 11.93989519 | -3.502753239 | 0.010814299 | 0.991835066 | -2.786044409 |
| **A_55_P2428967** | Lrrc2 | -0.1689151 | -0.284248151 | -0.053582049 | 9.070186523 | -3.500449567 | 0.010846801 | 0.991835066 | -2.787650237 |
| **A_55_P1958349** | Nipsnap3b | -0.156936735 | -0.264421936 | -0.049451533 | 8.754456023 | -3.489675684 | 0.011000225 | 0.991835066 | -2.795173825 |
| **A_55_P2180415** | Cd72 | -0.205244862 | -0.346355755 | -0.064133969 | 7.96175206 | -3.476329117 | 0.011193564 | 0.991835066 | -2.804524578 |
| **A_65_P01319** | Pde4b | -0.337082987 | -0.568908797 | -0.105257177 | 9.784846081 | -3.475235019 | 0.011209576 | 0.991835066 | -2.805292619 |
| **A_51_P185660** | Ccl9 | -0.149741309 | -0.252768378 | -0.046714239 | 8.380459198 | -3.473756799 | 0.01123125 | 0.991835066 | -2.806330668 |
| **A_51_P261379** | Spats2 | 0.225599192 | 0.070253417 | 0.380944968 | 9.55520933 | 3.47094095 | 0.011272661 | 0.991835066 | -2.80830919 |
| **A_55_P2120807** | Dld | -0.241804591 | -0.408609107 | -0.075000075 | 11.92056769 | -3.46470187 | 0.011365008 | 0.991835066 | -2.812698379 |
| **A_55_P1961761** | Dcc | 0.765640722 | 0.236262003 | 1.295019441 | 12.15265528 | 3.456747663 | 0.011483936 | 0.991835066 | -2.818304898 |
| **A_52_P408757** | Fcgr3 | -0.157785531 | -0.267118292 | -0.048452771 | 7.989859616 | -3.449260503 | 0.011597116 | 0.991835066 | -2.823593218 |
| **A_55_P2018557** | NA | -0.155531173 | -0.263613607 | -0.047448739 | 11.94791796 | -3.439311119 | 0.011749394 | 0.991835066 | -2.830637157 |
| **A_55_P2081105** | NA | -0.191306863 | -0.324439697 | -0.058174029 | 8.176537198 | -3.434428709 | 0.011824913 | 0.991835066 | -2.834100682 |

| | | | | | | | | | |
|---|---|---|---|---|---|---|---|---|---|
| **A_51_P241995** | Col5a3 | -0.261653376 | -0.443794593 | -0.079512159 | 9.002983619 | -3.433422429 | 0.011840543 | 0.991835066 | -2.834815088 |
| **A_55_P1955479** | NA | -0.245144099 | -0.416303165 | -0.073985032 | 10.44101467 | -3.423187328 | 0.0120008 | 0.991835066 | -2.842092428 |
| **A_55_P2148001** | 6820431F2 | 0.312037917 | 0.09408955 | 0.529986285 | 10.13645613 | 3.421865081 | 0.012021674 | 0.991835066 | -2.843034023 |
| **A_55_P2015599** | NA | 0.156813716 | 0.047241591 | 0.266385842 | 11.87629111 | 3.42052756 | 0.01204283 | 0.991835066 | -2.843986834 |
| **A_55_P2056325** | Anxa3 | -0.150242211 | -0.25525088 | -0.045233542 | 8.932746272 | -3.419605005 | 0.012057445 | 0.991835066 | -2.844644233 |
| **A_51_P336833** | Fabp4 | -0.22846365 | -0.3883653 | -0.068562001 | 15.17288102 | -3.414863128 | 0.012132873 | 0.991835066 | -2.848025778 |
| **A_55_P1985788** | Usp2 | -0.161942211 | -0.275365944 | -0.048518478 | 12.16281322 | -3.412441857 | 0.012171584 | 0.991835066 | -2.849754094 |
| **A_51_P461040** | Crct1 | 0.346926386 | 0.103866719 | 0.589986053 | 7.979249938 | 3.411407232 | 0.012188167 | 0.991835066 | -2.850492955 |
| **A_66_P135165** | Nfib | 0.178306249 | 0.053016562 | 0.303595936 | 9.184799451 | 3.401420866 | 0.012349489 | 0.991835066 | -2.857635023 |
| **A_55_P2027157** | NA | -0.306288426 | -0.521536677 | -0.091040175 | 9.773551142 | -3.400948751 | 0.012357172 | 0.991835066 | -2.857973141 |
| **A_55_P2103698** | Isg15 | -0.819794737 | -1.396181859 | -0.243407616 | 10.00798608 | -3.399382152 | 0.012382706 | 0.991835066 | -2.859095405 |
| **A_51_P400543** | Aif1 | -0.256029047 | -0.436468221 | -0.075589873 | 8.946900115 | -3.391310404 | 0.012515176 | 0.991835066 | -2.864885161 |
| **A_55_P2413458** | Tfcp2l1 | 0.153668497 | 0.045226024 | 0.262110971 | 7.991898171 | 3.386839069 | 0.012589218 | 0.991835066 | -2.86809772 |
| **A_55_P2079561** | Lilra6 | -0.165008795 | -0.281660794 | -0.048356796 | 7.953815859 | -3.380835497 | 0.012689381 | 0.991835066 | -2.872417136 |
| **A_51_P179258** | Kif26b | 0.178029567 | 0.051711282 | 0.304347852 | 9.902932557 | 3.368488322 | 0.012898102 | 0.991835066 | -2.881322139 |
| **A_55_P2023707** | Camta1 | -0.184702151 | -0.315855877 | -0.053548425 | 10.97997481 | -3.365894137 | 0.012942426 | 0.991835066 | -2.883196794 |
| **A_55_P2150058** | Zfp593 | 0.206466269 | 0.059558915 | 0.353373622 | 9.313765804 | 3.359036377 | 0.013060394 | 0.991835066 | -2.888158621 |
| **A_55_P2180491** | Rrbp1 | 0.289300389 | 0.083378792 | 0.495221986 | 9.848054016 | 3.357811035 | 0.013081595 | 0.991835066 | -2.889046139 |
| **A_55_P2156731** | H2-Eb1 | -0.169957393 | -0.290982927 | -0.048931858 | 8.596064572 | -3.356388182 | 0.01310626 | 0.991835066 | -2.890077072 |
| **A_55_P2037608** | Acp5 | -0.404259657 | -0.692520673 | -0.115998641 | 9.581363561 | -3.351842653 | 0.013185393 | 0.991835066 | -2.893373127 |
| **A_51_P436386** | Qrich1 | -0.163622834 | -0.280308058 | -0.04693761 | 10.93752034 | -3.351484219 | 0.013191655 | 0.991835066 | -2.893633202 |
| **A_65_P11603** | Ldb3 | -0.232960221 | -0.399586072 | -0.066334371 | 13.85179575 | -3.341554307 | 0.01336642 | 0.991835066 | -2.900847897 |
| **A_55_P2159890** | Zbtb7a | -0.223034993 | -0.382632201 | -0.063437785 | 9.807153938 | -3.340079857 | 0.013392583 | 0.991835066 | -2.901920771 |
| **A_51_P254425** | Ahrr | -0.211074216 | -0.362131264 | -0.060017169 | 8.892025598 | -3.339668237 | 0.013399897 | 0.991835066 | -2.902220357 |
| **A_55_P1973560** | NA | -0.178155021 | -0.3056841 | -0.050625941 | 12.36908612 | -3.338858192 | 0.013414303 | 0.991835066 | -2.902810021 |
| **A_66_P119968** | Clec4a1 | -0.283546174 | -0.486858822 | -0.080233525 | 8.58546208 | -3.33325489 | 0.013514412 | 0.991835066 | -2.906892288 |
| **A_55_P2058117** | Slc35d1 | 0.215547421 | 0.060682071 | 0.370412771 | 8.860966679 | 3.326577955 | 0.013634758 | 0.991835066 | -2.911764521 |
| **A_51_P100828** | Prdx3 | -0.177055389 | -0.304275017 | -0.049835761 | 12.18647349 | -3.326321018 | 0.013639412 | 0.991835066 | -2.920455185 |
| **A_52_P186937** | Cmpk2 | -0.17737421 | -0.304935118 | -0.049813302 | 8.335464444 | -3.323395319 | 0.01369253 | 0.991835066 | -2.9140899 |
| **A_52_P114722** | Ptpn6 | -0.175910699 | -0.302507942 | -0.049313457 | 8.207607546 | -3.321063217 | 0.013735031 | 0.991835066 | -2.915795059 |
| **A_66_P122087** | Zfp622 | 0.209722796 | 0.058297304 | 0.361148288 | 13.11203759 | 3.310211761 | 0.013934676 | 0.991835066 | -2.923742848 |
| **A_51_P359570** | Ifit3 | -0.223081271 | -0.384294893 | -0.061867648 | 8.015718886 | -3.307276498 | 0.013989216 | 0.991835066 | -2.925896515 |
| **A_55_P1994275** | Gar1 | 0.212674426 | 0.058888617 | 0.366460234 | 12.25730548 | 3.305279128 | 0.014026461 | 0.991835066 | -2.927362962 |
| **A_55_P2081616** | Timeless | 0.22075356 | 0.061075897 | 0.380431222 | 10.84447496 | 3.304248341 | 0.014045724 | 0.991835066 | -2.928120051 |
| **A_55_P2094412** | Ssbp1 | 0.216110707 | 0.059303159 | 0.372918256 | 9.749352024 | 3.293960989 | 0.014239546 | 0.991835066 | -2.935686876 |
| **A_55_P1998697** | NA | -0.208293436 | -0.35950872 | -0.057078152 | 13.10404092 | -3.292221355 | 0.014272606 | 0.991835066 | -2.936968436 |
| **A_52_P219473** | Cdc6 | 0.195434565 | 0.053372531 | 0.337496598 | 8.59862434 | 3.288005096 | 0.014353078 | 0.991835066 | -2.94007686 |
| **A_55_P2053943** | Olfr1368 | -0.137806161 | -0.237992727 | -0.037619595 | 8.468502895 | -3.287519028 | 0.014362387 | 0.991835066 | -2.940435429 |
| **A_55_P2149951** | Prx | 0.17169374 | 0.046676021 | 0.296711459 | 8.876205968 | 3.28240391 | 0.014460742 | 0.991835066 | -2.944211513 |
| **A_52_P223333** | Mib1 | 0.141183645 | 0.038367316 | 0.243999974 | 8.847782809 | 3.281945978 | 0.014469582 | 0.991835066 | -2.944549809 |
| **A_51_P399614** | Zkscan14 | -0.252957845 | -0.437793684 | -0.068122005 | 8.614186835 | -3.270928876 | 0.014684043 | 0.991835066 | -2.952700599 |
| **A_55_P2165560** | Plekhg5 | 0.232134341 | 0.062425264 | 0.401843418 | 10.48341468 | 3.269214774 | 0.014717718 | 0.991835066 | -2.953970804 |
| **A_55_P1976504** | NA | -0.35472658 | -0.614736833 | -0.094716328 | 10.55986108 | -3.260711621 | 0.014886003 | 0.991835066 | -2.960280109 |
| **A_55_P2066230** | Hck | -0.248105402 | -0.430153993 | -0.066056811 | 8.870935952 | -3.25730203 | 0.014954064 | 0.991835066 | -2.962813842 |
| **A_55_P2026562** | Cdkn1b | -0.186840463 | -0.323965747 | -0.049715179 | 11.69007535 | -3.256585808 | 0.014968404 | 0.991835066 | -2.963346358 |
| **A_52_P99888** | Cxcl16 | -0.139070896 | -0.241507045 | -0.036634748 | 8.529917857 | -3.244831467 | 0.015205871 | 0.991835066 | -2.972099612 |
| **A_66_P115116** | Wwtr1 | -0.157911176 | -0.274263279 | -0.041559072 | 11.09621769 | -3.243752876 | 0.015227864 | 0.991835066 | -2.972904123 |
| **A_55_P2056342** | Mgst3 | -0.289170271 | -0.50247947 | -0.075861073 | 12.82966847 | -3.24006103 | 0.015303402 | 0.991835066 | -2.975659493 |

| | | | | | | | | | |
|---|---|---|---|---|---|---|---|---|---|
| **A_55_P2167416** | NA | 0.160540973 | 0.041951031 | 0.279130914 | 8.40988314 | 3.235542926 | 0.015396397 | 0.991835066 | -2.979035022 |
| **A_66_P116311** | Kif5b | -0.176761741 | -0.307424494 | -0.046098988 | 9.535176838 | -3.233296999 | 0.015442851 | 0.991835066 | -2.980714409 |
| **A_52_P222725** | Lims1 | -0.181703274 | -0.316039125 | -0.047367422 | 11.90698575 | -3.232808292 | 0.015452979 | 0.991835066 | -2.981079964 |
| **A_55_P2223851** | NA | 0.29891964 | 0.077682081 | 0.520157198 | 9.089846917 | 3.229272731 | 0.015526465 | 0.991835066 | -2.983725917 |
| **A_51_P437289** | Smco4 | 0.232059961 | 0.060121864 | 0.403998057 | 9.408260427 | 3.225798479 | 0.015599044 | 0.991835066 | -2.986328276 |
| **A_51_P237754** | H2-T23 | -0.239393096 | -0.416813918 | -0.061972274 | 9.629539866 | -3.224899362 | 0.015617887 | 0.991835066 | -2.987002121 |
| **A_55_P2351193** | Snora47 | -0.589621855 | -1.026619096 | -0.152624614 | 8.511774581 | -3.224809096 | 0.01561978 | 0.991835066 | -2.987069779 |
| **A_51_P224064** | Klhl38 | -0.380339933 | -0.662646766 | -0.0980331 | 8.748022568 | -3.220028146 | 0.015720398 | 0.991835066 | -2.990655502 |
| **A_51_P447545** | Igfbp1 | -0.596308582 | -1.039158229 | -0.153458936 | 8.470381162 | -3.218280473 | 0.015757354 | 0.991835066 | -2.991967332 |
| **A_55_P2175275** | Hmbox1 | -0.231440525 | -0.403408004 | -0.059473046 | 9.136477594 | -3.216638179 | 0.015792166 | 0.991835066 | -2.993200584 |
| **A_55_P2181738** | Lyz1 | -0.473813709 | -0.825909936 | -0.121717482 | 10.0918076 | -3.216291352 | 0.015799528 | 0.991835066 | -2.993461092 |
| **A_52_P108346** | Myc | 0.195255832 | 0.050077196 | 0.340434467 | 9.428541309 | 3.21447784 | 0.015838086 | 0.991835066 | -2.994823624 |
| **A_55_P1971229** | Slc25a36 | 0.14146243 | 0.036237733 | 0.246687128 | 9.084693005 | 3.213161554 | 0.015866135 | 0.991835066 | -2.995812965 |
| **A_51_P219789** | H2-Q2 | -0.22268643 | -0.388499932 | -0.056872927 | 9.827491173 | -3.209837087 | 0.015937216 | 0.991835066 | -2.998313133 |
| **A_55_P1997400** | Slc7a4 | 0.18506848 | 0.047081022 | 0.323055939 | 9.583287758 | 3.205545726 | 0.016029477 | 0.991835066 | -3.001543515 |
| **A_52_P10017** | Tmem64 | -0.148216072 | -0.25874412 | -0.037688023 | 9.121943385 | -3.205028754 | 0.016040631 | 0.991835066 | -3.001932907 |
| **A_52_P356119** | Hus1 | -0.139834684 | -0.244120882 | -0.035548486 | 8.886141426 | -3.204772487 | 0.016046163 | 0.991835066 | -3.002125949 |
| **A_52_P318073** | NA | -0.210847781 | -0.368252287 | -0.053443276 | 9.034813798 | -3.201555311 | 0.016115785 | 0.991835066 | -3.004550452 |
| **A_55_P2024555** | Plpp1 | -0.147680546 | -0.257968593 | -0.0373925 | 11.59590331 | -3.200397909 | 0.016140912 | 0.991835066 | -3.005423158 |
| **A_55_P2093770** | NA | -0.335088208 | -0.586641868 | -0.083534547 | 12.40306898 | -3.183740304 | 0.016507251 | 0.991835066 | -3.018011107 |
| **A_51_P200776** | Grsf1 | -0.18684086 | -0.327133044 | -0.046548676 | 11.11115091 | -3.183079706 | 0.016521962 | 0.991835066 | -3.018511381 |
| **A_55_P2062727** | Nasp | 0.200904577 | 0.049969589 | 0.351839564 | 10.81778905 | 3.181332326 | 0.016560944 | 0.991835066 | -3.019835076 |
| **A_55_P2027640** | NA | 0.203725355 | 0.050587229 | 0.35686348 | 17.11029838 | 3.179588287 | 0.016599949 | 0.991835066 | -3.021156807 |
| **A_65_P12993** | Kif1b | -0.19342729 | -0.338864253 | -0.047990327 | 9.825369964 | -3.178718598 | 0.016619437 | 0.991835066 | -3.021816118 |
| **A_65_P05238** | Knop1 | 0.181246587 | 0.04490375 | 0.317589424 | 9.789176063 | 3.177215285 | 0.016653181 | 0.991835066 | -3.022956113 |
| **A_55_P1999439** | Ino80e | 0.14759928 | 0.036561729 | 0.258636832 | 9.788469756 | 3.177045952 | 0.016656986 | 0.991835066 | -3.023084548 |
| **A_51_P235705** | Tsfm | -0.17015478 | -0.298544002 | -0.041765558 | 10.14309503 | -3.167559046 | 0.016871693 | 0.991835066 | -3.030288695 |
| **A_55_P2002712** | Mllt10 | 0.134655009 | 0.032964972 | 0.236345045 | 9.772926673 | 3.164850795 | 0.01693353 | 0.991835066 | -3.032348358 |
| **A_55_P2060996** | NA | -0.172781175 | -0.303364079 | -0.042198271 | 8.699511409 | -3.1624177 | 0.016989293 | 0.991835066 | -3.034199926 |
| **A_55_P1983523** | Cd300ld | -0.138124222 | -0.242516112 | -0.033732332 | 7.984751428 | -3.162366611 | 0.016990466 | 0.991835066 | -3.034238816 |
| **A_55_P1989504** | Tmem164 | -0.184330417 | -0.323866444 | -0.04479439 | 13.03199597 | -3.157328609 | 0.017106566 | 0.991835066 | -3.038076247 |
| **A_55_P2094925** | Srgn | -0.236390637 | -0.41535411 | -0.057427163 | 9.365433271 | -3.157003057 | 0.017114098 | 0.991835066 | -3.038324382 |
| **A_55_P1991718** | Hmmr | -0.229572253 | -0.403424749 | -0.055719757 | 10.5874203 | -3.156077068 | 0.01713554 | 0.991835066 | -3.039030275 |
| **A_51_P485240** | Adipor1 | -0.303103458 | -0.532751569 | -0.073455347 | 11.37385104 | -3.15454804 | 0.017171009 | 0.991835066 | -3.040196221 |
| **A_52_P591153** | Pld4 | -0.136479448 | -0.240117813 | -0.032841083 | 7.851444431 | -3.147428231 | 0.017337211 | 0.991835066 | -3.045631083 |
| **A_55_P2028491** | Hmgn2 | 0.175578292 | 0.042215503 | 0.30894108 | 10.60056643 | 3.146627142 | 0.017356019 | 0.991835066 | -3.046243179 |
| **A_51_P214251** | Rnls | -0.16995143 | -0.29921517 | -0.040687689 | 8.81165489 | -3.142369391 | 0.017456353 | 0.991835066 | -3.049498436 |
| **A_51_P465350** | Ly86 | -0.423374099 | -0.745496584 | -0.101251614 | 9.292201791 | -3.141321432 | 0.017481144 | 0.991835066 | -3.050300167 |
| **A_66_P124806** | Tlr4 | -0.135260726 | -0.23826144 | -0.032260012 | 8.66806763 | -3.138633531 | 0.017544902 | 0.991835066 | -3.052357451 |
| **A_52_P597860** | Wasf2 | 0.19319817 | 0.046035896 | 0.340360444 | 9.65379901 | 3.137730573 | 0.017566376 | 0.991835066 | -3.053048864 |
| **A_55_P1995318** | NA | -0.178032053 | -0.313651852 | -0.042412255 | 8.71372544 | -3.137503634 | 0.017571778 | 0.991835066 | -3.05322266 |
| **A_55_P2007991** | Tuba3b | -0.194908726 | -0.343741885 | -0.046075567 | 8.531337897 | -3.129973888 | 0.017752016 | 0.991835066 | -3.058994532 |
| **A_52_P91235** | Ccni | -0.162167434 | -0.286093967 | -0.038240901 | 13.5239981 | -3.127580371 | 0.017809725 | 0.991835066 | -3.060831463 |
| **A_55_P2139913** | Ssb | 0.171493893 | 0.040272221 | 0.302715565 | 11.93502749 | 3.123577065 | 0.017906696 | 0.991835066 | -3.063906212 |
| **A_55_P2139407** | Myl12b | -0.203879422 | -0.359918174 | -0.04784067 | 12.36577262 | -3.122842324 | 0.017924555 | 0.991835066 | -3.064470853 |
| **A_51_P175555** | Pelo | 0.173700115 | 0.04061953 | 0.3067807 | 12.84944387 | 3.119568566 | 0.018004362 | 0.991835066 | -3.066987916 |
| **A_51_P458384** | Slc38a2 | -0.249077083 | -0.440154508 | -0.057999659 | 13.34308489 | -3.115541386 | 0.018103058 | 0.991835066 | -3.070086971 |
| **A_55_P2070869** | Lcn2 | -0.195308927 | -0.345204261 | -0.045413593 | 8.424763991 | -3.114175698 | 0.01813666 | 0.991835066 | -3.071138594 |

| | | | | | | | | | |
|---|---|---|---|---|---|---|---|---|---|
| **A_51_P451151** | Ube2c | -0.182442155 | -0.322508943 | -0.042375367 | 12.56468868 | -3.11314361 | 0.018162098 | 0.991835066 | -3.071933564 |
| **A_51_P466288** | Frmd5 | -0.291861829 | -0.516282284 | -0.067441374 | 11.86399541 | -3.108307623 | 0.018281802 | 0.991835066 | -3.075661115 |
| **A_52_P507382** | Unc93b1 | -0.24152791 | -0.428013572 | -0.055042248 | 8.789858124 | -3.095501425 | 0.018602898 | 0.991835066 | -3.08555286 |
| **A_55_P2070005** | NA | -0.195954784 | -0.347338855 | -0.044570714 | 12.90763999 | -3.093747216 | 0.018647351 | 0.991835066 | -3.086910191 |
| **A_52_P654965** | Eif3j1 | 0.206176793 | 0.046889002 | 0.365464584 | 11.54566423 | 3.093616124 | 0.018650677 | 0.991835066 | -3.087011648 |
| **A_55_P2108201** | Gnptab | -0.245932422 | -0.435957559 | -0.055907286 | 10.29057723 | -3.093241775 | 0.01866018 | 0.991835066 | -3.087301385 |
| **A_52_P600318** | Scamp2 | 0.136342355 | 0.030891499 | 0.24179321 | 10.67879718 | 3.090222962 | 0.018737003 | 0.991835066 | -3.089638816 |
| **A_51_P418725** | Plekhf1 | -0.326992071 | -0.580224374 | -0.073759768 | 9.480348288 | -3.086222615 | 0.018839326 | 0.991835066 | -3.092738814 |
| **A_51_P476261** | Zfp120 | -0.136591571 | -0.242434652 | -0.03074849 | 8.867419589 | -3.084399041 | 0.018886169 | 0.991835066 | -3.094152934 |
| **A_55_P2170569** | Ggps1 | 0.221018713 | 0.049575213 | 0.392462212 | 11.25131762 | 3.081180593 | 0.018969149 | 0.991835066 | -3.096650219 |
| **A_55_P1955715** | Gm14326 | 0.217067661 | 0.048613624 | 0.385521698 | 9.981530944 | 3.079802273 | 0.019004804 | 0.991835066 | -3.097720277 |
| **A_52_P252258** | Mapk12 | -0.212568806 | -0.377940244 | -0.047197367 | 9.929579884 | -3.072190633 | 0.019203007 | 0.991835066 | -3.10363583 |
| **A_55_P1956819** | Gm4832 | -0.185541537 | -0.329893272 | -0.041189802 | 15.19702806 | -3.072049953 | 0.019206691 | 0.991835066 | -3.103745262 |
| **A_55_P1999314** | Dhx33 | 0.193117161 | 0.042757615 | 0.343476707 | 10.65458208 | 3.069721642 | 0.019267773 | 0.991835066 | -3.105556935 |
| **A_55_P2179314** | Macrod2 | -0.135376632 | -0.240838862 | -0.029914401 | 8.447538206 | -3.068003741 | 0.019312975 | 0.991835066 | -3.10689428 |
| **A_55_P2158011** | Knstrn | -0.237927689 | -0.423379379 | -0.052475998 | 9.37100482 | -3.066361271 | 0.019356298 | 0.991835066 | -3.108173407 |
| **A_52_P425839** | Retnlg | -0.248793041 | -0.442735281 | -0.054850801 | 8.086396383 | -3.066019788 | 0.019365318 | 0.991835066 | -3.10843941 |
| **A_51_P121891** | Rac2 | -0.229955294 | -0.40933409 | -0.050576499 | 8.39482691 | -3.063948361 | 0.019420132 | 0.991835066 | -3.110053432 |
| **A_51_P258372** | Igsf6 | -0.184675412 | -0.328788329 | -0.040562495 | 8.186224012 | -3.062776451 | 0.019451216 | 0.991835066 | -3.110966912 |
| **A_52_P93467** | Sdc4 | 0.206341401 | 0.045252289 | 0.367430512 | 8.28289389 | 3.06146515 | 0.019486061 | 0.991835066 | -3.111989342 |
| **A_51_P371485** | Abraxas2 | 0.156618971 | 0.034203187 | 0.279034755 | 9.521062368 | 3.057849309 | 0.01958249 | 0.991835066 | -3.114810259 |
| **A_55_P2053810** | NA | 0.176376248 | 0.038431538 | 0.314320958 | 8.200433389 | 3.055935492 | 0.019633735 | 0.991835066 | -3.116304297 |
| **A_55_P2179604** | Vash1 | 0.246147274 | 0.053431186 | 0.438863361 | 14.09819153 | 3.052714521 | 0.019720306 | 0.991835066 | -3.118820281 |
| **A_55_P2109554** | Serinc3 | -0.211115823 | -0.376460231 | -0.045771414 | 11.26938186 | -3.051689913 | 0.01974793 | 0.991835066 | -3.119621025 |
| **A_55_P2026445** | Tfam | -0.16804639 | -0.299820904 | -0.036271877 | 10.17100572 | -3.047943412 | 0.019849292 | 0.991835066 | -3.122550586 |
| **A_66_P118752** | Rabif | -0.153826682 | -0.274503521 | -0.033149844 | 8.986661334 | -3.046610136 | 0.019885498 | 0.991835066 | -3.123593751 |
| **A_55_P2401646** | Trdn | -0.189136224 | -0.337630159 | -0.040642288 | 11.5284681 | -3.044213638 | 0.019950754 | 0.991835066 | -3.125469601 |
| **A_55_P2179498** | Ppp1r8 | 0.155372514 | 0.033365501 | 0.277379527 | 9.751375818 | 3.043676777 | 0.019965405 | 0.991835066 | -3.125889969 |
| **A_55_P2173737** | Hspa4l | -0.151590145 | -0.270649732 | -0.032530558 | 10.77704339 | -3.04309652 | 0.019981252 | 0.991835066 | -3.126344377 |
| **A_66_P124011** | Spint2 | -0.20816549 | -0.371698475 | -0.044632504 | 10.29280239 | -3.042373278 | 0.020001023 | 0.991835066 | -3.126910843 |
| **A_51_P382369** | Zfp608 | 0.143905285 | 0.030586106 | 0.257224464 | 10.21884794 | 3.035166058 | 0.020199193 | 0.991835066 | -3.132560951 |
| **A_52_P75384** | B230219D2 | 0.244680074 | 0.051734392 | 0.437625757 | 12.17853436 | 3.030907423 | 0.020317274 | 0.991835066 | -3.135903934 |
| **A_55_P2026635** | Asnsd1 | -0.138837484 | -0.248377554 | -0.029297414 | 12.45087684 | -3.029303969 | 0.020361925 | 0.991835066 | -3.137163478 |
| **A_55_P2067308** | NA | 0.241488252 | 0.050800203 | 0.432176302 | 12.86181045 | 3.026785641 | 0.020432265 | 0.991835066 | -3.139142612 |
| **A_55_P2042953** | Acd | 0.155450726 | 0.03242191 | 0.278479542 | 10.85932343 | 3.019917254 | 0.02062543 | 0.991835066 | -3.144546245 |
| **A_51_P330133** | Pspc1 | 0.172471647 | 0.035970389 | 0.308972904 | 9.582543831 | 3.019883501 | 0.020626384 | 0.991835066 | -3.144572821 |
| **A_55_P2095909** | Lamtor3 | -0.158256961 | -0.283636734 | -0.032877188 | 8.625554633 | -3.016785787 | 0.020714145 | 0.991835066 | -3.147012718 |
| **A_55_P2030160** | Tcf7l2 | 0.150878183 | 0.031307225 | 0.270449141 | 9.038314463 | 3.015850923 | 0.020740709 | 0.991835066 | -3.147749398 |
| **A_51_P359272** | Alcam | 0.235847248 | 0.048578904 | 0.423115593 | 8.69966906 | 3.010062934 | 0.020905987 | 0.991835066 | -3.152313882 |
| **A_51_P142813** | Aspn | -0.295493084 | -0.530238309 | -0.060747858 | 9.64523929 | -3.008566979 | 0.020948933 | 0.991835066 | -3.153494591 |
| **A_55_P1961650** | Gm2066 | 0.156017639 | 0.032055864 | 0.279979415 | 8.055472859 | 3.008119228 | 0.020961805 | 0.991835066 | -3.153848065 |
| **A_52_P387009** | Egln3 | -0.21114444 | -0.379044739 | -0.04324414 | 11.02918637 | -3.005642414 | 0.021033164 | 0.991835066 | -3.155804018 |
| **A_52_P52618** | Csf2rb | -0.189835929 | -0.340972531 | -0.038699328 | 8.256847376 | -3.002049803 | 0.021137131 | 0.991835066 | -3.158643081 |
| **A_55_P1957459** | Lilrb4a | -0.249832076 | -0.448885309 | -0.050778844 | 8.419178983 | -2.999770953 | 0.021203364 | 0.991835066 | -3.160445143 |
| **A_52_P535484** | Gvin1 | -0.28310343 | -0.508732148 | -0.057474712 | 8.688736478 | -2.998885522 | 0.021229158 | 0.991835066 | -3.161145573 |
| **A_51_P268734** | Kptn | -0.164247032 | -0.295210841 | -0.033283222 | 11.39992103 | -2.997473575 | 0.021270359 | 0.991835066 | -3.162262799 |
| **A_55_P2053838** | Tnfaip3 | -0.190928934 | -0.343222363 | -0.038635505 | 8.691501521 | -2.996399496 | 0.021301759 | 0.991835066 | -3.163112921 |
| **A_66_P121459** | Cenpa | -0.249923277 | -0.449376257 | -0.050470297 | 10.91098303 | -2.994851614 | 0.021347096 | 0.991835066 | -3.164338416 |

| | | | | | | | | | |
|---|---|---|---|---|---|---|---|---|---|
| **A_55_P1965416** | Atp9a | 0.152019546 | 0.0306867 | 0.273352392 | 13.59575547 | 2.994540412 | 0.021356224 | 0.991835066 | -3.164584854 |
| **A_55_P2035167** | Pcdh7 | -0.177099259 | -0.31845133 | -0.035747188 | 9.276162835 | -2.994495753 | 0.021357534 | 0.991835066 | -3.16462022 |
| **A_55_P2075769** | Ube2n | 0.178852839 | 0.035969286 | 0.321736391 | 10.59426684 | 2.991732307 | 0.021438774 | 0.991835066 | -3.166809349 |
| **A_55_P1987439** | Pigt | 0.503810231 | 0.100564449 | 0.907056012 | 12.1019361 | 2.986113301 | 0.021604983 | 0.991835066 | -3.171264786 |
| **A_55_P2185605** | Cd48 | -0.157808516 | -0.284118452 | -0.03149858 | 8.099303616 | -2.986084222 | 0.021605846 | 0.991835066 | -3.171287858 |
| **A_65_P12530** | NA | -0.142003494 | -0.255690551 | -0.028316436 | 8.730185542 | -2.985362914 | 0.021627283 | 0.991835066 | -3.171860212 |
| **A_52_P349298** | Rprd1b | 0.231475525 | 0.046080426 | 0.416870624 | 8.823248954 | 2.984117779 | 0.02166434 | 0.991835066 | -3.172848437 |
| **A_51_P187901** | Nop56 | 0.238729121 | 0.047333127 | 0.430125114 | 10.07689954 | 2.981135305 | 0.021753379 | 0.991835066 | -3.17521666 |
| **A_55_P2067221** | Iqsec3 | -0.51913859 | -0.935383205 | -0.102893975 | 10.77371969 | -2.980875364 | 0.021761158 | 0.991835066 | -3.17542314 |
| **A_55_P1958160** | Sgce | -0.208599554 | -0.376064806 | -0.041134302 | 10.6651651 | -2.977130005 | 0.021873569 | 0.991835066 | -3.178399548 |
| **A_55_P2133993** | NA | -0.252934893 | -0.456017238 | -0.049852548 | 10.85473292 | -2.976772933 | 0.021884318 | 0.991835066 | -3.178683441 |
| **A_52_P52156** | Isoc2a | 0.211049232 | 0.041592971 | 0.380505494 | 8.521187932 | 2.976701511 | 0.021886469 | 0.991835066 | -3.178740228 |
| **A_55_P2183750** | Ncoa3 | 0.211243659 | 0.041503113 | 0.380984204 | 11.20781403 | 2.974453745 | 0.021954272 | 0.991835066 | -3.180527881 |
| **A_55_P2089488** | Coq10b | 0.163116359 | 0.032018318 | 0.2942144 | 8.921311145 | 2.973791046 | 0.021974305 | 0.991835066 | -3.181055098 |
| **A_52_P16563** | Lsm14b | 0.19499639 | 0.037993422 | 0.351999357 | 11.37006615 | 2.968437043 | 0.022136869 | 0.991835066 | -3.185317395 |
| **A_55_P1980611** | Trmt2b | -0.140092013 | -0.252961987 | -0.027222039 | 12.62923067 | -2.966498157 | 0.022196056 | 0.991835066 | -3.186862189 |
| **A_66_P114102** | Zfp36l2 | -0.175684928 | -0.31734992 | -0.034019936 | 11.82476837 | -2.964019772 | 0.022271957 | 0.991835066 | -3.188837795 |
| **A_55_P2169356** | Gm1966 | -0.212275055 | -0.383484238 | -0.041065873 | 8.271972845 | -2.963336993 | 0.022292916 | 0.991835066 | -3.189382252 |
| **A_51_P338485** | Aldh6a1 | -0.173441591 | -0.313389076 | -0.033494105 | 12.24836247 | -2.962083392 | 0.022331452 | 0.991835066 | -3.190382108 |
| **A_55_P2181191** | Btg1 | 0.260358891 | 0.050153463 | 0.47056432 | 8.313743459 | 2.960312902 | 0.022385998 | 0.991835066 | -3.1917947 |
| **A_52_P460703** | Rassf3 | -0.203584753 | -0.367991875 | -0.039177631 | 9.227418495 | -2.959605017 | 0.022407846 | 0.991835066 | -3.192359644 |
| **A_55_P2171116** | Lgals3 | -0.569256414 | -1.029142453 | -0.109370376 | 10.04467066 | -2.958467753 | 0.022442995 | 0.991835066 | -3.19326745 |
| **A_55_P2003309** | NA | 0.200705229 | 0.038461922 | 0.362948536 | 13.60518711 | 2.956657582 | 0.022499063 | 0.991835066 | -3.194712867 |
| **A_55_P1969431** | Etfrf1 | -0.176685287 | -0.319732774 | -0.033637799 | 9.613364277 | -2.952087896 | 0.022641267 | 0.991835066 | -3.198364325 |
| **A_55_P2091196** | Hrct1 | -0.148339017 | -0.268503417 | -0.028174616 | 8.816871064 | -2.950453029 | 0.022692375 | 0.991835066 | -3.199671579 |
| **A_51_P359237** | Tmem9 | 0.133566104 | 0.025228862 | 0.241903346 | 11.66064133 | 2.946643635 | 0.022811939 | 0.991835066 | -3.202719432 |
| **A_51_P143470** | Timm44 | -0.121517289 | -0.220101216 | -0.022933363 | 12.29424301 | -2.946056651 | 0.022830423 | 0.991835066 | -3.203189299 |
| **A_52_P590625** | Nlrp10 | -0.146583236 | -0.265506966 | -0.027659506 | 9.282427644 | -2.945946894 | 0.02283388 | 0.991835066 | -3.203277163 |
| **A_55_P2073489** | NA | -0.218387401 | -0.395654991 | -0.041119811 | 9.095585071 | -2.944471615 | 0.022880412 | 0.991835066 | -3.204458385 |
| **A_66_P113868** | Cdh3 | 0.134442325 | 0.025202068 | 0.243682583 | 9.549033427 | 2.941456512 | 0.022975826 | 0.991835066 | -3.2068737 |
| **A_66_P137495** | Btaf1 | 0.188400196 | 0.035234902 | 0.341565491 | 9.30669961 | 2.939883454 | 0.023025775 | 0.991835066 | -3.208134468 |
| **A_52_P99411** | Itm2b | -0.196817131 | -0.357026852 | -0.03660741 | 13.43995644 | -2.9361833 | 0.02314372 | 0.991835066 | -3.211101765 |
| **A_51_P369803** | Psmb9 | -0.142167142 | -0.257972116 | -0.026362169 | 8.050104728 | -2.934142178 | 0.023209057 | 0.991835066 | -3.212739649 |
| **A_52_P18081** | Pacc1 | -0.208793161 | -0.378903253 | -0.03868307 | 11.05690697 | -2.933562361 | 0.023227653 | 0.991835066 | -3.213205052 |
| **A_52_P620793** | Eif2a | -0.2026207 | -0.367854395 | -0.037387005 | 9.17151993 | -2.930854974 | 0.023314696 | 0.991835066 | -3.215378977 |
| **A_55_P1981829** | Rhox8 | 0.869575007 | 0.159585383 | 1.57956463 | 12.36924299 | 2.927279419 | 0.023430182 | 0.991835066 | -3.218251975 |
| **A_55_P1960117** | Dipk2a | 0.244638517 | 0.044856582 | 0.444420452 | 9.638892473 | 2.92669693 | 0.023449054 | 0.991835066 | -3.218720223 |
| **A_51_P351194** | Cnfn | 0.390334372 | 0.070803689 | 0.709865055 | 8.151459806 | 2.919667264 | 0.02367808 | 0.991835066 | -3.224375869 |
| **A_55_P2158946** | Lama2 | -0.315006403 | -0.57311219 | -0.056900616 | 11.82197616 | -2.916962002 | 0.023766851 | 0.991835066 | -3.226554657 |
| **A_55_P2092156** | NA | 0.138341992 | 0.024902444 | 0.251781539 | 13.83695679 | 2.914732281 | 0.023840283 | 0.991835066 | -3.228351409 |
| **A_52_P404005** | NA | -0.27176251 | -0.494830409 | -0.048694611 | 11.39942619 | -2.911800396 | 0.023937208 | 0.991835066 | -3.230715294 |
| **A_55_P2051414** | Axl | -0.211528983 | -0.385212361 | -0.037845606 | 10.99791169 | -2.910856249 | 0.02396851 | 0.991835066 | -3.231476848 |
| **A_66_P125071** | Ssb | 0.150004483 | 0.026797798 | 0.273211167 | 10.44214285 | 2.909906922 | 0.024000027 | 0.991835066 | -3.232242736 |
| **A_55_P1986596** | Cacna1h | 0.218846254 | 0.03898904 | 0.398703468 | 10.58951422 | 2.908174004 | 0.024057673 | 0.991835066 | -3.233641206 |
| **A_51_P106799** | Pparg | -0.158714637 | -0.289156167 | -0.028273107 | 9.29445515 | -2.908105974 | 0.024059939 | 0.991835066 | -3.245498273 |
| **A_65_P10450** | Tmpo | -0.196904914 | -0.35883971 | -0.034970118 | 9.308687778 | -2.906200065 | 0.024123516 | 0.991835066 | -3.235234816 |
| **A_55_P1961760** | NA | 0.710352818 | 0.125810719 | 1.294894916 | 12.1792205 | 2.904473678 | 0.024181259 | 0.991835066 | -3.236629125 |
| **A_51_P375146** | Cd36 | -0.25749935 | -0.469476776 | -0.045521925 | 12.24436533 | -2.903325019 | 0.024219759 | 0.991835066 | -3.237557121 |

| | | | | | | | | | |
|---|---|---|---|---|---|---|---|---|---|
| **A_55_P1970578** | Aamdc | -0.238266789 | -0.434444108 | -0.042089469 | 10.96641332 | -2.902845003 | 0.024235868 | 0.991835066 | -3.237944991 |
| **A_51_P508191** | Vps26c | -0.121821817 | -0.222190651 | -0.021452984 | 11.20921754 | -2.900917182 | 0.024300677 | 0.991835066 | -3.239503142 |
| **A_55_P2018181** | A730008H2 | 0.138093403 | 0.024284986 | 0.25190182 | 8.529315648 | 2.900064664 | 0.024329396 | 0.991835066 | -3.24019239 |
| **A_55_P2018116** | Pabpc4l | 0.130507685 | 0.022927405 | 0.238087964 | 8.316226042 | 2.899429479 | 0.024350816 | 0.991835066 | -3.240706009 |
| **A_55_P2433218** | Bloc1s6 | 0.167925717 | 0.029478111 | 0.306373324 | 10.8625386 | 2.898951117 | 0.024366962 | 0.991835066 | -3.241092865 |
| **A_55_P1965378** | Uqcrq | -0.228966228 | -0.417867926 | -0.040064529 | 16.55414992 | -2.896974677 | 0.02443379 | 0.991835066 | -3.242691653 |
| **A_52_P23225** | Gpc3 | 0.172106273 | 0.030088883 | 0.314123662 | 12.23314679 | 2.896438466 | 0.024451954 | 0.991835066 | -3.243125522 |
| **A_55_P2111855** | Gale | 0.292571908 | 0.051138975 | 0.534004842 | 9.870314794 | 2.896311344 | 0.024456263 | 0.991835066 | -3.243228389 |
| **A_55_P2010788** | Slc6a9 | 0.197597537 | 0.034536239 | 0.360658835 | 9.958176153 | 2.896274785 | 0.024457502 | 0.991835066 | -3.243257973 |
| **A_51_P443569** | Pwwp3a | 0.164692802 | 0.028710805 | 0.300674799 | 9.836588405 | 2.894691841 | 0.024511221 | 0.991835066 | -3.244539131 |
| **A_55_P2059134** | Smarca5 | 0.167887291 | 0.02916902 | 0.306605563 | 11.05542137 | 2.89263266 | 0.024581289 | 0.991835066 | -3.246206378 |
| **A_55_P2086659** | Rad54l | 0.130090491 | 0.022591022 | 0.237589959 | 8.522177731 | 2.892333504 | 0.024591487 | 0.991835066 | -3.246448655 |
| **A_55_P1973838** | Slc6a9 | 0.205002518 | 0.035367465 | 0.374637571 | 10.03994162 | 2.88836936 | 0.024727036 | 0.991835066 | -3.24966055 |
| **A_55_P2085546** | Dnaja4 | -0.24470869 | -0.447246354 | -0.042171027 | 11.46722613 | -2.887704364 | 0.024749852 | 0.991835066 | -3.250199619 |
| **A_51_P224593** | Arl8a | -0.147441263 | -0.269601893 | -0.025280634 | 10.49376332 | -2.884675158 | 0.02485407 | 0.991835066 | -3.252656164 |
| **A_52_P462125** | Smarca5-p | 0.219634018 | 0.037560044 | 0.401707993 | 10.45261452 | 2.883107708 | 0.024908179 | 0.991835066 | -3.253927912 |
| **A_55_P2207055** | NA | -0.193729492 | -0.354442615 | -0.033016369 | 8.867662539 | -2.881068156 | 0.024978774 | 0.991835066 | -3.255583332 |
| **A_55_P2363030** | Cmtm4 | 0.183217275 | 0.031180997 | 0.335253553 | 9.759339461 | 2.880237643 | 0.025007581 | 0.991835066 | -3.256257629 |
| **A_51_P369862** | Nceh1 | -0.248364812 | -0.454700295 | -0.042029329 | 11.21256705 | -2.876903398 | 0.025123589 | 0.991835066 | -3.258965911 |
| **A_52_P70854** | Tob2 | -0.347792991 | -0.63673246 | -0.058853522 | 9.845416917 | -2.876888881 | 0.025124095 | 0.991835066 | -3.258977706 |
| **A_55_P1966660** | LOC54734 | -0.152461386 | -0.279140006 | -0.025782767 | 8.271565293 | -2.876508582 | 0.025137364 | 0.991835066 | -3.259286731 |
| **A_52_P55190** | BC005537 | 0.19095284 | 0.032141679 | 0.349764001 | 9.433683449 | 2.873784738 | 0.025232613 | 0.991835066 | -3.261500805 |
| **A_55_P2065866** | Cygb | -0.23289887 | -0.426757607 | -0.039040132 | 12.28933133 | -2.871383055 | 0.025316916 | 0.991835066 | -3.263454064 |
| **A_55_P2019833** | NA | 0.558787383 | 0.093587462 | 1.023987303 | 11.85719903 | 2.870886955 | 0.025334367 | 0.991835066 | -3.263857659 |
| **A_51_P131358** | Selplg | -0.129868299 | -0.238116932 | -0.021619665 | 8.00159153 | -2.867410434 | 0.025457017 | 0.991835066 | -3.26668711 |
| **A_55_P1976664** | Gm16430 | 0.150959767 | 0.025108863 | 0.276810671 | 8.118066841 | 2.866909675 | 0.025474735 | 0.991835066 | -3.267094836 |
| **A_55_P1976694** | 11-Sep | 0.149884858 | 0.02490598 | 0.274863736 | 8.848228279 | 2.86635694 | 0.025494308 | 0.991835066 | -3.26754493 |
| **A_55_P2060454** | Nsd3 | 0.140479376 | 0.023253161 | 0.257705592 | 10.21940816 | 2.864157914 | 0.025572335 | 0.991835066 | -3.269336122 |
| **A_55_P2175381** | NA | 0.181975376 | 0.030041893 | 0.333908859 | 16.54960447 | 2.862650081 | 0.025625983 | 0.991835066 | -3.270564787 |
| **A_52_P553316** | Snrk | -0.214133177 | -0.393002712 | -0.035263642 | 11.33747311 | -2.861255991 | 0.025675691 | 0.991835066 | -3.271701112 |
| **A_55_P2066533** | NA | -0.144998325 | -0.26634015 | -0.0236565 | 15.65528314 | -2.856022311 | 0.025863218 | 0.991835066 | -3.275970043 |
| **A_55_P2121081** | Cul2 | -0.150803843 | -0.277135753 | -0.024471932 | 9.490558741 | -2.853044045 | 0.02597058 | 0.991835066 | -3.278401389 |
| **A_55_P2008879** | Cltc | -0.158374962 | -0.291095266 | -0.025654658 | 9.868832999 | -2.852057577 | 0.026006245 | 0.991835066 | -3.279207036 |
| **A_65_P03719** | Fuca2 | -0.26114957 | -0.480152538 | -0.042146602 | 9.101231465 | -2.850023536 | 0.026079948 | 0.991835066 | -3.280868755 |
| **A_55_P2086954** | NA | 0.811395746 | 0.130411709 | 1.492379782 | 12.12179057 | 2.847770108 | 0.02616186 | 0.991835066 | -3.282710522 |
| **A_55_P2107750** | Hectd1 | -0.160491765 | -0.29527388 | -0.025709651 | 9.261941868 | -2.845965428 | 0.026227656 | 0.991835066 | -3.284186138 |
| **A_52_P361993** | Zc2hc1a | -0.147312652 | -0.271060927 | -0.023564378 | 9.460883985 | -2.845181848 | 0.026256279 | 0.991835066 | -3.284827012 |
| **A_55_P1986158** | Klhdc4 | 0.151595349 | 0.024218059 | 0.27897264 | 11.22351094 | 2.844480776 | 0.026281916 | 0.991835066 | -3.285400493 |
| **A_55_P1952379** | Fkbp5 | 0.192696642 | 0.03058174 | 0.354811544 | 10.23791909 | 2.840928749 | 0.026412216 | 0.991835066 | -3.288307343 |
| **A_55_P2152926** | Mybbp1a | 0.207899001 | 0.032970423 | 0.38282758 | 12.53247184 | 2.840539084 | 0.026426552 | 0.991835066 | -3.28862636 |
| **A_51_P284608** | Cd74 | -0.121061606 | -0.222956904 | -0.019166309 | 7.934269661 | -2.839627825 | 0.02646011 | 0.991835066 | -3.289372505 |
| **A_52_P302041** | NA | -0.170566114 | -0.314140631 | -0.026991596 | 10.97532688 | -2.839386556 | 0.026469002 | 0.991835066 | -3.289570081 |
| **A_51_P112355** | Igtp | -0.219755857 | -0.404805624 | -0.034706091 | 9.271332765 | -2.838317994 | 0.026508425 | 0.991835066 | -3.290445248 |
| **A_55_P2028847** | NA | 0.649942408 | 0.102436639 | 1.197448176 | 12.29300705 | 2.837234997 | 0.026548443 | 0.991835066 | -3.291332432 |
| **A_66_P136813** | Smim41 | -0.164717772 | -0.303723743 | -0.025711802 | 9.475726772 | -2.832149273 | 0.026737225 | 0.991835066 | -3.295501266 |
| **A_55_P2187225** | NA | 0.244510059 | 0.037956324 | 0.451063794 | 9.952110582 | 2.829259673 | 0.02684512 | 0.991835066 | -3.297871845 |
| **A_55_P2048209** | Ganab | 0.184373652 | 0.028615722 | 0.340131581 | 11.93863305 | 2.829162048 | 0.026848774 | 0.991835066 | -3.297951959 |
| **A_55_P2039603** | Drosha | 0.136323513 | 0.021119364 | 0.251527661 | 11.83271266 | 2.828210845 | 0.026884397 | 0.991835066 | -3.298732631 |

| | | | | | | | | | |
|---|---|---|---|---|---|---|---|---|---|
| **A_55_P2021285** | Cpped1 | -0.128033437 | -0.236370619 | -0.019696255 | 12.95310697 | -2.824587299 | 0.02702056 | 0.991835066 | -3.301707939 |
| **A_51_P408946** | Ccne1 | 0.288632708 | 0.043825731 | 0.533439686 | 11.22992962 | 2.817934447 | 0.027272463 | 0.991835066 | -3.307176336 |
| **A_55_P2172233** | NA | 0.250657376 | 0.037979341 | 0.46333541 | 15.68868866 | 2.816871156 | 0.027312953 | 0.991835066 | -3.308051006 |
| **A_55_P1975090** | NA | -0.141187765 | -0.261030989 | -0.021344542 | 12.3038682 | -2.815741068 | 0.027356057 | 0.991835066 | -3.30898083 |
| **A_55_P2143499** | Pgbd5 | 0.149711012 | 0.022560555 | 0.27686147 | 8.693603637 | 2.814135155 | 0.027417433 | 0.991835066 | -3.310302523 |
| **A_55_P1997766** | Acp5 | -0.405935824 | -0.750831465 | -0.061040184 | 9.459737893 | -2.813058785 | 0.027458653 | 0.991835066 | -3.311188634 |
| **A_52_P440709** | Ryr2 | -0.373710708 | -0.691306232 | -0.056115184 | 12.23052574 | -2.812356045 | 0.0274856 | 0.991835066 | -3.311767261 |
| **A_55_P2102628** | Usp4 | 0.128857787 | 0.01912464 | 0.238590935 | 11.20831932 | 2.806609328 | 0.027707012 | 0.991835066 | -3.316502113 |
| **A_51_P372992** | Vsir | -0.230128925 | -0.426203946 | -0.034053905 | 8.768308842 | -2.805162813 | 0.027763041 | 0.991835066 | -3.317694793 |
| **A_55_P2076797** | Ddx5 | 0.323488279 | 0.047718553 | 0.599258005 | 10.57087268 | 2.803632552 | 0.027822445 | 0.991835066 | -3.3189569 |
| **A_51_P361448** | Scara5 | -0.155725438 | -0.288534527 | -0.022916349 | 8.174629478 | -2.802469509 | 0.027867683 | 0.991835066 | -3.319916397 |
| **A_51_P152826** | Golt1b | 0.172577949 | 0.025313253 | 0.319842644 | 10.33321625 | 2.8008883 | 0.027929311 | 0.991835066 | -3.321221235 |
| **A_51_P497402** | Tonsl | 0.183071824 | 0.026528051 | 0.339615597 | 9.314143528 | 2.795083862 | 0.02815678 | 0.991835066 | -3.326014683 |
| **A_55_P2245920** | Lrrc8a | 0.134894467 | 0.019527122 | 0.250261812 | 13.11350667 | 2.794604607 | 0.028175649 | 0.991835066 | -3.326410712 |
| **A_55_P2074291** | Fbxo6 | -0.169650131 | -0.314822552 | -0.024477711 | 11.5718972 | -2.793053252 | 0.02823682 | 0.991835066 | -3.327692922 |
| **A_51_P507280** | Patl1 | -0.125079955 | -0.232134146 | -0.018025764 | 9.674393505 | -2.792499943 | 0.028258671 | 0.991835066 | -3.328150333 |
| **A_55_P2152304** | Atp5h | -0.147959231 | -0.274730642 | -0.021187821 | 16.76850755 | -2.789522559 | 0.02837656 | 0.991835066 | -3.330612548 |
| **A_55_P2086423** | Recql4 | 0.113154862 | 0.016181763 | 0.210127961 | 8.44916155 | 2.788888013 | 0.028401752 | 0.991835066 | -3.331137489 |
| **A_52_P398925** | Stfa2l1 | -0.5000986 | -0.929009816 | -0.071187384 | 8.937293005 | -2.78674588 | 0.028486972 | 0.991835066 | -3.332910098 |
| **A_66_P140858** | NA | -0.295983875 | -0.549846811 | -0.042120939 | 12.1315289 | -2.786620867 | 0.028491953 | 0.991835066 | -3.333013568 |
| **A_55_P1967168** | Lrp4 | 0.203922578 | 0.02888245 | 0.378962705 | 8.630919181 | 2.784433325 | 0.028579274 | 0.991835066 | -3.334824574 |
| **A_55_P2424988** | C78228 | -0.12873542 | -0.239392807 | -0.018078033 | 7.849592404 | -2.780524807 | 0.028735997 | 0.991835066 | -3.338062277 |
| **A_52_P442676** | Cux1 | -0.139547375 | -0.259546711 | -0.019548039 | 12.10814583 | -2.779405815 | 0.028781034 | 0.991835066 | -3.338989678 |
| **A_55_P2182433** | Atg10 | -0.162084709 | -0.301494996 | -0.022674423 | 9.315484034 | -2.778794039 | 0.028805687 | 0.991835066 | -3.339496792 |
| **A_55_P2185273** | Umps | 0.243523902 | 0.034062596 | 0.452985208 | 11.33700637 | 2.778733646 | 0.028808122 | 0.991835066 | -3.339546857 |
| **A_52_P153327** | 2-Sep | 0.150705042 | 0.021029677 | 0.280380406 | 8.551153634 | 2.777662254 | 0.028851356 | 0.991835066 | -3.340435116 |
| **A_51_P279308** | Acbd5 | -0.144632103 | -0.269131241 | -0.020132964 | 9.307687144 | -2.776562709 | 0.028895796 | 0.991835066 | -3.341346911 |
| **A_55_P2012641** | NA | 0.133396834 | 0.01849363 | 0.248300037 | 16.85658276 | 2.774741417 | 0.028969567 | 0.991835066 | -3.342857645 |
| **A_51_P103780** | Sft2d2 | -0.122632404 | -0.228385385 | -0.016879422 | 9.987714235 | -2.771543846 | 0.029099566 | 0.991835066 | -3.355622257 |
| **A_55_P2145557** | NA | -0.146551815 | -0.273021444 | -0.020082186 | 12.06007902 | -2.769581109 | 0.029179667 | 0.991835066 | -3.347140974 |
| **A_55_P1967591** | NA | 0.738342965 | 0.101063126 | 1.375622805 | 11.8212113 | 2.769090146 | 0.02919974 | 0.991835066 | -3.347548724 |
| **A_55_P2046907** | Wdr83os | 0.133638711 | 0.018288425 | 0.248988997 | 11.72775983 | 2.768998601 | 0.029203484 | 0.991835066 | -3.347624758 |
| **A_51_P457528** | Ccnb2 | -0.186022064 | -0.34661391 | -0.025430219 | 12.79320419 | -2.768535536 | 0.029222432 | 0.991835066 | -3.348009382 |
| **A_51_P168695** | Mast4 | -0.165792267 | -0.309002271 | -0.022582264 | 9.977847189 | -2.766941953 | 0.02928774 | 0.991835066 | -3.349333284 |
| **A_55_P1952537** | Rrp7a | 0.132937307 | 0.018104063 | 0.247770551 | 11.41196176 | 2.766867576 | 0.029290792 | 0.991835066 | -3.349395084 |
| **A_55_P2149896** | Lrrc75a | 0.209879384 | 0.028494533 | 0.391264235 | 12.18687297 | 2.765526986 | 0.029345857 | 0.991835066 | -3.350509141 |
| **A_51_P167843** | Timm17a | -0.144364571 | -0.269358623 | -0.019370518 | 13.7415909 | -2.76045332 | 0.029555251 | 0.991835066 | -3.354728095 |
| **A_55_P2064321** | Lama5 | 0.187053683 | 0.025062522 | 0.349044844 | 12.03533926 | 2.759841096 | 0.029580625 | 0.991835066 | -3.355237464 |
| **A_52_P159256** | Lpp | 0.150544287 | 0.020156877 | 0.280931698 | 9.329757037 | 2.759546722 | 0.029592833 | 0.991835066 | -3.355482404 |
| **A_55_P2070766** | NA | 0.602217361 | 0.080056613 | 1.124378108 | 12.40336227 | 2.756501177 | 0.029719452 | 0.991835066 | -3.358017337 |
| **A_51_P342871** | S100a10 | -0.148704635 | -0.277773062 | -0.019636208 | 12.65217859 | -2.753680956 | 0.029837213 | 0.991835066 | -3.360366054 |
| **A_55_P2086258** | Pan3 | 0.222718794 | 0.029264806 | 0.416172782 | 10.41139095 | 2.751619074 | 0.029923621 | 0.991835066 | -3.362084026 |
| **A_52_P665202** | NA | 0.231165192 | 0.030329805 | 0.432000579 | 14.27406892 | 2.751004715 | 0.029949418 | 0.991835066 | -3.362596045 |
| **A_55_P2062642** | Magt1 | 0.127074837 | 0.016670172 | 0.237479502 | 10.20953549 | 2.750940962 | 0.029952096 | 0.991835066 | -3.362649181 |
| **A_51_P341789** | Sugp1 | 0.12657565 | 0.01659377 | 0.236557531 | 11.40397176 | 2.750667903 | 0.02996357 | 0.991835066 | -3.362876777 |
| **A_51_P466673** | Srsf7 | 0.12226019 | 0.015998069 | 0.228522312 | 9.798334078 | 2.74989251 | 0.029996179 | 0.991835066 | -3.363523134 |
| **A_55_P2014807** | Rpl4 | -0.149615011 | -0.279733366 | -0.019496657 | 15.31153375 | -2.748183578 | 0.030068179 | 0.991835066 | -3.364948016 |
| **A_55_P2021821** | Prmt1 | 0.24745703 | 0.032222245 | 0.462691816 | 13.23406347 | 2.747871798 | 0.030081335 | 0.991835066 | -3.365208024 |

| | | | | | | | | | |
|---|---|---|---|---|---|---|---|---|---|
| **A_55_P2111302** | Cp | -0.171109241 | -0.320145998 | -0.022072484 | 8.295227669 | -2.744032218 | 0.030243844 | 0.991835066 | -3.368411306 |
| **A_52_P358076** | Ppp2ca | -0.137052626 | -0.256435115 | -0.017670137 | 12.42465264 | -2.743821553 | 0.030252787 | 0.991835066 | -3.368587128 |
| **A_55_P2063166** | Ogfod1 | -0.14237201 | -0.26639276 | -0.01835126 | 12.71272804 | -2.74371751 | 0.030257205 | 0.991835066 | -3.368673966 |
| **A_55_P2076048** | Nuf2 | -0.222905495 | -0.417104591 | -0.028706399 | 9.940744665 | -2.743359372 | 0.030272417 | 0.991835066 | -3.368972891 |
| **A_55_P2030225** | NA | 0.118639406 | 0.015237106 | 0.222041706 | 15.81199676 | 2.742255378 | 0.030319361 | 0.991835066 | -3.369894485 |
| **A_55_P2051727** | Rnf8 | 0.14223439 | 0.018251847 | 0.266216933 | 11.2873346 | 2.741910079 | 0.03033406 | 0.991835066 | -3.370182774 |
| **A_52_P570652** | Flywch1 | 0.133946317 | 0.017158312 | 0.250734323 | 8.860466191 | 2.741205961 | 0.030364056 | 0.991835066 | -3.370770699 |
| **A_55_P2140076** | NA | 0.231156902 | 0.029603743 | 0.432710062 | 16.51473408 | 2.741109516 | 0.030368167 | 0.991835066 | -3.370851235 |
| **A_55_P2000107** | Atox1 | -0.128032662 | -0.239730355 | -0.016334968 | 14.53485728 | -2.739590821 | 0.030432981 | 0.991835066 | -3.37211961 |
| **A_66_P120507** | Plcd3 | -0.192394977 | -0.36037233 | -0.024417624 | 10.75441166 | -2.737487395 | 0.03052299 | 0.991835066 | -3.373876943 |
| **A_55_P2148744** | NA | -0.217751629 | -0.407892354 | -0.027610905 | 11.32125003 | -2.737129848 | 0.030538318 | 0.991835066 | -3.37417573 |
| **A_52_P655285** | Zfp462 | 0.135836507 | 0.017211286 | 0.254461728 | 9.272219471 | 2.736834925 | 0.030550967 | 0.991835066 | -3.3744222 |
| **A_51_P169693** | Bst2 | -0.327086943 | -0.612839644 | -0.041334242 | 10.36522047 | -2.735785202 | 0.030596035 | 0.991835066 | -3.375299577 |
| **A_55_P2052535** | Gm14393 | 0.159529721 | 0.020061856 | 0.298997586 | 8.857394159 | 2.733861982 | 0.030678787 | 0.991835066 | -3.376907492 |
| **A_55_P1998942** | Oas1a | -0.132265454 | -0.247944353 | -0.016586554 | 7.908034051 | -2.732759488 | 0.030726332 | 0.991835066 | -3.3778295 |
| **A_52_P580533** | Spata33 | -0.173764939 | -0.325817687 | -0.021712191 | 8.630706704 | -2.73134783 | 0.030787322 | 0.991835066 | -3.379010342 |
| **A_51_P358722** | Lancl3 | -0.169357122 | -0.317611321 | -0.021102922 | 8.862048568 | -2.730269964 | 0.030833977 | 0.991835066 | -3.379912182 |
| **A_55_P2068247** | NA | 0.481791267 | 0.059958161 | 0.903624373 | 11.42480499 | 2.729778417 | 0.030855278 | 0.991835066 | -3.380323516 |
| **A_55_P1971498** | Exoc1 | 0.209721743 | 0.026077711 | 0.393365775 | 11.38260911 | 2.729453953 | 0.030869347 | 0.991835066 | -3.380595053 |
| **A_52_P634822** | Nap1l4 | 0.187673931 | 0.023328062 | 0.3520198 | 8.877947043 | 2.729318951 | 0.030875202 | 0.991835066 | -3.380708038 |
| **A_55_P1954331** | Cavin4 | -0.317811268 | -0.596393175 | -0.039229361 | 11.38793555 | -2.726625656 | 0.030992271 | 0.991835066 | -3.382962701 |
| **A_55_P1968483** | Gtpbp6 | 0.12380223 | 0.015148987 | 0.232455473 | 10.07558 | 2.723296358 | 0.031137631 | 0.991835066 | -3.38575137 |
| **A_66_P129484** | Tmed9 | 0.159379789 | 0.019468468 | 0.299291109 | 8.815387458 | 2.72263563 | 0.031166564 | 0.991835066 | -3.386305014 |
| **A_55_P2151466** | Casc3 | 0.244880803 | 0.029862638 | 0.459898968 | 11.39428249 | 2.72200377 | 0.03119426 | 0.991835066 | -3.386834532 |
| **A_51_P149455** | Acadl | -0.173207026 | -0.325375248 | -0.021038803 | 13.59000572 | -2.720512147 | 0.031259744 | 0.991835066 | -3.388084809 |
| **A_55_P2067682** | Atp5l | -0.172358884 | -0.324069616 | -0.020648152 | 14.24923899 | -2.715354285 | 0.031487297 | 0.991835066 | -3.392410818 |
| **A_55_P1969181** | Paox | 0.115829967 | 0.013861274 | 0.21779866 | 10.56013052 | 2.714958613 | 0.031504825 | 0.991835066 | -3.392742849 |
| **A_51_P294555** | Ifitm6 | -0.188616831 | -0.354693239 | -0.022540423 | 7.990641322 | -2.714448663 | 0.031527431 | 0.991835066 | -3.393170813 |
| **A_55_P1956847** | Nolc1 | 0.20162667 | 0.025228213 | 0.378025128 | 10.87768533 | 2.839359245 | 0.031619032 | 0.991835066 | -3.454025493 |
| **A_55_P1985273** | Lmcd1 | -0.224965076 | -0.423344378 | -0.026585774 | 10.0134728 | -2.710365602 | 0.031709047 | 0.991835066 | -3.396598901 |
| **A_55_P1972991** | Pcsk6 | -0.204128297 | -0.384262031 | -0.023994564 | 10.20093403 | -2.708428 | 0.031795618 | 0.991835066 | -3.398226601 |
| **A_51_P339503** | Cct4 | -0.166736978 | -0.313982418 | -0.019491538 | 11.89794453 | -2.706444989 | 0.031884474 | 0.991835066 | -3.399893053 |
| **A_51_P478610** | Asb6 | 0.123186932 | 0.014378999 | 0.231994865 | 9.89248143 | 2.705909156 | 0.031908529 | 0.991835066 | -3.400343453 |
| **A_55_P2027022** | Prdm1 | 0.18360372 | 0.021403489 | 0.345803952 | 8.272643603 | 2.705447675 | 0.031929262 | 0.991835066 | -3.400731392 |
| **A_52_P315345** | Lats2 | -0.13653683 | -0.257175528 | -0.015898132 | 10.62902545 | -2.705031379 | 0.031947977 | 0.991835066 | -3.401081374 |
| **A_55_P2041397** | Ezh2 | 0.180216686 | 0.020857057 | 0.339576316 | 10.48809457 | 2.702874131 | 0.032045141 | 0.991835066 | -3.40289542 |
| **A_55_P2166985** | Dedd2 | -0.224828839 | -0.423664272 | -0.025993407 | 10.58472754 | -2.702510391 | 0.032061555 | 0.991835066 | -3.403201362 |
| **A_55_P2041509** | 1110004F1 | -0.123252515 | -0.23237426 | -0.014130771 | 11.49070696 | -2.699563969 | 0.032194839 | 0.991835066 | -3.405680364 |
| **A_51_P133612** | Cdt1 | 0.247852363 | 0.02837654 | 0.467328186 | 12.63191967 | 2.699078461 | 0.032216858 | 0.991835066 | -3.406088981 |
| **A_55_P2093196** | NA | 0.14657587 | 0.016753145 | 0.276398595 | 15.78827161 | 2.698490489 | 0.032243544 | 0.991835066 | -3.406583882 |
| **A_55_P1963579** | NA | -0.208633775 | -0.39343463 | -0.023832919 | 9.739976524 | -2.698297128 | 0.032252325 | 0.991835066 | -3.406746648 |
| **A_55_P2129316** | NA | -0.178125533 | -0.33595542 | -0.020295646 | 8.024046103 | -2.697404417 | 0.032292899 | 0.991835066 | -3.407498181 |
| **A_55_P2032538** | Pde3b | -0.152716879 | -0.28805361 | -0.017380149 | 10.05209844 | -2.696997185 | 0.032311426 | 0.991835066 | -3.407841052 |
| **A_65_P19250** | Ap2m1 | 0.197988151 | 0.022440865 | 0.373535437 | 9.042148235 | 2.695592338 | 0.032375423 | 0.991835066 | -3.409024067 |
| **A_55_P2124273** | Myo1f | -0.114470687 | -0.216071446 | -0.012869928 | 7.949837894 | -2.692814691 | 0.032502352 | 0.991835066 | -3.411364008 |
| **A_51_P193146** | Ms4a6c | -0.11163893 | -0.210763765 | -0.012514095 | 7.899904012 | -2.691797089 | 0.032548983 | 0.991835066 | -3.412221553 |
| **A_55_P2060642** | Spidr | 0.257728869 | 0.028816398 | 0.48664134 | 9.647449152 | 2.69093215 | 0.032588675 | 0.991835066 | -3.412950573 |
| **A_55_P2061620** | Chst1 | -0.286042635 | -0.54016972 | -0.031915549 | 10.38920701 | -2.690227122 | 0.032621065 | 0.991835066 | -3.413544896 |

| | | | | | | | | | |
|---|---|---|---|---|---|---|---|---|---|
| **A_55_P2139748** | Phf14 | 0.129853738 | 0.014418877 | 0.245288599 | 8.5205871 | 2.688602531 | 0.032695832 | 0.991835066 | -3.414914682 |
| **A_51_P206268** | Seh1l | -0.169932258 | -0.321069599 | -0.018794916 | 10.83412936 | -2.687281614 | 0.032756756 | 0.991835066 | -3.416028722 |
| **A_55_P2048406** | NA | -0.164739046 | -0.311502517 | -0.017975576 | 13.10845333 | -2.682796357 | 0.032964519 | 0.991835066 | -3.419813516 |
| **A_52_P556140** | Vapb | 0.25146978 | 0.027437503 | 0.475502057 | 12.52745703 | 2.682775558 | 0.032965485 | 0.991835066 | -3.419831073 |
| **A_55_P2070241** | Preb | 0.146728703 | 0.015982384 | 0.277475021 | 11.031928 | 2.682222124 | 0.032991218 | 0.991835066 | -3.420298293 |
| **A_55_P1971159** | NA | 0.25408655 | 0.027506105 | 0.480666996 | 9.656097737 | 2.680207283 | 0.033085078 | 0.991835066 | -3.421999657 |
| **A_55_P2019068** | Flii | 0.132917203 | 0.014309342 | 0.251525063 | 8.762574148 | 2.678408734 | 0.0331691 | 0.991835066 | -3.423518905 |
| **A_66_P102260** | Irs2 | -0.210632165 | -0.398619932 | -0.022644398 | 10.70351607 | -2.677960918 | 0.033190055 | 0.991835066 | -3.423897255 |
| **A_55_P2085127** | Ramac | -0.208690949 | -0.395249757 | -0.022132141 | 10.2732181 | -2.673603416 | 0.033394689 | 0.991835066 | -3.427580415 |
| **A_52_P425092** | NA | -0.310556529 | -0.588211342 | -0.032901716 | 8.282627851 | -2.673280866 | 0.033409889 | 0.991835066 | -3.427853165 |
| **A_55_P2089358** | Hspe1 | 0.135541993 | 0.014320406 | 0.256763581 | 13.92433264 | 2.672409659 | 0.03345098 | 0.991835066 | -3.428589938 |
| **A_51_P479052** | Tars | 0.198700011 | 0.020951653 | 0.37644837 | 11.51802015 | 2.671784497 | 0.033480499 | 0.991835066 | -3.429118704 |
| **A_52_P585233** | Ccpg1 | -0.190744983 | -0.361419877 | -0.020070088 | 9.601647666 | -2.671115153 | 0.033512135 | 0.991835066 | -3.429684905 |
| **A_52_P153848** | Prpf4b | -0.161978698 | -0.307012208 | -0.016945188 | 8.778838597 | -2.669307949 | 0.033597707 | 0.991835066 | -3.431213969 |
| **A_55_P2069012** | Pidd1 | 0.183288045 | 0.019114838 | 0.347461253 | 9.185331919 | 2.668338911 | 0.033643686 | 0.991835066 | -3.432034069 |
| **A_52_P663757** | Tnik | -0.169072872 | -0.320521742 | -0.017624003 | 9.46407552 | -2.66819169 | 0.033650677 | 0.991835066 | -3.432158675 |
| **A_51_P246924** | Tppp3 | -0.224231126 | -0.425125565 | -0.023336687 | 9.165740115 | -2.66770078 | 0.033674 | 0.991835066 | -3.432574199 |
| **A_55_P1960456** | Larp1 | 0.128672756 | 0.013383246 | 0.243962266 | 12.79134163 | 2.66750931 | 0.033683101 | 0.991835066 | -3.432736276 |
| **A_55_P2027701** | NA | -0.20964658 | -0.397629206 | -0.021663955 | 15.43855781 | -2.665503181 | 0.033778616 | 0.991835066 | -3.434434774 |
| **A_55_P1956063** | Pbld1 | -0.237379664 | -0.450437608 | -0.024321721 | 16.74843843 | -2.662900335 | 0.033902965 | 0.991835066 | -3.436639393 |
| **A_52_P476029** | Nol4l | 0.243712297 | 0.02485965 | 0.462564945 | 10.51031198 | 2.661550869 | 0.033967624 | 0.991835066 | -3.437782798 |
| **A_51_P223709** | Pask | 0.262596796 | 0.026755942 | 0.498437651 | 9.523104419 | 2.661212266 | 0.033983868 | 0.991835066 | -3.438069739 |
| **A_52_P218379** | Kif26a | -0.13405188 | -0.254451436 | -0.013652324 | 10.43924855 | -2.661075247 | 0.033990444 | 0.991835066 | -3.438185858 |
| **A_55_P2141786** | Pard6g | 0.126322137 | 0.012828433 | 0.239815841 | 11.0858527 | 2.660215536 | 0.034031733 | 0.991835066 | -3.438914497 |
| **A_51_P476851** | Fance | 0.194285778 | 0.019661411 | 0.368910144 | 11.50486509 | 2.659165073 | 0.034082255 | 0.991835066 | -3.439804957 |
| **A_51_P124254** | Col4a1 | -0.168370723 | -0.319786768 | -0.016954677 | 13.26796877 | -2.657686861 | 0.034153483 | 0.991835066 | -3.459457988 |
| **A_55_P2232988** | Scube2 | 0.169417463 | 0.017025411 | 0.321809516 | 11.06262108 | 2.657082187 | 0.034182664 | 0.991835066 | -3.441571076 |
| **A_55_P2015585** | Zcchc24 | -0.134346065 | -0.2553029 | -0.013389229 | 12.65545908 | -2.654627972 | 0.034301372 | 0.991835066 | -3.443652888 |
| **A_55_P2022773** | Glcci1 | 0.226955115 | 0.022616318 | 0.431293912 | 9.713654216 | 2.654595017 | 0.034302969 | 0.991835066 | -3.443680849 |
| **A_55_P2066017** | NA | -0.515926334 | -0.980608231 | -0.051244436 | 8.807234328 | -2.65363431 | 0.034349558 | 0.991835066 | -3.444496029 |
| **A_52_P175952** | Map3k7 | 0.11073904 | 0.010914109 | 0.210563972 | 8.852968803 | 2.651373225 | 0.03445947 | 0.991835066 | -3.446415152 |
| **A_55_P2113459** | Washc1 | -0.168705309 | -0.320848526 | -0.016562092 | 11.74457091 | -2.650240497 | 0.03451467 | 0.991835066 | -3.447376856 |
| **A_51_P191611** | Cat | -0.128317775 | -0.244126273 | -0.012509278 | 8.465573697 | -2.648228931 | 0.034612928 | 0.991835066 | -3.44908518 |
| **A_52_P574668** | Nt5e | -0.181358874 | -0.345065037 | -0.01765271 | 9.198696375 | -2.647786177 | 0.034634594 | 0.991835066 | -3.44946127 |
| **A_52_P313080** | Hnrnpab | 0.130351403 | 0.012656991 | 0.248045816 | 12.75220959 | 2.647091776 | 0.034668603 | 0.991835066 | -3.450051177 |
| **A_55_P2280821** | Dlc1 | -0.174226072 | -0.331581517 | -0.016870626 | 10.25646747 | -2.646308677 | 0.034706999 | 0.991835066 | -3.450716521 |
| **A_55_P1965076** | NA | -0.185009588 | -0.352106143 | -0.017913034 | 12.80154675 | -2.646280482 | 0.034708382 | 0.991835066 | -3.450740478 |
| **A_55_P2106593** | NA | -0.133524352 | -0.254219428 | -0.012829276 | 9.636646486 | -2.644113283 | 0.034814878 | 0.991835066 | -3.452582277 |
| **A_55_P1978511** | H2-Q7 | -0.23898469 | -0.455038914 | -0.022930466 | 9.826715602 | -2.643726047 | 0.034833942 | 0.991835066 | -3.452911444 |
| **A_52_P313102** | Yipf7 | -0.270627381 | -0.515317233 | -0.02593753 | 10.45235285 | -2.643412319 | 0.034849396 | 0.991835066 | -3.453178142 |
| **A_51_P353008** | Minpp1 | -0.132629646 | -0.252625576 | -0.012633717 | 10.12896982 | -2.641698372 | 0.034933949 | 0.991835066 | -3.454635418 |
| **A_51_P194273** | Sgcg | -0.460006781 | -0.876351551 | -0.043662012 | 11.26966678 | -2.640707221 | 0.034982943 | 0.991835066 | -3.455478338 |
| **A_55_P2082841** | NA | 0.647071439 | 0.060978404 | 1.233164475 | 11.97210753 | 2.638729076 | 0.035080942 | 0.991835066 | -3.457161079 |
| **A_55_P2008722** | Zfp984 | 0.209791653 | 0.019548544 | 0.400034761 | 10.14429871 | 2.63565404 | 0.035233852 | 0.991835066 | -3.459778057 |
| **A_55_P1953899** | Mccc1 | -0.157520002 | -0.300603881 | -0.014436123 | 12.44955482 | -2.631201662 | 0.035456493 | 0.991835066 | -3.46356968 |
| **A_51_P501018** | Nek2 | -0.200276591 | -0.382204726 | -0.018348457 | 8.817708925 | -2.631112738 | 0.035460954 | 0.991835066 | -3.463645438 |
| **A_52_P367294** | Fsd1l | 0.132254952 | 0.012028441 | 0.252481463 | 9.143485759 | 2.629183099 | 0.035557916 | 0.991835066 | -3.46528964 |
| **A_55_P1964559** | Smarca5-p | 0.17739866 | 0.01612295 | 0.338674371 | 9.8753502 | 2.628999512 | 0.035567155 | 0.991835066 | -3.465446098 |

| | | | | | | | | | |
|---|---|---|---|---|---|---|---|---|---|
| **A_55_P2122709** | Arhgef3 | -0.180987723 | -0.345555685 | -0.01641976 | 9.833021328 | -2.628530109 | 0.035590791 | 0.991835066 | -3.465846162 |
| **A_55_P2036377** | NA | 0.144365838 | 0.013074206 | 0.27565747 | 16.13957848 | 2.628067544 | 0.035614098 | 0.991835066 | -3.466240429 |
| **A_51_P267278** | Slc15a2 | -0.234699798 | -0.448410504 | -0.020989092 | 7.911942278 | -2.624796078 | 0.035779393 | 0.991835066 | -3.469029755 |
| **A_66_P139703** | NA | 0.263600405 | 0.023559471 | 0.503641338 | 8.985728764 | 2.624640936 | 0.035787251 | 0.991835066 | -3.469162072 |
| **A_51_P379710** | Rpn1 | 0.13927249 | 0.01239776 | 0.266147221 | 12.97364618 | 2.623610377 | 0.0358395 | 0.991835066 | -3.470041098 |
| **A_55_P2167025** | Nectin2 | 0.242258366 | 0.021483287 | 0.463033445 | 10.66826281 | 2.622635085 | 0.03588902 | 0.991835066 | -3.470873127 |
| **A_55_P2063146** | NA | -0.186176431 | -0.355845569 | -0.016507294 | 11.59332273 | -2.622593509 | 0.035891133 | 0.991835066 | -3.470908598 |
| **A_55_P2045268** | Tmem59 | -0.191480035 | -0.366146173 | -0.016813897 | 12.23368128 | -2.620136475 | 0.036016214 | 0.991835066 | -3.473005341 |
| **A_55_P2128626** | Gm4349 | 0.134839231 | 0.011834208 | 0.257844253 | 8.397322495 | 2.620007618 | 0.036022786 | 0.991835066 | -3.473115327 |
| **A_51_P101858** | Dlg1 | 0.128851054 | 0.011290381 | 0.246411728 | 8.412557224 | 2.619600369 | 0.036043566 | 0.991835066 | -3.473462951 |
| **A_55_P2014987** | Castor2 | -0.202221453 | -0.386725261 | -0.017717646 | 11.47438916 | -2.61957614 | 0.036044803 | 0.991835066 | -3.473483634 |
| **A_55_P2075631** | Slc20a2 | -0.13823404 | -0.26441074 | -0.012057339 | 11.97087228 | -2.618454106 | 0.036102122 | 0.991835066 | -3.474441521 |
| **A_55_P2016346** | Eif3d | 0.107821167 | 0.009342196 | 0.206300138 | 13.86183547 | 2.616794745 | 0.036187066 | 0.991835066 | -3.475858462 |
| **A_55_P1962384** | Churc1 | -0.165857541 | -0.317493911 | -0.014221172 | 12.60278996 | -2.614213042 | 0.036319641 | 0.991835066 | -3.478063793 |
| **A_52_P270164** | E2f6 | 0.164716532 | 0.01402723 | 0.315405834 | 11.56400231 | 2.612545722 | 0.036405531 | 0.991835066 | -3.479488558 |
| **A_55_P2036673** | Ddx52 | 0.144719815 | 0.0123089 | 0.277130729 | 10.39231975 | 2.612241623 | 0.036421219 | 0.991835066 | -3.479748461 |
| **A_55_P2019520** | Cma2 | -0.175772242 | -0.33660627 | -0.014938214 | 8.402620194 | -2.612049998 | 0.036431109 | 0.991835066 | -3.479912244 |
| **A_66_P105689** | Trim34a | -0.127502171 | -0.244175434 | -0.010828907 | 8.447176521 | -2.611892901 | 0.036439218 | 0.991835066 | -3.480046518 |
| **A_55_P2148402** | Cplx2 | 0.231945091 | 0.019618674 | 0.444271509 | 13.11344785 | 2.610900285 | 0.036490503 | 0.991835066 | -3.480895017 |
| **A_51_P116665** | 2610005L0 | 0.257242632 | 0.021491545 | 0.492993718 | 9.724880359 | 2.607944672 | 0.036643654 | 0.991835066 | -3.483422349 |
| **A_55_P2162344** | Lrsam1 | -0.142750763 | -0.273577168 | -0.011924358 | 9.981187946 | -2.607907369 | 0.036645591 | 0.991835066 | -3.483454254 |
| **A_52_P286928** | NA | 0.219128412 | 0.018258841 | 0.419997984 | 9.322037642 | 2.607316007 | 0.036676316 | 0.991835066 | -3.483960079 |
| **A_51_P317673** | Zfp764 | -0.147002517 | -0.281765013 | -0.012240021 | 10.91460275 | -2.607143018 | 0.036685309 | 0.991835066 | -3.484108056 |
| **A_52_P68702** | Frmd4b | 0.110910475 | 0.009101242 | 0.212719707 | 8.289280313 | 2.603721522 | 0.036863654 | 0.991835066 | -3.487035729 |
| **A_55_P2028961** | Idi1 | 0.265988704 | 0.021725605 | 0.510251803 | 11.39423322 | 2.602642171 | 0.036920103 | 0.991835066 | -3.487959646 |
| **A_51_P492595** | Fbxo42 | 0.137883527 | 0.011209653 | 0.264557401 | 9.355947134 | 2.601563689 | 0.036976597 | 0.991835066 | -3.488882987 |
| **A_51_P199104** | Casq1 | -0.212417377 | -0.40761547 | -0.017219283 | 9.647833802 | -2.600899683 | 0.037011424 | 0.991835066 | -3.489451557 |
| **A_51_P443359** | Trappc2l | -0.138892705 | -0.266609269 | -0.011176141 | 12.06457798 | -2.59920983 | 0.037100212 | 0.991835066 | -3.490898817 |
| **A_55_P2148519** | Teddm2 | -0.177168051 | -0.340140545 | -0.014195558 | 8.412358124 | -2.598245774 | 0.037150966 | 0.991835066 | -3.491724657 |
| **A_55_P2172146** | Pds5a | 0.151270515 | 0.012069788 | 0.290471242 | 9.004809105 | 2.597298795 | 0.037200891 | 0.991835066 | -3.492535998 |
| **A_55_P2037408** | Lrrc57 | -0.108491742 | -0.208347165 | -0.00863632 | 9.178705731 | -2.596774056 | 0.037228585 | 0.991835066 | -3.492985633 |
| **A_52_P571371** | Tmem144 | -0.123303575 | -0.236797012 | -0.009810137 | 9.307422556 | -2.596653786 | 0.037234936 | 0.991835066 | -3.493088694 |
| **A_55_P2060863** | NA | 0.153277719 | 0.012145553 | 0.294409885 | 12.02245752 | 2.59574576 | 0.037282919 | 0.991835066 | -3.493866864 |
| **A_55_P2062255** | Syne3 | 0.169210536 | 0.013390587 | 0.325030485 | 8.296316973 | 2.595454839 | 0.037298306 | 0.991835066 | -3.494116207 |
| **A_55_P2349265** | Ptk2 | 0.146253606 | 0.011510337 | 0.280996876 | 9.215270427 | 2.594230947 | 0.037363111 | 0.991835066 | -3.495165309 |
| **A_52_P212686** | Lrrk1 | 0.106367652 | 0.008368431 | 0.204366873 | 8.255747266 | 2.594155952 | 0.037367086 | 0.991835066 | -3.4952296 |
| **A_52_P295104** | Smim5 | -0.232458586 | -0.446640307 | -0.018276864 | 10.97164405 | -2.594014019 | 0.037374609 | 0.991835066 | -3.495351278 |
| **A_55_P1982613** | Hnrnpa3 | 0.203325571 | 0.015976512 | 0.390674631 | 13.96108083 | 2.59387842 | 0.037381799 | 0.991835066 | -3.495467529 |
| **A_55_P2018994** | Elk3 | 0.128440549 | 0.010073385 | 0.246807713 | 12.01239696 | 2.593462927 | 0.037403837 | 0.991835066 | -3.495823752 |
| **A_55_P1991802** | Kat14 | 0.111386815 | 0.008642833 | 0.214130798 | 10.74989439 | 2.591114011 | 0.037528684 | 0.991835066 | -3.497838061 |
| **A_55_P1986743** | NA | 0.165709114 | 0.012853368 | 0.31856486 | 11.08830781 | 2.591037857 | 0.037532739 | 0.991835066 | -3.49790338 |
| **A_55_P2005081** | Rpl10 | 0.115082918 | 0.00891469 | 0.221251145 | 17.33235303 | 2.590749541 | 0.037548095 | 0.991835066 | -3.498150682 |
| **A_55_P2154536** | Hagh | -0.198512078 | -0.381907478 | -0.015116677 | 11.38821114 | -2.587066706 | 0.037744825 | 0.991835066 | -3.501310655 |
| **A_55_P2062797** | 4921524J1 | -0.157168033 | -0.302504092 | -0.011831974 | 9.508231555 | -2.584639469 | 0.037875072 | 0.991835066 | -3.503394339 |
| **A_55_P1991239** | NA | 0.110892527 | 0.008315182 | 0.213469872 | 8.798982748 | 2.583806341 | 0.037919887 | 0.991835066 | -3.504109738 |
| **A_55_P2094641** | NA | -0.224870696 | -0.432941177 | -0.016800215 | 11.79611912 | -2.583042344 | 0.037961031 | 0.991835066 | -3.504765859 |
| **A_55_P2011659** | Anapc5 | -0.134842879 | -0.259615998 | -0.01006976 | 11.62561307 | -2.582950704 | 0.03796597 | 0.991835066 | -3.504844566 |
| **A_55_P2101906** | Rpl10a | 0.126649149 | 0.009439066 | 0.243859232 | 16.31091946 | 2.582536307 | 0.037988309 | 0.991835066 | -3.50520049 |

| | | | | | | | | | |
|---|---|---|---|---|---|---|---|---|---|
| **A_52_P75465** | Gdpd1 | -0.226315731 | -0.435938701 | -0.016692761 | 10.57903473 | -2.580387943 | 0.038104345 | 0.991835066 | -3.507046101 |
| **A_55_P2048656** | Atp5l | -0.141692718 | -0.272968908 | -0.010416528 | 15.90131157 | -2.579708894 | 0.038141098 | 0.991835066 | -3.507629592 |
| **A_55_P2144556** | Flrt3 | 0.295022965 | 0.021623933 | 0.568421997 | 9.767196549 | 2.57909882 | 0.03817415 | 0.991835066 | -3.508153869 |
| **A_55_P1963062** | Rps16 | 0.137728313 | 0.010064239 | 0.265392386 | 17.10673151 | 2.578479372 | 0.03820774 | 0.991835066 | -3.508686256 |
| **A_55_P1990859** | Rpl6 | 0.113215433 | 0.008255894 | 0.218174972 | 15.71210256 | 2.578058988 | 0.038230553 | 0.991835066 | -3.509047587 |
| **A_55_P2180071** | Zfand5 | -0.114014284 | -0.219740635 | -0.008287932 | 12.74130021 | -2.577419731 | 0.038265272 | 0.991835066 | -3.509597092 |
| **A_51_P158018** | Bend5 | 0.128280344 | 0.009298744 | 0.247261945 | 9.954814563 | 2.576851811 | 0.038296143 | 0.991835066 | -3.510085323 |
| **A_55_P1956807** | Gm14391 | 0.184756881 | 0.0132595 | 0.356254262 | 9.772675007 | 2.574851946 | 0.038405061 | 0.991835066 | -3.511804935 |
| **A_51_P500344** | Mat2a | 0.316882393 | 0.022711663 | 0.611053122 | 13.65420645 | 2.574588256 | 0.038419446 | 0.991835066 | -3.512031715 |
| **A_52_P183038** | Impdh1 | 0.141934713 | 0.010162019 | 0.273707407 | 10.17648077 | 2.574378102 | 0.038430915 | 0.991835066 | -3.512212459 |
| **A_65_P09523** | Scrn3 | 0.119611444 | 0.008481277 | 0.230741611 | 9.457477037 | 2.572467519 | 0.038535346 | 0.991835066 | -3.513855948 |
| **A_55_P2046378** | Eif4g1 | 0.125685571 | 0.00875403 | 0.242617111 | 13.29676338 | 2.568992767 | 0.038726033 | 0.991835066 | -3.516846242 |
| **A_55_P2443568** | Ube4a | 0.151096978 | 0.010517371 | 0.291676586 | 9.762265321 | 2.568872711 | 0.03873264 | 0.991835066 | -3.51694959 |
| **A_55_P2132147** | NA | -0.253925096 | -0.490219059 | -0.017631133 | 15.6644521 | -2.568396868 | 0.038758834 | 0.991835066 | -3.517359228 |
| **A_55_P2020532** | Selenov | 0.130525025 | 0.00902592 | 0.25202413 | 8.319293796 | 2.567614606 | 0.038801937 | 0.991835066 | -3.518032722 |
| **A_51_P197175** | Pigk | -0.116903854 | -0.225804665 | -0.008003043 | 8.549048636 | -2.565705758 | 0.038907325 | 0.991835066 | -3.519676512 |
| **A_51_P249313** | Herc3 | -0.161091095 | -0.311164343 | -0.011017847 | 11.87749636 | -2.565531688 | 0.03891695 | 0.991835066 | -3.519826436 |
| **A_55_P2319468** | NA | 0.312524329 | 0.021322298 | 0.603726359 | 8.669158054 | 2.565066121 | 0.038942706 | 0.991835066 | -3.520227443 |
| **A_51_P519276** | Ndufb7 | -0.459698163 | -0.888049751 | -0.031346575 | 15.85544321 | -2.564965448 | 0.038948278 | 0.991835066 | -3.52031416 |
| **A_55_P2065732** | Medag | -0.176883272 | -0.34189599 | -0.011870555 | 13.43965059 | -2.561996196 | 0.039112986 | 0.991835066 | -3.522872414 |
| **A_55_P2096834** | Fbxl19 | 0.156089785 | 0.01046901 | 0.301710561 | 10.54278813 | 2.561888802 | 0.039118957 | 0.991835066 | -3.522964966 |
| **A_52_P423814** | Cox8b | -0.263892129 | -0.510102062 | -0.017682196 | 16.00950189 | -2.561710214 | 0.039128888 | 0.991835066 | -3.523118875 |
| **A_52_P371718** | Tgs1 | 0.13297196 | 0.00889106 | 0.25705286 | 10.00239383 | 2.561322523 | 0.039150456 | 0.991835066 | -3.523453008 |
| **A_52_P117999** | Selenot | 0.106561325 | 0.007114813 | 0.206007837 | 8.991715651 | 2.561056683 | 0.039165253 | 0.991835066 | -3.523682135 |
| **A_55_P2063316** | Mgll | -0.200951389 | -0.38850057 | -0.013402208 | 12.20059961 | -2.560854905 | 0.039176488 | 0.991835066 | -3.523856053 |
| **A_55_P2313861** | Zbed6 | 0.114999306 | 0.007612212 | 0.2223864 | 8.493939003 | 2.559483077 | 0.03925296 | 0.991835066 | -3.525038622 |
| **A_51_P199135** | Cd83 | -0.159423382 | -0.308496396 | -0.010350369 | 8.442743912 | -2.556007485 | 0.039447402 | 0.991835066 | -3.528035874 |
| **A_66_P135391** | Igfbp2 | 0.251344261 | 0.016244451 | 0.486444071 | 9.298711509 | 2.555205459 | 0.039492414 | 0.991835066 | -3.528727753 |
| **A_51_P259879** | Fkrp | -0.218537302 | -0.423091856 | -0.013982748 | 10.05748435 | -2.553439409 | 0.039591719 | 0.991835066 | -3.530251572 |
| **A_55_P2046363** | Mapkap1 | -0.132381634 | -0.25633953 | -0.008423737 | 9.444419634 | -2.552481897 | 0.039645669 | 0.991835066 | -3.53107793 |
| **A_55_P2068248** | NA | 0.661049248 | 0.041695701 | 1.280402796 | 11.8107435 | 2.550963934 | 0.039731353 | 0.991835066 | -3.532388228 |
| **A_66_P109562** | Gm12271 | -0.15461874 | -0.299591565 | -0.009645915 | 11.15606513 | -2.549087007 | 0.039837568 | 0.991835066 | -3.534008816 |
| **A_55_P2171788** | NA | 0.347478523 | 0.021560289 | 0.673396757 | 12.09624997 | 2.548170247 | 0.039889554 | 0.991835066 | -3.534800546 |
| **A_52_P422494** | Cd300lf | -0.122572275 | -0.237579804 | -0.007564746 | 7.851381091 | -2.547270734 | 0.039940631 | 0.991835066 | -3.535577491 |
| **A_55_P1976237** | Dnajb3 | 0.119761418 | 0.00733866 | 0.232184177 | 8.677777618 | 2.546078714 | 0.040008423 | 0.991835066 | -3.536607255 |
| **A_55_P1961265** | NA | 0.319804713 | 0.019431039 | 0.620178388 | 9.054021116 | 2.544673838 | 0.040088474 | 0.991835066 | -3.537821149 |
| **A_52_P37894** | Cox7a2 | -0.128157636 | -0.248605735 | -0.007709537 | 15.95137618 | -2.543042806 | 0.040181622 | 0.991835066 | -3.539230789 |
| **A_55_P1966086** | NA | 0.213624703 | 0.01271159 | 0.414537817 | 9.228923326 | 2.541278848 | 0.040282614 | 0.991835066 | -3.540755717 |
| **A_55_P2142808** | NA | -0.277920853 | -0.539439173 | -0.016402534 | 14.80312069 | -2.539967446 | 0.040357868 | 0.991835066 | -3.541889686 |
| **A_55_P2008327** | Eno1b | 0.187614555 | 0.011060668 | 0.364168441 | 15.09594758 | 2.539793327 | 0.04036787 | 0.991835066 | -3.542040264 |
| **A_51_P186476** | Slc11a1 | -0.14164873 | -0.274973549 | -0.00832391 | 8.168820023 | -2.539281287 | 0.040397301 | 0.991835066 | -3.5424831 |
| **A_55_P2174203** | Lims2 | -0.208902429 | -0.405820416 | -0.011984441 | 11.07928048 | -2.535521119 | 0.040614108 | 0.991835066 | -3.545736144 |
| **A_55_P2201676** | Cyp2r1 | -0.182106243 | -0.353873306 | -0.01033918 | 9.756630652 | -2.533926874 | 0.040706397 | 0.991835066 | -3.54711595 |
| **A_51_P422223** | 1700006J1 | -0.124601165 | -0.242183708 | -0.007018622 | 8.123331704 | -2.532727034 | 0.040775997 | 0.991835066 | -3.548154625 |
| **A_55_P1995127** | NA | 0.177685068 | 0.010001103 | 0.345369033 | 13.07828003 | 2.532611242 | 0.040782721 | 0.991835066 | -3.548254874 |
| **A_51_P435068** | Acadsb | -0.129950281 | -0.25265581 | -0.007244752 | 9.887113254 | -2.531175298 | 0.040866195 | 0.991835066 | -3.549498215 |
| **A_51_P339540** | Cdkn1c | 0.284608616 | 0.01550879 | 0.553708441 | 14.10147555 | 2.527806112 | 0.041062751 | 0.991835066 | -3.552416571 |
| **A_52_P432969** | Pcdh19 | -0.166605371 | -0.32414123 | -0.009069512 | 9.286426496 | -2.527660282 | 0.041071281 | 0.991835066 | -3.552542922 |

| | | | | | | | | | |
|---|---|---|---|---|---|---|---|---|---|
| **A_55_P2139625** | Gtf2a2 | -0.165848783 | -0.322673328 | -0.009024237 | 11.59035175 | -2.527594391 | 0.041075136 | 0.991835066 | -3.552600011 |
| **A_55_P2124976** | Grhl1 | 0.213522412 | 0.011569803 | 0.415475021 | 8.059777031 | 2.526987721 | 0.041110644 | 0.991835066 | -3.553125681 |
| **A_55_P1985673** | Zbtb37 | 0.112665628 | 0.006094329 | 0.219236926 | 8.420976313 | 2.526738608 | 0.041125234 | 0.991835066 | -3.553341547 |
| **A_55_P2044380** | NA | 0.119800101 | 0.006470588 | 0.233129614 | 8.273106478 | 2.52652322 | 0.041137853 | 0.991835066 | -3.553528196 |
| **A_51_P395293** | Ccz1 | -0.137419605 | -0.267448581 | -0.007390629 | 11.60080398 | -2.525908921 | 0.041173865 | 0.991835066 | -3.554060563 |
| **A_55_P1962035** | Coa5 | -0.114062709 | -0.222028327 | -0.006097092 | 10.3491748 | -2.525034657 | 0.041225174 | 0.991835066 | -3.554818308 |
| **A_51_P386810** | Gmppb | 0.167534405 | 0.008906362 | 0.326162448 | 9.896233152 | 2.524254704 | 0.041271004 | 0.991835066 | -3.555494397 |
| **A_55_P2001558** | Zfp128 | -0.112676951 | -0.219448832 | -0.005905069 | 8.900488279 | -2.522245293 | 0.041389322 | 0.991835066 | -3.557236592 |
| **A_52_P686785** | Lyve1 | 0.230669943 | 0.012077459 | 0.449262426 | 9.531471289 | 2.522115176 | 0.041396996 | 0.991835066 | -3.557349424 |
| **A_55_P1995055** | Prr7 | 0.164520014 | 0.008331271 | 0.320708757 | 8.61751474 | 2.517550213 | 0.041667155 | 0.991835066 | -3.561309384 |
| **A_55_P2096422** | Inhbb | 0.176758488 | 0.009400123 | 0.344116854 | 8.737044283 | 2.623614378 | 0.041684708 | 0.991835066 | -3.599618655 |
| **A_55_P2162503** | Tbx20 | 0.157167213 | 0.00794013 | 0.306394296 | 8.843512707 | 2.517233108 | 0.04168599 | 0.991835066 | -3.561584564 |
| **A_51_P411264** | Txndc16 | -0.141467476 | -0.275806694 | -0.007128258 | 9.555811964 | -2.516882387 | 0.041706831 | 0.991835066 | -3.561888931 |
| **A_51_P493105** | Mettl15 | -0.15256984 | -0.297453318 | -0.007686362 | 9.753321409 | -2.516859437 | 0.041708196 | 0.991835066 | -3.561908849 |
| **A_51_P288103** | Psmd6 | -0.108515078 | -0.211737062 | -0.005293093 | 12.32445818 | -2.512621156 | 0.041960923 | 0.991835066 | -3.565588255 |
| **A_55_P2146384** | Filip1 | -0.191190616 | -0.373130193 | -0.009251038 | 9.737607943 | -2.511588703 | 0.042022728 | 0.991835066 | -3.566484919 |
| **A_51_P327530** | Ciao2a | -0.141818088 | -0.277020039 | -0.006616138 | 12.06375175 | -2.507020022 | 0.042297362 | 0.991835066 | -3.570454396 |
| **A_55_P1957018** | Caskin2 | 0.149011415 | 0.006802232 | 0.291220597 | 10.80603421 | 2.504384639 | 0.042456629 | 0.991835066 | -3.572745368 |
| **A_55_P2170536** | Zfp949 | -0.147908412 | -0.289209815 | -0.006607008 | 8.58051737 | -2.501816942 | 0.042612405 | 0.991835066 | -3.574978361 |
| **A_55_P1953851** | Stk39 | -0.123897431 | -0.242341882 | -0.00545298 | 12.86472867 | -2.500096173 | 0.042717132 | 0.991835066 | -3.576475301 |
| **A_55_P2066081** | Prmt6 | 0.143714628 | 0.006320225 | 0.28110903 | 8.355128031 | 2.500006089 | 0.042722622 | 0.991835066 | -3.576553678 |
| **A_55_P2078831** | NA | 0.151396588 | 0.006656452 | 0.296136725 | 9.585578992 | 2.499978331 | 0.042724314 | 0.991835066 | -3.576577829 |
| **A_55_P2159665** | Zfp534 | 0.138957713 | 0.005996049 | 0.271919377 | 10.91583626 | 2.497844209 | 0.042854593 | 0.991835066 | -3.578434916 |
| **A_55_P2032388** | Trim24 | -0.144945571 | -0.28364926 | -0.006241883 | 10.17646043 | -2.497618324 | 0.042868406 | 0.991835066 | -3.578631512 |
| **A_55_P2015715** | Rab4a | 0.131874707 | 0.005509561 | 0.258239853 | 9.339861018 | 2.494269276 | 0.043073752 | 0.991835066 | -3.581547086 |
| **A_55_P2055854** | Ppara | -0.120318796 | -0.235616688 | -0.005020903 | 11.46531188 | -2.49414237 | 0.043081554 | 0.991835066 | -3.581657594 |
| **A_52_P76034** | Rcc2 | 0.195568285 | 0.008112566 | 0.383024005 | 11.12485327 | 2.493497084 | 0.043121244 | 0.991835066 | -3.582219535 |
| **A_55_P2012955** | NA | 0.119427908 | 0.004850741 | 0.234005075 | 15.93187183 | 2.491247513 | 0.043259908 | 0.991835066 | -3.584178964 |
| **A_55_P1981836** | Ccm2l | 0.214132706 | 0.008545476 | 0.419719936 | 10.76861056 | 2.48940755 | 0.043373668 | 0.991835066 | -3.585782093 |
| **A_51_P223498** | Slc39a10 | -0.122296699 | -0.239719026 | -0.004874373 | 8.507273035 | -2.489276772 | 0.043381766 | 0.991835066 | -3.585896055 |
| **A_51_P246727** | Mlxip | -0.145888852 | -0.285978688 | -0.005799016 | 8.549860305 | -2.488998371 | 0.043399009 | 0.991835066 | -3.586138663 |
| **A_51_P235801** | Zfp36l1 | -0.243996257 | -0.478306112 | -0.009686402 | 13.02325893 | -2.488867293 | 0.04340713 | 0.991835066 | -3.586252892 |
| **A_55_P2063992** | Rps5 | 0.13936956 | 0.005493313 | 0.273245807 | 17.35447376 | 2.488132665 | 0.043452673 | 0.991835066 | -3.586893132 |
| **A_51_P227345** | Dpep1 | -0.185565034 | -0.363832544 | -0.007297524 | 9.062242389 | -2.487900911 | 0.043467051 | 0.991835066 | -3.587095123 |
| **A_55_P2029750** | Ndufs5 | -0.176424055 | -0.345944197 | -0.006903913 | 15.66173679 | -2.487399975 | 0.043498146 | 0.991835066 | -3.587531751 |
| **A_55_P2062777** | Mfap5 | -0.257395239 | -0.504773125 | -0.010017354 | 11.29199943 | -2.486845296 | 0.043532603 | 0.991835066 | -3.588015259 |
| **A_55_P2102454** | NA | 0.223706004 | 0.008609576 | 0.438802432 | 10.74286974 | 2.485727828 | 0.043602109 | 0.991835066 | -3.588989463 |
| **A_51_P175974** | NA | -0.114333854 | -0.224401358 | -0.004266351 | 9.521099444 | -2.482703526 | 0.043790795 | 0.991835066 | -3.591626825 |
| **A_55_P1995092** | H2-Q10 | -0.1061698 | -0.208409394 | -0.003930207 | 7.880261621 | -2.481938511 | 0.043838659 | 0.991835066 | -3.592294144 |
| **A_55_P1978597** | Trit1 | 0.197684952 | 0.007272578 | 0.388097325 | 10.13076437 | 2.48134743 | 0.043875677 | 0.991835066 | -3.592809791 |
| **A_52_P260629** | Cpne3 | -0.131750448 | -0.258757404 | -0.004743493 | 9.260644071 | -2.479326529 | 0.044002488 | 0.991835066 | -3.594573116 |
| **A_55_P2023732** | Limch1 | 0.136570684 | 0.004904577 | 0.268236792 | 11.07541727 | 2.479091868 | 0.044017238 | 0.991835066 | -3.594777902 |
| **A_55_P1955015** | Myo1g | -0.270563471 | -0.531434183 | -0.00969276 | 8.492819205 | -2.478865554 | 0.044031468 | 0.991835066 | -3.594975409 |
| **A_66_P119518** | Tuba8 | -0.182984501 | -0.359627114 | -0.006341888 | 8.009991318 | -2.475870686 | 0.044220224 | 0.991835066 | -3.597589677 |
| **A_51_P482711** | Dhcr24 | 0.329937708 | 0.011421105 | 0.648454311 | 11.82693805 | 2.475762668 | 0.044227047 | 0.991835066 | -3.597683988 |
| **A_55_P2167620** | NA | 0.302954399 | 0.010466161 | 0.595442638 | 9.916702417 | 2.475585835 | 0.044238221 | 0.991835066 | -3.597838386 |
| **A_51_P301215** | Knop1 | 0.155113034 | 0.005331569 | 0.3048945 | 10.91800218 | 2.475137622 | 0.044266554 | 0.991835066 | -3.59822975 |
| **A_55_P1971330** | NA | -0.158752744 | -0.312380688 | -0.005124799 | 9.582789041 | -2.46979071 | 0.04460601 | 0.991835066 | -3.602900416 |

| | | | | | | | | | |
|---|---|---|---|---|---|---|---|---|---|
| **A_55_P2159432** | Fndc4 | 0.210962902 | 0.006718236 | 0.415207568 | 10.63666734 | 2.468678299 | 0.044676972 | 0.991835066 | -3.603872581 |
| **A_51_P486239** | Clec3b | -0.281953158 | -0.554934827 | -0.00897149 | 8.527586263 | -2.46861073 | 0.044681286 | 0.991835066 | -3.603931636 |
| **A_55_P1999213** | Tdrd7 | 0.111594476 | 0.003546804 | 0.219642149 | 10.56108133 | 2.46851866 | 0.044687165 | 0.991835066 | -3.604012107 |
| **A_66_P100349** | Esyt2 | -0.117092708 | -0.230481148 | -0.003704267 | 9.754783171 | -2.468142322 | 0.044711204 | 0.991835066 | -3.60434104 |
| **A_55_P2448065** | Sfxn2 | 0.134183833 | 0.004243852 | 0.264123814 | 8.734552885 | 2.46812145 | 0.044712538 | 0.991835066 | -3.604359284 |
| **A_55_P2083401** | Acp1 | 0.168758234 | 0.00531425 | 0.332202219 | 11.12819072 | 2.467772743 | 0.044734824 | 0.991835066 | -3.604664083 |
| **A_55_P2326517** | Ube2i | 0.335545846 | 0.010554477 | 0.660537215 | 9.751494561 | 2.467681879 | 0.044740634 | 0.991835066 | -3.604743509 |
| **A_55_P2032453** | Mrpl23 | -0.164616552 | -0.324090762 | -0.005142343 | 14.36247606 | -2.467130802 | 0.044775883 | 0.991835066 | -3.605225234 |
| **A_55_P1983488** | Lsr | 0.344656697 | 0.010725783 | 0.678587611 | 9.881081482 | 2.466830035 | 0.044795133 | 0.991835066 | -3.605488165 |
| **A_51_P257550** | Marcksl1 | -0.150062792 | -0.295468332 | -0.004657253 | 12.36777337 | -2.46661407 | 0.044808961 | 0.991835066 | -3.60567697 |
| **A_51_P190111** | Mcm5 | 0.166184865 | 0.005154961 | 0.327214768 | 11.46064441 | 2.46657354 | 0.044811557 | 0.991835066 | -3.605712403 |
| **A_55_P2108067** | Dgcr8 | 0.190024686 | 0.005757679 | 0.374291692 | 10.02369838 | 2.464742617 | 0.044928973 | 0.991835066 | -3.607313301 |
| **A_65_P19395** | H2-D1 | -0.230378188 | -0.453778362 | -0.006978013 | 10.36791024 | -2.464716551 | 0.044930647 | 0.991835066 | -3.607336095 |
| **A_55_P2062981** | Abca9 | -0.121609368 | -0.239560741 | -0.003657994 | 8.505885809 | -2.464184156 | 0.044964851 | 0.991835066 | -3.607801683 |
| **A_55_P1983733** | Aldh1l1 | -0.160343109 | -0.315959352 | -0.004726865 | 9.069922378 | -2.462660271 | 0.045062902 | 0.991835066 | -3.609134536 |
| **A_55_P2067842** | 5-Mar | -0.139822251 | -0.27552997 | -0.004114533 | 10.49779595 | -2.462526274 | 0.045071534 | 0.991835066 | -3.609251748 |
| **A_52_P370022** | NA | -0.125754393 | -0.247883906 | -0.003624879 | 14.49819735 | -2.461000315 | 0.04516996 | 0.991835066 | -3.610586725 |
| **A_52_P627925** | Usp36 | 0.148504608 | 0.004261437 | 0.292747779 | 8.903607824 | 2.460672409 | 0.045191139 | 0.991835066 | -3.610873629 |
| **A_55_P2147126** | Zfp712 | -0.130146894 | -0.256564043 | -0.003729745 | 8.626852155 | -2.46057694 | 0.045197307 | 0.991835066 | -3.610957163 |
| **A_55_P1961965** | Plagl1 | 0.099855981 | 0.002860853 | 0.196851108 | 8.479543365 | 2.460556239 | 0.045198645 | 0.991835066 | -3.610975276 |
| **A_55_P2026291** | Smarce1 | 0.157851368 | 0.004451564 | 0.311251172 | 12.33513393 | 2.459419872 | 0.045272135 | 0.991835066 | -3.611969667 |
| **A_55_P2067262** | Cdv3 | 0.125953115 | 0.00354348 | 0.248362751 | 8.436311737 | 2.459248643 | 0.045283219 | 0.991835066 | -3.612119516 |
| **A_55_P2021398** | NA | 0.724797193 | 0.019830748 | 1.429763637 | 12.3844865 | 2.45729439 | 0.045409924 | 0.991835066 | -3.613830016 |
| **A_55_P2061025** | Hspa4 | 0.412673027 | 0.011245687 | 0.814100368 | 10.92832036 | 2.457017603 | 0.045427899 | 0.991835066 | -3.614072317 |
| **A_55_P2115257** | Fer | -0.101061667 | -0.199466822 | -0.002656513 | 10.0024232 | -2.454583126 | 0.045586319 | 0.991835066 | -3.616203874 |
| **A_55_P2007964** | Cx3cr1 | -0.188461624 | -0.371995979 | -0.004927268 | 9.892665292 | -2.454226774 | 0.045609556 | 0.991835066 | -3.616515945 |
| **A_55_P2200628** | 4733401D( | -0.183427224 | -0.362164476 | -0.004689973 | 9.529192546 | -2.452775789 | 0.045704299 | 0.991835066 | -3.617786788 |
| **A_55_P1992838** | Socs2 | -0.514871056 | -1.016763319 | -0.012978793 | 9.556122894 | -2.451868134 | 0.045763668 | 0.991835066 | -3.618581887 |
| **A_52_P124173** | Zdhhc8 | -0.129320971 | -0.255535959 | -0.003105984 | 9.383227666 | -2.448878063 | 0.045959813 | 0.991835066 | -3.621201866 |
| **A_55_P1965050** | Cks2 | -0.110410957 | -0.218215414 | -0.002606501 | 10.82177727 | -2.447848822 | 0.046027531 | 0.991835066 | -3.622103962 |
| **A_55_P2111459** | Trpv2 | -0.129551752 | -0.256140703 | -0.002962802 | 9.027786264 | -2.446000965 | 0.046149368 | 0.991835066 | -3.623723868 |
| **A_55_P2180805** | Pex3 | -0.121838235 | -0.241009714 | -0.002666756 | 9.607027783 | -2.443545338 | 0.046311796 | 0.991835066 | -3.625877202 |
| **A_55_P1953231** | Map2k5 | -0.122328899 | -0.242019984 | -0.002637813 | 12.13881423 | -2.442735212 | 0.046365512 | 0.991835066 | -3.626587758 |
| **A_52_P380369** | Phf11d | -0.128377713 | -0.253989056 | -0.002766371 | 8.002934537 | -2.442698754 | 0.046367931 | 0.991835066 | -3.626619737 |
| **A_52_P194680** | NA | -0.210174274 | -0.415929577 | -0.004418971 | 9.610260908 | -2.44139275 | 0.046454666 | 0.991835066 | -3.627765393 |
| **A_55_P1993493** | Efr3a | -0.149344219 | -0.295565326 | -0.003123112 | 10.78040191 | -2.441110729 | 0.046473418 | 0.991835066 | -3.628012814 |
| **A_51_P421664** | Tmem123 | -0.194979425 | -0.385993873 | -0.003964977 | 11.01538289 | -2.439673446 | 0.046569106 | 0.991835066 | -3.629273914 |
| **A_51_P175424** | Car14 | -0.221571596 | -0.438651885 | -0.004491307 | 11.82148521 | -2.439511256 | 0.046579917 | 0.991835066 | -3.629416238 |
| **A_52_P579640** | Rrs1 | 0.233208041 | 0.004704177 | 0.461711905 | 10.55338988 | 2.439265675 | 0.046596291 | 0.991835066 | -3.629631745 |
| **A_55_P2139181** | Cald1 | 0.151231812 | 0.003005811 | 0.299457813 | 9.592041331 | 2.438528844 | 0.046645454 | 0.991835066 | -3.630278385 |
| **A_55_P2262593** | 4930429F2 | -0.161177503 | -0.319205569 | -0.003149436 | 10.53532598 | -2.437694788 | 0.04670117 | 0.991835066 | -3.631010427 |
| **A_51_P115326** | Surf4 | 0.127594135 | 0.002363626 | 0.252824644 | 13.386762 | 2.435172317 | 0.046870093 | 0.991835066 | -3.633224872 |
| **A_55_P2184478** | NA | 0.123876555 | 0.0022724 | 0.24548071 | 16.39514336 | 2.434724563 | 0.046900143 | 0.991835066 | -3.633618028 |
| **A_55_P2025078** | Csf1r | -0.395523744 | -0.783926504 | -0.007120984 | 11.47337127 | -2.433881249 | 0.046956796 | 0.991835066 | -3.634358574 |
| **A_55_P2022277** | Nfe2l1 | 0.169219598 | 0.002946395 | 0.335492801 | 8.800895649 | 2.432414196 | 0.04705552 | 0.991835066 | -3.635647049 |
| **A_55_P2166773** | Pitpna | -0.147013915 | -0.29157006 | -0.002457771 | 12.88768609 | -2.430698087 | 0.047171276 | 0.991835066 | -3.637154583 |
| **A_55_P1982420** | Fam8a1 | 0.144221215 | 0.002402622 | 0.286039809 | 10.17651977 | 2.430553072 | 0.047181071 | 0.991835066 | -3.637281989 |
| **A_55_P1962139** | Mff | -0.129508224 | -0.256861348 | -0.002155101 | 12.15681662 | -2.430507023 | 0.047184182 | 0.991835066 | -3.637322447 |

| | | | | | | | | | |
|---|---|---|---|---|---|---|---|---|---|
| **A_51_P137388** | Zadh2 | -0.167511002 | -0.332283833 | -0.00273817 | 11.88098579 | -2.429779494 | 0.047233358 | 0.991835066 | -3.637961673 |
| **A_51_P184100** | Capn7 | -0.114851038 | -0.227842067 | -0.001860008 | 9.409638164 | -2.429405939 | 0.047258629 | 0.991835066 | -3.638289912 |
| **A_65_P17218** | Mndal | -0.198469216 | -0.393863181 | -0.003075252 | 8.323366813 | -2.427678331 | 0.047375681 | 0.991835066 | -3.639808156 |
| **A_55_P2432388** | Prr14l | 0.130245034 | 0.001969674 | 0.258520394 | 9.927209774 | 2.426761308 | 0.047437935 | 0.991835066 | -3.640614189 |
| **A_55_P2116802** | NA | 0.128201008 | 0.001898821 | 0.254503195 | 10.74934029 | 2.425993888 | 0.047490098 | 0.991835066 | -3.6412888 |
| **A_52_P257547** | Dele1 | -0.122817471 | -0.243835157 | -0.001799785 | 11.49517034 | -2.425606998 | 0.047516418 | 0.991835066 | -3.641628928 |
| **A_55_P1990170** | NA | -0.123835305 | -0.245868942 | -0.001801668 | 14.29272463 | -2.425347956 | 0.047534049 | 0.991835066 | -3.64185667 |
| **A_55_P2076777** | Mdfi | 0.138816722 | 0.00195673 | 0.275676715 | 12.82571313 | 2.424233266 | 0.047609995 | 0.991835066 | -3.642836759 |
| **A_55_P1987770** | Psenen | -0.155918156 | -0.309654461 | -0.00218185 | 13.19520127 | -2.423981945 | 0.047627135 | 0.991835066 | -3.643057753 |
| **A_55_P2089840** | Eif2d | -0.147214001 | -0.292421693 | -0.002006308 | 9.412381064 | -2.423084862 | 0.047688369 | 0.991835066 | -3.643846643 |
| **A_55_P1978351** | Apobec2 | -0.19512967 | -0.387688031 | -0.002571309 | 10.23650998 | -2.42197726 | 0.047764084 | 0.991835066 | -3.64482079 |
| **A_55_P1994887** | Zfpm1 | 0.128680864 | 0.001684759 | 0.25567697 | 12.10721667 | 2.421768905 | 0.047778341 | 0.991835066 | -3.645004056 |
| **A_55_P2174836** | Vldlr | -0.163556603 | -0.3249783 | -0.002134907 | 9.295051266 | -2.421671929 | 0.047784979 | 0.991835066 | -3.645089357 |
| **A_51_P178735** | Hibch | -0.130892632 | -0.260089959 | -0.001695306 | 11.5564418 | -2.421423797 | 0.047801966 | 0.991835066 | -3.645307619 |
| **A_55_P1994817** | Taf15 | 0.210183426 | 0.002709676 | 0.417657176 | 9.712199792 | 2.421276807 | 0.047812031 | 0.991835066 | -3.645436919 |
| **A_55_P2156016** | NA | 0.119703381 | 0.001476945 | 0.237929818 | 8.51282839 | 2.41991968 | 0.04790507 | 0.991835066 | -3.646630828 |
| **A_55_P2135331** | Evl | 0.166334288 | 0.001994683 | 0.330673893 | 13.55214536 | 2.419071346 | 0.047963323 | 0.991835066 | -3.647377243 |
| **A_55_P2381538** | NA | 0.288318134 | 0.0034511 | 0.573185168 | 9.911057378 | 2.419016872 | 0.047967067 | 0.991835066 | -3.647425175 |
| **A_55_P1993483** | NA | 0.211250197 | 0.002493237 | 0.420007157 | 10.062975 | 2.418606917 | 0.047995246 | 0.991835066 | -3.647785911 |
| **A_55_P2048176** | NA | 0.147537878 | 0.001725508 | 0.293350247 | 8.112592347 | 2.418345217 | 0.048013243 | 0.991835066 | -3.648016201 |
| **A_66_P108178** | Lin52 | -0.108349072 | -0.215434606 | -0.001263538 | 9.405569169 | -2.418262951 | 0.048018902 | 0.991835066 | -3.648088595 |
| **A_55_P2180551** | Sinhcaf | 0.139260201 | 0.001581209 | 0.276939192 | 8.959455009 | 2.417511077 | 0.048070655 | 0.991835066 | -3.648750277 |
| **A_55_P2083419** | Pkp4 | -0.154283696 | -0.306833597 | -0.001733796 | 10.33078081 | -2.417225894 | 0.0480903 | 0.991835066 | -3.649001268 |
| **A_55_P2016252** | Ctnnal1 | 0.187102608 | 0.002099111 | 0.372106106 | 9.111659023 | 2.417180245 | 0.048093445 | 0.991835066 | -3.649041444 |
| **A_55_P2010936** | Fbxo17 | 0.21366775 | 0.00231446 | 0.42502104 | 8.985960169 | 2.416234587 | 0.04815865 | 0.991835066 | -3.649873794 |
| **A_55_P2174905** | Cuta | -0.199569572 | -0.396993436 | -0.002145708 | 12.79693245 | -2.416038276 | 0.048172197 | 0.991835066 | -3.650046595 |
| **A_51_P243857** | Copb2 | -0.136339543 | -0.271256659 | -0.001422427 | 12.38074315 | -2.415260165 | 0.048225933 | 0.991835066 | -3.650731567 |
| **A_51_P412338** | Vps41 | -0.16709662 | -0.332486078 | -0.001707162 | 10.05336803 | -2.414732201 | 0.04826243 | 0.991835066 | -3.651196374 |
| **A_51_P403636** | Smad7 | -0.266701156 | -0.530874494 | -0.002527818 | 10.65959614 | -2.412931801 | 0.0483871 | 0.991835066 | -3.652781641 |
| **A_55_P2051791** | Cbr4 | -0.138862462 | -0.276476356 | -0.001248568 | 9.076241264 | -2.411746791 | 0.048469338 | 0.991835066 | -3.653825253 |
| **A_51_P513449** | Pogz | 0.136970318 | 0.001209282 | 0.272731354 | 9.981917707 | 2.411351117 | 0.04849683 | 0.991835066 | -3.65417375 |
| **A_55_P2115990** | 2010001K2 | 0.145934651 | 0.001198309 | 0.290670993 | 8.595939531 | 2.409849738 | 0.048601293 | 0.991835066 | -3.655496277 |
| **A_55_P2003398** | Rmdn1 | -0.116496197 | -0.232074623 | -0.000917771 | 9.836467142 | -2.409040516 | 0.048657693 | 0.991835066 | -3.656209207 |
| **A_51_P112627** | St6galnac2 | -0.159711709 | -0.318208777 | -0.001214642 | 8.820813865 | -2.408378043 | 0.048703915 | 0.991835066 | -3.656792905 |
| **A_55_P2038458** | Lancl2 | -0.10295427 | -0.20513781 | -0.00077073 | 8.93633731 | -2.408089106 | 0.04872409 | 0.991835066 | -3.657047499 |
| **A_52_P432685** | Ctdsp2 | 0.125110964 | 0.000848708 | 0.24937322 | 8.897747801 | 2.406385876 | 0.048843188 | 0.991835066 | -3.658548481 |
| **A_51_P172688** | Stard3nl | -0.133403991 | -0.265930628 | -0.000877353 | 10.02081374 | -2.405884506 | 0.048878303 | 0.991835066 | -3.658990378 |
| **A_55_P2004781** | Ass1 | 0.202106443 | 0.001194092 | 0.403018795 | 9.294332735 | 2.40426678 | 0.048991785 | 0.991835066 | -3.660416401 |
| **A_52_P481686** | Wtip | 0.11307196 | 0.000579426 | 0.225564494 | 12.23077062 | 2.402372523 | 0.049125011 | 0.991835066 | -3.662086559 |
| **A_51_P483118** | Hmga1 | 0.169529175 | 0.000804382 | 0.338253968 | 14.17174763 | 2.401456233 | 0.049189589 | 0.991835066 | -3.662894592 |
| **A_52_P572702** | Gxylt1 | 0.16074931 | 0.000741889 | 0.320756732 | 10.24847339 | 2.401143541 | 0.049211647 | 0.991835066 | -3.663170362 |
| **A_55_P2099358** | Cars | 0.32520781 | 0.001499869 | 0.648915752 | 11.07686025 | 2.401135925 | 0.049212185 | 0.991835066 | -3.663177079 |
| **A_55_P1968231** | Rbpms | -0.135173793 | -0.269729101 | -0.000618485 | 10.08491702 | -2.40104775 | 0.049218407 | 0.991835066 | -3.663254844 |
| **A_55_P2110758** | B430306N( | -0.099834257 | -0.199222053 | -0.000446461 | 7.899317883 | -2.400798233 | 0.049236018 | 0.991835066 | -3.663474909 |
| **A_66_P105736** | Net1 | 0.110882579 | 0.000434857 | 0.221330301 | 9.055887897 | 2.399472005 | 0.049329734 | 0.991835066 | -3.66464471 |
| **A_55_P2164615** | NA | 0.162136642 | 0.000623955 | 0.323649329 | 15.63142747 | 2.39929508 | 0.04934225 | 0.991835066 | -3.664800782 |
| **A_55_P1983070** | Uhrf1bp1 | 0.151173159 | 0.000530125 | 0.301816192 | 10.32841322 | 2.398472633 | 0.049400475 | 0.991835066 | -3.665526337 |
| **A_55_P2016462** | Cxcl10 | -0.14502322 | -0.289685502 | -0.000360938 | 8.125643648 | -2.396025113 | 0.049574166 | 0.991835066 | -3.667685961 |

| | | | | | | | | | |
|---|---|---|---|---|---|---|---|---|---|
| **A_51_P496845** | Capza2 | -0.152161194 | -0.303958155 | -0.000364233 | 11.30074825 | -2.395796706 | 0.049590407 | 0.991835066 | -3.667887534 |
| **A_55_P1984243** | Naip2 | -0.098703075 | -0.197180836 | -0.000225315 | 7.825428915 | -2.39553021 | 0.049609364 | 0.991835066 | -3.66812273 |
| **A_55_P1964568** | Megf10 | -0.128286349 | -0.256309423 | -0.000263275 | 8.306700697 | -2.394976887 | 0.049648747 | 0.991835066 | -3.668611089 |
| **A_55_P2027906** | Nsa2 | 0.161731429 | 0.000312121 | 0.323150738 | 13.04691943 | 2.394683236 | 0.049669661 | 0.991835066 | -3.668870277 |
| **A_55_P2029902** | Gab3 | 0.687807 | 0.001181663 | 1.374432336 | 11.86551152 | 2.39417504 | 0.049705876 | 0.991835066 | -3.669318854 |
| **A_55_P2079928** | Ccdc68 | -0.102857548 | -0.20554059 | -0.000174506 | 8.502683818 | -2.394123635 | 0.049709541 | 0.991835066 | -3.66936423 |
| **A_55_P2078285** | Uck1 | 0.109749623 | 0.000177071 | 0.219322174 | 12.1819106 | 2.393924195 | 0.049723762 | 0.991835066 | -3.669540281 |
| **A_52_P239707** | Nfyb | -0.11271663 | -0.225314727 | -0.000118532 | 9.085815561 | -2.392577828 | 0.049819875 | 0.991835066 | -3.670728873 |
| **A_51_P240253** | Rrad | -0.328466848 | -0.657049034 | 0.000115339 | 10.35514591 | -2.389222849 | 0.050060211 | 0.991835066 | -3.673691562 |
| **A_51_P319379** | Tmx4 | -0.162731757 | -0.325674572 | 0.000211057 | 10.52767716 | -2.386966003 | 0.050222554 | 0.991835066 | -3.675685214 |
| **A_51_P252199** | Mmgt1 | -0.139339552 | -0.278886107 | 0.000207002 | 10.11022876 | -2.386516411 | 0.05025496 | 0.991835066 | -3.67608244 |
| **A_51_P203182** | Apobr | -0.100593673 | -0.201364411 | 0.000177065 | 7.819014133 | -2.385862222 | 0.050302151 | 0.991835066 | -3.676660473 |
| **A_55_P1962641** | Rnf26 | 0.116520874 | -0.000238141 | 0.233279889 | 12.71640339 | 2.385187053 | 0.050350903 | 0.991835066 | -3.677257092 |
| **A_51_P344399** | Rilpl2 | -0.194832584 | -0.390069078 | 0.00040391 | 8.808699848 | -2.385117192 | 0.05035595 | 0.991835066 | -3.677318829 |
| **A_55_P2004930** | Hddc2 | 0.113220143 | -0.000250157 | 0.226690444 | 11.86051004 | 2.384792673 | 0.050379403 | 0.991835066 | -3.677605613 |
| **A_51_P159711** | Car7 | -0.177994417 | -0.35638601 | 0.000397176 | 8.989813349 | -2.384740505 | 0.050383174 | 0.991835066 | -3.677651716 |
| **A_55_P2008692** | Eef1a1 | -0.220357095 | -0.441256152 | 0.000541963 | 15.70105658 | -2.384197932 | 0.050422413 | 0.991835066 | -3.678131228 |
| **A_66_P127458** | Zfp652 | 0.13594904 | -0.000388866 | 0.272286946 | 11.40145536 | 2.383244828 | 0.050491418 | 0.991835066 | -3.678973634 |
| **A_55_P2130930** | Nhp2 | 0.217875595 | -0.000649018 | 0.436400208 | 13.23606249 | 2.382963331 | 0.050511817 | 0.991835066 | -3.679222455 |
| **A_51_P127756** | Ddx20 | 0.137266111 | -0.000501314 | 0.275033536 | 10.23657947 | 2.381364744 | 0.050627822 | 0.991835066 | -3.683281983 |
| **A_55_P1959763** | Dach1 | -0.202285249 | -0.405319856 | 0.000749357 | 10.21916354 | -2.3812406 | 0.050636842 | 0.991835066 | -3.680745401 |
| **A_55_P2049266** | Mettl6 | 0.123884005 | -0.000482246 | 0.248250256 | 9.641998734 | 2.380794032 | 0.050669303 | 0.991835066 | -3.681140233 |
| **A_55_P2106834** | Ndufs3 | -0.164636633 | -0.329966741 | 0.000693476 | 15.20331804 | -2.380036716 | 0.050724401 | 0.991835066 | -3.681809861 |
| **A_55_P1990525** | Ube2v1 | 0.147208237 | -0.00062518 | 0.295041653 | 13.95978435 | 2.379954367 | 0.050730396 | 0.991835066 | -3.681882679 |
| **A_51_P368713** | Tmbim4 | -0.210439478 | -0.421773994 | 0.000895039 | 11.01152289 | -2.379939475 | 0.050731481 | 0.991835066 | -3.681895848 |
| **A_55_P1975341** | Nasp | 0.217592798 | -0.000995039 | 0.436180634 | 12.3114896 | 2.379181953 | 0.050786664 | 0.991835066 | -3.682565727 |
| **A_52_P536494** | Mycn | 0.359552512 | -0.001654245 | 0.72075927 | 10.11212131 | 2.379115866 | 0.050791481 | 0.991835066 | -3.682624171 |
| **A_55_P2065303** | NA | -0.142829801 | -0.286398175 | 0.000738572 | 15.40235918 | -2.377766389 | 0.050889949 | 0.991835066 | -3.683817682 |
| **A_55_P2168664** | Atp5k | -0.167874181 | -0.336699607 | 0.000951244 | 16.33149504 | -2.376595041 | 0.050975579 | 0.991835066 | -3.684853809 |
| **A_51_P217498** | Slc2a4 | -0.162689825 | -0.326313549 | 0.000933898 | 12.98188183 | -2.376420303 | 0.050988366 | 0.991835066 | -3.685008388 |
| **A_55_P2052116** | NA | -0.148565109 | -0.297992419 | 0.0008622 | 10.59087078 | -2.376271081 | 0.050999288 | 0.991835066 | -3.685140397 |
| **A_51_P307316** | Syde1 | 0.141053649 | -0.000873738 | 0.282981035 | 10.30719973 | 2.37534804 | 0.051066903 | 0.991835066 | -3.685957017 |
| **A_55_P2036526** | Mbd2 | -0.148886208 | -0.298698491 | 0.000926075 | 9.748664859 | -2.375287477 | 0.051071342 | 0.991835066 | -3.686010601 |
| **A_55_P2135025** | NA | 0.142494763 | -0.000909935 | 0.285899461 | 10.78618107 | 2.374896324 | 0.051100026 | 0.991835066 | -3.686356686 |
| **A_55_P2158152** | NA | -0.097886817 | -0.196403391 | 0.000629757 | 15.52876116 | -2.374783575 | 0.051108297 | 0.991835066 | -3.686456448 |
| **A_51_P130427** | Psmd3 | 0.094514031 | -0.000619083 | 0.189647145 | 14.35312553 | 2.374508374 | 0.05112849 | 0.991835066 | -3.686699954 |
| **A_55_P2148002** | 2610005L0 | 0.22293577 | -0.001551569 | 0.44742311 | 9.168116868 | 2.373542632 | 0.05119942 | 0.991835066 | -3.687554536 |
| **A_55_P2109454** | Gtf3c5 | 0.124362323 | -0.000886458 | 0.249611103 | 11.0918535 | 2.373145964 | 0.051228583 | 0.991835066 | -3.687905575 |
| **A_51_P228171** | Cenpp | -0.234381883 | -0.470460108 | 0.001696343 | 10.41659959 | -2.372887991 | 0.051247558 | 0.991835066 | -3.688133882 |
| **A_55_P1955841** | Ctsa | 0.12922641 | -0.001146045 | 0.259598864 | 14.20196111 | 2.369051867 | 0.05153058 | 0.991835066 | -3.691529699 |
| **A_55_P1981830** | NA | 0.661556978 | -0.005979287 | 1.329093243 | 10.47895476 | 2.368653438 | 0.051560067 | 0.991835066 | -3.691882486 |
| **A_51_P281593** | Plpp5 | 0.166596528 | -0.00153322 | 0.334726276 | 9.89189712 | 2.368266204 | 0.051588743 | 0.991835066 | -3.692225376 |
| **A_51_P491976** | Fbxw17 | 0.184910496 | -0.001821203 | 0.371642194 | 10.46094846 | 2.366751426 | 0.051701073 | 0.991835066 | -3.693566843 |
| **A_55_P2134852** | Cnot2 | -0.114023707 | -0.229240225 | 0.001192811 | 10.84336817 | -2.365318032 | 0.051807601 | 0.991835066 | -3.694836459 |
| **A_55_P2076906** | NA | -0.154111809 | -0.309883826 | 0.001660207 | 12.48791377 | -2.364588698 | 0.05186189 | 0.991835066 | -3.695482543 |
| **A_55_P2026380** | Pdcd2l | -0.123269809 | -0.247965102 | 0.001425483 | 11.05248345 | -2.362739258 | 0.051999819 | 0.991835066 | -3.697121128 |
| **A_52_P167511** | NA | -0.175680455 | -0.353396414 | 0.002035504 | 8.916266429 | -2.362686781 | 0.052003738 | 0.991835066 | -3.697167627 |
| **A_66_P139460** | Mia3 | 0.110230136 | -0.001315887 | 0.221776159 | 8.745977862 | 2.361866716 | 0.052065024 | 0.991835066 | -3.697894315 |

| | | | | | | | | | |
|---|---|---|---|---|---|---|---|---|---|
| **A_55_P2015989** | Rpl22 | 0.160145395 | -0.002021277 | 0.322312066 | 15.42291297 | 2.360271616 | 0.052184441 | 0.991835066 | -3.699307987 |
| **A_51_P331328** | Gpihbp1 | -0.255976977 | -0.51519434 | 0.003240387 | 12.40739875 | -2.360184472 | 0.052190974 | 0.991835066 | -3.699385228 |
| **A_55_P1963483** | Vps16 | 0.14145483 | -0.001793533 | 0.284703192 | 11.70679123 | 2.360137175 | 0.052194519 | 0.991835066 | -3.69942715 |
| **A_52_P615952** | Qk | -0.127554231 | -0.256755322 | 0.00164686 | 11.06481932 | -2.359596922 | 0.052235037 | 0.991835066 | -3.699906021 |
| **A_51_P147373** | Dpp3 | 0.125883052 | -0.001741824 | 0.253507929 | 11.92608334 | 2.357442246 | 0.052396957 | 0.991835066 | -3.701816195 |
| **A_52_P434052** | Wdr37 | -0.1249912 | -0.251810993 | 0.001828594 | 9.030397454 | -2.355599899 | 0.052535815 | 0.991835066 | -3.70344986 |
| **A_51_P315904** | Gadd45g | 0.242811363 | -0.003730998 | 0.489353724 | 12.55535941 | 2.353892297 | 0.052664855 | 0.991835066 | -3.704964355 |
| **A_66_P116804** | Smchd1 | 0.169080455 | -0.002640513 | 0.340801422 | 10.75134905 | 2.353310393 | 0.052708903 | 0.991835066 | -3.705480522 |
| **A_52_P572284** | Laptm4a | -0.11939845 | -0.24070902 | 0.001912121 | 11.88368795 | -2.35238919 | 0.052778712 | 0.991835066 | -3.706297728 |
| **A_55_P2125208** | Arid5a | 0.198765208 | -0.003239665 | 0.40077008 | 10.71032331 | 2.351731058 | 0.052828643 | 0.991835066 | -3.706881616 |
| **A_55_P2430472** | Dido1 | 0.196344227 | -0.003237351 | 0.395925805 | 9.194243218 | 2.351293357 | 0.052861878 | 0.991835066 | -3.707269964 |
| **A_55_P2033932** | Zzz3 | 0.22309304 | -0.003773827 | 0.449959907 | 9.927880578 | 2.350304219 | 0.052937062 | 0.991835066 | -3.708147643 |
| **A_55_P2048769** | Zfp454 | 0.114177324 | -0.00193436 | 0.230289009 | 8.883564375 | 2.35024462 | 0.052941596 | 0.991835066 | -3.70820053 |
| **A_55_P1952256** | Ccnb1 | -0.171273809 | -0.345466501 | 0.002918883 | 8.53316836 | -2.350012418 | 0.052959263 | 0.991835066 | -3.708406582 |
| **A_52_P283802** | Tapt1 | -0.110127031 | -0.222242186 | 0.001988124 | 9.090473586 | -2.347679148 | 0.053137127 | 0.991835066 | -3.710477396 |
| **A_55_P2070277** | Mfn1 | -0.186833111 | -0.377256189 | 0.003589967 | 11.73129294 | -2.345002971 | 0.053341887 | 0.991835066 | -3.712853225 |
| **A_55_P2032081** | Dbp | -0.188732848 | -0.381126542 | 0.003660845 | 11.05540192 | -2.344583985 | 0.053374018 | 0.991835066 | -3.713225254 |
| **A_55_P2162965** | Ptms | -0.142641081 | -0.288063247 | 0.002781086 | 14.2474068 | -2.34435373 | 0.053391684 | 0.991835066 | -3.713429711 |
| **A_52_P288873** | Chmp4b | 0.116126658 | -0.00226722 | 0.234520537 | 13.0343707 | 2.344292575 | 0.053396377 | 0.991835066 | -3.713484014 |
| **A_52_P621588** | Ifnlr1 | -0.101073547 | -0.204166508 | 0.002019414 | 8.842428456 | -2.34324461 | 0.053476864 | 0.991835066 | -3.714414638 |
| **A_52_P96748** | Ttc30b | -0.155154714 | -0.313411447 | 0.003102019 | 9.264022228 | -2.343213771 | 0.053479235 | 0.991835066 | -3.714442025 |
| **A_55_P2049256** | Mapkapk5 | 0.112183196 | -0.002298444 | 0.226664836 | 10.65014924 | 2.342076628 | 0.053566716 | 0.991835066 | -3.71545197 |
| **A_52_P135707** | Creb3 | -0.098994638 | -0.200027502 | 0.002038226 | 10.99684669 | -2.341844966 | 0.053584556 | 0.991835066 | -3.715657734 |
| **A_51_P151484** | Atp1b1 | 0.21544075 | -0.004468415 | 0.435349916 | 13.7986619 | 2.341497263 | 0.053611343 | 0.991835066 | -3.715966578 |
| **A_55_P2042958** | Rad50 | 0.112493199 | -0.002337165 | 0.227323564 | 9.795569271 | 2.341416408 | 0.053617574 | 0.991835066 | -3.716038398 |
| **A_55_P2019585** | Klhl17 | 0.184071484 | -0.003828289 | 0.371971256 | 10.10273281 | 2.341366453 | 0.053621425 | 0.991835066 | -3.716082772 |
| **A_55_P2022404** | Gtf2h5 | -0.107912928 | -0.218091376 | 0.00226552 | 11.73116379 | -2.340916692 | 0.053656102 | 0.991835066 | -3.716482291 |
| **A_55_P2098935** | Sh2d3c | 0.140952338 | -0.00297166 | 0.284876336 | 11.45159079 | 2.340713185 | 0.0536718 | 0.991835066 | -3.716663071 |
| **A_51_P479808** | Mapk6 | 0.138515817 | -0.002925688 | 0.279957322 | 8.801674757 | 2.340623886 | 0.05367869 | 0.991835066 | -3.716742399 |
| **A_51_P244824** | Dapp1 | -0.146160848 | -0.295419994 | 0.003098297 | 8.655055677 | -2.340449301 | 0.053692163 | 0.991835066 | -3.716897492 |
| **A_52_P467389** | Slc15a3 | -0.09811586 | -0.198321154 | 0.002089435 | 7.884992839 | -2.340225339 | 0.053709451 | 0.991835066 | -3.717096454 |
| **A_55_P2148935** | NA | 0.67929704 | -0.014752411 | 1.37334649 | 11.84801524 | 2.339259703 | 0.053784057 | 0.991835066 | -3.717954356 |
| **A_55_P1984481** | Mfn2 | -0.114573222 | -0.231697584 | 0.00255114 | 10.9756283 | -2.338002761 | 0.05388133 | 0.991835066 | -3.719071203 |
| **A_55_P1962699** | NA | 0.535510339 | -0.011965484 | 1.082986161 | 11.91574617 | 2.337825262 | 0.05389508 | 0.991835066 | -3.719228931 |
| **A_55_P2016119** | Sgcb | -0.117201338 | -0.237033194 | 0.002630517 | 10.78786458 | -2.337595804 | 0.053912862 | 0.991835066 | -3.719432836 |
| **A_51_P109369** | Fbxo32 | -0.111664364 | -0.225845527 | 0.002516799 | 8.895695369 | -2.337379686 | 0.053929615 | 0.991835066 | -3.724418883 |
| **A_52_P325237** | Rgs6 | -0.144206353 | -0.291668781 | 0.003256074 | 8.712497517 | -2.337287558 | 0.053936758 | 0.991835066 | -3.719706762 |
| **A_66_P103271** | Irf2bp2 | -0.11654515 | -0.235734969 | 0.00264467 | 12.74461298 | -2.337029392 | 0.053956781 | 0.991835066 | -3.71993619 |
| **A_55_P2027836** | Tnfrsf10b | 0.18367491 | -0.004219045 | 0.371568864 | 9.20328041 | 2.336394422 | 0.054006059 | 0.991835066 | -3.72050051 |
| **A_55_P2112922** | Impdh2 | 0.212734938 | -0.004967348 | 0.430437224 | 12.76590198 | 2.335527382 | 0.054073423 | 0.991835066 | -3.721271142 |
| **A_55_P1977548** | Mapk1 | -0.128175025 | -0.259416632 | 0.003066581 | 10.88137563 | -2.334215811 | 0.054175487 | 0.991835066 | -3.722437017 |
| **A_55_P1998442** | Sema3f | 0.246995364 | -0.005966186 | 0.499956914 | 10.65240487 | 2.333691372 | 0.054216354 | 0.991835066 | -3.722903247 |
| **A_52_P650108** | Uri1 | 0.141307155 | -0.003456943 | 0.286071253 | 9.244256128 | 2.332987525 | 0.05427125 | 0.991835066 | -3.723529014 |
| **A_51_P387302** | Zmym2 | -0.109757268 | -0.222226489 | 0.002711953 | 10.2295451 | -2.332430612 | 0.054314727 | 0.991835066 | -3.72402418 |
| **A_52_P95740** | Smim19 | -0.163220729 | -0.330474788 | 0.00403333 | 10.65541975 | -2.332425488 | 0.054315127 | 0.991835066 | -3.724028737 |
| **A_52_P502267** | Epc2 | 0.182985605 | -0.004628836 | 0.370600047 | 9.867063067 | 2.331094042 | 0.054419215 | 0.991835066 | -3.725212687 |
| **A_52_P629895** | Adh1 | 0.29318106 | -0.007419365 | 0.593781486 | 8.487402384 | 2.331070735 | 0.054421039 | 0.991835066 | -3.725233414 |
| **A_55_P1980180** | Tardbp | 0.130691005 | -0.003325926 | 0.264707935 | 13.58959925 | 2.330747146 | 0.054446368 | 0.991835066 | -3.725521183 |

| | | | | | | | | | |
|---|---|---|---|---|---|---|---|---|---|
| **A_52_P681310** | Plaur | 0.118148488 | -0.003007875 | 0.23930485 | 9.024098148 | 2.330725209 | 0.054448086 | 0.991835066 | -3.725540692 |
| **A_55_P2098880** | Galnt10 | 0.176970653 | -0.004611112 | 0.358552417 | 11.08576294 | 2.329368255 | 0.054554437 | 0.991835066 | -3.726747552 |
| **A_55_P2028193** | Mgat1 | 0.12999499 | -0.003430105 | 0.263420085 | 13.06334672 | 2.328617865 | 0.05461334 | 0.991835066 | -3.727415019 |
| **A_55_P2183172** | Rdx | 0.158945722 | -0.004363391 | 0.322254834 | 12.23466835 | 2.326202708 | 0.054803365 | 0.991835066 | -3.729563654 |
| **A_55_P2123673** | Trnp1 | -0.125855996 | -0.255167839 | 0.003455847 | 9.474675588 | -2.326187625 | 0.054804554 | 0.991835066 | -3.729577074 |
| **A_55_P1995447** | NA | -0.177629353 | -0.360138373 | 0.004879667 | 13.13779484 | -2.326159726 | 0.054806753 | 0.991835066 | -3.729601897 |
| **A_52_P891775** | Cdr2l | 0.148141657 | -0.004073669 | 0.300356983 | 10.62241449 | 2.326097681 | 0.054811644 | 0.991835066 | -3.729657104 |
| **A_55_P2012206** | Ndufb11 | -0.129222994 | -0.262023575 | 0.003577586 | 15.85187434 | -2.325674648 | 0.054845003 | 0.991835066 | -3.730033515 |
| **A_52_P620748** | Ncl | 0.203973683 | -0.005682664 | 0.413630029 | 9.230382779 | 2.325279997 | 0.054876143 | 0.991835066 | -3.73038469 |
| **A_55_P2093291** | Coil | 0.12851611 | -0.003599566 | 0.260631786 | 9.574009043 | 2.324943236 | 0.05490273 | 0.991835066 | -3.730684363 |
| **A_55_P2047620** | NA | 0.370536797 | -0.01043587 | 0.751509463 | 13.25564921 | 2.324591563 | 0.054930507 | 0.991835066 | -3.730997318 |
| **A_66_P125212** | Mapk1ip1l | 0.219480888 | -0.006304368 | 0.445266144 | 11.2148089 | 2.32332659 | 0.055030544 | 0.991835066 | -3.732123118 |
| **A_55_P1959606** | Tom1l1 | 0.131112475 | -0.00377816 | 0.26600311 | 11.28596397 | 2.323118421 | 0.055047024 | 0.991835066 | -3.7323084 |
| **A_52_P305851** | 2410002F2 | 0.134057679 | -0.003882183 | 0.27199754 | 9.574533334 | 2.322795847 | 0.055072572 | 0.991835066 | -3.732595515 |
| **A_55_P1985882** | Fastkd1 | -0.118816582 | -0.241080484 | 0.00344732 | 9.795672891 | -2.322672264 | 0.055082363 | 0.991835066 | -3.732705516 |
| **A_51_P251205** | Psd | 0.388382791 | -0.011647372 | 0.788412954 | 9.181680356 | 2.320472207 | 0.055256961 | 0.991835066 | -3.73466403 |
| **A_55_P2172299** | Timm17b | -0.109657601 | -0.222618519 | 0.003303318 | 13.04318127 | -2.320169201 | 0.055281052 | 0.991835066 | -3.734933805 |
| **A_51_P103850** | Fbn2 | 0.175170219 | -0.005326427 | 0.355666866 | 10.7309997 | 2.319531465 | 0.055331792 | 0.991835066 | -3.735501629 |
| **A_66_P136834** | Usp7 | 0.117214526 | -0.003575387 | 0.238004439 | 8.647210645 | 2.319315878 | 0.055348956 | 0.991835066 | -3.735693591 |
| **A_55_P2077816** | Hmgxb3 | 0.142622497 | -0.00438318 | 0.289628174 | 11.36210727 | 2.318798772 | 0.055390146 | 0.991835066 | -3.736154049 |
| **A_51_P151038** | Pmpcb | -0.161679753 | -0.32844448 | 0.005084974 | 13.34811982 | -2.317184275 | 0.055518953 | 0.991835066 | -3.737591843 |
| **A_55_P2038046** | NA | -0.106526717 | -0.216454514 | 0.003401079 | 10.45935087 | -2.316115182 | 0.055604415 | 0.991835066 | -3.738544063 |
| **A_52_P199614** | Wdfy4 | -0.221654246 | -0.450660852 | 0.007352359 | 8.571123864 | -2.313327812 | 0.05582787 | 0.991835066 | -3.741027226 |
| **A_51_P497692** | Mageh1 | 0.1124104 | -0.003759189 | 0.228579989 | 9.603623719 | 2.31272063 | 0.055876667 | 0.991835066 | -3.74156824 |
| **A_52_P700056** | Wfdc17 | -0.116286057 | -0.236692164 | 0.004120049 | 7.853300731 | -2.308278809 | 0.056234978 | 0.991835066 | -3.745527056 |
| **A_52_P343627** | Rbp7 | -0.20849566 | -0.424388906 | 0.007397586 | 11.95571381 | -2.308166302 | 0.056244084 | 0.991835066 | -3.745627354 |
| **A_55_P2067181** | Usp21 | -0.11555806 | -0.235237681 | 0.004121561 | 10.03395371 | -2.307752175 | 0.056277616 | 0.991835066 | -3.745996547 |
| **A_55_P2023235** | Fen1 | 0.205204544 | -0.00732476 | 0.417733848 | 12.17310043 | 2.307689031 | 0.056282731 | 0.991835066 | -3.746052841 |
| **A_55_P2056236** | Pdlim5 | -0.222754897 | -0.45354151 | 0.008031717 | 11.93341944 | -2.306884111 | 0.05634797 | 0.991835066 | -3.746770475 |
| **A_55_P1967478** | Hbb-y | 1.262255944 | -0.045810101 | 2.570321989 | 9.791619161 | 2.30635887 | 0.056390583 | 0.991835066 | -3.747238792 |
| **A_55_P2108655** | Mtch2 | -0.105923261 | -0.215701426 | 0.003854905 | 13.9811681 | -2.306133824 | 0.056408851 | 0.991835066 | -3.747439455 |
| **A_51_P246653** | Clec7a | -0.172330102 | -0.350960416 | 0.006300211 | 8.097085543 | -2.305765402 | 0.05643877 | 0.991835066 | -3.747767971 |
| **A_55_P1961039** | Pnpla2 | -0.200583733 | -0.408606567 | 0.0074391 | 12.29977259 | -2.304590856 | 0.056534264 | 0.991835066 | -3.748815379 |
| **A_51_P346923** | P3h1 | 0.116980119 | -0.004364548 | 0.238324786 | 8.447384911 | 2.304095604 | 0.056574579 | 0.991835066 | -3.749257061 |
| **A_55_P2327027** | Ano6 | 0.140044596 | -0.005252175 | 0.285341367 | 8.317063308 | 2.303666063 | 0.056609569 | 0.991835066 | -3.749640158 |
| **A_65_P19780** | Klf9 | -0.463893279 | -0.945201818 | 0.01741526 | 9.492838683 | -2.303581841 | 0.056616432 | 0.991835066 | -3.749715275 |
| **A_52_P429774** | Cnst | -0.132140559 | -0.269322299 | 0.005041181 | 9.798616506 | -2.30223135 | 0.056726599 | 0.991835066 | -3.750919866 |
| **A_51_P149714** | Ms4a6d | -0.105607046 | -0.215265918 | 0.004051825 | 8.046819803 | -2.301750551 | 0.056765873 | 0.991835066 | -3.751348763 |
| **A_52_P263251** | 3110040N1 | -0.114688359 | -0.233812992 | 0.004436273 | 9.103880347 | -2.301054467 | 0.056822783 | 0.991835066 | -3.751969742 |
| **A_55_P2043466** | Eif5 | 0.125527193 | -0.004887705 | 0.255942092 | 12.56157894 | 2.300486783 | 0.056869238 | 0.991835066 | -3.752476207 |
| **A_55_P2107207** | Gpr137b-p | -0.133161554 | -0.271529778 | 0.00520667 | 8.847999374 | -2.300125964 | 0.056898785 | 0.991835066 | -3.752798131 |
| **A_55_P2074473** | Gtf2f1 | 0.109155518 | -0.004312903 | 0.22262394 | 13.08868537 | 2.299216221 | 0.056973352 | 0.991835066 | -3.75360986 |
| **A_51_P284988** | Nrbf2 | -0.115155096 | -0.234879257 | 0.004569064 | 10.05430876 | -2.298849267 | 0.057003458 | 0.991835066 | -3.753937301 |
| **A_51_P360492** | Mcm6 | 0.157992961 | -0.006279366 | 0.322265288 | 14.06597453 | 2.298700875 | 0.057015637 | 0.991835066 | -3.754069717 |
| **A_55_P2056186** | Siva1 | 0.166241327 | -0.006626213 | 0.339108866 | 12.70942687 | 2.298447977 | 0.057036399 | 0.991835066 | -3.754295393 |
| **A_51_P119544** | Aga | -0.102951026 | -0.210037728 | 0.004135677 | 9.228471274 | -2.297757872 | 0.057093095 | 0.991835066 | -3.759386712 |
| **A_55_P1969710** | NA | 0.180737158 | -0.00726483 | 0.368739146 | 13.9441691 | 2.297704308 | 0.057097498 | 0.991835066 | -3.754959049 |
| **A_55_P2098801** | Gm15393 | -0.128655247 | -0.26250946 | 0.005198967 | 16.87534526 | -2.297230575 | 0.057136454 | 0.991835066 | -3.755381838 |

| | | | | | | | | | |
|---|---|---|---|---|---|---|---|---|---|
| **A_55_P2047489** | Calm1 | 0.120699056 | -0.004947883 | 0.246345996 | 14.64676952 | 2.295942954 | 0.057242475 | 0.991835066 | -3.756531095 |
| **A_55_P2169714** | Gcn1 | 0.100746353 | -0.004132053 | 0.20562476 | 13.09961927 | 2.295896923 | 0.057246269 | 0.991835066 | -3.756572182 |
| **A_55_P1967335** | NA | 0.178925549 | -0.007432718 | 0.365283815 | 10.36835593 | 2.294736524 | 0.057341993 | 0.991835066 | -3.757608021 |
| **A_51_P174016** | Slc39a7 | 0.172580437 | -0.007288712 | 0.352449585 | 13.44499056 | 2.293211007 | 0.057468087 | 0.991835066 | -3.75896997 |
| **A_55_P2047598** | Arhgdia | 0.110599443 | -0.00469429 | 0.225893177 | 14.72501003 | 2.292748249 | 0.057506393 | 0.991835066 | -3.759383152 |
| **A_55_P2058992** | Vamp4 | -0.11074141 | -0.226307003 | 0.004824184 | 8.732250797 | -2.29029078 | 0.057710254 | 0.991835066 | -3.76157767 |
| **A_51_P451106** | Brca2 | 0.190675519 | -0.008314658 | 0.389665696 | 10.1157583 | 2.290194839 | 0.057718227 | 0.991835066 | -3.761663356 |
| **A_55_P2102936** | Pnrc1 | -0.284361468 | -0.581206386 | 0.012483451 | 10.28274782 | -2.289550674 | 0.057771794 | 0.991835066 | -3.76223869 |
| **A_51_P424256** | Gm6211 | 0.121772545 | -0.005385996 | 0.248931085 | 12.62650001 | 2.288827055 | 0.057832029 | 0.991835066 | -3.762885032 |
| **A_52_P653966** | Atf7ip | 0.099242567 | -0.004397673 | 0.202882807 | 10.31024416 | 2.288646476 | 0.057847071 | 0.991835066 | -3.763046333 |
| **A_55_P2205156** | Aamdc | -0.222962543 | -0.455880796 | 0.00995571 | 9.589756606 | -2.287902527 | 0.057909081 | 0.991835066 | -3.763710895 |
| **A_55_P2075268** | Tmem216 | 0.107642027 | -0.00488425 | 0.220168305 | 9.702401369 | 2.286320173 | 0.058041201 | 0.991835066 | -3.765124556 |
| **A_55_P2020458** | NA | -0.136451173 | -0.279176103 | 0.006273758 | 11.69945322 | -2.285001899 | 0.058151507 | 0.991835066 | -3.766302462 |
| **A_55_P2021065** | E130308A1 | 0.185067785 | -0.008665815 | 0.378801385 | 9.260166484 | 2.283152977 | 0.058306576 | 0.991835066 | -3.767954768 |
| **A_55_P2089310** | 2900097C1 | 0.125055465 | -0.005891433 | 0.256002364 | 8.887540502 | 2.282530523 | 0.058358876 | 0.991835066 | -3.768511098 |
| **A_55_P2040888** | Dhx15 | -0.107597612 | -0.220265815 | 0.005070591 | 9.544331633 | -2.282497952 | 0.058361614 | 0.991835066 | -3.76854021 |
| **A_51_P405606** | Ndrg1 | -0.134280098 | -0.274903768 | 0.006343572 | 11.69973034 | -2.282245471 | 0.058382843 | 0.991835066 | -3.76876588 |
| **A_55_P2047778** | Krtap4-8 | -0.136703036 | -0.279871268 | 0.006465195 | 10.03353209 | -2.282131324 | 0.058392443 | 0.991835066 | -3.768867908 |
| **A_55_P2072468** | Sirt6 | 0.179730404 | -0.008537508 | 0.367998316 | 9.678668628 | 2.281678115 | 0.058430576 | 0.991835066 | -3.769273009 |
| **A_55_P2040115** | Cdv3 | 0.207645228 | -0.009956293 | 0.425246748 | 13.40167829 | 2.280705237 | 0.058512518 | 0.991835066 | -3.770142678 |
| **A_51_P518592** | Atxn7 | 0.151402555 | -0.007278278 | 0.310083387 | 9.138863112 | 2.280435877 | 0.058535226 | 0.991835066 | -3.770383476 |
| **A_55_P2092776** | Apoo | -0.18049021 | -0.369730963 | 0.008750542 | 12.37454796 | -2.279544725 | 0.058610418 | 0.991835066 | -3.771180182 |
| **A_55_P2046198** | Slc25a25 | 0.097112939 | -0.004725424 | 0.198951302 | 8.51205025 | 2.27916003 | 0.058642908 | 0.991835066 | -3.771524127 |
| **A_55_P2360661** | Cbx5 | 0.160848159 | -0.007922709 | 0.329619026 | 10.69311704 | 2.277863753 | 0.058752521 | 0.991835066 | -3.772683188 |
| **A_55_P1983754** | Pcp4l1 | -0.416640822 | -0.853837209 | 0.020555565 | 12.21846436 | -2.277688807 | 0.05876733 | 0.991835066 | -3.772839626 |
| **A_55_P2049211** | Pisd-ps1 | 0.136754259 | -0.006759948 | 0.280268465 | 10.96125148 | 2.277482759 | 0.058784777 | 0.991835066 | -3.77302388 |
| **A_52_P246889** | Chmp3 | -0.162305883 | -0.332635787 | 0.008024021 | 8.596754347 | -2.277469095 | 0.058785934 | 0.991835066 | -3.773036099 |
| **A_51_P292566** | Xpot | 0.142085953 | -0.007041843 | 0.291213748 | 10.32143525 | 2.277202633 | 0.058808505 | 0.991835066 | -3.773274382 |
| **A_52_P58283** | Kcna5 | -0.134624448 | -0.275927017 | 0.00667812 | 9.645770433 | -2.277104769 | 0.058816797 | 0.991835066 | -3.773361898 |
| **A_55_P1959061** | NA | -0.154028402 | -0.315703931 | 0.007647127 | 11.42512427 | -2.277013748 | 0.058824511 | 0.991835066 | -3.773443296 |
| **A_55_P2091985** | Gpr157 | -0.169491451 | -0.347442615 | 0.008459713 | 10.91953893 | -2.276439413 | 0.058873204 | 0.991835066 | -3.773956924 |
| **A_51_P470715** | Cish | -0.53761634 | -1.102235484 | 0.027002805 | 12.25745485 | -2.275757551 | 0.058931067 | 0.991835066 | -3.77456675 |
| **A_55_P1970229** | Csnk1d | 0.142519981 | -0.007207361 | 0.292247323 | 13.04597463 | 2.275012433 | 0.058994365 | 0.991835066 | -3.775233196 |
| **A_52_P211418** | G2e3 | -0.111144368 | -0.227931822 | 0.005643086 | 8.390785019 | -2.274575745 | 0.059031494 | 0.991835066 | -3.775623799 |
| **A_55_P1998506** | Aplf | 0.103139199 | -0.005263034 | 0.211541432 | 8.350053201 | 2.274021979 | 0.059078612 | 0.991835066 | -3.776119148 |
| **A_51_P246224** | Tram2 | 0.142570582 | -0.007301379 | 0.292442544 | 9.302873273 | 2.273624102 | 0.059112489 | 0.991835066 | -3.776475069 |
| **A_51_P470311** | Cby1 | -0.264986982 | -0.543594412 | 0.013620448 | 9.244558053 | -2.273217427 | 0.059147136 | 0.991835066 | -3.776838874 |
| **A_55_P2126951** | Zfp467 | -0.230042826 | -0.472046794 | 0.011961142 | 10.40432539 | -2.271932058 | 0.059256781 | 0.991835066 | -3.777988839 |
| **A_51_P160439** | Crybg3 | -0.126963553 | -0.26055101 | 0.006623903 | 12.45916778 | -2.271551141 | 0.059289313 | 0.991835066 | -3.778329656 |
| **A_55_P2003251** | Magel2 | -0.181576564 | -0.3726275 | 0.009474371 | 8.665822447 | -2.271536707 | 0.059290547 | 0.991835066 | -3.778342571 |
| **A_51_P448032** | Cox6b1 | -0.15422243 | -0.316608772 | 0.008163911 | 13.8699059 | -2.269902366 | 0.05943034 | 0.991835066 | -3.779805003 |
| **A_51_P120717** | Lmnb1 | 0.344642917 | -0.01831103 | 0.707596863 | 11.7781748 | 2.269483169 | 0.059466251 | 0.991835066 | -3.784160713 |
| **A_55_P1957583** | Gps2 | -0.119430724 | -0.245215294 | 0.006353847 | 9.93289157 | -2.269330897 | 0.0594793 | 0.991835066 | -3.780316416 |
| **A_66_P124788** | Tbcb | -0.181801081 | -0.373278605 | 0.009676443 | 11.39995438 | -2.269278461 | 0.059483795 | 0.991835066 | -3.780363343 |
| **A_55_P2029106** | Bmf | -0.215080653 | -0.441613716 | 0.011452411 | 9.503926182 | -2.269231898 | 0.059487786 | 0.991835066 | -3.780405014 |
| **A_52_P8227** | Lypla1 | -0.185814135 | -0.38158155 | 0.00995328 | 10.24099824 | -2.268545389 | 0.059546665 | 0.991835066 | -3.78101942 |
| **A_66_P125209** | Kif2c | -0.118001557 | -0.242332683 | 0.006329569 | 8.469779094 | -2.268386231 | 0.059560324 | 0.991835066 | -3.781161867 |
| **A_51_P365019** | Gclc | 0.163719081 | -0.008799139 | 0.336237301 | 9.488776145 | 2.268158825 | 0.059579845 | 0.991835066 | -3.781365401 |

| | | | | | | | | | |
|---|---|---|---|---|---|---|---|---|---|
| **A_52_P504068** | Cdk8 | 0.166760216 | -0.008973572 | 0.342494004 | 8.548934969 | 2.268017027 | 0.059592021 | 0.991835066 | -3.781492315 |
| **A_52_P354823** | Irf8 | -0.141261059 | -0.290189438 | 0.007667319 | 9.818910732 | -2.267013625 | 0.059678253 | 0.991835066 | -3.782390447 |
| **A_51_P265348** | Ldb3 | -0.292603802 | -0.601139815 | 0.01593221 | 11.44382691 | -2.266643582 | 0.059710086 | 0.991835066 | -3.782721689 |
| **A_55_P1980566** | Ndufb11 | -0.185740207 | -0.38162112 | 0.010140706 | 16.1264987 | -2.266328909 | 0.05973717 | 0.991835066 | -3.783003376 |
| **A_52_P490770** | Zfp26 | 0.120951357 | -0.006610412 | 0.248513126 | 8.638772769 | 2.266205787 | 0.059747771 | 0.991835066 | -3.783113593 |
| **A_55_P2048409** | Apool | -0.112548882 | -0.231321643 | 0.006223879 | 11.41726469 | -2.264818824 | 0.059867317 | 0.991835066 | -3.784355278 |
| **A_51_P159814** | Smpd4 | 0.128175154 | -0.007105115 | 0.263455424 | 9.785826929 | 2.264532316 | 0.059892042 | 0.991835066 | -3.784611796 |
| **A_51_P162098** | Ndufaf4 | 0.159024811 | -0.008838029 | 0.32688765 | 11.56914405 | 2.264224339 | 0.059918632 | 0.991835066 | -3.784887543 |
| **A_55_P1966332** | Plac9a | -0.275482347 | -0.566302112 | 0.015337418 | 12.31993964 | -2.264013368 | 0.059936854 | 0.991835066 | -3.78507644 |
| **A_55_P1959639** | NA | 0.188216479 | -0.010525372 | 0.38695833 | 13.24117768 | 2.263484092 | 0.059982592 | 0.991835066 | -3.785550355 |
| **A_52_P658122** | Ets2 | -0.125632798 | -0.258296153 | 0.007030557 | 12.14546784 | -2.263399363 | 0.059989917 | 0.991835066 | -3.785626224 |
| **A_55_P2046398** | Eif4a1 | 0.126555448 | -0.007083996 | 0.260194892 | 14.74644268 | 2.263368754 | 0.059992564 | 0.991835066 | -3.785653632 |
| **A_51_P397548** | Mtrf1l | 0.106189614 | -0.005998121 | 0.218377349 | 9.840654041 | 2.262277067 | 0.060087033 | 0.991835066 | -3.786631216 |
| **A_52_P515282** | 1110051M2 | -0.136164787 | -0.280142214 | 0.00781264 | 9.108026066 | -2.260369999 | 0.060252424 | 0.991835066 | -3.788339193 |
| **A_55_P2095727** | Pml | -0.188225437 | -0.38730904 | 0.010858167 | 10.94763136 | -2.259706074 | 0.060310112 | 0.991835066 | -3.788933877 |
| **A_55_P1998825** | Rpl9 | 0.135707215 | -0.007855085 | 0.279269514 | 16.94059557 | 2.259288357 | 0.060346437 | 0.991835066 | -3.78930805 |
| **A_66_P116807** | Selenot | -0.138701658 | -0.285448634 | 0.008045318 | 12.32704125 | -2.25902806 | 0.060369083 | 0.991835066 | -3.78954122 |
| **A_55_P2078269** | NA | -0.196105539 | -0.4036535 | 0.011442422 | 13.55313234 | -2.258294219 | 0.060432976 | 0.991835066 | -3.790198612 |
| **A_55_P2428968** | Lrrc2 | -0.126725639 | -0.260852505 | 0.007401227 | 9.753295854 | -2.258176295 | 0.060443249 | 0.991835066 | -3.790304255 |
| **A_55_P2032079** | Dbp | -0.209826208 | -0.431966476 | 0.01231406 | 11.20415656 | -2.257571809 | 0.06049594 | 0.991835066 | -3.790845807 |
| **A_52_P276792** | Foxp1 | 0.109128236 | -0.006421565 | 0.224678037 | 10.64096757 | 2.25723651 | 0.060525187 | 0.991835066 | -3.79114621 |
| **A_55_P1974855** | Zfp868 | 0.115429433 | -0.006843748 | 0.237702615 | 8.935472716 | 2.256287737 | 0.060608024 | 0.991835066 | -3.791996292 |
| **A_51_P268167** | Tasor | 0.114685364 | -0.006819303 | 0.236190031 | 9.111097181 | 2.255922475 | 0.060639945 | 0.991835066 | -3.792323579 |
| **A_51_P148355** | Zfp710 | 0.152427495 | -0.009080909 | 0.313935898 | 11.01461621 | 2.255679125 | 0.060661222 | 0.991835066 | -3.792541634 |
| **A_55_P2074718** | Fh1 | -0.103480029 | -0.213125795 | 0.006165738 | 14.85650617 | -2.255660871 | 0.060662818 | 0.991835066 | -3.792557991 |
| **A_52_P140356** | Calm3 | 0.109869051 | -0.006558853 | 0.226296954 | 11.27338897 | 2.255420001 | 0.060683886 | 0.991835066 | -3.79277383 |
| **A_55_P2055123** | Frrs1 | 0.139787513 | -0.008351207 | 0.287926232 | 8.531632788 | 2.255323905 | 0.060692294 | 0.991835066 | -3.792859941 |
| **A_55_P2099560** | Apobec1 | -0.170690594 | -0.351642661 | 0.010261473 | 8.519095241 | -2.254525615 | 0.06076218 | 0.991835066 | -3.793575312 |
| **A_51_P286737** | Ccl2 | 0.281814129 | -0.017097681 | 0.58072594 | 9.146745463 | 2.253350873 | 0.060865173 | 0.991835066 | -3.794628128 |
| **A_52_P17146** | Bmpr1a | -0.178064657 | -0.367024659 | 0.010895345 | 11.81005636 | -2.252251967 | 0.060961678 | 0.991835066 | -3.795613079 |
| **A_55_P1974645** | Entpd2 | -0.210509454 | -0.433952706 | 0.012933798 | 8.799002483 | -2.251715373 | 0.061008857 | 0.991835066 | -3.796094065 |
| **A_66_P139618** | Stfa2 | -0.565863568 | -1.166499784 | 0.034772648 | 9.44674687 | -2.2516939 | 0.061010746 | 0.991835066 | -3.796113313 |
| **A_51_P472217** | Sapcd2 | 0.104296404 | -0.006444606 | 0.215037414 | 10.53457099 | 2.250971459 | 0.061074327 | 0.991835066 | -3.796760924 |
| **A_65_P05358** | NA | -0.176545264 | -0.364040406 | 0.010949878 | 14.6395198 | -2.250480141 | 0.061117606 | 0.991835066 | -3.797201376 |
| **A_51_P165165** | Hibadh | -0.111068608 | -0.229049459 | 0.006912243 | 12.72763561 | -2.250033261 | 0.061156998 | 0.991835066 | -3.797602007 |
| **A_55_P2145696** | P3h4 | 0.167499421 | -0.010432139 | 0.345430982 | 11.63078507 | 2.249932325 | 0.061165899 | 0.991835066 | -3.7976925 |
| **A_55_P2108404** | NA | 0.125226753 | -0.007838265 | 0.258291772 | 11.40851069 | 2.249273963 | 0.061223988 | 0.991835066 | -3.79828276 |
| **A_55_P2042312** | NA | 0.110209118 | -0.006932614 | 0.22735085 | 15.94736107 | 2.248614557 | 0.061282226 | 0.991835066 | -3.798873992 |
| **A_51_P155763** | NA | 0.126579374 | -0.00796363 | 0.261122378 | 8.627141543 | 2.248593523 | 0.061284084 | 0.991835066 | -3.798892852 |
| **A_55_P2166743** | Sp3os | -0.130543581 | -0.26930497 | 0.008217808 | 12.05525305 | -2.248516175 | 0.061290919 | 0.991835066 | -3.798962206 |
| **A_51_P200477** | Hspa4 | 0.171866539 | -0.010947354 | 0.354680431 | 12.07626646 | 2.246938924 | 0.061430469 | 0.991835066 | -3.800376542 |
| **A_55_P1995126** | NA | -0.096795664 | -0.199772586 | 0.006181259 | 16.8124081 | -2.246596754 | 0.061460785 | 0.991835066 | -3.800683395 |
| **A_51_P301636** | Kazn | 0.14023897 | -0.008998967 | 0.289476907 | 8.835775072 | 2.245942379 | 0.061518806 | 0.991835066 | -3.801270255 |
| **A_51_P475076** | Vbp1 | -0.105063235 | -0.216904272 | 0.006777801 | 10.58022094 | -2.245219054 | 0.061583006 | 0.991835066 | -3.801918991 |
| **A_51_P350453** | Pdk4 | -0.236433108 | -0.488232403 | 0.015366186 | 10.65080743 | -2.244207015 | 0.061672945 | 0.991835066 | -3.802826739 |
| **A_55_P2080333** | NA | -0.115342253 | -0.238191831 | 0.007507324 | 8.312884262 | -2.244005391 | 0.061690879 | 0.991835066 | -3.803007595 |
| **A_52_P121491** | Msrb3 | 0.163600568 | -0.010659234 | 0.337860371 | 10.24755637 | 2.243864989 | 0.06170337 | 0.991835066 | -3.803133537 |
| **A_55_P2032960** | Brap | 0.106808199 | -0.007004348 | 0.220620746 | 11.33622462 | 2.242970609 | 0.061783005 | 0.991835066 | -3.803935842 |

| | | | | | | | | | |
|---|---|---|---|---|---|---|---|---|---|
| **A_55_P2156583** | Cdk5r1 | -0.161983939 | -0.334741644 | 0.010773766 | 8.731898776 | -2.241009317 | 0.061958001 | 0.991835066 | -3.805695442 |
| **A_51_P361201** | Atp6v0d1 | -0.329851765 | -0.681747084 | 0.022043554 | 10.57911739 | -2.240342691 | 0.062017596 | 0.991835066 | -3.806293584 |
| **A_51_P243930** | Qrsl1 | -0.134512725 | -0.278040289 | 0.009014838 | 9.538201106 | -2.239944156 | 0.062053252 | 0.991835066 | -3.806651192 |
| **A_51_P215475** | Ptprb | -0.133259533 | -0.27546214 | 0.008943073 | 11.22202897 | -2.239751644 | 0.062070483 | 0.991835066 | -3.806823939 |
| **A_52_P86158** | Mtx2 | -0.171816061 | -0.355215594 | 0.011583473 | 12.50974247 | -2.239106048 | 0.062128304 | 0.991835066 | -3.807403273 |
| **A_55_P2003158** | Wwp1 | 0.110000483 | -0.007428804 | 0.22742977 | 8.257594198 | 2.238861873 | 0.062150187 | 0.991835066 | -3.807622396 |
| **A_66_P139333** | Atf6 | -0.12327747 | -0.254909884 | 0.008354944 | 8.745301999 | -2.238360318 | 0.062195161 | 0.991835066 | -3.808072505 |
| **A_55_P1987245** | Ywhaz | 0.175898295 | -0.01193382 | 0.36373041 | 9.561638078 | 2.23821042 | 0.062208609 | 0.991835066 | -3.808207032 |
| **A_51_P324934** | Mcm3 | 0.183713543 | -0.012508696 | 0.379935782 | 12.22900719 | 2.23770112 | 0.062254321 | 0.991835066 | -3.808664118 |
| **A_51_P240723** | 1700022A2 | -0.253432292 | -0.52415312 | 0.017288535 | 8.881643388 | -2.237429784 | 0.062278689 | 0.991835066 | -3.808907644 |
| **A_52_P156190** | Ednra | -0.147954 | -0.306006116 | 0.010098116 | 12.20487046 | -2.237358239 | 0.062285116 | 0.991835066 | -3.808971857 |
| **A_55_P2162573** | Art3 | -0.308304184 | -0.637889473 | 0.021281106 | 9.414642143 | -2.235737086 | 0.062430924 | 0.991835066 | -3.810426983 |
| **A_55_P2045837** | NA | 0.119580388 | -0.008267007 | 0.247427784 | 8.64363644 | 2.235513042 | 0.062451102 | 0.991835066 | -3.810628097 |
| **A_55_P2145800** | NA | 0.161554505 | -0.011314855 | 0.334423866 | 8.299130182 | 2.233624589 | 0.062621447 | 0.991835066 | -3.812323436 |
| **A_52_P286098** | Epb41l4b | 0.115502579 | -0.00812932 | 0.239134478 | 8.276730502 | 2.232905137 | 0.062686468 | 0.991835066 | -3.812969389 |
| **A_51_P382688** | Zmynd19 | 0.165891851 | -0.011806706 | 0.343590408 | 9.885602821 | 2.231260535 | 0.062835358 | 0.991835066 | -3.814446125 |
| **A_51_P186487** | NA | -0.171983731 | -0.356285147 | 0.012317684 | 8.978291396 | -2.230323339 | 0.062920366 | 0.991835066 | -3.815287753 |
| **A_51_P128648** | Uqcrc2 | -0.136980383 | -0.283858407 | 0.00989764 | 15.53321733 | -2.229003185 | 0.063040309 | 0.991835066 | -3.818200872 |
| **A_55_P1992571** | NA | 0.209393977 | -0.015169264 | 0.433957218 | 12.67694363 | 2.22861295 | 0.063075808 | 0.991835066 | -3.8168239 |
| **A_52_P590474** | Ccnl1 | 0.153831581 | -0.011232322 | 0.318895483 | 8.847001207 | 2.227422104 | 0.063184264 | 0.991835066 | -3.817893562 |
| **A_55_P2161535** | Ddx23 | 0.143823436 | -0.010510918 | 0.29815779 | 8.367510848 | 2.227287013 | 0.063196579 | 0.991835066 | -3.818014913 |
| **A_55_P2159830** | Yaf2 | 0.14508142 | -0.010642324 | 0.300805163 | 9.395677422 | 2.22672248 | 0.06324807 | 0.991835066 | -3.81852204 |
| **A_55_P2157102** | NA | -0.103623788 | -0.214862417 | 0.00761484 | 13.57947254 | -2.226450129 | 0.063272927 | 0.991835066 | -3.818766705 |
| **A_55_P1959953** | Helz2 | -0.195688146 | -0.405791999 | 0.014415706 | 9.735527029 | -2.226074204 | 0.063307252 | 0.991835066 | -3.819104424 |
| **A_52_P430262** | Rbm15 | 0.176266697 | -0.013046138 | 0.365579532 | 9.89528346 | 2.225355193 | 0.063372958 | 0.991835066 | -3.819750391 |
| **A_55_P2180869** | Ocel1 | -0.206440493 | -0.428164601 | 0.015283616 | 9.920216162 | -2.225312983 | 0.063376817 | 0.991835066 | -3.819788313 |
| **A_52_P127024** | Tnks | 0.097628882 | -0.007246499 | 0.202504263 | 9.096969261 | 2.224917418 | 0.063412997 | 0.991835066 | -3.82014371 |
| **A_55_P2106525** | Nmnat3 | 0.155124005 | -0.011603811 | 0.321851822 | 10.75205431 | 2.22371988 | 0.063522656 | 0.991835066 | -3.821219714 |
| **A_55_P2021187** | Malat1 | 0.39170978 | -0.029431832 | 0.812851391 | 8.816638141 | 2.223030351 | 0.063585884 | 0.991835066 | -3.821839314 |
| **A_55_P2154595** | Mrpl39 | -0.124637546 | -0.25864953 | 0.009374438 | 12.24812354 | -2.222871645 | 0.063600446 | 0.991835066 | -3.821981929 |
| **A_52_P632093** | Gtf3c3 | -0.101286294 | -0.210225546 | 0.007652957 | 9.437762579 | -2.222160526 | 0.063665735 | 0.991835066 | -3.822620974 |
| **A_55_P2080248** | Cyp2j8 | -0.107445433 | -0.223020314 | 0.008129447 | 8.458162236 | -2.221946732 | 0.063685378 | 0.991835066 | -3.822813106 |
| **A_51_P234308** | Rpa2 | 0.123339069 | -0.009394118 | 0.256072256 | 9.882023555 | 2.220906496 | 0.063781037 | 0.991835066 | -3.823747994 |
| **A_51_P427828** | Mlip | -0.133738854 | -0.277667016 | 0.010189308 | 11.17197028 | -2.220858813 | 0.063785426 | 0.991835066 | -3.82379085 |
| **A_51_P189959** | Oxsr1 | -0.181362178 | -0.376598724 | 0.013874368 | 9.128365284 | -2.220213505 | 0.063844846 | 0.991835066 | -3.824370848 |
| **A_51_P228658** | Isyna1 | 0.144525743 | -0.011186972 | 0.300238458 | 12.07474677 | 2.218351006 | 0.06401666 | 0.991835066 | -3.82604502 |
| **A_55_P1999972** | NA | 0.196332613 | -0.015197502 | 0.407862728 | 8.289889681 | 2.218346455 | 0.064017081 | 0.991835066 | -3.82604911 |
| **A_51_P469285** | Nrp1 | -0.150533391 | -0.312742557 | 0.011675774 | 10.45262007 | -2.218025777 | 0.064046711 | 0.991835066 | -3.826337388 |
| **A_66_P121196** | Katnb1 | 0.106975536 | -0.008303377 | 0.222254449 | 10.73751415 | 2.217909044 | 0.0640575 | 0.991835066 | -3.826442329 |
| **A_55_P1980746** | Dram2 | -0.13221536 | -0.274725343 | 0.010294624 | 10.14882526 | -2.217408736 | 0.064103763 | 0.991835066 | -3.826892108 |
| **A_55_P1958464** | NA | 0.407137969 | -0.031719805 | 0.845995743 | 10.12081283 | 2.217312688 | 0.064112648 | 0.991835066 | -3.826978457 |
| **A_52_P51078** | Ctsh | -0.172642706 | -0.358940915 | 0.013655503 | 11.73202311 | -2.214872275 | 0.064338831 | 0.991835066 | -3.829172672 |
| **A_51_P386562** | Ebf2 | -0.112952705 | -0.234840423 | 0.008935013 | 8.849129183 | -2.214857663 | 0.064340187 | 0.991835066 | -3.829185811 |
| **A_55_P2035812** | Rnf208 | 0.243863099 | -0.019322402 | 0.507048599 | 8.526635551 | 2.214589624 | 0.06436508 | 0.991835066 | -3.829426836 |
| **A_66_P112855** | AI413582 | -0.099629906 | -0.207155169 | 0.007895357 | 10.94104914 | -2.214564523 | 0.064367411 | 0.991835066 | -3.829449407 |
| **A_55_P2010152** | Sell | -0.107949908 | -0.224460734 | 0.008560918 | 7.756211223 | -2.214446166 | 0.064378406 | 0.991835066 | -3.829555837 |
| **A_55_P2095196** | Akr1b10 | -0.180499325 | -0.375471194 | 0.014472544 | 12.60239904 | -2.212650194 | 0.064545483 | 0.991835066 | -3.831170954 |
| **A_52_P359257** | NA | -0.142099337 | -0.295699391 | 0.011500717 | 8.557567757 | -2.211107289 | 0.064689368 | 0.991835066 | -3.832558672 |

| | | | | | | | | | |
|---|---|---|---|---|---|---|---|---|---|
| **A_55_P2100963** | Hmces | 0.138095718 | -0.011198477 | 0.287389913 | 10.7831808 | 2.21078457 | 0.064719505 | 0.991835066 | -3.832848953 |
| **A_55_P2161585** | Coq10a | -0.227888099 | -0.474282864 | 0.018506667 | 9.512959393 | -2.210544694 | 0.064741915 | 0.991835066 | -3.833064723 |
| **A_55_P2074226** | Fbxo8 | -0.149202852 | -0.310591625 | 0.012185921 | 8.690382482 | -2.20959632 | 0.064830592 | 0.991835066 | -3.83391783 |
| **A_51_P324303** | Mylip | -0.11889546 | -0.247525966 | 0.009735046 | 9.43067169 | -2.209176556 | 0.06486988 | 0.991835066 | -3.834295448 |
| **A_55_P2185548** | Cdk8 | 0.19283273 | -0.015946686 | 0.401612146 | 10.62064321 | 2.207507579 | 0.065026332 | 0.991835066 | -3.835796975 |
| **A_55_P2175752** | Mfap1b | 0.21558127 | -0.017890309 | 0.449052849 | 9.506473744 | 2.206917699 | 0.06508172 | 0.991835066 | -3.836327719 |
| **A_55_P2023772** | Fam107b | -0.161256433 | -0.336087917 | 0.013575051 | 10.67478077 | -2.204481903 | 0.06531094 | 0.991835066 | -3.838519581 |
| **A_55_P2156101** | Skp1a | -0.105348476 | -0.219616629 | 0.008919676 | 13.44602281 | -2.203495594 | 0.065403989 | 0.991835066 | -3.839407234 |
| **A_55_P2033041** | Sirpb1b | -0.089401724 | -0.186374849 | 0.007571402 | 7.796967611 | -2.203452193 | 0.065408086 | 0.991835066 | -3.839446294 |
| **A_55_P2012734** | Fndc10 | 0.159260562 | -0.013550665 | 0.332071789 | 9.545214623 | 2.202649637 | 0.065483905 | 0.991835066 | -3.840168626 |
| **A_55_P1994229** | Hrc | -0.191030731 | -0.398332289 | 0.016270826 | 13.89695552 | -2.202469008 | 0.065500981 | 0.991835066 | -3.840331205 |
| **A_51_P220896** | Chmp2b | -0.129072209 | -0.269211416 | 0.011066997 | 10.05569153 | -2.201315148 | 0.065610172 | 0.991835066 | -3.841369816 |
| **A_55_P2072035** | Thy1 | -0.189841386 | -0.395981656 | 0.016298884 | 10.63493391 | -2.201086897 | 0.065631794 | 0.991835066 | -3.84157528 |
| **A_66_P104980** | Tceal5 | 0.110707171 | -0.009513607 | 0.230927949 | 12.1592513 | 2.200925547 | 0.065647082 | 0.991835066 | -3.841720523 |
| **A_51_P180724** | Mlh1 | 0.116850003 | -0.010146174 | 0.243846179 | 10.25895028 | 2.199111315 | 0.065819237 | 0.991835066 | -3.843353781 |
| **A_51_P214796** | Suz12 | 0.127377784 | -0.011102573 | 0.26585814 | 10.44089377 | 2.198440151 | 0.065883041 | 0.991835066 | -3.843958051 |
| **A_55_P2160331** | Bicd2 | -0.111764455 | -0.233278836 | 0.009749926 | 11.89660609 | -2.198290883 | 0.065897239 | 0.991835066 | -3.844092446 |
| **A_66_P116998** | Tro | 0.209960454 | -0.018333495 | 0.438254404 | 11.42230226 | 2.198124236 | 0.065913095 | 0.991835066 | -3.844242491 |
| **A_55_P2004811** | NA | 0.144765664 | -0.012690802 | 0.302222131 | 14.7680643 | 2.197425698 | 0.065979598 | 0.991835066 | -3.844871454 |
| **A_66_P102607** | Btrc | 0.098014117 | -0.008593246 | 0.20462148 | 8.894620676 | 2.197407296 | 0.065981351 | 0.991835066 | -3.844888024 |
| **A_55_P2015670** | Itga6 | 0.18815517 | -0.016551683 | 0.392862022 | 14.07329798 | 2.196812073 | 0.066038073 | 0.991835066 | -3.845423989 |
| **A_51_P509098** | Asah1 | -0.108663691 | -0.226928659 | 0.009601277 | 9.240398397 | -2.196025935 | 0.066113065 | 0.991835066 | -3.8461319 |
| **A_55_P1984886** | Hcst | -0.115850645 | -0.241951878 | 0.010250588 | 7.982642966 | -2.195777117 | 0.066136819 | 0.991835066 | -3.846355967 |
| **A_55_P2002819** | Rabgap1l | -0.09080951 | -0.189654589 | 0.008035569 | 10.63097629 | -2.195762752 | 0.06613819 | 0.991835066 | -3.846368903 |
| **A_55_P1973553** | Bbs12 | 0.134559172 | -0.011921747 | 0.281040092 | 8.877475796 | 2.195540146 | 0.066159449 | 0.991835066 | -3.84656937 |
| **A_55_P1958314** | Mospd1 | -0.194276865 | -0.405799054 | 0.017245323 | 11.96997347 | -2.195200982 | 0.066191853 | 0.991835066 | -3.846874808 |
| **A_55_P1955237** | NA | 0.108825141 | -0.009699361 | 0.227349644 | 16.5483524 | 2.194472945 | 0.066261463 | 0.991835066 | -3.847530477 |
| **A_51_P235117** | Zc3h11a | 0.166148698 | -0.014811723 | 0.347109119 | 13.1456796 | 2.194433766 | 0.066265212 | 0.991835066 | -3.847565763 |
| **A_52_P160518** | Sfmbt1 | 0.130385472 | -0.011704951 | 0.272475894 | 8.744261111 | 2.193176224 | 0.066385632 | 0.991835066 | -3.84869839 |
| **A_51_P447189** | 1700030C1 | 0.105800607 | -0.009509163 | 0.221110378 | 8.0277639 | 2.192962408 | 0.066406129 | 0.991835066 | -3.848890977 |
| **A_55_P2044439** | Mfng | 0.146384923 | -0.013161751 | 0.305931597 | 10.45117381 | 2.192894435 | 0.066412646 | 0.991835066 | -3.848952202 |
| **A_55_P2045549** | Dpy19l1 | 0.124758695 | -0.011217567 | 0.260734957 | 9.847459653 | 2.192890048 | 0.066413067 | 0.991835066 | -3.848956154 |
| **A_55_P2141585** | NA | -0.128209014 | -0.267980861 | 0.011562833 | 11.17978412 | -2.192340402 | 0.066465792 | 0.991835066 | -3.849451246 |
| **A_55_P2174967** | NA | 0.157514007 | -0.014208036 | 0.329236049 | 9.537560054 | 2.19231152 | 0.066468564 | 0.991835066 | -3.849477261 |
| **A_52_P582907** | Entpd4 | 0.120564276 | -0.010974758 | 0.25210331 | 8.65465248 | 2.190650656 | 0.066628148 | 0.991835066 | -3.850973404 |
| **A_55_P2082057** | NA | 0.112331125 | -0.01023333 | 0.234895581 | 16.45373746 | 2.19050728 | 0.066641942 | 0.991835066 | -3.851102569 |
| **A_51_P403273** | Tax1bp1 | -0.130754021 | -0.273423617 | 0.011915574 | 10.49700596 | -2.190447032 | 0.06664774 | 0.991835066 | -3.851156846 |
| **A_55_P2129187** | NA | 0.125014737 | -0.011401307 | 0.261430782 | 15.16283413 | 2.190306501 | 0.066661265 | 0.991835066 | -3.85128345 |
| **A_55_P1969058** | Epha7 | 0.099062553 | -0.009044535 | 0.20716964 | 8.469360029 | 2.190102691 | 0.066680884 | 0.991835066 | -3.851467063 |
| **A_55_P2094520** | Fam171a1 | -0.119569009 | -0.250067369 | 0.010929351 | 11.38030479 | -2.189892049 | 0.066701168 | 0.991835066 | -3.851656835 |
| **A_55_P2050426** | Ankrd50 | 0.159807648 | -0.014614085 | 0.334229381 | 10.1781735 | 2.18980829 | 0.066709235 | 0.991835066 | -3.851732296 |
| **A_52_P641892** | Hacd3 | 0.19118399 | -0.017487111 | 0.399855091 | 12.91734704 | 2.189769211 | 0.066712999 | 0.991835066 | -3.851767504 |
| **A_55_P2060021** | Otub2 | 0.190091156 | -0.01744869 | 0.397631003 | 10.05027241 | 2.189119923 | 0.066775572 | 0.991835066 | -3.852352484 |
| **A_52_P176619** | Atat1 | 0.133433062 | -0.012265584 | 0.279131709 | 8.941335499 | 2.188855379 | 0.066801084 | 0.991835066 | -3.852590834 |
| **A_55_P1954896** | Rpl17 | 0.124996032 | -0.011527417 | 0.261519481 | 14.85329934 | 2.188255899 | 0.066858932 | 0.991835066 | -3.853130975 |
| **A_51_P286865** | Wdr33 | -0.101895742 | -0.213211909 | 0.009420424 | 10.57746843 | -2.187796528 | 0.066903295 | 0.991835066 | -3.853544891 |
| **A_51_P183630** | Bcor | -0.092781028 | -0.194163964 | 0.008601908 | 10.42637635 | -2.187275297 | 0.066953667 | 0.991835066 | -3.854014563 |
| **A_55_P2055134** | NA | -0.200570712 | -0.41975363 | 0.018612206 | 11.09406543 | -2.187106567 | 0.066969982 | 0.991835066 | -3.854166606 |

| | | | | | | | | | |
|---|---|---|---|---|---|---|---|---|---|
| **A_55_P2050192** | NA | -0.181941635 | -0.38077055 | 0.016887279 | 15.05717225 | -2.187064971 | 0.066974004 | 0.991835066 | -3.854204089 |
| **A_55_P2149020** | Rpl35 | 0.114344547 | -0.010648098 | 0.239337192 | 17.43471938 | 2.186452928 | 0.06703322 | 0.991835066 | -3.85475562 |
| **A_55_P2079967** | Map3k5 | -0.160804593 | -0.336597041 | 0.014987854 | 9.536957585 | -2.186287992 | 0.067049187 | 0.991835066 | -3.854904254 |
| **A_51_P215489** | Slc37a1 | -0.127788897 | -0.26753103 | 0.011953236 | 8.346077808 | -2.185621157 | 0.06711378 | 0.991835066 | -3.855505195 |
| **A_55_P2067960** | Cplx2 | 0.154980799 | -0.014553704 | 0.324515302 | 13.76068282 | 2.184886754 | 0.067184991 | 0.991835066 | -3.856167061 |
| **A_55_P2013765** | Raly | 0.178759499 | -0.016795157 | 0.374314155 | 13.57057523 | 2.184792019 | 0.067194183 | 0.991835066 | -3.856252441 |
| **A_55_P1980125** | Srsf3 | -0.114139795 | -0.239055266 | 0.010775677 | 12.06215128 | -2.183886117 | 0.067282141 | 0.991835066 | -3.857068921 |
| **A_51_P205779** | Cd5l | -0.088887602 | -0.186175782 | 0.008400578 | 7.937436872 | -2.183686266 | 0.067301561 | 0.991835066 | -3.857249052 |
| **A_51_P401907** | Cstdc4 | -0.434086837 | -0.909198449 | 0.041024775 | 8.954905093 | -2.183685571 | 0.067301629 | 0.991835066 | -3.857249678 |
| **A_55_P2100305** | Lrba | -0.123855657 | -0.259517617 | 0.011806302 | 10.83604004 | -2.182061036 | 0.067459701 | 0.991835066 | -3.858714007 |
| **A_51_P337918** | Aldh4a1 | -0.117267039 | -0.245736998 | 0.01120292 | 11.20394936 | -2.181642097 | 0.067500526 | 0.991835066 | -3.859091659 |
| **A_52_P175376** | Tfcp2l1 | 0.15969517 | -0.015314488 | 0.334704829 | 8.191616764 | 2.180915792 | 0.067571363 | 0.991835066 | -3.859746412 |
| **A_51_P130757** | Acer3 | -0.094725406 | -0.198555361 | 0.00910455 | 8.486747832 | -2.180484163 | 0.067613495 | 0.991835066 | -3.860135535 |
| **A_55_P1989328** | Rps12 | 0.162565998 | -0.0156979 | 0.340829897 | 16.93904705 | 2.179593219 | 0.067700547 | 0.991835066 | -3.860938779 |
| **A_65_P19286** | Alg10b | 0.111866026 | -0.010815046 | 0.234547097 | 8.743302881 | 2.179364041 | 0.067722958 | 0.991835066 | -3.861145405 |
| **A_51_P517870** | Nfatc1 | 0.109094902 | -0.010580551 | 0.228770356 | 9.965247092 | 2.178755554 | 0.067782496 | 0.991835066 | -3.861694033 |
| **A_55_P2343648** | 1700120C1 | -0.222699124 | -0.467057505 | 0.021659257 | 9.012592936 | -2.178213278 | 0.067835601 | 0.991835066 | -3.862182982 |
| **A_51_P221651** | Coq8a | -0.418192075 | -0.877165757 | 0.040781607 | 10.50559726 | -2.177695468 | 0.06788635 | 0.991835066 | -3.86264989 |
| **A_55_P1953400** | Rbfox1 | -0.223464087 | -0.468825767 | 0.021897593 | 9.671242742 | -2.176757921 | 0.067978332 | 0.991835066 | -3.863495315 |
| **A_55_P2112622** | Smad4 | -0.118935884 | -0.249550763 | 0.011678995 | 12.18162642 | -2.176353235 | 0.068018075 | 0.991835066 | -3.863860253 |
| **A_55_P2080562** | Arhgap26 | -0.142086466 | -0.298126623 | 0.01395369 | 8.869335222 | -2.176333604 | 0.068020003 | 0.991835066 | -3.863877957 |
| **A_51_P401668** | Laptm5 | -0.130620612 | -0.274165191 | 0.012923968 | 8.31767758 | -2.174873725 | 0.068163572 | 0.991835066 | -3.865194539 |
| **A_51_P400752** | NA | -0.150401167 | -0.315688719 | 0.014886385 | 9.265627444 | -2.174804334 | 0.068170404 | 0.991835066 | -3.865257121 |
| **A_55_P1978726** | NA | 0.24768606 | -0.024634663 | 0.520006783 | 9.868333866 | 2.173852162 | 0.068264218 | 0.991835066 | -3.866115906 |
| **A_55_P2083654** | Prmt2 | -0.110998667 | -0.233051711 | 0.011054377 | 12.10276395 | -2.173593265 | 0.068289749 | 0.991835066 | -3.86634942 |
| **A_55_P2052563** | Id1 | -0.143586885 | -0.301513453 | 0.014339683 | 12.4953636 | -2.173044941 | 0.068343854 | 0.991835066 | -3.866844001 |
| **A_55_P2025611** | Psme2 | -0.220877443 | -0.463850964 | 0.022096078 | 11.51300271 | -2.172708943 | 0.068377029 | 0.991835066 | -3.867147075 |
| **A_52_P66199** | Fbxo11 | -0.118820151 | -0.249528671 | 0.011888369 | 9.33576846 | -2.17267784 | 0.0683801 | 0.991835066 | -3.86717513 |
| **A_55_P2150048** | Sypl | -0.121185565 | -0.254536065 | 0.012164935 | 10.89735475 | -2.172027774 | 0.068444334 | 0.991835066 | -3.867761518 |
| **A_52_P106463** | Jak1 | -0.133855399 | -0.281168314 | 0.013457515 | 11.57504337 | -2.171721875 | 0.068474582 | 0.991835066 | -3.868037461 |
| **A_55_P1985623** | Abcc3 | -0.107562155 | -0.226038287 | 0.010913977 | 8.20242348 | -2.169890217 | 0.06865598 | 0.991835066 | -3.869689871 |
| **A_55_P2037313** | Gm12027 | -0.138101731 | -0.290227869 | 0.014024407 | 12.39624369 | -2.169723609 | 0.068672505 | 0.991835066 | -3.869840185 |
| **A_55_P1974288** | Rela | 0.110217108 | -0.011304546 | 0.231738763 | 13.00532112 | 2.167726418 | 0.068870899 | 0.991835066 | -3.871642181 |
| **A_51_P209518** | Galnt1 | 0.116836086 | -0.012001817 | 0.245673989 | 9.46642566 | 2.167417051 | 0.068901683 | 0.991835066 | -3.871921333 |
| **A_52_P182659** | Cs | -0.114170766 | -0.24007191 | 0.011730378 | 16.59532183 | -2.167376556 | 0.068905713 | 0.991835066 | -3.871957873 |
| **A_51_P389714** | Phlpp2 | 0.144291131 | -0.01483225 | 0.303414512 | 8.913304374 | 2.16727874 | 0.06891545 | 0.991835066 | -3.872046138 |
| **A_55_P2091506** | Srp54c | -0.116782854 | -0.24558236 | 0.012016652 | 12.17229598 | -2.167075395 | 0.068935695 | 0.991835066 | -3.872229628 |
| **A_55_P1956097** | Cyb5d2 | -0.115660365 | -0.243232017 | 0.011911286 | 9.706459194 | -2.166903218 | 0.068952842 | 0.991835066 | -3.872384995 |
| **A_65_P17866** | Vma21 | -0.127087684 | -0.267282923 | 0.013107556 | 8.909582176 | -2.166602948 | 0.068982756 | 0.991835066 | -3.872655954 |
| **A_55_P2111523** | Higd1b | -0.251194703 | -0.528325303 | 0.025935898 | 10.68700272 | -2.166382437 | 0.069004732 | 0.991835066 | -3.872854943 |
| **A_55_P2125006** | Gpatch1 | 0.158686781 | -0.016396984 | 0.333770547 | 9.977908215 | 2.1662272 | 0.069020208 | 0.991835066 | -3.872995031 |
| **A_55_P2169059** | Strada | 0.120086075 | -0.012413033 | 0.252585184 | 10.6755888 | 2.166151495 | 0.069027756 | 0.991835066 | -3.873063348 |
| **A_55_P2058923** | Bud23 | 0.141505954 | -0.014658052 | 0.29766996 | 10.62830977 | 2.165722986 | 0.069070497 | 0.991835066 | -3.873450049 |
| **A_55_P1962831** | Ssr1 | 0.156891992 | -0.016283594 | 0.330067578 | 12.4435345 | 2.165325764 | 0.06911014 | 0.991835066 | -3.873808525 |
| **A_51_P214747** | Parp12 | -0.087903816 | -0.184946682 | 0.009139051 | 8.559308185 | -2.164976784 | 0.069144988 | 0.991835066 | -3.874123473 |
| **A_55_P1955722** | Med15 | 0.095842773 | -0.009999036 | 0.201684582 | 10.38362093 | 2.164269048 | 0.069215715 | 0.991835066 | -3.874762212 |
| **A_55_P2139753** | Jade2 | -0.160706473 | -0.338201109 | 0.016788162 | 9.293191605 | -2.164000072 | 0.069242614 | 0.991835066 | -3.875004974 |
| **A_55_P2094901** | Phyhd1 | 0.226923393 | -0.023895552 | 0.477742337 | 16.48391602 | 2.162360323 | 0.069406827 | 0.991835066 | -3.876485006 |

| | | | | | | | | | |
|---|---|---|---|---|---|---|---|---|---|
| **A_51_P324228** | Satb1 | -0.22981689 | -0.484019492 | 0.024385713 | 10.04299685 | -2.160782645 | 0.069565195 | 0.991835066 | -3.877909163 |
| **A_55_P2031481** | Fam131a | -0.122065691 | -0.257121574 | 0.012990192 | 10.363282 | -2.160176505 | 0.069626136 | 0.991835066 | -3.878456359 |
| **A_55_P1982683** | Cib1 | -0.170831331 | -0.359864886 | 0.018202223 | 12.23227389 | -2.159920455 | 0.069651896 | 0.991835066 | -3.878687516 |
| **A_55_P1967677** | Cped1 | -0.122549831 | -0.258158606 | 0.013058944 | 7.978874228 | -2.159902046 | 0.069653748 | 0.991835066 | -3.878704135 |
| **A_52_P141161** | Hnrnpf | 0.151042372 | -0.016152824 | 0.318237569 | 9.539643054 | 2.159156565 | 0.069728804 | 0.991835066 | -3.879377164 |
| **A_55_P2095709** | NA | 0.092056588 | -0.009864222 | 0.193977398 | 15.55087256 | 2.158743992 | 0.069770377 | 0.991835066 | -3.879749653 |
| **A_55_P1966583** | Bpifc | -0.305006793 | -0.643168207 | 0.033154621 | 11.64600064 | -2.15573113 | 0.070074729 | 0.991835066 | -3.882470096 |
| **A_55_P2095345** | Rara | -0.090052259 | -0.189896466 | 0.009791948 | 12.88352363 | -2.155663027 | 0.070081624 | 0.991835066 | -3.882531596 |
| **A_51_P222467** | Abcg1 | -0.186749825 | -0.393883562 | 0.020383913 | 10.30194017 | -2.154857193 | 0.070163263 | 0.991835066 | -3.883259309 |
| **A_55_P1976764** | Polr3e | 0.211486049 | -0.023098572 | 0.44607067 | 9.625320976 | 2.154722368 | 0.070176932 | 0.991835066 | -3.883381068 |
| **A_55_P2081615** | Timeless | 0.119216889 | -0.013051232 | 0.251485011 | 8.358773935 | 2.154228323 | 0.070227041 | 0.991835066 | -3.883827241 |
| **A_55_P1973178** | Haus5 | 0.126471346 | -0.013858463 | 0.266801154 | 9.965984476 | 2.154027976 | 0.070247371 | 0.991835066 | -3.884008178 |
| **A_51_P459944** | Tcf21 | -0.243575337 | -0.513869112 | 0.026718437 | 9.471126918 | -2.153805103 | 0.070269995 | 0.991835066 | -3.884209462 |
| **A_55_P1969750** | Stim1 | -0.202632721 | -0.427502026 | 0.022236584 | 10.74088865 | -2.153716481 | 0.070278993 | 0.991835066 | -3.884289501 |
| **A_55_P2049429** | Phtf2 | -0.136619786 | -0.288236514 | 0.014996943 | 12.30724037 | -2.153652408 | 0.070285499 | 0.991835066 | -3.884347368 |
| **A_55_P2157551** | NA | 0.092074156 | -0.010140015 | 0.194288327 | 9.074765001 | 2.152959038 | 0.070355945 | 0.991835066 | -3.884973597 |
| **A_51_P234728** | Lpar6 | -0.113822519 | -0.240271157 | 0.012626118 | 11.44197766 | -2.151409945 | 0.070513589 | 0.991835066 | -3.886372786 |
| **A_55_P2020200** | Snx6 | -0.124001478 | -0.261823278 | 0.013820322 | 8.681318745 | -2.150394188 | 0.070617152 | 0.991835066 | -3.887290322 |
| **A_51_P263591** | Pank1 | -0.140538789 | -0.29674714 | 0.015669562 | 9.589604596 | -2.150310077 | 0.070625734 | 0.991835066 | -3.887366302 |
| **A_55_P1953341** | Wfdc2 | 0.138264951 | -0.015420528 | 0.291950431 | 7.969292379 | 2.150247256 | 0.070632145 | 0.991835066 | -3.887423051 |
| **A_52_P344376** | Eif4a2 | 0.124992376 | -0.013964093 | 0.263948845 | 9.191092948 | 2.149878346 | 0.070669804 | 0.991835066 | -3.887756306 |
| **A_55_P2163319** | Peg3 | 0.101653912 | -0.011356801 | 0.214664625 | 8.04935483 | 2.149876996 | 0.070669942 | 0.991835066 | -3.887757525 |
| **A_55_P2056260** | NA | 0.110695198 | -0.01239688 | 0.233787276 | 14.94652535 | 2.149353313 | 0.070723435 | 0.991835066 | -3.888230608 |
| **A_51_P432199** | Sap30 | -0.144105455 | -0.304349482 | 0.016138571 | 10.28595048 | -2.149352788 | 0.070723488 | 0.991835066 | -3.888231082 |
| **A_52_P422859** | Hspg2 | -0.113147551 | -0.238971674 | 0.012676572 | 13.92910586 | -2.149267034 | 0.070732252 | 0.991835066 | -3.888308551 |
| **A_55_P2109439** | Lsmem1 | -0.154737522 | -0.326812486 | 0.017337442 | 10.19898525 | -2.149250728 | 0.070733918 | 0.991835066 | -3.888323282 |
| **A_52_P219904** | Afmid | -0.149697728 | -0.316220231 | 0.016824774 | 8.911245634 | -2.148579429 | 0.07080256 | 0.991835066 | -3.888929742 |
| **A_55_P1958374** | NA | -0.143574133 | -0.303291644 | 0.016143378 | 12.10620574 | -2.148487349 | 0.070811981 | 0.991835066 | -3.88901293 |
| **A_55_P2107402** | NA | 0.250459752 | -0.028323155 | 0.52924266 | 8.572386293 | 2.147241714 | 0.070939545 | 0.991835066 | -3.890138326 |
| **A_55_P2361547** | 2610201A1 | 0.199016763 | -0.022605134 | 0.42063866 | 8.835975747 | 2.146278733 | 0.071038322 | 0.991835066 | -3.891008411 |
| **A_51_P108459** | Gpr65 | -0.106953952 | -0.226132847 | 0.012224944 | 7.950984815 | -2.144897829 | 0.071180209 | 0.991835066 | -3.897772788 |
| **A_55_P2057321** | Myh7b | -0.143440898 | -0.303348397 | 0.0164666 | 14.68370396 | -2.143943315 | 0.071278453 | 0.991835066 | -3.893118746 |
| **A_52_P679101** | Tjp2 | 0.118666925 | -0.013680323 | 0.251014173 | 11.706364 | 2.143008568 | 0.071374794 | 0.991835066 | -3.893963487 |
| **A_55_P2074796** | Cdkn2c | -0.125599184 | -0.265681424 | 0.014483055 | 10.05600152 | -2.142954133 | 0.071380408 | 0.991835066 | -3.894012681 |
| **A_66_P131708** | Znhit1 | -0.136678327 | -0.289147617 | 0.015790963 | 13.55839902 | -2.142527522 | 0.071424425 | 0.991835066 | -3.894398231 |
| **A_51_P466759** | Mau2 | -0.10050794 | -0.212649642 | 0.011633762 | 8.842833163 | -2.142112913 | 0.071467229 | 0.991835066 | -3.894772944 |
| **A_55_P1987156** | Crhr2 | -0.160489836 | -0.339557245 | 0.018577573 | 10.69690691 | -2.142101842 | 0.071468373 | 0.991835066 | -3.89478295 |
| **A_51_P237585** | Btnl9 | -0.160302189 | -0.339190059 | 0.018585681 | 9.096649342 | -2.141744659 | 0.07150527 | 0.991835066 | -3.895105769 |
| **A_55_P2134690** | Ankrd26 | 0.091499962 | -0.010617121 | 0.193617045 | 8.442731863 | 2.141566906 | 0.071523639 | 0.991835066 | -3.895266424 |
| **A_51_P317941** | Perp | -0.149925387 | -0.317294245 | 0.017443471 | 14.54165512 | -2.140965449 | 0.071585829 | 0.991835066 | -3.895810039 |
| **A_55_P2148400** | NA | 0.155557855 | -0.01812921 | 0.32924492 | 13.45714118 | 2.140590537 | 0.071624622 | 0.991835066 | -3.896148905 |
| **A_55_P2174935** | Mn1 | 0.095913307 | -0.011180128 | 0.203006742 | 9.457835548 | 2.140548873 | 0.071628935 | 0.991835066 | -3.896186563 |
| **A_55_P2171623** | Ptbp1 | 0.160070227 | -0.018677857 | 0.338818311 | 9.08353012 | 2.140317975 | 0.071652838 | 0.991835066 | -3.896395266 |
| **A_55_P2006662** | Zfp967 | 0.093795841 | -0.010969418 | 0.1985611 | 8.399179618 | 2.139811034 | 0.071705348 | 0.991835066 | -3.896853486 |
| **A_51_P138923** | Peg12 | 0.112849644 | -0.013200352 | 0.23889964 | 9.040324993 | 2.13976702 | 0.071709908 | 0.991835066 | -3.896893271 |
| **A_55_P1999309** | Zfp668 | 0.113611701 | -0.01330191 | 0.240525312 | 10.0668876 | 2.139557661 | 0.071731607 | 0.991835066 | -3.897082514 |
| **A_55_P2017789** | Il4ra | -0.114044553 | -0.241473902 | 0.013384795 | 8.078477995 | -2.139016915 | 0.071787681 | 0.991835066 | -3.897571313 |
| **A_52_P24160** | Mrpl51 | -0.135294889 | -0.286500221 | 0.015910443 | 14.86948757 | -2.138569731 | 0.071834086 | 0.991835066 | -3.897975549 |

| | | | | | | | | | |
|---|---|---|---|---|---|---|---|---|---|
| **A_55_P2083967** | Trmt5 | -0.112389893 | -0.238021852 | 0.013242066 | 9.12656199 | -2.138140582 | 0.071878649 | 0.991835066 | -3.898363493 |
| **A_55_P2035023** | Zfp653 | 0.19582709 | -0.023076721 | 0.414730901 | 9.571132188 | 2.138102786 | 0.071882575 | 0.991835066 | -3.898397661 |
| **A_55_P2153797** | Slc38a6 | 0.132885909 | -0.015696287 | 0.281468104 | 9.068197856 | 2.137574659 | 0.071937456 | 0.991835066 | -3.898875093 |
| **A_55_P2057065** | Gtf3c5 | 0.142324624 | -0.016822164 | 0.301471411 | 11.10709493 | 2.137427045 | 0.071952804 | 0.991835066 | -3.89900854 |
| **A_52_P325161** | Zbtb10 | -0.099933951 | -0.211724568 | 0.011856666 | 8.858193459 | -2.136568579 | 0.072042122 | 0.991835066 | -3.89978464 |
| **A_51_P155196** | Abtb2 | -0.105284108 | -0.223074092 | 0.012505876 | 11.61317997 | -2.136306648 | 0.072069397 | 0.991835066 | -3.900021448 |
| **A_55_P2131379** | Ybx1 | -0.133503341 | -0.283048769 | 0.016042087 | 11.90385262 | -2.133674301 | 0.072344081 | 0.991835066 | -3.902401508 |
| **A_51_P493016** | Cmya5 | -0.282937058 | -0.599883427 | 0.034009311 | 11.84646787 | -2.133600898 | 0.072351756 | 0.991835066 | -3.902467881 |
| **A_66_P116635** | Igf2os | 0.285993084 | -0.034601987 | 0.606588156 | 11.10141264 | 2.132101237 | 0.072508734 | 0.991835066 | -3.903823977 |
| **A_52_P376106** | Slc30a9 | -0.112676781 | -0.238991718 | 0.013638156 | 9.814338987 | -2.132008116 | 0.072518492 | 0.991835066 | -3.903908187 |
| **A_55_P2030183** | NA | -0.353589567 | -0.752233851 | 0.045054716 | 13.46259238 | -2.20333763 | 0.072552168 | 0.991835066 | -3.905189213 |
| **A_55_P1985001** | NA | 0.290537836 | -0.035369718 | 0.61644539 | 9.591152721 | 2.130675947 | 0.072658244 | 0.991835066 | -3.905112929 |
| **A_55_P2105542** | Pde2a | 0.11239843 | -0.013702677 | 0.238499538 | 12.14543142 | 2.130347627 | 0.072692729 | 0.991835066 | -3.905409857 |
| **A_51_P172251** | Ifrd2 | 0.154343677 | -0.018821292 | 0.327508646 | 11.17406268 | 2.130286108 | 0.072699192 | 0.991835066 | -3.905465494 |
| **A_55_P2051047** | Rps10 | 0.129696162 | -0.015827581 | 0.275219905 | 17.25451834 | 2.130111808 | 0.072717508 | 0.991835066 | -3.905623132 |
| **A_51_P286488** | Ier3 | -0.170502887 | -0.361892625 | 0.020886851 | 10.10997543 | -2.129228261 | 0.072810423 | 0.991835066 | -3.906422239 |
| **A_55_P2157616** | NA | 0.115701531 | -0.014198154 | 0.245601216 | 16.79268903 | 2.128825866 | 0.072852779 | 0.991835066 | -3.90678619 |
| **A_55_P2348676** | B130052P1 | 0.148730015 | -0.018278601 | 0.315738632 | 8.9510612 | 2.128476581 | 0.072889565 | 0.991835066 | -3.907102112 |
| **A_55_P2150908** | Atp5c1 | -0.096179869 | -0.204195606 | 0.011835869 | 13.92742591 | -2.128169805 | 0.072921889 | 0.991835066 | -3.907379589 |
| **A_55_P2151143** | Dynlt1c | -0.123255584 | -0.261734163 | 0.015222995 | 14.31619357 | -2.127321542 | 0.073011344 | 0.991835066 | -3.908146866 |
| **A_55_P2126790** | 1500009L1 | -0.100191075 | -0.212900832 | 0.012518681 | 8.209791381 | -2.124597465 | 0.073299364 | 0.991835066 | -3.910611106 |
| **A_52_P123864** | Rab21 | -0.142122405 | -0.3020113 | 0.01776649 | 9.309753575 | -2.124483587 | 0.07331143 | 0.991835066 | -3.910714129 |
| **A_55_P2106975** | Clcn5 | 0.120618531 | -0.015090525 | 0.256327587 | 8.949287776 | 2.124292615 | 0.073331668 | 0.991835066 | -3.9108869 |
| **A_66_P102879** | Thra | 0.150976024 | -0.018920859 | 0.320872907 | 8.725278572 | 2.123888456 | 0.073374516 | 0.991835066 | -3.911252545 |
| **A_55_P1966432** | Gstm1 | -0.253423265 | -0.538672904 | 0.031826375 | 9.905105867 | -2.123393624 | 0.073427012 | 0.991835066 | -3.911700234 |
| **A_55_P2389528** | D130061D1 | 0.132302668 | -0.016638486 | 0.281243822 | 8.349056952 | 2.123063678 | 0.073462037 | 0.991835066 | -3.911998752 |
| **A_55_P2119802** | NA | 0.197955404 | -0.024974938 | 0.420885746 | 15.4557743 | 2.122302634 | 0.073542888 | 0.991835066 | -3.912687324 |
| **A_52_P194250** | Ets1 | -0.206596472 | -0.439266581 | 0.026073636 | 10.37118064 | -2.122225077 | 0.073551132 | 0.991835066 | -3.912757498 |
| **A_52_P359502** | Pdxk | 0.175394983 | -0.022243178 | 0.373033145 | 10.96657379 | 2.121072413 | 0.073673772 | 0.991835066 | -3.913800457 |
| **A_52_P168338** | Fundc2 | -0.162114987 | -0.34480551 | 0.020575536 | 8.950926103 | -2.120880885 | 0.07369417 | 0.991835066 | -3.913973762 |
| **A_55_P2135631** | Rps18 | 0.124202456 | -0.015764083 | 0.264168995 | 14.5967499 | 2.120875091 | 0.073694787 | 0.991835066 | -3.913979005 |
| **A_55_P1957762** | Tsr1 | 0.104427198 | -0.013259847 | 0.222114243 | 9.354304195 | 2.120772583 | 0.073705707 | 0.991835066 | -3.91407176 |
| **A_51_P214127** | Cpa3 | -0.503703581 | -1.072051616 | 0.064644453 | 9.193242704 | -2.118213875 | 0.073978801 | 0.991835066 | -3.916387205 |
| **A_55_P2041470** | NA | -0.117704211 | -0.250515493 | 0.015107072 | 9.182708606 | -2.118196084 | 0.073980703 | 0.991835066 | -3.916403306 |
| **A_55_P1952269** | Xylt2 | 0.099956741 | -0.012839398 | 0.212752879 | 10.62789421 | 2.118005019 | 0.074001137 | 0.991835066 | -3.916576219 |
| **A_51_P327121** | Was | -0.109731955 | -0.233561264 | 0.014097354 | 8.275058634 | -2.117965109 | 0.074005406 | 0.991835066 | -3.916612337 |
| **A_55_P2139430** | NA | 0.666761372 | -0.085967448 | 1.419490191 | 11.39435247 | 2.117098282 | 0.074098188 | 0.991835066 | -3.917396833 |
| **A_55_P1954302** | Esrrb | -0.142950836 | -0.304367402 | 0.018465729 | 10.42187435 | -2.116643568 | 0.074146906 | 0.991835066 | -3.917808372 |
| **A_55_P2129706** | Srsf4 | 0.134529054 | -0.0174312 | 0.286489307 | 11.2858844 | 2.11590035 | 0.074226603 | 0.991835066 | -3.918481044 |
| **A_55_P1962084** | H2ac18 | -0.196486034 | -0.418448382 | 0.025476315 | 11.46361947 | -2.115736155 | 0.074244222 | 0.991835066 | -3.918629656 |
| **A_55_P2091836** | Fnip1 | -0.143371251 | -0.305351616 | 0.018609114 | 10.78700488 | -2.115479565 | 0.074271763 | 0.991835066 | -3.918861898 |
| **A_55_P2121618** | Tox4 | -0.670675627 | -1.428447525 | 0.087096271 | 9.527839031 | -2.115354509 | 0.07428519 | 0.991835066 | -3.918975088 |
| **A_51_P136460** | Zmat2 | 0.151264633 | -0.019716242 | 0.322245508 | 11.80436487 | 2.114457663 | 0.074381552 | 0.991835066 | -3.919786862 |
| **A_55_P2180445** | NA | 0.44086892 | -0.057659911 | 0.939397751 | 10.56823835 | 2.113626942 | 0.074470922 | 0.991835066 | -3.920538816 |
| **A_55_P2067976** | Gmeb1 | 0.115261536 | -0.015192047 | 0.245715118 | 9.889279485 | 2.111725787 | 0.074675857 | 0.991835066 | -3.922259827 |
| **A_52_P577729** | Rnf169 | 0.148090822 | -0.019611265 | 0.315792908 | 10.4515513 | 2.11056537 | 0.074801223 | 0.991835066 | -3.923310368 |
| **A_52_P10041** | Akr1b3 | -0.167558597 | -0.357327106 | 0.022209912 | 13.22172994 | -2.11033646 | 0.074825978 | 0.991835066 | -3.923517611 |
| **A_55_P2101806** | Esf1 | 0.125272619 | -0.016605143 | 0.26715038 | 10.54839059 | 2.11033286 | 0.074826367 | 0.991835066 | -3.92352087 |

| | | | | | | | | | |
|---|---|---|---|---|---|---|---|---|---|
| **A_52_P131672** | Rbm7 | -0.117008852 | -0.249530772 | 0.015513068 | 10.16577587 | -2.110280231 | 0.07483206 | 0.991835066 | -3.923568518 |
| **A_66_P110909** | Ppp1r12a | 0.171361291 | -0.022846984 | 0.365569566 | 12.91593968 | 2.10889097 | 0.07498249 | 0.991835066 | -3.924826334 |
| **A_55_P2067722** | Camsap3 | 0.101178595 | -0.013496051 | 0.21585324 | 9.268746042 | 2.10877561 | 0.074994995 | 0.991835066 | -3.924930783 |
| **A_55_P1973723** | Etfdh | -0.094519451 | -0.201653091 | 0.012614189 | 13.67666314 | -2.108649821 | 0.075008633 | 0.991835066 | -3.925044675 |
| **A_55_P2102424** | Msto1 | 0.178511572 | -0.023830474 | 0.380853618 | 11.03825358 | 2.108576535 | 0.075016579 | 0.991835066 | -3.92511103 |
| **A_55_P1964724** | NA | -0.093628599 | -0.199778825 | 0.012521627 | 8.512042226 | -2.10812683 | 0.075065361 | 0.991835066 | -3.925518211 |
| **A_55_P1953489** | Lrrc8b | 0.13952496 | -0.018714721 | 0.297764641 | 9.396362041 | 2.107393523 | 0.075144975 | 0.991835066 | -3.926182195 |
| **A_55_P2083481** | Lpin1 | -0.178354153 | -0.380643135 | 0.02393483 | 8.583854502 | -2.107269727 | 0.075158423 | 0.991835066 | -3.92629429 |
| **A_66_P117022** | Atp6v1e1 | -0.140578963 | -0.300056761 | 0.018898835 | 12.16354954 | -2.106828754 | 0.075206349 | 0.991835066 | -3.92669359 |
| **A_51_P238734** | Mfsd11 | -0.150705876 | -0.321681607 | 0.020269854 | 9.118939966 | -2.106710456 | 0.07521921 | 0.991835066 | -3.92680071 |
| **A_52_P82209** | Wnk1 | 0.188664622 | -0.025375417 | 0.402704661 | 10.48440949 | 2.106709146 | 0.075219353 | 0.991835066 | -3.926801896 |
| **A_55_P1973995** | NA | 0.47466133 | -0.063853994 | 1.013176655 | 10.16419725 | 2.106662275 | 0.07522445 | 0.991835066 | -3.926844339 |
| **A_55_P1985569** | Tomm6 | 0.109289204 | -0.014719735 | 0.233298144 | 13.96372406 | 2.106363905 | 0.075256902 | 0.991835066 | -3.927114519 |
| **A_51_P234174** | Pms2 | 0.123679676 | -0.016686099 | 0.26404545 | 9.277434303 | 2.105941217 | 0.0753029 | 0.991835066 | -3.927497278 |
| **A_51_P270184** | Nhlrc2 | -0.100891398 | -0.215430924 | 0.013648129 | 10.91275827 | -2.105270417 | 0.075375956 | 0.991835066 | -3.928104728 |
| **A_52_P263870** | Cdc23 | 0.102545326 | -0.013892372 | 0.218983025 | 8.464371087 | 2.104899625 | 0.075416369 | 0.991835066 | -3.928440512 |
| **A_55_P2148478** | Klhl25 | 0.113700078 | -0.015476208 | 0.242876364 | 10.66795077 | 2.103715962 | 0.075545523 | 0.991835066 | -3.929512457 |
| **A_52_P589568** | Foxo6 | 0.216058445 | -0.029421782 | 0.461538672 | 9.441702782 | 2.103603395 | 0.075557817 | 0.991835066 | -3.929614402 |
| **A_55_P2058962** | Mcm10 | 0.2246467 | -0.030610216 | 0.479903616 | 9.669335506 | 2.103447408 | 0.075574857 | 0.991835066 | -3.929755672 |
| **A_55_P2199118** | Bend4 | 0.209090376 | -0.028514907 | 0.44669566 | 11.20719357 | 2.103231527 | 0.075598446 | 0.991835066 | -3.929951187 |
| **A_55_P2133716** | Tax1bp3 | -0.098683943 | -0.21084732 | 0.013479434 | 14.14833053 | -2.102831859 | 0.075642136 | 0.991835066 | -3.930313154 |
| **A_55_P2076278** | NA | 0.11393492 | -0.015576586 | 0.243446425 | 17.08025907 | 2.102604704 | 0.07566698 | 0.991835066 | -3.930518886 |
| **A_55_P2083128** | Sec16a | 0.111520524 | -0.015263725 | 0.238304773 | 10.6369323 | 2.102319086 | 0.075698228 | 0.991835066 | -3.930777569 |
| **A_55_P2032362** | NA | 0.108181101 | -0.014834147 | 0.231196349 | 8.332982793 | 2.101849335 | 0.075749651 | 0.991835066 | -3.931203028 |
| **A_52_P97670** | Atad2b | 0.136439174 | -0.018723595 | 0.291601942 | 10.07258011 | 2.10165145 | 0.075771323 | 0.991835066 | -3.931382258 |
| **A_51_P252677** | Card19 | -0.160999412 | -0.344126956 | 0.022128132 | 13.23317281 | -2.101259793 | 0.075814236 | 0.991835066 | -3.931736997 |
| **A_55_P1992234** | Slx | 0.103802209 | -0.014283479 | 0.221887896 | 8.012602852 | 2.100963294 | 0.075846739 | 0.991835066 | -3.932005552 |
| **A_51_P392005** | Car8 | 0.111605088 | -0.015390293 | 0.238600469 | 8.49085141 | 2.100415431 | 0.075906834 | 0.991835066 | -3.932501791 |
| **A_51_P450021** | Higd2a | -0.216929325 | -0.46380484 | 0.029946189 | 13.83635296 | -2.100145479 | 0.075936462 | 0.991835066 | -3.932746311 |
| **A_52_P116264** | Adhfe1 | -0.151687298 | -0.324452259 | 0.021077663 | 8.43701847 | -2.098469601 | 0.076120658 | 0.991835066 | -3.934264371 |
| **A_51_P419286** | Batf3 | -0.174782685 | -0.373865664 | 0.024300295 | 8.248855712 | -2.098328148 | 0.076136225 | 0.991835066 | -3.934392509 |
| **A_55_P2027999** | Hk1 | 0.102069837 | -0.014198795 | 0.218338469 | 14.75393635 | 2.098186029 | 0.076151869 | 0.991835066 | -3.934521251 |
| **A_55_P2117959** | Cyp4b1 | -0.157747008 | -0.337535477 | 0.022041461 | 8.246798136 | -2.097048283 | 0.076277226 | 0.991835066 | -3.935551935 |
| **A_55_P2049147** | NA | 0.132107331 | -0.018466659 | 0.28268132 | 14.80856292 | 2.096940418 | 0.076289121 | 0.991835066 | -3.935649652 |
| **A_52_P304128** | Mmp14 | 0.135344049 | -0.01892716 | 0.289615258 | 9.157186931 | 2.096830926 | 0.076301198 | 0.991835066 | -3.935748845 |
| **A_52_P621064** | NA | 0.138983246 | -0.019501993 | 0.297468484 | 9.043511685 | 2.095958908 | 0.076397448 | 0.991835066 | -3.93653885 |
| **A_51_P260035** | Sar1b | -0.114797747 | -0.245715429 | 0.016119936 | 11.28660837 | -2.095772744 | 0.076418012 | 0.991835066 | -3.93670751 |
| **A_55_P2152976** | NA | 0.084085833 | -0.011845825 | 0.180017491 | 14.25865485 | 2.094932395 | 0.076510906 | 0.991835066 | -3.937468861 |
| **A_55_P2171143** | Hnrnpc | 0.145363492 | -0.020482267 | 0.311209251 | 13.62140219 | 2.094884628 | 0.07651619 | 0.991835066 | -3.937512138 |
| **A_51_P391380** | Mindy2 | -0.133992624 | -0.286915263 | 0.018930015 | 10.09939919 | -2.09420041 | 0.076591914 | 0.991835066 | -3.938132057 |
| **A_66_P120260** | Taf10 | 0.111873085 | -0.015815027 | 0.239561198 | 13.14751064 | 2.094036665 | 0.076610048 | 0.991835066 | -3.938280418 |
| **A_55_P2114908** | Nek3 | 0.119018614 | -0.016879827 | 0.254917056 | 9.077234147 | 2.093194319 | 0.076703398 | 0.991835066 | -3.939043637 |
| **A_55_P1996862** | Psme2 | -0.137740178 | -0.295020347 | 0.019539991 | 14.12114678 | -2.093128079 | 0.076710744 | 0.991835066 | -3.939103655 |
| **A_55_P2094966** | H3c14 | 0.24115016 | -0.034229815 | 0.516530134 | 11.79756158 | 2.092976399 | 0.076727567 | 0.991835066 | -3.939241091 |
| **A_52_P141338** | Acot10 | -0.130972512 | -0.280536466 | 0.018591442 | 8.508733645 | -2.092966852 | 0.076728626 | 0.991835066 | -3.939249741 |
| **A_51_P254583** | Foxred1 | -0.155019189 | -0.332111686 | 0.022073307 | 11.38068388 | -2.092157783 | 0.076818426 | 0.991835066 | -3.939982844 |
| **A_51_P302458** | Plscr3 | 0.101834742 | -0.014545146 | 0.21821463 | 11.31212939 | 2.091352129 | 0.076907951 | 0.991835066 | -3.940712877 |
| **A_51_P333622** | 2010300C0 | 0.226848455 | -0.032515807 | 0.486212717 | 9.274647808 | 2.090426128 | 0.07701098 | 0.991835066 | -3.941551995 |

| | | | | | | | | | |
|---|---|---|---|---|---|---|---|---|---|
| **A_55_P1996941** | Ube2c | -0.138681907 | -0.297272893 | 0.019909079 | 11.82395609 | -2.090019985 | 0.077056211 | 0.991835066 | -3.941920042 |
| **A_55_P2148896** | NA | 0.17190664 | -0.02470307 | 0.36851635 | 9.23237855 | 2.089761967 | 0.07708496 | 0.991835066 | -3.94215386 |
| **A_66_P131788** | Ythdf2 | 0.11221977 | -0.016214627 | 0.240654166 | 8.454041548 | 2.088320525 | 0.077245768 | 0.991835066 | -3.94346016 |
| **A_66_P116126** | Sdhb | -0.146406967 | -0.313990615 | 0.021176681 | 16.42253003 | -2.088042033 | 0.077276875 | 0.991835066 | -3.943712551 |
| **A_51_P349878** | Txndc17 | 0.132091309 | -0.019106669 | 0.283289287 | 13.99754924 | 2.088033171 | 0.077277865 | 0.991835066 | -3.943720583 |
| **A_52_P653902** | Eny2 | 0.119534985 | -0.017331094 | 0.256401065 | 10.34642163 | 2.087412783 | 0.077347209 | 0.991835066 | -3.944282839 |
| **A_52_P470316** | Laptm4b | -0.136211948 | -0.292189183 | 0.019765287 | 10.84097316 | -2.087195442 | 0.077371516 | 0.991835066 | -3.944479818 |
| **A_51_P206251** | Ahcyl1 | -0.118985807 | -0.255251869 | 0.017280255 | 9.766933092 | -2.08697183 | 0.077396534 | 0.991835066 | -3.944682482 |
| **A_55_P2073977** | NA | -0.152400606 | -0.326945358 | 0.022144146 | 14.49218356 | -2.086839409 | 0.077411352 | 0.991835066 | -3.944802499 |
| **A_52_P454994** | Pi4ka | 0.138185879 | -0.02011689 | 0.296488648 | 13.2172472 | 2.086336163 | 0.077467695 | 0.991835066 | -3.945258611 |
| **A_55_P2028655** | Slc31a1 | 0.137058171 | -0.019967103 | 0.294083445 | 11.25296897 | 2.086145063 | 0.077489101 | 0.991835066 | -3.945431815 |
| **A_52_P418665** | Ttpal | 0.14840663 | -0.021705795 | 0.318519055 | 10.60024399 | 2.085097655 | 0.077606531 | 0.991835066 | -3.946381162 |
| **A_51_P423091** | Flot1 | -0.215724133 | -0.46313219 | 0.031683924 | 11.88921603 | -2.083982299 | 0.077731775 | 0.991835066 | -3.947392141 |
| **A_55_P2010341** | Rpl11 | 0.110661028 | -0.016258061 | 0.237580117 | 15.6508877 | 2.083900056 | 0.077741018 | 0.991835066 | -3.947466689 |
| **A_51_P321341** | Sult1a1 | -0.231371133 | -0.496787163 | 0.034044898 | 10.19398821 | -2.083488731 | 0.077787263 | 0.991835066 | -3.947839534 |
| **A_51_P172155** | Hal | 0.116197525 | -0.017204466 | 0.249599516 | 7.725069769 | 2.081822497 | 0.077974878 | 0.991835066 | -3.949349956 |
| **A_55_P2140745** | NA | -0.137961444 | -0.296350631 | 0.020427742 | 8.18761904 | -2.081811185 | 0.077976154 | 0.991835066 | -3.949360211 |
| **A_55_P2183378** | Snx11 | 0.138001038 | -0.020443517 | 0.296445593 | 10.76316521 | 2.081680946 | 0.077990838 | 0.991835066 | -3.949478276 |
| **A_52_P238242** | Sertad2 | -0.094706925 | -0.203458867 | 0.014045016 | 9.990334193 | -2.081391863 | 0.078023441 | 0.991835066 | -3.949740338 |
| **A_51_P161362** | Gnl2 | 0.128876424 | -0.019177562 | 0.276930411 | 11.08992888 | 2.08047502 | 0.078126934 | 0.991835066 | -3.950571504 |
| **A_51_P473576** | Apex1 | 0.104148797 | -0.015532205 | 0.223829798 | 11.23866796 | 2.07987949 | 0.078194231 | 0.991835066 | -3.9511114 |
| **A_55_P2008407** | Ppp1r14c | -0.263382191 | -0.566099894 | 0.039335513 | 12.36714593 | -2.079494223 | 0.078237798 | 0.991835066 | -3.951460682 |
| **A_52_P218537** | Hnrnpc | 0.125652621 | -0.018808741 | 0.270113983 | 11.37209417 | 2.078877884 | 0.078307547 | 0.991835066 | -3.952019465 |
| **A_55_P2146127** | Chmp4b | 0.127063445 | -0.019031452 | 0.273158341 | 12.07990777 | 2.078713865 | 0.078326119 | 0.991835066 | -3.952168169 |
| **A_55_P2043987** | Luzp1 | 0.121309773 | -0.018172377 | 0.260791922 | 12.33997166 | 2.078673554 | 0.078330684 | 0.991835066 | -3.952204717 |
| **A_55_P1966649** | Glmn | 0.130075122 | -0.019505989 | 0.279656232 | 9.584056519 | 2.078387974 | 0.078363033 | 0.991835066 | -3.952463635 |
| **A_52_P11833** | NA | -0.158785432 | -0.341422149 | 0.023851286 | 11.25648389 | -2.077933733 | 0.078414514 | 0.991835066 | -3.952875473 |
| **A_55_P1965233** | Sh3tc1 | 0.209380764 | -0.031479564 | 0.450241092 | 10.07604963 | 2.077689475 | 0.078442211 | 0.991835066 | -3.953096934 |
| **A_55_P1954598** | NA | 0.117703299 | -0.017702638 | 0.253109235 | 11.47749077 | 2.077591032 | 0.078453376 | 0.991835066 | -3.953186189 |
| **A_55_P2028571** | Sgsh | 0.185018642 | -0.027957077 | 0.39799436 | 9.677890865 | 2.07632115 | 0.078597549 | 0.991835066 | -3.954337586 |
| **A_55_P2127702** | Racgap1 | -0.108952054 | -0.234377274 | 0.016473167 | 11.1870454 | -2.076154555 | 0.078616483 | 0.991835066 | -3.954488641 |
| **A_51_P171999** | Apoe | -0.191378323 | -0.411794427 | 0.029037781 | 13.62328009 | -2.075193293 | 0.07872582 | 0.991835066 | -3.956181157 |
| **A_51_P400585** | Api5 | -0.106494196 | -0.229152568 | 0.016164176 | 10.4232018 | -2.075094483 | 0.078737068 | 0.991835066 | -3.955449854 |
| **A_55_P2028721** | Rnf130 | -0.105641462 | -0.227364592 | 0.016081667 | 10.80570402 | -2.074294551 | 0.078828184 | 0.991835066 | -3.956175213 |
| **A_51_P150087** | Abhd10 | -0.085969016 | -0.185041602 | 0.01310357 | 9.107692543 | -2.073946684 | 0.078867841 | 0.991835066 | -3.956490658 |
| **A_51_P208361** | Ak3 | -0.160396289 | -0.345243037 | 0.02445046 | 12.23049567 | -2.073918238 | 0.078871085 | 0.991835066 | -3.956516453 |
| **A_51_P440892** | Dph2 | 0.130733127 | -0.019969659 | 0.281435913 | 9.880988719 | 2.073354203 | 0.07893543 | 0.991835066 | -3.957027927 |
| **A_55_P2079009** | Slco2b1 | -0.090504516 | -0.194835879 | 0.013826846 | 8.338420066 | -2.073311251 | 0.078940332 | 0.991835066 | -3.957066877 |
| **A_51_P423578** | Slfn2 | -0.103555921 | -0.222942563 | 0.015830722 | 7.896651415 | -2.07313855 | 0.078960045 | 0.991835066 | -3.957223486 |
| **A_55_P2027587** | Supt4b | -0.179058367 | -0.385524299 | 0.027407566 | 10.1171516 | -2.07279021 | 0.078999823 | 0.991835066 | -3.957539374 |
| **A_51_P231320** | Mmp8 | -0.113431766 | -0.244243499 | 0.017379968 | 7.850632836 | -2.072512334 | 0.079031568 | 0.991835066 | -3.957791364 |
| **A_51_P502798** | Nxpe4 | -0.096477348 | -0.207746075 | 0.01479138 | 8.421680691 | -2.072341704 | 0.079051068 | 0.991835066 | -3.957946101 |
| **A_55_P2083449** | Spryd7 | -0.108505508 | -0.233666859 | 0.016655842 | 9.130842787 | -2.072004427 | 0.079089626 | 0.991835066 | -3.958251965 |
| **A_55_P2166995** | NA | 0.099785986 | -0.015339055 | 0.214911028 | 15.3026196 | 2.071614235 | 0.079134257 | 0.991835066 | -3.95860582 |
| **A_51_P116906** | Rapgef3 | 0.141449292 | -0.021745585 | 0.304644168 | 11.36648752 | 2.071588017 | 0.079137257 | 0.991835066 | -3.958629597 |
| **A_55_P2025174** | Rps27a | 0.146250643 | -0.022492624 | 0.31499391 | 15.56067467 | 2.07147866 | 0.07914977 | 0.991835066 | -3.958728771 |
| **A_55_P2137206** | Tgfbr1 | 0.209975044 | -0.032332326 | 0.452282414 | 10.39411382 | 2.071143496 | 0.079188134 | 0.991835066 | -3.959032729 |
| **A_51_P261443** | Mlx | -0.115716644 | -0.249316889 | 0.0178836 | 9.334628295 | -2.070130433 | 0.079304208 | 0.991835066 | -3.959951491 |

| | | | | | | | | | |
|---|---|---|---|---|---|---|---|---|---|
| **A_65_P15518** | NA | 0.199895008 | -0.030932688 | 0.430722703 | 8.539460896 | 2.069775129 | 0.079344958 | 0.991835066 | -3.96027373 |
| **A_51_P216593** | Wfs1 | 0.107132154 | -0.016592426 | 0.230856734 | 10.75930727 | 2.069535973 | 0.079372398 | 0.991835066 | -3.960490631 |
| **A_51_P114353** | Ctu1 | -0.086744943 | -0.186940816 | 0.01345093 | 8.546722661 | -2.069204735 | 0.07941042 | 0.991835066 | -3.96079105 |
| **A_55_P2110497** | Ddc | 0.088048267 | -0.013655584 | 0.189752119 | 13.51914567 | 2.069152715 | 0.079416393 | 0.991835066 | -3.960838231 |
| **A_51_P338443** | Angptl4 | -0.32014736 | -0.690044947 | 0.049750227 | 10.57540432 | -2.068604946 | 0.079479315 | 0.991835066 | -3.961335043 |
| **A_55_P1967443** | NA | 0.441728595 | -0.068671404 | 0.952128594 | 9.406268325 | 2.068492649 | 0.07949222 | 0.991835066 | -3.961436895 |
| **A_51_P229105** | Cetn3 | -0.130695405 | -0.28177295 | 0.02038214 | 11.89762119 | -2.067614321 | 0.079593233 | 0.991835066 | -3.962233539 |
| **A_51_P260504** | Arhgef4 | 0.171308876 | -0.026741434 | 0.369359186 | 8.487253365 | 2.067347449 | 0.079623951 | 0.991835066 | -3.962475597 |
| **A_55_P2087352** | NA | -0.279325955 | -0.602261669 | 0.043609759 | 15.1562091 | -2.067304014 | 0.079628951 | 0.991835066 | -3.962514994 |
| **A_55_P2133170** | Luc7l2 | 0.147069383 | -0.023025917 | 0.317164682 | 9.241057532 | 2.066517512 | 0.079719553 | 0.991835066 | -3.963228381 |
| **A_51_P341571** | Klhdc2 | -0.126719742 | -0.273333874 | 0.01989439 | 11.65038339 | -2.065749134 | 0.079808167 | 0.991835066 | -3.96392535 |
| **A_55_P2026109** | Rpap1 | 0.094060989 | -0.014773853 | 0.202895832 | 11.3939627 | 2.06562139 | 0.079822909 | 0.991835066 | -3.964041223 |
| **A_55_P2048607** | Hp | -0.403382761 | -0.87065395 | 0.063888428 | 8.629866765 | -2.063276644 | 0.08009398 | 0.991835066 | -3.966168185 |
| **A_52_P90363** | Ifi27l2a | -0.720632223 | -1.555463787 | 0.11419934 | 10.0196312 | -2.063117437 | 0.080112419 | 0.991835066 | -3.966312611 |
| **A_66_P119017** | Smarca1 | 0.110004673 | -0.017465188 | 0.237474535 | 10.12444897 | 2.062589274 | 0.08017362 | 0.991835066 | -3.966791744 |
| **A_51_P509229** | Tnrc6a | 0.196835732 | -0.031265199 | 0.424936663 | 9.709310963 | 2.06246228 | 0.080188342 | 0.991835066 | -3.966906951 |
| **A_51_P460954** | Ccl6 | -0.115869284 | -0.250204719 | 0.01846615 | 8.32287222 | -2.061516774 | 0.08029804 | 0.991835066 | -3.967764709 |
| **A_55_P2064676** | Gas2l3 | -0.117191826 | -0.253084465 | 0.018700813 | 9.371788845 | -2.061154384 | 0.080340124 | 0.991835066 | -3.968093476 |
| **A_51_P244558** | Rab3gap2 | -0.09250653 | -0.199776158 | 0.014763097 | 9.522154045 | -2.061126995 | 0.080343306 | 0.991835066 | -3.968118324 |
| **A_52_P497188** | Prrg1 | -0.1039464 | -0.224532944 | 0.016640144 | 9.082392373 | -2.060249118 | 0.080445349 | 0.991835066 | -3.968914766 |
| **A_55_P1972263** | Nop56 | 0.236419685 | -0.03787046 | 0.51070983 | 11.55405853 | 2.060072774 | 0.080465863 | 0.991835066 | -3.969074754 |
| **A_51_P265571** | Adm | -0.297558389 | -0.642842403 | 0.047725624 | 9.813004961 | -2.059704231 | 0.080508751 | 0.991835066 | -3.969409118 |
| **A_55_P2093689** | Pold3 | 0.146139019 | -0.023451677 | 0.315729714 | 11.23009303 | 2.059554545 | 0.080526177 | 0.991835066 | -3.969544924 |
| **A_52_P355398** | Dars | 0.183591096 | -0.029520384 | 0.396702577 | 9.44592102 | 2.058988406 | 0.080592119 | 0.991835066 | -3.970058571 |
| **A_51_P514961** | Tiparp | -0.125446594 | -0.271070422 | 0.020177235 | 10.14972542 | -2.05890145 | 0.080602252 | 0.991835066 | -3.970137464 |
| **A_55_P2347976** | Mir17hg | 0.291515279 | -0.046890358 | 0.629920916 | 8.439857578 | 2.058888678 | 0.080603741 | 0.991835066 | -3.970149052 |
| **A_66_P115053** | Anp32b | 0.113387553 | -0.01827232 | 0.245047425 | 13.00982769 | 2.058358821 | 0.080665515 | 0.991835066 | -3.970629792 |
| **A_55_P1974577** | Nfam1 | -0.129060321 | -0.278938756 | 0.020818114 | 8.189739721 | -2.058082227 | 0.080697781 | 0.991835066 | -3.970880748 |
| **A_55_P2143516** | NA | 0.622468422 | -0.100695134 | 1.345631978 | 10.5046786 | 2.05726352 | 0.080793362 | 0.991835066 | -3.971623584 |
| **A_55_P2134616** | Med12l | 0.223054664 | -0.036114522 | 0.482223851 | 8.648513703 | 2.057013187 | 0.08082261 | 0.991835066 | -3.971850722 |
| **A_55_P2335903** | NA | 0.184869418 | -0.030101825 | 0.399840661 | 10.55367281 | 2.055388105 | 0.081012738 | 0.991835066 | -3.973325275 |
| **A_55_P2145754** | Atp5pb | -0.099442299 | -0.215099511 | 0.016214913 | 14.09062507 | -2.054979862 | 0.08106057 | 0.991835066 | -3.973695716 |
| **A_55_P2064171** | NA | 0.424652537 | -0.069273496 | 0.91857857 | 11.75959698 | 2.05485385 | 0.081075341 | 0.991835066 | -3.97381006 |
| **A_55_P2135027** | NA | 0.140816748 | -0.023094692 | 0.304728187 | 9.874381984 | 2.053308373 | 0.081256709 | 0.991835066 | -3.975212477 |
| **A_52_P377299** | Zbtb11 | 0.149364232 | -0.024598508 | 0.323326971 | 8.573046551 | 2.052104644 | 0.081398254 | 0.991835066 | -3.976304829 |
| **A_55_P2074246** | Zfp184 | 0.11946615 | -0.019684132 | 0.258616432 | 8.505781218 | 2.051964818 | 0.081414712 | 0.991835066 | -3.97643172 |
| **A_55_P2140626** | Arl5b | 0.142668707 | -0.023511361 | 0.308848776 | 9.523901946 | 2.051912914 | 0.081420822 | 0.991835066 | -3.976478823 |
| **A_51_P345699** | Tnnt3 | -0.131695721 | -0.285125292 | 0.021733849 | 8.239069265 | -2.051500974 | 0.081469331 | 0.991835066 | -3.976852659 |
| **A_52_P232314** | NA | 0.19251359 | -0.033416812 | 0.418443992 | 9.099842154 | 2.116673475 | 0.081497561 | 0.991835066 | -3.971190119 |
| **A_51_P490305** | Ifi30 | -0.092931268 | -0.201223995 | 0.015361458 | 9.395059675 | -2.05102856 | 0.081524997 | 0.991835066 | -3.977281381 |
| **A_55_P2138739** | NA | 0.441145615 | -0.073136252 | 0.955427482 | 9.368635852 | 2.050170061 | 0.081626254 | 0.991835066 | -3.978060497 |
| **A_55_P2122849** | NA | 0.203297834 | -0.033719504 | 0.440315172 | 9.535244004 | 2.050037325 | 0.081641921 | 0.991835066 | -3.978180961 |
| **A_52_P58181** | Suclg2 | -0.099361183 | -0.215225107 | 0.016502742 | 10.90585182 | -2.049640287 | 0.081688802 | 0.991835066 | -3.978541294 |
| **A_55_P2375121** | AI225934 | 0.26085423 | -0.043414455 | 0.565122915 | 8.783807927 | 2.0490368 | 0.081760111 | 0.991835066 | -3.979088999 |
| **A_66_P101942** | Gm9706 | -0.232111717 | -0.502928265 | 0.038704832 | 8.287909144 | -2.048476553 | 0.081826367 | 0.991835066 | -3.97959747 |
| **A_55_P2096010** | NA | 0.128582704 | -0.021441847 | 0.278607255 | 12.58532382 | 2.048468784 | 0.081827286 | 0.991835066 | -3.979604521 |
| **A_55_P1998213** | NA | 0.127024538 | -0.02121945 | 0.275268526 | 16.32375468 | 2.047951501 | 0.081888509 | 0.991835066 | -3.980074006 |
| **A_66_P138829** | Ndufs8 | -0.286739383 | -0.621439165 | 0.047960399 | 12.79095617 | -2.047580803 | 0.08193241 | 0.991835066 | -3.980410455 |

| | | | | | | | | | |
|---|---|---|---|---|---|---|---|---|---|
| **A_51_P331207** | Tmem218 | -0.127960515 | -0.277395422 | 0.021474391 | 10.51156943 | -2.046600408 | 0.082048633 | 0.991835066 | -3.981300289 |
| **A_55_P2022201** | NA | 0.183163934 | -0.030775017 | 0.397102886 | 8.633693689 | 2.046252549 | 0.08208991 | 0.991835066 | -3.981616022 |
| **A_55_P2077548** | Git1 | 0.093325249 | -0.015688698 | 0.202339197 | 11.56784915 | 2.046097024 | 0.082108371 | 0.991835066 | -3.981757185 |
| **A_55_P1999790** | Pkdcc | 0.228063092 | -0.03835289 | 0.494479075 | 12.87460926 | 2.045991692 | 0.082120876 | 0.991835066 | -3.98185279 |
| **A_52_P232830** | Ets1 | 0.200485039 | -0.033874946 | 0.434845023 | 10.10616281 | 2.044596627 | 0.082286686 | 0.991835066 | -3.983119057 |
| **A_55_P2022663** | NA | 0.114269207 | -0.019311172 | 0.247849586 | 15.49529116 | 2.044540293 | 0.082293388 | 0.991835066 | -3.983170191 |
| **A_55_P1967761** | Foxc1 | 0.251014216 | -0.042498212 | 0.544526643 | 10.19069731 | 2.044000298 | 0.082357664 | 0.991835066 | -3.983660346 |
| **A_55_P2072213** | NA | 0.190275511 | -0.032216951 | 0.412767973 | 11.54595339 | 2.0439804 | 0.082360033 | 0.991835066 | -3.983678407 |
| **A_52_P491244** | Zfp287 | 0.104615687 | -0.017737963 | 0.226969336 | 8.702244116 | 2.043567634 | 0.0824092 | 0.991835066 | -3.984053082 |
| **A_51_P208680** | Chtf18 | 0.177550731 | -0.030109438 | 0.385210899 | 9.936839609 | 2.043517652 | 0.082415156 | 0.991835066 | -3.984098452 |
| **A_55_P1994339** | Dhrs7c | -0.362356518 | -0.786297967 | 0.06158493 | 12.31465176 | -2.042863415 | 0.082493151 | 0.991835066 | -3.984692323 |
| **A_51_P346826** | Rragc | -0.186378017 | -0.404443937 | 0.031687902 | 11.05789895 | -2.042753782 | 0.082506228 | 0.991835066 | -3.984791841 |
| **A_55_P2143070** | Ass1 | 0.154244672 | -0.026261364 | 0.334750709 | 8.552402546 | 2.042337792 | 0.082555867 | 0.991835066 | -3.985169454 |
| **A_51_P210082** | Ercc4 | 0.094368663 | -0.016095543 | 0.204832869 | 9.851086238 | 2.041810154 | 0.082618871 | 0.991835066 | -3.985648423 |
| **A_55_P2182423** | NA | -0.149335145 | -0.324172029 | 0.02550174 | 16.659508 | -2.041446962 | 0.082662268 | 0.991835066 | -3.985978118 |
| **A_55_P1981296** | Nup85 | 0.16055956 | -0.027419232 | 0.348538351 | 12.16676404 | 2.041439192 | 0.082663196 | 0.991835066 | -3.985985171 |
| **A_55_P2005680** | Ipo11 | 0.16810658 | -0.02871928 | 0.364932439 | 12.14882865 | 2.041322802 | 0.082677108 | 0.991835066 | -3.986090827 |
| **A_55_P2136752** | Ermn | 0.591232764 | -0.101170047 | 1.283635576 | 11.23142353 | 2.040839272 | 0.082734928 | 0.991835066 | -3.986529767 |
| **A_51_P232371** | Stab1 | 0.158041111 | -0.027049225 | 0.343131446 | 11.44888387 | 2.040776585 | 0.082742427 | 0.991835066 | -3.986586675 |
| **A_51_P495780** | Plin4 | -0.318007942 | -0.690614715 | 0.054598832 | 10.65500636 | -2.039841169 | 0.082854409 | 0.991835066 | -3.987435849 |
| **A_51_P416509** | H1f1 | 0.251228317 | -0.043169408 | 0.545626042 | 9.586496068 | 2.039591866 | 0.082884279 | 0.991835066 | -3.98766217 |
| **A_51_P317191** | Eepd1 | -0.107246125 | -0.233028092 | 0.018535841 | 8.729934415 | -2.037850696 | 0.083093198 | 0.991835066 | -3.989242877 |
| **A_55_P2143041** | Phgdh | 0.367185161 | -0.063548892 | 0.797919214 | 10.08779079 | 2.037441024 | 0.083142429 | 0.991835066 | -3.989614805 |
| **A_55_P1985495** | Gm12005 | 0.120126596 | -0.020857246 | 0.261110439 | 14.29318746 | 2.036474426 | 0.083258705 | 0.991835066 | -3.990492365 |
| **A_55_P2178203** | Derl2 | -0.153078687 | -0.332785778 | 0.026628404 | 9.942226985 | -2.035910334 | 0.083326636 | 0.991835066 | -3.991004506 |
| **A_55_P2018611** | Igfbp7 | -0.09062562 | -0.197044474 | 0.015793234 | 7.890420867 | -2.035361447 | 0.08339279 | 0.991835066 | -3.991502849 |
| **A_55_P2162765** | Dnajc24 | -0.095224662 | -0.207120871 | 0.016671547 | 10.6787739 | -2.033963703 | 0.083561486 | 0.991835066 | -3.992771914 |
| **A_55_P1955483** | Grb14 | -0.234527039 | -0.510163606 | 0.041109528 | 12.27142145 | -2.033598539 | 0.083605615 | 0.991835066 | -3.993103468 |
| **A_52_P670026** | Rsad2 | -0.308186646 | -0.670397057 | 0.054023766 | 8.578633366 | -2.033583544 | 0.083607427 | 0.991835066 | -3.993117083 |
| **A_55_P1976108** | Rps27a | 0.145473893 | -0.025520062 | 0.316467849 | 16.81357902 | 2.033356069 | 0.083634929 | 0.991835066 | -3.993323623 |
| **A_52_P108089** | Mosmo | 0.119962065 | -0.021062062 | 0.260986192 | 10.96390102 | 2.033104232 | 0.083665387 | 0.991835066 | -3.993552283 |
| **A_55_P2074536** | Ppm1b | -0.095900991 | -0.20866446 | 0.016862478 | 12.83474884 | -2.032655599 | 0.083719673 | 0.991835066 | -3.993959632 |
| **A_55_P2345116** | NA | 0.199311475 | -0.035078026 | 0.433700976 | 8.681847286 | 2.032372365 | 0.083753963 | 0.991835066 | -3.994216805 |
| **A_55_P2110925** | Flrt3 | 0.241932877 | -0.042617631 | 0.526483386 | 8.934536334 | 2.032098041 | 0.083787188 | 0.991835066 | -3.99446589 |
| **A_55_P2363442** | Zfp867 | -0.119362702 | -0.259781826 | 0.021056422 | 9.128719899 | -2.03166227 | 0.083839994 | 0.991835066 | -3.994861571 |
| **A_55_P1996952** | NA | 0.172966466 | -0.030556247 | 0.37648918 | 9.342217389 | 2.031225595 | 0.083892942 | 0.991835066 | -3.995258076 |
| **A_52_P123485** | Tcf25 | 0.117044765 | -0.020683041 | 0.254772571 | 13.94391171 | 2.031138305 | 0.08390353 | 0.991835066 | -3.995337337 |
| **A_55_P2087726** | Cops9 | -0.085681592 | -0.18653651 | 0.015173325 | 15.45204211 | -2.030484054 | 0.083982933 | 0.991835066 | -3.995931414 |
| **A_55_P2065465** | Mettl1 | 0.21549168 | -0.038274127 | 0.469257487 | 9.657140527 | 2.029581686 | 0.084092572 | 0.991835066 | -3.996750801 |
| **A_51_P481398** | Kif11 | 0.100461789 | -0.0178477 | 0.218771279 | 9.51061906 | 2.02950657 | 0.084101705 | 0.991835066 | -3.99681901 |
| **A_51_P285271** | Ccdc61 | -0.106279041 | -0.231504701 | 0.018946619 | 10.20567183 | -2.028445905 | 0.084230773 | 0.991835066 | -3.997782163 |
| **A_55_P2159264** | Lifr | -0.103475924 | -0.225424737 | 0.018472889 | 8.446817389 | -2.028013619 | 0.084283433 | 0.991835066 | -3.998174714 |
| **A_55_P2119032** | Rasl2-9 | -0.134561177 | -0.29316157 | 0.024039217 | 10.59617002 | -2.0277978 | 0.084309735 | 0.991835066 | -3.998370696 |
| **A_51_P131942** | Dph5 | 0.138039926 | -0.024729401 | 0.300809253 | 10.07221223 | 2.026941823 | 0.084414137 | 0.991835066 | -3.999148008 |
| **A_51_P199899** | Fto | -0.108444078 | -0.236338901 | 0.019450746 | 10.07223772 | -2.026571838 | 0.084459304 | 0.991835066 | -3.999483995 |
| **A_55_P2013986** | Ddx31 | 0.133499373 | -0.024052225 | 0.291050972 | 9.144005761 | 2.025188929 | 0.084628337 | 0.991835066 | -4.000739855 |
| **A_55_P2026214** | NA | -0.111515246 | -0.243186003 | 0.020155511 | 12.16831197 | -2.024202926 | 0.084749062 | 0.991835066 | -4.001635297 |
| **A_55_P2085485** | Pidd1 | 0.097713458 | -0.017669857 | 0.213096773 | 8.236029116 | 2.024046586 | 0.08476822 | 0.991835066 | -4.00177728 |

| | | | | | | | | | |
|---|---|---|---|---|---|---|---|---|---|
| **A_52_P680710** | Kpna3 | -0.11895441 | -0.259443955 | 0.021535136 | 10.19421134 | -2.023697848 | 0.08481097 | 0.991835066 | -4.002093993 |
| **A_55_P2037439** | Depdc1a | -0.110768711 | -0.241605645 | 0.020068223 | 8.444532021 | -2.023465832 | 0.084839424 | 0.991835066 | -4.002304705 |
| **A_55_P2173373** | Zfp62 | 0.115967731 | -0.021013421 | 0.252948883 | 9.007165789 | 2.023417385 | 0.084845366 | 0.991835066 | -4.002348703 |
| **A_51_P100289** | Srf | -0.120521524 | -0.262888576 | 0.021845528 | 12.32672815 | -2.02331851 | 0.084857495 | 0.991835066 | -4.002438499 |
| **A_55_P1962305** | Plac8 | -0.428727652 | -0.935190121 | 0.077734816 | 8.81323185 | -2.02322117 | 0.084869438 | 0.991835066 | -4.002526902 |
| **A_51_P280492** | Ndufa9 | -0.11976281 | -0.26127368 | 0.02174806 | 13.91511916 | -2.022745803 | 0.084927785 | 0.991835066 | -4.002958624 |
| **A_51_P299287** | Impa2 | 0.146767433 | -0.026678172 | 0.320213039 | 12.87667413 | 2.022439465 | 0.084965407 | 0.991835066 | -4.003236839 |
| **A_55_P2091201** | Thoc2 | 0.108905576 | -0.019806058 | 0.237617209 | 8.614614544 | 2.022280743 | 0.084984906 | 0.991835066 | -4.00338099 |
| **A_55_P2066180** | Hid1 | 0.092517029 | -0.016856052 | 0.201890111 | 9.581326284 | 2.021716996 | 0.0850542 | 0.991835066 | -4.003892988 |
| **A_55_P2057459** | Mall | -0.160137109 | -0.349464675 | 0.029190458 | 10.15379948 | -2.021562911 | 0.085073149 | 0.991835066 | -4.00403293 |
| **A_52_P306305** | NA | -0.139294134 | -0.305287476 | 0.026699208 | 13.65284723 | -2.084535558 | 0.085098597 | 0.991835066 | -3.995866956 |
| **A_55_P2183735** | Pisd | 0.108482844 | -0.019882625 | 0.236848314 | 10.03176981 | 2.019863321 | 0.085282443 | 0.991835066 | -4.00557655 |
| **A_51_P169920** | Noc2l | 0.128150012 | -0.02356209 | 0.279862115 | 10.72153832 | 2.018866295 | 0.085405459 | 0.991835066 | -4.006482105 |
| **A_66_P125701** | NA | 0.102818372 | -0.018909093 | 0.224545837 | 8.226640377 | 2.018790618 | 0.085414803 | 0.991835066 | -4.00655084 |
| **A_55_P2126950** | Zfp467 | -0.179956031 | -0.393101051 | 0.033188989 | 11.48780837 | -2.017903293 | 0.085524444 | 0.991835066 | -4.007356774 |
| **A_51_P432851** | Dhrs11 | -0.215263878 | -0.470279148 | 0.039751393 | 9.141826759 | -2.017502606 | 0.085574001 | 0.991835066 | -4.007720713 |
| **A_55_P1976682** | 4932431P2 | -0.393213875 | -0.859060757 | 0.072633008 | 12.14297904 | -2.017412804 | 0.085585111 | 0.991835066 | -4.007802279 |
| **A_55_P2024012** | Rpl5 | 0.091784835 | -0.017022078 | 0.200591748 | 14.632903 | 2.01615342 | 0.085741077 | 0.991835066 | -4.008946181 |
| **A_51_P140871** | Ankhd1 | 0.124214256 | -0.023043703 | 0.271472215 | 11.44712161 | 2.01605231 | 0.08575361 | 0.991835066 | -4.00903802 |
| **A_55_P1960231** | Txn2 | -0.11185338 | -0.244484374 | 0.020777615 | 14.52765799 | -2.015641155 | 0.085804598 | 0.991835066 | -4.00941148 |
| **A_55_P2083121** | Vill | 0.16098263 | -0.030000569 | 0.351965828 | 9.681011074 | 2.014619289 | 0.085931449 | 0.991835066 | -4.010339673 |
| **A_55_P2095271** | Pkn3 | 0.153908216 | -0.028688246 | 0.336504679 | 9.741949321 | 2.014552444 | 0.085939754 | 0.991835066 | -4.01040039 |
| **A_55_P1999785** | NA | 0.102287026 | -0.01908897 | 0.223663022 | 10.81044604 | 2.014173501 | 0.085986847 | 0.991835066 | -4.010744601 |
| **A_66_P117495** | Capza1 | 0.122832853 | -0.02300839 | 0.268674097 | 9.410174953 | 2.012997864 | 0.086133112 | 0.991835066 | -4.0118125 |
| **A_55_P1956728** | NA | 0.126816618 | -0.023763025 | 0.277396262 | 13.48557313 | 2.012885332 | 0.086147126 | 0.991835066 | -4.01191472 |
| **A_55_P2002466** | Zfp747 | -0.088517079 | -0.193631751 | 0.016597592 | 9.92242422 | -2.012671376 | 0.086173776 | 0.991835066 | -4.012109071 |
| **A_55_P1990490** | Recql | 0.125041425 | -0.02344911 | 0.273531961 | 8.628196895 | 2.012631544 | 0.086178738 | 0.991835066 | -4.012145253 |
| **A_51_P411917** | Gata6 | -0.141512843 | -0.309585766 | 0.026560081 | 13.01496489 | -2.012367209 | 0.086211676 | 0.991835066 | -4.012385369 |
| **A_51_P225771** | NA | 0.104040867 | -0.019580795 | 0.227662528 | 12.57532921 | 2.011492953 | 0.086320705 | 0.991835066 | -4.013179529 |
| **A_55_P1985895** | Thumpd1 | 0.14066048 | -0.026481387 | 0.307802346 | 11.5105815 | 2.011388576 | 0.086333731 | 0.991835066 | -4.013274344 |
| **A_52_P651948** | Fam229b | -0.096193037 | -0.210563518 | 0.018177444 | 8.488397036 | -2.010197922 | 0.086482461 | 0.991835066 | -4.014355937 |
| **A_51_P112762** | Slc5a3 | 0.095135101 | -0.017985927 | 0.208256129 | 8.116466285 | 2.010048672 | 0.086501123 | 0.991835066 | -4.016046584 |
| **A_66_P132855** | Baz1b | 0.101301964 | -0.01917604 | 0.221779969 | 8.624553363 | 2.009644473 | 0.086551682 | 0.991835066 | -4.014858699 |
| **A_55_P2012630** | NA | 0.108241609 | -0.020494381 | 0.2369776 | 14.75282499 | 2.009571177 | 0.086560853 | 0.991835066 | -4.014925283 |
| **A_55_P2014511** | Amd2 | 0.217460185 | -0.04125226 | 0.47617263 | 10.36261602 | 2.008961274 | 0.086637206 | 0.991835066 | -4.015479335 |
| **A_55_P1999576** | Josd1 | 0.142888094 | -0.028517367 | 0.314293556 | 13.24133556 | 2.070801818 | 0.086687103 | 0.991835066 | -4.006442555 |
| **A_51_P248638** | Myoz2 | -0.203674903 | -0.44605314 | 0.038703335 | 13.283959 | -2.008413016 | 0.0867059 | 0.991835066 | -4.015977391 |
| **A_55_P2003673** | Mtmr12 | 0.117543801 | -0.022352292 | 0.257439894 | 10.02518815 | 2.00818295 | 0.086734742 | 0.991835066 | -4.016186392 |
| **A_55_P2179020** | Mrpl15 | 0.181124823 | -0.034484148 | 0.396733793 | 9.797952937 | 2.007799214 | 0.086782869 | 0.991835066 | -4.016534996 |
| **A_55_P2092831** | NA | 0.48605641 | -0.092588752 | 1.064701572 | 12.54356234 | 2.007629095 | 0.086804214 | 0.991835066 | -4.01668954 |
| **A_51_P127464** | Ppp1r37 | 0.247404472 | -0.047146795 | 0.54195574 | 9.461578556 | 2.007501059 | 0.086820282 | 0.991835066 | -4.016805855 |
| **A_51_P425080** | Hdgf | -0.105987313 | -0.232192455 | 0.020217829 | 13.08865951 | -2.00717836 | 0.086860793 | 0.991835066 | -4.017099013 |
| **A_55_P2042685** | NA | -0.100419159 | -0.220032353 | 0.019194034 | 10.11779503 | -2.006534486 | 0.086941678 | 0.991835066 | -4.017683948 |
| **A_52_P641758** | Ggct | 0.175328791 | -0.03355618 | 0.384213761 | 9.419012196 | 2.006111999 | 0.086994793 | 0.991835066 | -4.018067764 |
| **A_55_P1954544** | Zfp446 | 0.092506945 | -0.01770992 | 0.202723811 | 8.968879901 | 2.006020726 | 0.087006273 | 0.991835066 | -4.018150683 |
| **A_55_P2181109** | Il10rb | -0.169837862 | -0.372222569 | 0.032546845 | 10.34457071 | -2.005699902 | 0.087046634 | 0.991835066 | -4.018442144 |
| **A_55_P2122643** | Dtymk | 0.125240971 | -0.024001074 | 0.274483016 | 12.17240132 | 2.00569257 | 0.087047556 | 0.991835066 | -4.018448805 |
| **A_51_P283292** | Gm14326 | 0.134649709 | -0.025805227 | 0.295104645 | 8.936039382 | 2.0056792 | 0.087049239 | 0.991835066 | -4.018460952 |

| | | | | | | | | | |
|---|---|---|---|---|---|---|---|---|---|
| **A_51_P342481** | Gpd2 | -0.083875097 | -0.183874686 | 0.016124493 | 8.766558745 | -2.004674886 | 0.087175709 | 0.991835066 | -4.019373356 |
| **A_51_P184353** | Nr2f6 | -0.13405502 | -0.293966581 | 0.025856541 | 12.77692734 | -2.003606134 | 0.087310495 | 0.991835066 | -4.020344318 |
| **A_52_P49321** | Adamts9 | -0.140037856 | -0.307115593 | 0.027039882 | 10.7467437 | -2.003253911 | 0.087354961 | 0.991835066 | -4.020664316 |
| **A_55_P1953087** | Mcm3 | 0.187345241 | -0.036234075 | 0.410924556 | 12.74812341 | 2.002719726 | 0.087422442 | 0.991835066 | -4.021149632 |
| **A_55_P1967481** | Hbb-y | 1.071635728 | -0.207317973 | 2.35058943 | 9.406920807 | 2.002633579 | 0.087433329 | 0.991835066 | -4.021227898 |
| **A_52_P550185** | Urm1 | -0.104188348 | -0.228558235 | 0.020181538 | 11.24276118 | -2.002225773 | 0.087484886 | 0.991835066 | -4.021598399 |
| **A_55_P1990629** | NA | -0.23912439 | -0.524612329 | 0.046363549 | 11.58157577 | -2.001913196 | 0.087524425 | 0.991835066 | -4.021882384 |
| **A_66_P140533** | NA | 0.154829128 | -0.030040773 | 0.339699029 | 8.63799455 | 2.001684336 | 0.087553385 | 0.991835066 | -4.02209031 |
| **A_55_P1971474** | NA | 0.083212403 | -0.016146056 | 0.182570862 | 11.07080739 | 2.001669388 | 0.087555277 | 0.991835066 | -4.022103891 |
| **A_66_P128055** | Spindoc | 0.135414762 | -0.026329054 | 0.297158578 | 8.672617423 | 2.001001695 | 0.087639823 | 0.991835066 | -4.022710516 |
| **A_55_P2207255** | Srpx | -0.154113664 | -0.338221058 | 0.029993729 | 9.678195896 | -2.000686538 | 0.087679758 | 0.991835066 | -4.02299685 |
| **A_55_P2187266** | Pop1 | 0.151453331 | -0.029503349 | 0.332410012 | 9.918341338 | 2.000383857 | 0.087718129 | 0.991835066 | -4.02327185 |
| **A_52_P14666** | NA | 0.094898353 | -0.018543426 | 0.208340131 | 8.434895682 | 1.999377407 | 0.087845837 | 0.991835066 | -4.024186266 |
| **A_52_P20966** | Hnrnpab | 0.095999441 | -0.018788993 | 0.210787875 | 13.7988958 | 1.998847704 | 0.087913125 | 0.991835066 | -4.024667535 |
| **A_51_P468329** | Psmb7 | -0.244445129 | -0.536804289 | 0.047914031 | 11.35623686 | -1.998360401 | 0.087975072 | 0.991835066 | -4.025110285 |
| **A_55_P2045188** | NA | -0.133766381 | -0.2937575 | 0.026224739 | 8.936144333 | -1.998297897 | 0.087983021 | 0.991835066 | -4.025167074 |
| **A_55_P2031989** | Rin2 | -0.14307098 | -0.314208985 | 0.028067024 | 10.03913559 | -1.998086212 | 0.088009947 | 0.991835066 | -4.025359406 |
| **A_55_P2012478** | Zfp618 | 0.115044792 | -0.022585499 | 0.252675084 | 10.59299728 | 1.997846273 | 0.088040476 | 0.991835066 | -4.02557741 |
| **A_51_P367125** | Pold1 | 0.173065398 | -0.034035074 | 0.38016587 | 11.09081921 | 1.997276951 | 0.088112958 | 0.991835066 | -4.026094686 |
| **A_55_P2105833** | Chchd5 | -0.086037111 | -0.189071937 | 0.016997714 | 9.875564038 | -1.995771944 | 0.088304851 | 0.991835066 | -4.027462129 |
| **A_51_P222522** | Mesd | 0.115938419 | -0.023035626 | 0.254912464 | 12.71018146 | 1.993897408 | 0.08854444 | 0.991835066 | -4.029165358 |
| **A_55_P2064230** | Tmx3 | -0.100276471 | -0.220477894 | 0.019924953 | 8.983277925 | -1.993877907 | 0.088546936 | 0.991835066 | -4.029183078 |
| **A_51_P397564** | Snrnp40 | 0.129893524 | -0.02586697 | 0.285654018 | 10.63125669 | 1.993146934 | 0.088640541 | 0.991835066 | -4.02984726 |
| **A_55_P2114776** | Greb1 | 0.094824449 | -0.018887333 | 0.20853623 | 8.110433135 | 1.993076623 | 0.08864955 | 0.991835066 | -4.029911147 |
| **A_52_P303928** | Rab2a | -0.161348335 | -0.35488004 | 0.032183369 | 11.53200584 | -1.992606307 | 0.088709834 | 0.991835066 | -4.030338492 |
| **A_55_P2063505** | NA | 0.475352827 | -0.094872752 | 1.045578406 | 11.93771226 | 1.992409107 | 0.088735123 | 0.991835066 | -4.030517675 |
| **A_51_P214825** | Orc5 | -0.090094882 | -0.198207198 | 0.018017433 | 8.391704131 | -1.991746609 | 0.088820135 | 0.991835066 | -4.031119649 |
| **A_51_P509211** | Rnf34 | -0.097029751 | -0.213496979 | 0.019437476 | 11.27095596 | -1.991179042 | 0.088893029 | 0.991835066 | -4.031635368 |
| **A_52_P387954** | Dctn5 | 0.115905592 | -0.023247669 | 0.255058852 | 11.26484177 | 1.990765626 | 0.088946162 | 0.991835066 | -4.03201102 |
| **A_55_P1974412** | Kat2a | 0.104171587 | -0.020897088 | 0.229240262 | 10.38555709 | 1.990718559 | 0.088952213 | 0.991835066 | -4.032053787 |
| **A_66_P103301** | Dicer1 | 0.113909389 | -0.022863073 | 0.250681851 | 10.52324403 | 1.99053579 | 0.088975715 | 0.991835066 | -4.032219862 |
| **A_55_P2069818** | Mapk10 | -0.325210577 | -0.715891048 | 0.065469894 | 9.422415786 | -1.989537325 | 0.089104213 | 0.991835066 | -4.033127129 |
| **A_55_P2153021** | Ttn | -0.134080744 | -0.295206186 | 0.027044698 | 14.99386797 | -1.988893007 | 0.089187232 | 0.991835066 | -4.033712601 |
| **A_55_P2004797** | Tacc2 | -0.11454921 | -0.252259811 | 0.023161391 | 12.20842491 | -1.988079998 | 0.089292096 | 0.991835066 | -4.034451363 |
| **A_55_P2050997** | Serinc4 | -0.111059645 | -0.244588697 | 0.022469408 | 8.738510297 | -1.987877623 | 0.089318218 | 0.991835066 | -4.034635257 |
| **A_55_P1970033** | Per1 | -0.19740911 | -0.436803034 | 0.041984814 | 12.34383044 | -2.048430423 | 0.089340276 | 0.991835066 | -4.023706228 |
| **A_55_P2029533** | Zfp57 | -0.095249156 | -0.209800613 | 0.019302301 | 7.987451328 | -1.987328451 | 0.089389141 | 0.991835066 | -4.035134281 |
| **A_55_P1971951** | Cd86 | -0.088148013 | -0.194178546 | 0.017882521 | 7.839869073 | -1.986967264 | 0.089435817 | 0.991835066 | -4.035462487 |
| **A_51_P502456** | H2-K2 | -0.17416023 | -0.383690416 | 0.035369955 | 9.536513474 | -1.986605003 | 0.089482657 | 0.991835066 | -4.03579167 |
| **A_55_P2151751** | Matr3 | -0.106625512 | -0.234932914 | 0.021681889 | 10.93283714 | -1.986179769 | 0.089537669 | 0.991835066 | -4.036178077 |
| **A_55_P1967659** | Lactb | -0.147510379 | -0.325039907 | 0.03001915 | 9.616991498 | -1.985917079 | 0.08957167 | 0.991835066 | -4.036416783 |
| **A_55_P2148708** | NA | 0.105039135 | -0.021419263 | 0.231497534 | 12.95986734 | 1.985238057 | 0.089659618 | 0.991835066 | -4.037033809 |
| **A_55_P2153251** | Plagl1 | 0.150613132 | -0.030761544 | 0.331987807 | 8.580256594 | 1.984702071 | 0.0897291 | 0.991835066 | -4.037520862 |
| **A_51_P252784** | Fam216a | 0.090491933 | -0.018511118 | 0.199494983 | 10.85661097 | 1.98417669 | 0.08979726 | 0.991835066 | -4.037998281 |
| **A_51_P107362** | Socs2 | -0.488848361 | -1.07786468 | 0.100167958 | 10.39135874 | -1.983608535 | 0.089871026 | 0.991835066 | -4.03851457 |
| **A_55_P2280237** | Zfp697 | 0.126207292 | -0.025879674 | 0.278294258 | 8.507186193 | 1.983360151 | 0.089903294 | 0.991835066 | -4.03874028 |
| **A_55_P2118734** | Rnmt | 0.123217243 | -0.025306261 | 0.271740747 | 9.690498953 | 1.982829781 | 0.089972233 | 0.991835066 | -4.039222238 |
| **A_55_P2087225** | Hnrnpa3 | 0.212325073 | -0.043690025 | 0.468340171 | 11.90954446 | 1.982187976 | 0.090055727 | 0.991835066 | -4.03980546 |

| | | | | | | | | | |
|---|---|---|---|---|---|---|---|---|---|
| **A_55_P2125963** | NA | 0.137863156 | -0.028397887 | 0.304124199 | 9.300471782 | 1.981832055 | 0.090102063 | 0.991835066 | -4.040128894 |
| **A_52_P201206** | Scrn1 | -0.131997502 | -0.291208237 | 0.027213233 | 9.323694457 | -1.981538415 | 0.090140309 | 0.991835066 | -4.040395734 |
| **A_52_P407796** | Mdh1 | -0.093728192 | -0.20678905 | 0.019332665 | 16.32478099 | -1.981376911 | 0.090161351 | 0.991835066 | -4.040542497 |
| **A_55_P2001168** | NA | 0.132178491 | -0.027322178 | 0.291679161 | 10.82982185 | 1.980648519 | 0.090256313 | 0.991835066 | -4.041204411 |
| **A_51_P187842** | Eif4e3 | -0.094883606 | -0.209400529 | 0.019633317 | 8.625666738 | -1.980298427 | 0.09030199 | 0.991835066 | -4.041522552 |
| **A_51_P189722** | Cabin1 | 0.11891229 | -0.024656908 | 0.262481489 | 9.724671982 | 1.979587031 | 0.090394878 | 0.991835066 | -4.042169024 |
| **A_52_P240453** | Polr3k | 0.14744323 | -0.03065361 | 0.32554007 | 10.69511017 | 1.97868998 | 0.090512143 | 0.991835066 | -4.042984214 |
| **A_55_P2102621** | Eaf2 | -0.15177789 | -0.335160743 | 0.031604963 | 11.36744954 | -1.978148629 | 0.090582983 | 0.991835066 | -4.043476165 |
| **A_55_P2116149** | NA | 0.598785165 | -0.124755635 | 1.322325965 | 12.31535728 | 1.977958332 | 0.090607898 | 0.991835066 | -4.043649098 |
| **A_55_P2000329** | Gskip | 0.105145513 | -0.02194872 | 0.232239745 | 8.640892916 | 1.97730668 | 0.090693267 | 0.991835066 | -4.044241288 |
| **A_51_P372702** | Il16 | -0.1222541 | -0.270049846 | 0.025541645 | 8.498269867 | -1.977018046 | 0.090731105 | 0.991835066 | -4.044503585 |
| **A_55_P2128695** | Nckap1 | -0.114344168 | -0.252597869 | 0.023909533 | 10.34621099 | -1.976725598 | 0.090769459 | 0.991835066 | -4.044769349 |
| **A_52_P197095** | Cdk17 | -0.091360182 | -0.201837401 | 0.019117037 | 8.478464044 | -1.976484228 | 0.090801126 | 0.991835066 | -4.044988695 |
| **A_55_P1955821** | Ccl21b | 0.311073889 | -0.065190048 | 0.687337827 | 12.91910576 | 1.975968858 | 0.090868778 | 0.991835066 | -4.045457042 |
| **A_52_P142208** | Akirin1 | 0.189288777 | -0.039689877 | 0.41826743 | 10.70769114 | 1.975781887 | 0.090893334 | 0.991835066 | -4.045626955 |
| **A_55_P2026125** | Srrm2 | 0.120421283 | -0.025309415 | 0.266151982 | 8.651251338 | 1.974973789 | 0.090999542 | 0.991835066 | -4.046361324 |
| **A_51_P387235** | Nampt | -0.088556292 | -0.195727264 | 0.01861468 | 9.753177989 | -1.974928543 | 0.091005492 | 0.991835066 | -4.046402442 |
| **A_66_P100365** | NA | 0.13343022 | -0.028080054 | 0.294940494 | 8.868256382 | 1.974527466 | 0.091058254 | 0.991835066 | -4.046766927 |
| **A_55_P2058657** | NA | 0.126399163 | -0.026665748 | 0.279464074 | 16.30755932 | 1.973684304 | 0.091169272 | 0.991835066 | -4.047533165 |
| **A_55_P1967683** | Bex4 | 0.120754139 | -0.026766872 | 0.268275151 | 13.17701853 | 2.03336477 | 0.091173857 | 0.991835066 | -4.035356648 |
| **A_51_P450123** | Mrpl36 | 0.093003993 | -0.019641611 | 0.205649597 | 12.6835471 | 1.973315286 | 0.091217902 | 0.991835066 | -4.047868519 |
| **A_55_P2028054** | Incenp | 0.147689544 | -0.031202909 | 0.326581998 | 12.75683183 | 1.973180719 | 0.091235642 | 0.991835066 | -4.04799081 |
| **A_55_P2049867** | Ccrl2 | -0.13126547 | -0.290294824 | 0.027763884 | 8.286047472 | -1.972796708 | 0.091286285 | 0.991835066 | -4.048339788 |
| **A_55_P2114569** | Zfp24 | 0.099564976 | -0.021061521 | 0.220191473 | 11.30009038 | 1.972754353 | 0.091291872 | 0.991835066 | -4.048378279 |
| **A_51_P348665** | Ramp1 | -0.307630226 | -0.680428234 | 0.065167783 | 11.69435444 | -1.972261755 | 0.09135688 | 0.991835066 | -4.04882594 |
| **A_55_P2018191** | Ccm2l | 0.08788212 | -0.018629174 | 0.194393415 | 9.103978455 | 1.972032173 | 0.091387194 | 0.991835066 | -4.049034578 |
| **A_66_P111534** | 5430431A1 | 0.348405101 | -0.074106728 | 0.77091693 | 12.91935414 | 1.970855417 | 0.091542727 | 0.991835066 | -4.050103988 |
| **A_66_P114333** | Tlr12 | -0.095645093 | -0.211665089 | 0.020374902 | 8.045683046 | -1.97033006 | 0.091612248 | 0.991835066 | -4.050581422 |
| **A_55_P2181071** | NA | 0.111201493 | -0.023694201 | 0.246097186 | 15.19045409 | 1.970251483 | 0.091622651 | 0.991835066 | -4.050652832 |
| **A_55_P2065631** | Hook3 | 0.127462097 | -0.027163271 | 0.282087465 | 10.68110716 | 1.970196055 | 0.09162999 | 0.991835066 | -4.050703204 |
| **A_52_P480064** | Tmem134 | -0.18008918 | -0.398632313 | 0.038453953 | 12.58923823 | -1.96951634 | 0.091720033 | 0.991835066 | -4.051320917 |
| **A_55_P1956842** | NA | 0.156974647 | -0.033524642 | 0.347473936 | 14.65540628 | 1.969451487 | 0.091728629 | 0.991835066 | -4.051379854 |
| **A_55_P2112445** | Tbx4 | 0.097289959 | -0.020799842 | 0.215379759 | 7.724252858 | 1.969086356 | 0.091777039 | 0.991835066 | -4.05171168 |
| **A_55_P2048518** | Zfpm1 | 0.095670825 | -0.020497733 | 0.211839382 | 12.08249026 | 1.968339705 | 0.091876111 | 0.991835066 | -4.052390226 |
| **A_55_P2154107** | Gcm1 | 0.385323648 | -0.082828968 | 0.853476264 | 10.96067139 | 1.967194678 | 0.092028249 | 0.991835066 | -4.053430812 |
| **A_66_P106654** | Camsap1 | 0.115228998 | -0.024788022 | 0.255246018 | 11.04256001 | 1.966935355 | 0.092062739 | 0.991835066 | -4.053666482 |
| **A_55_P2135633** | NA | 0.102362567 | -0.022036578 | 0.226761712 | 15.96485492 | 1.966676387 | 0.092097195 | 0.991835066 | -4.05390183 |
| **A_55_P2023747** | Mfsd10 | 0.125524696 | -0.027118095 | 0.278167488 | 10.77343469 | 1.965450054 | 0.092260533 | 0.991835066 | -4.05501631 |
| **A_55_P1968763** | Shc4 | 0.298256715 | -0.064491795 | 0.661005225 | 10.4154805 | 1.965141041 | 0.092301737 | 0.991835066 | -4.055297139 |
| **A_51_P393426** | Pros1 | -0.092657708 | -0.205377042 | 0.020061626 | 9.13882227 | -1.964682017 | 0.092362976 | 0.991835066 | -4.055714296 |
| **A_55_P2150757** | Gzmm | -0.121641578 | -0.269640659 | 0.026357503 | 8.873610246 | -1.964410099 | 0.092399272 | 0.991835066 | -4.055961413 |
| **A_55_P2417569** | Prrx2 | 0.224071896 | -0.048554562 | 0.496698354 | 10.34671776 | 1.964393644 | 0.092401469 | 0.991835066 | -4.055976367 |
| **A_52_P282987** | Yeats2 | 0.102982617 | -0.022333346 | 0.228298581 | 9.922082262 | 1.964113855 | 0.092438831 | 0.991835066 | -4.056230638 |
| **A_55_P1961049** | Cep295 | 0.194378498 | -0.042155721 | 0.430912717 | 9.959041507 | 1.964099008 | 0.092440815 | 0.991835066 | -4.056244131 |
| **A_52_P178998** | Fam168b | 0.19541753 | -0.042383068 | 0.433218129 | 12.3988225 | 1.964082422 | 0.09244303 | 0.991835066 | -4.056259204 |
| **A_51_P303180** | Fam114a1 | 0.174759114 | -0.037913366 | 0.387431593 | 9.932248006 | 1.963982757 | 0.092456343 | 0.991835066 | -4.056349778 |
| **A_55_P2131288** | Ank2 | -0.113324699 | -0.251275821 | 0.024626423 | 11.4987998 | -1.96339856 | 0.092534418 | 0.991835066 | -4.056880693 |
| **A_66_P105830** | 1700028B0 | -0.199859388 | -0.44319684 | 0.043478064 | 16.03063452 | -1.963020022 | 0.092585043 | 0.991835066 | -4.057224706 |

| | | | | | | | | | |
|---|---|---|---|---|---|---|---|---|---|
| **A_51_P474459** | Socs3 | -0.22102562 | -0.490162337 | 0.048111097 | 9.206260706 | -1.962812429 | 0.092612817 | 0.991835066 | -4.057413365 |
| **A_52_P202725** | Rragc | 0.161675715 | -0.035201308 | 0.358552737 | 8.947365607 | 1.962722447 | 0.092624859 | 0.991835066 | -4.057495141 |
| **A_55_P2078680** | Tpm1 | 0.121945716 | -0.026599429 | 0.270490861 | 8.42684157 | 1.962082293 | 0.09271057 | 0.991835066 | -4.058076909 |
| **A_52_P425706** | Rnf168 | 0.125588977 | -0.027397586 | 0.278575541 | 10.92405215 | 1.962037791 | 0.092716532 | 0.991835066 | -4.058117352 |
| **A_51_P454993** | Tmcc2 | 0.096314889 | -0.021012642 | 0.21364242 | 8.144062129 | 1.962016384 | 0.092719399 | 0.991835066 | -4.058136806 |
| **A_55_P2114269** | Fam177a2 | -0.161981417 | -0.359317253 | 0.035354419 | 16.44837767 | -1.961861593 | 0.092740138 | 0.991835066 | -4.058277479 |
| **A_52_P45738** | Trim35 | 0.138886849 | -0.030316522 | 0.30809022 | 8.567918967 | 1.961829439 | 0.092744447 | 0.991835066 | -4.058306701 |
| **A_55_P1962092** | Mpc1 | -0.147431198 | -0.327088036 | 0.032225639 | 13.58865767 | -1.961348541 | 0.09280891 | 0.991835066 | -4.058743738 |
| **A_55_P2462377** | Nsf | 0.079408375 | -0.017359305 | 0.176176055 | 10.45970218 | 1.961304899 | 0.092814762 | 0.991835066 | -4.0587834 |
| **A_55_P2032997** | Srp19 | 0.156427336 | -0.034215581 | 0.347070254 | 11.45680269 | 1.961106173 | 0.092841415 | 0.991835066 | -4.058964001 |
| **A_51_P506787** | Nap1l1 | 0.195166176 | -0.042711969 | 0.43304432 | 10.90395175 | 1.960916685 | 0.092866837 | 0.991835066 | -4.059136206 |
| **A_55_P2048279** | Tlr13 | -0.199921732 | -0.443606613 | 0.04376315 | 8.64408208 | -1.960832748 | 0.0928781 | 0.991835066 | -4.059212488 |
| **A_55_P2186929** | NA | 0.373984211 | -0.081948831 | 0.829917253 | 10.05003229 | 1.96047511 | 0.092926104 | 0.991835066 | -4.059537507 |
| **A_55_P2127825** | Bcas3 | -0.137711346 | -0.305623705 | 0.030201014 | 9.241401594 | -1.960181066 | 0.092965591 | 0.991835066 | -4.059804733 |
| **A_55_P1975742** | Dap3 | -0.237718583 | -0.527585119 | 0.052147953 | 12.01326564 | -1.960081746 | 0.092978933 | 0.991835066 | -4.059894994 |
| **A_55_P2079520** | Mpp1 | -0.161247033 | -0.357874874 | 0.035380808 | 9.224316163 | -1.959999014 | 0.092990047 | 0.991835066 | -4.059970181 |
| **A_55_P2154173** | NA | 0.204192634 | -0.044875196 | 0.453260465 | 14.92906986 | 1.959438181 | 0.093065427 | 0.991835066 | -4.060479862 |
| **A_55_P1998872** | Slitrk5 | 0.171411243 | -0.037822853 | 0.38064534 | 8.911201081 | 1.95801484 | 0.093257004 | 0.991835066 | -4.061773384 |
| **A_55_P2065370** | NA | -0.117729388 | -0.261480421 | 0.026021646 | 12.80345837 | -1.957415588 | 0.093337778 | 0.991835066 | -4.06231798 |
| **A_55_P1993758** | Nsa2 | 0.088088094 | -0.019499314 | 0.195675503 | 11.22001545 | 1.956883187 | 0.093409599 | 0.991835066 | -4.062801821 |
| **A_55_P1992834** | Socs2 | -0.435601632 | -0.96765016 | 0.096446897 | 9.940620651 | -1.956804254 | 0.093420252 | 0.991835066 | -4.062873555 |
| **A_55_P2005828** | Hbb-bh1 | 0.187556532 | -0.041579362 | 0.416692426 | 8.000775885 | 1.956357407 | 0.093480581 | 0.991835066 | -4.063279645 |
| **A_52_P88091** | Dsg2 | 0.103968339 | -0.023060744 | 0.230997423 | 8.103327964 | 1.956172166 | 0.093505601 | 0.991835066 | -4.06344799 |
| **A_51_P463428** | Pik3ip1 | -0.319571515 | -0.710120354 | 0.070977324 | 10.88552981 | -1.955698232 | 0.093569646 | 0.991835066 | -4.063878696 |
| **A_51_P183853** | Polr2m | 0.09148523 | -0.020335823 | 0.203306283 | 14.41633235 | 1.955404182 | 0.093609404 | 0.991835066 | -4.064145925 |
| **A_55_P2340448** | B230114P1 | -0.221700139 | -0.49270403 | 0.049303753 | 8.501087345 | -1.955237732 | 0.093631917 | 0.991835066 | -4.064297192 |
| **A_55_P2112230** | Pcyt2 | 0.10852546 | -0.024147804 | 0.241198725 | 12.71595991 | 1.955047688 | 0.093657628 | 0.991835066 | -4.064469902 |
| **A_55_P1962811** | Anp32b | 0.178372048 | -0.039732737 | 0.396476833 | 13.41094437 | 1.954657802 | 0.093710397 | 0.991835066 | -4.064824226 |
| **A_55_P2110875** | 2700046G0 | 0.109496858 | -0.024399129 | 0.243392845 | 9.577302548 | 1.954534003 | 0.093727158 | 0.991835066 | -4.064936733 |
| **A_52_P472324** | Slpi | -0.155815388 | -0.346362575 | 0.0347318 | 8.336714231 | -1.954415661 | 0.093743184 | 0.991835066 | -4.06504428 |
| **A_51_P490257** | Fbxo4 | -0.086014079 | -0.191204796 | 0.019176638 | 10.38660838 | -1.95434512 | 0.093752738 | 0.991835066 | -4.065108387 |
| **A_55_P2031167** | Efna1 | -0.107164594 | -0.238261966 | 0.023932779 | 11.35554086 | -1.953738648 | 0.093834915 | 0.991835066 | -4.06565954 |
| **A_55_P2021659** | Abcb8 | -0.095760172 | -0.212942928 | 0.021422585 | 11.67674441 | -1.953126349 | 0.093917954 | 0.991835066 | -4.066215987 |
| **A_55_P2082806** | Trib1 | 0.146745393 | -0.03283334 | 0.326324125 | 9.255360375 | 1.953074027 | 0.093925053 | 0.991835066 | -4.066263536 |
| **A_55_P1968370** | Gprc5c | 0.143118879 | -0.032054101 | 0.318291859 | 9.516367413 | 1.95271535 | 0.093973734 | 0.991835066 | -4.066589496 |
| **A_52_P328078** | Atp5b | -0.223155825 | -0.496349148 | 0.050037498 | 17.13365752 | -1.952303261 | 0.094029694 | 0.991835066 | -4.066963994 |
| **A_52_P200617** | Iars | 0.191432869 | -0.043024468 | 0.425890206 | 11.19298261 | 1.9514697 | 0.09414299 | 0.991835066 | -4.067721517 |
| **A_51_P181400** | Efcab2 | -0.143234762 | -0.318666039 | 0.032196515 | 9.440724196 | -1.951419044 | 0.09414988 | 0.991835066 | -4.067767552 |
| **A_55_P1958246** | Tcof1 | 0.132450676 | -0.02980349 | 0.294704842 | 11.4141173 | 1.951045764 | 0.094200663 | 0.991835066 | -4.06810678 |
| **A_55_P2159710** | Acvr1 | 0.123307914 | -0.027746608 | 0.274362436 | 12.33968961 | 1.951040802 | 0.094201338 | 0.991835066 | -4.068111289 |
| **A_51_P374900** | P2ry13 | -0.079274464 | -0.176409017 | 0.017860088 | 7.722669059 | -1.950602183 | 0.094261045 | 0.991835066 | -4.068509896 |
| **A_55_P1970775** | Hdac1 | 0.109014414 | -0.024564599 | 0.242593426 | 13.06425307 | 1.950539842 | 0.094269534 | 0.991835066 | -4.068566549 |
| **A_52_P489729** | Sec61a1 | 0.165351349 | -0.03727941 | 0.367982107 | 13.28160898 | 1.950345281 | 0.094296033 | 0.991835066 | -4.068743362 |
| **A_55_P2007738** | Mir22hg | 0.090003817 | -0.02032078 | 0.200328414 | 7.942056371 | 1.949834313 | 0.094365661 | 0.991835066 | -4.069207716 |
| **A_55_P2115442** | Clec2d | 0.122134649 | -0.027580316 | 0.271849615 | 8.440712083 | 1.949767414 | 0.094374781 | 0.991835066 | -4.069268511 |
| **A_55_P2039572** | NA | 0.100372659 | -0.022667461 | 0.223412778 | 15.89437369 | 1.949745 | 0.094377837 | 0.991835066 | -4.069288881 |
| **A_52_P22324** | Pln | -0.268579436 | -0.597824099 | 0.060665227 | 13.49894854 | -1.949679143 | 0.094386816 | 0.991835066 | -4.069348729 |
| **A_55_P1992403** | Simc1 | 0.118633252 | -0.026800998 | 0.264067502 | 8.914113104 | 1.949615058 | 0.094395554 | 0.991835066 | -4.069406968 |

| | | | | | | | | | |
|---|---|---|---|---|---|---|---|---|---|
| **A_51_P512591** | Cdc42se1 | -0.118026074 | -0.262746879 | 0.026694732 | 9.966631526 | -1.949198741 | 0.09445234 | 0.991835066 | -4.069785304 |
| **A_52_P265002** | Pik3ca | -0.093424741 | -0.20804641 | 0.021196928 | 8.684350466 | -1.948068881 | 0.094606623 | 0.991835066 | -4.070812084 |
| **A_52_P581435** | Pdha1 | -0.114170685 | -0.254280881 | 0.025939511 | 12.46520906 | -1.947574139 | 0.094674259 | 0.991835066 | -4.071261687 |
| **A_66_P126552** | Prr14l | 0.135036039 | -0.030735307 | 0.300807384 | 8.817213764 | 1.946925619 | 0.09476299 | 0.991835066 | -4.071851036 |
| **A_52_P400999** | Arhgap31 | -0.132901171 | -0.296059146 | 0.030256804 | 12.10885138 | -1.946837192 | 0.094775095 | 0.991835066 | -4.071931395 |
| **A_55_P2052185** | Lamtor1 | -0.124395603 | -0.277129744 | 0.028338537 | 12.35955995 | -1.946605908 | 0.094806763 | 0.991835066 | -4.072141577 |
| **A_52_P539414** | Gtf2h3 | -0.143067214 | -0.318747989 | 0.032613561 | 9.519705377 | -1.946368263 | 0.094839313 | 0.991835066 | -4.072357538 |
| **A_55_P1976709** | NA | -0.083786391 | -0.186682168 | 0.019109387 | 13.08736706 | -1.946189219 | 0.094863844 | 0.991835066 | -4.072520246 |
| **A_55_P2113165** | Camkk1 | 0.113213966 | -0.025855173 | 0.252283106 | 8.582556085 | 1.94571116 | 0.094929373 | 0.991835066 | -4.072954684 |
| **A_55_P1956707** | Rpl36a | 0.111192422 | -0.025409211 | 0.247794055 | 14.82767717 | 1.945487435 | 0.094960056 | 0.991835066 | -4.073157995 |
| **A_52_P271377** | Efr3a | 0.091594882 | -0.020982551 | 0.204172315 | 8.499474002 | 1.944594252 | 0.095082647 | 0.991835066 | -4.073969673 |
| **A_55_P2159026** | Rpl22l1 | 0.187845506 | -0.043153321 | 0.418844333 | 14.33849187 | 1.943569916 | 0.095223432 | 0.991835066 | -4.074900533 |
| **A_55_P2142028** | Clip4 | -0.117590128 | -0.262194249 | 0.027013992 | 11.68234 | -1.943566163 | 0.095223948 | 0.991835066 | -4.074903944 |
| **A_52_P254149** | Irgq | -0.143337784 | -0.319616919 | 0.032941351 | 9.870556136 | -1.943430016 | 0.095242676 | 0.991835066 | -4.075027665 |
| **A_55_P2173664** | Frmpd2 | -0.142526151 | -0.317829589 | 0.032777286 | 13.05868517 | -1.943181017 | 0.095276936 | 0.991835066 | -4.075253941 |
| **A_52_P125350** | Anp32e | 0.132460595 | -0.030514251 | 0.295435442 | 10.8368851 | 1.942563641 | 0.095361935 | 0.991835066 | -4.075814974 |
| **A_55_P2129407** | NA | 0.273463859 | -0.063018434 | 0.609946153 | 8.612605245 | 1.942436612 | 0.095379433 | 0.991835066 | -4.075930409 |
| **A_52_P198898** | Samd5 | 0.114028101 | -0.026296982 | 0.254353185 | 9.117924497 | 1.942163176 | 0.09541711 | 0.991835066 | -4.076178889 |
| **A_55_P2057913** | Hmgb1-ps7 | -0.32573213 | -0.726588261 | 0.075124002 | 12.95441242 | -1.942142984 | 0.095419893 | 0.991835066 | -4.076197239 |
| **A_55_P2181655** | Tns3 | 0.109156892 | -0.025202441 | 0.243516225 | 8.255798051 | 1.941746159 | 0.095474599 | 0.991835066 | -4.076557846 |
| **A_55_P2075731** | NA | 0.557084904 | -0.128761151 | 1.24293096 | 12.04969941 | 1.941350169 | 0.095529221 | 0.991835066 | -4.076917693 |
| **A_52_P15388** | Ltf | -0.1713408 | -0.382315363 | 0.039633763 | 8.059078791 | -1.941063877 | 0.095568731 | 0.991835066 | -4.077177854 |
| **A_55_P1976864** | Fbxl20 | 0.307071068 | -0.071167563 | 0.685309699 | 9.90920899 | 1.940359268 | 0.095666039 | 0.991835066 | -4.077818148 |
| **A_55_P2080895** | Rab7 | 0.127755088 | -0.029618032 | 0.285128209 | 10.09558209 | 1.940245939 | 0.095681699 | 0.991835066 | -4.077921132 |
| **A_51_P247873** | Ndufb8 | -0.176833155 | -0.394663282 | 0.040996973 | 16.28058676 | -1.940237445 | 0.095682873 | 0.991835066 | -4.077928851 |
| **A_55_P2077009** | Acap3 | 0.161768949 | -0.037560334 | 0.361098233 | 10.4198972 | 1.939693864 | 0.095758022 | 0.991835066 | -4.078422813 |
| **A_55_P2028576** | NA | 0.300682923 | -0.069844741 | 0.671210586 | 11.85102588 | 1.939533378 | 0.09578022 | 0.991835066 | -4.078568649 |
| **A_51_P479583** | Obsl1 | 0.182735205 | -0.042460788 | 0.407931198 | 12.58406577 | 1.939414773 | 0.095796629 | 0.991835066 | -4.078676427 |
| **A_55_P2034400** | Cdk8 | 0.199547617 | -0.046443916 | 0.44553915 | 10.23038479 | 1.938811195 | 0.095880174 | 0.991835066 | -4.079224904 |
| **A_51_P114878** | Elp1 | 0.130943667 | -0.0304805 | 0.292367834 | 10.79926291 | 1.93876458 | 0.09588663 | 0.991835066 | -4.079306518 |
| **A_51_P512119** | AF067063 | -0.158388893 | -0.353680188 | 0.036902402 | 8.941011981 | -1.93843379 | 0.09593245 | 0.991835066 | -4.079567855 |
| **A_52_P157880** | Gm1947 | 0.115494556 | -0.026936462 | 0.257925574 | 9.142611087 | 1.938054866 | 0.095984965 | 0.991835066 | -4.079912186 |
| **A_55_P2158404** | Cmpk2 | -0.091055974 | -0.203357369 | 0.021245422 | 8.038930448 | -1.937904733 | 0.096005779 | 0.991835066 | -4.080048612 |
| **A_55_P2095488** | Syncrip | 0.171030221 | -0.039948349 | 0.38200879 | 9.62009343 | 1.937508631 | 0.096060717 | 0.991835066 | -4.08040855 |
| **A_55_P2184128** | Ndufa3 | -0.131030002 | -0.292695185 | 0.030635181 | 16.26686832 | -1.937150583 | 0.096110403 | 0.991835066 | -4.080733907 |
| **A_55_P2052281** | Rnf208 | 0.234313702 | -0.054875827 | 0.523503232 | 8.700672354 | 1.936530108 | 0.096196566 | 0.991835066 | -4.081297729 |
| **A_55_P1966790** | Impdh2 | 0.152356252 | -0.035722559 | 0.340435064 | 11.97161428 | 1.936107832 | 0.096255249 | 0.991835066 | -4.081681446 |
| **A_51_P371876** | Zmynd8 | 0.13637962 | -0.032032562 | 0.304791802 | 10.28216798 | 1.935464037 | 0.096344785 | 0.991835066 | -4.082266453 |
| **A_55_P2114333** | NA | 0.108472165 | -0.025480046 | 0.242424376 | 14.93310187 | 1.93543038 | 0.096349468 | 0.991835066 | -4.082297036 |
| **A_55_P2014259** | Zfp82 | -0.151778535 | -0.339290899 | 0.035733829 | 9.336123021 | -1.934592862 | 0.096466074 | 0.991835066 | -4.08305807 |
| **A_51_P230987** | Pi4k2b | 0.10038787 | -0.023646976 | 0.224422717 | 8.227944389 | 1.934401677 | 0.096492712 | 0.991835066 | -4.083231795 |
| **A_51_P484254** | Pcca | -0.126280617 | -0.282318791 | 0.029757556 | 9.373032245 | -1.934260535 | 0.096512382 | 0.991835066 | -4.083360046 |
| **A_55_P2058613** | NA | -0.080831732 | -0.180720327 | 0.019056864 | 8.182154971 | -1.934083008 | 0.096537128 | 0.991835066 | -4.08352136 |
| **A_66_P125437** | Rpl30 | 0.144146875 | -0.033998916 | 0.322292665 | 10.18120279 | 1.93392131 | 0.096559673 | 0.991835066 | -4.08366829 |
| **A_55_P2043782** | Trpm1 | 0.241873495 | -0.057116348 | 0.540863338 | 8.654969863 | 1.93348576 | 0.096620427 | 0.991835066 | -4.08406406 |
| **A_51_P448814** | Zfp90 | 0.136695129 | -0.032282446 | 0.305672704 | 10.1005531 | 1.933450679 | 0.096625322 | 0.991835066 | -4.084095936 |
| **A_52_P494372** | Zxdc | -0.116074705 | -0.259563058 | 0.027413649 | 8.650013678 | -1.933437188 | 0.096627205 | 0.991835066 | -4.084108195 |
| **A_66_P117913** | NA | 0.097310282 | -0.022983074 | 0.217603638 | 15.7453555 | 1.933420065 | 0.096629594 | 0.991835066 | -4.084123754 |

| | | | | | | | | | |
|---|---|---|---|---|---|---|---|---|---|
| **A_55_P2419514** | Ccny | 0.181645548 | -0.042927056 | 0.406218152 | 8.81189765 | 1.933201467 | 0.096660103 | 0.991835066 | -4.084322385 |
| **A_51_P330213** | Asf1b | 0.117705745 | -0.027835657 | 0.263247146 | 10.94800141 | 1.932948299 | 0.096695449 | 0.991835066 | -4.084552429 |
| **A_55_P2060404** | Pus3 | 0.126091158 | -0.029846915 | 0.282029232 | 9.835194625 | 1.932598337 | 0.096744329 | 0.991835066 | -4.084870425 |
| **A_52_P552547** | Lppos | -0.177111185 | -0.396148602 | 0.041926232 | 9.470561351 | -1.932577022 | 0.096747307 | 0.991835066 | -4.084889793 |
| **A_51_P245525** | NA | -0.103087821 | -0.230600448 | 0.024424806 | 16.03215678 | -1.932249932 | 0.096793016 | 0.991835066 | -4.085187004 |
| **A_52_P375312** | Jaml | -0.088459688 | -0.197886888 | 0.020967512 | 7.751793756 | -1.932098432 | 0.096814194 | 0.991835066 | -4.085324665 |
| **A_51_P477440** | Sec23ip | 0.082630175 | -0.019619279 | 0.184879628 | 10.20729075 | 1.931464847 | 0.096902814 | 0.991835066 | -4.08590037 |
| **A_52_P269461** | Ddhd2 | 0.093236103 | -0.022148985 | 0.208621191 | 8.190945019 | 1.931272509 | 0.096929732 | 0.991835066 | -4.086075137 |
| **A_51_P151902** | Spon1 | 0.167197375 | -0.039940718 | 0.374335468 | 9.568733791 | 1.929206038 | 0.097219404 | 0.991835066 | -4.087952798 |
| **A_55_P1974593** | NA | -0.155498459 | -0.348208489 | 0.037211571 | 15.51303709 | -1.928549996 | 0.097311545 | 0.991835066 | -4.088548889 |
| **A_51_P161874** | Med30 | -0.116493294 | -0.260959118 | 0.027972531 | 11.08995523 | -1.927280544 | 0.097490083 | 0.991835066 | -4.089702321 |
| **A_51_P149325** | Psmd8 | -0.16423616 | -0.368083188 | 0.039610869 | 11.79469323 | -1.925633042 | 0.097722272 | 0.991835066 | -4.091199225 |
| **A_55_P1981241** | Grasp | 0.096259526 | -0.023220432 | 0.215739485 | 10.25937998 | 1.925563258 | 0.097732119 | 0.991835066 | -4.091262629 |
| **A_55_P2095933** | Zranb1 | -0.120159311 | -0.269306639 | 0.028988017 | 9.806004024 | -1.925533524 | 0.097736315 | 0.991835066 | -4.091289645 |
| **A_55_P1962771** | Cyfip2 | -0.134747129 | -0.302041202 | 0.032546943 | 10.03187325 | -1.925076979 | 0.097800763 | 0.991835066 | -4.09170445 |
| **A_55_P2010487** | Unk | 0.17886017 | -0.043240574 | 0.400960914 | 10.71168077 | 1.924743038 | 0.09784793 | 0.991835066 | -4.09200786 |
| **A_51_P461475** | Atn1 | -0.105671153 | -0.236922503 | 0.025580197 | 10.12699944 | -1.924251352 | 0.097917419 | 0.991835066 | -4.092454589 |
| **A_55_P2027969** | Myl6 | 0.150529428 | -0.036476391 | 0.337535248 | 15.78598436 | 1.923868671 | 0.097971535 | 0.991835066 | -4.092802278 |
| **A_52_P561650** | Vwa1 | -0.168151568 | -0.377056203 | 0.040753067 | 9.231167674 | -1.923809114 | 0.09797996 | 0.991835066 | -4.092856389 |
| **A_51_P138538** | Eif1b | -0.136351984 | -0.305775262 | 0.033071294 | 12.31454292 | -1.923523582 | 0.098020361 | 0.991835066 | -4.093115811 |
| **A_52_P346458** | Tollip | -0.096412236 | -0.216232043 | 0.023407572 | 11.03555925 | -1.923147825 | 0.098073554 | 0.991835066 | -4.093457206 |
| **A_55_P2030899** | NA | -0.103038834 | -0.231100048 | 0.02502238 | 11.40378188 | -1.923058304 | 0.098086231 | 0.991835066 | -4.093538539 |
| **A_66_P123527** | Ndufb4 | -0.117578145 | -0.263773387 | 0.028617097 | 15.09823868 | -1.922217371 | 0.098205392 | 0.991835066 | -4.094302563 |
| **A_51_P297586** | Birc6 | 0.096783721 | -0.023588067 | 0.21715551 | 9.979091086 | 1.921705069 | 0.098278056 | 0.991835066 | -4.094768006 |
| **A_55_P2081482** | Atp5g2 | -0.111263122 | -0.249648031 | 0.027121786 | 16.33536429 | -1.921638295 | 0.098287531 | 0.991835066 | -4.094828672 |
| **A_55_P2151138** | Dynlt1c | -0.139844765 | -0.313779053 | 0.034089522 | 13.66145209 | -1.92163166 | 0.098288472 | 0.991835066 | -4.0948347 |
| **A_55_P2080850** | Ptpn2 | 0.104321207 | -0.025441653 | 0.234084066 | 10.32087828 | 1.921459902 | 0.098312848 | 0.991835066 | -4.094990747 |
| **A_55_P2038298** | Rpl31 | 0.142948747 | -0.03488501 | 0.320782505 | 16.07433339 | 1.921211965 | 0.098348046 | 0.991835066 | -4.095216005 |
| **A_55_P1973848** | Sema4b | 0.139622748 | -0.034078217 | 0.313323714 | 9.54827737 | 1.921157996 | 0.09835571 | 0.991835066 | -4.095265037 |
| **A_51_P296416** | Orc6 | 0.12004959 | -0.029332589 | 0.26943177 | 11.06453992 | 1.920750808 | 0.098413547 | 0.991835066 | -4.095634975 |
| **A_55_P2360501** | NA | 0.211621707 | -0.051778938 | 0.475022351 | 8.658208657 | 1.920226735 | 0.098488036 | 0.991835066 | -4.096111102 |
| **A_51_P269078** | Habp4 | -0.179578885 | -0.403100625 | 0.043942856 | 10.26031816 | -1.920191892 | 0.09849299 | 0.991835066 | -4.096142758 |
| **A_55_P2175579** | Olfr1466 | -0.20026444 | -0.449555571 | 0.049026692 | 8.583492798 | -1.92002173 | 0.098517189 | 0.991835066 | -4.096297352 |
| **A_55_P1975045** | Sgcg | -0.248047044 | -0.556822638 | 0.060728551 | 10.72462679 | -1.919995545 | 0.098520914 | 0.991835066 | -4.09632114 |
| **A_66_P109720** | Jpt2 | 0.103813905 | -0.02542139 | 0.233049199 | 8.068849727 | 1.919921723 | 0.098531414 | 0.991835066 | -4.096388209 |
| **A_51_P165293** | Ckmt2 | -0.400485126 | -0.899153584 | 0.098183332 | 12.83259446 | -1.919480145 | 0.098594248 | 0.991835066 | -4.096789383 |
| **A_55_P2048867** | NA | -0.23189097 | -0.520686458 | 0.056904518 | 8.159150968 | -1.919121917 | 0.098645251 | 0.991835066 | -4.097114832 |
| **A_55_P2060243** | Itgbl1 | -0.145273881 | -0.326228245 | 0.035680483 | 8.08871889 | -1.918790726 | 0.098692427 | 0.991835066 | -4.097415716 |
| **A_51_P451346** | Klf6 | 0.11879709 | -0.029177953 | 0.266772133 | 12.72731684 | 1.918785643 | 0.098693151 | 0.991835066 | -4.097420334 |
| **A_52_P429876** | Tbx20 | 0.155893809 | -0.038302451 | 0.350090068 | 12.20429368 | 1.918656101 | 0.09871161 | 0.991835066 | -4.097538021 |
| **A_55_P1961983** | Uqcr11 | -0.153090164 | -0.343842957 | 0.037662628 | 16.708343 | -1.918163036 | 0.098781899 | 0.991835066 | -4.097985964 |
| **A_51_P198076** | Mien1 | -0.119383184 | -0.268186431 | 0.029420064 | 12.31010875 | -1.917519899 | 0.098873656 | 0.991835066 | -4.098570237 |
| **A_55_P2098771** | Taf13 | 0.09326022 | -0.022986892 | 0.209507332 | 10.27168728 | 1.917447115 | 0.098884045 | 0.991835066 | -4.09863636 |
| **A_66_P109183** | Apold1 | -0.184121138 | -0.413633824 | 0.045391549 | 10.38442351 | -1.917370698 | 0.098894954 | 0.991835066 | -4.098705782 |
| **A_55_P2121856** | Ier5l | -0.140990922 | -0.316740762 | 0.034758918 | 11.69314976 | -1.917367419 | 0.098895422 | 0.991835066 | -4.098708761 |
| **A_52_P287992** | Tbcel | 0.093134695 | -0.02300225 | 0.209271641 | 8.203292234 | 1.916682731 | 0.09899322 | 0.991835066 | -4.099330775 |
| **A_55_P2030756** | Nedd4l | 0.145157976 | -0.035866256 | 0.326182207 | 9.832176448 | 1.916519857 | 0.099016499 | 0.991835066 | -4.099478739 |
| **A_55_P1966538** | Ctu2 | 0.133586549 | -0.033021248 | 0.300194346 | 10.71486628 | 1.916357541 | 0.099039702 | 0.991835066 | -4.099626196 |

| | | | | | | | | | |
|---|---|---|---|---|---|---|---|---|---|
| **A_55_P1955243** | Rps24 | 0.119613477 | -0.029585949 | 0.268812904 | 17.00837842 | 1.91611731 | 0.099074055 | 0.991835066 | -4.099844435 |
| **A_51_P105927** | Rasl12 | -0.105129996 | -0.23627521 | 0.026015219 | 9.734545166 | -1.915946289 | 0.099098517 | 0.991835066 | -4.099999798 |
| **A_55_P2002903** | Smoc2 | -0.244985461 | -0.550650366 | 0.060679443 | 11.90889657 | -1.91559576 | 0.099148674 | 0.991835066 | -4.100318234 |
| **A_51_P276063** | Phyh | -0.12552033 | -0.282153377 | 0.031112717 | 13.4083335 | -1.915313233 | 0.099189119 | 0.991835066 | -4.100574893 |
| **A_55_P2092661** | Nup210 | 0.210183611 | -0.052107456 | 0.472474678 | 10.53074624 | 1.915245637 | 0.099198799 | 0.991835066 | -4.1006363 |
| **A_52_P79385** | Magt1 | 0.108039098 | -0.026793997 | 0.242872194 | 9.499631589 | 1.91510936 | 0.099218315 | 0.991835066 | -4.100760099 |
| **A_55_P1994608** | Itsn1 | -0.09142845 | -0.205531512 | 0.022674613 | 9.269785466 | -1.915107636 | 0.099218562 | 0.991835066 | -4.100761665 |
| **A_55_P1991688** | Rad51ap1 | 0.234255527 | -0.058109953 | 0.526621007 | 10.08558075 | 1.915018112 | 0.099231385 | 0.991835066 | -4.100842992 |
| **A_55_P1967138** | Cenpx | -0.089202781 | -0.200533643 | 0.022128081 | 11.59041989 | -1.915014014 | 0.099231972 | 0.991835066 | -4.100846714 |
| **A_55_P2158697** | H4c9 | 0.168680091 | -0.041863976 | 0.379224158 | 9.04430496 | 1.914828801 | 0.099258506 | 0.991835066 | -4.101014968 |
| **A_55_P1981274** | NA | 0.117839974 | -0.029259587 | 0.264939534 | 17.16883582 | 1.914654402 | 0.099283497 | 0.991835066 | -4.101173397 |
| **A_66_P119429** | NA | 0.111045682 | -0.02757396 | 0.249665324 | 13.85594227 | 1.914635176 | 0.099286253 | 0.991835066 | -4.101190862 |
| **A_55_P2122314** | Emc1 | 0.085704602 | -0.02132116 | 0.192730365 | 10.75311446 | 1.913925132 | 0.099388069 | 0.991835066 | -4.101835882 |
| **A_55_P2037777** | NA | 0.114141563 | -0.028404561 | 0.256687686 | 8.854166259 | 1.913804341 | 0.0994054 | 0.991835066 | -4.10194561 |
| **A_55_P1955309** | Zfp800 | 0.113279537 | -0.028212608 | 0.254771681 | 8.429968865 | 1.913499127 | 0.099449204 | 0.991835066 | -4.10222287 |
| **A_51_P134812** | Chac1 | 0.334687117 | -0.083421868 | 0.752796102 | 8.033341507 | 1.913192315 | 0.099493258 | 0.991835066 | -4.10250158 |
| **A_55_P1992769** | Pafah1b1 | -0.142786281 | -0.321203321 | 0.035630759 | 12.64133393 | -1.912754728 | 0.099556122 | 0.991835066 | -4.102899084 |
| **A_52_P327664** | Gbp5 | -0.08342259 | -0.187686089 | 0.020840909 | 7.879246584 | -1.912319729 | 0.099618652 | 0.991835066 | -4.103294234 |
| **A_52_P78257** | Plekha2 | -0.178242309 | -0.401014964 | 0.044530347 | 10.126453 | -1.912308912 | 0.099620208 | 0.991835066 | -4.10330406 |
| **A_52_P181468** | Kmt5a | 0.221649943 | -0.055383826 | 0.498683712 | 10.1312854 | 1.912247252 | 0.099629074 | 0.991835066 | -4.103360071 |
| **A_52_P179178** | Bend3 | 0.12132264 | -0.03034474 | 0.27299002 | 10.93730277 | 1.911871938 | 0.099683062 | 0.991835066 | -4.103701001 |
| **A_52_P314129** | Pkia | -0.117617761 | -0.26466325 | 0.029427729 | 9.98552678 | -1.911746621 | 0.099701095 | 0.991835066 | -4.103814836 |
| **A_51_P342271** | Setd6 | 0.116272498 | -0.029097439 | 0.261642434 | 9.919386104 | 1.911663875 | 0.099713004 | 0.991835066 | -4.103890001 |
| **A_52_P209311** | Snx12 | 0.093764622 | -0.023466552 | 0.210995796 | 11.77066744 | 1.911635223 | 0.099717127 | 0.991835066 | -4.103916028 |
| **A_66_P128702** | Nup43 | 0.156525428 | -0.039181687 | 0.352232543 | 9.949064213 | 1.911557728 | 0.099728282 | 0.991835066 | -4.103986423 |
| **A_51_P407165** | Abhd5 | -0.107746738 | -0.242506708 | 0.027013232 | 9.028758028 | -1.910963353 | 0.099813878 | 0.991835066 | -4.104526336 |
| **A_52_P452689** | Atf3 | 0.342985423 | -0.086078787 | 0.772049633 | 8.985629543 | 1.910568026 | 0.099870849 | 0.991835066 | -4.104885436 |
| **A_51_P461005** | Nrk | 0.102922036 | -0.025943028 | 0.231787099 | 10.01899861 | 1.908896166 | 0.100112136 | 0.991835066 | -4.106404063 |
| **A_55_P2157200** | NA | -0.126947276 | -0.285903777 | 0.032009225 | 11.03575761 | -1.908772747 | 0.100129971 | 0.991835066 | -4.106516168 |
| **A_55_P2014867** | Iws1 | 0.11578733 | -0.029203925 | 0.260778585 | 8.990741191 | 1.908659085 | 0.100146398 | 0.991835066 | -4.10661941 |
| **A_66_P112065** | Sertad2 | 0.271507684 | -0.068491823 | 0.611507191 | 10.16903292 | 1.908591431 | 0.100156178 | 0.991835066 | -4.106680862 |
| **A_55_P1997991** | Ecpas | 0.092823834 | -0.023416223 | 0.20906389 | 8.682055947 | 1.90859078 | 0.100156272 | 0.991835066 | -4.106681453 |
| **A_52_P612693** | Polb | -0.113245305 | -0.255069104 | 0.028578494 | 9.882030175 | -1.908447528 | 0.100176982 | 0.991835066 | -4.106811573 |
| **A_51_P453909** | Cyp2f2 | 0.156204844 | -0.039503455 | 0.351913144 | 7.754564715 | 1.90763107 | 0.100295098 | 0.991835066 | -4.107553174 |
| **A_66_P126425** | Eif1 | 0.108691653 | -0.027494341 | 0.244877646 | 14.80769267 | 1.907536605 | 0.100308773 | 0.991835066 | -4.107638977 |
| **A_51_P118650** | Wdr6 | 0.113458271 | -0.02875786 | 0.255674402 | 9.821009521 | 1.906761765 | 0.10042101 | 0.991835066 | -4.10818624 |
| **A_55_P2127804** | Map3k7 | 0.124104842 | -0.031474352 | 0.279684035 | 8.586523915 | 1.906541846 | 0.100452889 | 0.991835066 | -4.108542513 |
| **A_55_P2096827** | Art3 | -0.304530287 | -0.686295531 | 0.077234957 | 10.6511003 | -1.906528214 | 0.100454865 | 0.991835066 | -4.108554896 |
| **A_55_P2079579** | Pira7 | -0.093700734 | -0.211168048 | 0.023766579 | 7.821634623 | -1.906492448 | 0.100460051 | 0.991835066 | -4.108587381 |
| **A_55_P1952220** | Pnpo | 0.298052883 | -0.075648552 | 0.671754319 | 11.90741992 | 1.906240506 | 0.100496585 | 0.991835066 | -4.108816216 |
| **A_55_P1977473** | Dab2 | 0.132901356 | -0.033733072 | 0.299535784 | 9.517229958 | 1.906223458 | 0.100499058 | 0.991835066 | -4.108831701 |
| **A_52_P387724** | Tcf7l1 | 0.181413053 | -0.046120341 | 0.408946447 | 9.231736237 | 1.905603407 | 0.100589031 | 0.991835066 | -4.109394878 |
| **A_52_P162298** | Ydjc | 0.184511939 | -0.046925296 | 0.415949173 | 9.425481103 | 1.905462356 | 0.10060951 | 0.991835066 | -4.10952299 |
| **A_51_P304397** | Cpm | -0.122369198 | -0.275934056 | 0.03119566 | 8.189126406 | -1.90453695 | 0.100743966 | 0.991835066 | -4.110363498 |
| **A_51_P176377** | Usp25 | -0.110155265 | -0.248508775 | 0.028198244 | 10.74492942 | -1.902936135 | 0.100976972 | 0.991835066 | -4.111817413 |
| **A_55_P1998545** | Ncdn | 0.111604194 | -0.028670042 | 0.251878431 | 11.23105557 | 1.901567455 | 0.10117661 | 0.991835066 | -4.113060455 |
| **A_55_P2003643** | Tstd2 | 0.113926576 | -0.029337736 | 0.257190888 | 9.483935747 | 1.900623781 | 0.101314481 | 0.991835066 | -4.113917481 |
| **A_55_P2014269** | Zfp771 | -0.112407967 | -0.253769708 | 0.028953774 | 12.33187489 | -1.900528305 | 0.101328441 | 0.991835066 | -4.11400419 |

| | | | | | | | | | |
|---|---|---|---|---|---|---|---|---|---|
| **A_51_P257762** | Kat2b | -0.105890312 | -0.239075369 | 0.027294745 | 8.654942064 | -1.900246138 | 0.101369707 | 0.991835066 | -4.114260444 |
| **A_55_P2041090** | Tyw5 | 0.213643345 | -0.055167679 | 0.48245437 | 9.981451015 | 1.899553045 | 0.10147114 | 0.991835066 | -4.114889881 |
| **A_55_P2010773** | Tbx2 | 0.106974969 | -0.027657845 | 0.241607782 | 8.947221352 | 1.899067403 | 0.101542272 | 0.991835066 | -4.115330914 |
| **A_52_P549166** | Arsj | -0.11185566 | -0.252680145 | 0.028968826 | 8.48197128 | -1.898405232 | 0.101639339 | 0.991835066 | -4.115932252 |
| **A_52_P227307** | Mllt1 | 0.17778475 | -0.046066122 | 0.401635621 | 11.08769785 | 1.89821258 | 0.101667597 | 0.991835066 | -4.116107205 |
| **A_65_P13527** | Osbpl8 | -0.195357583 | -0.441376734 | 0.050661568 | 12.14824751 | -1.897887611 | 0.101715281 | 0.991835066 | -4.116402314 |
| **A_55_P2011672** | NA | -0.118909346 | -0.268674025 | 0.030855333 | 13.26977692 | -1.897648282 | 0.101750412 | 0.991835066 | -4.116619651 |
| **A_55_P2059686** | Rcc1 | 0.178774742 | -0.046397541 | 0.403947025 | 10.60839409 | 1.897581171 | 0.101760265 | 0.991835066 | -4.116680596 |
| **A_51_P406557** | Myorg | -0.099687362 | -0.225292794 | 0.02591807 | 8.330409869 | -1.896884179 | 0.101862655 | 0.991835066 | -4.117313533 |
| **A_55_P2130610** | Smc3 | 0.138032232 | -0.035899828 | 0.311964293 | 12.14994641 | 1.896749603 | 0.101882436 | 0.991835066 | -4.11743574 |
| **A_51_P314111** | Psip1 | 0.105214321 | -0.027383746 | 0.237812389 | 10.61242781 | 1.896473571 | 0.101923022 | 0.991835066 | -4.117686401 |
| **A_55_P2143139** | Myadml2 | 0.099470449 | -0.025896016 | 0.224836913 | 11.0929248 | 1.896364559 | 0.101939054 | 0.991835066 | -4.117785393 |
| **A_55_P2115281** | Chchd2-ps | -0.092157627 | -0.208314207 | 0.023998952 | 16.01668778 | -1.896254411 | 0.101955256 | 0.991835066 | -4.117885416 |
| **A_51_P391764** | Cbx6 | 0.217936725 | -0.056787398 | 0.492660847 | 12.9985899 | 1.896019316 | 0.101989846 | 0.991835066 | -4.1180989 |
| **A_51_P505868** | Lhfp | -0.120892669 | -0.273313897 | 0.031528558 | 9.688654874 | -1.895673957 | 0.10204068 | 0.991835066 | -4.11841251 |
| **A_66_P127255** | Ing3 | 0.116611171 | -0.030421187 | 0.263643529 | 10.47131285 | 1.895554899 | 0.10205821 | 0.991835066 | -4.118520622 |
| **A_55_P1969078** | Adar | -0.110805896 | -0.250534127 | 0.028922335 | 9.039181471 | -1.895343106 | 0.102089401 | 0.991835066 | -4.118712943 |
| **A_55_P2049687** | Efna2 | 0.083625152 | -0.021832922 | 0.189083226 | 8.504966263 | 1.895248753 | 0.1021033 | 0.991835066 | -4.11879862 |
| **A_55_P2293013** | Ces2a | -0.082948202 | -0.187562518 | 0.021666114 | 7.714319855 | -1.895068822 | 0.10212981 | 0.991835066 | -4.118962007 |
| **A_52_P654043** | Msto1 | 0.156616702 | -0.040917809 | 0.354151214 | 12.01307647 | 1.894978231 | 0.102143159 | 0.991835066 | -4.119044269 |
| **A_66_P106231** | Nccrp1 | 0.099157088 | -0.02591623 | 0.224230406 | 7.988028198 | 1.894821149 | 0.102166311 | 0.991835066 | -4.119186906 |
| **A_52_P302395** | Lnpep | -0.157001899 | -0.355058566 | 0.041054767 | 9.309136114 | -1.894630716 | 0.102194386 | 0.991835066 | -4.119359827 |
| **A_51_P304510** | BC017158 | 0.145118524 | -0.037997483 | 0.32823453 | 10.38370324 | 1.894112089 | 0.102270883 | 0.991835066 | -4.119830759 |
| **A_55_P2033120** | Srxn1 | 0.133145882 | -0.034869588 | 0.301161351 | 9.743024501 | 1.894033257 | 0.102282515 | 0.991835066 | -4.119902341 |
| **A_51_P417858** | Zfp825 | -0.096944202 | -0.219312082 | 0.025423678 | 8.486044729 | -1.893492271 | 0.102362379 | 0.991835066 | -4.120393568 |
| **A_51_P257675** | Tspyl4 | -0.108683235 | -0.245897929 | 0.028531459 | 10.23407921 | -1.893089158 | 0.102421928 | 0.991835066 | -4.120759598 |
| **A_51_P204053** | Hba-x | 0.380553182 | -0.100037365 | 0.861143728 | 8.429096487 | 1.892558297 | 0.102500401 | 0.991835066 | -4.12124162 |
| **A_55_P2118391** | Lrp6 | 0.102376454 | -0.0269277 | 0.231680609 | 11.85389308 | 1.892329402 | 0.102534255 | 0.991835066 | -4.121449455 |
| **A_55_P2117689** | Rmnd1 | -0.115053133 | -0.260374886 | 0.030268619 | 11.87873047 | -1.892243212 | 0.102547005 | 0.991835066 | -4.121527714 |
| **A_55_P2090025** | Mest | 0.248855689 | -0.065485044 | 0.563196422 | 13.56593465 | 1.892152098 | 0.102560486 | 0.991835066 | -4.121610444 |
| **A_55_P1972699** | Rars2 | 0.106213192 | -0.027958267 | 0.240384651 | 10.94512735 | 1.892027534 | 0.102578918 | 0.991835066 | -4.121723546 |
| **A_55_P2105152** | Trim59 | -0.104441682 | -0.236384832 | 0.027501467 | 11.14479072 | -1.891891143 | 0.102599105 | 0.991835066 | -4.121847386 |
| **A_55_P1967149** | Irx3 | 0.150135556 | -0.039556086 | 0.339827197 | 10.74985462 | 1.891666154 | 0.102632412 | 0.991835066 | -4.12205167 |
| **A_52_P475450** | Rbm48 | 0.090787097 | -0.02392585 | 0.205500045 | 8.325152216 | 1.891563064 | 0.102647677 | 0.991835066 | -4.122145273 |
| **A_55_P2147831** | Gimap4 | -0.215375775 | -0.487527673 | 0.056776123 | 10.638128 | -1.891448921 | 0.102664581 | 0.991835066 | -4.122248911 |
| **A_66_P104221** | NA | 0.121557461 | -0.032051522 | 0.275166444 | 8.440130593 | 1.89135974 | 0.102677791 | 0.991835066 | -4.122329885 |
| **A_52_P531651** | Parvg | -0.078305981 | -0.177290945 | 0.020678984 | 7.783301362 | -1.890753157 | 0.102767682 | 0.991835066 | -4.122880636 |
| **A_52_P646034** | Nfatc2ip | 0.094742623 | -0.025019608 | 0.214504854 | 8.955292789 | 1.890752396 | 0.102767794 | 0.991835066 | -4.122881327 |
| **A_52_P674165** | Rsbn1l | -0.088816655 | -0.201106961 | 0.023473651 | 8.588959962 | -1.890432948 | 0.102815165 | 0.991835066 | -4.123171369 |
| **A_51_P278427** | 3830406C1 | -0.103796644 | -0.235050044 | 0.027456757 | 9.20854996 | -1.890087357 | 0.102866437 | 0.991835066 | -4.123485144 |
| **A_55_P2082708** | Pwp2 | 0.117116324 | -0.030984009 | 0.265216658 | 10.53813401 | 1.890037976 | 0.102873765 | 0.991835066 | -4.123529979 |
| **A_51_P232705** | Commd9 | -0.109855862 | -0.248798111 | 0.029086386 | 9.187645753 | -1.889722562 | 0.102920584 | 0.991835066 | -4.123816352 |
| **A_51_P386358** | Wwp2 | 0.114815608 | -0.030416165 | 0.260047382 | 10.97494044 | 1.88950664 | 0.102952648 | 0.991835066 | -4.124012393 |
| **A_52_P384574** | Stard4 | 0.095854574 | -0.025398931 | 0.217108079 | 9.532054712 | 1.889416365 | 0.102966056 | 0.991835066 | -4.124094355 |
| **A_55_P2115136** | Vsir | -0.112094547 | -0.253909961 | 0.029720868 | 8.089229068 | -1.889166249 | 0.103003214 | 0.991835066 | -4.124321439 |
| **A_51_P108226** | Wfdc21 | -0.189124368 | -0.428395582 | 0.050146846 | 7.835338469 | -1.889148807 | 0.103005805 | 0.991835066 | -4.123819374 |
| **A_55_P2182806** | NA | 0.121582354 | -0.032255033 | 0.275419741 | 9.940741151 | 1.888938361 | 0.103037081 | 0.991835066 | -4.12452834 |
| **A_55_P2021721** | NA | 0.37268677 | -0.098888454 | 0.844261995 | 10.01469613 | 1.888870258 | 0.103047204 | 0.991835066 | -4.124590171 |

| | | | | | | | | | |
|---|---|---|---|---|---|---|---|---|---|
| **A_66_P106212** | Secisbp2 | -0.099860522 | -0.226225671 | 0.026504627 | 9.684916741 | -1.888755099 | 0.103064324 | 0.991835066 | -4.124694724 |
| **A_52_P22617** | NA | -0.113774631 | -0.257760582 | 0.030211321 | 8.434995755 | -1.888575913 | 0.103090968 | 0.991835066 | -4.124857407 |
| **A_55_P2110713** | Anxa2 | 0.113777342 | -0.030230813 | 0.257785498 | 15.67575477 | 1.888329726 | 0.103127585 | 0.991835066 | -4.125080919 |
| **A_51_P369971** | Fra10ac1 | 0.124216533 | -0.033004529 | 0.281437594 | 11.29929722 | 1.888329643 | 0.103127598 | 0.991835066 | -4.125080994 |
| **A_51_P416869** | Nedd8 | -0.105927457 | -0.240056737 | 0.028201823 | 13.47057928 | -1.887530966 | 0.10324648 | 0.991835066 | -4.125806097 |
| **A_55_P2177539** | Sod1 | 0.093225919 | -0.024856769 | 0.211308606 | 15.14779934 | 1.886946446 | 0.10333357 | 0.991835066 | -4.126336762 |
| **A_51_P425048** | H2-Q2 | -0.098433471 | -0.223150601 | 0.026283659 | 8.177169751 | -1.886365402 | 0.103420213 | 0.991835066 | -4.126864262 |
| **A_55_P2019019** | Rpl36a | 0.1056608 | -0.028238935 | 0.239560536 | 15.04104838 | 1.886007036 | 0.103473687 | 0.991835066 | -4.127189599 |
| **A_51_P291417** | Thbd | 0.097434617 | -0.026044765 | 0.220913999 | 10.78535249 | 1.885940417 | 0.103483631 | 0.991835066 | -4.127250077 |
| **A_55_P1959738** | Zfp511 | 0.123808376 | -0.033097054 | 0.280713807 | 9.677584579 | 1.885910964 | 0.103488027 | 0.991835066 | -4.127276816 |
| **A_55_P2163143** | Cdk4 | 0.143687964 | -0.038413171 | 0.325789099 | 13.87892993 | 1.885892225 | 0.103490825 | 0.991835066 | -4.127293827 |
| **A_55_P1956517** | Polr2l | 0.082864516 | -0.022164364 | 0.187893396 | 14.84843075 | 1.885684351 | 0.10352186 | 0.991835066 | -4.127482541 |
| **A_55_P2166745** | Sp3os | -0.13466547 | -0.3053792 | 0.036048261 | 11.96109192 | -1.885371467 | 0.103568591 | 0.991835066 | -4.127766583 |
| **A_55_P2417212** | NA | 0.17455195 | -0.046758824 | 0.395862724 | 9.375817694 | 1.88508649 | 0.103611172 | 0.991835066 | -4.128025288 |
| **A_55_P2071486** | Suv39h1 | 0.16562093 | -0.044382139 | 0.375623999 | 8.898138908 | 1.88494512 | 0.103632301 | 0.991835066 | -4.128153625 |
| **A_51_P480233** | Nmral1 | 0.111777348 | -0.029957752 | 0.253512448 | 10.16815117 | 1.884887863 | 0.103640861 | 0.991835066 | -4.128205604 |
| **A_51_P347823** | Ipo4 | 0.135954222 | -0.036476056 | 0.3083845 | 10.68927918 | 1.88446599 | 0.103703945 | 0.991835066 | -4.128588578 |
| **A_55_P2044252** | Zfp512b | 0.224363835 | -0.060234806 | 0.508962476 | 10.41343546 | 1.884209394 | 0.103742334 | 0.991835066 | -4.128821513 |
| **A_52_P81783** | Gspt1 | 0.121588847 | -0.032728451 | 0.275906145 | 9.861100091 | 1.883164519 | 0.103898797 | 0.991835066 | -4.12977002 |
| **A_55_P2033510** | Ppm1m | 0.113993398 | -0.030687965 | 0.25867476 | 10.23361552 | 1.883112385 | 0.10390661 | 0.991835066 | -4.129817344 |
| **A_55_P2174258** | Pin4 | -0.08341791 | -0.189315779 | 0.022479958 | 11.9015556 | -1.882700422 | 0.103968367 | 0.991835066 | -4.130191304 |
| **A_52_P476400** | NA | 0.198124823 | -0.053491998 | 0.449741643 | 10.39762252 | 1.88195118 | 0.104080778 | 0.991835066 | -4.130871416 |
| **A_51_P250123** | Zfp763 | -0.098612973 | -0.223855005 | 0.026629058 | 8.417307878 | -1.881885005 | 0.104090712 | 0.991835066 | -4.130931485 |
| **A_55_P1988760** | Sestd1 | -0.137461918 | -0.312153929 | 0.037230093 | 9.271335961 | -1.880695507 | 0.104269437 | 0.991835066 | -4.132011199 |
| **A_55_P1990548** | Tbc1d22a | -0.080141807 | -0.182008988 | 0.021725375 | 11.75362949 | -1.880329552 | 0.104324482 | 0.991835066 | -4.13234337 |
| **A_55_P1979463** | Prdx5 | -0.175006978 | -0.397510132 | 0.047496177 | 13.41133582 | -1.879872192 | 0.104393317 | 0.991835066 | -4.132758503 |
| **A_55_P2090708** | Fam207a | 0.105900746 | -0.028798405 | 0.240599898 | 12.04263507 | 1.879071443 | 0.104513941 | 0.991835066 | -4.133485307 |
| **A_55_P2080758** | Slc8a1 | 0.138849195 | -0.037758366 | 0.315456757 | 10.57032586 | 1.879071073 | 0.104513997 | 0.991835066 | -4.133485643 |
| **A_55_P2022074** | Klf10 | -0.184425826 | -0.419027031 | 0.050175378 | 10.51275421 | -1.878886878 | 0.104541763 | 0.991835066 | -4.133652826 |
| **A_55_P2144391** | NA | -0.110359827 | -0.250785212 | 0.030065558 | 10.33394475 | -1.878341352 | 0.104624039 | 0.991835066 | -4.134147962 |
| **A_55_P2117649** | Slfn9 | 0.119509315 | -0.032686163 | 0.271704793 | 8.808850138 | 1.876761734 | 0.104862636 | 0.991835066 | -4.135581623 |
| **A_51_P105380** | Csta2 | -0.22577459 | -0.513299767 | 0.061750587 | 8.344230997 | -1.876758168 | 0.104863176 | 0.991835066 | -4.135039264 |
| **A_51_P272876** | Tent5a | 0.130192437 | -0.035614284 | 0.295999157 | 10.53259963 | 1.876690943 | 0.104873342 | 0.991835066 | -4.135645871 |
| **A_51_P393768** | Osbp | -0.127427648 | -0.289721763 | 0.034866468 | 9.541528299 | -1.876592712 | 0.104888198 | 0.991835066 | -4.135735023 |
| **A_52_P661071** | Snhg3 | 0.180904414 | -0.049499145 | 0.411307972 | 10.84800468 | 1.87658877 | 0.104888795 | 0.991835066 | -4.135738601 |
| **A_51_P351285** | Pcsk7 | -0.106140859 | -0.241366801 | 0.029085083 | 9.597229057 | -1.875995164 | 0.104978617 | 0.991835066 | -4.136277336 |
| **A_55_P1958419** | NA | 0.121469179 | -0.033298134 | 0.276236492 | 11.55732691 | 1.875840841 | 0.105001981 | 0.991835066 | -4.136417392 |
| **A_55_P2032192** | Tpm1 | -0.211125813 | -0.48308082 | 0.060829193 | 9.276205318 | -1.928463517 | 0.105053331 | 0.991835066 | -4.116948927 |
| **A_51_P314763** | Rabggtb | -0.088333735 | -0.20090317 | 0.0242357 | 10.67673617 | -1.875492109 | 0.105054796 | 0.991835066 | -4.136733882 |
| **A_55_P2001143** | Ehmt2 | 0.125371438 | -0.034466381 | 0.285209257 | 13.0505561 | 1.874684524 | 0.105177204 | 0.991835066 | -4.137466788 |
| **A_51_P406638** | Rfc1 | 0.140721847 | -0.038708887 | 0.320152581 | 11.57110502 | 1.874449846 | 0.105212801 | 0.991835066 | -4.137679761 |
| **A_55_P2165544** | Aebp2 | 0.113233646 | -0.03118635 | 0.257653643 | 9.167739134 | 1.873946961 | 0.105289121 | 0.991835066 | -4.13813613 |
| **A_55_P2072032** | NA | 0.103422157 | -0.028498716 | 0.235343031 | 15.74341379 | 1.873739473 | 0.105320626 | 0.991835066 | -4.138324423 |
| **A_55_P1996086** | Gm14326 | 0.090170118 | -0.024851629 | 0.205191865 | 8.759359475 | 1.873664424 | 0.105332024 | 0.991835066 | -4.138392529 |
| **A_55_P1960479** | NA | 0.567586945 | -0.15651041 | 1.291684301 | 12.12553448 | 1.873460621 | 0.105362981 | 0.991835066 | -4.138577477 |
| **A_51_P322473** | Perm1 | -0.235699921 | -0.536405564 | 0.065005722 | 10.77833959 | -1.873384795 | 0.105374502 | 0.991835066 | -4.138646287 |
| **A_52_P275354** | Osbpl7 | 0.123016767 | -0.033956493 | 0.279990027 | 9.725294276 | 1.873043073 | 0.105426435 | 0.991835066 | -4.138956391 |
| **A_55_P2023912** | NA | 0.466350734 | -0.128733589 | 1.061435056 | 10.65390294 | 1.873023765 | 0.10542937 | 0.991835066 | -4.138973911 |

| | | | | | | | | | |
|---|---|---|---|---|---|---|---|---|---|
| **A_51_P327751** | Ifit1 | -0.075167854 | -0.171101811 | 0.020766103 | 7.734228937 | -1.872703075 | 0.105478131 | 0.991835066 | -4.139264925 |
| **A_55_P2006479** | 3300002I08 | 0.113003941 | -0.03122628 | 0.257234162 | 8.502143803 | 1.87260619 | 0.105492866 | 0.991835066 | -4.139352844 |
| **A_52_P602091** | Csf1r | -0.102448941 | -0.233285489 | 0.028387606 | 8.024144113 | -1.871490084 | 0.105662765 | 0.991835066 | -4.140365638 |
| **A_55_P1979403** | Tigar | -0.141646364 | -0.32266175 | 0.039369022 | 10.50907277 | -1.870247451 | 0.10585224 | 0.991835066 | -4.141493203 |
| **A_55_P1963489** | NA | -0.132376907 | -0.301572058 | 0.036818243 | 8.698942345 | -1.869964889 | 0.105895371 | 0.991835066 | -4.141749592 |
| **A_55_P2007022** | Adgre5 | -0.088803582 | -0.202331397 | 0.024724232 | 12.62036319 | -1.869551102 | 0.105958563 | 0.991835066 | -4.142125048 |
| **A_55_P2053320** | Zfr2 | 0.141533255 | -0.039415916 | 0.322482425 | 9.055547571 | 1.869437841 | 0.105975867 | 0.991835066 | -4.142227816 |
| **A_51_P437786** | Rbm28 | 0.159957606 | -0.044577415 | 0.364492628 | 11.77731823 | 1.86915944 | 0.106018411 | 0.991835066 | -4.142480423 |
| **A_55_P2017769** | B020031M | 0.343710328 | -0.095885259 | 0.783305916 | 8.842669555 | 1.868737886 | 0.106082863 | 0.991835066 | -4.142862914 |
| **A_52_P453884** | Foxo1 | 0.110249446 | -0.030789604 | 0.251288496 | 10.65965141 | 1.868298116 | 0.106150141 | 0.991835066 | -4.143261929 |
| **A_55_P2114218** | Zcchc7 | 0.225325679 | -0.062966672 | 0.513618031 | 9.943319324 | 1.868042281 | 0.106189299 | 0.991835066 | -4.143494052 |
| **A_51_P420547** | Clic5 | -0.208043505 | -0.474243158 | 0.058156149 | 11.48087018 | -1.867909398 | 0.106209643 | 0.991835066 | -4.143614617 |
| **A_55_P2015734** | Syvn1 | 0.166976068 | -0.046700915 | 0.380653051 | 11.89932815 | 1.867693549 | 0.106242698 | 0.991835066 | -4.143810457 |
| **A_55_P1962404** | Ugt1a6b | -0.143944636 | -0.328157428 | 0.040268156 | 8.429148448 | -1.867604157 | 0.10625639 | 0.991835066 | -4.143891562 |
| **A_55_P2177488** | H2bc21 | -0.105923782 | -0.241505844 | 0.029658279 | 8.589545169 | -1.867241016 | 0.106312031 | 0.991835066 | -4.144221036 |
| **A_55_P1999419** | Bag5 | 0.100300758 | -0.02811755 | 0.228719066 | 10.03528142 | 1.866751056 | 0.106387148 | 0.991835066 | -4.144665565 |
| **A_51_P503542** | Zfp414 | 0.142323213 | -0.039899738 | 0.324546165 | 10.82220542 | 1.866731241 | 0.106390187 | 0.991835066 | -4.144683542 |
| **A_55_P2058757** | Cap1 | 0.084742994 | -0.023771674 | 0.193257663 | 10.97514545 | 1.866484945 | 0.106427968 | 0.991835066 | -4.144906997 |
| **A_51_P367060** | Ifrd1 | 0.323449437 | -0.090760784 | 0.737659659 | 10.97635796 | 1.866357003 | 0.106447599 | 0.991835066 | -4.145023073 |
| **A_51_P479043** | Nif3l1 | 0.112034313 | -0.031458543 | 0.255527168 | 11.29426569 | 1.866078497 | 0.106490344 | 0.991835066 | -4.145275747 |
| **A_51_P117618** | Ethe1 | -0.23559153 | -0.537408517 | 0.066225457 | 10.37262881 | -1.865628317 | 0.106559474 | 0.991835066 | -4.145684168 |
| **A_51_P376347** | Hebp1 | 0.194035709 | -0.054570409 | 0.442641828 | 10.13597825 | 1.865430107 | 0.106589925 | 0.991835066 | -4.14586399 |
| **A_52_P649296** | Nras | 0.112581838 | -0.031672934 | 0.25683661 | 9.759303309 | 1.865293936 | 0.10661085 | 0.991835066 | -4.145987527 |
| **A_55_P2036514** | Dis3 | 0.145044932 | -0.04081974 | 0.330909603 | 10.45505566 | 1.865154624 | 0.106632262 | 0.991835066 | -4.146113913 |
| **A_55_P1966327** | NA | -0.139112312 | -0.317393186 | 0.039168561 | 8.760088317 | -1.864961836 | 0.106661899 | 0.991835066 | -4.146288812 |
| **A_51_P508580** | Tcf15 | -0.148055585 | -0.337800963 | 0.041689792 | 10.53615172 | -1.864930813 | 0.106666669 | 0.991835066 | -4.146316957 |
| **A_66_P133928** | Fundc2 | -0.123127189 | -0.280971465 | 0.034717086 | 13.47888899 | -1.864379253 | 0.10675151 | 0.991835066 | -4.146817331 |
| **A_55_P1998186** | NA | 0.1066969 | -0.030133585 | 0.243527384 | 12.88945146 | 1.863708851 | 0.10685472 | 0.991835066 | -4.147425504 |
| **A_52_P31814** | 1110038B1 | 0.17996395 | -0.050826007 | 0.410753907 | 11.09444789 | 1.863707461 | 0.106854934 | 0.991835066 | -4.147426765 |
| **A_51_P329251** | Chchd1 | 0.112054914 | -0.031657657 | 0.255767484 | 14.44879435 | 1.863568149 | 0.106876393 | 0.991835066 | -4.147553144 |
| **A_55_P2061874** | Prelid1 | 0.106722689 | -0.030240006 | 0.243685384 | 14.06970448 | 1.862359835 | 0.107062698 | 0.991835066 | -4.148649254 |
| **A_51_P275454** | Trim30a | -0.098811676 | -0.225629209 | 0.028005857 | 7.982003575 | -1.86225049 | 0.107079573 | 0.991835066 | -4.148748443 |
| **A_55_P2112737** | Actb | 0.129375228 | -0.036675493 | 0.295425949 | 16.33581838 | 1.862170728 | 0.107091885 | 0.991835066 | -4.148820797 |
| **A_51_P389885** | Spic | -0.07402732 | -0.16906965 | 0.021015011 | 7.774579818 | -1.861590174 | 0.107181535 | 0.991835066 | -4.149347419 |
| **A_65_P20104** | Mical2 | 0.131021606 | -0.0372424 | 0.299285613 | 8.651724908 | 1.861061935 | 0.10726317 | 0.991835066 | -4.149826576 |
| **A_52_P213181** | Cul4b | 0.126986909 | -0.036096758 | 0.290070576 | 9.324871344 | 1.861048176 | 0.107265297 | 0.991835066 | -4.149839057 |
| **A_55_P2036615** | Lrif1 | -0.12064457 | -0.275599579 | 0.03431044 | 9.434507062 | -1.860849672 | 0.107295991 | 0.991835066 | -4.150019114 |
| **A_55_P2171493** | BC030867 | 0.158903934 | -0.045201636 | 0.363009503 | 9.690754419 | 1.860753844 | 0.107310811 | 0.991835066 | -4.150106036 |
| **A_51_P196973** | Chaf1a | 0.178965669 | -0.050912847 | 0.408844186 | 9.978983342 | 1.86071764 | 0.107316411 | 0.991835066 | -4.150138875 |
| **A_55_P2118684** | Idh2 | -0.091124221 | -0.208182615 | 0.025934173 | 17.01664997 | -1.86054594 | 0.107342972 | 0.991835066 | -4.150294617 |
| **A_52_P208222** | Pde1c | -0.181111379 | -0.413768659 | 0.0515459 | 8.95561578 | -1.860536631 | 0.107344412 | 0.991835066 | -4.150303062 |
| **A_52_P423128** | Arglu1 | 0.187713133 | -0.053451744 | 0.42887801 | 11.24495082 | 1.860328896 | 0.107376556 | 0.991835066 | -4.150491488 |
| **A_55_P2128153** | Col6a1 | -0.155235307 | -0.354724009 | 0.044253394 | 13.21709752 | -1.859864632 | 0.107448429 | 0.991835066 | -4.150912595 |
| **A_55_P1976212** | Selplg | -0.096440214 | -0.220373145 | 0.027492717 | 8.439533903 | -1.85986138 | 0.107448932 | 0.991835066 | -4.150915544 |
| **A_55_P2013680** | NA | 0.159803039 | -0.045609828 | 0.365215905 | 16.37955439 | 1.859373005 | 0.107524588 | 0.991835066 | -4.151358512 |
| **A_55_P2254968** | BB212172 | 0.13347938 | -0.038142628 | 0.305101387 | 8.817974822 | 1.858875633 | 0.107601692 | 0.991835066 | -4.151809632 |
| **A_66_P122086** | 9030619P0 | -0.101073459 | -0.231084861 | 0.028937944 | 7.870169029 | -1.858081744 | 0.107724874 | 0.991835066 | -4.152529675 |
| **A_55_P2012615** | NA | 0.113406057 | -0.032505858 | 0.259317971 | 14.75854368 | 1.857610371 | 0.107798079 | 0.991835066 | -4.152957193 |

| | | | | | | | | | |
|---|---|---|---|---|---|---|---|---|---|
| **A_55_P2288117** | NA | 0.108377709 | -0.031075318 | 0.247830736 | 8.984813404 | 1.857467197 | 0.107820323 | 0.991835066 | -4.153087045 |
| **A_55_P2099064** | Nsd2 | 0.09768256 | -0.028073793 | 0.223438914 | 8.425627081 | 1.85650546 | 0.107969863 | 0.991835066 | -4.153959272 |
| **A_51_P225427** | Pkp2 | -0.10873696 | -0.248732005 | 0.031258086 | 13.556004 | -1.856408942 | 0.107984881 | 0.991835066 | -4.154046806 |
| **A_55_P2159575** | Reep6 | 0.157155649 | -0.045216398 | 0.359527696 | 8.718442046 | 1.856045434 | 0.108041463 | 0.991835066 | -4.154376472 |
| **A_55_P2085781** | Smim26 | -0.097372693 | -0.222788442 | 0.028043056 | 13.11010473 | -1.855642195 | 0.108104263 | 0.991835066 | -4.154742165 |
| **A_55_P2062310** | NA | 0.097737254 | -0.028163214 | 0.223637722 | 14.58215836 | 1.855418655 | 0.108139092 | 0.991835066 | -4.154944887 |
| **A_55_P1965674** | Alg2 | 0.148221859 | -0.042744639 | 0.339188358 | 10.76941506 | 1.855086664 | 0.108190839 | 0.991835066 | -4.155245958 |
| **A_51_P453167** | Naxd | -0.087920274 | -0.201209147 | 0.025368599 | 12.0606516 | -1.854859037 | 0.108226333 | 0.991835066 | -4.155452384 |
| **A_55_P1966204** | Cxcl12 | 0.18564307 | -0.053644535 | 0.424930675 | 10.05964605 | 1.854247364 | 0.108321767 | 0.991835066 | -4.156007074 |
| **A_55_P1965571** | Rpl8 | 0.106965571 | -0.03096748 | 0.244898622 | 14.92653169 | 1.853466772 | 0.108443676 | 0.991835066 | -4.156714925 |
| **A_55_P2229645** | NA | 0.134073007 | -0.038823926 | 0.306969941 | 8.988899907 | 1.853374544 | 0.108458089 | 0.991835066 | -4.156798558 |
| **A_65_P07627** | Dach1 | -0.130279704 | -0.298306256 | 0.037746848 | 8.729442061 | -1.853138926 | 0.108494917 | 0.991835066 | -4.157012214 |
| **A_55_P1985288** | Bhlha9 | 0.17465929 | -0.05060611 | 0.39992469 | 7.860890756 | 1.8531319 | 0.108496016 | 0.991835066 | -4.157018585 |
| **A_55_P1972481** | NA | -0.14905887 | -0.341383072 | 0.043265333 | 9.518244676 | -1.852392504 | 0.108611668 | 0.991835066 | -4.15768905 |
| **A_51_P362176** | Fgg | -0.699312867 | -1.601624766 | 0.202999032 | 8.994391661 | -1.852353912 | 0.108617708 | 0.991835066 | -4.157724043 |
| **A_55_P2121222** | Clcc1 | 0.087783373 | -0.025502366 | 0.201069111 | 11.31339059 | 1.85202206 | 0.108669657 | 0.991835066 | -4.158024951 |
| **A_52_P517224** | Trim63 | -0.226695325 | -0.519345317 | 0.065954668 | 12.64063799 | -1.85141244 | 0.108765151 | 0.991835066 | -4.158577713 |
| **A_52_P220485** | Uhrf2 | 0.089119437 | -0.025950703 | 0.204189577 | 9.092660684 | 1.851053311 | 0.108821445 | 0.991835066 | -4.15890334 |
| **A_55_P1984932** | Zc3h18 | 0.095535989 | -0.02782256 | 0.218894538 | 11.54604736 | 1.851001984 | 0.108829493 | 0.991835066 | -4.158949879 |
| **A_55_P2183955** | Olfr97 | -0.099572137 | -0.228151279 | 0.029007006 | 8.550966769 | -1.850872206 | 0.108849844 | 0.991835066 | -4.159067549 |
| **A_51_P266774** | Mfn2 | -0.126930734 | -0.290838581 | 0.036977113 | 13.15780312 | -1.850871111 | 0.108850016 | 0.991835066 | -4.159068542 |
| **A_55_P2047290** | Ankrd40 | -0.197704145 | -0.453127499 | 0.05771921 | 10.19874305 | -1.849968366 | 0.108991684 | 0.991835066 | -4.159887046 |
| **A_55_P2094613** | NA | -0.102406695 | -0.234735364 | 0.029921974 | 8.848514761 | -1.849624372 | 0.109045715 | 0.991835066 | -4.160198931 |
| **A_55_P2427900** | Unc13c | 0.176348465 | -0.051606645 | 0.404303574 | 8.678559496 | 1.848976897 | 0.109147484 | 0.991835066 | -4.160785957 |
| **A_52_P624149** | Hsp90b1 | 0.104853625 | -0.030692534 | 0.240399785 | 14.45499101 | 1.848865703 | 0.10916497 | 0.991835066 | -4.160886769 |
| **A_55_P2026945** | Zranb2 | 0.099358008 | -0.029144853 | 0.227860869 | 10.79297779 | 1.84798828 | 0.109303051 | 0.991835066 | -4.161682245 |
| **A_51_P428483** | Fgb | -0.739339436 | -1.695959402 | 0.217280531 | 9.079415642 | -1.847198481 | 0.109427488 | 0.991835066 | -4.162398254 |
| **A_55_P2024358** | Rpl34 | 0.126978036 | -0.037324601 | 0.291280672 | 15.78341351 | 1.847111887 | 0.10944114 | 0.991835066 | -4.162476757 |
| **A_51_P309184** | Pxmp2 | -0.208486667 | -0.478270046 | 0.061296712 | 9.997878716 | -1.847022686 | 0.109455205 | 0.991835066 | -4.162557622 |
| **A_55_P2075627** | Apc2 | -0.078221672 | -0.179444949 | 0.023001604 | 9.206352497 | -1.846952971 | 0.109466198 | 0.991835066 | -4.162620822 |
| **A_55_P2124120** | Ywhae | -0.099821072 | -0.228999733 | 0.029357588 | 15.09966478 | -1.846888122 | 0.109476425 | 0.991835066 | -4.162679611 |
| **A_66_P113578** | Washc1 | -0.094378894 | -0.216540479 | 0.027782692 | 11.43067698 | -1.846500178 | 0.109537625 | 0.991835066 | -4.163031296 |
| **A_55_P2057212** | Tpr | 0.146400238 | -0.0431493 | 0.335949776 | 11.2443737 | 1.845985071 | 0.109618937 | 0.991835066 | -4.16349825 |
| **A_55_P2174743** | Akap7 | 0.143035073 | -0.042228552 | 0.328298698 | 9.679029847 | 1.845276774 | 0.109730841 | 0.991835066 | -4.164140317 |
| **A_55_P2077613** | Fam102a | -0.137311051 | -0.315232048 | 0.040609946 | 11.52530752 | -1.844537206 | 0.109847804 | 0.991835066 | -4.164810708 |
| **A_51_P223929** | Ash1l | -0.114639256 | -0.26323159 | 0.033953077 | 9.835282442 | -1.843937047 | 0.109942809 | 0.991835066 | -4.165354714 |
| **A_52_P359621** | Luzp1 | 0.134240523 | -0.039768246 | 0.308249292 | 11.24952493 | 1.843833211 | 0.109959254 | 0.991835066 | -4.165448833 |
| **A_52_P105840** | Fnbp1 | 0.09269227 | -0.02749231 | 0.212876849 | 8.234474743 | 1.843333438 | 0.110038441 | 0.991835066 | -4.165901832 |
| **A_55_P2008061** | Itpr2 | -0.112137768 | -0.257535655 | 0.033260118 | 11.95978396 | -1.843329394 | 0.110039082 | 0.991835066 | -4.165905498 |
| **A_55_P2050932** | Tlr7 | -0.161198047 | -0.37021717 | 0.047821075 | 8.34981249 | -1.843244253 | 0.110052577 | 0.991835066 | -4.165982669 |
| **A_55_P2162452** | Mpdu1 | 0.170638731 | -0.05064441 | 0.391921873 | 11.942743 | 1.843055516 | 0.1100825 | 0.991835066 | -4.166153739 |
| **A_51_P429212** | Npl | 0.091206732 | -0.027071456 | 0.20948492 | 8.042712733 | 1.843025591 | 0.110087245 | 0.991835066 | -4.166180863 |
| **A_55_P2112085** | Vwa8 | -0.109223894 | -0.250891825 | 0.032444036 | 9.578856171 | -1.842702564 | 0.110138479 | 0.991835066 | -4.166473648 |
| **A_55_P2065050** | NA | -0.130843228 | -0.30057344 | 0.038886985 | 9.071240523 | -1.842473399 | 0.11017484 | 0.991835066 | -4.166681356 |
| **A_51_P257541** | Plppr1 | -0.137869131 | -0.316751939 | 0.041013676 | 8.556665215 | -1.842076107 | 0.110237904 | 0.991835066 | -4.167041443 |
| **A_55_P2425235** | NA | 0.139957214 | -0.041652048 | 0.321566476 | 9.374253414 | 1.841901615 | 0.110265613 | 0.991835066 | -4.167199593 |
| **A_51_P140819** | Wdr36 | -0.08172186 | -0.187781914 | 0.024338194 | 9.247262651 | -1.84160096 | 0.110313373 | 0.991835066 | -4.167472086 |
| **A_51_P182131** | 5330417C2 | -0.084137654 | -0.193342826 | 0.025067518 | 7.801928869 | -1.841434706 | 0.110339792 | 0.991835066 | -4.167622767 |

| | | | | | | | | | |
|---|---|---|---|---|---|---|---|---|---|
| **A_52_P490272** | Nup98 | 0.124814116 | -0.037292176 | 0.286920408 | 8.860641704 | 1.840233639 | 0.11053083 | 0.991835066 | -4.168711287 |
| **A_52_P561936** | Sptssa | -0.131801797 | -0.302991592 | 0.039387998 | 11.44262167 | -1.840147312 | 0.110544574 | 0.991835066 | -4.168789523 |
| **A_52_P462472** | Nexmif | 0.08318348 | -0.024875783 | 0.191242743 | 8.125988318 | 1.839857617 | 0.110590705 | 0.991835066 | -4.169052061 |
| **A_55_P1962926** | Snapc3 | -0.096395282 | -0.221630566 | 0.028840002 | 8.702556896 | -1.839662716 | 0.110621753 | 0.991835066 | -4.16922869 |
| **A_52_P362981** | NA | -0.157383932 | -0.361941972 | 0.047174108 | 10.62869678 | -1.838878222 | 0.110746806 | 0.991835066 | -4.169939621 |
| **A_55_P2088385** | Tnnt1 | 0.235441887 | -0.070580944 | 0.541464718 | 13.68986105 | 1.838819215 | 0.110756218 | 0.991835066 | -4.169993093 |
| **A_55_P2154684** | Smarca4 | 0.094659825 | -0.028377849 | 0.217697499 | 13.619075 | 1.838809403 | 0.110757783 | 0.991835066 | -4.170001985 |
| **A_55_P2111005** | Pafah1b1 | -0.09946481 | -0.228750178 | 0.029820559 | 12.12372636 | -1.838777627 | 0.110762852 | 0.991835066 | -4.170030781 |
| **A_51_P429682** | Ebp | -0.136864624 | -0.314777466 | 0.041048217 | 12.25330334 | -1.83862451 | 0.110787279 | 0.991835066 | -4.170169536 |
| **A_55_P2056285** | Sf3b5 | 0.090667864 | -0.027270153 | 0.208605881 | 13.15303511 | 1.837421082 | 0.110979449 | 0.991835066 | -4.171260051 |
| **A_51_P309740** | Usp40 | 0.1098674 | -0.033065246 | 0.252800046 | 9.227239573 | 1.837158155 | 0.111021478 | 0.991835066 | -4.171498301 |
| **A_55_P2159666** | NA | 0.16889747 | -0.050845811 | 0.38864075 | 9.342334529 | 1.837031792 | 0.111041683 | 0.991835066 | -4.171612802 |
| **A_55_P2132631** | NA | -0.101588072 | -0.233781492 | 0.030605347 | 12.74429744 | -1.836716022 | 0.111092188 | 0.991835066 | -4.17189893 |
| **A_55_P2003824** | Asgr1 | 0.08508411 | -0.025658651 | 0.19582687 | 7.860313889 | 1.836294135 | 0.111159701 | 0.991835066 | -4.172281204 |
| **A_51_P168632** | Prim1 | 0.159048024 | -0.047980115 | 0.366076164 | 11.90330786 | 1.836149475 | 0.111182859 | 0.991835066 | -4.172412281 |
| **A_65_P07196** | COX3 | -0.121741632 | -0.28020992 | 0.036726657 | 17.25806856 | -1.836140386 | 0.111184315 | 0.991835066 | -4.172420516 |
| **A_51_P293069** | Flvcr1 | 0.113653128 | -0.034287204 | 0.261593459 | 8.558382588 | 1.83613216 | 0.111185632 | 0.991835066 | -4.172427969 |
| **A_51_P440443** | Exosc5 | 0.119063683 | -0.035944836 | 0.274072202 | 11.45621111 | 1.835831757 | 0.111233739 | 0.991835066 | -4.17270016 |
| **A_66_P126492** | Brk1 | -0.131322465 | -0.302312232 | 0.039667301 | 12.34637157 | -1.835599959 | 0.111270874 | 0.991835066 | -4.172910185 |
| **A_55_P2107357** | Gm15421 | 0.211015218 | -0.063789391 | 0.485819827 | 14.09380491 | 1.835265483 | 0.111324479 | 0.991835066 | -4.173213242 |
| **A_51_P433127** | Rdm1 | -0.13115872 | -0.302001162 | 0.039683722 | 12.67008805 | -1.834892097 | 0.11138435 | 0.991835066 | -4.173551546 |
| **A_55_P1963061** | Rps16 | 0.121978993 | -0.036933326 | 0.280891312 | 16.56187966 | 1.834579815 | 0.111434447 | 0.991835066 | -4.173834482 |
| **A_55_P1970806** | Mif-ps6 | 0.227711292 | -0.069009034 | 0.524431619 | 16.0734432 | 1.834198787 | 0.111495603 | 0.991835066 | -4.174179699 |
| **A_55_P2150053** | Yipf5 | -0.116208623 | -0.267684833 | 0.035267587 | 10.02870501 | -1.833593479 | 0.111592822 | 0.991835066 | -4.174728104 |
| **A_55_P2062707** | Mthfsl | -0.085051826 | -0.195943988 | 0.025840337 | 8.997045369 | -1.833124328 | 0.11166823 | 0.991835066 | -4.175153139 |
| **A_55_P2033287** | NA | 0.104350763 | -0.031718004 | 0.24041953 | 14.99624835 | 1.832931818 | 0.111699187 | 0.991835066 | -4.175327544 |
| **A_55_P2126391** | Srl | -0.156409342 | -0.360466324 | 0.04764764 | 13.54943937 | -1.831978459 | 0.111852618 | 0.991835066 | -4.176191218 |
| **A_55_P2015687** | Phf11d | -0.146050107 | -0.336642061 | 0.044541848 | 8.603266179 | -1.83149799 | 0.11193002 | 0.991835066 | -4.176626472 |
| **A_51_P507023** | Nt5dc3 | -0.134068283 | -0.309028233 | 0.040891667 | 11.91844518 | -1.831456186 | 0.111936757 | 0.991835066 | -4.176664342 |
| **A_55_P1990785** | Zranb3 | 0.118870558 | -0.036278154 | 0.27401927 | 9.020661568 | 1.831197805 | 0.111978406 | 0.991835066 | -4.176898403 |
| **A_52_P462350** | Dr1 | -0.091603102 | -0.211216026 | 0.028009821 | 9.395732977 | -1.830379782 | 0.112110363 | 0.991835066 | -4.177639409 |
| **A_55_P2099700** | Klc4 | -0.221006429 | -0.509676637 | 0.067663778 | 11.46325334 | -1.829835613 | 0.112198228 | 0.991835066 | -4.178132328 |
| **A_51_P282394** | Mrps28 | -0.106250607 | -0.245066369 | 0.032565154 | 13.23342539 | -1.829370928 | 0.112273313 | 0.991835066 | -4.178553238 |
| **A_51_P318381** | Pgf | -0.099506802 | -0.22953979 | 0.030526187 | 8.220283688 | -1.828977465 | 0.112336927 | 0.991835066 | -4.178909627 |
| **A_55_P2043702** | Tfg | 0.1211199 | -0.037170244 | 0.279410044 | 13.12819253 | 1.828819155 | 0.112362532 | 0.991835066 | -4.179053019 |
| **A_51_P240801** | Tmem173 | 0.125049566 | -0.038399881 | 0.288499013 | 9.919717698 | 1.828554317 | 0.112405379 | 0.991835066 | -4.179292896 |
| **A_55_P2147971** | Ranbp1 | 0.18019106 | -0.055367635 | 0.415749755 | 12.36441501 | 1.828282207 | 0.11244942 | 0.991835066 | -4.179539358 |
| **A_55_P2012498** | Cdca7 | 0.202281158 | -0.062169992 | 0.466732308 | 10.88412898 | 1.828180632 | 0.112465864 | 0.991835066 | -4.179631357 |
| **A_55_P2182392** | Adat3 | 0.095642826 | -0.029422127 | 0.22070778 | 8.701788715 | 1.827788369 | 0.11252939 | 0.991835066 | -4.179986638 |
| **A_52_P134924** | Wdr20 | -0.077735591 | -0.179402693 | 0.023931512 | 8.980548079 | -1.827462984 | 0.112582112 | 0.991835066 | -4.18028134 |
| **A_51_P296608** | Gadd45a | -0.092619721 | -0.213821404 | 0.028581963 | 8.302648289 | -1.826433848 | 0.112749019 | 0.991835066 | -4.181213398 |
| **A_55_P2014169** | Ppa2 | -0.080830409 | -0.186607642 | 0.024946824 | 13.13803227 | -1.82638237 | 0.112757374 | 0.991835066 | -4.181260019 |
| **A_55_P1972128** | Ccndbp1 | -0.163295972 | -0.377003528 | 0.050411585 | 9.450993059 | -1.826268906 | 0.112775792 | 0.991835066 | -4.181362777 |
| **A_55_P2166384** | Cnot1 | 0.121924268 | -0.037664191 | 0.281512727 | 11.57158612 | 1.825987526 | 0.112821479 | 0.991835066 | -4.181617604 |
| **A_66_P137943** | Tctn2 | -0.126603626 | -0.292352683 | 0.03914543 | 9.834088014 | -1.825594054 | 0.112885397 | 0.991835066 | -4.181973938 |
| **A_66_P112394** | NA | -0.17624453 | -0.406995328 | 0.054506269 | 12.12950338 | -1.825498861 | 0.112900866 | 0.991835066 | -4.182060145 |
| **A_55_P2145521** | Stk38l | 0.165870098 | -0.051335202 | 0.383075399 | 10.09471628 | 1.82518468 | 0.112951935 | 0.991835066 | -4.182344666 |
| **A_52_P215106** | Pik3r6 | -0.078737435 | -0.181851946 | 0.024377076 | 10.98165388 | -1.825032523 | 0.112976676 | 0.991835066 | -4.182482456 |

| | | | | | | | | | |
|---|---|---|---|---|---|---|---|---|---|
| **A_55_P2061645** | NA | 0.172699035 | -0.053468585 | 0.398866656 | 8.882827949 | 1.825024151 | 0.112978037 | 0.991835066 | -4.182490038 |
| **A_55_P2008591** | Kras | 0.080430217 | -0.024914898 | 0.185775331 | 7.916025273 | 1.824794531 | 0.113015384 | 0.991835066 | -4.182697976 |
| **A_55_P2004084** | Iqce | 0.125266357 | -0.038864354 | 0.289397068 | 9.121782444 | 1.824121362 | 0.113124941 | 0.991835066 | -4.183307565 |
| **A_55_P1960916** | Egln3 | -0.137589927 | -0.317885338 | 0.042705483 | 9.810396213 | -1.823942325 | 0.113154096 | 0.991835066 | -4.183469688 |
| **A_51_P205481** | Mtmr6 | -0.08918436 | -0.206067444 | 0.027698724 | 10.91225475 | -1.823669647 | 0.113198515 | 0.991835066 | -4.183716604 |
| **A_52_P629451** | Gon4l | 0.086466348 | -0.026858896 | 0.199791592 | 8.700993739 | 1.823600015 | 0.11320986 | 0.991835066 | -4.183779656 |
| **A_55_P1972923** | Polr1d | 0.131051531 | -0.040755888 | 0.302858951 | 11.81172858 | 1.82309507 | 0.113292167 | 0.991835066 | -4.184236883 |
| **A_55_P2068172** | Nufip2 | 0.239549722 | -0.074508126 | 0.55360757 | 9.928688412 | 1.823035612 | 0.113301862 | 0.991835066 | -4.184290721 |
| **A_55_P2046262** | Phgdh | 0.382129213 | -0.119201476 | 0.883459902 | 10.71780876 | 1.821776442 | 0.113507379 | 0.991835066 | -4.185430838 |
| **A_51_P153053** | Smpdl3a | -0.103278449 | -0.238796877 | 0.032239979 | 11.83604944 | -1.821463545 | 0.113558505 | 0.991835066 | -4.185714139 |
| **A_55_P2013598** | Ccnq | 0.086118994 | -0.026899062 | 0.199137051 | 10.48765271 | 1.82121093 | 0.113599797 | 0.991835066 | -4.185942856 |
| **A_55_P2145307** | Ints11 | 0.097126184 | -0.030343888 | 0.224596255 | 9.53772262 | 1.821114391 | 0.113615581 | 0.991835066 | -4.186030262 |
| **A_55_P2045163** | Ehmt1 | -0.101719242 | -0.235232046 | 0.031793561 | 11.79110322 | -1.820913574 | 0.113648421 | 0.991835066 | -4.186212078 |
| **A_55_P1953391** | Rbfox1 | -0.139982056 | -0.323720291 | 0.043756178 | 8.69574298 | -1.820882666 | 0.113653477 | 0.991835066 | -4.186240061 |
| **A_51_P173100** | Pttg1ip | -0.124284351 | -0.287457483 | 0.038888781 | 11.69182337 | -1.820442354 | 0.113725518 | 0.991835066 | -4.186638704 |
| **A_55_P1996927** | 4933409K0 | 0.213001695 | -0.066703512 | 0.492706903 | 9.806210838 | 1.820084876 | 0.113784039 | 0.991835066 | -4.186962345 |
| **A_55_P2016902** | Entr1 | 0.098776495 | -0.030938255 | 0.228491246 | 9.65249741 | 1.820008351 | 0.113796571 | 0.991835066 | -4.187031626 |
| **A_55_P2098100** | Arhgef9 | 0.100793147 | -0.031609215 | 0.233195509 | 8.625217923 | 1.819467922 | 0.113885107 | 0.991835066 | -4.187520886 |
| **A_52_P394539** | Tex2 | -0.087998994 | -0.203626469 | 0.02762848 | 11.08630244 | -1.81897111 | 0.113966556 | 0.991835066 | -4.187970646 |
| **A_51_P157255** | Sdc2 | -0.143608276 | -0.332329254 | 0.045112703 | 10.67104499 | -1.818730795 | 0.114005975 | 0.991835066 | -4.188188196 |
| **A_52_P340669** | Bhlha15 | 0.078499715 | -0.024702589 | 0.181702018 | 7.698863838 | 1.817974634 | 0.114130093 | 0.991835066 | -4.188872711 |
| **A_55_P2096187** | Hax1 | 0.0847762 | -0.026678941 | 0.196231342 | 12.0197187 | 1.817954346 | 0.114133425 | 0.991835066 | -4.188891076 |
| **A_55_P2040011** | Itga7 | -0.192105057 | -0.444673226 | 0.060463113 | 10.8124394 | -1.817897165 | 0.114142817 | 0.991835066 | -4.188942837 |
| **A_51_P295442** | Nek7 | -0.111363455 | -0.257911501 | 0.035184592 | 11.33535763 | -1.816233969 | 0.11441631 | 0.991835066 | -4.190448334 |
| **A_51_P427964** | Lmod3 | -0.13251106 | -0.306948832 | 0.041926712 | 12.8938682 | -1.815602324 | 0.114520344 | 0.991835066 | -4.191020051 |
| **A_55_P2071466** | Ncf1 | -0.073100843 | -0.169419867 | 0.023218181 | 7.864857358 | -1.813925492 | 0.114796967 | 0.991835066 | -4.19253769 |
| **A_55_P2086811** | Sp140 | -0.097928566 | -0.226974486 | 0.031117354 | 9.012126738 | -1.813736729 | 0.114828148 | 0.991835066 | -4.192708524 |
| **A_52_P262676** | NA | 0.090708909 | -0.028845757 | 0.210263575 | 9.606859797 | 1.81339554 | 0.114884527 | 0.991835066 | -4.1930173 |
| **A_51_P440852** | Mtrf1 | -0.08413912 | -0.195043534 | 0.026765293 | 9.400046157 | -1.813252441 | 0.114908181 | 0.991835066 | -4.193146803 |
| **A_55_P1959515** | Cep135 | 0.103779149 | -0.033031071 | 0.240589369 | 8.598581661 | 1.813012065 | 0.114947926 | 0.991835066 | -4.193364338 |
| **A_55_P1996988** | Tent4b | 0.110570248 | -0.035204691 | 0.256345187 | 10.53577038 | 1.812861176 | 0.114972882 | 0.991835066 | -4.193500888 |
| **A_51_P413545** | Txndc11 | 0.118905712 | -0.037938387 | 0.275749812 | 11.13822343 | 1.811939389 | 0.115125449 | 0.991835066 | -4.194335049 |
| **A_51_P365468** | Clpb | -0.167290635 | -0.388152469 | 0.0535712 | 11.24860988 | -1.810339742 | 0.115390678 | 0.991835066 | -4.195782526 |
| **A_55_P1988728** | Fbxl19 | 0.120857383 | -0.038722043 | 0.280436809 | 9.355354863 | 1.81011188 | 0.115428507 | 0.991835066 | -4.1959887 |
| **A_51_P332602** | Surf1 | -0.130255334 | -0.302252549 | 0.041741881 | 10.7817173 | -1.810019423 | 0.11544386 | 0.991835066 | -4.196072356 |
| **A_52_P330395** | Farp1 | 0.122200949 | -0.03916713 | 0.283569028 | 11.60335593 | 1.80994793 | 0.115455733 | 0.991835066 | -4.196137044 |
| **A_55_P2412214** | Mhrt | 0.140297981 | -0.045013246 | 0.325609208 | 8.575464422 | 1.809500972 | 0.115529987 | 0.991835066 | -4.19654145 |
| **A_55_P2091206** | NA | 0.20502149 | -0.065792897 | 0.475835877 | 9.830788822 | 1.809409164 | 0.115545245 | 0.991835066 | -4.196624516 |
| **A_66_P125734** | Tnip2 | 0.132228131 | -0.042438493 | 0.306894756 | 8.772147125 | 1.809352002 | 0.115554746 | 0.991835066 | -4.196676235 |
| **A_51_P395921** | Uchl4 | -0.110071485 | -0.255561992 | 0.035419022 | 10.60459452 | -1.808211812 | 0.115744417 | 0.991835066 | -4.197707816 |
| **A_65_P15875** | Chpf2 | 0.10761438 | -0.034633544 | 0.249862304 | 9.546757981 | 1.808146032 | 0.115755369 | 0.991835066 | -4.197767328 |
| **A_55_P2145656** | NA | -0.139781543 | -0.32455964 | 0.044996554 | 14.1726594 | -1.808041823 | 0.115772721 | 0.991835066 | -4.197861607 |
| **A_55_P2125491** | Tnfrsf4 | 0.110266394 | -0.035520719 | 0.256053507 | 8.246788936 | 1.807728344 | 0.115824934 | 0.991835066 | -4.19814521 |
| **A_55_P1990217** | Zfp13 | 0.135068969 | -0.043542495 | 0.313680433 | 8.581700916 | 1.807404619 | 0.115878877 | 0.991835066 | -4.198438077 |
| **A_55_P1997141** | Mybl2 | 0.175490623 | -0.056582137 | 0.407563383 | 10.2580566 | 1.807335924 | 0.115890327 | 0.991835066 | -4.198500223 |
| **A_52_P621567** | Tcf20 | 0.157841643 | -0.050904366 | 0.366587651 | 10.84856143 | 1.807226325 | 0.115908597 | 0.991835066 | -4.198599373 |
| **A_55_P1977478** | Zfp629 | 0.11715663 | -0.037801759 | 0.272115018 | 12.04298899 | 1.807011479 | 0.11594442 | 0.991835066 | -4.198793735 |
| **A_51_P179131** | Vasp | -0.249416563 | -0.579375716 | 0.08054259 | 10.11003201 | -1.8066509 | 0.116004566 | 0.991835066 | -4.199119928 |

| | | | | | | | | | |
|---|---|---|---|---|---|---|---|---|---|
| **A_51_P498388** | Sbk1 | -0.144633063 | -0.335991552 | 0.046725425 | 9.241572236 | -1.806462645 | 0.116035979 | 0.991835066 | -4.199290228 |
| **A_55_P2061094** | Fnta | 0.117327891 | -0.037946052 | 0.272601835 | 8.502666906 | 1.805975337 | 0.116117334 | 0.991835066 | -4.199731048 |
| **A_52_P632423** | NA | 0.08920708 | -0.028948508 | 0.207362668 | 8.514872404 | 1.804488794 | 0.116365848 | 0.991835066 | -4.2010757 |
| **A_55_P2159390** | NA | -0.384001587 | -0.892627207 | 0.124624034 | 13.12038001 | -1.80444612 | 0.11637299 | 0.991835066 | -4.201114299 |
| **A_55_P1999641** | Frem1 | 0.222979315 | -0.072436223 | 0.518394852 | 9.352925595 | 1.80401596 | 0.116445003 | 0.991835066 | -4.201503375 |
| **A_51_P112308** | Tcim | -0.132607235 | -0.308309481 | 0.04309501 | 10.28523807 | -1.803844275 | 0.116473756 | 0.991835066 | -4.200950143 |
| **A_51_P286034** | Ccdc117 | -0.126706787 | -0.294591362 | 0.041177788 | 8.86836349 | -1.803840839 | 0.116474332 | 0.991835066 | -4.201661768 |
| **A_55_P1963868** | Plekho2 | 0.099200744 | -0.032279579 | 0.230681067 | 11.19819407 | 1.803280549 | 0.116568217 | 0.991835066 | -4.202168526 |
| **A_51_P261001** | Mtf2 | 0.153489629 | -0.049988464 | 0.356967723 | 10.44774221 | 1.802895317 | 0.116632812 | 0.991835066 | -4.20251694 |
| **A_66_P130759** | 4631405J1 | -0.199133589 | -0.463157646 | 0.064890468 | 8.739388392 | -1.802644769 | 0.116674841 | 0.991835066 | -4.202743538 |
| **A_55_P1991605** | Aplp1 | 0.224381108 | -0.073173378 | 0.521935594 | 8.863386659 | 1.802307615 | 0.116731422 | 0.991835066 | -4.203048458 |
| **A_55_P1965402** | Uqcrb | -0.13784385 | -0.320734156 | 0.045046455 | 15.11920347 | -1.801382103 | 0.116886878 | 0.991835066 | -4.203885452 |
| **A_51_P409429** | Aars | 0.144193101 | -0.047142432 | 0.335528634 | 11.64031217 | 1.801183603 | 0.116920245 | 0.991835066 | -4.204064962 |
| **A_51_P421182** | Sys1 | -0.104023839 | -0.242109359 | 0.034061681 | 8.397162552 | -1.800503085 | 0.117034709 | 0.991835066 | -4.204680354 |
| **A_51_P259902** | Arl6ip5 | 0.281149993 | -0.092100725 | 0.654400711 | 11.82326041 | 1.80030695 | 0.11706772 | 0.991835066 | -4.204857714 |
| **A_55_P2031496** | Rufy3 | 0.159593246 | -0.052320079 | 0.371506571 | 8.79613974 | 1.799970445 | 0.117124376 | 0.991835066 | -4.205162002 |
| **A_52_P638895** | Vegfa | -0.212466981 | -0.494594796 | 0.069660834 | 11.37961469 | -1.7999261 | 0.117131844 | 0.991835066 | -4.205202101 |
| **A_55_P2100824** | LOC67668 | -0.128860456 | -0.30000464 | 0.042283727 | 8.671519881 | -1.799561334 | 0.117193292 | 0.991835066 | -4.205531936 |
| **A_51_P162437** | Bcr | 0.112892995 | -0.037064567 | 0.262850557 | 11.26530333 | 1.799317295 | 0.11723442 | 0.991835066 | -4.205752601 |
| **A_55_P1953920** | Me3 | -0.261055991 | -0.607849516 | 0.085737534 | 9.726598272 | -1.799168407 | 0.117259519 | 0.991835066 | -4.205887226 |
| **A_55_P1980521** | Ccdc85a | -0.129381801 | -0.301326927 | 0.042563325 | 8.058136339 | -1.798425519 | 0.11738483 | 0.991835066 | -4.206558933 |
| **A_55_P2053541** | NA | 0.154356054 | -0.050786184 | 0.359498292 | 14.07604762 | 1.798364455 | 0.117395137 | 0.991835066 | -4.206614144 |
| **A_55_P1989765** | NA | 0.347678604 | -0.11440403 | 0.809761238 | 9.013202899 | 1.798321971 | 0.117402307 | 0.991835066 | -4.206652556 |
| **A_55_P1984416** | NA | 0.410582401 | -0.135148372 | 0.956313173 | 10.19196184 | 1.798171122 | 0.117427772 | 0.991835066 | -4.206788946 |
| **A_52_P100252** | Fasn | 0.078299657 | -0.025835169 | 0.182434484 | 8.49357138 | 1.797103109 | 0.117608217 | 0.991835066 | -4.207754551 |
| **A_55_P1991783** | NA | 0.623405668 | -0.2060404 | 1.452851736 | 12.20485098 | 1.796353177 | 0.117735081 | 0.991835066 | -4.208432535 |
| **A_55_P2147220** | Ak2 | 0.309657873 | -0.102359308 | 0.721675054 | 11.00534236 | 1.796287852 | 0.117746138 | 0.991835066 | -4.208491592 |
| **A_55_P1954400** | NA | -0.133142843 | -0.310347774 | 0.044062088 | 11.94694286 | -1.79577184 | 0.117833516 | 0.991835066 | -4.208958076 |
| **A_51_P380401** | Cars2 | -0.124086124 | -0.289243548 | 0.0410713 | 10.4213662 | -1.79570194 | 0.117845357 | 0.991835066 | -4.209021266 |
| **A_55_P1964332** | NA | 0.175528517 | -0.058147635 | 0.40920467 | 10.30783744 | 1.795322291 | 0.11790969 | 0.991835066 | -4.209364465 |
| **A_55_P2017600** | Crmp1 | 0.141117762 | -0.046850754 | 0.329086279 | 9.046224954 | 1.794343974 | 0.118075626 | 0.991835066 | -4.210248812 |
| **A_55_P2125496** | Gm10731 | -0.099437235 | -0.231887853 | 0.033013383 | 12.56616636 | -1.794337709 | 0.11807669 | 0.991835066 | -4.210254475 |
| **A_52_P299358** | Lclat1 | -0.096157762 | -0.224258328 | 0.031942804 | 11.5820072 | -1.794082589 | 0.118119999 | 0.991835066 | -4.21048508 |
| **A_55_P2007640** | Eprs | 0.1894446 | -0.062976217 | 0.441865417 | 10.7359707 | 1.793767678 | 0.11817348 | 0.991835066 | -4.210769726 |
| **A_66_P121369** | Gars | 0.13509859 | -0.044915889 | 0.315113069 | 15.01962433 | 1.793711162 | 0.118183081 | 0.991835066 | -4.210820811 |
| **A_55_P2149440** | Ehmt1 | 0.107625655 | -0.03579523 | 0.251046539 | 11.01152805 | 1.793546091 | 0.118211126 | 0.991835066 | -4.210970014 |
| **A_55_P2039354** | NA | 0.1745525 | -0.058091856 | 0.407196857 | 8.403447764 | 1.793257617 | 0.118260153 | 0.991835066 | -4.211230755 |
| **A_51_P104527** | Atrip | 0.092276056 | -0.030728843 | 0.215280955 | 9.860387503 | 1.792981255 | 0.11830714 | 0.991835066 | -4.211480543 |
| **A_66_P135700** | Cyp2d11 | -0.11904342 | -0.277769643 | 0.039682803 | 9.415386297 | -1.79252757 | 0.118384314 | 0.991835066 | -4.211890593 |
| **A_55_P2101920** | Eva1c | 0.142680228 | -0.04757998 | 0.332940435 | 9.495074892 | 1.792358827 | 0.11841303 | 0.991835066 | -4.212043104 |
| **A_55_P2170732** | Fndc10 | 0.153097625 | -0.051098704 | 0.357293954 | 9.398881146 | 1.791965544 | 0.118479985 | 0.991835066 | -4.212398548 |
| **A_51_P338998** | Prim2 | 0.122811889 | -0.040999895 | 0.286623674 | 9.645669437 | 1.791861359 | 0.118497728 | 0.991835066 | -4.212492707 |
| **A_55_P1961235** | Rpl23 | 0.083618599 | -0.027942278 | 0.195179475 | 16.06862829 | 1.791431061 | 0.118571037 | 0.991835066 | -4.212881591 |
| **A_51_P456208** | Tff3 | -0.180903185 | -0.422357479 | 0.060551109 | 7.947179417 | -1.790690016 | 0.11869739 | 0.991835066 | -4.213551288 |
| **A_52_P545810** | Lrrfip1 | 0.248248325 | -0.083119834 | 0.579616485 | 9.906470913 | 1.790542708 | 0.118722522 | 0.991835066 | -4.213684409 |
| **A_52_P263518** | Gng2 | -0.121509308 | -0.283707416 | 0.0406888 | 12.42760295 | -1.790494102 | 0.118730816 | 0.991835066 | -4.213728334 |
| **A_55_P2015299** | NA | -0.108298117 | -0.252918542 | 0.036322307 | 15.18554037 | -1.789783116 | 0.118852199 | 0.991835066 | -4.214370825 |
| **A_51_P441387** | Kansl1l | -0.127634953 | -0.298079747 | 0.042809841 | 8.636916167 | -1.78976089 | 0.118855995 | 0.991835066 | -4.214390909 |

| | | | | | | | | | |
|---|---|---|---|---|---|---|---|---|---|
| **A_55_P2158102** | Camp | -0.214075593 | -0.500013905 | 0.071862719 | 8.287766875 | -1.789385606 | 0.118920116 | 0.991835066 | -4.214730026 |
| **A_55_P2098140** | Tmed2 | 0.102328366 | -0.034362353 | 0.239019085 | 13.91118283 | 1.789229893 | 0.118946731 | 0.991835066 | -4.21487073 |
| **A_55_P2021099** | Tmem181a | -0.132623499 | -0.309802847 | 0.044555849 | 10.21252268 | -1.789025432 | 0.118981687 | 0.991835066 | -4.215055481 |
| **A_55_P2151868** | Mest | 0.282797066 | -0.095008011 | 0.660602142 | 10.44440484 | 1.789024312 | 0.118981878 | 0.991835066 | -4.215056493 |
| **A_52_P136782** | Rgs5 | -0.263966401 | -0.616628266 | 0.088695465 | 9.658862937 | -1.78895445 | 0.118993824 | 0.991835066 | -4.21511962 |
| **A_55_P1966124** | Erh | 0.136206488 | -0.045790612 | 0.318203588 | 14.57197133 | 1.788720393 | 0.119033856 | 0.991835066 | -4.215331109 |
| **A_66_P106065** | Ttc3 | 0.074955635 | -0.025202266 | 0.175113537 | 9.365186132 | 1.788661692 | 0.119043898 | 0.991835066 | -4.21538415 |
| **A_55_P2131353** | 1110059G1 | -0.103699232 | -0.242292875 | 0.034894411 | 9.308077855 | -1.788304051 | 0.119105097 | 0.991835066 | -4.2157073 |
| **A_55_P2171273** | Mier3 | 0.087163299 | -0.0293384 | 0.203664998 | 8.67126984 | 1.788177108 | 0.119126826 | 0.991835066 | -4.215821999 |
| **A_51_P377508** | Srebf2 | 0.196889692 | -0.066459739 | 0.460239123 | 8.553786145 | 1.786897855 | 0.119346017 | 0.991835066 | -4.216977805 |
| **A_55_P2042525** | Apoo | -0.088976033 | -0.208015503 | 0.030063437 | 12.98828354 | -1.786451323 | 0.119422619 | 0.991835066 | -4.217381223 |
| **A_55_P2113359** | Zfp523 | 0.133587389 | -0.045156152 | 0.31233093 | 11.08638943 | 1.786258205 | 0.119455763 | 0.991835066 | -4.217555691 |
| **A_55_P2171897** | Notch1 | 0.177406339 | -0.05999052 | 0.414803199 | 8.435337124 | 1.786089829 | 0.119484668 | 0.991835066 | -4.217707804 |
| **A_55_P1975380** | Nol11 | 0.115521766 | -0.039066334 | 0.270109866 | 10.50295187 | 1.786063485 | 0.119489191 | 0.991835066 | -4.217731604 |
| **A_51_P504815** | Stfa3 | -0.389849533 | -0.911594128 | 0.131895063 | 8.907871985 | -1.785863213 | 0.119523581 | 0.991835066 | -4.21791253 |
| **A_55_P2039611** | Eed | 0.146339972 | -0.049533048 | 0.342212992 | 11.72891339 | 1.785654702 | 0.119559396 | 0.991835066 | -4.218100895 |
| **A_55_P2095047** | Mat2a | 0.162202892 | -0.05490831 | 0.379314093 | 9.083164101 | 1.785605411 | 0.119567864 | 0.991835066 | -4.218145424 |
| **A_55_P1953846** | Abca8b | -0.111374709 | -0.26045957 | 0.037710152 | 8.344460243 | -1.785509512 | 0.119584341 | 0.991835066 | -4.218232057 |
| **A_55_P1976007** | Psme4 | -0.119761621 | -0.280092109 | 0.040568867 | 9.690530134 | -1.785297861 | 0.119620714 | 0.991835066 | -4.218423255 |
| **A_55_P2195157** | Dbn1 | -0.111972231 | -0.261891435 | 0.037946972 | 10.10898699 | -1.785098555 | 0.119654975 | 0.991835066 | -4.2186033 |
| **A_55_P1989469** | NA | 0.124992241 | -0.04244367 | 0.292428153 | 14.87910745 | 1.78420018 | 0.119809523 | 0.991835066 | -4.219414818 |
| **A_55_P2128663** | Uggt1 | 0.079812969 | -0.027111893 | 0.186737832 | 8.447365339 | 1.784037173 | 0.119837586 | 0.991835066 | -4.219562059 |
| **A_55_P2047768** | Serpinb6c | -0.119702962 | -0.280185726 | 0.040779803 | 13.0837047 | -1.782730237 | 0.120062816 | 0.991835066 | -4.220742531 |
| **A_52_P594049** | Actr5 | 0.10685 | -0.036408487 | 0.250108486 | 9.596723036 | 1.782638566 | 0.120078629 | 0.991835066 | -4.220825328 |
| **A_55_P2165234** | Slurp2 | -0.095263124 | -0.223010102 | 0.032483854 | 8.260448295 | -1.782310305 | 0.120135271 | 0.991835066 | -4.221121807 |
| **A_66_P113505** | Fgd4 | 0.272426667 | -0.092905982 | 0.637759317 | 10.29898342 | 1.782256731 | 0.120144518 | 0.991835066 | -4.221170193 |
| **A_55_P2066007** | Crocc | 0.259210085 | -0.088476239 | 0.606896409 | 9.825427739 | 1.781859344 | 0.120213127 | 0.991835066 | -4.221529093 |
| **A_51_P343356** | Isoc2b | 0.079484434 | -0.027152864 | 0.186121733 | 8.551571315 | 1.781484652 | 0.120277853 | 0.991835066 | -4.221867488 |
| **A_55_P2034027** | Nos3 | 0.096460255 | -0.032993632 | 0.225914142 | 12.35128302 | 1.780911936 | 0.120376851 | 0.991835066 | -4.222384705 |
| **A_55_P2037544** | NA | 0.091442013 | -0.031333793 | 0.214217819 | 15.70430645 | 1.780090647 | 0.120518954 | 0.991835066 | -4.223126369 |
| **A_55_P1969566** | Atp9b | 0.081488979 | -0.027925763 | 0.190903721 | 8.180049551 | 1.780049861 | 0.120526015 | 0.991835066 | -4.2231632 |
| **A_55_P1998461** | Rpl17 | 0.144384736 | -0.04949939 | 0.338268862 | 10.15789652 | 1.779869503 | 0.120557245 | 0.991835066 | -4.223326066 |
| **A_55_P2081373** | Glis1 | -0.088629436 | -0.20768309 | 0.030424219 | 8.048445464 | -1.779280363 | 0.120659312 | 0.991835066 | -4.223858053 |
| **A_55_P2042988** | Aoah | -0.078263374 | -0.183414112 | 0.026887365 | 7.83057799 | -1.778915711 | 0.120722529 | 0.991835066 | -4.224187318 |
| **A_55_P2002460** | Gm2399 | 0.135463418 | -0.0465931 | 0.317519937 | 8.142963246 | 1.778381488 | 0.1208152 | 0.991835066 | -4.224669683 |
| **A_52_P398989** | Cytip | -0.074922434 | -0.175617631 | 0.025772762 | 7.805935187 | -1.778329597 | 0.120824205 | 0.991835066 | -4.224716536 |
| **A_52_P367675** | Acin1 | 0.146605557 | -0.050446301 | 0.343657415 | 8.823776026 | 1.778193552 | 0.120847818 | 0.991835066 | -4.224839371 |
| **A_51_P472829** | Aif1l | 0.214498679 | -0.07387562 | 0.502872977 | 10.77011494 | 1.777776671 | 0.1209202 | 0.991835066 | -4.225215766 |
| **A_52_P339791** | Hdgf | 0.156072163 | -0.05376146 | 0.365905787 | 8.949861546 | 1.777704213 | 0.120932785 | 0.991835066 | -4.225281186 |
| **A_55_P2096867** | Gap43 | 0.261075695 | -0.090156651 | 0.61230804 | 10.07061329 | 1.776564874 | 0.121130841 | 0.991835066 | -4.226309812 |
| **A_66_P102733** | Ccdc28b | -0.126603839 | -0.29692904 | 0.043721362 | 12.52667474 | -1.776548613 | 0.121133671 | 0.991835066 | -4.226324493 |
| **A_51_P506674** | Rpp38 | 0.102444724 | -0.035392781 | 0.240282229 | 9.78952245 | 1.776361389 | 0.121166247 | 0.991835066 | -4.226493515 |
| **A_55_P2126850** | Nudt11 | 0.109504985 | -0.037854908 | 0.256864877 | 9.287926025 | 1.776084917 | 0.121214368 | 0.991835066 | -4.226743105 |
| **A_55_P2111148** | Gemin5 | -0.113141826 | -0.265501467 | 0.039217816 | 9.612119006 | -1.774852935 | 0.121429024 | 0.991835066 | -4.227855234 |
| **A_55_P1971579** | Acox3 | 0.141231408 | -0.049020771 | 0.331483586 | 8.520458608 | 1.77423353 | 0.121537085 | 0.991835066 | -4.228414341 |
| **A_52_P167382** | Tmem8 | 0.080509454 | -0.027990837 | 0.189009745 | 7.72840645 | 1.773475163 | 0.121669515 | 0.991835066 | -4.229098847 |
| **A_55_P2437253** | Nae1 | -0.097013729 | -0.227792039 | 0.033764581 | 9.118197167 | -1.772991325 | 0.121754079 | 0.991835066 | -4.229535542 |
| **A_65_P08521** | Ssrp1 | 0.151905665 | -0.052917593 | 0.356728922 | 9.418841591 | 1.772571788 | 0.12182745 | 0.991835066 | -4.229914188 |

| | | | | | | | | | |
|---|---|---|---|---|---|---|---|---|---|
| **A_55_P2061386** | Chd3 | 0.124549932 | -0.043389798 | 0.292489663 | 11.99607482 | 1.772552781 | 0.121830775 | 0.991835066 | -4.229931342 |
| **A_55_P2005691** | NA | -0.13701239 | -0.321839213 | 0.047814434 | 15.61568174 | -1.771756253 | 0.121970197 | 0.991835066 | -4.2306502 |
| **A_51_P452629** | Tlr2 | -0.079833244 | -0.187545066 | 0.027878578 | 8.291084333 | -1.771452613 | 0.122023386 | 0.991835066 | -4.225935689 |
| **A_51_P317181** | Pcdhb21 | 0.163888422 | -0.057322815 | 0.385099658 | 8.894415533 | 1.770721341 | 0.122151576 | 0.991835066 | -4.231584132 |
| **A_51_P257065** | Prpf40b | 0.120253661 | -0.042072917 | 0.282580238 | 9.033035367 | 1.770589178 | 0.122174757 | 0.991835066 | -4.231703394 |
| **A_55_P2144636** | Ccdc163 | 0.120762036 | -0.042261857 | 0.283785929 | 9.484179025 | 1.77046889 | 0.12219586 | 0.991835066 | -4.231811939 |
| **A_55_P2123234** | NA | -0.10175894 | -0.239136738 | 0.035618859 | 9.147667383 | -1.770374526 | 0.122212417 | 0.991835066 | -4.231897091 |
| **A_55_P2096043** | Acot11 | 0.107678873 | -0.037694038 | 0.253051785 | 10.51411063 | 1.770337813 | 0.122218859 | 0.991835066 | -4.231930219 |
| **A_55_P1976805** | Isca2 | -0.196409327 | -0.461642842 | 0.068824187 | 10.41600926 | -1.769876004 | 0.122299922 | 0.991835066 | -4.232346931 |
| **A_55_P2066996** | Cav2 | -0.09474318 | -0.222694782 | 0.033208422 | 8.162516523 | -1.769747729 | 0.122322448 | 0.991835066 | -4.232462677 |
| **A_55_P2077964** | Vps8 | -0.173173584 | -0.407048942 | 0.060701774 | 8.787482308 | -1.769727149 | 0.122326062 | 0.991835066 | -4.232481247 |
| **A_55_P1966337** | Polr2h | 0.103295323 | -0.036213914 | 0.24280456 | 10.56465871 | 1.769647738 | 0.12234001 | 0.991835066 | -4.232552901 |
| **A_51_P496751** | Nop58 | 0.191086486 | -0.067016741 | 0.449189713 | 11.88562473 | 1.769480057 | 0.122369466 | 0.991835066 | -4.232704201 |
| **A_51_P332201** | Cst3 | -0.13243091 | -0.311310961 | 0.04644914 | 14.24241369 | -1.769443041 | 0.122375969 | 0.991835066 | -4.232737601 |
| **A_55_P2182006** | Cacng6 | -0.172260609 | -0.404946565 | 0.060425347 | 8.706224035 | -1.769395587 | 0.122384307 | 0.991835066 | -4.232780419 |
| **A_55_P2163812** | 4-Sep | -0.166858651 | -0.392272457 | 0.058555154 | 8.999563663 | -1.769201707 | 0.122418378 | 0.991835066 | -4.232955355 |
| **A_52_P671769** | Arhgap18 | -0.120283751 | -0.282784578 | 0.042217077 | 9.723483679 | -1.769133138 | 0.12243043 | 0.991835066 | -4.233017224 |
| **A_51_P142046** | Oxld1 | -0.203800744 | -0.479174402 | 0.071572914 | 11.69494788 | -1.768856097 | 0.122479135 | 0.991835066 | -4.233267189 |
| **A_51_P303725** | Tmem38a | -0.261076783 | -0.613888014 | 0.091734447 | 12.98782555 | -1.768621844 | 0.122520332 | 0.991835066 | -4.233478545 |
| **A_55_P2019690** | Slc43a1 | 0.15953749 | -0.056108646 | 0.375183627 | 8.555673286 | 1.768195195 | 0.1225954 | 0.991835066 | -4.23386348 |
| **A_55_P1973965** | Ppp1r13b | -0.101786314 | -0.239373384 | 0.035800757 | 11.45367183 | -1.768157284 | 0.122602072 | 0.991835066 | -4.233897684 |
| **A_55_P2110935** | Mlip | -0.144896142 | -0.340758323 | 0.050966039 | 10.29105314 | -1.768134788 | 0.122606032 | 0.991835066 | -4.23391798 |
| **A_51_P107782** | Kdsr | 0.101909903 | -0.035864783 | 0.239684588 | 8.96662818 | 1.767893466 | 0.122648515 | 0.991835066 | -4.234135701 |
| **A_55_P2099019** | NA | 0.085197931 | -0.029997641 | 0.200393502 | 10.58077727 | 1.767674902 | 0.122687004 | 0.991835066 | -4.234332886 |
| **A_55_P2065567** | Chac2 | -0.104094861 | -0.244841028 | 0.036651307 | 9.222400317 | -1.767672655 | 0.1226874 | 0.991835066 | -4.234334913 |
| **A_55_P1956302** | H2-Q4 | 0.226971 | -0.079972162 | 0.533914162 | 13.32225477 | 1.767345834 | 0.122744975 | 0.991835066 | -4.234629759 |
| **A_51_P171200** | Golm1 | 0.216613885 | -0.076377778 | 0.509605548 | 12.08142175 | 1.767014691 | 0.122803338 | 0.991835066 | -4.234928498 |
| **A_51_P167292** | Chil3 | -0.121225795 | -0.285209981 | 0.042758392 | 7.963161849 | -1.766860262 | 0.122830565 | 0.991835066 | -4.235067812 |
| **A_55_P2047155** | Meis2 | 0.115254056 | -0.040657169 | 0.271165282 | 11.02739374 | 1.766802343 | 0.122840778 | 0.991835066 | -4.235120062 |
| **A_55_P2032695** | Tut4 | 0.145867021 | -0.051501802 | 0.343235844 | 11.66467743 | 1.766394464 | 0.122912723 | 0.991835066 | -4.23548801 |
| **A_55_P2030334** | D0Kist3 | 0.150641196 | -0.053210422 | 0.354492814 | 8.442954343 | 1.766195297 | 0.122947869 | 0.991835066 | -4.235667675 |
| **A_55_P1978321** | Fbxo8 | -0.225893919 | -0.531636279 | 0.079848442 | 10.58921714 | -1.765867273 | 0.123005774 | 0.991835066 | -4.235963572 |
| **A_52_P587738** | P2ry2 | -0.136312778 | -0.32083684 | 0.048211284 | 8.930428615 | -1.765601524 | 0.123052705 | 0.991835066 | -4.236203289 |
| **A_55_P1957613** | Ak2 | 0.157692693 | -0.055778312 | 0.371163699 | 13.35585552 | 1.765557261 | 0.123060523 | 0.991835066 | -4.236243215 |
| **A_52_P450934** | Paqr9 | -0.121386592 | -0.285729002 | 0.042955817 | 8.253611627 | -1.765347484 | 0.123097584 | 0.991835066 | -4.236432438 |
| **A_55_P2174953** | Akr1b3 | -0.206812985 | -0.486926941 | 0.07330097 | 10.1145858 | -1.764624031 | 0.123225478 | 0.991835066 | -4.237084982 |
| **A_55_P1965194** | Fbxl5 | -0.080101566 | -0.188625785 | 0.028422653 | 8.750332128 | -1.764101092 | 0.123318004 | 0.991835066 | -4.237556642 |
| **A_55_P2001233** | Ifi208 | 0.103735413 | -0.036852654 | 0.244323479 | 10.32667266 | 1.763549741 | 0.12341563 | 0.991835066 | -4.238053907 |
| **A_55_P2131707** | Lpcat1 | 0.106278705 | -0.037769226 | 0.250326637 | 7.985400304 | 1.763389946 | 0.123443938 | 0.991835066 | -4.238198023 |
| **A_55_P2063251** | Dusp3 | 0.09019878 | -0.03205759 | 0.212455149 | 11.33528924 | 1.763349098 | 0.123451176 | 0.991835066 | -4.238234862 |
| **A_52_P596592** | Rassf9 | -0.124340548 | -0.292875348 | 0.044194253 | 8.282819775 | -1.763324807 | 0.12345548 | 0.991835066 | -4.23825677 |
| **A_55_P2008691** | NA | -0.165108309 | -0.388941603 | 0.058724984 | 14.94153088 | -1.763004325 | 0.123512278 | 0.991835066 | -4.238545798 |
| **A_55_P2168124** | NA | 0.150323564 | -0.053473141 | 0.354120269 | 13.5865465 | 1.762946116 | 0.123522597 | 0.991835066 | -4.238598293 |
| **A_55_P2140751** | NA | -0.279168253 | -0.657671703 | 0.099335198 | 12.25008589 | -1.762809239 | 0.123546865 | 0.991835066 | -4.238721734 |
| **A_66_P132832** | Myzap | -0.182571093 | -0.430158241 | 0.065016054 | 10.28346477 | -1.762434768 | 0.123613282 | 0.991835066 | -4.239059437 |
| **A_55_P2106901** | Nkain4 | 0.276607543 | -0.098566327 | 0.651781413 | 10.34342182 | 1.762140644 | 0.123665472 | 0.991835066 | -4.239324676 |
| **A_55_P2181271** | Nhsl2 | 0.12418109 | -0.044271998 | 0.292634179 | 10.20336564 | 1.761917718 | 0.123705043 | 0.991835066 | -4.239525705 |
| **A_55_P2015092** | E430024I0 | 0.117149088 | -0.041771163 | 0.276069339 | 9.080316258 | 1.761849476 | 0.123717159 | 0.991835066 | -4.239587243 |

| | | | | | | | | | |
|---|---|---|---|---|---|---|---|---|---|
| **A_55_P2039444** | Cmss1 | -0.268891669 | -0.633741007 | 0.095957669 | 11.65180715 | -1.761460531 | 0.123786235 | 0.991835066 | -4.239937972 |
| **A_51_P199624** | Sfswap | 0.112626129 | -0.040248077 | 0.265500335 | 8.935990172 | 1.760816398 | 0.123900713 | 0.991835066 | -4.240518791 |
| **A_55_P2378827** | NA | 0.231874372 | -0.082882859 | 0.546631602 | 8.879280504 | 1.760703256 | 0.123920832 | 0.991835066 | -4.24062081 |
| **A_65_P15689** | Mmut | -0.123359901 | -0.290824307 | 0.044104505 | 11.40529515 | -1.760599725 | 0.123939244 | 0.991835066 | -4.24071416 |
| **A_55_P2140151** | Flcn | -0.174052351 | -0.410364846 | 0.062260143 | 11.26282409 | -1.76036345 | 0.123981275 | 0.991835066 | -4.2409272 |
| **A_52_P215539** | C030006K1 | 0.136079324 | -0.048688844 | 0.320847492 | 9.770136534 | 1.760249062 | 0.124001628 | 0.991835066 | -4.241030337 |
| **A_55_P2052634** | Rpl6 | 0.098405632 | -0.035233908 | 0.232045172 | 15.75370705 | 1.759924812 | 0.124059339 | 0.991835066 | -4.241322691 |
| **A_51_P491095** | Gzf1 | -0.106551321 | -0.251305812 | 0.038203171 | 8.961257606 | -1.759283875 | 0.124173491 | 0.991835066 | -4.241900557 |
| **A_55_P2102385** | Etl4 | -0.087687358 | -0.2068443 | 0.031469585 | 9.548317203 | -1.758841749 | 0.124252294 | 0.991835066 | -4.242299158 |
| **A_55_P2010778** | Il11ra1 | 0.146246891 | -0.052520475 | 0.345014258 | 13.70492876 | 1.758533686 | 0.12430723 | 0.991835066 | -4.242576885 |
| **A_51_P249286** | Rgs16 | 0.140878945 | -0.050633064 | 0.332390955 | 8.725660276 | 1.758163299 | 0.124373311 | 0.991835066 | -4.242910789 |
| **A_52_P210338** | Ndufb6 | -0.138471634 | -0.326730005 | 0.049786737 | 14.30446198 | -1.757986976 | 0.124404781 | 0.991835066 | -4.243069741 |
| **A_51_P229599** | Etnk1 | 0.089126656 | -0.032082116 | 0.210335427 | 8.555279968 | 1.757448851 | 0.124500872 | 0.991835066 | -4.243554836 |
| **A_51_P275101** | Chst11 | 0.125283799 | -0.045099578 | 0.295667175 | 8.826669954 | 1.757425097 | 0.124505115 | 0.991835066 | -4.243576249 |
| **A_55_P1966102** | Nme5 | -0.130790013 | -0.308677789 | 0.047097763 | 8.781517859 | -1.757266415 | 0.124533465 | 0.991835066 | -4.243719289 |
| **A_55_P2001948** | Cab39l | 0.14519492 | -0.052305367 | 0.342695207 | 10.2575839 | 1.757085209 | 0.124565847 | 0.991835066 | -4.243882631 |
| **A_55_P2281624** | 2610034E0 | 0.128621826 | -0.046338976 | 0.303582628 | 8.26647255 | 1.757045635 | 0.12457292 | 0.991835066 | -4.243918303 |
| **A_51_P465809** | Slc30a9 | 0.169226019 | -0.060993392 | 0.39944543 | 9.14739404 | 1.756848579 | 0.124608145 | 0.991835066 | -4.244095929 |
| **A_55_P2028600** | Myh9 | 0.107104775 | -0.038611518 | 0.252821068 | 13.33964249 | 1.756749544 | 0.124625852 | 0.991835066 | -4.244185197 |
| **A_51_P222092** | Igbp1 | -0.110743766 | -0.261419768 | 0.039932236 | 10.00683678 | -1.756646329 | 0.124644308 | 0.991835066 | -4.244278233 |
| **A_55_P2125049** | Gm11127 | -0.140545585 | -0.331785528 | 0.050694359 | 9.217806686 | -1.756498293 | 0.124670785 | 0.991835066 | -4.244411668 |
| **A_55_P2068330** | Cryzl1 | -0.095676648 | -0.225915119 | 0.034561823 | 9.21065221 | -1.755803038 | 0.124795203 | 0.991835066 | -4.245038328 |
| **A_51_P109840** | Vtn | -0.287439396 | -0.678786755 | 0.103907962 | 8.307259525 | -1.755468406 | 0.124855129 | 0.991835066 | -4.242488797 |
| **A_55_P2014189** | NA | -0.14363756 | -0.339218558 | 0.051943438 | 9.491436804 | -1.755296526 | 0.12488592 | 0.991835066 | -4.245494844 |
| **A_51_P342773** | Vipr2 | -0.073929394 | -0.174593819 | 0.02673503 | 7.923178191 | -1.755295618 | 0.124886083 | 0.991835066 | -4.245495661 |
| **A_55_P2166421** | Rpusd3 | 0.107076779 | -0.038765654 | 0.252919212 | 9.391557468 | 1.754771331 | 0.124980051 | 0.991835066 | -4.245968177 |
| **A_51_P211325** | Nudt2 | -0.110289059 | -0.260528464 | 0.039950346 | 9.636053789 | -1.754517524 | 0.125025565 | 0.991835066 | -4.246196914 |
| **A_51_P237668** | Bex2 | 0.266362127 | -0.096504326 | 0.62922858 | 9.098715625 | 1.754424917 | 0.125042175 | 0.991835066 | -4.246280372 |
| **A_51_P291388** | Pafah1b1 | -0.129970438 | -0.307119907 | 0.047179032 | 12.65602324 | -1.753532653 | 0.125202329 | 0.991835066 | -4.247084457 |
| **A_51_P116651** | Dpt | -0.211269985 | -0.499421678 | 0.076881708 | 9.091608246 | -1.752369794 | 0.125411347 | 0.991835066 | -4.248132302 |
| **A_55_P2109033** | Hmgcs2 | 0.099866506 | -0.036370682 | 0.236103693 | 7.935761071 | 1.751996836 | 0.125478455 | 0.991835066 | -4.248468351 |
| **A_51_P125135** | Cdca5 | 0.11883036 | -0.043279395 | 0.280940116 | 10.98913287 | 1.751972943 | 0.125482755 | 0.991835066 | -4.248489879 |
| **A_51_P122246** | Creld2 | 0.151244877 | -0.055086078 | 0.357575832 | 12.29095973 | 1.751964967 | 0.125484191 | 0.991835066 | -4.243403768 |
| **A_52_P305685** | NA | 0.106119042 | -0.038701165 | 0.250939248 | 8.929364916 | 1.751351384 | 0.125594676 | 0.991835066 | -4.249049901 |
| **A_55_P2186287** | Sub1 | 0.104925039 | -0.038295938 | 0.248146016 | 14.68278616 | 1.750981834 | 0.125661265 | 0.991835066 | -4.24938285 |
| **A_55_P2247852** | Ap2s1 | -0.191760699 | -0.453527179 | 0.070005782 | 11.93050735 | -1.750873225 | 0.125680841 | 0.991835066 | -4.2494807 |
| **A_55_P2139713** | Phf13 | 0.09409868 | -0.034390653 | 0.222588013 | 8.030021636 | 1.750352786 | 0.12577469 | 0.991835066 | -4.249949572 |
| **A_55_P2124619** | Smurf2 | 0.148123474 | -0.054146101 | 0.350393049 | 9.749741775 | 1.75025956 | 0.125791508 | 0.991835066 | -4.250033557 |
| **A_55_P2001963** | NA | -0.1349207 | -0.319171012 | 0.049329612 | 17.16662922 | -1.750166975 | 0.125808213 | 0.991835066 | -4.250116966 |
| **A_51_P352381** | Dhrs4 | -0.291456714 | -0.68959722 | 0.106683793 | 9.039183536 | -1.749632477 | 0.125904691 | 0.991835066 | -4.250598474 |
| **A_51_P356642** | Krt19 | 0.440056286 | -0.161215172 | 1.041327745 | 10.19128368 | 1.749229418 | 0.125977492 | 0.991835066 | -4.250961558 |
| **A_55_P2000833** | E2f8 | 0.190367699 | -0.06975735 | 0.450492749 | 10.61001922 | 1.749122466 | 0.125996816 | 0.991835066 | -4.251057902 |
| **A_51_P126177** | Map1lc3b | -0.144955534 | -0.343070164 | 0.053159096 | 12.47466096 | -1.748748623 | 0.126064386 | 0.991835066 | -4.251394653 |
| **A_55_P2092310** | Phgdh | 0.411382273 | -0.15089342 | 0.973657965 | 10.49273685 | 1.748660083 | 0.126080394 | 0.991835066 | -4.251474407 |
| **A_55_P2053923** | Rab17 | -0.197576467 | -0.467687801 | 0.072534868 | 8.965941906 | -1.748241955 | 0.126156018 | 0.991835066 | -4.251851034 |
| **A_55_P2059164** | H3f3b | 0.100521442 | -0.036925104 | 0.237967989 | 12.89202142 | 1.747970145 | 0.126205202 | 0.991835066 | -4.252095859 |
| **A_65_P06872** | Fam210a | -0.115034731 | -0.272334517 | 0.042265055 | 9.321502172 | -1.74787344 | 0.126222705 | 0.991835066 | -4.252182962 |
| **A_65_P11773** | Adi1 | -0.115387479 | -0.273257979 | 0.042483022 | 9.913592499 | -1.746895114 | 0.126399909 | 0.991835066 | -4.253064103 |

| | | | | | | | | | |
|---|---|---|---|---|---|---|---|---|---|
| **A_52_P199928** | NA | -0.174623392 | -0.413739085 | 0.064492301 | 12.97434713 | -1.74543417 | 0.126664974 | 0.991835066 | -4.25437978 |
| **A_55_P2046338** | Mapk14 | 0.094027745 | -0.034806698 | 0.222862188 | 11.04196652 | 1.744348152 | 0.126862361 | 0.991835066 | -4.255357702 |
| **A_51_P210395** | Glrx | -0.189467598 | -0.449193422 | 0.070258226 | 10.39663693 | -1.743528088 | 0.127011606 | 0.991835066 | -4.256096078 |
| **A_55_P1980190** | NA | -0.096935986 | -0.229821079 | 0.035949107 | 11.74648358 | -1.743483728 | 0.127019684 | 0.991835066 | -4.256136017 |
| **A_55_P2151855** | Acp1 | 0.115802053 | -0.042963818 | 0.274567924 | 11.42815422 | 1.743284384 | 0.12705599 | 0.991835066 | -4.256315496 |
| **A_51_P292754** | Yaf2 | 0.13010592 | -0.048275392 | 0.308487232 | 11.30736383 | 1.743238615 | 0.127064328 | 0.991835066 | -4.256356703 |
| **A_55_P1974243** | Dbh | -0.218942466 | -0.519153659 | 0.081268727 | 9.656523055 | -1.743059683 | 0.127096927 | 0.991835066 | -4.2565178 |
| **A_55_P2149293** | Agfg2 | 0.134134801 | -0.049798907 | 0.318068509 | 8.041482175 | 1.742967446 | 0.127113735 | 0.991835066 | -4.256600843 |
| **A_51_P231738** | Dnajc4 | -0.145749141 | -0.345678941 | 0.054180659 | 11.89835924 | -1.742358848 | 0.127224689 | 0.991835066 | -4.257148755 |
| **A_51_P123297** | Ppp1r7 | 0.102109186 | -0.037970074 | 0.242188446 | 8.452830905 | 1.742208419 | 0.127252128 | 0.991835066 | -4.257284179 |
| **A_55_P1963414** | NA | -0.079133304 | -0.187696975 | 0.029430368 | 15.94150968 | -1.742143433 | 0.127263984 | 0.991835066 | -4.257342683 |
| **A_51_P134062** | Stxbp2 | 0.085234364 | -0.03170156 | 0.202170288 | 9.231292267 | 1.742111324 | 0.127269842 | 0.991835066 | -4.257371589 |
| **A_55_P2055063** | Mcts1 | 0.127671474 | -0.047489643 | 0.302832591 | 11.19673856 | 1.742068784 | 0.127277604 | 0.991835066 | -4.257409885 |
| **A_51_P271370** | Wsb1 | 0.110427032 | -0.041114655 | 0.26196872 | 8.565359974 | 1.741616035 | 0.127360238 | 0.991835066 | -4.257817458 |
| **A_55_P2027225** | Fau | 0.096537322 | -0.035945903 | 0.229020548 | 17.29036608 | 1.741580239 | 0.127366774 | 0.991835066 | -4.257849682 |
| **A_55_P1978441** | Kif3a | -0.091477346 | -0.217021752 | 0.034067059 | 10.47995051 | -1.74150741 | 0.127380071 | 0.991835066 | -4.257915242 |
| **A_52_P482925** | Serbp1 | 0.099211077 | -0.036949475 | 0.23537163 | 13.93239888 | 1.741478 | 0.127385442 | 0.991835066 | -4.257941717 |
| **A_55_P2124701** | NA | 0.128625181 | -0.047912312 | 0.305162675 | 12.6755135 | 1.74139854 | 0.127399953 | 0.991835066 | -4.258013246 |
| **A_55_P1960972** | Lrrc59 | 0.146964337 | -0.054778269 | 0.348706942 | 8.448897633 | 1.741098998 | 0.127454669 | 0.991835066 | -4.258282885 |
| **A_55_P2060349** | Lzts2 | -0.116267009 | -0.275881186 | 0.043347167 | 12.10741878 | -1.740981563 | 0.127476127 | 0.991835066 | -4.258388595 |
| **A_51_P343309** | Cmas | 0.139742011 | -0.052140345 | 0.331624366 | 10.7212802 | 1.740608417 | 0.12754433 | 0.991835066 | -4.258724477 |
| **A_55_P1955457** | Grb10 | 0.131978376 | -0.049277102 | 0.313233855 | 12.43068354 | 1.740286582 | 0.127603184 | 0.991835066 | -4.259014162 |
| **A_55_P1993049** | Gm4076 | -0.233768211 | -0.554850055 | 0.087313632 | 14.03971429 | -1.740118556 | 0.12763392 | 0.991835066 | -4.2591654 |
| **A_55_P2033420** | Sppl2a | -0.099716533 | -0.236690642 | 0.037257576 | 10.36284833 | -1.73995421 | 0.127663991 | 0.991835066 | -4.259313324 |
| **A_55_P1999841** | Cox7c | -0.11066836 | -0.262717572 | 0.041380851 | 15.15777282 | -1.739596145 | 0.12772953 | 0.991835066 | -4.259635601 |
| **A_51_P119429** | Nckap1l | -0.088574724 | -0.210277823 | 0.033128375 | 8.269313247 | -1.73947145 | 0.127752361 | 0.991835066 | -4.257649555 |
| **A_55_P1968035** | 1810009A1 | 0.084711511 | -0.031761066 | 0.201184088 | 12.06863407 | 1.738312598 | 0.127964732 | 0.991835066 | -4.260790773 |
| **A_55_P2023783** | Leng8 | 0.205089261 | -0.076950462 | 0.487128984 | 9.101951786 | 1.737967988 | 0.12802795 | 0.991835066 | -4.261100893 |
| **A_55_P2155438** | Rps27 | 0.083132003 | -0.031191924 | 0.19745593 | 16.53847975 | 1.737961862 | 0.128029074 | 0.991835066 | -4.261106406 |
| **A_55_P1956672** | Rab34 | 0.156017958 | -0.058542331 | 0.370578247 | 11.88026984 | 1.737938388 | 0.128033382 | 0.991835066 | -4.26112753 |
| **A_51_P115715** | Asb2 | -0.209714208 | -0.498119361 | 0.078690945 | 12.425744 | -1.737936765 | 0.12803368 | 0.991835066 | -4.261128991 |
| **A_51_P288138** | Fpr2 | -0.086941454 | -0.206511121 | 0.032628212 | 7.790388564 | -1.737860906 | 0.1280476 | 0.991835066 | -4.261197256 |
| **A_66_P122758** | Tsr1 | 0.122451079 | -0.045958207 | 0.290860364 | 9.364998427 | 1.737823691 | 0.12805443 | 0.991835066 | -4.261230746 |
| **A_52_P316405** | Lamtor4 | -0.153025818 | -0.363494232 | 0.057442596 | 12.81620235 | -1.73774847 | 0.128068236 | 0.991835066 | -4.261298435 |
| **A_55_P2045903** | NA | 0.110631087 | -0.041587472 | 0.262849647 | 8.587815053 | 1.737075544 | 0.128191807 | 0.991835066 | -4.261903973 |
| **A_51_P148105** | Rad51 | 0.11363836 | -0.042733742 | 0.270010462 | 9.980227164 | 1.736900005 | 0.128224061 | 0.991835066 | -4.262061927 |
| **A_55_P2405784** | BC023202 | 0.101337313 | -0.038111489 | 0.240786115 | 8.905347571 | 1.736855661 | 0.12823221 | 0.991835066 | -4.262101828 |
| **A_52_P487686** | Cstdc5 | -0.584807803 | -1.389623477 | 0.22000787 | 10.47781276 | -1.736704245 | 0.128260039 | 0.991835066 | -4.262238073 |
| **A_51_P320481** | Mcee | -0.114506606 | -0.272113645 | 0.043100432 | 11.9884236 | -1.736457142 | 0.128305467 | 0.991835066 | -4.262460413 |
| **A_51_P321150** | Lyz2 | -0.106075081 | -0.252083372 | 0.03993321 | 8.02018678 | -1.736380849 | 0.128319496 | 0.991835066 | -4.26252906 |
| **A_65_P10357** | Srsf6 | 0.175406574 | -0.066128052 | 0.416941199 | 11.14198783 | 1.735703742 | 0.128444069 | 0.991835066 | -4.263138282 |
| **A_55_P2120080** | NA | 0.100709431 | -0.03796947 | 0.239388333 | 8.626266532 | 1.735676896 | 0.128449011 | 0.991835066 | -4.263162436 |
| **A_55_P1999760** | Lrrc51 | 0.123034692 | -0.046432572 | 0.292501956 | 8.722402859 | 1.735205446 | 0.128535819 | 0.991835066 | -4.263586596 |
| **A_52_P602996** | Cdip1 | -0.136578582 | -0.324713831 | 0.051556667 | 8.860073801 | -1.735088218 | 0.128557413 | 0.991835066 | -4.263692062 |
| **A_66_P132888** | Dhfr | 0.131087636 | -0.049500673 | 0.311675945 | 9.308077126 | 1.734927104 | 0.128587097 | 0.991835066 | -4.263837009 |
| **A_51_P118603** | Wdr82 | -0.083674093 | -0.198964665 | 0.031616478 | 11.2771431 | -1.734628015 | 0.128642219 | 0.991835066 | -4.256771294 |
| **A_66_P112853** | Eif1a | 0.131335762 | -0.049819344 | 0.312490868 | 10.81565906 | 1.732772516 | 0.128984691 | 0.991835066 | -4.265775189 |
| **A_55_P1960999** | Pigr | 0.458824783 | -0.174160068 | 1.091809634 | 12.76675106 | 1.732457876 | 0.129042851 | 0.991835066 | -4.266058193 |

| | | | | | | | | | |
|---|---|---|---|---|---|---|---|---|---|
| **A_55_P2055682** | Atp5o | -0.160339553 | -0.381581831 | 0.060902724 | 15.06672124 | -1.732134775 | 0.129102602 | 0.991835066 | -4.266348799 |
| **A_55_P2012221** | Armcx2 | 0.117788024 | -0.044741516 | 0.280317564 | 12.96177967 | 1.732119941 | 0.129105346 | 0.991835066 | -4.266362141 |
| **A_52_P220229** | Plekhb2 | -0.141223526 | -0.336101277 | 0.053654226 | 9.779268433 | -1.732024064 | 0.129123081 | 0.991835066 | -4.266448374 |
| **A_55_P2158107** | Ddx31 | 0.12960778 | -0.049244988 | 0.308460548 | 9.36315041 | 1.731986638 | 0.129130005 | 0.991835066 | -4.266482035 |
| **A_51_P314186** | Syne1 | -0.136907992 | -0.325910022 | 0.052094039 | 9.317519768 | -1.731296545 | 0.129257737 | 0.991835066 | -4.267102685 |
| **A_55_P2009201** | NA | -0.105355931 | -0.250812178 | 0.040100316 | 11.80108124 | -1.731154163 | 0.129284107 | 0.991835066 | -4.267230735 |
| **A_55_P2169621** | Prorsd1 | -0.116980627 | -0.278494564 | 0.044533311 | 14.57152101 | -1.731063785 | 0.129300847 | 0.991835066 | -4.267312014 |
| **A_55_P2083093** | Haus7 | -0.155593442 | -0.370492601 | 0.059305717 | 9.009364103 | -1.730476498 | 0.129409681 | 0.991835066 | -4.26784016 |
| **A_55_P1957733** | Cysrt1 | 0.117997167 | -0.044988754 | 0.280983089 | 7.775145201 | 1.730336715 | 0.129435598 | 0.991835066 | -4.267965862 |
| **A_55_P2014997** | Amz2 | -0.103269242 | -0.245914014 | 0.039375529 | 10.53942341 | -1.730311384 | 0.129440295 | 0.991835066 | -4.267988641 |
| **A_55_P1977633** | Irf2bpl | -0.122077127 | -0.290749635 | 0.046595381 | 12.34776418 | -1.729812893 | 0.129532763 | 0.991835066 | -4.268436902 |
| **A_52_P480301** | Sppl3 | -0.114325103 | -0.272300481 | 0.043650275 | 8.843620793 | -1.729662344 | 0.129560701 | 0.991835066 | -4.268572277 |
| **A_51_P502614** | Dusp6 | 0.19186048 | -0.073256375 | 0.456977335 | 11.73953266 | 1.729646368 | 0.129563667 | 0.991835066 | -4.268586643 |
| **A_55_P1960331** | Srbd1 | 0.098251016 | -0.037520522 | 0.234022554 | 10.02496004 | 1.729567215 | 0.129578358 | 0.991835066 | -4.268657818 |
| **A_51_P267354** | Lrfn3 | 0.093230126 | -0.035627609 | 0.222087862 | 8.840297608 | 1.729238552 | 0.129639379 | 0.991835066 | -4.268953345 |
| **A_55_P1952618** | Ear2 | -0.072203355 | -0.172006993 | 0.027600283 | 7.837768622 | -1.7291001 | 0.129665093 | 0.991835066 | -4.269077835 |
| **A_55_P2169888** | Prag1 | 0.079955413 | -0.030595856 | 0.190506682 | 9.134296003 | 1.728595081 | 0.129758928 | 0.991835066 | -4.269531914 |
| **A_52_P161707** | Cand1 | 0.113519656 | -0.043452941 | 0.270492253 | 9.50883709 | 1.728448153 | 0.12978624 | 0.991835066 | -4.269664018 |
| **A_55_P2032598** | NA | 0.086567132 | -0.033145575 | 0.206279839 | 15.43905236 | 1.728311072 | 0.129811727 | 0.991835066 | -4.269787266 |
| **A_66_P113749** | Il2rg | -0.132903862 | -0.316782907 | 0.050975184 | 8.583810968 | -1.727485822 | 0.129965263 | 0.991835066 | -4.270529208 |
| **A_55_P2176145** | Rgs17 | -0.075913027 | -0.180973874 | 0.02914782 | 8.151767413 | -1.726969002 | 0.130061504 | 0.991835066 | -4.270993824 |
| **A_55_P2083841** | Atp13a1 | 0.147054397 | -0.056484931 | 0.350593725 | 11.8659168 | 1.726787165 | 0.130095382 | 0.991835066 | -4.271157289 |
| **A_55_P2058433** | Cyp2c68 | -0.090098978 | -0.214841018 | 0.034643062 | 7.746533447 | -1.726299534 | 0.130186273 | 0.991835066 | -4.271595636 |
| **A_55_P1995745** | Gm14308 | 0.120711528 | -0.046479209 | 0.287902264 | 8.783491091 | 1.725621994 | 0.130312664 | 0.991835066 | -4.272204664 |
| **A_55_P1999240** | NA | -0.084463394 | -0.201459714 | 0.032532925 | 9.503824093 | -1.725462248 | 0.13034248 | 0.991835066 | -4.272348251 |
| **A_52_P139097** | NA | -0.121510877 | -0.289831469 | 0.046809714 | 11.24971355 | -1.725389063 | 0.130356142 | 0.991835066 | -4.272414032 |
| **A_52_P31543** | Btg2 | 0.149738813 | -0.057777242 | 0.357254869 | 9.566596497 | 1.724613641 | 0.130500982 | 0.991835066 | -4.273110979 |
| **A_51_P297968** | Pdia6 | 0.109239082 | -0.042152198 | 0.260630361 | 9.99632788 | 1.724591781 | 0.130505067 | 0.991835066 | -4.273130626 |
| **A_51_P495269** | Lor | 0.546653932 | -0.210982262 | 1.304290126 | 8.407106356 | 1.724490851 | 0.130523932 | 0.991835066 | -4.273221337 |
| **A_51_P345467** | Pi4k2a | -0.08203763 | -0.195751512 | 0.031676251 | 8.956839533 | -1.724283834 | 0.130562633 | 0.991835066 | -4.273407393 |
| **A_52_P151320** | Tnfaip8l1 | -0.131593829 | -0.314033848 | 0.050846189 | 10.27488439 | -1.723949545 | 0.13062515 | 0.991835066 | -4.273707825 |
| **A_51_P348617** | Borcs8 | -0.157986997 | -0.377027884 | 0.06105389 | 12.12388878 | -1.723873079 | 0.130639454 | 0.991835066 | -4.273776545 |
| **A_51_P142515** | Antxr2 | -0.096049085 | -0.229220794 | 0.037122623 | 12.43038014 | -1.72381397 | 0.130650512 | 0.991835066 | -4.273829666 |
| **A_52_P566348** | Nras | -0.083222958 | -0.198625037 | 0.03217912 | 10.31497592 | -1.723608597 | 0.130688941 | 0.991835066 | -4.274014231 |
| **A_55_P1997791** | Fbl | 0.13804421 | -0.053425832 | 0.329514252 | 12.18606511 | 1.723163538 | 0.130772257 | 0.991835066 | -4.274414185 |
| **A_52_P558939** | Rab3gap1 | -0.161352616 | -0.385181711 | 0.062476479 | 9.082089663 | -1.72293385 | 0.130815275 | 0.991835066 | -4.274620588 |
| **A_55_P1966189** | Il4ra | -0.088456943 | -0.211189673 | 0.034275788 | 7.919813829 | -1.722585003 | 0.130880636 | 0.991835066 | -4.274934062 |
| **A_55_P1967539** | Hunk | 0.162036388 | -0.062945061 | 0.387017838 | 10.98136591 | 1.721372963 | 0.131107971 | 0.991835066 | -4.276023115 |
| **A_55_P2121760** | NA | 0.185759859 | -0.07217931 | 0.443699028 | 8.272886625 | 1.721249031 | 0.131131237 | 0.991835066 | -4.276134465 |
| **A_55_P2017066** | NA | -0.179063718 | -0.427715707 | 0.06958827 | 11.84907968 | -1.721174068 | 0.131145312 | 0.991835066 | -4.276201816 |
| **A_52_P228236** | Tfrc | -0.19221685 | -0.459246642 | 0.074812942 | 10.45427848 | -1.720445302 | 0.131282221 | 0.991835066 | -4.274624067 |
| **A_55_P2081164** | Cd300lf | -0.102394252 | -0.244662662 | 0.039874158 | 7.916187952 | -1.720189258 | 0.131330355 | 0.991835066 | -4.277086582 |
| **A_51_P495379** | Fhod1 | 0.110254413 | -0.042949387 | 0.263458213 | 9.418850477 | 1.720028236 | 0.131360634 | 0.991835066 | -4.277231239 |
| **A_51_P160413** | Yod1 | -0.097307548 | -0.232534541 | 0.037919445 | 8.295228537 | -1.719856728 | 0.131392893 | 0.991835066 | -4.277385312 |
| **A_55_P2153904** | Gm6132 | -0.116352259 | -0.278075565 | 0.045371047 | 12.60347078 | -1.719536269 | 0.131453187 | 0.991835066 | -4.277673188 |
| **A_55_P2006668** | Pi4kb | 0.088950356 | -0.034702802 | 0.212603514 | 8.73720293 | 1.719299797 | 0.131497697 | 0.991835066 | -4.277885611 |
| **A_51_P334308** | Tspan14 | 0.105162122 | -0.041039492 | 0.251363736 | 11.07285827 | 1.719160026 | 0.131524012 | 0.991835066 | -4.278011165 |
| **A_55_P1968718** | Plekha6 | 0.137744519 | -0.053790787 | 0.329279824 | 8.718612911 | 1.718836702 | 0.131584903 | 0.991835066 | -4.278301594 |

| | | | | | | | | | |
|---|---|---|---|---|---|---|---|---|---|
| **A_55_P2078204** | Prr5 | 0.199632993 | -0.078061912 | 0.477327897 | 9.403647389 | 1.718199305 | 0.131705024 | 0.991835066 | -4.278874115 |
| **A_65_P10433** | Rpp14 | -0.105130267 | -0.251374593 | 0.04111406 | 9.635925511 | -1.718137322 | 0.13171671 | 0.991835066 | -4.278929788 |
| **A_55_P2156944** | Rbis | 0.197698519 | -0.077325092 | 0.472722131 | 12.99753337 | 1.718076782 | 0.131728126 | 0.991835066 | -4.278984164 |
| **A_55_P2124085** | NA | 0.099840204 | -0.039065727 | 0.238746135 | 10.30444362 | 1.717883878 | 0.131764506 | 0.991835066 | -4.279157425 |
| **A_51_P157042** | Ccn2 | 0.487694441 | -0.191058503 | 1.166447384 | 12.87895535 | 1.717296209 | 0.131875396 | 0.991835066 | -4.279685229 |
| **A_55_P2093543** | Snapc4 | 0.193194608 | -0.075710212 | 0.462099427 | 10.13627031 | 1.717139375 | 0.131905005 | 0.991835066 | -4.279826082 |
| **A_55_P2006280** | Ogdh | -0.092953859 | -0.222339639 | 0.036431921 | 14.63727028 | -1.717077935 | 0.131916606 | 0.991835066 | -4.279881261 |
| **A_55_P2077522** | NA | 0.137352355 | -0.053847251 | 0.32855196 | 9.465039661 | 1.716952374 | 0.131940317 | 0.991835066 | -4.279994025 |
| **A_52_P98625** | Sowahc | -0.078075848 | -0.186763405 | 0.030611708 | 8.032854066 | -1.716904033 | 0.131949447 | 0.991835066 | -4.280037439 |
| **A_51_P416191** | Hectd1 | -0.100947719 | -0.24152351 | 0.039628072 | 10.96207426 | -1.71630753 | 0.132062155 | 0.991835066 | -4.280573124 |
| **A_55_P2084758** | Agfg1 | -0.091043959 | -0.217838911 | 0.035750993 | 10.09651756 | -1.716162081 | 0.132089651 | 0.991835066 | -4.280703739 |
| **A_55_P1991377** | NA | -0.099917827 | -0.239072186 | 0.039236532 | 15.4026244 | -1.716150216 | 0.132091895 | 0.991835066 | -4.280714394 |
| **A_66_P123735** | NA | 0.218748579 | -0.085958724 | 0.523455883 | 9.623826729 | 1.715819144 | 0.132154503 | 0.991835066 | -4.281011692 |
| **A_52_P350537** | Mtmr11 | -0.072362379 | -0.173176181 | 0.028451422 | 8.11944832 | -1.715544462 | 0.13220647 | 0.991835066 | -4.281258345 |
| **A_52_P273821** | Abhd5 | -0.097861588 | -0.234200458 | 0.038477281 | 8.283109293 | -1.715543378 | 0.132206675 | 0.991835066 | -4.281259319 |
| **A_55_P2027240** | Engase | 0.097108164 | -0.038182362 | 0.232398689 | 9.261903471 | 1.71552673 | 0.132209825 | 0.991835066 | -4.281274268 |
| **A_55_P2109613** | Ahr | 0.088252312 | -0.03471915 | 0.211223774 | 8.976518961 | 1.715263659 | 0.132259614 | 0.991835066 | -4.281510488 |
| **A_55_P1953809** | Plaat3 | -0.101463195 | -0.242891765 | 0.039965374 | 10.58516931 | -1.714669881 | 0.13237206 | 0.991835066 | -4.282043637 |
| **A_55_P1985015** | Pard3 | 0.12374778 | -0.048760531 | 0.29625609 | 10.00920161 | 1.714496196 | 0.132404968 | 0.991835066 | -4.282199582 |
| **A_55_P2086664** | Tspan33 | 0.2157024 | -0.085001471 | 0.516406272 | 9.535734806 | 1.714451053 | 0.132413523 | 0.991835066 | -4.282240114 |
| **A_51_P307082** | Hey2 | -0.161835032 | -0.387460754 | 0.063790691 | 9.591029943 | -1.71432461 | 0.132437487 | 0.991835066 | -4.282353639 |
| **A_51_P347452** | Htatsf1 | -0.190341248 | -0.455736083 | 0.075053588 | 12.15147875 | -1.714152977 | 0.132470022 | 0.991835066 | -4.282507736 |
| **A_55_P2114498** | Gm8801 | -0.138683446 | -0.332081499 | 0.054714606 | 9.778710429 | -1.713884937 | 0.132520847 | 0.991835066 | -4.282748384 |
| **A_55_P2011235** | Rpl13 | 0.075954867 | -0.029977767 | 0.181887502 | 16.07948614 | 1.71370068 | 0.132555796 | 0.991835066 | -4.282913806 |
| **A_52_P113700** | Sfxn1 | 0.175034452 | -0.069099981 | 0.419168885 | 10.35178416 | 1.713577041 | 0.132579253 | 0.991835066 | -4.283024806 |
| **A_55_P2075884** | Senp8 | -0.096994154 | -0.232305117 | 0.038316809 | 8.564514321 | -1.713253819 | 0.132640592 | 0.991835066 | -4.283314978 |
| **A_55_P2119155** | Set | 0.150442719 | -0.059467934 | 0.360353373 | 10.93026306 | 1.7129545 | 0.132697419 | 0.991835066 | -4.283583684 |
| **A_66_P139167** | Safb | 0.142950239 | -0.056523932 | 0.34242441 | 11.39641438 | 1.71280274 | 0.132726241 | 0.991835066 | -4.283719918 |
| **A_51_P366161** | Mpg | 0.08032281 | -0.031792112 | 0.192437731 | 10.35862791 | 1.712318735 | 0.1328182 | 0.991835066 | -4.284154396 |
| **A_55_P1992079** | Ptprf | 0.144192654 | -0.057088204 | 0.345473512 | 10.56416538 | 1.712181471 | 0.132844291 | 0.991835066 | -4.284277609 |
| **A_55_P2172747** | Mfn2 | -0.157917994 | -0.378364408 | 0.062528419 | 11.33789244 | -1.712133851 | 0.132853343 | 0.991835066 | -4.284320354 |
| **A_51_P126525** | D230025D1 | -0.088564921 | -0.212229244 | 0.035099401 | 9.293005807 | -1.71169528 | 0.132936744 | 0.991835066 | -4.281122191 |
| **A_52_P411601** | Ep300 | 0.123350414 | -0.048885348 | 0.295586176 | 8.917897265 | 1.71169512 | 0.132936775 | 0.991835066 | -4.284714165 |
| **A_51_P282179** | Mtor | -0.094663422 | -0.226843477 | 0.037516634 | 9.169070661 | -1.711691131 | 0.132937534 | 0.991835066 | -4.284717745 |
| **A_51_P461894** | Tnnc1 | -0.246485497 | -0.590686682 | 0.097715687 | 15.01676482 | -1.711544294 | 0.132965468 | 0.991835066 | -4.284849543 |
| **A_51_P170911** | Ptpn9 | -0.099074 | -0.237454848 | 0.039306848 | 8.486209159 | -1.711168764 | 0.133036935 | 0.991835066 | -4.285186603 |
| **A_55_P1958919** | Olfr1347 | 0.084398194 | -0.033516323 | 0.202312711 | 8.036520432 | 1.710704548 | 0.133125331 | 0.991835066 | -4.285603245 |
| **A_55_P1959500** | Ces2e | -0.120687597 | -0.28931143 | 0.047936237 | 7.844115829 | -1.710617112 | 0.133141986 | 0.991835066 | -4.285681718 |
| **A_51_P256827** | S100a8 | -0.571186484 | -1.369268767 | 0.226895799 | 10.13345694 | -1.710564225 | 0.133152062 | 0.991835066 | -4.285729184 |
| **A_55_P2026535** | Erc2 | -0.089816713 | -0.215333261 | 0.035699836 | 8.585312094 | -1.71027245 | 0.133207661 | 0.991835066 | -4.285991044 |
| **A_55_P1998801** | NA | 0.150994789 | -0.06008072 | 0.362070299 | 10.04644101 | 1.709752497 | 0.133306795 | 0.991835066 | -4.286457665 |
| **A_52_P445253** | Dst | 0.112931842 | -0.044968174 | 0.270831858 | 8.804821876 | 1.709398706 | 0.133374289 | 0.991835066 | -4.286775155 |
| **A_55_P2013840** | Krt42 | 0.240073132 | -0.09560929 | 0.575755554 | 8.645102217 | 1.709322823 | 0.133388769 | 0.991835066 | -4.28684325 |
| **A_51_P265869** | Hspa9 | -0.080569122 | -0.193231483 | 0.03209324 | 13.24112282 | -1.709223721 | 0.133407683 | 0.991835066 | -4.28693218 |
| **A_55_P2179106** | Ppig | 0.159845468 | -0.063678378 | 0.383369315 | 9.513195499 | 1.709171324 | 0.133417684 | 0.991835066 | -4.286979199 |
| **A_52_P591166** | Dpysl2 | 0.131252245 | -0.052305789 | 0.314810279 | 12.3490579 | 1.709001622 | 0.133450081 | 0.991835066 | -4.287131479 |
| **A_52_P641477** | Wars2 | 0.076799204 | -0.030616646 | 0.184215054 | 8.758760063 | 1.708824579 | 0.133483886 | 0.991835066 | -4.287290346 |
| **A_66_P126138** | Ncoa1 | -0.143020269 | -0.343062587 | 0.057022049 | 9.820555704 | -1.708774859 | 0.133493382 | 0.991835066 | -4.28733496 |

| | | | | | | | | | |
|---|---|---|---|---|---|---|---|---|---|
| **A_55_P2151506** | Nudt9 | -0.084460285 | -0.202623834 | 0.033703265 | 12.39021802 | -1.708355087 | 0.133573574 | 0.991835066 | -4.287711619 |
| **A_52_P589699** | Polr3a | 0.149115903 | -0.059516222 | 0.357748028 | 8.708269529 | 1.708251906 | 0.133593292 | 0.991835066 | -4.2878042 |
| **A_52_P478629** | Arcn1 | 0.137910351 | -0.055067459 | 0.33088816 | 8.732289677 | 1.708042301 | 0.133633357 | 0.991835066 | -4.28799227 |
| **A_51_P450100** | Rab5if | 0.125959312 | -0.050297301 | 0.302215925 | 13.25652227 | 1.708024085 | 0.13363684 | 0.991835066 | -4.288008614 |
| **A_55_P2108708** | Kcne3 | -0.238286405 | -0.571730536 | 0.095157725 | 8.939874557 | -1.70798999 | 0.133643358 | 0.991835066 | -4.288039205 |
| **A_55_P2027521** | NA | 0.285974936 | -0.114317926 | 0.686267799 | 10.61444973 | 1.707494281 | 0.133738163 | 0.991835066 | -4.288483963 |
| **A_52_P551526** | NA | 0.121634418 | -0.04865403 | 0.291922866 | 16.16633907 | 1.707184366 | 0.133797468 | 0.991835066 | -4.288762013 |
| **A_66_P118256** | Strn3 | -0.116448847 | -0.279481227 | 0.046583532 | 11.68007879 | -1.707145194 | 0.133804965 | 0.991835066 | -4.288797157 |
| **A_55_P2020706** | NA | -0.140540659 | -0.337305345 | 0.056224027 | 11.60089881 | -1.707119648 | 0.133809855 | 0.991835066 | -4.288820075 |
| **A_51_P262533** | Cxxc1 | -0.079602247 | -0.191107793 | 0.0319033 | 9.64308795 | -1.706231624 | 0.133979939 | 0.991835066 | -4.289616735 |
| **A_55_P2041457** | NA | 0.462260968 | -0.185396954 | 1.10991889 | 11.24877721 | 1.705888631 | 0.134045688 | 0.991835066 | -4.289924419 |
| **A_55_P2001718** | Pex1 | 0.088278585 | -0.035454614 | 0.212011783 | 8.624307943 | 1.705211504 | 0.134175578 | 0.991835066 | -4.290531806 |
| **A_51_P310398** | Tk2 | -0.237024055 | -0.569403287 | 0.095355177 | 8.486448843 | -1.704384892 | 0.134334305 | 0.991835066 | -4.291273224 |
| **A_52_P100199** | Abhd10 | -0.098611952 | -0.236961724 | 0.03973782 | 8.623003829 | -1.703571007 | 0.134490764 | 0.991835066 | -4.29200316 |
| **A_51_P335770** | Afap1 | -0.104348087 | -0.250774613 | 0.042078439 | 11.86591729 | -1.703232222 | 0.134555943 | 0.991835066 | -4.292306983 |
| **A_55_P2058942** | Aldh3b1 | -0.094234898 | -0.22650454 | 0.038034743 | 8.441403863 | -1.702788553 | 0.134641345 | 0.991835066 | -4.292704848 |
| **A_55_P2032966** | Hmgcs1 | 0.160416784 | -0.06476464 | 0.385598208 | 12.7121315 | 1.702653898 | 0.134667275 | 0.991835066 | -4.292825598 |
| **A_66_P139250** | Kctd3 | 0.115424165 | -0.046614386 | 0.277462715 | 11.19944508 | 1.702501579 | 0.134696612 | 0.991835066 | -4.292962185 |
| **A_55_P2135311** | Nol3 | -0.136640427 | -0.328500217 | 0.055219363 | 9.022602187 | -1.702175663 | 0.134759406 | 0.991835066 | -4.293254433 |
| **A_51_P420415** | Srd5a1 | 0.082644565 | -0.033405984 | 0.198695114 | 9.100858005 | 1.702065345 | 0.134780667 | 0.991835066 | -4.293353353 |
| **A_55_P1996212** | Zbtb2 | 0.106194978 | -0.042929471 | 0.255319428 | 9.651764176 | 1.702018434 | 0.134789709 | 0.991835066 | -4.293395416 |
| **A_55_P2162988** | Tmem179 | 0.075300277 | -0.030468698 | 0.181069252 | 7.803324598 | 1.701560569 | 0.13487799 | 0.991835066 | -4.293805957 |
| **A_51_P106859** | Lsm10 | -0.076202916 | -0.183250939 | 0.030845107 | 11.27005405 | -1.701382932 | 0.134912256 | 0.991835066 | -4.291532595 |
| **A_55_P2136556** | Pkn2 | 0.095428386 | -0.038634934 | 0.229491706 | 9.396865454 | 1.701283704 | 0.1349314 | 0.991835066 | -4.294054196 |
| **A_52_P86928** | Cog2 | -0.082893663 | -0.199351824 | 0.033564498 | 8.677254261 | -1.701220218 | 0.13494365 | 0.991835066 | -4.294111117 |
| **A_66_P115323** | Utp4 | 0.096284959 | -0.039034796 | 0.231604715 | 10.40288098 | 1.700616468 | 0.135060198 | 0.991835066 | -4.294652414 |
| **A_51_P367310** | Chaf1b | 0.136281349 | -0.055267951 | 0.327830649 | 11.33183871 | 1.700454389 | 0.135091503 | 0.991835066 | -4.294797721 |
| **A_55_P2171091** | Sap30bp | 0.075624007 | -0.030677556 | 0.18192557 | 9.551919297 | 1.700314135 | 0.135118597 | 0.991835066 | -4.294923459 |
| **A_55_P1968381** | Cast | -0.08332495 | -0.200451666 | 0.033801766 | 9.891156822 | -1.700310465 | 0.135119306 | 0.991835066 | -4.294926749 |
| **A_51_P158210** | Mcm2 | 0.163034061 | -0.066147529 | 0.392215651 | 11.51395246 | 1.7002303 | 0.135134795 | 0.991835066 | -4.294998617 |
| **A_52_P69194** | Rimkla | 0.081250491 | -0.032976238 | 0.195477221 | 10.42001833 | 1.700072281 | 0.135165331 | 0.991835066 | -4.295140278 |
| **A_55_P1971179** | Cd1d1 | -0.155542343 | -0.374249624 | 0.063164938 | 8.800763684 | -1.699787099 | 0.135220457 | 0.991835066 | -4.295395933 |
| **A_51_P405565** | Msh6 | 0.17766455 | -0.072220586 | 0.427549686 | 11.96323444 | 1.699297778 | 0.135315093 | 0.991835066 | -4.29583457 |
| **A_51_P142896** | Cd59a | -0.104712899 | -0.252025909 | 0.042600112 | 10.08583189 | -1.698901535 | 0.135391774 | 0.991835066 | -4.296189754 |
| **A_55_P2145833** | Dmpk | -0.250086139 | -0.60204182 | 0.101869542 | 11.68746555 | -1.698285786 | 0.135511016 | 0.991835066 | -4.296741667 |
| **A_52_P525837** | Trmt6 | 0.088438915 | -0.036038357 | 0.212916187 | 8.823545448 | 1.698096923 | 0.135547611 | 0.991835066 | -4.296910943 |
| **A_51_P375406** | Rprd1a | -0.133435467 | -0.328043668 | 0.061172734 | 11.28777398 | -1.803166193 | 0.135606162 | 0.991835066 | -4.258719679 |
| **A_52_P533146** | Ddit3 | -0.109865431 | -0.264530481 | 0.044799619 | 9.049147176 | -1.697766698 | 0.135611618 | 0.991835066 | -4.297206911 |
| **A_55_P2152009** | Tesc | 0.098527699 | -0.040182442 | 0.237237841 | 11.25649614 | 1.697693392 | 0.13562583 | 0.991835066 | -4.297272611 |
| **A_55_P1968606** | Synrg | -0.08104941 | -0.19518767 | 0.033088849 | 16.59712918 | -1.697179379 | 0.135725528 | 0.991835066 | -4.297733276 |
| **A_55_P2169804** | Nanos3 | -0.100396264 | -0.241827665 | 0.041035137 | 9.595890931 | -1.696605387 | 0.135836941 | 0.991835066 | -4.298247664 |
| **A_52_P244803** | D630033O | -0.158850652 | -0.382638267 | 0.064936964 | 8.740450044 | -1.696532113 | 0.13585117 | 0.991835066 | -4.298313327 |
| **A_55_P2099540** | H2afj | 0.083890624 | -0.034312255 | 0.202093503 | 8.755644018 | 1.696268124 | 0.135902446 | 0.991835066 | -4.298549891 |
| **A_55_P2055199** | Airn | 0.322231297 | -0.131869366 | 0.776331961 | 11.020493 | 1.695995577 | 0.135955403 | 0.991835066 | -4.298794115 |
| **A_51_P126626** | Zfp503 | 0.153874023 | -0.063012698 | 0.370760745 | 8.381409175 | 1.695670552 | 0.136018582 | 0.991835066 | -4.296592996 |
| **A_51_P481676** | Zfp787 | -0.079695671 | -0.192040525 | 0.032649182 | 10.36544393 | -1.695472241 | 0.136057145 | 0.991835066 | -4.299263047 |
| **A_55_P1996698** | Gm16381 | -0.137132341 | -0.330457289 | 0.056192607 | 13.10152146 | -1.695356833 | 0.136079591 | 0.991835066 | -4.299366454 |
| **A_55_P2074636** | Adap1 | -0.080315544 | -0.193570768 | 0.032939681 | 10.94428623 | -1.694925022 | 0.136163607 | 0.991835066 | -4.29975335 |

| | | | | | | | | | |
|---|---|---|---|---|---|---|---|---|---|
| **A_52_P471502** | Dcaf13 | -0.106735578 | -0.257261118 | 0.043789961 | 9.529378397 | -1.694759776 | 0.136195771 | 0.991835066 | -4.299901402 |
| **A_66_P115996** | Kyat1 | -0.168384409 | -0.405973075 | 0.069204258 | 12.34483936 | -1.693890331 | 0.136365124 | 0.991835066 | -4.300680339 |
| **A_55_P2235911** | 0610009E0 | 0.152330901 | -0.062635677 | 0.367297479 | 9.271353867 | 1.693659878 | 0.136410046 | 0.991835066 | -4.30088679 |
| **A_55_P2168823** | NA | 0.353382148 | -0.145310612 | 0.852074907 | 11.17416566 | 1.693638335 | 0.136414246 | 0.991835066 | -4.300906089 |
| **A_55_P2032600** | Rps24 | 0.077611609 | -0.031926113 | 0.18714933 | 16.96534315 | 1.693448967 | 0.136451171 | 0.991835066 | -4.301075729 |
| **A_55_P2107025** | Flad1 | 0.087415703 | -0.035994964 | 0.21082637 | 8.97044872 | 1.692956844 | 0.136547175 | 0.991835066 | -4.301516566 |
| **A_55_P2120577** | Cited2 | -0.109776255 | -0.264782891 | 0.045230381 | 10.25753361 | -1.692650339 | 0.136607001 | 0.991835066 | -4.301791118 |
| **A_66_P115061** | Smc6 | 0.132323144 | -0.054538177 | 0.319184465 | 10.58283597 | 1.692487732 | 0.13663875 | 0.991835066 | -4.301936769 |
| **A_55_P2093794** | Usp3 | -0.116624738 | -0.281337822 | 0.048088345 | 10.14178346 | -1.692278032 | 0.136679704 | 0.991835066 | -4.302124598 |
| **A_52_P604515** | Zfp319 | 0.088898841 | -0.036656061 | 0.214453742 | 8.857117078 | 1.692277411 | 0.136679825 | 0.991835066 | -4.302125154 |
| **A_55_P2085979** | Igf1 | -0.143584259 | -0.34637638 | 0.059207863 | 9.235193187 | -1.692251409 | 0.136684904 | 0.991835066 | -4.302148444 |
| **A_55_P1955612** | Nup93 | 0.134621724 | -0.055531717 | 0.324775164 | 11.8752477 | 1.692076884 | 0.136718999 | 0.991835066 | -4.302304764 |
| **A_51_P430973** | Paqr7 | -0.128236402 | -0.309381106 | 0.052908303 | 9.675227063 | -1.691978393 | 0.136738243 | 0.991835066 | -4.302392978 |
| **A_55_P1975456** | Nfyb | -0.103017813 | -0.248543137 | 0.042507512 | 9.203370096 | -1.691931906 | 0.136747327 | 0.991835066 | -4.302434615 |
| **A_51_P324410** | Vti1a | 0.134306156 | -0.055463917 | 0.32407623 | 9.757602443 | 1.691520741 | 0.136827699 | 0.991835066 | -4.30280287 |
| **A_55_P2039436** | Ilkap | -0.084861032 | -0.204767262 | 0.035045198 | 12.09249832 | -1.691514375 | 0.136828943 | 0.991835066 | -4.302808571 |
| **A_55_P1953252** | Ctnnd1 | -0.101103275 | -0.24398189 | 0.04177534 | 13.17911267 | -1.691247327 | 0.136881169 | 0.991835066 | -4.30304774 |
| **A_51_P275949** | Loxl2 | 0.169082651 | -0.069884938 | 0.408050239 | 10.99068997 | 1.691099571 | 0.136910073 | 0.991835066 | -4.303180068 |
| **A_55_P2238079** | A830054O( | 0.129861397 | -0.053737033 | 0.313459827 | 8.357703971 | 1.690519709 | 0.137023562 | 0.991835066 | -4.30369936 |
| **A_51_P297165** | Sf3a3 | 0.14456576 | -0.059825739 | 0.348957259 | 12.68267436 | 1.690486661 | 0.137030032 | 0.991835066 | -4.303728955 |
| **A_55_P2003414** | Kng1 | -0.614537626 | -1.48348115 | 0.254405898 | 9.05856852 | -1.69030883 | 0.137064856 | 0.991835066 | -4.303888203 |
| **A_55_P2313033** | Ptprb | 0.071674847 | -0.029677661 | 0.173027355 | 8.80526693 | 1.69021287 | 0.137083651 | 0.991835066 | -4.303974134 |
| **A_66_P125257** | Cep120 | -0.080278523 | -0.193808814 | 0.033251768 | 9.263167719 | -1.690039115 | 0.13711769 | 0.991835066 | -4.304129727 |
| **A_66_P122415** | Snhg6 | 0.196751108 | -0.081509407 | 0.475011623 | 12.66483843 | 1.689953417 | 0.137134481 | 0.991835066 | -4.304206466 |
| **A_52_P502577** | S1pr3 | -0.172878506 | -0.417431603 | 0.071674591 | 10.29141039 | -1.68957302 | 0.137209037 | 0.991835066 | -4.304547087 |
| **A_51_P359603** | Itgb7 | -0.072373184 | -0.17475589 | 0.030009521 | 7.841092133 | -1.689507843 | 0.137221815 | 0.991835066 | -4.304605448 |
| **A_55_P2032678** | Lox | 0.14102676 | -0.058564267 | 0.340617787 | 8.147427558 | 1.688766665 | 0.137367207 | 0.991835066 | -4.305269079 |
| **A_55_P2040362** | Bola2 | 0.117802039 | -0.048949423 | 0.2845535 | 13.40867599 | 1.688465885 | 0.137426251 | 0.991835066 | -4.305538372 |
| **A_52_P308507** | Chd6 | -0.102446423 | -0.247465267 | 0.042572422 | 8.739911402 | -1.688423888 | 0.137434497 | 0.991835066 | -4.305575972 |
| **A_55_P2137476** | Mpv17 | -0.119914097 | -0.289717769 | 0.049889576 | 11.70897328 | -1.687843961 | 0.137548412 | 0.991835066 | -4.306095164 |
| **A_52_P603421** | Ccdc189 | -0.142292376 | -0.343811231 | 0.05922648 | 8.192639318 | -1.687621593 | 0.137592116 | 0.991835066 | -4.306294233 |
| **A_55_P1968433** | Gpat3 | -0.094925732 | -0.229372233 | 0.039520769 | 11.03194403 | -1.687499233 | 0.13761617 | 0.991835066 | -4.306403771 |
| **A_55_P1962901** | Osr1 | 0.224747209 | -0.093579612 | 0.543074031 | 8.595120759 | 1.687447256 | 0.137626389 | 0.991835066 | -4.306450301 |
| **A_51_P300936** | Rbbp8 | 0.101204206 | -0.042148883 | 0.244557294 | 10.67601126 | 1.687332372 | 0.137648978 | 0.991835066 | -4.306553144 |
| **A_55_P1962946** | Syk | -0.138615281 | -0.334978489 | 0.057747927 | 9.855808021 | -1.687174967 | 0.137679934 | 0.991835066 | -4.306694049 |
| **A_51_P272250** | Snf8 | -0.156459895 | -0.378205375 | 0.065285584 | 10.94222194 | -1.686387575 | 0.137834887 | 0.991835066 | -4.307398866 |
| **A_51_P189916** | Nol12 | 0.143909764 | -0.06016481 | 0.347984337 | 11.04553332 | 1.685429124 | 0.138023726 | 0.991835066 | -4.308256716 |
| **A_55_P1983095** | Adam19 | -0.102817817 | -0.248646169 | 0.043010536 | 12.01091067 | -1.685138269 | 0.13808108 | 0.991835066 | -4.308517023 |
| **A_55_P2074942** | Slc6a4 | 0.107348619 | -0.044927112 | 0.25962435 | 7.892591328 | 1.684902995 | 0.138127491 | 0.991835066 | -4.30872758 |
| **A_55_P2007273** | Pole | 0.14360637 | -0.060105572 | 0.347318313 | 10.83117317 | 1.684869804 | 0.13813404 | 0.991835066 | -4.308757283 |
| **A_55_P2072945** | NA | 0.089381687 | -0.037454394 | 0.216217768 | 16.41115137 | 1.684282228 | 0.138250017 | 0.991835066 | -4.309283104 |
| **A_51_P474701** | Fbp1 | -0.233343593 | -0.56450815 | 0.097820965 | 8.215174762 | -1.684073963 | 0.138291147 | 0.991835066 | -4.309469471 |
| **A_52_P583973** | Lrpap1 | -0.127191519 | -0.307742919 | 0.05335988 | 11.43524215 | -1.683706652 | 0.138363715 | 0.991835066 | -4.30979815 |
| **A_55_P1989981** | Fam129b | 0.122945165 | -0.051619167 | 0.297509498 | 13.04015033 | 1.683313767 | 0.138441376 | 0.991835066 | -4.310149698 |
| **A_55_P1988070** | Eef1g | 0.171595661 | -0.072085018 | 0.41527634 | 11.19834789 | 1.68303961 | 0.138495593 | 0.991835066 | -4.310395001 |
| **A_55_P2078271** | Rbis | 0.158470634 | -0.066571694 | 0.383512962 | 12.85077541 | 1.683037201 | 0.13849607 | 0.991835066 | -4.310397156 |
| **A_55_P2105858** | Atf5 | 0.116019705 | -0.048773396 | 0.280812806 | 12.59801022 | 1.682681281 | 0.138566486 | 0.991835066 | -4.310715603 |
| **A_51_P458866** | F11r | -0.141754373 | -0.343123707 | 0.05961496 | 11.68061495 | -1.682489127 | 0.138604516 | 0.991835066 | -4.310887521 |

| | | | | | | | | | |
|---|---|---|---|---|---|---|---|---|---|
| **A_52_P456059** | Pfdn1 | -0.122645082 | -0.29691892 | 0.051628755 | 12.21058155 | -1.682004203 | 0.138700535 | 0.991835066 | -4.311321359 |
| **A_55_P2185489** | Zfp865 | -0.092173552 | -0.223166415 | 0.038819312 | 8.82754122 | -1.681774715 | 0.138745997 | 0.991835066 | -4.311526662 |
| **A_55_P2024858** | Fars2 | -0.125902694 | -0.304835209 | 0.05302982 | 12.66606947 | -1.681724656 | 0.138755916 | 0.991835066 | -4.311571445 |
| **A_51_P298666** | Scaf4 | 0.191023461 | -0.080467984 | 0.462514905 | 9.882129412 | 1.681665805 | 0.138767578 | 0.991835066 | -4.311624093 |
| **A_55_P2060737** | Trmt10b | 0.078763033 | -0.033184606 | 0.190710672 | 8.756366416 | 1.681576485 | 0.138785279 | 0.991835066 | -4.311703997 |
| **A_52_P356780** | Pdzd8 | -0.125646522 | -0.304252444 | 0.052959399 | 9.195932514 | -1.681371768 | 0.138825857 | 0.991835066 | -4.311887131 |
| **A_55_P2045482** | NA | 0.12866856 | -0.054285492 | 0.311622611 | 13.12159676 | 1.680890959 | 0.138921205 | 0.991835066 | -4.312317231 |
| **A_55_P1960924** | Egln2 | 0.102303741 | -0.043218545 | 0.247826027 | 13.10743836 | 1.680239306 | 0.139050533 | 0.991835066 | -4.312900117 |
| **A_55_P1973402** | Tada2a | 0.179933956 | -0.076016381 | 0.435884292 | 9.903539421 | 1.68022157 | 0.139054054 | 0.991835066 | -4.312915981 |
| **A_55_P2171047** | Oaz1 | -0.10367355 | -0.251153487 | 0.043806387 | 16.08978078 | -1.68013492 | 0.13907126 | 0.991835066 | -4.312993483 |
| **A_52_P160970** | Akap10 | -0.073914239 | -0.179101853 | 0.031273375 | 9.380156522 | -1.679471496 | 0.139203059 | 0.991835066 | -4.313586844 |
| **A_52_P79639** | Vat1 | -0.143124719 | -0.346822332 | 0.060572894 | 9.174560014 | -1.679336934 | 0.139229806 | 0.991835066 | -4.313707189 |
| **A_55_P2002117** | Kbtbd4 | 0.111224173 | -0.04718818 | 0.269636526 | 10.88924613 | 1.678105546 | 0.1394748 | 0.991835066 | -4.314808389 |
| **A_55_P2015530** | NA | 0.096402469 | -0.040903494 | 0.233708431 | 8.482419136 | 1.678061568 | 0.139483557 | 0.991835066 | -4.314847715 |
| **A_51_P118765** | Eef1a1 | -0.125276617 | -0.303710483 | 0.053157248 | 16.17019494 | -1.678038289 | 0.139488193 | 0.991835066 | -4.314868531 |
| **A_55_P2065671** | Ccnb1 | -0.148088795 | -0.359024356 | 0.062846767 | 12.12251218 | -1.677959703 | 0.139503843 | 0.991835066 | -4.314938802 |
| **A_51_P163252** | Schip1 | -0.097840291 | -0.237202669 | 0.041522087 | 8.628081921 | -1.677958908 | 0.139504002 | 0.991835066 | -4.314939513 |
| **A_55_P2048110** | Tmem97 | 0.137910749 | -0.058531021 | 0.33435252 | 10.70035992 | 1.677928344 | 0.139510089 | 0.991835066 | -4.314966843 |
| **A_55_P1991693** | Pprc1 | 0.174384441 | -0.074022155 | 0.422791037 | 8.568528291 | 1.677852359 | 0.139525224 | 0.991835066 | -4.315034787 |
| **A_51_P382928** | Cstf3 | -0.081853086 | -0.19851905 | 0.034812878 | 10.82383388 | -1.676872401 | 0.139720553 | 0.991835066 | -4.315910996 |
| **A_51_P194498** | Wfdc1 | -0.117488596 | -0.285024527 | 0.050047335 | 8.686186681 | -1.676088255 | 0.13987704 | 0.991835066 | -4.316612048 |
| **A_51_P175580** | Trp53inp1 | -0.209474486 | -0.508196032 | 0.089247061 | 9.718663251 | -1.67599885 | 0.139894892 | 0.991835066 | -4.316691975 |
| **A_55_P2133840** | Sipa1 | 0.13438599 | -0.057276897 | 0.326048878 | 11.69062798 | 1.675811249 | 0.139932359 | 0.991835066 | -4.316859685 |
| **A_51_P254986** | Serf1 | -0.084772272 | -0.205692872 | 0.036148327 | 13.6469851 | -1.675570343 | 0.139980487 | 0.991835066 | -4.317075043 |
| **A_55_P2093143** | Casd1 | 0.072922389 | -0.031123957 | 0.176968736 | 8.062769744 | 1.675109443 | 0.140072608 | 0.991835066 | -4.317487045 |
| **A_51_P466221** | Amhr2 | 0.192808292 | -0.082294112 | 0.467910697 | 9.645031723 | 1.675098903 | 0.140074715 | 0.991835066 | -4.317496467 |
| **A_55_P1958434** | Got1 | -0.133309257 | -0.323531521 | 0.056913006 | 14.54884426 | -1.674974097 | 0.14009967 | 0.991835066 | -4.317608028 |
| **A_55_P2118906** | Atp5j2 | -0.084697856 | -0.205556224 | 0.036160512 | 16.38016171 | -1.674961489 | 0.140102192 | 0.991835066 | -4.317619298 |
| **A_51_P384113** | Acad12 | -0.114071692 | -0.276858114 | 0.04871473 | 9.141212702 | -1.674822697 | 0.140129949 | 0.991835066 | -4.317743358 |
| **A_55_P2002557** | Srebf1 | -0.076875151 | -0.186582653 | 0.032832351 | 8.541371887 | -1.674783934 | 0.140137702 | 0.991835066 | -4.317778007 |
| **A_55_P2017804** | Cox5b | -0.193196808 | -0.46897954 | 0.082585923 | 16.16134915 | -1.67433367 | 0.140227793 | 0.991835066 | -4.318180464 |
| **A_55_P2169963** | Zfp982 | 0.111853765 | -0.047860627 | 0.271568158 | 8.505246366 | 1.673846721 | 0.140325285 | 0.991835066 | -4.318615685 |
| **A_55_P2157966** | Map1a | -0.071356476 | -0.17325087 | 0.030537919 | 8.546527076 | -1.673756327 | 0.14034339 | 0.991835066 | -4.318696474 |
| **A_51_P311785** | Man2b2 | -0.079960026 | -0.194148942 | 0.034228891 | 8.371551076 | -1.673624807 | 0.140369735 | 0.991835066 | -4.318814017 |
| **A_51_P411345** | Mogat2 | 0.087901961 | -0.037641274 | 0.213445196 | 7.96136207 | 1.673456327 | 0.140403492 | 0.991835066 | -4.31896459 |
| **A_66_P132493** | Siglece | -0.091966777 | -0.223327216 | 0.039393663 | 9.583140052 | -1.673306529 | 0.140433512 | 0.991835066 | -4.319098463 |
| **A_51_P370470** | Cyp20a1 | -0.084819678 | -0.206011052 | 0.036371696 | 8.797672882 | -1.672761573 | 0.140542774 | 0.991835066 | -4.319585467 |
| **A_55_P1956827** | Ogfr | 0.146764219 | -0.062971891 | 0.356500328 | 13.32793388 | 1.672461434 | 0.140602985 | 0.991835066 | -4.319853674 |
| **A_55_P2298029** | Nsd3 | 0.102045304 | -0.043789877 | 0.247880485 | 8.72232503 | 1.672398821 | 0.140615549 | 0.991835066 | -4.319909624 |
| **A_55_P1997145** | Nlrx1 | -0.070787359 | -0.171984377 | 0.030409658 | 8.71727428 | -1.671849311 | 0.14072586 | 0.991835066 | -4.320400642 |
| **A_51_P183051** | Upb1 | 0.151930253 | -0.065335563 | 0.36919607 | 8.729427033 | 1.671329172 | 0.140830351 | 0.991835066 | -4.320865386 |
| **A_51_P259029** | Dusp26 | 0.11427565 | -0.049153057 | 0.277704357 | 8.962040457 | 1.671223329 | 0.140851622 | 0.991835066 | -4.320959952 |
| **A_52_P207614** | NA | 0.130068647 | -0.055973048 | 0.316110341 | 10.43788752 | 1.670980829 | 0.14090037 | 0.991835066 | -4.321176612 |
| **A_52_P222096** | 4931428F0 | 0.088316622 | -0.038026381 | 0.214659625 | 8.759327561 | 1.670707368 | 0.140955361 | 0.991835066 | -4.321420925 |
| **A_55_P2121456** | Fam43a | -0.125016645 | -0.303880408 | 0.053847118 | 9.953446657 | -1.670531266 | 0.140990785 | 0.991835066 | -4.321578254 |
| **A_55_P2106358** | NA | 0.189730815 | -0.081831472 | 0.461293103 | 8.332447303 | 1.669850333 | 0.141127836 | 0.991835066 | -4.32218656 |
| **A_55_P2083149** | NA | 0.194119033 | -0.083782399 | 0.472020465 | 9.287530821 | 1.66950017 | 0.141198363 | 0.991835066 | -4.322499355 |
| **A_55_P2036459** | Med24 | -0.107048799 | -0.260346161 | 0.046248563 | 8.915843417 | -1.668999662 | 0.141299229 | 0.991835066 | -4.322946427 |

| | | | | | | | | | |
|---|---|---|---|---|---|---|---|---|---|
| **A_66_P134690** | Spin1 | 0.109795239 | -0.047467027 | 0.267057504 | 8.40734094 | 1.668660981 | 0.141367521 | 0.991835066 | -4.323248934 |
| **A_55_P2065360** | Siva1 | 0.12114782 | -0.0524343 | 0.29472994 | 12.37881682 | 1.668091035 | 0.141482516 | 0.991835066 | -4.323757975 |
| **A_52_P95910** | Ugcg | -0.083836886 | -0.20396599 | 0.036292217 | 9.000704218 | -1.667999971 | 0.141500898 | 0.991835066 | -4.323839304 |
| **A_55_P2337138** | Ago2 | 0.103767746 | -0.04498037 | 0.252515863 | 11.71884131 | 1.667324153 | 0.141637386 | 0.991835066 | -4.32444285 |
| **A_52_P379337** | Rtn4 | -0.134683099 | -0.32777064 | 0.058404442 | 10.94776943 | -1.667124295 | 0.141677773 | 0.991835066 | -4.324621325 |
| **A_66_P128313** | Mtmr2 | 0.110559168 | -0.047955709 | 0.269074044 | 10.52910639 | 1.666993349 | 0.141704241 | 0.991835066 | -4.324738259 |
| **A_52_P569549** | Lig4 | 0.104490861 | -0.045331202 | 0.254312925 | 9.67210137 | 1.666908138 | 0.141721466 | 0.991835066 | -4.32481435 |
| **A_51_P296878** | Triqk | -0.091294104 | -0.222207489 | 0.03961928 | 9.210852925 | -1.666739829 | 0.141755497 | 0.991835066 | -4.324964644 |
| **A_55_P1957204** | NA | -0.090839161 | -0.221138482 | 0.039460161 | 17.28015437 | -1.666249725 | 0.141854634 | 0.991835066 | -4.325402271 |
| **A_55_P2058270** | Gm6507 | 0.108197926 | -0.047002449 | 0.2633983 | 7.925138498 | 1.666231354 | 0.141858352 | 0.991835066 | -4.325418675 |
| **A_51_P462448** | Gpx4 | -0.114506549 | -0.278763587 | 0.049750489 | 14.65596169 | -1.666155275 | 0.141873747 | 0.991835066 | -4.325486606 |
| **A_55_P1963712** | Cyb5b | 0.177031772 | -0.076984826 | 0.431048369 | 11.23585641 | 1.665705631 | 0.14196477 | 0.991835066 | -4.325888076 |
| **A_55_P1984976** | Wnt5b | 0.114330216 | -0.049742649 | 0.278403081 | 10.22979765 | 1.665456889 | 0.142015148 | 0.991835066 | -4.326110159 |
| **A_55_P2068988** | NA | -0.122727262 | -0.298879262 | 0.053424739 | 10.15334097 | -1.665185411 | 0.142070149 | 0.991835066 | -4.326352533 |
| **A_51_P206824** | Hjv | -0.152596796 | -0.3716511 | 0.066457508 | 8.323885418 | -1.664955988 | 0.142116646 | 0.991835066 | -4.326557354 |
| **A_55_P2116924** | Dbi | -0.083564361 | -0.203525021 | 0.036396298 | 15.15100636 | -1.664912391 | 0.142125484 | 0.991835066 | -4.326596275 |
| **A_55_P2032723** | Dnajc10 | 0.098968338 | -0.043144273 | 0.241080949 | 8.688178165 | 1.664457802 | 0.142217663 | 0.991835066 | -4.327002097 |
| **A_66_P107353** | NA | 0.121337295 | -0.052918352 | 0.295592943 | 15.8148342 | 1.664242393 | 0.142261362 | 0.991835066 | -4.32719439 |
| **A_55_P2136209** | Ublcp1 | 0.092482273 | -0.040336704 | 0.22530125 | 9.494009794 | 1.664207591 | 0.142268424 | 0.991835066 | -4.327225456 |
| **A_51_P107234** | Abcf1 | 0.091688471 | -0.040021906 | 0.223398847 | 12.16333698 | 1.66381053 | 0.142349011 | 0.991835066 | -4.327579891 |
| **A_51_P282424** | Eapp | -0.103476334 | -0.252148401 | 0.045195733 | 10.05511184 | -1.663492267 | 0.142413637 | 0.991835066 | -4.327863974 |
| **A_55_P2100526** | Ptges3 | 0.168125774 | -0.073468982 | 0.409720531 | 11.99825519 | 1.663243847 | 0.1424641 | 0.991835066 | -4.328085707 |
| **A_55_P2043627** | Fam89a | -0.105363483 | -0.25680383 | 0.046076864 | 8.305219798 | -1.662867535 | 0.142540575 | 0.991835066 | -4.32842158 |
| **A_51_P341010** | Ercc8 | 0.166531892 | -0.072858604 | 0.405922387 | 9.997685315 | 1.662645431 | 0.14258573 | 0.991835066 | -4.328619808 |
| **A_51_P423509** | Faap100 | -0.096109322 | -0.234272582 | 0.042053938 | 8.632620055 | -1.66257817 | 0.142599407 | 0.991835066 | -4.328679838 |
| **A_55_P2113160** | Mal | -0.125879766 | -0.306845265 | 0.055085732 | 9.665894495 | -1.662529182 | 0.142609369 | 0.991835066 | -4.328723558 |
| **A_52_P303891** | Nr1d2 | -0.117797898 | -0.287150594 | 0.051554799 | 8.890424893 | -1.662472831 | 0.142620829 | 0.991835066 | -4.32877385 |
| **A_51_P169745** | Tuba1a | -0.092460697 | -0.225399608 | 0.040478214 | 15.1885884 | -1.662318273 | 0.142652267 | 0.991835066 | -4.328911787 |
| **A_55_P2167035** | Zfand1 | -0.102535403 | -0.249977333 | 0.044906526 | 9.151955316 | -1.662118458 | 0.14269292 | 0.991835066 | -4.329090111 |
| **A_55_P2447879** | Asxl2 | 0.085310615 | -0.03737359 | 0.207994819 | 9.061589121 | 1.661971429 | 0.14272284 | 0.991835066 | -4.329221322 |
| **A_55_P1968295** | Pacsin3 | -0.132959511 | -0.324175143 | 0.058256121 | 12.13653382 | -1.661900993 | 0.142737176 | 0.991835066 | -4.32928418 |
| **A_55_P2122605** | Cbr2 | 0.325672116 | -0.142780602 | 0.794124834 | 8.603280801 | 1.661590286 | 0.14280043 | 0.991835066 | -4.32956145 |
| **A_52_P408657** | Mycbp | -0.082456793 | -0.201070379 | 0.036156792 | 8.220853451 | -1.661503036 | 0.142818197 | 0.991835066 | -4.329639309 |
| **A_55_P2048937** | Kif5c | 0.110403207 | -0.04846716 | 0.269273574 | 8.224297396 | 1.660916974 | 0.142937595 | 0.991835066 | -4.330162267 |
| **A_55_P2101944** | NA | 0.244432547 | -0.112564761 | 0.601429856 | 12.4203973 | 1.700831308 | 0.142937814 | 0.991835066 | -4.295569872 |
| **A_51_P415395** | C2cd4b | -0.108041644 | -0.263547821 | 0.047464533 | 8.411098828 | -1.660552732 | 0.143011849 | 0.991835066 | -4.330487269 |
| **A_55_P1952016** | R3hdm2 | -0.084756399 | -0.20679626 | 0.037283463 | 11.79720865 | -1.659892342 | 0.14314657 | 0.991835066 | -4.331076476 |
| **A_66_P108685** | Cideb | -0.078907487 | -0.192528478 | 0.034713505 | 8.974244131 | -1.659849719 | 0.143155269 | 0.991835066 | -4.331114503 |
| **A_51_P496432** | Acsl1 | -0.118913096 | -0.290181771 | 0.052355579 | 13.9880804 | -1.659437421 | 0.143239444 | 0.991835066 | -4.331482332 |
| **A_66_P118513** | 11-Sep | 0.12031284 | -0.052972816 | 0.293598496 | 12.84971589 | 1.65942831 | 0.143241305 | 0.991835066 | -4.331490461 |
| **A_55_P1987839** | Erbin | -0.132363682 | -0.323011484 | 0.058284119 | 10.99674644 | -1.659381226 | 0.14325092 | 0.991835066 | -4.331532465 |
| **A_55_P1990324** | Ptprc | -0.071334862 | -0.174125664 | 0.03145594 | 7.787128611 | -1.65865744 | 0.143398813 | 0.991835066 | -4.332178134 |
| **A_55_P2136657** | Thbs3 | -0.212211287 | -0.518035813 | 0.09361324 | 10.68671553 | -1.658461138 | 0.143438949 | 0.991835066 | -4.332353239 |
| **A_55_P2054310** | M6pr | -0.124192727 | -0.303205832 | 0.054820377 | 10.15635316 | -1.658137241 | 0.143505196 | 0.991835066 | -4.332642151 |
| **A_55_P2031112** | Dhps | 0.096794571 | -0.042755188 | 0.23634433 | 10.85991756 | 1.657795825 | 0.143575058 | 0.991835066 | -4.332946676 |
| **A_55_P2116111** | Prag1 | 0.158058011 | -0.069841671 | 0.385957694 | 11.56016149 | 1.65760835 | 0.143613433 | 0.991835066 | -4.333113889 |
| **A_55_P2151822** | Plxnd1 | 0.120417392 | -0.053220896 | 0.294055681 | 13.88620007 | 1.65749739 | 0.143636151 | 0.991835066 | -4.333212855 |
| **A_52_P171033** | Mcmbp | 0.176531991 | -0.078051563 | 0.431115546 | 9.12732034 | 1.657304106 | 0.143675732 | 0.991835066 | -4.333385241 |

| | | | | | | | | | |
|---|---|---|---|---|---|---|---|---|---|
| **A_51_P111462** | Arl15 | 0.118773622 | -0.05252745 | 0.290074695 | 10.39565744 | 1.65717759 | 0.143701645 | 0.991835066 | -4.333498076 |
| **A_55_P2022509** | Mgat5b | -0.11950572 | -0.291865821 | 0.052854382 | 9.179746923 | -1.657147177 | 0.143707875 | 0.991835066 | -4.3335252 |
| **A_51_P465292** | Hnmt | -0.132185616 | -0.322856153 | 0.05848492 | 8.541915019 | -1.656951302 | 0.143748005 | 0.991835066 | -4.33369989 |
| **A_51_P446315** | Cdc37l1 | -0.119366772 | -0.291560975 | 0.05282743 | 11.16274344 | -1.656815151 | 0.143775906 | 0.991835066 | -4.333821312 |
| **A_55_P2088640** | Trip4 | 0.103624665 | -0.045882248 | 0.253131578 | 9.552622624 | 1.656574599 | 0.143825213 | 0.991835066 | -4.334035836 |
| **A_55_P2105512** | Nop56 | 0.144928793 | -0.064173253 | 0.354030839 | 12.19926031 | 1.656553726 | 0.143829492 | 0.991835066 | -4.334054451 |
| **A_55_P2108738** | Nck1 | 0.095959316 | -0.042524894 | 0.234443527 | 10.649645 | 1.656136076 | 0.14391514 | 0.991835066 | -4.334426894 |
| **A_55_P1958961** | 2010001K2 | 0.089879514 | -0.039895286 | 0.219654314 | 8.476399437 | 1.655310545 | 0.144084575 | 0.991835066 | -4.335163009 |
| **A_55_P2178808** | Zbtb5 | 0.08849647 | -0.03932351 | 0.216316449 | 9.160650292 | 1.65476503 | 0.144196641 | 0.991835066 | -4.335649393 |
| **A_52_P316437** | 1810013L2 | 0.073836247 | -0.032814332 | 0.180486825 | 8.320611657 | 1.654685752 | 0.144212935 | 0.991835066 | -4.335720075 |
| **A_52_P563340** | Smim4 | -0.141529056 | -0.345958062 | 0.062899949 | 12.71265821 | -1.65467318 | 0.144215519 | 0.991835066 | -4.335731283 |
| **A_51_P193336** | Nucb2 | -0.08032255 | -0.196357842 | 0.035712742 | 8.844075882 | -1.654460948 | 0.144259146 | 0.991835066 | -4.335920499 |
| **A_55_P2099890** | Atxn7 | -0.16478471 | -0.402848839 | 0.073279419 | 12.28383375 | -1.654367852 | 0.144278287 | 0.991835066 | -4.336003497 |
| **A_55_P2030721** | Ankle1 | 0.110352417 | -0.049076799 | 0.269781634 | 8.398465349 | 1.65433353 | 0.144285344 | 0.991835066 | -4.336034096 |
| **A_55_P1969615** | Pnck | 0.131168948 | -0.058349618 | 0.320687515 | 9.686568037 | 1.654201483 | 0.144312499 | 0.991835066 | -4.336151817 |
| **A_51_P470724** | Eif2s2 | 0.143250442 | -0.063750236 | 0.35025112 | 10.18527363 | 1.653991739 | 0.144355642 | 0.991835066 | -4.336338803 |
| **A_51_P318551** | Nle1 | 0.071407827 | -0.031797245 | 0.174612898 | 8.468038379 | 1.653689274 | 0.144417879 | 0.991835066 | -4.33660844 |
| **A_55_P1983284** | Flot2 | -0.12529861 | -0.306415179 | 0.055817959 | 12.71051029 | -1.653473367 | 0.144462321 | 0.991835066 | -4.336800906 |
| **A_55_P2121945** | Cnot6l | -0.149174282 | -0.368070861 | 0.069722297 | 12.04603702 | -1.692864246 | 0.144484477 | 0.991835066 | -4.301825801 |
| **A_51_P270364** | Mmaa | -0.097996161 | -0.239657406 | 0.043665085 | 11.06093462 | -1.653358901 | 0.144485887 | 0.991835066 | -4.336902943 |
| **A_55_P2144716** | Hecw2 | -0.120085788 | -0.293703187 | 0.053531611 | 9.785442308 | -1.653131878 | 0.144532638 | 0.991835066 | -4.33710531 |
| **A_55_P2096121** | NA | 0.155259801 | -0.069267146 | 0.379786748 | 13.86922163 | 1.652721539 | 0.144617176 | 0.991835066 | -4.337471068 |
| **A_52_P520495** | Vcam1 | 0.126501261 | -0.056456928 | 0.309459451 | 8.727235268 | 1.652540585 | 0.144654471 | 0.991835066 | -4.337632356 |
| **A_55_P2244112** | Amotl1 | 0.07595571 | -0.033900336 | 0.185811756 | 8.372237544 | 1.652515709 | 0.144659598 | 0.991835066 | -4.337654528 |
| **A_51_P326229** | Ddx25 | 0.165314767 | -0.073924624 | 0.404554159 | 8.798322879 | 1.651536183 | 0.144861644 | 0.991835066 | -4.338527529 |
| **A_55_P2084611** | B4galt6 | -0.086203785 | -0.210969489 | 0.038561918 | 9.380417412 | -1.651354257 | 0.144899199 | 0.991835066 | -4.338689658 |
| **A_55_P1997429** | Gm13698 | 0.121724368 | -0.054508034 | 0.297956771 | 9.200142798 | 1.650824475 | 0.145008614 | 0.991835066 | -4.339161766 |
| **A_55_P2094706** | Sox17 | -0.12628074 | -0.309142348 | 0.056580868 | 9.243717804 | -1.650531117 | 0.145069235 | 0.991835066 | -4.339423174 |
| **A_55_P2010003** | Zfp113 | -0.080808737 | -0.197837171 | 0.036219697 | 8.773710225 | -1.65035 | 0.145106674 | 0.991835066 | -4.33958456 |
| **A_55_P2452384** | Mga | -0.083823897 | -0.20522078 | 0.037572986 | 9.151843068 | -1.650324875 | 0.145111868 | 0.991835066 | -4.339606947 |
| **A_55_P1975445** | NA | 0.093260093 | -0.041850061 | 0.228370248 | 13.09166576 | 1.649745632 | 0.145231669 | 0.991835066 | -4.340123058 |
| **A_52_P375493** | Tfdp1 | -0.084612957 | -0.207208829 | 0.037982915 | 9.711304039 | -1.649567745 | 0.145268479 | 0.991835066 | -4.340281549 |
| **A_51_P414396** | Mmrn2 | 0.123631762 | -0.055509273 | 0.302772797 | 13.49656781 | 1.649468831 | 0.145288951 | 0.991835066 | -4.340369677 |
| **A_51_P279127** | Fam173a | 0.106005814 | -0.047615836 | 0.259627464 | 12.290399 | 1.649249614 | 0.145334331 | 0.991835066 | -4.340564984 |
| **A_51_P486512** | Letmd1 | 0.135174392 | -0.060747903 | 0.331096687 | 9.3657365 | 1.648996363 | 0.145386774 | 0.991835066 | -4.340790605 |
| **A_52_P78770** | Chpt1 | -0.085063609 | -0.208364369 | 0.038237151 | 8.372649688 | -1.648872915 | 0.145412343 | 0.991835066 | -4.340900582 |
| **A_55_P1982818** | Odc1 | -0.104656408 | -0.25639512 | 0.047082305 | 9.115412131 | -1.648460559 | 0.145497785 | 0.991835066 | -4.341267928 |
| **A_55_P1996613** | NA | -0.192147388 | -0.470746927 | 0.086452151 | 14.50150076 | -1.648402349 | 0.14550985 | 0.991835066 | -4.341319782 |
| **A_52_P505234** | Stt3a | 0.070806111 | -0.03195444 | 0.173566661 | 8.581858585 | 1.646847749 | 0.145832422 | 0.991835066 | -4.342704494 |
| **A_55_P1980831** | Arpc2 | -0.095177328 | -0.233356806 | 0.04300215 | 15.24287748 | -1.646262527 | 0.145954028 | 0.991835066 | -4.343225687 |
| **A_55_P1966843** | Gm12070 | -0.090739152 | -0.222490063 | 0.041011758 | 16.6765074 | -1.646077291 | 0.145992539 | 0.991835066 | -4.343390647 |
| **A_55_P2175550** | Atf2 | -0.100770964 | -0.247196784 | 0.045654857 | 8.725207086 | -1.644852193 | 0.14624748 | 0.991835066 | -4.344481544 |
| **A_55_P2152468** | NA | -0.119501114 | -0.293194605 | 0.054192378 | 9.765466462 | -1.644362411 | 0.146349521 | 0.991835066 | -4.344917622 |
| **A_52_P1029978** | Lasp1 | 0.116989823 | -0.05305926 | 0.287038906 | 12.48329513 | 1.644307067 | 0.146361056 | 0.991835066 | -4.344966896 |
| **A_55_P2246650** | 9330159M0 | -0.100133074 | -0.245695735 | 0.045429587 | 8.923512273 | -1.644132055 | 0.146397537 | 0.991835066 | -4.34512271 |
| **A_55_P2100241** | Rn18s | 0.143730445 | -0.065225956 | 0.352686846 | 17.41312927 | 1.644001554 | 0.146424745 | 0.991835066 | -4.345238893 |
| **A_52_P90684** | Klhl31 | -0.254089406 | -0.623622812 | 0.115443999 | 11.63448663 | -1.643395096 | 0.146551248 | 0.991835066 | -4.345778787 |
| **A_55_P2133115** | NA | -0.081421245 | -0.199842878 | 0.037000388 | 14.43096441 | -1.643296098 | 0.146571909 | 0.991835066 | -4.345866914 |

| | | | | | | | | | |
|---|---|---|---|---|---|---|---|---|---|
| **A_51_P290819** | Hint3 | -0.153369177 | -0.37644328 | 0.069704927 | 11.88936406 | -1.643228889 | 0.146585936 | 0.991835066 | -4.345926742 |
| **A_55_P1984675** | Abhd6 | 0.076539871 | -0.034805336 | 0.187885079 | 8.664005523 | 1.64295372 | 0.146643382 | 0.991835066 | -4.346171689 |
| **A_55_P2021854** | NA | 0.111662042 | -0.050783566 | 0.274107649 | 8.500144925 | 1.64288333 | 0.14665808 | 0.991835066 | -4.346234346 |
| **A_55_P2164139** | NA | -0.114916651 | -0.282099466 | 0.052266165 | 13.5293901 | -1.642859631 | 0.146663029 | 0.991835066 | -4.346255441 |
| **A_51_P195573** | Dnmt1 | 0.106921468 | -0.048647487 | 0.262490422 | 10.01160297 | 1.642672972 | 0.146702014 | 0.991835066 | -4.346421591 |
| **A_51_P251717** | Hikeshi | -0.1027289 | -0.252207188 | 0.046749387 | 10.48302137 | -1.642569139 | 0.146723704 | 0.991835066 | -4.346514013 |
| **A_55_P2001203** | Dnm2 | 0.070835867 | -0.032266009 | 0.173937743 | 12.89678964 | 1.642085545 | 0.146824765 | 0.991835066 | -4.346944447 |
| **A_51_P278887** | Hint2 | -0.09674015 | -0.237618718 | 0.044138417 | 13.3861413 | -1.641235734 | 0.147002517 | 0.991835066 | -4.347700768 |
| **A_55_P2007966** | Tmem215 | -0.135818145 | -0.333608661 | 0.06197237 | 10.90716307 | -1.641199838 | 0.147010029 | 0.991835066 | -4.347732714 |
| **A_55_P1966987** | Limk1 | 0.124944808 | -0.057017254 | 0.30690687 | 10.31886929 | 1.641143277 | 0.147021868 | 0.991835066 | -4.347783049 |
| **A_55_P2169276** | 4732491K2 | -0.099793689 | -0.245127299 | 0.045539921 | 8.7985897 | -1.641141951 | 0.147022145 | 0.991835066 | -4.347784229 |
| **A_55_P2044202** | Vps29 | 0.110834007 | -0.050583659 | 0.272251673 | 11.55181089 | 1.641085111 | 0.147034043 | 0.991835066 | -4.347834812 |
| **A_66_P108059** | Ttc39c | -0.079715364 | -0.195842168 | 0.03641144 | 8.05016026 | -1.640660391 | 0.147122974 | 0.991835066 | -4.348212766 |
| **A_52_P595225** | Vps35 | -0.114499119 | -0.281359342 | 0.052361105 | 8.865673189 | -1.640055155 | 0.147249791 | 0.991835066 | -4.348751322 |
| **A_55_P2185722** | Fermt2 | -0.073923264 | -0.181656129 | 0.033809601 | 10.36084587 | -1.639993239 | 0.14726277 | 0.991835066 | -4.348806414 |
| **A_52_P33453** | NA | -0.122333891 | -0.300642735 | 0.055974953 | 14.285226 | -1.639770384 | 0.147309496 | 0.991835066 | -4.349004703 |
| **A_55_P2110513** | Lmo7 | 0.122107743 | -0.055873419 | 0.300088905 | 13.29148243 | 1.639752492 | 0.147313248 | 0.991835066 | -4.349020622 |
| **A_65_P06061** | Rpl37 | 0.101983156 | -0.046708232 | 0.250674544 | 17.57359699 | 1.639274804 | 0.147413452 | 0.991835066 | -4.349445632 |
| **A_51_P157976** | Mmab | 0.075025854 | -0.034364893 | 0.184416601 | 8.986739252 | 1.639228478 | 0.147423174 | 0.991835066 | -4.349486848 |
| **A_52_P160955** | Sox11 | 0.189990865 | -0.087083646 | 0.467065376 | 9.059773298 | 1.638872914 | 0.147497807 | 0.991835066 | -4.34980318 |
| **A_55_P2121364** | Tdg | 0.086407782 | -0.039610558 | 0.212426122 | 8.260922105 | 1.638808602 | 0.14751131 | 0.991835066 | -4.349860395 |
| **A_55_P2092526** | Tgif1 | -0.187586884 | -0.461219247 | 0.086045479 | 10.20100585 | -1.638491307 | 0.147577946 | 0.991835066 | -4.350142665 |
| **A_55_P1993168** | Ppargc1b | -0.092131646 | -0.226561034 | 0.042297742 | 9.44302791 | -1.638037123 | 0.147673381 | 0.991835066 | -4.350546691 |
| **A_66_P129730** | Pdia3 | 0.153314525 | -0.070432024 | 0.377061073 | 12.19393554 | 1.637706562 | 0.147742875 | 0.991835066 | -4.350840731 |
| **A_51_P164030** | Tcp1 | -0.123996566 | -0.304976467 | 0.056983336 | 12.68506039 | -1.637526897 | 0.14778066 | 0.991835066 | -4.35100054 |
| **A_51_P190845** | Atp6v1b2 | -0.088613113 | -0.217967292 | 0.040741065 | 8.733961658 | -1.637293979 | 0.147829657 | 0.991835066 | -4.35120771 |
| **A_55_P2066559** | Endou | -0.109075051 | -0.268327645 | 0.050177544 | 8.176206054 | -1.636997587 | 0.14789203 | 0.991835066 | -4.351471329 |
| **A_55_P1961913** | Pygm | -0.134449139 | -0.330791923 | 0.061893645 | 13.03879057 | -1.636636422 | 0.147968066 | 0.991835066 | -4.351792543 |
| **A_55_P2088375** | Tnnt3 | -0.109990349 | -0.270672624 | 0.050691927 | 8.40322809 | -1.636046851 | 0.148092269 | 0.991835066 | -4.352316863 |
| **A_52_P341449** | Pgm3 | 0.126352226 | -0.058242717 | 0.31094717 | 9.357055491 | 1.635958303 | 0.148110931 | 0.991835066 | -4.352395607 |
| **A_55_P2109382** | Adora2a | -0.238815333 | -0.587719226 | 0.11008856 | 9.644737524 | -1.635933043 | 0.148116255 | 0.991835066 | -4.35241807 |
| **A_51_P295192** | Nfkbia | -0.123444403 | -0.303798981 | 0.056910174 | 12.0576861 | -1.635887249 | 0.148125908 | 0.991835066 | -4.352458793 |
| **A_52_P234354** | Hic2 | -0.10744249 | -0.264422822 | 0.049537842 | 9.823101546 | -1.63583672 | 0.14813656 | 0.991835066 | -4.352503726 |
| **A_51_P414448** | Naa15 | -0.103559086 | -0.25488468 | 0.047766508 | 10.21592741 | -1.635629577 | 0.148180233 | 0.991835066 | -4.352687928 |
| **A_55_P2085335** | Mia | 0.213264242 | -0.098373737 | 0.524902221 | 8.08530362 | 1.635598853 | 0.148186712 | 0.991835066 | -4.352715249 |
| **A_55_P2103351** | Ndufa5 | -0.131093153 | -0.322765235 | 0.060578929 | 14.87590263 | -1.634670705 | 0.148382558 | 0.991835066 | -4.353540531 |
| **A_51_P394190** | Lmo4 | 0.110713105 | -0.051165463 | 0.272591674 | 12.72373218 | 1.634627533 | 0.148391674 | 0.991835066 | -4.353578916 |
| **A_55_P2429480** | Atad2 | 0.120262028 | -0.055618626 | 0.296142681 | 8.770822697 | 1.634254104 | 0.148470543 | 0.991835066 | -4.353910925 |
| **A_52_P257812** | Lpl | -0.302225115 | -0.744224338 | 0.139774107 | 11.10541258 | -1.634248815 | 0.14847166 | 0.991835066 | -4.353915627 |
| **A_55_P1956457** | 6030458C1 | 0.120531393 | -0.055787819 | 0.296850605 | 9.156056378 | 1.633840552 | 0.148557933 | 0.991835066 | -4.354278586 |
| **A_52_P298088** | Itgb1bp2 | -0.287096135 | -0.707123542 | 0.132931271 | 11.42057241 | -1.633649372 | 0.148598348 | 0.991835066 | -4.354448544 |
| **A_52_P434820** | Gnpda2 | -0.082929349 | -0.204261712 | 0.038403014 | 8.7994818 | -1.633581215 | 0.14861276 | 0.991835066 | -4.354509135 |
| **A_51_P459320** | Gkap1 | 0.099004457 | -0.045857328 | 0.243866242 | 11.54254337 | 1.633465795 | 0.148637167 | 0.991835066 | -4.354611739 |
| **A_51_P332355** | Kif18a | -0.103143824 | -0.254069991 | 0.047782342 | 8.477180569 | -1.633382207 | 0.148654845 | 0.991835066 | -4.354686044 |
| **A_66_P106289** | Vps54 | 0.138327353 | -0.064093686 | 0.340748392 | 9.268592583 | 1.633283407 | 0.148675743 | 0.991835066 | -4.354773872 |
| **A_51_P463440** | Elovl6 | 0.18392773 | -0.085259125 | 0.453114586 | 9.388360671 | 1.633061329 | 0.148722726 | 0.991835066 | -4.354971282 |
| **A_55_P1979743** | NA | -0.093871226 | -0.231261578 | 0.043519127 | 16.22017875 | -1.632996983 | 0.148736342 | 0.991835066 | -4.355028479 |
| **A_55_P1955517** | NA | 0.159896998 | -0.074174943 | 0.393968938 | 10.24393638 | 1.632676291 | 0.148804219 | 0.991835066 | -4.355313535 |

| | | | | | | | | | |
|---|---|---|---|---|---|---|---|---|---|
| **A_55_P2063594** | Gm5801 | 0.099227231 | -0.046044511 | 0.244498972 | 11.34243011 | 1.632521315 | 0.148837032 | 0.991835066 | -4.355451285 |
| **A_55_P2021266** | Hpse | 0.170493745 | -0.07911963 | 0.420107119 | 8.160310801 | 1.632486995 | 0.148844299 | 0.991835066 | -4.355481791 |
| **A_55_P2079020** | Snhg7os | -0.112950558 | -0.278360432 | 0.052459316 | 9.374365705 | -1.632059849 | 0.148934776 | 0.991835066 | -4.35586144 |
| **A_55_P2114288** | Mrpl24 | -0.09462115 | -0.233203189 | 0.043960889 | 13.01322305 | -1.631888223 | 0.148971145 | 0.991835066 | -4.356013976 |
| **A_55_P2186621** | Rbmxl1 | 0.124147814 | -0.057694158 | 0.305989786 | 12.93198287 | 1.631751706 | 0.149000079 | 0.991835066 | -4.356135305 |
| **A_55_P2169729** | NA | 0.176946955 | -0.082246263 | 0.436140173 | 13.2945095 | 1.631655964 | 0.149020374 | 0.991835066 | -4.356220394 |
| **A_55_P2123101** | Cbarp | -0.127055468 | -0.313233841 | 0.059122905 | 13.89908333 | -1.631072481 | 0.149144118 | 0.991835066 | -4.35673893 |
| **A_55_P2014665** | Racgap1 | -0.100214074 | -0.247069143 | 0.046640994 | 10.37554368 | -1.630981039 | 0.149163519 | 0.991835066 | -4.35682019 |
| **A_55_P1998072** | NA | -0.082014215 | -0.202206116 | 0.038177686 | 8.057005522 | -1.630883965 | 0.149184118 | 0.991835066 | -4.356906453 |
| **A_55_P2076631** | Gypc | -0.140834916 | -0.347287374 | 0.065617543 | 12.97117447 | -1.630419689 | 0.149282674 | 0.991835066 | -4.357319008 |
| **A_52_P104824** | Diaph3 | 0.129094703 | -0.060159362 | 0.318348768 | 9.7027589 | 1.630318056 | 0.149304257 | 0.991835066 | -4.357409315 |
| **A_55_P2165719** | Pcyt1a | -0.130974816 | -0.323012227 | 0.061062594 | 12.18423082 | -1.630088144 | 0.149353092 | 0.991835066 | -4.357613601 |
| **A_55_P1972892** | NA | 0.163853622 | -0.076445977 | 0.404153221 | 13.3720952 | 1.629716764 | 0.149432007 | 0.991835066 | -4.357943573 |
| **A_55_P2090996** | Ebna1bp2 | 0.075526382 | -0.035256546 | 0.186309309 | 12.57120223 | 1.629427246 | 0.149493555 | 0.991835066 | -4.358200798 |
| **A_55_P1985945** | NA | -0.085294402 | -0.210407377 | 0.039818573 | 14.30932361 | -1.629398486 | 0.149499671 | 0.991835066 | -4.35822635 |
| **A_52_P625624** | Stau2 | -0.112945496 | -0.278627786 | 0.052736793 | 9.14773369 | -1.629303399 | 0.149519891 | 0.991835066 | -4.358310827 |
| **A_55_P2029612** | Dhx35 | 0.145942699 | -0.068146144 | 0.360031542 | 10.60167573 | 1.629286547 | 0.149523475 | 0.991835066 | -4.3583258 |
| **A_55_P2015304** | NA | -0.288724761 | -0.712271351 | 0.134821829 | 13.24916898 | -1.62926592 | 0.149527861 | 0.991835066 | -4.358344124 |
| **A_51_P253633** | Mrps9 | -0.093630061 | -0.231002867 | 0.043742745 | 11.2264608 | -1.629009702 | 0.149582362 | 0.991835066 | -4.35857175 |
| **A_55_P2141654** | NA | 0.085060941 | -0.039752572 | 0.209874454 | 15.85880804 | 1.62883731 | 0.149619042 | 0.991835066 | -4.358724898 |
| **A_51_P173086** | Eif3l | 0.104335494 | -0.048763315 | 0.257434303 | 14.60986759 | 1.628806136 | 0.149625675 | 0.991835066 | -4.358752592 |
| **A_55_P1996872** | Polr2k | 0.118667238 | -0.055465969 | 0.292800444 | 12.47824506 | 1.628764777 | 0.149634477 | 0.991835066 | -4.358789333 |
| **A_55_P2005838** | Ticrr | 0.136242126 | -0.063690559 | 0.33617481 | 9.244943991 | 1.628683682 | 0.149651736 | 0.991835066 | -4.358861375 |
| **A_51_P111259** | Cdk2ap2 | 0.109853918 | -0.051393162 | 0.271100998 | 11.04252847 | 1.628294012 | 0.149734695 | 0.991835066 | -4.359207525 |
| **A_55_P1969356** | Ccm2l | 0.105151195 | -0.049207105 | 0.259509495 | 9.289754678 | 1.62814605 | 0.149766207 | 0.991835066 | -4.359338958 |
| **A_55_P2093418** | Zmpste24 | 0.159112461 | -0.074497545 | 0.392722468 | 11.06866128 | 1.627878116 | 0.149823285 | 0.991835066 | -4.359576952 |
| **A_55_P1984961** | NA | -0.11244573 | -0.277569082 | 0.052677622 | 11.46626659 | -1.627584726 | 0.14988581 | 0.991835066 | -4.359837547 |
| **A_55_P2029471** | NA | 0.115511683 | -0.054114602 | 0.285137968 | 13.49236488 | 1.62757831 | 0.149887178 | 0.991835066 | -4.359843245 |
| **A_66_P100620** | Ensa | -0.133528768 | -0.329663053 | 0.062605516 | 11.45509442 | -1.627160753 | 0.149976207 | 0.991835066 | -4.360214109 |
| **A_55_P1954965** | Npm1 | 0.137440785 | -0.064467702 | 0.339349271 | 14.27075011 | 1.626934936 | 0.150024375 | 0.991835066 | -4.360414664 |
| **A_55_P2151071** | Cox7a2l | -0.100612359 | -0.248434919 | 0.047210201 | 15.60460899 | -1.626745996 | 0.150064688 | 0.991835066 | -4.360582462 |
| **A_51_P130095** | Fcgr2b | -0.067369028 | -0.166390032 | 0.031651976 | 7.746883704 | -1.626080678 | 0.150206726 | 0.991835066 | -4.359340654 |
| **A_55_P1998962** | Oas1b | -0.099351036 | -0.245484781 | 0.046782709 | 13.34168703 | -1.624916379 | 0.150455594 | 0.991835066 | -4.362207112 |
| **A_55_P2153541** | Sik3 | -0.120144192 | -0.296870563 | 0.056582178 | 12.13837288 | -1.624839827 | 0.15047197 | 0.991835066 | -4.362275078 |
| **A_55_P2029051** | NA | -0.082018572 | -0.202665603 | 0.03862846 | 8.008329279 | -1.624817894 | 0.150476663 | 0.991835066 | -4.362294551 |
| **A_55_P2172334** | Nup155 | 0.162096053 | -0.076344811 | 0.400536917 | 10.30278719 | 1.62480365 | 0.15047971 | 0.991835066 | -4.362307197 |
| **A_55_P2270412** | C230096K1 | 0.283341724 | -0.133479049 | 0.700162496 | 9.865800423 | 1.624689261 | 0.150504185 | 0.991835066 | -4.362408755 |
| **A_66_P140613** | NA | 0.168590426 | -0.07943839 | 0.416619243 | 9.222671105 | 1.624575503 | 0.150528529 | 0.991835066 | -4.362509752 |
| **A_55_P1982227** | NA | 0.410504442 | -0.193473125 | 1.014482009 | 12.32856845 | 1.624449388 | 0.150555522 | 0.991835066 | -4.362621716 |
| **A_55_P2050857** | D11Wsu47 | 0.115235697 | -0.054334934 | 0.284806328 | 10.16911331 | 1.624222524 | 0.150604089 | 0.991835066 | -4.362823121 |
| **A_52_P639229** | Fgd4 | -0.096036271 | -0.237364104 | 0.045291563 | 8.665319033 | -1.624114778 | 0.150627161 | 0.991835066 | -4.362918773 |
| **A_65_P11718** | Nol8 | -0.088633691 | -0.219081172 | 0.04181379 | 8.628055473 | -1.623948574 | 0.150662757 | 0.991835066 | -4.363066318 |
| **A_52_P599264** | Mdfic | 0.07388406 | -0.034858878 | 0.182626999 | 9.747064985 | 1.623898283 | 0.150673529 | 0.991835066 | -4.363110963 |
| **A_55_P1958447** | Prepl | -0.098253714 | -0.24287214 | 0.046364712 | 8.586453984 | -1.623807252 | 0.15069303 | 0.991835066 | -4.363191772 |
| **A_55_P2187261** | NA | 0.094940902 | -0.044825232 | 0.234707036 | 8.533843786 | 1.623530808 | 0.150752264 | 0.991835066 | -4.363437168 |
| **A_52_P655803** | Chpt1 | -0.128920162 | -0.318760834 | 0.060920509 | 9.047669611 | -1.623082943 | 0.150848276 | 0.991835066 | -4.36383471 |
| **A_51_P465740** | Lysmd4 | -0.069943821 | -0.172943806 | 0.033056165 | 9.406784516 | -1.623010469 | 0.150863818 | 0.991835066 | -4.36389904 |
| **A_55_P2024366** | Raver2 | 0.092698324 | -0.043810288 | 0.229206937 | 9.334584328 | 1.623009132 | 0.150864105 | 0.991835066 | -4.363900226 |

| | | | | | | | | | |
|---|---|---|---|---|---|---|---|---|---|
| **A_55_P2047986** | Ankrd23 | -0.136637253 | -0.337900935 | 0.06462643 | 9.103167561 | -1.622605107 | 0.150950777 | 0.991835066 | -4.364258828 |
| **A_55_P2102698** | NA | 0.095519656 | -0.045183235 | 0.236222548 | 13.14090013 | 1.622552884 | 0.150961983 | 0.991835066 | -4.364305178 |
| **A_55_P2005585** | Trps1 | -0.092128633 | -0.227843114 | 0.043585848 | 8.917041291 | -1.622473342 | 0.150979053 | 0.991835066 | -4.364375775 |
| **A_55_P2006579** | Rbbp6 | 0.103513476 | -0.049003104 | 0.256030057 | 9.633711785 | 1.622142361 | 0.151050103 | 0.991835066 | -4.364669523 |
| **A_55_P2060793** | NA | -0.2649931 | -0.655455997 | 0.125469796 | 13.32061452 | -1.622048841 | 0.151070184 | 0.991835066 | -4.364752521 |
| **A_51_P377526** | Sts | -0.140706739 | -0.348059646 | 0.066646168 | 9.633748439 | -1.621862014 | 0.151110307 | 0.991835066 | -4.364918322 |
| **A_52_P469956** | Zc3h6 | -0.10527204 | -0.260418361 | 0.049874282 | 9.338830501 | -1.62173798 | 0.151136951 | 0.991835066 | -4.365028395 |
| **A_55_P2010038** | Tnfsf9 | -0.145153083 | -0.359074731 | 0.068768564 | 8.395530527 | -1.621737888 | 0.151136971 | 0.991835066 | -4.365028477 |
| **A_52_P59206** | Cst6 | -0.081148622 | -0.200768752 | 0.038471509 | 8.067793011 | -1.621384472 | 0.151212912 | 0.991835066 | -4.365342102 |
| **A_52_P652316** | Mapre2 | -0.09208032 | -0.227921415 | 0.043760775 | 11.54414329 | -1.620111036 | 0.151486843 | 0.991835066 | -4.366472022 |
| **A_55_P2049647** | H2-K1 | -0.13127505 | -0.324939966 | 0.062389866 | 9.388554131 | -1.620094597 | 0.151490382 | 0.991835066 | -4.366486607 |
| **A_51_P483329** | Sdad1 | 0.114160174 | -0.054303261 | 0.282623608 | 11.02613573 | 1.619638542 | 0.151588601 | 0.991835066 | -4.366891212 |
| **A_51_P104897** | Itpr3 | 0.128713933 | -0.06123574 | 0.318663605 | 10.65183987 | 1.619556646 | 0.151606245 | 0.991835066 | -4.366963865 |
| **A_55_P2050902** | Fetub | -0.095597375 | -0.236741865 | 0.045547115 | 7.812501815 | -1.618792451 | 0.151770978 | 0.991835066 | -4.367641776 |
| **A_51_P385952** | Man2a2 | 0.086590162 | -0.041268528 | 0.214448853 | 9.210491181 | 1.618629438 | 0.15180614 | 0.991835066 | -4.367786373 |
| **A_55_P2033255** | NA | -0.093069917 | -0.230509391 | 0.044369558 | 14.894333 | -1.618478642 | 0.151838673 | 0.991835066 | -4.36792013 |
| **A_55_P2122987** | Dlx3 | -0.149433869 | -0.37014005 | 0.071272312 | 11.49136051 | -1.618242776 | 0.151889572 | 0.991835066 | -4.368129339 |
| **A_52_P281209** | Kdm4a | 0.097889219 | -0.046700443 | 0.242478881 | 10.8329592 | 1.618105201 | 0.151919268 | 0.991835066 | -4.368251362 |
| **A_55_P1996456** | Srrm1 | 0.079240924 | -0.037804768 | 0.196286616 | 9.258769795 | 1.618092073 | 0.151922102 | 0.991835066 | -4.368263005 |
| **A_51_P161354** | Sesn2 | 0.107290588 | -0.05119982 | 0.265780996 | 9.400695986 | 1.617959977 | 0.151950621 | 0.991835066 | -4.368380167 |
| **A_55_P2137521** | Nmral1 | 0.081714722 | -0.039012385 | 0.202441829 | 8.603650911 | 1.617724815 | 0.152001404 | 0.991835066 | -4.368588735 |
| **A_51_P224311** | Popdc2 | -0.128972291 | -0.319527518 | 0.061582936 | 14.53385303 | -1.617650447 | 0.152017467 | 0.991835066 | -4.36865469 |
| **A_55_P1953980** | Sdhb | -0.138440834 | -0.343028408 | 0.066146741 | 14.67864535 | -1.617313027 | 0.152090367 | 0.991835066 | -4.368953936 |
| **A_55_P2094352** | Zfp523 | 0.164134522 | -0.078461881 | 0.406730924 | 8.451814008 | 1.617054693 | 0.152146203 | 0.991835066 | -4.369183033 |
| **A_51_P214197** | Stk17b | -0.08594071 | -0.212968569 | 0.041087149 | 9.306072697 | -1.616996548 | 0.152158773 | 0.991835066 | -4.369234596 |
| **A_52_P672453** | Prpf3 | 0.086496595 | -0.041362602 | 0.214355792 | 10.58097143 | 1.616873973 | 0.152185275 | 0.991835066 | -4.369343295 |
| **A_55_P2051313** | Gstk1 | -0.209740692 | -0.519823932 | 0.100342549 | 10.9033178 | -1.616640798 | 0.152235702 | 0.991835066 | -4.369550066 |
| **A_55_P1974098** | Gm12063 | -0.117324967 | -0.290781344 | 0.056131409 | 10.23050993 | -1.616625053 | 0.152239107 | 0.991835066 | -4.369564027 |
| **A_55_P2054362** | NA | -0.145539635 | -0.360738721 | 0.069659451 | 9.574656493 | -1.616404281 | 0.152286867 | 0.991835066 | -4.369759794 |
| **A_51_P348325** | Poc1b | 0.123807729 | -0.059269296 | 0.306884755 | 9.822136422 | 1.61630398 | 0.15230857 | 0.991835066 | -4.369848732 |
| **A_55_P2069721** | Hspa8 | 0.105832382 | -0.050680595 | 0.262345359 | 15.25407756 | 1.616133936 | 0.152345371 | 0.991835066 | -4.369999509 |
| **A_51_P387220** | Ddx1 | -0.083512508 | -0.207024619 | 0.039999602 | 11.28273966 | -1.616036319 | 0.1523665 | 0.991835066 | -4.370086062 |
| **A_51_P205215** | Klhl30 | -0.115354252 | -0.286000984 | 0.055292481 | 12.78612547 | -1.61564061 | 0.152452182 | 0.991835066 | -4.370436915 |
| **A_51_P153486** | Dnajb1 | 0.112588304 | -0.053972049 | 0.279148658 | 13.1358996 | 1.615588592 | 0.152463449 | 0.991835066 | -4.370483034 |
| **A_52_P588881** | Iqgap3 | -0.104589574 | -0.259346025 | 0.050166877 | 10.12088724 | -1.615283526 | 0.152529539 | 0.991835066 | -4.370753502 |
| **A_51_P412914** | Efs | 0.104160675 | -0.04999252 | 0.258313869 | 10.73186591 | 1.614954859 | 0.152600773 | 0.991835066 | -4.371044879 |
| **A_55_P2068673** | Stmn2 | 0.086422805 | -0.041480818 | 0.214326429 | 8.233050887 | 1.614933507 | 0.152605402 | 0.991835066 | -4.371063809 |
| **A_55_P1986998** | Enoph1 | 0.153107326 | -0.073536971 | 0.379751624 | 10.77343544 | 1.614582752 | 0.152681459 | 0.991835066 | -4.371374751 |
| **A_55_P2076639** | Vps25 | -0.166854952 | -0.413860367 | 0.080150464 | 9.473110236 | -1.614513782 | 0.152696419 | 0.991835066 | -4.371435891 |
| **A_55_P1958245** | Tcof1 | 0.137835897 | -0.066213068 | 0.341884861 | 10.61605969 | 1.61449637 | 0.152700195 | 0.991835066 | -4.371451326 |
| **A_55_P2050226** | Ackr4 | -0.113798134 | -0.282306068 | 0.054709801 | 8.743512593 | -1.614075762 | 0.152791457 | 0.991835066 | -4.371824165 |
| **A_55_P2130975** | Ncoa1 | -0.09315876 | -0.231105186 | 0.044787666 | 11.38579323 | -1.614070053 | 0.152792696 | 0.991835066 | -4.371829226 |
| **A_55_P2050988** | NA | 0.158480039 | -0.076219749 | 0.393179826 | 9.380099436 | 1.613879124 | 0.152834141 | 0.991835066 | -4.371998464 |
| **A_66_P122158** | Pisd-ps3 | 0.077346191 | -0.037222873 | 0.191915256 | 7.810396314 | 1.613543577 | 0.152907002 | 0.991835066 | -4.372295876 |
| **A_52_P614731** | Gng12 | -0.126207873 | -0.313160157 | 0.060744412 | 9.685548966 | -1.613484515 | 0.15291983 | 0.991835066 | -4.372348224 |
| **A_55_P1991985** | Ptprs | 0.130416515 | -0.06280607 | 0.3236391 | 13.5258928 | 1.61318374 | 0.152985174 | 0.991835066 | -4.372614801 |
| **A_51_P389895** | Slc35f5 | -0.101049204 | -0.250763845 | 0.048665437 | 11.65820185 | -1.613161152 | 0.152990082 | 0.991835066 | -4.372634821 |
| **A_55_P1964347** | NA | 0.098202403 | -0.047370285 | 0.243775091 | 8.695201227 | 1.612320397 | 0.153172882 | 0.991835066 | -4.373379915 |

| | | | | | | | | | |
|---|---|---|---|---|---|---|---|---|---|
| **A_55_P2015430** | Gpn2 | 0.094413832 | -0.045555307 | 0.234382972 | 9.648382888 | 1.612176061 | 0.153204285 | 0.991835066 | -4.373507818 |
| **A_55_P1957319** | Krr1 | 0.077590955 | -0.037476416 | 0.192658326 | 9.943063897 | 1.611640004 | 0.153320966 | 0.991835066 | -4.373982821 |
| **A_55_P2041894** | NA | 0.087745917 | -0.042385967 | 0.217877801 | 16.45695269 | 1.61158171 | 0.15333366 | 0.991835066 | -4.374034473 |
| **A_55_P2144526** | Ripor2 | 0.185868798 | -0.089789531 | 0.461527128 | 8.788718004 | 1.611552668 | 0.153339984 | 0.991835066 | -4.374060206 |
| **A_55_P2010966** | Zfp688 | 0.081694128 | -0.03947406 | 0.202862315 | 9.213091263 | 1.611429689 | 0.153366767 | 0.991835066 | -4.374169172 |
| **A_55_P2062598** | E2f7 | 0.118174252 | -0.057131666 | 0.29348017 | 10.32883212 | 1.611147924 | 0.153428148 | 0.991835066 | -4.374418821 |
| **A_51_P393897** | Mpi | -0.089454194 | -0.22216134 | 0.043252951 | 10.95760652 | -1.611074165 | 0.15344422 | 0.991835066 | -4.374484171 |
| **A_51_P342906** | Sh3bp1 | 0.139847951 | -0.067624491 | 0.347320392 | 8.674725499 | 1.611034428 | 0.153452879 | 0.991835066 | -4.374519378 |
| **A_51_P481221** | Bace2 | 0.138468232 | -0.06700597 | 0.343942434 | 9.033530645 | 1.610652965 | 0.153536028 | 0.991835066 | -4.374857338 |
| **A_55_P1959485** | NA | -0.102323286 | -0.254167412 | 0.04952084 | 12.10707716 | -1.610592282 | 0.153549259 | 0.991835066 | -4.374911099 |
| **A_55_P2144364** | NA | 0.105823598 | -0.051263851 | 0.262911047 | 8.02365087 | 1.610090065 | 0.153658803 | 0.991835066 | -4.375356008 |
| **A_55_P2121846** | NA | -0.080476989 | -0.199940053 | 0.038986074 | 15.68082826 | -1.610079072 | 0.153661202 | 0.991835066 | -4.375365746 |
| **A_66_P109296** | NA | -0.185845056 | -0.461743881 | 0.09005377 | 10.67771941 | -1.609942231 | 0.153691063 | 0.991835066 | -4.375486966 |
| **A_55_P2113210** | Cyth4 | -0.142575577 | -0.354281374 | 0.06913022 | 8.699882461 | -1.609613185 | 0.153762888 | 0.991835066 | -4.375778439 |
| **A_55_P2014326** | NA | 0.095742815 | -0.04642703 | 0.23791266 | 9.368967174 | 1.609562463 | 0.153773963 | 0.991835066 | -4.375823368 |
| **A_55_P2094292** | Erp27 | -0.098029873 | -0.243626847 | 0.0475671 | 9.265554077 | -1.609219275 | 0.153848914 | 0.991835066 | -4.376127349 |
| **A_55_P2093535** | Rps8 | 0.102973324 | -0.049979811 | 0.25592646 | 16.48279154 | 1.609072011 | 0.153881087 | 0.991835066 | -4.376257785 |
| **A_55_P1982404** | Gpm6b | -0.106420728 | -0.264517632 | 0.051676176 | 9.502097913 | -1.608836804 | 0.153932485 | 0.991835066 | -4.376466108 |
| **A_55_P2011967** | Akirin2 | -0.108706096 | -0.270225859 | 0.052813666 | 12.2115709 | -1.608560372 | 0.153992912 | 0.991835066 | -4.376710933 |
| **A_51_P321374** | Plpp7 | -0.187842147 | -0.466953472 | 0.091269178 | 9.954502299 | -1.608513523 | 0.154003156 | 0.991835066 | -4.376752424 |
| **A_55_P1985422** | NA | 0.426393158 | -0.207324137 | 1.060110452 | 12.6474566 | 1.608139798 | 0.154084892 | 0.991835066 | -4.377083399 |
| **A_55_P2028710** | Rnf170 | 0.121418693 | -0.059139675 | 0.30197706 | 10.28706098 | 1.607226429 | 0.154284823 | 0.991835066 | -4.377892207 |
| **A_51_P450296** | Thap3 | 0.093918528 | -0.045774259 | 0.233611316 | 8.805718519 | 1.606891035 | 0.1543583 | 0.991835066 | -4.378189177 |
| **A_55_P2170515** | NA | -0.066492366 | -0.165426951 | 0.03244222 | 7.726518964 | -1.606322637 | 0.154482897 | 0.991835066 | -4.37869242 |
| **A_55_P2404484** | 1700001L0 | 0.156704809 | -0.07646229 | 0.389871908 | 8.118021105 | 1.606290855 | 0.154489867 | 0.991835066 | -4.378720558 |
| **A_52_P578043** | NA | -0.182636415 | -0.454491827 | 0.089218997 | 9.802012118 | -1.605678246 | 0.154624267 | 0.991835066 | -4.379262891 |
| **A_55_P2011061** | Tspan12 | -0.102266315 | -0.256925942 | 0.052393311 | 11.30420225 | -1.642565895 | 0.15463311 | 0.991835066 | -4.341276675 |
| **A_52_P139747** | Ndufa1 | -0.224875261 | -0.559617853 | 0.109867331 | 15.92045892 | -1.605609162 | 0.15463943 | 0.991835066 | -4.379324047 |
| **A_55_P1953894** | Mccc2 | -0.083496387 | -0.207807679 | 0.040814905 | 9.879579807 | -1.605337072 | 0.154699164 | 0.991835066 | -4.379564904 |
| **A_55_P2153570** | Zfp655 | -0.149846328 | -0.373003188 | 0.073310532 | 8.192868972 | -1.604888977 | 0.154797586 | 0.991835066 | -4.379961541 |
| **A_55_P2132902** | Gprasp1 | 0.119066026 | -0.058284789 | 0.296416842 | 11.12243453 | 1.604588973 | 0.154863513 | 0.991835066 | -4.380227078 |
| **A_55_P2095000** | NA | 0.115926068 | -0.056790581 | 0.288642716 | 16.15251789 | 1.604190848 | 0.154951043 | 0.991835066 | -4.380579442 |
| **A_55_P1956843** | NA | 0.106208799 | -0.052084303 | 0.2645019 | 13.81225672 | 1.603642804 | 0.15507161 | 0.991835066 | -4.381064458 |
| **A_52_P588539** | Snapin | -0.130711779 | -0.325563779 | 0.06414022 | 8.808256224 | -1.603315504 | 0.155143656 | 0.991835066 | -4.381354097 |
| **A_51_P497317** | Tcp11l2 | -0.181834378 | -0.452995259 | 0.089326502 | 10.1453215 | -1.602721611 | 0.155274465 | 0.991835066 | -4.381879613 |
| **A_55_P2068882** | Ccng2 | -0.142639835 | -0.355394863 | 0.070115193 | 8.517208158 | -1.602397019 | 0.155346002 | 0.991835066 | -4.382166813 |
| **A_55_P1986660** | Anapc2 | 0.132386079 | -0.065077722 | 0.32984988 | 13.0918523 | 1.602374256 | 0.15535102 | 0.991835066 | -4.382186953 |
| **A_55_P1994477** | Prrc2c | 0.110473274 | -0.054384251 | 0.275330799 | 9.236105878 | 1.601612982 | 0.155518925 | 0.991835066 | -4.382860469 |
| **A_51_P205385** | Uox | -0.105624447 | -0.263270005 | 0.05202111 | 7.847465452 | -1.601370577 | 0.155572425 | 0.991835066 | -4.383074913 |
| **A_52_P599238** | Rspry1 | -0.111849047 | -0.278803525 | 0.055105432 | 9.314515116 | -1.601191759 | 0.155611902 | 0.991835066 | -4.383233099 |
| **A_55_P1982628** | Tma16 | 0.153314138 | -0.075567566 | 0.382195841 | 9.50765188 | 1.600959186 | 0.155663261 | 0.991835066 | -4.383438832 |
| **A_51_P220723** | B230118H( | -0.082397968 | -0.205426393 | 0.040630457 | 13.52802904 | -1.600737695 | 0.155712187 | 0.991835066 | -4.383634753 |
| **A_55_P2016968** | NA | -0.075845057 | -0.189105707 | 0.037415592 | 15.531185 | -1.600506228 | 0.155763332 | 0.991835066 | -4.383839492 |
| **A_55_P2403453** | BC037032 | -0.121477755 | -0.302887233 | 0.059931724 | 9.530546985 | -1.600464016 | 0.15577266 | 0.991835066 | -4.383876829 |
| **A_51_P333949** | Ints7 | 0.08375536 | -0.041325193 | 0.208835913 | 8.813974839 | 1.600412552 | 0.155784035 | 0.991835066 | -4.383922349 |
| **A_55_P1983868** | NA | -0.243873799 | -0.60811081 | 0.120363212 | 10.64793164 | -1.600258722 | 0.155818038 | 0.991835066 | -4.38405841 |
| **A_51_P155152** | Ank | -0.095563685 | -0.238311244 | 0.047183873 | 10.51376452 | -1.600049182 | 0.155864367 | 0.991835066 | -4.384243739 |
| **A_51_P297105** | Ucp2 | -0.078063355 | -0.194673295 | 0.038546585 | 9.058194394 | -1.600002913 | 0.155874599 | 0.991835066 | -4.384284662 |

| | | | | | | | | | |
|---|---|---|---|---|---|---|---|---|---|
| **A_55_P1973868** | Sema3b | 0.118630988 | -0.05859148 | 0.295853456 | 11.1082332 | 1.599884015 | 0.155900894 | 0.991835066 | -4.384389819 |
| **A_55_P1996087** | Gm14326 | 0.123589149 | -0.061044621 | 0.308222919 | 11.13712984 | 1.59984658 | 0.155909174 | 0.991835066 | -4.384422927 |
| **A_51_P239984** | Exo1 | 0.211582678 | -0.104510662 | 0.527676017 | 9.903178755 | 1.59983022 | 0.155912793 | 0.991835066 | -4.384437396 |
| **A_55_P1957098** | Mtfr1l | 0.110111676 | -0.054408352 | 0.274631705 | 12.85129404 | 1.599645432 | 0.155953672 | 0.991835066 | -4.384600823 |
| **A_55_P2025463** | Brpf3 | 0.159281912 | -0.078709752 | 0.397273576 | 9.164803838 | 1.599609034 | 0.155961725 | 0.991835066 | -4.384633013 |
| **A_55_P2077497** | NA | 0.512949289 | -0.254072755 | 1.279971333 | 12.16543986 | 1.598364109 | 0.156237402 | 0.991835066 | -4.385733893 |
| **A_66_P121495** | Psat1 | 0.106125398 | -0.052566692 | 0.264817488 | 8.245310548 | 1.598354785 | 0.156239469 | 0.991835066 | -4.385742137 |
| **A_66_P121059** | Mirg | 0.209399071 | -0.103795611 | 0.522593754 | 9.098740461 | 1.597973246 | 0.156324049 | 0.991835066 | -4.386079484 |
| **A_55_P2052824** | Lrrc14b | -0.154932265 | -0.386700995 | 0.076836465 | 11.06705843 | -1.597703404 | 0.156383895 | 0.991835066 | -4.38631806 |
| **A_55_P2015113** | Atg5 | -0.075892356 | -0.18943589 | 0.037651178 | 10.19301849 | -1.597514324 | 0.156425842 | 0.991835066 | -4.386485224 |
| **A_66_P101620** | Cfap97 | 0.111032966 | -0.055087601 | 0.277153532 | 9.032136363 | 1.597488235 | 0.15643163 | 0.991835066 | -4.38650829 |
| **A_55_P2061268** | Rnpc3 | 0.130206828 | -0.064638145 | 0.325051802 | 8.363582703 | 1.597179353 | 0.15650018 | 0.991835066 | -4.386781359 |
| **A_52_P483129** | Utp23 | 0.119021962 | -0.059140155 | 0.297184078 | 9.180980375 | 1.596690981 | 0.15660862 | 0.991835066 | -4.387213079 |
| **A_51_P363914** | Cct2 | 0.110794089 | -0.055067838 | 0.276656016 | 13.62340546 | 1.596537109 | 0.156642801 | 0.991835066 | -4.387349094 |
| **A_55_P2009449** | Pnma2 | 0.309214477 | -0.153755988 | 0.772184942 | 9.440567653 | 1.596304231 | 0.156694546 | 0.991835066 | -4.387554941 |
| **A_55_P2044197** | Vps29 | 0.071494324 | -0.03555036 | 0.178539009 | 11.90824746 | 1.59630396 | 0.156694606 | 0.991835066 | -4.387555181 |
| **A_55_P1983683** | Rtl5 | 0.091800045 | -0.045664311 | 0.229264401 | 8.404308728 | 1.596106712 | 0.156738447 | 0.991835066 | -4.387729528 |
| **A_55_P1994000** | Snx16 | 0.104004316 | -0.051760123 | 0.259768755 | 9.69198919 | 1.59585041 | 0.15679543 | 0.991835066 | -4.387956062 |
| **A_55_P1988113** | Cyth2 | -0.127926288 | -0.319550877 | 0.0636983 | 10.82554526 | -1.595576736 | 0.156856296 | 0.991835066 | -4.388197942 |
| **A_55_P2058871** | Otud5 | 0.103179627 | -0.051414872 | 0.257774126 | 11.17403796 | 1.595177626 | 0.1569451 | 0.991835066 | -4.388550665 |
| **A_55_P1984496** | Usp48 | 0.191402609 | -0.095635204 | 0.478440422 | 9.954431658 | 1.593741472 | 0.157265041 | 0.991835066 | -4.38981971 |
| **A_55_P1986351** | Tbc1d32 | 0.11276979 | -0.056346142 | 0.281885722 | 8.540405912 | 1.593739663 | 0.157265445 | 0.991835066 | -4.389821307 |
| **A_55_P2101757** | Dscc1 | 0.114081079 | -0.057046785 | 0.285208943 | 8.97528425 | 1.593316385 | 0.157359858 | 0.991835066 | -4.390195276 |
| **A_52_P242251** | Utp25 | 0.073946297 | -0.036982884 | 0.184875479 | 8.614755249 | 1.5932347 | 0.157378083 | 0.991835066 | -4.390267442 |
| **A_55_P2109263** | Ppp1r14b | 0.122782727 | -0.061411096 | 0.30697655 | 13.87046638 | 1.593203846 | 0.157384968 | 0.991835066 | -4.390294701 |
| **A_55_P2103145** | Ppp1r2 | -0.141174914 | -0.353037699 | 0.070687872 | 9.203856674 | -1.59261934 | 0.15751545 | 0.991835066 | -4.390811063 |
| **A_55_P1991505** | Gamt | -0.100849263 | -0.25219874 | 0.050500214 | 10.95768962 | -1.592578818 | 0.1575245 | 0.991835066 | -4.390846859 |
| **A_55_P2188464** | Uhrf1bp1l | 0.083184172 | -0.041668003 | 0.208036346 | 10.70644297 | 1.592405678 | 0.157563172 | 0.991835066 | -4.390999802 |
| **A_51_P234140** | Cdkl1 | 0.075767538 | -0.037964415 | 0.189499491 | 9.164020668 | 1.592244698 | 0.157599136 | 0.991835066 | -4.391142001 |
| **A_52_P89717** | NA | -0.081491261 | -0.203846733 | 0.04086421 | 8.361436944 | -1.591830337 | 0.157691743 | 0.991835066 | -4.391508001 |
| **A_52_P354373** | Lockd | -0.17073541 | -0.427096413 | 0.085625593 | 11.67191934 | -1.591771671 | 0.157704858 | 0.991835066 | -4.391559818 |
| **A_55_P2266295** | Fzd3 | 0.080772428 | -0.040511693 | 0.202056549 | 8.322330139 | 1.591726046 | 0.157715059 | 0.991835066 | -4.391600116 |
| **A_55_P2046374** | NA | 0.080876903 | -0.040577466 | 0.202331272 | 8.43706413 | 1.591550785 | 0.15775425 | 0.991835066 | -4.391754912 |
| **A_55_P2030736** | Scmh1 | -0.120311303 | -0.300996562 | 0.060373956 | 9.462754433 | -1.591449424 | 0.157776919 | 0.991835066 | -4.391844435 |
| **A_52_P13815** | Laptm5 | -0.090765282 | -0.227084176 | 0.045553611 | 8.106890461 | -1.591376147 | 0.15779331 | 0.991835066 | -4.391909153 |
| **A_52_P162099** | Ckap2 | -0.131097426 | -0.328030965 | 0.065836113 | 9.398104237 | -1.591049211 | 0.157866458 | 0.991835066 | -4.392197892 |
| **A_55_P1954820** | D1Ertd622e | 0.137389935 | -0.06899728 | 0.343777151 | 10.00627102 | 1.591040592 | 0.157868386 | 0.991835066 | -4.392205503 |
| **A_52_P473975** | Cpsf4 | 0.080387607 | -0.04037808 | 0.201153294 | 9.783937066 | 1.590943208 | 0.157890181 | 0.991835066 | -4.392291506 |
| **A_52_P219266** | Frmd8 | 0.088272729 | -0.044350298 | 0.220895756 | 11.49700231 | 1.590804281 | 0.157921278 | 0.991835066 | -4.392414196 |
| **A_55_P2091751** | NA | -0.17259342 | -0.431919852 | 0.086733011 | 12.72458102 | -1.590693781 | 0.157946017 | 0.991835066 | -4.392511778 |
| **A_55_P2075294** | NA | -0.084866063 | -0.212390231 | 0.042658105 | 8.120004409 | -1.590562319 | 0.157975452 | 0.991835066 | -4.39262787 |
| **A_51_P502119** | F11 | -0.115510253 | -0.289123538 | 0.058103033 | 7.8093843 | -1.590181553 | 0.158060739 | 0.991835066 | -4.392964103 |
| **A_55_P2073308** | Bin1 | 0.096224126 | -0.048405809 | 0.240854062 | 10.456351 | 1.590138367 | 0.158070415 | 0.991835066 | -4.393002236 |
| **A_51_P141535** | Oat | -0.140580204 | -0.351881668 | 0.070721261 | 12.14073799 | -1.590123273 | 0.158073797 | 0.991835066 | -4.393015564 |
| **A_55_P2153937** | NA | -0.084465097 | -0.211454582 | 0.042524388 | 8.499285791 | -1.589712743 | 0.158165806 | 0.991835066 | -4.393378053 |
| **A_55_P2048164** | Msantd3 | 0.087996535 | -0.044323637 | 0.220316708 | 9.792757854 | 1.589456497 | 0.158223261 | 0.991835066 | -4.3936043 |
| **A_55_P1968028** | Tdgf1 | 0.167908896 | -0.084618326 | 0.420436117 | 8.497041799 | 1.589185659 | 0.15828401 | 0.991835066 | -4.393843419 |
| **A_55_P2018949** | Fbxo7 | 0.115917751 | -0.058419969 | 0.290255472 | 11.37393295 | 1.589160332 | 0.158289692 | 0.991835066 | -4.393865779 |

| | | | | | | | | | |
|---|---|---|---|---|---|---|---|---|---|
| **A_55_P2065819** | Fbxo22 | 0.11004459 | -0.055499419 | 0.2755886 | 11.24803144 | 1.588782189 | 0.158374549 | 0.991835066 | -4.394199618 |
| **A_55_P1970062** | Igsf9b | -0.212873835 | -0.533164299 | 0.107416629 | 9.764133604 | -1.588500693 | 0.158437745 | 0.991835066 | -4.39444812 |
| **A_55_P1972182** | Siah1b | 0.147100082 | -0.074257756 | 0.368457919 | 10.64300954 | 1.588280276 | 0.158487245 | 0.991835066 | -4.394642693 |
| **A_52_P572045** | Pfkp | 0.12908883 | -0.065176616 | 0.323354276 | 8.979483749 | 1.588189199 | 0.158507703 | 0.991835066 | -4.394723089 |
| **A_52_P142191** | Aph1b | -0.085594583 | -0.21444545 | 0.043256283 | 8.282439775 | -1.587698636 | 0.158617937 | 0.991835066 | -4.395156101 |
| **A_55_P2120919** | NA | 0.067466898 | -0.034098932 | 0.169032728 | 7.930577417 | 1.587640812 | 0.158630935 | 0.991835066 | -4.395207139 |
| **A_52_P253915** | Cds2 | -0.153733193 | -0.385174764 | 0.077708378 | 9.305487745 | -1.587579242 | 0.158644776 | 0.991835066 | -4.395261483 |
| **A_51_P336874** | Atp6v0e | -0.084146028 | -0.210831308 | 0.042539253 | 13.89843101 | -1.587510455 | 0.158660242 | 0.991835066 | -4.395322197 |
| **A_55_P2062148** | Arl1 | 0.079806833 | -0.040352689 | 0.199966355 | 10.58797664 | 1.587416968 | 0.158681262 | 0.991835066 | -4.395404709 |
| **A_55_P2008181** | Il3ra | 0.197175167 | -0.099749778 | 0.494100112 | 9.808005522 | 1.587137908 | 0.158744025 | 0.991835066 | -4.395651004 |
| **A_52_P578562** | Slc41a1 | -0.085898342 | -0.215257534 | 0.04346085 | 8.75475904 | -1.587071966 | 0.158758859 | 0.991835066 | -4.395709202 |
| **A_52_P162957** | Frat2 | -0.108334163 | -0.271558685 | 0.054890359 | 8.525102429 | -1.586314003 | 0.158929463 | 0.991835066 | -4.396378103 |
| **A_51_P513163** | Tra2a | 0.180391408 | -0.091429571 | 0.452212388 | 12.33184127 | 1.586141791 | 0.158968248 | 0.991835066 | -4.396530069 |
| **A_55_P2086885** | Slc45a4 | 0.085162978 | -0.043178054 | 0.21350401 | 9.473826875 | 1.58596809 | 0.159007378 | 0.991835066 | -4.396683343 |
| **A_52_P390114** | Eif4b | 0.093755399 | -0.047546937 | 0.235057735 | 8.468611499 | 1.585827988 | 0.159038946 | 0.991835066 | -4.396806966 |
| **A_55_P2036633** | Ssrp1 | 0.159792257 | -0.081053172 | 0.400637686 | 13.07787168 | 1.585719824 | 0.159063321 | 0.991835066 | -4.396902406 |
| **A_55_P2408848** | Lgr4 | -0.086878507 | -0.217851784 | 0.044094769 | 8.435009749 | -1.58539977 | 0.159135467 | 0.991835066 | -4.397184799 |
| **A_55_P2021886** | Mrpl2 | -0.075636826 | -0.189679572 | 0.038405921 | 13.9934333 | -1.585166041 | 0.159188174 | 0.991835066 | -4.397391016 |
| **A_51_P298190** | Sbf2 | -0.083100554 | -0.2084016 | 0.042200493 | 10.05248329 | -1.585106157 | 0.15920168 | 0.991835066 | -4.397443849 |
| **A_55_P1996171** | Pcgf5 | -0.169346784 | -0.424711186 | 0.086017619 | 12.16507496 | -1.584987084 | 0.15922854 | 0.991835066 | -4.397548902 |
| **A_52_P360650** | Abcb7 | -0.099891496 | -0.250540884 | 0.050757892 | 8.973304777 | -1.584784733 | 0.159274194 | 0.991835066 | -4.397727422 |
| **A_55_P2006422** | Gm6900 | -0.14166635 | -0.355335099 | 0.0720024 | 15.56736205 | -1.584655372 | 0.159303387 | 0.991835066 | -4.397841544 |
| **A_52_P323820** | Ufm1 | 0.081568128 | -0.041458337 | 0.204594594 | 8.046771032 | 1.584641703 | 0.159306472 | 0.991835066 | -4.397853603 |
| **A_55_P2042723** | Gm2769 | 0.07218196 | -0.036688736 | 0.181052656 | 8.870634209 | 1.584626087 | 0.159309997 | 0.991835066 | -4.397867379 |
| **A_55_P2105271** | Pdlim7 | 0.173646507 | -0.088337701 | 0.435630714 | 11.56200843 | 1.584163746 | 0.159414378 | 0.991835066 | -4.398275236 |
| **A_51_P247637** | Rnf144a | -0.130607376 | -0.327666655 | 0.066451903 | 9.073795846 | -1.584090346 | 0.159430955 | 0.991835066 | -4.398339984 |
| **A_55_P2329923** | Camkk2 | 0.134661653 | -0.068516347 | 0.337839652 | 11.28502205 | 1.584077383 | 0.159433883 | 0.991835066 | -4.398351419 |
| **A_51_P267494** | Cdc42ep3 | 0.147958914 | -0.075291254 | 0.371209082 | 10.81077554 | 1.584012019 | 0.159448647 | 0.991835066 | -4.398409076 |
| **A_52_P451073** | Tnfrsf21 | -0.066501126 | -0.166852633 | 0.033850382 | 8.04898523 | -1.583850648 | 0.159485101 | 0.991835066 | -4.398551419 |
| **A_51_P364786** | U2af1 | 0.104709481 | -0.053309437 | 0.262728399 | 8.875272179 | 1.583747909 | 0.159508315 | 0.991835066 | -4.398642042 |
| **A_51_P492765** | Wdr75 | 0.16299061 | -0.082987856 | 0.408969077 | 10.51286127 | 1.583706246 | 0.159517729 | 0.991835066 | -4.398678791 |
| **A_51_P189269** | Mbnl2 | -0.115716315 | -0.290358707 | 0.058926077 | 9.818540654 | -1.583631228 | 0.159534682 | 0.991835066 | -4.39874496 |
| **A_51_P356762** | Mcm4 | 0.117723361 | -0.059951422 | 0.295398143 | 12.42000727 | 1.583601817 | 0.159541329 | 0.991835066 | -4.398770902 |
| **A_55_P2007228** | Prkaca | 0.141977179 | -0.072341401 | 0.356295758 | 9.105709015 | 1.583316916 | 0.15960573 | 0.991835066 | -4.399022189 |
| **A_55_P2051962** | Sec61b | 0.133343676 | -0.067974885 | 0.334662237 | 13.14201399 | 1.58306132 | 0.159663527 | 0.991835066 | -4.399247618 |
| **A_51_P225948** | Snx33 | -0.084582645 | -0.212343571 | 0.043178281 | 9.271282314 | -1.582312809 | 0.159832899 | 0.991835066 | -4.399907729 |
| **A_66_P117289** | Cdk13 | 0.083981589 | -0.042903512 | 0.210866689 | 8.774211901 | 1.581912981 | 0.159923439 | 0.991835066 | -4.400260302 |
| **A_55_P1999289** | Ovol2 | -0.12854231 | -0.322759773 | 0.065675152 | 11.39077224 | -1.581856042 | 0.159936337 | 0.991835066 | -4.40031051 |
| **A_55_P2005720** | NA | 0.165812102 | -0.084749798 | 0.416374002 | 10.13990178 | 1.581649775 | 0.159983068 | 0.991835066 | -4.400492387 |
| **A_55_P1984422** | Nme2 | -0.138726224 | -0.348478297 | 0.071025849 | 15.3763075 | -1.580743612 | 0.160188517 | 0.991835066 | -4.401291328 |
| **A_55_P2054887** | Trappc1 | -0.086463881 | -0.217259648 | 0.044331885 | 13.11515783 | -1.57997484 | 0.160363009 | 0.991835066 | -4.401969036 |
| **A_55_P2135098** | Rplp0 | 0.106410522 | -0.054572448 | 0.267393491 | 17.25592099 | 1.57984242 | 0.160393083 | 0.991835066 | -4.402085762 |
| **A_51_P300297** | Srek1ip1 | -0.076627715 | -0.19255626 | 0.03930083 | 9.692598748 | -1.579809139 | 0.160400643 | 0.991835066 | -4.402115098 |
| **A_51_P142421** | Rspo1 | 0.441976063 | -0.226687078 | 1.110639204 | 9.613867611 | 1.579794137 | 0.16040405 | 0.991835066 | -4.402128322 |
| **A_51_P256682** | Kpna1 | -0.086337721 | -0.216968007 | 0.044292565 | 9.946495899 | -1.579668058 | 0.160432691 | 0.991835066 | -4.402239454 |
| **A_55_P2083489** | Fam129a | -0.076603351 | -0.192582495 | 0.039375793 | 9.807211868 | -1.578617821 | 0.160671452 | 0.991835066 | -4.403165093 |
| **A_66_P134785** | 2610044O1 | -0.089592622 | -0.225256916 | 0.046071672 | 8.887517365 | -1.578395451 | 0.160722048 | 0.991835066 | -4.40336106 |
| **A_55_P1964459** | NA | -0.104357112 | -0.262393274 | 0.05367905 | 8.554341953 | -1.578246053 | 0.160756049 | 0.991835066 | -4.403492715 |

| | | | | | | | | | |
|---|---|---|---|---|---|---|---|---|---|
| **A_55_P2014357** | Pdpk1 | -0.172326817 | -0.433301235 | 0.088647601 | 10.48430739 | -1.578207346 | 0.16076486 | 0.991835066 | -4.403526824 |
| **A_55_P2049071** | NA | 0.070736887 | -0.036393339 | 0.177867114 | 13.40958566 | 1.578131012 | 0.160782236 | 0.991835066 | -4.403594091 |
| **A_51_P514675** | Zcchc10 | 0.126337302 | -0.065011129 | 0.317685733 | 9.826393385 | 1.578032069 | 0.160804761 | 0.991835066 | -4.403681279 |
| **A_55_P2089362** | Wfdc13 | -0.115588769 | -0.29070694 | 0.059529403 | 8.452542122 | -1.577587857 | 0.160905926 | 0.991835066 | -4.404072701 |
| **A_55_P1987775** | Pla2g4d | 0.083458111 | -0.043000953 | 0.209917174 | 7.740485699 | 1.577348731 | 0.16096041 | 0.991835066 | -4.404283396 |
| **A_51_P508485** | Tmtc4 | -0.120451312 | -0.303031253 | 0.062128629 | 10.23631773 | -1.576767301 | 0.161092956 | 0.991835066 | -4.40479566 |
| **A_55_P2052749** | Pcm1 | 0.146613385 | -0.075638283 | 0.368865052 | 11.0385647 | 1.576658819 | 0.161117698 | 0.991835066 | -4.404891233 |
| **A_55_P2103115** | Tor1aip2 | -0.086811043 | -0.218420196 | 0.04479811 | 10.8089596 | -1.576514654 | 0.161150583 | 0.991835066 | -4.405018238 |
| **A_51_P327778** | Pfdn2 | -0.101682564 | -0.25584095 | 0.052475823 | 10.70284991 | -1.576479996 | 0.16115849 | 0.991835066 | -4.405048769 |
| **A_55_P2022231** | Gm826 | 0.136940867 | -0.070682834 | 0.344564568 | 9.361474011 | 1.576395829 | 0.161177693 | 0.991835066 | -4.405122917 |
| **A_52_P987071** | Gm6803 | -0.153182687 | -0.385448249 | 0.079082874 | 12.41450919 | -1.576282288 | 0.161203601 | 0.991835066 | -4.405222938 |
| **A_52_P197627** | Umps | 0.099774555 | -0.051518166 | 0.251067276 | 9.099636306 | 1.576198456 | 0.161222733 | 0.991835066 | -4.405296787 |
| **A_55_P2038262** | Phf11b | -0.075488455 | -0.189971718 | 0.038994808 | 7.88120393 | -1.575969004 | 0.161275108 | 0.991835066 | -4.40549891 |
| **A_55_P2175301** | NA | 0.084515516 | -0.043674041 | 0.212705073 | 15.52292522 | 1.575770384 | 0.161320458 | 0.991835066 | -4.405673867 |
| **A_55_P1958097** | Eno1 | 0.131843009 | -0.068153324 | 0.331839342 | 15.22633416 | 1.57559359 | 0.161360834 | 0.991835066 | -4.405829593 |
| **A_51_P270733** | Syngr1 | -0.256824719 | -0.646455126 | 0.132805689 | 10.39445553 | -1.575408233 | 0.161403177 | 0.991835066 | -4.405992856 |
| **A_55_P2090209** | Tpm3 | 0.080184418 | -0.041514517 | 0.201883353 | 14.3257312 | 1.574752611 | 0.161553027 | 0.991835066 | -4.406570288 |
| **A_66_P119771** | Nme8 | -0.125210254 | -0.315288429 | 0.064867921 | 8.410864536 | -1.574406143 | 0.16163227 | 0.991835066 | -4.406875408 |
| **A_55_P2252184** | NA | 0.196912534 | -0.102020751 | 0.495845818 | 8.97100684 | 1.57437512 | 0.161639367 | 0.991835066 | -4.406902729 |
| **A_55_P2040775** | NA | 0.139270854 | -0.0721804 | 0.350722109 | 13.10310709 | 1.57419709 | 0.1616801 | 0.991835066 | -4.407059504 |
| **A_52_P494115** | Gatad2b | 0.074852449 | -0.038796935 | 0.188501834 | 8.993290405 | 1.574157054 | 0.161689262 | 0.991835066 | -4.40709476 |
| **A_51_P253207** | NA | -0.197081498 | -0.49633115 | 0.102168155 | 8.653535216 | -1.574060177 | 0.161711433 | 0.991835066 | -4.407180069 |
| **A_55_P2270137** | B930036G( | 0.118898402 | -0.061684811 | 0.299481615 | 11.4712638 | 1.573648653 | 0.161805644 | 0.991835066 | -4.407542436 |
| **A_55_P2107347** | Ro60 | -0.116820111 | -0.294279976 | 0.060639755 | 9.248947954 | -1.573354534 | 0.161873009 | 0.991835066 | -4.407801406 |
| **A_66_P114769** | NA | 0.10016365 | -0.051995694 | 0.252322993 | 10.9558569 | 1.573332983 | 0.161877946 | 0.991835066 | -4.407820381 |
| **A_55_P1969273** | Zfp758 | -0.072153989 | -0.18179821 | 0.037490233 | 8.700545405 | -1.57283703 | 0.161991602 | 0.991835066 | -4.408257034 |
| **A_52_P1042732** | Akap5 | 0.109031273 | -0.056656736 | 0.274719282 | 8.990160925 | 1.572784195 | 0.162003714 | 0.991835066 | -4.408303549 |
| **A_55_P1960989** | Nlgn2 | 0.174207347 | -0.090604243 | 0.439018938 | 10.40366897 | 1.572311571 | 0.1621121 | 0.991835066 | -4.408719622 |
| **A_55_P1968143** | Rbm25 | 0.195468113 | -0.101669095 | 0.492605321 | 10.20819984 | 1.572273212 | 0.1621209 | 0.991835066 | -4.40875339 |
| **A_55_P2016667** | Gstm7 | -0.110744116 | -0.27909182 | 0.057603588 | 8.326392813 | -1.572253594 | 0.1621254 | 0.991835066 | -4.408770659 |
| **A_51_P438821** | Pycard | -0.135226863 | -0.340800107 | 0.070346381 | 11.32797683 | -1.572191762 | 0.162139586 | 0.991835066 | -4.40882509 |
| **A_55_P1956502** | Hnrnph1 | 0.260276938 | -0.135510003 | 0.656063879 | 13.53846902 | 1.571749608 | 0.162241062 | 0.991835066 | -4.409214299 |
| **A_55_P1979330** | Dapp1 | -0.109929855 | -0.27713251 | 0.057272799 | 9.570834349 | -1.571381445 | 0.162325601 | 0.991835066 | -4.409538354 |
| **A_51_P497821** | Zfp790 | -0.087797453 | -0.221355438 | 0.045760532 | 8.321707225 | -1.571162816 | 0.162375823 | 0.991835066 | -4.40973078 |
| **A_55_P2048848** | Smn1 | 0.074492869 | -0.03887737 | 0.187863108 | 10.66931262 | 1.570452373 | 0.162539122 | 0.991835066 | -4.410356023 |
| **A_52_P10732** | Map6 | 0.077933655 | -0.04069324 | 0.196560549 | 8.61835198 | 1.570185685 | 0.162600461 | 0.991835066 | -4.410590709 |
| **A_55_P2186572** | Lypla2 | 0.091608371 | -0.047854942 | 0.231071684 | 12.81041535 | 1.569944553 | 0.16265594 | 0.991835066 | -4.410802895 |
| **A_55_P2255628** | 9430013L1 | 0.196129992 | -0.102503109 | 0.494763094 | 8.925142222 | 1.569694724 | 0.162713439 | 0.991835066 | -4.411022724 |
| **A_52_P337126** | Bves | -0.102447694 | -0.258447247 | 0.053551859 | 14.72126862 | -1.569596302 | 0.162736097 | 0.991835066 | -4.411109325 |
| **A_52_P606774** | Stk4 | 0.115490706 | -0.060388865 | 0.291370277 | 10.11037457 | 1.569425738 | 0.162775369 | 0.991835066 | -4.4112594 |
| **A_51_P370380** | Rdh14 | -0.127322599 | -0.321266793 | 0.066621595 | 11.76970754 | -1.569053831 | 0.16286103 | 0.991835066 | -4.411586615 |
| **A_55_P2093286** | Apod | -0.07315994 | -0.184610258 | 0.038290378 | 8.004790543 | -1.568921305 | 0.162891565 | 0.991835066 | -4.41170321 |
| **A_55_P2061809** | Ndufc2 | -0.105245384 | -0.265617496 | 0.055126728 | 15.59220213 | -1.568495724 | 0.162989658 | 0.991835066 | -4.412077613 |
| **A_55_P2085664** | Meaf6 | 0.179155546 | -0.093857697 | 0.452168789 | 11.29213223 | 1.568395821 | 0.163012693 | 0.991835066 | -4.412165499 |
| **A_55_P2022297** | Ptrh2 | 0.067628116 | -0.035439777 | 0.170696009 | 9.507835611 | 1.568241799 | 0.163048211 | 0.991835066 | -4.412300989 |
| **A_51_P514922** | Ccar2 | 0.079575922 | -0.041704249 | 0.200856093 | 9.764250879 | 1.568198415 | 0.163058217 | 0.991835066 | -4.412339153 |
| **A_52_P77245** | Snx8 | 0.100637355 | -0.052774639 | 0.254049349 | 10.69746488 | 1.567866324 | 0.16313483 | 0.991835066 | -4.412631273 |
| **A_52_P362033** | Polr2e | -0.226710967 | -0.572473974 | 0.119052039 | 10.34873027 | -1.567123187 | 0.16330639 | 0.991835066 | -4.413284904 |

| | | | | | | | | | |
|---|---|---|---|---|---|---|---|---|---|
| **A_66_P127024** | Mbd3 | 0.075336345 | -0.039568215 | 0.190240905 | 13.47185743 | 1.567026773 | 0.163328661 | 0.991835066 | -4.413369699 |
| **A_55_P2125466** | Ube2l3 | -0.082656637 | -0.208730176 | 0.043416902 | 12.5089972 | -1.56697807 | 0.163339912 | 0.991835066 | -4.413412532 |
| **A_55_P2151156** | Farsa | 0.130606269 | -0.068607349 | 0.329819888 | 10.89777348 | 1.566946368 | 0.163347235 | 0.991835066 | -4.413440413 |
| **A_51_P423290** | Mmrn1 | 0.185146589 | -0.097284919 | 0.467578096 | 8.898494275 | 1.566793291 | 0.163382604 | 0.991835066 | -4.413575038 |
| **A_52_P17112** | Znrf1 | 0.144027654 | -0.07577296 | 0.363828269 | 10.19194971 | 1.566123902 | 0.16353735 | 0.991835066 | -4.414163692 |
| **A_55_P2040774** | NA | 0.081952218 | -0.043116742 | 0.207021177 | 13.58950649 | 1.56610294 | 0.163542198 | 0.991835066 | -4.414182125 |
| **A_55_P1998471** | S100a9 | -0.617407866 | -1.559668039 | 0.324852307 | 11.214591 | -1.566067423 | 0.163550413 | 0.991835066 | -4.414213356 |
| **A_55_P2366273** | Amn1 | 0.130169336 | -0.068523054 | 0.328861726 | 8.427939026 | 1.565801074 | 0.163612028 | 0.991835066 | -4.414447559 |
| **A_55_P2002113** | Rpl36 | 0.074101939 | -0.039019405 | 0.187223282 | 16.64498146 | 1.565648078 | 0.163647431 | 0.991835066 | -4.414582085 |
| **A_55_P2125643** | NA | 0.120516859 | -0.063463373 | 0.30449709 | 14.85438513 | 1.56561789 | 0.163654418 | 0.991835066 | -4.414608628 |
| **A_55_P2176430** | 0610010K1 | -0.179290554 | -0.453006982 | 0.094425874 | 10.94385297 | -1.565545438 | 0.163671186 | 0.991835066 | -4.414672332 |
| **A_55_P2159807** | Rfc4 | 0.096269269 | -0.050722459 | 0.243260997 | 11.80488319 | 1.565322795 | 0.163722725 | 0.991835066 | -4.414868086 |
| **A_55_P1965075** | NA | -0.072613572 | -0.183499329 | 0.038272184 | 15.50053679 | -1.565132722 | 0.163766737 | 0.991835066 | -4.415035197 |
| **A_51_P262878** | Tmed9 | 0.131955277 | -0.069562693 | 0.333473246 | 14.21316155 | 1.565028011 | 0.163790987 | 0.991835066 | -4.415127257 |
| **A_52_P367060** | Atxn2l | 0.099784927 | -0.052606949 | 0.252176804 | 12.41647506 | 1.564992503 | 0.163799211 | 0.991835066 | -4.415158474 |
| **A_55_P1979432** | Prdx2 | -0.079353448 | -0.200561749 | 0.041854854 | 14.92726918 | -1.564741382 | 0.163857386 | 0.991835066 | -4.415379244 |
| **A_55_P2063970** | Ube2l3 | -0.077748739 | -0.196519 | 0.041021521 | 12.83794601 | -1.564569205 | 0.163897284 | 0.991835066 | -4.415530606 |
| **A_55_P1995007** | Arhgap11a | -0.107862395 | -0.272678343 | 0.056953553 | 9.473210611 | -1.564155616 | 0.163993159 | 0.991835066 | -4.415894175 |
| **A_66_P120125** | Nrep | -0.130162266 | -0.329053118 | 0.068728586 | 11.35494561 | -1.564153698 | 0.163993604 | 0.991835066 | -4.415895861 |
| **A_51_P307243** | Ppp5c | -0.098361207 | -0.248664531 | 0.051942117 | 10.66524823 | -1.564099566 | 0.164006156 | 0.991835066 | -4.415943444 |
| **A_52_P565940** | Nsd1 | 0.126551193 | -0.066841061 | 0.319943447 | 10.12075947 | 1.56399839 | 0.16402962 | 0.991835066 | -4.416032378 |
| **A_55_P2040838** | Gm14548 | -0.066050101 | -0.167026823 | 0.034926621 | 7.790568646 | -1.563368471 | 0.164175774 | 0.991835066 | -4.416586046 |
| **A_65_P13910** | Ndufb2 | -0.157582796 | -0.398505684 | 0.083340093 | 15.66857623 | -1.563291158 | 0.164193721 | 0.991835066 | -4.416653996 |
| **A_52_P597461** | Skil | 0.077693334 | -0.041148305 | 0.196534974 | 8.093574244 | 1.562515223 | 0.164373939 | 0.991835066 | -4.417335907 |
| **A_55_P2097244** | Bptf | 0.082032124 | -0.043460538 | 0.207524786 | 12.36376838 | 1.562337134 | 0.164415327 | 0.991835066 | -4.417492403 |
| **A_55_P2010292** | Serpina1e | -0.640830092 | -1.621233209 | 0.339573026 | 9.203918389 | -1.562238532 | 0.164438247 | 0.991835066 | -4.417579047 |
| **A_55_P1997872** | Apip | -0.089622691 | -0.226739954 | 0.047494571 | 11.58256065 | -1.562194048 | 0.164448588 | 0.991835066 | -4.417618136 |
| **A_55_P2005898** | NA | -0.225249045 | -0.569888306 | 0.119390216 | 13.03939944 | -1.56209463 | 0.164471702 | 0.991835066 | -4.417705494 |
| **A_55_P2171086** | NA | 0.138129901 | -0.073305941 | 0.349565742 | 8.93479802 | 1.561414551 | 0.164629895 | 0.991835066 | -4.418303041 |
| **A_55_P2093232** | Myh7 | 0.067626167 | -0.035890763 | 0.171143097 | 18.26978331 | 1.561394054 | 0.164634665 | 0.991835066 | -4.418321049 |
| **A_55_P2069579** | NA | 0.075036208 | -0.039828403 | 0.189900819 | 8.197593527 | 1.561326623 | 0.164650358 | 0.991835066 | -4.418380292 |
| **A_55_P2108868** | Nufip1 | 0.081551118 | -0.043326285 | 0.206428521 | 8.164484068 | 1.56082853 | 0.164766323 | 0.991835066 | -4.418817883 |
| **A_51_P173285** | Nkx2-5 | 0.096586086 | -0.051320027 | 0.244492199 | 13.03898958 | 1.560765215 | 0.164781069 | 0.991835066 | -4.418873504 |
| **A_51_P175871** | Got2 | -0.129115084 | -0.326842617 | 0.068612449 | 14.1664045 | -1.560698337 | 0.164796647 | 0.991835066 | -4.418932255 |
| **A_55_P2111429** | Btf3l4 | -0.087175773 | -0.220690919 | 0.046339373 | 11.29421737 | -1.560538204 | 0.164833951 | 0.991835066 | -4.419072925 |
| **A_55_P1992655** | Hdac6 | -0.089342686 | -0.226189891 | 0.047504519 | 11.49432038 | -1.560386576 | 0.164869281 | 0.991835066 | -4.419206121 |
| **A_55_P2025765** | Adam8 | -0.120577356 | -0.305272424 | 0.064117712 | 8.411902674 | -1.560341252 | 0.164879843 | 0.991835066 | -4.419245934 |
| **A_55_P2026420** | Pou6f1 | -0.078589371 | -0.19897377 | 0.041795028 | 8.625276584 | -1.560280708 | 0.164893953 | 0.991835066 | -4.419299116 |
| **A_55_P2057720** | Zfp937 | 0.073484432 | -0.039113392 | 0.186082255 | 8.743734381 | 1.559819973 | 0.165001365 | 0.991835066 | -4.419703811 |
| **A_55_P2033906** | Mrpl22 | -0.084404007 | -0.213792122 | 0.044984109 | 12.93521517 | -1.559113776 | 0.165166128 | 0.991835066 | -4.420324045 |
| **A_52_P634090** | Jag1 | 0.117960105 | -0.062886543 | 0.298806753 | 9.65810869 | 1.558955863 | 0.165202991 | 0.991835066 | -4.420462724 |
| **A_55_P2079568** | NA | -0.111308388 | -0.281988642 | 0.059371867 | 8.540745408 | -1.558668446 | 0.165270106 | 0.991835066 | -4.420715125 |
| **A_51_P453246** | Uqcc3 | -0.084412235 | -0.213857079 | 0.045032609 | 13.05992884 | -1.558582426 | 0.165290198 | 0.991835066 | -4.420790662 |
| **A_55_P1954486** | Mapt | 0.07505423 | -0.04004734 | 0.1901558 | 8.034543419 | 1.558486543 | 0.165312596 | 0.991835066 | -4.420874858 |
| **A_51_P489153** | Crot | -0.079636179 | -0.201772284 | 0.042499927 | 10.00024488 | -1.558387572 | 0.165335718 | 0.991835066 | -4.420961766 |
| **A_55_P2017640** | Phf7 | 0.078872787 | -0.042097762 | 0.199843336 | 9.908784673 | 1.558320076 | 0.165351489 | 0.991835066 | -4.421021033 |
| **A_66_P111426** | Lce1a2 | 0.275720246 | -0.147211148 | 0.698651641 | 8.63731114 | 1.558144984 | 0.165392406 | 0.991835066 | -4.421174777 |
| **A_51_P493720** | Tfam | -0.095705076 | -0.242514252 | 0.0511041 | 11.07713773 | -1.558084121 | 0.165406631 | 0.991835066 | -4.421228218 |

| | | | | | | | | | |
|---|---|---|---|---|---|---|---|---|---|
| **A_55_P2160032** | Atp5pb | -0.09490468 | -0.240509258 | 0.050699897 | 15.06069839 | -1.557835997 | 0.165464636 | 0.991835066 | -4.421446079 |
| **A_52_P163849** | Ivns1abp | 0.097254181 | -0.051965934 | 0.246474297 | 9.120778494 | 1.557722323 | 0.165491216 | 0.991835066 | -4.421545884 |
| **A_55_P1953998** | NA | 0.08825331 | -0.047168673 | 0.223675293 | 11.63814878 | 1.557582163 | 0.165523994 | 0.991835066 | -4.421668941 |
| **A_52_P672740** | Rala | -0.087076285 | -0.220694175 | 0.046541605 | 9.030160638 | -1.557558669 | 0.165529489 | 0.991835066 | -4.421689568 |
| **A_55_P2117146** | Pa2g4 | 0.145046062 | -0.07753526 | 0.367627383 | 11.14297536 | 1.557493909 | 0.165544637 | 0.991835066 | -4.421746425 |
| **A_52_P48777** | Copg2 | 0.108716568 | -0.058175661 | 0.275608797 | 10.81286 | 1.556928795 | 0.165676875 | 0.991835066 | -4.422242543 |
| **A_55_P1961302** | NA | 0.259361806 | -0.138895754 | 0.657619367 | 11.21508601 | 1.55650717 | 0.1657756 | 0.991835066 | -4.422612657 |
| **A_55_P2053958** | Rabep2 | 0.102212181 | -0.054751087 | 0.259175449 | 10.74875206 | 1.556373241 | 0.165806972 | 0.991835066 | -4.422730217 |
| **A_51_P160713** | Alb | -0.907682 | -2.3017424 | 0.486378401 | 10.06824325 | -1.556185142 | 0.165851041 | 0.991835066 | -4.422895321 |
| **A_55_P2156126** | Sema6c | 0.132599117 | -0.071070558 | 0.336268792 | 9.2475574 | 1.55604945 | 0.165882839 | 0.991835066 | -4.423014423 |
| **A_51_P491504** | Aldh5a1 | -0.159341791 | -0.404096811 | 0.085413229 | 12.14214706 | -1.555991491 | 0.165896423 | 0.991835066 | -4.423065294 |
| **A_66_P102062** | NA | -0.163909641 | -0.415750456 | 0.087931174 | 9.398514526 | -1.555562678 | 0.165996955 | 0.991835066 | -4.423441651 |
| **A_55_P2077079** | Rab29 | -0.119501661 | -0.303126266 | 0.064122943 | 9.723074065 | -1.555436198 | 0.166026618 | 0.991835066 | -4.423552654 |
| **A_55_P2100700** | Pygo2 | 0.083784516 | -0.045004703 | 0.212573736 | 12.42297345 | 1.554867504 | 0.166160054 | 0.991835066 | -4.424051723 |
| **A_55_P1984721** | Ube2i | 0.082211443 | -0.04417346 | 0.208596347 | 12.4513224 | 1.554698585 | 0.166199708 | 0.991835066 | -4.424199951 |
| **A_55_P1977144** | Cyp2c67 | -0.206054405 | -0.522852704 | 0.110743894 | 7.891657013 | -1.554562526 | 0.166231654 | 0.991835066 | -4.424319341 |
| **A_51_P321886** | Cmtm3 | 0.108103461 | -0.058156031 | 0.274362953 | 11.49084461 | 1.554040313 | 0.16635432 | 0.991835066 | -4.424777547 |
| **A_55_P1980316** | Pcbp2 | 0.10907889 | -0.058721006 | 0.276878785 | 14.10726067 | 1.553667762 | 0.166441883 | 0.991835066 | -4.425104408 |
| **A_52_P525183** | Acot2 | -0.110276976 | -0.279930063 | 0.059376111 | 8.918907145 | -1.55357497 | 0.166463699 | 0.991835066 | -4.425185817 |
| **A_51_P347961** | Npc2 | -0.130593287 | -0.331507194 | 0.07032062 | 12.07305847 | -1.553531228 | 0.166473984 | 0.991835066 | -4.425224192 |
| **A_55_P1986993** | Brd8 | 0.097225515 | -0.052412279 | 0.24686331 | 8.485607273 | 1.55291644 | 0.166618601 | 0.991835066 | -4.425763518 |
| **A_55_P2069306** | Ptpn18 | -0.127159403 | -0.322891761 | 0.068572955 | 10.78333409 | -1.552726569 | 0.166663287 | 0.991835066 | -4.42593007 |
| **A_55_P2010298** | Serpina1d | -0.536211825 | -1.361791615 | 0.289367966 | 8.714141615 | -1.552338633 | 0.166754624 | 0.991835066 | -4.426270345 |
| **A_51_P487547** | Ccdc91 | -0.128250138 | -0.325741395 | 0.069241119 | 9.716280247 | -1.552097856 | 0.166811337 | 0.991835066 | -4.426481528 |
| **A_51_P148612** | Cox7a1 | -0.184441121 | -0.468472332 | 0.099590091 | 16.54547169 | -1.552032525 | 0.166826728 | 0.991835066 | -4.426538827 |
| **A_55_P2069392** | NA | -0.131672969 | -0.334550178 | 0.071204239 | 14.23915728 | -1.551216803 | 0.167019013 | 0.991835066 | -4.427254208 |
| **A_51_P288505** | Tradd | 0.104756465 | -0.056658574 | 0.266171504 | 10.29797372 | 1.551122049 | 0.167041362 | 0.991835066 | -4.427337299 |
| **A_55_P2065239** | NA | 0.081195476 | -0.043924046 | 0.206314997 | 15.99157585 | 1.551014599 | 0.167066709 | 0.991835066 | -4.427431522 |
| **A_55_P1967504** | Abhd11 | -0.159762423 | -0.406017065 | 0.086492218 | 11.38559378 | -1.550598456 | 0.167164908 | 0.991835066 | -4.427796419 |
| **A_51_P133137** | Kif20a | -0.142629554 | -0.362480699 | 0.07722159 | 11.87142268 | -1.550564821 | 0.167172848 | 0.991835066 | -4.427825911 |
| **A_55_P1986406** | Ddx54 | 0.119060831 | -0.064469688 | 0.30259135 | 11.39221628 | 1.550492785 | 0.167189853 | 0.991835066 | -4.427889073 |
| **A_66_P138673** | Whamm | 0.072288983 | -0.039159383 | 0.183737348 | 8.188951464 | 1.550270713 | 0.167242286 | 0.991835066 | -4.428083782 |
| **A_52_P409769** | Mrvi1 | -0.09111645 | -0.231614197 | 0.049381297 | 8.951530768 | -1.550017364 | 0.167302122 | 0.991835066 | -4.428305905 |
| **A_55_P2063471** | NA | 0.092367453 | -0.050073575 | 0.234808481 | 16.89157083 | 1.549861901 | 0.167338849 | 0.991835066 | -4.428442201 |
| **A_55_P1955346** | NA | 0.0700817 | -0.038007707 | 0.178171107 | 14.46459703 | 1.54963931 | 0.167391448 | 0.991835066 | -4.428637342 |
| **A_51_P205106** | Mlf1 | -0.182923569 | -0.465094591 | 0.099247453 | 13.62832999 | -1.549410118 | 0.167445623 | 0.991835066 | -4.428838261 |
| **A_51_P240453** | Nusap1 | -0.106632765 | -0.271123579 | 0.057858049 | 10.3789305 | -1.549380738 | 0.167452569 | 0.991835066 | -4.428864017 |
| **A_55_P2176350** | 0610010F0 | 0.107981948 | -0.058601363 | 0.274565258 | 9.546501608 | 1.549276026 | 0.167477326 | 0.991835066 | -4.428955808 |
| **A_52_P607683** | Ppm1e | -0.085582466 | -0.217619489 | 0.046454558 | 9.866658475 | -1.549166875 | 0.167503136 | 0.991835066 | -4.429051489 |
| **A_51_P464911** | Msrb2 | -0.232736537 | -0.591812826 | 0.126339753 | 12.16723765 | -1.549126814 | 0.167512611 | 0.991835066 | -4.429086606 |
| **A_51_P192089** | Pinx1 | 0.254072453 | -0.13796489 | 0.646109797 | 10.04730473 | 1.548956693 | 0.167552848 | 0.991835066 | -4.429235727 |
| **A_51_P463846** | Gbp7 | -0.073888529 | -0.187902466 | 0.040125409 | 7.940935727 | -1.54891722 | 0.167562185 | 0.991835066 | -4.429270327 |
| **A_55_P2419854** | NA | 0.102485623 | -0.055751192 | 0.260722438 | 8.492348639 | 1.547977144 | 0.167784707 | 0.991835066 | -4.430094269 |
| **A_51_P446510** | Emp3 | -0.120471639 | -0.306494425 | 0.065551148 | 10.03578273 | -1.547846197 | 0.167815725 | 0.991835066 | -4.430209027 |
| **A_55_P1964525** | NA | 0.085799777 | -0.046702745 | 0.218302299 | 16.71733435 | 1.547644284 | 0.167863563 | 0.991835066 | -4.430385973 |
| **A_55_P2125551** | Mmadhc | 0.083189834 | -0.045292155 | 0.211671823 | 13.25863325 | 1.547523094 | 0.167892282 | 0.991835066 | -4.430492174 |
| **A_55_P1988048** | Emcn | -0.131220833 | -0.333963454 | 0.071521788 | 9.560691996 | -1.546916475 | 0.168036104 | 0.991835066 | -4.431023728 |
| **A_55_P2043767** | NA | 0.084186191 | -0.045893119 | 0.214265501 | 15.85283658 | 1.546827083 | 0.168057307 | 0.991835066 | -4.431102054 |

| | | | | | | | | | |
|---|---|---|---|---|---|---|---|---|---|
| **A_51_P359333** | Fh1 | -0.105959192 | -0.269682288 | 0.057763904 | 14.68954738 | -1.546813028 | 0.168060641 | 0.991835066 | -4.431114369 |
| **A_65_P19458** | Rock2 | -0.080120765 | -0.203931353 | 0.043689824 | 9.880600933 | -1.546665617 | 0.168095612 | 0.991835066 | -4.431243527 |
| **A_51_P235123** | Nfkbie | -0.087822726 | -0.223535327 | 0.047889876 | 8.219460069 | -1.546663611 | 0.168096088 | 0.991835066 | -4.431245284 |
| **A_52_P284495** | Ankrd33b | -0.097317582 | -0.247703777 | 0.053068613 | 8.234933679 | -1.546651512 | 0.168098959 | 0.991835066 | -4.431255885 |
| **A_51_P244969** | Rmnd5b | -0.092018212 | -0.234218142 | 0.050181718 | 9.026833069 | -1.546619703 | 0.168106506 | 0.991835066 | -4.431283754 |
| **A_55_P2020143** | Slc39a3 | 0.06517476 | -0.035547601 | 0.165897122 | 9.252925623 | 1.546545406 | 0.168124136 | 0.991835066 | -4.43134885 |
| **A_55_P1980411** | Zkscan1 | 0.095742902 | -0.052248077 | 0.243733881 | 10.89702099 | 1.546252719 | 0.168193602 | 0.991835066 | -4.431605279 |
| **A_52_P335259** | Cdkal1 | -0.115633632 | -0.294376315 | 0.063109051 | 10.54899696 | -1.546197715 | 0.16820666 | 0.991835066 | -4.431653468 |
| **A_55_P2156785** | Ppfia3 | -0.097392675 | -0.247945544 | 0.053160193 | 9.348465291 | -1.546131372 | 0.168222411 | 0.991835066 | -4.43171159 |
| **A_52_P35961** | NA | 0.084262292 | -0.045997279 | 0.214521863 | 8.494329178 | 1.546082833 | 0.168233936 | 0.991835066 | -4.431754113 |
| **A_51_P435922** | Rsph9 | -0.074826969 | -0.190535175 | 0.040881237 | 9.255776702 | -1.545621404 | 0.16834353 | 0.991835066 | -4.432158338 |
| **A_55_P1956130** | LOC68395 | -0.127659762 | -0.325106953 | 0.069787428 | 9.413182248 | -1.545297866 | 0.168420414 | 0.991835066 | -4.432441745 |
| **A_51_P323521** | Snx21 | -0.138919272 | -0.353803926 | 0.075965382 | 12.12149863 | -1.545134285 | 0.168459299 | 0.991835066 | -4.432585029 |
| **A_66_P110088** | NA | 0.096779641 | -0.052929661 | 0.246488943 | 9.101955398 | 1.545056463 | 0.168477801 | 0.991835066 | -4.432653193 |
| **A_55_P2052425** | Setd4 | 0.155483372 | -0.085063216 | 0.39602996 | 9.168491311 | 1.544876907 | 0.168520497 | 0.991835066 | -4.432810463 |
| **A_51_P143190** | Lyl1 | -0.095201177 | -0.242490196 | 0.052087843 | 11.01616251 | -1.544831364 | 0.168531328 | 0.991835066 | -4.432850352 |
| **A_51_P354077** | Svil | -0.125373522 | -0.319354334 | 0.068607291 | 12.8700397 | -1.544742811 | 0.16855239 | 0.991835066 | -4.432927911 |
| **A_55_P2320085** | 4833415N1 | -0.076395004 | -0.194616091 | 0.041826083 | 8.198450271 | -1.544468808 | 0.168617575 | 0.991835066 | -4.433167888 |
| **A_55_P2108389** | D830030K2 | -0.116147349 | -0.295897916 | 0.063603217 | 9.76019139 | -1.544358658 | 0.168643786 | 0.991835066 | -4.433264356 |
| **A_55_P2048498** | NA | -0.071080723 | -0.181100827 | 0.038939382 | 8.123736284 | -1.544147965 | 0.168693933 | 0.991835066 | -4.433448873 |
| **A_55_P2024391** | NA | -0.112127351 | -0.285688157 | 0.061433456 | 9.669660422 | -1.544077284 | 0.168710758 | 0.991835066 | -4.43351077 |
| **A_55_P2273705** | NA | 0.074537672 | -0.040857651 | 0.189932994 | 9.705909953 | 1.543820311 | 0.168771945 | 0.991835066 | -4.433735804 |
| **A_52_P316933** | Sh3bgrl2 | 0.120073327 | -0.06588725 | 0.306033905 | 9.040163818 | 1.543244691 | 0.168909077 | 0.991835066 | -4.434239837 |
| **A_51_P320444** | Rock2 | 0.094435489 | -0.051833653 | 0.240704632 | 8.956800919 | 1.5430914 | 0.168945613 | 0.991835066 | -4.434374056 |
| **A_52_P161237** | Cpa1 | -0.240074187 | -0.611927342 | 0.131778968 | 10.54053993 | -1.543061118 | 0.168952831 | 0.991835066 | -4.434400569 |
| **A_52_P295313** | Wdr46 | 0.129446133 | -0.07108144 | 0.329973706 | 11.35112745 | 1.542851459 | 0.169002817 | 0.991835066 | -4.434584134 |
| **A_51_P484526** | Wif1 | 0.109795869 | -0.060378012 | 0.279969751 | 10.67507757 | 1.542063402 | 0.169190822 | 0.991835066 | -4.435274043 |
| **A_55_P2072906** | Gm6788 | 0.113296177 | -0.062321643 | 0.288913996 | 13.28453798 | 1.541898567 | 0.169230171 | 0.991835066 | -4.435418335 |
| **A_55_P2081630** | Gpa33 | -0.112121922 | -0.285940714 | 0.06169687 | 9.954337968 | -1.54171088 | 0.169274986 | 0.991835066 | -4.435582626 |
| **A_51_P361426** | Arl8b | -0.119850821 | -0.305657838 | 0.065956197 | 11.27792232 | -1.541657967 | 0.169287622 | 0.991835066 | -4.435628941 |
| **A_51_P387868** | Tekt4 | 0.099647353 | -0.054866951 | 0.254161658 | 8.373672146 | 1.541367542 | 0.169356993 | 0.991835066 | -4.435883148 |
| **A_51_P466685** | Fbxl22 | -0.075301797 | -0.192085766 | 0.041482172 | 9.245831651 | -1.541101504 | 0.169420563 | 0.991835066 | -4.436115998 |
| **A_51_P107090** | BC031181 | -0.068863554 | -0.175687416 | 0.037960309 | 10.42896024 | -1.540743287 | 0.169506194 | 0.991835066 | -4.436429507 |
| **A_55_P2051124** | Scaf1 | -0.080210725 | -0.204637649 | 0.044216199 | 12.90127956 | -1.540732377 | 0.169508803 | 0.991835066 | -4.436439055 |
| **A_65_P05396** | Rad18 | 0.068960042 | -0.038017879 | 0.175937964 | 8.484270709 | 1.540680189 | 0.169521282 | 0.991835066 | -4.436484727 |
| **A_55_P2075592** | Clstn1 | -0.06384119 | -0.162879161 | 0.03519678 | 8.252619481 | -1.540665561 | 0.16952478 | 0.991835066 | -4.43649753 |
| **A_65_P15675** | Ipo9 | 0.073541341 | -0.040618263 | 0.187700945 | 9.683370481 | 1.53967204 | 0.169762512 | 0.991835066 | -4.437366925 |
| **A_51_P251736** | Reep5 | -0.168510318 | -0.430105196 | 0.093084559 | 10.60071733 | -1.539594663 | 0.16978104 | 0.991835066 | -4.437434628 |
| **A_55_P2012394** | Sfxn2 | 0.183870737 | -0.101569923 | 0.469311397 | 10.18492273 | 1.539592943 | 0.169781452 | 0.991835066 | -4.437436132 |
| **A_52_P476431** | Evc | -0.0760915 | -0.194218368 | 0.042035368 | 8.117277472 | -1.539559893 | 0.169789366 | 0.991835066 | -4.437465051 |
| **A_55_P2084303** | Nid1 | 0.097426016 | -0.053833269 | 0.2486853 | 8.096729926 | 1.539437398 | 0.169818703 | 0.991835066 | -4.437572227 |
| **A_55_P2094721** | Sox11 | 0.191144244 | -0.105620464 | 0.487908951 | 9.461675793 | 1.539423472 | 0.169822038 | 0.991835066 | -4.437584412 |
| **A_55_P2120931** | Dleu2 | 0.088652314 | -0.048994749 | 0.226299376 | 8.262343923 | 1.53933186 | 0.169843982 | 0.991835066 | -4.437664566 |
| **A_55_P1958623** | Ncln | 0.09016756 | -0.049900871 | 0.23023599 | 11.63185779 | 1.538576827 | 0.170024935 | 0.991835066 | -4.438325112 |
| **A_55_P1956171** | NA | -0.113123256 | -0.288893548 | 0.062647037 | 14.43371271 | -1.538209723 | 0.17011298 | 0.991835066 | -4.43864624 |
| **A_55_P2112947** | Srp9 | -0.082648333 | -0.211087783 | 0.045791116 | 9.703650108 | -1.537959139 | 0.170173104 | 0.991835066 | -4.438865428 |
| **A_55_P2103206** | NA | 0.1266711 | -0.070211258 | 0.323553458 | 8.711897453 | 1.537729239 | 0.170228283 | 0.991835066 | -4.439066514 |
| **A_51_P337944** | Bmp2k | -0.081252749 | -0.20754247 | 0.045036972 | 10.00750446 | -1.537726824 | 0.170228862 | 0.991835066 | -4.439068626 |

| | | | | | | | | | |
|---|---|---|---|---|---|---|---|---|---|
| **A_55_P2132671** | NA | 0.099376151 | -0.055084175 | 0.253836478 | 9.601422598 | 1.537709708 | 0.170232971 | 0.991835066 | -4.439083597 |
| **A_55_P2008740** | Fcgr1 | -0.064262155 | -0.164152446 | 0.035628136 | 7.909483215 | -1.537592098 | 0.170261206 | 0.991835066 | -4.439186462 |
| **A_55_P2020256** | Sik2 | -0.084029052 | -0.214693711 | 0.046635608 | 9.029511552 | -1.537023314 | 0.170397815 | 0.991835066 | -4.439683906 |
| **A_51_P473259** | Dpyd | -0.070337553 | -0.179722149 | 0.039047042 | 8.059334485 | -1.536880944 | 0.170432025 | 0.991835066 | -4.439808411 |
| **A_51_P103975** | Mpc2 | -0.092961029 | -0.237538332 | 0.051616273 | 14.16889205 | -1.53677377 | 0.170457782 | 0.991835066 | -4.439902134 |
| **A_66_P111437** | Ube2j2 | 0.090824849 | -0.050435887 | 0.232085585 | 13.97780455 | 1.536711539 | 0.170472739 | 0.991835066 | -4.439956553 |
| **A_55_P2005873** | Zfp280d | 0.100522162 | -0.055844519 | 0.256888843 | 9.368722593 | 1.536479379 | 0.170528551 | 0.991835066 | -4.440159565 |
| **A_51_P143296** | Myh8 | -0.166644172 | -0.425891904 | 0.092603561 | 15.47324697 | -1.536329235 | 0.170564655 | 0.991835066 | -4.440290855 |
| **A_55_P2018479** | Carnmt1 | -0.094670189 | -0.242010161 | 0.052669782 | 11.57240528 | -1.535683779 | 0.170719943 | 0.991835066 | -4.440855209 |
| **A_51_P173735** | Dock11 | -0.1327932 | -0.339477504 | 0.073891104 | 9.919315005 | -1.535597769 | 0.170740645 | 0.991835066 | -4.440930406 |
| **A_55_P2327958** | Mtx3 | 0.206451971 | -0.114898961 | 0.527802903 | 9.019607419 | 1.535495693 | 0.170765218 | 0.991835066 | -4.441019648 |
| **A_52_P595717** | Smug1 | -0.086027696 | -0.219965615 | 0.047910223 | 8.786542041 | -1.535125469 | 0.17085437 | 0.991835066 | -4.441343309 |
| **A_52_P526265** | Wdr73 | 0.113267542 | -0.063125575 | 0.289660658 | 11.07087649 | 1.534733504 | 0.170948804 | 0.991835066 | -4.441685951 |
| **A_55_P1954758** | Prickle1 | -0.080263648 | -0.20529072 | 0.044763424 | 12.88639933 | -1.534348332 | 0.171041648 | 0.991835066 | -4.442022628 |
| **A_55_P2157023** | Sox17 | -0.11725018 | -0.299901937 | 0.065401577 | 9.8519127 | -1.53425941 | 0.171063089 | 0.991835066 | -4.442100351 |
| **A_51_P508029** | Pcyox1 | -0.078667601 | -0.201215975 | 0.043880773 | 9.241670516 | -1.534254776 | 0.171064206 | 0.991835066 | -4.442104401 |
| **A_51_P509997** | Cox6a2 | -0.184885711 | -0.472953539 | 0.103182118 | 17.30070132 | -1.533973019 | 0.17113216 | 0.991835066 | -4.442350662 |
| **A_52_P574190** | Psmd12 | -0.079455717 | -0.203293292 | 0.044381858 | 13.50464484 | -1.533493164 | 0.17124795 | 0.991835066 | -4.442770035 |
| **A_51_P169516** | Ppp1r3d | -0.082031826 | -0.209890024 | 0.045826372 | 11.99539098 | -1.533426392 | 0.171264067 | 0.991835066 | -4.442828387 |
| **A_55_P2126870** | Nsf | 0.108147671 | -0.060419665 | 0.276715007 | 8.27420974 | 1.533390898 | 0.171272636 | 0.991835066 | -4.442859405 |
| **A_55_P2084666** | H2ac10 | 0.098570946 | -0.055074256 | 0.252216148 | 17.00064184 | 1.533342081 | 0.171284421 | 0.991835066 | -4.442902066 |
| **A_55_P2052775** | Morc2a | -0.076380573 | -0.195438334 | 0.042677188 | 10.66669504 | -1.533325412 | 0.171288445 | 0.991835066 | -4.442916633 |
| **A_55_P1980931** | NA | -0.226441772 | -0.579453732 | 0.126570188 | 12.61449024 | -1.533120384 | 0.171337952 | 0.991835066 | -4.443095801 |
| **A_55_P1996223** | Sf3b4 | 0.132582598 | -0.074109754 | 0.33927495 | 11.39672684 | 1.533102706 | 0.171342221 | 0.991835066 | -4.443111248 |
| **A_52_P407280** | Adamtsl5 | 0.117781837 | -0.065855332 | 0.301419006 | 8.514421796 | 1.53294604 | 0.171380059 | 0.991835066 | -4.443248148 |
| **A_51_P407879** | Fam193b | 0.203262845 | -0.113673131 | 0.52019882 | 9.977767748 | 1.53283565 | 0.171406725 | 0.991835066 | -4.443344608 |
| **A_55_P1965757** | Eme2 | 0.196404422 | -0.109859346 | 0.502668191 | 8.922928675 | 1.532726874 | 0.171433006 | 0.991835066 | -4.443439655 |
| **A_52_P246749** | Ghitm | -0.14470624 | -0.370467761 | 0.081055282 | 13.10327162 | -1.531956619 | 0.171619206 | 0.991835066 | -4.444112637 |
| **A_52_P127643** | NA | -0.156835431 | -0.401577208 | 0.087906346 | 9.591993584 | -1.531599459 | 0.171705608 | 0.991835066 | -4.444424657 |
| **A_55_P1995902** | Zfp992 | 0.136326622 | -0.076444692 | 0.349097936 | 9.105576348 | 1.531357996 | 0.171764045 | 0.991835066 | -4.44463559 |
| **A_55_P2113857** | Serpina1e | -0.975437131 | -2.498059439 | 0.547185177 | 10.37418052 | -1.531144672 | 0.171815687 | 0.991835066 | -4.444821934 |
| **A_55_P2045036** | NA | -0.084169188 | -0.215560771 | 0.047222395 | 9.690921492 | -1.531068868 | 0.171834041 | 0.991835066 | -4.444888148 |
| **A_51_P155755** | Pld6 | 0.102375804 | -0.057467182 | 0.262218789 | 8.617348926 | 1.530780333 | 0.17190392 | 0.991835066 | -4.445140173 |
| **A_55_P2044385** | Fgfbp3 | 0.316578502 | -0.177728542 | 0.810885547 | 9.115783153 | 1.530712938 | 0.171920245 | 0.991835066 | -4.445199039 |
| **A_51_P418357** | Mov10l1 | -0.316738484 | -0.811394135 | 0.177917166 | 9.966317747 | -1.530407171 | 0.171994333 | 0.991835066 | -4.445466096 |
| **A_52_P399584** | Ckap2l | -0.097354108 | -0.249405172 | 0.054696956 | 9.029103387 | -1.53029074 | 0.172022552 | 0.991835066 | -4.445567783 |
| **A_55_P1963873** | NA | -0.076209462 | -0.195236924 | 0.042818001 | 13.63990369 | -1.530279822 | 0.172025198 | 0.991835066 | -4.445577318 |
| **A_55_P1992487** | NA | 0.07255024 | -0.04077456 | 0.185875041 | 10.52869741 | 1.530111304 | 0.172066049 | 0.991835066 | -4.445724491 |
| **A_51_P348456** | Rest | 0.134302963 | -0.075489402 | 0.344095328 | 9.044572636 | 1.53004797 | 0.172081405 | 0.991835066 | -4.445779801 |
| **A_52_P437230** | Rbx1 | -0.073923135 | -0.189401525 | 0.041555255 | 14.43550605 | -1.529990687 | 0.172095294 | 0.991835066 | -4.445829827 |
| **A_55_P1990944** | Casp1 | -0.065061906 | -0.166728348 | 0.036604536 | 7.954860543 | -1.529531014 | 0.172206789 | 0.991835066 | -4.446231242 |
| **A_52_P84901** | Slc44a1 | -0.079588655 | -0.2039617 | 0.04478439 | 9.398054467 | -1.529445587 | 0.172227517 | 0.991835066 | -4.446305838 |
| **A_55_P2058348** | Pacc1 | -0.063480139 | -0.162686799 | 0.035726521 | 8.624535745 | -1.529347493 | 0.172251321 | 0.991835066 | -4.446391493 |
| **A_55_P2129000** | Cyyr1 | 0.125827105 | -0.070818454 | 0.322472664 | 9.024108429 | 1.52932291 | 0.172257287 | 0.991835066 | -4.446412959 |
| **A_55_P2011747** | Lsg1 | 0.067085471 | -0.037762517 | 0.171933459 | 10.87394303 | 1.529246531 | 0.172275824 | 0.991835066 | -4.446479651 |
| **A_51_P503261** | Lrrc8c | -0.090127379 | -0.230994501 | 0.050739743 | 9.059110655 | -1.52917163 | 0.172294004 | 0.991835066 | -4.446545052 |
| **A_51_P414706** | Ddt | -0.104366491 | -0.267500338 | 0.058767357 | 12.22157252 | -1.529065656 | 0.17231973 | 0.991835066 | -4.446637582 |
| **A_51_P144319** | Efna5 | 0.1008483 | -0.056819969 | 0.25851657 | 10.87246758 | 1.52873924 | 0.172398991 | 0.991835066 | -4.44692258 |

| | | | | | | | | | |
|---|---|---|---|---|---|---|---|---|---|
| **A_52_P87892** | Abraxas2 | -0.116610801 | -0.298925241 | 0.065703638 | 8.870283976 | -1.528716123 | 0.172404606 | 0.991835066 | -4.446942763 |
| **A_55_P2022921** | Nsa2 | 0.110052724 | -0.062009275 | 0.282114723 | 12.23376219 | 1.528709497 | 0.172406215 | 0.991835066 | -4.446948547 |
| **A_55_P1988699** | Cacna2d1 | -0.119598127 | -0.306682893 | 0.067486638 | 11.01428467 | -1.52790055 | 0.172602799 | 0.991835066 | -4.447654762 |
| **A_51_P406008** | Hgh1 | 0.0806578 | -0.045526 | 0.2068416 | 8.569388447 | 1.527748632 | 0.17263974 | 0.991835066 | -4.447787374 |
| **A_55_P2181288** | Ppp1r2 | -0.080999568 | -0.207718177 | 0.04571904 | 10.13187644 | -1.527747004 | 0.172640136 | 0.991835066 | -4.447788796 |
| **A_55_P2168312** | NA | 0.0776906 | -0.043867832 | 0.199249033 | 14.82452802 | 1.527539747 | 0.172690545 | 0.991835066 | -4.447969708 |
| **A_66_P126094** | Pycrl | 0.126509052 | -0.071443849 | 0.324461954 | 9.565047293 | 1.527456544 | 0.172710786 | 0.991835066 | -4.448042332 |
| **A_52_P281702** | Igfbp5 | 0.329802802 | -0.186296905 | 0.845902509 | 12.75570738 | 1.527319373 | 0.17274416 | 0.991835066 | -4.448162061 |
| **A_66_P119164** | Zfp354c | 0.158141932 | -0.089336034 | 0.405619898 | 9.596551019 | 1.527283415 | 0.17275291 | 0.991835066 | -4.448193446 |
| **A_55_P2028064** | Kdm4b | -0.063165662 | -0.162019613 | 0.035688288 | 11.09809832 | -1.527200847 | 0.172773002 | 0.991835066 | -4.448265512 |
| **A_55_P2052062** | Cd200 | 0.106541858 | -0.060238768 | 0.273322484 | 9.897934559 | 1.526805791 | 0.172869168 | 0.991835066 | -4.448610308 |
| **A_51_P231630** | Zfp239 | 0.063581818 | -0.035961513 | 0.16312515 | 7.879210876 | 1.526616334 | 0.172915304 | 0.991835066 | -4.448775651 |
| **A_55_P2062787** | Mfap3 | 0.077478188 | -0.043829274 | 0.198785651 | 10.99176718 | 1.526514982 | 0.17293999 | 0.991835066 | -4.448864101 |
| **A_55_P2036764** | Prr15l | 0.064966767 | -0.03675589 | 0.166689423 | 7.709441971 | 1.526450385 | 0.172955725 | 0.991835066 | -4.448920473 |
| **A_55_P2118362** | Tmem121 | 0.103863332 | -0.058772209 | 0.266498873 | 8.308456977 | 1.526356307 | 0.172978643 | 0.991835066 | -4.449002572 |
| **A_51_P493857** | Katna1 | 0.105857355 | -0.059940837 | 0.271655548 | 10.17850566 | 1.525985411 | 0.173069026 | 0.991835066 | -4.449326226 |
| **A_55_P2084980** | Dcaf12l2 | -0.11054413 | -0.283704041 | 0.062615781 | 9.816498197 | -1.525799487 | 0.17311435 | 0.991835066 | -4.449488458 |
| **A_51_P314277** | Parp1 | -0.120020058 | -0.308079593 | 0.068039477 | 9.700618605 | -1.525343324 | 0.173225598 | 0.991835066 | -4.44988647 |
| **A_55_P2135701** | BC024978 | 0.072750931 | -0.041250394 | 0.186752255 | 8.380264251 | 1.525238608 | 0.173251145 | 0.991835066 | -4.449977832 |
| **A_55_P2059105** | Gm9920 | 0.205973942 | -0.116793625 | 0.52874151 | 9.470508624 | 1.525216603 | 0.173256514 | 0.991835066 | -4.44999703 |
| **A_65_P15503** | Map3k2 | 0.111961859 | -0.063494697 | 0.287418415 | 9.51371713 | 1.525139725 | 0.173275272 | 0.991835066 | -4.450064102 |
| **A_55_P1967617** | Med16 | -0.186485835 | -0.478747726 | 0.105776057 | 16.36872059 | -1.525045464 | 0.173298275 | 0.991835066 | -4.450146339 |
| **A_55_P1955869** | NA | -0.077971159 | -0.200191648 | 0.044249329 | 12.42116341 | -1.524751637 | 0.173369995 | 0.991835066 | -4.450402673 |
| **A_55_P1970537** | Bcl7b | 0.098156937 | -0.055710339 | 0.252024214 | 10.45502434 | 1.524698118 | 0.173383062 | 0.991835066 | -4.450449362 |
| **A_52_P367774** | Zfp821 | 0.079968645 | -0.045395367 | 0.205332658 | 9.655078968 | 1.524600249 | 0.173406958 | 0.991835066 | -4.450534738 |
| **A_51_P481693** | Ero1l | -0.082348737 | -0.211457574 | 0.0467601 | 8.183552448 | -1.524439197 | 0.173446289 | 0.991835066 | -4.450675229 |
| **A_52_P460537** | Lpar2 | 0.066839454 | -0.037975438 | 0.171654346 | 7.851567797 | 1.524119559 | 0.173524373 | 0.991835066 | -4.450954047 |
| **A_52_P508991** | Fmo1 | -0.091399829 | -0.234730094 | 0.051930436 | 8.323810905 | -1.524111048 | 0.173526452 | 0.991835066 | -4.45096147 |
| **A_55_P2018747** | Cox8a | -0.065965572 | -0.16942488 | 0.037493736 | 16.58240102 | -1.523901502 | 0.17357766 | 0.991835066 | -4.451144245 |
| **A_66_P121745** | Arhgef28 | 0.102058157 | -0.05801635 | 0.262132663 | 8.205142397 | 1.52382355 | 0.173596713 | 0.991835066 | -4.451212236 |
| **A_51_P297679** | Hcls1 | -0.130823771 | -0.336016994 | 0.074369452 | 9.514062924 | -1.523816892 | 0.17359834 | 0.991835066 | -4.451218043 |
| **A_51_P429983** | Supt5 | 0.096693395 | -0.054970776 | 0.248357565 | 11.94847014 | 1.523782378 | 0.173606777 | 0.991835066 | -4.451248146 |
| **A_51_P396570** | Plod2 | 0.137238511 | -0.078027013 | 0.352504035 | 9.410029628 | 1.523739227 | 0.173617325 | 0.991835066 | -4.451285782 |
| **A_55_P2160077** | NA | 0.089208403 | -0.050725737 | 0.229142544 | 17.18808106 | 1.523671044 | 0.173633994 | 0.991835066 | -4.451345251 |
| **A_51_P394802** | Fam111a | 0.116587104 | -0.066316294 | 0.299490501 | 9.933943057 | 1.52348391 | 0.17367975 | 0.991835066 | -4.451508462 |
| **A_55_P2139878** | NA | -0.095955904 | -0.246576143 | 0.054664334 | 8.453948519 | -1.522640945 | 0.173886002 | 0.991835066 | -4.452243589 |
| **A_55_P2128526** | Pfn2 | 0.072790025 | -0.041469582 | 0.187049633 | 10.93890387 | 1.522608587 | 0.173893923 | 0.991835066 | -4.452271805 |
| **A_66_P111877** | Snx13 | -0.077814223 | -0.199970261 | 0.044341814 | 10.2332583 | -1.522485568 | 0.173924043 | 0.991835066 | -4.452379075 |
| **A_55_P2042526** | Apoo | -0.097158217 | -0.249686098 | 0.055369665 | 12.71437453 | -1.522437345 | 0.173935851 | 0.991835066 | -4.452421124 |
| **A_55_P1985764** | NA | 0.088841817 | -0.05065174 | 0.228335374 | 15.14186203 | 1.522202447 | 0.17399338 | 0.991835066 | -4.45262594 |
| **A_55_P2064150** | NA | -0.130582823 | -0.335627039 | 0.074461393 | 15.70923905 | -1.522115693 | 0.174014631 | 0.991835066 | -4.452701582 |
| **A_55_P1967538** | Hunk | 0.129572849 | -0.073885833 | 0.333031532 | 10.9805984 | 1.522113063 | 0.174015276 | 0.991835066 | -4.452703875 |
| **A_55_P2118630** | Lasp1 | 0.067857909 | -0.038694837 | 0.174410655 | 7.889562193 | 1.522106209 | 0.174016955 | 0.991835066 | -4.452709851 |
| **A_55_P2091116** | Kbtbd12 | -0.13668073 | -0.351355372 | 0.077993913 | 11.04358872 | -1.521723237 | 0.174110798 | 0.991835066 | -4.453043751 |
| **A_55_P2077866** | Nktr | 0.185136032 | -0.105718125 | 0.475990189 | 10.81959314 | 1.521334833 | 0.174206019 | 0.991835066 | -4.453382361 |
| **A_55_P2117400** | Cacng7 | 0.150030077 | -0.085708202 | 0.385768357 | 9.382954114 | 1.521098565 | 0.174263966 | 0.991835066 | -4.453588325 |
| **A_55_P2143306** | Rest | 0.121469389 | -0.069394228 | 0.312333006 | 10.39708734 | 1.521082707 | 0.174267857 | 0.991835066 | -4.453602149 |
| **A_51_P475378** | Echdc3 | -0.113477331 | -0.291793754 | 0.064839093 | 9.987583136 | -1.520991887 | 0.174290136 | 0.991835066 | -4.453681318 |

| | | | | | | | | | |
|---|---|---|---|---|---|---|---|---|---|
| **A_55_P2134387** | Olfr544 | 0.286784949 | -0.163939887 | 0.737509785 | 11.29866984 | 1.520736598 | 0.174352777 | 0.991835066 | -4.453903847 |
| **A_55_P2055087** | Prkar2b | 0.101303031 | -0.05793722 | 0.260543282 | 10.11272132 | 1.520473019 | 0.174417475 | 0.991835066 | -4.454133591 |
| **A_55_P2045812** | Sigmar1 | 0.149830064 | -0.085691423 | 0.38535155 | 11.98161421 | 1.520468974 | 0.174418467 | 0.991835066 | -4.454137116 |
| **A_52_P266530** | Ide | 0.29881262 | -0.171022711 | 0.768647951 | 12.07901788 | 1.520065827 | 0.174517466 | 0.991835066 | -4.454488487 |
| **A_55_P1953972** | Pdhb | -0.118294364 | -0.304304989 | 0.06771626 | 15.34133743 | -1.519971467 | 0.174540644 | 0.991835066 | -4.454570724 |
| **A_55_P2130219** | Tpr | 0.126928157 | -0.072721521 | 0.326577835 | 9.342283661 | 1.519492258 | 0.174658402 | 0.991835066 | -4.454988341 |
| **A_55_P2101666** | Vamp5 | 0.24862004 | -0.142489732 | 0.639729811 | 10.27314117 | 1.519310702 | 0.174703036 | 0.991835066 | -4.455146551 |
| **A_55_P2026218** | Igsf8 | 0.075618114 | -0.043345029 | 0.194581256 | 10.43048463 | 1.519226563 | 0.174723724 | 0.991835066 | -4.45521987 |
| **A_55_P2056876** | Akap13 | 0.144031133 | -0.082574522 | 0.370636789 | 9.141070472 | 1.519129389 | 0.17474762 | 0.991835066 | -4.455304544 |
| **A_55_P2000077** | Nrf1 | 0.087648723 | -0.050257968 | 0.225555415 | 9.873905603 | 1.519040623 | 0.174769452 | 0.991835066 | -4.455381891 |
| **A_55_P2017267** | Nanp | 0.080378979 | -0.04609767 | 0.206855629 | 8.955222796 | 1.518942266 | 0.174793645 | 0.991835066 | -4.455467594 |
| **A_51_P318830** | Syt10 | -0.124284035 | -0.319853834 | 0.071285764 | 7.8302944 | -1.518877291 | 0.174809628 | 0.991835066 | -4.455524208 |
| **A_55_P2088525** | Myh10 | 0.142938677 | -0.082000766 | 0.36787812 | 11.40255734 | 1.518774425 | 0.174834936 | 0.991835066 | -4.455613836 |
| **A_55_P2079973** | NA | -0.198360868 | -0.510527153 | 0.113805417 | 14.82395757 | -1.518724979 | 0.174847102 | 0.991835066 | -4.455656918 |
| **A_55_P1965620** | Bud31 | 0.114715286 | -0.065826444 | 0.295257017 | 12.54892486 | 1.518632971 | 0.174869742 | 0.991835066 | -4.455737083 |
| **A_55_P2179726** | Mettl18 | 0.070499511 | -0.040466697 | 0.181465719 | 9.03953124 | 1.518463971 | 0.174911335 | 0.991835066 | -4.455884325 |
| **A_55_P2335648** | Eprn | -0.135032551 | -0.347603273 | 0.07753817 | 9.548301581 | -1.518253043 | 0.17496326 | 0.991835066 | -4.456068091 |
| **A_55_P2010301** | Serpina1c | -0.989239034 | -2.54654463 | 0.568066561 | 10.58868896 | -1.518226379 | 0.174969825 | 0.991835066 | -4.456091321 |
| **A_51_P279606** | Socs1 | -0.081003023 | -0.208522015 | 0.046515969 | 8.676257862 | -1.518222731 | 0.174970723 | 0.991835066 | -4.456094499 |
| **A_55_P2011802** | Pdk2 | -0.21320944 | -0.548930594 | 0.122511714 | 11.78398657 | -1.51787796 | 0.17505563 | 0.991835066 | -4.456394852 |
| **A_55_P2023953** | Arpc3 | 0.12922897 | -0.07425808 | 0.332716021 | 13.30799121 | 1.517861832 | 0.175059603 | 0.991835066 | -4.456408903 |
| **A_55_P2020306** | Aasdh | 0.186591404 | -0.107274855 | 0.480457662 | 9.573245186 | 1.517578063 | 0.175129517 | 0.991835066 | -4.456656096 |
| **A_55_P2133895** | Hnrnph2 | -0.110007047 | -0.283270814 | 0.06325672 | 10.35182867 | -1.517476198 | 0.17515462 | 0.991835066 | -4.456744828 |
| **A_51_P264495** | Pgam2 | -0.141842123 | -0.365247848 | 0.081563602 | 15.29441483 | -1.517469805 | 0.175156196 | 0.991835066 | -4.456750396 |
| **A_55_P2123484** | Gm11627 | -0.093941266 | -0.241904131 | 0.0540216 | 9.729964762 | -1.517444467 | 0.17516244 | 0.991835066 | -4.456772467 |
| **A_55_P2086866** | NA | 0.082874405 | -0.047661804 | 0.213410614 | 13.19349515 | 1.51739469 | 0.175174709 | 0.991835066 | -4.456815826 |
| **A_51_P492446** | Ldah | 0.077010003 | -0.044292583 | 0.198312589 | 11.1957879 | 1.517351544 | 0.175185344 | 0.991835066 | -4.456853408 |
| **A_55_P2056212** | Gpr3 | -0.157200073 | -0.404850052 | 0.090449906 | 9.610037946 | -1.517132741 | 0.175239285 | 0.991835066 | -4.45704399 |
| **A_55_P2125229** | Arhgef2 | 0.084217774 | -0.048475346 | 0.216910894 | 12.2873356 | 1.51692631 | 0.17529019 | 0.991835066 | -4.457223788 |
| **A_51_P351896** | Gask1b | -0.132269611 | -0.34068723 | 0.076148008 | 10.35935524 | -1.516822562 | 0.17531578 | 0.991835066 | -4.457314148 |
| **A_51_P512629** | Ttll5 | -0.077728081 | -0.200207803 | 0.044751641 | 9.205580595 | -1.516781026 | 0.175326025 | 0.991835066 | -4.457350323 |
| **A_55_P2069989** | Klhl12 | 0.079090041 | -0.045551905 | 0.203731986 | 10.96439647 | 1.516584846 | 0.175374424 | 0.991835066 | -4.45752118 |
| **A_66_P136857** | Phrf1 | 0.081439795 | -0.046913445 | 0.209793034 | 10.32147188 | 1.516487958 | 0.175398332 | 0.991835066 | -4.457605559 |
| **A_51_P153013** | Map2k3os | -0.068382392 | -0.17615785 | 0.039393065 | 8.552221772 | -1.516469034 | 0.175403002 | 0.991835066 | -4.45762204 |
| **A_51_P388847** | Stap2 | 0.080511364 | -0.046402802 | 0.207425531 | 9.293930292 | 1.516199041 | 0.175469641 | 0.991835066 | -4.457857164 |
| **A_55_P2337213** | Odr4 | -0.065126164 | -0.167795505 | 0.037543177 | 9.733885613 | -1.516086071 | 0.175497532 | 0.991835066 | -4.457955542 |
| **A_66_P112024** | NA | -0.153637429 | -0.39590095 | 0.088626092 | 11.77814567 | -1.515717055 | 0.175588663 | 0.991835066 | -4.458276873 |
| **A_55_P2067518** | Slc13a3 | -0.077304448 | -0.199218636 | 0.044609739 | 7.831183953 | -1.515511965 | 0.17563933 | 0.991835066 | -4.458455449 |
| **A_51_P225083** | Snx17 | 0.109860836 | -0.063400024 | 0.283121697 | 13.19880339 | 1.515484739 | 0.175646058 | 0.991835066 | -4.458479155 |
| **A_52_P617327** | Rcan1 | -0.173095847 | -0.446159617 | 0.099967924 | 10.44559722 | -1.515066507 | 0.175749427 | 0.991835066 | -4.458843294 |
| **A_55_P1952882** | Cyp4f18 | -0.083317681 | -0.214759758 | 0.048124395 | 8.297716971 | -1.514997425 | 0.175766506 | 0.991835066 | -4.458903438 |
| **A_55_P2212908** | Pon2 | 0.131507904 | -0.0759869 | 0.339002708 | 8.137820862 | 1.514794648 | 0.175816649 | 0.991835066 | -4.459079975 |
| **A_51_P343323** | Cox6c | -0.147518777 | -0.380304945 | 0.08526739 | 16.43336038 | -1.51460458 | 0.175863661 | 0.991835066 | -4.459245439 |
| **A_66_P109049** | Brd9 | 0.118569141 | -0.068548228 | 0.305686511 | 9.722474023 | 1.514491023 | 0.175891754 | 0.991835066 | -4.459344294 |
| **A_55_P1986983** | Gne | -0.100704079 | -0.259638996 | 0.058230839 | 10.97612574 | -1.514386996 | 0.175917493 | 0.991835066 | -4.45943485 |
| **A_52_P592951** | Ostc | -0.09729162 | -0.250858703 | 0.056275463 | 11.67005872 | -1.514211128 | 0.175961015 | 0.991835066 | -4.459587941 |
| **A_52_P665675** | Abca1 | -0.148788423 | -0.383639392 | 0.086062546 | 9.833137367 | -1.514209327 | 0.175961461 | 0.991835066 | -4.459589509 |
| **A_55_P1964477** | Srsf1 | 0.169637184 | -0.098127069 | 0.437401436 | 10.36120925 | 1.514180293 | 0.175968647 | 0.991835066 | -4.459614782 |

| | | | | | | | | | |
|---|---|---|---|---|---|---|---|---|---|
| **A_55_P1977792** | Folr1 | 0.216180632 | -0.125089853 | 0.557451116 | 8.466340844 | 1.514004566 | 0.176012146 | 0.991835066 | -4.459767742 |
| **A_51_P519385** | Sacm1l | -0.120893333 | -0.311785454 | 0.069998788 | 10.24246596 | -1.513643081 | 0.176101658 | 0.991835066 | -4.460082378 |
| **A_55_P2129856** | Mtfmt | 0.092614908 | -0.053643696 | 0.238873512 | 10.15692261 | 1.513451847 | 0.176149029 | 0.991835066 | -4.460248818 |
| **A_55_P2163659** | Rspo3 | 0.204628426 | -0.11852585 | 0.527782701 | 8.776281161 | 1.513439935 | 0.176151981 | 0.991835066 | -4.460259185 |
| **A_55_P2073526** | H2-Ke6 | -0.105078727 | -0.271054919 | 0.060897464 | 11.73650442 | -1.513136616 | 0.176227141 | 0.991835066 | -4.460523163 |
| **A_55_P2039110** | NA | -0.081896112 | -0.211267829 | 0.047475604 | 15.79444728 | -1.512979625 | 0.176266055 | 0.991835066 | -4.460659786 |
| **A_52_P2732** | Mfn1 | -0.138293502 | -0.356862561 | 0.080275556 | 12.18563999 | -1.512245239 | 0.176448191 | 0.991835066 | -4.461298829 |
| **A_55_P2230968** | NA | 0.171503654 | -0.099583781 | 0.44259109 | 8.740640918 | 1.512074263 | 0.17649062 | 0.991835066 | -4.461447593 |
| **A_55_P2143311** | Pdxk | 0.087918182 | -0.051057533 | 0.226893898 | 8.434043065 | 1.511989979 | 0.17651154 | 0.991835066 | -4.461520926 |
| **A_55_P1962602** | NA | 0.149759699 | -0.091158244 | 0.390677642 | 12.71666085 | 1.54416236 | 0.176519443 | 0.991835066 | -4.418079323 |
| **A_51_P459070** | Aplf | 0.10152604 | -0.058973146 | 0.262025226 | 8.838706668 | 1.511867545 | 0.176541931 | 0.991835066 | -4.461627449 |
| **A_66_P138976** | Lpin2 | 0.07093732 | -0.041225568 | 0.183100209 | 8.280753383 | 1.511592496 | 0.176610225 | 0.991835066 | -4.461866745 |
| **A_52_P385489** | Ppp1r3a | -0.116039526 | -0.299566771 | 0.06748772 | 9.113501094 | -1.511174203 | 0.176714132 | 0.991835066 | -4.462230638 |
| **A_55_P2175451** | NA | -0.082258402 | -0.212367195 | 0.047850391 | 11.59305523 | -1.511063631 | 0.176741608 | 0.991835066 | -4.462326825 |
| **A_65_P11193** | Flad1 | -0.14871211 | -0.383935368 | 0.086511148 | 10.49989754 | -1.511037373 | 0.176748134 | 0.991835066 | -4.462349666 |
| **A_51_P518051** | Dusp4 | 0.082939589 | -0.048256393 | 0.214135571 | 8.228788078 | 1.510951335 | 0.176769517 | 0.991835066 | -4.462424508 |
| **A_55_P2132472** | Prpf4b | 0.074250814 | -0.04321242 | 0.191714048 | 8.515158338 | 1.510804947 | 0.176805904 | 0.991835066 | -4.462551844 |
| **A_55_P2000133** | Cdh2 | -0.100435122 | -0.259338377 | 0.058468132 | 14.23204445 | -1.510643381 | 0.176846073 | 0.991835066 | -4.462692379 |
| **A_51_P125467** | Igf2r | 0.083130439 | -0.048430344 | 0.214691221 | 14.46459412 | 1.510228831 | 0.176949176 | 0.991835066 | -4.456498788 |
| **A_55_P2063041** | Prelid3b | -0.089751579 | -0.231800859 | 0.052297701 | 10.30500723 | -1.510122558 | 0.176975617 | 0.991835066 | -4.463145371 |
| **A_52_P136914** | Nudt7 | -0.060295215 | -0.15572446 | 0.03513403 | 8.988823594 | -1.510116628 | 0.176977092 | 0.991835066 | -4.463150528 |
| **A_55_P2129354** | Pilra | -0.106210792 | -0.274317814 | 0.06189623 | 8.289381404 | -1.51005208 | 0.176993153 | 0.991835066 | -4.463206666 |
| **A_55_P2066897** | A230045G1 | 0.109299677 | -0.063738156 | 0.28233751 | 8.911712972 | 1.509687091 | 0.177083997 | 0.991835066 | -4.463524085 |
| **A_51_P341918** | Tsc22d1 | -0.14943606 | -0.386099719 | 0.087227599 | 12.11305452 | -1.509151941 | 0.17721727 | 0.991835066 | -4.463989442 |
| **A_52_P93902** | Alg5 | -0.111841366 | -0.288972524 | 0.065289792 | 10.20765777 | -1.509095181 | 0.177231411 | 0.991835066 | -4.464038797 |
| **A_51_P161021** | Ifit2 | -0.102103887 | -0.263824832 | 0.059617058 | 8.671490156 | -1.50898575 | 0.177258677 | 0.991835066 | -4.464133948 |
| **A_55_P2007919** | Akr1c19 | -0.076219383 | -0.196982947 | 0.04454418 | 8.165757862 | -1.508476826 | 0.177385532 | 0.991835066 | -4.464576435 |
| **A_55_P2103856** | Hgs | 0.102497728 | -0.059902902 | 0.264898358 | 12.12881229 | 1.508466473 | 0.177388113 | 0.991835066 | -4.464585436 |
| **A_66_P107680** | Med8 | 0.083755644 | -0.048958391 | 0.21646968 | 8.360365088 | 1.508364702 | 0.177413491 | 0.991835066 | -4.464673915 |
| **A_55_P2061520** | NA | 0.098940985 | -0.057884989 | 0.25576696 | 8.42268516 | 1.507882045 | 0.177533893 | 0.991835066 | -4.465093509 |
| **A_55_P2197638** | 1110046J0 | -0.072505636 | -0.187431023 | 0.042419751 | 8.355116766 | -1.507873548 | 0.177536013 | 0.991835066 | -4.465100896 |
| **A_51_P417854** | Rab20 | -0.16420857 | -0.424522589 | 0.09610545 | 8.707598565 | -1.507673817 | 0.17758586 | 0.991835066 | -4.465274517 |
| **A_55_P2006876** | NA | -0.126801855 | -0.327821112 | 0.074217402 | 15.30662691 | -1.507638001 | 0.1775948 | 0.991835066 | -4.465305651 |
| **A_51_P243914** | Skap2 | -0.085359654 | -0.220754436 | 0.050035127 | 9.588799355 | -1.506814727 | 0.17780041 | 0.991835066 | -4.46602122 |
| **A_52_P631547** | Cytl1 | 0.116446479 | -0.068264955 | 0.301157912 | 9.267040035 | 1.506751781 | 0.177816139 | 0.991835066 | -4.466075926 |
| **A_55_P2183433** | Rab30 | -0.094436942 | -0.244239634 | 0.055365749 | 8.567432103 | -1.506716112 | 0.177825053 | 0.991835066 | -4.466106925 |
| **A_55_P2058467** | Gm4415 | 0.319160542 | -0.187213452 | 0.825534536 | 12.30709164 | 1.506422984 | 0.177898323 | 0.991835066 | -4.466361668 |
| **A_51_P465148** | Ctsb | -0.121045737 | -0.313107807 | 0.071016334 | 13.44647939 | -1.506319241 | 0.177924261 | 0.991835066 | -4.466451822 |
| **A_52_P456750** | Aph1b | -0.082870934 | -0.214386823 | 0.048644956 | 9.298794908 | -1.506028316 | 0.177997017 | 0.991835066 | -4.466704629 |
| **A_55_P2159070** | 2810025M1 | -0.125294104 | -0.324137105 | 0.073548898 | 12.49544051 | -1.506015553 | 0.17800021 | 0.991835066 | -4.46671572 |
| **A_55_P2064901** | NA | 0.0824599 | -0.048425567 | 0.213345368 | 15.19179999 | 1.505776481 | 0.17806002 | 0.991835066 | -4.466923456 |
| **A_55_P1974028** | Stab2 | 0.151859104 | -0.089254071 | 0.392972278 | 8.622758821 | 1.505320667 | 0.178174104 | 0.991835066 | -4.467319494 |
| **A_55_P2109429** | Rfng | 0.150319242 | -0.088352829 | 0.388991313 | 11.67082577 | 1.505296692 | 0.178180107 | 0.991835066 | -4.467340325 |
| **A_55_P2022364** | Nxn | 0.11300994 | -0.066439161 | 0.292459041 | 12.63607581 | 1.505166311 | 0.178212753 | 0.991835066 | -4.4674536 |
| **A_51_P155873** | Ppp1r3g | -0.196280602 | -0.507968184 | 0.115406979 | 10.14441183 | -1.505105752 | 0.178227918 | 0.991835066 | -4.467506212 |
| **A_55_P1981879** | Ercc6l2 | 0.121023141 | -0.071158643 | 0.313204925 | 10.47116032 | 1.505099921 | 0.178229379 | 0.991835066 | -4.467511278 |
| **A_52_P467930** | NA | -0.111458162 | -0.288454207 | 0.065537883 | 9.822536811 | -1.50507259 | 0.178236223 | 0.991835066 | -4.467535022 |
| **A_55_P2003133** | Ngrn | -0.068749476 | -0.177924512 | 0.04042556 | 12.26833588 | -1.505064742 | 0.178238189 | 0.991835066 | -4.467541841 |

| | | | | | | | | | |
|---|---|---|---|---|---|---|---|---|---|
| **A_55_P2183055** | E4f1 | 0.186625136 | -0.109762075 | 0.483012346 | 10.05823456 | 1.504942163 | 0.178268891 | 0.991835066 | -4.467648332 |
| **A_55_P2100155** | Mapt | 0.06454071 | -0.037992227 | 0.167073646 | 7.847465986 | 1.50445594 | 0.17839072 | 0.991835066 | -4.468070714 |
| **A_55_P2052696** | Synm | 0.116998843 | -0.068885235 | 0.302882921 | 8.681729596 | 1.504348679 | 0.178417606 | 0.991835066 | -4.468163886 |
| **A_66_P100249** | Snhg12 | 0.190886549 | -0.112390019 | 0.494163117 | 10.33195308 | 1.504338613 | 0.178420129 | 0.991835066 | -4.46817263 |
| **A_55_P1954604** | NA | 0.094365337 | -0.055590757 | 0.24432143 | 8.728312482 | 1.504033488 | 0.178496633 | 0.991835066 | -4.468437663 |
| **A_55_P2050448** | Slc25a3 | -0.099550746 | -0.257771369 | 0.058669877 | 12.55245879 | -1.503801659 | 0.178554779 | 0.991835066 | -4.468639019 |
| **A_51_P142187** | Abhd11 | -0.094841133 | -0.245577164 | 0.055894899 | 11.50726302 | -1.503795526 | 0.178556318 | 0.991835066 | -4.468644346 |
| **A_65_P15809** | Gm4759 | 0.206847991 | -0.121930538 | 0.535626519 | 9.474767574 | 1.503685428 | 0.178583938 | 0.991835066 | -4.468739968 |
| **A_55_P1974587** | Vangl1 | 0.071891612 | -0.042385419 | 0.186168643 | 8.730058957 | 1.503586453 | 0.178608772 | 0.991835066 | -4.468825927 |
| **A_51_P245796** | Ddit4 | -0.231996334 | -0.60077538 | 0.136782712 | 12.23557251 | -1.503571269 | 0.178612582 | 0.991835066 | -4.468839115 |
| **A_52_P408690** | Sft2d2 | -0.115453343 | -0.298992937 | 0.068086252 | 8.759157798 | -1.503439224 | 0.178645718 | 0.991835066 | -4.468953792 |
| **A_51_P254045** | Traip | 0.16461994 | -0.097090966 | 0.426330846 | 9.493751316 | 1.503383399 | 0.178659729 | 0.991835066 | -4.469002274 |
| **A_55_P2034953** | NA | 0.114495266 | -0.067533257 | 0.296523788 | 10.22981843 | 1.503340014 | 0.178670619 | 0.991835066 | -4.469039952 |
| **A_55_P2008918** | Cbll1 | 0.096302998 | -0.056817281 | 0.249423277 | 9.115259795 | 1.50319814 | 0.178706233 | 0.991835066 | -4.46916316 |
| **A_52_P10781** | Lman1 | 0.081837733 | -0.048296692 | 0.211972157 | 9.445293008 | 1.503039958 | 0.178745948 | 0.991835066 | -4.469300525 |
| **A_55_P2046812** | Sparc | 0.109604172 | -0.064687547 | 0.283895891 | 17.16100033 | 1.503001675 | 0.178755562 | 0.991835066 | -4.46933377 |
| **A_55_P2169839** | Rufy3 | 0.126126499 | -0.074450449 | 0.326703448 | 10.25677391 | 1.502915127 | 0.178777296 | 0.991835066 | -4.469408926 |
| **A_51_P490509** | Bub1b | -0.096677833 | -0.250446808 | 0.057091142 | 10.33394018 | -1.502682813 | 0.178835648 | 0.991835066 | -4.469610654 |
| **A_52_P274555** | Snx14 | -0.090198265 | -0.233685767 | 0.053289237 | 9.817530103 | -1.50242652 | 0.178900044 | 0.991835066 | -4.469833192 |
| **A_55_P2069485** | Ptpn13 | 0.113839258 | -0.067266431 | 0.294944946 | 9.851481239 | 1.502342994 | 0.178921035 | 0.991835066 | -4.469905715 |
| **A_55_P1996953** | NA | 0.117569796 | -0.069493585 | 0.304633176 | 8.930103653 | 1.502159738 | 0.178967097 | 0.991835066 | -4.470064825 |
| **A_55_P1990866** | Mylk3 | 0.147310947 | -0.087132 | 0.381753893 | 9.348307484 | 1.501782302 | 0.179062002 | 0.991835066 | -4.470392509 |
| **A_55_P1977071** | Fam110a | -0.134769716 | -0.3492665 | 0.079727069 | 9.404889796 | -1.501691279 | 0.179084897 | 0.991835066 | -4.47047153 |
| **A_55_P2058933** | Kif1b | -0.140929432 | -0.365270908 | 0.083412044 | 11.84612263 | -1.501416765 | 0.17915396 | 0.991835066 | -4.470709838 |
| **A_55_P2084533** | Zbtb40 | 0.101558406 | -0.060171294 | 0.263288106 | 10.53319907 | 1.500842875 | 0.179298419 | 0.991835066 | -4.47120799 |
| **A_55_P2014645** | Knop1 | 0.086293386 | -0.051149672 | 0.223736445 | 9.528204881 | 1.500596165 | 0.179360554 | 0.991835066 | -4.471422121 |
| **A_55_P2033215** | Cabp5 | 0.389167481 | -0.230926736 | 1.009261697 | 11.24232227 | 1.499988725 | 0.179513623 | 0.991835066 | -4.471949298 |
| **A_51_P408989** | L3hypdh | 0.063723456 | -0.03781809 | 0.165265001 | 9.789954933 | 1.499908214 | 0.17953392 | 0.991835066 | -4.472019166 |
| **A_51_P478172** | Tcrb-J | -0.430297538 | -1.116092795 | 0.255497719 | 7.898747959 | -1.499627917 | 0.179604601 | 0.991835066 | -4.472262399 |
| **A_55_P1980863** | NA | -0.084014567 | -0.217953719 | 0.049924585 | 8.465013501 | -1.499188292 | 0.179715509 | 0.991835066 | -4.472643861 |
| **A_55_P2038007** | Csrp1 | 0.137717169 | -0.081893381 | 0.357327718 | 15.78572889 | 1.498801157 | 0.179813227 | 0.991835066 | -4.472979747 |
| **A_55_P2105371** | Dlk1 | -0.219136622 | -0.568655072 | 0.130381828 | 11.24878851 | -1.498490483 | 0.179891681 | 0.991835066 | -4.473249273 |
| **A_51_P120093** | Snx10 | -0.075079273 | -0.194837398 | 0.044678852 | 8.821681083 | -1.498387707 | 0.179917642 | 0.991835066 | -4.473338433 |
| **A_55_P1984366** | Cdadc1 | -0.123953095 | -0.321674671 | 0.073768482 | 9.374056623 | -1.498347135 | 0.179927891 | 0.991835066 | -4.473373629 |
| **A_55_P2083879** | Vsig2 | 0.166068177 | -0.098863748 | 0.431000103 | 8.755248013 | 1.498170551 | 0.179972505 | 0.991835066 | -4.473526812 |
| **A_55_P2060184** | Lamtor3 | -0.072527452 | -0.188233794 | 0.04317889 | 12.45418542 | -1.498146868 | 0.17997849 | 0.991835066 | -4.473547356 |
| **A_51_P143468** | Klhl26 | 0.084087855 | -0.050066352 | 0.218242062 | 8.744462909 | 1.498090705 | 0.179992682 | 0.991835066 | -4.473596075 |
| **A_55_P1953593** | Ece2 | 0.127977976 | -0.07620234 | 0.332158292 | 10.39863219 | 1.498064449 | 0.179999317 | 0.991835066 | -4.473618851 |
| **A_51_P368009** | E2f2 | 0.177622721 | -0.105780885 | 0.461026328 | 10.51018476 | 1.497967113 | 0.180023917 | 0.991835066 | -4.473703282 |
| **A_55_P1954768** | NA | -0.0806095 | -0.209232329 | 0.04801333 | 8.840954297 | -1.497880955 | 0.180045694 | 0.991835066 | -4.473778017 |
| **A_52_P207964** | Mrps18b | -0.137923867 | -0.358031748 | 0.082184014 | 9.002717978 | -1.497659085 | 0.180101786 | 0.991835066 | -4.473970465 |
| **A_55_P2034481** | Erdr1 | 0.374587011 | -0.223288992 | 0.972463014 | 12.62195074 | 1.497444461 | 0.180156061 | 0.991835066 | -4.474156617 |
| **A_55_P2175942** | NA | -0.128275071 | -0.333023253 | 0.07647311 | 14.84873681 | -1.497377639 | 0.180172962 | 0.991835066 | -4.474214574 |
| **A_55_P2147487** | Cyth3 | 0.158183237 | -0.094309059 | 0.410675533 | 9.757588115 | 1.49734356 | 0.180181582 | 0.991835066 | -4.47424413 |
| **A_55_P1957378** | Unc119 | 0.079275091 | -0.047273574 | 0.205823756 | 11.82649743 | 1.497229289 | 0.180210489 | 0.991835066 | -4.474343237 |
| **A_55_P1975510** | H6pd | 0.063614805 | -0.037939587 | 0.165169197 | 10.4225129 | 1.497161394 | 0.180227667 | 0.991835066 | -4.47440212 |
| **A_66_P118759** | Safb2 | 0.182025979 | -0.108637049 | 0.472689008 | 11.53731563 | 1.49676188 | 0.180328774 | 0.991835066 | -4.474748591 |
| **A_55_P2080603** | Hspa8 | -0.220860621 | -0.573600439 | 0.131879197 | 15.89291334 | -1.496486955 | 0.180398381 | 0.991835066 | -4.474986997 |

| | | | | | | | | | |
|---|---|---|---|---|---|---|---|---|---|
| **A_55_P2094881** | Fgfr2 | -0.074683117 | -0.193988063 | 0.04462183 | 8.528594376 | -1.496143039 | 0.18048549 | 0.991835066 | -4.475285209 |
| **A_51_P313157** | Arhgef5 | 0.088300206 | -0.052763475 | 0.229363888 | 9.244896641 | 1.496082824 | 0.180500746 | 0.991835066 | -4.475337419 |
| **A_55_P2088416** | Lamtor5 | -0.064149733 | -0.166661225 | 0.038361758 | 12.61626368 | -1.495655025 | 0.180609164 | 0.991835066 | -4.47570833 |
| **A_55_P1957227** | Las1l | -0.097956343 | -0.254541558 | 0.058628871 | 9.913421969 | -1.495171274 | 0.180731834 | 0.991835066 | -4.47612771 |
| **A_55_P2052420** | Setd5 | 0.11273932 | -0.067478377 | 0.292957017 | 12.81999016 | 1.495158068 | 0.180735184 | 0.991835066 | -4.476139158 |
| **A_55_P2019751** | Paox | 0.072854249 | -0.043631706 | 0.189340203 | 10.53599324 | 1.494825351 | 0.1808196 | 0.991835066 | -4.476427574 |
| **A_52_P214612** | NA | -0.080919317 | -0.210307446 | 0.048468813 | 9.620750921 | -1.49474429 | 0.180840172 | 0.991835066 | -4.476497839 |
| **A_51_P379976** | Arl4a | -0.090289832 | -0.23470054 | 0.054120875 | 10.1431656 | -1.49433712 | 0.180943539 | 0.991835066 | -4.47685076 |
| **A_55_P2143436** | NA | -0.07693721 | -0.200047454 | 0.046173033 | 7.931779655 | -1.493658718 | 0.181115882 | 0.991835066 | -4.477438703 |
| **A_55_P2004333** | Rpl24 | -0.071531959 | -0.185994744 | 0.042930827 | 15.70659692 | -1.493636574 | 0.18112151 | 0.991835066 | -4.477457893 |
| **A_55_P1996674** | Itih3 | -0.230930347 | -0.600469978 | 0.138609284 | 8.227148685 | -1.49358217 | 0.181135338 | 0.991835066 | -4.477505039 |
| **A_55_P2078325** | Smco3 | 0.066791641 | -0.040098524 | 0.173681806 | 7.709497256 | 1.493459673 | 0.181166477 | 0.991835066 | -4.477611191 |
| **A_55_P1997614** | Gm9693 | -0.222208264 | -0.577859489 | 0.133442962 | 11.39427476 | -1.493292998 | 0.181208854 | 0.991835066 | -4.477755621 |
| **A_55_P1988498** | Naa20 | -0.066572412 | -0.173123667 | 0.039978843 | 12.03344784 | -1.493292403 | 0.181209005 | 0.991835066 | -4.477756136 |
| **A_51_P451075** | Atp2a2 | -0.213831038 | -0.556076928 | 0.128414852 | 15.40199987 | -1.493281327 | 0.181211821 | 0.991835066 | -4.477765734 |
| **A_51_P159610** | Ccdc120 | 0.095500521 | -0.05735847 | 0.248359513 | 7.824379072 | 1.4932203 | 0.18122734 | 0.991835066 | -4.477818615 |
| **A_51_P308814** | Toe1 | 0.082125367 | -0.049332355 | 0.213583088 | 8.854974344 | 1.493139392 | 0.181247915 | 0.991835066 | -4.477888722 |
| **A_66_P126563** | Ptges3l | -0.11035983 | -0.287032785 | 0.066313124 | 11.91764238 | -1.492966572 | 0.181291872 | 0.991835066 | -4.478038466 |
| **A_55_P2072908** | NA | -0.086303171 | -0.224465787 | 0.051859444 | 14.07029004 | -1.49295027 | 0.181296019 | 0.991835066 | -4.478052591 |
| **A_55_P1999648** | Plin2 | -0.096376943 | -0.250671278 | 0.057917392 | 11.31889301 | -1.492905431 | 0.181307426 | 0.991835066 | -4.478091443 |
| **A_51_P325485** | Utp3 | 0.088211514 | -0.053020081 | 0.229443109 | 12.12028088 | 1.492803153 | 0.181333447 | 0.991835066 | -4.47818006 |
| **A_55_P2155616** | Gm9457 | 0.067690035 | -0.040687581 | 0.176067651 | 14.84207516 | 1.492774732 | 0.181340679 | 0.991835066 | -4.478204685 |
| **A_51_P503937** | Calm1 | -0.085986057 | -0.223668164 | 0.05169605 | 10.08137788 | -1.492655762 | 0.181370952 | 0.991835066 | -4.478307762 |
| **A_55_P2081378** | NA | 0.099881848 | -0.060097121 | 0.259860818 | 15.51796205 | 1.492219835 | 0.181481919 | 0.991835066 | -4.478685431 |
| **A_51_P304859** | Zfand6 | -0.092947752 | -0.24182342 | 0.055927916 | 9.16649737 | -1.492190594 | 0.181489365 | 0.991835066 | -4.478710762 |
| **A_55_P2082658** | NA | 0.084558734 | -0.050897323 | 0.220014792 | 8.311923135 | 1.492001207 | 0.181537595 | 0.991835066 | -4.478874826 |
| **A_52_P434841** | Coa4 | -0.139456453 | -0.362887405 | 0.0839745 | 9.834555655 | -1.491778724 | 0.181594269 | 0.991835066 | -4.479067553 |
| **A_52_P18816** | Gosr2 | 0.081398389 | -0.049038461 | 0.211835238 | 8.719080928 | 1.491504746 | 0.181664082 | 0.991835066 | -4.479304874 |
| **A_55_P2164189** | Sfpq | 0.166162411 | -0.100127441 | 0.432452263 | 10.73294614 | 1.491376521 | 0.181696764 | 0.991835066 | -4.479415937 |
| **A_55_P2110537** | Slfn9 | 0.153742996 | -0.092659983 | 0.400145976 | 9.66587486 | 1.491277683 | 0.18172196 | 0.991835066 | -4.479501545 |
| **A_51_P466378** | Itgal | -0.066101538 | -0.172052717 | 0.03984964 | 8.013065633 | -1.491127944 | 0.181760137 | 0.991835066 | -4.479631236 |
| **A_55_P2138422** | Obox1 | -0.089627261 | -0.233302829 | 0.054048307 | 7.848293272 | -1.490961173 | 0.181802665 | 0.991835066 | -4.479775674 |
| **A_55_P2011702** | Ppargc1a | -0.07858937 | -0.20458402 | 0.04740528 | 9.259239437 | -1.490804976 | 0.181842505 | 0.991835066 | -4.47991095 |
| **A_52_P516409** | Col4a6 | 0.185405426 | -0.111901009 | 0.48271186 | 9.929487944 | 1.490483811 | 0.181924448 | 0.991835066 | -4.480189081 |
| **A_55_P1975400** | Flnc | -0.069964494 | -0.182166209 | 0.04223722 | 14.2698209 | -1.490346802 | 0.181959415 | 0.991835066 | -4.480307726 |
| **A_52_P513177** | Sla | -0.067156463 | -0.174867448 | 0.040554522 | 7.81374787 | -1.490173891 | 0.182003553 | 0.991835066 | -4.480457456 |
| **A_52_P37681** | Fkbp9 | -0.087070678 | -0.22672899 | 0.052587635 | 12.28717861 | -1.490096061 | 0.182023424 | 0.991835066 | -4.480524849 |
| **A_55_P2106922** | Arfgap3 | 0.074839622 | -0.045208919 | 0.194888164 | 10.81856177 | 1.48999164 | 0.182050086 | 0.991835066 | -4.480615267 |
| **A_51_P490296** | Slx4 | 0.12999713 | -0.078567955 | 0.338562215 | 10.00774911 | 1.489708485 | 0.182122405 | 0.991835066 | -4.480860437 |
| **A_51_P379750** | Myoz1 | -0.0933546 | -0.243162773 | 0.056453574 | 9.454747274 | -1.489393122 | 0.18220298 | 0.991835066 | -4.481133478 |
| **A_66_P128942** | Mipep | -0.089462898 | -0.233030632 | 0.054104836 | 8.52926641 | -1.489344788 | 0.182215332 | 0.991835066 | -4.481175323 |
| **A_51_P444437** | Ptgr1 | -0.105144807 | -0.273897516 | 0.063607902 | 9.985561933 | -1.489176615 | 0.182258316 | 0.991835066 | -4.481320916 |
| **A_55_P2247608** | 4931428AC | -0.074936345 | -0.195232205 | 0.045359515 | 8.766205042 | -1.488850041 | 0.182341814 | 0.991835066 | -4.481603628 |
| **A_51_P504442** | Sf3a1 | 0.086317505 | -0.052254158 | 0.224889169 | 11.00363382 | 1.488790477 | 0.182357046 | 0.991835066 | -4.48165519 |
| **A_65_P03442** | Pgk1 | 0.132493873 | -0.080238642 | 0.345226388 | 15.42738258 | 1.488576138 | 0.182411871 | 0.991835066 | -4.481840727 |
| **A_55_P2089565** | Tbx22 | 0.199561188 | -0.120881254 | 0.520003629 | 8.554729615 | 1.488453186 | 0.182443327 | 0.991835066 | -4.481947154 |
| **A_55_P2094237** | NA | 0.09132073 | -0.055347233 | 0.237988693 | 16.26657811 | 1.488138144 | 0.18252395 | 0.991835066 | -4.482219838 |
| **A_51_P173022** | Nsmce1 | -0.101195798 | -0.26375159 | 0.061359994 | 11.60016188 | -1.48788431 | 0.182588933 | 0.991835066 | -4.482439529 |

| | | | | | | | | | |
|---|---|---|---|---|---|---|---|---|---|
| **A_55_P2165414** | Serpina1a | -0.999064243 | -2.60420894 | 0.606080454 | 10.58350417 | -1.487607501 | 0.182659822 | 0.991835066 | -4.482679092 |
| **A_51_P134972** | Shkbp1 | 0.070142655 | -0.042569019 | 0.18285433 | 10.25711611 | 1.487381702 | 0.182717666 | 0.991835066 | -4.482874496 |
| **A_55_P2107477** | Rab5c | 0.081511182 | -0.049483312 | 0.212505675 | 12.25780412 | 1.487213374 | 0.182760799 | 0.991835066 | -4.48302016 |
| **A_51_P311905** | Elp6 | 0.089623589 | -0.054455658 | 0.233702836 | 8.473414241 | 1.486722906 | 0.18288653 | 0.991835066 | -4.483444556 |
| **A_51_P349727** | Slc25a45 | -0.063645484 | -0.165964098 | 0.038673129 | 8.12364312 | -1.486695692 | 0.182893508 | 0.991835066 | -4.483468102 |
| **A_51_P490795** | Mxd1 | -0.106043889 | -0.276540153 | 0.064452376 | 9.062352221 | -1.486551328 | 0.182930532 | 0.991835066 | -4.483593009 |
| **A_55_P2018274** | NA | -0.076892155 | -0.200523746 | 0.046739436 | 12.82158503 | -1.486489024 | 0.182946513 | 0.991835066 | -4.483646914 |
| **A_51_P472481** | Dync2li1 | -0.070523423 | -0.183932553 | 0.042885707 | 8.23629928 | -1.486259002 | 0.183005524 | 0.991835066 | -4.483845922 |
| **A_55_P2419483** | 4732460I02 | 0.201389584 | -0.122487362 | 0.52526653 | 8.502503526 | 1.486161823 | 0.18303046 | 0.991835066 | -4.483929996 |
| **A_55_P2145543** | Eef2k | 0.107537009 | -0.065415156 | 0.280489175 | 8.429578998 | 1.486076213 | 0.18305243 | 0.991835066 | -4.484004059 |
| **A_52_P1197913** | Gadd45b | -0.134975106 | -0.352077216 | 0.082127005 | 10.59110536 | -1.485931411 | 0.183089596 | 0.991835066 | -4.484129327 |
| **A_55_P2099232** | Bdp1 | 0.08900678 | -0.054169235 | 0.232182795 | 10.14979815 | 1.485805463 | 0.183121928 | 0.991835066 | -4.484238281 |
| **A_51_P314521** | Smim11 | -0.08336282 | -0.217474467 | 0.050748827 | 12.94895091 | -1.485644965 | 0.183163137 | 0.991835066 | -4.484377119 |
| **A_51_P286373** | Ttll12 | 0.100576803 | -0.061236404 | 0.26239001 | 10.08896329 | 1.485569572 | 0.183182498 | 0.991835066 | -4.484442336 |
| **A_55_P2062437** | Begain | 0.100635952 | -0.061309983 | 0.262581887 | 8.303047449 | 1.48522497 | 0.183271014 | 0.991835066 | -4.484740408 |
| **A_55_P2008538** | Chmp4b | 0.616789756 | -0.375834552 | 1.609414065 | 12.55789702 | 1.485119424 | 0.183298133 | 0.991835066 | -4.484831699 |
| **A_52_P633752** | Apc | 0.060934178 | -0.037169503 | 0.15903786 | 8.100458209 | 1.484515671 | 0.183453333 | 0.991835066 | -4.485353863 |
| **A_55_P1962224** | Afap1l2 | -0.101516399 | -0.264959765 | 0.061926967 | 8.406823754 | -1.484492601 | 0.183459265 | 0.991835066 | -4.485373814 |
| **A_55_P2027831** | Cryba4 | -0.096024477 | -0.250643857 | 0.058594903 | 8.45745897 | -1.484318688 | 0.183503994 | 0.991835066 | -4.485524211 |
| **A_55_P2031979** | Sspn | -0.087503943 | -0.228419116 | 0.053411229 | 13.74597755 | -1.484154113 | 0.183546331 | 0.991835066 | -4.485666527 |
| **A_51_P483159** | Gchfr | -0.126163608 | -0.329462582 | 0.077135366 | 9.181232832 | -1.483228444 | 0.183784625 | 0.991835066 | -4.486466898 |
| **A_52_P192426** | Tnfrsf1a | -0.093397084 | -0.243925366 | 0.057131199 | 12.04977342 | -1.482942604 | 0.183858266 | 0.991835066 | -4.486714012 |
| **A_55_P2107035** | Mcmbp | 0.075013696 | -0.045895565 | 0.195922956 | 11.5546499 | 1.482825788 | 0.183888369 | 0.991835066 | -4.486814998 |
| **A_51_P108266** | Actn2 | -0.164524955 | -0.429731343 | 0.100681433 | 14.78020758 | -1.482712446 | 0.183917581 | 0.991835066 | -4.486912977 |
| **A_55_P2125149** | Tulp1 | -0.26537985 | -0.693259218 | 0.162499518 | 9.57377265 | -1.482366976 | 0.184006646 | 0.991835066 | -4.487211605 |
| **A_55_P2088440** | Arrdc2 | 0.087838285 | -0.053854859 | 0.229531428 | 8.949334426 | 1.481644944 | 0.18419292 | 0.991835066 | -4.487835661 |
| **A_55_P2022658** | Cdk11b | 0.131915796 | -0.080892994 | 0.344724586 | 12.22637072 | 1.481550206 | 0.184217374 | 0.991835066 | -4.487917536 |
| **A_52_P366932** | Gna11 | -0.128550826 | -0.335990716 | 0.078889064 | 10.78358421 | -1.481125058 | 0.18432715 | 0.991835066 | -4.488284937 |
| **A_55_P1994294** | NA | -0.138959868 | -0.363230995 | 0.085311258 | 10.64418955 | -1.480898052 | 0.184385789 | 0.991835066 | -4.488481095 |
| **A_52_P545643** | Tmem263 | 0.08858907 | -0.054387593 | 0.231565733 | 10.92841217 | 1.48089449 | 0.184386709 | 0.991835066 | -4.488484172 |
| **A_52_P123400** | Dnaaf2 | 0.096646454 | -0.059349344 | 0.252642251 | 8.987925054 | 1.480751415 | 0.184423677 | 0.991835066 | -4.488607799 |
| **A_55_P1968729** | Rpl10 | 0.145283613 | -0.089233627 | 0.379800854 | 16.5739956 | 1.480645154 | 0.184451136 | 0.991835066 | -4.488699613 |
| **A_55_P1980651** | Ubr5 | 0.103968882 | -0.063864404 | 0.271802167 | 12.10052537 | 1.480588624 | 0.184465746 | 0.991835066 | -4.488748456 |
| **A_55_P2119877** | Pnisr | 0.109182831 | -0.067077416 | 0.285443078 | 9.216784868 | 1.48050238 | 0.184488038 | 0.991835066 | -4.488822973 |
| **A_55_P1985565** | NA | 0.08122021 | -0.049907708 | 0.212348128 | 15.67775778 | 1.480396585 | 0.184515386 | 0.991835066 | -4.488914379 |
| **A_51_P153423** | Fndc1 | -0.202749332 | -0.530116831 | 0.124618167 | 11.21966359 | -1.480242958 | 0.184555105 | 0.991835066 | -4.489047108 |
| **A_51_P384879** | Mdh2 | -0.104551872 | -0.273379362 | 0.064275618 | 17.69402449 | -1.480122917 | 0.184586146 | 0.991835066 | -4.489150816 |
| **A_55_P2338535** | Slc35a1 | 0.077478982 | -0.047679313 | 0.202637277 | 8.840411469 | 1.479562783 | 0.184731054 | 0.991835066 | -4.489634701 |
| **A_55_P2050592** | NA | -0.060554752 | -0.158383373 | 0.037273868 | 8.354487595 | -1.479419825 | 0.184768054 | 0.991835066 | -4.489758189 |
| **A_55_P2023081** | Dnaja2 | -0.075811468 | -0.198335283 | 0.046712347 | 10.6734859 | -1.478847962 | 0.18491613 | 0.991835066 | -4.490252124 |
| **A_55_P2008639** | NA | -0.094366366 | -0.246924523 | 0.058191791 | 12.76211865 | -1.47839651 | 0.185033104 | 0.991835066 | -4.49064201 |
| **A_52_P322141** | Ccdc88b | -0.113252908 | -0.296371257 | 0.069865441 | 8.439510242 | -1.478177644 | 0.185089838 | 0.991835066 | -4.490831015 |
| **A_55_P1963900** | Mib2 | 0.097479003 | -0.060151076 | 0.255109082 | 10.97311433 | 1.478022751 | 0.185129999 | 0.991835066 | -4.490964768 |
| **A_55_P2399688** | Glipr2 | -0.088890398 | -0.23263505 | 0.054854254 | 8.502179398 | -1.477992695 | 0.185137793 | 0.991835066 | -4.490990722 |
| **A_55_P1998396** | Eif2ak4 | 0.143384457 | -0.088497535 | 0.375266448 | 9.889430062 | 1.477897064 | 0.185162593 | 0.991835066 | -4.491073299 |
| **A_55_P2168504** | Mcmbp | 0.089718045 | -0.055378382 | 0.234814472 | 9.948514011 | 1.477856325 | 0.185173159 | 0.991835066 | -4.491108475 |
| **A_55_P1996907** | BC067074 | 0.102046799 | -0.06301499 | 0.267108589 | 8.458273802 | 1.47761731 | 0.185235159 | 0.991835066 | -4.491314853 |
| **A_55_P2359807** | NA | 0.139540378 | -0.086268091 | 0.365348848 | 9.195069146 | 1.476960231 | 0.185405703 | 0.991835066 | -4.491882149 |

| | | | | | | | | | |
|---|---|---|---|---|---|---|---|---|---|
| **A_55_P1966982** | Gsr | 0.144855744 | -0.089569929 | 0.379281418 | 11.75330672 | 1.476861205 | 0.185431418 | 0.991835066 | -4.491967636 |
| **A_52_P495372** | Twnk | -0.09217607 | -0.241362498 | 0.057010358 | 8.953903242 | -1.476719483 | 0.185468225 | 0.991835066 | -4.492089978 |
| **A_55_P2000027** | Spink2 | -0.095021051 | -0.24885598 | 0.058813879 | 8.582442626 | -1.476297908 | 0.185577753 | 0.991835066 | -4.492453883 |
| **A_55_P2060938** | NA | -0.171306825 | -0.448684513 | 0.106070863 | 13.05423557 | -1.476088084 | 0.185632289 | 0.991835066 | -4.49263499 |
| **A_51_P476711** | Mtrex | 0.100505408 | -0.062246503 | 0.263257318 | 11.26155867 | 1.475952788 | 0.185667462 | 0.991835066 | -4.492751764 |
| **A_55_P1980548** | NA | -0.08994775 | -0.235647963 | 0.055752463 | 8.44663736 | -1.475500128 | 0.185785185 | 0.991835066 | -4.49314243 |
| **A_51_P381218** | Gulp1 | 0.143565576 | -0.089023769 | 0.376154922 | 9.737661417 | 1.475263625 | 0.185846719 | 0.991835066 | -4.493346526 |
| **A_55_P2039180** | Zswim8 | 0.097679621 | -0.060573269 | 0.255932511 | 12.02725494 | 1.475235823 | 0.185853954 | 0.991835066 | -4.493370518 |
| **A_55_P2171348** | Grm2 | -0.101015122 | -0.264699233 | 0.062668989 | 9.216327996 | -1.474989741 | 0.185918002 | 0.991835066 | -4.493582866 |
| **A_51_P229911** | Adcy4 | -0.09436232 | -0.247267679 | 0.058543039 | 10.49295071 | -1.474976276 | 0.185921507 | 0.991835066 | -4.493594485 |
| **A_52_P503208** | NA | -0.177732018 | -0.465744077 | 0.11028004 | 12.07386256 | -1.474905291 | 0.185939986 | 0.991835066 | -4.493655737 |
| **A_52_P494514** | Insr | -0.075853644 | -0.198782035 | 0.047074747 | 9.511898542 | -1.474800873 | 0.185967173 | 0.991835066 | -4.493745836 |
| **A_51_P285077** | Hhatl | -0.218613818 | -0.572919898 | 0.135692263 | 9.612205083 | -1.474715127 | 0.1859895 | 0.991835066 | -4.493819822 |
| **A_55_P2178608** | NA | -0.10892522 | -0.285460913 | 0.067610473 | 8.993162423 | -1.474704658 | 0.185992226 | 0.991835066 | -4.493828855 |
| **A_55_P1952324** | NA | -0.155431038 | -0.407359269 | 0.096497194 | 11.40357886 | -1.474585778 | 0.186023186 | 0.991835066 | -4.493931427 |
| **A_55_P2045114** | Tmem242 | -0.157793423 | -0.41356413 | 0.097977284 | 11.07919788 | -1.474508316 | 0.186043362 | 0.991835066 | -4.493998262 |
| **A_51_P512561** | Zfp68 | -0.099900653 | -0.261858307 | 0.062057002 | 11.22787446 | -1.474266441 | 0.186106373 | 0.991835066 | -4.494206945 |
| **A_52_P220783** | Tmem229b | -0.093302606 | -0.244568979 | 0.057963768 | 9.469654716 | -1.474213931 | 0.186120056 | 0.991835066 | -4.494252248 |
| **A_52_P200265** | Mocs1 | 0.085678453 | -0.05325912 | 0.224616026 | 9.245890094 | 1.473876318 | 0.186208047 | 0.991835066 | -4.494543507 |
| **A_55_P1981494** | H2-Bl | -0.077633363 | -0.203543623 | 0.048276897 | 8.030845441 | -1.473657002 | 0.186265227 | 0.991835066 | -4.494732699 |
| **A_52_P238816** | Ppt2 | -0.09029315 | -0.236751638 | 0.056165338 | 10.15000713 | -1.473497457 | 0.186306834 | 0.991835066 | -4.494870324 |
| **A_55_P2176953** | Usp51 | 0.074816401 | -0.046539017 | 0.196171819 | 8.081557318 | 1.473488589 | 0.186309147 | 0.991835066 | -4.494877974 |
| **A_66_P134809** | Gas5 | 0.151877354 | -0.094503852 | 0.398258561 | 12.51268954 | 1.473311497 | 0.18635534 | 0.991835066 | -4.495030728 |
| **A_55_P2104978** | Edc4 | 0.095663001 | -0.059610615 | 0.250936617 | 11.65889294 | 1.472500548 | 0.186567003 | 0.991835066 | -4.495730147 |
| **A_55_P2089125** | Zfp976 | 0.090230086 | -0.056226067 | 0.236686239 | 11.56919255 | 1.472491788 | 0.186569291 | 0.991835066 | -4.495737702 |
| **A_55_P2111907** | Noxo1 | 0.062674324 | -0.039064409 | 0.164413056 | 7.998586029 | 1.472354767 | 0.186605077 | 0.991835066 | -4.495855865 |
| **A_55_P2123802** | NA | -0.100402558 | -0.263452643 | 0.062647527 | 9.539117033 | -1.471746054 | 0.186764129 | 0.991835066 | -4.496380756 |
| **A_66_P137383** | Ddx55 | 0.067145406 | -0.041898221 | 0.176189034 | 8.354577086 | 1.471719846 | 0.18677098 | 0.991835066 | -4.496403354 |
| **A_51_P498631** | Gsdme | 0.093230926 | -0.058175656 | 0.244637508 | 8.363939787 | 1.471717232 | 0.186771663 | 0.991835066 | -4.496405607 |
| **A_52_P89425** | Pcnt | 0.093405133 | -0.05832363 | 0.245133895 | 11.1146967 | 1.47133633 | 0.186871256 | 0.991835066 | -4.496734016 |
| **A_51_P123805** | Sbno1 | -0.086144249 | -0.226117909 | 0.053829411 | 8.885320256 | -1.47092017 | 0.186980124 | 0.991835066 | -4.497092791 |
| **A_55_P1997696** | Plbd1 | -0.150581229 | -0.395320169 | 0.094157711 | 10.12649471 | -1.470540181 | 0.18707958 | 0.991835066 | -4.497420352 |
| **A_55_P1956948** | Rpl27 | 0.080691946 | -0.050457122 | 0.211841013 | 16.14199951 | 1.470530761 | 0.187082046 | 0.991835066 | -4.497428472 |
| **A_55_P2131128** | Helq | 0.094035437 | -0.058822432 | 0.246893306 | 8.484651692 | 1.470323436 | 0.187136331 | 0.991835066 | -4.497607178 |
| **A_55_P1991151** | Rpl13 | -0.075870185 | -0.199219684 | 0.047479314 | 14.80964061 | -1.47008648 | 0.187198392 | 0.991835066 | -4.497811414 |
| **A_66_P126999** | Fis1 | -0.067267664 | -0.176636967 | 0.04210164 | 13.82425233 | -1.470009127 | 0.187218656 | 0.991835066 | -4.497878084 |
| **A_51_P447258** | Alox12b | 0.103257284 | -0.064633911 | 0.271148479 | 7.887936699 | 1.469947783 | 0.187234727 | 0.991835066 | -4.497930955 |
| **A_55_P2028661** | Dusp12 | 0.125395533 | -0.078495479 | 0.329286545 | 10.98469593 | 1.469918026 | 0.187242523 | 0.991835066 | -4.497956602 |
| **A_55_P2089118** | Zfp976 | 0.190087024 | -0.118993289 | 0.499167337 | 8.979852487 | 1.469908359 | 0.187245056 | 0.991835066 | -4.497964933 |
| **A_55_P1966438** | Gstm2 | -0.174756096 | -0.458949414 | 0.109437222 | 8.965391693 | -1.469696309 | 0.187300622 | 0.991835066 | -4.498147685 |
| **A_55_P2003005** | Capza1 | -0.099642407 | -0.261690854 | 0.06240604 | 12.77332329 | -1.469631561 | 0.187317592 | 0.991835066 | -4.498203486 |
| **A_55_P2065529** | Mok | -0.080445415 | -0.211274713 | 0.050383882 | 8.868370774 | -1.469621242 | 0.187320297 | 0.991835066 | -4.498212378 |
| **A_55_P2117243** | Npm1 | 0.186882434 | -0.117222394 | 0.490987263 | 11.7386312 | 1.468771709 | 0.187543084 | 0.991835066 | -4.49894443 |
| **A_51_P313483** | Dyrk1a | -0.075635131 | -0.198720023 | 0.047449761 | 10.15398217 | -1.468682596 | 0.187566468 | 0.991835066 | -4.499021212 |
| **A_52_P209514** | Cacnb2 | -0.139445643 | -0.366406773 | 0.087515486 | 8.643720462 | -1.468461615 | 0.187624465 | 0.991835066 | -4.499211605 |
| **A_51_P287198** | Krt23 | 0.070859402 | -0.044478931 | 0.186197735 | 7.804746546 | 1.468361353 | 0.187650785 | 0.991835066 | -4.499297986 |
| **A_55_P2010641** | Slc38a5 | 0.154922312 | -0.097274698 | 0.407119321 | 11.49098269 | 1.468193065 | 0.18769497 | 0.991835066 | -4.499442969 |
| **A_55_P1997544** | NA | 0.172025914 | -0.108024926 | 0.452076753 | 9.70524591 | 1.468135455 | 0.187710097 | 0.991835066 | -4.4994926 |

| | | | | | | | | | |
|---|---|---|---|---|---|---|---|---|---|
| **A_55_P2171303** | Bin3 | 0.074408297 | -0.046754557 | 0.195571151 | 8.192434176 | 1.467780121 | 0.18780343 | 0.991835066 | -4.499798704 |
| **A_55_P1952638** | Fgd6 | 0.083493127 | -0.052477225 | 0.219463479 | 8.492803441 | 1.467626815 | 0.187843711 | 0.991835066 | -4.499930763 |
| **A_55_P2170419** | NA | 0.089401621 | -0.05619256 | 0.234995803 | 15.76585115 | 1.467609478 | 0.187848266 | 0.991835066 | -4.499945696 |
| **A_55_P2029592** | Dus2 | 0.092877718 | -0.058392149 | 0.244147584 | 9.741918665 | 1.467466662 | 0.187885799 | 0.991835066 | -4.500068714 |
| **A_51_P200561** | Shtn1 | -0.074988991 | -0.197140974 | 0.047162993 | 8.2079324 | -1.467256756 | 0.187940975 | 0.991835066 | -4.500249513 |
| **A_52_P676076** | Rbx1 | 0.095001605 | -0.059757052 | 0.249760262 | 12.13497016 | 1.467185823 | 0.187959623 | 0.991835066 | -4.500310607 |
| **A_55_P2042973** | 1010001N( | 0.226126755 | -0.142250413 | 0.594503923 | 10.04430252 | 1.46712926 | 0.187974495 | 0.991835066 | -4.500359324 |
| **A_55_P1972927** | Elk3 | 0.080446846 | -0.050607792 | 0.211501484 | 11.2951291 | 1.467120414 | 0.187976821 | 0.991835066 | -4.500366943 |
| **A_51_P181851** | Eml3 | -0.113737551 | -0.299030216 | 0.071555114 | 9.696625604 | -1.467083314 | 0.187986577 | 0.991835066 | -4.500398897 |
| **A_51_P472888** | Enkd1 | 0.099614083 | -0.062684123 | 0.261912289 | 9.786312503 | 1.466952852 | 0.188020885 | 0.991835066 | -4.500511259 |
| **A_52_P278354** | Bmp7 | 0.176603645 | -0.111160048 | 0.464367339 | 12.22006764 | 1.466806405 | 0.188059404 | 0.991835066 | -4.500637384 |
| **A_55_P1990046** | NA | 0.104127253 | -0.06556911 | 0.273823616 | 8.367594471 | 1.466563963 | 0.188123188 | 0.991835066 | -4.500846174 |
| **A_55_P1991160** | Mrrf | -0.071804058 | -0.188832726 | 0.045224611 | 9.378718873 | -1.466445261 | 0.188154424 | 0.991835066 | -4.500948394 |
| **A_55_P1993390** | NA | 0.089199839 | -0.056198495 | 0.234598173 | 14.52680326 | 1.466269404 | 0.188200709 | 0.991835066 | -4.501099829 |
| **A_52_P102559** | Dars2 | -0.075825771 | -0.199431138 | 0.047779597 | 9.76852302 | -1.466184541 | 0.188223049 | 0.991835066 | -4.501172904 |
| **A_51_P184284** | Dld | -0.131168008 | -0.34499216 | 0.082656143 | 13.02145215 | -1.466156398 | 0.188230457 | 0.991835066 | -4.501197138 |
| **A_55_P2083988** | Lmo2 | 0.144218752 | -0.090906986 | 0.379344489 | 10.41366912 | 1.465988937 | 0.188274549 | 0.991835066 | -4.501341332 |
| **A_51_P258409** | Hey1 | -0.180073012 | -0.473659185 | 0.11351316 | 11.08329927 | -1.465960154 | 0.188282128 | 0.991835066 | -4.501366116 |
| **A_51_P224564** | Ppm1f | 0.087361198 | -0.05507716 | 0.229799555 | 11.73503309 | 1.46588788 | 0.188301161 | 0.991835066 | -4.501428347 |
| **A_52_P581440** | Wasf2 | 0.123160818 | -0.077703488 | 0.324025123 | 12.48604656 | 1.465476734 | 0.188409467 | 0.991835066 | -4.501782337 |
| **A_51_P282227** | As3mt | -0.074888514 | -0.197027516 | 0.047250487 | 10.66493214 | -1.465446546 | 0.188417421 | 0.991835066 | -4.501808327 |
| **A_55_P2095118** | Dancr | 0.281126116 | -0.177379672 | 0.739631904 | 9.545983922 | 1.465431431 | 0.188421404 | 0.991835066 | -4.50182134 |
| **A_52_P433828** | Tctn3 | -0.08167807 | -0.214895022 | 0.051538883 | 8.674324547 | -1.465396344 | 0.18843065 | 0.991835066 | -4.501851547 |
| **A_55_P2022539** | NA | 0.140937257 | -0.088942439 | 0.370816953 | 8.727209973 | 1.465326264 | 0.188449118 | 0.991835066 | -4.50191188 |
| **A_55_P2058783** | Hilpda | -0.10775265 | -0.28350849 | 0.06800319 | 10.14691342 | -1.465302622 | 0.188455349 | 0.991835066 | -4.501932234 |
| **A_55_P2097729** | Gtpbp4 | 0.102904007 | -0.064950515 | 0.27075853 | 8.95622536 | 1.465238672 | 0.188472204 | 0.991835066 | -4.501987289 |
| **A_51_P521161** | Caml | 0.081641216 | -0.051574143 | 0.214856574 | 10.5908761 | 1.464752663 | 0.188600343 | 0.991835066 | -4.502405666 |
| **A_51_P518621** | H3c1 | 0.099808581 | -0.063158022 | 0.262775184 | 8.207998659 | 1.463788729 | 0.188854724 | 0.991835066 | -4.503235315 |
| **A_55_P1969993** | NA | 0.083624379 | -0.052942592 | 0.220191349 | 17.25958182 | 1.463512251 | 0.188927743 | 0.991835066 | -4.503473242 |
| **A_55_P2148385** | NA | 0.089506108 | -0.056680418 | 0.235692635 | 8.879070864 | 1.463371051 | 0.188965045 | 0.991835066 | -4.503594747 |
| **A_55_P1982334** | Llph | 0.091991528 | -0.058259152 | 0.242242208 | 10.32275338 | 1.463324073 | 0.188977457 | 0.991835066 | -4.503635171 |
| **A_55_P2024406** | Napepld | -0.12847898 | -0.338329553 | 0.081371592 | 9.741166822 | -1.46329219 | 0.188985881 | 0.991835066 | -4.503662607 |
| **A_55_P2014229** | Sema4a | -0.078292084 | -0.206176891 | 0.049592722 | 8.607218445 | -1.46321464 | 0.189006373 | 0.991835066 | -4.503729337 |
| **A_55_P1955676** | Kat14 | 0.065411736 | -0.041450012 | 0.172273485 | 10.07559369 | 1.462993956 | 0.189064698 | 0.991835066 | -4.503919223 |
| **A_52_P584078** | Pop1 | 0.170617206 | -0.108117176 | 0.449351587 | 10.10001863 | 1.462990193 | 0.189065693 | 0.991835066 | -4.503922462 |
| **A_51_P185135** | Gpr108 | -0.100486838 | -0.264656855 | 0.06368318 | 10.34301145 | -1.462933105 | 0.189080783 | 0.991835066 | -4.503971581 |
| **A_55_P1988328** | NA | 0.123285719 | -0.078134885 | 0.324706322 | 15.89280931 | 1.462911356 | 0.189086533 | 0.991835066 | -4.503990294 |
| **A_55_P1960145** | Clip4 | -0.077821272 | -0.205029091 | 0.049386547 | 9.778167758 | -1.462155798 | 0.189286365 | 0.991835066 | -4.504640322 |
| **A_52_P175157** | NA | -0.136815027 | -0.360473467 | 0.086843414 | 11.26316378 | -1.462034564 | 0.189318447 | 0.991835066 | -4.504744612 |
| **A_52_P180565** | 2310061I0 | -0.13438228 | -0.354091579 | 0.08532702 | 10.79155831 | -1.461849614 | 0.1893674 | 0.991835066 | -4.504903708 |
| **A_51_P192800** | Grn | -0.154337803 | -0.406684047 | 0.09800844 | 11.91307623 | -1.46178871 | 0.189383523 | 0.991835066 | -4.504956096 |
| **A_55_P2152049** | Xrcc3 | 0.105951463 | -0.067307786 | 0.279210712 | 8.673000505 | 1.461570141 | 0.189441393 | 0.991835066 | -4.505144097 |
| **A_51_P300709** | Srm | 0.172831896 | -0.109827351 | 0.455491142 | 11.23123296 | 1.461402443 | 0.189485806 | 0.991835066 | -4.505288337 |
| **A_52_P254174** | Nr3c1 | 0.104902238 | -0.066661253 | 0.276465729 | 8.85006815 | 1.461399686 | 0.189486536 | 0.991835066 | -4.505290708 |
| **A_52_P184149** | Mthfd2 | 0.310556018 | -0.197397614 | 0.818509651 | 9.307244553 | 1.461251641 | 0.189525752 | 0.991835066 | -4.505418038 |
| **A_55_P2035892** | Prkcsh | 0.095725742 | -0.060860335 | 0.252311819 | 13.95418966 | 1.461116119 | 0.189561657 | 0.991835066 | -4.505534593 |
| **A_55_P2058497** | Tmem50a | -0.142281739 | -0.37502495 | 0.090461471 | 11.29408433 | -1.461104496 | 0.189564736 | 0.991835066 | -4.505544589 |
| **A_55_P2049282** | Ly6h | 0.107167845 | -0.068150326 | 0.282486016 | 9.03256554 | 1.46098817 | 0.189595561 | 0.991835066 | -4.505644631 |

| | | | | | | | | | |
|---|---|---|---|---|---|---|---|---|---|
| **A_51_P127915** | Rnasek | -0.165983248 | -0.43754601 | 0.105579514 | 10.64432171 | -1.460841758 | 0.189634364 | 0.991835066 | -4.491439483 |
| **A_55_P2099526** | H2afv | -0.092768875 | -0.244551369 | 0.059013619 | 14.3988864 | -1.460796568 | 0.189646342 | 0.991835066 | -4.505809406 |
| **A_55_P2178658** | NA | -0.134031428 | -0.353339823 | 0.085276968 | 13.16969596 | -1.460698281 | 0.189672396 | 0.991835066 | -4.505893928 |
| **A_55_P2038299** | Rpl31 | 0.082531664 | -0.052581672 | 0.217645 | 15.18459252 | 1.45992826 | 0.18987663 | 0.991835066 | -4.506556041 |
| **A_55_P1996088** | NA | 0.114506901 | -0.072964537 | 0.301978338 | 12.15468954 | 1.45984142 | 0.189899675 | 0.991835066 | -4.506630704 |
| **A_52_P625683** | Pxmp4 | -0.130294657 | -0.343620814 | 0.083031501 | 9.522018696 | -1.459794181 | 0.189912212 | 0.991835066 | -4.506671318 |
| **A_55_P2425437** | D030036P1 | 0.15387197 | -0.098066716 | 0.405810655 | 9.831828578 | 1.459734211 | 0.189928129 | 0.991835066 | -4.506722877 |
| **A_55_P2076176** | NA | -0.082186863 | -0.216781267 | 0.052407542 | 15.0505993 | -1.459434237 | 0.190007766 | 0.991835066 | -4.506980768 |
| **A_55_P2029740** | Gm10240 | 0.130504117 | -0.083247446 | 0.344255681 | 15.09881563 | 1.459230993 | 0.19006174 | 0.991835066 | -4.507155487 |
| **A_55_P2081483** | Atp5g2 | -0.065436718 | -0.172615407 | 0.041741972 | 16.9028239 | -1.459224786 | 0.190063388 | 0.991835066 | -4.507160823 |
| **A_51_P387608** | Hif1a | -0.086858229 | -0.229146524 | 0.055430066 | 10.51394748 | -1.45898534 | 0.190126994 | 0.991835066 | -4.507366653 |
| **A_55_P2004213** | Gprasp2 | 0.168801278 | -0.107728694 | 0.445331251 | 10.91446787 | 1.458957541 | 0.19013438 | 0.991835066 | -4.507390549 |
| **A_55_P1952202** | NA | -0.132676403 | -0.350036799 | 0.084683994 | 8.997876733 | -1.458889513 | 0.190152455 | 0.991835066 | -4.507449024 |
| **A_55_P1978676** | Mospd3 | -0.100995844 | -0.266462433 | 0.064470746 | 10.31599959 | -1.458822049 | 0.190170382 | 0.991835066 | -4.507507013 |
| **A_52_P91618** | Lysmd3 | -0.075828523 | -0.200072515 | 0.048415468 | 8.984988001 | -1.458701191 | 0.190202501 | 0.991835066 | -4.507610894 |
| **A_51_P165060** | Slc22a5 | -0.084506367 | -0.22299653 | 0.053983797 | 8.369247655 | -1.458410002 | 0.190279906 | 0.991835066 | -4.507861169 |
| **A_51_P129435** | Arid2 | 0.134773186 | -0.086129531 | 0.355675903 | 8.688072935 | 1.458181456 | 0.190340679 | 0.991835066 | -4.508057591 |
| **A_55_P2186558** | Tmem242 | -0.197700689 | -0.521774844 | 0.126373466 | 10.57433065 | -1.45805168 | 0.190375196 | 0.991835066 | -4.50816912 |
| **A_55_P2236818** | A430103D1 | 0.084984398 | -0.05434199 | 0.224310786 | 8.694791842 | 1.457857095 | 0.190426961 | 0.991835066 | -4.50833634 |
| **A_55_P2108486** | NA | 0.233214868 | -0.14912736 | 0.615557095 | 11.31525928 | 1.457850869 | 0.190428618 | 0.991835066 | -4.50834169 |
| **A_66_P137157** | Arl6ip4 | 0.097877287 | -0.062588238 | 0.258342812 | 13.07425704 | 1.457838157 | 0.190432 | 0.991835066 | -4.508352614 |
| **A_55_P1952744** | Timm8a1 | 0.129264331 | -0.082661837 | 0.341190499 | 11.92276595 | 1.457817802 | 0.190437416 | 0.991835066 | -4.508370106 |
| **A_52_P676063** | Tmem126b | -0.123115475 | -0.324976388 | 0.078745438 | 8.829090257 | -1.457704663 | 0.190467521 | 0.991835066 | -4.508467329 |
| **A_55_P1968276** | Tomm22 | -0.089851536 | -0.237174193 | 0.057471122 | 13.09336022 | -1.457689726 | 0.190471496 | 0.991835066 | -4.508480164 |
| **A_55_P1989813** | Gcm1 | 0.115257136 | -0.073760304 | 0.304274576 | 9.911073967 | 1.457387632 | 0.190551904 | 0.991835066 | -4.508739748 |
| **A_55_P2076994** | Defa-rs10 | 0.069367724 | -0.044414653 | 0.1831501 | 8.170048293 | 1.457107444 | 0.190626509 | 0.991835066 | -4.50898049 |
| **A_51_P295610** | Cyc1 | -0.199938692 | -0.527900476 | 0.128023092 | 15.05889545 | -1.457077793 | 0.190634405 | 0.991835066 | -4.509005965 |
| **A_52_P151853** | Tpd52 | 0.118513328 | -0.075917345 | 0.312944 | 9.575302955 | 1.456838954 | 0.190698024 | 0.991835066 | -4.509211165 |
| **A_51_P361830** | Lgi2 | 0.137685746 | -0.088212591 | 0.363584084 | 8.711297211 | 1.456750183 | 0.190721674 | 0.991835066 | -4.50928743 |
| **A_52_P660922** | Dock1 | 0.112150839 | -0.071854389 | 0.296156066 | 11.52572886 | 1.456738162 | 0.190724877 | 0.991835066 | -4.509297757 |
| **A_51_P130079** | Lrmp | -0.070208567 | -0.185416495 | 0.044999362 | 7.89988055 | -1.456521407 | 0.190782637 | 0.991835066 | -4.509483969 |
| **A_55_P2039523** | Phf10 | 0.083202111 | -0.053351777 | 0.219755999 | 11.13666141 | 1.456261636 | 0.190851881 | 0.991835066 | -4.509707121 |
| **A_52_P269158** | Pid1 | -0.134066912 | -0.354109857 | 0.085976032 | 8.378275286 | -1.456207599 | 0.190866288 | 0.991835066 | -4.509753539 |
| **A_55_P2013038** | Serpinb6c | -0.127653742 | -0.337180624 | 0.081873139 | 11.77934356 | -1.456139335 | 0.190884489 | 0.991835066 | -4.509812177 |
| **A_55_P1999805** | NA | 0.120922261 | -0.077577949 | 0.319422471 | 10.81391772 | 1.455976682 | 0.190927863 | 0.991835066 | -4.50995189 |
| **A_55_P2089710** | Ednrb | 0.109736074 | -0.070430785 | 0.289902932 | 8.654026059 | 1.455739425 | 0.190991149 | 0.991835066 | -4.510155675 |
| **A_66_P129873** | Npm1 | 0.159750479 | -0.10254535 | 0.422046308 | 14.61735607 | 1.455659894 | 0.191012367 | 0.991835066 | -4.510223983 |
| **A_55_P2050040** | Rpl36al | 0.07608955 | -0.048875412 | 0.201054512 | 14.37572317 | 1.455277748 | 0.19111435 | 0.991835066 | -4.510552185 |
| **A_51_P117226** | NA | -0.096577491 | -0.255194514 | 0.062039532 | 9.650646038 | -1.455242125 | 0.191123859 | 0.991835066 | -4.496084573 |
| **A_51_P419959** | Eaf1 | 0.098442819 | -0.063246358 | 0.260131997 | 9.258905687 | 1.455164941 | 0.191144464 | 0.991835066 | -4.510649062 |
| **A_55_P2006713** | NA | 0.432357704 | -0.277814049 | 1.142529456 | 16.52816837 | 1.45508693 | 0.191165292 | 0.991835066 | -4.510716055 |
| **A_55_P2149763** | Tapbp | -0.14923907 | -0.394452103 | 0.095973963 | 10.56240025 | -1.454615186 | 0.191291285 | 0.991835066 | -4.511121145 |
| **A_51_P277444** | Rnf19a | -0.078836097 | -0.208382124 | 0.050709931 | 10.56304045 | -1.454488002 | 0.191325266 | 0.991835066 | -4.511230351 |
| **A_55_P2092750** | Car9 | 0.120776104 | -0.077699861 | 0.319252069 | 8.336702425 | 1.454394508 | 0.191350249 | 0.991835066 | -4.511310626 |
| **A_55_P1973770** | Unc5b | 0.136645729 | -0.087984737 | 0.361276195 | 12.47859135 | 1.453906692 | 0.19148065 | 0.991835066 | -4.511729445 |
| **A_51_P306066** | Eif4g2 | -0.092937501 | -0.245721473 | 0.059846471 | 11.84889944 | -1.453859128 | 0.191493369 | 0.991835066 | -4.511770279 |
| **A_51_P336599** | Kcne3 | -0.228942256 | -0.605324098 | 0.147439585 | 9.192963396 | -1.453805907 | 0.191507602 | 0.991835066 | -4.511815968 |
| **A_51_P346453** | Noc4l | 0.07597613 | -0.04893638 | 0.200888641 | 9.350608339 | 1.453718662 | 0.191530935 | 0.991835066 | -4.511890866 |

| | | | | | | | | | |
|---|---|---|---|---|---|---|---|---|---|
| **A_55_P2060255** | NA | -0.124811939 | -0.330015592 | 0.080391714 | 14.81402058 | -1.453718025 | 0.191531106 | 0.991835066 | -4.511891413 |
| **A_51_P141610** | Cables2 | 0.066753987 | -0.043019386 | 0.176527359 | 9.149372508 | 1.453413976 | 0.191612443 | 0.991835066 | -4.512152419 |
| **A_55_P2089462** | Ncoa5 | 0.098292837 | -0.063377456 | 0.259963129 | 13.32862264 | 1.453117644 | 0.191691746 | 0.991835066 | -4.512406781 |
| **A_55_P1966573** | Gemin4 | 0.163134377 | -0.105227721 | 0.431496476 | 10.50792932 | 1.452892368 | 0.191752054 | 0.991835066 | -4.512600138 |
| **A_55_P2091191** | Slc28a2 | -0.169590365 | -0.44861204 | 0.10943131 | 8.366026991 | -1.452688057 | 0.191806764 | 0.991835066 | -4.512775492 |
| **A_55_P1952182** | Vmn2r111 | 0.115629505 | -0.074616913 | 0.305875924 | 7.805325785 | 1.452651078 | 0.191816668 | 0.991835066 | -4.512807229 |
| **A_51_P447866** | Sash3 | -0.076193987 | -0.201576884 | 0.049188909 | 7.892399006 | -1.452417706 | 0.19187918 | 0.991835066 | -4.513007511 |
| **A_55_P2004099** | NA | -0.118980014 | -0.314813766 | 0.076853737 | 8.291965879 | -1.452096925 | 0.191965136 | 0.991835066 | -4.51328279 |
| **A_55_P1977910** | NA | 0.178003526 | -0.115128816 | 0.471135867 | 10.1517256 | 1.451356159 | 0.192163765 | 0.991835066 | -4.513918397 |
| **A_55_P2260648** | 6330418K0 | 0.082034709 | -0.053061378 | 0.217130797 | 8.936415654 | 1.451322755 | 0.192172726 | 0.991835066 | -4.513947056 |
| **A_55_P1961168** | NA | -0.106249855 | -0.281248338 | 0.068748628 | 12.04256202 | -1.451119554 | 0.192227247 | 0.991835066 | -4.514121388 |
| **A_55_P2179834** | Gatad2a | 0.137097929 | -0.088764242 | 0.362960101 | 11.59399172 | 1.450763192 | 0.192322897 | 0.991835066 | -4.514427101 |
| **A_66_P106509** | NA | 0.096740781 | -0.062677515 | 0.256159076 | 12.18724156 | 1.450375849 | 0.192426912 | 0.991835066 | -4.51475936 |
| **A_55_P1968993** | NA | 0.159464851 | -0.10340514 | 0.422334841 | 10.77659352 | 1.449883453 | 0.19255921 | 0.991835066 | -4.515181686 |
| **A_52_P195922** | Exosc6 | 0.178875845 | -0.116009147 | 0.473760837 | 11.61916569 | 1.449800221 | 0.192581581 | 0.991835066 | -4.515253069 |
| **A_55_P1962234** | Bad | -0.091609756 | -0.242638163 | 0.059418652 | 12.31957245 | -1.449746984 | 0.192595892 | 0.991835066 | -4.515298726 |
| **A_55_P2093920** | Pzp | -0.334847412 | -0.886923503 | 0.21722868 | 8.563806773 | -1.449629901 | 0.192627368 | 0.991835066 | -4.515399137 |
| **A_55_P2013138** | NA | 0.104095423 | -0.067540479 | 0.275731326 | 11.53771805 | 1.449548096 | 0.192649362 | 0.991835066 | -4.515469291 |
| **A_55_P2024625** | Ccl21a | 0.339019574 | -0.220008182 | 0.89804733 | 12.45068188 | 1.449440976 | 0.192678167 | 0.991835066 | -4.515561152 |
| **A_51_P484111** | Matn2 | -0.208454351 | -0.552281776 | 0.135373075 | 10.6105821 | -1.449037355 | 0.192786735 | 0.991835066 | -4.515907259 |
| **A_55_P1971025** | Paqr6 | 0.079750887 | -0.051826568 | 0.211328343 | 8.227091154 | 1.448649001 | 0.19289125 | 0.991835066 | -4.516240241 |
| **A_55_P2292435** | Slc8a1 | 0.168440282 | -0.109510066 | 0.446390631 | 8.661003475 | 1.448397843 | 0.192958869 | 0.991835066 | -4.516455571 |
| **A_55_P1981499** | Rpl10a | 0.102060956 | -0.066368037 | 0.27048995 | 15.83473075 | 1.448277928 | 0.192991161 | 0.991835066 | -4.516558375 |
| **A_55_P2115871** | Inpp5j | -0.138929637 | -0.368216333 | 0.090357059 | 11.39896525 | -1.448188773 | 0.193015173 | 0.991835066 | -4.516634807 |
| **A_51_P438200** | Sil1 | 0.078078748 | -0.050789114 | 0.206946609 | 11.9805257 | 1.448095989 | 0.193040166 | 0.991835066 | -4.516714347 |
| **A_55_P2011712** | NA | 0.070738614 | -0.046029787 | 0.187507016 | 8.875811655 | 1.447905923 | 0.193091371 | 0.991835066 | -4.516877279 |
| **A_55_P2157423** | NA | 0.117543435 | -0.076494389 | 0.311581259 | 8.463596345 | 1.447841814 | 0.193108645 | 0.991835066 | -4.516932233 |
| **A_55_P2085672** | NA | 0.080992595 | -0.05271067 | 0.214695861 | 15.21544417 | 1.447812878 | 0.193116443 | 0.991835066 | -4.516957037 |
| **A_51_P356283** | Fbxo31 | 0.079960969 | -0.052039882 | 0.211961821 | 11.11539204 | 1.447806259 | 0.193118226 | 0.991835066 | -4.516962711 |
| **A_55_P2006869** | Fyb | -0.069880792 | -0.185251593 | 0.04549001 | 7.895687061 | -1.447674883 | 0.193153632 | 0.991835066 | -4.517075324 |
| **A_51_P236324** | Gtf2e1 | 0.109190863 | -0.071172818 | 0.289554544 | 10.07745228 | 1.446926061 | 0.19335555 | 0.991835066 | -4.517717128 |
| **A_66_P128909** | Numbl | 0.092640906 | -0.060386113 | 0.245667926 | 9.485049572 | 1.446917627 | 0.193357825 | 0.991835066 | -4.517724356 |
| **A_66_P125831** | Pnn | 0.146447395 | -0.095513594 | 0.388408385 | 11.12365784 | 1.446589912 | 0.193446254 | 0.991835066 | -4.518005196 |
| **A_55_P1993807** | Nudt10 | 0.083775272 | -0.054664676 | 0.222215219 | 8.505097998 | 1.446317203 | 0.193519868 | 0.991835066 | -4.518238881 |
| **A_55_P2056926** | Mtss2 | 0.10066745 | -0.065728392 | 0.267063292 | 10.3361575 | 1.445958172 | 0.193616823 | 0.991835066 | -4.51854651 |
| **A_55_P2040971** | Tom1 | -0.128568915 | -0.341143091 | 0.08400526 | 12.59358846 | -1.445554965 | 0.193725759 | 0.991835066 | -4.518891958 |
| **A_55_P2247633** | Gpatch2 | 0.095882753 | -0.062656764 | 0.254422269 | 8.899573759 | 1.445480034 | 0.19374601 | 0.991835066 | -4.51895615 |
| **A_66_P108152** | Cbr1 | -0.139760017 | -0.370877787 | 0.091357753 | 11.3868271 | -1.445302453 | 0.19379401 | 0.991835066 | -4.519108279 |
| **A_66_P137048** | Men1 | 0.096870361 | -0.063327489 | 0.257068211 | 11.40400923 | 1.445251295 | 0.19380784 | 0.991835066 | -4.519152103 |
| **A_55_P2029310** | Arf4 | 0.135275047 | -0.08845556 | 0.359005654 | 10.93526015 | 1.445111721 | 0.193845577 | 0.991835066 | -4.519271665 |
| **A_52_P31011** | Cep162 | 0.106556878 | -0.069702151 | 0.282815908 | 8.462617467 | 1.444904844 | 0.193901523 | 0.991835066 | -4.519448873 |
| **A_55_P2141395** | Ly6g6c | -0.083947535 | -0.22286588 | 0.054970809 | 8.356166952 | -1.444300241 | 0.19406511 | 0.991835066 | -4.519966715 |
| **A_66_P116771** | Vps8 | -0.155209058 | -0.41207915 | 0.101661034 | 11.08810107 | -1.444151164 | 0.194105465 | 0.991835066 | -4.520094386 |
| **A_52_P443167** | Naa35 | -0.05832616 | -0.154867627 | 0.038215306 | 12.22818526 | -1.443971522 | 0.194154104 | 0.991835066 | -4.520248228 |
| **A_55_P2048398** | Kmt5b | 0.067421791 | -0.044214348 | 0.17905793 | 8.493055658 | 1.443459516 | 0.194292793 | 0.991835066 | -4.520686661 |
| **A_55_P2262560** | 5730471H1 | 0.072232336 | -0.047374549 | 0.191839221 | 8.746327318 | 1.44339306 | 0.194310801 | 0.991835066 | -4.520743563 |
| **A_51_P388737** | Slc11a2 | 0.064248049 | -0.042142337 | 0.170638435 | 8.403036449 | 1.443333507 | 0.194326939 | 0.991835066 | -4.520794553 |
| **A_55_P2081169** | NA | 0.079739261 | -0.052316871 | 0.211795394 | 15.40311022 | 1.443187526 | 0.194366505 | 0.991835066 | -4.520919544 |

| | | | | | | | | | |
|---|---|---|---|---|---|---|---|---|---|
| **A_55_P2161923** | Rabgap1 | 0.072287619 | -0.047436231 | 0.192011469 | 8.671198016 | 1.443086556 | 0.194393875 | 0.991835066 | -4.521005991 |
| **A_55_P2036220** | Chadl | 0.095181414 | -0.062462157 | 0.252824984 | 8.773176006 | 1.443062102 | 0.194400504 | 0.991835066 | -4.521026929 |
| **A_55_P2031382** | Crebzf | 0.070718975 | -0.046433841 | 0.18787179 | 8.397284465 | 1.442754231 | 0.194483983 | 0.991835066 | -4.521290505 |
| **A_51_P172085** | Arhgdig | 0.226241929 | -0.148596316 | 0.601080175 | 8.305865365 | 1.442574769 | 0.194532659 | 0.991835066 | -4.521444138 |
| **A_51_P282523** | Adgra2 | 0.160506486 | -0.105439117 | 0.426452089 | 10.88332978 | 1.442477022 | 0.194559176 | 0.991835066 | -4.521527813 |
| **A_55_P2006698** | Bmp1 | -0.100261644 | -0.266467337 | 0.065944049 | 10.7245262 | -1.441776884 | 0.194749206 | 0.991835066 | -4.5221271 |
| **A_55_P2176792** | Sh3gl3 | 0.137541364 | -0.090477035 | 0.365559762 | 8.735030662 | 1.441692257 | 0.194772187 | 0.991835066 | -4.52219953 |
| **A_55_P2007756** | Krt16 | 0.061925198 | -0.040748712 | 0.164599107 | 8.245506245 | 1.441505938 | 0.194822791 | 0.991835066 | -4.52235899 |
| **A_51_P412595** | Eef1b2 | 0.09545809 | -0.062824497 | 0.253740677 | 14.82905439 | 1.441413988 | 0.194847769 | 0.991835066 | -4.522437681 |
| **A_51_P310896** | Pfkl | 0.131182309 | -0.086376229 | 0.348740848 | 11.10784479 | 1.441146965 | 0.194920322 | 0.991835066 | -4.522666192 |
| **A_55_P2283207** | Maz | 0.196828694 | -0.129713905 | 0.523371292 | 8.30210353 | 1.440647395 | 0.195056125 | 0.991835066 | -4.523093666 |
| **A_52_P251615** | Arhgef40 | 0.168018085 | -0.11077906 | 0.446815231 | 10.88816264 | 1.440379198 | 0.195129067 | 0.991835066 | -4.523323135 |
| **A_55_P2076635** | Vps25 | -0.163671956 | -0.43528901 | 0.107945099 | 9.488749319 | -1.440211812 | 0.195174604 | 0.991835066 | -4.523466344 |
| **A_51_P372456** | Wdr86 | 0.141011071 | -0.093046981 | 0.375069124 | 9.27617609 | 1.439921306 | 0.195253658 | 0.991835066 | -4.523714872 |
| **A_66_P125226** | NA | 0.089628066 | -0.059145872 | 0.238402004 | 12.67257712 | 1.43988 | 0.195264901 | 0.991835066 | -4.523750208 |
| **A_51_P246387** | Nktr | 0.161665098 | -0.106838006 | 0.430168202 | 9.722967218 | 1.439050684 | 0.19549075 | 0.991835066 | -4.52445958 |
| **A_52_P671812** | NA | 0.120988651 | -0.079988867 | 0.32196617 | 12.47164481 | 1.438819409 | 0.195553776 | 0.991835066 | -4.524657378 |
| **A_51_P205390** | Cabcoco1 | -0.1154541 | -0.307244173 | 0.076335973 | 9.853123798 | -1.438773294 | 0.195566345 | 0.991835066 | -4.524696817 |
| **A_55_P1980817** | Gm6548 | -0.095219845 | -0.253398131 | 0.062958441 | 15.40542724 | -1.438764584 | 0.195568719 | 0.991835066 | -4.524704266 |
| **A_55_P2031939** | Aqp4 | -0.116744191 | -0.310725363 | 0.07723698 | 9.847779871 | -1.438417092 | 0.195663457 | 0.991835066 | -4.525001433 |
| **A_55_P2121252** | Amd2 | 0.10585689 | -0.07006122 | 0.281774999 | 9.38220998 | 1.438194791 | 0.195724085 | 0.991835066 | -4.525191525 |
| **A_55_P2062039** | Tsen34 | -0.109376182 | -0.291204447 | 0.072452084 | 13.24850124 | -1.437707355 | 0.195857084 | 0.991835066 | -4.5256083 |
| **A_55_P1979728** | Atf4 | 0.164813066 | -0.122432905 | 0.452059036 | 14.11380549 | 1.508909898 | 0.195987869 | 0.991835066 | -4.440414062 |
| **A_52_P467046** | Wdr48 | 0.097461537 | -0.064617704 | 0.259540777 | 9.357777471 | 1.437192674 | 0.195997605 | 0.991835066 | -4.526048313 |
| **A_55_P2040371** | Rnaset2a | -0.094873398 | -0.252653286 | 0.062906489 | 13.8289763 | -1.437149503 | 0.196009396 | 0.991835066 | -4.526085218 |
| **A_55_P2016618** | Neurl1a | 0.131690855 | -0.087365336 | 0.350747046 | 8.822512594 | 1.436842677 | 0.196093215 | 0.991835066 | -4.5263475 |
| **A_51_P392090** | Tmem19 | -0.064128623 | -0.170801287 | 0.04254404 | 10.16325451 | -1.43683835 | 0.196094398 | 0.991835066 | -4.526351198 |
| **A_55_P2394490** | NA | -0.06639571 | -0.176884348 | 0.044092929 | 8.479266336 | -1.436254912 | 0.196253873 | 0.991835066 | -4.526849877 |
| **A_51_P516705** | Ern1 | 0.069649473 | -0.046254152 | 0.185553097 | 8.217990309 | 1.436249692 | 0.1962553 | 0.991835066 | -4.526854338 |
| **A_51_P218975** | Kcns3 | 0.06535341 | -0.043408825 | 0.174115646 | 7.728760471 | 1.436148208 | 0.196283052 | 0.991835066 | -4.526941071 |
| **A_55_P1987938** | NA | 0.182131585 | -0.121007185 | 0.485270355 | 11.28976117 | 1.435994958 | 0.196324966 | 0.991835066 | -4.527072041 |
| **A_52_P135873** | NA | 0.253961044 | -0.168793351 | 0.676715438 | 10.3803812 | 1.435780677 | 0.196383585 | 0.991835066 | -4.52725516 |
| **A_51_P101137** | Samd14 | 0.073365539 | -0.04880097 | 0.195532048 | 9.680842405 | 1.435321136 | 0.196509353 | 0.991835066 | -4.527647839 |
| **A_55_P1979645** | S100a1 | -0.251178293 | -0.669438451 | 0.167081866 | 11.77575502 | -1.435306789 | 0.19651328 | 0.991835066 | -4.527660098 |
| **A_55_P2055557** | Sdsl | 0.10966018 | -0.072969565 | 0.292289925 | 11.59064996 | 1.435114568 | 0.19656591 | 0.991835066 | -4.527824336 |
| **A_55_P2075000** | NA | -0.107256933 | -0.285914246 | 0.07140038 | 11.86580453 | -1.434873811 | 0.196631847 | 0.991835066 | -4.528030034 |
| **A_66_P139393** | Psmd10 | -0.081759718 | -0.21795983 | 0.054440394 | 9.626752706 | -1.434732884 | 0.196670452 | 0.991835066 | -4.528150432 |
| **A_55_P2021476** | NA | 0.238751213 | -0.159065327 | 0.636567753 | 13.44849949 | 1.434405308 | 0.196760214 | 0.991835066 | -4.528430275 |
| **A_55_P2089304** | 2900097C1 | 0.098744754 | -0.065813386 | 0.263302895 | 9.37165983 | 1.434180441 | 0.196821854 | 0.991835066 | -4.52862236 |
| **A_55_P2048323** | Cdk10 | 0.135566244 | -0.090372105 | 0.361504593 | 10.62133448 | 1.434071304 | 0.196851776 | 0.991835066 | -4.528715583 |
| **A_52_P628067** | Cdca3 | -0.130979504 | -0.349294354 | 0.087335346 | 12.33413951 | -1.433934114 | 0.196889396 | 0.991835066 | -4.528832766 |
| **A_51_P217382** | Kif1c | 0.063964612 | -0.042655029 | 0.170584252 | 12.27706503 | 1.433876297 | 0.196905252 | 0.991835066 | -4.528882149 |
| **A_51_P326030** | Zfp959 | 0.061598436 | -0.041100592 | 0.164297464 | 8.31084099 | 1.433548809 | 0.196995087 | 0.991835066 | -4.529161853 |
| **A_52_P585652** | Fndc3b | -0.127756702 | -0.340763698 | 0.085250293 | 10.17693701 | -1.433504164 | 0.197007336 | 0.991835066 | -4.529199982 |
| **A_55_P2077746** | Mphosph8 | 0.070071847 | -0.046762967 | 0.186906661 | 10.91830068 | 1.433442999 | 0.19702412 | 0.991835066 | -4.52925222 |
| **A_55_P2102816** | Rpl36a | 0.085024106 | -0.056742435 | 0.226790647 | 14.73329621 | 1.433433221 | 0.197026803 | 0.991835066 | -4.52926057 |
| **A_55_P1990864** | NA | 0.144224696 | -0.096321206 | 0.384770598 | 9.493861192 | 1.43301522 | 0.197141538 | 0.991835066 | -4.529617535 |
| **A_55_P2133255** | Mad2l1 | -0.109394616 | -0.291874575 | 0.073085344 | 11.34311479 | -1.432814289 | 0.197196711 | 0.991835066 | -4.529789112 |

| | | | | | | | | | |
|---|---|---|---|---|---|---|---|---|---|
| **A_55_P2041017** | Gm15365 | 0.133715883 | -0.089395714 | 0.35682748 | 12.97905987 | 1.432418707 | 0.197305376 | 0.991835066 | -4.530126879 |
| **A_55_P1993609** | Slx4ip | 0.070046186 | -0.046846662 | 0.186939033 | 8.842846467 | 1.432206646 | 0.19736365 | 0.991835066 | -4.530307931 |
| **A_55_P2040121** | Cdv3 | 0.12209668 | -0.081709351 | 0.32590271 | 10.28228057 | 1.431844827 | 0.197463113 | 0.991835066 | -4.53061682 |
| **A_55_P1989102** | Hmgn1 | 0.066356007 | -0.044415124 | 0.177127138 | 11.08855819 | 1.431735477 | 0.197493182 | 0.991835066 | -4.530710168 |
| **A_55_P2172924** | NA | 0.070615297 | -0.047278996 | 0.18850959 | 8.492164552 | 1.431578414 | 0.197536379 | 0.991835066 | -4.530844242 |
| **A_55_P1959973** | Dcaf12l2 | 0.233179077 | -0.156197131 | 0.622555284 | 12.15747436 | 1.431295478 | 0.197614215 | 0.991835066 | -4.53108575 |
| **A_51_P379478** | Nckap5 | -0.08254332 | -0.22038065 | 0.055294009 | 8.816209362 | -1.431278724 | 0.197618825 | 0.991835066 | -4.531100051 |
| **A_55_P2113633** | Bzw1 | 0.086611327 | -0.05802628 | 0.231248934 | 10.61889114 | 1.431207484 | 0.197638429 | 0.991835066 | -4.531160857 |
| **A_55_P1995572** | Meis1 | 0.079407231 | -0.055736076 | 0.214550537 | 11.9908177 | 1.459595983 | 0.197659164 | 0.991835066 | -4.483419839 |
| **A_55_P2090718** | Pop5 | 0.067275552 | -0.04507894 | 0.179630044 | 12.09581388 | 1.4311197 | 0.197662586 | 0.991835066 | -4.531235782 |
| **A_55_P2173702** | Ebpl | 0.069589389 | -0.046642375 | 0.185821154 | 10.33552772 | 1.430959446 | 0.197706695 | 0.991835066 | -4.531372557 |
| **A_55_P1975185** | Sqle | 0.266405949 | -0.178560948 | 0.711372846 | 10.98804249 | 1.430952928 | 0.197708489 | 0.991835066 | -4.53137812 |
| **A_55_P2180301** | Clcn5 | 0.114641949 | -0.076842295 | 0.306126193 | 9.012301235 | 1.430934153 | 0.197713657 | 0.991835066 | -4.531394144 |
| **A_55_P2086687** | Nfib | 0.109019025 | -0.073077491 | 0.291115541 | 9.03809046 | 1.430901664 | 0.197722601 | 0.991835066 | -4.531421872 |
| **A_55_P2045225** | NA | -0.091256777 | -0.243724518 | 0.061210963 | 11.85003784 | -1.43052778 | 0.197825551 | 0.991835066 | -4.531740951 |
| **A_55_P2163493** | Tgm3 | -0.179526911 | -0.479549815 | 0.120495993 | 10.9685653 | -1.430158858 | 0.197927182 | 0.991835066 | -4.532055765 |
| **A_51_P279571** | Syap1 | -0.079930171 | -0.213523133 | 0.053662791 | 10.91806813 | -1.430000853 | 0.197970724 | 0.991835066 | -4.532190587 |
| **A_51_P422124** | Fam126a | -0.092214287 | -0.246400822 | 0.061972247 | 10.93607547 | -1.429423437 | 0.198129918 | 0.991835066 | -4.532683233 |
| **A_55_P2090345** | BC085271 | 0.117775995 | -0.079219559 | 0.314771549 | 11.39700124 | 1.428925178 | 0.198267381 | 0.991835066 | -4.533108283 |
| **A_51_P181002** | Vps50 | -0.093556981 | -0.250111395 | 0.062997433 | 8.930555695 | -1.4283019 | 0.198439456 | 0.991835066 | -4.533639903 |
| **A_55_P2127265** | NA | -0.16802587 | -0.449305967 | 0.113254226 | 16.39512446 | -1.427730653 | 0.198597285 | 0.991835066 | -4.534127067 |
| **A_55_P2083406** | Myh3 | -0.41956408 | -1.121984863 | 0.282856704 | 8.235882295 | -1.427611635 | 0.198630183 | 0.991835066 | -4.534228557 |
| **A_55_P1965478** | Kcnd1 | -0.067375795 | -0.180212712 | 0.045461122 | 9.191799585 | -1.427124381 | 0.198764916 | 0.991835066 | -4.53464402 |
| **A_55_P2153923** | Slc17a5 | 0.067616487 | -0.045628167 | 0.180861141 | 8.055200841 | 1.427065884 | 0.198781096 | 0.991835066 | -4.534693894 |
| **A_55_P2240116** | 2310005E1 | 0.077538781 | -0.05236219 | 0.207439752 | 7.931728424 | 1.426644289 | 0.198897749 | 0.991835066 | -4.535053322 |
| **A_51_P105709** | Trip13 | 0.103087302 | -0.069615755 | 0.275790359 | 10.57879488 | 1.426639622 | 0.19889904 | 0.991835066 | -4.535057301 |
| **A_55_P1953683** | Nsmaf | 0.129201622 | -0.087253245 | 0.345656489 | 11.80249259 | 1.426624712 | 0.198903167 | 0.991835066 | -4.535070012 |
| **A_55_P2024337** | Tmem53 | -0.136213655 | -0.364445849 | 0.092018539 | 11.51020794 | -1.426437911 | 0.198954874 | 0.991835066 | -4.535229254 |
| **A_51_P413866** | Cfb | -0.118243692 | -0.316398023 | 0.079910639 | 8.021865571 | -1.426210225 | 0.199017915 | 0.991835066 | -4.535423339 |
| **A_51_P324814** | Krt18 | 0.307047946 | -0.207606153 | 0.821702044 | 10.91020888 | 1.425935537 | 0.199093994 | 0.991835066 | -4.535657473 |
| **A_51_P173678** | Slc10a6 | -0.062644681 | -0.167648852 | 0.04235949 | 8.016711866 | -1.42589249 | 0.199105919 | 0.991835066 | -4.535694163 |
| **A_55_P1973683** | Morn4 | 0.102538795 | -0.069346472 | 0.274424062 | 10.14048481 | 1.425800256 | 0.199131472 | 0.991835066 | -4.535772776 |
| **A_51_P481159** | Cbr3 | -0.061989407 | -0.165906946 | 0.041928132 | 8.51521303 | -1.425731544 | 0.19915051 | 0.991835066 | -4.535831339 |
| **A_55_P2136902** | Cisd3 | -0.125119005 | -0.334911402 | 0.084673391 | 11.84880197 | -1.425419423 | 0.199237011 | 0.991835066 | -4.536097345 |
| **A_55_P1970027** | Itch | -0.070021583 | -0.187439742 | 0.047396577 | 10.3973956 | -1.425298364 | 0.19927057 | 0.991835066 | -4.536200512 |
| **A_55_P1970002** | Sh3kbp1 | -0.106652812 | -0.285501681 | 0.072196057 | 10.3596265 | -1.425263763 | 0.199280163 | 0.991835066 | -4.536229999 |
| **A_52_P35064** | Tm7sf2 | -0.084764896 | -0.226919419 | 0.057389627 | 8.910333697 | -1.425162813 | 0.199308152 | 0.991835066 | -4.536316025 |
| **A_51_P298486** | Cog6 | -0.112211593 | -0.300453774 | 0.076030589 | 10.60410413 | -1.424721281 | 0.199430615 | 0.991835066 | -4.536692257 |
| **A_55_P2143204** | Pbxip1 | -0.118277038 | -0.316728166 | 0.080174091 | 11.23981317 | -1.424478822 | 0.199497892 | 0.991835066 | -4.53689884 |
| **A_55_P1959985** | Alas1 | -0.096845535 | -0.259338327 | 0.065647258 | 9.673551496 | -1.424474344 | 0.199499134 | 0.991835066 | -4.536902655 |
| **A_55_P1959703** | Zfp646 | -0.064210877 | -0.171951789 | 0.043530035 | 15.24635052 | -1.424416793 | 0.199515107 | 0.991835066 | -4.536951688 |
| **A_51_P440210** | Prkg1 | 0.122555289 | -0.08314108 | 0.328251658 | 9.043071798 | 1.424015007 | 0.199626647 | 0.991835066 | -4.537293986 |
| **A_51_P205820** | Klf11 | -0.104881252 | -0.280961845 | 0.071199341 | 8.607084366 | -1.423624664 | 0.199735065 | 0.991835066 | -4.537626501 |
| **A_55_P2030805** | Actr10 | -0.074467112 | -0.199493618 | 0.050559395 | 13.31059807 | -1.423546128 | 0.199756884 | 0.991835066 | -4.537693398 |
| **A_55_P2075479** | Rpl37a | 0.096081142 | -0.065235233 | 0.257397516 | 17.29147992 | 1.423537244 | 0.199759353 | 0.991835066 | -4.537700965 |
| **A_55_P2062543** | Kifc5b | 0.171292937 | -0.116326307 | 0.458912181 | 10.86160852 | 1.423412081 | 0.199794132 | 0.991835066 | -4.537807576 |
| **A_55_P2060111** | NA | -0.091342984 | -0.244742822 | 0.062056855 | 14.86487545 | -1.423178661 | 0.199859007 | 0.991835066 | -4.538006387 |
| **A_51_P123077** | Nubp1 | 0.071302652 | -0.048447316 | 0.191052621 | 11.02010899 | 1.423113075 | 0.199877238 | 0.991835066 | -4.538062246 |

| | | | | | | | | | |
|---|---|---|---|---|---|---|---|---|---|
| **A_55_P2006940** | Sfmbt1 | 0.103720344 | -0.070481096 | 0.277921783 | 9.235012709 | 1.423053868 | 0.199893698 | 0.991835066 | -4.538112672 |
| **A_66_P105711** | Ap4s1 | -0.104802777 | -0.280837615 | 0.071232062 | 9.147715417 | -1.422929216 | 0.199928356 | 0.991835066 | -4.538218834 |
| **A_65_P02543** | Srp72 | -0.075401342 | -0.202052298 | 0.051249615 | 9.089215604 | -1.422917535 | 0.199931604 | 0.991835066 | -4.538228781 |
| **A_55_P2143953** | Pank2 | -0.088361221 | -0.23678364 | 0.060061198 | 9.716590325 | -1.422890029 | 0.199939253 | 0.991835066 | -4.538252207 |
| **A_55_P2093173** | NA | -0.199456719 | -0.534489536 | 0.135576097 | 11.75710012 | -1.422887143 | 0.199940055 | 0.991835066 | -4.538254664 |
| **A_55_P2123716** | Tug1 | 0.121482248 | -0.082615948 | 0.325580445 | 9.16265864 | 1.422599939 | 0.200019934 | 0.991835066 | -4.538499249 |
| **A_51_P134551** | Coq9 | -0.181929771 | -0.487651473 | 0.12379193 | 11.46207718 | -1.422285025 | 0.200107552 | 0.991835066 | -4.538767409 |
| **A_55_P2176963** | Hsph1 | 0.072401644 | -0.049270661 | 0.194073948 | 10.48629617 | 1.422216861 | 0.200126522 | 0.991835066 | -4.53882545 |
| **A_55_P2021689** | Chrdl2 | 0.063230075 | -0.043034817 | 0.169494966 | 7.722408148 | 1.4221422 | 0.200147302 | 0.991835066 | -4.538889023 |
| **A_55_P1995503** | Gm5643 | 0.07778876 | -0.053009555 | 0.208587075 | 13.5672252 | 1.421424612 | 0.200347122 | 0.991835066 | -4.539499966 |
| **A_66_P124017** | Map7d3 | -0.096714531 | -0.259366658 | 0.065937596 | 8.799487286 | -1.421153909 | 0.200422548 | 0.991835066 | -4.539730407 |
| **A_66_P135694** | NA | 0.060874752 | -0.0415251 | 0.163274604 | 15.8940997 | 1.420846008 | 0.20050837 | 0.991835066 | -4.539992493 |
| **A_51_P239750** | Inhba | 0.17405529 | -0.118759437 | 0.466870018 | 10.11623507 | 1.420703482 | 0.200548109 | 0.991835066 | -4.540113805 |
| **A_51_P336622** | Dexi | -0.07522342 | -0.201777489 | 0.051330649 | 12.74417415 | -1.420646718 | 0.200563937 | 0.991835066 | -4.540162118 |
| **A_52_P230688** | Rep15 | 0.078743059 | -0.053736633 | 0.211222751 | 8.640008029 | 1.420600961 | 0.200576697 | 0.991835066 | -4.540201063 |
| **A_51_P145785** | Ambp | -0.579409538 | -1.554241233 | 0.395422158 | 9.24723525 | -1.420578151 | 0.200583058 | 0.991835066 | -4.540220476 |
| **A_55_P2219765** | Wnt5a | 0.100451446 | -0.068580198 | 0.269483089 | 9.011264548 | 1.420356325 | 0.20064493 | 0.991835066 | -4.540409269 |
| **A_55_P1999172** | Nat10 | 0.134166514 | -0.091641082 | 0.35997411 | 10.65360971 | 1.420086243 | 0.200720284 | 0.991835066 | -4.540639115 |
| **A_55_P2042655** | NA | -0.086739122 | -0.232725729 | 0.059247485 | 11.67524854 | -1.420074536 | 0.200723551 | 0.991835066 | -4.540649077 |
| **A_55_P2033586** | Mmab | 0.06269213 | -0.04282313 | 0.16820739 | 8.511973363 | 1.420060627 | 0.200727433 | 0.991835066 | -4.540660914 |
| **A_51_P116755** | Cap2 | -0.090541 | -0.242994736 | 0.061912736 | 12.13801788 | -1.419437742 | 0.200901324 | 0.991835066 | -4.535783174 |
| **A_55_P2123502** | Jam2 | -0.147628216 | -0.396302592 | 0.10104616 | 9.551703521 | -1.418885878 | 0.201055502 | 0.991835066 | -4.541660451 |
| **A_55_P1995652** | Rexo4 | 0.107333254 | -0.073504922 | 0.288171431 | 11.81911689 | 1.418578297 | 0.201141479 | 0.991835066 | -4.541922104 |
| **A_51_P248381** | Zdhhc4 | -0.107497631 | -0.288634401 | 0.073639138 | 8.617141836 | -1.418408772 | 0.20118888 | 0.991835066 | -4.542066307 |
| **A_51_P297963** | Recql | 0.06938045 | -0.047544023 | 0.186304923 | 8.392050896 | 1.418210922 | 0.201244214 | 0.991835066 | -4.542234594 |
| **A_51_P462692** | Ptbp1 | 0.116139415 | -0.079642422 | 0.311921253 | 11.82355941 | 1.417804553 | 0.201357908 | 0.991835066 | -4.542580216 |
| **A_51_P363749** | Irf6 | 0.182697399 | -0.125291724 | 0.490686521 | 9.236701716 | 1.417771094 | 0.201367272 | 0.991835066 | -4.542608672 |
| **A_51_P365980** | 2310022B0 | -0.085064371 | -0.22847423 | 0.058345488 | 12.32586805 | -1.417678718 | 0.201393126 | 0.991835066 | -4.542687233 |
| **A_55_P2068152** | Anapc15 | -0.062195099 | -0.167063874 | 0.042673676 | 11.45882725 | -1.417487061 | 0.201446777 | 0.991835066 | -4.54285022 |
| **A_51_P307325** | Rbbp4 | 0.077092266 | -0.052923461 | 0.207107993 | 13.32037403 | 1.417176866 | 0.201533637 | 0.991835066 | -4.543113997 |
| **A_52_P676198** | Fam210b | -0.084240076 | -0.226321108 | 0.057840957 | 8.944340148 | -1.417071543 | 0.201563137 | 0.991835066 | -4.543203553 |
| **A_51_P430327** | Gmpr2 | -0.113315536 | -0.304450992 | 0.077819921 | 9.26954087 | -1.416959152 | 0.201594621 | 0.991835066 | -4.543299117 |
| **A_55_P2073647** | Elp2 | 0.096068113 | -0.065975945 | 0.258112172 | 11.15775195 | 1.416952467 | 0.201596494 | 0.991835066 | -4.543304801 |
| **A_51_P348433** | Rasal1 | 0.098919543 | -0.067939674 | 0.265778759 | 7.718282838 | 1.416905975 | 0.201609519 | 0.991835066 | -4.543344332 |
| **A_51_P323610** | Selenos | -0.10064571 | -0.270472154 | 0.069180735 | 11.15885863 | -1.416442935 | 0.201739285 | 0.991835066 | -4.54373801 |
| **A_51_P415607** | Fkbp7 | -0.093315798 | -0.250792471 | 0.064160874 | 11.21018824 | -1.416276599 | 0.201785919 | 0.991835066 | -4.543879417 |
| **A_55_P1953919** | Me3 | -0.209086055 | -0.561970431 | 0.14379832 | 10.1144077 | -1.416125592 | 0.201828263 | 0.991835066 | -4.544007787 |
| **A_51_P146753** | Csf2rb2 | -0.081758037 | -0.219772675 | 0.0562566 | 8.144551115 | -1.415840856 | 0.201908129 | 0.991835066 | -4.544249824 |
| **A_55_P2125564** | Lrrc58 | -0.11898883 | -0.319865502 | 0.081887842 | 11.67832635 | -1.41574756 | 0.201934304 | 0.991835066 | -4.544329126 |
| **A_51_P194377** | Pcbp4 | 0.099960887 | -0.068795869 | 0.268717643 | 12.07295998 | 1.415722277 | 0.201941398 | 0.991835066 | -4.544350616 |
| **A_55_P2178378** | Gas5 | 0.237603048 | -0.163562209 | 0.638768305 | 8.729128097 | 1.415591107 | 0.201978205 | 0.991835066 | -4.544462105 |
| **A_55_P2076448** | Tmem255a | -0.14150295 | -0.380416132 | 0.097410233 | 11.24183403 | -1.415580303 | 0.201981237 | 0.991835066 | -4.544471289 |
| **A_55_P2122841** | NA | -0.085386768 | -0.229609019 | 0.058835484 | 14.72969535 | -1.415035827 | 0.202134087 | 0.991835066 | -4.544934031 |
| **A_51_P296866** | Msi1 | 0.088142684 | -0.060754343 | 0.237039711 | 8.498897848 | 1.414846666 | 0.202187214 | 0.991835066 | -4.545094779 |
| **A_55_P2087865** | Tinf2 | 0.085798098 | -0.059170162 | 0.230766358 | 8.955603381 | 1.414535546 | 0.202274621 | 0.991835066 | -4.545359151 |
| **A_55_P2173098** | Commd1 | -0.081393017 | -0.218943947 | 0.056157912 | 12.52178031 | -1.414271374 | 0.202348866 | 0.991835066 | -4.54558361 |
| **A_51_P118168** | Zbtb43 | 0.066048973 | -0.045585106 | 0.177683051 | 9.932045362 | 1.414094417 | 0.202398613 | 0.991835066 | -4.545733957 |
| **A_55_P2099530** | NA | -0.087615799 | -0.235716963 | 0.060485364 | 12.70617161 | -1.413946869 | 0.2024401 | 0.991835066 | -4.545859311 |

| | | | | | | | | | |
|---|---|---|---|---|---|---|---|---|---|
| **A_52_P565051** | Cpne3 | 0.099824728 | -0.068976811 | 0.268626267 | 8.664115665 | 1.413418805 | 0.202588644 | 0.991835066 | -4.546307904 |
| **A_55_P1972841** | Rps28 | 0.090203597 | -0.062353654 | 0.242760848 | 17.09231914 | 1.413188627 | 0.202653424 | 0.991835066 | -4.54650342 |
| **A_55_P2131340** | Churc1 | -0.152178095 | -0.40956208 | 0.10520589 | 11.90669562 | -1.41312232 | 0.202672088 | 0.991835066 | -4.54655974 |
| **A_51_P114826** | Cdh13 | -0.103826058 | -0.279463444 | 0.071811328 | 12.50618462 | -1.412858062 | 0.202746488 | 0.991835066 | -4.546784186 |
| **A_55_P2122618** | Asb11 | -0.14742013 | -0.396827701 | 0.101987441 | 11.85059715 | -1.412720636 | 0.202785189 | 0.991835066 | -4.546900901 |
| **A_55_P2170737** | Igf2bp2 | 0.101617523 | -0.07038734 | 0.273622386 | 9.570797259 | 1.412007529 | 0.202986117 | 0.991835066 | -4.547506468 |
| **A_55_P2063598** | Tomm5 | 0.090759164 | -0.062893399 | 0.244411727 | 14.44644301 | 1.411756548 | 0.203056877 | 0.991835066 | -4.547719571 |
| **A_51_P324572** | Ctnna1 | -0.104384851 | -0.281114047 | 0.072344346 | 14.93260639 | -1.411686635 | 0.203076591 | 0.991835066 | -4.547778931 |
| **A_52_P226127** | Dido1 | 0.153776877 | -0.106621157 | 0.414174911 | 8.591870853 | 1.411440153 | 0.203146111 | 0.991835066 | -4.547988196 |
| **A_51_P494775** | Rcor1 | 0.073446271 | -0.050967812 | 0.197860354 | 11.8629488 | 1.410942578 | 0.203286515 | 0.991835066 | -4.548410597 |
| **A_52_P491073** | Nek8 | 0.124286667 | -0.086263636 | 0.334836969 | 9.513889188 | 1.410840127 | 0.203315435 | 0.991835066 | -4.548497563 |
| **A_55_P1960837** | NA | 0.163988012 | -0.113830853 | 0.441806876 | 13.49638129 | 1.410780672 | 0.20333222 | 0.991835066 | -4.54854803 |
| **A_52_P330694** | Tle1 | 0.085755815 | -0.059544173 | 0.231055803 | 9.026647118 | 1.410610562 | 0.20338025 | 0.991835066 | -4.548692419 |
| **A_52_P365011** | Fes | -0.092490449 | -0.249215499 | 0.0642346 | 9.172285149 | -1.410482185 | 0.203416505 | 0.991835066 | -4.548801382 |
| **A_55_P2005475** | Sult1a1 | -0.153719272 | -0.414212892 | 0.106774348 | 9.934791257 | -1.410393701 | 0.203441496 | 0.991835066 | -4.548876482 |
| **A_52_P555235** | Rgs19 | 0.079426531 | -0.055183256 | 0.214036317 | 9.158665948 | 1.410256429 | 0.203480273 | 0.991835066 | -4.548992987 |
| **A_51_P124535** | Mest | 0.189036567 | -0.131393376 | 0.50946651 | 15.31677673 | 1.410008922 | 0.203550206 | 0.991835066 | -4.546496324 |
| **A_52_P539632** | Plxna3 | 0.089181928 | -0.06199024 | 0.240354096 | 8.403532028 | 1.40998388 | 0.203557283 | 0.991835066 | -4.54922429 |
| **A_66_P108692** | NA | 0.082325697 | -0.057242324 | 0.221893717 | 14.64359065 | 1.409803635 | 0.203608227 | 0.991835066 | -4.549377249 |
| **A_51_P332932** | NA | 0.073918828 | -0.051397543 | 0.199235199 | 8.154491756 | 1.409796389 | 0.203610275 | 0.991835066 | -4.549383397 |
| **A_55_P2004452** | Tceal8 | 0.122167299 | -0.08498042 | 0.329315019 | 10.29647247 | 1.409561241 | 0.203676753 | 0.991835066 | -4.549582935 |
| **A_52_P15212** | Ctbp2 | -0.07782452 | -0.212371621 | 0.056722582 | 11.44208417 | -1.436842791 | 0.203740564 | 0.991835066 | -4.500856899 |
| **A_55_P2056014** | NA | 0.242889115 | -0.169057595 | 0.654835824 | 9.430313595 | 1.409211393 | 0.203775695 | 0.991835066 | -4.549879779 |
| **A_52_P253179** | Igfbp3 | 0.154924914 | -0.10784443 | 0.417694258 | 9.551716031 | 1.409145052 | 0.203794462 | 0.991835066 | -4.549936066 |
| **A_52_P537466** | Zfp330 | 0.118423041 | -0.082451764 | 0.319297846 | 9.844951826 | 1.409028814 | 0.203827348 | 0.991835066 | -4.550034685 |
| **A_55_P2046728** | C920009B1 | 0.251083935 | -0.174853291 | 0.67702116 | 11.75026832 | 1.408907431 | 0.203861695 | 0.991835066 | -4.550137665 |
| **A_55_P2161219** | Thsd7a | -0.079168271 | -0.213515303 | 0.055178761 | 9.493062027 | -1.4084201 | 0.203999643 | 0.991835066 | -4.550551079 |
| **A_55_P2408588** | Arntl | -0.089960504 | -0.24266018 | 0.062739171 | 9.089691081 | -1.408065634 | 0.204100034 | 0.991835066 | -4.550851746 |
| **A_55_P2155062** | Naa40 | 0.111271099 | -0.077620956 | 0.300163153 | 10.21656464 | 1.407919479 | 0.20414144 | 0.991835066 | -4.550975709 |
| **A_51_P451458** | Mamdc2 | -0.09571505 | -0.258207444 | 0.066777343 | 8.110301154 | -1.407849817 | 0.204161179 | 0.991835066 | -4.551034792 |
| **A_66_P114381** | Ypel2 | -0.085618119 | -0.230980252 | 0.059744014 | 9.461909104 | -1.407743488 | 0.20419131 | 0.991835066 | -4.551124972 |
| **A_55_P1976112** | C130074G1 | -0.078615713 | -0.212098807 | 0.054867381 | 9.64985696 | -1.407642034 | 0.204220063 | 0.991835066 | -4.551211014 |
| **A_55_P2006261** | Krt15 | 0.127500671 | -0.088991101 | 0.343992443 | 7.832231044 | 1.407603074 | 0.204231106 | 0.991835066 | -4.551244055 |
| **A_55_P2046709** | NA | -0.931868461 | -2.514240585 | 0.650503662 | 10.43660452 | -1.407521775 | 0.20425415 | 0.991835066 | -4.551313001 |
| **A_51_P472621** | Zfp719 | 0.071476298 | -0.049896418 | 0.192849013 | 8.347308479 | 1.407505543 | 0.204258752 | 0.991835066 | -4.551326767 |
| **A_55_P2279035** | 9030419F2 | 0.153777122 | -0.107356771 | 0.414911015 | 9.040007438 | 1.407465043 | 0.204270233 | 0.991835066 | -4.551361113 |
| **A_55_P2074591** | Ppm1h | -0.074007652 | -0.199693963 | 0.051678658 | 8.666450797 | -1.40733595 | 0.204306833 | 0.991835066 | -4.551470586 |
| **A_51_P185757** | Casz1 | -0.061437822 | -0.165805407 | 0.042929763 | 11.18008346 | -1.406952101 | 0.204415695 | 0.991835066 | -4.551796076 |
| **A_52_P515769** | Pcdh12 | -0.100331362 | -0.270791459 | 0.070128735 | 10.85822795 | -1.406770028 | 0.20446735 | 0.991835066 | -4.551950455 |
| **A_51_P226429** | Rhobtb2 | -0.096600824 | -0.260759111 | 0.067557463 | 10.00017838 | -1.406459239 | 0.20455555 | 0.991835066 | -4.552213953 |
| **A_55_P1992445** | Vdac1 | -0.085779292 | -0.231572061 | 0.060013478 | 15.73139076 | -1.406227554 | 0.204621323 | 0.991835066 | -4.552410369 |
| **A_51_P336391** | Tmem18 | 0.125234839 | -0.087621017 | 0.338090696 | 10.30699954 | 1.406205177 | 0.204627676 | 0.991835066 | -4.552429339 |
| **A_55_P2017694** | Gprc5c | 0.098323403 | -0.06881848 | 0.265465286 | 9.789868857 | 1.405985178 | 0.204690151 | 0.991835066 | -4.552615835 |
| **A_55_P2079560** | Lilra6 | -0.078157229 | -0.211031397 | 0.054716939 | 7.779670414 | -1.405845924 | 0.204729705 | 0.991835066 | -4.552733877 |
| **A_51_P487501** | NA | -0.098877387 | -0.266979497 | 0.069224724 | 10.51248567 | -1.405830454 | 0.204734099 | 0.991835066 | -4.552746989 |
| **A_55_P2107686** | Ttll1 | -0.096087241 | -0.259449349 | 0.067274867 | 8.598908095 | -1.405799965 | 0.20474276 | 0.991835066 | -4.552772833 |
| **A_51_P433430** | Phc1 | -0.070672974 | -0.190877058 | 0.04953111 | 9.058339924 | -1.405216621 | 0.204908538 | 0.991835066 | -4.55326726 |
| **A_55_P2019964** | Lancl1 | -0.068240317 | -0.184310722 | 0.047830087 | 11.82392358 | -1.405169365 | 0.204921973 | 0.991835066 | -4.553307309 |

| | | | | | | | | | |
|---|---|---|---|---|---|---|---|---|---|
| **A_55_P1985012** | Pard3 | 0.101952459 | -0.071460294 | 0.275365212 | 8.494148699 | 1.405160082 | 0.204924612 | 0.991835066 | -4.553315177 |
| **A_52_P293120** | Prrg3 | -0.078027602 | -0.210756387 | 0.054701183 | 9.206264361 | -1.405051599 | 0.204955457 | 0.991835066 | -4.553407114 |
| **A_51_P478688** | Samd10 | -0.121766464 | -0.328912669 | 0.08537974 | 10.62454506 | -1.404946699 | 0.204985287 | 0.991835066 | -4.553496011 |
| **A_51_P114005** | Gstm7 | -0.099033329 | -0.267507416 | 0.069440759 | 8.269847324 | -1.40493877 | 0.204987542 | 0.991835066 | -4.545423559 |
| **A_55_P2031999** | Dglucy | -0.137928087 | -0.372659663 | 0.096803489 | 9.954289153 | -1.404398413 | 0.205141264 | 0.991835066 | -4.553960614 |
| **A_52_P480351** | Loxl2 | 0.134688675 | -0.094606087 | 0.363983436 | 13.13361702 | 1.403932016 | 0.205274029 | 0.991835066 | -4.55435577 |
| **A_51_P345902** | Dohh | -0.081672495 | -0.220713193 | 0.057368204 | 12.659208 | -1.403922104 | 0.205276851 | 0.991835066 | -4.554364168 |
| **A_52_P541270** | Crebl2 | -0.076156888 | -0.205813433 | 0.053499657 | 8.378133104 | -1.40386024 | 0.205294467 | 0.991835066 | -4.554416578 |
| **A_55_P1960298** | Tmem80 | 0.118365639 | -0.083151881 | 0.31988316 | 9.251601603 | 1.403854081 | 0.205296221 | 0.991835066 | -4.554421796 |
| **A_55_P2069554** | Brix1 | 0.09204432 | -0.064676138 | 0.248764778 | 11.51520026 | 1.403719824 | 0.205334457 | 0.991835066 | -4.554535534 |
| **A_66_P116299** | Pias1 | 0.097560724 | -0.068560573 | 0.263682021 | 10.37196623 | 1.403650012 | 0.205354342 | 0.991835066 | -4.554594674 |
| **A_66_P111798** | Znrf2 | -0.082869894 | -0.223981475 | 0.058241687 | 9.076089775 | -1.403599669 | 0.205368682 | 0.991835066 | -4.554637321 |
| **A_55_P2008347** | Prelid1 | 0.090839597 | -0.063843833 | 0.245523027 | 14.48957462 | 1.403590876 | 0.205371187 | 0.991835066 | -4.554644769 |
| **A_55_P2117726** | Stk33 | -0.123356062 | -0.333415105 | 0.086702982 | 7.713107103 | -1.403551152 | 0.205382503 | 0.991835066 | -4.55467842 |
| **A_55_P1986252** | NA | 0.073193448 | -0.051485599 | 0.197872495 | 16.16832393 | 1.403097543 | 0.205511763 | 0.991835066 | -4.55506265 |
| **A_55_P1955791** | Fopnl | -0.073532861 | -0.198800263 | 0.051734542 | 12.61764321 | -1.402983364 | 0.205544311 | 0.991835066 | -4.555159357 |
| **A_55_P2103352** | Ndufb5 | -0.101887904 | -0.275467235 | 0.071691427 | 14.67573188 | -1.402922726 | 0.205561599 | 0.991835066 | -4.555210716 |
| **A_55_P2149719** | NA | -0.132412919 | -0.358010025 | 0.093184188 | 16.48320088 | -1.402832974 | 0.205587188 | 0.991835066 | -4.55528673 |
| **A_51_P255565** | Smarcad1 | -0.065864178 | -0.178111313 | 0.046382957 | 9.355463909 | -1.402436302 | 0.205700321 | 0.991835066 | -4.555622666 |
| **A_55_P2004927** | Dnajc25 | -0.088123597 | -0.238321043 | 0.062073849 | 8.8464445 | -1.402293104 | 0.205741175 | 0.991835066 | -4.555743929 |
| **A_66_P119999** | Atg7 | 0.06044164 | -0.042580007 | 0.163463287 | 9.840532923 | 1.402222345 | 0.205761366 | 0.991835066 | -4.555803848 |
| **A_52_P121960** | NA | -0.084087926 | -0.227422882 | 0.05924703 | 16.37948698 | -1.402137668 | 0.20578553 | 0.991835066 | -4.55587555 |
| **A_55_P2029711** | Cdh2 | -0.105196882 | -0.284543562 | 0.074149798 | 14.35348653 | -1.401905236 | 0.205851871 | 0.991835066 | -4.556072359 |
| **A_66_P121480** | Sardh | -0.101139648 | -0.273582299 | 0.071303003 | 9.960076455 | -1.401799427 | 0.205882077 | 0.991835066 | -4.556161948 |
| **A_52_P657240** | Cdc40 | 0.076302535 | -0.053807831 | 0.206412902 | 8.400349196 | 1.401639087 | 0.205927859 | 0.991835066 | -4.556297702 |
| **A_51_P446477** | Cldnd1 | 0.085268006 | -0.060151447 | 0.230687459 | 11.68103645 | 1.401434267 | 0.205986355 | 0.991835066 | -4.556471107 |
| **A_55_P1962315** | NA | 0.059990806 | -0.042333593 | 0.162315205 | 16.03281104 | 1.401246784 | 0.206039912 | 0.991835066 | -4.556629826 |
| **A_55_P2131238** | Ttc39a | 0.072802068 | -0.051385525 | 0.196989662 | 8.480573957 | 1.401117761 | 0.206076776 | 0.991835066 | -4.556739048 |
| **A_51_P154469** | Ccndbp1 | -0.160668233 | -0.434741361 | 0.113404895 | 11.29860291 | -1.401111486 | 0.206078569 | 0.991835066 | -4.55674436 |
| **A_55_P1952950** | Cic | 0.148370901 | -0.104753784 | 0.401495586 | 14.1805106 | 1.400952353 | 0.206124045 | 0.991835066 | -4.556879066 |
| **A_55_P1990663** | Ddx5 | 0.138222577 | -0.097595449 | 0.374040603 | 12.09816358 | 1.400912847 | 0.206135336 | 0.991835066 | -4.556912507 |
| **A_51_P328400** | Cops2 | -0.082697431 | -0.223791352 | 0.058396491 | 10.9974298 | -1.400853902 | 0.206152184 | 0.991835066 | -4.556962402 |
| **A_55_P2016540** | Cry2 | -0.083842391 | -0.22692146 | 0.059236678 | 10.87159952 | -1.400543751 | 0.206240854 | 0.991835066 | -4.55722492 |
| **A_51_P407904** | Trappc4 | 0.076317985 | -0.053922355 | 0.206558325 | 12.42855686 | 1.400523838 | 0.206246548 | 0.991835066 | -4.557241774 |
| **A_51_P134142** | Cyp2c70 | -0.062973287 | -0.17045599 | 0.044509416 | 7.904384538 | -1.400318794 | 0.206305189 | 0.991835066 | -4.557415314 |
| **A_51_P190254** | NA | -0.139169473 | -0.376775789 | 0.098436842 | 9.803139379 | -1.399893947 | 0.206426739 | 0.991835066 | -4.557774852 |
| **A_52_P305890** | Yipf5 | -0.063296982 | -0.171375336 | 0.044781372 | 11.39923788 | -1.399759472 | 0.206465226 | 0.991835066 | -4.557888646 |
| **A_51_P424781** | Sf3b6 | -0.074877166 | -0.202735828 | 0.052981496 | 12.1944952 | -1.399678775 | 0.206488325 | 0.991835066 | -4.55795693 |
| **A_55_P2115995** | Sms | 0.114752912 | -0.081231506 | 0.31073733 | 11.10137318 | 1.399430403 | 0.206559434 | 0.991835066 | -4.558167089 |
| **A_52_P515036** | Htatip2 | -0.108161835 | -0.29289613 | 0.07657246 | 8.113360808 | -1.399379962 | 0.206573878 | 0.991835066 | -4.558209768 |
| **A_51_P483351** | Lix1l | -0.104375812 | -0.282653665 | 0.073902041 | 8.991160796 | -1.399302485 | 0.206596065 | 0.991835066 | -4.558275321 |
| **A_52_P302544** | Col8a2 | 0.131012156 | -0.092792064 | 0.354816375 | 8.598616051 | 1.399111915 | 0.206650649 | 0.991835066 | -4.558436554 |
| **A_55_P2224582** | Fam219b | 0.071086829 | -0.050354529 | 0.192528187 | 8.362926944 | 1.399044914 | 0.206669843 | 0.991835066 | -4.558493239 |
| **A_55_P2059931** | Prom1 | 0.12598507 | -0.089255836 | 0.341225975 | 9.069795884 | 1.398953898 | 0.206695919 | 0.991835066 | -4.55857024 |
| **A_55_P2016682** | Alkbh7 | -0.123087786 | -0.333407057 | 0.087231485 | 13.35020406 | -1.398765858 | 0.206749801 | 0.991835066 | -4.558729317 |
| **A_55_P2173968** | Ctdspl | 0.08777768 | -0.062252274 | 0.237807634 | 11.98600623 | 1.398347957 | 0.206869594 | 0.991835066 | -4.559082821 |
| **A_52_P111031** | Pcdh17 | -0.183532985 | -0.497246114 | 0.130180144 | 10.2276552 | -1.398268477 | 0.206892385 | 0.991835066 | -4.559150048 |
| **A_55_P2035038** | NA | 0.407219101 | -0.288891335 | 1.103329537 | 12.13195176 | 1.39816726 | 0.206921411 | 0.991835066 | -4.55923566 |

| | | | | | | | | | |
|---|---|---|---|---|---|---|---|---|---|
| **A_66_P101835** | Cd38 | -0.132378031 | -0.358731349 | 0.093975287 | 9.218063486 | -1.397777947 | 0.207033091 | 0.991835066 | -4.559564925 |
| **A_55_P2010312** | Serpina1a | -0.53364302 | -1.446285479 | 0.378999439 | 8.899324235 | -1.397524069 | 0.207105948 | 0.991835066 | -4.559779626 |
| **A_55_P1982902** | Tceal3 | 0.195903551 | -0.139183863 | 0.530990966 | 9.015016003 | 1.397311793 | 0.207166884 | 0.991835066 | -4.559959133 |
| **A_55_P2049061** | NA | -0.090542103 | -0.245430343 | 0.064346137 | 10.44406466 | -1.397144304 | 0.207214975 | 0.991835066 | -4.560100758 |
| **A_52_P578436** | NA | 0.061550983 | -0.043784422 | 0.166886388 | 7.817944448 | 1.396592663 | 0.207373438 | 0.991835066 | -4.560567167 |
| **A_55_P2015941** | NA | -0.11134642 | -0.301912742 | 0.079219902 | 9.096025016 | -1.396494526 | 0.20740164 | 0.991835066 | -4.560650133 |
| **A_55_P2063465** | Thbs3 | -0.210741659 | -0.571426392 | 0.149943075 | 10.37918917 | -1.396470498 | 0.207408546 | 0.991835066 | -4.560670447 |
| **A_55_P2114779** | Greb1 | 0.075540823 | -0.053755082 | 0.204836727 | 7.98418788 | 1.396387889 | 0.207432288 | 0.991835066 | -4.560740283 |
| **A_55_P2001023** | Lrrc19 | -0.118640806 | -0.321708647 | 0.084427036 | 11.66057763 | -1.396375006 | 0.207435991 | 0.991835066 | -4.560751174 |
| **A_55_P2086475** | NA | 0.111419384 | -0.079288918 | 0.302127685 | 11.19006852 | 1.39636928 | 0.207437637 | 0.991835066 | -4.560756014 |
| **A_55_P1968188** | Btf3 | 0.082106166 | -0.058429145 | 0.222641476 | 14.11383745 | 1.396366577 | 0.207438414 | 0.991835066 | -4.5607583 |
| **A_55_P2122619** | Asb11 | -0.106008257 | -0.28745891 | 0.075442397 | 11.99123041 | -1.396337136 | 0.207446877 | 0.991835066 | -4.560783188 |
| **A_55_P2074154** | Dhx57 | 0.14355455 | -0.102174416 | 0.389283515 | 8.66464574 | 1.396271076 | 0.207465866 | 0.991835066 | -4.560839033 |
| **A_55_P1968600** | NA | 0.08989991 | -0.064007424 | 0.243807243 | 8.788337161 | 1.396076039 | 0.20752194 | 0.991835066 | -4.561003902 |
| **A_52_P182520** | Timm9 | 0.190749708 | -0.135817452 | 0.517316868 | 9.312454113 | 1.396048495 | 0.207529861 | 0.991835066 | -4.561027185 |
| **A_51_P303620** | Whrn | 0.075482637 | -0.05375941 | 0.204724683 | 8.971094909 | 1.395893766 | 0.207574357 | 0.991835066 | -4.561157973 |
| **A_55_P2047932** | Cyb5r4 | -0.096285214 | -0.261163641 | 0.068593212 | 9.707621758 | -1.395741203 | 0.207618239 | 0.991835066 | -4.561286925 |
| **A_51_P452798** | Zfp524 | -0.112890406 | -0.306259868 | 0.080479056 | 10.25076168 | -1.395334326 | 0.207735311 | 0.991835066 | -4.561630806 |
| **A_55_P1984541** | NA | -0.143584433 | -0.389573052 | 0.102404187 | 8.687673272 | -1.395087585 | 0.207806335 | 0.991835066 | -4.561839324 |
| **A_55_P2116993** | Hnrnpdl | 0.152246328 | -0.108583783 | 0.413076439 | 12.22144015 | 1.395077171 | 0.207809333 | 0.991835066 | -4.561848125 |
| **A_55_P1964213** | Tprkb | 0.136637923 | -0.097471871 | 0.370747718 | 9.041225722 | 1.394956937 | 0.207843951 | 0.991835066 | -4.561949727 |
| **A_55_P2102578** | NA | 0.0852991 | -0.060860993 | 0.231459193 | 14.60109552 | 1.394841208 | 0.207877276 | 0.991835066 | -4.56204752 |
| **A_52_P428496** | Rhoa | 0.139520333 | -0.099549749 | 0.378590415 | 10.90362319 | 1.394830408 | 0.207880386 | 0.991835066 | -4.562056646 |
| **A_55_P2077228** | Cenpl | -0.100987914 | -0.274042153 | 0.072066325 | 9.141097627 | -1.394749744 | 0.207903617 | 0.991835066 | -4.562124806 |
| **A_55_P1968355** | Tle1 | 0.118656379 | -0.084706377 | 0.322019135 | 10.46548362 | 1.394533025 | 0.207966044 | 0.991835066 | -4.562307923 |
| **A_51_P218774** | Rgs10 | -0.104131984 | -0.282612528 | 0.074348561 | 9.741637304 | -1.394448221 | 0.207990477 | 0.991835066 | -4.562379576 |
| **A_51_P318640** | Rps3 | 0.089251378 | -0.063726429 | 0.242229185 | 14.6395411 | 1.394426516 | 0.20799673 | 0.991835066 | -4.562397914 |
| **A_66_P102090** | Pkmyt1 | 0.110075183 | -0.078620583 | 0.298770949 | 11.03818718 | 1.394236322 | 0.208051537 | 0.991835066 | -4.562558605 |
| **A_55_P2045224** | NA | -0.076225635 | -0.206903675 | 0.054452405 | 12.05580451 | -1.394143801 | 0.208078203 | 0.991835066 | -4.56263677 |
| **A_55_P2293351** | Slc2a4rg-p | 0.270076671 | -0.192998033 | 0.733151375 | 9.101297698 | 1.39394342 | 0.208135966 | 0.991835066 | -4.562806052 |
| **A_52_P594917** | Pip4k2c | -0.117158569 | -0.318056451 | 0.083739313 | 10.72201118 | -1.393823656 | 0.208170497 | 0.991835066 | -4.562907225 |
| **A_55_P2130178** | Fn1 | 0.215239213 | -0.153896737 | 0.584375163 | 15.18017561 | 1.393619406 | 0.208229399 | 0.991835066 | -4.563079761 |
| **A_55_P2133637** | Etfa | -0.168802618 | -0.458322938 | 0.120717702 | 10.63242793 | -1.393507338 | 0.208261724 | 0.991835066 | -4.563174423 |
| **A_52_P683306** | Fam13b | 0.059150931 | -0.042314121 | 0.160615983 | 8.126054839 | 1.393330788 | 0.208312657 | 0.991835066 | -4.563323547 |
| **A_51_P329966** | D6Wsu163 | 0.065783596 | -0.04706422 | 0.178631412 | 9.106120832 | 1.393264543 | 0.208331771 | 0.991835066 | -4.563379499 |
| **A_66_P123938** | Smim13 | 0.06719942 | -0.048106843 | 0.182505683 | 11.39658124 | 1.39290584 | 0.208435295 | 0.991835066 | -4.563682448 |
| **A_51_P466829** | Pheta1 | 0.080871615 | -0.057956207 | 0.219699437 | 11.57450836 | 1.392286908 | 0.208614034 | 0.991835066 | -4.564205107 |
| **A_55_P2027812** | Arxes2 | 0.12159849 | -0.087227414 | 0.330424393 | 10.58103582 | 1.391723446 | 0.208776872 | 0.991835066 | -4.56468084 |
| **A_66_P134878** | D11Wsu47 | 0.071808293 | -0.051514876 | 0.195131461 | 10.80427223 | 1.391678955 | 0.208789735 | 0.991835066 | -4.564718402 |
| **A_55_P2077879** | Kat6b | 0.099287869 | -0.071231427 | 0.269807164 | 8.739014176 | 1.391655658 | 0.20879647 | 0.991835066 | -4.564738069 |
| **A_55_P2056241** | Pdlim5 | 0.080347412 | -0.05764789 | 0.218342714 | 9.68913889 | 1.391607389 | 0.208810426 | 0.991835066 | -4.564778818 |
| **A_52_P442756** | Btf3l4 | -0.095105584 | -0.258463264 | 0.068252097 | 11.22142168 | -1.391475581 | 0.20884854 | 0.991835066 | -4.56489009 |
| **A_55_P2081349** | Focad | 0.109712367 | -0.078740617 | 0.298165352 | 10.07376638 | 1.391431074 | 0.208861411 | 0.991835066 | -4.564927661 |
| **A_52_P577748** | Lpxn | -0.062939978 | -0.171083017 | 0.045203061 | 7.909900703 | -1.39103209 | 0.208976827 | 0.991835066 | -4.56526445 |
| **A_55_P2075429** | Ttll4 | 0.082829899 | -0.059494687 | 0.225154486 | 8.34844024 | 1.390965424 | 0.208996117 | 0.991835066 | -4.56532072 |
| **A_51_P185108** | Dpy30 | 0.070811715 | -0.05092554 | 0.19254897 | 11.42114746 | 1.390243068 | 0.209205238 | 0.991835066 | -4.56593036 |
| **A_55_P1969197** | NA | -0.063618716 | -0.172995256 | 0.045757824 | 16.69886025 | -1.390176197 | 0.209224607 | 0.991835066 | -4.56598679 |
| **A_52_P277139** | Ankrd17 | -0.069851624 | -0.189950207 | 0.050246959 | 11.23929519 | -1.390105482 | 0.20924509 | 0.991835066 | -4.566046464 |

| | | | | | | | | | |
|---|---|---|---|---|---|---|---|---|---|
| **A_55_P2034175** | Usp28 | -0.097611914 | -0.26545162 | 0.070227791 | 12.9861486 | -1.390007855 | 0.209273372 | 0.991835066 | -4.566128843 |
| **A_55_P1983544** | Tmed5 | 0.0651217 | -0.046897768 | 0.177141169 | 9.220688585 | 1.389444985 | 0.2094365 | 0.991835066 | -4.566603763 |
| **A_51_P386539** | Rnf125 | -0.069261305 | -0.188406051 | 0.049883442 | 8.500171173 | -1.389392349 | 0.20945176 | 0.991835066 | -4.566648169 |
| **A_65_P15245** | Nrp2 | -0.067277281 | -0.18301454 | 0.048459978 | 8.845673219 | -1.389326664 | 0.209470805 | 0.991835066 | -4.566703585 |
| **A_52_P509185** | Zfp318 | 0.072938701 | -0.052565196 | 0.198442598 | 9.331320766 | 1.38902462 | 0.209558401 | 0.991835066 | -4.566958392 |
| **A_55_P2057446** | Ccdc134 | 0.111002991 | -0.080005112 | 0.302011093 | 9.664531431 | 1.388967301 | 0.209575028 | 0.991835066 | -4.567006744 |
| **A_52_P211223** | Cdca2 | 0.066243675 | -0.047759252 | 0.180246602 | 8.949729278 | 1.388793095 | 0.209625569 | 0.991835066 | -4.567153694 |
| **A_55_P2028345** | Ccser2 | -0.08444723 | -0.229799912 | 0.060905452 | 12.59512227 | -1.388581868 | 0.209686864 | 0.991835066 | -4.567331862 |
| **A_55_P2076001** | Rbm12 | 0.088792317 | -0.064070991 | 0.241655625 | 11.06571945 | 1.388293433 | 0.20977059 | 0.991835066 | -4.567575135 |
| **A_66_P106536** | Atp5mpl | -0.104260857 | -0.283757592 | 0.075235879 | 16.38146915 | -1.38826977 | 0.20977746 | 0.991835066 | -4.567595092 |
| **A_51_P466148** | Bckdha | -0.204890132 | -0.557642456 | 0.147862191 | 12.00268426 | -1.38822638 | 0.209790058 | 0.991835066 | -4.567631687 |
| **A_51_P515452** | Fkbpl | -0.10035541 | -0.273138086 | 0.072427265 | 10.50018316 | -1.388192612 | 0.209799863 | 0.991835066 | -4.567660166 |
| **A_55_P2129614** | Itprip | -0.121447706 | -0.330549341 | 0.08765393 | 12.6014182 | -1.38816477 | 0.209807947 | 0.991835066 | -4.567683646 |
| **A_51_P377452** | Ncf4 | -0.186732787 | -0.508257536 | 0.134791962 | 8.87238789 | -1.388082581 | 0.209831814 | 0.991835066 | -4.56775296 |
| **A_55_P2139673** | Rpl11 | 0.079642183 | -0.057532111 | 0.216816476 | 15.28197138 | 1.387648768 | 0.209957829 | 0.991835066 | -4.568118789 |
| **A_55_P2134790** | Qsox1 | 0.10823219 | -0.078210615 | 0.294674995 | 11.2503986 | 1.387458337 | 0.210013167 | 0.991835066 | -4.568279361 |
| **A_55_P2063336** | Tedc1 | 0.11576246 | -0.083678957 | 0.315203876 | 10.03145087 | 1.387271706 | 0.210067414 | 0.991835066 | -4.568436721 |
| **A_55_P2031045** | Stat5b | -0.069514821 | -0.189293846 | 0.050264203 | 10.59098159 | -1.387093614 | 0.21011919 | 0.991835066 | -4.568586874 |
| **A_66_P127956** | Zfp947 | 0.09887122 | -0.071528356 | 0.269270796 | 11.9841384 | 1.386789404 | 0.210207659 | 0.991835066 | -4.56884334 |
| **A_51_P311904** | Ptk2b | -0.095717485 | -0.260694671 | 0.0692597 | 8.659732174 | -1.386680867 | 0.210239231 | 0.991835066 | -4.568934837 |
| **A_55_P2176838** | Mitf | -0.061993216 | -0.168852426 | 0.044865993 | 12.74822665 | -1.386568544 | 0.210271909 | 0.991835066 | -4.569029522 |
| **A_55_P2068265** | Mif | 0.170572333 | -0.12351958 | 0.464664246 | 15.99855701 | 1.386227912 | 0.210371037 | 0.991835066 | -4.569316649 |
| **A_52_P660032** | Cpsf2 | 0.118563539 | -0.085865735 | 0.322992812 | 10.2788777 | 1.386172252 | 0.210387238 | 0.991835066 | -4.569363563 |
| **A_51_P182572** | Phactr1 | 0.087272816 | -0.063236962 | 0.237782595 | 9.200076188 | 1.385872915 | 0.210474389 | 0.991835066 | -4.569615852 |
| **A_55_P2121613** | Sox4 | -0.244083598 | -0.665035678 | 0.176868482 | 11.57538145 | -1.385846307 | 0.210482137 | 0.991835066 | -4.569638277 |
| **A_51_P161037** | Cep170 | -0.084081947 | -0.229097193 | 0.060933299 | 8.578608518 | -1.385792567 | 0.210497788 | 0.991835066 | -4.569683568 |
| **A_55_P2168663** | Atp5k | -0.074816293 | -0.203858015 | 0.05422543 | 16.1719253 | -1.385718978 | 0.21051922 | 0.991835066 | -4.569745585 |
| **A_55_P2134427** | NA | -0.135365148 | -0.368865681 | 0.098135386 | 9.779675444 | -1.385568864 | 0.210562945 | 0.991835066 | -4.569872092 |
| **A_55_P2025730** | Hif3a | 0.129006787 | -0.093693162 | 0.351706735 | 9.046584515 | 1.384527462 | 0.210866511 | 0.991835066 | -4.570749562 |
| **A_55_P2138436** | NA | 0.069372095 | -0.050392922 | 0.189137113 | 15.57240668 | 1.38440756 | 0.210901487 | 0.991835066 | -4.570850572 |
| **A_55_P2007384** | Sgca | -0.172522207 | -0.470386465 | 0.12534205 | 10.02201845 | -1.384317615 | 0.210927727 | 0.991835066 | -4.570926343 |
| **A_51_P341379** | Acad9 | -0.127657975 | -0.348073236 | 0.092757286 | 11.40510399 | -1.384252833 | 0.210946629 | 0.991835066 | -4.570980915 |
| **A_55_P1955034** | Myo1c | 0.10473341 | -0.076112027 | 0.285578847 | 12.12224129 | 1.384161678 | 0.210973228 | 0.991835066 | -4.571057702 |
| **A_51_P101460** | Dsp | -0.069855996 | -0.190480206 | 0.050768214 | 13.04649959 | -1.384134647 | 0.210981116 | 0.991835066 | -4.568253911 |
| **A_55_P2004876** | Srpx2 | -0.115855775 | -0.315941197 | 0.084229648 | 8.245711038 | -1.383921219 | 0.211043408 | 0.991835066 | -4.571260249 |
| **A_55_P2029191** | Pde4a | -0.083252697 | -0.227052745 | 0.060547352 | 11.07910571 | -1.383720605 | 0.211101976 | 0.991835066 | -4.571429222 |
| **A_65_P02958** | Rnf13 | -0.115285318 | -0.314420321 | 0.083849686 | 11.00633359 | -1.383679566 | 0.211113958 | 0.991835066 | -4.571463786 |
| **A_55_P2096367** | NA | -0.072717412 | -0.198376832 | 0.052942008 | 13.69978574 | -1.383096539 | 0.211284258 | 0.991835066 | -4.571954795 |
| **A_51_P404236** | Atp6v0b | -0.15324148 | -0.418079101 | 0.111596141 | 10.48040972 | -1.382947817 | 0.211327719 | 0.991835066 | -4.572080031 |
| **A_55_P1958857** | Nek6 | 0.080344976 | -0.05852197 | 0.219211922 | 9.47185454 | 1.382830579 | 0.211361984 | 0.991835066 | -4.572178751 |
| **A_52_P652513** | Lman1 | 0.153627656 | -0.111963314 | 0.419218625 | 12.63884328 | 1.382500293 | 0.211458546 | 0.991835066 | -4.572456849 |
| **A_55_P1978795** | Pja2 | -0.094702657 | -0.258457234 | 0.069051919 | 12.56895703 | -1.382222158 | 0.211539891 | 0.991835066 | -4.572691014 |
| **A_55_P2104769** | Utf1 | -0.143140615 | -0.390654098 | 0.104372869 | 9.686375142 | -1.38220719 | 0.211544269 | 0.991835066 | -4.572703615 |
| **A_51_P346523** | Zcchc7 | 0.231506408 | -0.168813866 | 0.631826683 | 9.469971276 | 1.38217987 | 0.211552261 | 0.991835066 | -4.572726615 |
| **A_66_P120249** | Nr1h3 | -0.109064101 | -0.297665068 | 0.079536865 | 8.722541902 | -1.382124111 | 0.211568573 | 0.991835066 | -4.572773556 |
| **A_55_P2128499** | Mgl2 | 0.071192119 | -0.05193041 | 0.194314649 | 12.54130961 | 1.381985619 | 0.211609093 | 0.991835066 | -4.572890144 |
| **A_51_P439612** | Dnajb2 | -0.12871691 | -0.35137342 | 0.0939396 | 10.43785393 | -1.38168595 | 0.211696793 | 0.991835066 | -4.5731424 |
| **A_55_P1988368** | Upp1 | -0.164546755 | -0.449199878 | 0.120106367 | 11.06952627 | -1.381600567 | 0.211721786 | 0.991835066 | -4.57321427 |

| | | | | | | | | | |
|---|---|---|---|---|---|---|---|---|---|
| **A_55_P2081133** | Gapdh | 0.096553616 | -0.07047866 | 0.263585892 | 17.22067224 | 1.381583939 | 0.211726654 | 0.991835066 | -4.573228266 |
| **A_55_P1985385** | Acp2 | -0.063713865 | -0.173941612 | 0.046513881 | 8.095977559 | -1.381504032 | 0.211750048 | 0.991835066 | -4.573295524 |
| **A_55_P1976908** | NA | -0.090544408 | -0.247202113 | 0.066113297 | 11.98619396 | -1.381398579 | 0.211780924 | 0.991835066 | -4.573384283 |
| **A_55_P2127439** | Kri1 | 0.059017715 | -0.043099935 | 0.161135365 | 11.85694912 | 1.381308591 | 0.211807275 | 0.991835066 | -4.573460022 |
| **A_55_P2051894** | Zfp598 | 0.100113036 | -0.07318218 | 0.273408252 | 9.225675345 | 1.380744083 | 0.211972647 | 0.991835066 | -4.5739351 |
| **A_55_P2245422** | NA | 0.24110308 | -0.176263142 | 0.658469302 | 9.041202959 | 1.380684957 | 0.211989975 | 0.991835066 | -4.573984855 |
| **A_66_P105032** | Gm13889 | 0.094772185 | -0.069285442 | 0.258829812 | 11.76395908 | 1.380681799 | 0.2119909 | 0.991835066 | -4.573987513 |
| **A_51_P421734** | Slc12a7 | -0.12621786 | -0.344739678 | 0.092303959 | 11.26587366 | -1.380495951 | 0.212045374 | 0.991835066 | -4.574143898 |
| **A_52_P477431** | Mnt | -0.159453899 | -0.435519891 | 0.116612092 | 11.48617678 | -1.380483966 | 0.212048887 | 0.991835066 | -4.574153982 |
| **A_55_P2068554** | NA | 0.138826263 | -0.101583475 | 0.379236 | 16.2625906 | 1.380157692 | 0.212144552 | 0.991835066 | -4.57442851 |
| **A_55_P2266880** | Kirrel | 0.115114113 | -0.084284211 | 0.314512437 | 10.02653304 | 1.379800189 | 0.212249418 | 0.991835066 | -4.574729283 |
| **A_51_P135517** | Coch | -0.069968382 | -0.191169672 | 0.051232908 | 7.757015455 | -1.379760545 | 0.212261049 | 0.991835066 | -4.574762633 |
| **A_52_P400224** | Ccs | 0.079367429 | -0.058123708 | 0.216858566 | 11.56472799 | 1.379674824 | 0.212286201 | 0.991835066 | -4.574834746 |
| **A_66_P125828** | NA | -0.164364908 | -0.44914386 | 0.120414045 | 9.876800901 | -1.379463913 | 0.212348099 | 0.991835066 | -4.575012168 |
| **A_55_P1962661** | Cyp2c67 | -0.058570554 | -0.160086845 | 0.042945738 | 7.792841965 | -1.378963325 | 0.212495074 | 0.991835066 | -4.575433223 |
| **A_55_P1957633** | Gpsm3 | -0.07689725 | -0.210195892 | 0.056401393 | 9.837422457 | -1.378777579 | 0.212549633 | 0.991835066 | -4.575589443 |
| **A_55_P2121372** | Trp53i13 | 0.101685715 | -0.074588519 | 0.27795995 | 11.89220984 | 1.378733232 | 0.212562661 | 0.991835066 | -4.575626738 |
| **A_55_P2141415** | Ppp6r3 | 0.090577939 | -0.066459131 | 0.247615009 | 8.558489398 | 1.378571782 | 0.212610096 | 0.991835066 | -4.575762515 |
| **A_55_P1993789** | Hmgn2 | -0.090231119 | -0.246712532 | 0.066250294 | 14.58587841 | -1.378169763 | 0.212728254 | 0.991835066 | -4.576100576 |
| **A_66_P107379** | Mdk | 0.165446867 | -0.121603638 | 0.452497372 | 11.42382685 | 1.377556323 | 0.212908663 | 0.991835066 | -4.576616345 |
| **A_55_P2018017** | Tnfsf10 | -0.070599106 | -0.193094844 | 0.051896633 | 7.871984754 | -1.377486498 | 0.212929207 | 0.991835066 | -4.576675047 |
| **A_55_P2162582** | Rab5b | 0.071929366 | -0.05288977 | 0.196748502 | 8.779874852 | 1.377317899 | 0.212978819 | 0.991835066 | -4.576816782 |
| **A_52_P635182** | Rras | -0.111072577 | -0.303817925 | 0.081672772 | 11.96777962 | -1.377311182 | 0.212980796 | 0.991835066 | -4.576822428 |
| **A_55_P2136020** | Uba2 | 0.085284398 | -0.062740339 | 0.233309134 | 12.58546904 | 1.377033235 | 0.213062608 | 0.991835066 | -4.577056072 |
| **A_55_P1964664** | Fus | 0.127400952 | -0.093756119 | 0.348558022 | 13.62305505 | 1.376832079 | 0.213121835 | 0.991835066 | -4.577225154 |
| **A_55_P2033610** | Fastkd3 | 0.098942644 | -0.072821444 | 0.270706732 | 9.887393083 | 1.376766455 | 0.21314116 | 0.991835066 | -4.577280311 |
| **A_55_P2010962** | Surf6 | 0.072583914 | -0.053428451 | 0.19859628 | 10.30581096 | 1.376690618 | 0.213163495 | 0.991835066 | -4.577344052 |
| **A_51_P135654** | Eif4ebp2 | -0.105783179 | -0.289477331 | 0.077910973 | 11.7608335 | -1.376354847 | 0.213262406 | 0.991835066 | -4.577626248 |
| **A_55_P1989333** | NA | -0.126514695 | -0.346229076 | 0.093199686 | 11.55091679 | -1.376231904 | 0.213298633 | 0.991835066 | -4.577729567 |
| **A_52_P50329** | Stimate | 0.073978777 | -0.054516959 | 0.202474513 | 8.953788891 | 1.376028922 | 0.213358457 | 0.991835066 | -4.577900142 |
| **A_52_P355075** | Rab40c | 0.065810472 | -0.048498537 | 0.180119481 | 10.88110802 | 1.376016618 | 0.213362083 | 0.991835066 | -4.577910481 |
| **A_51_P330630** | Usp3 | -0.079076609 | -0.216429468 | 0.058276251 | 8.775106427 | -1.376003263 | 0.21336602 | 0.991835066 | -4.577921703 |
| **A_55_P2065537** | NA | 0.083301519 | -0.061535727 | 0.228138766 | 9.674921439 | 1.374617271 | 0.213774917 | 0.991835066 | -4.579086122 |
| **A_55_P2172121** | Mxd4 | 1.228853684 | -0.90817262 | 3.365879989 | 8.172341801 | 1.374356625 | 0.213851891 | 0.991835066 | -4.579305045 |
| **A_55_P2081488** | Pglyrp1 | -0.065412624 | -0.179177751 | 0.048352502 | 7.852380809 | -1.37423673 | 0.213887307 | 0.991835066 | -4.579405742 |
| **A_51_P472329** | Meak7 | -0.097875211 | -0.268102735 | 0.072352313 | 9.042653016 | -1.3742067 | 0.213896178 | 0.991835066 | -4.579430962 |
| **A_55_P2010906** | Rbm45 | 0.14466429 | -0.106975554 | 0.396304133 | 9.801009275 | 1.374013706 | 0.2139532 | 0.991835066 | -4.579593045 |
| **A_55_P2095213** | Dusp22 | 0.086350086 | -0.063853862 | 0.236554034 | 8.532449305 | 1.374012101 | 0.213953674 | 0.991835066 | -4.579594393 |
| **A_55_P2174288** | Snap29 | 0.103885047 | -0.076821792 | 0.284591885 | 9.734732875 | 1.374002691 | 0.213956455 | 0.991835066 | -4.579602295 |
| **A_55_P2058157** | Ppp4r3b | 0.083604116 | -0.06183765 | 0.229045882 | 9.259029081 | 1.373876365 | 0.213993787 | 0.991835066 | -4.579708382 |
| **A_51_P278994** | Swap70 | -0.058235305 | -0.159583725 | 0.043113115 | 9.508235015 | -1.373341371 | 0.214151952 | 0.991835066 | -4.580157616 |
| **A_51_P399960** | Mier2 | 0.081000352 | -0.060016213 | 0.222016918 | 8.752643218 | 1.37285891 | 0.214294676 | 0.991835066 | -4.580562676 |
| **A_51_P156434** | Slc25a33 | -0.086338472 | -0.236654716 | 0.063977772 | 8.082404277 | -1.372800962 | 0.214311824 | 0.991835066 | -4.580611324 |
| **A_52_P409055** | Dcaf8 | -0.093067622 | -0.255112013 | 0.068976769 | 12.12697514 | -1.372694038 | 0.214343469 | 0.991835066 | -4.580701085 |
| **A_51_P185292** | 4930581F2 | -0.126388419 | -0.346456746 | 0.093679908 | 7.827345586 | -1.37264702 | 0.214357385 | 0.991835066 | -4.580740554 |
| **A_55_P1969477** | Abcb1a | -0.09854848 | -0.270145484 | 0.073048524 | 8.258952988 | -1.372616961 | 0.214366283 | 0.991835066 | -4.580765787 |
| **A_66_P130882** | Eif3e | -0.082880851 | -0.227207157 | 0.061445454 | 10.92002121 | -1.372517348 | 0.21439577 | 0.991835066 | -4.580849406 |
| **A_66_P112886** | Ubl3 | -0.085861422 | -0.235379157 | 0.063656314 | 10.64586001 | -1.37250677 | 0.214398902 | 0.991835066 | -4.580858285 |

| | | | | | | | | | |
|---|---|---|---|---|---|---|---|---|---|
| **A_55_P1994773** | Gm16381 | -0.10615471 | -0.291012978 | 0.078703559 | 9.561154523 | -1.372491042 | 0.214403558 | 0.991835066 | -4.580871487 |
| **A_55_P1962010** | Clec1b | 0.132041043 | -0.097907611 | 0.361989697 | 8.243139333 | 1.37242053 | 0.214424434 | 0.991835066 | -4.580930675 |
| **A_55_P2139886** | Sft2d1 | -0.066608737 | -0.182637118 | 0.049419645 | 7.882211742 | -1.372069452 | 0.214528401 | 0.991835066 | -4.581225354 |
| **A_51_P238523** | Shisa4 | -0.084755286 | -0.232403725 | 0.062893152 | 9.375315542 | -1.371977754 | 0.214555563 | 0.991835066 | -4.581302315 |
| **A_51_P439085** | Hilpda | -0.134963062 | -0.370078337 | 0.100152214 | 9.947389065 | -1.371965556 | 0.214559177 | 0.991835066 | -4.581312553 |
| **A_51_P305437** | Rcn1 | 0.15628514 | -0.115981574 | 0.428551854 | 12.9278275 | 1.371930998 | 0.214569414 | 0.991835066 | -4.581341557 |
| **A_51_P293269** | Pde7a | 0.087394763 | -0.0648698 | 0.239659326 | 8.401731241 | 1.371815488 | 0.214603637 | 0.991835066 | -4.581438499 |
| **A_51_P295625** | Arf4 | -0.200449389 | -0.549685423 | 0.148786644 | 11.24243317 | -1.37181271 | 0.21460446 | 0.991835066 | -4.58144083 |
| **A_66_P124952** | Hspe1 | 0.098826537 | -0.073361375 | 0.271014449 | 14.3343058 | 1.371766051 | 0.214618285 | 0.991835066 | -4.581479988 |
| **A_55_P2106459** | Zfhx3 | 0.170442031 | -0.126595514 | 0.467479576 | 9.556279154 | 1.371432653 | 0.214717095 | 0.991835066 | -4.581759772 |
| **A_55_P2117741** | Nck2 | 0.105178488 | -0.07813311 | 0.288490085 | 11.87675912 | 1.371343055 | 0.214743656 | 0.991835066 | -4.581834956 |
| **A_66_P131398** | Ror2 | 0.113934331 | -0.084677711 | 0.312546374 | 11.23289229 | 1.371065371 | 0.214825994 | 0.991835066 | -4.582067957 |
| **A_55_P1968977** | Stk38l | -0.107550176 | -0.295035181 | 0.07993483 | 10.30850661 | -1.371051337 | 0.214830155 | 0.991835066 | -4.582079731 |
| **A_55_P2070823** | Ino80d | 0.107659636 | -0.080033801 | 0.295353073 | 10.24763011 | 1.370922646 | 0.214868325 | 0.991835066 | -4.582187707 |
| **A_55_P1964573** | Arhgef6 | -0.064248144 | -0.176259743 | 0.047763455 | 11.44956596 | -1.370902977 | 0.214874159 | 0.991835066 | -4.582204209 |
| **A_52_P59681** | Rida | 0.095455304 | -0.070980145 | 0.261890754 | 9.097461076 | 1.370766132 | 0.214914754 | 0.991835066 | -4.582319021 |
| **A_55_P2138174** | Psd3 | -0.104831044 | -0.287640888 | 0.0779788 | 8.465969128 | -1.370564454 | 0.214974594 | 0.991835066 | -4.582488218 |
| **A_55_P2073482** | NA | -0.091443378 | -0.250919286 | 0.068032531 | 11.66430683 | -1.370459822 | 0.215005646 | 0.991835066 | -4.582575995 |
| **A_55_P2133165** | Wwc1 | 0.070398481 | -0.05238455 | 0.193181512 | 8.278047639 | 1.370358099 | 0.215035838 | 0.991835066 | -4.582661328 |
| **A_55_P2098802** | Uqcrq | -0.098878162 | -0.271336109 | 0.073579785 | 14.88348599 | -1.370333594 | 0.215043112 | 0.991835066 | -4.582681885 |
| **A_55_P1998354** | Wbp4 | 0.072188302 | -0.0537445 | 0.198121104 | 8.761164923 | 1.370052132 | 0.215126674 | 0.991835066 | -4.582917982 |
| **A_55_P1975370** | Apoc1 | -0.258384513 | -0.70921813 | 0.192449104 | 9.975094951 | -1.369806805 | 0.215199531 | 0.991835066 | -4.583123752 |
| **A_55_P1968568** | Rapgef5 | 0.084281981 | -0.062791582 | 0.231355544 | 9.26208595 | 1.369648899 | 0.215246438 | 0.991835066 | -4.583256189 |
| **A_55_P2425661** | Ncbp3 | -0.154659403 | -0.424563062 | 0.115244257 | 10.89812319 | -1.369546201 | 0.21527695 | 0.991835066 | -4.583342319 |
| **A_55_P1976033** | Cnih2 | 0.107824573 | -0.080351824 | 0.296000969 | 8.6743539 | 1.369499036 | 0.215290964 | 0.991835066 | -4.583381874 |
| **A_55_P2028570** | Sgsh | -0.129523763 | -0.355574778 | 0.096527251 | 13.64597503 | -1.369468749 | 0.215299963 | 0.991835066 | -4.583407274 |
| **A_55_P1982096** | Drap1 | -0.061013436 | -0.167504831 | 0.045477958 | 14.44288656 | -1.369367773 | 0.21532997 | 0.991835066 | -4.583491955 |
| **A_55_P2003793** | Tspan3 | 0.094073365 | -0.070150223 | 0.258296954 | 14.30590921 | 1.369116097 | 0.215404776 | 0.991835066 | -4.583703006 |
| **A_52_P560728** | Serhl | -0.108958232 | -0.299178557 | 0.081262093 | 10.61709425 | -1.369027776 | 0.215431034 | 0.991835066 | -4.583777067 |
| **A_55_P1999715** | Tcrb-J | -0.471761033 | -1.295459642 | 0.351937576 | 8.001727839 | -1.36887208 | 0.215477328 | 0.991835066 | -4.583907618 |
| **A_51_P480928** | Zfyve16 | -0.08366413 | -0.229775159 | 0.062446898 | 9.235618644 | -1.368565021 | 0.215568654 | 0.991835066 | -4.58416507 |
| **A_51_P498720** | Cacna2d1 | -0.073536099 | -0.201974601 | 0.054902402 | 10.46076033 | -1.368404502 | 0.21561641 | 0.991835066 | -4.584299648 |
| **A_55_P2119302** | Cspp1 | 0.14158353 | -0.105710016 | 0.388877076 | 8.337753068 | 1.368387465 | 0.215621479 | 0.991835066 | -4.584313931 |
| **A_55_P1971639** | Abitram | 0.111258211 | -0.083070329 | 0.305586751 | 9.681809531 | 1.368373376 | 0.215625672 | 0.991835066 | -4.584325743 |
| **A_51_P154534** | Bloc1s5 | -0.083974834 | -0.230703366 | 0.062753699 | 10.76952873 | -1.367866491 | 0.215776542 | 0.991835066 | -4.58475066 |
| **A_51_P312497** | Atp7a | 0.098169135 | -0.073363648 | 0.269701918 | 10.27316137 | 1.36784524 | 0.215782869 | 0.991835066 | -4.584768473 |
| **A_55_P1976109** | NA | 0.118371645 | -0.088482309 | 0.325225599 | 14.06583379 | 1.367706743 | 0.215824109 | 0.991835066 | -4.584884562 |
| **A_51_P456721** | Azgp1 | -0.062225131 | -0.170963849 | 0.046513587 | 7.813531282 | -1.367699681 | 0.215826212 | 0.991835066 | -4.584890481 |
| **A_65_P20371** | Bclaf1 | 0.103064219 | -0.077045026 | 0.283173465 | 10.79927475 | 1.367669124 | 0.215835312 | 0.991835066 | -4.584916093 |
| **A_51_P513392** | Azi2 | -0.102461036 | -0.281562708 | 0.076640635 | 10.05074219 | -1.367313926 | 0.215941115 | 0.991835066 | -4.585213795 |
| **A_55_P2133096** | Tgds | -0.062581794 | -0.171991806 | 0.046828218 | 8.303059893 | -1.367099343 | 0.216005056 | 0.991835066 | -4.585393627 |
| **A_55_P1966209** | Fez2 | -0.071563618 | -0.196682771 | 0.053555535 | 11.87015567 | -1.367028675 | 0.216026117 | 0.991835066 | -4.585452848 |
| **A_51_P174275** | Colec11 | -0.122442578 | -0.336530942 | 0.091645786 | 10.617125 | -1.366937109 | 0.216053409 | 0.991835066 | -4.585529581 |
| **A_55_P2059025** | U2af1 | 0.081853789 | -0.06127512 | 0.224982698 | 13.34882497 | 1.366849063 | 0.216079654 | 0.991835066 | -4.58560336 |
| **A_55_P2001148** | Pdlim5 | -0.172775809 | -0.474899868 | 0.12934825 | 10.83035137 | -1.366805623 | 0.216092605 | 0.991835066 | -4.585639762 |
| **A_55_P2096127** | NA | 0.487797207 | -0.365450966 | 1.34104538 | 11.9148539 | 1.366384966 | 0.216218044 | 0.991835066 | -4.585992229 |
| **A_55_P2000793** | Rbm34 | 0.084255253 | -0.063125722 | 0.231636228 | 9.088928758 | 1.36635859 | 0.216225912 | 0.991835066 | -4.586014327 |
| **A_52_P66313** | Psmf1 | -0.097240972 | -0.267404517 | 0.072922574 | 8.684200255 | -1.365815059 | 0.216388094 | 0.991835066 | -4.586469679 |

| | | | | | | | | | |
|---|---|---|---|---|---|---|---|---|---|
| **A_55_P2071097** | Adrm1 | 0.1255557 | -0.094229555 | 0.345340956 | 13.52201301 | 1.36535949 | 0.216524113 | 0.991835066 | -4.586851281 |
| **A_55_P2050298** | NA | 0.068879226 | -0.051700779 | 0.189459232 | 14.72532806 | 1.365281151 | 0.21654751 | 0.991835066 | -4.586916895 |
| **A_55_P1992049** | Gucy1a1 | 0.211035547 | -0.158438628 | 0.580509722 | 11.63986274 | 1.365150896 | 0.216586417 | 0.991835066 | -4.587025988 |
| **A_55_P2103429** | Psmc3 | -0.096798238 | -0.266270526 | 0.07267405 | 15.38491313 | -1.365142195 | 0.216589017 | 0.991835066 | -4.587033275 |
| **A_51_P359549** | Fem1c | 0.077479546 | -0.058179652 | 0.213138744 | 9.19184088 | 1.365044956 | 0.216618067 | 0.991835066 | -4.587114714 |
| **A_51_P480982** | Glo1 | -0.112439977 | -0.309358293 | 0.084478339 | 11.96103655 | -1.364720661 | 0.216714975 | 0.991835066 | -4.587386296 |
| **A_52_P491544** | Nedd9 | -0.129172157 | -0.355398104 | 0.097053791 | 9.166596363 | -1.364695082 | 0.21672262 | 0.991835066 | -4.587407716 |
| **A_51_P271725** | Fundc1 | -0.074984262 | -0.206323904 | 0.05635538 | 10.85204407 | -1.364531063 | 0.21677165 | 0.991835066 | -4.587545063 |
| **A_55_P2005465** | Zfp422 | -0.077880414 | -0.2143087 | 0.058547873 | 10.15012902 | -1.364372499 | 0.216819058 | 0.991835066 | -4.587677835 |
| **A_52_P364140** | Itga5 | -0.119491838 | -0.328827078 | 0.089843401 | 9.395800063 | -1.364284773 | 0.216845291 | 0.991835066 | -4.587751289 |
| **A_55_P1978023** | NA | -0.104333357 | -0.287117667 | 0.078450952 | 9.371148996 | -1.364248241 | 0.216856216 | 0.991835066 | -4.587781877 |
| **A_55_P2236291** | Plpp1 | -0.075265047 | -0.207131439 | 0.056601346 | 8.229794636 | -1.364169519 | 0.216879759 | 0.991835066 | -4.587847789 |
| **A_55_P2151011** | NA | -0.061578475 | -0.169493155 | 0.046336205 | 7.954619605 | -1.363821514 | 0.216983866 | 0.991835066 | -4.588139147 |
| **A_51_P417074** | Arhgap8 | 0.152043229 | -0.114415413 | 0.41850187 | 8.950750498 | 1.363786559 | 0.216994325 | 0.991835066 | -4.58816841 |
| **A_55_P2023937** | Impg1 | -0.13481502 | -0.371096287 | 0.101466248 | 9.789305469 | -1.363697737 | 0.217020905 | 0.991835066 | -4.588242767 |
| **A_51_P151835** | Tmtc3 | -0.110194891 | -0.303330342 | 0.08294056 | 9.441251769 | -1.363667828 | 0.217029856 | 0.991835066 | -4.588267806 |
| **A_55_P1979356** | Morf4l1 | 0.109043813 | -0.08213362 | 0.300221247 | 14.14513908 | 1.363243815 | 0.217156784 | 0.991835066 | -4.58862274 |
| **A_55_P2091861** | Hipk2 | 0.064814491 | -0.048863592 | 0.178492574 | 8.228068516 | 1.362713336 | 0.217315676 | 0.991835066 | -4.589066728 |
| **A_55_P2091551** | Arhgap9 | -0.062467539 | -0.172030054 | 0.047094977 | 8.332853442 | -1.362704005 | 0.217318471 | 0.991835066 | -4.589074536 |
| **A_51_P386270** | Cyp2d12 | -0.0641127 | -0.176571193 | 0.048345793 | 7.861590977 | -1.362576647 | 0.217356634 | 0.991835066 | -4.589181119 |
| **A_55_P2130090** | Cnn3 | 0.112446201 | -0.08479935 | 0.309691753 | 13.38142588 | 1.36253198 | 0.21737002 | 0.991835066 | -4.589218498 |
| **A_52_P161495** | Bcl6 | 0.295120809 | -0.222578985 | 0.812820603 | 8.74348675 | 1.362482626 | 0.217384812 | 0.991835066 | -4.589259799 |
| **A_55_P2058794** | Cmip | 0.133141847 | -0.10047154 | 0.366755235 | 11.82776266 | 1.362153291 | 0.217483535 | 0.991835066 | -4.589535381 |
| **A_52_P539310** | Serp2 | 0.114895262 | -0.086724928 | 0.316515451 | 9.479876661 | 1.36200039 | 0.217529384 | 0.991835066 | -4.589663317 |
| **A_55_P2099418** | Neil2 | 0.078779049 | -0.059475184 | 0.217033282 | 8.230743142 | 1.361888108 | 0.217563058 | 0.991835066 | -4.589757262 |
| **A_55_P2044592** | NA | -0.075009362 | -0.206659197 | 0.056640474 | 12.82749241 | -1.36177163 | 0.217597995 | 0.991835066 | -4.589854714 |
| **A_55_P2162324** | Bmx | 0.111624682 | -0.084380554 | 0.307629918 | 9.365350035 | 1.361136544 | 0.217788575 | 0.991835066 | -4.590386001 |
| **A_55_P2090055** | Phrf1 | 0.073333829 | -0.055442083 | 0.202109742 | 10.62537208 | 1.361064975 | 0.217810061 | 0.991835066 | -4.590445866 |
| **A_55_P2138177** | Psd3 | -0.137351311 | -0.378548095 | 0.103845473 | 9.060617531 | -1.361038558 | 0.217817992 | 0.991835066 | -4.590467963 |
| **A_55_P2044684** | Rsph1 | 0.064788677 | -0.048984364 | 0.178561719 | 8.105613098 | 1.361033695 | 0.217819452 | 0.991835066 | -4.59047203 |
| **A_55_P2066384** | Akt1 | 0.07918413 | -0.059869835 | 0.218238096 | 14.24651599 | 1.361018112 | 0.217824131 | 0.991835066 | -4.590485065 |
| **A_52_P321318** | NA | -0.095563619 | -0.26338878 | 0.072261541 | 16.45207734 | -1.360957779 | 0.217842247 | 0.991835066 | -4.59053553 |
| **A_55_P2063999** | Gpatch8 | 0.091739469 | -0.06937145 | 0.252850387 | 10.45526232 | 1.360944385 | 0.217846268 | 0.991835066 | -4.590546733 |
| **A_55_P1975670** | Prps1l3 | 0.114488755 | -0.086581663 | 0.315559174 | 10.90560162 | 1.360892389 | 0.217861882 | 0.991835066 | -4.590590223 |
| **A_55_P2034042** | Gga3 | 0.066653888 | -0.050411163 | 0.18371894 | 10.8850471 | 1.360840922 | 0.217877337 | 0.991835066 | -4.59063327 |
| **A_55_P2140449** | B3galt4 | -0.082390067 | -0.227121649 | 0.062341515 | 8.494370109 | -1.360569338 | 0.217958911 | 0.991835066 | -4.590860414 |
| **A_52_P574214** | Rrp1b | 0.168102735 | -0.127270636 | 0.463476105 | 8.9722024 | 1.360230698 | 0.218060663 | 0.991835066 | -4.591143613 |
| **A_55_P2082156** | Rab11fip3 | -0.087427585 | -0.241075066 | 0.066219895 | 10.65815263 | -1.359978907 | 0.218136346 | 0.991835066 | -4.591354161 |
| **A_55_P2082425** | Mtln | 0.073807651 | -0.055907528 | 0.20352283 | 12.14074288 | 1.35993991 | 0.21814807 | 0.991835066 | -4.591386769 |
| **A_55_P2143042** | Gm8096 | 0.207052618 | -0.157036385 | 0.571141621 | 8.574432016 | 1.359196656 | 0.218371625 | 0.991835066 | -4.592008177 |
| **A_55_P2440032** | Usp24 | 0.110472129 | -0.083804142 | 0.3047484 | 8.46533132 | 1.359070845 | 0.218409487 | 0.991835066 | -4.59211335 |
| **A_52_P190973** | Vcl | -0.082551104 | -0.227731343 | 0.062629135 | 11.34254743 | -1.35901581 | 0.218426051 | 0.991835066 | -4.592159355 |
| **A_52_P157170** | Rnf157 | 0.081615596 | -0.061941936 | 0.225173127 | 8.61478908 | 1.358802399 | 0.218490292 | 0.991835066 | -4.592337743 |
| **A_55_P2100928** | Ptgds | -0.184484349 | -0.509032324 | 0.140063626 | 14.54401341 | -1.358594199 | 0.218552981 | 0.991835066 | -4.592511764 |
| **A_55_P2348582** | NA | -0.102370657 | -0.282464723 | 0.077723409 | 8.804409018 | -1.358580006 | 0.218557255 | 0.991835066 | -4.592523626 |
| **A_52_P205797** | Vgll4 | -0.142571145 | -0.393434024 | 0.108291735 | 8.528124359 | -1.358327103 | 0.218633427 | 0.991835066 | -4.592734996 |
| **A_55_P1963017** | Stfa1 | -0.471858283 | -1.30213236 | 0.358415794 | 10.09149622 | -1.358311061 | 0.218638259 | 0.991835066 | -4.592748403 |
| **A_52_P677891** | Fam174b | -0.215457819 | -0.594628725 | 0.163713087 | 10.66005371 | -1.358114497 | 0.21869748 | 0.991835066 | -4.592912672 |

| | | | | | | | | | |
|---|---|---|---|---|---|---|---|---|---|
| **A_55_P2124346** | Ivns1abp | 0.155658615 | -0.118279608 | 0.429596838 | 12.77837065 | 1.358093463 | 0.218703818 | 0.991835066 | -4.59293025 |
| **A_51_P353173** | Fam98a | 0.094435888 | -0.071782478 | 0.260654254 | 11.34368434 | 1.357898132 | 0.218762682 | 0.991835066 | -4.593093479 |
| **A_52_P656867** | Fam98b | 0.09316636 | -0.070818292 | 0.257151012 | 10.49060765 | 1.357891456 | 0.218764694 | 0.991835066 | -4.593099057 |
| **A_52_P745** | Acox3 | -0.10231849 | -0.282430185 | 0.077793205 | 9.665767242 | -1.357754785 | 0.21880589 | 0.991835066 | -4.59321326 |
| **A_55_P2058127** | Pde4dip | 0.197237543 | -0.150038918 | 0.544514003 | 9.046172742 | 1.357448521 | 0.21889823 | 0.991835066 | -4.593469158 |
| **A_55_P1985850** | Timp1 | 0.162674112 | -0.123763669 | 0.449111892 | 10.72585075 | 1.357366969 | 0.218922824 | 0.991835066 | -4.593537295 |
| **A_55_P2097508** | Mcc | 0.152218035 | -0.115814334 | 0.420250403 | 11.35135839 | 1.357337972 | 0.218931569 | 0.991835066 | -4.593561521 |
| **A_66_P106861** | B3glct | 0.102774934 | -0.078225813 | 0.283775681 | 10.0680297 | 1.357112878 | 0.218999467 | 0.991835066 | -4.593749576 |
| **A_52_P651034** | Tmx1 | -0.107148059 | -0.295880125 | 0.081584007 | 10.18406145 | -1.356899698 | 0.219063788 | 0.991835066 | -4.593927664 |
| **A_66_P123526** | 4930504D1 | 0.091668759 | -0.069801974 | 0.253139491 | 7.847454379 | 1.356865082 | 0.219074234 | 0.991835066 | -4.59395658 |
| **A_55_P2102533** | Fuca2 | -0.091681022 | -0.253223347 | 0.069861304 | 8.919279453 | -1.356445176 | 0.219200984 | 0.991835066 | -4.594307327 |
| **A_55_P2135039** | Hyi | 0.099535671 | -0.075924491 | 0.274995834 | 10.50748353 | 1.355842849 | 0.219382912 | 0.991835066 | -4.594810367 |
| **A_55_P2007601** | Sftpd | 0.085124059 | -0.064973774 | 0.235221893 | 7.692449853 | 1.355461023 | 0.219498308 | 0.991835066 | -4.595129204 |
| **A_55_P2016089** | Uggt1 | 0.100494937 | -0.076720867 | 0.277710741 | 11.45456443 | 1.355348139 | 0.219532435 | 0.991835066 | -4.595223458 |
| **A_52_P543824** | Srsf6 | 0.12038465 | -0.091920089 | 0.332689389 | 9.583185735 | 1.355253567 | 0.219561029 | 0.991835066 | -4.59530242 |
| **A_55_P2174816** | Zfp266 | 0.063302479 | -0.048335959 | 0.174940916 | 8.726471576 | 1.355239646 | 0.219565238 | 0.991835066 | -4.595314043 |
| **A_52_P199776** | Snhg7 | 0.127101755 | -0.097065569 | 0.351269078 | 8.580503251 | 1.355153124 | 0.219591401 | 0.991835066 | -4.595386281 |
| **A_51_P150678** | Tnfaip8l2 | -0.063380382 | -0.175168653 | 0.04840789 | 7.855304589 | -1.355088761 | 0.219610866 | 0.991835066 | -4.595440018 |
| **A_52_P335064** | Mustn1 | -0.090040515 | -0.248852295 | 0.068771265 | 8.175314612 | -1.3550783 | 0.21961403 | 0.991835066 | -4.595448751 |
| **A_51_P283876** | Bphl | -0.154950838 | -0.428269197 | 0.118367522 | 11.63634075 | -1.354984275 | 0.219642468 | 0.991835066 | -4.59552725 |
| **A_55_P2017677** | Cap2 | -0.120989937 | -0.334416828 | 0.092436954 | 10.88821122 | -1.354906247 | 0.21966607 | 0.991835066 | -4.595592391 |
| **A_55_P2115127** | Mphosph1 | 0.095494921 | -0.07297484 | 0.263964682 | 8.586194155 | 1.354775851 | 0.219705518 | 0.991835066 | -4.595701247 |
| **A_55_P1980956** | Osbp | 0.076787261 | -0.058681208 | 0.21225573 | 8.298598608 | 1.354752891 | 0.219712465 | 0.991835066 | -4.595720415 |
| **A_51_P192964** | Clec12b | 0.09894271 | -0.075648776 | 0.273534196 | 7.711166284 | 1.354471505 | 0.219797614 | 0.991835066 | -4.595955303 |
| **A_55_P2002210** | Babam2 | 0.115628714 | -0.088407385 | 0.319664813 | 12.22575461 | 1.354465093 | 0.219799554 | 0.991835066 | -4.595960655 |
| **A_51_P343900** | Xrcc1 | 0.079316239 | -0.060672422 | 0.2193049 | 11.69343512 | 1.354186213 | 0.219883975 | 0.991835066 | -4.596193431 |
| **A_66_P109577** | Plcb3 | 0.118471794 | -0.090696147 | 0.327639734 | 10.00656838 | 1.353720406 | 0.220025045 | 0.991835066 | -4.596582185 |
| **A_55_P1993293** | Ndrg3 | 0.08335285 | -0.063827074 | 0.230532774 | 10.18810439 | 1.353570911 | 0.220070337 | 0.991835066 | -4.596706938 |
| **A_55_P2139901** | Tsr3 | 0.087070528 | -0.066677346 | 0.240818403 | 11.19683953 | 1.353540304 | 0.22007961 | 0.991835066 | -4.596732479 |
| **A_55_P2157814** | Gmds | 0.103233922 | -0.079058888 | 0.285526732 | 10.78631656 | 1.353511713 | 0.220088274 | 0.991835066 | -4.596756338 |
| **A_55_P2001438** | Coq2 | -0.090663935 | -0.250772522 | 0.069444653 | 11.56797315 | -1.353409027 | 0.220119391 | 0.991835066 | -4.596842024 |
| **A_55_P2018559** | Morf4l1 | -0.066847299 | -0.18491179 | 0.051217193 | 11.75024782 | -1.353236462 | 0.220171693 | 0.991835066 | -4.596986014 |
| **A_51_P165435** | Cox4i1 | -0.214646125 | -0.593812589 | 0.164520339 | 17.02642035 | -1.353013927 | 0.220239156 | 0.991835066 | -4.597171689 |
| **A_55_P2085716** | Atg12 | 0.075517389 | -0.057895308 | 0.208930087 | 9.625688217 | 1.352878938 | 0.220280087 | 0.991835066 | -4.597284312 |
| **A_65_P01783** | Med28 | -0.130284106 | -0.360453369 | 0.099885158 | 9.774237351 | -1.352861197 | 0.220285467 | 0.991835066 | -4.597299113 |
| **A_55_P2086770** | Uevld | -0.064883183 | -0.179534987 | 0.049768622 | 8.435865555 | -1.352571974 | 0.220373191 | 0.991835066 | -4.597540398 |
| **A_55_P1991964** | Iqsec2 | 0.0735197 | -0.056414505 | 0.203453906 | 9.335605029 | 1.352350805 | 0.220440294 | 0.991835066 | -4.597724894 |
| **A_55_P2011490** | NA | -0.093270897 | -0.258153618 | 0.071611824 | 12.35077773 | -1.352010742 | 0.220543506 | 0.991835066 | -4.598008543 |
| **A_55_P2000643** | NA | -0.068630257 | -0.189957305 | 0.05269679 | 11.11914948 | -1.351970231 | 0.220555804 | 0.991835066 | -4.598042332 |
| **A_55_P2169659** | Vwf | 0.12806092 | -0.098345985 | 0.354467825 | 8.293975042 | 1.351873583 | 0.220585146 | 0.991835066 | -4.59812294 |
| **A_51_P316801** | Wdr60 | 0.095765611 | -0.0735663 | 0.265097521 | 8.432359211 | 1.351698732 | 0.22063824 | 0.991835066 | -4.598268766 |
| **A_55_P2156340** | NA | -0.083331582 | -0.230707207 | 0.064044043 | 16.32069525 | -1.351428581 | 0.220720295 | 0.991835066 | -4.598494058 |
| **A_66_P114451** | Adgrg6 | 0.155050988 | -0.119170503 | 0.429272478 | 9.232796951 | 1.351394609 | 0.220730615 | 0.991835066 | -4.598522387 |
| **A_55_P1980080** | Rbms2 | -0.08802701 | -0.243754064 | 0.067700045 | 11.6554078 | -1.35101762 | 0.22084517 | 0.991835066 | -4.598836739 |
| **A_65_P07569** | Rock1 | -0.081787312 | -0.226484606 | 0.062909983 | 9.179842972 | -1.350935625 | 0.220870092 | 0.991835066 | -4.598905105 |
| **A_55_P2186615** | Acp1 | -0.060786988 | -0.168339679 | 0.046765703 | 10.40480854 | -1.350823083 | 0.220904304 | 0.991835066 | -4.598998938 |
| **A_55_P2178862** | U2surp | -0.072299795 | -0.200245343 | 0.055645753 | 8.942370968 | -1.350582194 | 0.220977547 | 0.991835066 | -4.599199771 |
| **A_52_P602898** | Nkrf | 0.06511712 | -0.050118242 | 0.180352483 | 8.304637291 | 1.350574503 | 0.220979886 | 0.991835066 | -4.599206183 |

| | | | | | | | | | |
|---|---|---|---|---|---|---|---|---|---|
| **A_55_P1981670** | 8-Sep | 0.075307947 | -0.057965254 | 0.208581148 | 8.162445706 | 1.350538943 | 0.2209907 | 0.991835066 | -4.599235829 |
| **A_55_P2124228** | Uaca | 0.057942452 | -0.044610005 | 0.16049491 | 12.40825906 | 1.350392231 | 0.221035321 | 0.991835066 | -4.599358135 |
| **A_55_P2044167** | NA | -0.060255967 | -0.16690593 | 0.046393996 | 8.069878869 | -1.350356645 | 0.221046146 | 0.991835066 | -4.599387801 |
| **A_55_P2046184** | Ccdc90b | -0.102783997 | -0.284740541 | 0.079172546 | 11.08507898 | -1.350103169 | 0.221123261 | 0.991835066 | -4.599599095 |
| **A_51_P128065** | Prcc | 0.087885927 | -0.067719367 | 0.243491221 | 10.96985293 | 1.349907785 | 0.221182719 | 0.991835066 | -4.599761953 |
| **A_55_P2136757** | Tomm34 | 0.107436752 | -0.082789469 | 0.297662973 | 11.56736567 | 1.349868999 | 0.221194523 | 0.991835066 | -4.599794281 |
| **A_55_P1977875** | Arglu1 | 0.115023454 | -0.088657424 | 0.318704332 | 11.00629523 | 1.349724955 | 0.221238369 | 0.991835066 | -4.599914338 |
| **A_55_P1958730** | NA | 0.103645394 | -0.079897713 | 0.287188502 | 10.82824984 | 1.34964969 | 0.221261282 | 0.991835066 | -4.599977067 |
| **A_52_P22781** | Zfp866 | 0.066140065 | -0.050990314 | 0.183270444 | 9.449771986 | 1.349597299 | 0.221277232 | 0.991835066 | -4.600020731 |
| **A_51_P157462** | Rgn | -0.099467546 | -0.2756228 | 0.076687708 | 8.149405107 | -1.349568506 | 0.221285999 | 0.991835066 | -4.600044727 |
| **A_55_P1965699** | NA | 0.112384331 | -0.086654694 | 0.311423356 | 7.701989042 | 1.349511722 | 0.221303289 | 0.991835066 | -4.600092051 |
| **A_52_P32864** | Prps2 | -0.077006901 | -0.213392085 | 0.059378283 | 9.250580242 | -1.349495949 | 0.221308092 | 0.991835066 | -4.600105197 |
| **A_55_P2146229** | NA | 1.526018194 | -1.176751497 | 4.228787885 | 9.229489937 | 1.34945934 | 0.221319239 | 0.991835066 | -4.600135706 |
| **A_51_P507942** | Atp13a2 | -0.113523167 | -0.314625933 | 0.0875796 | 11.80912833 | -1.349197673 | 0.221398934 | 0.991835066 | -4.600353765 |
| **A_55_P2121082** | Cul2 | -0.067069531 | -0.185898893 | 0.051759831 | 10.66247688 | -1.348995921 | 0.221460397 | 0.991835066 | -4.600521882 |
| **A_55_P2006525** | Adamtsl4 | -0.13307296 | -0.368877747 | 0.102731828 | 9.600857572 | -1.34879619 | 0.22152126 | 0.991835066 | -4.600688304 |
| **A_51_P384618** | Dhdds | 0.087833788 | -0.067826622 | 0.243494198 | 10.34840045 | 1.348629258 | 0.22157214 | 0.991835066 | -4.600827388 |
| **A_65_P20641** | NA | 0.058685125 | -0.045344314 | 0.162714565 | 8.106156508 | 1.348282539 | 0.22167785 | 0.991835066 | -4.601116245 |
| **A_51_P165082** | Tmem191c | 0.100026207 | -0.077307877 | 0.277360292 | 8.463619062 | 1.34812672 | 0.221725372 | 0.991835066 | -4.601246049 |
| **A_52_P522222** | NA | -0.078280085 | -0.21710829 | 0.06054812 | 8.736817591 | -1.347667367 | 0.221865519 | 0.991835066 | -4.601628672 |
| **A_51_P121031** | 1-Mar | -0.057355741 | -0.159076049 | 0.044364567 | 7.795214104 | -1.347653856 | 0.221869642 | 0.991835066 | -4.594284734 |
| **A_55_P1964108** | B230118H( | -0.106170162 | -0.294569911 | 0.082229587 | 11.69495744 | -1.34688741 | 0.22210366 | 0.991835066 | -4.602278216 |
| **A_55_P2150361** | NA | 0.068622304 | -0.053150565 | 0.190395173 | 12.76354753 | 1.346864445 | 0.222110675 | 0.991835066 | -4.60229734 |
| **A_52_P351882** | Impad1 | -0.083808355 | -0.232530399 | 0.064913689 | 12.78843784 | -1.34685581 | 0.222113313 | 0.991835066 | -4.602304529 |
| **A_55_P1976242** | Arhgef7 | 0.106087903 | -0.082201859 | 0.294377664 | 14.24549952 | 1.346630013 | 0.222182299 | 0.991835066 | -4.60249254 |
| **A_51_P501550** | NA | 0.078859566 | -0.061145816 | 0.218864949 | 8.1797682 | 1.346228512 | 0.222305014 | 0.991835066 | -4.602826816 |
| **A_55_P2172239** | NA | 0.144373911 | -0.111976085 | 0.400723908 | 16.26139101 | 1.346060372 | 0.222356423 | 0.991835066 | -4.602966791 |
| **A_51_P494006** | Scaf8 | 0.075802779 | -0.058810339 | 0.210415897 | 11.03152369 | 1.345881664 | 0.222411074 | 0.991835066 | -4.603115555 |
| **A_52_P158710** | Dcbld1 | 0.066108261 | -0.051324525 | 0.183541047 | 8.712505874 | 1.345474587 | 0.222535607 | 0.991835066 | -4.603454391 |
| **A_51_P438711** | Tbx18 | 0.07447987 | -0.057836213 | 0.206795953 | 9.271826068 | 1.345350387 | 0.222573615 | 0.991835066 | -4.603557762 |
| **A_55_P2295671** | NA | -0.062124829 | -0.172501603 | 0.048251946 | 8.44256392 | -1.345230296 | 0.22261037 | 0.991835066 | -4.603657709 |
| **A_55_P1996046** | Zfp931 | 0.077204919 | -0.05996623 | 0.214376069 | 9.354269235 | 1.345213841 | 0.222615407 | 0.991835066 | -4.603671403 |
| **A_52_P275408** | Ppp4c | -0.087782794 | -0.243785198 | 0.068219611 | 10.28288552 | -1.344891466 | 0.222714103 | 0.991835066 | -4.603939682 |
| **A_51_P174696** | Casq2 | -0.110350939 | -0.306473836 | 0.085771958 | 13.03729812 | -1.344797415 | 0.222742905 | 0.991835066 | -4.604017946 |
| **A_55_P2049289** | NA | -0.080009107 | -0.222218225 | 0.062200011 | 16.26959433 | -1.344686708 | 0.222776811 | 0.991835066 | -4.604110067 |
| **A_51_P455671** | Psmd5 | -0.089942946 | -0.24991239 | 0.070026497 | 9.584854483 | -1.343814147 | 0.223044209 | 0.991835066 | -4.604836018 |
| **A_51_P108901** | Ccdc86 | 0.14033569 | -0.109325075 | 0.389996455 | 10.37789272 | 1.343466897 | 0.223150704 | 0.991835066 | -4.605124865 |
| **A_55_P2022585** | Pira6 | -0.065243243 | -0.181312702 | 0.050826216 | 8.058359249 | -1.343466096 | 0.223150949 | 0.991835066 | -4.605125531 |
| **A_51_P181523** | Tmem214 | 0.098521497 | -0.076759425 | 0.273802418 | 10.68307387 | 1.343400439 | 0.22317109 | 0.991835066 | -4.605180142 |
| **A_51_P137094** | Sall2 | 0.097873336 | -0.076261408 | 0.272008081 | 9.442282878 | 1.343346637 | 0.223187595 | 0.991835066 | -4.605224891 |
| **A_55_P2001683** | Atp5g1 | -0.109128551 | -0.30329732 | 0.085040219 | 16.71394845 | -1.343284927 | 0.223206528 | 0.991835066 | -4.605276217 |
| **A_51_P355301** | Cyp3a11 | 0.070950201 | -0.055306496 | 0.197206898 | 7.762656531 | 1.343099975 | 0.22326328 | 0.991835066 | -4.605430039 |
| **A_55_P2126469** | NA | -0.057501158 | -0.159835359 | 0.044833044 | 15.78844716 | -1.342965686 | 0.223304494 | 0.991835066 | -4.605541721 |
| **A_55_P2193419** | NA | 0.132920033 | -0.103672102 | 0.369512169 | 8.412855222 | 1.342762702 | 0.223366804 | 0.991835066 | -4.605710522 |
| **A_55_P2113061** | NA | 0.091853631 | -0.071663567 | 0.255370829 | 8.653217242 | 1.342585729 | 0.223421142 | 0.991835066 | -4.605857684 |
| **A_55_P2030761** | NA | -0.125075087 | -0.347741271 | 0.097591096 | 12.51985346 | -1.342535204 | 0.223436657 | 0.991835066 | -4.605899697 |
| **A_55_P1954850** | Pars2 | 0.067670652 | -0.052811823 | 0.188153127 | 10.28263193 | 1.342411341 | 0.223474697 | 0.991835066 | -4.606002688 |
| **A_55_P2100149** | Mapt | 0.098663734 | -0.077038257 | 0.274365725 | 8.196477178 | 1.342115825 | 0.223565478 | 0.991835066 | -4.606248391 |

| | | | | | | | | | |
|---|---|---|---|---|---|---|---|---|---|
| **A_55_P2011922** | 9-Mar | -0.087682838 | -0.243844546 | 0.06847887 | 9.16987175 | -1.341989695 | 0.223604234 | 0.991835066 | -4.606353254 |
| **A_55_P2177911** | Lepr | -0.059919835 | -0.166661198 | 0.046821529 | 7.870572414 | -1.341673969 | 0.223701274 | 0.991835066 | -4.606615723 |
| **A_66_P128631** | Cinp | 0.085417947 | -0.066754679 | 0.237590572 | 8.873861358 | 1.341595912 | 0.223725271 | 0.991835066 | -4.606680609 |
| **A_51_P363719** | Stx7 | -0.092309091 | -0.256763973 | 0.072145791 | 10.58008377 | -1.34154991 | 0.223739414 | 0.991835066 | -4.606718848 |
| **A_55_P2005859** | Fn3k | -0.205322977 | -0.571199351 | 0.160553397 | 9.637286567 | -1.341257977 | 0.223829188 | 0.991835066 | -4.606961504 |
| **A_66_P140531** | Selenok | 0.091945056 | -0.071924871 | 0.255814982 | 12.15953408 | 1.341029255 | 0.223899545 | 0.991835066 | -4.607151603 |
| **A_52_P55717** | Adnp | -0.068421139 | -0.190390573 | 0.053548295 | 8.819970667 | -1.34075191 | 0.223984886 | 0.991835066 | -4.607382095 |
| **A_55_P2083559** | Hrk | -0.065962027 | -0.183554434 | 0.051630381 | 7.786200338 | -1.340676021 | 0.224008242 | 0.991835066 | -4.607445159 |
| **A_55_P2112120** | Tnk2 | -0.072674619 | -0.202254749 | 0.056905512 | 8.192412949 | -1.340458831 | 0.224075099 | 0.991835066 | -4.607625639 |
| **A_55_P2141084** | Tenm4 | 0.131234416 | -0.102816721 | 0.365285554 | 11.26054414 | 1.340127503 | 0.224177125 | 0.991835066 | -4.60790094 |
| **A_51_P130110** | Idh3b | -0.174253856 | -0.48502995 | 0.136522238 | 13.26205488 | -1.340120731 | 0.224179211 | 0.991835066 | -4.607906567 |
| **A_55_P2019699** | Samhd1 | 0.081574723 | -0.063945623 | 0.22709507 | 10.47675544 | 1.339803233 | 0.224277017 | 0.991835066 | -4.608170347 |
| **A_55_P1973886** | Socs4 | 0.082881603 | -0.065027756 | 0.230790961 | 8.625162175 | 1.339280728 | 0.224438058 | 0.991835066 | -4.608604391 |
| **A_51_P159565** | Arhgef9 | 0.108987852 | -0.085545281 | 0.303520984 | 9.068648201 | 1.339040291 | 0.224512198 | 0.991835066 | -4.608804096 |
| **A_51_P158388** | Cdk19 | -0.05554863 | -0.154720174 | 0.043622913 | 9.114978742 | -1.338737456 | 0.224605608 | 0.991835066 | -4.609055605 |
| **A_51_P295312** | Gabarap | -0.259171929 | -0.721885285 | 0.203541427 | 12.35068033 | -1.338705531 | 0.224615458 | 0.991835066 | -4.609082118 |
| **A_55_P2090415** | Nat9 | -0.055023015 | -0.153262171 | 0.04321614 | 8.052458194 | -1.338655721 | 0.224630826 | 0.991835066 | -4.609123483 |
| **A_55_P2121521** | H3c11 | 0.1468108 | -0.115330594 | 0.408952195 | 8.564052888 | 1.338540554 | 0.224666362 | 0.991835066 | -4.609219123 |
| **A_55_P2116794** | NA | 0.090223376 | -0.070904948 | 0.251351701 | 11.02578801 | 1.338308747 | 0.224737904 | 0.991835066 | -4.609411613 |
| **A_52_P561377** | Fam160b1 | 0.089542997 | -0.070371068 | 0.249457061 | 11.58659437 | 1.338301903 | 0.224740017 | 0.991835066 | -4.609417295 |
| **A_51_P138772** | Ddx27 | 0.090812819 | -0.071369765 | 0.252995403 | 10.23643444 | 1.338295675 | 0.224741939 | 0.991835066 | -4.609422467 |
| **A_55_P2002319** | Ap5s1 | -0.115816079 | -0.322655532 | 0.091023374 | 11.91142013 | -1.338272674 | 0.224749039 | 0.991835066 | -4.609441566 |
| **A_55_P2022158** | Gc | -0.330728193 | -0.9214237 | 0.259967314 | 8.294876401 | -1.338186621 | 0.224775604 | 0.991835066 | -4.609513019 |
| **A_55_P1974892** | Zbtb18 | 0.106605238 | -0.083816745 | 0.297027221 | 10.21201989 | 1.338044606 | 0.22481945 | 0.991835066 | -4.609630935 |
| **A_55_P2138714** | Sgcd | 0.096319119 | -0.075757804 | 0.268396042 | 8.762078043 | 1.337824055 | 0.224887559 | 0.991835066 | -4.609814049 |
| **A_51_P306067** | Lpp | -0.086655851 | -0.241541593 | 0.068229892 | 11.07357505 | -1.337197574 | 0.225081123 | 0.991835066 | -4.610334115 |
| **A_55_P2054240** | Med8 | 0.101143309 | -0.079655103 | 0.281941721 | 12.5327465 | 1.337062405 | 0.225122906 | 0.991835066 | -4.61044631 |
| **A_51_P191469** | Rnf31 | -0.08971791 | -0.250097303 | 0.070661482 | 10.20713418 | -1.337025589 | 0.225134287 | 0.991835066 | -4.610476868 |
| **A_52_P505827** | Cyb5r4 | 0.135394952 | -0.106698609 | 0.377488514 | 9.186717793 | 1.33668282 | 0.225240277 | 0.991835066 | -4.610761352 |
| **A_55_P2021540** | Myocd | -0.140628215 | -0.392120154 | 0.110863723 | 11.98787266 | -1.336464812 | 0.225307712 | 0.991835066 | -4.610942273 |
| **A_52_P164161** | Cyp51 | 0.146541633 | -0.115572389 | 0.408655656 | 8.887258807 | 1.336225958 | 0.225381615 | 0.991835066 | -4.611140479 |
| **A_55_P2033660** | Crls1 | 0.098089631 | -0.077382167 | 0.27356143 | 9.959210309 | 1.336056756 | 0.22543398 | 0.991835066 | -4.611280877 |
| **A_55_P2169879** | NA | -0.05840334 | -0.162886594 | 0.046079914 | 13.94394604 | -1.335980531 | 0.225457574 | 0.991835066 | -4.611344122 |
| **A_51_P203474** | Tsg101 | -0.07574362 | -0.211252662 | 0.059765422 | 10.90374462 | -1.335939877 | 0.225470158 | 0.991835066 | -4.611377854 |
| **A_55_P1958597** | Slc27a3 | 0.15185373 | -0.119861106 | 0.423568565 | 9.254237833 | 1.335737885 | 0.225532695 | 0.991835066 | -4.611545442 |
| **A_55_P2034893** | Socs7 | 0.080175135 | -0.063296142 | 0.223646412 | 11.34185427 | 1.33562294 | 0.225568288 | 0.991835066 | -4.611640805 |
| **A_66_P120572** | Gm14488 | 0.054854779 | -0.043311745 | 0.153021303 | 8.139938042 | 1.335550112 | 0.225590842 | 0.991835066 | -4.611701224 |
| **A_52_P663526** | Nmrk1 | -0.055330529 | -0.154373101 | 0.043712043 | 7.953748234 | -1.33521759 | 0.225693847 | 0.991835066 | -4.611977069 |
| **A_52_P550843** | Desi2 | 0.058243043 | -0.046027755 | 0.162513841 | 10.64259632 | 1.335028357 | 0.225752484 | 0.991835066 | -4.612134034 |
| **A_55_P2016976** | Zfp729a | -0.061595016 | -0.171874831 | 0.0486848 | 8.837579766 | -1.334930551 | 0.225782796 | 0.991835066 | -4.612215159 |
| **A_55_P1970380** | NA | 0.081742149 | -0.064671615 | 0.228155913 | 8.638258898 | 1.334360804 | 0.225959443 | 0.991835066 | -4.612687679 |
| **A_51_P165330** | Mrpl33 | -0.072499938 | -0.202363325 | 0.05736345 | 15.12761442 | -1.334320129 | 0.225972059 | 0.991835066 | -4.61272141 |
| **A_55_P1985079** | Mcrip1 | 0.095733065 | -0.075779433 | 0.267245562 | 13.42711555 | 1.334059881 | 0.226052792 | 0.991835066 | -4.612937214 |
| **A_52_P428446** | Tsn | 0.077694784 | -0.061524114 | 0.216913682 | 12.1044099 | 1.33383713 | 0.226121913 | 0.991835066 | -4.613121909 |
| **A_55_P2088615** | Fbln5 | 0.89137384 | -0.706065978 | 2.488813658 | 12.20146282 | 1.333658112 | 0.226177477 | 0.991835066 | -4.613270333 |
| **A_55_P2082241** | NA | -0.072874622 | -0.203475996 | 0.057726753 | 13.89655867 | -1.333637184 | 0.226183973 | 0.991835066 | -4.613287684 |
| **A_52_P446457** | Ly6g6f | -0.068844098 | -0.192230384 | 0.054542188 | 7.716388639 | -1.33354893 | 0.226211371 | 0.991835066 | -4.613360852 |
| **A_55_P2050799** | Nup35 | 0.103404471 | -0.081933683 | 0.288742624 | 10.45708436 | 1.333471126 | 0.226235527 | 0.991835066 | -4.613425355 |

| | | | | | | | | | |
|---|---|---|---|---|---|---|---|---|---|
| **A_55_P2081690** | Tex38 | -0.065792722 | -0.183739377 | 0.052153934 | 8.023472454 | -1.333218574 | 0.226313953 | 0.991835066 | -4.613634718 |
| **A_51_P106428** | Nsmce4a | 0.093613267 | -0.074279862 | 0.261506397 | 12.91195095 | 1.332642353 | 0.226492979 | 0.991835066 | -4.604608623 |
| **A_52_P336825** | Matr3 | -0.090926027 | -0.254013167 | 0.072161114 | 11.4461865 | -1.332531943 | 0.226527296 | 0.991835066 | -4.61420384 |
| **A_51_P144770** | 2410002F2 | 0.141348701 | -0.112193182 | 0.394890584 | 11.92793439 | 1.332450987 | 0.226552462 | 0.991835066 | -4.614270932 |
| **A_52_P128068** | Kazn | 0.054906711 | -0.043594628 | 0.153408049 | 8.113748208 | 1.332270547 | 0.226608561 | 0.991835066 | -4.614420466 |
| **A_51_P171772** | Bcl11b | 0.084353027 | -0.067015871 | 0.235721925 | 8.493584006 | 1.331904715 | 0.226722337 | 0.991835066 | -4.61472361 |
| **A_51_P520139** | Ccpg1os | -0.156649448 | -0.437913426 | 0.124614529 | 9.630594576 | -1.331140471 | 0.226960185 | 0.991835066 | -4.615356776 |
| **A_51_P128667** | Lynx1 | -0.197381052 | -0.551782501 | 0.157020397 | 10.92632494 | -1.331125806 | 0.226964751 | 0.991835066 | -4.615368924 |
| **A_55_P1952788** | NA | 0.097237554 | -0.077391481 | 0.271866589 | 9.20996453 | 1.330842627 | 0.227052939 | 0.991835066 | -4.615603491 |
| **A_55_P2040044** | NA | 0.095161782 | -0.075786213 | 0.266109777 | 8.449509935 | 1.330477968 | 0.227166546 | 0.991835066 | -4.615905519 |
| **A_55_P2019214** | NA | -0.104227879 | -0.291467172 | 0.083011413 | 8.106346858 | -1.33044229 | 0.227177664 | 0.991835066 | -4.615935066 |
| **A_51_P246773** | Sesn3 | 0.118196957 | -0.094145802 | 0.330539715 | 9.404808781 | 1.330386939 | 0.227194913 | 0.991835066 | -4.615980907 |
| **A_55_P1994204** | Ythdc1 | -0.120825114 | -0.337898019 | 0.09624779 | 12.06579494 | -1.330334121 | 0.227211374 | 0.991835066 | -4.616024649 |
| **A_55_P2139889** | Zfp783 | 0.087493261 | -0.069698025 | 0.244684546 | 8.16893735 | 1.330317394 | 0.227216588 | 0.991835066 | -4.616038502 |
| **A_55_P2077608** | Hdac1 | -0.104203321 | -0.291459305 | 0.083052664 | 9.551249012 | -1.330010242 | 0.227312336 | 0.991835066 | -4.616292857 |
| **A_55_P2060966** | Mov10 | 0.109642848 | -0.087400507 | 0.306686203 | 10.30339219 | 1.329926517 | 0.227338442 | 0.991835066 | -4.616362185 |
| **A_55_P2141938** | 1810058I2 | -0.104473101 | -0.292251911 | 0.083305709 | 8.912546619 | -1.329740923 | 0.227396321 | 0.991835066 | -4.61651586 |
| **A_55_P2154977** | Rpap3 | 0.084269793 | -0.067206616 | 0.235746201 | 11.8363866 | 1.329646079 | 0.227425903 | 0.991835066 | -4.616594389 |
| **A_55_P1996573** | Ttc13 | 0.113614982 | -0.090636054 | 0.317866017 | 11.77761199 | 1.329475899 | 0.227478992 | 0.991835066 | -4.616735288 |
| **A_55_P2082653** | Zfp711 | 0.109628266 | -0.087472606 | 0.306729139 | 8.862037243 | 1.32936161 | 0.227514652 | 0.991835066 | -4.616829907 |
| **A_51_P239654** | Nr4a1 | -0.109095997 | -0.305260225 | 0.087068232 | 10.97626927 | -1.329223872 | 0.227557634 | 0.991835066 | -4.616943936 |
| **A_51_P513959** | NA | -0.131225694 | -0.367182443 | 0.104731056 | 9.413779727 | -1.329216136 | 0.227560048 | 0.991835066 | -4.61695034 |
| **A_52_P471143** | Pusl1 | 0.08414874 | -0.067165588 | 0.235463068 | 9.23489124 | 1.329158262 | 0.227578111 | 0.991835066 | -4.616998251 |
| **A_55_P2060459** | Nsd3 | 0.057394714 | -0.045812916 | 0.160602345 | 8.239285007 | 1.329135391 | 0.227585249 | 0.991835066 | -4.617017183 |
| **A_66_P120882** | Nfkbib | -0.126632661 | -0.354355739 | 0.101090417 | 8.987518157 | -1.329069892 | 0.227605693 | 0.991835066 | -4.617071404 |
| **A_55_P2128451** | Sfpq | 0.095225752 | -0.076021351 | 0.266472855 | 12.27477625 | 1.329046911 | 0.227612867 | 0.991835066 | -4.617090428 |
| **A_55_P1967633** | Lsm5 | -0.124533143 | -0.348500943 | 0.099434657 | 9.85725642 | -1.328949562 | 0.227643256 | 0.991835066 | -4.617171012 |
| **A_52_P541833** | Vps37b | -0.101174593 | -0.283138481 | 0.080789295 | 11.34697651 | -1.328909451 | 0.227655779 | 0.991835066 | -4.617204215 |
| **A_51_P388801** | Tbcc | 0.135994171 | -0.108633276 | 0.380621618 | 11.86969855 | 1.328691769 | 0.227723749 | 0.991835066 | -4.617384396 |
| **A_55_P2106414** | Haus7 | -0.114967145 | -0.321817452 | 0.091883163 | 8.948992294 | -1.328393393 | 0.227816944 | 0.991835066 | -4.617631348 |
| **A_51_P468780** | Hmg20a | 0.083784114 | -0.066975185 | 0.234543413 | 11.08609872 | 1.328271046 | 0.227855168 | 0.991835066 | -4.617732603 |
| **A_55_P2005808** | Abhd18 | -0.100109006 | -0.280255527 | 0.080037515 | 10.56511964 | -1.328178372 | 0.227884125 | 0.991835066 | -4.617809296 |
| **A_52_P505501** | Nsmce2 | -0.06279046 | -0.175837611 | 0.05025669 | 8.81196144 | -1.327526437 | 0.228087921 | 0.991835066 | -4.618348748 |
| **A_66_P138915** | Fyn | -0.077893035 | -0.218132271 | 0.0623462 | 8.911934784 | -1.327511294 | 0.228092657 | 0.991835066 | -4.618361277 |
| **A_55_P2092909** | Rassf1 | 0.115704426 | -0.092611342 | 0.324020193 | 11.69806772 | 1.327507428 | 0.228093866 | 0.991835066 | -4.618364475 |
| **A_52_P515057** | Slc25a24 | 0.109042283 | -0.087298338 | 0.305382905 | 10.8074757 | 1.327375835 | 0.228135023 | 0.991835066 | -4.618473349 |
| **A_55_P2103884** | Supt4b | -0.137349389 | -0.38466063 | 0.109961852 | 10.99177252 | -1.327370031 | 0.228136838 | 0.991835066 | -4.618478151 |
| **A_55_P1989986** | Fam149b | 0.076325606 | -0.061123804 | 0.213775015 | 9.610832867 | 1.327200425 | 0.228189894 | 0.991835066 | -4.618618467 |
| **A_51_P121466** | Tab1 | 0.074338398 | -0.059532962 | 0.208209758 | 11.79078591 | 1.327194747 | 0.22819167 | 0.991835066 | -4.618623164 |
| **A_55_P2058152** | Ppp4r3b | 0.075036658 | -0.060111698 | 0.210185014 | 9.082825536 | 1.327002827 | 0.228251719 | 0.991835066 | -4.618781929 |
| **A_51_P389531** | Prmt1 | 0.172981266 | -0.138583342 | 0.484545874 | 10.65319042 | 1.326966883 | 0.228262967 | 0.991835066 | -4.618811663 |
| **A_66_P138999** | NA | 0.29753962 | -0.238484794 | 0.833564033 | 8.500171428 | 1.326689725 | 0.228349715 | 0.991835066 | -4.619040922 |
| **A_52_P235241** | Fntb | -0.054482888 | -0.152674985 | 0.043709208 | 8.015815959 | -1.326150223 | 0.228518656 | 0.991835066 | -4.619487123 |
| **A_55_P2167530** | NA | 0.216957452 | -0.174138616 | 0.60805352 | 8.214788357 | 1.325867892 | 0.22860711 | 0.991835066 | -4.619720595 |
| **A_55_P2045085** | NA | 0.073637342 | -0.059109474 | 0.206384157 | 13.99822932 | 1.325815596 | 0.228623498 | 0.991835066 | -4.619763839 |
| **A_55_P2068892** | Il6ra | -0.106102964 | -0.297383263 | 0.085177335 | 8.495317908 | -1.325764567 | 0.228639489 | 0.991835066 | -4.619806034 |
| **A_55_P2019397** | Mtch1 | 0.096496749 | -0.077503807 | 0.270497305 | 15.69909266 | 1.325473892 | 0.2287306 | 0.991835066 | -4.620046374 |
| **A_52_P407871** | Lsm12 | 0.111202277 | -0.089375859 | 0.311780414 | 11.0390567 | 1.325071224 | 0.228856868 | 0.991835066 | -4.620379276 |

| | | | | | | | | | |
|---|---|---|---|---|---|---|---|---|---|
| **A_51_P341025** | Eif3k | -0.096797477 | -0.271396404 | 0.07780145 | 11.42065096 | -1.325047964 | 0.228864163 | 0.991835066 | -4.620398505 |
| **A_66_P115539** | Gm11240 | -0.10186145 | -0.285602083 | 0.081879183 | 10.52513757 | -1.32499359 | 0.228881219 | 0.991835066 | -4.620443454 |
| **A_55_P2029417** | 2610318N( | 0.125251978 | -0.100689741 | 0.351193697 | 9.482069246 | 1.324943307 | 0.228896992 | 0.991835066 | -4.62048502 |
| **A_55_P2061804** | NA | -0.082085007 | -0.230159622 | 0.065989608 | 9.709923567 | -1.324928244 | 0.228901718 | 0.991835066 | -4.620497473 |
| **A_55_P1973451** | Stra6 | -0.813237832 | -2.280543119 | 0.654067454 | 7.322545539 | -1.324665496 | 0.228984156 | 0.991835066 | -4.620714661 |
| **A_51_P348119** | Ube2q1 | 0.086643234 | -0.069688219 | 0.242974687 | 10.48926287 | 1.324638653 | 0.22899258 | 0.991835066 | -4.620736848 |
| **A_55_P2234361** | Rnf150 | -0.067836202 | -0.190233912 | 0.054561508 | 10.28199289 | -1.324638473 | 0.228992636 | 0.991835066 | -4.620736997 |
| **A_55_P2051874** | Psmc2 | 0.081222254 | -0.06533341 | 0.227777919 | 13.27171422 | 1.324590274 | 0.229007763 | 0.991835066 | -4.620776836 |
| **A_51_P344566** | Plk1 | -0.110384273 | -0.309584712 | 0.088816165 | 12.10846751 | -1.324420958 | 0.229060905 | 0.991835066 | -4.62091678 |
| **A_52_P279152** | Vamp8 | -0.125967147 | -0.353295087 | 0.101360794 | 11.68512899 | -1.32438303 | 0.229072811 | 0.991835066 | -4.620948127 |
| **A_55_P2045765** | Rnf224 | -0.058100585 | -0.162958244 | 0.046757075 | 7.937678509 | -1.324309439 | 0.229095914 | 0.991835066 | -4.621008949 |
| **A_55_P2160594** | NA | 0.096190133 | -0.077425919 | 0.269806185 | 15.93943272 | 1.324188408 | 0.229133913 | 0.991835066 | -4.621108975 |
| **A_55_P2136353** | NA | 0.107980939 | -0.0869349 | 0.302896778 | 9.387325152 | 1.324064379 | 0.229172859 | 0.991835066 | -4.621211474 |
| **A_55_P1999633** | Cit | 0.07662322 | -0.061702037 | 0.214948477 | 8.195010586 | 1.323939212 | 0.229212169 | 0.991835066 | -4.62131491 |
| **A_55_P1962339** | Trim41 | 0.153238189 | -0.123420687 | 0.429897065 | 13.02228167 | 1.323827916 | 0.229247127 | 0.991835066 | -4.621406879 |
| **A_55_P1970929** | Hnf1a | 0.056414931 | -0.045456999 | 0.158286861 | 7.690035129 | 1.323575311 | 0.229326489 | 0.991835066 | -4.621615606 |
| **A_55_P1976310** | Tbx20 | 0.130499649 | -0.105172536 | 0.366171835 | 8.911619068 | 1.32345795 | 0.229363368 | 0.991835066 | -4.621712574 |
| **A_51_P351394** | Vps45 | 0.066352352 | -0.053493508 | 0.186198211 | 9.544082244 | 1.323251571 | 0.229428234 | 0.991835066 | -4.621883085 |
| **A_51_P390078** | NA | -0.068306681 | -0.191694656 | 0.055081295 | 9.765416797 | -1.32312073 | 0.229469366 | 0.991835066 | -4.621991179 |
| **A_52_P80593** | NA | 0.088534667 | -0.071426174 | 0.248495508 | 11.53834438 | 1.322844551 | 0.229556209 | 0.991835066 | -4.622219329 |
| **A_66_P125777** | Slc43a2 | -0.074787543 | -0.209923456 | 0.060348369 | 8.345227367 | -1.322719089 | 0.229595669 | 0.991835066 | -4.622322965 |
| **A_55_P2158350** | Gm15427 | 0.060829002 | -0.049100102 | 0.170758106 | 16.05376987 | 1.322534879 | 0.229653617 | 0.991835066 | -4.622475121 |
| **A_55_P2113208** | Snrpa1 | 0.085036157 | -0.068676234 | 0.238748549 | 11.68815186 | 1.322220475 | 0.229752552 | 0.991835066 | -4.622734794 |
| **A_55_P2183473** | Cldn3 | 0.124814834 | -0.100805413 | 0.350435081 | 8.206480338 | 1.32220034 | 0.229758889 | 0.991835066 | -4.622751423 |
| **A_51_P368483** | 1700123O2 | 0.060518585 | -0.048878418 | 0.169915587 | 10.06359712 | 1.322185746 | 0.229763483 | 0.991835066 | -4.622763477 |
| **A_55_P2032445** | Slc25a37 | 0.177005169 | -0.142984286 | 0.496994625 | 9.629851324 | 1.322085112 | 0.229795159 | 0.991835066 | -4.622846585 |
| **A_51_P295967** | Proc | -0.120602387 | -0.338662325 | 0.097457551 | 7.91560873 | -1.321871232 | 0.229862493 | 0.991835066 | -4.62302321 |
| **A_55_P2178317** | Gm9199 | -0.077038548 | -0.216341898 | 0.062264802 | 8.726690513 | -1.321769297 | 0.229894591 | 0.991835066 | -4.623107384 |
| **A_51_P112223** | Gsta4 | -0.0939891 | -0.263966603 | 0.075988403 | 9.675882304 | -1.321585236 | 0.229952559 | 0.991835066 | -4.623259368 |
| **A_51_P487999** | Sgo1 | 0.097969457 | -0.079250265 | 0.275189179 | 10.32297596 | 1.321258466 | 0.230055503 | 0.991835066 | -4.623529166 |
| **A_52_P177847** | Tril | -0.064294835 | -0.180614183 | 0.052024514 | 7.862615666 | -1.321092589 | 0.230107776 | 0.991835066 | -4.623666112 |
| **A_55_P1967568** | Chtop | 0.071218947 | -0.057641903 | 0.200079798 | 11.69957771 | 1.320941815 | 0.230155298 | 0.991835066 | -4.623790581 |
| **A_51_P406429** | Pdk1 | 0.138912303 | -0.112430854 | 0.39025546 | 10.87573149 | 1.320939048 | 0.23015617 | 0.991835066 | -4.623792865 |
| **A_51_P248705** | Cse1l | -0.126991421 | -0.356778985 | 0.102796144 | 10.29821943 | -1.32086062 | 0.230180893 | 0.991835066 | -4.623857608 |
| **A_55_P2061045** | Rpl29 | -0.070885877 | -0.199159607 | 0.057387853 | 15.08685915 | -1.320781946 | 0.230205696 | 0.991835066 | -4.623922552 |
| **A_52_P302345** | Cyp4v3 | -0.062257617 | -0.174918297 | 0.050403063 | 8.038216159 | -1.320776267 | 0.230207487 | 0.991835066 | -4.62392724 |
| **A_55_P2004741** | Cyb5rl | 0.058062495 | -0.047006807 | 0.163131797 | 8.712511923 | 1.320775425 | 0.230207752 | 0.991835066 | -4.623927935 |
| **A_55_P2078257** | NA | 0.09825828 | -0.079571531 | 0.27608809 | 8.378239778 | 1.320607384 | 0.230260738 | 0.991835066 | -4.624066644 |
| **A_52_P358093** | Fbxo10 | -0.066235223 | -0.186113489 | 0.053643043 | 11.07149515 | -1.32055861 | 0.230276119 | 0.991835066 | -4.624106902 |
| **A_52_P86693** | Ifi27 | -0.113553091 | -0.319084528 | 0.091978345 | 11.56773254 | -1.320473945 | 0.23030282 | 0.991835066 | -4.624176784 |
| **A_55_P2048041** | Slc44a1 | 0.118247425 | -0.095792277 | 0.332287127 | 8.631247555 | 1.320402953 | 0.230325212 | 0.991835066 | -4.624235379 |
| **A_66_P101703** | Krtap5-5 | -0.107402136 | -0.301859152 | 0.087054879 | 8.313265756 | -1.320074482 | 0.230428839 | 0.991835066 | -4.624506471 |
| **A_51_P179498** | Fam92a | -0.07074103 | -0.198838166 | 0.057356107 | 10.96387226 | -1.31990018 | 0.230483845 | 0.991835066 | -4.624650314 |
| **A_52_P637730** | Ubl4a | 0.08381901 | -0.06799344 | 0.235631461 | 12.25375331 | 1.319605971 | 0.230576718 | 0.991835066 | -4.624893088 |
| **A_55_P2001072** | NA | -0.123490056 | -0.34716537 | 0.100185259 | 14.17957722 | -1.319541527 | 0.230597065 | 0.991835066 | -4.624946263 |
| **A_55_P2091212** | NA | 0.060881812 | -0.049420291 | 0.171183915 | 16.74908235 | 1.319206881 | 0.230702751 | 0.991835066 | -4.625222369 |
| **A_55_P2179398** | Alg1 | -0.061435409 | -0.172742138 | 0.04987132 | 10.39456793 | -1.319187312 | 0.230708933 | 0.991835066 | -4.625238515 |
| **A_55_P2137586** | Nphp4 | 0.103046493 | -0.08367181 | 0.289764797 | 8.356680694 | 1.319032382 | 0.230757877 | 0.991835066 | -4.625366331 |

| | | | | | | | | | |
|---|---|---|---|---|---|---|---|---|---|
| **A_55_P1971448** | Snrpd1 | 0.07863079 | -0.063866954 | 0.221128534 | 13.48791668 | 1.318845078 | 0.230817061 | 0.991835066 | -4.625520847 |
| **A_51_P489138** | Sptb | -0.165784648 | -0.466246481 | 0.134677185 | 9.872420883 | -1.318755041 | 0.230845515 | 0.991835066 | -4.62559512 |
| **A_55_P2054998** | NA | 0.080057359 | -0.06506437 | 0.225179088 | 10.29395067 | 1.318493357 | 0.230928233 | 0.991835066 | -4.625810971 |
| **A_51_P426633** | Dnttip2 | 0.081772762 | -0.066458985 | 0.230004508 | 11.93595795 | 1.318489183 | 0.230929552 | 0.991835066 | -4.625814414 |
| **A_55_P2006975** | NA | 0.091404603 | -0.074298289 | 0.257107494 | 9.271977704 | 1.318399746 | 0.230957829 | 0.991835066 | -4.625888182 |
| **A_55_P1959263** | Dnpep | 0.122480343 | -0.099583678 | 0.344544363 | 13.06253341 | 1.318248622 | 0.231005617 | 0.991835066 | -4.626012825 |
| **A_51_P424338** | Nqo1 | -0.090738233 | -0.255283013 | 0.073806547 | 8.651494694 | -1.317999788 | 0.23108432 | 0.991835066 | -4.626218043 |
| **A_51_P343913** | Parp4 | 0.101348774 | -0.0825194 | 0.285216949 | 9.095756599 | 1.317410344 | 0.231270848 | 0.991835066 | -4.626704096 |
| **A_55_P2161170** | Srp68 | 0.087700656 | -0.07143284 | 0.246834151 | 11.75770898 | 1.317195899 | 0.231338741 | 0.991835066 | -4.626880901 |
| **A_55_P1990067** | NA | -0.10678186 | -0.300571504 | 0.087007785 | 13.17379507 | -1.316970498 | 0.231410122 | 0.991835066 | -4.627066726 |
| **A_55_P2030433** | Gpi1 | 0.098217961 | -0.080045818 | 0.27648174 | 16.52638487 | 1.316851909 | 0.231447685 | 0.991835066 | -4.627164487 |
| **A_52_P57526** | Sin3b | 0.084213944 | -0.068635305 | 0.237063192 | 14.4286698 | 1.316830357 | 0.231454512 | 0.991835066 | -4.627182253 |
| **A_55_P1963878** | NA | -0.104001264 | -0.292779252 | 0.084776724 | 11.82837708 | -1.316728991 | 0.231486625 | 0.991835066 | -4.627265813 |
| **A_55_P2129437** | Plpp2 | 0.13265389 | -0.108156753 | 0.373464532 | 10.76554618 | 1.31659877 | 0.231527884 | 0.991835066 | -4.627373153 |
| **A_55_P2115255** | Fer | -0.077264064 | -0.21755727 | 0.063029142 | 8.921720412 | -1.316285329 | 0.231627223 | 0.991835066 | -4.627631502 |
| **A_52_P56397** | Aunip | 0.188034744 | -0.153431837 | 0.529501325 | 9.129558765 | 1.316130728 | 0.231676234 | 0.991835066 | -4.627758919 |
| **A_52_P118310** | Chn1 | -0.086825451 | -0.244500991 | 0.070850089 | 9.167838694 | -1.316108979 | 0.231683129 | 0.991835066 | -4.627776844 |
| **A_55_P2002376** | Srm | 0.192458588 | -0.15707478 | 0.541991955 | 11.00401836 | 1.316005747 | 0.231715861 | 0.991835066 | -4.62786192 |
| **A_55_P1999923** | Nyap1 | 0.105208879 | -0.085893231 | 0.29631099 | 8.882264995 | 1.315818667 | 0.23177519 | 0.991835066 | -4.628016089 |
| **A_55_P2379480** | NA | 0.179116237 | -0.146239394 | 0.504471868 | 8.667927014 | 1.315787517 | 0.23178507 | 0.991835066 | -4.628041759 |
| **A_55_P2138416** | NA | 0.080426121 | -0.065704818 | 0.22655706 | 13.18860569 | 1.315418916 | 0.231902008 | 0.991835066 | -4.628345484 |
| **A_51_P244504** | LTO1 | 0.249216504 | -0.203636163 | 0.702069172 | 10.02105197 | 1.315312667 | 0.231935725 | 0.991835066 | -4.628433025 |
| **A_55_P1960197** | P2ry14 | -0.054508094 | -0.15356109 | 0.044544902 | 7.941415899 | -1.315232434 | 0.231961188 | 0.991835066 | -4.628499129 |
| **A_55_P2092073** | Taf11 | 0.055008714 | -0.044957464 | 0.154974892 | 9.55646023 | 1.315187079 | 0.231975584 | 0.991835066 | -4.628536496 |
| **A_55_P1955532** | Tfr2 | -0.060877454 | -0.171514438 | 0.04975953 | 7.770035485 | -1.315119697 | 0.231996972 | 0.991835066 | -4.62859201 |
| **A_55_P1999334** | Inpp5b | 0.079112491 | -0.064669159 | 0.222894141 | 10.36888244 | 1.315075628 | 0.232010961 | 0.991835066 | -4.628628316 |
| **A_55_P1995623** | Pomp | 0.090508236 | -0.073986468 | 0.25500294 | 13.58915171 | 1.315059232 | 0.232016166 | 0.991835066 | -4.628641824 |
| **A_55_P1981090** | Thap7 | 0.089280433 | -0.072994431 | 0.251555297 | 9.719462266 | 1.314964915 | 0.232046109 | 0.991835066 | -4.628719524 |
| **A_55_P2095177** | Usp5 | 0.061230676 | -0.050069545 | 0.172530897 | 13.42609947 | 1.31486801 | 0.232076878 | 0.991835066 | -4.628799355 |
| **A_55_P2065991** | S100a11 | -0.18994541 | -0.535263993 | 0.155373172 | 11.96358262 | -1.314673737 | 0.232138572 | 0.991835066 | -4.628959388 |
| **A_55_P2160322** | Anapc15 | -0.074940459 | -0.211188044 | 0.061307126 | 8.601498263 | -1.314609206 | 0.232159068 | 0.991835066 | -4.629012544 |
| **A_55_P2058928** | Ralgds | 0.058880254 | -0.048172099 | 0.165932606 | 10.51268589 | 1.314566587 | 0.232172605 | 0.991835066 | -4.629047649 |
| **A_52_P220810** | Trib2 | -0.134280623 | -0.378449738 | 0.109888492 | 10.71284042 | -1.314412714 | 0.232221486 | 0.991835066 | -4.629174391 |
| **A_52_P240152** | Snx27 | 0.089266783 | -0.073060472 | 0.251594038 | 10.33178326 | 1.31433954 | 0.232244735 | 0.991835066 | -4.62923466 |
| **A_55_P1978471** | NA | -0.143775764 | -0.405243705 | 0.117692176 | 11.57181546 | -1.314245115 | 0.232274738 | 0.991835066 | -4.62931243 |
| **A_55_P2087985** | NA | 0.133167141 | -0.109034251 | 0.375368533 | 11.46584696 | 1.314103501 | 0.232319742 | 0.991835066 | -4.629429061 |
| **A_55_P1992074** | Ptpra | 0.075017865 | -0.06142571 | 0.21146144 | 12.56311007 | 1.314076787 | 0.232328232 | 0.991835066 | -4.629451062 |
| **A_51_P334104** | Dcn | -0.190056015 | -0.535738022 | 0.155625992 | 14.83625907 | -1.314056311 | 0.23233474 | 0.991835066 | -4.629467925 |
| **A_55_P2089655** | Eif2s2 | 0.101668371 | -0.083289874 | 0.286626616 | 12.97115398 | 1.313775927 | 0.232423871 | 0.991835066 | -4.629698824 |
| **A_52_P5901** | Saal1 | 0.073320596 | -0.060095559 | 0.206736751 | 8.668371109 | 1.313489778 | 0.232514866 | 0.991835066 | -4.629934447 |
| **A_51_P501844** | Cyp26b1 | 0.219763355 | -0.180153408 | 0.619680119 | 8.986929308 | 1.313393313 | 0.232545549 | 0.991835066 | -4.630013874 |
| **A_52_P456977** | Deaf1 | 0.097019597 | -0.079560229 | 0.273599422 | 8.463805902 | 1.31318984 | 0.23261028 | 0.991835066 | -4.6301814 |
| **A_51_P316951** | Wipf3 | 0.184554734 | -0.15140122 | 0.520510688 | 10.58692573 | 1.312961467 | 0.232682951 | 0.991835066 | -4.630369413 |
| **A_51_P310254** | Fam118a | -0.060680707 | -0.17114235 | 0.049780936 | 8.499988967 | -1.312950235 | 0.232686525 | 0.991835066 | -4.630378659 |
| **A_51_P392546** | Haus3 | 0.115993216 | -0.09524613 | 0.327232561 | 10.67924976 | 1.312402075 | 0.23286104 | 0.991835066 | -4.630829879 |
| **A_51_P317317** | Gatd1 | -0.077071023 | -0.217430719 | 0.063288673 | 8.252500161 | -1.312374661 | 0.23286977 | 0.991835066 | -4.630852443 |
| **A_65_P20699** | Tlk2 | -0.090882704 | -0.256402571 | 0.074637162 | 9.213179401 | -1.312321509 | 0.232886698 | 0.991835066 | -4.63089619 |
| **A_52_P232580** | Dyrk3 | 0.075982241 | -0.062428173 | 0.214392654 | 8.228689368 | 1.312056277 | 0.232971188 | 0.991835066 | -4.631114479 |

| | | | | | | | | | |
|---|---|---|---|---|---|---|---|---|---|
| **A_55_P2089820** | Adk | -0.103098535 | -0.290922232 | 0.084725161 | 9.676961036 | -1.311931754 | 0.233010864 | 0.991835066 | -4.631216957 |
| **A_52_P35377** | Plagl2 | 0.305003835 | -0.250681974 | 0.860689643 | 9.561212144 | 1.311852859 | 0.233036005 | 0.991835066 | -4.631281882 |
| **A_55_P2056286** | Sf3b5 | 0.071319271 | -0.05862617 | 0.201264713 | 13.15023993 | 1.311761807 | 0.233065023 | 0.991835066 | -4.631356809 |
| **A_55_P1966508** | Hoxc6 | 0.054946853 | -0.045177476 | 0.155071182 | 7.807746214 | 1.311633013 | 0.233106075 | 0.991835066 | -4.63146279 |
| **A_55_P2041320** | NA | -0.074418875 | -0.210044849 | 0.061207099 | 13.37618009 | -1.31144283 | 0.233166704 | 0.991835066 | -4.631619277 |
| **A_51_P385635** | Mrpl53 | -0.081323847 | -0.229540206 | 0.066892512 | 13.60651047 | -1.311387093 | 0.233184476 | 0.991835066 | -4.631665136 |
| **A_55_P2011440** | Ralgapb | 0.065942226 | -0.054251856 | 0.186136308 | 10.98822574 | 1.311262528 | 0.233224197 | 0.991835066 | -4.631767625 |
| **A_55_P2082986** | Hmgb1 | -0.193549052 | -0.546376176 | 0.159278073 | 13.04032639 | -1.311107237 | 0.233273725 | 0.991835066 | -4.631895386 |
| **A_55_P2024431** | Gaa | 0.111046884 | -0.091388467 | 0.313482235 | 13.79326506 | 1.311079881 | 0.233282451 | 0.991835066 | -4.631917892 |
| **A_66_P140599** | NA | -0.070614493 | -0.199344047 | 0.058115061 | 8.897345584 | -1.311066478 | 0.233286726 | 0.991835066 | -4.631928918 |
| **A_55_P2054128** | NA | -0.126185926 | -0.356250151 | 0.103878299 | 14.75338817 | -1.31090422 | 0.233338487 | 0.991835066 | -4.632062402 |
| **A_55_P1998194** | Snrpn | 0.076863086 | -0.063276652 | 0.217002825 | 8.500953821 | 1.310888185 | 0.233343604 | 0.991835066 | -4.632075594 |
| **A_55_P1990134** | Cox18 | 0.085531868 | -0.070428405 | 0.241492142 | 9.563032678 | 1.310759768 | 0.233384578 | 0.991835066 | -4.632181232 |
| **A_55_P2115176** | Pdhx | -0.06777788 | -0.191366518 | 0.055810758 | 10.87082832 | -1.310746074 | 0.233388947 | 0.991835066 | -4.632192496 |
| **A_55_P1980237** | Acad11 | -0.093286052 | -0.263404617 | 0.076832513 | 10.88719258 | -1.310611981 | 0.23343174 | 0.991835066 | -4.632302798 |
| **A_52_P167278** | Mthfd1l | 0.252543059 | -0.208069269 | 0.713155388 | 10.06605076 | 1.310415471 | 0.233494464 | 0.991835066 | -4.632464433 |
| **A_55_P2032740** | Iars2 | 0.09488372 | -0.078185056 | 0.267952496 | 8.783538706 | 1.310334307 | 0.233520375 | 0.991835066 | -4.63253119 |
| **A_55_P1996653** | D030056L2 | 0.094327863 | -0.077729609 | 0.266385336 | 11.7769744 | 1.310314634 | 0.233526656 | 0.991835066 | -4.632547371 |
| **A_55_P1978608** | NA | -0.100956809 | -0.285126215 | 0.083212598 | 8.999108572 | -1.310168812 | 0.233573216 | 0.991835066 | -4.632667302 |
| **A_51_P165451** | Pbld2 | -0.065340721 | -0.18454157 | 0.053860128 | 7.768402401 | -1.310127927 | 0.233586272 | 0.991835066 | -4.632700927 |
| **A_51_P107433** | Mrpl34 | -0.066893609 | -0.189009817 | 0.055222599 | 13.00150952 | -1.309243572 | 0.233868829 | 0.991835066 | -4.630263733 |
| **A_51_P293753** | Cul7 | 0.09223701 | -0.07616916 | 0.26064318 | 13.23689467 | 1.309050343 | 0.233930607 | 0.991835066 | -4.633586983 |
| **A_51_P161636** | Kng2 | -0.147534374 | -0.416911498 | 0.12184275 | 7.916038436 | -1.309006005 | 0.233944785 | 0.991835066 | -4.633623433 |
| **A_51_P463860** | Fam161b | -0.093246811 | -0.263514492 | 0.077020869 | 8.760351809 | -1.308913362 | 0.23397441 | 0.991835066 | -4.633699593 |
| **A_55_P2150737** | NA | 0.167167703 | -0.138089745 | 0.472425152 | 9.629928539 | 1.308866158 | 0.233989507 | 0.991835066 | -4.633738398 |
| **A_55_P2035872** | Mical3 | -0.105589008 | -0.298408943 | 0.087230927 | 10.61973019 | -1.308807903 | 0.234008139 | 0.991835066 | -4.633786286 |
| **A_55_P2289765** | Fam160a2 | 0.139391478 | -0.115235006 | 0.394017961 | 9.075755602 | 1.308403753 | 0.234137434 | 0.991835066 | -4.634118488 |
| **A_55_P2029726** | NA | 0.060242981 | -0.049820885 | 0.170306847 | 14.95509091 | 1.308189997 | 0.234205845 | 0.991835066 | -4.634294172 |
| **A_55_P2115068** | Psmb5 | -0.085098562 | -0.240579284 | 0.070382159 | 13.3087456 | -1.308141755 | 0.234221286 | 0.991835066 | -4.634333819 |
| **A_55_P2010539** | NA | -0.060551538 | -0.171208123 | 0.050105046 | 8.224630906 | -1.307847333 | 0.234315547 | 0.991835066 | -4.634575775 |
| **A_55_P1976958** | NA | 0.112998428 | -0.093512777 | 0.319509633 | 9.961843356 | 1.307789701 | 0.234334002 | 0.991835066 | -4.634623134 |
| **A_55_P2034998** | Tjp1 | -0.071961569 | -0.203482871 | 0.059559733 | 11.80432233 | -1.307716655 | 0.234357394 | 0.991835066 | -4.634683158 |
| **A_51_P189814** | Cldn5 | -0.165811021 | -0.468908998 | 0.137286956 | 11.48979009 | -1.307493346 | 0.234428921 | 0.991835066 | -4.634866648 |
| **A_55_P2173073** | Zfp931 | 0.06414197 | -0.053129031 | 0.181412971 | 8.727437687 | 1.307256457 | 0.234504818 | 0.991835066 | -4.63506128 |
| **A_55_P2076846** | Tead2 | 0.081471866 | -0.067485372 | 0.230429104 | 11.66852459 | 1.307239565 | 0.234510232 | 0.991835066 | -4.635075159 |
| **A_55_P1998164** | Rab28 | -0.084881028 | -0.240098883 | 0.070336827 | 8.53993493 | -1.307007519 | 0.234584599 | 0.991835066 | -4.635265795 |
| **A_51_P260169** | Gstm5 | -0.224999729 | -0.636448617 | 0.18644916 | 11.62195317 | -1.306998933 | 0.234587351 | 0.991835066 | -4.635272848 |
| **A_55_P2069885** | Slc50a1 | -0.136650212 | -0.386548193 | 0.113247768 | 9.188031006 | -1.306943151 | 0.234605232 | 0.991835066 | -4.635318673 |
| **A_55_P1952990** | Myl6b | -0.084307878 | -0.238498439 | 0.069882684 | 9.191626484 | -1.306831213 | 0.234641117 | 0.991835066 | -4.635410628 |
| **A_55_P2001920** | Flna | 0.160249054 | -0.13284946 | 0.453347569 | 13.73421987 | 1.30674543 | 0.234668621 | 0.991835066 | -4.635481094 |
| **A_52_P441044** | Rfxap | -0.083354872 | -0.235825021 | 0.069115276 | 8.571078024 | -1.306638048 | 0.234703053 | 0.991835066 | -4.6355693 |
| **A_55_P2175291** | NA | 0.080197012 | -0.066498713 | 0.226892738 | 14.75934196 | 1.306621688 | 0.234708299 | 0.991835066 | -4.635582738 |
| **A_51_P235878** | Guk1 | -0.08511519 | -0.240838234 | 0.070607854 | 11.36676962 | -1.306361346 | 0.234791799 | 0.991835066 | -4.635796573 |
| **A_55_P1964068** | Slc25a38 | -0.090110951 | -0.254995431 | 0.074773529 | 10.31970378 | -1.306191717 | 0.234846218 | 0.991835066 | -4.635935889 |
| **A_51_P255657** | Misp3 | -0.059328861 | -0.167890149 | 0.049232427 | 7.831110443 | -1.306171358 | 0.23485275 | 0.991835066 | -4.63595261 |
| **A_55_P2172440** | Scarf2 | 0.251653888 | -0.208855064 | 0.712162841 | 11.60376576 | 1.306094797 | 0.234877316 | 0.991835066 | -4.636015486 |
| **A_55_P2186110** | R3hdm2 | -0.082949908 | -0.234760879 | 0.068861063 | 12.44669417 | -1.305935959 | 0.23492829 | 0.991835066 | -4.636145927 |
| **A_55_P2003891** | Txlna | 0.124146421 | -0.103123571 | 0.351416414 | 10.51552118 | 1.30557324 | 0.235044727 | 0.991835066 | -4.636443774 |

| | | | | | | | | | |
|---|---|---|---|---|---|---|---|---|---|
| **A_52_P185907** | Crabp1 | 0.064507965 | -0.05358514 | 0.182601071 | 7.76728918 | 1.305563296 | 0.23504792 | 0.991835066 | -4.636451938 |
| **A_55_P2050602** | Ncam1 | -0.104934846 | -0.297098584 | 0.087228892 | 11.39841509 | -1.305140975 | 0.235183556 | 0.991835066 | -4.636798676 |
| **A_55_P2008884** | Cltc | 0.110780309 | -0.092103318 | 0.313663937 | 13.5488936 | 1.30504265 | 0.235215145 | 0.991835066 | -4.636879397 |
| **A_55_P2346639** | Sec14l3 | 0.078287404 | -0.065096404 | 0.221671211 | 7.676607372 | 1.30497116 | 0.235238114 | 0.991835066 | -4.636938084 |
| **A_55_P2001608** | Klhl4 | 0.096602866 | -0.080347261 | 0.273552994 | 9.096829944 | 1.304812969 | 0.235288948 | 0.991835066 | -4.637067943 |
| **A_51_P237695** | Gnai3 | -0.068979668 | -0.195355662 | 0.057396327 | 11.50614701 | -1.304564761 | 0.235368728 | 0.991835066 | -4.637271681 |
| **A_55_P2126109** | Usp16 | 0.122921306 | -0.102302325 | 0.348144936 | 11.71929456 | 1.304434696 | 0.235410543 | 0.991835066 | -4.637378436 |
| **A_51_P109421** | Eif2d | -0.068938306 | -0.195293633 | 0.057417021 | 9.336660497 | -1.303995781 | 0.2355517 | 0.991835066 | -4.627829819 |
| **A_51_P195374** | Baiap2l1 | -0.090767442 | -0.257136001 | 0.075601116 | 8.738268062 | -1.303971128 | 0.235559631 | 0.991835066 | -4.637758883 |
| **A_55_P2112997** | Ddx18 | 0.099160035 | -0.082641618 | 0.280961687 | 10.64179633 | 1.303610875 | 0.235675549 | 0.991835066 | -4.638054497 |
| **A_66_P108567** | Xrcc6 | 0.074937767 | -0.06247627 | 0.212351803 | 11.93933361 | 1.303403195 | 0.235742396 | 0.991835066 | -4.638224896 |
| **A_55_P2035043** | Gm2436 | -0.06086991 | -0.172489605 | 0.050749785 | 10.75472224 | -1.303379714 | 0.235749955 | 0.991835066 | -4.638244161 |
| **A_52_P660242** | Isy1 | -0.063998857 | -0.181363855 | 0.053366141 | 10.47747336 | -1.303295076 | 0.235777204 | 0.991835066 | -4.638313602 |
| **A_51_P480328** | Adgrl4 | -0.126037962 | -0.357199529 | 0.105123605 | 8.515067614 | -1.303151399 | 0.235823465 | 0.991835066 | -4.638431475 |
| **A_52_P27332** | Ppp2r5c | -0.08178293 | -0.231783474 | 0.068217614 | 11.4088167 | -1.30310366 | 0.235838839 | 0.991835066 | -4.638470639 |
| **A_55_P2142296** | Ywhaq | 0.064118706 | -0.053488626 | 0.181726038 | 13.14342153 | 1.303045201 | 0.235857665 | 0.991835066 | -4.638518596 |
| **A_55_P2083213** | Purb | -0.068789512 | -0.194992563 | 0.05741354 | 9.565608094 | -1.302751257 | 0.235952347 | 0.991835066 | -4.638759722 |
| **A_52_P80944** | Zfp36 | -0.063724205 | -0.180635757 | 0.053187347 | 7.904152031 | -1.302735157 | 0.235957534 | 0.991835066 | -4.638772929 |
| **A_55_P1954271** | Col26a1 | 0.082860551 | -0.069164322 | 0.234885424 | 10.41774053 | 1.302693661 | 0.235970904 | 0.991835066 | -4.638806966 |
| **A_55_P1962503** | Nup62cl | -0.08544291 | -0.24220972 | 0.0713239 | 7.7049073 | -1.302659895 | 0.235981783 | 0.991835066 | -4.638834662 |
| **A_55_P2013928** | Dhrs13 | 0.112319858 | -0.093769034 | 0.318408749 | 9.549042346 | 1.302600059 | 0.236001063 | 0.991835066 | -4.638883741 |
| **A_52_P182331** | NA | 0.068144402 | -0.056905956 | 0.193194759 | 15.28324754 | 1.302429961 | 0.236055878 | 0.991835066 | -4.639023256 |
| **A_55_P2044468** | Marf1 | 0.085604726 | -0.071488154 | 0.242697606 | 8.474745836 | 1.302417953 | 0.236059748 | 0.991835066 | -4.639033105 |
| **A_55_P2127243** | Tpsb2 | -0.093345229 | -0.264653212 | 0.077962754 | 8.637675047 | -1.302337825 | 0.236085574 | 0.991835066 | -4.639098822 |
| **A_55_P2122688** | Zmym3 | 0.117303444 | -0.097992227 | 0.332599115 | 11.76006427 | 1.302220707 | 0.236123327 | 0.991835066 | -4.639194875 |
| **A_55_P2051914** | Zfp568 | 0.121002981 | -0.101101508 | 0.343107471 | 11.02314446 | 1.302110574 | 0.236158833 | 0.991835066 | -4.639285194 |
| **A_52_P64356** | Sparcl1 | -0.106476576 | -0.301923234 | 0.088970082 | 15.17262827 | -1.302071879 | 0.23617131 | 0.991835066 | -4.639316927 |
| **A_51_P424532** | Vnn1 | 0.070441477 | -0.058862619 | 0.199745573 | 7.985984629 | 1.302042928 | 0.236180644 | 0.991835066 | -4.639340668 |
| **A_55_P2013078** | Thoc5 | 0.060879668 | -0.050887047 | 0.172646383 | 9.942346832 | 1.301873899 | 0.236235152 | 0.991835066 | -4.639479278 |
| **A_55_P2163093** | Nxf7 | -0.099167044 | -0.281232232 | 0.082898144 | 7.677011993 | -1.301815939 | 0.236253844 | 0.991835066 | -4.639526805 |
| **A_55_P2153621** | Ahnak | -0.108778536 | -0.308502677 | 0.090945605 | 10.33229462 | -1.301732598 | 0.236280726 | 0.991835066 | -4.639595144 |
| **A_55_P2015632** | Gm5766 | 0.116011835 | -0.097025873 | 0.329049544 | 15.50584402 | 1.301532289 | 0.236345345 | 0.991835066 | -4.639759385 |
| **A_51_P250058** | Epas1 | -0.083300501 | -0.23628303 | 0.069682028 | 12.94117679 | -1.301412306 | 0.236384059 | 0.991835066 | -4.639857758 |
| **A_55_P2082609** | NA | 0.108993961 | -0.091222732 | 0.309210654 | 11.50727885 | 1.30110182 | 0.236484265 | 0.991835066 | -4.640112303 |
| **A_52_P115579** | Fbxo45 | -0.058217473 | -0.165169687 | 0.048734741 | 8.43367067 | -1.300986234 | 0.236521579 | 0.991835066 | -4.640207056 |
| **A_55_P2058107** | Pigyl | 0.077965007 | -0.065304544 | 0.221234559 | 11.78820404 | 1.300633554 | 0.236635465 | 0.991835066 | -4.640496148 |
| **A_55_P1984622** | Smarcc1 | 0.088203945 | -0.073888565 | 0.250296455 | 8.472486829 | 1.300571384 | 0.236655546 | 0.991835066 | -4.640547105 |
| **A_55_P2042071** | Slc25a3 | -0.077385165 | -0.219598316 | 0.064827985 | 15.13560609 | -1.300550106 | 0.236662419 | 0.991835066 | -4.640564544 |
| **A_51_P381618** | Pla1a | -0.120072953 | -0.340800325 | 0.100654419 | 9.886050832 | -1.300163984 | 0.23678717 | 0.991835066 | -4.640880996 |
| **A_52_P628117** | Abcc5 | 0.1073318 | -0.09001979 | 0.30468339 | 8.247504534 | 1.299861009 | 0.236885098 | 0.991835066 | -4.641129272 |
| **A_55_P2106961** | 2900097C1 | 0.069534722 | -0.058324669 | 0.197394113 | 8.49425367 | 1.299805065 | 0.236903185 | 0.991835066 | -4.641175113 |
| **A_55_P2038452** | NA | -0.058999873 | -0.167489874 | 0.049490129 | 8.104810626 | -1.299781917 | 0.236910669 | 0.991835066 | -4.641194081 |
| **A_66_P109673** | NA | 0.0778197 | -0.065327614 | 0.220967015 | 10.74813659 | 1.29931808 | 0.237060674 | 0.991835066 | -4.641574115 |
| **A_55_P2050353** | Prpmp5 | -0.081100068 | -0.230306075 | 0.068105939 | 8.457506719 | -1.299104364 | 0.237129817 | 0.991835066 | -4.641749196 |
| **A_51_P163778** | 2810025M1 | -0.090879256 | -0.258088678 | 0.076330165 | 12.70787593 | -1.299011974 | 0.237159714 | 0.991835066 | -4.64182488 |
| **A_55_P2035593** | Hsf1 | 0.073517391 | -0.061749402 | 0.208784184 | 10.57887058 | 1.298996635 | 0.237164677 | 0.991835066 | -4.641837445 |
| **A_55_P2037902** | Brd3 | 0.069252404 | -0.05816813 | 0.196672938 | 8.357121324 | 1.298986282 | 0.237168028 | 0.991835066 | -4.641845926 |
| **A_55_P2176492** | Tollip | 0.057217462 | -0.048078913 | 0.162513837 | 8.244851336 | 1.298746234 | 0.237245723 | 0.991835066 | -4.642042556 |

| | | | | | | | | | |
|---|---|---|---|---|---|---|---|---|---|
| **A_55_P1973352** | NA | -0.15821302 | -0.44940187 | 0.13297583 | 10.5892452 | -1.298603624 | 0.237291891 | 0.991835066 | -4.642159363 |
| **A_55_P1968129** | Eloc | 0.09964286 | -0.083779834 | 0.283065554 | 10.66787608 | 1.29838129 | 0.237363884 | 0.991835066 | -4.642341457 |
| **A_55_P2113698** | Nup205 | 0.134523247 | -0.113179215 | 0.38222571 | 11.30861823 | 1.298004356 | 0.237485982 | 0.991835066 | -4.642650137 |
| **A_66_P124776** | NA | -0.075753703 | -0.215244422 | 0.063737015 | 8.465904774 | -1.297979072 | 0.237494175 | 0.991835066 | -4.642670842 |
| **A_51_P421140** | Tubb6 | -0.083666071 | -0.237755744 | 0.070423602 | 13.63591617 | -1.297731875 | 0.23757428 | 0.991835066 | -4.642873253 |
| **A_55_P2024933** | Zfp706 | -0.055012019 | -0.156334338 | 0.046310301 | 12.95772145 | -1.297662006 | 0.237596925 | 0.991835066 | -4.64293046 |
| **A_55_P2037101** | Exoc2 | -0.094336871 | -0.268126164 | 0.079452421 | 10.96285938 | -1.297381156 | 0.237687972 | 0.991835066 | -4.643160399 |
| **A_55_P1967928** | Srfbp1 | 0.082155085 | -0.069242177 | 0.233552347 | 9.413451657 | 1.296956948 | 0.237825551 | 0.991835066 | -4.643507665 |
| **A_52_P546635** | Zeb1 | -0.091261714 | -0.259461453 | 0.076938025 | 11.53922881 | -1.296798309 | 0.237877018 | 0.991835066 | -4.643637516 |
| **A_55_P1960281** | Med7 | -0.074275094 | -0.211185357 | 0.062635169 | 9.631556384 | -1.29663081 | 0.237931371 | 0.991835066 | -4.643774612 |
| **A_51_P245090** | Aqp3 | 0.059435454 | -0.050130096 | 0.169001003 | 7.849931655 | 1.2965244 | 0.237965906 | 0.991835066 | -4.643861703 |
| **A_55_P1983135** | NA | -0.112983682 | -0.321286918 | 0.095319554 | 8.527119821 | -1.296369601 | 0.238016153 | 0.991835066 | -4.643988392 |
| **A_51_P160482** | Vsig2 | 0.095150384 | -0.080279877 | 0.270580645 | 7.790898412 | 1.296328794 | 0.238029401 | 0.991835066 | -4.644021787 |
| **A_55_P2082271** | NA | -0.077819968 | -0.221347908 | 0.065707971 | 15.23806645 | -1.295876847 | 0.238176162 | 0.991835066 | -4.644391617 |
| **A_51_P192162** | NA | -0.085155137 | -0.24224179 | 0.071931515 | 9.211242167 | -1.295629124 | 0.238256639 | 0.991835066 | -4.644594305 |
| **A_55_P2409306** | Vps35l | -0.061125817 | -0.173907427 | 0.051655792 | 8.374460818 | -1.295375044 | 0.238339205 | 0.991835066 | -4.644802174 |
| **A_55_P2075200** | Tor3a | -0.114997712 | -0.327180619 | 0.097185195 | 8.919459735 | -1.295352408 | 0.238346562 | 0.991835066 | -4.644820693 |
| **A_55_P1984841** | NA | -0.091700771 | -0.260921174 | 0.077519633 | 11.09678145 | -1.295177796 | 0.238403321 | 0.991835066 | -4.644963534 |
| **A_51_P324651** | Adgrl1 | -0.087855149 | -0.249993516 | 0.074283218 | 9.432232775 | -1.295061985 | 0.238440972 | 0.991835066 | -4.645058269 |
| **A_55_P1987979** | Plrg1 | 0.053926813 | -0.045610652 | 0.153464278 | 11.56782399 | 1.294873396 | 0.238502296 | 0.991835066 | -4.645212529 |
| **A_52_P485766** | Zdhhc17 | -0.114837582 | -0.326855327 | 0.097180163 | 9.010423068 | -1.294556353 | 0.238605419 | 0.991835066 | -4.645471836 |
| **A_55_P2052578** | Bclaf3 | 0.122684228 | -0.103844358 | 0.349212814 | 9.013720556 | 1.294418917 | 0.238650135 | 0.991835066 | -4.645584235 |
| **A_55_P2136334** | NA | -0.061438791 | -0.174925842 | 0.05204826 | 11.66623523 | -1.293914213 | 0.238814406 | 0.991835066 | -4.645996945 |
| **A_51_P343350** | Amn | 0.077728825 | -0.065898582 | 0.221356232 | 8.054411688 | 1.293462719 | 0.238961442 | 0.991835066 | -4.64636608 |
| **A_51_P199567** | Med26 | 0.114966731 | -0.097547899 | 0.327481361 | 9.864594011 | 1.292982008 | 0.239118079 | 0.991835066 | -4.646759036 |
| **A_55_P2085157** | Eef1g | 0.103428169 | -0.087785751 | 0.29464209 | 12.75498171 | 1.292791431 | 0.239180203 | 0.991835066 | -4.646914803 |
| **A_66_P116678** | Rps8 | 0.112670602 | -0.095636299 | 0.320977503 | 8.985181659 | 1.292754596 | 0.239192212 | 0.991835066 | -4.646944909 |
| **A_55_P2129925** | Mybl2 | 0.144807579 | -0.122924954 | 0.412540112 | 9.324465421 | 1.292704556 | 0.239208527 | 0.991835066 | -4.646985807 |
| **A_55_P2057821** | Golga1 | 0.086893257 | -0.073782354 | 0.247568869 | 9.943368532 | 1.292543742 | 0.239260964 | 0.991835066 | -4.647117235 |
| **A_55_P2121608** | NA | -0.172296786 | -0.490973484 | 0.146379913 | 13.35638246 | -1.292218633 | 0.239367006 | 0.991835066 | -4.647382912 |
| **A_66_P125067** | Snrpd1 | 0.097616611 | -0.083000427 | 0.278233648 | 10.97017587 | 1.291737127 | 0.239524134 | 0.991835066 | -4.647776338 |
| **A_51_P209327** | Apln | -0.156710838 | -0.446759927 | 0.133338251 | 9.912511473 | -1.291328269 | 0.239657626 | 0.991835066 | -4.648110351 |
| **A_51_P469968** | H2-M3 | -0.05518639 | -0.157331998 | 0.046959217 | 8.008213492 | -1.291282974 | 0.239672419 | 0.991835066 | -4.648147351 |
| **A_52_P504478** | Adprh | -0.074124682 | -0.211326303 | 0.06307694 | 12.53740359 | -1.291257123 | 0.239680862 | 0.991835066 | -4.648168468 |
| **A_55_P2126063** | Mrps21 | -0.079463654 | -0.226553138 | 0.067625829 | 14.27440652 | -1.291207507 | 0.239697068 | 0.991835066 | -4.648208998 |
| **A_51_P398467** | Bsdc1 | -0.0954827 | -0.272224489 | 0.08125909 | 10.10615806 | -1.291203142 | 0.239698494 | 0.991835066 | -4.648212563 |
| **A_55_P1991351** | S100a13 | -0.20639878 | -0.588501794 | 0.175704235 | 12.44811172 | -1.291028392 | 0.239755578 | 0.991835066 | -4.648355301 |
| **A_55_P2047375** | NA | -0.071027786 | -0.202539538 | 0.060483966 | 16.28500685 | -1.290841287 | 0.239816711 | 0.991835066 | -4.648508121 |
| **A_51_P105408** | Alg14 | -0.07017206 | -0.200100301 | 0.059756181 | 10.36314945 | -1.290832222 | 0.239819673 | 0.991835066 | -4.635724961 |
| **A_55_P2299524** | Scube1 | 0.140364839 | -0.119546603 | 0.400276282 | 8.5601607 | 1.290749797 | 0.239846609 | 0.991835066 | -4.648582843 |
| **A_55_P1993404** | NA | -0.056830258 | -0.162071127 | 0.048410611 | 11.3964244 | -1.290637661 | 0.239883258 | 0.991835066 | -4.648674424 |
| **A_55_P1974650** | Npdc1 | -0.067335734 | -0.192054249 | 0.05738278 | 11.26212013 | -1.290398365 | 0.239961482 | 0.991835066 | -4.648869842 |
| **A_51_P493558** | Rnpep | -0.080019951 | -0.228266064 | 0.068226163 | 10.61645739 | -1.290102136 | 0.240058348 | 0.991835066 | -4.649111729 |
| **A_51_P339200** | Abhd16a | -0.104791325 | -0.298937002 | 0.089354352 | 8.629785157 | -1.290050582 | 0.24007521 | 0.991835066 | -4.649153823 |
| **A_51_P389386** | Nsmce2 | -0.093557217 | -0.26691244 | 0.079798006 | 10.51642426 | -1.289880553 | 0.240130828 | 0.991835066 | -4.649292647 |
| **A_55_P2080774** | Qdpr | -0.124631048 | -0.355570515 | 0.106308418 | 10.48479051 | -1.289844103 | 0.240142752 | 0.991835066 | -4.649322406 |
| **A_51_P416439** | Zdhhc6 | 0.089631591 | -0.076484616 | 0.255747799 | 10.3877696 | 1.289609521 | 0.240219508 | 0.991835066 | -4.649513919 |
| **A_55_P2072841** | NA | 0.118495919 | -0.101116981 | 0.338108819 | 9.335778356 | 1.289598976 | 0.240222959 | 0.991835066 | -4.649522527 |

| | | | | | | | | | |
|---|---|---|---|---|---|---|---|---|---|
| **A_52_P172014** | Ramp1 | -0.124613484 | -0.355569727 | 0.10634276 | 8.460621653 | -1.289568637 | 0.240232887 | 0.991835066 | -4.649547294 |
| **A_55_P2073709** | Tfdp2 | 2.514078303 | -2.145480277 | 7.173636882 | 7.857151733 | 1.289564759 | 0.240234156 | 0.991835066 | -4.64955046 |
| **A_55_P2064595** | NA | 0.090829302 | -0.077524966 | 0.25918357 | 8.556954946 | 1.289469219 | 0.240265425 | 0.991835066 | -4.649628453 |
| **A_51_P371255** | Rwdd2b | 0.106563945 | -0.09100503 | 0.304132921 | 9.190294784 | 1.289141753 | 0.240372626 | 0.991835066 | -4.649895754 |
| **A_55_P2041121** | Rn4.5s | 0.103075123 | -0.088043239 | 0.294193485 | 17.1363733 | 1.289022743 | 0.240411596 | 0.991835066 | -4.649992891 |
| **A_55_P2103668** | Zfp316 | -0.103222819 | -0.29464902 | 0.088203382 | 10.48809035 | -1.288793883 | 0.240486552 | 0.991835066 | -4.650179676 |
| **A_55_P1976017** | Cnih4 | -0.079344238 | -0.226490223 | 0.067801746 | 13.62530383 | -1.288772064 | 0.240493699 | 0.991835066 | -4.650197483 |
| **A_51_P288618** | Dusp11 | 0.093540617 | -0.079939217 | 0.267020451 | 8.563833484 | 1.288725331 | 0.240509008 | 0.991835066 | -4.650235621 |
| **A_55_P2140835** | NA | 0.12540818 | -0.107319799 | 0.358136158 | 8.317437007 | 1.287912621 | 0.240775372 | 0.991835066 | -4.650898773 |
| **A_55_P2104810** | Mrpl27 | -0.089674123 | -0.256119592 | 0.076771346 | 12.50236234 | -1.287669153 | 0.240855218 | 0.991835066 | -4.651097398 |
| **A_55_P2108773** | Tedc1 | 0.127963364 | -0.109564319 | 0.365491047 | 9.258272241 | 1.287598757 | 0.240878309 | 0.991835066 | -4.651154825 |
| **A_51_P465772** | Ruvbl1 | 0.076327023 | -0.065390896 | 0.218044942 | 12.40292769 | 1.287249368 | 0.240992942 | 0.991835066 | -4.651439823 |
| **A_55_P2031403** | NA | -0.074815766 | -0.213744224 | 0.064112692 | 9.101552056 | -1.287096306 | 0.241043176 | 0.991835066 | -4.651564665 |
| **A_55_P2045407** | Ppp4r1 | 0.066063785 | -0.056627865 | 0.188755435 | 11.77873499 | 1.286937856 | 0.241095188 | 0.991835066 | -4.651693895 |
| **A_55_P1963002** | Ppfia4 | 0.098190953 | -0.084166669 | 0.280548574 | 9.37727258 | 1.28693522 | 0.241096054 | 0.991835066 | -4.651696044 |
| **A_55_P1975475** | Mdm2 | 0.076142839 | -0.065273969 | 0.217559647 | 12.09245528 | 1.286877381 | 0.241115042 | 0.991835066 | -4.651743215 |
| **A_55_P2083023** | Sac3d1 | 0.134357622 | -0.115199385 | 0.38391463 | 11.58295295 | 1.286772208 | 0.241149573 | 0.991835066 | -4.651828986 |
| **A_51_P491051** | Ints9 | -0.08941965 | -0.255516573 | 0.076677273 | 11.19907849 | -1.286709505 | 0.241170163 | 0.991835066 | -4.65188012 |
| **A_55_P2141287** | Erlin1 | 0.122144985 | -0.10474431 | 0.34903428 | 10.01011201 | 1.286680644 | 0.24117964 | 0.991835066 | -4.651903656 |
| **A_51_P209372** | Msmo1 | 0.231801549 | -0.198854009 | 0.662457107 | 11.92324072 | 1.286457401 | 0.24125296 | 0.991835066 | -4.652085699 |
| **A_55_P1985514** | Dlg3 | 0.153234945 | -0.131568004 | 0.438037893 | 10.31860704 | 1.285945216 | 0.241421249 | 0.991835066 | -4.652503302 |
| **A_55_P2037141** | Jsrp1 | 0.164920927 | -0.141660446 | 0.471502301 | 10.99235078 | 1.285698492 | 0.241502352 | 0.991835066 | -4.652704435 |
| **A_51_P284946** | Rnd3 | 0.109488284 | -0.09406857 | 0.313045139 | 10.35680379 | 1.285556151 | 0.241549153 | 0.991835066 | -4.652820467 |
| **A_55_P1977741** | NA | -0.128677974 | -0.36792047 | 0.110564522 | 8.974500858 | -1.285508709 | 0.241564754 | 0.991835066 | -4.652859138 |
| **A_55_P1981994** | Krt17 | 0.069255642 | -0.059517909 | 0.198029192 | 7.830675701 | 1.285397994 | 0.241601164 | 0.991835066 | -4.652949383 |
| **A_52_P561020** | Slc25a32 | 0.073877071 | -0.063490553 | 0.211244695 | 8.537749544 | 1.285388509 | 0.241604283 | 0.991835066 | -4.652957114 |
| **A_55_P2171648** | Ppp1r7 | -0.108242963 | -0.309579695 | 0.093093768 | 11.67147395 | -1.284948709 | 0.241748966 | 0.991835066 | -4.653315562 |
| **A_51_P305320** | B4gat1 | -0.067881896 | -0.194167269 | 0.058403477 | 11.68782351 | -1.284724616 | 0.241822716 | 0.991835066 | -4.65349818 |
| **A_51_P375693** | Tmem135 | -0.063178423 | -0.180718299 | 0.054361452 | 10.25393534 | -1.284673275 | 0.241839615 | 0.991835066 | -4.653540016 |
| **A_51_P140901** | Pmm2 | 0.071073021 | -0.061163257 | 0.2033093 | 8.908129462 | 1.284586313 | 0.241868242 | 0.991835066 | -4.653610878 |
| **A_55_P2268221** | Arhgap10 | 0.060313888 | -0.051911872 | 0.172539648 | 8.202760245 | 1.284499394 | 0.241896857 | 0.991835066 | -4.653681703 |
| **A_55_P2148268** | Xxylt1 | 0.096656092 | -0.083215995 | 0.276528179 | 8.869656936 | 1.284323977 | 0.241954616 | 0.991835066 | -4.653824631 |
| **A_51_P349192** | NA | -0.075644823 | -0.216438244 | 0.065148598 | 12.436291 | -1.284121092 | 0.242021435 | 0.991835066 | -4.653989929 |
| **A_52_P599055** | Atg14 | -0.080821713 | -0.231251463 | 0.069608037 | 9.234314685 | -1.284113607 | 0.242023901 | 0.991835066 | -4.653996028 |
| **A_52_P82247** | Cycs | -0.089950773 | -0.257390406 | 0.07748886 | 14.79722883 | -1.283972639 | 0.242070337 | 0.991835066 | -4.654110871 |
| **A_55_P1959828** | Slc35g1 | 0.125108784 | -0.107796376 | 0.358013944 | 8.417546093 | 1.283860465 | 0.242107294 | 0.991835066 | -4.654202253 |
| **A_55_P2030859** | NA | 0.057068211 | -0.049180925 | 0.163317347 | 15.9465965 | 1.283742697 | 0.2421461 | 0.991835066 | -4.654298188 |
| **A_51_P136337** | Galm | -0.06095768 | -0.174465135 | 0.052549776 | 8.118229681 | -1.283551127 | 0.242209235 | 0.991835066 | -4.654454233 |
| **A_55_P2114680** | Car4 | 0.102354964 | -0.088246828 | 0.292956756 | 9.926397557 | 1.283485785 | 0.242230773 | 0.991835066 | -4.654507456 |
| **A_52_P171791** | Smg1 | 0.104705225 | -0.090314432 | 0.299724882 | 9.812803604 | 1.283214027 | 0.242320367 | 0.991835066 | -4.654728794 |
| **A_52_P142531** | Phc3 | 0.078202209 | -0.067461001 | 0.223865419 | 9.301535861 | 1.283152511 | 0.242340651 | 0.991835066 | -4.654778894 |
| **A_55_P2025902** | NA | -0.055875422 | -0.159965386 | 0.048214541 | 9.800094009 | -1.282983569 | 0.242396368 | 0.991835066 | -4.654916478 |
| **A_51_P221014** | Cd47 | -0.10707082 | -0.306575886 | 0.092434245 | 11.15940919 | -1.282703666 | 0.242488702 | 0.991835066 | -4.655144408 |
| **A_55_P1957168** | Ube2s | 0.235217521 | -0.203161632 | 0.673596674 | 13.74891194 | 1.282415943 | 0.242583648 | 0.991835066 | -4.655378681 |
| **A_55_P2169149** | Washc3 | -0.081475092 | -0.233322285 | 0.070372102 | 10.60843301 | -1.282410961 | 0.242585292 | 0.991835066 | -4.655382737 |
| **A_52_P409746** | Serpinc1 | -0.243786443 | -0.698301539 | 0.210728654 | 8.246370657 | -1.281947885 | 0.242738172 | 0.991835066 | -4.655759733 |
| **A_55_P1994042** | Zbp1 | -0.070911299 | -0.203134949 | 0.061312352 | 7.937369096 | -1.28178572 | 0.24279173 | 0.991835066 | -4.655891739 |
| **A_55_P2078675** | NA | 0.065703967 | -0.05682149 | 0.188229423 | 8.421892929 | 1.28166461 | 0.242831734 | 0.991835066 | -4.655990319 |

| | | | | | | | | | |
|---|---|---|---|---|---|---|---|---|---|
| **A_51_P247694** | Adgrg3 | 0.082735567 | -0.07157088 | 0.237042014 | 8.465934036 | 1.28149616 | 0.242887386 | 0.991835066 | -4.656127426 |
| **A_52_P683211** | NA | 0.076807321 | -0.066445456 | 0.220060099 | 8.271489876 | 1.281470758 | 0.242895779 | 0.991835066 | -4.656148101 |
| **A_55_P1966035** | Apoa2 | -0.329323506 | -0.943570823 | 0.28492381 | 9.136790245 | -1.281411434 | 0.242915381 | 0.991835066 | -4.656196384 |
| **A_55_P1954418** | NA | -0.096983981 | -0.27788298 | 0.083915019 | 11.4991241 | -1.281365345 | 0.242930611 | 0.991835066 | -4.656233895 |
| **A_55_P2053157** | Zc3h3 | 0.111164902 | -0.096219945 | 0.31854975 | 10.50890565 | 1.281149474 | 0.243001956 | 0.991835066 | -4.656409578 |
| **A_51_P428578** | Retreg1 | -0.226172847 | -0.64813512 | 0.195789426 | 10.00809434 | -1.281079184 | 0.243025191 | 0.991835066 | -4.656466779 |
| **A_52_P541161** | Rgs18 | -0.066397969 | -0.190281629 | 0.057485691 | 7.782257153 | -1.281002276 | 0.243050616 | 0.991835066 | -4.656529364 |
| **A_55_P2111399** | Pcp2 | -0.063021593 | -0.180608342 | 0.054565155 | 8.029994005 | -1.280973454 | 0.243060145 | 0.991835066 | -4.656552818 |
| **A_55_P1959648** | Hspg2 | 0.109286669 | -0.094634798 | 0.313208136 | 9.28119173 | 1.280894544 | 0.243086234 | 0.991835066 | -4.65661703 |
| **A_55_P1956938** | Rpl24 | 0.091036731 | -0.078842302 | 0.260915764 | 16.6678046 | 1.280813828 | 0.243112924 | 0.991835066 | -4.656682709 |
| **A_51_P160372** | Wbp1 | -0.079889527 | -0.228971187 | 0.069192134 | 13.38130452 | -1.280780657 | 0.243123893 | 0.991835066 | -4.656709701 |
| **A_51_P495485** | Pip4p2 | -0.058881789 | -0.168779598 | 0.05101602 | 11.69356614 | -1.280563436 | 0.243195734 | 0.991835066 | -4.656886443 |
| **A_55_P1968908** | Hypk | -0.077369873 | -0.221807675 | 0.06706793 | 11.63282015 | -1.280265789 | 0.243294204 | 0.991835066 | -4.657128601 |
| **A_55_P2012780** | Rnf167 | -0.080349231 | -0.230375336 | 0.069676875 | 10.79627852 | -1.280041413 | 0.243368457 | 0.991835066 | -4.65731113 |
| **A_51_P323299** | Trim28 | 0.097570914 | -0.084625238 | 0.279767066 | 11.70352268 | 1.279942047 | 0.243401347 | 0.991835066 | -4.657391959 |
| **A_55_P2073248** | Slc25a34 | 0.138167333 | -0.119882426 | 0.396217091 | 8.539473067 | 1.279708482 | 0.243478671 | 0.991835066 | -4.65758194 |
| **A_55_P2081318** | Gnas | -0.132011245 | -0.378568058 | 0.114545568 | 8.319615094 | -1.279684917 | 0.243486474 | 0.991835066 | -4.657601107 |
| **A_51_P503866** | Edem1 | -0.072353706 | -0.207504501 | 0.06279709 | 9.100415247 | -1.279532446 | 0.243536964 | 0.991835066 | -4.657725115 |
| **A_55_P2056140** | Flad1 | -0.120615751 | -0.345936686 | 0.104705183 | 10.68228556 | -1.279415521 | 0.243575689 | 0.991835066 | -4.65782021 |
| **A_55_P2143233** | Rasgrp2 | -0.113691894 | -0.326089814 | 0.098706027 | 10.28767729 | -1.27934705 | 0.243598369 | 0.991835066 | -4.657875894 |
| **A_55_P2010008** | Zfp113 | -0.067211526 | -0.19278125 | 0.058358198 | 8.829603129 | -1.279286893 | 0.243618296 | 0.991835066 | -4.657924816 |
| **A_52_P601385** | Cd99l2 | 0.064458211 | -0.056006752 | 0.184923174 | 11.71247438 | 1.278870669 | 0.243756213 | 0.991835066 | -4.658263275 |
| **A_55_P2016249** | Ppid | -0.095942907 | -0.275292439 | 0.083406624 | 9.441757014 | -1.278561909 | 0.243858565 | 0.991835066 | -4.658514314 |
| **A_55_P1984333** | Emd | 0.087324402 | -0.075916284 | 0.250565089 | 8.793280478 | 1.27854595 | 0.243863856 | 0.991835066 | -4.658527289 |
| **A_55_P1980854** | Ankrd24 | 0.119062876 | -0.103524899 | 0.341650651 | 11.11133561 | 1.27845131 | 0.243895237 | 0.991835066 | -4.65860423 |
| **A_51_P288891** | Tbc1d15 | -0.09187992 | -0.2636801 | 0.07992026 | 9.485082336 | -1.278221518 | 0.243971446 | 0.991835066 | -4.658791037 |
| **A_55_P2131785** | Kiz | -0.109106699 | -0.313123006 | 0.094909609 | 10.93553103 | -1.278190731 | 0.243981657 | 0.991835066 | -4.658816064 |
| **A_55_P1976351** | Gpcpd1 | 0.10060338 | -0.087551221 | 0.288757981 | 8.684337703 | 1.277929403 | 0.244068353 | 0.991835066 | -4.659028485 |
| **A_52_P168271** | Spg20 | 0.073737468 | -0.064190951 | 0.211665888 | 8.230571899 | 1.277743249 | 0.244130126 | 0.991835066 | -4.659179788 |
| **A_55_P2045946** | Tepsin | 0.076145651 | -0.066293327 | 0.21858463 | 9.950944334 | 1.27768968 | 0.244147905 | 0.991835066 | -4.659223327 |
| **A_55_P2076757** | Znfx1 | -0.068507565 | -0.196675097 | 0.059659968 | 10.99201425 | -1.277525675 | 0.244202343 | 0.991835066 | -4.659356616 |
| **A_55_P2082733** | Cybb | -0.077251694 | -0.221798524 | 0.067295137 | 7.779905806 | -1.277346047 | 0.244261978 | 0.991835066 | -4.659502593 |
| **A_55_P2020378** | Ehd4 | -0.091413517 | -0.262481756 | 0.079654722 | 11.1388944 | -1.277174288 | 0.244319013 | 0.991835066 | -4.659642165 |
| **A_55_P1980636** | Aurka | -0.065690932 | -0.188622934 | 0.057241069 | 11.71542957 | -1.277172634 | 0.244319563 | 0.991835066 | -4.659643509 |
| **A_55_P2013273** | NA | 0.135586534 | -0.118185185 | 0.389358252 | 14.03683645 | 1.27697522 | 0.244385131 | 0.991835066 | -4.659803919 |
| **A_52_P365768** | Nsun2 | 0.079569472 | -0.069361017 | 0.228499961 | 11.33686701 | 1.27694442 | 0.244395362 | 0.991835066 | -4.659828944 |
| **A_51_P223458** | Polr3d | 0.07764121 | -0.067707987 | 0.222990407 | 9.014899459 | 1.276699808 | 0.244476631 | 0.991835066 | -4.660027685 |
| **A_55_P2116235** | NA | 0.105414955 | -0.091940798 | 0.302770709 | 13.77340938 | 1.276619783 | 0.244503224 | 0.991835066 | -4.660092699 |
| **A_51_P419226** | S100a14 | 0.077618162 | -0.067701062 | 0.222937387 | 7.789653919 | 1.276584061 | 0.244515095 | 0.991835066 | -4.66012172 |
| **A_66_P106661** | Slc7a1 | 0.089957436 | -0.078467402 | 0.258382275 | 12.54665579 | 1.276556563 | 0.244524233 | 0.991835066 | -4.660144059 |
| **A_55_P2066440** | Myh3 | -0.497581499 | -1.429362128 | 0.43419913 | 8.520311511 | -1.276320307 | 0.244602761 | 0.991835066 | -4.660335983 |
| **A_51_P184062** | Hbs1l | 0.079308604 | -0.069250818 | 0.227868026 | 11.12943224 | 1.275937013 | 0.244730209 | 0.991835066 | -4.660647317 |
| **A_51_P185939** | Gper1 | -0.064039635 | -0.184000057 | 0.055920788 | 8.094303596 | -1.275909848 | 0.244739244 | 0.991835066 | -4.660669381 |
| **A_55_P2048759** | Mgarp | 0.375163634 | -0.327693415 | 1.078020683 | 10.1440758 | 1.275742023 | 0.244795067 | 0.991835066 | -4.660805682 |
| **A_55_P2011551** | Anapc5 | -0.069863761 | -0.20077997 | 0.061052448 | 10.95736082 | -1.275462441 | 0.244888088 | 0.991835066 | -4.66103273 |
| **A_52_P433864** | Naa16 | 0.099564784 | -0.087009883 | 0.28613945 | 8.451878153 | 1.275446404 | 0.244893425 | 0.991835066 | -4.661045753 |
| **A_55_P2000280** | Mthfd1l | 0.155095909 | -0.1355783 | 0.445770118 | 8.684998718 | 1.275272441 | 0.244951321 | 0.991835066 | -4.661187015 |
| **A_55_P2142728** | NA | -0.111334573 | -0.320033085 | 0.097363939 | 11.84382587 | -1.275028314 | 0.245032589 | 0.991835066 | -4.661385236 |

| | | | | | | | | | |
|---|---|---|---|---|---|---|---|---|---|
| **A_51_P286357** | Adgre4 | -0.075888629 | -0.218152819 | 0.066375562 | 7.769591924 | -1.27494145 | 0.245061511 | 0.991835066 | -4.661455761 |
| **A_55_P2075195** | Cd302 | -0.096735702 | -0.27811162 | 0.084640217 | 9.90689338 | -1.274724382 | 0.245133798 | 0.991835066 | -4.66163199 |
| **A_55_P1996716** | Cachd1 | 0.09732446 | -0.085205186 | 0.279854105 | 9.830513262 | 1.274376408 | 0.245249717 | 0.991835066 | -4.661914466 |
| **A_52_P441974** | Evc2 | 0.107799533 | -0.094381617 | 0.309980684 | 10.15154731 | 1.274340098 | 0.245261816 | 0.991835066 | -4.661943939 |
| **A_55_P2033462** | Rps23-ps1 | 0.07848375 | -0.068733044 | 0.225700543 | 16.44018854 | 1.274182165 | 0.245314446 | 0.991835066 | -4.662072131 |
| **A_51_P221248** | Tspan32 | -0.053510005 | -0.153883029 | 0.046863018 | 7.752384248 | -1.27416925 | 0.24531875 | 0.991835066 | -4.662082613 |
| **A_55_P2015217** | Alg11 | 0.09443393 | -0.082711403 | 0.271579262 | 8.942382995 | 1.274111632 | 0.245337953 | 0.991835066 | -4.662129378 |
| **A_52_P99531** | Ippk | 0.112869472 | -0.098871137 | 0.32461008 | 9.58987426 | 1.274035317 | 0.24536339 | 0.991835066 | -4.662191317 |
| **A_51_P509760** | Actr1a | -0.053287682 | -0.153256179 | 0.046680815 | 12.93147707 | -1.274009892 | 0.245371865 | 0.991835066 | -4.662211953 |
| **A_55_P2114839** | NA | -0.084994773 | -0.244453895 | 0.074464349 | 14.69716173 | -1.273948824 | 0.245392222 | 0.991835066 | -4.662261516 |
| **A_55_P2147141** | Spata2 | 0.060402362 | -0.052937149 | 0.173741872 | 11.53746499 | 1.273742729 | 0.245460935 | 0.991835066 | -4.662428773 |
| **A_51_P216702** | Eogt | 0.07741517 | -0.067865915 | 0.222696256 | 9.356759789 | 1.273579706 | 0.245515299 | 0.991835066 | -4.662561065 |
| **A_55_P2386592** | Cep63 | 0.075585972 | -0.066303257 | 0.217475201 | 11.38163772 | 1.273212538 | 0.245637779 | 0.991835066 | -4.662858992 |
| **A_55_P1998578** | Ifitm2 | -0.196975157 | -0.566736442 | 0.172786128 | 11.98355365 | -1.273207387 | 0.245639498 | 0.991835066 | -4.662863171 |
| **A_51_P199608** | Ttll1 | -0.093515155 | -0.269065211 | 0.082034902 | 9.869417345 | -1.273181023 | 0.245648294 | 0.991835066 | -4.662884562 |
| **A_52_P657338** | Mrps17 | -0.079912832 | -0.22993783 | 0.070112165 | 12.37777155 | -1.27309856 | 0.245675811 | 0.991835066 | -4.662951467 |
| **A_55_P1964628** | NA | 0.130760444 | -0.114845088 | 0.376365976 | 12.91009787 | 1.272469478 | 0.245885811 | 0.991835066 | -4.663461794 |
| **A_55_P1968438** | Arxes1 | 0.148700054 | -0.130688431 | 0.428088539 | 9.714507204 | 1.272072181 | 0.246018517 | 0.991835066 | -4.663784029 |
| **A_52_P562467** | Sync | -0.060139316 | -0.173180886 | 0.052902255 | 8.466194144 | -1.271538258 | 0.246196957 | 0.991835066 | -4.664216999 |
| **A_55_P2183288** | Cryzl1 | -1.722476708 | -4.960394615 | 1.5154412 | 9.609376592 | -1.271442301 | 0.246229038 | 0.991835066 | -4.664294802 |
| **A_55_P2114809** | Gtf2e2 | -0.068968461 | -0.198647281 | 0.060710358 | 11.21034328 | -1.271131908 | 0.246332836 | 0.991835066 | -4.664546457 |
| **A_55_P1959595** | Fmo5 | -0.083276638 | -0.239878578 | 0.073325301 | 8.998701469 | -1.270969654 | 0.246387111 | 0.991835066 | -4.664677995 |
| **A_55_P2089506** | NA | -0.107515151 | -0.309707085 | 0.094676783 | 13.5008842 | -1.270910525 | 0.246406892 | 0.991835066 | -4.664725928 |
| **A_51_P337675** | Cd53 | -0.058465704 | -0.168431465 | 0.051500057 | 7.733466852 | -1.270728685 | 0.246467734 | 0.991835066 | -4.66487333 |
| **A_51_P440743** | Celsr1 | 0.083993546 | -0.074005511 | 0.241992602 | 11.57964671 | 1.270575721 | 0.246518925 | 0.991835066 | -4.664997317 |
| **A_52_P18596** | NA | 0.170561913 | -0.150285356 | 0.491409183 | 11.9455282 | 1.270553169 | 0.246526473 | 0.991835066 | -4.665015596 |
| **A_66_P102719** | Spop | -0.086240187 | -0.248469 | 0.075988626 | 11.65359818 | -1.270547278 | 0.246528445 | 0.991835066 | -4.665020372 |
| **A_55_P2059432** | Crabp2 | 0.094324894 | -0.083157432 | 0.27180722 | 8.621179407 | 1.270224091 | 0.246636636 | 0.991835066 | -4.665282308 |
| **A_52_P652913** | Akip1 | -0.11708532 | -0.33739626 | 0.10322562 | 10.25080933 | -1.270209968 | 0.246641365 | 0.991835066 | -4.665293754 |
| **A_55_P2418745** | D230034L2 | 0.188405384 | -0.166103687 | 0.542914454 | 8.324212309 | 1.270208718 | 0.246641783 | 0.991835066 | -4.665294767 |
| **A_52_P667913** | Pcdhga9 | 0.102840508 | -0.090686028 | 0.296367043 | 9.130093319 | 1.270085101 | 0.246683178 | 0.991835066 | -4.665394947 |
| **A_55_P1997977** | Rab2b | 0.113944602 | -0.100554796 | 0.328444 | 9.999547956 | 1.269628935 | 0.246835981 | 0.991835066 | -4.665764586 |
| **A_52_P344978** | NA | 0.059484359 | -0.052520317 | 0.171489035 | 8.806396603 | 1.269333559 | 0.246934967 | 0.991835066 | -4.666003899 |
| **A_55_P2019719** | Oas2 | -0.061354102 | -0.17688411 | 0.054175905 | 7.835410305 | -1.269281461 | 0.24695243 | 0.991835066 | -4.666046106 |
| **A_51_P233160** | Lysmd2 | 0.124842992 | -0.110254874 | 0.359940858 | 10.03254823 | 1.269184071 | 0.246985077 | 0.991835066 | -4.666125004 |
| **A_55_P2144746** | Apip | -0.091507895 | -0.263844152 | 0.080828362 | 11.04332945 | -1.269085968 | 0.247017966 | 0.991835066 | -4.666204476 |
| **A_55_P2186928** | NA | 0.118272295 | -0.104469246 | 0.341013837 | 8.279364899 | 1.269085664 | 0.247018068 | 0.991835066 | -4.666204722 |
| **A_55_P2049553** | NA | -0.165033281 | -0.475853261 | 0.145786699 | 10.80678508 | -1.269029561 | 0.247036879 | 0.991835066 | -4.66625017 |
| **A_55_P2065731** | Medag | -0.085337714 | -0.246107732 | 0.075432304 | 16.52071392 | -1.268659507 | 0.247160984 | 0.991835066 | -4.666549913 |
| **A_55_P2168383** | Myo18b | -0.0862127 | -0.248635667 | 0.076210268 | 13.53174219 | -1.268624043 | 0.247172881 | 0.991835066 | -4.666578637 |
| **A_52_P408530** | St3gal2 | -0.069081796 | -0.199286617 | 0.061123025 | 10.75517909 | -1.268077181 | 0.247356388 | 0.991835066 | -4.667021511 |
| **A_52_P24783** | Pds5b | -1.291815065 | -3.726880826 | 1.143250695 | 8.934395403 | -1.267940235 | 0.247402362 | 0.991835066 | -4.667132402 |
| **A_55_P2101739** | NA | -0.093835138 | -0.270734827 | 0.083064551 | 13.13066335 | -1.26779069 | 0.247452572 | 0.991835066 | -4.667253487 |
| **A_52_P268231** | NA | 0.071732083 | -0.063505192 | 0.206969358 | 9.165110393 | 1.267728231 | 0.247473546 | 0.991835066 | -4.667304058 |
| **A_52_P540434** | Ppp1cc | -0.088771012 | -0.25614272 | 0.078600697 | 11.08080914 | -1.267646764 | 0.247500905 | 0.991835066 | -4.667370017 |
| **A_51_P322994** | Mrpl22 | -0.204988649 | -0.591561538 | 0.18158424 | 10.21135915 | -1.267382052 | 0.24758982 | 0.991835066 | -4.667584324 |
| **A_55_P1961863** | Pcdhga9 | 0.095669087 | -0.084832661 | 0.276170835 | 13.29563548 | 1.266774609 | 0.247793962 | 0.991835066 | -4.668076017 |
| **A_55_P2305294** | Ptpn21 | 0.132292737 | -0.117338589 | 0.381924063 | 9.062028216 | 1.266619151 | 0.24784623 | 0.991835066 | -4.668201833 |

| | | | | | | | | | |
|---|---|---|---|---|---|---|---|---|---|
| **A_55_P1953311** | Stx2 | 0.070799435 | -0.062814663 | 0.204413533 | 10.11154451 | 1.266445899 | 0.247904491 | 0.991835066 | -4.668342042 |
| **A_55_P2116059** | Crybg1 | -0.07489992 | -0.21626378 | 0.066463939 | 7.978221929 | -1.266345159 | 0.247938374 | 0.991835066 | -4.668423565 |
| **A_55_P1985890** | Tiparp | -0.092480581 | -0.267094634 | 0.082133471 | 9.841334059 | -1.265844883 | 0.248106695 | 0.991835066 | -4.668828358 |
| **A_55_P1969776** | Pdhb | -0.085849371 | -0.248000463 | 0.07630172 | 14.97889381 | -1.26539576 | 0.248257889 | 0.991835066 | -4.669191694 |
| **A_55_P2113524** | Nsd2 | 0.05953536 | -0.052918261 | 0.17198898 | 8.273988387 | 1.265350001 | 0.248273298 | 0.991835066 | -4.66922871 |
| **A_51_P462918** | Ehhadh | -0.063037718 | -0.182145057 | 0.056069621 | 8.316981895 | -1.264943394 | 0.248410255 | 0.991835066 | -4.66955759 |
| **A_51_P141926** | Fxyd4 | 0.088310169 | -0.078549085 | 0.255169423 | 7.689944102 | 1.264938908 | 0.248411767 | 0.991835066 | -4.669561218 |
| **A_51_P292368** | Tmco6 | 0.062845413 | -0.055911856 | 0.181602682 | 8.152882477 | 1.264801909 | 0.248457927 | 0.991835066 | -4.669672017 |
| **A_55_P1988951** | NA | 0.110800502 | -0.098577253 | 0.320178257 | 14.79628103 | 1.264795529 | 0.248460077 | 0.991835066 | -4.669677177 |
| **A_66_P117477** | Dlk1 | -0.101924008 | -0.294537907 | 0.090689891 | 14.47020872 | -1.264730526 | 0.248481982 | 0.991835066 | -4.669729746 |
| **A_55_P2056393** | Stk11ip | 0.070321819 | -0.062580934 | 0.203224571 | 10.02733618 | 1.264635157 | 0.248514122 | 0.991835066 | -4.66980687 |
| **A_51_P136781** | Pnliprp1 | 0.060104424 | -0.05349275 | 0.173701598 | 7.852131451 | 1.264585055 | 0.248531009 | 0.991835066 | -4.669847387 |
| **A_66_P135804** | NA | 0.072526099 | -0.064549848 | 0.209602046 | 13.79159901 | 1.264568023 | 0.24853675 | 0.991835066 | -4.66986116 |
| **A_55_P2007630** | Sez6l2 | -0.166993657 | -0.482660399 | 0.148673086 | 9.306573568 | -1.264387746 | 0.24859752 | 0.991835066 | -4.670006937 |
| **A_52_P486253** | Polr2b | 0.12066048 | -0.107442295 | 0.348763256 | 11.29584711 | 1.264281002 | 0.248633509 | 0.991835066 | -4.67009325 |
| **A_51_P401964** | Aph1a | 0.077621756 | -0.069129732 | 0.224373244 | 11.8751158 | 1.264183404 | 0.248666418 | 0.991835066 | -4.670172163 |
| **A_55_P1971804** | Hdac10 | 0.056628111 | -0.050457979 | 0.163714201 | 9.215331456 | 1.26388671 | 0.248766484 | 0.991835066 | -4.670412038 |
| **A_55_P2062495** | Smyd1 | 0.143590753 | -0.127980296 | 0.415161803 | 8.971255499 | 1.263723715 | 0.248821472 | 0.991835066 | -4.670543806 |
| **A_52_P411331** | Vac14 | -0.084564889 | -0.244509014 | 0.075379236 | 8.420218833 | -1.263661996 | 0.248842296 | 0.991835066 | -4.670593699 |
| **A_55_P2171976** | NA | 0.087711961 | -0.078208032 | 0.253631953 | 15.09654779 | 1.263482499 | 0.248902868 | 0.991835066 | -4.670738794 |
| **A_52_P536293** | NA | -0.075892067 | -0.219463617 | 0.067679483 | 14.4450817 | -1.263389094 | 0.248934393 | 0.991835066 | -4.670814294 |
| **A_55_P1972322** | Btg3 | 0.099828712 | -0.089054143 | 0.288711567 | 12.66988161 | 1.263199837 | 0.248998279 | 0.991835066 | -4.670967263 |
| **A_51_P505521** | NA | 0.095779607 | -0.085461792 | 0.277021007 | 9.672906028 | 1.263062314 | 0.24904471 | 0.991835066 | -4.671078409 |
| **A_51_P297694** | Prpf6 | 0.055236356 | -0.049288176 | 0.159760888 | 11.68785397 | 1.263036557 | 0.249053408 | 0.991835066 | -4.671099226 |
| **A_55_P1996419** | NA | 0.08105507 | -0.072368136 | 0.234478277 | 15.50589139 | 1.262694426 | 0.249168956 | 0.991835066 | -4.671375711 |
| **A_55_P1975947** | Fam234b | -0.120223309 | -0.347819723 | 0.107373105 | 11.00454894 | -1.26250293 | 0.249233651 | 0.991835066 | -4.671530447 |
| **A_55_P2436715** | Paics | -0.077317096 | -0.223695885 | 0.069061693 | 11.89934962 | -1.262427709 | 0.249259067 | 0.991835066 | -4.671591225 |
| **A_66_P124179** | Atp6v0d2 | -0.052513847 | -0.151941188 | 0.046913493 | 7.674825199 | -1.262342328 | 0.249287919 | 0.991835066 | -4.67166021 |
| **A_55_P2104327** | Pxdn | -0.0755313 | -0.21858601 | 0.067523411 | 13.80166846 | -1.261926116 | 0.249428608 | 0.991835066 | -4.671996465 |
| **A_55_P1973858** | NA | -0.086360941 | -0.249945591 | 0.077223709 | 9.961690264 | -1.261780904 | 0.249477709 | 0.991835066 | -4.672113768 |
| **A_55_P2157788** | NA | -0.211179143 | -0.6112014 | 0.188843113 | 13.32932704 | -1.261757806 | 0.24948552 | 0.991835066 | -4.672132425 |
| **A_55_P2127131** | Trmt112 | -0.084345327 | -0.244128146 | 0.075437492 | 11.37079346 | -1.261653445 | 0.249520814 | 0.991835066 | -4.672216724 |
| **A_55_P1969276** | Hhip | 0.054166025 | -0.048450138 | 0.156782188 | 7.721452165 | 1.261596066 | 0.249540221 | 0.991835066 | -4.672263071 |
| **A_55_P2034555** | Pdcd5 | 0.100075724 | -0.08951628 | 0.289667727 | 14.60865388 | 1.261588891 | 0.249542648 | 0.991835066 | -4.672268866 |
| **A_55_P1996623** | Dnajb12 | -0.081611517 | -0.23624732 | 0.073024285 | 9.594162251 | -1.261393339 | 0.249608799 | 0.991835066 | -4.672426812 |
| **A_51_P459368** | Appbp2 | -0.065686852 | -0.190177894 | 0.05880419 | 12.23219529 | -1.261099859 | 0.249708104 | 0.991835066 | -4.67266383 |
| **A_55_P2036883** | H2bc18 | 0.210740647 | -0.188674888 | 0.610156183 | 11.28110632 | 1.26105053 | 0.249724799 | 0.991835066 | -4.672703665 |
| **A_55_P1957900** | Dlg1 | 0.070012813 | -0.062693019 | 0.202718644 | 8.901041438 | 1.260946473 | 0.249760019 | 0.991835066 | -4.672787695 |
| **A_51_P467410** | Npm1 | 0.147873732 | -0.132414595 | 0.42816206 | 15.1720617 | 1.260942128 | 0.24976149 | 0.991835066 | -4.672791204 |
| **A_55_P2097578** | NA | -0.083065916 | -0.240612408 | 0.074480575 | 11.61926303 | -1.260152936 | 0.25002875 | 0.991835066 | -4.67342839 |
| **A_55_P1972275** | Wars | 0.071107057 | -0.063773616 | 0.205987729 | 12.16938298 | 1.260004546 | 0.25007903 | 0.991835066 | -4.673548177 |
| **A_55_P1952828** | NA | -0.069625543 | -0.201720673 | 0.062469588 | 8.58868718 | -1.259768997 | 0.250158861 | 0.991835066 | -4.673738307 |
| **A_51_P110759** | Slc1a1 | -0.111481924 | -0.32299054 | 0.100026692 | 8.552601655 | -1.259753361 | 0.250164161 | 0.991835066 | -4.663229304 |
| **A_51_P450527** | Tagln | 0.462563461 | -0.415267547 | 1.340394468 | 13.04052214 | 1.259416962 | 0.250278211 | 0.991835066 | -4.674022431 |
| **A_55_P2079912** | NA | 0.086105197 | -0.077305132 | 0.249515526 | 9.338747035 | 1.259386386 | 0.250288579 | 0.991835066 | -4.674047106 |
| **A_55_P2162394** | Pdp1 | -0.080147614 | -0.232278527 | 0.0719833 | 8.991573488 | -1.259163875 | 0.250364045 | 0.991835066 | -4.67422667 |
| **A_52_P157274** | Abi1 | -0.063442734 | -0.183886846 | 0.057001377 | 8.313175234 | -1.258941215 | 0.250439581 | 0.991835066 | -4.674406338 |
| **A_52_P466090** | Samhd1 | -0.083397208 | -0.241729804 | 0.074935388 | 8.908616169 | -1.258897329 | 0.250454471 | 0.991835066 | -4.674441749 |

| | | | | | | | | | |
|---|---|---|---|---|---|---|---|---|---|
| **A_51_P342387** | Apoh | -0.286694689 | -0.831014721 | 0.257625343 | 8.289112954 | -1.258851387 | 0.25047006 | 0.991835066 | -4.674478818 |
| **A_55_P2086746** | Fam3a | -0.069502477 | -0.201461156 | 0.062456203 | 8.130988811 | -1.258842664 | 0.25047302 | 0.991835066 | -4.674485856 |
| **A_55_P2161818** | NA | 0.069791547 | -0.062726316 | 0.202309411 | 16.18408896 | 1.258744352 | 0.250506381 | 0.991835066 | -4.674565177 |
| **A_52_P686136** | Prrc2b | -0.083131304 | -0.241099357 | 0.074836749 | 12.95146185 | -1.257779347 | 0.250834056 | 0.991835066 | -4.675343613 |
| **A_55_P1998688** | Gm2012 | 0.077070741 | -0.069468178 | 0.22360966 | 8.067404429 | 1.257030123 | 0.251088715 | 0.991835066 | -4.675947781 |
| **A_51_P362538** | Syde2 | -0.090191103 | -0.261702064 | 0.081319859 | 9.57492855 | -1.256842755 | 0.251152436 | 0.991835066 | -4.676098845 |
| **A_55_P2038928** | NA | 0.084794345 | -0.076457135 | 0.246045825 | 11.67772328 | 1.256817772 | 0.251160933 | 0.991835066 | -4.676118987 |
| **A_51_P262515** | Phf11a | -0.0834273 | -0.242131174 | 0.075276573 | 8.054532335 | -1.256405404 | 0.251301226 | 0.991835066 | -4.676451413 |
| **A_55_P2117319** | Ntan1 | 0.075174897 | -0.0678446 | 0.218194394 | 12.10770583 | 1.25628082 | 0.251343624 | 0.991835066 | -4.676551835 |
| **A_55_P2084965** | Klk13 | 0.07130163 | -0.064351762 | 0.206955022 | 7.739411784 | 1.256255375 | 0.251352284 | 0.991835066 | -4.676572344 |
| **A_51_P391159** | Ang | -0.062179012 | -0.180483145 | 0.056125121 | 8.198542873 | -1.256183343 | 0.251376802 | 0.991835066 | -4.676630403 |
| **A_55_P2012979** | Itgb2l | -0.053556976 | -0.155473679 | 0.048359728 | 7.672913082 | -1.25597157 | 0.251448894 | 0.991835066 | -4.676801085 |
| **A_55_P2039454** | Gm5088 | 0.135316186 | -0.122185107 | 0.392817479 | 8.767912007 | 1.255970577 | 0.251449232 | 0.991835066 | -4.676801885 |
| **A_55_P2068233** | Phactr1 | 0.066939881 | -0.060448596 | 0.194328359 | 8.634862659 | 1.255925636 | 0.251464534 | 0.991835066 | -4.676838105 |
| **A_55_P2129217** | NA | -0.149757356 | -0.434767663 | 0.135252951 | 14.94483692 | -1.255846997 | 0.251491311 | 0.991835066 | -4.676901481 |
| **A_55_P2087622** | Il4i1 | -0.060119055 | -0.17453961 | 0.054301501 | 8.068569354 | -1.255790593 | 0.251510518 | 0.991835066 | -4.676946936 |
| **A_55_P2277825** | E430010N( | 0.059556616 | -0.053798304 | 0.172911535 | 9.34258126 | 1.255737225 | 0.251528692 | 0.991835066 | -4.676989943 |
| **A_52_P437850** | Fam13c | 0.285342299 | -0.258437675 | 0.829122273 | 9.141361682 | 1.254157498 | 0.25206718 | 0.991835066 | -4.678262583 |
| **A_52_P467140** | Snd1 | 0.098564881 | -0.089272527 | 0.286402288 | 10.20920062 | 1.25414932 | 0.252069971 | 0.991835066 | -4.67826917 |
| **A_55_P2166543** | Pam16 | -0.069330081 | -0.20145527 | 0.062795108 | 13.86110459 | -1.254137688 | 0.252073939 | 0.991835066 | -4.678278537 |
| **A_55_P2073329** | NA | -0.084735204 | -0.246219764 | 0.076749355 | 10.83395542 | -1.254128421 | 0.252077101 | 0.991835066 | -4.678286 |
| **A_55_P2078710** | Morc4 | 0.140033632 | -0.126870488 | 0.406937752 | 10.57996163 | 1.253967292 | 0.252132084 | 0.991835066 | -4.678415761 |
| **A_52_P49406** | Pde12 | 0.095232546 | -0.086287712 | 0.276752804 | 8.975874666 | 1.253918841 | 0.252148619 | 0.991835066 | -4.678454778 |
| **A_55_P2208071** | Nbeal1 | -0.108797313 | -0.316183043 | 0.098588417 | 9.585466202 | -1.253858221 | 0.252169308 | 0.991835066 | -4.678503593 |
| **A_55_P2016316** | Xrn2 | 0.114456063 | -0.103725907 | 0.332638033 | 10.24250631 | 1.253802341 | 0.252188381 | 0.991835066 | -4.67854859 |
| **A_51_P266143** | Adpgk | 0.10418001 | -0.094447856 | 0.302807876 | 8.832345604 | 1.253583742 | 0.252263005 | 0.991835066 | -4.678724608 |
| **A_52_P486279** | Popdc3 | -0.14635648 | -0.425435292 | 0.132722333 | 9.683134427 | -1.253413077 | 0.252321279 | 0.991835066 | -4.678862018 |
| **A_55_P2098911** | Lrrcc1 | 0.099719249 | -0.090443943 | 0.28988244 | 9.194621316 | 1.253319144 | 0.252353357 | 0.991835066 | -4.678937643 |
| **A_51_P167426** | Herc1 | 0.059876312 | -0.054341969 | 0.174094592 | 8.044626533 | 1.252935038 | 0.252484567 | 0.991835066 | -4.679246858 |
| **A_55_P1983162** | Pstpip2 | -0.144127862 | -0.419083445 | 0.130827721 | 8.078998011 | -1.252836895 | 0.252518102 | 0.991835066 | -4.679325858 |
| **A_52_P432780** | Mrpl50 | -0.132139704 | -0.38427638 | 0.119996971 | 9.197157625 | -1.25258279 | 0.252604946 | 0.991835066 | -4.679530384 |
| **A_55_P2140955** | NA | 0.060453648 | -0.05490493 | 0.175812227 | 15.94778727 | 1.252511586 | 0.252629286 | 0.991835066 | -4.679587692 |
| **A_55_P2070039** | Zfyve9 | 0.079505903 | -0.072235192 | 0.231246997 | 8.506893604 | 1.252291094 | 0.25270467 | 0.991835066 | -4.679765142 |
| **A_55_P2086949** | Cntf | -0.06060452 | -0.176278372 | 0.055069332 | 8.503705327 | -1.252215136 | 0.252730643 | 0.991835066 | -4.679826269 |
| **A_55_P2081168** | Rpl7 | -0.073066235 | -0.212530374 | 0.066397905 | 16.34872325 | -1.25217004 | 0.252746065 | 0.991835066 | -4.679862559 |
| **A_55_P2015666** | NA | 0.066086406 | -0.060078433 | 0.192251244 | 16.66194715 | 1.251938307 | 0.252825324 | 0.991835066 | -4.680049029 |
| **A_55_P2015700** | NA | -0.073031023 | -0.212469936 | 0.066407889 | 10.46433136 | -1.251793037 | 0.252875021 | 0.991835066 | -4.680165917 |
| **A_55_P1990457** | Rpl15 | 0.054935099 | -0.049964986 | 0.159835185 | 15.63545861 | 1.251650869 | 0.252923665 | 0.991835066 | -4.680280301 |
| **A_55_P2023181** | Lsm2 | 0.087572166 | -0.079668246 | 0.254812578 | 12.82166689 | 1.251509056 | 0.252972196 | 0.991835066 | -4.680394393 |
| **A_55_P2090324** | Zfp956 | -0.084610599 | -0.246209909 | 0.076988711 | 9.043958373 | -1.251394957 | 0.253011248 | 0.991835066 | -4.680486185 |
| **A_55_P2095688** | Sh3bp5l | -0.079008893 | -0.229919553 | 0.071901767 | 11.84958525 | -1.251310794 | 0.253040057 | 0.991835066 | -4.680553889 |
| **A_52_P96782** | Wasl | 0.088770377 | -0.080789127 | 0.258329881 | 9.780139877 | 1.251281604 | 0.25305005 | 0.991835066 | -4.680577371 |
| **A_55_P2031786** | Fcsk | 0.103856613 | -0.094524613 | 0.302237839 | 10.57929676 | 1.25124604 | 0.253062225 | 0.991835066 | -4.68060598 |
| **A_55_P2179296** | Mad2l1 | -0.136755259 | -0.397992661 | 0.124482142 | 8.048623483 | -1.2511743 | 0.253086786 | 0.991835066 | -4.680663688 |
| **A_55_P1991920** | Dhx32 | -0.076484171 | -0.22259587 | 0.069627529 | 10.61338615 | -1.251110593 | 0.253108599 | 0.991835066 | -4.680714933 |
| **A_51_P311038** | Adamts15 | -0.075491964 | -0.219755157 | 0.068771229 | 9.744154461 | -1.250703363 | 0.253248069 | 0.991835066 | -4.681042473 |
| **A_55_P2086198** | NA | -0.084192468 | -0.245088796 | 0.076703859 | 9.719337064 | -1.250651317 | 0.253265898 | 0.991835066 | -4.681084329 |
| **A_55_P1998651** | Entpd5 | -0.099284933 | -0.289025618 | 0.090455752 | 11.83948585 | -1.250639141 | 0.25327007 | 0.991835066 | -4.681094122 |

| | | | | | | | | | |
|---|---|---|---|---|---|---|---|---|---|
| **A_55_P2078238** | NA | 0.07603388 | -0.069283787 | 0.221351546 | 8.382838508 | 1.250540808 | 0.25330376 | 0.991835066 | -4.681173202 |
| **A_51_P227392** | Rhou | 0.194046319 | -0.176850905 | 0.564943543 | 10.75880428 | 1.250434531 | 0.253340175 | 0.991835066 | -4.681258667 |
| **A_55_P2016478** | Ddx39b | 0.131211224 | -0.119687594 | 0.382110042 | 9.402107981 | 1.249917948 | 0.253517247 | 0.991835066 | -4.681674039 |
| **A_51_P323435** | Gtf2h2 | -0.053933295 | -0.157069106 | 0.049202516 | 8.592433137 | -1.249846273 | 0.253541824 | 0.991835066 | -4.681731664 |
| **A_55_P1962736** | Sec23b | 0.065002898 | -0.059313875 | 0.189319672 | 11.17595495 | 1.249718288 | 0.253585714 | 0.991835066 | -4.681834557 |
| **A_55_P2047275** | Tspan7 | -0.08954039 | -0.260792259 | 0.08171148 | 12.49105876 | -1.249662657 | 0.253604794 | 0.991835066 | -4.68187928 |
| **A_51_P383399** | Aldh1a7 | 0.062288093 | -0.056856085 | 0.18143227 | 7.943719035 | 1.249514615 | 0.253655574 | 0.991835066 | -4.681998289 |
| **A_51_P238383** | B4galt4 | -0.076214368 | -0.222015975 | 0.069587238 | 10.20717343 | -1.249348726 | 0.253712486 | 0.991835066 | -4.682131636 |
| **A_52_P93837** | Mme | 0.105646507 | -0.096473852 | 0.307766867 | 7.841022972 | 1.249263966 | 0.253741569 | 0.991835066 | -4.682199765 |
| **A_55_P2108483** | NA | -0.099493647 | -0.289937747 | 0.090950454 | 8.68060901 | -1.248639181 | 0.253956036 | 0.991835066 | -4.682701893 |
| **A_66_P113884** | NA | -0.067274937 | -0.196056139 | 0.061506264 | 7.997270669 | -1.248561567 | 0.253982689 | 0.991835066 | -4.68276426 |
| **A_55_P2119257** | Serpine1 | 0.29615595 | -0.271005382 | 0.863317283 | 9.601481567 | 1.248024126 | 0.254167315 | 0.991835066 | -4.683196076 |
| **A_55_P1995992** | NA | 0.099772575 | -0.091312501 | 0.290857651 | 9.297706318 | 1.247939538 | 0.254196384 | 0.991835066 | -4.683264031 |
| **A_55_P2135271** | Zfp358 | -0.079328128 | -0.231259881 | 0.072603626 | 11.51254278 | -1.24792299 | 0.254202071 | 0.991835066 | -4.683277325 |
| **A_52_P196105** | Ttyh3 | 0.097937441 | -0.089650566 | 0.285525448 | 11.86489684 | 1.247822509 | 0.254236606 | 0.991835066 | -4.683358044 |
| **A_51_P190437** | Ccdc181 | 0.086956389 | -0.079601162 | 0.25351394 | 10.47877381 | 1.247803802 | 0.254243036 | 0.991835066 | -4.683373072 |
| **A_55_P2146136** | Gla | 0.209161917 | -0.19154601 | 0.609869844 | 15.22195034 | 1.247566807 | 0.254324508 | 0.991835066 | -4.683563443 |
| **A_55_P1962586** | NA | -0.066661763 | -0.194384745 | 0.061061219 | 8.47316584 | -1.247431986 | 0.254370865 | 0.991835066 | -4.683671732 |
| **A_52_P402705** | Kcnj11 | -0.119734657 | -0.349188473 | 0.10971916 | 8.722865679 | -1.247193158 | 0.254453003 | 0.991835066 | -4.683863546 |
| **A_52_P370935** | Glcci1 | 0.083802232 | -0.076805088 | 0.244409552 | 10.07083463 | 1.247094555 | 0.254486921 | 0.991835066 | -4.683942734 |
| **A_55_P1957228** | Sgk3 | -0.075812332 | -0.221137254 | 0.06951259 | 13.11315755 | -1.24683473 | 0.254576316 | 0.991835066 | -4.684151383 |
| **A_55_P1971963** | Tmem176b | -0.110042422 | -0.320987324 | 0.10090248 | 13.57859902 | -1.246809895 | 0.254584862 | 0.991835066 | -4.684171325 |
| **A_52_P331762** | Lmo1 | 0.064365855 | -0.059039258 | 0.187770969 | 8.154231435 | 1.246612628 | 0.254652754 | 0.991835066 | -4.684329721 |
| **A_55_P2003638** | Stxbp6 | 0.130151317 | -0.11942479 | 0.379727423 | 9.289900755 | 1.246392116 | 0.254728664 | 0.991835066 | -4.684506767 |
| **A_51_P424854** | Ahsg | -0.616562339 | -1.798957633 | 0.565832956 | 11.48076881 | -1.246302405 | 0.254759552 | 0.991835066 | -4.684578791 |
| **A_55_P2135035** | NA | 0.130256382 | -0.119560811 | 0.380073574 | 8.623319526 | 1.246194467 | 0.25479672 | 0.991835066 | -4.684665443 |
| **A_55_P2002572** | Ephx2 | -0.098236906 | -0.286655456 | 0.090181644 | 11.86579417 | -1.246120818 | 0.254822083 | 0.991835066 | -4.684724567 |
| **A_55_P2054342** | 2810408I1 | 0.163259273 | -0.149892704 | 0.47641125 | 9.752114488 | 1.246039564 | 0.254850068 | 0.991835066 | -4.684789793 |
| **A_55_P2170509** | Yipf4 | -0.064645873 | -0.18865031 | 0.059358565 | 11.28668255 | -1.245984696 | 0.254868967 | 0.991835066 | -4.684833837 |
| **A_55_P1974533** | Eif4e | 0.094663835 | -0.086944789 | 0.276272459 | 9.366233769 | 1.245824189 | 0.254924259 | 0.991835066 | -4.684962675 |
| **A_52_P399054** | Tmed2 | 0.084357041 | -0.077507917 | 0.246221999 | 8.777366143 | 1.245597224 | 0.255002462 | 0.991835066 | -4.685144843 |
| **A_51_P407193** | Clp1 | -0.053520917 | -0.156250609 | 0.049208775 | 8.046195771 | -1.245193065 | 0.25514177 | 0.991835066 | -4.685469191 |
| **A_51_P130544** | Gas5 | 0.10038399 | -0.092306054 | 0.293074035 | 8.343254657 | 1.245128893 | 0.255163896 | 0.991835066 | -4.685520686 |
| **A_51_P382970** | Itga9 | -0.149812666 | -0.437399145 | 0.137773813 | 10.89303993 | -1.245056907 | 0.255188717 | 0.991835066 | -4.68557845 |
| **A_55_P2115732** | Kif1c | -0.103402203 | -0.301931334 | 0.095126927 | 8.735326981 | -1.244843298 | 0.255262383 | 0.991835066 | -4.685749845 |
| **A_51_P105124** | P2ry6 | -0.07328946 | -0.21402492 | 0.067446 | 8.789235935 | -1.244649643 | 0.255329184 | 0.991835066 | -4.682460696 |
| **A_52_P18765** | Hsbp1l1 | 0.076204677 | -0.070138191 | 0.222547546 | 8.400483661 | 1.24456962 | 0.255356792 | 0.991835066 | -4.685969418 |
| **A_55_P1973698** | Htra2 | 0.061363518 | -0.056482832 | 0.179209869 | 10.57149492 | 1.244523922 | 0.255372559 | 0.991835066 | -4.686006079 |
| **A_55_P2052939** | Dyrk2 | 0.053542955 | -0.04933995 | 0.15642586 | 8.787043399 | 1.243850688 | 0.25560494 | 0.991835066 | -4.686546102 |
| **A_55_P2033665** | NA | -0.113110469 | -0.330456435 | 0.104235498 | 8.351004576 | -1.243828056 | 0.255612755 | 0.991835066 | -4.686564254 |
| **A_55_P2064442** | Pabpn1 | 0.070644938 | -0.065145413 | 0.206435288 | 13.0905195 | 1.243429795 | 0.255750312 | 0.991835066 | -4.686883639 |
| **A_52_P88793** | Zfp933 | 0.078922509 | -0.072779776 | 0.230624794 | 9.235350426 | 1.243420125 | 0.255753653 | 0.991835066 | -4.686891393 |
| **A_51_P337125** | Inpp5d | -0.071371697 | -0.208564149 | 0.065820756 | 9.091092161 | -1.243383022 | 0.255766472 | 0.991835066 | -4.686921145 |
| **A_55_P2120433** | Wdr12 | 0.088471977 | -0.081606444 | 0.258550399 | 8.706845803 | 1.243270561 | 0.255805329 | 0.991835066 | -4.687011322 |
| **A_51_P239673** | Hprt | 0.111246617 | -0.102655458 | 0.325148692 | 13.25412976 | 1.243028103 | 0.255889119 | 0.991835066 | -4.687205724 |
| **A_55_P1990127** | Dolpp1 | 0.102534709 | -0.094621705 | 0.299691124 | 10.76946034 | 1.242994265 | 0.255900815 | 0.991835066 | -4.687232853 |
| **A_51_P160673** | Kcne1l | 0.14802426 | -0.13669544 | 0.43274396 | 9.725919374 | 1.242580443 | 0.256043886 | 0.991835066 | -4.687564604 |
| **A_55_P1955672** | NA | 0.071032879 | -0.065608922 | 0.20767468 | 15.07193055 | 1.242467319 | 0.256083009 | 0.991835066 | -4.687655284 |

| | | | | | | | | | |
|---|---|---|---|---|---|---|---|---|---|
| **A_55_P2068125** | Myct1 | -0.152256052 | -0.445157737 | 0.140645633 | 8.594570151 | -1.242401098 | 0.256105913 | 0.991835066 | -4.687708364 |
| **A_55_P2055284** | Chrm2 | -0.090463109 | -0.264492097 | 0.083565879 | 8.845041297 | -1.242393149 | 0.256108663 | 0.991835066 | -4.687714735 |
| **A_55_P1975420** | Dhx40 | -0.07084064 | -0.207125936 | 0.065444656 | 10.87274201 | -1.242346112 | 0.256124933 | 0.991835066 | -4.687752437 |
| **A_66_P138567** | NA | 0.087157658 | -0.080542258 | 0.254857574 | 10.11086835 | 1.242172299 | 0.256185062 | 0.991835066 | -4.687891748 |
| **A_52_P256279** | Zbtb11 | 0.094303739 | -0.087149366 | 0.275756845 | 8.808287339 | 1.242148848 | 0.256193176 | 0.991835066 | -4.687910544 |
| **A_55_P2168761** | Sf1 | 0.114940125 | -0.106222826 | 0.336103075 | 12.71750301 | 1.242133917 | 0.256198342 | 0.991835066 | -4.68792251 |
| **A_51_P265406** | Ric8a | 0.056522141 | -0.052237439 | 0.165281721 | 11.37513475 | 1.242110454 | 0.25620646 | 0.991835066 | -4.687941315 |
| **A_52_P484118** | Urb2 | 0.091622439 | -0.084696992 | 0.267941871 | 8.660057553 | 1.241969143 | 0.256255358 | 0.991835066 | -4.688054566 |
| **A_55_P2059957** | Snx7 | -0.07448104 | -0.217822505 | 0.068860424 | 8.915663952 | -1.241889711 | 0.256282848 | 0.991835066 | -4.688118223 |
| **A_51_P350817** | Cnn1 | 0.232953599 | -0.215433011 | 0.681340209 | 13.66397385 | 1.241726421 | 0.256339367 | 0.991835066 | -4.688249076 |
| **A_52_P707503** | Tmem212 | -0.050408738 | -0.147440284 | 0.046622809 | 7.655746394 | -1.241658024 | 0.256363044 | 0.991835066 | -4.688303884 |
| **A_55_P2230638** | Ece1 | 0.07061565 | -0.065347481 | 0.206578781 | 9.149616148 | 1.241334821 | 0.256474953 | 0.991835066 | -4.688562851 |
| **A_51_P482410** | Ceacam15 | -0.057574518 | -0.168456059 | 0.053307024 | 7.765401911 | -1.241024017 | 0.256582608 | 0.991835066 | -4.688811851 |
| **A_55_P2110081** | Cisd3 | -0.116205007 | -0.340102838 | 0.107692824 | 12.95087303 | -1.240463776 | 0.256776761 | 0.991835066 | -4.689260609 |
| **A_55_P2056639** | Bcar1 | -0.072569542 | -0.212394561 | 0.067255478 | 11.59578647 | -1.240448176 | 0.256782169 | 0.991835066 | -4.689273102 |
| **A_55_P1971614** | NA | -0.178505524 | -0.522453843 | 0.165442794 | 10.08862445 | -1.240416696 | 0.256793083 | 0.991835066 | -4.689298315 |
| **A_52_P478745** | Paxbp1 | 0.13203853 | -0.122439604 | 0.386516664 | 8.660421165 | 1.24010752 | 0.256900288 | 0.991835066 | -4.689545918 |
| **A_55_P1973548** | NA | 0.064324892 | -0.059649045 | 0.188298829 | 14.85910914 | 1.240103137 | 0.256901808 | 0.991835066 | -4.689549428 |
| **A_55_P1997936** | Hsd17b7 | -0.069770242 | -0.204258538 | 0.064718055 | 9.055254183 | -1.239923435 | 0.256964137 | 0.991835066 | -4.689693328 |
| **A_55_P1966644** | Trpc6 | -0.052651665 | -0.154160434 | 0.048857104 | 7.816490977 | -1.239703085 | 0.257040582 | 0.991835066 | -4.689869762 |
| **A_52_P300730** | Hmga2 | 0.237935355 | -0.220805267 | 0.696675977 | 9.038689587 | 1.239655216 | 0.257057192 | 0.991835066 | -4.689908089 |
| **A_51_P104418** | Dusp10 | -0.064561359 | -0.189088082 | 0.059965364 | 8.71742169 | -1.239136742 | 0.257237151 | 0.991835066 | -4.690323161 |
| **A_55_P1967880** | Chtf8 | 0.223822038 | -0.207895698 | 0.655539775 | 13.84552772 | 1.23911635 | 0.257244232 | 0.991835066 | -4.690339484 |
| **A_55_P2032128** | Dnase1l1 | -0.080132375 | -0.238146147 | 0.077881396 | 8.356512482 | -1.259737994 | 0.257294137 | 0.991835066 | -4.633662729 |
| **A_51_P225832** | 2700097O( | 0.128838366 | -0.11973609 | 0.377412822 | 10.07779272 | 1.238790431 | 0.257357414 | 0.991835066 | -4.690600355 |
| **A_52_P190405** | Ifnar2 | -0.071203449 | -0.208594953 | 0.066188054 | 10.92394668 | -1.238654801 | 0.257404527 | 0.991835066 | -4.690708906 |
| **A_51_P436817** | Cbarp | 0.105739428 | -0.09833719 | 0.309816047 | 8.792410615 | 1.238376897 | 0.257501084 | 0.991835066 | -4.690931305 |
| **A_55_P2114148** | NA | 0.098829264 | -0.09195923 | 0.289617758 | 9.990911253 | 1.238062344 | 0.257610413 | 0.991835066 | -4.691183004 |
| **A_55_P2045203** | Chp1 | -0.080490713 | -0.235897879 | 0.074916454 | 12.08481164 | -1.237895152 | 0.257668539 | 0.991835066 | -4.691316773 |
| **A_52_P357133** | Selenom | -0.090443551 | -0.265104614 | 0.084217512 | 12.08893247 | -1.237629454 | 0.257760936 | 0.991835066 | -4.691529339 |
| **A_55_P1954643** | 2310081J2 | 0.058614678 | -0.054593677 | 0.171823034 | 8.166117158 | 1.237476717 | 0.257814063 | 0.991835066 | -4.691651522 |
| **A_52_P532227** | S1pr1 | 0.130303413 | -0.121365452 | 0.381972277 | 11.28960887 | 1.237472148 | 0.257815653 | 0.991835066 | -4.691655177 |
| **A_55_P2186648** | Bard1 | 0.086027037 | -0.080148827 | 0.252202901 | 9.774305437 | 1.237303244 | 0.257874415 | 0.991835066 | -4.691790284 |
| **A_51_P285770** | Fem1a | -0.106125554 | -0.311187825 | 0.098936716 | 13.46937782 | -1.236924929 | 0.258006073 | 0.991835066 | -4.692092865 |
| **A_51_P224771** | Tmem184b | -0.053977508 | -0.158284657 | 0.050329641 | 12.57031075 | -1.236823951 | 0.258041224 | 0.991835066 | -4.69217362 |
| **A_51_P474468** | Utp11 | 0.088343604 | -0.082381735 | 0.259068943 | 12.169949 | 1.236762365 | 0.258062665 | 0.991835066 | -4.692222871 |
| **A_51_P142989** | Zfp429 | -0.077490701 | -0.227264737 | 0.072283334 | 8.707655902 | -1.23657993 | 0.258126187 | 0.991835066 | -4.692368758 |
| **A_55_P2067583** | Cd68 | -0.052131359 | -0.152892042 | 0.048629324 | 7.721307507 | -1.236565358 | 0.258131262 | 0.991835066 | -4.692380411 |
| **A_51_P270891** | 1700056E2 | -0.079131071 | -0.232086281 | 0.073824139 | 9.511695731 | -1.236493682 | 0.258156222 | 0.991835066 | -4.692437724 |
| **A_51_P487933** | Psmd7 | -0.06333745 | -0.185769907 | 0.059095007 | 12.33434912 | -1.236440278 | 0.258174822 | 0.991835066 | -4.692480426 |
| **A_55_P1966444** | NA | -0.061986721 | -0.181840063 | 0.057866621 | 15.58224582 | -1.236111504 | 0.25828935 | 0.991835066 | -4.692743293 |
| **A_55_P1986526** | Cdh26 | 0.112433232 | -0.105003754 | 0.329870219 | 8.570214241 | 1.235863222 | 0.258375868 | 0.991835066 | -4.69294178 |
| **A_55_P2274592** | Wnt7b | 0.066590368 | -0.062193972 | 0.195374708 | 7.718058679 | 1.235826471 | 0.258388676 | 0.991835066 | -4.692971158 |
| **A_51_P383629** | Vps4a | -0.066520796 | -0.195193773 | 0.062152182 | 8.691161467 | -1.235603751 | 0.258466311 | 0.991835066 | -4.693149189 |
| **A_51_P184806** | Elmod2 | 0.078227027 | -0.073105362 | 0.229559415 | 9.357473621 | 1.235475307 | 0.258511093 | 0.991835066 | -4.693251853 |
| **A_55_P1960451** | Thsd7a | -0.066344641 | -0.194695122 | 0.062005841 | 8.68931725 | -1.235428103 | 0.258527552 | 0.991835066 | -4.693289582 |
| **A_55_P1982384** | NA | -0.072914724 | -0.213975832 | 0.068146384 | 11.67549363 | -1.23542697 | 0.258527947 | 0.991835066 | -4.693290488 |
| **A_51_P512992** | Entpd3 | -0.213469906 | -0.626483083 | 0.199543271 | 7.765079707 | -1.235326857 | 0.258562858 | 0.991835066 | -4.693370502 |

| | | | | | | | | | |
|---|---|---|---|---|---|---|---|---|---|
| **A_52_P267824** | Ces2g | 0.063791707 | -0.059632415 | 0.187215829 | 8.063346433 | 1.235302462 | 0.258571365 | 0.991835066 | -4.693389998 |
| **A_55_P1964218** | Tprkb | 0.100128193 | -0.093657988 | 0.293914374 | 13.58773217 | 1.234931045 | 0.258700921 | 0.991835066 | -4.693686817 |
| **A_55_P2001238** | Luc7l3 | -0.172185333 | -0.505444661 | 0.161073994 | 12.16370596 | -1.234874931 | 0.2587205 | 0.991835066 | -4.693731657 |
| **A_55_P1990879** | Banp | 0.056935145 | -0.053300747 | 0.167171038 | 8.317779136 | 1.234430213 | 0.258875708 | 0.991835066 | -4.694086987 |
| **A_55_P1955276** | Specc1 | 0.069298005 | -0.064882932 | 0.203478943 | 8.052441572 | 1.234352053 | 0.258902994 | 0.991835066 | -4.694149429 |
| **A_51_P411770** | Nip7 | 0.098090825 | -0.091898686 | 0.288080336 | 11.6938269 | 1.233979358 | 0.25903314 | 0.991835066 | -4.694447153 |
| **A_55_P1980601** | Oscp1 | -0.066576634 | -0.195534204 | 0.062380936 | 9.031174533 | -1.233911828 | 0.259056727 | 0.991835066 | -4.694501093 |
| **A_55_P2171196** | Gm3411 | 0.265348175 | -0.248715553 | 0.779411903 | 9.399815854 | 1.23369634 | 0.259132007 | 0.991835066 | -4.694673207 |
| **A_55_P2106424** | NA | -0.056118732 | -0.164843052 | 0.052605589 | 16.88151081 | -1.233645217 | 0.25914987 | 0.991835066 | -4.694714038 |
| **A_52_P438036** | Lpar4 | 0.083716489 | -0.078620665 | 0.246053643 | 10.0014815 | 1.232543374 | 0.25953511 | 0.991835066 | -4.695593839 |
| **A_55_P2076772** | Hspa5 | 0.08244892 | -0.077455179 | 0.242353019 | 14.65620966 | 1.232351244 | 0.259602335 | 0.991835066 | -4.69574721 |
| **A_51_P390967** | Slc43a3 | -0.131154647 | -0.385528612 | 0.123219317 | 10.87059617 | -1.232310526 | 0.259616584 | 0.991835066 | -4.695779713 |
| **A_55_P2059085** | Zfp457 | 0.068687071 | -0.064551394 | 0.201925535 | 9.314843188 | 1.23212426 | 0.259681774 | 0.991835066 | -4.695928388 |
| **A_51_P502872** | 2200002D( | -0.158227959 | -0.46521089 | 0.148754973 | 9.352699095 | -1.231907583 | 0.259757626 | 0.991835066 | -4.696101323 |
| **A_55_P2034740** | Exosc2 | 0.087693832 | -0.082445836 | 0.257833499 | 17.66781438 | 1.231891894 | 0.259763119 | 0.991835066 | -4.696113844 |
| **A_55_P1959218** | Usp51 | 0.069461072 | -0.065322472 | 0.204244615 | 8.274071776 | 1.231724955 | 0.259821573 | 0.991835066 | -4.69624707 |
| **A_51_P215066** | Emc4 | -0.081778259 | -0.240472408 | 0.076915891 | 12.54828478 | -1.231646496 | 0.25984905 | 0.991835066 | -4.696309681 |
| **A_51_P493270** | Mrps5 | -0.128261645 | -0.3771903 | 0.120667011 | 9.944639701 | -1.231490434 | 0.25990371 | 0.991835066 | -4.696434216 |
| **A_52_P504361** | Naglu | -0.106182224 | -0.312266725 | 0.099902277 | 9.276828512 | -1.231446696 | 0.259919031 | 0.991835066 | -4.696469116 |
| **A_55_P2038422** | Adamts17 | 0.123148262 | -0.115883849 | 0.362180373 | 9.572010529 | 1.23134903 | 0.259953245 | 0.991835066 | -4.696547045 |
| **A_51_P172853** | Cd14 | -0.057479465 | -0.169060622 | 0.054101692 | 7.870857518 | -1.231206759 | 0.260003092 | 0.991835066 | -4.69666056 |
| **A_55_P2151951** | NA | 0.053236428 | -0.050145815 | 0.156618672 | 13.56991936 | 1.230756362 | 0.26016095 | 0.991835066 | -4.697019877 |
| **A_51_P278868** | H2-DMb1 | -0.090287993 | -0.265639202 | 0.085063216 | 7.914267995 | -1.230638131 | 0.260202402 | 0.991835066 | -4.697114188 |
| **A_55_P2087567** | Olfr373 | -0.055570609 | -0.163512904 | 0.052371685 | 8.57571119 | -1.230446248 | 0.260269689 | 0.991835066 | -4.69726724 |
| **A_51_P118539** | 2700049A( | 0.084295071 | -0.079474716 | 0.248064857 | 9.967243428 | 1.230205116 | 0.260354266 | 0.991835066 | -4.697459558 |
| **A_55_P2275279** | Mid2 | 0.079884617 | -0.075332863 | 0.235102098 | 8.625665005 | 1.230075197 | 0.260399845 | 0.991835066 | -4.697563167 |
| **A_55_P1970700** | Rpsa | -0.073933353 | -0.217604062 | 0.069737357 | 16.10445743 | -1.229932558 | 0.260449894 | 0.991835066 | -4.697676915 |
| **A_55_P2104583** | C87436 | 0.052062925 | -0.049112259 | 0.15323811 | 9.271971941 | 1.2298827 | 0.26046739 | 0.991835066 | -4.697716673 |
| **A_55_P2052495** | Tmx2 | -0.103796426 | -0.305560642 | 0.097967791 | 12.66159829 | -1.229553373 | 0.260582983 | 0.991835066 | -4.697979265 |
| **A_55_P2052475** | NA | -0.074132181 | -0.218234205 | 0.069969842 | 15.27034503 | -1.229548973 | 0.260584528 | 0.991835066 | -4.697982772 |
| **A_55_P2426549** | NA | 0.145101849 | -0.136988099 | 0.427191797 | 8.616529814 | 1.229403568 | 0.260635579 | 0.991835066 | -4.6980987 |
| **A_55_P2155574** | NA | 0.073768841 | -0.06966826 | 0.217205942 | 13.68530219 | 1.229194456 | 0.260709012 | 0.991835066 | -4.698265408 |
| **A_55_P2177386** | Cradd | -0.104050806 | -0.306390641 | 0.098289028 | 9.751980795 | -1.2290603 | 0.260756132 | 0.991835066 | -4.698372352 |
| **A_55_P2143968** | Gps1 | 0.052076241 | -0.049199294 | 0.153351775 | 12.78062018 | 1.228978294 | 0.260784939 | 0.991835066 | -4.698437721 |
| **A_52_P389347** | Snap29 | 0.079631693 | -0.075243379 | 0.234506765 | 10.85262334 | 1.228891555 | 0.260815412 | 0.991835066 | -4.69850686 |
| **A_55_P2055107** | Agpat1 | 0.147828229 | -0.139704874 | 0.435361332 | 11.62384201 | 1.228792794 | 0.260850112 | 0.991835066 | -4.698585579 |
| **A_55_P2049979** | Dbi | -0.05651347 | -0.166439792 | 0.053412853 | 14.62096376 | -1.228738328 | 0.26086925 | 0.991835066 | -4.69862899 |
| **A_51_P378856** | Pfkp | 0.160302145 | -0.151541281 | 0.47214557 | 11.21688542 | 1.228603853 | 0.260916508 | 0.991835066 | -4.698736168 |
| **A_51_P396708** | Med21 | -0.077065568 | -0.227020822 | 0.072889685 | 10.95493793 | -1.228309562 | 0.261019953 | 0.991835066 | -4.698970698 |
| **A_55_P1979620** | Eif3h | -0.108620169 | -0.320038784 | 0.102798447 | 13.24555672 | -1.227937833 | 0.261150669 | 0.991835066 | -4.699266901 |
| **A_55_P2080592** | Spata31d1 | -0.102257169 | -0.301343056 | 0.096828717 | 8.237405803 | -1.227615676 | 0.261263998 | 0.991835066 | -4.699523566 |
| **A_52_P487615** | Otulinl | -0.059251343 | -0.174618035 | 0.056115349 | 7.703401048 | -1.227515238 | 0.261299339 | 0.991835066 | -4.699603578 |
| **A_55_P2059010** | Rbp1 | 0.113097432 | -0.107120084 | 0.333314947 | 13.68658796 | 1.22746754 | 0.261316124 | 0.991835066 | -4.699641576 |
| **A_55_P1967648** | Flcn | -0.1251925 | -0.368990228 | 0.118605228 | 10.93134812 | -1.227319936 | 0.261368071 | 0.991835066 | -4.699759153 |
| **A_55_P2168781** | NA | 0.130847529 | -0.123993802 | 0.38568886 | 14.17160843 | 1.227170181 | 0.261420785 | 0.991835066 | -4.699878437 |
| **A_52_P203560** | Fzd10 | 0.102220477 | -0.096891403 | 0.301332357 | 8.312877877 | 1.227014977 | 0.261475426 | 0.991835066 | -4.700002054 |
| **A_52_P517063** | Immp2l | -0.083062059 | -0.244861183 | 0.078737066 | 8.475935102 | -1.226974835 | 0.26148956 | 0.991835066 | -4.700034025 |
| **A_51_P245631** | Rftn2 | -0.088166788 | -0.259911272 | 0.083577697 | 9.385155656 | -1.226962677 | 0.261493841 | 0.991835066 | -4.700043708 |

| | | | | | | | | | |
|---|---|---|---|---|---|---|---|---|---|
| **A_55_P2004511** | Cd300lf | 0.130063837 | -0.123357463 | 0.383485138 | 8.624154938 | 1.226655412 | 0.261602052 | 0.991835066 | -4.700288409 |
| **A_55_P1983768** | Birc5 | -0.13482488 | -0.397531292 | 0.127881532 | 10.32752495 | -1.226615655 | 0.261616056 | 0.991835066 | -4.700320068 |
| **A_55_P2121141** | Tbc1d22b | 0.101300869 | -0.09609273 | 0.298694468 | 9.603846392 | 1.226561245 | 0.261635223 | 0.991835066 | -4.700363396 |
| **A_55_P2022251** | Cdkn2d | -0.085232725 | -0.251321407 | 0.080855958 | 13.1826687 | -1.226522345 | 0.261648927 | 0.991835066 | -4.700394371 |
| **A_52_P429534** | Elavl1 | 0.071046028 | -0.067404822 | 0.209496879 | 8.899465236 | 1.226459778 | 0.261670969 | 0.991835066 | -4.700444192 |
| **A_55_P2155996** | NA | 0.063920856 | -0.0606585 | 0.188500211 | 15.12363002 | 1.226325142 | 0.261718407 | 0.991835066 | -4.700551394 |
| **A_51_P155073** | Nudt14 | -0.104354183 | -0.30776387 | 0.099055503 | 8.909787753 | -1.226160622 | 0.261776385 | 0.991835066 | -4.700682384 |
| **A_55_P1952915** | Sh3bgr | -0.082825475 | -0.244355326 | 0.078704376 | 15.14122218 | -1.225519634 | 0.262002377 | 0.991835066 | -4.701192647 |
| **A_51_P157083** | Gas1 | -0.114028509 | -0.336458708 | 0.108401689 | 9.758227875 | -1.225261621 | 0.262093391 | 0.991835066 | -4.701398002 |
| **A_55_P2370394** | Ppp2r5e | 0.059735015 | -0.056802539 | 0.176272569 | 8.595634168 | 1.225101895 | 0.262149748 | 0.991835066 | -4.701525117 |
| **A_55_P1968943** | NA | -0.131186906 | -0.387136551 | 0.12476274 | 15.96704994 | -1.225025382 | 0.262176748 | 0.991835066 | -4.701586006 |
| **A_51_P335480** | 1810055G0 | 0.09051115 | -0.086085126 | 0.267107425 | 9.132574677 | 1.224981909 | 0.26219209 | 0.991835066 | -4.701620601 |
| **A_51_P327904** | Yipf1 | -0.105975885 | -0.312774335 | 0.100822566 | 8.876634565 | -1.224810506 | 0.262252586 | 0.991835066 | -4.701756992 |
| **A_55_P2082778** | NA | -0.131234073 | -0.387344698 | 0.124876552 | 14.16590385 | -1.224695566 | 0.262293162 | 0.991835066 | -4.70184845 |
| **A_55_P2077055** | Itih4 | -0.189571452 | -0.55954651 | 0.180403606 | 8.320690203 | -1.224643334 | 0.262311602 | 0.991835066 | -4.701890009 |
| **A_52_P451834** | Stk35 | 0.115658582 | -0.11008284 | 0.341400004 | 9.674682762 | 1.224547789 | 0.262345336 | 0.991835066 | -4.701966027 |
| **A_55_P2046153** | NA | 0.104669388 | -0.09967602 | 0.309014796 | 14.60792974 | 1.224232582 | 0.262456653 | 0.991835066 | -4.702216796 |
| **A_55_P2118570** | Cd37 | -0.052599737 | -0.155303552 | 0.050104078 | 8.260405966 | -1.22406965 | 0.262514209 | 0.991835066 | -4.702346407 |
| **A_55_P1955227** | Rabac1 | -0.068885192 | -0.203392416 | 0.065622032 | 14.15095434 | -1.224022482 | 0.262530874 | 0.991835066 | -4.702383927 |
| **A_55_P2047325** | Mbtps1 | 0.083893636 | -0.07996417 | 0.247751442 | 12.82019067 | 1.223688885 | 0.262648758 | 0.991835066 | -4.702649267 |
| **A_55_P2008870** | NA | -1.670337967 | -4.932830656 | 1.592154722 | 12.02008426 | -1.223668944 | 0.262655806 | 0.991835066 | -4.702665126 |
| **A_66_P116326** | NA | -0.094039285 | -0.277730074 | 0.089651504 | 9.051549345 | -1.223576338 | 0.262688539 | 0.991835066 | -4.702738776 |
| **A_55_P2131506** | NA | 0.063521417 | -0.060569705 | 0.18761254 | 8.191095811 | 1.223456688 | 0.262730837 | 0.991835066 | -4.702833931 |
| **A_55_P1959800** | NA | -0.067521467 | -0.199430138 | 0.064387204 | 8.906455241 | -1.223425865 | 0.262741734 | 0.991835066 | -4.702858442 |
| **A_52_P574653** | Bid | 0.071004817 | -0.067757093 | 0.209766727 | 8.277510279 | 1.223000614 | 0.262892117 | 0.991835066 | -4.703196591 |
| **A_51_P473219** | Siae | -0.08894349 | -0.262763938 | 0.084876957 | 9.217690365 | -1.222988681 | 0.262896338 | 0.991835066 | -4.703206079 |
| **A_52_P303077** | Rogdi | -0.061120912 | -0.180589963 | 0.058348139 | 8.566896007 | -1.222766541 | 0.262974924 | 0.991835066 | -4.703382692 |
| **A_55_P2022342** | Stim1 | -0.087006061 | -0.257089271 | 0.083077148 | 10.16409935 | -1.222636053 | 0.263021097 | 0.991835066 | -4.703486431 |
| **A_51_P426270** | Mgp | 0.207501748 | -0.198133196 | 0.613136692 | 14.19271668 | 1.222631359 | 0.263022758 | 0.991835066 | -4.703490162 |
| **A_55_P1975210** | Fubp3 | -0.084789228 | -0.250550292 | 0.080971837 | 9.196793261 | -1.222551841 | 0.263050898 | 0.991835066 | -4.703553376 |
| **A_66_P132037** | Jmjd4 | 0.101928073 | -0.09736444 | 0.301220586 | 8.73416617 | 1.222396125 | 0.263106011 | 0.991835066 | -4.703677158 |
| **A_51_P477456** | Cnot10 | 0.054259633 | -0.051837834 | 0.160357099 | 9.45507142 | 1.222308887 | 0.263136892 | 0.991835066 | -4.703746502 |
| **A_51_P114177** | Eci3 | -0.064607729 | -0.190946193 | 0.061730734 | 11.56747764 | -1.222244294 | 0.263159759 | 0.991835066 | -4.696584876 |
| **A_55_P2067086** | Togaram1 | 0.150933067 | -0.144264089 | 0.446130224 | 10.83560369 | 1.222028574 | 0.263236139 | 0.991835066 | -4.7039693 |
| **A_55_P2055763** | Gm5878 | 0.160820372 | -0.153801634 | 0.475442378 | 11.41147232 | 1.221690228 | 0.263355976 | 0.991835066 | -4.704238188 |
| **A_51_P279639** | Col6a2 | -0.144940744 | -0.4285015 | 0.138620011 | 10.82271531 | -1.221668833 | 0.263363555 | 0.991835066 | -4.704255189 |
| **A_55_P2001773** | Copg2 | 0.06381865 | -0.061039172 | 0.188676473 | 10.92360508 | 1.221633665 | 0.263376014 | 0.991835066 | -4.704283135 |
| **A_55_P2013948** | Gsn | -0.071557798 | -0.211562787 | 0.068447191 | 14.39846133 | -1.221581902 | 0.263394353 | 0.991835066 | -4.704324268 |
| **A_51_P314285** | Tmem86a | -0.126443047 | -0.373899883 | 0.121013789 | 9.257214672 | -1.221250144 | 0.263511917 | 0.991835066 | -4.704587871 |
| **A_55_P2071389** | Erh | 0.075491789 | -0.07231193 | 0.223295507 | 13.88285741 | 1.220740873 | 0.263692471 | 0.991835066 | -4.704992446 |
| **A_55_P2093949** | Zfp428 | 0.106670272 | -0.102205886 | 0.315546429 | 8.557070318 | 1.220572735 | 0.263752105 | 0.991835066 | -4.705126 |
| **A_52_P229783** | Ptgr2 | -0.125310848 | -0.370696001 | 0.120074305 | 10.6735742 | -1.220532983 | 0.263766206 | 0.991835066 | -4.705157573 |
| **A_51_P350073** | Ndel1 | 0.069137999 | -0.066289951 | 0.204565949 | 12.08441287 | 1.220162394 | 0.26389769 | 0.991835066 | -4.705451896 |
| **A_51_P437021** | Trak1 | -0.067305752 | -0.199148706 | 0.064537202 | 11.46924554 | -1.220125173 | 0.263910899 | 0.991835066 | -4.705481454 |
| **A_52_P675617** | Agbl3 | 0.098266637 | -0.094252501 | 0.290785775 | 8.623415459 | 1.219947991 | 0.263973786 | 0.991835066 | -4.705622153 |
| **A_51_P193173** | Slc25a25 | 0.087161456 | -0.083635641 | 0.257958554 | 9.238543871 | 1.219700281 | 0.264061726 | 0.991835066 | -4.705818841 |
| **A_55_P2152467** | NA | -0.061606279 | -0.182337533 | 0.059124974 | 8.768950789 | -1.21959154 | 0.264100338 | 0.991835066 | -4.705905178 |
| **A_52_P534583** | Ahsp | 0.312501112 | -0.299952612 | 0.924954837 | 11.38993089 | 1.219515767 | 0.264127247 | 0.991835066 | -4.705965336 |

| | | | | | | | | | |
|---|---|---|---|---|---|---|---|---|---|
| **A_55_P2348799** | 1700025G( | 0.069272809 | -0.066498026 | 0.205043643 | 9.014083687 | 1.219454054 | 0.264149164 | 0.991835066 | -4.70601433 |
| **A_55_P2068260** | H3c8 | 0.062995844 | -0.060476328 | 0.186468017 | 8.102577174 | 1.219416153 | 0.264162626 | 0.991835066 | -4.70604442 |
| **A_55_P2010758** | Npm3 | 0.060202028 | -0.057808296 | 0.178212352 | 8.157336529 | 1.219271018 | 0.264214179 | 0.991835066 | -4.706159637 |
| **A_51_P213153** | Tmem208 | 0.074790035 | -0.071818042 | 0.221398113 | 12.29045151 | 1.2192562 | 0.264219443 | 0.991835066 | -4.7061714 |
| **A_55_P1961385** | NA | -0.072170916 | -0.213685888 | 0.069344055 | 8.23051154 | -1.218902485 | 0.264345123 | 0.991835066 | -4.706452168 |
| **A_55_P2286503** | A930001C( | 0.083260882 | -0.080016479 | 0.246538243 | 8.470744414 | 1.218776768 | 0.264389805 | 0.991835066 | -4.706551949 |
| **A_55_P2110245** | Fignl1 | 0.16803273 | -0.161487401 | 0.497552861 | 11.75205146 | 1.218768057 | 0.264392901 | 0.991835066 | -4.706558862 |
| **A_55_P1953157** | Mettl18 | 0.075721979 | -0.072812975 | 0.224256934 | 9.449336218 | 1.218435161 | 0.264511249 | 0.991835066 | -4.706823051 |
| **A_55_P2007377** | Elf2 | -0.078331433 | -0.232007842 | 0.075344976 | 10.85512207 | -1.218254434 | 0.264575518 | 0.991835066 | -4.706966462 |
| **A_52_P355084** | Metrnl | -0.107241527 | -0.317667874 | 0.10318482 | 10.76551776 | -1.218069325 | 0.264641359 | 0.991835066 | -4.707113338 |
| **A_52_P648304** | Pigs | 0.061636294 | -0.059305784 | 0.182578371 | 11.46504407 | 1.21805872 | 0.264645132 | 0.991835066 | -4.707121753 |
| **A_66_P119191** | Inpp5e | -0.087959638 | -0.260558585 | 0.084639308 | 9.160420363 | -1.218020023 | 0.264658898 | 0.991835066 | -4.707152455 |
| **A_55_P2013465** | L2hgdh | 0.074478913 | -0.071668779 | 0.220626606 | 8.419148922 | 1.218009009 | 0.264662816 | 0.991835066 | -4.707161193 |
| **A_51_P137611** | Card14 | 0.059961069 | -0.057706604 | 0.177628743 | 7.925440685 | 1.217927216 | 0.264691916 | 0.991835066 | -4.707226088 |
| **A_51_P482029** | Mcoln1 | -0.056372348 | -0.167000136 | 0.05425544 | 11.87967015 | -1.217898306 | 0.264702201 | 0.991835066 | -4.707249024 |
| **A_51_P434171** | Myo19 | 0.139225425 | -0.133999029 | 0.41244988 | 8.775333015 | 1.217890148 | 0.264705104 | 0.991835066 | -4.707255496 |
| **A_55_P2115141** | Lcor | 0.054809211 | -0.052756513 | 0.162374935 | 8.012650803 | 1.217835919 | 0.264724399 | 0.991835066 | -4.707298519 |
| **A_52_P2706** | Mypn | -0.073154184 | -0.216740482 | 0.070432114 | 8.457623726 | -1.217685967 | 0.26477776 | 0.991835066 | -4.707417479 |
| **A_55_P1959748** | Asns | 0.440371843 | -0.424027388 | 1.304771074 | 10.88594586 | 1.217627094 | 0.264798713 | 0.991835066 | -4.707464183 |
| **A_66_P129852** | Fxn | -0.167017981 | -0.49494987 | 0.160913909 | 11.22049217 | -1.217275018 | 0.264924044 | 0.991835066 | -4.707743455 |
| **A_55_P2061452** | NA | 0.063105624 | -0.06082711 | 0.187038357 | 14.95932059 | 1.217001651 | 0.265021392 | 0.991835066 | -4.707960265 |
| **A_55_P2001510** | NA | 0.069044817 | -0.06657503 | 0.204664664 | 17.28237176 | 1.216793732 | 0.265095454 | 0.991835066 | -4.708125151 |
| **A_55_P1977589** | Nfu1 | -0.064697987 | -0.191840058 | 0.062444084 | 12.6181936 | -1.216215732 | 0.265301433 | 0.991835066 | -4.708583443 |
| **A_55_P1954718** | Cyb561 | 0.104747625 | -0.101131632 | 0.310626883 | 9.821302579 | 1.21602002 | 0.265371208 | 0.991835066 | -4.708738595 |
| **A_51_P272123** | Ndufa10 | -0.14996222 | -0.444770803 | 0.144846363 | 13.3808001 | -1.215768453 | 0.265460921 | 0.991835066 | -4.708938008 |
| **A_55_P2091631** | Coa4 | -0.085300109 | -0.252993813 | 0.082393595 | 9.993623578 | -1.215743511 | 0.265469817 | 0.991835066 | -4.708957778 |
| **A_52_P263095** | Ahnak | -0.054842345 | -0.162665412 | 0.052980722 | 13.94101604 | -1.215663754 | 0.265498265 | 0.991835066 | -4.709020995 |
| **A_55_P1952127** | Psma3 | 0.079560997 | -0.076889401 | 0.236011396 | 14.03434287 | 1.21543763 | 0.265578936 | 0.991835066 | -4.709200214 |
| **A_55_P2078335** | Mapk4 | 0.086158075 | -0.083277123 | 0.255593274 | 7.7644116 | 1.215350338 | 0.265610083 | 0.991835066 | -4.709269394 |
| **A_51_P365578** | Ppp2r1a | -0.118604015 | -0.351861623 | 0.114653594 | 10.39742459 | -1.215269791 | 0.265638827 | 0.991835066 | -4.709333226 |
| **A_55_P2181029** | NA | 0.131233824 | -0.126864107 | 0.389331755 | 8.428182632 | 1.215263326 | 0.265641134 | 0.991835066 | -4.70933835 |
| **A_51_P398164** | Klhl2 | 0.123149227 | -0.119085052 | 0.365383506 | 9.254004377 | 1.215080974 | 0.265706217 | 0.991835066 | -4.709482852 |
| **A_66_P122877** | NA | 0.061834022 | -0.059819979 | 0.183488024 | 16.13303288 | 1.21481524 | 0.265801084 | 0.991835066 | -4.709693409 |
| **A_52_P597726** | Ccdc84 | -0.05379123 | -0.159632119 | 0.05204966 | 8.861314396 | -1.214694662 | 0.26584414 | 0.991835066 | -4.709788942 |
| **A_66_P127036** | NA | 0.072554461 | -0.070209384 | 0.215318306 | 15.23776656 | 1.214660795 | 0.265856234 | 0.991835066 | -4.709815774 |
| **A_55_P2039190** | Gfra4 | 0.096934113 | -0.093827107 | 0.287695333 | 8.436477306 | 1.214494859 | 0.265915499 | 0.991835066 | -4.709947234 |
| **A_55_P2001553** | Igdcc4 | 0.15109068 | -0.146258196 | 0.448439557 | 10.05317161 | 1.214452427 | 0.265930655 | 0.991835066 | -4.709980848 |
| **A_51_P272106** | Cirbp | -0.055789133 | -0.165636448 | 0.054058182 | 12.03057526 | -1.21386195 | 0.266141646 | 0.991835066 | -4.710448559 |
| **A_66_P112862** | Gm17821 | 0.20553334 | -0.199256615 | 0.610323296 | 12.30980509 | 1.213561207 | 0.266249163 | 0.991835066 | -4.710686729 |
| **A_55_P2030638** | Dusp23 | 0.083975115 | -0.081414781 | 0.249365012 | 10.04451043 | 1.213530695 | 0.266260073 | 0.991835066 | -4.71071089 |
| **A_51_P443364** | Cbll1 | 0.07401288 | -0.071762805 | 0.219788565 | 10.23656466 | 1.213476436 | 0.266279476 | 0.991835066 | -4.710753857 |
| **A_51_P237383** | Rnase4 | -0.131168632 | -0.389528033 | 0.127190769 | 8.873254039 | -1.213430348 | 0.266295958 | 0.991835066 | -4.710790351 |
| **A_55_P1957593** | Ino80b | -0.169532619 | -0.503543154 | 0.164477915 | 9.771491628 | -1.213115747 | 0.266408485 | 0.991835066 | -4.711039448 |
| **A_51_P487105** | Bud31 | -0.105345584 | -0.312918757 | 0.102227589 | 10.33848951 | -1.21298169 | 0.266456448 | 0.991835066 | -4.711145582 |
| **A_51_P335081** | Gnb1 | -0.084899166 | -0.252198886 | 0.082400554 | 11.48825565 | -1.212878628 | 0.266493326 | 0.991835066 | -4.711227173 |
| **A_51_P341664** | Zfp87 | -0.07679987 | -0.228165405 | 0.074565666 | 8.930157183 | -1.212669942 | 0.266568012 | 0.991835066 | -4.711392372 |
| **A_55_P1974671** | Sft2d1 | -0.057468375 | -0.170750136 | 0.055813386 | 9.044124942 | -1.2124897 | 0.266632533 | 0.991835066 | -4.711535042 |
| **A_55_P2011172** | NA | -0.063378827 | -0.188318114 | 0.06156046 | 15.48672848 | -1.212423386 | 0.266656275 | 0.991835066 | -4.71158753 |

| | | | | | | | | | |
|---|---|---|---|---|---|---|---|---|---|
| **A_52_P201232** | Xpo5 | 0.094182059 | -0.091492206 | 0.279856324 | 9.085680931 | 1.212343253 | 0.266684966 | 0.991835066 | -4.711650953 |
| **A_55_P2127782** | NA | -0.083538885 | -0.248245945 | 0.081168174 | 15.9527284 | -1.212231581 | 0.266724955 | 0.991835066 | -4.711739336 |
| **A_55_P1986552** | Mcam | -0.084989256 | -0.252556679 | 0.082578166 | 13.12066608 | -1.212225941 | 0.266726974 | 0.991835066 | -4.7117438 |
| **A_55_P2129286** | Zc3hc1 | -0.108894707 | -0.323598907 | 0.105809493 | 9.987442398 | -1.212203024 | 0.266735181 | 0.991835066 | -4.711761937 |
| **A_52_P276794** | Tox4 | -0.135129365 | -0.40158502 | 0.13132629 | 9.18812291 | -1.212087372 | 0.266776602 | 0.991835066 | -4.711853464 |
| **A_55_P2067772** | Cplane1 | 0.054017845 | -0.052507683 | 0.160543373 | 8.843351735 | 1.211972295 | 0.266817821 | 0.991835066 | -4.711944531 |
| **A_55_P2037677** | Cttn | -0.066477433 | -0.197583123 | 0.064628257 | 12.24102452 | -1.211886174 | 0.266848673 | 0.991835066 | -4.71201268 |
| **A_51_P252410** | Cope | -0.123563078 | -0.367301367 | 0.120175211 | 12.09416938 | -1.21164137 | 0.266936387 | 0.991835066 | -4.712206386 |
| **A_66_P111572** | Fam122b | -0.117578531 | -0.3495268 | 0.114369738 | 9.827361941 | -1.211563065 | 0.266964448 | 0.991835066 | -4.712268341 |
| **A_52_P222709** | Pigu | -0.099122692 | -0.29469639 | 0.096451007 | 10.78082237 | -1.211355931 | 0.267038691 | 0.991835066 | -4.712432218 |
| **A_55_P2020592** | Trim3 | -0.070123733 | -0.208483929 | 0.068236462 | 8.31212639 | -1.211331452 | 0.267047466 | 0.991835066 | -4.712451584 |
| **A_51_P156243** | Eps15 | -0.078561036 | -0.233573675 | 0.076451602 | 8.671229013 | -1.211293055 | 0.267061231 | 0.991835066 | -4.712481961 |
| **A_55_P2125972** | Gorasp1 | 0.07121646 | -0.069316658 | 0.211749579 | 9.772623653 | 1.211185972 | 0.267099622 | 0.991835066 | -4.712566673 |
| **A_55_P2110507** | NA | 0.054293662 | -0.052848855 | 0.161436179 | 7.969528864 | 1.211145786 | 0.26711403 | 0.991835066 | -4.712598462 |
| **A_51_P312485** | Fpr1 | -0.063863596 | -0.189908429 | 0.062181238 | 7.79495126 | -1.210981332 | 0.267173001 | 0.991835066 | -4.712728549 |
| **A_51_P513062** | B630019K( | 0.082152056 | -0.080021053 | 0.244325164 | 8.476140077 | 1.210733965 | 0.267261725 | 0.991835066 | -4.712924205 |
| **A_55_P2097820** | NA | -0.065330479 | -0.194299735 | 0.063638776 | 12.48174424 | -1.210706247 | 0.267271668 | 0.991835066 | -4.712946128 |
| **A_55_P2046145** | Gm10693 | -0.051168169 | -0.152205132 | 0.049868794 | 7.752352031 | -1.210399477 | 0.267381736 | 0.991835066 | -4.713188737 |
| **A_55_P2006340** | Uba52 | 0.113368479 | -0.110491611 | 0.337228569 | 17.34341014 | 1.21038847 | 0.267385686 | 0.991835066 | -4.713197441 |
| **A_55_P2145804** | Aen | 0.147576096 | -0.143832727 | 0.43898492 | 12.41239484 | 1.210381992 | 0.267388011 | 0.991835066 | -4.713202564 |
| **A_51_P267314** | Tbc1d10c | -0.073309396 | -0.218083809 | 0.071465017 | 8.203177188 | -1.210255206 | 0.267433514 | 0.991835066 | -4.713302822 |
| **A_55_P1984601** | Vapb | 0.116436226 | -0.113514442 | 0.346386894 | 9.995497112 | 1.210215129 | 0.267447899 | 0.991835066 | -4.713334513 |
| **A_55_P1997421** | Csta3 | -0.055713304 | -0.165765519 | 0.054338911 | 7.849255835 | -1.209955122 | 0.267541238 | 0.991835066 | -4.713540098 |
| **A_55_P2130399** | Rilp | -0.153202793 | -0.455835196 | 0.14942961 | 10.96770388 | -1.209930398 | 0.267550115 | 0.991835066 | -4.713559645 |
| **A_55_P2084122** | Dus4l | 0.11364232 | -0.110844806 | 0.338129445 | 9.113272615 | 1.209923142 | 0.267552721 | 0.991835066 | -4.713565382 |
| **A_55_P2020577** | Pcolce | -0.116924836 | -0.347899362 | 0.11404969 | 14.60775461 | -1.209906524 | 0.267558688 | 0.991835066 | -4.713578521 |
| **A_55_P2016237** | Hand2 | 0.069624071 | -0.067930015 | 0.207178156 | 10.77604801 | 1.209748378 | 0.267615477 | 0.991835066 | -4.713703551 |
| **A_66_P107818** | Gdf3 | -0.050329691 | -0.149768214 | 0.049108832 | 7.733503609 | -1.209702919 | 0.267631803 | 0.991835066 | -4.713739489 |
| **A_55_P1971759** | Zfp688 | 0.063189823 | -0.061663414 | 0.18804306 | 9.374211324 | 1.209640906 | 0.267654075 | 0.991835066 | -4.713788513 |
| **A_55_P2010586** | Gm6252 | 0.261894086 | -0.255653541 | 0.779441712 | 11.25313992 | 1.209440486 | 0.267726069 | 0.991835066 | -4.713946946 |
| **A_55_P1996946** | Cdc20 | -0.094475638 | -0.281253784 | 0.092302507 | 12.6923838 | -1.208934882 | 0.267907761 | 0.991835066 | -4.714346564 |
| **A_52_P432153** | Yipf6 | -0.06335919 | -0.188646603 | 0.061928224 | 8.273189098 | -1.208679904 | 0.267999428 | 0.991835066 | -4.71454806 |
| **A_51_P349546** | Cd109 | -0.091905494 | -0.273683153 | 0.089872164 | 8.879218188 | -1.208398292 | 0.268100702 | 0.991835066 | -4.714770577 |
| **A_55_P1967905** | Dcbld1 | 0.069798786 | -0.068259374 | 0.207856947 | 8.848560107 | 1.208356048 | 0.268115897 | 0.991835066 | -4.714803953 |
| **A_55_P2009329** | Gm8709 | -0.278024634 | -0.828008184 | 0.271958915 | 11.47086789 | -1.208210794 | 0.268168149 | 0.991835066 | -4.714918713 |
| **A_55_P2157388** | Skint9 | -0.056159963 | -0.167266715 | 0.054946789 | 8.202463811 | -1.208079437 | 0.268215409 | 0.991835066 | -4.715022488 |
| **A_66_P118696** | Nup107 | 0.07678129 | -0.075124772 | 0.228687353 | 11.86253132 | 1.208062579 | 0.268221474 | 0.991835066 | -4.715035805 |
| **A_52_P527500** | Helb | 0.062216368 | -0.060884486 | 0.185317223 | 9.86712419 | 1.207960473 | 0.268258216 | 0.991835066 | -4.715116466 |
| **A_66_P118772** | Tmem136 | 0.092455395 | -0.090480557 | 0.275391346 | 10.08496371 | 1.207931554 | 0.268268623 | 0.991835066 | -4.71513931 |
| **A_55_P2066299** | Gpr137b | -0.067536796 | -0.201168776 | 0.066095184 | 9.313821108 | -1.207922816 | 0.268271768 | 0.991835066 | -4.715146213 |
| **A_55_P2053551** | Mast4 | 0.099991993 | -0.097924456 | 0.297908442 | 9.88103058 | 1.207514812 | 0.268418632 | 0.991835066 | -4.715468484 |
| **A_55_P2077343** | Ahdc1 | 0.082641165 | -0.080950208 | 0.246232537 | 9.86199956 | 1.207383301 | 0.268465985 | 0.991835066 | -4.715572348 |
| **A_51_P119401** | Tug1 | -0.061219326 | -0.182423993 | 0.05998534 | 9.244163976 | -1.207197533 | 0.268532887 | 0.991835066 | -4.715719053 |
| **A_55_P2176908** | Tdpoz8 | -0.09542406 | -0.284358366 | 0.093510246 | 10.12026196 | -1.207135992 | 0.268555053 | 0.991835066 | -4.71576765 |
| **A_55_P1978136** | Tnip2 | 0.068336324 | -0.066991603 | 0.203664252 | 9.17067546 | 1.206905637 | 0.268638037 | 0.991835066 | -4.715949545 |
| **A_55_P2169135** | NA | -0.238368255 | -0.710450916 | 0.233714406 | 13.47797643 | -1.206811665 | 0.268671896 | 0.991835066 | -4.716023743 |
| **A_55_P2232754** | Smarcal1 | 0.049622359 | -0.048658082 | 0.1479028 | 9.39659046 | 1.206755932 | 0.268691979 | 0.991835066 | -4.716067746 |
| **A_55_P2094686** | Smim10l1 | -0.078997263 | -0.235459571 | 0.077465045 | 9.126581451 | -1.206733703 | 0.26869999 | 0.991835066 | -4.716085297 |

| | | | | | | | | | |
|---|---|---|---|---|---|---|---|---|---|
| **A_55_P2058861** | Mvk | 0.081065595 | -0.079514702 | 0.241645891 | 11.62227502 | 1.206572576 | 0.268758059 | 0.991835066 | -4.716212507 |
| **A_55_P1994492** | Usp31 | 0.079724674 | -0.078204943 | 0.237654291 | 9.13494862 | 1.20653049 | 0.268773229 | 0.991835066 | -4.716245732 |
| **A_55_P2194958** | Nfia | 0.076597304 | -0.075141569 | 0.228336177 | 8.996909769 | 1.20649565 | 0.268785787 | 0.991835066 | -4.716273237 |
| **A_52_P311245** | Scarb2 | -0.106517661 | -0.317556344 | 0.104521022 | 9.29378916 | -1.206337103 | 0.268842942 | 0.991835066 | -4.716398397 |
| **A_51_P409010** | Comp | -0.149852428 | -0.446768706 | 0.147063851 | 8.712177715 | -1.206254391 | 0.268872763 | 0.991835066 | -4.716463687 |
| **A_55_P1975690** | Bicc1 | 0.185576705 | -0.182129514 | 0.553282925 | 12.27044607 | 1.206234141 | 0.268880065 | 0.991835066 | -4.716479672 |
| **A_55_P2107362** | Gjc1 | -0.083650727 | -0.249467881 | 0.082166426 | 10.98434445 | -1.205728145 | 0.269062563 | 0.991835066 | -4.716879038 |
| **A_55_P2051716** | Tbx1 | 0.145210976 | -0.142643401 | 0.433065353 | 8.182049409 | 1.205690221 | 0.269076245 | 0.991835066 | -4.716908966 |
| **A_51_P498212** | Ube2j1 | 0.085297802 | -0.08379545 | 0.254391053 | 8.378681768 | 1.205648461 | 0.269091312 | 0.991835066 | -4.716941922 |
| **A_55_P1976898** | Pnp2 | 0.055744859 | -0.054798744 | 0.166288461 | 7.884470217 | 1.205258873 | 0.26923191 | 0.991835066 | -4.717249339 |
| **A_52_P342896** | Mrps33 | 0.082777938 | -0.081395973 | 0.246951848 | 10.03665778 | 1.205090304 | 0.269292764 | 0.991835066 | -4.717382338 |
| **A_55_P2136413** | NA | -0.077200012 | -0.230312194 | 0.07591217 | 11.0658496 | -1.205082432 | 0.269295606 | 0.991835066 | -4.717388549 |
| **A_55_P2408415** | Kcnq1ot1 | 0.213663882 | -0.210122279 | 0.637450043 | 8.78059798 | 1.205017839 | 0.269318928 | 0.991835066 | -4.717439509 |
| **A_52_P350554** | Kcnb1 | -0.102175556 | -0.304855443 | 0.100504331 | 8.717298092 | -1.204884694 | 0.269367006 | 0.991835066 | -4.717544548 |
| **A_51_P496934** | Hps3 | -0.053495913 | -0.15961549 | 0.052623663 | 8.401603715 | -1.204853469 | 0.269378282 | 0.991835066 | -4.717569181 |
| **A_55_P2123421** | Dnal4 | -0.099928451 | -0.29820007 | 0.098343168 | 8.731646734 | -1.204585793 | 0.269474964 | 0.991835066 | -4.717780331 |
| **A_51_P332652** | Pqlc3 | -0.074070179 | -0.221044626 | 0.072904269 | 8.141418208 | -1.204510771 | 0.269502066 | 0.991835066 | -4.717839506 |
| **A_55_P2194064** | NA | 0.141344889 | -0.139152948 | 0.421842727 | 8.60075031 | 1.204369432 | 0.269553133 | 0.991835066 | -4.717950984 |
| **A_55_P2049597** | Tpp2 | -0.080180666 | -0.239314826 | 0.078953494 | 10.57098741 | -1.204246456 | 0.269597571 | 0.991835066 | -4.718047974 |
| **A_55_P1971897** | Mdk | 0.177148582 | -0.174452269 | 0.528749434 | 11.8721989 | 1.20419521 | 0.269616091 | 0.991835066 | -4.718088389 |
| **A_51_P511236** | Pik3r1 | 0.071909385 | -0.07082548 | 0.214644251 | 8.767240469 | 1.204105773 | 0.269648416 | 0.991835066 | -4.718158922 |
| **A_55_P2068862** | Pigv | -0.06370928 | -0.190211088 | 0.062792529 | 10.14489042 | -1.203691223 | 0.269798287 | 0.991835066 | -4.718485812 |
| **A_55_P1962483** | NA | -0.12675617 | -0.378472644 | 0.124960304 | 11.85009609 | -1.20355683 | 0.269846889 | 0.991835066 | -4.718591774 |
| **A_55_P2031337** | Trmt2a | 0.104900882 | -0.103504223 | 0.313305987 | 10.66008071 | 1.203039593 | 0.270034012 | 0.991835066 | -4.718999529 |
| **A_55_P2067700** | Tas2r113 | -0.141579919 | -0.42293759 | 0.139777752 | 8.274289183 | -1.202685383 | 0.270162219 | 0.991835066 | -4.71927871 |
| **A_51_P110035** | Pik3r4 | 0.070389014 | -0.069494683 | 0.21027271 | 9.722429613 | 1.202671203 | 0.270167353 | 0.991835066 | -4.711908857 |
| **A_52_P385615** | Rnpc3 | 0.103532715 | -0.102228349 | 0.309293778 | 8.879380824 | 1.202606473 | 0.270190788 | 0.991835066 | -4.719340899 |
| **A_55_P2056293** | Sf3b4 | 0.075661362 | -0.074725014 | 0.226047738 | 8.353182922 | 1.2024715 | 0.27023966 | 0.991835066 | -4.719447267 |
| **A_51_P161086** | Ppic | 0.071131151 | -0.070255462 | 0.212517765 | 12.98560862 | 1.202432422 | 0.270253811 | 0.991835066 | -4.719478062 |
| **A_55_P2032714** | Fhit | -0.063413291 | -0.189460183 | 0.0626336 | 8.13670934 | -1.202423031 | 0.270257212 | 0.991835066 | -4.719485462 |
| **A_55_P1995074** | Ubald2 | -0.122970222 | -0.367482536 | 0.121542092 | 14.14863483 | -1.202010753 | 0.270406547 | 0.991835066 | -4.719810318 |
| **A_52_P535255** | Syf2 | -0.16059645 | -0.479996953 | 0.158804054 | 10.19088352 | -1.20173712 | 0.2705057 | 0.991835066 | -4.720025896 |
| **A_51_P262208** | Itgb2 | -0.059852641 | -0.178896439 | 0.059191157 | 8.044812759 | -1.20167127 | 0.270529567 | 0.991835066 | -4.720077771 |
| **A_55_P1977533** | Eno3 | -0.147484551 | -0.440904439 | 0.145935338 | 15.55000025 | -1.20134049 | 0.270649478 | 0.991835066 | -4.720338328 |
| **A_52_P106789** | Rbm18 | -0.082598556 | -0.246928969 | 0.081731858 | 12.03738319 | -1.201333638 | 0.270651963 | 0.991835066 | -4.720343726 |
| **A_55_P2054927** | Fam136a | 0.0586052 | -0.058040914 | 0.175251314 | 12.25833074 | 1.20081197 | 0.270841166 | 0.991835066 | -4.720754568 |
| **A_55_P2041060** | Rbp3 | -0.100358124 | -0.30013036 | 0.099414111 | 10.64180894 | -1.20067796 | 0.270889788 | 0.991835066 | -4.720860092 |
| **A_55_P1980311** | Pcbp2 | 0.099006863 | -0.098085569 | 0.296099295 | 10.15364168 | 1.200616989 | 0.270911913 | 0.991835066 | -4.720908101 |
| **A_55_P2095384** | Glrx2 | -0.0665505 | -0.199048597 | 0.065947596 | 11.77304385 | -1.200468639 | 0.27096575 | 0.991835066 | -4.721024907 |
| **A_55_P2004137** | Gnai2 | -0.130330085 | -0.389927944 | 0.129267774 | 14.11501597 | -1.199921136 | 0.271164522 | 0.991835066 | -4.721455927 |
| **A_55_P2079761** | Ggps1 | 0.074906598 | -0.07431554 | 0.224128736 | 8.969756583 | 1.199764339 | 0.27122147 | 0.991835066 | -4.721579346 |
| **A_51_P465582** | Hdhd3 | 0.107108563 | -0.106279327 | 0.320496454 | 9.041239967 | 1.199674852 | 0.271253976 | 0.991835066 | -4.721649779 |
| **A_51_P170946** | Hnrnpm | 0.102241538 | -0.101453363 | 0.305936438 | 13.7677082 | 1.199654947 | 0.271261207 | 0.991835066 | -4.721665445 |
| **A_52_P351021** | NA | 0.073981209 | -0.073423907 | 0.221386324 | 10.29760717 | 1.199549014 | 0.271299692 | 0.991835066 | -4.721748819 |
| **A_52_P52128** | Leo1 | 0.118073102 | -0.117228085 | 0.353374288 | 10.98105411 | 1.199322514 | 0.271381994 | 0.991835066 | -4.721927069 |
| **A_66_P133202** | Nfyc | 0.089626466 | -0.089033683 | 0.268286616 | 13.2178536 | 1.198995942 | 0.271500696 | 0.991835066 | -4.722184043 |
| **A_51_P251587** | Psmd9 | -0.065671405 | -0.196582049 | 0.065239238 | 9.17361098 | -1.198975983 | 0.271507952 | 0.991835066 | -4.722199747 |
| **A_55_P1999823** | Zmynd19 | 0.09195703 | -0.09138293 | 0.275296991 | 8.485873233 | 1.198772953 | 0.271581773 | 0.991835066 | -4.722359488 |

| | | | | | | | | | |
|---|---|---|---|---|---|---|---|---|---|
| **A_55_P2016838** | NA | 0.051877228 | -0.051557017 | 0.155311473 | 14.48556525 | 1.198730477 | 0.271597219 | 0.991835066 | -4.722392905 |
| **A_55_P2064240** | Hemk1 | -0.060237541 | -0.180343608 | 0.059868525 | 8.552604431 | -1.198702546 | 0.271607377 | 0.991835066 | -4.72241488 |
| **A_55_P1980986** | Lca5l | 0.091888451 | -0.091346822 | 0.275123724 | 9.540059933 | 1.198563323 | 0.271658011 | 0.991835066 | -4.722524407 |
| **A_55_P2153296** | Pycrl | 0.062326034 | -0.061965302 | 0.186617369 | 8.503020683 | 1.198499253 | 0.271681316 | 0.991835066 | -4.722574807 |
| **A_51_P443293** | Decr2 | 0.076565293 | -0.076128169 | 0.229258756 | 10.00326582 | 1.198451988 | 0.271698509 | 0.991835066 | -4.722611988 |
| **A_55_P2039324** | Ccnf | 0.090409105 | -0.089898741 | 0.27071695 | 10.577652 | 1.198413458 | 0.271712525 | 0.991835066 | -4.722642297 |
| **A_52_P340073** | Efnb2 | 0.062598019 | -0.062260302 | 0.18745634 | 10.27265063 | 1.198263231 | 0.27176718 | 0.991835066 | -4.722760464 |
| **A_55_P2012349** | Fermt2 | -0.080900501 | -0.242265571 | 0.08046457 | 12.08807176 | -1.198259305 | 0.271768609 | 0.991835066 | -4.722763552 |
| **A_55_P2129691** | AB124611 | -0.155011559 | -0.464222239 | 0.154199122 | 11.31213673 | -1.198170792 | 0.271800816 | 0.991835066 | -4.722833173 |
| **A_55_P2050652** | Ikbkg | -0.049972823 | -0.1496579 | 0.049712254 | 8.462982836 | -1.198154621 | 0.271806701 | 0.991835066 | -4.722845892 |
| **A_55_P2117843** | Coasy | 0.082855487 | -0.082424602 | 0.248135576 | 11.7512431 | 1.19814635 | 0.27180971 | 0.991835066 | -4.722852397 |
| **A_55_P2152305** | Atp5h | -0.085581721 | -0.25634402 | 0.085180578 | 16.98052377 | -1.197838192 | 0.271921867 | 0.991835066 | -4.723094755 |
| **A_55_P2076866** | Megf6 | 0.173001871 | -0.172239243 | 0.518242986 | 11.25450602 | 1.197670696 | 0.271982846 | 0.991835066 | -4.723226472 |
| **A_52_P233335** | Pphln1 | 0.092765239 | -0.092377431 | 0.27790791 | 10.32213537 | 1.197534069 | 0.272032594 | 0.991835066 | -4.723333906 |
| **A_52_P630632** | Rcbtb2 | 0.060031158 | -0.059788639 | 0.179850955 | 10.49552764 | 1.197449681 | 0.272063326 | 0.991835066 | -4.72340026 |
| **A_51_P431870** | Map1s | 0.224005148 | -0.223104905 | 0.671115201 | 10.54387311 | 1.197437064 | 0.27206792 | 0.991835066 | -4.723410181 |
| **A_55_P2115582** | Slc20a1 | 0.14261342 | -0.142089597 | 0.427316438 | 11.24266439 | 1.197229631 | 0.272143474 | 0.991835066 | -4.723573273 |
| **A_55_P2316335** | Gfm2 | 0.069686726 | -0.069455081 | 0.208828534 | 8.143225627 | 1.197020406 | 0.272219699 | 0.991835066 | -4.723737759 |
| **A_51_P402435** | Trap1 | -0.126169079 | -0.378091375 | 0.125753216 | 11.17494709 | -1.197003613 | 0.272225817 | 0.991835066 | -4.723750961 |
| **A_55_P1971347** | Mta3 | 0.061429627 | -0.061266969 | 0.184126222 | 7.959252021 | 1.196615142 | 0.272367395 | 0.991835066 | -4.72405632 |
| **A_52_P177293** | Snx24 | 0.080610963 | -0.080401673 | 0.241623598 | 8.502243052 | 1.196584247 | 0.272378657 | 0.991835066 | -4.724080603 |
| **A_55_P2008452** | Wtap | 0.107817788 | -0.107543968 | 0.323179543 | 11.22755393 | 1.196550316 | 0.272391026 | 0.991835066 | -4.724107271 |
| **A_52_P635338** | Fes | -0.05894145 | -0.176680026 | 0.058797127 | 10.22255139 | -1.196495764 | 0.272410914 | 0.991835066 | -4.724150147 |
| **A_55_P2054261** | C2cd4b | -0.065360366 | -0.195936728 | 0.065215997 | 8.453263356 | -1.196352158 | 0.272463274 | 0.991835066 | -4.72426301 |
| **A_52_P404461** | Tmem60 | -0.063596784 | -0.190651332 | 0.063457764 | 10.82674331 | -1.196338474 | 0.272468264 | 0.991835066 | -4.724273764 |
| **A_51_P209930** | Rtn2 | -0.195361643 | -0.585674497 | 0.194951212 | 10.20080869 | -1.196287531 | 0.27248684 | 0.991835066 | -4.724313799 |
| **A_55_P2022569** | Pak3 | -0.086467094 | -0.259221506 | 0.086287318 | 9.113946754 | -1.196274508 | 0.272491589 | 0.991835066 | -4.724324034 |
| **A_55_P2016326** | NA | 0.067295895 | -0.067190148 | 0.201781937 | 9.944985914 | 1.195970561 | 0.272602448 | 0.991835066 | -4.724562879 |
| **A_55_P2180846** | Wdsub1 | -0.080716527 | -0.242037075 | 0.080604021 | 8.969061504 | -1.195864323 | 0.272641206 | 0.991835066 | -4.724646354 |
| **A_66_P140067** | Wdyhv1 | 0.091570252 | -0.09146173 | 0.274602234 | 9.19054268 | 1.195739454 | 0.272686766 | 0.991835066 | -4.724744464 |
| **A_55_P2147186** | A430005L1 | 0.072502587 | -0.072419277 | 0.217424452 | 11.88824911 | 1.195717882 | 0.272694637 | 0.991835066 | -4.724761413 |
| **A_66_P109002** | Kcng2 | -0.112202936 | -0.336480013 | 0.112074141 | 9.705070859 | -1.195717171 | 0.272694897 | 0.991835066 | -4.724761971 |
| **A_55_P2247235** | Rab26os | 0.079807174 | -0.0797751 | 0.239389448 | 8.169251573 | 1.195271087 | 0.272857713 | 0.991835066 | -4.725112411 |
| **A_55_P2158351** | Rpl18a | 0.057709903 | -0.057706158 | 0.173125964 | 15.80457448 | 1.195069686 | 0.27293125 | 0.991835066 | -4.725270607 |
| **A_51_P405985** | Fig4 | -0.084403876 | -0.253215471 | 0.084407719 | 10.92440483 | -1.1950037 | 0.272955347 | 0.991835066 | -4.725322434 |
| **A_55_P1958220** | Crebl2 | -0.059014316 | -0.177056406 | 0.059027775 | 7.876260275 | -1.194894653 | 0.272995173 | 0.991835066 | -4.725408079 |
| **A_55_P2001903** | Smim4 | -0.105202529 | -0.315632524 | 0.105227467 | 12.81447185 | -1.194889277 | 0.272997136 | 0.991835066 | -4.725412302 |
| **A_52_P278295** | Pear1 | 0.067389356 | -0.06740919 | 0.202187901 | 9.107178389 | 1.194855069 | 0.27300963 | 0.991835066 | -4.725439168 |
| **A_52_P646755** | Tex30 | 0.07804913 | -0.078077849 | 0.234176108 | 10.49076792 | 1.194811083 | 0.273025697 | 0.991835066 | -4.725473712 |
| **A_55_P2258567** | D930043N1 | 0.156055542 | -0.156127248 | 0.468238332 | 8.787159694 | 1.194756417 | 0.273045666 | 0.991835066 | -4.725516644 |
| **A_51_P391367** | Tfap4 | -0.057781743 | -0.173374723 | 0.057811238 | 8.024154931 | -1.194725978 | 0.273056785 | 0.991835066 | -4.725540548 |
| **A_55_P2161465** | Gm10516 | -0.076164401 | -0.22853254 | 0.076203737 | 8.622259947 | -1.194722389 | 0.273058096 | 0.991835066 | -4.725543367 |
| **A_66_P118592** | Atrx | -0.057456603 | -0.172419134 | 0.057505929 | 9.089635214 | -1.19451816 | 0.273132711 | 0.991835066 | -4.725703744 |
| **A_55_P1968133** | Ttc4 | 0.069805501 | -0.069886067 | 0.209497068 | 11.30544322 | 1.194341682 | 0.273197202 | 0.991835066 | -4.725842317 |
| **A_65_P13713** | Ank3 | -0.051205404 | -0.153689284 | 0.051278477 | 9.416467877 | -1.194178821 | 0.273256728 | 0.991835066 | -4.725970188 |
| **A_51_P181538** | Brwd1 | 0.095005805 | -0.095143752 | 0.285155361 | 8.95768641 | 1.194163949 | 0.273262164 | 0.991835066 | -4.725981864 |
| **A_55_P2204491** | AI131754 | -0.04888648 | -0.14673631 | 0.04896335 | 7.904726992 | -1.194092097 | 0.27328843 | 0.991835066 | -4.726038275 |
| **A_55_P1955706** | Spg7 | -0.063034841 | -0.18920401 | 0.063134327 | 13.63915831 | -1.194088606 | 0.273289706 | 0.991835066 | -4.726041017 |

| | | | | | | | | | |
|---|---|---|---|---|---|---|---|---|---|
| **A_51_P140237** | Fhl2 | -0.226796081 | -0.680783337 | 0.227191175 | 15.92340134 | -1.1939909 | 0.273325427 | 0.991835066 | -4.726117723 |
| **A_51_P452153** | Mzb1 | -0.051335041 | -0.154131891 | 0.05146181 | 7.800579024 | -1.193557195 | 0.273484034 | 0.991835066 | -4.71899961 |
| **A_55_P2181222** | Ifi208 | -0.079423533 | -0.238486381 | 0.079639315 | 8.600081674 | -1.193409751 | 0.273537972 | 0.991835066 | -4.726573899 |
| **A_55_P2134712** | NA | 0.179918088 | -0.180443411 | 0.540279586 | 11.63115581 | 1.193288829 | 0.273582215 | 0.991835066 | -4.726668803 |
| **A_55_P2009507** | Ocrl | 0.103736352 | -0.104045166 | 0.31151787 | 11.26260382 | 1.193254797 | 0.273594667 | 0.991835066 | -4.726695511 |
| **A_55_P2063126** | Patj | 0.087482958 | -0.087745206 | 0.262711122 | 9.433154918 | 1.193242409 | 0.2735992 | 0.991835066 | -4.726705233 |
| **A_55_P2160712** | NA | -0.054736529 | -0.16437925 | 0.054906192 | 8.744770335 | -1.193181688 | 0.27362142 | 0.991835066 | -4.726752886 |
| **A_55_P1952901** | Gm833 | -0.092563442 | -0.27799998 | 0.092873097 | 8.455819447 | -1.193035355 | 0.273674975 | 0.991835066 | -4.72686772 |
| **A_65_P10673** | Tmem44 | 0.068530176 | -0.068784232 | 0.205844584 | 9.686036529 | 1.192819888 | 0.273753847 | 0.991835066 | -4.727036794 |
| **A_52_P244463** | D16Ertd472 | 0.05532017 | -0.055527603 | 0.166167943 | 8.636404014 | 1.1927946 | 0.273763105 | 0.991835066 | -4.727056635 |
| **A_55_P1953630** | E330016L1 | -0.074764719 | -0.224625943 | 0.075096506 | 7.665070948 | -1.19238515 | 0.273913042 | 0.991835066 | -4.727377875 |
| **A_55_P2066269** | Aak1 | 0.076762733 | -0.077119872 | 0.230645339 | 8.630642938 | 1.192257411 | 0.273959833 | 0.991835066 | -4.727478081 |
| **A_51_P510106** | B3gntl1 | -0.075861151 | -0.227940047 | 0.076217745 | 8.002286004 | -1.192228796 | 0.273970316 | 0.991835066 | -4.727500528 |
| **A_51_P369628** | Ptov1 | -0.145056954 | -0.435871405 | 0.145757496 | 12.17038779 | -1.1921522 | 0.273998378 | 0.991835066 | -4.727560612 |
| **A_51_P509140** | U2af1l4 | -0.174961417 | -0.525736864 | 0.175814029 | 10.48118345 | -1.192126205 | 0.274007902 | 0.991835066 | -4.727581002 |
| **A_55_P1984497** | Usp48 | 0.117727098 | -0.118303477 | 0.353757673 | 9.965173402 | 1.192112679 | 0.274012858 | 0.991835066 | -4.727591612 |
| **A_55_P2102050** | Cc2d1b | 0.091440797 | -0.091893965 | 0.274775559 | 11.53336919 | 1.192077017 | 0.274025924 | 0.991835066 | -4.727619584 |
| **A_51_P192751** | Mrpl14 | -0.109690356 | -0.329631297 | 0.110250584 | 14.11606583 | -1.191986952 | 0.274058926 | 0.991835066 | -4.727690229 |
| **A_51_P148060** | Ptgfrn | 0.195013161 | -0.196039236 | 0.586065558 | 9.782994626 | 1.191895287 | 0.274092517 | 0.991835066 | -4.727762124 |
| **A_55_P2124461** | NA | 0.204366714 | -0.205459787 | 0.614193215 | 13.88593529 | 1.191843567 | 0.274111472 | 0.991835066 | -4.727802688 |
| **A_51_P173564** | Ahsa1 | 0.048233904 | -0.048497818 | 0.144965627 | 13.75823693 | 1.191770489 | 0.274138256 | 0.991835066 | -4.727860002 |
| **A_55_P1969845** | Lrpprc | 0.072649 | -0.073053127 | 0.218351127 | 13.82941814 | 1.191716307 | 0.274158116 | 0.991835066 | -4.727902495 |
| **A_55_P2014282** | NA | 0.070019913 | -0.0704095 | 0.210449326 | 16.96338807 | 1.191715583 | 0.274158382 | 0.991835066 | -4.727903063 |
| **A_66_P111285** | Pdp2 | -0.070609966 | -0.212243291 | 0.071023359 | 9.007769285 | -1.191542899 | 0.274221686 | 0.991835066 | -4.728038485 |
| **A_55_P1952779** | Dcp1b | -0.180737963 | -0.54329014 | 0.181814214 | 11.08095507 | -1.191483407 | 0.274243498 | 0.991835066 | -4.728085138 |
| **A_55_P1961009** | Apbb1ip | -0.064203059 | -0.193046611 | 0.064640493 | 8.158765081 | -1.190973679 | 0.274430444 | 0.991835066 | -4.728484804 |
| **A_51_P296249** | Foxs1 | -0.066481752 | -0.199904379 | 0.066940874 | 9.220436012 | -1.190918669 | 0.274450625 | 0.991835066 | -4.728527931 |
| **A_55_P1952676** | Gtf2a2 | -0.060007585 | -0.180441674 | 0.060426503 | 11.92219818 | -1.19087411 | 0.274466974 | 0.991835066 | -4.728562863 |
| **A_55_P1995210** | Gm11849 | 0.072414048 | -0.072926699 | 0.217754794 | 16.31175331 | 1.190815748 | 0.274488388 | 0.991835066 | -4.728608616 |
| **A_55_P1959838** | NA | -0.057059753 | -0.171585143 | 0.057465636 | 14.84917854 | -1.190795663 | 0.274495758 | 0.991835066 | -4.728624361 |
| **A_52_P286141** | Cisd2 | -0.063587222 | -0.191248868 | 0.064074424 | 10.3822205 | -1.190470245 | 0.274615187 | 0.991835066 | -4.728879445 |
| **A_66_P120901** | Atxn10 | -0.056372774 | -0.169567405 | 0.056821858 | 12.22447447 | -1.19028978 | 0.274681437 | 0.991835066 | -4.729020889 |
| **A_52_P522864** | Zfp442 | 0.069860606 | -0.070425213 | 0.210146426 | 11.48112657 | 1.190221277 | 0.274706588 | 0.991835066 | -4.729074577 |
| **A_55_P2044091** | Tdg | 0.256254926 | -0.258340221 | 0.770850072 | 8.94676397 | 1.190188278 | 0.274718705 | 0.991835066 | -4.729100438 |
| **A_55_P2122718** | Gpr19 | 0.092340634 | -0.093115487 | 0.277796756 | 9.631521417 | 1.190037956 | 0.274773906 | 0.991835066 | -4.729218243 |
| **A_55_P1969645** | Sdcbp | -0.060107106 | -0.180859284 | 0.060645072 | 10.79644947 | -1.189706884 | 0.274895514 | 0.991835066 | -4.729477669 |
| **A_52_P179583** | Adnp2 | 0.059013199 | -0.059559223 | 0.177585622 | 8.482840791 | 1.189527804 | 0.274961313 | 0.991835066 | -4.729617979 |
| **A_51_P267740** | Zc3h15 | 0.069492512 | -0.070155014 | 0.209140037 | 9.917354816 | 1.189361555 | 0.275022408 | 0.991835066 | -4.729748226 |
| **A_55_P2060854** | NA | -0.149738274 | -0.450662409 | 0.151185861 | 12.36184119 | -1.18928224 | 0.27505156 | 0.991835066 | -4.72981036 |
| **A_52_P21550** | Gcnt1 | -0.061207874 | -0.184219818 | 0.061804071 | 9.936358469 | -1.189238996 | 0.275067455 | 0.991835066 | -4.729844238 |
| **A_55_P2093483** | NA | 0.061323737 | -0.061945764 | 0.184593239 | 16.15338925 | 1.189000686 | 0.275155065 | 0.991835066 | -4.730030911 |
| **A_55_P2110362** | Zfp637 | -0.116459127 | -0.350564371 | 0.117646118 | 8.990910486 | -1.1889717 | 0.275165723 | 0.991835066 | -4.730053615 |
| **A_66_P122173** | Mrtfb | 0.103411543 | -0.104465782 | 0.311288868 | 11.24865719 | 1.188970369 | 0.275166212 | 0.991835066 | -4.730054658 |
| **A_52_P420308** | NA | 0.112111729 | -0.113358955 | 0.337582413 | 10.63389083 | 1.188420407 | 0.275368491 | 0.991835066 | -4.730485374 |
| **A_52_P674759** | Btbd3 | 0.078067456 | -0.078971462 | 0.235106375 | 9.322239992 | 1.188151627 | 0.275467396 | 0.991835066 | -4.730695837 |
| **A_51_P426276** | Pdk2 | -0.17525165 | -0.527837424 | 0.177334123 | 9.483940089 | -1.187972708 | 0.27553325 | 0.991835066 | -4.730835921 |
| **A_51_P331003** | Sema4g | 0.050469194 | -0.051077646 | 0.152016035 | 7.853231479 | 1.187870478 | 0.275570884 | 0.991835066 | -4.730915956 |
| **A_55_P2032668** | NA | -0.053642817 | -0.161575707 | 0.054290072 | 13.34521348 | -1.187864506 | 0.275573082 | 0.991835066 | -4.730920632 |

| | | | | | | | | | |
|---|---|---|---|---|---|---|---|---|---|
| **A_55_P1954835** | NA | 0.08431339 | -0.085343427 | 0.253970206 | 9.07723755 | 1.187775517 | 0.275605846 | 0.991835066 | -4.730990298 |
| **A_52_P508089** | Psap | -0.239685102 | -0.722076842 | 0.242706638 | 11.47959246 | -1.187545641 | 0.275690494 | 0.991835066 | -4.731170246 |
| **A_51_P221510** | Fam81a | 0.098984268 | -0.100251294 | 0.29821983 | 9.657100836 | 1.187431181 | 0.27573265 | 0.991835066 | -4.731259839 |
| **A_55_P2167512** | NA | 0.093255187 | -0.094462322 | 0.280972696 | 12.11193736 | 1.187346144 | 0.275763973 | 0.991835066 | -4.731326398 |
| **A_55_P2137023** | NA | -0.112704403 | -0.339573606 | 0.1141648 | 8.855364375 | -1.187338284 | 0.275766869 | 0.991835066 | -4.73133255 |
| **A_55_P2031357** | Zfpm2 | -0.104826021 | -0.315840998 | 0.106188957 | 10.23880959 | -1.187312257 | 0.275776456 | 0.991835066 | -4.731352921 |
| **A_55_P1983988** | Enox2 | -0.079308818 | -0.238992638 | 0.080375002 | 8.670898102 | -1.187051874 | 0.27587239 | 0.991835066 | -4.731556705 |
| **A_51_P328133** | Gle1 | -0.074429256 | -0.224305554 | 0.075447041 | 8.313925061 | -1.186915646 | 0.275922593 | 0.991835066 | -4.731663312 |
| **A_51_P330615** | Zfp119b | -0.070042082 | -0.211096686 | 0.071012523 | 9.933041053 | -1.186809215 | 0.27596182 | 0.991835066 | -4.731746596 |
| **A_55_P2131875** | Poldip3 | 0.115760466 | -0.117367002 | 0.348887934 | 12.47672262 | 1.186795664 | 0.275966814 | 0.991835066 | -4.7317572 |
| **A_55_P2169039** | Olfr558 | -0.110222906 | -0.332230136 | 0.111784324 | 8.531668334 | -1.186626032 | 0.276029345 | 0.991835066 | -4.731889931 |
| **A_55_P1972426** | Pdzd11 | -0.086434539 | -0.260557689 | 0.087688611 | 9.871166085 | -1.186424031 | 0.276103825 | 0.991835066 | -4.732047976 |
| **A_55_P2079142** | Fam124b | 0.051867351 | -0.052629115 | 0.156363817 | 8.186078656 | 1.186319311 | 0.276142442 | 0.991835066 | -4.732129902 |
| **A_55_P2018701** | Kxd1 | -0.165218737 | -0.498105225 | 0.167667752 | 10.85109046 | -1.186239173 | 0.276171998 | 0.991835066 | -4.732192595 |
| **A_55_P2103626** | Slc27a4 | -0.056129438 | -0.169237951 | 0.056979076 | 8.087851193 | -1.186054179 | 0.276240235 | 0.991835066 | -4.732337309 |
| **A_51_P341465** | Csf2ra | 0.183678414 | -0.186461748 | 0.553818576 | 11.3889871 | 1.18604466 | 0.276243747 | 0.991835066 | -4.732344755 |
| **A_52_P173442** | Wscd2 | 0.372687957 | -0.378370465 | 1.123746379 | 9.326914474 | 1.1859893 | 0.27626417 | 0.991835066 | -4.732388059 |
| **A_52_P98614** | Slc6a2 | 0.051773296 | -0.052566423 | 0.156113015 | 7.652324457 | 1.185947002 | 0.276279775 | 0.991835066 | -4.732421144 |
| **A_55_P1959393** | Hhat | 0.071320545 | -0.072434278 | 0.215075368 | 9.270726988 | 1.18577246 | 0.276344179 | 0.991835066 | -4.732557664 |
| **A_51_P104077** | Usp13 | -0.082334493 | -0.248300574 | 0.083631587 | 10.08443109 | -1.185691244 | 0.27637415 | 0.991835066 | -4.732621184 |
| **A_55_P2004119** | NA | -0.125525933 | -0.378686809 | 0.127634944 | 14.17136166 | -1.185075445 | 0.276601492 | 0.991835066 | -4.733102731 |
| **A_55_P2029235** | Btn1a1 | -0.120741789 | -0.364256504 | 0.122772925 | 12.44981668 | -1.18506325 | 0.276605995 | 0.991835066 | -4.733112266 |
| **A_52_P598714** | Afg3l1 | 0.059780491 | -0.060842651 | 0.180403633 | 10.018377 | 1.184507927 | 0.276811148 | 0.991835066 | -4.733546402 |
| **A_51_P267239** | Litaf | 0.055740945 | -0.056733728 | 0.168215619 | 12.04136446 | 1.184482703 | 0.276820469 | 0.991835066 | -4.733566119 |
| **A_55_P2027052** | Prdm9 | 0.060600753 | -0.06168161 | 0.182883117 | 7.797992565 | 1.184467999 | 0.276825903 | 0.991835066 | -4.733577612 |
| **A_55_P1966420** | Scrn2 | 0.057578548 | -0.058608828 | 0.173765924 | 10.95739526 | 1.184434084 | 0.276838437 | 0.991835066 | -4.733604122 |
| **A_52_P399717** | Ccdc50 | 0.062832232 | -0.063956668 | 0.189621132 | 8.686810399 | 1.184432692 | 0.276838952 | 0.991835066 | -4.73360521 |
| **A_55_P2259125** | D7Ertd143 | 0.078734213 | -0.080145211 | 0.237613637 | 8.307621561 | 1.184417916 | 0.276844413 | 0.991835066 | -4.73361676 |
| **A_55_P1967855** | Kctd20 | -0.075409883 | -0.227593201 | 0.076773435 | 10.22210482 | -1.184323514 | 0.276879304 | 0.991835066 | -4.733690547 |
| **A_55_P2040996** | Klf7 | 0.069821074 | -0.071092379 | 0.210734527 | 10.70250651 | 1.184249472 | 0.276906672 | 0.991835066 | -4.733748417 |
| **A_55_P1953132** | Camkv | 0.059016614 | -0.060125671 | 0.178158898 | 14.67974837 | 1.183906745 | 0.277033387 | 0.991835066 | -4.734016264 |
| **A_51_P475875** | Lrwd1 | -0.072012156 | -0.217391787 | 0.073367476 | 9.727937467 | -1.183890076 | 0.277039551 | 0.991835066 | -4.734029291 |
| **A_51_P342549** | Hand1 | 0.108562245 | -0.110609367 | 0.327733856 | 8.396218241 | 1.183868993 | 0.277047347 | 0.991835066 | -4.734045766 |
| **A_52_P255841** | Dbr1 | 0.092105318 | -0.093862763 | 0.2780734 | 8.93491819 | 1.183737562 | 0.277095956 | 0.991835066 | -4.734148469 |
| **A_55_P2348815** | Casd1 | -0.067036563 | -0.202407264 | 0.068334138 | 9.911577299 | -1.183576118 | 0.277155673 | 0.991835066 | -4.734274615 |
| **A_51_P436068** | Gpr182 | 0.141969515 | -0.144727393 | 0.428666423 | 10.93193507 | 1.183535316 | 0.277170768 | 0.991835066 | -4.734306496 |
| **A_52_P189132** | Thap1 | 0.081908872 | -0.083523113 | 0.247340856 | 10.35548471 | 1.183370109 | 0.277231892 | 0.991835066 | -4.734435571 |
| **A_55_P2022444** | Pdss2 | -0.068160884 | -0.205835004 | 0.069513236 | 8.985408839 | -1.183292303 | 0.277260683 | 0.991835066 | -4.734496356 |
| **A_55_P1958394** | Slit2 | 0.195583979 | -0.199465892 | 0.59063385 | 10.16101904 | 1.183288069 | 0.277262249 | 0.991835066 | -4.734499664 |
| **A_55_P2137314** | Kctd21 | -0.050985266 | -0.153971579 | 0.052001047 | 8.343026061 | -1.183244008 | 0.277278555 | 0.991835066 | -4.734534086 |
| **A_52_P295067** | Snn | 0.074116628 | -0.075596608 | 0.223829864 | 10.83510969 | 1.183217512 | 0.27728836 | 0.991835066 | -4.734554785 |
| **A_55_P2253284** | NA | 0.050411979 | -0.051451321 | 0.15227528 | 7.714744735 | 1.182837645 | 0.277428972 | 0.991835066 | -4.734851514 |
| **A_55_P2101246** | Aldh6a1 | -0.094898479 | -0.28667707 | 0.096880113 | 8.342407415 | -1.182682739 | 0.27748633 | 0.991835066 | -4.734972503 |
| **A_52_P306537** | Emc1 | 0.051472091 | -0.05254804 | 0.155492222 | 8.323236459 | 1.182669902 | 0.277491083 | 0.991835066 | -4.734982529 |
| **A_51_P301998** | Fmo2 | 0.175444399 | -0.179123319 | 0.530012117 | 8.392301034 | 1.182631514 | 0.277505299 | 0.991835066 | -4.73501251 |
| **A_55_P2140953** | NA | 0.068343871 | -0.069825077 | 0.206512819 | 14.96837289 | 1.182219877 | 0.277657774 | 0.991835066 | -4.735333966 |
| **A_55_P2106956** | Gm5523 | -0.09521162 | -0.287750248 | 0.097327008 | 17.6116026 | -1.181901315 | 0.277775821 | 0.991835066 | -4.735582696 |
| **A_55_P2098071** | Trim25 | -0.100947623 | -0.30983127 | 0.107936025 | 12.09944736 | -1.200490708 | 0.277847053 | 0.991835066 | -4.676611852 |

| | | | | | | | | | |
|---|---|---|---|---|---|---|---|---|---|
| **A_55_P2082186** | NA | -0.056059557 | -0.169448635 | 0.057329521 | 13.76662763 | -1.18164649 | 0.27787028 | 0.991835066 | -4.735781635 |
| **A_55_P2028259** | Rhbdf2 | 0.06076869 | -0.062145371 | 0.183682752 | 9.01787357 | 1.181646142 | 0.277870409 | 0.991835066 | -4.735781906 |
| **A_51_P377760** | Rnasel | -0.16118123 | -0.487221696 | 0.164859236 | 9.364745735 | -1.181549969 | 0.277906066 | 0.991835066 | -4.735856981 |
| **A_52_P252322** | Pdcd11 | 0.07546836 | -0.077203433 | 0.228140154 | 8.430550116 | 1.181449709 | 0.277943242 | 0.991835066 | -4.735935243 |
| **A_66_P113659** | Ripply1 | -0.08510044 | -0.257273478 | 0.087072598 | 7.732561214 | -1.181342409 | 0.277983033 | 0.991835066 | -4.736018996 |
| **A_55_P2011991** | St7 | 0.064252123 | -0.065769634 | 0.19427388 | 10.68128506 | 1.181083456 | 0.278079083 | 0.991835066 | -4.736221105 |
| **A_55_P2161595** | Hpca | 0.064924639 | -0.066471644 | 0.196320922 | 7.717065018 | 1.180961108 | 0.278124474 | 0.991835066 | -4.736316587 |
| **A_52_P456640** | Fgr | -0.047628236 | -0.144025774 | 0.048769303 | 7.782267393 | -1.180885206 | 0.278152636 | 0.991835066 | -4.73637582 |
| **A_55_P2097612** | Zfp131 | 0.105595528 | -0.108203896 | 0.319394952 | 10.33917418 | 1.180451445 | 0.278313624 | 0.991835066 | -4.736714278 |
| **A_52_P37757** | Eps8l1 | 0.056502568 | -0.057964742 | 0.170969878 | 7.781282605 | 1.179765914 | 0.278568213 | 0.991835066 | -4.73724905 |
| **A_55_P2013188** | Dnajb6 | -0.062723626 | -0.189805867 | 0.064358615 | 11.90703397 | -1.179656119 | 0.278609007 | 0.991835066 | -4.737334684 |
| **A_52_P452019** | Usp31 | 0.058756877 | -0.060294558 | 0.177808311 | 8.473727804 | 1.179595755 | 0.278631437 | 0.991835066 | -4.737381762 |
| **A_55_P2009915** | Stxbp3 | 0.061911281 | -0.063534047 | 0.187356609 | 8.218459654 | 1.17957193 | 0.27864029 | 0.991835066 | -4.737400344 |
| **A_52_P24951** | Ado | 0.084878664 | -0.087116811 | 0.256874139 | 10.10202349 | 1.179480181 | 0.278674386 | 0.991835066 | -4.737471896 |
| **A_55_P2093634** | Ddi2 | -0.053752986 | -0.162683827 | 0.055177855 | 9.220884367 | -1.17939931 | 0.278704442 | 0.991835066 | -4.737534963 |
| **A_55_P2192633** | E330017L1 | 0.070216414 | -0.072090766 | 0.212523593 | 9.505136717 | 1.179290952 | 0.278744718 | 0.991835066 | -4.737619461 |
| **A_55_P2027609** | Scgb3a2 | 0.10711092 | -0.110006216 | 0.324228056 | 7.738480824 | 1.179094956 | 0.278817581 | 0.991835066 | -4.737772289 |
| **A_55_P2062652** | Zfp941 | 0.095355526 | -0.097945648 | 0.288656699 | 7.92723066 | 1.179018194 | 0.278846123 | 0.991835066 | -4.737832141 |
| **A_55_P2172956** | Cnih2 | 0.100488255 | -0.103290094 | 0.304266605 | 8.6316404 | 1.178599897 | 0.279001695 | 0.991835066 | -4.738158251 |
| **A_55_P2132865** | Sumo2 | -0.056897808 | -0.172312307 | 0.05851669 | 12.90383471 | -1.178268594 | 0.279124965 | 0.991835066 | -4.738416494 |
| **A_55_P1962011** | Clec1b | 0.222864381 | -0.22923531 | 0.674964071 | 9.232093775 | 1.178190687 | 0.279153958 | 0.991835066 | -4.738477215 |
| **A_55_P1990210** | Scpep1 | -0.102661261 | -0.310944677 | 0.105622154 | 11.62404196 | -1.178042712 | 0.279209036 | 0.991835066 | -4.738592541 |
| **A_51_P401527** | Mrm3 | -0.089641232 | -0.271549195 | 0.09226673 | 9.257851956 | -1.177782894 | 0.279305764 | 0.991835066 | -4.738795014 |
| **A_55_P2073557** | Slc27a5 | -0.084703544 | -0.256627283 | 0.087220196 | 7.835362876 | -1.177537821 | 0.279397029 | 0.991835066 | -4.738985974 |
| **A_51_P337771** | Pcdhb17 | 0.080732694 | -0.083141443 | 0.24460683 | 8.466076093 | 1.17746541 | 0.279423999 | 0.991835066 | -4.739042392 |
| **A_55_P2127959** | Zfp36l3 | -0.106411012 | -0.322445313 | 0.109623289 | 8.442490299 | -1.177261641 | 0.279499908 | 0.991835066 | -4.739201147 |
| **A_55_P2113733** | NA | 0.102757984 | -0.105870564 | 0.311386533 | 10.42598611 | 1.177201949 | 0.279522147 | 0.991835066 | -4.739247648 |
| **A_51_P191463** | H2-M2 | -0.063902257 | -0.193702932 | 0.065898419 | 8.440127968 | -1.176652915 | 0.279726774 | 0.991835066 | -4.728566638 |
| **A_55_P2045642** | Stmn4 | 0.208587864 | -0.215184874 | 0.632360602 | 9.352260424 | 1.176427462 | 0.279810838 | 0.991835066 | -4.739850887 |
| **A_51_P413539** | Nek6 | 0.071767728 | -0.074046066 | 0.217581522 | 10.19797572 | 1.17635856 | 0.279836534 | 0.991835066 | -4.739904544 |
| **A_55_P2015178** | Nap1l1 | 0.074563812 | -0.076971057 | 0.226098682 | 13.14265578 | 1.176046944 | 0.279952768 | 0.991835066 | -4.740147188 |
| **A_51_P143712** | Tdh | 0.052532515 | -0.054244128 | 0.159309159 | 7.726036115 | 1.175874744 | 0.280017017 | 0.991835066 | -4.740281259 |
| **A_55_P1988358** | Lrrc47 | 0.068589766 | -0.070842571 | 0.208022102 | 11.83941161 | 1.17572282 | 0.280073711 | 0.991835066 | -4.740399535 |
| **A_51_P198775** | BC055324 | 0.123479568 | -0.127549113 | 0.374508249 | 10.68543545 | 1.175657693 | 0.280098018 | 0.991835066 | -4.740450235 |
| **A_55_P2111416** | Rcc2 | 0.091393746 | -0.094410131 | 0.277197624 | 9.812285762 | 1.175630485 | 0.280108173 | 0.991835066 | -4.740471415 |
| **A_55_P1999541** | Ccdc42 | 0.057426529 | -0.059353187 | 0.174206245 | 8.274775167 | 1.175315023 | 0.280225939 | 0.991835066 | -4.740716971 |
| **A_55_P1990121** | Aqp5 | 0.063444313 | -0.065582015 | 0.192470641 | 7.753429162 | 1.175231692 | 0.280257054 | 0.991835066 | -4.740781829 |
| **A_55_P2097017** | Eid1 | 0.075340282 | -0.077905366 | 0.228585931 | 13.50230946 | 1.175028022 | 0.280333116 | 0.991835066 | -4.74094034 |
| **A_51_P406365** | Maf1 | -0.09603361 | -0.291385697 | 0.099318477 | 12.3054957 | -1.174936327 | 0.280367366 | 0.991835066 | -4.741011699 |
| **A_55_P2071329** | Rhog | -0.111340573 | -0.337882295 | 0.115201148 | 11.44857729 | -1.174665974 | 0.280468367 | 0.991835066 | -4.741222075 |
| **A_52_P316474** | St3gal4 | -0.112296945 | -0.34080603 | 0.11621214 | 8.789489506 | -1.174555657 | 0.28050959 | 0.991835066 | -4.741307911 |
| **A_55_P2055419** | Arid4a | 0.061141002 | -0.063325753 | 0.185607756 | 8.611766115 | 1.17405466 | 0.280696862 | 0.991835066 | -4.741697673 |
| **A_52_P64687** | Camk2n1 | -0.098967904 | -0.300442117 | 0.102506309 | 8.974800233 | -1.174043087 | 0.280701189 | 0.991835066 | -4.741706676 |
| **A_51_P418809** | Airn | 0.086809649 | -0.089973657 | 0.263592955 | 8.842569655 | 1.17364264 | 0.280850954 | 0.991835066 | -4.742018145 |
| **A_55_P1976734** | Cacna2d2 | 0.283793192 | -0.294264993 | 0.861851377 | 11.58223051 | 1.173382347 | 0.280948338 | 0.991835066 | -4.74222057 |
| **A_51_P410927** | NA | 0.091944216 | -0.095345565 | 0.279233998 | 8.715631989 | 1.173328082 | 0.280968644 | 0.991835066 | -4.742262768 |
| **A_52_P213696** | Fndc5 | -0.132277857 | -0.401795449 | 0.137239736 | 12.91278399 | -1.173030118 | 0.281080163 | 0.991835066 | -4.742494453 |
| **A_55_P2250249** | Taok1 | 0.108498189 | -0.1126042 | 0.329600578 | 13.49781259 | 1.172838427 | 0.281151927 | 0.991835066 | -4.742643487 |

| | | | | | | | | | |
|---|---|---|---|---|---|---|---|---|---|
| **A_55_P2082625** | NA | -0.165892071 | -0.503979019 | 0.172194878 | 11.28592384 | -1.172752468 | 0.281184113 | 0.991835066 | -4.742710314 |
| **A_51_P497113** | Igf2bp1 | 0.085415671 | -0.088714225 | 0.259545568 | 8.817915901 | 1.172393358 | 0.281318608 | 0.991835066 | -4.742989463 |
| **A_55_P2053202** | Itsn2 | -0.106534751 | -0.323745543 | 0.11067604 | 10.18461424 | -1.172246732 | 0.281373539 | 0.991835066 | -4.743103428 |
| **A_51_P183102** | Rnf139 | 0.065192885 | -0.067733561 | 0.198119332 | 10.46382527 | 1.172189802 | 0.281394868 | 0.991835066 | -4.743147674 |
| **A_55_P2028394** | Atg13 | -0.069831287 | -0.212216359 | 0.072553785 | 11.48505883 | -1.172181108 | 0.281398126 | 0.991835066 | -4.743154432 |
| **A_51_P405912** | Lmcd1 | -0.110259977 | -0.335150536 | 0.114630581 | 10.17587967 | -1.171806243 | 0.281538613 | 0.991835066 | -4.743445749 |
| **A_55_P2048563** | Cdk2ap1 | 0.062450427 | -0.064974512 | 0.189875365 | 13.71870866 | 1.171359244 | 0.281706211 | 0.991835066 | -4.743793056 |
| **A_55_P2115846** | Gm10836 | -0.048641605 | -0.147910699 | 0.05062749 | 7.711054965 | -1.171124231 | 0.28179436 | 0.991835066 | -4.743975625 |
| **A_55_P2083411** | Cyp4f18 | -0.059867079 | -0.182063818 | 0.062329659 | 8.26840976 | -1.170947942 | 0.281860498 | 0.991835066 | -4.744112562 |
| **A_55_P2168899** | NA | -0.120633732 | -0.366872772 | 0.125605309 | 8.441980529 | -1.170903177 | 0.281877294 | 0.991835066 | -4.744147333 |
| **A_55_P2062444** | Memo1 | -0.060646954 | -0.184449295 | 0.063155387 | 11.54252361 | -1.170817663 | 0.281909383 | 0.991835066 | -4.744213752 |
| **A_55_P2184264** | Timm23 | -0.074559866 | -0.226766789 | 0.077647056 | 12.33657304 | -1.170792278 | 0.281918909 | 0.991835066 | -4.744233469 |
| **A_55_P2035932** | Fgf7 | 0.06136303 | -0.063921599 | 0.186647659 | 7.844271585 | 1.170625921 | 0.281981344 | 0.991835066 | -4.744362671 |
| **A_55_P2077068** | Malt1 | 0.170415846 | -0.18561057 | 0.526442261 | 8.969236201 | 1.189035724 | 0.281982642 | 0.991835066 | -4.684811497 |
| **A_51_P226417** | Rraga | 0.069044653 | -0.071943268 | 0.210032574 | 11.45238455 | 1.170461885 | 0.282042918 | 0.991835066 | -4.74449006 |
| **A_66_P121636** | Ablim3 | -0.178379947 | -0.542727516 | 0.185967622 | 10.93744595 | -1.170143935 | 0.282162301 | 0.991835066 | -4.74473695 |
| **A_55_P2171413** | Me1 | 0.086599588 | -0.090328035 | 0.26352721 | 9.686916027 | 1.169847671 | 0.282273579 | 0.991835066 | -4.744966967 |
| **A_55_P2139211** | NA | -0.077591093 | -0.236119398 | 0.080937211 | 14.0757403 | -1.169806927 | 0.282288885 | 0.991835066 | -4.744998597 |
| **A_55_P2031466** | Lrrc32 | -0.064100751 | -0.19509449 | 0.066892989 | 10.71392457 | -1.169557851 | 0.282382472 | 0.991835066 | -4.745191949 |
| **A_52_P167189** | Isca1 | -0.139752424 | -0.425345033 | 0.145840184 | 11.97377432 | -1.169557342 | 0.282382664 | 0.991835066 | -4.745192344 |
| **A_51_P247249** | Alox5 | -0.131119628 | -0.399101008 | 0.136861752 | 8.894829779 | -1.16942459 | 0.282432554 | 0.991835066 | -4.745295387 |
| **A_51_P229602** | Plxna2 | -0.098985661 | -0.301294484 | 0.103323162 | 10.99431266 | -1.169409443 | 0.282438247 | 0.991835066 | -4.745307144 |
| **A_51_P248122** | Bbc3 | -0.080125924 | -0.243891181 | 0.083639334 | 9.949712488 | -1.169392782 | 0.282444509 | 0.991835066 | -4.745320075 |
| **A_55_P2084433** | NA | -0.085186907 | -0.259299041 | 0.088925227 | 13.56431394 | -1.169372686 | 0.282452062 | 0.991835066 | -4.745335673 |
| **A_51_P105068** | Lypd6b | 0.053506275 | -0.055860519 | 0.162873069 | 7.875487285 | 1.169306514 | 0.282476935 | 0.991835066 | -4.741629575 |
| **A_55_P2115968** | NA | -0.054311052 | -0.165334156 | 0.056712052 | 9.192996629 | -1.169187009 | 0.282521859 | 0.991835066 | -4.745479782 |
| **A_55_P2110177** | NA | 0.06853296 | -0.071574858 | 0.208640778 | 13.35219632 | 1.169085442 | 0.282560044 | 0.991835066 | -4.745558606 |
| **A_51_P212038** | Atp6v0e2 | -0.149996448 | -0.456652845 | 0.156659949 | 9.974869962 | -1.169063439 | 0.282568317 | 0.991835066 | -4.745575682 |
| **A_65_P07561** | Tpd52l2 | -0.107145366 | -0.326202287 | 0.111911555 | 9.111739822 | -1.169029699 | 0.282581004 | 0.991835066 | -4.745601865 |
| **A_51_P375078** | Lmbrd2 | 0.068634508 | -0.071700496 | 0.208969512 | 8.489694617 | 1.168922307 | 0.282621386 | 0.991835066 | -4.745685203 |
| **A_55_P1959174** | Pmepa1 | 0.092768962 | -0.096926172 | 0.282464096 | 10.33037475 | 1.168841542 | 0.28265176 | 0.991835066 | -4.745747876 |
| **A_66_P121446** | Gm7969 | 0.054455143 | -0.056899821 | 0.165810107 | 7.87978258 | 1.16879529 | 0.282669155 | 0.991835066 | -4.745783764 |
| **A_52_P435670** | Npepl1 | 0.075492477 | -0.078923052 | 0.229908006 | 10.54870252 | 1.168481485 | 0.2827872 | 0.991835066 | -4.746027243 |
| **A_55_P1975650** | Fbxo33 | 0.059659119 | -0.062415906 | 0.181734143 | 8.77276199 | 1.168043841 | 0.282951899 | 0.991835066 | -4.746366747 |
| **A_55_P1978972** | Rp2 | -0.051526802 | -0.156963614 | 0.053910009 | 8.151717496 | -1.16801941 | 0.282961096 | 0.991835066 | -4.746385697 |
| **A_55_P1981259** | Fkbp14 | 0.086134475 | -0.090164004 | 0.262432954 | 10.74029207 | 1.167716939 | 0.283074975 | 0.991835066 | -4.746620297 |
| **A_55_P2145879** | Tpt1 | 0.065984079 | -0.069077298 | 0.201045456 | 17.66605075 | 1.167661917 | 0.283095695 | 0.991835066 | -4.746662968 |
| **A_55_P2045886** | Stk3 | 0.084555359 | -0.08853121 | 0.257641928 | 9.776976038 | 1.167580685 | 0.283126287 | 0.991835066 | -4.746725965 |
| **A_52_P417654** | Tcea1 | -0.078645096 | -0.239644798 | 0.082354606 | 8.977408845 | -1.167496824 | 0.283157872 | 0.991835066 | -4.746790998 |
| **A_51_P430383** | Dnajc30 | -0.122719101 | -0.373945887 | 0.128507685 | 11.45453724 | -1.167495878 | 0.283158228 | 0.991835066 | -4.746791731 |
| **A_55_P1962284** | Klhl24 | -0.116825886 | -0.355990152 | 0.12233838 | 8.591880606 | -1.167486652 | 0.283161703 | 0.991835066 | -4.746798886 |
| **A_55_P2353004** | Pitpnm2os | -0.098067296 | -0.298858473 | 0.102723882 | 8.224266946 | -1.167316717 | 0.283225717 | 0.991835066 | -4.746930659 |
| **A_55_P2190152** | 4921509J1 | -0.062650851 | -0.190930371 | 0.065628668 | 8.466055635 | -1.167290053 | 0.283235762 | 0.991835066 | -4.746951334 |
| **A_52_P592111** | Brf1 | -0.066094851 | -0.201427367 | 0.069237665 | 8.934072604 | -1.167278818 | 0.283239994 | 0.991835066 | -4.746960046 |
| **A_51_P453043** | Aacs | 0.089698171 | -0.093965257 | 0.273361598 | 9.594777517 | 1.167266534 | 0.283244622 | 0.991835066 | -4.746969571 |
| **A_55_P2172430** | Zxdb | 0.097555805 | -0.102200439 | 0.297312048 | 9.004752258 | 1.167244629 | 0.283252875 | 0.991835066 | -4.746986555 |
| **A_55_P2212027** | Utrn | 0.100304089 | -0.105106209 | 0.305714386 | 9.135857844 | 1.16709325 | 0.283309913 | 0.991835066 | -4.747103927 |
| **A_51_P382152** | Procr | 0.069481236 | -0.072826247 | 0.211788719 | 9.023362626 | 1.166941088 | 0.283367256 | 0.991835066 | -4.747221897 |

| | | | | | | | | | |
|---|---|---|---|---|---|---|---|---|---|
| **A_55_P2088325** | Ctdsp2 | 0.074954803 | -0.078575353 | 0.228484959 | 9.833960654 | 1.166849668 | 0.283401713 | 0.991835066 | -4.74729277 |
| **A_55_P2199040** | NA | -0.055621834 | -0.169553651 | 0.058309982 | 8.320196219 | -1.16683492 | 0.283407272 | 0.991835066 | -4.747304203 |
| **A_65_P18433** | Ankrd28 | 0.091129261 | -0.095556086 | 0.277814607 | 9.88523326 | 1.166693424 | 0.28346061 | 0.991835066 | -4.747413891 |
| **A_51_P279038** | Ppargc1a | -0.126658723 | -0.386129302 | 0.132811856 | 12.74635462 | -1.166691719 | 0.283461253 | 0.991835066 | -4.747415213 |
| **A_55_P2135023** | NA | -0.14988591 | -0.456981666 | 0.157209846 | 13.01464772 | -1.166530576 | 0.283522008 | 0.991835066 | -4.747540121 |
| **A_52_P96552** | Vkorc1l1 | -0.047427129 | -0.144602675 | 0.049748418 | 9.061568182 | -1.166484499 | 0.283539382 | 0.991835066 | -4.747575836 |
| **A_51_P263965** | Hmox1 | 0.174147129 | -0.18267338 | 0.530967638 | 9.124120918 | 1.166475558 | 0.283542753 | 0.991835066 | -4.747582766 |
| **A_55_P2139587** | NA | 0.047683425 | -0.050023511 | 0.145390362 | 7.760323958 | 1.166409864 | 0.283567526 | 0.991835066 | -4.747633685 |
| **A_55_P2041961** | Bmp2 | 0.193176935 | -0.202661443 | 0.589015314 | 9.039119561 | 1.1663973 | 0.283572264 | 0.991835066 | -4.747643423 |
| **A_55_P1974189** | Ptges | 0.048869539 | -0.051277837 | 0.149016916 | 7.769349961 | 1.166293358 | 0.283611465 | 0.991835066 | -4.747723982 |
| **A_55_P2187206** | NA | -0.073427459 | -0.22390332 | 0.077048402 | 8.558287772 | -1.166274541 | 0.283618562 | 0.991835066 | -4.747738566 |
| **A_55_P1957584** | Gps2 | -0.103812164 | -0.316611611 | 0.108987283 | 9.988267336 | -1.165968669 | 0.283733946 | 0.991835066 | -4.747975604 |
| **A_55_P2052897** | Arhgap30 | -0.050137824 | -0.15294029 | 0.052664641 | 7.772383926 | -1.165657825 | 0.283851246 | 0.991835066 | -4.748216461 |
| **A_55_P1985498** | Gm12005 | 0.053758452 | -0.056480456 | 0.163997359 | 14.18942093 | 1.165523364 | 0.283901998 | 0.991835066 | -4.748320636 |
| **A_55_P2074438** | Gtf2a1 | 0.069481265 | -0.073022344 | 0.211984874 | 10.51160508 | 1.165335525 | 0.283972912 | 0.991835066 | -4.748466157 |
| **A_55_P1999878** | Sec24b | 0.093504452 | -0.098277958 | 0.285286862 | 11.43005177 | 1.165286323 | 0.283991489 | 0.991835066 | -4.748504271 |
| **A_51_P468140** | Serpind1 | -0.079202154 | -0.241654746 | 0.083250439 | 7.919582152 | -1.165250982 | 0.284004833 | 0.991835066 | -4.748531649 |
| **A_55_P2117681** | Rnf115 | 0.059035357 | -0.062054406 | 0.180125121 | 9.919843338 | 1.16523601 | 0.284010487 | 0.991835066 | -4.748543246 |
| **A_52_P264902** | Dnajc3 | 0.079617091 | -0.083688882 | 0.242923064 | 9.356989802 | 1.165234596 | 0.284011021 | 0.991835066 | -4.748544342 |
| **A_55_P2069515** | Dennd4a | 0.053293209 | -0.056018945 | 0.162605363 | 8.474982534 | 1.165232402 | 0.284011849 | 0.991835066 | -4.748546041 |
| **A_55_P2121437** | NA | -0.124826722 | -0.380866725 | 0.131213281 | 14.52503233 | -1.165222532 | 0.284015576 | 0.991835066 | -4.748553686 |
| **A_55_P2154049** | Myo18a | 0.079666306 | -0.083750502 | 0.243083114 | 8.318482201 | 1.165164087 | 0.284037646 | 0.991835066 | -4.748598959 |
| **A_55_P2183854** | Sctr | 0.065898543 | -0.069294401 | 0.201091487 | 7.905828476 | 1.165013404 | 0.284094554 | 0.991835066 | -4.748715675 |
| **A_55_P2141699** | Rnf149 | 0.11201818 | -0.117795904 | 0.341832264 | 10.28662609 | 1.164986799 | 0.284104602 | 0.991835066 | -4.748736282 |
| **A_51_P415809** | Tusc1 | 0.052359695 | -0.055060913 | 0.159780304 | 9.290850927 | 1.164980446 | 0.284107002 | 0.991835066 | -4.748741202 |
| **A_55_P2015934** | NA | 0.06166744 | -0.06485742 | 0.188192301 | 16.50609751 | 1.16490145 | 0.284136841 | 0.991835066 | -4.748802387 |
| **A_51_P222386** | Tbc1d10a | 0.08369191 | -0.088034396 | 0.255418216 | 8.948620453 | 1.164811862 | 0.284170684 | 0.991835066 | -4.748871772 |
| **A_52_P257728** | Cnnm4 | 0.074904176 | -0.078808044 | 0.228616395 | 9.172509876 | 1.164680403 | 0.28422035 | 0.991835066 | -4.74897358 |
| **A_51_P478003** | Papolg | -0.05819921 | -0.177653001 | 0.061254581 | 9.291975832 | -1.164464579 | 0.284301906 | 0.991835066 | -4.749140712 |
| **A_52_P194242** | Sel1l | -0.083292294 | -0.254262079 | 0.087677491 | 9.529354595 | -1.164379609 | 0.28433402 | 0.991835066 | -4.749206507 |
| **A_51_P294807** | Stom | -0.170843374 | -0.521544082 | 0.179857333 | 9.987954274 | -1.164315372 | 0.2843583 | 0.991835066 | -4.749256246 |
| **A_55_P1961210** | Trp53inp2 | -0.150559512 | -0.459665892 | 0.158546868 | 14.69067543 | -1.164151123 | 0.28442039 | 0.991835066 | -4.749383418 |
| **A_55_P2002553** | Stx16 | 0.076925368 | -0.081056546 | 0.234907281 | 8.783611906 | 1.163781217 | 0.284560265 | 0.991835066 | -4.749669784 |
| **A_55_P2117033** | Mtmr3 | 0.066070107 | -0.069644213 | 0.201784427 | 9.433511859 | 1.163559153 | 0.284644263 | 0.991835066 | -4.749841674 |
| **A_55_P2070769** | Dus1l | 0.102060736 | -0.107584752 | 0.311706225 | 11.55467828 | 1.163542654 | 0.284650505 | 0.991835066 | -4.749854444 |
| **A_55_P2301312** | Wdr78 | 0.077347684 | -0.08156047 | 0.236255837 | 8.373268087 | 1.16334965 | 0.284723529 | 0.991835066 | -4.750003823 |
| **A_51_P313761** | Shmt2 | 0.148551984 | -0.156705749 | 0.453809717 | 11.44540006 | 1.163110331 | 0.284814098 | 0.991835066 | -4.750189029 |
| **A_51_P479408** | Hyal1 | 0.101247713 | -0.106829001 | 0.309324427 | 8.240244389 | 1.162976321 | 0.284864824 | 0.991835066 | -4.750292728 |
| **A_51_P348397** | Hexim1 | 0.07716074 | -0.081419753 | 0.235741234 | 9.13847504 | 1.162935837 | 0.28488015 | 0.991835066 | -4.750324054 |
| **A_55_P1967533** | Gm5069 | -0.189996561 | -0.580520123 | 0.200527001 | 14.09921025 | -1.162806983 | 0.284928934 | 0.991835066 | -4.750423756 |
| **A_51_P512783** | Rmdn1 | -0.080970908 | -0.247407513 | 0.085465696 | 11.47278055 | -1.162757896 | 0.28494752 | 0.991835066 | -4.750461736 |
| **A_55_P2009267** | Gm16060 | -0.090559435 | -0.276717805 | 0.095598936 | 14.69185084 | -1.162680172 | 0.284976951 | 0.991835066 | -4.750521871 |
| **A_55_P2044704** | NA | -0.079238884 | -0.242144724 | 0.083666956 | 11.69242043 | -1.162547828 | 0.28502707 | 0.991835066 | -4.750624259 |
| **A_51_P437717** | Glmp | -0.124173082 | -0.379473271 | 0.131127108 | 10.19708964 | -1.162479905 | 0.285052796 | 0.991835066 | -4.750676806 |
| **A_55_P2335695** | Ndor1 | 0.215999042 | -0.228100389 | 0.660098472 | 8.18560276 | 1.162467287 | 0.285057575 | 0.991835066 | -4.750686568 |
| **A_55_P2159740** | Mfsd2b | -0.115907864 | -0.354222802 | 0.122407074 | 15.53067943 | -1.162440595 | 0.285067685 | 0.991835066 | -4.750707217 |
| **A_52_P631568** | Thoc6 | 0.108445957 | -0.114528228 | 0.331420142 | 9.91247729 | 1.16243295 | 0.285070581 | 0.991835066 | -4.750713131 |
| **A_55_P2058851** | Fancf | 0.099288792 | -0.10489184 | 0.303469424 | 9.526033049 | 1.162237316 | 0.285144692 | 0.991835066 | -4.750864465 |

| | | | | | | | | | |
|---|---|---|---|---|---|---|---|---|---|
| **A_51_P327828** | Ppib | -0.109560228 | -0.334885144 | 0.115764687 | 13.41758599 | -1.162124998 | 0.285187247 | 0.991835066 | -4.750951343 |
| **A_51_P320401** | Rab24 | 0.055033068 | -0.058173893 | 0.168240029 | 10.63604044 | 1.161875846 | 0.285281667 | 0.991835066 | -4.751144044 |
| **A_55_P2025468** | Tst | -0.093150764 | -0.284773065 | 0.098471538 | 9.164883834 | -1.161848495 | 0.285292033 | 0.991835066 | -4.751165197 |
| **A_55_P2158478** | Nkiras2 | -0.0527551 | -0.161280353 | 0.055770154 | 12.73609915 | -1.161830495 | 0.285298856 | 0.991835066 | -4.751179118 |
| **A_51_P516615** | Ndufb10 | -0.104271381 | -0.318777566 | 0.110234803 | 16.28701233 | -1.161808211 | 0.285307302 | 0.991835066 | -4.751196351 |
| **A_55_P2085551** | Dennd4b | -0.055030005 | -0.16823747 | 0.05817746 | 10.74838881 | -1.161806019 | 0.285308133 | 0.991835066 | -4.751198046 |
| **A_55_P2003548** | NA | 0.081822726 | -0.086516471 | 0.250161924 | 16.82016442 | 1.161710263 | 0.285344431 | 0.991835066 | -4.751272098 |
| **A_52_P536735** | Hpcal1 | 0.066312818 | -0.070119125 | 0.202744761 | 8.968878266 | 1.1616908 | 0.285351809 | 0.991835066 | -4.751287149 |
| **A_52_P413855** | Slc35a3 | -0.063812113 | -0.1951013 | 0.067477074 | 8.338181228 | -1.161671403 | 0.285359163 | 0.991835066 | -4.751302149 |
| **A_66_P120745** | Armc1 | -0.060755763 | -0.185757064 | 0.064245538 | 12.07454886 | -1.161668135 | 0.285360402 | 0.991835066 | -4.751304676 |
| **A_55_P2036854** | Pnkd | -0.078984379 | -0.241503306 | 0.083534549 | 13.20878656 | -1.161572683 | 0.28539659 | 0.991835066 | -4.751378489 |
| **A_51_P363525** | Fbrsl1 | 0.0754546 | -0.079835753 | 0.230744953 | 10.58845585 | 1.161315913 | 0.285493956 | 0.991835066 | -4.751577029 |
| **A_51_P233815** | Tmem126a | -0.064662348 | -0.197763415 | 0.068438719 | 11.38580179 | -1.161125241 | 0.285566277 | 0.991835066 | -4.751724445 |
| **A_52_P159429** | AU021092 | -0.054656408 | -0.167244059 | 0.057931244 | 7.879603204 | -1.160271049 | 0.285890453 | 0.991835066 | -4.752384689 |
| **A_55_P2107155** | Rasd2 | 0.074378449 | -0.078845698 | 0.227602596 | 8.322556074 | 1.160189783 | 0.285921311 | 0.991835066 | -4.752447488 |
| **A_55_P2162855** | Nipa2 | -0.069695987 | -0.213284398 | 0.073892425 | 9.649012946 | -1.160105571 | 0.28595329 | 0.991835066 | -4.752512562 |
| **A_55_P2336943** | Dph6 | 0.066115186 | -0.070104635 | 0.202335008 | 9.218670757 | 1.160032222 | 0.285981146 | 0.991835066 | -4.752569239 |
| **A_55_P2029021** | NA | 0.158606492 | -0.168242461 | 0.485455445 | 15.05143254 | 1.159799705 | 0.286069466 | 0.991835066 | -4.752748895 |
| **A_55_P2049383** | Erp29 | 0.066675398 | -0.070732231 | 0.204083026 | 13.9235577 | 1.159748724 | 0.286088833 | 0.991835066 | -4.752788283 |
| **A_51_P141521** | Sap130 | 0.053993178 | -0.057279278 | 0.165265634 | 11.11276798 | 1.15973923 | 0.28609244 | 0.991835066 | -4.752795617 |
| **A_52_P97192** | Ergic1 | 0.115171151 | -0.122235108 | 0.35257741 | 10.06924736 | 1.159473097 | 0.286193563 | 0.991835066 | -4.753001215 |
| **A_55_P1982107** | NA | 0.068479681 | -0.072717403 | 0.209676764 | 15.75081351 | 1.159164664 | 0.286310797 | 0.991835066 | -4.753239458 |
| **A_52_P39418** | Ccpg1 | -0.059893109 | -0.183396334 | 0.063610116 | 8.07159748 | -1.159064739 | 0.286348786 | 0.991835066 | -4.753316635 |
| **A_51_P462271** | Acan | 0.224359424 | -0.238310762 | 0.687029611 | 8.275672131 | 1.158995991 | 0.286374925 | 0.991835066 | -4.75336973 |
| **A_55_P2089530** | Fkbp10 | 0.095393318 | -0.101326743 | 0.292113379 | 13.43265424 | 1.158986662 | 0.286378472 | 0.991835066 | -4.753376935 |
| **A_52_P328497** | Cdk10 | 0.091480822 | -0.097201908 | 0.280163553 | 9.747745047 | 1.158796138 | 0.286450924 | 0.991835066 | -4.753524071 |
| **A_55_P2039125** | Fam219a | -0.05730427 | -0.175496929 | 0.06088839 | 8.01465074 | -1.158792325 | 0.286452374 | 0.991835066 | -4.753527016 |
| **A_52_P330214** | Fanca | 0.145166058 | -0.154287138 | 0.444619253 | 9.635699388 | 1.158631315 | 0.286513614 | 0.991835066 | -4.753651347 |
| **A_51_P504354** | Nrarp | -0.107862442 | -0.330384173 | 0.114659289 | 9.082483602 | -1.158529108 | 0.286552494 | 0.991835066 | -4.753730267 |
| **A_51_P126835** | Arl14ep | 0.058772832 | -0.062489214 | 0.180034879 | 8.616111113 | 1.158406163 | 0.286599268 | 0.991835066 | -4.746071936 |
| **A_55_P2137910** | NA | 0.100192448 | -0.106530814 | 0.30691571 | 13.51563415 | 1.158389925 | 0.286605447 | 0.991835066 | -4.753837732 |
| **A_51_P459661** | Lipa | 0.067914424 | -0.072230141 | 0.208058988 | 9.699291327 | 1.158230224 | 0.286666216 | 0.991835066 | -4.753961029 |
| **A_55_P2074776** | Gm8894 | 0.108445435 | -0.115364123 | 0.332254993 | 15.81559365 | 1.158088578 | 0.286720124 | 0.991835066 | -4.754070379 |
| **A_52_P546090** | Ssr4 | -0.060296511 | -0.184746895 | 0.064153874 | 12.8062008 | -1.157990698 | 0.28675738 | 0.991835066 | -4.754145937 |
| **A_55_P2166884** | NA | 0.079442776 | -0.084525293 | 0.243410845 | 13.38668093 | 1.157988534 | 0.286758204 | 0.991835066 | -4.754147608 |
| **A_55_P2027771** | Zfp260 | 0.080733712 | -0.085909056 | 0.247376479 | 12.14119457 | 1.157917405 | 0.286785281 | 0.991835066 | -4.754202513 |
| **A_52_P196568** | Meg3 | 0.087185336 | -0.092802676 | 0.267173348 | 7.975399052 | 1.157734561 | 0.286854894 | 0.991835066 | -4.754343645 |
| **A_55_P2240478** | NA | -0.128416404 | -0.393543135 | 0.136710327 | 8.4232766 | -1.157646918 | 0.286888267 | 0.991835066 | -4.75441129 |
| **A_55_P2096967** | Gjb1 | -0.059151366 | -0.181282451 | 0.062979719 | 8.591111393 | -1.157571147 | 0.286917121 | 0.991835066 | -4.754469769 |
| **A_55_P2139763** | Trim56 | -0.119114419 | -0.365097407 | 0.12686857 | 10.56172349 | -1.157359803 | 0.286997617 | 0.991835066 | -4.75463287 |
| **A_55_P2012819** | Sec61a2 | 0.114251578 | -0.121689613 | 0.350192769 | 10.12449767 | 1.157357613 | 0.286998451 | 0.991835066 | -4.75463456 |
| **A_52_P431872** | Ptcd3 | 0.088438731 | -0.094200384 | 0.271077847 | 13.24544382 | 1.157331681 | 0.28700833 | 0.991835066 | -4.754654572 |
| **A_51_P475613** | Fahd2a | -0.104474497 | -0.320230613 | 0.111281618 | 10.59875547 | -1.157327597 | 0.287009885 | 0.991835066 | -4.754657724 |
| **A_51_P376149** | Eef1akmt1 | -0.086818163 | -0.266115363 | 0.092479036 | 11.55199898 | -1.157300711 | 0.287020127 | 0.991835066 | -4.754678471 |
| **A_55_P2183777** | NA | -0.086257742 | -0.264422001 | 0.091906517 | 9.95246548 | -1.15714193 | 0.287080619 | 0.991835066 | -4.754800993 |
| **A_55_P1995165** | Mdc1 | 0.102631444 | -0.109373265 | 0.314636154 | 10.60911372 | 1.15702852 | 0.287123833 | 0.991835066 | -4.754888499 |
| **A_55_P2052848** | Mif | 0.076739007 | -0.081780526 | 0.235258541 | 16.82606074 | 1.157024415 | 0.287125397 | 0.991835066 | -4.754891666 |
| **A_51_P380432** | Scx | -0.159965414 | -0.490464753 | 0.170533925 | 10.67729866 | -1.156816915 | 0.287204476 | 0.991835066 | -4.755051759 |

| | | | | | | | | | |
|---|---|---|---|---|---|---|---|---|---|
| **A_52_P202142** | Sv2a | -0.06355706 | -0.194872926 | 0.067758806 | 8.869844916 | -1.156793212 | 0.287213511 | 0.991835066 | -4.755070045 |
| **A_55_P2017684** | Macf1 | -0.054754025 | -0.167886406 | 0.058378356 | 12.72129909 | -1.156746663 | 0.287231254 | 0.991835066 | -4.755105957 |
| **A_51_P484919** | Polrmt | 0.065439586 | -0.069782226 | 0.200661397 | 10.38309101 | 1.156652562 | 0.287267125 | 0.991835066 | -4.755178551 |
| **A_55_P2007464** | Rps2 | 0.063144912 | -0.067349564 | 0.193639388 | 16.88741799 | 1.156525911 | 0.28731541 | 0.991835066 | -4.755276251 |
| **A_55_P1955861** | Cux2 | 0.069374244 | -0.073998413 | 0.212746901 | 8.104591925 | 1.156487816 | 0.287329935 | 0.991835066 | -4.755305636 |
| **A_55_P1973809** | Hbb-bt | 0.309502694 | -0.330213784 | 0.949219173 | 17.44537472 | 1.156341276 | 0.287385813 | 0.991835066 | -4.755418668 |
| **A_55_P2108883** | NA | 0.088335709 | -0.094258274 | 0.270929691 | 8.980679349 | 1.156269231 | 0.287413289 | 0.991835066 | -4.755474236 |
| **A_55_P2045481** | NA | 0.110265574 | -0.117683513 | 0.338214661 | 12.26961203 | 1.156142107 | 0.287461775 | 0.991835066 | -4.755572282 |
| **A_55_P2131279** | Atp6v1a | -0.092542934 | -0.283901016 | 0.098815148 | 10.11631021 | -1.155860942 | 0.287569038 | 0.991835066 | -4.755789112 |
| **A_55_P2032278** | NA | 0.080150666 | -0.085583584 | 0.245884915 | 8.79372838 | 1.155856716 | 0.28757065 | 0.991835066 | -4.75579237 |
| **A_51_P412640** | Ccdc51 | -0.079220802 | -0.24303271 | 0.084591107 | 9.720661721 | -1.155853773 | 0.287571773 | 0.991835066 | -4.755794641 |
| **A_55_P1965945** | NA | 0.057716168 | -0.06164175 | 0.177074086 | 16.27258211 | 1.155727331 | 0.287620021 | 0.991835066 | -4.75589214 |
| **A_55_P1980051** | Il17d | -0.049317654 | -0.151307851 | 0.052672543 | 9.457806333 | -1.155721279 | 0.287622331 | 0.991835066 | -4.755896807 |
| **A_55_P2014506** | Amd2 | 0.098163171 | -0.104847126 | 0.301173469 | 12.16108245 | 1.155685447 | 0.287636005 | 0.991835066 | -4.755924436 |
| **A_55_P2032034** | Tbp | 0.073350931 | -0.078358601 | 0.225060462 | 9.022870564 | 1.155584993 | 0.287674343 | 0.991835066 | -4.756001891 |
| **A_51_P446106** | Lrrc38 | -0.05552602 | -0.170382544 | 0.059330504 | 8.485107052 | -1.155446945 | 0.287727036 | 0.991835066 | -4.756108326 |
| **A_51_P450573** | Tgfbr2 | -0.085441884 | -0.262191848 | 0.09130808 | 11.82665751 | -1.155368745 | 0.287756888 | 0.991835066 | -4.756168615 |
| **A_55_P2105820** | Tbc1d31 | 0.105157727 | -0.112403121 | 0.322718574 | 10.1431388 | 1.155232978 | 0.287808723 | 0.991835066 | -4.756273281 |
| **A_55_P2096762** | Arhgdig | 0.141360516 | -0.151127682 | 0.433848713 | 8.081200358 | 1.155124799 | 0.28785003 | 0.991835066 | -4.756356674 |
| **A_51_P211934** | Larp7 | -0.065192894 | -0.200093688 | 0.069707901 | 11.25968058 | -1.155034307 | 0.287884587 | 0.991835066 | -4.756426428 |
| **A_55_P2035400** | Ripk4 | -0.060759499 | -0.186520835 | 0.065001838 | 8.934531816 | -1.154718622 | 0.288005169 | 0.991835066 | -4.756669746 |
| **A_51_P345747** | E130309D( | 0.059531824 | -0.063698816 | 0.182762464 | 8.915496139 | 1.154621435 | 0.2880423 | 0.991835066 | -4.756744646 |
| **A_55_P2056496** | Tk1 | 0.085244424 | -0.091262415 | 0.261751264 | 11.73772666 | 1.154286397 | 0.288170334 | 0.991835066 | -4.757002827 |
| **A_52_P274238** | Maea | -0.068057559 | -0.208984031 | 0.072868913 | 12.05746739 | -1.154231501 | 0.288191317 | 0.991835066 | -4.757045126 |
| **A_55_P2009518** | NA | 0.05096403 | -0.054578512 | 0.156506572 | 15.5615937 | 1.154105061 | 0.288239651 | 0.991835066 | -4.757142547 |
| **A_51_P296036** | Nrbp2 | -0.111969721 | -0.343903871 | 0.11996443 | 10.22997582 | -1.153838502 | 0.28834157 | 0.991835066 | -4.75734791 |
| **A_55_P2163198** | NA | 0.094627073 | -0.101404682 | 0.290658827 | 11.08776911 | 1.153713862 | 0.288389237 | 0.991835066 | -4.757443926 |
| **A_55_P1982245** | Phactr2 | 0.078848419 | -0.084513582 | 0.24221042 | 8.500158127 | 1.153588917 | 0.288437026 | 0.991835066 | -4.75754017 |
| **A_52_P235479** | Usp38 | 0.081539125 | -0.087449073 | 0.250527323 | 9.64939002 | 1.153237628 | 0.288571425 | 0.991835066 | -4.757810737 |
| **A_55_P2140516** | Pex13 | -0.084955311 | -0.261044288 | 0.091133666 | 11.28591162 | -1.153101392 | 0.288623562 | 0.991835066 | -4.757915654 |
| **A_55_P2052759** | Scd3 | 0.138270416 | -0.148344792 | 0.424885624 | 8.813105522 | 1.153026185 | 0.288652346 | 0.991835066 | -4.757973569 |
| **A_55_P2182810** | Arl16 | 0.053601685 | -0.057507831 | 0.164711201 | 8.839376147 | 1.15301862 | 0.288655241 | 0.991835066 | -4.757979395 |
| **A_52_P323830** | Zfp830 | 0.064198685 | -0.068883684 | 0.197281053 | 9.117457541 | 1.152961332 | 0.28867717 | 0.991835066 | -4.75802351 |
| **A_51_P468418** | Ice2 | 0.071898282 | -0.077158043 | 0.220954606 | 8.496529077 | 1.152861765 | 0.288715284 | 0.991835066 | -4.758100178 |
| **A_55_P2046838** | Amfr | -0.064003603 | -0.196708689 | 0.068701482 | 11.11317495 | -1.152725743 | 0.28876736 | 0.991835066 | -4.758204912 |
| **A_52_P334670** | Clock | -0.063630054 | -0.195564728 | 0.068304621 | 9.12235997 | -1.152689852 | 0.288781102 | 0.991835066 | -4.758232546 |
| **A_66_P123448** | Pigx | 0.06900999 | -0.074081956 | 0.212101935 | 10.98324457 | 1.152672431 | 0.288787772 | 0.991835066 | -4.758245959 |
| **A_55_P2072706** | Chsy1 | 0.105376831 | -0.113137049 | 0.32389071 | 12.23662928 | 1.152591038 | 0.288818939 | 0.991835066 | -4.758308625 |
| **A_55_P1969523** | Sumo2 | -0.10631079 | -0.326768028 | 0.114146448 | 11.27095257 | -1.152556215 | 0.288832275 | 0.991835066 | -4.758335435 |
| **A_55_P2038303** | NA | 0.103586223 | -0.11123572 | 0.318408165 | 10.04593341 | 1.152477592 | 0.288862385 | 0.991835066 | -4.758395964 |
| **A_52_P206881** | Prpf18 | 0.072620962 | -0.078001271 | 0.223243195 | 10.54040478 | 1.152343746 | 0.288913649 | 0.991835066 | -4.758499003 |
| **A_55_P2100770** | Usp48 | 0.112748524 | -0.121163265 | 0.346660312 | 9.317973012 | 1.152040871 | 0.289029682 | 0.991835066 | -4.75873214 |
| **A_52_P87964** | Pla2g12a | 0.095378095 | -0.102498317 | 0.293254507 | 10.6162911 | 1.152029898 | 0.289033886 | 0.991835066 | -4.758740586 |
| **A_51_P325836** | Hpx | -0.089815087 | -0.27616133 | 0.096531156 | 7.859546141 | -1.151961028 | 0.289060276 | 0.991835066 | -4.758793593 |
| **A_55_P2054332** | Ephb3 | -0.116376064 | -0.357835635 | 0.125083507 | 13.4253681 | -1.151936059 | 0.289069845 | 0.991835066 | -4.758812811 |
| **A_55_P2313163** | D10Bwg10 | 0.214363002 | -0.230408337 | 0.659134341 | 10.87294979 | 1.151919603 | 0.289076151 | 0.991835066 | -4.758825476 |
| **A_51_P262721** | 0610009L1 | -0.093270273 | -0.286810679 | 0.100270133 | 11.98916449 | -1.151809704 | 0.289118269 | 0.991835066 | -4.758910057 |
| **A_55_P1970596** | NA | 0.092062961 | -0.098975033 | 0.283100956 | 15.31485438 | 1.151792706 | 0.289124784 | 0.991835066 | -4.758923139 |

| | | | | | | | | | |
|---|---|---|---|---|---|---|---|---|---|
| **A_52_P447284** | Clic6 | 0.077630948 | -0.083460731 | 0.238722627 | 8.956905997 | 1.151783655 | 0.289128253 | 0.991835066 | -4.758930104 |
| **A_51_P329490** | Slc25a40 | 0.077237288 | -0.083073307 | 0.237547883 | 8.650039763 | 1.151526463 | 0.289226843 | 0.991835066 | -4.759128025 |
| **A_51_P411271** | Nfe2l2 | -0.071332606 | -0.219396388 | 0.076731176 | 11.47565203 | -1.151458753 | 0.289252803 | 0.991835066 | -4.759180127 |
| **A_51_P319592** | Usp42 | 0.094751926 | -0.101943849 | 0.2914477 | 10.0979018 | 1.151336164 | 0.289299808 | 0.991835066 | -4.759274452 |
| **A_55_P2000553** | Slc6a8 | -0.069340862 | -0.213286981 | 0.074605257 | 12.72303562 | -1.151326251 | 0.28930361 | 0.991835066 | -4.75928208 |
| **A_55_P2133280** | Ap4b1 | -0.104865315 | -0.322617194 | 0.112886563 | 10.04437692 | -1.151009981 | 0.289424912 | 0.991835066 | -4.759525406 |
| **A_55_P2083794** | Tmem159 | -0.061693328 | -0.189803004 | 0.066416347 | 9.950823411 | -1.150973706 | 0.289438827 | 0.991835066 | -4.759553312 |
| **A_55_P1968053** | NA | 0.06650593 | -0.071627265 | 0.204639124 | 15.31966638 | 1.150724732 | 0.289534352 | 0.991835066 | -4.759744832 |
| **A_51_P232901** | Cnp | -0.094439322 | -0.290615627 | 0.101736984 | 8.864425522 | -1.150576344 | 0.289591296 | 0.991835066 | -4.759858966 |
| **A_55_P2064038** | NA | -0.067830549 | -0.208736576 | 0.073075478 | 8.133196102 | -1.150548411 | 0.289602017 | 0.991835066 | -4.759880451 |
| **A_51_P221031** | Slc16a12 | 0.079394694 | -0.085562173 | 0.244351562 | 8.324546748 | 1.150350573 | 0.289677956 | 0.991835066 | -4.760032606 |
| **A_51_P391805** | Dkkl1 | -0.056731963 | -0.174606338 | 0.061142413 | 7.971855927 | -1.150316995 | 0.289690846 | 0.991835066 | -4.760058428 |
| **A_66_P138046** | Gm12298 | -0.049018037 | -0.150874739 | 0.052838666 | 7.924845552 | -1.150205477 | 0.28973366 | 0.991835066 | -4.760144187 |
| **A_55_P1987988** | Atg2b | 0.053133733 | -0.057275935 | 0.163543401 | 8.522683025 | 1.150197331 | 0.289736788 | 0.991835066 | -4.760150452 |
| **A_55_P2025687** | Muc4 | 0.047258956 | -0.050943364 | 0.145461276 | 7.668441078 | 1.150195078 | 0.289737653 | 0.991835066 | -4.760152185 |
| **A_55_P2091230** | NA | 0.1180967 | -0.127337985 | 0.363531385 | 11.58167544 | 1.150034731 | 0.289799224 | 0.991835066 | -4.760275485 |
| **A_55_P2031966** | Fbxo39 | 0.05526405 | -0.059596403 | 0.170124502 | 8.15063808 | 1.149956245 | 0.289829365 | 0.991835066 | -4.760335834 |
| **A_66_P128608** | Fut11 | 0.053440928 | -0.057640251 | 0.164522108 | 8.850589508 | 1.149853849 | 0.289868693 | 0.991835066 | -4.760414565 |
| **A_51_P302149** | Cfl2 | -0.075004178 | -0.230946049 | 0.080937692 | 10.54855381 | -1.149560551 | 0.289981365 | 0.991835066 | -4.760640054 |
| **A_66_P121361** | C330021F2 | 0.059864048 | -0.06460033 | 0.184328427 | 8.66988558 | 1.149556019 | 0.289983106 | 0.991835066 | -4.760643537 |
| **A_51_P396811** | Fam122a | -0.081229806 | -0.250120381 | 0.087660768 | 9.281620525 | -1.149526896 | 0.289994296 | 0.991835066 | -4.760665926 |
| **A_51_P442719** | Usp6nl | -0.051149963 | -0.157502356 | 0.055202429 | 8.455227789 | -1.149495292 | 0.29000644 | 0.991835066 | -4.760690221 |
| **A_51_P114062** | Ncs1 | 0.073748436 | -0.079609404 | 0.227106275 | 9.695230866 | 1.149359694 | 0.290058547 | 0.991835066 | -4.760794455 |
| **A_55_P1968703** | Gfra2 | -0.104508633 | -0.321832391 | 0.112815125 | 9.712011769 | -1.149354745 | 0.290060449 | 0.991835066 | -4.760798259 |
| **A_52_P594768** | Aprt | 0.102439934 | -0.110593399 | 0.315473267 | 13.11558801 | 1.149293263 | 0.290084077 | 0.991835066 | -4.760845518 |
| **A_55_P1994539** | Cldn12 | -0.067571958 | -0.208111284 | 0.072967367 | 10.37658957 | -1.149152781 | 0.290138073 | 0.991835066 | -4.760953496 |
| **A_55_P2024709** | 4933427D1 | 0.144351503 | -0.155886739 | 0.444589745 | 9.959334724 | 1.149117486 | 0.290151641 | 0.991835066 | -4.760980623 |
| **A_52_P437795** | Cdh11 | 0.136802417 | -0.147745286 | 0.421350121 | 9.89340699 | 1.149073527 | 0.290168539 | 0.991835066 | -4.761014409 |
| **A_55_P2070347** | Rps2 | 0.0555497 | -0.060009821 | 0.171109222 | 16.04628759 | 1.148907641 | 0.290232316 | 0.991835066 | -4.761141898 |
| **A_55_P2020636** | NA | -0.084911915 | -0.261563355 | 0.091739526 | 8.615050571 | -1.148842738 | 0.290257272 | 0.991835066 | -4.761191775 |
| **A_51_P213030** | Macrod1 | -0.139378482 | -0.429373591 | 0.150616627 | 12.73141832 | -1.148720019 | 0.290304463 | 0.991835066 | -4.761286078 |
| **A_55_P2140843** | NA | 0.059698229 | -0.064522207 | 0.183918666 | 8.939660362 | 1.148623066 | 0.290341751 | 0.991835066 | -4.761360579 |
| **A_52_P284348** | Smim20 | -0.082743611 | -0.254927479 | 0.089440257 | 12.73745452 | -1.148553271 | 0.290368597 | 0.991835066 | -4.761414208 |
| **A_51_P454196** | Sh2d4a | -0.065236658 | -0.200991917 | 0.070518602 | 8.803057313 | -1.148534831 | 0.290375689 | 0.991835066 | -4.761428376 |
| **A_52_P423247** | Pde4b | -0.101682467 | -0.31334296 | 0.109978025 | 8.764813239 | -1.148194349 | 0.290506682 | 0.991835066 | -4.761689967 |
| **A_51_P494125** | Alpl | 0.061235594 | -0.066257188 | 0.188728376 | 8.221166559 | 1.147961881 | 0.290596146 | 0.991835066 | -4.761868546 |
| **A_55_P2007001** | Ctnnbip1 | 0.076254665 | -0.082525462 | 0.235034793 | 10.84978997 | 1.147834848 | 0.290645045 | 0.991835066 | -4.761966123 |
| **A_51_P286563** | Gna13 | -0.050076962 | -0.154380937 | 0.054227012 | 10.67386472 | -1.147482983 | 0.290780522 | 0.991835066 | -4.762236364 |
| **A_55_P2046918** | NA | -0.068591598 | -0.211459549 | 0.074276353 | 15.49585258 | -1.14748029 | 0.290781559 | 0.991835066 | -4.762238432 |
| **A_55_P1968399** | Rdh7 | -0.071154184 | -0.219377941 | 0.077069573 | 7.758309946 | -1.147339003 | 0.290835973 | 0.991835066 | -4.762346931 |
| **A_51_P507801** | F13a1 | 0.171180633 | -0.185446883 | 0.527808149 | 9.569618369 | 1.147225818 | 0.29087957 | 0.991835066 | -4.762433844 |
| **A_51_P151775** | Mief1 | 0.07280173 | -0.078876328 | 0.224479787 | 9.410853147 | 1.147170767 | 0.290900777 | 0.991835066 | -4.762476115 |
| **A_66_P108345** | Sgcb | -0.080975288 | -0.2497024 | 0.087751823 | 10.89698542 | -1.147035249 | 0.290952987 | 0.991835066 | -4.762580167 |
| **A_51_P324871** | Cyb561a3 | -0.07316808 | -0.225646411 | 0.079310252 | 10.47397758 | -1.146892353 | 0.291008047 | 0.991835066 | -4.762689878 |
| **A_55_P1993744** | Myo10 | 0.069332746 | -0.075197724 | 0.213863217 | 8.251686112 | 1.146537116 | 0.291144965 | 0.991835066 | -4.762962581 |
| **A_66_P111660** | Mt1 | 0.090278624 | -0.097963021 | 0.278520268 | 16.36364565 | 1.146247377 | 0.291256678 | 0.991835066 | -4.763184969 |
| **A_51_P362881** | Fbxo38 | 0.086040711 | -0.093392764 | 0.265474185 | 11.11286945 | 1.14606607 | 0.291326601 | 0.991835066 | -4.763324114 |
| **A_51_P193863** | Gtf3c4 | 0.057646373 | -0.062573414 | 0.17786616 | 8.503223422 | 1.146054225 | 0.29133117 | 0.991835066 | -4.763333204 |

| | | | | | | | | | |
|---|---|---|---|---|---|---|---|---|---|
| **A_51_P494361** | Mbnl2 | -0.059827565 | -0.184605049 | 0.064949918 | 12.22354918 | -1.145972616 | 0.291362648 | 0.991835066 | -4.763395831 |
| **A_65_P12104** | Tmtc1 | -0.114039637 | -0.351905163 | 0.123825889 | 9.603128368 | -1.145865001 | 0.291404162 | 0.991835066 | -4.76347841 |
| **A_55_P1993594** | Dcp1b | -0.196757326 | -0.60735756 | 0.213842909 | 11.48398989 | -1.145304192 | 0.291620582 | 0.991835066 | -4.763908683 |
| **A_51_P337269** | Aldob | -0.201219536 | -0.621167281 | 0.218728208 | 8.196522983 | -1.145207078 | 0.291658072 | 0.991835066 | -4.76398318 |
| **A_51_P325318** | Serinc2 | 0.071668388 | -0.077910219 | 0.221246994 | 7.947303858 | 1.145162932 | 0.291675117 | 0.991835066 | -4.764017044 |
| **A_55_P2151056** | Rarg | -0.100712155 | -0.310917215 | 0.109492904 | 9.960524447 | -1.145111714 | 0.291694891 | 0.991835066 | -4.764056332 |
| **A_66_P104473** | Polg2 | 0.060347536 | -0.065614336 | 0.186309408 | 9.082986893 | 1.145063489 | 0.291713512 | 0.991835066 | -4.764093322 |
| **A_55_P2221365** | NA | 0.113617026 | -0.123535721 | 0.350769773 | 8.597711155 | 1.145049837 | 0.291718783 | 0.991835066 | -4.764103794 |
| **A_51_P115159** | Fam162a | 0.133376479 | -0.145044171 | 0.411797128 | 15.37846036 | 1.144951094 | 0.291756913 | 0.991835066 | -4.764179531 |
| **A_66_P111747** | NA | 0.11177139 | -0.121550326 | 0.345093105 | 15.15251108 | 1.144944993 | 0.29175927 | 0.991835066 | -4.76418421 |
| **A_66_P133244** | Vcl | 0.082482841 | -0.089709259 | 0.254674942 | 8.465483236 | 1.144878819 | 0.291784825 | 0.991835066 | -4.764234964 |
| **A_51_P488510** | Appl1 | -0.051526748 | -0.159098495 | 0.056045 | 8.975246632 | -1.144836949 | 0.291800996 | 0.991835066 | -4.764267077 |
| **A_55_P1973159** | Serpinf1 | -0.126007557 | -0.389127013 | 0.137111899 | 10.30337984 | -1.144597417 | 0.291893521 | 0.991835066 | -4.764450775 |
| **A_51_P440865** | Fam110b | -0.05362695 | -0.165610659 | 0.058356759 | 8.854354462 | -1.144556879 | 0.291909183 | 0.991835066 | -4.764481861 |
| **A_55_P2049158** | NA | -0.168937181 | -0.521750561 | 0.183876198 | 12.02969088 | -1.144430253 | 0.291958107 | 0.991835066 | -4.76457896 |
| **A_52_P412046** | NA | 0.050052772 | -0.054481242 | 0.154586786 | 14.55859006 | 1.144404721 | 0.291967973 | 0.991835066 | -4.764598538 |
| **A_55_P2101216** | Aldh3a1 | 0.065690292 | -0.071502593 | 0.202883176 | 8.688300816 | 1.144402338 | 0.291968894 | 0.991835066 | -4.764600366 |
| **A_51_P144160** | Colec10 | 0.104196868 | -0.113434246 | 0.321827981 | 7.858169618 | 1.144307679 | 0.292005472 | 0.991835066 | -4.764672947 |
| **A_55_P2084626** | Rab5b | -0.049005381 | -0.151364626 | 0.053353865 | 11.50251707 | -1.144262917 | 0.292022771 | 0.991835066 | -4.764707267 |
| **A_55_P1998416** | Ifi47 | -0.073738552 | -0.22776386 | 0.080286756 | 8.440512169 | -1.144225582 | 0.2920372 | 0.991835066 | -4.764735893 |
| **A_51_P110841** | Mrps35 | -0.079172685 | -0.244550973 | 0.086205602 | 11.500535 | -1.144210731 | 0.29204294 | 0.991835066 | -4.760889007 |
| **A_55_P2012729** | Kcnh2 | -0.097659362 | -0.301663004 | 0.106344281 | 10.31287237 | -1.144155605 | 0.292064246 | 0.991835066 | -4.764789545 |
| **A_52_P518997** | Epha2 | 0.125188779 | -0.136332577 | 0.386710136 | 9.432710473 | 1.144108934 | 0.292082285 | 0.991835066 | -4.764825326 |
| **A_51_P397768** | Csnk2a1 | 0.075853182 | -0.082623187 | 0.23432955 | 9.070923378 | 1.14397998 | 0.292132134 | 0.991835066 | -4.764924188 |
| **A_55_P1960208** | Vcan | 0.157491816 | -0.171548274 | 0.486531905 | 9.529142326 | 1.143979673 | 0.292132253 | 0.991835066 | -4.764924423 |
| **A_51_P464710** | Adap2 | -0.063982264 | -0.197658207 | 0.069693679 | 7.942918344 | -1.143972218 | 0.292135135 | 0.991835066 | -4.764930139 |
| **A_55_P2168431** | Amer1 | 0.075144695 | -0.081852922 | 0.232142311 | 10.23801713 | 1.143969375 | 0.292136234 | 0.991835066 | -4.764932318 |
| **A_55_P2047748** | NA | 0.148906547 | -0.162213198 | 0.460026293 | 10.23688254 | 1.143919204 | 0.29215563 | 0.991835066 | -4.76497078 |
| **A_51_P484753** | Saraf | -0.063200762 | -0.195262031 | 0.068860506 | 12.37266638 | -1.14381552 | 0.292195718 | 0.991835066 | -4.765050262 |
| **A_55_P2015007** | Zfp46 | -0.133694722 | -0.413082358 | 0.145692913 | 12.54894342 | -1.143710777 | 0.29223622 | 0.991835066 | -4.765130551 |
| **A_52_P89477** | Bcl9 | 0.086750136 | -0.094551978 | 0.268052251 | 9.229362838 | 1.143606007 | 0.292276737 | 0.991835066 | -4.765210858 |
| **A_55_P2425938** | 2810468N( | 0.0641404 | -0.069914553 | 0.198195353 | 8.892561873 | 1.143557297 | 0.292295576 | 0.991835066 | -4.765248192 |
| **A_55_P2056070** | Hfe | -0.098282159 | -0.303712602 | 0.107148284 | 9.62902286 | -1.143454842 | 0.292335205 | 0.991835066 | -4.765326718 |
| **A_55_P2114838** | NA | 0.070435058 | -0.076830564 | 0.21770068 | 14.70804519 | 1.143132665 | 0.292459848 | 0.991835066 | -4.765573621 |
| **A_55_P1996469** | Zfp592 | 0.068099129 | -0.074295722 | 0.210493981 | 11.20427355 | 1.143026781 | 0.292500822 | 0.991835066 | -4.765654757 |
| **A_52_P327615** | Rab18 | -0.055577604 | -0.171806261 | 0.060651053 | 11.70071155 | -1.1428671 | 0.292562623 | 0.991835066 | -4.765777109 |
| **A_55_P1955078** | Igflr1 | 0.270396872 | -0.295140639 | 0.835934382 | 9.214104184 | 1.14274513 | 0.292609837 | 0.991835066 | -4.765870559 |
| **A_55_P2057681** | Cdyl2 | 0.059445576 | -0.064887326 | 0.183778477 | 8.323774171 | 1.142727296 | 0.29261674 | 0.991835066 | -4.765884223 |
| **A_51_P520639** | Mslnl | -0.049026532 | -0.151567696 | 0.053514633 | 7.682835454 | -1.142725858 | 0.292617297 | 0.991835066 | -4.765885324 |
| **A_51_P284466** | Tsc22d4 | -0.095113606 | -0.294073743 | 0.103846532 | 10.80401148 | -1.142577601 | 0.292674695 | 0.991835066 | -4.765998906 |
| **A_51_P134262** | 1700052K1 | -0.05973836 | -0.184714889 | 0.065238169 | 9.732563067 | -1.142441505 | 0.292727393 | 0.991835066 | -4.766103163 |
| **A_52_P550049** | Nfya | -0.074108861 | -0.229159532 | 0.080941809 | 8.859947591 | -1.142366934 | 0.292756272 | 0.991835066 | -4.766160286 |
| **A_51_P254656** | Hdc | -0.162621018 | -0.502880251 | 0.177638215 | 9.131472158 | -1.142288725 | 0.292786561 | 0.991835066 | -4.766220193 |
| **A_55_P2150224** | NA | -0.162678718 | -0.50318357 | 0.177826135 | 14.05694296 | -1.141869749 | 0.292948871 | 0.991835066 | -4.766541084 |
| **A_55_P2039289** | Hspb6 | -0.127106972 | -0.39315655 | 0.138942606 | 13.09762039 | -1.141868073 | 0.29294952 | 0.991835066 | -4.766542367 |
| **A_66_P128431** | Fam122b | 0.07232301 | -0.079067723 | 0.223713742 | 8.344470961 | 1.141790258 | 0.292979674 | 0.991835066 | -4.766601958 |
| **A_55_P2008735** | Cdc123 | -0.055928226 | -0.173009897 | 0.061153446 | 8.179090779 | -1.141698053 | 0.293015407 | 0.991835066 | -4.766672565 |
| **A_55_P2003789** | Taf4 | -0.083652724 | -0.258885936 | 0.091580488 | 10.15251121 | -1.14096625 | 0.293299137 | 0.991835066 | -4.767232837 |

| | | | | | | | | | |
|---|---|---|---|---|---|---|---|---|---|
| **A_51_P319732** | Mrps31 | -0.090225845 | -0.27922989 | 0.0987782 | 12.35645081 | -1.140956244 | 0.293303019 | 0.991835066 | -4.767240497 |
| **A_51_P386280** | Olfr187 | -0.071212679 | -0.220400241 | 0.077974882 | 8.117829619 | -1.140863914 | 0.293338833 | 0.991835066 | -4.76731117 |
| **A_55_P1970905** | Lin28b | 0.064230169 | -0.070330196 | 0.198790535 | 8.190318152 | 1.140856557 | 0.293341687 | 0.991835066 | -4.767316801 |
| **A_55_P2011146** | Reep1 | 0.071859807 | -0.078685166 | 0.22240478 | 10.54911749 | 1.14085098 | 0.29334385 | 0.991835066 | -4.76732107 |
| **A_66_P116360** | NA | -0.086549545 | -0.267871932 | 0.094772843 | 8.526503168 | -1.140834094 | 0.293350401 | 0.991835066 | -4.767333995 |
| **A_66_P108019** | Heatr1 | 0.133983694 | -0.146720038 | 0.414687425 | 11.1664825 | 1.14080888 | 0.293360182 | 0.991835066 | -4.767353294 |
| **A_55_P2043892** | Srd5a3 | -0.076645856 | -0.237228655 | 0.083936944 | 8.901607432 | -1.14077182 | 0.293374559 | 0.991835066 | -4.76738166 |
| **A_55_P2158030** | Glrx3 | 0.079597263 | -0.087188117 | 0.246382643 | 13.70566832 | 1.140641819 | 0.293424997 | 0.991835066 | -4.767481158 |
| **A_55_P1956482** | NA | -0.119530601 | -0.370142277 | 0.131081075 | 10.44129588 | -1.139952973 | 0.293692376 | 0.991835066 | -4.768008268 |
| **A_55_P2124736** | Col14a1 | -0.104979663 | -0.325103594 | 0.115144269 | 12.02739638 | -1.139848272 | 0.293733034 | 0.991835066 | -4.76808837 |
| **A_55_P2102715** | NA | 0.056202214 | -0.061677086 | 0.174081515 | 15.32862513 | 1.139528019 | 0.293857425 | 0.991835066 | -4.768333354 |
| **A_55_P1969650** | Rasgrp1 | -0.059390843 | -0.183963305 | 0.065181619 | 7.730808607 | -1.139479653 | 0.293876214 | 0.991835066 | -4.768370349 |
| **A_55_P2090484** | Pde10a | -0.065222825 | -0.202070513 | 0.071624863 | 7.756577585 | -1.13912471 | 0.294014138 | 0.991835066 | -4.768641819 |
| **A_51_P293901** | Dhrs1 | -0.100432257 | -0.311182185 | 0.110317671 | 9.1124338 | -1.138976914 | 0.294071584 | 0.991835066 | -4.768754843 |
| **A_51_P505696** | Inpp5k | -0.09941916 | -0.308100917 | 0.109262596 | 9.74561991 | -1.138661769 | 0.294194108 | 0.991835066 | -4.768995814 |
| **A_55_P2052959** | Ctbp1 | 0.064364495 | -0.070745283 | 0.199474273 | 14.3010819 | 1.138593547 | 0.294220637 | 0.991835066 | -4.769047974 |
| **A_52_P109403** | Cebpz | 0.099014831 | -0.108831005 | 0.306860667 | 9.37771005 | 1.13859181 | 0.294221313 | 0.991835066 | -4.769049302 |
| **A_51_P401987** | Tmem37 | 0.088203303 | -0.096960133 | 0.273366739 | 10.34754396 | 1.138514983 | 0.294251191 | 0.991835066 | -4.769108039 |
| **A_55_P2071137** | Asap1 | 0.114706601 | -0.126098107 | 0.355511309 | 9.940473227 | 1.138498776 | 0.294257494 | 0.991835066 | -4.769120429 |
| **A_55_P1990261** | Chchd6 | -0.125535014 | -0.389072607 | 0.13800258 | 12.04050099 | -1.138495795 | 0.294258653 | 0.991835066 | -4.769122708 |
| **A_52_P464437** | Trp53bp1 | -0.058428343 | -0.181088102 | 0.064231415 | 9.646852727 | -1.138493617 | 0.2942595 | 0.991835066 | -4.769124373 |
| **A_65_P05040** | Ston2 | 0.054213065 | -0.059601587 | 0.168027718 | 7.830419218 | 1.138452508 | 0.294275489 | 0.991835066 | -4.769155802 |
| **A_55_P2112295** | Hmg20b | 0.061131601 | -0.067217308 | 0.18948051 | 13.27348672 | 1.138368111 | 0.294308316 | 0.991835066 | -4.769220321 |
| **A_55_P1994175** | Ppil3 | -0.073574119 | -0.228073445 | 0.080925206 | 9.348231406 | -1.138171267 | 0.294384892 | 0.991835066 | -4.769370794 |
| **A_55_P2041521** | Doxl2 | -0.256360933 | -0.794850803 | 0.282128937 | 8.022871834 | -1.137845872 | 0.294511513 | 0.991835066 | -4.769619501 |
| **A_55_P2079425** | Spred1 | 0.070702068 | -0.077809297 | 0.219213433 | 8.842493406 | 1.137840953 | 0.294513427 | 0.991835066 | -4.76962326 |
| **A_55_P1982951** | Dcaf10 | 0.065028433 | -0.071572347 | 0.201629213 | 8.30986385 | 1.137782479 | 0.294536186 | 0.991835066 | -4.769667949 |
| **A_55_P2006226** | NA | 0.080955225 | -0.089124025 | 0.251034475 | 16.31460701 | 1.137634319 | 0.294593859 | 0.991835066 | -4.769781174 |
| **A_51_P518600** | Atp6v1c2 | 0.047482219 | -0.052278789 | 0.147243227 | 7.637712198 | 1.137573088 | 0.294617696 | 0.991835066 | -4.769827965 |
| **A_55_P2035159** | Pcdh1 | 0.079275542 | -0.087291072 | 0.245842157 | 9.677744752 | 1.137523544 | 0.294636985 | 0.991835066 | -4.769865824 |
| **A_66_P106606** | 2900097C1 | 0.068759403 | -0.07571299 | 0.213231797 | 10.92722067 | 1.137512985 | 0.294641096 | 0.991835066 | -4.769873892 |
| **A_55_P2031501** | Hnrnpa3 | 0.070629531 | -0.077772604 | 0.219031665 | 14.92673668 | 1.137510213 | 0.294642175 | 0.991835066 | -4.769876011 |
| **A_55_P1987396** | Tbl1xr1 | 0.109316325 | -0.12048206 | 0.33911471 | 8.756566482 | 1.136965233 | 0.294854423 | 0.991835066 | -4.77029239 |
| **A_55_P2105436** | Foxp1 | 0.146836774 | -0.161891335 | 0.455564882 | 8.792698359 | 1.136757409 | 0.294935395 | 0.991835066 | -4.770451142 |
| **A_55_P2049577** | Aggf1 | 0.089327944 | -0.098493469 | 0.277149357 | 12.22789143 | 1.13671441 | 0.294952151 | 0.991835066 | -4.770483987 |
| **A_55_P2078780** | Etv4 | 0.175246874 | -0.193280267 | 0.543774015 | 9.291159439 | 1.136553629 | 0.295014809 | 0.991835066 | -4.770606789 |
| **A_51_P357207** | Zfp518b | 0.057518328 | -0.063452498 | 0.178489155 | 8.464577442 | 1.136409196 | 0.295071107 | 0.991835066 | -4.770717098 |
| **A_55_P1978860** | Trp53bp2 | 0.128806875 | -0.142112678 | 0.399726429 | 9.459454752 | 1.136338774 | 0.295098559 | 0.991835066 | -4.770770879 |
| **A_55_P2121426** | NA | 0.057750566 | -0.063722291 | 0.179223423 | 14.03864413 | 1.136282008 | 0.29512069 | 0.991835066 | -4.770814229 |
| **A_55_P1981217** | Aatf | -0.07545118 | -0.234193715 | 0.083291356 | 8.767089004 | -1.136009212 | 0.29522706 | 0.991835066 | -4.771022537 |
| **A_52_P494380** | Dgkq | 0.141410495 | -0.156189153 | 0.439010143 | 8.941972824 | 1.135686237 | 0.295353037 | 0.991835066 | -4.771269124 |
| **A_55_P1992989** | Mrps26 | 0.086205861 | -0.095224238 | 0.26763596 | 13.23595169 | 1.135629304 | 0.295375248 | 0.991835066 | -4.771312588 |
| **A_51_P298178** | Dctn3 | -0.069449541 | -0.215641611 | 0.076742529 | 13.68783445 | -1.135415184 | 0.295458796 | 0.991835066 | -4.771476039 |
| **A_52_P401484** | Inha | 0.073605078 | -0.081377728 | 0.228587884 | 8.407519129 | 1.135098085 | 0.295582561 | 0.991835066 | -4.771718067 |
| **A_55_P2316612** | 6720407P1 | 0.120443165 | -0.133162661 | 0.374048992 | 9.955313621 | 1.135094621 | 0.295583914 | 0.991835066 | -4.771720711 |
| **A_52_P420466** | H2ac4 | 0.117133907 | -0.129526744 | 0.363794558 | 15.20626291 | 1.134989619 | 0.295624906 | 0.991835066 | -4.771800846 |
| **A_55_P2109544** | Trnt1 | 0.061050274 | -0.067509538 | 0.189610085 | 8.778179947 | 1.134988657 | 0.295625282 | 0.991835066 | -4.771801581 |
| **A_51_P221428** | Cbx6 | 0.08652109 | -0.095718463 | 0.268760644 | 11.65381269 | 1.13471938 | 0.295730428 | 0.991835066 | -4.772007066 |

| | | | | | | | | | |
|---|---|---|---|---|---|---|---|---|---|
| **A_52_P530280** | Nsa2 | 0.107093076 | -0.118521829 | 0.332707981 | 10.79985372 | 1.134495394 | 0.295817913 | 0.991835066 | -4.772177969 |
| **A_51_P485458** | Txlna | 0.064864309 | -0.071791997 | 0.201520614 | 8.41285247 | 1.134449716 | 0.295835756 | 0.991835066 | -4.772212819 |
| **A_55_P1972222** | NA | 0.075782124 | -0.08393364 | 0.235497887 | 14.81348012 | 1.134039339 | 0.295996106 | 0.991835066 | -4.772525882 |
| **A_55_P2156409** | Rpl26 | 0.127998603 | -0.141807668 | 0.397804873 | 13.9146536 | 1.133867537 | 0.296063256 | 0.991835066 | -4.772656924 |
| **A_55_P2005570** | Celf2 | 0.09491517 | -0.105163701 | 0.294994041 | 12.31109515 | 1.133818483 | 0.296082432 | 0.991835066 | -4.772694338 |
| **A_55_P2024654** | Trrap | -0.069636376 | -0.216429415 | 0.077156662 | 10.15235081 | -1.133808832 | 0.296086205 | 0.991835066 | -4.772701699 |
| **A_52_P495565** | Efnb3 | -0.234306981 | -0.728273272 | 0.25965931 | 11.85865024 | -1.133697129 | 0.296129875 | 0.991835066 | -4.772786892 |
| **A_51_P315391** | Txlnb | -0.073729818 | -0.229183501 | 0.081723866 | 11.78843087 | -1.133577651 | 0.29617659 | 0.991835066 | -4.772878009 |
| **A_55_P2112693** | Sipa1l1 | 0.076774812 | -0.085102415 | 0.238652039 | 9.550865328 | 1.133553798 | 0.296185917 | 0.991835066 | -4.7728962 |
| **A_52_P25812** | Prdm2 | 0.063809965 | -0.070743677 | 0.198363607 | 8.279630614 | 1.133449514 | 0.296226698 | 0.991835066 | -4.772975724 |
| **A_66_P103490** | Srsf1 | 0.072231422 | -0.080080613 | 0.224543456 | 10.77138956 | 1.133446631 | 0.296227826 | 0.991835066 | -4.772977923 |
| **A_66_P107734** | NA | 0.088622962 | -0.098277554 | 0.275523479 | 12.26885082 | 1.133300014 | 0.296285169 | 0.991835066 | -4.773089722 |
| **A_51_P459091** | Ybey | -0.062193222 | -0.193360879 | 0.068974435 | 8.816092968 | -1.13324922 | 0.296305037 | 0.991835066 | -4.773128452 |
| **A_55_P2451183** | Pdpk1 | 0.051697188 | -0.057342075 | 0.16073645 | 10.18085448 | 1.133164986 | 0.296337987 | 0.991835066 | -4.773192677 |
| **A_52_P830312** | Iba57 | 0.075910765 | -0.084247691 | 0.23606922 | 9.50238516 | 1.132824482 | 0.296471217 | 0.991835066 | -4.77345227 |
| **A_52_P12877** | Hspa8 | 0.334641611 | -0.371510884 | 1.040794107 | 11.88661092 | 1.132636563 | 0.296544765 | 0.991835066 | -4.773595517 |
| **A_55_P2011852** | Cdk2 | -0.052172267 | -0.162274735 | 0.057930201 | 9.162094924 | -1.132535402 | 0.296584364 | 0.991835066 | -4.773672623 |
| **A_55_P2131672** | Mical2 | 0.17686617 | -0.196388229 | 0.550120569 | 9.055246853 | 1.132528055 | 0.29658724 | 0.991835066 | -4.773678223 |
| **A_55_P1984861** | NA | 0.0579042 | -0.06430209 | 0.180110491 | 14.65724831 | 1.132467217 | 0.296611057 | 0.991835066 | -4.773724593 |
| **A_55_P2167050** | Paip2 | -0.090759268 | -0.282335627 | 0.10081709 | 12.08087321 | -1.132291389 | 0.2966799 | 0.991835066 | -4.773858599 |
| **A_52_P412452** | Cntn6 | 0.056148477 | -0.062413576 | 0.17471053 | 7.765371177 | 1.131882649 | 0.296839987 | 0.991835066 | -4.774170069 |
| **A_55_P2034345** | Terf2 | 0.052400752 | -0.058254792 | 0.163056296 | 10.40649146 | 1.131809865 | 0.296868501 | 0.991835066 | -4.774225526 |
| **A_55_P2068167** | Nufip2 | 0.109658513 | -0.121986387 | 0.341303413 | 11.42700196 | 1.131432738 | 0.297016281 | 0.991835066 | -4.774512837 |
| **A_55_P2209767** | 2610002J2 | 0.118346981 | -0.131698889 | 0.368392851 | 9.251803795 | 1.131218841 | 0.297100126 | 0.991835066 | -4.774675769 |
| **A_51_P413785** | Commd3 | -0.081833481 | -0.254772166 | 0.091105205 | 12.23619692 | -1.130961974 | 0.297200839 | 0.991835066 | -4.774871408 |
| **A_55_P2136962** | NA | 0.108988201 | -0.121341734 | 0.339318135 | 10.86073527 | 1.130936522 | 0.29721082 | 0.991835066 | -4.774890792 |
| **A_52_P116372** | Ppp1r15b | 0.073518946 | -0.081859777 | 0.228897669 | 8.287384975 | 1.130880873 | 0.297232643 | 0.991835066 | -4.774933172 |
| **A_55_P2114995** | NA | 0.082666667 | -0.09206751 | 0.257400845 | 14.17662119 | 1.130737258 | 0.29728897 | 0.991835066 | -4.775042539 |
| **A_51_P245156** | Gdf2 | -0.068463987 | -0.213179131 | 0.076251157 | 7.842079208 | -1.13072589 | 0.297293429 | 0.991835066 | -4.775051196 |
| **A_55_P2056871** | Akap13 | 0.060695367 | -0.067610595 | 0.189001329 | 11.69686285 | 1.130623051 | 0.297333769 | 0.991835066 | -4.775129505 |
| **A_55_P2035539** | D3Ertd751( | -0.059571191 | -0.185513295 | 0.066370913 | 11.13498123 | -1.130510168 | 0.297378054 | 0.991835066 | -4.775215458 |
| **A_55_P2088296** | Fam160b2 | -0.099384298 | -0.309582483 | 0.110813887 | 13.13599686 | -1.130050743 | 0.297558347 | 0.991835066 | -4.775565228 |
| **A_55_P2079103** | Pan2 | 0.124667085 | -0.139006251 | 0.38834042 | 10.95493691 | 1.130042356 | 0.297561639 | 0.991835066 | -4.775571612 |
| **A_55_P2089507** | NA | 0.061407974 | -0.068480969 | 0.191296917 | 15.07232006 | 1.129956488 | 0.297595347 | 0.991835066 | -4.775636976 |
| **A_55_P2106096** | Fam171a1 | -0.062229222 | -0.193885349 | 0.069426905 | 9.72684493 | -1.129698183 | 0.297696764 | 0.991835066 | -4.775833583 |
| **A_55_P2130965** | NA | 0.126007305 | -0.140656159 | 0.392670769 | 13.21958907 | 1.129383243 | 0.297820457 | 0.991835066 | -4.776073262 |
| **A_52_P1026777** | Ecscr | 0.06380631 | -0.071241386 | 0.198854005 | 11.45737547 | 1.129238252 | 0.297877416 | 0.991835066 | -4.776183591 |
| **A_55_P2033075** | NA | 0.070505336 | -0.082325951 | 0.223336622 | 15.93635607 | 1.145979547 | 0.298007537 | 0.991835066 | -4.715303825 |
| **A_52_P599624** | Pfas | 0.055716779 | -0.062246628 | 0.173680186 | 8.465154077 | 1.128880129 | 0.298018142 | 0.991835066 | -4.776456067 |
| **A_51_P383140** | Slk | -0.090886641 | -0.283325826 | 0.101552545 | 9.416135598 | -1.12879655 | 0.298050993 | 0.991835066 | -4.77651965 |
| **A_52_P544555** | Hsbp1 | -0.102430399 | -0.319317196 | 0.114456398 | 12.33098997 | -1.128768506 | 0.298062016 | 0.991835066 | -4.776540984 |
| **A_51_P327564** | Glb1 | -0.06884095 | -0.214643002 | 0.076961102 | 10.34738647 | -1.128476064 | 0.298176987 | 0.991835066 | -4.776763437 |
| **A_55_P2278604** | A330009N2 | -0.066695265 | -0.207962665 | 0.074572134 | 8.743498493 | -1.128397687 | 0.298207807 | 0.991835066 | -4.77682305 |
| **A_51_P461884** | Abl2 | 0.103316401 | -0.115624977 | 0.32225778 | 10.2211182 | 1.12784795 | 0.298424049 | 0.991835066 | -4.777241111 |
| **A_66_P140408** | D8Ertd738( | -0.08863254 | -0.276488336 | 0.099223255 | 10.8992148 | -1.127658844 | 0.298498465 | 0.991835066 | -4.777384895 |
| **A_51_P277718** | Cenpc1 | -0.12694209 | -0.396006547 | 0.142122368 | 9.814222233 | -1.127608766 | 0.298518174 | 0.991835066 | -4.777422968 |
| **A_55_P2134012** | Oxa1l | -0.063109409 | -0.196885925 | 0.070667106 | 13.5025219 | -1.127517698 | 0.298554018 | 0.991835066 | -4.777492203 |
| **A_52_P134666** | Srpr | -0.098838789 | -0.308370637 | 0.11069306 | 10.20316618 | -1.127421991 | 0.298591691 | 0.991835066 | -4.777564961 |

| | | | | | | | | | |
|---|---|---|---|---|---|---|---|---|---|
| **A_52_P162509** | Ahcyl2 | 0.063517907 | -0.071140664 | 0.198176478 | 8.177043143 | 1.127382548 | 0.298607219 | 0.991835066 | -4.777594945 |
| **A_52_P608132** | Snx32 | -0.064813473 | -0.202220437 | 0.072593491 | 8.700045979 | -1.127367945 | 0.298612968 | 0.991835066 | -4.777606047 |
| **A_51_P430620** | Slc25a12 | -0.061463962 | -0.191789881 | 0.068861957 | 12.46528522 | -1.127194564 | 0.298681231 | 0.991835066 | -4.777737841 |
| **A_55_P1994355** | Ptges3 | 0.113035505 | -0.126768309 | 0.352839319 | 13.37216423 | 1.126595273 | 0.298917281 | 0.991835066 | -4.778193299 |
| **A_55_P1992617** | Clec1b | 0.190134779 | -0.213289612 | 0.59355917 | 8.978204342 | 1.126441246 | 0.298977974 | 0.991835066 | -4.778310336 |
| **A_51_P249608** | Trdn | -0.084017208 | -0.262292811 | 0.094258396 | 12.78392346 | -1.126381377 | 0.299001568 | 0.991835066 | -4.778355824 |
| **A_55_P2081510** | Atp5j2 | -0.107639182 | -0.336048665 | 0.120770301 | 15.83903521 | -1.126329325 | 0.299022082 | 0.991835066 | -4.778395372 |
| **A_55_P2125561** | Mrps36 | -0.102956691 | -0.321460216 | 0.115546834 | 13.71709942 | -1.126173389 | 0.299083546 | 0.991835066 | -4.778513843 |
| **A_66_P125862** | A930005H1 | -0.126634845 | -0.395423357 | 0.142153668 | 9.593256554 | -1.126034379 | 0.299138346 | 0.991835066 | -4.778619447 |
| **A_66_P131530** | Rpp30 | -0.057931569 | -0.180900301 | 0.065037163 | 8.439012493 | -1.125977538 | 0.299160757 | 0.991835066 | -4.778662626 |
| **A_51_P217430** | Riox1 | 0.098343876 | -0.11042306 | 0.307110812 | 9.425056051 | 1.125886822 | 0.299196526 | 0.991835066 | -4.778731535 |
| **A_51_P449935** | Ftsj3 | 0.100702546 | -0.113084248 | 0.31448934 | 9.292752325 | 1.125819351 | 0.299223131 | 0.991835066 | -4.778782785 |
| **A_55_P1997486** | Snu13 | -0.091149285 | -0.284669868 | 0.102371297 | 12.06484573 | -1.125732591 | 0.299257346 | 0.991835066 | -4.778848684 |
| **A_51_P433870** | Vegfc | 0.086187777 | -0.101260225 | 0.273635779 | 10.63234099 | 1.142173324 | 0.299461081 | 0.991835066 | -4.717973444 |
| **A_51_P137029** | Plce1 | -0.071166791 | -0.222347485 | 0.080013903 | 8.77299931 | -1.125097558 | 0.299507877 | 0.991835066 | -4.779330935 |
| **A_55_P2079272** | Kars | 0.057714614 | -0.064896375 | 0.180325603 | 13.06621886 | 1.12503371 | 0.299533076 | 0.991835066 | -4.779379413 |
| **A_51_P400016** | Kalrn | -0.064355613 | -0.201100803 | 0.072389576 | 11.20461089 | -1.124821239 | 0.299616944 | 0.991835066 | -4.779540725 |
| **A_55_P2040860** | Sun3 | -0.070056164 | -0.218951104 | 0.078838776 | 8.511662782 | -1.124541646 | 0.299727336 | 0.991835066 | -4.77975297 |
| **A_55_P2095325** | AI182371 | -0.047455375 | -0.148350506 | 0.053439756 | 7.653046333 | -1.124150181 | 0.299881955 | 0.991835066 | -4.780050089 |
| **A_55_P1990235** | Bin3 | 0.060641535 | -0.068330886 | 0.189613956 | 10.5564829 | 1.123783016 | 0.300027036 | 0.991835066 | -4.780328708 |
| **A_55_P2219657** | Tmf1 | 0.062994164 | -0.071014187 | 0.197002514 | 8.516596865 | 1.123511662 | 0.300134296 | 0.991835066 | -4.780534589 |
| **A_51_P233855** | Adprhl1 | -0.236524078 | -0.739762609 | 0.266714454 | 13.37876809 | -1.123338397 | 0.3002028 | 0.991835066 | -4.780666032 |
| **A_51_P465128** | Chst15 | -0.055091156 | -0.172308982 | 0.062126671 | 8.323042961 | -1.123304116 | 0.300216355 | 0.991835066 | -4.780692038 |
| **A_55_P2115057** | Tuba1a | 0.056551722 | -0.063773971 | 0.176877416 | 14.19507233 | 1.123302161 | 0.300217128 | 0.991835066 | -4.78069352 |
| **A_52_P628162** | Zfp592 | 0.062600437 | -0.070635202 | 0.195836077 | 8.449728386 | 1.122964655 | 0.30035061 | 0.991835066 | -4.780949522 |
| **A_66_P106090** | Rpa3 | 0.084547247 | -0.095402468 | 0.264496963 | 11.97934698 | 1.122942299 | 0.300359454 | 0.991835066 | -4.780966478 |
| **A_55_P2253289** | NA | 0.168673689 | -0.190334168 | 0.527681547 | 9.592488813 | 1.122929581 | 0.300364485 | 0.991835066 | -4.780976124 |
| **A_55_P2007334** | Mepce | 0.079184776 | -0.089370619 | 0.247740171 | 11.77772161 | 1.122814902 | 0.300409853 | 0.991835066 | -4.781063098 |
| **A_55_P1968878** | Rhpn2 | 0.114239384 | -0.128936034 | 0.357414802 | 9.436846899 | 1.122807525 | 0.300412772 | 0.991835066 | -4.781068692 |
| **A_55_P2024376** | NA | -0.070651369 | -0.221061803 | 0.079759064 | 15.77370861 | -1.122669056 | 0.300467559 | 0.991835066 | -4.781173701 |
| **A_51_P340402** | Eefsec | 0.064174685 | -0.072456276 | 0.200805646 | 9.285467152 | 1.122596678 | 0.3004962 | 0.991835066 | -4.781228586 |
| **A_55_P2126014** | NA | -0.103416277 | -0.323596534 | 0.11676398 | 15.92156642 | -1.122586095 | 0.300500388 | 0.991835066 | -4.781236611 |
| **A_55_P2112617** | Smad4 | -0.069102681 | -0.216236946 | 0.078031583 | 10.45298056 | -1.122509977 | 0.300530511 | 0.991835066 | -4.78129433 |
| **A_55_P1960083** | NA | 0.077192106 | -0.087185712 | 0.241569923 | 14.23402647 | 1.122377132 | 0.30058309 | 0.991835066 | -4.781395058 |
| **A_51_P444052** | Scp2 | -0.094075961 | -0.29449892 | 0.106346998 | 11.31302861 | -1.121864294 | 0.300786139 | 0.991835066 | -4.781783847 |
| **A_55_P2174601** | Plekha7 | 0.111721076 | -0.126329442 | 0.349771594 | 11.05812581 | 1.121695844 | 0.300852858 | 0.991835066 | -4.781911527 |
| **A_51_P262144** | Ddx19b | 0.081957815 | -0.092686652 | 0.256602283 | 9.170772351 | 1.121617233 | 0.300883999 | 0.991835066 | -4.781971109 |
| **A_51_P487918** | Rinl | -0.061678198 | -0.193118579 | 0.069762183 | 8.320920226 | -1.121532856 | 0.300917426 | 0.991835066 | -4.782035058 |
| **A_66_P111689** | NA | 0.068081883 | -0.077010928 | 0.213174693 | 9.452623375 | 1.121488426 | 0.300935029 | 0.991835066 | -4.78206873 |
| **A_55_P2164070** | Grip1 | 0.098409427 | -0.111329658 | 0.308148513 | 8.213160839 | 1.121415273 | 0.300964014 | 0.991835066 | -4.782124169 |
| **A_55_P2069221** | Prr11 | -0.067632957 | -0.211798524 | 0.076532611 | 8.408279304 | -1.121259048 | 0.301025921 | 0.991835066 | -4.782242557 |
| **A_52_P468343** | Bdh1 | -0.076291801 | -0.238916139 | 0.086332537 | 7.78754223 | -1.121247422 | 0.301030528 | 0.991835066 | -4.782251366 |
| **A_52_P89567** | Rhob | -0.088539087 | -0.27733385 | 0.100255677 | 8.70221875 | -1.120867371 | 0.301181176 | 0.991835066 | -4.782539328 |
| **A_55_P2026785** | Dcaf1 | 0.089796266 | -0.101684588 | 0.28127712 | 10.01781572 | 1.12083596 | 0.301193629 | 0.991835066 | -4.782563124 |
| **A_55_P2088825** | Foxp1 | 0.070148399 | -0.079448836 | 0.219745634 | 12.31091855 | 1.120736021 | 0.301233256 | 0.991835066 | -4.782638837 |
| **A_55_P2013780** | NA | -0.154447085 | -0.483850134 | 0.174955963 | 13.16065441 | -1.120627397 | 0.301276331 | 0.991835066 | -4.782721124 |
| **A_51_P487678** | Dtnbp1 | -0.090018124 | -0.282010875 | 0.101974627 | 10.65313697 | -1.120609393 | 0.301283471 | 0.991835066 | -4.782734762 |
| **A_51_P320980** | Sfxn4 | -0.082458773 | -0.258354698 | 0.093437151 | 8.710516787 | -1.120444179 | 0.301348997 | 0.991835066 | -4.782859908 |

| | | | | | | | | | |
|---|---|---|---|---|---|---|---|---|---|
| **A_51_P414837** | Faf1 | -0.055268313 | -0.173164448 | 0.062627821 | 10.54415818 | -1.120432703 | 0.301353549 | 0.991835066 | -4.782868601 |
| **A_52_P448466** | BC004004 | -0.079242343 | -0.248284027 | 0.089799341 | 8.690806536 | -1.120398786 | 0.301367003 | 0.991835066 | -4.782894291 |
| **A_52_P520466** | Kif18b | -0.049882342 | -0.156294004 | 0.05652932 | 8.36214313 | -1.120383598 | 0.301373028 | 0.991835066 | -4.782905795 |
| **A_52_P603654** | Stx8 | -0.096653549 | -0.302876828 | 0.109569731 | 9.722545665 | -1.120183696 | 0.301452333 | 0.991835066 | -4.783057197 |
| **A_55_P2051322** | Efhd2 | -0.069316545 | -0.217218072 | 0.078584982 | 11.52486958 | -1.120142772 | 0.301468571 | 0.991835066 | -4.78308819 |
| **A_55_P1988481** | NA | -0.08809284 | -0.2760625 | 0.099876819 | 9.263002668 | -1.120113395 | 0.301480227 | 0.991835066 | -4.783110439 |
| **A_55_P1966804** | Fdps | 0.170900499 | -0.193828349 | 0.535629347 | 12.6025186 | 1.119908004 | 0.301561734 | 0.991835066 | -4.783265976 |
| **A_55_P1985070** | Robo1 | 0.192103146 | -0.217973619 | 0.602179911 | 10.34009657 | 1.119640107 | 0.301668073 | 0.991835066 | -4.783468822 |
| **A_55_P2009895** | Gm5113 | -0.053146918 | -0.16660643 | 0.060312594 | 8.775669867 | -1.119557252 | 0.301700968 | 0.991835066 | -4.783531553 |
| **A_52_P124472** | Kcnd2 | -0.082986647 | -0.260165367 | 0.094192073 | 9.087640515 | -1.119452805 | 0.301742439 | 0.991835066 | -4.783610627 |
| **A_51_P200610** | Zbtb12 | 0.092101128 | -0.104541933 | 0.28874419 | 10.36756028 | 1.119426171 | 0.301753015 | 0.991835066 | -4.78363079 |
| **A_55_P2081981** | NA | -0.090891892 | -0.284977246 | 0.103193461 | 8.517655991 | -1.119287144 | 0.301808226 | 0.991835066 | -4.783736035 |
| **A_55_P1993864** | Sh2b3 | 0.056418112 | -0.064059805 | 0.176896028 | 9.454085651 | 1.119232286 | 0.301830013 | 0.991835066 | -4.783777562 |
| **A_55_P2008056** | Cideb | -0.04827504 | -0.151399554 | 0.054849474 | 7.884082996 | -1.118844827 | 0.301983934 | 0.991835066 | -4.784070825 |
| **A_52_P497534** | Wipi1 | -0.093240702 | -0.292445869 | 0.105964466 | 10.68103141 | -1.118701099 | 0.302041048 | 0.991835066 | -4.784179597 |
| **A_55_P1995633** | NA | -0.162087287 | -0.50844204 | 0.184267467 | 12.33505205 | -1.118502432 | 0.302120007 | 0.991835066 | -4.784329931 |
| **A_55_P2051254** | Pvt1 | 0.069434262 | -0.078947282 | 0.217815806 | 8.227745722 | 1.118415224 | 0.302154672 | 0.991835066 | -4.784395917 |
| **A_55_P1990755** | Ppp3r1 | 0.105597121 | -0.120103606 | 0.331297847 | 11.76613566 | 1.118222563 | 0.302231268 | 0.991835066 | -4.784541685 |
| **A_55_P1995996** | Zfp984 | 0.067064491 | -0.076296797 | 0.21042578 | 9.241086985 | 1.118072256 | 0.302291036 | 0.991835066 | -4.784655398 |
| **A_52_P504268** | B3galnt1 | 0.118948821 | -0.135363909 | 0.373261551 | 9.425847333 | 1.11789541 | 0.30236137 | 0.991835066 | -4.784789177 |
| **A_51_P187665** | Rmi1 | -0.060326919 | -0.189319743 | 0.068665905 | 9.868972905 | -1.117775857 | 0.302408925 | 0.991835066 | -4.784879608 |
| **A_55_P2144285** | Nnt | 0.112089956 | -0.12760936 | 0.351789272 | 14.02693531 | 1.117658271 | 0.302455703 | 0.991835066 | -4.784968545 |
| **A_51_P329158** | Zc3h10 | -0.077256601 | -0.242468334 | 0.087955131 | 8.961775538 | -1.117644917 | 0.302461017 | 0.991835066 | -4.784978646 |
| **A_55_P2028362** | NA | -0.086080916 | -0.270166204 | 0.098004372 | 8.550758476 | -1.117627117 | 0.302468098 | 0.991835066 | -4.784992108 |
| **A_55_P2075794** | Tti2 | 0.066907153 | -0.076205963 | 0.210020269 | 11.20987552 | 1.117383469 | 0.30256505 | 0.991835066 | -4.785176374 |
| **A_52_P5652** | Sae1 | 0.086718184 | -0.098807181 | 0.272243549 | 9.726133477 | 1.117161637 | 0.302653343 | 0.991835066 | -4.78534412 |
| **A_55_P1964174** | Nme1 | 0.120166601 | -0.1369502 | 0.377283402 | 11.54030769 | 1.117023869 | 0.302708188 | 0.991835066 | -4.785448289 |
| **A_52_P3845** | Rnf217 | -0.051451496 | -0.161544549 | 0.058641557 | 8.904903324 | -1.116984705 | 0.30272378 | 0.991835066 | -4.7854779 |
| **A_55_P2023424** | Trip10 | 0.082353001 | -0.093892124 | 0.258598125 | 11.66008976 | 1.116789821 | 0.30280138 | 0.991835066 | -4.785625239 |
| **A_55_P2122504** | Zpr1 | 0.067788848 | -0.077297861 | 0.212875558 | 11.35567304 | 1.11670833 | 0.302833833 | 0.991835066 | -4.785686844 |
| **A_55_P2145908** | Phb | 0.082154945 | -0.093704468 | 0.258014357 | 14.0407889 | 1.116547548 | 0.302897872 | 0.991835066 | -4.785808384 |
| **A_55_P2087384** | Gps1 | 0.065827893 | -0.07509037 | 0.206746155 | 13.40397002 | 1.116482204 | 0.302923901 | 0.991835066 | -4.785857776 |
| **A_55_P2088490** | Acad10 | -0.104143277 | -0.327108704 | 0.118822151 | 10.78413104 | -1.116356339 | 0.302974044 | 0.991835066 | -4.785952911 |
| **A_51_P510849** | Tmem50b | 0.095104003 | -0.108533023 | 0.298741028 | 12.57526577 | 1.116223558 | 0.303026949 | 0.991835066 | -4.786053266 |
| **A_55_P2064053** | NA | 0.090159361 | -0.1029049 | 0.283223622 | 10.10278132 | 1.116138452 | 0.303060863 | 0.991835066 | -4.786117586 |
| **A_55_P2041270** | Arhgef1 | 0.106401052 | -0.121475548 | 0.334277653 | 11.36034226 | 1.115977206 | 0.303125126 | 0.991835066 | -4.786239439 |
| **A_51_P265219** | Ppm1k | -0.137399762 | -0.431691562 | 0.156892039 | 10.06885338 | -1.115878604 | 0.303164428 | 0.991835066 | -4.786313948 |
| **A_55_P2077872** | Deaf1 | 0.084191536 | -0.0961364 | 0.264519472 | 9.943339856 | 1.115872451 | 0.303166881 | 0.991835066 | -4.786318597 |
| **A_55_P2087182** | Car4 | -0.053429498 | -0.167877493 | 0.061018498 | 7.761418749 | -1.115788887 | 0.303200192 | 0.991835066 | -4.786381739 |
| **A_55_P2136289** | NA | -0.048949625 | -0.153810073 | 0.055910822 | 8.782159524 | -1.115698365 | 0.303236282 | 0.991835066 | -4.786450135 |
| **A_55_P2102569** | Cox7a2l | -0.150338722 | -0.472419862 | 0.171742418 | 12.16022696 | -1.11561589 | 0.303269165 | 0.991835066 | -4.786512449 |
| **A_51_P109939** | Naxe | -0.050326023 | -0.158145711 | 0.057493665 | 13.03896377 | -1.115587581 | 0.303280453 | 0.991835066 | -4.786533837 |
| **A_55_P2043337** | Colq | 0.163168538 | -0.186455491 | 0.512792568 | 10.17920468 | 1.115435036 | 0.303341285 | 0.991835066 | -4.786649082 |
| **A_66_P114974** | NA | -0.144191232 | -0.453236261 | 0.164853797 | 13.05301292 | -1.115131859 | 0.303462214 | 0.991835066 | -4.7868781 |
| **A_52_P286002** | Obsl1 | 0.072035812 | -0.082371249 | 0.226442872 | 8.756563066 | 1.115039954 | 0.30349888 | 0.991835066 | -4.786947517 |
| **A_55_P2093221** | Rnf111 | 0.094553746 | -0.108152179 | 0.29725967 | 11.3759089 | 1.114862811 | 0.303569564 | 0.991835066 | -4.787081306 |
| **A_51_P312310** | Mtbp | 0.113156641 | -0.129438691 | 0.355751972 | 10.44525045 | 1.114825105 | 0.303584611 | 0.991835066 | -4.787109782 |
| **A_51_P157524** | E2f4 | 0.087343617 | -0.099918977 | 0.274606211 | 8.313248795 | 1.114780255 | 0.30360251 | 0.991835066 | -4.787143653 |

| | | | | | | | | | |
|---|---|---|---|---|---|---|---|---|---|
| **A_55_P1974681** | Nsun4 | -0.072410308 | -0.227669625 | 0.082849009 | 9.834192076 | -1.114684227 | 0.303640836 | 0.991835066 | -4.787216171 |
| **A_55_P1960873** | Mbd3 | 0.143690952 | -0.164416804 | 0.451798708 | 9.670848538 | 1.114643333 | 0.303657158 | 0.991835066 | -4.787247051 |
| **A_55_P2075237** | Ulk3 | 0.094005742 | -0.107571875 | 0.295583359 | 9.684612854 | 1.114605564 | 0.303672234 | 0.991835066 | -4.787275572 |
| **A_55_P2080308** | Gm12538 | 0.074589593 | -0.085437375 | 0.234616561 | 8.434173683 | 1.114023093 | 0.30390481 | 0.991835066 | -4.787715341 |
| **A_51_P350332** | Rbpms | -0.069829728 | -0.219671778 | 0.080012323 | 10.69065987 | -1.11382195 | 0.303985159 | 0.991835066 | -4.787867173 |
| **A_55_P1960311** | Dnase1l1 | -0.084421069 | -0.265579895 | 0.096737757 | 9.609354773 | -1.113782734 | 0.304000826 | 0.991835066 | -4.787896773 |
| **A_51_P172752** | Gpatch4 | 0.141643191 | -0.1623288 | 0.445615183 | 11.2455259 | 1.113707814 | 0.30403076 | 0.991835066 | -4.787953321 |
| **A_55_P2033500** | Ltbp4 | -0.136468365 | -0.429375645 | 0.156438916 | 11.60801189 | -1.113553156 | 0.304092559 | 0.991835066 | -4.788070046 |
| **A_52_P159885** | Ddx49 | 0.11616561 | -0.133200813 | 0.365532033 | 9.141139066 | 1.113393632 | 0.304156313 | 0.991835066 | -4.788190434 |
| **A_55_P2028365** | Lrrc36 | -0.062157802 | -0.195597009 | 0.071281405 | 7.93420163 | -1.11332338 | 0.304184394 | 0.991835066 | -4.788243448 |
| **A_51_P280437** | Slc12a6 | -0.078894109 | -0.248275767 | 0.090487549 | 9.616251848 | -1.113236223 | 0.304219234 | 0.991835066 | -4.788309216 |
| **A_55_P2153587** | Afg3l1 | -0.080636667 | -0.25376433 | 0.092490997 | 11.41424596 | -1.113205211 | 0.304231631 | 0.991835066 | -4.788332617 |
| **A_55_P2176300** | Slc33a1 | -0.052953744 | -0.166647926 | 0.060740438 | 8.269352852 | -1.113185558 | 0.304239488 | 0.991835066 | -4.788347445 |
| **A_52_P36680** | Pggt1b | 0.061316477 | -0.070338443 | 0.192971398 | 8.696775486 | 1.113138576 | 0.304258271 | 0.991835066 | -4.788382896 |
| **A_55_P2048671** | Rpl27a | 0.065184197 | -0.07480906 | 0.205177454 | 16.20614387 | 1.112869744 | 0.304365765 | 0.991835066 | -4.788585724 |
| **A_55_P2145136** | NA | 0.070344179 | -0.080752775 | 0.221441132 | 8.300575698 | 1.112708962 | 0.30443007 | 0.991835066 | -4.788707017 |
| **A_55_P2162204** | Kctd15 | 0.133670162 | -0.153502817 | 0.420843141 | 9.985559987 | 1.112500036 | 0.304513647 | 0.991835066 | -4.788864614 |
| **A_55_P2014075** | Ercc2 | 0.062127954 | -0.071367832 | 0.195623739 | 10.38881596 | 1.112317136 | 0.304586828 | 0.991835066 | -4.789002565 |
| **A_51_P245405** | Ppp3cc | 0.048486922 | -0.055701827 | 0.152675671 | 10.06073275 | 1.11227691 | 0.304602925 | 0.991835066 | -4.789032903 |
| **A_55_P2121275** | Gm4907 | 0.077150229 | -0.088665199 | 0.242965657 | 8.39121956 | 1.112042582 | 0.304696708 | 0.991835066 | -4.78920962 |
| **A_55_P2024075** | Prkacb | -0.085288595 | -0.268608959 | 0.098031769 | 12.82526382 | -1.11196055 | 0.304729545 | 0.991835066 | -4.789271479 |
| **A_51_P292057** | Tbl1xr1 | 0.053313245 | -0.06128487 | 0.16791136 | 8.500728665 | 1.111902674 | 0.304752713 | 0.991835066 | -4.78931512 |
| **A_51_P120295** | Cyp2d26 | -0.231835597 | -0.730364033 | 0.266692839 | 8.317534102 | -1.111474023 | 0.304924356 | 0.991835066 | -4.789638302 |
| **A_51_P154780** | Vav1 | -0.04745535 | -0.149503061 | 0.054592361 | 7.792789937 | -1.111452862 | 0.304932832 | 0.991835066 | -4.789654255 |
| **A_52_P581390** | Kif1c | 0.098696288 | -0.113569282 | 0.310961858 | 8.497366867 | 1.111297652 | 0.304995003 | 0.991835066 | -4.789771256 |
| **A_55_P2011436** | NA | -0.048589915 | -0.153094208 | 0.055914378 | 15.14786261 | -1.111273968 | 0.30500449 | 0.991835066 | -4.789789109 |
| **A_55_P1953475** | NA | -0.081004638 | -0.255252689 | 0.093243413 | 16.15709081 | -1.111094735 | 0.305076299 | 0.991835066 | -4.789924206 |
| **A_55_P2058746** | Mterf1a | 0.074608651 | -0.085888287 | 0.235105589 | 8.76482217 | 1.111044797 | 0.305096309 | 0.991835066 | -4.789961844 |
| **A_55_P2110567** | Proca1 | -0.053383624 | -0.168233371 | 0.061466123 | 8.293584188 | -1.110931144 | 0.305141853 | 0.991835066 | -4.790047501 |
| **A_52_P306357** | NA | 0.074719716 | -0.086056897 | 0.235496329 | 7.704000765 | 1.11076316 | 0.305209179 | 0.991835066 | -4.790174096 |
| **A_51_P114634** | Amz1 | -0.045847075 | -0.144507817 | 0.052813667 | 8.195157114 | -1.110647874 | 0.305255391 | 0.991835066 | -4.790260971 |
| **A_51_P424959** | Bcl6b | -0.082542343 | -0.260179553 | 0.095094867 | 9.59630679 | -1.110585454 | 0.305280415 | 0.991835066 | -4.790308006 |
| **A_66_P130813** | Samd4 | -0.101121421 | -0.318753538 | 0.116510695 | 10.47984695 | -1.110527484 | 0.305303656 | 0.991835066 | -4.790351686 |
| **A_51_P504065** | Spa17 | -0.06256762 | -0.197234012 | 0.072098772 | 9.462641372 | -1.110451368 | 0.305334174 | 0.991835066 | -4.790409037 |
| **A_55_P1989673** | Slco2a1 | 0.083074843 | -0.095740124 | 0.261889809 | 10.07385772 | 1.11038809 | 0.305359547 | 0.991835066 | -4.790456713 |
| **A_55_P2083997** | Zfp664 | 0.055991575 | -0.064544088 | 0.176527239 | 12.18798901 | 1.110238428 | 0.305419564 | 0.991835066 | -4.790569469 |
| **A_52_P28806** | Foxm1 | -0.063661027 | -0.200722601 | 0.073400547 | 9.337044478 | -1.110112665 | 0.305470005 | 0.991835066 | -4.790664212 |
| **A_55_P1985304** | Krtap5-3 | 0.052898103 | -0.061010238 | 0.166806444 | 8.424890759 | 1.109925182 | 0.305545213 | 0.991835066 | -4.790805439 |
| **A_51_P165098** | Gga2 | 0.10783128 | -0.12438031 | 0.340042871 | 11.84249857 | 1.109864602 | 0.305569518 | 0.991835066 | -4.790851069 |
| **A_52_P352362** | NA | 0.236511301 | -0.27292547 | 0.745948072 | 9.420259922 | 1.109610968 | 0.305671293 | 0.991835066 | -4.791042099 |
| **A_52_P38639** | Fermt3 | -0.06156946 | -0.194190725 | 0.071051805 | 7.958179484 | -1.10958687 | 0.305680964 | 0.991835066 | -4.791060247 |
| **A_55_P1968969** | NA | 0.115429917 | -0.133222446 | 0.364082281 | 14.52836667 | 1.109519464 | 0.305708018 | 0.991835066 | -4.79111101 |
| **A_65_P13209** | Ctsd | 0.166866188 | -0.1926404 | 0.526372775 | 9.509506048 | 1.1093552 | 0.305773953 | 0.991835066 | -4.791234708 |
| **A_55_P2183894** | Gm5617 | -0.076893308 | -0.242559845 | 0.08877323 | 11.13264135 | -1.109335422 | 0.305781893 | 0.991835066 | -4.791249602 |
| **A_51_P352687** | Rnf216 | 0.122529936 | -0.141468836 | 0.386528707 | 10.60543442 | 1.109301069 | 0.305795683 | 0.991835066 | -4.791275469 |
| **A_51_P259571** | Angptl6 | 0.074698548 | -0.086270103 | 0.2356672 | 9.30395022 | 1.109123705 | 0.305866894 | 0.991835066 | -4.791409016 |
| **A_55_P2076941** | Lrr1 | 0.064024578 | -0.073950637 | 0.201999794 | 8.783149983 | 1.109059323 | 0.305892747 | 0.991835066 | -4.79145749 |
| **A_55_P1965333** | Naa60 | 0.070672051 | -0.081637318 | 0.222981421 | 11.44321187 | 1.108996584 | 0.305917941 | 0.991835066 | -4.791504725 |

| | | | | | | | | | |
|---|---|---|---|---|---|---|---|---|---|
| **A_55_P2067905** | Scfd2 | 0.069240905 | -0.080000584 | 0.218482394 | 10.50901539 | 1.108874239 | 0.305967077 | 0.991835066 | -4.791596832 |
| **A_55_P1983999** | Desi1 | -0.094199711 | -0.29724557 | 0.108846149 | 8.696383785 | -1.108828965 | 0.305985261 | 0.991835066 | -4.791630915 |
| **A_55_P2021054** | Elp5 | 0.078474348 | -0.090684978 | 0.247633673 | 11.74887261 | 1.108768556 | 0.306009526 | 0.991835066 | -4.79167639 |
| **A_55_P1999154** | Gm4987 | 0.091170376 | -0.105406725 | 0.287747477 | 13.97548923 | 1.108485336 | 0.306123309 | 0.991835066 | -4.791889575 |
| **A_55_P2008197** | Esr2 | 0.201192301 | -0.232673928 | 0.635058529 | 10.25989991 | 1.108318653 | 0.30619029 | 0.991835066 | -4.792015026 |
| **A_51_P222657** | Tspo | -0.164128811 | -0.518089719 | 0.189832096 | 11.7400947 | -1.108252339 | 0.306216941 | 0.991835066 | -4.792064933 |
| **A_52_P575178** | Cdon | 0.12834526 | -0.148467513 | 0.405158033 | 10.83567482 | 1.108160943 | 0.306253676 | 0.991835066 | -4.792133712 |
| **A_55_P1968841** | NA | 0.096148159 | -0.111237821 | 0.30353414 | 11.8164728 | 1.10807897 | 0.306286627 | 0.991835066 | -4.792195399 |
| **A_66_P105764** | Tor1b | 0.050579902 | -0.058538377 | 0.159698181 | 9.501701502 | 1.107872059 | 0.306369811 | 0.991835066 | -4.79235109 |
| **A_52_P466887** | Hnrnpd | 0.070756601 | -0.081900128 | 0.223413329 | 12.30013634 | 1.107796884 | 0.306400038 | 0.991835066 | -4.792407652 |
| **A_55_P2054883** | Trappc1 | -0.113597063 | -0.358683164 | 0.131489038 | 9.896721631 | -1.107790283 | 0.306402693 | 0.991835066 | -4.792412619 |
| **A_55_P2000813** | Cops2 | 0.085595015 | -0.099105238 | 0.270295268 | 10.01586883 | 1.107618281 | 0.306471863 | 0.991835066 | -4.792542024 |
| **A_51_P152747** | Nrip1 | 0.068202504 | -0.078973625 | 0.215378632 | 8.900334341 | 1.107572276 | 0.306490366 | 0.991835066 | -4.792576633 |
| **A_52_P88695** | Dgka | -0.071553924 | -0.225969984 | 0.082862135 | 9.313618586 | -1.107516295 | 0.306512882 | 0.991835066 | -4.792618747 |
| **A_51_P370510** | Clba1 | -0.073823739 | -0.233142855 | 0.085495378 | 8.793531028 | -1.10748353 | 0.306526061 | 0.991835066 | -4.792643395 |
| **A_55_P2186085** | Atp11c | 0.096558291 | -0.111859744 | 0.304976325 | 8.963610157 | 1.107295172 | 0.306601835 | 0.991835066 | -4.792785082 |
| **A_55_P2048434** | Hsp90ab1 | 0.080063676 | -0.092751386 | 0.252878739 | 16.33696983 | 1.10729431 | 0.306602182 | 0.991835066 | -4.79278573 |
| **A_52_P587948** | Zfp11 | -0.05180662 | -0.163634988 | 0.060021748 | 8.989804853 | -1.107241625 | 0.306623379 | 0.991835066 | -4.792825358 |
| **A_55_P1958912** | NA | -0.74420004 | -2.351031505 | 0.862631425 | 9.74906456 | -1.106951247 | 0.30674023 | 0.991835066 | -4.793043752 |
| **A_51_P166244** | Osbpl1a | -0.062042342 | -0.196019594 | 0.07193491 | 12.51851785 | -1.10679261 | 0.306804083 | 0.991835066 | -4.793163048 |
| **A_52_P118560** | Kynu | 0.072009022 | -0.083515658 | 0.227533703 | 7.674006374 | 1.106615451 | 0.306875404 | 0.991835066 | -4.793296262 |
| **A_55_P2227335** | Ptpre | 0.052893632 | -0.061348118 | 0.167135381 | 8.393464395 | 1.106592379 | 0.306884693 | 0.991835066 | -4.79331361 |
| **A_52_P501721** | Ccdc127 | -0.065738086 | -0.207729056 | 0.076252884 | 8.715218567 | -1.106535782 | 0.306907481 | 0.991835066 | -4.793356164 |
| **A_55_P2009022** | NA | -0.064568646 | -0.204042322 | 0.074905031 | 8.843752777 | -1.106467239 | 0.306935081 | 0.991835066 | -4.7934077 |
| **A_55_P2053783** | Ppp2r3a | -0.092935431 | -0.293697371 | 0.107826509 | 8.587209088 | -1.106392103 | 0.306965338 | 0.991835066 | -4.793464189 |
| **A_55_P2011137** | Armc10 | 0.069208816 | -0.080303088 | 0.21872072 | 8.914823833 | 1.106355701 | 0.306979998 | 0.991835066 | -4.793491556 |
| **A_55_P1959753** | NA | 0.158785863 | -0.184349085 | 0.501920812 | 8.784565547 | 1.106002257 | 0.307122368 | 0.991835066 | -4.793757251 |
| **A_55_P2131607** | Trnau1ap | -0.089303429 | -0.282325038 | 0.10371818 | 10.50384103 | -1.105786636 | 0.307209248 | 0.991835066 | -4.793919315 |
| **A_55_P2159485** | NA | 0.148864993 | -0.172948399 | 0.470678385 | 9.457558078 | 1.105598908 | 0.307284906 | 0.991835066 | -4.794060399 |
| **A_66_P129469** | NA | -0.049986554 | -0.158059717 | 0.058086609 | 7.767146632 | -1.105463651 | 0.307339427 | 0.991835066 | -4.79416204 |
| **A_51_P322265** | Grap | -0.088023048 | -0.278353033 | 0.102306936 | 8.698216879 | -1.105346203 | 0.307386775 | 0.991835066 | -4.794250293 |
| **A_55_P2025103** | Snx3 | -0.097601361 | -0.308668267 | 0.113465545 | 11.4891211 | -1.105210144 | 0.307441634 | 0.991835066 | -4.794352523 |
| **A_51_P438701** | Atpaf2 | -0.070808974 | -0.22394477 | 0.082326822 | 10.00861398 | -1.105148696 | 0.307466412 | 0.991835066 | -4.79439869 |
| **A_55_P2112105** | NA | -0.172679887 | -0.54616333 | 0.200803556 | 13.49330262 | -1.105043905 | 0.307508672 | 0.991835066 | -4.794477419 |
| **A_55_P2071406** | NA | 0.059724147 | -0.069455432 | 0.188903726 | 8.713994145 | 1.105007495 | 0.307523356 | 0.991835066 | -4.794504772 |
| **A_55_P2008896** | NA | -0.072839964 | -0.230423247 | 0.084743318 | 12.70523242 | -1.104761966 | 0.307622395 | 0.991835066 | -4.794689215 |
| **A_55_P2016069** | Adamtsl1 | 0.074834762 | -0.087076507 | 0.23674603 | 8.469791482 | 1.104677321 | 0.307656544 | 0.991835066 | -4.794752796 |
| **A_55_P1992884** | NA | 0.046957732 | -0.054643743 | 0.148559206 | 7.8960196 | 1.104628466 | 0.307676256 | 0.991835066 | -4.794789491 |
| **A_52_P214891** | Stx6 | -0.059867527 | -0.189428823 | 0.06969377 | 10.16223161 | -1.104396859 | 0.307769716 | 0.991835066 | -4.794963441 |
| **A_51_P166023** | Hip1 | -0.05657631 | -0.179021271 | 0.06586865 | 8.684390564 | -1.104340089 | 0.307792628 | 0.991835066 | -4.795006075 |
| **A_52_P78373** | Sorbs3 | 0.063667297 | -0.074143413 | 0.201478007 | 8.286563199 | 1.104186859 | 0.307854477 | 0.991835066 | -4.795121143 |
| **A_55_P2038514** | NA | -0.050317136 | -0.159239265 | 0.058604992 | 7.701481856 | -1.104101317 | 0.307889009 | 0.991835066 | -4.795185377 |
| **A_52_P317393** | Adgrg1 | -0.082807939 | -0.262066332 | 0.096450454 | 9.008450281 | -1.10408271 | 0.307896521 | 0.991835066 | -4.795199349 |
| **A_51_P116616** | Rbm14 | 0.081512006 | -0.094989668 | 0.258013681 | 11.97586904 | 1.103778384 | 0.308019402 | 0.991835066 | -4.783439809 |
| **A_55_P1970414** | Mccc1 | 0.081875815 | -0.095423675 | 0.259175304 | 10.27536676 | 1.103715854 | 0.308044656 | 0.991835066 | -4.795474787 |
| **A_55_P1979067** | A430073D2 | 0.243266687 | -0.283548171 | 0.770081545 | 13.237591 | 1.103656075 | 0.3080688 | 0.991835066 | -4.795519664 |
| **A_51_P255304** | Cav3 | -0.108472618 | -0.343414096 | 0.12646886 | 14.27016855 | -1.103492936 | 0.308134697 | 0.991835066 | -4.795642128 |
| **A_55_P2371801** | A430046D1 | -0.055252519 | -0.174945982 | 0.064440944 | 8.492754307 | -1.103292799 | 0.308215556 | 0.991835066 | -4.795792351 |

| | | | | | | | | | |
|---|---|---|---|---|---|---|---|---|---|
| **A_55_P2136732** | Cd99 | -0.139281877 | -0.441078312 | 0.162514557 | 13.27252543 | -1.103035881 | 0.30831938 | 0.991835066 | -4.79598517 |
| **A_55_P2054456** | Adprm | 0.09354079 | -0.109156324 | 0.296237905 | 10.50857903 | 1.10296721 | 0.308347136 | 0.991835066 | -4.796036703 |
| **A_51_P503131** | Wnt11 | 0.093460865 | -0.109068834 | 0.295990564 | 8.555617287 | 1.102935745 | 0.308359854 | 0.991835066 | -4.796060315 |
| **A_55_P2133632** | Sipa1l2 | -0.08624112 | -0.273156778 | 0.100674537 | 13.50073607 | -1.102751966 | 0.308434147 | 0.991835066 | -4.796198219 |
| **A_55_P2147816** | NA | -0.105635749 | -0.33458882 | 0.123317323 | 17.0993852 | -1.102741128 | 0.308438529 | 0.991835066 | -4.796206351 |
| **A_55_P2133427** | Vkorc1 | 0.059837394 | -0.069864087 | 0.189538876 | 10.83082002 | 1.102647937 | 0.308476208 | 0.991835066 | -4.796276274 |
| **A_51_P174681** | Bzw2 | -0.072557464 | -0.22984268 | 0.084727751 | 12.79429947 | -1.102562784 | 0.30851064 | 0.991835066 | -4.796340163 |
| **A_51_P238183** | Etnk2 | 0.077162274 | -0.090106846 | 0.244431395 | 8.236492605 | 1.102550221 | 0.30851572 | 0.991835066 | -4.796349588 |
| **A_51_P249867** | Snrnp48 | -0.060083423 | -0.19034397 | 0.070177123 | 9.589368149 | -1.102429704 | 0.308564458 | 0.991835066 | -4.796440004 |
| **A_52_P120740** | 0610030E2 | 0.04791233 | -0.055971397 | 0.151796056 | 8.173269114 | 1.102323079 | 0.308607583 | 0.991835066 | -4.796519992 |
| **A_51_P517120** | Zfp930 | 0.068733522 | -0.080299149 | 0.217766193 | 10.99532831 | 1.102290955 | 0.308620576 | 0.991835066 | -4.79654409 |
| **A_52_P630493** | Dnajb6 | 0.099280199 | -0.115986013 | 0.314546411 | 8.957567562 | 1.102290088 | 0.308620927 | 0.991835066 | -4.796544741 |
| **A_55_P2128229** | NA | 0.15760964 | -0.184157149 | 0.499376428 | 8.54708662 | 1.102204174 | 0.30865568 | 0.991835066 | -4.796609188 |
| **A_55_P1982451** | Hmgn5 | -0.074492903 | -0.236048881 | 0.087063074 | 11.01148586 | -1.102049243 | 0.30871836 | 0.991835066 | -4.796725399 |
| **A_55_P2086835** | NA | -0.156987156 | -0.497503923 | 0.18352961 | 15.59449067 | -1.101881152 | 0.308786375 | 0.991835066 | -4.796851471 |
| **A_51_P494686** | Sumf1 | -0.053066124 | -0.16822688 | 0.062094631 | 10.15310009 | -1.101341482 | 0.309004826 | 0.991835066 | -4.797256156 |
| **A_55_P2069787** | NA | -0.121066779 | -0.383834396 | 0.141700837 | 12.86084171 | -1.101190053 | 0.309066146 | 0.991835066 | -4.797369688 |
| **A_55_P2000658** | Gm15645 | -0.102373108 | -0.324584132 | 0.119837916 | 9.896985508 | -1.101106739 | 0.309099887 | 0.991835066 | -4.797432147 |
| **A_51_P194249** | Stmn4 | 0.157743701 | -0.184661063 | 0.500148466 | 9.053527408 | 1.101086304 | 0.309108164 | 0.991835066 | -4.797447467 |
| **A_55_P1974783** | Tsnax | -0.05398098 | -0.171158531 | 0.06319657 | 12.39577668 | -1.101046055 | 0.309124465 | 0.991835066 | -4.797477639 |
| **A_52_P576720** | Calm4 | 0.062788674 | -0.073514803 | 0.199092152 | 7.70668636 | 1.100990346 | 0.30914703 | 0.991835066 | -4.797519401 |
| **A_51_P143031** | Eps8l3 | -0.069162416 | -0.219331803 | 0.08100697 | 7.728096308 | -1.100773297 | 0.309234957 | 0.991835066 | -4.797682095 |
| **A_55_P1963631** | Yae1d1 | 0.072557054 | -0.084988716 | 0.230102824 | 10.68669201 | 1.100733102 | 0.309251243 | 0.991835066 | -4.797712223 |
| **A_55_P2131579** | NA | -0.061655 | -0.195534223 | 0.072224223 | 14.62147763 | -1.100688049 | 0.309269497 | 0.991835066 | -4.79774599 |
| **A_55_P1989076** | Slc22a23 | 0.093483824 | -0.109519939 | 0.296487586 | 9.560828493 | 1.100630425 | 0.309292846 | 0.991835066 | -4.797789179 |
| **A_55_P2037584** | Trf | -0.48684497 | -1.544145422 | 0.570455482 | 11.71329545 | -1.100528773 | 0.309334039 | 0.991835066 | -4.797865362 |
| **A_55_P2173952** | Myh6 | -0.095139387 | -0.301767647 | 0.111488872 | 17.50900959 | -1.100473943 | 0.30935626 | 0.991835066 | -4.797906453 |
| **A_55_P2076984** | Farp1 | 0.068894722 | -0.080734603 | 0.218524048 | 8.789870545 | 1.100470404 | 0.309357694 | 0.991835066 | -4.797909105 |
| **A_52_P404363** | Cdc42bpb | -0.049530023 | -0.15710566 | 0.058045613 | 8.536809878 | -1.100433343 | 0.309372715 | 0.991835066 | -4.797936879 |
| **A_55_P2135220** | NA | -0.188641749 | -0.598386876 | 0.221103378 | 8.714514152 | -1.100355829 | 0.309404132 | 0.991835066 | -4.797994966 |
| **A_55_P1986680** | NA | 0.072668539 | -0.085218911 | 0.230555989 | 8.206793182 | 1.10003867 | 0.309532709 | 0.991835066 | -4.798232613 |
| **A_51_P472274** | Sox18 | 0.046419127 | -0.054440054 | 0.147278307 | 10.68116259 | 1.099994882 | 0.309550464 | 0.991835066 | -4.79826542 |
| **A_55_P2071893** | NA | -0.076325248 | -0.24221382 | 0.089563325 | 11.59053241 | -1.099666222 | 0.309683755 | 0.991835066 | -4.798511635 |
| **A_55_P1970935** | Usp7 | 0.081423566 | -0.095550142 | 0.258397273 | 11.71466876 | 1.099639924 | 0.309694423 | 0.991835066 | -4.798531334 |
| **A_55_P2065926** | Akap8l | -0.101421249 | -0.32189509 | 0.119052592 | 9.526093369 | -1.099464014 | 0.309765786 | 0.991835066 | -4.798663097 |
| **A_55_P2018631** | NA | -0.056724105 | -0.180048238 | 0.066600028 | 12.83050664 | -1.099331602 | 0.309819512 | 0.991835066 | -4.79876227 |
| **A_55_P2138222** | NA | -0.051315504 | -0.162893636 | 0.060262629 | 8.928189918 | -1.099204862 | 0.309870944 | 0.991835066 | -4.798857188 |
| **A_55_P2152114** | Zfyve19 | -0.074754652 | -0.237371484 | 0.08786218 | 9.503209215 | -1.098706922 | 0.310073077 | 0.991835066 | -4.799230041 |
| **A_52_P512427** | Mkrn2os | -0.051160049 | -0.162451884 | 0.060131786 | 7.901627931 | -1.09869407 | 0.310078296 | 0.991835066 | -4.799239663 |
| **A_51_P415905** | Pola1 | 0.123602344 | -0.145313521 | 0.392518209 | 9.945502987 | 1.098548955 | 0.310137225 | 0.991835066 | -4.799348304 |
| **A_55_P2164629** | Ptn | 0.10534149 | -0.123846031 | 0.334529011 | 12.45829402 | 1.098544417 | 0.310139068 | 0.991835066 | -4.799351701 |
| **A_55_P2170134** | Eif2b4 | 0.054029361 | -0.063522146 | 0.171580869 | 12.51311948 | 1.098527067 | 0.310146114 | 0.991835066 | -4.799364689 |
| **A_55_P2123727** | Grb2 | -0.096484895 | -0.306418562 | 0.113448771 | 8.550814342 | -1.098465374 | 0.31017117 | 0.991835066 | -4.799410873 |
| **A_55_P2069277** | Mtf1 | 0.048866871 | -0.057472311 | 0.155206053 | 8.074018314 | 1.098323688 | 0.310228721 | 0.991835066 | -4.799516934 |
| **A_55_P2066697** | Trim47 | -0.0931671 | -0.295925295 | 0.109591094 | 12.94569509 | -1.098229981 | 0.310266788 | 0.991835066 | -4.799587075 |
| **A_52_P217437** | Htra3 | -0.047759514 | -0.15174242 | 0.056223392 | 8.063169574 | -1.097759188 | 0.310458098 | 0.991835066 | -4.799939415 |
| **A_52_P965502** | Ssr1 | 0.066354456 | -0.078116389 | 0.210825301 | 8.771900469 | 1.097738787 | 0.31046639 | 0.991835066 | -4.799954681 |
| **A_55_P2140031** | Fam129b | 0.067146684 | -0.079060049 | 0.213353416 | 13.00237389 | 1.097656185 | 0.310499967 | 0.991835066 | -4.80001649 |

| | | | | | | | | | |
|---|---|---|---|---|---|---|---|---|---|
| **A_55_P2142580** | NA | -0.052633448 | -0.167246397 | 0.0619795 | 10.99251459 | -1.097582742 | 0.310529823 | 0.991835066 | -4.800071444 |
| **A_51_P106144** | Zfp64 | 0.067652377 | -0.079680263 | 0.214985017 | 10.06435386 | 1.097471428 | 0.310575079 | 0.991835066 | -4.788092718 |
| **A_51_P441687** | Lrrc10 | -0.149186084 | -0.474124715 | 0.175752547 | 12.28369014 | -1.097327091 | 0.310633769 | 0.991835066 | -4.800262716 |
| **A_55_P2005433** | NA | 0.048741542 | -0.057432156 | 0.154915241 | 8.056498097 | 1.09721429 | 0.310679642 | 0.991835066 | -4.800347102 |
| **A_55_P2125774** | Ola1 | 0.102130829 | -0.120361066 | 0.324622724 | 11.70015457 | 1.097114093 | 0.310720394 | 0.991835066 | -4.800422055 |
| **A_55_P1973213** | Trmt61a | 0.146678321 | -0.172905091 | 0.466261733 | 9.834236951 | 1.096960108 | 0.310783031 | 0.991835066 | -4.800537237 |
| **A_55_P2002752** | Zswim6 | -0.054533607 | -0.173366528 | 0.064299314 | 8.361774142 | -1.096823086 | 0.310838776 | 0.991835066 | -4.800639722 |
| **A_55_P2003443** | Lsm7 | 0.113167882 | -0.133443196 | 0.35977896 | 14.22633533 | 1.096780544 | 0.310856085 | 0.991835066 | -4.800671539 |
| **A_55_P1956734** | NA | 0.068624115 | -0.080986389 | 0.21823462 | 10.21142409 | 1.096285835 | 0.311057428 | 0.991835066 | -4.801041481 |
| **A_52_P609868** | Timd4 | 0.066049116 | -0.077951038 | 0.21004927 | 7.707122225 | 1.096259033 | 0.31106834 | 0.991835066 | -4.801061521 |
| **A_55_P2126448** | Snhg16 | 0.203885365 | -0.240706541 | 0.648477272 | 9.731572052 | 1.096058246 | 0.311150091 | 0.991835066 | -4.801211638 |
| **A_55_P2075996** | Cpne1 | 0.099085186 | -0.117012505 | 0.315182877 | 11.12814911 | 1.095891941 | 0.311217816 | 0.991835066 | -4.801335963 |
| **A_55_P1961715** | Nme5 | -0.053596935 | -0.170490783 | 0.063296913 | 7.907876089 | -1.095865945 | 0.311228404 | 0.991835066 | -4.801355395 |
| **A_52_P154880** | Ing3 | -0.060223728 | -0.191574168 | 0.071126712 | 8.934042277 | -1.09583518 | 0.311240934 | 0.991835066 | -4.801378393 |
| **A_55_P2078460** | NA | -0.066147551 | -0.21041932 | 0.078124217 | 10.46669788 | -1.095825868 | 0.311244727 | 0.991835066 | -4.801385354 |
| **A_51_P299805** | Slc46a3 | -0.062314941 | -0.198241268 | 0.073611386 | 9.356465032 | -1.095715329 | 0.311289752 | 0.991835066 | -4.80146798 |
| **A_52_P650453** | Spink4 | -0.063026412 | -0.200536394 | 0.07448357 | 8.634185783 | -1.095462437 | 0.311392781 | 0.991835066 | -4.801656994 |
| **A_55_P1978506** | H2-Q6 | -0.110733589 | -0.352376327 | 0.130909149 | 9.113970816 | -1.095253776 | 0.311477811 | 0.991835066 | -4.80181293 |
| **A_55_P1990440** | R3hdm1 | 0.065691277 | -0.077675351 | 0.209057906 | 10.87805558 | 1.095137793 | 0.311525083 | 0.991835066 | -4.801899598 |
| **A_55_P2013655** | Cd4 | -0.083221911 | -0.264848041 | 0.09840422 | 8.982397595 | -1.095137078 | 0.311525374 | 0.991835066 | -4.801900132 |
| **A_51_P292877** | Snrnp200 | -0.069848561 | -0.222289044 | 0.082591923 | 10.7958029 | -1.095131513 | 0.311527643 | 0.991835066 | -4.80190429 |
| **A_52_P200359** | Aco2 | -0.090930209 | -0.289393023 | 0.107532604 | 16.16877414 | -1.095060665 | 0.311556522 | 0.991835066 | -4.801957229 |
| **A_55_P2156394** | Btbd16 | 0.130106634 | -0.153869484 | 0.414082752 | 9.050663933 | 1.095031863 | 0.311568263 | 0.991835066 | -4.801978749 |
| **A_51_P392147** | Xab2 | 0.087345373 | -0.103299309 | 0.277990055 | 9.634624049 | 1.095025775 | 0.311570744 | 0.991835066 | -4.801983298 |
| **A_55_P2296118** | 9630010G1 | 0.316821674 | -0.374745808 | 1.008389156 | 10.22233887 | 1.09493781 | 0.311606605 | 0.991835066 | -4.802049022 |
| **A_55_P2124115** | Invs | 0.08257296 | -0.097706299 | 0.262852218 | 9.297836814 | 1.094715379 | 0.311697297 | 0.991835066 | -4.802215199 |
| **A_55_P2084512** | NA | 0.058000917 | -0.068633991 | 0.184635826 | 8.500036317 | 1.094688496 | 0.31170826 | 0.991835066 | -4.802235282 |
| **A_55_P2027212** | NA | -0.111046779 | -0.353559571 | 0.131466012 | 8.214525336 | -1.094410999 | 0.311821439 | 0.991835066 | -4.802442567 |
| **A_55_P2007525** | Smu1 | -0.082438424 | -0.262479441 | 0.097602593 | 9.493833935 | -1.094378006 | 0.311834897 | 0.991835066 | -4.802467211 |
| **A_52_P463346** | Serinc3 | -0.075313279 | -0.239834668 | 0.089208111 | 10.72019846 | -1.094103277 | 0.311946984 | 0.991835066 | -4.802672394 |
| **A_55_P2019814** | Rnf20 | 0.096288817 | -0.114056555 | 0.306634189 | 11.26834059 | 1.094087413 | 0.311953457 | 0.991835066 | -4.802684241 |
| **A_51_P328300** | Pdia4 | 0.099841876 | -0.118290176 | 0.317973928 | 9.567896272 | 1.09396236 | 0.312004489 | 0.991835066 | -4.802777626 |
| **A_55_P2075343** | Arfrp1 | -0.067894978 | -0.21624661 | 0.080456655 | 8.673081126 | -1.093841643 | 0.312053758 | 0.991835066 | -4.802867767 |
| **A_51_P156934** | Dnajc15 | -0.07645716 | -0.243528819 | 0.090614499 | 14.10863689 | -1.093766225 | 0.312084543 | 0.991835066 | -4.802924081 |
| **A_51_P176505** | Rps19bp1 | 0.067733838 | -0.080307058 | 0.215774734 | 11.25050781 | 1.093536069 | 0.312178503 | 0.991835066 | -4.803095918 |
| **A_51_P406835** | Mei1 | 0.044316305 | -0.052557105 | 0.141189715 | 7.723158555 | 1.093372356 | 0.312245352 | 0.991835066 | -4.803218135 |
| **A_55_P2052490** | Ushbp1 | -0.068175384 | -0.217235892 | 0.080885124 | 8.41945915 | -1.093135827 | 0.312341955 | 0.991835066 | -4.803394691 |
| **A_52_P625171** | Asph | 0.113902257 | -0.135172033 | 0.362976546 | 9.690529829 | 1.092980872 | 0.312405255 | 0.991835066 | -4.803510345 |
| **A_55_P2060445** | Apopt1 | -0.11684236 | -0.372408784 | 0.138724064 | 13.79844224 | -1.092711856 | 0.312515174 | 0.991835066 | -4.803711107 |
| **A_55_P2024155** | Zbtb16 | -0.124943895 | -0.398280369 | 0.148392579 | 10.0994654 | -1.092512932 | 0.312596474 | 0.991835066 | -4.803859541 |
| **A_51_P499698** | Asprv1 | 0.084325935 | -0.100152355 | 0.268804224 | 7.792330035 | 1.092509023 | 0.312598072 | 0.991835066 | -4.803862457 |
| **A_52_P237159** | Spty2d1 | 0.069285564 | -0.082312545 | 0.220883673 | 9.233034833 | 1.092340676 | 0.312666889 | 0.991835066 | -4.803988062 |
| **A_55_P2075359** | Eif4a3l1 | 0.078192302 | -0.092898457 | 0.249283061 | 12.32198332 | 1.092311681 | 0.312678743 | 0.991835066 | -4.804009694 |
| **A_52_P238044** | Psmc3ip | 0.081241951 | -0.09654009 | 0.259023991 | 9.35898915 | 1.092198532 | 0.312725004 | 0.991835066 | -4.804094108 |
| **A_51_P283016** | Slc1a6 | 0.129424707 | -0.153821103 | 0.412670517 | 8.08021622 | 1.092101061 | 0.31276486 | 0.991835066 | -4.80416682 |
| **A_55_P2085186** | NA | -0.06508027 | -0.207521365 | 0.077360826 | 9.547534159 | -1.092001338 | 0.312805641 | 0.991835066 | -4.804241209 |
| **A_55_P2069949** | Dynlrb1 | -0.061444587 | -0.19592927 | 0.073040096 | 14.18689841 | -1.091993216 | 0.312808963 | 0.991835066 | -4.804247267 |
| **A_55_P2249055** | NA | 0.090215195 | -0.107258483 | 0.287688873 | 9.955287908 | 1.091891816 | 0.312850435 | 0.991835066 | -4.804322903 |

| | | | | | | | | | |
|---|---|---|---|---|---|---|---|---|---|
| **A_51_P365008** | Tns1 | -0.109323571 | -0.348695485 | 0.130048342 | 13.83447559 | -1.091565374 | 0.312983976 | 0.991835066 | -4.804566369 |
| **A_52_P318438** | Nab1 | -0.058513829 | -0.186635326 | 0.069607668 | 10.15383816 | -1.091555051 | 0.3129882 | 0.991835066 | -4.804574068 |
| **A_51_P206445** | Zfp773 | -0.050930041 | -0.162447007 | 0.060586924 | 8.059320635 | -1.091546436 | 0.312991725 | 0.991835066 | -4.804580492 |
| **A_51_P457331** | Hmgn5 | -0.087288618 | -0.278425319 | 0.103848083 | 10.75710028 | -1.091497295 | 0.313011832 | 0.991835066 | -4.804617138 |
| **A_52_P655948** | Mosmo | 0.100901606 | -0.12004689 | 0.321850102 | 9.931061057 | 1.091480952 | 0.31301852 | 0.991835066 | -4.804629326 |
| **A_51_P336721** | Tipin | 0.088533309 | -0.105335812 | 0.282402429 | 11.68933549 | 1.091458398 | 0.313027749 | 0.991835066 | -4.804646144 |
| **A_55_P2034420** | NA | -0.070177398 | -0.223852968 | 0.083498173 | 9.640680688 | -1.091444256 | 0.313033536 | 0.991835066 | -4.80465669 |
| **A_55_P2034067** | Myom3 | -0.057997465 | -0.185012959 | 0.069018028 | 8.231736812 | -1.091343449 | 0.313074789 | 0.991835066 | -4.80473186 |
| **A_55_P1953439** | Lemd3 | 0.078764127 | -0.093763847 | 0.2512921 | 9.578677423 | 1.091133962 | 0.313160531 | 0.991835066 | -4.804888057 |
| **A_55_P2175284** | Ei24 | -0.085563325 | -0.273000542 | 0.101873891 | 9.347382469 | -1.091040719 | 0.313198701 | 0.991835066 | -4.804957574 |
| **A_55_P1981739** | Donson | 0.094639196 | -0.112708464 | 0.301986857 | 10.09258429 | 1.090890188 | 0.313260331 | 0.991835066 | -4.805069796 |
| **A_55_P2145109** | Rpn2 | 0.100661282 | -0.119893269 | 0.321215834 | 13.64201497 | 1.090826214 | 0.313286526 | 0.991835066 | -4.805117486 |
| **A_55_P1969625** | Hnrnpr | 0.063497888 | -0.075632135 | 0.202627911 | 12.18175881 | 1.090806099 | 0.313294763 | 0.991835066 | -4.805132481 |
| **A_51_P489413** | Luc7l | 0.134170626 | -0.159823889 | 0.428165141 | 9.434099636 | 1.090755349 | 0.313315545 | 0.991835066 | -4.805170311 |
| **A_52_P59579** | Tmbim6 | -0.102602751 | -0.327433537 | 0.122228034 | 11.40386254 | -1.090717702 | 0.313330962 | 0.991835066 | -4.805198374 |
| **A_55_P2037398** | NA | 0.063427444 | -0.07558277 | 0.202437658 | 15.08612651 | 1.090535049 | 0.31340577 | 0.991835066 | -4.805334517 |
| **A_52_P399934** | Dusp2 | -0.089160896 | -0.284610261 | 0.106288468 | 10.46383768 | -1.090308244 | 0.313498681 | 0.991835066 | -4.805503549 |
| **A_51_P448946** | Sertad2 | -0.102066598 | -0.325816356 | 0.12168316 | 11.69118553 | -1.090260295 | 0.313518327 | 0.991835066 | -4.805539282 |
| **A_66_P129419** | Cox7a2l | -0.075022402 | -0.239511692 | 0.089466888 | 15.38501397 | -1.090090297 | 0.313587985 | 0.991835066 | -4.805665961 |
| **A_55_P2110355** | Zfp637 | -0.14871518 | -0.474786672 | 0.177356312 | 9.927170253 | -1.090063012 | 0.313599167 | 0.991835066 | -4.805686292 |
| **A_55_P2146249** | NA | -0.04947989 | -0.157998151 | 0.05903837 | 13.71757098 | -1.08977047 | 0.313719072 | 0.991835066 | -4.805904256 |
| **A_51_P201338** | Mtss1 | -0.048310201 | -0.154275951 | 0.057655549 | 9.81568549 | -1.08963855 | 0.313773155 | 0.991835066 | -4.806002534 |
| **A_55_P2130890** | Parva | 0.094114345 | -0.112342598 | 0.300571288 | 10.75381028 | 1.089520642 | 0.3138215 | 0.991835066 | -4.806090367 |
| **A_51_P302520** | Myom1 | -0.105667599 | -0.337478931 | 0.126143733 | 13.32440282 | -1.089472592 | 0.313841203 | 0.991835066 | -4.806126159 |
| **A_52_P44030** | Exoc3 | -0.095428467 | -0.304780766 | 0.113923832 | 9.156508479 | -1.089455125 | 0.313848366 | 0.991835066 | -4.80613917 |
| **A_55_P1969904** | Abcc6 | -0.046692284 | -0.149150518 | 0.055765949 | 7.616242171 | -1.089199391 | 0.31395325 | 0.991835066 | -4.806329647 |
| **A_55_P2038247** | Tyro3 | 0.097764669 | -0.116773802 | 0.31230314 | 10.48050863 | 1.089145463 | 0.313975371 | 0.991835066 | -4.806369811 |
| **A_51_P372473** | Auh | -0.081939803 | -0.261762496 | 0.09788289 | 11.92424712 | -1.089079417 | 0.314002465 | 0.991835066 | -4.806418998 |
| **A_52_P668715** | Nutf2 | 0.066538599 | -0.079503844 | 0.212581042 | 9.154235409 | 1.088939358 | 0.314059926 | 0.991835066 | -4.806523299 |
| **A_55_P2141724** | Gm13275 | -0.065709539 | -0.209933489 | 0.07851441 | 8.448390359 | -1.088930523 | 0.314063551 | 0.991835066 | -4.806529878 |
| **A_51_P122740** | Slc37a3 | 0.0614697 | -0.07345251 | 0.19639191 | 9.70299952 | 1.088896947 | 0.314077328 | 0.991835066 | -4.80655488 |
| **A_55_P2031100** | Ahcy | 0.058788496 | -0.07026361 | 0.187840601 | 8.000816642 | 1.088770596 | 0.314129176 | 0.991835066 | -4.806648964 |
| **A_55_P2130535** | Dnmt3b | 0.107454221 | -0.128433493 | 0.343341935 | 9.330742309 | 1.088747801 | 0.31413853 | 0.991835066 | -4.806665937 |
| **A_55_P2092286** | NA | 0.053873572 | -0.064399658 | 0.172146802 | 8.124010219 | 1.088675496 | 0.314168204 | 0.991835066 | -4.806719773 |
| **A_51_P453948** | Drg1 | -0.0544408 | -0.173980357 | 0.065098757 | 10.67113437 | -1.088483846 | 0.314246868 | 0.991835066 | -4.806862459 |
| **A_51_P193185** | Mb | -0.251433425 | -0.803629148 | 0.300762299 | 15.8533821 | -1.088276132 | 0.314332144 | 0.991835066 | -4.807017088 |
| **A_52_P674357** | Thtpa | -0.0832638 | -0.266129096 | 0.099601497 | 9.981615402 | -1.088263499 | 0.314337331 | 0.991835066 | -4.807026492 |
| **A_55_P1960775** | Rps15a | 0.056288514 | -0.06733975 | 0.179916779 | 15.74584974 | 1.088206078 | 0.314360908 | 0.991835066 | -4.807069234 |
| **A_52_P22331** | Akap10 | 0.088952442 | -0.106433747 | 0.284338631 | 8.408625037 | 1.088110856 | 0.31440001 | 0.991835066 | -4.807140112 |
| **A_55_P2149575** | Sdr39u1 | -0.075331236 | -0.240812451 | 0.09014998 | 8.627819398 | -1.088016601 | 0.314438719 | 0.991835066 | -4.807210266 |
| **A_51_P314264** | Ssr2 | -0.100410399 | -0.32100974 | 0.120188942 | 11.24324103 | -1.087886567 | 0.314492128 | 0.991835066 | -4.807307045 |
| **A_52_P585124** | Cxcr4 | 0.088385522 | -0.105797115 | 0.282568159 | 8.352959524 | 1.087877181 | 0.314495983 | 0.991835066 | -4.80731403 |
| **A_51_P311990** | P4hb | 0.065754792 | -0.078722241 | 0.210231824 | 12.44343065 | 1.087771628 | 0.314539343 | 0.991835066 | -4.807392583 |
| **A_55_P1997741** | Slc9a8 | 0.061896002 | -0.074117887 | 0.197909891 | 11.29573193 | 1.087648268 | 0.314590024 | 0.991835066 | -4.807484381 |
| **A_52_P579252** | NA | 0.059826064 | -0.071641607 | 0.191293735 | 13.67867933 | 1.087628538 | 0.31459813 | 0.991835066 | -4.807499063 |
| **A_55_P2076303** | 8-Mar | 0.092965507 | -0.111389521 | 0.297320536 | 11.57560784 | 1.087290635 | 0.31473699 | 0.991835066 | -4.807750479 |
| **A_51_P193490** | Fbxo21 | 0.084087065 | -0.10076316 | 0.268937291 | 12.18424393 | 1.087222279 | 0.314765087 | 0.991835066 | -4.807801334 |
| **A_55_P2054743** | Sh3bgrl2 | 0.045244162 | -0.05422556 | 0.144713883 | 7.911477231 | 1.087128238 | 0.314803744 | 0.991835066 | -4.807871294 |

| | | | | | | | | | |
|---|---|---|---|---|---|---|---|---|---|
| **A_55_P2024011** | NA | 0.08602867 | -0.103149404 | 0.275206743 | 13.653602 | 1.08687986 | 0.314905863 | 0.991835066 | -4.808056053 |
| **A_51_P454666** | Arl6ip4 | 0.083132145 | -0.099698621 | 0.265962911 | 10.46622006 | 1.086747982 | 0.314960095 | 0.991835066 | -4.808154141 |
| **A_52_P355057** | Wdr4 | 0.074983411 | -0.08993303 | 0.239899852 | 10.41065585 | 1.086701758 | 0.314979105 | 0.991835066 | -4.80818852 |
| **A_55_P2005470** | Mfap4 | 0.335990385 | -0.403299586 | 1.075280356 | 12.40668754 | 1.086228434 | 0.315173821 | 0.991835066 | -4.808540501 |
| **A_51_P394394** | Tspan2 | -0.079615874 | -0.254817486 | 0.095585737 | 11.46180616 | -1.086102228 | 0.315225756 | 0.991835066 | -4.808634336 |
| **A_55_P2077826** | Ppp1cc | -0.068515189 | -0.219301 | 0.082270622 | 13.17643102 | -1.086014227 | 0.315261973 | 0.991835066 | -4.808699762 |
| **A_66_P134728** | Psmg1 | 0.083014976 | -0.099684115 | 0.265714067 | 12.88689557 | 1.085998416 | 0.315268481 | 0.991835066 | -4.808711517 |
| **A_52_P555672** | Vrk1 | -0.069063363 | -0.221072788 | 0.082946062 | 8.707154479 | -1.085891258 | 0.315312588 | 0.991835066 | -4.80879118 |
| **A_55_P2057523** | Ppm1b | -0.083517373 | -0.267446133 | 0.100411386 | 9.760713555 | -1.0852663 | 0.315569925 | 0.991835066 | -4.809255691 |
| **A_55_P2004501** | Immp1l | -0.066822753 | -0.213992898 | 0.080347391 | 12.48037417 | -1.085209988 | 0.315593121 | 0.991835066 | -4.809297538 |
| **A_55_P2062262** | Aspscr1 | -0.121631113 | -0.389577525 | 0.146315299 | 9.466835649 | -1.084940366 | 0.315704201 | 0.991835066 | -4.809497882 |
| **A_51_P196972** | Slc4a1 | 0.284353628 | -0.342166245 | 0.910873501 | 10.58050874 | 1.084758483 | 0.315779153 | 0.991835066 | -4.809633015 |
| **A_55_P2108938** | Hps5 | 0.105542379 | -0.127001037 | 0.338085795 | 9.380201831 | 1.084755754 | 0.315780277 | 0.991835066 | -4.809635042 |
| **A_55_P1995152** | NA | -0.160462237 | -0.514056432 | 0.193131958 | 16.75354754 | -1.084618099 | 0.315837012 | 0.991835066 | -4.809737304 |
| **A_55_P2055477** | Scaf11 | 0.080351067 | -0.096719451 | 0.257421586 | 10.33206907 | 1.084562345 | 0.315859994 | 0.991835066 | -4.809778722 |
| **A_52_P509710** | Zwint | 0.076350225 | -0.091903718 | 0.244604169 | 10.49879605 | 1.084561551 | 0.315860322 | 0.991835066 | -4.809779312 |
| **A_55_P2085156** | NA | 0.081798823 | -0.098487117 | 0.262084763 | 13.34188134 | 1.084412031 | 0.31592196 | 0.991835066 | -4.809890377 |
| **A_55_P2039699** | Camp | -0.236161122 | -0.75666642 | 0.284344177 | 8.739085246 | -1.084407171 | 0.315923964 | 0.991835066 | -4.809893987 |
| **A_55_P2158458** | Iqcc | 0.059391766 | -0.071516609 | 0.190300142 | 9.193618284 | 1.08434615 | 0.315949122 | 0.991835066 | -4.809939311 |
| **A_52_P533792** | Sucla2 | -0.051471999 | -0.164931173 | 0.061987175 | 14.11438586 | -1.084277763 | 0.31597732 | 0.991835066 | -4.809990104 |
| **A_52_P209484** | Tmem88 | -0.112869005 | -0.361677935 | 0.135939926 | 12.38758351 | -1.08422112 | 0.316000676 | 0.991835066 | -4.810032174 |
| **A_51_P511899** | Mthfsd | 0.065033348 | -0.078331954 | 0.20839865 | 8.166099509 | 1.084179504 | 0.316017837 | 0.991835066 | -4.810063082 |
| **A_66_P129508** | Hyls1 | -0.053821245 | -0.172470907 | 0.064828417 | 8.446222474 | -1.084167451 | 0.316022808 | 0.991835066 | -4.810072034 |
| **A_52_P649064** | NA | -0.085829239 | -0.275065998 | 0.103407519 | 14.96654075 | -1.084023996 | 0.316081971 | 0.991835066 | -4.81017857 |
| **A_51_P407683** | Ints3 | 0.072208277 | -0.08704133 | 0.231457884 | 9.925312439 | 1.083721637 | 0.316206697 | 0.991835066 | -4.810403088 |
| **A_55_P2144512** | NA | 0.192639341 | -0.232268068 | 0.61754675 | 9.509747172 | 1.083577087 | 0.31626634 | 0.991835066 | -4.810510411 |
| **A_55_P2095342** | Rara | -0.074364942 | -0.238415063 | 0.089685178 | 10.26982391 | -1.083429918 | 0.316327073 | 0.991835066 | -4.81061967 |
| **A_55_P2040907** | Gtf2i | 0.082493182 | -0.099508047 | 0.264494411 | 8.429924404 | 1.083310286 | 0.316376449 | 0.991835066 | -4.810708478 |
| **A_51_P159896** | Dnajc1 | 0.058179732 | -0.070244091 | 0.186603556 | 10.42410807 | 1.082767611 | 0.316600506 | 0.991835066 | -4.811111254 |
| **A_51_P323878** | Coro7 | 0.073306064 | -0.08852273 | 0.235134857 | 10.70571881 | 1.082662854 | 0.316643773 | 0.991835066 | -4.811188991 |
| **A_51_P314019** | Banf1 | -0.198903418 | -0.63801594 | 0.240209105 | 12.21516655 | -1.082618777 | 0.316661979 | 0.991835066 | -4.811221698 |
| **A_52_P36928** | Morn2 | -0.084121703 | -0.269839111 | 0.101595706 | 9.265561361 | -1.082591399 | 0.316673288 | 0.991835066 | -4.811242013 |
| **A_55_P1970867** | Gm5779 | -0.125447731 | -0.402444449 | 0.151548988 | 13.51498002 | -1.082423763 | 0.31674254 | 0.991835066 | -4.811366397 |
| **A_55_P1964309** | Mlf2 | 0.053607151 | -0.064767649 | 0.17198195 | 14.10021609 | 1.082362158 | 0.316767993 | 0.991835066 | -4.811412104 |
| **A_55_P1971674** | Golga7b | -0.091655552 | -0.294078511 | 0.110767406 | 9.256135603 | -1.08220153 | 0.316834365 | 0.991835066 | -4.811531272 |
| **A_52_P434974** | Ubqln1 | -0.061103334 | -0.196077948 | 0.07387128 | 10.97460169 | -1.081986757 | 0.316923129 | 0.991835066 | -4.811690593 |
| **A_51_P161554** | Arid3b | -0.056779216 | -0.182210754 | 0.068652323 | 8.205686588 | -1.081911585 | 0.316954201 | 0.991835066 | -4.811746351 |
| **A_55_P2005335** | Kcng2 | -0.097284731 | -0.312198962 | 0.1176295 | 10.43683097 | -1.08190379 | 0.316957424 | 0.991835066 | -4.811752133 |
| **A_51_P106488** | Speg | 0.093698112 | -0.113333338 | 0.300729562 | 11.32393767 | 1.081692081 | 0.317044948 | 0.991835066 | -4.811909155 |
| **A_51_P378789** | Cxcl13 | 0.115412603 | -0.139620712 | 0.370445919 | 7.762801063 | 1.081596948 | 0.317084284 | 0.991835066 | -4.811979708 |
| **A_52_P185485** | Arpc4 | -0.127493933 | -0.409224786 | 0.15423692 | 10.0663626 | -1.081593926 | 0.317085534 | 0.991835066 | -4.811981949 |
| **A_51_P301259** | Araf | 0.066454058 | -0.080414106 | 0.213322222 | 10.79415902 | 1.081441355 | 0.317148629 | 0.991835066 | -4.812095091 |
| **A_55_P2000439** | Ptprz1 | 0.240284459 | -0.290890773 | 0.771459692 | 8.536497517 | 1.0811775 | 0.317257768 | 0.991835066 | -4.812290735 |
| **A_52_P429450** | Ngp | -0.063076925 | -0.202525131 | 0.076371281 | 7.846262009 | -1.081102113 | 0.317288956 | 0.991835066 | -4.812346627 |
| **A_51_P218953** | Zfp536 | 0.121202828 | -0.146872226 | 0.389277881 | 9.560056806 | 1.080601294 | 0.317496211 | 0.991835066 | -4.812717879 |
| **A_55_P2141013** | Siglech | 0.158975562 | -0.192662395 | 0.510613519 | 14.22217871 | 1.080547228 | 0.317518593 | 0.991835066 | -4.812757952 |
| **A_55_P1982568** | Blcap | -0.103481804 | -0.332377306 | 0.125413698 | 9.731850534 | -1.080527601 | 0.317526718 | 0.991835066 | -4.812772499 |
| **A_55_P2169829** | Armh4 | -0.059344034 | -0.190612461 | 0.071924394 | 9.686701278 | -1.080502838 | 0.317536969 | 0.991835066 | -4.812790852 |

| | | | | | | | | | |
|---|---|---|---|---|---|---|---|---|---|
| **A_52_P221024** | NA | -0.144717067 | -0.464841481 | 0.175407347 | 13.95343634 | -1.080463468 | 0.317553268 | 0.991835066 | -4.81282003 |
| **A_55_P1983959** | Ftl2-ps | -0.100135556 | -0.321644547 | 0.121373434 | 13.27552482 | -1.080453522 | 0.317557386 | 0.991835066 | -4.812827401 |
| **A_51_P333274** | Gzmb | -0.045321102 | -0.145580643 | 0.05493844 | 7.674391456 | -1.080398259 | 0.317580266 | 0.991835066 | -4.812868358 |
| **A_55_P1988872** | NA | 0.066195938 | -0.080267072 | 0.212658948 | 9.677667194 | 1.080220757 | 0.317653764 | 0.991835066 | -4.812999898 |
| **A_55_P2066778** | AB124611 | -0.048084661 | -0.154493678 | 0.058324355 | 7.725785886 | -1.080033599 | 0.317731276 | 0.991835066 | -4.813138579 |
| **A_55_P2233271** | Zfp626 | 0.059796675 | -0.072624333 | 0.192217684 | 8.951131768 | 1.079267949 | 0.31804853 | 0.991835066 | -4.813705761 |
| **A_52_P684378** | Gpx1 | -0.111081218 | -0.357093931 | 0.134931495 | 14.1023306 | -1.079175842 | 0.318086712 | 0.991835066 | -4.813773976 |
| **A_51_P443618** | Mettl26 | -0.05147593 | -0.165490947 | 0.062539087 | 10.58303717 | -1.07907412 | 0.318128885 | 0.991835066 | -4.813849308 |
| **A_51_P138895** | Ccdc102a | 0.062336566 | -0.075779329 | 0.200452461 | 11.16452906 | 1.078719036 | 0.318276135 | 0.991835066 | -4.814112236 |
| **A_51_P184910** | Phpt1 | -0.068673298 | -0.22087882 | 0.083532224 | 14.11647737 | -1.078367092 | 0.318422137 | 0.991835066 | -4.814372788 |
| **A_51_P457492** | Tmem223 | -0.098636465 | -0.317255643 | 0.119982714 | 13.90725797 | -1.078346598 | 0.31843064 | 0.991835066 | -4.814387958 |
| **A_55_P1975732** | Sipa1l3 | 0.073865041 | -0.089872039 | 0.23760212 | 9.001164195 | 1.078204235 | 0.318489715 | 0.991835066 | -4.814493336 |
| **A_52_P464129** | Gon7 | -0.061649508 | -0.198324802 | 0.075025786 | 9.793730921 | -1.078074391 | 0.318543604 | 0.991835066 | -4.81458944 |
| **A_51_P329413** | Pomgnt1 | -0.087345833 | -0.280989967 | 0.106298301 | 10.80021972 | -1.078070037 | 0.318545411 | 0.991835066 | -4.814592662 |
| **A_52_P456134** | Dgat1 | -0.063197533 | -0.203311671 | 0.076916605 | 9.112770443 | -1.078021187 | 0.318565686 | 0.991835066 | -4.814628817 |
| **A_52_P130912** | Pcdh7 | -0.073984638 | -0.238032445 | 0.090063168 | 8.660410175 | -1.077904437 | 0.318614149 | 0.991835066 | -4.81471522 |
| **A_55_P2101441** | Gpbar1 | -0.054802187 | -0.176329742 | 0.066725369 | 7.95727322 | -1.077785302 | 0.318663608 | 0.991835066 | -4.814803384 |
| **A_55_P2153862** | Irak1 | 0.062564797 | -0.076191242 | 0.201320837 | 9.156896848 | 1.077673686 | 0.318709951 | 0.991835066 | -4.814885977 |
| **A_55_P2049080** | Fam160a2 | 0.071352258 | -0.086898705 | 0.229603222 | 11.04194816 | 1.077632034 | 0.318727246 | 0.991835066 | -4.814916797 |
| **A_51_P501757** | NA | -0.04655831 | -0.149822219 | 0.056705598 | 8.750344696 | -1.077600499 | 0.318740341 | 0.991835066 | -4.814940131 |
| **A_55_P2088018** | Fhod3 | -0.112100472 | -0.360770448 | 0.136569503 | 13.56700668 | -1.07744032 | 0.318806862 | 0.991835066 | -4.815058645 |
| **A_51_P376789** | Pgghg | 0.0794932 | -0.096846431 | 0.255832831 | 9.694972126 | 1.0774303 | 0.318811024 | 0.991835066 | -4.815066059 |
| **A_55_P2254779** | NA | -0.092076187 | -0.296333466 | 0.112181093 | 8.625023921 | -1.077404819 | 0.318821607 | 0.991835066 | -4.815084911 |
| **A_55_P2081223** | Sptbn1 | 0.091947947 | -0.112035378 | 0.295931272 | 15.01242107 | 1.077349224 | 0.318844699 | 0.991835066 | -4.815126042 |
| **A_51_P161946** | Epop | 0.081068516 | -0.098782952 | 0.260919984 | 8.549967319 | 1.077326567 | 0.31885411 | 0.991835066 | -4.815142803 |
| **A_55_P2107203** | Gpr137b | -0.060897769 | -0.196012655 | 0.074217118 | 8.574969994 | -1.077227201 | 0.318895387 | 0.991835066 | -4.815216313 |
| **A_55_P2157610** | Rpusd3 | 0.076602458 | -0.09339307 | 0.246597986 | 9.943663187 | 1.076996618 | 0.318991187 | 0.991835066 | -4.815386879 |
| **A_55_P2003966** | Rybp | 0.081556044 | -0.099440811 | 0.2625529 | 11.49073396 | 1.076946812 | 0.319011883 | 0.991835066 | -4.815423718 |
| **A_51_P144249** | Ubr1 | -0.04747589 | -0.152856829 | 0.057905049 | 8.283770409 | -1.076763156 | 0.319088208 | 0.991835066 | -4.815559552 |
| **A_55_P2024640** | Eral1 | 0.077743023 | -0.094878086 | 0.250364132 | 10.27745545 | 1.076407352 | 0.319236116 | 0.991835066 | -4.815822667 |
| **A_55_P2081571** | Raver1 | -0.105233277 | -0.338930673 | 0.12846412 | 13.00592581 | -1.076238073 | 0.319306506 | 0.991835066 | -4.815947829 |
| **A_66_P106716** | Akap8 | 0.098704057 | -0.12059387 | 0.318001985 | 9.982281136 | 1.075745678 | 0.319511323 | 0.991835066 | -4.81631183 |
| **A_55_P2020962** | NA | -0.060207656 | -0.193976591 | 0.073561278 | 16.83652266 | -1.075735709 | 0.319515471 | 0.991835066 | -4.816319198 |
| **A_66_P102395** | Elac2 | 0.063318185 | -0.077369786 | 0.204006156 | 8.694464413 | 1.075673886 | 0.319541195 | 0.991835066 | -4.816364893 |
| **A_52_P229044** | Slc20a2 | -0.101384639 | -0.326689819 | 0.123920541 | 9.222080071 | -1.075499254 | 0.319613866 | 0.991835066 | -4.816493958 |
| **A_51_P400463** | Agmat | -0.063195447 | -0.203635373 | 0.077244478 | 9.054148772 | -1.075484941 | 0.319619823 | 0.991835066 | -4.816504536 |
| **A_51_P417251** | 6330403K0 | 0.163342186 | -0.199756229 | 0.526440601 | 9.932023075 | 1.075184864 | 0.319744731 | 0.991835066 | -4.816726283 |
| **A_55_P1963772** | Atp5j | -0.081911335 | -0.264002736 | 0.100180066 | 15.33814062 | -1.075136734 | 0.319764769 | 0.991835066 | -4.816761846 |
| **A_52_P621418** | NA | -0.08356553 | -0.2693569 | 0.10222584 | 13.37800442 | -1.075005696 | 0.319819328 | 0.991835066 | -4.816858664 |
| **A_55_P2171993** | Mrgbp | 0.046621923 | -0.057068871 | 0.150312717 | 8.574607201 | 1.074630382 | 0.319975638 | 0.991835066 | -4.817135926 |
| **A_52_P641216** | Galk2 | 0.08167409 | -0.100000327 | 0.263348507 | 11.03569533 | 1.074483274 | 0.320036922 | 0.991835066 | -4.817244586 |
| **A_65_P13402** | Rnf14 | 0.05650405 | -0.069183052 | 0.182191153 | 8.747559057 | 1.074479162 | 0.320038635 | 0.991835066 | -4.817247623 |
| **A_55_P2009994** | NA | -0.106292184 | -0.342736916 | 0.130152547 | 11.33727695 | -1.074436671 | 0.320056338 | 0.991835066 | -4.817279006 |
| **A_55_P2325803** | Tnfrsf11a | -0.075348412 | -0.242963372 | 0.092266549 | 9.032685956 | -1.074411024 | 0.320067024 | 0.991835066 | -4.817297949 |
| **A_55_P2169586** | Ccdc80 | -0.118704332 | -0.382780153 | 0.14537149 | 11.53897627 | -1.074353145 | 0.320091141 | 0.991835066 | -4.817340696 |
| **A_52_P274496** | Tspan18 | 0.078615451 | -0.096278854 | 0.253509757 | 13.25437836 | 1.074339087 | 0.320096998 | 0.991835066 | -4.817351079 |
| **A_55_P2115151** | Pxylp1 | 0.078853129 | -0.096571053 | 0.254277311 | 10.76250849 | 1.074332232 | 0.320099855 | 0.991835066 | -4.817356142 |
| **A_51_P446558** | Timm29 | -0.048859849 | -0.157569883 | 0.059850186 | 8.310050531 | -1.074216 | 0.320148291 | 0.991835066 | -4.817441981 |

| | | | | | | | | | |
|---|---|---|---|---|---|---|---|---|---|
| **A_55_P1962918** | Ifi211 | -0.084539941 | -0.272636901 | 0.103557019 | 8.110621056 | -1.074210264 | 0.320150681 | 0.991835066 | -4.817446217 |
| **A_51_P289889** | Npnt | 0.048607949 | -0.059564915 | 0.156780812 | 7.756360257 | 1.073984715 | 0.320244688 | 0.991835066 | -4.817612771 |
| **A_55_P2339601** | NA | -0.057971105 | -0.186988471 | 0.071046261 | 7.968427178 | -1.073921514 | 0.320271034 | 0.991835066 | -4.817659438 |
| **A_52_P544043** | Pcsk5 | 0.061151757 | -0.074948849 | 0.197252363 | 9.128700128 | 1.073885581 | 0.320286014 | 0.991835066 | -4.81768597 |
| **A_55_P2037897** | Brd3 | 0.103330048 | -0.126731259 | 0.333391355 | 11.27342021 | 1.073475607 | 0.320456963 | 0.991835066 | -4.817988639 |
| **A_55_P2163729** | Tvp23a | 0.077397727 | -0.094995853 | 0.249791308 | 8.378655473 | 1.073040839 | 0.320638332 | 0.991835066 | -4.818309536 |
| **A_55_P2034275** | Arfgap2 | 0.063667936 | -0.078160522 | 0.205496395 | 11.87620799 | 1.072917978 | 0.3206896 | 0.991835066 | -4.818400204 |
| **A_55_P1957219** | Ptma | 0.110074863 | -0.135134652 | 0.355284379 | 14.25120672 | 1.072901786 | 0.320696357 | 0.991835066 | -4.818412153 |
| **A_55_P1968493** | Telo2 | 0.082263814 | -0.101018654 | 0.265546282 | 10.21931675 | 1.072746361 | 0.320761224 | 0.991835066 | -4.818526841 |
| **A_55_P1985094** | Wrap73 | 0.077556029 | -0.095261783 | 0.250373841 | 10.17532461 | 1.072596056 | 0.320823965 | 0.991835066 | -4.818637742 |
| **A_52_P278497** | Eif4ebp3 | 0.066514565 | -0.081710285 | 0.214739414 | 10.16954568 | 1.072518682 | 0.320856266 | 0.991835066 | -4.818694827 |
| **A_55_P2037585** | Trf | -0.442352589 | -1.428150446 | 0.543445269 | 10.69651685 | -1.072481564 | 0.320871762 | 0.991835066 | -4.818722212 |
| **A_55_P2149288** | Cmya5 | 0.109822632 | -0.134938865 | 0.354584128 | 8.600816176 | 1.072402652 | 0.32090471 | 0.991835066 | -4.818780429 |
| **A_51_P393067** | Ap4m1 | 0.065629099 | -0.080640092 | 0.21189829 | 8.557620091 | 1.072389898 | 0.320910035 | 0.991835066 | -4.818789837 |
| **A_55_P2019118** | NA | 0.053856461 | -0.06618416 | 0.173897082 | 8.265442707 | 1.072305935 | 0.320945095 | 0.991835066 | -4.818851777 |
| **A_51_P382925** | Fadd | -0.079926505 | -0.2580853 | 0.098232291 | 9.629197397 | -1.072241682 | 0.320971926 | 0.991835066 | -4.818899174 |
| **A_55_P2150179** | C1s2 | -0.111379927 | -0.359668419 | 0.136908564 | 8.217984155 | -1.072159684 | 0.321006171 | 0.991835066 | -4.818959659 |
| **A_51_P280532** | Supt16 | 0.059890478 | -0.073633447 | 0.193414402 | 8.226755945 | 1.07203218 | 0.321059426 | 0.991835066 | -4.819053706 |
| **A_55_P2027327** | Usp1 | 0.090843516 | -0.111691401 | 0.293378433 | 11.04211344 | 1.072020677 | 0.321064231 | 0.991835066 | -4.81906219 |
| **A_55_P2009543** | NA | 0.076817146 | -0.094456629 | 0.248090922 | 16.43531193 | 1.071954695 | 0.321091793 | 0.991835066 | -4.819110855 |
| **A_51_P393862** | Spryd4 | 0.072334293 | -0.092924167 | 0.237592753 | 10.63293837 | 1.087295749 | 0.321097097 | 0.991835066 | -4.75596266 |
| **A_52_P465121** | NA | -0.077800687 | -0.251302196 | 0.095700821 | 10.85367937 | -1.071739686 | 0.321181619 | 0.991835066 | -4.819269421 |
| **A_55_P2098578** | Tubd1 | -0.064643307 | -0.208814974 | 0.079528361 | 9.205203507 | -1.07164952 | 0.321219295 | 0.991835066 | -4.819335912 |
| **A_51_P195240** | Gpatch11 | 0.067023712 | -0.082468474 | 0.216515898 | 10.79163479 | 1.071566477 | 0.321253998 | 0.991835066 | -4.819397146 |
| **A_51_P134301** | Pfdn5 | -0.047251672 | -0.152695577 | 0.058192232 | 12.98389532 | -1.071037891 | 0.321474958 | 0.991835066 | -4.81978685 |
| **A_55_P2146185** | Rs1 | 0.112562007 | -0.138679276 | 0.363803291 | 13.07407391 | 1.070803934 | 0.321572796 | 0.991835066 | -4.819959298 |
| **A_51_P110931** | Znrd1as | -0.055060306 | -0.177967448 | 0.067846836 | 9.759334795 | -1.070706981 | 0.321613348 | 0.991835066 | -4.820030755 |
| **A_55_P2032238** | Zbtb8os | 0.057898236 | -0.071419334 | 0.187215807 | 11.03452051 | 1.07008168 | 0.321874986 | 0.991835066 | -4.820491521 |
| **A_55_P1967643** | Plb1 | 0.088388343 | -0.109048981 | 0.285825668 | 8.295973173 | 1.06997805 | 0.321918364 | 0.991835066 | -4.820567867 |
| **A_55_P2169849** | Thoc6 | 0.086663811 | -0.106953483 | 0.280281105 | 9.801627881 | 1.069800435 | 0.321992721 | 0.991835066 | -4.820698708 |
| **A_51_P338600** | Tmem14c | -0.078356073 | -0.253413184 | 0.096701038 | 12.24988123 | -1.069798627 | 0.321993478 | 0.991835066 | -4.82070004 |
| **A_52_P93933** | Mcl1 | -0.059733456 | -0.19319304 | 0.073726129 | 10.86100308 | -1.069736969 | 0.322019294 | 0.991835066 | -4.820745457 |
| **A_55_P2044932** | Gpr84 | 0.042937081 | -0.052996458 | 0.138870619 | 7.712005243 | 1.069722626 | 0.322025299 | 0.991835066 | -4.820756022 |
| **A_55_P2064074** | Vps37d | 0.083587187 | -0.103173435 | 0.270347809 | 9.444480013 | 1.069703777 | 0.322033191 | 0.991835066 | -4.820769905 |
| **A_51_P405423** | Smcr8 | -0.059863148 | -0.193618877 | 0.073892582 | 9.83150952 | -1.069685939 | 0.322040661 | 0.991835066 | -4.820783044 |
| **A_52_P384901** | Wdr43 | 0.087911322 | -0.108526016 | 0.284348659 | 9.556009583 | 1.069620955 | 0.322067873 | 0.991835066 | -4.820830909 |
| **A_55_P2134082** | Clip4 | -0.061480847 | -0.198860206 | 0.075898513 | 10.41136329 | -1.069615 | 0.322070366 | 0.991835066 | -4.820835295 |
| **A_66_P108276** | Gsg1l | -0.073359036 | -0.237323507 | 0.090605435 | 9.414838393 | -1.069333059 | 0.32218845 | 0.991835066 | -4.821042938 |
| **A_55_P1962781** | NA | 0.09597786 | -0.118556124 | 0.310511843 | 8.302909169 | 1.069261909 | 0.322218255 | 0.991835066 | -4.821095333 |
| **A_51_P314153** | Nr2c2ap | 0.152717019 | -0.188669913 | 0.494103952 | 9.061042208 | 1.069177176 | 0.322253753 | 0.991835066 | -4.821157728 |
| **A_55_P1984113** | Nup62 | 0.059118049 | -0.073037457 | 0.191273554 | 13.52948456 | 1.069163098 | 0.322259651 | 0.991835066 | -4.821168095 |
| **A_51_P143721** | Vipas39 | -0.086000946 | -0.278270988 | 0.106269095 | 10.55944941 | -1.069056707 | 0.322304227 | 0.991835066 | -4.821246433 |
| **A_55_P1968085** | NA | 0.124386544 | -0.153727524 | 0.402500611 | 8.525547208 | 1.068955374 | 0.322346689 | 0.991835066 | -4.821321042 |
| **A_55_P2020911** | Rps6ka6 | 0.101223195 | -0.125110249 | 0.32755664 | 8.830051222 | 1.068908283 | 0.322366423 | 0.991835066 | -4.821355713 |
| **A_51_P189208** | 4933417G( | -0.097061738 | -0.314094305 | 0.119970829 | 9.317201699 | -1.068888216 | 0.322374832 | 0.991835066 | -4.821370486 |
| **A_55_P2157008** | Ear10 | -0.043898219 | -0.142081403 | 0.054284966 | 7.766618314 | -1.068609212 | 0.322491775 | 0.991835066 | -4.821575881 |
| **A_52_P192307** | Rbak | -0.061398532 | -0.198727186 | 0.075930121 | 9.010711604 | -1.068577335 | 0.322505138 | 0.991835066 | -4.821599346 |
| **A_52_P176300** | Med1 | 0.074568292 | -0.09222461 | 0.241361195 | 9.615958115 | 1.068527643 | 0.322525971 | 0.991835066 | -4.821635923 |

| | | | | | | | | | |
|---|---|---|---|---|---|---|---|---|---|
| **A_52_P56394** | Psmg4 | 0.094155838 | -0.116471906 | 0.304783582 | 11.17387011 | 1.068417053 | 0.322572337 | 0.991835066 | -4.821717323 |
| **A_55_P1961894** | Fbxw11 | -0.059475248 | -0.192522722 | 0.073572227 | 12.67168521 | -1.068411997 | 0.322574457 | 0.991835066 | -4.821721044 |
| **A_55_P2040110** | Cdv3 | -0.093888513 | -0.303939944 | 0.116162918 | 9.146026609 | -1.06830669 | 0.322618614 | 0.991835066 | -4.82179855 |
| **A_52_P48689** | Bzw1 | -0.064400289 | -0.208484362 | 0.079683784 | 9.669702346 | -1.068269848 | 0.322634063 | 0.991835066 | -4.821825665 |
| **A_66_P119170** | Phf21a | 0.093745975 | -0.116077414 | 0.303569363 | 9.063620441 | 1.067844131 | 0.322812629 | 0.991835066 | -4.822138939 |
| **A_55_P1992084** | Ptprg | -0.060538813 | -0.196086823 | 0.075009197 | 8.930580918 | -1.067455771 | 0.322975595 | 0.991835066 | -4.822424655 |
| **A_55_P2316300** | Neo1 | 0.056839875 | -0.070437948 | 0.184117699 | 10.79586968 | 1.067356524 | 0.323017253 | 0.991835066 | -4.822497661 |
| **A_52_P681016** | NA | 0.047997875 | -0.059487518 | 0.155483267 | 7.753806669 | 1.067288158 | 0.323045951 | 0.991835066 | -4.822547948 |
| **A_52_P613977** | NA | -0.078382484 | -0.253929591 | 0.097164622 | 12.80602773 | -1.067172149 | 0.323094653 | 0.991835066 | -4.822633275 |
| **A_55_P1953459** | Yap1 | 0.059667473 | -0.073975081 | 0.193310027 | 12.85049579 | 1.06709236 | 0.323128152 | 0.991835066 | -4.822691958 |
| **A_51_P139108** | Cpxm1 | 0.110357193 | -0.136821321 | 0.357535707 | 9.423453011 | 1.067085111 | 0.323131196 | 0.991835066 | -4.82269729 |
| **A_51_P242201** | Naaa | -0.077667994 | -0.251642088 | 0.096306101 | 8.907668775 | -1.067005444 | 0.323164648 | 0.991835066 | -4.82275588 |
| **A_51_P337089** | Gins1 | 0.069482543 | -0.086159666 | 0.225124753 | 10.57485901 | 1.06698288 | 0.323174123 | 0.991835066 | -4.822772475 |
| **A_55_P2026530** | Erc2 | 0.088394217 | -0.109610513 | 0.286398948 | 8.331289223 | 1.066982807 | 0.323174153 | 0.991835066 | -4.822772528 |
| **A_51_P431554** | Ten1 | -0.090040466 | -0.291783523 | 0.111702592 | 10.1702046 | -1.066714664 | 0.323286768 | 0.991835066 | -4.822969709 |
| **A_51_P127297** | Hsd11b1 | 0.059677105 | -0.074041457 | 0.193395667 | 8.104448122 | 1.066657967 | 0.323310584 | 0.991835066 | -4.797771372 |
| **A_55_P2024600** | Zfp658 | 0.206175452 | -0.255818723 | 0.668169626 | 13.62656962 | 1.066619668 | 0.323326672 | 0.991835066 | -4.823039558 |
| **A_52_P633410** | Gabarapl1 | -0.09376352 | -0.303872051 | 0.116345011 | 13.41506995 | -1.06659452 | 0.323337236 | 0.991835066 | -4.823058048 |
| **A_55_P2060707** | Il15ra | -0.074492447 | -0.241471272 | 0.092486378 | 11.52704505 | -1.066252276 | 0.323481036 | 0.991835066 | -4.823309657 |
| **A_55_P1996852** | NA | -0.084202131 | -0.272959476 | 0.104555215 | 10.1969308 | -1.066174651 | 0.323513658 | 0.991835066 | -4.823366718 |
| **A_55_P2072110** | 3-Sep | -0.192730212 | -0.624794092 | 0.239333668 | 13.51650595 | -1.066131981 | 0.323531592 | 0.991835066 | -4.823398083 |
| **A_52_P523480** | Rfx1 | 0.062581784 | -0.077745568 | 0.202909137 | 9.422786048 | 1.065895778 | 0.323630879 | 0.991835066 | -4.823571692 |
| **A_55_P2183551** | NA | -0.04449267 | -0.144258914 | 0.055273573 | 14.80250834 | -1.065893924 | 0.323631659 | 0.991835066 | -4.823573055 |
| **A_55_P1969002** | Tbx3 | 0.118375 | -0.147060237 | 0.383810237 | 10.44814486 | 1.065885486 | 0.323635206 | 0.991835066 | -4.823579256 |
| **A_52_P112888** | Atp6v1h | -0.048651688 | -0.157755591 | 0.060452216 | 8.939519051 | -1.065778009 | 0.323680392 | 0.991835066 | -4.823658244 |
| **A_55_P2043684** | Ubac1 | 0.053599278 | -0.066620283 | 0.17381884 | 12.72577369 | 1.065596862 | 0.323756563 | 0.991835066 | -4.823791362 |
| **A_55_P2009375** | Cdk18 | 0.165138305 | -0.205280031 | 0.535556641 | 9.548767485 | 1.065527047 | 0.323785923 | 0.991835066 | -4.823842662 |
| **A_55_P2151398** | Tmem116 | 0.048856322 | -0.060735779 | 0.158448423 | 7.884396153 | 1.065493121 | 0.323800191 | 0.991835066 | -4.82386759 |
| **A_51_P309158** | Snx20 | -0.088505652 | -0.287042433 | 0.11003113 | 9.006931949 | -1.065464929 | 0.323812048 | 0.991835066 | -4.823888305 |
| **A_55_P1972799** | Eif4enif1 | 0.063011261 | -0.078352955 | 0.204375476 | 10.66642187 | 1.06533897 | 0.323865029 | 0.991835066 | -4.823980852 |
| **A_51_P214319** | Cox11 | 0.064216335 | -0.079851834 | 0.208284504 | 11.96984452 | 1.065336024 | 0.323866268 | 0.991835066 | -4.823983016 |
| **A_55_P2016842** | Me1 | 0.076553163 | -0.095227879 | 0.248334205 | 9.31697728 | 1.065116319 | 0.323958696 | 0.991835066 | -4.824144425 |
| **A_55_P1992329** | Gmpr | -0.250146537 | -0.811484701 | 0.311191628 | 10.30137892 | -1.065072217 | 0.323977252 | 0.991835066 | -4.824176823 |
| **A_55_P1962303** | Hba-a1 | 0.281370778 | -0.350123306 | 0.912864862 | 17.85068726 | 1.064924545 | 0.324039392 | 0.991835066 | -4.824285298 |
| **A_55_P2175314** | NA | 0.057007317 | -0.070966693 | 0.184981327 | 15.12308293 | 1.064677203 | 0.324143494 | 0.991835066 | -4.824466966 |
| **A_51_P426283** | Dhx30 | 0.070494427 | -0.087762826 | 0.228751679 | 13.07810046 | 1.064633906 | 0.324161719 | 0.991835066 | -4.824498763 |
| **A_55_P2083764** | NA | 0.083120604 | -0.10349381 | 0.269735017 | 8.759153683 | 1.064566107 | 0.32419026 | 0.991835066 | -4.824548554 |
| **A_55_P2083624** | Wbp2 | -0.091950173 | -0.298394539 | 0.114494193 | 13.40847231 | -1.064531818 | 0.324204696 | 0.991835066 | -4.824573735 |
| **A_55_P2045066** | Eif5a | -0.079379513 | -0.257605026 | 0.098846001 | 12.69119572 | -1.064504953 | 0.324216006 | 0.991835066 | -4.824593464 |
| **A_66_P104479** | NA | 0.051288139 | -0.063869043 | 0.166445321 | 16.65671956 | 1.064473967 | 0.324229052 | 0.991835066 | -4.824616219 |
| **A_51_P140347** | Slc4a9 | 0.097834977 | -0.12185711 | 0.317527065 | 8.163455085 | 1.064360786 | 0.324276707 | 0.991835066 | -4.824699329 |
| **A_55_P2072776** | Fpgs | 0.118134323 | -0.147157079 | 0.383425725 | 9.938550304 | 1.064295078 | 0.324304376 | 0.991835066 | -4.824747577 |
| **A_55_P2093241** | Zc3h4 | 0.073060746 | -0.091027723 | 0.237149214 | 10.95767534 | 1.064180191 | 0.324352758 | 0.991835066 | -4.824831931 |
| **A_55_P2083063** | Mob1b | -0.050443769 | -0.163737938 | 0.0628504 | 8.633595175 | -1.064165319 | 0.324359021 | 0.991835066 | -4.824842851 |
| **A_55_P2136678** | NA | 0.079571454 | -0.099193876 | 0.258336785 | 8.547981675 | 1.063856701 | 0.32448902 | 0.991835066 | -4.82506942 |
| **A_55_P1992640** | Hdac3 | 0.086648207 | -0.108045961 | 0.281342374 | 9.537023737 | 1.063691697 | 0.324558541 | 0.991835066 | -4.825190539 |
| **A_55_P2126827** | NA | 0.11969021 | -0.149251458 | 0.388631878 | 13.34309023 | 1.063676754 | 0.324564838 | 0.991835066 | -4.825201508 |
| **A_66_P101473** | Fbxl3 | -0.077888621 | -0.252903711 | 0.097126468 | 9.26905816 | -1.063671819 | 0.324566918 | 0.991835066 | -4.82520513 |

| | | | | | | | | | |
|---|---|---|---|---|---|---|---|---|---|
| **A_55_P1960896** | Fndc3c1 | 0.063011794 | -0.078596732 | 0.20462032 | 7.81272887 | 1.063509989 | 0.324635115 | 0.991835066 | -4.825323907 |
| **A_51_P461429** | Cyp7b1 | 0.050297159 | -0.06274253 | 0.163336847 | 8.147127136 | 1.063461156 | 0.324655697 | 0.991835066 | -4.825359746 |
| **A_52_P421897** | NA | 0.06440455 | -0.080347922 | 0.209157021 | 8.149449834 | 1.06340743 | 0.324678341 | 0.991835066 | -4.825399175 |
| **A_52_P471395** | Ifngr2 | -0.088146353 | -0.286279891 | 0.109987186 | 12.64869168 | -1.063299192 | 0.324723965 | 0.991835066 | -4.825478607 |
| **A_55_P2001063** | Fcf1 | 0.078740849 | -0.098260054 | 0.255741751 | 11.91405091 | 1.063245963 | 0.324746404 | 0.991835066 | -4.825517667 |
| **A_55_P2155397** | Rps8 | 0.053765022 | -0.067100012 | 0.174630057 | 17.45595968 | 1.063183635 | 0.32477268 | 0.991835066 | -4.825563404 |
| **A_55_P1983209** | Rcbtb1 | 0.097775504 | -0.122065235 | 0.317616244 | 9.53859749 | 1.062994509 | 0.324852422 | 0.991835066 | -4.825702173 |
| **A_55_P1972097** | NA | 0.072461485 | -0.09051528 | 0.235438249 | 16.07239372 | 1.06265103 | 0.324997286 | 0.991835066 | -4.825954161 |
| **A_55_P2130970** | Parp10 | -0.082830288 | -0.269132904 | 0.103472328 | 11.24226747 | -1.062623342 | 0.325008966 | 0.991835066 | -4.825974471 |
| **A_51_P438805** | Txnip | -0.193379763 | -0.628524802 | 0.241765275 | 12.19793514 | -1.062150655 | 0.325208413 | 0.991835066 | -4.82632116 |
| **A_55_P2035286** | Uhrf1 | 0.123891276 | -0.154966777 | 0.40274933 | 12.38112774 | 1.061858549 | 0.325331715 | 0.991835066 | -4.826535355 |
| **A_66_P123683** | Nabp1 | -0.052208312 | -0.169740169 | 0.065323545 | 8.157474669 | -1.0616789 | 0.325407566 | 0.991835066 | -4.826667069 |
| **A_51_P308590** | Ppif | -0.160410054 | -0.521606337 | 0.200786229 | 9.916423569 | -1.061444877 | 0.325506396 | 0.991835066 | -4.826838629 |
| **A_51_P430082** | Tst | -0.088398154 | -0.287461276 | 0.110664967 | 10.0135259 | -1.061357077 | 0.325543481 | 0.991835066 | -4.826902988 |
| **A_55_P2123199** | Akirin1 | 0.086923919 | -0.108834345 | 0.282682183 | 8.932535034 | 1.061275955 | 0.325577748 | 0.991835066 | -4.826962449 |
| **A_52_P467690** | Sptbn1 | 0.090909131 | -0.113896029 | 0.295714291 | 11.29615595 | 1.060903164 | 0.325735258 | 0.991835066 | -4.827235662 |
| **A_55_P1956973** | Tln1 | 0.08225565 | -0.103078377 | 0.267589676 | 14.02234722 | 1.060766283 | 0.325793108 | 0.991835066 | -4.827335964 |
| **A_55_P2040046** | Swi5 | 0.150991099 | -0.189307989 | 0.491290186 | 14.94003089 | 1.060473188 | 0.325917007 | 0.991835066 | -4.82755071 |
| **A_55_P2085412** | Ankle1 | 0.094393478 | -0.118352098 | 0.307139055 | 8.501963655 | 1.060450946 | 0.32592641 | 0.991835066 | -4.827567005 |
| **A_52_P109304** | Tbl2 | -0.06348774 | -0.206588668 | 0.079613187 | 8.99588666 | -1.060367859 | 0.325961541 | 0.991835066 | -4.827627874 |
| **A_51_P247481** | Adck1 | -0.058947063 | -0.191820356 | 0.07392623 | 10.07110474 | -1.06031182 | 0.325985237 | 0.991835066 | -4.827668926 |
| **A_55_P2083464** | Lpin3 | 0.097368613 | -0.122128265 | 0.316865492 | 8.151811428 | 1.060229217 | 0.326020168 | 0.991835066 | -4.827729435 |
| **A_55_P2292737** | 2310002F0 | -0.097298671 | -0.316639831 | 0.122042489 | 12.75437183 | -1.060219785 | 0.326024157 | 0.991835066 | -4.827736344 |
| **A_55_P1972448** | NA | -0.090014597 | -0.292937873 | 0.112908679 | 9.206967773 | -1.060205884 | 0.326030035 | 0.991835066 | -4.827746526 |
| **A_55_P1970670** | Scp2 | -0.07013529 | -0.228246209 | 0.087975629 | 12.5569355 | -1.060190402 | 0.326036583 | 0.991835066 | -4.827757867 |
| **A_51_P163378** | Zwint | 0.085209909 | -0.106893161 | 0.277312978 | 13.3635047 | 1.060144167 | 0.326056137 | 0.991835066 | -4.827791734 |
| **A_55_P2097478** | Gda | -0.072841409 | -0.237096367 | 0.09141355 | 8.064369101 | -1.05990997 | 0.326155199 | 0.991835066 | -4.827963265 |
| **A_55_P2185461** | Rpl9 | 0.067826561 | -0.085120334 | 0.220773456 | 16.35199296 | 1.059908227 | 0.326155936 | 0.991835066 | -4.827964542 |
| **A_55_P2398534** | Gid8 | 0.112307944 | -0.14094582 | 0.365561709 | 11.17185404 | 1.059897092 | 0.326160647 | 0.991835066 | -4.827972697 |
| **A_51_P246166** | Wfdc18 | 0.094460425 | -0.11854951 | 0.307470359 | 9.508617965 | 1.059886026 | 0.326165328 | 0.991835066 | -4.827980801 |
| **A_55_P2181371** | NA | -0.05856259 | -0.190624911 | 0.07349973 | 8.524199915 | -1.05986484 | 0.326174291 | 0.991835066 | -4.827996317 |
| **A_51_P291682** | Tmed4 | -0.052540294 | -0.171023995 | 0.065943407 | 9.607569904 | -1.05984662 | 0.326181999 | 0.991835066 | -4.828009661 |
| **A_55_P2037883** | Ino80 | 0.086990718 | -0.109255817 | 0.283237252 | 8.907681364 | 1.059448987 | 0.326350259 | 0.991835066 | -4.828300832 |
| **A_55_P1960566** | Ybx3 | 0.065406432 | -0.082162977 | 0.21297584 | 14.26074579 | 1.059334828 | 0.326398578 | 0.991835066 | -4.828384414 |
| **A_51_P347240** | Lrr1 | 0.065254379 | -0.081995229 | 0.212503987 | 8.586335753 | 1.059167507 | 0.32646941 | 0.991835066 | -4.828506908 |
| **A_55_P2150299** | NA | -0.095067485 | -0.309625561 | 0.11949059 | 9.132710871 | -1.059000764 | 0.326540009 | 0.991835066 | -4.828628967 |
| **A_55_P1980421** | Wdr81 | -0.084153745 | -0.27409662 | 0.105789131 | 10.33987746 | -1.058911269 | 0.326577906 | 0.991835066 | -4.828694474 |
| **A_55_P1982156** | Nelfcd | 0.058216571 | -0.073211616 | 0.189644758 | 12.65565453 | 1.058686158 | 0.326673247 | 0.991835066 | -4.828859232 |
| **A_55_P2050488** | Cep170b | 0.06997108 | -0.088010579 | 0.227952738 | 13.75304977 | 1.058573552 | 0.326720947 | 0.991835066 | -4.828941639 |
| **A_51_P239766** | Plcd1 | 0.097136234 | -0.122194212 | 0.316466679 | 10.8965345 | 1.058501488 | 0.326751477 | 0.991835066 | -4.828994374 |
| **A_55_P1953143** | Wt1 | 0.107870489 | -0.135725338 | 0.351466316 | 11.04227751 | 1.058380755 | 0.326802629 | 0.991835066 | -4.82908272 |
| **A_51_P353592** | Commd4 | -0.064076253 | -0.208788717 | 0.080636211 | 10.56954063 | -1.058279303 | 0.326845618 | 0.991835066 | -4.829156952 |
| **A_55_P1988678** | Znhit6 | 0.097927778 | -0.123236408 | 0.319091965 | 11.25331555 | 1.05827913 | 0.326845691 | 0.991835066 | -4.829157078 |
| **A_55_P2234744** | Spice1 | 0.056924276 | -0.071640065 | 0.185488617 | 9.293550481 | 1.058244745 | 0.326860263 | 0.991835066 | -4.829182237 |
| **A_51_P329441** | Bsg | -0.158228569 | -0.515695989 | 0.199238852 | 11.87616654 | -1.057931541 | 0.326993011 | 0.991835066 | -4.829411372 |
| **A_51_P375987** | Fign | -0.1110909 | -0.362189416 | 0.140007615 | 10.28171622 | -1.057410147 | 0.327214096 | 0.991835066 | -4.829792724 |
| **A_55_P2244677** | 6720422M2 | 0.194168928 | -0.244732858 | 0.633070715 | 9.758218896 | 1.057356689 | 0.32723677 | 0.991835066 | -4.829831817 |
| **A_55_P2026233** | Uba7 | -0.092102653 | -0.300334639 | 0.116129332 | 9.962631032 | -1.057143234 | 0.32732732 | 0.991835066 | -4.8299879 |

| | | | | | | | | | |
|---|---|---|---|---|---|---|---|---|---|
| **A_52_P265877** | Aldh9a1 | 0.060364025 | -0.076117693 | 0.196845742 | 9.737398188 | 1.057092137 | 0.327348999 | 0.991835066 | -4.830025261 |
| **A_55_P2133943** | NA | 0.048250505 | -0.060856849 | 0.157357859 | 8.280815554 | 1.056956147 | 0.327406701 | 0.991835066 | -4.830124687 |
| **A_51_P283266** | Zfp874b | -0.063033612 | -0.205587795 | 0.079520571 | 8.304338273 | -1.056820826 | 0.327464128 | 0.991835066 | -4.830223615 |
| **A_55_P2135860** | Prss36 | 0.079000774 | -0.099682335 | 0.257683883 | 9.361588092 | 1.05671283 | 0.327509964 | 0.991835066 | -4.830302562 |
| **A_52_P111963** | 1810030O( | -0.050545014 | -0.164894218 | 0.06380419 | 9.617593666 | -1.056463039 | 0.327616001 | 0.991835066 | -4.830485144 |
| **A_51_P462746** | Tex10 | 0.070028118 | -0.0884103 | 0.228466536 | 9.846773029 | 1.056382242 | 0.327650306 | 0.991835066 | -4.830544196 |
| **A_52_P679205** | Slc35f6 | -0.065401542 | -0.213465141 | 0.082662057 | 9.569180937 | -1.055720173 | 0.327931514 | 0.991835066 | -4.831027973 |
| **A_51_P208801** | Ndufab1 | 0.058441203 | -0.073867062 | 0.190749468 | 10.58067919 | 1.055701903 | 0.327939276 | 0.991835066 | -4.83104132 |
| **A_51_P163444** | Carhsp1 | -0.051286702 | -0.167432383 | 0.06485898 | 11.90162696 | -1.055384801 | 0.328074034 | 0.991835066 | -4.831272958 |
| **A_52_P455404** | Itpa | 0.057332681 | -0.072540051 | 0.187205414 | 12.87156925 | 1.055099474 | 0.328195326 | 0.991835066 | -4.831481348 |
| **A_52_P20357** | Aqr | 0.046538542 | -0.058897158 | 0.151974241 | 9.545004265 | 1.054955683 | 0.328256465 | 0.991835066 | -4.831586353 |
| **A_51_P462102** | Adgra3 | -0.044153799 | -0.14421534 | 0.055907742 | 11.04764425 | -1.054654039 | 0.328384751 | 0.991835066 | -4.831806602 |
| **A_55_P1970695** | NA | 0.067470373 | -0.085433848 | 0.220374593 | 9.977319717 | 1.054636429 | 0.328392242 | 0.991835066 | -4.831819459 |
| **A_55_P2095578** | NA | 0.077121232 | -0.097659008 | 0.251901473 | 8.54672831 | 1.054607268 | 0.328404647 | 0.991835066 | -4.831840749 |
| **A_51_P131025** | Ngdn | 0.065773139 | -0.083305928 | 0.214852206 | 13.25555672 | 1.054486531 | 0.328456009 | 0.991835066 | -4.831928894 |
| **A_55_P1987201** | Capn2 | -0.059187054 | -0.19334581 | 0.074971702 | 12.89669428 | -1.054427769 | 0.328481009 | 0.991835066 | -4.83197179 |
| **A_55_P2360427** | BC005561 | 0.122614954 | -0.155321536 | 0.400551443 | 8.695681147 | 1.054403899 | 0.328491165 | 0.991835066 | -4.831989215 |
| **A_55_P2131481** | NA | -0.146997332 | -0.480225913 | 0.186231248 | 14.08304412 | -1.054329462 | 0.328522836 | 0.991835066 | -4.832043552 |
| **A_51_P519837** | Cherp | 0.093692546 | -0.118732343 | 0.306117435 | 11.3489249 | 1.054165436 | 0.328592636 | 0.991835066 | -4.832163278 |
| **A_55_P2151986** | Gngt2 | -0.075000487 | -0.245049947 | 0.095048974 | 11.47741372 | -1.054139183 | 0.328603809 | 0.991835066 | -4.83218244 |
| **A_52_P234958** | Dvl1 | 0.059376334 | -0.07530216 | 0.194054827 | 10.20891805 | 1.053717662 | 0.328783241 | 0.991835066 | -4.832490058 |
| **A_52_P282838** | Spin2c | 0.06290699 | -0.079800275 | 0.205614254 | 8.313601906 | 1.053566497 | 0.328847608 | 0.991835066 | -4.832600357 |
| **A_55_P2024675** | Shld2 | -0.050193297 | -0.164064753 | 0.06367816 | 8.896419015 | -1.053513193 | 0.328870308 | 0.991835066 | -4.832639249 |
| **A_55_P2112330** | Anapc7 | -0.067642065 | -0.221120936 | 0.085836807 | 12.07442111 | -1.053361376 | 0.328934966 | 0.991835066 | -4.832750009 |
| **A_52_P270145** | Zfp329 | 0.078837555 | -0.100054245 | 0.257729355 | 10.30899247 | 1.053299422 | 0.328961355 | 0.991835066 | -4.832795206 |
| **A_55_P1972187** | Slc25a14 | 0.070367105 | -0.089350552 | 0.230084763 | 9.792311886 | 1.052993973 | 0.329091484 | 0.991835066 | -4.833018013 |
| **A_51_P235139** | Klhl40 | -0.127416885 | -0.416653031 | 0.161819261 | 8.707891939 | -1.052891331 | 0.329135221 | 0.991835066 | -4.833092875 |
| **A_55_P2159595** | NA | -0.160502189 | -0.52487563 | 0.203871251 | 10.15876397 | -1.052793949 | 0.329176722 | 0.991835066 | -4.833163897 |
| **A_55_P2059347** | Ormdl3 | 0.055722147 | -0.070787908 | 0.182232202 | 12.67260544 | 1.052717713 | 0.329209214 | 0.991835066 | -4.833219494 |
| **A_55_P2043122** | Arsg | 0.073700371 | -0.093676899 | 0.241077641 | 8.673557978 | 1.052403608 | 0.329343112 | 0.991835066 | -4.833448535 |
| **A_66_P131645** | Ahnak2 | 0.052142244 | -0.066287262 | 0.17057175 | 8.148727741 | 1.052298454 | 0.329387947 | 0.991835066 | -4.833525202 |
| **A_55_P1959152** | Gm10324 | -0.077736835 | -0.2543266 | 0.098852929 | 8.520082011 | -1.052132562 | 0.32945869 | 0.991835066 | -4.833646144 |
| **A_52_P54856** | Ubald1 | -0.092755082 | -0.303511245 | 0.118001081 | 9.483684426 | -1.051880885 | 0.329566037 | 0.991835066 | -4.833829602 |
| **A_55_P2105457** | Zfp7 | 0.045958884 | -0.05847567 | 0.150393438 | 7.978708664 | 1.051802968 | 0.329599277 | 0.991835066 | -4.833886393 |
| **A_55_P2008016** | Armc3 | -0.048944386 | -0.160164882 | 0.062276109 | 7.778888836 | -1.051785536 | 0.329606714 | 0.991835066 | -4.833899099 |
| **A_55_P1995233** | Lonp2 | 0.060642872 | -0.07720849 | 0.198494234 | 11.80235168 | 1.051423871 | 0.329761039 | 0.991835066 | -4.834162669 |
| **A_51_P437240** | Emp2 | -0.061494447 | -0.20128999 | 0.078301097 | 11.88783802 | -1.051360609 | 0.329788039 | 0.991835066 | -4.834208766 |
| **A_55_P1976376** | Gimap4 | -0.095756962 | -0.313443355 | 0.121929431 | 8.553317219 | -1.051352155 | 0.329791648 | 0.991835066 | -4.834214927 |
| **A_55_P2076805** | Ddx5 | 0.133330132 | -0.169791577 | 0.436451841 | 8.948487138 | 1.051284834 | 0.329820382 | 0.991835066 | -4.83426398 |
| **A_55_P2136340** | NA | -0.117792099 | -0.385666319 | 0.150082121 | 14.09561288 | -1.05097981 | 0.329950602 | 0.991835066 | -4.834486209 |
| **A_51_P116932** | Lad1 | -0.052058751 | -0.170455259 | 0.066337757 | 7.922350803 | -1.050906264 | 0.329982006 | 0.991835066 | -4.834539786 |
| **A_55_P2074843** | Ccdc122 | 0.056905864 | -0.072516043 | 0.186327772 | 7.818011661 | 1.050892663 | 0.329987814 | 0.991835066 | -4.834549694 |
| **A_51_P436483** | Kin | 0.066986942 | -0.085363139 | 0.219337023 | 10.14600689 | 1.050888391 | 0.329989638 | 0.991835066 | -4.834552805 |
| **A_55_P2065364** | Tdrkh | 0.087539654 | -0.111567081 | 0.286646388 | 8.035031912 | 1.050819218 | 0.330019178 | 0.991835066 | -4.834603194 |
| **A_51_P280893** | Pla2g1b | 0.045242371 | -0.057665154 | 0.148149896 | 7.6772875 | 1.050769257 | 0.330040515 | 0.991835066 | -4.834639587 |
| **A_55_P2098428** | Bloc1s4 | -0.050929336 | -0.166772694 | 0.064914022 | 9.221778144 | -1.050765989 | 0.330041911 | 0.991835066 | -4.834641967 |
| **A_55_P2122789** | Homer1 | 0.065395731 | -0.083358762 | 0.214150225 | 9.015671909 | 1.05072349 | 0.330060061 | 0.991835066 | -4.834672923 |
| **A_55_P2150786** | Mul1 | 0.379706514 | -0.484071306 | 1.243484335 | 10.88899158 | 1.050642905 | 0.33009448 | 0.991835066 | -4.834731618 |

| | | | | | | | | | |
|---|---|---|---|---|---|---|---|---|---|
| **A_55_P2003931** | NA | 0.054063623 | -0.068924593 | 0.177051839 | 7.872453634 | 1.050632368 | 0.330098981 | 0.991835066 | -4.834739293 |
| **A_55_P2006281** | Ogdh | -0.061046712 | -0.199939126 | 0.077845702 | 14.44427619 | -1.050492331 | 0.330158801 | 0.991835066 | -4.834841284 |
| **A_55_P2168048** | Gramd4 | -0.071782599 | -0.235114714 | 0.091549517 | 8.536728011 | -1.050404854 | 0.330196172 | 0.991835066 | -4.83490499 |
| **A_51_P393598** | Atp6v1d | -0.078917708 | -0.258492119 | 0.100656702 | 13.39938114 | -1.050362355 | 0.33021433 | 0.991835066 | -4.83493594 |
| **A_51_P125260** | Acaa2 | -0.128463208 | -0.420782711 | 0.163856296 | 15.92725466 | -1.050340476 | 0.330223678 | 0.991835066 | -4.834951872 |
| **A_55_P2116496** | Tuba1b | -0.110873827 | -0.363180716 | 0.141433062 | 14.51852262 | -1.050289595 | 0.330245419 | 0.991835066 | -4.834988924 |
| **A_51_P233603** | Psma7 | -0.056749765 | -0.185915017 | 0.072415487 | 14.00349469 | -1.050092379 | 0.330329695 | 0.991835066 | -4.835132527 |
| **A_52_P40954** | Dubr | -0.067870793 | -0.222358404 | 0.086616819 | 9.163252002 | -1.050021996 | 0.330359777 | 0.991835066 | -4.835183773 |
| **A_55_P2139385** | Gm12070 | -0.04974595 | -0.162994699 | 0.0635028 | 16.85962192 | -1.049864971 | 0.330426896 | 0.991835066 | -4.835298094 |
| **A_55_P2152245** | Rpl19 | 0.054332079 | -0.06938682 | 0.178050978 | 15.47703226 | 1.049613502 | 0.330534407 | 0.991835066 | -4.835481151 |
| **A_55_P1978875** | Kmt2b | 0.072317583 | -0.092370396 | 0.237005563 | 10.51907264 | 1.04952101 | 0.330573958 | 0.991835066 | -4.835548474 |
| **A_55_P2086203** | Kcnq1 | -0.117383741 | -0.384714392 | 0.14994691 | 12.12794574 | -1.049465877 | 0.330597535 | 0.991835066 | -4.835588603 |
| **A_52_P123354** | Tusc2 | -0.073452143 | -0.240733023 | 0.093828738 | 9.389202034 | -1.049463394 | 0.330598597 | 0.991835066 | -4.83559041 |
| **A_52_P273594** | Zc3h14 | -0.064525792 | -0.211479221 | 0.082427637 | 10.57071881 | -1.049452414 | 0.330603293 | 0.991835066 | -4.835598402 |
| **A_55_P2118302** | Fam110a | -0.052297871 | -0.171403773 | 0.06680803 | 11.07160184 | -1.049445439 | 0.330606275 | 0.991835066 | -4.835603478 |
| **A_55_P1968450** | Gm5766 | -0.068050172 | -0.223037537 | 0.086937194 | 15.05277711 | -1.049402421 | 0.330624673 | 0.991835066 | -4.835634787 |
| **A_55_P2117525** | Rbm3 | -0.049678867 | -0.162831452 | 0.063473718 | 12.57414788 | -1.049340256 | 0.330651261 | 0.991835066 | -4.83568003 |
| **A_55_P2153292** | Tubb4b | 0.07686202 | -0.098242953 | 0.251966993 | 14.54951702 | 1.04911343 | 0.330748288 | 0.991835066 | -4.835845098 |
| **A_55_P2091494** | Tango2 | -0.078704687 | -0.258008538 | 0.100599164 | 11.53686719 | -1.049107787 | 0.330750702 | 0.991835066 | -4.835849204 |
| **A_55_P2142393** | Cacna1a | 0.050878931 | -0.065033258 | 0.16679112 | 8.597636796 | 1.049102692 | 0.330752882 | 0.991835066 | -4.835852912 |
| **A_55_P2058493** | NA | -0.064037846 | -0.210010159 | 0.081934466 | 15.80959775 | -1.048516726 | 0.331003643 | 0.991835066 | -4.83627923 |
| **A_55_P2017764** | Cep89 | -0.078990401 | -0.259048278 | 0.101067476 | 11.1748665 | -1.048506982 | 0.331007815 | 0.991835066 | -4.836286318 |
| **A_51_P346445** | Hspb7 | 0.147460527 | -0.18875046 | 0.483671513 | 12.09933165 | 1.04826965 | 0.331109424 | 0.991835066 | -4.823831753 |
| **A_51_P300207** | Rsl24d1 | -0.098939947 | -0.324573307 | 0.126693414 | 9.436806399 | -1.048039117 | 0.331208146 | 0.991835066 | -4.836626601 |
| **A_55_P2053077** | Chuk | 0.131325602 | -0.168187677 | 0.43083888 | 10.67133727 | 1.047954555 | 0.331244365 | 0.991835066 | -4.836688094 |
| **A_52_P373893** | NA | -0.085109236 | -0.279234793 | 0.109016322 | 10.90114713 | -1.047859626 | 0.331285027 | 0.991835066 | -4.836757122 |
| **A_55_P1957277** | NA | 0.067531804 | -0.08652383 | 0.221587438 | 9.615229618 | 1.047707127 | 0.331350358 | 0.991835066 | -4.836868003 |
| **A_55_P1957003** | Timm10 | 0.072638455 | -0.093090841 | 0.238367751 | 13.32769655 | 1.047554058 | 0.331415943 | 0.991835066 | -4.836979289 |
| **A_52_P355095** | Dstyk | 0.101897383 | -0.130590176 | 0.334384942 | 8.350594052 | 1.04754441 | 0.331420077 | 0.991835066 | -4.836986303 |
| **A_55_P1997837** | Osbp2 | -0.091228452 | -0.29939088 | 0.116933977 | 8.381532988 | -1.04745914 | 0.331456617 | 0.991835066 | -4.837048292 |
| **A_55_P2000853** | Mvb12b | 0.043648412 | -0.055988918 | 0.143285742 | 11.98669376 | 1.047021259 | 0.331644311 | 0.991835066 | -4.837366569 |
| **A_55_P2097309** | Nudt4 | -0.073650662 | -0.241775683 | 0.09447436 | 11.3573484 | -1.047016278 | 0.331646447 | 0.991835066 | -4.837370189 |
| **A_55_P2062108** | Apold1 | 0.16981066 | -0.217869951 | 0.55749127 | 8.799021775 | 1.046887468 | 0.331701676 | 0.991835066 | -4.837463799 |
| **A_52_P628590** | Pvr | 0.076361038 | -0.097973616 | 0.250695692 | 9.121865953 | 1.046880789 | 0.33170454 | 0.991835066 | -4.837468653 |
| **A_55_P2137527** | Fam183b | 0.086043903 | -0.110424583 | 0.282512389 | 8.399467628 | 1.046734009 | 0.331767484 | 0.991835066 | -4.837575314 |
| **A_52_P7183** | Tmem248 | 0.090711644 | -0.116521599 | 0.297944887 | 10.30227747 | 1.046195254 | 0.331998602 | 0.991835066 | -4.837966728 |
| **A_52_P155805** | 2010204K1 | 0.08822699 | -0.113338141 | 0.289792122 | 8.587387862 | 1.046152965 | 0.332016749 | 0.991835066 | -4.837997447 |
| **A_55_P2373987** | A730009E1 | 0.111405498 | -0.143136812 | 0.365947807 | 8.434798176 | 1.046058025 | 0.332057491 | 0.991835066 | -4.838066407 |
| **A_51_P505823** | Endod1 | 0.085681975 | -0.11009212 | 0.28145607 | 9.634779367 | 1.046028159 | 0.332070309 | 0.991835066 | -4.8380881 |
| **A_66_P112436** | Prkar2b | -0.047678478 | -0.156628683 | 0.061271727 | 8.103657358 | -1.045932034 | 0.332111566 | 0.991835066 | -4.838157915 |
| **A_51_P238448** | Ccnd3 | 0.124024801 | -0.159400315 | 0.407449916 | 12.68459772 | 1.045873931 | 0.332136505 | 0.991835066 | -4.838200114 |
| **A_55_P2336173** | Mirg | 0.152908607 | -0.19658082 | 0.502398034 | 8.921605447 | 1.045699794 | 0.33221126 | 0.991835066 | -4.838326576 |
| **A_52_P272534** | NA | 0.077900565 | -0.10016953 | 0.25597066 | 11.57087726 | 1.045583571 | 0.332261161 | 0.991835066 | -4.838410972 |
| **A_55_P1960748** | Arl13b | -0.05823825 | -0.191411906 | 0.074935405 | 8.807641428 | -1.045199347 | 0.332426169 | 0.991835066 | -4.838689936 |
| **A_55_P1962603** | Cplx2 | 0.111105808 | -0.142987706 | 0.365199322 | 13.51399399 | 1.045086688 | 0.332474564 | 0.991835066 | -4.838771719 |
| **A_55_P2011678** | Pdzk1ip1 | 0.10008229 | -0.128814995 | 0.328979575 | 8.185785676 | 1.045022699 | 0.332502055 | 0.991835066 | -4.838818168 |
| **A_66_P107973** | Prpf38b | 0.064394219 | -0.082890173 | 0.211678611 | 8.60404735 | 1.044959087 | 0.332529385 | 0.991835066 | -4.838864342 |
| **A_55_P2028798** | Nqo2 | -0.085140906 | -0.279922001 | 0.109640188 | 11.1634785 | -1.044721661 | 0.332631408 | 0.991835066 | -4.839036667 |

| | | | | | | | | | |
|---|---|---|---|---|---|---|---|---|---|
| **A_52_P201551** | Nav1 | 0.059790668 | -0.077005784 | 0.196587119 | 10.50321383 | 1.044642534 | 0.332665415 | 0.991835066 | -4.839094092 |
| **A_55_P2009783** | Sfr1 | 0.084121107 | -0.108362076 | 0.27660429 | 14.20874956 | 1.044530972 | 0.332713366 | 0.991835066 | -4.839175051 |
| **A_51_P374305** | Rchy1 | -0.058509168 | -0.192393574 | 0.075375237 | 11.20763383 | -1.044487059 | 0.332732242 | 0.991835066 | -4.839206917 |
| **A_51_P488673** | Rack1 | -0.06456166 | -0.212319851 | 0.083196532 | 16.71454783 | -1.044316765 | 0.332805451 | 0.991835066 | -4.839330484 |
| **A_55_P2016670** | Alpk2 | -0.122722556 | -0.403591649 | 0.158146536 | 8.728753011 | -1.044310331 | 0.332808218 | 0.991835066 | -4.839335152 |
| **A_51_P158552** | Zscan12 | 0.061253254 | -0.078934919 | 0.201441428 | 10.31702387 | 1.044303954 | 0.332810959 | 0.991835066 | -4.839339779 |
| **A_55_P1954306** | Esrra | -0.052970726 | -0.174204955 | 0.068263504 | 13.54266862 | -1.044286822 | 0.332818325 | 0.991835066 | -4.839352209 |
| **A_55_P1995737** | Gm14288 | 0.059406338 | -0.07656907 | 0.195381746 | 12.06280092 | 1.044194844 | 0.332857873 | 0.991835066 | -4.839418943 |
| **A_55_P2001714** | Blvra | -0.111231046 | -0.365854821 | 0.14339273 | 9.10685142 | -1.044085822 | 0.332904754 | 0.991835066 | -4.839498038 |
| **A_55_P2184811** | Eif2s1 | 0.084045718 | -0.108349968 | 0.276441404 | 9.383275111 | 1.044069463 | 0.33291179 | 0.991835066 | -4.839509906 |
| **A_51_P294113** | Chd2 | -0.059987682 | -0.197355891 | 0.077380527 | 8.738091373 | -1.043722332 | 0.333061099 | 0.991835066 | -4.83976171 |
| **A_55_P2060752** | Usp12 | 0.065316263 | -0.084267197 | 0.214899722 | 11.00783953 | 1.043630798 | 0.333100479 | 0.991835066 | -4.839828099 |
| **A_51_P162162** | Inmt | 0.043457135 | -0.056087398 | 0.143001669 | 7.67121298 | 1.043404749 | 0.333197746 | 0.991835066 | -4.839992033 |
| **A_52_P204618** | Rnf222 | 0.044898394 | -0.057952395 | 0.147749183 | 7.707031146 | 1.043355504 | 0.333218939 | 0.991835066 | -4.840027744 |
| **A_55_P2123381** | Fga | -0.063736345 | -0.209758906 | 0.082286216 | 7.826992642 | -1.043221011 | 0.333276824 | 0.991835066 | -4.840125267 |
| **A_55_P1967105** | Adgrb2 | 0.125555578 | -0.162133681 | 0.413244836 | 10.27321931 | 1.043089313 | 0.333333513 | 0.991835066 | -4.840220756 |
| **A_55_P1967867** | Xpa | -0.058421276 | -0.192295185 | 0.075452633 | 9.2802504 | -1.042999805 | 0.333372047 | 0.991835066 | -4.84028565 |
| **A_55_P2077884** | Kat6b | 0.11452137 | -0.147909172 | 0.376951911 | 8.473855062 | 1.042992749 | 0.333375084 | 0.991835066 | -4.840290766 |
| **A_51_P320614** | Txnrd1 | -0.06843626 | -0.225278689 | 0.088406169 | 11.50782036 | -1.042874004 | 0.333426211 | 0.991835066 | -4.840376852 |
| **A_55_P2156464** | Nt5c3b | -0.104810764 | -0.34504112 | 0.135419591 | 9.81522651 | -1.04276666 | 0.333472433 | 0.991835066 | -4.840454666 |
| **A_51_P144349** | Dtx4 | -0.084395698 | -0.277923493 | 0.109132097 | 9.283511485 | -1.042284056 | 0.333680306 | 0.991835066 | -4.840804448 |
| **A_55_P2044593** | 1810037I17 | -0.067865941 | -0.2234999 | 0.087768018 | 13.00759598 | -1.042213382 | 0.333710757 | 0.991835066 | -4.840855663 |
| **A_51_P272283** | Cmbl | -0.160332791 | -0.528038537 | 0.207372954 | 10.38277042 | -1.042152009 | 0.333737201 | 0.991835066 | -4.840900136 |
| **A_52_P498355** | Lyrm4 | 0.076700864 | -0.099233276 | 0.252635005 | 10.69309401 | 1.041979719 | 0.333811447 | 0.991835066 | -4.841024974 |
| **A_51_P490678** | Trim68 | -0.071215339 | -0.234597652 | 0.092166973 | 8.620443226 | -1.041783899 | 0.333895849 | 0.991835066 | -4.841166844 |
| **A_51_P374772** | Arfgef1 | -0.116105821 | -0.382486641 | 0.150274999 | 10.25837832 | -1.041741999 | 0.333913911 | 0.991835066 | -4.841197198 |
| **A_51_P356931** | Orc1 | 0.049158008 | -0.063654482 | 0.161970499 | 7.860580791 | 1.0414687 | 0.334031741 | 0.991835066 | -4.841395169 |
| **A_55_P1989215** | Entpd8 | -0.052774975 | -0.173901062 | 0.068351112 | 7.779638516 | -1.041356621 | 0.334080073 | 0.991835066 | -4.841476346 |
| **A_52_P2947** | Ndufaf2 | -0.057447343 | -0.189309278 | 0.074414593 | 11.75944563 | -1.041261065 | 0.334121283 | 0.991835066 | -4.841545552 |
| **A_55_P1982494** | H2al1m | -0.062131831 | -0.204752343 | 0.080488681 | 7.763421239 | -1.041217104 | 0.334140244 | 0.991835066 | -4.84157739 |
| **A_55_P2021530** | Clic4 | 0.060719377 | -0.078667882 | 0.200106637 | 8.676300936 | 1.041150141 | 0.334169127 | 0.991835066 | -4.841625883 |
| **A_55_P2108689** | NA | -0.108461834 | -0.357463589 | 0.140539922 | 15.86369789 | -1.041078948 | 0.334199837 | 0.991835066 | -4.841677438 |
| **A_55_P2151897** | NA | 0.04893462 | -0.063407783 | 0.161277023 | 14.98013858 | 1.041074111 | 0.334201924 | 0.991835066 | -4.841680941 |
| **A_55_P2084032** | AW146154 | -0.066404644 | -0.218857292 | 0.086048004 | 9.448934416 | -1.041052455 | 0.334211265 | 0.991835066 | -4.841696622 |
| **A_51_P267454** | Slc25a11 | -0.058135995 | -0.191612403 | 0.075340413 | 15.03021914 | -1.040997602 | 0.334234929 | 0.991835066 | -4.841736343 |
| **A_55_P2119441** | Vma21 | -0.07807459 | -0.257330168 | 0.101180989 | 10.70163933 | -1.04098905 | 0.334238619 | 0.991835066 | -4.841742536 |
| **A_51_P317542** | Slc24a2 | 0.063429267 | -0.082227196 | 0.20908573 | 9.0925795 | 1.040804276 | 0.33431834 | 0.991835066 | -4.841876323 |
| **A_55_P2082979** | Irak2 | 0.221602788 | -0.287278195 | 0.730483771 | 11.40160611 | 1.040802032 | 0.334319308 | 0.991835066 | -4.841877947 |
| **A_55_P2052799** | Cask | 0.045708636 | -0.059264403 | 0.150681675 | 10.16705631 | 1.040709751 | 0.334359129 | 0.991835066 | -4.841944759 |
| **A_55_P2121701** | Mir670hg | -0.093070852 | -0.30681612 | 0.120674415 | 8.888910021 | -1.040701821 | 0.334362552 | 0.991835066 | -4.8419505 |
| **A_55_P2155002** | Svbp | -0.087731444 | -0.289224441 | 0.113761553 | 12.51108838 | -1.040649438 | 0.334385158 | 0.991835066 | -4.841988423 |
| **A_55_P2004152** | NA | -0.061930633 | -0.204173694 | 0.080312428 | 12.39796012 | -1.040599373 | 0.334406765 | 0.991835066 | -4.842024667 |
| **A_51_P500550** | Mfsd1 | -0.050828489 | -0.167585295 | 0.065928316 | 8.82473825 | -1.040480943 | 0.334457881 | 0.991835066 | -4.842110399 |
| **A_51_P280401** | Tcta | -0.097693474 | -0.322117304 | 0.126730356 | 8.256804333 | -1.040412869 | 0.334487265 | 0.991835066 | -4.842159675 |
| **A_51_P162955** | Serpina7 | 0.044551103 | -0.057809028 | 0.146911235 | 7.664960835 | 1.040247687 | 0.334558575 | 0.991835066 | -4.842279236 |
| **A_55_P2467801** | Srp54a | 0.063563517 | -0.082507203 | 0.209634237 | 10.14237355 | 1.040049194 | 0.334644283 | 0.991835066 | -4.842422891 |
| **A_51_P388696** | Ndufs4 | -0.075855118 | -0.250211205 | 0.098500969 | 13.97734566 | -1.039816979 | 0.334744572 | 0.991835066 | -4.842590929 |
| **A_52_P69998** | Kdelr2 | 0.073017182 | -0.094836193 | 0.240870556 | 10.15354061 | 1.039690613 | 0.334799158 | 0.991835066 | -4.842682362 |

| | | | | | | | | | |
|---|---|---|---|---|---|---|---|---|---|
| **A_55_P2029022** | Rpl34 | 0.059426238 | -0.077188106 | 0.196040582 | 16.75969904 | 1.039659371 | 0.334812655 | 0.991835066 | -4.842704966 |
| **A_51_P468240** | Dennd4a | -0.065799392 | -0.217073526 | 0.085474743 | 8.271625674 | -1.039600153 | 0.334838238 | 0.991835066 | -4.84274781 |
| **A_51_P290139** | Dlst | -0.0922313 | -0.304323908 | 0.119861307 | 13.61047431 | -1.039350266 | 0.334946211 | 0.991835066 | -4.842928586 |
| **A_55_P2262287** | 5430434G1 | 0.131690259 | -0.171143293 | 0.434523812 | 9.074579885 | 1.039342755 | 0.334949457 | 0.991835066 | -4.842934019 |
| **A_66_P136801** | Peg13 | 0.073042556 | -0.094927654 | 0.241012766 | 10.98167833 | 1.03932848 | 0.334955626 | 0.991835066 | -4.842944346 |
| **A_55_P2048224** | NA | 0.071293902 | -0.092690759 | 0.235278564 | 8.43931898 | 1.039102263 | 0.335053397 | 0.991835066 | -4.843107972 |
| **A_55_P2027117** | Rgs3 | 0.107060648 | -0.139212054 | 0.353333349 | 11.12549717 | 1.039017169 | 0.335090181 | 0.991835066 | -4.843169515 |
| **A_55_P2155312** | Dock3 | -0.068390931 | -0.225738945 | 0.088957084 | 9.547152557 | -1.038834531 | 0.335169141 | 0.991835066 | -4.843301596 |
| **A_52_P193925** | Sulf2 | 0.061997702 | -0.080662909 | 0.204658312 | 12.34667169 | 1.038677309 | 0.335237125 | 0.991835066 | -4.843415286 |
| **A_51_P198411** | Grb2 | -0.067201907 | -0.221845031 | 0.087441216 | 12.07643977 | -1.038628221 | 0.335258353 | 0.991835066 | -4.84345078 |
| **A_55_P2144850** | Psmb5 | 0.070475658 | -0.091703361 | 0.232654676 | 15.76196301 | 1.038612633 | 0.335265095 | 0.991835066 | -4.84346205 |
| **A_51_P189272** | Tmcc3 | -0.077064544 | -0.254425027 | 0.100295939 | 10.68960239 | -1.038500914 | 0.335313413 | 0.991835066 | -4.843542826 |
| **A_55_P1976839** | Spata5 | 0.071010644 | -0.092425209 | 0.234446498 | 9.102134161 | 1.038449187 | 0.335335786 | 0.991835066 | -4.843580224 |
| **A_55_P2084571** | NA | 0.058324943 | -0.075928358 | 0.192578244 | 15.98717493 | 1.038337361 | 0.335384159 | 0.991835066 | -4.843661068 |
| **A_55_P2141709** | Gabarapl2 | -0.183915756 | -0.607268906 | 0.239437394 | 12.55186468 | -1.038305786 | 0.335397818 | 0.991835066 | -4.843683895 |
| **A_52_P658034** | Tspyl1 | -0.055318191 | -0.182670835 | 0.072034454 | 11.88086349 | -1.038171564 | 0.335455887 | 0.991835066 | -4.843780921 |
| **A_55_P2098471** | Rps19 | 0.055244694 | -0.071959858 | 0.182449246 | 15.04902159 | 1.037999277 | 0.335530437 | 0.991835066 | -4.843905452 |
| **A_51_P372550** | Cgref1 | 0.068643798 | -0.089435891 | 0.226723486 | 8.938703755 | 1.037849458 | 0.335595275 | 0.991835066 | -4.844013733 |
| **A_51_P256957** | Ankmy2 | -0.062864117 | -0.207644875 | 0.081916641 | 9.206622915 | -1.037769984 | 0.335629674 | 0.991835066 | -4.844071168 |
| **A_51_P501815** | Smdt1 | -0.085354487 | -0.281946219 | 0.111237244 | 15.47718504 | -1.037696237 | 0.335661596 | 0.991835066 | -4.844124462 |
| **A_55_P2132812** | NA | 0.181402701 | -0.236443029 | 0.599248432 | 11.156011 | 1.037616606 | 0.335696068 | 0.991835066 | -4.844182005 |
| **A_51_P132459** | Rpl27a | 0.076529656 | -0.099757467 | 0.252816779 | 12.84033618 | 1.037572147 | 0.335715315 | 0.991835066 | -4.84421413 |
| **A_55_P2053459** | Timd2 | 0.07273116 | -0.094819599 | 0.240281919 | 7.725171012 | 1.03748839 | 0.335751578 | 0.991835066 | -4.844274651 |
| **A_52_P600625** | Rad23b | 0.072586306 | -0.094660076 | 0.239832689 | 9.430510535 | 1.037306493 | 0.335830342 | 0.991835066 | -4.844406073 |
| **A_55_P2001155** | Rpl31 | 0.070733145 | -0.092247287 | 0.233713576 | 14.99778578 | 1.037281509 | 0.335841161 | 0.991835066 | -4.844424123 |
| **A_55_P2073841** | NA | 0.062937708 | -0.082080908 | 0.207956325 | 14.23468338 | 1.037280706 | 0.335841509 | 0.991835066 | -4.844424703 |
| **A_52_P493091** | Fmr1 | 0.060262805 | -0.078603032 | 0.199128642 | 8.543749373 | 1.037201312 | 0.335875893 | 0.991835066 | -4.84448206 |
| **A_55_P2051405** | Tet2 | 0.07168175 | -0.09356474 | 0.23692824 | 8.988738942 | 1.036777322 | 0.336059562 | 0.991835066 | -4.844788319 |
| **A_55_P1960371** | NA | 0.052835751 | -0.068991059 | 0.17466256 | 14.87438189 | 1.03655928 | 0.336154048 | 0.991835066 | -4.844945785 |
| **A_66_P115443** | Snx4 | -0.050331483 | -0.166394129 | 0.065731163 | 8.776722587 | -1.03646918 | 0.336193097 | 0.991835066 | -4.845010848 |
| **A_55_P1963309** | G3bp1 | 0.123653981 | -0.161553192 | 0.408861155 | 12.84375475 | 1.036231503 | 0.336296125 | 0.991835066 | -4.84518246 |
| **A_51_P219721** | Ola1 | 0.066171077 | -0.08646784 | 0.218809994 | 11.71123474 | 1.036124775 | 0.336342397 | 0.991835066 | -4.845259514 |
| **A_55_P2034705** | Nmi | -0.100202544 | -0.331346304 | 0.130941215 | 9.860969731 | -1.036109624 | 0.336348966 | 0.991835066 | -4.845270452 |
| **A_51_P459108** | Insl6 | -0.065403532 | -0.216278319 | 0.085471254 | 8.839656484 | -1.036080898 | 0.336361422 | 0.991835066 | -4.84529119 |
| **A_55_P2054744** | Sh3bgrl2 | -0.072093068 | -0.238407635 | 0.094221498 | 7.944564302 | -1.036030055 | 0.336383468 | 0.991835066 | -4.845327894 |
| **A_52_P222230** | Gm40812 | 0.080233242 | -0.104882067 | 0.26534855 | 8.945678839 | 1.03590788 | 0.336436449 | 0.991835066 | -4.845416089 |
| **A_55_P2136533** | Kmt2d | 0.052697235 | -0.068914738 | 0.174309207 | 8.473480232 | 1.035668166 | 0.336540419 | 0.991835066 | -4.845589112 |
| **A_55_P2307621** | 5730526G1 | 0.125853386 | -0.16466415 | 0.416370923 | 8.85913194 | 1.035384562 | 0.336663459 | 0.991835066 | -4.845793781 |
| **A_55_P2045127** | NA | -0.050760882 | -0.167969706 | 0.066447943 | 7.968259192 | -1.035089681 | 0.336791429 | 0.991835066 | -4.846006551 |
| **A_52_P8685** | Tmem183a | 0.077592342 | -0.101603098 | 0.256787782 | 9.38151496 | 1.034906318 | 0.336871023 | 0.991835066 | -4.846138836 |
| **A_51_P265576** | Rbm18 | 0.057477724 | -0.075285807 | 0.190241256 | 8.491106061 | 1.034736814 | 0.336944614 | 0.991835066 | -4.846261109 |
| **A_55_P1956179** | Git2 | -0.114393765 | -0.378637506 | 0.149849976 | 11.08637326 | -1.034681725 | 0.336968535 | 0.991835066 | -4.846300846 |
| **A_55_P2092963** | Traf2 | 0.078496099 | -0.102826578 | 0.259818775 | 12.27684517 | 1.034677688 | 0.336970288 | 0.991835066 | -4.846303758 |
| **A_55_P2025712** | Immt | -0.103294588 | -0.341915491 | 0.135326315 | 11.82098294 | -1.034613679 | 0.336998082 | 0.991835066 | -4.846349926 |
| **A_55_P1984283** | NA | -0.059280331 | -0.196236737 | 0.077676075 | 16.50595748 | -1.034516453 | 0.337040305 | 0.991835066 | -4.84642005 |
| **A_65_P19319** | Soga1 | 0.077793188 | -0.101952365 | 0.257538741 | 10.83465485 | 1.034409614 | 0.337086707 | 0.991835066 | -4.846497102 |
| **A_55_P2046593** | Gm15226 | -0.10387707 | -0.343918226 | 0.136164085 | 9.663131142 | -1.034291883 | 0.337137846 | 0.991835066 | -4.846582004 |
| **A_51_P383032** | Clec4d | -0.042612413 | -0.14109483 | 0.055870004 | 7.90921535 | -1.034157201 | 0.337196354 | 0.991835066 | -4.846679122 |

| | | | | | | | | | |
|---|---|---|---|---|---|---|---|---|---|
| **A_55_P2021351** | NA | 0.068684527 | -0.090066672 | 0.227435726 | 16.30153397 | 1.034072603 | 0.33723311 | 0.991835066 | -4.846740121 |
| **A_51_P363630** | Snrpe | 0.074051978 | -0.097108559 | 0.245212515 | 14.39778723 | 1.034051466 | 0.337242294 | 0.991835066 | -4.846755361 |
| **A_55_P1984009** | Rsph3b | -0.09249159 | -0.306300956 | 0.121317777 | 13.48530449 | -1.033914555 | 0.337301786 | 0.991835066 | -4.846854072 |
| **A_55_P2185347** | NA | -0.042853706 | -0.141937717 | 0.056230304 | 7.768323977 | -1.033698634 | 0.337395627 | 0.991835066 | -4.847009731 |
| **A_55_P1999800** | Gm14066 | -0.047244279 | -0.156481999 | 0.061993441 | 8.072734916 | -1.033679089 | 0.337404123 | 0.991835066 | -4.84702382 |
| **A_55_P2098558** | Slc39a1 | 0.086755024 | -0.113893634 | 0.287403681 | 13.40534768 | 1.033397739 | 0.337526434 | 0.991835066 | -4.847226613 |
| **A_55_P2068734** | Ccl27a | -0.08530575 | -0.282607439 | 0.111995938 | 10.9786953 | -1.033371875 | 0.337537679 | 0.991835066 | -4.847245253 |
| **A_55_P2017075** | Zfp54 | -0.042329559 | -0.140240742 | 0.055581624 | 8.064161002 | -1.03328607 | 0.337574989 | 0.991835066 | -4.847307092 |
| **A_55_P1967069** | Lpar6 | -0.093099027 | -0.308452616 | 0.122254562 | 11.00570217 | -1.033242262 | 0.337594039 | 0.991835066 | -4.847338663 |
| **A_51_P465281** | Lgals1 | -0.103699339 | -0.343587837 | 0.13618916 | 14.66904617 | -1.033179293 | 0.337621423 | 0.991835066 | -4.847384041 |
| **A_55_P2122329** | Nfic | -0.079245238 | -0.262566766 | 0.104076289 | 9.012291502 | -1.033162994 | 0.337628511 | 0.991835066 | -4.847395787 |
| **A_55_P2178247** | Fam20b | 0.071408047 | -0.093784178 | 0.236600271 | 9.13464358 | 1.033157855 | 0.337630746 | 0.991835066 | -4.84739949 |
| **A_52_P458697** | Higd1a | 0.136100487 | -0.178817033 | 0.451018006 | 13.39868737 | 1.032932609 | 0.337728716 | 0.991835066 | -4.847561794 |
| **A_51_P392861** | Med25 | -0.143680042 | -0.476148629 | 0.188788545 | 10.80506731 | -1.032892114 | 0.337746332 | 0.991835066 | -4.847590971 |
| **A_55_P2159094** | P3h1 | 0.09017602 | -0.118525653 | 0.298877692 | 9.184321344 | 1.03270021 | 0.337829822 | 0.991835066 | -4.847729229 |
| **A_51_P132718** | 2310010J1 | 0.102135142 | -0.134260066 | 0.33853035 | 10.96701065 | 1.032632196 | 0.337859416 | 0.991835066 | -4.847778227 |
| **A_51_P124568** | Mpp1 | -0.079410312 | -0.263216849 | 0.104396225 | 10.44756824 | -1.032583262 | 0.337880709 | 0.991835066 | -4.839357527 |
| **A_55_P2048101** | Gm6762 | -0.074206849 | -0.245993779 | 0.097580082 | 8.059252158 | -1.032435672 | 0.337944938 | 0.991835066 | -4.84791979 |
| **A_55_P2140212** | NA | 0.242592807 | -0.319007522 | 0.804193136 | 14.41608783 | 1.032427821 | 0.337948355 | 0.991835066 | -4.847925444 |
| **A_55_P1968738** | Nfkb1 | -0.056976507 | -0.1888773 | 0.074924286 | 8.802374327 | -1.03242271 | 0.33795058 | 0.991835066 | -4.847929126 |
| **A_55_P2000909** | Atp5l | -0.059404699 | -0.196947324 | 0.078137927 | 16.55250238 | -1.032268367 | 0.338017759 | 0.991835066 | -4.848040292 |
| **A_51_P170959** | NA | -0.066635898 | -0.220922992 | 0.087651196 | 7.806564184 | -1.03225689 | 0.338022755 | 0.991835066 | -4.848048558 |
| **A_52_P399095** | Mettl24 | -0.049591791 | -0.164424069 | 0.065240488 | 8.093940404 | -1.032178812 | 0.338056744 | 0.991835066 | -4.848104789 |
| **A_55_P1977747** | Picalm | -0.087931267 | -0.291553615 | 0.115691081 | 13.10238319 | -1.03211246 | 0.33808563 | 0.991835066 | -4.848152574 |
| **A_51_P426195** | Nppb | -0.290742312 | -0.964085743 | 0.38260112 | 13.01486934 | -1.032002487 | 0.338133511 | 0.991835066 | -4.848231768 |
| **A_55_P2168628** | Sac3d1 | 0.083685366 | -0.110174794 | 0.277545526 | 12.21908408 | 1.031739565 | 0.338248006 | 0.991835066 | -4.848421082 |
| **A_55_P2004920** | NA | 0.060554836 | -0.079725235 | 0.200834908 | 9.477474323 | 1.031720328 | 0.338256384 | 0.991835066 | -4.848434932 |
| **A_52_P622442** | Pdcl3 | -0.069274297 | -0.229755756 | 0.091207162 | 10.97329822 | -1.03170705 | 0.338262167 | 0.991835066 | -4.848444492 |
| **A_55_P2147191** | NA | 0.059832159 | -0.078775681 | 0.198439998 | 16.63607946 | 1.031706128 | 0.338262569 | 0.991835066 | -4.848445155 |
| **A_51_P107020** | Kif5a | 0.07931963 | -0.104498574 | 0.263137833 | 8.052838608 | 1.031338649 | 0.338422653 | 0.991835066 | -4.839950701 |
| **A_55_P1953485** | Copz1 | 0.082469376 | -0.108648923 | 0.273587675 | 13.00091843 | 1.031334559 | 0.338424435 | 0.991835066 | -4.848712642 |
| **A_55_P2145389** | Flrt3 | 0.089263165 | -0.11761871 | 0.29614504 | 8.617752173 | 1.031238147 | 0.338466444 | 0.991835066 | -4.848782036 |
| **A_55_P2021471** | Pwwp3a | 0.049003613 | -0.06458792 | 0.162595147 | 8.012200341 | 1.031077413 | 0.338536491 | 0.991835066 | -4.84889772 |
| **A_55_P2173623** | Samd1 | 0.081558795 | -0.112270667 | 0.275388257 | 12.41295494 | 1.045245148 | 0.338549507 | 0.991835066 | -4.784396954 |
| **A_52_P290725** | Arfgef2 | -0.07390425 | -0.245272726 | 0.097464227 | 8.800722559 | -1.030736392 | 0.338685142 | 0.991835066 | -4.849143121 |
| **A_51_P481644** | Mbip | 0.077798984 | -0.102609328 | 0.258207296 | 9.03324998 | 1.030686324 | 0.338706971 | 0.991835066 | -4.849179147 |
| **A_55_P1979178** | NA | 0.103784548 | -0.136889636 | 0.344458732 | 13.3916369 | 1.030652644 | 0.338721656 | 0.991835066 | -4.849203379 |
| **A_51_P351413** | Phospho2 | -0.083694924 | -0.277807686 | 0.110417837 | 8.479468499 | -1.030514637 | 0.338781833 | 0.991835066 | -4.84930267 |
| **A_52_P154501** | Ube2q2 | 0.060637655 | -0.080013836 | 0.201289147 | 8.69643979 | 1.030403184 | 0.338830438 | 0.991835066 | -4.84938285 |
| **A_55_P1959550** | Carmil2 | -0.04280302 | -0.142097146 | 0.056491106 | 8.235432807 | -1.030291193 | 0.338879283 | 0.991835066 | -4.849463412 |
| **A_51_P111612** | Arrdc4 | 0.080412982 | -0.106236902 | 0.267062866 | 9.009360166 | 1.02969256 | 0.339140472 | 0.991835066 | -4.849893946 |
| **A_66_P111430** | 2410006H1 | 0.087995772 | -0.116268637 | 0.292260181 | 11.49125038 | 1.029623003 | 0.339170831 | 0.991835066 | -4.849943961 |
| **A_55_P1979027** | Tmem65 | 0.087657099 | -0.11582633 | 0.291140528 | 10.18194656 | 1.029596786 | 0.339182274 | 0.991835066 | -4.849962811 |
| **A_52_P52263** | Ilrun | 0.066478222 | -0.087845605 | 0.220802048 | 8.377952665 | 1.029569199 | 0.339194316 | 0.991835066 | -4.849982647 |
| **A_51_P453111** | Hexb | -0.075077957 | -0.249417313 | 0.0992614 | 10.56641835 | -1.029262472 | 0.339328222 | 0.991835066 | -4.850203165 |
| **A_52_P462203** | Dclre1c | 0.044245349 | -0.058527786 | 0.147018484 | 8.049414148 | 1.02895682 | 0.339461701 | 0.991835066 | -4.850422867 |
| **A_51_P212023** | Sart3 | 0.063608834 | -0.084148525 | 0.211366194 | 12.40015234 | 1.028910144 | 0.339482088 | 0.991835066 | -4.850456414 |
| **A_55_P2006236** | Ugdh | 0.10011809 | -0.132474341 | 0.332710522 | 10.98263891 | 1.02878852 | 0.339535216 | 0.991835066 | -4.850543823 |

| | | | | | | | | | |
|---|---|---|---|---|---|---|---|---|---|
| **A_55_P2078459** | Kpna2 | 0.058538601 | -0.077471167 | 0.194548369 | 12.71271165 | 1.028682547 | 0.339581512 | 0.991835066 | -4.850619979 |
| **A_52_P647740** | Klhl41 | -0.155252091 | -0.51601808 | 0.205513899 | 8.880567094 | -1.028539562 | 0.339643986 | 0.991835066 | -4.850722725 |
| **A_55_P1963300** | Vsig10 | -0.060172244 | -0.200023458 | 0.079678969 | 9.117560685 | -1.028345619 | 0.339728739 | 0.991835066 | -4.850862073 |
| **A_55_P2131088** | Vwa5a | -0.080958842 | -0.269170197 | 0.107252512 | 9.619338617 | -1.028081636 | 0.339844126 | 0.991835066 | -4.851051717 |
| **A_55_P2050207** | Ankrd27 | 0.092309554 | -0.122323329 | 0.306942438 | 10.53636021 | 1.027920497 | 0.339914576 | 0.991835066 | -4.851167463 |
| **A_55_P1981306** | Hdgfl3 | 0.072920943 | -0.096638317 | 0.242480203 | 10.61554361 | 1.02787404 | 0.339934889 | 0.991835066 | -4.851200831 |
| **A_51_P409039** | Echs1 | -0.080651536 | -0.268202234 | 0.106899162 | 13.81907018 | -1.027786925 | 0.339972982 | 0.991835066 | -4.851263399 |
| **A_51_P135340** | Panx1 | 0.069598434 | -0.0923076 | 0.231504467 | 10.68780894 | 1.027414199 | 0.340136003 | 0.991835066 | -4.851531061 |
| **A_55_P2177712** | NA | -0.04640083 | -0.154354992 | 0.061553333 | 8.537608482 | -1.027295732 | 0.340187831 | 0.991835066 | -4.851616122 |
| **A_55_P2185543** | Tle2 | 0.080588799 | -0.106910162 | 0.268087761 | 10.47446165 | 1.027270817 | 0.340198732 | 0.991835066 | -4.851634011 |
| **A_52_P313217** | Adgrd1 | 0.117832374 | -0.156319883 | 0.391984631 | 8.203683394 | 1.027263681 | 0.340201854 | 0.991835066 | -4.851639134 |
| **A_55_P2083153** | 1700001J0 | -0.051890577 | -0.1726382 | 0.068857045 | 7.752231254 | -1.027114939 | 0.340266937 | 0.991835066 | -4.851745921 |
| **A_55_P2099105** | Rhobtb1 | -0.096725536 | -0.32180287 | 0.128351799 | 9.034795609 | -1.027113675 | 0.34026749 | 0.991835066 | -4.851746828 |
| **A_51_P288916** | Tmtc2 | 0.105008051 | -0.139342897 | 0.349359 | 8.929013326 | 1.027111764 | 0.340268327 | 0.991835066 | -4.8517482 |
| **A_55_P2057912** | NA | -0.180773082 | -0.601451029 | 0.239904866 | 12.13766876 | -1.027053691 | 0.34029374 | 0.991835066 | -4.85178989 |
| **A_52_P586141** | Adcy7 | -0.087998971 | -0.292782127 | 0.116784186 | 8.661018814 | -1.02705214 | 0.340294419 | 0.991835066 | -4.851791004 |
| **A_52_P258218** | Proser1 | 0.059834186 | -0.079422101 | 0.199090472 | 8.474004063 | 1.026936781 | 0.340344905 | 0.991835066 | -4.851873814 |
| **A_52_P527625** | Colec12 | 0.159207321 | -0.211422543 | 0.529837185 | 8.940559882 | 1.026672093 | 0.340460767 | 0.991835066 | -4.852063796 |
| **A_55_P2120466** | Snx11 | 0.058822503 | -0.078138315 | 0.195783322 | 8.774513495 | 1.026493707 | 0.34053887 | 0.991835066 | -4.852191817 |
| **A_55_P2021413** | Celf4 | 0.119791773 | -0.159140016 | 0.398723562 | 9.170759735 | 1.026450741 | 0.340557684 | 0.991835066 | -4.85222265 |
| **A_55_P1979814** | Abcc8 | 0.162845367 | -0.216379793 | 0.542070527 | 11.91262663 | 1.026330881 | 0.340610172 | 0.991835066 | -4.852308659 |
| **A_51_P224227** | Rlf | -0.123263272 | -0.410314462 | 0.163787919 | 10.37020215 | -1.026321605 | 0.340614235 | 0.991835066 | -4.852315315 |
| **A_52_P552550** | Agtrap | -0.065590009 | -0.218364714 | 0.087184696 | 8.664318554 | -1.026113423 | 0.340705417 | 0.991835066 | -4.852464685 |
| **A_52_P1020860** | AW112010 | -0.046464135 | -0.154693585 | 0.061765314 | 7.792335491 | -1.026080761 | 0.340719724 | 0.991835066 | -4.852488118 |
| **A_51_P446012** | Phf13 | 0.075710768 | -0.10067831 | 0.252099845 | 11.38953528 | 1.025876527 | 0.340809199 | 0.991835066 | -4.852634633 |
| **A_52_P5891** | Rflnb | -0.071298563 | -0.237488001 | 0.094890875 | 8.187927905 | -1.025383896 | 0.341025097 | 0.991835066 | -4.852987963 |
| **A_52_P155124** | Zfp229 | -0.05099805 | -0.169885938 | 0.067889838 | 8.285649002 | -1.025238928 | 0.341088651 | 0.991835066 | -4.853091918 |
| **A_55_P2358679** | NA | 0.07712229 | -0.102676773 | 0.256921354 | 10.73544705 | 1.025183545 | 0.341112933 | 0.991835066 | -4.85313163 |
| **A_55_P2024679** | NA | 0.05353891 | -0.071308335 | 0.178386155 | 9.278856528 | 1.024942947 | 0.341218437 | 0.991835066 | -4.853304133 |
| **A_55_P1988373** | Cenpu | 0.064358541 | -0.085727954 | 0.214445035 | 8.850799435 | 1.024881623 | 0.341245332 | 0.991835066 | -4.853348097 |
| **A_52_P12806** | Gm5069 | -0.063230966 | -0.210728954 | 0.084267023 | 16.10667059 | -1.024596455 | 0.341370422 | 0.991835066 | -4.853552513 |
| **A_51_P483922** | Tm9sf4 | -0.066465589 | -0.221579393 | 0.088648214 | 9.392329956 | -1.024131078 | 0.341574639 | 0.991835066 | -4.85388603 |
| **A_55_P1970159** | Rnf11 | -0.063279413 | -0.21096788 | 0.084409055 | 11.27029722 | -1.024059018 | 0.341606269 | 0.991835066 | -4.853937663 |
| **A_51_P276479** | 4930486L2 | 0.055647083 | -0.074257299 | 0.185551464 | 8.026684278 | 1.023829721 | 0.341706933 | 0.991835066 | -4.854101948 |
| **A_55_P2030752** | Nedd4l | -0.089988953 | -0.300067312 | 0.120089406 | 11.9496517 | -1.023804456 | 0.341718025 | 0.991835066 | -4.854120048 |
| **A_52_P378084** | NA | -0.044781724 | -0.149341236 | 0.059777789 | 8.919492815 | -1.02363798 | 0.341791127 | 0.991835066 | -4.854239306 |
| **A_55_P2192492** | Slmap | -0.076004857 | -0.253473911 | 0.101464196 | 9.137358219 | -1.023594274 | 0.34181032 | 0.991835066 | -4.854270613 |
| **A_55_P1978052** | Gatb | -0.065074235 | -0.217070321 | 0.086921852 | 8.767338544 | -1.023259523 | 0.341957356 | 0.991835066 | -4.854510374 |
| **A_52_P253004** | Ralb | 0.076632315 | -0.102371414 | 0.255636044 | 12.83564029 | 1.023196392 | 0.341985091 | 0.991835066 | -4.854555585 |
| **A_51_P436201** | Gart | 0.089958228 | -0.120191507 | 0.300107963 | 9.452086391 | 1.023107283 | 0.342024242 | 0.991835066 | -4.854619396 |
| **A_55_P2073099** | Trex1 | -0.092161086 | -0.307460551 | 0.123138378 | 10.83054964 | -1.02308983 | 0.342031911 | 0.991835066 | -4.854631894 |
| **A_52_P360330** | Map1b | 0.061698153 | -0.082440632 | 0.205836939 | 8.917204332 | 1.023058433 | 0.342045706 | 0.991835066 | -4.854654377 |
| **A_51_P283499** | Drd4 | -0.045881017 | -0.153071118 | 0.061309084 | 7.698198171 | -1.023027932 | 0.342059109 | 0.991835066 | -4.854676218 |
| **A_51_P163261** | Gm6985 | -0.065240717 | -0.217679599 | 0.087198165 | 8.681460829 | -1.022897468 | 0.342116441 | 0.991835066 | -4.854769633 |
| **A_51_P304126** | Necap1 | 0.065097171 | -0.087008681 | 0.217203024 | 10.55870766 | 1.022881504 | 0.342123457 | 0.991835066 | -4.854781063 |
| **A_55_P2145720** | Thoc2 | 0.077340653 | -0.10339602 | 0.258077325 | 8.81335092 | 1.022752816 | 0.342180017 | 0.991835066 | -4.854873199 |
| **A_55_P1958140** | B9d1 | 0.048859387 | -0.06532061 | 0.163039383 | 9.933677135 | 1.02274442 | 0.342183707 | 0.991835066 | -4.854879209 |
| **A_55_P2184189** | Ncf2 | -0.056216429 | -0.187614051 | 0.075181192 | 8.373016171 | -1.02255078 | 0.342268829 | 0.991835066 | -4.855017832 |

| | | | | | | | | | |
|---|---|---|---|---|---|---|---|---|---|
| **A_51_P193686** | 1700012B0 | 0.042673991 | -0.057083235 | 0.142431217 | 7.664695585 | 1.022416917 | 0.342327683 | 0.991835066 | -4.855113653 |
| **A_55_P2182705** | Kpna2 | 0.061756992 | -0.08261029 | 0.206124273 | 13.04832163 | 1.022413286 | 0.342329279 | 0.991835066 | -4.855116251 |
| **A_51_P212754** | Tgfbi | 0.192540713 | -0.257578533 | 0.642659959 | 11.47290253 | 1.022360649 | 0.342352424 | 0.991835066 | -4.855153928 |
| **A_66_P108792** | Rad1 | 0.076576085 | -0.102449616 | 0.255601785 | 10.04810584 | 1.022320117 | 0.342370247 | 0.991835066 | -4.855182938 |
| **A_51_P246816** | Napa | 0.055686195 | -0.074512524 | 0.185884914 | 9.977969399 | 1.022233161 | 0.342408486 | 0.991835066 | -4.855245174 |
| **A_55_P2170210** | Ap1s3 | 0.230411998 | -0.3083203 | 0.769144297 | 9.007834382 | 1.022212551 | 0.34241755 | 0.991835066 | -4.855259924 |
| **A_55_P2102901** | Fbxl4 | -0.055182475 | -0.18420839 | 0.07384344 | 8.33613819 | -1.022194069 | 0.342425678 | 0.991835066 | -4.855273151 |
| **A_55_P2091472** | Bnip3 | 0.162329115 | -0.21722471 | 0.541882939 | 13.46860856 | 1.0221913 | 0.342426896 | 0.991835066 | -4.855275133 |
| **A_51_P249793** | Dusp28 | -0.097108087 | -0.324175886 | 0.129959712 | 9.833784687 | -1.022136696 | 0.342450911 | 0.991835066 | -4.85531421 |
| **A_52_P681** | Cnot4 | 0.058473285 | -0.078288941 | 0.19523551 | 9.226161804 | 1.02188133 | 0.34256324 | 0.991835066 | -4.855496949 |
| **A_51_P362738** | Dusp18 | -0.058775759 | -0.196248758 | 0.07869724 | 8.495677597 | -1.021856634 | 0.342574104 | 0.991835066 | -4.85551462 |
| **A_51_P432520** | Wrnip1 | -0.047137972 | -0.157397154 | 0.063121209 | 10.8356446 | -1.021798509 | 0.342599676 | 0.991835066 | -4.855556208 |
| **A_51_P260300** | Ikbkg | -0.046160068 | -0.154149961 | 0.061829824 | 8.177473404 | -1.021627248 | 0.342675032 | 0.991835066 | -4.855678738 |
| **A_52_P491861** | Stag1 | 0.064036592 | -0.085793512 | 0.213866696 | 9.670687013 | 1.021499744 | 0.342731143 | 0.991835066 | -4.855769953 |
| **A_52_P79187** | Ankrd12 | 0.055567819 | -0.074451696 | 0.185587334 | 8.901424491 | 1.021466062 | 0.342745967 | 0.991835066 | -4.855794048 |
| **A_55_P2133845** | Eid2b | -0.041889553 | -0.13990647 | 0.056127363 | 8.391961784 | -1.021442264 | 0.342756441 | 0.991835066 | -4.855811072 |
| **A_55_P1953419** | Rab13 | -0.099496512 | -0.332341235 | 0.13334821 | 8.446879329 | -1.021293554 | 0.342821897 | 0.991835066 | -4.855917444 |
| **A_52_P241484** | Rcl1 | 0.047316719 | -0.063421857 | 0.158055295 | 8.635781057 | 1.021232956 | 0.342848573 | 0.991835066 | -4.855960787 |
| **A_52_P577388** | Epdr1 | 0.068828353 | -0.092258597 | 0.229915303 | 11.2667657 | 1.021212573 | 0.342857546 | 0.991835066 | -4.855975366 |
| **A_51_P418737** | Rbm33 | -0.055459744 | -0.185272843 | 0.074353355 | 8.595972124 | -1.021100467 | 0.342906901 | 0.991835066 | -4.856055544 |
| **A_55_P2114915** | Ube2i | 0.080350976 | -0.107756034 | 0.268457986 | 13.955414 | 1.020928459 | 0.342982639 | 0.991835066 | -4.856178554 |
| **A_52_P172405** | Washc4 | -0.052486524 | -0.175369009 | 0.07039596 | 8.443670049 | -1.020861822 | 0.343011984 | 0.991835066 | -4.856226205 |
| **A_55_P2067226** | Arpc5l | 0.048308813 | -0.064796514 | 0.161414139 | 10.94363716 | 1.020827681 | 0.343027019 | 0.991835066 | -4.856250618 |
| **A_55_P2085295** | Inf2 | 0.070067531 | -0.093986102 | 0.234121165 | 10.55652913 | 1.020798669 | 0.343039797 | 0.991835066 | -4.856271363 |
| **A_66_P123643** | Ptbp2 | -0.050103658 | -0.167442475 | 0.067235159 | 9.490340076 | -1.020556045 | 0.343146664 | 0.991835066 | -4.856444836 |
| **A_51_P394728** | Kti12 | 0.065267807 | -0.08759795 | 0.218133563 | 9.494587557 | 1.020464591 | 0.343186954 | 0.991835066 | -4.856510217 |
| **A_55_P2105808** | Wdr62 | -0.218373386 | -0.729877335 | 0.293130564 | 10.53931117 | -1.020375091 | 0.343226386 | 0.991835066 | -4.856574198 |
| **A_55_P2157068** | Cdhr2 | -0.072208211 | -0.241353087 | 0.096936665 | 9.224336686 | -1.020321111 | 0.343250171 | 0.991835066 | -4.856612785 |
| **A_55_P2249536** | Nucks1 | 0.058148416 | -0.078073818 | 0.194370649 | 10.6747898 | 1.020232188 | 0.343289354 | 0.991835066 | -4.856676348 |
| **A_51_P286946** | Lhpp | -0.06002041 | -0.20063181 | 0.08059099 | 11.95471378 | -1.020205257 | 0.343301222 | 0.991835066 | -4.856695598 |
| **A_55_P1979305** | Gle1 | -0.065394286 | -0.218611882 | 0.087823311 | 8.987928419 | -1.02009422 | 0.343350156 | 0.991835066 | -4.856774961 |
| **A_51_P353882** | Zfp580 | 0.062248298 | -0.083669189 | 0.208165786 | 10.3074032 | 1.0195987 | 0.343568601 | 0.991835066 | -4.857129064 |
| **A_52_P68477** | Zfp809 | -0.046252078 | -0.15467359 | 0.062169434 | 9.194989378 | -1.019588481 | 0.343573107 | 0.991835066 | -4.857136365 |
| **A_55_P2000390** | Arpc5 | 0.098648562 | -0.132681389 | 0.329978513 | 13.45716819 | 1.019220205 | 0.343735531 | 0.991835066 | -4.857399465 |
| **A_55_P2094100** | BC085271 | 0.134532321 | -0.180949253 | 0.450013895 | 10.34328983 | 1.019205522 | 0.343742008 | 0.991835066 | -4.857409953 |
| **A_51_P190765** | Rtca | 0.066295512 | -0.089207953 | 0.221798978 | 10.83995416 | 1.018950744 | 0.343854411 | 0.991835066 | -4.85759193 |
| **A_55_P2063151** | NA | 0.087017479 | -0.117100916 | 0.291135874 | 10.31061227 | 1.018904511 | 0.343874812 | 0.991835066 | -4.85762495 |
| **A_66_P121117** | Serpina1d | -0.116932502 | -0.391278931 | 0.157413927 | 7.887565436 | -1.018697084 | 0.343966351 | 0.991835066 | -4.85777308 |
| **A_55_P2114118** | Cadm1 | 0.049482609 | -0.066623464 | 0.165588683 | 8.117595366 | 1.018607309 | 0.344005975 | 0.991835066 | -4.857837185 |
| **A_51_P337195** | Pipox | 0.077021252 | -0.103705033 | 0.257747537 | 8.683070455 | 1.018587597 | 0.344014676 | 0.991835066 | -4.85785126 |
| **A_51_P220135** | Nfatc1 | 0.110230057 | -0.148441282 | 0.368901395 | 9.187248277 | 1.018499579 | 0.344053529 | 0.991835066 | -4.857914107 |
| **A_55_P2155216** | Ankrd34a | 0.097762069 | -0.131696803 | 0.327220941 | 10.6441158 | 1.018297467 | 0.344142759 | 0.991835066 | -4.858058404 |
| **A_55_P2066518** | Akt2 | 0.046126315 | -0.06214789 | 0.15440052 | 12.84347953 | 1.01819953 | 0.344186004 | 0.991835066 | -4.85812832 |
| **A_55_P1975903** | Dnmt3b | 0.045320113 | -0.061067481 | 0.151707707 | 8.316742661 | 1.018143822 | 0.344210604 | 0.991835066 | -4.858168087 |
| **A_66_P122699** | NA | 0.052576913 | -0.070864374 | 0.1760182 | 8.598222232 | 1.017990613 | 0.344278266 | 0.991835066 | -4.858277447 |
| **A_52_P94845** | Nicn1 | -0.071557434 | -0.239564603 | 0.096449735 | 8.98140925 | -1.017972568 | 0.344286236 | 0.991835066 | -4.858290327 |
| **A_65_P11306** | NA | -0.06404046 | -0.214413162 | 0.086332242 | 8.999991109 | -1.017875289 | 0.344329204 | 0.991835066 | -4.858359758 |
| **A_55_P1953053** | NA | 0.053581789 | -0.072243179 | 0.179406758 | 11.23591377 | 1.017793131 | 0.344365497 | 0.991835066 | -4.858418394 |

| | | | | | | | | | |
|---|---|---|---|---|---|---|---|---|---|
| **A_55_P2128944** | Bcl2l13 | -0.080834792 | -0.270670159 | 0.109000575 | 11.10314732 | -1.017724736 | 0.344395712 | 0.991835066 | -4.858467204 |
| **A_55_P2250424** | 9530029O1 | 0.209658633 | -0.282722819 | 0.702040086 | 9.804243339 | 1.017700991 | 0.344406202 | 0.991835066 | -4.858484149 |
| **A_55_P2289800** | Fryl | -0.062597759 | -0.209642124 | 0.084446607 | 11.08256546 | -1.017465111 | 0.344510426 | 0.991835066 | -4.858652468 |
| **A_52_P477819** | Hmox2 | -0.074224522 | -0.248612261 | 0.100163217 | 12.11794198 | -1.017280205 | 0.344592145 | 0.991835066 | -4.858784395 |
| **A_52_P531662** | Ttc9c | -0.084329486 | -0.282498624 | 0.113839652 | 9.354489545 | -1.017074034 | 0.344683279 | 0.991835066 | -4.858931475 |
| **A_51_P174720** | Ccdc43 | -0.061244486 | -0.205169889 | 0.082680917 | 8.446769939 | -1.017041491 | 0.344697666 | 0.991835066 | -4.85895469 |
| **A_55_P2051229** | Gpr132 | -0.062922854 | -0.210794225 | 0.084948517 | 7.772798445 | -1.017029251 | 0.344703078 | 0.991835066 | -4.858963421 |
| **A_51_P506748** | Arhgap35 | -0.043329696 | -0.14517521 | 0.058515819 | 13.36196507 | -1.016840571 | 0.344786501 | 0.991835066 | -4.859098003 |
| **A_51_P285446** | Lig1 | 0.061855298 | -0.083569434 | 0.20728003 | 12.23778108 | 1.016594516 | 0.344895316 | 0.991835066 | -4.859273484 |
| **A_55_P2339650** | Hnrnpl | 0.083077421 | -0.112274148 | 0.278428991 | 9.107722755 | 1.016424759 | 0.344970405 | 0.991835066 | -4.859394536 |
| **A_51_P210877** | Pcnp | 0.081743894 | -0.110496314 | 0.273984101 | 11.88958539 | 1.01629602 | 0.34502736 | 0.991835066 | -4.85948633 |
| **A_51_P358906** | U2af2 | 0.086313587 | -0.116766594 | 0.289393768 | 8.84694465 | 1.015829346 | 0.345233878 | 0.991835066 | -4.859819013 |
| **A_55_P2176917** | NA | -0.126326016 | -0.423587057 | 0.170935026 | 10.45745557 | -1.015696454 | 0.345292705 | 0.991835066 | -4.859913732 |
| **A_51_P459340** | Pacrgl | 0.051764283 | -0.070059925 | 0.17358849 | 8.860512518 | 1.01556035 | 0.345352962 | 0.991835066 | -4.860010731 |
| **A_51_P483557** | Ube2b | -0.094930382 | -0.318364385 | 0.128503621 | 10.70748335 | -1.015465316 | 0.345395041 | 0.991835066 | -4.860078455 |
| **A_55_P1963585** | Mbd5 | 0.189153841 | -0.256073938 | 0.63438162 | 9.278538442 | 1.01541142 | 0.345418907 | 0.991835066 | -4.860116861 |
| **A_51_P361951** | Uqcrfs1 | -0.074414786 | -0.249586533 | 0.100756961 | 15.41021254 | -1.015323202 | 0.345457974 | 0.991835066 | -4.860179722 |
| **A_55_P2305525** | C81189 | 0.108126431 | -0.146418815 | 0.362671678 | 8.978326485 | 1.015257043 | 0.345487274 | 0.991835066 | -4.860226862 |
| **A_52_P429364** | Ccdc69 | 0.09392229 | -0.127193006 | 0.315037587 | 9.788168659 | 1.015217321 | 0.345504867 | 0.991835066 | -4.860255164 |
| **A_52_P410636** | Trp73 | -0.04076223 | -0.136728727 | 0.055204268 | 8.060301924 | -1.015190208 | 0.345516876 | 0.991835066 | -4.860274482 |
| **A_55_P2120906** | NA | 0.072632306 | -0.098369641 | 0.243634253 | 8.866284093 | 1.015167975 | 0.345526724 | 0.991835066 | -4.860290323 |
| **A_55_P1973387** | Srrm1 | 0.133059918 | -0.180225166 | 0.446345003 | 12.45938816 | 1.015118321 | 0.345548717 | 0.991835066 | -4.860325699 |
| **A_55_P1980401** | Cbx7 | -0.051407622 | -0.172454177 | 0.069638932 | 7.665851359 | -1.015042482 | 0.345582312 | 0.991835066 | -4.86037973 |
| **A_55_P2140453** | Myh6 | -0.214180849 | -0.718568993 | 0.290207296 | 11.25559396 | -1.014903843 | 0.345643732 | 0.991835066 | -4.860478494 |
| **A_55_P2130855** | Nap1l1 | 0.067209581 | -0.091094003 | 0.225513166 | 10.95097686 | 1.014727839 | 0.345721718 | 0.991835066 | -4.860603863 |
| **A_52_P381806** | Terf2ip | -0.059934376 | -0.201140086 | 0.081271333 | 8.957135994 | -1.014455184 | 0.345842556 | 0.991835066 | -4.86079805 |
| **A_55_P2044369** | Hs2st1 | -0.163199947 | -0.547795628 | 0.221395733 | 9.948022859 | -1.014202658 | 0.345954503 | 0.991835066 | -4.860977871 |
| **A_55_P1994224** | Ythdf1 | 0.073750626 | -0.100091822 | 0.247593075 | 12.59666545 | 1.013955782 | 0.346063973 | 0.991835066 | -4.861153641 |
| **A_55_P2062190** | Gstm1 | -0.094830135 | -0.318360635 | 0.128700365 | 11.57407538 | -1.013955069 | 0.34606429 | 0.991835066 | -4.861154149 |
| **A_51_P212560** | Cog7 | 0.073884122 | -0.100287107 | 0.248055352 | 9.597262896 | 1.013873641 | 0.346100402 | 0.991835066 | -4.861212117 |
| **A_51_P158772** | Elof1 | -0.082295136 | -0.276301135 | 0.111710863 | 13.27028219 | -1.013837007 | 0.346116651 | 0.991835066 | -4.861238196 |
| **A_55_P1962304** | Plac8 | -0.045797022 | -0.153765495 | 0.062171451 | 7.766409754 | -1.01379328 | 0.346136045 | 0.991835066 | -4.861269323 |
| **A_52_P542290** | Pdxdc1 | 0.053164786 | -0.072180468 | 0.178510041 | 9.326632156 | 1.013737024 | 0.346160998 | 0.991835066 | -4.861309368 |
| **A_55_P2042156** | Aatk | 0.04863486 | -0.066041507 | 0.163311228 | 8.354604425 | 1.013637984 | 0.346204932 | 0.991835066 | -4.861379864 |
| **A_55_P2069246** | Ddx39b | 0.104449236 | -0.141841956 | 0.350740428 | 14.13230715 | 1.013597471 | 0.346222905 | 0.991835066 | -4.8614087 |
| **A_55_P2017082** | Hnrnpa2b1 | 0.186755184 | -0.253645774 | 0.627156142 | 11.73273904 | 1.013522848 | 0.346256011 | 0.991835066 | -4.861461811 |
| **A_55_P2075358** | NA | 0.068106597 | -0.092505515 | 0.228718708 | 12.39996777 | 1.01349129 | 0.346270013 | 0.991835066 | -4.861484272 |
| **A_51_P289239** | F8 | -0.043793331 | -0.147088417 | 0.059501755 | 7.90589795 | -1.013298629 | 0.346355502 | 0.991835066 | -4.861621381 |
| **A_52_P574697** | Khdrbs1 | 0.048352745 | -0.065706737 | 0.162412226 | 12.32953837 | 1.013208608 | 0.346395452 | 0.991835066 | -4.861685439 |
| **A_55_P2058737** | Mex3d | -0.072747953 | -0.244353869 | 0.098857962 | 10.50004117 | -1.013205778 | 0.346396708 | 0.991835066 | -4.861687453 |
| **A_55_P1977761** | Ttc14 | 0.084919334 | -0.1154494 | 0.285288069 | 8.3341685 | 1.01294475 | 0.346512571 | 0.991835066 | -4.861873178 |
| **A_55_P1957875** | NA | -0.05406493 | -0.181633027 | 0.073503166 | 10.07737457 | -1.01293763 | 0.346515732 | 0.991835066 | -4.861878244 |
| **A_52_P367565** | Mrps2 | -0.054374757 | -0.18267533 | 0.073925817 | 10.62430886 | -1.012926335 | 0.346520746 | 0.991835066 | -4.86188628 |
| **A_51_P310030** | Naca | 0.050633198 | -0.068851429 | 0.170117824 | 15.01261196 | 1.012820441 | 0.346567759 | 0.991835066 | -4.861961615 |
| **A_55_P2026669** | Atp5l | -0.061037159 | -0.205074981 | 0.083000662 | 15.62505805 | -1.012807485 | 0.346573512 | 0.991835066 | -4.861970832 |
| **A_52_P19016** | Clasrp | 0.167181911 | -0.227364599 | 0.561728421 | 11.74921927 | 1.012745244 | 0.346601147 | 0.991835066 | -4.862015108 |
| **A_55_P2347529** | 3010003L2 | 0.070056643 | -0.095286537 | 0.235399822 | 9.088529419 | 1.012679854 | 0.346630182 | 0.991835066 | -4.862061624 |
| **A_66_P117786** | Crxos | 0.051816038 | -0.07050048 | 0.174132556 | 8.16458694 | 1.012484125 | 0.346717104 | 0.991835066 | -4.862200843 |

| | | | | | | | | | |
|---|---|---|---|---|---|---|---|---|---|
| **A_52_P597634** | Fzd1 | 0.102494264 | -0.139459817 | 0.344448344 | 11.15528221 | 1.012455025 | 0.346730028 | 0.991835066 | -4.86222154 |
| **A_55_P2019254** | NA | -0.123397832 | -0.414738002 | 0.167942337 | 12.72341907 | -1.012316452 | 0.34679158 | 0.991835066 | -4.862320093 |
| **A_55_P2099378** | Tfe3 | 0.058192559 | -0.079201825 | 0.195586943 | 11.22193196 | 1.012296204 | 0.346800574 | 0.991835066 | -4.862334492 |
| **A_55_P1980551** | Dennd1a | -0.048204607 | -0.162027931 | 0.065618716 | 9.167852092 | -1.012200202 | 0.346843222 | 0.991835066 | -4.862402762 |
| **A_55_P2108206** | Rrp7a | 0.061537694 | -0.083773688 | 0.206849076 | 10.7964714 | 1.012163602 | 0.346859483 | 0.991835066 | -4.862428789 |
| **A_51_P421846** | Ech1 | -0.109130217 | -0.366867098 | 0.148606665 | 12.07264277 | -1.011993167 | 0.34693521 | 0.991835066 | -4.862549977 |
| **A_55_P2137887** | Mpdz | 0.057952064 | -0.078920058 | 0.194824187 | 8.584655497 | 1.01195929 | 0.346950264 | 0.991835066 | -4.862574065 |
| **A_55_P2106827** | NA | -0.103410677 | -0.34767464 | 0.140853287 | 11.42057792 | -1.011847615 | 0.346999892 | 0.991835066 | -4.862653463 |
| **A_51_P197321** | Clta | 0.056843944 | -0.077439628 | 0.191127516 | 13.42095166 | 1.011743559 | 0.347046139 | 0.991835066 | -4.862727438 |
| **A_51_P274947** | Atad1 | -0.092357078 | -0.310590876 | 0.125876719 | 12.06856442 | -1.011480022 | 0.347163288 | 0.991835066 | -4.862914772 |
| **A_66_P127118** | Trmt112 | -0.062154748 | -0.209050233 | 0.084740738 | 8.177071162 | -1.011288318 | 0.347248525 | 0.991835066 | -4.863051022 |
| **A_55_P1998823** | NA | -0.133945952 | -0.450584111 | 0.182692207 | 10.4841802 | -1.01105661 | 0.347351571 | 0.991835066 | -4.863215684 |
| **A_55_P2104790** | NA | -0.075404454 | -0.253659272 | 0.102850364 | 13.48580074 | -1.011031894 | 0.347362565 | 0.991835066 | -4.863233246 |
| **A_51_P251357** | Ctps | 0.094549237 | -0.128981605 | 0.318080079 | 13.3680903 | 1.010950067 | 0.347398962 | 0.991835066 | -4.863291389 |
| **A_55_P2068451** | NA | 0.107491413 | -0.146667128 | 0.361649954 | 14.12589948 | 1.010830172 | 0.347452297 | 0.991835066 | -4.863376575 |
| **A_55_P2187053** | Tpm1 | -0.057229505 | -0.192547206 | 0.078088196 | 9.090224888 | -1.010821593 | 0.347456114 | 0.991835066 | -4.863382671 |
| **A_55_P2026270** | Cfi | 0.115410068 | -0.157516998 | 0.388337135 | 8.676125411 | 1.010662666 | 0.347526823 | 0.991835066 | -4.863495579 |
| **A_51_P464029** | Fads3 | 0.063189308 | -0.086246489 | 0.212625105 | 12.00873122 | 1.010643736 | 0.347535246 | 0.991835066 | -4.863509027 |
| **A_55_P2009940** | Xlr | 0.068598864 | -0.093646726 | 0.230844455 | 8.041005654 | 1.010539177 | 0.347581773 | 0.991835066 | -4.863583302 |
| **A_51_P115626** | Shank3 | 0.061230603 | -0.083599763 | 0.206060969 | 8.70080525 | 1.010457471 | 0.347618134 | 0.991835066 | -4.86364134 |
| **A_66_P115034** | Bend5 | 0.053494555 | -0.073055285 | 0.180044395 | 8.941835497 | 1.010315719 | 0.347681225 | 0.991835066 | -4.863742024 |
| **A_55_P1994564** | Sap30bp | 0.085113544 | -0.116252014 | 0.286479101 | 8.937154121 | 1.010235475 | 0.347716943 | 0.991835066 | -4.863799015 |
| **A_55_P2126368** | Atad5 | 0.110639743 | -0.151121203 | 0.372400689 | 8.965316738 | 1.010218785 | 0.347724373 | 0.991835066 | -4.863810868 |
| **A_51_P209527** | Bcl10 | -0.061721161 | -0.207753548 | 0.084311227 | 10.18402593 | -1.010168987 | 0.347746541 | 0.991835066 | -4.863846234 |
| **A_66_P133993** | Gm5093 | 0.074585252 | -0.101888415 | 0.251058919 | 9.357892086 | 1.010141427 | 0.34775881 | 0.991835066 | -4.863865807 |
| **A_55_P2085905** | Anpep | -0.163354443 | -0.549886209 | 0.223177323 | 10.89639973 | -1.010077953 | 0.347787069 | 0.991835066 | -4.863910882 |
| **A_55_P2121096** | Fbn2 | 0.112058493 | -0.159882482 | 0.383999468 | 12.06588961 | 1.023616502 | 0.347826414 | 0.991835066 | -4.798777256 |
| **A_55_P2025829** | NA | 0.057845763 | -0.079044949 | 0.194736475 | 14.45983252 | 1.009965881 | 0.347836968 | 0.991835066 | -4.863990466 |
| **A_55_P2444515** | NA | 0.110095165 | -0.150470868 | 0.370661198 | 10.06378801 | 1.009856298 | 0.347885764 | 0.991835066 | -4.864068276 |
| **A_55_P2084496** | Ssbp3 | 0.066380386 | -0.090756352 | 0.223517125 | 10.95481648 | 1.009650755 | 0.347977305 | 0.991835066 | -4.864214209 |
| **A_51_P378051** | Aoc3 | -0.070275967 | -0.23664948 | 0.096097547 | 10.84248256 | -1.00955916 | 0.348018104 | 0.991835066 | -4.864279234 |
| **A_52_P537566** | Cenpt | -0.043993407 | -0.148153513 | 0.060166699 | 9.231850447 | -1.009474422 | 0.348055852 | 0.991835066 | -4.864339387 |
| **A_55_P2144931** | NA | 0.059033825 | -0.080786123 | 0.198853772 | 14.48716577 | 1.009115597 | 0.348215732 | 0.991835066 | -4.864594072 |
| **A_55_P2020443** | Hnrnpa3 | 0.113091737 | -0.154775446 | 0.380958919 | 9.821468045 | 1.009068144 | 0.348236879 | 0.991835066 | -4.864627748 |
| **A_51_P208020** | Ulk3 | 0.104889166 | -0.143617499 | 0.35339583 | 9.413078869 | 1.008792219 | 0.348359868 | 0.991835066 | -4.864823547 |
| **A_51_P170619** | Ddx54 | 0.066546533 | -0.0911376 | 0.224230665 | 11.72480152 | 1.008664117 | 0.348416978 | 0.991835066 | -4.864914436 |
| **A_55_P2024290** | Fam149a | 0.045587121 | -0.062442889 | 0.153617132 | 8.115441567 | 1.008571943 | 0.348458076 | 0.991835066 | -4.864979831 |
| **A_55_P2057112** | Prl8a2 | -0.184038337 | -0.620189819 | 0.252113145 | 8.019544622 | -1.0085097 | 0.34848583 | 0.991835066 | -4.865023987 |
| **A_55_P1974967** | H4c14 | 0.04377594 | -0.059978962 | 0.147530843 | 7.866381196 | 1.00840732 | 0.348531486 | 0.991835066 | -4.865096614 |
| **A_55_P2166128** | NA | 0.113744953 | -0.155847087 | 0.383336993 | 8.813317207 | 1.008403168 | 0.348533337 | 0.991835066 | -4.865099559 |
| **A_51_P244213** | Hapln3 | 0.058887341 | -0.080705627 | 0.198480309 | 8.397125895 | 1.00824838 | 0.348602373 | 0.991835066 | -4.865209354 |
| **A_51_P141164** | Ddx50 | -0.044488181 | -0.149953305 | 0.060976943 | 9.806811887 | -1.008195875 | 0.348625792 | 0.991835066 | -4.865246595 |
| **A_55_P2127020** | Nup54 | 0.061829312 | -0.084749522 | 0.208408146 | 10.22301302 | 1.008166555 | 0.348638871 | 0.991835066 | -4.865267391 |
| **A_55_P2141860** | Aen | 0.101357685 | -0.13893746 | 0.341652831 | 12.61070998 | 1.008139939 | 0.348650744 | 0.991835066 | -4.865286268 |
| **A_55_P2084332** | Pigp | -0.047001793 | -0.15844958 | 0.064445994 | 7.978353597 | -1.007980454 | 0.348721893 | 0.991835066 | -4.865399376 |
| **A_52_P330540** | Mis18a | 0.100418812 | -0.137709984 | 0.338547608 | 9.493332228 | 1.007888049 | 0.348763122 | 0.991835066 | -4.865464904 |
| **A_55_P2025974** | Ruvbl2 | 0.061015266 | -0.083684788 | 0.20571532 | 11.42486144 | 1.007810663 | 0.348797653 | 0.991835066 | -4.865519779 |
| **A_52_P537571** | Trap1a | 0.0723351 | -0.099232532 | 0.243902732 | 7.717057237 | 1.007680518 | 0.348855732 | 0.991835066 | -4.865612059 |

| | | | | | | | | | |
|---|---|---|---|---|---|---|---|---|---|
| **A_51_P214083** | 2310039H0 | 0.080129163 | -0.109924877 | 0.270183203 | 11.41443976 | 1.007679989 | 0.348855968 | 0.991835066 | -4.865612434 |
| **A_51_P328652** | Acot7 | 0.060005051 | -0.082317724 | 0.202327826 | 11.55965005 | 1.007679764 | 0.348856068 | 0.991835066 | -4.865612594 |
| **A_55_P1989573** | Tmem175 | 0.062584111 | -0.085859825 | 0.211028046 | 9.593387948 | 1.007652432 | 0.348868266 | 0.991835066 | -4.865631973 |
| **A_55_P2044207** | Bphl | -0.080167425 | -0.270324001 | 0.109989152 | 8.433050469 | -1.007617528 | 0.348883845 | 0.991835066 | -4.865656721 |
| **A_55_P2052936** | NA | 0.058235208 | -0.079899526 | 0.196369942 | 15.94459342 | 1.007608604 | 0.348887827 | 0.991835066 | -4.865663047 |
| **A_51_P120066** | 9330151L1 | 0.070436976 | -0.096642997 | 0.237516949 | 8.908959473 | 1.007593692 | 0.348894483 | 0.991835066 | -4.86567362 |
| **A_51_P477833** | Klhl21 | -0.070663836 | -0.238282262 | 0.096954591 | 11.13261157 | -1.007591696 | 0.348895374 | 0.991835066 | -4.865675035 |
| **A_55_P1998097** | Tmem214 | 0.098700914 | -0.135426231 | 0.332828058 | 9.165877763 | 1.007577677 | 0.348901631 | 0.991835066 | -4.865684974 |
| **A_66_P132846** | Exosc1 | 0.053136609 | -0.07292604 | 0.179199258 | 8.839051222 | 1.007433849 | 0.348965831 | 0.991835066 | -4.865786941 |
| **A_55_P2028788** | Slc39a11 | 0.106214353 | -0.145793913 | 0.358222619 | 11.07263828 | 1.007343418 | 0.349006202 | 0.991835066 | -4.865851047 |
| **A_66_P134428** | Sfrp1 | -0.056221776 | -0.189642819 | 0.077199268 | 8.973334671 | -1.007138868 | 0.349097531 | 0.991835066 | -4.865996037 |
| **A_55_P2092953** | Traf5 | -0.09459334 | -0.319077214 | 0.129890534 | 9.589651682 | -1.007127709 | 0.349102514 | 0.991835066 | -4.866003946 |
| **A_51_P281582** | Lrrc42 | -0.057837023 | -0.195117576 | 0.079443529 | 8.532816095 | -1.006945689 | 0.3491838 | 0.991835066 | -4.86613295 |
| **A_55_P2020831** | NA | -0.054227867 | -0.182975453 | 0.074519718 | 12.23926973 | -1.006682606 | 0.349301314 | 0.991835066 | -4.866319378 |
| **A_55_P2036894** | Med29 | 0.075676996 | -0.104003421 | 0.255357413 | 11.24696556 | 1.006635561 | 0.349322331 | 0.991835066 | -4.866352712 |
| **A_52_P503334** | Erlin2 | -0.046804445 | -0.157943421 | 0.064334531 | 9.892064821 | -1.006537226 | 0.349366265 | 0.991835066 | -4.866422384 |
| **A_55_P2060663** | NA | 0.110546662 | -0.152009659 | 0.373102983 | 15.36988441 | 1.006311157 | 0.349467285 | 0.991835066 | -4.866582542 |
| **A_51_P195483** | Ube2q2 | -0.063256213 | -0.213494533 | 0.086982108 | 9.647407927 | -1.006309556 | 0.349468001 | 0.991835066 | -4.866583676 |
| **A_55_P2047698** | Pop7 | 0.043234172 | -0.059454783 | 0.145923126 | 12.63290538 | 1.006265413 | 0.349487729 | 0.991835066 | -4.866614947 |
| **A_55_P2036559** | Fbxl12os | 0.139631083 | -0.192021843 | 0.471284009 | 11.25083584 | 1.006253504 | 0.349493051 | 0.991835066 | -4.866623382 |
| **A_52_P106259** | Egfr | -0.068305624 | -0.230594667 | 0.093983419 | 8.458749919 | -1.005949999 | 0.349628718 | 0.991835066 | -4.866838354 |
| **A_66_P110251** | Tmem91 | 0.062377001 | -0.085831226 | 0.210585229 | 8.588314176 | 1.005915064 | 0.349644337 | 0.991835066 | -4.866863096 |
| **A_51_P386526** | Rcc1l | 0.054369444 | -0.074843297 | 0.183582184 | 9.098620686 | 1.005677386 | 0.349750611 | 0.991835066 | -4.867031409 |
| **A_55_P2170076** | Taf9-ps | 0.064863675 | -0.08929906 | 0.21902641 | 12.6952758 | 1.00561392 | 0.349778993 | 0.991835066 | -4.867076349 |
| **A_55_P1990919** | Msrb1 | -0.095124785 | -0.321277391 | 0.131027821 | 12.60591853 | -1.005312824 | 0.349913669 | 0.991835066 | -4.867289526 |
| **A_55_P2035946** | Penk | 0.254726647 | -0.350952372 | 0.860405666 | 8.892333636 | 1.005173386 | 0.349976051 | 0.991835066 | -4.867388233 |
| **A_55_P2144456** | Kmt5b | 0.0808554 | -0.111418528 | 0.273129327 | 8.32167667 | 1.005073361 | 0.350020807 | 0.991835066 | -4.867459036 |
| **A_55_P1953758** | Serinc2 | 0.111059325 | -0.153043098 | 0.375161747 | 8.198288118 | 1.005059508 | 0.350027005 | 0.991835066 | -4.867468841 |
| **A_55_P2068570** | NA | -0.04550222 | -0.153708846 | 0.062704406 | 7.950399551 | -1.005050453 | 0.350031057 | 0.991835066 | -4.867475251 |
| **A_51_P455897** | Pimreg | -0.106113931 | -0.358482901 | 0.146255038 | 10.12491425 | -1.004952612 | 0.35007484 | 0.991835066 | -4.867544501 |
| **A_55_P2122066** | Ppp2r3d | -0.076202108 | -0.257448874 | 0.105044658 | 8.462816756 | -1.004860678 | 0.350115984 | 0.991835066 | -4.867609567 |
| **A_55_P2131925** | Kprp | 0.045732182 | -0.063044814 | 0.154509179 | 7.710191739 | 1.004833243 | 0.350128263 | 0.991835066 | -4.867628983 |
| **A_55_P2244142** | Klf13 | -0.114458463 | -0.386720848 | 0.157803921 | 10.42822281 | -1.004776338 | 0.350153732 | 0.991835066 | -4.867669254 |
| **A_51_P185247** | Gdf10 | -0.068303 | -0.230799947 | 0.094193946 | 8.578782439 | -1.004624369 | 0.350221758 | 0.991835066 | -4.867776795 |
| **A_55_P2061690** | Atp5d | -0.099098899 | -0.334870886 | 0.136673089 | 15.06180911 | -1.004582839 | 0.35024035 | 0.991835066 | -4.867806182 |
| **A_66_P138765** | Trappc10 | 0.068256579 | -0.094146662 | 0.230659821 | 9.681471528 | 1.004520855 | 0.3502681 | 0.991835066 | -4.86785004 |
| **A_55_P2046877** | Foxq1 | 0.048064223 | -0.066296428 | 0.162424875 | 7.759566218 | 1.004510411 | 0.350272776 | 0.991835066 | -4.86785743 |
| **A_51_P286142** | Znrd1 | -0.052480024 | -0.17736849 | 0.072408441 | 11.13500606 | -1.004340163 | 0.350349005 | 0.991835066 | -4.867977884 |
| **A_66_P130914** | NA | 0.056011158 | -0.077305353 | 0.189327669 | 15.08869064 | 1.00415266 | 0.350432975 | 0.991835066 | -4.868110531 |
| **A_66_P105244** | Tor1aip1 | 0.055628285 | -0.07680158 | 0.188058149 | 8.444273057 | 1.003965676 | 0.350516728 | 0.991835066 | -4.868242794 |
| **A_51_P452779** | Pygl | 0.071318032 | -0.098484852 | 0.241120916 | 13.09070856 | 1.003837515 | 0.350574143 | 0.991835066 | -4.86833344 |
| **A_55_P1975695** | Snx18 | 0.089102714 | -0.123046379 | 0.301251808 | 11.10091021 | 1.003827031 | 0.35057884 | 0.991835066 | -4.868340855 |
| **A_66_P103092** | NA | -0.090153475 | -0.304845557 | 0.124538606 | 11.14009578 | -1.003634493 | 0.350665109 | 0.991835066 | -4.868477017 |
| **A_52_P288251** | Tmem204 | -0.086140136 | -0.291292084 | 0.119011811 | 9.338499167 | -1.003550065 | 0.350702943 | 0.991835066 | -4.868536719 |
| **A_51_P403334** | Cuedc2 | 0.093010187 | -0.128520663 | 0.314541037 | 13.34219899 | 1.003472408 | 0.350737746 | 0.991835066 | -4.86859163 |
| **A_55_P1972230** | NA | -0.080884488 | -0.273552246 | 0.111783271 | 11.81974652 | -1.00337974 | 0.35077928 | 0.991835066 | -4.868657151 |
| **A_52_P5213** | Chd1l | 0.055067921 | -0.076112955 | 0.186248797 | 9.2831098 | 1.003314952 | 0.350808321 | 0.991835066 | -4.868702958 |
| **A_55_P2112569** | Ern1 | 0.040296962 | -0.055706079 | 0.136300002 | 7.708753691 | 1.003220617 | 0.350850608 | 0.991835066 | -4.868769651 |

| | | | | | | | | | |
|---|---|---|---|---|---|---|---|---|---|
| **A_55_P2068104** | NA | 0.097306434 | -0.134527759 | 0.329140626 | 9.464131109 | 1.003166914 | 0.350874683 | 0.991835066 | -4.868807616 |
| **A_51_P435198** | Cryz | -0.066115881 | -0.223658285 | 0.091426524 | 10.78228378 | -1.003038148 | 0.350932415 | 0.991835066 | -4.868898642 |
| **A_55_P1978311** | NA | 0.084466091 | -0.116811838 | 0.285744019 | 14.24332882 | 1.002987158 | 0.350955278 | 0.991835066 | -4.868934685 |
| **A_55_P1979833** | Cited1 | 0.067741205 | -0.093693944 | 0.229176354 | 14.83098481 | 1.002914589 | 0.350987819 | 0.991835066 | -4.868985979 |
| **A_52_P206925** | Parp16 | -0.049595341 | -0.167788297 | 0.068597615 | 8.497972787 | -1.002901818 | 0.350993546 | 0.991835066 | -4.868995006 |
| **A_55_P2171378** | Pdzk1ip1 | 0.044264891 | -0.06122535 | 0.149755132 | 7.698470149 | 1.002896803 | 0.350995795 | 0.991835066 | -4.868998551 |
| **A_55_P1953578** | Top3a | 0.057115713 | -0.079001762 | 0.193233187 | 9.189514648 | 1.00288434 | 0.351001384 | 0.991835066 | -4.86900736 |
| **A_55_P2206461** | A930006K( | 0.088954468 | -0.123057126 | 0.300966062 | 8.158680916 | 1.002806836 | 0.351036142 | 0.991835066 | -4.869062138 |
| **A_55_P1999102** | Pi16 | 0.194894583 | -0.269615117 | 0.659404283 | 9.830452672 | 1.002799512 | 0.351039427 | 0.991835066 | -4.869067315 |
| **A_55_P2070721** | Rapgef1 | 0.05021177 | -0.069463886 | 0.169887427 | 10.86699599 | 1.002787355 | 0.351044879 | 0.991835066 | -4.869075907 |
| **A_51_P419439** | Gnmt | -0.064031661 | -0.216685354 | 0.088622032 | 8.101674546 | -1.00252817 | 0.351161135 | 0.991835066 | -4.869259073 |
| **A_55_P2368625** | Sp4 | -0.062387476 | -0.211133217 | 0.086358265 | 8.847816888 | -1.002448361 | 0.351196939 | 0.991835066 | -4.869315467 |
| **A_55_P2017850** | Rpsa | 0.079400809 | -0.109909721 | 0.268711339 | 13.23164562 | 1.002442078 | 0.351199758 | 0.991835066 | -4.869319907 |
| **A_55_P2017055** | Kctd12b | 0.161796835 | -0.223967216 | 0.547560887 | 9.72711934 | 1.002437722 | 0.351201712 | 0.991835066 | -4.869322985 |
| **A_55_P2046703** | Atp5g3 | -0.058852503 | -0.199240766 | 0.08153576 | 16.75296753 | -1.001943584 | 0.35142346 | 0.991835066 | -4.869672085 |
| **A_55_P2160648** | Gm6900 | 0.059166956 | -0.082001589 | 0.200335501 | 15.48130998 | 1.001729399 | 0.351519611 | 0.991835066 | -4.869823367 |
| **A_55_P2129271** | Tead2 | 0.094126657 | -0.130470052 | 0.318723367 | 12.84052683 | 1.001655499 | 0.351552791 | 0.991835066 | -4.869875559 |
| **A_52_P23308** | Fam241a | -0.066893024 | -0.226529913 | 0.092743865 | 8.831075184 | -1.001513268 | 0.351616656 | 0.991835066 | -4.869976003 |
| **A_55_P2084812** | Sec31a | -0.060961798 | -0.206455846 | 0.08453225 | 9.840776057 | -1.001432477 | 0.351652938 | 0.991835066 | -4.870033053 |
| **A_55_P2129319** | Mre11a | 0.068116059 | -0.094476939 | 0.230709056 | 9.440350775 | 1.001282918 | 0.351720109 | 0.991835066 | -4.870138656 |
| **A_55_P2259500** | D130012P( | 0.100675317 | -0.139671506 | 0.34102214 | 8.513060374 | 1.001137552 | 0.351785407 | 0.991835066 | -4.870241288 |
| **A_55_P1986833** | Tulp4 | 0.057100224 | -0.07923899 | 0.193439439 | 10.56807888 | 1.000981749 | 0.351855404 | 0.991835066 | -4.870351278 |
| **A_55_P2031671** | Gstm6 | -0.05059929 | -0.171426724 | 0.070228143 | 8.405036564 | -1.000893819 | 0.351894913 | 0.991835066 | -4.870413348 |
| **A_55_P2179030** | Ostf1 | -0.06610404 | -0.223979688 | 0.091771608 | 11.28996561 | -1.00074168 | 0.35196328 | 0.991835066 | -4.870520734 |
| **A_52_P665386** | Ube2m | -0.141566654 | -0.479707852 | 0.196574543 | 10.99699663 | -1.000626532 | 0.352015031 | 0.991835066 | -4.870602003 |
| **A_51_P255193** | Faf2 | -0.092104638 | -0.312137603 | 0.127928328 | 9.941714162 | -1.000467255 | 0.352086626 | 0.991835066 | -4.870714407 |
| **A_55_P2006677** | Gm7969 | 0.048326783 | -0.067124427 | 0.163777993 | 7.861597587 | 1.00045724 | 0.352091128 | 0.991835066 | -4.870721475 |
| **A_55_P2088360** | NA | 0.078988658 | -0.109719201 | 0.267696518 | 14.22314826 | 1.000423491 | 0.352106299 | 0.991835066 | -4.870745291 |
| **A_55_P2172196** | Thnsl1 | 0.073333833 | -0.101866579 | 0.248534245 | 8.070492167 | 1.000410851 | 0.352111982 | 0.991835066 | -4.87075421 |
| **A_55_P2030126** | Tfpt | -0.067696637 | -0.229440889 | 0.094047615 | 9.502225307 | -1.000339397 | 0.352144105 | 0.991835066 | -4.87080463 |
| **A_55_P2150294** | Synj2bp | 0.109720105 | -0.152439208 | 0.371879418 | 11.3401704 | 1.000299511 | 0.352162038 | 0.991835066 | -4.870832774 |
| **A_55_P1974582** | Eif4h | 0.055199548 | -0.07671816 | 0.187117256 | 14.36687032 | 1.000095696 | 0.352253684 | 0.991835066 | -4.870976576 |
| **A_55_P2054352** | Rbm6 | 0.059863821 | -0.083205881 | 0.202933523 | 10.22037977 | 1.000059622 | 0.352269907 | 0.991835066 | -4.871002027 |
| **A_51_P501069** | Spta1 | -0.048283012 | -0.163677201 | 0.067111178 | 8.092689552 | -1.000045 | 0.352276483 | 0.991835066 | -4.871012343 |
| **A_55_P2140107** | Cyp21a1 | 0.04537619 | -0.063072636 | 0.153825017 | 7.916130086 | 1.000028342 | 0.352283974 | 0.991835066 | -4.871024094 |
| **A_55_P1974477** | Msh3 | 0.090385266 | -0.125649324 | 0.306419856 | 10.4947599 | 0.99996196 | 0.352313829 | 0.991835066 | -4.871070924 |
| **A_55_P2070936** | Tra2b | -0.052840695 | -0.179142188 | 0.073460799 | 8.192500669 | -0.999929007 | 0.35232865 | 0.991835066 | -4.87109417 |
| **A_55_P2099269** | Ift80 | -0.083867984 | -0.28440201 | 0.116666042 | 9.177412584 | -0.999579324 | 0.352485956 | 0.991835066 | -4.871340818 |
| **A_51_P354706** | Lefty1 | -0.048268986 | -0.163755734 | 0.067217762 | 7.999624546 | -0.998953236 | 0.35276774 | 0.991835066 | -4.871782284 |
| **A_55_P1957393** | NA | -0.117714644 | -0.399417816 | 0.163988528 | 15.51257272 | -0.998729524 | 0.352868468 | 0.991835066 | -4.871939983 |
| **A_55_P2090519** | Ppdpf | -0.087571975 | -0.29714324 | 0.12199929 | 13.27605914 | -0.998717231 | 0.352874004 | 0.991835066 | -4.871948648 |
| **A_55_P1960491** | Snta1 | -0.079609322 | -0.270129497 | 0.110910854 | 12.61609717 | -0.998693177 | 0.352884836 | 0.991835066 | -4.871965603 |
| **A_55_P2034663** | Csf2rb2 | -0.040431348 | -0.137192913 | 0.056330217 | 7.772695453 | -0.998675672 | 0.352892719 | 0.991835066 | -4.871977941 |
| **A_55_P2022840** | Csnk2a1 | 0.045852183 | -0.063891939 | 0.155596304 | 11.68070322 | 0.998591531 | 0.352930613 | 0.991835066 | -4.872037245 |
| **A_55_P2119430** | Usp10 | 0.102829507 | -0.143300543 | 0.348959557 | 11.78292597 | 0.998532597 | 0.352957156 | 0.991835066 | -4.872078782 |
| **A_51_P491595** | AW549877 | -0.109757678 | -0.372490805 | 0.152975448 | 9.660578639 | -0.998456644 | 0.352991367 | 0.991835066 | -4.87213231 |
| **A_55_P1991792** | Slc25a46 | 0.045469678 | -0.063375543 | 0.154314899 | 12.14460444 | 0.998439249 | 0.352999202 | 0.991835066 | -4.872144569 |
| **A_51_P260871** | Sdhd | -0.066814687 | -0.226778804 | 0.093149429 | 15.07514691 | -0.998294097 | 0.353064589 | 0.991835066 | -4.872246857 |

| | | | | | | | | | |
|---|---|---|---|---|---|---|---|---|---|
| **A_55_P1991134** | Fkbp1a | 0.05660805 | -0.07893033 | 0.192146429 | 14.59909247 | 0.99821717 | 0.353099247 | 0.991835066 | -4.872301064 |
| **A_51_P399985** | Myo9b | 0.082218605 | -0.114691031 | 0.279128241 | 12.03515701 | 0.997958009 | 0.353216025 | 0.991835066 | -4.87248366 |
| **A_51_P224468** | Maip1 | -0.068511288 | -0.232617424 | 0.095594848 | 11.08054378 | -0.99780677 | 0.353284188 | 0.991835066 | -4.872590204 |
| **A_55_P1969882** | Morf4l2 | 0.096602674 | -0.134866968 | 0.328072316 | 14.02635402 | 0.997480103 | 0.35343145 | 0.991835066 | -4.872820294 |
| **A_55_P1972915** | NA | -0.083149075 | -0.282417271 | 0.116119122 | 13.66412727 | -0.997306302 | 0.353509819 | 0.991835066 | -4.872942692 |
| **A_55_P2141479** | Rnu2-10 | 0.11969106 | -0.167169381 | 0.406551501 | 10.59622164 | 0.997241134 | 0.353539208 | 0.991835066 | -4.872988583 |
| **A_66_P125161** | 1700057H1 | -0.042589815 | -0.144720706 | 0.059541077 | 7.967365413 | -0.996684629 | 0.353790251 | 0.991835066 | -4.873380385 |
| **A_55_P2022263** | NA | 0.073488555 | -0.102743767 | 0.249720876 | 17.0028664 | 0.996651389 | 0.353805251 | 0.991835066 | -4.873403783 |
| **A_55_P2077558** | Sod3 | -0.093983236 | -0.319375832 | 0.13140936 | 8.786407879 | -0.996597704 | 0.353829476 | 0.991835066 | -4.873441571 |
| **A_55_P2109585** | Plekha7 | 0.087019965 | -0.121680439 | 0.295720369 | 9.141624656 | 0.996562975 | 0.353845149 | 0.991835066 | -4.873466016 |
| **A_51_P432511** | Afg1l | -0.094347603 | -0.320723163 | 0.132027958 | 8.809067394 | -0.996117256 | 0.354046341 | 0.991835066 | -4.87377969 |
| **A_55_P2102155** | Rgs20 | -0.088584341 | -0.301181447 | 0.124012765 | 11.21377903 | -0.99588397 | 0.354151679 | 0.991835066 | -4.873943827 |
| **A_52_P464629** | Nme7 | -0.044340691 | -0.15078698 | 0.062105598 | 8.323927866 | -0.99559123 | 0.354283898 | 0.991835066 | -4.87414976 |
| **A_52_P426513** | Sh3bgrl | -0.077999067 | -0.265298375 | 0.109300241 | 10.15148977 | -0.995319172 | 0.35440681 | 0.991835066 | -4.874341108 |
| **A_52_P463183** | Asap1 | -0.054098138 | -0.184004205 | 0.075807929 | 8.652382411 | -0.995318363 | 0.354407176 | 0.991835066 | -4.874341677 |
| **A_55_P2159870** | Tead4 | 0.047274963 | -0.066246987 | 0.160796913 | 7.961823725 | 0.995314857 | 0.35440876 | 0.991835066 | -4.874344143 |
| **A_55_P2013833** | Ddx3x | 0.14461246 | -0.202677242 | 0.491902163 | 12.41736277 | 0.995228814 | 0.35444764 | 0.991835066 | -4.874404653 |
| **A_55_P2325729** | Arhgap1 | -0.105528221 | -0.358961782 | 0.147905341 | 10.51087912 | -0.995207458 | 0.35445729 | 0.991835066 | -4.87441967 |
| **A_55_P2155931** | Tnfsfm13 | -0.061889857 | -0.21061976 | 0.086840046 | 8.662069885 | -0.994558457 | 0.354750668 | 0.991835066 | -4.874875961 |
| **A_55_P2043668** | Zfp617 | -0.077916384 | -0.265175933 | 0.109343166 | 9.849065315 | -0.994475171 | 0.35478833 | 0.991835066 | -4.874934501 |
| **A_55_P1959869** | Ppp1r2 | -0.06410115 | -0.218166634 | 0.089964334 | 11.45980652 | -0.994419433 | 0.354813537 | 0.991835066 | -4.874973677 |
| **A_52_P289692** | NA | 0.103591859 | -0.145432123 | 0.352615841 | 9.823255939 | 0.994245387 | 0.354892257 | 0.991835066 | -4.875095998 |
| **A_52_P204331** | D630039A( | -0.042224196 | -0.14372853 | 0.059280138 | 7.74678906 | -0.994227873 | 0.354900179 | 0.991835066 | -4.875108306 |
| **A_55_P2069251** | Prr18 | 0.042726277 | -0.0599965 | 0.145449055 | 7.778320917 | 0.994116852 | 0.354950401 | 0.991835066 | -4.875186324 |
| **A_55_P2043287** | Tle5 | -0.109364072 | -0.372375845 | 0.1536477 | 11.74895612 | -0.993822026 | 0.355083796 | 0.991835066 | -4.87539348 |
| **A_55_P1971709** | Dnajc8 | -0.118300643 | -0.402835367 | 0.166234082 | 9.747026482 | -0.993712974 | 0.355133148 | 0.991835066 | -4.875470093 |
| **A_55_P2094049** | Hectd4 | -0.049728326 | -0.169349635 | 0.069892983 | 11.5615572 | -0.99358362 | 0.355191693 | 0.991835066 | -4.875560962 |
| **A_51_P163233** | Khdc4 | -0.060183203 | -0.204985658 | 0.084619252 | 8.988534712 | -0.993364199 | 0.35529102 | 0.991835066 | -4.875715084 |
| **A_51_P156158** | Phlpp1 | -0.070830084 | -0.241254975 | 0.099594807 | 9.115295159 | -0.99333071 | 0.355306182 | 0.991835066 | -4.875738605 |
| **A_55_P1964408** | Cryab | -0.089700022 | -0.305552701 | 0.126152656 | 15.71292363 | -0.993217221 | 0.355357566 | 0.991835066 | -4.87581831 |
| **A_55_P2007496** | NA | 0.130331537 | -0.18337266 | 0.444035734 | 9.079648968 | 0.992975012 | 0.35546725 | 0.991835066 | -4.875988395 |
| **A_55_P2039771** | Itgav | 0.04261622 | -0.059964092 | 0.145196533 | 8.030797922 | 0.992933227 | 0.355486175 | 0.991835066 | -4.876017735 |
| **A_55_P2185840** | Nnmt | -0.060544793 | -0.206297018 | 0.085207432 | 8.433541792 | -0.992820507 | 0.355537231 | 0.991835066 | -4.876096877 |
| **A_51_P426739** | Gpt | -0.09194083 | -0.313278688 | 0.129397029 | 8.857732562 | -0.992800182 | 0.355546438 | 0.991835066 | -4.876111148 |
| **A_51_P269134** | Sftpb | -0.066006855 | -0.22491872 | 0.09290501 | 7.687865484 | -0.992754464 | 0.355567148 | 0.991835066 | -4.876143245 |
| **A_51_P362066** | Chil1 | -0.04195767 | -0.142985599 | 0.059070259 | 7.765805859 | -0.992610915 | 0.355632181 | 0.991835066 | -4.876244021 |
| **A_51_P248304** | Vps26b | -0.05174297 | -0.176348159 | 0.07286222 | 9.202774896 | -0.992485916 | 0.355688818 | 0.991835066 | -4.876331766 |
| **A_55_P1970154** | Rnf10 | 0.090950915 | -0.128107928 | 0.310009758 | 11.49631908 | 0.992328389 | 0.355760202 | 0.991835066 | -4.876442335 |
| **A_55_P2040863** | Cnn3 | 0.067908923 | -0.095666437 | 0.231484283 | 12.40277372 | 0.992243109 | 0.355798853 | 0.991835066 | -4.876502188 |
| **A_52_P430934** | Dcun1d1 | 0.065404736 | -0.092145459 | 0.222954931 | 9.347988172 | 0.992200367 | 0.355818225 | 0.991835066 | -4.876532185 |
| **A_55_P2008889** | Tmsb15b2 | -0.051105963 | -0.174220356 | 0.07200843 | 9.053281064 | -0.99213754 | 0.355846702 | 0.991835066 | -4.876576275 |
| **A_55_P1956989** | BC004004 | -0.073589492 | -0.250887179 | 0.103708194 | 13.47227286 | -0.992023294 | 0.355898491 | 0.991835066 | -4.876656448 |
| **A_55_P1994898** | NA | -0.12660552 | -0.431714227 | 0.178503187 | 13.99690074 | -0.991761333 | 0.356017262 | 0.991835066 | -4.876840255 |
| **A_55_P2014500** | Phf24 | 0.088234338 | -0.124432553 | 0.300901229 | 9.14450501 | 0.991623663 | 0.356079692 | 0.991835066 | -4.876936839 |
| **A_55_P2068822** | Cdk20 | -0.063707543 | -0.21728606 | 0.089870974 | 9.021052442 | -0.99144703 | 0.356159805 | 0.991835066 | -4.877060745 |
| **A_66_P125496** | NA | -0.101099128 | -0.344893205 | 0.142694949 | 12.50034682 | -0.99113632 | 0.356300762 | 0.991835066 | -4.877278669 |
| **A_55_P2123933** | D2hgdh | -0.059618941 | -0.203403792 | 0.084165911 | 8.987353341 | -0.991015055 | 0.356355787 | 0.991835066 | -4.877363709 |
| **A_51_P425824** | C1qtnf12 | 0.096164855 | -0.135805851 | 0.328135561 | 11.79068144 | 0.990814535 | 0.356446789 | 0.991835066 | -4.877504313 |

| | | | | | | | | | |
|---|---|---|---|---|---|---|---|---|---|
| **A_55_P2038207** | Zfp687 | 0.070477194 | -0.09955291 | 0.240507297 | 9.970492746 | 0.990676622 | 0.356509389 | 0.991835066 | -4.877601006 |
| **A_55_P2101526** | L1td1 | 0.117308364 | -0.165732455 | 0.400349182 | 8.77637426 | 0.99057882 | 0.356553788 | 0.991835066 | -4.877669571 |
| **A_55_P2088630** | Ap3s2 | 0.084980391 | -0.120104179 | 0.290064962 | 10.44917417 | 0.99036406 | 0.356651295 | 0.991835066 | -4.877820115 |
| **A_55_P2053444** | Map4 | -0.086632825 | -0.29577192 | 0.122506271 | 12.53211876 | -0.990048298 | 0.356794699 | 0.991835066 | -4.878041421 |
| **A_52_P217646** | Snapc3 | -0.08336943 | -0.284640892 | 0.117902032 | 10.48453059 | -0.989996734 | 0.356818121 | 0.991835066 | -4.878077556 |
| **A_55_P2004168** | NA | 0.13623576 | -0.192704653 | 0.465176172 | 9.814064372 | 0.989881065 | 0.356870666 | 0.991835066 | -4.878158609 |
| **A_55_P1986088** | Prr14 | 0.056298524 | -0.079635565 | 0.192232612 | 11.57762873 | 0.98986908 | 0.356876111 | 0.991835066 | -4.878167007 |
| **A_55_P1963124** | NA | -0.045302525 | -0.154688353 | 0.064083303 | 14.60491458 | -0.989852488 | 0.356883649 | 0.991835066 | -4.878178633 |
| **A_55_P2187038** | Tpm1 | 0.295195649 | -0.417748947 | 1.008140245 | 17.64041902 | 0.989608241 | 0.356994627 | 0.991835066 | -4.878349762 |
| **A_51_P134452** | Lpar3 | -0.091457326 | -0.312351401 | 0.129436748 | 10.79224869 | -0.98956327 | 0.357015063 | 0.991835066 | -4.878381267 |
| **A_51_P227004** | Cks1b | 0.093210958 | -0.131934011 | 0.318355927 | 12.86682233 | 0.989495575 | 0.357045827 | 0.991835066 | -4.878428691 |
| **A_55_P2027107** | Rgs3 | -0.095297451 | -0.325487334 | 0.134892432 | 9.070035659 | -0.989473537 | 0.357055843 | 0.991835066 | -4.878444128 |
| **A_66_P105801** | Igf1r | 0.117789723 | -0.166737595 | 0.402317041 | 11.82853604 | 0.989447058 | 0.357067878 | 0.991835066 | -4.878462678 |
| **A_55_P2107957** | Abraxas1 | 0.071650306 | -0.10144436 | 0.244744971 | 8.517114006 | 0.98933528 | 0.357118683 | 0.991835066 | -4.878540975 |
| **A_55_P2045682** | Rassf5 | 0.112420138 | -0.159189336 | 0.384029613 | 9.750304735 | 0.98925518 | 0.357155093 | 0.991835066 | -4.87859708 |
| **A_55_P1964302** | Timm8a1 | 0.098649063 | -0.139691777 | 0.336989903 | 9.476213231 | 0.989244471 | 0.357159961 | 0.991835066 | -4.878604581 |
| **A_51_P241943** | Sap30l | -0.075526588 | -0.258005369 | 0.106952192 | 13.02628894 | -0.989228525 | 0.35716721 | 0.991835066 | -4.878615749 |
| **A_55_P1997061** | NA | 0.092039001 | -0.13033852 | 0.314416521 | 15.11872518 | 0.989213747 | 0.357173928 | 0.991835066 | -4.878626099 |
| **A_51_P481788** | Mapk1 | 0.077217117 | -0.109366404 | 0.263800637 | 10.79480593 | 0.989121013 | 0.357216087 | 0.991835066 | -4.864788091 |
| **A_55_P1955239** | Rps6 | 0.064822904 | -0.091822129 | 0.221467937 | 14.53194078 | 0.989056236 | 0.357245538 | 0.991835066 | -4.878736413 |
| **A_55_P2101231** | Aldh7a1 | 0.040986665 | -0.058073156 | 0.140046486 | 12.91050857 | 0.9889041 | 0.357314715 | 0.991835066 | -4.878842952 |
| **A_55_P2110235** | Trip12 | 0.088401966 | -0.125273308 | 0.30207724 | 10.79306795 | 0.988818965 | 0.357353431 | 0.991835066 | -4.878902566 |
| **A_55_P2111478** | Polq | 0.088445292 | -0.125340519 | 0.302231102 | 8.51726201 | 0.988792069 | 0.357365662 | 0.991835066 | -4.878921398 |
| **A_55_P2096257** | Myl6 | 0.077256914 | -0.10952654 | 0.264040369 | 14.30488693 | 0.988571505 | 0.357465983 | 0.991835066 | -4.879075824 |
| **A_51_P373619** | Mbl1 | -0.040522788 | -0.138496517 | 0.057450941 | 7.678336123 | -0.98855039 | 0.357475589 | 0.991835066 | -4.879090606 |
| **A_55_P1953133** | NA | 0.055846549 | -0.079203299 | 0.190896398 | 14.38224873 | 0.988351389 | 0.357566123 | 0.991835066 | -4.879229913 |
| **A_55_P2184314** | NA | -0.095884553 | -0.32777623 | 0.136007124 | 11.25357289 | -0.988263191 | 0.357606253 | 0.991835066 | -4.879291649 |
| **A_55_P1970105** | Ltbp2 | 0.574982047 | -0.816038991 | 1.966003084 | 10.42278039 | 0.987938064 | 0.357754219 | 0.991835066 | -4.879519194 |
| **A_52_P430274** | Dnaaf5 | 0.079750689 | -0.113199695 | 0.272701072 | 10.20024273 | 0.987865749 | 0.357787136 | 0.991835066 | -4.879569799 |
| **A_52_P458870** | Slc25a16 | -0.051701228 | -0.17683066 | 0.073428204 | 8.628535251 | -0.987530499 | 0.35793977 | 0.991835066 | -4.879804365 |
| **A_55_P2055369** | NA | -0.137255267 | -0.469471138 | 0.194960603 | 8.866017501 | -0.987456055 | 0.35797367 | 0.991835066 | -4.879856444 |
| **A_55_P2064292** | Col7a1 | 0.064149275 | -0.091120644 | 0.219419194 | 8.235640153 | 0.98744646 | 0.357978039 | 0.991835066 | -4.879863156 |
| **A_55_P2065449** | Kdm5b | 0.075155632 | -0.10679724 | 0.257108505 | 12.24745618 | 0.98721501 | 0.358083453 | 0.991835066 | -4.880025056 |
| **A_66_P117695** | Zpr1 | 0.050371007 | -0.071585032 | 0.172327046 | 12.12973329 | 0.987157509 | 0.358109645 | 0.991835066 | -4.880065274 |
| **A_52_P424826** | Txnl4b | 0.054515487 | -0.077484101 | 0.186515075 | 10.51539763 | 0.987089327 | 0.358140705 | 0.991835066 | -4.880112961 |
| **A_52_P644534** | Dhrsx | -0.081751277 | -0.279699011 | 0.116196456 | 8.931695597 | -0.987081803 | 0.358144132 | 0.991835066 | -4.880118222 |
| **A_55_P2180764** | Gls | -0.07379824 | -0.252521657 | 0.104925177 | 10.4291097 | -0.986901199 | 0.358226416 | 0.991835066 | -4.880244526 |
| **A_55_P2058864** | Mvk | 0.056111826 | -0.079796764 | 0.192020417 | 9.329222936 | 0.986771568 | 0.358285485 | 0.991835066 | -4.880335173 |
| **A_55_P2077994** | Ube2v2 | -0.076912129 | -0.263203216 | 0.109378959 | 9.192084155 | -0.986760791 | 0.358290396 | 0.991835066 | -4.880342709 |
| **A_55_P2049262** | NA | 0.251374383 | -0.357515877 | 0.860264644 | 10.24645629 | 0.986713621 | 0.358311892 | 0.991835066 | -4.880375691 |
| **A_55_P1967076** | Rpl11 | 0.056614492 | -0.08052473 | 0.193753714 | 16.69682912 | 0.986677134 | 0.358328521 | 0.991835066 | -4.880401203 |
| **A_55_P2029630** | Mpdu1 | 0.091847007 | -0.130668271 | 0.314362284 | 9.007963965 | 0.986539105 | 0.35839143 | 0.991835066 | -4.880497707 |
| **A_51_P268697** | Slc1a3 | 0.049211558 | -0.070029368 | 0.168452484 | 8.085377248 | 0.986395094 | 0.358457076 | 0.991835066 | -4.880598384 |
| **A_55_P1973930** | Krt79 | 0.050136147 | -0.071380096 | 0.17165239 | 8.665228553 | 0.986110891 | 0.358586654 | 0.991835066 | -4.880797039 |
| **A_55_P2373121** | Dcaf17 | 0.081515912 | -0.116066652 | 0.279098475 | 8.751185405 | 0.986059013 | 0.358610311 | 0.991835066 | -4.880833297 |
| **A_55_P2039463** | Timmdc1 | -0.19068842 | -0.652924517 | 0.271547677 | 11.6123034 | -0.985983381 | 0.358644802 | 0.991835066 | -4.880886154 |
| **A_52_P124734** | Cybrd1 | 0.138589959 | -0.197384134 | 0.474564052 | 8.477612998 | 0.985905091 | 0.358680508 | 0.991835066 | -4.880940867 |
| **A_66_P119155** | NA | -0.055995299 | -0.191746276 | 0.079755678 | 12.95118871 | -0.985865657 | 0.358698494 | 0.991835066 | -4.880968424 |

| | | | | | | | | | |
|---|---|---|---|---|---|---|---|---|---|
| **A_65_P11455** | Pphln1 | 0.074458543 | -0.106054924 | 0.25497201 | 8.542619494 | 0.985857304 | 0.358702304 | 0.991835066 | -4.880974262 |
| **A_52_P313856** | Slc25a26 | -0.052609283 | -0.180164506 | 0.07494594 | 8.790645497 | -0.98576472 | 0.358744535 | 0.991835066 | -4.881038957 |
| **A_51_P496715** | Pomk | -0.070111217 | -0.240104163 | 0.09988173 | 9.111202341 | -0.985747611 | 0.358752339 | 0.991835066 | -4.881050912 |
| **A_52_P208613** | Fancb | 0.077493227 | -0.110415534 | 0.265401988 | 8.75855155 | 0.985657084 | 0.358793636 | 0.991835066 | -4.881114166 |
| **A_51_P422208** | Dpagt1 | 0.055758531 | -0.079461163 | 0.190978226 | 11.83511967 | 0.985554184 | 0.358840581 | 0.991835066 | -4.88118606 |
| **A_55_P2064876** | Fam89b | -0.063723045 | -0.218303079 | 0.090856989 | 12.81799206 | -0.985263178 | 0.358973372 | 0.991835066 | -4.881389354 |
| **A_66_P114862** | Ankrd11 | -0.050521988 | -0.173087938 | 0.072043963 | 10.69443134 | -0.985189382 | 0.359007052 | 0.991835066 | -4.881440901 |
| **A_55_P2034453** | NA | -0.045263056 | -0.155091301 | 0.064565189 | 8.178590873 | -0.985006196 | 0.359090669 | 0.991835066 | -4.881568846 |
| **A_52_P547447** | Sirt2 | -0.119624616 | -0.409897991 | 0.170648759 | 10.60658908 | -0.98496883 | 0.359107727 | 0.991835066 | -4.881594941 |
| **A_51_P279163** | Plcg2 | 0.067158445 | -0.095810412 | 0.230127302 | 10.28120028 | 0.984929495 | 0.359125684 | 0.991835066 | -4.881622412 |
| **A_55_P2064766** | NA | 0.062442094 | -0.089096077 | 0.213980265 | 11.79840188 | 0.984837433 | 0.359167715 | 0.991835066 | -4.881686702 |
| **A_51_P491987** | Ripk3 | 0.095897547 | -0.136837195 | 0.328632288 | 9.646243975 | 0.984816715 | 0.359177175 | 0.991835066 | -4.88170117 |
| **A_51_P149373** | Fbxl20 | 0.10743472 | -0.153346841 | 0.368216281 | 10.07052243 | 0.984638713 | 0.359258454 | 0.991835066 | -4.881825462 |
| **A_55_P2078623** | Stxbp3 | -0.07049481 | -0.241610968 | 0.100621348 | 9.938563692 | -0.984634969 | 0.359260164 | 0.991835066 | -4.881828077 |
| **A_51_P152845** | Trim24 | -0.056659565 | -0.194205319 | 0.08088619 | 11.19979152 | -0.984544103 | 0.359301661 | 0.991835066 | -4.881891519 |
| **A_66_P140886** | Inafm1 | 0.055039087 | -0.078588524 | 0.188666698 | 10.63848632 | 0.984428437 | 0.359354489 | 0.991835066 | -4.881972271 |
| **A_51_P516637** | Bmp5 | 0.040836143 | -0.058333016 | 0.140005302 | 7.963250209 | 0.984186078 | 0.359465202 | 0.991835066 | -4.882141454 |
| **A_51_P429366** | Hes6 | -0.078210187 | -0.268159759 | 0.111739384 | 11.3545423 | -0.984088462 | 0.359509802 | 0.991835066 | -4.882209588 |
| **A_55_P2170681** | Cdca8 | -0.098923755 | -0.339189758 | 0.141342247 | 9.105808391 | -0.98405054 | 0.359527129 | 0.991835066 | -4.882236056 |
| **A_65_P11853** | Arhgap28 | -0.053371554 | -0.18300418 | 0.076261073 | 9.070529598 | -0.984021662 | 0.359540325 | 0.991835066 | -4.882256211 |
| **A_55_P2108451** | Galns | -0.111059751 | -0.380836505 | 0.158717004 | 10.61385628 | -0.983923427 | 0.359585214 | 0.991835066 | -4.88232477 |
| **A_51_P230934** | Cab39 | 0.099629081 | -0.142431185 | 0.341689347 | 10.92456227 | 0.983720564 | 0.359677929 | 0.991835066 | -4.882466335 |
| **A_55_P2123962** | Micos10 | 0.085409025 | -0.122109293 | 0.292927343 | 15.6781561 | 0.983685921 | 0.359693764 | 0.991835066 | -4.882490508 |
| **A_55_P1965154** | Spc25 | -0.095749387 | -0.328397774 | 0.136899 | 13.66974367 | -0.983660177 | 0.359705532 | 0.991835066 | -4.882508471 |
| **A_51_P417469** | Rnaseh2c | 0.047578128 | -0.068025628 | 0.163181885 | 13.76812749 | 0.983658931 | 0.359706101 | 0.991835066 | -4.88250934 |
| **A_55_P2075048** | Atic | 0.120888063 | -0.172897656 | 0.414673782 | 10.18081251 | 0.983471709 | 0.359791689 | 0.991835066 | -4.882639968 |
| **A_52_P366525** | Coq10b | -0.051676458 | -0.177280535 | 0.073927619 | 8.517367832 | -0.983327379 | 0.35985768 | 0.991835066 | -4.882740657 |
| **A_51_P148314** | Fam117a | -0.084034573 | -0.288313702 | 0.120244556 | 8.637080046 | -0.983202858 | 0.359914622 | 0.991835066 | -4.882827519 |
| **A_52_P118958** | Pex19 | -0.135492233 | -0.464867929 | 0.193883463 | 10.21343065 | -0.983177619 | 0.359926164 | 0.991835066 | -4.882845124 |
| **A_51_P487175** | Acsm3 | -0.047181983 | -0.161928858 | 0.067564892 | 7.679396822 | -0.982753174 | 0.360120313 | 0.991835066 | -4.883141143 |
| **A_51_P416419** | Calr | 0.114127401 | -0.163447501 | 0.391702303 | 12.20704836 | 0.982695267 | 0.360146807 | 0.991835066 | -4.883181522 |
| **A_51_P142153** | Filip1l | 0.088934887 | -0.127398051 | 0.305267825 | 10.85597303 | 0.982559009 | 0.360209155 | 0.991835066 | -4.88327653 |
| **A_55_P1963031** | Ampd2 | 0.049339828 | -0.070692675 | 0.169372331 | 8.942366063 | 0.982444215 | 0.360261688 | 0.991835066 | -4.883356565 |
| **A_55_P2125823** | NA | -0.086310974 | -0.296294974 | 0.123673026 | 13.92801494 | -0.982401341 | 0.360281309 | 0.991835066 | -4.883386456 |
| **A_55_P1976356** | Fgf18 | 0.109433825 | -0.156818165 | 0.375685814 | 9.94139603 | 0.982353616 | 0.360303153 | 0.991835066 | -4.883419727 |
| **A_51_P202964** | Csnk2a2 | 0.052849591 | -0.075733437 | 0.181432618 | 10.15649754 | 0.982351948 | 0.360303916 | 0.991835066 | -4.88342089 |
| **A_55_P2143015** | Sema6c | -0.067911806 | -0.233156967 | 0.097333354 | 8.582351187 | -0.982258205 | 0.360346824 | 0.991835066 | -4.883486239 |
| **A_55_P2061134** | NA | -0.095185553 | -0.326813948 | 0.136442842 | 14.77440329 | -0.982173864 | 0.360385432 | 0.991835066 | -4.883545031 |
| **A_55_P2102529** | Fuca2 | -0.083790467 | -0.287695067 | 0.120114133 | 8.849629013 | -0.982147511 | 0.360397495 | 0.991835066 | -4.883563399 |
| **A_55_P2058591** | NA | -0.044159663 | -0.15162485 | 0.063305524 | 7.92808467 | -0.982125724 | 0.360407469 | 0.991835066 | -4.883578586 |
| **A_55_P2064776** | 4930412O1 | 0.045567667 | -0.06532714 | 0.156462475 | 7.744866498 | 0.982097759 | 0.360420272 | 0.991835066 | -4.883598078 |
| **A_55_P2014337** | Bach1 | -0.085452158 | -0.293440763 | 0.122536447 | 10.12791873 | -0.981957346 | 0.360484559 | 0.991835066 | -4.883695943 |
| **A_51_P349825** | 1700021F0 | -0.058683887 | -0.201523729 | 0.084155955 | 13.32124371 | -0.981925732 | 0.360499034 | 0.991835066 | -4.883717976 |
| **A_51_P375431** | Phip | 0.088126512 | -0.126397361 | 0.302650385 | 8.886666038 | 0.981838563 | 0.36053895 | 0.991835066 | -4.883778725 |
| **A_55_P2266178** | Il2rb | -0.04319654 | -0.148348782 | 0.061955701 | 7.752641924 | -0.981837378 | 0.360539492 | 0.991835066 | -4.883779551 |
| **A_55_P2092364** | Ermard | 0.082084768 | -0.117740063 | 0.281909599 | 9.622624365 | 0.981798249 | 0.360557411 | 0.991835066 | -4.883806819 |
| **A_55_P1957747** | Inip | 0.117822805 | -0.169013683 | 0.404659294 | 9.606456114 | 0.981757198 | 0.36057621 | 0.991835066 | -4.883835426 |
| **A_55_P2333126** | Oga | 0.115702636 | -0.165980619 | 0.397385891 | 9.041778542 | 0.981728404 | 0.360589397 | 0.991835066 | -4.88385549 |

| | | | | | | | | | |
|---|---|---|---|---|---|---|---|---|---|
| **A_66_P113059** | Itgb1 | 0.071424872 | -0.102467692 | 0.245317437 | 11.48837598 | 0.981697294 | 0.360603645 | 0.991835066 | -4.883877169 |
| **A_51_P445683** | 4931406C0 | -0.07702353 | -0.264580845 | 0.110533784 | 10.13546755 | -0.981518626 | 0.360685481 | 0.991835066 | -4.884001661 |
| **A_55_P2011937** | Fignl1 | 0.111760787 | -0.160410761 | 0.383932336 | 11.68265279 | 0.981422162 | 0.36072967 | 0.991835066 | -4.884068869 |
| **A_55_P2150108** | Pgm2l1 | 0.072755048 | -0.104446202 | 0.249956298 | 8.998899486 | 0.981308328 | 0.360781822 | 0.991835066 | -4.884148173 |
| **A_55_P2083806** | Sp8 | -0.057236762 | -0.196655575 | 0.082182051 | 7.908611314 | -0.981211899 | 0.360826005 | 0.991835066 | -4.884215346 |
| **A_55_P2018345** | Amz2 | -0.066675954 | -0.229126478 | 0.095774569 | 9.974464464 | -0.980973463 | 0.360935272 | 0.991835066 | -4.884381425 |
| **A_51_P394515** | Tkt | 0.184288381 | -0.264755854 | 0.633332616 | 12.18792249 | 0.980884704 | 0.360975954 | 0.991835066 | -4.884443242 |
| **A_55_P2134042** | NA | -0.050831789 | -0.174721614 | 0.073058036 | 16.20387979 | -0.980638376 | 0.361088874 | 0.991835066 | -4.884614778 |
| **A_55_P2072801** | Lypd5 | 0.046009135 | -0.066128424 | 0.158146694 | 8.205857773 | 0.980623063 | 0.361095894 | 0.991835066 | -4.88462544 |
| **A_51_P289881** | Txndc12 | -0.081263189 | -0.279347043 | 0.116820665 | 10.78942361 | -0.980514262 | 0.36114578 | 0.991835066 | -4.884701197 |
| **A_55_P2081123** | Srcin1 | 0.040284887 | -0.057915666 | 0.138485441 | 7.663335804 | 0.980476863 | 0.361162929 | 0.991835066 | -4.884727236 |
| **A_66_P119034** | Pla2g7 | -0.146176786 | -0.50253367 | 0.210180098 | 9.185855541 | -0.980397937 | 0.361199121 | 0.991835066 | -4.884782185 |
| **A_55_P2184551** | NA | 0.166905686 | -0.239991872 | 0.573803243 | 11.88645236 | 0.980381667 | 0.361206582 | 0.991835066 | -4.884793513 |
| **A_65_P16059** | Tgfbr3 | -0.065489786 | -0.225147716 | 0.094168143 | 11.09084438 | -0.980374963 | 0.361209656 | 0.991835066 | -4.88479818 |
| **A_52_P559498** | Clk3 | 0.063540526 | -0.091380224 | 0.218461276 | 9.453547241 | 0.980280463 | 0.361252995 | 0.991835066 | -4.884863967 |
| **A_51_P466490** | Armc5 | 0.065381523 | -0.094030468 | 0.224793513 | 10.19015663 | 0.980264282 | 0.361260416 | 0.991835066 | -4.884875232 |
| **A_55_P2001583** | Tspan18 | 0.086122836 | -0.123878651 | 0.296124323 | 8.715832741 | 0.980178305 | 0.36129985 | 0.991835066 | -4.884935082 |
| **A_55_P2040168** | Pmp22 | 0.060092341 | -0.08647572 | 0.206660402 | 9.574014034 | 0.979916149 | 0.361420111 | 0.991835066 | -4.885117551 |
| **A_51_P464588** | Dnajc28 | -0.089306056 | -0.307181892 | 0.12856978 | 8.428420116 | -0.979672635 | 0.361531848 | 0.991835066 | -4.885287016 |
| **A_55_P1982201** | Sgsm2 | 0.083201907 | -0.119782618 | 0.286186432 | 10.08853523 | 0.979669265 | 0.361533394 | 0.991835066 | -4.885289361 |
| **A_55_P2003688** | Tmem143 | -0.085405819 | -0.293795617 | 0.122983978 | 11.59195626 | -0.979535414 | 0.361594823 | 0.991835066 | -4.885382496 |
| **A_55_P2022211** | Plxdc2 | 0.117253675 | -0.168870781 | 0.403378132 | 11.11967205 | 0.979446269 | 0.36163574 | 0.991835066 | -4.885444521 |
| **A_52_P541175** | Marveld1 | 0.104097375 | -0.149949033 | 0.358143783 | 9.294399579 | 0.979345319 | 0.361682079 | 0.991835066 | -4.885514754 |
| **A_52_P289256** | Arid1b | 0.089944425 | -0.129608436 | 0.309497285 | 10.13553219 | 0.979138847 | 0.361776871 | 0.991835066 | -4.885658386 |
| **A_55_P1981479** | Irgm1 | -0.043441616 | -0.14948237 | 0.062599138 | 7.984774257 | -0.97913438 | 0.361778922 | 0.991835066 | -4.885661493 |
| **A_66_P139799** | Cenpb | 0.091636186 | -0.132119379 | 0.31539175 | 8.750146744 | 0.978818774 | 0.361923854 | 0.991835066 | -4.885881003 |
| **A_55_P2108845** | NA | 0.065239392 | -0.094121444 | 0.224600228 | 8.154449683 | 0.978447295 | 0.362094503 | 0.991835066 | -4.886139313 |
| **A_55_P2002578** | Ephx1 | 0.055326287 | -0.079819888 | 0.190472463 | 10.95911588 | 0.978446088 | 0.362095058 | 0.991835066 | -4.886140153 |
| **A_51_P196444** | Foxc2 | 0.091306006 | -0.131762165 | 0.314374177 | 9.24942595 | 0.978297339 | 0.362163407 | 0.991835066 | -4.886243568 |
| **A_55_P2104846** | Rsl24d1 | 0.049849429 | -0.071968142 | 0.171667 | 8.544989336 | 0.978046237 | 0.36227881 | 0.991835066 | -4.886418117 |
| **A_55_P2041131** | NA | 0.045507578 | -0.065725269 | 0.156740425 | 13.33164617 | 0.977821993 | 0.362381893 | 0.991835066 | -4.886573971 |
| **A_55_P2008258** | Larp1b | -0.060075307 | -0.20692529 | 0.086774676 | 11.46101811 | -0.977757673 | 0.362411465 | 0.991835066 | -4.886618671 |
| **A_66_P106418** | Prkacb | -0.072523379 | -0.249823948 | 0.10477719 | 9.883662181 | -0.977635654 | 0.362467569 | 0.991835066 | -4.886703462 |
| **A_51_P155323** | Hc | -0.07305336 | -0.251699846 | 0.105593125 | 7.881018387 | -0.977360656 | 0.362594037 | 0.991835066 | -4.886894534 |
| **A_55_P2125811** | Esam | -0.060819806 | -0.20958455 | 0.087944939 | 11.35154655 | -0.977134036 | 0.362698283 | 0.991835066 | -4.887051965 |
| **A_51_P206405** | Ptprz1 | 0.205926962 | -0.297776217 | 0.709630142 | 8.539248672 | 0.977119439 | 0.362704999 | 0.991835066 | -4.887062105 |
| **A_55_P1963639** | Sycn | 0.047047352 | -0.068055339 | 0.162150044 | 7.829612368 | 0.976919645 | 0.362796925 | 0.991835066 | -4.887200878 |
| **A_55_P1976490** | Lman2l | 0.085830504 | -0.124166829 | 0.295827836 | 11.02827776 | 0.976870546 | 0.362819518 | 0.991835066 | -4.887234978 |
| **A_55_P1960738** | Gm4470 | 0.080389169 | -0.116355507 | 0.277133844 | 8.687015078 | 0.976570679 | 0.362957529 | 0.991835066 | -4.887443217 |
| **A_55_P1989061** | Tsc22d3 | 0.072438427 | -0.104851348 | 0.249728203 | 12.81987912 | 0.976549938 | 0.362967077 | 0.991835066 | -4.88745762 |
| **A_55_P2077033** | Ube2e1 | 0.068832287 | -0.099655114 | 0.237319688 | 11.45604676 | 0.976413786 | 0.363029754 | 0.991835066 | -4.887552153 |
| **A_51_P326393** | Gapvd1 | -0.070661038 | -0.243631492 | 0.102309415 | 10.57434788 | -0.976376291 | 0.363047017 | 0.991835066 | -4.887578185 |
| **A_55_P2077133** | Oxsr1 | 0.054297341 | -0.078620952 | 0.187215633 | 8.633762767 | 0.976344173 | 0.363061804 | 0.991835066 | -4.887600483 |
| **A_55_P1970841** | Nr2c2 | 0.095798628 | -0.138742671 | 0.330339927 | 9.80646023 | 0.976223133 | 0.363117536 | 0.991835066 | -4.887684513 |
| **A_51_P371331** | Mpp5 | -0.063201582 | -0.21794702 | 0.091543856 | 9.184776433 | -0.976155997 | 0.36314845 | 0.991835066 | -4.887731118 |
| **A_55_P2077628** | NA | -0.057304262 | -0.197618342 | 0.083009819 | 8.218493321 | -0.976101094 | 0.363173734 | 0.991835066 | -4.887769229 |
| **A_55_P1988158** | Esd | 0.050894197 | -0.073729813 | 0.175518208 | 13.58930037 | 0.976058119 | 0.363193525 | 0.991835066 | -4.887799059 |
| **A_55_P2075230** | BC051226 | 0.071122995 | -0.103046418 | 0.245292408 | 8.602912054 | 0.975994301 | 0.363222917 | 0.991835066 | -4.887843355 |

| | | | | | | | | | |
|---|---|---|---|---|---|---|---|---|---|
| **A_55_P1962966** | NA | 0.074420953 | -0.10785839 | 0.256700296 | 12.74038351 | 0.975813685 | 0.36330611 | 0.991835066 | -4.887968711 |
| **A_52_P354744** | Slc2a3 | 0.102415151 | -0.148443958 | 0.353274261 | 9.217482503 | 0.975761024 | 0.363330369 | 0.991835066 | -4.888005257 |
| **A_51_P511386** | Zfp267 | 0.041008531 | -0.05944012 | 0.141457182 | 8.457210286 | 0.975751519 | 0.363334748 | 0.991835066 | -4.888011853 |
| **A_55_P1955931** | Slc25a42 | 0.092957713 | -0.134779067 | 0.320694493 | 10.25588293 | 0.975576624 | 0.363415325 | 0.991835066 | -4.888133219 |
| **A_65_P18948** | Ppm1h | -0.045795881 | -0.158003626 | 0.066411863 | 8.844144136 | -0.975467309 | 0.363465696 | 0.991835066 | -4.888209068 |
| **A_52_P105138** | Frs2 | 0.097124845 | -0.140878739 | 0.335128429 | 11.05799344 | 0.975339863 | 0.363524427 | 0.991835066 | -4.888297491 |
| **A_55_P2106033** | Myo18b | -0.106415831 | -0.367191702 | 0.15436004 | 11.9139374 | -0.975321886 | 0.363532713 | 0.991835066 | -4.888309963 |
| **A_52_P419162** | Cyhr1 | -0.098782085 | -0.340875868 | 0.143311697 | 9.310840758 | -0.97522244 | 0.363578547 | 0.991835066 | -4.888378953 |
| **A_51_P260658** | Amdhd2 | 0.041028356 | -0.059525053 | 0.141581766 | 11.03174077 | 0.975206195 | 0.363586035 | 0.991835066 | -4.888390222 |
| **A_51_P114297** | Kifc3 | -0.066410933 | -0.229174244 | 0.096352379 | 9.25393121 | -0.97519664 | 0.363590439 | 0.991835066 | -4.888396851 |
| **A_55_P1998596** | Nupl2 | -0.05626098 | -0.194161828 | 0.081639867 | 9.980766305 | -0.975100749 | 0.36363464 | 0.991835066 | -4.888463368 |
| **A_51_P288026** | Haus6 | 0.056265506 | -0.081651947 | 0.19418296 | 8.448070621 | 0.975061787 | 0.363652601 | 0.991835066 | -4.888490395 |
| **A_51_P243304** | Galk1 | 0.17950755 | -0.260507925 | 0.619523025 | 11.98281847 | 0.975043299 | 0.363661123 | 0.991835066 | -4.888503218 |
| **A_51_P100174** | Mns1 | -0.050746535 | -0.175149009 | 0.073655939 | 9.661678812 | -0.97495935 | 0.363699826 | 0.991835066 | -4.888561447 |
| **A_55_P2117711** | Rnf185 | 0.063814921 | -0.092624617 | 0.220254459 | 11.42278689 | 0.97495561 | 0.36370155 | 0.991835066 | -4.88856404 |
| **A_55_P2091153** | BC051142 | -0.071651261 | -0.247327415 | 0.104024892 | 8.659096908 | -0.974810408 | 0.363768499 | 0.991835066 | -4.888664746 |
| **A_55_P2085181** | Chaf1b | 0.107794873 | -0.156513675 | 0.372103421 | 11.80437326 | 0.974756253 | 0.36379347 | 0.991835066 | -4.888702302 |
| **A_51_P312437** | Dhrs7 | -0.171697749 | -0.592810985 | 0.249415487 | 13.70129733 | -0.97448429 | 0.363918897 | 0.991835066 | -4.888890889 |
| **A_55_P1980302** | Psenen | -0.058890282 | -0.203332219 | 0.085551655 | 12.56694885 | -0.974449778 | 0.363934816 | 0.991835066 | -4.888914819 |
| **A_51_P362054** | Atp5e | -0.116817376 | -0.403340717 | 0.169705965 | 16.43657184 | -0.974443298 | 0.363937805 | 0.991835066 | -4.888919312 |
| **A_55_P2073253** | NA | 0.056498372 | -0.082078421 | 0.195075165 | 9.669295035 | 0.974438783 | 0.363939888 | 0.991835066 | -4.888922442 |
| **A_55_P2024463** | Tbkbp1 | 0.049677863 | -0.072178957 | 0.171534682 | 8.780446238 | 0.974366168 | 0.363973385 | 0.991835066 | -4.888972788 |
| **A_66_P138780** | Elob | -0.079499589 | -0.27456362 | 0.115564443 | 13.64636692 | -0.974084917 | 0.364103145 | 0.991835066 | -4.889167761 |
| **A_55_P2018387** | Zkscan17 | -0.044567484 | -0.153943343 | 0.064808374 | 11.70988277 | -0.973880747 | 0.364197365 | 0.991835066 | -4.889309276 |
| **A_51_P168792** | Xylb | 0.057091152 | -0.083028041 | 0.197210345 | 8.209398307 | 0.973823641 | 0.364223722 | 0.991835066 | -4.889348854 |
| **A_52_P649332** | Sdf4 | 0.269026884 | -0.391323906 | 0.929377673 | 10.88592985 | 0.97371108 | 0.364275677 | 0.991835066 | -4.889426861 |
| **A_55_P2181162** | NA | 0.045193209 | -0.065742155 | 0.156128572 | 15.34707217 | 0.973671137 | 0.364294116 | 0.991835066 | -4.88945454 |
| **A_55_P1993940** | Sh3bgrl3 | -0.11785584 | -0.407165143 | 0.171453464 | 10.84169503 | -0.973638723 | 0.364309079 | 0.991835066 | -4.889477002 |
| **A_51_P328001** | Sftpc | 0.239060791 | -0.347828103 | 0.825949685 | 8.07538364 | 0.973557469 | 0.36434659 | 0.991835066 | -4.889533306 |
| **A_55_P1991109** | Gatd3a | 0.064142686 | -0.093335469 | 0.221620841 | 14.34497188 | 0.973500001 | 0.364373122 | 0.991835066 | -4.889573126 |
| **A_55_P1973818** | Mmel1 | 0.047066788 | -0.068490419 | 0.162623995 | 7.786822307 | 0.973479154 | 0.364382747 | 0.991835066 | -4.88958757 |
| **A_55_P2039274** | Hspb3 | -0.078082014 | -0.269789171 | 0.113625142 | 11.69609417 | -0.973468304 | 0.364387757 | 0.991835066 | -4.889595088 |
| **A_55_P2258261** | 1810008I18 | -0.049494436 | -0.171032714 | 0.072043843 | 7.720359469 | -0.973312788 | 0.364459566 | 0.991835066 | -4.889702835 |
| **A_55_P2104876** | NA | -0.071017451 | -0.245430595 | 0.103395693 | 8.534316658 | -0.973184097 | 0.364518997 | 0.991835066 | -4.889791988 |
| **A_55_P2005941** | Plcxd1 | -0.071561265 | -0.247320721 | 0.104198191 | 10.44334546 | -0.973124579 | 0.364546485 | 0.991835066 | -4.889833217 |
| **A_55_P2127688** | 4933434E2 | 0.058411829 | -0.085092571 | 0.201916228 | 12.87048771 | 0.972847393 | 0.364674526 | 0.991835066 | -4.890025208 |
| **A_55_P2180884** | Psmd10 | 0.055078279 | -0.080236587 | 0.190393144 | 9.166644814 | 0.972845743 | 0.364675289 | 0.991835066 | -4.890026351 |
| **A_52_P520408** | Serp1 | 0.08668431 | -0.126334903 | 0.299703523 | 10.31582805 | 0.97259236 | 0.364792364 | 0.991835066 | -4.890201821 |
| **A_55_P2014249** | Sema3a | 0.085927086 | -0.125232082 | 0.297086255 | 8.615740457 | 0.972588825 | 0.364793998 | 0.991835066 | -4.890204269 |
| **A_52_P174915** | Gja1 | -0.079921671 | -0.276373914 | 0.116530573 | 11.34787559 | -0.972336733 | 0.364910507 | 0.991835066 | -4.885878528 |
| **A_52_P727235** | Trabd2b | -0.144653768 | -0.500251943 | 0.210944407 | 8.735587297 | -0.972253153 | 0.364949141 | 0.991835066 | -4.890436677 |
| **A_55_P1976470** | 2410018L1 | 0.072149934 | -0.105279723 | 0.249579591 | 8.942271183 | 0.971893904 | 0.365115238 | 0.991835066 | -4.89068535 |
| **A_51_P483324** | Ptpn22 | -0.042159707 | -0.145885559 | 0.061566144 | 7.791523451 | -0.971448344 | 0.36532132 | 0.991835066 | -4.89099368 |
| **A_55_P2171158** | Hmgn1 | 0.107677371 | -0.157279609 | 0.37263435 | 14.48224147 | 0.971310784 | 0.365384964 | 0.991835066 | -4.891088853 |
| **A_55_P2148912** | S100a16 | -0.084036306 | -0.290853353 | 0.122780742 | 12.03624021 | -0.971157683 | 0.365455807 | 0.991835066 | -4.891194767 |
| **A_52_P50478** | Rnf123 | -0.066287284 | -0.229442185 | 0.096867617 | 9.61093179 | -0.971044726 | 0.365508082 | 0.991835066 | -4.891272903 |
| **A_55_P2138306** | NA | 0.058787987 | -0.085919712 | 0.203495686 | 10.57878423 | 0.970970603 | 0.365542388 | 0.991835066 | -4.891324173 |
| **A_51_P435817** | Zbtb8a | -0.046849241 | -0.162170327 | 0.068471844 | 8.693807147 | -0.970963649 | 0.365545606 | 0.991835066 | -4.891328982 |

| | | | | | | | | | |
|---|---|---|---|---|---|---|---|---|---|
| **A_55_P2376423** | Igkv4-72 | -0.051850764 | -0.179524086 | 0.075822559 | 8.344456846 | -0.970653284 | 0.365689279 | 0.991835066 | -4.891543627 |
| **A_55_P2159216** | NA | -0.089338919 | -0.309329758 | 0.13065192 | 11.35893006 | -0.970611045 | 0.365708836 | 0.991835066 | -4.891572836 |
| **A_52_P394395** | Abcd3 | -0.067015139 | -0.232069209 | 0.098038931 | 12.09329549 | -0.970411239 | 0.365801356 | 0.991835066 | -4.891710991 |
| **A_55_P2038697** | Zfp235 | 0.043662441 | -0.063882401 | 0.151207284 | 8.592400243 | 0.970348099 | 0.365830597 | 0.991835066 | -4.891754645 |
| **A_55_P2033849** | Sphkap | -0.077553291 | -0.268580883 | 0.113474302 | 8.847005983 | -0.970316149 | 0.365845394 | 0.991835066 | -4.891776733 |
| **A_55_P1961140** | Mettl3 | 0.069594367 | -0.101844634 | 0.241033369 | 10.52666524 | 0.970227536 | 0.365886436 | 0.991835066 | -4.891837994 |
| **A_55_P1963364** | NA | 0.086188096 | -0.126159966 | 0.298536158 | 8.816296871 | 0.970081263 | 0.365954191 | 0.991835066 | -4.891939109 |
| **A_51_P254299** | Dlk2 | -0.048672557 | -0.168605263 | 0.07126015 | 8.37235143 | -0.969964092 | 0.366008474 | 0.991835066 | -4.892020098 |
| **A_55_P2092501** | Med1 | 0.0628878 | -0.09209008 | 0.217865679 | 11.19272067 | 0.969852784 | 0.366060045 | 0.991835066 | -4.892097029 |
| **A_51_P394788** | Actn4 | -0.118355943 | -0.410072045 | 0.173360159 | 13.1590554 | -0.969703141 | 0.366129387 | 0.991835066 | -4.892200446 |
| **A_52_P319123** | Vps13b | 0.066349178 | -0.097197691 | 0.229896047 | 9.302113224 | 0.969621961 | 0.366167009 | 0.991835066 | -4.892256544 |
| **A_55_P2093553** | NA | 0.048007146 | -0.07033398 | 0.166348272 | 7.802170554 | 0.969570348 | 0.36619093 | 0.991835066 | -4.892292209 |
| **A_55_P2326727** | 9430085M1 | -0.088428848 | -0.306437954 | 0.129580258 | 10.5712116 | -0.969456807 | 0.366243557 | 0.991835066 | -4.892370662 |
| **A_51_P414115** | Naprt | -0.086904522 | -0.301157611 | 0.127348568 | 10.12364277 | -0.969447761 | 0.366247749 | 0.991835066 | -4.892376911 |
| **A_55_P2175880** | Mgst1 | -0.104670334 | -0.362816455 | 0.153475788 | 10.47023077 | -0.969096746 | 0.366410484 | 0.991835066 | -4.892619408 |
| **A_51_P448203** | Cnbp | 0.092778104 | -0.136053833 | 0.32161004 | 13.13812043 | 0.969031705 | 0.366440644 | 0.991835066 | -4.892664335 |
| **A_51_P138859** | Tmem68 | -0.069350735 | -0.240418777 | 0.101717307 | 9.79632535 | -0.968927573 | 0.366488935 | 0.991835066 | -4.89273626 |
| **A_55_P1971962** | Tmem176b | -0.108526921 | -0.376247306 | 0.159193464 | 8.982370713 | -0.968869254 | 0.366515982 | 0.991835066 | -4.892776538 |
| **A_51_P396818** | Rex1bd | -0.135296225 | -0.469149572 | 0.198557122 | 10.75115406 | -0.968587982 | 0.366646452 | 0.991835066 | -4.892970779 |
| **A_55_P2035205** | Nifk | 0.082040106 | -0.120405389 | 0.284485602 | 11.75068852 | 0.968561563 | 0.366658709 | 0.991835066 | -4.892989022 |
| **A_55_P1993813** | NA | 0.075257984 | -0.110453083 | 0.26096905 | 16.25109233 | 0.968554197 | 0.366662126 | 0.991835066 | -4.892994108 |
| **A_55_P2146650** | NA | 0.052894252 | -0.077643679 | 0.183432184 | 8.450033421 | 0.968458213 | 0.366706659 | 0.991835066 | -4.893060382 |
| **A_55_P2109922** | NA | 0.052811333 | -0.077523148 | 0.183145815 | 8.74117003 | 0.968449401 | 0.366710748 | 0.991835066 | -4.893066467 |
| **A_65_P19832** | Ttr | -0.330270912 | -1.14536363 | 0.484821805 | 8.894359506 | -0.96843939 | 0.366715392 | 0.991835066 | -4.893073379 |
| **A_55_P2062627** | Tmem238 | 0.05846679 | -0.085840305 | 0.202773885 | 8.140276395 | 0.968346304 | 0.366758586 | 0.991835066 | -4.893137647 |
| **A_55_P1956627** | Gpr160 | -0.043014115 | -0.14919102 | 0.063162789 | 7.770959136 | -0.968255712 | 0.366800625 | 0.991835066 | -4.893200189 |
| **A_55_P2098683** | St7l | 0.05361218 | -0.078728229 | 0.18595259 | 10.0649922 | 0.968233552 | 0.366810909 | 0.991835066 | -4.893215488 |
| **A_55_P2042062** | Mier1 | 0.052247845 | -0.076736508 | 0.181232198 | 8.011770068 | 0.968145179 | 0.366851923 | 0.991835066 | -4.893276493 |
| **A_55_P2009677** | Zmiz1 | -0.073754263 | -0.255836114 | 0.108327587 | 12.6843734 | -0.968120918 | 0.366863183 | 0.991835066 | -4.89329324 |
| **A_66_P124497** | Imp3 | 0.050655618 | -0.074406795 | 0.175718031 | 13.31660525 | 0.968077101 | 0.366883521 | 0.991835066 | -4.893323486 |
| **A_52_P212214** | Pomt2 | -0.045344908 | -0.157295993 | 0.066606176 | 8.489609914 | -0.968075786 | 0.366884131 | 0.991835066 | -4.893324394 |
| **A_55_P2159865** | Zfp59 | -0.045169542 | -0.156733495 | 0.066394411 | 8.747688179 | -0.96767812 | 0.367068745 | 0.991835066 | -4.89359885 |
| **A_55_P2042272** | Pbrm1 | 0.099576119 | -0.146381182 | 0.34553342 | 8.529023416 | 0.967619494 | 0.367095968 | 0.991835066 | -4.893639306 |
| **A_55_P2000090** | Arf1 | 0.050769806 | -0.074643685 | 0.176183297 | 14.4586407 | 0.967543226 | 0.367131385 | 0.991835066 | -4.893691932 |
| **A_55_P2024649** | Rpe | 0.077370404 | -0.113767715 | 0.268508522 | 9.32847753 | 0.967468177 | 0.367166239 | 0.991835066 | -4.893743716 |
| **A_55_P2018549** | Gcsh | 0.061697393 | -0.090723047 | 0.214117832 | 12.18934917 | 0.96745937 | 0.367170329 | 0.991835066 | -4.893749792 |
| **A_51_P464394** | Klb | -0.039863497 | -0.138348155 | 0.058621162 | 7.682812676 | -0.967421959 | 0.367187704 | 0.991835066 | -4.893775604 |
| **A_52_P207335** | Tmem246 | -0.087668272 | -0.304292394 | 0.128955849 | 12.62357599 | -0.967263426 | 0.367261341 | 0.991835066 | -4.893884977 |
| **A_55_P1984640** | Ube2f | 0.094263034 | -0.138661778 | 0.327187846 | 11.17502774 | 0.967241212 | 0.36727166 | 0.991835066 | -4.893900302 |
| **A_51_P487228** | B9d2 | -0.056137801 | -0.194885956 | 0.082610354 | 11.81392688 | -0.967024138 | 0.367372508 | 0.991835066 | -4.894050041 |
| **A_55_P1972605** | Adora3 | -0.049112977 | -0.170504814 | 0.072278861 | 7.835009768 | -0.966976464 | 0.367394659 | 0.991835066 | -4.894082924 |
| **A_55_P1966109** | Epn2 | 0.057145197 | -0.084119968 | 0.198410363 | 9.814244757 | 0.966838167 | 0.367458924 | 0.991835066 | -4.894178307 |
| **A_55_P1987730** | Prxl2a | 0.075955667 | -0.111809992 | 0.263721326 | 11.74732825 | 0.966836743 | 0.367459585 | 0.991835066 | -4.894179289 |
| **A_55_P1970846** | Pemt | -0.062926695 | -0.218531344 | 0.092677954 | 8.118793633 | -0.96654368 | 0.367595796 | 0.991835066 | -4.894381383 |
| **A_55_P2039038** | Zfp664 | 0.092841905 | -0.136775107 | 0.322458917 | 10.77214485 | 0.966382626 | 0.367670667 | 0.991835066 | -4.894492427 |
| **A_51_P129464** | Scd2 | 0.1123395 | -0.165508334 | 0.390187334 | 14.19483317 | 0.966350339 | 0.367685679 | 0.991835066 | -4.894514686 |
| **A_52_P238468** | Gipr | 0.168338503 | -0.248130121 | 0.584807127 | 9.822038829 | 0.966073801 | 0.367814268 | 0.991835066 | -4.89470532 |
| **A_55_P1981002** | NA | 0.080080913 | -0.118043225 | 0.27820505 | 9.463790997 | 0.966052565 | 0.367824144 | 0.991835066 | -4.894719958 |

| | | | | | | | | | |
|---|---|---|---|---|---|---|---|---|---|
| **A_55_P1968928** | Def6 | -0.066226927 | -0.230077816 | 0.097623962 | 9.876395803 | -0.966039611 | 0.367830169 | 0.991835066 | -4.894728887 |
| **A_51_P176365** | Gimap5 | -0.076472971 | -0.265703128 | 0.112757185 | 9.026084134 | -0.96588795 | 0.367900708 | 0.991835066 | -4.894833417 |
| **A_66_P104312** | Pdcl | -0.049915625 | -0.17343134 | 0.07360009 | 9.74263525 | -0.965880575 | 0.367904139 | 0.991835066 | -4.8948385 |
| **A_55_P2054942** | Iffo2 | 0.062369942 | -0.091979727 | 0.216719612 | 8.929317818 | 0.965781257 | 0.367950338 | 0.991835066 | -4.894906947 |
| **A_51_P380986** | Epc1 | 0.072562903 | -0.10703759 | 0.252163396 | 10.26539182 | 0.965642241 | 0.368015012 | 0.991835066 | -4.895002746 |
| **A_51_P357573** | Cald1 | 0.077535627 | -0.114424573 | 0.269495827 | 9.828163679 | 0.965382102 | 0.368136059 | 0.991835066 | -4.895181986 |
| **A_55_P1977318** | Wbp11 | -0.050593012 | -0.175851147 | 0.074665122 | 12.14414139 | -0.965369848 | 0.368141762 | 0.991835066 | -4.895190429 |
| **A_55_P2052685** | Dguok | -0.053315595 | -0.185318839 | 0.078687648 | 10.71921275 | -0.965336643 | 0.368157215 | 0.991835066 | -4.895213305 |
| **A_52_P342159** | Nfatc4 | 0.082060394 | -0.121129418 | 0.285250207 | 9.420300713 | 0.965252204 | 0.368196514 | 0.991835066 | -4.895271476 |
| **A_55_P2361437** | Pitpnc1 | -0.077679337 | -0.270043564 | 0.114684889 | 9.555223164 | -0.965140043 | 0.36824872 | 0.991835066 | -4.89534874 |
| **A_66_P109941** | Mamdc4 | 0.106944321 | -0.157981151 | 0.371869792 | 8.257419235 | 0.96481299 | 0.368400983 | 0.991835066 | -4.895574 |
| **A_55_P2046842** | Tmem253 | -0.040529579 | -0.140937371 | 0.059878212 | 8.007253361 | -0.964747835 | 0.368431322 | 0.991835066 | -4.89561887 |
| **A_55_P2227967** | NA | -0.047450067 | -0.165008724 | 0.070108589 | 9.169367435 | -0.964697939 | 0.368454558 | 0.991835066 | -4.895653231 |
| **A_55_P2067146** | NA | 0.049066797 | -0.072527506 | 0.170661101 | 8.092649076 | 0.964458655 | 0.368566001 | 0.991835066 | -4.895817993 |
| **A_55_P2025228** | Akt2 | -0.070172964 | -0.244076322 | 0.103730394 | 12.64452165 | -0.96443061 | 0.368579064 | 0.991835066 | -4.895837301 |
| **A_55_P2025626** | NA | 0.050848233 | -0.075178587 | 0.176875054 | 8.471579271 | 0.964321883 | 0.368629713 | 0.991835066 | -4.895912156 |
| **A_55_P2040341** | NA | 0.105859077 | -0.156520781 | 0.368238936 | 8.941607441 | 0.964287958 | 0.368645517 | 0.991835066 | -4.895935511 |
| **A_55_P2427685** | Agl | -0.059980238 | -0.208690604 | 0.088730128 | 13.90972628 | -0.963997868 | 0.368780681 | 0.991835066 | -4.896135195 |
| **A_55_P1977766** | Ttc14 | 0.129078706 | -0.190958912 | 0.449116323 | 8.533169449 | 0.963968194 | 0.368794509 | 0.991835066 | -4.896155619 |
| **A_52_P605556** | Nxnl2 | 0.056004907 | -0.082870828 | 0.194880642 | 7.986407622 | 0.963848651 | 0.368850222 | 0.991835066 | -4.896237893 |
| **A_51_P121228** | Ascc2 | 0.040679262 | -0.060207983 | 0.141566506 | 10.6664055 | 0.963709045 | 0.368915292 | 0.991835066 | -4.896333966 |
| **A_55_P1970164** | Rnf11 | -0.064335667 | -0.22389957 | 0.095228236 | 10.76629907 | -0.963665452 | 0.368935613 | 0.991835066 | -4.896363964 |
| **A_55_P2088103** | Ash2l | 0.056997773 | -0.08437465 | 0.198370196 | 11.32555404 | 0.96361227 | 0.368960404 | 0.991835066 | -4.896400558 |
| **A_55_P1993128** | Gas8 | 0.050968287 | -0.075460793 | 0.177397368 | 10.03074219 | 0.96352324 | 0.36900191 | 0.991835066 | -4.896461817 |
| **A_55_P1966159** | Dlg5 | 0.055749827 | -0.082540957 | 0.19404061 | 11.01531798 | 0.963517078 | 0.369004783 | 0.991835066 | -4.896466057 |
| **A_55_P2124096** | NA | -0.138836093 | -0.483253997 | 0.205581812 | 15.6906438 | -0.963442488 | 0.369039559 | 0.991835066 | -4.896517377 |
| **A_55_P2064233** | Ipo8 | 0.088623481 | -0.131261537 | 0.3085085 | 10.068886 | 0.963301639 | 0.369105235 | 0.991835066 | -4.896614278 |
| **A_51_P405167** | Maf | 0.073579115 | -0.108984315 | 0.256142546 | 8.756780735 | 0.963274149 | 0.369118055 | 0.991835066 | -4.89663319 |
| **A_51_P388478** | Efnb1 | 0.058166954 | -0.08615628 | 0.202490187 | 11.83250056 | 0.96327259 | 0.369118781 | 0.991835066 | -4.896634262 |
| **A_55_P2128704** | NA | 0.070075589 | -0.103830246 | 0.243981425 | 14.77688814 | 0.963078606 | 0.369209251 | 0.991835066 | -4.896767699 |
| **A_66_P106113** | Rhoj | 0.054081355 | -0.080155614 | 0.188318325 | 11.23332513 | 0.962907478 | 0.369289075 | 0.991835066 | -4.8968854 |
| **A_51_P414305** | Acot12 | -0.041672161 | -0.145108258 | 0.061763936 | 7.618836379 | -0.962904084 | 0.369290658 | 0.991835066 | -4.896887734 |
| **A_51_P278103** | Stambp | -0.058169581 | -0.202598559 | 0.086259396 | 10.03805639 | -0.962610804 | 0.369427492 | 0.991835066 | -4.897089415 |
| **A_55_P2231747** | NA | 0.244427931 | -0.362528272 | 0.851384134 | 9.450626667 | 0.962504147 | 0.369477264 | 0.991835066 | -4.897162751 |
| **A_66_P139267** | Rbpj | -0.052849934 | -0.184091843 | 0.078391976 | 10.39259546 | -0.962456344 | 0.369499574 | 0.991835066 | -4.897195617 |
| **A_51_P250807** | Spata2l | -0.071227654 | -0.248110169 | 0.105654861 | 8.319455959 | -0.962438236 | 0.369508025 | 0.991835066 | -4.897208066 |
| **A_52_P650387** | Ccnjl | 0.073874866 | -0.109617791 | 0.257367523 | 10.22709942 | 0.962248295 | 0.36959668 | 0.991835066 | -4.897338646 |
| **A_55_P2002854** | Aifm2 | -0.068411118 | -0.238350236 | 0.101528 | 8.530838086 | -0.962149282 | 0.369642901 | 0.991835066 | -4.897406708 |
| **A_66_P129263** | Lekr1 | -0.045758282 | -0.159437334 | 0.06792077 | 8.448713261 | -0.96205167 | 0.369688473 | 0.991835066 | -4.897473802 |
| **A_52_P309737** | Cdk12 | 0.04946839 | -0.073436884 | 0.172373664 | 8.584583665 | 0.961980764 | 0.369721579 | 0.991835066 | -4.897522536 |
| **A_51_P427674** | Cpt1a | 0.060493151 | -0.089810823 | 0.210797125 | 10.33595484 | 0.961933113 | 0.369743828 | 0.991835066 | -4.897555286 |
| **A_55_P2004806** | Hexim2 | 0.130729544 | -0.194128217 | 0.455587305 | 13.07109035 | 0.96181076 | 0.369800963 | 0.991835066 | -4.897639372 |
| **A_55_P2182586** | Esrp1 | 0.043661889 | -0.064852065 | 0.152175843 | 7.900823696 | 0.961669987 | 0.369866707 | 0.991835066 | -4.897736108 |
| **A_55_P1996583** | Ppdpf | -0.121688008 | -0.424145159 | 0.180769143 | 11.70136663 | -0.961596908 | 0.369900841 | 0.991835066 | -4.897786323 |
| **A_55_P2026119** | Dlec1 | -0.053952983 | -0.188055985 | 0.08015002 | 9.11602151 | -0.961581476 | 0.369908049 | 0.991835066 | -4.897796926 |
| **A_55_P1988970** | Mkks | -0.080305321 | -0.279949358 | 0.119338716 | 11.83548477 | -0.96138449 | 0.37000007 | 0.991835066 | -4.897932266 |
| **A_52_P763938** | NA | -0.039610926 | -0.138105214 | 0.058883363 | 8.247635494 | -0.961198476 | 0.370086981 | 0.991835066 | -4.89806005 |
| **A_51_P275350** | Pten | -0.075343822 | -0.262757896 | 0.112070252 | 10.88466374 | -0.96084775 | 0.370250893 | 0.991835066 | -4.898300938 |

| | | | | | | | | | |
|---|---|---|---|---|---|---|---|---|---|
| **A_51_P302167** | Cobl | -0.091917748 | -0.320575093 | 0.136739598 | 8.969417058 | -0.960778658 | 0.37028319 | 0.991835066 | -4.898348386 |
| **A_52_P282700** | Zbtb33 | -0.044287202 | -0.15446801 | 0.065893606 | 8.942296861 | -0.960685918 | 0.370326545 | 0.991835066 | -4.898412069 |
| **A_55_P2137203** | B4galt3 | -0.070245467 | -0.245020468 | 0.104529533 | 11.88723107 | -0.960612266 | 0.370360979 | 0.991835066 | -4.898462642 |
| **A_51_P302566** | Maob | -0.086208849 | -0.300704496 | 0.128286798 | 9.395755801 | -0.960599812 | 0.370366801 | 0.991835066 | -4.898471193 |
| **A_55_P2069955** | NA | 0.041019793 | -0.061054675 | 0.143094261 | 17.4955568 | 0.960473684 | 0.370425775 | 0.991835066 | -4.898557791 |
| **A_52_P429749** | Tmem115 | -0.093597514 | -0.326564148 | 0.13936912 | 9.448114503 | -0.960239841 | 0.370535133 | 0.991835066 | -4.898718324 |
| **A_51_P453088** | Coa6 | -0.082215944 | -0.286862178 | 0.12243029 | 11.3549828 | -0.960199386 | 0.370554054 | 0.991835066 | -4.898746094 |
| **A_52_P144285** | Fam72a | 0.05089575 | -0.075791365 | 0.177582865 | 8.912980172 | 0.96019226 | 0.370557387 | 0.991835066 | -4.898750986 |
| **A_52_P16877** | Tmcc3 | -0.051676485 | -0.180312864 | 0.076959894 | 7.981876835 | -0.960148242 | 0.370577976 | 0.991835066 | -4.8987812 |
| **A_55_P2013591** | NA | -0.062207862 | -0.217065201 | 0.092649478 | 10.53268646 | -0.960113577 | 0.370594191 | 0.991835066 | -4.898804993 |
| **A_66_P104422** | Ckap4 | 0.10738157 | -0.159942546 | 0.374705685 | 10.75780118 | 0.960065231 | 0.370616806 | 0.991835066 | -4.898838177 |
| **A_52_P377576** | NA | 0.102291686 | -0.152373565 | 0.356956937 | 8.224136135 | 0.960018896 | 0.370638481 | 0.991835066 | -4.898869979 |
| **A_55_P1955970** | Cbfa2t2 | 0.046037675 | -0.0685983 | 0.16067365 | 11.18894239 | 0.959846052 | 0.370719346 | 0.991835066 | -4.8989886 |
| **A_55_P1993640** | Gm6994 | -0.082450007 | -0.287769003 | 0.122868989 | 12.50917014 | -0.959777793 | 0.370751284 | 0.991835066 | -4.899035442 |
| **A_51_P363103** | Blvra | -0.07901705 | -0.275804009 | 0.117769909 | 10.18134137 | -0.959695882 | 0.370789613 | 0.991835066 | -4.899091649 |
| **A_52_P479500** | Npepps | -0.067831155 | -0.236763754 | 0.101101444 | 12.00188417 | -0.959676545 | 0.370798662 | 0.991835066 | -4.899104918 |
| **A_66_P117578** | Mtmr7 | 0.043206596 | -0.064416599 | 0.15082979 | 8.07475177 | 0.959518392 | 0.370872678 | 0.991835066 | -4.899213431 |
| **A_55_P1969530** | Zfp146 | 0.090106824 | -0.134355982 | 0.314569629 | 9.330713619 | 0.959450176 | 0.370904606 | 0.991835066 | -4.899260233 |
| **A_51_P111757** | Asb1 | -0.050911472 | -0.177764825 | 0.075941881 | 9.54021235 | -0.959230183 | 0.371007588 | 0.991835066 | -4.899411148 |
| **A_55_P2154387** | Bmp4 | 0.251103456 | -0.37459177 | 0.876798682 | 10.21951215 | 0.959177496 | 0.371032255 | 0.991835066 | -4.899447288 |
| **A_55_P2055404** | Slc15a4 | -0.057160633 | -0.199712874 | 0.085391607 | 11.83471189 | -0.958367591 | 0.371411592 | 0.991835066 | -4.90000266 |
| **A_52_P636343** | Wipi2 | 0.097876586 | -0.146228756 | 0.341981929 | 8.937827292 | 0.958320245 | 0.371433777 | 0.991835066 | -4.900035116 |
| **A_52_P514317** | Slc35a3 | 0.052212016 | -0.078046892 | 0.182470925 | 9.01244109 | 0.958014676 | 0.371576981 | 0.991835066 | -4.900244562 |
| **A_55_P2151952** | NA | 0.053696456 | -0.080300775 | 0.187693688 | 13.26434791 | 0.957764936 | 0.371694052 | 0.991835066 | -4.900415706 |
| **A_51_P131653** | Tsacc | -0.053780459 | -0.188009238 | 0.080448319 | 9.935990161 | -0.957608518 | 0.371767392 | 0.991835066 | -4.900522883 |
| **A_55_P1977422** | NA | 0.040854601 | -0.061121181 | 0.142830383 | 16.39514445 | 0.95753148 | 0.371803516 | 0.991835066 | -4.900575664 |
| **A_51_P389664** | Scrt2 | -0.049039708 | -0.171479554 | 0.073400138 | 7.895961661 | -0.957269528 | 0.37192637 | 0.991835066 | -4.900755114 |
| **A_51_P514449** | Nr5a2 | -0.06898386 | -0.241220197 | 0.103252478 | 8.50714792 | -0.95726425 | 0.371928846 | 0.991835066 | -4.90075873 |
| **A_51_P493649** | Sult1e1 | -0.088576036 | -0.309764982 | 0.132612911 | 7.793256109 | -0.957110216 | 0.372001102 | 0.991835066 | -4.900864235 |
| **A_55_P1960960** | Zdhhc9 | 0.100283982 | -0.150143722 | 0.350711687 | 11.86269594 | 0.957102239 | 0.372004844 | 0.991835066 | -4.900869698 |
| **A_55_P1956497** | Serpinf2 | -0.163404628 | -0.571468065 | 0.244658809 | 8.137106929 | -0.957074623 | 0.3720178 | 0.991835066 | -4.900888612 |
| **A_55_P1954393** | Susd4 | 0.118428265 | -0.177444859 | 0.414301389 | 9.672313406 | 0.956663011 | 0.372210943 | 0.991835066 | -4.901170479 |
| **A_55_P1996578** | Fgfr3 | 0.074066912 | -0.111026737 | 0.259160561 | 8.970235369 | 0.956405036 | 0.372332034 | 0.991835066 | -4.901347095 |
| **A_55_P2125322** | NA | 0.053276033 | -0.079862508 | 0.186414574 | 15.68733762 | 0.956394821 | 0.37233683 | 0.991835066 | -4.901354088 |
| **A_51_P512061** | Doxl1 | -0.045805843 | -0.160292658 | 0.068680972 | 7.810019195 | -0.956256798 | 0.372401629 | 0.991835066 | -4.901448569 |
| **A_55_P2125820** | Gm14288 | 0.062551587 | -0.093822763 | 0.218925937 | 10.32956528 | 0.956052949 | 0.372497349 | 0.991835066 | -4.90158809 |
| **A_55_P2155834** | NA | 0.05233473 | -0.078506633 | 0.183176094 | 16.60140178 | 0.955991568 | 0.372526174 | 0.991835066 | -4.901630098 |
| **A_51_P254262** | Nxph3 | 0.084180777 | -0.126329465 | 0.294691019 | 8.177257565 | 0.955759957 | 0.37263496 | 0.991835066 | -4.90178859 |
| **A_66_P112573** | Isg20 | -0.347977339 | -1.218180313 | 0.522225636 | 10.74978331 | -0.955739491 | 0.372644573 | 0.991835066 | -4.901802593 |
| **A_52_P382914** | Trip6 | 0.097495033 | -0.152736548 | 0.347726614 | 11.93561095 | 0.967848809 | 0.372700739 | 0.991835066 | -4.835035503 |
| **A_55_P1976926** | Tmem104 | 0.044526916 | -0.066861282 | 0.155915114 | 7.969959124 | 0.955416132 | 0.372796495 | 0.991835066 | -4.90202382 |
| **A_55_P2060284** | Itpripl2 | -0.074175485 | -0.259732806 | 0.111381836 | 8.404202583 | -0.955413628 | 0.372797671 | 0.991835066 | -4.902025533 |
| **A_66_P130777** | Ephb4 | 0.071826667 | -0.107894644 | 0.251547979 | 8.940193406 | 0.955202104 | 0.372897076 | 0.991835066 | -4.90217022 |
| **A_52_P451888** | Tlk2 | -0.06011665 | -0.210542697 | 0.090309397 | 8.498904423 | -0.955170416 | 0.372911969 | 0.991835066 | -4.902191893 |
| **A_52_P383653** | Cpne8 | -0.048797135 | -0.170958654 | 0.073364383 | 8.029584255 | -0.954704646 | 0.373130934 | 0.991835066 | -4.902510405 |
| **A_55_P2009064** | Psmc1 | -0.065031712 | -0.227841856 | 0.097778431 | 12.0971398 | -0.954669091 | 0.373147654 | 0.991835066 | -4.902534714 |
| **A_66_P137654** | Smpd2 | -0.109827209 | -0.384817583 | 0.165163166 | 8.674450289 | -0.954556382 | 0.373200656 | 0.991835066 | -4.902611771 |
| **A_55_P2113768** | NA | -0.114137066 | -0.399930816 | 0.171656683 | 11.46740301 | -0.954515777 | 0.373219753 | 0.991835066 | -4.902639531 |

| | | | | | | | | | |
|---|---|---|---|---|---|---|---|---|---|
| **A_51_P184969** | Rilpl1 | -0.077987341 | -0.273266779 | 0.117292096 | 10.69716289 | -0.954501757 | 0.373226346 | 0.991835066 | -4.902649114 |
| **A_51_P211491** | Gusb | -0.048535222 | -0.170086482 | 0.073016037 | 10.67123915 | -0.954347834 | 0.373298744 | 0.991835066 | -4.879202472 |
| **A_51_P308681** | Mcur1 | -0.082880827 | -0.29045032 | 0.124688665 | 9.865699178 | -0.954332438 | 0.373305985 | 0.991835066 | -4.902764858 |
| **A_52_P84096** | Pgd | 0.051855006 | -0.078015457 | 0.181725469 | 11.40443271 | 0.954309902 | 0.373316586 | 0.991835066 | -4.902780262 |
| **A_55_P2156186** | BC028528 | -0.108593404 | -0.380595658 | 0.16340885 | 9.69895724 | -0.954201455 | 0.373367602 | 0.991835066 | -4.902854386 |
| **A_55_P2131068** | NA | -0.104471178 | -0.366182717 | 0.157240362 | 15.71542927 | -0.954075513 | 0.373426855 | 0.991835066 | -4.902940461 |
| **A_52_P445360** | Krt20 | 0.071200588 | -0.107169072 | 0.249570247 | 8.188530931 | 0.954051296 | 0.373438249 | 0.991835066 | -4.90295701 |
| **A_55_P1993268** | Ap1s1 | 0.050869367 | -0.076582068 | 0.178320803 | 13.305242 | 0.953939291 | 0.373490953 | 0.991835066 | -4.903033552 |
| **A_51_P417891** | Trim10 | 0.051868911 | -0.078102097 | 0.181839919 | 7.878793147 | 0.953827361 | 0.373543626 | 0.991835066 | -4.903110036 |
| **A_55_P2199717** | 9330102E0 | 0.043006001 | -0.064791742 | 0.150803743 | 8.202881066 | 0.953517187 | 0.373689621 | 0.991835066 | -4.903321951 |
| **A_51_P405089** | NA | 0.040018919 | -0.06029872 | 0.140336558 | 8.37320122 | 0.953448373 | 0.373722017 | 0.991835066 | -4.903368959 |
| **A_55_P2430367** | Zbtb8b | 0.096718624 | -0.145748104 | 0.339185352 | 9.238708627 | 0.953382312 | 0.373753119 | 0.991835066 | -4.903414085 |
| **A_55_P2072193** | Firre | 0.050841512 | -0.076616943 | 0.178299967 | 8.104057731 | 0.953364421 | 0.373761543 | 0.991835066 | -4.903426306 |
| **A_55_P2350022** | Akap6 | -0.077062047 | -0.270265351 | 0.116141256 | 10.51143451 | -0.953312149 | 0.373786154 | 0.991835066 | -4.90346201 |
| **A_51_P202033** | Wls | -0.077995148 | -0.273574538 | 0.117584241 | 11.51787173 | -0.953133282 | 0.373870382 | 0.991835066 | -4.903584175 |
| **A_51_P317272** | Bod1 | -0.059579882 | -0.208987895 | 0.089828132 | 12.76703319 | -0.953092116 | 0.373889768 | 0.991835066 | -4.903612288 |
| **A_66_P140742** | Azin2 | -0.101276386 | -0.355254599 | 0.152701826 | 8.060801773 | -0.953061368 | 0.373904249 | 0.991835066 | -4.903633287 |
| **A_52_P485450** | Vta1 | 0.08237082 | -0.124200825 | 0.288942465 | 9.75375922 | 0.953041504 | 0.373913605 | 0.991835066 | -4.903646852 |
| **A_55_P2454896** | Amn1 | -0.053185348 | -0.186566201 | 0.080195506 | 9.864117933 | -0.953032351 | 0.373917915 | 0.991835066 | -4.903653102 |
| **A_51_P114693** | Parm1 | 0.053127993 | -0.08011509 | 0.186371077 | 12.87308769 | 0.952988962 | 0.373938351 | 0.991835066 | -4.89002285 |
| **A_66_P133328** | Tpd52 | 0.081059984 | -0.122236624 | 0.284356592 | 11.1878832 | 0.952983791 | 0.373940787 | 0.991835066 | -4.903686264 |
| **A_55_P1994913** | Ppp2cb | 0.072955117 | -0.110088169 | 0.255998403 | 12.61180379 | 0.952601121 | 0.374121058 | 0.991835066 | -4.903947543 |
| **A_66_P111216** | 4632427E1 | 0.072674577 | -0.109670515 | 0.255019669 | 8.414642444 | 0.952571464 | 0.374135032 | 0.991835066 | -4.903967789 |
| **A_52_P504342** | Coa7 | -0.070930644 | -0.248913431 | 0.107052144 | 9.16407171 | -0.952500097 | 0.37416866 | 0.991835066 | -4.904016507 |
| **A_51_P139400** | Taf1 | 0.044164682 | -0.066657816 | 0.154987179 | 8.739768889 | 0.952480963 | 0.374177677 | 0.991835066 | -4.904029569 |
| **A_51_P251352** | Slc25a13 | -0.098829885 | -0.346865581 | 0.149205812 | 11.79212686 | -0.952320719 | 0.374253194 | 0.991835066 | -4.904138949 |
| **A_51_P435321** | Ap1s2 | -0.047013655 | -0.165006665 | 0.070979355 | 8.799250839 | -0.952306769 | 0.374259769 | 0.991835066 | -4.904148471 |
| **A_51_P154753** | Klc3 | 0.054629445 | -0.082486952 | 0.191745842 | 8.152256066 | 0.952240231 | 0.37429113 | 0.991835066 | -4.904193885 |
| **A_55_P1956083** | Gpr68 | 0.15065913 | -0.227517083 | 0.528835343 | 9.955083411 | 0.952160978 | 0.374328486 | 0.991835066 | -4.904247974 |
| **A_55_P2181386** | Adm2 | 0.055967452 | -0.084519175 | 0.196454078 | 7.707819936 | 0.952159449 | 0.374329207 | 0.991835066 | -4.904249017 |
| **A_51_P211968** | Lcmt1 | -0.068570264 | -0.240741669 | 0.10360114 | 10.07191856 | -0.951883795 | 0.374459161 | 0.991835066 | -4.904437124 |
| **A_51_P486618** | Akr1b10 | -0.084627912 | -0.297124035 | 0.127868211 | 13.18973931 | -0.951857082 | 0.374471756 | 0.991835066 | -4.90445535 |
| **A_55_P1981544** | Lactb | -0.071385115 | -0.25063858 | 0.10786835 | 9.890871397 | -0.95180775 | 0.374495018 | 0.991835066 | -4.90448901 |
| **A_55_P2111355** | Egfl7 | 0.173456469 | -0.262230233 | 0.609143171 | 13.92636196 | 0.95153623 | 0.374623066 | 0.991835066 | -4.904674248 |
| **A_51_P329332** | Slc19a2 | 0.079017773 | -0.119534399 | 0.277569945 | 9.514464739 | 0.951172476 | 0.374794664 | 0.991835066 | -4.904922352 |
| **A_55_P2129034** | Rheb | 0.104047939 | -0.157441917 | 0.365537795 | 11.19052652 | 0.951015881 | 0.374868555 | 0.991835066 | -4.905029141 |
| **A_55_P2024260** | Vps33b | 0.070108993 | -0.106091253 | 0.24630924 | 9.199719587 | 0.950990878 | 0.374880354 | 0.991835066 | -4.90504619 |
| **A_52_P466853** | Bmyc | 0.057068597 | -0.08635894 | 0.200496135 | 9.730168628 | 0.950985266 | 0.374883002 | 0.991835066 | -4.905050017 |
| **A_52_P8324** | Tmem178 | 0.062210674 | -0.094201138 | 0.218622487 | 8.522634857 | 0.950614626 | 0.375057942 | 0.991835066 | -4.905302717 |
| **A_66_P127567** | Nxf1 | 0.082366464 | -0.124787882 | 0.28952081 | 12.50884875 | 0.950310453 | 0.375201556 | 0.991835066 | -4.90551005 |
| **A_52_P279687** | Lrp8 | 0.040296442 | -0.061074568 | 0.141667451 | 7.64247713 | 0.950084124 | 0.375308444 | 0.991835066 | -4.905664292 |
| **A_65_P16952** | Ywhaq | -0.069365482 | -0.243872395 | 0.105141431 | 12.19991473 | -0.95003566 | 0.375331335 | 0.991835066 | -4.905697316 |
| **A_55_P1959379** | BC080695 | 0.048427503 | -0.073404877 | 0.170259883 | 8.132751467 | 0.950032543 | 0.375332807 | 0.991835066 | -4.90569944 |
| **A_55_P2035662** | Aebp1 | 0.187363774 | -0.284071734 | 0.658799283 | 12.73343102 | 0.949888147 | 0.375401016 | 0.991835066 | -4.905797829 |
| **A_55_P2025675** | Dnah8 | 0.051580431 | -0.078218132 | 0.181378994 | 8.334981878 | 0.949782617 | 0.375450872 | 0.991835066 | -4.905869728 |
| **A_51_P412835** | Daxx | 0.047392175 | -0.071923388 | 0.166707738 | 11.35390728 | 0.949333219 | 0.375663239 | 0.991835066 | -4.906175848 |
| **A_55_P2102574** | Prorsd1 | 0.058205095 | -0.08836063 | 0.204770819 | 9.96919992 | 0.949156254 | 0.37574689 | 0.991835066 | -4.906296365 |
| **A_51_P508349** | Cdc16 | 0.0833141 | -0.126482003 | 0.293110202 | 12.48081339 | 0.949139879 | 0.375754631 | 0.991835066 | -4.906307516 |

| | | | | | | | | | |
|---|---|---|---|---|---|---|---|---|---|
| **A_55_P1990066** | NA | -0.070034793 | -0.24639897 | 0.106329385 | 9.313045047 | -0.949101376 | 0.375772834 | 0.991835066 | -4.906333735 |
| **A_55_P2078516** | NA | 0.080628978 | -0.122484256 | 0.283742212 | 9.362592897 | 0.948772457 | 0.37592836 | 0.991835066 | -4.906557686 |
| **A_51_P176086** | Ffar2 | -0.043174482 | -0.151941124 | 0.065592159 | 7.782906772 | -0.948725454 | 0.375950589 | 0.991835066 | -4.906589685 |
| **A_55_P2028977** | Tcf20 | 0.064532192 | -0.098042961 | 0.227107345 | 8.162862402 | 0.948705408 | 0.37596007 | 0.991835066 | -4.906603331 |
| **A_51_P123879** | Tecrl | -0.063079582 | -0.22201181 | 0.095852646 | 10.38853296 | -0.948606226 | 0.37600698 | 0.991835066 | -4.906670848 |
| **A_51_P491667** | Derl3 | 0.052845846 | -0.080310889 | 0.186002581 | 7.905253179 | 0.948542619 | 0.376037066 | 0.991835066 | -4.906714144 |
| **A_52_P144925** | Taf1d | 0.077150117 | -0.117311961 | 0.271612194 | 9.413144498 | 0.948223684 | 0.376187952 | 0.991835066 | -4.90693121 |
| **A_55_P1965989** | Zfp329 | 0.092865758 | -0.141259088 | 0.326990603 | 9.234272424 | 0.948019423 | 0.37628461 | 0.991835066 | -4.907070202 |
| **A_55_P1955726** | Ptpn5 | 0.039660974 | -0.060338323 | 0.139660271 | 7.650601703 | 0.947928449 | 0.376327666 | 0.991835066 | -4.9071321 |
| **A_51_P208377** | Trappc5 | -0.094966789 | -0.334418596 | 0.144485018 | 11.78702305 | -0.947900535 | 0.376340878 | 0.991835066 | -4.907151092 |
| **A_51_P249930** | Tceal1 | -0.068202469 | -0.240177755 | 0.103772817 | 9.679358846 | -0.947857799 | 0.376361106 | 0.991835066 | -4.907180167 |
| **A_55_P1976207** | Arf5 | -0.086534941 | -0.304748818 | 0.131678936 | 11.98915399 | -0.947803413 | 0.376386849 | 0.991835066 | -4.907217167 |
| **A_52_P17098** | Mpp7 | 0.044504742 | -0.067722838 | 0.156732322 | 8.635637869 | 0.947798079 | 0.376389374 | 0.991835066 | -4.907220796 |
| **A_52_P141112** | Csnk1g3 | -0.059106898 | -0.208173163 | 0.089959368 | 8.229968039 | -0.947693551 | 0.376438856 | 0.991835066 | -4.907291904 |
| **A_55_P2106800** | Bglap | -0.057851547 | -0.203758498 | 0.088055404 | 7.7003962 | -0.947650344 | 0.376459311 | 0.991835066 | -4.907321295 |
| **A_66_P100780** | Katnbl1 | 0.064705323 | -0.098501205 | 0.227911851 | 9.665140364 | 0.947570688 | 0.376497023 | 0.991835066 | -4.907375478 |
| **A_55_P1999269** | Grk2 | 0.093129144 | -0.141800696 | 0.328058984 | 11.33964222 | 0.947450569 | 0.376553898 | 0.991835066 | -4.907457178 |
| **A_55_P2128853** | Ear12 | -0.04118535 | -0.145097054 | 0.062726354 | 7.708066909 | -0.947299764 | 0.376625312 | 0.991835066 | -4.90755974 |
| **A_55_P2061319** | Mthfs | -0.051058526 | -0.179893643 | 0.07777659 | 8.272444161 | -0.947203194 | 0.376671048 | 0.991835066 | -4.90762541 |
| **A_55_P2179069** | NA | -0.061237415 | -0.215762336 | 0.093287507 | 8.199997116 | -0.947168942 | 0.376687271 | 0.991835066 | -4.907648702 |
| **A_52_P129428** | Fzd2 | 0.046016081 | -0.070124014 | 0.162156176 | 8.888132341 | 0.946970793 | 0.376781132 | 0.991835066 | -4.907783431 |
| **A_55_P2003129** | Nemp1 | 0.056329426 | -0.085842705 | 0.198501557 | 8.346880698 | 0.9469564 | 0.376787951 | 0.991835066 | -4.907793217 |
| **A_51_P518246** | Fibp | -0.083788641 | -0.29526674 | 0.127689458 | 10.49323679 | -0.946953995 | 0.37678909 | 0.991835066 | -4.907794853 |
| **A_52_P131836** | Bysl | 0.104640433 | -0.159497366 | 0.368778232 | 11.19715898 | 0.946843291 | 0.376841538 | 0.991835066 | -4.907870116 |
| **A_66_P123294** | NA | -0.074627097 | -0.263010086 | 0.113755891 | 8.462865176 | -0.946812535 | 0.37685611 | 0.991835066 | -4.907891024 |
| **A_51_P402193** | Map3k1 | 0.097415849 | -0.148588143 | 0.343419841 | 11.0106675 | 0.94644765 | 0.377029025 | 0.991835066 | -4.908139045 |
| **A_55_P2153067** | 2610028H2 | 0.074203283 | -0.113184424 | 0.261590989 | 17.12910732 | 0.946435789 | 0.377034647 | 0.991835066 | -4.908147106 |
| **A_55_P2102002** | Noc3l | 0.096638769 | -0.147430686 | 0.340708225 | 9.132995393 | 0.946339767 | 0.377080161 | 0.991835066 | -4.908212362 |
| **A_55_P2009953** | Cd300lg | -0.166307854 | -0.58644934 | 0.253833631 | 10.27948317 | -0.946076656 | 0.377204898 | 0.991835066 | -4.90839115 |
| **A_55_P2023484** | Osbpl5 | -0.045278315 | -0.159665084 | 0.069108453 | 12.57371688 | -0.946070726 | 0.377207709 | 0.991835066 | -4.908395179 |
| **A_51_P510891** | Afp | -0.73759958 | -2.601276158 | 1.126076999 | 11.34553108 | -0.945930537 | 0.377274184 | 0.991835066 | -4.908490425 |
| **A_52_P227445** | Parp2 | -0.052698383 | -0.185850947 | 0.080454182 | 8.648109995 | -0.945925396 | 0.377276621 | 0.991835066 | -4.908493917 |
| **A_66_P101407** | Acat3 | 0.043926943 | -0.067063451 | 0.154917338 | 7.889998427 | 0.945920679 | 0.377278858 | 0.991835066 | -4.908497122 |
| **A_52_P329917** | Prrg2 | -0.057449469 | -0.202612717 | 0.087713779 | 9.860747031 | -0.945885292 | 0.377295639 | 0.991835066 | -4.908521162 |
| **A_55_P2079064** | NA | 0.168816487 | -0.257759473 | 0.595392448 | 9.269245764 | 0.945861645 | 0.377306854 | 0.991835066 | -4.908537227 |
| **A_55_P2032272** | Habp2 | -0.048306432 | -0.170372639 | 0.073759776 | 7.83068871 | -0.945842089 | 0.377316128 | 0.991835066 | -4.908550512 |
| **A_51_P191782** | Olfml3 | -0.092090536 | -0.324802155 | 0.140621083 | 9.432131687 | -0.945814713 | 0.377329111 | 0.991835066 | -4.90856911 |
| **A_52_P255849** | NA | 0.05144007 | -0.078550083 | 0.181430223 | 8.416611862 | 0.945802006 | 0.377335138 | 0.991835066 | -4.908577742 |
| **A_55_P1958133** | NA | -0.188652966 | -0.665403511 | 0.288097578 | 11.44331659 | -0.945761373 | 0.377354409 | 0.991835066 | -4.908605343 |
| **A_55_P2087013** | Hipk1 | 0.046541528 | -0.071079816 | 0.164162871 | 8.018136312 | 0.945722298 | 0.377372942 | 0.991835066 | -4.908631886 |
| **A_52_P640386** | Usp53 | 0.04088821 | -0.062453763 | 0.144230183 | 7.750151855 | 0.945650119 | 0.377407177 | 0.991835066 | -4.908680914 |
| **A_66_P107255** | Nom1 | -0.044418635 | -0.156690565 | 0.067853295 | 8.864394152 | -0.945590618 | 0.377435402 | 0.991835066 | -4.908721328 |
| **A_55_P2028243** | Dnajc19 | -0.107166421 | -0.37805947 | 0.163726627 | 10.35800082 | -0.945518434 | 0.377469644 | 0.991835066 | -4.908770355 |
| **A_55_P1978047** | Aven | -0.063519248 | -0.224104396 | 0.0970659 | 8.385248886 | -0.945385864 | 0.377532539 | 0.991835066 | -4.908860388 |
| **A_55_P2125743** | NA | 0.059163612 | -0.09042445 | 0.208751674 | 16.18477137 | 0.945293949 | 0.37757615 | 0.991835066 | -4.908922805 |
| **A_51_P286748** | Frzb | -0.05460496 | -0.192675314 | 0.083465394 | 8.454584292 | -0.945237162 | 0.377603096 | 0.991835066 | -4.908961367 |
| **A_55_P2057792** | NA | 0.08956937 | -0.136910735 | 0.316049474 | 12.07184022 | 0.945232388 | 0.377605362 | 0.991835066 | -4.908964609 |
| **A_55_P2030524** | Vldlr | -0.078243099 | -0.276089095 | 0.119602898 | 13.17865027 | -0.945209131 | 0.377616398 | 0.991835066 | -4.9089804 |

| | | | | | | | | | |
|---|---|---|---|---|---|---|---|---|---|
| **A_55_P2415197** | 2810013P0 | -0.056293351 | -0.198694152 | 0.086107451 | 8.854263569 | -0.944830265 | 0.377796216 | 0.991835066 | -4.90923762 |
| **A_55_P1982872** | Naa10 | -0.102633814 | -0.362267446 | 0.156999818 | 11.51179446 | -0.944797317 | 0.377811857 | 0.991835066 | -4.909259986 |
| **A_66_P106133** | Magi3 | 0.066252839 | -0.101354741 | 0.233860419 | 9.925843011 | 0.944756683 | 0.377831147 | 0.991835066 | -4.909287568 |
| **A_52_P279184** | Srp54a | 0.067509497 | -0.103312505 | 0.238331499 | 9.231857277 | 0.944561411 | 0.37792386 | 0.991835066 | -4.909420108 |
| **A_52_P630867** | Abcc4 | 0.064283979 | -0.09838904 | 0.226956998 | 8.778877941 | 0.944487803 | 0.377958812 | 0.991835066 | -4.909470064 |
| **A_66_P102237** | Nemf | 0.063372776 | -0.096995238 | 0.22374079 | 10.79785384 | 0.94448293 | 0.377961126 | 0.991835066 | -4.90947337 |
| **A_55_P2113081** | Tmem132a | 0.05146117 | -0.078765947 | 0.181688287 | 13.4241349 | 0.94446824 | 0.377968102 | 0.991835066 | -4.90948334 |
| **A_52_P437084** | NA | 0.089276248 | -0.136657056 | 0.315209553 | 14.71765226 | 0.944419203 | 0.377991389 | 0.991835066 | -4.909516618 |
| **A_66_P134542** | Anln | -0.102527297 | -0.362111735 | 0.15705714 | 12.25755584 | -0.943995642 | 0.378192577 | 0.991835066 | -4.909804011 |
| **A_55_P2129348** | Pilra | -0.058134082 | -0.20532222 | 0.089054056 | 8.364155553 | -0.943989444 | 0.378195521 | 0.991835066 | -4.909808216 |
| **A_51_P500424** | Dnttip1 | -0.049688869 | -0.175510262 | 0.076132523 | 11.7217183 | -0.943873424 | 0.378250644 | 0.991835066 | -4.909886921 |
| **A_55_P1998066** | Tmem121 | 0.070689392 | -0.108345861 | 0.249724645 | 8.318613881 | 0.943680161 | 0.37834248 | 0.991835066 | -4.910018012 |
| **A_55_P1992054** | Armc10 | 0.064120385 | -0.098293292 | 0.226534062 | 11.31099256 | 0.943588533 | 0.378386027 | 0.991835066 | -4.910080156 |
| **A_55_P1975989** | Wdr63 | 0.043998772 | -0.06745122 | 0.155448764 | 7.709867241 | 0.943560269 | 0.37839946 | 0.991835066 | -4.910099324 |
| **A_51_P463765** | Timp3 | -0.060087496 | -0.212304501 | 0.09212951 | 12.86263291 | -0.943474276 | 0.378440333 | 0.991835066 | -4.910157643 |
| **A_51_P371091** | Rcsd1 | -0.059021407 | -0.208539598 | 0.090496784 | 12.74901654 | -0.943462526 | 0.378445918 | 0.991835066 | -4.910165611 |
| **A_55_P2027698** | Ndufa5 | -0.067081541 | -0.237023872 | 0.102860791 | 16.54670598 | -0.94343197 | 0.378460442 | 0.991835066 | -4.910186332 |
| **A_51_P242603** | E430018J2 | -0.060684985 | -0.21442418 | 0.09305421 | 8.513898456 | -0.943421521 | 0.378465409 | 0.991835066 | -4.910193418 |
| **A_55_P1980721** | NA | -0.10634504 | -0.375762683 | 0.163072603 | 14.27496601 | -0.943409706 | 0.378471026 | 0.991835066 | -4.91020143 |
| **A_51_P144783** | Cct5 | 0.047468458 | -0.072814748 | 0.167751664 | 14.86756855 | 0.943211878 | 0.378565073 | 0.991835066 | -4.91033557 |
| **A_52_P223571** | Mlec | 0.050608773 | -0.077631994 | 0.178849541 | 8.514558052 | 0.943210951 | 0.378565514 | 0.991835066 | -4.910336199 |
| **A_55_P2159869** | Zfp59 | -0.03980016 | -0.140665387 | 0.061065067 | 9.124525102 | -0.943088569 | 0.378623703 | 0.991835066 | -4.910419172 |
| **A_51_P153170** | Cyb5r3 | -0.128732431 | -0.454986265 | 0.197521404 | 11.47444061 | -0.943064674 | 0.378635065 | 0.991835066 | -4.910435372 |
| **A_51_P497061** | Ppp2r2d | 0.051072305 | -0.078365475 | 0.180510085 | 12.16656126 | 0.943047432 | 0.378643264 | 0.991835066 | -4.910447061 |
| **A_52_P322535** | Mplkip | 0.046471146 | -0.071314233 | 0.164256525 | 8.277412403 | 0.942977066 | 0.378676725 | 0.991835066 | -4.910494763 |
| **A_51_P468762** | Alkbh6 | 0.0442851 | -0.067977185 | 0.156547385 | 11.22282172 | 0.942828898 | 0.378747192 | 0.991835066 | -4.910595201 |
| **A_55_P2181419** | Tcf4 | 0.044047918 | -0.067634666 | 0.155730501 | 8.580719798 | 0.942646944 | 0.37883374 | 0.991835066 | -4.910718527 |
| **A_55_P2261882** | D230040J2 | 0.070285742 | -0.107931325 | 0.248502809 | 8.354040258 | 0.94259921 | 0.378856448 | 0.991835066 | -4.910750877 |
| **A_55_P2180425** | Uqcc1 | -0.097298512 | -0.344037929 | 0.149440904 | 10.4275345 | -0.942490103 | 0.378908355 | 0.991835066 | -4.910824817 |
| **A_66_P124052** | Bok | 0.115686591 | -0.177810747 | 0.409183929 | 10.1222088 | 0.942080443 | 0.379103298 | 0.991835066 | -4.911102384 |
| **A_55_P2089035** | Pole | -0.04716664 | -0.1668291 | 0.072495819 | 8.047914356 | -0.942076453 | 0.379105198 | 0.991835066 | -4.911105087 |
| **A_51_P419246** | Mirt1 | -0.043271586 | -0.153055017 | 0.066511845 | 7.781163607 | -0.942052582 | 0.379116559 | 0.991835066 | -4.911121258 |
| **A_51_P159673** | Snx1 | -0.045895986 | -0.162400112 | 0.070608139 | 12.13514359 | -0.941548151 | 0.379356712 | 0.991835066 | -4.911462916 |
| **A_55_P2020463** | Rufy2 | 0.096471125 | -0.148464593 | 0.341406844 | 9.757062772 | 0.941356997 | 0.379447748 | 0.991835066 | -4.911592353 |
| **A_52_P27271** | Dhdds | 0.05676814 | -0.087398756 | 0.200935035 | 9.564102227 | 0.941127031 | 0.37955729 | 0.991835066 | -4.911748048 |
| **A_51_P315646** | Nfs1 | -0.062965907 | -0.222918187 | 0.096986373 | 11.40256497 | -0.940858167 | 0.379685392 | 0.991835066 | -4.911930044 |
| **A_55_P1961089** | Npb | 0.066189503 | -0.101973394 | 0.2343524 | 10.1619172 | 0.940736669 | 0.379743291 | 0.991835066 | -4.912012275 |
| **A_51_P257684** | Stau1 | -0.066131291 | -0.234147638 | 0.101885056 | 9.028833454 | -0.940729139 | 0.37974688 | 0.991835066 | -4.912017372 |
| **A_55_P1991911** | NA | -0.123493706 | -0.437265837 | 0.190278426 | 13.74266629 | -0.940674965 | 0.379772698 | 0.991835066 | -4.912054034 |
| **A_51_P185259** | Gnb5 | -0.068370872 | -0.242098717 | 0.105356974 | 8.433991738 | -0.940612653 | 0.379802397 | 0.991835066 | -4.912096203 |
| **A_55_P1987326** | Prl3a1 | -0.385869974 | -1.366426004 | 0.594686056 | 8.294958845 | -0.940540938 | 0.379836579 | 0.991835066 | -4.912144732 |
| **A_55_P2088535** | Snrpd2 | 0.052483405 | -0.080885639 | 0.185852448 | 14.29482175 | 0.940537457 | 0.379838238 | 0.991835066 | -4.912147088 |
| **A_51_P171107** | Tmem35a | -0.197178321 | -0.698248823 | 0.30389218 | 10.93061346 | -0.940523089 | 0.379845087 | 0.991835066 | -4.912156811 |
| **A_51_P397925** | Smap2 | -0.088319383 | -0.312765984 | 0.136127217 | 9.305174735 | -0.940485555 | 0.379862979 | 0.991835066 | -4.912182208 |
| **A_52_P481182** | Stard5 | -0.045339077 | -0.160562839 | 0.069884686 | 8.170604774 | -0.940458747 | 0.379875758 | 0.991835066 | -4.912200348 |
| **A_55_P2092869** | Anxa9 | 0.053038714 | -0.081813267 | 0.187890695 | 7.942389173 | 0.94003665 | 0.380077011 | 0.991835066 | -4.912485911 |
| **A_51_P165087** | Snai1 | -0.067718621 | -0.239917842 | 0.104480601 | 9.215090764 | -0.939909526 | 0.380137639 | 0.991835066 | -4.912571897 |
| **A_55_P1962274** | Ccnt1 | 0.052476836 | -0.080966087 | 0.185919759 | 9.004823234 | 0.939899084 | 0.380142619 | 0.991835066 | -4.91257896 |

| | | | | | | | | | |
|---|---|---|---|---|---|---|---|---|---|
| **A_66_P108468** | 2610528A1 | -0.054479024 | -0.193019454 | 0.084061405 | 8.216746879 | -0.93985731 | 0.380162543 | 0.991835066 | -4.912607214 |
| **A_51_P150430** | Mkrn2 | -0.09726746 | -0.344654148 | 0.150119228 | 9.681518038 | -0.939724135 | 0.380226068 | 0.991835066 | -4.912697281 |
| **A_55_P2045770** | Gmfb | 0.072639984 | -0.112110524 | 0.257390493 | 12.19143753 | 0.939721648 | 0.380227254 | 0.991835066 | -4.912698963 |
| **A_51_P438692** | Vti1b | 0.04445137 | -0.068611281 | 0.157514021 | 12.77859326 | 0.939669478 | 0.380252142 | 0.991835066 | -4.912734243 |
| **A_55_P1965655** | Mtmr14 | -0.055712315 | -0.197420057 | 0.085995428 | 8.51056105 | -0.939651378 | 0.380260777 | 0.991835066 | -4.912746483 |
| **A_55_P2173982** | Rrm2 | 0.062272749 | -0.096144905 | 0.220690402 | 13.65520265 | 0.939514728 | 0.380325972 | 0.991835066 | -4.912838886 |
| **A_51_P203878** | Dynll2 | -0.088031086 | -0.31203118 | 0.135969009 | 11.3324199 | -0.93928414 | 0.380436004 | 0.991835066 | -4.912994788 |
| **A_51_P352283** | Cyb561d1 | 0.068133694 | -0.105254081 | 0.241521469 | 9.030102807 | 0.93918813 | 0.380481825 | 0.991835066 | -4.913059694 |
| **A_66_P132510** | Scamp3 | 0.043330496 | -0.066938339 | 0.153599331 | 12.68507154 | 0.939182537 | 0.380484494 | 0.991835066 | -4.913063474 |
| **A_52_P76402** | Ago1 | 0.045095971 | -0.069680127 | 0.159872069 | 8.864163765 | 0.939064487 | 0.38054084 | 0.991835066 | -4.913143273 |
| **A_55_P1979605** | Eif3m | 0.044879463 | -0.069351697 | 0.159110624 | 12.93094777 | 0.939014286 | 0.380564803 | 0.991835066 | -4.913177205 |
| **A_55_P2131173** | Sphk2 | -0.064611869 | -0.229072536 | 0.099848798 | 10.70275989 | -0.938986593 | 0.380578023 | 0.991835066 | -4.913195923 |
| **A_55_P2109932** | Tmem63b | 0.057671733 | -0.093018174 | 0.20836164 | 12.21001312 | 0.950705456 | 0.380626281 | 0.991835066 | -4.845931802 |
| **A_55_P1998631** | Capg | 0.054427387 | -0.084126865 | 0.192981639 | 11.13807257 | 0.9388728 | 0.380632347 | 0.991835066 | -4.913272832 |
| **A_66_P128694** | NA | -0.101328126 | -0.359291761 | 0.156635509 | 9.231770565 | -0.93881637 | 0.380659288 | 0.991835066 | -4.913310969 |
| **A_51_P481569** | Ddx42 | 0.060816105 | -0.094022942 | 0.215655152 | 11.43242966 | 0.938744154 | 0.380693768 | 0.991835066 | -4.913359773 |
| **A_51_P365694** | Rala | 0.081918902 | -0.126648909 | 0.290486713 | 12.42412206 | 0.938741404 | 0.380695082 | 0.991835066 | -4.913361631 |
| **A_51_P217047** | Gng10 | -0.081136391 | -0.287750769 | 0.125477987 | 9.805832329 | -0.938564831 | 0.380779398 | 0.991835066 | -4.913480948 |
| **A_55_P2038318** | NA | -0.039056257 | -0.138535905 | 0.060423392 | 7.694120697 | -0.938351402 | 0.380881334 | 0.991835066 | -4.913625148 |
| **A_55_P2153141** | NA | 0.068294182 | -0.105662837 | 0.242251202 | 11.47250133 | 0.938319809 | 0.380896425 | 0.991835066 | -4.913646492 |
| **A_55_P2008681** | Ktn1 | -0.071462572 | -0.253505016 | 0.110579871 | 12.63277054 | -0.938242538 | 0.380933337 | 0.991835066 | -4.913698692 |
| **A_55_P2146560** | H2-Ab1 | 0.328642268 | -0.508624751 | 1.165909288 | 8.819525008 | 0.938141973 | 0.380981379 | 0.991835066 | -4.913766623 |
| **A_55_P2133195** | Gm4951 | -0.046217533 | -0.163978384 | 0.071543318 | 7.771508918 | -0.938026175 | 0.381036705 | 0.991835066 | -4.91384484 |
| **A_55_P1952547** | Sh3rf1 | 0.048042065 | -0.074381169 | 0.170465299 | 8.167640645 | 0.937922488 | 0.38108625 | 0.991835066 | -4.913914869 |
| **A_52_P101399** | Tmem130 | 0.090857139 | -0.140842272 | 0.322556551 | 8.336966449 | 0.937223695 | 0.38142028 | 0.991835066 | -4.914386687 |
| **A_52_P671661** | Mak16 | 0.056981461 | -0.088345093 | 0.202308015 | 8.679747055 | 0.937125456 | 0.381467257 | 0.991835066 | -4.914452998 |
| **A_55_P2143000** | NA | -0.046964713 | -0.166746295 | 0.07281687 | 8.012430037 | -0.937110394 | 0.38147446 | 0.991835066 | -4.914463165 |
| **A_55_P2015541** | Hif3a | 0.067635031 | -0.104873549 | 0.24014361 | 12.68159723 | 0.937065878 | 0.381495749 | 0.991835066 | -4.91449321 |
| **A_55_P2085425** | Ophn1 | 0.06645945 | -0.103087684 | 0.236006584 | 8.92400811 | 0.936861562 | 0.381593471 | 0.991835066 | -4.9146311 |
| **A_55_P2099363** | Stac2 | 0.051715685 | -0.080245423 | 0.183676794 | 8.107065715 | 0.936667524 | 0.381686295 | 0.991835066 | -4.914762034 |
| **A_51_P500541** | Flywch2 | -0.045267018 | -0.160776851 | 0.070242816 | 8.381154937 | -0.936638608 | 0.381700129 | 0.991835066 | -4.914781544 |
| **A_55_P2074499** | Kcp | -0.068224458 | -0.242357526 | 0.10590861 | 9.675559198 | -0.936414166 | 0.381807522 | 0.991835066 | -4.914932967 |
| **A_55_P1970642** | NA | -0.047510066 | -0.168774408 | 0.073754277 | 9.290088031 | -0.936400523 | 0.381814051 | 0.991835066 | -4.914942171 |
| **A_52_P496260** | Exosc8 | -0.08223183 | -0.292185878 | 0.127722218 | 11.00123127 | -0.93610558 | 0.381955214 | 0.991835066 | -4.915141118 |
| **A_51_P241769** | Rhd | -0.108718456 | -0.386338574 | 0.168901661 | 10.06063626 | -0.935969022 | 0.382020586 | 0.991835066 | -4.915233215 |
| **A_55_P2009427** | Nrsn2 | -0.071606836 | -0.254487537 | 0.111273864 | 9.369502222 | -0.935827371 | 0.382088404 | 0.991835066 | -4.915328737 |
| **A_51_P163106** | Bdh1 | 0.130607231 | -0.202982652 | 0.464197113 | 9.391844831 | 0.935757859 | 0.382121688 | 0.991835066 | -4.915375608 |
| **A_52_P287683** | Pik3c3 | -0.053642286 | -0.190701581 | 0.08341701 | 8.316500624 | -0.935422713 | 0.382282194 | 0.991835066 | -4.915601561 |
| **A_51_P340668** | Bcl9l | 0.091655574 | -0.142537532 | 0.32584868 | 8.291467345 | 0.935392552 | 0.382296641 | 0.991835066 | -4.915621893 |
| **A_55_P2136522** | Tmem128 | -0.04830545 | -0.171733557 | 0.075122656 | 10.47985106 | -0.935386723 | 0.382299434 | 0.991835066 | -4.915625822 |
| **A_52_P515497** | Atp1a1 | 0.082518813 | -0.12833783 | 0.293375456 | 15.11698215 | 0.935351433 | 0.382316338 | 0.991835066 | -4.91564961 |
| **A_55_P1990500** | Masp2 | 0.041932897 | -0.065226019 | 0.149091812 | 7.698962663 | 0.935267166 | 0.382356706 | 0.991835066 | -4.915706411 |
| **A_66_P104527** | Tbl1x | 0.074521322 | -0.115917902 | 0.264960545 | 11.64051256 | 0.935261979 | 0.38235919 | 0.991835066 | -4.915709907 |
| **A_51_P398260** | Tppp | -0.067047457 | -0.23839059 | 0.104295676 | 8.704456925 | -0.935243586 | 0.382368002 | 0.991835066 | -4.915722304 |
| **A_55_P2009993** | Tomm20 | -0.076494108 | -0.271981237 | 0.11899302 | 12.08701305 | -0.935231123 | 0.382373972 | 0.991835066 | -4.915730704 |
| **A_51_P438619** | Sobp | -0.102767641 | -0.365401604 | 0.159866322 | 11.12394629 | -0.935221822 | 0.382378429 | 0.991835066 | -4.915736974 |
| **A_52_P29151** | Ankrd44 | 0.089696958 | -0.139592903 | 0.318986819 | 9.082693662 | 0.934979302 | 0.382494629 | 0.991835066 | -4.915900419 |
| **A_55_P2003473** | Zc4h2 | 0.049356956 | -0.076821727 | 0.175535638 | 8.398372365 | 0.934913665 | 0.382526082 | 0.991835066 | -4.915944649 |

| | | | | | | | | | |
|---|---|---|---|---|---|---|---|---|---|
| **A_55_P2093679** | Zfp599 | -0.05566758 | -0.197991779 | 0.086656618 | 8.513051105 | -0.934830193 | 0.382566086 | 0.991835066 | -4.916000895 |
| **A_51_P386983** | Tnni3 | -0.075724383 | -0.26935399 | 0.117905224 | 16.12036639 | -0.934701868 | 0.382627591 | 0.991835066 | -4.916087358 |
| **A_51_P145511** | Prl2a1 | -0.366203345 | -1.302909643 | 0.570502952 | 8.166097838 | -0.934389608 | 0.382777285 | 0.991835066 | -4.916297717 |
| **A_55_P2121142** | Tbc1d22b | -0.039444919 | -0.140342556 | 0.061452719 | 9.595600455 | -0.934370681 | 0.38278636 | 0.991835066 | -4.916310466 |
| **A_51_P117995** | Pfkm | -0.08053072 | -0.286541905 | 0.125480466 | 13.60955917 | -0.934286148 | 0.382826892 | 0.991835066 | -4.916367403 |
| **A_55_P2054420** | Tcf4 | -0.062942346 | -0.223978398 | 0.098093705 | 12.57768908 | -0.934176516 | 0.382879465 | 0.991835066 | -4.916441241 |
| **A_55_P2177023** | Mafg | 0.06118884 | -0.09536492 | 0.2177426 | 9.433038951 | 0.93415265 | 0.38289091 | 0.991835066 | -4.916457315 |
| **A_55_P2088965** | Scarb1 | -0.048723515 | -0.173386109 | 0.075939079 | 8.068448296 | -0.934139167 | 0.382897376 | 0.991835066 | -4.916466394 |
| **A_51_P140803** | Slco1b2 | 0.041284923 | -0.064345657 | 0.146915503 | 7.670692471 | 0.93413781 | 0.382898027 | 0.991835066 | -4.916467309 |
| **A_55_P1962466** | NA | 0.046276535 | -0.072151682 | 0.164704752 | 16.81504408 | 0.933930965 | 0.382997234 | 0.991835066 | -4.916606599 |
| **A_55_P2186210** | NA | 0.070053829 | -0.109226366 | 0.249334025 | 13.99455017 | 0.93391789 | 0.383003506 | 0.991835066 | -4.916615403 |
| **A_55_P2038489** | NA | 0.117171281 | -0.182708252 | 0.417050813 | 8.201946967 | 0.933863678 | 0.383029511 | 0.991835066 | -4.916651906 |
| **A_55_P2251181** | 2900076A1 | 0.074077568 | -0.115523922 | 0.263679058 | 8.211085709 | 0.933800502 | 0.383059818 | 0.991835066 | -4.916694443 |
| **A_51_P201721** | Crtap | 0.047810805 | -0.074571021 | 0.170192631 | 12.35199127 | 0.933723436 | 0.383096789 | 0.991835066 | -4.916746329 |
| **A_55_P2118103** | Zfp68 | -0.059502989 | -0.211862218 | 0.09285624 | 9.155639256 | -0.933424399 | 0.383240277 | 0.991835066 | -4.916947633 |
| **A_55_P2336558** | Chn2 | -0.040122201 | -0.142857447 | 0.062613045 | 8.114561396 | -0.933414227 | 0.383245159 | 0.991835066 | -4.91695448 |
| **A_55_P1990894** | Psmd4 | 0.041669577 | -0.065038029 | 0.148377183 | 14.09346443 | 0.933324884 | 0.383288037 | 0.991835066 | -4.917014615 |
| **A_55_P2127884** | Btbd1 | 0.068507085 | -0.106937235 | 0.243951406 | 8.945849695 | 0.933265709 | 0.383316438 | 0.991835066 | -4.917054441 |
| **A_55_P2171373** | Eloc | 0.067205899 | -0.104912165 | 0.239323962 | 13.33479891 | 0.933232971 | 0.383332152 | 0.991835066 | -4.917076474 |
| **A_55_P2031021** | Bex1 | 0.155005349 | -0.241972487 | 0.551983184 | 14.38265818 | 0.933231859 | 0.383332686 | 0.991835066 | -4.917077222 |
| **A_51_P483013** | Arel1 | -0.041827954 | -0.148955323 | 0.065299415 | 10.92333265 | -0.933201252 | 0.383347377 | 0.991835066 | -4.917097821 |
| **A_55_P2143431** | Nfib | 0.057647332 | -0.090013389 | 0.205308053 | 12.85915065 | 0.933089623 | 0.383400962 | 0.991835066 | -4.917172942 |
| **A_55_P2165160** | Arhgef1 | 0.042954378 | -0.06707609 | 0.152984846 | 8.236321761 | 0.933047194 | 0.383421331 | 0.991835066 | -4.917201494 |
| **A_55_P2373852** | 2310058N2 | 0.067164374 | -0.10490855 | 0.239237297 | 8.607801338 | 0.932901011 | 0.383491514 | 0.991835066 | -4.917299856 |
| **A_51_P516870** | Itm2a | -0.107486592 | -0.382868553 | 0.167895369 | 13.04843448 | -0.932884631 | 0.383499379 | 0.991835066 | -4.917310877 |
| **A_55_P2166262** | Rnf215 | 0.090900967 | -0.142024235 | 0.323826169 | 10.66260921 | 0.932741188 | 0.383568257 | 0.991835066 | -4.917407385 |
| **A_55_P2063312** | Mgll | -0.156459818 | -0.557377252 | 0.244457617 | 9.193960426 | -0.932732285 | 0.383572532 | 0.991835066 | -4.917413373 |
| **A_52_P472660** | Smurf2 | -0.045236898 | -0.161209221 | 0.070735426 | 9.472497989 | -0.932282621 | 0.383788514 | 0.991835066 | -4.917715834 |
| **A_55_P2083426** | NA | -0.14778901 | -0.526928805 | 0.231350786 | 13.36457644 | -0.931648093 | 0.384093446 | 0.991835066 | -4.918142466 |
| **A_51_P321965** | Ssna1 | -0.072317932 | -0.257887486 | 0.113251621 | 11.08254575 | -0.931426118 | 0.384200163 | 0.991835066 | -4.918291665 |
| **A_55_P1971769** | NA | -0.045197901 | -0.16117711 | 0.070781308 | 14.96732779 | -0.931423644 | 0.384201353 | 0.991835066 | -4.918293328 |
| **A_51_P253547** | Ctsl | 0.092048306 | -0.144201518 | 0.328298131 | 13.47682283 | 0.931222456 | 0.384298096 | 0.991835066 | -4.918428534 |
| **A_66_P128586** | Cwc22 | 0.061901509 | -0.096981363 | 0.220784382 | 9.544001898 | 0.931179245 | 0.384318876 | 0.991835066 | -4.91845757 |
| **A_55_P2037186** | Alkbh2 | 0.074953195 | -0.117441322 | 0.267347711 | 9.672933897 | 0.931122007 | 0.384346404 | 0.991835066 | -4.918496031 |
| **A_55_P1982291** | Clca3a1 | 0.129827322 | -0.203544949 | 0.463199594 | 10.48332235 | 0.930777248 | 0.384512243 | 0.991835066 | -4.918727656 |
| **A_55_P2153031** | Ttn | -0.109533149 | -0.390801892 | 0.171735595 | 10.88535936 | -0.9307504 | 0.38452516 | 0.991835066 | -4.918745691 |
| **A_52_P271155** | Mtpap | 0.048308607 | -0.075744172 | 0.172361386 | 10.35081741 | 0.930737371 | 0.384531429 | 0.991835066 | -4.918754443 |
| **A_55_P2197847** | Lhx1os | -0.065863273 | -0.235015219 | 0.103288672 | 8.334296073 | -0.930626567 | 0.384584742 | 0.991835066 | -4.918828872 |
| **A_55_P2027496** | Adck5 | 0.087575138 | -0.137345266 | 0.312495541 | 11.35806716 | 0.930595838 | 0.384599528 | 0.991835066 | -4.918849511 |
| **A_55_P2024575** | Sec24c | 0.09515741 | -0.149238932 | 0.339553751 | 13.53802466 | 0.930587133 | 0.384603717 | 0.991835066 | -4.918855358 |
| **A_51_P418056** | Sc5d | 0.078328799 | -0.122914006 | 0.279571604 | 9.816422727 | 0.930272615 | 0.384755084 | 0.991835066 | -4.919066584 |
| **A_52_P264905** | Cab39l | -0.056531605 | -0.201789321 | 0.088726112 | 10.30492326 | -0.930167655 | 0.384805608 | 0.991835066 | -4.919137062 |
| **A_51_P116407** | Pald1 | 0.059267451 | -0.093024382 | 0.211559285 | 8.871216828 | 0.93014096 | 0.384818458 | 0.991835066 | -4.919154986 |
| **A_55_P2066116** | Bcl3 | -0.044684532 | -0.159543493 | 0.07017443 | 8.887924094 | -0.929825514 | 0.384970336 | 0.991835066 | -4.919366763 |
| **A_55_P1953728** | Nes | 0.080556544 | -0.126590313 | 0.287703401 | 11.21306514 | 0.929461937 | 0.385145443 | 0.991835066 | -4.91961079 |
| **A_52_P515094** | Tmem176a | -0.106269247 | -0.379560188 | 0.167021694 | 9.148886262 | -0.929376098 | 0.385186794 | 0.991835066 | -4.919668394 |
| **A_51_P442964** | Knl1 | 0.059664747 | -0.093789771 | 0.213119266 | 8.483004469 | 0.92928143 | 0.385232402 | 0.991835066 | -4.919731919 |
| **A_55_P2128646** | Gmnn | 0.105284887 | -0.165504583 | 0.376074358 | 10.36557157 | 0.929273163 | 0.385236385 | 0.991835066 | -4.919737465 |

| | | | | | | | | | |
|---|---|---|---|---|---|---|---|---|---|
| **A_55_P2132756** | Cacna1g | -0.135255946 | -0.483195453 | 0.212683561 | 9.259585203 | -0.92909849 | 0.385320548 | 0.991835066 | -4.919854663 |
| **A_55_P1952690** | Rpl12 | 0.090336463 | -0.142052877 | 0.322725803 | 14.73096189 | 0.929086206 | 0.385326467 | 0.991835066 | -4.919862904 |
| **A_52_P583155** | Ykt6 | -0.093043644 | -0.332410884 | 0.146323597 | 9.728586552 | -0.929032974 | 0.385352119 | 0.991835066 | -4.919898617 |
| **A_55_P2026475** | NA | 0.069955477 | -0.110019152 | 0.249930105 | 11.71107559 | 0.929008241 | 0.385364038 | 0.991835066 | -4.919915209 |
| **A_52_P195018** | Arap3 | 0.058204954 | -0.091567306 | 0.207977213 | 12.02420063 | 0.928833135 | 0.385448431 | 0.991835066 | -4.920032673 |
| **A_66_P118307** | NA | -0.118372983 | -0.422993453 | 0.186247488 | 14.36383577 | -0.928758152 | 0.385484574 | 0.991835066 | -4.920082969 |
| **A_55_P1996643** | Polr2j | 0.04062011 | -0.063967828 | 0.145208049 | 12.54944496 | 0.928257847 | 0.385725791 | 0.991835066 | -4.920418477 |
| **A_55_P2078670** | Cbs | 0.058213251 | -0.091699334 | 0.208125836 | 8.276249262 | 0.928095983 | 0.385803857 | 0.991835066 | -4.920526997 |
| **A_55_P2185148** | Mettl7a3 | -0.049389281 | -0.176589748 | 0.077811186 | 8.529189668 | -0.928011016 | 0.385844841 | 0.991835066 | -4.920583957 |
| **A_51_P349961** | Gc | -0.091483259 | -0.327122159 | 0.14415564 | 7.953242448 | -0.927905557 | 0.385895713 | 0.991835066 | -4.920654648 |
| **A_51_P175771** | Flii | -0.06554723 | -0.234403504 | 0.103309044 | 11.84097509 | -0.92778271 | 0.38595498 | 0.991835066 | -4.920736989 |
| **A_52_P647906** | NA | -0.038556121 | -0.137892791 | 0.060780549 | 12.23582041 | -0.927668635 | 0.386010021 | 0.991835066 | -4.920813444 |
| **A_51_P391926** | Ubxn6 | -0.064989377 | -0.232440909 | 0.102462154 | 12.36339026 | -0.927603512 | 0.386041446 | 0.991835066 | -4.920857087 |
| **A_51_P387123** | Oasl2 | -0.139194488 | -0.497918723 | 0.219529746 | 8.567844562 | -0.927407182 | 0.386136194 | 0.991835066 | -4.920988647 |
| **A_55_P2182159** | Ywhaq | 0.060932382 | -0.096100241 | 0.217965006 | 13.30178287 | 0.927400671 | 0.386139336 | 0.991835066 | -4.920993009 |
| **A_51_P114910** | Cstb | 0.05447965 | -0.085945755 | 0.194905055 | 11.21856055 | 0.927251948 | 0.386211122 | 0.991835066 | -4.921092654 |
| **A_51_P136792** | Calcoco1 | -0.09620059 | -0.344167333 | 0.151766152 | 8.819822757 | -0.92724272 | 0.386215576 | 0.991835066 | -4.921098837 |
| **A_52_P231737** | Actl6a | 0.060611294 | -0.09567408 | 0.216896667 | 10.08898841 | 0.926924478 | 0.386369221 | 0.991835066 | -4.921312022 |
| **A_51_P352264** | Sirt7 | 0.084550209 | -0.133473694 | 0.302574113 | 10.37544295 | 0.926871885 | 0.386394617 | 0.991835066 | -4.921347248 |
| **A_55_P2057906** | Cetn2 | -0.05623941 | -0.201261591 | 0.088782772 | 12.0252406 | -0.926862797 | 0.386399005 | 0.991835066 | -4.921353335 |
| **A_66_P127722** | NA | -0.105732365 | -0.378419355 | 0.166954624 | 12.29043435 | -0.926728806 | 0.386463712 | 0.991835066 | -4.921443074 |
| **A_52_P523330** | Tmem147 | -0.092297528 | -0.330339718 | 0.145744663 | 11.61975248 | -0.926713017 | 0.386471338 | 0.991835066 | -4.921453647 |
| **A_55_P2124016** | Nipsnap1 | 0.085804008 | -0.135523207 | 0.307131222 | 9.473568723 | 0.926577795 | 0.386536649 | 0.991835066 | -4.9215442 |
| **A_51_P520857** | Selenok-ps | -0.051348339 | -0.183835955 | 0.081139277 | 12.40343646 | -0.92631831 | 0.386662001 | 0.991835066 | -4.921717941 |
| **A_55_P2061294** | NA | -0.148945517 | -0.533280879 | 0.235389845 | 15.7578214 | -0.926245741 | 0.386697063 | 0.991835066 | -4.921766524 |
| **A_66_P102299** | Ascc1 | -0.077703679 | -0.278214317 | 0.12280696 | 8.76421496 | -0.926218162 | 0.386710389 | 0.991835066 | -4.921784986 |
| **A_52_P12001** | Erp44 | 0.05993514 | -0.094738619 | 0.214608899 | 10.03446714 | 0.926134399 | 0.386750863 | 0.991835066 | -4.921841059 |
| **A_66_P102163** | NA | -0.054266549 | -0.194349477 | 0.085816379 | 8.59307264 | -0.925883038 | 0.386872341 | 0.991835066 | -4.922009304 |
| **A_55_P2006768** | NA | -0.038614242 | -0.138295131 | 0.061066647 | 12.89091034 | -0.925858766 | 0.386884073 | 0.991835066 | -4.922025549 |
| **A_52_P117408** | Tg | -0.111600977 | -0.399764991 | 0.176563037 | 8.740197124 | -0.925629919 | 0.386994698 | 0.991835066 | -4.922178693 |
| **A_52_P464003** | Atg3 | -0.051114705 | -0.183101876 | 0.080872465 | 10.23513216 | -0.925599849 | 0.387009236 | 0.991835066 | -4.922198814 |
| **A_55_P2028883** | Rnft2 | 0.068110699 | -0.107773581 | 0.24399498 | 8.418606043 | 0.925544797 | 0.387035852 | 0.991835066 | -4.92223565 |
| **A_55_P2436438** | Dusp22 | -0.045854894 | -0.164275149 | 0.072565361 | 8.333610593 | -0.925483826 | 0.387065332 | 0.991835066 | -4.922276445 |
| **A_52_P478256** | Samm50 | -0.12045042 | -0.431543008 | 0.190642168 | 11.64769193 | -0.92539636 | 0.387107625 | 0.991835066 | -4.922334963 |
| **A_55_P2013534** | Slc25a19 | -0.042191294 | -0.151161672 | 0.066779083 | 11.42023866 | -0.92538728 | 0.387112016 | 0.991835066 | -4.922341038 |
| **A_51_P207153** | 1110008P1 | -0.101366772 | -0.36317413 | 0.160440586 | 11.01414534 | -0.925385949 | 0.38711266 | 0.991835066 | -4.922341928 |
| **A_52_P321627** | NA | 0.049957891 | -0.079086904 | 0.179002686 | 14.23566541 | 0.925279068 | 0.387164346 | 0.991835066 | -4.922413431 |
| **A_55_P2103880** | Gtf2ird1 | 0.047629249 | -0.075406519 | 0.170665018 | 10.44183372 | 0.925233785 | 0.387186246 | 0.991835066 | -4.922443723 |
| **A_51_P494481** | Snx30 | 0.043962682 | -0.06960642 | 0.157531784 | 8.145394102 | 0.925194663 | 0.387205167 | 0.991835066 | -4.922469893 |
| **A_55_P2183597** | Tbc1d2 | -0.060782669 | -0.217809089 | 0.096243751 | 8.451149527 | -0.925158557 | 0.38722263 | 0.991835066 | -4.922494045 |
| **A_55_P2103284** | Twf2 | -0.117915802 | -0.422544169 | 0.186712565 | 10.28442584 | -0.925147114 | 0.387228165 | 0.991835066 | -4.922501699 |
| **A_55_P2050268** | Nfkbid | 0.046326369 | -0.073358505 | 0.166011243 | 7.978601821 | 0.925120116 | 0.387241223 | 0.991835066 | -4.922519758 |
| **A_52_P625431** | Hnrnpa0 | 0.176743217 | -0.279901277 | 0.633387712 | 9.012566888 | 0.92506801 | 0.387266427 | 0.991835066 | -4.922554609 |
| **A_51_P415247** | NA | 0.056637125 | -0.089709993 | 0.202984243 | 8.63245921 | 0.924966829 | 0.387315371 | 0.991835066 | -4.922622282 |
| **A_55_P2015485** | Cfap45 | 0.054336941 | -0.086069739 | 0.19474362 | 8.699956493 | 0.92494636 | 0.387325274 | 0.991835066 | -4.922635972 |
| **A_55_P2009285** | Fbxo25 | 0.084650327 | -0.13415354 | 0.303454194 | 10.12863864 | 0.924661507 | 0.387463094 | 0.991835066 | -4.922826459 |
| **A_52_P577384** | Il18bp | -0.044608025 | -0.160014601 | 0.070798551 | 7.887507154 | -0.923828958 | 0.387866119 | 0.991835066 | -4.923382965 |
| **A_55_P2176731** | Prdx6 | -0.080238584 | -0.287862359 | 0.127385192 | 11.51762693 | -0.923666731 | 0.387944687 | 0.991835066 | -4.923491362 |

| | | | | | | | | | |
|---|---|---|---|---|---|---|---|---|---|
| **A_52_P177021** | Pts | -0.080076972 | -0.287325659 | 0.127171716 | 10.15954249 | -0.92347466 | 0.388037725 | 0.991835066 | -4.923619683 |
| **A_55_P2302413** | BC002189 | 0.199638631 | -0.317077569 | 0.716354832 | 11.18584435 | 0.923425021 | 0.388061772 | 0.991835066 | -4.923652842 |
| **A_52_P484405** | Twf1 | 0.047221075 | -0.075003389 | 0.16944554 | 10.4183087 | 0.923393602 | 0.388076994 | 0.991835066 | -4.92367383 |
| **A_52_P86517** | Cbwd1 | 0.07326652 | -0.116430531 | 0.262963571 | 9.480076484 | 0.923111404 | 0.38821373 | 0.991835066 | -4.923862319 |
| **A_55_P1985219** | Pnpla6 | -0.058063787 | -0.208402046 | 0.092274472 | 9.318877205 | -0.923091966 | 0.388223149 | 0.991835066 | -4.923875301 |
| **A_55_P2056995** | Gcgr | 0.05861192 | -0.093149096 | 0.210372935 | 8.766963234 | 0.923070465 | 0.388233569 | 0.991835066 | -4.92388966 |
| **A_66_P105564** | Gjc2 | 0.079137329 | -0.125774269 | 0.284048928 | 10.43998822 | 0.923047352 | 0.38824477 | 0.991835066 | -4.923905095 |
| **A_55_P2170256** | Ptgr2 | -0.076999351 | -0.276380516 | 0.122381814 | 9.90252519 | -0.92302203 | 0.388257042 | 0.991835066 | -4.923922005 |
| **A_51_P469252** | Zfp647 | -0.052077005 | -0.186938545 | 0.082784534 | 8.926170982 | -0.922926297 | 0.388303441 | 0.991835066 | -4.923985935 |
| **A_55_P2112170** | Crtc1 | 0.093612511 | -0.148843119 | 0.336068142 | 9.799184359 | 0.92280673 | 0.388361396 | 0.991835066 | -4.924065773 |
| **A_55_P2072631** | Prr5l | 0.068857269 | -0.109483638 | 0.247198175 | 8.407519881 | 0.922800782 | 0.38836428 | 0.991835066 | -4.924069745 |
| **A_51_P463087** | Cenpm | -0.058038186 | -0.208362712 | 0.092286341 | 8.71938766 | -0.922769254 | 0.388379563 | 0.991835066 | -4.924090796 |
| **A_55_P2178553** | Apoa2 | -0.115218547 | -0.413669629 | 0.183232535 | 10.61587807 | -0.92269543 | 0.388415351 | 0.991835066 | -4.924140085 |
| **A_55_P2033055** | Pdp1 | -0.056620545 | -0.203286764 | 0.090045674 | 10.19821318 | -0.9226842 | 0.388420796 | 0.991835066 | -4.924147583 |
| **A_52_P535012** | NA | 0.059841492 | -0.095186355 | 0.214869339 | 15.021842 | 0.922575314 | 0.388473586 | 0.991835066 | -4.924220276 |
| **A_55_P2060607** | NA | 0.048781678 | -0.077633071 | 0.175196426 | 15.14717667 | 0.922291315 | 0.3886113 | 0.991835066 | -4.924409848 |
| **A_51_P464918** | Mefv | -0.036808674 | -0.132209581 | 0.058592232 | 7.712591516 | -0.922161119 | 0.388674446 | 0.991835066 | -4.924496742 |
| **A_55_P1968523** | Btbd1 | -0.074506611 | -0.267640888 | 0.118627667 | 12.64220394 | -0.922029 | 0.388738532 | 0.991835066 | -4.924584909 |
| **A_66_P103780** | Gm3646 | -0.080431596 | -0.28893696 | 0.128073768 | 9.640898341 | -0.921973817 | 0.388765302 | 0.991835066 | -4.924621732 |
| **A_55_P2044547** | NA | 0.063383937 | -0.100954514 | 0.227722388 | 8.828921881 | 0.921826423 | 0.388836811 | 0.991835066 | -4.924720079 |
| **A_51_P227445** | NA | 0.080753638 | -0.128658321 | 0.290165596 | 8.496966221 | 0.921657894 | 0.388918586 | 0.991835066 | -4.924832513 |
| **A_52_P8903** | Tmem251 | -0.049851543 | -0.17914076 | 0.079437673 | 9.189074451 | -0.921563865 | 0.388964217 | 0.991835066 | -4.924895239 |
| **A_52_P652212** | Psmd14 | -0.083298263 | -0.299346979 | 0.132750453 | 9.831504972 | -0.921495857 | 0.388997223 | 0.991835066 | -4.924940604 |
| **A_51_P129866** | Atxn1l | 0.07407269 | -0.118060259 | 0.26620564 | 10.80989804 | 0.921436478 | 0.389026043 | 0.991835066 | -4.92498021 |
| **A_55_P2041050** | Zmym5 | 0.0810008 | -0.129150624 | 0.291152224 | 9.519530454 | 0.921225827 | 0.389128295 | 0.991835066 | -4.925120703 |
| **A_55_P2473316** | Mapre1 | -0.070227844 | -0.252441753 | 0.111986066 | 11.75956366 | -0.921163964 | 0.389158329 | 0.991835066 | -4.925161958 |
| **A_52_P456077** | Commd2 | 0.057802509 | -0.092177809 | 0.207782826 | 10.41915447 | 0.921131326 | 0.389174174 | 0.991835066 | -4.925183722 |
| **A_55_P2005982** | NA | -0.056892033 | -0.204527379 | 0.090743313 | 9.783698721 | -0.921022502 | 0.389227012 | 0.991835066 | -4.925256288 |
| **A_55_P2197338** | Tnfsf13os | 0.044553964 | -0.071068727 | 0.160176656 | 8.251305531 | 0.920984686 | 0.389245374 | 0.991835066 | -4.925281503 |
| **A_55_P2151021** | NA | 0.066366767 | -0.105911177 | 0.238644711 | 13.77663568 | 0.920725374 | 0.389371304 | 0.991835066 | -4.925454386 |
| **A_55_P1997747** | Cox16 | -0.056830686 | -0.204357397 | 0.090696024 | 12.14824579 | -0.920706853 | 0.3893803 | 0.991835066 | -4.925466733 |
| **A_52_P681391** | G0s2 | -0.122445351 | -0.440301741 | 0.19541104 | 10.68958575 | -0.920704965 | 0.389381217 | 0.991835066 | -4.925467992 |
| **A_55_P1955244** | NA | -0.087235263 | -0.313706609 | 0.139236083 | 12.50620308 | -0.920635986 | 0.389414721 | 0.991835066 | -4.925513973 |
| **A_55_P2111277** | NA | 0.052035986 | -0.083071936 | 0.187143908 | 9.161163561 | 0.920517623 | 0.389472216 | 0.991835066 | -4.925592869 |
| **A_65_P19623** | Ssu72 | 0.041456385 | -0.066191397 | 0.149104168 | 12.14157255 | 0.920439985 | 0.389509933 | 0.991835066 | -4.925644615 |
| **A_51_P151820** | Urb1 | 0.060840033 | -0.097159192 | 0.218839258 | 8.807140746 | 0.920330076 | 0.389563332 | 0.991835066 | -4.925717865 |
| **A_55_P2249897** | NA | 0.067024985 | -0.107056405 | 0.241106374 | 8.70219706 | 0.920223908 | 0.389614919 | 0.991835066 | -4.925788616 |
| **A_55_P2156490** | Gm3146 | 0.047217243 | -0.07542306 | 0.169857546 | 8.228569443 | 0.920187952 | 0.389632391 | 0.991835066 | -4.925812576 |
| **A_55_P2087374** | NA | -0.040159564 | -0.14449621 | 0.064177082 | 14.93449947 | -0.919943704 | 0.389751093 | 0.991835066 | -4.925975317 |
| **A_52_P229210** | NA | -0.121353577 | -0.436670254 | 0.1939631 | 13.66359203 | -0.919845255 | 0.389798947 | 0.991835066 | -4.926040905 |
| **A_51_P492410** | Pmvk | 0.04966056 | -0.079376365 | 0.178697486 | 11.30023776 | 0.919828247 | 0.389807214 | 0.991835066 | -4.926052236 |
| **A_55_P2101585** | Cd55 | 0.103222953 | -0.165013595 | 0.371459501 | 9.267610642 | 0.919745054 | 0.389847656 | 0.991835066 | -4.926107654 |
| **A_55_P2097126** | Cnep1r1 | -0.062294419 | -0.224193242 | 0.099604404 | 10.15259721 | -0.919633069 | 0.389902099 | 0.991835066 | -4.926182248 |
| **A_51_P247359** | Ptprcap | -0.062206153 | -0.223967045 | 0.09955474 | 8.372989893 | -0.919113068 | 0.390154979 | 0.991835066 | -4.926528538 |
| **A_51_P370561** | Dcakd | 0.075296978 | -0.12051415 | 0.271108106 | 9.837621772 | 0.919071522 | 0.390175189 | 0.991835066 | -4.926556199 |
| **A_51_P398723** | Flt1 | 0.082356139 | -0.131845558 | 0.296557835 | 13.1200862 | 0.918929521 | 0.390244269 | 0.991835066 | -4.926650736 |
| **A_55_P2037454** | Etv5 | 0.13986672 | -0.223926621 | 0.50366006 | 11.19639348 | 0.918901111 | 0.390258091 | 0.991835066 | -4.926669648 |
| **A_52_P642207** | Mkrn1 | -0.169030861 | -0.608746679 | 0.270684957 | 10.654192 | -0.918762047 | 0.390325753 | 0.991835066 | -4.926762218 |

| | | | | | | | | | |
|---|---|---|---|---|---|---|---|---|---|
| **A_55_P2156068** | NA | 0.053486604 | -0.085655984 | 0.192629193 | 8.830208197 | 0.918743081 | 0.390334982 | 0.991835066 | -4.926774842 |
| **A_55_P1987984** | NA | 0.05427146 | -0.086914226 | 0.195457147 | 8.829148207 | 0.918734381 | 0.390339215 | 0.991835066 | -4.926780633 |
| **A_55_P1980446** | Sbno1 | 0.04307926 | -0.069002951 | 0.15516147 | 10.69755526 | 0.918630109 | 0.390389956 | 0.991835066 | -4.926850035 |
| **A_55_P2079855** | Prl6a1 | -0.0379982 | -0.136880888 | 0.060884488 | 7.664684599 | -0.918442338 | 0.390481343 | 0.991835066 | -4.926974999 |
| **A_55_P2297485** | Ago2 | 0.061001519 | -0.097749428 | 0.219752465 | 11.86340829 | 0.918403344 | 0.390500323 | 0.991835066 | -4.927000947 |
| **A_55_P2035981** | Mindy1 | -0.072439501 | -0.26098466 | 0.116105658 | 11.66981791 | -0.918267468 | 0.390566465 | 0.991835066 | -4.92709136 |
| **A_55_P2115320** | Metap1d | -0.051336893 | -0.184964768 | 0.082290982 | 8.455348334 | -0.918209225 | 0.390594819 | 0.991835066 | -4.927130113 |
| **A_55_P2132711** | Tmem248 | 0.055932699 | -0.089673941 | 0.201539338 | 9.54251197 | 0.918107902 | 0.390644149 | 0.991835066 | -4.927197524 |
| **A_55_P2177721** | Adra1b | -0.104589999 | -0.376890362 | 0.167710365 | 10.42579937 | -0.918017728 | 0.390688055 | 0.991835066 | -4.927257514 |
| **A_51_P302651** | Topbp1 | 0.105656106 | -0.16942723 | 0.380739442 | 10.32246158 | 0.917993173 | 0.390700012 | 0.991835066 | -4.927273849 |
| **A_55_P2102664** | NA | -0.079785693 | -0.287522861 | 0.127951475 | 8.208622309 | -0.917951951 | 0.390720085 | 0.991835066 | -4.927301271 |
| **A_52_P236448** | Ngfr | -0.070034145 | -0.252398824 | 0.112330534 | 8.937713744 | -0.917863787 | 0.390763019 | 0.991835066 | -4.903847868 |
| **A_51_P346641** | NA | -0.053987459 | -0.194579311 | 0.086604393 | 9.214323088 | -0.917786921 | 0.390800454 | 0.991835066 | -4.927411043 |
| **A_55_P2104761** | NA | 0.076462527 | -0.122659755 | 0.275584809 | 14.56247312 | 0.917778579 | 0.390804517 | 0.991835066 | -4.927416592 |
| **A_51_P461031** | Ostm1 | -0.050701359 | -0.182747218 | 0.081344499 | 9.905876496 | -0.917706803 | 0.390839475 | 0.991835066 | -4.92746433 |
| **A_51_P161691** | Coq7 | -0.090776565 | -0.327233467 | 0.145680337 | 12.30102515 | -0.917552413 | 0.390914679 | 0.991835066 | -4.927567006 |
| **A_55_P2091446** | Trim33 | 0.068483112 | -0.109936559 | 0.246902783 | 9.42453901 | 0.917381308 | 0.390998038 | 0.991835066 | -4.927680784 |
| **A_55_P2016852** | Zyx | 0.076223177 | -0.122364016 | 0.27481037 | 13.56433897 | 0.917370861 | 0.391003128 | 0.991835066 | -4.92768773 |
| **A_55_P2035613** | Tmbim1 | 0.092821301 | -0.149042353 | 0.334684954 | 11.55278498 | 0.917246734 | 0.391063608 | 0.991835066 | -4.92777026 |
| **A_55_P2340101** | NA | 0.070771006 | -0.113646264 | 0.255188276 | 10.4352389 | 0.917197609 | 0.391087547 | 0.991835066 | -4.92780292 |
| **A_51_P333869** | Nsmce3 | -0.066207262 | -0.238735882 | 0.106321359 | 9.373795689 | -0.917177958 | 0.391097123 | 0.991835066 | -4.927815984 |
| **A_51_P367285** | Rnf135 | -0.057761764 | -0.208297626 | 0.092774098 | 8.710698947 | -0.917085039 | 0.391142405 | 0.991835066 | -4.927877756 |
| **A_55_P1965114** | Spsb3 | 0.067013109 | -0.107646936 | 0.241673155 | 12.0531179 | 0.917012666 | 0.391177677 | 0.991835066 | -4.927925865 |
| **A_51_P224575** | Gnpat | -0.052858811 | -0.190632914 | 0.084915291 | 13.3508576 | -0.916978036 | 0.391194556 | 0.991835066 | -4.927948884 |
| **A_51_P229676** | Plscr1 | 0.064711566 | -0.103979562 | 0.233402693 | 9.496833806 | 0.916851076 | 0.39125644 | 0.991835066 | -4.928033272 |
| **A_55_P2048454** | Rps2 | -0.060764106 | -0.219251158 | 0.097722946 | 15.12383336 | -0.916352272 | 0.391499643 | 0.991835066 | -4.928364734 |
| **A_51_P155458** | Dok7 | 0.067815889 | -0.109128502 | 0.244760279 | 11.2770324 | 0.916017539 | 0.391662914 | 0.991835066 | -4.928587098 |
| **A_51_P380309** | Ncam1 | 0.093867668 | -0.151062307 | 0.338797643 | 12.1434328 | 0.915974163 | 0.391684075 | 0.991835066 | -4.928615908 |
| **A_55_P1964078** | NA | -0.068602144 | -0.247647671 | 0.110443383 | 12.72274114 | -0.915763535 | 0.391786841 | 0.991835066 | -4.928755794 |
| **A_51_P140731** | Cops5 | 0.055096419 | -0.088724306 | 0.198917145 | 13.36624753 | 0.915611066 | 0.391861245 | 0.991835066 | -4.92885704 |
| **A_55_P2129685** | Pkig | -0.141473827 | -0.510864847 | 0.227917193 | 9.5713553 | -0.91537469 | 0.391976615 | 0.991835066 | -4.92901398 |
| **A_55_P2109874** | Rpl23a | -0.0534275 | -0.192947625 | 0.086092625 | 15.90011184 | -0.915244495 | 0.392040171 | 0.991835066 | -4.92910041 |
| **A_52_P283055** | Slc5a3 | 0.048334781 | -0.077910327 | 0.174579889 | 8.327188616 | 0.915070029 | 0.392125351 | 0.991835066 | -4.929216215 |
| **A_55_P2096204** | Kdm1b | 0.067816259 | -0.109318509 | 0.244951026 | 9.953395447 | 0.91503804 | 0.392140971 | 0.991835066 | -4.929237447 |
| **A_55_P1971594** | Pqbp1 | -0.053966229 | -0.194926251 | 0.086993793 | 10.05389483 | -0.915029819 | 0.392144985 | 0.991835066 | -4.929242903 |
| **A_55_P2049025** | NA | 0.049342002 | -0.079559563 | 0.178243567 | 14.42311277 | 0.914887525 | 0.39221447 | 0.991835066 | -4.929337339 |
| **A_51_P224517** | Jade1 | -0.056142421 | -0.202828549 | 0.090543706 | 9.74526087 | -0.914768552 | 0.392272575 | 0.991835066 | -4.92941629 |
| **A_55_P2122997** | Tet1 | -0.076805278 | -0.277486357 | 0.123875801 | 10.34174289 | -0.914731788 | 0.392290531 | 0.991835066 | -4.929440684 |
| **A_52_P123944** | Mospd2 | -0.041183411 | -0.148809192 | 0.066442371 | 8.192445546 | -0.914566151 | 0.392371439 | 0.991835066 | -4.929550586 |
| **A_65_P19112** | Zfp397 | -0.039993203 | -0.144515074 | 0.064528668 | 8.599279216 | -0.914509335 | 0.392399194 | 0.991835066 | -4.929588281 |
| **A_51_P440568** | Stk19 | 0.065767467 | -0.106130736 | 0.237665671 | 8.764054963 | 0.914426731 | 0.39243955 | 0.991835066 | -4.929643081 |
| **A_66_P124925** | Pdcd5 | 0.067183624 | -0.108449149 | 0.242816396 | 13.51126828 | 0.914254274 | 0.392523813 | 0.991835066 | -4.92975748 |
| **A_55_P2037957** | NA | 0.049999221 | -0.080713502 | 0.180711945 | 14.38456835 | 0.914227981 | 0.392536661 | 0.991835066 | -4.92977492 |
| **A_51_P294515** | Gde1 | -0.084405894 | -0.30507264 | 0.136260853 | 9.559744293 | -0.914207991 | 0.39254643 | 0.991835066 | -4.92978818 |
| **A_55_P2134004** | Gstm2 | -0.092229862 | -0.333379067 | 0.148919343 | 11.53138987 | -0.914102413 | 0.392598024 | 0.991835066 | -4.929858204 |
| **A_55_P2160750** | Col25a1 | 0.048777805 | -0.078763558 | 0.176319168 | 7.903987727 | 0.914071837 | 0.392612968 | 0.991835066 | -4.929878483 |
| **A_51_P317695** | Trpc3 | 0.099058602 | -0.16003805 | 0.358155253 | 9.720058307 | 0.913775533 | 0.3927578 | 0.991835066 | -4.930074972 |
| **A_55_P2039105** | Rbck1 | -0.046563735 | -0.168359151 | 0.075231681 | 11.67755351 | -0.913747067 | 0.392771716 | 0.991835066 | -4.930093846 |

| | | | | | | | | | |
|---|---|---|---|---|---|---|---|---|---|
| **A_55_P1993976** | Aimp2 | 0.097877756 | -0.158157811 | 0.353913323 | 13.29383494 | 0.913677301 | 0.392805824 | 0.991835066 | -4.930140103 |
| **A_55_P1964832** | Reck | 0.069640051 | -0.112561484 | 0.251841585 | 8.361173706 | 0.913516045 | 0.392884669 | 0.991835066 | -4.930247009 |
| **A_51_P279437** | Mfsd2a | 0.045978628 | -0.074320985 | 0.166278242 | 7.746817823 | 0.91348393 | 0.392900373 | 0.991835066 | -4.930268299 |
| **A_55_P2129578** | Pkp3 | -0.10421493 | -0.376891522 | 0.168461663 | 11.37534052 | -0.913463531 | 0.392910348 | 0.991835066 | -4.930281822 |
| **A_55_P2081405** | BC052688 | -0.054977879 | -0.198844308 | 0.08888855 | 12.11352499 | -0.913350873 | 0.392965442 | 0.991835066 | -4.930356499 |
| **A_52_P125363** | Rbp2 | -0.037693322 | -0.136333916 | 0.060947271 | 7.724662013 | -0.913309285 | 0.392985781 | 0.991835066 | -4.930384065 |
| **A_51_P214343** | Sephs1 | -0.039013085 | -0.141138977 | 0.063112807 | 9.165048276 | -0.913026889 | 0.393123912 | 0.991835066 | -4.93057122 |
| **A_55_P2015681** | AU015836 | 0.045999851 | -0.074427211 | 0.166426913 | 7.682276869 | 0.912938383 | 0.393167211 | 0.991835066 | -4.930629869 |
| **A_51_P487360** | Hpcal1 | -0.065481005 | -0.236942441 | 0.105980431 | 9.992773822 | -0.912762967 | 0.39325304 | 0.991835066 | -4.930746095 |
| **A_51_P473240** | Zfp426 | -0.04978885 | -0.180175562 | 0.080597862 | 10.47617484 | -0.912657637 | 0.393304583 | 0.991835066 | -4.930815877 |
| **A_52_P585028** | Cnep1r1 | -0.059846332 | -0.216588628 | 0.096895965 | 10.90336161 | -0.912557965 | 0.393353362 | 0.991835066 | -4.930881905 |
| **A_55_P1961127** | H19 | 0.068444806 | -0.110866391 | 0.247756004 | 17.54042301 | 0.912309545 | 0.393474958 | 0.991835066 | -4.931046449 |
| **A_55_P1989013** | Eif6 | -0.123159297 | -0.445860294 | 0.1995417 | 11.20399703 | -0.912170505 | 0.393543026 | 0.991835066 | -4.93113853 |
| **A_51_P366672** | Slc36a2 | -0.044745324 | -0.162008885 | 0.072518236 | 8.38586672 | -0.911997646 | 0.393627664 | 0.991835066 | -4.931252994 |
| **A_55_P2144210** | Snx5 | 0.057048056 | -0.092478028 | 0.206574139 | 12.40955073 | 0.911870195 | 0.393690078 | 0.991835066 | -4.93133738 |
| **A_55_P2080804** | Slirp | 0.077543213 | -0.125722793 | 0.28080922 | 13.14417732 | 0.911776032 | 0.393736194 | 0.991835066 | -4.93139972 |
| **A_55_P1969831** | Ptpmt1 | -0.069077076 | -0.250170507 | 0.112016355 | 12.7300531 | -0.911675706 | 0.393785334 | 0.991835066 | -4.931466136 |
| **A_55_P2172022** | B4galt1 | 0.074672708 | -0.121185546 | 0.270530963 | 8.890988202 | 0.911232402 | 0.394002518 | 0.991835066 | -4.931759538 |
| **A_55_P2161600** | Diaph2 | -0.049213035 | -0.178335664 | 0.079909594 | 9.788561046 | -0.910934015 | 0.394148756 | 0.991835066 | -4.931956971 |
| **A_52_P403443** | Tns3 | 0.069384064 | -0.112666482 | 0.25143461 | 11.50510894 | 0.910912962 | 0.394159075 | 0.991835066 | -4.931970899 |
| **A_66_P137265** | Gm9933 | -0.040718487 | -0.147580623 | 0.066143649 | 8.418458 | -0.91070331 | 0.39426185 | 0.991835066 | -4.932109588 |
| **A_52_P644674** | Trappc6b | -0.069344865 | -0.251336701 | 0.11264697 | 10.79794981 | -0.910692029 | 0.394267381 | 0.991835066 | -4.932117049 |
| **A_66_P139493** | Tmem250- | 0.100489307 | -0.163245279 | 0.364223893 | 11.36562902 | 0.910671816 | 0.394277291 | 0.991835066 | -4.93213042 |
| **A_51_P129149** | Gatad2a | 0.109962634 | -0.178652588 | 0.398577856 | 10.38965379 | 0.910615493 | 0.394304906 | 0.991835066 | -4.932167674 |
| **A_55_P2120843** | NA | -0.047382753 | -0.171767256 | 0.077001749 | 8.89079276 | -0.910464785 | 0.394378804 | 0.991835066 | -4.932267349 |
| **A_51_P452875** | Pigyl | 0.066266109 | -0.107706299 | 0.240238516 | 10.11020656 | 0.910374798 | 0.394422933 | 0.991835066 | -4.932326859 |
| **A_66_P133112** | Gm14635 | -0.041239851 | -0.149516501 | 0.067036799 | 8.076501528 | -0.910314395 | 0.394452557 | 0.991835066 | -4.932366803 |
| **A_51_P429246** | Fbxl6 | -0.071588713 | -0.259551306 | 0.11637388 | 8.886303962 | -0.910295212 | 0.394461965 | 0.991835066 | -4.932379487 |
| **A_55_P2143336** | Rpp21 | -0.091240888 | -0.330834273 | 0.148352497 | 9.510689786 | -0.910172717 | 0.394522046 | 0.991835066 | -4.932460484 |
| **A_52_P706060** | Mex3a | 0.079449038 | -0.129183134 | 0.288081211 | 10.58759464 | 0.910157382 | 0.394529568 | 0.991835066 | -4.932470624 |
| **A_51_P431785** | Myom2 | -0.140218818 | -0.508435422 | 0.227997785 | 14.66405977 | -0.910148101 | 0.394534121 | 0.991835066 | -4.93247676 |
| **A_55_P2203478** | Scn5a | 0.080226149 | -0.130456476 | 0.290908775 | 8.450088524 | 0.910115178 | 0.39455027 | 0.991835066 | -4.932498527 |
| **A_51_P513224** | Cep164 | 0.056836132 | -0.092436383 | 0.206108647 | 8.831025601 | 0.910025989 | 0.394594022 | 0.991835066 | -4.932557494 |
| **A_52_P203981** | Srek1ip1 | -0.077977661 | -0.282791757 | 0.126836435 | 10.46319721 | -0.909954117 | 0.394629282 | 0.991835066 | -4.932605008 |
| **A_51_P374101** | Pard6b | 0.042824456 | -0.069658227 | 0.15530714 | 7.790855908 | 0.90994537 | 0.394633574 | 0.991835066 | -4.932610791 |
| **A_55_P2140390** | NA | -0.127703199 | -0.46315766 | 0.207751261 | 13.25796506 | -0.909865795 | 0.394672616 | 0.991835066 | -4.932663394 |
| **A_55_P2077048** | Itih5 | -0.133763308 | -0.48515145 | 0.217624833 | 9.812766456 | -0.909827444 | 0.394691433 | 0.991835066 | -4.932688745 |
| **A_55_P2064686** | Lrrc61 | -0.050260834 | -0.182300766 | 0.081779098 | 9.438748003 | -0.909774016 | 0.394717649 | 0.991835066 | -4.93272406 |
| **A_66_P124555** | Rps15 | -0.132557693 | -0.480801982 | 0.215686596 | 14.02361807 | -0.909766763 | 0.394721208 | 0.991835066 | -4.932728854 |
| **A_52_P533779** | Ankrd40 | -0.053705644 | -0.194831669 | 0.08742038 | 8.667517146 | -0.909540321 | 0.394832333 | 0.991835066 | -4.932878515 |
| **A_51_P295215** | Alkbh5 | -0.066441641 | -0.241037576 | 0.108154294 | 12.01232314 | -0.909526491 | 0.394839121 | 0.991835066 | -4.932887654 |
| **A_52_P383862** | Cmc1 | -0.080119911 | -0.290692204 | 0.130452381 | 11.38392132 | -0.909386214 | 0.394907973 | 0.991835066 | -4.932980352 |
| **A_55_P2146620** | Atf2 | 0.046591252 | -0.07588328 | 0.169065785 | 8.410457718 | 0.909217377 | 0.394990857 | 0.991835066 | -4.933091909 |
| **A_55_P2073294** | Hadhb | 0.047107009 | -0.076747739 | 0.170961758 | 14.19111147 | 0.909037931 | 0.395078962 | 0.991835066 | -4.93321046 |
| **A_51_P486068** | Zfp280b | 0.076017467 | -0.123860313 | 0.275895248 | 9.420942374 | 0.908987709 | 0.395103623 | 0.991835066 | -4.933243636 |
| **A_52_P630571** | Epg5 | -0.043307989 | -0.157190532 | 0.070574555 | 8.731660409 | -0.908908129 | 0.395142702 | 0.991835066 | -4.933296203 |
| **A_66_P124674** | Ubxn2a | 0.064167538 | -0.104575957 | 0.232911033 | 8.692893677 | 0.908861004 | 0.395165845 | 0.991835066 | -4.93332733 |
| **A_55_P2090817** | Macrod2 | 0.044037359 | -0.071770497 | 0.159845215 | 9.474530186 | 0.908850342 | 0.395171081 | 0.991835066 | -4.933334373 |

| | | | | | | | | | |
|---|---|---|---|---|---|---|---|---|---|
| **A_55_P1975793** | Srd5a3 | 0.045755026 | -0.074580346 | 0.166090397 | 9.475391709 | 0.908771362 | 0.39520987 | 0.991835066 | -4.933386537 |
| **A_55_P1970788** | C5ar2 | 0.046337581 | -0.075533748 | 0.16820891 | 8.118329463 | 0.908742719 | 0.395223939 | 0.991835066 | -4.933405455 |
| **A_51_P417839** | Hdac4 | -0.049988358 | -0.18147119 | 0.081494474 | 8.893713272 | -0.908675786 | 0.395256815 | 0.991835066 | -4.93344966 |
| **A_55_P2102897** | Gm12657 | -0.071495983 | -0.259555129 | 0.116563163 | 13.98700898 | -0.908649341 | 0.395269804 | 0.991835066 | -4.933467124 |
| **A_52_P821** | Fli1 | 0.06439593 | -0.105059103 | 0.233850963 | 10.65296728 | 0.90826605 | 0.395458112 | 0.991835066 | -4.933720211 |
| **A_52_P222350** | Flnb | -0.040551468 | -0.147262781 | 0.066159844 | 10.4454493 | -0.908249688 | 0.395466152 | 0.991835066 | -4.933731013 |
| **A_52_P590535** | Fbln2 | 0.180390623 | -0.294317983 | 0.65509923 | 12.8799132 | 0.908230299 | 0.395475679 | 0.991835066 | -4.933743813 |
| **A_51_P439824** | Gipc1 | -0.080778691 | -0.293377905 | 0.131820524 | 10.60156212 | -0.908122185 | 0.395528809 | 0.991835066 | -4.933815185 |
| **A_55_P1959056** | Pom121 | 0.040671451 | -0.066370974 | 0.147713876 | 12.36235819 | 0.908119201 | 0.395530275 | 0.991835066 | -4.933817155 |
| **A_55_P2064995** | Capzb | -0.130771739 | -0.474967548 | 0.21342407 | 9.957397753 | -0.908066082 | 0.395556381 | 0.991835066 | -4.93385222 |
| **A_52_P479269** | Sdc1 | 0.061543617 | -0.100464184 | 0.223551418 | 9.14125752 | 0.907938062 | 0.395619303 | 0.991835066 | -4.933936721 |
| **A_55_P2038757** | Fblim1 | 0.059415481 | -0.096991497 | 0.21582246 | 11.306227 | 0.907930542 | 0.395622999 | 0.991835066 | -4.933941684 |
| **A_52_P629037** | Gm6987 | 0.062227003 | -0.101593849 | 0.226047855 | 11.31664957 | 0.907859903 | 0.395657722 | 0.991835066 | -4.933988307 |
| **A_51_P397375** | Gatb | -0.079430463 | -0.288545401 | 0.129684475 | 10.46569476 | -0.907843879 | 0.395665599 | 0.991835066 | -4.933998882 |
| **A_55_P2083814** | Zzz3 | 0.070333164 | -0.114845721 | 0.255512048 | 11.18733062 | 0.907774173 | 0.395699865 | 0.991835066 | -4.934044885 |
| **A_55_P2001073** | Arid2 | 0.049006108 | -0.080030459 | 0.178042675 | 9.367675477 | 0.9077088 | 0.395732004 | 0.991835066 | -4.934088027 |
| **A_52_P162486** | Bgn | 0.139874769 | -0.228436532 | 0.50818607 | 9.474095955 | 0.907681469 | 0.395745441 | 0.991835066 | -4.934106063 |
| **A_55_P2020203** | Sfrp5 | -0.05606456 | -0.203710874 | 0.091581754 | 9.375365063 | -0.907559159 | 0.395805577 | 0.991835066 | -4.934186771 |
| **A_55_P2106058** | Gm11110 | 0.057983057 | -0.09473616 | 0.210702275 | 7.918256128 | 0.907437147 | 0.395865574 | 0.991835066 | -4.934267275 |
| **A_55_P1961034** | Gm13272 | -0.098942598 | -0.359591099 | 0.161705903 | 8.761297064 | -0.907271375 | 0.3959471 | 0.991835066 | -4.934376639 |
| **A_55_P2070940** | Krtap20-2 | -0.07956764 | -0.289206498 | 0.130071217 | 8.750039639 | -0.907138974 | 0.396012223 | 0.991835066 | -4.934463977 |
| **A_55_P2035454** | Oaz1 | -0.057800741 | -0.210104114 | 0.094502632 | 16.19626893 | -0.907053736 | 0.396054153 | 0.991835066 | -4.934520199 |
| **A_55_P2159506** | Gm5458 | 0.038587247 | -0.063093113 | 0.140267607 | 7.691651769 | 0.907017891 | 0.396071787 | 0.991835066 | -4.934543841 |
| **A_52_P46310** | Dcaf7 | -0.054461007 | -0.198018434 | 0.089096419 | 8.752603156 | -0.906711532 | 0.396222521 | 0.991835066 | -4.934745876 |
| **A_55_P2092046** | NA | -0.179900578 | -0.654276652 | 0.294475496 | 11.00366648 | -0.906397951 | 0.396376853 | 0.991835066 | -4.934952623 |
| **A_55_P1967983** | Use1 | -0.04708212 | -0.171255607 | 0.077091368 | 12.42238052 | -0.906225459 | 0.396461766 | 0.991835066 | -4.935066327 |
| **A_55_P2070661** | Taf7l | 0.116482901 | -0.190748591 | 0.423714394 | 10.18372788 | 0.906161447 | 0.396493281 | 0.991835066 | -4.935108518 |
| **A_66_P134534** | Atl2 | -0.040239344 | -0.146377619 | 0.065898931 | 8.349555828 | -0.906124764 | 0.396511342 | 0.991835066 | -4.935132697 |
| **A_52_P393589** | Clmp | -0.098126357 | -0.356972355 | 0.160719641 | 8.765253269 | -0.906052475 | 0.396546935 | 0.991835066 | -4.935180339 |
| **A_55_P2006808** | Ntrk3 | 0.165845765 | -0.271660787 | 0.603352317 | 8.226780861 | 0.906001583 | 0.396571994 | 0.991835066 | -4.935213879 |
| **A_55_P2160323** | NA | -0.057288219 | -0.208417835 | 0.093841398 | 8.252655911 | -0.905993062 | 0.39657619 | 0.991835066 | -4.935219495 |
| **A_55_P2186352** | Ttc12 | -0.064598273 | -0.235031734 | 0.105835188 | 9.987878761 | -0.905889401 | 0.396627237 | 0.991835066 | -4.935287806 |
| **A_55_P2011674** | Rpl17 | -0.036711923 | -0.133592401 | 0.060168555 | 15.51830919 | -0.905690879 | 0.396725011 | 0.991835066 | -4.935418615 |
| **A_55_P2035018** | Clec4n | -0.07033048 | -0.255951985 | 0.115291026 | 8.452465541 | -0.905574994 | 0.396782094 | 0.991835066 | -4.935494963 |
| **A_55_P2005853** | Nacc2 | 0.086762549 | -0.14223452 | 0.315759618 | 10.40037869 | 0.905548074 | 0.396795355 | 0.991835066 | -4.935512698 |
| **A_55_P2060567** | Ddx39 | 0.066820502 | -0.109549058 | 0.243190062 | 12.6934099 | 0.90551414 | 0.396812072 | 0.991835066 | -4.935535053 |
| **A_55_P2077635** | NA | -0.086627943 | -0.315279658 | 0.142023773 | 11.15391608 | -0.905508787 | 0.396814709 | 0.991835066 | -4.935538579 |
| **A_55_P2069022** | Vstm4 | -0.041229109 | -0.150065972 | 0.067607753 | 8.068095415 | -0.905392871 | 0.396871817 | 0.991835066 | -4.935614937 |
| **A_55_P2160451** | Stk26 | 0.057311511 | -0.094001268 | 0.20862429 | 7.93465898 | 0.905264277 | 0.396935178 | 0.991835066 | -4.935699638 |
| **A_55_P1965846** | Sp2 | 0.085466602 | -0.140224693 | 0.311157897 | 9.596254258 | 0.905087905 | 0.397022092 | 0.991835066 | -4.935815795 |
| **A_51_P187716** | Zc3h11a | -0.086510809 | -0.314971054 | 0.141949435 | 12.54695895 | -0.905042285 | 0.397044575 | 0.991835066 | -4.935845837 |
| **A_55_P2185214** | NA | 0.058591919 | -0.096141383 | 0.213325221 | 16.43317483 | 0.905030179 | 0.397050542 | 0.991835066 | -4.935853809 |
| **A_52_P524895** | Eif3a | -0.052094708 | -0.189675006 | 0.08548559 | 11.51024679 | -0.904995655 | 0.397067558 | 0.991835066 | -4.935876544 |
| **A_52_P150114** | NA | -0.0468437 | -0.170561557 | 0.076874157 | 7.895385739 | -0.904956986 | 0.397086617 | 0.991835066 | -4.935902007 |
| **A_55_P2096777** | Tprn | 0.068595731 | -0.112610106 | 0.249801569 | 8.328283593 | 0.904761346 | 0.397183055 | 0.991835066 | -4.936030821 |
| **A_52_P283910** | Hnrnpdl | 0.099841963 | -0.163911373 | 0.363595299 | 10.79723015 | 0.904741023 | 0.397193074 | 0.991835066 | -4.936044201 |
| **A_52_P218955** | D3Ertd751e | 0.067788437 | -0.11131074 | 0.246887614 | 8.792553474 | 0.904630369 | 0.397247627 | 0.991835066 | -4.936117048 |
| **A_55_P2043612** | Prdm6 | 0.148144734 | -0.243319772 | 0.53960924 | 8.223583041 | 0.904488313 | 0.397317672 | 0.991835066 | -4.93621056 |

| | | | | | | | | | |
|---|---|---|---|---|---|---|---|---|---|
| **A_52_P5549** | Fam133b | 0.073830874 | -0.121296773 | 0.26895852 | 9.613087175 | 0.904332904 | 0.39739431 | 0.991835066 | -4.936312849 |
| **A_55_P2142008** | 4933434E2 | 0.105244436 | -0.172946357 | 0.383435229 | 8.592092991 | 0.90420213 | 0.397458809 | 0.991835066 | -4.936398913 |
| **A_52_P239536** | Ppp1r9a | 0.126327298 | -0.207605755 | 0.460260352 | 10.08174703 | 0.904163418 | 0.397477903 | 0.991835066 | -4.936424389 |
| **A_55_P2020373** | Chd4 | -0.044606457 | -0.16252964 | 0.073316726 | 13.54852391 | -0.904081681 | 0.397518222 | 0.991835066 | -4.936478175 |
| **A_52_P129443** | Wdpcp | -0.059754508 | -0.217757369 | 0.098248353 | 8.923346817 | -0.903888494 | 0.397613527 | 0.991835066 | -4.936605286 |
| **A_52_P248604** | Cdh5 | 0.056802979 | -0.093403441 | 0.207009399 | 12.47147781 | 0.903840405 | 0.397637254 | 0.991835066 | -4.936636924 |
| **A_55_P2035265** | Trpm7 | 0.09655292 | -0.158797672 | 0.351903512 | 10.89697608 | 0.903727874 | 0.397692779 | 0.991835066 | -4.936710954 |
| **A_55_P2162782** | Ahnak | 0.113688136 | -0.18700512 | 0.414381392 | 8.954938684 | 0.903650703 | 0.397730861 | 0.991835066 | -4.936761718 |
| **A_55_P1992475** | NA | 0.059384282 | -0.097686242 | 0.216454806 | 8.968180415 | 0.903620234 | 0.397745897 | 0.991835066 | -4.936781761 |
| **A_51_P100573** | Map3k3 | 0.076238817 | -0.125448934 | 0.277926568 | 8.632650337 | 0.903453401 | 0.397828236 | 0.991835066 | -4.898139492 |
| **A_66_P138137** | Pnp | 0.067157402 | -0.110521319 | 0.244836123 | 10.8615297 | 0.903374026 | 0.397867415 | 0.991835066 | -4.936943695 |
| **A_55_P1989865** | Thrap3 | 0.088631859 | -0.145865014 | 0.323128731 | 9.574400867 | 0.90336224 | 0.397873233 | 0.991835066 | -4.936951446 |
| **A_51_P400375** | Erg28 | 0.099605751 | -0.164003486 | 0.363214989 | 12.24040251 | 0.90309393 | 0.398005693 | 0.991835066 | -4.93712788 |
| **A_66_P110742** | Gm9949 | -0.040667878 | -0.148307077 | 0.066971321 | 7.736623328 | -0.903005058 | 0.398049574 | 0.991835066 | -4.937186311 |
| **A_55_P1959818** | Tmem40 | -0.098912422 | -0.360742025 | 0.16291718 | 8.24599499 | -0.902903264 | 0.39809984 | 0.991835066 | -4.937253233 |
| **A_51_P300655** | Tbccd1 | 0.063389199 | -0.104414726 | 0.231193124 | 8.976341934 | 0.902863883 | 0.398119288 | 0.991835066 | -4.937279122 |
| **A_55_P2156288** | NA | 0.069920835 | -0.115177773 | 0.255019444 | 8.737334685 | 0.90284373 | 0.398129241 | 0.991835066 | -4.937292371 |
| **A_51_P348022** | Nudt21 | -0.065990818 | -0.240704789 | 0.108723153 | 10.89103523 | -0.902744827 | 0.398178087 | 0.991835066 | -4.937357384 |
| **A_55_P1999384** | NA | 0.075014309 | -0.123600088 | 0.273628707 | 10.75741333 | 0.902698084 | 0.398201174 | 0.991835066 | -4.937388109 |
| **A_55_P2005920** | NA | 0.042275483 | -0.069667096 | 0.154218062 | 14.47373566 | 0.902614691 | 0.398242365 | 0.991835066 | -4.937442921 |
| **A_51_P357422** | Coprs | -0.054988786 | -0.200620601 | 0.090643028 | 10.0973201 | -0.902458014 | 0.398319763 | 0.991835066 | -4.937545891 |
| **A_55_P2106175** | Gch1 | -0.074899727 | -0.273273777 | 0.123474323 | 10.32975869 | -0.902411266 | 0.398342858 | 0.991835066 | -4.937576612 |
| **A_51_P148196** | Sdf2 | -0.066027756 | -0.240941188 | 0.108885676 | 13.33272239 | -0.902220122 | 0.398437302 | 0.991835066 | -4.937702212 |
| **A_51_P233027** | Rmdn3 | -0.041582841 | -0.151741183 | 0.068575501 | 8.47178305 | -0.902206391 | 0.398444087 | 0.991835066 | -4.937711234 |
| **A_51_P351970** | Hells | 0.064749736 | -0.106799938 | 0.23629941 | 8.333016732 | 0.902105309 | 0.398494039 | 0.991835066 | -4.937777646 |
| **A_55_P2172688** | Srbd1 | 0.051868183 | -0.085555384 | 0.18929175 | 8.888299895 | 0.902088085 | 0.398502551 | 0.991835066 | -4.937788962 |
| **A_55_P2084631** | Hist1h2an | 0.064441759 | -0.106389248 | 0.235272766 | 16.88505679 | 0.90159152 | 0.39874801 | 0.991835066 | -4.938115129 |
| **A_55_P2090782** | NA | -0.097718694 | -0.356803273 | 0.161365886 | 12.25885117 | -0.901457421 | 0.398814317 | 0.991835066 | -4.938203189 |
| **A_55_P1954461** | Ptdss1 | 0.055390996 | -0.09148338 | 0.202265372 | 8.379431407 | 0.901368283 | 0.398858397 | 0.991835066 | -4.93826172 |
| **A_55_P2143885** | Etl4 | -0.063798311 | -0.232970676 | 0.105374055 | 9.355847173 | -0.901340506 | 0.398872133 | 0.991835066 | -4.938279957 |
| **A_52_P1092823** | Irx1 | 0.076832637 | -0.126922596 | 0.280587871 | 10.70891105 | 0.901251708 | 0.39891605 | 0.991835066 | -4.938338259 |
| **A_51_P277006** | Chst8 | 0.038836656 | -0.064167259 | 0.141840572 | 7.813057036 | 0.9011503 | 0.398966207 | 0.991835066 | -4.938404835 |
| **A_55_P2410626** | A430061O | 0.080610398 | -0.133245949 | 0.294466744 | 8.340309225 | 0.900903043 | 0.399088521 | 0.991835066 | -4.93856714 |
| **A_51_P107808** | Ttc19 | -0.039648839 | -0.144845791 | 0.065548113 | 8.240842906 | -0.900816739 | 0.399131221 | 0.991835066 | -4.928934246 |
| **A_55_P2026794** | Unc50 | 0.047052372 | -0.077817236 | 0.171921979 | 11.09319333 | 0.900604065 | 0.399236459 | 0.991835066 | -4.938763352 |
| **A_55_P2055638** | Gsn | -0.072974254 | -0.266642521 | 0.120694012 | 13.67301318 | -0.900575927 | 0.399250384 | 0.991835066 | -4.938781816 |
| **A_55_P2093974** | Mphosph1( | 0.09086371 | -0.150294143 | 0.332021564 | 8.820195753 | 0.90053001 | 0.399273108 | 0.991835066 | -4.938811945 |
| **A_55_P2015495** | Abat | -0.045097394 | -0.164789366 | 0.074594577 | 11.49340231 | -0.900524557 | 0.399275807 | 0.991835066 | -4.938815524 |
| **A_55_P2028511** | NA | 0.092769073 | -0.15351426 | 0.339052407 | 11.68939214 | 0.900279433 | 0.399397136 | 0.991835066 | -4.938976348 |
| **A_55_P2135261** | Ccdc174 | 0.055393977 | -0.091701455 | 0.202489409 | 10.25626402 | 0.90006214 | 0.399504713 | 0.991835066 | -4.939118886 |
| **A_55_P1956418** | Efr3b | 0.096072557 | -0.159116552 | 0.351261665 | 10.69397861 | 0.899800739 | 0.399634156 | 0.991835066 | -4.939290325 |
| **A_51_P325592** | 2410004B1 | -0.051109406 | -0.186873939 | 0.084655128 | 10.76654294 | -0.899753683 | 0.399657461 | 0.991835066 | -4.939321183 |
| **A_52_P641367** | Desi1 | -0.05051766 | -0.184714188 | 0.083678869 | 8.843903122 | -0.899727663 | 0.399670347 | 0.991835066 | -4.939338245 |
| **A_55_P2059864** | Igsf1 | -0.088608816 | -0.32400048 | 0.146782848 | 10.11743692 | -0.899694338 | 0.399686853 | 0.991835066 | -4.939360098 |
| **A_52_P194316** | Atp13a3 | 0.058582677 | -0.097069519 | 0.214234872 | 8.431434231 | 0.899545412 | 0.399760621 | 0.991835066 | -4.939457746 |
| **A_52_P222967** | Adck2 | -0.044573277 | -0.163007945 | 0.073861391 | 8.735874692 | -0.899507624 | 0.399779339 | 0.991835066 | -4.939482521 |
| **A_51_P200083** | Grina | -0.074211785 | -0.271418121 | 0.122994551 | 10.47555554 | -0.899417112 | 0.399824179 | 0.991835066 | -4.939541861 |
| **A_55_P1964359** | Cnot8 | -0.061243145 | -0.224001686 | 0.101515396 | 11.90190966 | -0.899337765 | 0.399863491 | 0.991835066 | -4.939593877 |

| | | | | | | | | | |
|---|---|---|---|---|---|---|---|---|---|
| **A_55_P2054842** | Clpb | -0.076458989 | -0.279667103 | 0.126749124 | 11.31177822 | -0.899283535 | 0.39989036 | 0.991835066 | -4.939629426 |
| **A_52_P14453** | Atxn1 | -0.042860909 | -0.156780298 | 0.07105848 | 8.430111321 | -0.899234301 | 0.399914756 | 0.991835066 | -4.939661699 |
| **A_52_P600531** | Bahd1 | -0.046495612 | -0.170100593 | 0.077109369 | 8.791908001 | -0.899052665 | 0.400004765 | 0.991835066 | -4.939780749 |
| **A_55_P2032294** | Zfp932 | 0.07214817 | -0.119655275 | 0.263951614 | 9.040630289 | 0.899038 | 0.400012033 | 0.991835066 | -4.93979036 |
| **A_52_P527842** | Reps1 | 0.059463801 | -0.098620261 | 0.217547864 | 9.110057493 | 0.899029027 | 0.400016479 | 0.991835066 | -4.939796241 |
| **A_55_P2076134** | NA | -0.075412226 | -0.275896324 | 0.125071872 | 12.25151595 | -0.899023332 | 0.400019302 | 0.991835066 | -4.939799973 |
| **A_51_P507053** | Slc38a1 | 0.049438496 | -0.081997518 | 0.18087451 | 12.98814953 | 0.899000637 | 0.40003055 | 0.991835066 | -4.939814847 |
| **A_55_P1987449** | Ep400 | 0.094796542 | -0.157232893 | 0.346825978 | 10.07312462 | 0.898980687 | 0.400040437 | 0.991835066 | -4.939827921 |
| **A_55_P2071191** | Prss8 | -0.041712976 | -0.152661605 | 0.069235653 | 7.704367439 | -0.898583345 | 0.400237404 | 0.991835066 | -4.940088276 |
| **A_55_P1965507** | Trnau1ap | 0.062771082 | -0.104212518 | 0.229754682 | 12.75766992 | 0.898452098 | 0.400302481 | 0.991835066 | -4.940174257 |
| **A_52_P134083** | Cyhr1 | -0.068653649 | -0.251292373 | 0.113985074 | 10.64591082 | -0.898421002 | 0.400317901 | 0.991835066 | -4.940194627 |
| **A_51_P268343** | Aagab | 0.055517885 | -0.092214369 | 0.20325014 | 11.93434509 | 0.898186915 | 0.400433991 | 0.991835066 | -4.940347952 |
| **A_55_P2015032** | Kazald1 | 0.18612634 | -0.309218979 | 0.681471659 | 9.477450293 | 0.898067349 | 0.400493298 | 0.991835066 | -4.940426256 |
| **A_55_P2012389** | Sfxn3 | 0.057375743 | -0.095387711 | 0.210139196 | 10.46775025 | 0.897672632 | 0.400689128 | 0.991835066 | -4.940684702 |
| **A_55_P2069768** | Kiss1 | -0.038167017 | -0.139826875 | 0.063492841 | 7.659052788 | -0.897321048 | 0.400863619 | 0.991835066 | -4.940914838 |
| **A_55_P2048660** | NA | 0.113154036 | -0.188277482 | 0.414585554 | 11.03052958 | 0.897202595 | 0.40092242 | 0.991835066 | -4.940992358 |
| **A_55_P2038573** | NA | -0.103256439 | -0.378374647 | 0.171861769 | 15.11007979 | -0.897029948 | 0.401008135 | 0.991835066 | -4.941105333 |
| **A_55_P1992004** | NA | -0.053997375 | -0.197873891 | 0.089879141 | 10.77134856 | -0.896998812 | 0.401023594 | 0.991835066 | -4.941125706 |
| **A_55_P1985428** | Atg16l2 | 0.101730557 | -0.169365041 | 0.372826155 | 9.349693364 | 0.896887744 | 0.401078745 | 0.991835066 | -4.941198375 |
| **A_55_P2110101** | Tm2d3 | 0.047879947 | -0.07973241 | 0.175492305 | 10.88369609 | 0.896747274 | 0.401148504 | 0.991835066 | -4.941290272 |
| **A_51_P428754** | Grik5 | 0.045012917 | -0.074958356 | 0.16498419 | 9.158178596 | 0.896745122 | 0.401149573 | 0.991835066 | -4.94129168 |
| **A_55_P1974273** | Ndufs7 | -0.061561802 | -0.225641877 | 0.102518272 | 15.08581612 | -0.896736014 | 0.401154096 | 0.991835066 | -4.941297638 |
| **A_51_P242733** | Tex261 | -0.053823815 | -0.197288841 | 0.089641211 | 12.00482789 | -0.896680182 | 0.401181826 | 0.991835066 | -4.941334161 |
| **A_66_P112164** | Rab29 | -0.057405074 | -0.210420957 | 0.09561081 | 10.47335861 | -0.896649885 | 0.401196874 | 0.991835066 | -4.941353979 |
| **A_52_P467266** | Mtf1 | -0.052567746 | -0.192695835 | 0.087560342 | 9.248635622 | -0.896609412 | 0.401216976 | 0.991835066 | -4.941380452 |
| **A_55_P2157922** | Ndufaf1 | 0.057595847 | -0.095936378 | 0.211128073 | 11.72001729 | 0.89660418 | 0.401219575 | 0.991835066 | -4.941383874 |
| **A_51_P419571** | Polr2d | -0.045907711 | -0.168283776 | 0.076468354 | 11.09880828 | -0.896599068 | 0.401222115 | 0.991835066 | -4.941387219 |
| **A_65_P15812** | Pum1 | 0.05367259 | -0.089431216 | 0.196776396 | 10.75638128 | 0.896417857 | 0.401312131 | 0.991835066 | -4.941505739 |
| **A_55_P2007495** | NA | 0.146449126 | -0.244036815 | 0.536935067 | 11.20119485 | 0.896376607 | 0.401332624 | 0.991835066 | -4.941532716 |
| **A_51_P356467** | Cldn4 | 0.039285933 | -0.065473076 | 0.144044941 | 7.687532394 | 0.896302938 | 0.401369225 | 0.991835066 | -4.941580892 |
| **A_51_P445841** | Deptor | -0.077045868 | -0.282498681 | 0.128406946 | 7.990139273 | -0.896285538 | 0.40137787 | 0.991835066 | -4.941592271 |
| **A_55_P2047626** | Cutal | -0.046006142 | -0.168701783 | 0.076689498 | 8.527507489 | -0.896181175 | 0.401429726 | 0.991835066 | -4.941660514 |
| **A_51_P273538** | Syce2 | 0.119920074 | -0.19995439 | 0.439794538 | 9.861250956 | 0.896027729 | 0.401505979 | 0.991835066 | -4.941760843 |
| **A_55_P1963533** | Pcdhac2 | 0.040735513 | -0.067978878 | 0.149449903 | 7.906427017 | 0.895561232 | 0.401737864 | 0.991835066 | -4.942065778 |
| **A_52_P625277** | Accs | 0.077489534 | -0.129346009 | 0.284325078 | 9.142325779 | 0.895420457 | 0.40180786 | 0.991835066 | -4.942157776 |
| **A_55_P2070084** | Zfp988 | -0.094997694 | -0.34861696 | 0.158621573 | 9.4995516 | -0.895240976 | 0.401897113 | 0.991835066 | -4.942275054 |
| **A_55_P2060483** | Rnf141 | 0.045703819 | -0.076338375 | 0.167746013 | 8.400671901 | 0.895058906 | 0.40198767 | 0.991835066 | -4.942394005 |
| **A_51_P365399** | Smg5 | 0.057924525 | -0.096801971 | 0.212651022 | 11.05604778 | 0.894760752 | 0.402135996 | 0.991835066 | -4.942588761 |
| **A_55_P1976012** | Cnih4 | -0.052840127 | -0.194031256 | 0.088351003 | 11.51776604 | -0.89446957 | 0.402280893 | 0.991835066 | -4.942778917 |
| **A_55_P2079251** | Ap2m1 | -0.045140465 | -0.16582748 | 0.075546549 | 11.88739235 | -0.893952861 | 0.402538112 | 0.991835066 | -4.943116243 |
| **A_55_P2147842** | Zglp1 | -0.062015765 | -0.227832297 | 0.103800767 | 9.208711763 | -0.893888621 | 0.402570099 | 0.991835066 | -4.943158172 |
| **A_55_P2026818** | Slc4a7 | 0.044329022 | -0.074220005 | 0.16287805 | 9.800972642 | 0.893715495 | 0.402656313 | 0.991835066 | -4.943271157 |
| **A_55_P2024704** | Cpe | 0.110807701 | -0.185558976 | 0.407174379 | 14.35952596 | 0.89361347 | 0.402707127 | 0.991835066 | -4.943337733 |
| **A_55_P2096933** | Cntln | 0.077638505 | -0.130043437 | 0.285320447 | 10.21677236 | 0.893485604 | 0.402770818 | 0.991835066 | -4.943421164 |
| **A_51_P371745** | Havcr1 | -0.038014904 | -0.139708354 | 0.063678546 | 7.648199296 | -0.893449587 | 0.402788759 | 0.991835066 | -4.943444663 |
| **A_51_P517051** | Castor1 | -0.057562019 | -0.211557182 | 0.096433143 | 7.817300183 | -0.893383803 | 0.402821531 | 0.991835066 | -4.943487582 |
| **A_52_P460791** | Ppy | 0.040565632 | -0.067976295 | 0.14910756 | 7.642179278 | 0.893243473 | 0.402891444 | 0.991835066 | -4.943579127 |
| **A_51_P114028** | Bcl7c | -0.135846116 | -0.499335348 | 0.227643116 | 10.02195844 | -0.893233103 | 0.402896612 | 0.991835066 | -4.943585892 |

| | | | | | | | | | |
|---|---|---|---|---|---|---|---|---|---|
| **A_55_P2111770** | Klhl13 | 0.071759777 | -0.120284583 | 0.263804137 | 12.6410342 | 0.893076489 | 0.402974649 | 0.991835066 | -4.943688048 |
| **A_55_P1992388** | NA | -0.054912762 | -0.201874389 | 0.092048865 | 14.05861557 | -0.893055546 | 0.402985086 | 0.991835066 | -4.943701707 |
| **A_52_P529360** | Rab11fip5 | 0.06173095 | -0.103484773 | 0.226946672 | 12.88524417 | 0.893019036 | 0.40300328 | 0.991835066 | -4.94372552 |
| **A_52_P347591** | Ndufa7 | -0.076705788 | -0.28202058 | 0.128609004 | 15.31652882 | -0.892929205 | 0.403048049 | 0.991835066 | -4.943784106 |
| **A_55_P2145257** | NA | 0.059705164 | -0.10010576 | 0.219516088 | 9.397454753 | 0.892924144 | 0.403050571 | 0.991835066 | -4.943787407 |
| **A_51_P464300** | Gdf1 | 0.080564815 | -0.135111672 | 0.296241303 | 9.422962952 | 0.892794994 | 0.403114943 | 0.991835066 | -4.943871628 |
| **A_55_P2169445** | Mfsd6 | 0.059018121 | -0.098981232 | 0.217017474 | 10.20864493 | 0.892769212 | 0.403127794 | 0.991835066 | -4.94388844 |
| **A_55_P2154097** | NA | 0.042539978 | -0.071348608 | 0.156428565 | 14.79183635 | 0.892742469 | 0.403141124 | 0.991835066 | -4.943905878 |
| **A_51_P146760** | Pcdhb20 | 0.066141271 | -0.110951071 | 0.243233614 | 8.482403258 | 0.892651393 | 0.403186525 | 0.991835066 | -4.943965263 |
| **A_52_P820196** | Cxcl15 | 0.037681053 | -0.063229022 | 0.138591128 | 7.678900733 | 0.892478233 | 0.403272854 | 0.991835066 | -4.944078157 |
| **A_55_P2021665** | Zfp638 | 0.075548057 | -0.126780994 | 0.277877107 | 10.59173187 | 0.892430053 | 0.403296877 | 0.991835066 | -4.944109566 |
| **A_55_P2075829** | NA | -0.049459788 | -0.181946062 | 0.083026486 | 8.799500131 | -0.892258091 | 0.403382627 | 0.991835066 | -4.944221659 |
| **A_55_P2060072** | Clk2 | 0.070979076 | -0.119154164 | 0.261112316 | 10.2765353 | 0.892239458 | 0.403391919 | 0.991835066 | -4.944233804 |
| **A_52_P504302** | Pmpca | -0.106016121 | -0.390019477 | 0.177987235 | 10.27109841 | -0.892190449 | 0.403416361 | 0.991835066 | -4.944265748 |
| **A_55_P2028532** | Ate1 | 0.039901423 | -0.066996342 | 0.146799189 | 8.610004357 | 0.892131536 | 0.403445742 | 0.991835066 | -4.944304144 |
| **A_55_P1992814** | Shq1 | 0.04465621 | -0.075001405 | 0.164313826 | 8.100311963 | 0.89197083 | 0.403525901 | 0.991835066 | -4.944408875 |
| **A_55_P1953377** | NA | -0.046758556 | -0.172053013 | 0.078535902 | 9.004163926 | -0.891945582 | 0.403538496 | 0.991835066 | -4.944425328 |
| **A_55_P2169391** | Atp6v1a | -0.08123965 | -0.299010607 | 0.136531306 | 11.2870382 | -0.891614697 | 0.403703578 | 0.991835066 | -4.944640916 |
| **A_55_P1962747** | H2-Ab1 | 0.32214826 | -0.541411092 | 1.185707611 | 8.882405605 | 0.891605483 | 0.403708176 | 0.991835066 | -4.944646919 |
| **A_55_P2120254** | Avpi1 | -0.053305348 | -0.196208349 | 0.089597654 | 12.62147849 | -0.891535336 | 0.40374318 | 0.991835066 | -4.944692615 |
| **A_55_P2017183** | NA | 0.073525623 | -0.123617187 | 0.270668433 | 10.53820268 | 0.891388241 | 0.403816589 | 0.991835066 | -4.94478843 |
| **A_65_P10399** | Gna13 | -0.051393925 | -0.189203265 | 0.086415415 | 8.172808942 | -0.891337678 | 0.403841825 | 0.991835066 | -4.944821363 |
| **A_55_P1959833** | Tmem266 | 0.070644262 | -0.118807812 | 0.260096335 | 9.139334152 | 0.891223563 | 0.403898784 | 0.991835066 | -4.944895685 |
| **A_55_P2027930** | Nfx1 | -0.04313235 | -0.158818896 | 0.072554196 | 8.243747052 | -0.891106068 | 0.403957437 | 0.991835066 | -4.9449722 |
| **A_55_P1980951** | 1110012L1 | -0.103213881 | -0.380058575 | 0.173630814 | 11.41902716 | -0.891068384 | 0.40397625 | 0.991835066 | -4.94499674 |
| **A_55_P2093720** | Dnajc5 | -0.071449792 | -0.263096685 | 0.120197101 | 8.658678142 | -0.891062807 | 0.403979034 | 0.991835066 | -4.945000371 |
| **A_55_P2032659** | Tox | -0.08139397 | -0.299723278 | 0.136935337 | 8.670912393 | -0.89102385 | 0.403998484 | 0.991835066 | -4.945025738 |
| **A_55_P2182867** | Sgcb | -0.048976577 | -0.180357053 | 0.0824039 | 10.50812759 | -0.890977477 | 0.404021636 | 0.991835066 | -4.945055934 |
| **A_55_P1978451** | Ttc1 | 0.057645621 | -0.096998063 | 0.212289306 | 11.0576048 | 0.890929355 | 0.404045663 | 0.991835066 | -4.945087267 |
| **A_55_P2090617** | Pilra | -0.042866658 | -0.157872687 | 0.072139371 | 7.837590679 | -0.890857308 | 0.404081637 | 0.991835066 | -4.945134175 |
| **A_52_P684138** | Dpf3 | 0.067543588 | -0.113669433 | 0.248756608 | 8.285594716 | 0.890848511 | 0.40408603 | 0.991835066 | -4.945139902 |
| **A_55_P2155421** | Sec61g | 0.098779931 | -0.166267268 | 0.36382713 | 14.74006763 | 0.890747539 | 0.404136451 | 0.991835066 | -4.945205638 |
| **A_51_P176352** | Ndrg2 | -0.107490403 | -0.395912909 | 0.180932104 | 13.59097587 | -0.890737375 | 0.404141528 | 0.991835066 | -4.945212255 |
| **A_51_P305628** | Atp8a2 | -0.049866799 | -0.18367717 | 0.083943571 | 8.503832869 | -0.890698771 | 0.404160806 | 0.991835066 | -4.945237386 |
| **A_55_P2051099** | Tnfrsf1a | -0.096374723 | -0.354992389 | 0.162242942 | 10.29428855 | -0.890664389 | 0.404177977 | 0.991835066 | -4.945259767 |
| **A_51_P252859** | Ccn1 | 0.145656632 | -0.245229265 | 0.53654253 | 9.874324924 | 0.890613747 | 0.404203269 | 0.991835066 | -4.945292732 |
| **A_51_P341736** | Mmp2 | -0.099637897 | -0.36703317 | 0.167757375 | 10.91625221 | -0.890594403 | 0.404212931 | 0.991835066 | -4.945305324 |
| **A_55_P1999408** | NA | 0.109891303 | -0.185035434 | 0.404818039 | 11.82308393 | 0.890550002 | 0.404235108 | 0.991835066 | -4.945334225 |
| **A_55_P2184816** | Eif2s1 | -0.071066165 | -0.26179909 | 0.119666759 | 11.92985366 | -0.890525471 | 0.40424736 | 0.991835066 | -4.945350192 |
| **A_55_P2084344** | NA | 0.089933472 | -0.151444713 | 0.331311657 | 13.09280537 | 0.890497033 | 0.404261565 | 0.991835066 | -4.945368702 |
| **A_51_P470989** | Paip1 | 0.05924466 | -0.099770504 | 0.218259824 | 9.404134192 | 0.890471049 | 0.404274544 | 0.991835066 | -4.945385614 |
| **A_55_P2039210** | Cpt1b | -0.056484877 | -0.208094719 | 0.095124965 | 11.88507262 | -0.890459001 | 0.404280562 | 0.991835066 | -4.945393455 |
| **A_51_P232411** | Rag1 | -0.084536352 | -0.311461151 | 0.142388446 | 7.702722069 | -0.89037033 | 0.404324856 | 0.991835066 | -4.945451164 |
| **A_51_P425696** | Nt5c2 | -0.115451348 | -0.425368525 | 0.19446583 | 10.02078554 | -0.890353544 | 0.404333242 | 0.991835066 | -4.945462089 |
| **A_51_P110395** | Setd2 | 0.049407723 | -0.083231928 | 0.182047374 | 11.46418479 | 0.890288161 | 0.404365906 | 0.991835066 | -4.940729198 |
| **A_51_P148037** | Sh3pxd2b | 0.063833989 | -0.107548488 | 0.235216467 | 9.194338671 | 0.890214576 | 0.40440267 | 0.991835066 | -4.945552524 |
| **A_52_P584335** | Folr1 | 0.03934986 | -0.066306965 | 0.145006685 | 7.742497953 | 0.890132727 | 0.404443566 | 0.991835066 | -4.945605782 |
| **A_55_P2046084** | Dcun1d4 | 0.076697109 | -0.129245824 | 0.282640042 | 9.361813317 | 0.890104985 | 0.404457428 | 0.991835066 | -4.945623834 |

| | | | | | | | | | |
|---|---|---|---|---|---|---|---|---|---|
| **A_55_P1994907** | Senp2 | 0.048414578 | -0.081616913 | 0.178446068 | 10.0842627 | 0.889890846 | 0.404564438 | 0.991835066 | -4.945763153 |
| **A_55_P2008704** | Gstp2 | -0.132246681 | -0.48745849 | 0.222965127 | 11.72375399 | -0.889828928 | 0.404595385 | 0.991835066 | -4.945803433 |
| **A_55_P2117130** | Zfp937 | 0.046513786 | -0.078422423 | 0.171449995 | 8.746971817 | 0.889820683 | 0.404599505 | 0.991835066 | -4.945808796 |
| **A_51_P211573** | Wdr47 | 0.052483054 | -0.088505658 | 0.193471767 | 9.122774983 | 0.889700609 | 0.404659523 | 0.991835066 | -4.945886902 |
| **A_55_P1984830** | Fgl1 | -0.048519639 | -0.17888282 | 0.081843542 | 8.125297467 | -0.889552826 | 0.404733399 | 0.991835066 | -4.945983022 |
| **A_55_P2142758** | Slc9a7 | -0.058730881 | -0.216531623 | 0.09906986 | 8.553093086 | -0.889542311 | 0.404738656 | 0.991835066 | -4.94598986 |
| **A_52_P529570** | Nsl1 | 0.069830404 | -0.117805501 | 0.257466309 | 9.807326631 | 0.889483183 | 0.404768217 | 0.991835066 | -4.946028314 |
| **A_51_P194609** | Prss34 | -0.045075922 | -0.16619657 | 0.076044727 | 7.827574584 | -0.889478718 | 0.404770449 | 0.991835066 | -4.946031218 |
| **A_55_P2253885** | AI503301 | 0.071072011 | -0.119904146 | 0.262048168 | 8.36014755 | 0.889464438 | 0.404777589 | 0.991835066 | -4.946040505 |
| **A_51_P425632** | Mad2l1bp | -0.068619308 | -0.253047881 | 0.115809264 | 12.02856383 | -0.889256943 | 0.40488134 | 0.991835066 | -4.946175432 |
| **A_51_P160344** | Cenpv | 0.057402695 | -0.096887458 | 0.211692848 | 11.59463007 | 0.889207684 | 0.404905974 | 0.991835066 | -4.94620746 |
| **A_51_P156857** | Smim3 | 0.071166024 | -0.120163625 | 0.262495674 | 9.392126395 | 0.888995499 | 0.405012095 | 0.991835066 | -4.946345408 |
| **A_55_P2097736** | Patz1 | 0.03909176 | -0.066014524 | 0.144198044 | 8.525414389 | 0.888926124 | 0.405046796 | 0.991835066 | -4.946390505 |
| **A_52_P354390** | Snrpn | -0.09915364 | -0.365757562 | 0.167450282 | 9.092099449 | -0.888896629 | 0.405061551 | 0.991835066 | -4.946409678 |
| **A_55_P2279575** | 5430400D1 | 0.062178602 | -0.105033102 | 0.229390306 | 8.448701085 | 0.888757778 | 0.405131012 | 0.991835066 | -4.946499929 |
| **A_55_P2091473** | Bnip3 | 0.114647075 | -0.193684764 | 0.422978914 | 14.31414087 | 0.888697047 | 0.405161397 | 0.991835066 | -4.9465394 |
| **A_51_P168395** | Ttc30a1 | 0.054832657 | -0.09263969 | 0.202305004 | 9.507003377 | 0.888664505 | 0.405177678 | 0.991835066 | -4.946560549 |
| **A_66_P115531** | Crebzf | 0.197653576 | -0.333951134 | 0.729258286 | 10.36272585 | 0.888638219 | 0.405190831 | 0.991835066 | -4.946577632 |
| **A_55_P2012146** | NA | -0.039314317 | -0.145078793 | 0.066450159 | 8.085994137 | -0.888423513 | 0.405298269 | 0.991835066 | -4.946717154 |
| **A_55_P2012599** | NA | 0.056038478 | -0.094733857 | 0.206810813 | 8.000908951 | 0.888328928 | 0.405345606 | 0.991835066 | -4.94677861 |
| **A_66_P100093** | Ccdc47 | -0.067107441 | -0.247663788 | 0.113448906 | 12.05481264 | -0.888315113 | 0.40535252 | 0.991835066 | -4.946787586 |
| **A_51_P108190** | Rpl23 | 0.059094284 | -0.099905828 | 0.218094396 | 11.61861017 | 0.888294916 | 0.405362629 | 0.991835066 | -4.946800708 |
| **A_55_P1956918** | Adamts5 | -0.037316134 | -0.137723005 | 0.063090737 | 8.12785351 | -0.888264579 | 0.405377813 | 0.991835066 | -4.946820418 |
| **A_55_P2255399** | F630048H1 | 0.097428763 | -0.164766187 | 0.359623713 | 8.419733374 | 0.888120715 | 0.405449825 | 0.991835066 | -4.946913879 |
| **A_55_P2046509** | Amot | 0.077077362 | -0.130419568 | 0.284574291 | 9.995665605 | 0.887818718 | 0.405601021 | 0.991835066 | -4.947110035 |
| **A_55_P1960178** | NA | 0.067296782 | -0.113899632 | 0.248493195 | 17.21592113 | 0.887674675 | 0.405673151 | 0.991835066 | -4.947203577 |
| **A_55_P2052892** | Xiap | 0.110214189 | -0.18655659 | 0.406984969 | 11.753711 | 0.887616785 | 0.405702143 | 0.991835066 | -4.947241169 |
| **A_55_P1986306** | Ltv1 | 0.110792986 | -0.18754237 | 0.409128342 | 11.6571213 | 0.887598736 | 0.405711182 | 0.991835066 | -4.947252889 |
| **A_55_P1967002** | NA | 0.14743195 | -0.249621918 | 0.544485817 | 9.505744098 | 0.887465155 | 0.405778086 | 0.991835066 | -4.947339622 |
| **A_55_P1999852** | Tchp | 0.046468677 | -0.078687341 | 0.171624695 | 8.9457948 | 0.887396489 | 0.405812481 | 0.991835066 | -4.947384203 |
| **A_52_P204689** | Map4k2 | 0.040384272 | -0.068388649 | 0.149157192 | 8.358345843 | 0.887361531 | 0.405829993 | 0.991835066 | -4.947406898 |
| **A_51_P121302** | Them4 | -0.054658517 | -0.201911598 | 0.092594565 | 9.184559691 | -0.887161287 | 0.405930311 | 0.991835066 | -4.93775628 |
| **A_52_P447944** | Epcam | -0.036785413 | -0.135903942 | 0.062333117 | 7.684145329 | -0.887012857 | 0.406004683 | 0.991835066 | -4.947633227 |
| **A_52_P84803** | NA | 0.085868043 | -0.145504837 | 0.317240922 | 16.12775403 | 0.887009445 | 0.406006392 | 0.991835066 | -4.947635441 |
| **A_51_P180452** | Crk | 0.057878377 | -0.098091991 | 0.213848745 | 11.46069926 | 0.886917821 | 0.406052307 | 0.991835066 | -4.947694905 |
| **A_55_P2181246** | NA | -0.086414562 | -0.319297484 | 0.14646836 | 9.895783485 | -0.886866852 | 0.40607785 | 0.991835066 | -4.947727982 |
| **A_55_P1956687** | Rab37 | -0.045848419 | -0.169420822 | 0.077723983 | 8.195975508 | -0.886772074 | 0.40612535 | 0.991835066 | -4.947789484 |
| **A_55_P2011387** | Tifa | -0.048428889 | -0.179000646 | 0.082142867 | 8.028848189 | -0.88647072 | 0.40627641 | 0.991835066 | -4.947985006 |
| **A_51_P155345** | Chmp2a | -0.138669533 | -0.512544797 | 0.235205732 | 11.56600276 | -0.886468791 | 0.406277377 | 0.991835066 | -4.947986257 |
| **A_55_P1962951** | Stx5a | 0.063966642 | -0.10849814 | 0.236431425 | 12.12817088 | 0.886466365 | 0.406278594 | 0.991835066 | -4.947987831 |
| **A_55_P1968723** | Noct | 0.053752043 | -0.091189141 | 0.198693227 | 11.08031042 | 0.886364403 | 0.406329714 | 0.991835066 | -4.948053974 |
| **A_55_P2103796** | Arhgap11a | -0.057338681 | -0.211978969 | 0.097301608 | 9.249858205 | -0.88620496 | 0.406409663 | 0.991835066 | -4.948157394 |
| **A_52_P488305** | Ndel1 | 0.055812389 | -0.094720127 | 0.206344906 | 8.380006089 | 0.886154453 | 0.406434991 | 0.991835066 | -4.948190152 |
| **A_51_P120729** | Polr2f | -0.148316535 | -0.548349708 | 0.251716639 | 12.0257881 | -0.886140721 | 0.406441877 | 0.991835066 | -4.948199058 |
| **A_55_P2168692** | Cep131 | 0.044888112 | -0.076183053 | 0.165959277 | 9.872715386 | 0.886134708 | 0.406444893 | 0.991835066 | -4.948202958 |
| **A_55_P1960735** | Gdf15 | 0.140443221 | -0.238370091 | 0.519256533 | 8.803073987 | 0.886103968 | 0.406460309 | 0.991835066 | -4.948222894 |
| **A_51_P176071** | Cmtr2 | 0.048179467 | -0.081782998 | 0.178141933 | 9.418731205 | 0.886039702 | 0.40649254 | 0.991835066 | -4.948264572 |
| **A_55_P2004732** | Ttc39a | 0.053737442 | -0.091227387 | 0.198702272 | 8.134461337 | 0.885979093 | 0.406522938 | 0.991835066 | -4.948303875 |

| | | | | | | | | | |
|---|---|---|---|---|---|---|---|---|---|
| **A_66_P126348** | Smim31 | 0.109209776 | -0.185438701 | 0.403858253 | 11.17367918 | 0.885862764 | 0.406581288 | 0.991835066 | -4.948379308 |
| **A_52_P634122** | Rexo1 | 0.073426237 | -0.124682861 | 0.271535335 | 8.845305338 | 0.885841422 | 0.406591993 | 0.991835066 | -4.948393146 |
| **A_55_P1973208** | Tax1bp3 | -0.116119961 | -0.429454106 | 0.197214183 | 9.726868548 | -0.885744147 | 0.406640791 | 0.991835066 | -4.948456216 |
| **A_55_P2072095** | Mucl3 | 0.053655408 | -0.091156193 | 0.198467009 | 7.720941138 | 0.885562621 | 0.406731865 | 0.991835066 | -4.948573899 |
| **A_55_P2131920** | Prrx1 | -0.074062531 | -0.27396741 | 0.125842348 | 9.867497123 | -0.885491279 | 0.406767663 | 0.991835066 | -4.948620144 |
| **A_55_P2065059** | Wnt2 | 0.040542972 | -0.068917579 | 0.150003524 | 7.747353923 | 0.885252346 | 0.406887569 | 0.991835066 | -4.948775008 |
| **A_65_P20433** | Parpbp | -0.069351654 | -0.256611817 | 0.117908509 | 8.882170696 | -0.885157506 | 0.406935171 | 0.991835066 | -4.948836469 |
| **A_55_P2054857** | Ube2v2 | 0.074228587 | -0.126220679 | 0.274677854 | 11.28900503 | 0.885066406 | 0.4069809 | 0.991835066 | -4.948895503 |
| **A_55_P2095854** | Rab3c | 0.165769585 | -0.281886785 | 0.613425954 | 11.78387712 | 0.885052867 | 0.406987697 | 0.991835066 | -4.948904276 |
| **A_55_P2047753** | Tec | -0.053597567 | -0.198377219 | 0.091182085 | 8.494296358 | -0.884803192 | 0.407113044 | 0.991835066 | -4.949066042 |
| **A_55_P2051384** | Ccdc88c | -0.096854413 | -0.3585094 | 0.164800574 | 10.28496852 | -0.884707133 | 0.407161278 | 0.991835066 | -4.949128271 |
| **A_55_P2427150** | C430002N1 | -0.036478803 | -0.135029712 | 0.062072106 | 7.66930066 | -0.884685844 | 0.407171968 | 0.991835066 | -4.949142062 |
| **A_51_P384091** | Rad21 | -0.07465342 | -0.276350019 | 0.127043178 | 13.65492532 | -0.884627158 | 0.407201439 | 0.991835066 | -4.949180076 |
| **A_51_P474454** | Rnaseh2b | -0.043771739 | -0.162139792 | 0.074596315 | 11.58551814 | -0.883829359 | 0.407602225 | 0.991835066 | -4.94969668 |
| **A_52_P449629** | Ankrd13c | -0.048659073 | -0.18024925 | 0.082931104 | 8.69775502 | -0.883790838 | 0.407621584 | 0.991835066 | -4.949721615 |
| **A_55_P2009042** | NA | 0.16380656 | -0.279233143 | 0.606846262 | 11.53476615 | 0.883685593 | 0.407674479 | 0.991835066 | -4.949789737 |
| **A_55_P2110978** | Atp6ap1 | 0.067311032 | -0.114743772 | 0.249365837 | 8.836723856 | 0.883676366 | 0.407679117 | 0.991835066 | -4.94979571 |
| **A_55_P1955382** | Rgma | -0.06203539 | -0.22982438 | 0.1057536 | 9.759800657 | -0.883659985 | 0.407687351 | 0.991835066 | -4.949806312 |
| **A_52_P106739** | Ddx46 | 0.054173174 | -0.0923587 | 0.200705049 | 10.55785635 | 0.883611403 | 0.40771177 | 0.991835066 | -4.949837755 |
| **A_55_P2072661** | Prex1 | 0.069173248 | -0.117948309 | 0.256294805 | 10.01605137 | 0.88353443 | 0.407750461 | 0.991835066 | -4.949887571 |
| **A_51_P416919** | Ube2j2 | -0.092310955 | -0.342126015 | 0.157504105 | 9.932955279 | -0.883168882 | 0.407934246 | 0.991835066 | -4.950124105 |
| **A_55_P2177105** | NA | -0.101578874 | -0.376529986 | 0.173372238 | 13.47248108 | -0.882992565 | 0.408022914 | 0.991835066 | -4.950238168 |
| **A_51_P290601** | Znhit2 | 0.08104035 | -0.138352023 | 0.300432724 | 10.68748091 | 0.882854056 | 0.408092579 | 0.991835066 | -4.950327762 |
| **A_55_P1974830** | Pde1c | -0.093095497 | -0.345126419 | 0.158935425 | 8.972733168 | -0.882844018 | 0.408097628 | 0.991835066 | -4.950334254 |
| **A_51_P378858** | Atg10 | 0.052786386 | -0.090130957 | 0.195703728 | 9.725365877 | 0.88276708 | 0.408136329 | 0.991835066 | -4.950384016 |
| **A_55_P1960621** | NA | 0.037104818 | -0.063365323 | 0.137574959 | 15.83152565 | 0.882678254 | 0.408181013 | 0.991835066 | -4.950441462 |
| **A_55_P2163263** | NA | 0.068113428 | -0.116356623 | 0.252583478 | 9.803540473 | 0.882502616 | 0.40826938 | 0.991835066 | -4.950555041 |
| **A_55_P2128821** | Selenok | -0.048761555 | -0.180824807 | 0.083301697 | 13.00627116 | -0.882479631 | 0.408280945 | 0.991835066 | -4.950569903 |
| **A_55_P2414479** | NA | -0.049552429 | -0.183779491 | 0.084674633 | 8.011879331 | -0.882335995 | 0.408353222 | 0.991835066 | -4.950662772 |
| **A_55_P2074080** | NA | -0.122069442 | -0.45273215 | 0.208593266 | 14.17245373 | -0.882329648 | 0.408356416 | 0.991835066 | -4.950666876 |
| **A_66_P118849** | NA | 0.039896983 | -0.06819682 | 0.147990786 | 8.141179439 | 0.882162096 | 0.40844074 | 0.991835066 | -4.950775194 |
| **A_65_P16680** | B4galt5 | 0.05469014 | -0.093506666 | 0.202886947 | 11.56005834 | 0.882021812 | 0.408511351 | 0.991835066 | -4.950865873 |
| **A_55_P2126537** | Fam53c | 0.060456462 | -0.103365888 | 0.224278812 | 11.88079027 | 0.882020561 | 0.408511981 | 0.991835066 | -4.950866682 |
| **A_55_P1982414** | NA | -0.084761764 | -0.314463375 | 0.144939847 | 8.34817728 | -0.881952261 | 0.408546362 | 0.991835066 | -4.950910827 |
| **A_55_P2045876** | NA | -0.06699138 | -0.2485396 | 0.11455684 | 8.982574312 | -0.881933953 | 0.408555579 | 0.991835066 | -4.950922659 |
| **A_51_P439452** | Insig2 | -0.057067026 | -0.211730817 | 0.097596765 | 10.76955404 | -0.881872342 | 0.408586595 | 0.991835066 | -4.950962478 |
| **A_55_P2032258** | Pdlim1 | -0.0574559 | -0.213195676 | 0.098283875 | 13.7038729 | -0.881747474 | 0.408649463 | 0.991835066 | -4.951043173 |
| **A_55_P2049313** | NA | -0.040713862 | -0.151079069 | 0.069651345 | 8.249303036 | -0.881696773 | 0.408674991 | 0.991835066 | -4.951075936 |
| **A_55_P1984361** | NA | 0.059223077 | -0.101325182 | 0.219771337 | 13.95997042 | 0.881646526 | 0.408700292 | 0.991835066 | -4.951108404 |
| **A_55_P1960576** | Kdm5b | -0.065729041 | -0.243920924 | 0.112462841 | 9.0025638 | -0.881614075 | 0.408716633 | 0.991835066 | -4.951129372 |
| **A_51_P239036** | Ubn2 | 0.070603416 | -0.120838247 | 0.262045079 | 10.90114832 | 0.881451434 | 0.408798539 | 0.991835066 | -4.951234453 |
| **A_55_P2184113** | Aurkaip1 | -0.049991232 | -0.185559546 | 0.085577082 | 14.71221336 | -0.881342628 | 0.40885334 | 0.991835066 | -4.951304745 |
| **A_52_P161400** | Cnpy4 | 0.040215684 | -0.068880482 | 0.149311851 | 9.020842231 | 0.881038941 | 0.409006324 | 0.991835066 | -4.9515009 |
| **A_55_P2369474** | Nr1h4 | -0.03994916 | -0.148341273 | 0.068442954 | 8.112918111 | -0.880884761 | 0.409084009 | 0.991835066 | -4.951600467 |
| **A_52_P354682** | Elovl7 | 0.070464015 | -0.120729083 | 0.261657113 | 8.509357875 | 0.880854763 | 0.409099125 | 0.991835066 | -4.951619838 |
| **A_55_P2154430** | Cd320 | 0.04730117 | -0.081052834 | 0.175655173 | 11.5403854 | 0.880788411 | 0.409132561 | 0.991835066 | -4.951662683 |
| **A_55_P2153431** | Ralgds | 0.044970246 | -0.077098982 | 0.167039473 | 9.365149936 | 0.880497643 | 0.409279109 | 0.991835066 | -4.951850408 |
| **A_55_P2110887** | NA | 0.068084108 | -0.116748424 | 0.252916641 | 11.15627908 | 0.880392777 | 0.409331972 | 0.991835066 | -4.9519181 |

| | | | | | | | | | |
|---|---|---|---|---|---|---|---|---|---|
| **A_55_P1989578** | NA | 0.146026521 | -0.250401692 | 0.542454734 | 9.105166247 | 0.880392464 | 0.409332129 | 0.991835066 | -4.951918302 |
| **A_51_P322972** | Hkdc1 | 0.039975125 | -0.068554939 | 0.148505189 | 7.73044481 | 0.880336894 | 0.409360144 | 0.991835066 | -4.951954171 |
| **A_55_P2029151** | Snap47 | 0.07012682 | -0.120272971 | 0.260526612 | 9.050526392 | 0.880292111 | 0.409382722 | 0.991835066 | -4.951983075 |
| **A_52_P141136** | Fggy | 0.04163509 | -0.07141633 | 0.154686511 | 8.102615765 | 0.880222811 | 0.409417662 | 0.991835066 | -4.952027803 |
| **A_55_P1974452** | Lime1 | 0.151495949 | -0.259927702 | 0.5629196 | 9.359379648 | 0.880077461 | 0.409490951 | 0.991835066 | -4.952121604 |
| **A_51_P340699** | Rasl11a | 0.049646796 | -0.085196606 | 0.184490199 | 8.050199585 | 0.879975659 | 0.409542289 | 0.991835066 | -4.952187297 |
| **A_55_P1971840** | Muc13 | 0.050723854 | -0.08705539 | 0.188503098 | 7.863346599 | 0.879908636 | 0.40957609 | 0.991835066 | -4.952230542 |
| **A_66_P106583** | NA | 0.0536567 | -0.092108763 | 0.199422162 | 13.83467605 | 0.879788851 | 0.409636506 | 0.991835066 | -4.952307827 |
| **A_52_P113916** | NA | -0.077023165 | -0.286275892 | 0.132229563 | 8.880943703 | -0.879750177 | 0.409656013 | 0.991835066 | -4.952332777 |
| **A_55_P2093727** | Gemin6 | 0.063254838 | -0.108624028 | 0.235133704 | 9.519268091 | 0.879590233 | 0.409736697 | 0.991835066 | -4.952435957 |
| **A_55_P2016681** | Alkbh7 | -0.058646266 | -0.218004085 | 0.100711554 | 11.29236854 | -0.879581562 | 0.409741071 | 0.991835066 | -4.95244155 |
| **A_51_P130727** | Fkbp11 | 0.050551115 | -0.086834015 | 0.187936244 | 7.948554296 | 0.879427706 | 0.409818696 | 0.991835066 | -4.952540788 |
| **A_55_P2000399** | Rhox4a | -0.111452794 | -0.414374567 | 0.191468979 | 7.831739772 | -0.879365864 | 0.409849899 | 0.991835066 | -4.952580673 |
| **A_51_P442053** | Ptdss1 | -0.065150036 | -0.242250585 | 0.111950512 | 9.016038821 | -0.879232815 | 0.409917038 | 0.991835066 | -4.952666476 |
| **A_51_P219465** | Tmem11 | 0.060059333 | -0.103213152 | 0.223331819 | 13.22362517 | 0.879177647 | 0.40994488 | 0.991835066 | -4.952702051 |
| **A_52_P379277** | Enpp3 | -0.052407147 | -0.194926098 | 0.090111803 | 8.467627455 | -0.878874848 | 0.410097715 | 0.991835066 | -4.952897281 |
| **A_55_P2009752** | Hlf | 0.093714389 | -0.161231343 | 0.348660121 | 9.740566196 | 0.87855239 | 0.41026052 | 0.991835066 | -4.953105132 |
| **A_55_P2074453** | Slc16a9 | 0.036544489 | -0.062906552 | 0.135995529 | 7.682861854 | 0.878257138 | 0.410409631 | 0.991835066 | -4.953295398 |
| **A_51_P454691** | 4930430F0 | 0.04601804 | -0.07921765 | 0.17125373 | 8.823879335 | 0.878231762 | 0.410422448 | 0.991835066 | -4.953311748 |
| **A_51_P103706** | Cyp2c29 | -0.04045778 | -0.15056675 | 0.069651189 | 7.702378843 | -0.878189999 | 0.410443544 | 0.991835066 | -4.953338657 |
| **A_55_P2324921** | NA | 0.108061365 | -0.186144886 | 0.402267617 | 8.442592069 | 0.877864902 | 0.410607783 | 0.991835066 | -4.953548089 |
| **A_51_P380178** | Id3 | -0.070501334 | -0.26245092 | 0.121448252 | 11.61872534 | -0.877847926 | 0.41061636 | 0.991835066 | -4.953559024 |
| **A_52_P513123** | Dna2 | 0.058748155 | -0.10122639 | 0.218722699 | 8.325109915 | 0.877712896 | 0.410684593 | 0.991835066 | -4.953645994 |
| **A_65_P12392** | Timp2 | -0.0868368 | -0.323305331 | 0.14963173 | 10.68255065 | -0.877686851 | 0.410697755 | 0.991835066 | -4.953662768 |
| **A_51_P137800** | Acp7 | -0.037643695 | -0.140167169 | 0.064879778 | 7.642092301 | -0.877562529 | 0.410760585 | 0.991835066 | -4.953742831 |
| **A_55_P1979669** | Zfp946 | 0.064887033 | -0.111842343 | 0.241616408 | 9.194899233 | 0.877522587 | 0.410780773 | 0.991835066 | -4.953768552 |
| **A_51_P375390** | Gemin7 | -0.0755996 | -0.281507543 | 0.130308343 | 11.61250411 | -0.877516986 | 0.410783604 | 0.991835066 | -4.953772159 |
| **A_52_P368262** | Bcl7c | 0.05188496 | -0.089478305 | 0.193248225 | 8.609780956 | 0.877231159 | 0.410928089 | 0.991835066 | -4.953956192 |
| **A_55_P2124026** | Synj2 | -0.049507801 | -0.184407198 | 0.085391595 | 10.11855735 | -0.877147773 | 0.410970248 | 0.991835066 | -4.954009873 |
| **A_52_P578790** | Dok3 | -0.037168258 | -0.138447514 | 0.064110999 | 7.972655063 | -0.877123672 | 0.410982433 | 0.991835066 | -4.954025388 |
| **A_55_P2045476** | Hmgxb4 | 0.080259551 | -0.138443568 | 0.298962671 | 9.664623085 | 0.877103577 | 0.410992594 | 0.991835066 | -4.954038323 |
| **A_52_P15197** | Dcun1d2 | -0.050811791 | -0.18928609 | 0.087662508 | 11.62149855 | -0.877009826 | 0.411039999 | 0.991835066 | -4.95409867 |
| **A_55_P2021565** | Ntf3 | 0.063280477 | -0.109222826 | 0.23578378 | 9.207454402 | 0.876761477 | 0.411165594 | 0.991835066 | -4.954258508 |
| **A_66_P112005** | Olfr1252 | 0.041822507 | -0.072187728 | 0.155832743 | 8.067111119 | 0.876749153 | 0.411171828 | 0.991835066 | -4.954266439 |
| **A_55_P2227154** | A530041M( | 0.091342011 | -0.157671355 | 0.340355377 | 8.806874376 | 0.876712184 | 0.411190526 | 0.991835066 | -4.954290229 |
| **A_51_P342716** | Ppp1cb | -0.044567281 | -0.166068505 | 0.076933944 | 13.36787819 | -0.876687092 | 0.411203218 | 0.991835066 | -4.954306376 |
| **A_66_P137556** | Tle1 | -0.053407367 | -0.199013353 | 0.092198619 | 8.173821752 | -0.876659754 | 0.411217046 | 0.991835066 | -4.954323967 |
| **A_55_P2014531** | Hes7 | -0.048019618 | -0.17894387 | 0.082904634 | 8.659103113 | -0.876612654 | 0.411240871 | 0.991835066 | -4.954354275 |
| **A_55_P2035439** | Uxt | -0.072167402 | -0.268937287 | 0.124602482 | 11.19609306 | -0.876580035 | 0.411257372 | 0.991835066 | -4.954375263 |
| **A_55_P2021149** | NA | 0.043189646 | -0.074599834 | 0.160979126 | 8.084840428 | 0.876359444 | 0.411368971 | 0.991835066 | -4.954517187 |
| **A_51_P511015** | Fzd9 | 0.123908523 | -0.214116774 | 0.46193382 | 9.544865087 | 0.876114986 | 0.411492672 | 0.991835066 | -4.954674435 |
| **A_51_P125446** | Lzic | -0.065003692 | -0.242371061 | 0.112363677 | 9.681196193 | -0.875938135 | 0.411582179 | 0.991835066 | -4.954788174 |
| **A_55_P2043996** | Cops7a | -0.041677441 | -0.155397841 | 0.072042958 | 8.463660119 | -0.875934848 | 0.411583843 | 0.991835066 | -4.954790288 |
| **A_51_P225592** | Tpm4 | 0.088830302 | -0.153567745 | 0.331228348 | 13.90749172 | 0.875873027 | 0.411615135 | 0.991835066 | -4.954830043 |
| **A_51_P495825** | Apof | -0.051712471 | -0.192830121 | 0.089405178 | 7.789106334 | -0.875836603 | 0.411633572 | 0.991835066 | -4.954853466 |
| **A_51_P114049** | Tmem109 | -0.043853346 | -0.163526292 | 0.0758196 | 13.74149409 | -0.875822065 | 0.411640931 | 0.991835066 | -4.954862814 |
| **A_52_P193236** | Tedc2 | 0.041237599 | -0.071330192 | 0.15380539 | 8.37111845 | 0.875564926 | 0.411771112 | 0.991835066 | -4.955028144 |
| **A_55_P1961094** | Tmem230 | -0.037910344 | -0.141402209 | 0.065581521 | 10.84524361 | -0.87550905 | 0.411799405 | 0.991835066 | -4.955064065 |

| | | | | | | | | | |
|---|---|---|---|---|---|---|---|---|---|
| **A_55_P2176729** | Prdx6 | -0.05860895 | -0.218618757 | 0.101400857 | 12.42911959 | -0.875440172 | 0.411834283 | 0.991835066 | -4.955108343 |
| **A_52_P197666** | Med12 | 0.055361832 | -0.095799678 | 0.206523342 | 10.6312191 | 0.875343204 | 0.411883387 | 0.991835066 | -4.955170673 |
| **A_55_P2125982** | Lrrc4b | -0.08857288 | -0.330416482 | 0.153270722 | 12.69332102 | -0.87533702 | 0.411886519 | 0.991835066 | -4.955174648 |
| **A_55_P2103952** | NA | 0.059464879 | -0.102941786 | 0.221871545 | 8.078628822 | 0.875116403 | 0.411998258 | 0.991835066 | -4.955316441 |
| **A_55_P1990299** | Ophn1 | 0.037400846 | -0.064773694 | 0.139575386 | 7.798172274 | 0.874878752 | 0.412118648 | 0.991835066 | -4.955469151 |
| **A_52_P480939** | Tnpo1 | 0.060718766 | -0.105189796 | 0.226627328 | 10.74115686 | 0.874708344 | 0.412204991 | 0.991835066 | -4.955578634 |
| **A_55_P1956593** | Plekha4 | 0.066971008 | -0.116098574 | 0.250040591 | 8.402443734 | 0.874338854 | 0.412392249 | 0.991835066 | -4.955815968 |
| **A_55_P2089477** | Sypl | -0.061834725 | -0.230867547 | 0.107198096 | 12.03753416 | -0.874320231 | 0.412401689 | 0.991835066 | -4.955827928 |
| **A_55_P2010111** | Zxdc | -0.078638797 | -0.293644632 | 0.136367039 | 9.003794595 | -0.874169692 | 0.412478002 | 0.991835066 | -4.955924601 |
| **A_52_P605517** | Phactr1 | 0.048248171 | -0.083692564 | 0.180188905 | 8.305158602 | 0.873999299 | 0.412564392 | 0.991835066 | -4.956034009 |
| **A_55_P1967414** | Asph | -0.040261025 | -0.150370972 | 0.069848923 | 8.60592932 | -0.873911397 | 0.412608964 | 0.991835066 | -4.956090444 |
| **A_66_P125110** | NA | 0.097871399 | -0.169812379 | 0.365555176 | 10.57764441 | 0.873862041 | 0.412633992 | 0.991835066 | -4.95612213 |
| **A_52_P981697** | Zfp874a | -0.041716162 | -0.155837188 | 0.072404865 | 8.049687233 | -0.873670772 | 0.412730994 | 0.991835066 | -4.956244909 |
| **A_55_P1995045** | Zc3hav1 | -0.047977822 | -0.179239151 | 0.083283507 | 8.238023689 | -0.873600484 | 0.412766645 | 0.991835066 | -4.956290024 |
| **A_55_P1955287** | NA | 0.086713055 | -0.15052458 | 0.32395069 | 16.28576432 | 0.873594789 | 0.412769533 | 0.991835066 | -4.956293679 |
| **A_55_P2149122** | Farp2 | 0.035844206 | -0.062229553 | 0.133917966 | 9.083357565 | 0.873524881 | 0.412804994 | 0.991835066 | -4.956338546 |
| **A_55_P2021515** | Rpl3 | 0.06609551 | -0.114762511 | 0.246953532 | 16.25902451 | 0.873460594 | 0.412837605 | 0.991835066 | -4.956379804 |
| **A_55_P1958867** | Gm9790 | 0.099246774 | -0.172343809 | 0.370837357 | 15.19923639 | 0.873395249 | 0.412870754 | 0.991835066 | -4.956421738 |
| **A_55_P2144597** | 9030025P2 | 0.051000217 | -0.08861367 | 0.190614104 | 8.395036456 | 0.873076987 | 0.413032238 | 0.991835066 | -4.956625946 |
| **A_55_P2047188** | Fgf1 | -0.108852112 | -0.406859506 | 0.189155283 | 11.27452935 | -0.873009461 | 0.413066506 | 0.991835066 | -4.956669266 |
| **A_51_P106259** | Camk1 | -0.079483376 | -0.297130387 | 0.138163636 | 8.707814764 | -0.872836153 | 0.413154466 | 0.991835066 | -4.956780436 |
| **A_51_P449824** | NA | 0.043997878 | -0.076492471 | 0.164488227 | 8.125914726 | 0.872747471 | 0.413199481 | 0.991835066 | -4.956837317 |
| **A_55_P1989798** | NA | 0.070678987 | -0.122894264 | 0.264252237 | 14.73267564 | 0.872678155 | 0.413234667 | 0.991835066 | -4.956881773 |
| **A_55_P2168189** | NA | 0.038862188 | -0.067586249 | 0.145310624 | 12.08648226 | 0.872563592 | 0.413292828 | 0.991835066 | -4.956955242 |
| **A_51_P187018** | Magohb | 0.083851088 | -0.145827702 | 0.313529878 | 10.75655536 | 0.872563302 | 0.413292975 | 0.991835066 | -4.956955428 |
| **A_51_P496804** | Adat2 | 0.05085951 | -0.088497536 | 0.190216556 | 8.80984883 | 0.872272884 | 0.413440439 | 0.991835066 | -4.957141642 |
| **A_51_P208697** | Ttl | 0.03509553 | -0.061085337 | 0.131276397 | 9.592644678 | 0.872111976 | 0.413522159 | 0.991835066 | -4.957244795 |
| **A_51_P487278** | Yipf3 | -0.08280868 | -0.309766252 | 0.144148892 | 11.00912023 | -0.872047852 | 0.413554729 | 0.991835066 | -4.957285899 |
| **A_55_P1956488** | Dmtn | 0.037994565 | -0.066185618 | 0.142174747 | 7.926035972 | 0.871656741 | 0.413753424 | 0.991835066 | -4.957536557 |
| **A_51_P172323** | Pak2 | 0.042619763 | -0.074260224 | 0.159499751 | 10.90455035 | 0.871525319 | 0.413820205 | 0.991835066 | -4.957620764 |
| **A_51_P210474** | Mrpl18 | 0.042661535 | -0.074339932 | 0.159663001 | 12.83238596 | 0.871473731 | 0.413846421 | 0.991835066 | -4.957653816 |
| **A_55_P2119420** | Ndufb2 | -0.066063616 | -0.24726265 | 0.115135418 | 14.92603555 | -0.871396072 | 0.413885889 | 0.991835066 | -4.95770357 |
| **A_55_P2003823** | Asgr1 | 0.044927289 | -0.078313955 | 0.168168533 | 7.68304842 | 0.871291097 | 0.413939242 | 0.991835066 | -4.957770817 |
| **A_52_P205789** | Unc119b | 0.049497136 | -0.086287584 | 0.185281857 | 8.71441159 | 0.871240993 | 0.41396471 | 0.991835066 | -4.957802912 |
| **A_55_P2028804** | Uchl3 | -0.03973815 | -0.148774243 | 0.069297942 | 7.981749327 | -0.871056851 | 0.414058318 | 0.991835066 | -4.957920857 |
| **A_55_P1980677** | Fam24b | 0.058195827 | -0.101488334 | 0.217879989 | 9.336255348 | 0.871042079 | 0.414065828 | 0.991835066 | -4.957930317 |
| **A_52_P368306** | Tmem100 | -0.059553059 | -0.223006729 | 0.103900612 | 8.410862853 | -0.870800211 | 0.414188806 | 0.991835066 | -4.958085205 |
| **A_55_P2000818** | Cab39l | -0.057301514 | -0.214640442 | 0.100037413 | 8.817454128 | -0.870440415 | 0.414371793 | 0.991835066 | -4.958315553 |
| **A_55_P2100108** | Zfp148 | 0.046891446 | -0.081868193 | 0.175651085 | 8.685849698 | 0.870408261 | 0.414388149 | 0.991835066 | -4.958336135 |
| **A_52_P644354** | Zfp106 | -0.06518025 | -0.244159795 | 0.113799295 | 11.11213412 | -0.870405759 | 0.414389422 | 0.991835066 | -4.958337737 |
| **A_52_P390944** | Chst3 | -0.047425181 | -0.177651788 | 0.082801426 | 8.314986017 | -0.870399039 | 0.414392841 | 0.991835066 | -4.958342038 |
| **A_51_P399889** | Ap4b1 | -0.036143658 | -0.135399868 | 0.063112551 | 10.8046761 | -0.870329201 | 0.414428368 | 0.991835066 | -4.95838674 |
| **A_51_P462299** | Aida | -0.050303018 | -0.188458655 | 0.087852619 | 9.521595513 | -0.870231032 | 0.41447831 | 0.991835066 | -4.958449572 |
| **A_55_P1958537** | Map6 | 0.111976421 | -0.195587978 | 0.419540819 | 8.975951771 | 0.870161072 | 0.414513905 | 0.991835066 | -4.958494345 |
| **A_55_P2047571** | Rps9 | 0.043575533 | -0.07614489 | 0.163295957 | 15.90377496 | 0.869928576 | 0.414632211 | 0.991835066 | -4.958643122 |
| **A_52_P259335** | Fkbp15 | -0.072763031 | -0.272674469 | 0.127148407 | 10.0485773 | -0.869925918 | 0.414633563 | 0.991835066 | -4.958644822 |
| **A_55_P2322039** | NA | -0.04141027 | -0.155198197 | 0.072377658 | 8.105514607 | -0.869803201 | 0.414696018 | 0.991835066 | -4.958723338 |
| **A_51_P109144** | Grtp1 | 0.039426737 | -0.068974662 | 0.147828136 | 9.173170347 | 0.869290804 | 0.414956868 | 0.991835066 | -4.959051086 |

| | | | | | | | | | |
|---|---|---|---|---|---|---|---|---|---|
| **A_55_P2062851** | 7-Mar | 0.072651037 | -0.127129364 | 0.272431438 | 11.62745258 | 0.86915667 | 0.415025173 | 0.991835066 | -4.95913686 |
| **A_55_P2123220** | Slc35e1 | -0.147084444 | -0.55161966 | 0.257450772 | 9.956059324 | -0.868999531 | 0.415105203 | 0.991835066 | -4.959237332 |
| **A_55_P2120652** | Snrpa1 | 0.076735849 | -0.134407036 | 0.287878733 | 11.88096333 | 0.868622315 | 0.415297362 | 0.991835066 | -4.959478462 |
| **A_51_P155142** | Cdca8 | -0.049262989 | -0.184823443 | 0.086297465 | 12.54221973 | -0.8685541 | 0.415332119 | 0.991835066 | -4.95952206 |
| **A_55_P2228918** | Rslcan18 | -0.050654588 | -0.190047576 | 0.0887384 | 8.203805882 | -0.868534333 | 0.415342191 | 0.991835066 | -4.959534693 |
| **A_51_P161890** | Fcgrt | -0.082021424 | -0.307732502 | 0.143689654 | 9.841564186 | -0.868527482 | 0.415345682 | 0.991835066 | -4.959539071 |
| **A_51_P442097** | Slc41a3 | -0.071635833 | -0.268818844 | 0.125547177 | 9.812712561 | -0.868300311 | 0.415461447 | 0.991835066 | -4.959684239 |
| **A_55_P2083499** | Shisal2a | -0.046127041 | -0.173133197 | 0.080879115 | 7.956741171 | -0.868040431 | 0.41559391 | 0.991835066 | -4.959850273 |
| **A_55_P2113920** | NA | 0.051493702 | -0.090303122 | 0.193290526 | 8.696797969 | 0.867954069 | 0.415637937 | 0.991835066 | -4.959905441 |
| **A_51_P415220** | Zmat3 | 0.071303438 | -0.125048662 | 0.267655539 | 9.968294065 | 0.867928709 | 0.415650865 | 0.991835066 | -4.95992164 |
| **A_55_P2097759** | NA | -0.059628481 | -0.223837379 | 0.104580416 | 13.49640103 | -0.86789302 | 0.41566906 | 0.991835066 | -4.959944436 |
| **A_55_P2103105** | Hacd1 | -0.071713977 | -0.269285747 | 0.125857793 | 14.36395792 | -0.867537083 | 0.415850559 | 0.991835066 | -4.960171754 |
| **A_55_P2263098** | A130078K2 | 0.081000293 | -0.142165099 | 0.304165685 | 8.389465836 | 0.867498784 | 0.415870091 | 0.991835066 | -4.960196208 |
| **A_52_P357611** | Neu3 | 0.052470345 | -0.092091792 | 0.197032483 | 8.856988187 | 0.867498018 | 0.415870482 | 0.991835066 | -4.960196697 |
| **A_52_P289893** | Prdm4 | 0.042618144 | -0.07482161 | 0.160057898 | 9.215897607 | 0.867338312 | 0.41595194 | 0.991835066 | -4.960298667 |
| **A_55_P2126627** | Phf20l1 | 0.080131471 | -0.140684871 | 0.300947814 | 8.736823484 | 0.867323347 | 0.415959574 | 0.991835066 | -4.960308221 |
| **A_55_P1978502** | H2-Q1 | -0.094412152 | -0.354654799 | 0.165830494 | 7.74452983 | -0.867078792 | 0.416084334 | 0.991835066 | -4.960464336 |
| **A_55_P2168267** | NA | 0.119385001 | -0.20979893 | 0.448568932 | 12.30181442 | 0.866802734 | 0.416225198 | 0.991835066 | -4.960640523 |
| **A_51_P418168** | Manf | -0.083546274 | -0.313952926 | 0.146860378 | 11.41541991 | -0.866644943 | 0.416305729 | 0.991835066 | -4.96074121 |
| **A_52_P399990** | B4galt2 | 0.046845915 | -0.082363469 | 0.1760553 | 9.742200422 | 0.866536385 | 0.416361141 | 0.991835066 | -4.960810473 |
| **A_51_P365912** | Ciao3 | -0.053513434 | -0.201113067 | 0.094086199 | 11.92658322 | -0.866536125 | 0.416361274 | 0.991835066 | -4.960810639 |
| **A_55_P2062015** | Acyp1 | -0.045987783 | -0.172841318 | 0.080865752 | 8.348468378 | -0.866461021 | 0.416399612 | 0.991835066 | -4.960858554 |
| **A_52_P47255** | Gmps | -0.061403657 | -0.230804831 | 0.107997517 | 9.350610458 | -0.866337179 | 0.416462836 | 0.991835066 | -4.960937557 |
| **A_55_P1988083** | Prc1 | -0.07039611 | -0.264606203 | 0.123813983 | 13.03283573 | -0.866335274 | 0.416463808 | 0.991835066 | -4.960938772 |
| **A_55_P2162811** | NA | 0.163846512 | -0.288297462 | 0.615990485 | 14.17495699 | 0.866103085 | 0.416582364 | 0.991835066 | -4.961086869 |
| **A_55_P2074811** | Zfp609 | -0.049064118 | -0.184487529 | 0.086359293 | 8.974602747 | -0.865923212 | 0.416674224 | 0.991835066 | -4.961201577 |
| **A_52_P24986** | Gpat4 | 0.056067008 | -0.098689967 | 0.210823982 | 12.18853766 | 0.865897089 | 0.416687566 | 0.991835066 | -4.961218235 |
| **A_55_P2030204** | Supt6 | 0.057154018 | -0.100603677 | 0.214911712 | 12.44433971 | 0.865895227 | 0.416688517 | 0.991835066 | -4.961219422 |
| **A_55_P1981455** | NA | -0.038105976 | -0.143291727 | 0.067079775 | 7.825892018 | -0.86585528 | 0.41670892 | 0.991835066 | -4.961244894 |
| **A_55_P1999637** | Gm15455 | 0.045845248 | -0.080703936 | 0.172394431 | 11.12618649 | 0.865852888 | 0.416710142 | 0.991835066 | -4.961246419 |
| **A_55_P1991124** | Tom1l2 | -0.07216489 | -0.271372827 | 0.127043048 | 12.22408744 | -0.865821655 | 0.416726095 | 0.991835066 | -4.961266334 |
| **A_55_P2134246** | Serpinb9 | -0.044195883 | -0.166209183 | 0.077817418 | 8.37157577 | -0.865732594 | 0.416771588 | 0.991835066 | -4.961323119 |
| **A_55_P1958833** | Gm9079 | 0.084660539 | -0.149145028 | 0.318466105 | 9.63616132 | 0.865436712 | 0.416922751 | 0.991835066 | -4.961511738 |
| **A_55_P2070686** | Wdfy1 | 0.035561352 | -0.062665163 | 0.133787867 | 8.515425353 | 0.865283979 | 0.417000797 | 0.991835066 | -4.961609085 |
| **A_55_P1970710** | Mllt6 | 0.081837187 | -0.144231506 | 0.307905881 | 12.59576084 | 0.865205763 | 0.417040769 | 0.991835066 | -4.961658932 |
| **A_55_P2049186** | Plvap | -0.057575018 | -0.216621713 | 0.101471677 | 8.66709703 | -0.865204091 | 0.417041623 | 0.991835066 | -4.961659997 |
| **A_55_P2154027** | Ptgis | 0.146540854 | -0.258276108 | 0.551357815 | 11.62991039 | 0.865185334 | 0.417051209 | 0.991835066 | -4.96167195 |
| **A_66_P113245** | Mfng | 0.087162977 | -0.153649127 | 0.327975082 | 11.30572025 | 0.865093155 | 0.417098321 | 0.991835066 | -4.961730691 |
| **A_55_P1958280** | Arrdc1 | 0.059471935 | -0.104879525 | 0.223823396 | 9.174505577 | 0.86486363 | 0.417215647 | 0.991835066 | -4.961876934 |
| **A_51_P443443** | 5830417I1( | 0.052326072 | -0.092288263 | 0.196940406 | 11.19821472 | 0.864800483 | 0.41724793 | 0.991835066 | -4.961917164 |
| **A_55_P2081388** | Mdm4 | 0.097920099 | -0.172746837 | 0.368587034 | 8.48521012 | 0.864660796 | 0.417319349 | 0.991835066 | -4.962006147 |
| **A_55_P1982449** | NA | -0.040072889 | -0.150878178 | 0.070732401 | 7.782089453 | -0.864369211 | 0.41746846 | 0.991835066 | -4.962191858 |
| **A_55_P2058646** | Nhlh1 | -0.041403702 | -0.155965491 | 0.073158087 | 8.055142617 | -0.863790697 | 0.417764414 | 0.991835066 | -4.962560177 |
| **A_55_P2135880** | Fancl | -0.047035863 | -0.177202402 | 0.083130676 | 10.73184744 | -0.863652216 | 0.417835281 | 0.991835066 | -4.962648316 |
| **A_51_P208410** | Ppp1r12c | -0.061483405 | -0.231633608 | 0.108666799 | 11.06577437 | -0.863643615 | 0.417839683 | 0.991835066 | -4.96265379 |
| **A_55_P2031242** | Sun1 | -0.040158789 | -0.151316261 | 0.070998684 | 8.213763782 | -0.863477591 | 0.417924657 | 0.991835066 | -4.962759445 |
| **A_51_P279854** | Ndufa11 | -0.088231185 | -0.332452561 | 0.155990191 | 13.27710803 | -0.863470633 | 0.417928218 | 0.991835066 | -4.962763872 |
| **A_55_P2137828** | Gprc5b | 0.076487995 | -0.135235546 | 0.288211535 | 9.912839626 | 0.863442178 | 0.417942783 | 0.991835066 | -4.962781979 |

| | | | | | | | | | |
|---|---|---|---|---|---|---|---|---|---|
| **A_55_P2078474** | Fam50a | 0.049754416 | -0.087976486 | 0.187485318 | 12.79984025 | 0.863394692 | 0.41796709 | 0.991835066 | -4.962812195 |
| **A_55_P1970901** | Gramd2 | 0.034428958 | -0.060887482 | 0.129745398 | 7.657920595 | 0.863306866 | 0.418012049 | 0.991835066 | -4.962868075 |
| **A_51_P370825** | Ccdc124 | -0.086621209 | -0.326451987 | 0.153209569 | 12.05985097 | -0.863233839 | 0.418049435 | 0.991835066 | -4.962914536 |
| **A_55_P1976574** | Tmod4 | -0.045343499 | -0.170919749 | 0.080232751 | 8.544168388 | -0.863011638 | 0.418163206 | 0.991835066 | -4.963055888 |
| **A_51_P468055** | Smyd1 | -0.070161369 | -0.264495558 | 0.124172819 | 11.39024706 | -0.862895052 | 0.418222909 | 0.991835066 | -4.963130043 |
| **A_52_P268279** | Fbxl17 | -0.051504583 | -0.19417445 | 0.091165284 | 10.31425005 | -0.862825058 | 0.418258755 | 0.991835066 | -4.963174559 |
| **A_51_P100787** | Snw1 | -0.063978201 | -0.241214342 | 0.11325794 | 11.03549903 | -0.862757751 | 0.418293227 | 0.991835066 | -4.963217363 |
| **A_55_P1994882** | Ccdc174 | 0.042396433 | -0.075069562 | 0.159862427 | 10.94918547 | 0.862633439 | 0.418356902 | 0.991835066 | -4.963296415 |
| **A_51_P235088** | Dusp27 | 0.104400475 | -0.184881061 | 0.393682012 | 9.881376236 | 0.862563135 | 0.418392915 | 0.991835066 | -4.963341118 |
| **A_51_P414653** | Plvap | 0.091150021 | -0.161424126 | 0.343724167 | 11.44072529 | 0.862535561 | 0.418407041 | 0.991835066 | -4.96335865 |
| **A_55_P1963080** | Rps12 | -0.057715238 | -0.217642813 | 0.102212336 | 16.60478149 | -0.8625341 | 0.418407789 | 0.991835066 | -4.963359579 |
| **A_51_P407866** | 1700020D0 | -0.041506486 | -0.156520825 | 0.073507853 | 8.357231327 | -0.862527824 | 0.418411004 | 0.991835066 | -4.96336357 |
| **A_52_P600535** | Zfp39 | 0.04450523 | -0.078821277 | 0.167831736 | 8.189344273 | 0.862509227 | 0.418420532 | 0.991835066 | -4.963375394 |
| **A_55_P2179299** | Ceacam13 | -0.368055762 | -1.38799464 | 0.651883117 | 8.348988357 | -0.862479153 | 0.418435939 | 0.991835066 | -4.963394515 |
| **A_55_P1986691** | Card6 | 0.073592919 | -0.130361333 | 0.277547172 | 8.599168167 | 0.86240725 | 0.418472776 | 0.991835066 | -4.963440229 |
| **A_55_P2042086** | Nrtn | -0.095699977 | -0.360927519 | 0.169527564 | 12.80363551 | -0.86238729 | 0.418483002 | 0.991835066 | -4.963452918 |
| **A_51_P482121** | Edn2 | -0.046563974 | -0.175639624 | 0.082511675 | 7.84069764 | -0.862213574 | 0.418572013 | 0.991835066 | -4.963563348 |
| **A_55_P2082989** | Rab7b | -0.081474614 | -0.307329079 | 0.144379852 | 8.23088789 | -0.862189561 | 0.418584319 | 0.991835066 | -4.963578612 |
| **A_55_P1999246** | Gm10318 | -0.073235377 | -0.276272972 | 0.129802218 | 11.36384847 | -0.862091961 | 0.418634335 | 0.991835066 | -4.963640647 |
| **A_52_P279557** | NA | -0.081155355 | -0.306185471 | 0.14387476 | 8.51182704 | -0.861957143 | 0.418703432 | 0.991835066 | -4.96372633 |
| **A_55_P2048368** | Fggy | 0.035794674 | -0.063461418 | 0.135050766 | 7.903074671 | 0.861926774 | 0.418718998 | 0.991835066 | -4.963745629 |
| **A_55_P2002884** | Zcchc24 | -0.058656948 | -0.221316535 | 0.104002639 | 12.90568024 | -0.861884217 | 0.418740811 | 0.991835066 | -4.963772673 |
| **A_51_P277795** | Resf1 | 0.065043696 | -0.11533022 | 0.245417612 | 9.52457373 | 0.861867711 | 0.418749272 | 0.991835066 | -4.963783162 |
| **A_51_P487818** | Fabp1 | -0.110254729 | -0.416074734 | 0.195565277 | 8.010058761 | -0.861668995 | 0.41885114 | 0.991835066 | -4.963909427 |
| **A_55_P2065726** | Snx29 | 0.047016795 | -0.083408226 | 0.177441815 | 8.239374811 | 0.861591167 | 0.418891043 | 0.991835066 | -4.963958873 |
| **A_51_P384640** | Mrpl10 | -0.075848957 | -0.28625699 | 0.134559076 | 10.0499252 | -0.861581625 | 0.418895935 | 0.991835066 | -4.963964935 |
| **A_55_P1962731** | Clcn4 | -0.082239846 | -0.310400532 | 0.145920841 | 12.924174 | -0.86149072 | 0.418942546 | 0.991835066 | -4.964022685 |
| **A_51_P239113** | Zfp35 | 0.043225796 | -0.07671198 | 0.163163572 | 10.67968045 | 0.861382696 | 0.418997939 | 0.991835066 | -4.964091304 |
| **A_52_P135822** | Myo18a | 0.042431512 | -0.075326011 | 0.160189034 | 14.18238021 | 0.861209826 | 0.419086595 | 0.991835066 | -4.964201101 |
| **A_55_P2078994** | Lrrc31 | -0.118076547 | -0.445802744 | 0.20964965 | 16.09229348 | -0.861115918 | 0.419134762 | 0.991835066 | -4.96426074 |
| **A_51_P209319** | Slc35b1 | 0.053614902 | -0.095213791 | 0.202443596 | 12.58523894 | 0.861009575 | 0.419189312 | 0.991835066 | -4.96432827 |
| **A_55_P2320283** | Hmga2-ps1 | -0.04191554 | -0.158274796 | 0.074443716 | 8.156239251 | -0.860960572 | 0.41921445 | 0.991835066 | -4.964359385 |
| **A_55_P2380806** | Gm2115 | 0.035779527 | -0.063551655 | 0.13511071 | 7.76922651 | 0.860910746 | 0.419240011 | 0.991835066 | -4.964391022 |
| **A_55_P2124941** | Brinp1 | 0.091941324 | -0.163308262 | 0.34719091 | 8.182371536 | 0.860904223 | 0.419243358 | 0.991835066 | -4.964395164 |
| **A_55_P2013296** | Prkag2 | -0.062497428 | -0.236018206 | 0.111023349 | 13.56828333 | -0.860834759 | 0.419278996 | 0.991835066 | -4.964439267 |
| **A_55_P2151082** | NA | 0.092105363 | -0.163683486 | 0.347894213 | 11.90891112 | 0.860622 | 0.419388165 | 0.991835066 | -4.964574334 |
| **A_55_P2235059** | 6230415J0 | 0.063489903 | -0.112857784 | 0.239837591 | 9.150119124 | 0.860486438 | 0.419457734 | 0.991835066 | -4.96466038 |
| **A_52_P274028** | Ssr1 | 0.103477571 | -0.18398381 | 0.390938952 | 9.211965841 | 0.86035136 | 0.419527063 | 0.991835066 | -4.964746109 |
| **A_51_P276939** | Gmip | 0.043928614 | -0.07810611 | 0.165963338 | 8.121901824 | 0.860346131 | 0.419529747 | 0.991835066 | -4.964749428 |
| **A_55_P2098539** | Itgb3bp | -0.065441185 | -0.24724075 | 0.116358379 | 10.8900775 | -0.860334721 | 0.419535603 | 0.991835066 | -4.964756669 |
| **A_55_P2186446** | NA | -0.092753631 | -0.350463266 | 0.164956004 | 16.05693645 | -0.860219723 | 0.419594634 | 0.991835066 | -4.964829645 |
| **A_51_P115666** | Fam83h | 0.047985475 | -0.08535398 | 0.181324931 | 8.164442428 | 0.860122396 | 0.419644598 | 0.991835066 | -4.964891402 |
| **A_55_P1993909** | Taz | 0.069070281 | -0.122898813 | 0.261039375 | 10.56798464 | 0.859941763 | 0.41973734 | 0.991835066 | -4.965006005 |
| **A_55_P1972118** | NA | 0.046622562 | -0.0829572 | 0.176202325 | 15.76766166 | 0.859939886 | 0.419738304 | 0.991835066 | -4.965007196 |
| **A_51_P466910** | Cacna1g | -0.121893317 | -0.460692376 | 0.216905742 | 9.477558353 | -0.859897788 | 0.419759921 | 0.991835066 | -4.965033903 |
| **A_51_P376057** | Bcorl1 | 0.041910518 | -0.074586803 | 0.158407839 | 8.758720528 | 0.859837188 | 0.419791039 | 0.991835066 | -4.965072345 |
| **A_55_P2295786** | Mindy2 | 0.040406283 | -0.071931482 | 0.152744049 | 8.123074428 | 0.859670959 | 0.419876407 | 0.991835066 | -4.965177784 |
| **A_55_P1965308** | Ccdc66 | -0.037790532 | -0.142868271 | 0.067287207 | 8.844222328 | -0.859570337 | 0.419928088 | 0.991835066 | -4.965241601 |

| | | | | | | | | | |
|---|---|---|---|---|---|---|---|---|---|
| **A_52_P450188** | Mtmr4 | 0.07034729 | -0.125325968 | 0.266020547 | 8.814387849 | 0.859260854 | 0.420087073 | 0.991835066 | -4.965437849 |
| **A_66_P133642** | NA | 0.06425604 | -0.114520102 | 0.243032183 | 9.715116621 | 0.859040282 | 0.420200409 | 0.991835066 | -4.965577685 |
| **A_51_P279183** | Tkfc | -0.042290799 | -0.159967348 | 0.07538575 | 8.908582572 | -0.858944486 | 0.420249639 | 0.991835066 | -4.965638409 |
| **A_66_P113760** | Acvr2a | -0.059939101 | -0.226725874 | 0.106847671 | 9.202650268 | -0.858929966 | 0.420257101 | 0.991835066 | -4.965647612 |
| **A_51_P234544** | Azin1 | 0.045993354 | -0.082040823 | 0.174027532 | 12.74819728 | 0.858575128 | 0.420439493 | 0.991835066 | -4.965872491 |
| **A_52_P662013** | Plg | -0.068182143 | -0.258002435 | 0.121638149 | 7.922866751 | -0.85849376 | 0.420481325 | 0.991835066 | -4.965924049 |
| **A_55_P2091762** | NA | 0.066558999 | -0.118748739 | 0.251866738 | 15.92344869 | 0.858464539 | 0.420496349 | 0.991835066 | -4.965942563 |
| **A_55_P2029420** | 2610318N( | 0.094373864 | -0.16839448 | 0.357142207 | 10.26895944 | 0.858396273 | 0.420531449 | 0.991835066 | -4.965985814 |
| **A_51_P405668** | 5-Mar | 0.063602909 | -0.113502566 | 0.240708384 | 11.13681698 | 0.858329674 | 0.420565694 | 0.991835066 | -4.966028008 |
| **A_55_P2035495** | Zbtb20 | -0.090660811 | -0.343111385 | 0.161789763 | 10.57884628 | -0.858326203 | 0.420567478 | 0.991835066 | -4.966030207 |
| **A_51_P245989** | Ccr2 | -0.040560243 | -0.153513152 | 0.072392667 | 7.81744919 | -0.858246922 | 0.420608247 | 0.991835066 | -4.966080431 |
| **A_51_P105017** | Rad52 | 0.036621681 | -0.06537686 | 0.138620223 | 8.393325153 | 0.858130714 | 0.420668009 | 0.991835066 | -4.961276544 |
| **A_55_P2019190** | NA | -0.04726863 | -0.178923091 | 0.084385832 | 15.12370183 | -0.85811711 | 0.420675006 | 0.991835066 | -4.966162659 |
| **A_55_P2063711** | Snu13 | 0.051397603 | -0.09178586 | 0.194581067 | 12.93792833 | 0.857944389 | 0.420763843 | 0.991835066 | -4.966272053 |
| **A_55_P2131348** | Loxl2 | 0.130316004 | -0.232731316 | 0.493363325 | 11.07457651 | 0.857913796 | 0.42077958 | 0.991835066 | -4.966291427 |
| **A_55_P2145982** | NA | 0.048882773 | -0.087310155 | 0.185075701 | 8.447603151 | 0.857848132 | 0.420813359 | 0.991835066 | -4.966333011 |
| **A_51_P263246** | Dusp8 | 0.086420106 | -0.154383668 | 0.32722388 | 10.16970809 | 0.857749826 | 0.420863932 | 0.991835066 | -4.966395261 |
| **A_55_P2017312** | Eapp | 0.047989862 | -0.08573614 | 0.181715864 | 11.02213026 | 0.857714547 | 0.420882083 | 0.991835066 | -4.966417599 |
| **A_52_P648715** | Triml1 | 0.040261226 | -0.071933847 | 0.1524563 | 7.67682569 | 0.857674191 | 0.420902846 | 0.991835066 | -4.966443152 |
| **A_52_P470401** | Tmem186 | -0.057816567 | -0.218961961 | 0.103328827 | 8.844047215 | -0.857518577 | 0.420982915 | 0.991835066 | -4.966541673 |
| **A_51_P415475** | Dok4 | -0.095807823 | -0.362843133 | 0.171227487 | 10.32954252 | -0.85751438 | 0.420985075 | 0.991835066 | -4.96654433 |
| **A_55_P2047882** | Zc3h7b | -0.062741212 | -0.237615993 | 0.112133569 | 10.45520418 | -0.857501428 | 0.42099174 | 0.991835066 | -4.96655253 |
| **A_51_P164207** | Mta2 | -0.106748638 | -0.40433179 | 0.190834514 | 11.28178312 | -0.857359839 | 0.421064604 | 0.991835066 | -4.966642159 |
| **A_51_P234864** | Sfxn5 | 0.071450966 | -0.127743394 | 0.270645326 | 8.253627712 | 0.857314559 | 0.421087908 | 0.991835066 | -4.966670821 |
| **A_55_P2035300** | Rhot1 | -0.068422646 | -0.259378102 | 0.122532811 | 11.73270545 | -0.85640052 | 0.421558528 | 0.991835066 | -4.967249144 |
| **A_55_P1979684** | Rhoc | -0.082588757 | -0.313082006 | 0.147904492 | 12.91905647 | -0.856390522 | 0.421563678 | 0.991835066 | -4.967255467 |
| **A_55_P1979281** | Rab1a | 0.046936752 | -0.08406356 | 0.177937063 | 13.2499248 | 0.856347104 | 0.421586043 | 0.991835066 | -4.967282926 |
| **A_55_P2138529** | Mrpl40 | -0.045570487 | -0.172762859 | 0.081621885 | 12.14556856 | -0.856311416 | 0.421604426 | 0.991835066 | -4.967305496 |
| **A_66_P130602** | Pdzd8 | -0.058710523 | -0.227083659 | 0.109662612 | 8.397606973 | -0.866184288 | 0.421640059 | 0.991835066 | -4.89779641 |
| **A_55_P2122555** | Csnk1e | -0.106072338 | -0.402199706 | 0.19005503 | 10.36565773 | -0.85611622 | 0.421704986 | 0.991835066 | -4.96742893 |
| **A_55_P2037156** | Pja1 | -0.042866008 | -0.16254215 | 0.076810134 | 12.14882341 | -0.856080476 | 0.421723402 | 0.991835066 | -4.96745153 |
| **A_55_P1990608** | Ube2j1 | -0.045966179 | -0.174299856 | 0.082367498 | 8.90011305 | -0.856065303 | 0.42173122 | 0.991835066 | -4.967461124 |
| **A_55_P2042823** | Sh3bp2 | 0.06311806 | -0.113124979 | 0.239361099 | 9.008423622 | 0.855954735 | 0.421788192 | 0.991835066 | -4.967531031 |
| **A_55_P2134251** | Hmgb1 | -0.073421477 | -0.278451526 | 0.131608571 | 12.66685896 | -0.855883662 | 0.421824817 | 0.991835066 | -4.967575962 |
| **A_55_P2156435** | NA | -0.073566253 | -0.27902481 | 0.131892303 | 11.44992288 | -0.855782771 | 0.421876811 | 0.991835066 | -4.96763974 |
| **A_55_P2172999** | Ptpn13 | 0.077918985 | -0.139745499 | 0.29558347 | 9.065598535 | 0.855588323 | 0.421977032 | 0.991835066 | -4.967762645 |
| **A_55_P2184741** | Katnal1 | -0.038609761 | -0.146465248 | 0.069245727 | 10.69920241 | -0.855586637 | 0.421977901 | 0.991835066 | -4.96776371 |
| **A_55_P2127204** | Amigo1 | -0.049877617 | -0.189254211 | 0.089498977 | 8.547679876 | -0.855312823 | 0.422119059 | 0.991835066 | -4.967936744 |
| **A_66_P137516** | NA | 0.047983518 | -0.086104914 | 0.182071951 | 13.70841638 | 0.855283131 | 0.422134367 | 0.991835066 | -4.967955504 |
| **A_51_P488768** | Slc25a44 | -0.03797278 | -0.144088294 | 0.068142734 | 8.271935353 | -0.855268836 | 0.422141738 | 0.991835066 | -4.967964537 |
| **A_55_P2038121** | Rbx1 | 0.042398843 | -0.076088737 | 0.160886424 | 14.15740247 | 0.855244544 | 0.422154263 | 0.991835066 | -4.967979885 |
| **A_55_P2182324** | NA | -0.054675231 | -0.207484403 | 0.098133941 | 8.955205125 | -0.855165828 | 0.422194852 | 0.991835066 | -4.968029618 |
| **A_55_P2107398** | NA | 0.111062528 | -0.19935546 | 0.421480515 | 11.86388683 | 0.855125402 | 0.422215699 | 0.991835066 | -4.968055158 |
| **A_55_P2035102** | Odc1 | 0.102515467 | -0.184026489 | 0.389057423 | 11.00019656 | 0.855087006 | 0.422235498 | 0.991835066 | -4.968079415 |
| **A_55_P2068459** | Hspa1a | -0.039835006 | -0.15118345 | 0.071513438 | 7.93504255 | -0.855046763 | 0.422256252 | 0.991835066 | -4.968104837 |
| **A_55_P2047345** | Sat2 | 0.061203189 | -0.10994427 | 0.232350649 | 9.297390287 | 0.854698081 | 0.422436097 | 0.991835066 | -4.968325072 |
| **A_55_P2388197** | 1010001B2 | -0.050214745 | -0.190637202 | 0.090207712 | 9.408183494 | -0.854680561 | 0.422445135 | 0.991835066 | -4.968336136 |
| **A_51_P403049** | Heatr5b | -0.093937363 | -0.356705977 | 0.168831251 | 9.434235874 | -0.854425114 | 0.422576929 | 0.991835066 | -4.968497437 |

| | | | | | | | | | |
|---|---|---|---|---|---|---|---|---|---|
| **A_51_P379775** | Nrm | -0.088334091 | -0.335436039 | 0.158767857 | 9.262525819 | -0.854400133 | 0.422589819 | 0.991835066 | -4.968513209 |
| **A_52_P522372** | Aard | 0.039399829 | -0.07085079 | 0.149650447 | 7.893393105 | 0.854126961 | 0.422730793 | 0.991835066 | -4.968685658 |
| **A_55_P1969522** | NA | -0.131631539 | -0.500051913 | 0.236788835 | 9.89717616 | -0.853936254 | 0.422829231 | 0.991835066 | -4.968806024 |
| **A_55_P2097474** | Creb5 | -0.060219437 | -0.228780075 | 0.108341202 | 14.13652152 | -0.853865866 | 0.422865567 | 0.991835066 | -4.968850445 |
| **A_51_P381360** | Ndufa13 | -0.084460103 | -0.32088453 | 0.151964325 | 17.35006914 | -0.853824063 | 0.422887148 | 0.991835066 | -4.968876825 |
| **A_55_P2132681** | Fzr1 | 0.067587499 | -0.121639735 | 0.256814733 | 9.844464917 | 0.85367363 | 0.422964817 | 0.991835066 | -4.968971748 |
| **A_51_P109835** | Uqcrc1 | -0.123317825 | -0.468606135 | 0.221970485 | 13.99672164 | -0.853597456 | 0.42300415 | 0.991835066 | -4.969019809 |
| **A_55_P2066414** | Dffa | -0.054144153 | -0.205753064 | 0.097464758 | 8.976682128 | -0.853563764 | 0.423021548 | 0.991835066 | -4.969041066 |
| **A_51_P469480** | Yes1 | -0.047876559 | -0.181960546 | 0.086207428 | 8.132322056 | -0.853404932 | 0.423103572 | 0.991835066 | -4.969141266 |
| **A_55_P2025090** | Gins3 | 0.050887846 | -0.091655006 | 0.193430697 | 12.51015515 | 0.853252867 | 0.423182112 | 0.991835066 | -4.969237183 |
| **A_52_P489778** | Ablim1 | -0.083052908 | -0.315696901 | 0.149591086 | 9.698413227 | -0.853241812 | 0.423187822 | 0.991835066 | -4.969244156 |
| **A_52_P503730** | Fchsd2 | -0.056389813 | -0.214354578 | 0.101574952 | 9.319490206 | -0.85319747 | 0.423210727 | 0.991835066 | -4.969272123 |
| **A_52_P631499** | NA | 0.067961658 | -0.122450068 | 0.258373383 | 10.8955298 | 0.85305966 | 0.423281917 | 0.991835066 | -4.969359034 |
| **A_55_P1974780** | NA | 0.191877718 | -0.345860387 | 0.729615823 | 9.82037784 | 0.852830776 | 0.423400174 | 0.991835066 | -4.969503359 |
| **A_55_P2041693** | Gm5834 | 0.037285714 | -0.067214889 | 0.141786318 | 7.931521231 | 0.852771745 | 0.423430678 | 0.991835066 | -4.969540577 |
| **A_55_P2140201** | Dupd1 | 0.035228398 | -0.063520496 | 0.133977292 | 7.745063486 | 0.852648017 | 0.423494617 | 0.991835066 | -4.969618578 |
| **A_66_P114463** | NA | -0.082697077 | -0.314563846 | 0.149169693 | 11.94998562 | -0.85243403 | 0.423605217 | 0.991835066 | -4.969753462 |
| **A_51_P439836** | NA | -0.106742148 | -0.406089994 | 0.192605698 | 11.10585842 | -0.852253773 | 0.423698399 | 0.991835066 | -4.969867065 |
| **A_51_P258766** | Smo | 0.078215824 | -0.141164896 | 0.297596544 | 11.34106468 | 0.852128908 | 0.423762956 | 0.991835066 | -4.969945747 |
| **A_55_P2178084** | Dnmt3a | 0.059111824 | -0.10670276 | 0.224926407 | 12.57529217 | 0.852041532 | 0.423808135 | 0.991835066 | -4.970000801 |
| **A_55_P2176894** | NA | -0.041971288 | -0.159737387 | 0.075794812 | 7.941192051 | -0.851806859 | 0.423929492 | 0.991835066 | -4.970148644 |
| **A_51_P111932** | Tor1aip2 | -0.055616049 | -0.21170987 | 0.100477772 | 8.897547406 | -0.851576271 | 0.424048762 | 0.991835066 | -4.970293884 |
| **A_55_P2175469** | Chd2 | 0.076126848 | -0.13755507 | 0.289808765 | 10.17614935 | 0.851489323 | 0.424093741 | 0.991835066 | -4.970348641 |
| **A_55_P2025695** | C330007P( | 0.081134096 | -0.146629394 | 0.308897586 | 10.4589278 | 0.851389767 | 0.424145247 | 0.991835066 | -4.970411334 |
| **A_55_P1993064** | Mbd6 | 0.103946974 | -0.187947143 | 0.395841091 | 11.22616857 | 0.851129496 | 0.424279921 | 0.991835066 | -4.970575208 |
| **A_55_P2086075** | Gm2058 | 0.075469849 | -0.13645957 | 0.287399269 | 8.856330219 | 0.851121119 | 0.424284256 | 0.991835066 | -4.970580482 |
| **A_52_P282500** | Kif21b | 0.074774819 | -0.135237247 | 0.284786884 | 9.246773854 | 0.850981766 | 0.424356376 | 0.991835066 | -4.970668206 |
| **A_55_P2177885** | Dact3 | -0.039402879 | -0.150081546 | 0.071275789 | 11.21274253 | -0.850889496 | 0.424404133 | 0.991835066 | -4.970726285 |
| **A_51_P337856** | Ubl7 | -0.074138548 | -0.282449446 | 0.134172351 | 10.13537527 | -0.850631016 | 0.424537939 | 0.991835066 | -4.97088896 |
| **A_51_P506755** | Tep1 | -0.050403401 | -0.192043609 | 0.091236807 | 9.634243292 | -0.850515862 | 0.424597561 | 0.991835066 | -4.97096142 |
| **A_51_P236412** | Cdkl2 | 0.044731296 | -0.080979371 | 0.170441964 | 7.894197061 | 0.850449413 | 0.424631968 | 0.991835066 | -4.97100323 |
| **A_55_P2012669** | NA | 0.041741306 | -0.075575319 | 0.159057932 | 16.90861889 | 0.850385028 | 0.424665308 | 0.991835066 | -4.971043739 |
| **A_55_P2056473** | Spc24 | -0.077999784 | -0.297244649 | 0.14124508 | 10.47961839 | -0.85030181 | 0.424708402 | 0.991835066 | -4.971096093 |
| **A_55_P2102322** | Yif1b | 0.066022494 | -0.119603149 | 0.251648137 | 13.56095937 | 0.850086433 | 0.424819952 | 0.991835066 | -4.971231573 |
| **A_52_P584188** | Frat1 | -0.053549392 | -0.20411334 | 0.097014557 | 8.57812633 | -0.850046489 | 0.424840642 | 0.991835066 | -4.971256697 |
| **A_55_P2002155** | Sumo1 | 0.091658642 | -0.166070343 | 0.349387628 | 13.38554915 | 0.850000712 | 0.424864355 | 0.991835066 | -4.971285488 |
| **A_55_P1994957** | NA | -0.051392266 | -0.195930289 | 0.093145756 | 14.90977476 | -0.849815787 | 0.424960156 | 0.991835066 | -4.971401784 |
| **A_51_P300337** | Csrp2 | -0.077147469 | -0.294122489 | 0.139827551 | 14.55457697 | -0.849808513 | 0.424963925 | 0.991835066 | -4.971406358 |
| **A_55_P1968245** | Ftl1 | -0.054220231 | -0.2067443 | 0.098303838 | 16.21290045 | -0.849634449 | 0.425054114 | 0.991835066 | -4.971515805 |
| **A_51_P450812** | Cog4 | 0.050104352 | -0.09085195 | 0.191060653 | 10.95840009 | 0.849571793 | 0.425086582 | 0.991835066 | -4.971555198 |
| **A_55_P1971522** | Kif16b | -0.053329114 | -0.203363053 | 0.096704825 | 9.984953046 | -0.849540311 | 0.425102897 | 0.991835066 | -4.97157499 |
| **A_51_P289697** | Nek4 | 0.045648783 | -0.082782747 | 0.174080313 | 8.767057898 | 0.84950645 | 0.425120445 | 0.991835066 | -4.971596277 |
| **A_52_P214630** | Sox9 | 0.054010441 | -0.097963884 | 0.205984766 | 9.117415994 | 0.849408555 | 0.42517118 | 0.991835066 | -4.971657817 |
| **A_55_P2382262** | A830039N2 | 0.063207148 | -0.11465195 | 0.241066247 | 8.686737018 | 0.849374546 | 0.425188806 | 0.991835066 | -4.971679195 |
| **A_51_P257258** | Cpox | 0.077441749 | -0.14047845 | 0.295361948 | 10.77235666 | 0.849350208 | 0.425201421 | 0.991835066 | -4.971694493 |
| **A_55_P2009091** | Mad1l1 | 0.052932333 | -0.096035258 | 0.201899924 | 9.63358393 | 0.84925551 | 0.425250506 | 0.991835066 | -4.971754015 |
| **A_55_P2144376** | NA | 0.061233573 | -0.111102963 | 0.233570109 | 8.061876337 | 0.849222266 | 0.425267738 | 0.991835066 | -4.971774909 |
| **A_55_P2147291** | NA | 0.080628308 | -0.146299359 | 0.307555975 | 15.66402559 | 0.849198522 | 0.425280046 | 0.991835066 | -4.971789832 |

| | | | | | | | | | |
|---|---|---|---|---|---|---|---|---|---|
| **A_55_P2131438** | H2ac19 | -0.058596977 | -0.22352267 | 0.106328716 | 16.4525271 | -0.849172709 | 0.425293428 | 0.991835066 | -4.971806055 |
| **A_52_P20048** | Tcf12 | -0.052687547 | -0.201013495 | 0.095638401 | 10.08272073 | -0.848984926 | 0.425390781 | 0.991835066 | -4.971924062 |
| **A_55_P1979616** | Tmem189 | 0.070552373 | -0.128093359 | 0.269198105 | 9.009548218 | 0.848870654 | 0.425450031 | 0.991835066 | -4.971995863 |
| **A_55_P2045916** | NA | -0.045890544 | -0.175102716 | 0.083321627 | 7.879572447 | -0.848846017 | 0.425462806 | 0.991835066 | -4.972011342 |
| **A_51_P487913** | NA | 0.040026843 | -0.072685339 | 0.152739025 | 7.92503639 | 0.848769195 | 0.425502643 | 0.991835066 | -4.972059608 |
| **A_55_P2091691** | Nudt3 | 0.048357889 | -0.087818637 | 0.184534416 | 14.10627426 | 0.848739112 | 0.425518243 | 0.991835066 | -4.972078507 |
| **A_55_P2005719** | NA | -0.09713243 | -0.37066222 | 0.176397361 | 14.50734463 | -0.848728434 | 0.425523781 | 0.991835066 | -4.972085215 |
| **A_55_P2138960** | Tpk1 | 0.039707207 | -0.072147459 | 0.151561873 | 8.220947393 | 0.848446319 | 0.425670102 | 0.991835066 | -4.972262425 |
| **A_66_P126523** | Phb2 | -0.106541769 | -0.406669098 | 0.19358556 | 13.09695355 | -0.848444607 | 0.425670991 | 0.991835066 | -4.972263501 |
| **A_55_P2060415** | NA | 0.065171555 | -0.118430556 | 0.248773667 | 16.08616985 | 0.848378288 | 0.425705393 | 0.991835066 | -4.972305152 |
| **A_51_P327874** | Pth1r | -0.112545405 | -0.4298354 | 0.20474459 | 10.93077976 | -0.847774839 | 0.426018518 | 0.991835066 | -4.972684037 |
| **A_55_P2145401** | H3c3 | 0.073298559 | -0.133366468 | 0.279963587 | 10.65906288 | 0.847691018 | 0.426062025 | 0.991835066 | -4.97273665 |
| **A_55_P2052330** | NA | -0.059867771 | -0.228683641 | 0.108948098 | 13.43862799 | -0.84759611 | 0.426111291 | 0.991835066 | -4.972796216 |
| **A_55_P2335529** | 4921531C2 | -0.043399004 | -0.165783391 | 0.078985383 | 8.637460569 | -0.847545224 | 0.426137707 | 0.991835066 | -4.972828151 |
| **A_52_P678895** | Dynlt3 | -0.072744179 | -0.277904717 | 0.132416359 | 10.16196955 | -0.847448957 | 0.426187684 | 0.991835066 | -4.972888563 |
| **A_51_P195153** | Gtse1 | 0.057701693 | -0.105061337 | 0.220464723 | 8.965950554 | 0.847309203 | 0.426260246 | 0.991835066 | -4.972976256 |
| **A_55_P2294184** | Cemip2 | 0.093502507 | -0.170277039 | 0.357282053 | 11.17309472 | 0.847210388 | 0.426311557 | 0.991835066 | -4.973038253 |
| **A_55_P2025038** | Cpe | 0.069434388 | -0.126447437 | 0.265316213 | 14.68145693 | 0.84720713 | 0.426313249 | 0.991835066 | -4.973040297 |
| **A_51_P488991** | Oaf | 0.06044634 | -0.110082164 | 0.230974843 | 9.790905837 | 0.847192609 | 0.42632079 | 0.991835066 | -4.973049407 |
| **A_51_P358225** | Serinc1 | -0.058148321 | -0.222208478 | 0.105911836 | 13.52635957 | -0.847116593 | 0.426360266 | 0.991835066 | -4.973097097 |
| **A_52_P599317** | Hs6st2 | 0.085894924 | -0.156451786 | 0.328241634 | 8.855475032 | 0.847109404 | 0.426363999 | 0.991835066 | -4.973101606 |
| **A_55_P1962026** | Pigg | 0.042717115 | -0.077827459 | 0.16326169 | 8.322708319 | 0.846960936 | 0.426441109 | 0.991835066 | -4.973194739 |
| **A_55_P1989544** | Cklf | -0.042379537 | -0.161996223 | 0.077237149 | 8.890675072 | -0.846785809 | 0.426532077 | 0.991835066 | -4.973304578 |
| **A_55_P2163961** | Kcnj5 | -0.131823692 | -0.503917344 | 0.240269961 | 12.06145505 | -0.846740515 | 0.426555607 | 0.991835066 | -4.973332984 |
| **A_55_P2134032** | NA | 0.142390888 | -0.259551957 | 0.544333732 | 9.31447572 | 0.846695064 | 0.426579219 | 0.991835066 | -4.973361486 |
| **A_51_P307220** | Ankrd54 | 0.046077494 | -0.083990679 | 0.176145666 | 8.140963689 | 0.846694897 | 0.426579307 | 0.991835066 | -4.973361591 |
| **A_51_P183971** | Barhl1 | -0.106959462 | -0.408906282 | 0.194987357 | 9.994527766 | -0.846638247 | 0.426608738 | 0.991835066 | -4.973397115 |
| **A_55_P2095019** | NA | 0.06565361 | -0.119693725 | 0.251000945 | 8.781898761 | 0.846606108 | 0.426625436 | 0.991835066 | -4.973417268 |
| **A_55_P1981550** | Park7 | -0.077410789 | -0.295950751 | 0.141129174 | 13.20770412 | -0.846602918 | 0.426627094 | 0.991835066 | -4.973419269 |
| **A_55_P1984168** | Casp4 | 0.080741314 | -0.147245681 | 0.308728309 | 8.563299119 | 0.84643745 | 0.426713071 | 0.991835066 | -4.973523018 |
| **A_55_P2144126** | NA | -0.055085036 | -0.210674896 | 0.100504823 | 14.53371353 | -0.84617752 | 0.426848157 | 0.991835066 | -4.973685963 |
| **A_51_P462428** | NA | -0.066400844 | -0.253958071 | 0.121156384 | 8.360044989 | -0.84615305 | 0.426860875 | 0.991835066 | -4.973701301 |
| **A_51_P372463** | Fxr1 | -0.051243855 | -0.19599354 | 0.093505831 | 12.49442149 | -0.846122598 | 0.426876704 | 0.991835066 | -4.973720388 |
| **A_55_P1994939** | Hmgb2 | 0.086160038 | -0.157260544 | 0.32958062 | 14.00141531 | 0.845975365 | 0.426953238 | 0.991835066 | -4.973812665 |
| **A_55_P1985463** | NA | -0.07262727 | -0.2778633 | 0.132608759 | 14.28487892 | -0.845775793 | 0.427056995 | 0.991835066 | -4.973937725 |
| **A_55_P2104617** | Kctd5 | -0.043144171 | -0.165068611 | 0.07878027 | 8.772701318 | -0.84574704 | 0.427071945 | 0.991835066 | -4.973955742 |
| **A_51_P394014** | Qprt | -0.054520913 | -0.208598344 | 0.099556517 | 8.215896833 | -0.845732906 | 0.427079294 | 0.991835066 | -4.973964597 |
| **A_55_P2160563** | Mrps17 | -0.044377928 | -0.169791742 | 0.081035886 | 11.85180202 | -0.84572813 | 0.427081778 | 0.991835066 | -4.97396759 |
| **A_51_P499195** | Nkg7 | 0.034746245 | -0.063450242 | 0.132942731 | 7.679613734 | 0.845709203 | 0.427091619 | 0.991835066 | -4.973979449 |
| **A_55_P2081005** | Cox7c | -0.038211875 | -0.146211336 | 0.069787586 | 8.506448862 | -0.845640731 | 0.427127224 | 0.991835066 | -4.974022349 |
| **A_55_P2137927** | Cd82 | 0.055232648 | -0.100887753 | 0.21135305 | 8.367677549 | 0.845561772 | 0.427168284 | 0.991835066 | -4.974071816 |
| **A_55_P2123362** | NA | 0.10417094 | -0.190364885 | 0.398706765 | 8.692169908 | 0.845313082 | 0.427297628 | 0.991835066 | -4.974227596 |
| **A_55_P2028837** | Tspan2 | 0.071766515 | -0.131181932 | 0.274714961 | 9.856349431 | 0.8451723 | 0.427370861 | 0.991835066 | -4.974315766 |
| **A_52_P659762** | Eid2 | -0.056006681 | -0.214398093 | 0.102384732 | 8.807492634 | -0.845117967 | 0.427399126 | 0.991835066 | -4.974349791 |
| **A_52_P504421** | Rnf138 | 0.047153093 | -0.086200609 | 0.180506796 | 9.19855807 | 0.845111954 | 0.427402254 | 0.991835066 | -4.974353557 |
| **A_55_P2000968** | Ppp2r3c | -0.059112534 | -0.226293626 | 0.108068559 | 10.4921772 | -0.845087247 | 0.427415108 | 0.991835066 | -4.974369029 |
| **A_55_P2160862** | Wrn | -0.045893091 | -0.175699353 | 0.083913171 | 8.092748877 | -0.84500795 | 0.427456365 | 0.991835066 | -4.974418683 |
| **A_51_P101228** | Gmeb2 | -0.039689006 | -0.151956623 | 0.07257861 | 8.166218099 | -0.844938026 | 0.427492747 | 0.991835066 | -4.957195464 |

| | | | | | | | | | |
|---|---|---|---|---|---|---|---|---|---|
| **A_52_P275627** | Kpnb1 | 0.059990729 | -0.109714412 | 0.229695869 | 11.39611504 | 0.844886309 | 0.427519657 | 0.991835066 | -4.974494846 |
| **A_51_P412225** | Tbc1d16 | -0.07377094 | -0.282482995 | 0.134941115 | 11.93240478 | -0.844786401 | 0.427571646 | 0.991835066 | -4.974557395 |
| **A_51_P503654** | Tmem205 | -0.062834936 | -0.240613604 | 0.114943732 | 9.409546093 | -0.844754786 | 0.427588099 | 0.991835066 | -4.974577187 |
| **A_51_P144512** | Rbm19 | 0.080822073 | -0.147865783 | 0.30950993 | 10.41083634 | 0.844687399 | 0.427623169 | 0.991835066 | -4.974619372 |
| **A_55_P1978696** | NA | 0.03724506 | -0.068141595 | 0.142631715 | 7.727834385 | 0.844679962 | 0.427627039 | 0.991835066 | -4.974624027 |
| **A_55_P2260094** | Tcf7l2 | 0.036559301 | -0.066910375 | 0.140028977 | 8.101104183 | 0.844488868 | 0.427726501 | 0.991835066 | -4.974743638 |
| **A_51_P476509** | Zbtb38 | 0.056888546 | -0.104126839 | 0.21790393 | 8.66007281 | 0.844435707 | 0.427754174 | 0.991835066 | -4.974776909 |
| **A_55_P2139665** | NA | 0.046077489 | -0.084367745 | 0.176522724 | 16.3890099 | 0.84424738 | 0.427852217 | 0.991835066 | -4.974894762 |
| **A_52_P27020** | Sgo2a | -0.059453207 | -0.227768631 | 0.108862216 | 8.744599786 | -0.844229466 | 0.427861543 | 0.991835066 | -4.974905971 |
| **A_55_P2007768** | Gm8709 | -0.067947723 | -0.260350166 | 0.12445472 | 16.6780309 | -0.84406027 | 0.427949642 | 0.991835066 | -4.975011835 |
| **A_52_P158211** | NA | 0.065395167 | -0.119797436 | 0.250587771 | 9.354364901 | 0.843978048 | 0.427992458 | 0.991835066 | -4.975063273 |
| **A_55_P1964154** | Asah1 | 0.039322348 | -0.072046861 | 0.150691557 | 8.353332362 | 0.843885335 | 0.428040742 | 0.991835066 | -4.975121271 |
| **A_51_P519592** | Srsf10 | 0.037260941 | -0.068309982 | 0.142831863 | 8.57559176 | 0.843565153 | 0.428207518 | 0.991835066 | -4.975321528 |
| **A_51_P227962** | Dynlrb2 | 0.06295071 | -0.115451591 | 0.241353011 | 7.776499999 | 0.843352848 | 0.428318129 | 0.991835066 | -4.975454283 |
| **A_55_P2085940** | NA | 0.081401617 | -0.149295133 | 0.312098367 | 12.11615531 | 0.84333609 | 0.428326861 | 0.991835066 | -4.97546476 |
| **A_66_P118600** | Lama1 | 0.169634985 | -0.311120966 | 0.650390936 | 8.262699945 | 0.843334541 | 0.428327668 | 0.991835066 | -4.975465729 |
| **A_65_P19138** | 7-Mar | 0.108336593 | -0.198709448 | 0.415382635 | 11.43751581 | 0.843297483 | 0.428346978 | 0.991835066 | -4.975488898 |
| **A_55_P2086978** | Col23a1 | 0.042600848 | -0.078137948 | 0.163339643 | 8.275074746 | 0.843296961 | 0.42834725 | 0.991835066 | -4.975489224 |
| **A_55_P1956799** | Smpx | -0.058308999 | -0.22364914 | 0.107031141 | 13.46461141 | -0.842881302 | 0.428563879 | 0.991835066 | -4.97574905 |
| **A_52_P549827** | Mgst1 | -0.105759648 | -0.405702275 | 0.194182979 | 10.56245299 | -0.842734818 | 0.428640241 | 0.991835066 | -4.975840593 |
| **A_51_P489229** | Eif3g | 0.0370286 | -0.068011051 | 0.142068252 | 14.3204321 | 0.842545094 | 0.428739158 | 0.991835066 | -4.975959141 |
| **A_55_P2037962** | Trpm6 | 0.039796915 | -0.0731325 | 0.15272633 | 7.878776072 | 0.842270245 | 0.428882487 | 0.991835066 | -4.976130843 |
| **A_55_P1963547** | Prdm16 | -0.081561691 | -0.313014538 | 0.149891156 | 9.816054488 | -0.842234109 | 0.428901334 | 0.991835066 | -4.976153414 |
| **A_55_P2046867** | Gata5 | -0.122288667 | -0.469357362 | 0.224780028 | 9.6316397 | -0.842131478 | 0.428954864 | 0.991835066 | -4.976217516 |
| **A_55_P1973526** | Fam216b | -0.037093863 | -0.142383371 | 0.068195644 | 7.930018485 | -0.842027174 | 0.429009273 | 0.991835066 | -4.976282658 |
| **A_52_P150236** | Apom | -0.156006021 | -0.598831649 | 0.286819607 | 8.196794345 | -0.84201096 | 0.429017731 | 0.991835066 | -4.976292783 |
| **A_51_P364984** | Gabpa | 0.062971657 | -0.115822956 | 0.24176627 | 8.793581588 | 0.841782371 | 0.429136989 | 0.991835066 | -4.976435521 |
| **A_52_P260818** | Card10 | 0.074079118 | -0.136254062 | 0.284412298 | 9.359224721 | 0.841777174 | 0.429139701 | 0.991835066 | -4.976438766 |
| **A_55_P2029558** | Adrb2 | 0.034159362 | -0.062848562 | 0.131167286 | 7.705430244 | 0.841611526 | 0.429226137 | 0.991835066 | -4.976542182 |
| **A_66_P110633** | Apol9b | -0.046449872 | -0.178371258 | 0.085471515 | 7.967289728 | -0.841547131 | 0.429259742 | 0.991835066 | -4.976582381 |
| **A_55_P1970954** | Pknox2 | -0.070065743 | -0.26909682 | 0.128965334 | 8.554842922 | -0.841383464 | 0.429345162 | 0.991835066 | -4.97668454 |
| **A_55_P2033997** | Ddo | -0.04494627 | -0.172638455 | 0.082745916 | 8.718697952 | -0.84127593 | 0.429401292 | 0.991835066 | -4.976751653 |
| **A_55_P2004153** | NA | -0.054619836 | -0.209796324 | 0.100556652 | 10.10945352 | -0.841266525 | 0.429406201 | 0.991835066 | -4.976757522 |
| **A_55_P2022499** | Pnisr | 0.15825747 | -0.291475618 | 0.607990559 | 9.498792442 | 0.8410436 | 0.42952258 | 0.991835066 | -4.976896631 |
| **A_66_P101107** | NA | 0.10252846 | -0.188837974 | 0.393894894 | 9.509606337 | 0.841034961 | 0.42952709 | 0.991835066 | -4.976902021 |
| **A_55_P1988754** | H4f16 | 0.13258663 | -0.244258318 | 0.509431578 | 13.87244618 | 0.840903511 | 0.429595726 | 0.991835066 | -4.976984035 |
| **A_51_P124345** | Cpne5 | 0.17298917 | -0.318746781 | 0.664725121 | 10.36085967 | 0.84080655 | 0.429646357 | 0.991835066 | -4.977044523 |
| **A_55_P2383902** | 8030453O2 | 0.129953288 | -0.23948911 | 0.499395686 | 8.198747597 | 0.840716691 | 0.429693285 | 0.991835066 | -4.977100577 |
| **A_55_P2027136** | H3c7 | 0.099994891 | -0.184340343 | 0.384330124 | 8.961746571 | 0.840535892 | 0.429787715 | 0.991835066 | -4.977213345 |
| **A_51_P320539** | Otulin | 0.062629257 | -0.115464208 | 0.240722723 | 10.80626733 | 0.840501338 | 0.429805764 | 0.991835066 | -4.977234895 |
| **A_66_P115389** | NA | -0.067696981 | -0.260206408 | 0.124812447 | 11.41456182 | -0.840478154 | 0.429817874 | 0.991835066 | -4.977249353 |
| **A_66_P116154** | Cstf2 | 0.092678456 | -0.170966026 | 0.356322939 | 9.364064001 | 0.840173999 | 0.429976775 | 0.991835066 | -4.97743901 |
| **A_55_P2067787** | Psg29 | -0.089022584 | -0.342283771 | 0.164238604 | 7.799783584 | -0.840118769 | 0.430005633 | 0.991835066 | -4.977473443 |
| **A_51_P483311** | Mpv17l2 | -0.11944318 | -0.459261913 | 0.220375553 | 9.44077294 | -0.84008489 | 0.430023336 | 0.991835066 | -4.977494564 |
| **A_51_P295582** | Cutc | 0.069107063 | -0.127507649 | 0.265721774 | 8.295185095 | 0.840070159 | 0.430031034 | 0.991835066 | -4.977503748 |
| **A_51_P179921** | Igdcc3 | -0.037986158 | -0.146062002 | 0.070089685 | 8.038385749 | -0.840051425 | 0.430040824 | 0.991835066 | -4.977515426 |
| **A_55_P1955528** | NA | 0.105365314 | -0.194569448 | 0.405300075 | 9.64706391 | 0.839614623 | 0.430269121 | 0.991835066 | -4.977787675 |
| **A_51_P470328** | Selenop | -0.061220278 | -0.235507546 | 0.113066991 | 13.73307261 | -0.839534917 | 0.430310789 | 0.991835066 | -4.977837342 |

| | | | | | | | | | |
|---|---|---|---|---|---|---|---|---|---|
| **A_52_P187132** | Ybx1 | -0.089413632 | -0.343973708 | 0.165146445 | 14.14962373 | -0.839503621 | 0.430327151 | 0.991835066 | -4.977856842 |
| **A_55_P1979595** | Selenon | 0.061522085 | -0.113639261 | 0.236683431 | 11.76217954 | 0.839463671 | 0.430348037 | 0.991835066 | -4.977881735 |
| **A_55_P2139640** | Fbh1 | -0.073872377 | -0.284250564 | 0.136505811 | 10.08338685 | -0.839248348 | 0.430460624 | 0.991835066 | -4.978015882 |
| **A_55_P2098321** | NA | -0.060484458 | -0.232827871 | 0.111858954 | 10.55058848 | -0.838799649 | 0.430695305 | 0.991835066 | -4.978295343 |
| **A_51_P297069** | Tmod1 | -0.077469193 | -0.298239371 | 0.143300985 | 11.53437265 | -0.838682842 | 0.430756414 | 0.991835066 | -4.978368075 |
| **A_55_P2099448** | Slc12a2 | 0.085107322 | -0.157432011 | 0.327646655 | 11.51463598 | 0.83867535 | 0.430760333 | 0.991835066 | -4.97837274 |
| **A_52_P177341** | Cox19 | -0.060022345 | -0.231078044 | 0.111033354 | 11.91321408 | -0.838657321 | 0.430769766 | 0.991835066 | -4.978383965 |
| **A_55_P2154252** | Gfpt2 | -0.051742631 | -0.199219755 | 0.095734493 | 8.964367131 | -0.838557757 | 0.43082186 | 0.991835066 | -4.978445953 |
| **A_51_P315634** | Nedd1 | -0.058700667 | -0.226031099 | 0.108629765 | 9.464217162 | -0.838450131 | 0.430878176 | 0.991835066 | -4.978512953 |
| **A_52_P605812** | Ptrh1 | 0.084527712 | -0.156438051 | 0.325493475 | 9.558807103 | 0.838403155 | 0.430902759 | 0.991835066 | -4.978542195 |
| **A_55_P2004833** | NA | -0.080042754 | -0.308229573 | 0.148144064 | 13.02948378 | -0.838379409 | 0.430915185 | 0.991835066 | -4.978556977 |
| **A_51_P387188** | Vrk3 | -0.078322505 | -0.301608473 | 0.144963463 | 10.93964525 | -0.838367183 | 0.430921583 | 0.991835066 | -4.978564586 |
| **A_51_P117477** | Slc27a1 | -0.046628317 | -0.179580012 | 0.086323378 | 8.905898593 | -0.838233459 | 0.430991569 | 0.991835066 | -4.978647819 |
| **A_55_P2119957** | Mlxipl | -0.079270458 | -0.305345037 | 0.146804122 | 8.796940404 | -0.838047753 | 0.431088773 | 0.991835066 | -4.978763389 |
| **A_51_P227108** | Bri3bp | 0.061253923 | -0.113438644 | 0.235946491 | 12.54539525 | 0.838047463 | 0.431088924 | 0.991835066 | -4.978763569 |
| **A_51_P381321** | Ddx17 | 0.079023901 | -0.146434497 | 0.304482298 | 14.03268451 | 0.837724427 | 0.431258049 | 0.991835066 | -4.978964558 |
| **A_55_P2026275** | Ppp1r1b | -0.071094277 | -0.273946548 | 0.131757994 | 11.28354667 | -0.837652521 | 0.431295701 | 0.991835066 | -4.979009289 |
| **A_55_P1999202** | Vmn1r217 | 0.070233359 | -0.130167337 | 0.270634055 | 8.486475108 | 0.837632164 | 0.431306361 | 0.991835066 | -4.979021952 |
| **A_55_P2168990** | NA | 0.076180673 | -0.141200749 | 0.293562095 | 14.21480076 | 0.837590059 | 0.43132841 | 0.991835066 | -4.979048143 |
| **A_55_P1959403** | Ccdc77 | -0.057111123 | -0.220097093 | 0.105874848 | 9.454624634 | -0.837489959 | 0.431380833 | 0.991835066 | -4.979110404 |
| **A_55_P2083009** | Catsperd | -0.055966388 | -0.215748052 | 0.103815275 | 8.514568425 | -0.837161941 | 0.431552649 | 0.991835066 | -4.97931439 |
| **A_66_P139196** | Baz1a | 0.061685976 | -0.114473973 | 0.237845925 | 8.532813886 | 0.836928579 | 0.431674914 | 0.991835066 | -4.979459474 |
| **A_55_P2457154** | G2e3 | -0.064332564 | -0.248088707 | 0.119423579 | 9.437835209 | -0.836754637 | 0.431766064 | 0.991835066 | -4.979567597 |
| **A_66_P123989** | Ikzf5 | 0.041715533 | -0.077459468 | 0.160890534 | 8.638913245 | 0.83660752 | 0.431843167 | 0.991835066 | -4.979659032 |
| **A_55_P2114110** | Cadm4 | -0.062815794 | -0.242279268 | 0.11664768 | 8.707415661 | -0.836569283 | 0.431863209 | 0.991835066 | -4.979682794 |
| **A_52_P604613** | S1pr2 | -0.05484527 | -0.211568589 | 0.101878049 | 9.079394937 | -0.836401281 | 0.431951273 | 0.991835066 | -4.979787191 |
| **A_55_P2020395** | 2-Mar | 0.039855217 | -0.074046608 | 0.153757041 | 8.0211873 | 0.836302945 | 0.432002825 | 0.991835066 | -4.97984829 |
| **A_55_P2026049** | NA | 0.036391642 | -0.067620283 | 0.140403566 | 7.986633779 | 0.836233665 | 0.432039147 | 0.991835066 | -4.979891332 |
| **A_55_P1971199** | Slc4a1ap | -0.037190698 | -0.143491932 | 0.069110536 | 10.86003882 | -0.836190356 | 0.432061855 | 0.991835066 | -4.979918238 |
| **A_55_P2370250** | Syn3 | 0.061296214 | -0.113925582 | 0.236518011 | 8.027238972 | 0.836093137 | 0.432112831 | 0.991835066 | -4.979978631 |
| **A_52_P487362** | Ppp4r4 | 0.038618801 | -0.071785144 | 0.149022745 | 7.719536085 | 0.836032811 | 0.432144465 | 0.991835066 | -4.980016104 |
| **A_55_P2028415** | Hacd2 | -0.056149314 | -0.216727968 | 0.10442934 | 10.36572167 | -0.835729577 | 0.432303499 | 0.991835066 | -4.980204431 |
| **A_51_P327232** | Lenep | 0.056106701 | -0.104381526 | 0.216594927 | 9.099090198 | 0.835565858 | 0.43238938 | 0.991835066 | -4.980306089 |
| **A_51_P302942** | Rasl10a | -0.043678257 | -0.168629296 | 0.081272781 | 7.780076782 | -0.83547713 | 0.432435929 | 0.991835066 | -4.980361177 |
| **A_51_P269687** | Pole2 | 0.061723907 | -0.114851828 | 0.238299642 | 8.857449694 | 0.835471264 | 0.432439007 | 0.991835066 | -4.980364819 |
| **A_55_P2039556** | Pak6 | -0.07298739 | -0.281788038 | 0.135813257 | 9.335721205 | -0.835458969 | 0.432445458 | 0.991835066 | -4.980372452 |
| **A_51_P113003** | Cenpq | -0.049282175 | -0.190269253 | 0.091704904 | 9.43519668 | -0.835448501 | 0.432450949 | 0.991835066 | -4.975431466 |
| **A_55_P2113912** | NA | -0.101180125 | -0.390675178 | 0.188314928 | 13.99511669 | -0.835339845 | 0.43250796 | 0.991835066 | -4.980446403 |
| **A_51_P214663** | Tpmt | -0.037935861 | -0.146505513 | 0.070633792 | 8.615558322 | -0.835123348 | 0.432621568 | 0.991835066 | -4.980580782 |
| **A_51_P173709** | Gprc5b | -0.054888227 | -0.21199642 | 0.102219966 | 10.13673008 | -0.835005818 | 0.432683251 | 0.991835066 | -4.980653722 |
| **A_52_P338110** | Chmp1b | -0.046226541 | -0.178542531 | 0.086089449 | 9.121990384 | -0.835003312 | 0.432684567 | 0.991835066 | -4.980655277 |
| **A_51_P162671** | Folr2 | -0.040378688 | -0.155961425 | 0.075204048 | 7.844900305 | -0.834965189 | 0.432704576 | 0.991835066 | -4.980678934 |
| **A_55_P2036180** | Mcf2l | -0.044628917 | -0.172399262 | 0.083141429 | 10.86146071 | -0.834824925 | 0.432778203 | 0.991835066 | -4.98076597 |
| **A_55_P2253746** | Nip7 | 0.060604462 | -0.112903756 | 0.23411268 | 9.873967069 | 0.834821605 | 0.432779946 | 0.991835066 | -4.98076803 |
| **A_55_P2175813** | Ythdf2 | 0.071212462 | -0.132705621 | 0.275130544 | 10.31772187 | 0.834659599 | 0.432864996 | 0.991835066 | -4.980868542 |
| **A_55_P2061135** | Rpl17 | 0.055080963 | -0.102650432 | 0.212812357 | 14.13626689 | 0.834627157 | 0.432882028 | 0.991835066 | -4.980888668 |
| **A_55_P2066409** | Gabpb1 | 0.076128639 | -0.141885352 | 0.29414263 | 10.57753829 | 0.834589337 | 0.432901885 | 0.991835066 | -4.980912129 |
| **A_55_P2057132** | Scn8a | -0.097302576 | -0.375961871 | 0.181356718 | 10.64368818 | -0.834564561 | 0.432914894 | 0.991835066 | -4.980927498 |

| | | | | | | | | | |
|---|---|---|---|---|---|---|---|---|---|
| **A_55_P2120055** | Fgfr4 | 0.09347209 | -0.174241657 | 0.361185837 | 8.048434226 | 0.834488607 | 0.432954776 | 0.991835066 | -4.980974613 |
| **A_51_P449048** | Cndp2 | -0.060111573 | -0.232299353 | 0.112076208 | 9.05617189 | -0.834381941 | 0.433010789 | 0.991835066 | -4.981040772 |
| **A_55_P1981964** | Tnfrsf13b | -0.046537775 | -0.179861415 | 0.086785866 | 8.071884117 | -0.834271832 | 0.433068614 | 0.991835066 | -4.981109061 |
| **A_51_P165475** | C1galt1 | 0.041463862 | -0.077354632 | 0.160282356 | 8.764043196 | 0.834055291 | 0.433182351 | 0.991835066 | -4.981243337 |
| **A_51_P402458** | Anxa7 | -0.0525746 | -0.203237821 | 0.098088622 | 8.738238605 | -0.834022673 | 0.433199486 | 0.991835066 | -4.981263562 |
| **A_55_P1952312** | NA | -0.12111406 | -0.468192791 | 0.225964672 | 13.03288246 | -0.834018516 | 0.433201669 | 0.991835066 | -4.981266139 |
| **A_55_P2146783** | Ropn1l | -0.037164404 | -0.143679858 | 0.06935105 | 8.19301349 | -0.833918639 | 0.433254138 | 0.991835066 | -4.981328062 |
| **A_55_P2171033** | NA | 0.084141862 | -0.157014061 | 0.325297785 | 8.910282658 | 0.833917941 | 0.433254505 | 0.991835066 | -4.981328494 |
| **A_55_P1957474** | Pus7 | 0.079079062 | -0.14757075 | 0.305728874 | 10.59651785 | 0.833902501 | 0.433262616 | 0.991835066 | -4.981338066 |
| **A_51_P123705** | Dus3l | 0.073074333 | -0.136377255 | 0.282525921 | 9.977730202 | 0.833854613 | 0.433287775 | 0.991835066 | -4.981367754 |
| **A_55_P1957927** | Nono | 0.088110654 | -0.164499442 | 0.34072075 | 14.05820859 | 0.833655951 | 0.433392159 | 0.991835066 | -4.981490898 |
| **A_52_P634447** | Hs1bp3 | -0.043034358 | -0.166413004 | 0.080344288 | 9.864612895 | -0.833651356 | 0.433394573 | 0.991835066 | -4.981493746 |
| **A_66_P124551** | Akap13 | -0.054096473 | -0.209208226 | 0.10101528 | 11.86943326 | -0.833553306 | 0.433446099 | 0.991835066 | -4.981554516 |
| **A_52_P175242** | Irs1 | -0.113217856 | -0.437856464 | 0.211420753 | 11.78410533 | -0.833535093 | 0.43345567 | 0.991835066 | -4.981565803 |
| **A_51_P420236** | Srrt | 0.075162349 | -0.140384613 | 0.290709311 | 8.778269555 | 0.83342701 | 0.433512474 | 0.991835066 | -4.981632784 |
| **A_55_P2056674** | Lrrk1 | 0.065489579 | -0.122353029 | 0.253332187 | 10.24137074 | 0.83327283 | 0.433593513 | 0.991835066 | -4.98172832 |
| **A_52_P312398** | Ssr3 | 0.084482159 | -0.157860673 | 0.32682499 | 9.996696469 | 0.833189825 | 0.433637146 | 0.991835066 | -4.981779747 |
| **A_51_P219028** | Wdr53 | 0.068711346 | -0.128414976 | 0.265837668 | 10.27490506 | 0.833092009 | 0.433688569 | 0.991835066 | -4.981840347 |
| **A_55_P2129920** | Chd7 | 0.055217387 | -0.103207657 | 0.213642432 | 11.81642063 | 0.83303097 | 0.43372066 | 0.991835066 | -4.981878159 |
| **A_51_P232858** | Rpl14 | 0.064439671 | -0.120470414 | 0.249349757 | 10.65780447 | 0.832917236 | 0.43378046 | 0.991835066 | -4.981948609 |
| **A_55_P1999601** | NA | 0.142355925 | -0.26619204 | 0.550903891 | 12.6998358 | 0.832801749 | 0.433841188 | 0.991835066 | -4.982020137 |
| **A_65_P07361** | Trim14 | -0.039887348 | -0.154372722 | 0.074598025 | 7.86135709 | -0.832710981 | 0.433888922 | 0.991835066 | -4.982076351 |
| **A_55_P1961458** | Speer1 | -0.084021183 | -0.325194424 | 0.157152058 | 8.025606557 | -0.832662113 | 0.433914622 | 0.991835066 | -4.982106613 |
| **A_55_P1986185** | NA | 0.042781871 | -0.080037854 | 0.165601596 | 14.01033617 | 0.832531716 | 0.433983205 | 0.991835066 | -4.982187357 |
| **A_55_P2053456** | Gm6548 | -0.065673702 | -0.254213786 | 0.122866382 | 16.15163349 | -0.832524334 | 0.433987088 | 0.991835066 | -4.982191927 |
| **A_51_P101660** | Ncbp3 | 0.042667039 | -0.079828808 | 0.165162887 | 8.666094387 | 0.832492392 | 0.43400389 | 0.991835066 | -4.977255222 |
| **A_55_P2141093** | Eya2 | -0.066879763 | -0.258894783 | 0.125135256 | 8.482267559 | -0.832470129 | 0.4340156 | 0.991835066 | -4.982225489 |
| **A_55_P2005754** | Manbal | -0.08860233 | -0.343057026 | 0.165852367 | 10.75336749 | -0.832230835 | 0.434141486 | 0.991835066 | -4.98237363 |
| **A_55_P2000668** | Stxbp2 | -0.072599187 | -0.281121249 | 0.135922876 | 9.307080255 | -0.832125585 | 0.434196863 | 0.991835066 | -4.982438778 |
| **A_55_P2166446** | NA | 0.036653594 | -0.068629403 | 0.141936591 | 9.237743989 | 0.832084554 | 0.434218453 | 0.991835066 | -4.982464174 |
| **A_51_P334942** | Aldh1a1 | 0.107384783 | -0.201104929 | 0.415874495 | 8.73225134 | 0.831976752 | 0.434275181 | 0.991835066 | -4.982530892 |
| **A_55_P2027230** | NA | -0.039657543 | -0.15360007 | 0.074284985 | 7.922026261 | -0.831857784 | 0.43433779 | 0.991835066 | -4.982604514 |
| **A_55_P2034670** | Fbxl12 | 0.06635792 | -0.124333276 | 0.257049117 | 11.33261894 | 0.831708719 | 0.434416247 | 0.991835066 | -4.982696749 |
| **A_55_P1988774** | Slc17a5 | 0.128791006 | -0.241337402 | 0.498919415 | 12.2581623 | 0.831653175 | 0.434445484 | 0.991835066 | -4.982731114 |
| **A_55_P1960941** | Wac | 0.043587525 | -0.081683415 | 0.168858466 | 11.57833448 | 0.831612492 | 0.4344669 | 0.991835066 | -4.982756283 |
| **A_52_P112958** | Ywhab | -0.046754126 | -0.181133947 | 0.087625696 | 11.39352777 | -0.831562721 | 0.4344931 | 0.991835066 | -4.982787074 |
| **A_55_P2123982** | Rhoh | -0.048748284 | -0.188861849 | 0.091365281 | 8.310558218 | -0.831549827 | 0.434499888 | 0.991835066 | -4.98279505 |
| **A_55_P1973779** | Pde4c | 0.141789254 | -0.265841268 | 0.549419775 | 14.20658662 | 0.831353549 | 0.434603225 | 0.991835066 | -4.982916462 |
| **A_55_P2107422** | Kif21a | 0.052025308 | -0.097543181 | 0.201593797 | 9.297856217 | 0.831349586 | 0.434605312 | 0.991835066 | -4.982918914 |
| **A_55_P2040357** | Bola1 | 0.054093463 | -0.101422408 | 0.209609335 | 9.451572903 | 0.831341006 | 0.434609829 | 0.991835066 | -4.98292422 |
| **A_55_P2172832** | Kcne1l | 0.042590295 | -0.079854975 | 0.165035566 | 8.049791468 | 0.831338262 | 0.434611274 | 0.991835066 | -4.982925917 |
| **A_52_P9703** | Cyp2a4 | 0.037872798 | -0.071017737 | 0.146763333 | 7.683362244 | 0.831278195 | 0.434642902 | 0.991835066 | -4.982963068 |
| **A_51_P464234** | Fkbp2 | -0.098490993 | -0.381714928 | 0.184732942 | 12.10988608 | -0.831142896 | 0.434714149 | 0.991835066 | -4.983046742 |
| **A_51_P327295** | Akr1a1 | -0.0844156 | -0.327231789 | 0.158400588 | 13.35682077 | -0.830910426 | 0.434836586 | 0.991835066 | -4.983190486 |
| **A_55_P2359436** | A930026B( | 0.094846465 | -0.177974862 | 0.367667793 | 8.681905898 | 0.830906136 | 0.434838845 | 0.991835066 | -4.983193138 |
| **A_66_P105175** | Bche | 0.051148289 | -0.096046086 | 0.198342664 | 11.11345952 | 0.830517967 | 0.43504334 | 0.991835066 | -4.983433086 |
| **A_51_P330369** | Ssx2ip | -0.034623201 | -0.134263176 | 0.065016775 | 10.09278344 | -0.830505923 | 0.435049686 | 0.991835066 | -4.98344053 |
| **A_55_P2067741** | Psg23 | -0.494316059 | -1.917000224 | 0.928368106 | 10.51410908 | -0.830434445 | 0.43508735 | 0.991835066 | -4.983484704 |

| | | | | | | | | | |
|---|---|---|---|---|---|---|---|---|---|
| **A_55_P2072284** | Dnaaf3 | -0.037427378 | -0.145161076 | 0.07030632 | 8.225026376 | -0.830322805 | 0.435146181 | 0.991835066 | -4.983553694 |
| **A_51_P440460** | Hip1r | -0.077055068 | -0.29886315 | 0.144753014 | 8.584472223 | -0.830296054 | 0.435160279 | 0.991835066 | -4.983570224 |
| **A_55_P2230688** | Pigm | 0.068889094 | -0.129428512 | 0.267206701 | 8.974905128 | 0.830229831 | 0.43519518 | 0.991835066 | -4.983611143 |
| **A_52_P281145** | Kank4 | -0.033364545 | -0.129415027 | 0.062685937 | 8.087713397 | -0.830223065 | 0.435198746 | 0.991835066 | -4.983615324 |
| **A_55_P2023777** | Tmx2 | -0.073290156 | -0.284321062 | 0.13774075 | 9.233162893 | -0.830058526 | 0.435285471 | 0.991835066 | -4.983716981 |
| **A_52_P514306** | Spata2 | -0.075482396 | -0.29282947 | 0.141864678 | 9.410562987 | -0.830043801 | 0.435293233 | 0.991835066 | -4.983726078 |
| **A_51_P464238** | Atf4 | 0.054142026 | -0.101770721 | 0.210054773 | 8.588497216 | 0.829969271 | 0.435332521 | 0.991835066 | -4.983772119 |
| **A_51_P153063** | Fabp12 | -0.03828789 | -0.148554677 | 0.071978896 | 7.936622157 | -0.829900168 | 0.43536895 | 0.991835066 | -4.983814805 |
| **A_51_P475291** | Moap1 | -0.051927441 | -0.201480219 | 0.097625338 | 8.763035152 | -0.829872869 | 0.435383342 | 0.991835066 | -4.983831667 |
| **A_66_P121288** | Tceal6 | 0.159447058 | -0.299819463 | 0.618713579 | 9.270697653 | 0.829775971 | 0.435434429 | 0.991835066 | -4.983891516 |
| **A_51_P121447** | 1110038F1 | 0.040319326 | -0.075830785 | 0.156469437 | 11.93030746 | 0.82966499 | 0.435492946 | 0.991835066 | -4.983960057 |
| **A_55_P2047621** | Cutal | -0.036986299 | -0.143582694 | 0.069610096 | 7.708528693 | -0.829292028 | 0.435689639 | 0.991835066 | -4.984190346 |
| **A_55_P2086087** | NA | 0.044023014 | -0.082868788 | 0.170914817 | 14.07628447 | 0.829192453 | 0.435742164 | 0.991835066 | -4.984251815 |
| **A_55_P2046807** | Sparc | 0.088219589 | -0.166109908 | 0.342549086 | 13.46702332 | 0.829043715 | 0.43582063 | 0.991835066 | -4.984343625 |
| **A_55_P2093650** | NA | -0.116820537 | -0.45372983 | 0.220088756 | 14.2989388 | -0.828734348 | 0.435983868 | 0.991835066 | -4.984534543 |
| **A_55_P1999902** | Pip5k1a | 0.072223047 | -0.136097422 | 0.280543515 | 12.07813285 | 0.828615388 | 0.436046649 | 0.991835066 | -4.984607941 |
| **A_52_P650028** | Dock10 | -0.063510746 | -0.246730205 | 0.119708713 | 8.314416689 | -0.828485187 | 0.43611537 | 0.991835066 | -4.984688267 |
| **A_51_P340704** | Prpf31 | 0.06271514 | -0.118214627 | 0.243644907 | 11.74292598 | 0.828459923 | 0.436128705 | 0.991835066 | -4.984703852 |
| **A_55_P1980214** | NA | 0.062433032 | -0.117701222 | 0.242567285 | 10.03815829 | 0.828375511 | 0.436173263 | 0.991835066 | -4.984755922 |
| **A_55_P2176724** | Fbxw7 | 0.046733098 | -0.088115404 | 0.181581599 | 8.827650182 | 0.828299837 | 0.436213211 | 0.991835066 | -4.984802599 |
| **A_52_P417825** | Osgin2 | 0.057267257 | -0.107980444 | 0.222514959 | 9.018120795 | 0.828285561 | 0.436220748 | 0.991835066 | -4.984811404 |
| **A_55_P1995924** | Zfp984 | 0.07013798 | -0.132250876 | 0.272526835 | 10.53772551 | 0.828277361 | 0.436225077 | 0.991835066 | -4.984816462 |
| **A_51_P125183** | Coq5 | -0.059314422 | -0.230471249 | 0.111842406 | 9.826460859 | -0.828276246 | 0.436225666 | 0.991835066 | -4.944012154 |
| **A_55_P2132132** | Gm5532 | 0.070695987 | -0.133310933 | 0.274702906 | 9.48285716 | 0.828245329 | 0.436241988 | 0.991835066 | -4.984836218 |
| **A_51_P502422** | Acot13 | -0.077425441 | -0.300856009 | 0.146005127 | 15.58022629 | -0.828228617 | 0.436250811 | 0.991835066 | -4.984846525 |
| **A_55_P2157705** | Uqcr10 | -0.062154408 | -0.241519221 | 0.117210405 | 15.53609656 | -0.828216388 | 0.436257267 | 0.991835066 | -4.984854067 |
| **A_55_P2072403** | Suv39h2 | -0.074315396 | -0.288795025 | 0.140164234 | 12.7419329 | -0.828136405 | 0.436299497 | 0.991835066 | -4.984903395 |
| **A_51_P434776** | Jmjd8 | 0.042348392 | -0.079895418 | 0.164592202 | 8.597890763 | 0.827978729 | 0.436382754 | 0.991835066 | -4.985000626 |
| **A_55_P2032222** | Tm9sf2 | -0.062103157 | -0.241383095 | 0.117176781 | 10.51942772 | -0.827925223 | 0.436411009 | 0.991835066 | -4.985033617 |
| **A_55_P1993423** | NA | 0.074485158 | -0.140545376 | 0.289515692 | 13.19601883 | 0.827901637 | 0.436423465 | 0.991835066 | -4.98504816 |
| **A_55_P2176325** | Prex1 | -0.05112427 | -0.198749612 | 0.096501071 | 8.760740267 | -0.827704545 | 0.436527558 | 0.991835066 | -4.985169669 |
| **A_55_P1996374** | Prmt9 | -0.042280649 | -0.164393251 | 0.079831954 | 10.78671763 | -0.827542461 | 0.436613175 | 0.991835066 | -4.985269579 |
| **A_55_P2071300** | Rnf126 | 0.050010296 | -0.094426807 | 0.194447398 | 10.90820315 | 0.82754151 | 0.436613678 | 0.991835066 | -4.985270165 |
| **A_55_P2038587** | NA | -0.045743233 | -0.177877677 | 0.086391212 | 7.846444856 | -0.827408428 | 0.436683984 | 0.991835066 | -4.985352186 |
| **A_55_P1975385** | Dctn1 | 0.046736064 | -0.088278726 | 0.181750855 | 13.62601094 | 0.827332189 | 0.436724265 | 0.991835066 | -4.98539917 |
| **A_51_P161265** | Ccl25 | 0.101265919 | -0.191304447 | 0.393836285 | 9.22290896 | 0.827260154 | 0.436762326 | 0.991835066 | -4.98544356 |
| **A_55_P2018457** | Hdac7 | 0.042306418 | -0.079927748 | 0.164540585 | 11.59660581 | 0.827223333 | 0.436781783 | 0.991835066 | -4.985466249 |
| **A_51_P405397** | Ecm1 | -0.085417829 | -0.332226215 | 0.161390557 | 11.82211159 | -0.827175661 | 0.436806974 | 0.991835066 | -4.985495623 |
| **A_55_P2156401** | NA | 0.066313175 | -0.125328282 | 0.257954633 | 14.52843767 | 0.827026624 | 0.436885734 | 0.991835066 | -4.985587446 |
| **A_55_P2083859** | NA | 0.060222874 | -0.113835325 | 0.234281073 | 13.35736029 | 0.826944051 | 0.436929376 | 0.991835066 | -4.985638315 |
| **A_55_P2076871** | Lef1 | 0.110532488 | -0.208953437 | 0.430018412 | 8.540670147 | 0.826889254 | 0.436958339 | 0.991835066 | -4.985672071 |
| **A_51_P344046** | Nt5m | 0.049204309 | -0.093041905 | 0.191450523 | 10.32967313 | 0.826744954 | 0.437034615 | 0.991835066 | -4.985760952 |
| **A_52_P110070** | 5730416F0 | 0.042321387 | -0.080075556 | 0.164718329 | 8.107464601 | 0.826415496 | 0.4372088 | 0.991835066 | -4.985963838 |
| **A_51_P104933** | Trub2 | 0.040428451 | -0.076498378 | 0.157355281 | 12.75454219 | 0.826384313 | 0.43722529 | 0.991835066 | -4.945532554 |
| **A_55_P1962419** | Ccar1 | 0.072291182 | -0.136794324 | 0.281376688 | 9.997846132 | 0.826362369 | 0.437236893 | 0.991835066 | -4.985996549 |
| **A_66_P117838** | Dhx29 | 0.035545577 | -0.067266459 | 0.138357613 | 9.025595079 | 0.826324718 | 0.437256804 | 0.991835066 | -4.98601973 |
| **A_55_P1970676** | Sh3yl1 | 0.044960894 | -0.085091945 | 0.175013733 | 8.457349249 | 0.826274277 | 0.437283478 | 0.991835066 | -4.986050784 |
| **A_55_P2137049** | AA467197 | -0.042553195 | -0.165674433 | 0.080568044 | 7.787856795 | -0.826053785 | 0.437400095 | 0.991835066 | -4.986186515 |

| | | | | | | | | | |
|---|---|---|---|---|---|---|---|---|---|
| **A_51_P513785** | Ctcf | 0.069753733 | -0.13208283 | 0.271590296 | 11.12744639 | 0.825993714 | 0.43743187 | 0.991835066 | -4.986223489 |
| **A_55_P2112991** | Slc4a3 | -0.104209611 | -0.405760297 | 0.197341075 | 11.09081456 | -0.82595538 | 0.437452148 | 0.991835066 | -4.986247083 |
| **A_55_P2124381** | NA | -0.0442891 | -0.172491006 | 0.083912806 | 12.76239502 | -0.82567951 | 0.437598097 | 0.991835066 | -4.986416849 |
| **A_51_P443872** | Slu7 | 0.059382232 | -0.112524865 | 0.231289329 | 9.667690831 | 0.825604104 | 0.437637997 | 0.991835066 | -4.986463246 |
| **A_51_P102860** | Herpud2 | -0.041073182 | -0.159982945 | 0.07783658 | 9.600058104 | -0.825562532 | 0.437659995 | 0.991835066 | -4.961462325 |
| **A_55_P2062714** | Zfp975 | 0.046663217 | -0.088452138 | 0.181778573 | 8.512542578 | 0.825427823 | 0.437731283 | 0.991835066 | -4.986571696 |
| **A_55_P1984981** | NA | -0.040896724 | -0.159338043 | 0.077544595 | 8.555455483 | -0.825266888 | 0.43781646 | 0.991835066 | -4.98667069 |
| **A_55_P2144436** | Rrp1 | 0.077332861 | -0.14669167 | 0.301357391 | 12.04773395 | 0.825044994 | 0.43793392 | 0.991835066 | -4.986807157 |
| **A_52_P297332** | Aida | 0.041828977 | -0.079359017 | 0.16301697 | 8.551046823 | 0.824948383 | 0.437985068 | 0.991835066 | -4.986866565 |
| **A_66_P131979** | Cdc6 | 0.082690284 | -0.156892237 | 0.322272805 | 8.825746991 | 0.824913641 | 0.438003462 | 0.991835066 | -4.986887927 |
| **A_55_P2102419** | Lime1 | 0.163240569 | -0.309738658 | 0.636219796 | 9.257611416 | 0.824888339 | 0.438016859 | 0.991835066 | -4.986903485 |
| **A_55_P2099520** | H2afv | -0.070880854 | -0.276267862 | 0.134506154 | 10.01242537 | -0.824831251 | 0.438047087 | 0.991835066 | -4.986938585 |
| **A_55_P2160381** | NA | 0.066209502 | -0.125748117 | 0.258167122 | 10.73266644 | 0.824373645 | 0.438289437 | 0.991835066 | -4.987219873 |
| **A_52_P102002** | Srrd | 0.058335674 | -0.110803316 | 0.227474663 | 8.629229574 | 0.824327174 | 0.438314053 | 0.991835066 | -4.987248432 |
| **A_51_P130660** | Mrps30 | -0.065291161 | -0.254640875 | 0.124058553 | 11.83920776 | -0.824135975 | 0.438415346 | 0.991835066 | -4.987365922 |
| **A_55_P2156776** | NA | -0.043654974 | -0.170308718 | 0.082998769 | 7.96885013 | -0.82380579 | 0.438590309 | 0.991835066 | -4.987568767 |
| **A_51_P446825** | Trmt9b | -0.053675727 | -0.209414967 | 0.102063513 | 8.734756533 | -0.82373784 | 0.438626321 | 0.991835066 | -4.987610503 |
| **A_55_P2040763** | Sec62 | 0.043572409 | -0.082869908 | 0.170014726 | 12.04618944 | 0.823622603 | 0.438687399 | 0.991835066 | -4.987681278 |
| **A_55_P2040993** | NA | 0.168149346 | -0.319848485 | 0.656147177 | 9.06974479 | 0.823543271 | 0.438729451 | 0.991835066 | -4.987729997 |
| **A_51_P393761** | Ndufs2 | -0.14940051 | -0.583020759 | 0.284219739 | 12.28157884 | -0.823477351 | 0.438764395 | 0.991835066 | -4.987770477 |
| **A_55_P1998023** | NA | -0.087187646 | -0.340284597 | 0.165909304 | 8.807511661 | -0.823336128 | 0.438839264 | 0.991835066 | -4.987857191 |
| **A_51_P262701** | Cpb2 | -0.044517425 | -0.173751102 | 0.084716252 | 7.875038054 | -0.823310137 | 0.438853044 | 0.991835066 | -4.987873148 |
| **A_55_P1997501** | Gm10778 | 0.043884305 | -0.083516393 | 0.171285003 | 8.93467018 | 0.823278074 | 0.438870044 | 0.991835066 | -4.987892833 |
| **A_55_P2005025** | Mbnl1 | -0.064638653 | -0.252320575 | 0.12304327 | 10.86405572 | -0.823150003 | 0.438937952 | 0.991835066 | -4.987971457 |
| **A_55_P2156345** | Zfp51 | -0.046351779 | -0.180956109 | 0.088252552 | 8.938752374 | -0.82303159 | 0.439000746 | 0.991835066 | -4.988044143 |
| **A_55_P2067533** | Foxa2 | 0.063424152 | -0.12076988 | 0.247618185 | 7.683934686 | 0.822978043 | 0.439029144 | 0.991835066 | -4.98807701 |
| **A_55_P2003517** | Zfp536 | 0.096161322 | -0.183115153 | 0.375437796 | 9.364092877 | 0.822953322 | 0.439042254 | 0.991835066 | -4.988092183 |
| **A_55_P2111735** | Ppp1r2 | -0.055212486 | -0.215636894 | 0.105211922 | 10.19057422 | -0.822575916 | 0.439242447 | 0.991835066 | -4.988323779 |
| **A_51_P393078** | Tigd2 | 0.045951156 | -0.087584154 | 0.179486465 | 9.489500741 | 0.822449902 | 0.439309304 | 0.991835066 | -4.98840109 |
| **A_55_P2229351** | NA | 0.04417077 | -0.084200534 | 0.172542075 | 8.134004257 | 0.822386835 | 0.439342767 | 0.991835066 | -4.988439778 |
| **A_55_P2167323** | Cort | -0.039002629 | -0.152360192 | 0.074354933 | 8.05534031 | -0.822342092 | 0.439366509 | 0.991835066 | -4.988467225 |
| **A_55_P2152566** | Sp110 | -0.055568661 | -0.217095484 | 0.105958162 | 8.775396572 | -0.822232074 | 0.439424891 | 0.991835066 | -4.988534708 |
| **A_55_P2048751** | Prox1 | 0.056300592 | -0.107381491 | 0.219982675 | 8.629206831 | 0.822092997 | 0.439498702 | 0.991835066 | -4.988620005 |
| **A_51_P322273** | Ptp4a3 | -0.146179907 | -0.571203031 | 0.278843216 | 11.38516671 | -0.822023541 | 0.439535566 | 0.991835066 | -4.988662599 |
| **A_51_P315666** | Nid2 | -0.046529424 | -0.181831627 | 0.08877278 | 11.3023759 | -0.821924518 | 0.439588128 | 0.991835066 | -4.98872332 |
| **A_55_P2419299** | NA | 0.104676357 | -0.19972613 | 0.409078843 | 8.548984422 | 0.821882125 | 0.439610632 | 0.991835066 | -4.988749314 |
| **A_52_P311104** | Brd7 | 0.054821881 | -0.104697273 | 0.214341035 | 13.30956333 | 0.821391543 | 0.439871109 | 0.991835066 | -4.989050045 |
| **A_55_P2051721** | Psg22 | -0.034470839 | -0.134784015 | 0.065842337 | 7.690204515 | -0.821302237 | 0.439918539 | 0.991835066 | -4.989104776 |
| **A_55_P2068134** | NA | -0.042920646 | -0.167823608 | 0.081982316 | 7.892627501 | -0.821301561 | 0.439918898 | 0.991835066 | -4.989105191 |
| **A_55_P2065834** | Dhx8 | 0.046788745 | -0.089400103 | 0.182977593 | 10.21181305 | 0.821124442 | 0.440012974 | 0.991835066 | -4.989213723 |
| **A_55_P1964348** | Gdpd2 | 0.085192468 | -0.162796874 | 0.333181809 | 8.311164248 | 0.821064578 | 0.440044774 | 0.991835066 | -4.989250403 |
| **A_55_P1955059** | Rpl38 | 0.070229563 | -0.134250845 | 0.274709972 | 16.32077269 | 0.820875692 | 0.440145122 | 0.991835066 | -4.98936612 |
| **A_51_P362209** | Commd10 | 0.088374937 | -0.169000722 | 0.345750595 | 10.28086899 | 0.820674194 | 0.440252187 | 0.991835066 | -4.989489541 |
| **A_51_P324923** | Man2a1 | 0.07919721 | -0.151457 | 0.309851421 | 11.14920152 | 0.820649349 | 0.44026539 | 0.991835066 | -4.989504758 |
| **A_51_P147714** | Sugp2 | -0.043121444 | -0.168715285 | 0.082472397 | 9.276144164 | -0.820604864 | 0.44028903 | 0.991835066 | -4.989532002 |
| **A_51_P374464** | Gstp1 | -0.082555897 | -0.323011523 | 0.157899729 | 11.98721076 | -0.820582573 | 0.440300876 | 0.991835066 | -4.989545653 |
| **A_52_P468068** | Tchh | 0.041851954 | -0.080048846 | 0.163752754 | 8.047888663 | 0.820575067 | 0.440304865 | 0.991835066 | -4.98955025 |
| **A_52_P504330** | Ankib1 | 0.060293652 | -0.115330991 | 0.235918295 | 9.019610067 | 0.820531518 | 0.440328009 | 0.991835066 | -4.989576919 |

| | | | | | | | | | |
|---|---|---|---|---|---|---|---|---|---|
| **A_55_P2137406** | Bax | 0.114375863 | -0.218857107 | 0.447608834 | 11.47543639 | 0.820343144 | 0.440428131 | 0.991835066 | -4.989692266 |
| **A_51_P240019** | Gyg | 0.076329349 | -0.146060264 | 0.298718963 | 15.70003576 | 0.820325462 | 0.440437529 | 0.991835066 | -4.989703091 |
| **A_55_P2149906** | NA | 0.075431715 | -0.1443862 | 0.295249631 | 13.49493695 | 0.820162732 | 0.440524035 | 0.991835066 | -4.989802718 |
| **A_55_P1960857** | Sorbs2 | 0.052518496 | -0.100558779 | 0.205595771 | 9.37655271 | 0.819994027 | 0.44061373 | 0.991835066 | -4.989905985 |
| **A_55_P1967144** | NA | -0.039836364 | -0.155957108 | 0.076284381 | 16.43249505 | -0.819934215 | 0.440645533 | 0.991835066 | -4.989942594 |
| **A_52_P657435** | Psmc5 | -0.062779731 | -0.245789848 | 0.120230385 | 12.62769415 | -0.819886034 | 0.440671153 | 0.991835066 | -4.989972082 |
| **A_55_P2035484** | Gucd1 | -0.079209138 | -0.310157664 | 0.151739387 | 11.39190666 | -0.819726975 | 0.440755739 | 0.991835066 | -4.990069421 |
| **A_55_P2038126** | Tpmt | 0.034877275 | -0.066825813 | 0.136580364 | 8.516671049 | 0.819629431 | 0.440807618 | 0.991835066 | -4.990129108 |
| **A_55_P2043723** | Rabac1 | -0.114874582 | -0.449891213 | 0.22014205 | 12.63858366 | -0.819533496 | 0.440858645 | 0.991835066 | -4.990187804 |
| **A_55_P2082826** | Fam160b1 | 0.060017351 | -0.11503285 | 0.235067552 | 11.15362969 | 0.819451663 | 0.440902174 | 0.991835066 | -4.990237869 |
| **A_55_P2053988** | Epas1 | 0.041339409 | -0.079248904 | 0.161927722 | 8.655233981 | 0.819347588 | 0.440957539 | 0.991835066 | -4.990301535 |
| **A_55_P1981366** | Lamc2 | 0.088465453 | -0.169601419 | 0.346532325 | 10.81685581 | 0.819314387 | 0.440975203 | 0.991835066 | -4.990321844 |
| **A_55_P2113837** | Tbc1d1 | 0.052421462 | -0.100506917 | 0.205349841 | 11.91966018 | 0.819275891 | 0.440995683 | 0.991835066 | -4.990345391 |
| **A_55_P2111985** | Prss35 | 0.117509709 | -0.225317173 | 0.460336591 | 8.322810724 | 0.819234086 | 0.441017926 | 0.991835066 | -4.990370961 |
| **A_55_P1996504** | Abcg2 | -0.05888144 | -0.230665111 | 0.112902231 | 12.22249006 | -0.819229676 | 0.441020272 | 0.991835066 | -4.990373658 |
| **A_55_P2155041** | Ssbp2 | 0.044603495 | -0.085531822 | 0.174738811 | 12.43234887 | 0.819186612 | 0.441043185 | 0.991835066 | -4.990399998 |
| **A_55_P2020338** | Scml4 | -0.046733189 | -0.183086083 | 0.089619704 | 8.434892285 | -0.819162748 | 0.441055882 | 0.991835066 | -4.990414593 |
| **A_51_P211822** | Neurl4 | 0.04934675 | -0.094632204 | 0.193325703 | 12.12799686 | 0.819159876 | 0.44105741 | 0.991835066 | -4.990416349 |
| **A_51_P193813** | Fga | -0.036040531 | -0.141201082 | 0.06912002 | 7.697882575 | -0.819119876 | 0.441078694 | 0.991835066 | -4.990440813 |
| **A_55_P2091364** | Cep164 | -0.054887259 | -0.215052867 | 0.10527835 | 11.13980131 | -0.819051871 | 0.441114881 | 0.991835066 | -4.990482402 |
| **A_51_P478895** | Ppp6r1 | 0.06035239 | -0.115767832 | 0.236472611 | 9.383531299 | 0.81901976 | 0.441131968 | 0.991835066 | -4.990502038 |
| **A_52_P126055** | Pon2 | -0.048469994 | -0.189927605 | 0.092987617 | 8.682740213 | -0.818946974 | 0.441170702 | 0.991835066 | -4.990546547 |
| **A_51_P409452** | Cldn11 | 0.087896893 | -0.1686819 | 0.344475686 | 8.487600923 | 0.818769959 | 0.441264914 | 0.991835066 | -4.990654779 |
| **A_55_P2122220** | Kcnk4 | 0.038648755 | -0.074175105 | 0.151472616 | 8.809616349 | 0.81873563 | 0.441283186 | 0.991835066 | -4.990675766 |
| **A_55_P2132024** | Tekt1 | 0.038898118 | -0.074695053 | 0.152491289 | 7.829690961 | 0.818437458 | 0.441441917 | 0.991835066 | -4.99085803 |
| **A_55_P2069875** | Trim44 | 0.102844907 | -0.197511919 | 0.403201733 | 13.38856513 | 0.818378887 | 0.441473101 | 0.991835066 | -4.990893827 |
| **A_55_P2054350** | Fbxo44 | -0.079719592 | -0.312542534 | 0.15310335 | 10.02982526 | -0.818367602 | 0.44147911 | 0.991835066 | -4.990900724 |
| **A_55_P1954860** | NA | -0.072025363 | -0.282388559 | 0.138337833 | 13.05352298 | -0.818323135 | 0.441502787 | 0.991835066 | -4.990927899 |
| **A_52_P180335** | Mrpl28 | -0.10546046 | -0.413526194 | 0.202605275 | 10.88699028 | -0.818192316 | 0.441572447 | 0.991835066 | -4.99100784 |
| **A_55_P2035344** | Tcf25 | 0.061151448 | -0.117486746 | 0.239789642 | 11.03406666 | 0.818166247 | 0.44158633 | 0.991835066 | -4.99102377 |
| **A_51_P253537** | Atf6b | 0.051533512 | -0.099037019 | 0.202104042 | 8.526059746 | 0.818010521 | 0.441669265 | 0.991835066 | -4.991118916 |
| **A_55_P1957909** | Mrpl49 | -0.063317593 | -0.248352342 | 0.121717156 | 10.8354773 | -0.817862382 | 0.44174817 | 0.991835066 | -4.991209414 |
| **A_55_P2000823** | Rbm39 | 0.119109997 | -0.228980114 | 0.467200109 | 13.19624078 | 0.817834941 | 0.441762787 | 0.991835066 | -4.991226176 |
| **A_55_P2066484** | Crtc3 | -0.054660985 | -0.214409576 | 0.105087605 | 9.188590687 | -0.817804604 | 0.441778947 | 0.991835066 | -4.991244707 |
| **A_55_P1980267** | Sidt2 | 0.035546972 | -0.068349605 | 0.139443549 | 11.24658802 | 0.817731081 | 0.441818114 | 0.991835066 | -4.991289615 |
| **A_66_P105074** | Enpp5 | 0.043876916 | -0.084379845 | 0.172133677 | 8.451213931 | 0.817645322 | 0.441863803 | 0.991835066 | -4.991341993 |
| **A_66_P100186** | Psg28 | -0.146917799 | -0.576457717 | 0.282622118 | 7.871368499 | -0.817485423 | 0.441948999 | 0.991835066 | -4.991439641 |
| **A_66_P100561** | Bnip2 | 0.074142364 | -0.142630349 | 0.290915077 | 13.30653728 | 0.817468348 | 0.441958097 | 0.991835066 | -4.991450068 |
| **A_52_P42834** | Usp25 | -0.058210356 | -0.228408312 | 0.111987601 | 8.96925252 | -0.817438415 | 0.441974048 | 0.991835066 | -4.991468346 |
| **A_55_P2113539** | NA | -0.09422211 | -0.369737679 | 0.181293459 | 8.910233005 | -0.817364577 | 0.442013395 | 0.991835066 | -4.991513431 |
| **A_51_P513541** | Rfk | -0.035349637 | -0.138728626 | 0.068029352 | 10.22305394 | -0.81726295 | 0.442067554 | 0.991835066 | -4.991575479 |
| **A_55_P1957373** | Pdcl | -0.036316702 | -0.142527523 | 0.069894118 | 9.854580934 | -0.817234657 | 0.442082633 | 0.991835066 | -4.991592752 |
| **A_55_P2001218** | Cd80 | 0.058718117 | -0.113031771 | 0.230468005 | 8.374196196 | 0.817118022 | 0.442144798 | 0.991835066 | -4.991663954 |
| **A_52_P558699** | Ppp4r3a | 0.060891029 | -0.117215231 | 0.238997288 | 9.795846706 | 0.817115145 | 0.442146332 | 0.991835066 | -4.99166571 |
| **A_51_P419389** | Bmpr2 | 0.070204077 | -0.135153999 | 0.275562154 | 9.832378 | 0.817070782 | 0.442169978 | 0.991835066 | -4.99169279 |
| **A_52_P40241** | Zfp746 | 0.059196914 | -0.113983782 | 0.23237761 | 9.278722251 | 0.816974907 | 0.442221085 | 0.991835066 | -4.99175131 |
| **A_55_P2006906** | Armcx3 | -0.052180812 | -0.204844054 | 0.100482431 | 9.044580173 | -0.816931197 | 0.442244387 | 0.991835066 | -4.991777988 |
| **A_55_P2101392** | Tmem237 | 0.055668519 | -0.107246674 | 0.218583712 | 8.186546778 | 0.816689955 | 0.442373006 | 0.991835066 | -4.991925207 |

| | | | | | | | | | |
|---|---|---|---|---|---|---|---|---|---|
| **A_55_P2078395** | Gm1943 | -0.041764425 | -0.163993034 | 0.080464185 | 8.272520016 | -0.816662788 | 0.442387491 | 0.991835066 | -4.991941783 |
| **A_55_P2002597** | NA | 0.064722209 | -0.124738004 | 0.254182422 | 9.846439253 | 0.816477912 | 0.442486079 | 0.991835066 | -4.99205458 |
| **A_55_P2215880** | Ephb2 | -0.041176747 | -0.161715319 | 0.079361825 | 8.147886381 | -0.816460396 | 0.44249542 | 0.991835066 | -4.992065266 |
| **A_55_P2044659** | NA | -0.040249488 | -0.158118144 | 0.077619169 | 7.734154617 | -0.8161522 | 0.442659807 | 0.991835066 | -4.992253255 |
| **A_66_P139974** | Arl6 | -0.053534021 | -0.210311421 | 0.103243379 | 9.518317579 | -0.816122851 | 0.442675464 | 0.991835066 | -4.992271154 |
| **A_52_P558859** | Rras2 | -0.059247177 | -0.232797857 | 0.114303502 | 9.548979206 | -0.815925447 | 0.442780781 | 0.991835066 | -4.992391533 |
| **A_52_P381418** | Slc35b4 | -0.037617777 | -0.147830459 | 0.072594906 | 8.296200679 | -0.81577555 | 0.442860765 | 0.991835066 | -4.992482926 |
| **A_55_P2038767** | Fam49a | 0.078375701 | -0.151253009 | 0.308004412 | 9.454707838 | 0.815763714 | 0.442867081 | 0.991835066 | -4.992490141 |
| **A_66_P123155** | Ddo | -0.07912608 | -0.310983803 | 0.152731643 | 10.30714115 | -0.815656341 | 0.442924381 | 0.991835066 | -4.992555599 |
| **A_55_P2062911** | Epha5 | -0.078533893 | -0.308670038 | 0.151602253 | 12.22591947 | -0.815607895 | 0.442950236 | 0.991835066 | -4.992585131 |
| **A_55_P2152901** | 2900040C0 | -0.092373713 | -0.363085166 | 0.17833774 | 10.32873589 | -0.815550585 | 0.442980824 | 0.991835066 | -4.992620065 |
| **A_55_P1963434** | Gpc4 | -0.057329703 | -0.225379749 | 0.110720343 | 9.117224335 | -0.815361473 | 0.443081766 | 0.991835066 | -4.992735324 |
| **A_55_P2117420** | NA | -0.08136557 | -0.319907394 | 0.157176254 | 15.70203903 | -0.8152396 | 0.443146827 | 0.991835066 | -4.992809593 |
| **A_55_P2055881** | NA | 0.059239159 | -0.114434424 | 0.232912741 | 16.15199332 | 0.815237696 | 0.443147843 | 0.991835066 | -4.992810753 |
| **A_55_P2072030** | NA | 0.049174494 | -0.095008499 | 0.193357487 | 15.37375817 | 0.815145234 | 0.443197208 | 0.991835066 | -4.992867093 |
| **A_51_P110341** | Scgb3a1 | -0.072595112 | -0.285455778 | 0.140265554 | 7.900515058 | -0.815119149 | 0.443211135 | 0.991835066 | -4.987874195 |
| **A_55_P1957754** | NA | 0.057308695 | -0.110764976 | 0.225382367 | 14.52681228 | 0.814948126 | 0.443302455 | 0.991835066 | -4.99298718 |
| **A_55_P2061525** | Cdc27 | -0.041958085 | -0.165013416 | 0.081097245 | 9.292579489 | -0.814937618 | 0.443308067 | 0.991835066 | -4.992993582 |
| **A_52_P254795** | Ndufc1 | -0.045771999 | -0.18002531 | 0.088481311 | 16.01462899 | -0.814861896 | 0.443348504 | 0.991835066 | -4.993039709 |
| **A_51_P262230** | Ggact | -0.057629118 | -0.226692283 | 0.111434047 | 9.47465642 | -0.814708244 | 0.443430566 | 0.991835066 | -4.993133298 |
| **A_55_P2181542** | Sult1d1 | -0.036994814 | -0.145530508 | 0.071540879 | 7.667942688 | -0.81466189 | 0.443455325 | 0.991835066 | -4.993161529 |
| **A_55_P1969660** | Pex26 | -0.065118478 | -0.256222075 | 0.125985118 | 10.20367698 | -0.814412656 | 0.443588462 | 0.991835066 | -4.993313301 |
| **A_55_P2033160** | Pomgnt1 | -0.037188851 | -0.146331063 | 0.07195336 | 12.52549851 | -0.81438384 | 0.443603858 | 0.991835066 | -4.993330846 |
| **A_55_P2000047** | 9930104L0 | 0.052954412 | -0.102473153 | 0.208381976 | 8.388793766 | 0.814297757 | 0.44364985 | 0.991835066 | -4.993383257 |
| **A_52_P346581** | Colgalt1 | -0.09640791 | -0.379399305 | 0.186583485 | 10.47048834 | -0.814232757 | 0.44368458 | 0.991835066 | -4.993422829 |
| **A_55_P2044359** | Rxrg | -0.061394445 | -0.241629656 | 0.118840766 | 9.743129582 | -0.814139026 | 0.443734665 | 0.991835066 | -4.993479888 |
| **A_51_P233928** | B4galnt1 | -0.040176327 | -0.158129657 | 0.077777003 | 7.804702146 | -0.814083877 | 0.443764136 | 0.991835066 | -4.993513457 |
| **A_66_P124659** | Mark1 | 0.051494374 | -0.099687906 | 0.202676654 | 9.336847259 | 0.814081758 | 0.443765268 | 0.991835066 | -4.993514748 |
| **A_55_P2031692** | Gstm6 | -0.047180337 | -0.185708518 | 0.091347844 | 9.476967249 | -0.814014313 | 0.443801312 | 0.991835066 | -4.993555799 |
| **A_55_P1961059** | Usp16 | 0.056595389 | -0.109593172 | 0.222783949 | 11.08113529 | 0.813933744 | 0.443844371 | 0.991835066 | -4.993604836 |
| **A_55_P2151546** | Adarb1 | 0.073541955 | -0.142420884 | 0.289504794 | 10.99234426 | 0.813889179 | 0.44386819 | 0.991835066 | -4.993631959 |
| **A_52_P657123** | Gm10270 | -0.056990686 | -0.224356293 | 0.110374922 | 10.30553958 | -0.813854547 | 0.443886701 | 0.991835066 | -4.993653035 |
| **A_55_P2082045** | Vps13a | 0.050773859 | -0.098344355 | 0.199892074 | 8.507877085 | 0.813801734 | 0.44391493 | 0.991835066 | -4.993685174 |
| **A_52_P110257** | Wdr83 | -0.064114501 | -0.252430878 | 0.124201875 | 9.767774418 | -0.813724351 | 0.443956295 | 0.991835066 | -4.993732262 |
| **A_51_P268843** | Rasip1 | 0.064026212 | -0.124036937 | 0.252089361 | 11.47485047 | 0.813697981 | 0.443970391 | 0.991835066 | -4.993748308 |
| **A_65_P04472** | Atp6v1c1 | -0.043273271 | -0.17042841 | 0.083881868 | 10.16165209 | -0.813382714 | 0.444138946 | 0.991835066 | -4.99394011 |
| **A_66_P135106** | Slco3a1 | 0.051035966 | -0.098950732 | 0.201022664 | 9.770476108 | 0.813266205 | 0.444201248 | 0.991835066 | -4.994010978 |
| **A_66_P114054** | Sco2 | -0.099993759 | -0.393936414 | 0.193948896 | 10.39936346 | -0.813054045 | 0.444314715 | 0.991835066 | -4.994140006 |
| **A_55_P2138070** | NA | 0.048688357 | -0.094539996 | 0.19191671 | 13.81227887 | 0.81246611 | 0.444629259 | 0.991835066 | -4.994497433 |
| **A_55_P2018507** | Gorasp2 | -0.072063592 | -0.28406277 | 0.139935586 | 9.010183986 | -0.812439182 | 0.444643669 | 0.991835066 | -4.994513798 |
| **A_55_P2190299** | Fggy | 0.054355574 | -0.105572928 | 0.214284076 | 8.52992329 | 0.812320375 | 0.444707251 | 0.991835066 | -4.994585999 |
| **A_66_P110970** | Tpd52l2 | -0.043773948 | -0.172571093 | 0.085023198 | 8.522806783 | -0.812304031 | 0.444715998 | 0.991835066 | -4.994595931 |
| **A_51_P164203** | Nme4 | -0.047996943 | -0.189236815 | 0.093242928 | 11.70629086 | -0.812204512 | 0.444769264 | 0.991835066 | -4.994656403 |
| **A_51_P369897** | Zfp410 | -0.054569816 | -0.215178603 | 0.106038971 | 9.685529422 | -0.81206785 | 0.444842416 | 0.991835066 | -4.994739436 |
| **A_55_P2169529** | NA | 0.051804629 | -0.100671228 | 0.204280487 | 9.640996597 | 0.812038499 | 0.444858128 | 0.991835066 | -4.994757267 |
| **A_55_P1973462** | Pex11b | 0.037294865 | -0.072479462 | 0.147069193 | 10.27347314 | 0.812002543 | 0.444877377 | 0.991835066 | -4.994779111 |
| **A_51_P345649** | Pdgfra | -0.087887234 | -0.346587854 | 0.170813386 | 10.16935509 | -0.811965278 | 0.444897327 | 0.991835066 | -4.994801749 |
| **A_55_P2038484** | 4933427D0 | -0.082787554 | -0.326503621 | 0.160928514 | 12.09733717 | -0.811876593 | 0.444944807 | 0.991835066 | -4.994855621 |

| | | | | | | | | | |
|---|---|---|---|---|---|---|---|---|---|
| **A_55_P2043662** | Zfp607a | 0.034568101 | -0.067197509 | 0.136333712 | 8.464549026 | 0.811864626 | 0.444951214 | 0.991835066 | -4.99486289 |
| **A_55_P2160682** | Set | 0.07271909 | -0.141369421 | 0.286807602 | 10.86751243 | 0.81182834 | 0.444970642 | 0.991835066 | -4.994884931 |
| **A_55_P1993517** | Tbc1d14 | -0.035867869 | -0.141488587 | 0.069752849 | 7.798922477 | -0.811644012 | 0.445069343 | 0.991835066 | -4.994996881 |
| **A_55_P1989276** | Romo1 | 0.050699381 | -0.098597363 | 0.199996125 | 14.60115965 | 0.811636284 | 0.445073481 | 0.991835066 | -4.995001574 |
| **A_51_P147064** | 1600014C2 | 0.126881032 | -0.246758228 | 0.500520291 | 8.708909317 | 0.811621103 | 0.445081611 | 0.991835066 | -4.995010794 |
| **A_55_P1980543** | Ube2i | 0.036317182 | -0.070641719 | 0.143276084 | 10.70836361 | 0.811529557 | 0.445130637 | 0.991835066 | -4.995066385 |
| **A_55_P1970920** | Timm17b | -0.041713902 | -0.164573236 | 0.081145431 | 13.15302256 | -0.811487432 | 0.445153198 | 0.991835066 | -4.995091964 |
| **A_51_P343517** | Ly6d | -0.04266375 | -0.168338235 | 0.083010736 | 7.733016781 | -0.811373912 | 0.445214 | 0.991835066 | -4.995160891 |
| **A_51_P404193** | Sp5 | 0.03463706 | -0.067420797 | 0.136694917 | 7.784106468 | 0.811154731 | 0.44533141 | 0.991835066 | -4.99529395 |
| **A_55_P2153291** | NA | -0.046067924 | -0.18180712 | 0.089671272 | 14.59923528 | -0.811152479 | 0.445332616 | 0.991835066 | -4.995295316 |
| **A_55_P1972817** | Trmt11 | -0.045697909 | -0.180359137 | 0.08896332 | 9.210504264 | -0.811078488 | 0.445372257 | 0.991835066 | -4.995340228 |
| **A_51_P506683** | Fbxw7 | -0.099790199 | -0.393886095 | 0.194305697 | 9.601240919 | -0.810976102 | 0.445427114 | 0.991835066 | -4.99540237 |
| **A_55_P1970299** | Mttp | 0.049523251 | -0.096442571 | 0.195489073 | 8.091589171 | 0.810899627 | 0.445468092 | 0.991835066 | -4.995448782 |
| **A_55_P2106844** | NA | -0.037877313 | -0.149522636 | 0.07376801 | 8.231453002 | -0.810863519 | 0.44548744 | 0.991835066 | -4.995470694 |
| **A_55_P2117375** | NA | 0.040884027 | -0.079632486 | 0.161400541 | 8.717989737 | 0.81080467 | 0.445518976 | 0.991835066 | -4.995506406 |
| **A_66_P101519** | Abcc9 | -0.101811397 | -0.402023878 | 0.198401083 | 11.92723717 | -0.810544359 | 0.445658488 | 0.991835066 | -4.995664346 |
| **A_51_P309733** | Cyp11a1 | -0.033590182 | -0.132647334 | 0.06546697 | 7.665194114 | -0.810467584 | 0.445699641 | 0.991835066 | -4.99571092 |
| **A_55_P1971194** | Timm23 | -0.071087393 | -0.28074448 | 0.138569694 | 13.74567201 | -0.810386457 | 0.44574313 | 0.991835066 | -4.995760131 |
| **A_66_P102046** | NA | -0.131349215 | -0.518740956 | 0.256042525 | 12.11169939 | -0.810375416 | 0.445749049 | 0.991835066 | -4.995766828 |
| **A_52_P37091** | Magi1 | 0.091562938 | -0.178511896 | 0.361637771 | 9.171017074 | 0.810297939 | 0.445790585 | 0.991835066 | -4.995813821 |
| **A_55_P1954907** | Rpl12 | 0.05683428 | -0.110819369 | 0.224487928 | 16.67169316 | 0.810226576 | 0.445828845 | 0.991835066 | -4.995857103 |
| **A_52_P251623** | Fam222a | -0.053532973 | -0.211458528 | 0.104392581 | 8.496699212 | -0.810173604 | 0.445857247 | 0.991835066 | -4.995889229 |
| **A_55_P1974788** | Cfl1 | -0.083563533 | -0.330124762 | 0.162997696 | 11.31477217 | -0.810030066 | 0.445934213 | 0.991835066 | -4.995976271 |
| **A_51_P426754** | Anxa5 | -0.076462715 | -0.302080737 | 0.149155307 | 13.64360293 | -0.810000073 | 0.445950296 | 0.991835066 | -4.995994457 |
| **A_55_P2165299** | Mybpc1 | 0.08765408 | -0.170989135 | 0.346297294 | 8.484041139 | 0.809990969 | 0.445955178 | 0.991835066 | -4.995999977 |
| **A_55_P2032768** | NA | -0.063966074 | -0.252729388 | 0.124797239 | 12.73481307 | -0.809918348 | 0.445994123 | 0.991835066 | -4.996044009 |
| **A_55_P2079089** | Fuca1 | -0.074529184 | -0.294480654 | 0.145422285 | 10.63402562 | -0.809857546 | 0.446026731 | 0.991835066 | -4.996080872 |
| **A_55_P2185712** | Copa | -0.057588269 | -0.22758329 | 0.112406752 | 11.68695847 | -0.809667963 | 0.446128416 | 0.991835066 | -4.9961958 |
| **A_55_P2001250** | Cpz | 0.119467082 | -0.233232132 | 0.472166295 | 9.445795138 | 0.809567185 | 0.446182476 | 0.991835066 | -4.996256884 |
| **A_55_P2124153** | Rusc2 | -0.05162481 | -0.204036576 | 0.100786955 | 11.00697566 | -0.809560124 | 0.446186264 | 0.991835066 | -4.996261163 |
| **A_55_P2139027** | Plec | -0.064726749 | -0.255931819 | 0.12647832 | 13.01872547 | -0.809083837 | 0.446441821 | 0.991835066 | -4.996549773 |
| **A_52_P657317** | Tmem160 | -0.16100671 | -0.636640024 | 0.314626604 | 12.05883651 | -0.80906021 | 0.446454502 | 0.991835066 | -4.996564086 |
| **A_55_P2150044** | Cdkl2 | -0.060142434 | -0.237818723 | 0.117533856 | 8.134556339 | -0.8090226 | 0.446474686 | 0.991835066 | -4.99658687 |
| **A_55_P1959064** | Oas1b | -0.033033886 | -0.130636504 | 0.064568732 | 8.085231558 | -0.808923266 | 0.446528001 | 0.991835066 | -4.996647042 |
| **A_55_P2182790** | NA | -0.069062925 | -0.273168691 | 0.135042842 | 15.62995257 | -0.808721192 | 0.446636472 | 0.991835066 | -4.996769432 |
| **A_51_P349008** | Ptges2 | -0.083899249 | -0.331853255 | 0.164054756 | 11.55059752 | -0.808716081 | 0.446639216 | 0.991835066 | -4.996772528 |
| **A_55_P2430221** | Shld2 | 0.038926671 | -0.076125313 | 0.153978655 | 8.370800758 | 0.808653152 | 0.446672999 | 0.991835066 | -4.996810637 |
| **A_55_P1954086** | Postn | -0.093277162 | -0.368985093 | 0.182430769 | 13.72843255 | -0.808602718 | 0.446700076 | 0.991835066 | -4.996841177 |
| **A_55_P2047310** | Pdgfb | -0.045237422 | -0.178953148 | 0.088478305 | 11.34923922 | -0.808582776 | 0.446710783 | 0.991835066 | -4.996853253 |
| **A_55_P2187034** | NA | -0.076877215 | -0.304116916 | 0.150362485 | 12.63596446 | -0.808579204 | 0.446712701 | 0.991835066 | -4.996855416 |
| **A_55_P1953402** | Mlana | -0.103615923 | -0.409893012 | 0.202661166 | 8.682749929 | -0.80857651 | 0.446714148 | 0.991835066 | -4.996857048 |
| **A_51_P468073** | Ggt1 | -0.033293471 | -0.131729875 | 0.065142934 | 7.68580536 | -0.808374233 | 0.446822759 | 0.991835066 | -4.99697952 |
| **A_66_P129800** | Rab8b | -0.051762454 | -0.204831412 | 0.101306503 | 10.39457924 | -0.808233541 | 0.446898314 | 0.991835066 | -4.997064691 |
| **A_51_P267194** | Faim | 0.055109446 | -0.1078605 | 0.218079393 | 9.834541418 | 0.808216398 | 0.446907521 | 0.991835066 | -4.997075067 |
| **A_55_P1991440** | Zscan21 | 0.061960857 | -0.121276886 | 0.245198599 | 11.75044177 | 0.80818654 | 0.446923557 | 0.991835066 | -4.997093141 |
| **A_55_P2324976** | 5033406O( | 0.069949589 | -0.136937542 | 0.27683672 | 8.47420127 | 0.808092032 | 0.446974317 | 0.991835066 | -4.997150344 |
| **A_55_P2046852** | Gata1 | -0.047742808 | -0.188960011 | 0.093474395 | 7.996205323 | -0.808033727 | 0.447005635 | 0.991835066 | -4.997185632 |
| **A_55_P2041011** | Slc2a13 | 0.091226829 | -0.178630489 | 0.361084146 | 8.983627477 | 0.807974233 | 0.447037593 | 0.991835066 | -4.997221638 |

| | | | | | | | | | |
|---|---|---|---|---|---|---|---|---|---|
| **A_51_P437978** | Agtr2 | -0.092216539 | -0.365013599 | 0.180580521 | 8.032407173 | -0.807938427 | 0.447056827 | 0.991835066 | -4.997243307 |
| **A_51_P324491** | Cmpk1 | -0.041822226 | -0.165574844 | 0.081930391 | 9.834140808 | -0.807721954 | 0.447173125 | 0.991835066 | -4.997374294 |
| **A_51_P130332** | Hspa12b | 0.033074569 | -0.064796422 | 0.13094556 | 10.40830667 | 0.807698618 | 0.447185663 | 0.991835066 | -4.997388412 |
| **A_55_P2040518** | NA | -0.076829118 | -0.304206015 | 0.150547779 | 10.46654727 | -0.807585747 | 0.447246312 | 0.991835066 | -4.997456698 |
| **A_55_P2047837** | NA | 0.054211632 | -0.106252674 | 0.214675938 | 8.008190055 | 0.807464006 | 0.447311733 | 0.991835066 | -4.997530342 |
| **A_55_P1993473** | Gm5935 | 0.04169953 | -0.081762582 | 0.165161642 | 7.915921579 | 0.807247285 | 0.447428211 | 0.991835066 | -4.99766142 |
| **A_51_P164630** | Fitm1 | -0.132988089 | -0.526737674 | 0.260761496 | 10.48629491 | -0.807238319 | 0.44743303 | 0.991835066 | -4.997666842 |
| **A_66_P118233** | Hnrnpa0 | 0.074468563 | -0.146038159 | 0.294975286 | 13.74936282 | 0.807161191 | 0.447474489 | 0.991835066 | -4.997713484 |
| **A_55_P2052097** | Ip6k2 | 0.05412011 | -0.106144694 | 0.214384913 | 8.467864741 | 0.80710427 | 0.447505087 | 0.991835066 | -4.997747903 |
| **A_55_P1954755** | NA | -0.078209732 | -0.30983406 | 0.153414597 | 15.88114842 | -0.807022708 | 0.447548934 | 0.991835066 | -4.99779722 |
| **A_51_P281024** | Rad9a | -0.048497114 | -0.192148249 | 0.095154021 | 9.022195334 | -0.806893036 | 0.44761865 | 0.991835066 | -4.997875619 |
| **A_52_P654618** | Znhit3 | -0.047392361 | -0.1877748 | 0.092990079 | 8.783850493 | -0.806872081 | 0.447629917 | 0.991835066 | -4.997888288 |
| **A_51_P134007** | Ncl | 0.144950065 | -0.284429703 | 0.574329834 | 12.40741427 | 0.806837305 | 0.447648616 | 0.991835066 | -4.997909311 |
| **A_51_P127681** | Clic4 | -0.04688978 | -0.185794919 | 0.092015359 | 14.17968368 | -0.8068058 | 0.447665556 | 0.991835066 | -4.967126936 |
| **A_52_P148212** | Mis18bp1 | -0.04330553 | -0.171595776 | 0.084984717 | 8.242035575 | -0.806786917 | 0.447675709 | 0.991835066 | -4.997939771 |
| **A_55_P1997926** | NA | -0.103493865 | -0.410107417 | 0.203119687 | 12.92492648 | -0.806737772 | 0.447702136 | 0.991835066 | -4.997969478 |
| **A_55_P2009857** | NA | 0.155567732 | -0.305342318 | 0.616477782 | 11.1118773 | 0.806700775 | 0.447722031 | 0.991835066 | -4.997991841 |
| **A_55_P2136786** | Zkscan1 | 0.06336953 | -0.12442713 | 0.25116619 | 13.0283942 | 0.80649514 | 0.447832621 | 0.991835066 | -4.998116124 |
| **A_55_P2112185** | Nhsl1 | -0.058019705 | -0.229968839 | 0.113929428 | 11.15042444 | -0.806463395 | 0.447849695 | 0.991835066 | -4.998135308 |
| **A_66_P140949** | NA | 0.052022715 | -0.102190515 | 0.206235946 | 12.5425197 | 0.80627002 | 0.447953713 | 0.991835066 | -4.998252155 |
| **A_55_P2025665** | Mrpl40 | -0.045648891 | -0.180990881 | 0.0896931 | 11.76766 | -0.806133185 | 0.448027327 | 0.991835066 | -4.998334824 |
| **A_55_P2090633** | Ptp4a1 | 0.053080145 | -0.104314519 | 0.21047481 | 9.239563798 | 0.806030042 | 0.448082822 | 0.991835066 | -4.998397132 |
| **A_55_P2021490** | Gm16386 | -0.044888633 | -0.178009047 | 0.08823178 | 8.241331161 | -0.805936558 | 0.448133124 | 0.991835066 | -4.998453599 |
| **A_51_P516452** | Eif3f | -0.067225493 | -0.266592495 | 0.132141509 | 12.57939144 | -0.805916133 | 0.448144115 | 0.991835066 | -4.998465936 |
| **A_51_P338262** | Tnnt2 | -0.077219019 | -0.306225437 | 0.151787398 | 17.30716959 | -0.805908546 | 0.448148197 | 0.991835066 | -4.998470519 |
| **A_52_P614960** | Skiv2l | -0.068950975 | -0.273500636 | 0.135598687 | 8.902723135 | -0.805658095 | 0.448282983 | 0.991835066 | -4.998621769 |
| **A_55_P2113155** | Scnm1 | 0.053966849 | -0.106160101 | 0.214093799 | 9.776607206 | 0.805511528 | 0.448361874 | 0.991835066 | -4.998710266 |
| **A_52_P526852** | Eri1 | 0.044212626 | -0.086995871 | 0.175421122 | 8.69707501 | 0.805366353 | 0.448440025 | 0.991835066 | -4.998797911 |
| **A_55_P2002642** | Deup1 | -0.038500268 | -0.152775609 | 0.075775073 | 8.022603268 | -0.805230773 | 0.448513021 | 0.991835066 | -4.998879752 |
| **A_65_P03031** | Ncapd3 | 0.053058983 | -0.104434754 | 0.210552721 | 8.502044705 | 0.80520186 | 0.448528588 | 0.991835066 | -4.998897203 |
| **A_55_P2113071** | Ago4 | 0.063031561 | -0.124122457 | 0.250185579 | 8.745061419 | 0.804948396 | 0.448665076 | 0.991835066 | -4.99905017 |
| **A_55_P2082185** | NA | 0.052309125 | -0.103021523 | 0.207639773 | 13.64818277 | 0.804876849 | 0.448703609 | 0.991835066 | -4.999093342 |
| **A_52_P541157** | Calu | -0.08100789 | -0.321591885 | 0.159576106 | 9.933739664 | -0.804766182 | 0.448763216 | 0.991835066 | -4.999160114 |
| **A_52_P315976** | Tpm2 | 0.102693141 | -0.202295277 | 0.407681559 | 10.7842818 | 0.804761555 | 0.448765708 | 0.991835066 | -4.999162906 |
| **A_52_P582309** | Nub1 | -0.0511882 | -0.20324127 | 0.10086487 | 10.3321293 | -0.80460699 | 0.448848967 | 0.991835066 | -4.999256151 |
| **A_55_P2005555** | Celf2 | -0.044079094 | -0.175019729 | 0.086861541 | 9.126784594 | -0.804576506 | 0.448865389 | 0.991835066 | -4.99927454 |
| **A_52_P665817** | Abhd13 | -0.053590685 | -0.212797664 | 0.105616294 | 8.449880965 | -0.804519066 | 0.448896334 | 0.991835066 | -4.999309188 |
| **A_55_P2077909** | Tpgs2 | 0.035756104 | -0.070479546 | 0.141991754 | 8.104149443 | 0.804431462 | 0.448943532 | 0.991835066 | -4.999362027 |
| **A_51_P120254** | Fbxl8 | -0.052098326 | -0.20691678 | 0.102720128 | 8.593552867 | -0.804285379 | 0.449022244 | 0.991835066 | -4.999450128 |
| **A_55_P2129234** | Calm2 | 0.072189906 | -0.142348031 | 0.286727844 | 14.09962016 | 0.804232298 | 0.449050848 | 0.991835066 | -4.999482138 |
| **A_51_P113182** | Lifr | -0.050580023 | -0.200903696 | 0.099743651 | 8.143982871 | -0.804193894 | 0.449071543 | 0.991835066 | -4.999505295 |
| **A_52_P553890** | Itgb3 | 0.099483877 | -0.196242393 | 0.395210147 | 10.04988646 | 0.804029399 | 0.449160194 | 0.991835066 | -4.999604477 |
| **A_52_P559975** | Cxcr2 | -0.0342628 | -0.136113733 | 0.067588132 | 7.710000745 | -0.804020235 | 0.449165133 | 0.991835066 | -4.999610002 |
| **A_51_P281806** | Cactin | 0.051515222 | -0.101649526 | 0.20467997 | 12.24517706 | 0.803870125 | 0.449246043 | 0.991835066 | -4.999700496 |
| **A_66_P138929** | Lox | 0.117149111 | -0.231163529 | 0.465461751 | 8.56033295 | 0.803857176 | 0.449253023 | 0.991835066 | -4.999708302 |
| **A_55_P2137186** | Rps25 | 0.054067686 | -0.10674055 | 0.214875921 | 15.42536584 | 0.803597588 | 0.449392969 | 0.991835066 | -4.999864762 |
| **A_55_P2159289** | NA | -0.102440446 | -0.407141735 | 0.202260843 | 8.700490477 | -0.803537783 | 0.449425214 | 0.991835066 | -4.999900802 |
| **A_55_P2057102** | Sft2d3 | 0.056694842 | -0.111980745 | 0.22537043 | 8.592251857 | 0.803341959 | 0.449530809 | 0.991835066 | -5.000018797 |

| | | | | | | | | | |
|---|---|---|---|---|---|---|---|---|---|
| **A_52_P217710** | Fzd6 | -0.045650309 | -0.181470893 | 0.090170275 | 8.48816207 | -0.80331756 | 0.449543967 | 0.991835066 | -5.000033498 |
| **A_52_P364232** | Pik3c2a | 0.039884286 | -0.078786398 | 0.158554971 | 8.315313983 | 0.803281027 | 0.44956367 | 0.991835066 | -5.000055508 |
| **A_55_P2220283** | Stk35os1 | -0.035252558 | -0.140147744 | 0.069642628 | 8.738251331 | -0.803237941 | 0.449586906 | 0.991835066 | -5.000081465 |
| **A_51_P293781** | Fkbp1b | 0.057674784 | -0.113950556 | 0.229300125 | 9.691633547 | 0.803181502 | 0.449617346 | 0.991835066 | -5.000115466 |
| **A_51_P132170** | Ccdc141 | 0.102795369 | -0.203100215 | 0.408690953 | 9.808251234 | 0.803173691 | 0.449621559 | 0.991835066 | -5.000120171 |
| **A_52_P552832** | Ndufa4 | -0.086545481 | -0.344097186 | 0.171006225 | 14.77616381 | -0.803136007 | 0.449641885 | 0.991835066 | -5.000142872 |
| **A_51_P330428** | Eif4ebp1 | -0.047017053 | -0.186937109 | 0.092903003 | 14.4124157 | -0.803127623 | 0.449646407 | 0.991835066 | -5.000147922 |
| **A_55_P2119985** | Baiap2 | 0.048208747 | -0.095259123 | 0.191676617 | 8.173749186 | 0.803119786 | 0.449650634 | 0.991835066 | -5.000152643 |
| **A_55_P2050688** | Zbed4 | 0.076550436 | -0.151269058 | 0.30436993 | 10.94319662 | 0.80309314 | 0.449665006 | 0.991835066 | -5.000168693 |
| **A_55_P2269459** | Prl8a6 | -0.633890181 | -2.520562989 | 1.252782626 | 9.063045397 | -0.803020379 | 0.449704254 | 0.991835066 | -5.00021252 |
| **A_55_P2037930** | NA | -0.039928336 | -0.1588007 | 0.078944029 | 7.989834684 | -0.802803842 | 0.449821071 | 0.991835066 | -5.00034293 |
| **A_55_P1990495** | Stat5a | -0.048372392 | -0.19239011 | 0.095645326 | 9.124475501 | -0.802769325 | 0.449839694 | 0.991835066 | -5.000363715 |
| **A_55_P1995195** | Fosl2 | -0.049160415 | -0.195536901 | 0.09721607 | 9.832228924 | -0.802700178 | 0.449877002 | 0.991835066 | -5.000405352 |
| **A_51_P498403** | Cnpy3 | 0.058744784 | -0.116189617 | 0.233679185 | 10.0899117 | 0.802607513 | 0.449927004 | 0.991835066 | -5.000461146 |
| **A_55_P2097161** | Ifitm7 | -0.070211744 | -0.279322225 | 0.138898737 | 12.92237073 | -0.802496402 | 0.449986963 | 0.991835066 | -5.00052804 |
| **A_55_P2033680** | NA | 0.069254433 | -0.137028749 | 0.275537616 | 9.52743817 | 0.802403638 | 0.450037027 | 0.991835066 | -5.000583882 |
| **A_51_P270519** | Cenps | 0.04772447 | -0.094431799 | 0.189880738 | 8.597734187 | 0.802387635 | 0.450045664 | 0.991835066 | -5.000593515 |
| **A_55_P2034858** | Pias4 | -0.074125451 | -0.294955985 | 0.146705083 | 8.596815071 | -0.802264097 | 0.450112342 | 0.991835066 | -5.000667874 |
| **A_55_P2091284** | Ltbp3 | 0.090865063 | -0.179848387 | 0.361578513 | 10.48723258 | 0.80222507 | 0.450133408 | 0.991835066 | -5.000691363 |
| **A_51_P507290** | Klf5 | 0.040844947 | -0.080851346 | 0.162541241 | 8.156510041 | 0.802176842 | 0.450159442 | 0.991835066 | -5.000720388 |
| **A_55_P2048779** | Lratd2 | 0.041341605 | -0.081835288 | 0.164518498 | 7.932006175 | 0.802171485 | 0.450162333 | 0.991835066 | -5.000723613 |
| **A_51_P267700** | Gkn3 | -0.054682441 | -0.217619153 | 0.108254271 | 8.664494446 | -0.80211766 | 0.45019139 | 0.991835066 | -5.000756005 |
| **A_51_P110471** | Ddah1 | 0.074904884 | -0.148321369 | 0.298131138 | 9.037042572 | 0.801999321 | 0.450255276 | 0.991835066 | -4.990853775 |
| **A_55_P2098057** | Sf3a2 | -0.068581674 | -0.273041968 | 0.13587862 | 12.2036187 | -0.801693265 | 0.450420534 | 0.991835066 | -5.00101135 |
| **A_55_P2045818** | NA | -0.055296959 | -0.220153227 | 0.109559309 | 14.57090567 | -0.801687135 | 0.450423844 | 0.991835066 | -5.001015037 |
| **A_52_P31490** | Vamp3 | -0.043580519 | -0.17353228 | 0.086371242 | 8.997584513 | -0.801529221 | 0.450509129 | 0.991835066 | -5.001110022 |
| **A_55_P1967820** | AI661453 | 0.049570578 | -0.098261067 | 0.197402223 | 8.332771638 | 0.801430202 | 0.450562611 | 0.991835066 | -5.001169574 |
| **A_55_P1995141** | Casz1 | -0.082041916 | -0.32677944 | 0.162695607 | 8.788276052 | -0.80120632 | 0.450683552 | 0.991835066 | -5.0013042 |
| **A_55_P1966372** | Srek1 | -0.06950039 | -0.276829523 | 0.137828744 | 13.08286621 | -0.801190956 | 0.450691853 | 0.991835066 | -5.001313438 |
| **A_55_P1971433** | Serpinb9d | -0.066281836 | -0.264018398 | 0.131454726 | 7.689813993 | -0.801155253 | 0.450711142 | 0.991835066 | -5.001334904 |
| **A_51_P104891** | Selenoi | 0.075347771 | -0.149510495 | 0.300206038 | 9.529277125 | 0.800885969 | 0.450856645 | 0.991835066 | -5.001496786 |
| **A_55_P2014319** | Armcx6 | 0.040827622 | -0.081029936 | 0.16268518 | 10.06030168 | 0.800775438 | 0.450916378 | 0.991835066 | -5.00156322 |
| **A_55_P2114785** | NA | 0.068605146 | -0.136173064 | 0.273383357 | 9.324881255 | 0.800722595 | 0.450944938 | 0.991835066 | -5.001594978 |
| **A_55_P2088625** | Fbln7 | -0.180668306 | -0.719993939 | 0.358657327 | 8.257358196 | -0.800645085 | 0.45098683 | 0.991835066 | -5.001641558 |
| **A_55_P2008576** | Ubr2 | 0.040618296 | -0.080637472 | 0.161874063 | 8.463055471 | 0.800623667 | 0.450998408 | 0.991835066 | -5.001654429 |
| **A_55_P1954925** | Endov | 0.071926069 | -0.142845918 | 0.286698057 | 15.84548179 | 0.800419802 | 0.451108609 | 0.991835066 | -5.001776925 |
| **A_52_P596541** | Kdelr1 | -0.07147254 | -0.284898675 | 0.141953596 | 9.965358704 | -0.800388327 | 0.451125625 | 0.991835066 | -5.001795835 |
| **A_51_P491227** | Suclg1 | -0.051030744 | -0.203426544 | 0.101365056 | 13.93489777 | -0.800328045 | 0.451158215 | 0.991835066 | -5.00183205 |
| **A_55_P1991778** | Fam219a | -0.073706018 | -0.293819642 | 0.146407606 | 12.92973198 | -0.800322741 | 0.451161083 | 0.991835066 | -5.001835236 |
| **A_55_P2183672** | Pebp1 | -0.068616231 | -0.27354333 | 0.136310869 | 16.57710911 | -0.800270107 | 0.45118954 | 0.991835066 | -5.001866855 |
| **A_55_P1955717** | Hccs | 0.147756807 | -0.293556664 | 0.589070277 | 16.23419133 | 0.800220081 | 0.451216588 | 0.991835066 | -5.001896906 |
| **A_55_P2410007** | NA | 0.089117571 | -0.177073887 | 0.355309029 | 9.17957083 | 0.800162799 | 0.451247561 | 0.991835066 | -5.001931314 |
| **A_55_P2018989** | Elk3 | 0.04235111 | -0.084153483 | 0.168855704 | 8.453180646 | 0.800143055 | 0.451258237 | 0.991835066 | -5.001943173 |
| **A_51_P359293** | Ubxn2b | -0.036434912 | -0.145273337 | 0.072403513 | 8.796061567 | -0.800100619 | 0.451281184 | 0.991835066 | -5.001968661 |
| **A_52_P361551** | Hspa14 | 0.073380625 | -0.145834909 | 0.292596159 | 12.6036431 | 0.800053835 | 0.451306483 | 0.991835066 | -5.00199676 |
| **A_55_P2153887** | Tatdn3 | 0.092722909 | -0.184279115 | 0.369724932 | 9.737069232 | 0.80004283 | 0.451312435 | 0.991835066 | -5.00200337 |
| **A_55_P2064457** | Gng7 | -0.047727109 | -0.190327712 | 0.094873494 | 8.182758003 | -0.799931684 | 0.451372543 | 0.991835066 | -5.002070119 |
| **A_55_P2121913** | Adi1 | -0.063211053 | -0.252079246 | 0.125657139 | 9.265286644 | -0.799914069 | 0.451382069 | 0.991835066 | -5.002080697 |

| | | | | | | | | | |
|---|---|---|---|---|---|---|---|---|---|
| **A_55_P2121980** | Plcb4 | 0.084748678 | -0.168478542 | 0.337975897 | 9.497641202 | 0.799892597 | 0.451393682 | 0.991835066 | -5.002093591 |
| **A_66_P101755** | Sp1 | -0.045561821 | -0.181718669 | 0.090595026 | 10.98493033 | -0.799780337 | 0.451454401 | 0.991835066 | -5.002161 |
| **A_55_P2097548** | NA | -0.032153843 | -0.128243709 | 0.063936024 | 16.83877156 | -0.799768736 | 0.451460676 | 0.991835066 | -5.002167965 |
| **A_55_P2225087** | 5430439M( | 0.046148579 | -0.091772438 | 0.184069595 | 9.056257057 | 0.799718264 | 0.451487977 | 0.991835066 | -5.002198269 |
| **A_55_P2069856** | Tnpo3 | 0.06376948 | -0.126819945 | 0.254358905 | 11.96666252 | 0.799692846 | 0.451501726 | 0.991835066 | -5.00221353 |
| **A_52_P370203** | Jarid2 | -0.037624584 | -0.150075908 | 0.07482674 | 10.72620396 | -0.79968006 | 0.451508643 | 0.991835066 | -5.002221206 |
| **A_55_P1979457** | Prdx4 | -0.063318224 | -0.252570533 | 0.125934085 | 12.55896215 | -0.79964398 | 0.45152816 | 0.991835066 | -5.002242868 |
| **A_51_P191909** | Rem2 | -0.032642114 | -0.130222976 | 0.064938748 | 7.701036763 | -0.799507893 | 0.451601782 | 0.991835066 | -5.002324563 |
| **A_55_P2139390** | NA | -0.056059768 | -0.223654876 | 0.111535339 | 16.26497273 | -0.799464332 | 0.451625349 | 0.991835066 | -5.00235071 |
| **A_52_P76196** | NA | -0.051116781 | -0.203948823 | 0.101715261 | 13.36376637 | -0.799389088 | 0.451666061 | 0.991835066 | -5.002395874 |
| **A_51_P466875** | Sbds | -0.073244096 | -0.292259828 | 0.145771636 | 11.31499427 | -0.799293799 | 0.451717621 | 0.991835066 | -5.002453064 |
| **A_66_P126293** | Itgb3 | -0.033193137 | -0.132463153 | 0.06607688 | 7.773970284 | -0.799170292 | 0.451784457 | 0.991835066 | -5.002527183 |
| **A_55_P2102624** | Eaf2 | -0.053772894 | -0.214599519 | 0.107053731 | 7.808295609 | -0.79912477 | 0.451809093 | 0.991835066 | -5.002554499 |
| **A_55_P2109486** | Milr1 | -0.042066417 | -0.167880905 | 0.083748071 | 17.06599433 | -0.79912369 | 0.451809677 | 0.991835066 | -5.002555147 |
| **A_55_P1982996** | Ramp1 | -0.087499849 | -0.349225317 | 0.174225619 | 9.847353029 | -0.799043555 | 0.451853048 | 0.991835066 | -5.002603231 |
| **A_55_P2025790** | Fanci | -0.064179771 | -0.256155934 | 0.127796393 | 9.193669766 | -0.799024294 | 0.451863472 | 0.991835066 | -5.002614787 |
| **A_52_P378914** | AU040320 | 0.064547729 | -0.128532792 | 0.25762825 | 8.398292364 | 0.79900894 | 0.451871782 | 0.991835066 | -5.002623999 |
| **A_51_P122425** | Ctf1 | 0.044749281 | -0.089177685 | 0.178676246 | 8.514521826 | 0.798596063 | 0.45209529 | 0.991835066 | -5.00287167 |
| **A_55_P2106426** | NA | -0.070757477 | -0.282550063 | 0.14103511 | 16.46073319 | -0.798492268 | 0.45215149 | 0.991835066 | -5.002933917 |
| **A_55_P2172058** | Trim55 | 0.108603498 | -0.216472658 | 0.433679654 | 9.990946534 | 0.798486963 | 0.452154363 | 0.991835066 | -5.002937099 |
| **A_51_P210928** | Tsta3 | 0.041283619 | -0.082307078 | 0.164874317 | 9.712243694 | 0.798364308 | 0.452220782 | 0.991835066 | -5.003010648 |
| **A_52_P30550** | Acp2 | -0.039565352 | -0.158025044 | 0.078894341 | 8.532653732 | -0.798276894 | 0.452268122 | 0.991835066 | -5.00306306 |
| **A_55_P1977681** | Vps13d | -0.042818635 | -0.17103473 | 0.08539746 | 9.726471375 | -0.79817736 | 0.45232203 | 0.991835066 | -5.003122734 |
| **A_55_P2091486** | Tmem219 | 0.066522708 | -0.13267798 | 0.265723397 | 10.26904456 | 0.798156804 | 0.452333163 | 0.991835066 | -5.003135057 |
| **A_52_P676956** | Tirap | -0.049330229 | -0.197050306 | 0.098389849 | 8.684031469 | -0.79814672 | 0.452338625 | 0.991835066 | -5.003141102 |
| **A_51_P179864** | Prkx | -0.045831749 | -0.183141935 | 0.091478436 | 10.53814862 | -0.797761018 | 0.45254757 | 0.991835066 | -5.00337228 |
| **A_52_P67212** | 2900052N( | -0.043689289 | -0.174581873 | 0.087203295 | 7.869290036 | -0.797754139 | 0.452551297 | 0.991835066 | -5.003376403 |
| **A_51_P155501** | Alg3 | -0.056996401 | -0.227767297 | 0.113774494 | 8.496728549 | -0.79770573 | 0.452577527 | 0.991835066 | -5.003405411 |
| **A_55_P1956308** | NA | -0.051683775 | -0.206538384 | 0.103170834 | 13.37096645 | -0.797699318 | 0.452581001 | 0.991835066 | -5.003409253 |
| **A_55_P2162422** | NA | 0.037481862 | -0.07483497 | 0.149798695 | 16.99440741 | 0.797600555 | 0.452634517 | 0.991835066 | -5.003468432 |
| **A_55_P2248320** | C430003N: | 0.123486769 | -0.246621948 | 0.493595485 | 8.902689407 | 0.797444092 | 0.452719308 | 0.991835066 | -5.003562172 |
| **A_65_P01641** | Rnf6 | -0.055282617 | -0.220989213 | 0.110423979 | 12.95838357 | -0.797366397 | 0.452761417 | 0.991835066 | -5.003608715 |
| **A_55_P2174821** | Rab8a | 0.077104011 | -0.154011888 | 0.30821991 | 10.38114156 | 0.797363372 | 0.452763056 | 0.991835066 | -5.003610527 |
| **A_55_P2104597** | Kctd9 | -0.053897931 | -0.215477609 | 0.107681748 | 12.49570137 | -0.797249917 | 0.452824551 | 0.991835066 | -5.003678486 |
| **A_55_P1992849** | Adrb3 | 0.05556388 | -0.111017244 | 0.222145004 | 7.834826095 | 0.797215822 | 0.452843032 | 0.991835066 | -5.003698907 |
| **A_55_P2074736** | Prkar1b | 0.046752656 | -0.093412466 | 0.186917777 | 8.983821521 | 0.797214996 | 0.45284348 | 0.991835066 | -5.003699402 |
| **A_51_P482051** | Cyp3a16 | -0.057116801 | -0.228395092 | 0.11416149 | 7.691102529 | -0.797022693 | 0.452947729 | 0.991835066 | -5.003814569 |
| **A_51_P141554** | Med22 | -0.034895904 | -0.139547305 | 0.069755496 | 10.47289238 | -0.796963713 | 0.452979705 | 0.991835066 | -5.003849887 |
| **A_52_P542970** | NA | -0.054602568 | -0.218364162 | 0.109159026 | 10.74543434 | -0.796911593 | 0.453007964 | 0.991835066 | -5.003881095 |
| **A_55_P2413359** | NA | 0.075661044 | -0.151259431 | 0.30258152 | 8.422354458 | 0.796907254 | 0.453010317 | 0.991835066 | -5.003883694 |
| **A_55_P2019929** | Ythdf3 | 0.036159461 | -0.072312296 | 0.144631218 | 8.643966513 | 0.796735927 | 0.453103217 | 0.991835066 | -5.003986269 |
| **A_66_P110798** | NA | 0.20388157 | -0.40773601 | 0.81549915 | 13.16837198 | 0.796722609 | 0.453110439 | 0.991835066 | -5.003994242 |
| **A_52_P645632** | Senp8 | -0.05095117 | -0.203803119 | 0.10190078 | 8.433185342 | -0.796695397 | 0.453125196 | 0.991835066 | -5.004010532 |
| **A_51_P163958** | Spr | 0.112231515 | -0.224536264 | 0.448999293 | 11.51279466 | 0.79651402 | 0.453223564 | 0.991835066 | -5.004119101 |
| **A_55_P1997291** | Fam53a | 0.085221837 | -0.170502095 | 0.340945768 | 12.34300953 | 0.796505262 | 0.453228314 | 0.991835066 | -5.004124343 |
| **A_55_P1984069** | NA | -0.069261452 | -0.277093357 | 0.138570452 | 14.854609 | -0.796505006 | 0.453228453 | 0.991835066 | -5.004124496 |
| **A_51_P161929** | Tspo2 | 0.222346989 | -0.444988989 | 0.889682966 | 9.915043244 | 0.796335076 | 0.453320627 | 0.991835066 | -5.004226196 |
| **A_55_P1981011** | Slc35c2 | 0.078394354 | -0.156893925 | 0.313682634 | 9.639351798 | 0.796331005 | 0.453322835 | 0.991835066 | -5.004228632 |

| | | | | | | | | | |
|---|---|---|---|---|---|---|---|---|---|
| **A_55_P2002577** | Ephx1 | 0.040449936 | -0.080954364 | 0.161854235 | 8.467738512 | 0.796329677 | 0.453323555 | 0.991835066 | -5.004229426 |
| **A_55_P1966194** | Plek | -0.107557518 | -0.430376306 | 0.215261269 | 8.636060433 | -0.796326382 | 0.453325343 | 0.991835066 | -5.004231398 |
| **A_55_P2091147** | H4c12 | 0.0897532 | -0.179648756 | 0.359155156 | 12.76155312 | 0.796266288 | 0.453357942 | 0.991835066 | -5.004267359 |
| **A_55_P2000758** | Gkap1 | 0.058487635 | -0.117105886 | 0.234081155 | 10.10156695 | 0.796094649 | 0.453451062 | 0.991835066 | -5.004370057 |
| **A_51_P436878** | Sertad1 | -0.090855938 | -0.363674006 | 0.181962131 | 10.3175554 | -0.795956469 | 0.453526039 | 0.991835066 | -5.004452723 |
| **A_55_P2163344** | NA | 0.057684153 | -0.115553017 | 0.230921323 | 9.190240971 | 0.795837813 | 0.453590429 | 0.991835066 | -5.004523699 |
| **A_55_P1982524** | Cyhr1 | -0.068864575 | -0.275680321 | 0.137951172 | 8.803193018 | -0.795832006 | 0.45359358 | 0.991835066 | -5.004527172 |
| **A_51_P326685** | Lrtm1 | 0.10479414 | -0.209935045 | 0.419523325 | 9.599275458 | 0.79580949 | 0.453605799 | 0.991835066 | -5.00454064 |
| **A_52_P647607** | Ubr4 | 0.045593904 | -0.0913633 | 0.182551109 | 8.467488064 | 0.795666426 | 0.453683444 | 0.991835066 | -5.004626203 |
| **A_55_P1992019** | Ptpru | 0.058812052 | -0.117854816 | 0.235478921 | 8.335275078 | 0.795646865 | 0.453694062 | 0.991835066 | -5.004637902 |
| **A_55_P1989928** | Lin52 | -0.052820072 | -0.211492247 | 0.105852103 | 9.002673276 | -0.795623036 | 0.453706996 | 0.991835066 | -5.004652152 |
| **A_55_P2133027** | Kirrel3 | -0.033518773 | -0.134212265 | 0.067174719 | 8.000921042 | -0.795601951 | 0.45371844 | 0.991835066 | -5.00466476 |
| **A_52_P516091** | Pla2g15 | -0.04607805 | -0.184503719 | 0.092347619 | 8.448536078 | -0.795585017 | 0.453727632 | 0.991835066 | -5.004674887 |
| **A_55_P2126695** | NA | 0.04127177 | -0.082720333 | 0.165263872 | 7.971545191 | 0.79555132 | 0.453745923 | 0.991835066 | -5.004695037 |
| **A_55_P1967025** | NA | 0.034201795 | -0.068574154 | 0.136977744 | 8.062318965 | 0.795365109 | 0.453847009 | 0.991835066 | -5.004806375 |
| **A_52_P582394** | Mrps11 | -0.095614964 | -0.382940141 | 0.191710212 | 9.694191637 | -0.795355552 | 0.453852198 | 0.991835066 | -5.004812089 |
| **A_55_P1990648** | Ndufaf4 | 0.070546185 | -0.141448978 | 0.282541348 | 11.12026933 | 0.795347122 | 0.453856774 | 0.991835066 | -5.004817129 |
| **A_55_P2142033** | Clip4 | -0.041950763 | -0.168093798 | 0.084192272 | 9.441663697 | -0.794850989 | 0.454126185 | 0.991835066 | -5.00511367 |
| **A_55_P2176688** | NA | 0.046707658 | -0.093744721 | 0.187160037 | 13.96908371 | 0.794818786 | 0.454143676 | 0.991835066 | -5.005132914 |
| **A_51_P271865** | Clec4f | -0.041143765 | -0.16486719 | 0.082579659 | 7.757345078 | -0.79480618 | 0.454150523 | 0.991835066 | -5.005140446 |
| **A_55_P2038273** | NA | 0.100709996 | -0.202146312 | 0.403566303 | 9.147461456 | 0.794776626 | 0.454166576 | 0.991835066 | -5.005158106 |
| **A_55_P2137309** | Kctd11 | 0.060204473 | -0.120878529 | 0.241287476 | 9.627057473 | 0.794621308 | 0.454250945 | 0.991835066 | -5.005250903 |
| **A_55_P2121484** | Smo | 0.099852686 | -0.200565342 | 0.400270715 | 10.1901803 | 0.794406692 | 0.454367543 | 0.991835066 | -5.005379108 |
| **A_55_P2133266** | Rtl8c | 0.059530469 | -0.119600428 | 0.238661366 | 14.04961459 | 0.794287876 | 0.454432103 | 0.991835066 | -5.005450072 |
| **A_55_P2069525** | Zmym6 | 0.094954477 | -0.190875543 | 0.380784497 | 9.215461481 | 0.793993119 | 0.454592291 | 0.991835066 | -5.005626085 |
| **A_52_P193194** | Slc35d2 | -0.035171637 | -0.141057363 | 0.070714088 | 7.980554988 | -0.793897264 | 0.454644392 | 0.991835066 | -5.005683313 |
| **A_55_P2095035** | NA | 0.032431229 | -0.065239137 | 0.130101596 | 8.149739426 | 0.793614743 | 0.454797979 | 0.991835066 | -5.005851956 |
| **A_55_P2398264** | Rnf130 | 0.041235513 | -0.082976574 | 0.1654476 | 8.682045287 | 0.793444717 | 0.454890428 | 0.991835066 | -5.005953425 |
| **A_55_P2050877** | Plekhg2 | 0.091226426 | -0.183664294 | 0.366117147 | 12.2298451 | 0.793176273 | 0.455036417 | 0.991835066 | -5.006113595 |
| **A_55_P2159522** | Col14a1 | -0.044182134 | -0.177321298 | 0.08895703 | 8.492075309 | -0.793140256 | 0.455056006 | 0.991835066 | -5.006135082 |
| **A_55_P2010116** | Rab27b | 0.047558116 | -0.095765769 | 0.190882002 | 8.280182724 | 0.793076721 | 0.455090565 | 0.991835066 | -5.006172983 |
| **A_51_P339713** | Mrps26 | -0.068428605 | -0.274650039 | 0.137792829 | 11.2211116 | -0.793072725 | 0.455092738 | 0.991835066 | -5.006175367 |
| **A_55_P2040276** | Prcd | -0.077970552 | -0.312949994 | 0.157008889 | 8.125388635 | -0.793066993 | 0.455095856 | 0.991835066 | -5.006178786 |
| **A_55_P2007389** | Fam189b | 0.059077761 | -0.11906579 | 0.237221312 | 10.05054015 | 0.792616402 | 0.455340998 | 0.991835066 | -5.006447512 |
| **A_66_P105075** | Mmp11 | -0.055928413 | -0.224586517 | 0.11272969 | 9.130119142 | -0.792564137 | 0.455369438 | 0.991835066 | -5.006478674 |
| **A_66_P128293** | Pheta2 | 0.055339036 | -0.111545077 | 0.222223149 | 8.796302232 | 0.792548278 | 0.455378068 | 0.991835066 | -5.00648813 |
| **A_55_P1960167** | Bcat2 | -0.068246574 | -0.274094607 | 0.13760146 | 10.62513449 | -0.792397803 | 0.455459958 | 0.991835066 | -5.006577838 |
| **A_55_P2402134** | NA | 0.189835009 | -0.382846068 | 0.762516085 | 9.656038499 | 0.792268895 | 0.45553012 | 0.991835066 | -5.006654679 |
| **A_55_P1992650** | Bloc1s1 | -0.07361195 | -0.29574009 | 0.14851619 | 13.95225858 | -0.792052327 | 0.45564801 | 0.991835066 | -5.006783751 |
| **A_55_P1980041** | Kpna6 | 0.065045229 | -0.131239164 | 0.261329621 | 9.064588006 | 0.792024851 | 0.455662968 | 0.991835066 | -5.006800124 |
| **A_55_P1989728** | C2cd2l | 0.04164055 | -0.084035726 | 0.167316825 | 11.28232684 | 0.791903542 | 0.455729014 | 0.991835066 | -5.006872409 |
| **A_51_P249848** | Dsg2 | 0.085589144 | -0.172735589 | 0.343913878 | 9.140937643 | 0.791884474 | 0.455739396 | 0.991835066 | -5.00688377 |
| **A_52_P512575** | Hopx | -0.074108999 | -0.297838772 | 0.149620774 | 13.55339415 | -0.791692074 | 0.455844162 | 0.991835066 | -5.006998396 |
| **A_55_P2134236** | Foxa2 | 0.042418677 | -0.08564596 | 0.170483315 | 7.694368417 | 0.791656953 | 0.455863288 | 0.991835066 | -5.007019317 |
| **A_51_P521176** | Rassf7 | 0.078473257 | -0.158464448 | 0.315410963 | 9.89624686 | 0.791583319 | 0.455903389 | 0.991835066 | -5.007063179 |
| **A_55_P2032849** | NA | 0.068127591 | -0.137623843 | 0.273879025 | 8.558101195 | 0.791387694 | 0.456009938 | 0.991835066 | -5.007179691 |
| **A_55_P1959228** | Slc4a4 | 0.058175313 | -0.117559336 | 0.233909962 | 9.003377727 | 0.791207623 | 0.456108031 | 0.991835066 | -5.00728692 |
| **A_51_P425737** | Tbc1d7 | -0.085036398 | -0.341913809 | 0.171841014 | 9.678631197 | -0.791203264 | 0.456110406 | 0.991835066 | -5.007289516 |

| | | | | | | | | | |
|---|---|---|---|---|---|---|---|---|---|
| **A_55_P2065330** | NA | 0.037707082 | -0.076215259 | 0.151629423 | 14.98402305 | 0.791085015 | 0.456174829 | 0.991835066 | -5.00735992 |
| **A_52_P33800** | Fndc3a | 0.036450593 | -0.073676851 | 0.146578037 | 8.161269038 | 0.791075931 | 0.456179778 | 0.991835066 | -5.007365328 |
| **A_55_P2144003** | NA | -0.066202246 | -0.266242027 | 0.133837535 | 15.28742585 | -0.790979965 | 0.456232067 | 0.991835066 | -5.007422459 |
| **A_55_P1976744** | NA | -0.088794565 | -0.357119419 | 0.179530289 | 9.961245037 | -0.790923746 | 0.456262701 | 0.991835066 | -5.007455925 |
| **A_55_P1982718** | Rtf2 | -0.049921462 | -0.200829148 | 0.100986225 | 10.26126506 | -0.790651435 | 0.456411104 | 0.991835066 | -5.007618 |
| **A_52_P13730** | Cbx2 | 0.049282011 | -0.099709948 | 0.198273969 | 9.335968096 | 0.790559792 | 0.456461055 | 0.991835066 | -5.007672534 |
| **A_55_P2009008** | NA | -0.053123 | -0.213733092 | 0.107487092 | 13.83466545 | -0.790530984 | 0.456476758 | 0.991835066 | -5.007689676 |
| **A_51_P260490** | Ralbp1 | -0.043688709 | -0.175820543 | 0.088443124 | 10.86174286 | -0.790261612 | 0.456623607 | 0.991835066 | -5.00784994 |
| **A_55_P2152592** | Agpat3 | -0.069904134 | -0.281326565 | 0.141518297 | 12.1721717 | -0.790243498 | 0.456633483 | 0.991835066 | -5.007860715 |
| **A_52_P606303** | Mrpl3 | 0.042954375 | -0.086962088 | 0.172870838 | 13.06580148 | 0.790227879 | 0.456641999 | 0.991835066 | -5.007870006 |
| **A_51_P511707** | Ntpcr | 0.043841611 | -0.088758385 | 0.176441608 | 10.51306647 | 0.79022748 | 0.456642217 | 0.991835066 | -5.007870244 |
| **A_55_P2060608** | NA | 0.045222484 | -0.091588009 | 0.182032977 | 14.23581161 | 0.790031003 | 0.456749351 | 0.991835066 | -5.007987107 |
| **A_51_P365656** | Tmem101 | 0.055638339 | -0.112686967 | 0.223963644 | 10.46387345 | 0.790012341 | 0.456759527 | 0.991835066 | -5.007998206 |
| **A_55_P2186672** | Rps14 | 0.061657926 | -0.124917206 | 0.248233058 | 17.57620585 | 0.789849389 | 0.456848396 | 0.991835066 | -5.00809511 |
| **A_52_P179729** | Txnl4a | -0.082888972 | -0.33371613 | 0.167938187 | 11.74609062 | -0.789825818 | 0.456861251 | 0.991835066 | -5.008109126 |
| **A_55_P2025283** | NA | 0.047409376 | -0.096095811 | 0.190914563 | 15.04665527 | 0.789597519 | 0.45698578 | 0.991835066 | -5.008244861 |
| **A_52_P560146** | NA | 0.05280059 | -0.107042611 | 0.212643792 | 8.610588906 | 0.789502922 | 0.457037386 | 0.991835066 | -5.008301095 |
| **A_55_P1964225** | Elac2 | 0.046928394 | -0.09516802 | 0.189024808 | 9.882423968 | 0.789335631 | 0.457128659 | 0.991835066 | -5.008400529 |
| **A_52_P381430** | Tbc1d4 | 0.056862366 | -0.115401123 | 0.229125855 | 8.435257287 | 0.788934267 | 0.457347694 | 0.991835066 | -5.008639025 |
| **A_55_P2118173** | Igf2 | 0.083651583 | -0.169776108 | 0.337079274 | 11.91924866 | 0.788913213 | 0.457359185 | 0.991835066 | -5.008651533 |
| **A_55_P2152771** | Lhfpl2 | -0.053801786 | -0.216797704 | 0.109194132 | 9.367227919 | -0.788912969 | 0.457359319 | 0.991835066 | -5.008651678 |
| **A_55_P1980341** | Romo1 | 0.051351137 | -0.104227798 | 0.206930072 | 14.58231388 | 0.78887538 | 0.457379836 | 0.991835066 | -5.008674008 |
| **A_51_P376445** | Rhox5 | -0.10818614 | -0.436190662 | 0.219818381 | 11.28817084 | -0.788317066 | 0.457684658 | 0.991835066 | -5.009005588 |
| **A_51_P249414** | Ess2 | 0.033973945 | -0.069054895 | 0.137002785 | 10.18627742 | 0.788127164 | 0.457788371 | 0.991835066 | -5.009118328 |
| **A_55_P2011727** | Neurl1b | -0.070848332 | -0.285727967 | 0.144031304 | 12.98257842 | -0.788031361 | 0.457840698 | 0.991835066 | -5.009175196 |
| **A_55_P2036026** | Robo4 | -0.057024087 | -0.230005255 | 0.115957081 | 12.15290672 | -0.787895553 | 0.457914885 | 0.991835066 | -5.009255801 |
| **A_55_P2184106** | Gpn2 | 0.055526529 | -0.112936037 | 0.223989096 | 8.496791394 | 0.787782356 | 0.457976726 | 0.991835066 | -5.009322978 |
| **A_55_P2024322** | Rps27 | 0.098799244 | -0.201007771 | 0.398606259 | 15.64023195 | 0.787627666 | 0.458061244 | 0.991835066 | -5.009414767 |
| **A_55_P2079054** | Ndufv1 | -0.087555937 | -0.353260622 | 0.178148748 | 15.89863502 | -0.787581526 | 0.458086456 | 0.991835066 | -5.009442142 |
| **A_55_P2015238** | Dctn4 | 0.059173679 | -0.120410966 | 0.238758324 | 11.68505659 | 0.787532534 | 0.458113226 | 0.991835066 | -5.009471208 |
| **A_55_P2001188** | Phf6 | 0.045602564 | -0.092805532 | 0.184010659 | 10.24593615 | 0.787475221 | 0.458144546 | 0.991835066 | -5.009505209 |
| **A_55_P2064243** | Rhot2 | -0.130715927 | -0.527486151 | 0.266054297 | 9.106286424 | -0.787405725 | 0.458182525 | 0.991835066 | -5.009546436 |
| **A_51_P364421** | Vamp7 | -0.097785655 | -0.394627568 | 0.199056258 | 9.249551219 | -0.787334096 | 0.458221672 | 0.991835066 | -5.009588924 |
| **A_52_P410765** | Sema7a | -0.044788767 | -0.180752462 | 0.091174928 | 10.9659058 | -0.787327244 | 0.458225417 | 0.991835066 | -5.009592988 |
| **A_52_P229972** | Slc22a1 | 0.249121157 | -0.507173958 | 1.005416271 | 10.20605904 | 0.787278605 | 0.458252001 | 0.991835066 | -5.009621838 |
| **A_55_P2056300** | Sf3b3 | 0.060506383 | -0.123189164 | 0.244201929 | 12.23949521 | 0.787248234 | 0.458268601 | 0.991835066 | -5.009639852 |
| **A_51_P120615** | Rabl3 | -0.048359268 | -0.19519026 | 0.098471724 | 8.57366721 | -0.787174678 | 0.458308806 | 0.991835066 | -4.994431091 |
| **A_55_P2064492** | Pcmtd2 | -0.093713881 | -0.378273422 | 0.19084566 | 10.34140804 | -0.787118107 | 0.45833973 | 0.991835066 | -5.009717025 |
| **A_55_P2096530** | Icmt | 0.046589287 | -0.094893528 | 0.188072103 | 9.350615356 | 0.787030391 | 0.45838768 | 0.991835066 | -5.00976904 |
| **A_55_P2059342** | Lamp1 | -0.040263155 | -0.162541583 | 0.082015274 | 14.84716663 | -0.786986142 | 0.45841187 | 0.991835066 | -5.009795278 |
| **A_55_P2119969** | Bdh2 | -0.052217178 | -0.210805376 | 0.10637102 | 8.628738688 | -0.786958201 | 0.458427146 | 0.991835066 | -5.009811846 |
| **A_51_P415225** | Zfp105 | 0.047510129 | -0.096803189 | 0.191823448 | 9.938504332 | 0.786844533 | 0.458489292 | 0.991835066 | -5.009879239 |
| **A_51_P241074** | Map2k1 | -0.060317171 | -0.243541555 | 0.122907213 | 10.21187362 | -0.786804478 | 0.458511194 | 0.991835066 | -5.009902986 |
| **A_52_P84275** | Derl2 | 0.048549271 | -0.098929595 | 0.196028136 | 9.411088977 | 0.786795841 | 0.458515916 | 0.991835066 | -5.009908107 |
| **A_52_P564951** | C2cd3 | 0.067424559 | -0.137407413 | 0.272256532 | 8.759344345 | 0.786736867 | 0.458548163 | 0.991835066 | -5.009943067 |
| **A_51_P441942** | Tial1 | 0.068625689 | -0.139855827 | 0.277107206 | 10.62081714 | 0.78673468 | 0.458549359 | 0.991835066 | -5.009944364 |
| **A_51_P424272** | Mt4 | -0.034152508 | -0.13790819 | 0.069603173 | 7.95388966 | -0.786719385 | 0.458557722 | 0.991835066 | -5.009953431 |
| **A_51_P472323** | Fbh1 | -0.065163971 | -0.26316118 | 0.132833238 | 11.97328908 | -0.786606637 | 0.458619378 | 0.991835066 | -5.010020263 |

| | | | | | | | | | |
|---|---|---|---|---|---|---|---|---|---|
| **A_55_P2033958** | Sltm | 0.084300586 | -0.171851181 | 0.340452352 | 11.293588 | 0.786579039 | 0.458634471 | 0.991835066 | -5.010036621 |
| **A_52_P920158** | Rbm24 | -0.064323047 | -0.259810231 | 0.131164137 | 13.49988281 | -0.786425252 | 0.458718579 | 0.991835066 | -5.010127765 |
| **A_66_P119418** | Rb1cc1 | -0.047750779 | -0.192883714 | 0.097382156 | 8.707681896 | -0.786363994 | 0.458752086 | 0.991835066 | -5.010164067 |
| **A_51_P421876** | Irf7 | -0.033596814 | -0.135711541 | 0.068517912 | 7.747174915 | -0.786355364 | 0.458756806 | 0.991835066 | -5.01016918 |
| **A_55_P2096486** | Bcs1l | -0.063302887 | -0.2557488 | 0.129143025 | 10.71454262 | -0.786183565 | 0.458850784 | 0.991835066 | -5.010270976 |
| **A_55_P2115003** | Prpf4 | 0.040887055 | -0.083431818 | 0.165205929 | 8.843900689 | 0.786063988 | 0.458916203 | 0.991835066 | -5.010341818 |
| **A_51_P165934** | Phb | -0.078661418 | -0.317837752 | 0.160514915 | 12.92201299 | -0.786054579 | 0.458921351 | 0.991835066 | -5.010347392 |
| **A_55_P2183208** | Prl2c1 | -0.061703728 | -0.24934921 | 0.125941754 | 9.45458523 | -0.785927385 | 0.458990945 | 0.991835066 | -5.010422737 |
| **A_55_P1994132** | Tmem184a | 0.04516308 | -0.09223829 | 0.18256445 | 8.168849419 | 0.785600272 | 0.459169958 | 0.991835066 | -5.010616463 |
| **A_55_P2033922** | Tsga8 | 0.035341211 | -0.072184111 | 0.142866532 | 7.941744584 | 0.785560803 | 0.459191561 | 0.991835066 | -5.010639833 |
| **A_55_P1985351** | Slc35f2 | 0.034997646 | -0.071490813 | 0.141486105 | 7.782964403 | 0.785498622 | 0.459225596 | 0.991835066 | -5.01067665 |
| **A_55_P2167451** | Cnnm2 | 0.070825033 | -0.144688799 | 0.286338865 | 9.68147463 | 0.785454023 | 0.459250009 | 0.991835066 | -5.010703055 |
| **A_55_P2009066** | Cfap74 | 0.068909653 | -0.140822687 | 0.278641992 | 13.66454456 | 0.785278653 | 0.459346012 | 0.991835066 | -5.010806873 |
| **A_52_P270035** | Tspyl2 | -0.048342928 | -0.195490799 | 0.098804943 | 8.629160477 | -0.785214123 | 0.459381341 | 0.991835066 | -5.01084507 |
| **A_51_P498257** | NA | -0.047722816 | -0.192991569 | 0.097545936 | 9.864468669 | -0.785168722 | 0.459406198 | 0.991835066 | -5.010871942 |
| **A_66_P123804** | Fbxo30 | 0.043538885 | -0.088998663 | 0.176076432 | 8.965000611 | 0.785140725 | 0.459421527 | 0.991835066 | -5.010888513 |
| **A_55_P1981674** | NA | 0.039905283 | -0.081586644 | 0.16139721 | 8.170223959 | 0.785040573 | 0.459476366 | 0.991835066 | -5.010947786 |
| **A_55_P2110718** | Npm1 | 0.107428283 | -0.219674813 | 0.434531379 | 11.60019827 | 0.784952024 | 0.459524855 | 0.991835066 | -5.011000187 |
| **A_66_P134412** | Mettl25 | 0.038892717 | -0.079540549 | 0.157325984 | 10.28177835 | 0.784880813 | 0.459563853 | 0.991835066 | -5.011042325 |
| **A_55_P2000102** | Mettl21a | -0.044370159 | -0.17949667 | 0.090756351 | 9.663524712 | -0.78480102 | 0.459607553 | 0.991835066 | -5.011089537 |
| **A_51_P142164** | Hspa13 | 0.035426088 | -0.072472786 | 0.143324961 | 7.849515416 | 0.784721258 | 0.45965124 | 0.991835066 | -5.011136727 |
| **A_51_P419971** | Tmem106b | -0.070088297 | -0.283565323 | 0.143388729 | 11.97443084 | -0.784699719 | 0.459663037 | 0.991835066 | -5.01114947 |
| **A_52_P496655** | Leprotl1 | -0.032950465 | -0.133317241 | 0.067416311 | 8.653067575 | -0.784658538 | 0.459685594 | 0.991835066 | -5.011173832 |
| **A_51_P399071** | Anp32a | 0.04371154 | -0.089446553 | 0.176869632 | 9.300973878 | 0.784580792 | 0.459728181 | 0.991835066 | -5.011219823 |
| **A_55_P1982084** | NA | 0.066310037 | -0.135715905 | 0.268335978 | 10.77099038 | 0.784478888 | 0.459784004 | 0.991835066 | -5.011280099 |
| **A_52_P611798** | Ddx5 | 0.056751764 | -0.116169638 | 0.229673167 | 12.5086126 | 0.78440391 | 0.459825081 | 0.991835066 | -5.011324445 |
| **A_55_P2051094** | Rorc | -0.102869431 | -0.416314692 | 0.210575829 | 9.212298447 | -0.784393097 | 0.459831005 | 0.991835066 | -5.01133084 |
| **A_55_P2105256** | Trim47 | -0.07354808 | -0.297654882 | 0.150558723 | 13.05482415 | -0.784378049 | 0.45983925 | 0.991835066 | -5.01133974 |
| **A_65_P18167** | Hnrnpul2 | -0.04182855 | -0.169300683 | 0.085643582 | 10.21137469 | -0.78427197 | 0.459897371 | 0.991835066 | -5.011402473 |
| **A_52_P146164** | Ube2n | -0.035461809 | -0.14353659 | 0.072612972 | 9.247212256 | -0.784233976 | 0.459918189 | 0.991835066 | -5.01142494 |
| **A_55_P2059904** | Chsy1 | 0.074308841 | -0.152169121 | 0.300786803 | 11.87704532 | 0.784194285 | 0.459939938 | 0.991835066 | -5.01144841 |
| **A_55_P1965772** | Atp2a3 | -0.053539877 | -0.216723025 | 0.10964327 | 8.255402841 | -0.784171758 | 0.459952282 | 0.991835066 | -5.01146173 |
| **A_51_P103594** | Fam162b | 0.032433762 | -0.066421935 | 0.131289459 | 7.745376967 | 0.784160127 | 0.459958655 | 0.991835066 | -5.001040896 |
| **A_55_P2046661** | Tnfrsf14 | -0.036926675 | -0.149483811 | 0.075630462 | 7.989470305 | -0.784108742 | 0.459986813 | 0.991835066 | -5.011498989 |
| **A_66_P117058** | Aasdh | 0.092389568 | -0.189252839 | 0.374031976 | 9.174886314 | 0.784032422 | 0.460028638 | 0.991835066 | -5.011544112 |
| **A_52_P427759** | NA | 0.037148426 | -0.076095676 | 0.150392527 | 12.68209423 | 0.784032302 | 0.460028704 | 0.991835066 | -5.011544183 |
| **A_51_P173961** | Pdrg1 | -0.051292708 | -0.207665061 | 0.105079645 | 13.12073137 | -0.783979651 | 0.460057559 | 0.991835066 | -5.01157531 |
| **A_55_P2035827** | Vma21 | 0.082194523 | -0.168457491 | 0.332846537 | 10.38540388 | 0.783755886 | 0.460180207 | 0.991835066 | -5.01170758 |
| **A_55_P2325378** | NA | 0.058041655 | -0.118963736 | 0.235047046 | 9.054751562 | 0.783722699 | 0.460198399 | 0.991835066 | -5.011727194 |
| **A_51_P301804** | St3gal1 | 0.06080045 | -0.124692872 | 0.246293772 | 10.05023673 | 0.783407359 | 0.460371283 | 0.991835066 | -5.011913539 |
| **A_51_P186053** | Rrp15 | 0.065085206 | -0.133484564 | 0.263654976 | 10.05575128 | 0.78339047 | 0.460380543 | 0.991835066 | -5.011923517 |
| **A_65_P19933** | Zdhhc23 | 0.053462276 | -0.109664809 | 0.216589361 | 8.213642134 | 0.783304278 | 0.460427807 | 0.991835066 | -5.01197444 |
| **A_51_P461578** | Fdxr | -0.064644899 | -0.261906781 | 0.132616983 | 10.05163822 | -0.78324967 | 0.460457752 | 0.991835066 | -5.0120067 |
| **A_55_P2020926** | Trip11 | 0.077757649 | -0.159518201 | 0.3150335 | 9.274489232 | 0.783246958 | 0.46045924 | 0.991835066 | -5.012008303 |
| **A_55_P2040878** | Ung | -0.077938746 | -0.315768966 | 0.159891473 | 13.29800523 | -0.783241178 | 0.460462409 | 0.991835066 | -5.012011717 |
| **A_51_P348804** | Amotl2 | -0.094635203 | -0.383476273 | 0.194205867 | 10.42449192 | -0.78307418 | 0.460553996 | 0.991835066 | -5.012110361 |
| **A_51_P423008** | Klhl7 | -0.082790056 | -0.335486009 | 0.169905897 | 9.434401091 | -0.78304915 | 0.460567724 | 0.991835066 | -5.012125144 |
| **A_51_P396774** | Sugt1 | 0.036100085 | -0.074087725 | 0.146287895 | 13.62503171 | 0.783039751 | 0.46057288 | 0.991835066 | -5.012130696 |

| | | | | | | | | | |
|---|---|---|---|---|---|---|---|---|---|
| **A_52_P292684** | Exosc4 | -0.099661822 | -0.403939296 | 0.204615653 | 10.98088161 | -0.782831245 | 0.460687251 | 0.991835066 | -5.012253831 |
| **A_55_P2249220** | A430102J1 | 0.106557332 | -0.218805988 | 0.431920653 | 8.891122861 | 0.782751448 | 0.460731028 | 0.991835066 | -5.012300949 |
| **A_55_P2105242** | Arhgap39 | -0.040571751 | -0.164459207 | 0.083315704 | 10.92231338 | -0.782718422 | 0.460749147 | 0.991835066 | -5.012320449 |
| **A_55_P2128181** | NA | -0.060753258 | -0.24632788 | 0.124821364 | 10.10801052 | -0.782456355 | 0.46089294 | 0.991835066 | -5.012475162 |
| **A_55_P2018825** | NA | -0.07094234 | -0.287684187 | 0.145799507 | 9.286285157 | -0.782297375 | 0.460980186 | 0.991835066 | -5.012568996 |
| **A_51_P233947** | Tbc1d10b | -0.033179965 | -0.134555481 | 0.068195551 | 12.28482684 | -0.782261534 | 0.460999856 | 0.991835066 | -5.012590149 |
| **A_55_P2105100** | Lrig1 | 0.055281102 | -0.113637887 | 0.22420009 | 10.39016301 | 0.782181157 | 0.461043971 | 0.991835066 | -5.012637582 |
| **A_51_P477419** | Nfic | -0.044951502 | -0.182307028 | 0.092404023 | 11.73246095 | -0.782180903 | 0.461044111 | 0.991835066 | -5.012637732 |
| **A_55_P2136501** | Midn | 0.041045462 | -0.084376338 | 0.166467262 | 13.13881876 | 0.782170175 | 0.461049999 | 0.991835066 | -5.012644062 |
| **A_55_P1999962** | Podn | 0.108463839 | -0.223198351 | 0.440126029 | 8.170101585 | 0.781624457 | 0.461349599 | 0.991835066 | -5.012966007 |
| **A_55_P2293381** | Cldn18 | -0.032300394 | -0.131069243 | 0.066468454 | 7.690907003 | -0.781622343 | 0.461350759 | 0.991835066 | -5.012967253 |
| **A_52_P584293** | Atrnl1 | 0.062933086 | -0.129523387 | 0.255389558 | 9.225784316 | 0.781547967 | 0.461391603 | 0.991835066 | -5.013011118 |
| **A_55_P2173892** | Isg20 | -0.223401613 | -0.906803205 | 0.459999979 | 9.336453611 | -0.781302927 | 0.461526182 | 0.991835066 | -5.01315561 |
| **A_52_P262219** | Fos | -0.060026902 | -0.243677397 | 0.123623593 | 8.130118623 | -0.781201306 | 0.461582001 | 0.991835066 | -5.008107466 |
| **A_55_P2011385** | NA | -0.059589804 | -0.24201722 | 0.122837612 | 8.193783856 | -0.780712231 | 0.461850713 | 0.991835066 | -5.013503778 |
| **A_55_P2098734** | Ttc21b | 0.056186977 | -0.115837256 | 0.228211209 | 9.742241735 | 0.780647849 | 0.461886093 | 0.991835066 | -5.013541713 |
| **A_55_P2093301** | NA | -0.032663219 | -0.13267176 | 0.067345322 | 8.015261565 | -0.780604443 | 0.461909948 | 0.991835066 | -5.013567287 |
| **A_51_P351697** | Mrpl35 | 0.037326065 | -0.076961305 | 0.151613436 | 9.075384479 | 0.78059022 | 0.461917765 | 0.991835066 | -5.013575667 |
| **A_51_P206134** | Akna | -0.033000713 | -0.13405594 | 0.068054515 | 8.136857447 | -0.780501367 | 0.4619666 | 0.991835066 | -5.013628015 |
| **A_55_P2106255** | Mbnl3 | 0.070185665 | -0.144747487 | 0.285118818 | 9.972706971 | 0.780466281 | 0.461985884 | 0.991835066 | -5.013648684 |
| **A_55_P2024595** | Atxn7l1 | 0.054887869 | -0.113215905 | 0.222991642 | 8.119755053 | 0.780383424 | 0.462031428 | 0.991835066 | -5.013697493 |
| **A_55_P2077143** | Mcidas | 0.112086337 | -0.231197944 | 0.455370617 | 12.28428382 | 0.780383165 | 0.462031571 | 0.991835066 | -5.013697646 |
| **A_55_P2008427** | Dnd1 | 0.057450066 | -0.118502794 | 0.233402925 | 8.740301451 | 0.780374974 | 0.462036073 | 0.991835066 | -5.01370247 |
| **A_55_P2157159** | NA | -0.038165138 | -0.155078246 | 0.07874797 | 11.82969593 | -0.780212245 | 0.46212553 | 0.991835066 | -5.013798318 |
| **A_55_P1973941** | Slc7a5 | 0.099824198 | -0.20618286 | 0.405831255 | 8.905920836 | 0.779674836 | 0.462421042 | 0.991835066 | -5.014114738 |
| **A_55_P2005315** | NA | -0.090638648 | -0.368499712 | 0.187222416 | 10.46443393 | -0.779641336 | 0.462439468 | 0.991835066 | -5.014134457 |
| **A_52_P238858** | Shisa9 | -0.034009424 | -0.138270769 | 0.070251921 | 7.846748848 | -0.779623799 | 0.462449114 | 0.991835066 | -5.01414478 |
| **A_51_P424232** | Mecp2 | -0.06101178 | -0.248136264 | 0.126112704 | 10.48025639 | -0.779277634 | 0.462639541 | 0.991835066 | -5.014348497 |
| **A_51_P495171** | Pbdc1 | 0.125689366 | -0.259820455 | 0.511199187 | 11.05572039 | 0.779241766 | 0.462659275 | 0.991835066 | -5.014369601 |
| **A_52_P127362** | Vmp1 | 0.057530772 | -0.118952821 | 0.234014365 | 11.43190578 | 0.779121154 | 0.462725639 | 0.991835066 | -5.014440561 |
| **A_55_P1968998** | Tmub2 | -0.051782169 | -0.210653138 | 0.107088801 | 10.26442377 | -0.779013211 | 0.462785038 | 0.991835066 | -5.014504061 |
| **A_55_P2415472** | 4931408D1 | 0.039647424 | -0.082016037 | 0.161310886 | 7.764387582 | 0.778868145 | 0.462864874 | 0.991835066 | -5.014589388 |
| **A_52_P552589** | Map4k1 | 0.039820658 | -0.082399693 | 0.162041009 | 8.44669425 | 0.778706927 | 0.462953609 | 0.991835066 | -5.014684201 |
| **A_55_P2114863** | Mgll | -0.062439135 | -0.254082905 | 0.129204635 | 10.81890177 | -0.778702028 | 0.462956306 | 0.991835066 | -5.014687082 |
| **A_51_P407999** | 1500011B0 | 0.047006895 | -0.09727687 | 0.19129066 | 9.21783031 | 0.778669624 | 0.462974143 | 0.991835066 | -5.014706137 |
| **A_51_P511250** | Tent5b | -0.081089296 | -0.330017274 | 0.167838682 | 8.381699292 | -0.778572304 | 0.463027715 | 0.991835066 | -5.014763362 |
| **A_51_P288558** | Cfap52 | 0.036948575 | -0.076499011 | 0.150396162 | 7.67722422 | 0.778415672 | 0.463113948 | 0.991835066 | -5.014855451 |
| **A_51_P381873** | Bub3 | -0.042296324 | -0.172189596 | 0.087596947 | 11.80028514 | -0.778260712 | 0.46319927 | 0.991835066 | -5.014946542 |
| **A_55_P2031898** | NA | -0.14575451 | -0.59337501 | 0.301865989 | 10.48254437 | -0.778253652 | 0.463203158 | 0.991835066 | -5.014950692 |
| **A_55_P2083844** | Dis3l | -0.039333513 | -0.160134997 | 0.08146797 | 10.67315656 | -0.778215015 | 0.463224433 | 0.991835066 | -5.014973401 |
| **A_55_P2273929** | Gm20558 | 0.101519789 | -0.210319377 | 0.413358955 | 9.475768571 | 0.778088828 | 0.463293924 | 0.991835066 | -5.015047565 |
| **A_55_P2025820** | NA | 0.098259517 | -0.203598989 | 0.400118023 | 8.828750415 | 0.778001331 | 0.463342112 | 0.991835066 | -5.015098984 |
| **A_55_P2140423** | Arfip2 | 0.051453586 | -0.106620343 | 0.209527515 | 9.278809277 | 0.777973011 | 0.46335771 | 0.991835066 | -5.015115625 |
| **A_52_P457411** | Ubl5 | -0.063815877 | -0.259909098 | 0.132277344 | 13.35906351 | -0.777813175 | 0.463445749 | 0.991835066 | -5.015209541 |
| **A_52_P558087** | Poc5 | -0.048640993 | -0.19811709 | 0.100835104 | 9.895650691 | -0.777749635 | 0.463480751 | 0.991835066 | -5.015246871 |
| **A_55_P2068486** | Arfgap1 | 0.057877179 | -0.120017804 | 0.235772163 | 9.956780127 | 0.777593794 | 0.463566605 | 0.991835066 | -5.015338418 |
| **A_51_P256224** | Zfp768 | 0.051497168 | -0.106800802 | 0.209795138 | 12.16872442 | 0.777529963 | 0.463601774 | 0.991835066 | -5.015375911 |
| **A_55_P2012399** | NA | -0.072706362 | -0.296221845 | 0.150809121 | 12.19508671 | -0.777452624 | 0.463644387 | 0.991835066 | -5.015421335 |

| | | | | | | | | | |
|---|---|---|---|---|---|---|---|---|---|
| **A_55_P2094626** | Ralgps1 | -0.077006836 | -0.313746188 | 0.159732515 | 11.07699122 | -0.77744193 | 0.46365028 | 0.991835066 | -5.015427616 |
| **A_55_P2172807** | NA | 0.083314692 | -0.172847824 | 0.339477209 | 7.989755111 | 0.777347395 | 0.463702372 | 0.991835066 | -5.015483134 |
| **A_55_P1987409** | Nbeal1 | 0.051168225 | -0.106165899 | 0.208502349 | 8.459488772 | 0.777296224 | 0.463730571 | 0.991835066 | -5.015513184 |
| **A_52_P280899** | Wdr33 | 0.057083085 | -0.118442778 | 0.232608947 | 9.797483226 | 0.777276346 | 0.463741525 | 0.991835066 | -5.015524856 |
| **A_55_P2044463** | NA | 0.062937184 | -0.13061658 | 0.256490947 | 8.957011441 | 0.77716783 | 0.463801329 | 0.991835066 | -5.015588574 |
| **A_52_P547491** | Lurap1 | 0.046422721 | -0.096353182 | 0.189198624 | 8.511613582 | 0.777114134 | 0.463830924 | 0.991835066 | -5.0156201 |
| **A_55_P1967736** | Nkd1 | 0.099791891 | -0.207149585 | 0.406733368 | 9.647196991 | 0.77704972 | 0.463866428 | 0.991835066 | -5.015657918 |
| **A_51_P327418** | Rsrc2 | -0.054057523 | -0.220345995 | 0.112230949 | 10.64904197 | -0.776967995 | 0.463911475 | 0.991835066 | -5.015705894 |
| **A_51_P428742** | Pus7l | 0.041932184 | -0.08705741 | 0.170921779 | 8.437045892 | 0.776965873 | 0.463912645 | 0.991835066 | -5.01570714 |
| **A_52_P436238** | Odc1 | 0.083750582 | -0.173899734 | 0.341400898 | 9.608605155 | 0.776902087 | 0.463947807 | 0.991835066 | -5.015744582 |
| **A_55_P2012960** | Gpx7 | -0.044013764 | -0.179433858 | 0.09140633 | 9.858679806 | -0.776809509 | 0.463998843 | 0.991835066 | -5.015798922 |
| **A_55_P2030373** | Atp1a2 | -0.051901548 | -0.211595152 | 0.107792055 | 8.657677968 | -0.77678696 | 0.464011275 | 0.991835066 | -5.015812156 |
| **A_52_P663742** | Ganc | 0.03802857 | -0.079015798 | 0.155072939 | 8.339006025 | 0.776548537 | 0.464142733 | 0.991835066 | -5.015952074 |
| **A_55_P1964757** | Stam2 | 0.040085836 | -0.083298329 | 0.16347 | 8.485540197 | 0.776498553 | 0.464170295 | 0.991835066 | -5.015981402 |
| **A_51_P103650** | Trim8 | 0.032195927 | -0.06691248 | 0.131304333 | 11.52794027 | 0.776425103 | 0.464210799 | 0.991835066 | -5.016024497 |
| **A_51_P105520** | Nomo1 | 0.053209345 | -0.110622459 | 0.21704115 | 12.7697438 | 0.776245032 | 0.464310111 | 0.991835066 | -5.016130135 |
| **A_51_P378667** | Wrap53 | 0.038704728 | -0.080473388 | 0.157882844 | 10.75338281 | 0.776205359 | 0.464331993 | 0.991835066 | -5.016153407 |
| **A_55_P2122425** | NA | 0.04988936 | -0.103741846 | 0.203520565 | 12.5739702 | 0.776135639 | 0.46437045 | 0.991835066 | -5.016194301 |
| **A_55_P2027412** | NA | 0.182478905 | -0.379468361 | 0.744426171 | 10.61106046 | 0.776115283 | 0.464381679 | 0.991835066 | -5.01620624 |
| **A_55_P2072443** | Kansl1 | 0.059527885 | -0.123912995 | 0.242968765 | 8.619635793 | 0.775592249 | 0.464670253 | 0.991835066 | -5.016512927 |
| **A_51_P158638** | Ggnbp1 | -0.067717391 | -0.276507172 | 0.14107239 | 9.814910787 | -0.775175632 | 0.464900203 | 0.991835066 | -5.0167571 |
| **A_51_P446583** | Zfp445 | 0.079443506 | -0.165536671 | 0.324423682 | 9.614003172 | 0.775062258 | 0.464962792 | 0.991835066 | -5.016823528 |
| **A_51_P208031** | Cnot6l | -0.036585426 | -0.149416789 | 0.076245936 | 9.08148516 | -0.774974516 | 0.465011236 | 0.991835066 | -5.016874933 |
| **A_55_P2095894** | 6430548M( | -0.053795623 | -0.219712255 | 0.112121009 | 9.97243113 | -0.774936555 | 0.465032195 | 0.991835066 | -5.016897172 |
| **A_52_P6828** | Xk | 0.055075718 | -0.114796042 | 0.224947477 | 8.418640633 | 0.774904373 | 0.465049964 | 0.991835066 | -5.016916024 |
| **A_52_P678056** | Eva1b | -0.071240912 | -0.29098669 | 0.148504866 | 11.89999852 | -0.774850759 | 0.465079569 | 0.991835066 | -5.01694743 |
| **A_51_P475249** | Rnf44 | -0.04883034 | -0.199454371 | 0.10179369 | 10.95158028 | -0.774826775 | 0.465092812 | 0.991835066 | -5.016961479 |
| **A_51_P369516** | Nudt16l1 | -0.069244953 | -0.282841404 | 0.144351498 | 12.49696236 | -0.774824282 | 0.465094189 | 0.991835066 | -5.016962939 |
| **A_51_P427768** | Diaph1 | 0.062598294 | -0.130496838 | 0.255693427 | 8.362428013 | 0.774819075 | 0.465097064 | 0.991835066 | -5.016965989 |
| **A_66_P121965** | 1110032F0 | 0.069999778 | -0.145930441 | 0.285929998 | 8.612641954 | 0.774804922 | 0.46510488 | 0.991835066 | -5.016974279 |
| **A_66_P133450** | Dnajb9 | 0.033457505 | -0.069754733 | 0.136669743 | 7.88093326 | 0.774767669 | 0.465125451 | 0.991835066 | -5.016996099 |
| **A_52_P474089** | Capn6 | 0.140135778 | -0.292287208 | 0.572558763 | 10.90027215 | 0.774549878 | 0.465245731 | 0.991835066 | -5.017123649 |
| **A_55_P2310898** | Kantr | 0.057544609 | -0.120024818 | 0.235114035 | 8.597115039 | 0.774543083 | 0.465249484 | 0.991835066 | -5.017127628 |
| **A_55_P2091681** | Nudt1 | 0.042157078 | -0.087940982 | 0.172255138 | 10.26811825 | 0.77447752 | 0.465285697 | 0.991835066 | -5.017166019 |
| **A_51_P476601** | Ly96 | -0.037368521 | -0.152705055 | 0.077968013 | 9.261107842 | -0.774369327 | 0.465345461 | 0.991835066 | -5.017229367 |
| **A_52_P530291** | Pim1 | 0.036124216 | -0.075438191 | 0.147686624 | 8.36684612 | 0.773908627 | 0.4656 | 0.991835066 | -5.017499034 |
| **A_52_P370402** | Kcmf1 | -0.052848169 | -0.216063561 | 0.110367224 | 11.9264694 | -0.773887727 | 0.465611549 | 0.991835066 | -5.017511264 |
| **A_55_P2047852** | Clcn3 | 0.046112156 | -0.096300382 | 0.188524695 | 11.03677698 | 0.773884833 | 0.465613149 | 0.991835066 | -5.017512958 |
| **A_51_P227077** | Mdh1b | 0.052720695 | -0.110104801 | 0.21554619 | 7.877934187 | 0.773869707 | 0.465621508 | 0.991835066 | -5.017521809 |
| **A_51_P282144** | Adam23 | 0.050046363 | -0.104549937 | 0.204642662 | 8.142208776 | 0.773717745 | 0.465705492 | 0.991835066 | -5.017610728 |
| **A_55_P2016079** | Erich4 | -0.035057934 | -0.143359898 | 0.073244029 | 7.814538162 | -0.773676 | 0.465728564 | 0.991835066 | -5.017635152 |
| **A_52_P381784** | Nsd2 | 0.082665384 | -0.172783585 | 0.338114354 | 10.55731163 | 0.773443628 | 0.465857012 | 0.991835066 | -5.017771088 |
| **A_55_P1955548** | Ezr | -0.065165255 | -0.266558202 | 0.136227691 | 12.17549493 | -0.773358703 | 0.465903962 | 0.991835066 | -5.017820761 |
| **A_55_P2138097** | NA | -0.047051519 | -0.192472921 | 0.098369883 | 14.77755758 | -0.773311476 | 0.465930072 | 0.991835066 | -5.017848383 |
| **A_55_P1985519** | NA | -0.047741034 | -0.195305569 | 0.099823501 | 16.50495485 | -0.77324828 | 0.465965013 | 0.991835066 | -5.017885341 |
| **A_66_P122155** | Wdr26 | -0.069217102 | -0.283169476 | 0.144735271 | 11.38377537 | -0.773224202 | 0.465978326 | 0.991835066 | -5.017899422 |
| **A_51_P431996** | Zfp367 | -0.05995682 | -0.245287379 | 0.12537374 | 10.62014457 | -0.773215735 | 0.465983007 | 0.991835066 | -5.017904374 |
| **A_55_P1954897** | NA | 0.042382659 | -0.088637486 | 0.173402804 | 14.21462445 | 0.773141976 | 0.466023792 | 0.991835066 | -5.017947506 |

| | | | | | | | | | |
|---|---|---|---|---|---|---|---|---|---|
| **A_55_P2013268** | NA | 0.075214046 | -0.157322735 | 0.307750827 | 7.735547228 | 0.773065738 | 0.466065949 | 0.991835066 | -5.017992084 |
| **A_51_P196895** | Ist1 | -0.038798346 | -0.158755325 | 0.081158634 | 9.732413989 | -0.773030836 | 0.46608525 | 0.991835066 | -5.018012491 |
| **A_55_P1964313** | Pla2g4b | -0.054271096 | -0.222072165 | 0.113529974 | 10.97586225 | -0.773006235 | 0.466098854 | 0.991835066 | -5.018026874 |
| **A_55_P1963374** | Klhdc7a | -0.04086166 | -0.167237121 | 0.085513802 | 9.948512549 | -0.772791579 | 0.466217574 | 0.991835066 | -5.018152363 |
| **A_55_P2025233** | Llph | 0.077958475 | -0.163160033 | 0.319076984 | 9.00429223 | 0.77275517 | 0.466237712 | 0.991835066 | -5.018173645 |
| **A_66_P101477** | Bglap3 | 0.031486751 | -0.065912618 | 0.128886119 | 7.652873325 | 0.77264649 | 0.466297829 | 0.991835066 | -5.018237166 |
| **A_55_P1966383** | NA | 0.140025756 | -0.293214292 | 0.573265804 | 10.67196581 | 0.772482168 | 0.466388735 | 0.991835066 | -5.018333197 |
| **A_55_P2142668** | NA | -0.091314511 | -0.373842183 | 0.191213161 | 8.188793756 | -0.772481234 | 0.466389251 | 0.991835066 | -5.018333743 |
| **A_55_P1991906** | Acadvl | -0.063119884 | -0.258414952 | 0.132175184 | 14.87727524 | -0.77247432 | 0.466393077 | 0.991835066 | -5.018337783 |
| **A_51_P363801** | Pgpep1 | -0.055542342 | -0.227404154 | 0.116319471 | 8.890452747 | -0.772420748 | 0.466422716 | 0.991835066 | -5.018369086 |
| **A_52_P207635** | Zfc3h1 | 0.086054444 | -0.180267899 | 0.352376787 | 9.210307454 | 0.772280079 | 0.46650055 | 0.991835066 | -5.018451276 |
| **A_55_P2030080** | Kdm6b | -0.070950197 | -0.290549118 | 0.148648725 | 11.28005953 | -0.772204863 | 0.466542172 | 0.991835066 | -5.018495218 |
| **A_52_P537852** | Fam120b | 0.044490614 | -0.093234696 | 0.182215923 | 8.265571091 | 0.772082616 | 0.466609824 | 0.991835066 | -5.018566629 |
| **A_55_P2120388** | Skint7 | -0.068234053 | -0.279477619 | 0.143009513 | 14.10836055 | -0.772016908 | 0.46664619 | 0.991835066 | -5.018605008 |
| **A_51_P174854** | Maged1 | 0.0439288 | -0.092097271 | 0.179954871 | 13.21855199 | 0.771856058 | 0.466735221 | 0.991835066 | -5.018698949 |
| **A_55_P1996816** | NA | 0.056988692 | -0.119561248 | 0.233538632 | 13.82673821 | 0.771489903 | 0.466937931 | 0.991835066 | -5.018912736 |
| **A_51_P286814** | Ncor2 | 0.059646453 | -0.125138689 | 0.244431595 | 12.29992877 | 0.771483617 | 0.466941412 | 0.991835066 | -5.018916406 |
| **A_55_P1953884** | Dpm1 | 0.03887166 | -0.0815664 | 0.15930972 | 10.81812301 | 0.771397925 | 0.466988861 | 0.991835066 | -5.018966427 |
| **A_66_P128440** | NA | -0.04699682 | -0.192626784 | 0.098633145 | 9.165276734 | -0.771306264 | 0.46703962 | 0.991835066 | -5.019019928 |
| **A_66_P119801** | Optn | -0.038037355 | -0.155907133 | 0.079832423 | 8.030950814 | -0.771288714 | 0.467049339 | 0.991835066 | -5.019030171 |
| **A_55_P1952720** | Kctd2 | 0.057090644 | -0.119902696 | 0.234083985 | 12.13039381 | 0.770933912 | 0.467245854 | 0.991835066 | -5.01923721 |
| **A_55_P2154235** | 1700008J0 | 0.118064591 | -0.247992138 | 0.484121319 | 8.406800192 | 0.770868685 | 0.467281987 | 0.991835066 | -5.019275264 |
| **A_52_P236398** | NA | 0.107184503 | -0.225144587 | 0.439513593 | 8.411591534 | 0.770855142 | 0.46728949 | 0.991835066 | -5.019283165 |
| **A_51_P224445** | Gpn3 | -0.035559855 | -0.145823962 | 0.074704252 | 8.886127475 | -0.770788011 | 0.46732668 | 0.991835066 | -5.019322327 |
| **A_55_P2066657** | Vps13a | -0.043605339 | -0.178899737 | 0.09168906 | 8.444597807 | -0.77031611 | 0.467588172 | 0.991835066 | -5.01959754 |
| **A_55_P2128869** | Ccdc80 | -0.039964027 | -0.163986599 | 0.084058545 | 8.919130525 | -0.770154121 | 0.467677958 | 0.991835066 | -5.019691981 |
| **A_52_P400857** | Zfp595 | 0.041549869 | -0.087405465 | 0.170505202 | 8.366263286 | 0.770086447 | 0.46771547 | 0.991835066 | -5.019731431 |
| **A_55_P2125456** | NA | 0.064618422 | -0.135989418 | 0.265226261 | 14.06699318 | 0.76987032 | 0.467835288 | 0.991835066 | -5.019857403 |
| **A_55_P2088741** | Mtif3 | -0.079038554 | -0.324442051 | 0.166364943 | 8.766213033 | -0.769781329 | 0.467884629 | 0.991835066 | -5.019909264 |
| **A_55_P2181508** | Gm10653 | 0.039847036 | -0.08388168 | 0.163575752 | 15.55614436 | 0.769723328 | 0.467916789 | 0.991835066 | -5.019943062 |
| **A_55_P2353462** | 9530036O1 | -0.033468971 | -0.137403629 | 0.070465687 | 7.817825276 | -0.769646145 | 0.467959589 | 0.991835066 | -5.019988036 |
| **A_55_P1973033** | Ltbp1 | 0.068875272 | -0.145113245 | 0.282863789 | 10.85755524 | 0.76927566 | 0.468165066 | 0.991835066 | -5.020203861 |
| **A_51_P188772** | Arhgef28 | 0.065478669 | -0.137971884 | 0.268929221 | 9.412364134 | 0.769219173 | 0.4681964 | 0.991835066 | -5.02023676 |
| **A_52_P109548** | Map7 | -0.038377716 | -0.157623892 | 0.080868461 | 8.16926092 | -0.769208001 | 0.468202597 | 0.991835066 | -5.020243267 |
| **A_55_P2023692** | Neurl1a | 0.08190032 | -0.172655146 | 0.336455786 | 8.841148801 | 0.768975149 | 0.468331779 | 0.991835066 | -5.020378863 |
| **A_51_P246705** | Nop14 | -0.043045393 | -0.176848671 | 0.090757885 | 8.426463907 | -0.768898579 | 0.468374264 | 0.991835066 | -5.020423444 |
| **A_55_P2149215** | Camk2a | 0.06217191 | -0.131099571 | 0.255443392 | 8.97909427 | 0.768839289 | 0.468407164 | 0.991835066 | -5.020457963 |
| **A_52_P682990** | Timm50 | -0.077485476 | -0.318366437 | 0.163395485 | 10.93006857 | -0.768824052 | 0.468415618 | 0.991835066 | -5.020466833 |
| **A_55_P1984123** | Kmt2c | 0.067869673 | -0.143137391 | 0.278876737 | 10.30594664 | 0.768754895 | 0.468453995 | 0.991835066 | -5.020507093 |
| **A_51_P375862** | Srpk3 | -0.086438585 | -0.355183232 | 0.182306061 | 10.69944198 | -0.768735543 | 0.468464734 | 0.991835066 | -5.020518358 |
| **A_51_P299216** | Wdr90 | 0.068356206 | -0.144173184 | 0.280885597 | 9.01742511 | 0.768719835 | 0.468473451 | 0.991835066 | -5.020527502 |
| **A_55_P1970389** | Srsf10 | 0.068706906 | -0.144914307 | 0.28232812 | 9.907208976 | 0.768714635 | 0.468476337 | 0.991835066 | -5.020530529 |
| **A_55_P2035554** | Strip1 | 0.049770072 | -0.104996173 | 0.204536316 | 12.5110295 | 0.768601373 | 0.468539195 | 0.991835066 | -5.020596455 |
| **A_51_P153042** | Psg16 | -0.676608178 | -2.780683309 | 1.427466952 | 9.344989946 | -0.768573015 | 0.468554934 | 0.991835066 | -5.02061296 |
| **A_51_P301809** | Slit3 | 0.070466482 | -0.148685549 | 0.289618512 | 8.493650748 | 0.768504157 | 0.468593152 | 0.991835066 | -5.020653035 |
| **A_51_P108489** | Cfap20 | -0.046145907 | -0.189668618 | 0.097376805 | 11.72401762 | -0.768460745 | 0.468617249 | 0.991835066 | -5.010512356 |
| **A_51_P353386** | Micos13 | -0.091810136 | -0.377389563 | 0.193769292 | 13.35607667 | -0.768374323 | 0.46866522 | 0.991835066 | -5.020728591 |
| **A_55_P2035717** | Pgap2 | 0.040222119 | -0.084905126 | 0.165349363 | 11.13483408 | 0.768284716 | 0.468714963 | 0.991835066 | -5.02078073 |

| | | | | | | | | | |
|---|---|---|---|---|---|---|---|---|---|
| **A_51_P464981** | Map1lc3a | -0.15310581 | -0.629425575 | 0.323213956 | 11.91079369 | -0.768249346 | 0.468734599 | 0.991835066 | -5.02080131 |
| **A_55_P2100973** | Irf6 | -0.041026609 | -0.168666732 | 0.086613515 | 8.899985418 | -0.768223409 | 0.468748998 | 0.991835066 | -5.0208164 |
| **A_55_P2126805** | Wdr81 | -0.045075425 | -0.185324065 | 0.095173214 | 10.65618196 | -0.768157557 | 0.468785559 | 0.991835066 | -5.020854712 |
| **A_55_P2150442** | NA | -0.056748684 | -0.233322564 | 0.119825196 | 13.93282878 | -0.768136613 | 0.468797187 | 0.991835066 | -5.020866896 |
| **A_51_P118132** | Skil | 0.056617897 | -0.119566665 | 0.23280246 | 9.968506234 | 0.76805977 | 0.468839853 | 0.991835066 | -5.015766107 |
| **A_55_P2023042** | Ift43 | 0.079268043 | -0.167486946 | 0.326023032 | 12.19975725 | 0.767788012 | 0.468990762 | 0.991835066 | -5.021069661 |
| **A_52_P168575** | Fam169b | -0.032963276 | -0.135576912 | 0.06965036 | 7.694540815 | -0.767775809 | 0.468997539 | 0.991835066 | -5.021076757 |
| **A_51_P374818** | Shd | 0.056775278 | -0.119988268 | 0.233538824 | 8.728879696 | 0.767671996 | 0.469055197 | 0.991835066 | -5.021137126 |
| **A_55_P1975843** | NA | -0.092638934 | -0.38107332 | 0.195795451 | 11.50536416 | -0.767636557 | 0.469074881 | 0.991835066 | -5.021157732 |
| **A_55_P2012011** | Tceal7 | 0.132346086 | -0.279893354 | 0.544585527 | 10.88733816 | 0.767309713 | 0.469256447 | 0.991835066 | -5.021347746 |
| **A_55_P1984825** | Bloc1s2 | 0.040872162 | -0.086458034 | 0.168202357 | 10.56653953 | 0.767194242 | 0.469320604 | 0.991835066 | -5.02141486 |
| **A_55_P2044710** | Asb10 | -0.109021128 | -0.448714936 | 0.230672681 | 9.036087887 | -0.767065006 | 0.469392416 | 0.991835066 | -5.021489966 |
| **A_51_P386304** | Ccnl2 | 0.121184572 | -0.256428318 | 0.498797462 | 10.65060319 | 0.76702524 | 0.469414514 | 0.991835066 | -5.021513074 |
| **A_55_P2185870** | Bcl7a | 0.059272138 | -0.125434003 | 0.243978278 | 9.1164567 | 0.766970021 | 0.469445201 | 0.991835066 | -5.02154516 |
| **A_55_P2012430** | NA | 0.052757675 | -0.111661796 | 0.217177147 | 8.444471152 | 0.76690494 | 0.46948137 | 0.991835066 | -5.021582975 |
| **A_55_P2142172** | Olfr1229 | 0.074231525 | -0.157123079 | 0.30558613 | 8.686668503 | 0.766865799 | 0.469503124 | 0.991835066 | -5.021605716 |
| **A_55_P1969396** | NA | 0.113562005 | -0.240375136 | 0.467499145 | 11.32873751 | 0.766859929 | 0.469506386 | 0.991835066 | -5.021609127 |
| **A_55_P2000062** | Irf1 | -0.066945501 | -0.275624001 | 0.141732998 | 10.01487192 | -0.766748307 | 0.469568427 | 0.991835066 | -5.021673974 |
| **A_51_P455241** | Rrp36 | -0.066035731 | -0.271878997 | 0.139807535 | 9.991017666 | -0.766745896 | 0.469569767 | 0.991835066 | -5.021675375 |
| **A_55_P2016967** | Gm8213 | -0.04750258 | -0.19560728 | 0.10060212 | 14.79437796 | -0.766580012 | 0.469661978 | 0.991835066 | -5.021771732 |
| **A_51_P515605** | Col3a1 | -0.125655166 | -0.517539666 | 0.266229334 | 14.88679237 | -0.766357468 | 0.469785705 | 0.991835066 | -5.021900976 |
| **A_55_P2077666** | Porcn | 0.041202171 | -0.087304467 | 0.169708808 | 8.668448279 | 0.766308548 | 0.469812905 | 0.991835066 | -5.021929383 |
| **A_55_P1966134** | Prl3c1 | -0.031567275 | -0.130051029 | 0.06691648 | 7.629747655 | -0.766093231 | 0.46993264 | 0.991835066 | -5.022054395 |
| **A_55_P2081560** | Phf20 | 0.057819255 | -0.122568326 | 0.238206836 | 9.765399347 | 0.766081526 | 0.469939149 | 0.991835066 | -5.02206119 |
| **A_52_P427240** | Ltn1 | -0.031854475 | -0.131243988 | 0.067535038 | 8.429227504 | -0.76601808 | 0.469974435 | 0.991835066 | -5.02209802 |
| **A_55_P2142251** | H3c15 | 0.061045405 | -0.129488608 | 0.251579418 | 8.22590636 | 0.765754568 | 0.470121008 | 0.991835066 | -5.022250964 |
| **A_55_P1990678** | NA | -0.042911238 | -0.176849976 | 0.091027501 | 8.328376203 | -0.765727013 | 0.470136336 | 0.991835066 | -5.022266955 |
| **A_66_P126313** | Rhof | -0.03623957 | -0.149359578 | 0.076880438 | 8.39100947 | -0.765689591 | 0.470157155 | 0.991835066 | -5.022288671 |
| **A_52_P595954** | Nelfa | 0.062242611 | -0.132055487 | 0.256540708 | 9.910507051 | 0.765646646 | 0.470181046 | 0.991835066 | -5.022313591 |
| **A_51_P317214** | Hpdl | 0.070818465 | -0.150302767 | 0.291939698 | 8.833256779 | 0.765464749 | 0.470282249 | 0.991835066 | -5.022419129 |
| **A_51_P341177** | Ddah2 | 0.057282982 | -0.121588489 | 0.236154452 | 13.42431649 | 0.765409184 | 0.470313167 | 0.991835066 | -5.022451365 |
| **A_55_P2166555** | NA | 0.147839002 | -0.313940258 | 0.609618262 | 11.39366907 | 0.765180212 | 0.470440588 | 0.991835066 | -5.02258418 |
| **A_52_P470956** | NA | 0.063133186 | -0.134115899 | 0.260382271 | 16.39652271 | 0.764983101 | 0.470550298 | 0.991835066 | -5.02269849 |
| **A_55_P2007303** | Snrpg | 0.051335935 | -0.109058042 | 0.211729913 | 14.0885563 | 0.764966745 | 0.470559402 | 0.991835066 | -5.022707974 |
| **A_55_P2032727** | NA | 0.034624799 | -0.073561868 | 0.142811466 | 15.05950851 | 0.764931692 | 0.470578915 | 0.991835066 | -5.022728299 |
| **A_51_P359137** | Doc2g | -0.079989882 | -0.329925905 | 0.16994614 | 11.04231093 | -0.764918804 | 0.470586089 | 0.991835066 | -5.022735772 |
| **A_51_P249360** | Suox | -0.043716751 | -0.180343476 | 0.092909974 | 10.87524577 | -0.764753288 | 0.470678232 | 0.991835066 | -5.022831734 |
| **A_66_P137605** | Cgas | -0.048500663 | -0.200096994 | 0.103095669 | 8.911442628 | -0.764659543 | 0.470730425 | 0.991835066 | -5.022886078 |
| **A_55_P1958808** | Stoml2 | -0.052278349 | -0.215686636 | 0.111129937 | 12.35651013 | -0.764639839 | 0.470741396 | 0.991835066 | -5.0228975 |
| **A_55_P2061341** | Ift74 | -0.040002947 | -0.165067033 | 0.08506114 | 10.03699455 | -0.764484181 | 0.47082807 | 0.991835066 | -5.022987721 |
| **A_51_P178815** | H1f8 | -0.071816627 | -0.296345092 | 0.152711839 | 8.114192146 | -0.764474009 | 0.470833734 | 0.991835066 | -5.022993616 |
| **A_55_P2186480** | NA | 0.075487036 | -0.160520481 | 0.311494553 | 9.856863329 | 0.764461589 | 0.47084065 | 0.991835066 | -5.023000814 |
| **A_55_P1966958** | Mef2d | -0.072451765 | -0.29902117 | 0.15411764 | 8.368605372 | -0.76428764 | 0.470937524 | 0.991835066 | -5.023101617 |
| **A_52_P372032** | Ddx21 | 0.08544246 | -0.181752157 | 0.352637077 | 9.877703737 | 0.76428471 | 0.470939156 | 0.991835066 | -5.023103315 |
| **A_55_P2099785** | Usp15 | 0.071472833 | -0.152089009 | 0.295034675 | 10.72227902 | 0.764103963 | 0.47103983 | 0.991835066 | -5.023208038 |
| **A_51_P469902** | Ggps1 | 0.046030832 | -0.097973383 | 0.190035046 | 9.378680657 | 0.76398134 | 0.471108138 | 0.991835066 | -5.023279073 |
| **A_51_P518893** | Ccin | -0.042513444 | -0.175528717 | 0.09050183 | 8.541602787 | -0.763895418 | 0.471156005 | 0.991835066 | -5.023328843 |
| **A_55_P2108850** | 2-Mar | -0.056884015 | -0.234875037 | 0.121107007 | 9.183294612 | -0.763838036 | 0.471187975 | 0.991835066 | -5.023362078 |

| | | | | | | | | | |
|---|---|---|---|---|---|---|---|---|---|
| **A_51_P309920** | Itga8 | 0.13240073 | -0.281929081 | 0.546730542 | 10.12864325 | 0.763753705 | 0.471234961 | 0.991835066 | -5.023410918 |
| **A_51_P502150** | Slc9a3r1 | 0.037653887 | -0.080202144 | 0.155509918 | 8.518107424 | 0.763602137 | 0.471319418 | 0.991835066 | -5.023498688 |
| **A_52_P14526** | Zyg11b | 0.046022214 | -0.098045054 | 0.190089482 | 9.259532753 | 0.763504012 | 0.471374101 | 0.991835066 | -5.023555503 |
| **A_51_P249118** | Abcb4 | -0.054111698 | -0.223515577 | 0.115292182 | 8.457617789 | -0.763443567 | 0.471407788 | 0.991835066 | -5.023590498 |
| **A_55_P1959313** | Rbm42 | 0.044432757 | -0.094689106 | 0.183554619 | 12.97042307 | 0.763338219 | 0.471466504 | 0.991835066 | -5.023651485 |
| **A_55_P2066799** | Sctr | 0.032147124 | -0.068513091 | 0.132807339 | 7.795294157 | 0.763296738 | 0.471489624 | 0.991835066 | -5.023675497 |
| **A_66_P107703** | Smox | -0.071119931 | -0.293917224 | 0.151677363 | 8.846100823 | -0.762940282 | 0.471688339 | 0.991835066 | -5.023881793 |
| **A_55_P2013218** | Paip1 | 0.037882536 | -0.080799125 | 0.156564197 | 8.102473259 | 0.762894629 | 0.471713793 | 0.991835066 | -5.023908209 |
| **A_55_P2128224** | NA | -0.063781816 | -0.263617296 | 0.136053665 | 16.27769266 | -0.762839918 | 0.471744299 | 0.991835066 | -5.023939864 |
| **A_55_P2042184** | NA | 0.04067797 | -0.086776416 | 0.168132356 | 8.035631417 | 0.762805151 | 0.471763686 | 0.991835066 | -5.023959979 |
| **A_55_P2001078** | Mmel1 | 0.031348086 | -0.066898084 | 0.129594256 | 7.776623705 | 0.762613582 | 0.471870516 | 0.991835066 | -5.024070802 |
| **A_66_P132663** | Uqcrb | -0.056190828 | -0.232315583 | 0.119933927 | 16.12956375 | -0.762525134 | 0.471919845 | 0.991835066 | -5.024121961 |
| **A_55_P2122604** | Cbr2 | 0.032279513 | -0.068902308 | 0.133461333 | 7.787586021 | 0.762489054 | 0.471939968 | 0.991835066 | -5.024142829 |
| **A_51_P180140** | H2bc1 | -0.085765148 | -0.354654548 | 0.183124252 | 11.20166616 | -0.762335762 | 0.472025473 | 0.991835066 | -5.02423148 |
| **A_55_P1998761** | Coq8b | -0.052382682 | -0.216642317 | 0.111876953 | 9.573350143 | -0.762194847 | 0.472104083 | 0.991835066 | -5.024312961 |
| **A_55_P2134785** | Zscan22 | 0.077171627 | -0.164878386 | 0.31922164 | 9.55080236 | 0.762011766 | 0.472206229 | 0.991835066 | -5.024418806 |
| **A_66_P130469** | Rce1 | 0.045039274 | -0.096227962 | 0.18630651 | 8.607546287 | 0.762007178 | 0.47220879 | 0.991835066 | -5.024421459 |
| **A_55_P2062469** | Col12a1 | 0.139010048 | -0.297105079 | 0.575125176 | 9.410372682 | 0.761823167 | 0.472311471 | 0.991835066 | -5.024527821 |
| **A_55_P2104791** | NA | -0.05594104 | -0.231464891 | 0.119582811 | 14.34211796 | -0.761734331 | 0.472361048 | 0.991835066 | -5.024579162 |
| **A_52_P401504** | Thbs4 | 0.08449263 | -0.180625026 | 0.349610285 | 11.55584469 | 0.761709387 | 0.472374969 | 0.991835066 | -5.024593577 |
| **A_55_P2025601** | Lsm6 | 0.064263348 | -0.13739062 | 0.265917315 | 14.30122683 | 0.761667994 | 0.472398071 | 0.991835066 | -5.024617498 |
| **A_52_P447196** | Col4a6 | 0.034002123 | -0.072716736 | 0.140720981 | 8.307398195 | 0.761507158 | 0.472487844 | 0.991835066 | -5.024710432 |
| **A_55_P2068701** | NA | 0.038773353 | -0.082924101 | 0.160470808 | 8.109519594 | 0.761484383 | 0.472500557 | 0.991835066 | -5.024723591 |
| **A_52_P548202** | NA | -0.031255 | -0.129371199 | 0.066861199 | 7.720349672 | -0.761356251 | 0.472572085 | 0.991835066 | -5.024797615 |
| **A_52_P554703** | Gprin3 | 0.048789147 | -0.104394254 | 0.201972548 | 8.009356116 | 0.761238337 | 0.472637915 | 0.991835066 | -5.024865728 |
| **A_55_P2057405** | Cpt1a | 0.059244155 | -0.126765402 | 0.245253713 | 8.47369154 | 0.761236117 | 0.472639155 | 0.991835066 | -5.02486701 |
| **A_55_P2055819** | Adam15 | 0.056626806 | -0.12116679 | 0.234420403 | 13.38750516 | 0.761228581 | 0.472643362 | 0.991835066 | -5.024871363 |
| **A_55_P1971872** | NA | 0.074196313 | -0.15877666 | 0.307169286 | 9.113320793 | 0.761177453 | 0.472671908 | 0.991835066 | -5.024900894 |
| **A_55_P2063795** | Bag6 | -0.044999568 | -0.186297456 | 0.096298319 | 13.40429734 | -0.761170259 | 0.472675925 | 0.991835066 | -5.024905049 |
| **A_55_P2042521** | Cop1 | -0.049211307 | -0.203736075 | 0.105313461 | 11.66210769 | -0.761159953 | 0.47268168 | 0.991835066 | -5.024911002 |
| **A_55_P2002133** | Ago2 | 0.065467046 | -0.140108494 | 0.271042586 | 8.915430452 | 0.761132801 | 0.47269684 | 0.991835066 | -5.024926684 |
| **A_51_P212068** | Aoc1 | -0.043247651 | -0.17906428 | 0.092568978 | 7.721992877 | -0.761059673 | 0.472737674 | 0.991835066 | -5.024968917 |
| **A_55_P2097657** | Rps11 | -0.042995875 | -0.178030534 | 0.092038784 | 14.98569565 | -0.761010536 | 0.472765112 | 0.991835066 | -5.024997293 |
| **A_55_P2124831** | Brdt | 0.040028834 | -0.08570482 | 0.165762489 | 8.062000157 | 0.760905171 | 0.472823952 | 0.991835066 | -5.025058134 |
| **A_51_P398235** | Dnajc11 | 0.046491445 | -0.099551611 | 0.192534502 | 9.38983123 | 0.760853888 | 0.472852593 | 0.991835066 | -5.004090881 |
| **A_55_P2148906** | Cltb | -0.051856857 | -0.214773569 | 0.111059855 | 10.11141947 | -0.760763535 | 0.472903056 | 0.991835066 | -5.025139911 |
| **A_52_P137378** | Prkaa2 | -0.066072228 | -0.273663253 | 0.141518798 | 12.23876896 | -0.760710668 | 0.472932585 | 0.991835066 | -5.025170431 |
| **A_51_P428134** | Lrig3 | 0.113455793 | -0.243019443 | 0.469931029 | 8.776135334 | 0.760687784 | 0.472945367 | 0.991835066 | -5.025183642 |
| **A_55_P2088218** | Uqcrb | -0.060186665 | -0.249360572 | 0.128987241 | 14.54224953 | -0.760410634 | 0.473100191 | 0.991835066 | -5.025343612 |
| **A_55_P2096395** | Cdc42bpa | 0.047734164 | -0.102300347 | 0.197768676 | 10.50146445 | 0.760409066 | 0.473101067 | 0.991835066 | -5.025344517 |
| **A_55_P2163928** | Rtl8c | -0.072859964 | -0.3018819 | 0.156161972 | 8.826990342 | -0.760363046 | 0.473126779 | 0.991835066 | -5.025371075 |
| **A_52_P678816** | Ctsm | -0.240378138 | -0.995993299 | 0.515237023 | 8.071107577 | -0.760332293 | 0.473143961 | 0.991835066 | -5.025388821 |
| **A_55_P2086511** | Ppid | 0.071994331 | -0.154317691 | 0.298306352 | 12.54105304 | 0.760325939 | 0.473147511 | 0.991835066 | -5.025392488 |
| **A_55_P2078940** | Nsfl1c | 0.047418824 | -0.101647442 | 0.196485089 | 13.32431391 | 0.760292199 | 0.473166363 | 0.991835066 | -5.025411958 |
| **A_51_P244531** | NA | 0.04038662 | -0.086584789 | 0.167358029 | 10.80284064 | 0.760222466 | 0.473205327 | 0.991835066 | -5.025452195 |
| **A_55_P2011220** | Armcx1 | 0.09266916 | -0.198691486 | 0.384029805 | 10.25008864 | 0.760174795 | 0.473231965 | 0.991835066 | -5.025479701 |
| **A_66_P118989** | Gm8350 | -0.040437748 | -0.167577875 | 0.086702379 | 7.9927886 | -0.760174771 | 0.473231978 | 0.991835066 | -5.025479714 |
| **A_51_P383270** | Fras1 | 0.160891186 | -0.344994312 | 0.666776684 | 10.51644216 | 0.760132243 | 0.473255743 | 0.991835066 | -5.025504251 |

| | | | | | | | | | |
|---|---|---|---|---|---|---|---|---|---|
| **A_52_P356204** | Nostrin | -0.055085606 | -0.228293443 | 0.118122231 | 8.395241557 | -0.760115736 | 0.473264967 | 0.991835066 | -5.025513775 |
| **A_55_P2117169** | Zmym1 | 0.052338838 | -0.112294362 | 0.216972038 | 9.9864926 | 0.759828869 | 0.473425294 | 0.991835066 | -5.025679253 |
| **A_51_P224983** | Cln8 | -0.042987383 | -0.178213938 | 0.092239173 | 10.50055717 | -0.759780514 | 0.473452323 | 0.991835066 | -5.025707142 |
| **A_51_P228193** | Ociad1 | 0.056612284 | -0.121493205 | 0.234717772 | 11.75914438 | 0.759700662 | 0.473496959 | 0.991835066 | -5.025753193 |
| **A_55_P2003658** | Gnpda1 | -0.030686891 | -0.127229716 | 0.065855935 | 8.336445492 | -0.759699804 | 0.473497439 | 0.991835066 | -5.025753688 |
| **A_55_P2062678** | Hmox2 | -0.065967936 | -0.273511679 | 0.141575808 | 10.34735608 | -0.75968295 | 0.473506861 | 0.991835066 | -5.025763407 |
| **A_66_P112816** | B4galt6 | -0.048886269 | -0.202738632 | 0.104966093 | 10.29851564 | -0.759437186 | 0.473644261 | 0.991835066 | -5.025905114 |
| **A_55_P2060183** | Lamtor3 | -0.05634398 | -0.233674081 | 0.120986121 | 13.01801982 | -0.759406292 | 0.473661534 | 0.991835066 | -5.025922925 |
| **A_55_P2087825** | Ncoa2 | -0.060936685 | -0.252767423 | 0.130894052 | 10.62017125 | -0.759223711 | 0.473763631 | 0.991835066 | -5.026028174 |
| **A_55_P2072453** | 1-Mar | -0.042435373 | -0.176030832 | 0.091160085 | 7.739213459 | -0.759181236 | 0.473787384 | 0.991835066 | -5.026052656 |
| **A_55_P2019650** | NA | 0.05042038 | -0.10834709 | 0.209187851 | 10.69687994 | 0.759020881 | 0.473877067 | 0.991835066 | -5.026145072 |
| **A_55_P2142527** | NA | -0.063576091 | -0.263772823 | 0.136620642 | 9.019314864 | -0.759007324 | 0.47388465 | 0.991835066 | -5.026152884 |
| **A_55_P2005868** | Bbs7 | -0.031634743 | -0.131250894 | 0.067981409 | 8.466820865 | -0.759003323 | 0.473886888 | 0.991835066 | -5.02615519 |
| **A_55_P2010386** | Tspan32 | -0.032605429 | -0.135283004 | 0.070072146 | 7.993269871 | -0.758967971 | 0.473906661 | 0.991835066 | -5.026175562 |
| **A_55_P2055042** | Uqcc2 | -0.053595157 | -0.222389409 | 0.115199096 | 14.22937339 | -0.758886844 | 0.47395204 | 0.991835066 | -5.026222309 |
| **A_55_P2171206** | Matn4 | -0.055830268 | -0.231680161 | 0.120019625 | 8.149480421 | -0.758816443 | 0.473991422 | 0.991835066 | -5.026262871 |
| **A_55_P1965836** | Crebbp | 0.056049246 | -0.120539704 | 0.232638196 | 11.60447205 | 0.758604444 | 0.474110026 | 0.991835066 | -5.026385001 |
| **A_55_P2141943** | NA | -0.040577777 | -0.16842819 | 0.087272636 | 8.108007595 | -0.758569271 | 0.474129705 | 0.991835066 | -5.026405261 |
| **A_51_P434758** | Fam83e | 0.049654389 | -0.106797382 | 0.20610616 | 7.71982288 | 0.758553632 | 0.474138456 | 0.991835066 | -5.026414269 |
| **A_55_P2178510** | Mc3r | -0.050977311 | -0.211600395 | 0.109645773 | 7.784099138 | -0.758539314 | 0.474146467 | 0.991835066 | -5.026422516 |
| **A_66_P116252** | Dusp16 | 0.059054206 | -0.127020846 | 0.245129258 | 10.88849041 | 0.758528348 | 0.474152603 | 0.991835066 | -5.026428832 |
| **A_55_P1976689** | Snrnp35 | 0.056366286 | -0.121274797 | 0.234007369 | 10.73267683 | 0.758376974 | 0.474237308 | 0.991835066 | -5.026516013 |
| **A_55_P2141889** | NA | 0.052190876 | -0.112298203 | 0.216679956 | 11.39880119 | 0.758344689 | 0.474255375 | 0.991835066 | -5.026534605 |
| **A_51_P429335** | Prss16 | -0.062627382 | -0.260019899 | 0.134765134 | 7.697702316 | -0.758302883 | 0.474278771 | 0.991835066 | -5.026558679 |
| **A_55_P2101133** | Tomm7 | -0.056406165 | -0.234198342 | 0.121386012 | 14.6355933 | -0.758268579 | 0.474297968 | 0.991835066 | -5.026578432 |
| **A_55_P1976152** | Vrk2 | 0.040739698 | -0.087751292 | 0.169230688 | 9.255096028 | 0.75779941 | 0.47456059 | 0.991835066 | -5.026848522 |
| **A_55_P2006250** | Myo5a | 0.103995744 | -0.224001777 | 0.431993266 | 9.473206272 | 0.757799192 | 0.474560712 | 0.991835066 | -5.026848648 |
| **A_52_P109319** | E2f5 | 0.057840861 | -0.124587442 | 0.240269164 | 8.147084275 | 0.757794872 | 0.474563131 | 0.991835066 | -5.026851134 |
| **A_51_P249215** | Ptger2 | -0.035039239 | -0.145561801 | 0.075483322 | 7.699672789 | -0.757727166 | 0.474601038 | 0.991835066 | -5.0268901 |
| **A_55_P1973699** | Htra2 | 0.065526402 | -0.141184211 | 0.272237015 | 12.25821893 | 0.757639622 | 0.474650056 | 0.991835066 | -5.026940478 |
| **A_55_P1959119** | NA | -0.048809545 | -0.20280301 | 0.10518392 | 12.81785428 | -0.757550515 | 0.474699952 | 0.991835066 | -5.026991751 |
| **A_51_P520849** | Sfrp2 | 0.36979649 | -0.797219488 | 1.536812469 | 11.45772095 | 0.757347359 | 0.474813724 | 0.991835066 | -5.027108632 |
| **A_52_P315030** | NA | -0.037913714 | -0.157586072 | 0.081758644 | 15.12840693 | -0.757201751 | 0.474895278 | 0.991835066 | -5.027192388 |
| **A_52_P249920** | Zscan29 | -0.038792771 | -0.161255279 | 0.083669737 | 8.482399494 | -0.757106167 | 0.47494882 | 0.991835066 | -5.027247363 |
| **A_51_P210603** | Cyp4f13 | 0.033695214 | -0.072690164 | 0.140080592 | 8.225508246 | 0.756999183 | 0.475008753 | 0.991835066 | -5.027308887 |
| **A_52_P72839** | Sec23a | -0.060273701 | -0.250588937 | 0.130041536 | 9.498276472 | -0.756943439 | 0.475039983 | 0.991835066 | -5.027340942 |
| **A_55_P1956064** | NA | -0.092873165 | -0.386130434 | 0.200384104 | 11.99419485 | -0.756921068 | 0.475052516 | 0.991835066 | -5.027353806 |
| **A_55_P2042615** | NA | 0.096554476 | -0.208345937 | 0.401454889 | 9.813329264 | 0.756873904 | 0.475078941 | 0.991835066 | -5.027380925 |
| **A_55_P2232023** | NA | 0.075826453 | -0.163643499 | 0.315296406 | 8.038356449 | 0.756796031 | 0.475122572 | 0.991835066 | -5.027425697 |
| **A_55_P2042729** | NA | -0.038397705 | -0.159669494 | 0.082874084 | 12.07610858 | -0.756753813 | 0.475146229 | 0.991835066 | -5.02744997 |
| **A_51_P353392** | Cript | -0.099495619 | -0.413792785 | 0.214801547 | 9.640775319 | -0.756610954 | 0.475226282 | 0.991835066 | -5.027532093 |
| **A_55_P1965391** | Klhl18 | 0.054292196 | -0.117219416 | 0.225803808 | 10.13055305 | 0.756576789 | 0.475245428 | 0.991835066 | -5.027551731 |
| **A_55_P2147846** | Npcd | 0.05030258 | -0.108618995 | 0.209224154 | 9.052554085 | 0.756513232 | 0.475281047 | 0.991835066 | -5.027588262 |
| **A_55_P2017845** | Ehbp1 | -0.037982181 | -0.157983414 | 0.082019051 | 9.210962363 | -0.756490238 | 0.475293934 | 0.991835066 | -5.027601478 |
| **A_66_P136998** | Tmem43 | -0.049773124 | -0.207075775 | 0.107529526 | 9.241179633 | -0.756254541 | 0.475426043 | 0.991835066 | -5.027736927 |
| **A_52_P335609** | Idnk | -0.058612399 | -0.243851579 | 0.126626781 | 10.01039859 | -0.756250688 | 0.475428203 | 0.991835066 | -5.027739141 |
| **A_55_P2151899** | NA | -0.101591621 | -0.422689022 | 0.21950578 | 12.58526299 | -0.756188786 | 0.475462904 | 0.991835066 | -5.027774708 |
| **A_55_P2071834** | Tln2 | -0.047196217 | -0.196370686 | 0.101978252 | 10.79327775 | -0.756174143 | 0.475471113 | 0.991835066 | -5.027783121 |

| | | | | | | | | | |
|---|---|---|---|---|---|---|---|---|---|
| **A_52_P284778** | Rae1 | -0.043450452 | -0.180797947 | 0.093897043 | 11.25018941 | -0.756105999 | 0.475509315 | 0.991835066 | -5.027822272 |
| **A_55_P2067747** | Psg21 | -0.404118786 | -1.681610494 | 0.873372922 | 8.485392176 | -0.75606665 | 0.475531375 | 0.991835066 | -5.027844878 |
| **A_51_P401343** | Cldn14 | -0.031713214 | -0.131991584 | 0.068565156 | 7.826963975 | -0.755861319 | 0.475646502 | 0.991835066 | -5.027962826 |
| **A_52_P71686** | Atp6v0d2 | -0.032668898 | -0.135986741 | 0.070648944 | 7.665149467 | -0.75573284 | 0.475718548 | 0.991835066 | -5.028036614 |
| **A_55_P2081829** | Rps4x | 0.048339439 | -0.104538989 | 0.201217867 | 15.11197879 | 0.755726288 | 0.475722223 | 0.991835066 | -5.028040377 |
| **A_55_P1962756** | Ttll2 | 0.030929675 | -0.066897154 | 0.128756504 | 7.664064538 | 0.755660139 | 0.47575932 | 0.991835066 | -5.028078363 |
| **A_55_P1996509** | Rpl29 | 0.055818256 | -0.120764073 | 0.232400586 | 16.79340243 | 0.755506412 | 0.475845539 | 0.991835066 | -5.028166632 |
| **A_51_P101196** | Psme1 | -0.078923237 | -0.328677733 | 0.170831259 | 13.12865686 | -0.755267339 | 0.475979646 | 0.991835066 | -5.028303878 |
| **A_55_P2017038** | Cbx3 | 0.108261334 | -0.234342155 | 0.450864824 | 11.71981039 | 0.755249986 | 0.475989381 | 0.991835066 | -5.028313839 |
| **A_55_P2122546** | Csnk1e | 0.054847193 | -0.118779301 | 0.228473687 | 11.45016177 | 0.755001026 | 0.476129064 | 0.991835066 | -5.028456721 |
| **A_55_P1962665** | Exoc3l2 | -0.035654181 | -0.148524455 | 0.077216093 | 8.027705199 | -0.754987945 | 0.476136404 | 0.991835066 | -5.028464227 |
| **A_55_P1989419** | AI987944 | -0.041063705 | -0.171063561 | 0.088936151 | 8.336842661 | -0.754960786 | 0.476151644 | 0.991835066 | -5.028479812 |
| **A_51_P160824** | Cspg4 | 0.078877319 | -0.170881321 | 0.328635958 | 11.50623607 | 0.754815398 | 0.476233232 | 0.991835066 | -5.028563231 |
| **A_55_P1974954** | NA | 0.03695481 | -0.080065772 | 0.153975393 | 15.48792706 | 0.754775603 | 0.476255566 | 0.991835066 | -5.028586063 |
| **A_55_P1953301** | Sord | -0.06794625 | -0.283142157 | 0.147249658 | 11.74622814 | -0.754641382 | 0.476330897 | 0.991835066 | -5.02866306 |
| **A_52_P109010** | Hipk3 | -0.045787288 | -0.190812061 | 0.099237486 | 13.58399631 | -0.754591409 | 0.476358947 | 0.991835066 | -5.028691726 |
| **A_55_P2052380** | Gbp6 | 0.058578153 | -0.126981026 | 0.244137333 | 8.160882211 | 0.754505421 | 0.476407214 | 0.991835066 | -5.028741045 |
| **A_52_P78123** | Dcaf12 | 0.035451493 | -0.076865115 | 0.147768101 | 9.905390703 | 0.754396528 | 0.476468343 | 0.991835066 | -5.028803496 |
| **A_52_P163515** | NA | 0.059051957 | -0.128047407 | 0.246151321 | 9.198785685 | 0.754346912 | 0.476496197 | 0.991835066 | -5.028831949 |
| **A_52_P32466** | Yif1a | -0.053351369 | -0.222407828 | 0.11570509 | 11.63039644 | -0.754263216 | 0.476543187 | 0.991835066 | -5.028879942 |
| **A_55_P2143337** | Rpp21 | -0.063171703 | -0.26340093 | 0.137057523 | 12.22220042 | -0.754057125 | 0.476658907 | 0.991835066 | -5.0289981 |
| **A_55_P2026748** | Zmat1 | 0.040796955 | -0.088515146 | 0.170109055 | 8.347082107 | 0.754045778 | 0.476665279 | 0.991835066 | -5.029004605 |
| **A_51_P360963** | Mbtd1 | -0.079455415 | -0.331331048 | 0.172420218 | 10.68971514 | -0.753956823 | 0.476715233 | 0.991835066 | -5.029055597 |
| **A_55_P1959600** | Tmem222 | 0.0433198 | -0.094010906 | 0.180650505 | 11.51484877 | 0.753924615 | 0.476733321 | 0.991835066 | -5.029074059 |
| **A_55_P2065149** | Kdm2b | -0.065570687 | -0.273447236 | 0.142305862 | 11.65939086 | -0.753899346 | 0.476747513 | 0.991835066 | -5.029088542 |
| **A_55_P2119864** | Cd2bp2 | -0.038124002 | -0.159008758 | 0.082760754 | 9.100112053 | -0.753765191 | 0.47682286 | 0.991835066 | -5.029165431 |
| **A_51_P336161** | Man2b1 | -0.050713234 | -0.211528754 | 0.110102286 | 10.73751621 | -0.753706881 | 0.476855612 | 0.991835066 | -5.029198847 |
| **A_55_P2160686** | Tsc22d1 | -0.063456451 | -0.264706679 | 0.137793776 | 11.02122404 | -0.753613262 | 0.4769082 | 0.991835066 | -5.029252493 |
| **A_55_P1968240** | Cbr4 | -0.045210307 | -0.188771652 | 0.098351038 | 11.18835629 | -0.752677741 | 0.47743392 | 0.991835066 | -5.029788281 |
| **A_55_P2024220** | NA | 0.038299486 | -0.083325218 | 0.159924189 | 8.346477384 | 0.75262784 | 0.477461973 | 0.991835066 | -5.029816844 |
| **A_55_P2048912** | Themis2 | -0.041753337 | -0.174349762 | 0.090843088 | 7.844059155 | -0.752607442 | 0.47747344 | 0.991835066 | -5.02982852 |
| **A_51_P265106** | Dlat | -0.040495712 | -0.169140671 | 0.088149247 | 11.53392537 | -0.752359477 | 0.477612859 | 0.991835066 | -5.029970434 |
| **A_55_P2082688** | Ak1 | -0.075848404 | -0.316833278 | 0.165136471 | 15.31378926 | -0.752256228 | 0.477670919 | 0.991835066 | -5.030029514 |
| **A_55_P2077664** | Gm5458 | 0.148176197 | -0.322675042 | 0.619027436 | 9.236717859 | 0.752148958 | 0.477731245 | 0.991835066 | -5.030090887 |
| **A_55_P1969885** | NA | 0.072986286 | -0.158956714 | 0.304929287 | 9.297686374 | 0.752088811 | 0.477765072 | 0.991835066 | -5.030125297 |
| **A_66_P136099** | Coasy | 0.044416173 | -0.096776521 | 0.185608868 | 12.07886337 | 0.751861845 | 0.477892736 | 0.991835066 | -5.030255123 |
| **A_52_P449417** | Vangl1 | -0.039643362 | -0.165665189 | 0.086378465 | 11.08376872 | -0.751854567 | 0.47789683 | 0.991835066 | -5.030259285 |
| **A_55_P2097178** | Idua | 0.074373958 | -0.162126773 | 0.310874689 | 9.389217812 | 0.751618633 | 0.478029563 | 0.991835066 | -5.030394206 |
| **A_55_P2071271** | Abracl | 0.104303192 | -0.227369468 | 0.435975853 | 12.84228325 | 0.751617803 | 0.47803003 | 0.991835066 | -5.030394681 |
| **A_51_P393958** | Fbxo5 | 0.04876682 | -0.106306867 | 0.203840506 | 10.70237685 | 0.751615028 | 0.478031591 | 0.991835066 | -5.030396267 |
| **A_51_P254234** | Chchd4 | 0.054167695 | -0.118089686 | 0.226425076 | 10.50306914 | 0.751573828 | 0.478054772 | 0.991835066 | -5.030419825 |
| **A_55_P2140118** | Qpct | 0.131961777 | -0.287710678 | 0.551634232 | 9.788699543 | 0.751530868 | 0.478078945 | 0.991835066 | -5.030444387 |
| **A_55_P1973583** | Hpn | -0.037129077 | -0.155261486 | 0.081003333 | 8.137364495 | -0.751197645 | 0.478266467 | 0.991835066 | -5.030634868 |
| **A_52_P112651** | Lman2 | 0.047150676 | -0.102872751 | 0.197174103 | 10.03547753 | 0.751169552 | 0.478282279 | 0.991835066 | -5.030650924 |
| **A_66_P136364** | NA | -0.088163922 | -0.368820027 | 0.192492183 | 14.94660392 | -0.750802206 | 0.478489066 | 0.991835066 | -5.030860827 |
| **A_52_P33382** | NA | -0.311620898 | -1.303618958 | 0.680377161 | 11.05091664 | -0.750801074 | 0.478489703 | 0.991835066 | -5.030861474 |
| **A_55_P2055213** | Abcf1 | 0.059642739 | -0.130225425 | 0.249510904 | 8.295906088 | 0.75078323 | 0.478499749 | 0.991835066 | -5.030871667 |
| **A_51_P259555** | Gpatch3 | 0.045732526 | -0.099863007 | 0.191328058 | 8.328166432 | 0.75073432 | 0.478527287 | 0.991835066 | -5.030899608 |

| | | | | | | | | | |
|---|---|---|---|---|---|---|---|---|---|
| **A_55_P2178036** | Serpina6 | 0.034225318 | -0.074742986 | 0.143193621 | 7.69624129 | 0.750682736 | 0.478556331 | 0.991835066 | -5.030929074 |
| **A_52_P168097** | Atp2a2 | -0.134934707 | -0.564583165 | 0.29471375 | 10.54820835 | -0.750618988 | 0.478592226 | 0.991835066 | -5.030965487 |
| **A_55_P2073219** | Hps4 | 0.052671768 | -0.115054683 | 0.220398219 | 11.05312119 | 0.750560099 | 0.478625386 | 0.991835066 | -5.030999122 |
| **A_55_P2003494** | NA | 0.078265978 | -0.171023336 | 0.327555291 | 13.51169823 | 0.750375223 | 0.4787295 | 0.991835066 | -5.031104703 |
| **A_52_P152952** | Leng1 | -0.095648141 | -0.400332084 | 0.209035802 | 9.628756094 | -0.750301992 | 0.478770745 | 0.991835066 | -5.031146518 |
| **A_52_P51029** | Ralgapa1 | -0.066364548 | -0.277800909 | 0.145071813 | 10.32372686 | -0.750180203 | 0.478839343 | 0.991835066 | -5.031216054 |
| **A_51_P417321** | Zfp236 | -0.041264485 | -0.172752838 | 0.090223867 | 8.871447705 | -0.750063933 | 0.478904839 | 0.991835066 | -5.03128243 |
| **A_55_P2073642** | 1600014C1 | -0.040884849 | -0.171197605 | 0.089427906 | 8.049361631 | -0.749867627 | 0.479015434 | 0.991835066 | -5.031394477 |
| **A_55_P2093014** | NA | -0.055691272 | -0.233206646 | 0.121824102 | 16.308882 | -0.749825656 | 0.479039082 | 0.991835066 | -5.031418431 |
| **A_52_P112178** | Srgap2 | -0.04430739 | -0.185553504 | 0.096938724 | 8.501322281 | -0.749736737 | 0.479089184 | 0.991835066 | -5.031469175 |
| **A_52_P40367** | Mrpl42 | -0.042904471 | -0.179691491 | 0.093882549 | 12.57663231 | -0.749664245 | 0.479130033 | 0.991835066 | -5.03151054 |
| **A_55_P2131153** | Smarcc1 | -0.050055218 | -0.209646345 | 0.109535909 | 11.52890463 | -0.749634818 | 0.479146616 | 0.991835066 | -5.031527331 |
| **A_55_P1994074** | Mrtfa | -0.045880303 | -0.192170934 | 0.100410328 | 8.870672233 | -0.749581566 | 0.479176625 | 0.991835066 | -5.031557714 |
| **A_55_P2157914** | NA | -0.030765318 | -0.128862801 | 0.067332165 | 7.822612071 | -0.749570824 | 0.479182679 | 0.991835066 | -5.031563843 |
| **A_51_P432802** | Rbm4 | -0.051241611 | -0.214647857 | 0.112164635 | 11.77854284 | -0.749485531 | 0.479230748 | 0.991835066 | -5.031612504 |
| **A_55_P2117710** | Snhg1 | 0.088907222 | -0.194683002 | 0.372497447 | 9.138804617 | 0.749298594 | 0.479336113 | 0.991835066 | -5.031719139 |
| **A_52_P595642** | Smim7 | -0.062960233 | -0.263804287 | 0.137883821 | 10.69909842 | -0.749232277 | 0.479373495 | 0.991835066 | -5.031756963 |
| **A_55_P2063163** | Agk | 0.042617846 | -0.093337385 | 0.178573078 | 10.70112601 | 0.74921197 | 0.479384943 | 0.991835066 | -5.031768545 |
| **A_55_P2097269** | NA | 0.049291058 | -0.107964083 | 0.206546199 | 11.32861088 | 0.749156278 | 0.479416338 | 0.991835066 | -5.031800306 |
| **A_51_P237752** | Cavin1 | -0.043032569 | -0.180333486 | 0.094268349 | 15.72280378 | -0.749088211 | 0.479454711 | 0.991835066 | -5.031839123 |
| **A_55_P2159585** | Pm20d1 | -0.033734996 | -0.141376574 | 0.073906581 | 8.042270119 | -0.749048169 | 0.479477286 | 0.991835066 | -5.031861956 |
| **A_51_P237575** | Akirin1 | 0.059481289 | -0.13036926 | 0.249331839 | 11.4927394 | 0.748820367 | 0.47960573 | 0.991835066 | -5.031991839 |
| **A_55_P2146535** | Cnr2 | 0.131337707 | -0.287898139 | 0.550573553 | 9.975046982 | 0.74875572 | 0.479642185 | 0.991835066 | -5.032028691 |
| **A_51_P409349** | Trim29 | 0.033729456 | -0.073961451 | 0.141420363 | 7.693436705 | 0.748582094 | 0.479740104 | 0.991835066 | -5.032127657 |
| **A_52_P310530** | Rbfox2 | 0.070773943 | -0.155193637 | 0.296741523 | 10.76696793 | 0.74857685 | 0.479743061 | 0.991835066 | -5.032130646 |
| **A_55_P2053516** | Cacfd1 | 0.067993008 | -0.149098602 | 0.285084618 | 12.24300008 | 0.748566431 | 0.479748937 | 0.991835066 | -5.032136584 |
| **A_66_P110343** | Snx10 | -0.038157414 | -0.159995247 | 0.08368042 | 8.127409648 | -0.748524285 | 0.479772708 | 0.991835066 | -5.032160603 |
| **A_51_P196925** | Cx3cl1 | 0.097813701 | -0.214578532 | 0.410205934 | 9.735473246 | 0.748356605 | 0.479867291 | 0.991835066 | -5.032256156 |
| **A_55_P1973347** | A130040M | 0.140991962 | -0.309308447 | 0.59129237 | 11.63508345 | 0.748343765 | 0.479874534 | 0.991835066 | -5.032263473 |
| **A_55_P2139894** | Cwc15 | 0.053554839 | -0.117508226 | 0.224617903 | 10.94631304 | 0.7482584 | 0.47992269 | 0.991835066 | -5.032312111 |
| **A_55_P2165869** | Cebpb | 0.074300407 | -0.163060697 | 0.31166151 | 10.39524523 | 0.7481536 | 0.479981815 | 0.991835066 | -5.032371816 |
| **A_52_P68714** | NA | 0.046151397 | -0.101319135 | 0.193621929 | 8.905032168 | 0.747977847 | 0.48008098 | 0.991835066 | -5.03247193 |
| **A_55_P2080756** | Olfr704 | -0.039763907 | -0.166846852 | 0.087319037 | 8.025402323 | -0.74784383 | 0.480156606 | 0.991835066 | -5.032548256 |
| **A_55_P1985554** | B4galt4 | -0.042263526 | -0.177366138 | 0.092839087 | 8.589861224 | -0.747671986 | 0.480253589 | 0.991835066 | -5.032646111 |
| **A_51_P248644** | Tdp1 | 0.045394975 | -0.099723913 | 0.190513862 | 10.28180824 | 0.747640764 | 0.480271211 | 0.991835066 | -5.032663887 |
| **A_55_P2016287** | Tpst1 | -0.031646096 | -0.132813713 | 0.069521522 | 8.022951372 | -0.747631769 | 0.480276288 | 0.991835066 | -5.032669009 |
| **A_55_P1963980** | Carmil1 | 0.031827665 | -0.069946808 | 0.133602138 | 7.841992093 | 0.747437777 | 0.480385791 | 0.991835066 | -5.032779449 |
| **A_51_P514647** | Paip2 | -0.091702742 | -0.384942565 | 0.201537081 | 12.73024113 | -0.747426522 | 0.480392144 | 0.991835066 | -5.032785855 |
| **A_55_P2096281** | NA | -0.061327162 | -0.257444752 | 0.134790428 | 16.30319768 | -0.74738685 | 0.48041454 | 0.991835066 | -5.032808438 |
| **A_55_P1963454** | Cenpq | 0.054922733 | -0.120733861 | 0.230579326 | 8.791698657 | 0.747303152 | 0.480461792 | 0.991835066 | -5.032856077 |
| **A_55_P2145034** | Swt1 | 0.052763744 | -0.115997058 | 0.221524547 | 9.201555119 | 0.747262447 | 0.480484774 | 0.991835066 | -5.032879244 |
| **A_52_P222073** | Ackr2 | -0.124676311 | -0.523452151 | 0.27409953 | 9.533969324 | -0.747247097 | 0.48049344 | 0.991835066 | -5.032887981 |
| **A_51_P237834** | Tmcc1 | -0.044373881 | -0.186311882 | 0.09756412 | 9.182693415 | -0.74720172 | 0.48051906 | 0.991835066 | -5.032913805 |
| **A_55_P2031676** | Gstm6 | -0.034689813 | -0.145673953 | 0.076294327 | 7.996546439 | -0.747050858 | 0.480604245 | 0.991835066 | -5.032999653 |
| **A_55_P1977699** | NA | 0.046646542 | -0.102623033 | 0.195916117 | 13.8481333 | 0.746891108 | 0.480694459 | 0.991835066 | -5.033090544 |
| **A_55_P2329680** | Dyrk2 | -0.048403526 | -0.203313369 | 0.106506318 | 10.54284169 | -0.746804822 | 0.480743191 | 0.991835066 | -5.033139631 |
| **A_55_P2002314** | Zmiz2 | 0.051985217 | -0.114391932 | 0.218362366 | 10.92434311 | 0.746784535 | 0.480754649 | 0.991835066 | -5.033151171 |
| **A_55_P2179221** | Man1a2 | 0.079793034 | -0.175642829 | 0.335228897 | 11.44249565 | 0.746607316 | 0.48085475 | 0.991835066 | -5.033251971 |

| | | | | | | | | | |
|---|---|---|---|---|---|---|---|---|---|
| **A_55_P2023797** | Cep112 | -0.049309896 | -0.207162817 | 0.108543024 | 9.041636727 | -0.746604497 | 0.480856343 | 0.991835066 | -5.033253574 |
| **A_51_P352738** | Mpv17l | -0.048194304 | -0.20250128 | 0.106112672 | 7.914338147 | -0.746481907 | 0.480925595 | 0.991835066 | -5.03332329 |
| **A_55_P2180944** | Tmco4 | -0.045895495 | -0.192843682 | 0.101052693 | 10.82786558 | -0.746474462 | 0.480929801 | 0.991835066 | -5.033327524 |
| **A_51_P331288** | Akr1b7 | -0.035623929 | -0.149699952 | 0.078452093 | 7.724398616 | -0.746374134 | 0.480986483 | 0.991835066 | -5.033384573 |
| **A_55_P2368355** | 2610319H1 | 0.188366503 | -0.414947375 | 0.791680382 | 11.4732364 | 0.74622448 | 0.481071041 | 0.991835066 | -5.033469658 |
| **A_55_P2024763** | Lama3 | -0.035764345 | -0.150326565 | 0.078797875 | 8.175320278 | -0.746135992 | 0.481121043 | 0.991835066 | -5.033519961 |
| **A_51_P467629** | Efcab10 | -0.036305414 | -0.152606668 | 0.079995839 | 7.659476862 | -0.746098441 | 0.481142263 | 0.991835066 | -5.033541306 |
| **A_66_P125634** | Wdr63 | -0.07990482 | -0.335906311 | 0.176096672 | 9.432952862 | -0.746001348 | 0.481197134 | 0.991835066 | -5.033596493 |
| **A_51_P307168** | Ddah1 | 0.072592739 | -0.159986145 | 0.305171624 | 8.827632731 | 0.745988331 | 0.481204491 | 0.991835066 | -5.033603892 |
| **A_51_P293607** | Mrps14 | 0.049467014 | -0.109054445 | 0.207988472 | 8.517171959 | 0.745824705 | 0.481296971 | 0.991835066 | -5.033696881 |
| **A_51_P268234** | Il34 | 0.035094279 | -0.077371015 | 0.147559573 | 7.817865724 | 0.745807827 | 0.481306511 | 0.991835066 | -5.033706472 |
| **A_55_P2091858** | Hipk2 | 0.079692576 | -0.175712058 | 0.335097211 | 9.701692614 | 0.745758524 | 0.48133438 | 0.991835066 | -5.033734488 |
| **A_52_P657360** | Tnni1 | -0.122181228 | -0.513763383 | 0.269400927 | 15.59705013 | -0.745745643 | 0.481341661 | 0.991835066 | -5.033741807 |
| **A_55_P2071601** | Ppara | 0.061854977 | -0.136387911 | 0.260097864 | 10.26716041 | 0.745737814 | 0.481346087 | 0.991835066 | -5.033746255 |
| **A_52_P93910** | Nrp2 | -0.058504602 | -0.246015976 | 0.129006771 | 11.68480553 | -0.745712719 | 0.481360273 | 0.991835066 | -5.033760514 |
| **A_52_P504787** | Epha7 | -0.034694239 | -0.145910129 | 0.076521652 | 9.033955001 | -0.74558927 | 0.481430059 | 0.991835066 | -5.033830652 |
| **A_51_P448489** | Mea1 | 0.097883846 | -0.215912846 | 0.411680537 | 11.15844423 | 0.745541452 | 0.481457093 | 0.991835066 | -5.033857817 |
| **A_55_P2025248** | Mxd1 | 0.04403735 | -0.097174761 | 0.185249462 | 8.863829333 | 0.745346752 | 0.481567178 | 0.991835066 | -5.033968412 |
| **A_52_P615401** | Tpra1 | -0.049053799 | -0.206375029 | 0.10826743 | 8.42996563 | -0.745237069 | 0.4816292 | 0.991835066 | -5.034030703 |
| **A_55_P2022885** | Zfp866 | -0.048149723 | -0.202585077 | 0.106285631 | 8.638280786 | -0.745171435 | 0.481666317 | 0.991835066 | -5.034067976 |
| **A_52_P528600** | Hhex | -0.064199591 | -0.270123427 | 0.141724245 | 9.304226623 | -0.745134678 | 0.481687105 | 0.991835066 | -5.034088848 |
| **A_55_P2218223** | Pacs2 | 0.036058419 | -0.079604816 | 0.151721655 | 8.783970279 | 0.745110153 | 0.481700975 | 0.991835066 | -5.034102774 |
| **A_55_P2107785** | Gm13105 | -0.126353391 | -0.531658801 | 0.278952018 | 10.65485324 | -0.745098408 | 0.481707617 | 0.991835066 | -5.034109443 |
| **A_55_P2103857** | Hgs | -0.052042175 | -0.223762904 | 0.119678554 | 12.02298035 | -0.752835103 | 0.481723401 | 0.991835066 | -4.962005358 |
| **A_55_P2180909** | Ccdc166 | 0.049964537 | -0.110323031 | 0.210252105 | 8.602859464 | 0.745025538 | 0.48174883 | 0.991835066 | -5.034150818 |
| **A_55_P2145706** | Thoc7 | -0.047917362 | -0.201663626 | 0.105828903 | 10.81960424 | -0.744899116 | 0.481820337 | 0.991835066 | -5.034222591 |
| **A_55_P2054331** | Gm9222 | -0.083509614 | -0.351472312 | 0.184453084 | 8.398761461 | -0.744854194 | 0.481845748 | 0.991835066 | -5.034248092 |
| **A_55_P1961255** | Ppp1cc | 0.047383746 | -0.104666864 | 0.199434356 | 14.42074754 | 0.744818335 | 0.481866032 | 0.991835066 | -5.034268448 |
| **A_55_P2031125** | Ube2l6 | -0.164054694 | -0.690516393 | 0.362407006 | 10.53456492 | -0.744785153 | 0.481884803 | 0.991835066 | -5.034287283 |
| **A_51_P184728** | Cnksr3 | 0.036120661 | -0.079811352 | 0.152052674 | 10.10716975 | 0.744665859 | 0.481952291 | 0.991835066 | -5.034354992 |
| **A_55_P2074771** | Exosc7 | 0.046271448 | -0.102243354 | 0.194786251 | 11.4880865 | 0.744650496 | 0.481960983 | 0.991835066 | -5.034363711 |
| **A_52_P543040** | Utp14a | 0.062643128 | -0.138419702 | 0.263705958 | 8.979295668 | 0.744647566 | 0.481962641 | 0.991835066 | -5.034365375 |
| **A_52_P24428** | Ankrd11 | 0.073089063 | -0.161507954 | 0.30768608 | 11.25066721 | 0.74462745 | 0.481974021 | 0.991835066 | -5.034376791 |
| **A_55_P2052016** | Crispld2 | 0.127705249 | -0.282245702 | 0.5376562 | 9.087923449 | 0.744536483 | 0.48202549 | 0.991835066 | -5.034428415 |
| **A_55_P2115119** | Rnaseh2a | -0.086640532 | -0.364848443 | 0.191567379 | 9.858342121 | -0.744321848 | 0.482146943 | 0.991835066 | -5.034550199 |
| **A_55_P1986441** | Gcdh | -0.042388269 | -0.178532745 | 0.093756208 | 9.793136672 | -0.744140228 | 0.48224973 | 0.991835066 | -5.034653229 |
| **A_55_P2146171** | Cox20 | -0.062010062 | -0.261185922 | 0.137165797 | 14.15975858 | -0.744105648 | 0.482269302 | 0.991835066 | -5.034672843 |
| **A_55_P1974134** | Mrpl58 | -0.07051279 | -0.297043676 | 0.156018096 | 13.40760213 | -0.743960034 | 0.482351725 | 0.991835066 | -5.034755429 |
| **A_55_P2127952** | NA | 0.03838889 | -0.084952543 | 0.161730322 | 8.023078913 | 0.74388482 | 0.482394302 | 0.991835066 | -5.034798083 |
| **A_55_P1996821** | Myo15b | -0.038512038 | -0.162302576 | 0.0852785 | 7.706940654 | -0.743563708 | 0.482576107 | 0.991835066 | -5.034980144 |
| **A_52_P386837** | Trmu | -0.033913642 | -0.142923978 | 0.075096693 | 9.787103376 | -0.743559784 | 0.482578329 | 0.991835066 | -5.034982369 |
| **A_52_P570240** | Kbtbd11 | 0.135265937 | -0.299540709 | 0.570072584 | 9.35191834 | 0.74353498 | 0.482592374 | 0.991835066 | -5.034996429 |
| **A_55_P2024953** | Tbc1d4 | 0.055897304 | -0.123790951 | 0.235585559 | 10.35049775 | 0.743498854 | 0.482612831 | 0.991835066 | -5.035016907 |
| **A_55_P1958230** | Dhodh | 0.051036641 | -0.113049205 | 0.215122487 | 10.87058976 | 0.74339579 | 0.482671196 | 0.991835066 | -5.035075323 |
| **A_55_P2050083** | Gak | 0.081874915 | -0.181387712 | 0.345137543 | 10.25358572 | 0.743311385 | 0.482718998 | 0.991835066 | -5.03512316 |
| **A_51_P368903** | Anapc13 | -0.079974868 | -0.337128838 | 0.177179102 | 15.36191414 | -0.743309071 | 0.482720309 | 0.991835066 | -5.035124471 |
| **A_55_P1968628** | NA | -0.033848501 | -0.142691705 | 0.074994703 | 8.192107637 | -0.743271119 | 0.482741804 | 0.991835066 | -5.035145979 |
| **A_55_P2161640** | Slc38a1 | 0.086554729 | -0.19177387 | 0.364883329 | 9.698569649 | 0.74326229 | 0.482746804 | 0.991835066 | -5.035150982 |

| | | | | | | | | | |
|---|---|---|---|---|---|---|---|---|---|
| **A_52_P55772** | Tbxa2r | 0.076061083 | -0.168527682 | 0.320649848 | 9.247494419 | 0.743250367 | 0.482753557 | 0.991835066 | -5.035157739 |
| **A_55_P2167447** | Cnnm3 | -0.069846901 | -0.294453335 | 0.154759534 | 12.28545263 | -0.743248564 | 0.482754578 | 0.991835066 | -5.03515876 |
| **A_55_P2115477** | Parva | 0.055713841 | -0.123455181 | 0.234882864 | 9.245967876 | 0.743206176 | 0.482778587 | 0.991835066 | -5.03518278 |
| **A_55_P2034390** | Ankhd1 | -0.033806117 | -0.142539146 | 0.074926912 | 8.774365516 | -0.7430926 | 0.482842919 | 0.991835066 | -5.035247134 |
| **A_55_P1962344** | Trim21 | -0.037485314 | -0.158070559 | 0.083099931 | 8.521000034 | -0.742978277 | 0.482907682 | 0.991835066 | -5.035311904 |
| **A_55_P2183688** | NA | 0.04839533 | -0.10730297 | 0.20409363 | 15.28491638 | 0.742897198 | 0.482953615 | 0.991835066 | -5.035357835 |
| **A_55_P1977968** | Epb41 | -0.258081733 | -1.149207612 | 0.633044145 | 13.33193305 | -0.761629621 | 0.483091717 | 0.991835066 | -4.840533775 |
| **A_55_P2094602** | Lrch1 | -0.039673234 | -0.167370874 | 0.088024405 | 9.109517628 | -0.742546867 | 0.48315212 | 0.991835066 | -5.035556247 |
| **A_55_P2030486** | Srsf2 | 0.050618095 | -0.112310922 | 0.213547112 | 9.540264122 | 0.742534247 | 0.483159272 | 0.991835066 | -5.035563393 |
| **A_51_P357744** | Jund | 0.071556033 | -0.15880678 | 0.301918847 | 13.5284694 | 0.742408636 | 0.48323046 | 0.991835066 | -5.035634515 |
| **A_55_P1980020** | Rsl1d1 | 0.072290296 | -0.160437848 | 0.30501844 | 12.85909894 | 0.742403873 | 0.483233159 | 0.991835066 | -5.035637211 |
| **A_55_P2164534** | Dtl | 0.126264989 | -0.280321604 | 0.532851581 | 10.10762403 | 0.742230888 | 0.483331208 | 0.991835066 | -5.03573514 |
| **A_51_P419925** | Arhgef40 | 0.044797034 | -0.099481822 | 0.18907589 | 8.939108696 | 0.742088504 | 0.483411921 | 0.991835066 | -5.035815731 |
| **A_55_P2029746** | NA | 0.183707108 | -0.408005254 | 0.77541947 | 14.76022231 | 0.742035102 | 0.483442196 | 0.991835066 | -5.035845954 |
| **A_66_P111714** | Utp20 | 0.038369007 | -0.085263356 | 0.162001369 | 8.424917466 | 0.741749936 | 0.483603883 | 0.991835066 | -5.036007315 |
| **A_55_P2099379** | Tfe3 | -0.091268795 | -0.385419318 | 0.202881727 | 8.888631066 | -0.74158652 | 0.483696555 | 0.991835066 | -5.036099761 |
| **A_52_P88983** | Dock5 | 0.073372169 | -0.163258821 | 0.310003158 | 8.903378156 | 0.741086442 | 0.48398022 | 0.991835066 | -5.036382559 |
| **A_52_P26953** | Ergic2 | -0.040172453 | -0.1697574 | 0.089412493 | 9.273801681 | -0.740939813 | 0.484063415 | 0.991835066 | -5.03646545 |
| **A_51_P103397** | Vwf | 0.082860972 | -0.184443454 | 0.350165399 | 11.90728231 | 0.74088876 | 0.484092384 | 0.991835066 | -5.036494308 |
| **A_55_P2039250** | Mboat2 | -0.047477404 | -0.200639786 | 0.105684979 | 10.63361704 | -0.740873363 | 0.484101121 | 0.991835066 | -5.036503011 |
| **A_55_P2100884** | Fjx1 | 0.071310404 | -0.158761544 | 0.301382352 | 8.671217587 | 0.740795538 | 0.484145284 | 0.991835066 | -5.036546997 |
| **A_65_P11504** | Zfp85 | -0.033373459 | -0.141070484 | 0.074323565 | 8.160674063 | -0.740639131 | 0.484234048 | 0.991835066 | -5.036635388 |
| **A_55_P2117081** | Trgv4 | -0.032082564 | -0.135617274 | 0.071452147 | 7.655921035 | -0.740614524 | 0.484248013 | 0.991835066 | -5.036649292 |
| **A_52_P117352** | Gja4 | 0.05458118 | -0.121612026 | 0.230774387 | 10.61059943 | 0.740394009 | 0.48437318 | 0.991835066 | -5.036773882 |
| **A_52_P393314** | P2rx7 | -0.035768449 | -0.15125571 | 0.079718813 | 8.093672851 | -0.740244438 | 0.484458091 | 0.991835066 | -5.036858372 |
| **A_51_P418317** | Mis12 | 0.048712295 | -0.108571343 | 0.205995934 | 8.163775001 | 0.740225734 | 0.48446871 | 0.991835066 | -5.036868937 |
| **A_66_P134149** | Snx30 | -0.04036116 | -0.170717251 | 0.089994932 | 8.615571346 | -0.740016559 | 0.484587476 | 0.991835066 | -5.036987071 |
| **A_55_P1957598** | Atad2b | 0.054986767 | -0.122614931 | 0.232588465 | 10.73049252 | 0.739980376 | 0.484608022 | 0.991835066 | -5.037007503 |
| **A_55_P2100008** | NA | 0.050528901 | -0.112687755 | 0.213745558 | 10.75254915 | 0.739919564 | 0.484642555 | 0.991835066 | -5.037041841 |
| **A_55_P2084002** | Aptx | -0.03219971 | -0.136221643 | 0.071822224 | 8.663202935 | -0.739837206 | 0.484689325 | 0.991835066 | -5.037088341 |
| **A_55_P2138792** | Fdx1 | -0.042078234 | -0.17802212 | 0.093865651 | 12.7737643 | -0.739787453 | 0.484717581 | 0.991835066 | -5.037116431 |
| **A_55_P2170225** | Sh3bgrl2 | 0.032033977 | -0.071465445 | 0.1355334 | 7.977996521 | 0.739745057 | 0.48474166 | 0.991835066 | -5.037140365 |
| **A_55_P2051566** | NA | -0.050041741 | -0.211756877 | 0.111673395 | 7.739320597 | -0.739589735 | 0.484829881 | 0.991835066 | -5.037228042 |
| **A_55_P2002517** | Fhl1 | 0.037351023 | -0.083364185 | 0.158066232 | 8.387796554 | 0.739519526 | 0.484869762 | 0.991835066 | -5.037267669 |
| **A_55_P2214625** | 4930519A1 | -0.041274631 | -0.174695537 | 0.092146274 | 8.005671595 | -0.739381278 | 0.484948298 | 0.991835066 | -5.037345689 |
| **A_55_P2125766** | Ola1 | 0.081768487 | -0.182560851 | 0.346097825 | 9.922721585 | 0.739349402 | 0.484966408 | 0.991835066 | -5.037363677 |
| **A_51_P139651** | Nos3 | 0.038222064 | -0.085400143 | 0.161844272 | 10.1222617 | 0.73896995 | 0.485182017 | 0.991835066 | -5.037577753 |
| **A_52_P83959** | Taf7 | -0.048881904 | -0.207037558 | 0.109273751 | 8.797208036 | -0.738707518 | 0.485331172 | 0.991835066 | -5.037725758 |
| **A_55_P2002703** | NA | -0.053390882 | -0.226137918 | 0.119356153 | 9.698366894 | -0.738695796 | 0.485337835 | 0.991835066 | -5.037732367 |
| **A_51_P240136** | Gng8 | 0.066315292 | -0.148324857 | 0.280955441 | 8.326343252 | 0.738434292 | 0.485486495 | 0.991835066 | -5.037879805 |
| **A_55_P2120526** | NA | 0.034274665 | -0.0766695 | 0.145218831 | 8.361126372 | 0.738376535 | 0.485519333 | 0.991835066 | -5.037912363 |
| **A_55_P1986706** | Pih1d1 | 0.042852271 | -0.095875556 | 0.181580098 | 12.21640455 | 0.73827709 | 0.485575875 | 0.991835066 | -5.037968416 |
| **A_51_P235548** | Mtmr9 | -0.039542906 | -0.167567387 | 0.088481575 | 8.570018736 | -0.738218107 | 0.485609414 | 0.991835066 | -5.038001659 |
| **A_55_P1993805** | Zfp947 | 0.058620942 | -0.131174469 | 0.248416354 | 10.85466783 | 0.738203699 | 0.485617607 | 0.991835066 | -5.038009779 |
| **A_55_P1971828** | Csnk2a1 | 0.058505014 | -0.130921336 | 0.247931365 | 9.000353936 | 0.738179242 | 0.485631515 | 0.991835066 | -5.038023563 |
| **A_52_P195798** | Dmac2l | -0.042096051 | -0.178400386 | 0.094208284 | 8.91279334 | -0.738143531 | 0.485651822 | 0.991835066 | -5.038043688 |
| **A_51_P369252** | Ddias | 0.0520627 | -0.116515039 | 0.220640439 | 8.629544616 | 0.738134655 | 0.48565687 | 0.991835066 | -5.03804869 |
| **A_55_P2024868** | Psg18 | -0.469020143 | -1.987981906 | 1.049941619 | 8.604843091 | -0.737995623 | 0.485735937 | 0.991835066 | -5.038127034 |

| | | | | | | | | | |
|---|---|---|---|---|---|---|---|---|---|
| **A_51_P102507** | Vps33a | -0.042903218 | -0.181858213 | 0.096051776 | 8.944392629 | -0.737946443 | 0.485763908 | 0.991835066 | -5.038154745 |
| **A_51_P362959** | Fbxo36 | -0.039283926 | -0.166523445 | 0.087955594 | 8.222698103 | -0.73790761 | 0.485785995 | 0.991835066 | -5.038176623 |
| **A_55_P2087984** | Dnaja1 | 0.045429666 | -0.101746508 | 0.192605839 | 13.76381135 | 0.737753304 | 0.485873765 | 0.991835066 | -5.038263552 |
| **A_52_P583458** | E2f3 | 0.055388271 | -0.124117376 | 0.234893918 | 10.52251165 | 0.737477581 | 0.486030625 | 0.991835066 | -5.038418846 |
| **A_55_P2028922** | Stoml2 | 0.037375799 | -0.083759781 | 0.158511378 | 11.87888878 | 0.737442038 | 0.486050848 | 0.991835066 | -5.038438861 |
| **A_55_P2169198** | NA | -0.031419154 | -0.133253249 | 0.070414942 | 7.926732992 | -0.737412348 | 0.486067741 | 0.991835066 | -5.038455579 |
| **A_55_P2156093** | NA | -0.103877313 | -0.440575174 | 0.232820547 | 15.04829906 | -0.737376823 | 0.486087955 | 0.991835066 | -5.038475583 |
| **A_55_P1970270** | B3gnt2 | -0.050997851 | -0.216314675 | 0.114318972 | 9.069674038 | -0.737299529 | 0.486131937 | 0.991835066 | -5.038519103 |
| **A_55_P2139611** | Acbd4 | 0.036075099 | -0.080878396 | 0.153028594 | 8.639692847 | 0.73723078 | 0.486171059 | 0.991835066 | -5.038557809 |
| **A_51_P159415** | Sin3a | -0.034312154 | -0.145616391 | 0.076992083 | 9.460351862 | -0.736792885 | 0.486420297 | 0.991835066 | -5.038804278 |
| **A_55_P2074055** | NA | -0.03802808 | -0.16138916 | 0.085333001 | 8.636061657 | -0.736775822 | 0.486430011 | 0.991835066 | -5.03881388 |
| **A_55_P2027259** | Zcchc2 | 0.03630672 | -0.081478784 | 0.154092224 | 9.456075796 | 0.736723131 | 0.486460006 | 0.991835066 | -5.038843528 |
| **A_51_P511949** | Setd7 | 0.064067841 | -0.143857466 | 0.271993147 | 11.42080907 | 0.736447631 | 0.486616866 | 0.991835066 | -5.038998521 |
| **A_55_P1998892** | Smox | 0.060561262 | -0.135987477 | 0.257110001 | 10.77839738 | 0.736433925 | 0.486624671 | 0.991835066 | -5.039006231 |
| **A_51_P279465** | Farsa | -0.058764319 | -0.249497572 | 0.131968933 | 8.757217136 | -0.736370576 | 0.486660744 | 0.991835066 | -5.039041863 |
| **A_55_P2063216** | Dusp14 | 0.03922514 | -0.088093761 | 0.16654404 | 7.99004503 | 0.736343998 | 0.486675879 | 0.991835066 | -5.039056811 |
| **A_51_P246543** | Ebf3 | -0.045299134 | -0.192340285 | 0.101742016 | 10.38960047 | -0.736309058 | 0.486695777 | 0.991835066 | -5.039076463 |
| **A_52_P138806** | Dlgap3 | 0.041102023 | -0.09231585 | 0.174519896 | 8.248298492 | 0.736305962 | 0.48669754 | 0.991835066 | -5.039078204 |
| **A_55_P2112240** | Cdk5rap2 | -0.052378291 | -0.222409677 | 0.117653095 | 10.44371535 | -0.736260261 | 0.486723567 | 0.991835066 | -5.039103907 |
| **A_55_P1997751** | Prpf8 | 0.033009989 | -0.074166064 | 0.140186041 | 14.56587809 | 0.736133791 | 0.486795596 | 0.991835066 | -5.039175026 |
| **A_52_P637353** | Zbtb43 | -0.032304259 | -0.137201389 | 0.072592872 | 9.471277782 | -0.736046586 | 0.486845266 | 0.991835066 | -5.03922406 |
| **A_55_P2004447** | Fgl1 | -0.031834481 | -0.13520869 | 0.071539729 | 7.740094592 | -0.736028617 | 0.486855501 | 0.991835066 | -5.039234163 |
| **A_55_P2001198** | NA | 0.032966861 | -0.074086311 | 0.140020034 | 8.304579734 | 0.736015896 | 0.486862748 | 0.991835066 | -5.039241316 |
| **A_55_P2118675** | Tnip1 | 0.042155203 | -0.094739537 | 0.179049943 | 11.64079266 | 0.73599279 | 0.486875909 | 0.991835066 | -5.039254307 |
| **A_55_P1973360** | NA | 0.04934923 | -0.11092768 | 0.209626141 | 14.56960792 | 0.735899576 | 0.486929009 | 0.991835066 | -5.039306711 |
| **A_51_P342707** | Pold2 | 0.051225441 | -0.115163627 | 0.217614508 | 9.732346959 | 0.735817389 | 0.486975829 | 0.991835066 | -5.039352912 |
| **A_55_P2136935** | Rgs7bp | 0.047895154 | -0.107686965 | 0.203477273 | 8.970804648 | 0.735768217 | 0.487003843 | 0.991835066 | -5.039380551 |
| **A_55_P2147427** | Prdx1 | -0.059112644 | -0.251182335 | 0.132957048 | 13.34691741 | -0.735581295 | 0.487110345 | 0.991835066 | -5.039485606 |
| **A_55_P2126780** | Pgam5 | 0.053668863 | -0.120726006 | 0.228063732 | 12.21702358 | 0.735525652 | 0.487142052 | 0.991835066 | -5.039516875 |
| **A_66_P126984** | Gm10523 | 0.039601357 | -0.089093819 | 0.168296534 | 8.310390586 | 0.735456404 | 0.487181513 | 0.991835066 | -5.039555785 |
| **A_55_P2014640** | Knop1 | 0.05814917 | -0.130840302 | 0.247138641 | 11.11510425 | 0.735385459 | 0.487221943 | 0.991835066 | -5.039595647 |
| **A_55_P2073915** | Nos1 | -0.032157032 | -0.136681684 | 0.07236762 | 7.992190706 | -0.73530304 | 0.487268915 | 0.991835066 | -5.039641952 |
| **A_55_P1977468** | Dab2ip | 0.050035576 | -0.112604962 | 0.212676114 | 13.23319274 | 0.735290975 | 0.487275791 | 0.991835066 | -5.03964873 |
| **A_66_P106760** | Adam32 | -0.043161184 | -0.183465421 | 0.097143052 | 8.3601989 | -0.735244362 | 0.487302358 | 0.991835066 | -5.039674916 |
| **A_55_P2005080** | NA | -0.047779464 | -0.203122272 | 0.107563345 | 16.29463322 | -0.735121713 | 0.487372267 | 0.991835066 | -5.039743811 |
| **A_51_P143805** | Tmem42 | -0.034909839 | -0.148412741 | 0.078593063 | 11.03965144 | -0.735106068 | 0.487381185 | 0.991835066 | -5.039752599 |
| **A_55_P1998380** | Traf6 | 0.085836118 | -0.193304425 | 0.364976661 | 8.667797432 | 0.734947437 | 0.487471613 | 0.991835066 | -5.03984169 |
| **A_55_P2175405** | Zfp467 | 0.054999374 | -0.123975709 | 0.233974457 | 8.965339031 | 0.734470415 | 0.487743611 | 0.991835066 | -5.040109505 |
| **A_51_P109335** | Copa | 0.0379194 | -0.085480335 | 0.161319135 | 12.14545462 | 0.73444007 | 0.487760917 | 0.991835066 | -5.011387152 |
| **A_55_P2088720** | NA | 0.164613842 | -0.371105198 | 0.700332883 | 13.41702467 | 0.734409696 | 0.48777824 | 0.991835066 | -5.040143584 |
| **A_55_P2143494** | Doc2b | 0.032005647 | -0.072170123 | 0.136181417 | 7.794864416 | 0.734292375 | 0.487845155 | 0.991835066 | -5.040209425 |
| **A_55_P1966734** | Cyp2u1 | -0.066477542 | -0.282861535 | 0.149906451 | 14.30657168 | -0.734275364 | 0.487854858 | 0.991835066 | -5.040218972 |
| **A_55_P2046408** | NA | -0.057734074 | -0.245677744 | 0.130209596 | 8.712096542 | -0.734198741 | 0.487898564 | 0.991835066 | -5.040261967 |
| **A_51_P408082** | Apoa1 | -0.071751028 | -0.305349568 | 0.161847513 | 7.87514712 | -0.734120133 | 0.487943406 | 0.991835066 | -5.040306074 |
| **A_55_P2059323** | Gm13315 | 0.095828058 | -0.216226796 | 0.407882913 | 12.97203085 | 0.733957441 | 0.488036221 | 0.991835066 | -5.040397347 |
| **A_55_P2162875** | NA | 0.048806631 | -0.110141739 | 0.207755002 | 9.709510564 | 0.73389155 | 0.488073815 | 0.991835066 | -5.040434308 |
| **A_51_P429276** | Tmod3 | -0.077033626 | -0.328011249 | 0.173943996 | 10.01359373 | -0.73359181 | 0.488244856 | 0.991835066 | -5.040602412 |
| **A_51_P290074** | Fabp7 | -0.05795005 | -0.246788402 | 0.130888302 | 9.659947929 | -0.733453774 | 0.488323637 | 0.991835066 | -5.040679809 |

| | | | | | | | | | |
|---|---|---|---|---|---|---|---|---|---|
| **A_55_P1988952** | NA | 0.032608687 | -0.07368009 | 0.138897463 | 14.36948614 | 0.733254996 | 0.488437099 | 0.991835066 | -5.040791242 |
| **A_55_P2069530** | NA | -0.12249151 | -0.521785937 | 0.276802917 | 10.8958607 | -0.733199015 | 0.488469057 | 0.991835066 | -5.04082262 |
| **A_55_P1989251** | Chfr | 0.035410345 | -0.080026259 | 0.150846949 | 10.54996789 | 0.733154914 | 0.488494233 | 0.991835066 | -5.040847338 |
| **A_51_P383001** | Meox2 | -0.089234411 | -0.380184293 | 0.201715471 | 10.61021343 | -0.733032632 | 0.488564046 | 0.991835066 | -5.040915869 |
| **A_66_P116373** | Pwwp2b | -0.037077792 | -0.157996137 | 0.083840554 | 9.787701373 | -0.732876502 | 0.488653194 | 0.991835066 | -5.041003356 |
| **A_51_P464576** | Psen1 | -0.041329788 | -0.17611559 | 0.093456015 | 9.208555895 | -0.732872049 | 0.488655736 | 0.991835066 | -5.041005851 |
| **A_55_P2070913** | Rpl14 | 0.047688951 | -0.107837484 | 0.203215387 | 16.72174623 | 0.732862813 | 0.48866101 | 0.991835066 | -5.041011026 |
| **A_52_P342620** | Csrnp2 | 0.042699015 | -0.096603974 | 0.182002004 | 8.275177372 | 0.732599391 | 0.488811445 | 0.991835066 | -5.041158597 |
| **A_55_P1989748** | NA | 0.038715357 | -0.087615746 | 0.16504646 | 8.14593341 | 0.732456965 | 0.488892795 | 0.991835066 | -5.041238368 |
| **A_55_P2094034** | Nrg2 | 0.037289911 | -0.084418087 | 0.15899791 | 7.878542841 | 0.732287064 | 0.488989849 | 0.991835066 | -5.04133351 |
| **A_55_P2049549** | Rab22a | -0.06898126 | -0.294169463 | 0.156206942 | 10.27999676 | -0.732140822 | 0.489073398 | 0.991835066 | -5.041415389 |
| **A_51_P375201** | Plk3 | 0.036230286 | -0.08208007 | 0.154540642 | 8.516577404 | 0.731910766 | 0.489204851 | 0.991835066 | -5.041544168 |
| **A_55_P2176565** | B3galt5 | -0.030124694 | -0.128499042 | 0.068249655 | 7.660420489 | -0.731896889 | 0.489212781 | 0.991835066 | -5.041551935 |
| **A_55_P2067757** | Psg27 | -0.263287774 | -1.123205637 | 0.596630089 | 8.154811128 | -0.731783907 | 0.489277348 | 0.991835066 | -5.041615167 |
| **A_55_P2086536** | Safb2 | 0.043946769 | -0.099597471 | 0.187491009 | 8.439999287 | 0.731729075 | 0.489308685 | 0.991835066 | -5.041645851 |
| **A_55_P2073010** | Gm4832 | -0.037699311 | -0.160841399 | 0.085442777 | 13.67085229 | -0.731705019 | 0.489322434 | 0.991835066 | -5.041659312 |
| **A_65_P01418** | Wnk2 | 0.076982994 | -0.174648401 | 0.328614388 | 10.51423005 | 0.731204918 | 0.489608316 | 0.991835066 | -5.041939078 |
| **A_55_P2169010** | Ndfip1 | -0.044824916 | -0.19136381 | 0.101713979 | 10.9148911 | -0.73109818 | 0.489669347 | 0.991835066 | -5.041998769 |
| **A_51_P467751** | Hgsnat | 0.03735156 | -0.084793782 | 0.159496902 | 10.4735279 | 0.730871405 | 0.48979903 | 0.991835066 | -5.042125566 |
| **A_51_P258698** | Phf8 | 0.032972894 | -0.074873626 | 0.140819413 | 10.53965774 | 0.730735256 | 0.489876899 | 0.991835066 | -5.042201675 |
| **A_52_P660693** | Dsel | 0.067984403 | -0.154432899 | 0.290401706 | 9.373236053 | 0.730549844 | 0.489982956 | 0.991835066 | -5.042305305 |
| **A_51_P254912** | Rfc2 | 0.038647435 | -0.087812317 | 0.165107188 | 10.88822313 | 0.730428123 | 0.49005259 | 0.991835066 | -5.042373325 |
| **A_51_P352594** | St5 | -0.060061669 | -0.256603297 | 0.136479959 | 9.124877962 | -0.730385224 | 0.490077133 | 0.991835066 | -5.042397296 |
| **A_55_P2112355** | NA | -0.057015206 | -0.243602656 | 0.129572245 | 13.85391511 | -0.730327066 | 0.490110407 | 0.991835066 | -5.042429791 |
| **A_55_P2120961** | NA | -0.050505251 | -0.215798434 | 0.114787932 | 10.6749461 | -0.730282212 | 0.490136072 | 0.991835066 | -5.042454852 |
| **A_55_P2143279** | Nkd2 | 0.082308596 | -0.18708279 | 0.351699982 | 9.728006076 | 0.73024841 | 0.490155412 | 0.991835066 | -5.042473736 |
| **A_55_P1991406** | Fbxo34 | 0.041045279 | -0.09331356 | 0.175404117 | 9.609760039 | 0.730139932 | 0.490217484 | 0.991835066 | -5.042534336 |
| **A_52_P407854** | Dda1 | 0.069616492 | -0.158304562 | 0.297537547 | 8.589253091 | 0.730023474 | 0.490284129 | 0.991835066 | -5.042599386 |
| **A_55_P2055869** | Dvl2 | 0.047550978 | -0.108141072 | 0.203243028 | 9.232385767 | 0.729965193 | 0.490317482 | 0.991835066 | -5.042631936 |
| **A_51_P368496** | Tmem98 | -0.060425869 | -0.258273299 | 0.137421561 | 11.51827699 | -0.729964309 | 0.490317988 | 0.991835066 | -5.04263243 |
| **A_55_P2036863** | Rabl2 | 0.062973273 | -0.143248098 | 0.269194644 | 10.36456891 | 0.729846832 | 0.490385225 | 0.991835066 | -5.042698037 |
| **A_55_P2168321** | NA | 0.053599169 | -0.121945441 | 0.229143779 | 8.267497613 | 0.729759386 | 0.490435278 | 0.991835066 | -5.042746866 |
| **A_51_P378558** | Rpap2 | -0.049763429 | -0.212747295 | 0.113220436 | 8.971240785 | -0.729751201 | 0.490439963 | 0.991835066 | -5.042751436 |
| **A_55_P2069714** | Sqstm1 | -0.092777702 | -0.396708865 | 0.211153461 | 9.483141543 | -0.729587713 | 0.490533551 | 0.991835066 | -5.042842714 |
| **A_52_P570543** | Sdk2 | 0.034463122 | -0.078458568 | 0.147384811 | 8.011921898 | 0.729434629 | 0.490621193 | 0.991835066 | -5.042928168 |
| **A_55_P1962289** | NA | 0.093663557 | -0.213260145 | 0.400587258 | 9.382899495 | 0.729372443 | 0.490656798 | 0.991835066 | -5.042962877 |
| **A_55_P2179796** | NA | -0.04646021 | -0.198705782 | 0.105785363 | 14.14609701 | -0.729366192 | 0.490660377 | 0.991835066 | -5.042966365 |
| **A_52_P448006** | Nit2 | -0.043323203 | -0.185289756 | 0.098643351 | 10.22071372 | -0.729362864 | 0.490662283 | 0.991835066 | -5.042968223 |
| **A_55_P2025155** | NA | -0.055985438 | -0.239496535 | 0.12752566 | 13.08745537 | -0.729158391 | 0.490779369 | 0.991835066 | -5.043082331 |
| **A_52_P229316** | Galnt4 | 0.0406727 | -0.092675391 | 0.174020791 | 9.046358592 | 0.728996311 | 0.490872194 | 0.991835066 | -5.043172763 |
| **A_55_P1958354** | Rps29 | 0.041660856 | -0.094964949 | 0.178286661 | 17.2279898 | 0.728793658 | 0.490988271 | 0.991835066 | -5.04328581 |
| **A_51_P475501** | Ndufa8 | -0.062107133 | -0.265811412 | 0.141597146 | 14.18543527 | -0.728702837 | 0.491040298 | 0.991835066 | -5.043336465 |
| **A_55_P2037111** | Ran | 0.081249534 | -0.185252251 | 0.34775132 | 11.21943921 | 0.728668323 | 0.49106007 | 0.991835066 | -5.043355713 |
| **A_55_P2156895** | Tmem39b | 0.037467212 | -0.085432007 | 0.160366431 | 8.938601086 | 0.728637281 | 0.491077854 | 0.991835066 | -5.043373025 |
| **A_51_P208121** | Klhl25 | 0.039136577 | -0.089266396 | 0.167539549 | 8.307300512 | 0.728478754 | 0.49116868 | 0.991835066 | -5.043461423 |
| **A_66_P102954** | Gm2155 | -0.030383977 | -0.130092923 | 0.069324969 | 7.897028858 | -0.728315623 | 0.491262156 | 0.991835066 | -5.043552373 |
| **A_66_P100565** | Ccdc107 | -0.047545248 | -0.203636207 | 0.108545712 | 12.96628853 | -0.728011932 | 0.491436205 | 0.991835066 | -5.043721644 |
| **A_52_P511381** | Tesk1 | 0.06039642 | -0.137886456 | 0.258679296 | 11.43631324 | 0.728006273 | 0.491439448 | 0.991835066 | -5.043724798 |

| | | | | | | | | | |
|---|---|---|---|---|---|---|---|---|---|
| **A_55_P2103902** | Mrpl4 | -0.09380319 | -0.401774104 | 0.214167725 | 11.52095289 | -0.727975956 | 0.491456826 | 0.991835066 | -5.043741692 |
| **A_55_P1981884** | Ercc6l2 | -0.063705798 | -0.272873788 | 0.145462191 | 9.123138014 | -0.727935456 | 0.49148004 | 0.991835066 | -5.043764261 |
| **A_51_P188795** | Akr7a5 | -0.08678527 | -0.371760665 | 0.198190125 | 9.55343248 | -0.72785989 | 0.491523357 | 0.991835066 | -5.043806368 |
| **A_55_P2167922** | Dmd | 0.057799303 | -0.132031004 | 0.247629611 | 8.830521716 | 0.727723139 | 0.491601754 | 0.991835066 | -5.043882558 |
| **A_55_P2065513** | Ebf1 | -0.048041985 | -0.205852746 | 0.109768776 | 9.391389734 | -0.727601289 | 0.491671614 | 0.991835066 | -5.043950437 |
| **A_55_P2169396** | NA | 0.040932563 | -0.093526401 | 0.175391527 | 15.56661023 | 0.727592669 | 0.491676557 | 0.991835066 | -5.043955239 |
| **A_55_P2044533** | NA | -0.086403108 | -0.370239762 | 0.197433546 | 8.628763177 | -0.72756202 | 0.49169413 | 0.991835066 | -5.043972311 |
| **A_55_P2117333** | NA | 0.043761969 | -0.100010624 | 0.187534562 | 9.689968853 | 0.727494783 | 0.491732683 | 0.991835066 | -5.04400976 |
| **A_55_P1991728** | Zfp956 | -0.049452328 | -0.211931416 | 0.113026761 | 8.947655625 | -0.727442037 | 0.491762929 | 0.991835066 | -5.044039138 |
| **A_52_P556290** | Arih1 | 0.067388647 | -0.154027878 | 0.288805173 | 9.626479747 | 0.72742101 | 0.491774987 | 0.991835066 | -5.044050848 |
| **A_55_P1987761** | Prkcz | 0.030465757 | -0.069649719 | 0.130581233 | 8.013459412 | 0.727310545 | 0.491838335 | 0.991835066 | -5.044112364 |
| **A_55_P2144781** | Erich5 | -0.07268406 | -0.311538147 | 0.166170027 | 9.190277306 | -0.727303426 | 0.491842418 | 0.991835066 | -5.044116328 |
| **A_55_P2083904** | Tmem185a | 0.038047588 | -0.086993575 | 0.163088752 | 8.466387059 | 0.727249213 | 0.491873509 | 0.991835066 | -5.044146516 |
| **A_55_P1965902** | Pi4k2a | -0.055802267 | -0.239193836 | 0.127589301 | 9.597915924 | -0.727246452 | 0.491875092 | 0.991835066 | -5.044148053 |
| **A_52_P295533** | Slc25a28 | 0.043580277 | -0.09965138 | 0.186811934 | 9.814856769 | 0.727210433 | 0.491895751 | 0.991835066 | -5.044168108 |
| **A_55_P2007656** | Cryab | -0.109796538 | -0.470676124 | 0.251083047 | 12.74016854 | -0.727169181 | 0.491919411 | 0.991835066 | -5.044191076 |
| **A_55_P1980165** | NA | -0.03591758 | -0.153995989 | 0.08216083 | 15.34910574 | -0.727018901 | 0.492005609 | 0.991835066 | -5.04427474 |
| **A_51_P485479** | Prrc2a | 0.055181491 | -0.126245623 | 0.236608604 | 10.00138702 | 0.726943019 | 0.492049138 | 0.991835066 | -5.044316979 |
| **A_52_P572447** | Agpat5 | -0.073496166 | -0.315174387 | 0.168182055 | 10.45517633 | -0.726835787 | 0.492110655 | 0.991835066 | -5.044376663 |
| **A_52_P399677** | Tprkb | -0.034165404 | -0.14651504 | 0.078184233 | 8.385633854 | -0.726815227 | 0.49212245 | 0.991835066 | -5.044388106 |
| **A_55_P1978103** | Ppip5k2 | -0.09339818 | -0.400536561 | 0.213740202 | 11.7863326 | -0.726797547 | 0.492132594 | 0.991835066 | -5.044397945 |
| **A_52_P337259** | Heyl | -0.062346741 | -0.267374575 | 0.142681093 | 9.975274777 | -0.726791882 | 0.492135844 | 0.991835066 | -5.044401098 |
| **A_55_P2121827** | Dgcr2 | 0.100273879 | -0.229498133 | 0.43004589 | 9.213467883 | 0.726746841 | 0.492161685 | 0.991835066 | -5.044426164 |
| **A_55_P2075013** | Tpt1 | 0.041433983 | -0.09483216 | 0.177700125 | 16.78220526 | 0.726737971 | 0.492166774 | 0.991835066 | -5.0444311 |
| **A_55_P2104332** | Pxdn | 0.078209658 | -0.179008714 | 0.33542803 | 10.23672487 | 0.726720704 | 0.492176682 | 0.991835066 | -5.044440709 |
| **A_55_P2072980** | Sptan1 | 0.11384347 | -0.26068618 | 0.48837312 | 9.049736605 | 0.72649236 | 0.492307706 | 0.991835066 | -5.044567762 |
| **A_52_P60353** | Greb1l | 0.050168524 | -0.114915709 | 0.215252758 | 8.262036983 | 0.726331471 | 0.492400039 | 0.991835066 | -5.044657264 |
| **A_55_P1968618** | NA | 0.033574508 | -0.076918554 | 0.14406757 | 7.880621933 | 0.726246048 | 0.492449067 | 0.991835066 | -5.044704777 |
| **A_55_P2127425** | Sys1 | 0.062603405 | -0.143429886 | 0.268636697 | 10.7070585 | 0.726222481 | 0.492462594 | 0.991835066 | -5.044717885 |
| **A_55_P2161077** | NA | 0.055963218 | -0.128227354 | 0.240153789 | 9.364580491 | 0.726180217 | 0.492486853 | 0.991835066 | -5.044741391 |
| **A_55_P2004588** | Nabp2 | 0.036862504 | -0.084483779 | 0.158208786 | 11.05403053 | 0.726051598 | 0.492560683 | 0.991835066 | -5.044812916 |
| **A_55_P2183493** | Spcs3 | 0.076056742 | -0.174333335 | 0.326446818 | 8.934012071 | 0.725988491 | 0.49259691 | 0.991835066 | -5.044848007 |
| **A_55_P2070576** | Ccl27a | -0.054932129 | -0.235805784 | 0.125941527 | 9.399068426 | -0.725872337 | 0.492663595 | 0.991835066 | -5.044912588 |
| **A_55_P2157872** | Srsf5 | 0.08627468 | -0.197869975 | 0.370419335 | 11.09116372 | 0.725693111 | 0.492766502 | 0.991835066 | -5.04501222 |
| **A_52_P546552** | Pelp1 | 0.062080275 | -0.142407249 | 0.266567799 | 9.963118638 | 0.725597783 | 0.492821242 | 0.991835066 | -5.045065204 |
| **A_51_P329356** | Mcts2 | -0.035122996 | -0.150817764 | 0.080571772 | 11.22686567 | -0.725582785 | 0.492829855 | 0.991835066 | -5.04507354 |
| **A_51_P115817** | Ralgps2 | -0.052571905 | -0.2258151 | 0.120671289 | 9.520670164 | -0.72528161 | 0.493002827 | 0.991835066 | -5.029463979 |
| **A_52_P424784** | Clstn2 | -0.041065383 | -0.176394418 | 0.094263651 | 8.310443752 | -0.725260507 | 0.493014949 | 0.991835066 | -5.045252622 |
| **A_66_P129747** | Dnajc24 | 0.031567197 | -0.072473849 | 0.135608242 | 10.37157204 | 0.725171021 | 0.493066352 | 0.991835066 | -5.045302336 |
| **A_55_P1997894** | NA | -0.041191424 | -0.176971025 | 0.094588176 | 15.26839279 | -0.725072468 | 0.493122967 | 0.991835066 | -5.045357081 |
| **A_55_P2153827** | Fgfr1op | 0.060846066 | -0.139759468 | 0.261451599 | 10.4899096 | 0.724934428 | 0.493202273 | 0.991835066 | -5.045433751 |
| **A_55_P2184449** | NA | 0.039498863 | -0.090736215 | 0.169733941 | 7.732666265 | 0.72487939 | 0.493233896 | 0.991835066 | -5.045464316 |
| **A_55_P2037797** | Tex44 | -0.085064741 | -0.365542206 | 0.195412724 | 9.338865718 | -0.724871027 | 0.493238701 | 0.991835066 | -5.045468961 |
| **A_55_P2034081** | Hrh1 | -0.031717334 | -0.136302447 | 0.072867779 | 7.945795737 | -0.724829631 | 0.493262487 | 0.991835066 | -5.045491949 |
| **A_55_P2018480** | Carnmt1 | -0.042792763 | -0.183920261 | 0.098334736 | 11.37625924 | -0.72471594 | 0.493327816 | 0.991835066 | -5.045555078 |
| **A_52_P81468** | Psma4 | -0.045864912 | -0.197129803 | 0.10539998 | 11.82807782 | -0.724688806 | 0.493343408 | 0.991835066 | -5.045570143 |
| **A_52_P387564** | Tmpo | -0.042787437 | -0.183923371 | 0.098348498 | 12.57850165 | -0.724582429 | 0.493404541 | 0.991835066 | -5.045629203 |
| **A_55_P2043117** | Ccdc138 | -0.053136034 | -0.228409153 | 0.122137085 | 13.96011779 | -0.724574344 | 0.493409187 | 0.991835066 | -5.045633691 |

| | | | | | | | | | |
|---|---|---|---|---|---|---|---|---|---|
| **A_51_P342082** | Cnpy2 | 0.045374009 | -0.104314771 | 0.19506279 | 12.56079736 | 0.724481066 | 0.493462797 | 0.991835066 | -5.045685472 |
| **A_51_P414243** | Pomgnt2 | 0.043348031 | -0.099710579 | 0.186406642 | 11.38992004 | 0.724209981 | 0.493618619 | 0.991835066 | -5.045835926 |
| **A_51_P135389** | Zfp474 | -0.029261808 | -0.125838176 | 0.067314561 | 7.632308786 | -0.724168132 | 0.493642677 | 0.991835066 | -5.045859148 |
| **A_51_P219385** | Six5 | 0.041988237 | -0.096604542 | 0.180581017 | 9.443520419 | 0.724096041 | 0.493684122 | 0.991835066 | -5.045899149 |
| **A_55_P2154709** | Pter | 0.070953226 | -0.163251908 | 0.30515836 | 9.536812107 | 0.72407719 | 0.49369496 | 0.991835066 | -5.045909609 |
| **A_55_P2078955** | Aqp8 | 0.034184436 | -0.078657566 | 0.147026438 | 7.710278999 | 0.724047017 | 0.493712308 | 0.991835066 | -5.045926349 |
| **A_55_P1968285** | Tle5 | -0.065274351 | -0.280781429 | 0.150232726 | 14.97494733 | -0.723919306 | 0.493785738 | 0.991835066 | -5.045997201 |
| **A_51_P176387** | Hook3 | -0.039434491 | -0.169651036 | 0.090782054 | 8.532393859 | -0.723801039 | 0.493853744 | 0.991835066 | -5.046062804 |
| **A_55_P1997931** | Map9 | -0.030560492 | -0.131479373 | 0.070358389 | 10.19054618 | -0.723764125 | 0.493874972 | 0.991835066 | -5.046083279 |
| **A_52_P284889** | Prkcz | 0.045020469 | -0.103719066 | 0.193760004 | 8.121500334 | 0.723423686 | 0.494070773 | 0.991835066 | -5.046272065 |
| **A_51_P135357** | Sike1 | -0.033689658 | -0.145009663 | 0.077630348 | 8.742436608 | -0.723323394 | 0.494128465 | 0.991835066 | -5.046327667 |
| **A_55_P2339379** | Sass6 | 0.034867766 | -0.080352838 | 0.15008837 | 8.288456492 | 0.723274422 | 0.494156637 | 0.991835066 | -5.046354815 |
| **A_66_P106814** | NA | -0.054433725 | -0.234324148 | 0.125456697 | 12.48880586 | -0.723217868 | 0.494189172 | 0.991835066 | -5.046386164 |
| **A_51_P314907** | Dbf4 | -0.055245122 | -0.237817815 | 0.127327571 | 8.707446229 | -0.7232147 | 0.494190995 | 0.991835066 | -5.04638792 |
| **A_55_P1957957** | Dhx34 | 0.048619928 | -0.112065686 | 0.209305543 | 9.757728048 | 0.723180069 | 0.494210919 | 0.991835066 | -5.046407116 |
| **A_55_P2015692** | Elob | -0.079562896 | -0.342539491 | 0.183413698 | 13.99057023 | -0.72310709 | 0.494252907 | 0.991835066 | -5.046447565 |
| **A_52_P252425** | 2700062C0 | 0.032724058 | -0.075456731 | 0.140904848 | 8.665086838 | 0.722979768 | 0.494326166 | 0.991835066 | -5.046518126 |
| **A_51_P201137** | Lbr | -0.046461331 | -0.20009435 | 0.107171688 | 9.97699899 | -0.722796794 | 0.49443146 | 0.991835066 | -5.046619512 |
| **A_55_P2067153** | NA | -0.074972582 | -0.322948832 | 0.173003668 | 15.22958808 | -0.722605914 | 0.494541319 | 0.991835066 | -5.046725256 |
| **A_52_P425860** | Iqsec1 | 0.035724387 | -0.08247016 | 0.153918934 | 10.91899895 | 0.722397908 | 0.494661053 | 0.991835066 | -5.046840461 |
| **A_55_P2072611** | 1-Mar | -0.032170151 | -0.13860887 | 0.074268568 | 7.738306745 | -0.722374803 | 0.494674354 | 0.991835066 | -5.046853257 |
| **A_55_P2162537** | Acrbp | 0.043481063 | -0.100389349 | 0.187351475 | 8.349526153 | 0.722333566 | 0.494698093 | 0.991835066 | -5.046876092 |
| **A_55_P1961391** | NA | 0.039401503 | -0.090988674 | 0.16979168 | 7.966303287 | 0.722232533 | 0.49475626 | 0.991835066 | -5.046932036 |
| **A_55_P2121662** | Mapk3 | -0.096818879 | -0.417294603 | 0.223656844 | 9.43434721 | -0.722061264 | 0.494854875 | 0.991835066 | -5.047026857 |
| **A_55_P2203051** | Gt(ROSA)2 | 0.099579539 | -0.230048122 | 0.429207201 | 10.48575485 | 0.722030587 | 0.494872539 | 0.991835066 | -5.047043839 |
| **A_55_P2184925** | Tmem185b | 0.050728058 | -0.117201408 | 0.218657525 | 10.44790762 | 0.721988805 | 0.4948966 | 0.991835066 | -5.047066968 |
| **A_55_P2074864** | NA | 0.047752438 | -0.110372728 | 0.205877605 | 12.85103931 | 0.72177808 | 0.495017956 | 0.991835066 | -5.047183598 |
| **A_51_P308844** | Nrn1 | -0.053394753 | -0.230216554 | 0.123427049 | 8.304080131 | -0.721725251 | 0.495048384 | 0.991835066 | -5.047212833 |
| **A_55_P2097869** | Frem2 | -0.053103273 | -0.228975958 | 0.122769413 | 9.530963726 | -0.721658988 | 0.49508655 | 0.991835066 | -5.0472495 |
| **A_51_P242076** | Ubr7 | -0.042030812 | -0.181244352 | 0.097182729 | 11.54987971 | -0.721598183 | 0.495121575 | 0.991835066 | -5.047283144 |
| **A_52_P589550** | Nr2c1 | -0.044810145 | -0.193276061 | 0.10365577 | 8.477190255 | -0.721371075 | 0.495252407 | 0.991835066 | -5.047408785 |
| **A_55_P1977583** | Anxa11 | 0.050232341 | -0.116211112 | 0.216675794 | 11.7982236 | 0.721316442 | 0.495283883 | 0.991835066 | -5.047439005 |
| **A_55_P2219903** | AW061147 | -0.051513834 | -0.22227192 | 0.119244252 | 10.79170941 | -0.721027328 | 0.495450475 | 0.991835066 | -5.047598892 |
| **A_55_P2151308** | G430095P1 | 0.043952634 | -0.101746254 | 0.189651523 | 8.930960633 | 0.721004216 | 0.495463795 | 0.991835066 | -5.047611672 |
| **A_51_P232474** | Zswim1 | 0.030953008 | -0.071686714 | 0.13359273 | 8.086414728 | 0.720769707 | 0.495598952 | 0.991835066 | -5.04774132 |
| **A_55_P1971938** | Atp2b2 | -0.050148735 | -0.216462196 | 0.116164725 | 8.100580919 | -0.720678751 | 0.49565138 | 0.991835066 | -5.047791596 |
| **A_55_P2055324** | Spint1 | -0.032516394 | -0.140395455 | 0.075362667 | 7.726115945 | -0.720401072 | 0.495811462 | 0.991835066 | -5.04794505 |
| **A_55_P2120303** | NA | -0.039565165 | -0.17084507 | 0.09171474 | 8.129000862 | -0.720317316 | 0.495859754 | 0.991835066 | -5.047991327 |
| **A_55_P2143075** | Ubc | -0.09356027 | -0.404051965 | 0.216931425 | 15.57911792 | -0.720195844 | 0.495929797 | 0.991835066 | -5.048058435 |
| **A_51_P321651** | Fam192a | 0.036744435 | -0.085199694 | 0.158688565 | 10.45762805 | 0.720177936 | 0.495940124 | 0.991835066 | -5.048068328 |
| **A_55_P2089233** | Pou2af1 | -0.028860337 | -0.124642844 | 0.06692217 | 7.677918554 | -0.720152258 | 0.495954931 | 0.991835066 | -5.048082512 |
| **A_52_P568805** | Ubxn1 | 0.073343619 | -0.170122602 | 0.31680984 | 13.30701079 | 0.720000429 | 0.496042492 | 0.991835066 | -5.048166374 |
| **A_51_P485899** | Srsf1 | -0.043264992 | -0.186896342 | 0.100366357 | 11.32279157 | -0.719940364 | 0.496077134 | 0.991835066 | -5.048199547 |
| **A_55_P2155942** | Ezh1 | -0.062704898 | -0.270890379 | 0.145480583 | 9.330097711 | -0.719880084 | 0.496111902 | 0.991835066 | -5.048232836 |
| **A_55_P2154054** | Myo18a | -0.047648534 | -0.205850546 | 0.110553478 | 12.33634308 | -0.719857728 | 0.496124797 | 0.991835066 | -5.048245181 |
| **A_55_P2108864** | Fer | -0.031305688 | -0.135249618 | 0.072638242 | 8.598210053 | -0.719835482 | 0.496137628 | 0.991835066 | -5.048257465 |
| **A_51_P461108** | Osbpl10 | 0.035540439 | -0.082486232 | 0.15356711 | 8.551251442 | 0.719700436 | 0.496215528 | 0.991835066 | -5.048332031 |
| **A_55_P1952725** | NA | -0.070458036 | -0.304492296 | 0.163576225 | 7.763506033 | -0.719548751 | 0.496303036 | 0.991835066 | -5.04841577 |

| | | | | | | | | | |
|---|---|---|---|---|---|---|---|---|---|
| **A_55_P1973339** | Mapk1 | 0.121832105 | -0.282928834 | 0.526593044 | 10.25940435 | 0.71940307 | 0.496387089 | 0.991835066 | -5.048496181 |
| **A_55_P1995507** | Aqp11 | -0.047317205 | -0.204530821 | 0.10989641 | 8.284021361 | -0.719346382 | 0.496419799 | 0.991835066 | -5.048527468 |
| **A_51_P464308** | Gnb4 | 0.055468413 | -0.128834115 | 0.23977094 | 9.497391547 | 0.719322389 | 0.496433644 | 0.991835066 | -5.048540709 |
| **A_55_P2003378** | Atrnl1 | 0.036584889 | -0.085021639 | 0.158191417 | 8.251239408 | 0.719041545 | 0.496595718 | 0.991835066 | -5.048695674 |
| **A_55_P2032936** | Cisd1 | -0.063789666 | -0.275867869 | 0.148288537 | 13.36839756 | -0.718891628 | 0.49668225 | 0.991835066 | -5.048778376 |
| **A_55_P2021465** | Phldb2 | 0.057199297 | -0.133016704 | 0.247415298 | 12.10348791 | 0.718708494 | 0.496787967 | 0.991835066 | -5.048879383 |
| **A_55_P2099141** | NA | -0.048771644 | -0.210969274 | 0.113425985 | 14.84208052 | -0.718674156 | 0.496807791 | 0.991835066 | -5.048898319 |
| **A_52_P377723** | Peg10 | 0.034632514 | -0.080561277 | 0.149826304 | 7.983170821 | 0.71856172 | 0.496872706 | 0.991835066 | -5.04896032 |
| **A_55_P2009875** | NA | -0.072712054 | -0.314595119 | 0.169171011 | 11.01908719 | -0.718472389 | 0.496924285 | 0.991835066 | -5.049009574 |
| **A_51_P216496** | Tfpi2 | -0.030067613 | -0.130099716 | 0.06996449 | 7.784967145 | -0.718403898 | 0.496963834 | 0.991835066 | -5.049047335 |
| **A_51_P346964** | Mrap | -0.076771191 | -0.332181899 | 0.178639516 | 11.49527786 | -0.718403292 | 0.496964183 | 0.991835066 | -5.049047669 |
| **A_52_P330984** | Lin9 | 0.042348668 | -0.098560502 | 0.183257838 | 9.424000553 | 0.718306221 | 0.497020239 | 0.991835066 | -5.049101181 |
| **A_51_P234714** | Ceacam14 | -0.150086598 | -0.649731969 | 0.349558773 | 7.894188137 | -0.717941698 | 0.497230775 | 0.991835066 | -5.049302078 |
| **A_55_P2057777** | Fgfr1 | 0.098550968 | -0.22957629 | 0.426678226 | 11.53596803 | 0.717840105 | 0.497289462 | 0.991835066 | -5.049358053 |
| **A_55_P2107528** | NA | 0.044786065 | -0.104330589 | 0.193902719 | 7.698894072 | 0.717837078 | 0.497291211 | 0.991835066 | -5.049359721 |
| **A_66_P131324** | Platr21 | 0.030941866 | -0.072093397 | 0.133977129 | 7.673322618 | 0.717744286 | 0.497344819 | 0.991835066 | -5.049410841 |
| **A_55_P1985005** | Ghrh | 0.046692899 | -0.108809218 | 0.202195016 | 7.783124904 | 0.717668141 | 0.497388812 | 0.991835066 | -5.049452786 |
| **A_52_P293556** | Wdr41 | -0.054983233 | -0.238161347 | 0.12819488 | 9.500322205 | -0.717407359 | 0.497539498 | 0.991835066 | -5.049596413 |
| **A_51_P132978** | Idh1 | 0.039399911 | -0.091863746 | 0.170663568 | 8.665835029 | 0.71739752 | 0.497545184 | 0.991835066 | -5.049601831 |
| **A_51_P506312** | Cdkn2aipnl | 0.038368757 | -0.089468856 | 0.166206369 | 11.32799915 | 0.717345221 | 0.497575408 | 0.991835066 | -5.04963063 |
| **A_52_P88818** | Clybl | -0.097750616 | -0.423656252 | 0.228155019 | 9.359568516 | -0.716863991 | 0.497853567 | 0.991835066 | -5.049895539 |
| **A_51_P165683** | Hirip3 | 0.038087034 | -0.088903257 | 0.165077326 | 10.98169317 | 0.716829337 | 0.497873602 | 0.991835066 | -5.04991461 |
| **A_55_P2329313** | C530014P2 | 0.194356361 | -0.453737973 | 0.842450695 | 9.863029949 | 0.716753243 | 0.497917596 | 0.991835066 | -5.049956483 |
| **A_66_P117251** | Snx6 | -0.04551816 | -0.197309065 | 0.106272745 | 11.99471824 | -0.716717617 | 0.497938195 | 0.991835066 | -5.049976087 |
| **A_55_P2107247** | Tssk5 | 0.202944098 | -0.473828389 | 0.879716584 | 8.270313519 | 0.716709007 | 0.497943173 | 0.991835066 | -5.049980824 |
| **A_55_P2017112** | Mpnd | -0.071043772 | -0.307963571 | 0.165876027 | 11.31889176 | -0.716694032 | 0.497951832 | 0.991835066 | -5.049989064 |
| **A_55_P2258832** | 5230400M( | 0.069045914 | -0.16130275 | 0.299394578 | 8.327514791 | 0.716409633 | 0.498116289 | 0.991835066 | -5.050145523 |
| **A_52_P546660** | 1700029I15 | 0.030144953 | -0.070435561 | 0.130725468 | 8.098761437 | 0.716324646 | 0.498165441 | 0.991835066 | -5.050192268 |
| **A_55_P2210213** | 4732423E2 | 0.144069471 | -0.336634614 | 0.624773557 | 9.915803599 | 0.71631374 | 0.498171749 | 0.991835066 | -5.050198266 |
| **A_55_P2214841** | Irs3 | -0.030880106 | -0.133941928 | 0.072181716 | 7.853837484 | -0.716127086 | 0.498279711 | 0.991835066 | -5.050300912 |
| **A_55_P2142218** | NA | 0.050779768 | -0.118722629 | 0.220282164 | 14.32790971 | 0.7160181 | 0.498342757 | 0.991835066 | -5.050360837 |
| **A_55_P2174506** | Stx17 | 0.037068339 | -0.086688343 | 0.160825021 | 8.400191162 | 0.715885564 | 0.498419433 | 0.991835066 | -5.0504337 |
| **A_52_P39157** | Bmi1 | -0.05648012 | -0.245045864 | 0.132085623 | 9.823698989 | -0.715882829 | 0.498421015 | 0.991835066 | -5.050435203 |
| **A_55_P2216996** | Tmem87a | -0.033465519 | -0.145197049 | 0.078266012 | 8.197741764 | -0.715864695 | 0.498431507 | 0.991835066 | -5.050445172 |
| **A_51_P115005** | Edn1 | 0.098694729 | -0.230818894 | 0.428208352 | 8.614347547 | 0.715862672 | 0.498432677 | 0.991835066 | -5.045157822 |
| **A_55_P2007413** | Agps | 0.054303233 | -0.127019244 | 0.235625709 | 11.4756576 | 0.715785958 | 0.498477063 | 0.991835066 | -5.050488452 |
| **A_51_P212801** | Ube2a | -0.042380345 | -0.183892052 | 0.099131362 | 10.65659147 | -0.715782786 | 0.498478898 | 0.991835066 | -5.050490196 |
| **A_52_P113672** | Chml | 0.061233223 | -0.14326152 | 0.265727965 | 10.17120641 | 0.715672125 | 0.49854293 | 0.991835066 | -5.050551018 |
| **A_55_P2053798** | NA | 0.039945594 | -0.093464314 | 0.173355502 | 14.41181948 | 0.715632294 | 0.498565978 | 0.991835066 | -5.050572907 |
| **A_55_P2111870** | Rogdi | -0.049839935 | -0.216306479 | 0.116626609 | 8.49731535 | -0.71558237 | 0.498594868 | 0.991835066 | -5.050600343 |
| **A_52_P650855** | Myo1d | -0.10965866 | -0.476044933 | 0.256727613 | 11.07430072 | -0.715340599 | 0.498734792 | 0.991835066 | -5.050733185 |
| **A_51_P519251** | Nupr1 | 0.155856811 | -0.364916466 | 0.676630088 | 9.338949251 | 0.715296711 | 0.498760195 | 0.991835066 | -5.050757296 |
| **A_55_P2174022** | Brd4 | -0.054833926 | -0.238056474 | 0.128388622 | 12.49950763 | -0.71528572 | 0.498766557 | 0.991835066 | -5.050763333 |
| **A_51_P333923** | Tspan1 | 0.030472005 | -0.071390863 | 0.132334873 | 7.684456436 | 0.714980615 | 0.49894318 | 0.991835066 | -5.050930912 |
| **A_55_P2140288** | Phka2 | -0.059175507 | -0.256991798 | 0.138640785 | 11.10111798 | -0.714972047 | 0.49894814 | 0.991835066 | -5.050935617 |
| **A_55_P2004571** | Pitpnm2 | 0.046471329 | -0.108900359 | 0.201843016 | 11.44407004 | 0.714862209 | 0.499011735 | 0.991835066 | -5.05099593 |
| **A_55_P2141955** | NA | 0.052862234 | -0.123883194 | 0.229607661 | 15.97093522 | 0.714836064 | 0.499026873 | 0.991835066 | -5.051010286 |
| **A_51_P121818** | Prkag1 | -0.058784631 | -0.255416598 | 0.137847335 | 11.53527542 | -0.714527274 | 0.499205691 | 0.991835066 | -5.051179801 |

| | | | | | | | | | |
|---|---|---|---|---|---|---|---|---|---|
| **A_51_P345046** | Metap2 | -0.03089706 | -0.134279621 | 0.072485501 | 11.95058489 | -0.714297287 | 0.499338901 | 0.991835066 | -5.051306016 |
| **A_55_P2055232** | Sdf4 | -0.093049434 | -0.404429644 | 0.218330776 | 9.899273227 | -0.714219762 | 0.49938381 | 0.991835066 | -5.051348554 |
| **A_55_P2007673** | Lrsam1 | -0.038444279 | -0.16709916 | 0.090210601 | 10.33022568 | -0.714191357 | 0.499400265 | 0.991835066 | -5.051364139 |
| **A_55_P2105002** | Pcna | 0.061346284 | -0.143964618 | 0.266657186 | 13.36918732 | 0.714143328 | 0.499428089 | 0.991835066 | -5.051390489 |
| **A_55_P2123831** | Thap6 | -0.047556442 | -0.206734796 | 0.111621911 | 8.848705859 | -0.714059628 | 0.49947658 | 0.991835066 | -5.051436407 |
| **A_55_P2018941** | Fmr1 | -0.062596964 | -0.272138444 | 0.146944515 | 10.04754719 | -0.713990444 | 0.499516664 | 0.991835066 | -5.051474359 |
| **A_66_P117189** | Rad54l2 | 0.033965095 | -0.079743888 | 0.147674079 | 8.995517587 | 0.713916125 | 0.499559725 | 0.991835066 | -5.051515123 |
| **A_55_P2091368** | Cep164 | 0.048638514 | -0.114199651 | 0.211476679 | 9.050692295 | 0.713893176 | 0.499573023 | 0.991835066 | -5.05152771 |
| **A_51_P216219** | Ptk2 | 0.050440033 | -0.118440266 | 0.219320332 | 10.52975017 | 0.713847604 | 0.499599429 | 0.991835066 | -5.051552704 |
| **A_55_P2033863** | Psmc1 | 0.040142547 | -0.094297467 | 0.174582561 | 14.15149569 | 0.713650387 | 0.499713717 | 0.991835066 | -5.051660854 |
| **A_51_P514319** | Slc13a4 | 0.051992172 | -0.122151152 | 0.226135496 | 8.245047554 | 0.713576048 | 0.499756801 | 0.991835066 | -5.051701613 |
| **A_51_P481482** | Dram1 | -0.030234464 | -0.131511103 | 0.071042175 | 7.742322573 | -0.713513386 | 0.49979312 | 0.991835066 | -5.051735968 |
| **A_55_P2442017** | Samd8 | 0.038910479 | -0.091437436 | 0.169258393 | 8.361814457 | 0.713463265 | 0.499822171 | 0.991835066 | -5.051763445 |
| **A_51_P118885** | Amacr | -0.041713374 | -0.181504425 | 0.098077677 | 10.81106218 | -0.713189728 | 0.499980737 | 0.991835066 | -5.041446531 |
| **A_55_P2058201** | Scgb2b20 | -0.078213645 | -0.340370694 | 0.183943403 | 8.364146582 | -0.713066644 | 0.500052099 | 0.991835066 | -5.051980823 |
| **A_55_P2065799** | Naa50 | 0.059085461 | -0.138974209 | 0.257145131 | 10.03356915 | 0.713006863 | 0.500086761 | 0.991835066 | -5.052013579 |
| **A_55_P2187043** | Tpm1 | -0.036668782 | -0.159598646 | 0.086261082 | 16.96435827 | -0.712932178 | 0.500130067 | 0.991835066 | -5.052054497 |
| **A_52_P75777** | NA | 0.044846061 | -0.10551214 | 0.195204263 | 8.170940306 | 0.7128634 | 0.50016995 | 0.991835066 | -5.052092178 |
| **A_55_P2102679** | NA | -0.032564287 | -0.141761031 | 0.076632456 | 8.162590796 | -0.712756236 | 0.500232096 | 0.991835066 | -5.052150881 |
| **A_51_P248444** | Poglut1 | -0.035381155 | -0.154044768 | 0.083282459 | 8.642515364 | -0.712629126 | 0.500305816 | 0.991835066 | -5.0522205 |
| **A_52_P652104** | Brd3 | 0.061370551 | -0.14447569 | 0.267216791 | 9.77743557 | 0.712567829 | 0.500341369 | 0.991835066 | -5.05225407 |
| **A_55_P2256686** | NA | 0.0692916 | -0.163124079 | 0.301707279 | 8.128348039 | 0.712564695 | 0.500343187 | 0.991835066 | -5.052255786 |
| **A_52_P496202** | Pcsk1n | -0.034359553 | -0.149619939 | 0.080900834 | 8.179665888 | -0.712486367 | 0.50038862 | 0.991835066 | -5.052298679 |
| **A_55_P2094242** | NA | 0.030927293 | -0.07283286 | 0.134687445 | 8.086731871 | 0.712394297 | 0.500442028 | 0.991835066 | -5.052349093 |
| **A_55_P2059055** | Zfp943 | 0.049853949 | -0.117410203 | 0.217118101 | 10.28756989 | 0.712370336 | 0.500455929 | 0.991835066 | -5.052362212 |
| **A_52_P578768** | Slc38a9 | 0.055604608 | -0.130971376 | 0.242180591 | 8.953451818 | 0.712302015 | 0.500495563 | 0.991835066 | -5.052399618 |
| **A_66_P133383** | NA | -0.042840205 | -0.186594412 | 0.100914001 | 7.788162516 | -0.712262554 | 0.500518456 | 0.991835066 | -5.05242122 |
| **A_55_P2010014** | Map7d1 | -0.057024349 | -0.248387232 | 0.134338535 | 13.50606066 | -0.712216056 | 0.500545433 | 0.991835066 | -5.052446675 |
| **A_55_P1985693** | Fhdc1 | 0.035107944 | -0.082710327 | 0.152926214 | 8.070569239 | 0.712199861 | 0.50055483 | 0.991835066 | -5.05245554 |
| **A_55_P2066727** | Serpinb6a | -0.067240687 | -0.292900982 | 0.158419608 | 13.01456101 | -0.712174015 | 0.500569825 | 0.991835066 | -5.052469688 |
| **A_55_P2075298** | Atp8a1 | -0.053366808 | -0.232477051 | 0.125743436 | 9.247331604 | -0.712131065 | 0.500594745 | 0.991835066 | -5.052493198 |
| **A_52_P181709** | Crkl | -0.048007198 | -0.209131156 | 0.113116761 | 8.862932696 | -0.71212358 | 0.500599088 | 0.991835066 | -5.052497295 |
| **A_55_P2106645** | Adcy5 | -0.042036769 | -0.183125691 | 0.099052153 | 9.789387101 | -0.712107478 | 0.500608431 | 0.991835066 | -5.052506108 |
| **A_55_P2069832** | Prpsap1 | -0.040152635 | -0.174920841 | 0.094615571 | 12.86179836 | -0.712091393 | 0.500617764 | 0.991835066 | -5.052514913 |
| **A_55_P2103055** | Gsr | 0.039827864 | -0.093945032 | 0.17360076 | 10.43742142 | 0.711587023 | 0.500910473 | 0.991835066 | -5.052790892 |
| **A_55_P1954921** | Fbxw5 | -0.030378693 | -0.132414276 | 0.07165689 | 12.55862955 | -0.711584641 | 0.500911856 | 0.991835066 | -5.052792196 |
| **A_55_P2410304** | Iigp1 | -0.034691753 | -0.151255547 | 0.081872041 | 8.168888764 | -0.711330952 | 0.501059126 | 0.991835066 | -5.052930948 |
| **A_55_P2002472** | Zfp777 | 0.037843583 | -0.089313754 | 0.165000919 | 10.41170682 | 0.711311707 | 0.501070299 | 0.991835066 | -5.052941472 |
| **A_55_P2007535** | Edf1 | -0.05163475 | -0.22513774 | 0.12186824 | 14.63507932 | -0.711285977 | 0.501085238 | 0.991835066 | -5.052955542 |
| **A_55_P2026761** | NA | 0.113877768 | -0.268854249 | 0.496609785 | 11.9835274 | 0.711137013 | 0.50117173 | 0.991835066 | -5.053036993 |
| **A_66_P135742** | Wdr1 | 0.060206639 | -0.142157869 | 0.262571147 | 12.11973769 | 0.71108116 | 0.501204162 | 0.991835066 | -5.053067529 |
| **A_52_P243249** | Mapk3 | -0.09621593 | -0.419632029 | 0.227200169 | 11.12609608 | -0.711040729 | 0.50122764 | 0.991835066 | -5.053089632 |
| **A_55_P2107182** | Angptl8 | -0.034378242 | -0.149990173 | 0.08123369 | 8.190965216 | -0.710706255 | 0.501421893 | 0.991835066 | -5.053272445 |
| **A_66_P128839** | Gm13262 | 0.031183565 | -0.073692566 | 0.136059695 | 7.997462488 | 0.710654052 | 0.501452216 | 0.991835066 | -5.053300972 |
| **A_55_P2133370** | NA | -0.03118506 | -0.136082119 | 0.073711998 | 7.664210715 | -0.710546349 | 0.501514779 | 0.991835066 | -5.05335982 |
| **A_55_P2024342** | Tmem33 | 0.059935872 | -0.141677942 | 0.261549687 | 10.02172708 | 0.710518969 | 0.501530685 | 0.991835066 | -5.05337478 |
| **A_55_P1975537** | Nutf2 | -0.044643355 | -0.194821695 | 0.105534984 | 11.78277174 | -0.710491138 | 0.501546853 | 0.991835066 | -5.053389985 |
| **A_52_P58145** | Aldh1a2 | -0.121418633 | -0.529886894 | 0.287049627 | 10.4808111 | -0.710454314 | 0.501568246 | 0.991835066 | -5.053410102 |

| | | | | | | | | | |
|---|---|---|---|---|---|---|---|---|---|
| **A_55_P2134800** | Cinp | -0.049910036 | -0.217889281 | 0.118069209 | 9.037544487 | -0.71013577 | 0.50175333 | 0.991835066 | -5.053584093 |
| **A_55_P1979221** | Txndc9 | 0.054799604 | -0.129650486 | 0.239249695 | 10.35383513 | 0.710080659 | 0.501785355 | 0.991835066 | -5.053614188 |
| **A_51_P468599** | Smchd1 | 0.051534341 | -0.121927756 | 0.224996438 | 11.318507 | 0.710070169 | 0.501791451 | 0.991835066 | -5.053619917 |
| **A_51_P260740** | Pcdh7 | -0.033356106 | -0.145644583 | 0.078932371 | 8.564632056 | -0.709985184 | 0.501840839 | 0.991835066 | -5.053666321 |
| **A_55_P2162890** | Zfp780b | -0.039310382 | -0.171645127 | 0.093024363 | 8.871768191 | -0.709974107 | 0.501847277 | 0.991835066 | -5.05367237 |
| **A_51_P165704** | Mcm7 | 0.052600657 | -0.124480804 | 0.229682119 | 12.68988532 | 0.709949091 | 0.501861816 | 0.991835066 | -5.053686028 |
| **A_51_P448391** | Nkiras1 | -0.049675019 | -0.216977378 | 0.117627339 | 9.080280462 | -0.70965148 | 0.502034801 | 0.991835066 | -5.053848492 |
| **A_55_P1953356** | Triobp | 0.082036891 | -0.194262179 | 0.358335961 | 11.32272381 | 0.709641336 | 0.502040698 | 0.991835066 | -5.053854029 |
| **A_55_P1952743** | NA | 0.086215766 | -0.204183437 | 0.376614969 | 11.99512436 | 0.709578428 | 0.502077269 | 0.991835066 | -5.053888362 |
| **A_55_P2106150** | Cenpk | 0.066559005 | -0.157631942 | 0.290749952 | 10.61330485 | 0.709574311 | 0.502079662 | 0.991835066 | -5.053890609 |
| **A_55_P2361647** | 4831440E1 | -0.029701082 | -0.129747983 | 0.07034582 | 8.464701045 | -0.709541429 | 0.502098778 | 0.991835066 | -5.053908554 |
| **A_66_P131110** | Zfp473 | 0.029434607 | -0.069759099 | 0.128628314 | 7.877605447 | 0.709223736 | 0.502283494 | 0.991835066 | -5.054081897 |
| **A_55_P2180854** | Mrgprg | -0.031415584 | -0.137307052 | 0.074475884 | 7.75412477 | -0.709076842 | 0.502368918 | 0.991835066 | -5.054162025 |
| **A_52_P130727** | Rtl6 | -0.05095703 | -0.222729081 | 0.12081502 | 9.255403027 | -0.709023683 | 0.502399834 | 0.991835066 | -5.05419102 |
| **A_52_P89267** | Castor2 | -0.058271629 | -0.254711996 | 0.138168739 | 8.642334498 | -0.708982555 | 0.502423754 | 0.991835066 | -5.05421345 |
| **A_52_P615312** | Mrps27 | -0.097114518 | -0.424499367 | 0.23027033 | 9.997080816 | -0.708981197 | 0.502424544 | 0.991835066 | -5.054214191 |
| **A_55_P2150309** | Lhfpl4 | -0.049348805 | -0.215715844 | 0.117018234 | 11.15104891 | -0.708954699 | 0.502439956 | 0.991835066 | -5.054228643 |
| **A_52_P59457** | Nprl2 | -0.056838826 | -0.248468132 | 0.13479048 | 9.818714674 | -0.70891196 | 0.502464814 | 0.991835066 | -5.05425195 |
| **A_55_P2130660** | Slitrk4 | 0.079049073 | -0.18748065 | 0.345578796 | 9.650904558 | 0.708859668 | 0.502495229 | 0.991835066 | -5.054280466 |
| **A_52_P459929** | Itga1 | -0.07065566 | -0.308889617 | 0.167578297 | 10.14594337 | -0.708846869 | 0.502502674 | 0.991835066 | -5.054287445 |
| **A_55_P2064567** | Anks1 | 0.042143201 | -0.099953592 | 0.184239993 | 8.950582382 | 0.708846784 | 0.502502723 | 0.991835066 | -5.054287491 |
| **A_55_P2008891** | Tmsb15b1 | 0.052433799 | -0.124367044 | 0.229234643 | 8.269355143 | 0.708820268 | 0.502518147 | 0.991835066 | -5.05430195 |
| **A_55_P2052924** | Selenow | 0.039459581 | -0.093601425 | 0.172520587 | 14.0284889 | 0.708778931 | 0.502542193 | 0.991835066 | -5.05432449 |
| **A_55_P2142200** | NA | -0.040078237 | -0.175269293 | 0.095112819 | 11.10407448 | -0.708548827 | 0.502676055 | 0.991835066 | -5.054449937 |
| **A_51_P349495** | Mboat1 | 0.044143212 | -0.104850413 | 0.193136837 | 8.435511138 | 0.708117579 | 0.502926995 | 0.991835066 | -5.054684953 |
| **A_51_P334789** | Nudt16 | -0.036841924 | -0.161224903 | 0.087541055 | 8.372850251 | -0.707930264 | 0.503036018 | 0.991835066 | -5.054786997 |
| **A_55_P2173183** | Cdk5r1 | -0.032140586 | -0.140658025 | 0.076376852 | 8.204300576 | -0.707886112 | 0.503061718 | 0.991835066 | -5.054811047 |
| **A_52_P639567** | Ate1 | -0.047763716 | -0.209102757 | 0.113575324 | 10.88572983 | -0.707567333 | 0.503247299 | 0.991835066 | -5.05498465 |
| **A_52_P84353** | Homez | 0.035834438 | -0.08521545 | 0.156884326 | 9.714576086 | 0.707530779 | 0.503268582 | 0.991835066 | -5.055004552 |
| **A_51_P382484** | Itgav | 0.043974938 | -0.104576683 | 0.19252656 | 8.863448434 | 0.707517156 | 0.503276514 | 0.991835066 | -5.05501197 |
| **A_55_P2072391** | Acp1 | 0.042982607 | -0.102239938 | 0.188205151 | 8.628327072 | 0.707404536 | 0.50334209 | 0.991835066 | -5.055073282 |
| **A_51_P108020** | Zmynd15 | -0.054978763 | -0.240775644 | 0.130818118 | 8.503056592 | -0.707238148 | 0.503438985 | 0.991835066 | -5.041427808 |
| **A_66_P135130** | Tmem161a | -0.078402058 | -0.343374187 | 0.186570071 | 9.871316919 | -0.707190472 | 0.50346675 | 0.991835066 | -5.0551898 |
| **A_55_P2118784** | Rcor2 | 0.07173756 | -0.170731965 | 0.314207085 | 11.96421633 | 0.707128875 | 0.503502625 | 0.991835066 | -5.055223323 |
| **A_55_P2039225** | Rap1gap | 0.083404642 | -0.198503035 | 0.365312319 | 9.576357903 | 0.707118908 | 0.503508431 | 0.991835066 | -5.055228747 |
| **A_52_P54261** | Tmem56 | -0.031861899 | -0.139561315 | 0.075837517 | 7.903187344 | -0.707078182 | 0.503532151 | 0.991835066 | -5.05525091 |
| **A_55_P2077213** | Cenpb | 0.045105935 | -0.107365966 | 0.197577836 | 11.26847147 | 0.707054691 | 0.503545833 | 0.991835066 | -5.055263692 |
| **A_55_P2177103** | Slc25a5 | -0.040263979 | -0.176368902 | 0.095840943 | 14.5053667 | -0.707053042 | 0.503546793 | 0.991835066 | -5.05526459 |
| **A_55_P2071799** | Slbp | -0.038077696 | -0.166803294 | 0.090647901 | 8.467723779 | -0.706992621 | 0.503581987 | 0.991835066 | -5.055297468 |
| **A_51_P273508** | Ube2g1 | -0.043379873 | -0.190054624 | 0.103294879 | 9.004066272 | -0.706874062 | 0.503651049 | 0.991835066 | -5.055361974 |
| **A_55_P2368710** | NA | 0.116496914 | -0.277467799 | 0.510461628 | 8.780861686 | 0.706750671 | 0.503722932 | 0.991835066 | -5.055429099 |
| **A_55_P2059179** | Slc48a1 | 0.059427582 | -0.141609928 | 0.260465092 | 10.47045341 | 0.706512899 | 0.503861468 | 0.991835066 | -5.055558422 |
| **A_55_P2431118** | Itsn1 | 0.040687152 | -0.096956816 | 0.178331119 | 10.77668607 | 0.706495234 | 0.503871762 | 0.991835066 | -5.055568029 |
| **A_55_P1976057** | Smurf2 | 0.05357986 | -0.127683076 | 0.234842796 | 8.634765854 | 0.70648297 | 0.503878908 | 0.991835066 | -5.055574698 |
| **A_51_P423880** | Smarcd3 | 0.038610364 | -0.09201059 | 0.169231317 | 11.71373846 | 0.70648049 | 0.503880353 | 0.991835066 | -5.055576046 |
| **A_55_P2056896** | Kif2a | -0.05471415 | -0.239884768 | 0.130456469 | 10.04097327 | -0.706214628 | 0.504035289 | 0.991835066 | -5.055720599 |
| **A_55_P2185203** | Snrnp40 | 0.057206558 | -0.136416461 | 0.250829576 | 8.950078124 | 0.706151622 | 0.504072011 | 0.991835066 | -5.05575485 |
| **A_55_P1994117** | Ifnl3 | -0.033258641 | -0.145887625 | 0.079370342 | 7.92387355 | -0.705770451 | 0.504294211 | 0.991835066 | -5.055962005 |

| | | | | | | | | | |
|---|---|---|---|---|---|---|---|---|---|
| **A_51_P343252** | Cd151 | -0.074332836 | -0.326139805 | 0.177474132 | 13.47664733 | -0.705540734 | 0.504428153 | 0.991835066 | -5.056086805 |
| **A_52_P240542** | Id2 | 0.092414937 | -0.220675135 | 0.405505008 | 10.92594113 | 0.705475613 | 0.504466127 | 0.991835066 | -5.056122177 |
| **A_51_P117794** | Bik | -0.061517229 | -0.269944337 | 0.14690988 | 8.890872426 | -0.705426371 | 0.504494844 | 0.991835066 | -5.056148923 |
| **A_55_P2066399** | Fcho2 | -0.028181251 | -0.123677714 | 0.067315212 | 8.032304606 | -0.705313363 | 0.504560749 | 0.991835066 | -5.056210297 |
| **A_66_P129893** | 1700047M1 | 0.028185979 | -0.067342869 | 0.123714827 | 7.801878464 | 0.70519255 | 0.504631214 | 0.991835066 | -5.056275902 |
| **A_51_P233788** | Dmap1 | 0.04099052 | -0.097946475 | 0.179927515 | 11.41362777 | 0.705138874 | 0.504662522 | 0.991835066 | -5.056305046 |
| **A_55_P2070045** | Golga7b | 0.030615712 | -0.073157227 | 0.13438865 | 7.72711076 | 0.705130305 | 0.504667521 | 0.991835066 | -5.056309698 |
| **A_55_P2169721** | Rps17 | 0.047135824 | -0.112635153 | 0.206906802 | 15.34823164 | 0.705118884 | 0.504674183 | 0.991835066 | -5.056315899 |
| **A_55_P1957918** | Asap2 | 0.046193681 | -0.110411869 | 0.202799231 | 9.686474837 | 0.704992596 | 0.504747851 | 0.991835066 | -5.056384461 |
| **A_52_P609972** | Eln | 0.413572528 | -0.988541492 | 1.815686547 | 10.57772634 | 0.704981114 | 0.50475455 | 0.991835066 | -5.056390694 |
| **A_55_P2009762** | Mapkapk3 | -0.091076183 | -0.399847227 | 0.217694861 | 10.04653538 | -0.704980957 | 0.504754641 | 0.991835066 | -5.056390779 |
| **A_55_P1993719** | Tmem216 | 0.046694271 | -0.111611403 | 0.204999944 | 8.2593693 | 0.704979111 | 0.504755718 | 0.991835066 | -5.056391781 |
| **A_51_P384673** | Lrrc59 | 0.06295616 | -0.150497413 | 0.276409733 | 10.80049569 | 0.704926652 | 0.504786322 | 0.991835066 | -5.056420257 |
| **A_55_P2108171** | Glud1 | -0.036953106 | -0.162252011 | 0.0883458 | 12.34495247 | -0.70487612 | 0.504815803 | 0.991835066 | -5.056447686 |
| **A_51_P427476** | Mxra8 | -0.069879299 | -0.306834277 | 0.167075678 | 8.964240691 | -0.704842101 | 0.50483565 | 0.991835066 | -5.056466151 |
| **A_51_P248857** | Inka1 | -0.064103614 | -0.281479646 | 0.153272417 | 8.368804479 | -0.704822878 | 0.504846865 | 0.991835066 | -5.056476584 |
| **A_55_P2067251** | Tmpo | 0.038097627 | -0.091126414 | 0.167321669 | 9.505223832 | 0.70463424 | 0.504956932 | 0.991835066 | -5.056578956 |
| **A_55_P1978481** | Nr1h2 | -0.092747308 | -0.407347309 | 0.221852693 | 11.45771166 | -0.704614742 | 0.504968309 | 0.991835066 | -5.056589536 |
| **A_55_P2036007** | Rai2 | -0.039127336 | -0.171856709 | 0.093602036 | 8.470609467 | -0.704567121 | 0.504996099 | 0.991835066 | -5.056615376 |
| **A_55_P2182222** | Rab3c | 0.14472979 | -0.346323271 | 0.635782851 | 10.96922838 | 0.704431296 | 0.505075364 | 0.991835066 | -5.056689067 |
| **A_51_P121275** | Cenpo | 0.042502655 | -0.101722732 | 0.186728042 | 8.807145497 | 0.704341823 | 0.505127583 | 0.991835066 | -5.056737604 |
| **A_55_P2051991** | Brsk1 | 0.078682262 | -0.188330263 | 0.345694788 | 9.070527472 | 0.704294563 | 0.505155167 | 0.991835066 | -5.056763238 |
| **A_52_P251183** | Tmem177 | -0.040939964 | -0.179882038 | 0.098002111 | 9.604957597 | -0.704243437 | 0.505185008 | 0.991835066 | -5.056790969 |
| **A_52_P139650** | Ska1 | -0.060013228 | -0.263695466 | 0.143669009 | 10.21194905 | -0.704211262 | 0.505203789 | 0.991835066 | -5.05680842 |
| **A_51_P104392** | Rpp25 | -0.057432719 | -0.252357813 | 0.137492374 | 9.327449439 | -0.704207683 | 0.505205878 | 0.991835066 | -5.056810361 |
| **A_55_P1954001** | Snrpc | -0.053299686 | -0.23420189 | 0.127602517 | 12.2589374 | -0.704190123 | 0.505216128 | 0.991835066 | -5.056819885 |
| **A_55_P1964852** | Kbtbd2 | 0.040553603 | -0.097124325 | 0.178231532 | 12.30599892 | 0.704002593 | 0.505325601 | 0.991835066 | -5.056921581 |
| **A_55_P2084345** | NA | -0.072286146 | -0.31770976 | 0.173137468 | 14.65987815 | -0.703959795 | 0.505350586 | 0.991835066 | -5.056944786 |
| **A_55_P2192577** | H2bfm | -0.048023981 | -0.211088614 | 0.115040653 | 7.970447017 | -0.703894396 | 0.505388768 | 0.991835066 | -5.056980245 |
| **A_55_P2084492** | NA | 0.045622133 | -0.109294284 | 0.200538549 | 8.301554857 | 0.703861605 | 0.505407914 | 0.991835066 | -5.056998023 |
| **A_52_P389230** | Smarcc2 | -0.05373057 | -0.236197425 | 0.128736286 | 8.611988252 | -0.703795666 | 0.505446414 | 0.991835066 | -5.05703377 |
| **A_55_P2160094** | Plscr2 | 0.059690486 | -0.143020865 | 0.262401837 | 8.593204642 | 0.703778797 | 0.505456264 | 0.991835066 | -5.057042914 |
| **A_52_P120925** | Metap1 | -0.032921204 | -0.144730578 | 0.07888817 | 8.460292798 | -0.703730915 | 0.505484222 | 0.991835066 | -5.057068869 |
| **A_55_P1953508** | Mob3c | -0.029157591 | -0.128194811 | 0.069879629 | 8.0419521 | -0.703659133 | 0.505526139 | 0.991835066 | -5.057107778 |
| **A_66_P111089** | Ube2f | -0.035127582 | -0.154462126 | 0.084206963 | 8.764270645 | -0.703543904 | 0.50559343 | 0.991835066 | -5.057170229 |
| **A_55_P1965313** | Mctp2 | 0.033796574 | -0.081021153 | 0.148614302 | 7.685883863 | 0.703514196 | 0.50561078 | 0.991835066 | -5.057186329 |
| **A_55_P2138095** | NA | -0.036398637 | -0.160073528 | 0.087276253 | 16.22809061 | -0.703416782 | 0.505667673 | 0.991835066 | -5.057239117 |
| **A_52_P484838** | Rfxank | -0.062302703 | -0.274090531 | 0.149485125 | 9.602311269 | -0.703096643 | 0.505854676 | 0.991835066 | -5.057412555 |
| **A_55_P2132942** | Adal | -0.040295126 | -0.17728136 | 0.096691107 | 9.691363538 | -0.703047598 | 0.505883328 | 0.991835066 | -5.057439119 |
| **A_55_P1993518** | Tbc1d14 | 0.041014191 | -0.098426026 | 0.180454408 | 10.39343572 | 0.702999849 | 0.505911225 | 0.991835066 | -5.057464981 |
| **A_55_P1970755** | Hdac9 | 0.04531719 | -0.108752858 | 0.199387237 | 10.36658506 | 0.702997666 | 0.5059125 | 0.991835066 | -5.057466163 |
| **A_52_P118461** | Ipmk | 0.060343014 | -0.144861174 | 0.265547202 | 9.729010362 | 0.702829384 | 0.506010824 | 0.991835066 | -5.057557294 |
| **A_51_P103757** | Kat8 | 0.035381094 | -0.084967396 | 0.155729585 | 11.00992707 | 0.702651119 | 0.506114994 | 0.991835066 | -5.057653812 |
| **A_55_P2121961** | Psmc3 | -0.060158442 | -0.264844071 | 0.144527188 | 13.76402145 | -0.702454756 | 0.506229756 | 0.991835066 | -5.057760106 |
| **A_55_P2002562** | Sec11a | -0.048406747 | -0.213119196 | 0.116305701 | 10.99538317 | -0.702406644 | 0.506257877 | 0.991835066 | -5.057786146 |
| **A_51_P316553** | Kdr | 0.040228553 | -0.096711277 | 0.177168382 | 12.67216044 | 0.702123896 | 0.506423162 | 0.991835066 | -5.036282137 |
| **A_52_P191537** | NA | 0.058098809 | -0.139682535 | 0.255880154 | 9.820231917 | 0.702087173 | 0.506444632 | 0.991835066 | -5.057959017 |
| **A_55_P1983858** | Scel | 0.032152152 | -0.077305648 | 0.141609953 | 7.695730028 | 0.702057144 | 0.506462188 | 0.991835066 | -5.057975263 |

| | | | | | | | | | |
|---|---|---|---|---|---|---|---|---|---|
| **A_55_P2112934** | C030034I2 | 0.03534765 | -0.084993134 | 0.155688434 | 8.481345177 | 0.702031887 | 0.506476954 | 0.991835066 | -5.057988927 |
| **A_55_P2285599** | 5730437C1 | 0.029391738 | -0.070681107 | 0.129464583 | 7.656590485 | 0.701969351 | 0.506513518 | 0.991835066 | -5.058022756 |
| **A_55_P2140328** | NA | -0.03298004 | -0.145307813 | 0.079347733 | 13.18791824 | -0.701735042 | 0.50665053 | 0.991835066 | -5.058149486 |
| **A_55_P2030802** | Rpl3 | 0.038059299 | -0.091574255 | 0.167692853 | 15.53422456 | 0.701701632 | 0.506670068 | 0.991835066 | -5.058167554 |
| **A_55_P2160043** | A330069E1 | -0.037377651 | -0.164707173 | 0.089951872 | 8.473138674 | -0.701603949 | 0.506727197 | 0.991835066 | -5.058220375 |
| **A_55_P2300421** | NA | -0.04173583 | -0.183923098 | 0.100451439 | 8.359056779 | -0.701548134 | 0.506759841 | 0.991835066 | -5.058250554 |
| **A_55_P1960683** | F2 | -0.120530782 | -0.531175741 | 0.290114176 | 8.091055772 | -0.701520896 | 0.506775772 | 0.991835066 | -5.058265281 |
| **A_55_P2040031** | NA | -0.044074074 | -0.194266217 | 0.106118069 | 12.46022811 | -0.70136666 | 0.506865988 | 0.991835066 | -5.058348661 |
| **A_51_P308948** | Tamm41 | 0.056216728 | -0.135359464 | 0.247792921 | 11.24194278 | 0.701347351 | 0.506877284 | 0.991835066 | -5.058359099 |
| **A_55_P2186562** | Hars2 | -0.036216815 | -0.159644626 | 0.087210996 | 10.35242579 | -0.701304076 | 0.506902598 | 0.991835066 | -5.058382491 |
| **A_55_P2168397** | NA | 0.031402107 | -0.075617722 | 0.138421936 | 15.30256788 | 0.701299718 | 0.506905148 | 0.991835066 | -5.058384846 |
| **A_51_P322989** | Ptrhd1 | -0.034034287 | -0.150030726 | 0.081962152 | 11.95933486 | -0.701263329 | 0.506926435 | 0.991835066 | -5.058404515 |
| **A_55_P2155826** | Rplp1 | 0.040581697 | -0.097737716 | 0.178901109 | 16.65076148 | 0.701223072 | 0.506949986 | 0.991835066 | -5.058426273 |
| **A_51_P241667** | Prkdc | 0.033463447 | -0.080657091 | 0.147583986 | 8.620462694 | 0.70083535 | 0.507176845 | 0.991835066 | -5.053307864 |
| **A_51_P451428** | Marveld2 | 0.039106986 | -0.094261439 | 0.172475411 | 7.84535885 | 0.700826403 | 0.507182081 | 0.991835066 | -5.05864061 |
| **A_55_P2117630** | Snrnp25 | -0.065758866 | -0.290024885 | 0.158507153 | 11.31763306 | -0.700809488 | 0.50719198 | 0.991835066 | -5.058649747 |
| **A_52_P576689** | Zfp944 | 0.039364792 | -0.094927872 | 0.173657455 | 8.800019349 | 0.700591396 | 0.507319619 | 0.991835066 | -5.058767547 |
| **A_55_P1994603** | Itsn1 | -0.062212224 | -0.274456442 | 0.150031993 | 13.971575 | -0.700565903 | 0.507334541 | 0.991835066 | -5.058781315 |
| **A_52_P333097** | NA | -0.033512776 | -0.14787399 | 0.080848438 | 7.697474978 | -0.700391364 | 0.507436707 | 0.991835066 | -5.058875565 |
| **A_55_P2041893** | NA | 0.032378921 | -0.078114625 | 0.142872466 | 15.00176148 | 0.700381379 | 0.507442552 | 0.991835066 | -5.058880956 |
| **A_55_P2157360** | Tagap1 | -0.039937543 | -0.176251748 | 0.096376662 | 8.45787689 | -0.700243939 | 0.507523012 | 0.991835066 | -5.058955159 |
| **A_55_P2132512** | Ngef | 0.055369062 | -0.133666937 | 0.244405061 | 7.892347323 | 0.700054387 | 0.507633994 | 0.991835066 | -5.059057476 |
| **A_51_P369200** | Tpx2 | -0.040843599 | -0.180288377 | 0.09860118 | 10.59118514 | -0.700052926 | 0.50763485 | 0.991835066 | -5.059058265 |
| **A_55_P1997225** | Agap2 | -0.038316509 | -0.169141134 | 0.092508116 | 8.160712935 | -0.70001213 | 0.507658737 | 0.991835066 | -5.059080282 |
| **A_66_P134040** | Vps25 | -0.085663604 | -0.37819511 | 0.206867902 | 9.897692789 | -0.699894898 | 0.507727387 | 0.991835066 | -5.059143548 |
| **A_55_P2134979** | NA | -0.117377353 | -0.518214665 | 0.283459958 | 13.39458356 | -0.699882775 | 0.507734486 | 0.991835066 | -5.05915009 |
| **A_55_P2001342** | Cacul1 | 0.037194809 | -0.089840591 | 0.164230209 | 8.771938803 | 0.699788353 | 0.507789782 | 0.991835066 | -5.059201039 |
| **A_55_P2054300** | Alg8 | 0.10618941 | -0.256545594 | 0.468924413 | 9.950036517 | 0.699682277 | 0.507851909 | 0.991835066 | -5.059258269 |
| **A_55_P2006930** | 2610005L0 | 0.054671683 | -0.132084659 | 0.241428024 | 10.90214766 | 0.699674771 | 0.507856305 | 0.991835066 | -5.059262318 |
| **A_55_P2067513** | Slc10a3 | 0.045517859 | -0.11002959 | 0.201065308 | 9.740559958 | 0.699403929 | 0.508014954 | 0.991835066 | -5.05940841 |
| **A_51_P163173** | Rbm12b1 | 0.049966532 | -0.12079351 | 0.220726574 | 8.593000036 | 0.699362092 | 0.508039463 | 0.991835066 | -5.059430973 |
| **A_55_P2061219** | Ces3a | -0.029334131 | -0.129603094 | 0.070934832 | 7.728641946 | -0.699223215 | 0.508120828 | 0.991835066 | -5.059505861 |
| **A_55_P2099338** | Akt1s1 | 0.105531791 | -0.255199375 | 0.466262957 | 8.532343063 | 0.699211845 | 0.508127489 | 0.991835066 | -5.059511992 |
| **A_55_P2286650** | 6720462K0 | 0.097204132 | -0.23512948 | 0.429537744 | 8.772865559 | 0.699068268 | 0.508211616 | 0.991835066 | -5.0595894 |
| **A_55_P2183668** | NA | 0.197139723 | -0.476885196 | 0.871164641 | 12.78953574 | 0.699048521 | 0.508223187 | 0.991835066 | -5.059600046 |
| **A_55_P1985749** | Rad23a | 0.035830285 | -0.086698559 | 0.15835913 | 8.061481759 | 0.698909691 | 0.508304543 | 0.991835066 | -5.059674881 |
| **A_55_P2131954** | Gm2590 | 0.084081731 | -0.203505761 | 0.371669222 | 8.554368788 | 0.698780506 | 0.508380254 | 0.991835066 | -5.059744506 |
| **A_55_P1974961** | Plekhg3 | -0.076534648 | -0.338348527 | 0.185279231 | 12.55543695 | -0.698673958 | 0.508442704 | 0.991835066 | -5.059801923 |
| **A_55_P2102126** | Lamp2 | -0.042646808 | -0.188539195 | 0.103245579 | 10.14769223 | -0.698655424 | 0.508453568 | 0.991835066 | -5.05981191 |
| **A_51_P496381** | Ptk7 | 0.11497116 | -0.278412491 | 0.50835481 | 9.852042557 | 0.698524653 | 0.508530223 | 0.991835066 | -5.059882369 |
| **A_55_P2116900** | NA | -0.039913185 | -0.17649895 | 0.09667258 | 9.323186949 | -0.698425488 | 0.508588356 | 0.991835066 | -5.059935791 |
| **A_55_P2239317** | Afap1l1 | -0.070552986 | -0.312000233 | 0.170894261 | 11.83115289 | -0.698396849 | 0.508605147 | 0.991835066 | -5.059951218 |
| **A_55_P2112599** | Lgals8 | 0.03002361 | -0.072734209 | 0.132781428 | 8.997835948 | 0.698324315 | 0.508647672 | 0.991835066 | -5.059990288 |
| **A_51_P427232** | Zbed5 | -0.031594931 | -0.139753142 | 0.076563279 | 8.127262587 | -0.698179445 | 0.508732613 | 0.991835066 | -5.060068312 |
| **A_55_P2178678** | P2rx5 | -0.064617412 | -0.285863805 | 0.15662898 | 9.101497838 | -0.698043516 | 0.50881232 | 0.991835066 | -5.060141509 |
| **A_55_P2093994** | Maged2 | 0.060992397 | -0.147843357 | 0.269828152 | 14.21863422 | 0.69803947 | 0.508814693 | 0.991835066 | -5.060143687 |
| **A_55_P2002449** | NA | 0.045709603 | -0.110812989 | 0.202232194 | 9.488894577 | 0.697974486 | 0.508852802 | 0.991835066 | -5.060178676 |
| **A_66_P101423** | Ogfod3 | 0.062602033 | -0.151808834 | 0.277012901 | 11.28187326 | 0.697831833 | 0.508936465 | 0.991835066 | -5.060255475 |

| | | | | | | | | | |
|---|---|---|---|---|---|---|---|---|---|
| **A_55_P2137376** | Dip2c | -0.047193774 | -0.20886078 | 0.114473233 | 10.99969679 | -0.697705972 | 0.509010287 | 0.991835066 | -5.060323222 |
| **A_55_P2056192** | NA | -0.046072593 | -0.203902708 | 0.111757523 | 11.81330272 | -0.697689053 | 0.509020212 | 0.991835066 | -5.060332328 |
| **A_55_P2022678** | C1qtnf1 | 0.049250398 | -0.119472527 | 0.217973324 | 8.46978005 | 0.697661542 | 0.509036349 | 0.991835066 | -5.060347135 |
| **A_52_P35048** | Serpinf1 | -0.046739914 | -0.20686528 | 0.113385453 | 8.03565904 | -0.697648879 | 0.509043777 | 0.991835066 | -5.060353951 |
| **A_55_P2274233** | Agap3 | -0.0592802 | -0.277434229 | 0.158873828 | 12.11452257 | -0.714614856 | 0.509096757 | 0.991835066 | -4.861373053 |
| **A_52_P134141** | Pyurf | -0.057885383 | -0.256266245 | 0.14049548 | 10.41466624 | -0.697394096 | 0.509193245 | 0.991835066 | -5.060491053 |
| **A_55_P1964867** | Ndufs6 | -0.051534471 | -0.228153318 | 0.125084375 | 16.70603748 | -0.697380685 | 0.509201114 | 0.991835066 | -5.060498268 |
| **A_52_P160936** | Gusb | 0.035417749 | -0.085968172 | 0.15680367 | 9.736645784 | 0.697367609 | 0.509208786 | 0.991835066 | -5.060505304 |
| **A_51_P138790** | Plekhf2 | -0.039702316 | -0.175805901 | 0.096401269 | 8.750680663 | -0.697196843 | 0.509308983 | 0.991835066 | -5.060597169 |
| **A_55_P2079388** | Snora74a | -0.03805958 | -0.168536437 | 0.092417276 | 7.995613676 | -0.697171525 | 0.509323839 | 0.991835066 | -5.060610788 |
| **A_55_P2073383** | NA | -0.034983202 | -0.154950921 | 0.084984516 | 14.60373042 | -0.696954292 | 0.509451322 | 0.991835066 | -5.06072762 |
| **A_55_P2106690** | Tal1 | -0.043446409 | -0.192489641 | 0.105596823 | 8.953187534 | -0.696707935 | 0.509595922 | 0.991835066 | -5.06086008 |
| **A_55_P1965448** | NA | 0.060136121 | -0.146169258 | 0.2664415 | 10.74591667 | 0.696681042 | 0.509611708 | 0.991835066 | -5.060874537 |
| **A_55_P2024041** | Psg19 | -0.061157682 | -0.271024403 | 0.148709038 | 7.789466154 | -0.696492713 | 0.509722268 | 0.991835066 | -5.060975768 |
| **A_55_P1954724** | A130040M | 0.168499044 | -0.409775686 | 0.746773774 | 13.5892283 | 0.696421802 | 0.509763901 | 0.991835066 | -5.061013878 |
| **A_55_P1989703** | Galnt11 | 0.03159974 | -0.076848816 | 0.140048295 | 9.739133528 | 0.696416198 | 0.509767192 | 0.991835066 | -5.06101689 |
| **A_52_P485376** | Ube2k | -0.039779009 | -0.176300525 | 0.096742507 | 12.83480874 | -0.696405171 | 0.509773666 | 0.991835066 | -5.061022816 |
| **A_55_P2050697** | Nfkbil1 | 0.061694011 | -0.150061686 | 0.273449707 | 10.89245406 | 0.696333088 | 0.50981599 | 0.991835066 | -5.061061552 |
| **A_55_P2034928** | BC147527 | -0.03218375 | -0.14267246 | 0.07830496 | 8.480417587 | -0.696190154 | 0.509899921 | 0.991835066 | -5.061138351 |
| **A_55_P2106608** | NA | 0.028538991 | -0.069448884 | 0.126526865 | 7.715295264 | 0.696106041 | 0.509949316 | 0.991835066 | -5.061183541 |
| **A_55_P2178578** | Tmprss4 | 0.029249014 | -0.071180972 | 0.129678999 | 7.663104679 | 0.696076473 | 0.509966681 | 0.991835066 | -5.061199425 |
| **A_52_P253230** | Ankrd46 | -0.034927577 | -0.154862989 | 0.085007836 | 11.01718486 | -0.696033514 | 0.50999191 | 0.991835066 | -5.061222501 |
| **A_55_P1978866** | NA | 0.126006528 | -0.306738583 | 0.558751639 | 12.57998186 | 0.695937129 | 0.510048519 | 0.991835066 | -5.061274273 |
| **A_55_P1979998** | Spaca9 | 0.037869057 | -0.092188017 | 0.167926131 | 7.951282964 | 0.69592052 | 0.510058275 | 0.991835066 | -5.061283194 |
| **A_55_P2000219** | Ubap2 | 0.031912013 | -0.077705339 | 0.141529366 | 12.79385431 | 0.695799362 | 0.51012944 | 0.991835066 | -5.061348262 |
| **A_51_P193446** | Prpf40a | 0.034998718 | -0.085224761 | 0.155222198 | 9.127599351 | 0.695780063 | 0.510140777 | 0.991835066 | -5.061358626 |
| **A_51_P447976** | Tent5c | 0.237477826 | -0.578286636 | 1.053242288 | 11.27530689 | 0.695772749 | 0.510145072 | 0.991835066 | -5.061362553 |
| **A_52_P112110** | Tmem82 | -0.029406943 | -0.130476124 | 0.071662238 | 7.759502288 | -0.695408944 | 0.510358806 | 0.991835066 | -5.061557875 |
| **A_51_P422685** | Zmat4 | -0.040107148 | -0.177955291 | 0.097740994 | 7.784865977 | -0.695392491 | 0.510368474 | 0.991835066 | -5.061566706 |
| **A_55_P1964076** | Dnajb6 | -0.057953786 | -0.257184622 | 0.14127705 | 9.058841009 | -0.695239418 | 0.510458422 | 0.991835066 | -5.061648862 |
| **A_55_P2064025** | A430073D2 | 0.135374256 | -0.330112312 | 0.600860824 | 12.62760966 | 0.695085234 | 0.510549033 | 0.991835066 | -5.061731598 |
| **A_55_P2021709** | Oxnad1 | -0.032844149 | -0.145819905 | 0.080131607 | 12.01919714 | -0.694835318 | 0.510695926 | 0.991835066 | -5.061865674 |
| **A_55_P2003483** | Gldc | 0.041034221 | -0.100133686 | 0.182202128 | 8.091347526 | 0.69473527 | 0.510754739 | 0.991835066 | -5.061919336 |
| **A_52_P225260** | Ywhaz | 0.055077179 | -0.134409241 | 0.244563599 | 9.045961389 | 0.694708687 | 0.510770367 | 0.991835066 | -5.061933594 |
| **A_55_P2020341** | Camk2g | -0.047674438 | -0.21170038 | 0.116351503 | 13.52840279 | -0.694675814 | 0.510789692 | 0.991835066 | -5.061951224 |
| **A_51_P472726** | Pdlim2 | 0.055937766 | -0.136582968 | 0.248458501 | 10.4209351 | 0.694443217 | 0.510926447 | 0.991835066 | -5.062075948 |
| **A_55_P1957967** | Dhx32 | -0.045365405 | -0.201521488 | 0.110790679 | 10.87280355 | -0.694344523 | 0.510984481 | 0.991835066 | -5.06212886 |
| **A_51_P320304** | Plod1 | -0.057323023 | -0.254658605 | 0.14001256 | 8.478512052 | -0.694277056 | 0.511024155 | 0.991835066 | -5.062165027 |
| **A_55_P2054098** | Gm9495 | 0.060974942 | -0.148933386 | 0.270883269 | 8.402233253 | 0.694273933 | 0.511025992 | 0.991835066 | -5.062166701 |
| **A_55_P2030428** | Sirt1 | 0.046487537 | -0.113550991 | 0.206526065 | 10.35280863 | 0.694258365 | 0.511035147 | 0.991835066 | -5.062175046 |
| **A_51_P239286** | Bcl2l12 | -0.040637022 | -0.180535224 | 0.099261179 | 9.713411452 | -0.694254781 | 0.511037255 | 0.991835066 | -5.062176967 |
| **A_55_P2067942** | D16Ertd47 | 0.04596804 | -0.112374833 | 0.204310912 | 9.515644551 | 0.693851606 | 0.51127439 | 0.991835066 | -5.06239303 |
| **A_55_P2051592** | Gfm2 | 0.042957853 | -0.105028622 | 0.190944329 | 10.36554365 | 0.693792619 | 0.51130909 | 0.991835066 | -5.062424633 |
| **A_55_P2123471** | Kpna2 | 0.067491835 | -0.165033922 | 0.300017593 | 14.00204295 | 0.693728129 | 0.511347029 | 0.991835066 | -5.062459181 |
| **A_55_P1990780** | Ptp4a1 | 0.047253091 | -0.115545353 | 0.210051534 | 12.98241922 | 0.69372781 | 0.511347217 | 0.991835066 | -5.062459352 |
| **A_55_P1992811** | Nudc | 0.081230808 | -0.198649662 | 0.361111279 | 10.20398746 | 0.693677025 | 0.511377094 | 0.991835066 | -5.062486556 |
| **A_52_P672803** | Ctsa | 0.040797782 | -0.099856602 | 0.181452167 | 9.054574933 | 0.693254045 | 0.511625987 | 0.991835066 | -5.062713073 |
| **A_55_P2037732** | Foxl2 | 0.097309603 | -0.238236263 | 0.432855468 | 14.04302779 | 0.693127197 | 0.511700643 | 0.991835066 | -5.062780982 |

| | | | | | | | | | |
|---|---|---|---|---|---|---|---|---|---|
| **A_55_P2383877** | 2810040C0 | 0.076942916 | -0.188374621 | 0.342260454 | 8.863370403 | 0.693125408 | 0.511701696 | 0.991835066 | -5.06278194 |
| **A_66_P113487** | Pacsin2 | -0.057220267 | -0.254550148 | 0.140109614 | 13.45848844 | -0.693052543 | 0.511744584 | 0.991835066 | -5.062820943 |
| **A_55_P2113774** | Yif1b | -0.114181217 | -0.50802298 | 0.279660546 | 10.90188674 | -0.692918303 | 0.511823602 | 0.991835066 | -5.062892791 |
| **A_55_P2017967** | Usf2 | -0.057937529 | -0.257815182 | 0.141940124 | 12.79671149 | -0.692795183 | 0.511896081 | 0.991835066 | -5.062958677 |
| **A_55_P2134862** | Cmip | 0.090775085 | -0.222390243 | 0.403940414 | 9.657411926 | 0.692790817 | 0.511898652 | 0.991835066 | -5.062961013 |
| **A_55_P2084739** | Gcnt1 | -0.02924176 | -0.130126405 | 0.071642885 | 8.058742823 | -0.692767615 | 0.511912312 | 0.991835066 | -5.062973429 |
| **A_51_P369636** | Hat1 | -0.035401579 | -0.157539103 | 0.086735945 | 10.35781872 | -0.692759763 | 0.511916934 | 0.991835066 | -5.06297763 |
| **A_55_P1979893** | Tef | 0.0674165 | -0.165190804 | 0.300023804 | 12.83269998 | 0.692710846 | 0.511945734 | 0.991835066 | -5.063003804 |
| **A_52_P366105** | Pip4p1 | -0.048575892 | -0.216193616 | 0.119041832 | 8.417666463 | -0.692643839 | 0.511985185 | 0.991835066 | -5.063039654 |
| **A_55_P2177639** | NA | -0.084244913 | -0.374995909 | 0.206506083 | 15.37405036 | -0.692518863 | 0.512058773 | 0.991835066 | -5.063106511 |
| **A_55_P1975743** | Dap3 | 0.04261141 | -0.10447344 | 0.189696261 | 13.77314349 | 0.692416005 | 0.512119342 | 0.991835066 | -5.063161529 |
| **A_52_P255804** | NA | -0.037045581 | -0.164932945 | 0.090841782 | 11.79391644 | -0.692337593 | 0.512165519 | 0.991835066 | -5.063203466 |
| **A_65_P18858** | Btf3 | -0.040708969 | -0.181259846 | 0.099841909 | 12.39403433 | -0.692254317 | 0.512214563 | 0.991835066 | -5.063248 |
| **A_55_P2097176** | Idua | 0.106587115 | -0.261503633 | 0.474677862 | 12.06275948 | 0.692084203 | 0.512314759 | 0.991835066 | -5.06333896 |
| **A_52_P313071** | Gosr1 | 0.064601199 | -0.15849954 | 0.287701938 | 10.09785839 | 0.69206789 | 0.512324368 | 0.991835066 | -5.063347682 |
| **A_51_P421958** | Sgms1 | 0.044355211 | -0.108834706 | 0.197545127 | 11.69784392 | 0.692027891 | 0.51234793 | 0.991835066 | -5.063369066 |
| **A_55_P2054777** | Maml1 | 0.048211201 | -0.118317199 | 0.214739602 | 10.56477228 | 0.691940538 | 0.512399387 | 0.991835066 | -5.063415763 |
| **A_55_P1977917** | Cln3 | 0.030448932 | -0.074749019 | 0.135646884 | 9.566500407 | 0.691789424 | 0.512488412 | 0.991835066 | -5.063496534 |
| **A_55_P2414924** | Luc7l | 0.059065871 | -0.145031373 | 0.263163116 | 8.436305634 | 0.691685397 | 0.512549703 | 0.991835066 | -5.063552129 |
| **A_52_P134639** | Tgoln2 | 0.031465919 | -0.077274414 | 0.140206251 | 8.072421277 | 0.691606218 | 0.512596357 | 0.991835066 | -5.063594439 |
| **A_55_P1960813** | NA | 0.03188084 | -0.078295492 | 0.142057171 | 14.41342119 | 0.691592975 | 0.51260416 | 0.991835066 | -5.063601515 |
| **A_55_P2089223** | Ak4 | 0.098054074 | -0.240842749 | 0.436950897 | 11.37191562 | 0.691524032 | 0.512644786 | 0.991835066 | -5.063638352 |
| **A_55_P1976494** | Bbof1 | 0.036829228 | -0.090463083 | 0.16412154 | 8.110539557 | 0.691511772 | 0.51265201 | 0.991835066 | -5.063644902 |
| **A_55_P2043762** | N6amt1 | 0.069552041 | -0.170842942 | 0.309947025 | 9.761043207 | 0.691502275 | 0.512657607 | 0.991835066 | -5.063649976 |
| **A_51_P419300** | Pdcd2l | -0.035152975 | -0.156675852 | 0.086369902 | 11.66384799 | -0.691374211 | 0.512733077 | 0.991835066 | -5.063718392 |
| **A_51_P339822** | Arhgap4 | -0.033127685 | -0.147666192 | 0.081410821 | 7.880849797 | -0.691271586 | 0.51279356 | 0.991835066 | -5.063773209 |
| **A_55_P1967693** | Wipi2 | -0.042985167 | -0.191632756 | 0.105662422 | 8.390382929 | -0.691146124 | 0.512867509 | 0.991835066 | -5.063840217 |
| **A_52_P528707** | Mrpl57 | -0.033732261 | -0.150394688 | 0.082930167 | 14.05442827 | -0.691072435 | 0.512910945 | 0.991835066 | -5.063879569 |
| **A_55_P2142116** | NA | -0.058083868 | -0.258981129 | 0.142813393 | 13.93834605 | -0.691020046 | 0.512941828 | 0.991835066 | -5.063907543 |
| **A_55_P2074381** | NA | 0.044804468 | -0.110184331 | 0.199793268 | 11.54462051 | 0.690923786 | 0.512998574 | 0.991835066 | -5.063958939 |
| **A_55_P2012047** | Gigyf1 | -0.077695324 | -0.346486815 | 0.191096167 | 10.07713853 | -0.690857534 | 0.513037633 | 0.991835066 | -5.06399431 |
| **A_55_P2097553** | C77080 | -0.087703222 | -0.391162701 | 0.215756256 | 9.920781323 | -0.690754899 | 0.513098146 | 0.991835066 | -5.064049099 |
| **A_55_P1990874** | Banp | 0.078808322 | -0.193879396 | 0.35149604 | 10.16354046 | 0.690741639 | 0.513105964 | 0.991835066 | -5.064056177 |
| **A_51_P198645** | Parvb | 0.064097467 | -0.157704424 | 0.285899359 | 9.140113754 | 0.690692527 | 0.513134922 | 0.991835066 | -5.064082392 |
| **A_51_P104125** | Fam76b | -0.066898989 | -0.298405128 | 0.16460715 | 9.758500628 | -0.690662977 | 0.513152346 | 0.991835066 | -5.025954007 |
| **A_55_P1956752** | NA | -0.045609456 | -0.203453489 | 0.112234578 | 8.525532383 | -0.690614754 | 0.513180781 | 0.991835066 | -5.064123902 |
| **A_55_P2043382** | Llgl2 | 0.040834252 | -0.10050121 | 0.182169714 | 9.935430484 | 0.690530064 | 0.513230723 | 0.991835066 | -5.064169099 |
| **A_55_P2081148** | Foxk2 | 0.05805599 | -0.142937522 | 0.259049501 | 10.67616946 | 0.690357629 | 0.513332415 | 0.991835066 | -5.064261109 |
| **A_55_P2282969** | Gm2a | 0.028205666 | -0.069455059 | 0.12586639 | 7.830918786 | 0.690280406 | 0.513377962 | 0.991835066 | -5.064302309 |
| **A_55_P2454004** | Smim8 | -0.03977547 | -0.177500163 | 0.097949223 | 11.92371312 | -0.690259898 | 0.513390058 | 0.991835066 | -5.064313249 |
| **A_55_P1961152** | Pou3f1 | -0.049418894 | -0.220534662 | 0.121696874 | 8.225349282 | -0.690259071 | 0.513390546 | 0.991835066 | -5.06431369 |
| **A_55_P2341890** | NA | 0.068864873 | -0.169594363 | 0.307324109 | 8.3840466 | 0.69022826 | 0.513408719 | 0.991835066 | -5.064330127 |
| **A_55_P2082529** | Sav1 | 0.050943413 | -0.125469647 | 0.227356472 | 11.45257374 | 0.690186459 | 0.513433375 | 0.991835066 | -5.064352425 |
| **A_55_P2106666** | Adcy9 | -0.086526133 | -0.386205523 | 0.213153257 | 10.15634105 | -0.690080175 | 0.513496071 | 0.991835066 | -5.064409117 |
| **A_55_P1974317** | Surf6 | 0.040550836 | -0.099906712 | 0.181008384 | 9.059201055 | 0.690023462 | 0.513529527 | 0.991835066 | -5.064439364 |
| **A_55_P2069176** | NA | 0.073923069 | -0.182155183 | 0.330001321 | 11.99863874 | 0.689948105 | 0.513573983 | 0.991835066 | -5.064479552 |
| **A_55_P2120545** | Wdtc1 | -0.062287333 | -0.27807621 | 0.153501545 | 8.753237008 | -0.689889934 | 0.513608303 | 0.991835066 | -5.064510572 |
| **A_66_P107943** | NA | -0.055874585 | -0.249501811 | 0.137752641 | 12.5873177 | -0.689694905 | 0.513723376 | 0.991835066 | -5.064614557 |

| | | | | | | | | | |
|---|---|---|---|---|---|---|---|---|---|
| **A_55_P2145449** | Cep68 | 0.044597457 | -0.109952619 | 0.199147534 | 8.677722984 | 0.689683771 | 0.513729946 | 0.991835066 | -5.064620493 |
| **A_55_P2054708** | NA | 0.045204473 | -0.11147207 | 0.201881017 | 7.826689544 | 0.689583028 | 0.513789395 | 0.991835066 | -5.064674197 |
| **A_55_P2087589** | Rps11 | -0.032238926 | -0.143978114 | 0.079500263 | 15.22612144 | -0.689579246 | 0.513791627 | 0.991835066 | -5.064676213 |
| **A_66_P121774** | Tmem59 | -0.042588614 | -0.190202341 | 0.105025114 | 14.75771362 | -0.689566073 | 0.5137994 | 0.991835066 | -5.064683234 |
| **A_52_P475870** | Tmem29 | 0.053301612 | -0.131447384 | 0.238050608 | 8.757728037 | 0.689552586 | 0.51380736 | 0.991835066 | -5.064690423 |
| **A_52_P98778** | Ang4 | -0.029568916 | -0.132058711 | 0.072920879 | 7.898437036 | -0.689547062 | 0.513810619 | 0.991835066 | -5.064693368 |
| **A_55_P2012789** | Impa1 | 0.044087475 | -0.108745119 | 0.196920069 | 10.76030936 | 0.689458893 | 0.513862653 | 0.991835066 | -5.064740361 |
| **A_52_P14456** | Srrm4 | 0.035194004 | -0.086829459 | 0.157217467 | 8.201643845 | 0.68934157 | 0.513931897 | 0.991835066 | -5.064802885 |
| **A_66_P120414** | Adcy5 | -0.075058141 | -0.335365541 | 0.18524926 | 10.94229831 | -0.689160568 | 0.514038737 | 0.991835066 | -5.064899328 |
| **A_55_P2088530** | Htt | 0.050900457 | -0.12565244 | 0.227453354 | 10.28112686 | 0.689058299 | 0.51409911 | 0.991835066 | -5.064953811 |
| **A_51_P424561** | Zfp93 | -0.060411495 | -0.26996421 | 0.14914122 | 9.136485169 | -0.689025705 | 0.514118352 | 0.991835066 | -5.064971174 |
| **A_55_P2148171** | Ncmap | -0.030617474 | -0.136828133 | 0.075593185 | 7.698074495 | -0.688985986 | 0.514141801 | 0.991835066 | -5.064992331 |
| **A_55_P1959633** | Hnf4a | -0.062654476 | -0.280002918 | 0.154693966 | 7.839919665 | -0.688976966 | 0.514147126 | 0.991835066 | -5.064997136 |
| **A_52_P677147** | Prmt5 | 0.057890855 | -0.142945759 | 0.258727469 | 10.98455341 | 0.688931759 | 0.514173816 | 0.991835066 | -5.065021215 |
| **A_55_P1982817** | NA | 0.057093327 | -0.141051379 | 0.255238034 | 9.980696228 | 0.688671346 | 0.514327581 | 0.991835066 | -5.065159896 |
| **A_52_P551909** | Arhgap5 | 0.040338545 | -0.099669196 | 0.180346287 | 9.228803581 | 0.688616328 | 0.514360071 | 0.991835066 | -5.065189189 |
| **A_55_P2072164** | NA | -0.034060844 | -0.152280988 | 0.0841593 | 8.032762385 | -0.68860957 | 0.514364062 | 0.991835066 | -5.065192787 |
| **A_55_P2015232** | NA | 0.072498309 | -0.179160355 | 0.324156974 | 9.478821234 | 0.688533574 | 0.514408943 | 0.991835066 | -5.065233247 |
| **A_51_P266640** | Nob1 | 0.05207379 | -0.128695158 | 0.232842738 | 8.47217195 | 0.688500864 | 0.514428261 | 0.991835066 | -5.065250661 |
| **A_55_P2185595** | Samd4b | -0.035136594 | -0.157120475 | 0.086847286 | 12.03872559 | -0.688440404 | 0.514463969 | 0.991835066 | -5.065282845 |
| **A_55_P2023391** | Grhl3 | -0.05824084 | -0.260451665 | 0.143969985 | 8.789347938 | -0.688386528 | 0.51449579 | 0.991835066 | -5.065311523 |
| **A_55_P2115458** | Pot1a | -0.087412632 | -0.390931074 | 0.21610581 | 10.77547245 | -0.688332454 | 0.514527729 | 0.991835066 | -5.065340304 |
| **A_52_P658437** | Espl1 | -0.039826518 | -0.178146671 | 0.098493634 | 8.894429915 | -0.688170439 | 0.514623432 | 0.991835066 | -5.065426527 |
| **A_55_P1973289** | Mapk9 | 0.05115425 | -0.126518016 | 0.228826517 | 11.12538581 | 0.688131142 | 0.514646647 | 0.991835066 | -5.065447438 |
| **A_51_P456957** | Pex11a | -0.03842505 | -0.171888099 | 0.095037999 | 10.23575703 | -0.688117388 | 0.514654772 | 0.991835066 | -5.060094061 |
| **A_55_P2024080** | Rpl7a | -0.047159007 | -0.21096959 | 0.116651575 | 15.08883364 | -0.688068747 | 0.514683508 | 0.991835066 | -5.065480638 |
| **A_55_P1991896** | Ceacam5 | -0.315913166 | -1.413291007 | 0.781464675 | 8.279409232 | -0.688051066 | 0.514693954 | 0.991835066 | -5.065490045 |
| **A_55_P2003719** | NA | 0.053842962 | -0.133190296 | 0.240876221 | 15.75720069 | 0.688048793 | 0.514695297 | 0.991835066 | -5.065491255 |
| **A_55_P2102250** | Dzip1 | -0.030361808 | -0.13583067 | 0.075107054 | 8.323095113 | -0.688038121 | 0.514701602 | 0.991835066 | -5.065496932 |
| **A_52_P380301** | Unc5c | 0.054534391 | -0.134909539 | 0.243978321 | 8.033213722 | 0.688016586 | 0.514714325 | 0.991835066 | -5.06550839 |
| **A_55_P2077618** | Csgalnact1 | -0.032472551 | -0.145308519 | 0.080363418 | 8.806406185 | -0.687825027 | 0.514827507 | 0.991835066 | -5.065610295 |
| **A_55_P1964842** | Opa1 | -0.047291656 | -0.211624133 | 0.117040821 | 11.17828569 | -0.687812796 | 0.514834734 | 0.991835066 | -5.065616801 |
| **A_55_P2062836** | Ttf2 | 0.047002829 | -0.116331477 | 0.210337136 | 10.80709594 | 0.687789785 | 0.514848331 | 0.991835066 | -5.06562904 |
| **A_66_P134394** | Pde6c | -0.028335717 | -0.126809856 | 0.070138422 | 7.71981406 | -0.687735034 | 0.514880685 | 0.991835066 | -5.065658161 |
| **A_55_P2095301** | NA | 0.029894608 | -0.074036225 | 0.133825441 | 7.80463241 | 0.68747608 | 0.515033723 | 0.991835066 | -5.065795866 |
| **A_52_P227391** | Kif15 | -0.035327609 | -0.158151165 | 0.087495948 | 8.152477283 | -0.687450937 | 0.515048584 | 0.991835066 | -5.065809234 |
| **A_55_P2041828** | Tubb3 | 0.042090338 | -0.104319073 | 0.18849975 | 7.953779097 | 0.68710412 | 0.515253598 | 0.991835066 | -5.06599359 |
| **A_55_P2087671** | Abhd12 | -0.051535752 | -0.230807074 | 0.127735571 | 8.646383535 | -0.687079395 | 0.515268216 | 0.991835066 | -5.06600673 |
| **A_55_P2030282** | Gm5796 | 0.165352548 | -0.409906034 | 0.74061113 | 11.10982646 | 0.687000286 | 0.515314988 | 0.991835066 | -5.066048769 |
| **A_55_P2091974** | Gpr137 | -0.057777786 | -0.258855192 | 0.143299621 | 10.51912398 | -0.68676278 | 0.515455426 | 0.991835066 | -5.066174958 |
| **A_55_P1983968** | Slc8b1 | 0.033975379 | -0.084267199 | 0.152217958 | 7.880305219 | 0.686751397 | 0.515462157 | 0.991835066 | -5.066181005 |
| **A_55_P2068154** | Anapc15 | -0.03602332 | -0.161411359 | 0.08936472 | 10.36944955 | -0.686652098 | 0.515520881 | 0.991835066 | -5.066233752 |
| **A_55_P2090995** | Ebna1bp2 | -0.048402515 | -0.216880208 | 0.120075178 | 9.498213789 | -0.686648787 | 0.515522839 | 0.991835066 | -5.066235511 |
| **A_55_P2082439** | Spout1 | 0.048594786 | -0.120556333 | 0.217745905 | 11.15215939 | 0.686631826 | 0.51553287 | 0.991835066 | -5.06624452 |
| **A_55_P2069460** | NA | -0.066787864 | -0.299275628 | 0.1656999 | 15.93387325 | -0.686604407 | 0.515549086 | 0.991835066 | -5.066259083 |
| **A_55_P1995698** | Thumpd1 | 0.047437684 | -0.117709555 | 0.212584924 | 10.63308832 | 0.686532805 | 0.515591434 | 0.991835066 | -5.066297111 |
| **A_51_P431329** | Car3 | 0.080775949 | -0.200517124 | 0.362069022 | 8.348885538 | 0.68632871 | 0.515712156 | 0.991835066 | -5.06640549 |
| **A_55_P1991016** | Basp1 | -0.038687244 | -0.173454285 | 0.096079798 | 8.528554926 | -0.686109175 | 0.515842031 | 0.991835066 | -5.066522037 |

| | | | | | | | | | |
|---|---|---|---|---|---|---|---|---|---|
| **A_55_P2145617** | Krt80 | 0.042882435 | -0.106507679 | 0.192272549 | 8.198547427 | 0.686067284 | 0.515866816 | 0.991835066 | -5.066544273 |
| **A_52_P185044** | Adipor2 | 0.036155538 | -0.089824463 | 0.16213554 | 13.40238901 | 0.685934037 | 0.515945657 | 0.991835066 | -5.066614993 |
| **A_51_P355413** | Nisch | -0.056002168 | -0.251169552 | 0.139165216 | 15.66576731 | -0.685814613 | 0.516016325 | 0.991835066 | -5.066678367 |
| **A_65_P02323** | Sec22b | 0.055235787 | -0.137304742 | 0.247776316 | 8.258974028 | 0.685657954 | 0.516109036 | 0.991835066 | -5.066761486 |
| **A_66_P130960** | Psmd11 | 0.055161098 | -0.137129258 | 0.247451454 | 10.21055172 | 0.685621663 | 0.516130515 | 0.991835066 | -5.066780739 |
| **A_55_P1955921** | NA | 0.032747139 | -0.081416344 | 0.146910622 | 14.5554728 | 0.6855755 | 0.516157837 | 0.991835066 | -5.066805227 |
| **A_51_P480241** | Elf3 | -0.028438208 | -0.127584628 | 0.070708212 | 7.66439687 | -0.685542397 | 0.51617743 | 0.991835066 | -5.066822787 |
| **A_55_P2024749** | Hspb11 | 0.047141868 | -0.117250118 | 0.211533855 | 10.7904225 | 0.685386078 | 0.516269958 | 0.991835066 | -5.066905699 |
| **A_55_P1988689** | Rbpjl | -0.028494239 | -0.127912487 | 0.070924009 | 7.727727223 | -0.685015018 | 0.51648964 | 0.991835066 | -5.067102445 |
| **A_55_P2404157** | Rbm27 | 0.040817542 | -0.101633876 | 0.18326896 | 8.364649992 | 0.68484014 | 0.516593194 | 0.991835066 | -5.06719514 |
| **A_51_P518586** | Emg1 | -0.072990891 | -0.327728026 | 0.181746244 | 10.26681707 | -0.684834356 | 0.51659662 | 0.991835066 | -5.067198205 |
| **A_55_P2088716** | Ss18l2 | 0.048981214 | -0.122031509 | 0.219993937 | 12.9791773 | 0.684558008 | 0.516760288 | 0.991835066 | -5.067344643 |
| **A_52_P316697** | Ost4 | 0.054530027 | -0.135867262 | 0.244927316 | 12.02132099 | 0.684516756 | 0.516784723 | 0.991835066 | -5.067366499 |
| **A_51_P443403** | Smoc1 | 0.057704966 | -0.143793327 | 0.259203259 | 8.696875197 | 0.684464534 | 0.516815656 | 0.991835066 | -5.067394164 |
| **A_51_P101545** | Hgfac | -0.036982062 | -0.166140585 | 0.092176461 | 7.841285232 | -0.68434829 | 0.516884517 | 0.991835066 | -5.067455741 |
| **A_66_P134294** | Uba1y | 0.050409657 | -0.125662919 | 0.226482233 | 8.564161146 | 0.684275762 | 0.516927485 | 0.991835066 | -5.067494156 |
| **A_51_P521171** | Hif1an | 0.042377929 | -0.105650069 | 0.190405926 | 8.444198808 | 0.684234542 | 0.516951905 | 0.991835066 | -5.067515987 |
| **A_51_P254646** | Jdp2 | 0.052960143 | -0.132051904 | 0.23797219 | 9.365468443 | 0.684160933 | 0.516995516 | 0.991835066 | -5.067554969 |
| **A_55_P2084741** | Zkscan6 | 0.0315956 | -0.078783553 | 0.141974752 | 9.559249161 | 0.684145824 | 0.517004469 | 0.991835066 | -5.06756297 |
| **A_51_P448664** | Tbxas1 | -0.040459278 | -0.181817379 | 0.100898823 | 8.085861384 | -0.684079475 | 0.517043781 | 0.991835066 | -5.067598104 |
| **A_66_P114784** | Pla2g7 | -0.02923774 | -0.131409449 | 0.072933969 | 7.797617717 | -0.683946731 | 0.517122439 | 0.991835066 | -5.067668387 |
| **A_55_P2016388** | Eif3k | -0.034827486 | -0.156538573 | 0.086883601 | 14.45591205 | -0.683913406 | 0.517142187 | 0.991835066 | -5.06768603 |
| **A_55_P2170335** | Bmi1 | -0.044477205 | -0.19991426 | 0.11095985 | 10.74604894 | -0.683899145 | 0.517150638 | 0.991835066 | -5.06769358 |
| **A_55_P2151273** | Elovl1 | 0.043084369 | -0.107501192 | 0.193669929 | 13.23414035 | 0.683825886 | 0.517194053 | 0.991835066 | -5.06773236 |
| **A_51_P192149** | Slc9a6 | -0.031402582 | -0.141162348 | 0.078357184 | 8.423475493 | -0.683803496 | 0.517207322 | 0.991835066 | -5.067744212 |
| **A_55_P2435504** | Cep295 | 0.047538776 | -0.118626721 | 0.213704273 | 9.370879807 | 0.683779817 | 0.517221355 | 0.991835066 | -5.067756746 |
| **A_52_P257625** | Esm1 | 0.110273277 | -0.275190452 | 0.495737006 | 8.454463752 | 0.683747727 | 0.517240373 | 0.991835066 | -5.067773731 |
| **A_51_P456465** | Cldn10 | -0.031157382 | -0.140111191 | 0.077796427 | 7.692636111 | -0.683482923 | 0.51739733 | 0.991835066 | -5.067913868 |
| **A_55_P2457153** | G2e3 | -0.043963027 | -0.197700052 | 0.109773998 | 9.339655459 | -0.683468099 | 0.517406118 | 0.991835066 | -5.067921712 |
| **A_55_P1970715** | NA | 0.031970355 | -0.079845141 | 0.14378585 | 8.290749971 | 0.683367926 | 0.517465502 | 0.991835066 | -5.067974711 |
| **A_52_P508974** | Igf2bp3 | 0.042642315 | -0.106539623 | 0.191824254 | 9.389071701 | 0.683177673 | 0.517578298 | 0.991835066 | -5.068075353 |
| **A_51_P357433** | Ap1g2 | -0.028275915 | -0.127197973 | 0.070646143 | 8.253962034 | -0.683176085 | 0.517579239 | 0.991835066 | -5.068076193 |
| **A_51_P443754** | Hivep2 | -0.038305191 | -0.172335426 | 0.095725043 | 8.579845902 | -0.683068078 | 0.517643281 | 0.991835066 | -5.068133317 |
| **A_51_P221625** | Stk24 | -0.065902719 | -0.296538006 | 0.164732567 | 9.837653253 | -0.682946548 | 0.517715348 | 0.991835066 | -5.068197584 |
| **A_55_P2096485** | Bcs1l | -0.033792791 | -0.152074136 | 0.084488555 | 11.32268728 | -0.682836827 | 0.517780417 | 0.991835066 | -5.068255598 |
| **A_55_P2043062** | Hs3st6 | 0.060680182 | -0.151725137 | 0.273085502 | 8.137490327 | 0.682795453 | 0.517804954 | 0.991835066 | -5.068277472 |
| **A_55_P2016851** | Lce1a2 | -0.044378939 | -0.199733283 | 0.110975404 | 11.169567 | -0.682751478 | 0.517831036 | 0.991835066 | -5.06830072 |
| **A_55_P2362601** | D4Ertd681 | 0.132347545 | -0.330975125 | 0.595670215 | 8.946302916 | 0.682718188 | 0.51785078 | 0.991835066 | -5.068318318 |
| **A_55_P2087647** | Grhpr | 0.0669133 | -0.167337102 | 0.301163702 | 8.957024266 | 0.682717814 | 0.517851002 | 0.991835066 | -5.068318516 |
| **A_51_P214985** | Zfp521 | 0.069764418 | -0.174501845 | 0.31403068 | 10.14441823 | 0.68262096 | 0.51790845 | 0.991835066 | -5.068369713 |
| **A_55_P2032845** | Rbm10 | 0.049899233 | -0.12483841 | 0.224636876 | 11.33719128 | 0.682521806 | 0.517967267 | 0.991835066 | -5.06842212 |
| **A_51_P333349** | Tmem120a | 0.083607099 | -0.209240059 | 0.376454257 | 10.40887299 | 0.682356407 | 0.518065387 | 0.991835066 | -5.068509525 |
| **A_55_P2061381** | Ghitm | -0.047480381 | -0.213848239 | 0.118887477 | 13.28124458 | -0.6821092 | 0.518212063 | 0.991835066 | -5.068640129 |
| **A_55_P1974467** | Ndufb6 | -0.03783491 | -0.170406488 | 0.094736669 | 14.59974179 | -0.682105273 | 0.518214393 | 0.991835066 | -5.068642203 |
| **A_52_P497424** | Aanat | -0.039612126 | -0.17846196 | 0.099237708 | 7.887249089 | -0.68185483 | 0.518363015 | 0.991835066 | -5.068774476 |
| **A_55_P2128263** | Phf21a | 0.065034164 | -0.162964009 | 0.293032337 | 11.04908457 | 0.681740864 | 0.518430656 | 0.991835066 | -5.068834654 |
| **A_55_P2175692** | Reep3 | -0.087676951 | -0.395076954 | 0.219723052 | 10.38805231 | -0.681695933 | 0.518457325 | 0.991835066 | -5.068858377 |
| **A_55_P2024515** | Crtc2 | 0.054991581 | -0.13781261 | 0.247795772 | 9.397408149 | 0.681693054 | 0.518459034 | 0.991835066 | -5.068859897 |

| | | | | | | | | | |
|---|---|---|---|---|---|---|---|---|---|
| **A_55_P1976224** | Ckb | -0.087287616 | -0.393368192 | 0.218792959 | 12.36679518 | -0.681594372 | 0.51851761 | 0.991835066 | -5.068911995 |
| **A_55_P1956882** | Cpt1b | -0.042532238 | -0.191707646 | 0.10664317 | 12.9316441 | -0.681443943 | 0.51860691 | 0.991835066 | -5.0689914 |
| **A_51_P472901** | Slc3a2 | 0.070581987 | -0.177052631 | 0.318216604 | 10.90344225 | 0.681226691 | 0.518735897 | 0.991835066 | -5.069106052 |
| **A_55_P1996817** | NA | 0.074464202 | -0.186831017 | 0.335759421 | 12.76693555 | 0.681122472 | 0.518797781 | 0.991835066 | -5.069161042 |
| **A_51_P402536** | Uvrag | -0.044553593 | -0.20090338 | 0.111796194 | 11.3364704 | -0.681074422 | 0.518826314 | 0.991835066 | -5.069186392 |
| **A_51_P486091** | Ift52 | 0.05313388 | -0.133353712 | 0.239621472 | 10.01255015 | 0.680974299 | 0.518885773 | 0.991835066 | -5.069239211 |
| **A_55_P2036967** | Cox17 | 0.030209561 | -0.075827622 | 0.136246745 | 7.817427092 | 0.680918863 | 0.518918696 | 0.991835066 | -5.069268453 |
| **A_51_P469661** | Stk38 | -0.054004677 | -0.24362849 | 0.135619137 | 10.18797901 | -0.680687264 | 0.519056254 | 0.991835066 | -5.069390597 |
| **A_66_P127432** | Tial1 | 0.0437403 | -0.109875551 | 0.197356152 | 11.41433283 | 0.680541887 | 0.519142613 | 0.991835066 | -5.069467251 |
| **A_55_P2133187** | NA | 0.079088638 | -0.198697911 | 0.356875186 | 9.173795185 | 0.680474752 | 0.519182496 | 0.991835066 | -5.069502644 |
| **A_55_P2074406** | Ldoc1 | 0.029169375 | -0.073285344 | 0.131624093 | 7.706481262 | 0.680462642 | 0.51918969 | 0.991835066 | -5.069509028 |
| **A_55_P2052831** | 0610009B2 | -0.065234324 | -0.294393831 | 0.163925183 | 10.00867016 | -0.680373545 | 0.519242625 | 0.991835066 | -5.069555996 |
| **A_55_P2021908** | NA | -0.148424051 | -0.669897959 | 0.373049858 | 12.49419476 | -0.680269233 | 0.519304603 | 0.991835066 | -5.069610977 |
| **A_55_P2074521** | Psg28 | -0.372175654 | -1.679788574 | 0.935437266 | 8.456884305 | -0.680264628 | 0.519307339 | 0.991835066 | -5.069613404 |
| **A_55_P2171406** | Tm7sf2 | -0.031213067 | -0.140892898 | 0.078466765 | 7.891357204 | -0.680172074 | 0.519362335 | 0.991835066 | -5.069662182 |
| **A_52_P521507** | Trio | -0.074494393 | -0.336290962 | 0.187302175 | 11.388943 | -0.680093728 | 0.519408892 | 0.991835066 | -5.069703467 |
| **A_52_P374669** | Kdm5a | -0.049009399 | -0.221246601 | 0.123227802 | 10.62416436 | -0.680082426 | 0.519415608 | 0.991835066 | -5.069709422 |
| **A_55_P2130256** | Polr2l | -0.056190134 | -0.253715899 | 0.14133563 | 9.145557599 | -0.679900639 | 0.519523644 | 0.991835066 | -5.0698052 |
| **A_51_P180001** | Ago2 | 0.053485784 | -0.134561487 | 0.241533055 | 11.63699552 | 0.679798906 | 0.519584111 | 0.991835066 | -5.069858791 |
| **A_52_P601980** | Kdm5b | 0.039665996 | -0.099805856 | 0.179137849 | 8.547225152 | 0.679737031 | 0.51962089 | 0.991835066 | -5.069891383 |
| **A_52_P366653** | Hnrnph3 | -0.039225696 | -0.177182178 | 0.098730786 | 8.596031316 | -0.679575446 | 0.519716943 | 0.991835066 | -5.069976482 |
| **A_55_P2143522** | Fstl3 | 0.108778878 | -0.273819272 | 0.491377028 | 10.45125266 | 0.679533454 | 0.519741908 | 0.991835066 | -5.069998595 |
| **A_51_P319453** | Zfp143 | -0.034191156 | -0.154468962 | 0.086086651 | 8.670704396 | -0.679418575 | 0.519810206 | 0.991835066 | -5.070059083 |
| **A_51_P352385** | Ism1 | 0.028418523 | -0.071553549 | 0.128390595 | 7.829653031 | 0.679410005 | 0.519815301 | 0.991835066 | -5.070063595 |
| **A_55_P1957468** | She | 0.036602598 | -0.092164563 | 0.16536976 | 10.78122627 | 0.67938496 | 0.519830192 | 0.991835066 | -5.070076781 |
| **A_51_P171616** | Wnt10a | 0.039209503 | -0.098762458 | 0.177181463 | 7.697250586 | 0.679218692 | 0.519929056 | 0.991835066 | -5.070164309 |
| **A_55_P1990609** | Ube2j1 | 0.044204997 | -0.111349948 | 0.199759942 | 11.43980523 | 0.67919844 | 0.519941099 | 0.991835066 | -5.070174969 |
| **A_52_P63905** | Ddc | -0.10464889 | -0.472922148 | 0.263624369 | 10.86768415 | -0.679162303 | 0.519962588 | 0.991835066 | -5.070193989 |
| **A_55_P2066041** | Hnrnpa1 | 0.059001577 | -0.148667747 | 0.266670901 | 9.18390999 | 0.679047888 | 0.520030629 | 0.991835066 | -5.070254206 |
| **A_55_P2099760** | NA | 0.079766298 | -0.200995437 | 0.360528033 | 8.756350391 | 0.679032643 | 0.520039696 | 0.991835066 | -5.070262229 |
| **A_55_P1955632** | Trp53i11 | -0.050558995 | -0.228534534 | 0.127416545 | 11.96937282 | -0.678964777 | 0.520080057 | 0.991835066 | -5.070297941 |
| **A_52_P348250** | Klhl29 | -0.032959074 | -0.148980319 | 0.083062172 | 7.832360993 | -0.678963775 | 0.520080654 | 0.991835066 | -5.070298469 |
| **A_55_P2280089** | 4930570B1 | 0.079960379 | -0.201533684 | 0.361454442 | 10.21643568 | 0.678913958 | 0.520110283 | 0.991835066 | -5.070324682 |
| **A_51_P419017** | Pde10a | 0.030216663 | -0.076161607 | 0.136594932 | 8.016548282 | 0.678895155 | 0.520121466 | 0.991835066 | -5.070334575 |
| **A_51_P264695** | Crym | 0.036957584 | -0.093160161 | 0.167075328 | 8.130034381 | 0.678853674 | 0.520146139 | 0.991835066 | -5.070356401 |
| **A_55_P1961853** | Zfand3 | 0.052921393 | -0.133467482 | 0.239310269 | 9.334918617 | 0.678610245 | 0.520290943 | 0.991835066 | -5.070484458 |
| **A_51_P387379** | Tshz3 | 0.037291578 | -0.094049229 | 0.168632384 | 9.075597195 | 0.678609935 | 0.520291127 | 0.991835066 | -5.070484621 |
| **A_55_P2040962** | Gm1604b | -0.061711912 | -0.279076935 | 0.155653111 | 10.04116071 | -0.678560337 | 0.520320634 | 0.991835066 | -5.070510708 |
| **A_55_P2001193** | Stx18 | 0.036166704 | -0.09122303 | 0.163556437 | 10.4745919 | 0.678552776 | 0.520325132 | 0.991835066 | -5.070514685 |
| **A_55_P2022595** | Gm10653 | -0.03553775 | -0.160719207 | 0.089643706 | 14.95337294 | -0.678514388 | 0.52034797 | 0.991835066 | -5.070534874 |
| **A_51_P113784** | Prpf19 | 0.096606671 | -0.24369336 | 0.436906702 | 9.782210868 | 0.678506889 | 0.520352432 | 0.991835066 | -5.070538818 |
| **A_55_P2343151** | Akap1 | -0.042555062 | -0.192459332 | 0.107349208 | 11.58267821 | -0.678494534 | 0.520359782 | 0.991835066 | -5.070545315 |
| **A_55_P2175767** | Rangrf | 0.065669592 | -0.165705417 | 0.297044601 | 10.59912536 | 0.678354955 | 0.52044283 | 0.991835066 | -5.070618714 |
| **A_55_P2122260** | Tmem87b | -0.0338076 | -0.15296058 | 0.08534538 | 8.295064068 | -0.678138754 | 0.520571483 | 0.991835066 | -5.07073238 |
| **A_51_P487690** | Ifi44 | -0.068720457 | -0.311088779 | 0.173647865 | 8.067667542 | -0.677671645 | 0.52084951 | 0.991835066 | -5.070977857 |
| **A_55_P2087805** | Hsd3b1 | -0.04554174 | -0.206161989 | 0.115078508 | 7.787334127 | -0.677670316 | 0.520850302 | 0.991835066 | -5.070978555 |
| **A_55_P2143657** | Gm5784 | 0.041070489 | -0.103829115 | 0.185970093 | 8.325541726 | 0.677441523 | 0.520986516 | 0.991835066 | -5.071098739 |
| **A_52_P518808** | Mmd | 0.043613738 | -0.110275705 | 0.197503181 | 13.23386447 | 0.677366348 | 0.521031277 | 0.991835066 | -5.071138221 |

| | | | | | | | | | |
|---|---|---|---|---|---|---|---|---|---|
| **A_51_P464387** | Hspb8 | -0.095199864 | -0.431124971 | 0.240725243 | 10.29907535 | -0.677334187 | 0.521050428 | 0.991835066 | -5.071155111 |
| **A_55_P2096535** | Icmt | 0.075370937 | -0.19059632 | 0.341338195 | 9.516879979 | 0.677305925 | 0.521067257 | 0.991835066 | -5.071169952 |
| **A_52_P67995** | Eif4a3 | -0.063780545 | -0.288956455 | 0.161395364 | 10.39726961 | -0.676979373 | 0.521261732 | 0.991835066 | -5.071341402 |
| **A_66_P127411** | Acat2 | -0.040322991 | -0.182684701 | 0.10203872 | 8.09598023 | -0.676968829 | 0.521268012 | 0.991835066 | -5.071346937 |
| **A_55_P2382070** | 4930593A0 | -0.053180637 | -0.240960952 | 0.134599678 | 8.399859537 | -0.676881438 | 0.521320065 | 0.991835066 | -5.071392807 |
| **A_52_P140497** | NA | -0.046425334 | -0.210358977 | 0.117508308 | 8.96816401 | -0.67685569 | 0.521335402 | 0.991835066 | -5.071406321 |
| **A_66_P104190** | Gsk3b | 0.06825439 | -0.172782202 | 0.309290982 | 10.42665283 | 0.676794382 | 0.521371922 | 0.991835066 | -5.071438497 |
| **A_55_P2459897** | A2m | 0.027794673 | -0.070366112 | 0.125955459 | 7.680292589 | 0.676756884 | 0.521394259 | 0.991835066 | -5.071458175 |
| **A_52_P447424** | NA | 0.054232258 | -0.137308121 | 0.245772637 | 9.241508428 | 0.676716051 | 0.521418584 | 0.991835066 | -5.071479603 |
| **A_55_P1954092** | NA | 0.123484554 | -0.312657441 | 0.559626548 | 14.31099632 | 0.676696397 | 0.521430293 | 0.991835066 | -5.071489917 |
| **A_52_P432919** | Rab3d | -0.040801223 | -0.184919963 | 0.103317517 | 8.874958333 | -0.676646529 | 0.521460002 | 0.991835066 | -5.071516084 |
| **A_52_P608322** | Maff | -0.068047211 | -0.308408964 | 0.172314542 | 9.661617226 | -0.67663444 | 0.521467204 | 0.991835066 | -5.071522427 |
| **A_55_P2140780** | Clip4 | -0.066605163 | -0.30188328 | 0.168672954 | 10.37880441 | -0.676605454 | 0.521484472 | 0.991835066 | -5.071537635 |
| **A_55_P2006504** | Eif4a2 | 0.070299815 | -0.17804018 | 0.31863981 | 12.78559557 | 0.67657609 | 0.521501967 | 0.991835066 | -5.071553041 |
| **A_51_P502054** | Gtf2b | -0.040305148 | -0.182701278 | 0.102090981 | 11.24816277 | -0.676505714 | 0.521543897 | 0.991835066 | -5.071589964 |
| **A_51_P140321** | Mocos | -0.03464101 | -0.15706567 | 0.08778365 | 7.993391542 | -0.676286578 | 0.521674472 | 0.991835066 | -5.07170491 |
| **A_52_P525317** | Gja5 | 0.097711011 | -0.247751677 | 0.443173699 | 11.26895813 | 0.676007465 | 0.521840816 | 0.991835066 | -5.071851273 |
| **A_55_P1964615** | Dnttip1 | -0.043094153 | -0.195485336 | 0.10929703 | 8.45743706 | -0.675876957 | 0.521918607 | 0.991835066 | -5.071919692 |
| **A_66_P102581** | Ube2z | -0.029724043 | -0.134836122 | 0.075388036 | 10.80933798 | -0.675871898 | 0.521921623 | 0.991835066 | -5.071922344 |
| **A_55_P1953387** | Fabp5 | -0.04342425 | -0.197010958 | 0.110162459 | 15.60798865 | -0.675752751 | 0.521992649 | 0.991835066 | -5.071984797 |
| **A_55_P2181444** | NA | -0.046937944 | -0.212987141 | 0.119111252 | 12.80116664 | -0.675610544 | 0.522077428 | 0.991835066 | -5.072059325 |
| **A_66_P106593** | Vhl | -0.039716075 | -0.180238287 | 0.100806138 | 10.81086609 | -0.67550796 | 0.522138592 | 0.991835066 | -5.072113079 |
| **A_55_P2065040** | NA | -0.033201608 | -0.150687071 | 0.084283854 | 13.66208911 | -0.675435874 | 0.522181574 | 0.991835066 | -5.072150849 |
| **A_55_P2101357** | Aftph | 0.062321029 | -0.158220965 | 0.282863024 | 10.70744118 | 0.675386619 | 0.522210945 | 0.991835066 | -5.072176654 |
| **A_52_P605834** | Tm9sf3 | -0.03126236 | -0.141902116 | 0.079377396 | 10.59325242 | -0.675335659 | 0.522241333 | 0.991835066 | -5.07220335 |
| **A_52_P330289** | Inpp4b | -0.043813979 | -0.198912771 | 0.111284812 | 8.740408786 | -0.675170436 | 0.522339865 | 0.991835066 | -5.072289895 |
| **A_51_P414548** | Casp7 | 0.048695224 | -0.123696919 | 0.221087367 | 9.837710565 | 0.675115428 | 0.522372673 | 0.991835066 | -5.072318705 |
| **A_52_P724940** | Ndufa12 | -0.049958592 | -0.226835495 | 0.126918311 | 15.67471717 | -0.675069048 | 0.522400335 | 0.991835066 | -5.072342994 |
| **A_52_P176737** | Lcorl | 0.044902675 | -0.114079463 | 0.203884814 | 8.663811878 | 0.67504545 | 0.52241441 | 0.991835066 | -5.072355351 |
| **A_55_P1993840** | Tmem132c | 0.054389683 | -0.138190817 | 0.246970184 | 7.99361724 | 0.675014886 | 0.52243264 | 0.991835066 | -5.072371357 |
| **A_55_P1962906** | Als2cl | 0.039798136 | -0.101118593 | 0.180714865 | 8.650057832 | 0.675008609 | 0.522436384 | 0.991835066 | -5.072374644 |
| **A_52_P412321** | Pcgf6 | 0.047988535 | -0.121930416 | 0.217907486 | 8.631090702 | 0.675001606 | 0.522440561 | 0.991835066 | -5.072378311 |
| **A_52_P656545** | Ogt | -0.036007297 | -0.163565569 | 0.091550974 | 8.339483227 | -0.674669426 | 0.52263872 | 0.991835066 | -5.072552218 |
| **A_55_P2099479** | H2afz | 0.053771659 | -0.136740482 | 0.244283799 | 13.96732581 | 0.674590015 | 0.522686099 | 0.991835066 | -5.072593782 |
| **A_51_P255441** | Mrpl43 | 0.058389366 | -0.148488076 | 0.265266808 | 13.31266057 | 0.674574243 | 0.52269551 | 0.991835066 | -5.072602037 |
| **A_55_P2140171** | NA | 0.060217623 | -0.153165738 | 0.273600984 | 11.22774658 | 0.674484836 | 0.522748857 | 0.991835066 | -5.072648827 |
| **A_55_P2017774** | NA | 0.057505377 | -0.14626708 | 0.261277834 | 17.64034496 | 0.674484707 | 0.522748933 | 0.991835066 | -5.072648894 |
| **A_55_P2046563** | Cym | -0.035471272 | -0.161200713 | 0.090258169 | 7.708066452 | -0.674293395 | 0.522863096 | 0.991835066 | -5.072748997 |
| **A_55_P1993599** | Oaz2 | -0.038566955 | -0.175277282 | 0.098143371 | 14.77309999 | -0.674253431 | 0.522886946 | 0.991835066 | -5.072769905 |
| **A_51_P453995** | Gm11961 | 0.038181486 | -0.097164027 | 0.173526998 | 9.657560475 | 0.674245557 | 0.522891645 | 0.991835066 | -5.072774024 |
| **A_55_P1991475** | Sesn1 | 0.048426374 | -0.123290444 | 0.220143191 | 11.0800588 | 0.674028483 | 0.523021204 | 0.991835066 | -5.072887571 |
| **A_55_P2049999** | Slc9a1 | 0.035553765 | -0.090621166 | 0.161728696 | 11.76125717 | 0.673475279 | 0.523351473 | 0.991835066 | -5.073176804 |
| **A_55_P2149615** | Bex3 | -0.071804025 | -0.326637268 | 0.183029218 | 11.90593015 | -0.673444545 | 0.523369825 | 0.991835066 | -5.073192866 |
| **A_51_P205278** | Tmem161b | -0.041910352 | -0.190663653 | 0.10684295 | 8.598080714 | -0.6733856 | 0.523405025 | 0.991835066 | -5.073223672 |
| **A_51_P194306** | Lrrc1 | -0.053497224 | -0.243379928 | 0.13638548 | 11.06883116 | -0.673371877 | 0.52341322 | 0.991835066 | -5.073230844 |
| **A_55_P2168307** | Gm13213 | 0.038272003 | -0.097603809 | 0.174147816 | 16.80916494 | 0.673206303 | 0.523512102 | 0.991835066 | -5.073317361 |
| **A_55_P2154132** | Tuba1b | 0.050851698 | -0.129708315 | 0.23141171 | 17.56100278 | 0.673120803 | 0.523563169 | 0.991835066 | -5.07336203 |
| **A_55_P2108678** | NA | -0.035046758 | -0.159519547 | 0.089426031 | 8.815752593 | -0.672949637 | 0.52366541 | 0.991835066 | -5.073451442 |

| | | | | | | | | | |
|---|---|---|---|---|---|---|---|---|---|
| **A_55_P1992739** | Dync1i2 | -0.046871998 | -0.213350457 | 0.119606462 | 13.55426047 | -0.672921721 | 0.523682086 | 0.991835066 | -5.073466022 |
| **A_52_P876147** | Crocc2 | -0.043512493 | -0.198081603 | 0.111056617 | 7.879705208 | -0.672822328 | 0.523741462 | 0.991835066 | -5.073517931 |
| **A_55_P2011912** | Ecm2 | -0.042115574 | -0.191723391 | 0.107492244 | 8.5430222 | -0.672817944 | 0.523744082 | 0.991835066 | -5.07352022 |
| **A_55_P2142146** | Glul | -0.031440186 | -0.143133247 | 0.080252874 | 11.32825991 | -0.672772224 | 0.523771396 | 0.991835066 | -5.073544096 |
| **A_52_P263201** | Tmem33 | -0.036022633 | -0.164016732 | 0.091971465 | 8.101148504 | -0.672658516 | 0.523839332 | 0.991835066 | -5.073603469 |
| **A_55_P2115713** | Nid1 | 0.051918957 | -0.132571952 | 0.236409866 | 8.386957682 | 0.672605046 | 0.52387128 | 0.991835066 | -5.073631386 |
| **A_55_P1952696** | NA | -0.055423899 | -0.252376758 | 0.141528961 | 12.68338589 | -0.672579945 | 0.523886278 | 0.991835066 | -5.073644491 |
| **A_55_P2062418** | Atp6v0a1 | -0.053862113 | -0.245313126 | 0.1375889 | 11.9401608 | -0.672411065 | 0.523987193 | 0.991835066 | -5.073732648 |
| **A_55_P2102738** | NA | 0.124995496 | -0.319455304 | 0.569446295 | 13.97500261 | 0.67217105 | 0.524130636 | 0.991835066 | -5.073857908 |
| **A_55_P1995774** | NA | 0.036121284 | -0.092323907 | 0.164566475 | 9.518031979 | 0.672131837 | 0.524154074 | 0.991835066 | -5.073878369 |
| **A_55_P2130695** | Armc4 | 0.028308764 | -0.072357697 | 0.128975225 | 7.644203196 | 0.672117547 | 0.524162616 | 0.991835066 | -5.073885826 |
| **A_51_P414637** | Col5a1 | -0.070281086 | -0.320236976 | 0.179674805 | 12.39254853 | -0.672023126 | 0.524219055 | 0.991835066 | -5.073935088 |
| **A_66_P120484** | Mbtd1 | -0.06763869 | -0.308212091 | 0.17293471 | 11.07554282 | -0.671980569 | 0.524244494 | 0.991835066 | -5.07395729 |
| **A_55_P2100209** | Kif2c | -0.030649371 | -0.139668141 | 0.078369399 | 8.374165788 | -0.671938334 | 0.524269741 | 0.991835066 | -5.073979323 |
| **A_55_P2035087** | Arhgef15 | -0.068413274 | -0.311773743 | 0.174947195 | 10.89088351 | -0.671892006 | 0.524297436 | 0.991835066 | -5.074003489 |
| **A_55_P2065646** | Zfp189 | -0.039672093 | -0.180859121 | 0.101514935 | 9.453900762 | -0.671582617 | 0.524482415 | 0.991835066 | -5.074164843 |
| **A_51_P436727** | Arrb1 | 0.040387147 | -0.103348485 | 0.184122778 | 10.81255447 | 0.671564702 | 0.524493127 | 0.991835066 | -5.074174184 |
| **A_52_P642488** | Kcnk1 | 0.030288651 | -0.077508039 | 0.138085342 | 7.776143919 | 0.671558169 | 0.524497034 | 0.991835066 | -5.07417759 |
| **A_55_P2128734** | Pbxip1 | -0.05986315 | -0.273006126 | 0.153279826 | 13.43134433 | -0.671270673 | 0.524668963 | 0.991835066 | -5.074327465 |
| **A_52_P307961** | Abca5 | -0.051594498 | -0.235301419 | 0.132112423 | 8.52003174 | -0.671254187 | 0.524678823 | 0.991835066 | -5.074336058 |
| **A_66_P124445** | Rmc1 | -0.038589564 | -0.176023846 | 0.098844718 | 8.768489097 | -0.67109488 | 0.524774109 | 0.991835066 | -5.074419082 |
| **A_55_P2074081** | NA | -0.096180893 | -0.438729455 | 0.246367669 | 14.18876632 | -0.67108231 | 0.524781628 | 0.991835066 | -5.074425632 |
| **A_55_P1953030** | Tnfsf12 | -0.035229726 | -0.160730743 | 0.090271292 | 8.108670082 | -0.670920632 | 0.524878344 | 0.991835066 | -5.074509873 |
| **A_55_P2077751** | NA | 0.058588362 | -0.150148703 | 0.267325426 | 8.64988791 | 0.670843034 | 0.524924767 | 0.991835066 | -5.074550299 |
| **A_51_P230624** | Ccdc71l | 0.03162843 | -0.081083465 | 0.144340325 | 8.714370496 | 0.670682568 | 0.525020774 | 0.991835066 | -5.074633884 |
| **A_55_P2118740** | NA | 0.050201516 | -0.128779217 | 0.229182248 | 12.04291083 | 0.670377891 | 0.525203095 | 0.991835066 | -5.074792541 |
| **A_55_P2164502** | NA | -0.038415871 | -0.175412187 | 0.098580444 | 8.781711301 | -0.670210049 | 0.52530355 | 0.991835066 | -5.074879917 |
| **A_55_P2100120** | Nme1 | 0.060245487 | -0.154617565 | 0.275108539 | 12.27192463 | 0.67014983 | 0.525339594 | 0.991835066 | -5.074911261 |
| **A_55_P2427377** | NA | 0.027933923 | -0.071694989 | 0.127562835 | 7.674345964 | 0.670124769 | 0.525354595 | 0.991835066 | -5.074924305 |
| **A_55_P2041355** | Gm4750 | -0.069235888 | -0.316190735 | 0.177718958 | 11.6411935 | -0.670074123 | 0.525384911 | 0.991835066 | -5.074950664 |
| **A_55_P2062133** | Etv3 | 0.042977278 | -0.110329727 | 0.196284282 | 8.621105001 | 0.670017328 | 0.52541891 | 0.991835066 | -5.074980221 |
| **A_51_P433026** | Plpp6 | -0.037752257 | -0.172422021 | 0.096917508 | 9.045216979 | -0.670011022 | 0.525422684 | 0.991835066 | -5.074983503 |
| **A_55_P2079713** | Gramd1c | 0.034006301 | -0.087310432 | 0.155323033 | 9.298506698 | 0.669958372 | 0.525454203 | 0.991835066 | -5.075010901 |
| **A_52_P405206** | Tm7sf3 | 0.040450656 | -0.103860622 | 0.184761934 | 11.56613471 | 0.669937716 | 0.525466569 | 0.991835066 | -5.075021649 |
| **A_51_P408649** | Shisa2 | 0.120156039 | -0.308559716 | 0.548871794 | 8.791989829 | 0.669861924 | 0.525511944 | 0.991835066 | -5.075061086 |
| **A_55_P1992910** | Gyg | 0.053160446 | -0.136516174 | 0.242837067 | 15.62719711 | 0.669859849 | 0.525513186 | 0.991835066 | -5.075062165 |
| **A_55_P2000618** | Gm12690 | -0.047903883 | -0.218826441 | 0.123018674 | 8.747478791 | -0.669854485 | 0.525516397 | 0.991835066 | -5.075064956 |
| **A_55_P2015753** | Enho | -0.038927561 | -0.177892279 | 0.100037158 | 9.386241203 | -0.669517249 | 0.525718324 | 0.991835066 | -5.075240382 |
| **A_52_P171724** | 2310009B1 | -0.037426483 | -0.171057732 | 0.096204766 | 10.55161926 | -0.669391387 | 0.525793699 | 0.991835066 | -5.075305835 |
| **A_52_P127465** | Ypel1 | -0.043262035 | -0.197756836 | 0.111232766 | 8.216260554 | -0.66927131 | 0.525865616 | 0.991835066 | -5.07536827 |
| **A_55_P2152836** | NA | 0.079906461 | -0.205456742 | 0.365269665 | 11.8723876 | 0.669257212 | 0.52587406 | 0.991835066 | -5.0753756 |
| **A_55_P2015936** | NA | 0.034126168 | -0.087749864 | 0.1560022 | 16.35181991 | 0.669234544 | 0.525887637 | 0.991835066 | -5.075387385 |
| **A_55_P2022104** | Cop1 | 0.029568014 | -0.076032003 | 0.135168031 | 8.732199954 | 0.669217512 | 0.525897839 | 0.991835066 | -5.075396239 |
| **A_52_P385206** | Harbi1 | -0.032675144 | -0.149373332 | 0.084023043 | 8.507676902 | -0.669210171 | 0.525902236 | 0.991835066 | -5.075400056 |
| **A_55_P1968448** | NA | -0.057838583 | -0.264446258 | 0.148769091 | 14.55544103 | -0.669083514 | 0.525978104 | 0.991835066 | -5.075465897 |
| **A_55_P2060922** | Unc5a | 0.041752931 | -0.107410212 | 0.190916075 | 8.868298068 | 0.669013032 | 0.526020326 | 0.991835066 | -5.075502531 |
| **A_55_P2019483** | Ndrg4 | 0.06451814 | -0.165990483 | 0.295026764 | 14.61100898 | 0.668965615 | 0.526048732 | 0.991835066 | -5.075527175 |
| **A_55_P2062213** | Tmem41a | -0.058355509 | -0.266857322 | 0.150146305 | 11.86922347 | -0.66893074 | 0.526069625 | 0.991835066 | -5.0755453 |

| | | | | | | | | | |
|---|---|---|---|---|---|---|---|---|---|
| **A_66_P117389** | Rcn2 | -0.042834812 | -0.195904882 | 0.110235258 | 10.24999416 | -0.668829959 | 0.526130005 | 0.991835066 | -5.075597673 |
| **A_52_P361081** | Arhgef16 | 0.027979426 | -0.072011868 | 0.127970721 | 7.836565675 | 0.668783803 | 0.526157659 | 0.991835066 | -5.075621656 |
| **A_55_P2112797** | NA | -0.110886519 | -0.507168952 | 0.285395914 | 10.93884499 | -0.668779667 | 0.526160137 | 0.991835066 | -5.075623805 |
| **A_52_P483301** | Mios | 0.040138202 | -0.103321212 | 0.183597616 | 9.069605197 | 0.668710269 | 0.526201719 | 0.991835066 | -5.075659862 |
| **A_55_P2162269** | Bri3 | -0.084688655 | -0.387406549 | 0.218029239 | 12.90545357 | -0.668646035 | 0.526240208 | 0.991835066 | -5.075693234 |
| **A_52_P576442** | Dap | -0.03933837 | -0.179988816 | 0.101312076 | 10.67134026 | -0.668473785 | 0.52634343 | 0.991835066 | -5.07578271 |
| **A_51_P475228** | Armc6 | 0.048408811 | -0.124674363 | 0.221491984 | 9.686745629 | 0.668465036 | 0.526348673 | 0.991835066 | -5.075787254 |
| **A_55_P2014635** | Ccdc82 | 0.073825931 | -0.190136969 | 0.337788832 | 9.569520182 | 0.668459615 | 0.526351922 | 0.991835066 | -5.075790069 |
| **A_52_P574720** | Oit3 | 0.072957038 | -0.187928173 | 0.333842249 | 8.819938639 | 0.668385262 | 0.526396483 | 0.991835066 | -5.075828686 |
| **A_55_P1994519** | Tpd52l2 | -0.034296802 | -0.156940672 | 0.088347068 | 8.443546938 | -0.668369941 | 0.526405666 | 0.991835066 | -5.075836643 |
| **A_55_P2151609** | Sorl1 | 0.060070168 | -0.154752552 | 0.274892888 | 8.561993628 | 0.668325094 | 0.526432544 | 0.991835066 | -5.075859932 |
| **A_55_P2057390** | Smap1 | 0.04040017 | -0.104083197 | 0.184883537 | 11.17261742 | 0.668304634 | 0.526444807 | 0.991835066 | -5.075870557 |
| **A_55_P1981210** | Pex26 | -0.027140306 | -0.124212802 | 0.06993219 | 7.800162761 | -0.668232627 | 0.526487967 | 0.991835066 | -5.075907948 |
| **A_52_P301223** | Brwd1 | 0.036146391 | -0.093168244 | 0.165461026 | 10.39907675 | 0.66807681 | 0.526581369 | 0.991835066 | -5.075988846 |
| **A_55_P2121225** | Cyp3a11 | -0.027216323 | -0.124595142 | 0.070162495 | 7.65090587 | -0.667996346 | 0.526629606 | 0.991835066 | -5.076030616 |
| **A_55_P2047225** | Uxs1 | -0.037789309 | -0.173033233 | 0.097454615 | 8.252440458 | -0.667821388 | 0.5267345 | 0.991835066 | -5.076121425 |
| **A_55_P2135967** | NA | 0.038523389 | -0.099348785 | 0.176395563 | 12.93844516 | 0.667816265 | 0.526737571 | 0.991835066 | -5.076124083 |
| **A_51_P391825** | Exosc10 | 0.061718724 | -0.159170681 | 0.28260813 | 10.65257186 | 0.667807339 | 0.526742923 | 0.991835066 | -5.076128716 |
| **A_55_P1960049** | 2810408A1 | 0.053446527 | -0.137849851 | 0.244742905 | 9.685000895 | 0.667762268 | 0.526769947 | 0.991835066 | -5.076152105 |
| **A_55_P2084696** | H2ac1 | -0.03878583 | -0.177623884 | 0.100052223 | 8.690848252 | -0.667688211 | 0.526814353 | 0.991835066 | -5.076190534 |
| **A_55_P1960496** | Ppp1r2 | -0.032416629 | -0.14846599 | 0.083632732 | 8.88260171 | -0.66762752 | 0.526850746 | 0.991835066 | -5.076222024 |
| **A_52_P447971** | NA | 0.045293779 | -0.116875448 | 0.207463006 | 10.45703044 | 0.667542996 | 0.526901433 | 0.991835066 | -5.076265877 |
| **A_51_P386389** | Actr6 | -0.042401071 | -0.194214326 | 0.109412184 | 10.06866743 | -0.667538422 | 0.526904176 | 0.991835066 | -5.07626825 |
| **A_51_P476209** | Fscn3 | 0.039977932 | -0.103168138 | 0.183124003 | 8.181104405 | 0.667498094 | 0.526928361 | 0.991835066 | -5.076289171 |
| **A_52_P565847** | AU018091 | -0.028743033 | -0.131678447 | 0.07419238 | 7.654482391 | -0.667385728 | 0.526995751 | 0.991835066 | -5.076347458 |
| **A_51_P493037** | Dclre1a | -0.034266384 | -0.15699721 | 0.088464441 | 8.218049527 | -0.667304042 | 0.527044745 | 0.991835066 | -5.076389825 |
| **A_55_P2040815** | Lrrc45 | 0.064037655 | -0.165345931 | 0.293421241 | 10.92096962 | 0.667240214 | 0.52708303 | 0.991835066 | -5.076422928 |
| **A_51_P421094** | NA | 0.031819697 | -0.082166557 | 0.145805951 | 10.31206659 | 0.667194862 | 0.527110234 | 0.991835066 | -5.076446446 |
| **A_51_P126563** | Otc | 0.028261461 | -0.072991214 | 0.129514135 | 7.659229655 | 0.667109666 | 0.52716134 | 0.991835066 | -5.049202292 |
| **A_52_P671543** | Pomc | 0.031464723 | -0.081277867 | 0.144207313 | 8.332581114 | 0.667029492 | 0.527209436 | 0.991835066 | -5.076532192 |
| **A_52_P369123** | Fam189a2 | -0.040600513 | -0.186092012 | 0.104890986 | 8.399080785 | -0.666964988 | 0.527248134 | 0.991835066 | -5.076565633 |
| **A_55_P2127557** | Abracl | -0.033792579 | -0.154910252 | 0.087325094 | 12.92380394 | -0.666842013 | 0.527321915 | 0.991835066 | -5.07662938 |
| **A_55_P2020050** | 6720475J1 | -0.086891169 | -0.398353004 | 0.224570666 | 9.931754383 | -0.66677596 | 0.527361548 | 0.991835066 | -5.076663616 |
| **A_51_P228817** | NA | 0.047079114 | -0.121684394 | 0.215842621 | 8.886886596 | 0.666743616 | 0.527380955 | 0.991835066 | -5.076680379 |
| **A_55_P1955827** | NA | -0.05533916 | -0.253775837 | 0.143097517 | 11.72803354 | -0.666530076 | 0.527509098 | 0.991835066 | -5.076791036 |
| **A_55_P2128501** | Krt8 | 0.095647401 | -0.247378895 | 0.438673697 | 8.169762173 | 0.666430541 | 0.527568834 | 0.991835066 | -5.076842604 |
| **A_66_P104309** | Myl2 | 0.03273261 | -0.084672552 | 0.150137773 | 17.80592 | 0.66635027 | 0.527617013 | 0.991835066 | -5.076884188 |
| **A_55_P1971619** | NA | 0.032174151 | -0.083237485 | 0.147585787 | 7.92208013 | 0.666295114 | 0.527650118 | 0.991835066 | -5.076912758 |
| **A_55_P1955289** | Slc18b1 | 0.035119023 | -0.090888073 | 0.161126118 | 8.901830378 | 0.666126255 | 0.527751479 | 0.991835066 | -5.077000214 |
| **A_52_P367791** | Mri1 | 0.037654104 | -0.097453061 | 0.17276127 | 10.4306206 | 0.66610558 | 0.527763891 | 0.991835066 | -5.077010921 |
| **A_55_P1967592** | Ccnk | -0.046903308 | -0.21521885 | 0.121412234 | 12.25133543 | -0.66602171 | 0.527814241 | 0.991835066 | -5.077054351 |
| **A_55_P2041539** | Pigx | -0.039444845 | -0.181072324 | 0.102182634 | 11.13499659 | -0.665659078 | 0.528031976 | 0.991835066 | -5.077242078 |
| **A_55_P1966863** | Mad2l2 | 0.038647235 | -0.100129166 | 0.177423636 | 11.19333116 | 0.665597897 | 0.528068716 | 0.991835066 | -5.077273742 |
| **A_55_P2142863** | Parp9 | -0.044886899 | -0.206123309 | 0.11634951 | 9.212155124 | -0.665373685 | 0.528203374 | 0.991835066 | -5.07738976 |
| **A_51_P444335** | Acp6 | 0.05302058 | -0.137504443 | 0.243545602 | 11.29199975 | 0.665122412 | 0.52835431 | 0.991835066 | -5.077519741 |
| **A_55_P2059019** | Mapk8ip3 | 0.094003587 | -0.2437913 | 0.431798474 | 12.13473982 | 0.665120735 | 0.528355317 | 0.991835066 | -5.077520608 |
| **A_55_P2089761** | NA | -0.057834854 | -0.265660902 | 0.149991195 | 10.35728288 | -0.665118144 | 0.528356874 | 0.991835066 | -5.077521949 |
| **A_55_P2104259** | Plin5 | -0.074495502 | -0.342197567 | 0.193206563 | 9.592841021 | -0.665100786 | 0.528367302 | 0.991835066 | -5.077530926 |

| | | | | | | | | | |
|---|---|---|---|---|---|---|---|---|---|
| **A_55_P2073734** | Bfar | 0.056590656 | -0.146785039 | 0.25996635 | 11.33383718 | 0.665050784 | 0.528397341 | 0.991835066 | -5.077556786 |
| **A_52_P175066** | Cd2ap | 0.099668889 | -0.258567246 | 0.457905024 | 10.61062354 | 0.664965877 | 0.528448352 | 0.991835066 | -5.077600695 |
| **A_55_P2088028** | Mtss1 | 0.028706693 | -0.074473171 | 0.131886557 | 8.904138465 | 0.664962795 | 0.528450204 | 0.991835066 | -5.077602288 |
| **A_55_P2002702** | Ddx18 | 0.041163534 | -0.106800659 | 0.189127727 | 10.29404501 | 0.664913514 | 0.528479813 | 0.991835066 | -5.077627771 |
| **A_51_P456914** | Bmt2 | -0.057573337 | -0.26456011 | 0.149413437 | 9.263421376 | -0.664795294 | 0.528550846 | 0.991835066 | -5.077688895 |
| **A_55_P1961320** | Tes | 0.049946828 | -0.129627366 | 0.229521023 | 9.119804387 | 0.664772616 | 0.528564473 | 0.991835066 | -5.077700619 |
| **A_55_P2115166** | AI661453 | -0.051457558 | -0.236500427 | 0.13358531 | 9.644619344 | -0.664639207 | 0.528644641 | 0.991835066 | -5.077769584 |
| **A_55_P2181114** | NA | -0.055014406 | -0.252884699 | 0.142855887 | 13.96716095 | -0.664515272 | 0.528719123 | 0.991835066 | -5.07783364 |
| **A_51_P124089** | Gtpbp2 | -0.069143455 | -0.317860256 | 0.179573346 | 9.558209212 | -0.664438954 | 0.528764991 | 0.991835066 | -5.07787308 |
| **A_52_P245766** | NA | -0.068647138 | -0.315697905 | 0.17840363 | 10.80412915 | -0.664118164 | 0.528957819 | 0.991835066 | -5.078038819 |
| **A_55_P1975110** | Pnpt1 | 0.067701377 | -0.175958214 | 0.311360968 | 10.68773952 | 0.664084163 | 0.52897826 | 0.991835066 | -5.078056382 |
| **A_51_P211341** | Eif3j1 | 0.03316499 | -0.086205329 | 0.152535308 | 9.858136916 | 0.664037557 | 0.529006279 | 0.991835066 | -5.078080455 |
| **A_51_P139030** | Slc38a3 | -0.031976248 | -0.147070645 | 0.083118149 | 8.128374358 | -0.664021975 | 0.529015647 | 0.991835066 | -5.078088503 |
| **A_55_P1957198** | Apob | -0.141785178 | -0.652126425 | 0.36855607 | 8.074252093 | -0.664017144 | 0.529018552 | 0.991835066 | -5.078090998 |
| **A_55_P2150633** | Gadd45gip | -0.047313213 | -0.217612323 | 0.122985896 | 12.54513081 | -0.664016999 | 0.529018639 | 0.991835066 | -5.078091073 |
| **A_51_P150120** | Mgst2 | -0.071687933 | -0.329731196 | 0.186355329 | 9.679453597 | -0.663991729 | 0.529033831 | 0.991835066 | -5.078104124 |
| **A_55_P2094964** | NA | 0.067401472 | -0.175246446 | 0.310049391 | 10.34484614 | 0.663898896 | 0.529089647 | 0.991835066 | -5.078152067 |
| **A_52_P161806** | Ovca2 | 0.04490967 | -0.116792291 | 0.206611632 | 9.911249625 | 0.663794596 | 0.529152361 | 0.991835066 | -5.078205925 |
| **A_51_P269084** | Chchd10 | -0.06255462 | -0.287826361 | 0.162717121 | 17.34562154 | -0.6636847 | 0.529218446 | 0.991835066 | -5.078262665 |
| **A_55_P2175206** | Rbbp6 | -0.029656159 | -0.136469566 | 0.077157247 | 8.965391061 | -0.663587617 | 0.529276829 | 0.991835066 | -5.078312783 |
| **A_55_P2282257** | NA | 0.089289889 | -0.232438813 | 0.41101859 | 11.0564632 | 0.663317734 | 0.529439153 | 0.991835066 | -5.078452074 |
| **A_55_P2064598** | Cnot6 | 0.048939698 | -0.132630555 | 0.230509952 | 12.93072203 | 0.669551116 | 0.529535275 | 0.991835066 | -5.004811704 |
| **A_66_P107584** | Fam133b | 0.037392196 | -0.097376182 | 0.172160575 | 8.356315287 | 0.663135237 | 0.529548935 | 0.991835066 | -5.078546237 |
| **A_55_P2162678** | Dnajc10 | 0.053204646 | -0.138581337 | 0.244990629 | 11.07863098 | 0.663043204 | 0.529604304 | 0.991835066 | -5.078593716 |
| **A_55_P2050408** | NA | 0.056551391 | -0.14732484 | 0.260427622 | 9.154970683 | 0.662957715 | 0.529655739 | 0.991835066 | -5.078637813 |
| **A_55_P2182939** | NA | -0.106853209 | -0.492122891 | 0.278416474 | 13.54652474 | -0.662875344 | 0.5297053 | 0.991835066 | -5.078680297 |
| **A_55_P2045792** | Pes1 | 0.047161709 | -0.122957437 | 0.217280854 | 11.08353921 | 0.662590907 | 0.529876465 | 0.991835066 | -5.078826966 |
| **A_55_P2008790** | Zfp30 | 0.208197889 | -0.54301742 | 0.959413198 | 9.25586837 | 0.662401069 | 0.529990723 | 0.991835066 | -5.078924826 |
| **A_52_P528963** | Eps15l1 | 0.039976943 | -0.104302967 | 0.184256852 | 8.898216386 | 0.662236093 | 0.53009003 | 0.991835066 | -5.079009851 |
| **A_55_P2018076** | Arap1 | 0.053444018 | -0.139439743 | 0.246327778 | 11.51576377 | 0.662235663 | 0.530090289 | 0.991835066 | -5.079010072 |
| **A_55_P2025443** | Oprm1 | 0.091631172 | -0.239086144 | 0.422348488 | 13.00408687 | 0.66220955 | 0.530106009 | 0.991835066 | -5.079023528 |
| **A_55_P2002093** | Dph7 | 0.030733665 | -0.080203834 | 0.141671163 | 8.741141411 | 0.662132816 | 0.530152203 | 0.991835066 | -5.079063068 |
| **A_55_P2011087** | Gnptg | 0.048065703 | -0.125533297 | 0.221664703 | 10.17023142 | 0.661754966 | 0.53037971 | 0.991835066 | -5.07925771 |
| **A_55_P2109875** | Rpl23a | 0.031134543 | -0.081322665 | 0.143591751 | 14.76603668 | 0.66170487 | 0.530409878 | 0.991835066 | -5.079283509 |
| **A_55_P2058494** | NA | -0.049503103 | -0.228336061 | 0.129329856 | 14.20022908 | -0.661597705 | 0.530474416 | 0.991835066 | -5.079338692 |
| **A_55_P2020461** | Hmgn2 | 0.050358241 | -0.13156596 | 0.232282442 | 10.61488784 | 0.661590421 | 0.530478803 | 0.991835066 | -5.079342443 |
| **A_55_P2176345** | Psma5 | -0.045267297 | -0.20880144 | 0.118266847 | 13.14872349 | -0.661584392 | 0.530482434 | 0.991835066 | -5.079345547 |
| **A_51_P316523** | Irf2 | -0.044275118 | -0.204226779 | 0.115676543 | 8.946794622 | -0.661576557 | 0.530487153 | 0.991835066 | -5.079349582 |
| **A_55_P2036638** | Ints11 | -0.039738401 | -0.183310944 | 0.103834143 | 9.974973511 | -0.661527834 | 0.530516497 | 0.991835066 | -5.079374667 |
| **A_55_P2162910** | Rtn1 | 0.075917775 | -0.198458829 | 0.350294379 | 9.633878782 | 0.661310663 | 0.530647307 | 0.991835066 | -5.079486464 |
| **A_55_P2049261** | NA | 0.141816647 | -0.370883534 | 0.654516827 | 9.154795468 | 0.661108703 | 0.530768972 | 0.991835066 | -5.079590403 |
| **A_51_P102503** | Metap1 | 0.041716311 | -0.109136971 | 0.192569593 | 11.0021256 | 0.660937307 | 0.530872239 | 0.991835066 | -5.079678591 |
| **A_51_P195100** | Vps4b | -0.042087278 | -0.194285201 | 0.110110644 | 10.9944376 | -0.660923588 | 0.530880505 | 0.991835066 | -5.079685649 |
| **A_51_P264053** | Pigf | 0.063167812 | -0.165320484 | 0.291656107 | 9.60336126 | 0.660755834 | 0.530981591 | 0.991835066 | -5.079771942 |
| **A_55_P2124791** | Col18a1 | 0.046975192 | -0.123022067 | 0.21697245 | 12.79664646 | 0.660443656 | 0.531169736 | 0.991835066 | -5.079932479 |
| **A_52_P238846** | Bpnt1 | -0.035802911 | -0.165380917 | 0.093775094 | 9.013873465 | -0.660383454 | 0.531206024 | 0.991835066 | -5.079963431 |
| **A_52_P602669** | Serpinb6d | -0.027580345 | -0.127444847 | 0.072284156 | 8.03870202 | -0.6600817 | 0.531387934 | 0.991835066 | -5.080118535 |
| **A_51_P137778** | NA | 0.031679378 | -0.083033866 | 0.146392622 | 7.803356399 | 0.660042974 | 0.531411283 | 0.991835066 | -5.080138436 |

| | | | | | | | | | |
|---|---|---|---|---|---|---|---|---|---|
| **A_55_P2081925** | Zfp53 | 0.059968589 | -0.157204524 | 0.277141702 | 8.782995856 | 0.659974119 | 0.531452799 | 0.991835066 | -5.080173818 |
| **A_55_P2065621** | Hook2 | 0.04307033 | -0.112927917 | 0.199068578 | 8.795537245 | 0.659884028 | 0.531507121 | 0.991835066 | -5.080220108 |
| **A_51_P331661** | Ubiad1 | -0.045572772 | -0.21064156 | 0.119496016 | 10.9675872 | -0.659856672 | 0.531523616 | 0.991835066 | -5.080234162 |
| **A_51_P357561** | Fbxw9 | -0.036568305 | -0.169052346 | 0.095915737 | 8.600917644 | -0.65970593 | 0.53161452 | 0.991835066 | -5.080311601 |
| **A_55_P2170787** | Braf | 0.029880446 | -0.078395523 | 0.138156414 | 8.03802417 | 0.65957491 | 0.531693538 | 0.991835066 | -5.080378895 |
| **A_51_P117739** | Vegfd | 0.117140959 | -0.307344504 | 0.541626421 | 8.999883524 | 0.659561178 | 0.53170182 | 0.991835066 | -5.063324777 |
| **A_55_P2012689** | Stk16 | -0.055414243 | -0.256236673 | 0.145408187 | 9.969186705 | -0.659505349 | 0.531735493 | 0.991835066 | -5.080414619 |
| **A_52_P126825** | Cramp1l | 0.065753646 | -0.172540386 | 0.304047678 | 10.52572661 | 0.65950153 | 0.531737797 | 0.991835066 | -5.08041658 |
| **A_55_P2020552** | Iscu | -0.058800969 | -0.271911339 | 0.154309401 | 12.15331892 | -0.659460874 | 0.531762319 | 0.991835066 | -5.080437457 |
| **A_51_P150763** | Ropn1l | -0.034969414 | -0.161728339 | 0.091789511 | 8.828712856 | -0.65935445 | 0.531826514 | 0.991835066 | -5.080492103 |
| **A_52_P460957** | Crem | -0.036825534 | -0.17033209 | 0.096681023 | 9.316724164 | -0.659258276 | 0.531884529 | 0.991835066 | -5.080541478 |
| **A_52_P181759** | Eif5a2 | -0.03477582 | -0.160860514 | 0.091308874 | 8.915839563 | -0.659210549 | 0.531913322 | 0.991835066 | -5.080565979 |
| **A_55_P2163446** | NA | -0.047787019 | -0.221053304 | 0.125479266 | 9.308580903 | -0.659181496 | 0.531930849 | 0.991835066 | -5.080580893 |
| **A_51_P214107** | Tbc1d20 | 0.069796086 | -0.183279754 | 0.322871926 | 9.516105783 | 0.659157979 | 0.531945037 | 0.991835066 | -5.080592964 |
| **A_55_P2056493** | NA | -0.060253142 | -0.278762891 | 0.158256606 | 9.273107524 | -0.659049473 | 0.532010502 | 0.991835066 | -5.080648657 |
| **A_55_P2030568** | Ifit1bl2 | -0.034962643 | -0.161760165 | 0.09183488 | 9.312952117 | -0.659026103 | 0.532024603 | 0.991835066 | -5.080660651 |
| **A_51_P223720** | Rtcb | 0.029696228 | -0.078017287 | 0.137409742 | 11.30786376 | 0.658931424 | 0.53208173 | 0.991835066 | -5.080709239 |
| **A_51_P151968** | Eif2b5 | -0.071329268 | -0.330136458 | 0.187477923 | 10.09161218 | -0.658719562 | 0.532209578 | 0.991835066 | -5.080817942 |
| **A_55_P1995442** | NA | 0.037027128 | -0.097341405 | 0.171395662 | 8.567821203 | 0.658614953 | 0.532272711 | 0.991835066 | -5.080871604 |
| **A_55_P2052690** | Synm | 0.035432033 | -0.093166789 | 0.164030856 | 8.045142928 | 0.658518857 | 0.532330711 | 0.991835066 | -5.080920894 |
| **A_55_P2182412** | Adat1 | 0.032439014 | -0.085297759 | 0.150175786 | 8.168866397 | 0.658513447 | 0.532333976 | 0.991835066 | -5.080923668 |
| **A_51_P371001** | Tm4sf4 | -0.026928852 | -0.124668591 | 0.070810886 | 7.945419334 | -0.658500039 | 0.532342069 | 0.991835066 | -5.080930545 |
| **A_55_P2003813** | Scn3b | 0.049583277 | -0.13044443 | 0.229610984 | 8.283526761 | 0.658271436 | 0.532480062 | 0.991835066 | -5.081047771 |
| **A_52_P680870** | Ctxn1 | -0.046123128 | -0.213730195 | 0.121483939 | 8.517392607 | -0.657711687 | 0.532818042 | 0.991835066 | -5.081334664 |
| **A_51_P137419** | Cst7 | -0.02685019 | -0.124426095 | 0.070725715 | 7.6775513 | -0.657678893 | 0.532837847 | 0.991835066 | -5.081351465 |
| **A_55_P2070239** | Preb | -0.052568606 | -0.243639262 | 0.13850205 | 8.881369505 | -0.657569404 | 0.532903975 | 0.991835066 | -5.081407556 |
| **A_55_P2163128** | NA | 0.031134701 | -0.082032991 | 0.144302393 | 8.377114644 | 0.65755393 | 0.532913321 | 0.991835066 | -5.081415482 |
| **A_51_P232399** | Acy3 | -0.061103175 | -0.28325822 | 0.16105187 | 10.14559835 | -0.657380365 | 0.533018159 | 0.991835066 | -5.081504381 |
| **A_52_P96847** | Smlr1 | 0.02919655 | -0.076954985 | 0.135348085 | 7.69474078 | 0.657376831 | 0.533020294 | 0.991835066 | -5.081506191 |
| **A_65_P16796** | Psd4 | 0.030462369 | -0.08031195 | 0.141236688 | 7.866413268 | 0.65725473 | 0.533094055 | 0.991835066 | -5.081568718 |
| **A_51_P418016** | Ccnj | 0.031448902 | -0.082924946 | 0.14582275 | 8.341324303 | 0.65718537 | 0.533135958 | 0.991835066 | -5.081604233 |
| **A_51_P180032** | Sem1 | -0.071879575 | -0.333305698 | 0.189546548 | 15.88845275 | -0.657151724 | 0.533156285 | 0.991835066 | -5.081621459 |
| **A_55_P2039479** | Twistnb | -0.042923938 | -0.19905328 | 0.113205404 | 8.388401944 | -0.657088948 | 0.533194213 | 0.991835066 | -5.081653598 |
| **A_55_P2062733** | NA | -0.072214839 | -0.334955654 | 0.190525975 | 16.00059806 | -0.656913275 | 0.53330036 | 0.991835066 | -5.081743523 |
| **A_55_P2037787** | NA | 0.027508727 | -0.072584096 | 0.12760155 | 7.900401868 | 0.656865858 | 0.533329013 | 0.991835066 | -5.081767791 |
| **A_51_P270213** | Ceacam11 | -0.657223881 | -3.048637565 | 1.734189802 | 9.717452924 | -0.65685235 | 0.533337176 | 0.991835066 | -5.081774704 |
| **A_55_P1993636** | Rab9 | -0.036531598 | -0.169490222 | 0.096427026 | 10.45217518 | -0.656691342 | 0.533434478 | 0.991835066 | -5.081857098 |
| **A_51_P327632** | Igsf3 | 0.059141355 | -0.156129358 | 0.274412067 | 11.4296426 | 0.656622033 | 0.533476367 | 0.991835066 | -5.081892561 |
| **A_52_P508700** | Zfp951 | 0.031478727 | -0.083143989 | 0.146101442 | 8.013474079 | 0.656380389 | 0.533622428 | 0.991835066 | -5.082016175 |
| **A_55_P2150901** | Foxo4 | 0.028108346 | -0.074243445 | 0.130460136 | 7.862705201 | 0.656370373 | 0.533628482 | 0.991835066 | -5.082021298 |
| **A_55_P2134148** | Mzt2 | -0.062209968 | -0.288753082 | 0.164333147 | 11.04778421 | -0.656323932 | 0.533656556 | 0.991835066 | -5.082045051 |
| **A_55_P1991901** | NA | -0.176555015 | -0.819690086 | 0.466580056 | 7.952104342 | -0.656125623 | 0.533776447 | 0.991835066 | -5.082146461 |
| **A_55_P2007470** | Pdgfa | -0.030695422 | -0.142509748 | 0.081118903 | 13.32387892 | -0.656123054 | 0.533778 | 0.991835066 | -5.082147775 |
| **A_66_P127160** | Eif2b3 | 0.041691526 | -0.110189219 | 0.193572271 | 8.857268134 | 0.656076082 | 0.5338064 | 0.991835066 | -5.082171791 |
| **A_55_P2064996** | Capzb | -0.097517791 | -0.452795211 | 0.257759628 | 11.05998501 | -0.656032541 | 0.533832727 | 0.991835066 | -5.082194052 |
| **A_55_P1957954** | Rsu1 | -0.040770364 | -0.189307348 | 0.107766619 | 11.54291943 | -0.656023091 | 0.533838441 | 0.991835066 | -5.082198883 |
| **A_55_P2090330** | Kcnmb4 | 0.103203637 | -0.272796287 | 0.479203561 | 9.760305425 | 0.656018937 | 0.533840952 | 0.991835066 | -5.082201007 |
| **A_66_P100601** | NA | -0.084044883 | -0.3902508 | 0.222161035 | 13.38974752 | -0.656004516 | 0.533849672 | 0.991835066 | -5.082208379 |

| | | | | | | | | | |
|---|---|---|---|---|---|---|---|---|---|
| **A_51_P131164** | Enkur | -0.045516871 | -0.211361496 | 0.120327755 | 8.351372531 | -0.655964184 | 0.533874059 | 0.991835066 | -5.082228998 |
| **A_55_P2119100** | Yipf1 | -0.040287349 | -0.187089336 | 0.106514638 | 12.25702092 | -0.655912467 | 0.533905332 | 0.991835066 | -5.082255435 |
| **A_51_P427812** | Tmem108 | -0.036789333 | -0.17084513 | 0.097266464 | 8.169231654 | -0.655911803 | 0.533905733 | 0.991835066 | -5.082255774 |
| **A_55_P2130905** | NA | 0.106077253 | -0.280491362 | 0.492645868 | 11.5883775 | 0.655850427 | 0.533942848 | 0.991835066 | -5.082287147 |
| **A_55_P1997951** | Htatsf1 | -0.048250439 | -0.224125517 | 0.127624638 | 9.70091332 | -0.655701386 | 0.534032982 | 0.991835066 | -5.082363319 |
| **A_51_P198137** | Ubxn8 | -0.043584508 | -0.202471412 | 0.115302396 | 8.546603647 | -0.655621487 | 0.534081305 | 0.991835066 | -5.082404147 |
| **A_55_P1964282** | Sbk3 | -0.308119268 | -1.431593584 | 0.815355047 | 9.050398535 | -0.655488146 | 0.534161956 | 0.991835066 | -5.082472276 |
| **A_55_P2160286** | Olfm1 | -0.078440349 | -0.364472619 | 0.20759192 | 8.906901283 | -0.655441021 | 0.534190462 | 0.991835066 | -5.082496351 |
| **A_55_P2108690** | Gm8709 | -0.117877197 | -0.547777243 | 0.312022849 | 13.5236864 | -0.655347187 | 0.534247224 | 0.991835066 | -5.082544284 |
| **A_55_P2192729** | 5830469G1 | 0.109433354 | -0.289720865 | 0.508587574 | 9.520554629 | 0.655266732 | 0.534295896 | 0.991835066 | -5.082585378 |
| **A_55_P1980336** | NA | 0.073332811 | -0.194169625 | 0.340835248 | 8.479692184 | 0.655208804 | 0.534330941 | 0.991835066 | -5.082614963 |
| **A_55_P2048249** | NA | 0.051555172 | -0.136508976 | 0.239619321 | 12.8851805 | 0.655202226 | 0.534334921 | 0.991835066 | -5.082618322 |
| **A_52_P101238** | B230354K1 | -0.02813553 | -0.130787616 | 0.074516557 | 9.238730718 | -0.655083176 | 0.53440695 | 0.991835066 | -5.082679117 |
| **A_55_P2007249** | Spdef | 0.029405899 | -0.077893474 | 0.136705272 | 7.811678983 | 0.655007713 | 0.53445261 | 0.991835066 | -5.082717648 |
| **A_55_P2021555** | Tbrg1 | 0.039797984 | -0.105465355 | 0.185061324 | 13.78828891 | 0.654808316 | 0.53457327 | 0.991835066 | -5.082819442 |
| **A_55_P2064557** | Furin | -0.030129065 | -0.140108454 | 0.079850324 | 11.63548027 | -0.654762028 | 0.534601283 | 0.991835066 | -5.082843068 |
| **A_55_P1988789** | Acss2 | -0.037107638 | -0.172565663 | 0.098350387 | 10.55368668 | -0.654738235 | 0.534615683 | 0.991835066 | -5.082855213 |
| **A_66_P132890** | NA | 0.048017347 | -0.127317087 | 0.22335178 | 8.565256277 | 0.654545852 | 0.53473212 | 0.991835066 | -5.082953392 |
| **A_51_P511482** | Ift57 | -0.03989356 | -0.185565349 | 0.105778228 | 10.65489288 | -0.654540432 | 0.534735402 | 0.991835066 | -5.082956158 |
| **A_51_P247799** | Casp8 | -0.034531519 | -0.160624538 | 0.091561501 | 9.104293729 | -0.654536341 | 0.534737877 | 0.991835066 | -5.082958245 |
| **A_51_P259214** | Slc39a6 | 0.058861496 | -0.15607848 | 0.273801473 | 10.35859555 | 0.65452047 | 0.534747484 | 0.991835066 | -5.082966343 |
| **A_55_P2002602** | NA | 0.0382361 | -0.101389003 | 0.177861203 | 16.09446576 | 0.654514414 | 0.53475115 | 0.991835066 | -5.082969433 |
| **A_51_P300434** | Dst | -0.048344517 | -0.224884124 | 0.128195089 | 9.780232567 | -0.654506862 | 0.534755721 | 0.991835066 | -5.082973287 |
| **A_55_P2055612** | Zfp800 | 0.043405175 | -0.115142043 | 0.201952392 | 8.136875171 | 0.654322741 | 0.534867177 | 0.991835066 | -5.083067223 |
| **A_55_P2053258** | Poc1a | -0.036894897 | -0.171682387 | 0.097892593 | 11.80027883 | -0.654223058 | 0.534927525 | 0.991835066 | -5.08311807 |
| **A_55_P2066782** | Fam104a | 0.062724582 | -0.166428451 | 0.291877614 | 12.14093388 | 0.654216204 | 0.534931674 | 0.991835066 | -5.083121565 |
| **A_51_P284823** | Polr1b | 0.043309306 | -0.114931797 | 0.201550409 | 9.234785063 | 0.65414052 | 0.534977497 | 0.991835066 | -5.083160166 |
| **A_55_P2029775** | Gm38396 | 0.035524914 | -0.094282388 | 0.165332216 | 8.365334776 | 0.654098331 | 0.535003041 | 0.991835066 | -5.083181682 |
| **A_55_P2044104** | Fgfr1op | 0.057411935 | -0.152420149 | 0.267244019 | 10.74781152 | 0.65394229 | 0.535097524 | 0.991835066 | -5.083261251 |
| **A_55_P2141301** | Synj2bp | -0.047602384 | -0.221611836 | 0.126407068 | 11.32909724 | -0.653830231 | 0.535165384 | 0.991835066 | -5.083318383 |
| **A_51_P488092** | Pde6d | -0.05237038 | -0.243810801 | 0.139070041 | 10.01311645 | -0.653824541 | 0.535168829 | 0.991835066 | -5.083321284 |
| **A_55_P2135601** | Polr2i | 0.048987334 | -0.130086334 | 0.228061001 | 9.780180792 | 0.653824526 | 0.535168839 | 0.991835066 | -5.083321291 |
| **A_66_P132249** | Akr1c13 | 0.042609252 | -0.113181704 | 0.198400207 | 8.120854283 | 0.653688432 | 0.53525126 | 0.991835066 | -5.083390665 |
| **A_66_P107585** | Taf2 | 0.034757522 | -0.092352551 | 0.161867596 | 10.53086588 | 0.653548729 | 0.535335875 | 0.991835066 | -5.083461866 |
| **A_55_P1983235** | Tmed10 | -0.034030917 | -0.158486415 | 0.090424582 | 13.9478662 | -0.653534757 | 0.535344338 | 0.991835066 | -5.083468986 |
| **A_51_P460710** | Tdrd3 | -0.043148092 | -0.20097111 | 0.114674926 | 10.42138642 | -0.653431979 | 0.535406595 | 0.991835066 | -5.083521359 |
| **A_51_P518298** | Rnf187 | 0.041010808 | -0.109011175 | 0.191032791 | 13.44281005 | 0.653360021 | 0.535450185 | 0.991835066 | -5.083558022 |
| **A_55_P2071802** | Slbp | -0.047432994 | -0.220984663 | 0.126118676 | 8.404337114 | -0.653222105 | 0.535533738 | 0.991835066 | -5.083628282 |
| **A_55_P2360266** | 4833411C0 | -0.026186832 | -0.122021177 | 0.069647513 | 7.699115992 | -0.653086806 | 0.535615712 | 0.991835066 | -5.083697197 |
| **A_55_P2008567** | Rps6 | 0.046291806 | -0.123124067 | 0.215707678 | 11.83676157 | 0.653069132 | 0.535626421 | 0.991835066 | -5.083706199 |
| **A_55_P1959989** | D430042O0 | -0.062308249 | -0.290353711 | 0.165737213 | 10.6240771 | -0.653030166 | 0.535650031 | 0.991835066 | -5.083726043 |
| **A_51_P133097** | Gm4013 | -0.05134088 | -0.239264777 | 0.136583018 | 8.090949132 | -0.652965785 | 0.535689043 | 0.991835066 | -5.083758829 |
| **A_52_P129697** | Stradb | -0.028222256 | -0.131537259 | 0.075092747 | 7.962353347 | -0.652886161 | 0.535737293 | 0.991835066 | -5.083799374 |
| **A_55_P2009952** | Me1 | 0.061213609 | -0.162881882 | 0.2853091 | 10.82263231 | 0.65286592 | 0.535749559 | 0.991835066 | -5.08380968 |
| **A_51_P390285** | Larp6 | 0.029756806 | -0.079214963 | 0.138728575 | 8.195263988 | 0.652651652 | 0.535879416 | 0.991835066 | -5.083918762 |
| **A_55_P2117164** | Tmem106b | 0.053219013 | -0.141679621 | 0.248117647 | 9.585677733 | 0.652630176 | 0.535892432 | 0.991835066 | -5.083929693 |
| **A_51_P297865** | Kctd6 | 0.035820737 | -0.095363534 | 0.167005009 | 8.649330582 | 0.652622265 | 0.535897227 | 0.991835066 | -5.08393372 |
| **A_55_P2043559** | NA | -0.04642613 | -0.216522699 | 0.123670439 | 9.042822796 | -0.652343087 | 0.536066455 | 0.991835066 | -5.084075796 |

| | | | | | | | | | |
|---|---|---|---|---|---|---|---|---|---|
| **A_55_P2029678** | Etfa | -0.073015188 | -0.340536072 | 0.194505695 | 14.49010976 | -0.652325943 | 0.536076849 | 0.991835066 | -5.084084519 |
| **A_55_P2250246** | Taok3 | 0.033326018 | -0.088794436 | 0.155446472 | 7.958183326 | 0.652235074 | 0.536131938 | 0.991835066 | -5.084130751 |
| **A_51_P351363** | Tyms-ps | 0.039092127 | -0.104162852 | 0.182347106 | 10.10934951 | 0.652211879 | 0.536146001 | 0.991835066 | -5.084142552 |
| **A_51_P175699** | Mtfp1 | -0.085669067 | -0.399664294 | 0.228326161 | 12.37790122 | -0.652093876 | 0.536217547 | 0.991835066 | -5.084202579 |
| **A_51_P204247** | C8a | -0.0260757 | -0.121651289 | 0.06949989 | 7.670241387 | -0.652075853 | 0.536228475 | 0.991835066 | -5.084211746 |
| **A_51_P396983** | Ssu2 | -0.038237341 | -0.178420145 | 0.101945462 | 8.321369098 | -0.65193167 | 0.536315903 | 0.991835066 | -5.084285077 |
| **A_55_P2018393** | NA | 0.032814136 | -0.087538209 | 0.153166481 | 15.37805504 | 0.651651729 | 0.536485676 | 0.991835066 | -5.084427413 |
| **A_55_P2073945** | Eif2b3 | 0.039806038 | -0.106196128 | 0.185808204 | 8.418865633 | 0.651626568 | 0.536500937 | 0.991835066 | -5.084440204 |
| **A_55_P2087399** | Tex22 | 0.027078007 | -0.072250807 | 0.12640682 | 7.688140281 | 0.651554244 | 0.536544805 | 0.991835066 | -5.084476968 |
| **A_55_P1954633** | NA | -0.044167285 | -0.206201294 | 0.117866724 | 12.13597224 | -0.651483861 | 0.536587498 | 0.991835066 | -5.084512741 |
| **A_52_P595824** | Cpeb4 | 0.034994431 | -0.093388814 | 0.163377676 | 8.001204845 | 0.651477948 | 0.536591085 | 0.991835066 | -5.084515747 |
| **A_55_P2038077** | Rpsa | 0.042665788 | -0.113872639 | 0.199204216 | 16.22020378 | 0.651430277 | 0.536620002 | 0.991835066 | -5.084539975 |
| **A_51_P146044** | Ccdc92 | 0.053092195 | -0.141714487 | 0.247898876 | 8.244056879 | 0.651382313 | 0.536649098 | 0.991835066 | -5.08456435 |
| **A_52_P82660** | Mcm9 | 0.034511266 | -0.092128775 | 0.161151307 | 8.702466025 | 0.651326847 | 0.536682747 | 0.991835066 | -5.084592536 |
| **A_52_P599162** | Tug1 | 0.030275903 | -0.080830983 | 0.14138279 | 8.719984829 | 0.651276287 | 0.53671342 | 0.991835066 | -5.084618226 |
| **A_55_P1992199** | Psg20 | -0.037080923 | -0.173185189 | 0.099023343 | 7.707718792 | -0.651160324 | 0.536783775 | 0.991835066 | -5.084677144 |
| **A_55_P2143219** | Rasgrp2 | -0.05665442 | -0.264620009 | 0.151311168 | 11.1199729 | -0.651105636 | 0.536816957 | 0.991835066 | -5.084704927 |
| **A_55_P2067016** | Phykpl | 0.028780142 | -0.07686538 | 0.134425663 | 8.655025149 | 0.651104907 | 0.536817399 | 0.991835066 | -5.084705297 |
| **A_55_P2008417** | Mnd1 | -0.045023578 | -0.210306333 | 0.120259178 | 10.05251884 | -0.651060862 | 0.536844124 | 0.991835066 | -5.084727671 |
| **A_55_P1988102** | Hmbs | -0.054280027 | -0.253550342 | 0.144990289 | 8.453435061 | -0.651038359 | 0.536857778 | 0.991835066 | -5.084739102 |
| **A_55_P2037912** | Cyp2d10 | -0.037232241 | -0.173943381 | 0.099478899 | 7.739969785 | -0.650915182 | 0.536932524 | 0.991835066 | -5.084801665 |
| **A_51_P335981** | Narf | -0.047401936 | -0.221464629 | 0.126660757 | 9.288901578 | -0.650877879 | 0.53695516 | 0.991835066 | -5.08482061 |
| **A_55_P2063266** | Piga | 0.064950997 | -0.173555082 | 0.303457076 | 9.536350624 | 0.650871869 | 0.536958808 | 0.991835066 | -5.084823662 |
| **A_55_P2081137** | NA | -0.126662595 | -0.591781416 | 0.338456226 | 12.68624207 | -0.650869019 | 0.536960538 | 0.991835066 | -5.08482511 |
| **A_55_P2040048** | Swi5 | 0.084467017 | -0.225707737 | 0.39464177 | 13.67919335 | 0.650863388 | 0.536963954 | 0.991835066 | -5.084827969 |
| **A_51_P362903** | Mrps22 | -0.043760338 | -0.204458415 | 0.116937739 | 11.73690793 | -0.650847321 | 0.536973705 | 0.991835066 | -5.084836128 |
| **A_55_P2412349** | AW111846 | -0.026553577 | -0.124073572 | 0.070966418 | 7.666334243 | -0.650786443 | 0.537010651 | 0.991835066 | -5.084867043 |
| **A_51_P357685** | Dhx16 | 0.039585826 | -0.105859199 | 0.185030851 | 10.58613836 | 0.650504004 | 0.537182077 | 0.991835066 | -5.085010436 |
| **A_55_P2018666** | Thrsp | -0.043028174 | -0.201151472 | 0.115095123 | 7.996724664 | -0.650378518 | 0.537258252 | 0.991835066 | -5.085074128 |
| **A_55_P2111444** | Trpv6 | -0.027247896 | -0.127403607 | 0.072907815 | 7.783740551 | -0.650229069 | 0.537348982 | 0.991835066 | -5.085149969 |
| **A_55_P2143095** | NA | -0.039373058 | -0.184106762 | 0.105360646 | 8.358512725 | -0.650187479 | 0.537374233 | 0.991835066 | -5.085171072 |
| **A_55_P2130448** | NA | 0.08076845 | -0.216146427 | 0.377683327 | 11.57910656 | 0.650158017 | 0.537392121 | 0.991835066 | -5.08518602 |
| **A_51_P282508** | Rhoc | 0.03889198 | -0.104087287 | 0.181871246 | 14.6507798 | 0.650123879 | 0.537412848 | 0.991835066 | -5.08520334 |
| **A_55_P1960375** | NA | 0.040929996 | -0.109582444 | 0.191442436 | 14.21820534 | 0.64994774 | 0.537519802 | 0.991835066 | -5.085292694 |
| **A_55_P2126489** | Atxn7l3b | 0.061194072 | -0.163874305 | 0.286262449 | 12.76104726 | 0.64983636 | 0.53758744 | 0.991835066 | -5.085349186 |
| **A_55_P2022116** | NA | 0.046252375 | -0.123862203 | 0.216366953 | 15.3635412 | 0.649832809 | 0.537589596 | 0.991835066 | -5.085350987 |
| **A_55_P2454784** | Ifnar1 | -0.030492802 | -0.142671246 | 0.081685642 | 8.253476539 | -0.649676339 | 0.537684625 | 0.991835066 | -5.085430334 |
| **A_55_P2016387** | Eif3k | 0.029446244 | -0.078988006 | 0.137880495 | 14.91953309 | 0.649041639 | 0.538070206 | 0.991835066 | -5.085752029 |
| **A_55_P2038237** | Rps3 | 0.046622556 | -0.125065043 | 0.218310156 | 10.31941843 | 0.649032262 | 0.538075904 | 0.991835066 | -5.08575678 |
| **A_55_P2040161** | Upf3a | 0.029384586 | -0.078824647 | 0.137593818 | 9.536327893 | 0.649029428 | 0.538077626 | 0.991835066 | -5.085758215 |
| **A_55_P2033864** | Plekha8 | -0.039790917 | -0.186324746 | 0.106742912 | 9.93263296 | -0.649015671 | 0.538085985 | 0.991835066 | -5.085765185 |
| **A_51_P295420** | Zc3h7a | 0.108367598 | -0.290727737 | 0.507462934 | 10.3051829 | 0.64898092 | 0.538107102 | 0.991835066 | -5.08578279 |
| **A_51_P436189** | Nudt22 | -0.08216096 | -0.384778362 | 0.220456442 | 10.08457054 | -0.648904429 | 0.538153584 | 0.991835066 | -5.085821538 |
| **A_55_P2005778** | Uap1 | 0.060486779 | -0.16230153 | 0.283275088 | 11.50295661 | 0.648899129 | 0.538156804 | 0.991835066 | -5.085824222 |
| **A_51_P268559** | Idh3a | -0.037716335 | -0.17664338 | 0.101210711 | 13.75834138 | -0.648861211 | 0.538179847 | 0.991835066 | -5.085843429 |
| **A_55_P2091247** | NA | -0.102913076 | -0.482018785 | 0.276192632 | 15.22230063 | -0.648812741 | 0.538209304 | 0.991835066 | -5.085867979 |
| **A_55_P1984690** | Ptprr | -0.026625263 | -0.124718965 | 0.071468438 | 7.824642284 | -0.648726922 | 0.53826146 | 0.991835066 | -5.085911442 |
| **A_51_P113274** | Hps6 | -0.072958945 | -0.341821754 | 0.195903864 | 10.20742481 | -0.648570138 | 0.538356754 | 0.991835066 | -5.085990834 |

| | | | | | | | | | |
|---|---|---|---|---|---|---|---|---|---|
| **A_55_P1963443** | Vps18 | -0.051041315 | -0.239146092 | 0.137063463 | 8.412010187 | -0.648531626 | 0.538380164 | 0.991835066 | -5.086010333 |
| **A_55_P2174198** | H4c9 | -0.070018607 | -0.328108871 | 0.188071657 | 9.612230578 | -0.6484119 | 0.538452942 | 0.991835066 | -5.086070945 |
| **A_55_P2162935** | Ntn1 | -0.043845375 | -0.205483597 | 0.117792848 | 10.84761577 | -0.648319151 | 0.538509326 | 0.991835066 | -5.086117893 |
| **A_66_P123432** | NA | -0.046659718 | -0.218687271 | 0.125367835 | 8.081349075 | -0.648265983 | 0.53854165 | 0.991835066 | -5.086144804 |
| **A_51_P368855** | Pfdn6 | -0.066130158 | -0.309967995 | 0.177707679 | 11.7943614 | -0.64819786 | 0.538583068 | 0.991835066 | -5.086179281 |
| **A_55_P1990810** | 4932412D2 | 0.028996342 | -0.077933485 | 0.13592617 | 7.928717537 | 0.648117105 | 0.538632168 | 0.991835066 | -5.086220147 |
| **A_55_P2150976** | NA | 0.070951186 | -0.190722847 | 0.332625219 | 15.88548457 | 0.648049472 | 0.538673292 | 0.991835066 | -5.08625437 |
| **A_51_P271503** | Il1r1 | -0.059940216 | -0.281026706 | 0.161146274 | 9.176141226 | -0.647985417 | 0.538712242 | 0.991835066 | -5.086286778 |
| **A_52_P69043** | Nkap | -0.032504344 | -0.152446908 | 0.087438219 | 9.443662566 | -0.647704955 | 0.538882804 | 0.991835066 | -5.086428649 |
| **A_51_P520106** | Adprhl2 | 0.038422324 | -0.103373981 | 0.18021863 | 9.171937259 | 0.647631332 | 0.538927582 | 0.991835066 | -5.086465881 |
| **A_51_P105589** | Eci1 | -0.059101659 | -0.277248543 | 0.159045224 | 13.98910347 | -0.647529849 | 0.538989311 | 0.991835066 | -5.086517198 |
| **A_55_P2204804** | Dclk1 | 0.064284204 | -0.173009416 | 0.301577824 | 8.533201691 | 0.647481469 | 0.53901874 | 0.991835066 | -5.08654166 |
| **A_55_P2167645** | NA | -0.085951728 | -0.403288726 | 0.23138527 | 8.334185789 | -0.647355785 | 0.539095197 | 0.991835066 | -5.086605201 |
| **A_55_P1988795** | Acss2 | -0.046620558 | -0.21874904 | 0.125507924 | 8.151491163 | -0.647342114 | 0.539103514 | 0.991835066 | -5.086612112 |
| **A_52_P638605** | Ap1m2 | 0.033153906 | -0.089262279 | 0.155570091 | 7.687176368 | 0.64729908 | 0.539129694 | 0.991835066 | -5.086633865 |
| **A_52_P190647** | Mxd3 | -0.053072055 | -0.24903417 | 0.14289006 | 8.792415675 | -0.647295994 | 0.539131572 | 0.991835066 | -5.086635425 |
| **A_55_P2008021** | Srpx2 | -0.080892961 | -0.379597571 | 0.217811648 | 10.16375534 | -0.647258767 | 0.53915422 | 0.991835066 | -5.086654242 |
| **A_55_P2179335** | Usp33 | 0.052753399 | -0.142063241 | 0.247570038 | 10.62644094 | 0.64719258 | 0.539194489 | 0.991835066 | -5.086687695 |
| **A_51_P208263** | NA | 0.033115979 | -0.089189222 | 0.155421179 | 14.65337489 | 0.647145301 | 0.539223255 | 0.991835066 | -5.08671159 |
| **A_52_P485891** | Dnah5 | 0.02659002 | -0.071615403 | 0.124795444 | 7.644095475 | 0.647131189 | 0.539231841 | 0.991835066 | -5.086718721 |
| **A_55_P1972490** | Gm4055 | 0.050101201 | -0.134948583 | 0.235150985 | 9.774808572 | 0.647095958 | 0.539253278 | 0.991835066 | -5.086736526 |
| **A_52_P480360** | Dut | 0.051690136 | -0.139237633 | 0.242617904 | 11.49826054 | 0.647064699 | 0.539272299 | 0.991835066 | -5.086752322 |
| **A_55_P2026710** | 4930445E1 | 0.026604411 | -0.071676002 | 0.124884825 | 7.699411792 | 0.646987386 | 0.539319343 | 0.991835066 | -5.086791388 |
| **A_55_P2092668** | Kyat3 | -0.033551014 | -0.157519932 | 0.090417904 | 8.490716992 | -0.646847602 | 0.539404407 | 0.991835066 | -5.086862011 |
| **A_55_P2122419** | Arl1 | 0.032651237 | -0.088021254 | 0.153323729 | 9.959516528 | 0.646696479 | 0.539496381 | 0.991835066 | -5.086938348 |
| **A_55_P1988433** | Stard7 | 0.038360713 | -0.1034408 | 0.180162227 | 13.718803 | 0.646569092 | 0.539573917 | 0.991835066 | -5.087002684 |
| **A_51_P158678** | Fam181b | 0.05154958 | -0.139012314 | 0.242111475 | 8.875889746 | 0.64654418 | 0.539589081 | 0.991835066 | -5.087015264 |
| **A_66_P135185** | Etv1 | -0.049816991 | -0.234069651 | 0.134435669 | 12.01977957 | -0.646208782 | 0.539793262 | 0.991835066 | -5.087184597 |
| **A_55_P2071846** | NA | 0.056272191 | -0.151861294 | 0.264405677 | 12.27866931 | 0.646191144 | 0.539804 | 0.991835066 | -5.087193499 |
| **A_55_P1968235** | Rab33b | 0.042538968 | -0.114831415 | 0.199909351 | 10.07989078 | 0.646060336 | 0.539883647 | 0.991835066 | -5.087259518 |
| **A_52_P324754** | Nupr1l | 0.027290599 | -0.073704096 | 0.128285294 | 8.193767481 | 0.645838074 | 0.540018994 | 0.991835066 | -5.087371669 |
| **A_52_P117313** | Azin1 | 0.043761349 | -0.118212848 | 0.205735545 | 8.52940485 | 0.645734512 | 0.540082065 | 0.991835066 | -5.087423914 |
| **A_51_P208603** | Dio2 | 0.038588738 | -0.104257385 | 0.18143486 | 9.151277292 | 0.645656083 | 0.540129833 | 0.991835066 | -5.087463474 |
| **A_55_P1956572** | Dcaf8 | -0.039502699 | -0.185740797 | 0.106735399 | 12.301878 | -0.645617612 | 0.540153266 | 0.991835066 | -5.087482878 |
| **A_55_P1957088** | Fam53b | 0.032163876 | -0.086907315 | 0.151235068 | 11.27317647 | 0.645610848 | 0.540157386 | 0.991835066 | -5.08748629 |
| **A_51_P515242** | Strip2 | -0.063081748 | -0.296633708 | 0.170470212 | 12.9897791 | -0.645549184 | 0.540194946 | 0.991835066 | -5.08751739 |
| **A_66_P128079** | Krt8 | 0.200664154 | -0.542289749 | 0.943618058 | 9.39702479 | 0.645530939 | 0.540206059 | 0.991835066 | -5.087526591 |
| **A_66_P111011** | Gata3 | 0.033412212 | -0.090311686 | 0.157136111 | 8.111814821 | 0.64544727 | 0.540257026 | 0.991835066 | -5.087568784 |
| **A_52_P346033** | Trmt1l | 0.03418833 | -0.092438116 | 0.160814776 | 8.695641209 | 0.645301389 | 0.540345897 | 0.991835066 | -5.087642338 |
| **A_52_P284958** | Ubtd2 | 0.049974785 | -0.135205405 | 0.235154975 | 10.04215401 | 0.645008652 | 0.540524259 | 0.991835066 | -5.087789895 |
| **A_51_P193716** | Supv3l1 | -0.052071209 | -0.245063261 | 0.140920843 | 9.704095012 | -0.644862865 | 0.5406131 | 0.991835066 | -5.087863359 |
| **A_51_P416137** | Slc31a2 | 0.06569303 | -0.177875801 | 0.309261862 | 9.750700541 | 0.644624363 | 0.540758458 | 0.991835066 | -5.087983513 |
| **A_55_P2161858** | Pdlim7 | 0.03848162 | -0.104203168 | 0.181166409 | 8.669357004 | 0.644591844 | 0.54077828 | 0.991835066 | -5.087999893 |
| **A_65_P09285** | 2610524H0 | 0.039875558 | -0.107991909 | 0.187743025 | 8.604894444 | 0.644530199 | 0.540815855 | 0.991835066 | -5.088030941 |
| **A_52_P73126** | Epb42 | -0.027857945 | -0.131177682 | 0.075461791 | 7.722955475 | -0.64442878 | 0.540877678 | 0.991835066 | -5.088082017 |
| **A_55_P2065334** | Gm14326 | 0.060040666 | -0.162654021 | 0.282735352 | 9.140172819 | 0.644384042 | 0.540904951 | 0.991835066 | -5.088104545 |
| **A_55_P2228327** | 4921529L0 | -0.047973511 | -0.225951657 | 0.130004635 | 8.765048015 | -0.644234469 | 0.540996138 | 0.991835066 | -5.088179855 |
| **A_55_P2106803** | Bglap2 | 0.028440738 | -0.077096259 | 0.133977735 | 7.768594505 | 0.644088073 | 0.541085398 | 0.991835066 | -5.08825355 |

| | | | | | | | | | |
|---|---|---|---|---|---|---|---|---|---|
| **A_55_P2147641** | Cstf1 | 0.041282607 | -0.111931172 | 0.194496386 | 11.68904602 | 0.643988958 | 0.541145835 | 0.991835066 | -5.088303437 |
| **A_55_P2042870** | Aoc2 | 0.087907452 | -0.238428776 | 0.414243681 | 14.62287761 | 0.64382752 | 0.541244284 | 0.991835066 | -5.088384678 |
| **A_55_P1984144** | NA | 0.030013443 | -0.081442031 | 0.141468917 | 15.18556425 | 0.643611126 | 0.541376263 | 0.991835066 | -5.088493547 |
| **A_51_P158073** | Shfl | 0.048077089 | -0.130462877 | 0.226617056 | 8.694615612 | 0.643593796 | 0.541386833 | 0.991835066 | -5.088502264 |
| **A_55_P1982593** | Rad51d | -0.039699797 | -0.187137779 | 0.107738185 | 8.612373912 | -0.643558515 | 0.541408353 | 0.991835066 | -5.088520011 |
| **A_55_P2087414** | Sema3e | 0.028292587 | -0.076800265 | 0.133385439 | 7.698577676 | 0.643440831 | 0.54148014 | 0.991835066 | -5.088579201 |
| **A_55_P2026823** | Uspl1 | 0.027579715 | -0.074876442 | 0.130035872 | 10.53081647 | 0.643370053 | 0.541523317 | 0.991835066 | -5.088614795 |
| **A_51_P102438** | Ugt2b36 | -0.027240595 | -0.128448711 | 0.073967522 | 7.660391874 | -0.64329529 | 0.541568928 | 0.991835066 | -5.08865239 |
| **A_51_P462972** | Ap5m1 | 0.058176407 | -0.157979705 | 0.274332519 | 9.530924857 | 0.643262905 | 0.541588685 | 0.991835066 | -5.088668673 |
| **A_66_P123186** | Mapk1ip1l | 0.042496473 | -0.115401109 | 0.200394055 | 8.643679256 | 0.643259991 | 0.541590463 | 0.991835066 | -5.088670139 |
| **A_51_P435251** | Cul4a | 0.029141751 | -0.079138349 | 0.137421851 | 11.01956227 | 0.643244568 | 0.541599873 | 0.991835066 | -5.088677893 |
| **A_55_P1970636** | Unc45a | 0.041353869 | -0.112320365 | 0.195028103 | 11.14381127 | 0.643167689 | 0.541646778 | 0.991835066 | -5.088716545 |
| **A_55_P1988260** | NA | 0.126006002 | -0.342250305 | 0.59426231 | 13.17544214 | 0.643156598 | 0.541653545 | 0.991835066 | -5.088722121 |
| **A_55_P1981681** | Syt8 | -0.027618099 | -0.130262211 | 0.075026012 | 7.705906857 | -0.643085743 | 0.541696778 | 0.991835066 | -5.088757741 |
| **A_52_P87503** | Hbq1b | 0.035148053 | -0.095495944 | 0.16579205 | 7.875018625 | 0.64301477 | 0.541740085 | 0.991835066 | -5.088793416 |
| **A_51_P267544** | Frg1 | -0.066700864 | -0.314663701 | 0.181261974 | 10.53589084 | -0.642915642 | 0.541800575 | 0.991835066 | -5.088843239 |
| **A_55_P1985159** | Mef2a | -0.052687938 | -0.24861253 | 0.143236654 | 10.12224384 | -0.642734159 | 0.541911331 | 0.991835066 | -5.088934436 |
| **A_55_P1969072** | Cbx3 | 0.05965258 | -0.162178727 | 0.281483886 | 13.87812213 | 0.642710692 | 0.541925653 | 0.991835066 | -5.088946226 |
| **A_55_P1983268** | 4930444P1 | -0.029275814 | -0.138157444 | 0.079605816 | 7.793789295 | -0.64263371 | 0.541972639 | 0.991835066 | -5.088984903 |
| **A_55_P2000943** | Nrip3 | 0.031935719 | -0.086882727 | 0.150754165 | 7.833654694 | 0.642394721 | 0.542118522 | 0.991835066 | -5.089104949 |
| **A_55_P1982737** | Dnajc5 | -0.089196043 | -0.421082175 | 0.242690089 | 14.61772427 | -0.642340956 | 0.542151343 | 0.991835066 | -5.08913195 |
| **A_55_P2126207** | Fthl17a | -0.318701704 | -1.504550939 | 0.867147532 | 8.483229801 | -0.642338628 | 0.542152765 | 0.991835066 | -5.089133119 |
| **A_55_P2140941** | Tinagl1 | -0.043175989 | -0.203940429 | 0.11758845 | 8.104987674 | -0.64189123 | 0.542425939 | 0.991835066 | -5.08935773 |
| **A_55_P1986103** | Suco | -0.033774434 | -0.15955263 | 0.092003762 | 8.195308801 | -0.641788381 | 0.542488749 | 0.991835066 | -5.089409346 |
| **A_51_P194853** | Sec14l4 | -0.030222655 | -0.142802529 | 0.082357219 | 7.694575373 | -0.641624575 | 0.542588795 | 0.991835066 | -5.089491539 |
| **A_55_P2043486** | Msx2 | 0.039542728 | -0.10775898 | 0.186844435 | 8.196582312 | 0.64160535 | 0.542600537 | 0.991835066 | -5.089501184 |
| **A_55_P2016193** | Prxl2c | -0.038312532 | -0.181035631 | 0.104410568 | 10.31746344 | -0.641587236 | 0.542611601 | 0.991835066 | -5.089510271 |
| **A_55_P1986113** | Ppil1 | 0.029346667 | -0.079980297 | 0.138673631 | 10.67327757 | 0.641564953 | 0.542625212 | 0.991835066 | -5.08952145 |
| **A_52_P513842** | Fubp1 | -0.030142889 | -0.142437325 | 0.082151547 | 8.50220248 | -0.641557766 | 0.542629602 | 0.991835066 | -5.089525056 |
| **A_51_P388374** | Ppil4 | 0.05154847 | -0.140504979 | 0.24360192 | 10.4931305 | 0.641509074 | 0.542659345 | 0.991835066 | -5.089549482 |
| **A_52_P367745** | Acads | 0.152279763 | -0.415437473 | 0.719996998 | 9.298297305 | 0.641090357 | 0.542915152 | 0.991835066 | -5.089759468 |
| **A_55_P2045622** | Setdb2 | 0.047363044 | -0.129223306 | 0.223949395 | 8.722518122 | 0.641049569 | 0.542940074 | 0.991835066 | -5.089779917 |
| **A_55_P1957213** | NA | 0.162538197 | -0.443541311 | 0.768617704 | 13.52208034 | 0.640965965 | 0.542991161 | 0.991835066 | -5.089821827 |
| **A_51_P427401** | Fam90a1b | -0.026738316 | -0.126450591 | 0.072973959 | 7.675069186 | -0.64090632 | 0.543027609 | 0.991835066 | -5.089851725 |
| **A_51_P147034** | Ica1l | -0.033010947 | -0.156118137 | 0.090096243 | 7.967944297 | -0.640890302 | 0.543037397 | 0.991835066 | -5.089859754 |
| **A_52_P222593** | Cebpzos | 0.045121633 | -0.123154469 | 0.213397735 | 11.55045452 | 0.640872293 | 0.543048403 | 0.991835066 | -5.08986878 |
| **A_55_P1999842** | NA | -0.096555454 | -0.456682189 | 0.263571281 | 11.93088677 | -0.640811917 | 0.5430853 | 0.991835066 | -5.08989904 |
| **A_55_P2156425** | Upk1a | -0.039311773 | -0.185948546 | 0.107325001 | 8.189746388 | -0.640750371 | 0.543122914 | 0.991835066 | -5.089929884 |
| **A_55_P2214487** | E330013P0 | 0.074718152 | -0.203993222 | 0.353429526 | 8.915586518 | 0.640738119 | 0.543130402 | 0.991835066 | -5.089936024 |
| **A_55_P1963389** | NA | 0.05007067 | -0.136703087 | 0.236844427 | 12.12967905 | 0.640732393 | 0.543133902 | 0.991835066 | -5.089938893 |
| **A_55_P1961567** | Ypel3 | -0.16161568 | -0.764545889 | 0.441314529 | 11.63414389 | -0.64065701 | 0.543179975 | 0.991835066 | -5.089976667 |
| **A_55_P2070174** | NA | -0.047206811 | -0.223379001 | 0.128965379 | 8.69087514 | -0.640437038 | 0.543314432 | 0.991835066 | -5.090086872 |
| **A_55_P2014555** | Lrp8 | -0.028817266 | -0.136386702 | 0.07875217 | 7.711039989 | -0.640284545 | 0.543407656 | 0.991835066 | -5.090163252 |
| **A_51_P195034** | Esco2 | -0.054153268 | -0.256300898 | 0.147994363 | 9.285310663 | -0.640272936 | 0.543414753 | 0.991835066 | -5.090169066 |
| **A_55_P2042918** | Trim37 | 0.045085834 | -0.123230187 | 0.213401856 | 9.291234396 | 0.640211963 | 0.543452031 | 0.991835066 | -5.090199601 |
| **A_51_P248044** | Lamb2 | -0.086584915 | -0.409873888 | 0.236704057 | 10.84333608 | -0.640118648 | 0.543509085 | 0.991835066 | -5.090246327 |
| **A_51_P429472** | Mindy3 | 0.043504644 | -0.118935613 | 0.2059449 | 9.85985309 | 0.640104796 | 0.543517554 | 0.991835066 | -5.090253263 |
| **A_51_P417720** | Itga11 | 0.109477423 | -0.299297639 | 0.518252484 | 8.53338943 | 0.640102178 | 0.543519155 | 0.991835066 | -5.090254574 |

| | | | | | | | | | |
|---|---|---|---|---|---|---|---|---|---|
| **A_55_P2018929** | Spns2 | 0.040780319 | -0.11149062 | 0.193051258 | 9.859604634 | 0.640092477 | 0.543525087 | 0.991835066 | -5.090259431 |
| **A_55_P2176530** | Ldah | 0.034457327 | -0.094207675 | 0.163122329 | 9.116979843 | 0.640074153 | 0.543536291 | 0.991835066 | -5.090268606 |
| **A_55_P2052145** | Ly6g6d | -0.041186112 | -0.195016752 | 0.112644527 | 8.405633655 | -0.639907331 | 0.5436383 | 0.991835066 | -5.090352121 |
| **A_55_P2033947** | Fnd3c2 | 0.032986822 | -0.090266138 | 0.156239782 | 7.687468155 | 0.639664503 | 0.543786807 | 0.991835066 | -5.090473654 |
| **A_52_P677956** | Pex11g | -0.042267751 | -0.200248739 | 0.115713236 | 10.4548138 | -0.639460098 | 0.543911835 | 0.991835066 | -5.090575926 |
| **A_55_P1984070** | NA | -0.070185383 | -0.332553464 | 0.192182698 | 14.65951315 | -0.639359037 | 0.543973657 | 0.991835066 | -5.09062648 |
| **A_55_P1958082** | Prpsap2 | 0.044984603 | -0.12321155 | 0.213180757 | 9.263436746 | 0.639229733 | 0.544052763 | 0.991835066 | -5.090691154 |
| **A_55_P1975667** | Rhox4g | 0.030567606 | -0.083727404 | 0.144862616 | 7.754258459 | 0.639209601 | 0.54406508 | 0.991835066 | -5.090701222 |
| **A_55_P2080264** | NA | -0.037978825 | -0.17999838 | 0.104040729 | 8.672458152 | -0.639149589 | 0.544101797 | 0.991835066 | -5.090731233 |
| **A_55_P2035122** | NA | -0.046022173 | -0.21814587 | 0.126101525 | 11.21453512 | -0.639051091 | 0.544162064 | 0.991835066 | -5.090780485 |
| **A_55_P2091245** | NA | -0.038073398 | -0.180488884 | 0.104342089 | 17.32149103 | -0.63895982 | 0.544217914 | 0.991835066 | -5.090826118 |
| **A_55_P2154817** | Ccdc28a | -0.058919693 | -0.279362833 | 0.161523447 | 9.628559515 | -0.638811933 | 0.544308414 | 0.991835066 | -5.090900045 |
| **A_55_P1980591** | Hace1 | 0.052719996 | -0.144575165 | 0.250015157 | 9.325613506 | 0.638657576 | 0.544402882 | 0.991835066 | -5.09097719 |
| **A_55_P2098120** | NA | 0.042290052 | -0.115974901 | 0.200555006 | 10.48731427 | 0.638649537 | 0.544407803 | 0.991835066 | -5.090981208 |
| **A_55_P2024530** | Siah2 | 0.053894899 | -0.147807244 | 0.255597041 | 9.718836231 | 0.638625535 | 0.544422494 | 0.991835066 | -5.090993202 |
| **A_55_P2138090** | U2af1l4 | -0.034238375 | -0.162382977 | 0.093906228 | 12.56805017 | -0.638589766 | 0.544444387 | 0.991835066 | -5.091011076 |
| **A_55_P2034250** | Zscan26 | 0.0774767 | -0.212509373 | 0.367462772 | 10.68141493 | 0.63856205 | 0.544461352 | 0.991835066 | -5.091024926 |
| **A_55_P1965659** | Havcr2 | -0.025965719 | -0.123165006 | 0.071233567 | 7.693770652 | -0.638478693 | 0.544512375 | 0.991835066 | -5.091066575 |
| **A_55_P1970860** | Pemt | -0.053027881 | -0.251595698 | 0.145539937 | 10.02414282 | -0.638270159 | 0.544640033 | 0.991835066 | -5.091170747 |
| **A_55_P1980501** | Znhit1 | -0.083281753 | -0.395232353 | 0.228668847 | 10.68246581 | -0.638077107 | 0.54475823 | 0.991835066 | -5.09126716 |
| **A_55_P1984173** | Pde4dip | 0.055952575 | -0.15363162 | 0.26553677 | 9.615462959 | 0.638073462 | 0.544760463 | 0.991835066 | -5.09126898 |
| **A_51_P509384** | Aldh8a1 | -0.05241291 | -0.248744822 | 0.143919002 | 8.131134087 | -0.638052642 | 0.544773211 | 0.991835066 | -5.091279377 |
| **A_51_P216550** | Trp63 | 0.026628842 | -0.073169436 | 0.126427121 | 7.675038939 | 0.637732235 | 0.54496942 | 0.991835066 | -5.091439332 |
| **A_66_P100789** | Kansl2 | 0.042830149 | -0.117699453 | 0.203359751 | 10.50430482 | 0.637681162 | 0.545000699 | 0.991835066 | -5.091464823 |
| **A_52_P321140** | Defb1 | -0.026861659 | -0.127541523 | 0.073818206 | 7.664400378 | -0.637674923 | 0.545004521 | 0.991835066 | -5.091467937 |
| **A_51_P400366** | Rhbg | 0.026345758 | -0.072407513 | 0.125099029 | 7.643745224 | 0.637629403 | 0.5450324 | 0.991835066 | -5.091490654 |
| **A_66_P140335** | Mbnl3 | 0.040916706 | -0.112466285 | 0.194299697 | 8.269078451 | 0.637576929 | 0.545064541 | 0.991835066 | -5.09151684 |
| **A_52_P559817** | Fbln7 | -0.184729257 | -0.877286985 | 0.507828472 | 8.353558904 | -0.637512692 | 0.545103887 | 0.991835066 | -5.091548894 |
| **A_55_P2158281** | NA | 0.082199009 | -0.225972055 | 0.390370073 | 16.43223913 | 0.637505383 | 0.545108364 | 0.991835066 | -5.091552541 |
| **A_55_P2166049** | Vmn1r65 | -0.03001584 | -0.142548624 | 0.082516944 | 7.729658731 | -0.637500567 | 0.545111314 | 0.991835066 | -5.091554944 |
| **A_55_P2016064** | NA | -0.06665235 | -0.316573085 | 0.183268385 | 8.856361691 | -0.637415043 | 0.545163703 | 0.991835066 | -5.091597615 |
| **A_55_P2100968** | Dnah7b | -0.038134603 | -0.181137029 | 0.104867823 | 8.432833427 | -0.637360229 | 0.545197281 | 0.991835066 | -5.091624961 |
| **A_55_P1963206** | NA | -0.03418184 | -0.162372769 | 0.094009089 | 8.477423531 | -0.637304925 | 0.545231161 | 0.991835066 | -5.09165255 |
| **A_55_P2019512** | Mocs2 | 0.04282535 | -0.117812905 | 0.203463605 | 10.84630626 | 0.637178442 | 0.545308651 | 0.991835066 | -5.091715639 |
| **A_55_P2040490** | NA | 0.058603728 | -0.161313438 | 0.278520895 | 9.139808584 | 0.636905861 | 0.545475671 | 0.991835066 | -5.091851564 |
| **A_55_P2007663** | Alkbh8 | -0.038120339 | -0.181178971 | 0.104938293 | 9.007243076 | -0.636871507 | 0.545496723 | 0.991835066 | -5.091868691 |
| **A_55_P2001347** | Rpl7a | -0.032223321 | -0.15315349 | 0.088706847 | 15.36407519 | -0.636861179 | 0.545503052 | 0.991835066 | -5.09187384 |
| **A_51_P122845** | Naa50 | 0.04446717 | -0.122416344 | 0.211350684 | 9.993658555 | 0.636847113 | 0.545511672 | 0.991835066 | -5.086393656 |
| **A_55_P2032608** | Dipk2a | 0.072820553 | -0.20048715 | 0.346128255 | 11.1245645 | 0.636811988 | 0.545533197 | 0.991835066 | -5.091898363 |
| **A_52_P375546** | Lrrc46 | 0.028478135 | -0.078433337 | 0.135389607 | 7.893188716 | 0.636643595 | 0.5456364 | 0.991835066 | -5.091982298 |
| **A_55_P1967553** | Tmem254a | -0.052300776 | -0.248655636 | 0.144054084 | 10.54362425 | -0.636613154 | 0.545655058 | 0.991835066 | -5.09199747 |
| **A_66_P129029** | Taf10 | 0.055033401 | -0.151591504 | 0.261658305 | 11.83112353 | 0.636579747 | 0.545675533 | 0.991835066 | -5.092014118 |
| **A_51_P485756** | Nts | 0.068762134 | -0.189429639 | 0.326953908 | 10.69832057 | 0.6365259 | 0.545708539 | 0.991835066 | -5.092040952 |
| **A_51_P116421** | Abcc5 | 0.079009767 | -0.217725671 | 0.375745205 | 9.66691385 | 0.636385827 | 0.545794401 | 0.991835066 | -5.075649715 |
| **A_51_P440790** | Dpp7 | 0.038322469 | -0.105617365 | 0.182262302 | 9.032182183 | 0.636328853 | 0.545829327 | 0.991835066 | -5.09213913 |
| **A_55_P2031442** | Pithd1 | -0.041725578 | -0.198473893 | 0.115022738 | 10.87614012 | -0.63622189 | 0.545894902 | 0.991835066 | -5.092192413 |
| **A_55_P1955437** | Cmtm5 | 0.033285967 | -0.091761351 | 0.158333286 | 7.938585114 | 0.636203317 | 0.545906289 | 0.991835066 | -5.092201665 |
| **A_55_P2063381** | Tmem74b | 0.097832806 | -0.269757586 | 0.465423199 | 9.461369242 | 0.636105999 | 0.545965956 | 0.991835066 | -5.092250136 |

| | | | | | | | | | |
|---|---|---|---|---|---|---|---|---|---|
| **A_66_P135882** | Rrp1 | -0.04106621 | -0.195371002 | 0.113238581 | 13.31575596 | -0.636083818 | 0.545979556 | 0.991835066 | -5.092261182 |
| **A_51_P302358** | Ltb | -0.046019014 | -0.218975048 | 0.12693702 | 7.909102343 | -0.63593206 | 0.546072609 | 0.991835066 | -5.092336753 |
| **A_51_P152990** | Grem2 | 0.02644993 | -0.072966361 | 0.125866221 | 7.676852757 | 0.635881369 | 0.546103694 | 0.991835066 | -5.092361992 |
| **A_51_P136303** | Cyp4f15 | 0.026300249 | -0.072567031 | 0.12516753 | 7.634986078 | 0.635793977 | 0.546157286 | 0.991835066 | -5.0924055 |
| **A_55_P2085445** | Acsm1 | -0.026094137 | -0.124200835 | 0.072012561 | 7.642908871 | -0.635701762 | 0.546213841 | 0.991835066 | -5.092451405 |
| **A_55_P2120866** | NA | 0.107911765 | -0.297862941 | 0.513686472 | 9.393699125 | 0.635613273 | 0.546268113 | 0.991835066 | -5.092495449 |
| **A_52_P114494** | Stat6 | -0.047458508 | -0.225933529 | 0.131016513 | 9.268810133 | -0.635544215 | 0.546310469 | 0.991835066 | -5.092529818 |
| **A_51_P226364** | Coa3 | 0.047112486 | -0.130118614 | 0.224343586 | 14.06369842 | 0.635338564 | 0.546436618 | 0.991835066 | -5.092632147 |
| **A_51_P411297** | Nup50 | 0.04265021 | -0.117835958 | 0.203136378 | 9.165865117 | 0.635173979 | 0.546537589 | 0.991835066 | -5.081857778 |
| **A_55_P2325663** | Cers3 | 0.030456881 | -0.084158265 | 0.145072026 | 7.85551121 | 0.635115253 | 0.54657362 | 0.991835066 | -5.092743232 |
| **A_55_P1994694** | Serinc2 | 0.028435093 | -0.078581915 | 0.135452102 | 7.774386701 | 0.63505448 | 0.546610908 | 0.991835066 | -5.092773458 |
| **A_55_P1970655** | Kdm3b | 0.030833187 | -0.085221342 | 0.146887715 | 8.369885264 | 0.634987911 | 0.546651754 | 0.991835066 | -5.092806564 |
| **A_52_P556082** | Trmt2b | -0.027607135 | -0.131521825 | 0.076307554 | 9.267916772 | -0.634970479 | 0.546662451 | 0.991835066 | -5.092815232 |
| **A_51_P391445** | Ifngr1 | 0.042239879 | -0.116768082 | 0.201247839 | 11.30162958 | 0.634911109 | 0.546698881 | 0.991835066 | -5.092844754 |
| **A_55_P2002968** | Coro2a | -0.027584953 | -0.131442128 | 0.076272222 | 7.711717361 | -0.634811627 | 0.546759929 | 0.991835066 | -5.092894217 |
| **A_55_P2141088** | NA | 0.083140512 | -0.229886128 | 0.396167151 | 10.54867047 | 0.634805274 | 0.546763828 | 0.991835066 | -5.092897375 |
| **A_51_P271425** | Lhfpl4 | 0.082240454 | -0.227477394 | 0.391958302 | 9.32285601 | 0.634641397 | 0.546864402 | 0.991835066 | -5.09297884 |
| **A_52_P42269** | Fgf10 | 0.030500659 | -0.084396293 | 0.145397611 | 7.688157766 | 0.63446819 | 0.546970714 | 0.991835066 | -5.06950387 |
| **A_55_P2093013** | NA | -0.059030726 | -0.281423568 | 0.163362116 | 15.63205481 | -0.634404785 | 0.547009634 | 0.991835066 | -5.09309643 |
| **A_55_P2036460** | Synb | 0.028118215 | -0.077818816 | 0.134055247 | 7.775251856 | 0.634379419 | 0.547025205 | 0.991835066 | -5.093109034 |
| **A_51_P386670** | Dse | 0.02901008 | -0.080294287 | 0.138314446 | 8.107352924 | 0.634337727 | 0.547050799 | 0.991835066 | -5.09312975 |
| **A_55_P2015519** | Mtln | 0.056544412 | -0.156536642 | 0.269625466 | 11.03858315 | 0.634240524 | 0.547110472 | 0.991835066 | -5.093178042 |
| **A_51_P507974** | Brf2 | 0.041607649 | -0.11518873 | 0.198404028 | 8.5949661 | 0.634229274 | 0.547117378 | 0.991835066 | -5.093183631 |
| **A_55_P2146175** | NA | -0.030471125 | -0.1453151 | 0.08437285 | 9.040316279 | -0.63414622 | 0.547168369 | 0.991835066 | -5.093224888 |
| **A_55_P2052252** | NA | 0.04558273 | -0.126220841 | 0.217386301 | 8.944583409 | 0.634128512 | 0.54717924 | 0.991835066 | -5.093233683 |
| **A_55_P2359610** | AI646519 | -0.042381286 | -0.20212076 | 0.117358188 | 9.823973756 | -0.634119355 | 0.547184863 | 0.991835066 | -5.093238232 |
| **A_55_P2155833** | NA | 0.031004209 | -0.085862725 | 0.147871144 | 15.8477086 | 0.634071364 | 0.547214328 | 0.991835066 | -5.093262069 |
| **A_55_P2074393** | NA | -0.064183944 | -0.306125254 | 0.177757366 | 8.764592613 | -0.634052916 | 0.547225655 | 0.991835066 | -5.093271231 |
| **A_55_P1977653** | Palld | -0.048496073 | -0.231319244 | 0.134327098 | 13.14804329 | -0.63399301 | 0.547262438 | 0.991835066 | -5.093300983 |
| **A_55_P1978089** | Dcaf17 | 0.038029536 | -0.105364354 | 0.181423425 | 8.920686275 | 0.633868998 | 0.547338588 | 0.991835066 | -5.093362565 |
| **A_55_P2172934** | Rgs7bp | 0.124015094 | -0.343671498 | 0.591701686 | 10.98478181 | 0.633765742 | 0.547401997 | 0.991835066 | -5.093413832 |
| **A_55_P2129972** | NA | 0.090780024 | -0.251577677 | 0.433137725 | 13.92244437 | 0.633751973 | 0.547410453 | 0.991835066 | -5.093420667 |
| **A_55_P1989071** | Oas1d | -0.028609558 | -0.136536983 | 0.079317868 | 7.738124404 | -0.633561029 | 0.547527724 | 0.991835066 | -5.093515451 |
| **A_51_P218653** | Jph2 | 0.052828193 | -0.146553379 | 0.252209765 | 8.585920848 | 0.633271394 | 0.547705637 | 0.991835066 | -5.093659178 |
| **A_55_P2003720** | NA | -0.044384087 | -0.211908583 | 0.12314041 | 14.73632865 | -0.633225069 | 0.547734097 | 0.991835066 | -5.093682161 |
| **A_55_P2073809** | AU015836 | -0.033379408 | -0.159385717 | 0.092626902 | 7.685958172 | -0.633133748 | 0.547790201 | 0.991835066 | -5.093727463 |
| **A_51_P284686** | Aak1 | -0.047441867 | -0.226539158 | 0.131655423 | 8.666622423 | -0.63311396 | 0.547802359 | 0.991835066 | -5.093737279 |
| **A_51_P302139** | Mastl | 0.030066497 | -0.083448848 | 0.143581842 | 8.624256389 | 0.633049094 | 0.547842213 | 0.991835066 | -5.093769453 |
| **A_55_P1996299** | Pofut1 | 0.03251421 | -0.090303744 | 0.155332165 | 11.85963933 | 0.632732997 | 0.548036451 | 0.991835066 | -5.0939262 |
| **A_55_P2178006** | NA | -0.034634164 | -0.165468375 | 0.096200046 | 8.557678374 | -0.632692271 | 0.54806148 | 0.991835066 | -5.093946391 |
| **A_55_P2052472** | NA | -0.076230738 | -0.364201924 | 0.211740448 | 15.7695284 | -0.632688911 | 0.548063545 | 0.991835066 | -5.093948056 |
| **A_55_P2128326** | Ubap2l | -0.044535997 | -0.212780937 | 0.123708942 | 9.450115248 | -0.632671547 | 0.548074217 | 0.991835066 | -5.093956665 |
| **A_55_P1981599** | Ccdc125 | -0.032137809 | -0.153553108 | 0.08927749 | 8.757858361 | -0.632633206 | 0.54809778 | 0.991835066 | -5.093975671 |
| **A_52_P467726** | Nsg1 | 0.063944675 | -0.177675021 | 0.305564371 | 9.567998724 | 0.632530082 | 0.548161162 | 0.991835066 | -5.094026787 |
| **A_55_P1994586** | Itsn1 | 0.033640683 | -0.09347718 | 0.160758546 | 8.149053348 | 0.632509944 | 0.54817354 | 0.991835066 | -5.094036768 |
| **A_52_P343641** | Siah1a | -0.036454168 | -0.174228747 | 0.101320412 | 8.427322127 | -0.632393249 | 0.548245269 | 0.991835066 | -5.094094601 |
| **A_55_P1988623** | Add3 | 0.058732514 | -0.163260739 | 0.280725767 | 11.77739424 | 0.632336056 | 0.548280426 | 0.991835066 | -5.094122941 |
| **A_55_P1972720** | Pmm1 | 0.052928252 | -0.147144744 | 0.253001248 | 11.01321644 | 0.632278201 | 0.548315992 | 0.991835066 | -5.094151608 |

| | | | | | | | | | |
|---|---|---|---|---|---|---|---|---|---|
| **A_55_P2114318** | NA | 0.126047813 | -0.350428094 | 0.602523721 | 11.1377951 | 0.632271349 | 0.548320204 | 0.991835066 | -5.094155003 |
| **A_51_P431737** | Cth | 0.029980223 | -0.083353337 | 0.143313783 | 7.858552589 | 0.632245077 | 0.548336355 | 0.991835066 | -5.09416802 |
| **A_55_P2034245** | Meox1 | 0.073782218 | -0.205184066 | 0.352748501 | 10.2675787 | 0.632133956 | 0.548404671 | 0.991835066 | -5.094223071 |
| **A_55_P2039532** | Pax8 | 0.025274054 | -0.070286416 | 0.120834524 | 7.667732148 | 0.632129079 | 0.548407669 | 0.991835066 | -5.094225487 |
| **A_55_P1953693** | Pex2 | -0.048936046 | -0.233961975 | 0.136089884 | 10.64136326 | -0.632128558 | 0.54840799 | 0.991835066 | -5.094225745 |
| **A_52_P516034** | Ptp4a1 | -0.039341855 | -0.188100844 | 0.109417133 | 9.122218557 | -0.632092669 | 0.548430055 | 0.991835066 | -5.094243523 |
| **A_55_P2092384** | Pik3r2 | 0.062055637 | -0.172708043 | 0.296819316 | 9.389593731 | 0.631770672 | 0.548628049 | 0.991835066 | -5.094402989 |
| **A_51_P342669** | Pgam1 | 0.047546062 | -0.132381738 | 0.227473861 | 15.07348182 | 0.631575701 | 0.548747956 | 0.991835066 | -5.094499512 |
| **A_51_P344947** | Rab1b | -0.043940969 | -0.210234273 | 0.122352335 | 12.63230559 | -0.631544564 | 0.548767107 | 0.991835066 | -5.094514925 |
| **A_55_P1965101** | Xlr3b | 0.067357524 | -0.187567464 | 0.322282512 | 8.175093634 | 0.631513793 | 0.548786033 | 0.991835066 | -5.094530156 |
| **A_55_P2022519** | Tmem108 | 0.046644901 | -0.129957983 | 0.223247784 | 9.737227623 | 0.631270527 | 0.548935671 | 0.991835066 | -5.094650543 |
| **A_51_P520936** | Bcar3 | -0.043070105 | -0.206165376 | 0.120025167 | 8.919073918 | -0.631166134 | 0.548999893 | 0.991835066 | -5.094702192 |
| **A_51_P197883** | Exo5 | -0.04131238 | -0.197752672 | 0.115127913 | 9.644740956 | -0.631161827 | 0.549002543 | 0.991835066 | -5.094704323 |
| **A_55_P2048794** | Cd82 | 0.062931994 | -0.175381255 | 0.301245243 | 11.51993881 | 0.631149779 | 0.549009955 | 0.991835066 | -5.094710284 |
| **A_51_P385773** | Eif2ak1 | -0.032970886 | -0.157826454 | 0.091884682 | 8.437549633 | -0.63114891 | 0.54901049 | 0.991835066 | -5.094710714 |
| **A_66_P137134** | NA | -0.032177114 | -0.154027315 | 0.089673087 | 8.573182247 | -0.631146199 | 0.549012158 | 0.991835066 | -5.094712055 |
| **A_55_P2015426** | Asnsd1 | -0.031471688 | -0.150657378 | 0.087714001 | 12.22993648 | -0.631109997 | 0.54903443 | 0.991835066 | -5.094729964 |
| **A_55_P2079860** | Asap3 | -0.03176711 | -0.152166589 | 0.088632369 | 8.137220859 | -0.630611997 | 0.549340872 | 0.991835066 | -5.094976236 |
| **A_55_P2145496** | Wdr45 | -0.067802984 | -0.324830397 | 0.189224429 | 10.47091039 | -0.630490424 | 0.549415698 | 0.991835066 | -5.095036332 |
| **A_55_P2184246** | Rps26 | 0.058614823 | -0.163611835 | 0.280841481 | 16.36101986 | 0.63040614 | 0.549467576 | 0.991835066 | -5.095077989 |
| **A_55_P2096226** | Loxl1 | 0.0652053 | -0.182189783 | 0.312600383 | 9.098344485 | 0.629942582 | 0.549752958 | 0.991835066 | -5.095307015 |
| **A_52_P198009** | Mrpl44 | 0.039782441 | -0.111171748 | 0.19073663 | 11.37286521 | 0.629876484 | 0.549793658 | 0.991835066 | -5.095339659 |
| **A_52_P186672** | NA | -0.042552807 | -0.204045092 | 0.118939477 | 8.911103143 | -0.629775225 | 0.549856011 | 0.991835066 | -5.095389664 |
| **A_66_P136788** | Ubald2 | -0.067299246 | -0.322713842 | 0.188115351 | 13.25834674 | -0.629757888 | 0.549866687 | 0.991835066 | -5.095398224 |
| **A_55_P2055523** | NA | -0.087920104 | -0.421609893 | 0.245769685 | 12.79316438 | -0.629730038 | 0.549883838 | 0.991835066 | -5.095411976 |
| **A_51_P130905** | Gcm1 | 0.046898858 | -0.131116526 | 0.224914243 | 7.693307638 | 0.629671252 | 0.54992004 | 0.991835066 | -5.095441002 |
| **A_55_P2370096** | NA | 0.045132431 | -0.126179058 | 0.216443919 | 8.321304938 | 0.629667628 | 0.549922272 | 0.991835066 | -5.095442791 |
| **A_51_P505493** | Elovl5 | 0.033020015 | -0.092318803 | 0.158358834 | 11.22547293 | 0.629652318 | 0.549931701 | 0.991835066 | -5.095450349 |
| **A_55_P2431862** | Srpk2 | 0.035228333 | -0.09850904 | 0.168965706 | 11.51082215 | 0.629576387 | 0.549978464 | 0.991835066 | -5.095487835 |
| **A_55_P1985793** | Dock6 | 0.081255153 | -0.227247432 | 0.389757739 | 12.26014918 | 0.629507977 | 0.550020598 | 0.991835066 | -5.095521605 |
| **A_51_P168203** | Aig1 | -0.040854039 | -0.195965522 | 0.114257445 | 8.62546865 | -0.629506439 | 0.550021545 | 0.991835066 | -5.095522364 |
| **A_55_P2005420** | NA | 0.048940941 | -0.13687652 | 0.234758402 | 9.340255248 | 0.629498831 | 0.550026231 | 0.991835066 | -5.09552612 |
| **A_65_P09031** | Cdc42se2 | 0.0278266 | -0.077837128 | 0.133490329 | 8.0861607 | 0.629424075 | 0.550072276 | 0.991835066 | -5.095563018 |
| **A_55_P1963634** | Yae1d1 | -0.043780052 | -0.210085807 | 0.122525702 | 10.04339125 | -0.629184666 | 0.550219752 | 0.991835066 | -5.095681161 |
| **A_52_P418934** | Sec22b | 0.044301959 | -0.124007784 | 0.212611702 | 12.46697163 | 0.629104518 | 0.550269129 | 0.991835066 | -5.095720704 |
| **A_51_P465273** | Mettl17 | -0.035477011 | -0.170279711 | 0.099325688 | 8.860310427 | -0.629010029 | 0.550327344 | 0.991835066 | -5.095767316 |
| **A_55_P2004201** | NA | -0.042118674 | -0.202211797 | 0.117974449 | 11.71369703 | -0.62879799 | 0.550457995 | 0.991835066 | -5.095871895 |
| **A_52_P622391** | Usp9x | 0.026990902 | -0.075608083 | 0.129589887 | 8.652164494 | 0.628757923 | 0.550482686 | 0.991835066 | -5.095891654 |
| **A_55_P1971513** | NA | -0.031757792 | -0.152478014 | 0.08896243 | 15.9516758 | -0.628752035 | 0.550486314 | 0.991835066 | -5.095894557 |
| **A_52_P354271** | 2310057M2 | 0.031346281 | -0.087824112 | 0.150516674 | 9.15044417 | 0.628675864 | 0.550533254 | 0.991835066 | -5.095932116 |
| **A_51_P400555** | Cts3 | -0.357833378 | -1.718298657 | 1.0026319 | 8.489333148 | -0.628640734 | 0.550554904 | 0.991835066 | -5.095949436 |
| **A_51_P108629** | Prl3b1 | -0.588493165 | -2.826186811 | 1.649200481 | 9.672804269 | -0.628564612 | 0.550601817 | 0.991835066 | -5.095986965 |
| **A_55_P2022724** | Inafm2 | -0.068425398 | -0.328641546 | 0.19179075 | 10.11969614 | -0.628481098 | 0.55065329 | 0.991835066 | -5.096028133 |
| **A_51_P434198** | Anapc4 | 0.031919555 | -0.089469031 | 0.15330814 | 13.07281661 | 0.62847514 | 0.550656962 | 0.991835066 | -5.09603107 |
| **A_55_P2024689** | Rsbn1 | -0.0394477 | -0.189466949 | 0.110571548 | 8.79946704 | -0.628468964 | 0.550660768 | 0.991835066 | -5.096034114 |
| **A_55_P2160716** | NA | 0.047241378 | -0.132459198 | 0.226941954 | 14.60539285 | 0.628321939 | 0.550751393 | 0.991835066 | -5.096106578 |
| **A_55_P2043147** | Akr1e1 | 0.046921962 | -0.131578776 | 0.225422701 | 10.30910201 | 0.628268491 | 0.550784339 | 0.991835066 | -5.096132917 |
| **A_55_P1972852** | Mcrs1 | 0.054006795 | -0.151468046 | 0.259481636 | 12.79855271 | 0.628201378 | 0.550825711 | 0.991835066 | -5.096165988 |

| | | | | | | | | | |
|---|---|---|---|---|---|---|---|---|---|
| **A_52_P533198** | Dnajc2 | 0.054889428 | -0.153959607 | 0.263738462 | 9.60461116 | 0.628152893 | 0.550855601 | 0.991835066 | -5.096189878 |
| **A_52_P358860** | Gss | 0.057356113 | -0.16095623 | 0.275668456 | 8.829258748 | 0.627929016 | 0.550993628 | 0.991835066 | -5.096300166 |
| **A_55_P2072041** | NA | 0.058496191 | -0.164161665 | 0.281154048 | 8.508730116 | 0.627911878 | 0.551004195 | 0.991835066 | -5.096308607 |
| **A_51_P251129** | Stard8 | 0.037120773 | -0.104217723 | 0.178459269 | 9.753372555 | 0.627719579 | 0.551122772 | 0.991835066 | -5.09640331 |
| **A_55_P2168549** | Ap5b1 | 0.043210647 | -0.121326934 | 0.207748228 | 9.244391363 | 0.627674944 | 0.551150297 | 0.991835066 | -5.096425288 |
| **A_55_P2032559** | NA | -0.076827037 | -0.369378865 | 0.215724791 | 13.41987669 | -0.627654143 | 0.551163125 | 0.991835066 | -5.09643553 |
| **A_55_P2038831** | NA | 0.039878305 | -0.112045023 | 0.191801634 | 14.80658722 | 0.627366548 | 0.551340501 | 0.991835066 | -5.096577105 |
| **A_55_P2088331** | Ctdsp2 | 0.030133794 | -0.084676647 | 0.144944236 | 10.80772935 | 0.627309073 | 0.551375952 | 0.991835066 | -5.096605391 |
| **A_55_P2095133** | Synpo2 | 0.120712535 | -0.339255936 | 0.580681006 | 8.63409708 | 0.627239554 | 0.551418835 | 0.991835066 | -5.096639603 |
| **A_55_P1961690** | Tctex1d2 | -0.04581055 | -0.2203742 | 0.1287531 | 10.48347169 | -0.627221332 | 0.551430076 | 0.991835066 | -5.096648569 |
| **A_55_P2083929** | Robo2 | 0.027784467 | -0.078090693 | 0.133659628 | 7.872772212 | 0.627215996 | 0.551433368 | 0.991835066 | -5.096651195 |
| **A_51_P464064** | Zc3h13 | -0.030937597 | -0.148866273 | 0.086991079 | 8.475785673 | -0.627012628 | 0.55155883 | 0.991835066 | -5.096751253 |
| **A_55_P1963737** | Aktip | -0.041674209 | -0.200534304 | 0.117185885 | 8.750640462 | -0.626991546 | 0.551571837 | 0.991835066 | -5.096761624 |
| **A_55_P2107288** | Tgfbr2 | 0.042813279 | -0.120423557 | 0.206050115 | 8.335559829 | 0.62685841 | 0.551653982 | 0.991835066 | -5.096827111 |
| **A_55_P2162136** | Hmgn2 | -0.035675066 | -0.171698328 | 0.100348196 | 12.85013972 | -0.626845811 | 0.551661756 | 0.991835066 | -5.096833308 |
| **A_51_P509999** | Prl8a9 | -0.532200315 | -2.561526951 | 1.497126321 | 9.159152985 | -0.626804785 | 0.551687071 | 0.991835066 | -5.096853484 |
| **A_55_P2166123** | Atp9a | 0.03344298 | -0.094117129 | 0.161003089 | 8.253521586 | 0.626612743 | 0.551805578 | 0.991835066 | -5.096947918 |
| **A_66_P124817** | Ing1 | -0.037251022 | -0.179364579 | 0.104862535 | 8.569843605 | -0.626486643 | 0.551883402 | 0.991835066 | -5.097009912 |
| **A_55_P2000289** | Mab21l2 | 0.033110341 | -0.09321959 | 0.159440273 | 8.058194161 | 0.626421316 | 0.551923722 | 0.991835066 | -5.097042024 |
| **A_55_P2023952** | Arpc3 | 0.060333626 | -0.169865524 | 0.290532776 | 9.629346709 | 0.626418889 | 0.55192522 | 0.991835066 | -5.097043217 |
| **A_55_P2097828** | Gmfg | 0.066634368 | -0.187670362 | 0.320939098 | 9.842368963 | 0.626257554 | 0.552024803 | 0.991835066 | -5.09712251 |
| **A_55_P2012037** | Lrrc8e | 0.058172245 | -0.16391457 | 0.28025906 | 8.814547164 | 0.626040145 | 0.552159016 | 0.991835066 | -5.097229336 |
| **A_51_P123047** | Smc1a | 0.061626964 | -0.173653831 | 0.296907758 | 13.34676159 | 0.626027518 | 0.552166811 | 0.991835066 | -5.097235539 |
| **A_55_P2236607** | Ccnc | 0.031835102 | -0.089746925 | 0.153417129 | 8.229955179 | 0.625815038 | 0.552298001 | 0.991835066 | -5.09733991 |
| **A_55_P2093704** | Meig1 | 0.025586743 | -0.072134795 | 0.123308281 | 7.655522719 | 0.625797532 | 0.552308811 | 0.991835066 | -5.097348508 |
| **A_52_P105765** | Cdc42bpg | 0.024872673 | -0.070123503 | 0.11986885 | 7.924041351 | 0.625785466 | 0.552316262 | 0.991835066 | -5.097354434 |
| **A_52_P16419** | Gpd1 | -0.084303084 | -0.406416202 | 0.237810033 | 8.884788953 | -0.625524301 | 0.55247754 | 0.991835066 | -5.097482672 |
| **A_51_P394921** | 1700020L2 | -0.043479682 | -0.209635783 | 0.122676419 | 8.98349953 | -0.625430703 | 0.552535347 | 0.991835066 | -5.097528619 |
| **A_55_P2038947** | Lig3 | -0.045226648 | -0.21809508 | 0.127641785 | 9.548119801 | -0.625299146 | 0.552616604 | 0.991835066 | -5.097593191 |
| **A_55_P2094986** | Atp1a3 | 0.033253499 | -0.093860247 | 0.160367246 | 8.163342619 | 0.625250384 | 0.552646724 | 0.991835066 | -5.097617122 |
| **A_55_P1964375** | Gm9731 | 0.038281998 | -0.108054331 | 0.184618326 | 11.02813767 | 0.625246932 | 0.552648856 | 0.991835066 | -5.097618816 |
| **A_51_P370286** | Med24 | 0.025736173 | -0.072649975 | 0.124122322 | 10.78349042 | 0.625200261 | 0.552677686 | 0.991835066 | -5.097641719 |
| **A_55_P2078266** | H3f3a | -0.066781638 | -0.32209603 | 0.188532754 | 13.51158113 | -0.625159596 | 0.552702805 | 0.991835066 | -5.097661673 |
| **A_66_P138319** | Acox2 | -0.037572701 | -0.181261118 | 0.106115717 | 7.82929614 | -0.624970881 | 0.55281939 | 0.991835066 | -5.097754262 |
| **A_55_P2092616** | Tbc1d25 | -0.045014011 | -0.217165301 | 0.127137279 | 9.71340663 | -0.624951859 | 0.552831143 | 0.991835066 | -5.097763594 |
| **A_55_P1990331** | Add1 | 0.050926946 | -0.143880697 | 0.245734588 | 8.737584637 | 0.624814028 | 0.552916303 | 0.991835066 | -5.0978312 |
| **A_66_P109595** | Ube4b | 0.038695084 | -0.109326355 | 0.186716523 | 11.24906371 | 0.624798976 | 0.552925603 | 0.991835066 | -5.097838583 |
| **A_52_P85864** | Eif6 | 0.060280152 | -0.17032382 | 0.290884123 | 11.44608559 | 0.624764992 | 0.552946602 | 0.991835066 | -5.097855249 |
| **A_55_P2266977** | Ahcyl2 | -0.03695224 | -0.178332019 | 0.104427538 | 9.765202287 | -0.624687206 | 0.552994668 | 0.991835066 | -5.097893396 |
| **A_55_P2021841** | Calm3 | 0.047893487 | -0.135348551 | 0.231135525 | 13.62559278 | 0.624684138 | 0.552996564 | 0.991835066 | -5.0978949 |
| **A_66_P102467** | Casz1 | 0.123365476 | -0.348636313 | 0.595367265 | 8.85423135 | 0.624682195 | 0.552997764 | 0.991835066 | -5.097895853 |
| **A_55_P2045086** | NA | 0.040343169 | -0.114024008 | 0.194710346 | 11.77961798 | 0.624631931 | 0.553028825 | 0.991835066 | -5.0979205 |
| **A_51_P290921** | Sytl2 | -0.043420697 | -0.209574459 | 0.122733065 | 8.517200539 | -0.624591032 | 0.553054099 | 0.991835066 | -5.097940553 |
| **A_51_P267346** | Topors | 0.045729403 | -0.129351098 | 0.220809904 | 10.91138276 | 0.624261981 | 0.55325747 | 0.991835066 | -5.098101853 |
| **A_55_P2011943** | Gramd1a | 0.073898598 | -0.209038868 | 0.356836065 | 10.79970223 | 0.624244714 | 0.553268142 | 0.991835066 | -5.098110315 |
| **A_52_P453624** | Arl6ip6 | 0.033858329 | -0.095797414 | 0.163514072 | 8.966276212 | 0.624141265 | 0.55333209 | 0.991835066 | -5.098161009 |
| **A_51_P261059** | Cpne2 | 0.034820568 | -0.098523331 | 0.168164467 | 8.337439648 | 0.624125371 | 0.553341914 | 0.991835066 | -5.098168797 |
| **A_52_P323658** | Zfp955a | -0.038655344 | -0.186692978 | 0.10938229 | 10.11688008 | -0.624089018 | 0.553364388 | 0.991835066 | -5.09818661 |

| | | | | | | | | | |
|---|---|---|---|---|---|---|---|---|---|
| **A_51_P285487** | Rnf113a1 | -0.03647443 | -0.176197771 | 0.10324891 | 8.755858133 | -0.623919689 | 0.553469071 | 0.991835066 | -5.098269566 |
| **A_55_P2061076** | Gm3219 | -0.025858685 | -0.124935052 | 0.073217681 | 7.850971986 | -0.623800191 | 0.553542955 | 0.991835066 | -5.098328097 |
| **A_55_P1983314** | Crat | 0.052493998 | -0.14868039 | 0.253668386 | 8.4818984 | 0.623657423 | 0.553631235 | 0.991835066 | -5.098398015 |
| **A_55_P2353079** | NA | -0.054788842 | -0.264762813 | 0.15518513 | 8.572342711 | -0.623642621 | 0.553640388 | 0.991835066 | -5.098405263 |
| **A_55_P2015410** | 0610010K1 | -0.030284506 | -0.146356336 | 0.085787324 | 13.76488772 | -0.623595246 | 0.553669684 | 0.991835066 | -5.09842846 |
| **A_55_P2165074** | Rnf31 | -0.057616785 | -0.278455221 | 0.163221651 | 9.742011329 | -0.62356753 | 0.553686824 | 0.991835066 | -5.098442031 |
| **A_52_P609200** | Serinc3 | -0.05603717 | -0.270827846 | 0.158753507 | 14.1534912 | -0.623548014 | 0.553698893 | 0.991835066 | -5.098451586 |
| **A_55_P2152946** | NA | -0.050930986 | -0.246177029 | 0.144315057 | 8.837048875 | -0.623460548 | 0.553752985 | 0.991835066 | -5.098494407 |
| **A_55_P1985025** | Pard3 | 0.042032951 | -0.119172469 | 0.203238372 | 9.918930808 | 0.623188425 | 0.553921296 | 0.991835066 | -5.098627598 |
| **A_55_P2126470** | Rpl7 | 0.028629063 | -0.081192788 | 0.138450913 | 14.90305034 | 0.623056607 | 0.554002838 | 0.991835066 | -5.098692099 |
| **A_55_P2037918** | NA | -0.052249085 | -0.252683715 | 0.148185546 | 8.576683945 | -0.62303875 | 0.554013885 | 0.991835066 | -5.098700836 |
| **A_52_P1004509** | BC065397 | 0.071843568 | -0.203816552 | 0.347503687 | 8.56901674 | 0.622906815 | 0.554095508 | 0.991835066 | -5.098765381 |
| **A_51_P354433** | Ppie | 0.059183002 | -0.167948213 | 0.286314217 | 8.469775482 | 0.622772316 | 0.554178725 | 0.991835066 | -5.098831168 |
| **A_52_P276525** | Ankrd9 | -0.063077706 | -0.305199477 | 0.179044065 | 9.788721037 | -0.622660308 | 0.554248031 | 0.991835066 | -5.098885944 |
| **A_66_P129111** | NA | 0.055325064 | -0.157067439 | 0.267717567 | 9.03237881 | 0.622575284 | 0.554300645 | 0.991835066 | -5.098927519 |
| **A_55_P1954886** | NA | -0.033813657 | -0.16362785 | 0.096000536 | 16.4779238 | -0.622556961 | 0.554311984 | 0.991835066 | -5.098936478 |
| **A_51_P160113** | Copb1 | 0.079071767 | -0.224570786 | 0.382714319 | 11.35417904 | 0.622397645 | 0.554410579 | 0.991835066 | -5.099014365 |
| **A_55_P2070799** | Tyw1 | 0.028465615 | -0.080862421 | 0.137793652 | 9.468035946 | 0.622297648 | 0.554472469 | 0.991835066 | -5.099063242 |
| **A_55_P2209298** | NA | 0.07379919 | -0.209649258 | 0.357247638 | 8.525309291 | 0.622281146 | 0.554482683 | 0.991835066 | -5.099071308 |
| **A_55_P2243431** | Gdap10 | -0.092319355 | -0.446911873 | 0.262273163 | 8.88164034 | -0.622260632 | 0.55449538 | 0.991835066 | -5.099081334 |
| **A_51_P160536** | Cfap161 | 0.032271917 | -0.091684023 | 0.156227856 | 8.017070077 | 0.622252355 | 0.554500503 | 0.991835066 | -5.099085379 |
| **A_51_P307644** | Gtf3a | -0.050758937 | -0.245734057 | 0.144216184 | 10.62628957 | -0.622217828 | 0.554521875 | 0.991835066 | -5.099102253 |
| **A_55_P2380132** | NA | 0.026460213 | -0.075184105 | 0.12810453 | 7.832102242 | 0.622184743 | 0.554542354 | 0.991835066 | -5.099118422 |
| **A_51_P278018** | Vps36 | 0.05446298 | -0.15477494 | 0.263700899 | 12.35110657 | 0.622114234 | 0.554585999 | 0.991835066 | -5.099152877 |
| **A_55_P2040748** | Scrib | 0.041342898 | -0.117490842 | 0.200176637 | 12.06713871 | 0.622110148 | 0.554588528 | 0.991835066 | -5.099154873 |
| **A_55_P2175356** | Uros | 0.027107115 | -0.077035378 | 0.131249607 | 7.834288195 | 0.622106093 | 0.554591038 | 0.991835066 | -5.099156854 |
| **A_65_P19149** | Nln | 0.044784158 | -0.127287285 | 0.216855602 | 9.589818965 | 0.622049216 | 0.554626247 | 0.991835066 | -5.099184645 |
| **A_55_P2053083** | Gm6098 | -0.033690959 | -0.163144605 | 0.095762688 | 10.55296938 | -0.622025539 | 0.554640904 | 0.991835066 | -5.099196213 |
| **A_52_P89099** | Nolc1 | 0.043324938 | -0.123170578 | 0.209820454 | 8.543692096 | 0.621934348 | 0.554697358 | 0.991835066 | -5.099240764 |
| **A_52_P134023** | Rtp3 | 0.026934706 | -0.07658037 | 0.130449782 | 7.714350404 | 0.621895999 | 0.5547211 | 0.991835066 | -5.099259498 |
| **A_55_P2131483** | Hmgb1-ps1 | -0.102287196 | -0.495423633 | 0.290849241 | 11.96514949 | -0.621852106 | 0.554748275 | 0.991835066 | -5.099280938 |
| **A_52_P229728** | Cdc42 | -0.056305138 | -0.272713738 | 0.160103462 | 12.21820442 | -0.621845715 | 0.554752232 | 0.991835066 | -5.09928406 |
| **A_66_P103398** | Ogfrl1 | 0.039812539 | -0.113208585 | 0.192833663 | 8.834179083 | 0.621838518 | 0.554756688 | 0.991835066 | -5.099287575 |
| **A_55_P2076560** | Nr5a1 | -0.028769793 | -0.139351296 | 0.08181171 | 8.048462663 | -0.621818127 | 0.554769313 | 0.991835066 | -5.099297536 |
| **A_55_P2241553** | 9330133O1 | 0.074953044 | -0.213190242 | 0.36309633 | 9.043047313 | 0.621712936 | 0.554834444 | 0.991835066 | -5.099348911 |
| **A_52_P424585** | Ctnnb1 | 0.02707513 | -0.077020666 | 0.131170925 | 12.58614785 | 0.62165079 | 0.554872925 | 0.991835066 | -5.09937926 |
| **A_51_P192860** | Hint1 | -0.038096053 | -0.184582342 | 0.108390236 | 16.88137692 | -0.621572991 | 0.5549211 | 0.991835066 | -5.099417249 |
| **A_51_P366024** | Ercc3 | -0.043527287 | -0.21090445 | 0.123849876 | 9.067158577 | -0.621547796 | 0.554936702 | 0.991835066 | -5.09942955 |
| **A_52_P110088** | Ndufaf3 | -0.039539699 | -0.19160098 | 0.112521582 | 10.61910335 | -0.621475256 | 0.554981623 | 0.991835066 | -5.099464966 |
| **A_51_P289414** | Spg11 | 0.033710376 | -0.09594089 | 0.163361643 | 10.03589294 | 0.621435373 | 0.555006323 | 0.991835066 | -5.099484437 |
| **A_51_P456366** | Pard6a | -0.037880561 | -0.183602722 | 0.1078416 | 8.856368014 | -0.621297961 | 0.555091427 | 0.991835066 | -5.099551512 |
| **A_51_P229886** | Fpgt | -0.045048616 | -0.218387982 | 0.12829075 | 9.01492568 | -0.621145554 | 0.555185827 | 0.991835066 | -5.099625892 |
| **A_52_P102248** | Mex3b | -0.05646855 | -0.273772863 | 0.160835763 | 10.13713654 | -0.621079826 | 0.555226541 | 0.991835066 | -5.099657965 |
| **A_55_P2092984** | NA | -0.044205242 | -0.21431817 | 0.125907686 | 14.08718854 | -0.621077202 | 0.555228166 | 0.991835066 | -5.099659245 |
| **A_51_P278550** | Tecr | -0.118117023 | -0.572700366 | 0.336466319 | 11.31591319 | -0.621023604 | 0.555261368 | 0.991835066 | -5.099685396 |
| **A_55_P2043862** | Stmn1 | -0.054877809 | -0.266080788 | 0.156325171 | 13.81901019 | -0.621020382 | 0.555263364 | 0.991835066 | -5.099686968 |
| **A_55_P2080794** | Taf9 | -0.039392427 | -0.191002596 | 0.112217742 | 12.73449166 | -0.621002772 | 0.555274273 | 0.991835066 | -5.09969556 |
| **A_55_P2002357** | Tnrc18 | 0.034668285 | -0.098760145 | 0.168096715 | 9.379977568 | 0.621002161 | 0.555274651 | 0.991835066 | -5.099695858 |

| | | | | | | | | | |
|---|---|---|---|---|---|---|---|---|---|
| **A_55_P2102469** | Gm6428 | 0.026452024 | -0.0753608 | 0.128264847 | 16.02999176 | 0.620962756 | 0.555299063 | 0.991835066 | -5.099715083 |
| **A_55_P2281095** | 2610002M( | 0.036439045 | -0.103840042 | 0.176718132 | 9.654626946 | 0.620845 | 0.555372016 | 0.991835066 | -5.099772527 |
| **A_51_P392928** | Mthfd1 | 0.051496034 | -0.146770673 | 0.249762742 | 11.05297248 | 0.620773436 | 0.555416354 | 0.991835066 | -5.099807432 |
| **A_55_P2177659** | Mast1 | -0.024770505 | -0.120157304 | 0.070616293 | 7.855278151 | -0.620662809 | 0.555484899 | 0.991835066 | -5.099861384 |
| **A_55_P2087186** | Ier3ip1 | 0.041193025 | -0.117475195 | 0.199861244 | 13.46194598 | 0.620501539 | 0.555584831 | 0.991835066 | -5.099940019 |
| **A_55_P2053374** | Sorbs1 | -0.050827072 | -0.246619227 | 0.144965084 | 12.68339387 | -0.620453064 | 0.555614872 | 0.991835066 | -5.099963653 |
| **A_51_P315085** | Mia3 | 0.031133803 | -0.0888181 | 0.151085705 | 11.88343826 | 0.620346253 | 0.555681066 | 0.991835066 | -5.100015721 |
| **A_51_P466162** | C1qbp | 0.048920523 | -0.139579043 | 0.23742009 | 14.17631613 | 0.620282991 | 0.555720274 | 0.991835066 | -5.100046556 |
| **A_55_P2181928** | Slc45a3 | -0.035489331 | -0.172240393 | 0.10126173 | 8.242018715 | -0.620263523 | 0.55573234 | 0.991835066 | -5.100056044 |
| **A_55_P1969914** | Tcf25 | 0.038227729 | -0.109081222 | 0.185536679 | 11.56313433 | 0.620238173 | 0.555748052 | 0.991835066 | -5.1000684 |
| **A_55_P2118609** | St6galnac1 | 0.143500003 | -0.409477669 | 0.696477674 | 9.886182376 | 0.620230967 | 0.555752518 | 0.991835066 | -5.100071911 |
| **A_55_P1979650** | S100a1 | -0.083899657 | -0.407277315 | 0.239478 | 11.99413815 | -0.62009654 | 0.555835841 | 0.991835066 | -5.10013742 |
| **A_55_P1990285** | Rplp2 | 0.049766449 | -0.142064111 | 0.241597009 | 17.09401715 | 0.620051829 | 0.555863556 | 0.991835066 | -5.100159206 |
| **A_55_P2143150** | Qk | -0.065659589 | -0.318786472 | 0.187467295 | 10.67526892 | -0.619967637 | 0.555915747 | 0.991835066 | -5.100200226 |
| **A_51_P352562** | Mrpl38 | -0.045057788 | -0.218790906 | 0.128675329 | 10.84331989 | -0.619863965 | 0.555980017 | 0.991835066 | -5.10025073 |
| **A_52_P357402** | Mlst8 | 0.040254653 | -0.114996795 | 0.1955061 | 10.34892923 | 0.6197115 | 0.556074544 | 0.991835066 | -5.10032499 |
| **A_51_P454103** | Manba | 0.032615594 | -0.09317902 | 0.158410207 | 11.14529061 | 0.619686981 | 0.556089746 | 0.991835066 | -5.100336931 |
| **A_55_P2010197** | Serpina10 | -0.036138727 | -0.175526267 | 0.103248813 | 7.84921706 | -0.61966652 | 0.556102433 | 0.991835066 | -5.100346895 |
| **A_55_P1985143** | NA | -0.067710926 | -0.32892884 | 0.193506988 | 10.7137887 | -0.619533687 | 0.556184799 | 0.991835066 | -5.100411576 |
| **A_55_P2155827** | NA | 0.037461359 | -0.107085613 | 0.182008331 | 15.28230014 | 0.619417774 | 0.556256679 | 0.991835066 | -5.100468009 |
| **A_55_P2044364** | Mms22l | -0.048471738 | -0.235527256 | 0.138583781 | 13.11380391 | -0.619337243 | 0.556306621 | 0.991835066 | -5.100507211 |
| **A_55_P2024021** | Atp5l | -0.051697731 | -0.251228227 | 0.147832766 | 16.01903597 | -0.619257577 | 0.55635603 | 0.991835066 | -5.100545987 |
| **A_55_P2118717** | Ninl | 0.031455193 | -0.089987679 | 0.152898065 | 8.080286284 | 0.619055315 | 0.556481484 | 0.991835066 | -5.100644415 |
| **A_51_P473888** | Il6st | -0.037445098 | -0.182026983 | 0.107136786 | 11.18725593 | -0.618999402 | 0.556516167 | 0.991835066 | -5.100671619 |
| **A_55_P2112125** | Tnk2 | 0.070156025 | -0.200753917 | 0.341065966 | 11.62152584 | 0.618940871 | 0.556552476 | 0.991835066 | -5.100700095 |
| **A_51_P420655** | Reep4 | 0.030192387 | -0.086397528 | 0.146782302 | 9.171366888 | 0.618935787 | 0.55655563 | 0.991835066 | -5.100702569 |
| **A_55_P2037513** | Pld3 | -0.036886146 | -0.179324941 | 0.10555265 | 10.37066556 | -0.618933679 | 0.556556938 | 0.991835066 | -5.100703594 |
| **A_55_P2141876** | Ldha | 0.086022904 | -0.246206044 | 0.418251853 | 12.40817885 | 0.618850523 | 0.556608525 | 0.991835066 | -5.100744046 |
| **A_55_P2043833** | Srgap2 | -0.036542226 | -0.17770838 | 0.104623928 | 8.642941287 | -0.618690639 | 0.55670772 | 0.991835066 | -5.10082181 |
| **A_55_P2130666** | Armc1 | -0.035873845 | -0.174474051 | 0.102726361 | 11.27501073 | -0.618618899 | 0.556752232 | 0.991835066 | -5.100856697 |
| **A_55_P2144796** | Gfer | -0.042808665 | -0.208208059 | 0.122590729 | 9.285989244 | -0.618595707 | 0.556766622 | 0.991835066 | -5.100867974 |
| **A_52_P28792** | Isg20l2 | 0.038335688 | -0.109796006 | 0.186467382 | 9.672857195 | 0.618535176 | 0.556804181 | 0.991835066 | -5.100897407 |
| **A_55_P2015022** | NA | 0.090213634 | -0.258445131 | 0.438872399 | 14.31911355 | 0.61841601 | 0.556878129 | 0.991835066 | -5.100955343 |
| **A_52_P517001** | Rpf1 | -0.049988211 | -0.243183866 | 0.143207445 | 9.978240389 | -0.618414078 | 0.556879328 | 0.991835066 | -5.100956282 |
| **A_55_P2102365** | Spag7 | -0.055012219 | -0.26764295 | 0.157618512 | 14.56244169 | -0.618361248 | 0.556912113 | 0.991835066 | -5.100981963 |
| **A_55_P2126890** | NA | 0.053429205 | -0.153095377 | 0.259953786 | 8.558959106 | 0.618323982 | 0.55693524 | 0.991835066 | -5.101000078 |
| **A_55_P2005908** | Lin9 | 0.048747315 | -0.139684313 | 0.237178943 | 9.869932365 | 0.618309664 | 0.556944126 | 0.991835066 | -5.101007037 |
| **A_51_P215995** | Zswim6 | 0.059658447 | -0.170961762 | 0.290278656 | 9.062036803 | 0.618277891 | 0.556963845 | 0.991835066 | -5.101022481 |
| **A_55_P2080703** | Cstf2 | 0.046992571 | -0.134676244 | 0.228661385 | 8.84066268 | 0.618241213 | 0.556986608 | 0.991835066 | -5.101040307 |
| **A_52_P377416** | Disp1 | -0.049572083 | -0.241221066 | 0.1420769 | 8.858298854 | -0.618215344 | 0.557002663 | 0.991835066 | -5.10105288 |
| **A_55_P2083894** | NA | 0.028036027 | -0.080378987 | 0.13645104 | 8.021820982 | 0.618067865 | 0.5570942 | 0.991835066 | -5.101124547 |
| **A_55_P2093231** | NA | 0.074584472 | -0.213856034 | 0.363024978 | 17.48702975 | 0.618018256 | 0.557124994 | 0.991835066 | -5.101148652 |
| **A_51_P243808** | Traf3ip3 | -0.026008164 | -0.126591941 | 0.074575612 | 7.758116571 | -0.61800345 | 0.557134185 | 0.991835066 | -5.101155846 |
| **A_55_P2136906** | Vpreb3 | -0.0284214 | -0.138374012 | 0.081531212 | 7.985007027 | -0.617801624 | 0.557259474 | 0.991835066 | -5.10125389 |
| **A_55_P2025143** | Tlk1 | 0.041935738 | -0.120312919 | 0.204184395 | 11.35699773 | 0.617749369 | 0.557291915 | 0.991835066 | -5.101279271 |
| **A_55_P2119962** | Shld1 | -0.033149753 | -0.161412582 | 0.095113076 | 9.978765871 | -0.617715656 | 0.557312846 | 0.991835066 | -5.101295644 |
| **A_66_P130647** | NA | -0.027395857 | -0.133400804 | 0.07860909 | 7.747930256 | -0.617686186 | 0.557331143 | 0.991835066 | -5.101309957 |
| **A_55_P2208260** | D230019N2 | 0.088479971 | -0.253883509 | 0.43084345 | 9.039453489 | 0.617684454 | 0.557332219 | 0.991835066 | -5.101310798 |

| | | | | | | | | | |
|---|---|---|---|---|---|---|---|---|---|
| **A_52_P114282** | Armcx5 | -0.057961667 | -0.282241533 | 0.1663182 | 9.430323173 | -0.617674549 | 0.557338368 | 0.991835066 | -5.101315608 |
| **A_55_P2133680** | NA | -0.095451551 | -0.464804109 | 0.273901007 | 13.10723031 | -0.617662181 | 0.557346047 | 0.991835066 | -5.101321614 |
| **A_55_P2087840** | Gpr50 | 0.026471055 | -0.075968505 | 0.128910615 | 7.655721924 | 0.617607664 | 0.557379896 | 0.991835066 | -5.101348088 |
| **A_55_P2125912** | Cct6a | -0.047940418 | -0.233471634 | 0.137590798 | 10.51207041 | -0.617581042 | 0.557396426 | 0.991835066 | -5.101361015 |
| **A_55_P2138816** | Mplkip | 0.033779956 | -0.096961679 | 0.16452159 | 9.072334242 | 0.617524647 | 0.557431443 | 0.991835066 | -5.101388398 |
| **A_55_P2011525** | Rnf214 | -0.032776689 | -0.159668921 | 0.094115543 | 9.144717761 | -0.617360979 | 0.557533076 | 0.991835066 | -5.101467855 |
| **A_55_P2112510** | NA | -0.044974595 | -0.219148196 | 0.129199005 | 14.43280638 | -0.617154741 | 0.55766116 | 0.991835066 | -5.101567953 |
| **A_55_P1952517** | Sult2a1 | -0.025160976 | -0.122643978 | 0.072322027 | 7.655218626 | -0.616889977 | 0.557825617 | 0.991835066 | -5.101696414 |
| **A_55_P2043454** | NA | -0.035286459 | -0.172007551 | 0.101434634 | 8.450185475 | -0.616853009 | 0.557848582 | 0.991835066 | -5.101714347 |
| **A_51_P429209** | Smpd1 | -0.045566465 | -0.22212526 | 0.13099233 | 13.14166148 | -0.616829469 | 0.557863206 | 0.991835066 | -5.101725766 |
| **A_55_P2132845** | NA | 0.029179972 | -0.083893986 | 0.142253931 | 15.70771298 | 0.616781605 | 0.55789294 | 0.991835066 | -5.101748982 |
| **A_52_P554451** | 1300017J0 | 0.030850772 | -0.088721409 | 0.150422952 | 7.757633831 | 0.616658911 | 0.557969167 | 0.991835066 | -5.101808487 |
| **A_51_P358243** | Tstd3 | -0.041393092 | -0.201874693 | 0.119088508 | 9.830544974 | -0.616469729 | 0.558086713 | 0.991835066 | -5.101900218 |
| **A_51_P179241** | Tssc4 | -0.063340558 | -0.308949307 | 0.182268191 | 11.21250872 | -0.616378079 | 0.558143663 | 0.991835066 | -5.101944649 |
| **A_66_P108770** | Oxct1 | -0.035897084 | -0.175093345 | 0.103299177 | 14.24814038 | -0.616368918 | 0.558149356 | 0.991835066 | -5.10194909 |
| **A_55_P1997866** | Mtr | -0.053762167 | -0.262245874 | 0.154721541 | 10.49055707 | -0.616330662 | 0.558173129 | 0.991835066 | -5.101967634 |
| **A_51_P196862** | Amdhd1 | -0.028219025 | -0.137651868 | 0.081213818 | 7.704746564 | -0.616316022 | 0.558182227 | 0.991835066 | -5.10197473 |
| **A_51_P366227** | Zfp799 | -0.041659743 | -0.203226928 | 0.119907443 | 8.601585552 | -0.616272169 | 0.55820948 | 0.991835066 | -5.101995985 |
| **A_52_P183152** | Txnl1 | 0.028230725 | -0.081273082 | 0.137734531 | 8.255005907 | 0.616171974 | 0.558271749 | 0.991835066 | -5.102044545 |
| **A_55_P2093764** | Eif2s3x | 0.125194995 | -0.360506476 | 0.610896466 | 9.404862913 | 0.616065206 | 0.558338108 | 0.991835066 | -5.102096283 |
| **A_51_P401876** | Kif1bp | -0.037339964 | -0.182211901 | 0.107531972 | 11.67164511 | -0.616025608 | 0.55836272 | 0.991835066 | -5.102115469 |
| **A_55_P1952420** | Odf2 | 0.03743835 | -0.107828772 | 0.182705472 | 8.504644267 | 0.615968498 | 0.558398218 | 0.991835066 | -5.102143139 |
| **A_51_P422893** | Tmem14a | 0.040018706 | -0.115263138 | 0.195300551 | 9.594345546 | 0.615958564 | 0.558404393 | 0.991835066 | -5.102147952 |
| **A_55_P1987746** | NA | 0.037131283 | -0.106999983 | 0.181262548 | 14.96960337 | 0.615730806 | 0.558545976 | 0.991835066 | -5.102258276 |
| **A_51_P361788** | Vapa | 0.040066794 | -0.115468261 | 0.195601849 | 13.57972751 | 0.615694727 | 0.558568406 | 0.991835066 | -5.10227575 |
| **A_55_P2146780** | Sdhc | 0.037159615 | -0.107096773 | 0.181416003 | 14.23451663 | 0.615666164 | 0.558586164 | 0.991835066 | -5.102289582 |
| **A_65_P02321** | Usp36 | 0.031012443 | -0.089380803 | 0.151405689 | 8.638661864 | 0.615662902 | 0.558588192 | 0.991835066 | -5.102291162 |
| **A_55_P2379599** | Asph | -0.028141344 | -0.137422782 | 0.081140094 | 10.00671607 | -0.615470953 | 0.558707536 | 0.991835066 | -5.102384105 |
| **A_55_P2025490** | Tnfrsf18 | 0.027149414 | -0.07828142 | 0.132580248 | 7.77214295 | 0.615462998 | 0.558712483 | 0.991835066 | -5.102387956 |
| **A_55_P2131766** | Rsrp1 | -0.127953685 | -0.624931841 | 0.369024472 | 13.1293329 | -0.615353433 | 0.558780612 | 0.991835066 | -5.102440996 |
| **A_55_P1964403** | Agfg2 | 0.04612177 | -0.133030371 | 0.225273911 | 8.469729054 | 0.615308756 | 0.558808395 | 0.991835066 | -5.102462622 |
| **A_55_P1962110** | NA | 0.035574323 | -0.10265365 | 0.173802295 | 14.86627131 | 0.615105817 | 0.558934603 | 0.991835066 | -5.102560837 |
| **A_55_P2010957** | Gm4971 | -0.076839801 | -0.37543583 | 0.221756229 | 12.22950267 | -0.61505129 | 0.558968517 | 0.991835066 | -5.102587221 |
| **A_52_P155708** | Creg1 | 0.05224028 | -0.150852367 | 0.255332928 | 8.068621007 | 0.614780991 | 0.559136649 | 0.991835066 | -5.102717982 |
| **A_55_P2169415** | Cgn | 0.078631233 | -0.227126343 | 0.384388808 | 8.132849471 | 0.614648734 | 0.559218927 | 0.991835066 | -5.102781945 |
| **A_52_P468041** | Alkbh5 | 0.037588595 | -0.108577882 | 0.183755072 | 10.56350458 | 0.614635229 | 0.559227329 | 0.991835066 | -5.102788476 |
| **A_55_P2065897** | NA | -0.032684147 | -0.159798905 | 0.094430611 | 12.75353707 | -0.614540218 | 0.559286441 | 0.991835066 | -5.102834418 |
| **A_55_P2048202** | Ss18 | 0.046083805 | -0.133160033 | 0.225327642 | 12.53683692 | 0.614487743 | 0.559319091 | 0.991835066 | -5.102859789 |
| **A_51_P372141** | Pnkd | -0.091057969 | -0.445236046 | 0.263120108 | 9.023806723 | -0.614476697 | 0.559325964 | 0.991835066 | -5.102865129 |
| **A_55_P1975285** | Lrig2 | -0.043482642 | -0.212630136 | 0.125664852 | 8.902917353 | -0.614411713 | 0.559366398 | 0.991835066 | -5.102896545 |
| **A_51_P361580** | Rab11b | 0.030588669 | -0.088418133 | 0.14959547 | 12.44534921 | 0.614324626 | 0.559420588 | 0.991835066 | -5.102938642 |
| **A_55_P2047509** | Ufd1 | 0.034961419 | -0.101075314 | 0.170998151 | 11.86234726 | 0.614245507 | 0.559469823 | 0.991835066 | -5.102976884 |
| **A_55_P1972842** | Klf3 | -0.039384654 | -0.192638692 | 0.113869385 | 11.49595694 | -0.614220395 | 0.55948545 | 0.991835066 | -5.10298902 |
| **A_55_P2122633** | Airn | 0.171115737 | -0.494810572 | 0.837042045 | 12.48433178 | 0.614147814 | 0.559530619 | 0.991835066 | -5.103024096 |
| **A_52_P639774** | Gart | 0.056994093 | -0.164814438 | 0.278802624 | 9.739820522 | 0.614130594 | 0.559541336 | 0.991835066 | -5.103032418 |
| **A_55_P2003410** | Kng1 | -0.033621604 | -0.16450226 | 0.097259053 | 7.713089322 | -0.613976983 | 0.559636941 | 0.991835066 | -5.103106639 |
| **A_51_P485594** | Tab3 | 0.031874651 | -0.092241541 | 0.155990843 | 9.47374933 | 0.613798937 | 0.559747766 | 0.991835066 | -5.103192647 |
| **A_52_P220879** | Tgm2 | 0.03909792 | -0.113165155 | 0.191360995 | 14.86796459 | 0.613717056 | 0.559798738 | 0.991835066 | -5.103232194 |

| | | | | | | | | | |
|---|---|---|---|---|---|---|---|---|---|
| **A_55_P2115507** | Egfl8 | -0.058040012 | -0.284087805 | 0.16800778 | 9.480923819 | -0.61367207 | 0.559826743 | 0.991835066 | -5.103253919 |
| **A_55_P2035504** | Nt5c3 | -0.051788346 | -0.259523967 | 0.155947275 | 11.80388882 | -0.619281659 | 0.559852524 | 0.991835066 | -5.028693683 |
| **A_51_P270666** | Uspl1 | 0.029886163 | -0.086526751 | 0.146299078 | 10.670616 | 0.6135898 | 0.559877961 | 0.991835066 | -5.103293647 |
| **A_55_P2110743** | Mettl7a2 | 0.101491019 | -0.293921876 | 0.496903913 | 13.08007698 | 0.613459528 | 0.559959068 | 0.991835066 | -5.103356545 |
| **A_55_P2382741** | NA | -0.047671744 | -0.233403925 | 0.138060436 | 8.428231008 | -0.613455434 | 0.559961617 | 0.991835066 | -5.103358521 |
| **A_51_P111164** | Rnd1 | 0.056838265 | -0.164608973 | 0.278285504 | 8.622380199 | 0.613450719 | 0.559964553 | 0.991835066 | -5.103360798 |
| **A_66_P136097** | Trim13 | -0.037250428 | -0.182395471 | 0.107894614 | 8.668440197 | -0.613392126 | 0.560001036 | 0.991835066 | -5.103389083 |
| **A_55_P2109377** | Camk2b | -0.050599991 | -0.247764384 | 0.146564402 | 9.135094822 | -0.613382083 | 0.560007289 | 0.991835066 | -5.103393932 |
| **A_55_P2021114** | Ier5 | 0.04208714 | -0.121935172 | 0.206109452 | 8.609119629 | 0.613275503 | 0.560073655 | 0.991835066 | -5.103445376 |
| **A_55_P2136385** | Rps3a1 | -0.031947844 | -0.156461255 | 0.092565568 | 15.18215315 | -0.613245755 | 0.560092179 | 0.991835066 | -5.103459734 |
| **A_55_P1959137** | Usp32 | 0.044949546 | -0.130251414 | 0.220150506 | 10.90504632 | 0.613194089 | 0.560124353 | 0.991835066 | -5.103484668 |
| **A_51_P417701** | Apaf1 | -0.028502367 | -0.139614236 | 0.082609502 | 8.521636627 | -0.613097581 | 0.560184455 | 0.991835066 | -5.103531239 |
| **A_55_P2005930** | NA | 0.050790899 | -0.147226842 | 0.24880864 | 13.14050757 | 0.613042992 | 0.560218452 | 0.991835066 | -5.103557579 |
| **A_51_P353095** | Zfyve27 | 0.049359446 | -0.143097908 | 0.241816801 | 10.30307241 | 0.612978017 | 0.56025892 | 0.991835066 | -5.103588928 |
| **A_55_P2027067** | Phc2 | -0.035052676 | -0.171735436 | 0.101630083 | 12.83285513 | -0.612938042 | 0.560283818 | 0.991835066 | -5.103608213 |
| **A_55_P1995205** | Top2a | 0.036669054 | -0.106324747 | 0.179662856 | 10.68852456 | 0.612902834 | 0.560305747 | 0.991835066 | -5.103625198 |
| **A_55_P2137773** | NA | 0.047734942 | -0.138426964 | 0.233896848 | 16.12208915 | 0.61285074 | 0.560338195 | 0.991835066 | -5.103650327 |
| **A_51_P367298** | NA | 0.026649716 | -0.077284894 | 0.130584326 | 8.061385405 | 0.612832128 | 0.560349788 | 0.991835066 | -5.103659305 |
| **A_55_P2009576** | Rnu12 | 0.040354452 | -0.117040624 | 0.197749527 | 12.65967051 | 0.612786856 | 0.560377988 | 0.991835066 | -5.103681141 |
| **A_51_P395555** | Vkorc1 | 0.046197739 | -0.133997008 | 0.226392485 | 12.52311176 | 0.612756218 | 0.560397073 | 0.991835066 | -5.103695918 |
| **A_55_P2177941** | Tmem208 | -0.047528786 | -0.232929339 | 0.137871767 | 10.72147885 | -0.612709799 | 0.560425989 | 0.991835066 | -5.103718305 |
| **A_66_P109175** | Farsb | -0.033708999 | -0.165224384 | 0.097806386 | 11.27578451 | -0.612602023 | 0.56049313 | 0.991835066 | -5.103770277 |
| **A_52_P420504** | Acta2 | 0.097479222 | -0.282870532 | 0.477828976 | 15.483197 | 0.612545067 | 0.560528613 | 0.991835066 | -5.064240513 |
| **A_55_P1960074** | Pafah1b3 | 0.086047045 | -0.249705232 | 0.421799323 | 12.24198078 | 0.612528256 | 0.560539087 | 0.991835066 | -5.103805846 |
| **A_55_P2013371** | Ttc29 | -0.024606289 | -0.120625233 | 0.071412655 | 7.657616096 | -0.612489051 | 0.560563512 | 0.991835066 | -5.103824747 |
| **A_55_P1975011** | Ufc1 | 0.038481213 | -0.111718455 | 0.188680881 | 13.20832535 | 0.612334759 | 0.560659646 | 0.991835066 | -5.103899126 |
| **A_51_P370615** | Neurog2 | -0.024686565 | -0.121044983 | 0.071671853 | 7.638599699 | -0.612322395 | 0.56066735 | 0.991835066 | -5.103905086 |
| **A_51_P108334** | Slc25a4 | -0.030184823 | -0.148021148 | 0.087651501 | 17.78242911 | -0.612235609 | 0.560721428 | 0.991835066 | -5.092601901 |
| **A_51_P493117** | Slc16a9 | -0.036223694 | -0.177638575 | 0.105191186 | 8.095824833 | -0.612218941 | 0.560731815 | 0.991835066 | -5.103954947 |
| **A_55_P1986863** | Ccdc126 | 0.051501536 | -0.149581551 | 0.252584623 | 9.599598984 | 0.612144247 | 0.560778361 | 0.991835066 | -5.103990942 |
| **A_55_P2080870** | Cmtm7 | 0.051817155 | -0.150566218 | 0.254200528 | 11.93787541 | 0.611938629 | 0.560906507 | 0.991835066 | -5.104090012 |
| **A_51_P258848** | Mfsd13b | 0.027395363 | -0.079603364 | 0.13439409 | 7.708366451 | 0.61193822 | 0.560906761 | 0.991835066 | -5.104090208 |
| **A_55_P1968108** | NA | -0.055788802 | -0.273718427 | 0.162140823 | 15.5658631 | -0.611842857 | 0.560966199 | 0.991835066 | -5.104136146 |
| **A_51_P142091** | Asb8 | -0.046929334 | -0.230283138 | 0.13642447 | 12.24400984 | -0.611735381 | 0.561033191 | 0.991835066 | -5.10418791 |
| **A_55_P2089219** | NA | 0.045906283 | -0.133453818 | 0.225266384 | 7.830278625 | 0.611723863 | 0.561040371 | 0.991835066 | -5.104193458 |
| **A_55_P2024525** | Prdm8 | 0.025088433 | -0.072954207 | 0.123131073 | 7.73746368 | 0.611600264 | 0.56111742 | 0.991835066 | -5.104252977 |
| **A_55_P1991960** | Nwd1 | 0.046656586 | -0.135674951 | 0.228988123 | 8.14869355 | 0.611589888 | 0.561123889 | 0.991835066 | -5.104257973 |
| **A_51_P103222** | Slc39a4 | 0.044483249 | -0.129446715 | 0.218413214 | 8.308361275 | 0.611267389 | 0.561324959 | 0.991835066 | -5.104413223 |
| **A_55_P2133524** | Nbr1 | -0.036011424 | -0.176832682 | 0.104809834 | 8.429719359 | -0.611196995 | 0.561368853 | 0.991835066 | -5.104447101 |
| **A_55_P2012984** | Ldha | -0.028467362 | -0.139797989 | 0.082863266 | 16.78840457 | -0.611141388 | 0.561403529 | 0.991835066 | -5.10447386 |
| **A_51_P490388** | Nudt8 | -0.074531479 | -0.366079059 | 0.217016101 | 10.98381524 | -0.610997497 | 0.561493262 | 0.991835066 | -5.104543093 |
| **A_55_P2133356** | NA | 0.028394541 | -0.082694502 | 0.139483584 | 15.45430993 | 0.610903707 | 0.561551756 | 0.991835066 | -5.104588212 |
| **A_52_P189957** | Dbnl | -0.080618822 | -0.396036481 | 0.234798836 | 9.210882987 | -0.610885163 | 0.561563322 | 0.991835066 | -5.104597133 |
| **A_55_P2025023** | NA | 0.049984472 | -0.145588746 | 0.245557691 | 15.80979258 | 0.610850397 | 0.561585006 | 0.991835066 | -5.104613855 |
| **A_52_P451644** | Ndn | -0.058262255 | -0.286243757 | 0.169719247 | 9.117224634 | -0.610796883 | 0.561618384 | 0.991835066 | -5.104639595 |
| **A_55_P2031432** | Usp47 | 0.038920349 | -0.113384863 | 0.191225561 | 8.45665691 | 0.610760715 | 0.561640943 | 0.991835066 | -5.10465699 |
| **A_55_P2010066** | Capn3 | -0.054941314 | -0.269941787 | 0.160059158 | 8.730744037 | -0.610757434 | 0.56164299 | 0.991835066 | -5.104658568 |
| **A_55_P2109857** | Rgs2 | 0.038669467 | -0.112673051 | 0.190011984 | 11.6720201 | 0.610683744 | 0.561688955 | 0.991835066 | -5.104694006 |

| | | | | | | | | | |
|---|---|---|---|---|---|---|---|---|---|
| **A_55_P2277171** | Gm10134 | 0.028610877 | -0.083368188 | 0.140589943 | 7.918158532 | 0.61066562 | 0.561700261 | 0.991835066 | -5.104702722 |
| **A_55_P2061979** | NA | 0.029278705 | -0.085334172 | 0.143891582 | 15.68700878 | 0.610558918 | 0.561766823 | 0.991835066 | -5.104754028 |
| **A_55_P2084448** | Cybc1 | 0.032862905 | -0.095780979 | 0.161506789 | 10.28104331 | 0.610556617 | 0.561768258 | 0.991835066 | -5.104755135 |
| **A_55_P2022604** | Exoc3l4 | 0.073195396 | -0.213410568 | 0.35980136 | 8.827819329 | 0.610390371 | 0.561871974 | 0.991835066 | -5.104835056 |
| **A_55_P2046699** | Tvp23b | 0.037755735 | -0.110094793 | 0.185606263 | 9.776775319 | 0.610336276 | 0.561905725 | 0.991835066 | -5.104861059 |
| **A_55_P2185359** | NA | -0.069942929 | -0.343915388 | 0.20402953 | 15.41549689 | -0.610163238 | 0.562013695 | 0.991835066 | -5.104944219 |
| **A_51_P392867** | Pex12 | -0.039506412 | -0.194256495 | 0.115243671 | 8.791224642 | -0.610162944 | 0.562013878 | 0.991835066 | -5.10494436 |
| **A_51_P118223** | Gm1943 | -0.027311606 | -0.134302089 | 0.079678877 | 8.245369021 | -0.610114326 | 0.562044216 | 0.991835066 | -5.093612706 |
| **A_55_P1973036** | NA | -0.027481581 | -0.13513997 | 0.080176808 | 7.799763188 | -0.610102723 | 0.562051457 | 0.991835066 | -5.104973298 |
| **A_51_P245546** | Synrg | 0.031767382 | -0.092704459 | 0.156239224 | 9.203884499 | 0.60998541 | 0.562124665 | 0.991835066 | -5.105029661 |
| **A_51_P137007** | Zw10 | 0.039034778 | -0.113941476 | 0.192011031 | 9.456188531 | 0.609869369 | 0.562197086 | 0.991835066 | -5.105085404 |
| **A_51_P323842** | Ciao2b | -0.045464455 | -0.223654287 | 0.132725376 | 14.31636087 | -0.609815146 | 0.562230929 | 0.991835066 | -5.105111448 |
| **A_51_P129317** | Mob4 | 0.034765969 | -0.101519502 | 0.171051441 | 11.38102413 | 0.609696799 | 0.562304797 | 0.991835066 | -5.07633507 |
| **A_55_P1995497** | Atad3a | 0.076333242 | -0.222926327 | 0.375592811 | 9.287925471 | 0.609641879 | 0.562339078 | 0.991835066 | -5.105194657 |
| **A_51_P385598** | Slc37a4 | -0.042594034 | -0.209620104 | 0.124432036 | 11.22707289 | -0.609499906 | 0.562427704 | 0.991835066 | -5.105262823 |
| **A_52_P409731** | Scamp5 | 0.025838166 | -0.075484425 | 0.127160757 | 8.055864476 | 0.609487118 | 0.562435688 | 0.991835066 | -5.105268962 |
| **A_55_P2107972** | Smad5 | 0.036832538 | -0.107622713 | 0.181287789 | 11.92612101 | 0.609407005 | 0.562485702 | 0.991835066 | -5.10530742 |
| **A_51_P424499** | Txn1 | 0.039706276 | -0.11605212 | 0.195464672 | 14.10829939 | 0.60927986 | 0.562565084 | 0.991835066 | -5.105368447 |
| **A_51_P279100** | Ptgs1 | -0.069573406 | -0.34250184 | 0.203355028 | 10.29809966 | -0.609261332 | 0.562576652 | 0.991835066 | -5.105377339 |
| **A_55_P2028159** | C9orf72 | -0.045121685 | -0.2221501 | 0.13190673 | 11.369191 | -0.609188166 | 0.562622336 | 0.991835066 | -5.10541245 |
| **A_55_P1977850** | Sall1 | 0.032912935 | -0.09624653 | 0.162072401 | 8.344756811 | 0.609045179 | 0.562711621 | 0.991835066 | -5.105481058 |
| **A_55_P2152387** | Wwox | -0.038281327 | -0.188530389 | 0.111967735 | 8.19524639 | -0.6089538 | 0.562768686 | 0.991835066 | -5.105524897 |
| **A_51_P204080** | Hk2 | -0.072407657 | -0.35660228 | 0.211786966 | 11.54683782 | -0.608944582 | 0.562774443 | 0.991835066 | -5.105529319 |
| **A_55_P2064811** | Nxt1 | 0.066877008 | -0.195665128 | 0.329419144 | 12.09454081 | 0.608817255 | 0.562853962 | 0.991835066 | -5.105590392 |
| **A_52_P353417** | Gm2a | -0.056582531 | -0.278749349 | 0.165584288 | 8.940395025 | -0.608712618 | 0.562919316 | 0.991835066 | -5.105640574 |
| **A_52_P141488** | Grk5 | -0.037678539 | -0.185723387 | 0.11036631 | 9.5398024 | -0.608288888 | 0.563184015 | 0.991835066 | -5.105843711 |
| **A_66_P117730** | Hapln1 | 0.045485299 | -0.133234316 | 0.224204914 | 8.190975655 | 0.608286205 | 0.563185692 | 0.991835066 | -5.105844997 |
| **A_55_P2010843** | Fahd1 | 0.081887854 | -0.239869234 | 0.403644942 | 10.29816592 | 0.608275746 | 0.563192226 | 0.991835066 | -5.105850009 |
| **A_52_P53906** | Ccnd2 | 0.07270732 | -0.212984545 | 0.358399185 | 12.07126243 | 0.608260194 | 0.563201943 | 0.991835066 | -5.105857462 |
| **A_51_P394115** | Aadac | -0.057261942 | -0.282282918 | 0.167759035 | 7.870290783 | -0.608208095 | 0.563234494 | 0.991835066 | -5.105882429 |
| **A_55_P2043982** | Mtm1 | -0.028416474 | -0.140086371 | 0.083253423 | 8.266746106 | -0.608195509 | 0.563242358 | 0.991835066 | -5.105888461 |
| **A_55_P2065751** | Krtap19-5 | 0.082872315 | -0.242901296 | 0.408645925 | 9.184913377 | 0.607998771 | 0.563365291 | 0.991835066 | -5.105982723 |
| **A_51_P206971** | Mnat1 | 0.029736027 | -0.08715823 | 0.146630283 | 10.03752656 | 0.607993444 | 0.563368619 | 0.991835066 | -5.105985274 |
| **A_66_P123824** | Sorl1 | 0.037135183 | -0.108862682 | 0.183133048 | 8.050207663 | 0.607922467 | 0.563412974 | 0.991835066 | -5.106019275 |
| **A_55_P2003246** | Asxl2 | 0.06302716 | -0.184795594 | 0.310849915 | 9.935539533 | 0.60784898 | 0.563458899 | 0.991835066 | -5.106054473 |
| **A_51_P345593** | Sptlc1 | -0.037124215 | -0.183097487 | 0.108849056 | 9.235424332 | -0.607845318 | 0.563461188 | 0.991835066 | -5.106056227 |
| **A_55_P2159299** | Ing1 | 0.028729631 | -0.08423722 | 0.141696482 | 11.03896632 | 0.607838436 | 0.563465489 | 0.991835066 | -5.106059524 |
| **A_55_P2163622** | Rps27 | 0.029381652 | -0.086162857 | 0.144926161 | 16.29612015 | 0.607765482 | 0.563511083 | 0.991835066 | -5.106094463 |
| **A_55_P2150831** | Cdh4 | -0.04669376 | -0.230352703 | 0.136965182 | 8.157139973 | -0.607653357 | 0.563581163 | 0.991835066 | -5.106148155 |
| **A_55_P2087561** | Eno1b | 0.097077598 | -0.284874995 | 0.479030192 | 11.32958858 | 0.607461408 | 0.563701147 | 0.991835066 | -5.106240053 |
| **A_51_P499816** | Pbld1 | 0.027166438 | -0.07973142 | 0.134064297 | 7.708295757 | 0.607397259 | 0.563741248 | 0.991835066 | -5.106270759 |
| **A_55_P1981415** | Gsk3b | 0.033819201 | -0.099274805 | 0.166913208 | 12.46527882 | 0.60731496 | 0.563792698 | 0.991835066 | -5.106310149 |
| **A_55_P2295445** | 2600002B0 | 0.103326652 | -0.303316662 | 0.509969966 | 10.24326828 | 0.607306393 | 0.563798054 | 0.991835066 | -5.106314249 |
| **A_55_P2040743** | Zwilch | 0.042238848 | -0.124009134 | 0.20848683 | 8.954976943 | 0.607246218 | 0.563835675 | 0.991835066 | -5.106343047 |
| **A_55_P2045234** | NA | 0.030797789 | -0.090458097 | 0.152053675 | 8.609424596 | 0.607051928 | 0.563957154 | 0.991835066 | -5.10643601 |
| **A_55_P1974283** | Rbmx | -0.044823693 | -0.221319751 | 0.131672366 | 10.70005288 | -0.606990304 | 0.563995688 | 0.991835066 | -5.106465491 |
| **A_55_P2036280** | Psen2 | 0.036226232 | -0.106417124 | 0.178869588 | 10.70397028 | 0.606988905 | 0.563996563 | 0.991835066 | -5.10646616 |
| **A_66_P104118** | NA | 0.033333213 | -0.097945909 | 0.164612335 | 8.262852584 | 0.606862986 | 0.564075305 | 0.991835066 | -5.10652639 |

| | | | | | | | | | |
|---|---|---|---|---|---|---|---|---|---|
| **A_55_P2166990** | Ppcs | -0.045766528 | -0.226104848 | 0.134571793 | 10.04244386 | -0.606553447 | 0.5642689 | 0.991835066 | -5.106674405 |
| **A_55_P1964896** | NA | 0.154810101 | -0.455238352 | 0.764858554 | 10.24250594 | 0.606518562 | 0.564290721 | 0.991835066 | -5.106691082 |
| **A_51_P249127** | Pkd1 | 0.052778055 | -0.155205944 | 0.260762055 | 10.3217903 | 0.606502492 | 0.564300773 | 0.991835066 | -5.106698764 |
| **A_51_P268903** | Mon1a | 0.036490961 | -0.107313127 | 0.180295049 | 9.351607336 | 0.606489381 | 0.564308974 | 0.991835066 | -5.106705032 |
| **A_55_P2182931** | Sim2 | -0.048657756 | -0.24040968 | 0.143094169 | 8.432944037 | -0.606486971 | 0.564310481 | 0.991835066 | -5.106706184 |
| **A_51_P144926** | Cops8 | -0.042001198 | -0.207520789 | 0.123518393 | 10.17820929 | -0.606486871 | 0.564310544 | 0.991835066 | -5.106706232 |
| **A_51_P461761** | Zcchc8 | 0.050794928 | -0.149403208 | 0.250993063 | 9.791323516 | 0.606414321 | 0.564355927 | 0.991835066 | -5.106740911 |
| **A_55_P2042426** | Rpl6 | 0.030753734 | -0.090515609 | 0.152023077 | 16.27160025 | 0.6061163 | 0.564542373 | 0.991835066 | -5.106883328 |
| **A_55_P1953608** | NA | 0.063536586 | -0.187052276 | 0.314125447 | 10.8042565 | 0.605998074 | 0.564616347 | 0.991835066 | -5.106939808 |
| **A_52_P318673** | Saa1 | 0.041467271 | -0.122095368 | 0.20502991 | 8.399898656 | 0.605941192 | 0.56465194 | 0.991835066 | -5.106966979 |
| **A_65_P06453** | Nup35 | 0.038451591 | -0.113222557 | 0.190125738 | 10.43293788 | 0.605915246 | 0.564668176 | 0.991835066 | -5.106979373 |
| **A_51_P114843** | H1f10 | 0.03311763 | -0.097523058 | 0.163758318 | 8.489665057 | 0.605884613 | 0.564687345 | 0.991835066 | -5.106994004 |
| **A_55_P2052555** | Dpysl5 | -0.058190515 | -0.287743957 | 0.171362928 | 11.23524318 | -0.605867312 | 0.564698171 | 0.991835066 | -5.107002267 |
| **A_52_P660385** | Trmt6 | -0.029927367 | -0.148009093 | 0.088154359 | 8.791295366 | -0.605752132 | 0.56477025 | 0.991835066 | -5.107057272 |
| **A_55_P2010611** | Slc30a5 | 0.02622232 | -0.077272492 | 0.129717132 | 10.24508719 | 0.605566251 | 0.564886587 | 0.991835066 | -5.107146024 |
| **A_55_P2167299** | NA | 0.032563115 | -0.095960594 | 0.161086825 | 15.10941787 | 0.605552534 | 0.564895173 | 0.991835066 | -5.107152572 |
| **A_55_P2279690** | 2900076A0 | -0.041293888 | -0.20430386 | 0.121716084 | 9.008137376 | -0.605453418 | 0.564957212 | 0.991835066 | -5.107199886 |
| **A_55_P2052240** | Crlf1 | -0.027160143 | -0.134428397 | 0.080108112 | 7.906147671 | -0.605159658 | 0.565141108 | 0.991835066 | -5.107340074 |
| **A_51_P309854** | Kcnn2 | -0.055488814 | -0.274697027 | 0.163719398 | 9.748460483 | -0.605003319 | 0.565238993 | 0.991835066 | -5.107414658 |
| **A_51_P312336** | Slc14a1 | -0.024987999 | -0.123703957 | 0.073727958 | 7.700149801 | -0.604997054 | 0.565242915 | 0.991835066 | -5.107417647 |
| **A_52_P194805** | Zfp706 | -0.043561345 | -0.215661545 | 0.128538854 | 13.27048507 | -0.604963317 | 0.565264039 | 0.991835066 | -5.107433739 |
| **A_55_P2154714** | Klhl9 | 0.05635989 | -0.166332108 | 0.279051888 | 13.46199527 | 0.604887565 | 0.565311473 | 0.991835066 | -5.10746987 |
| **A_55_P2035677** | Srp19 | 0.052381298 | -0.154611328 | 0.259373925 | 11.03281117 | 0.604826087 | 0.56534997 | 0.991835066 | -5.10749919 |
| **A_55_P2045693** | NA | -0.068063561 | -0.337102168 | 0.200975047 | 8.612079024 | -0.604657147 | 0.565455767 | 0.991835066 | -5.107579746 |
| **A_55_P2051899** | Atp2c1 | 0.040331774 | -0.119124651 | 0.199788199 | 10.20945987 | 0.604525236 | 0.565538384 | 0.991835066 | -5.107642632 |
| **A_55_P2064928** | Nebl | 0.041131377 | -0.121536887 | 0.203799642 | 8.688148902 | 0.604337512 | 0.565655969 | 0.991835066 | -5.107732106 |
| **A_55_P1967534** | NA | 0.049150706 | -0.145253045 | 0.243554456 | 16.27409211 | 0.604274476 | 0.565695456 | 0.991835066 | -5.107762145 |
| **A_55_P2116853** | Brpf1 | 0.03313291 | -0.097963754 | 0.164229573 | 11.18417559 | 0.604055813 | 0.565832445 | 0.991835066 | -5.107866325 |
| **A_55_P2137431** | Rtel1 | 0.058630749 | -0.173383025 | 0.290644524 | 10.24209223 | 0.603977567 | 0.565881469 | 0.991835066 | -5.107903596 |
| **A_51_P333099** | Mrps15 | -0.040625005 | -0.201395109 | 0.120145099 | 11.70045391 | -0.603944828 | 0.565901982 | 0.991835066 | -5.10791919 |
| **A_55_P2131193** | NA | 0.045343922 | -0.134139549 | 0.224827392 | 14.65309598 | 0.6038148 | 0.565983457 | 0.991835066 | -5.107981115 |
| **A_52_P423364** | Fbxl7 | -0.049393734 | -0.244934163 | 0.146146695 | 10.26625634 | -0.603732323 | 0.566035141 | 0.991835066 | -5.108020389 |
| **A_55_P1967332** | NA | -0.057651219 | -0.285898645 | 0.170596208 | 15.38715614 | -0.603686875 | 0.566063622 | 0.991835066 | -5.108042028 |
| **A_55_P1993708** | Lrp12 | 0.030316133 | -0.089712149 | 0.150344415 | 8.81747764 | 0.603669664 | 0.566074408 | 0.991835066 | -5.108050222 |
| **A_51_P486001** | Mob3a | -0.038825437 | -0.192550782 | 0.114899907 | 8.365056887 | -0.603642788 | 0.566091251 | 0.991835066 | -5.108063017 |
| **A_51_P234692** | Neat1 | 0.115455235 | -0.341691917 | 0.572602387 | 8.646590611 | 0.603624338 | 0.566102813 | 0.991835066 | -5.108071801 |
| **A_55_P2048650** | Dennd4a | -0.046552345 | -0.230903984 | 0.137799295 | 14.56660548 | -0.603536705 | 0.566157735 | 0.991835066 | -5.108113518 |
| **A_55_P2066553** | Oacyl | 0.094355483 | -0.279304866 | 0.468015832 | 11.67843045 | 0.603530553 | 0.56616159 | 0.991835066 | -5.108116446 |
| **A_51_P184197** | Smarcd2 | 0.049877163 | -0.147748527 | 0.247502853 | 8.832542412 | 0.603208533 | 0.566363435 | 0.991835066 | -5.108269694 |
| **A_55_P2052210** | Gdi2 | 0.031450157 | -0.093200044 | 0.156100358 | 13.94256592 | 0.603030071 | 0.566475315 | 0.991835066 | -5.108354593 |
| **A_51_P268193** | Slc7a10 | -0.037367182 | -0.18547002 | 0.110735655 | 7.805583503 | -0.603026095 | 0.566477808 | 0.991835066 | -5.108356484 |
| **A_52_P144818** | Ppp1r1a | -0.036361207 | -0.180479934 | 0.107757519 | 8.861879 | -0.60301347 | 0.566485723 | 0.991835066 | -5.10836249 |
| **A_66_P112399** | BC029722 | -0.037366062 | -0.18546828 | 0.110736155 | 10.83445536 | -0.603010543 | 0.566487559 | 0.991835066 | -5.108363882 |
| **A_52_P28057** | Gm14124 | 0.031627861 | -0.093796895 | 0.157052616 | 8.248779177 | 0.602692359 | 0.566687067 | 0.991835066 | -5.108515192 |
| **A_55_P1976097** | NA | -0.044742815 | -0.222184539 | 0.132698909 | 10.4727374 | -0.602665997 | 0.566703598 | 0.991835066 | -5.108527725 |
| **A_51_P200517** | Rasgef1a | -0.026415501 | -0.131180115 | 0.078349112 | 8.004291893 | -0.602633647 | 0.566723886 | 0.991835066 | -5.108543104 |
| **A_55_P2025954** | Acly | 0.033141956 | -0.098302836 | 0.164586749 | 12.47252122 | 0.602620479 | 0.566732143 | 0.991835066 | -5.108549364 |
| **A_55_P2069083** | Galnt12 | -0.061483126 | -0.305402919 | 0.182436668 | 8.56351849 | -0.602445863 | 0.566841654 | 0.991835066 | -5.108632363 |

| | | | | | | | | | |
|---|---|---|---|---|---|---|---|---|---|
| **A_55_P2424042** | Ttn | -0.092289037 | -0.458448028 | 0.273869955 | 9.826888411 | -0.60240635 | 0.566866437 | 0.991835066 | -5.108651142 |
| **A_55_P2003115** | Rnf7 | 0.039570315 | -0.117437644 | 0.196578274 | 12.73126017 | 0.602361173 | 0.566894773 | 0.991835066 | -5.108672611 |
| **A_55_P2154690** | Smarca4 | 0.070382936 | -0.208902184 | 0.349668057 | 9.333803945 | 0.602321986 | 0.566919352 | 0.991835066 | -5.108691232 |
| **A_65_P08768** | Fbln1 | -0.060338756 | -0.299768145 | 0.179090633 | 10.42610625 | -0.602321028 | 0.566919953 | 0.991835066 | -5.108691688 |
| **A_55_P2060672** | Hoxa3 | 0.073259717 | -0.217497603 | 0.364017037 | 8.250641548 | 0.602204106 | 0.566993294 | 0.991835066 | -5.108747242 |
| **A_55_P1974114** | NA | -0.028411446 | -0.141185593 | 0.084362702 | 15.09954575 | -0.602133669 | 0.567037479 | 0.991835066 | -5.108780705 |
| **A_52_P36218** | Upf2 | -0.03569007 | -0.177373771 | 0.105993632 | 8.870733422 | -0.602055645 | 0.567086426 | 0.991835066 | -5.108817768 |
| **A_51_P453401** | Cipc | 0.033337356 | -0.099022945 | 0.165697657 | 11.77021386 | 0.601980659 | 0.56713347 | 0.991835066 | -5.108853384 |
| **A_52_P473953** | Ctdspl | 0.035943878 | -0.106772546 | 0.178660302 | 9.668933891 | 0.601949569 | 0.567152975 | 0.991835066 | -5.10886815 |
| **A_51_P443723** | Slc35c1 | 0.042781863 | -0.127123353 | 0.212687079 | 10.46873937 | 0.601813762 | 0.567238184 | 0.991835066 | -5.108932642 |
| **A_52_P277854** | Snrpb | -0.081654376 | -0.405955781 | 0.242647029 | 11.04908385 | -0.601782795 | 0.567257615 | 0.991835066 | -5.108947346 |
| **A_55_P2122429** | Cym | 0.039420999 | -0.117150647 | 0.195992645 | 8.374635675 | 0.601760447 | 0.567271637 | 0.991835066 | -5.108957957 |
| **A_51_P423549** | Shbg | -0.026336851 | -0.130948559 | 0.078274856 | 7.679101816 | -0.601717566 | 0.567298544 | 0.991835066 | -5.108978315 |
| **A_51_P466591** | 1600012P1 | -0.470264312 | -2.338286043 | 1.397757418 | 8.949783071 | -0.601685063 | 0.56731894 | 0.991835066 | -5.108993746 |
| **A_51_P290207** | Insig1 | 0.036694369 | -0.109104394 | 0.182493131 | 8.184183455 | 0.601526433 | 0.567418486 | 0.991835066 | -5.109069046 |
| **A_66_P131091** | Rabl6 | -0.038079908 | -0.189393949 | 0.113234133 | 12.16918483 | -0.60148637 | 0.567443628 | 0.991835066 | -5.109088061 |
| **A_55_P2051949** | Elmod3 | -0.057767657 | -0.287333005 | 0.17179769 | 10.71498829 | -0.601433419 | 0.567476861 | 0.991835066 | -5.109113191 |
| **A_52_P372151** | Ctnnal1 | 0.055175836 | -0.164099365 | 0.274451037 | 11.18039001 | 0.601407081 | 0.56749339 | 0.991835066 | -5.10912569 |
| **A_55_P2174401** | NA | -0.057065511 | -0.283895704 | 0.169764682 | 8.228891414 | -0.601287231 | 0.567568614 | 0.991835066 | -5.10918256 |
| **A_55_P2060188** | NA | 0.057098961 | -0.169885219 | 0.284083141 | 8.961955474 | 0.601231528 | 0.567603577 | 0.991835066 | -5.109208989 |
| **A_55_P2114994** | NA | 0.055795926 | -0.16603015 | 0.277622002 | 14.50477412 | 0.601172391 | 0.567640697 | 0.991835066 | -5.109237044 |
| **A_66_P140658** | Olfr787 | -0.039872559 | -0.198410545 | 0.118665427 | 7.885016632 | -0.601104397 | 0.567683379 | 0.991835066 | -5.109269298 |
| **A_52_P251699** | Prodh2 | -0.027321804 | -0.135971087 | 0.08132748 | 7.696324973 | -0.601023748 | 0.567734007 | 0.991835066 | -5.109307551 |
| **A_55_P1977330** | Zbtb1 | 0.034424495 | -0.102549104 | 0.171398094 | 8.515403917 | 0.600675395 | 0.56795272 | 0.991835066 | -5.10947273 |
| **A_52_P66371** | Nlrx1 | -0.045408372 | -0.226094262 | 0.135277517 | 8.640763876 | -0.600649098 | 0.567969233 | 0.991835066 | -5.109485196 |
| **A_55_P2152547** | Myl3 | -0.043799996 | -0.218097083 | 0.130497091 | 18.19309307 | -0.600610715 | 0.567993334 | 0.991835066 | -5.10950339 |
| **A_55_P2073940** | Abcc5 | -0.027800832 | -0.138444779 | 0.082843115 | 8.233310132 | -0.600536302 | 0.568040063 | 0.991835066 | -5.109538661 |
| **A_66_P134704** | Gm4952 | -0.024726546 | -0.123152427 | 0.073699335 | 7.682819329 | -0.600431232 | 0.568106047 | 0.991835066 | -5.109588456 |
| **A_55_P2235970** | C130068B( | 0.074579918 | -0.222362726 | 0.371522563 | 8.498481537 | 0.600286344 | 0.568197043 | 0.991835066 | -5.109657109 |
| **A_55_P2167769** | Arid1a | 0.048229687 | -0.143809143 | 0.240268516 | 11.68870973 | 0.600253254 | 0.568217827 | 0.991835066 | -5.109672786 |
| **A_52_P137765** | Lmna | -0.05081285 | -0.253148449 | 0.151522749 | 12.49863552 | -0.600219895 | 0.56823878 | 0.991835066 | -5.10968859 |
| **A_51_P344008** | Slc25a20 | -0.09896468 | -0.493073872 | 0.295144513 | 9.061019279 | -0.600167937 | 0.568271415 | 0.991835066 | -5.109713203 |
| **A_51_P212679** | Cwf19l1 | 0.033583438 | -0.100157046 | 0.167323922 | 8.66206061 | 0.60016601 | 0.568272626 | 0.991835066 | -5.109714116 |
| **A_55_P2079430** | NA | 0.045261491 | -0.134985246 | 0.225508228 | 9.845226304 | 0.600164884 | 0.568273333 | 0.991835066 | -5.109714649 |
| **A_55_P1963491** | Ptma | 0.049108448 | -0.146483772 | 0.244700668 | 13.04223102 | 0.600086377 | 0.568322647 | 0.991835066 | -5.109751836 |
| **A_55_P2161347** | Acmsd | -0.027007042 | -0.134577023 | 0.080562938 | 7.68781395 | -0.600060538 | 0.568338878 | 0.991835066 | -5.109764075 |
| **A_55_P2002815** | Rabgap1l | -0.03187642 | -0.158854341 | 0.095101501 | 8.805204447 | -0.599998912 | 0.56837759 | 0.991835066 | -5.109793261 |
| **A_52_P559808** | Arhgef12 | -0.048991553 | -0.244171902 | 0.146188795 | 12.14691987 | -0.599921261 | 0.568426372 | 0.991835066 | -5.109830034 |
| **A_55_P1962380** | NA | -0.067536545 | -0.336602795 | 0.201529706 | 13.01134568 | -0.599913648 | 0.568431155 | 0.991835066 | -5.109833639 |
| **A_51_P454309** | Zfp772 | 0.051022476 | -0.152297006 | 0.254341957 | 8.365119709 | 0.599779568 | 0.568515392 | 0.991835066 | -5.109897123 |
| **A_55_P2176248** | NA | 0.034857775 | -0.104049012 | 0.173764562 | 15.75240387 | 0.599770817 | 0.56852089 | 0.991835066 | -5.109901266 |
| **A_51_P149699** | P3h3 | 0.075178294 | -0.224423138 | 0.374779726 | 10.50313175 | 0.599732678 | 0.568544853 | 0.991835066 | -5.109919322 |
| **A_51_P461877** | Thbs2 | 0.056153491 | -0.167699294 | 0.280006276 | 9.434638147 | 0.599547218 | 0.568661385 | 0.991835066 | -5.110007108 |
| **A_51_P109449** | Ltb4r1 | -0.031627482 | -0.157756426 | 0.094501462 | 7.922152633 | -0.599320301 | 0.568803987 | 0.991835066 | -5.110114485 |
| **A_51_P499169** | Tram1 | 0.057690322 | -0.172392542 | 0.287773186 | 9.903808234 | 0.599277291 | 0.568831018 | 0.991835066 | -5.110134833 |
| **A_51_P210560** | Pptc7 | -0.043170591 | -0.215416375 | 0.129075193 | 12.78093124 | -0.599029942 | 0.568986488 | 0.991835066 | -5.110251832 |
| **A_55_P2011132** | Armc10 | -0.040208278 | -0.200654052 | 0.120237496 | 10.97583973 | -0.598957939 | 0.56903175 | 0.991835066 | -5.110285881 |
| **A_51_P412508** | Irx4 | -0.075767943 | -0.378147238 | 0.226611352 | 11.62053704 | -0.598883819 | 0.569078345 | 0.991835066 | -5.110320929 |

| | | | | | | | | | |
|---|---|---|---|---|---|---|---|---|---|
| **A_55_P2132781** | Slc16a2 | -0.034769689 | -0.173548141 | 0.104008763 | 10.82478334 | -0.598808419 | 0.569125747 | 0.991835066 | -5.110356578 |
| **A_55_P2036629** | NA | -0.054842246 | -0.273768983 | 0.164084492 | 11.04832363 | -0.598722469 | 0.569179784 | 0.991835066 | -5.11039721 |
| **A_55_P2000127** | Tgfb1i1 | -0.035978839 | -0.179605893 | 0.107648214 | 12.76166667 | -0.59871485 | 0.569184574 | 0.991835066 | -5.110400812 |
| **A_51_P125842** | Tmem51 | -0.075044918 | -0.374703664 | 0.224613828 | 11.57537034 | -0.598554172 | 0.569285602 | 0.991835066 | -5.110476756 |
| **A_55_P2164258** | Prl7b1 | -0.086895813 | -0.433894394 | 0.260102768 | 7.840606201 | -0.598522227 | 0.569305689 | 0.991835066 | -5.110491853 |
| **A_52_P426768** | Cited4 | -0.111267762 | -0.555712452 | 0.333176928 | 11.43693608 | -0.598357536 | 0.569409253 | 0.991835066 | -5.110569673 |
| **A_52_P134381** | Psmd12 | 0.098148615 | -0.294065424 | 0.490362654 | 10.86743993 | 0.598095 | 0.569574368 | 0.991835066 | -5.110693687 |
| **A_55_P2019113** | Apol7b | 0.032241401 | -0.096609604 | 0.161092406 | 8.135826507 | 0.598046878 | 0.569604636 | 0.991835066 | -5.110716413 |
| **A_55_P2080119** | NA | -0.044800057 | -0.223851685 | 0.134251571 | 14.47515427 | -0.598011349 | 0.569626985 | 0.991835066 | -5.110733191 |
| **A_55_P2131433** | NA | -0.028348642 | -0.141654556 | 0.084957272 | 7.785476488 | -0.59798297 | 0.569644836 | 0.991835066 | -5.110746592 |
| **A_66_P126300** | Mrpl47 | -0.060059992 | -0.300119494 | 0.179999511 | 11.87094432 | -0.597964631 | 0.569656371 | 0.991835066 | -5.110755252 |
| **A_55_P2071496** | Fcf1 | -0.032575371 | -0.16278717 | 0.097636427 | 11.8553177 | -0.597927007 | 0.569680038 | 0.991835066 | -5.110773017 |
| **A_51_P116813** | Cyp17a1 | -0.150508325 | -0.752191099 | 0.451174449 | 10.97089111 | -0.597863551 | 0.569719956 | 0.991835066 | -5.110802978 |
| **A_55_P2147591** | Med27 | -0.039058096 | -0.195237643 | 0.117121451 | 8.455022531 | -0.597717605 | 0.569811772 | 0.991835066 | -5.110871875 |
| **A_52_P191975** | Rtn3 | 0.04723041 | -0.141656673 | 0.236117493 | 10.2493639 | 0.597624765 | 0.569870182 | 0.991835066 | -5.110915694 |
| **A_51_P335306** | Ppm1a | -0.059787723 | -0.298906898 | 0.179331452 | 9.62958192 | -0.597594708 | 0.569889093 | 0.991835066 | -5.110929879 |
| **A_55_P1987944** | Rai1 | 0.049299427 | -0.147876803 | 0.246475658 | 11.27863877 | 0.597580541 | 0.569898007 | 0.991835066 | -5.110936565 |
| **A_55_P2055537** | Slc16a6 | -0.093384241 | -0.466880927 | 0.280112446 | 9.869515138 | -0.597579886 | 0.569898419 | 0.991835066 | -5.110936874 |
| **A_51_P461779** | Ppp2r2c | 0.025450579 | -0.076355702 | 0.127256859 | 7.65661132 | 0.597492176 | 0.569953607 | 0.991835066 | -5.110978264 |
| **A_51_P386344** | Mtmr10 | -0.027511579 | -0.137565142 | 0.082541985 | 8.617139446 | -0.597476095 | 0.569963726 | 0.991835066 | -5.110985852 |
| **A_55_P2149363** | 6430548M( | -0.04321791 | -0.216113727 | 0.129677907 | 9.073401929 | -0.597431898 | 0.569991537 | 0.991835066 | -5.111006706 |
| **A_55_P2023076** | Arpc1b | 0.061565829 | -0.184748645 | 0.307880302 | 12.4847155 | 0.59739135 | 0.570017052 | 0.991835066 | -5.111025838 |
| **A_51_P484054** | F7 | -0.025402234 | -0.127054726 | 0.076250259 | 7.72035186 | -0.597259419 | 0.570100075 | 0.991835066 | -5.111088076 |
| **A_55_P2397240** | B430316J0 | 0.086412957 | -0.259401627 | 0.43222754 | 8.600203462 | 0.597234234 | 0.570115925 | 0.991835066 | -5.111099956 |
| **A_51_P472879** | NA | -0.028107025 | -0.140592137 | 0.084378088 | 7.862648945 | -0.597212599 | 0.570129541 | 0.991835066 | -5.111110161 |
| **A_55_P2170109** | Bivm | 0.055788014 | -0.167510338 | 0.279086366 | 10.11822849 | 0.597123987 | 0.57018531 | 0.991835066 | -5.111151954 |
| **A_65_P05152** | Flrt2 | 0.048883986 | -0.146783029 | 0.244551 | 9.337837412 | 0.597115193 | 0.570190844 | 0.991835066 | -5.111156101 |
| **A_55_P2041082** | NA | -0.028366396 | -0.141940607 | 0.085207814 | 8.088225166 | -0.596943971 | 0.570298615 | 0.991835066 | -5.111236841 |
| **A_66_P126332** | Zfp703 | 0.047398582 | -0.142427093 | 0.237224257 | 11.54884852 | 0.596787243 | 0.570397273 | 0.991835066 | -5.111310728 |
| **A_55_P1978316** | Adamts14 | -0.036398793 | -0.182234708 | 0.109437122 | 9.770429161 | -0.596529088 | 0.570559799 | 0.991835066 | -5.111432395 |
| **A_55_P2059986** | Chst14 | 0.044912735 | -0.135057407 | 0.224882877 | 9.920645967 | 0.596455677 | 0.570606021 | 0.991835066 | -5.111466985 |
| **A_55_P1987331** | Prl3d1 | 0.025872568 | -0.077821181 | 0.129566317 | 7.864307175 | 0.596342963 | 0.570676995 | 0.991835066 | -5.111520086 |
| **A_55_P2111790** | Gem | 0.051446722 | -0.154793616 | 0.25768706 | 8.700272809 | 0.596201722 | 0.570765938 | 0.991835066 | -5.111586615 |
| **A_55_P2020153** | Cycs | -0.047755926 | -0.239217763 | 0.143705911 | 16.23607701 | -0.596148123 | 0.570799693 | 0.991835066 | -5.111611858 |
| **A_51_P349023** | Lsm3 | -0.062744559 | -0.314315152 | 0.188826033 | 11.37160869 | -0.596108524 | 0.570824632 | 0.991835066 | -5.111630506 |
| **A_51_P455807** | Ehd4 | -0.037570987 | -0.188235612 | 0.113093638 | 14.05101882 | -0.596005737 | 0.570889369 | 0.991835066 | -5.111678907 |
| **A_52_P418489** | Tinagl1 | -0.060563298 | -0.303432688 | 0.182306093 | 9.17692803 | -0.595999458 | 0.570893323 | 0.991835066 | -5.111681863 |
| **A_51_P375187** | Rpf2 | 0.061436375 | -0.184940904 | 0.307813654 | 11.2174787 | 0.595983262 | 0.570903524 | 0.991835066 | -5.111689489 |
| **A_52_P298548** | Nbn | -0.047112219 | -0.236050702 | 0.141826264 | 8.983796765 | -0.595967082 | 0.570913715 | 0.991835066 | -5.111697107 |
| **A_51_P162176** | Trappc3 | -0.138213479 | -0.69251124 | 0.416084282 | 9.637668706 | -0.595959032 | 0.570918786 | 0.991835066 | -5.111700897 |
| **A_51_P306047** | Sec13 | 0.035156341 | -0.105848677 | 0.176161359 | 10.44962491 | 0.595906649 | 0.57095178 | 0.991835066 | -5.111725558 |
| **A_55_P2050236** | Hikeshi | 0.03846405 | -0.115813154 | 0.192741254 | 11.69253024 | 0.595884901 | 0.570965478 | 0.991835066 | -5.111735797 |
| **A_55_P2312783** | D030029J2 | -0.045185995 | -0.226471333 | 0.136099343 | 10.76676676 | -0.595731138 | 0.571062337 | 0.991835066 | -5.111808174 |
| **A_55_P2057786** | Rft1 | -0.032396531 | -0.162407708 | 0.097614647 | 9.121411935 | -0.59556195 | 0.571168923 | 0.991835066 | -5.111887794 |
| **A_55_P2080048** | NA | 0.027220094 | -0.08203806 | 0.136478248 | 14.84276406 | 0.595449445 | 0.571239805 | 0.991835066 | -5.111940728 |
| **A_55_P2141851** | Pls3 | 0.052684794 | -0.158844448 | 0.264214037 | 11.98263642 | 0.595283721 | 0.571344228 | 0.991835066 | -5.112018685 |
| **A_51_P346668** | Irf5 | -0.027327245 | -0.137047521 | 0.082393032 | 7.909767126 | -0.595275599 | 0.571349346 | 0.991835066 | -5.112022505 |
| **A_52_P169901** | Slc25a51 | -0.039181005 | -0.196520294 | 0.118158283 | 10.99371413 | -0.595178901 | 0.571410281 | 0.991835066 | -5.112067983 |

| | | | | | | | | | |
|---|---|---|---|---|---|---|---|---|---|
| **A_51_P467837** | Ap3m2 | 0.024094527 | -0.072666943 | 0.120855997 | 8.153439458 | 0.595148131 | 0.571429672 | 0.991835066 | -5.112082454 |
| **A_55_P2068566** | NA | 0.045365689 | -0.136848508 | 0.227579886 | 14.17245955 | 0.595051336 | 0.571490673 | 0.991835066 | -5.112127969 |
| **A_55_P2019890** | Rasef | 0.027617121 | -0.083308889 | 0.138543131 | 7.792921622 | 0.595050938 | 0.571490924 | 0.991835066 | -5.112128156 |
| **A_52_P565496** | Prl8a1 | -0.047272262 | -0.237147806 | 0.142603282 | 7.662049107 | -0.595040446 | 0.571497536 | 0.991835066 | -5.112133089 |
| **A_52_P188678** | Nectin2 | 0.055374364 | -0.167051829 | 0.277800558 | 10.5001277 | 0.595020539 | 0.571510082 | 0.991835066 | -5.112142449 |
| **A_51_P468294** | Polg | -0.057553091 | -0.288736103 | 0.173629922 | 9.442185602 | -0.595006711 | 0.571518797 | 0.991835066 | -5.11214895 |
| **A_55_P2043531** | NA | -0.108118259 | -0.542497421 | 0.326260903 | 13.32668104 | -0.594893459 | 0.571590176 | 0.991835066 | -5.112202192 |
| **A_55_P2033521** | Med6 | -0.047151776 | -0.236617841 | 0.142314288 | 9.848747382 | -0.594806567 | 0.571644945 | 0.991835066 | -5.112243035 |
| **A_55_P2045911** | Spata32 | -0.023979154 | -0.120355323 | 0.072397014 | 7.914182267 | -0.594666314 | 0.571733355 | 0.991835066 | -5.11230895 |
| **A_55_P2055632** | Rtf2 | -0.052005492 | -0.261027541 | 0.157016556 | 8.489647597 | -0.594656602 | 0.571739477 | 0.991835066 | -5.112313514 |
| **A_52_P609334** | Klhdc8b | 0.063188679 | -0.190782071 | 0.317159429 | 10.30058109 | 0.594654495 | 0.571740805 | 0.991835066 | -5.112314504 |
| **A_55_P2016877** | Nsd1 | -0.029182592 | -0.146480253 | 0.088115068 | 12.26271696 | -0.594625666 | 0.571758979 | 0.991835066 | -5.112328051 |
| **A_55_P2051400** | Prl3d2 | -0.025545122 | -0.128253215 | 0.077162971 | 7.633391934 | -0.594446056 | 0.571872212 | 0.991835066 | -5.112412438 |
| **A_51_P363400** | Prg2 | -0.026556994 | -0.133357776 | 0.080243788 | 7.687746833 | -0.594310795 | 0.571957495 | 0.991835066 | -5.112475974 |
| **A_55_P2064272** | NA | -0.028220335 | -0.141740112 | 0.085299442 | 7.992739317 | -0.594155017 | 0.572055723 | 0.991835066 | -5.112549131 |
| **A_52_P362111** | Fam3c | 0.037116546 | -0.112199809 | 0.186432901 | 10.95193145 | 0.59411334 | 0.572082004 | 0.991835066 | -5.112568701 |
| **A_51_P270206** | Cep83 | -0.047329996 | -0.237759352 | 0.143099361 | 10.12633586 | -0.594034539 | 0.572131699 | 0.991835066 | -5.112605699 |
| **A_51_P487389** | Drc3 | -0.026282265 | -0.132032165 | 0.079467634 | 7.925446376 | -0.594007548 | 0.572148721 | 0.991835066 | -5.112618371 |
| **A_55_P2100088** | Ccdc92b | 0.030144151 | -0.091145598 | 0.151433899 | 7.908794729 | 0.594002244 | 0.572152065 | 0.991835066 | -5.112620861 |
| **A_55_P2086825** | Rgl3 | 0.027241519 | -0.082373186 | 0.136856224 | 8.587089398 | 0.593979737 | 0.57216626 | 0.991835066 | -5.112631427 |
| **A_55_P1978905** | NA | -0.028115066 | -0.141246224 | 0.085016092 | 7.846942607 | -0.593972054 | 0.572171105 | 0.991835066 | -5.112635034 |
| **A_55_P2141038** | Gnpnat1 | 0.026881023 | -0.081343026 | 0.135105071 | 8.28553028 | 0.593650918 | 0.572373657 | 0.992030702 | -5.112785754 |
| **A_51_P370252** | Capn10 | -0.10477732 | -0.526629462 | 0.317074823 | 10.32218994 | -0.593630433 | 0.572386579 | 0.992030702 | -5.112795366 |
| **A_52_P49601** | Fth1 | -0.067637643 | -0.340081756 | 0.20480647 | 15.96293629 | -0.593362602 | 0.572555544 | 0.992145958 | -5.112921011 |
| **A_55_P2085536** | Smarcal1 | 0.046636799 | -0.141228927 | 0.234502525 | 11.04940889 | 0.593321809 | 0.572581282 | 0.992145958 | -5.112940143 |
| **A_55_P1972406** | Tor1b | 0.055725378 | -0.168767054 | 0.28021781 | 9.75880998 | 0.593281015 | 0.57260702 | 0.992145958 | -5.112959275 |
| **A_55_P2104689** | NA | 0.078924619 | -0.239121474 | 0.396970712 | 8.839392165 | 0.593104967 | 0.572718105 | 0.992208682 | -5.113041825 |
| **A_55_P2104612** | NA | 0.031132831 | -0.094333948 | 0.156599611 | 7.902985973 | 0.593060502 | 0.572746164 | 0.992208682 | -5.113062672 |
| **A_52_P215176** | Opn3 | -0.033586173 | -0.16897247 | 0.101800124 | 9.345074569 | -0.592918425 | 0.572835824 | 0.992208682 | -5.113129273 |
| **A_55_P1980621** | NA | -0.089778994 | -0.451691521 | 0.272133534 | 13.52441136 | -0.59289836 | 0.572848488 | 0.992208682 | -5.113138678 |
| **A_66_P106611** | Gna14 | 0.02883743 | -0.087438031 | 0.145112891 | 7.769954018 | 0.592758264 | 0.572936908 | 0.992227618 | -5.113204335 |
| **A_55_P2096465** | NA | -0.057416229 | -0.288986059 | 0.1741536 | 13.18080853 | -0.59260024 | 0.573036652 | 0.992227618 | -5.113278377 |
| **A_55_P2149540** | Srf | 0.059909234 | -0.181756748 | 0.301575217 | 11.22838313 | 0.59249867 | 0.573100768 | 0.992227618 | -5.11332596 |
| **A_51_P271439** | Garem1 | -0.043253696 | -0.217750318 | 0.131242927 | 9.323221017 | -0.592441304 | 0.573136983 | 0.992227618 | -5.11335283 |
| **A_52_P494126** | Usp46 | -0.04561862 | -0.229668358 | 0.138431119 | 9.462179027 | -0.592401392 | 0.57316218 | 0.992227618 | -5.113371525 |
| **A_51_P220806** | Gdf9 | -0.035247961 | -0.17745891 | 0.106962988 | 8.863120143 | -0.592393239 | 0.573167327 | 0.992227618 | -5.113375343 |
| **A_55_P2154982** | Cd8b1 | -0.029908011 | -0.150592155 | 0.090776132 | 7.840548063 | -0.592306444 | 0.573222123 | 0.992233639 | -5.113415991 |
| **A_55_P2038242** | Tchhl1 | 0.026190754 | -0.079509551 | 0.131891058 | 7.689918068 | 0.592217026 | 0.573278579 | 0.992234063 | -5.113457863 |
| **A_55_P2047559** | Ccp110 | 0.082638163 | -0.250928381 | 0.416204708 | 9.13217638 | 0.59211669 | 0.573341932 | 0.992234063 | -5.11350484 |
| **A_55_P2132014** | Tekt2 | 0.032373523 | -0.098313284 | 0.163060331 | 7.854308826 | 0.592062226 | 0.573376322 | 0.992234063 | -5.113530337 |
| **A_55_P2025702** | C330007P( | 0.049245369 | -0.149641835 | 0.248132573 | 8.34316773 | 0.591790085 | 0.573548181 | 0.992280687 | -5.113657709 |
| **A_55_P2005972** | NA | -0.074715023 | -0.376468177 | 0.22703813 | 9.489553568 | -0.591786768 | 0.573550276 | 0.992280687 | -5.113659261 |
| **A_55_P1966928** | Lnx1 | -0.029107011 | -0.1466643 | 0.088450278 | 8.025867409 | -0.591775764 | 0.573557226 | 0.992280687 | -5.113664411 |
| **A_55_P1972277** | Samd11 | -0.024783909 | -0.124910958 | 0.07534314 | 7.689206711 | -0.591599126 | 0.573668792 | 0.992304853 | -5.113747055 |
| **A_55_P2129469** | NA | -0.036438326 | -0.183692061 | 0.11081541 | 7.950510821 | -0.591427106 | 0.573777452 | 0.992304853 | -5.113827517 |
| **A_55_P2071236** | Aqp5 | 0.024578432 | -0.074755727 | 0.123912592 | 7.677946177 | 0.591377352 | 0.573808882 | 0.992304853 | -5.113850785 |
| **A_51_P408703** | Rnf138rt1 | -0.030083693 | -0.151711956 | 0.091544571 | 7.710644087 | -0.591160992 | 0.573945573 | 0.992304853 | -5.113951952 |
| **A_55_P2028015** | Pawr | 0.042643728 | -0.129864951 | 0.215152407 | 8.605749974 | 0.590817495 | 0.574162626 | 0.992304853 | -5.114112499 |

| | | | | | | | | | |
|---|---|---|---|---|---|---|---|---|---|
| **A_52_P203948** | Pla2g4e | -0.040130042 | -0.202477639 | 0.122217555 | 7.899150262 | -0.590789658 | 0.574180218 | 0.992304853 | -5.114125506 |
| **A_55_P2035138** | NA | -0.057221051 | -0.28873264 | 0.174290538 | 15.19695792 | -0.590734355 | 0.574215168 | 0.992304853 | -5.114151345 |
| **A_55_P2158701** | H4c4 | 0.061943647 | -0.188684561 | 0.312571854 | 14.16435921 | 0.590712217 | 0.574229159 | 0.992304853 | -5.114161688 |
| **A_66_P109395** | Rpia | -0.042563818 | -0.214846102 | 0.129718466 | 9.111548426 | -0.590485293 | 0.574372586 | 0.992304853 | -5.114267689 |
| **A_52_P163766** | Atp6v0a2 | 0.050270475 | -0.15322052 | 0.253761471 | 9.434055028 | 0.590441573 | 0.574400222 | 0.992304853 | -5.114288107 |
| **A_55_P2162651** | NA | -0.060635476 | -0.306220573 | 0.18494962 | 14.73573655 | -0.590111282 | 0.574609025 | 0.992304853 | -5.114442319 |
| **A_55_P2143195** | Dyrk1a | -0.030165948 | -0.152348393 | 0.092016496 | 9.440001545 | -0.59008871 | 0.574623296 | 0.992304853 | -5.114452855 |
| **A_51_P331021** | Ttc32 | -0.043666853 | -0.220552868 | 0.133219163 | 9.519834694 | -0.590021074 | 0.574666061 | 0.992304853 | -5.114484424 |
| **A_66_P111301** | Mplkip | 0.03350047 | -0.102205841 | 0.169206782 | 9.812999054 | 0.590010837 | 0.574672533 | 0.992304853 | -5.114489202 |
| **A_51_P326191** | Serpina3g | -0.041833652 | -0.211339363 | 0.12767206 | 8.018051244 | -0.589862208 | 0.574766514 | 0.992304853 | -5.114558561 |
| **A_51_P276235** | Pnpla7 | -0.029720809 | -0.150220242 | 0.090778625 | 8.906123885 | -0.589501276 | 0.574994776 | 0.992304853 | -5.114726931 |
| **A_52_P373798** | Zdhhc3 | 0.03193504 | -0.097542454 | 0.161412534 | 8.237037182 | 0.589497969 | 0.574996867 | 0.992304853 | -5.114728473 |
| **A_55_P2016675** | Ogfr | -0.058141804 | -0.29387235 | 0.177588742 | 10.77848136 | -0.589497238 | 0.57499733 | 0.992304853 | -5.114728814 |
| **A_55_P2055463** | Snapin | 0.033152219 | -0.101268331 | 0.167572769 | 11.71513079 | 0.589462349 | 0.575019397 | 0.992304853 | -5.114745085 |
| **A_51_P414889** | Ifi35 | -0.050918754 | -0.257407433 | 0.155569926 | 9.200734391 | -0.589373561 | 0.575075558 | 0.992304853 | -5.114786487 |
| **A_55_P1961160** | NA | -0.038484934 | -0.194565207 | 0.117595338 | 15.32143272 | -0.589320933 | 0.575108849 | 0.992304853 | -5.114811025 |
| **A_66_P100232** | Paxbp1 | 0.078703925 | -0.240528326 | 0.397936176 | 8.997009198 | 0.589248876 | 0.575154431 | 0.992304853 | -5.114844619 |
| **A_55_P2096305** | Rc3h1 | 0.053565697 | -0.163728708 | 0.270860102 | 10.91203325 | 0.589179119 | 0.57519856 | 0.992304853 | -5.114877137 |
| **A_55_P2157408** | NA | 0.026807806 | -0.081942587 | 0.1355582 | 7.932315899 | 0.589168572 | 0.575205232 | 0.992304853 | -5.114882053 |
| **A_52_P569348** | Dbt | 0.035672778 | -0.109045089 | 0.180390645 | 9.754264581 | 0.58914733 | 0.575218671 | 0.992304853 | -5.114891955 |
| **A_55_P2173313** | Plcb1 | 0.024636166 | -0.075309211 | 0.124581543 | 8.073805803 | 0.589141397 | 0.575222424 | 0.992304853 | -5.11489472 |
| **A_55_P2047614** | Zfp809 | -0.029494468 | -0.149173195 | 0.090184259 | 8.164204042 | -0.589023661 | 0.575296912 | 0.992304853 | -5.114949594 |
| **A_55_P2096340** | Rnf170 | 0.03115803 | -0.0952756 | 0.15759166 | 8.535301577 | 0.589001663 | 0.57531083 | 0.992304853 | -5.114959845 |
| **A_51_P185846** | Dnal4 | -0.032736724 | -0.165579207 | 0.10010576 | 11.01788436 | -0.588989237 | 0.575318692 | 0.992304853 | -5.114965636 |
| **A_51_P304125** | Slc29a2 | 0.050072352 | -0.153143652 | 0.253288355 | 9.635572247 | 0.588910389 | 0.57536858 | 0.992304853 | -5.115002377 |
| **A_55_P2038597** | NA | 0.061966899 | -0.189532915 | 0.313466713 | 12.31739003 | 0.588886001 | 0.575384012 | 0.992304853 | -5.115013741 |
| **A_55_P2145005** | NA | -0.034437271 | -0.174236394 | 0.105361853 | 8.583447474 | -0.588753371 | 0.575467936 | 0.992304853 | -5.115075532 |
| **A_55_P2005410** | NA | -0.039620896 | -0.200493663 | 0.121251871 | 13.75595716 | -0.588641523 | 0.575538716 | 0.992304853 | -5.115127631 |
| **A_51_P319070** | Retsat | -0.068664443 | -0.34747964 | 0.210150755 | 9.671843643 | -0.58860587 | 0.57556128 | 0.992304853 | -5.115144237 |
| **A_66_P119247** | Gatad1 | -0.044818822 | -0.226829132 | 0.137191488 | 11.49671857 | -0.588536738 | 0.575605032 | 0.992304853 | -5.115176433 |
| **A_51_P357735** | Inhbe | 0.031249403 | -0.095661732 | 0.158160538 | 7.859305728 | 0.588506318 | 0.575624284 | 0.992304853 | -5.115190599 |
| **A_51_P502348** | Tfip11 | 0.032374734 | -0.099113859 | 0.163863328 | 9.939771964 | 0.588473981 | 0.575644751 | 0.992304853 | -5.115205658 |
| **A_52_P235278** | Pms1 | 0.039000468 | -0.119402382 | 0.197403318 | 8.646849016 | 0.588458663 | 0.575654446 | 0.992304853 | -5.115212791 |
| **A_55_P2020072** | NA | 0.036479517 | -0.111698348 | 0.184657381 | 12.62118785 | 0.588402997 | 0.575689678 | 0.992304853 | -5.11523871 |
| **A_55_P2074281** | Nr2f1 | 0.17081856 | -0.523064367 | 0.864701486 | 10.79141917 | 0.588380115 | 0.575704162 | 0.992304853 | -5.115249364 |
| **A_55_P2098061** | Tchp | -0.029003149 | -0.146820804 | 0.088814506 | 8.590363672 | -0.588361049 | 0.57571623 | 0.992304853 | -5.11525824 |
| **A_52_P563450** | Brix1 | 0.040639582 | -0.124479289 | 0.205758452 | 9.157946559 | 0.588249614 | 0.575786766 | 0.992304853 | -5.115310117 |
| **A_51_P242414** | Micall1 | 0.045821769 | -0.140384088 | 0.232027626 | 12.35247311 | 0.58814938 | 0.575850217 | 0.992304853 | -5.115356772 |
| **A_55_P2176292** | NA | -0.04827108 | -0.244470243 | 0.147928083 | 14.07446296 | -0.588029345 | 0.575926208 | 0.992304853 | -5.115412635 |
| **A_55_P2038687** | Tmco1 | -0.031644129 | -0.160272242 | 0.096983985 | 12.44749589 | -0.587985169 | 0.575954176 | 0.992304853 | -5.115433191 |
| **A_52_P35304** | Chkb | -0.036643976 | -0.185600007 | 0.112312055 | 10.14070872 | -0.58796792 | 0.575965097 | 0.992304853 | -5.115441217 |
| **A_55_P1955427** | Cux1 | -0.04188266 | -0.212173596 | 0.128408277 | 9.554022091 | -0.587830139 | 0.576052332 | 0.992304853 | -5.115505321 |
| **A_55_P2122200** | Prkar2b | 0.030919842 | -0.094798631 | 0.156638316 | 7.92960941 | 0.587823989 | 0.576056227 | 0.992304853 | -5.115508182 |
| **A_52_P488427** | Sec14l2 | -0.02621621 | -0.132845511 | 0.080413091 | 7.889968059 | -0.587627996 | 0.576180333 | 0.992304853 | -5.115599345 |
| **A_55_P2046744** | Cspg5 | -0.030572409 | -0.154922512 | 0.093777693 | 8.18040244 | -0.587614698 | 0.576188754 | 0.992304853 | -5.115605529 |
| **A_55_P2044109** | NA | -0.032019818 | -0.162293555 | 0.098253919 | 16.11719602 | -0.58745029 | 0.576292873 | 0.992304853 | -5.115681979 |
| **A_52_P127682** | Dagla | 0.044227404 | -0.135779764 | 0.224234572 | 8.459254501 | 0.587233442 | 0.576430219 | 0.992304853 | -5.115782785 |
| **A_51_P410650** | Dpp9 | -0.03061656 | -0.155231261 | 0.093998142 | 9.213784798 | -0.587213783 | 0.576442672 | 0.992304853 | -5.115791922 |

| | | | | | | | | | |
|---|---|---|---|---|---|---|---|---|---|
| **A_51_P201785** | Morn5 | 0.02677254 | -0.082198065 | 0.135743144 | 7.865252885 | 0.587204453 | 0.576448582 | 0.992304853 | -5.115796258 |
| **A_51_P162471** | Dnajc9 | 0.045762817 | -0.140513052 | 0.232038685 | 12.50595002 | 0.587171928 | 0.576469184 | 0.992304853 | -5.115811375 |
| **A_55_P2021358** | Klhl15 | 0.027862715 | -0.085619792 | 0.141345223 | 8.032624606 | 0.586818299 | 0.576693214 | 0.992304853 | -5.115975681 |
| **A_55_P2101473** | NA | 0.059097651 | -0.181631847 | 0.299827149 | 15.29292073 | 0.586745871 | 0.576739105 | 0.992304853 | -5.116009323 |
| **A_55_P2026437** | Gm4802 | -0.036528777 | -0.185353203 | 0.112295648 | 16.30915562 | -0.58663781 | 0.576807576 | 0.992304853 | -5.116059508 |
| **A_66_P115378** | Orc2 | -0.03066579 | -0.155611234 | 0.094279654 | 8.114914669 | -0.586601092 | 0.576830844 | 0.992304853 | -5.116076559 |
| **A_55_P2153631** | NA | -0.076789831 | -0.3896865 | 0.236106839 | 8.806215603 | -0.58655927 | 0.576857346 | 0.992304853 | -5.116095979 |
| **A_55_P1998411** | Chchd3 | 0.043615416 | -0.134177631 | 0.221408463 | 13.30201222 | 0.586319557 | 0.577009262 | 0.992304853 | -5.116207264 |
| **A_51_P453856** | Csnk1a1 | 0.030600788 | -0.094150491 | 0.155352068 | 12.72801573 | 0.586268739 | 0.57704147 | 0.992304853 | -5.116230851 |
| **A_55_P2097656** | Rps11 | 0.039222575 | -0.120698145 | 0.199143295 | 15.18861105 | 0.586192827 | 0.577089586 | 0.992304853 | -5.116266082 |
| **A_55_P2102182** | NA | -0.027458231 | -0.139480841 | 0.08456438 | 7.9330508 | -0.585835917 | 0.577315836 | 0.992304853 | -5.11643167 |
| **A_51_P177518** | Emc6 | 0.035854281 | -0.110427541 | 0.182136103 | 11.89343759 | 0.585814057 | 0.577329696 | 0.992304853 | -5.11644181 |
| **A_55_P2156274** | Plekha2 | -0.039165207 | -0.199025034 | 0.12069462 | 8.661158617 | -0.585558409 | 0.577491789 | 0.992304853 | -5.116560359 |
| **A_55_P2275054** | Nrap | -0.049185136 | -0.249956789 | 0.151586518 | 12.58727461 | -0.585518487 | 0.577517103 | 0.992304853 | -5.116578868 |
| **A_51_P258435** | Zdhhc3 | -0.030008191 | -0.15251883 | 0.092502447 | 9.710852064 | -0.585430232 | 0.577573069 | 0.992304853 | -5.116619781 |
| **A_55_P2095153** | Pdf | -0.035939806 | -0.182715659 | 0.110836048 | 10.86977312 | -0.585234935 | 0.577696925 | 0.992304853 | -5.116710297 |
| **A_55_P2142873** | Mink1 | 0.056295041 | -0.173620975 | 0.286211057 | 10.33615061 | 0.585207722 | 0.577714184 | 0.992304853 | -5.116722907 |
| **A_52_P327402** | Cds1 | 0.023659289 | -0.072989568 | 0.120308146 | 7.703850302 | 0.585078453 | 0.577796176 | 0.992304853 | -5.116782804 |
| **A_66_P102510** | Fam114a2 | 0.03055727 | -0.094276502 | 0.155391042 | 8.846458122 | 0.585048124 | 0.577815413 | 0.992304853 | -5.116796854 |
| **A_51_P337897** | Prrc1 | 0.032177831 | -0.099291645 | 0.163647307 | 8.374046833 | 0.584979929 | 0.577858671 | 0.992304853 | -5.116828446 |
| **A_51_P155843** | Igsf10 | -0.038540973 | -0.196065823 | 0.118983878 | 9.249224175 | -0.584766822 | 0.577993861 | 0.992304853 | -5.116927149 |
| **A_55_P2060604** | Gm5797 | 0.131523504 | -0.406076092 | 0.669123101 | 10.19611021 | 0.584727568 | 0.578018765 | 0.992304853 | -5.116945326 |
| **A_55_P2012759** | Bicdl1 | 0.072367758 | -0.223442291 | 0.368177808 | 8.909079513 | 0.584711084 | 0.578029223 | 0.992304853 | -5.116952959 |
| **A_55_P1999958** | NA | 0.025476917 | -0.078696913 | 0.129650747 | 7.932405455 | 0.584517306 | 0.578152172 | 0.992304853 | -5.117042675 |
| **A_55_P1993475** | Gm5168 | 0.031408146 | -0.097062013 | 0.159878304 | 7.887432888 | 0.584317868 | 0.578278729 | 0.992304853 | -5.117134983 |
| **A_55_P2107447** | Rab37 | -0.026819674 | -0.13655567 | 0.082916322 | 8.129947095 | -0.584135388 | 0.578394538 | 0.992304853 | -5.117219419 |
| **A_55_P1975685** | Nup37 | 0.063759955 | -0.197138141 | 0.324658052 | 11.04145111 | 0.584098681 | 0.578417836 | 0.992304853 | -5.117236401 |
| **A_55_P1963764** | 1-Sep | -0.031922517 | -0.162581061 | 0.098736026 | 8.094209831 | -0.583940307 | 0.578518359 | 0.992304853 | -5.11730966 |
| **A_66_P109708** | Il1f6 | 0.024749944 | -0.076611338 | 0.126111225 | 7.641409133 | 0.583594585 | 0.578737833 | 0.992304853 | -5.117469519 |
| **A_52_P648601** | Dusp9 | 0.043805746 | -0.13560424 | 0.223215733 | 7.860043867 | 0.583570867 | 0.578752892 | 0.992304853 | -5.117480483 |
| **A_55_P2104472** | NA | 0.045772886 | -0.141757199 | 0.233302971 | 13.21718355 | 0.583373202 | 0.578878399 | 0.992304853 | -5.117571841 |
| **A_52_P260994** | Fgd2 | -0.025416877 | -0.129562161 | 0.078728407 | 7.820391076 | -0.583299642 | 0.57892511 | 0.992304853 | -5.117605833 |
| **A_66_P118195** | Ciao1 | -0.050407488 | -0.2569816 | 0.156166625 | 10.73005269 | -0.583214466 | 0.578979199 | 0.992304853 | -5.117645188 |
| **A_55_P2071532** | NA | -0.05353996 | -0.272953897 | 0.165873976 | 16.26891577 | -0.583207325 | 0.578983734 | 0.992304853 | -5.117648487 |
| **A_66_P111840** | NA | 0.04387561 | -0.135975355 | 0.223726576 | 14.10222993 | 0.583068431 | 0.579071943 | 0.992304853 | -5.11771265 |
| **A_55_P1981794** | Sgf29 | -0.04380277 | -0.223371219 | 0.135765679 | 8.423250723 | -0.583016269 | 0.579105072 | 0.992304853 | -5.117736743 |
| **A_52_P88773** | Tor2a | 0.053361083 | -0.165407509 | 0.272129676 | 10.06892725 | 0.582973479 | 0.57913225 | 0.992304853 | -5.117756506 |
| **A_55_P2070331** | NA | 0.024615099 | -0.076320127 | 0.125550325 | 7.660409862 | 0.582864978 | 0.579201167 | 0.992304853 | -5.117806612 |
| **A_55_P1963344** | Upf3b | 0.067611221 | -0.209724044 | 0.344946486 | 9.113851115 | 0.582670211 | 0.579324889 | 0.992304853 | -5.117896536 |
| **A_51_P386870** | Sprr2f | 0.029299698 | -0.094520632 | 0.153120028 | 7.678944563 | 0.587811844 | 0.579372763 | 0.992304853 | -5.042854949 |
| **A_51_P488175** | Phf20l1 | 0.049784408 | -0.154461524 | 0.254030339 | 9.055506342 | 0.582571266 | 0.579387748 | 0.992304853 | -5.117942208 |
| **A_51_P149422** | Gng13 | 0.147357796 | -0.457227789 | 0.75194338 | 8.4148042 | 0.582538269 | 0.579408712 | 0.992304853 | -5.117957439 |
| **A_55_P2097022** | Eid1 | 0.034238155 | -0.106260134 | 0.174736444 | 8.25814228 | 0.582436323 | 0.579473483 | 0.992304853 | -5.118004487 |
| **A_55_P2022693** | Tcerg1 | 0.047681199 | -0.147990944 | 0.243353342 | 9.19492796 | 0.582407954 | 0.579491508 | 0.992304853 | -5.118017578 |
| **A_55_P1971854** | Vcpkmt | 0.055990348 | -0.173830158 | 0.285810854 | 10.32405507 | 0.582282215 | 0.579571403 | 0.992304853 | -5.118075595 |
| **A_55_P2157838** | NA | 0.078980033 | -0.24522562 | 0.403185686 | 12.29324768 | 0.582245124 | 0.579594972 | 0.992304853 | -5.118092707 |
| **A_52_P594756** | Asb4 | 0.072816313 | -0.226104069 | 0.371736696 | 10.99253977 | 0.582213524 | 0.579615052 | 0.992304853 | -5.118107285 |
| **A_55_P2152246** | Rpl19 | 0.051594411 | -0.160241022 | 0.263429844 | 16.77132895 | 0.58212089 | 0.579673919 | 0.992304853 | -5.118150015 |

| | | | | | | | | | |
|---|---|---|---|---|---|---|---|---|---|
| **A_55_P2037528** | Acsf3 | -0.048686229 | -0.248689361 | 0.151316903 | 10.76831651 | -0.581806373 | 0.579873812 | 0.992304853 | -5.118295052 |
| **A_55_P2162920** | NA | -0.060216936 | -0.30759137 | 0.187157497 | 15.77060714 | -0.581799007 | 0.579878494 | 0.992304853 | -5.118298448 |
| **A_55_P2081283** | Dlx4 | -0.025622092 | -0.130884607 | 0.079640423 | 7.680301371 | -0.581768198 | 0.579898078 | 0.992304853 | -5.118312651 |
| **A_51_P212419** | Lrrc41 | -0.0407774 | -0.208322721 | 0.126767921 | 9.438558115 | -0.581696377 | 0.579943731 | 0.992304853 | -5.118345759 |
| **A_55_P1999818** | NA | 0.100622906 | -0.312844882 | 0.514090694 | 12.81847014 | 0.581653448 | 0.579971019 | 0.992304853 | -5.118365547 |
| **A_52_P337427** | Izumo4 | -0.034655394 | -0.177059659 | 0.107748871 | 7.961974902 | -0.581643632 | 0.579977259 | 0.992304853 | -5.118370071 |
| **A_51_P477941** | Fbxw4 | -0.0298793 | -0.152710489 | 0.092951889 | 10.17231586 | -0.581394463 | 0.580135666 | 0.992304853 | -5.118484897 |
| **A_55_P2046245** | Hes1 | -0.061733102 | -0.315516634 | 0.19205043 | 8.851110038 | -0.581384964 | 0.580141705 | 0.992304853 | -5.118489273 |
| **A_51_P109541** | E130308A1 | 0.039976919 | -0.124377696 | 0.204331534 | 9.926339117 | 0.581348487 | 0.580164898 | 0.992304853 | -5.118506079 |
| **A_55_P2030775** | Fnbp1 | -0.035526943 | -0.18158907 | 0.110535185 | 10.44080964 | -0.581338847 | 0.580171026 | 0.992304853 | -5.11851052 |
| **A_55_P2096599** | Tsc22d4 | -0.07624776 | -0.389760449 | 0.237264929 | 8.809761445 | -0.581274269 | 0.580212087 | 0.992304853 | -5.118540271 |
| **A_52_P436628** | Vegfb | -0.065753707 | -0.336195935 | 0.204688522 | 12.28179486 | -0.58110534 | 0.580319504 | 0.992304853 | -5.11861808 |
| **A_55_P2146520** | Carns1 | -0.064563655 | -0.330114256 | 0.200986945 | 10.26628174 | -0.58109877 | 0.580323682 | 0.992304853 | -5.118621106 |
| **A_55_P1972084** | Napb | 0.050960034 | -0.158651217 | 0.260571286 | 8.532950345 | 0.58106438 | 0.580345551 | 0.992304853 | -5.118636943 |
| **A_52_P317923** | Zfp948 | 0.056782249 | -0.176790336 | 0.290354834 | 9.227577083 | 0.581031738 | 0.580366309 | 0.992304853 | -5.118651975 |
| **A_55_P1963144** | Mtmr14 | 0.03583461 | -0.111580429 | 0.183249648 | 10.00718031 | 0.580991823 | 0.580391693 | 0.992304853 | -5.118670355 |
| **A_55_P2403325** | NA | 0.057819962 | -0.180066318 | 0.295706243 | 8.410910034 | 0.580921621 | 0.580436339 | 0.992304853 | -5.118702679 |
| **A_55_P2011352** | NA | -0.056195002 | -0.287411363 | 0.175021359 | 12.02708491 | -0.580882457 | 0.580461247 | 0.992304853 | -5.11872071 |
| **A_55_P2094445** | Ms4a10 | -0.02513417 | -0.128567784 | 0.078299444 | 7.712571667 | -0.580780434 | 0.580526135 | 0.992304853 | -5.118767677 |
| **A_55_P2098300** | Chmp6 | -0.05962379 | -0.305003651 | 0.185756071 | 8.507940682 | -0.580750773 | 0.580545001 | 0.992304853 | -5.11878133 |
| **A_51_P273755** | Exoc7 | -0.066217873 | -0.338751971 | 0.206316225 | 9.611123657 | -0.580715629 | 0.580567354 | 0.992304853 | -5.118797506 |
| **A_55_P2063553** | Gm4234 | -0.023423265 | -0.119839064 | 0.072992533 | 7.679089208 | -0.580641888 | 0.580614259 | 0.992304853 | -5.118831446 |
| **A_55_P2067031** | Agxt2 | 0.024562375 | -0.076558059 | 0.125682808 | 7.723450113 | 0.580551252 | 0.580671913 | 0.992304853 | -5.118873156 |
| **A_55_P1963150** | Creg1 | -0.114599139 | -0.586545844 | 0.357347566 | 10.39024381 | -0.580360076 | 0.580793534 | 0.992304853 | -5.118961115 |
| **A_55_P2027077** | Shc2 | -0.033004981 | -0.16894312 | 0.102933159 | 8.194572409 | -0.580292948 | 0.580836242 | 0.992304853 | -5.118991994 |
| **A_52_P392509** | Gin1 | -0.03120541 | -0.159735418 | 0.097324598 | 8.666636928 | -0.580275849 | 0.580847121 | 0.992304853 | -5.118999859 |
| **A_55_P2064984** | Cenpi | -0.04386912 | -0.224601603 | 0.136863362 | 9.542676929 | -0.58013871 | 0.580934379 | 0.992304853 | -5.119062932 |
| **A_55_P2130792** | Tuft1 | -0.043240169 | -0.221391164 | 0.134910826 | 8.194106551 | -0.580107211 | 0.580954421 | 0.992304853 | -5.119077417 |
| **A_55_P2154208** | NA | -0.040680095 | -0.208305146 | 0.126944957 | 10.52571154 | -0.580032283 | 0.5810021 | 0.992304853 | -5.119111871 |
| **A_55_P1970507** | Pef1 | 0.044112827 | -0.137657065 | 0.225882719 | 10.99790208 | 0.580032159 | 0.581002179 | 0.992304853 | -5.119111928 |
| **A_51_P509971** | Plekho1 | -0.048475361 | -0.248261224 | 0.151310502 | 13.16469591 | -0.579916452 | 0.581075811 | 0.992304853 | -5.119165125 |
| **A_55_P2327518** | 1500032P0 | 0.091998239 | -0.287239184 | 0.471235663 | 8.632808947 | 0.579798998 | 0.581150559 | 0.992304853 | -5.119219116 |
| **A_55_P2019312** | Car12 | -0.026389978 | -0.135181723 | 0.082401767 | 7.883135654 | -0.579765301 | 0.581172006 | 0.992304853 | -5.119234604 |
| **A_52_P451574** | Xpr1 | 0.045752141 | -0.142868699 | 0.234372982 | 11.79525691 | 0.579736818 | 0.581190134 | 0.992304853 | -5.119247695 |
| **A_51_P418935** | Neurl2 | -0.048707973 | -0.249523631 | 0.152107684 | 12.39332447 | -0.579711106 | 0.581206498 | 0.992304853 | -5.119259512 |
| **A_51_P349015** | Pacrg | -0.024670287 | -0.126385686 | 0.077045113 | 7.877757547 | -0.579691079 | 0.581219245 | 0.992304853 | -5.119268716 |
| **A_52_P417437** | Gab1 | -0.03722006 | -0.190683514 | 0.116243394 | 9.57660858 | -0.57967055 | 0.581232311 | 0.992304853 | -5.11927815 |
| **A_51_P234847** | 1700001L1 | 0.026296712 | -0.082165855 | 0.134759278 | 7.741530176 | 0.579469653 | 0.581360187 | 0.992304853 | -5.119370458 |
| **A_66_P126842** | Frg2f1 | 0.030286634 | -0.09465522 | 0.155228488 | 8.408660463 | 0.57936492 | 0.581426859 | 0.992304853 | -5.119418569 |
| **A_52_P557129** | Slc12a5 | 0.023672943 | -0.073996412 | 0.121342298 | 8.00751509 | 0.579299383 | 0.581468582 | 0.992304853 | -5.119448671 |
| **A_55_P1990970** | Casp9 | 0.041057773 | -0.128387715 | 0.210503261 | 11.04929714 | 0.579127935 | 0.581577738 | 0.992304853 | -5.119527405 |
| **A_55_P1993348** | NA | -0.033667518 | -0.172632838 | 0.105297801 | 15.77464966 | -0.579046988 | 0.581629278 | 0.992304853 | -5.119564571 |
| **A_51_P431903** | Pdcd6 | -0.038418084 | -0.197004852 | 0.120168683 | 12.58525497 | -0.578999106 | 0.581659766 | 0.992304853 | -5.119586553 |
| **A_55_P2004637** | NA | 0.047278975 | -0.147907563 | 0.242465512 | 10.34367791 | 0.578931687 | 0.581702697 | 0.992304853 | -5.119617503 |
| **A_55_P2022805** | Rubcn | -0.038674764 | -0.198355331 | 0.121005802 | 13.04453278 | -0.578874934 | 0.581738838 | 0.992304853 | -5.119643553 |
| **A_51_P299632** | Spef1 | 0.040989121 | -0.128253371 | 0.210231614 | 11.10935402 | 0.578853053 | 0.581752771 | 0.992304853 | -5.119653596 |
| **A_66_P100021** | Ncaph2 | -0.078675071 | -0.403632608 | 0.246282466 | 10.47565228 | -0.578654934 | 0.581878944 | 0.992304853 | -5.119744515 |
| **A_52_P584415** | Cars2 | -0.032924517 | -0.168925894 | 0.10307686 | 8.587598007 | -0.578609066 | 0.581908158 | 0.992304853 | -5.11976556 |

| | | | | | | | | | |
|---|---|---|---|---|---|---|---|---|---|
| **A_55_P2115704** | Aip | -0.091270725 | -0.468325413 | 0.285783963 | 9.521472615 | -0.57854386 | 0.58194969 | 0.992304853 | -5.119795476 |
| **A_66_P112356** | Agt | -0.024119243 | -0.123775511 | 0.075537025 | 7.827565157 | -0.578453137 | 0.582007476 | 0.992304853 | -5.119837094 |
| **A_55_P2007088** | Rnf128 | 0.049325535 | -0.160734816 | 0.259385885 | 12.64634463 | 0.583303881 | 0.582202367 | 0.992304853 | -5.04483246 |
| **A_51_P442206** | Ankrd42 | 0.024327961 | -0.076247576 | 0.124903499 | 7.845891414 | 0.578125984 | 0.582215887 | 0.992304853 | -5.119987122 |
| **A_55_P2161390** | NA | -0.049969466 | -0.256615422 | 0.156676491 | 9.284306861 | -0.577945554 | 0.582330848 | 0.992304853 | -5.120069833 |
| **A_55_P2012523** | Twf1 | -0.050244406 | -0.258096499 | 0.157607687 | 10.48869261 | -0.577753313 | 0.582453347 | 0.992304853 | -5.120157933 |
| **A_55_P2090914** | Tpst2 | 0.032015996 | -0.100447041 | 0.164479032 | 10.96652879 | 0.577672162 | 0.582505063 | 0.992304853 | -5.120195115 |
| **A_55_P2002723** | Inpp5a | -0.046665906 | -0.239807161 | 0.146475349 | 12.60089027 | -0.577475793 | 0.582630215 | 0.992304853 | -5.12028507 |
| **A_51_P344376** | Gpr162 | -0.047101326 | -0.24206661 | 0.147863959 | 9.978164938 | -0.577410895 | 0.58267158 | 0.992304853 | -5.120314793 |
| **A_52_P131548** | Ajuba | 0.047098859 | -0.147875855 | 0.242073574 | 9.318165585 | 0.577352738 | 0.58270865 | 0.992304853 | -5.120341426 |
| **A_55_P1956718** | Rab38 | 0.031702501 | -0.09954981 | 0.162954812 | 8.022912155 | 0.577292216 | 0.582747229 | 0.992304853 | -5.12036914 |
| **A_55_P2063376** | Add2 | 0.024299116 | -0.076307877 | 0.124906109 | 7.822748067 | 0.57725996 | 0.58276779 | 0.992304853 | -5.120383909 |
| **A_52_P206492** | Pop4 | 0.030841789 | -0.096856561 | 0.158540139 | 11.67687174 | 0.577249293 | 0.58277459 | 0.992304853 | -5.120388793 |
| **A_55_P2121195** | Lcorl | -0.039057842 | -0.200788823 | 0.122673139 | 8.974652366 | -0.577197118 | 0.58280785 | 0.992304853 | -5.120412681 |
| **A_55_P2001159** | Oser1 | 0.043052577 | -0.135221993 | 0.221327148 | 11.55307429 | 0.577190123 | 0.58281231 | 0.992304853 | -5.120415884 |
| **A_55_P2061170** | Tbc1d2 | -0.029442412 | -0.151364651 | 0.092479826 | 8.13230789 | -0.577164478 | 0.582828658 | 0.992304853 | -5.120427625 |
| **A_51_P501248** | Sphk1 | -0.040447926 | -0.207953449 | 0.127057597 | 9.26506734 | -0.577133464 | 0.582848429 | 0.992304853 | -5.120441822 |
| **A_51_P420400** | Lef1 | 0.040234956 | -0.126390392 | 0.206860304 | 8.030683622 | 0.577127272 | 0.582852377 | 0.992304853 | -5.120444657 |
| **A_55_P2084797** | Aff3 | -0.08226776 | -0.423013876 | 0.258478357 | 9.763743099 | -0.577042614 | 0.582906349 | 0.992304853 | -5.120483409 |
| **A_55_P1955998** | Nlrp1a | 0.04531625 | -0.142389258 | 0.233021759 | 11.62467886 | 0.577013642 | 0.58292482 | 0.992304853 | -5.12049667 |
| **A_55_P2062034** | Tsen34 | 0.049814218 | -0.156531223 | 0.256159659 | 14.39576287 | 0.57698905 | 0.582940499 | 0.992304853 | -5.120507925 |
| **A_55_P1993851** | Gm10021 | 0.116373019 | -0.365712915 | 0.598458953 | 9.128210158 | 0.5769484 | 0.582966416 | 0.992304853 | -5.120526529 |
| **A_52_P509906** | Ift172 | -0.035156005 | -0.180795799 | 0.110483789 | 9.236114609 | -0.576937271 | 0.582973512 | 0.992304853 | -5.120531622 |
| **A_55_P2000384** | Pcnx3 | 0.042356866 | -0.133162856 | 0.217876588 | 11.38691145 | 0.5767758 | 0.583076469 | 0.992304853 | -5.12060551 |
| **A_52_P502771** | Rad54b | 0.041157136 | -0.129404262 | 0.211718534 | 8.528923905 | 0.576731311 | 0.583104839 | 0.992304853 | -5.120625865 |
| **A_55_P2051039** | Gm11937 | -0.03730461 | -0.191948457 | 0.117339236 | 9.081356012 | -0.576552682 | 0.583218752 | 0.992304853 | -5.120707576 |
| **A_51_P455906** | Pold4 | -0.052633064 | -0.270825614 | 0.165559486 | 8.843811415 | -0.576537908 | 0.583228174 | 0.992304853 | -5.120714334 |
| **A_55_P2130877** | Parl | -0.048693547 | -0.250583306 | 0.153196212 | 10.88986091 | -0.57645612 | 0.583280335 | 0.992304853 | -5.120751738 |
| **A_55_P2056542** | Nin | 0.059796387 | -0.188147179 | 0.307739953 | 9.834453547 | 0.576409636 | 0.583309982 | 0.992304853 | -5.120772995 |
| **A_55_P2078008** | Ddx46 | 0.049879209 | -0.15695587 | 0.256714288 | 10.07057855 | 0.576374144 | 0.583332619 | 0.992304853 | -5.120789224 |
| **A_55_P2057946** | Hsp90aa1 | 0.048689415 | -0.153232989 | 0.25061182 | 14.0373258 | 0.576314018 | 0.583370969 | 0.992304853 | -5.120816716 |
| **A_51_P485391** | Parn | 0.028052373 | -0.088286386 | 0.144391132 | 9.162467513 | 0.576307556 | 0.583375091 | 0.992304853 | -5.12081967 |
| **A_66_P106375** | Strap | -0.048967887 | -0.252051395 | 0.154115621 | 10.13132899 | -0.576296311 | 0.583382264 | 0.992304853 | -5.120824812 |
| **A_51_P104747** | Gtf2h4 | 0.05337825 | -0.168028534 | 0.274785034 | 10.73417515 | 0.57621232 | 0.583435839 | 0.992304853 | -5.12086321 |
| **A_55_P2068772** | Ncbp2 | -0.035682121 | -0.183740171 | 0.112375928 | 9.193860063 | -0.576007019 | 0.583566805 | 0.992304853 | -5.120957046 |
| **A_51_P401504** | Col9a2 | 0.04889299 | -0.154001166 | 0.251787146 | 8.899608499 | 0.575951869 | 0.583601989 | 0.992304853 | -5.120982249 |
| **A_51_P333034** | Ier3ip1 | -0.035592985 | -0.183326445 | 0.112140475 | 12.0745419 | -0.575830514 | 0.583679415 | 0.992304853 | -5.121037698 |
| **A_52_P233441** | Gata2 | 0.03019345 | -0.095137027 | 0.155523927 | 8.991220414 | 0.575791402 | 0.583704369 | 0.992304853 | -5.121055566 |
| **A_52_P301374** | Aaas | -0.046925365 | -0.241716054 | 0.147865324 | 9.30752241 | -0.575769425 | 0.583718392 | 0.992304853 | -5.121065606 |
| **A_55_P2087385** | Gps1 | -0.030190761 | -0.155613118 | 0.095231597 | 13.06815916 | -0.57531835 | 0.584006245 | 0.992304853 | -5.1212716 |
| **A_52_P144297** | Tspyl3 | 0.029255032 | -0.092281633 | 0.150791697 | 9.200637798 | 0.575310619 | 0.58401118 | 0.992304853 | -5.12127513 |
| **A_55_P1981775** | NA | -0.035720311 | -0.184121806 | 0.112681183 | 12.72342898 | -0.575289028 | 0.58402496 | 0.992304853 | -5.121284986 |
| **A_55_P2143517** | Fstl1 | 0.073435573 | -0.231719905 | 0.37859105 | 14.79652969 | 0.575167646 | 0.584102436 | 0.992304853 | -5.121340391 |
| **A_55_P2006911** | Pik3c3 | -0.034153039 | -0.17608576 | 0.107779681 | 8.339751123 | -0.57511668 | 0.584134968 | 0.992304853 | -5.121363652 |
| **A_52_P423174** | Acaa1b | -0.031488652 | -0.162353311 | 0.099376008 | 9.555106706 | -0.57509662 | 0.584147773 | 0.992304853 | -5.121372807 |
| **A_55_P1984751** | Ppp1r14b | 0.047241168 | -0.149122568 | 0.243604905 | 12.78924317 | 0.575000831 | 0.584208919 | 0.992304853 | -5.121416518 |
| **A_51_P172532** | Nit1 | 0.037857565 | -0.119539225 | 0.195254356 | 11.17967403 | 0.574865097 | 0.584295571 | 0.992304853 | -5.121478446 |
| **A_55_P1983739** | NA | -0.056682261 | -0.292408856 | 0.179044335 | 11.49273888 | -0.57470862 | 0.584395475 | 0.992304853 | -5.121549823 |

| | | | | | | | | | |
|---|---|---|---|---|---|---|---|---|---|
| **A_55_P2179448** | NA | -0.06087792 | -0.314053785 | 0.192297946 | 9.314947907 | -0.574707191 | 0.584396387 | 0.992304853 | -5.121550474 |
| **A_51_P310548** | Osgep | -0.0461103 | -0.237883255 | 0.145662655 | 11.96077913 | -0.574671578 | 0.584419126 | 0.992304853 | -5.121566717 |
| **A_55_P2017929** | Clu | 0.055535577 | -0.175478109 | 0.286549264 | 13.70906829 | 0.574569689 | 0.584484184 | 0.992304853 | -5.121613181 |
| **A_52_P38157** | Lypla1 | 0.033425607 | -0.105625772 | 0.172476985 | 8.498283228 | 0.574530559 | 0.584509171 | 0.992304853 | -5.121631024 |
| **A_55_P2008895** | Tmsb15b1 | -0.025760179 | -0.132940501 | 0.081420142 | 8.354610575 | -0.574437731 | 0.584568448 | 0.992304853 | -5.121673347 |
| **A_55_P2177633** | NA | 0.024307474 | -0.076831135 | 0.125446083 | 7.658248879 | 0.574423207 | 0.584577723 | 0.992304853 | -5.121679968 |
| **A_55_P2021991** | Zmym4 | -0.027433991 | -0.141593109 | 0.086725128 | 8.974989765 | -0.574364398 | 0.584615279 | 0.992304853 | -5.121706777 |
| **A_51_P430563** | Mtpn | 0.037424787 | -0.118312376 | 0.19316195 | 9.068653518 | 0.574349447 | 0.584624828 | 0.992304853 | -5.121713593 |
| **A_51_P406334** | Dnlz | -0.064560751 | -0.33325864 | 0.204137137 | 10.66119555 | -0.574266462 | 0.584677826 | 0.992304853 | -5.121751419 |
| **A_51_P492087** | Pole3 | 0.038225703 | -0.120871089 | 0.197322496 | 9.52975426 | 0.574252898 | 0.584686489 | 0.992304853 | -5.121757601 |
| **A_51_P378825** | Actr8 | -0.033220717 | -0.171490312 | 0.105048877 | 11.01513609 | -0.574237363 | 0.584696411 | 0.992304853 | -5.121764681 |
| **A_52_P574945** | 1500026H1 | -0.025410967 | -0.131175798 | 0.080353865 | 8.265390117 | -0.574234178 | 0.584698445 | 0.992304853 | -5.121766133 |
| **A_55_P2144641** | Sdk1 | 0.031689631 | -0.100212224 | 0.163591486 | 10.02716201 | 0.574216164 | 0.58470995 | 0.992304853 | -5.121774342 |
| **A_55_P2041075** | Ccnd3 | 0.05848475 | -0.184951914 | 0.301921414 | 14.75288425 | 0.574203431 | 0.584718082 | 0.992304853 | -5.121780146 |
| **A_55_P2095010** | NA | 0.029802471 | -0.094276649 | 0.153881591 | 7.947327415 | 0.574067156 | 0.584805123 | 0.992304853 | -5.121842245 |
| **A_55_P2044953** | Lst1 | -0.059837685 | -0.308972241 | 0.189296871 | 10.32627772 | -0.574050298 | 0.584815891 | 0.992304853 | -5.121849926 |
| **A_55_P2123491** | B3gnt5 | -0.06867995 | -0.354649612 | 0.217289711 | 8.814397326 | -0.574009585 | 0.584841897 | 0.992304853 | -5.121868476 |
| **A_52_P127206** | Gm9 | -0.024540187 | -0.126751429 | 0.077671055 | 7.653940895 | -0.57383672 | 0.584952323 | 0.992304853 | -5.121947223 |
| **A_52_P552879** | Cracr2b | 0.041334526 | -0.13085275 | 0.213521801 | 8.962614718 | 0.573747804 | 0.585009128 | 0.992304853 | -5.12198772 |
| **A_51_P507851** | Gcc1 | 0.031976279 | -0.10122789 | 0.165180447 | 8.652524276 | 0.573745423 | 0.585010649 | 0.992304853 | -5.121988804 |
| **A_55_P2078173** | NA | -0.033614775 | -0.173669532 | 0.106439981 | 13.93673738 | -0.573642717 | 0.585076267 | 0.992304853 | -5.122035575 |
| **A_55_P2062058** | Dbndd2 | 0.032898467 | -0.104181019 | 0.169977953 | 8.314160682 | 0.573604204 | 0.585100874 | 0.992304853 | -5.122053111 |
| **A_55_P1982030** | H3f3b | -0.045497952 | -0.235079549 | 0.144083645 | 11.09141297 | -0.573594266 | 0.585107223 | 0.992304853 | -5.122057636 |
| **A_55_P1979986** | Crcp | 0.035821904 | -0.113446097 | 0.185089906 | 10.29417219 | 0.573576149 | 0.585118799 | 0.992304853 | -5.122065885 |
| **A_55_P2200618** | NA | 0.137000057 | -0.43393034 | 0.707930453 | 9.785501522 | 0.573517552 | 0.58515624 | 0.992304853 | -5.122092563 |
| **A_66_P100220** | NA | -0.03344518 | -0.172899712 | 0.106009353 | 8.129341072 | -0.573205081 | 0.585355917 | 0.992304853 | -5.122234784 |
| **A_55_P2065268** | NA | 0.051229281 | -0.16241137 | 0.264869933 | 9.169164789 | 0.573117277 | 0.585412034 | 0.992304853 | -5.122274736 |
| **A_55_P2057686** | Cdc42ep5 | 0.044257428 | -0.140323306 | 0.228838162 | 9.522862444 | 0.573071666 | 0.585441185 | 0.992304853 | -5.122295487 |
| **A_51_P334979** | Apoc2 | -0.152722352 | -0.789784117 | 0.484339412 | 8.68136503 | -0.572967774 | 0.585507589 | 0.992304853 | -5.122342749 |
| **A_51_P188073** | Dnajb11 | -0.040758022 | -0.210776893 | 0.129260849 | 12.07072895 | -0.572961057 | 0.585511883 | 0.992304853 | -5.122345804 |
| **A_51_P495077** | Ift22 | -0.03914393 | -0.202465108 | 0.124177248 | 8.813419139 | -0.572836993 | 0.585591186 | 0.992304853 | -5.122402232 |
| **A_52_P86073** | Mrpl1 | -0.036440691 | -0.188492962 | 0.11561158 | 9.590094958 | -0.572799759 | 0.585614988 | 0.992304853 | -5.122419165 |
| **A_51_P284486** | Gstm2 | -0.049023498 | -0.253595296 | 0.1555483 | 12.27990059 | -0.572753385 | 0.585644633 | 0.992304853 | -5.122440253 |
| **A_52_P596008** | Zeb2 | -0.062888237 | -0.325339678 | 0.199563205 | 10.15377077 | -0.57270317 | 0.585676734 | 0.992304853 | -5.122463086 |
| **A_52_P554267** | Ilf3 | 0.059445156 | -0.188747654 | 0.307637965 | 10.06664693 | 0.572448478 | 0.58583957 | 0.992304853 | -5.12257887 |
| **A_51_P328968** | Zfp322a | 0.027876057 | -0.088557665 | 0.144309778 | 11.65148933 | 0.572218229 | 0.5859868 | 0.992304853 | -5.122683502 |
| **A_55_P1955279** | Specc1 | 0.073074621 | -0.232152778 | 0.378302019 | 10.58367539 | 0.572205708 | 0.585994807 | 0.992304853 | -5.122689191 |
| **A_55_P2000117** | Ceacam13 | -0.202018537 | -1.045939364 | 0.64190229 | 8.109637034 | -0.572135175 | 0.586039914 | 0.992304853 | -5.122721236 |
| **A_55_P2130725** | Rhox8 | 0.030001393 | -0.095359144 | 0.15536193 | 8.132589447 | 0.571991676 | 0.586131688 | 0.992304853 | -5.122786419 |
| **A_55_P2115567** | Slc26a1 | -0.035570142 | -0.184231708 | 0.113091423 | 7.927090933 | -0.571868312 | 0.586210592 | 0.992304853 | -5.122842445 |
| **A_55_P2028773** | Slc22a17 | 0.054677124 | -0.173875749 | 0.283229998 | 11.98455866 | 0.571778885 | 0.586267794 | 0.992304853 | -5.122883052 |
| **A_55_P1975714** | Tor1aip1 | -0.030697338 | -0.159018204 | 0.097623527 | 9.837731947 | -0.571758425 | 0.586280881 | 0.992304853 | -5.122892342 |
| **A_55_P2025959** | Chd9 | 0.056463041 | -0.179568482 | 0.292494563 | 9.725707918 | 0.571746331 | 0.586288617 | 0.992304853 | -5.122897832 |
| **A_55_P1998384** | Wsb2 | -0.03128251 | -0.162078276 | 0.099513256 | 11.39895448 | -0.571632666 | 0.586361329 | 0.992304853 | -5.122949434 |
| **A_52_P664404** | Zfp286 | 0.029063588 | -0.092454778 | 0.150581955 | 8.912459038 | 0.571631878 | 0.586361833 | 0.992304853 | -5.122949792 |
| **A_55_P1959624** | NA | -0.052761999 | -0.273407221 | 0.167883222 | 9.063295413 | -0.571525812 | 0.586429688 | 0.992304853 | -5.122997936 |
| **A_55_P2053929** | NA | -0.033334582 | -0.172745738 | 0.106076574 | 12.25078741 | -0.571487349 | 0.586454295 | 0.992304853 | -5.123015393 |
| **A_55_P2003199** | Setd1b | 0.055978288 | -0.178134678 | 0.290091254 | 8.626349708 | 0.571482952 | 0.586457108 | 0.992304853 | -5.123017388 |

| | | | | | | | | | |
|---|---|---|---|---|---|---|---|---|---|
| **A_55_P2077053** | Itih4 | -0.02296607 | -0.119056033 | 0.073123894 | 7.659858003 | -0.571238914 | 0.586613251 | 0.992304853 | -5.123128121 |
| **A_55_P2148348** | NA | 0.032598466 | -0.10381716 | 0.169014092 | 14.01021197 | 0.57113948 | 0.586676879 | 0.992304853 | -5.123173228 |
| **A_55_P2076489** | Rac1 | -0.027554836 | -0.142911137 | 0.087801466 | 12.65525149 | -0.570907349 | 0.586825434 | 0.992304853 | -5.123278503 |
| **A_66_P116888** | Arid1a | 0.03201262 | -0.102017423 | 0.166042663 | 9.230229754 | 0.570858129 | 0.586856936 | 0.992304853 | -5.12330082 |
| **A_51_P371119** | Cops4 | 0.028501483 | -0.090828877 | 0.147831843 | 12.50251477 | 0.570854776 | 0.586859083 | 0.992304853 | -5.12330234 |
| **A_55_P2077535** | Zfp513 | 0.032226713 | -0.102702248 | 0.167155673 | 10.5511314 | 0.570847321 | 0.586863853 | 0.992304853 | -5.12330572 |
| **A_55_P2102579** | NA | 0.040074354 | -0.127792157 | 0.207940865 | 14.72183496 | 0.570573499 | 0.587039124 | 0.992304853 | -5.123429843 |
| **A_66_P129622** | Phldb3 | -0.029970421 | -0.155517503 | 0.095576661 | 7.908367919 | -0.570552161 | 0.587052784 | 0.992304853 | -5.123439513 |
| **A_51_P149621** | Stt3b | 0.044451956 | -0.14180727 | 0.230711182 | 9.583894737 | 0.570403539 | 0.587147929 | 0.992304853 | -5.123506859 |
| **A_52_P11441** | Rab6b | -0.068995681 | -0.35809842 | 0.220107058 | 10.09933977 | -0.570399102 | 0.58715077 | 0.992304853 | -5.123508869 |
| **A_52_P577662** | Ednrb | -0.054230371 | -0.281504106 | 0.173043365 | 9.62838065 | -0.570298797 | 0.587214989 | 0.992304853 | -5.123554312 |
| **A_51_P481920** | Ccna2 | -0.04760152 | -0.247109102 | 0.151906062 | 12.26894916 | -0.570256898 | 0.587241815 | 0.992304853 | -5.123573292 |
| **A_51_P204740** | Cd34 | -0.039214369 | -0.203584981 | 0.125156243 | 14.49556104 | -0.570203912 | 0.587275741 | 0.992304853 | -5.123597292 |
| **A_55_P2026315** | Gys1 | -0.066619181 | -0.34586235 | 0.212623989 | 10.60611674 | -0.570198224 | 0.587279383 | 0.992304853 | -5.123599869 |
| **A_55_P1969367** | Col4a5 | -0.059664169 | -0.309755304 | 0.190426965 | 11.5383179 | -0.570196353 | 0.587280582 | 0.992304853 | -5.123600716 |
| **A_55_P1972648** | Pheta2 | 0.027360243 | -0.087325259 | 0.142045744 | 8.51915337 | 0.570191266 | 0.587283838 | 0.992304853 | -5.12360302 |
| **A_52_P1077588** | Heatr6 | 0.046241034 | -0.14759793 | 0.240079999 | 8.959321876 | 0.570158485 | 0.587304828 | 0.992304853 | -5.123617868 |
| **A_55_P2081278** | Mrpl55 | 0.030463768 | -0.097271564 | 0.158199099 | 14.10773291 | 0.570008997 | 0.587400552 | 0.992304853 | -5.123685565 |
| **A_51_P405227** | Nradd | -0.025466408 | -0.132305516 | 0.081372699 | 8.373046983 | -0.569700467 | 0.587598144 | 0.992304853 | -5.123825236 |
| **A_52_P556337** | Txlng | 0.033622539 | -0.107441902 | 0.17468698 | 8.20118762 | 0.569668345 | 0.587618719 | 0.992304853 | -5.123839773 |
| **A_55_P2114799** | NA | 0.037022271 | -0.118341485 | 0.192386028 | 8.393824768 | 0.569537701 | 0.587702401 | 0.992304853 | -5.123898893 |
| **A_55_P2068877** | Ibtk | 0.044893643 | -0.14352139 | 0.233308677 | 11.14523112 | 0.569479942 | 0.5877394 | 0.992304853 | -5.123925026 |
| **A_55_P1975396** | Ighmbp2 | 0.035952075 | -0.114936072 | 0.186840222 | 8.893593218 | 0.569479331 | 0.587739791 | 0.992304853 | -5.123925303 |
| **A_55_P1954021** | Scarb1 | 0.033940801 | -0.108531579 | 0.176413182 | 11.1383336 | 0.569377821 | 0.587804819 | 0.992304853 | -5.123971226 |
| **A_55_P2065244** | Rbm26 | -0.043370875 | -0.225435366 | 0.138693615 | 9.519228718 | -0.569353597 | 0.587820337 | 0.992304853 | -5.123982184 |
| **A_51_P501107** | Cul5 | 0.033800217 | -0.108097002 | 0.175697436 | 9.214190791 | 0.569317764 | 0.587843293 | 0.992304853 | -5.123998392 |
| **A_55_P2129219** | NA | -0.064883316 | -0.337326202 | 0.20755957 | 13.43038874 | -0.569202368 | 0.587917225 | 0.992304853 | -5.124050584 |
| **A_52_P317653** | Car1 | 0.025425363 | -0.081351422 | 0.132202148 | 7.946089641 | 0.569114239 | 0.587973689 | 0.992304853 | -5.124090437 |
| **A_55_P2103412** | Golt1a | -0.043680573 | -0.227134539 | 0.139773393 | 8.66087866 | -0.569076109 | 0.587998121 | 0.992304853 | -5.124107678 |
| **A_52_P432570** | Chdh | -0.022605237 | -0.117582346 | 0.072371872 | 7.642295379 | -0.568851949 | 0.58814176 | 0.992304853 | -5.124209015 |
| **A_52_P496924** | Mgat4a | -0.026187007 | -0.136222561 | 0.083848547 | 7.98386101 | -0.568803103 | 0.588173063 | 0.992304853 | -5.124231092 |
| **A_55_P2019463** | Tnks1bp1 | 0.045542955 | -0.145856475 | 0.236942385 | 10.05037083 | 0.568708474 | 0.588233708 | 0.992304853 | -5.124273858 |
| **A_52_P140789** | Hspa13 | 0.058005913 | -0.185851538 | 0.301863364 | 10.33287628 | 0.568519506 | 0.588354823 | 0.992304853 | -5.124359239 |
| **A_55_P2039598** | Rpl7a | -0.06297752 | -0.327864108 | 0.201909068 | 13.08783574 | -0.568243817 | 0.588531546 | 0.992304853 | -5.124483759 |
| **A_51_P261351** | Mrpl49 | -0.038878102 | -0.202413478 | 0.124657274 | 10.7759114 | -0.568201627 | 0.588558594 | 0.992304853 | -5.12450281 |
| **A_55_P2024327** | Kmt5c | 0.047217675 | -0.151411433 | 0.245846783 | 13.08755998 | 0.56816024 | 0.588585127 | 0.992304853 | -5.124521497 |
| **A_55_P2056521** | Rpl21 | -0.031358416 | -0.163289554 | 0.100572721 | 8.474724217 | -0.568088434 | 0.588631164 | 0.992304853 | -5.124553917 |
| **A_51_P198434** | H2-K1 | -0.070977847 | -0.369619232 | 0.227663538 | 11.16410007 | -0.568043979 | 0.588659666 | 0.992304853 | -5.124573986 |
| **A_52_P522255** | Pitx1 | 0.025616358 | -0.082200118 | 0.133432834 | 7.687912317 | 0.567860139 | 0.588777543 | 0.992304853 | -5.124656965 |
| **A_55_P2094262** | Tph1 | -0.037484036 | -0.195252417 | 0.120284344 | 7.781267661 | -0.567852462 | 0.588782466 | 0.992304853 | -5.124660429 |
| **A_55_P1978241** | Artn | 0.025274093 | -0.081116223 | 0.131664409 | 7.883018423 | 0.567783296 | 0.588826818 | 0.992304853 | -5.124691642 |
| **A_55_P2058240** | Ipo5 | 0.068490531 | -0.219823891 | 0.356804953 | 10.8471586 | 0.567771123 | 0.588834624 | 0.992304853 | -5.124697135 |
| **A_55_P2116054** | Caprin1 | 0.03009694 | -0.096612423 | 0.156806304 | 8.874085615 | 0.567705062 | 0.588876988 | 0.992304853 | -5.124726943 |
| **A_51_P439299** | Relt | 0.028950895 | -0.092988932 | 0.150890723 | 7.929598704 | 0.567447328 | 0.589042284 | 0.992304853 | -5.124843207 |
| **A_66_P130887** | Pcdh18 | 0.062442277 | -0.200585141 | 0.325469695 | 10.46602834 | 0.567396749 | 0.589074726 | 0.992304853 | -5.124866018 |
| **A_51_P333071** | Anapc16 | -0.051777199 | -0.269903307 | 0.166348909 | 11.32183837 | -0.567335594 | 0.589113952 | 0.992304853 | -5.124893596 |
| **A_55_P2104653** | Vma21 | -0.034550443 | -0.180126257 | 0.111025372 | 8.56505032 | -0.567248716 | 0.589169681 | 0.992304853 | -5.12493277 |
| **A_55_P2077555** | C330021F2 | -0.032782768 | -0.170912207 | 0.105346671 | 8.505894412 | -0.567242159 | 0.589173887 | 0.992304853 | -5.124935726 |

| | | | | | | | | | |
|---|---|---|---|---|---|---|---|---|---|
| **A_51_P235726** | Icam4 | 0.025084603 | -0.080615168 | 0.130784373 | 8.118688141 | 0.567207957 | 0.589195827 | 0.992304853 | -5.124951146 |
| **A_55_P2044947** | NA | 0.03123566 | -0.100390204 | 0.162861523 | 8.276424307 | 0.56717696 | 0.589215712 | 0.992304853 | -5.124965121 |
| **A_51_P211998** | Sgms2 | -0.025672257 | -0.133874422 | 0.082529908 | 7.682439111 | -0.567070729 | 0.589283862 | 0.992304853 | -5.125013008 |
| **A_55_P2120993** | Zfp282 | 0.042635916 | -0.137079216 | 0.222351048 | 9.244401063 | 0.567022223 | 0.589314981 | 0.992304853 | -5.125034871 |
| **A_55_P2179791** | NA | 0.024109 | -0.077518143 | 0.125736144 | 8.022551219 | 0.566994202 | 0.589332959 | 0.992304853 | -5.125047501 |
| **A_55_P2007560** | NA | -0.034608305 | -0.180500138 | 0.111283528 | 8.566749081 | -0.566967907 | 0.589349829 | 0.992304853 | -5.125059351 |
| **A_55_P2068607** | Rnf19b | 0.027833887 | -0.08953089 | 0.145198664 | 9.966556816 | 0.566820072 | 0.589444682 | 0.992304853 | -5.125125969 |
| **A_55_P1969820** | Slc36a1 | 0.029161558 | -0.093823743 | 0.15214686 | 8.158464241 | 0.566717534 | 0.589510477 | 0.992304853 | -5.125172166 |
| **A_51_P517094** | Sh3gl1 | 0.041856026 | -0.134701568 | 0.21841362 | 10.17438249 | 0.566605417 | 0.589582424 | 0.992304853 | -5.125222671 |
| **A_55_P1984730** | Sgpl1 | 0.023840013 | -0.076729773 | 0.124409799 | 10.39706388 | 0.566562858 | 0.589609736 | 0.992304853 | -5.12524184 |
| **A_55_P2078073** | Pqlc2 | 0.050725993 | -0.163263449 | 0.264715436 | 9.021717269 | 0.566561874 | 0.589610367 | 0.992304853 | -5.125242283 |
| **A_55_P2009732** | Prr9 | -0.044062291 | -0.230031061 | 0.141906479 | 8.930669325 | -0.566286474 | 0.58978712 | 0.992304853 | -5.125366294 |
| **A_55_P1986711** | NA | -0.045399975 | -0.237024633 | 0.146224682 | 8.637153641 | -0.566256704 | 0.589806228 | 0.992304853 | -5.125379696 |
| **A_55_P2011031** | Cdk5rap1 | 0.085856399 | -0.276568182 | 0.44828098 | 8.031657119 | 0.566192557 | 0.589847403 | 0.992304853 | -5.125408572 |
| **A_51_P333159** | Rpa1 | -0.044830703 | -0.234082819 | 0.144421413 | 12.04572715 | -0.5661662 | 0.589864322 | 0.992304853 | -5.125420436 |
| **A_51_P122660** | Liph | -0.024085641 | -0.125786342 | 0.07761506 | 7.673598899 | -0.566035152 | 0.589948446 | 0.992304853 | -5.108536347 |
| **A_55_P2210575** | A830073O2 | 0.097034416 | -0.312760789 | 0.506829621 | 12.71920758 | 0.565936957 | 0.590011485 | 0.992304853 | -5.125523603 |
| **A_55_P2240468** | 1110019D1 | 0.036058917 | -0.116251659 | 0.188369493 | 8.448440878 | 0.565837537 | 0.590075314 | 0.992304853 | -5.125568333 |
| **A_55_P2040559** | NA | -0.048489505 | -0.253312563 | 0.156333553 | 15.64368983 | -0.565819669 | 0.590086787 | 0.992304853 | -5.125576372 |
| **A_55_P2106489** | Ankrd52 | 0.029185381 | -0.094108536 | 0.152479298 | 8.880166037 | 0.565760794 | 0.590124588 | 0.992304853 | -5.125602857 |
| **A_51_P246471** | Pbx2 | 0.030277231 | -0.097634187 | 0.158188649 | 10.60414317 | 0.565738815 | 0.5901387 | 0.992304853 | -5.125612743 |
| **A_55_P2131677** | NA | -0.039910207 | -0.208582341 | 0.128761927 | 8.595425593 | -0.56552235 | 0.590277696 | 0.992304853 | -5.125710096 |
| **A_65_P17827** | Cenpl | -0.046812116 | -0.244664837 | 0.151040605 | 9.073969935 | -0.565490581 | 0.590298098 | 0.992304853 | -5.125724381 |
| **A_66_P106789** | NA | 0.061519458 | -0.198537975 | 0.32157689 | 12.64881594 | 0.565395516 | 0.590359148 | 0.992304853 | -5.125767122 |
| **A_55_P2041626** | Ceacam12 | -0.479099817 | -2.504529459 | 1.546329824 | 9.070516723 | -0.565350755 | 0.590387895 | 0.992304853 | -5.125787245 |
| **A_55_P2096652** | Kyat1 | -0.065144221 | -0.340582323 | 0.210293881 | 11.34943218 | -0.565276606 | 0.590435516 | 0.992304853 | -5.125820576 |
| **A_52_P622434** | Nop16 | -0.062340028 | -0.32592588 | 0.201245824 | 9.382615712 | -0.565267518 | 0.590441354 | 0.992304853 | -5.125824661 |
| **A_55_P1954986** | Elk1 | -0.046028617 | -0.240656382 | 0.148599148 | 17.61113464 | -0.565239187 | 0.590459549 | 0.992304853 | -5.125837395 |
| **A_51_P220993** | Pfn1 | 0.051012217 | -0.164723268 | 0.266747701 | 11.18308613 | 0.565147412 | 0.590518496 | 0.992304853 | -5.125878641 |
| **A_55_P2149993** | NA | -0.02525544 | -0.132063213 | 0.081552333 | 7.776057356 | -0.565146722 | 0.590518939 | 0.992304853 | -5.125878951 |
| **A_52_P306351** | Asph | -0.042143213 | -0.220381809 | 0.136095384 | 12.39366064 | -0.565112636 | 0.590540834 | 0.992304853 | -5.12589427 |
| **A_55_P2112637** | NA | -0.024176833 | -0.126447029 | 0.078093364 | 7.653734283 | -0.565014308 | 0.590603994 | 0.992304853 | -5.125938452 |
| **A_55_P2033435** | Stra6l | -0.024605667 | -0.128697277 | 0.079485943 | 7.695707508 | -0.564974111 | 0.590629816 | 0.992304853 | -5.125956512 |
| **A_55_P1988866** | NA | 0.046257887 | -0.149440538 | 0.241956312 | 14.64942633 | 0.56494685 | 0.590647327 | 0.992304853 | -5.125968759 |
| **A_55_P2059640** | Abhd14b | -0.057318184 | -0.299838176 | 0.185201807 | 9.370504828 | -0.564877156 | 0.5906921 | 0.992304853 | -5.126000068 |
| **A_52_P507214** | Mmp9 | -0.025540851 | -0.133632844 | 0.082551142 | 7.666324564 | -0.564743151 | 0.590778191 | 0.992304853 | -5.114499135 |
| **A_55_P2058165** | Morn4 | 0.040060169 | -0.129508282 | 0.20962862 | 9.632132699 | 0.564646779 | 0.590840109 | 0.992304853 | -5.126103537 |
| **A_52_P676271** | Tuba4a | -0.039517659 | -0.206798874 | 0.127763556 | 14.1491383 | -0.564615984 | 0.590859895 | 0.992304853 | -5.126117365 |
| **A_55_P2028444** | Ube2j2 | 0.022830302 | -0.073823005 | 0.119483609 | 12.63884646 | 0.564552162 | 0.590900904 | 0.992304853 | -5.126146022 |
| **A_51_P229925** | Chst7 | -0.046161191 | -0.241593714 | 0.149271332 | 10.09310298 | -0.564532953 | 0.590913246 | 0.992304853 | -5.126154646 |
| **A_55_P1970120** | Vopp1 | 0.047641855 | -0.154078291 | 0.249362 | 12.54589384 | 0.564479949 | 0.590947305 | 0.992304853 | -5.126178442 |
| **A_52_P402663** | Ninl | 0.03089979 | -0.099946992 | 0.161746572 | 7.912903246 | 0.564418984 | 0.590986481 | 0.992304853 | -5.12620581 |
| **A_51_P507899** | Ttc8 | 0.060859052 | -0.196855042 | 0.318573146 | 9.208969314 | 0.564411875 | 0.590991049 | 0.992304853 | -5.126209 |
| **A_55_P1959006** | Imp4 | 0.035240744 | -0.113993045 | 0.184474533 | 11.02493961 | 0.564400032 | 0.590998659 | 0.992304853 | -5.126214316 |
| **A_52_P623775** | Eri2 | -0.02812835 | -0.14727293 | 0.09101623 | 8.413595379 | -0.564259778 | 0.591088792 | 0.992304853 | -5.126277266 |
| **A_55_P2045136** | Ly6a2 | -0.023720879 | -0.12420814 | 0.076766383 | 7.653304623 | -0.56419456 | 0.591130706 | 0.992304853 | -5.126306533 |
| **A_52_P275069** | Epp13 | -0.043417662 | -0.227415445 | 0.140580122 | 7.95039321 | -0.563979048 | 0.591269224 | 0.992304853 | -5.126403223 |
| **A_55_P2183032** | Zfp787 | 0.026820264 | -0.086847881 | 0.14048841 | 9.178077526 | 0.563940663 | 0.591293897 | 0.992304853 | -5.126420441 |

| | | | | | | | | | |
|---|---|---|---|---|---|---|---|---|---|
| **A_51_P320852** | Cd9 | 0.027621686 | -0.089456392 | 0.144699763 | 11.80886073 | 0.563876154 | 0.591335364 | 0.992304853 | -5.126449375 |
| **A_51_P414126** | Rab19 | 0.022908302 | -0.074205474 | 0.120022079 | 7.911163775 | 0.563794971 | 0.591387551 | 0.992304853 | -5.126485784 |
| **A_55_P2320263** | NA | 0.083410768 | -0.270210011 | 0.437031546 | 11.94497822 | 0.563758983 | 0.591410687 | 0.992304853 | -5.126501922 |
| **A_55_P2117043** | Mtmr1 | 0.046389475 | -0.150305297 | 0.243084247 | 9.842342941 | 0.563684082 | 0.591458839 | 0.992304853 | -5.126535507 |
| **A_52_P132165** | Hsd17b11 | -0.061856881 | -0.324146229 | 0.200432466 | 10.19242044 | -0.56365907 | 0.591474919 | 0.992304853 | -5.126546722 |
| **A_55_P2100550** | Mtif2 | 0.074243688 | -0.240629428 | 0.389116803 | 14.08107787 | 0.563550822 | 0.591544515 | 0.992304853 | -5.126595251 |
| **A_52_P164709** | Poc1a | -0.037317028 | -0.195589716 | 0.12095566 | 10.71126824 | -0.563521127 | 0.591563607 | 0.992304853 | -5.126608562 |
| **A_55_P2074924** | Ppp1r13l | -0.040759101 | -0.213669995 | 0.132151793 | 11.77515818 | -0.563392904 | 0.591646053 | 0.992304853 | -5.126666033 |
| **A_51_P212420** | Lama4 | -0.028476246 | -0.149284244 | 0.092331751 | 12.79505338 | -0.563373204 | 0.59165872 | 0.992304853 | -5.126674862 |
| **A_51_P234462** | Gpn1 | 0.039520553 | -0.128144475 | 0.207185581 | 10.25415784 | 0.563364734 | 0.591664166 | 0.992304853 | -5.126678657 |
| **A_55_P1984770** | Fam220a | -0.072075873 | -0.378056025 | 0.233904278 | 12.19941711 | -0.562996623 | 0.591900897 | 0.992304853 | -5.126843577 |
| **A_55_P1959628** | Pfdn4 | 0.044178162 | -0.143416003 | 0.231772328 | 9.662621108 | 0.5628562 | 0.591991217 | 0.992304853 | -5.126906464 |
| **A_55_P2052769** | Mast2 | -0.03787413 | -0.198699905 | 0.122951646 | 11.78772227 | -0.562854498 | 0.591992312 | 0.992304853 | -5.126907226 |
| **A_65_P01834** | Lima1 | 0.025733705 | -0.083540015 | 0.135007426 | 8.585697438 | 0.56285396 | 0.591992658 | 0.992304853 | -5.126907467 |
| **A_55_P2165839** | Cebpe | -0.040663009 | -0.213369576 | 0.132043558 | 7.80568183 | -0.562729646 | 0.592072623 | 0.992304853 | -5.126963128 |
| **A_55_P1964902** | NA | 0.119554919 | -0.388234943 | 0.627344781 | 9.917519494 | 0.562720267 | 0.592078656 | 0.992304853 | -5.126967326 |
| **A_51_P182257** | Catsperz | -0.025509767 | -0.133895091 | 0.082875557 | 8.015327372 | -0.56252929 | 0.592201515 | 0.992304853 | -5.127052813 |
| **A_55_P2187121** | Gpr137b-p | -0.035432659 | -0.185999753 | 0.115134435 | 9.611874734 | -0.562448552 | 0.592253459 | 0.992304853 | -5.127088945 |
| **A_52_P171212** | Irs1 | -0.060638158 | -0.318417073 | 0.197140757 | 8.697336339 | -0.562221877 | 0.592399309 | 0.992304853 | -5.127190364 |
| **A_55_P1986486** | Gtf2ird1 | 0.043619168 | -0.141872089 | 0.229110425 | 11.90027689 | 0.562034621 | 0.592519811 | 0.992304853 | -5.12727412 |
| **A_51_P253527** | Rffl | 0.02360856 | -0.076787262 | 0.124004382 | 7.821848217 | 0.562034527 | 0.592519872 | 0.992304853 | -5.127274162 |
| **A_52_P1128182** | Tmem185a | -0.027548515 | -0.144709129 | 0.089612099 | 9.187378708 | -0.561986246 | 0.592550944 | 0.992304853 | -5.127295752 |
| **A_55_P2133057** | Pdia2 | 0.023800022 | -0.077420364 | 0.125020407 | 7.779388234 | 0.561976942 | 0.592556931 | 0.992304853 | -5.127299913 |
| **A_55_P1990870** | Mylk3 | -0.055072207 | -0.289296099 | 0.179151684 | 12.35689806 | -0.561966496 | 0.592563654 | 0.992304853 | -5.127304584 |
| **A_65_P06611** | Wtap | 0.034790345 | -0.113191555 | 0.182772246 | 10.25556423 | 0.561900309 | 0.592606252 | 0.992304853 | -5.127334179 |
| **A_55_P2081353** | NA | 0.042678884 | -0.138890897 | 0.224248665 | 13.85602683 | 0.561795971 | 0.592673407 | 0.992304853 | -5.127380826 |
| **A_51_P323812** | Slc6a12 | -0.027983167 | -0.147037551 | 0.091071216 | 7.681251555 | -0.561772677 | 0.5926884 | 0.992304853 | -5.12739124 |
| **A_66_P128446** | Ttpa | 0.024985497 | -0.081324288 | 0.131295282 | 7.714251664 | 0.561725173 | 0.592718976 | 0.992304853 | -5.127412474 |
| **A_55_P2010020** | NA | 0.026045481 | -0.084779453 | 0.136870416 | 15.47888739 | 0.561699497 | 0.592735503 | 0.992304853 | -5.127423951 |
| **A_55_P2074762** | Lect2 | 0.023594699 | -0.076802163 | 0.123991561 | 7.712170267 | 0.561698727 | 0.592735999 | 0.992304853 | -5.127424295 |
| **A_55_P1995135** | Casz1 | -0.056371204 | -0.296240462 | 0.183498053 | 10.40417013 | -0.561683745 | 0.592745643 | 0.992304853 | -5.127430992 |
| **A_55_P2082276** | NA | 0.029812115 | -0.097060836 | 0.156685065 | 10.45122288 | 0.56160747 | 0.592794741 | 0.992304853 | -5.127465081 |
| **A_55_P2137286** | Gm10845 | 0.051051493 | -0.166212409 | 0.268315395 | 11.39747208 | 0.561603761 | 0.592797129 | 0.992304853 | -5.127466739 |
| **A_55_P2068199** | NA | -0.03595576 | -0.188983886 | 0.117072366 | 8.350117879 | -0.56157316 | 0.592816828 | 0.992304853 | -5.127480415 |
| **A_55_P2001891** | Prps1 | 0.052310638 | -0.170623479 | 0.275244756 | 11.67982053 | 0.560818866 | 0.593302506 | 0.992470753 | -5.127817297 |
| **A_51_P186547** | Pah | -0.027985964 | -0.147283824 | 0.091311897 | 7.667090651 | -0.560682169 | 0.593390547 | 0.992470753 | -5.127878306 |
| **A_55_P1960238** | Slc2a6 | -0.025074441 | -0.131964329 | 0.081815447 | 7.701900056 | -0.560665422 | 0.593401334 | 0.992470753 | -5.127885779 |
| **A_51_P417747** | Kap | -0.034741306 | -0.182873408 | 0.113390797 | 7.736756772 | -0.560539319 | 0.593482559 | 0.992470753 | -5.127942047 |
| **A_52_P384314** | Tmem198b | 0.037210834 | -0.121456684 | 0.195878352 | 10.37926584 | 0.560519216 | 0.593495509 | 0.992470753 | -5.127951016 |
| **A_52_P799815** | Tmem171 | -0.051284141 | -0.269960707 | 0.167392426 | 8.175288255 | -0.560518522 | 0.593495956 | 0.992470753 | -5.127951325 |
| **A_51_P146505** | Atp11b | -0.032767605 | -0.172502683 | 0.106967473 | 8.887424103 | -0.560464857 | 0.593530525 | 0.992470753 | -5.127975267 |
| **A_66_P108672** | Mtg1 | 0.032713496 | -0.106791219 | 0.172218211 | 11.31900833 | 0.560463327 | 0.59353151 | 0.992470753 | -5.127975949 |
| **A_55_P2049592** | NA | 0.116940977 | -0.381956965 | 0.615838919 | 9.320577016 | 0.560227133 | 0.593683671 | 0.992470753 | -5.128081296 |
| **A_55_P2021216** | Crlf3 | 0.068523127 | -0.22389224 | 0.360938494 | 11.27767053 | 0.560074903 | 0.593781753 | 0.992470753 | -5.128149173 |
| **A_55_P2052881** | NA | 0.082618006 | -0.269956636 | 0.435192647 | 10.2128955 | 0.560057692 | 0.593792843 | 0.992470753 | -5.128156846 |
| **A_55_P2110037** | Akap7 | 0.040503614 | -0.132350402 | 0.21335763 | 10.00079529 | 0.560045658 | 0.593800597 | 0.992470753 | -5.128162211 |
| **A_55_P2062622** | Nono | 0.047681358 | -0.155833955 | 0.25119667 | 13.65528194 | 0.559964705 | 0.593852759 | 0.992470753 | -5.128198298 |
| **A_52_P559919** | Eif2ak2 | -0.042866996 | -0.225859263 | 0.140125271 | 8.849929726 | -0.55988579 | 0.593903611 | 0.992470753 | -5.128233473 |

| | | | | | | | | | |
|---|---|---|---|---|---|---|---|---|---|
| **A_52_P3412** | Med17 | 0.025512374 | -0.083397359 | 0.134422106 | 9.077104472 | 0.55987788 | 0.593908708 | 0.992470753 | -5.128236998 |
| **A_55_P2034320** | NA | -0.040774231 | -0.214884449 | 0.133335986 | 12.72993065 | -0.559719782 | 0.594010593 | 0.992470753 | -5.128307453 |
| **A_51_P244061** | Snapc5 | -0.074459662 | -0.392414666 | 0.243495342 | 11.01192242 | -0.559711885 | 0.594015683 | 0.992470753 | -5.128310971 |
| **A_55_P1963578** | NA | -0.060733021 | -0.320081444 | 0.198615402 | 15.36682317 | -0.55969368 | 0.594027415 | 0.992470753 | -5.128319083 |
| **A_55_P2076592** | NA | -0.027206919 | -0.14339571 | 0.088981871 | 8.094565593 | -0.559660004 | 0.594049119 | 0.992470753 | -5.128334087 |
| **A_55_P2170309** | Ppfibp2 | -0.032429052 | -0.170943964 | 0.106085859 | 8.202587813 | -0.559560257 | 0.594113407 | 0.992470753 | -5.128378525 |
| **A_55_P2029528** | NA | -0.024522896 | -0.12926858 | 0.080222788 | 7.868596297 | -0.559557535 | 0.594115162 | 0.992470753 | -5.128379737 |
| **A_55_P2019349** | Txnrd2 | 0.028264747 | -0.092505696 | 0.14903519 | 7.7420939 | 0.559362787 | 0.594240691 | 0.992470753 | -5.128466478 |
| **A_55_P2141878** | Ldha | 0.095247565 | -0.311755306 | 0.502250436 | 13.51116109 | 0.559326687 | 0.594263962 | 0.992470753 | -5.128482553 |
| **A_55_P2159895** | Zfp52 | -0.036174276 | -0.1907992 | 0.118450648 | 8.547758161 | -0.559151483 | 0.594376909 | 0.992470753 | -5.128560561 |
| **A_51_P234359** | Sct | -0.166793388 | -0.879744513 | 0.546157737 | 10.08051597 | -0.559149838 | 0.594377969 | 0.992470753 | -5.128561294 |
| **A_55_P1957855** | NA | 0.043304222 | -0.14181581 | 0.228424255 | 10.01688493 | 0.559095449 | 0.594413034 | 0.992470753 | -5.128585505 |
| **A_55_P2137591** | Rnf220 | -0.029048849 | -0.153252393 | 0.095154694 | 8.962667054 | -0.55899005 | 0.594480989 | 0.992470753 | -5.128632419 |
| **A_55_P2161865** | Pdlim7 | -0.072128535 | -0.380572959 | 0.236315889 | 9.68681322 | -0.558906707 | 0.594534727 | 0.992470753 | -5.12866951 |
| **A_55_P2067453** | Tspan17 | 0.027469099 | -0.090013019 | 0.144951217 | 9.312815871 | 0.558832653 | 0.594582477 | 0.992470753 | -5.128702462 |
| **A_66_P123014** | Twistnb | 0.027829945 | -0.091220338 | 0.146880228 | 9.977918364 | 0.558715922 | 0.594657751 | 0.992470753 | -5.128754397 |
| **A_51_P145772** | Rnf113a2 | 0.029236622 | -0.095834408 | 0.154307652 | 9.470566526 | 0.558701198 | 0.594667246 | 0.992470753 | -5.128760947 |
| **A_55_P2131274** | Zfp663 | 0.094331943 | -0.309212414 | 0.4978763 | 13.44964244 | 0.558697377 | 0.59466971 | 0.992470753 | -5.128762647 |
| **A_66_P111021** | Nlrp6 | -0.024646705 | -0.130084227 | 0.080790817 | 7.752299521 | -0.558692456 | 0.594672883 | 0.992470753 | -5.128764836 |
| **A_55_P2159259** | Lifr | -0.047707945 | -0.251905188 | 0.156489297 | 10.92786428 | -0.558405869 | 0.594857713 | 0.992470753 | -5.128892297 |
| **A_55_P1962054** | NA | -0.032770992 | -0.173045668 | 0.107503684 | 15.58883617 | -0.558366619 | 0.594883029 | 0.992470753 | -5.128909749 |
| **A_55_P1975095** | Timmdc1 | 0.048098294 | -0.157792254 | 0.253988842 | 10.40426772 | 0.558344698 | 0.594897168 | 0.992470753 | -5.128919496 |
| **A_55_P1956862** | Egfr | -0.02485283 | -0.131247447 | 0.081541786 | 7.763601066 | -0.558297044 | 0.594927906 | 0.992470753 | -5.128940682 |
| **A_55_P1981301** | Nup88 | 0.030244976 | -0.099252673 | 0.159742625 | 11.97293325 | 0.558213694 | 0.594981671 | 0.992470753 | -5.128977736 |
| **A_55_P1978166** | Zfp945 | -0.028681115 | -0.151492741 | 0.094130512 | 8.449755275 | -0.558168954 | 0.595010532 | 0.992470753 | -5.128997623 |
| **A_51_P413916** | Pgrmc2 | 0.044427107 | -0.145824212 | 0.234678427 | 10.88052102 | 0.558122452 | 0.59504053 | 0.992470753 | -5.129018292 |
| **A_51_P207412** | Slf1 | 0.053135136 | -0.174408016 | 0.280678288 | 9.939212166 | 0.558119453 | 0.595042464 | 0.992470753 | -5.129019624 |
| **A_51_P303160** | Arg1 | -0.024427788 | -0.129058932 | 0.080203356 | 7.728157009 | -0.557997558 | 0.595121103 | 0.992470753 | -5.129073796 |
| **A_55_P2001068** | NA | -0.0996224 | -0.526662988 | 0.327418188 | 13.37472695 | -0.557566892 | 0.595398984 | 0.992470753 | -5.129265103 |
| **A_51_P418420** | Ddx3x | 0.062992714 | -0.207056434 | 0.333041862 | 10.38327681 | 0.557515107 | 0.595432403 | 0.992470753 | -5.129288098 |
| **A_55_P2183338** | Marveld3 | -0.027217992 | -0.143901451 | 0.089465467 | 7.803458882 | -0.557514183 | 0.595432999 | 0.992470753 | -5.129288508 |
| **A_52_P215170** | Acot4 | 0.023529586 | -0.077349827 | 0.124409 | 7.718730602 | 0.557469197 | 0.595462031 | 0.992470753 | -5.129308483 |
| **A_52_P636830** | G3bp2 | 0.044375754 | -0.145911017 | 0.234662526 | 11.0118817 | 0.557373458 | 0.595523819 | 0.992470753 | -5.129350987 |
| **A_55_P2061724** | NA | 0.030458486 | -0.100155666 | 0.161072639 | 8.518916097 | 0.557348984 | 0.595539614 | 0.992470753 | -5.129361851 |
| **A_55_P2033845** | NA | 0.029561238 | -0.097217829 | 0.156340305 | 15.68175614 | 0.557293783 | 0.595575242 | 0.992470753 | -5.129386354 |
| **A_55_P2016004** | Cd22 | -0.05767566 | -0.30511015 | 0.189758831 | 8.536337597 | -0.557110659 | 0.595693443 | 0.992470753 | -5.129467625 |
| **A_55_P2146254** | Ifitm1 | -0.037879385 | -0.2004269 | 0.12466813 | 8.589739568 | -0.556969895 | 0.59578431 | 0.992470753 | -5.129530081 |
| **A_55_P2070296** | NA | 0.03484988 | -0.114710831 | 0.184410591 | 9.354973585 | 0.55692011 | 0.595816449 | 0.992470753 | -5.129552166 |
| **A_51_P494675** | Cotl1 | -0.046216247 | -0.244574057 | 0.152141564 | 12.93784111 | -0.55687087 | 0.595848238 | 0.992470753 | -5.129574009 |
| **A_51_P315673** | Usp11 | 0.040136558 | -0.132167286 | 0.212440402 | 10.0905568 | 0.55674239 | 0.595931188 | 0.992470753 | -5.129630993 |
| **A_51_P168630** | Klf1 | 0.025050188 | -0.082489242 | 0.132589617 | 7.734763849 | 0.556739951 | 0.595932763 | 0.992470753 | -5.129632075 |
| **A_55_P2355345** | A130019P1 | 0.062109879 | -0.204566808 | 0.328786567 | 8.43734223 | 0.556653273 | 0.595988728 | 0.992470753 | -5.129670512 |
| **A_55_P2137706** | Tmpo | 0.033844788 | -0.111477592 | 0.179167168 | 11.97857566 | 0.556632328 | 0.596002251 | 0.992470753 | -5.129679799 |
| **A_55_P2183237** | Ccdc191 | -0.064812522 | -0.343311289 | 0.213686246 | 9.07208812 | -0.556217658 | 0.596270035 | 0.992470753 | -5.129863604 |
| **A_55_P2172001** | Mrgpra2a | -0.023365084 | -0.123799961 | 0.077069793 | 7.702231054 | -0.556021942 | 0.596396446 | 0.992470753 | -5.129950314 |
| **A_55_P1987302** | Eny2 | 0.042049575 | -0.138726879 | 0.222826029 | 12.37913623 | 0.555941221 | 0.596448588 | 0.992470753 | -5.129986068 |
| **A_51_P349341** | Npc1 | -0.028051374 | -0.148680718 | 0.092577971 | 8.393158926 | -0.555789451 | 0.596546631 | 0.992470753 | -5.130053281 |
| **A_55_P2038347** | Acot3 | -0.028414549 | -0.150642682 | 0.093813584 | 8.038322117 | -0.555621086 | 0.596655404 | 0.992470753 | -5.130127825 |

| | | | | | | | | | |
|---|---|---|---|---|---|---|---|---|---|
| **A_51_P143232** | Wdr59 | -0.031015134 | -0.164435832 | 0.102405563 | 8.744243603 | -0.555596617 | 0.596671213 | 0.992470753 | -5.130138656 |
| **A_55_P2155901** | Snhg10 | 0.030447114 | -0.100545109 | 0.161439336 | 8.686148956 | 0.555532858 | 0.596712409 | 0.992470753 | -5.130166879 |
| **A_55_P2049072** | NA | 0.024515612 | -0.08096909 | 0.130000314 | 12.9811494 | 0.555472287 | 0.596751547 | 0.992470753 | -5.130193688 |
| **A_51_P338317** | Rhoq | -0.033430252 | -0.177274033 | 0.110413529 | 12.3855986 | -0.555466269 | 0.596755435 | 0.992470753 | -5.130196352 |
| **A_55_P2119011** | Osbpl9 | 0.024793942 | -0.08193484 | 0.131522725 | 8.643146578 | 0.55523031 | 0.596907911 | 0.992470753 | -5.130300762 |
| **A_55_P1989125** | Rps17 | -0.024971512 | -0.132471867 | 0.082528843 | 15.69527344 | -0.555193121 | 0.596931945 | 0.992470753 | -5.130317215 |
| **A_66_P139116** | Rpl8 | 0.03966243 | -0.131084138 | 0.210408999 | 16.57963675 | 0.555183401 | 0.596938226 | 0.992470753 | -5.130321515 |
| **A_51_P125368** | Hars | -0.039806085 | -0.21117127 | 0.131559101 | 9.773122746 | -0.555182796 | 0.596938617 | 0.992470753 | -5.130321782 |
| **A_55_P2152292** | Map2k2 | 0.088880041 | -0.293759261 | 0.471519343 | 10.20438696 | 0.555167205 | 0.596948693 | 0.992470753 | -5.130328679 |
| **A_51_P339444** | Rab9 | 0.051345376 | -0.169731546 | 0.272422298 | 11.70559413 | 0.555094675 | 0.596995569 | 0.992470753 | -5.130360762 |
| **A_55_P2136743** | Slc36a4 | 0.03746798 | -0.123866795 | 0.198802755 | 8.378279766 | 0.555061903 | 0.597016749 | 0.992470753 | -5.130375257 |
| **A_52_P680495** | Pdcd10 | 0.037604921 | -0.12436659 | 0.199576433 | 10.87962942 | 0.554900581 | 0.597121018 | 0.992470753 | -5.1304466 |
| **A_51_P173906** | Mto1 | -0.028461465 | -0.151063916 | 0.094140986 | 8.712336292 | -0.55483932 | 0.597160616 | 0.992470753 | -5.130473687 |
| **A_55_P2079619** | Rnf43 | 0.031299584 | -0.103530043 | 0.166129212 | 7.933263023 | 0.554833109 | 0.597164632 | 0.992470753 | -5.130476433 |
| **A_55_P2112687** | Sipa1l1 | 0.036981201 | -0.122338678 | 0.196301079 | 9.809462596 | 0.554779205 | 0.597199476 | 0.992470753 | -5.130500264 |
| **A_55_P1958350** | Nipsnap3b | 0.056564808 | -0.187187791 | 0.300317408 | 10.82467662 | 0.554633627 | 0.597293585 | 0.992470753 | -5.130564616 |
| **A_55_P2165028** | NA | -0.028958432 | -0.153811195 | 0.095894332 | 14.92940283 | -0.554352502 | 0.597475342 | 0.992470753 | -5.130688842 |
| **A_55_P2021011** | Sfi1 | 0.02257789 | -0.074779601 | 0.119935381 | 7.802553857 | 0.554272217 | 0.597527255 | 0.992470753 | -5.130724309 |
| **A_52_P381665** | Aff1 | -0.038545599 | -0.204796457 | 0.12770526 | 10.89537523 | -0.554140676 | 0.597612316 | 0.992470753 | -5.130782409 |
| **A_55_P2040425** | Guf1 | 0.032139826 | -0.106484729 | 0.170764382 | 8.916929292 | 0.554131057 | 0.597618536 | 0.992470753 | -5.130786657 |
| **A_55_P2067332** | NA | -0.026389612 | -0.140214813 | 0.087435589 | 8.396085049 | -0.554119857 | 0.597625779 | 0.992470753 | -5.130791604 |
| **A_55_P1962801** | Max | 0.032922084 | -0.109138909 | 0.174983076 | 11.00798693 | 0.553887549 | 0.597776019 | 0.992470753 | -5.130894178 |
| **A_55_P2140646** | Creg1 | -0.114160522 | -0.606775902 | 0.378454859 | 12.34576441 | -0.553881819 | 0.597779726 | 0.992470753 | -5.130896708 |
| **A_55_P2126552** | Zdhhc13 | -0.038697578 | -0.205690391 | 0.128295235 | 10.10992164 | -0.553853794 | 0.597797852 | 0.992470753 | -5.130909079 |
| **A_52_P24631** | Azin1 | -0.050624414 | -0.269123403 | 0.167874574 | 11.25492913 | -0.553757619 | 0.597860058 | 0.992470753 | -5.130951531 |
| **A_51_P144581** | Pgrmc1 | 0.041060489 | -0.136163447 | 0.218284425 | 11.83440132 | 0.553746344 | 0.597867351 | 0.992470753 | -5.130956507 |
| **A_55_P2358002** | 5-Mar | 0.041037838 | -0.136161057 | 0.218236733 | 8.806046326 | 0.553519082 | 0.598014361 | 0.992470753 | -5.131056794 |
| **A_55_P2108644** | NA | -0.038915466 | -0.206999388 | 0.129168456 | 12.71965533 | -0.553356729 | 0.598119396 | 0.992470753 | -5.131128415 |
| **A_52_P686091** | Phf2 | 0.027062701 | -0.089854571 | 0.143979973 | 8.92804498 | 0.55322475 | 0.598204788 | 0.992470753 | -5.131186623 |
| **A_51_P521374** | Nudt13 | -0.033980117 | -0.180788968 | 0.112828733 | 10.54008056 | -0.553199485 | 0.598221135 | 0.992470753 | -5.131197765 |
| **A_51_P506045** | Cpq | 0.026444272 | -0.08782138 | 0.140709925 | 11.08032324 | 0.553127248 | 0.598267877 | 0.992470753 | -5.131229618 |
| **A_55_P2103596** | Slc30a4 | 0.037536967 | -0.124704479 | 0.199778414 | 11.0462471 | 0.552976284 | 0.598365566 | 0.992470753 | -5.131296173 |
| **A_55_P2083609** | Dkk3 | 0.237791481 | -0.790493989 | 1.266076951 | 12.44388059 | 0.552702876 | 0.598542512 | 0.992470753 | -5.13141667 |
| **A_52_P77332** | Ireb2 | 0.025918749 | -0.086177636 | 0.138015133 | 8.033587202 | 0.55262631 | 0.59859207 | 0.992470753 | -5.131450404 |
| **A_55_P2179483** | NA | 0.049271632 | -0.163838442 | 0.262381705 | 14.33644186 | 0.552588826 | 0.598616332 | 0.992470753 | -5.131466918 |
| **A_55_P2135632** | Rps18 | 0.031422608 | -0.104488935 | 0.16733415 | 15.35160347 | 0.552579816 | 0.598622165 | 0.992470753 | -5.131470887 |
| **A_51_P418908** | Larp1 | 0.038334446 | -0.127497548 | 0.204166439 | 9.026919312 | 0.552497094 | 0.598675711 | 0.992470753 | -5.131507327 |
| **A_51_P499891** | Krit1 | -0.045330021 | -0.241437166 | 0.150777125 | 8.700088423 | -0.552461009 | 0.59869907 | 0.992470753 | -5.131523221 |
| **A_55_P1987023** | NA | 0.037666434 | -0.125293007 | 0.200625875 | 8.821929134 | 0.552438724 | 0.598713496 | 0.992470753 | -5.131533037 |
| **A_66_P124958** | Ccdc47 | 0.051341681 | -0.170784474 | 0.273467836 | 9.292542956 | 0.552432876 | 0.598717282 | 0.992470753 | -5.131535612 |
| **A_55_P2022861** | Elovl1 | -0.051073966 | -0.272046875 | 0.169898942 | 9.484750391 | -0.552420372 | 0.598725376 | 0.992470753 | -5.131541119 |
| **A_66_P121053** | Sftpd | 0.023730782 | -0.078996039 | 0.126457603 | 7.752951785 | 0.552124904 | 0.598916665 | 0.992470753 | -5.131671219 |
| **A_55_P2009567** | Ptbp3 | -0.025657428 | -0.136727423 | 0.085412567 | 9.3226875 | -0.552109851 | 0.598926411 | 0.992470753 | -5.131677846 |
| **A_52_P200925** | NA | 0.061998099 | -0.206392889 | 0.330389087 | 15.57347477 | 0.552102328 | 0.598931282 | 0.992470753 | -5.131681158 |
| **A_55_P1992045** | Msi2 | 0.065356619 | -0.217577758 | 0.348290996 | 13.11746375 | 0.552093954 | 0.598936704 | 0.992470753 | -5.131684844 |
| **A_51_P425680** | Ivd | -0.03886693 | -0.207136761 | 0.129402902 | 11.77122698 | -0.552055965 | 0.598961302 | 0.992470753 | -5.131701566 |
| **A_51_P234627** | Nubpl | 0.039649647 | -0.132013088 | 0.211312383 | 9.847261134 | 0.552042399 | 0.598970085 | 0.992470753 | -5.131707537 |
| **A_55_P2064507** | Msra | 0.046039456 | -0.153316495 | 0.245395407 | 11.72962222 | 0.551963184 | 0.599021379 | 0.992470753 | -5.131742402 |

| | | | | | | | | | |
|---|---|---|---|---|---|---|---|---|---|
| **A_66_P104442** | Cog6 | 0.040085951 | -0.133496724 | 0.213668626 | 8.672891353 | 0.55194391 | 0.599033859 | 0.992470753 | -5.131750884 |
| **A_52_P15490** | Nectin3 | -0.047048209 | -0.250790153 | 0.156693734 | 8.56703602 | -0.551914478 | 0.599052918 | 0.992470753 | -5.131763836 |
| **A_66_P114782** | Nr2f1 | 0.151924475 | -0.506156446 | 0.810005395 | 9.689291436 | 0.551769355 | 0.599146896 | 0.992470753 | -5.131827692 |
| **A_55_P2038162** | Nt5c | -0.033948397 | -0.181006716 | 0.113109923 | 12.14135086 | -0.551745502 | 0.599162344 | 0.992470753 | -5.131838186 |
| **A_55_P2081178** | Msl3l2 | 0.036226991 | -0.120715945 | 0.193169927 | 8.896893149 | 0.551695732 | 0.599194576 | 0.992470753 | -5.131860082 |
| **A_55_P2040583** | Map6d1 | 0.053021026 | -0.176678192 | 0.282720243 | 11.01255446 | 0.551693339 | 0.599196126 | 0.992470753 | -5.131861135 |
| **A_52_P198473** | NA | -0.044299055 | -0.23622134 | 0.147623231 | 8.783241995 | -0.551668495 | 0.599212216 | 0.992470753 | -5.131872063 |
| **A_65_P04980** | Ilf3 | 0.037167822 | -0.123860872 | 0.198196515 | 10.66643096 | 0.551661877 | 0.599216502 | 0.992470753 | -5.131874974 |
| **A_52_P423810** | Mt1 | -0.024881612 | -0.132692008 | 0.082928785 | 7.940182791 | -0.551603475 | 0.599254327 | 0.992470753 | -5.131900664 |
| **A_51_P521052** | Ly6k | 0.023316435 | -0.077740384 | 0.124373253 | 7.763300722 | 0.55144938 | 0.599354135 | 0.992470753 | -5.131968433 |
| **A_55_P1985453** | NA | -0.029440871 | -0.157048404 | 0.098166663 | 7.774215175 | -0.55142121 | 0.599372382 | 0.992470753 | -5.13198082 |
| **A_55_P2029061** | Tnfrsf9 | -0.061409317 | -0.327593633 | 0.204774998 | 11.06468145 | -0.551392609 | 0.599390908 | 0.992470753 | -5.131993396 |
| **A_55_P1980826** | Kcnrg | -0.02231815 | -0.119060192 | 0.074423891 | 7.798327945 | -0.551381363 | 0.599398193 | 0.992470753 | -5.131998341 |
| **A_52_P440729** | Atf7ip | -0.05936434 | -0.316767867 | 0.198039187 | 10.68248545 | -0.551214056 | 0.599506573 | 0.992470753 | -5.132071894 |
| **A_55_P2255449** | NA | 0.064576981 | -0.215450422 | 0.344604384 | 9.868161471 | 0.551170972 | 0.599534484 | 0.992470753 | -5.132090832 |
| **A_51_P156438** | Slc25a33 | -0.067527775 | -0.360506323 | 0.225450772 | 9.766660351 | -0.55087841 | 0.599724035 | 0.992470753 | -5.132219394 |
| **A_51_P341130** | Iqgap1 | 0.041868809 | -0.139799593 | 0.223537212 | 10.87995442 | 0.550833503 | 0.599753134 | 0.992470753 | -5.132239122 |
| **A_55_P2172989** | Ptpn23 | -0.032797872 | -0.175156735 | 0.109560991 | 11.20715073 | -0.550643209 | 0.599876447 | 0.992470753 | -5.132322706 |
| **A_55_P2114402** | Notum | 0.080850552 | -0.270176737 | 0.43187784 | 9.987927951 | 0.550492289 | 0.599974255 | 0.992470753 | -5.132388976 |
| **A_51_P275591** | Zfp292 | -0.033234861 | -0.177563742 | 0.11109402 | 9.958819608 | -0.550363661 | 0.600057623 | 0.992470753 | -5.132445445 |
| **A_55_P2097206** | Camk4 | -0.024227312 | -0.129456874 | 0.081002251 | 7.965992078 | -0.550270958 | 0.600117711 | 0.992470753 | -5.132486136 |
| **A_51_P164495** | Dda1 | -0.027192778 | -0.145322043 | 0.090936486 | 11.45427519 | -0.550180526 | 0.600176331 | 0.992470753 | -5.132525824 |
| **A_55_P1993858** | Rmrp | 0.171226025 | -0.572655486 | 0.915107537 | 10.26167788 | 0.550142433 | 0.600201024 | 0.992470753 | -5.13254254 |
| **A_55_P2025273** | Trabd | -0.051434698 | -0.274913457 | 0.17204406 | 9.929813154 | -0.550084086 | 0.600238848 | 0.992470753 | -5.132568141 |
| **A_55_P2144110** | Mink1 | 0.050729436 | -0.169688165 | 0.271147037 | 10.31751926 | 0.550076248 | 0.600243929 | 0.992470753 | -5.132571581 |
| **A_52_P411430** | Gm5914 | 0.024518354 | -0.082069194 | 0.131105901 | 8.622169267 | 0.549786371 | 0.600431865 | 0.992470753 | -5.132698738 |
| **A_55_P1970825** | NA | -0.022954809 | -0.122751003 | 0.076841385 | 8.050212946 | -0.54975455 | 0.600452497 | 0.992470753 | -5.132712693 |
| **A_55_P2391065** | Cep128 | -0.046104391 | -0.246577932 | 0.15436915 | 8.668337918 | -0.549660284 | 0.600513621 | 0.992470753 | -5.132754029 |
| **A_52_P517896** | Man2c1 | 0.049155624 | -0.164610128 | 0.262921375 | 11.51774837 | 0.549596828 | 0.600554769 | 0.992470753 | -5.132781851 |
| **A_55_P2151672** | NA | -0.028804662 | -0.154071173 | 0.096461848 | 8.762153726 | -0.549587621 | 0.60056074 | 0.992470753 | -5.132785887 |
| **A_55_P2048348** | Aig1 | 0.04941177 | -0.165500467 | 0.264324008 | 10.58030079 | 0.549513545 | 0.600608776 | 0.992470753 | -5.132818361 |
| **A_52_P291971** | Stau2 | 0.028548894 | -0.095634407 | 0.152732196 | 9.452415068 | 0.549458914 | 0.600644205 | 0.992470753 | -5.132842309 |
| **A_55_P2127303** | Spsb2 | -0.040550462 | -0.216968272 | 0.135867347 | 9.777839993 | -0.549366935 | 0.600703856 | 0.992470753 | -5.132882622 |
| **A_55_P2279957** | 5730414N1 | 0.032010177 | -0.107265009 | 0.171285363 | 8.565289507 | 0.549317531 | 0.600735898 | 0.992470753 | -5.132904273 |
| **A_52_P51429** | Dennd1c | -0.025339618 | -0.1356461 | 0.084966865 | 7.800362645 | -0.549045272 | 0.600912491 | 0.992470753 | -5.133023558 |
| **A_52_P667477** | Fyco1 | -0.057241578 | -0.306484899 | 0.192001743 | 10.42159388 | -0.548905015 | 0.601003477 | 0.992470753 | -5.133084987 |
| **A_55_P1959576** | Slitrk4 | 0.050470372 | -0.169305385 | 0.27024613 | 8.992213332 | 0.548865402 | 0.601029175 | 0.992470753 | -5.133102335 |
| **A_55_P1971244** | Ndufa4l2 | 0.055966616 | -0.187762858 | 0.299696089 | 11.73183458 | 0.548820252 | 0.601058467 | 0.992470753 | -5.133122106 |
| **A_55_P1958379** | Slit1 | -0.049207636 | -0.263517804 | 0.165102532 | 8.216433319 | -0.548780736 | 0.601084104 | 0.992470753 | -5.133139408 |
| **A_66_P101631** | Zfp27 | 0.02808353 | -0.094279058 | 0.150446118 | 8.428152172 | 0.548544887 | 0.601237129 | 0.992470753 | -5.133242653 |
| **A_55_P1991530** | Cep126 | -0.037098717 | -0.198753316 | 0.124555881 | 8.277049713 | -0.54850421 | 0.601263523 | 0.992470753 | -5.133260456 |
| **A_55_P1971076** | Atp11a | 0.04600546 | -0.154470401 | 0.24648132 | 11.35142067 | 0.548474476 | 0.601282817 | 0.992470753 | -5.133273468 |
| **A_52_P200458** | Fut4 | 0.02393591 | -0.08039425 | 0.128266069 | 7.861827778 | 0.548339073 | 0.601370684 | 0.992470753 | -5.133332718 |
| **A_55_P1971124** | Mbd1 | -0.026818959 | -0.143721079 | 0.090083162 | 9.65963817 | -0.548313143 | 0.601387512 | 0.992470753 | -5.133344063 |
| **A_51_P281700** | Khsrp | -0.079053829 | -0.423708901 | 0.265601242 | 11.8248189 | -0.548210527 | 0.601454108 | 0.992470753 | -5.133388955 |
| **A_55_P2169514** | Fgf23 | 0.08721822 | -0.29303784 | 0.467474279 | 12.16229119 | 0.548201484 | 0.601459977 | 0.992470753 | -5.13339291 |
| **A_55_P1960641** | Trpt1 | -0.032158829 | -0.172373029 | 0.108055372 | 10.87522295 | -0.548172641 | 0.601478696 | 0.992470753 | -5.133405527 |
| **A_55_P2180744** | Clstn3 | 0.025863725 | -0.086905298 | 0.138632747 | 8.104653222 | 0.548163841 | 0.601484408 | 0.992470753 | -5.133409376 |

| | | | | | | | | | |
|---|---|---|---|---|---|---|---|---|---|
| **A_55_P2088571** | Repin1 | 0.034179466 | -0.114857981 | 0.183216913 | 10.98540308 | 0.548124232 | 0.601510115 | 0.992470753 | -5.133426701 |
| **A_55_P2014690** | NA | -0.041176755 | -0.220736852 | 0.138383343 | 8.279599834 | -0.548089417 | 0.601532712 | 0.992470753 | -5.133441928 |
| **A_55_P1998081** | Trmt1 | 0.03378328 | -0.11359486 | 0.18116142 | 12.73542044 | 0.54787045 | 0.601674842 | 0.992470753 | -5.133537677 |
| **A_55_P1987534** | Tnrc6b | 0.053615093 | -0.180328963 | 0.287559149 | 9.192535842 | 0.547752261 | 0.601751566 | 0.992470753 | -5.133589343 |
| **A_55_P2006118** | Rbp4 | -0.1898789 | -1.018687149 | 0.638929348 | 9.058833897 | -0.547560076 | 0.601876336 | 0.992470753 | -5.133673337 |
| **A_51_P133562** | Serpina6 | 0.055827273 | -0.18787941 | 0.299533955 | 8.036024115 | 0.547505024 | 0.60191208 | 0.992470753 | -5.133697393 |
| **A_55_P2172624** | Gm1123 | -0.026924277 | -0.14447458 | 0.090626026 | 7.87380763 | -0.547431051 | 0.60196011 | 0.992470753 | -5.133729712 |
| **A_55_P1989653** | Slco4a1 | 0.022244787 | -0.074876682 | 0.119366256 | 7.77748856 | 0.547421863 | 0.601966076 | 0.992470753 | -5.133733726 |
| **A_55_P2026156** | NA | -0.043699146 | -0.234495176 | 0.147096883 | 14.12729741 | -0.54741003 | 0.60197376 | 0.992470753 | -5.133738896 |
| **A_52_P654108** | Dync1li2 | 0.02351278 | -0.079147274 | 0.126172834 | 8.649119543 | 0.547408605 | 0.601974684 | 0.992470753 | -5.133739518 |
| **A_55_P2060253** | Gm12657 | -0.046002406 | -0.246868557 | 0.154863746 | 14.58612822 | -0.54737243 | 0.601998174 | 0.992470753 | -5.133755322 |
| **A_55_P2147566** | Tac2 | 0.027619909 | -0.093002508 | 0.148242326 | 7.751559693 | 0.547272146 | 0.602063293 | 0.992470753 | -5.133799126 |
| **A_55_P2144511** | NA | 0.132596081 | -0.446602017 | 0.711794179 | 9.489181268 | 0.547157923 | 0.602137469 | 0.992470753 | -5.133849011 |
| **A_55_P2071581** | NA | 0.026733461 | -0.090048678 | 0.1435156 | 8.126929632 | 0.547126688 | 0.602157754 | 0.992470753 | -5.133862651 |
| **A_66_P115467** | Sfta2 | 0.03238902 | -0.113411997 | 0.178190036 | 7.928823786 | 0.551828849 | 0.602187572 | 0.992470753 | -5.058276342 |
| **A_51_P256282** | Ppp6c | -0.046449294 | -0.249383388 | 0.1564848 | 11.23145223 | -0.547057825 | 0.602202476 | 0.992470753 | -5.13389272 |
| **A_51_P196673** | Khdc1b | 0.022442173 | -0.07561038 | 0.120494726 | 7.639318752 | 0.547035034 | 0.602217278 | 0.992470753 | -5.133902671 |
| **A_52_P587648** | Phf3 | 0.053587022 | -0.180592415 | 0.287766458 | 10.81333383 | 0.546915202 | 0.602295108 | 0.992470753 | -5.133954984 |
| **A_55_P2089293** | Rnf121 | 0.045737998 | -0.154148253 | 0.245624248 | 9.479953462 | 0.546894253 | 0.602308714 | 0.992470753 | -5.133964129 |
| **A_51_P324704** | Mrpl16 | -0.035922464 | -0.192925199 | 0.121080271 | 12.54299244 | -0.546849766 | 0.60233761 | 0.992470753 | -5.133983547 |
| **A_51_P505229** | Gtpbp8 | -0.027436309 | -0.147354426 | 0.092481808 | 9.197958014 | -0.546827079 | 0.602352346 | 0.992470753 | -5.133993449 |
| **A_51_P156274** | Ltbr | -0.041146387 | -0.221036801 | 0.138744027 | 11.72481612 | -0.546679536 | 0.602448186 | 0.992470753 | -5.134057837 |
| **A_55_P2269497** | Pdp1 | -0.034196262 | -0.183732509 | 0.115339985 | 9.99201552 | -0.54656433 | 0.602523027 | 0.992470753 | -5.134108103 |
| **A_55_P2114789** | Gtf2a2 | 0.025070064 | -0.084580161 | 0.13472029 | 8.872924853 | 0.546455813 | 0.602593527 | 0.992470753 | -5.134155442 |
| **A_55_P2416494** | 8430426J0 | 0.023403705 | -0.078961064 | 0.125768473 | 7.752989147 | 0.546440942 | 0.602603189 | 0.992470753 | -5.134161928 |
| **A_52_P1172945** | Ankrd34b | 0.022789152 | -0.076920347 | 0.122498652 | 7.774545207 | 0.546261724 | 0.602719632 | 0.992470753 | -5.134240088 |
| **A_55_P1999364** | Krt85 | -0.059318107 | -0.318930872 | 0.200294658 | 10.53444191 | -0.546097732 | 0.602826194 | 0.992470753 | -5.134311588 |
| **A_55_P1965827** | Ubb | -0.030903096 | -0.166184544 | 0.104378351 | 16.67059071 | -0.54597516 | 0.602905848 | 0.992470753 | -5.134365016 |
| **A_55_P2005213** | Ces2c | -0.024752523 | -0.1331216 | 0.083616555 | 7.729064994 | -0.545912734 | 0.602946418 | 0.992470753 | -5.134392223 |
| **A_55_P2415372** | Mta1 | -0.040439041 | -0.217487183 | 0.136609102 | 8.820277887 | -0.545906922 | 0.602950195 | 0.992470753 | -5.134394756 |
| **A_52_P339742** | Cyb5r3 | 0.037499369 | -0.126689678 | 0.201688415 | 8.565471286 | 0.545869598 | 0.602974453 | 0.992470753 | -5.134411022 |
| **A_55_P2047788** | Katnal1 | -0.046673954 | -0.251041626 | 0.157693719 | 8.853413507 | -0.545847749 | 0.602988653 | 0.992470753 | -5.134420543 |
| **A_55_P1953103** | Nudt7 | -0.054368379 | -0.29243625 | 0.183699492 | 10.59279771 | -0.545826638 | 0.603002373 | 0.992470753 | -5.134429742 |
| **A_55_P1954356** | Ttc23 | -0.022400461 | -0.120487487 | 0.075686565 | 9.661153033 | -0.545826382 | 0.60300254 | 0.992470753 | -5.134429853 |
| **A_55_P2022149** | NA | 0.037740073 | -0.127524202 | 0.203004348 | 8.312005299 | 0.545799183 | 0.603020218 | 0.992470753 | -5.134441705 |
| **A_51_P185693** | Slc2a2 | -0.023303343 | -0.125377812 | 0.078771126 | 7.698196417 | -0.545645068 | 0.603120389 | 0.992470753 | -5.134508848 |
| **A_55_P2076787** | Synpo | -0.063075071 | -0.339363649 | 0.213213507 | 10.66387498 | -0.545637172 | 0.603125522 | 0.992470753 | -5.134512288 |
| **A_52_P683441** | Capn5 | 0.031619941 | -0.106885798 | 0.17012568 | 8.529908539 | 0.545635249 | 0.603126771 | 0.992470753 | -5.134513126 |
| **A_55_P2005285** | Klhl22 | -0.040235105 | -0.216483557 | 0.136013346 | 11.69172625 | -0.545618343 | 0.603137761 | 0.992470753 | -5.13452049 |
| **A_55_P2034570** | Ddx19a | 0.03058878 | -0.103454228 | 0.164631788 | 8.605907149 | 0.545415051 | 0.603269912 | 0.992470753 | -5.134609029 |
| **A_55_P2056774** | Hsd17b14 | 0.0227104 | -0.076819261 | 0.122240061 | 7.856024483 | 0.545357621 | 0.603307248 | 0.992470753 | -5.134634036 |
| **A_51_P507509** | Gimap1 | 0.02997269 | -0.101395 | 0.16134038 | 9.937342904 | 0.54531356 | 0.603335893 | 0.992470753 | -5.13465322 |
| **A_55_P2005903** | Lhx6 | 0.07296525 | -0.246844677 | 0.392775177 | 12.20847793 | 0.545297198 | 0.603346531 | 0.992470753 | -5.134660343 |
| **A_55_P2090924** | Tbc1d8b | 0.041000724 | -0.138739571 | 0.220741018 | 8.544546916 | 0.545199194 | 0.603410249 | 0.992470753 | -5.134703008 |
| **A_51_P259975** | Aspa | 0.02324791 | -0.078754282 | 0.125250101 | 7.675957473 | 0.544732814 | 0.603713524 | 0.992470753 | -5.134905945 |
| **A_51_P401730** | Ints14 | 0.029196675 | -0.098912936 | 0.157306285 | 10.63848151 | 0.544704316 | 0.603732058 | 0.992470753 | -5.13491834 |
| **A_55_P2090953** | NA | 0.024445201 | -0.082821816 | 0.131712218 | 17.08847327 | 0.544673869 | 0.60375186 | 0.992470753 | -5.134931583 |
| **A_52_P479961** | NA | 0.031470831 | -0.106670861 | 0.169612522 | 15.36383565 | 0.544493336 | 0.603869283 | 0.992470753 | -5.13501009 |

| | | | | | | | | | |
|---|---|---|---|---|---|---|---|---|---|
| **A_51_P408363** | Cfp | -0.068126785 | -0.367204865 | 0.230951295 | 9.041688036 | -0.544430494 | 0.603910159 | 0.992470753 | -5.135037412 |
| **A_55_P1978151** | NA | -0.039642309 | -0.213702116 | 0.134417498 | 16.67519995 | -0.544339159 | 0.603969572 | 0.992470753 | -5.135077117 |
| **A_55_P2222870** | Mylk | -0.035011792 | -0.188767455 | 0.11874387 | 8.193219835 | -0.544242381 | 0.60403253 | 0.992470753 | -5.135119182 |
| **A_55_P2072985** | Sptan1 | 0.034274211 | -0.116295992 | 0.184844415 | 13.40524489 | 0.544048436 | 0.604158709 | 0.992470753 | -5.13520346 |
| **A_55_P2387284** | NA | 0.097712328 | -0.331596711 | 0.527021367 | 12.78340083 | 0.543986921 | 0.604198733 | 0.992470753 | -5.135230186 |
| **A_51_P308298** | Myl9 | 0.067359965 | -0.22863611 | 0.36335604 | 17.22675453 | 0.543907482 | 0.604250422 | 0.992470753 | -5.135264695 |
| **A_51_P483373** | Txndc5 | -0.076468766 | -0.412541653 | 0.259604122 | 9.677011275 | -0.54382571 | 0.604303632 | 0.992470753 | -5.135300213 |
| **A_52_P305995** | Ammecr1 | 0.092272998 | -0.313355277 | 0.497901273 | 10.64764006 | 0.543695254 | 0.604388525 | 0.992470753 | -5.135356866 |
| **A_55_P1984635** | Parp4 | -0.026814789 | -0.144708988 | 0.091079411 | 10.30362986 | -0.543614553 | 0.604441044 | 0.992470753 | -5.135391907 |
| **A_52_P16752** | Aox3 | -0.021787168 | -0.117583075 | 0.074008738 | 7.627067909 | -0.543579372 | 0.60446394 | 0.992470753 | -5.135407181 |
| **A_55_P2020054** | 2610035D1 | -0.02610018 | -0.140863651 | 0.088663291 | 8.38237898 | -0.543561846 | 0.604475346 | 0.992470753 | -5.13541479 |
| **A_52_P551743** | Aff4 | 0.053828291 | -0.18288132 | 0.290537903 | 9.832473443 | 0.543505364 | 0.604512106 | 0.992470753 | -5.135439309 |
| **A_55_P1988531** | Myh11 | -0.03941686 | -0.212776685 | 0.133942966 | 7.9846033 | -0.543428852 | 0.604561904 | 0.992470753 | -5.13547252 |
| **A_55_P2014284** | Serpina11 | -0.028054875 | -0.151460472 | 0.095350723 | 8.048570621 | -0.543353671 | 0.604610839 | 0.992470753 | -5.13550515 |
| **A_51_P103718** | Ip6k1 | 0.037059491 | -0.125975937 | 0.200094919 | 10.33469571 | 0.543283599 | 0.60465645 | 0.992470753 | -5.135535558 |
| **A_55_P2070054** | Senp3 | 0.028709666 | -0.097615446 | 0.155034777 | 11.55863374 | 0.543184759 | 0.604720789 | 0.992470753 | -5.135578445 |
| **A_55_P2168185** | NA | 0.026255289 | -0.089272492 | 0.14178307 | 16.5839498 | 0.543174656 | 0.604727366 | 0.992470753 | -5.135582829 |
| **A_52_P649224** | Sash1 | -0.047766574 | -0.257955344 | 0.162422196 | 11.0955275 | -0.543154918 | 0.604740215 | 0.992470753 | -5.135591392 |
| **A_51_P383218** | Tmem161a | 0.040450027 | -0.137552283 | 0.218452337 | 9.089388729 | 0.543128145 | 0.604757644 | 0.992470753 | -5.135603007 |
| **A_55_P2178486** | Spin1 | 0.037231706 | -0.126622712 | 0.201086123 | 8.433507294 | 0.543080127 | 0.604788904 | 0.992470753 | -5.135623838 |
| **A_52_P56658** | Emc3 | -0.041988213 | -0.226792878 | 0.142816452 | 9.907194723 | -0.543029712 | 0.604821725 | 0.992470753 | -5.135645706 |
| **A_55_P2127179** | NA | 0.023339819 | -0.079392429 | 0.126072066 | 7.719807131 | 0.542999995 | 0.604841071 | 0.992470753 | -5.135658596 |
| **A_55_P1973829** | NA | -0.029174349 | -0.157589445 | 0.099240746 | 9.430373223 | -0.542993006 | 0.604845622 | 0.992470753 | -5.135661627 |
| **A_52_P350664** | Pygb | -0.075593311 | -0.408394192 | 0.25720757 | 11.30808556 | -0.542885239 | 0.604915785 | 0.992470753 | -5.135708365 |
| **A_55_P2055875** | Josd2 | 0.056532283 | -0.192432353 | 0.305496919 | 10.03212632 | 0.542710216 | 0.605029745 | 0.992470753 | -5.135784253 |
| **A_51_P377948** | Gimap9 | 0.022361989 | -0.076119761 | 0.12084374 | 8.03193341 | 0.542704983 | 0.605033152 | 0.992470753 | -5.135786521 |
| **A_55_P1993762** | NA | 0.036256197 | -0.1234447 | 0.195957094 | 8.436200948 | 0.542605287 | 0.605098072 | 0.992470753 | -5.135829738 |
| **A_55_P2183015** | P2rx4 | 0.038984387 | -0.132753871 | 0.210722644 | 10.75114786 | 0.542541277 | 0.605139756 | 0.992470753 | -5.135857482 |
| **A_51_P300572** | Slc30a1 | 0.030570655 | -0.1041385 | 0.165279809 | 9.607023441 | 0.542396356 | 0.605234134 | 0.992470753 | -5.135920284 |
| **A_51_P228276** | Nipsnap2 | 0.080176824 | -0.273219724 | 0.433573372 | 11.05152971 | 0.542245153 | 0.605332614 | 0.992470753 | -5.135985793 |
| **A_52_P402319** | Med20 | -0.033080194 | -0.178898049 | 0.112737662 | 9.294134446 | -0.542208684 | 0.605356367 | 0.992470753 | -5.136001591 |
| **A_55_P2000148** | Dnaic1 | -0.03335141 | -0.180380836 | 0.113678016 | 8.332861866 | -0.542149507 | 0.605394913 | 0.992470753 | -5.136027223 |
| **A_51_P391727** | Slc3a1 | 0.027500431 | -0.093761913 | 0.148762776 | 7.766755481 | 0.542029196 | 0.605473283 | 0.992470753 | -5.136079328 |
| **A_55_P2074005** | Tctn1 | -0.027391084 | -0.148191183 | 0.093409016 | 8.305063168 | -0.541939805 | 0.605531515 | 0.992470753 | -5.136118035 |
| **A_52_P271** | Dnajc18 | -0.035840994 | -0.193924552 | 0.122242564 | 10.15601998 | -0.541879193 | 0.605571001 | 0.992470753 | -5.136144278 |
| **A_55_P2319035** | AW011956 | 0.022846032 | -0.077950904 | 0.123642968 | 7.739140054 | 0.541717148 | 0.605676575 | 0.992470753 | -5.136214423 |
| **A_52_P296632** | NA | -0.070698155 | -0.382632865 | 0.241236554 | 10.04290265 | -0.541693362 | 0.605692072 | 0.992470753 | -5.136224718 |
| **A_55_P2010196** | Serpina10 | 0.023201477 | -0.07920068 | 0.125603634 | 7.649344617 | 0.541521446 | 0.605804089 | 0.992470753 | -5.136299113 |
| **A_55_P2157992** | NA | -0.0735796 | -0.398368444 | 0.251209244 | 16.53544794 | -0.541458841 | 0.605844885 | 0.992470753 | -5.136326199 |
| **A_55_P2119030** | Rasl2-9 | -0.036208926 | -0.196053808 | 0.123635956 | 13.56833978 | -0.541409709 | 0.605876901 | 0.992470753 | -5.136347455 |
| **A_51_P238786** | Mcl1 | -0.040520579 | -0.21943332 | 0.138392163 | 12.83732013 | -0.541306822 | 0.605943951 | 0.992470753 | -5.13639196 |
| **A_66_P101538** | Lrrk2 | 0.036746566 | -0.125504785 | 0.198997916 | 8.377314058 | 0.54129943 | 0.605948768 | 0.992470753 | -5.136395157 |
| **A_52_P417620** | Pacs1 | -0.028956976 | -0.156815627 | 0.098901674 | 8.639624811 | -0.541292771 | 0.605953108 | 0.992470753 | -5.136398038 |
| **A_55_P2106434** | Zfp114 | 0.023613914 | -0.080657362 | 0.127885191 | 8.411326685 | 0.541268097 | 0.605969188 | 0.992470753 | -5.136408709 |
| **A_55_P1962651** | NA | 0.053451646 | -0.182595461 | 0.289498754 | 8.380301545 | 0.541217132 | 0.606002403 | 0.992470753 | -5.136430751 |
| **A_52_P558577** | Zfp263 | 0.033586763 | -0.114739031 | 0.181912558 | 10.18870659 | 0.541203507 | 0.606011283 | 0.992470753 | -5.136436643 |
| **A_55_P2067752** | Psg26 | -0.260036436 | -1.408792262 | 0.88871939 | 8.288473395 | -0.541022853 | 0.606129028 | 0.992470753 | -5.136514755 |
| **A_51_P397296** | Marveld3 | 0.024027024 | -0.082117401 | 0.130171449 | 7.681896098 | 0.541018259 | 0.606132022 | 0.992470753 | -5.136516741 |

| | | | | | | | | | |
|---|---|---|---|---|---|---|---|---|---|
| **A_51_P482425** | Lins1 | 0.024205226 | -0.082759767 | 0.131170219 | 8.683597602 | 0.540849714 | 0.606241888 | 0.992470753 | -5.136589597 |
| **A_55_P1984655** | Smtnl2 | 0.028110221 | -0.09613081 | 0.152351251 | 13.07365449 | 0.540764714 | 0.606297298 | 0.992470753 | -5.136626331 |
| **A_51_P272806** | Ctsr | -0.098113401 | -0.531798011 | 0.335571209 | 7.872372427 | -0.540708816 | 0.606333739 | 0.992470753 | -5.136650486 |
| **A_51_P269173** | Alg9 | -0.03040823 | -0.164822713 | 0.104006252 | 9.161794667 | -0.540697318 | 0.606341235 | 0.992470753 | -5.136655454 |
| **A_51_P322285** | Tecpr2 | -0.026898813 | -0.145803895 | 0.092006269 | 8.5618851 | -0.540681906 | 0.606351283 | 0.992470753 | -5.136662114 |
| **A_55_P1979014** | NA | 0.027806879 | -0.095130134 | 0.150743891 | 8.263521573 | 0.540603332 | 0.606402509 | 0.992470753 | -5.136696062 |
| **A_55_P2349148** | 5830408B1 | 0.030682132 | -0.105008926 | 0.16637319 | 9.310628613 | 0.54043497 | 0.606512282 | 0.992470753 | -5.136768788 |
| **A_55_P2036907** | Mycbpap | 0.032047461 | -0.10968645 | 0.173781372 | 8.505388322 | 0.540416989 | 0.606524007 | 0.992470753 | -5.136776554 |
| **A_52_P450835** | Cdpf1 | 0.030122221 | -0.103107633 | 0.163352075 | 11.36949534 | 0.540374151 | 0.606551939 | 0.992470753 | -5.136795055 |
| **A_55_P2050439** | Dlgap5 | -0.026567762 | -0.144161857 | 0.091026332 | 8.55312288 | -0.539981148 | 0.60680823 | 0.992470753 | -5.136964723 |
| **A_55_P2105321** | Acin1 | 0.083753392 | -0.286983706 | 0.454490489 | 11.93440753 | 0.539939985 | 0.606835078 | 0.992470753 | -5.136982487 |
| **A_55_P2181753** | Bdh1 | 0.046241944 | -0.158469073 | 0.250952961 | 8.70927707 | 0.539888405 | 0.60686872 | 0.992470753 | -5.137004746 |
| **A_65_P20249** | Prkcb | -0.037555653 | -0.203825286 | 0.12871398 | 8.12217871 | -0.539848019 | 0.606895062 | 0.992470753 | -5.137022172 |
| **A_55_P1996431** | NA | 0.021686408 | -0.074325614 | 0.11769843 | 7.674982627 | 0.539847552 | 0.606895367 | 0.992470753 | -5.137022374 |
| **A_52_P244064** | Ube2g2 | 0.038564411 | -0.132173541 | 0.209302363 | 10.90202786 | 0.539840878 | 0.606899721 | 0.992470753 | -5.137025254 |
| **A_55_P2007457** | Muc5b | 0.02746534 | -0.094157656 | 0.149088336 | 7.832770249 | 0.539732313 | 0.606970536 | 0.992470753 | -5.137072093 |
| **A_51_P236829** | Zswim7 | 0.046022244 | -0.157780188 | 0.249824676 | 10.17317624 | 0.539718819 | 0.606979338 | 0.992470753 | -5.137077915 |
| **A_51_P144264** | Klf2 | -0.046095906 | -0.250256561 | 0.158064749 | 11.92799929 | -0.539634165 | 0.607034561 | 0.992470753 | -5.137114431 |
| **A_55_P1961888** | Twistnb | 0.051588997 | -0.176930307 | 0.280108302 | 10.34353993 | 0.539564445 | 0.607080043 | 0.992470753 | -5.137144502 |
| **A_51_P113722** | Fbxl14 | 0.026476773 | -0.090831175 | 0.143784721 | 10.58033977 | 0.539444471 | 0.607158314 | 0.992470753 | -5.137196239 |
| **A_55_P2152364** | NA | -0.022670965 | -0.123164127 | 0.077822197 | 7.786991611 | -0.539190996 | 0.607323699 | 0.992470753 | -5.137305515 |
| **A_52_P475886** | Rc3h1 | -0.038884529 | -0.211268745 | 0.133499687 | 8.606705435 | -0.539123765 | 0.607367569 | 0.992470753 | -5.137334491 |
| **A_55_P2126644** | NA | 0.023170055 | -0.079556734 | 0.125896845 | 10.47513693 | 0.539079092 | 0.607396721 | 0.992470753 | -5.137353743 |
| **A_55_P2049717** | Agrp | 0.026692336 | -0.091678906 | 0.145063577 | 7.770118814 | 0.538951277 | 0.607480131 | 0.992470753 | -5.137408817 |
| **A_51_P285413** | Rbbp6 | 0.02794733 | -0.096012842 | 0.151907501 | 8.434073733 | 0.538849247 | 0.607546719 | 0.992470753 | -5.137452773 |
| **A_66_P134775** | Pter | -0.034247035 | -0.186152757 | 0.117658686 | 8.67242323 | -0.538837712 | 0.607554247 | 0.992470753 | -5.137457741 |
| **A_52_P671062** | Uba6 | 0.023444199 | -0.08054915 | 0.127437549 | 8.037149146 | 0.538814118 | 0.607569646 | 0.992470753 | -5.137467905 |
| **A_55_P2425190** | Nhsl2 | 0.035251289 | -0.121127095 | 0.191629673 | 8.923719546 | 0.538774977 | 0.607595192 | 0.992470753 | -5.137484764 |
| **A_51_P489903** | Ogg1 | 0.041724205 | -0.143374514 | 0.226822925 | 9.341804653 | 0.538758077 | 0.607606223 | 0.992470753 | -5.137492043 |
| **A_55_P2136378** | Dazap2 | 0.046018863 | -0.158135808 | 0.250173534 | 13.90824354 | 0.538748033 | 0.607612779 | 0.992470753 | -5.137496369 |
| **A_55_P2014570** | Zfp961 | 0.036498633 | -0.125449902 | 0.198447169 | 9.998178386 | 0.538652539 | 0.607675109 | 0.992470753 | -5.137537495 |
| **A_55_P1954258** | Zfp652 | 0.025724705 | -0.088435599 | 0.13988501 | 8.52248746 | 0.538572806 | 0.607727154 | 0.992470753 | -5.137571829 |
| **A_55_P1963084** | Rps13 | 0.025660738 | -0.088254358 | 0.139575834 | 16.00433548 | 0.538390019 | 0.607846476 | 0.992470753 | -5.13765052 |
| **A_55_P2158376** | NA | -0.062924997 | -0.342307963 | 0.216457969 | 13.62181816 | -0.538309957 | 0.607898744 | 0.992470753 | -5.13768498 |
| **A_51_P196726** | Itih2 | -0.037005305 | -0.201334122 | 0.127323512 | 7.842444871 | -0.538219454 | 0.607957831 | 0.992470753 | -5.137723929 |
| **A_55_P2016623** | S100a11-p | -0.059250098 | -0.322470073 | 0.203969877 | 10.55478273 | -0.537996394 | 0.608103476 | 0.992470753 | -5.137819899 |
| **A_55_P2243828** | NA | 0.101275229 | -0.34866261 | 0.551213067 | 8.869027556 | 0.537972217 | 0.608119263 | 0.992470753 | -5.137830299 |
| **A_51_P511315** | Pstpip1 | -0.024146426 | -0.131432269 | 0.083139417 | 7.843865524 | -0.537922335 | 0.608151837 | 0.992470753 | -5.137851755 |
| **A_51_P478311** | Nmd3 | -0.041782481 | -0.227434349 | 0.143869387 | 8.897094235 | -0.53790308 | 0.60816441 | 0.992470753 | -5.137860036 |
| **A_55_P2185821** | Cyp3a13 | 0.022406071 | -0.077162023 | 0.121974166 | 7.658207514 | 0.537841926 | 0.608204346 | 0.992470753 | -5.137886337 |
| **A_55_P2045802** | Nsmf | 0.04785994 | -0.164850051 | 0.26056993 | 11.97670856 | 0.537766063 | 0.608253889 | 0.992470753 | -5.13791896 |
| **A_55_P2116889** | Hax1 | -0.033855818 | -0.184352044 | 0.116640408 | 10.09111337 | -0.537671269 | 0.608315797 | 0.992470753 | -5.137959718 |
| **A_55_P2116650** | A130040M | 0.073387942 | -0.252847405 | 0.39962329 | 11.03332076 | 0.537653933 | 0.608327119 | 0.992470753 | -5.137967171 |
| **A_51_P386899** | Flvcr2 | -0.039011976 | -0.212448594 | 0.134424643 | 10.14806294 | -0.53760869 | 0.608356669 | 0.992470753 | -5.137986621 |
| **A_51_P502964** | Tmem63a | -0.045060723 | -0.24541783 | 0.155296383 | 10.15936451 | -0.537529794 | 0.608408199 | 0.992470753 | -5.138020535 |
| **A_55_P2125311** | Rab3b | 0.043282544 | -0.149196069 | 0.235761157 | 8.42710644 | 0.537451686 | 0.608459218 | 0.992470753 | -5.138054106 |
| **A_55_P2045040** | F7 | 0.023199293 | -0.079971126 | 0.126369711 | 7.662406235 | 0.537438393 | 0.608467901 | 0.992470753 | -5.138059819 |
| **A_55_P2064333** | Lama3 | 0.032492142 | -0.112011525 | 0.17699581 | 7.971811937 | 0.537413546 | 0.608484131 | 0.992470753 | -5.138070497 |

| | | | | | | | | | |
|---|---|---|---|---|---|---|---|---|---|
| **A_66_P102202** | Gm648 | 0.02928878 | -0.10097886 | 0.159556419 | 7.705368076 | 0.537370555 | 0.608512214 | 0.992470753 | -5.138088972 |
| **A_55_P2018944** | Snx27 | -0.040265657 | -0.219416985 | 0.138885671 | 9.388976707 | -0.537185016 | 0.608633419 | 0.992470753 | -5.138168689 |
| **A_51_P182471** | Celf2 | 0.063292593 | -0.218323317 | 0.344908504 | 10.9988997 | 0.537161448 | 0.608648816 | 0.992470753 | -5.138178813 |
| **A_55_P1955857** | Ap5z1 | 0.079570307 | -0.274477753 | 0.433618368 | 11.74935973 | 0.53715293 | 0.608654381 | 0.992470753 | -5.138182472 |
| **A_55_P2028310** | Trappc1 | -0.047383243 | -0.258226956 | 0.16346047 | 8.78452068 | -0.537122393 | 0.608674331 | 0.992470753 | -5.138195589 |
| **A_55_P2081585** | Dennd5b | 0.053668246 | -0.185208601 | 0.292545093 | 10.50478683 | 0.536973033 | 0.608771915 | 0.992475747 | -5.138259737 |
| **A_51_P284966** | Dolk | -0.041520826 | -0.22633391 | 0.143292258 | 11.15195949 | -0.536960575 | 0.608780055 | 0.992475747 | -5.138265087 |
| **A_55_P2056348** | NA | -0.034495303 | -0.188091038 | 0.119100431 | 13.41122583 | -0.536772116 | 0.608903198 | 0.992592811 | -5.138346003 |
| **A_55_P2101776** | Adora1 | 0.037052639 | -0.127957033 | 0.202062311 | 10.46380602 | 0.536684281 | 0.608960596 | 0.992602691 | -5.138383707 |
| **A_51_P346871** | Ift20 | -0.051488297 | -0.280861325 | 0.177884732 | 11.64307245 | -0.536506898 | 0.60907652 | 0.992707959 | -5.138459833 |
| **A_55_P2128042** | NA | -0.042384667 | -0.231275604 | 0.14650627 | 10.00801435 | -0.536298752 | 0.609212564 | 0.992771882 | -5.138549133 |
| **A_55_P2168243** | NA | 0.084330108 | -0.291525741 | 0.460185957 | 12.33840448 | 0.536253916 | 0.609241871 | 0.992771882 | -5.138568365 |
| **A_52_P49042** | Arfgef3 | 0.023172073 | -0.080153148 | 0.126497295 | 7.810231738 | 0.536003568 | 0.609405524 | 0.992771882 | -5.138675721 |
| **A_55_P2279807** | NA | 0.161085815 | -0.557231103 | 0.879402733 | 9.718522164 | 0.535982161 | 0.609419519 | 0.992771882 | -5.138684899 |
| **A_55_P2034415** | Lrrc8d | -0.041964418 | -0.229094108 | 0.145165273 | 9.326953469 | -0.535978827 | 0.609421699 | 0.992771882 | -5.138686328 |
| **A_52_P444457** | Dnlz | 0.0435876 | -0.150795127 | 0.237970328 | 9.289437232 | 0.535937841 | 0.609448494 | 0.992771882 | -5.138703899 |
| **A_55_P1970424** | Fam221a | 0.023014409 | -0.079650691 | 0.125679509 | 7.994489756 | 0.535779544 | 0.60955199 | 0.992771882 | -5.138771751 |
| **A_52_P239023** | Zfp955b | 0.046258032 | -0.160107436 | 0.2526235 | 8.608646534 | 0.535746391 | 0.609573667 | 0.992771882 | -5.138785959 |
| **A_55_P2114133** | Garnl3 | 0.05410877 | -0.187283149 | 0.295500689 | 9.113733979 | 0.53573999 | 0.609577853 | 0.992771882 | -5.138788702 |
| **A_51_P279623** | Pagr1a | 0.029845386 | -0.103346754 | 0.163037527 | 13.30462047 | 0.535559514 | 0.609695864 | 0.992812475 | -5.138866033 |
| **A_51_P484289** | Slc7a6 | 0.037675916 | -0.13046644 | 0.205818271 | 8.378293255 | 0.535544819 | 0.609705473 | 0.992812475 | -5.138872329 |
| **A_55_P1978466** | Pnma5 | 0.022195098 | -0.076881889 | 0.121272084 | 7.625264297 | 0.535418539 | 0.609788055 | 0.992846343 | -5.138926423 |
| **A_55_P2143802** | NA | -0.02918037 | -0.159471593 | 0.101110853 | 8.158985305 | -0.535284617 | 0.60987564 | 0.992846343 | -5.138983778 |
| **A_52_P512817** | Shisa5 | -0.026051398 | -0.142393793 | 0.090290997 | 12.87047616 | -0.535182826 | 0.609942217 | 0.992846343 | -5.139027364 |
| **A_51_P123676** | Synpo | 0.05129564 | -0.177785319 | 0.280376599 | 8.999038669 | 0.535180883 | 0.609943488 | 0.992846343 | -5.139028196 |
| **A_55_P1954436** | NA | 0.026522892 | -0.091938945 | 0.144984728 | 7.866380028 | 0.535120442 | 0.609983021 | 0.992846343 | -5.139054072 |
| **A_55_P2079981** | NA | 0.031661694 | -0.109777635 | 0.173101023 | 8.574878751 | 0.535023789 | 0.610046243 | 0.992865666 | -5.139095447 |
| **A_55_P2150025** | Otub1 | 0.036533171 | -0.126704113 | 0.199770455 | 9.709988833 | 0.534905594 | 0.610123561 | 0.992907924 | -5.139146034 |
| **A_55_P1975535** | Nutf2 | 0.031942932 | -0.11083815 | 0.174724014 | 13.01031382 | 0.534703766 | 0.6102556 | 0.993039221 | -5.139232392 |
| **A_55_P2090798** | Mtus1 | -0.030353081 | -0.166152608 | 0.105446445 | 11.51075832 | -0.534212024 | 0.610577371 | 0.993420565 | -5.139442677 |
| **A_52_P258557** | Slc26a9 | 0.023018693 | -0.079975057 | 0.126012443 | 7.679054795 | 0.534169291 | 0.610605338 | 0.993420565 | -5.139460943 |
| **A_55_P2116604** | NA | -0.049089158 | -0.268755713 | 0.170577397 | 13.84431384 | -0.534110084 | 0.610644087 | 0.993420565 | -5.139486248 |
| **A_55_P2059318** | Dgkz | -0.056835172 | -0.31126353 | 0.197593186 | 10.89121826 | -0.533901073 | 0.61078089 | 0.993432408 | -5.13957556 |
| **A_51_P215887** | Aldh18a1 | 0.032804004 | -0.114082884 | 0.179690892 | 8.971451554 | 0.533768524 | 0.610867655 | 0.993432408 | -5.139632183 |
| **A_55_P2106429** | Herc2 | -0.031822228 | -0.174313278 | 0.110668822 | 10.11294242 | -0.533767499 | 0.610868326 | 0.993432408 | -5.139632621 |
| **A_55_P2182458** | Kcnh4 | 0.02386165 | -0.082999229 | 0.13072253 | 8.128166115 | 0.533692211 | 0.610917612 | 0.993432408 | -5.139664778 |
| **A_55_P1986566** | Gm10318 | 0.038116764 | -0.132595873 | 0.208829401 | 9.842127041 | 0.533653649 | 0.610942857 | 0.993432408 | -5.139681246 |
| **A_52_P145767** | Vcpip1 | -0.030365062 | -0.166368759 | 0.105638635 | 9.374640133 | -0.533620609 | 0.610964487 | 0.993432408 | -5.139695356 |
| **A_55_P2015177** | NA | -0.038050049 | -0.208497149 | 0.132397051 | 11.48079039 | -0.533549522 | 0.611011027 | 0.993432408 | -5.13972571 |
| **A_55_P1999078** | NA | -0.026084441 | -0.142992076 | 0.090823194 | 8.770146971 | -0.533270782 | 0.611193533 | 0.993477227 | -5.1398447 |
| **A_52_P659312** | Spsb4 | 0.070078824 | -0.244040047 | 0.384197695 | 11.55861831 | 0.533214448 | 0.611230421 | 0.993477227 | -5.13986874 |
| **A_55_P2007699** | Nanog | -0.074869237 | -0.410574104 | 0.260835629 | 9.402915743 | -0.533033991 | 0.611348596 | 0.993477227 | -5.139945738 |
| **A_55_P2129029** | Smarcb1 | 0.047067107 | -0.164004194 | 0.258138409 | 11.60272986 | 0.532963483 | 0.611394773 | 0.993477227 | -5.139975815 |
| **A_51_P478952** | N4bp2l1 | 0.025211865 | -0.087873742 | 0.138297471 | 7.798357347 | 0.532852205 | 0.611467654 | 0.993477227 | -5.140023278 |
| **A_55_P1987904** | Chpf2 | 0.051848892 | -0.180724053 | 0.284421837 | 8.302655886 | 0.532830924 | 0.611481592 | 0.993477227 | -5.140032354 |
| **A_52_P646783** | Fbxl22 | -0.092842271 | -0.509310893 | 0.32362635 | 12.0440756 | -0.53281029 | 0.611495107 | 0.993477227 | -5.140041153 |
| **A_55_P2078765** | Kmt2e | 0.030035088 | -0.104704908 | 0.164775084 | 10.55581421 | 0.532772149 | 0.611520089 | 0.993477227 | -5.140057418 |
| **A_55_P2033010** | Mrrf | -0.040846054 | -0.224085397 | 0.142393288 | 10.89564124 | -0.532770925 | 0.611520891 | 0.993477227 | -5.14005794 |

| | | | | | | | | | |
|---|---|---|---|---|---|---|---|---|---|
| **A_55_P2001970** | Zswim9 | 0.039277569 | -0.136944921 | 0.21550006 | 9.617066434 | 0.532711904 | 0.61155955 | 0.993477227 | -5.140083107 |
| **A_51_P308961** | Hmgcl | -0.047578799 | -0.261172032 | 0.166014434 | 12.06298329 | -0.532396406 | 0.611766228 | 0.993477227 | -5.140217593 |
| **A_51_P190805** | Ap1m1 | 0.031920469 | -0.111395583 | 0.175236521 | 12.12383634 | 0.532333212 | 0.61180763 | 0.993477227 | -5.140244523 |
| **A_55_P1985559** | Gatad1 | -0.034322438 | -0.188433044 | 0.119788168 | 9.391738471 | -0.532297877 | 0.61183078 | 0.993477227 | -5.140259579 |
| **A_55_P1988246** | NA | 0.047579829 | -0.166087414 | 0.261247071 | 10.54061992 | 0.53222352 | 0.611879499 | 0.993477227 | -5.140291259 |
| **A_55_P2067463** | 1700110I07 | -0.054278015 | -0.298106409 | 0.189550378 | 11.46317488 | -0.532045551 | 0.611996112 | 0.993477227 | -5.140367068 |
| **A_55_P2009604** | NA | 0.021918131 | -0.076572193 | 0.120408455 | 8.143794088 | 0.531886638 | 0.612100249 | 0.993477227 | -5.140434741 |
| **A_51_P139184** | Fank1 | 0.022739566 | -0.079469557 | 0.124948688 | 7.6999269 | 0.531742824 | 0.6121945 | 0.993477227 | -5.140495968 |
| **A_55_P2055137** | NA | -0.048133577 | -0.264502213 | 0.168235058 | 12.94195519 | -0.531695474 | 0.612225533 | 0.993477227 | -5.140516123 |
| **A_55_P2020080** | NA | 0.050610797 | -0.176894138 | 0.278115731 | 9.438913687 | 0.531693664 | 0.61222672 | 0.993477227 | -5.140516894 |
| **A_55_P2023441** | Elp4 | 0.033470143 | -0.117005622 | 0.183945908 | 9.18135142 | 0.531618568 | 0.61227594 | 0.993477227 | -5.140548856 |
| **A_55_P2462358** | Fam118b | 0.022732184 | -0.079479943 | 0.12494431 | 11.19729372 | 0.531554577 | 0.612317883 | 0.993477227 | -5.140576089 |
| **A_55_P2026325** | NA | -0.051293198 | -0.282013183 | 0.179426787 | 16.42081547 | -0.531353684 | 0.61244957 | 0.993477227 | -5.140661565 |
| **A_55_P2079746** | Pls3 | 0.059484827 | -0.208152365 | 0.327122019 | 12.04657305 | 0.531213211 | 0.61254166 | 0.993477227 | -5.140721316 |
| **A_55_P2133901** | NA | 0.035628575 | -0.12471179 | 0.195968939 | 15.50086816 | 0.531085832 | 0.612625173 | 0.993477227 | -5.140775486 |
| **A_51_P351481** | Ccnl1 | 0.038124593 | -0.133538151 | 0.209787337 | 9.085927229 | 0.530809027 | 0.612806674 | 0.993477227 | -5.14089316 |
| **A_55_P2470474** | 9530082P2 | -0.023257298 | -0.128008243 | 0.081493648 | 7.932045865 | -0.530652765 | 0.612909148 | 0.993477227 | -5.140959565 |
| **A_51_P200667** | Clmn | 0.042786706 | -0.149983174 | 0.235556586 | 7.991794979 | 0.530491962 | 0.61301461 | 0.993477227 | -5.141027882 |
| **A_55_P2146555** | NA | 0.030769085 | -0.107861537 | 0.169399708 | 9.917957932 | 0.530474537 | 0.613026039 | 0.993477227 | -5.141035283 |
| **A_55_P2129477** | NA | -0.042730652 | -0.235280407 | 0.149819103 | 8.767490872 | -0.530402648 | 0.613073191 | 0.993477227 | -5.141065818 |
| **A_52_P345247** | Tfb1m | -0.024429213 | -0.134536913 | 0.085678486 | 10.24583613 | -0.530274721 | 0.613157102 | 0.993477227 | -5.141120146 |
| **A_52_P163939** | Cln6 | -0.047524041 | -0.261781998 | 0.166733917 | 8.913790299 | -0.530133843 | 0.613249516 | 0.993477227 | -5.14117996 |
| **A_51_P373901** | Rsph1 | 0.022735559 | -0.079785696 | 0.125256815 | 7.747824167 | 0.5300305 | 0.613317313 | 0.993477227 | -5.141223829 |
| **A_55_P2084338** | 2310009B1 | 0.028419674 | -0.09976184 | 0.156601189 | 9.769742504 | 0.529910877 | 0.613395795 | 0.993477227 | -5.141274599 |
| **A_51_P376711** | Gm15498 | 0.024641664 | -0.086528342 | 0.135811671 | 7.965741621 | 0.5297751 | 0.613484881 | 0.993477227 | -5.141332212 |
| **A_66_P103792** | AI606181 | -0.067772592 | -0.373530237 | 0.237985054 | 9.920480797 | -0.52976822 | 0.613489396 | 0.993477227 | -5.141335131 |
| **A_55_P2117903** | Stard5 | 0.039930005 | -0.140237251 | 0.220097262 | 8.745346874 | 0.529703248 | 0.613532028 | 0.993477227 | -5.141362695 |
| **A_55_P1958921** | Ankrd29 | 0.040627947 | -0.142689111 | 0.223945005 | 9.303183733 | 0.529701412 | 0.613533233 | 0.993477227 | -5.141363474 |
| **A_51_P370717** | Gsc | 0.023205092 | -0.081499965 | 0.12791015 | 7.689290969 | 0.529693658 | 0.613538321 | 0.993477227 | -5.141366764 |
| **A_51_P282268** | Snapc1 | 0.040977243 | -0.143935791 | 0.225890277 | 9.066648042 | 0.529644347 | 0.613570678 | 0.993477227 | -5.141387681 |
| **A_51_P294778** | Zfp954 | -0.035025348 | -0.19309571 | 0.123045014 | 9.313669539 | -0.529591665 | 0.613605249 | 0.993477227 | -5.141410027 |
| **A_55_P1990081** | Snrpg | 0.041658617 | -0.146364559 | 0.229681793 | 14.33942804 | 0.529544661 | 0.613636094 | 0.993477227 | -5.141429962 |
| **A_55_P2032318** | 4930522L1 | 0.029742615 | -0.10450049 | 0.163985721 | 8.2570962 | 0.529536984 | 0.613641133 | 0.993477227 | -5.141433218 |
| **A_55_P2105694** | Prr29 | -0.03105549 | -0.171248659 | 0.109137679 | 8.563859243 | -0.529444773 | 0.613701647 | 0.993477227 | -5.141472322 |
| **A_55_P1959758** | Top3b | -0.035111286 | -0.193614597 | 0.123392024 | 10.09040528 | -0.529440958 | 0.613704151 | 0.993477227 | -5.14147394 |
| **A_55_P2031631** | Igf1 | -0.036826321 | -0.2031014 | 0.129448758 | 9.753258259 | -0.529346818 | 0.613765934 | 0.993477227 | -5.141513855 |
| **A_52_P19532** | Fgd3 | -0.022222701 | -0.122578074 | 0.078132672 | 7.683968812 | -0.52925546 | 0.613825895 | 0.993477227 | -5.141552585 |
| **A_55_P1982668** | Nadk | 0.050807133 | -0.178633948 | 0.280248213 | 10.61913182 | 0.52925216 | 0.613828061 | 0.993477227 | -5.141553984 |
| **A_55_P2020602** | Pacsin1 | -0.047198973 | -0.260380474 | 0.165982528 | 11.68688074 | -0.529166288 | 0.613884426 | 0.993477227 | -5.141590382 |
| **A_55_P1982578** | Kdm3a | -0.052746858 | -0.291019373 | 0.185525658 | 11.11954739 | -0.529092708 | 0.613932723 | 0.993477227 | -5.141621566 |
| **A_55_P1964638** | Cxadr | 0.051006418 | -0.179404431 | 0.281417267 | 12.17839419 | 0.529091807 | 0.613933315 | 0.993477227 | -5.141621948 |
| **A_55_P2083396** | Radil | -0.040560871 | -0.22381743 | 0.142695689 | 8.538014746 | -0.529001462 | 0.61399262 | 0.993477227 | -5.141660232 |
| **A_55_P1993733** | Gm2824 | -0.024525478 | -0.135359846 | 0.08630889 | 8.364615714 | -0.52887394 | 0.614076334 | 0.993477227 | -5.141714259 |
| **A_51_P392303** | Lrrn3 | -0.04629167 | -0.255566546 | 0.162983206 | 9.215390253 | -0.528682442 | 0.614202059 | 0.993477227 | -5.14179537 |
| **A_51_P236013** | Rnf40 | 0.045491022 | -0.160167218 | 0.251149263 | 10.29321108 | 0.528674928 | 0.614206993 | 0.993477227 | -5.141798552 |
| **A_51_P202014** | Taf12 | -0.058412367 | -0.322497823 | 0.205673089 | 10.21958308 | -0.528651478 | 0.61422239 | 0.993477227 | -5.141808483 |
| **A_51_P216922** | Ddx59 | -0.05104801 | -0.281866478 | 0.179770459 | 8.910929949 | -0.528588111 | 0.614263996 | 0.993477227 | -5.141835315 |
| **A_55_P2058601** | Gm7097 | -0.022304078 | -0.12317354 | 0.078565384 | 7.814294301 | -0.528486262 | 0.614330873 | 0.993477227 | -5.141878436 |

| | | | | | | | | | |
|---|---|---|---|---|---|---|---|---|---|
| **A_55_P1966274** | Pdzd4 | 0.028832734 | -0.101593246 | 0.159258714 | 8.662196833 | 0.528361109 | 0.614413057 | 0.993477227 | -5.141931414 |
| **A_51_P108252** | Gpsm2 | -0.03930658 | -0.217111599 | 0.13849844 | 9.976768419 | -0.528360532 | 0.614413436 | 0.993477227 | -5.141931658 |
| **A_55_P2147276** | Orai1 | 0.054595852 | -0.192387294 | 0.301578999 | 9.014402493 | 0.528325368 | 0.614436528 | 0.993477227 | -5.141946541 |
| **A_55_P2029187** | Pde4a | -0.034530098 | -0.190750158 | 0.121689963 | 9.616964929 | -0.528287259 | 0.614461554 | 0.993477227 | -5.141962669 |
| **A_55_P1994997** | Arhgap11a | 0.035131914 | -0.123823202 | 0.19408703 | 8.35590785 | 0.528246266 | 0.614488476 | 0.993477227 | -5.141980017 |
| **A_55_P2034260** | Ctdsp1 | -0.060226796 | -0.332812241 | 0.212358649 | 11.97249192 | -0.528075758 | 0.614600461 | 0.993477227 | -5.142052162 |
| **A_52_P51548** | Pard3 | -0.023084119 | -0.12756993 | 0.081401692 | 10.30508051 | -0.528037933 | 0.614625304 | 0.993477227 | -5.142068163 |
| **A_55_P2071477** | Prl6a1 | -0.021307449 | -0.11779599 | 0.075181093 | 7.640549669 | -0.527794476 | 0.614785223 | 0.993477227 | -5.142171131 |
| **A_55_P2026863** | Trim32 | 0.034574963 | -0.122000825 | 0.191150751 | 11.69907652 | 0.52777188 | 0.614800066 | 0.993477227 | -5.142180685 |
| **A_51_P438149** | Mapre2 | -0.047857822 | -0.26459054 | 0.168874896 | 12.75922309 | -0.527761353 | 0.614806981 | 0.993477227 | -5.142185136 |
| **A_55_P1993549** | Npnt | 0.023991726 | -0.084685621 | 0.132669074 | 7.73577781 | 0.527632576 | 0.614891581 | 0.993477227 | -5.142239581 |
| **A_55_P2030702** | NA | -0.03107834 | -0.171888437 | 0.109731758 | 9.91990724 | -0.52751297 | 0.614970162 | 0.993477227 | -5.142290137 |
| **A_52_P609109** | Pitpnc1 | -0.046025805 | -0.254597656 | 0.162546047 | 9.783373797 | -0.527417852 | 0.615032658 | 0.993477227 | -5.142330335 |
| **A_66_P129795** | Ccdc59 | 0.043153402 | -0.152453207 | 0.238760011 | 11.57530748 | 0.527279207 | 0.615123758 | 0.993477227 | -5.142388917 |
| **A_55_P2076174** | NA | -0.052945139 | -0.293001531 | 0.187111253 | 14.08451925 | -0.527135123 | 0.615218441 | 0.993477227 | -5.142449782 |
| **A_52_P282035** | Rnf5 | -0.042785976 | -0.236810316 | 0.151238365 | 11.18134458 | -0.527053081 | 0.615272357 | 0.993477227 | -5.142484432 |
| **A_51_P175567** | Dact1 | -0.058517866 | -0.32391595 | 0.206880218 | 9.046422906 | -0.52698691 | 0.615315846 | 0.993477227 | -5.142512376 |
| **A_51_P167360** | Ptpn7 | -0.021712312 | -0.120189495 | 0.076764871 | 7.717208315 | -0.526962351 | 0.615331986 | 0.993477227 | -5.142522746 |
| **A_51_P388002** | Snrpb2 | -0.045403714 | -0.251353417 | 0.160545989 | 11.83662737 | -0.526913516 | 0.615364082 | 0.993477227 | -5.142543366 |
| **A_55_P2142724** | NA | -0.032761835 | -0.181453696 | 0.115930027 | 8.091531513 | -0.526611269 | 0.615562751 | 0.993477227 | -5.142670947 |
| **A_52_P213889** | Tmc7 | 0.035163982 | -0.124477887 | 0.194805851 | 8.747036108 | 0.526453938 | 0.61566618 | 0.993477227 | -5.142737332 |
| **A_52_P10254** | Psmc4 | -0.030042363 | -0.166433809 | 0.106349082 | 12.46302855 | -0.526448746 | 0.615669593 | 0.993477227 | -5.142739522 |
| **A_66_P126080** | Rpl39 | 0.042979735 | -0.152152547 | 0.238112017 | 9.200077953 | 0.526433772 | 0.615679437 | 0.993477227 | -5.14274584 |
| **A_55_P2005743** | Ubxn11 | 0.023622118 | -0.083665133 | 0.130909369 | 7.975571535 | 0.526235149 | 0.615810025 | 0.993477227 | -5.142829619 |
| **A_65_P07062** | Serf2 | 0.032630039 | -0.115583675 | 0.180843753 | 12.44273061 | 0.526184849 | 0.615843099 | 0.993477227 | -5.142850831 |
| **A_51_P273433** | Prkg2 | 0.021890814 | -0.077544849 | 0.121326478 | 7.756391759 | 0.526173381 | 0.615850639 | 0.993477227 | -5.142855667 |
| **A_51_P193563** | Cdk7 | -0.027632614 | -0.153175815 | 0.097910586 | 8.976647937 | -0.526063191 | 0.615923095 | 0.993477227 | -5.142902128 |
| **A_51_P355040** | NA | -0.029356143 | -0.162738716 | 0.104026429 | 8.444600306 | -0.526028218 | 0.615946092 | 0.993477227 | -5.142916872 |
| **A_52_P326214** | Cttn | 0.024728342 | -0.087659983 | 0.137116668 | 8.391346928 | 0.525875498 | 0.616046523 | 0.993477227 | -5.142981247 |
| **A_55_P2102449** | Nxf7 | 0.026918082 | -0.095455 | 0.149291163 | 7.690686603 | 0.525735546 | 0.616138566 | 0.993477227 | -5.143040226 |
| **A_52_P180826** | Ndufaf7 | 0.023481928 | -0.083309064 | 0.13027292 | 11.24406045 | 0.525543011 | 0.616265202 | 0.993477227 | -5.143121341 |
| **A_51_P247184** | Npr3 | 0.042632838 | -0.151282152 | 0.236547828 | 9.92150861 | 0.525462821 | 0.61631795 | 0.993477227 | -5.143155117 |
| **A_55_P2121603** | 2810047C2 | -0.025865959 | -0.143517293 | 0.091785376 | 8.267484155 | -0.52546144 | 0.616318859 | 0.993477227 | -5.143155699 |
| **A_51_P396917** | Zfyve21 | -0.04288591 | -0.23797426 | 0.152202441 | 10.73465062 | -0.525402849 | 0.616357401 | 0.993477227 | -5.143180375 |
| **A_51_P270899** | Zfp61 | 0.037420537 | -0.132833961 | 0.207675035 | 9.624906149 | 0.525315913 | 0.616414591 | 0.993477227 | -5.143216983 |
| **A_51_P193794** | Lrp1 | 0.052852485 | -0.187634447 | 0.293339417 | 10.20058715 | 0.525270561 | 0.616444426 | 0.993477227 | -5.143236078 |
| **A_52_P588559** | Ube2d3 | -0.034466082 | -0.19132217 | 0.122390005 | 12.66687562 | -0.525169719 | 0.61651077 | 0.993477227 | -5.143278533 |
| **A_55_P2062480** | NA | 0.035212367 | -0.125098816 | 0.19552355 | 13.62204177 | 0.524977306 | 0.616637368 | 0.993477227 | -5.143359518 |
| **A_51_P406020** | Abhd4 | -0.039474836 | -0.219208819 | 0.140259146 | 14.93614866 | -0.524927433 | 0.616670184 | 0.993477227 | -5.143380505 |
| **A_52_P439159** | Thap12 | 0.02984635 | -0.106071053 | 0.165763754 | 9.818632152 | 0.524838031 | 0.616729013 | 0.993477227 | -5.143418122 |
| **A_55_P2173358** | NA | -0.025134461 | -0.139607255 | 0.089338334 | 7.871564676 | -0.524778962 | 0.616767883 | 0.993477227 | -5.143442972 |
| **A_51_P286321** | Trir | -0.027427311 | -0.152369358 | 0.097514736 | 12.59307867 | -0.524667001 | 0.616841563 | 0.993477227 | -5.143490067 |
| **A_55_P2015258** | Trim13 | -0.027390349 | -0.152197135 | 0.097416437 | 9.035598283 | -0.52452778 | 0.616933189 | 0.993477227 | -5.143548616 |
| **A_55_P1991600** | Aplp2 | 0.046493764 | -0.165419269 | 0.258406797 | 13.62675985 | 0.524380063 | 0.617030414 | 0.993477227 | -5.143610723 |
| **A_55_P1996837** | Baz2b | -0.054100338 | -0.300689668 | 0.192488992 | 10.69010162 | -0.524366369 | 0.617039428 | 0.993477227 | -5.14361648 |
| **A_55_P2115225** | Fap | -0.060070112 | -0.333879163 | 0.213738938 | 9.833182626 | -0.524348195 | 0.61705139 | 0.993477227 | -5.14362412 |
| **A_55_P2375905** | 1110035E0 | 0.051949924 | -0.184884268 | 0.288784116 | 8.497940911 | 0.524263531 | 0.61710712 | 0.993477227 | -5.143659708 |
| **A_55_P1960216** | Cd151 | -0.077351973 | -0.430082856 | 0.275378909 | 10.09258897 | -0.524127617 | 0.61719659 | 0.993477227 | -5.143716827 |

| | | | | | | | | | |
|---|---|---|---|---|---|---|---|---|---|
| **A_55_P2153382** | Ermap | 0.021963807 | -0.078195799 | 0.122123412 | 7.67366629 | 0.524112046 | 0.617206841 | 0.993477227 | -5.14372337 |
| **A_66_P121249** | Ddi2 | 0.021834204 | -0.077737175 | 0.121405582 | 8.967724478 | 0.524097359 | 0.61721651 | 0.993477227 | -5.143729542 |
| **A_55_P2059382** | Arl6 | -0.038484229 | -0.214016097 | 0.137047639 | 9.748133687 | -0.524005626 | 0.617276901 | 0.993477227 | -5.143768084 |
| **A_55_P2069350** | Psph | 0.039430025 | -0.140421014 | 0.219281065 | 10.5067268 | 0.523990288 | 0.617286999 | 0.993477227 | -5.143774527 |
| **A_55_P2088070** | Spata5 | 0.040193269 | -0.143159578 | 0.223546116 | 9.543979933 | 0.52393185 | 0.617325473 | 0.993477227 | -5.143799077 |
| **A_51_P439626** | Map2k2 | 0.040846333 | -0.145489723 | 0.22718239 | 11.44753922 | 0.523920398 | 0.617333013 | 0.993477227 | -5.143803887 |
| **A_55_P1958103** | NA | -0.061018268 | -0.339426429 | 0.217389892 | 15.45419369 | -0.523825998 | 0.617395166 | 0.993477227 | -5.143843538 |
| **A_51_P201661** | Xrcc5 | -0.029871955 | -0.166177272 | 0.106433363 | 9.947810975 | -0.523793346 | 0.617416665 | 0.993477227 | -5.143857251 |
| **A_51_P460332** | Apoc4 | -0.029310502 | -0.163059217 | 0.104438214 | 7.737855624 | -0.523772586 | 0.617430335 | 0.993477227 | -5.14386597 |
| **A_55_P2272007** | Lrrfip2 | 0.038120064 | -0.135854884 | 0.212095012 | 8.794796808 | 0.523692117 | 0.61748332 | 0.993477227 | -5.143899761 |
| **A_55_P1997375** | NA | -0.023394967 | -0.130166915 | 0.08337698 | 7.823775221 | -0.523690157 | 0.617484611 | 0.993477227 | -5.143900584 |
| **A_55_P2116833** | 9430038I0' | 0.039735989 | -0.141621384 | 0.221093363 | 9.67025645 | 0.523670303 | 0.617497685 | 0.993477227 | -5.14390892 |
| **A_55_P2124751** | Col16a1 | 0.085732479 | -0.305607192 | 0.47707215 | 9.98630699 | 0.523601206 | 0.617543185 | 0.993477227 | -5.143937931 |
| **A_52_P579876** | Suds3 | 0.07168095 | -0.255558215 | 0.398920114 | 9.98528514 | 0.523537274 | 0.617585286 | 0.993477227 | -5.14396477 |
| **A_52_P429265** | Ppp6r2 | 0.024720155 | -0.088149436 | 0.137589745 | 8.398731785 | 0.523459839 | 0.617636281 | 0.993477227 | -5.143997274 |
| **A_55_P1958480** | NA | 0.127752706 | -0.455616422 | 0.711121834 | 11.84226153 | 0.523402507 | 0.617674038 | 0.993477227 | -5.144021337 |
| **A_55_P2104572** | Tmem268 | 0.0238863 | -0.085193585 | 0.132966184 | 8.480070319 | 0.523375437 | 0.617691866 | 0.993477227 | -5.144032697 |
| **A_55_P1975026** | Gm9104 | 0.031583313 | -0.112651245 | 0.17581787 | 15.49733222 | 0.523356334 | 0.617704447 | 0.993477227 | -5.144040714 |
| **A_51_P173384** | Tada1 | 0.026967594 | -0.096207113 | 0.150142302 | 9.826038369 | 0.523274772 | 0.617758166 | 0.993477227 | -5.14407494 |
| **A_51_P423743** | Cldn6 | -0.02230818 | -0.124209285 | 0.079592925 | 7.618630581 | -0.523232091 | 0.617786278 | 0.993477227 | -5.144092848 |
| **A_52_P154101** | Calca | 0.078106702 | -0.278696792 | 0.434910197 | 8.217991364 | 0.523200722 | 0.617806939 | 0.993477227 | -5.144106009 |
| **A_52_P587831** | Ddx56 | 0.032992675 | -0.117727381 | 0.183712731 | 10.09170828 | 0.523185397 | 0.617817033 | 0.993477227 | -5.144112439 |
| **A_51_P520966** | Icosl | 0.022851761 | -0.081545556 | 0.127249078 | 8.254281632 | 0.523165948 | 0.617829844 | 0.993477227 | -5.144120598 |
| **A_52_P644690** | Flt3l | -0.028341658 | -0.157840499 | 0.101157184 | 8.107152087 | -0.52308046 | 0.617886154 | 0.993477227 | -5.144156459 |
| **A_66_P123787** | Lars2 | -0.025504239 | -0.142045846 | 0.091037368 | 11.33844736 | -0.523046738 | 0.617908367 | 0.993477227 | -5.144170603 |
| **A_51_P451052** | Pgm5 | 0.035995929 | -0.128514186 | 0.200506044 | 12.10626544 | 0.52296173 | 0.617964365 | 0.993477227 | -5.144206256 |
| **A_55_P2148046** | NA | -0.026659539 | -0.148540179 | 0.095221101 | 7.761057468 | -0.522789724 | 0.618077681 | 0.993477227 | -5.144278379 |
| **A_55_P2107499** | Galnt18 | -0.042759721 | -0.238254414 | 0.152734973 | 9.659003329 | -0.522768028 | 0.618091975 | 0.993477227 | -5.144287475 |
| **A_51_P176783** | Atxn7l3b | 0.031332931 | -0.111925682 | 0.174591544 | 11.61607249 | 0.522744435 | 0.618107518 | 0.993477227 | -5.144297366 |
| **A_51_P262340** | Rbm3 | 0.039675022 | -0.141728651 | 0.221078695 | 13.96176918 | 0.522733381 | 0.618114802 | 0.993477227 | -5.144302 |
| **A_55_P2054913** | Olfr406 | 0.026262701 | -0.093834097 | 0.1463595 | 7.74927333 | 0.522657394 | 0.618164866 | 0.993477227 | -5.144333852 |
| **A_55_P2157048** | Mllt3 | 0.02737819 | -0.097820977 | 0.152577358 | 8.747127942 | 0.522651775 | 0.618168568 | 0.993477227 | -5.144336207 |
| **A_66_P116412** | Flt3 | 0.022072886 | -0.078867343 | 0.123013116 | 7.708829828 | 0.522641599 | 0.618175273 | 0.993477227 | -5.144340472 |
| **A_55_P2133225** | Mmgt2 | -0.023238382 | -0.129510157 | 0.083033393 | 11.02612473 | -0.522633308 | 0.618180736 | 0.993477227 | -5.144343948 |
| **A_55_P2062070** | Pdzd2 | -0.069695228 | -0.388685389 | 0.249294932 | 9.045840281 | -0.522197623 | 0.618467836 | 0.993515734 | -5.14452649 |
| **A_51_P394833** | Tshz1 | 0.028604851 | -0.102329461 | 0.159539163 | 11.02306434 | 0.522150089 | 0.618499164 | 0.993515734 | -5.144546397 |
| **A_55_P2149515** | Rsrc1 | 0.037522714 | -0.13424075 | 0.209286178 | 9.512074051 | 0.522122706 | 0.618517211 | 0.993515734 | -5.144557864 |
| **A_66_P120305** | Zmynd11 | -0.038270765 | -0.213473337 | 0.136931808 | 11.40890264 | -0.522078483 | 0.618546357 | 0.993515734 | -5.144576382 |
| **A_55_P2145149** | Cpn2 | 0.021013419 | -0.075186981 | 0.11721382 | 7.66735052 | 0.522070288 | 0.618551759 | 0.993515734 | -5.144579814 |
| **A_55_P2067505** | Slc16a3 | 0.10136741 | -0.362728077 | 0.565462898 | 13.12274313 | 0.522035621 | 0.618574608 | 0.993515734 | -5.14459433 |
| **A_51_P474053** | Erv3 | -0.025654033 | -0.143122459 | 0.091814394 | 7.748326355 | -0.521967694 | 0.61861938 | 0.993515734 | -5.144622769 |
| **A_52_P297009** | Itpk1 | 0.023246352 | -0.083198254 | 0.129690957 | 8.104488394 | 0.52196367 | 0.618622032 | 0.993515734 | -5.144624454 |
| **A_66_P134383** | NA | -0.043450024 | -0.242440545 | 0.155540498 | 10.89386919 | -0.52187532 | 0.618680268 | 0.993515734 | -5.144661438 |
| **A_55_P2069974** | Kctd1 | 0.030099146 | -0.107762947 | 0.16796124 | 8.91142023 | 0.521817261 | 0.618718539 | 0.993515734 | -5.14468574 |
| **A_55_P2135759** | NA | -0.036609364 | -0.20432847 | 0.131109741 | 16.23361454 | -0.521697533 | 0.618797466 | 0.993559957 | -5.144735847 |
| **A_55_P2148243** | Wdr12 | 0.039829271 | -0.142693264 | 0.222351805 | 10.65543963 | 0.521548855 | 0.618895483 | 0.993562293 | -5.144798054 |
| **A_51_P485862** | Eef1a2 | -0.103787267 | -0.579413779 | 0.371839246 | 10.35426727 | -0.521539435 | 0.618901694 | 0.993562293 | -5.144801995 |
| **A_66_P124239** | Gm6557 | -0.023621201 | -0.131908797 | 0.084666396 | 8.569208236 | -0.521353611 | 0.619024214 | 0.993619256 | -5.144879721 |

| | | | | | | | | | |
|---|---|---|---|---|---|---|---|---|---|
| **A_52_P592909** | Dgat2 | 0.028815361 | -0.103310772 | 0.160941494 | 11.81601317 | 0.521248085 | 0.619093796 | 0.993619256 | -5.144923849 |
| **A_52_P67200** | Stt3b | 0.048504203 | -0.173937782 | 0.270946188 | 11.85541292 | 0.521160803 | 0.619151352 | 0.993619256 | -5.144960342 |
| **A_55_P2140592** | Dmkn | 0.191020448 | -0.685348447 | 1.067389343 | 8.930619596 | 0.520957191 | 0.61928563 | 0.993619256 | -5.145045451 |
| **A_52_P463143** | Cdc37 | 0.025606154 | -0.091898308 | 0.143110617 | 14.83413401 | 0.520833764 | 0.619367035 | 0.993619256 | -5.145097029 |
| **A_55_P2040678** | Susd2 | 0.049235181 | -0.176821643 | 0.275292006 | 8.404268083 | 0.520555514 | 0.619550574 | 0.993619256 | -5.145213263 |
| **A_55_P1971419** | Serpinb9g | -0.065359215 | -0.365472067 | 0.234753638 | 7.803379098 | -0.52051274 | 0.619578791 | 0.993619256 | -5.145231127 |
| **A_52_P63709** | Fam49b | -0.023946684 | -0.133925395 | 0.086032026 | 8.605130809 | -0.520410318 | 0.619646359 | 0.993619256 | -5.145273895 |
| **A_52_P573255** | Cdc42ep1 | -0.023356976 | -0.130629652 | 0.0839157 | 12.55279128 | -0.520399209 | 0.619653688 | 0.993619256 | -5.145278533 |
| **A_55_P1968153** | Pcgf2 | 0.038503019 | -0.138368515 | 0.215374552 | 9.081082656 | 0.520290588 | 0.619725351 | 0.993619256 | -5.14532388 |
| **A_55_P2121491** | Bbs9 | -0.049027195 | -0.274244358 | 0.176189969 | 9.485194506 | -0.52028906 | 0.619726359 | 0.993619256 | -5.145324518 |
| **A_51_P384946** | Ndufa2 | 0.036916163 | -0.132667931 | 0.206500256 | 15.949483 | 0.520284118 | 0.61972962 | 0.993619256 | -5.145326581 |
| **A_55_P2107647** | NA | -0.036183928 | -0.202438156 | 0.130070299 | 10.54996212 | -0.520178202 | 0.619799502 | 0.993619256 | -5.14537079 |
| **A_55_P2019058** | Acaca | 0.045038746 | -0.16190749 | 0.251984982 | 10.5772793 | 0.520161125 | 0.61981077 | 0.993619256 | -5.145377917 |
| **A_51_P263220** | Taf5 | 0.032024658 | -0.115174453 | 0.179223769 | 8.545122188 | 0.519982159 | 0.619928861 | 0.993619256 | -5.145452597 |
| **A_51_P128463** | Grrp1 | -0.043807145 | -0.245173384 | 0.157559093 | 10.40935334 | -0.519956998 | 0.619945464 | 0.993619256 | -5.120812581 |
| **A_55_P2005300** | Ranbp3 | 0.028303368 | -0.101838399 | 0.158445135 | 12.51007901 | 0.519793147 | 0.620053594 | 0.993619256 | -5.145531444 |
| **A_55_P2014066** | NA | 0.02140221 | -0.07703954 | 0.11984396 | 7.872150235 | 0.519623072 | 0.620165842 | 0.993619256 | -5.145602369 |
| **A_55_P2024061** | Gtpbp10 | 0.025265113 | -0.090951033 | 0.141481258 | 9.673249533 | 0.519593736 | 0.620185205 | 0.993619256 | -5.145614601 |
| **A_51_P234044** | 1190005I0( | -0.032364677 | -0.181240675 | 0.116511322 | 9.648671123 | -0.519583937 | 0.620191673 | 0.993619256 | -5.145618686 |
| **A_55_P2148288** | Tmem79 | -0.022931389 | -0.128444581 | 0.082581804 | 7.950605684 | -0.519436809 | 0.620288787 | 0.993619256 | -5.14568002 |
| **A_52_P226929** | Cflar | 0.03358712 | -0.12096853 | 0.18814277 | 8.341097431 | 0.519394104 | 0.620316977 | 0.993619256 | -5.145697821 |
| **A_55_P1957344** | Hnrnpa3 | 0.027444032 | -0.098862486 | 0.15375055 | 12.97023167 | 0.519315499 | 0.620368866 | 0.993619256 | -5.14573058 |
| **A_52_P203691** | Arl5c | -0.026992507 | -0.151222732 | 0.097237718 | 8.015220378 | -0.519308078 | 0.620373765 | 0.993619256 | -5.145733673 |
| **A_55_P2043798** | Mrto4 | 0.04227871 | -0.152333159 | 0.236890578 | 11.1156714 | 0.519232098 | 0.620423923 | 0.993619256 | -5.145765334 |
| **A_55_P1997895** | Rpl35a | 0.03911408 | -0.140945982 | 0.219174142 | 14.43135746 | 0.519188245 | 0.620452874 | 0.993619256 | -5.145783606 |
| **A_55_P2169539** | NA | -0.035761772 | -0.200394706 | 0.128871163 | 8.846912881 | -0.519172213 | 0.620463459 | 0.993619256 | -5.145790286 |
| **A_55_P1981155** | Phldb1 | -0.046408766 | -0.260164756 | 0.167347224 | 12.34968593 | -0.518908589 | 0.620637515 | 0.993619256 | -5.145900096 |
| **A_55_P2050628** | Cyp4a31 | -0.028400693 | -0.159258093 | 0.102456706 | 7.88129423 | -0.518728116 | 0.620756686 | 0.993619256 | -5.145975241 |
| **A_55_P2064841** | Prss53 | 0.053495491 | -0.19311808 | 0.300109061 | 11.97909735 | 0.518452934 | 0.62093842 | 0.993619256 | -5.146089775 |
| **A_55_P2002846** | Cystm1 | -0.056812931 | -0.318850496 | 0.205224634 | 9.32409611 | -0.518194466 | 0.621109142 | 0.993619256 | -5.146197303 |
| **A_52_P382886** | Gjb2 | 0.029903749 | -0.108022349 | 0.167829847 | 7.720835744 | 0.518189157 | 0.621112649 | 0.993619256 | -5.146199511 |
| **A_51_P114854** | Clns1a | -0.038458151 | -0.215871291 | 0.138954988 | 9.850436688 | -0.518097809 | 0.621172993 | 0.993619256 | -5.121570996 |
| **A_55_P1981804** | Gp1ba | -0.028472191 | -0.159822891 | 0.102878509 | 8.004287392 | -0.518080957 | 0.621184125 | 0.993619256 | -5.14624451 |
| **A_51_P131800** | Cyba | -0.103036798 | -0.578377275 | 0.37230368 | 9.737125244 | -0.518079832 | 0.621184868 | 0.993619256 | -5.146244978 |
| **A_55_P2145606** | Kmo | -0.022757536 | -0.127802896 | 0.082287825 | 7.673327331 | -0.517794566 | 0.621373333 | 0.993619256 | -5.146363575 |
| **A_52_P515826** | Med13 | 0.021661173 | -0.078330145 | 0.121652491 | 7.917866427 | 0.517760374 | 0.621395924 | 0.993619256 | -5.146377786 |
| **A_51_P301964** | Arl6ip1 | 0.02795425 | -0.101117673 | 0.157026173 | 11.25455289 | 0.517636859 | 0.621477537 | 0.993619256 | -5.146429115 |
| **A_55_P2052290** | Psat1 | 0.077946929 | -0.281959595 | 0.437853453 | 12.05533693 | 0.51762879 | 0.621482869 | 0.993619256 | -5.146432468 |
| **A_55_P2111293** | 3110009E1 | -0.041019423 | -0.23043614 | 0.148397294 | 9.13575706 | -0.517583443 | 0.621512834 | 0.993619256 | -5.14645131 |
| **A_55_P1959238** | Nagk | 0.043521771 | -0.157507382 | 0.244550923 | 11.28892975 | 0.517436006 | 0.621610263 | 0.993619256 | -5.14651256 |
| **A_51_P293688** | Rab32 | -0.037432229 | -0.210337452 | 0.135472994 | 8.60151304 | -0.517424166 | 0.621618088 | 0.993619256 | -5.146517478 |
| **A_51_P509223** | Arfgap1 | 0.03899397 | -0.141126312 | 0.219114251 | 8.81476821 | 0.517420896 | 0.621620249 | 0.993619256 | -5.146518836 |
| **A_55_P1992421** | Hmgb3 | 0.032410745 | -0.117316858 | 0.182138348 | 12.71915325 | 0.517364087 | 0.621657792 | 0.993619256 | -5.146542431 |
| **A_55_P2086682** | Nfib | -0.028615712 | -0.160813972 | 0.103582548 | 13.88757399 | -0.517354159 | 0.621664354 | 0.993619256 | -5.146546555 |
| **A_55_P2354531** | 2610029K1 | -0.030445728 | -0.171113201 | 0.110221746 | 8.790312967 | -0.517299193 | 0.62170068 | 0.993619256 | -5.146569382 |
| **A_55_P2083028** | NA | 0.066397096 | -0.240417948 | 0.37321214 | 11.5448575 | 0.517227451 | 0.621748096 | 0.993619256 | -5.146599173 |
| **A_51_P501922** | Plxna1 | -0.039272635 | -0.220773108 | 0.142227838 | 9.071482845 | -0.517155814 | 0.621795444 | 0.993619256 | -5.146628917 |
| **A_55_P2023446** | NA | -0.033052441 | -0.185829362 | 0.11972448 | 11.83590618 | -0.51707664 | 0.621847775 | 0.993619256 | -5.146661786 |

| | | | | | | | | | |
|---|---|---|---|---|---|---|---|---|---|
| **A_55_P2048478** | Olfml1 | -0.034632997 | -0.194716447 | 0.125450454 | 8.514228098 | -0.517074077 | 0.62184947 | 0.993619256 | -5.14666285 |
| **A_51_P292757** | Acbd4 | -0.072416827 | -0.407156946 | 0.262323293 | 10.18692462 | -0.517059897 | 0.621858842 | 0.993619256 | -5.146668736 |
| **A_51_P330090** | Trappc9 | -0.028022032 | -0.157551768 | 0.101507703 | 11.40539199 | -0.517058028 | 0.621860078 | 0.993619256 | -5.146669512 |
| **A_55_P2135031** | NA | -0.028621923 | -0.1609399 | 0.103696053 | 7.86766129 | -0.51699828 | 0.621899572 | 0.993619256 | -5.146694313 |
| **A_55_P2157586** | NA | -0.021557687 | -0.121225826 | 0.078110452 | 7.87767794 | -0.516957634 | 0.62192644 | 0.993619256 | -5.146711183 |
| **A_55_P2099952** | Car1 | 0.020952801 | -0.075920347 | 0.117825948 | 7.661290589 | 0.516949123 | 0.621932066 | 0.993619256 | -5.146714715 |
| **A_55_P2167107** | Cdk14 | 0.032978876 | -0.119501258 | 0.18545901 | 10.88318215 | 0.516929973 | 0.621944724 | 0.993619256 | -5.146722662 |
| **A_55_P2103501** | Ankrd29 | 0.031936741 | -0.115727325 | 0.179600806 | 8.564230143 | 0.516921868 | 0.621950082 | 0.993619256 | -5.146726026 |
| **A_51_P374869** | Abca7 | 0.044916287 | -0.162786271 | 0.252618845 | 10.5075772 | 0.516857874 | 0.621992386 | 0.993619256 | -5.146752583 |
| **A_51_P234253** | Sdcbp2 | 0.045204303 | -0.163857319 | 0.254265925 | 8.493622817 | 0.516790586 | 0.622036869 | 0.993619256 | -5.146780503 |
| **A_55_P2000938** | Ahcy | 0.035899239 | -0.130128837 | 0.201927316 | 9.790118846 | 0.516788498 | 0.622038249 | 0.993619256 | -5.146781369 |
| **A_52_P126266** | Prkab2 | 0.038337865 | -0.139013946 | 0.215689676 | 9.732994751 | 0.516655945 | 0.622125883 | 0.993619256 | -5.14683636 |
| **A_55_P2107367** | NA | 0.051222909 | -0.185770269 | 0.288216086 | 11.11210788 | 0.516579924 | 0.622176145 | 0.993619256 | -5.146867893 |
| **A_52_P654437** | NA | -0.09930144 | -0.55879441 | 0.360191529 | 13.69098921 | -0.516518414 | 0.622216815 | 0.993619256 | -5.146893404 |
| **A_52_P234910** | Gm15032 | -0.035084045 | -0.197444585 | 0.127276494 | 12.69566263 | -0.516461928 | 0.622254164 | 0.993619256 | -5.146916828 |
| **A_51_P283473** | Fibin | 0.100305029 | -0.363927304 | 0.564537361 | 9.840373745 | 0.516412153 | 0.622287076 | 0.993619256 | -5.146937467 |
| **A_51_P290685** | Ppp6r3 | 0.041928034 | -0.152157984 | 0.236014051 | 8.939585052 | 0.516320513 | 0.622347674 | 0.993619256 | -5.146975462 |
| **A_55_P1962354** | Lexm | -0.025578701 | -0.143994521 | 0.092837119 | 7.652874495 | -0.516271199 | 0.622380285 | 0.993619256 | -5.146995905 |
| **A_55_P2052470** | Uqcrh | -0.029749364 | -0.167543395 | 0.108044666 | 16.11567525 | -0.516007981 | 0.622554361 | 0.993727868 | -5.147104993 |
| **A_55_P2013336** | Melk | 0.037615728 | -0.136650974 | 0.211882431 | 10.1063698 | 0.515898414 | 0.62262683 | 0.993727868 | -5.147150387 |
| **A_55_P2023114** | Hsd3b2 | -0.021646774 | -0.121933418 | 0.078639869 | 7.669399338 | -0.515892508 | 0.622630737 | 0.993727868 | -5.147152834 |
| **A_55_P2322139** | C530042K1 | 0.071943244 | -0.261382922 | 0.40526941 | 8.702276224 | 0.515857492 | 0.622653898 | 0.993727868 | -5.147167339 |
| **A_51_P473953** | Arhgef26 | -0.029355758 | -0.165398465 | 0.106686949 | 8.156452618 | -0.515735666 | 0.622734482 | 0.993774448 | -5.147217799 |
| **A_52_P370473** | Msl1 | 0.03277348 | -0.119132087 | 0.184679047 | 12.1811062 | 0.515653536 | 0.622788812 | 0.993779127 | -5.14725181 |
| **A_51_P240986** | Plekhg6 | 0.022276383 | -0.081014983 | 0.12556775 | 7.732470754 | 0.515453855 | 0.622920914 | 0.993793378 | -5.147334481 |
| **A_55_P2145531** | Mpst | 0.033086128 | -0.120343991 | 0.186516247 | 11.53553128 | 0.515400048 | 0.622956513 | 0.993793378 | -5.147356754 |
| **A_52_P474765** | Denr | -0.056004939 | -0.315798686 | 0.203788808 | 10.56568075 | -0.515236673 | 0.62306461 | 0.993793378 | -5.147424365 |
| **A_55_P2112967** | Diaph1 | 0.03156037 | -0.114855069 | 0.177975809 | 8.330973506 | 0.515186347 | 0.62309791 | 0.993793378 | -5.147445189 |
| **A_51_P454217** | Nfkb2 | -0.047098735 | -0.265614641 | 0.171417171 | 8.75694445 | -0.515151914 | 0.623120695 | 0.993793378 | -5.147459435 |
| **A_51_P406188** | Asb10 | -0.06836405 | -0.385588608 | 0.248860507 | 9.730918658 | -0.51507458 | 0.623171869 | 0.993793378 | -5.147491428 |
| **A_51_P148494** | Rnft1 | -0.026336862 | -0.148550908 | 0.095877184 | 9.073921451 | -0.515053137 | 0.623186059 | 0.993793378 | -5.147500298 |
| **A_55_P2003338** | Clint1 | -0.030910617 | -0.174391508 | 0.112570274 | 10.23155837 | -0.514899823 | 0.623287519 | 0.993793378 | -5.147563708 |
| **A_52_P72587** | Prkcq | 0.026705953 | -0.097276328 | 0.150688233 | 7.854091585 | 0.514822583 | 0.623338638 | 0.993793378 | -5.147595648 |
| **A_55_P1990735** | Ift122 | -0.032220534 | -0.181865391 | 0.117424323 | 9.744642919 | -0.51461219 | 0.623477893 | 0.993793378 | -5.147682626 |
| **A_55_P1987459** | Pigt | -0.022916961 | -0.129367583 | 0.083533661 | 8.100476163 | -0.514538594 | 0.623526608 | 0.993793378 | -5.147713044 |
| **A_51_P125648** | Vwce | 0.022069641 | -0.080447817 | 0.1245871 | 7.859630936 | 0.514525118 | 0.623535529 | 0.993793378 | -5.141966969 |
| **A_55_P1999324** | Nudt6 | -0.028946563 | -0.163425471 | 0.105532345 | 10.0838422 | -0.514460415 | 0.62357836 | 0.993793378 | -5.147745352 |
| **A_66_P122362** | Aldh16a1 | 0.033618389 | -0.122565968 | 0.189802746 | 11.83337296 | 0.514456312 | 0.623581076 | 0.993793378 | -5.147747048 |
| **A_55_P2127497** | Trim35 | 0.030860007 | -0.112517239 | 0.174237254 | 13.91231098 | 0.514428379 | 0.623599567 | 0.993793378 | -5.14775859 |
| **A_55_P2077671** | NA | -0.090420084 | -0.510647009 | 0.329806841 | 13.20366167 | -0.514268783 | 0.623705222 | 0.993793378 | -5.147824525 |
| **A_55_P2017645** | Tap2 | -0.094276871 | -0.532587043 | 0.344033301 | 9.357343731 | -0.51408241 | 0.623828616 | 0.993793378 | -5.147901501 |
| **A_51_P332141** | Clcn2 | -0.04422402 | -0.249893326 | 0.161445285 | 8.308800324 | -0.513922785 | 0.623934312 | 0.993793378 | -5.147967409 |
| **A_51_P254805** | Kif4 | -0.023882283 | -0.134968531 | 0.087203964 | 9.647589472 | -0.513836183 | 0.623991659 | 0.993793378 | -5.148003158 |
| **A_55_P1995892** | NA | 0.028298934 | -0.103349387 | 0.159947256 | 9.219594773 | 0.513764253 | 0.624039292 | 0.993793378 | -5.148032847 |
| **A_55_P2049757** | NA | -0.030396142 | -0.171800994 | 0.111008711 | 14.54986045 | -0.513763535 | 0.624039768 | 0.993793378 | -5.148033143 |
| **A_55_P2086692** | Nfib | 0.028084051 | -0.102572835 | 0.158740937 | 11.85282497 | 0.513731961 | 0.624060678 | 0.993793378 | -5.148046174 |
| **A_55_P2069052** | Sacs | 0.049046368 | -0.179138762 | 0.277231497 | 8.698469355 | 0.513722564 | 0.624066901 | 0.993793378 | -5.148050052 |
| **A_55_P1964228** | Elac1 | 0.02684149 | -0.098038254 | 0.151721234 | 8.274609418 | 0.513716778 | 0.624070733 | 0.993793378 | -5.14805244 |

| | | | | | | | | | |
|---|---|---|---|---|---|---|---|---|---|
| **A_55_P1984386** | Depdc5 | 0.029624033 | -0.108237072 | 0.167485137 | 8.878077406 | 0.513584084 | 0.624158613 | 0.993793378 | -5.148107195 |
| **A_51_P265806** | Clca3a2 | 0.076819616 | -0.280680092 | 0.434319325 | 9.988127143 | 0.513577011 | 0.624163298 | 0.993793378 | -5.148110113 |
| **A_55_P2022114** | NA | 0.028958573 | -0.105857971 | 0.163775117 | 8.175545573 | 0.513384913 | 0.624290533 | 0.993793378 | -5.148189356 |
| **A_52_P285470** | Lrp2 | 0.063292481 | -0.231408984 | 0.357993945 | 8.066639863 | 0.51330909 | 0.624340758 | 0.993793378 | -5.148220627 |
| **A_55_P2104119** | NA | 0.035440002 | -0.129624504 | 0.200504509 | 8.773824207 | 0.513155724 | 0.624442353 | 0.993793378 | -5.148283864 |
| **A_55_P2021049** | Ugp2 | -0.030898969 | -0.174865645 | 0.113067706 | 12.72669535 | -0.512969034 | 0.624566035 | 0.993793378 | -5.14836082 |
| **A_51_P494825** | Mfhas1 | -0.033432254 | -0.18921354 | 0.122349032 | 12.38433841 | -0.512931656 | 0.6245908 | 0.993793378 | -5.148376224 |
| **A_55_P2106394** | Knstrn | -0.028137004 | -0.15930266 | 0.103028652 | 12.73653092 | -0.512704166 | 0.624741533 | 0.993793378 | -5.148469958 |
| **A_51_P223443** | Cd81 | 0.029434226 | -0.107849512 | 0.166717963 | 15.37709805 | 0.512439565 | 0.62491688 | 0.993793378 | -5.148578935 |
| **A_55_P2039379** | Slc27a6 | 0.0215725 | -0.079066297 | 0.122211296 | 7.736153296 | 0.512323372 | 0.624993888 | 0.993793378 | -5.148626773 |
| **A_52_P220573** | Serpini1 | -0.031606883 | -0.179105031 | 0.115891264 | 8.563249131 | -0.512158329 | 0.62510328 | 0.993793378 | -5.148694707 |
| **A_51_P455946** | Rac3 | 0.030506338 | -0.111867815 | 0.172880492 | 8.293606861 | 0.512115664 | 0.62513156 | 0.993793378 | -5.148712265 |
| **A_51_P275642** | NA | -0.021513107 | -0.121955758 | 0.078929543 | 7.928370018 | -0.511910589 | 0.625267504 | 0.993793378 | -5.148796643 |
| **A_51_P181341** | Necab2 | 0.021740472 | -0.079777494 | 0.123258438 | 7.736793129 | 0.511841145 | 0.625313541 | 0.993793378 | -5.148825209 |
| **A_55_P2102540** | Olfml3 | -0.029621345 | -0.167966574 | 0.108723883 | 7.956258383 | -0.511740429 | 0.625380314 | 0.993793378 | -5.148866632 |
| **A_55_P2061428** | Fads6 | 0.022456381 | -0.082436291 | 0.127349052 | 7.778595195 | 0.511686257 | 0.625416231 | 0.993793378 | -5.14888891 |
| **A_51_P283649** | Brat1 | 0.041176427 | -0.15120042 | 0.233553274 | 10.31883866 | 0.511569908 | 0.625493375 | 0.993793378 | -5.148936748 |
| **A_65_P08971** | F3 | 0.021693921 | -0.079666762 | 0.123054605 | 7.790189981 | 0.511537714 | 0.625514721 | 0.993793378 | -5.148949984 |
| **A_66_P122559** | Myct1 | 0.038811855 | -0.142550606 | 0.220174316 | 9.028072206 | 0.511477025 | 0.625554964 | 0.993793378 | -5.148974932 |
| **A_51_P112445** | Wdr3 | 0.034139974 | -0.125402805 | 0.193682753 | 9.588705637 | 0.51144056 | 0.625579143 | 0.993793378 | -5.125532028 |
| **A_55_P2131522** | Arfrp1 | 0.031691962 | -0.116425946 | 0.17980987 | 11.74160987 | 0.511388182 | 0.625613876 | 0.993793378 | -5.149011449 |
| **A_55_P2257341** | 2410004I0 | 0.023015223 | -0.084570427 | 0.130600873 | 7.755326679 | 0.511293149 | 0.625676898 | 0.993793378 | -5.149050504 |
| **A_51_P252363** | Stx12 | -0.041025192 | -0.232832442 | 0.150782058 | 11.52928513 | -0.51120458 | 0.625735635 | 0.993793378 | -5.149086896 |
| **A_52_P175679** | Gm13023 | -0.025654763 | -0.145620742 | 0.094311216 | 8.015812853 | -0.511115478 | 0.625794729 | 0.993793378 | -5.149123502 |
| **A_55_P2048011** | Larp4b | 0.038158534 | -0.140291657 | 0.216608724 | 12.68216788 | 0.511074005 | 0.625822235 | 0.993793378 | -5.149140538 |
| **A_55_P2013143** | Esrp2 | 0.020941819 | -0.077002981 | 0.118886618 | 7.668637388 | 0.511025004 | 0.625854736 | 0.993793378 | -5.149160665 |
| **A_55_P2086500** | Dennd5a | -0.039132372 | -0.222217012 | 0.143952268 | 13.49036243 | -0.510849995 | 0.625970819 | 0.993793378 | -5.149232536 |
| **A_55_P2151116** | Micu1 | -0.027147323 | -0.158069369 | 0.103774724 | 11.46767149 | -0.515087925 | 0.626010549 | 0.993793378 | -5.073149714 |
| **A_51_P434426** | Rwdd4a | 0.049263862 | -0.181251092 | 0.279778816 | 9.77670813 | 0.510785413 | 0.626013659 | 0.993793378 | -5.149259052 |
| **A_55_P2001998** | Zfp503 | 0.028386153 | -0.104443842 | 0.161216149 | 8.350217584 | 0.510763103 | 0.626028458 | 0.993793378 | -5.149268211 |
| **A_55_P2110290** | Tmem254a | -0.041124326 | -0.233579337 | 0.151330686 | 9.389893875 | -0.510715099 | 0.626060303 | 0.993793378 | -5.149287918 |
| **A_55_P1993880** | Rps24 | -0.043760356 | -0.248589623 | 0.161068911 | 14.66331758 | -0.510620173 | 0.626123277 | 0.993793378 | -5.149326883 |
| **A_52_P508317** | Erlec1 | 0.078149047 | -0.287905206 | 0.4442033 | 9.051216111 | 0.510255108 | 0.626365493 | 0.993793378 | -5.14947667 |
| **A_55_P2154228** | Ncapd3 | 0.044017219 | -0.162200877 | 0.250235314 | 10.40755134 | 0.510158301 | 0.626429731 | 0.993793378 | -5.149516374 |
| **A_55_P1970474** | Tmem67 | -0.023572129 | -0.13402384 | 0.086879582 | 8.161882465 | -0.510076704 | 0.626483879 | 0.993793378 | -5.149549835 |
| **A_52_P65719** | Eme2 | 0.036784319 | -0.135587379 | 0.209156016 | 9.012496464 | 0.510041939 | 0.626506951 | 0.993793378 | -5.14956409 |
| **A_55_P1954006** | Ldhb | -0.046346435 | -0.263604936 | 0.170912066 | 15.90629834 | -0.509857354 | 0.626629454 | 0.993793378 | -5.14963976 |
| **A_51_P335460** | Scin | 0.032783512 | -0.120903209 | 0.186470232 | 8.631840822 | 0.509833376 | 0.626645369 | 0.993793378 | -5.149649588 |
| **A_55_P2176391** | NA | -0.039310827 | -0.223686117 | 0.145064464 | 11.63595029 | -0.509587293 | 0.626808709 | 0.993793378 | -5.149750427 |
| **A_52_P669964** | Cchcr1 | -0.053457529 | -0.304192172 | 0.197277114 | 8.869309267 | -0.509569785 | 0.62682033 | 0.993793378 | -5.149757599 |
| **A_51_P211506** | Muc20 | -0.025119072 | -0.142963689 | 0.092725544 | 7.983533921 | -0.509451661 | 0.626898745 | 0.993793378 | -5.149805987 |
| **A_55_P1970017** | Zer1 | 0.043766715 | -0.161567819 | 0.249101249 | 12.18805412 | 0.509437708 | 0.626908008 | 0.993793378 | -5.149811702 |
| **A_55_P2036952** | NA | -0.023727114 | -0.13505392 | 0.087599691 | 8.337850511 | -0.509394568 | 0.626936648 | 0.993793378 | -5.14982937 |
| **A_55_P2060592** | Hoxa1 | 0.045514717 | -0.168062261 | 0.259091696 | 8.059624722 | 0.509338547 | 0.626973839 | 0.993793378 | -5.149852312 |
| **A_55_P1965592** | NA | -0.025040496 | -0.142550001 | 0.092469008 | 8.739863065 | -0.509306326 | 0.626995231 | 0.993793378 | -5.149865506 |
| **A_55_P1961395** | Pdpn | 0.09201474 | -0.339861177 | 0.523890657 | 10.88199854 | 0.509222455 | 0.627050915 | 0.993793378 | -5.149899847 |
| **A_55_P1981780** | Cul4b | 0.034156105 | -0.126158994 | 0.194471203 | 9.75952818 | 0.509217172 | 0.627054422 | 0.993793378 | -5.14990201 |
| **A_55_P2180406** | Cdkn2aipnl | -0.031354881 | -0.178591287 | 0.115881526 | 8.840367579 | -0.508978073 | 0.627213181 | 0.993793378 | -5.149999881 |

| | | | | | | | | | |
|---|---|---|---|---|---|---|---|---|---|
| **A_55_P2070179** | Ing4 | -0.062139212 | -0.353937123 | 0.2296587 | 11.09532133 | -0.508970596 | 0.627218146 | 0.993793378 | -5.150002941 |
| **A_55_P1996553** | Ttc17 | 0.031123837 | -0.115048684 | 0.177296358 | 10.72718277 | 0.508904775 | 0.627261855 | 0.993793378 | -5.150029876 |
| **A_55_P2154416** | Car15 | 0.046776156 | -0.172922494 | 0.266474806 | 8.688020504 | 0.508869328 | 0.627285394 | 0.993793378 | -5.15004438 |
| **A_55_P2118799** | Rmi1 | 0.032359558 | -0.119628391 | 0.184347507 | 8.658135075 | 0.508864981 | 0.627288281 | 0.993793378 | -5.150046158 |
| **A_55_P2093665** | NA | 0.027277945 | -0.100844093 | 0.155399983 | 8.594984901 | 0.50885839 | 0.627292658 | 0.993793378 | -5.150048855 |
| **A_55_P2090142** | Kcna7 | -0.029094017 | -0.165751515 | 0.10756348 | 8.677275229 | -0.508837796 | 0.627306334 | 0.993793378 | -5.150057281 |
| **A_52_P331901** | Trp53rkb | 0.03316513 | -0.122616674 | 0.188946934 | 8.340965295 | 0.508831636 | 0.627310424 | 0.993793378 | -5.150059801 |
| **A_52_P552017** | Rbsn | 0.027454251 | -0.101525712 | 0.156434215 | 9.15028525 | 0.508740706 | 0.627370812 | 0.993793378 | -5.150097001 |
| **A_55_P1991585** | Blvrb | -0.037722242 | -0.215090872 | 0.139646387 | 10.70994454 | -0.508311369 | 0.627655979 | 0.993793378 | -5.150272564 |
| **A_51_P149623** | Gabrd | 0.020432687 | -0.075643193 | 0.116508568 | 7.718490872 | 0.508300162 | 0.627663423 | 0.993793378 | -5.150277144 |
| **A_55_P2103944** | Mrpl30 | -0.032250168 | -0.183932979 | 0.119432643 | 13.42860138 | -0.508164992 | 0.627753218 | 0.993793378 | -5.150332388 |
| **A_66_P135176** | Trak2 | -0.055991504 | -0.319343348 | 0.20736034 | 9.63326306 | -0.508153477 | 0.627760868 | 0.993793378 | -5.150337094 |
| **A_51_P299062** | Kel | 0.052202532 | -0.193383428 | 0.297788491 | 8.055505089 | 0.508039129 | 0.627836837 | 0.993793378 | -5.150383817 |
| **A_55_P2196027** | NA | -0.031390949 | -0.179070343 | 0.116288445 | 9.726923173 | -0.508035055 | 0.627839543 | 0.993793378 | -5.150385481 |
| **A_52_P486702** | G3bp1 | -0.03210872 | -0.183181593 | 0.118964154 | 9.632358574 | -0.507978849 | 0.627876886 | 0.993793378 | -5.150408444 |
| **A_55_P2060962** | Tm2d3 | 0.056961902 | -0.211188386 | 0.32511219 | 10.78107184 | 0.507709564 | 0.628055815 | 0.993793378 | -5.150518425 |
| **A_55_P2286493** | 4933405E2 | -0.054945663 | -0.313646647 | 0.203755321 | 8.95391543 | -0.507626715 | 0.62811087 | 0.993793378 | -5.150552251 |
| **A_55_P2042993** | Arl6ip6 | 0.045629895 | -0.169220933 | 0.260480724 | 13.71753996 | 0.507599951 | 0.628128656 | 0.993793378 | -5.150563177 |
| **A_55_P2030284** | Trem1 | -0.025021025 | -0.146477369 | 0.096435319 | 9.117397629 | -0.511743152 | 0.628205087 | 0.993793378 | -5.074459072 |
| **A_55_P2319370** | Plekha3 | 0.04854266 | -0.180101433 | 0.277186754 | 10.48288942 | 0.507426003 | 0.628244258 | 0.993793378 | -5.150634179 |
| **A_55_P2084646** | Csrnp3 | -0.03281679 | -0.187390879 | 0.121757299 | 8.111786224 | -0.507421114 | 0.628247507 | 0.993793378 | -5.150636174 |
| **A_52_P625270** | 1700024G1 | 0.021719166 | -0.080610654 | 0.124048986 | 7.685934855 | 0.507282723 | 0.628339488 | 0.993793378 | -5.150692646 |
| **A_55_P2037424** | NA | -0.056424578 | -0.322307971 | 0.209458814 | 15.91459554 | -0.507208172 | 0.628389041 | 0.993793378 | -5.150723061 |
| **A_51_P499776** | Denr | 0.024646071 | -0.091516067 | 0.140808209 | 8.451233065 | 0.507098386 | 0.628462017 | 0.993793378 | -5.150767845 |
| **A_55_P2068289** | Slc17a2 | -0.02064519 | -0.117956231 | 0.076665851 | 7.645890265 | -0.507067636 | 0.628482458 | 0.993793378 | -5.150780387 |
| **A_52_P323852** | Tnfrsf22 | -0.023775923 | -0.135856243 | 0.088304398 | 7.798143069 | -0.507010729 | 0.628520287 | 0.993793378 | -5.150803595 |
| **A_66_P108822** | NA | 0.027397549 | -0.10175782 | 0.156552918 | 15.49898917 | 0.507000491 | 0.628527093 | 0.993793378 | -5.15080777 |
| **A_55_P2020743** | Abcc10 | 0.038946284 | -0.144691893 | 0.222584462 | 9.17549619 | 0.506888206 | 0.62860174 | 0.993793378 | -5.150853556 |
| **A_51_P437478** | Zfp566 | 0.038411116 | -0.142713314 | 0.219535545 | 9.288406517 | 0.506861172 | 0.628619712 | 0.993793378 | -5.150864578 |
| **A_52_P1055590** | Bnip1 | -0.03009115 | -0.172019508 | 0.111837208 | 10.56764522 | -0.506732473 | 0.628705278 | 0.993793378 | -5.150917043 |
| **A_55_P1974442** | Sumf2 | -0.029980446 | -0.171417662 | 0.111456769 | 9.081354324 | -0.506621402 | 0.628779128 | 0.993793378 | -5.150962312 |
| **A_55_P2067112** | Gm4924 | 0.061454635 | -0.228508625 | 0.351417894 | 10.32710751 | 0.506548228 | 0.628827783 | 0.993793378 | -5.150992131 |
| **A_55_P2037478** | Kifap3 | -0.029513829 | -0.168815857 | 0.1097882 | 11.21680939 | -0.506380814 | 0.628939109 | 0.993793378 | -5.151060338 |
| **A_51_P181722** | Rbks | -0.030307808 | -0.173368871 | 0.112753254 | 9.084638928 | -0.506339977 | 0.628966266 | 0.993793378 | -5.151076972 |
| **A_52_P357484** | Nfrkb | -0.025707793 | -0.147107505 | 0.095691919 | 8.497769542 | -0.506123231 | 0.629110416 | 0.993793378 | -5.151165241 |
| **A_55_P2084706** | Acaca | 0.04530032 | -0.168626611 | 0.25922725 | 10.71913101 | 0.506110024 | 0.629119199 | 0.993793378 | -5.151170618 |
| **A_52_P95759** | Sh3rf3 | -0.024600062 | -0.140796444 | 0.09159632 | 7.781788698 | -0.506002577 | 0.629190665 | 0.993793378 | -5.151214362 |
| **A_52_P83623** | Rapgef6 | 0.027885173 | -0.103853409 | 0.159623755 | 8.934874877 | 0.505905599 | 0.629255171 | 0.993793378 | -5.151253836 |
| **A_55_P1977812** | Cd244a | 0.05809503 | -0.216403992 | 0.332594052 | 8.780683981 | 0.505833175 | 0.629303347 | 0.993793378 | -5.151283311 |
| **A_51_P351238** | Plgrkt | -0.035411694 | -0.202734349 | 0.131910961 | 10.74682091 | -0.50582593 | 0.629308166 | 0.993793378 | -5.151286259 |
| **A_55_P2185651** | NA | 0.031543058 | -0.117501112 | 0.180587228 | 8.679701794 | 0.505822255 | 0.629310611 | 0.993793378 | -5.151287754 |
| **A_52_P579531** | Pdlim3 | 0.055847563 | -0.208093604 | 0.319788731 | 12.85356982 | 0.505715457 | 0.629381657 | 0.993793378 | -5.151331212 |
| **A_55_P2060369** | Akap9 | 0.050049878 | -0.1864949 | 0.286594657 | 10.6120284 | 0.505706799 | 0.629387416 | 0.993793378 | -5.151334734 |
| **A_55_P1960708** | Setd1a | 0.033363531 | -0.124327535 | 0.191054597 | 10.09131787 | 0.505677987 | 0.629406584 | 0.993793378 | -5.151346456 |
| **A_52_P122696** | Arl2 | -0.081255055 | -0.465330223 | 0.302820113 | 9.681854384 | -0.505642176 | 0.629430408 | 0.993793378 | -5.151361025 |
| **A_55_P2012051** | Sox1 | 0.031497392 | -0.117385552 | 0.180380336 | 8.223723644 | 0.505636921 | 0.629433904 | 0.993793378 | -5.151363163 |
| **A_55_P2106078** | F10 | -0.030377625 | -0.173973376 | 0.113218126 | 17.39613634 | -0.505616643 | 0.629447395 | 0.993793378 | -5.151371412 |
| **A_55_P2002632** | Rap1b | 0.039234014 | -0.146270725 | 0.224738752 | 12.49670282 | 0.505494997 | 0.629528327 | 0.993793378 | -5.151420892 |

| | | | | | | | | | |
|---|---|---|---|---|---|---|---|---|---|
| **A_55_P2086099** | Gm2061 | 0.022887311 | -0.085344798 | 0.131119419 | 7.733208674 | 0.505414592 | 0.629581825 | 0.993793378 | -5.151453591 |
| **A_55_P2065971** | Setd3 | 0.025261297 | -0.094217588 | 0.144740181 | 11.39016809 | 0.505328292 | 0.629639247 | 0.993793378 | -5.151488682 |
| **A_55_P2186619** | Rbmxl1 | 0.038967839 | -0.145346011 | 0.223281689 | 12.79452176 | 0.505309523 | 0.629651735 | 0.993793378 | -5.151496313 |
| **A_55_P2066096** | Zfand3 | -0.031029378 | -0.177800564 | 0.115741807 | 12.29547823 | -0.505290813 | 0.629664186 | 0.993793378 | -5.15150392 |
| **A_51_P113195** | Upk1b | 0.08822653 | -0.329131432 | 0.505584492 | 8.359191714 | 0.505242211 | 0.629696526 | 0.993793378 | -5.151523679 |
| **A_51_P169880** | Zbtb3 | 0.024404781 | -0.091087391 | 0.139896954 | 8.590153224 | 0.505046658 | 0.62982666 | 0.993793378 | -5.151603162 |
| **A_65_P08022** | Hnrnpk | 0.022254655 | -0.083071894 | 0.127581204 | 10.80946343 | 0.505000897 | 0.629857114 | 0.993793378 | -5.151621757 |
| **A_55_P1956195** | Sap18b | -0.027691366 | -0.158776023 | 0.103393292 | 13.02202769 | -0.504895669 | 0.629927147 | 0.993793378 | -5.151664512 |
| **A_52_P651248** | Zdhhc2 | 0.052925396 | -0.197642077 | 0.303492868 | 9.624431029 | 0.504833949 | 0.629968226 | 0.993793378 | -5.151689586 |
| **A_51_P453010** | Mphosph6 | 0.043832918 | -0.163721184 | 0.25138702 | 9.996487242 | 0.504752166 | 0.63002266 | 0.993793378 | -5.151722806 |
| **A_55_P1955393** | 7-Sep | -0.024826266 | -0.142415906 | 0.092763375 | 12.96611963 | -0.504604901 | 0.630120686 | 0.993793378 | -5.151782612 |
| **A_55_P1970866** | Rplp0 | 0.039297707 | -0.146846389 | 0.225441802 | 16.26851464 | 0.504576564 | 0.630139549 | 0.993793378 | -5.151794118 |
| **A_51_P386612** | Pir | -0.02657568 | -0.152463519 | 0.099312158 | 8.339528448 | -0.504556427 | 0.630152953 | 0.993793378 | -5.151802294 |
| **A_55_P1959683** | Xlr3b | 0.038342506 | -0.143300265 | 0.219985277 | 7.916210644 | 0.504512006 | 0.630182524 | 0.993793378 | -5.15182033 |
| **A_52_P311297** | Als2 | -0.02386513 | -0.136934108 | 0.089203849 | 8.100043547 | -0.50446317 | 0.630215035 | 0.993793378 | -5.151840156 |
| **A_55_P1987043** | Tpp1 | 0.028378191 | -0.106090054 | 0.162846436 | 9.456635437 | 0.504398869 | 0.630257842 | 0.993793378 | -5.151866258 |
| **A_66_P111385** | Klhl28 | 0.036906708 | -0.138000651 | 0.211814067 | 10.25571699 | 0.504320199 | 0.630310217 | 0.993793378 | -5.151898188 |
| **A_51_P479321** | Acss1 | -0.031098785 | -0.178487015 | 0.116289445 | 11.40784517 | -0.504300908 | 0.630323061 | 0.993793378 | -5.151906017 |
| **A_55_P2013586** | Prss8 | -0.021358506 | -0.122616126 | 0.079899113 | 7.695339375 | -0.504141326 | 0.630429312 | 0.993793378 | -5.151970773 |
| **A_51_P480499** | Emc8 | 0.062949595 | -0.235487645 | 0.361386835 | 10.57972512 | 0.504137562 | 0.630431818 | 0.993793378 | -5.1519723 |
| **A_51_P175758** | Atg101 | -0.045729193 | -0.262531789 | 0.171073404 | 11.49047867 | -0.504124944 | 0.63044022 | 0.993793378 | -5.151977419 |
| **A_55_P2133205** | Nmt2 | 0.046066379 | -0.17233781 | 0.264470567 | 8.719636617 | 0.504118043 | 0.630444815 | 0.993793378 | -5.151980219 |
| **A_66_P116632** | Mafg | 0.035843265 | -0.134099248 | 0.205785779 | 11.43658415 | 0.504097635 | 0.630458403 | 0.993793378 | -5.151988498 |
| **A_52_P217875** | H3f3b | -0.024211584 | -0.139020041 | 0.090596873 | 8.123797376 | -0.504032408 | 0.630501836 | 0.993793378 | -5.152014958 |
| **A_55_P2008687** | Nlk | -0.030044147 | -0.172536879 | 0.112448584 | 10.67821675 | -0.503937071 | 0.63056532 | 0.993793378 | -5.152053628 |
| **A_55_P1970740** | Mrps10 | 0.060130912 | -0.225214043 | 0.345475867 | 11.64389246 | 0.503659144 | 0.63075041 | 0.993793378 | -5.152166319 |
| **A_65_P03022** | Cdc14b | -0.033568626 | -0.192885801 | 0.125748549 | 9.284217502 | -0.503593479 | 0.630794145 | 0.993793378 | -5.152192936 |
| **A_55_P2021029** | Tigd3 | -0.024709173 | -0.142001209 | 0.092582863 | 8.154003074 | -0.503499239 | 0.630856914 | 0.993793378 | -5.15223113 |
| **A_66_P131389** | Ugt2b38 | -0.022111308 | -0.127074516 | 0.082851901 | 7.700391751 | -0.50348491 | 0.630866458 | 0.993793378 | -5.152236937 |
| **A_55_P1962982** | NA | 0.036853365 | -0.138117568 | 0.211824299 | 10.49547806 | 0.503408302 | 0.630917487 | 0.993793378 | -5.152267979 |
| **A_55_P2064578** | Prkci | 0.053113004 | -0.199084833 | 0.30531084 | 11.68072268 | 0.503348336 | 0.630957431 | 0.993793378 | -5.152292275 |
| **A_65_P12977** | Dnaja2 | -0.040765115 | -0.234338078 | 0.152807848 | 11.89871451 | -0.503330339 | 0.63096942 | 0.993793378 | -5.152299567 |
| **A_51_P513244** | Zfp949 | -0.026195268 | -0.150584444 | 0.098193907 | 8.237992962 | -0.503326039 | 0.630972284 | 0.993793378 | -5.152301309 |
| **A_55_P2133405** | Dmkn | 0.197567337 | -0.740872865 | 1.136007539 | 9.0673561 | 0.5031734 | 0.631073968 | 0.993793378 | -5.152363138 |
| **A_52_P645159** | Cpsf6 | 0.038701166 | -0.145143246 | 0.222545578 | 8.984311681 | 0.503132941 | 0.631100922 | 0.993793378 | -5.152379524 |
| **A_52_P214752** | NA | -0.030174724 | -0.173531914 | 0.113182466 | 9.106830794 | -0.50307526 | 0.63113935 | 0.993793378 | -5.152402883 |
| **A_52_P38964** | Sap25 | 0.044106893 | -0.165464296 | 0.253678083 | 8.393856193 | 0.503018577 | 0.631177115 | 0.993793378 | -5.152425835 |
| **A_51_P217395** | Cyp2d34 | 0.021018546 | -0.078858681 | 0.120895773 | 7.631862773 | 0.50297375 | 0.631206982 | 0.993793378 | -5.152443985 |
| **A_51_P397920** | Ilrun | -0.031410038 | -0.180672183 | 0.117852107 | 11.3415815 | -0.502953597 | 0.631220409 | 0.993793378 | -5.152452144 |
| **A_55_P1971328** | Lage3 | 0.035377228 | -0.132753937 | 0.203508392 | 11.9234846 | 0.502903557 | 0.63125375 | 0.993793378 | -5.152472402 |
| **A_52_P664699** | Cul1 | 0.026656535 | -0.100039265 | 0.153352334 | 8.970303295 | 0.502864069 | 0.631280061 | 0.993793378 | -5.152488387 |
| **A_51_P257885** | Mmd2 | 0.021189709 | -0.079531031 | 0.121910448 | 7.648649294 | 0.502823085 | 0.631307369 | 0.993793378 | -5.152504976 |
| **A_55_P1986296** | Tagln2 | 0.040296857 | -0.151499575 | 0.23209329 | 15.24517365 | 0.502157304 | 0.63175108 | 0.993793378 | -5.152774295 |
| **A_55_P2000454** | Tspan18 | 0.054191214 | -0.203745059 | 0.312127487 | 10.36710606 | 0.502140895 | 0.631762018 | 0.993793378 | -5.152780929 |
| **A_55_P2065874** | D430042O( | 0.039048398 | -0.146816468 | 0.224913265 | 9.06883954 | 0.50212871 | 0.631770141 | 0.993793378 | -5.152785855 |
| **A_52_P272054** | NA | -0.104922313 | -0.604364686 | 0.39452006 | 12.08036408 | -0.502101598 | 0.631788213 | 0.993793378 | -5.152796815 |
| **A_52_P266320** | Akap17b | -0.021994962 | -0.126698591 | 0.082708667 | 8.494573551 | -0.502077337 | 0.631804385 | 0.993793378 | -5.152806622 |
| **A_55_P1972530** | Efl1 | 0.040375602 | -0.151830233 | 0.232581437 | 10.1336095 | 0.502066885 | 0.631811353 | 0.993793378 | -5.152810846 |

| | | | | | | | | | |
|---|---|---|---|---|---|---|---|---|---|
| **A_51_P311362** | 4930449E0 | -0.024263744 | -0.139803518 | 0.09127603 | 7.785220762 | -0.501921072 | 0.631908557 | 0.993793378 | -5.152869777 |
| **A_55_P2009861** | NA | 0.100524869 | -0.378162757 | 0.579212496 | 11.8747263 | 0.501915316 | 0.631912394 | 0.993793378 | -5.152872103 |
| **A_51_P379069** | Dkk1 | 0.022319116 | -0.083963868 | 0.128602101 | 7.659050081 | 0.501905999 | 0.631918605 | 0.993793378 | -5.152875868 |
| **A_51_P256945** | Opa3 | 0.117316295 | -0.441341605 | 0.675974196 | 9.152125716 | 0.501905006 | 0.631919268 | 0.993793378 | -5.15287627 |
| **A_66_P109368** | Prex2 | 0.023720282 | -0.089282205 | 0.136722768 | 7.97262131 | 0.501696385 | 0.632058357 | 0.993793378 | -5.152960555 |
| **A_55_P2117202** | Mtx1 | 0.028736513 | -0.108174023 | 0.165647049 | 11.92763269 | 0.501656365 | 0.63208504 | 0.993793378 | -5.15297672 |
| **A_55_P1975415** | Cpped1 | -0.036818315 | -0.212246298 | 0.138609668 | 12.70221945 | -0.501619226 | 0.632109804 | 0.993793378 | -5.15299172 |
| **A_51_P331385** | Paip2b | -0.02839517 | -0.163723605 | 0.106933264 | 9.192638694 | -0.501492628 | 0.632194219 | 0.993793378 | -5.153042845 |
| **A_51_P441974** | Pip5k1a | -0.028154985 | -0.16233994 | 0.106029969 | 10.13150215 | -0.501488078 | 0.632197253 | 0.993793378 | -5.153044682 |
| **A_51_P288447** | Nup133 | 0.025923529 | -0.09763097 | 0.149478028 | 11.33029681 | 0.501469695 | 0.632209511 | 0.993793378 | -5.153052104 |
| **A_55_P2076360** | NA | -0.022582842 | -0.130226815 | 0.085061131 | 8.014731066 | -0.501415792 | 0.632245456 | 0.993793378 | -5.153073868 |
| **A_55_P2007500** | Tmem151b | 0.037210312 | -0.140162041 | 0.214582664 | 8.417408696 | 0.501402521 | 0.632254305 | 0.993793378 | -5.153079226 |
| **A_55_P2108248** | Art4 | 0.054743471 | -0.206237344 | 0.315724285 | 11.8206639 | 0.5013406 | 0.632295599 | 0.993793378 | -5.153104223 |
| **A_51_P409988** | Arl13b | -0.032714857 | -0.188726893 | 0.123297179 | 8.642216517 | -0.501182686 | 0.632400913 | 0.993793378 | -5.15316796 |
| **A_55_P1994348** | Golim4 | 0.050440463 | -0.190162271 | 0.291043196 | 11.00089553 | 0.501057579 | 0.632484353 | 0.993793378 | -5.153218443 |
| **A_55_P1959303** | Rbm47 | -0.020829033 | -0.120202658 | 0.078544592 | 7.762813783 | -0.500964681 | 0.632546317 | 0.993793378 | -5.153255921 |
| **A_51_P209183** | Cxcl14 | 0.025055728 | -0.094485119 | 0.144596574 | 7.775294982 | 0.500956282 | 0.632551919 | 0.993793378 | -5.153259309 |
| **A_55_P1999847** | Nutf2 | 0.043919644 | -0.165676897 | 0.253516185 | 13.7996483 | 0.5008225 | 0.632641157 | 0.993793378 | -5.15331327 |
| **A_55_P2132517** | Gmppa | 0.056583281 | -0.213471686 | 0.326638247 | 9.609912907 | 0.500777821 | 0.632670962 | 0.993793378 | -5.153331288 |
| **A_52_P228318** | Smu1 | -0.02278323 | -0.131520789 | 0.085954329 | 12.90378211 | -0.500777541 | 0.632671149 | 0.993793378 | -5.153331401 |
| **A_55_P2083566** | NA | 0.03623001 | -0.136763422 | 0.209223441 | 15.79699557 | 0.50055058 | 0.632822561 | 0.993793378 | -5.153422908 |
| **A_51_P278540** | Myg1 | 0.046917517 | -0.177165977 | 0.271001011 | 9.86570599 | 0.500419568 | 0.632909971 | 0.993793378 | -5.153475712 |
| **A_55_P2100065** | Fsd1l | 0.042107659 | -0.15900585 | 0.243221167 | 9.115809319 | 0.500413461 | 0.632914046 | 0.993793378 | -5.153478173 |
| **A_55_P2137461** | NA | 0.033890551 | -0.128033552 | 0.195814654 | 15.59463586 | 0.500237519 | 0.633031444 | 0.993793378 | -5.153549067 |
| **A_51_P407004** | Prkrip1 | -0.038649506 | -0.22331563 | 0.146016617 | 10.40892901 | -0.500225528 | 0.633039446 | 0.993793378 | -5.153553898 |
| **A_55_P2211319** | Fgf18 | 0.05332174 | -0.201470991 | 0.308114471 | 10.76961261 | 0.5001801 | 0.63306976 | 0.993793378 | -5.153572198 |
| **A_55_P2198946** | Gm10768 | 0.020467095 | -0.077333533 | 0.118267724 | 7.653567278 | 0.500176983 | 0.63307184 | 0.993793378 | -5.153573454 |
| **A_51_P392687** | Vim | -0.037792043 | -0.218392887 | 0.142808802 | 15.41167198 | -0.500137849 | 0.633097954 | 0.993793378 | -5.153589217 |
| **A_55_P2014315** | NA | 0.046026289 | -0.173927789 | 0.265980367 | 8.526672493 | 0.500130194 | 0.633103063 | 0.993793378 | -5.153592301 |
| **A_55_P2012439** | Tnfrsf19 | 0.027972479 | -0.105768722 | 0.161713679 | 9.862905554 | 0.499890478 | 0.633263042 | 0.993793378 | -5.153688835 |
| **A_55_P2006392** | NA | 0.030055124 | -0.113652328 | 0.173762575 | 16.32363384 | 0.499859979 | 0.633283397 | 0.993793378 | -5.153701114 |
| **A_55_P1996683** | Itih1 | 0.020937164 | -0.079218902 | 0.121093229 | 7.751058653 | 0.499631403 | 0.633435965 | 0.993793378 | -5.153793119 |
| **A_55_P1978226** | Prkn | -0.046246948 | -0.267506101 | 0.175012206 | 9.08196811 | -0.499563799 | 0.633481092 | 0.993793378 | -5.153820322 |
| **A_51_P456870** | Foxj1 | -0.023479043 | -0.135816011 | 0.088857926 | 7.727838324 | -0.499536033 | 0.633499627 | 0.993793378 | -5.153831495 |
| **A_55_P2118520** | Col1a1 | 0.051606791 | -0.19533249 | 0.298546072 | 14.65684678 | 0.499488858 | 0.633531119 | 0.993793378 | -5.153850475 |
| **A_52_P445622** | 2610002M( | -0.030469617 | -0.176273279 | 0.115334044 | 8.550274127 | -0.499468039 | 0.633545017 | 0.993793378 | -5.153858851 |
| **A_55_P1993463** | NA | 0.100905463 | -0.381985282 | 0.583796209 | 9.568671958 | 0.499430351 | 0.633570177 | 0.993793378 | -5.153874013 |
| **A_52_P80444** | Dpm2 | 0.038635552 | -0.146274385 | 0.22354549 | 11.17039384 | 0.499385591 | 0.633600058 | 0.993793378 | -5.153892018 |
| **A_51_P257951** | Retnla | -0.020801235 | -0.12036008 | 0.07875761 | 7.650639945 | -0.49936534 | 0.633613578 | 0.993793378 | -5.153900164 |
| **A_55_P2136763** | NA | -0.034059165 | -0.197099117 | 0.128980786 | 14.68979127 | -0.499285664 | 0.633666771 | 0.993793378 | -5.15393221 |
| **A_55_P2008651** | Qdpr | 0.030446117 | -0.115350992 | 0.176243226 | 12.47290287 | 0.499105243 | 0.633787234 | 0.993793378 | -5.154004759 |
| **A_52_P679152** | Clk1 | 0.090665364 | -0.343577846 | 0.524908574 | 9.592636841 | 0.499019485 | 0.633844496 | 0.993793378 | -5.154039234 |
| **A_51_P473919** | Mycbp2 | 0.030222276 | -0.114536145 | 0.174980697 | 9.47317773 | 0.49899071 | 0.63386371 | 0.993793378 | -5.154050801 |
| **A_55_P2095732** | Fbrsl1 | 0.038375069 | -0.145453062 | 0.2222032 | 9.291808491 | 0.498937711 | 0.6338991 | 0.993793378 | -5.154072103 |
| **A_51_P517787** | Clptm1l | 0.036680031 | -0.139080856 | 0.212440918 | 10.69532276 | 0.498788683 | 0.63399862 | 0.993793378 | -5.154131992 |
| **A_55_P2279685** | 2900076A0 | -0.051543676 | -0.298553215 | 0.195465863 | 8.31906377 | -0.498736089 | 0.634033744 | 0.993793378 | -5.154153124 |
| **A_51_P288876** | Tmem45a | -0.069049032 | -0.400026459 | 0.261928394 | 9.400187179 | -0.498618461 | 0.634112304 | 0.993793378 | -5.154200379 |
| **A_52_P257523** | 1110065P2 | -0.057936433 | -0.335660539 | 0.219787673 | 12.00717206 | -0.498594298 | 0.634128442 | 0.993793378 | -5.154210085 |

| | | | | | | | | | |
|---|---|---|---|---|---|---|---|---|---|
| **A_52_P421713** | 9930012K1 | 0.025203231 | -0.095617618 | 0.14602408 | 7.968962943 | 0.498566929 | 0.634146722 | 0.993793378 | -5.154221078 |
| **A_55_P2042486** | Dpysl3 | 0.040467856 | -0.153533964 | 0.234469677 | 9.247104903 | 0.498555518 | 0.634154343 | 0.993793378 | -5.154225661 |
| **A_52_P676461** | Lpgat1 | -0.024497605 | -0.14194143 | 0.092946221 | 9.02802206 | -0.498542932 | 0.63416275 | 0.993793378 | -5.154230716 |
| **A_55_P1996329** | Kat5 | -0.041915055 | -0.242875896 | 0.159045785 | 9.194657135 | -0.498502953 | 0.634189452 | 0.993793378 | -5.154246772 |
| **A_55_P2041700** | Hoxa11os | -0.021594411 | -0.12513991 | 0.081951089 | 7.951543544 | -0.498447311 | 0.634226618 | 0.993793378 | -5.154269117 |
| **A_51_P232439** | Ice1 | -0.045953829 | -0.266364666 | 0.174457008 | 10.3264339 | -0.498308035 | 0.634319651 | 0.993793378 | -5.154325038 |
| **A_51_P462546** | Gmppa | 0.037796918 | -0.14350074 | 0.219094576 | 9.570787141 | 0.498279852 | 0.634338477 | 0.993793378 | -5.154336351 |
| **A_55_P2177910** | Lepr | 0.072810714 | -0.276511311 | 0.42213274 | 10.70997433 | 0.498171013 | 0.634411185 | 0.993793378 | -5.154380039 |
| **A_51_P157214** | Mfsd14a | -0.028921707 | -0.167687468 | 0.109844054 | 10.45683118 | -0.498139216 | 0.634432427 | 0.993793378 | -5.154392801 |
| **A_51_P517695** | Ly6f | 0.052811357 | -0.200591804 | 0.306214517 | 7.729301541 | 0.498109048 | 0.634452581 | 0.993793378 | -5.154404908 |
| **A_55_P2052479** | NA | 0.023651949 | -0.08984081 | 0.137144707 | 14.83641965 | 0.498090096 | 0.634465242 | 0.993793378 | -5.154412514 |
| **A_55_P2006093** | Nebl | 0.057012898 | -0.216640461 | 0.330666257 | 11.55474701 | 0.4979451 | 0.634562115 | 0.993798578 | -5.154470693 |
| **A_55_P1982733** | Dnajc5 | 0.04548821 | -0.172874622 | 0.263851043 | 9.351394027 | 0.497885256 | 0.6346021 | 0.993798578 | -5.154494701 |
| **A_51_P394814** | Svep1 | 0.046511707 | -0.176778213 | 0.269801627 | 9.294700583 | 0.497854337 | 0.634622759 | 0.993798578 | -5.154507104 |
| **A_55_P2167112** | Kank2 | 0.03319122 | -0.126177422 | 0.192559862 | 12.27004125 | 0.497770863 | 0.634678534 | 0.993805431 | -5.154540585 |
| **A_51_P260548** | Glg1 | 0.035951957 | -0.136721317 | 0.208625231 | 13.23015009 | 0.497629988 | 0.63477267 | 0.993809927 | -5.154597078 |
| **A_55_P2086631** | NA | -0.041424453 | -0.240411365 | 0.157562459 | 14.45592372 | -0.49755535 | 0.634822548 | 0.993809927 | -5.154627002 |
| **A_55_P2152872** | Fgfrl1 | 0.029018609 | -0.110397454 | 0.168434672 | 11.31752708 | 0.497476891 | 0.634874982 | 0.993809927 | -5.154658455 |
| **A_55_P2085974** | Igf1 | 0.06441382 | -0.245065033 | 0.373892672 | 10.66370802 | 0.497458902 | 0.634887004 | 0.993809927 | -5.154665666 |
| **A_55_P2050483** | Abcc12 | 0.033771062 | -0.128599378 | 0.196141502 | 7.719805132 | 0.49710357 | 0.6351245 | 0.993878996 | -5.15480805 |
| **A_55_P2120278** | Ube2l3 | -0.025777299 | -0.149717741 | 0.098163143 | 13.43823228 | -0.497088254 | 0.635134738 | 0.993878996 | -5.154814186 |
| **A_51_P136699** | Ttc7b | -0.043799991 | -0.25446533 | 0.166865349 | 10.85532843 | -0.496924104 | 0.635244469 | 0.993878996 | -5.154879928 |
| **A_55_P1955502** | Tasor2 | 0.02874737 | -0.109542275 | 0.167037016 | 9.290827624 | 0.49684119 | 0.635299899 | 0.993878996 | -5.154913128 |
| **A_55_P2091359** | Padi2 | 0.025339637 | -0.096564698 | 0.147243973 | 7.969026678 | 0.496810054 | 0.635320715 | 0.993878996 | -5.154925595 |
| **A_55_P2046499** | Vmn1r70 | 0.020720742 | -0.07897074 | 0.120412223 | 8.207552892 | 0.496771169 | 0.635346712 | 0.993878996 | -5.154941162 |
| **A_55_P1971298** | Lonp1 | -0.026453835 | -0.153729252 | 0.100821583 | 13.78777845 | -0.496767572 | 0.635349116 | 0.993878996 | -5.154942602 |
| **A_55_P2034230** | Polr2c | -0.028539562 | -0.165889831 | 0.108810708 | 10.79160475 | -0.496623097 | 0.635445712 | 0.993878996 | -5.155000432 |
| **A_55_P2172375** | NA | -0.022933628 | -0.133322684 | 0.087455428 | 8.212629614 | -0.496541875 | 0.63550002 | 0.993878996 | -5.155032937 |
| **A_51_P485405** | Gtsf1 | 0.125594567 | -0.479166077 | 0.730355211 | 11.96484488 | 0.496359643 | 0.635621875 | 0.993878996 | -5.155105848 |
| **A_55_P2105843** | NA | 0.097903768 | -0.373640956 | 0.569448492 | 12.68921502 | 0.496233008 | 0.635706562 | 0.993878996 | -5.1551565 |
| **A_55_P2332731** | D130062J2 | 0.085238588 | -0.325370406 | 0.495847582 | 8.613593581 | 0.496154486 | 0.635759075 | 0.993878996 | -5.155187902 |
| **A_52_P618187** | Mrs2 | 0.026641008 | -0.101728331 | 0.155010348 | 9.11926647 | 0.496019191 | 0.635849563 | 0.993878996 | -5.155241997 |
| **A_55_P2028894** | Gata4 | -0.026180197 | -0.152336284 | 0.09997589 | 12.33311627 | -0.495991047 | 0.635868387 | 0.993878996 | -5.155253248 |
| **A_51_P485161** | Cplx2 | -0.029448036 | -0.171383068 | 0.112486997 | 10.39742772 | -0.495879167 | 0.635943221 | 0.993878996 | -5.15529797 |
| **A_55_P2155504** | Lck | -0.079740739 | -0.464099799 | 0.304618322 | 8.412585765 | -0.495852222 | 0.635961244 | 0.993878996 | -5.155308739 |
| **A_55_P2005675** | NA | -0.02220672 | -0.129250353 | 0.084836913 | 15.09049045 | -0.495829889 | 0.635976183 | 0.993878996 | -5.155317664 |
| **A_55_P2034227** | NA | 0.051171032 | -0.195513289 | 0.297855353 | 13.29145439 | 0.495783148 | 0.636007449 | 0.993878996 | -5.155336343 |
| **A_55_P2055168** | Gm2016 | 0.024760396 | -0.094607682 | 0.144128475 | 8.633716559 | 0.495768039 | 0.636017556 | 0.993878996 | -5.155342381 |
| **A_55_P2171746** | NA | -0.031825174 | -0.185263034 | 0.121612686 | 11.60157586 | -0.495732493 | 0.636041334 | 0.993878996 | -5.155356584 |
| **A_51_P436719** | Eftud2 | -0.050035683 | -0.291283378 | 0.191212011 | 9.664534618 | -0.495707849 | 0.63605782 | 0.993878996 | -5.155366431 |
| **A_55_P2156293** | Pcyox1 | 0.025819865 | -0.098672516 | 0.150312246 | 10.83884247 | 0.495701606 | 0.636061996 | 0.993878996 | -5.155368926 |
| **A_51_P290576** | Plk2 | 0.0489659 | -0.187207119 | 0.285138918 | 10.05045692 | 0.495533011 | 0.636174785 | 0.993901575 | -5.155436277 |
| **A_55_P2072373** | Msn | 0.040842237 | -0.156235774 | 0.237920249 | 13.96714411 | 0.495313864 | 0.636321408 | 0.993901575 | -5.155523793 |
| **A_55_P2164111** | Nploc4 | -0.022411184 | -0.13055569 | 0.085733322 | 10.73737137 | -0.495301299 | 0.636329815 | 0.993901575 | -5.15552881 |
| **A_55_P2076713** | Immp1l | -0.028278742 | -0.164766886 | 0.108209402 | 10.99082393 | -0.495192762 | 0.63640244 | 0.993901575 | -5.155572139 |
| **A_52_P647919** | Usf2 | -0.036350154 | -0.211936156 | 0.139235848 | 10.38092967 | -0.494795226 | 0.636668479 | 0.993901575 | -5.155730769 |
| **A_55_P1966719** | NA | 0.079588867 | -0.30499507 | 0.464172804 | 10.22229588 | 0.494618453 | 0.636786798 | 0.993901575 | -5.15580127 |
| **A_55_P2046348** | Itpripl2 | 0.041277133 | -0.158209163 | 0.240763429 | 11.80257871 | 0.494544741 | 0.636836138 | 0.993901575 | -5.155830661 |

| | | | | | | | | | |
|---|---|---|---|---|---|---|---|---|---|
| **A_51_P376857** | Det1 | 0.022151741 | -0.084935539 | 0.12923902 | 9.067689166 | 0.49440073 | 0.636932541 | 0.993901575 | -5.155888071 |
| **A_51_P157270** | Plekhd1 | 0.021223246 | -0.081378645 | 0.123825138 | 7.745949267 | 0.494385336 | 0.636942846 | 0.993901575 | -5.155894207 |
| **A_55_P1994258** | Gapdh | -0.086485554 | -0.504608879 | 0.33163777 | 13.52108949 | -0.494365677 | 0.636956007 | 0.993901575 | -5.155902042 |
| **A_55_P2164724** | Gnaq | 0.036270971 | -0.139119142 | 0.211661085 | 10.66601789 | 0.494268811 | 0.637020856 | 0.993901575 | -5.155940647 |
| **A_55_P2106250** | Trappc6a | 0.042077169 | -0.161483557 | 0.245637895 | 13.0133237 | 0.49403948 | 0.637174399 | 0.993901575 | -5.156032016 |
| **A_51_P519108** | Dzank1 | 0.037893465 | -0.14544857 | 0.2212355 | 8.334583399 | 0.493982322 | 0.637212671 | 0.993901575 | -5.156054783 |
| **A_51_P122085** | Usp20 | 0.028694719 | -0.110156611 | 0.167546048 | 12.40467969 | 0.49392506 | 0.637251014 | 0.993901575 | -5.150286685 |
| **A_51_P462047** | Prpf38a | 0.025271028 | -0.097061859 | 0.147603914 | 10.95940006 | 0.493729201 | 0.637382171 | 0.993901575 | -5.156155575 |
| **A_55_P2090535** | Thrap3 | 0.026195047 | -0.100631101 | 0.153021196 | 8.342719287 | 0.493650424 | 0.637434928 | 0.993901575 | -5.156186934 |
| **A_55_P1965368** | Ctc1 | 0.031564622 | -0.121260227 | 0.184389471 | 12.34354348 | 0.493646145 | 0.637437794 | 0.993901575 | -5.156188637 |
| **A_55_P2045414** | Hipk4 | -0.030200167 | -0.176435082 | 0.116034748 | 8.751974616 | -0.493591185 | 0.637474602 | 0.993901575 | -5.156210513 |
| **A_51_P221337** | Ranbp10 | 0.03994941 | -0.153520501 | 0.233419322 | 10.18733949 | 0.493521493 | 0.637521278 | 0.993901575 | -5.156238248 |
| **A_55_P2057803** | Stum | 0.022048724 | -0.084738979 | 0.128836426 | 7.723521268 | 0.493482035 | 0.637547706 | 0.993901575 | -5.15625395 |
| **A_55_P2056739** | Col16a1 | 0.081533102 | -0.313486829 | 0.476553034 | 11.89326035 | 0.493314738 | 0.637659762 | 0.993901575 | -5.156320511 |
| **A_51_P141818** | Mrpl32 | -0.026662004 | -0.155842836 | 0.102518828 | 11.85765307 | -0.493291738 | 0.637675169 | 0.993901575 | -5.156329661 |
| **A_66_P126485** | Dipk1a | -0.025797633 | -0.150792844 | 0.099197578 | 8.85845538 | -0.493282402 | 0.637681422 | 0.993901575 | -5.156333374 |
| **A_55_P1958887** | DXBay18 | -0.03223486 | -0.188442649 | 0.123972929 | 7.780441483 | -0.493210407 | 0.637729649 | 0.993901575 | -5.15636201 |
| **A_55_P1964368** | Kctd14 | -0.030744196 | -0.179758054 | 0.118269661 | 7.852576914 | -0.493112052 | 0.637795536 | 0.993901575 | -5.156401125 |
| **A_55_P1956647** | Sfmbt2 | 0.026971719 | -0.103785069 | 0.157728506 | 8.019210224 | 0.493007488 | 0.637865586 | 0.993901575 | -5.156442701 |
| **A_52_P536481** | Mogs | 0.045544511 | -0.175297315 | 0.266386337 | 12.45433094 | 0.492905711 | 0.637933774 | 0.993901575 | -5.156483161 |
| **A_51_P455338** | Rnase2a | -0.02011622 | -0.117687712 | 0.077455273 | 7.754248074 | -0.492756718 | 0.638033601 | 0.993901575 | -5.156542377 |
| **A_55_P2009360** | Psg-ps1 | -0.184856103 | -1.081512282 | 0.711800075 | 8.176206467 | -0.49273905 | 0.63804544 | 0.993901575 | -5.156549398 |
| **A_51_P416585** | Hdlbp | -0.023749453 | -0.13895995 | 0.091461044 | 11.82396006 | -0.492686537 | 0.638080627 | 0.993901575 | -5.156570265 |
| **A_55_P2064063** | Casp14 | -0.037858877 | -0.221523429 | 0.145805675 | 7.766273073 | -0.492664778 | 0.638095206 | 0.993901575 | -5.15657891 |
| **A_51_P170758** | Grwd1 | 0.068137136 | -0.262447305 | 0.398721576 | 10.13859249 | 0.492618363 | 0.638126308 | 0.993901575 | -5.156597351 |
| **A_51_P407074** | Tspan31 | -0.032339271 | -0.189251006 | 0.124572465 | 11.54654707 | -0.492588115 | 0.638146577 | 0.993901575 | -5.156609368 |
| **A_52_P574759** | Rbpjl | 0.023232557 | -0.089501702 | 0.135966815 | 7.842342046 | 0.492549892 | 0.638172191 | 0.993901575 | -5.156624552 |
| **A_51_P262670** | Rbm38 | -0.047171076 | -0.276071749 | 0.181729596 | 9.234011222 | -0.492535854 | 0.638181598 | 0.993901575 | -5.156630129 |
| **A_55_P2128668** | Ccnb1 | 0.034346084 | -0.132340028 | 0.201032197 | 9.078099238 | 0.492478128 | 0.638220282 | 0.993901575 | -5.156653058 |
| **A_55_P2049086** | Map3k7 | 0.025343166 | -0.097657738 | 0.148344069 | 10.98548068 | 0.49244949 | 0.638239474 | 0.993901575 | -5.156664432 |
| **A_55_P2009572** | Ptbp3 | 0.023213224 | -0.089456708 | 0.135883155 | 8.001713538 | 0.492420993 | 0.638258572 | 0.993901575 | -5.15667575 |
| **A_55_P1959994** | Unc5cl | 0.020007247 | -0.077106158 | 0.117120652 | 7.655103312 | 0.492399146 | 0.638273213 | 0.993901575 | -5.156684426 |
| **A_51_P483511** | Dock8 | -0.022868512 | -0.133881758 | 0.088144734 | 7.874321711 | -0.492348069 | 0.638307444 | 0.993901575 | -5.156704709 |
| **A_52_P482983** | Cd164 | 0.052941671 | -0.2040909 | 0.309974242 | 14.13657705 | 0.492287281 | 0.638348183 | 0.993901575 | -5.156728846 |
| **A_55_P2041624** | Ceacam12 | -0.421004033 | -2.465742632 | 1.623734566 | 9.117219756 | -0.492104791 | 0.638470497 | 0.993901575 | -5.156801291 |
| **A_51_P321734** | Bhmt2 | -0.026163164 | -0.153244296 | 0.100917968 | 7.71224439 | -0.492060299 | 0.638500319 | 0.993901575 | -5.15681895 |
| **A_51_P421984** | Timm22 | -0.033218454 | -0.194574793 | 0.128137885 | 12.07384539 | -0.492042391 | 0.638512323 | 0.993901575 | -5.156826057 |
| **A_55_P1981992** | NA | -0.03344557 | -0.195934103 | 0.129042963 | 9.006925203 | -0.49195459 | 0.638571178 | 0.993901575 | -5.156860899 |
| **A_55_P2012899** | Vsig4 | 0.043366909 | -0.167343887 | 0.254077704 | 7.852798976 | 0.491904518 | 0.638604743 | 0.993901575 | -5.156880767 |
| **A_55_P2068177** | NA | -0.025669748 | -0.150401708 | 0.099062212 | 11.93521111 | -0.491873 | 0.638625872 | 0.993901575 | -5.156893272 |
| **A_55_P2016166** | Wfdc2 | -0.020406302 | -0.11958316 | 0.078770555 | 7.74101815 | -0.491771212 | 0.638694109 | 0.993901575 | -5.156933651 |
| **A_55_P2047305** | Adcy5 | -0.083552462 | -0.489693231 | 0.322588306 | 10.95449567 | -0.49169048 | 0.638748233 | 0.993901575 | -5.156965673 |
| **A_52_P120803** | Ankrd1 | -0.103385526 | -0.605936018 | 0.399164966 | 14.50232929 | -0.491687503 | 0.638750228 | 0.993901575 | -5.156966853 |
| **A_51_P181891** | Amer2 | 0.023608769 | -0.091158287 | 0.138375824 | 7.835964145 | 0.491660398 | 0.638768401 | 0.993901575 | -5.156977603 |
| **A_55_P1980556** | NA | -0.040203903 | -0.235693506 | 0.155285701 | 8.53699885 | -0.491534132 | 0.638853058 | 0.993901575 | -5.157027672 |
| **A_51_P504522** | Derl1 | -0.035893101 | -0.210427614 | 0.138641412 | 10.90730691 | -0.491517285 | 0.638864354 | 0.993901575 | -5.157034352 |
| **A_51_P316949** | Ccdc6 | -0.025301166 | -0.148347995 | 0.097745663 | 9.713011786 | -0.491449892 | 0.638909541 | 0.993901575 | -5.15706107 |
| **A_52_P413947** | Mthfr | 0.057952746 | -0.223920753 | 0.339826244 | 9.756131356 | 0.491392931 | 0.638947735 | 0.993901575 | -5.15708365 |

| | | | | | | | | | |
|---|---|---|---|---|---|---|---|---|---|
| **A_55_P2004347** | NA | 0.071825941 | -0.277548112 | 0.421199995 | 12.66992326 | 0.491360013 | 0.638969808 | 0.993901575 | -5.157096698 |
| **A_51_P483220** | Impact | 0.052640798 | -0.203463152 | 0.308744748 | 9.880615929 | 0.491264427 | 0.639033905 | 0.993901575 | -5.157134582 |
| **A_55_P1968863** | Synpo2l | -0.044910039 | -0.263418809 | 0.173598731 | 12.56267312 | -0.491228654 | 0.639057894 | 0.993901575 | -5.157148758 |
| **A_55_P2109752** | Nans | 0.033160168 | -0.128331848 | 0.194652183 | 12.48547361 | 0.49076637 | 0.639367941 | 0.993984331 | -5.157331868 |
| **A_51_P429295** | Pus10 | -0.028317324 | -0.16623714 | 0.109602492 | 8.958551689 | -0.490721027 | 0.639398357 | 0.993984331 | -5.15734982 |
| **A_52_P500488** | Lars | 0.035257882 | -0.136570688 | 0.207086453 | 12.10748987 | 0.490422039 | 0.63959893 | 0.993984331 | -5.157468155 |
| **A_55_P2126855** | E230025N2 | 0.023218928 | -0.08995005 | 0.136387905 | 7.713793883 | 0.490370013 | 0.639633835 | 0.993984331 | -5.157488739 |
| **A_55_P1990086** | Mks1 | 0.035286881 | -0.136702682 | 0.207276443 | 8.966245042 | 0.490365955 | 0.639636557 | 0.993984331 | -5.157490345 |
| **A_55_P2127934** | Palm3 | 0.041404324 | -0.160440749 | 0.243249396 | 7.955341758 | 0.490271531 | 0.639699909 | 0.993984331 | -5.157527699 |
| **A_51_P222590** | Arf6 | 0.045978193 | -0.178221113 | 0.2701775 | 10.68135639 | 0.490147475 | 0.639783147 | 0.993984331 | -5.157576765 |
| **A_51_P172344** | Foxn3 | -0.035409769 | -0.208083285 | 0.137263746 | 10.19816674 | -0.490124597 | 0.639798498 | 0.993984331 | -5.157585813 |
| **A_52_P88829** | Ctsq | 0.079703318 | -0.30898881 | 0.468395446 | 7.867853967 | 0.490094457 | 0.639818723 | 0.993984331 | -5.157597731 |
| **A_52_P221776** | Kif12 | 0.019761061 | -0.076612456 | 0.116134577 | 7.660647097 | 0.490074014 | 0.63983244 | 0.993984331 | -5.157605815 |
| **A_55_P2106180** | Dnah2 | -0.023538626 | -0.138343207 | 0.091265955 | 7.765909111 | -0.490039428 | 0.639855648 | 0.993984331 | -5.157619491 |
| **A_55_P2138071** | NA | -0.053868814 | -0.316624717 | 0.208887088 | 12.72990324 | -0.489997732 | 0.639883628 | 0.993984331 | -5.157635976 |
| **A_52_P97572** | Aplnr | -0.085123872 | -0.500376907 | 0.330129163 | 8.897787403 | -0.489945404 | 0.639918744 | 0.993984331 | -5.157656664 |
| **A_51_P278489** | Itfg2 | 0.0248165 | -0.096245289 | 0.14587829 | 9.145036456 | 0.489939643 | 0.63992261 | 0.993984331 | -5.157658941 |
| **A_55_P2090070** | Myh14 | 0.029961601 | -0.11622211 | 0.176145311 | 8.359234563 | 0.48986359 | 0.639973648 | 0.993984331 | -5.157689005 |
| **A_55_P1975021** | Rad18 | 0.030664415 | -0.118951953 | 0.180280783 | 9.0253092 | 0.489851798 | 0.639981561 | 0.993984331 | -5.157693665 |
| **A_51_P258620** | Rab4b | -0.058657038 | -0.344971656 | 0.22765758 | 8.579990145 | -0.489649975 | 0.640117013 | 0.993984331 | -5.157773422 |
| **A_55_P2143261** | Rpain | 0.048944737 | -0.189963317 | 0.28785279 | 10.91192348 | 0.4896484 | 0.64011807 | 0.993984331 | -5.157774045 |
| **A_52_P191468** | Ubfd1 | 0.038461069 | -0.149288227 | 0.226210365 | 9.874528001 | 0.489612124 | 0.640142418 | 0.993984331 | -5.157788377 |
| **A_51_P211305** | Alpk3 | 0.047039001 | -0.182593433 | 0.276671435 | 11.35056004 | 0.489591641 | 0.640156166 | 0.993984331 | -5.157796469 |
| **A_55_P2095311** | NA | -0.03081137 | -0.181240442 | 0.119617701 | 7.739760355 | -0.489540214 | 0.640190684 | 0.993984331 | -5.157816785 |
| **A_55_P2018210** | NA | 0.021381313 | -0.083024213 | 0.12578684 | 16.74795652 | 0.48946317 | 0.640242398 | 0.993984806 | -5.157847218 |
| **A_51_P159612** | Hebp2 | 0.036232487 | -0.14075712 | 0.213222094 | 9.038461671 | 0.489282311 | 0.640363805 | 0.994087192 | -5.15791864 |
| **A_55_P1982111** | Homer1 | -0.032591727 | -0.191834631 | 0.126651177 | 10.07927248 | -0.48916617 | 0.640441774 | 0.994087192 | -5.157964491 |
| **A_55_P2038747** | Ano1 | 0.12088468 | -0.469837679 | 0.711607038 | 10.05814727 | 0.48909924 | 0.640486709 | 0.994087192 | -5.157990911 |
| **A_55_P1962299** | Hba-a2 | 0.038040094 | -0.147864372 | 0.223944559 | 17.65185583 | 0.489058588 | 0.640514002 | 0.994087192 | -5.158006955 |
| **A_55_P2101910** | Rpl13a | 0.046655208 | -0.181427177 | 0.274737593 | 12.47523398 | 0.488897159 | 0.640622389 | 0.994170468 | -5.158070657 |
| **A_52_P116134** | Aldh2 | -0.094383255 | -0.5559215 | 0.367154989 | 10.14570738 | -0.488760826 | 0.640713934 | 0.994170468 | -5.158124441 |
| **A_55_P1965467** | 2300009A0 | 0.042077508 | -0.163692195 | 0.24784721 | 11.4558976 | 0.488739802 | 0.640728051 | 0.994170468 | -5.158132734 |
| **A_55_P2166538** | Mrps16 | 0.040154715 | -0.156245781 | 0.236555211 | 13.56815858 | 0.488655845 | 0.64078443 | 0.994170468 | -5.158165847 |
| **A_55_P1971094** | Esx1 | -0.020364006 | -0.119986007 | 0.079257996 | 7.638143305 | -0.488559065 | 0.640849424 | 0.994170468 | -5.158204012 |
| **A_52_P356068** | Proz | 0.022223226 | -0.086503063 | 0.130949516 | 7.690698068 | 0.488519242 | 0.640876168 | 0.994170468 | -5.158219713 |
| **A_55_P2096657** | NA | -0.028425856 | -0.16753711 | 0.110685399 | 16.54345127 | -0.48838286 | 0.640967764 | 0.99423279 | -5.158273479 |
| **A_55_P1997282** | Fam135a | -0.036794343 | -0.216903508 | 0.143314823 | 9.222945193 | -0.488263619 | 0.641047854 | 0.994277255 | -5.158320475 |
| **A_51_P136845** | Usp19 | 0.027639113 | -0.107686898 | 0.162965124 | 10.03737286 | 0.488148497 | 0.641125182 | 0.994302964 | -5.158365839 |
| **A_55_P2345853** | NA | 0.047309106 | -0.184354426 | 0.278972638 | 9.064218513 | 0.488085827 | 0.64116728 | 0.994302964 | -5.158390529 |
| **A_51_P116581** | Ccdc22 | -0.025436687 | -0.150112236 | 0.099238862 | 10.83578595 | -0.487627718 | 0.641475051 | 0.994392933 | -5.158570927 |
| **A_55_P2001593** | Rnf180 | -0.02428914 | -0.143355142 | 0.094776862 | 8.593501886 | -0.487566103 | 0.641516451 | 0.994392933 | -5.158595179 |
| **A_55_P2093822** | Ube2d2a | 0.022144448 | -0.086415245 | 0.13070414 | 12.53138602 | 0.48753453 | 0.641537667 | 0.994392933 | -5.158607605 |
| **A_51_P229893** | Cts6 | 0.06442619 | -0.251422987 | 0.380275367 | 7.813737985 | 0.487519319 | 0.641547888 | 0.994392933 | -5.158613591 |
| **A_51_P314679** | Rrn3 | 0.040824008 | -0.159367517 | 0.241015533 | 11.66895849 | 0.487392772 | 0.641632925 | 0.994392933 | -5.158663386 |
| **A_55_P2036813** | H2bu2 | 0.080215076 | -0.313199662 | 0.473629813 | 9.99898803 | 0.487320302 | 0.641681625 | 0.994392933 | -5.158691897 |
| **A_55_P2054511** | NA | 0.031268772 | -0.122151855 | 0.184689398 | 9.009188822 | 0.487120267 | 0.641816062 | 0.994392933 | -5.158770575 |
| **A_55_P1993121** | NA | 0.023663032 | -0.092452869 | 0.139778933 | 16.07397008 | 0.487066015 | 0.641852525 | 0.994392933 | -5.158791908 |
| **A_51_P355943** | Mvd | -0.051688874 | -0.305376619 | 0.201998871 | 8.969517474 | -0.486975057 | 0.64191366 | 0.994392933 | -5.15882767 |

| | | | | | | | | | |
|---|---|---|---|---|---|---|---|---|---|
| **A_55_P2102335** | Zfp423 | 0.031116653 | -0.121623468 | 0.183856773 | 8.646144094 | 0.486910203 | 0.641957253 | 0.994392933 | -5.158853166 |
| **A_55_P2037425** | Gapdh | -0.050615397 | -0.299147278 | 0.197916483 | 16.53165915 | -0.486754164 | 0.642062143 | 0.994392933 | -5.158914494 |
| **A_55_P2016486** | Tmem131 | -0.030546571 | -0.180539515 | 0.119446374 | 11.08262742 | -0.486744176 | 0.642068857 | 0.994392933 | -5.158918419 |
| **A_55_P1996202** | NA | -0.031513436 | -0.186259141 | 0.12323227 | 11.55259715 | -0.486727941 | 0.642079771 | 0.994392933 | -5.158924799 |
| **A_55_P2048840** | Ppil2 | -0.049851524 | -0.294659244 | 0.194956196 | 9.522186393 | -0.486701249 | 0.642097715 | 0.994392933 | -5.158935288 |
| **A_55_P1989296** | Tnxb | -0.083831091 | -0.495653097 | 0.327990914 | 10.70610854 | -0.486524486 | 0.64221655 | 0.994392933 | -5.159004733 |
| **A_52_P613808** | Zbtb7a | 0.046169503 | -0.180740749 | 0.273079755 | 8.319499936 | 0.486306657 | 0.642363008 | 0.994392933 | -5.15909028 |
| **A_55_P2058290** | Cluh | 0.046713395 | -0.182895887 | 0.276322677 | 9.479719147 | 0.486251687 | 0.64239997 | 0.994392933 | -5.159111863 |
| **A_55_P2076989** | NA | -0.042342381 | -0.25048046 | 0.165795699 | 9.596203765 | -0.486220046 | 0.642421246 | 0.994392933 | -5.159124285 |
| **A_55_P2159852** | NA | -0.053125036 | -0.314327758 | 0.208077687 | 9.646251641 | -0.486105648 | 0.642498172 | 0.994392933 | -5.159169191 |
| **A_51_P483379** | Rnaseh1 | 0.021986312 | -0.086124901 | 0.130097525 | 9.081593126 | 0.486061007 | 0.642528193 | 0.994392933 | -5.159186712 |
| **A_52_P604629** | Csrnp1 | -0.063109306 | -0.373436982 | 0.247218371 | 10.00875932 | -0.486051205 | 0.642534784 | 0.994392933 | -5.159190559 |
| **A_52_P498193** | Aldh1l2 | -0.047800515 | -0.28285904 | 0.187258009 | 9.323494982 | -0.486032943 | 0.642547065 | 0.994392933 | -5.159197725 |
| **A_52_P520564** | Wasl | -0.028991798 | -0.171583108 | 0.113599512 | 8.593375227 | -0.485949589 | 0.642603122 | 0.994392933 | -5.159230435 |
| **A_55_P1969131** | Cidec | 0.02438222 | -0.095571703 | 0.144336144 | 7.675377825 | 0.485811648 | 0.642695894 | 0.994392933 | -5.159284554 |
| **A_55_P1974542** | Gxylt2 | -0.047297239 | -0.280063495 | 0.185469017 | 10.96440698 | -0.48565169 | 0.642803482 | 0.994392933 | -5.159347293 |
| **A_55_P1961451** | Adamts7 | -0.048458861 | -0.286970542 | 0.190052821 | 8.700109449 | -0.485593289 | 0.642842765 | 0.994392933 | -5.159370194 |
| **A_55_P1980119** | Cptp | -0.030816381 | -0.18250268 | 0.120869918 | 9.663175216 | -0.485561692 | 0.642864019 | 0.994392933 | -5.159382584 |
| **A_52_P246165** | Ptprk | -0.034866176 | -0.206612263 | 0.136879911 | 9.164902447 | -0.485206489 | 0.643102974 | 0.994392933 | -5.159521811 |
| **A_51_P246903** | Rps27l | 0.028235972 | -0.110868708 | 0.167340653 | 14.17636331 | 0.48514341 | 0.643145414 | 0.994392933 | -5.159546527 |
| **A_52_P208521** | Khdc1a | -0.020239523 | -0.119971367 | 0.07949232 | 7.618637019 | -0.485037784 | 0.643216484 | 0.994392933 | -5.159587906 |
| **A_66_P105965** | Drp2 | -0.020380671 | -0.120813332 | 0.08005199 | 7.786176677 | -0.485012173 | 0.643233716 | 0.994392933 | -5.159597937 |
| **A_55_P2032916** | Slc6a20a | -0.182426847 | -1.081642137 | 0.716788444 | 10.30238083 | -0.484879922 | 0.643322706 | 0.994392933 | -5.159649733 |
| **A_55_P1970583** | Ttc38 | -0.037345392 | -0.221444833 | 0.146754049 | 9.137599875 | -0.484834691 | 0.643353143 | 0.994392933 | -5.159667444 |
| **A_51_P419117** | Rab15 | 0.035872937 | -0.140990175 | 0.212736048 | 9.361023336 | 0.484773424 | 0.643394372 | 0.994392933 | -5.159691433 |
| **A_52_P48569** | Slc38a4 | 0.056137654 | -0.220694593 | 0.332969901 | 11.26034242 | 0.484670642 | 0.643463541 | 0.994392933 | -5.15973167 |
| **A_51_P136355** | Gng11 | -0.029440911 | -0.174624609 | 0.115742787 | 14.07759 | -0.484665966 | 0.643466688 | 0.994392933 | -5.1597335 |
| **A_55_P2012974** | Sdha | -0.04725721 | -0.280321152 | 0.185806731 | 14.74957218 | -0.484620885 | 0.643497028 | 0.994392933 | -5.159751146 |
| **A_55_P2043008** | NA | -0.024351487 | -0.144469164 | 0.095766191 | 8.797492357 | -0.484537825 | 0.643552928 | 0.994392933 | -5.159783653 |
| **A_51_P505795** | Tapbpl | 0.023077188 | -0.090778368 | 0.136932745 | 8.10688879 | 0.484437548 | 0.643620421 | 0.994392933 | -5.159822892 |
| **A_55_P2122898** | Rnf25 | 0.058180177 | -0.228924547 | 0.345284901 | 10.33361465 | 0.484332744 | 0.643690963 | 0.994392933 | -5.159863894 |
| **A_52_P623751** | Fem1b | 0.033548587 | -0.132015859 | 0.199113033 | 12.31087296 | 0.484302029 | 0.643711638 | 0.994392933 | -5.15987591 |
| **A_55_P2147886** | Fam118a | 0.074866127 | -0.294657674 | 0.444389928 | 11.05231418 | 0.484230435 | 0.64375983 | 0.994392933 | -5.159903913 |
| **A_52_P103550** | Tfdp1 | -0.030379497 | -0.180364684 | 0.11960569 | 9.323038843 | -0.48410698 | 0.643842936 | 0.994392933 | -5.159952193 |
| **A_55_P2059154** | Dtx3l | -0.034540043 | -0.205092854 | 0.136012768 | 9.481075232 | -0.484030943 | 0.643894125 | 0.994392933 | -5.159981923 |
| **A_52_P496870** | 2310011J0 | -0.056261355 | -0.334096871 | 0.22157416 | 8.617702346 | -0.48398462 | 0.643925311 | 0.994392933 | -5.160000033 |
| **A_65_P08291** | Gpalpp1 | -0.032263453 | -0.191597099 | 0.127070194 | 11.32174131 | -0.483963352 | 0.643939629 | 0.994392933 | -5.160008347 |
| **A_55_P2201822** | Naf1 | -0.066995592 | -0.397913786 | 0.263922603 | 10.89275651 | -0.483876704 | 0.643997966 | 0.994392933 | -5.160042217 |
| **A_55_P2041634** | Vps51 | 0.031497562 | -0.124087139 | 0.187082263 | 10.33439077 | 0.483859395 | 0.64400962 | 0.994392933 | -5.160048982 |
| **A_52_P446120** | Commd6 | 0.025193389 | -0.099256572 | 0.149643351 | 10.56462624 | 0.483839105 | 0.644023281 | 0.994392933 | -5.160056912 |
| **A_55_P2075213** | Kbtbd11 | 0.091911106 | -0.362371091 | 0.546193304 | 8.524705939 | 0.483561157 | 0.644210434 | 0.994392933 | -5.160165513 |
| **A_55_P2027436** | Bri3 | -0.02659604 | -0.158053615 | 0.104861535 | 15.33792153 | -0.483549007 | 0.644218616 | 0.994392933 | -5.160170259 |
| **A_55_P1977593** | NA | 0.032885523 | -0.129678959 | 0.195450005 | 11.5686415 | 0.483490814 | 0.644257803 | 0.994392933 | -5.160192989 |
| **A_52_P208727** | Zmym4 | -0.026316322 | -0.156409453 | 0.103776809 | 9.507436608 | -0.483481606 | 0.644264004 | 0.994392933 | -5.160196585 |
| **A_55_P1956652** | Sfmbt2 | 0.022472239 | -0.088634635 | 0.133579114 | 7.851116752 | 0.483408798 | 0.644313035 | 0.994392933 | -5.160225019 |
| **A_55_P2026624** | NA | -0.049417599 | -0.293761674 | 0.194926477 | 13.41994774 | -0.483380311 | 0.64433222 | 0.994392933 | -5.160236144 |
| **A_55_P2046453** | Aarsd1 | 0.028714341 | -0.113273309 | 0.170701992 | 12.66148678 | 0.483345208 | 0.644355861 | 0.994392933 | -5.16024985 |
| **A_51_P410949** | Polr3g | 0.043048779 | -0.169855185 | 0.255952743 | 9.057695491 | 0.483265977 | 0.644409221 | 0.994392933 | -5.160280785 |

| | | | | | | | | | |
|---|---|---|---|---|---|---|---|---|---|
| **A_55_P2078645** | Mettl5 | -0.026100122 | -0.155193723 | 0.102993479 | 10.28341987 | -0.483222292 | 0.644438643 | 0.994392933 | -5.160297839 |
| **A_55_P1993365** | NA | -0.029387618 | -0.174773326 | 0.115998089 | 8.279839968 | -0.483116431 | 0.644509944 | 0.994392933 | -5.16033916 |
| **A_55_P1967315** | Ppp1r9b | -0.043107167 | -0.25639355 | 0.170179217 | 11.28040564 | -0.483053774 | 0.644552147 | 0.994392933 | -5.160363613 |
| **A_51_P191520** | Stard10 | -0.098345646 | -0.585038513 | 0.388347221 | 9.093037663 | -0.482957915 | 0.644616717 | 0.994392933 | -5.160401019 |
| **A_52_P438082** | Cebpg | -0.032592113 | -0.193910177 | 0.128725952 | 8.615148315 | -0.482879359 | 0.644669634 | 0.994392933 | -5.160431667 |
| **A_52_P72497** | Acbd3 | 0.033112607 | -0.130789638 | 0.197014852 | 10.84923671 | 0.482855974 | 0.644685387 | 0.994392933 | -5.16044079 |
| **A_51_P489337** | Becn1 | -0.033208316 | -0.197603382 | 0.13118675 | 9.070493851 | -0.482799937 | 0.644723137 | 0.994392933 | -5.160462649 |
| **A_65_P11092** | Tspan15 | -0.029932153 | -0.178137265 | 0.118272959 | 8.072521162 | -0.482707343 | 0.644785516 | 0.994392933 | -5.160498763 |
| **A_51_P394339** | Eif2b1 | -0.047027454 | -0.279882785 | 0.185827878 | 10.89114661 | -0.482696791 | 0.644792624 | 0.994392933 | -5.160502878 |
| **A_55_P2029252** | NA | 0.074924886 | -0.296065799 | 0.445915571 | 10.42286907 | 0.482694351 | 0.644794268 | 0.994392933 | -5.160503829 |
| **A_55_P2003492** | NA | -0.089951236 | -0.535625853 | 0.355723381 | 12.94773086 | -0.482390078 | 0.644999275 | 0.994392933 | -5.160622457 |
| **A_52_P32903** | Zzef1 | -0.030918269 | -0.184145464 | 0.122308925 | 9.980072218 | -0.482268009 | 0.645081529 | 0.994392933 | -5.160670028 |
| **A_55_P2034372** | Eif4g3 | -0.026509521 | -0.15792593 | 0.104906888 | 11.01114634 | -0.482126955 | 0.645176583 | 0.994392933 | -5.160724985 |
| **A_55_P2087872** | NA | -0.06069421 | -0.361592936 | 0.240204516 | 15.55348659 | -0.482098798 | 0.645195558 | 0.994392933 | -5.160735954 |
| **A_51_P370163** | Ctr9 | -0.033328113 | -0.19866817 | 0.132011943 | 9.199459643 | -0.481772246 | 0.645415647 | 0.994392933 | -5.16086312 |
| **A_52_P493519** | Igll1 | 0.020366551 | -0.080676849 | 0.121409951 | 7.643853882 | 0.481746607 | 0.645432929 | 0.994392933 | -5.160873101 |
| **A_55_P2085766** | NA | -0.020781225 | -0.123925566 | 0.082363117 | 7.768845918 | -0.481542764 | 0.645570336 | 0.994392933 | -5.160952438 |
| **A_55_P1974877** | Bcl2l1 | 0.047426856 | -0.188023459 | 0.282877172 | 12.70872934 | 0.481431159 | 0.645645573 | 0.994392933 | -5.160995863 |
| **A_55_P2090557** | Lrrc75a | 0.02492175 | -0.098809757 | 0.148653257 | 12.29364339 | 0.481401416 | 0.645665625 | 0.994392933 | -5.161007434 |
| **A_55_P2106235** | Syngr2 | -0.024561684 | -0.146513921 | 0.097390552 | 13.11457841 | -0.481368319 | 0.645687938 | 0.994392933 | -5.161020309 |
| **A_55_P2145709** | NA | -0.048738992 | -0.290862718 | 0.193384735 | 9.919922593 | -0.481114363 | 0.645859162 | 0.994392933 | -5.161119075 |
| **A_55_P2051849** | Snap25 | 0.084564001 | -0.33555602 | 0.504684023 | 11.79186961 | 0.481084404 | 0.645879363 | 0.994392933 | -5.161130723 |
| **A_55_P2032912** | NA | -0.029253681 | -0.174618097 | 0.116110735 | 13.51290884 | -0.480985017 | 0.64594638 | 0.994392933 | -5.161169361 |
| **A_66_P130911** | Proser2 | -0.028892316 | -0.172556956 | 0.114772325 | 8.336242522 | -0.480664 | 0.646162867 | 0.994392933 | -5.161294108 |
| **A_55_P2072000** | Pcmtd1 | -0.028642375 | -0.171098425 | 0.113813675 | 9.261110966 | -0.480548538 | 0.646240741 | 0.994392933 | -5.161338958 |
| **A_55_P2157770** | Neu2 | -0.020288127 | -0.121196713 | 0.080620459 | 7.865862847 | -0.480532721 | 0.646251409 | 0.994392933 | -5.161345101 |
| **A_55_P1978628** | Hnrnpa2b1 | 0.024693953 | -0.098145224 | 0.14753313 | 8.066301478 | 0.480466207 | 0.646296272 | 0.994392933 | -5.161370933 |
| **A_55_P1958007** | Ddx24 | -0.04397226 | -0.262771112 | 0.174826592 | 12.8986274 | -0.480333502 | 0.646385786 | 0.994392933 | -5.161422461 |
| **A_55_P1956812** | Fam83g | 0.039283868 | -0.156194307 | 0.234762043 | 8.011996237 | 0.480313839 | 0.64639905 | 0.994392933 | -5.161430094 |
| **A_55_P2041070** | Tle3 | 0.031124349 | -0.123829695 | 0.186078394 | 10.6436226 | 0.480072135 | 0.646562105 | 0.994392933 | -5.161523908 |
| **A_55_P2051429** | NA | 0.058078478 | -0.231089341 | 0.347246297 | 8.590389392 | 0.480036654 | 0.646586043 | 0.994392933 | -5.161537676 |
| **A_52_P644972** | Mzt1 | 0.030488667 | -0.121354206 | 0.182331539 | 10.25236636 | 0.479902657 | 0.646676448 | 0.994392933 | -5.161589662 |
| **A_51_P265545** | Maco1 | -0.024491416 | -0.146471954 | 0.097489123 | 10.13150394 | -0.479879808 | 0.646691865 | 0.994392933 | -5.161598526 |
| **A_55_P2057519** | Rpl17 | 0.045634455 | -0.181664545 | 0.272933456 | 13.64719588 | 0.47984887 | 0.64671274 | 0.994392933 | -5.161610527 |
| **A_55_P2111153** | NA | -0.028175911 | -0.168555625 | 0.112203802 | 12.00483689 | -0.479714399 | 0.646803475 | 0.994392933 | -5.161662678 |
| **A_52_P27122** | Ago4 | -0.024348472 | -0.145664066 | 0.096967121 | 8.02971285 | -0.47969393 | 0.646817287 | 0.994392933 | -5.161670616 |
| **A_55_P2091323** | Wdr26 | 0.026228858 | -0.104465525 | 0.156923241 | 9.807342507 | 0.479657891 | 0.646841605 | 0.994392933 | -5.16168459 |
| **A_55_P2027102** | Rgs3 | 0.03261517 | -0.129920618 | 0.195150959 | 11.97495963 | 0.479600669 | 0.646880219 | 0.994392933 | -5.161706776 |
| **A_55_P2013756** | NA | -0.048657607 | -0.291185208 | 0.193869994 | 8.068855377 | -0.479511149 | 0.646940631 | 0.994392933 | -5.16174148 |
| **A_55_P2109782** | Psg17 | -0.078420919 | -0.469397839 | 0.312556002 | 7.869741728 | -0.479391066 | 0.647021671 | 0.994392933 | -5.161788022 |
| **A_55_P2112111** | Ube2b | -0.041178516 | -0.246496561 | 0.164139529 | 13.39700349 | -0.479349966 | 0.647049409 | 0.994392933 | -5.161803949 |
| **A_55_P2084408** | Uba1 | 0.027983294 | -0.111553054 | 0.167519643 | 8.683626588 | 0.479314552 | 0.647073311 | 0.994392933 | -5.161817672 |
| **A_55_P2084308** | Nid1 | 0.0569789 | -0.227177703 | 0.341135504 | 13.27698569 | 0.479253665 | 0.647114406 | 0.994392933 | -5.161841264 |
| **A_55_P2032708** | Hmgb1 | 0.045119003 | -0.179893177 | 0.270131184 | 13.54393075 | 0.479250531 | 0.647116521 | 0.994392933 | -5.161842478 |
| **A_55_P2253348** | Galnt7 | -0.052878044 | -0.316688752 | 0.210932663 | 10.34683502 | -0.479062412 | 0.647243497 | 0.994392933 | -5.161915349 |
| **A_51_P479914** | Pik3cb | -0.040066238 | -0.239975689 | 0.159843212 | 9.045695139 | -0.479020804 | 0.647271584 | 0.994392933 | -5.161931463 |
| **A_52_P337679** | Canx | 0.024340415 | -0.100858601 | 0.14953943 | 13.72126454 | 0.482941145 | 0.647275542 | 0.994392933 | -5.08542035 |
| **A_55_P2080902** | NA | 0.021842275 | -0.087141742 | 0.130826291 | 7.941797804 | 0.479009565 | 0.64727917 | 0.994392933 | -5.161935815 |

| | | | | | | | | | |
|---|---|---|---|---|---|---|---|---|---|
| **A_66_P125392** | Atp8b4 | -0.01951171 | -0.116929923 | 0.077906503 | 7.788458397 | -0.478700976 | 0.647487495 | 0.994392933 | -5.162055284 |
| **A_55_P1966169** | Ppdpf | -0.035282631 | -0.211442561 | 0.140877299 | 13.11017871 | -0.47869949 | 0.647488498 | 0.994392933 | -5.162055859 |
| **A_55_P2133568** | Snhg5 | 0.029084381 | -0.116132203 | 0.174300966 | 8.09858738 | 0.478688225 | 0.647496103 | 0.994392933 | -5.162060219 |
| **A_55_P1967672** | Tas2r109 | -0.023271516 | -0.139475957 | 0.092932924 | 8.716999203 | -0.478642308 | 0.647527104 | 0.994392933 | -5.162077989 |
| **A_55_P2015964** | Rif1 | 0.038835593 | -0.155091678 | 0.232762865 | 9.393788817 | 0.478630303 | 0.64753521 | 0.994392933 | -5.162082635 |
| **A_55_P2002689** | Hmgb1 | 0.037147446 | -0.148352951 | 0.222647843 | 12.14283671 | 0.478622655 | 0.647540374 | 0.994392933 | -5.162085595 |
| **A_55_P2128225** | Rpl18 | 0.021049043 | -0.084075201 | 0.126173286 | 15.18571594 | 0.478562426 | 0.647581039 | 0.994392933 | -5.1621089 |
| **A_52_P243412** | Ulk2 | -0.0408718 | -0.245012073 | 0.163268474 | 9.141122979 | -0.478524525 | 0.64760663 | 0.994392933 | -5.162123565 |
| **A_51_P296057** | Upf1 | 0.035978419 | -0.143764296 | 0.215721134 | 9.917989201 | 0.478409629 | 0.647684211 | 0.994392933 | -5.162168013 |
| **A_55_P1958255** | Rnf114 | -0.05449852 | -0.326784608 | 0.217787569 | 8.932555578 | -0.478374899 | 0.647707662 | 0.994392933 | -5.162181446 |
| **A_55_P2071536** | NA | 0.038522393 | -0.154031621 | 0.231076407 | 8.889349278 | 0.478156228 | 0.647855331 | 0.994392933 | -5.162266009 |
| **A_52_P242973** | Sec11a | 0.023250078 | -0.09299175 | 0.139491905 | 12.7166849 | 0.47804757 | 0.647928714 | 0.994392933 | -5.162308014 |
| **A_51_P256202** | Ctsz | -0.039400314 | -0.236409797 | 0.157609168 | 11.90602044 | -0.47799317 | 0.647965455 | 0.994392933 | -5.162329041 |
| **A_51_P291839** | Cox14 | -0.023617376 | -0.141721444 | 0.094486691 | 14.57098358 | -0.477942806 | 0.647999471 | 0.994392933 | -5.162348506 |
| **A_55_P2031032** | NA | -0.037831734 | -0.227026623 | 0.151363156 | 10.62352038 | -0.47792085 | 0.6480143 | 0.994392933 | -5.162356992 |
| **A_52_P322096** | Patl2 | 0.023019397 | -0.092103994 | 0.138142789 | 7.678051324 | 0.477902719 | 0.648026547 | 0.994392933 | -5.162363998 |
| **A_55_P1961014** | Selenbp1 | 0.036022577 | -0.14413551 | 0.216180664 | 13.38236211 | 0.477892432 | 0.648033495 | 0.994392933 | -5.162367974 |
| **A_51_P332799** | Clptm1 | -0.048637029 | -0.29190502 | 0.194630963 | 12.0092868 | -0.477849569 | 0.648062446 | 0.994392933 | -5.162384536 |
| **A_55_P1989858** | Thrap3 | 0.040581632 | -0.162396741 | 0.243560005 | 11.23037403 | 0.477847011 | 0.648064174 | 0.994392933 | -5.162385525 |
| **A_55_P2076190** | NA | 0.070714373 | -0.28311208 | 0.424540826 | 15.18180905 | 0.477668418 | 0.64818481 | 0.994392933 | -5.162454521 |
| **A_55_P2035877** | Faap20 | -0.027050402 | -0.162422806 | 0.108322002 | 11.07170732 | -0.477587233 | 0.648239653 | 0.994392933 | -5.162485878 |
| **A_52_P137691** | Trappc5 | -0.04375597 | -0.262777569 | 0.175265629 | 8.769004113 | -0.47748475 | 0.648308886 | 0.994392933 | -5.162525454 |
| **A_55_P2074831** | Fam168a | -0.037361403 | -0.224395997 | 0.149673191 | 11.23819407 | -0.477430733 | 0.64834538 | 0.994392933 | -5.16254631 |
| **A_52_P472302** | Fxyd6 | 0.055414734 | -0.222108066 | 0.332937534 | 12.79755088 | 0.477238771 | 0.648475076 | 0.994392933 | -5.162620411 |
| **A_55_P2023449** | NA | 0.101136272 | -0.405446957 | 0.607719502 | 10.21820033 | 0.47716136 | 0.648527381 | 0.994392933 | -5.162650286 |
| **A_55_P2006148** | Aldh3b2 | -0.025663625 | -0.154213565 | 0.102886315 | 7.678173323 | -0.477150368 | 0.648534808 | 0.994392933 | -5.162654527 |
| **A_55_P2124090** | Ambra1 | 0.028432823 | -0.114003641 | 0.170869288 | 10.40375408 | 0.47709837 | 0.648569944 | 0.994392933 | -5.162674592 |
| **A_51_P223036** | Fbxo30 | 0.03122343 | -0.125211944 | 0.187658804 | 9.675928891 | 0.477039976 | 0.648609402 | 0.994392933 | -5.162697121 |
| **A_55_P1972039** | Nox4 | -0.022364437 | -0.134425753 | 0.089696879 | 8.059864748 | -0.47699232 | 0.648641605 | 0.994392933 | -5.162715506 |
| **A_51_P513032** | Trps1 | -0.038953944 | -0.234180783 | 0.156272895 | 9.28489422 | -0.476893102 | 0.648708654 | 0.994392933 | -5.162753777 |
| **A_55_P1965554** | Orc3 | 0.033396485 | -0.134000008 | 0.200792978 | 11.53313228 | 0.476829963 | 0.648751324 | 0.994392933 | -5.162778128 |
| **A_55_P2039196** | Gfra4 | 0.08222033 | -0.329958091 | 0.49439875 | 13.03430709 | 0.476763606 | 0.64879617 | 0.994392933 | -5.162803717 |
| **A_55_P2066301** | Gpr137b-p | -0.026141162 | -0.157213448 | 0.104931123 | 8.504269185 | -0.476675856 | 0.648855475 | 0.994392933 | -5.16283755 |
| **A_51_P185713** | Ptdss2 | -0.048807884 | -0.293543882 | 0.195928113 | 8.574927115 | -0.476651825 | 0.648871718 | 0.994392933 | -5.162846815 |
| **A_51_P315682** | Igf2bp2 | 0.055554517 | -0.223017254 | 0.334126287 | 10.30707463 | 0.476641007 | 0.648879029 | 0.994392933 | -5.162850985 |
| **A_52_P42380** | Tmem106c | -0.02832155 | -0.170413947 | 0.113770848 | 8.474840875 | -0.476381955 | 0.64905413 | 0.994392933 | -5.162950826 |
| **A_55_P2116165** | Pfkfb3 | -0.076676311 | -0.461460681 | 0.308108059 | 9.763667702 | -0.476269665 | 0.649130038 | 0.994392933 | -5.162994088 |
| **A_55_P2169332** | NA | -0.034489777 | -0.207584381 | 0.138604828 | 15.52180501 | -0.476229159 | 0.649157421 | 0.994392933 | -5.163009692 |
| **A_55_P2065231** | Gstm3 | -0.081627986 | -0.491383253 | 0.32812728 | 9.028927513 | -0.476127945 | 0.649225845 | 0.994392933 | -5.163048676 |
| **A_55_P1980326** | Pcbp1 | 0.032963059 | -0.132562368 | 0.198488486 | 14.37617849 | 0.475961613 | 0.649338302 | 0.994392933 | -5.163112724 |
| **A_55_P1982076** | Emsy | 0.04411546 | -0.177420264 | 0.265651184 | 12.74806049 | 0.475944346 | 0.649349977 | 0.994392933 | -5.163119372 |
| **A_51_P199098** | Tmem107 | 0.024316315 | -0.097803976 | 0.146436606 | 10.6450137 | 0.475903683 | 0.64937747 | 0.994392933 | -5.163135026 |
| **A_55_P2003043** | P4ha1 | -0.033411233 | -0.201233058 | 0.134410592 | 8.319791847 | -0.475831508 | 0.649426273 | 0.994392933 | -5.163162809 |
| **A_55_P2016732** | Slc30a2 | -0.019370261 | -0.116683587 | 0.077943065 | 7.804144537 | -0.475742871 | 0.649486207 | 0.994392933 | -5.163196922 |
| **A_55_P2006615** | Rbbp9 | 0.052623266 | -0.211824656 | 0.317071188 | 10.83358671 | 0.475605392 | 0.649579175 | 0.994392933 | -5.163249823 |
| **A_52_P582858** | Dgkd | -0.031406084 | -0.189339003 | 0.126526835 | 10.45945978 | -0.475280788 | 0.649798708 | 0.994392933 | -5.163374672 |
| **A_55_P2224830** | D230044B1 | 0.066182802 | -0.266669701 | 0.399035305 | 8.879160224 | 0.475228476 | 0.649834091 | 0.994392933 | -5.163394784 |
| **A_55_P2112474** | NA | 0.02028634 | -0.08174315 | 0.12231583 | 16.06001275 | 0.475211693 | 0.649845443 | 0.994392933 | -5.163401237 |

| | | | | | | | | | |
|---|---|---|---|---|---|---|---|---|---|
| **A_52_P413584** | NA | -0.028796124 | -0.173645514 | 0.116053266 | 8.345126323 | -0.475145367 | 0.649890305 | 0.994392933 | -5.163426734 |
| **A_52_P257426** | Eps8l2 | -0.021625676 | -0.130424187 | 0.087172835 | 7.726413617 | -0.475068098 | 0.649942573 | 0.994392933 | -5.163456434 |
| **A_51_P434240** | Mrpl11 | 0.029666757 | -0.119611454 | 0.178944967 | 13.40623976 | 0.474988159 | 0.649996649 | 0.994392933 | -5.163487155 |
| **A_55_P2041708** | Gas2 | -0.030160583 | -0.18197747 | 0.121656304 | 8.644153299 | -0.474819764 | 0.650110568 | 0.994392933 | -5.163551856 |
| **A_51_P115655** | Iqcd | -0.021505734 | -0.129766357 | 0.086754889 | 9.426493192 | -0.474780509 | 0.650137126 | 0.994392933 | -5.163566935 |
| **A_55_P2022987** | Psmd2 | 0.035099141 | -0.141597405 | 0.211795687 | 13.65463253 | 0.474763761 | 0.650148457 | 0.994392933 | -5.163573368 |
| **A_52_P400677** | AW209491 | -0.026362928 | -0.159097164 | 0.106371307 | 9.685524737 | -0.474700655 | 0.650191152 | 0.994392933 | -5.163597606 |
| **A_55_P2031471** | Rufy4 | 0.030393433 | -0.122690511 | 0.183477376 | 8.286760868 | 0.474525161 | 0.650309893 | 0.994392933 | -5.163664997 |
| **A_55_P2424459** | Atg9a | -0.035821722 | -0.216280876 | 0.144637433 | 8.430796437 | -0.474434943 | 0.650370939 | 0.994392933 | -5.163699632 |
| **A_51_P296429** | Tmem170 | 0.022448385 | -0.090643275 | 0.135540045 | 7.814460992 | 0.47442072 | 0.650380563 | 0.994392933 | -5.163705092 |
| **A_55_P1999197** | Clasp1 | -0.037187124 | -0.224546298 | 0.15017205 | 13.16279643 | -0.474380425 | 0.65040783 | 0.994392933 | -5.163720559 |
| **A_51_P348548** | Ecsit | -0.040238905 | -0.24301127 | 0.162533461 | 13.78857464 | -0.474292784 | 0.650467138 | 0.994392933 | -5.163754195 |
| **A_55_P1962359** | Trim11 | 0.0667027 | -0.269441168 | 0.402846568 | 12.21260636 | 0.474271852 | 0.650481303 | 0.994392933 | -5.163762228 |
| **A_52_P131559** | Srprb | -0.035815354 | -0.216328814 | 0.144698106 | 8.515555144 | -0.474207906 | 0.650524578 | 0.994392933 | -5.163786765 |
| **A_55_P1954116** | C8b | 0.022274636 | -0.090018331 | 0.134567603 | 7.720936383 | 0.474096978 | 0.650599651 | 0.994392933 | -5.163829324 |
| **A_51_P469951** | Srgap3 | 0.031474223 | -0.127223401 | 0.190171846 | 8.190124819 | 0.47401679 | 0.650653922 | 0.994392933 | -5.163860084 |
| **A_51_P368543** | Abhd8 | -0.067847623 | -0.409951231 | 0.274255986 | 9.632963762 | -0.474008481 | 0.650659546 | 0.994392933 | -5.16386327 |
| **A_55_P2162160** | Anp32a | 0.032574136 | -0.131687847 | 0.196836119 | 12.8609246 | 0.47396359 | 0.650689929 | 0.994392933 | -5.163880488 |
| **A_51_P346575** | Rhag | 0.020022114 | -0.080947162 | 0.12099139 | 7.702914704 | 0.473947044 | 0.650701129 | 0.994392933 | -5.163886833 |
| **A_55_P1998389** | NA | -0.043097421 | -0.260463636 | 0.174268795 | 16.73050466 | -0.473879987 | 0.650746517 | 0.994392933 | -5.163912549 |
| **A_55_P2074656** | Padi2 | 0.030789676 | -0.124507294 | 0.186086646 | 8.052165199 | 0.473861333 | 0.650759143 | 0.994392933 | -5.163919702 |
| **A_55_P2145480** | NA | -0.032317907 | -0.195323037 | 0.130687223 | 10.81983454 | -0.47386113 | 0.650759281 | 0.994392933 | -5.16391978 |
| **A_51_P232142** | Thumpd3 | 0.026677141 | -0.107888292 | 0.161242575 | 10.90690153 | 0.473821656 | 0.650786 | 0.994392933 | -5.163934915 |
| **A_55_P2170514** | Sfn | 0.03201533 | -0.129510243 | 0.193540903 | 7.86181775 | 0.473724481 | 0.650851778 | 0.994392933 | -5.16397217 |
| **A_52_P376214** | Cstf3 | 0.048914392 | -0.197908867 | 0.295737651 | 10.16100664 | 0.473652366 | 0.650900595 | 0.994392933 | -5.163999813 |
| **A_52_P397204** | Csk | 0.022011413 | -0.08909139 | 0.133114215 | 8.890475039 | 0.473513144 | 0.650994845 | 0.994392933 | -5.164053168 |
| **A_55_P1995243** | Cobl | 0.042756628 | -0.173059083 | 0.258572339 | 8.685457263 | 0.473510402 | 0.650996702 | 0.994392933 | -5.164054219 |
| **A_51_P198387** | Uckl1 | 0.045603825 | -0.184587459 | 0.27579511 | 10.24959238 | 0.473501686 | 0.651002602 | 0.994392933 | -5.164057558 |
| **A_55_P2082434** | Btbd11 | 0.021131867 | -0.085544117 | 0.127807851 | 8.104451612 | 0.473456781 | 0.651033004 | 0.994392933 | -5.164074764 |
| **A_51_P483261** | Pomt1 | -0.026886136 | -0.162614991 | 0.10884272 | 8.820455736 | -0.473440418 | 0.651044082 | 0.994392933 | -5.164081033 |
| **A_55_P1969746** | Pitpnm1 | 0.036617291 | -0.148322653 | 0.221557234 | 10.95153775 | 0.473221663 | 0.651192193 | 0.994392933 | -5.164164828 |
| **A_52_P103977** | Ints8 | -0.025535046 | -0.154611331 | 0.10354124 | 8.373483696 | -0.472823781 | 0.651461629 | 0.994392933 | -5.164317146 |
| **A_52_P37498** | Dpcd | -0.046733346 | -0.282973094 | 0.189506402 | 10.43187487 | -0.472806064 | 0.651473628 | 0.994392933 | -5.164323926 |
| **A_55_P2396375** | 1810073O( | -0.022420323 | -0.13576091 | 0.090920265 | 8.235955328 | -0.472787006 | 0.651486535 | 0.994392933 | -5.164331219 |
| **A_52_P18807** | Eif3c | -0.046066679 | -0.278962538 | 0.186829179 | 13.68605122 | -0.472752977 | 0.651509582 | 0.994392933 | -5.16434424 |
| **A_51_P479865** | Glb1l | -0.024013124 | -0.145443376 | 0.097417128 | 8.146474425 | -0.472640463 | 0.651585786 | 0.994392933 | -5.164387285 |
| **A_55_P1953650** | Zdhhc18 | 0.02057368 | -0.083465027 | 0.124612388 | 12.2832719 | 0.472635321 | 0.65158927 | 0.994392933 | -5.164389252 |
| **A_66_P133204** | NA | 0.023718068 | -0.096265526 | 0.143701661 | 9.337243223 | 0.472461659 | 0.651706898 | 0.994392933 | -5.164455673 |
| **A_52_P445474** | Lmtk2 | -0.024101494 | -0.146026631 | 0.097823642 | 8.529256355 | -0.472454348 | 0.651711851 | 0.994392933 | -5.164458468 |
| **A_66_P113008** | Pla2g2e | 0.019345275 | -0.078522889 | 0.11721344 | 7.735516091 | 0.472435588 | 0.651724558 | 0.994392933 | -5.164465642 |
| **A_52_P99807** | Dpy19l3 | 0.026548315 | -0.107802924 | 0.160899553 | 8.258999118 | 0.47228529 | 0.651826371 | 0.994392933 | -5.164523106 |
| **A_55_P2065519** | NA | 0.036105314 | -0.146635883 | 0.218846511 | 10.43417659 | 0.472219366 | 0.651871032 | 0.994392933 | -5.164548305 |
| **A_51_P495242** | Lat | -0.027033955 | -0.163875123 | 0.109807212 | 7.944165247 | -0.472173875 | 0.651901851 | 0.994392933 | -5.164565692 |
| **A_55_P2134122** | NA | 0.026011001 | -0.105659103 | 0.157681105 | 8.100133317 | 0.472148942 | 0.651918742 | 0.994392933 | -5.164575221 |
| **A_55_P1954476** | NA | 0.023686029 | -0.096221858 | 0.143593917 | 8.127244962 | 0.472121352 | 0.651937434 | 0.994392933 | -5.164585765 |
| **A_55_P2162404** | NA | -0.02097708 | -0.127174087 | 0.085219926 | 8.414681254 | -0.472108586 | 0.651946083 | 0.994392933 | -5.164590644 |
| **A_55_P2059556** | NA | -0.041424721 | -0.251155162 | 0.16830572 | 15.80641541 | -0.472070926 | 0.651971598 | 0.994392933 | -5.164605035 |
| **A_51_P303749** | Depdc1b | -0.032243675 | -0.195507608 | 0.131020259 | 9.367380983 | -0.472023269 | 0.652003887 | 0.994392933 | -5.164623244 |

| | | | | | | | | | |
|---|---|---|---|---|---|---|---|---|---|
| **A_51_P159284** | Bud13 | 0.026290736 | -0.106846468 | 0.15942794 | 8.409360924 | 0.471967881 | 0.652041415 | 0.994392933 | -5.164644406 |
| **A_55_P1966760** | NA | -0.041979756 | -0.261151452 | 0.177191941 | 17.53633895 | -0.475797972 | 0.652052421 | 0.994392933 | -5.088051059 |
| **A_51_P449911** | NA | 0.046567286 | -0.189295257 | 0.282429829 | 12.38044304 | 0.471879468 | 0.65210132 | 0.994392933 | -5.16467818 |
| **A_52_P179068** | Gucy1b1 | 0.033200901 | -0.134966497 | 0.201368299 | 10.65260967 | 0.471864386 | 0.65211154 | 0.994392933 | -5.164683941 |
| **A_55_P2108334** | Med10 | -0.046570106 | -0.282565316 | 0.189425103 | 10.77092408 | -0.471642761 | 0.652261718 | 0.994392933 | -5.164768575 |
| **A_55_P1977538** | Apba3 | -0.041488672 | -0.251778959 | 0.168801616 | 10.28849273 | -0.471540989 | 0.652330688 | 0.994392933 | -5.164807428 |
| **A_55_P2045026** | Mgrn1 | -0.040930337 | -0.24842896 | 0.166568287 | 14.08672934 | -0.471453896 | 0.652389712 | 0.994392933 | -5.16484067 |
| **A_55_P2044005** | NA | -0.037510274 | -0.227723274 | 0.152702725 | 9.286520229 | -0.471323592 | 0.652478027 | 0.994392933 | -5.164890396 |
| **A_52_P138598** | Pum2 | 0.039481058 | -0.160776486 | 0.239738603 | 8.668581682 | 0.471204068 | 0.65255904 | 0.994392933 | -5.164935996 |
| **A_52_P358349** | Mtdh | -0.045105904 | -0.27392939 | 0.183717583 | 11.77764059 | -0.471131264 | 0.65260839 | 0.994392933 | -5.164963767 |
| **A_55_P1994917** | Wdr5 | 0.030725148 | -0.125190356 | 0.186640652 | 11.16802972 | 0.470992309 | 0.652702583 | 0.994392933 | -5.165016759 |
| **A_55_P2079041** | NA | -0.03754778 | -0.22810793 | 0.15301237 | 15.84502205 | -0.470935371 | 0.652741182 | 0.994392933 | -5.165038469 |
| **A_55_P2161958** | Gm10324 | 0.024289845 | -0.098990107 | 0.147569798 | 9.442518419 | 0.470913804 | 0.652755803 | 0.994392933 | -5.165046692 |
| **A_52_P670812** | Snord123 | 0.053169121 | -0.216713086 | 0.323051329 | 11.15534773 | 0.470862778 | 0.652790395 | 0.994392933 | -5.165066145 |
| **A_55_P2083484** | Mettl21a | -0.029888884 | -0.181604376 | 0.121826607 | 8.382604335 | -0.470856864 | 0.652794404 | 0.994392933 | -5.1650684 |
| **A_51_P128147** | Chmp1a | -0.036317102 | -0.220703406 | 0.148069203 | 10.9264586 | -0.470751434 | 0.652865883 | 0.994392933 | -5.165108587 |
| **A_51_P112557** | Polr1c | 0.036019456 | -0.146858218 | 0.218897129 | 11.00486506 | 0.470744862 | 0.652870338 | 0.994392933 | -5.147600867 |
| **A_52_P244702** | Tcf7 | -0.029406509 | -0.178742801 | 0.119929782 | 7.822154493 | -0.470638275 | 0.652942604 | 0.994392933 | -5.165151711 |
| **A_55_P2139937** | Lrrc56 | -0.036795425 | -0.223695024 | 0.150104175 | 8.994510827 | -0.470537869 | 0.653010684 | 0.994392933 | -5.165189967 |
| **A_51_P518163** | Rrp9 | 0.058888782 | -0.240235826 | 0.358013391 | 9.284280733 | 0.470532432 | 0.653014371 | 0.994392933 | -5.165192039 |
| **A_55_P1971249** | Eya3 | 0.032968969 | -0.134521514 | 0.200459453 | 11.95523164 | 0.470461804 | 0.653062262 | 0.994392933 | -5.165218944 |
| **A_51_P194004** | Glyctk | -0.020627798 | -0.125423015 | 0.084167419 | 7.754169412 | -0.47045766 | 0.653065072 | 0.994392933 | -5.165220523 |
| **A_66_P140307** | NA | -0.052021128 | -0.316331692 | 0.212289437 | 15.23025639 | -0.470407645 | 0.653098987 | 0.994392933 | -5.165239573 |
| **A_52_P178904** | Seh1l | -0.027585489 | -0.167757274 | 0.112586296 | 8.439367899 | -0.470358737 | 0.653132152 | 0.994392933 | -5.165258201 |
| **A_51_P175232** | Fbxl4 | -0.045466347 | -0.276502137 | 0.185569442 | 9.42204447 | -0.470348686 | 0.653138968 | 0.994392933 | -5.165262028 |
| **A_55_P2108784** | Arhgap22 | -0.03744566 | -0.227782246 | 0.152890927 | 8.353203227 | -0.47020619 | 0.653235602 | 0.994392933 | -5.165316289 |
| **A_55_P2143030** | Timd2 | 0.019445289 | -0.079397466 | 0.118288044 | 7.656536454 | 0.470195745 | 0.653242686 | 0.994392933 | -5.165320265 |
| **A_51_P303906** | Actr2 | 0.050570705 | -0.206553124 | 0.307694533 | 12.03544076 | 0.470073547 | 0.65332556 | 0.994392933 | -5.165366783 |
| **A_55_P2124586** | Xlr4a | 0.06088715 | -0.248709885 | 0.370484184 | 7.956960615 | 0.470043426 | 0.653345989 | 0.994392933 | -5.165378248 |
| **A_55_P1979397** | Palm | 0.022765094 | -0.093009806 | 0.138539994 | 11.68420631 | 0.469963532 | 0.653400177 | 0.994392933 | -5.165408654 |
| **A_55_P2091756** | Nxnl1 | -0.035264478 | -0.214612662 | 0.144083706 | 8.286610008 | -0.469947783 | 0.653410859 | 0.994392933 | -5.165414647 |
| **A_55_P2070546** | 1700037C1 | 0.019741562 | -0.080678597 | 0.120161722 | 9.669409138 | 0.469861375 | 0.653469468 | 0.994392933 | -5.165447526 |
| **A_51_P163694** | Rufy1 | -0.030677648 | -0.186736658 | 0.125381363 | 10.21441187 | -0.469831723 | 0.653489582 | 0.994392933 | -5.165458808 |
| **A_51_P517012** | Tysnd1 | 0.037255939 | -0.152284642 | 0.226796521 | 11.1742972 | 0.469788563 | 0.653518857 | 0.994392933 | -5.165475227 |
| **A_55_P2002864** | Aifm1 | 0.031125655 | -0.127272188 | 0.189523497 | 12.95890235 | 0.469654368 | 0.653609889 | 0.994392933 | -5.165526271 |
| **A_55_P2340206** | E230006M | 0.025710106 | -0.10514964 | 0.156569852 | 8.930848253 | 0.469577117 | 0.653662295 | 0.994392933 | -5.165555649 |
| **A_55_P1987222** | NA | -0.026721202 | -0.1627437 | 0.109301296 | 8.347262892 | -0.469520304 | 0.653700837 | 0.994392933 | -5.165577251 |
| **A_55_P2178044** | Inppl1 | 0.048723244 | -0.199333594 | 0.296780082 | 13.25749947 | 0.469455166 | 0.653745029 | 0.994392933 | -5.165602016 |
| **A_52_P612382** | Cdc25b | 0.062653208 | -0.256341162 | 0.381647578 | 10.77142409 | 0.469428473 | 0.653763139 | 0.994392933 | -5.165612164 |
| **A_55_P2013043** | Serpinb6b | -0.042177128 | -0.256921216 | 0.17256696 | 10.37594255 | -0.469423603 | 0.653766443 | 0.994392933 | -5.165614016 |
| **A_55_P1960023** | NA | 0.024708925 | -0.101126641 | 0.150544491 | 8.805592697 | 0.469309749 | 0.65384369 | 0.994392933 | -5.165657292 |
| **A_55_P2092730** | Wdfy1 | 0.061867763 | -0.2532162 | 0.376951725 | 8.913352055 | 0.469296423 | 0.653852731 | 0.994392933 | -5.165662357 |
| **A_51_P225506** | Pnrc2 | -0.040680527 | -0.247905946 | 0.166544892 | 12.51743853 | -0.469194243 | 0.653922063 | 0.994392933 | -5.165701187 |
| **A_55_P1993054** | Psip1 | 0.030207778 | -0.123686656 | 0.184102213 | 9.747881017 | 0.469142743 | 0.653957008 | 0.994392933 | -5.165720755 |
| **A_51_P155174** | Zfp672 | -0.058619011 | -0.357310583 | 0.24007256 | 10.11350721 | -0.469055955 | 0.6540159 | 0.994392933 | -5.165753726 |
| **A_52_P176999** | Klhl31 | -0.037282064 | -0.227346891 | 0.152782764 | 8.577643625 | -0.468821283 | 0.654175154 | 0.994392933 | -5.165842852 |
| **A_51_P354272** | Cluh | 0.031892075 | -0.13074722 | 0.194531371 | 13.60966211 | 0.468669219 | 0.654278359 | 0.994392933 | -5.165900582 |
| **A_55_P2136662** | 8030462N1 | 0.030661589 | -0.125705933 | 0.187029112 | 8.397903218 | 0.4686593 | 0.654285092 | 0.994392933 | -5.165904347 |

| | | | | | | | | | |
|---|---|---|---|---|---|---|---|---|---|
| **A_52_P72237** | Actg1 | 0.045393818 | -0.186106375 | 0.276894011 | 11.36179562 | 0.468656331 | 0.654287107 | 0.994392933 | -5.165905474 |
| **A_52_P612803** | Ccng1 | -0.032984408 | -0.201209883 | 0.135241068 | 11.76587671 | -0.468625651 | 0.65430793 | 0.994392933 | -5.165917119 |
| **A_55_P2057916** | Pex5 | -0.030084355 | -0.183545652 | 0.123376942 | 9.66237131 | -0.468544644 | 0.654362914 | 0.994392933 | -5.165947863 |
| **A_55_P2153600** | Mfsd4b3-p | -0.019703527 | -0.120215651 | 0.080808597 | 7.928911565 | -0.468527035 | 0.654374866 | 0.994392933 | -5.165954546 |
| **A_52_P320261** | Cpa2 | 0.041286765 | -0.169339345 | 0.251912876 | 8.479643803 | 0.468498047 | 0.654394542 | 0.994392933 | -5.165965546 |
| **A_51_P348183** | Tmem141 | -0.040136219 | -0.24489386 | 0.164621423 | 11.61810795 | -0.46849555 | 0.654396238 | 0.994392933 | -5.165966493 |
| **A_51_P149349** | Nfxl1 | -0.02674256 | -0.163227011 | 0.10974189 | 8.188847944 | -0.468305159 | 0.654525478 | 0.994392933 | -5.166038725 |
| **A_55_P2155499** | Tmem9 | 0.045757228 | -0.187801211 | 0.279315667 | 8.841776804 | 0.468245133 | 0.654566228 | 0.994392933 | -5.166061493 |
| **A_55_P2007155** | Akr1c14 | -0.026045575 | -0.159001553 | 0.106910403 | 8.203817761 | -0.468204101 | 0.654594084 | 0.994392933 | -5.166077054 |
| **A_52_P407692** | H2afj | 0.026756406 | -0.109835675 | 0.163348486 | 16.81888254 | 0.468178412 | 0.654611523 | 0.994392933 | -5.166086796 |
| **A_55_P2098598** | Btg1 | -0.044451913 | -0.271445731 | 0.182541905 | 13.23001404 | -0.468042787 | 0.654703602 | 0.994392933 | -5.166138221 |
| **A_55_P2143025** | Sema3c | -0.073990204 | -0.45182711 | 0.303846702 | 9.699397171 | -0.4680357 | 0.654708414 | 0.994392933 | -5.166140908 |
| **A_55_P1957267** | Rab9 | 0.031021184 | -0.127397364 | 0.189439732 | 11.28430032 | 0.468016838 | 0.65472122 | 0.994392933 | -5.166148059 |
| **A_55_P2014304** | Kank1 | 0.05054426 | -0.207598893 | 0.308687413 | 12.25098953 | 0.467972532 | 0.654751302 | 0.994392933 | -5.166164855 |
| **A_55_P1974362** | Hp1bp3 | 0.048758331 | -0.200355788 | 0.297872451 | 13.20579563 | 0.467799361 | 0.654868885 | 0.994454555 | -5.166230487 |
| **A_55_P1962876** | Adam11 | -0.036125951 | -0.220714243 | 0.148462341 | 8.841406471 | -0.467761281 | 0.654894743 | 0.994454555 | -5.166244917 |
| **A_51_P346132** | Rhox9 | -0.140735786 | -0.860619054 | 0.579147481 | 8.057847615 | -0.467252461 | 0.6552403 | 0.99471946 | -5.166437619 |
| **A_55_P2042217** | Fam161a | -0.019167117 | -0.117258268 | 0.078924034 | 7.737597095 | -0.46702066 | 0.655397753 | 0.99471946 | -5.166525343 |
| **A_55_P2142451** | Echdc2 | -0.039484565 | -0.241596635 | 0.162627504 | 10.78623978 | -0.466921901 | 0.655464843 | 0.99471946 | -5.166562706 |
| **A_55_P2156718** | NA | -0.022193361 | -0.135835564 | 0.091448842 | 7.721035401 | -0.466758862 | 0.655575607 | 0.99471946 | -5.166624371 |
| **A_55_P2225082** | Mettl4 | 0.037077527 | -0.15281468 | 0.226969734 | 8.548061614 | 0.466673086 | 0.655633884 | 0.99471946 | -5.166656806 |
| **A_52_P239052** | Zfp148 | 0.023286479 | -0.095987181 | 0.14256014 | 11.53276678 | 0.466625447 | 0.655666252 | 0.99471946 | -5.166674818 |
| **A_55_P2164265** | Prl7c1 | -0.025044994 | -0.153336727 | 0.103246738 | 7.693304726 | -0.466585671 | 0.655693277 | 0.99471946 | -5.166689855 |
| **A_55_P2088095** | NA | -0.029796204 | -0.182426682 | 0.122834275 | 10.64601646 | -0.466582884 | 0.655695171 | 0.99471946 | -5.166690909 |
| **A_52_P232637** | Dhh | 0.03226125 | -0.133001117 | 0.197523617 | 9.059645972 | 0.466569512 | 0.655704257 | 0.99471946 | -5.166695964 |
| **A_55_P2170834** | NA | 0.08152648 | -0.336197769 | 0.499250729 | 9.939312779 | 0.466464005 | 0.655775947 | 0.99471946 | -5.166735844 |
| **A_55_P1997105** | Large2 | -0.028606028 | -0.175195142 | 0.117983087 | 8.511423828 | -0.466406894 | 0.655814755 | 0.99471946 | -5.166757427 |
| **A_55_P2035657** | Psmb6 | -0.036248491 | -0.222054565 | 0.149557583 | 15.07674247 | -0.46627181 | 0.655906551 | 0.99471946 | -5.166808469 |
| **A_51_P112237** | Angel1 | 0.02348809 | -0.096948103 | 0.143924282 | 9.221236702 | 0.466122221 | 0.656008211 | 0.99471946 | -5.149329785 |
| **A_51_P492830** | Cenph | 0.034013401 | -0.140404292 | 0.208431095 | 9.471743772 | 0.466088787 | 0.656030934 | 0.99471946 | -5.166877603 |
| **A_55_P2148431** | Slc4a3 | 0.024893232 | -0.102806435 | 0.152592898 | 8.519881697 | 0.465908517 | 0.656153457 | 0.99471946 | -5.166945673 |
| **A_51_P411757** | Ankrd13b | 0.022616999 | -0.093412446 | 0.138646444 | 11.56345233 | 0.465881961 | 0.656171507 | 0.99471946 | -5.166955699 |
| **A_55_P2108143** | NA | 0.034314479 | -0.141750385 | 0.210379343 | 10.62950693 | 0.465815402 | 0.656216748 | 0.99471946 | -5.166980824 |
| **A_55_P2001976** | Rhox12 | 0.019128426 | -0.079020573 | 0.117277425 | 7.669810926 | 0.465803226 | 0.656225025 | 0.99471946 | -5.16698542 |
| **A_55_P2136154** | Heca | -0.030961171 | -0.189885446 | 0.127963105 | 10.00555202 | -0.465624975 | 0.656346194 | 0.99471946 | -5.167052689 |
| **A_52_P339722** | Csnk1a1 | -0.047086369 | -0.288937191 | 0.194764454 | 11.95911942 | -0.465325403 | 0.656549858 | 0.99471946 | -5.16716569 |
| **A_51_P501897** | Pitx2 | 0.117315245 | -0.485330533 | 0.719961023 | 8.6661948 | 0.465266159 | 0.656590139 | 0.99471946 | -5.16718803 |
| **A_55_P1990964** | Osbpl11 | -0.029808218 | -0.18295168 | 0.123335245 | 10.95082139 | -0.46520747 | 0.656630044 | 0.99471946 | -5.167210158 |
| **A_55_P2042753** | Cltrn | 0.019076638 | -0.078942988 | 0.117096263 | 7.66328646 | 0.465155254 | 0.656665549 | 0.99471946 | -5.167229842 |
| **A_55_P2005229** | NA | 0.025124596 | -0.10399292 | 0.154242111 | 10.69898943 | 0.465075059 | 0.656720079 | 0.99471946 | -5.167260071 |
| **A_51_P331752** | Ccl11 | 0.02455357 | -0.101641631 | 0.15074877 | 7.72277351 | 0.465029958 | 0.656750748 | 0.99471946 | -5.167277069 |
| **A_55_P2123331** | Cog2 | -0.022215377 | -0.136400174 | 0.09196942 | 8.122492893 | -0.465001693 | 0.656769969 | 0.99471946 | -5.167287722 |
| **A_51_P278266** | Gltpd2 | -0.021421311 | -0.131539228 | 0.088696605 | 7.744848292 | -0.464940305 | 0.656811714 | 0.99471946 | -5.167310855 |
| **A_66_P112006** | H2-T10 | -0.029841004 | -0.183256725 | 0.123574717 | 8.092066834 | -0.464892667 | 0.65684411 | 0.99471946 | -5.167328804 |
| **A_66_P132448** | Gnpnat1 | 0.040621094 | -0.168235137 | 0.249477325 | 8.795057149 | 0.464850514 | 0.656872777 | 0.99471946 | -5.167344686 |
| **A_51_P319879** | Acadm | -0.055321983 | -0.339800466 | 0.2291565 | 13.92773908 | -0.464790721 | 0.656913441 | 0.99471946 | -5.167367211 |
| **A_55_P2041584** | NA | -0.030525017 | -0.187508657 | 0.126458622 | 7.876783195 | -0.464740648 | 0.656947496 | 0.99471946 | -5.167386073 |
| **A_55_P2124523** | Gm13620 | -0.020134606 | -0.123685151 | 0.083415939 | 7.829033563 | -0.464729114 | 0.65695534 | 0.99471946 | -5.167390417 |

| | | | | | | | | | |
|---|---|---|---|---|---|---|---|---|---|
| A_51_P460734 | Prkcd | -0.033626903 | -0.206605068 | 0.139351263 | 9.872760812 | -0.464627288 | 0.657024595 | 0.99471946 | -5.167428766 |
| A_55_P2314496 | Secisbp2l | -0.020049753 | -0.123219415 | 0.083119909 | 8.672544295 | -0.464479078 | 0.657125404 | 0.99471946 | -5.16748457 |
| A_55_P1999666 | Cntrl | 0.075833395 | -0.314495324 | 0.466162114 | 9.71284652 | 0.464343237 | 0.657217807 | 0.99471946 | -5.167535703 |
| A_51_P209782 | Cyp2c23 | -0.032367899 | -0.204155553 | 0.139419756 | 7.728958127 | -0.46804726 | 0.657256398 | 0.99471946 | -5.090865646 |
| A_51_P257156 | Ddrgk1 | -0.041644741 | -0.256039716 | 0.172750233 | 9.770435255 | -0.464252983 | 0.657279203 | 0.99471946 | -5.167569669 |
| A_51_P174961 | F10 | -0.023780429 | -0.146222962 | 0.098662104 | 8.082330218 | -0.464190783 | 0.657321517 | 0.99471946 | -5.167593073 |
| A_55_P1959718 | Cobll1 | -0.027263392 | -0.167687525 | 0.113160742 | 11.20620565 | -0.464031283 | 0.65743003 | 0.99471946 | -5.167653075 |
| A_51_P176265 | Trip11 | -0.025909486 | -0.159381632 | 0.107562661 | 8.637050089 | -0.463956515 | 0.6574809 | 0.99471946 | -5.167681196 |
| A_52_P24439 | Sftpa1 | -0.018699507 | -0.115031718 | 0.077632705 | 7.640651034 | -0.463946343 | 0.657487821 | 0.99471946 | -5.167685021 |
| A_55_P2118037 | Sirpa | -0.027216457 | -0.167451382 | 0.113018468 | 9.488968449 | -0.46385745 | 0.657548304 | 0.99471946 | -5.167718448 |
| A_55_P1981709 | Rreb1 | 0.077918692 | -0.323602679 | 0.479440064 | 9.617595563 | 0.463812149 | 0.657579128 | 0.99471946 | -5.167735481 |
| A_51_P268094 | Serpine2 | -0.029474271 | -0.181395397 | 0.122446856 | 8.624737574 | -0.463696726 | 0.657657669 | 0.99471946 | -5.167778871 |
| A_55_P1987514 | NA | 0.027796636 | -0.115488221 | 0.171081493 | 16.70015128 | 0.463661551 | 0.657681605 | 0.99471946 | -5.167792093 |
| A_51_P416059 | Slc49a4 | 0.035106361 | -0.145904786 | 0.216117508 | 8.611971244 | 0.463542573 | 0.657762571 | 0.99471946 | -5.167836806 |
| A_55_P2422243 | 2900072G1 | 0.059560385 | -0.247558261 | 0.36667903 | 10.05619713 | 0.463511424 | 0.657783769 | 0.99471946 | -5.16784851 |
| A_51_P177432 | NA | 0.020302416 | -0.084405578 | 0.125010409 | 8.009585983 | 0.463422389 | 0.657844362 | 0.99471946 | -5.167881962 |
| A_55_P2057142 | Cks1brt | -0.039366557 | -0.242421393 | 0.163688279 | 9.199346798 | -0.463364997 | 0.657883422 | 0.99471946 | -5.167903521 |
| A_52_P487024 | Smg7 | 0.028516503 | -0.118613476 | 0.175646483 | 9.422284215 | 0.463238055 | 0.657969821 | 0.99471946 | -5.167951199 |
| A_51_P194479 | 1110059G1 | -0.034385638 | -0.211805023 | 0.143033748 | 9.736630837 | -0.463217701 | 0.657983675 | 0.99471946 | -5.167958843 |
| A_55_P1975155 | Galnt6 | -0.073454291 | -0.452527013 | 0.30561843 | 8.297586355 | -0.463130915 | 0.658042747 | 0.99471946 | -5.16799143 |
| A_55_P2002426 | Rps6kb2 | -0.03427719 | -0.211208855 | 0.142654475 | 8.630756466 | -0.463029629 | 0.658111692 | 0.99471946 | -5.168029456 |
| A_55_P2326582 | 5730408K0 | 0.019360206 | -0.080585788 | 0.119306201 | 7.769670292 | 0.462970932 | 0.658151648 | 0.99471946 | -5.168051488 |
| A_55_P2087375 | Gm4987 | -0.026110749 | -0.160934045 | 0.108712548 | 15.96723702 | -0.462874773 | 0.658217108 | 0.99471946 | -5.168087577 |
| A_66_P132969 | NA | -0.046711517 | -0.287909609 | 0.194486574 | 14.4947105 | -0.462870218 | 0.658220209 | 0.99471946 | -5.168089286 |
| A_55_P2112892 | Dab1 | -0.036521561 | -0.225203307 | 0.152160186 | 8.621629346 | -0.462624439 | 0.658387538 | 0.99471946 | -5.168181497 |
| A_55_P2017636 | Thbs1 | 0.108745469 | -0.453133641 | 0.670624579 | 10.41600226 | 0.46256995 | 0.658424638 | 0.99471946 | -5.168201934 |
| A_66_P111773 | Gatad2a | -0.059711392 | -0.368239601 | 0.248816818 | 9.870571471 | -0.462563593 | 0.658428966 | 0.99471946 | -5.168204318 |
| A_55_P2157019 | Phb2 | -0.070230493 | -0.433218809 | 0.292757822 | 11.2950402 | -0.46242596 | 0.65852268 | 0.99471946 | -5.168255928 |
| A_55_P2308835 | E130110O2 | -0.02063936 | -0.127334562 | 0.086055841 | 8.322918943 | -0.462338942 | 0.658581934 | 0.99471946 | -5.168288551 |
| A_52_P222354 | Vps13c | -0.021564839 | -0.133052299 | 0.089922621 | 8.38224074 | -0.462305783 | 0.658604514 | 0.99471946 | -5.168300981 |
| A_55_P2078845 | Qk | -0.03290101 | -0.203037316 | 0.137235296 | 12.71513011 | -0.462190874 | 0.658682765 | 0.99471946 | -5.168344049 |
| A_55_P2058903 | Sgpp1 | 0.030137434 | -0.125719911 | 0.185994778 | 11.62842869 | 0.462155499 | 0.658706856 | 0.99471946 | -5.168357306 |
| A_51_P205740 | Ube3b | 0.031067004 | -0.129611452 | 0.191745459 | 12.31312901 | 0.462115834 | 0.658733869 | 0.99471946 | -5.168372169 |
| A_55_P2011390 | Tead1 | -0.031024616 | -0.191603009 | 0.129553778 | 8.693587206 | -0.461772891 | 0.658967445 | 0.99471946 | -5.168500627 |
| A_55_P2096055 | NA | 0.063409661 | -0.264793674 | 0.391612995 | 13.14717291 | 0.461765596 | 0.658972413 | 0.99471946 | -5.168503358 |
| A_51_P314339 | Chtop | 0.02182058 | -0.09112583 | 0.13476699 | 11.30881491 | 0.461745845 | 0.658985867 | 0.99471946 | -5.168510754 |
| A_55_P2064257 | Dnaic2 | -0.022844983 | -0.141107122 | 0.095417157 | 7.71867416 | -0.461694 | 0.659021183 | 0.99471946 | -5.168530165 |
| A_55_P2023118 | Prss28 | 0.020493982 | -0.085610287 | 0.126598251 | 7.875417862 | 0.461639139 | 0.659058553 | 0.99471946 | -5.168550703 |
| A_55_P2041843 | Nde1 | -0.025913028 | -0.160124872 | 0.108298816 | 8.307413053 | -0.461462541 | 0.659178857 | 0.99471946 | -5.1686168 |
| A_55_P2058791 | 4833420G1 | 0.022692667 | -0.09486966 | 0.140254995 | 8.518559468 | 0.461345727 | 0.65925844 | 0.99471946 | -5.168660509 |
| A_55_P2088912 | Gapdh | -0.060920064 | -0.376646332 | 0.254806204 | 15.08466159 | -0.46116758 | 0.659379818 | 0.99471946 | -5.168727147 |
| A_51_P240594 | Zfp7 | 0.026356355 | -0.110249415 | 0.162962124 | 8.732037636 | 0.461132184 | 0.659403936 | 0.99471946 | -5.168740384 |
| A_51_P202498 | Sar1a | -0.034369216 | -0.212522388 | 0.143783955 | 10.60140615 | -0.461089469 | 0.659433041 | 0.99471946 | -5.168756358 |
| A_55_P2063396 | Zfand2a | 0.028325443 | -0.11851584 | 0.175166727 | 10.57172016 | 0.461039014 | 0.659467421 | 0.99471946 | -5.168775224 |
| A_51_P176474 | Kndc1 | 0.019167251 | -0.080199391 | 0.118533893 | 7.67237151 | 0.461029108 | 0.659474172 | 0.99471946 | -5.168778928 |
| A_51_P420100 | Emc2 | -0.043210923 | -0.26730965 | 0.180887804 | 11.51990899 | -0.460853922 | 0.659593551 | 0.99471946 | -5.168844417 |
| A_55_P1968789 | Rasa3 | 0.036599206 | -0.153241432 | 0.226439844 | 11.99043648 | 0.460777869 | 0.65964538 | 0.99471946 | -5.16887284 |
| A_55_P2103682 | NA | -0.04439788 | -0.274785718 | 0.185989958 | 15.65588116 | -0.460587149 | 0.659775362 | 0.99471946 | -5.168944099 |

| | | | | | | | | | |
|---|---|---|---|---|---|---|---|---|---|
| **A_52_P9437** | Psmb3 | 0.024653955 | -0.103299145 | 0.152607054 | 14.53856351 | 0.460516205 | 0.659823717 | 0.99471946 | -5.168970599 |
| **A_55_P2010332** | Ewsr1 | 0.032889294 | -0.137822314 | 0.203600901 | 12.18970622 | 0.460469242 | 0.659855726 | 0.99471946 | -5.168988139 |
| **A_55_P2061456** | Nfyc | 0.01975754 | -0.08281092 | 0.122326001 | 9.830807982 | 0.46039243 | 0.659908083 | 0.99471946 | -5.169016824 |
| **A_55_P2017362** | Prr36 | -0.022222151 | -0.137589135 | 0.093144833 | 7.761912276 | -0.460377075 | 0.65991855 | 0.99471946 | -5.169022558 |
| **A_55_P2121185** | Omp | -0.019062805 | -0.118031763 | 0.079906154 | 7.726274778 | -0.460359306 | 0.659930662 | 0.99471946 | -5.169029193 |
| **A_52_P454295** | Ttn | -0.071043822 | -0.440142742 | 0.298055099 | 13.38912166 | -0.460036905 | 0.660150444 | 0.99471946 | -5.169149536 |
| **A_65_P20174** | Jade1 | -0.035692246 | -0.221180882 | 0.149796391 | 8.563523435 | -0.459902424 | 0.660242131 | 0.99471946 | -5.169199711 |
| **A_55_P1979699** | Spop | -0.058932045 | -0.365210914 | 0.247346824 | 10.20025332 | -0.459879035 | 0.660258078 | 0.99471946 | -5.169208436 |
| **A_55_P2008755** | NA | -0.029347888 | -0.181883903 | 0.123188127 | 13.52817383 | -0.459847248 | 0.660279751 | 0.99471946 | -5.169220293 |
| **A_51_P183995** | Paf1 | 0.023929947 | -0.100460445 | 0.14832034 | 12.49335272 | 0.459794779 | 0.660315526 | 0.99471946 | -5.169239863 |
| **A_55_P1969857** | Ap2m1 | 0.022691914 | -0.095279065 | 0.140662893 | 13.51315549 | 0.459732357 | 0.660358089 | 0.99471946 | -5.169263143 |
| **A_55_P2113984** | Mospd2 | -0.020042936 | -0.124260723 | 0.08417485 | 8.164490366 | -0.459651452 | 0.660413256 | 0.99471946 | -5.169293312 |
| **A_51_P293339** | Tob1 | 0.057052963 | -0.239667222 | 0.353773147 | 10.5043312 | 0.459557907 | 0.660477046 | 0.99471946 | -5.169328188 |
| **A_51_P451516** | Arhgef25 | 0.075559827 | -0.317453631 | 0.468573285 | 11.66189548 | 0.459507565 | 0.660511376 | 0.99471946 | -5.169346954 |
| **A_55_P2183404** | Ptn | 0.056572202 | -0.237681595 | 0.350825999 | 11.0156533 | 0.459504894 | 0.660513197 | 0.99471946 | -5.16934795 |
| **A_52_P533083** | Dnajc16 | -0.025436873 | -0.157763103 | 0.106889357 | 9.037936677 | -0.45943801 | 0.66055881 | 0.99471946 | -5.16937288 |
| **A_51_P428364** | Psma6 | -0.041357446 | -0.256545263 | 0.173830371 | 12.11572633 | -0.459351526 | 0.66061779 | 0.99471946 | -5.169405109 |
| **A_55_P2186787** | NA | 0.023147197 | -0.097309129 | 0.143603523 | 16.96342577 | 0.459280408 | 0.660666294 | 0.99471946 | -5.169431609 |
| **A_55_P2186025** | NA | -0.023758167 | -0.147432355 | 0.099916022 | 16.55762183 | -0.459137733 | 0.660763605 | 0.99471946 | -5.16948476 |
| **A_55_P2069425** | Ttll3 | 0.030208506 | -0.127049075 | 0.187466086 | 8.17117626 | 0.459120605 | 0.660775288 | 0.99471946 | -5.169491139 |
| **A_55_P2034517** | NA | 0.02231161 | -0.093922018 | 0.138545238 | 7.679235942 | 0.458783979 | 0.661004914 | 0.99471946 | -5.169616478 |
| **A_55_P2003981** | NA | -0.018868574 | -0.117176444 | 0.079439295 | 7.658258305 | -0.45873295 | 0.661039726 | 0.99471946 | -5.169635471 |
| **A_55_P2045741** | Fyb | -0.019327933 | -0.120072321 | 0.081416456 | 7.815394074 | -0.458536244 | 0.661173929 | 0.99471946 | -5.169708666 |
| **A_51_P242166** | Lap3 | 0.031410371 | -0.132352429 | 0.19517317 | 12.02976078 | 0.458423572 | 0.661250805 | 0.99471946 | -5.169750578 |
| **A_55_P2100895** | Arnt | 0.025077863 | -0.105684995 | 0.155840721 | 10.73385826 | 0.458369022 | 0.661288026 | 0.99471946 | -5.169770866 |
| **A_55_P2021119** | Sun2 | -0.034032141 | -0.211495449 | 0.143431168 | 12.75727935 | -0.458342181 | 0.661306342 | 0.99471946 | -5.169780848 |
| **A_51_P225224** | Htra1 | -0.040519033 | -0.251829525 | 0.17079146 | 12.39210581 | -0.458297129 | 0.661337083 | 0.99471946 | -5.169797602 |
| **A_51_P255329** | Riok2 | 0.026767911 | -0.112884776 | 0.166420598 | 10.57163954 | 0.458114793 | 0.661461509 | 0.99471946 | -5.16986539 |
| **A_52_P219966** | Spg21 | -0.022918295 | -0.142498044 | 0.096661455 | 8.415970541 | -0.458072054 | 0.661490676 | 0.99471946 | -5.169881277 |
| **A_52_P350111** | Cyth1 | -0.030323558 | -0.188544079 | 0.127896963 | 10.82345238 | -0.458064335 | 0.661495944 | 0.99471946 | -5.169884146 |
| **A_52_P68028** | Ace | 0.033257322 | -0.140275544 | 0.206790187 | 8.083663885 | 0.458051877 | 0.661504446 | 0.99471946 | -5.169888776 |
| **A_52_P56634** | Map3k11 | 0.061301469 | -0.258573675 | 0.381176612 | 9.601910641 | 0.458035899 | 0.66151535 | 0.99471946 | -5.169894714 |
| **A_55_P1972019** | 4632415L0 | -0.036683195 | -0.228135048 | 0.154768657 | 8.45249873 | -0.457948583 | 0.661574941 | 0.99471946 | -5.169927163 |
| **A_55_P2068812** | Pkm | 0.029787992 | -0.125680516 | 0.185256499 | 16.08664942 | 0.457939313 | 0.661581268 | 0.99471946 | -5.169930607 |
| **A_51_P212183** | Dmac1 | 0.035140164 | -0.148270218 | 0.218550547 | 12.6695912 | 0.457919357 | 0.661594887 | 0.99471946 | -5.169938022 |
| **A_51_P140368** | Cmc2 | -0.027922445 | -0.173695435 | 0.117850545 | 12.80006601 | -0.457810254 | 0.661669352 | 0.99471946 | -5.169978557 |
| **A_55_P1968365** | NA | -0.019670049 | -0.122376967 | 0.083036868 | 8.084279235 | -0.457735809 | 0.661720165 | 0.99471946 | -5.17000621 |
| **A_51_P320888** | Cln5 | -0.030587366 | -0.190305227 | 0.129130494 | 10.82715501 | -0.457717728 | 0.661732506 | 0.99471946 | -5.170012925 |
| **A_52_P302463** | Arl5a | -0.031481566 | -0.195896898 | 0.132933766 | 9.082884438 | -0.457639125 | 0.661786159 | 0.99471946 | -5.170042118 |
| **A_55_P2046393** | Rcn3 | 0.062955524 | -0.265875663 | 0.391786711 | 13.85291722 | 0.4575831 | 0.661824402 | 0.99471946 | -5.170062922 |
| **A_52_P604195** | Mbp | 0.119754676 | -0.505868124 | 0.745377476 | 9.16841951 | 0.457497836 | 0.661882606 | 0.99471946 | -5.170094579 |
| **A_55_P2122830** | Zfp185 | 0.067789517 | -0.286401631 | 0.421980665 | 8.479829064 | 0.457439821 | 0.66192221 | 0.99471946 | -5.170116115 |
| **A_51_P256066** | Tiam2 | -0.038437245 | -0.239289026 | 0.162414535 | 9.038449344 | -0.457388984 | 0.661956916 | 0.99471946 | -5.170134986 |
| **A_55_P1967720** | Polr2i | 0.025235564 | -0.106638414 | 0.157109542 | 13.15548917 | 0.457365122 | 0.661973206 | 0.99471946 | -5.170143842 |
| **A_55_P2126269** | Nmb | -0.029377925 | -0.182924593 | 0.124168743 | 9.522027279 | -0.457288049 | 0.662025824 | 0.99471946 | -5.170172446 |
| **A_52_P411003** | Dlgap5 | -0.025612503 | -0.159544382 | 0.108319376 | 8.290229294 | -0.457064186 | 0.662178668 | 0.99471946 | -5.170255501 |
| **A_55_P2033520** | Med6 | -0.021474715 | -0.133771747 | 0.090822316 | 9.891130017 | -0.457054801 | 0.662185077 | 0.99471946 | -5.170258982 |
| **A_51_P214487** | 2-Sep | 0.024659142 | -0.10429476 | 0.153613045 | 14.05880694 | 0.457038315 | 0.662196333 | 0.99471946 | -5.170265097 |

| | | | | | | | | | |
|---|---|---|---|---|---|---|---|---|---|
| **A_55_P2129696** | Smad9 | 0.03395798 | -0.143752665 | 0.211668626 | 8.082465821 | 0.456706865 | 0.662422668 | 0.99471946 | -5.170387993 |
| **A_55_P1959076** | Zfp930 | 0.0227512 | -0.096348836 | 0.141851236 | 8.514262399 | 0.456563872 | 0.662520325 | 0.99471946 | -5.170440987 |
| **A_55_P2073795** | Mecr | -0.032716359 | -0.203990346 | 0.138557628 | 8.528031813 | -0.456544054 | 0.662533861 | 0.99471946 | -5.170448331 |
| **A_55_P1967231** | Stbd1 | -0.053296562 | -0.332316863 | 0.22572374 | 12.75204966 | -0.456533362 | 0.662541163 | 0.99471946 | -5.170452292 |
| **A_55_P1998848** | NA | -0.024455702 | -0.152500068 | 0.103588663 | 14.56384837 | -0.456487401 | 0.662572554 | 0.99471946 | -5.170469321 |
| **A_55_P2026492** | NA | 0.018396394 | -0.077938577 | 0.114731365 | 7.657599418 | 0.456412847 | 0.662623475 | 0.99471946 | -5.170496941 |
| **A_51_P135732** | Ccdc136 | 0.031169192 | -0.132089005 | 0.19442739 | 10.03136995 | 0.456309683 | 0.66269394 | 0.99471946 | -5.170535153 |
| **A_51_P283344** | ccdc198 | -0.03125507 | -0.194982987 | 0.132472847 | 7.712539618 | -0.456254197 | 0.662731841 | 0.99471946 | -5.170555702 |
| **A_55_P1965892** | Gnpda1 | 0.035970437 | -0.152482194 | 0.224423068 | 10.99410513 | 0.456197228 | 0.662770756 | 0.99471946 | -5.170576797 |
| **A_51_P302856** | Ugt2b37 | -0.019856099 | -0.123885589 | 0.084173391 | 7.80169418 | -0.456190873 | 0.662775097 | 0.99471946 | -5.17057915 |
| **A_52_P46419** | Grk6 | -0.02935391 | -0.183167449 | 0.124459629 | 9.128785052 | -0.45612148 | 0.6628225 | 0.99471946 | -5.170604843 |
| **A_51_P249051** | Pycr2 | 0.04372597 | -0.185396841 | 0.272848781 | 11.74304292 | 0.456121202 | 0.66282269 | 0.99471946 | -5.170604946 |
| **A_55_P2130565** | Prdm11 | 0.025224976 | -0.106953827 | 0.157403779 | 8.197215476 | 0.456118912 | 0.662824254 | 0.99471946 | -5.170605793 |
| **A_55_P1958137** | NA | -0.041762481 | -0.260630934 | 0.177105972 | 8.837710803 | -0.456049789 | 0.662871476 | 0.99471946 | -5.170631383 |
| **A_55_P2087923** | Otud3 | 0.035483421 | -0.150500176 | 0.221467018 | 9.616043376 | 0.455994891 | 0.662908979 | 0.99471946 | -5.170651703 |
| **A_55_P2151461** | NA | -0.031969501 | -0.199538786 | 0.135599783 | 12.53315253 | -0.455985023 | 0.662915721 | 0.99471946 | -5.170655355 |
| **A_55_P2045859** | NA | 0.049154805 | -0.208501056 | 0.306810667 | 10.45793594 | 0.455968758 | 0.662926833 | 0.99471946 | -5.170661375 |
| **A_51_P389636** | Kcnn4 | 0.020840345 | -0.088419896 | 0.130100585 | 7.753139199 | 0.455881409 | 0.662986509 | 0.99471946 | -5.1706937 |
| **A_55_P1977245** | Slc22a13 | 0.019552741 | -0.082976392 | 0.122081873 | 8.175505868 | 0.455794931 | 0.663045593 | 0.99471946 | -5.170725698 |
| **A_55_P2156062** | NA | -0.022317631 | -0.139349534 | 0.094714272 | 8.535800349 | -0.455777578 | 0.663057449 | 0.99471946 | -5.170732118 |
| **A_55_P2017729** | Gm2138 | 0.028596464 | -0.121381712 | 0.178574641 | 8.055532473 | 0.455715084 | 0.663100149 | 0.99471946 | -5.170755237 |
| **A_52_P232648** | Slc38a1 | 0.025142229 | -0.10672858 | 0.157013039 | 8.03627333 | 0.455684483 | 0.663121058 | 0.99471946 | -5.170766557 |
| **A_51_P222475** | Ehbp1l1 | -0.036519422 | -0.22808555 | 0.155046707 | 12.50337241 | -0.455632091 | 0.663156856 | 0.99471946 | -5.170785935 |
| **A_55_P2035424** | Hpgd | -0.033222516 | -0.207497767 | 0.141052736 | 8.019140924 | -0.455623305 | 0.66316286 | 0.99471946 | -5.170789185 |
| **A_55_P2046488** | Nbeal2 | 0.029075313 | -0.123446224 | 0.18159685 | 8.321096826 | 0.455619555 | 0.663165422 | 0.99471946 | -5.170790571 |
| **A_65_P16753** | Bex1 | 0.070003647 | -0.297277627 | 0.437284921 | 12.86677779 | 0.455544715 | 0.663216561 | 0.99471946 | -5.170818248 |
| **A_55_P2097448** | Rbpms | 0.042713739 | -0.181389938 | 0.266817415 | 11.58957733 | 0.455541282 | 0.663218907 | 0.99471946 | -5.170819518 |
| **A_55_P2099061** | NA | -0.03954525 | -0.247087434 | 0.167996933 | 13.87941516 | -0.45540425 | 0.663312548 | 0.99471946 | -5.170870183 |
| **A_55_P2074317** | Ap2b1 | 0.050260584 | -0.213535877 | 0.314057045 | 9.813958227 | 0.455373441 | 0.663333602 | 0.99471946 | -5.170881572 |
| **A_52_P519882** | Etf1 | 0.048351326 | -0.205466831 | 0.302169482 | 10.08566931 | 0.455297046 | 0.66338581 | 0.99471946 | -5.17090981 |
| **A_55_P2059650** | Atpaf2 | -0.024869397 | -0.155444567 | 0.105705773 | 10.69076775 | -0.455212094 | 0.663443869 | 0.99471946 | -5.170941206 |
| **A_51_P427934** | Megf8 | -0.057025072 | -0.356433514 | 0.242383369 | 10.31252777 | -0.455209099 | 0.663445915 | 0.99471946 | -5.170942313 |
| **A_55_P2408330** | Rbm33 | 0.045395966 | -0.192977287 | 0.283769219 | 8.392736002 | 0.455165009 | 0.663476049 | 0.99471946 | -5.170958605 |
| **A_55_P2030647** | Dusp13 | -0.018800265 | -0.117532253 | 0.079931722 | 7.665580438 | -0.4551088 | 0.663514466 | 0.99471946 | -5.170979373 |
| **A_51_P440399** | Bcap31 | -0.027601699 | -0.172568831 | 0.117365434 | 12.14274189 | -0.455067057 | 0.663542997 | 0.99471946 | -5.170994795 |
| **A_52_P4698** | Rfwd3 | 0.023287777 | -0.099051961 | 0.145627515 | 11.70950245 | 0.454956239 | 0.663618742 | 0.99471946 | -5.17103573 |
| **A_55_P1967266** | Plxnb2 | -0.028158386 | -0.176093096 | 0.119776324 | 13.0091526 | -0.454932338 | 0.663635079 | 0.99471946 | -5.171044558 |
| **A_55_P2013011** | NA | -0.023298206 | -0.145699663 | 0.099103252 | 7.955813945 | -0.454930459 | 0.663636364 | 0.99471946 | -5.171045252 |
| **A_55_P1967500** | Nell1 | 0.02343009 | -0.099673911 | 0.146534091 | 7.712806961 | 0.45489475 | 0.663660772 | 0.99471946 | -5.171058439 |
| **A_55_P2032818** | Trim2 | -0.023055774 | -0.144248056 | 0.098136507 | 10.22857091 | -0.454688407 | 0.663801826 | 0.994753976 | -5.171134626 |
| **A_52_P609024** | Cenpw | -0.039846587 | -0.249310994 | 0.16961782 | 11.38995313 | -0.454663429 | 0.663818901 | 0.994753976 | -5.171143847 |
| **A_66_P122099** | Zc3h8 | 0.034772958 | -0.148151323 | 0.21769724 | 8.511189689 | 0.454338371 | 0.66404114 | 0.994753976 | -5.171263795 |
| **A_52_P69020** | Slc24a5 | 0.019109303 | -0.081430097 | 0.119648702 | 8.106956522 | 0.454273799 | 0.664085291 | 0.994753976 | -5.171287613 |
| **A_51_P413698** | Mrpl20 | -0.041355288 | -0.258999695 | 0.176289118 | 13.63988998 | -0.454143053 | 0.664174694 | 0.994753976 | -5.17133583 |
| **A_55_P2059090** | Tagap | -0.032003768 | -0.200463642 | 0.136456106 | 9.189495065 | -0.454060553 | 0.664231109 | 0.994753976 | -5.171366248 |
| **A_51_P145433** | Lin7c | 0.023512527 | -0.100287642 | 0.147312697 | 8.240412823 | 0.453928246 | 0.664321589 | 0.994753976 | -5.17141502 |
| **A_55_P2092864** | Emp2 | 0.040088996 | -0.17099505 | 0.251173041 | 10.89441281 | 0.453919561 | 0.664327529 | 0.994753976 | -5.171418221 |
| **A_51_P364210** | Ms4a3 | -0.021003418 | -0.131610147 | 0.089603312 | 7.663718629 | -0.453855447 | 0.664371376 | 0.994753976 | -5.17144185 |

| | | | | | | | | | |
|---|---|---|---|---|---|---|---|---|---|
| **A_55_P1988033** | Atp6v1g1 | -0.021889806 | -0.137174649 | 0.093395036 | 13.26287257 | -0.453814995 | 0.664399043 | 0.994753976 | -5.171456757 |
| **A_52_P39247** | Hbp1 | 0.057909782 | -0.247092849 | 0.362912413 | 11.87397678 | 0.453792671 | 0.66441431 | 0.994753976 | -5.171464983 |
| **A_51_P329928** | Phlda3 | 0.06329628 | -0.270103775 | 0.396696335 | 12.63828351 | 0.453755238 | 0.664439912 | 0.994753976 | -5.171478776 |
| **A_52_P418515** | Cfap299 | -0.02358393 | -0.147857381 | 0.100689522 | 7.7096392 | -0.453572742 | 0.664564735 | 0.994753976 | -5.171546003 |
| **A_51_P512157** | Slc30a6 | -0.024071859 | -0.150916862 | 0.102773144 | 8.807913729 | -0.453571128 | 0.66456584 | 0.994753976 | -5.171546598 |
| **A_55_P2055299** | NA | -0.020103403 | -0.126062917 | 0.08585611 | 8.315195927 | -0.453459766 | 0.664642015 | 0.994753976 | -5.171587609 |
| **A_55_P1968982** | Pbx3 | -0.029559298 | -0.185417925 | 0.126299328 | 10.51563695 | -0.453286109 | 0.664760809 | 0.994753976 | -5.171651542 |
| **A_55_P2008437** | Cdkn2a | -0.023104472 | -0.14493194 | 0.098722995 | 7.757562541 | -0.453273126 | 0.664769691 | 0.994753976 | -5.171656321 |
| **A_55_P2404434** | NA | -0.030354548 | -0.190424762 | 0.129715666 | 9.043342307 | -0.453233887 | 0.664796535 | 0.994753976 | -5.171670764 |
| **A_52_P459564** | Itga2b | 0.041509764 | -0.1774357 | 0.260455228 | 7.895885863 | 0.453130658 | 0.664867159 | 0.994753976 | -5.171708754 |
| **A_55_P2090794** | NA | -0.025731367 | -0.161462208 | 0.109999474 | 9.009333279 | -0.45309936 | 0.664888572 | 0.994753976 | -5.17172027 |
| **A_55_P2075474** | Gid8 | 0.035434121 | -0.15147901 | 0.222347252 | 9.394635592 | 0.453096793 | 0.664890329 | 0.994753976 | -5.171721215 |
| **A_66_P140711** | Rps10 | -0.023484814 | -0.147377699 | 0.100408071 | 16.02243851 | -0.453053913 | 0.664919666 | 0.994753976 | -5.171736992 |
| **A_51_P365952** | Trdmt1 | 0.027602715 | -0.118033488 | 0.173238917 | 10.09237788 | 0.452993096 | 0.664961277 | 0.994753976 | -5.171759367 |
| **A_55_P2081050** | Kcnj13 | -0.023017114 | -0.144472746 | 0.098438518 | 7.800185154 | -0.452941746 | 0.664996412 | 0.994753976 | -5.171778256 |
| **A_52_P626756** | Trnt1 | -0.035967012 | -0.225760419 | 0.153826395 | 9.923455197 | -0.452931335 | 0.665003535 | 0.994753976 | -5.171782085 |
| **A_52_P28651** | Nectin1 | 0.026831231 | -0.114761998 | 0.168424461 | 9.402723875 | 0.45290514 | 0.665021458 | 0.994753976 | -5.17179172 |
| **A_55_P2110027** | NA | -0.04913283 | -0.308459086 | 0.210193427 | 13.29138759 | -0.452829194 | 0.665073425 | 0.994754751 | -5.171819652 |
| **A_52_P13897** | Hook1 | 0.018655511 | -0.079838072 | 0.117149094 | 7.959803762 | 0.452697765 | 0.66516336 | 0.994812312 | -5.171867979 |
| **A_55_P2086387** | 1700030K0 | 0.025993519 | -0.111309237 | 0.163296275 | 9.120412213 | 0.452475385 | 0.665315546 | 0.994962959 | -5.171949719 |
| **A_51_P310649** | Arsa | 0.026784458 | -0.114739708 | 0.168308625 | 10.9532485 | 0.452336251 | 0.665410771 | 0.994971635 | -5.172000842 |
| **A_55_P1982747** | Slc8a3 | 0.07183635 | -0.3077503 | 0.451423001 | 12.07315058 | 0.452316533 | 0.665424267 | 0.994971635 | -5.172008086 |
| **A_51_P220260** | Phf1 | 0.033756964 | -0.144719804 | 0.212233732 | 8.621011154 | 0.452054522 | 0.665603611 | 0.995075473 | -5.172104314 |
| **A_55_P2140383** | Mis18a | -0.042844738 | -0.26937453 | 0.183685055 | 8.991224328 | -0.452044608 | 0.665610398 | 0.995075473 | -5.172107954 |
| **A_66_P126504** | Trabd2b | -0.086973404 | -0.547004819 | 0.373058011 | 9.673898382 | -0.451864381 | 0.665733775 | 0.995075473 | -5.172174115 |
| **A_55_P2065966** | Ctdp1 | 0.026695816 | -0.114507392 | 0.167899024 | 9.614555316 | 0.451864024 | 0.66573402 | 0.995075473 | -5.172174246 |
| **A_51_P488730** | Zfp612 | 0.041861448 | -0.179624127 | 0.263347022 | 9.425095602 | 0.451728956 | 0.665826491 | 0.995075473 | -5.172223813 |
| **A_55_P2073507** | 2610528J1 | 0.020392997 | -0.087521685 | 0.128307679 | 7.740228797 | 0.451657937 | 0.665875114 | 0.995075473 | -5.17224987 |
| **A_66_P132070** | Abra | 0.025702869 | -0.110326291 | 0.161732029 | 8.602166343 | 0.451604973 | 0.665911378 | 0.995075473 | -5.1722693 |
| **A_55_P2002160** | Sumo3 | 0.046599699 | -0.200052408 | 0.293251806 | 9.350964537 | 0.451551626 | 0.665947905 | 0.995075473 | -5.172288868 |
| **A_51_P288341** | Hpd | -0.019717388 | -0.124084516 | 0.08464974 | 7.668072207 | -0.451538491 | 0.665956898 | 0.995075473 | -5.172293686 |
| **A_55_P2157695** | Tsc1 | 0.055249372 | -0.237345502 | 0.347844245 | 10.37962138 | 0.451304605 | 0.666117052 | 0.995136929 | -5.172379451 |
| **A_52_P493620** | Fgfr1op2 | 0.041361034 | -0.177683387 | 0.260405454 | 10.02311903 | 0.451303102 | 0.666118081 | 0.995136929 | -5.172380002 |
| **A_51_P185509** | Mapre1 | -0.024121077 | -0.151913694 | 0.103671539 | 9.870142315 | -0.451128303 | 0.666237788 | 0.995136929 | -5.172444073 |
| **A_55_P2167999** | Ldlr | 0.072738989 | -0.312669104 | 0.458147083 | 10.48144123 | 0.451082068 | 0.666269452 | 0.995136929 | -5.172461016 |
| **A_55_P2169069** | Gm10405 | -0.048478734 | -0.305351794 | 0.208394326 | 11.85595876 | -0.45106782 | 0.666279211 | 0.995136929 | -5.172466237 |
| **A_55_P2068496** | NA | -0.039311375 | -0.247676963 | 0.169054213 | 11.17624317 | -0.450921944 | 0.666379121 | 0.995136929 | -5.172519682 |
| **A_51_P209736** | Atoh8 | -0.031046633 | -0.195615268 | 0.133522001 | 8.268479467 | -0.450896205 | 0.66639675 | 0.995136929 | -5.172529111 |
| **A_51_P498023** | Scyl1 | -0.021873776 | -0.137842212 | 0.094094661 | 12.32054217 | -0.450809524 | 0.666456122 | 0.995136929 | -5.172560859 |
| **A_51_P215097** | Klf16 | 0.040514864 | -0.174306743 | 0.25533647 | 8.593039793 | 0.45076019 | 0.666489915 | 0.995136929 | -5.172578926 |
| **A_55_P2136880** | Ppp1r15a | -0.048347204 | -0.304790171 | 0.208095763 | 11.21946649 | -0.450598459 | 0.666600702 | 0.995136929 | -5.172638142 |
| **A_55_P2139814** | Fgf13 | 0.021442993 | -0.092312885 | 0.135198871 | 8.443347619 | 0.450526856 | 0.666649754 | 0.995136929 | -5.172664352 |
| **A_51_P454008** | Lbp | 0.018986425 | -0.081749384 | 0.119722234 | 8.595057019 | 0.450472676 | 0.666686871 | 0.995136929 | -5.172684182 |
| **A_55_P2092760** | Rhbdd3 | -0.026223688 | -0.165371328 | 0.112923951 | 11.68393795 | -0.450429749 | 0.66671628 | 0.995136929 | -5.172699892 |
| **A_55_P1971425** | Serpinb9f | -0.064544636 | -0.407105604 | 0.278016333 | 7.768296672 | -0.450330548 | 0.666784243 | 0.995136929 | -5.172736191 |
| **A_55_P1963807** | Actg2 | -0.115283524 | -0.727334063 | 0.496767015 | 9.770532685 | -0.450183 | 0.666885336 | 0.995136929 | -5.172790167 |
| **A_51_P518926** | Borcs6 | 0.038926022 | -0.167773681 | 0.245625724 | 9.407020428 | 0.4501003 | 0.666942002 | 0.995136929 | -5.172820414 |
| **A_51_P205099** | Dazap1 | -0.027874685 | -0.175939144 | 0.120189773 | 11.69889413 | -0.449954175 | 0.667042132 | 0.995136929 | -5.172873844 |

| | | | | | | | | | |
|---|---|---|---|---|---|---|---|---|---|
| **A_55_P1957159** | NA | -0.026971916 | -0.170252094 | 0.116308262 | 15.1744686 | -0.449919504 | 0.667065891 | 0.995136929 | -5.172886519 |
| **A_55_P2041045** | Csnk1g2 | -0.027819778 | -0.175616177 | 0.119976622 | 12.61389533 | -0.449882332 | 0.667091364 | 0.995136929 | -5.172900107 |
| **A_51_P430357** | Lrp10 | -0.043417149 | -0.27417776 | 0.187343461 | 10.52738092 | -0.449685372 | 0.667226344 | 0.995136929 | -5.172972089 |
| **A_52_P617636** | Klf12 | -0.01934816 | -0.122217433 | 0.083521113 | 8.329636644 | -0.449534607 | 0.667329675 | 0.995136929 | -5.173027169 |
| **A_66_P137660** | Fam166b | -0.021382759 | -0.135077121 | 0.092311603 | 7.870828845 | -0.449504394 | 0.667350383 | 0.995136929 | -5.173038205 |
| **A_55_P2057994** | Mif4gd | 0.052784965 | -0.227926132 | 0.333496062 | 10.0188958 | 0.449427651 | 0.667402985 | 0.995136929 | -5.173066233 |
| **A_55_P2111292** | 3110009E1 | -0.035060801 | -0.221528177 | 0.151406576 | 9.741331316 | -0.449394861 | 0.667425461 | 0.995136929 | -5.173078208 |
| **A_51_P346104** | Sgta | 0.030603336 | -0.132163231 | 0.193369902 | 12.29432456 | 0.449378921 | 0.667436386 | 0.995136929 | -5.173084029 |
| **A_51_P171697** | Pebp1 | -0.037095235 | -0.23439397 | 0.1602035 | 14.16236767 | -0.449368844 | 0.667443294 | 0.995136929 | -5.173087708 |
| **A_55_P2031011** | S100pbp | 0.020328242 | -0.087838669 | 0.128495154 | 8.144165563 | 0.449173921 | 0.667576912 | 0.995136929 | -5.173158871 |
| **A_55_P2158547** | Cyp2c54 | -0.019392849 | -0.122644882 | 0.083859183 | 7.671699915 | -0.448902628 | 0.667762904 | 0.995136929 | -5.173257867 |
| **A_55_P1989594** | Mynn | -0.030272655 | -0.191463653 | 0.130918344 | 9.218215684 | -0.448868213 | 0.667786499 | 0.995136929 | -5.173270421 |
| **A_55_P2161545** | Ddx21 | 0.036998857 | -0.160025805 | 0.234023519 | 9.642152349 | 0.448824802 | 0.667816264 | 0.995136929 | -5.173286256 |
| **A_51_P333111** | Aox1 | -0.027174166 | -0.171897226 | 0.117548893 | 8.092506068 | -0.44877394 | 0.667851138 | 0.995136929 | -5.173304807 |
| **A_52_P638794** | Stk31 | -0.018027018 | -0.114041527 | 0.077987492 | 7.657296507 | -0.448741417 | 0.667873438 | 0.995136929 | -5.173316668 |
| **A_55_P2054491** | Slc31a2 | 0.02733366 | -0.118252638 | 0.172919958 | 8.14655887 | 0.448731352 | 0.667880339 | 0.995136929 | -5.173320338 |
| **A_55_P2068723** | Ccl27a | -0.035775586 | -0.226336611 | 0.15478544 | 11.08491835 | -0.448705925 | 0.667897774 | 0.995136929 | -5.17332961 |
| **A_52_P114620** | Elmo3 | 0.023866516 | -0.103291533 | 0.151024566 | 8.368938919 | 0.448594876 | 0.667973921 | 0.995136929 | -5.1733701 |
| **A_55_P2010622** | Sec16b | 0.031085777 | -0.134584609 | 0.196756162 | 8.203719748 | 0.448462333 | 0.668064812 | 0.995136929 | -5.173418415 |
| **A_55_P2076782** | 1700028P1 | 0.018947515 | -0.082032899 | 0.119927929 | 7.647072067 | 0.448460543 | 0.66806604 | 0.995136929 | -5.173419068 |
| **A_51_P520927** | Ccdc97 | 0.025475822 | -0.110309972 | 0.161261617 | 10.79197054 | 0.44841797 | 0.668095235 | 0.995136929 | -5.173434583 |
| **A_66_P124146** | Ttc16 | -0.019530045 | -0.123686919 | 0.08462683 | 8.272557493 | -0.448151062 | 0.66827829 | 0.995136929 | -5.173531828 |
| **A_55_P1982454** | Eps8 | -0.031855846 | -0.201784325 | 0.138072633 | 9.939920262 | -0.448055799 | 0.66834363 | 0.995136929 | -5.173566523 |
| **A_55_P2097840** | NA | 0.033288153 | -0.144379475 | 0.210955781 | 9.357368013 | 0.447806644 | 0.668514539 | 0.995136929 | -5.173657234 |
| **A_55_P2409570** | Luc7l2 | 0.066483538 | -0.288435051 | 0.421402128 | 14.3486788 | 0.447707645 | 0.668582453 | 0.995136929 | -5.173693263 |
| **A_52_P348720** | Rhobtb3 | -0.035518922 | -0.225153862 | 0.154116019 | 10.32752819 | -0.447662326 | 0.668613544 | 0.995136929 | -5.173709754 |
| **A_55_P2081116** | Fam89a | 0.018470996 | -0.080170545 | 0.117112538 | 7.696473054 | 0.447547983 | 0.668691991 | 0.995136929 | -5.173751355 |
| **A_55_P2019934** | Ythdf3 | -0.021179853 | -0.134318575 | 0.091958869 | 10.8216014 | -0.447425575 | 0.668775976 | 0.995136929 | -5.17379588 |
| **A_52_P424272** | Gatc | -0.023062015 | -0.146272957 | 0.100148926 | 8.654662578 | -0.447359957 | 0.668820998 | 0.995136929 | -5.173819743 |
| **A_66_P124715** | Ppard | -0.027507733 | -0.174486802 | 0.119471336 | 10.36474724 | -0.447309828 | 0.668855395 | 0.995136929 | -5.173837971 |
| **A_51_P352968** | Marcks | 0.038807865 | -0.168640352 | 0.246256083 | 13.65649882 | 0.447114936 | 0.66898913 | 0.995136929 | -5.17390882 |
| **A_51_P256093** | Map2k6 | 0.045749288 | -0.19880447 | 0.290303046 | 9.834402427 | 0.447114887 | 0.668989163 | 0.995136929 | -5.173908838 |
| **A_51_P164939** | Tmem150a | 0.026386227 | -0.11467149 | 0.167443945 | 8.233815281 | 0.447084462 | 0.669010043 | 0.995136929 | -5.173919896 |
| **A_55_P2052172** | NA | -0.033425348 | -0.21215605 | 0.145305354 | 11.03774235 | -0.446977753 | 0.669083273 | 0.995136929 | -5.173958674 |
| **A_55_P2090406** | H13 | -0.042329497 | -0.268725564 | 0.18406657 | 9.279176636 | -0.446872228 | 0.669155694 | 0.995136929 | -5.173997013 |
| **A_51_P349691** | Ano10 | -0.023202505 | -0.147308043 | 0.100903032 | 9.395332072 | -0.446840835 | 0.66917724 | 0.995136929 | -5.174008417 |
| **A_52_P639402** | Kcnk3 | -0.088167137 | -0.55979827 | 0.383463997 | 9.918436916 | -0.446800245 | 0.669205099 | 0.995136929 | -5.17402316 |
| **A_55_P1959338** | Spindoc | -0.032679077 | -0.207525282 | 0.142167127 | 10.26187684 | -0.44670695 | 0.669269132 | 0.995136929 | -5.174057044 |
| **A_55_P2151442** | C330007P( | 0.032655052 | -0.142081166 | 0.20739127 | 10.36340695 | 0.446659506 | 0.669301697 | 0.995136929 | -5.174074272 |
| **A_55_P2107745** | Yy1 | 0.031544884 | -0.137255845 | 0.200345613 | 10.50732794 | 0.446646302 | 0.66931076 | 0.995136929 | -5.174079067 |
| **A_55_P2054714** | NA | -0.03595915 | -0.228390008 | 0.156471708 | 9.660876803 | -0.446625821 | 0.669324818 | 0.995136929 | -5.174086503 |
| **A_52_P80806** | Syncrip | 0.041227842 | -0.179402355 | 0.261858039 | 10.27538206 | 0.446616524 | 0.6693312 | 0.995136929 | -5.174089879 |
| **A_55_P2246900** | NA | -0.028717421 | -0.182403247 | 0.124968405 | 8.678178312 | -0.446602093 | 0.669341105 | 0.995136929 | -5.174095119 |
| **A_55_P2068366** | Gm9292 | 0.018199858 | -0.079217898 | 0.115617614 | 8.127871935 | 0.446518039 | 0.669398802 | 0.995136929 | -5.174125634 |
| **A_55_P1978122** | Mutyh | -0.05293204 | -0.336260209 | 0.23039613 | 8.788104242 | -0.446517006 | 0.669399511 | 0.995136929 | -5.174126009 |
| **A_55_P1959425** | Slc16a3 | 0.089003132 | -0.387412535 | 0.565418799 | 13.2376647 | 0.446507117 | 0.6694063 | 0.995136929 | -5.174129599 |
| **A_51_P117130** | Naga | 0.02915493 | -0.126923138 | 0.185232998 | 8.100030865 | 0.446456611 | 0.669440969 | 0.995136929 | -5.174147931 |
| **A_55_P2177953** | Aqp11 | -0.022099977 | -0.140436594 | 0.096236639 | 7.886849344 | -0.44635645 | 0.669509728 | 0.995136929 | -5.174184282 |

| | | | | | | | | | |
|---|---|---|---|---|---|---|---|---|---|
| **A_51_P305547** | Snai2 | 0.041592548 | -0.181194853 | 0.264379949 | 9.608489594 | 0.446204588 | 0.669613984 | 0.995136929 | -5.174239382 |
| **A_55_P2088272** | Golga3 | 0.018771278 | -0.081794601 | 0.119337157 | 11.36455132 | 0.446120644 | 0.669671617 | 0.995136929 | -5.174269832 |
| **A_51_P180974** | Cavin3 | -0.040382104 | -0.256736267 | 0.175972058 | 11.88580613 | -0.446100619 | 0.669685365 | 0.995136929 | -5.174277095 |
| **A_52_P502141** | Hectd2 | -0.041895621 | -0.266389585 | 0.182598344 | 8.853949609 | -0.446039265 | 0.66972749 | 0.995136929 | -5.174299347 |
| **A_55_P2079344** | Zfp362 | 0.030355548 | -0.132344398 | 0.193055493 | 9.972923919 | 0.445922921 | 0.669807374 | 0.995136929 | -5.174341534 |
| **A_55_P2115318** | Fmr1nb | -0.026558044 | -0.168925014 | 0.115808925 | 7.849229264 | -0.445857404 | 0.669852362 | 0.995136929 | -5.174365286 |
| **A_55_P2294037** | NA | -0.025620265 | -0.162962204 | 0.111721673 | 8.816808998 | -0.445850837 | 0.669856871 | 0.995136929 | -5.174367667 |
| **A_51_P163578** | Ugt2b35 | 0.018422513 | -0.080340762 | 0.117185787 | 7.6307393 | 0.445823048 | 0.669875953 | 0.995136929 | -5.17437774 |
| **A_55_P2090880** | Stk40 | -0.040992906 | -0.260768194 | 0.178782382 | 11.65293088 | -0.445798889 | 0.669892543 | 0.995136929 | -5.174386497 |
| **A_51_P324287** | Kif23 | 0.038726516 | -0.168951054 | 0.246404086 | 9.706913744 | 0.445684944 | 0.669970789 | 0.995136929 | -5.174427793 |
| **A_55_P1987483** | Pgm2 | 0.030742577 | -0.134121914 | 0.195607068 | 15.04488088 | 0.445679105 | 0.669974799 | 0.995136929 | -5.17442991 |
| **A_51_P160083** | 1500035N2 | -0.03678956 | -0.234102708 | 0.160523587 | 9.308113135 | -0.44563337 | 0.670006207 | 0.995136929 | -5.174446482 |
| **A_55_P2131979** | 1-Mar | 0.052954831 | -0.231063262 | 0.336972923 | 11.70944225 | 0.445624142 | 0.670012544 | 0.995136929 | -5.174449826 |
| **A_55_P2007510** | Pef1 | -0.034247626 | -0.218028312 | 0.149533059 | 11.97850612 | -0.44538926 | 0.670173857 | 0.995221492 | -5.174534911 |
| **A_55_P2038987** | Gas7 | 0.029934325 | -0.130721853 | 0.190590502 | 8.633239328 | 0.445329197 | 0.67021511 | 0.995221492 | -5.174556662 |
| **A_51_P118763** | Ahctf1 | 0.033369202 | -0.145755629 | 0.212494032 | 9.508027581 | 0.445245107 | 0.670272868 | 0.995221492 | -5.162556053 |
| **A_51_P215038** | Tmem59l | -0.021973512 | -0.139927601 | 0.095980578 | 7.925417243 | -0.445241466 | 0.670275369 | 0.995221492 | -5.174588428 |
| **A_51_P174415** | Retreg2 | -0.027754629 | -0.176806887 | 0.121297629 | 9.640758454 | -0.44504713 | 0.67040886 | 0.995343263 | -5.174658772 |
| **A_55_P2042773** | Ctdnep1 | 0.049665521 | -0.217171126 | 0.316502169 | 11.9338487 | 0.444855185 | 0.670540722 | 0.995363133 | -5.174728224 |
| **A_55_P1987121** | Rtraf | -0.025741307 | -0.168356756 | 0.116874141 | 14.27141185 | -0.448364458 | 0.670567579 | 0.995363133 | -5.097825324 |
| **A_55_P2119463** | Ggh | -0.031878925 | -0.203174145 | 0.139416295 | 11.22433194 | -0.44480284 | 0.670576683 | 0.995363133 | -5.174747159 |
| **A_55_P2093595** | Mmd | -0.032072709 | -0.204446173 | 0.140300755 | 12.13091946 | -0.444707413 | 0.670642245 | 0.995384034 | -5.174781673 |
| **A_52_P272854** | NA | -0.050208581 | -0.320109787 | 0.219692626 | 8.052847727 | -0.444613098 | 0.670707047 | 0.995403803 | -5.174815778 |
| **A_55_P2124319** | NA | 0.018990811 | -0.083218813 | 0.121200436 | 7.756821294 | 0.444079632 | 0.671073634 | 0.995570006 | -5.175008558 |
| **A_52_P189005** | Gk5 | 0.01993906 | -0.087424918 | 0.127303038 | 7.994728142 | 0.44386941 | 0.671218121 | 0.995570006 | -5.175084468 |
| **A_65_P01065** | Phf14 | -0.025246302 | -0.161224873 | 0.110732269 | 11.3407245 | -0.443748022 | 0.671301559 | 0.995570006 | -5.175128286 |
| **A_55_P2008422** | Mpc1 | -0.02534526 | -0.161911806 | 0.111221286 | 14.36734491 | -0.443569374 | 0.671424364 | 0.995570006 | -5.175192752 |
| **A_52_P325527** | Edil3 | 0.030472437 | -0.133722905 | 0.19466778 | 7.973810524 | 0.443563183 | 0.67142862 | 0.995570006 | -5.175194985 |
| **A_55_P2127139** | H3c4 | 0.036447054 | -0.159948289 | 0.232842398 | 13.89754977 | 0.443547748 | 0.671439231 | 0.995570006 | -5.175200554 |
| **A_55_P2101336** | Ramp2 | 0.038744263 | -0.170036086 | 0.247524612 | 13.88323445 | 0.443533999 | 0.671448683 | 0.995570006 | -5.175205514 |
| **A_55_P2120354** | Cib2 | -0.035250889 | -0.225216278 | 0.1547145 | 9.610425743 | -0.443511336 | 0.671464263 | 0.995570006 | -5.17521369 |
| **A_51_P369371** | Ubtd1 | -0.04952641 | -0.316528467 | 0.217475647 | 9.472807757 | -0.443334341 | 0.671585946 | 0.995570006 | -5.175277528 |
| **A_55_P2413924** | Gm10404 | 0.019015433 | -0.083503823 | 0.12153469 | 8.684827532 | 0.443312436 | 0.671601006 | 0.995570006 | -5.175285427 |
| **A_55_P1999022** | Zfp760 | 0.037659954 | -0.165392824 | 0.240712732 | 8.60463233 | 0.443281879 | 0.671622015 | 0.995570006 | -5.175296445 |
| **A_55_P1974695** | NA | 0.039435125 | -0.173218929 | 0.252089179 | 8.490196763 | 0.443219322 | 0.671665027 | 0.995570006 | -5.175319 |
| **A_55_P1967341** | Gm9104 | -0.039007508 | -0.249362277 | 0.171347262 | 13.12856238 | -0.443205326 | 0.67167465 | 0.995570006 | -5.175324046 |
| **A_66_P133270** | Mfsd1 | -0.028202572 | -0.180374548 | 0.123969405 | 8.510799182 | -0.442958627 | 0.671844283 | 0.995570006 | -5.175412961 |
| **A_51_P435339** | Epor | -0.021358195 | -0.136614297 | 0.093897908 | 8.187108285 | -0.442904143 | 0.671881749 | 0.995570006 | -5.175432592 |
| **A_55_P1994194** | Heph | 0.024507925 | -0.107759654 | 0.156775505 | 10.41390397 | 0.442855739 | 0.671915036 | 0.995570006 | -5.17545003 |
| **A_55_P2364315** | D030002E0 | 0.043810784 | -0.192646532 | 0.280268101 | 8.598487161 | 0.442830377 | 0.671932477 | 0.995570006 | -5.175459167 |
| **A_55_P2041723** | Mid1ip1 | 0.030944534 | -0.13610778 | 0.197996848 | 11.18252923 | 0.442731665 | 0.672000363 | 0.995570006 | -5.175494722 |
| **A_55_P2019233** | Gsto2 | 0.028370097 | -0.124791695 | 0.18153189 | 8.45908046 | 0.442710192 | 0.67201513 | 0.995570006 | -5.175502455 |
| **A_51_P308254** | Surf2 | -0.027188895 | -0.173996248 | 0.119618458 | 8.908791781 | -0.442642264 | 0.672061848 | 0.995570006 | -5.175526917 |
| **A_55_P2077623** | NA | -0.019886622 | -0.127291748 | 0.087518504 | 7.693659042 | -0.442532468 | 0.672137363 | 0.995570006 | -5.175566449 |
| **A_51_P220162** | Notch3 | 0.042661509 | -0.187748711 | 0.273071729 | 11.02208861 | 0.442530907 | 0.672138437 | 0.995570006 | -5.175567011 |
| **A_55_P1963718** | Lias | -0.026783352 | -0.171464898 | 0.117898193 | 11.31355226 | -0.442446664 | 0.672196381 | 0.995570006 | -5.175597336 |
| **A_55_P2159475** | Sbf1 | 0.038055338 | -0.167640336 | 0.243751013 | 12.04137629 | 0.442180472 | 0.672379487 | 0.995570006 | -5.175693123 |
| **A_51_P367162** | Psmb1 | -0.026399985 | -0.169097582 | 0.116297611 | 12.60577859 | -0.442177008 | 0.67238187 | 0.995570006 | -5.175694369 |

| | | | | | | | | | |
|---|---|---|---|---|---|---|---|---|---|
| **A_51_P485421** | NA | -0.018591032 | -0.119111773 | 0.08192971 | 7.695658601 | -0.442035284 | 0.672479367 | 0.995570006 | -5.162799527 |
| **A_55_P2102515** | Daam1 | -0.043340988 | -0.27770335 | 0.191021373 | 10.21661846 | -0.44199777 | 0.672505176 | 0.995570006 | -5.175758835 |
| **A_51_P479786** | Smtn | -0.053004629 | -0.339704421 | 0.233695163 | 9.758403149 | -0.441871055 | 0.672592356 | 0.995570006 | -5.175804396 |
| **A_55_P2044282** | Mpp6 | 0.029787113 | -0.131363368 | 0.190937595 | 13.05668327 | 0.44177989 | 0.672655081 | 0.995570006 | -5.175837168 |
| **A_51_P509489** | Kras | -0.02116223 | -0.135662194 | 0.093337735 | 9.530630573 | -0.441738448 | 0.672683596 | 0.995570006 | -5.175852063 |
| **A_55_P2151209** | Krtap19-4 | 0.063836544 | -0.281610491 | 0.409283579 | 11.29666992 | 0.441669113 | 0.672731304 | 0.995570006 | -5.175876981 |
| **A_51_P517430** | Cd1d1 | -0.047448136 | -0.304245872 | 0.209349601 | 10.98941438 | -0.441608164 | 0.672773243 | 0.995570006 | -5.175898882 |
| **A_55_P2117425** | Syt5 | -0.020382866 | -0.130742965 | 0.089977234 | 8.478605682 | -0.44143045 | 0.672895536 | 0.995570006 | -5.175962724 |
| **A_55_P2112360** | Arhgap17 | -0.033268546 | -0.213441981 | 0.146904889 | 11.35135414 | -0.441318564 | 0.672972535 | 0.995570006 | -5.176002906 |
| **A_55_P2053718** | Pink1 | -0.043192598 | -0.277130963 | 0.190745767 | 12.41804361 | -0.441282809 | 0.672997142 | 0.995570006 | -5.176015744 |
| **A_55_P1999294** | Gpbp1 | -0.028064719 | -0.180084576 | 0.123955138 | 11.66462314 | -0.441234546 | 0.673030358 | 0.995570006 | -5.176033073 |
| **A_55_P2064538** | Zfp985 | 0.024222484 | -0.107009006 | 0.155453973 | 8.330861026 | 0.441153516 | 0.673086127 | 0.995570006 | -5.176062162 |
| **A_51_P411355** | Taf1c | 0.022863789 | -0.101019609 | 0.146747187 | 8.442726401 | 0.441107284 | 0.673117947 | 0.995570006 | -5.176078757 |
| **A_51_P184949** | F12 | -0.024529899 | -0.157444056 | 0.108384257 | 7.728871698 | -0.441096551 | 0.673125335 | 0.995570006 | -5.17608261 |
| **A_65_P17700** | Apobec3 | 0.048048375 | -0.212357984 | 0.308454734 | 9.406427851 | 0.440997627 | 0.673193424 | 0.995570006 | -5.176118112 |
| **A_55_P2140002** | Tnfaip8 | -0.042635144 | -0.273713065 | 0.188442777 | 10.6651009 | -0.440979518 | 0.673205889 | 0.995570006 | -5.17612461 |
| **A_51_P120461** | Riox2 | 0.023910974 | -0.105693231 | 0.15351518 | 10.33646383 | 0.440947935 | 0.673227629 | 0.995570006 | -5.152580068 |
| **A_55_P2054409** | Pira2 | -0.01783294 | -0.114497214 | 0.078831335 | 7.695981261 | -0.440926372 | 0.673242471 | 0.995570006 | -5.17614368 |
| **A_55_P2043682** | Ubac1 | -0.036090656 | -0.231721875 | 0.159540563 | 12.30672526 | -0.440926039 | 0.673242701 | 0.995570006 | -5.176143799 |
| **A_55_P2099750** | NA | -0.036179488 | -0.232325277 | 0.159966301 | 10.63704113 | -0.440851738 | 0.673293846 | 0.995570006 | -5.176170456 |
| **A_51_P280085** | Zfp655 | -0.042149918 | -0.270688826 | 0.186388991 | 11.50982792 | -0.440804191 | 0.673326576 | 0.995570006 | -5.176187512 |
| **A_51_P428715** | Mrpl13 | 0.022326296 | -0.098769293 | 0.143421886 | 13.05673718 | 0.440653771 | 0.673430128 | 0.995570006 | -5.176241459 |
| **A_51_P201480** | Stat3 | -0.033376983 | -0.214433418 | 0.147679452 | 11.96164854 | -0.440597718 | 0.673468717 | 0.995570006 | -5.146525799 |
| **A_52_P45834** | Pafah1b2 | 0.020657439 | -0.09141808 | 0.132732958 | 9.104214067 | 0.44052935 | 0.673515786 | 0.995570006 | -5.176286069 |
| **A_55_P2002782** | Vps26a | -0.027657978 | -0.177740227 | 0.122424271 | 11.27176789 | -0.440453667 | 0.673567893 | 0.995570006 | -5.176313199 |
| **A_52_P547234** | Tomm70a | 0.034008037 | -0.150549007 | 0.218565082 | 10.87868284 | 0.440412942 | 0.673595932 | 0.995570006 | -5.176327795 |
| **A_51_P398191** | Auts2 | 0.026831887 | -0.118866085 | 0.172529859 | 9.894408256 | 0.440156225 | 0.673772698 | 0.995570006 | -5.17641978 |
| **A_52_P523445** | Cdhr5 | 0.018762379 | -0.08316904 | 0.120693798 | 7.711036555 | 0.43993546 | 0.673924726 | 0.995570006 | -5.176498842 |
| **A_55_P1983177** | Acsl4 | 0.043853685 | -0.194404743 | 0.282112114 | 11.36624267 | 0.439913162 | 0.673940083 | 0.995570006 | -5.176506826 |
| **A_55_P2096737** | Elmod3 | -0.028690281 | -0.184611106 | 0.127230544 | 10.67693185 | -0.439784392 | 0.674028767 | 0.995570006 | -5.176552922 |
| **A_51_P358574** | Pdzd3 | -0.019983794 | -0.128594424 | 0.088626837 | 7.663167181 | -0.439758996 | 0.674046258 | 0.995570006 | -5.176562012 |
| **A_55_P2063937** | D630033A( | 0.027431077 | -0.121663928 | 0.176526082 | 8.374633806 | 0.439732839 | 0.674064274 | 0.995570006 | -5.176571374 |
| **A_52_P558368** | Zdhhc1 | -0.04145498 | -0.266800049 | 0.183890089 | 9.374954094 | -0.439681086 | 0.674099919 | 0.995570006 | -5.176589895 |
| **A_51_P202430** | Pdap1 | 0.03740567 | -0.165953644 | 0.240764984 | 13.68927964 | 0.439625126 | 0.674138463 | 0.995570006 | -5.176609919 |
| **A_51_P441782** | Micu2 | -0.029831747 | -0.1921004 | 0.132436905 | 11.53044355 | -0.439393062 | 0.674298314 | 0.995570006 | -5.176692934 |
| **A_52_P249588** | Cnot11 | 0.024586905 | -0.109196429 | 0.158370239 | 9.101412001 | 0.439249195 | 0.674397422 | 0.995570006 | -5.176744378 |
| **A_51_P367866** | Egr1 | 0.052143384 | -0.231641076 | 0.335927843 | 9.864445618 | 0.439156924 | 0.67446099 | 0.995570006 | -5.176777364 |
| **A_55_P2173005** | Dnase2a | 0.070770146 | -0.314480797 | 0.45602109 | 12.0545509 | 0.43905155 | 0.674533589 | 0.995570006 | -5.176815027 |
| **A_65_P15295** | NA | -0.017766979 | -0.114504942 | 0.078970984 | 7.662550802 | -0.438960838 | 0.674596089 | 0.995570006 | -5.176847442 |
| **A_51_P399853** | Zfp704 | -0.022665424 | -0.146176815 | 0.100845966 | 10.07599741 | -0.438597322 | 0.674846577 | 0.995570006 | -5.17697728 |
| **A_52_P322421** | Mpzl2 | -0.03578183 | -0.230800455 | 0.159236796 | 8.130449041 | -0.438526241 | 0.674895562 | 0.995570006 | -5.177002656 |
| **A_55_P2070406** | NA | 0.02397328 | -0.106686298 | 0.154632858 | 8.474340849 | 0.438525995 | 0.674895732 | 0.995570006 | -5.177002744 |
| **A_52_P458000** | AI467606 | -0.023897956 | -0.154216417 | 0.106420505 | 7.819312658 | -0.438292405 | 0.675056721 | 0.995570006 | -5.177086111 |
| **A_55_P1975322** | NA | 0.032604739 | -0.145260903 | 0.210470381 | 10.41970721 | 0.438124757 | 0.675172275 | 0.995570006 | -5.177145917 |
| **A_55_P1968833** | Rpgr | -0.040371568 | -0.26061316 | 0.179870024 | 9.51414746 | -0.438112269 | 0.675180883 | 0.995570006 | -5.177150371 |
| **A_51_P111952** | Nudt19 | 0.021136614 | -0.094199975 | 0.136473204 | 13.2413123 | 0.438003364 | 0.675255952 | 0.995570006 | -5.17718921 |
| **A_55_P2037294** | NA | 0.023058627 | -0.102797128 | 0.148914381 | 8.234407779 | 0.437894503 | 0.675330996 | 0.995570006 | -5.177228024 |
| **A_55_P2007733** | Epn1 | 0.036711252 | -0.163784119 | 0.237206624 | 14.50825654 | 0.43762687 | 0.675515506 | 0.995570006 | -5.177323409 |

| | | | | | | | | | |
|---|---|---|---|---|---|---|---|---|---|
| **A_52_P11174** | Hip1 | -0.036563723 | -0.23631443 | 0.163186984 | 10.28474489 | -0.437493109 | 0.675607732 | 0.995570006 | -5.177371061 |
| **A_55_P1964509** | Lcn12 | 0.025329937 | -0.113063793 | 0.163723666 | 8.996755076 | 0.437448387 | 0.675638569 | 0.995570006 | -5.17738699 |
| **A_52_P456271** | Tmem167 | 0.032395874 | -0.144644349 | 0.209436096 | 11.66754159 | 0.437347736 | 0.675707971 | 0.995570006 | -5.177422835 |
| **A_55_P2178127** | 1-Sep | 0.021428959 | -0.095687687 | 0.138545605 | 9.250912082 | 0.437312194 | 0.67573248 | 0.995570006 | -5.177435491 |
| **A_51_P150394** | Vps28 | -0.029123515 | -0.188314449 | 0.130067419 | 13.90472522 | -0.437254812 | 0.675772049 | 0.995570006 | -5.177455921 |
| **A_55_P2083649** | Alas1 | -0.028930679 | -0.192009459 | 0.134148102 | 13.80012261 | -0.440685047 | 0.675797665 | 0.995570006 | -5.100467016 |
| **A_55_P1952502** | Mapk7 | 0.040904007 | -0.18271991 | 0.264527923 | 11.18593823 | 0.437176427 | 0.675826103 | 0.995570006 | -5.177483825 |
| **A_52_P484956** | Nnat | -0.026996997 | -0.174595718 | 0.120601724 | 8.061636391 | -0.437161585 | 0.675836338 | 0.995570006 | -5.177489108 |
| **A_52_P269016** | Nek9 | 0.021500724 | -0.096056589 | 0.139058036 | 12.1328993 | 0.437131958 | 0.67585677 | 0.995570006 | -5.177499654 |
| **A_55_P2085170** | 6-Sep | 0.029282116 | -0.130823003 | 0.189387234 | 9.403076527 | 0.43712573 | 0.675861064 | 0.995570006 | -5.17750187 |
| **A_55_P1972018** | H4c3 | 0.072311927 | -0.323104178 | 0.467728032 | 10.3217972 | 0.437083803 | 0.675889979 | 0.995570006 | -5.177516792 |
| **A_55_P2120141** | NA | 0.13643096 | -0.609816845 | 0.882678765 | 12.21610753 | 0.436957301 | 0.675977223 | 0.995570006 | -5.177561807 |
| **A_51_P514412** | Gsk3b | 0.043188524 | -0.193062311 | 0.27943936 | 8.975138045 | 0.436922234 | 0.676001408 | 0.995570006 | -5.177574283 |
| **A_55_P2179582** | Nmur2 | -0.018150727 | -0.11744198 | 0.081140525 | 7.651804831 | -0.436910188 | 0.676009716 | 0.995570006 | -5.177578569 |
| **A_55_P2027147** | Mrps18c | 0.027938041 | -0.124947885 | 0.180823967 | 12.50414922 | 0.436754682 | 0.676116973 | 0.995570006 | -5.177633882 |
| **A_55_P1954365** | Ttc28 | 0.038241431 | -0.17104555 | 0.247528412 | 10.78851025 | 0.436717962 | 0.676142301 | 0.995570006 | -5.177646941 |
| **A_55_P2146374** | Nmnat1 | 0.041780179 | -0.186955227 | 0.270515586 | 9.520245252 | 0.436562106 | 0.676249809 | 0.995570006 | -5.177702356 |
| **A_55_P2117028** | Mtmr3 | -0.036253144 | -0.234765043 | 0.162258756 | 11.51264895 | -0.43648393 | 0.676303738 | 0.995570006 | -5.177730145 |
| **A_66_P126603** | NA | 0.031121448 | -0.139295952 | 0.201538848 | 13.2678632 | 0.436470596 | 0.676312936 | 0.995570006 | -5.177734884 |
| **A_55_P2073855** | Mrpl27 | -0.01924963 | -0.124659936 | 0.086160676 | 12.81376026 | -0.436464016 | 0.676317476 | 0.995570006 | -5.177737223 |
| **A_51_P353252** | Mal2 | -0.044564193 | -0.288822985 | 0.199694599 | 7.975564571 | -0.436058719 | 0.676597098 | 0.995570006 | -5.177881212 |
| **A_55_P2152567** | Plekha5 | -0.024623037 | -0.159591691 | 0.110345618 | 8.054625984 | -0.436031462 | 0.676615905 | 0.995570006 | -5.177890891 |
| **A_55_P2100050** | NA | -0.043744004 | -0.283531454 | 0.196043446 | 10.41505742 | -0.436014788 | 0.67662741 | 0.995570006 | -5.177896812 |
| **A_55_P2006792** | Ntrk2 | 0.044680523 | -0.200286644 | 0.289647689 | 8.772196783 | 0.435932752 | 0.676684017 | 0.995570006 | -5.177925939 |
| **A_55_P1957424** | Sulf1 | -0.044054434 | -0.285592784 | 0.197483915 | 8.58919857 | -0.435925892 | 0.67668875 | 0.995570006 | -5.177928374 |
| **A_55_P2003096** | Pla2g6 | -0.027052914 | -0.175421582 | 0.121315754 | 8.860860162 | -0.435793739 | 0.676779943 | 0.995570006 | -5.177975283 |
| **A_51_P506417** | Krt14 | 0.034492624 | -0.154709763 | 0.223695012 | 8.037692051 | 0.435721266 | 0.676829956 | 0.995570006 | -5.178001003 |
| **A_55_P1963777** | Gtf3c6 | 0.035623369 | -0.159842154 | 0.231088892 | 12.08826294 | 0.435586044 | 0.676923276 | 0.995570006 | -5.178048981 |
| **A_55_P1967342** | NA | -0.034231025 | -0.222067911 | 0.153605862 | 14.7875715 | -0.43556016 | 0.67694114 | 0.995570006 | -5.178058163 |
| **A_51_P228295** | Mpzl1 | 0.021784941 | -0.097770523 | 0.141340406 | 12.72062399 | 0.435507961 | 0.676977166 | 0.995570006 | -5.178076679 |
| **A_55_P2085460** | 5530601H( | -0.044378294 | -0.287996165 | 0.199239576 | 8.666864017 | -0.435382126 | 0.677064017 | 0.995570006 | -5.178121307 |
| **A_65_P09090** | Arih2 | 0.020810642 | -0.093468754 | 0.135090038 | 11.18781533 | 0.435237867 | 0.67716359 | 0.995570006 | -5.178172453 |
| **A_55_P1955562** | Extl3 | 0.042853415 | -0.192603533 | 0.278310364 | 13.1183776 | 0.434993794 | 0.677332075 | 0.995570006 | -5.178258953 |
| **A_51_P462385** | G6pc | -0.017925041 | -0.116441409 | 0.080591327 | 7.720498991 | -0.43487145 | 0.677416537 | 0.995570006 | -5.178302295 |
| **A_55_P1958840** | Itga6 | 0.059193407 | -0.266136176 | 0.384522989 | 13.18996312 | 0.434869461 | 0.67741791 | 0.995570006 | -5.178303 |
| **A_55_P2097391** | Tent5a | 0.018333508 | -0.082432583 | 0.119099598 | 8.074889167 | 0.434850819 | 0.677430781 | 0.995570006 | -5.178309603 |
| **A_55_P2014882** | Dhrs9 | -0.018836806 | -0.122380975 | 0.084707363 | 7.650308231 | -0.434801215 | 0.677465027 | 0.995570006 | -5.178327171 |
| **A_52_P325443** | Arhgap29 | 0.026785972 | -0.120490746 | 0.17406269 | 10.33354324 | 0.434692792 | 0.677539885 | 0.995570006 | -5.178365566 |
| **A_51_P203955** | Gbp2 | -0.042560502 | -0.276586162 | 0.191465159 | 8.607043983 | -0.434662716 | 0.677560651 | 0.995570006 | -5.178376215 |
| **A_55_P2168736** | Relb | -0.026807633 | -0.17424963 | 0.120634364 | 8.493908817 | -0.434556644 | 0.677633891 | 0.995570006 | -5.178413766 |
| **A_55_P2149049** | Rpl32 | 0.021190973 | -0.095359809 | 0.137741755 | 16.77230387 | 0.434555089 | 0.677634965 | 0.995570006 | -5.178414317 |
| **A_66_P139946** | Sfta3-ps | 0.017795352 | -0.080084976 | 0.11567568 | 7.662910397 | 0.434530534 | 0.67765192 | 0.995570006 | -5.178423009 |
| **A_55_P2009266** | Gm16060 | -0.030193954 | -0.196287912 | 0.135900005 | 14.38383802 | -0.434485496 | 0.67768302 | 0.995570006 | -5.17843895 |
| **A_52_P58041** | Arpc5 | 0.044055976 | -0.19833763 | 0.286449581 | 10.09990099 | 0.434402983 | 0.677739997 | 0.995570006 | -5.17846815 |
| **A_55_P2219059** | Lmnb2 | -0.031744634 | -0.206402261 | 0.142912993 | 9.066369001 | -0.434402082 | 0.67774062 | 0.995570006 | -5.178468469 |
| **A_51_P472932** | Atp8b1 | -0.018406599 | -0.119698585 | 0.082885386 | 7.845728683 | -0.434317779 | 0.677798836 | 0.995570006 | -5.178498298 |
| **A_55_P2129658** | Smad6 | 0.06702186 | -0.301903851 | 0.43594757 | 10.58092697 | 0.434196866 | 0.677882337 | 0.995570006 | -5.178541071 |
| **A_55_P2107432** | Slc18a1 | -0.019465389 | -0.126631192 | 0.087700414 | 7.849382369 | -0.434126203 | 0.677931139 | 0.995570006 | -5.178566063 |

| | | | | | | | | | |
|---|---|---|---|---|---|---|---|---|---|
| **A_55_P2147001** | Wdr55 | 0.031496871 | -0.141920897 | 0.204914639 | 10.66973315 | 0.434093174 | 0.677953951 | 0.995570006 | -5.178577743 |
| **A_55_P2113026** | Foxo4 | -0.018945136 | -0.123258576 | 0.085368304 | 7.818735396 | -0.434076823 | 0.677965243 | 0.995570006 | -5.178583525 |
| **A_51_P271984** | Tmem45b | -0.017394681 | -0.113193561 | 0.0784042 | 7.650565297 | -0.433975448 | 0.67803526 | 0.995570006 | -5.178619369 |
| **A_51_P182303** | Col1a2 | 0.076568311 | -0.345172415 | 0.498309038 | 15.79599014 | 0.433922988 | 0.678071494 | 0.995570006 | -5.178637914 |
| **A_55_P1956284** | NA | 0.029492889 | -0.132972686 | 0.191958463 | 8.645411527 | 0.433875465 | 0.678104319 | 0.995570006 | -5.178654713 |
| **A_55_P1985591** | Rtl8a | -0.053930438 | -0.351056642 | 0.243195765 | 9.587180045 | -0.433812568 | 0.678147764 | 0.995570006 | -5.178676943 |
| **A_66_P139310** | Cog1 | 0.029990774 | -0.135248314 | 0.195229862 | 10.65944048 | 0.433794472 | 0.678160263 | 0.995570006 | -5.178683338 |
| **A_52_P445387** | Clk4 | 0.077067647 | -0.347567414 | 0.501702707 | 9.89963361 | 0.433775862 | 0.678173119 | 0.995570006 | -5.178689915 |
| **A_55_P1964038** | Gm10032 | 0.035588563 | -0.160511115 | 0.231688241 | 8.497171641 | 0.43375321 | 0.678188765 | 0.995570006 | -5.178697919 |
| **A_51_P120830** | Mmp10 | 0.019629824 | -0.088550678 | 0.127810327 | 7.66627808 | 0.43368715 | 0.678234398 | 0.995570006 | -5.178721261 |
| **A_55_P2027323** | Golga7 | 0.028541744 | -0.128806804 | 0.185890291 | 12.20310283 | 0.433537725 | 0.678337622 | 0.995570006 | -5.178774048 |
| **A_55_P1971903** | Pla2r1 | -0.019397751 | -0.126339198 | 0.087543696 | 7.826533354 | -0.433525312 | 0.678346197 | 0.995570006 | -5.178778432 |
| **A_55_P2340593** | 1810019D2 | -0.019368844 | -0.126163845 | 0.087426156 | 7.873145001 | -0.433472865 | 0.678382431 | 0.995570006 | -5.178796955 |
| **A_55_P1971329** | Lage3 | -0.044385066 | -0.289161219 | 0.200391086 | 10.61026115 | -0.43338802 | 0.678441048 | 0.995570006 | -5.178826916 |
| **A_55_P2073699** | Bnip3l | 0.03634768 | -0.164121205 | 0.236816564 | 11.59814444 | 0.433350047 | 0.678467283 | 0.995570006 | -5.178840323 |
| **A_55_P2158072** | NA | -0.019776587 | -0.128863714 | 0.089310539 | 8.021569849 | -0.433298293 | 0.67850304 | 0.995570006 | -5.178858595 |
| **A_51_P233153** | Cadps2 | -0.018289125 | -0.119182185 | 0.082603934 | 7.79553414 | -0.433252196 | 0.678534889 | 0.995570006 | -5.178874867 |
| **A_55_P2069979** | Large1 | -0.032540733 | -0.21206389 | 0.146982424 | 11.5832133 | -0.433227467 | 0.678551975 | 0.995570006 | -5.178883596 |
| **A_55_P2174582** | Urah | -0.019392156 | -0.12638969 | 0.087605378 | 7.761587977 | -0.433173079 | 0.678589555 | 0.995570006 | -5.178902792 |
| **A_51_P504314** | Prpf38b | 0.028915325 | -0.130644592 | 0.188475241 | 9.350540687 | 0.433125153 | 0.67862267 | 0.995570006 | -5.178919706 |
| **A_55_P2426272** | Scaper | -0.023571753 | -0.153658311 | 0.106514805 | 8.474122662 | -0.43308046 | 0.678653552 | 0.995570006 | -5.178935477 |
| **A_52_P26626** | Fam92b | -0.025203656 | -0.164340608 | 0.113933295 | 7.746273236 | -0.432942479 | 0.678748898 | 0.995570006 | -5.178984157 |
| **A_51_P352303** | Homer2 | 0.049935697 | -0.225890909 | 0.325762303 | 9.718631716 | 0.432697207 | 0.678918399 | 0.995570006 | -5.179070655 |
| **A_55_P2076524** | 1700028I16 | -0.079756184 | -0.52037876 | 0.360866393 | 9.719289803 | -0.432620157 | 0.678971651 | 0.995570006 | -5.179097818 |
| **A_55_P2132237** | Fastkd3 | -0.019154518 | -0.125010218 | 0.086701181 | 9.060567589 | -0.432480094 | 0.679068458 | 0.995570006 | -5.179147185 |
| **A_66_P130241** | Slc25a1 | 0.03176356 | -0.143789688 | 0.207316807 | 9.033200358 | 0.432443558 | 0.679093712 | 0.995570006 | -5.17916006 |
| **A_52_P18068** | Txlng | -0.028708253 | -0.187386596 | 0.129970091 | 8.402687482 | -0.432412492 | 0.679115185 | 0.995570006 | -5.179171006 |
| **A_52_P482124** | Fam32a | -0.030378851 | -0.198362261 | 0.137604559 | 9.825849164 | -0.432229177 | 0.679241899 | 0.995570006 | -5.179235585 |
| **A_52_P372165** | Rpsa | 0.033710367 | -0.152706248 | 0.220126982 | 8.386051712 | 0.432203216 | 0.679259845 | 0.995570006 | -5.179244728 |
| **A_52_P455540** | Ric8b | 0.028159971 | -0.127577311 | 0.183897254 | 10.05626999 | 0.432164163 | 0.679286843 | 0.995570006 | -5.179258482 |
| **A_51_P508444** | St13 | 0.029157094 | -0.132110103 | 0.190424291 | 12.29550516 | 0.432122949 | 0.679315334 | 0.995570006 | -5.179272995 |
| **A_55_P2160198** | NA | 0.033017476 | -0.149679656 | 0.215714609 | 10.15135938 | 0.431937867 | 0.679443289 | 0.995570006 | -5.179338156 |
| **A_51_P158876** | Dctpp1 | -0.03691442 | -0.241247494 | 0.167418655 | 13.88796995 | -0.431783963 | 0.679549697 | 0.995570006 | -5.17939232 |
| **A_52_P639551** | Tinf2 | -0.023760871 | -0.15533305 | 0.107811309 | 9.526484567 | -0.431625818 | 0.679659046 | 0.995570006 | -5.179447958 |
| **A_51_P148311** | Chic2 | 0.029629758 | -0.134466121 | 0.193725637 | 10.06538861 | 0.431558386 | 0.679705674 | 0.995570006 | -5.179471676 |
| **A_55_P2029121** | Pex11g | 0.024458369 | -0.111084297 | 0.160001035 | 8.43129452 | 0.431281278 | 0.679897306 | 0.995570006 | -5.179569107 |
| **A_51_P340816** | Ipp | -0.021398529 | -0.140043204 | 0.097246145 | 9.181583677 | -0.431067033 | 0.680045484 | 0.995570006 | -5.179644396 |
| **A_55_P1963576** | Gm11595 | -0.02000246 | -0.130922821 | 0.0909179 | 8.154248397 | -0.431003977 | 0.680089098 | 0.995570006 | -5.179666548 |
| **A_55_P2044297** | Ciao2a | -0.026548363 | -0.173803399 | 0.120706674 | 13.28136613 | -0.430900221 | 0.680160866 | 0.995570006 | -5.179702991 |
| **A_52_P410685** | Krt7 | 0.091125635 | -0.414497719 | 0.596748989 | 8.699926486 | 0.430747312 | 0.680266639 | 0.995570006 | -5.179756685 |
| **A_55_P2073607** | Commd5 | -0.02230843 | -0.14609856 | 0.1014817 | 11.15684627 | -0.430717103 | 0.680287537 | 0.995570006 | -5.179767291 |
| **A_52_P742248** | Fam117b | -0.038595815 | -0.252780108 | 0.175588478 | 10.62060795 | -0.430686967 | 0.680308385 | 0.995570006 | -5.17977787 |
| **A_55_P2060278** | Fam45a | 0.042201944 | -0.192078468 | 0.276482357 | 9.811696084 | 0.430532172 | 0.680415473 | 0.995570006 | -5.179832201 |
| **A_55_P2061021** | Smg6 | -0.026063827 | -0.170888225 | 0.118760571 | 11.47306713 | -0.430135792 | 0.680689731 | 0.995570006 | -5.179971241 |
| **A_55_P2107854** | Hnrnpk | -0.039849055 | -0.261271955 | 0.181573846 | 11.25801573 | -0.430134839 | 0.68069039 | 0.995570006 | -5.179971575 |
| **A_52_P413395** | Sln | 0.449524078 | -2.048363676 | 2.947411833 | 13.52382608 | 0.43011954 | 0.680700977 | 0.995570006 | -5.17997694 |
| **A_55_P1984546** | Mmp1a | -0.017469221 | -0.114544839 | 0.079606398 | 7.657934839 | -0.430103025 | 0.680712405 | 0.995570006 | -5.17998273 |
| **A_52_P141583** | Macroh2a1 | 0.041777597 | -0.190402308 | 0.273957502 | 9.874003303 | 0.430058918 | 0.680742927 | 0.995570006 | -5.179998193 |

| | | | | | | | | | |
|---|---|---|---|---|---|---|---|---|---|
| **A_66_P126686** | NA | -0.024870023 | -0.163130847 | 0.1133908 | 15.08735235 | -0.429918549 | 0.680840065 | 0.995570006 | -5.180047394 |
| **A_52_P168567** | Cebpa | -0.018107061 | -0.118805814 | 0.082591691 | 7.836042436 | -0.429766948 | 0.680944984 | 0.995570006 | -5.180100516 |
| **A_52_P494686** | Kansl1l | 0.02320065 | -0.105825858 | 0.152227159 | 8.425266404 | 0.42976431 | 0.680946809 | 0.995570006 | -5.18010144 |
| **A_66_P111217** | NA | -0.041774763 | -0.274101214 | 0.190551689 | 15.87263302 | -0.429758488 | 0.680950838 | 0.995570006 | -5.180103479 |
| **A_51_P344878** | Thop1 | 0.042047243 | -0.191985003 | 0.276079489 | 9.222145495 | 0.429408815 | 0.681192867 | 0.995570006 | -5.180225937 |
| **A_55_P2325698** | 2010002M( | -0.023005926 | -0.15107051 | 0.105058657 | 9.074483372 | -0.429358253 | 0.681227867 | 0.995570006 | -5.180243637 |
| **A_55_P2093889** | Arhgef19 | -0.021000643 | -0.137929569 | 0.095928282 | 7.997909997 | -0.42925936 | 0.681296325 | 0.995570006 | -5.180278249 |
| **A_52_P641849** | Khnyn | 0.019136675 | -0.087425586 | 0.125698936 | 8.12707347 | 0.429212326 | 0.681328885 | 0.995570006 | -5.180294708 |
| **A_55_P1975305** | NA | 0.021673934 | -0.099035251 | 0.14238312 | 11.61656403 | 0.429147479 | 0.681373778 | 0.995570006 | -5.180317398 |
| **A_55_P2028496** | NA | -0.034750674 | -0.22828916 | 0.158787812 | 13.99582713 | -0.429145955 | 0.681374833 | 0.995570006 | -5.180317931 |
| **A_55_P1985040** | Gpbp1l1 | -0.025549249 | -0.167844732 | 0.116746235 | 8.663193101 | -0.429137191 | 0.681380901 | 0.995570006 | -5.180320998 |
| **A_66_P140452** | Map1a | 0.029591697 | -0.135225266 | 0.194408659 | 8.647093553 | 0.429118358 | 0.681393939 | 0.995570006 | -5.180327587 |
| **A_51_P446131** | Gipc2 | -0.018141506 | -0.119194111 | 0.0829111 | 7.970685647 | -0.429076721 | 0.681422765 | 0.995570006 | -5.180342153 |
| **A_55_P1965363** | Serpinb6e | -0.018106941 | -0.118972732 | 0.08275885 | 7.735790059 | -0.429052386 | 0.681439613 | 0.995570006 | -5.180350665 |
| **A_55_P2001871** | Uros | 0.022417859 | -0.102505596 | 0.147341314 | 10.96774488 | 0.428903194 | 0.681542907 | 0.995570006 | -5.180402845 |
| **A_55_P2164841** | NA | -0.022720257 | -0.149355012 | 0.103914497 | 10.20804372 | -0.428814502 | 0.681604317 | 0.995570006 | -5.180433856 |
| **A_52_P507498** | Plxnc1 | -0.034150397 | -0.224537219 | 0.156236425 | 8.985210357 | -0.428714332 | 0.681673677 | 0.995570006 | -5.180468874 |
| **A_55_P2175341** | NA | 0.030190628 | -0.138132778 | 0.198514034 | 9.497467133 | 0.428683502 | 0.681695026 | 0.995570006 | -5.18047965 |
| **A_55_P2012406** | NA | -0.033504276 | -0.220312577 | 0.153304025 | 16.61205504 | -0.428660237 | 0.681711135 | 0.995570006 | -5.180487782 |
| **A_66_P124420** | NA | 0.017301572 | -0.079185375 | 0.11378852 | 8.047625317 | 0.428574314 | 0.681770635 | 0.995570006 | -5.180517809 |
| **A_52_P428745** | Camk2d | 0.027437444 | -0.125575845 | 0.180450733 | 12.90164845 | 0.428571843 | 0.681772346 | 0.995570006 | -5.180518673 |
| **A_52_P562593** | Tpgs1 | 0.072934552 | -0.333821606 | 0.479690709 | 9.699241654 | 0.428556725 | 0.681782815 | 0.995570006 | -5.180523956 |
| **A_51_P516362** | Vps11 | -0.03604112 | -0.23704482 | 0.164962579 | 9.867844074 | -0.428551841 | 0.681786198 | 0.995570006 | -5.180525662 |
| **A_52_P645862** | Agtr1a | -0.033155715 | -0.218114156 | 0.151802726 | 10.62648046 | -0.428443321 | 0.681861349 | 0.995570006 | -5.180563577 |
| **A_55_P2043277** | Siglecf | 0.018734407 | -0.085805431 | 0.123274245 | 7.742141711 | 0.428318919 | 0.681947505 | 0.995570006 | -5.18060703 |
| **A_55_P2011151** | Tmprss3 | -0.035600731 | -0.234271918 | 0.163070456 | 10.65786839 | -0.428285297 | 0.681970791 | 0.995570006 | -5.180618772 |
| **A_55_P2126333** | Cyp2a12 | -0.020225381 | -0.133114744 | 0.092663983 | 7.693880459 | -0.428206061 | 0.68202567 | 0.995570006 | -5.18064644 |
| **A_55_P2010223** | Gabarap | -0.055373834 | -0.364458833 | 0.253711166 | 13.97631639 | -0.428189286 | 0.682037288 | 0.995570006 | -5.180652297 |
| **A_55_P1973448** | Ybx2 | -0.021284319 | -0.140116904 | 0.097548265 | 8.486871841 | -0.428088299 | 0.682107236 | 0.995570006 | -5.180687553 |
| **A_55_P2101431** | Mrps25 | 0.048092223 | -0.220432792 | 0.316617239 | 8.921632025 | 0.428054668 | 0.68213053 | 0.995570006 | -5.180699292 |
| **A_55_P2060722** | Uso1 | 0.028863934 | -0.132326544 | 0.190054412 | 11.16098294 | 0.427981768 | 0.682181026 | 0.995570006 | -5.180724735 |
| **A_51_P130808** | Tpbpb | -0.415190875 | -2.734141472 | 1.903759723 | 9.717611124 | -0.427922809 | 0.682221867 | 0.995570006 | -5.18074531 |
| **A_55_P1994047** | NA | 0.057111251 | -0.261917267 | 0.37613977 | 13.68751072 | 0.42785962 | 0.682265639 | 0.995570006 | -5.180767358 |
| **A_51_P371174** | Bag3 | -0.025078796 | -0.165197044 | 0.115039452 | 12.43643556 | -0.427780632 | 0.682320357 | 0.995570006 | -5.180794914 |
| **A_55_P1992470** | NA | -0.063682672 | -0.419496274 | 0.292130929 | 15.79165069 | -0.42776758 | 0.682329399 | 0.995570006 | -5.180799467 |
| **A_52_P133436** | Plekhm1 | 0.025862885 | -0.118653608 | 0.170379379 | 9.157885872 | 0.427728993 | 0.682356131 | 0.995570006 | -5.180812927 |
| **A_55_P2131964** | Pdia2 | 0.021639268 | -0.099280126 | 0.142558663 | 7.760315765 | 0.427716241 | 0.682364965 | 0.995570006 | -5.180817375 |
| **A_55_P2178064** | NA | 0.018298111 | -0.083960239 | 0.12055646 | 7.828814677 | 0.427677697 | 0.682391668 | 0.995570006 | -5.180830818 |
| **A_55_P2184567** | NA | -0.019241999 | -0.126800564 | 0.088316567 | 7.994890588 | -0.427576979 | 0.682461446 | 0.995570006 | -5.18086594 |
| **A_55_P2185900** | Nrg4 | -0.018312097 | -0.120689203 | 0.084065009 | 7.782951918 | -0.427508114 | 0.682509157 | 0.995570006 | -5.18088995 |
| **A_51_P103209** | Ttc7 | 0.036577154 | -0.167917665 | 0.241071973 | 9.053452894 | 0.427500609 | 0.682514358 | 0.995570006 | -5.180892567 |
| **A_55_P2027213** | Krtap20-2 | 0.031266021 | -0.14355858 | 0.206090622 | 8.328564813 | 0.427443979 | 0.682553594 | 0.995570006 | -5.180912308 |
| **A_55_P1985464** | Gm12271 | -0.031531712 | -0.207873824 | 0.1448104 | 14.0886016 | -0.427366669 | 0.682607161 | 0.995570006 | -5.180939255 |
| **A_55_P2096630** | Ssxb9 | 0.023981783 | -0.11015748 | 0.158121046 | 7.877907702 | 0.427301765 | 0.682652132 | 0.995570006 | -5.180961873 |
| **A_55_P1981814** | Myo3b | 0.035468733 | -0.16297602 | 0.233913487 | 9.184400744 | 0.427184209 | 0.682733591 | 0.995570006 | -5.181002833 |
| **A_51_P130994** | NA | -0.030441269 | -0.200785193 | 0.139902656 | 11.27633402 | -0.427115403 | 0.682781271 | 0.995570006 | -5.181026802 |
| **A_51_P267796** | Ddr2 | -0.046344536 | -0.305769969 | 0.213080897 | 9.233882948 | -0.426967794 | 0.682883564 | 0.995570006 | -5.181078211 |
| **A_55_P2114898** | Gdap2 | -0.034071579 | -0.224827418 | 0.15668426 | 9.519682372 | -0.426897444 | 0.682932319 | 0.995570006 | -5.181102706 |

| | | | | | | | | | |
|---|---|---|---|---|---|---|---|---|---|
| **A_52_P515247** | Arhgdib | 0.024126678 | -0.11096802 | 0.159221377 | 11.39279018 | 0.426843193 | 0.682969918 | 0.995570006 | -5.181121593 |
| **A_51_P106952** | Rbm26 | 0.044824803 | -0.206166838 | 0.295816444 | 10.31330118 | 0.426843098 | 0.682969984 | 0.995570006 | -5.16902774 |
| **A_51_P182296** | Washc3 | 0.023099399 | -0.106259131 | 0.15245793 | 10.92891255 | 0.426790497 | 0.68300644 | 0.995570006 | -5.181139937 |
| **A_55_P1983149** | Rps29 | 0.022276879 | -0.102489002 | 0.147042761 | 17.3538066 | 0.426744218 | 0.683038516 | 0.995570006 | -5.181156045 |
| **A_55_P2086510** | Ppid | 0.031926314 | -0.146903293 | 0.210755921 | 12.46238483 | 0.426695922 | 0.68307199 | 0.995570006 | -5.181172854 |
| **A_55_P2051825** | Ephx3 | -0.051014063 | -0.336769718 | 0.234741592 | 8.329387562 | -0.426681892 | 0.683081715 | 0.995570006 | -5.181177736 |
| **A_55_P1955891** | NA | 0.048797261 | -0.224600199 | 0.322194721 | 12.2603357 | 0.426589441 | 0.683145796 | 0.995570006 | -5.181209906 |
| **A_55_P1997661** | Smim15 | 0.033404489 | -0.15377814 | 0.220587117 | 10.72534376 | 0.426528857 | 0.68318779 | 0.995570006 | -5.181230984 |
| **A_55_P2230913** | Etv6 | -0.020652254 | -0.136378402 | 0.095073894 | 9.335855903 | -0.426525594 | 0.683190052 | 0.995570006 | -5.181232119 |
| **A_51_P480861** | Pga5 | -0.022731561 | -0.150136797 | 0.104673676 | 7.726088401 | -0.426433298 | 0.68325403 | 0.995570006 | -5.181264224 |
| **A_55_P1958088** | Lyn | -0.019292874 | -0.127478492 | 0.088892744 | 8.27672296 | -0.426222653 | 0.683400056 | 0.995570006 | -5.181337472 |
| **A_55_P2164221** | Tpt1 | 0.025866453 | -0.119186276 | 0.170919182 | 17.75906456 | 0.426206544 | 0.683411225 | 0.995570006 | -5.181343072 |
| **A_51_P228865** | Zfp72 | -0.022058012 | -0.145754755 | 0.101638731 | 8.617795273 | -0.426203727 | 0.683413178 | 0.995570006 | -5.181344052 |
| **A_52_P619248** | Prl2c5 | -0.064609741 | -0.426931603 | 0.297712122 | 7.921831825 | -0.426199161 | 0.683416343 | 0.995570006 | -5.181345639 |
| **A_55_P2055291** | Gpn2 | 0.027924687 | -0.128684543 | 0.184533918 | 8.971869437 | 0.426167273 | 0.68343845 | 0.995570006 | -5.181356724 |
| **A_52_P87843** | Aldh1a3 | -0.029101659 | -0.19232682 | 0.134123502 | 8.105168104 | -0.426127705 | 0.683465883 | 0.995570006 | -5.181370478 |
| **A_55_P1999182** | Cdk12 | 0.020938099 | -0.096501275 | 0.138377472 | 8.692958899 | 0.426120716 | 0.683470728 | 0.995570006 | -5.181372907 |
| **A_55_P2104542** | NA | -0.021491704 | -0.142043497 | 0.099060089 | 8.29316323 | -0.426094866 | 0.68348865 | 0.995570006 | -5.181381891 |
| **A_51_P336635** | Xpo1 | -0.033895009 | -0.224049489 | 0.15625947 | 11.00741742 | -0.426028185 | 0.683534881 | 0.995570006 | -5.181405065 |
| **A_55_P1986466** | Retreg2 | -0.027472545 | -0.181603053 | 0.126657964 | 8.436839196 | -0.426009624 | 0.68354775 | 0.995570006 | -5.181411515 |
| **A_55_P2033282** | Cept1 | 0.039309364 | -0.181247175 | 0.259865903 | 11.43429953 | 0.425976077 | 0.683571011 | 0.995570006 | -5.181423172 |
| **A_55_P1979575** | Shroom2 | 0.0545516 | -0.251550108 | 0.360653307 | 10.78254626 | 0.425942397 | 0.683594363 | 0.995570006 | -5.181434874 |
| **A_55_P1998299** | Itgb4 | -0.024471768 | -0.161809717 | 0.112866181 | 8.371706525 | -0.425876739 | 0.683639888 | 0.995570006 | -5.181457684 |
| **A_55_P2008176** | Zscan12 | -0.020201724 | -0.133579825 | 0.093176377 | 9.125600172 | -0.425861511 | 0.683650447 | 0.995570006 | -5.181462974 |
| **A_55_P2125241** | NA | 0.029030341 | -0.133910446 | 0.191971128 | 8.268845603 | 0.425825302 | 0.683675555 | 0.995570006 | -5.181475552 |
| **A_55_P2136765** | Uqcr10 | 0.035181545 | -0.162286443 | 0.232649532 | 14.60229246 | 0.425821255 | 0.68367836 | 0.995570006 | -5.181476958 |
| **A_55_P2317341** | NA | 0.040100185 | -0.184990238 | 0.265190608 | 8.620716044 | 0.425792974 | 0.683697971 | 0.995570006 | -5.181486781 |
| **A_55_P1993019** | NA | -0.023413452 | -0.154869751 | 0.108042847 | 10.65267427 | -0.425689741 | 0.683769556 | 0.995570006 | -5.181522632 |
| **A_55_P2155921** | Hace1 | -0.019903588 | -0.131678839 | 0.091871663 | 8.13089477 | -0.425593368 | 0.683836387 | 0.995570006 | -5.181556094 |
| **A_51_P113906** | Gkn2 | 0.085878171 | -0.396466824 | 0.568223167 | 7.847357403 | 0.425533881 | 0.683877641 | 0.995570006 | -5.163832583 |
| **A_52_P528463** | Mfsd3 | 0.02394709 | -0.110605732 | 0.158499913 | 9.556511557 | 0.425372166 | 0.683989794 | 0.995570006 | -5.181632871 |
| **A_51_P446978** | Cfdp1 | 0.030551624 | -0.14114175 | 0.202244998 | 12.67881659 | 0.42529463 | 0.68404357 | 0.995570006 | -5.181659774 |
| **A_55_P2043434** | Atmin | 0.022068792 | -0.101981422 | 0.146119006 | 10.71339532 | 0.425196982 | 0.684111298 | 0.995570006 | -5.181693649 |
| **A_51_P124798** | Tmem192 | -0.035997965 | -0.238352082 | 0.166356152 | 9.029064719 | -0.425182161 | 0.684121578 | 0.995570006 | -5.175788084 |
| **A_55_P2081555** | Phf23 | 0.027664097 | -0.12789237 | 0.183220564 | 10.04799018 | 0.425047592 | 0.68421492 | 0.995570006 | -5.18174546 |
| **A_55_P1985298** | Krtap5-2 | -0.019519522 | -0.129286317 | 0.090247274 | 8.125937897 | -0.42501799 | 0.684235454 | 0.995570006 | -5.181755724 |
| **A_55_P2009206** | Sumo2 | -0.070220373 | -0.465100398 | 0.324659652 | 9.632646656 | -0.425017783 | 0.684235597 | 0.995570006 | -5.181755796 |
| **A_55_P1961423** | Gsta3 | -0.027921426 | -0.184979958 | 0.129137106 | 7.976120737 | -0.424898492 | 0.684318348 | 0.995570006 | -5.181797153 |
| **A_55_P2064043** | Cd44 | 0.041985995 | -0.194210766 | 0.278182756 | 9.615997101 | 0.424853935 | 0.684349258 | 0.995570006 | -5.181812597 |
| **A_51_P108935** | Iars2 | -0.041977172 | -0.278165414 | 0.194211069 | 10.17114339 | -0.424779978 | 0.684400564 | 0.995570006 | -5.18183823 |
| **A_55_P2020128** | Dhrs3 | 0.048880888 | -0.226329527 | 0.324091304 | 11.42661545 | 0.424505533 | 0.684590973 | 0.995570006 | -5.181933311 |
| **A_55_P2030209** | Cir1 | 0.042606668 | -0.1972829 | 0.282496237 | 9.407864824 | 0.424497705 | 0.684596404 | 0.995570006 | -5.181936022 |
| **A_55_P2108754** | Ubfd1 | -0.042394307 | -0.281118794 | 0.19633018 | 11.67453038 | -0.42444332 | 0.684634139 | 0.995570006 | -5.181954857 |
| **A_55_P1952905** | Dnm1l | 0.041500382 | -0.192250684 | 0.275251448 | 12.25533992 | 0.424333802 | 0.684710131 | 0.995570006 | -5.181992778 |
| **A_51_P128336** | Serinc5 | -0.035730392 | -0.237015251 | 0.165554467 | 10.27237604 | -0.424263636 | 0.68475882 | 0.995570006 | -5.182017069 |
| **A_55_P2091348** | NA | 0.032661007 | -0.151361954 | 0.216683967 | 9.59389942 | 0.424196113 | 0.684805676 | 0.995570006 | -5.182040441 |
| **A_51_P291460** | Gm6498 | 0.0417427 | -0.193482565 | 0.276967965 | 16.10114668 | 0.424136551 | 0.684847009 | 0.995570006 | -5.182061055 |
| **A_51_P246339** | Rfc5 | -0.03466032 | -0.230002046 | 0.160681405 | 11.53921765 | -0.42407892 | 0.684887003 | 0.995570006 | -5.182080998 |

| | | | | | | | | | |
|---|---|---|---|---|---|---|---|---|---|
| **A_52_P580582** | Nppa | 0.220428542 | -1.021902966 | 1.46276005 | 16.58090181 | 0.424071865 | 0.684891899 | 0.995570006 | -5.146147305 |
| **A_55_P2015118** | NA | -0.052595837 | -0.34905505 | 0.243863375 | 11.17030775 | -0.424028994 | 0.684921651 | 0.995570006 | -5.182098272 |
| **A_55_P2107855** | Hnrnpk | -0.032437803 | -0.21528974 | 0.150414135 | 10.39241327 | -0.423995252 | 0.684945068 | 0.995570006 | -5.182109946 |
| **A_65_P17492** | Med29 | -0.036105152 | -0.239635626 | 0.167425321 | 10.52478815 | -0.423983419 | 0.68495328 | 0.995570006 | -5.18211404 |
| **A_55_P2311780** | 4930554H2 | -0.017114974 | -0.113602927 | 0.079372979 | 7.628073951 | -0.423947705 | 0.684978067 | 0.995570006 | -5.182126394 |
| **A_55_P2060498** | Zfp553 | -0.031585493 | -0.209660304 | 0.146489317 | 10.03179156 | -0.423930153 | 0.684990248 | 0.995570006 | -5.182132466 |
| **A_55_P2080708** | Cstf2 | 0.047479975 | -0.220221496 | 0.315181446 | 9.117527059 | 0.423905309 | 0.68500749 | 0.995570006 | -5.182141059 |
| **A_52_P147803** | Syne4 | 0.021953532 | -0.101829718 | 0.145736781 | 8.034946498 | 0.423888513 | 0.685019148 | 0.995570006 | -5.182146869 |
| **A_51_P404077** | Fzd2 | -0.042345297 | -0.281156266 | 0.196465672 | 11.46942731 | -0.423799114 | 0.685081195 | 0.995570006 | -5.182177787 |
| **A_55_P2182452** | Tnfsf15 | -0.017365354 | -0.115317687 | 0.080586978 | 7.76874122 | -0.423719062 | 0.685136758 | 0.995570006 | -5.182205467 |
| **A_55_P2087528** | Tmem131l | 0.019334763 | -0.089732527 | 0.128402052 | 11.20451997 | 0.423695116 | 0.685153379 | 0.995570006 | -5.182213746 |
| **A_55_P2098688** | Mcrs1 | -0.043735325 | -0.2905546 | 0.20308395 | 9.991811503 | -0.423508781 | 0.685282721 | 0.995570006 | -5.182278154 |
| **A_55_P1971184** | Ube2l3 | -0.026581795 | -0.176598355 | 0.123434764 | 12.88310575 | -0.423500805 | 0.685288258 | 0.995570006 | -5.18228091 |
| **A_66_P125401** | NA | 0.037322757 | -0.173455161 | 0.248100675 | 9.059445467 | 0.423211771 | 0.68548891 | 0.995570006 | -5.182380763 |
| **A_55_P2089804** | Atf7 | 0.018186931 | -0.084530931 | 0.120904793 | 7.885303021 | 0.423177509 | 0.685512697 | 0.995570006 | -5.182392596 |
| **A_55_P1976278** | Prkcb | -0.032590952 | -0.21670202 | 0.151520117 | 8.422086552 | -0.423083683 | 0.685577839 | 0.995570006 | -5.182424994 |
| **A_55_P2120348** | Pdcd7 | -0.029300264 | -0.194825554 | 0.136225026 | 10.08100974 | -0.423073976 | 0.685584579 | 0.995570006 | -5.182428345 |
| **A_51_P407675** | Zrsr1 | -0.048588816 | -0.323081607 | 0.225903976 | 9.012621841 | -0.423072213 | 0.685585803 | 0.995570006 | -5.182428954 |
| **A_51_P375533** | Polr3b | 0.021905276 | -0.101871594 | 0.145682146 | 8.99767297 | 0.422978571 | 0.685650821 | 0.995570006 | -5.182461281 |
| **A_66_P119643** | Atrx | 0.023450228 | -0.109062128 | 0.155962583 | 8.651671318 | 0.422960514 | 0.685663359 | 0.995570006 | -5.182467513 |
| **A_51_P101573** | Klc4 | -0.046848624 | -0.311605025 | 0.217907777 | 10.51784784 | -0.422921247 | 0.685690624 | 0.995570006 | -5.182481067 |
| **A_51_P278034** | Ufsp1 | -0.03629188 | -0.241400967 | 0.168817208 | 9.30676272 | -0.422896113 | 0.685708076 | 0.995570006 | -5.182489741 |
| **A_55_P1993836** | Dusp7 | 0.031554629 | -0.146805022 | 0.209914281 | 9.819424438 | 0.422839547 | 0.685747354 | 0.995570006 | -5.182509262 |
| **A_52_P509744** | Nup160 | 0.031010884 | -0.144282069 | 0.206303838 | 9.073631317 | 0.422823217 | 0.685758694 | 0.995570006 | -5.182514897 |
| **A_55_P2023572** | NA | -0.019461351 | -0.129510949 | 0.090588247 | 8.182353442 | -0.422662438 | 0.685870342 | 0.995570006 | -5.182570366 |
| **A_55_P2184385** | NA | -0.022595679 | -0.150389803 | 0.105198445 | 7.809560781 | -0.422594307 | 0.685917656 | 0.995570006 | -5.182593866 |
| **A_55_P2091626** | NA | 0.024449043 | -0.113837502 | 0.162735587 | 15.54814135 | 0.422562613 | 0.685939666 | 0.995570006 | -5.182604797 |
| **A_51_P148675** | Dag1 | -0.043268324 | -0.288034555 | 0.201497908 | 12.38430118 | -0.42250096 | 0.685982484 | 0.995570006 | -5.182626057 |
| **A_66_P134674** | Far1 | 0.046641455 | -0.217210102 | 0.310493012 | 9.073490201 | 0.422494985 | 0.685986633 | 0.995570006 | -5.182628117 |
| **A_55_P2129523** | NA | -0.017640252 | -0.117464043 | 0.082183539 | 7.717903726 | -0.422357162 | 0.686082354 | 0.995570006 | -5.182675634 |
| **A_52_P304720** | Crlf1 | -0.017371084 | -0.115680071 | 0.080937902 | 7.654693927 | -0.422321155 | 0.686107363 | 0.995570006 | -5.182688045 |
| **A_66_P103198** | NA | 0.028304268 | -0.131910754 | 0.188519291 | 8.268490162 | 0.422238498 | 0.686164774 | 0.995570006 | -5.182716533 |
| **A_55_P2161690** | Tomm40 | -0.08362347 | -0.557016771 | 0.389769831 | 9.567470357 | -0.422197065 | 0.686193553 | 0.995570006 | -5.182730811 |
| **A_51_P510764** | Agmo | -0.020513025 | -0.13674787 | 0.095721821 | 7.701961608 | -0.421796037 | 0.686472133 | 0.995570006 | -5.182868939 |
| **A_51_P296528** | Cers4 | -0.032530391 | -0.216865982 | 0.151805201 | 11.60628056 | -0.42178314 | 0.686481092 | 0.995570006 | -5.182873379 |
| **A_55_P1960411** | Lrrc23 | 0.02239049 | -0.104495618 | 0.149276598 | 7.80556096 | 0.421753465 | 0.686501709 | 0.995570006 | -5.182883595 |
| **A_55_P2004427** | Fbxl15 | 0.027088147 | -0.126462684 | 0.180638978 | 9.717212895 | 0.421634621 | 0.686584278 | 0.995570006 | -5.182924502 |
| **A_52_P247943** | Parl | -0.030904686 | -0.206124789 | 0.144315417 | 9.01045931 | -0.421550424 | 0.686642777 | 0.995570006 | -5.182953476 |
| **A_55_P2000515** | Peli1 | 0.041796061 | -0.195241423 | 0.278833546 | 11.31631849 | 0.421431953 | 0.686725094 | 0.995570006 | -5.182994236 |
| **A_52_P88808** | Vps37a | -0.033682733 | -0.224739995 | 0.15737453 | 9.432476288 | -0.421359604 | 0.686775367 | 0.995570006 | -5.183019122 |
| **A_52_P570266** | Psmb10 | -0.033545873 | -0.223854723 | 0.156762977 | 13.58088045 | -0.421297853 | 0.686818276 | 0.995570006 | -5.18304036 |
| **A_55_P2037538** | NA | -0.057518066 | -0.383829107 | 0.268792974 | 9.292877421 | -0.421290476 | 0.686823403 | 0.995570006 | -5.183042897 |
| **A_55_P2092103** | NA | 0.01982796 | -0.092727277 | 0.132383197 | 8.244586381 | 0.421038162 | 0.686998745 | 0.995570006 | -5.183129643 |
| **A_51_P213359** | Has2 | 0.023725973 | -0.110968318 | 0.158420263 | 8.413182397 | 0.42100182 | 0.687024003 | 0.995570006 | -5.183142133 |
| **A_55_P2331804** | AU015791 | 0.018799752 | -0.087953183 | 0.125552686 | 7.783978855 | 0.420902425 | 0.687093083 | 0.995570006 | -5.183176289 |
| **A_55_P2132099** | NA | -0.019854789 | -0.132622843 | 0.092913265 | 7.834304104 | -0.420812198 | 0.687155794 | 0.995570006 | -5.183207288 |
| **A_55_P2109997** | Lrig1 | 0.023611226 | -0.110551744 | 0.157774197 | 9.846566434 | 0.420624928 | 0.687285962 | 0.995570006 | -5.183271608 |
| **A_52_P551011** | Smim1 | 0.041139607 | -0.192631738 | 0.274910952 | 8.976016423 | 0.420608452 | 0.687297415 | 0.995570006 | -5.183277266 |

| | | | | | | | | | |
|---|---|---|---|---|---|---|---|---|---|
| **A_55_P2128493** | Rap1a | -0.034182056 | -0.22850855 | 0.160144437 | 9.257026303 | -0.420412195 | 0.687433843 | 0.995570006 | -5.183344641 |
| **A_55_P2184339** | Mdga1 | -0.04978054 | -0.33279763 | 0.233236549 | 9.782141536 | -0.420393584 | 0.687446781 | 0.995570006 | -5.183351029 |
| **A_52_P616591** | Eri3 | -0.029591275 | -0.197826715 | 0.138644165 | 8.703591651 | -0.420392851 | 0.687447291 | 0.995570006 | -5.18335128 |
| **A_66_P126153** | 1700011M( | 0.017224513 | -0.080706632 | 0.115155659 | 7.644951969 | 0.420373429 | 0.687460793 | 0.995570006 | -5.183357946 |
| **A_52_P296913** | Cnnm1 | -0.020657122 | -0.138105537 | 0.096791294 | 7.847238615 | -0.420370066 | 0.68746313 | 0.995570006 | -5.1833591 |
| **A_52_P681488** | Stx17 | 0.020565573 | -0.096378185 | 0.137509331 | 8.881151488 | 0.420313069 | 0.687502755 | 0.995570006 | -5.18337866 |
| **A_51_P209280** | Rab31 | -0.021936716 | -0.146703527 | 0.102830096 | 9.533644661 | -0.420224782 | 0.687564135 | 0.995570006 | -5.183408953 |
| **A_52_P182592** | Ube2w | 0.034394271 | -0.161252863 | 0.230041406 | 9.300675763 | 0.420166821 | 0.687604433 | 0.995570006 | -5.183428837 |
| **A_55_P2081437** | NA | 0.019411996 | -0.091016949 | 0.129840941 | 8.494995106 | 0.420142303 | 0.687621479 | 0.995570006 | -5.183437248 |
| **A_51_P336060** | Lcat | -0.017058684 | -0.11411973 | 0.080002361 | 7.654722105 | -0.420058423 | 0.6876798 | 0.995570006 | -5.183466018 |
| **A_51_P132081** | Rsl1 | -0.020814647 | -0.139275056 | 0.097645763 | 8.269707078 | -0.419957114 | 0.687750242 | 0.995570006 | -5.183500758 |
| **A_51_P240614** | Tm4sf1 | 0.035086073 | -0.164663555 | 0.234835702 | 13.67988766 | 0.419814966 | 0.687849085 | 0.995570006 | -5.18354949 |
| **A_55_P1960936** | NA | 0.077803748 | -0.365533307 | 0.521140803 | 14.22162247 | 0.419445577 | 0.688105973 | 0.995570006 | -5.183676054 |
| **A_55_P1975665** | Rhox4e | 0.017539358 | -0.082425157 | 0.117503874 | 7.677083576 | 0.419350305 | 0.688172236 | 0.995570006 | -5.183708681 |
| **A_55_P1961873** | Akt3 | 0.024993348 | -0.117489411 | 0.167476107 | 9.812228777 | 0.419248237 | 0.688243229 | 0.995570006 | -5.183743626 |
| **A_52_P476754** | Ubash3b | 0.043092603 | -0.202585665 | 0.28877087 | 10.71676936 | 0.419223017 | 0.688260771 | 0.995570006 | -5.18375226 |
| **A_55_P2106079** | NA | 0.025381606 | -0.119353075 | 0.170116288 | 8.011071079 | 0.419136641 | 0.688320853 | 0.995570006 | -5.183781825 |
| **A_51_P323620** | Thyn1 | 0.046327241 | -0.217854478 | 0.310508961 | 13.45367678 | 0.419124267 | 0.68832946 | 0.995570006 | -5.18378606 |
| **A_51_P269792** | Rad51b | -0.020755047 | -0.139118862 | 0.097608768 | 8.071857219 | -0.419096373 | 0.688348864 | 0.995570006 | -5.183795607 |
| **A_51_P214423** | Tead1 | -0.026419917 | -0.17709476 | 0.124254925 | 12.18923197 | -0.419082803 | 0.688358303 | 0.995570006 | -5.18380025 |
| **A_55_P1980426** | Nnat | -0.021125383 | -0.141627235 | 0.099376468 | 7.824460648 | -0.419005778 | 0.688411885 | 0.995570006 | -5.183826607 |
| **A_51_P292447** | Klhl36 | 0.023910829 | -0.112497309 | 0.160318967 | 8.574141389 | 0.418951243 | 0.688449822 | 0.995570006 | -5.183845265 |
| **A_55_P1965728** | Abca17 | -0.019731954 | -0.13230311 | 0.092839202 | 7.947113596 | -0.41894026 | 0.688457462 | 0.995570006 | -5.183849023 |
| **A_55_P2142590** | Mark3 | -0.030323 | -0.203332054 | 0.142686055 | 11.03793813 | -0.418902028 | 0.688484059 | 0.995570006 | -5.183862102 |
| **A_55_P2160732** | Tfcp2 | 0.027856103 | -0.131133318 | 0.186845524 | 9.085644006 | 0.418756215 | 0.688585502 | 0.995570006 | -5.183911973 |
| **A_55_P2043207** | Magef1 | 0.02464465 | -0.116056394 | 0.165345694 | 9.40403183 | 0.418633976 | 0.688670549 | 0.995570006 | -5.183953768 |
| **A_66_P125962** | Hipk1 | 0.023991478 | -0.113078976 | 0.161061932 | 8.187949127 | 0.418333158 | 0.688879862 | 0.995570006 | -5.184056574 |
| **A_55_P1998811** | NA | 0.094151359 | -0.44379514 | 0.632097858 | 11.093678 | 0.418308451 | 0.688897055 | 0.995570006 | -5.184065015 |
| **A_51_P275435** | Mab21l4 | -0.017595841 | -0.118133072 | 0.08294139 | 7.626613574 | -0.418304208 | 0.688900008 | 0.995570006 | -5.184066465 |
| **A_51_P467448** | Pif1 | -0.023321664 | -0.156594185 | 0.109950858 | 10.08463635 | -0.418242384 | 0.688943029 | 0.995570006 | -5.184087583 |
| **A_52_P264229** | Ing5 | 0.030438087 | -0.14350297 | 0.204379144 | 9.174150039 | 0.418238858 | 0.688945484 | 0.995570006 | -5.184088788 |
| **A_51_P477682** | Prss12 | 0.119938408 | -0.565516259 | 0.805393075 | 9.427694826 | 0.418204474 | 0.688969411 | 0.995570006 | -5.184100532 |
| **A_55_P2060101** | Tprgl | -0.031424172 | -0.211023454 | 0.14817511 | 13.61106129 | -0.418184929 | 0.688983013 | 0.995570006 | -5.184107207 |
| **A_55_P2073579** | Chchd5 | -0.025727974 | -0.172819172 | 0.121363225 | 9.60965765 | -0.41804981 | 0.689077046 | 0.995570006 | -5.184153347 |
| **A_51_P225525** | Gm9079 | -0.024996584 | -0.167938122 | 0.117944954 | 12.32611948 | -0.41795675 | 0.689141813 | 0.995570006 | -5.184185117 |
| **A_55_P1964433** | Zmynd12 | 0.016797332 | -0.079324998 | 0.112919662 | 7.805718409 | 0.417662178 | 0.689346845 | 0.995570006 | -5.184285637 |
| **A_55_P2131393** | Eef1akmt2 | 0.03736607 | -0.176530434 | 0.251262574 | 10.10968508 | 0.41752537 | 0.689442078 | 0.995570006 | -5.184332299 |
| **A_66_P119350** | 2310040G2 | -0.069529436 | -0.467559322 | 0.328500449 | 10.04635708 | -0.417505461 | 0.689455938 | 0.995570006 | -5.184339088 |
| **A_55_P2171185** | NA | -0.057191623 | -0.384830532 | 0.270447285 | 12.67319129 | -0.417201717 | 0.689667399 | 0.995570006 | -5.184442632 |
| **A_51_P284793** | Ints12 | -0.021104504 | -0.142010913 | 0.099801906 | 8.580639831 | -0.417191019 | 0.689674847 | 0.995570006 | -5.184446277 |
| **A_55_P2077313** | Ddr1 | -0.019521484 | -0.131427426 | 0.092384459 | 11.62849731 | -0.416935432 | 0.689852807 | 0.995570006 | -5.184533349 |
| **A_55_P2116664** | Hook2 | 0.030075884 | -0.14233372 | 0.202485488 | 9.18399603 | 0.416932817 | 0.689854627 | 0.995570006 | -5.184534239 |
| **A_55_P1968304** | NA | 0.043497611 | -0.205856382 | 0.292851603 | 12.53865899 | 0.416925262 | 0.689859888 | 0.995570006 | -5.184536812 |
| **A_55_P1961620** | Vax2 | -0.069905944 | -0.470807657 | 0.330995769 | 9.612609139 | -0.416759325 | 0.689975438 | 0.995570006 | -5.184593314 |
| **A_55_P2010076** | Miga2 | 0.038995936 | -0.184646676 | 0.262638548 | 11.93923232 | 0.416748387 | 0.689983055 | 0.995570006 | -5.184597038 |
| **A_55_P2017724** | NA | -0.090169947 | -0.607509945 | 0.427170051 | 8.828386802 | -0.416576618 | 0.690102676 | 0.995570006 | -5.184655502 |
| **A_55_P2096807** | Rbms1 | -0.025820661 | -0.173992288 | 0.122350966 | 10.25482803 | -0.416496581 | 0.690158418 | 0.995570006 | -5.184682736 |
| **A_55_P1998912** | Txnrd2 | -0.033144931 | -0.22334744 | 0.157057577 | 12.29831797 | -0.416495215 | 0.690159369 | 0.995570006 | -5.184683201 |

| | | | | | | | | | |
|---|---|---|---|---|---|---|---|---|---|
| **A_55_P1983016** | NA | 0.032856562 | -0.155736701 | 0.221449824 | 8.424015291 | 0.416394586 | 0.690229455 | 0.995570006 | -5.184717435 |
| **A_55_P2108424** | NA | 0.029925266 | -0.141911963 | 0.201762495 | 10.98260771 | 0.416226657 | 0.690346421 | 0.995570006 | -5.184774547 |
| **A_55_P2079158** | Saxo2 | -0.017497788 | -0.118013292 | 0.083017716 | 7.698084992 | -0.416063132 | 0.690460329 | 0.995570006 | -5.18483014 |
| **A_55_P1957291** | Rpl39 | 0.029666382 | -0.14077054 | 0.200103305 | 16.036752 | 0.416016003 | 0.690493159 | 0.995570006 | -5.184846159 |
| **A_52_P554650** | Gzmd | 0.017339711 | -0.082282654 | 0.116962076 | 7.666671964 | 0.416000775 | 0.690503767 | 0.995570006 | -5.184851334 |
| **A_51_P110301** | C3 | -0.19044962 | -1.285124005 | 0.904224765 | 11.17656952 | -0.415818959 | 0.69063043 | 0.995570006 | -5.184913113 |
| **A_55_P2133661** | NA | 0.020867892 | -0.099080897 | 0.140816681 | 14.36083015 | 0.41580704 | 0.690638733 | 0.995570006 | -5.184917162 |
| **A_52_P90289** | Oaz2 | -0.038255974 | -0.258161657 | 0.181649708 | 9.436805556 | -0.415787998 | 0.690652 | 0.995570006 | -5.184923631 |
| **A_55_P2159234** | Tatdn2 | 0.026193351 | -0.124411594 | 0.176798295 | 8.358796747 | 0.415681752 | 0.690726023 | 0.995570006 | -5.184959718 |
| **A_52_P243599** | Hsd17b12 | 0.04329305 | -0.205677024 | 0.292263124 | 9.860734085 | 0.415604424 | 0.6907799 | 0.995570006 | -5.184985977 |
| **A_65_P20683** | NA | -0.025466556 | -0.171920974 | 0.120987862 | 8.484220017 | -0.415601281 | 0.69078209 | 0.995570006 | -5.184987045 |
| **A_66_P139664** | Ppp2r5c | -0.020399012 | -0.137734181 | 0.096936156 | 8.48555987 | -0.41551822 | 0.690839964 | 0.995570006 | -5.185015246 |
| **A_51_P223524** | Ric1 | -0.025072181 | -0.16931777 | 0.119173409 | 9.072416873 | -0.415430806 | 0.690900874 | 0.995570006 | -5.185044919 |
| **A_55_P2071858** | Mgmt | 0.058010478 | -0.275948814 | 0.39196977 | 10.82543193 | 0.415166253 | 0.691085227 | 0.995570006 | -5.185134688 |
| **A_55_P1961099** | Atg4c | 0.039341081 | -0.187234474 | 0.265916636 | 8.850645815 | 0.414994526 | 0.691204907 | 0.995570006 | -5.18519293 |
| **A_66_P117659** | Nfatc3 | 0.02853743 | -0.135869176 | 0.192944036 | 9.052102885 | 0.41486302 | 0.691296563 | 0.995570006 | -5.185237516 |
| **A_51_P112677** | Mrpl50 | 0.020913898 | -0.099612116 | 0.141439912 | 11.78677188 | 0.414727971 | 0.691390694 | 0.995570006 | -5.179355886 |
| **A_55_P1998601** | Slc17a9 | 0.022012817 | -0.104884766 | 0.1489104 | 7.787032003 | 0.414602019 | 0.69147849 | 0.995570006 | -5.185325966 |
| **A_55_P2158653** | Cldn7 | -0.018303098 | -0.123816067 | 0.087209871 | 7.994468892 | -0.414598661 | 0.69148083 | 0.995570006 | -5.185327104 |
| **A_51_P370678** | Gfi1b | 0.017058213 | -0.081284177 | 0.115400603 | 7.759919613 | 0.414573845 | 0.691498129 | 0.995570006 | -5.185335511 |
| **A_66_P118165** | NA | 0.074328467 | -0.354185461 | 0.502842394 | 9.4612775 | 0.414571424 | 0.691499817 | 0.995570006 | -5.185336331 |
| **A_55_P2186395** | NA | -0.035381085 | -0.239407603 | 0.168645434 | 8.98608422 | -0.414470528 | 0.691570152 | 0.995570006 | -5.185370508 |
| **A_55_P1973854** | Sema6b | -0.02773915 | -0.187722961 | 0.132244661 | 8.347947054 | -0.414406203 | 0.691614995 | 0.995570006 | -5.185392292 |
| **A_51_P384148** | Ryk | -0.024754683 | -0.167641271 | 0.118131905 | 10.98518321 | -0.414071212 | 0.691848549 | 0.995570006 | -5.185505691 |
| **A_55_P1991500** | Stn1 | 0.027600807 | -0.131717014 | 0.186918627 | 11.23371814 | 0.414063122 | 0.69185419 | 0.995570006 | -5.185508428 |
| **A_55_P1983448** | S100a4 | -0.051278873 | -0.347298477 | 0.244740731 | 8.838851656 | -0.414025537 | 0.691880396 | 0.995570006 | -5.185521145 |
| **A_55_P1988183** | Cdkn2b | 0.021672141 | -0.103448005 | 0.146792288 | 7.925456176 | 0.413984153 | 0.691909251 | 0.995570006 | -5.185535147 |
| **A_51_P380750** | Cbfa2t3 | 0.037175089 | -0.17747089 | 0.251821069 | 9.835114568 | 0.413940951 | 0.691939376 | 0.995570006 | -5.185549762 |
| **A_51_P371882** | Ccdc71 | 0.090977693 | -0.434515439 | 0.616470824 | 11.33155947 | 0.413787156 | 0.692046619 | 0.995570006 | -5.18560178 |
| **A_55_P2109658** | NA | -0.027762167 | -0.18812219 | 0.132597855 | 11.21244405 | -0.413777041 | 0.692053673 | 0.995570006 | -5.1856052 |
| **A_55_P2105362** | Tmed5 | 0.019799645 | -0.094567918 | 0.134167207 | 8.629223622 | 0.413774441 | 0.692055486 | 0.995570006 | -5.185606079 |
| **A_55_P2108032** | Tmem255b | -0.030461097 | -0.206436968 | 0.145514774 | 8.500458867 | -0.413715265 | 0.692096752 | 0.995570006 | -5.185626089 |
| **A_55_P1989539** | Cklf | 0.033747146 | -0.161252938 | 0.22874723 | 8.401893921 | 0.413629382 | 0.692156645 | 0.995570006 | -5.185655124 |
| **A_55_P1972634** | Cmc4 | 0.027831826 | -0.133022055 | 0.188685707 | 9.811602247 | 0.413541682 | 0.692217808 | 0.995570006 | -5.185684768 |
| **A_55_P2095365** | Meaf6 | -0.026466684 | -0.179431071 | 0.126497703 | 8.549292859 | -0.413540771 | 0.692218443 | 0.995570006 | -5.185685076 |
| **A_51_P469789** | Agxt | 0.016452071 | -0.078635822 | 0.111539963 | 7.636928349 | 0.413527575 | 0.692227647 | 0.995570006 | -5.185689536 |
| **A_51_P276943** | Gpc1 | 0.027532865 | -0.131600143 | 0.186665872 | 14.33342316 | 0.41352356 | 0.692230446 | 0.995570006 | -5.185690892 |
| **A_52_P98232** | Hira | 0.036276764 | -0.173431892 | 0.245985419 | 8.68580608 | 0.413448396 | 0.692282869 | 0.995570006 | -5.185716293 |
| **A_55_P2021350** | NA | 0.020136985 | -0.096275524 | 0.136549494 | 15.70501664 | 0.413431852 | 0.692294407 | 0.995570006 | -5.185721883 |
| **A_55_P2116391** | Tesk2 | -0.023827676 | -0.161586871 | 0.113931518 | 8.189866925 | -0.413399768 | 0.692316784 | 0.995570006 | -5.185732724 |
| **A_55_P1986421** | Tcp11 | 0.016510356 | -0.078950595 | 0.111971307 | 7.793399671 | 0.413370806 | 0.692336985 | 0.995570006 | -5.185742509 |
| **A_55_P2037922** | Rpl7a | 0.02917301 | -0.139561862 | 0.197907882 | 14.06550862 | 0.41322399 | 0.692439389 | 0.995570006 | -5.185792102 |
| **A_55_P2088725** | Usp14 | -0.03470651 | -0.235676388 | 0.166263368 | 12.35278946 | -0.412751925 | 0.692768701 | 0.995570006 | -5.18595145 |
| **A_55_P2056615** | NA | -0.028825801 | -0.195756891 | 0.13810529 | 13.18029559 | -0.412717876 | 0.692792456 | 0.995570006 | -5.185962937 |
| **A_66_P120347** | Sstr5 | -0.039748199 | -0.269941475 | 0.190445077 | 11.01939398 | -0.412699511 | 0.692805269 | 0.995570006 | -5.185969132 |
| **A_55_P2009943** | Spen | 0.03687553 | -0.176700253 | 0.250451313 | 11.0565583 | 0.412662873 | 0.692830832 | 0.995570006 | -5.185981491 |
| **A_55_P1975682** | Rhox2e | -0.017669456 | -0.120030738 | 0.084691826 | 7.803349089 | -0.412569 | 0.692896328 | 0.995570006 | -5.186013151 |
| **A_51_P328883** | Zfp810 | -0.02632389 | -0.17886613 | 0.126218351 | 8.799286134 | -0.412447878 | 0.692980842 | 0.995570006 | -5.186053992 |

| | | | | | | | | | |
|---|---|---|---|---|---|---|---|---|---|
| **A_55_P2013601** | Ldb2 | 0.030814832 | -0.147789236 | 0.209418899 | 10.04977311 | 0.412361004 | 0.693041461 | 0.995570006 | -5.186083278 |
| **A_51_P492298** | Aqp2 | -0.045741354 | -0.31106814 | 0.219585432 | 8.364114145 | -0.412037789 | 0.693267018 | 0.995570006 | -5.186192186 |
| **A_66_P133269** | Hesx1 | 0.017637446 | -0.084700385 | 0.119975278 | 7.68539538 | 0.411915969 | 0.693352039 | 0.995570006 | -5.186233213 |
| **A_51_P367780** | Adamtsl2 | 0.031530324 | -0.151431095 | 0.214491742 | 8.366951704 | 0.411886963 | 0.693372284 | 0.995570006 | -5.18624298 |
| **A_52_P421344** | Paqr5 | 0.017225404 | -0.082753573 | 0.117204382 | 7.671160891 | 0.411784379 | 0.693443885 | 0.995570006 | -5.186277517 |
| **A_55_P2174632** | Rpl7 | -0.019082106 | -0.129838648 | 0.091674436 | 15.90973002 | -0.41178075 | 0.693446417 | 0.995570006 | -5.186278738 |
| **A_55_P2070131** | NA | 0.041709113 | -0.200397574 | 0.2838158 | 14.56092093 | 0.411749712 | 0.693468081 | 0.995570006 | -5.186289186 |
| **A_66_P139568** | Helz | 0.022813445 | -0.10969146 | 0.15531835 | 10.61916719 | 0.4114983 | 0.693643577 | 0.995570006 | -5.186373789 |
| **A_55_P2073218** | Hps4 | 0.020954783 | -0.100773727 | 0.142683294 | 10.75591927 | 0.411433832 | 0.693688582 | 0.995570006 | -5.186395476 |
| **A_55_P2052485** | Ushbp1 | -0.028998661 | -0.197467599 | 0.139470277 | 10.77504112 | -0.411402797 | 0.693710247 | 0.995570006 | -5.186405914 |
| **A_51_P287232** | Qars | -0.021654999 | -0.147476445 | 0.104166447 | 12.33260959 | -0.411351068 | 0.69374636 | 0.995570006 | -5.186423312 |
| **A_51_P192130** | Stk10 | -0.036914485 | -0.251525683 | 0.177696713 | 9.542263998 | -0.411105769 | 0.69391762 | 0.995570006 | -5.186505783 |
| **A_55_P1986371** | NA | 0.019579961 | -0.094264842 | 0.133424763 | 14.92178206 | 0.411062388 | 0.693947909 | 0.995570006 | -5.186520363 |
| **A_52_P137829** | NA | 0.036602817 | -0.176223644 | 0.249429278 | 9.364913926 | 0.411053185 | 0.693954334 | 0.995570006 | -5.186523456 |
| **A_55_P2067952** | NA | -0.043472739 | -0.296321073 | 0.209375594 | 11.25903847 | -0.410928291 | 0.694041541 | 0.995570006 | -5.186565424 |
| **A_55_P2146002** | Pank4 | 0.038928435 | -0.187566703 | 0.265423573 | 11.52464122 | 0.410787476 | 0.69413987 | 0.995570006 | -5.186612727 |
| **A_55_P2159555** | NA | -0.027484655 | -0.18740736 | 0.132438049 | 10.44700334 | -0.410761092 | 0.694158294 | 0.995570006 | -5.186621588 |
| **A_55_P2020035** | NA | 0.133759745 | -0.644724895 | 0.912244386 | 12.02872758 | 0.410661999 | 0.694227494 | 0.995570006 | -5.186654865 |
| **A_55_P2037338** | Plcd3 | -0.031885924 | -0.217462759 | 0.15369091 | 11.0830503 | -0.410661869 | 0.694227585 | 0.995570006 | -5.186654908 |
| **A_52_P456279** | Cct8 | 0.027649422 | -0.13330326 | 0.188602105 | 12.99281414 | 0.410579231 | 0.694285297 | 0.995570006 | -5.186682654 |
| **A_52_P183181** | Adar | 0.029156653 | -0.140600827 | 0.198914133 | 9.44103556 | 0.410504463 | 0.694337513 | 0.995570006 | -5.186707752 |
| **A_66_P127175** | Ankrd16 | 0.093903197 | -0.452834338 | 0.640640732 | 10.80972191 | 0.410497598 | 0.694342308 | 0.995570006 | -5.186710056 |
| **A_55_P2016262** | Isl2 | 0.019825716 | -0.095617728 | 0.13526916 | 7.695287858 | 0.410458014 | 0.694369953 | 0.995570006 | -5.186723342 |
| **A_55_P1967978** | Nfatc2 | -0.027692692 | -0.188959161 | 0.133573778 | 10.78215782 | -0.410421615 | 0.694395376 | 0.995570006 | -5.186735557 |
| **A_52_P537545** | Smpd3 | 0.081079051 | -0.391117757 | 0.553275859 | 8.681592132 | 0.410388084 | 0.694418794 | 0.995570006 | -5.18674681 |
| **A_55_P2173438** | Arhgap4 | 0.018204136 | -0.087832614 | 0.124240886 | 7.826586294 | 0.410320108 | 0.694466272 | 0.995570006 | -5.186769618 |
| **A_66_P100657** | Xpa | -0.019663514 | -0.134212452 | 0.094885424 | 10.62549453 | -0.410278912 | 0.694495046 | 0.995570006 | -5.186783439 |
| **A_55_P2149500** | Kifc2 | 0.046800715 | -0.225865139 | 0.319466569 | 8.515349352 | 0.41023326 | 0.694526933 | 0.995570006 | -5.186798754 |
| **A_51_P418820** | Tfap2c | -0.03092261 | -0.211151501 | 0.149306281 | 7.771890605 | -0.410072704 | 0.694639084 | 0.995570006 | -5.186852602 |
| **A_55_P2161342** | Pkib | 0.029075845 | -0.140390222 | 0.198541912 | 10.72297153 | 0.410070686 | 0.694640493 | 0.995570006 | -5.186853278 |
| **A_55_P2100628** | Gm12216 | -0.035130008 | -0.239883423 | 0.169623407 | 10.3747625 | -0.410068325 | 0.694642142 | 0.995570006 | -5.18685407 |
| **A_51_P239924** | Polr2c | -0.019181906 | -0.130986084 | 0.092622272 | 12.79831008 | -0.410055712 | 0.694650953 | 0.995570006 | -5.186858299 |
| **A_51_P445662** | Hsd17b4 | 0.033377525 | -0.161187656 | 0.227942707 | 12.48208967 | 0.410013488 | 0.694680449 | 0.995570006 | -5.186872457 |
| **A_55_P2033780** | Fabp3 | 0.02337838 | -0.112927398 | 0.159684159 | 16.51326754 | 0.409929602 | 0.694739049 | 0.995570006 | -5.186900579 |
| **A_55_P2069415** | Fxr1 | -0.036620276 | -0.250133701 | 0.176893148 | 11.46149171 | -0.409926091 | 0.694741501 | 0.995570006 | -5.186901756 |
| **A_55_P1955778** | Adamts4 | 0.022506102 | -0.108750662 | 0.153762867 | 8.710692401 | 0.409814887 | 0.694819189 | 0.995570006 | -5.186939028 |
| **A_55_P2054739** | Polr3c | -0.060368782 | -0.412450965 | 0.2917134 | 9.400746708 | -0.409805233 | 0.694825934 | 0.995570006 | -5.186942263 |
| **A_55_P1954212** | Snrpd2 | -0.028198884 | -0.192679284 | 0.136281515 | 14.07365641 | -0.409757495 | 0.694859285 | 0.995570006 | -5.18695826 |
| **A_52_P167411** | Tnk1 | 0.017703848 | -0.085582834 | 0.120990531 | 7.673450689 | 0.409668413 | 0.694921522 | 0.995570006 | -5.186988107 |
| **A_55_P2166506** | NA | -0.018664952 | -0.127565055 | 0.09023515 | 14.75861262 | -0.409645068 | 0.694937833 | 0.995570006 | -5.186995927 |
| **A_55_P1969987** | Fam204a | -0.028201778 | -0.192758392 | 0.136354837 | 8.572729447 | -0.409609736 | 0.694962519 | 0.995570006 | -5.187007763 |
| **A_51_P230098** | Pbk | -0.035621787 | -0.243480494 | 0.172236919 | 10.38842027 | -0.409596858 | 0.694971516 | 0.995570006 | -5.187012076 |
| **A_55_P2094197** | Dnase1l3 | -0.016809023 | -0.114912079 | 0.081294032 | 7.650673417 | -0.409514302 | 0.695029199 | 0.995570006 | -5.187039726 |
| **A_51_P390377** | Rcor3 | -0.04081127 | -0.279007239 | 0.1973847 | 10.5363194 | -0.409500872 | 0.695038583 | 0.995570006 | -5.187044224 |
| **A_55_P2137840** | Etfbkmt | 0.042107606 | -0.203868452 | 0.288083664 | 9.204871948 | 0.409144622 | 0.695287522 | 0.995570006 | -5.187163478 |
| **A_55_P2136817** | Coro2b | 0.04984479 | -0.241359646 | 0.341049227 | 10.5238656 | 0.409101356 | 0.695317759 | 0.995570006 | -5.187177954 |
| **A_55_P1953866** | Sema4c | -0.032024002 | -0.219128276 | 0.155080271 | 13.74506307 | -0.409073205 | 0.695337432 | 0.995570006 | -5.187187373 |
| **A_66_P112593** | Sec61b | -0.02396862 | -0.164020345 | 0.116083105 | 10.34594384 | -0.409038045 | 0.695362004 | 0.995570006 | -5.187199135 |

| | | | | | | | | | |
|---|---|---|---|---|---|---|---|---|---|
| **A_55_P2099390** | Ranbp9 | 0.030952096 | -0.149998449 | 0.211902642 | 8.570919802 | 0.408826751 | 0.695509679 | 0.995570006 | -5.187269802 |
| **A_55_P2134597** | NA | -0.042811387 | -0.293146596 | 0.207523822 | 8.988396992 | -0.408739393 | 0.695570737 | 0.995570006 | -5.187299009 |
| **A_55_P2056315** | Tbca | 0.051780154 | -0.25100129 | 0.354561598 | 13.54655661 | 0.408736302 | 0.695572898 | 0.995570006 | -5.187300042 |
| **A_55_P2097116** | Cnep1r1 | 0.023715816 | -0.114964767 | 0.162396399 | 8.187495293 | 0.408725326 | 0.69558057 | 0.995570006 | -5.187303711 |
| **A_52_P163795** | Tubb5 | 0.030255946 | -0.14669075 | 0.207202641 | 15.86873738 | 0.408674374 | 0.695616184 | 0.995570006 | -5.187320743 |
| **A_52_P589169** | Aopep | -0.019676696 | -0.134783411 | 0.095430019 | 9.326934275 | -0.408564513 | 0.695692978 | 0.995570006 | -5.187357459 |
| **A_55_P2127659** | Mtmr2 | 0.036656485 | -0.177829708 | 0.251142679 | 8.658795097 | 0.408470418 | 0.695758753 | 0.995570006 | -5.187388899 |
| **A_51_P317443** | Cd3eap | -0.038364779 | -0.262879128 | 0.18614957 | 9.579892611 | -0.408411286 | 0.69580009 | 0.995570006 | -5.187408653 |
| **A_55_P1986813** | Ibsp | 0.033688915 | -0.163468134 | 0.230845965 | 7.709349146 | 0.40839823 | 0.695809217 | 0.995570006 | -5.187413015 |
| **A_55_P1978576** | Dtx2 | 0.030976326 | -0.150324917 | 0.212277569 | 11.70257619 | 0.408355354 | 0.695839191 | 0.995570006 | -5.187427336 |
| **A_55_P2085225** | NA | 0.028297373 | -0.13735596 | 0.193950706 | 11.427826 | 0.408277148 | 0.695893865 | 0.995570006 | -5.187453455 |
| **A_55_P2423513** | B230369F2 | 0.037969001 | -0.184307217 | 0.260245219 | 10.92121758 | 0.408267965 | 0.695900285 | 0.995570006 | -5.187456522 |
| **A_52_P249424** | Vegfa | 0.031730614 | -0.154112325 | 0.217573553 | 10.12404589 | 0.408076458 | 0.696034177 | 0.995570006 | -5.187520459 |
| **A_51_P519791** | Ska3 | -0.023190199 | -0.15902124 | 0.112640842 | 8.392879451 | -0.408051127 | 0.696051888 | 0.995570006 | -5.187528914 |
| **A_52_P65237** | Zbtb7c | 0.018936624 | -0.092008667 | 0.129881916 | 7.751182601 | 0.407946134 | 0.696125299 | 0.995570006 | -5.187563954 |
| **A_66_P134481** | Skp2 | -0.025232162 | -0.173084709 | 0.122620385 | 8.567667907 | -0.407882229 | 0.696169984 | 0.995570006 | -5.187585277 |
| **A_51_P447988** | Atp6ap1 | -0.046603369 | -0.319734579 | 0.226527842 | 9.806606033 | -0.407807412 | 0.6962223 | 0.995570006 | -5.187610237 |
| **A_65_P12533** | Naa10 | 0.042231803 | -0.20531 | 0.289773606 | 11.42181428 | 0.407755857 | 0.696258351 | 0.995570006 | -5.187627434 |
| **A_55_P2069470** | 1110017D1 | -0.01852076 | -0.127088829 | 0.090047308 | 7.970275123 | -0.407723582 | 0.69628092 | 0.995570006 | -5.187638198 |
| **A_51_P507787** | Sergef | -0.028618421 | -0.196416276 | 0.139179433 | 8.860588842 | -0.407632124 | 0.696344877 | 0.995570006 | -5.187668698 |
| **A_52_P817257** | Smim22 | 0.01968863 | -0.095812245 | 0.135189505 | 8.594841017 | 0.407417191 | 0.696495192 | 0.995570006 | -5.18774035 |
| **A_51_P463497** | Rnh1 | -0.057816066 | -0.397073752 | 0.28144162 | 9.890147455 | -0.407312721 | 0.696568259 | 0.995570006 | -5.187775164 |
| **A_51_P270286** | Dync1li1 | -0.02420326 | -0.166276163 | 0.117869644 | 9.364291975 | -0.407166216 | 0.696670731 | 0.995570006 | -5.187823972 |
| **A_55_P2170963** | Gspt1 | -0.029980279 | -0.206012391 | 0.146051833 | 12.15557825 | -0.407054815 | 0.696748655 | 0.995570006 | -5.187861074 |
| **A_52_P93066** | Klhl42 | -0.018951708 | -0.130236892 | 0.092333476 | 8.537661992 | -0.407024116 | 0.69677013 | 0.995570006 | -5.187871297 |
| **A_55_P1988228** | Aspm | -0.039022528 | -0.268188986 | 0.19014393 | 10.59307599 | -0.406980387 | 0.696800719 | 0.995570006 | -5.187885857 |
| **A_55_P1957439** | NA | -0.018489538 | -0.1271226 | 0.090143525 | 8.275230324 | -0.406792715 | 0.696932007 | 0.995570006 | -5.187948329 |
| **A_51_P505617** | Il18r1 | 0.050399307 | -0.245746861 | 0.346545474 | 7.973625541 | 0.406750014 | 0.69696188 | 0.995570006 | -5.187962539 |
| **A_66_P104713** | Lrp5 | -0.019044795 | -0.130967786 | 0.092878197 | 9.561139874 | -0.406692454 | 0.697002149 | 0.995570006 | -5.187981693 |
| **A_55_P2181888** | Tmem54 | -0.025053551 | -0.172325026 | 0.122217923 | 7.961689551 | -0.4065929 | 0.697071801 | 0.995570006 | -5.188014813 |
| **A_52_P560628** | 1300002E1 | -0.017417109 | -0.119808314 | 0.084974097 | 8.850520981 | -0.406558025 | 0.697096202 | 0.995570006 | -5.188026414 |
| **A_55_P2209308** | 9530083O1 | -0.039295262 | -0.270310343 | 0.191719819 | 10.72273415 | -0.406545342 | 0.697105076 | 0.995570006 | -5.188030633 |
| **A_55_P1959923** | Cth | 0.019195374 | -0.093663382 | 0.13205413 | 7.797176979 | 0.406509268 | 0.697130315 | 0.995570006 | -5.188042631 |
| **A_51_P424878** | Prxl2b | -0.036971403 | -0.254378008 | 0.180435203 | 12.82834424 | -0.406445503 | 0.697174932 | 0.995570006 | -5.188063837 |
| **A_51_P395743** | Spopl | 0.018736031 | -0.091470417 | 0.128942479 | 8.648643683 | 0.406330779 | 0.697255206 | 0.995570006 | -5.188101982 |
| **A_55_P1992720** | NA | 0.016680292 | -0.081443652 | 0.114804236 | 7.630504146 | 0.406291537 | 0.697282665 | 0.995570006 | -5.188115028 |
| **A_55_P1981291** | Spink8 | 0.016597788 | -0.08104921 | 0.114244786 | 7.662450063 | 0.406256618 | 0.6973071 | 0.995570006 | -5.188126635 |
| **A_55_P2136622** | Camta2 | 0.02360314 | -0.115257937 | 0.162464217 | 8.536984781 | 0.40625469 | 0.697308449 | 0.995570006 | -5.188127276 |
| **A_55_P2030269** | Dnah6 | 0.019244672 | -0.093975088 | 0.132464432 | 7.94024592 | 0.406253788 | 0.697309081 | 0.995570006 | -5.188127576 |
| **A_52_P225570** | Parva | -0.034180261 | -0.235289316 | 0.166928793 | 10.39167943 | -0.406212126 | 0.697338234 | 0.995570006 | -5.188141423 |
| **A_55_P2145691** | Hlcs | 0.021879241 | -0.106854011 | 0.150612493 | 8.577272984 | 0.406210031 | 0.6973397 | 0.995570006 | -5.18814212 |
| **A_55_P2129786** | Slc38a10 | 0.038493224 | -0.188056109 | 0.265042557 | 8.542138871 | 0.406097796 | 0.697418241 | 0.995570006 | -5.188179417 |
| **A_51_P199168** | Cidea | -0.043842311 | -0.301923289 | 0.214238666 | 11.62463148 | -0.406019207 | 0.697473239 | 0.995570006 | -5.188205528 |
| **A_55_P2003266** | Fam107a | 0.050709388 | -0.247833503 | 0.349252279 | 10.69720705 | 0.405967032 | 0.697509754 | 0.995570006 | -5.18822286 |
| **A_55_P2005345** | Epb41 | -0.018752253 | -0.129154645 | 0.091650139 | 7.834186351 | -0.405960799 | 0.697514116 | 0.995570006 | -5.18822493 |
| **A_51_P272586** | Dym | -0.029400051 | -0.202525246 | 0.143725145 | 10.03490992 | -0.40587933 | 0.697571134 | 0.995570006 | -5.188251989 |
| **A_55_P2075303** | Zfp609 | -0.026020889 | -0.179307465 | 0.127265688 | 8.848460423 | -0.405720667 | 0.697682183 | 0.995570006 | -5.188304672 |
| **A_52_P228031** | Tmem179b | -0.017244419 | -0.118843422 | 0.084354584 | 9.817972443 | -0.405665671 | 0.697720677 | 0.995570006 | -5.188322929 |

| | | | | | | | | | |
|---|---|---|---|---|---|---|---|---|---|
| **A_55_P2088690** | NA | 0.053349578 | -0.261022701 | 0.367721856 | 9.205358816 | 0.405598065 | 0.697767999 | 0.995570006 | -5.188345368 |
| **A_55_P2048022** | Gm6578 | 0.021697189 | -0.106161812 | 0.149556191 | 11.70899265 | 0.405584453 | 0.697777526 | 0.995570006 | -5.188349886 |
| **A_55_P2007802** | Smarca2 | 0.04131658 | -0.202226709 | 0.28485987 | 9.871523256 | 0.4054687 | 0.697858553 | 0.995570006 | -5.188388296 |
| **A_55_P2171715** | Mrps33 | 0.031916384 | -0.15624266 | 0.220075427 | 13.47488473 | 0.405413043 | 0.697897514 | 0.995570006 | -5.188406762 |
| **A_55_P2000603** | Slc50a1 | 0.039017272 | -0.191021197 | 0.26905574 | 10.3248427 | 0.405383027 | 0.697918526 | 0.995570006 | -5.188416719 |
| **A_55_P2086265** | Gm8580 | 0.026275418 | -0.128653278 | 0.181204113 | 16.93758903 | 0.405346942 | 0.697943787 | 0.995570006 | -5.188428689 |
| **A_51_P159503** | Rnf213 | -0.043259745 | -0.298359172 | 0.211839682 | 10.36817454 | -0.405306535 | 0.697972074 | 0.995570006 | -5.188442091 |
| **A_55_P2066827** | Hexim2 | -0.02073648 | -0.143032075 | 0.101559116 | 8.46414914 | -0.405259633 | 0.698004909 | 0.995570006 | -5.188457646 |
| **A_55_P1958678** | Brd2 | 0.041788031 | -0.204684447 | 0.288260508 | 12.26924122 | 0.405221617 | 0.698031524 | 0.995570006 | -5.188470253 |
| **A_51_P430259** | Cfap126 | -0.028069923 | -0.193631992 | 0.137492146 | 9.675151833 | -0.405218729 | 0.698033545 | 0.995570006 | -5.18847121 |
| **A_52_P73475** | Fam78a | -0.040881767 | -0.282055656 | 0.200292121 | 11.1462999 | -0.405143159 | 0.698086452 | 0.995570006 | -5.188496267 |
| **A_55_P1982883** | Gys1 | -0.027607375 | -0.190499916 | 0.135285167 | 13.00638889 | -0.405072764 | 0.698135738 | 0.995570006 | -5.188519604 |
| **A_55_P2040245** | Piezo2 | 0.025501412 | -0.125031518 | 0.176034341 | 8.473670856 | 0.404894468 | 0.698260574 | 0.995570006 | -5.188578694 |
| **A_55_P2000798** | Ccdc144b | 0.034723466 | -0.170269939 | 0.239716871 | 8.129161606 | 0.404848294 | 0.698292906 | 0.995570006 | -5.188593993 |
| **A_55_P2123841** | Zfhx4 | -0.021095263 | -0.145680273 | 0.103489747 | 7.96997233 | -0.404695422 | 0.698399952 | 0.995570006 | -5.188644632 |
| **A_55_P2005753** | Manbal | 0.026799263 | -0.131631558 | 0.185230084 | 10.42433218 | 0.404289356 | 0.698684329 | 0.995570006 | -5.188779056 |
| **A_55_P1961461** | Bnip3l | 0.025111184 | -0.123349984 | 0.173572351 | 12.50647259 | 0.404262493 | 0.698703143 | 0.995570006 | -5.188787944 |
| **A_51_P123017** | Gmcl1 | -0.022629818 | -0.156423319 | 0.111163683 | 10.29874353 | -0.404254794 | 0.698708535 | 0.995570006 | -5.15875177 |
| **A_55_P2060107** | Pkm | 0.033475082 | -0.164504023 | 0.231454187 | 14.57171619 | 0.404121008 | 0.698802242 | 0.995570006 | -5.188834749 |
| **A_51_P455866** | Elf5 | 0.016962806 | -0.083379473 | 0.117305085 | 7.672855944 | 0.404038606 | 0.69885996 | 0.995570006 | -5.188862001 |
| **A_55_P2161702** | Nrip2 | -0.020218847 | -0.139825186 | 0.099387493 | 7.900014031 | -0.404027858 | 0.698867489 | 0.995570006 | -5.188865555 |
| **A_51_P288257** | Cdk5rap3 | 0.024057231 | -0.118277368 | 0.166391829 | 11.20093641 | 0.403965506 | 0.698911166 | 0.995570006 | -5.188886173 |
| **A_55_P1952877** | Snrpf | -0.026249433 | -0.181557612 | 0.129058746 | 12.76151037 | -0.403956625 | 0.698917387 | 0.995570006 | -5.188889109 |
| **A_55_P2034003** | Gtf3c1 | 0.025052026 | -0.123173509 | 0.173277561 | 10.42422444 | 0.403951255 | 0.698921148 | 0.995570006 | -5.188890884 |
| **A_51_P374137** | Bco1 | -0.01706293 | -0.118067052 | 0.083941193 | 7.646003325 | -0.403760314 | 0.699054907 | 0.995570006 | -5.188954 |
| **A_55_P2091350** | NA | -0.046593361 | -0.32252213 | 0.229335408 | 15.8086466 | -0.403586089 | 0.699176966 | 0.995570006 | -5.189011565 |
| **A_55_P2108599** | Tlcd1 | -0.02770645 | -0.191813939 | 0.136401038 | 8.301093429 | -0.4035168 | 0.699225511 | 0.995570006 | -5.189034453 |
| **A_51_P349073** | Endog | 0.044686038 | -0.219996231 | 0.309368307 | 10.84908595 | 0.403511705 | 0.699229081 | 0.995570006 | -5.189036136 |
| **A_52_P180842** | Exoc3 | 0.022087855 | -0.108742248 | 0.152917958 | 11.02490977 | 0.403510641 | 0.699229827 | 0.995570006 | -5.189036487 |
| **A_51_P499061** | Ube2o | 0.067234901 | -0.331106982 | 0.465576784 | 12.23775479 | 0.403411179 | 0.699299515 | 0.995570006 | -5.189069334 |
| **A_51_P428505** | Krtcap2 | 0.021264965 | -0.104756501 | 0.147286431 | 13.54993743 | 0.403300978 | 0.699376731 | 0.995570006 | -5.189105719 |
| **A_55_P1996518** | Arhgap42 | -0.016194705 | -0.112213418 | 0.079824007 | 8.009017011 | -0.403112531 | 0.699508781 | 0.995570006 | -5.189167916 |
| **A_55_P2006554** | Eif4e2 | -0.03387999 | -0.234768699 | 0.167008719 | 11.2202299 | -0.403085225 | 0.699527916 | 0.995570006 | -5.189176926 |
| **A_55_P1976978** | Zfp707 | -0.019506279 | -0.135170871 | 0.096158314 | 8.641535193 | -0.40307246 | 0.699536862 | 0.995570006 | -5.189181138 |
| **A_55_P2013396** | Gm14305 | 0.02151699 | -0.106093215 | 0.149127194 | 8.460398562 | 0.403000178 | 0.699587516 | 0.995570006 | -5.189204985 |
| **A_55_P2070352** | Csf3r | -0.070275641 | -0.487180646 | 0.346629364 | 14.43887876 | -0.402881047 | 0.699671005 | 0.995570006 | -5.18924428 |
| **A_51_P167527** | Lum | -0.044581942 | -0.309171336 | 0.220007452 | 12.54043502 | -0.402713031 | 0.69978876 | 0.995570006 | -5.189299681 |
| **A_51_P443693** | Rasl10b | -0.024076433 | -0.166982836 | 0.118829971 | 8.462333623 | -0.402670288 | 0.699818719 | 0.995570006 | -5.189313772 |
| **A_55_P2133365** | 1110017D1 | 0.017613694 | -0.086950741 | 0.122178129 | 7.706438309 | 0.402601681 | 0.699866806 | 0.995570006 | -5.189336385 |
| **A_51_P368591** | Tle6 | 0.020445327 | -0.100931738 | 0.141822392 | 11.15455454 | 0.402593313 | 0.699872672 | 0.995570006 | -5.189339143 |
| **A_52_P336958** | Jcad | -0.033886455 | -0.235154117 | 0.167381207 | 11.83874591 | -0.402403054 | 0.700006034 | 0.995570006 | -5.189401834 |
| **A_55_P2296088** | Sp3 | -0.017702617 | -0.122852131 | 0.087446897 | 8.363267572 | -0.402382734 | 0.700020279 | 0.995570006 | -5.189408529 |
| **A_52_P559545** | Cercam | -0.020045517 | -0.139126156 | 0.099035121 | 8.388078771 | -0.402332623 | 0.700055406 | 0.995570006 | -5.189425035 |
| **A_55_P2163404** | NA | -0.020286578 | -0.14083641 | 0.100263254 | 14.39567632 | -0.402208568 | 0.700142372 | 0.995570006 | -5.18946589 |
| **A_55_P1970785** | NA | -0.07474012 | -0.518902051 | 0.369421811 | 10.98889223 | -0.402181039 | 0.700161672 | 0.995570006 | -5.189474955 |
| **A_51_P204845** | Celf1 | 0.055798946 | -0.276037013 | 0.387634905 | 10.62221612 | 0.401894146 | 0.700362813 | 0.995570006 | -5.189569386 |
| **A_55_P2112005** | Tff1 | 0.017207176 | -0.085161752 | 0.119576104 | 7.660464553 | 0.401745082 | 0.700467332 | 0.995570006 | -5.189618426 |
| **A_55_P2197998** | D4Ertd58e | -0.020785087 | -0.144449838 | 0.102879663 | 7.95538068 | -0.40171224 | 0.700490361 | 0.995570006 | -5.189629228 |

| | | | | | | | | | |
|---|---|---|---|---|---|---|---|---|---|
| **A_51_P418259** | Pccb | -0.03486794 | -0.242446428 | 0.172710548 | 10.24064219 | -0.401469984 | 0.700660241 | 0.995570006 | -5.189708884 |
| **A_52_P299363** | NA | 0.019006798 | -0.094163966 | 0.132177563 | 8.821334401 | 0.401405992 | 0.700705118 | 0.995570006 | -5.189729917 |
| **A_55_P2161695** | Poglut2 | 0.033555565 | -0.166246225 | 0.233357355 | 9.249328537 | 0.401397174 | 0.700711302 | 0.995570006 | -5.189732816 |
| **A_55_P1968488** | Cblif | -0.017929873 | -0.1247059 | 0.088846155 | 7.669486277 | -0.401340125 | 0.700751311 | 0.995570006 | -5.189751564 |
| **A_55_P1957413** | Lsp1 | -0.035678027 | -0.248157724 | 0.17680167 | 12.94318684 | -0.401321587 | 0.700764313 | 0.995570006 | -5.189757656 |
| **A_52_P257502** | Igfbp4 | -0.045830826 | -0.318876103 | 0.22721445 | 9.145672247 | -0.401173422 | 0.700868228 | 0.995570006 | -5.189806334 |
| **A_55_P1965298** | Gm5640 | 0.021644797 | -0.107342572 | 0.150632167 | 7.887837627 | 0.401065653 | 0.700943817 | 0.995570006 | -5.189841731 |
| **A_52_P658974** | NA | -0.058169461 | -0.404937485 | 0.288598563 | 13.34547874 | -0.400926839 | 0.701041185 | 0.995570006 | -5.18988731 |
| **A_55_P2115245** | Zfyve19 | 0.023047746 | -0.114413289 | 0.160508781 | 11.39615035 | 0.40073566 | 0.701175294 | 0.995570006 | -5.18995006 |
| **A_51_P203675** | Trem3 | -0.016121619 | -0.112285648 | 0.080042411 | 7.675316476 | -0.400686883 | 0.701209512 | 0.995570006 | -5.189966065 |
| **A_66_P110817** | Mterf3 | 0.026945381 | -0.133787357 | 0.187678118 | 9.271820861 | 0.40067211 | 0.701219876 | 0.995570006 | -5.189970912 |
| **A_52_P454183** | Olfml2b | 0.021530385 | -0.106953731 | 0.150014501 | 9.114144739 | 0.400508267 | 0.701334821 | 0.995570006 | -5.190024659 |
| **A_51_P360880** | Zfp266 | 0.022622879 | -0.112426944 | 0.157672703 | 9.407728625 | 0.400371345 | 0.701430887 | 0.995570006 | -5.190069559 |
| **A_51_P372312** | Mrpl54 | -0.036699412 | -0.25579134 | 0.182392517 | 13.08304342 | -0.400351867 | 0.701444553 | 0.995570006 | -5.190075945 |
| **A_55_P2028214** | NA | -0.022622071 | -0.157681656 | 0.112437515 | 7.818793152 | -0.400328099 | 0.701461229 | 0.995570006 | -5.190083737 |
| **A_55_P2130388** | Mical1 | 0.02661281 | -0.1322871 | 0.18551272 | 12.02564293 | 0.400291363 | 0.701487005 | 0.995570006 | -5.19009578 |
| **A_51_P229633** | Rprd1b | 0.021266383 | -0.105724288 | 0.148257054 | 11.29761296 | 0.400249642 | 0.701516279 | 0.995570006 | -5.190109456 |
| **A_51_P363947** | Cdkn1a | 0.030076977 | -0.149540257 | 0.209694212 | 10.07401271 | 0.400216799 | 0.701539324 | 0.995570006 | -5.159150302 |
| **A_51_P422369** | Odf3b | -0.018848395 | -0.131413845 | 0.093717055 | 7.833081626 | -0.400201203 | 0.701550268 | 0.995570006 | -5.190125332 |
| **A_55_P2039284** | Hspb1 | 0.017076697 | -0.084935423 | 0.119088817 | 16.69242145 | 0.400093251 | 0.701626017 | 0.995570006 | -5.190160707 |
| **A_55_P2019014** | Tmem120b | 0.028332564 | -0.140924619 | 0.197589748 | 9.704500668 | 0.400080983 | 0.701634626 | 0.995570006 | -5.190164727 |
| **A_66_P110607** | Sart1 | 0.025326013 | -0.125970779 | 0.176622806 | 9.838595377 | 0.400079445 | 0.701635705 | 0.995570006 | -5.190165231 |
| **A_52_P566390** | Zfyve26 | -0.03063982 | -0.21371468 | 0.152435041 | 9.011555675 | -0.400006111 | 0.701687166 | 0.995570006 | -5.190189257 |
| **A_51_P258078** | Nop2 | 0.050191335 | -0.24976161 | 0.350144281 | 11.86105533 | 0.399930707 | 0.701740081 | 0.995570006 | -5.190213956 |
| **A_55_P2091145** | H4c12 | 0.034367586 | -0.171036297 | 0.239771468 | 15.37135382 | 0.399898254 | 0.701762855 | 0.995570006 | -5.190224585 |
| **A_55_P2076533** | Pmvk | 0.020787335 | -0.103489888 | 0.145064558 | 12.68267634 | 0.399775718 | 0.701848851 | 0.995570006 | -5.19026471 |
| **A_55_P2037081** | Zfp979 | -0.036593465 | -0.255400158 | 0.182213227 | 9.135097729 | -0.399716492 | 0.701890417 | 0.995570006 | -5.1902841 |
| **A_55_P2144666** | Prpsap1 | -0.024199385 | -0.168901088 | 0.120502319 | 12.60085061 | -0.399705211 | 0.701898334 | 0.995570006 | -5.190287793 |
| **A_66_P115406** | Pabpc1 | 0.042160761 | -0.209984932 | 0.294306454 | 14.65582561 | 0.399637305 | 0.701945995 | 0.995570006 | -5.190310021 |
| **A_55_P2184526** | NA | -0.021944085 | -0.153188424 | 0.109300255 | 8.018387929 | -0.3996189 | 0.701958912 | 0.995570006 | -5.190316045 |
| **A_55_P2004096** | Asb8 | -0.038579722 | -0.269328322 | 0.192168879 | 8.630485944 | -0.399603375 | 0.701969809 | 0.995570006 | -5.190321126 |
| **A_51_P441898** | Krt25 | 0.017155642 | -0.085465622 | 0.119776906 | 7.718755907 | 0.399556994 | 0.702002362 | 0.995570006 | -5.190336304 |
| **A_52_P278154** | Eif1ad | 0.019409002 | -0.096695357 | 0.135513361 | 10.68462572 | 0.399543259 | 0.702012002 | 0.995570006 | -5.190340799 |
| **A_55_P2092983** | NA | -0.018988538 | -0.132581603 | 0.094604527 | 14.50393957 | -0.399529496 | 0.702021663 | 0.995570006 | -5.190345302 |
| **A_52_P635271** | Btbd6 | 0.034840304 | -0.173731573 | 0.24341218 | 9.359963396 | 0.399241165 | 0.702224054 | 0.995570006 | -5.190439619 |
| **A_51_P385697** | Mgat3 | -0.027812444 | -0.194338486 | 0.138713598 | 8.155470928 | -0.399177571 | 0.702268696 | 0.995570006 | -5.190460412 |
| **A_51_P282616** | Maats1 | 0.018446464 | -0.092016764 | 0.128909691 | 7.736493825 | 0.399120951 | 0.702308444 | 0.995570006 | -5.190478923 |
| **A_51_P286665** | Rbl1 | 0.040130092 | -0.200189745 | 0.280449929 | 9.624179903 | 0.399107295 | 0.702318031 | 0.995570006 | -5.190483387 |
| **A_51_P479818** | Lonrf3 | -0.018083004 | -0.126383314 | 0.090217307 | 7.863109454 | -0.399070843 | 0.702343621 | 0.995570006 | -5.190495303 |
| **A_55_P2027737** | Il1rl1 | -0.038672421 | -0.270370005 | 0.193025164 | 8.319155276 | -0.398922914 | 0.702447477 | 0.995570006 | -5.190543647 |
| **A_52_P77227** | Rnf212 | -0.017211768 | -0.120352208 | 0.085928672 | 7.82477406 | -0.398846366 | 0.702501221 | 0.995570006 | -5.190568658 |
| **A_52_P303176** | 1810037I17 | -0.024502938 | -0.171363383 | 0.122357507 | 9.823205821 | -0.398769971 | 0.702554859 | 0.995570006 | -5.190593613 |
| **A_55_P2164909** | Gatd3a | 0.027257024 | -0.136116927 | 0.190630975 | 14.16639019 | 0.398753729 | 0.702566263 | 0.995570006 | -5.190598918 |
| **A_51_P519756** | Rusc1 | 0.031377822 | -0.156739285 | 0.219494929 | 9.84126596 | 0.398660892 | 0.702631449 | 0.995570006 | -5.190629238 |
| **A_55_P2096266** | Sftpb | 0.017924795 | -0.089543484 | 0.125393074 | 7.701909018 | 0.398641989 | 0.702644722 | 0.995570006 | -5.19063541 |
| **A_55_P2315921** | 5330431K0 | 0.071068289 | -0.355209214 | 0.497345792 | 8.998614706 | 0.398467202 | 0.702767458 | 0.995570006 | -5.190692473 |
| **A_51_P487308** | Pstk | -0.026656048 | -0.186570336 | 0.13325824 | 11.78082058 | -0.398398437 | 0.702815747 | 0.995570006 | -5.190714916 |
| **A_55_P2022181** | Arvcf | -0.023376977 | -0.163658941 | 0.116904987 | 9.040838515 | -0.398286557 | 0.702894316 | 0.995570006 | -5.190751423 |

| | | | | | | | | | |
|---|---|---|---|---|---|---|---|---|---|
| **A_55_P2079599** | NA | 0.033836563 | -0.169234968 | 0.236908093 | 12.43424767 | 0.398241326 | 0.702926082 | 0.995570006 | -5.19076618 |
| **A_55_P2084686** | Pdik1l | -0.038459694 | -0.26931809 | 0.192398703 | 9.151244664 | -0.39817068 | 0.702975697 | 0.995570006 | -5.190789225 |
| **A_55_P2030938** | Trim59 | -0.026692393 | -0.186918977 | 0.133534192 | 11.1200333 | -0.398164064 | 0.702980344 | 0.995570006 | -5.190791383 |
| **A_51_P399175** | Ppm1j | -0.041986619 | -0.294048312 | 0.210075075 | 9.862496257 | -0.398119254 | 0.703011815 | 0.995570006 | -5.190805998 |
| **A_55_P1970234** | Dclk2 | 0.025397034 | -0.127105943 | 0.17790001 | 8.463272032 | 0.398028168 | 0.70307579 | 0.995570006 | -5.190835701 |
| **A_52_P193233** | Trnau1ap | -0.046918547 | -0.328733939 | 0.234896845 | 10.65953282 | -0.397913777 | 0.703156136 | 0.995570006 | -5.190872995 |
| **A_55_P2091671** | Psmg1 | -0.026830385 | -0.187999536 | 0.134338766 | 11.29314949 | -0.397881844 | 0.703178566 | 0.995570006 | -5.190883403 |
| **A_52_P74524** | NA | -0.023801845 | -0.166784351 | 0.119180662 | 8.417790641 | -0.397866018 | 0.703189682 | 0.995570006 | -5.190888562 |
| **A_55_P2009910** | Stxbp3 | 0.033324371 | -0.166861889 | 0.233510631 | 10.05492038 | 0.397865997 | 0.703189697 | 0.995570006 | -5.190888569 |
| **A_51_P470851** | Cplx3 | -0.027147684 | -0.190270535 | 0.135975167 | 7.923032963 | -0.397765509 | 0.703260283 | 0.995570006 | -5.190921318 |
| **A_51_P291078** | Sel1l3 | 0.020758381 | -0.103976401 | 0.145493163 | 8.112655716 | 0.397754444 | 0.703268056 | 0.995570006 | -5.190924924 |
| **A_55_P2110915** | Flrt1 | 0.052032744 | -0.26066528 | 0.364730768 | 8.659551384 | 0.3977047 | 0.703302999 | 0.995570006 | -5.190941132 |
| **A_51_P396752** | Arl2bp | -0.020388178 | -0.142923586 | 0.10214723 | 13.10797735 | -0.397672863 | 0.703325363 | 0.995570006 | -5.190951505 |
| **A_66_P108815** | Pgls | 0.04371457 | -0.219035194 | 0.306464334 | 11.04587513 | 0.397642696 | 0.703346555 | 0.995570006 | -5.190961332 |
| **A_51_P358112** | Fads1 | -0.018388092 | -0.128936594 | 0.09216041 | 8.902530028 | -0.397551086 | 0.703410911 | 0.995570006 | -5.190991173 |
| **A_55_P2053848** | NA | 0.022854624 | -0.114587615 | 0.160296864 | 12.17800332 | 0.397432152 | 0.703494467 | 0.995570006 | -5.191029904 |
| **A_51_P157982** | Tcaf1 | 0.02342576 | -0.117452877 | 0.164304397 | 9.108542307 | 0.397427287 | 0.703497884 | 0.995570006 | -5.191031488 |
| **A_51_P123795** | Zfp637 | 0.056091485 | -0.281414288 | 0.393597258 | 12.34769538 | 0.397214291 | 0.703647533 | 0.995570006 | -5.179322324 |
| **A_66_P102628** | 1700016J1 | 0.017072369 | -0.08566714 | 0.119811878 | 7.895495058 | 0.397159942 | 0.70368572 | 0.995570006 | -5.191118508 |
| **A_66_P135842** | Fip1l1 | -0.02237173 | -0.157030207 | 0.112286747 | 10.97450462 | -0.397077249 | 0.703743824 | 0.995570006 | -5.191145413 |
| **A_55_P1985768** | Ttyh2 | -0.036069026 | -0.253177673 | 0.181039621 | 9.893316996 | -0.397069406 | 0.703749335 | 0.995570006 | -5.191147965 |
| **A_55_P2052744** | NA | 0.022515483 | -0.113021556 | 0.158052522 | 12.15484268 | 0.397038301 | 0.703771192 | 0.995570006 | -5.191158084 |
| **A_55_P2111713** | Spint2 | 0.066976476 | -0.336248964 | 0.470201916 | 9.640458237 | 0.396993594 | 0.703802607 | 0.995570006 | -5.191172626 |
| **A_55_P2083919** | Robo2 | 0.03451904 | -0.173319074 | 0.242357154 | 8.271690449 | 0.396956252 | 0.703828847 | 0.995570006 | -5.191184772 |
| **A_55_P1953382** | Prl8a8 | -0.305589968 | -2.145622169 | 1.534442233 | 8.897443594 | -0.396938114 | 0.703841592 | 0.995570006 | -5.191190671 |
| **A_55_P2013029** | NA | 0.020052239 | -0.100691849 | 0.140796326 | 8.491264604 | 0.396922873 | 0.703852302 | 0.995570006 | -5.191195627 |
| **A_55_P2068391** | Gm9222 | -0.025537804 | -0.179333791 | 0.128258184 | 9.114884783 | -0.396869449 | 0.703889845 | 0.995570006 | -5.191213001 |
| **A_52_P305073** | Pias1 | -0.016573759 | -0.116397019 | 0.0832495 | 9.685383644 | -0.396824444 | 0.703921472 | 0.995570006 | -5.191227634 |
| **A_55_P1964960** | Il33 | 0.054768937 | -0.27529709 | 0.384834964 | 8.562457902 | 0.396590783 | 0.704085684 | 0.995570006 | -5.191303585 |
| **A_66_P135534** | Zfp950 | 0.043116887 | -0.216819932 | 0.303053707 | 8.548538342 | 0.396450284 | 0.704184432 | 0.995570006 | -5.191349233 |
| **A_55_P2173224** | Kif2a | -0.020352311 | -0.143066164 | 0.102361543 | 8.619905636 | -0.396395998 | 0.704222588 | 0.995570006 | -5.191366867 |
| **A_55_P2000224** | Cd63 | 0.019001429 | -0.095575336 | 0.133578195 | 15.0493419 | 0.396368238 | 0.7042421 | 0.995570006 | -5.191375883 |
| **A_55_P1991214** | Lair1 | -0.030796682 | -0.216604113 | 0.155010749 | 8.215215589 | -0.396141171 | 0.70440171 | 0.995570006 | -5.191449611 |
| **A_55_P2007771** | Catsper2 | -0.017262866 | -0.121477157 | 0.086951426 | 7.95494716 | -0.395908423 | 0.70456533 | 0.995570006 | -5.191525142 |
| **A_55_P2088131** | Zfp975 | 0.052976777 | -0.266933491 | 0.372887045 | 11.74349541 | 0.39579152 | 0.704647519 | 0.995570006 | -5.191563064 |
| **A_55_P1996998** | Polr2a | -0.018009149 | -0.126793521 | 0.090775223 | 8.954642195 | -0.395672444 | 0.704731239 | 0.995570006 | -5.19160168 |
| **A_51_P390276** | Cwc25 | 0.021041979 | -0.106069453 | 0.148153412 | 8.867246754 | 0.395649942 | 0.704747061 | 0.995570006 | -5.191608976 |
| **A_55_P2033308** | Six2 | 0.019302396 | -0.097306115 | 0.135910908 | 7.819327291 | 0.395630814 | 0.70476051 | 0.995570006 | -5.191615177 |
| **A_55_P2155546** | Nlrc3 | -0.020423411 | -0.143825227 | 0.102978405 | 8.704939013 | -0.395563187 | 0.70480806 | 0.995570006 | -5.191637101 |
| **A_51_P198675** | Ttc36 | 0.016704534 | -0.084230007 | 0.117639075 | 7.663779338 | 0.395552099 | 0.704815857 | 0.995570006 | -5.191640695 |
| **A_51_P205545** | Creld1 | 0.031637005 | -0.15953397 | 0.22280798 | 9.945164204 | 0.395532835 | 0.704829402 | 0.995570006 | -5.19164694 |
| **A_51_P388973** | Zhx1 | -0.027505779 | -0.193716883 | 0.138705324 | 8.798943713 | -0.395524191 | 0.70483548 | 0.995570006 | -5.191649742 |
| **A_52_P422755** | Lsm6 | -0.017197463 | -0.12113083 | 0.086735904 | 8.516364091 | -0.395474533 | 0.704870397 | 0.995570006 | -5.191665837 |
| **A_55_P2090205** | Tpm3 | -0.02984209 | -0.210228892 | 0.150544713 | 8.48514981 | -0.395397215 | 0.704924766 | 0.995570006 | -5.191690893 |
| **A_55_P2038358** | Acot1 | 0.046780486 | -0.236095713 | 0.329656685 | 11.35331228 | 0.395255074 | 0.705024721 | 0.995570006 | -5.191736944 |
| **A_52_P32369** | Hmgn3 | 0.03781121 | -0.190830051 | 0.266452472 | 10.92478802 | 0.395252933 | 0.705026227 | 0.995570006 | -5.191737637 |
| **A_55_P2103603** | NA | 0.01606332 | -0.08108686 | 0.113213499 | 7.781075488 | 0.395185343 | 0.705073759 | 0.995570006 | -5.191759529 |
| **A_55_P2105220** | Hscb | 0.025691763 | -0.129742508 | 0.181126034 | 11.50036513 | 0.395053814 | 0.70516626 | 0.995570006 | -5.191802121 |

| | | | | | | | | | |
|---|---|---|---|---|---|---|---|---|---|
| **A_52_P610987** | Slc28a3 | -0.016620318 | -0.117183779 | 0.083943143 | 7.796430558 | -0.395010147 | 0.705196971 | 0.995570006 | -5.191816259 |
| **A_55_P1957624** | Insig2 | 0.019873543 | -0.100383399 | 0.140130484 | 9.490536538 | 0.394979243 | 0.705218707 | 0.995570006 | -5.191826263 |
| **A_55_P1974228** | NA | 0.024466817 | -0.123588694 | 0.172522327 | 14.89143479 | 0.394968103 | 0.705226542 | 0.995570006 | -5.191829869 |
| **A_51_P146407** | Fam104a | -0.024438133 | -0.172333892 | 0.123457627 | 9.431596436 | -0.394931184 | 0.705252508 | 0.995570006 | -5.19184182 |
| **A_55_P2087404** | Mapre3 | -0.03647729 | -0.257285902 | 0.184331322 | 10.30606396 | -0.39483504 | 0.705320131 | 0.995570006 | -5.191872936 |
| **A_55_P1998188** | Cdk4 | 0.02834237 | -0.143223519 | 0.199908258 | 12.94062042 | 0.39483382 | 0.705320989 | 0.995570006 | -5.19187333 |
| **A_55_P2071691** | Kank3 | 0.044223242 | -0.223560934 | 0.312007419 | 11.14352802 | 0.394706975 | 0.70541021 | 0.995570006 | -5.191914371 |
| **A_55_P1994052** | Asxl1 | 0.023242332 | -0.117504764 | 0.163989429 | 12.34216424 | 0.394683883 | 0.705426453 | 0.995570006 | -5.191921841 |
| **A_55_P2248651** | Aak1 | -0.025333487 | -0.178754039 | 0.128087065 | 11.40687001 | -0.394657688 | 0.705444879 | 0.995570006 | -5.191930315 |
| **A_52_P487599** | Thnsl2 | -0.018498093 | -0.130553636 | 0.09355745 | 8.657826515 | -0.394550637 | 0.705520183 | 0.995570006 | -5.191964938 |
| **A_55_P2280868** | 0610005C1 | 0.015741518 | -0.079668415 | 0.111151451 | 7.679035691 | 0.394332128 | 0.705673901 | 0.995570006 | -5.192035581 |
| **A_55_P2044788** | NA | -0.032944232 | -0.23266786 | 0.166779396 | 10.36153661 | -0.394238533 | 0.705739749 | 0.995570006 | -5.192065829 |
| **A_51_P378381** | Crnde | 0.030798181 | -0.155918719 | 0.217515081 | 9.048092626 | 0.39423082 | 0.705745176 | 0.995570006 | -5.192068321 |
| **A_51_P407774** | Slc22a21 | 0.024872444 | -0.125939266 | 0.175684154 | 8.107001263 | 0.394178138 | 0.705782241 | 0.995570006 | -5.192085344 |
| **A_51_P474169** | Proser2 | -0.033110493 | -0.233894893 | 0.167673908 | 10.68141625 | -0.394134822 | 0.705812717 | 0.995570006 | -5.192099338 |
| **A_55_P2128597** | Nr1h3 | -0.024486581 | -0.172979863 | 0.124006701 | 8.495900776 | -0.394121813 | 0.705821869 | 0.995570006 | -5.19210354 |
| **A_52_P592101** | Washc2 | -0.032733819 | -0.231335371 | 0.165867733 | 9.433589281 | -0.393933731 | 0.705954207 | 0.995570006 | -5.192164287 |
| **A_52_P73559** | NA | 0.021944434 | -0.111338881 | 0.155227749 | 8.520879837 | 0.393511777 | 0.706251142 | 0.995570006 | -5.19230047 |
| **A_55_P2091105** | NA | 0.029213963 | -0.148237846 | 0.206665772 | 16.14070023 | 0.393476844 | 0.706275727 | 0.995570006 | -5.192311738 |
| **A_55_P1972997** | A530016L2 | 0.020607896 | -0.104590982 | 0.145806774 | 8.484595769 | 0.393407242 | 0.706324713 | 0.995570006 | -5.192334186 |
| **A_55_P1997898** | Tfeb | -0.044902217 | -0.317739059 | 0.227934625 | 9.082441337 | -0.393345241 | 0.70636835 | 0.995570006 | -5.19235418 |
| **A_55_P2073377** | Mki67 | 0.026983179 | -0.137033129 | 0.190999487 | 12.76772394 | 0.39320154 | 0.706469495 | 0.995570006 | -5.192400509 |
| **A_55_P2029344** | Rpl23 | 0.019343526 | -0.098284233 | 0.136971286 | 16.68388409 | 0.393038378 | 0.706584344 | 0.995570006 | -5.192453092 |
| **A_55_P1992889** | Serping1 | -0.051672727 | -0.36590549 | 0.262560036 | 10.87195631 | -0.393023981 | 0.706594479 | 0.995570006 | -5.192457731 |
| **A_55_P2121408** | Tpm2 | 0.052386401 | -0.266205606 | 0.370978408 | 11.10855365 | 0.393000242 | 0.706611189 | 0.995570006 | -5.19246538 |
| **A_55_P2060330** | Cyren | -0.02216911 | -0.157006443 | 0.112668222 | 10.36055089 | -0.392959005 | 0.706640218 | 0.995570006 | -5.192478665 |
| **A_52_P366572** | Slain2 | -0.025786924 | -0.182628753 | 0.131054904 | 9.155878929 | -0.392958585 | 0.706640514 | 0.995570006 | -5.1924788 |
| **A_51_P187082** | G6pdx | 0.028572217 | -0.14529812 | 0.202442554 | 9.966064856 | 0.392760293 | 0.706780107 | 0.995570006 | -5.192542665 |
| **A_52_P30877** | Gm5176 | -0.036853573 | -0.268190697 | 0.19448355 | 13.06849715 | -0.395732229 | 0.706809339 | 0.995570006 | -5.115088471 |
| **A_55_P2062513** | Pdzrn3 | -0.054155115 | -0.383802464 | 0.275492234 | 12.01066057 | -0.392644056 | 0.706861942 | 0.995570006 | -5.192580088 |
| **A_55_P2087265** | NA | -0.020417464 | -0.144741262 | 0.103906335 | 8.308637584 | -0.392515355 | 0.706952556 | 0.995570006 | -5.192621512 |
| **A_51_P503303** | C1galt1c1 | 0.028227004 | -0.143674498 | 0.200128506 | 10.32941375 | 0.392458958 | 0.706992265 | 0.995570006 | -5.19263966 |
| **A_52_P538470** | Spats2l | -0.022041845 | -0.156289644 | 0.112205953 | 8.641310721 | -0.392418898 | 0.707020472 | 0.995570006 | -5.192652549 |
| **A_55_P2151907** | NA | -0.024850918 | -0.176213065 | 0.126511229 | 14.98866385 | -0.392404778 | 0.707030414 | 0.995570006 | -5.192657092 |
| **A_55_P2001589** | Klhl8 | -0.016678218 | -0.118269519 | 0.084913083 | 10.09229937 | -0.392375843 | 0.707050788 | 0.995570006 | -5.192666401 |
| **A_55_P2144055** | NA | 0.01812717 | -0.092290656 | 0.128544997 | 15.33504055 | 0.392373758 | 0.707052256 | 0.995570006 | -5.192667072 |
| **A_55_P1960646** | Ube2w | 0.019181757 | -0.097682744 | 0.136046259 | 8.668765504 | 0.392296937 | 0.707106349 | 0.995570006 | -5.192691783 |
| **A_55_P2015877** | Clps | -0.016413265 | -0.116417841 | 0.08359131 | 7.68925271 | -0.392269239 | 0.707125853 | 0.995570006 | -5.192700691 |
| **A_55_P1973362** | Kcnj12 | -0.033073448 | -0.234594224 | 0.168447328 | 9.799937712 | -0.392255265 | 0.707135693 | 0.995570006 | -5.192705185 |
| **A_55_P2152427** | Zwilch | -0.049576954 | -0.351676668 | 0.252522759 | 10.0207742 | -0.392228062 | 0.707154848 | 0.995570006 | -5.192713934 |
| **A_55_P2034300** | Tmem40 | 0.032371555 | -0.164899365 | 0.229642475 | 8.305096748 | 0.39220184 | 0.707173313 | 0.995570006 | -5.192722366 |
| **A_52_P207303** | Bmp3 | 0.023663636 | -0.120609243 | 0.167936516 | 7.717218148 | 0.392017913 | 0.707302837 | 0.995570006 | -5.192781498 |
| **A_51_P497724** | Apol7a | 0.021167045 | -0.107885153 | 0.150219244 | 7.741135464 | 0.392016154 | 0.707304076 | 0.995570006 | -5.192782063 |
| **A_52_P668984** | Tmem243 | 0.031338867 | -0.159737588 | 0.222415323 | 8.121925119 | 0.391999265 | 0.70731597 | 0.995570006 | -5.192787492 |
| **A_55_P2189571** | NA | -0.02436757 | -0.172980411 | 0.124245271 | 10.34900675 | -0.391890759 | 0.707392387 | 0.995570006 | -5.192822363 |
| **A_52_P305279** | Spata13 | 0.029232002 | -0.149076237 | 0.207540241 | 9.718698353 | 0.39182873 | 0.707436073 | 0.995570006 | -5.192842293 |
| **A_55_P1993629** | NA | 0.030577638 | -0.155965571 | 0.217120846 | 16.92150206 | 0.391772205 | 0.707475885 | 0.995570006 | -5.192860452 |
| **A_55_P2114963** | Exosc10 | -0.04145138 | -0.294338239 | 0.211435479 | 9.423511185 | -0.391761596 | 0.707483356 | 0.995570006 | -5.19286386 |

| | | | | | | | | | |
|---|---|---|---|---|---|---|---|---|---|
| **A_51_P218518** | Fiz1 | -0.03535614 | -0.251062271 | 0.180349992 | 8.534927061 | -0.391752235 | 0.70748995 | 0.995570006 | -5.192866867 |
| **A_52_P495104** | NA | -0.045178532 | -0.320841048 | 0.230483984 | 11.5165228 | -0.391708984 | 0.707520413 | 0.995570006 | -5.192880759 |
| **A_52_P474636** | Stk4 | 0.021201412 | -0.108195666 | 0.150598489 | 8.066362603 | 0.391606093 | 0.707592885 | 0.995570006 | -5.192913802 |
| **A_55_P2025760** | Adam9 | -0.052580986 | -0.373503509 | 0.268341537 | 11.95245164 | -0.391595469 | 0.707600368 | 0.995570006 | -5.192917214 |
| **A_65_P15215** | Mrpl15 | -0.03697727 | -0.262690678 | 0.188736138 | 11.7889211 | -0.391549447 | 0.707632785 | 0.995570006 | -5.19293199 |
| **A_55_P2424767** | Hecw1 | -0.044381186 | -0.315389 | 0.226626628 | 11.99167581 | -0.391404867 | 0.707734628 | 0.995570006 | -5.192978402 |
| **A_51_P433091** | Purb | -0.029690771 | -0.211010514 | 0.151628973 | 12.61646265 | -0.391368173 | 0.707760477 | 0.995570006 | -5.192990178 |
| **A_52_P97699** | D430019H | 0.037643561 | -0.192253556 | 0.267540678 | 8.65624761 | 0.391350871 | 0.707772665 | 0.995570006 | -5.192995731 |
| **A_55_P2105888** | G6pc3 | -0.037899603 | -0.26937362 | 0.193574414 | 11.17234786 | -0.39132856 | 0.707788383 | 0.995570006 | -5.19300289 |
| **A_51_P314161** | Qtrt1 | -0.025660101 | -0.182386651 | 0.131066448 | 8.357685422 | -0.391313582 | 0.707798934 | 0.995570006 | -5.193007697 |
| **A_51_P350048** | Gstt2 | -0.018818901 | -0.133768493 | 0.09613069 | 8.2959634 | -0.391287489 | 0.707817316 | 0.995570006 | -5.193016069 |
| **A_55_P2076248** | NA | -0.017531153 | -0.124637561 | 0.089575255 | 8.376157639 | -0.391204782 | 0.707875581 | 0.995570006 | -5.193042604 |
| **A_51_P365964** | Tor1a | 0.020994527 | -0.107302656 | 0.14929171 | 11.51798016 | 0.391109273 | 0.707942868 | 0.995570006 | -5.193073239 |
| **A_51_P363338** | Prkd1 | 0.021700459 | -0.110926066 | 0.154326984 | 8.671553064 | 0.391063835 | 0.707974881 | 0.995570006 | -5.193087812 |
| **A_55_P2023136** | Rnf214 | 0.031947885 | -0.163588545 | 0.227484315 | 8.948362564 | 0.390502269 | 0.708370575 | 0.995570006 | -5.193267777 |
| **A_55_P2173333** | Plcb4 | 0.032797278 | -0.167974431 | 0.233568987 | 10.85721215 | 0.39043111 | 0.708420721 | 0.995570006 | -5.193290564 |
| **A_55_P2023942** | Ccdc77 | -0.026325059 | -0.187567583 | 0.134917464 | 9.249543152 | -0.390210458 | 0.70857623 | 0.995570006 | -5.193361198 |
| **A_52_P525070** | Vamp4 | 0.021086444 | -0.108094864 | 0.150267752 | 8.677785798 | 0.390133106 | 0.708630749 | 0.995570006 | -5.19338595 |
| **A_51_P389864** | B3gnt9 | 0.033609142 | -0.172294775 | 0.239513059 | 9.091367296 | 0.390123354 | 0.708637622 | 0.995570006 | -5.193389071 |
| **A_55_P2095603** | Ccdc65 | 0.016325916 | -0.083696619 | 0.116348451 | 7.810866286 | 0.390111581 | 0.70864592 | 0.995570006 | -5.193392838 |
| **A_66_P122927** | NA | -0.083587832 | -0.595862079 | 0.428686415 | 8.862008827 | -0.389986586 | 0.708734023 | 0.995570006 | -5.193432824 |
| **A_55_P2027870** | Abhd17a | -0.039724716 | -0.283230526 | 0.203781093 | 10.76481951 | -0.389906623 | 0.708790388 | 0.995570006 | -5.193458398 |
| **A_51_P292073** | Haghl | -0.030497341 | -0.217446081 | 0.156451399 | 11.3633086 | -0.389895808 | 0.708798012 | 0.995570006 | -5.193461857 |
| **A_51_P208769** | Metrn | 0.025512957 | -0.13089133 | 0.181917243 | 9.043411247 | 0.389871304 | 0.708815284 | 0.995570006 | -5.193469692 |
| **A_51_P453351** | Lpgat1 | -0.022065362 | -0.157347068 | 0.113216344 | 11.68007883 | -0.389835264 | 0.70884069 | 0.995570006 | -5.193481216 |
| **A_52_P367760** | Calml4 | -0.017753041 | -0.126654027 | 0.091147944 | 7.874977201 | -0.389627936 | 0.708986846 | 0.995570006 | -5.19354749 |
| **A_66_P136899** | Nadk | 0.024036704 | -0.123445606 | 0.171519014 | 9.867542237 | 0.389532875 | 0.709053863 | 0.995570006 | -5.193577866 |
| **A_51_P225793** | Prr5l | -0.038567038 | -0.275258036 | 0.198123959 | 9.608151569 | -0.389442802 | 0.709117367 | 0.995570006 | -5.193606642 |
| **A_55_P2078895** | Gin1 | -0.02047318 | -0.146157574 | 0.105211214 | 8.286795372 | -0.389325708 | 0.709199925 | 0.995570006 | -5.19364404 |
| **A_51_P276599** | AU040320 | -0.029484823 | -0.210544561 | 0.151574916 | 10.72292359 | -0.389211588 | 0.709280391 | 0.995570006 | -5.193680479 |
| **A_55_P2008081** | Rhoa | 0.026118203 | -0.134336163 | 0.186572568 | 13.09403644 | 0.389045938 | 0.709397196 | 0.995570006 | -5.193733353 |
| **A_51_P234788** | Cxxc5 | -0.02392201 | -0.170961347 | 0.123117327 | 10.66562863 | -0.38884209 | 0.709540949 | 0.995570006 | -5.19379839 |
| **A_55_P1963184** | NA | 0.025331426 | -0.130373513 | 0.181036364 | 8.248465011 | 0.388835918 | 0.709545301 | 0.995570006 | -5.193800359 |
| **A_55_P2146749** | Rps13 | 0.033821289 | -0.174091166 | 0.241733744 | 16.37189175 | 0.388793304 | 0.709575354 | 0.995570006 | -5.193813951 |
| **A_52_P57308** | Cand1 | -0.0188936 | -0.135066231 | 0.097279031 | 12.68618969 | -0.38870491 | 0.709637694 | 0.995570006 | -5.193842139 |
| **A_51_P368285** | Rflna | 0.021846669 | -0.11249063 | 0.156183969 | 7.912562146 | 0.388684972 | 0.709651756 | 0.995570006 | -5.193848497 |
| **A_55_P2229137** | Tmem116 | 0.018421731 | -0.094896314 | 0.131739775 | 8.01134382 | 0.388544249 | 0.709751007 | 0.995570006 | -5.193893358 |
| **A_55_P2094852** | Rabep1 | -0.023012879 | -0.164574672 | 0.118548914 | 9.943895517 | -0.388538477 | 0.709755079 | 0.995570006 | -5.193895198 |
| **A_55_P2166282** | NA | -0.041470094 | -0.296584697 | 0.21364451 | 10.58312232 | -0.388515928 | 0.709770983 | 0.995570006 | -5.193902385 |
| **A_66_P134739** | Clpp | -0.037019587 | -0.264823264 | 0.19078409 | 10.29024947 | -0.388400668 | 0.70985228 | 0.995570006 | -5.193939115 |
| **A_51_P436447** | Zfp871 | -0.021064915 | -0.150739707 | 0.108609877 | 9.092889886 | -0.388251627 | 0.709957412 | 0.995570006 | -5.193986595 |
| **A_65_P06446** | Tfrc | 0.045502533 | -0.234615651 | 0.325620717 | 9.924757426 | 0.388242791 | 0.709963644 | 0.995570006 | -5.193989409 |
| **A_55_P1981099** | NA | 0.050202859 | -0.258886276 | 0.359291994 | 16.08416853 | 0.388198492 | 0.709994894 | 0.995570006 | -5.194003518 |
| **A_51_P221144** | Vwa8 | 0.01984395 | -0.102348435 | 0.142036335 | 10.33789854 | 0.388144216 | 0.710033182 | 0.995570006 | -5.194020802 |
| **A_55_P1997275** | Xpnpep1 | 0.018922993 | -0.097627819 | 0.135473805 | 13.34927131 | 0.388046395 | 0.710102191 | 0.995570006 | -5.194051948 |
| **A_52_P589119** | Rhno1 | -0.030634996 | -0.219338219 | 0.158068228 | 8.399144894 | -0.388014217 | 0.710124891 | 0.995570006 | -5.194062191 |
| **A_55_P2102135** | Rdh12 | -0.041048856 | -0.29391582 | 0.211818107 | 10.37455919 | -0.387987829 | 0.710143508 | 0.995570006 | -5.194070591 |
| **A_66_P132062** | Gm5608 | 0.015886837 | -0.081988205 | 0.113761878 | 7.799809499 | 0.387948976 | 0.710170918 | 0.995570006 | -5.194082958 |

| | | | | | | | | | |
|---|---|---|---|---|---|---|---|---|---|
| **A_55_P1973521** | Rnf7 | 0.025532066 | -0.131795018 | 0.182859151 | 10.54571042 | 0.387874834 | 0.710223226 | 0.995570006 | -5.194106553 |
| **A_55_P2032147** | Wnt9a | 0.036343457 | -0.187750203 | 0.260437118 | 8.760880577 | 0.387619663 | 0.710403265 | 0.995570006 | -5.194187729 |
| **A_55_P2105286** | Lrrc56 | -0.033807597 | -0.242337906 | 0.174722713 | 9.087752297 | -0.387484421 | 0.710498695 | 0.995570006 | -5.194230732 |
| **A_51_P488718** | Zfand2b | 0.02578149 | -0.133248469 | 0.18481145 | 10.67771732 | 0.387470105 | 0.710508797 | 0.995570006 | -5.194235283 |
| **A_52_P230167** | Setbp1 | 0.029514021 | -0.152563269 | 0.211591311 | 8.964554483 | 0.387419726 | 0.710544347 | 0.995570006 | -5.194251298 |
| **A_51_P460890** | Rgl2 | 0.035274758 | -0.182473389 | 0.253022905 | 10.54203872 | 0.387185164 | 0.710709878 | 0.995570006 | -5.194325837 |
| **A_51_P479769** | Ampd2 | 0.035887221 | -0.185746834 | 0.257521276 | 10.95999883 | 0.387001336 | 0.710839616 | 0.995570006 | -5.194384223 |
| **A_55_P1976984** | NA | 0.033661716 | -0.174230401 | 0.241553833 | 15.90552172 | 0.386996791 | 0.710842825 | 0.995570006 | -5.194385667 |
| **A_55_P1971430** | Serpinb9e | -0.049634466 | -0.356198393 | 0.256929461 | 7.800997638 | -0.386964776 | 0.710865421 | 0.995570006 | -5.194395832 |
| **A_52_P609120** | Pdxp | 0.074230133 | -0.384296353 | 0.532756619 | 10.17375112 | 0.386923356 | 0.710894655 | 0.995570006 | -5.194408983 |
| **A_55_P2039842** | Frmpd1 | -0.030417508 | -0.218333513 | 0.157498498 | 8.567862185 | -0.386873503 | 0.710929843 | 0.995570006 | -5.19442481 |
| **A_55_P2107667** | Enah | -0.027395963 | -0.196676444 | 0.141884518 | 8.526489059 | -0.386802096 | 0.710980245 | 0.995570006 | -5.194447475 |
| **A_55_P1991618** | Ppp1r16a | 0.026914435 | -0.139404218 | 0.193233088 | 11.07969132 | 0.386770585 | 0.711002487 | 0.995570006 | -5.194457476 |
| **A_55_P2183549** | NA | 0.036256828 | -0.18784781 | 0.260361466 | 10.88660723 | 0.386676782 | 0.711068701 | 0.995570006 | -5.194487243 |
| **A_52_P485850** | Mgam | 0.015392431 | -0.079761817 | 0.110546679 | 7.644648617 | 0.386623425 | 0.711106365 | 0.995570006 | -5.194504171 |
| **A_55_P1971278** | Kdm2a | 0.024364872 | -0.126318105 | 0.175047849 | 9.348258471 | 0.386464029 | 0.711218888 | 0.995570006 | -5.19455473 |
| **A_52_P588483** | Fbln1 | 0.056060499 | -0.290700291 | 0.402821289 | 10.14985849 | 0.386399101 | 0.711264725 | 0.995570006 | -5.194575319 |
| **A_55_P2007404** | Vbp1 | -0.031287497 | -0.224826084 | 0.16225109 | 9.927361754 | -0.386377993 | 0.711279626 | 0.995570006 | -5.194582012 |
| **A_55_P1979402** | Tigar | -0.024446181 | -0.175668169 | 0.126775806 | 9.937887702 | -0.386371619 | 0.711284127 | 0.995570006 | -5.194584033 |
| **A_55_P1955747** | Gm12070 | 0.029079661 | -0.150807255 | 0.208966577 | 16.46427255 | 0.386365997 | 0.711288096 | 0.995570006 | -5.194585815 |
| **A_52_P669922** | Hamp | -0.016335348 | -0.117387525 | 0.084716829 | 7.643463615 | -0.386359723 | 0.711292525 | 0.995570006 | -5.194587804 |
| **A_55_P1956762** | Sgms1 | 0.043180726 | -0.22403711 | 0.310398562 | 8.844065361 | 0.386218994 | 0.711391881 | 0.995570006 | -5.194632414 |
| **A_55_P2041988** | NA | -0.017349754 | -0.124721209 | 0.090021701 | 7.912633435 | -0.386201201 | 0.711404444 | 0.995570006 | -5.194638054 |
| **A_51_P460048** | Cnrip1 | -0.027951488 | -0.200953603 | 0.145050628 | 9.53043096 | -0.386155876 | 0.711436445 | 0.995570006 | -5.194652417 |
| **A_51_P170696** | Haus1 | -0.026711146 | -0.192041562 | 0.13861927 | 12.43203061 | -0.386143646 | 0.71144508 | 0.995570006 | -5.194656293 |
| **A_52_P297075** | Heatr5a | 0.035555226 | -0.184718647 | 0.255829099 | 10.3995904 | 0.385788776 | 0.711695657 | 0.995570006 | -5.194768695 |
| **A_55_P1968774** | Ptk2b | -0.026385642 | -0.189948481 | 0.137177197 | 8.353779836 | -0.385560165 | 0.711857102 | 0.995570006 | -5.194841054 |
| **A_55_P2038187** | BC051019 | -0.032897328 | -0.236860339 | 0.171065683 | 10.70512993 | -0.385494638 | 0.71190338 | 0.995570006 | -5.194861787 |
| **A_52_P304858** | Gnb1 | 0.052605102 | -0.273645358 | 0.378855561 | 10.25986984 | 0.385377064 | 0.71198642 | 0.995570006 | -5.19489898 |
| **A_51_P404463** | Ecrg4 | 0.080164221 | -0.41707388 | 0.577402323 | 9.581171401 | 0.385323336 | 0.712024368 | 0.995570006 | -5.194915972 |
| **A_55_P2150906** | Atp5c1 | -0.057667498 | -0.415405404 | 0.300070409 | 14.15081926 | -0.385278946 | 0.71205572 | 0.995570006 | -5.194930009 |
| **A_51_P167374** | Gpatch1 | 0.028732918 | -0.149564754 | 0.207030591 | 9.322520539 | 0.38516179 | 0.712138473 | 0.995570006 | -5.19496705 |
| **A_51_P363556** | Abi2 | -0.027771415 | -0.200205439 | 0.144662609 | 10.3245901 | -0.384932141 | 0.712300694 | 0.995570006 | -5.195039625 |
| **A_51_P511375** | Ranbp2 | 0.030040624 | -0.156486204 | 0.216567452 | 10.25079893 | 0.384925583 | 0.712305327 | 0.995570006 | -5.195041698 |
| **A_55_P2304507** | Noxa1 | -0.016139989 | -0.116366328 | 0.084086349 | 7.898815611 | -0.384884581 | 0.712334292 | 0.995570006 | -5.195054651 |
| **A_55_P2005833** | Adamts8 | 0.116324358 | -0.606672735 | 0.839321451 | 10.48269671 | 0.384541527 | 0.712576658 | 0.995570006 | -5.195162977 |
| **A_51_P190886** | Ankrd13a | -0.021606246 | -0.155984156 | 0.112771665 | 9.976838325 | -0.384291308 | 0.712753459 | 0.995570006 | -5.195241932 |
| **A_55_P2015767** | Yipf6 | 0.024561666 | -0.128210686 | 0.177334018 | 8.547697075 | 0.384257355 | 0.712777451 | 0.995570006 | -5.195252642 |
| **A_51_P312121** | Xdh | -0.019719135 | -0.142415512 | 0.102977243 | 7.859912844 | -0.384118523 | 0.712875557 | 0.995570006 | -5.195296424 |
| **A_55_P2098226** | C1rl | 0.021069957 | -0.110072661 | 0.152212575 | 8.068063574 | 0.383997978 | 0.712960746 | 0.995570006 | -5.195334428 |
| **A_55_P2074206** | Dlgap4 | -0.035699779 | -0.257921093 | 0.186521536 | 11.79968714 | -0.383962616 | 0.712985736 | 0.995570006 | -5.195345574 |
| **A_55_P2170978** | Ankrd13d | 0.020419189 | -0.106715027 | 0.147553405 | 8.127359054 | 0.383870883 | 0.713050567 | 0.995570006 | -5.195374484 |
| **A_55_P2160128** | Pls1 | -0.015578501 | -0.112604809 | 0.081447806 | 7.667244424 | -0.383747271 | 0.713137933 | 0.995570006 | -5.19541343 |
| **A_55_P1998987** | NA | 0.039730739 | -0.207754964 | 0.287216442 | 8.65208417 | 0.383694573 | 0.713175179 | 0.995570006 | -5.19543003 |
| **A_55_P2393483** | 6430584L0 | 0.019834719 | -0.103720045 | 0.143389483 | 8.382803292 | 0.38368577 | 0.713181401 | 0.995570006 | -5.195432803 |
| **A_51_P455997** | Meg3 | 0.173942094 | -0.909669678 | 1.257553866 | 10.49897171 | 0.383654337 | 0.713203618 | 0.995570006 | -5.195442703 |
| **A_66_P105307** | Adnp | -0.030701338 | -0.221970403 | 0.160567726 | 12.42190159 | -0.383638077 | 0.713215112 | 0.995570006 | -5.195447824 |
| **A_55_P2183692** | Rgs19 | 0.031855243 | -0.166695356 | 0.230405842 | 9.076987524 | 0.383458925 | 0.713341745 | 0.995570006 | -5.195504233 |

| | | | | | | | | | |
|---|---|---|---|---|---|---|---|---|---|
| **A_51_P260639** | Elp3 | 0.016572795 | -0.086781822 | 0.119927413 | 10.52942746 | 0.383243687 | 0.713493899 | 0.995570006 | -5.195571972 |
| **A_55_P2004752** | Prl2c5 | -0.10791719 | -0.7811432 | 0.56530882 | 8.03795409 | -0.383123572 | 0.713578815 | 0.995570006 | -5.195609758 |
| **A_55_P2073084** | Gtf3c2 | 0.031420782 | -0.164601659 | 0.227443223 | 10.60347463 | 0.383107214 | 0.71359038 | 0.995570006 | -5.195614903 |
| **A_55_P2008991** | Syp | -0.032521239 | -0.235424711 | 0.170382234 | 10.24086892 | -0.383077577 | 0.713611333 | 0.995570006 | -5.195624224 |
| **A_55_P2016962** | NA | -0.015550099 | -0.112579129 | 0.08147893 | 7.664860597 | -0.383036902 | 0.71364009 | 0.995570006 | -5.195637016 |
| **A_55_P2179453** | Lig3 | -0.025241671 | -0.182748602 | 0.13226526 | 8.225844084 | -0.38302539 | 0.713648229 | 0.995570006 | -5.195640636 |
| **A_55_P2130576** | NA | 0.026454281 | -0.138636567 | 0.191545129 | 14.64575058 | 0.382985288 | 0.713676582 | 0.995570006 | -5.195653246 |
| **A_55_P2128169** | Trappc13 | 0.021989888 | -0.115255846 | 0.159235622 | 9.251207066 | 0.382942259 | 0.713707005 | 0.995570006 | -5.195666775 |
| **A_55_P1966377** | 4930502E1 | 0.015534926 | -0.081427901 | 0.112497753 | 7.655043105 | 0.382924409 | 0.713719625 | 0.995570006 | -5.195672387 |
| **A_51_P222283** | Mlycd | -0.037436198 | -0.271112145 | 0.19623975 | 12.0608188 | -0.382901308 | 0.713735959 | 0.995570006 | -5.195679649 |
| **A_55_P1960431** | Ubn1 | 0.032635414 | -0.171074401 | 0.236345229 | 11.83981692 | 0.382900829 | 0.713736297 | 0.995570006 | -5.1956798 |
| **A_51_P165504** | Twist2 | 0.055241214 | -0.289583311 | 0.40006574 | 8.603842122 | 0.382890157 | 0.713743843 | 0.995570006 | -5.195683155 |
| **A_55_P2154782** | NA | -0.018696662 | -0.135407942 | 0.098014619 | 8.53684566 | -0.382877963 | 0.713752464 | 0.995570006 | -5.195686988 |
| **A_52_P326657** | Fam167b | 0.028566833 | -0.149799026 | 0.206932693 | 8.661038195 | 0.382789048 | 0.713815334 | 0.995570006 | -5.195714935 |
| **A_66_P118368** | Catip | -0.026928654 | -0.195078682 | 0.141221374 | 8.220547805 | -0.382760257 | 0.713835691 | 0.995570006 | -5.195723983 |
| **A_52_P613241** | Icam1 | -0.016399258 | -0.11880949 | 0.086010973 | 7.791444303 | -0.382727784 | 0.713858652 | 0.995570006 | -5.195734187 |
| **A_51_P122723** | Grpel1 | -0.024643587 | -0.178548268 | 0.129261095 | 13.55636801 | -0.382702429 | 0.713876581 | 0.995570006 | -5.195742154 |
| **A_55_P2014214** | NA | -0.029064812 | -0.210678722 | 0.152549099 | 14.24410165 | -0.382496562 | 0.714022158 | 0.995570006 | -5.195806824 |
| **A_55_P1953414** | Rab12 | -0.030100305 | -0.218189533 | 0.157988923 | 12.45996245 | -0.382486496 | 0.714029276 | 0.995570006 | -5.195809985 |
| **A_55_P2182975** | P2rx2 | 0.017892188 | -0.093932641 | 0.129717017 | 7.774206077 | 0.382414494 | 0.714080195 | 0.995570006 | -5.195832595 |
| **A_55_P2029763** | Peg10 | -0.01581397 | -0.114650815 | 0.083022875 | 7.745320533 | -0.382411699 | 0.714082171 | 0.995570006 | -5.195833472 |
| **A_55_P2037368** | Pdxdc1 | -0.022616559 | -0.163994609 | 0.118761492 | 9.626508256 | -0.382343464 | 0.714130427 | 0.995570006 | -5.195854895 |
| **A_55_P2412369** | Ubxn7 | 0.021178409 | -0.111212327 | 0.153569144 | 8.685396004 | 0.382335706 | 0.714135914 | 0.995570006 | -5.195857331 |
| **A_55_P2213348** | 5330421C1 | -0.063371246 | -0.459519488 | 0.332776997 | 9.253753981 | -0.382334636 | 0.714136671 | 0.995570006 | -5.195857667 |
| **A_55_P2053398** | NA | -0.025970243 | -0.18834592 | 0.136405434 | 17.36627173 | -0.382264677 | 0.714186148 | 0.995570006 | -5.195879627 |
| **A_55_P2062054** | Dbndd2 | 0.019885566 | -0.104457925 | 0.144229058 | 12.11972604 | 0.382229357 | 0.714211128 | 0.995570006 | -5.195890712 |
| **A_55_P2287260** | Snap23 | 0.056188184 | -0.295176526 | 0.407552894 | 12.42025459 | 0.382204669 | 0.714228588 | 0.995570006 | -5.19589846 |
| **A_51_P129803** | Cacybp | 0.016399876 | -0.086197123 | 0.118996875 | 12.17797423 | 0.382045456 | 0.714341197 | 0.995570006 | -5.195948415 |
| **A_55_P2099840** | NA | 0.019125735 | -0.100525565 | 0.138777036 | 7.885095662 | 0.38204089 | 0.714344427 | 0.995570006 | -5.195949848 |
| **A_51_P247614** | Smim10l2a | 0.021217941 | -0.111555832 | 0.153991714 | 8.304620997 | 0.381944339 | 0.71441272 | 0.995570006 | -5.195980132 |
| **A_51_P292030** | Nus1 | 0.032740101 | -0.172221917 | 0.237702119 | 9.96718506 | 0.381782275 | 0.714527358 | 0.995570006 | -5.196030949 |
| **A_55_P2076543** | NA | -0.024208625 | -0.175815011 | 0.127397761 | 9.392099464 | -0.381646918 | 0.71462311 | 0.995570006 | -5.196073376 |
| **A_51_P370050** | Nagpa | -0.025297933 | -0.183796325 | 0.13320046 | 8.52902159 | -0.38147783 | 0.714742732 | 0.995570006 | -5.196126355 |
| **A_52_P83557** | Arfgef2 | -0.019215607 | -0.139618661 | 0.101187447 | 8.387432767 | -0.381439566 | 0.714769803 | 0.995570006 | -5.196138341 |
| **A_52_P575668** | NA | -0.022458558 | -0.163191626 | 0.11827451 | 8.704653235 | -0.381412431 | 0.714789001 | 0.995570006 | -5.19614684 |
| **A_55_P1998374** | Ppfibp1 | 0.045703581 | -0.240706902 | 0.332114063 | 10.86151219 | 0.381391008 | 0.714804158 | 0.995570006 | -5.19615355 |
| **A_55_P2058718** | Ephb4 | 0.072319388 | -0.381620728 | 0.526259504 | 10.48113099 | 0.38077227 | 0.715241977 | 0.995570006 | -5.196347186 |
| **A_66_P115004** | Gabrp | 0.015237055 | -0.080417257 | 0.110891366 | 7.700664432 | 0.380719927 | 0.71527902 | 0.995570006 | -5.196363553 |
| **A_55_P2009121** | Agrn | 0.029496849 | -0.155713829 | 0.214707527 | 13.1398261 | 0.380643779 | 0.715332911 | 0.995570006 | -5.19638736 |
| **A_55_P2042487** | Dpysl3 | 0.019486384 | -0.102928141 | 0.141900909 | 14.41245968 | 0.380458628 | 0.715463955 | 0.995570006 | -5.196445228 |
| **A_55_P2166639** | Cdk4 | -0.023912328 | -0.174147281 | 0.126322625 | 11.72701973 | -0.38041708 | 0.715493362 | 0.995570006 | -5.19645821 |
| **A_55_P1985360** | Man1c1 | 0.020230283 | -0.106906933 | 0.147367499 | 9.456167665 | 0.380310568 | 0.715568753 | 0.995570006 | -5.196491483 |
| **A_55_P2027012** | Serpinb8 | 0.035502657 | -0.187634146 | 0.258639459 | 11.51665694 | 0.38027588 | 0.715593306 | 0.995570006 | -5.196502318 |
| **A_55_P2101182** | Pcmt1 | -0.034816281 | -0.253760759 | 0.184128197 | 11.42353082 | -0.38006468 | 0.71574281 | 0.995570006 | -5.196568264 |
| **A_51_P250551** | Dhx38 | 0.023044991 | -0.121910316 | 0.168000298 | 8.823901705 | 0.379971964 | 0.715808446 | 0.995570006 | -5.196597203 |
| **A_52_P229997** | Cbx8 | -0.021356899 | -0.155723482 | 0.113009684 | 8.352709662 | -0.379888423 | 0.715867588 | 0.995570006 | -5.196623273 |
| **A_51_P267063** | Ugt3a2 | 0.015514695 | -0.082110033 | 0.113139422 | 7.68997792 | 0.379832858 | 0.715906927 | 0.995570006 | -5.196640609 |
| **A_52_P540219** | Timp2 | -0.045449113 | -0.34055615 | 0.249657924 | 14.05871737 | -0.382571837 | 0.716011558 | 0.995570006 | -5.119093285 |

| | | | | | | | | | |
|---|---|---|---|---|---|---|---|---|---|
| **A_55_P2157927** | Tmem241 | 0.021338286 | -0.11298937 | 0.155665942 | 9.013260276 | 0.379667327 | 0.716024123 | 0.995570006 | -5.196692242 |
| **A_52_P937** | Spcs2 | 0.020190632 | -0.106946066 | 0.147327329 | 11.17936453 | 0.379566708 | 0.716095366 | 0.995570006 | -5.196723616 |
| **A_51_P503625** | Gsta3 | -0.057368163 | -0.418778637 | 0.304042312 | 8.221513131 | -0.379384285 | 0.716224537 | 0.995570006 | -5.196780478 |
| **A_55_P2073885** | Bbs5 | 0.027653887 | -0.146673734 | 0.201981509 | 8.834389667 | 0.379139573 | 0.71639783 | 0.995570006 | -5.196856715 |
| **A_51_P160907** | Morc3 | 0.041053632 | -0.21782752 | 0.299934783 | 9.617315507 | 0.379018388 | 0.716483654 | 0.995570006 | -5.196894452 |
| **A_51_P197378** | Atg12 | 0.026767196 | -0.142060749 | 0.195595141 | 11.33776661 | 0.37893758 | 0.716540886 | 0.995570006 | -5.196919609 |
| **A_51_P211786** | Chst13 | -0.015508281 | -0.113366692 | 0.082350131 | 7.659179387 | -0.378769167 | 0.716660169 | 0.995570006 | -5.196972023 |
| **A_51_P334281** | Gucy1b2 | -0.019287214 | -0.140997348 | 0.10242292 | 8.21527679 | -0.378749347 | 0.716674207 | 0.995570006 | -5.19697819 |
| **A_52_P175086** | Usp48 | -0.019314402 | -0.141204017 | 0.102575214 | 8.989217785 | -0.378724745 | 0.716691633 | 0.995570006 | -5.196985845 |
| **A_51_P462516** | Senp7 | -0.02358458 | -0.172430282 | 0.125261122 | 8.539818905 | -0.378704953 | 0.716705652 | 0.995570006 | -5.196992002 |
| **A_55_P2000022** | Ccdc151 | 0.019958463 | -0.106016481 | 0.145933406 | 7.774311746 | 0.378662279 | 0.716735879 | 0.995570006 | -5.197005277 |
| **A_52_P440962** | Cdc73 | -0.020645453 | -0.150969683 | 0.109678778 | 8.476480771 | -0.378624203 | 0.716762849 | 0.995570006 | -5.197017121 |
| **A_55_P2089253** | Atg4c | 0.029776307 | -0.158195791 | 0.217748406 | 9.294333404 | 0.378605206 | 0.716776306 | 0.995570006 | -5.19702303 |
| **A_55_P2010861** | Mrps23 | -0.02372492 | -0.173543514 | 0.126093674 | 13.83201266 | -0.378484568 | 0.716861762 | 0.995570006 | -5.197060546 |
| **A_51_P275953** | Dcps | 0.030871698 | -0.164092649 | 0.225836046 | 10.91491245 | 0.378455179 | 0.716882581 | 0.995570006 | -5.197069684 |
| **A_51_P441914** | Hsd17b2 | -0.01629163 | -0.119269127 | 0.086685867 | 7.67666131 | -0.37812147 | 0.717118996 | 0.995570006 | -5.197173395 |
| **A_55_P2017944** | Etfb | -0.022623978 | -0.165633147 | 0.120385192 | 16.07805502 | -0.378106558 | 0.717129561 | 0.995570006 | -5.197178027 |
| **A_51_P473940** | Slc22a15 | 0.018042756 | -0.096026859 | 0.13211237 | 8.338860623 | 0.378043719 | 0.717174083 | 0.995570006 | -5.197197546 |
| **A_52_P575771** | Mtfr1 | 0.022828684 | -0.121516561 | 0.167173928 | 10.33938119 | 0.377996275 | 0.717207698 | 0.995570006 | -5.197212281 |
| **A_55_P2185667** | NA | -0.016392708 | -0.120047638 | 0.087262223 | 7.772127764 | -0.37798091 | 0.717218585 | 0.995570006 | -5.197217052 |
| **A_55_P1957419** | Ensa | 0.032051628 | -0.170686398 | 0.234789654 | 9.054034844 | 0.377853989 | 0.717308516 | 0.995570006 | -5.197256461 |
| **A_55_P1953503** | Lrrc4b | -0.030284997 | -0.221887864 | 0.16131787 | 12.47304032 | -0.377776264 | 0.71736359 | 0.995570006 | -5.197280587 |
| **A_52_P431894** | Pknox1 | -0.026224904 | -0.192147483 | 0.139697676 | 9.377049118 | -0.377761367 | 0.717374147 | 0.995570006 | -5.197285211 |
| **A_51_P335900** | Cox5a | 0.022508518 | -0.119901392 | 0.164918429 | 17.10617967 | 0.377759878 | 0.717375201 | 0.995570006 | -5.197285673 |
| **A_55_P2168426** | Pced1b | 0.019366988 | -0.103198773 | 0.141932749 | 8.438527529 | 0.377660915 | 0.717445329 | 0.995570006 | -5.197316385 |
| **A_55_P2168058** | Glod4 | -0.036830678 | -0.270045057 | 0.1963837 | 11.66931735 | -0.377453563 | 0.717592272 | 0.995570006 | -5.197380708 |
| **A_55_P2153743** | Slc26a6 | 0.021619809 | -0.115335679 | 0.158575297 | 8.947293584 | 0.377295425 | 0.717704348 | 0.995570006 | -5.197429742 |
| **A_51_P161830** | Enpep | -0.041723755 | -0.306047882 | 0.222600372 | 8.796224797 | -0.377272986 | 0.717720251 | 0.995570006 | -5.197436698 |
| **A_51_P443475** | Bccip | -0.020399409 | -0.149654383 | 0.108855565 | 11.52943785 | -0.37720675 | 0.717767197 | 0.995570006 | -5.197457229 |
| **A_55_P2082570** | NA | -0.024145256 | -0.177184424 | 0.128893912 | 8.3014561 | -0.377084215 | 0.717854049 | 0.995570006 | -5.197495202 |
| **A_55_P2015363** | C1ra | 0.02691904 | -0.14370226 | 0.19754034 | 8.146913741 | 0.377081697 | 0.717855834 | 0.995570006 | -5.197495983 |
| **A_55_P2040367** | Bola3 | 0.020128238 | -0.107460396 | 0.147716872 | 12.75443772 | 0.377053434 | 0.717875867 | 0.995570006 | -5.197504739 |
| **A_55_P1964985** | Gm10240 | 0.032779008 | -0.175001929 | 0.240559946 | 15.70461182 | 0.377050259 | 0.717878117 | 0.995570006 | -5.197505723 |
| **A_55_P2177233** | Abhd5 | -0.01863606 | -0.136775099 | 0.099502979 | 8.994632899 | -0.377024695 | 0.717896237 | 0.995570006 | -5.197513643 |
| **A_55_P1991516** | E4f1 | 0.032268068 | -0.172314512 | 0.236850648 | 9.425047715 | 0.376975778 | 0.717930911 | 0.995570006 | -5.197528796 |
| **A_55_P1968043** | Jmjd1c | 0.046931289 | -0.260034777 | 0.353897355 | 11.15485831 | 0.379786276 | 0.717966207 | 0.995570006 | -5.119924754 |
| **A_51_P511511** | Stk33 | -0.017333044 | -0.127273001 | 0.092606913 | 7.746087801 | -0.376815192 | 0.718044745 | 0.995570006 | -5.19757853 |
| **A_52_P165654** | Star | 0.016545623 | -0.088418712 | 0.121509958 | 7.75060152 | 0.376747602 | 0.718092659 | 0.995570006 | -5.197599456 |
| **A_55_P2019577** | 1500011B0 | 0.017457238 | -0.093292261 | 0.128206737 | 8.265360845 | 0.376741011 | 0.718097332 | 0.995570006 | -5.197601497 |
| **A_55_P2167840** | Gm14151 | 0.018144199 | -0.096997058 | 0.133285455 | 7.873021625 | 0.376630909 | 0.718175387 | 0.995570006 | -5.197635577 |
| **A_55_P1974640** | NA | 0.020633826 | -0.110320179 | 0.151587832 | 7.854627131 | 0.376591157 | 0.718203569 | 0.995570006 | -5.19764788 |
| **A_51_P103509** | Cyb561d2 | -0.025051017 | -0.184094689 | 0.133992656 | 10.05419309 | -0.376459354 | 0.718297014 | 0.995570006 | -5.197688661 |
| **A_55_P1985804** | Thumpd2 | 0.020003663 | -0.107035055 | 0.147042381 | 8.50894954 | 0.376341888 | 0.718380299 | 0.995570006 | -5.197724995 |
| **A_55_P1982049** | Icam2 | 0.027593283 | -0.147661414 | 0.202847981 | 14.09497926 | 0.376307476 | 0.718404698 | 0.995570006 | -5.197735637 |
| **A_55_P1964093** | Rangrf | -0.036460136 | -0.268046252 | 0.195125981 | 11.62044344 | -0.376283257 | 0.718421871 | 0.995570006 | -5.197743126 |
| **A_55_P1981704** | Ccdc113 | 0.029867463 | -0.15996499 | 0.219699915 | 9.148478086 | 0.376042562 | 0.718592544 | 0.995570006 | -5.197817532 |
| **A_52_P572197** | Pank3 | 0.01638702 | -0.087779673 | 0.120553714 | 8.422770881 | 0.375993424 | 0.718627389 | 0.995570006 | -5.197832716 |
| **A_55_P1953063** | Hspe1 | 0.034491716 | -0.18480275 | 0.253786182 | 11.40482392 | 0.375920718 | 0.718678949 | 0.995570006 | -5.19785518 |

| | | | | | | | | | |
|---|---|---|---|---|---|---|---|---|---|
| **A_51_P286106** | NA | -0.019991628 | -0.147101807 | 0.10711855 | 9.745362022 | -0.375904022 | 0.718690789 | 0.995570006 | -5.197860338 |
| **A_55_P2108837** | Tuba1c | 0.030813918 | -0.165145834 | 0.22677367 | 14.67156453 | 0.375828032 | 0.718744679 | 0.995570006 | -5.197883811 |
| **A_51_P512002** | Rer1 | -0.020540698 | -0.151182742 | 0.110101346 | 12.72159331 | -0.375786661 | 0.718774019 | 0.995570006 | -5.197896589 |
| **A_55_P2060303** | Mras | -0.018314842 | -0.134831315 | 0.098201631 | 9.509615858 | -0.375685974 | 0.718845428 | 0.995570006 | -5.197927681 |
| **A_55_P2098995** | Cfhr2 | 0.023763207 | -0.127458042 | 0.174984456 | 9.37355811 | 0.375579054 | 0.71892126 | 0.995570006 | -5.197960688 |
| **A_52_P59228** | Zswim3 | 0.019755355 | -0.105996347 | 0.145507056 | 8.156905927 | 0.375474198 | 0.718995632 | 0.995570006 | -5.197993051 |
| **A_55_P2141311** | AU022252 | 0.040839495 | -0.219133381 | 0.30081237 | 9.342190629 | 0.375458078 | 0.719007065 | 0.995570006 | -5.197998025 |
| **A_55_P2038504** | 6-Mar | -0.022495305 | -0.165725292 | 0.120734683 | 9.53389627 | -0.375376479 | 0.719064944 | 0.995570006 | -5.198023202 |
| **A_55_P1996818** | NA | -0.033032361 | -0.243393299 | 0.177328577 | 14.12352319 | -0.375304394 | 0.719116076 | 0.995570006 | -5.198045439 |
| **A_51_P262101** | Ptcd2 | -0.029514494 | -0.217477609 | 0.158448622 | 10.58132955 | -0.375294185 | 0.719123318 | 0.995570006 | -5.198048588 |
| **A_55_P1966089** | NA | -0.026688941 | -0.196667775 | 0.143289894 | 8.711108793 | -0.375271535 | 0.719139385 | 0.995570006 | -5.198055575 |
| **A_55_P2109712** | Csnk1g2 | 0.041208787 | -0.221282579 | 0.303700153 | 8.898375716 | 0.375218238 | 0.719177192 | 0.995570006 | -5.198072012 |
| **A_55_P2023762** | Epha4 | -0.060685342 | -0.447397801 | 0.326027118 | 8.826586171 | -0.375063471 | 0.719286982 | 0.995570006 | -5.198119732 |
| **A_55_P2044381** | NA | 0.023423112 | -0.125857675 | 0.1727039 | 8.20076229 | 0.375016013 | 0.719320651 | 0.995570006 | -5.198134361 |
| **A_51_P269320** | Adsl | 0.036113013 | -0.194082034 | 0.26630806 | 9.909070454 | 0.374953041 | 0.719365325 | 0.995570006 | -5.19815377 |
| **A_55_P2037712** | Smc4 | -0.034759335 | -0.256479236 | 0.186960566 | 12.43575317 | -0.37469329 | 0.719549615 | 0.995570006 | -5.198233796 |
| **A_55_P2081805** | Slc46a1 | 0.026045098 | -0.140135432 | 0.192225629 | 8.691962245 | 0.374588977 | 0.719623629 | 0.995570006 | -5.198265918 |
| **A_55_P1953931** | Fam193a | -0.026643597 | -0.196679141 | 0.143391946 | 10.45216073 | -0.374509018 | 0.719680366 | 0.995570006 | -5.198290535 |
| **A_55_P2180292** | Dmrtc1c2 | -0.019951405 | -0.147287497 | 0.107384686 | 7.765464032 | -0.374482134 | 0.719699443 | 0.995570006 | -5.198298811 |
| **A_55_P2099650** | Slc24a3 | 0.035649844 | -0.19190192 | 0.263201608 | 9.870785516 | 0.374443726 | 0.719726697 | 0.995570006 | -5.198310633 |
| **A_55_P2104894** | Col5a2 | -0.046248374 | -0.341499368 | 0.24900262 | 10.2247818 | -0.374381375 | 0.719770942 | 0.995570006 | -5.198329823 |
| **A_52_P493810** | Srpk1 | 0.03020101 | -0.162663449 | 0.22306547 | 9.000874893 | 0.374264297 | 0.719854024 | 0.995570006 | -5.198365847 |
| **A_55_P2151463** | NA | 0.028117516 | -0.151529499 | 0.207764532 | 10.26590491 | 0.374081371 | 0.719983844 | 0.995570006 | -5.198422111 |
| **A_55_P2186282** | Agxt2 | 0.015758017 | -0.084935618 | 0.116451652 | 7.70831155 | 0.374031932 | 0.720018931 | 0.995570006 | -5.198437313 |
| **A_55_P2144115** | 1700125H2 | 0.016868346 | -0.090934529 | 0.124671221 | 7.734422351 | 0.373982508 | 0.720054009 | 0.995570006 | -5.198452509 |
| **A_55_P2010048** | NA | 0.016453892 | -0.088716225 | 0.121624008 | 8.986578633 | 0.373925785 | 0.720094268 | 0.995570006 | -5.198469946 |
| **A_55_P1987993** | NA | -0.036371397 | -0.268891717 | 0.196148922 | 14.13514536 | -0.373859316 | 0.720141446 | 0.995570006 | -5.198490376 |
| **A_55_P2042481** | Gm10653 | 0.026462325 | -0.142732181 | 0.19565683 | 15.2788184 | 0.37380996 | 0.720176477 | 0.995570006 | -5.198505544 |
| **A_55_P2138150** | Pias2 | -0.026368509 | -0.194979343 | 0.142242324 | 9.628003305 | -0.373774127 | 0.720201912 | 0.995570006 | -5.198516555 |
| **A_52_P501875** | Ccdc173 | 0.023257896 | -0.125480519 | 0.17199631 | 8.065829975 | 0.373728659 | 0.720234185 | 0.995570006 | -5.198530525 |
| **A_55_P2039439** | Ppp1ca | -0.02394716 | -0.177102305 | 0.129207984 | 15.29717742 | -0.373707286 | 0.720249356 | 0.995570006 | -5.198537091 |
| **A_51_P110672** | Mst1r | 0.050367828 | -0.271787069 | 0.372522725 | 8.396827225 | 0.373678076 | 0.720270091 | 0.995570006 | -5.198546065 |
| **A_52_P679105** | Prss23 | 0.055498814 | -0.299649719 | 0.410647348 | 9.633746408 | 0.373493296 | 0.720401257 | 0.995570006 | -5.198602814 |
| **A_65_P17602** | Pgm5 | -0.020412041 | -0.15107197 | 0.110247887 | 10.78527545 | -0.37338181 | 0.720480402 | 0.995570006 | -5.198637041 |
| **A_52_P231170** | Megf9 | 0.01648708 | -0.089101751 | 0.122075911 | 8.15328171 | 0.373194201 | 0.720613595 | 0.995570006 | -5.198694617 |
| **A_55_P1974984** | NA | 0.037489995 | -0.202642895 | 0.277622884 | 11.4460347 | 0.373140907 | 0.720651432 | 0.995570006 | -5.198710967 |
| **A_55_P2170813** | Pkig | -0.025873367 | -0.191634161 | 0.139887427 | 14.44612655 | -0.373061357 | 0.720707913 | 0.995570006 | -5.198735369 |
| **A_51_P232708** | Npff | 0.023617551 | -0.127692189 | 0.174927292 | 8.624729375 | 0.37305865 | 0.720709835 | 0.995570006 | -5.198736199 |
| **A_51_P173622** | Ptpn11 | 0.035173725 | -0.190173297 | 0.260520748 | 14.09328972 | 0.373057415 | 0.720710711 | 0.995570006 | -5.198736577 |
| **A_55_P2114187** | NA | -0.014890723 | -0.110292896 | 0.080511449 | 7.699475476 | -0.373049666 | 0.720716213 | 0.995570006 | -5.198738954 |
| **A_55_P2058812** | Fancl | 0.019854635 | -0.107374628 | 0.147083898 | 9.554690788 | 0.372978692 | 0.720766607 | 0.995570006 | -5.19876072 |
| **A_55_P2177198** | H2bc3 | 0.029328512 | -0.158611838 | 0.217268862 | 8.120732067 | 0.372974495 | 0.720769587 | 0.995570006 | -5.198762007 |
| **A_51_P108659** | Pon1 | -0.015378248 | -0.113949213 | 0.083192717 | 7.657078275 | -0.37287819 | 0.720837969 | 0.995570006 | -5.186528282 |
| **A_55_P2101503** | NA | 0.020338552 | -0.110027982 | 0.150705086 | 8.221399333 | 0.372874806 | 0.720840372 | 0.995570006 | -5.198792573 |
| **A_51_P164835** | Ppl | 0.071457435 | -0.386613808 | 0.529528677 | 8.408706093 | 0.372840881 | 0.720864461 | 0.995570006 | -5.198802973 |
| **A_51_P394471** | Tcf3 | -0.018187265 | -0.13480381 | 0.098429279 | 10.34784798 | -0.372748894 | 0.72092978 | 0.995570006 | -5.198831167 |
| **A_55_P2082361** | Ppt1 | 0.018097156 | -0.097951462 | 0.134145775 | 8.403109089 | 0.372717253 | 0.720952249 | 0.995570006 | -5.198840864 |
| **A_51_P372156** | 4930563E2 | 0.018770481 | -0.101657341 | 0.139198303 | 7.839580933 | 0.372526958 | 0.721087386 | 0.995570006 | -5.198899164 |

| | | | | | | | | | |
|---|---|---|---|---|---|---|---|---|---|
| **A_51_P517495** | Cox8a | -0.019413941 | -0.143977429 | 0.105149548 | 16.83424256 | -0.372504968 | 0.721103003 | 0.995570006 | -5.1989059 |
| **A_55_P2423586** | Eya1 | -0.037265741 | -0.276411026 | 0.201879545 | 13.32652026 | -0.372440642 | 0.721148686 | 0.995570006 | -5.1989256 |
| **A_52_P66130** | Myo5b | -0.041690111 | -0.30926987 | 0.225889647 | 8.192666535 | -0.372382213 | 0.721190183 | 0.995570006 | -5.198943491 |
| **A_55_P2048259** | Golph3l | -0.033075209 | -0.245363138 | 0.179212719 | 11.30883655 | -0.372380076 | 0.721191702 | 0.995570006 | -5.198944146 |
| **A_51_P491635** | Ly6g6e | -0.020435344 | -0.151596601 | 0.110725914 | 7.87718966 | -0.372379279 | 0.721192267 | 0.995570006 | -5.19894439 |
| **A_55_P2059606** | Stat2 | -0.03281506 | -0.243434364 | 0.177804244 | 9.641535028 | -0.372378131 | 0.721193083 | 0.995570006 | -5.198944741 |
| **A_51_P437608** | Tulp3 | -0.018744363 | -0.139099418 | 0.101610692 | 8.763279484 | -0.372233527 | 0.721295786 | 0.995570006 | -5.198989008 |
| **A_55_P2018403** | Sycp1 | 0.016090864 | -0.087231176 | 0.119412905 | 7.647842069 | 0.372216426 | 0.721307933 | 0.995570006 | -5.198994242 |
| **A_51_P465161** | Hoga1 | 0.01592422 | -0.086344872 | 0.118193311 | 7.795926795 | 0.372154181 | 0.721352144 | 0.995570006 | -5.199013291 |
| **A_52_P261322** | Tanc1 | 0.023753317 | -0.128815157 | 0.176321792 | 8.461387114 | 0.372107653 | 0.721385192 | 0.995570006 | -5.199027528 |
| **A_51_P167105** | B3galt4 | -0.028912175 | -0.214617726 | 0.156793375 | 8.732907765 | -0.37210458 | 0.721387375 | 0.995570006 | -5.199028468 |
| **A_51_P290626** | Gulo | -0.015744653 | -0.116905957 | 0.085416651 | 7.773447096 | -0.371987041 | 0.721470864 | 0.995570006 | -5.199064426 |
| **A_55_P1969441** | Eef1akmt2 | -0.026738314 | -0.198607487 | 0.145130858 | 10.16671746 | -0.371830638 | 0.721581967 | 0.995570006 | -5.199112256 |
| **A_55_P1979694** | NA | -0.041437998 | -0.307831888 | 0.224955892 | 15.78617151 | -0.371777957 | 0.72161939 | 0.995570006 | -5.199128362 |
| **A_55_P2098450** | Snrnp200 | -0.016538518 | -0.122897022 | 0.089819986 | 12.56090778 | -0.371649465 | 0.721710673 | 0.995570006 | -5.199167638 |
| **A_55_P2132651** | Ccn4 | 0.050071084 | -0.271977863 | 0.37212003 | 9.782550968 | 0.371598745 | 0.721746707 | 0.995570006 | -5.199183137 |
| **A_55_P2181084** | Rpl17 | 0.018079188 | -0.098248893 | 0.134407269 | 14.66720339 | 0.371452682 | 0.721850481 | 0.995570006 | -5.199227762 |
| **A_55_P1978830** | Camk2d | -0.039529757 | -0.293890845 | 0.214831332 | 10.00484001 | -0.371434806 | 0.721863181 | 0.995570006 | -5.199233222 |
| **A_55_P2036733** | Stk36 | -0.016199011 | -0.120472191 | 0.08807417 | 7.990825234 | -0.371300046 | 0.721958931 | 0.995570006 | -5.199274377 |
| **A_51_P487048** | Rhox6 | -0.077884766 | -0.579260778 | 0.423491245 | 7.923343 | -0.371277047 | 0.721975273 | 0.995570006 | -5.199281399 |
| **A_52_P6451** | Sec24a | -0.028268066 | -0.210246268 | 0.153710136 | 9.05837663 | -0.371266582 | 0.721982709 | 0.995570006 | -5.199284594 |
| **A_51_P328089** | Crb3 | -0.015844865 | -0.117881291 | 0.086191561 | 7.675191334 | -0.371144001 | 0.722069811 | 0.995570006 | -5.199322014 |
| **A_52_P642626** | Rbm20 | 0.033026347 | -0.179692265 | 0.245744958 | 9.440711077 | 0.371077119 | 0.722117337 | 0.995570006 | -5.199342426 |
| **A_55_P2198364** | Tsen54 | -0.029843348 | -0.222085181 | 0.162398484 | 8.625726345 | -0.371029793 | 0.722150967 | 0.995570006 | -5.199356867 |
| **A_55_P2058628** | Hnrnpa3 | 0.094625161 | -0.515033126 | 0.704283449 | 10.11570316 | 0.37096188 | 0.722199228 | 0.995570006 | -5.199377588 |
| **A_55_P2043837** | Cmtr1 | -0.019982556 | -0.14874893 | 0.108783817 | 12.33019355 | -0.370900752 | 0.722242669 | 0.995570006 | -5.199396235 |
| **A_55_P2262753** | 9530053J1 | -0.03986306 | -0.296829593 | 0.217103473 | 8.747840222 | -0.370768816 | 0.722336432 | 0.995570006 | -5.199436472 |
| **A_52_P175028** | NA | -0.024780262 | -0.184521982 | 0.134961459 | 10.16542619 | -0.370763233 | 0.7223404 | 0.995570006 | -5.199438175 |
| **A_55_P2052081** | Eif1ad | 0.020263357 | -0.110374043 | 0.150900756 | 11.81721165 | 0.370725957 | 0.722366893 | 0.995570006 | -5.199449541 |
| **A_55_P1993012** | NA | -0.056983105 | -0.424354427 | 0.310388217 | 10.73093694 | -0.370723395 | 0.722368713 | 0.995570006 | -5.199450322 |
| **A_52_P401311** | NA | -0.060290819 | -0.449256871 | 0.328675233 | 12.39743806 | -0.370466222 | 0.722551499 | 0.995570006 | -5.199528706 |
| **A_55_P2128303** | Ids | 0.017291849 | -0.0942749 | 0.128858598 | 8.606811256 | 0.370438216 | 0.722571405 | 0.995570006 | -5.199537238 |
| **A_52_P359819** | Ntf5 | 0.019138009 | -0.104342988 | 0.142619005 | 7.985817579 | 0.370429661 | 0.722577486 | 0.995570006 | -5.199539845 |
| **A_55_P2115691** | NA | 0.023253459 | -0.126785062 | 0.173291979 | 11.65724815 | 0.370419565 | 0.722584662 | 0.995570006 | -5.19954292 |
| **A_55_P2134877** | Eml1 | -0.031471091 | -0.234582501 | 0.171640319 | 8.611853897 | -0.370328051 | 0.722649712 | 0.995570006 | -5.199570797 |
| **A_51_P487813** | Lxn | -0.02682873 | -0.199980031 | 0.146322571 | 12.04539627 | -0.370325392 | 0.722651602 | 0.995570006 | -5.199571607 |
| **A_55_P2044557** | Rpl29 | 0.019330084 | -0.105441745 | 0.144101913 | 16.80206675 | 0.370276662 | 0.722686241 | 0.995570006 | -5.199586449 |
| **A_51_P174434** | Cenpj | -0.034052414 | -0.253870177 | 0.185765348 | 9.47038504 | -0.37024931 | 0.722705684 | 0.995570006 | -5.199594778 |
| **A_55_P2040723** | Dnali1 | 0.015633428 | -0.085317074 | 0.11658393 | 7.693481775 | 0.370130494 | 0.722790146 | 0.995570006 | -5.199630954 |
| **A_66_P130601** | Foxp2 | -0.016032426 | -0.119565756 | 0.087500903 | 7.817744337 | -0.370107772 | 0.722806299 | 0.995570006 | -5.199637871 |
| **A_55_P1992216** | Chchd2 | -0.024776207 | -0.184805747 | 0.135253333 | 17.47393091 | -0.370035847 | 0.722857431 | 0.995570006 | -5.199659764 |
| **A_55_P2109589** | Slc1a4 | -0.029564447 | -0.22053562 | 0.161406727 | 10.37977575 | -0.370007962 | 0.722877255 | 0.995570006 | -5.199668251 |
| **A_51_P343833** | Traf1 | -0.019151587 | -0.142862567 | 0.104559392 | 8.089867765 | -0.370003354 | 0.722880531 | 0.995570006 | -5.199669653 |
| **A_52_P256817** | Zik1 | 0.021784759 | -0.118960269 | 0.162529786 | 8.70540236 | 0.3699379 | 0.722927064 | 0.995570006 | -5.199689571 |
| **A_55_P1955417** | Atad3a | -0.036651659 | -0.27349511 | 0.200191792 | 13.81318373 | -0.369863427 | 0.722980011 | 0.995570006 | -5.199712229 |
| **A_51_P448618** | Slc16a10 | -0.018769979 | -0.140092422 | 0.102552464 | 8.706850729 | -0.369770091 | 0.723046371 | 0.995570006 | -5.19974062 |
| **A_55_P2026025** | NA | 0.018272984 | -0.099838897 | 0.136384865 | 16.3079488 | 0.369764336 | 0.723050463 | 0.995570006 | -5.199742371 |
| **A_55_P2151061** | NA | 0.053203379 | -0.290704986 | 0.397111745 | 9.083224841 | 0.369747809 | 0.723062214 | 0.995570006 | -5.199747397 |

| | | | | | | | | | |
|---|---|---|---|---|---|---|---|---|---|
| **A_55_P2032853** | Entpd1 | -0.016519219 | -0.123328877 | 0.090290438 | 8.173219272 | -0.369647801 | 0.723133321 | 0.995570006 | -5.199777809 |
| **A_55_P2054619** | NA | -0.02475485 | -0.184830196 | 0.135320495 | 15.44998876 | -0.369611087 | 0.723159426 | 0.995570006 | -5.199788971 |
| **A_51_P215922** | Casp6 | -0.021532126 | -0.160768634 | 0.117704382 | 10.61280207 | -0.369609326 | 0.723160678 | 0.995570006 | -5.199789506 |
| **A_55_P2024580** | Dtnbp1 | -0.017901722 | -0.133662631 | 0.097859187 | 13.20078106 | -0.369608547 | 0.723161232 | 0.995570006 | -5.199789743 |
| **A_51_P424810** | Ncapg2 | 0.034093937 | -0.186385388 | 0.254573262 | 10.13195179 | 0.369588474 | 0.723175504 | 0.995570006 | -5.199795846 |
| **A_52_P149391** | Usp40 | -0.029830361 | -0.222738999 | 0.163078278 | 9.307744278 | -0.369586381 | 0.723176993 | 0.995570006 | -5.199796482 |
| **A_55_P1979242** | Kcne2 | 0.019463392 | -0.106527917 | 0.145454701 | 8.091858324 | 0.36922158 | 0.723436404 | 0.995570006 | -5.199907331 |
| **A_51_P261107** | Ogt | -0.047676678 | -0.356320613 | 0.260967257 | 9.530518391 | -0.369196326 | 0.723454363 | 0.995570006 | -5.199915 |
| **A_55_P2001494** | Kitl | 0.035495809 | -0.194309355 | 0.265300974 | 9.004687638 | 0.369170025 | 0.723473067 | 0.995570006 | -5.199922988 |
| **A_55_P2088401** | H2-T9 | -0.017489652 | -0.130720449 | 0.095741145 | 8.038986375 | -0.369169433 | 0.723473489 | 0.995570006 | -5.199923168 |
| **A_51_P109258** | Cys1 | -0.028072823 | -0.209835892 | 0.153690247 | 8.402791727 | -0.369138691 | 0.723495351 | 0.995570006 | -5.199932503 |
| **A_51_P317640** | Tgfb2 | 0.043504523 | -0.23821441 | 0.325223457 | 10.61854878 | 0.369085949 | 0.72353286 | 0.995570006 | -5.199948517 |
| **A_55_P2029643** | NA | -0.025794673 | -0.192850799 | 0.141261453 | 12.13526758 | -0.369042818 | 0.723563536 | 0.995570006 | -5.199961612 |
| **A_55_P2069217** | Prr13 | -0.024554258 | -0.183629024 | 0.134520509 | 8.372390698 | -0.368922078 | 0.723649409 | 0.995570006 | -5.19999826 |
| **A_55_P2152962** | Mtr | -0.02017557 | -0.150885698 | 0.110534557 | 9.54668634 | -0.368914492 | 0.723654805 | 0.995570006 | -5.200000562 |
| **A_55_P1998159** | Deaf1 | 0.019062799 | -0.104459821 | 0.142585419 | 8.41673007 | 0.368849587 | 0.723700969 | 0.995570006 | -5.200020258 |
| **A_51_P367563** | Rsph3a | -0.05031535 | -0.376409195 | 0.275778496 | 9.074705427 | -0.368779715 | 0.723750667 | 0.995570006 | -5.200041457 |
| **A_66_P139295** | Tomm40 | 0.029055927 | -0.159343694 | 0.217455548 | 12.96159483 | 0.36860722 | 0.723873365 | 0.995570006 | -5.200093776 |
| **A_55_P2109257** | Nipal3 | -0.040801881 | -0.305372016 | 0.223768255 | 9.280428616 | -0.368594199 | 0.723882627 | 0.995570006 | -5.200097724 |
| **A_51_P495232** | Gpx8 | 0.030869358 | -0.169326208 | 0.231064925 | 12.11178594 | 0.368538004 | 0.723922601 | 0.995570006 | -5.200114763 |
| **A_51_P155522** | Psmb4 | -0.018335277 | -0.13729618 | 0.100625627 | 14.53210423 | -0.368376864 | 0.724037233 | 0.995570006 | -5.200163608 |
| **A_51_P424518** | Nin | 0.020483722 | -0.11242155 | 0.153388994 | 8.776327474 | 0.368362827 | 0.724047219 | 0.995570006 | -5.200167862 |
| **A_55_P2175502** | M1ap | 0.016506284 | -0.090597593 | 0.123610162 | 7.897303684 | 0.368343711 | 0.724060818 | 0.995570006 | -5.200173655 |
| **A_55_P2049572** | Pld2 | 0.035818678 | -0.196622255 | 0.268259611 | 11.18713786 | 0.368303696 | 0.724089286 | 0.995570006 | -5.20018578 |
| **A_51_P371750** | Marco | 0.019546832 | -0.107325452 | 0.146419117 | 8.127351548 | 0.368229654 | 0.724141963 | 0.995570006 | -5.200208213 |
| **A_52_P598835** | Ccdc71l | -0.022326643 | -0.167269948 | 0.122616663 | 9.279083631 | -0.368158128 | 0.72419285 | 0.995570006 | -5.200229879 |
| **A_55_P2014978** | Dvl3 | 0.061652253 | -0.338602548 | 0.461907053 | 12.0192323 | 0.368147226 | 0.724200607 | 0.995570006 | -5.200233181 |
| **A_55_P2004881** | Ldlrad4 | -0.027187466 | -0.203777427 | 0.149402496 | 8.354399285 | -0.367969518 | 0.724327045 | 0.995570006 | -5.200286993 |
| **A_55_P2004230** | Emc6 | 0.035576434 | -0.195582783 | 0.26673565 | 11.80100064 | 0.367841165 | 0.724418374 | 0.995570006 | -5.200325844 |
| **A_55_P2060158** | Ermn | 0.023965549 | -0.131773461 | 0.179704558 | 8.429380005 | 0.367789312 | 0.724455272 | 0.995570006 | -5.200341536 |
| **A_55_P1990092** | Snrpa | 0.027227461 | -0.149733982 | 0.204188904 | 11.63136355 | 0.367737245 | 0.724492322 | 0.995570006 | -5.20035729 |
| **A_55_P2151437** | NA | -0.025621066 | -0.192211683 | 0.140969551 | 8.379842507 | -0.367583311 | 0.724601864 | 0.995570006 | -5.200403855 |
| **A_55_P2079757** | NA | -0.027997409 | -0.210061753 | 0.154066935 | 9.238590596 | -0.367537853 | 0.724634214 | 0.995570006 | -5.200417603 |
| **A_55_P2126281** | NA | 0.016784034 | -0.092362265 | 0.125930332 | 8.190668568 | 0.367533102 | 0.724637595 | 0.995570006 | -5.200419039 |
| **A_55_P2013760** | Slc39a14 | 0.018906123 | -0.10404076 | 0.141853005 | 9.49607474 | 0.367531089 | 0.724639028 | 0.995570006 | -5.200419648 |
| **A_51_P415519** | Tmem39a | 0.03494499 | -0.192457569 | 0.26234755 | 10.02467391 | 0.36728121 | 0.724816865 | 0.995570006 | -5.200495187 |
| **A_51_P466886** | Scoc | 0.028782504 | -0.158533865 | 0.216098873 | 9.003853231 | 0.367250145 | 0.724838975 | 0.995570006 | -5.200504574 |
| **A_51_P156263** | Zfp511 | 0.020362314 | -0.112161167 | 0.152885795 | 10.40092858 | 0.367234466 | 0.724850134 | 0.995570006 | -5.200509312 |
| **A_52_P72434** | Khk | -0.023109824 | -0.17353238 | 0.127312732 | 9.712148061 | -0.367191658 | 0.724880603 | 0.995570006 | -5.200522247 |
| **A_55_P2163138** | Tm4sf5 | 0.037860901 | -0.208612668 | 0.28433447 | 8.409038228 | 0.367138323 | 0.724918564 | 0.995570006 | -5.20053836 |
| **A_66_P117543** | NA | 0.021457942 | -0.118326761 | 0.161242646 | 8.118332889 | 0.366891424 | 0.725094309 | 0.995570006 | -5.200612922 |
| **A_55_P2158522** | Pbx1 | 0.067023113 | -0.369598883 | 0.50364511 | 13.53487417 | 0.366883448 | 0.725099986 | 0.995570006 | -5.20061533 |
| **A_55_P2141884** | Tagln2 | -0.039189324 | -0.294516828 | 0.216138179 | 10.52158556 | -0.366842216 | 0.725129337 | 0.995570006 | -5.200627777 |
| **A_55_P2013058** | Shb | 0.025517202 | -0.140780828 | 0.191815233 | 12.01082481 | 0.366737301 | 0.725204024 | 0.995570006 | -5.200659442 |
| **A_52_P551903** | Slc33a1 | -0.017937322 | -0.134840656 | 0.098966012 | 8.162748934 | -0.366724428 | 0.725213188 | 0.995570006 | -5.200663327 |
| **A_55_P2094362** | Erp29 | 0.043968552 | -0.242679478 | 0.330616582 | 9.383679413 | 0.366608335 | 0.725295836 | 0.995570006 | -5.200698355 |
| **A_51_P377376** | Gnl3 | 0.035777496 | -0.19757873 | 0.269133722 | 9.377852712 | 0.366437304 | 0.725417601 | 0.995570006 | -5.200749939 |
| **A_55_P2149001** | NA | 0.025507374 | -0.140909283 | 0.191924031 | 13.01904299 | 0.366334724 | 0.725490638 | 0.995570006 | -5.200780868 |

| | | | | | | | | | |
|---|---|---|---|---|---|---|---|---|---|
| **A_55_P2134487** | NA | 0.024001713 | -0.132637848 | 0.180641274 | 11.24034184 | 0.366226625 | 0.725567606 | 0.995570006 | -5.200813451 |
| **A_51_P418526** | Sfxn1 | 0.028556151 | -0.157807312 | 0.214919614 | 9.70566118 | 0.366225033 | 0.72556874 | 0.995570006 | -5.200813931 |
| **A_55_P2022094** | Mta3 | -0.01867414 | -0.140569384 | 0.103221103 | 10.33919574 | -0.366153332 | 0.725619795 | 0.995570006 | -5.200835538 |
| **A_51_P395473** | Tns2 | 0.038512192 | -0.212977457 | 0.29000184 | 9.591164355 | 0.366005196 | 0.72572528 | 0.995570006 | -5.200880166 |
| **A_55_P2177513** | Chrna2 | 0.016165047 | -0.089401581 | 0.121731675 | 7.975477373 | 0.365981777 | 0.725741957 | 0.995570006 | -5.20088722 |
| **A_52_P37702** | NA | -0.017294139 | -0.130235835 | 0.095647556 | 7.853643298 | -0.365976996 | 0.725745361 | 0.995570006 | -5.200888659 |
| **A_52_P639101** | Lrrc14 | -0.024376533 | -0.183583329 | 0.134830263 | 8.376667442 | -0.36594808 | 0.725765953 | 0.995570006 | -5.200897368 |
| **A_55_P2049141** | Cops7b | -0.037076793 | -0.279235379 | 0.205081794 | 9.04211306 | -0.365941293 | 0.725770786 | 0.995570006 | -5.200899412 |
| **A_55_P2094611** | NA | 0.028527599 | -0.157796365 | 0.214851564 | 8.25357972 | 0.365936426 | 0.725774252 | 0.995570006 | -5.200900878 |
| **A_55_P1992555** | Gys2 | 0.018129982 | -0.100291543 | 0.136551507 | 7.772979051 | 0.365911327 | 0.725792126 | 0.995570006 | -5.200908436 |
| **A_55_P2279140** | F830014O1 | 0.035420909 | -0.19630238 | 0.267144199 | 8.1421994 | 0.365341623 | 0.726197878 | 0.995570006 | -5.201079866 |
| **A_51_P440047** | Lgalsl | 0.020075936 | -0.111269906 | 0.151421779 | 10.87747682 | 0.365315927 | 0.726216181 | 0.995570006 | -5.201087592 |
| **A_51_P183894** | Fbxo15 | 0.015044071 | -0.08339043 | 0.113478571 | 7.652433967 | 0.36528106 | 0.726241017 | 0.995570006 | -5.201098075 |
| **A_52_P85983** | Mrpl45 | -0.018048471 | -0.13616857 | 0.100071629 | 12.67976204 | -0.365195766 | 0.726301774 | 0.995570006 | -5.201123715 |
| **A_55_P2004037** | NA | -0.015386385 | -0.116093059 | 0.08532029 | 7.798179176 | -0.365163582 | 0.726324701 | 0.995570006 | -5.201133388 |
| **A_55_P2096297** | NA | 0.036458154 | -0.20217435 | 0.275090658 | 9.128757171 | 0.365152443 | 0.726332636 | 0.995570006 | -5.201136736 |
| **A_55_P2146050** | Dennd5b | 0.038450534 | -0.213254309 | 0.290155377 | 11.94009382 | 0.365106812 | 0.726365141 | 0.995570006 | -5.201150449 |
| **A_51_P336509** | Zfpm2 | -0.034893457 | -0.263339439 | 0.193552524 | 10.88070999 | -0.365064509 | 0.726395277 | 0.995570006 | -5.20116316 |
| **A_51_P493700** | Cep350 | -0.020956012 | -0.158197919 | 0.116285894 | 9.64186241 | -0.364948042 | 0.726478248 | 0.995570006 | -5.20119815 |
| **A_55_P2066543** | Orai2 | 0.017731974 | -0.098412496 | 0.133876444 | 8.270374708 | 0.364894811 | 0.726516171 | 0.995570006 | -5.201214138 |
| **A_55_P2108255** | Lgmn | 0.01722121 | -0.095604582 | 0.130047003 | 12.32184748 | 0.364808046 | 0.726577986 | 0.995570006 | -5.201240194 |
| **A_55_P2048655** | NA | 0.028077079 | -0.156007602 | 0.212161761 | 16.54457272 | 0.364538508 | 0.72677003 | 0.995570006 | -5.2013211 |
| **A_55_P2227321** | Ptprd | -0.019239817 | -0.145434629 | 0.106954995 | 8.593404635 | -0.364391778 | 0.726874583 | 0.995570006 | -5.201365119 |
| **A_55_P2099695** | Msantd4 | -0.020042275 | -0.151609778 | 0.111525229 | 11.53270239 | -0.364088962 | 0.727090377 | 0.995570006 | -5.201455911 |
| **A_55_P1964812** | Tfap2a | -0.024991108 | -0.1890835 | 0.139101285 | 7.791000535 | -0.364004027 | 0.727150908 | 0.995570006 | -5.201481364 |
| **A_52_P252737** | Rangap1 | 0.025956461 | -0.144485969 | 0.196398891 | 11.27770896 | 0.36397947 | 0.72716841 | 0.995570006 | -5.201488722 |
| **A_52_P282905** | Ces1b | -0.014742597 | -0.111549831 | 0.082064638 | 7.643874691 | -0.36397814 | 0.727169358 | 0.995570006 | -5.20148912 |
| **A_51_P177984** | Fam234a | -0.030160487 | -0.22821812 | 0.167897146 | 11.64990142 | -0.36396188 | 0.727180946 | 0.995570006 | -5.201493992 |
| **A_55_P1978465** | H2-Q8 | 0.018310569 | -0.102068612 | 0.13868975 | 8.082857089 | 0.363546185 | 0.727477239 | 0.995570006 | -5.20161847 |
| **A_52_P289213** | Notch2 | -0.026006852 | -0.197023588 | 0.145009885 | 10.4508165 | -0.363461401 | 0.727537676 | 0.995570006 | -5.201643842 |
| **A_55_P2159850** | NA | 0.057785922 | -0.322239968 | 0.437811812 | 12.93102686 | 0.363427674 | 0.727561718 | 0.995570006 | -5.201653933 |
| **A_51_P139069** | Wdr45 | -0.027285927 | -0.206788047 | 0.152216193 | 10.29293487 | -0.36331076 | 0.727645064 | 0.995570006 | -5.201688907 |
| **A_55_P2110032** | Akap7 | 0.022813967 | -0.127370462 | 0.172998397 | 9.441473696 | 0.363065547 | 0.727819883 | 0.995570006 | -5.201762226 |
| **A_55_P2045258** | Ehd1 | 0.050346002 | -0.281373063 | 0.382065066 | 9.75659121 | 0.362746882 | 0.728047095 | 0.995570006 | -5.201857438 |
| **A_55_P1989663** | Slco3a1 | 0.038033773 | -0.212755168 | 0.288822714 | 9.682549786 | 0.362468408 | 0.728245675 | 0.995570006 | -5.201940576 |
| **A_55_P1998947** | Trim16 | 0.0161543 | -0.090370206 | 0.122678805 | 8.238880762 | 0.362449698 | 0.728259017 | 0.995570006 | -5.201946159 |
| **A_55_P2091039** | NA | -0.017764248 | -0.134949488 | 0.099420992 | 8.461599145 | -0.362312274 | 0.728357024 | 0.995570006 | -5.201987163 |
| **A_66_P131604** | Otud4 | 0.038947541 | -0.217992344 | 0.295887426 | 12.12216311 | 0.362291087 | 0.728372134 | 0.995570006 | -5.201993483 |
| **A_51_P277745** | Abcc2 | 0.015589594 | -0.087279562 | 0.11845875 | 7.653588308 | 0.362208597 | 0.728430966 | 0.995570006 | -5.202018087 |
| **A_55_P2050727** | Fcrl5 | -0.024322993 | -0.184827862 | 0.136181876 | 10.75121961 | -0.362191231 | 0.728443352 | 0.995570006 | -5.202023266 |
| **A_55_P2049365** | NA | -0.019046039 | -0.144745333 | 0.106653255 | 12.34466311 | -0.362143728 | 0.728477232 | 0.995570006 | -5.202037432 |
| **A_52_P494930** | Fnip2 | 0.029033141 | -0.162584627 | 0.220650909 | 9.316498971 | 0.362132393 | 0.728485317 | 0.995570006 | -5.202040812 |
| **A_55_P2005135** | Tardbp | 0.03102176 | -0.17372417 | 0.235767691 | 9.245445333 | 0.362126487 | 0.728489529 | 0.995570006 | -5.202042572 |
| **A_51_P310821** | Hoxa5 | -0.018202254 | -0.138340329 | 0.101935821 | 7.727917464 | -0.36212093 | 0.728493493 | 0.995570006 | -5.20204423 |
| **A_55_P2043451** | NA | -0.018946127 | -0.144015197 | 0.106122942 | 12.19486696 | -0.362059268 | 0.728537473 | 0.995570006 | -5.202062614 |
| **A_55_P2109222** | Rps3 | -0.02497204 | -0.189826003 | 0.139881924 | 15.13702725 | -0.362046003 | 0.728546935 | 0.995570006 | -5.202066568 |
| **A_55_P1965219** | Syt6 | -0.018019372 | -0.13700526 | 0.100966515 | 8.16275302 | -0.361953968 | 0.728612581 | 0.995570006 | -5.202094001 |
| **A_55_P1953043** | E130311K1 | -0.021978976 | -0.167153573 | 0.123195621 | 8.996929414 | -0.361847813 | 0.728688301 | 0.995570006 | -5.202125635 |

| | | | | | | | | | |
|---|---|---|---|---|---|---|---|---|---|
| **A_52_P568781** | Sec63 | 0.018577286 | -0.104156183 | 0.141310754 | 8.540751698 | 0.361766527 | 0.728746286 | 0.995570006 | -5.202149852 |
| **A_55_P2044653** | Cyp2b10 | 0.015773432 | -0.088450656 | 0.11999752 | 7.671766248 | 0.361715597 | 0.728782617 | 0.995570006 | -5.202165022 |
| **A_55_P1975120** | NA | -0.018666692 | -0.142012137 | 0.104678753 | 8.912466748 | -0.361704047 | 0.728790856 | 0.995570006 | -5.202168462 |
| **A_55_P2184414** | Zfp330 | 0.014900102 | -0.08361773 | 0.113417934 | 10.89777447 | 0.361479383 | 0.72895113 | 0.995570006 | -5.202235357 |
| **A_52_P597775** | Gprc5a | -0.019006752 | -0.144784027 | 0.106770523 | 9.228297289 | -0.361172653 | 0.729169974 | 0.995570006 | -5.202326623 |
| **A_55_P2071384** | Crh | -0.014859186 | -0.113213078 | 0.083494705 | 7.655425086 | -0.361087635 | 0.729230636 | 0.995570006 | -5.202351906 |
| **A_52_P113830** | 1600025M1 | -0.034962444 | -0.26640876 | 0.196483872 | 7.836614016 | -0.361044421 | 0.729261472 | 0.995570006 | -5.202364755 |
| **A_55_P2061991** | Pnldc1 | -0.039528473 | -0.301208555 | 0.22215161 | 8.30025428 | -0.361034328 | 0.729268673 | 0.995570006 | -5.202367756 |
| **A_52_P504743** | Cdyl | -0.025139302 | -0.191623618 | 0.141345014 | 8.817414318 | -0.360901776 | 0.729363259 | 0.995570006 | -5.202407159 |
| **A_55_P2106745** | Trim63 | -0.075266513 | -0.573750704 | 0.423217679 | 9.882686274 | -0.360877276 | 0.729380742 | 0.995570006 | -5.202414441 |
| **A_55_P2008898** | NA | 0.028633064 | -0.161113458 | 0.218379586 | 13.27869368 | 0.360664285 | 0.72953274 | 0.995570006 | -5.202477722 |
| **A_52_P582150** | Gosr2 | 0.018769627 | -0.105613664 | 0.143152918 | 11.25421852 | 0.360663942 | 0.729532985 | 0.995570006 | -5.202477824 |
| **A_55_P2045952** | Mysm1 | 0.027916401 | -0.157232798 | 0.213065599 | 8.828695993 | 0.360368417 | 0.729743903 | 0.995570006 | -5.202565569 |
| **A_51_P350666** | Peli2 | 0.016740631 | -0.094292498 | 0.127773761 | 8.448489559 | 0.360353198 | 0.729754766 | 0.995570006 | -5.202570085 |
| **A_65_P13270** | Csnk1g1 | 0.018893778 | -0.106420729 | 0.144208286 | 8.779646944 | 0.360351719 | 0.729755822 | 0.995570006 | -5.202570525 |
| **A_55_P2011169** | NA | 0.017294651 | -0.097417627 | 0.132006929 | 8.501961543 | 0.36033881 | 0.729765035 | 0.995570006 | -5.202574356 |
| **A_55_P2169218** | NA | 0.087792425 | -0.49454182 | 0.67012667 | 10.27346914 | 0.360324545 | 0.729775217 | 0.995570006 | -5.202578589 |
| **A_55_P2157902** | Igsf10 | -0.024441886 | -0.186647732 | 0.137763959 | 8.486147502 | -0.360144969 | 0.729903396 | 0.995570006 | -5.202631867 |
| **A_52_P682745** | Dock4 | 0.030937706 | -0.174439417 | 0.236314828 | 8.931224311 | 0.360035371 | 0.729981631 | 0.995570006 | -5.202664371 |
| **A_66_P116195** | Psmd13 | -0.049268551 | -0.376366441 | 0.277829339 | 12.23043836 | -0.359998907 | 0.730007661 | 0.995570006 | -5.202675184 |
| **A_55_P2065716** | Glud1 | 0.032536175 | -0.183482539 | 0.248554889 | 11.82410368 | 0.359984876 | 0.730017677 | 0.995570006 | -5.202679344 |
| **A_55_P2065327** | Rpl27a | -0.019289627 | -0.147366209 | 0.108786955 | 15.33302883 | -0.359967452 | 0.730030116 | 0.995570006 | -5.20268451 |
| **A_52_P549977** | Fam32a | -0.031615305 | -0.241546759 | 0.17831615 | 8.855754276 | -0.359939069 | 0.730050377 | 0.995570006 | -5.202692925 |
| **A_55_P2085806** | Paxx | -0.025227767 | -0.192767223 | 0.142311689 | 11.01948419 | -0.359890885 | 0.730084775 | 0.995570006 | -5.202707208 |
| **A_66_P100932** | Tmem256 | -0.034929316 | -0.266927311 | 0.197068679 | 13.59733187 | -0.359844593 | 0.730117823 | 0.995570006 | -5.202720929 |
| **A_55_P2120189** | Gas2l3 | -0.027987465 | -0.213902732 | 0.157927803 | 9.790412354 | -0.359797079 | 0.730151743 | 0.995570006 | -5.202735011 |
| **A_55_P2068849** | Nudc | -0.028905431 | -0.220960747 | 0.163149886 | 11.1971283 | -0.359718059 | 0.730208158 | 0.995570006 | -5.202758426 |
| **A_55_P2147310** | Mrtfa | 0.024394783 | -0.137713654 | 0.186503219 | 10.9716025 | 0.3596669 | 0.730244683 | 0.995570006 | -5.202773582 |
| **A_55_P2184897** | Lbr | 0.028149143 | -0.158955824 | 0.215254111 | 8.216594009 | 0.359574594 | 0.730310586 | 0.995570006 | -5.202800924 |
| **A_55_P1998797** | NA | -0.035311195 | -0.270047414 | 0.199425023 | 11.1501952 | -0.359535227 | 0.730338694 | 0.995570006 | -5.202812583 |
| **A_55_P1980416** | Zkscan1 | -0.024766551 | -0.189421021 | 0.139887919 | 11.31645794 | -0.359501851 | 0.730362524 | 0.995570006 | -5.202822466 |
| **A_55_P1960326** | NA | -0.024119441 | -0.184537186 | 0.136298304 | 11.2102531 | -0.359355225 | 0.730467218 | 0.995570006 | -5.202865876 |
| **A_55_P2186027** | Gapdh | 0.036565097 | -0.206641253 | 0.279771448 | 17.53298449 | 0.359336186 | 0.730480813 | 0.995570006 | -5.202871512 |
| **A_52_P621617** | Dnajb4 | -0.017157797 | -0.131305935 | 0.09699034 | 8.80289785 | -0.359254188 | 0.730539365 | 0.995570006 | -5.202895779 |
| **A_55_P2033917** | Fut10 | -0.018757458 | -0.143554284 | 0.106039368 | 9.544840511 | -0.359235771 | 0.730552516 | 0.995570006 | -5.202901229 |
| **A_55_P1998957** | Oas1c | -0.022404518 | -0.171509826 | 0.126700789 | 8.229869332 | -0.359129958 | 0.730628076 | 0.995570006 | -5.202932536 |
| **A_52_P614472** | Rpusd1 | 0.023052117 | -0.130394097 | 0.17649833 | 8.501170439 | 0.359057302 | 0.730679961 | 0.995570006 | -5.202954027 |
| **A_55_P2038525** | C3 | -0.133303275 | -1.020691138 | 0.754084588 | 10.29876981 | -0.359034736 | 0.730696077 | 0.995570006 | -5.202960701 |
| **A_51_P479902** | Mars | 0.033681844 | -0.190555842 | 0.257919529 | 9.80280185 | 0.359001603 | 0.730719739 | 0.995570006 | -5.2029705 |
| **A_55_P2191002** | 2610027K0 | -0.015364493 | -0.117659448 | 0.086930461 | 8.08273318 | -0.358982408 | 0.730733447 | 0.995570006 | -5.202976176 |
| **A_55_P2138961** | Tpk1 | 0.019498915 | -0.110354158 | 0.149351988 | 8.868986582 | 0.358894952 | 0.730795905 | 0.995570006 | -5.203002034 |
| **A_51_P172054** | Gas6 | 0.032683051 | -0.184981822 | 0.250347924 | 14.22473743 | 0.358875142 | 0.730810052 | 0.995570006 | -5.203007891 |
| **A_52_P300416** | Dcp1a | 0.016963155 | -0.096029442 | 0.129955753 | 8.919497609 | 0.358811025 | 0.730855844 | 0.995570006 | -5.203026844 |
| **A_52_P430169** | Dip2b | -0.023432715 | -0.179522987 | 0.132657557 | 9.942147484 | -0.358802864 | 0.730861673 | 0.995570006 | -5.203029256 |
| **A_51_P120398** | Asb17 | -0.017685295 | -0.135533165 | 0.100162575 | 7.749556332 | -0.358673838 | 0.730953826 | 0.995570006 | -5.139099908 |
| **A_55_P1978615** | Rpl17 | -0.023675934 | -0.181460208 | 0.134108341 | 15.10715288 | -0.35863488 | 0.730981651 | 0.995570006 | -5.203078895 |
| **A_55_P1987291** | Plppr5 | 0.051148396 | -0.289736202 | 0.392032994 | 8.609300549 | 0.358619393 | 0.730992713 | 0.995570006 | -5.203083471 |
| **A_55_P2025840** | Spred2 | -0.018959654 | -0.145334031 | 0.107414724 | 8.821451064 | -0.358575415 | 0.731024125 | 0.995570006 | -5.203096462 |

| | | | | | | | | | |
|---|---|---|---|---|---|---|---|---|---|
| **A_51_P204402** | Shcbp1 | -0.037781876 | -0.289634286 | 0.214070534 | 10.45013226 | -0.358547369 | 0.731044158 | 0.995570006 | -5.203104746 |
| **A_55_P1973563** | Inava | 0.015772355 | -0.089368312 | 0.120913022 | 7.742915023 | 0.3585378 | 0.731050992 | 0.995570006 | -5.203107573 |
| **A_51_P244856** | Atp2a1 | -0.059844231 | -0.458784719 | 0.339096256 | 9.090433793 | -0.358528193 | 0.731057855 | 0.995570006 | -5.20311041 |
| **A_55_P2344598** | E330037I1! | 0.017435251 | -0.098845414 | 0.133715915 | 7.908541419 | 0.358368494 | 0.731171929 | 0.995570006 | -5.203157567 |
| **A_55_P2275057** | Nrap | 0.069520999 | -0.394153091 | 0.53319509 | 8.831977567 | 0.358354043 | 0.731182252 | 0.995570006 | -5.203161834 |
| **A_55_P2138878** | Lrfn4 | 0.029085708 | -0.164904808 | 0.223076223 | 11.29770046 | 0.35835071 | 0.731184632 | 0.995570006 | -5.203162818 |
| **A_66_P122110** | Mitd1 | 0.015447151 | -0.0876162 | 0.118510502 | 8.437967714 | 0.358222832 | 0.731275982 | 0.995570006 | -5.203200562 |
| **A_55_P2179027** | Gem | 0.019225717 | -0.10907575 | 0.147527184 | 8.102890785 | 0.358145959 | 0.731330899 | 0.995570006 | -5.203223246 |
| **A_51_P315595** | NA | -0.026851469 | -0.206044166 | 0.152341228 | 17.41892405 | -0.358143341 | 0.73133277 | 0.995570006 | -5.203224019 |
| **A_52_P245415** | AW554918 | 0.019909689 | -0.112959646 | 0.152779025 | 8.982003363 | 0.358136721 | 0.731337499 | 0.995570006 | -5.203225972 |
| **A_55_P2010931** | Cnn2 | 0.05028403 | -0.285480369 | 0.38604843 | 10.39331964 | 0.357935328 | 0.731481379 | 0.995570006 | -5.203285376 |
| **A_55_P1974554** | Eif4b | 0.018380293 | -0.104367382 | 0.141127969 | 11.49995386 | 0.357888949 | 0.731514515 | 0.995570006 | -5.203299052 |
| **A_55_P2119145** | NA | -0.015160073 | -0.116405765 | 0.086085619 | 7.681517554 | -0.357877066 | 0.731523006 | 0.995570006 | -5.203302556 |
| **A_51_P291501** | Ccno | 0.015978878 | -0.090758965 | 0.12271672 | 7.771375627 | 0.357797243 | 0.731580038 | 0.995570006 | -5.203326088 |
| **A_52_P101982** | Tmem129 | -0.019652205 | -0.150957498 | 0.111653088 | 10.78739737 | -0.357715849 | 0.731638194 | 0.995570006 | -5.203350079 |
| **A_55_P1972719** | Pmm1 | -0.026082059 | -0.200406805 | 0.148242687 | 8.44127995 | -0.357595433 | 0.731724235 | 0.995570006 | -5.203385562 |
| **A_51_P273556** | Fam83d | -0.019968623 | -0.153456248 | 0.113519002 | 9.577566088 | -0.357533099 | 0.731768777 | 0.995570006 | -5.203403925 |
| **A_51_P348652** | Spast | 0.041697986 | -0.237153748 | 0.320549721 | 11.38000797 | 0.357396969 | 0.731866054 | 0.995570006 | -5.203444018 |
| **A_55_P1955876** | Ifitm7 | -0.022353112 | -0.171844216 | 0.127137992 | 8.900501057 | -0.357381266 | 0.731877276 | 0.995570006 | -5.203448642 |
| **A_55_P2006133** | Tmc6 | -0.037039743 | -0.284800283 | 0.210720796 | 8.436056782 | -0.357309829 | 0.731928326 | 0.995570006 | -5.203469675 |
| **A_55_P2072288** | Tmem167b | -0.017062474 | -0.131218642 | 0.097093693 | 9.90643939 | -0.357233162 | 0.731983116 | 0.995570006 | -5.203492243 |
| **A_55_P2049473** | Sh3bp5 | 0.022370626 | -0.127355544 | 0.172096795 | 8.823767916 | 0.357099753 | 0.73207846 | 0.995570006 | -5.203531503 |
| **A_52_P32081** | Crbn | -0.021678864 | -0.16677767 | 0.123419942 | 9.052746459 | -0.357093397 | 0.732083003 | 0.995570006 | -5.203533373 |
| **A_51_P230977** | Cdc42 | -0.020055212 | -0.154294762 | 0.114184338 | 13.83263786 | -0.35707209 | 0.732098231 | 0.995570006 | -5.203539642 |
| **A_51_P477676** | Mcph1 | -0.017332265 | -0.133347539 | 0.098683008 | 8.737546629 | -0.357066657 | 0.732102114 | 0.995570006 | -5.20354124 |
| **A_51_P373890** | Guca1b | -0.029911164 | -0.2301375 | 0.170315172 | 8.738554251 | -0.357043595 | 0.732118597 | 0.995570006 | -5.203548025 |
| **A_52_P273169** | Sh3d19 | 0.023635072 | -0.134579249 | 0.181849393 | 8.350931441 | 0.357042789 | 0.732119173 | 0.995570006 | -5.203548262 |
| **A_55_P2037343** | H2-Q10 | -0.02374519 | -0.182704199 | 0.135213819 | 7.908818189 | -0.357025829 | 0.732131294 | 0.995570006 | -5.203553251 |
| **A_51_P117881** | Leap2 | -0.017264119 | -0.132848378 | 0.09832014 | 7.698090483 | -0.35698902 | 0.732157603 | 0.995570006 | -5.203564079 |
| **A_55_P2075636** | Epc2 | 0.031117335 | -0.177223902 | 0.239458571 | 9.27390227 | 0.356973752 | 0.732168515 | 0.995570006 | -5.20356857 |
| **A_55_P2019909** | Tmem35b | 0.019762507 | -0.112558567 | 0.152083582 | 9.127962447 | 0.356962138 | 0.732176816 | 0.995570006 | -5.203571986 |
| **A_55_P2102857** | Krtcap3 | 0.020950084 | -0.119350654 | 0.161250821 | 8.073549155 | 0.356890462 | 0.732228046 | 0.995570006 | -5.203593066 |
| **A_51_P243122** | Sec14l3 | 0.017838454 | -0.101630291 | 0.137307198 | 7.714991316 | 0.356871639 | 0.7322415 | 0.995570006 | -5.203598601 |
| **A_55_P2079659** | NA | 0.014242234 | -0.081142431 | 0.109626899 | 7.650724116 | 0.356868892 | 0.732243463 | 0.995570006 | -5.203599409 |
| **A_51_P156800** | Mob2 | 0.030050511 | -0.171216699 | 0.231317721 | 10.06364051 | 0.356851859 | 0.732255638 | 0.995570006 | -5.203604417 |
| **A_55_P2067777** | Cplane1 | 0.023732219 | -0.135345717 | 0.182810155 | 9.628055935 | 0.356564029 | 0.732461381 | 0.995570006 | -5.203689017 |
| **A_51_P357877** | Fbxo46 | 0.017818528 | -0.101620931 | 0.137257987 | 9.088874389 | 0.356560415 | 0.732463965 | 0.995570006 | -5.203690079 |
| **A_55_P1987914** | Map4k4 | 0.033987016 | -0.193878503 | 0.261852534 | 10.26313149 | 0.35648688 | 0.732516532 | 0.995570006 | -5.203711682 |
| **A_52_P675887** | NA | 0.016586727 | -0.094639008 | 0.127812463 | 15.44405558 | 0.356422041 | 0.732562884 | 0.995570006 | -5.203730727 |
| **A_66_P123582** | Mog | -0.023418006 | -0.180467315 | 0.133631303 | 8.26011567 | -0.356387958 | 0.73258725 | 0.995570006 | -5.203740737 |
| **A_51_P291749** | Pecr | -0.023214943 | -0.178904414 | 0.132474527 | 10.87821979 | -0.356383443 | 0.732590478 | 0.995570006 | -5.203742063 |
| **A_55_P2118810** | NA | 0.052407347 | -0.299075329 | 0.403890024 | 13.08869824 | 0.356366921 | 0.73260229 | 0.995570006 | -5.203746915 |
| **A_51_P484158** | Steap1 | 0.017903621 | -0.102198425 | 0.138005668 | 7.991943734 | 0.356286699 | 0.732659642 | 0.995570006 | -5.20377047 |
| **A_55_P1974845** | Pde1a | -0.020454662 | -0.157721338 | 0.116812014 | 9.063384967 | -0.356152769 | 0.732755395 | 0.995570006 | -5.203809784 |
| **A_51_P423518** | Amph | -0.018625638 | -0.143619222 | 0.106367946 | 7.89560838 | -0.356149694 | 0.732757593 | 0.995570006 | -5.203810686 |
| **A_55_P2009582** | Nkg7 | -0.022022725 | -0.169816219 | 0.12577077 | 9.172154888 | -0.35614337 | 0.732762115 | 0.995570006 | -5.203812542 |
| **A_55_P2162369** | Cacfd1 | 0.019374053 | -0.11065173 | 0.149399836 | 8.550486964 | 0.356123092 | 0.732776613 | 0.995570006 | -5.203818493 |
| **A_55_P2154943** | NA | -0.01581662 | -0.122003309 | 0.090370069 | 7.665593854 | -0.356002247 | 0.732863016 | 0.995570006 | -5.203853951 |

| | | | | | | | | | |
|---|---|---|---|---|---|---|---|---|---|
| **A_55_P2148116** | Tsr2 | 0.027879825 | -0.159349348 | 0.215108998 | 8.626906888 | 0.35589809 | 0.732937491 | 0.995570006 | -5.203884504 |
| **A_55_P2127174** | Zfp628 | 0.023732502 | -0.135729105 | 0.183194109 | 17.63477105 | 0.355710367 | 0.733071726 | 0.995570006 | -5.203939547 |
| **A_52_P297457** | 0610030E2 | -0.022191751 | -0.171301433 | 0.126917932 | 9.084220185 | -0.355708997 | 0.733072706 | 0.995570006 | -5.203939949 |
| **A_55_P2037982** | Agap1 | -0.02778663 | -0.214496854 | 0.158923595 | 10.79890462 | -0.355694298 | 0.733083217 | 0.995570006 | -5.203944257 |
| **A_55_P2057469** | Caskin1 | -0.015025658 | -0.115990538 | 0.085939221 | 7.697652443 | -0.35569053 | 0.733085912 | 0.995570006 | -5.203945362 |
| **A_55_P2003803** | Scml2 | -0.029722405 | -0.229474445 | 0.170029635 | 8.993902928 | -0.355632839 | 0.733127167 | 0.995570006 | -5.203962271 |
| **A_55_P2106569** | NA | 0.044656334 | -0.255531838 | 0.344844505 | 16.58795386 | 0.355548311 | 0.733187616 | 0.995570006 | -5.203987042 |
| **A_51_P178828** | Mbl2 | -0.015626547 | -0.120689297 | 0.089436203 | 7.710881185 | -0.355486719 | 0.733231663 | 0.995570006 | -5.204005088 |
| **A_55_P1983262** | Gm4026 | -0.027007043 | -0.208588102 | 0.154574017 | 8.5893287 | -0.355480364 | 0.733236208 | 0.995570006 | -5.20400695 |
| **A_55_P2090060** | Mab21l3 | -0.015915099 | -0.122937062 | 0.091106865 | 7.891081868 | -0.35542302 | 0.733277219 | 0.995570006 | -5.204023748 |
| **A_51_P289987** | Psmg3 | 0.023767468 | -0.136064461 | 0.183599396 | 10.86451291 | 0.355409065 | 0.733287199 | 0.995570006 | -5.204027835 |
| **A_52_P91274** | 1700018G( | 0.015192629 | -0.086980292 | 0.117365551 | 7.635843503 | 0.355390867 | 0.733300214 | 0.995570006 | -5.204033165 |
| **A_55_P2091120** | Kbtbd12 | -0.049089399 | -0.379227389 | 0.281048591 | 10.11122969 | -0.355386843 | 0.733303092 | 0.995570006 | -5.204034344 |
| **A_55_P2090194** | NA | 0.022328513 | -0.127836181 | 0.172493207 | 14.18218134 | 0.355386638 | 0.733303239 | 0.995570006 | -5.204034404 |
| **A_55_P2052754** | Pcm1 | 0.026802185 | -0.153460552 | 0.207064922 | 9.473787863 | 0.355363949 | 0.733319465 | 0.995570006 | -5.204041049 |
| **A_55_P2107901** | Dlk2 | -0.021129862 | -0.163255922 | 0.120996197 | 8.151424948 | -0.355330171 | 0.733343623 | 0.995570006 | -5.204050941 |
| **A_55_P1970910** | Stard13 | 0.025976264 | -0.148759342 | 0.200711869 | 9.137792361 | 0.355307525 | 0.73335982 | 0.995570006 | -5.204057573 |
| **A_51_P189733** | Sult2a8 | 0.025249854 | -0.144635053 | 0.19513476 | 8.72412303 | 0.355232915 | 0.733413182 | 0.995570006 | -5.204079418 |
| **A_52_P244193** | Cd24a | -0.046909029 | -0.362657799 | 0.268839741 | 11.20101081 | -0.355078116 | 0.733523903 | 0.995570006 | -5.204124729 |
| **A_55_P2113598** | Ikzf3 | 0.016160003 | -0.092628429 | 0.124948436 | 7.826170752 | 0.355032298 | 0.733556676 | 0.995570006 | -5.204138137 |
| **A_55_P2079927** | Ccdc68 | -0.015554439 | -0.12027878 | 0.089169901 | 7.828953265 | -0.354989783 | 0.733587087 | 0.995570006 | -5.204150576 |
| **A_51_P297336** | Chmp7 | -0.031454584 | -0.243277588 | 0.18036842 | 11.19089313 | -0.354911408 | 0.73364315 | 0.995570006 | -5.204173504 |
| **A_55_P1961414** | NA | -0.034570624 | -0.267416364 | 0.198275116 | 15.04705254 | -0.354852654 | 0.733685178 | 0.995570006 | -5.204190689 |
| **A_52_P149577** | Rab25 | -0.015560328 | -0.120399498 | 0.089278841 | 7.660335142 | -0.354735226 | 0.733769182 | 0.995570006 | -5.204225028 |
| **A_52_P556099** | Oip5 | -0.026927449 | -0.208420453 | 0.154565556 | 8.537228573 | -0.354604672 | 0.733862579 | 0.995570006 | -5.204263192 |
| **A_55_P2029687** | Hmox1 | -0.018547495 | -0.143577753 | 0.106482763 | 7.957810237 | -0.354551458 | 0.733900649 | 0.995570006 | -5.204278744 |
| **A_52_P106620** | Tnfrsf11b | 0.024600487 | -0.141239479 | 0.190440453 | 7.837492758 | 0.354538687 | 0.733909786 | 0.995570006 | -5.204282476 |
| **A_55_P2121076** | Cul2 | -0.020809096 | -0.161119161 | 0.119500969 | 8.680928756 | -0.354465128 | 0.733962413 | 0.995570006 | -5.204303969 |
| **A_65_P15412** | Usp13 | -0.048641542 | -0.376836839 | 0.279553754 | 11.09061385 | -0.354229003 | 0.734131357 | 0.995570006 | -5.204372934 |
| **A_51_P461724** | Atp6ap2 | 0.022242121 | -0.127836339 | 0.172320581 | 9.911594833 | 0.354215016 | 0.734141365 | 0.995570006 | -5.204377018 |
| **A_55_P2102210** | NA | -0.026593049 | -0.206050833 | 0.152864735 | 11.42172511 | -0.354172606 | 0.73417171 | 0.995570006 | -5.2043894 |
| **A_55_P1972157** | Espnl | 0.016382102 | -0.094204444 | 0.126968649 | 8.459710592 | 0.354059677 | 0.734252517 | 0.995570006 | -5.204422363 |
| **A_55_P2005330** | Egln2 | 0.041164676 | -0.236880922 | 0.319210275 | 10.34280105 | 0.353848869 | 0.734403371 | 0.995570006 | -5.204483869 |
| **A_55_P1977787** | NA | 0.078623918 | -0.452467263 | 0.609715099 | 8.701421875 | 0.353830059 | 0.734416832 | 0.995570006 | -5.204489355 |
| **A_55_P2008006** | Ap1g1 | -0.018514325 | -0.143620329 | 0.106591679 | 9.708646293 | -0.353703094 | 0.734507695 | 0.995570006 | -5.20452638 |
| **A_52_P73848** | Clint1 | 0.020271425 | -0.116732394 | 0.157275244 | 8.330494578 | 0.353639477 | 0.734553225 | 0.995570006 | -5.204544927 |
| **A_55_P2092826** | Anxa1 | 0.028243627 | -0.16269465 | 0.219181904 | 11.78613127 | 0.353538408 | 0.73462556 | 0.995570006 | -5.204574386 |
| **A_55_P2279977** | NA | -0.023907492 | -0.185670694 | 0.137855709 | 8.238240916 | -0.353234748 | 0.734842908 | 0.995570006 | -5.204662847 |
| **A_51_P221949** | Csnk1g3 | 0.036616361 | -0.211143728 | 0.284376451 | 11.30442478 | 0.353226247 | 0.734848993 | 0.995570006 | -5.204665322 |
| **A_55_P1961335** | Ctsk | 0.020664409 | -0.119171558 | 0.160500377 | 10.44235341 | 0.353193935 | 0.734872122 | 0.995570006 | -5.204674731 |
| **A_55_P1972234** | NA | 0.01625115 | -0.093727707 | 0.126230006 | 15.82921685 | 0.353170173 | 0.734889131 | 0.995570006 | -5.204681649 |
| **A_51_P140350** | Nadsyn1 | -0.017025408 | -0.132262347 | 0.098211532 | 9.508041642 | -0.353114003 | 0.73492934 | 0.995570006 | -5.204698002 |
| **A_55_P2157478** | NA | 0.021202559 | -0.122318161 | 0.164723279 | 8.614697154 | 0.353087875 | 0.734948043 | 0.995570006 | -5.204705607 |
| **A_55_P2025855** | Hnrnpul1 | 0.044411001 | -0.256252092 | 0.345074093 | 9.545732409 | 0.353036469 | 0.734984842 | 0.995570006 | -5.20472057 |
| **A_55_P2096887** | Steap3 | -0.022250274 | -0.172888446 | 0.128387897 | 8.737510758 | -0.353028255 | 0.734990722 | 0.995570006 | -5.20472296 |
| **A_55_P2182077** | NA | -0.035164349 | -0.273291724 | 0.202963026 | 16.63955742 | -0.352941223 | 0.735053026 | 0.995570006 | -5.204748286 |
| **A_55_P2218334** | 9430011C2 | 0.05510588 | -0.318314174 | 0.428525934 | 9.091295917 | 0.352703231 | 0.73522341 | 0.995570006 | -5.204817512 |
| **A_52_P523712** | Tbx5 | 0.060865558 | -0.35167889 | 0.473410005 | 10.86250363 | 0.352622476 | 0.735281228 | 0.995570006 | -5.204840991 |

| | | | | | | | | | |
|---|---|---|---|---|---|---|---|---|---|
| **A_51_P519364** | Fam83f | -0.036416651 | -0.283258599 | 0.210425297 | 8.818879388 | -0.352606387 | 0.735292748 | 0.995570006 | -5.204845669 |
| **A_55_P2051521** | NA | 0.0172062 | -0.099462077 | 0.133874478 | 8.497301257 | 0.352485554 | 0.735379264 | 0.995570006 | -5.204880789 |
| **A_55_P2163098** | Akr1c18 | -0.019705209 | -0.153350849 | 0.11394043 | 7.740164065 | -0.352399586 | 0.73544082 | 0.995570006 | -5.20490577 |
| **A_55_P2035315** | Rasgef1b | 0.035051754 | -0.2027308 | 0.272834307 | 11.07265533 | 0.352321297 | 0.735496878 | 0.995570006 | -5.204928513 |
| **A_55_P1994388** | Lsm14a | -0.023697622 | -0.184502113 | 0.137106869 | 10.99540675 | -0.352221384 | 0.735568424 | 0.995570006 | -5.204957532 |
| **A_55_P2085835** | 5330417C2 | -0.015652772 | -0.121875563 | 0.090570019 | 8.151577137 | -0.352194584 | 0.735587616 | 0.995570006 | -5.204965314 |
| **A_55_P2114993** | Actg1 | -0.025382269 | -0.197643056 | 0.146878518 | 15.25657844 | -0.352170643 | 0.73560476 | 0.995570006 | -5.204972266 |
| **A_55_P1999829** | Thoc7 | 0.019426692 | -0.112439117 | 0.1512925 | 13.46636094 | 0.352107908 | 0.735649686 | 0.995570006 | -5.20499048 |
| **A_55_P1962279** | Plac1 | 0.024024569 | -0.139059223 | 0.18710836 | 8.670506389 | 0.352090195 | 0.73566237 | 0.995570006 | -5.204995622 |
| **A_55_P2047361** | Ppih | 0.025658326 | -0.148540314 | 0.199856967 | 9.347809648 | 0.352040551 | 0.735697922 | 0.995570006 | -5.205010033 |
| **A_52_P550147** | NA | 0.042108352 | -0.24378464 | 0.328001343 | 8.683741698 | 0.352025289 | 0.735708852 | 0.995570006 | -5.205014463 |
| **A_55_P2030046** | Barx2 | -0.029414822 | -0.229181784 | 0.17035214 | 8.547257341 | -0.351926275 | 0.735779762 | 0.995570006 | -5.205043197 |
| **A_55_P2003513** | Hsph1 | 0.019995121 | -0.11580101 | 0.155791251 | 9.498758773 | 0.351921474 | 0.735783201 | 0.995570006 | -5.205044591 |
| **A_51_P221632** | 1700037H0 | 0.035612702 | -0.206266992 | 0.277492396 | 12.33194692 | 0.351896256 | 0.735801261 | 0.995570006 | -5.205051908 |
| **A_55_P2108579** | Phkg2 | 0.022281888 | -0.129083381 | 0.173647157 | 10.58675205 | 0.351831632 | 0.735847545 | 0.995570006 | -5.205070656 |
| **A_55_P2100864** | Tma7 | 0.025409426 | -0.147204417 | 0.198023269 | 14.42962017 | 0.351826349 | 0.735851329 | 0.995570006 | -5.205072189 |
| **A_51_P363642** | 1700088E0 | 0.022697297 | -0.131542172 | 0.176936767 | 8.398745624 | 0.351712458 | 0.735932899 | 0.995570006 | -5.205105222 |
| **A_52_P227937** | Cystm1 | -0.025544569 | -0.199135028 | 0.14804589 | 10.06457024 | -0.351707684 | 0.735936319 | 0.995570006 | -5.205106607 |
| **A_55_P2006499** | Esrrg | -0.028178979 | -0.219679764 | 0.163321806 | 9.729928034 | -0.351693078 | 0.735946781 | 0.995570006 | -5.205110843 |
| **A_51_P199866** | Pin1 | -0.027959044 | -0.217987334 | 0.162069246 | 11.90658918 | -0.351652076 | 0.735976148 | 0.995570006 | -5.205122732 |
| **A_55_P2018482** | Adh5 | 0.020616105 | -0.119507393 | 0.160739603 | 8.66083866 | 0.351645276 | 0.735981018 | 0.995570006 | -5.205124703 |
| **A_52_P305182** | Ccdc32 | 0.024882559 | -0.144266242 | 0.194031361 | 10.31842532 | 0.351588978 | 0.736021342 | 0.995570006 | -5.205141026 |
| **A_51_P372895** | Zfp433 | -0.021855576 | -0.170458423 | 0.126747272 | 9.346552683 | -0.351515315 | 0.736074106 | 0.995570006 | -5.205162379 |
| **A_55_P2032558** | NA | 0.01921484 | -0.111476393 | 0.149906074 | 14.57450034 | 0.351398133 | 0.736158045 | 0.995570006 | -5.205196338 |
| **A_55_P2035972** | Mark3 | 0.02818058 | -0.163551366 | 0.219912526 | 10.87582373 | 0.351289021 | 0.736236206 | 0.995570006 | -5.205227949 |
| **A_55_P2174576** | Nova1 | -0.02252624 | -0.175848113 | 0.130795633 | 7.934204083 | -0.351150852 | 0.736335188 | 0.995570006 | -5.205267964 |
| **A_52_P294510** | Fgl1 | 0.014534258 | -0.084414257 | 0.113482773 | 7.699139297 | 0.351069192 | 0.73639369 | 0.995570006 | -5.205291607 |
| **A_55_P2127258** | Dok7 | 0.014834705 | -0.08617851 | 0.11584792 | 8.608337582 | 0.351002216 | 0.736441674 | 0.995570006 | -5.205310995 |
| **A_55_P1970597** | Actg1 | 0.035863573 | -0.208352433 | 0.280079579 | 16.63050493 | 0.350985002 | 0.736454006 | 0.995570006 | -5.205315977 |
| **A_55_P2071028** | NA | -0.02491356 | -0.194574247 | 0.144747128 | 11.2821471 | -0.350964908 | 0.736468403 | 0.995570006 | -5.205321792 |
| **A_55_P2078048** | BC004004 | -0.018149432 | -0.141756998 | 0.105458134 | 12.79385299 | -0.350935354 | 0.736489577 | 0.995570006 | -5.205330346 |
| **A_55_P2043367** | Apol9a | -0.017579522 | -0.13734696 | 0.102187917 | 8.314047451 | -0.350814411 | 0.736576229 | 0.995570006 | -5.205365339 |
| **A_52_P510877** | Bcl2l1 | 0.016784812 | -0.097577372 | 0.131146997 | 7.937012507 | 0.350786747 | 0.73659605 | 0.995570006 | -5.205373342 |
| **A_51_P345340** | Upk3bl | 0.015882163 | -0.092353451 | 0.124117776 | 7.731778867 | 0.350710355 | 0.736650786 | 0.995570006 | -5.205395438 |
| **A_55_P1991075** | Lmf1 | -0.034405778 | -0.268902488 | 0.200090932 | 12.02502897 | -0.350674161 | 0.73667672 | 0.995570006 | -5.205405905 |
| **A_55_P1955577** | Ccm2 | -0.017276294 | -0.135058687 | 0.1005061 | 12.40433326 | -0.350573701 | 0.736748705 | 0.995570006 | -5.205434953 |
| **A_55_P2182736** | Limd2 | 0.028578886 | -0.166276023 | 0.223433794 | 8.703448185 | 0.350544434 | 0.736769676 | 0.995570006 | -5.205443413 |
| **A_55_P2122588** | Ubr2 | 0.023228365 | -0.13521803 | 0.181674759 | 9.164638167 | 0.35038492 | 0.736883983 | 0.995570006 | -5.205489516 |
| **A_55_P2100715** | Auh | 0.024534653 | -0.142833362 | 0.191902667 | 8.366086522 | 0.350361664 | 0.736900649 | 0.995570006 | -5.205496236 |
| **A_55_P2123002** | Tet1 | -0.030490546 | -0.238499386 | 0.177518295 | 8.825268177 | -0.350342268 | 0.736914548 | 0.995570006 | -5.20550184 |
| **A_55_P2081055** | 1700056E2 | -0.024628066 | -0.192644552 | 0.14338842 | 9.04235713 | -0.350338237 | 0.736917437 | 0.995570006 | -5.205503004 |
| **A_55_P2280581** | A130088B0 | 0.036689311 | -0.213618442 | 0.286997063 | 8.449459051 | 0.350327627 | 0.736925041 | 0.995570006 | -5.20550607 |
| **A_51_P403760** | Braf | -0.03436563 | -0.269066983 | 0.200335722 | 9.221401676 | -0.349959555 | 0.737188832 | 0.995570006 | -5.205612357 |
| **A_51_P417016** | Srsf3 | 0.027441478 | -0.160010441 | 0.214893397 | 8.993002928 | 0.349886137 | 0.737241454 | 0.995570006 | -5.205633545 |
| **A_51_P389751** | Relb | -0.02822666 | -0.221057439 | 0.16460412 | 8.887031353 | -0.349858365 | 0.73726136 | 0.995570006 | -5.205641558 |
| **A_55_P1952359** | Anapc5 | 0.03165042 | -0.184615705 | 0.247916545 | 14.78702807 | 0.349784136 | 0.737314565 | 0.995570006 | -5.205662974 |
| **A_55_P2018492** | Pgap3 | -0.021123239 | -0.165471544 | 0.123225066 | 10.67137135 | -0.34975019 | 0.737338897 | 0.995570006 | -5.205672767 |
| **A_51_P255387** | Med31 | 0.023079938 | -0.134740616 | 0.180900492 | 11.06331147 | 0.34952658 | 0.737499186 | 0.995570006 | -5.205737249 |

| | | | | | | | | | |
|---|---|---|---|---|---|---|---|---|---|
| **A_55_P1993503** | Adora2b | -0.016889557 | -0.132414965 | 0.098635852 | 7.896323511 | -0.349421695 | 0.737574376 | 0.995570006 | -5.205767481 |
| **A_52_P145349** | Stradb | -0.024527855 | -0.192328002 | 0.143272291 | 9.599084875 | -0.34936257 | 0.737616762 | 0.995570006 | -5.205784519 |
| **A_51_P359046** | Slurp1 | 0.114349111 | -0.668001475 | 0.896699697 | 8.743514712 | 0.349333723 | 0.737637443 | 0.995570006 | -5.205792831 |
| **A_55_P2330545** | NA | 0.021956236 | -0.128275496 | 0.172187967 | 9.305313721 | 0.349305432 | 0.737657725 | 0.995570006 | -5.205800982 |
| **A_51_P321743** | NA | 0.038218993 | -0.22334035 | 0.299778336 | 8.543771172 | 0.349235302 | 0.737708003 | 0.995570006 | -5.205821185 |
| **A_55_P2108988** | Kcnip3 | 0.023295504 | -0.136139655 | 0.182730664 | 8.719985567 | 0.349218422 | 0.737720106 | 0.995570006 | -5.205826047 |
| **A_52_P374292** | Tefm | -0.022536754 | -0.176780408 | 0.1317069 | 8.547253275 | -0.349215237 | 0.737722389 | 0.995570006 | -5.205826965 |
| **A_55_P2103097** | Ccr8 | -0.016494339 | -0.129393188 | 0.09640451 | 7.8020319 | -0.349184162 | 0.737744669 | 0.995570006 | -5.205835915 |
| **A_55_P1961084** | Map3k1 | 0.027154814 | -0.15876759 | 0.213077219 | 11.26207891 | 0.349079419 | 0.737819766 | 0.995570006 | -5.205866078 |
| **A_55_P1982122** | Homer1 | 0.031350499 | -0.183330127 | 0.246031126 | 9.028757239 | 0.34902838 | 0.737856361 | 0.995570006 | -5.205880773 |
| **A_51_P141836** | Ube3c | 0.022702756 | -0.132821044 | 0.178226557 | 10.34822937 | 0.348891881 | 0.737954233 | 0.995570006 | -5.205920062 |
| **A_55_P2023488** | NA | 0.051597237 | -0.301974454 | 0.405168928 | 11.38333885 | 0.348785235 | 0.738030704 | 0.995570006 | -5.205950748 |
| **A_51_P290018** | NA | 0.055331308 | -0.323864725 | 0.434527341 | 12.29392021 | 0.34875166 | 0.73805478 | 0.995570006 | -5.205960407 |
| **A_55_P2138130** | NA | -0.017133854 | -0.134642652 | 0.100374944 | 12.49399556 | -0.348492806 | 0.738240407 | 0.995570006 | -5.206034845 |
| **A_55_P2027278** | Gas2l1 | 0.018872967 | -0.110574318 | 0.148320253 | 11.36373655 | 0.348462762 | 0.738261953 | 0.995570006 | -5.206043482 |
| **A_52_P506529** | Rabgap1l | 0.035260815 | -0.20661034 | 0.27713197 | 9.134446196 | 0.348431489 | 0.738284381 | 0.995570006 | -5.206052471 |
| **A_51_P240693** | Tecpr1 | 0.027741541 | -0.162570973 | 0.218054054 | 11.14814954 | 0.348395364 | 0.738310289 | 0.995570006 | -5.206062853 |
| **A_55_P2170094** | Lrrc27 | 0.016359349 | -0.095895769 | 0.128614467 | 8.870874236 | 0.348312452 | 0.738369753 | 0.995570006 | -5.206086678 |
| **A_51_P345792** | Mfsd13a | 0.018903158 | -0.11081432 | 0.148620635 | 9.004057369 | 0.348293199 | 0.738383561 | 0.995570006 | -5.20609221 |
| **A_55_P2116744** | Xirp1 | -0.036007608 | -0.283154644 | 0.211139429 | 11.31291366 | -0.348215416 | 0.738439348 | 0.995570006 | -5.206114556 |
| **A_51_P510722** | Rita1 | -0.022926615 | -0.18030045 | 0.134447219 | 8.666261482 | -0.348190207 | 0.738457429 | 0.995570006 | -5.206121797 |
| **A_55_P2078776** | Kmt2e | 0.023638278 | -0.138622853 | 0.185899409 | 10.4583776 | 0.348185328 | 0.738460928 | 0.995570006 | -5.206123198 |
| **A_55_P2243566** | Redrum | 0.02562888 | -0.150383651 | 0.201641411 | 7.94174439 | 0.348012763 | 0.738584703 | 0.995570006 | -5.206172751 |
| **A_55_P2009187** | Iffo1 | 0.023365641 | -0.137105605 | 0.183836888 | 8.815833311 | 0.348008309 | 0.738587897 | 0.995570006 | -5.20617403 |
| **A_51_P377403** | Mettl14 | 0.016260786 | -0.095445555 | 0.127967128 | 9.023309632 | 0.347914756 | 0.738655003 | 0.995570006 | -5.206200884 |
| **A_55_P2039416** | Sgk1 | 0.02230512 | -0.130948242 | 0.175558482 | 11.24185083 | 0.347859358 | 0.738694741 | 0.995570006 | -5.206216783 |
| **A_55_P2002150** | Pgs1 | -0.019267941 | -0.151665378 | 0.113129495 | 12.01535149 | -0.347828267 | 0.738717044 | 0.995570006 | -5.206225705 |
| **A_51_P166740** | Kcnj4 | 0.096138588 | -0.564497716 | 0.756774893 | 9.351513159 | 0.3478119 | 0.738728785 | 0.995570006 | -5.206230401 |
| **A_52_P103391** | B4galt1 | 0.021555509 | -0.126618712 | 0.16972973 | 13.03533749 | 0.347692046 | 0.738814763 | 0.995570006 | -5.206264785 |
| **A_52_P373973** | Mex3c | 0.023244576 | -0.136541074 | 0.183030225 | 11.25963879 | 0.347690625 | 0.738815783 | 0.995570006 | -5.206265193 |
| **A_55_P2015337** | NA | -0.038017104 | -0.299357874 | 0.223323666 | 16.16701602 | -0.347681029 | 0.738822666 | 0.995570006 | -5.206267945 |
| **A_55_P2386236** | Ceacam2 | -0.023443315 | -0.184627865 | 0.137741235 | 8.287963684 | -0.347619989 | 0.738866456 | 0.995570006 | -5.206285452 |
| **A_51_P154997** | Brms1 | -0.028891754 | -0.22755065 | 0.169767142 | 8.82433036 | -0.347596202 | 0.73888352 | 0.995570006 | -5.206292273 |
| **A_55_P2171463** | Nptn | 0.037079601 | -0.217888518 | 0.292047721 | 13.06435175 | 0.347582823 | 0.738893118 | 0.995570006 | -5.20629611 |
| **A_52_P250555** | Dynll1 | 0.026181756 | -0.153873004 | 0.206236515 | 8.798399287 | 0.347538796 | 0.738924704 | 0.995570006 | -5.206308734 |
| **A_51_P430929** | Fam20a | -0.021690312 | -0.170860216 | 0.127479593 | 9.358880909 | -0.347531134 | 0.738930201 | 0.995570006 | -5.206310931 |
| **A_55_P1974005** | Tnfrsf23 | -0.01609932 | -0.126822728 | 0.094624088 | 7.692410294 | -0.347517932 | 0.738939673 | 0.995570006 | -5.206314716 |
| **A_55_P2423646** | Nf2 | 0.028603186 | -0.168121492 | 0.225327865 | 9.492588337 | 0.347507916 | 0.738946858 | 0.995570006 | -5.206317587 |
| **A_52_P15008** | Mapkapk2 | -0.030570102 | -0.240837837 | 0.179697633 | 11.80113669 | -0.347482856 | 0.738964837 | 0.995570006 | -5.206324771 |
| **A_55_P1981050** | Gemin8 | 0.016482623 | -0.096914404 | 0.129879649 | 8.148186624 | 0.347403177 | 0.739022003 | 0.995570006 | -5.206347611 |
| **A_51_P158044** | NA | 0.027003396 | -0.158776101 | 0.212782893 | 9.086709721 | 0.34739994 | 0.739024325 | 0.995570006 | -5.206348538 |
| **A_55_P2088750** | Mtif3 | -0.021619347 | -0.170358683 | 0.127119989 | 8.662108119 | -0.347396844 | 0.739026546 | 0.995570006 | -5.206349426 |
| **A_66_P128434** | Kit | 0.02891896 | -0.170076122 | 0.227914041 | 9.534435576 | 0.347335727 | 0.739070396 | 0.995570006 | -5.20636694 |
| **A_55_P2112424** | Madd | -0.032828059 | -0.258786208 | 0.19313009 | 13.70709524 | -0.347237269 | 0.739141039 | 0.995570006 | -5.206395151 |
| **A_55_P2075894** | NA | 0.022618131 | -0.133089624 | 0.178325886 | 12.01645709 | 0.347180726 | 0.739181609 | 0.995570006 | -5.206411348 |
| **A_55_P2160204** | NA | -0.051093888 | -0.403148254 | 0.300960479 | 13.09837188 | -0.346871281 | 0.739403655 | 0.995570006 | -5.206499945 |
| **A_52_P15300** | Hoxb2 | -0.017196265 | -0.135697348 | 0.101304817 | 8.167094027 | -0.346833431 | 0.739430817 | 0.995570006 | -5.206510777 |
| **A_51_P196207** | Capsl | 0.016034389 | -0.094469973 | 0.126538751 | 7.694787839 | 0.346802431 | 0.739453063 | 0.995570006 | -5.206519648 |

| | | | | | | | | | |
|---|---|---|---|---|---|---|---|---|---|
| **A_55_P1994032** | Xbp1 | -0.022855446 | -0.180381625 | 0.134670733 | 13.07972181 | -0.346773656 | 0.739473713 | 0.995570006 | -5.206527881 |
| **A_51_P495131** | Prl7c1 | 0.014019758 | -0.082658141 | 0.110697656 | 7.653129822 | 0.346595119 | 0.739601842 | 0.995570006 | -5.20657895 |
| **A_52_P382876** | Bloc1s3 | 0.023932368 | -0.141121104 | 0.188985839 | 8.196510863 | 0.346553381 | 0.739631797 | 0.995570006 | -5.206590885 |
| **A_55_P1992099** | Ptprj | -0.016235431 | -0.128248165 | 0.095777302 | 8.363484053 | -0.346422085 | 0.73972603 | 0.995570006 | -5.206628421 |
| **A_55_P1966878** | Rnaseh2a | 0.020889649 | -0.123274634 | 0.165053932 | 9.14590302 | 0.34632401 | 0.739796423 | 0.995570006 | -5.206656451 |
| **A_52_P465886** | Gramd3 | 0.025108031 | -0.148193766 | 0.198409828 | 8.906835611 | 0.346273073 | 0.739832984 | 0.995570006 | -5.206671006 |
| **A_51_P328014** | Cops3 | -0.035809645 | -0.283047635 | 0.211428346 | 11.24491331 | -0.346173597 | 0.739904387 | 0.995570006 | -5.206699424 |
| **A_55_P2030030** | Adssl1 | -0.022755596 | -0.179872433 | 0.134361241 | 13.95381772 | -0.346158198 | 0.73991544 | 0.995570006 | -5.206703822 |
| **A_51_P101006** | Ak8 | 0.015071456 | -0.088998638 | 0.11914155 | 7.754988979 | 0.346129322 | 0.739936168 | 0.995570006 | -5.194373149 |
| **A_51_P169047** | Tdp2 | 0.028832184 | -0.170295085 | 0.227959453 | 9.388937243 | 0.346063608 | 0.739983339 | 0.995570006 | -5.206730836 |
| **A_51_P292357** | Rps3a1 | 0.023793206 | -0.140612782 | 0.188199195 | 14.22635455 | 0.34589515 | 0.740104268 | 0.995570006 | -5.206778929 |
| **A_51_P308048** | Cmtm8 | -0.029479289 | -0.233184896 | 0.174226317 | 10.36246781 | -0.345878177 | 0.740116453 | 0.995570006 | -5.206783773 |
| **A_51_P475580** | Ugp2 | 0.019642503 | -0.116217412 | 0.155502417 | 11.45919482 | 0.345552956 | 0.740349939 | 0.995570006 | -5.206876552 |
| **A_55_P2089080** | Ssbp2 | 0.022043816 | -0.130455624 | 0.174543257 | 11.55963385 | 0.345483782 | 0.740399605 | 0.995570006 | -5.206896275 |
| **A_55_P2118931** | Foxj3 | 0.016424646 | -0.097219961 | 0.130069252 | 11.97535406 | 0.345427027 | 0.740440356 | 0.995570006 | -5.206912455 |
| **A_55_P2092611** | Slc45a3 | 0.031631787 | -0.187273798 | 0.250537373 | 8.983613081 | 0.345363171 | 0.740486206 | 0.995570006 | -5.206930655 |
| **A_51_P196395** | Cfap298 | 0.020480008 | -0.121270328 | 0.162230344 | 11.9295784 | 0.345314765 | 0.740520963 | 0.995570006 | -5.20694445 |
| **A_55_P2146525** | Camta2 | -0.026566703 | -0.210505621 | 0.157372214 | 11.79403218 | -0.345201897 | 0.740602009 | 0.995570006 | -5.206976609 |
| **A_55_P2109397** | Scn10a | -0.042122934 | -0.333790136 | 0.249544267 | 8.99604087 | -0.345175652 | 0.740620855 | 0.995570006 | -5.206984085 |
| **A_52_P391718** | Rpl22 | 0.016954727 | -0.100451351 | 0.134360805 | 11.91598507 | 0.345151169 | 0.740638436 | 0.995570006 | -5.206991059 |
| **A_55_P2108609** | Srgap3 | 0.014779772 | -0.087597545 | 0.117157089 | 7.704895775 | 0.345042928 | 0.740716164 | 0.995570006 | -5.207021885 |
| **A_55_P2142156** | Gm16441 | -0.022774662 | -0.180548982 | 0.134999658 | 8.502334193 | -0.345004501 | 0.74074376 | 0.995570006 | -5.207032826 |
| **A_51_P468464** | Gdpd5 | 0.033652625 | -0.199503098 | 0.266808348 | 10.44578957 | 0.344970532 | 0.740768154 | 0.995570006 | -5.207042497 |
| **A_52_P463235** | Ankrd33b | -0.01789748 | -0.141922382 | 0.106127422 | 8.147212994 | -0.344899158 | 0.740819411 | 0.995570006 | -5.207062815 |
| **A_55_P2002859** | Aifm2 | -0.025265951 | -0.200357725 | 0.149825823 | 8.411719201 | -0.344888756 | 0.740826881 | 0.995570006 | -5.207065776 |
| **A_55_P2078403** | Rgs19 | 0.027440503 | -0.162736337 | 0.217617344 | 10.44260172 | 0.3448606 | 0.740847103 | 0.995570006 | -5.207073789 |
| **A_55_P2036693** | Cd40 | 0.017958303 | -0.106510011 | 0.142426616 | 8.310351914 | 0.344838395 | 0.74086305 | 0.995570006 | -5.207080109 |
| **A_52_P488612** | Btbd9 | 0.043875495 | -0.260334521 | 0.348085511 | 8.838639948 | 0.344712991 | 0.740953114 | 0.995570006 | -5.207115791 |
| **A_55_P1989534** | Cklf | 0.023069714 | -0.136892705 | 0.183032132 | 9.084105463 | 0.344693722 | 0.740966953 | 0.995570006 | -5.207121273 |
| **A_55_P1967613** | NA | -0.017965286 | -0.142554652 | 0.10662408 | 7.998128385 | -0.344637311 | 0.741007469 | 0.995570006 | -5.207137319 |
| **A_55_P2156002** | Sumo2 | 0.027218293 | -0.161576945 | 0.216013532 | 14.86266805 | 0.344571209 | 0.741054947 | 0.995570006 | -5.207156119 |
| **A_55_P2094469** | Kdr | -0.063085807 | -0.5007295 | 0.374557886 | 11.2713868 | -0.344524508 | 0.74108849 | 0.995570006 | -5.207169398 |
| **A_51_P170641** | Dcaf4 | 0.023850968 | -0.141627501 | 0.189329436 | 10.04876119 | 0.34448764 | 0.741114971 | 0.995570006 | -5.207179881 |
| **A_55_P1966528** | Pmf1 | -0.029421834 | -0.233564137 | 0.174720469 | 10.66236033 | -0.344465607 | 0.741130797 | 0.995570006 | -5.207186145 |
| **A_51_P384718** | Efna4 | 0.015822009 | -0.09397192 | 0.125615939 | 8.139775966 | 0.344423234 | 0.741161233 | 0.995570006 | -5.207198191 |
| **A_55_P2115315** | Fmr1nb | -0.015001753 | -0.119109465 | 0.08910596 | 7.676075783 | -0.344404032 | 0.741175026 | 0.995570006 | -5.207203649 |
| **A_55_P1968664** | NA | -0.026416531 | -0.209740944 | 0.156907882 | 12.8452102 | -0.344401166 | 0.741177084 | 0.995570006 | -5.207204463 |
| **A_51_P513530** | Spag5 | 0.02418358 | -0.143657999 | 0.192025158 | 9.941773779 | 0.344373848 | 0.741196707 | 0.995570006 | -5.207212228 |
| **A_55_P2091846** | Tut4 | 0.02833156 | -0.168298766 | 0.224961886 | 9.609794614 | 0.344373022 | 0.7411973 | 0.995570006 | -5.207212463 |
| **A_52_P547612** | Tmem30b | 0.014151898 | -0.084068506 | 0.112372302 | 7.655823931 | 0.344367456 | 0.741201298 | 0.995570006 | -5.207214045 |
| **A_51_P338397** | Kctd10 | -0.03000869 | -0.238322384 | 0.178305004 | 8.964070525 | -0.344301053 | 0.741248997 | 0.995570006 | -5.207232916 |
| **A_52_P442710** | Ndnf | -0.015226393 | -0.121018418 | 0.090565631 | 7.782003497 | -0.343995888 | 0.741468219 | 0.995570006 | -5.207319596 |
| **A_55_P1978618** | Grcc10 | -0.018142709 | -0.144264559 | 0.10797914 | 14.12785015 | -0.343811933 | 0.741600379 | 0.995570006 | -5.207371812 |
| **A_52_P512749** | Ears2 | -0.038676437 | -0.307594938 | 0.230242063 | 11.12458782 | -0.34374383 | 0.741649309 | 0.995570006 | -5.207391136 |
| **A_52_P406036** | Cdkn2aip | 0.02003924 | -0.119311354 | 0.159389835 | 8.275582936 | 0.343701606 | 0.741679647 | 0.995570006 | -5.207403116 |
| **A_52_P69707** | Ergic1 | 0.036917725 | -0.219851749 | 0.2936872 | 9.512115632 | 0.343637597 | 0.741725638 | 0.995570006 | -5.207421273 |
| **A_52_P652193** | Sar1a | 0.023641982 | -0.140825156 | 0.188109119 | 13.25445823 | 0.343568923 | 0.741774982 | 0.995570006 | -5.20744075 |
| **A_52_P883557** | Slc30a10 | -0.014968421 | -0.119098303 | 0.089161462 | 7.651048739 | -0.343565647 | 0.741777335 | 0.995570006 | -5.207441679 |

| | | | | | | | | | |
|---|---|---|---|---|---|---|---|---|---|
| **A_55_P1982559** | 1700007K1 | 0.014218053 | -0.08469785 | 0.113133955 | 7.661300815 | 0.343544602 | 0.741792457 | 0.995570006 | -5.207447647 |
| **A_55_P2140195** | Gm15517 | -0.063002293 | -0.501419142 | 0.375414555 | 10.1488689 | -0.343461652 | 0.741852061 | 0.995570006 | -5.207471166 |
| **A_51_P221677** | Trim62 | 0.01615208 | -0.096260641 | 0.1285648 | 8.347707765 | 0.343417261 | 0.741883959 | 0.995570006 | -5.20748375 |
| **A_55_P1975087** | NA | -0.021738967 | -0.173057741 | 0.129579807 | 14.15871067 | -0.343364366 | 0.741921968 | 0.995570006 | -5.207498743 |
| **A_55_P2179216** | Man1a2 | -0.021174049 | -0.168601667 | 0.126253569 | 9.068428412 | -0.343268695 | 0.741990717 | 0.995570006 | -5.207525855 |
| **A_52_P413492** | Ldlrap1 | -0.025521418 | -0.203244397 | 0.152201562 | 9.009348213 | -0.343218225 | 0.742026986 | 0.995570006 | -5.207540154 |
| **A_66_P126320** | Mrpl46 | -0.027930487 | -0.222432588 | 0.166571614 | 10.90310445 | -0.343212696 | 0.74203096 | 0.995570006 | -5.207541721 |
| **A_55_P2467415** | B3galt6 | 0.031137044 | -0.185712022 | 0.247986109 | 8.638148961 | 0.343185517 | 0.742050492 | 0.995570006 | -5.207549421 |
| **A_55_P2027653** | Arhgap25 | -0.02269588 | -0.180774331 | 0.135382571 | 8.8856296 | -0.343149591 | 0.742076309 | 0.995570006 | -5.207559597 |
| **A_55_P2000930** | Ccnyl1 | 0.023660857 | -0.14116754 | 0.188489253 | 10.04149063 | 0.343089609 | 0.742119416 | 0.995570006 | -5.207576586 |
| **A_55_P2007871** | Ptpdc1 | 0.024079826 | -0.143732923 | 0.191892574 | 9.988889566 | 0.342955298 | 0.742215943 | 0.995570006 | -5.207614617 |
| **A_51_P381749** | Prelid3b | 0.029321254 | -0.175024389 | 0.233666898 | 13.32885441 | 0.342946434 | 0.742222314 | 0.995570006 | -5.207617126 |
| **A_51_P294020** | Ticam1 | -0.029226094 | -0.233108572 | 0.174656384 | 9.442818904 | -0.342609976 | 0.742464144 | 0.995570006 | -5.207712332 |
| **A_55_P2164469** | Prcd | -0.020025982 | -0.159758998 | 0.119707035 | 8.132092681 | -0.342534177 | 0.742518629 | 0.995570006 | -5.207733768 |
| **A_55_P2005655** | Ntsr1 | 0.040496518 | -0.242142714 | 0.32313575 | 8.516768457 | 0.34244779 | 0.742580727 | 0.995570006 | -5.207758192 |
| **A_55_P2084251** | Kcna4 | -0.041244918 | -0.329109634 | 0.246619798 | 11.50354173 | -0.342445249 | 0.742582554 | 0.995570006 | -5.207758911 |
| **A_55_P2139473** | Ilk | -0.02532293 | -0.20214576 | 0.151499899 | 12.83743199 | -0.342282548 | 0.742699514 | 0.995570006 | -5.207804896 |
| **A_66_P119174** | Cmtm6 | 0.023900861 | -0.143054471 | 0.190856194 | 11.81588697 | 0.342154604 | 0.742791494 | 0.995570006 | -5.207841043 |
| **A_55_P2103963** | Bhmt | -0.01541961 | -0.123141233 | 0.092302014 | 7.693824392 | -0.342120909 | 0.742815719 | 0.995570006 | -5.20785056 |
| **A_55_P2167005** | NA | 0.023606371 | -0.141380886 | 0.188593627 | 16.54221925 | 0.341969958 | 0.742924246 | 0.995570006 | -5.207893186 |
| **A_66_P127969** | Bcat1 | 0.013891403 | -0.083201378 | 0.110984185 | 7.821998231 | 0.341954489 | 0.742935368 | 0.995570006 | -5.207897553 |
| **A_55_P2077263** | Cenpk | 0.029653252 | -0.177685922 | 0.236992426 | 9.846936074 | 0.341822067 | 0.743030579 | 0.995570006 | -5.207934931 |
| **A_52_P87210** | Dcun1d5 | 0.026131881 | -0.156643056 | 0.208906819 | 11.25190344 | 0.341714311 | 0.74310806 | 0.995570006 | -5.207965336 |
| **A_55_P2010704** | Selenoh | 0.022287097 | -0.133614169 | 0.178188363 | 13.43648565 | 0.34167484 | 0.743136441 | 0.995570006 | -5.207976471 |
| **A_66_P109315** | Snapc2 | 0.020679285 | -0.123984504 | 0.165343074 | 12.0278502 | 0.341652665 | 0.743152387 | 0.995570006 | -5.207982726 |
| **A_51_P446085** | Map10 | -0.017960026 | -0.143622945 | 0.107702893 | 9.482690194 | -0.341592986 | 0.7431953 | 0.995570006 | -5.207999559 |
| **A_55_P2184541** | Cdadc1 | -0.026858716 | -0.214833732 | 0.161116299 | 10.88887108 | -0.341502788 | 0.743260161 | 0.995570006 | -5.208024994 |
| **A_51_P422751** | Clca1 | 0.01667954 | -0.100230938 | 0.133590019 | 7.670488295 | 0.340988531 | 0.743630004 | 0.995570006 | -5.208169887 |
| **A_52_P480402** | Avl9 | -0.020058233 | -0.160658675 | 0.120542209 | 8.794188318 | -0.34096918 | 0.743643922 | 0.995570006 | -5.208175336 |
| **A_55_P2144766** | Pigc | -0.026348438 | -0.211130824 | 0.158433949 | 8.618515905 | -0.340803016 | 0.743763441 | 0.995570006 | -5.208222106 |
| **A_55_P2180201** | Mrpl10 | 0.026092121 | -0.156899887 | 0.20908413 | 9.256243172 | 0.340789651 | 0.743773054 | 0.995570006 | -5.208225866 |
| **A_55_P1955975** | Afg3l2 | -0.019757267 | -0.158322991 | 0.118808457 | 11.96905404 | -0.340784778 | 0.743776559 | 0.995570006 | -5.208227237 |
| **A_55_P2163857** | Trim12c | -0.030654165 | -0.245709639 | 0.184401309 | 8.143503366 | -0.340681164 | 0.743851091 | 0.995570006 | -5.208256389 |
| **A_55_P2170414** | Zfp180 | 0.042055161 | -0.25314356 | 0.337253882 | 9.49634628 | 0.340497527 | 0.743983192 | 0.995570006 | -5.208308034 |
| **A_55_P2287910** | Rragd | -0.024456819 | -0.196169724 | 0.147256086 | 8.694862904 | -0.340413025 | 0.744043983 | 0.995570006 | -5.20833179 |
| **A_55_P2092958** | Traf2 | 0.017882164 | -0.107700893 | 0.143465222 | 8.496228851 | 0.340328372 | 0.744104884 | 0.995570006 | -5.208355583 |
| **A_55_P2143693** | Sytl1 | 0.044030019 | -0.265259531 | 0.353319569 | 8.226000783 | 0.340245786 | 0.7441643 | 0.995570006 | -5.208378789 |
| **A_52_P482897** | Areg | -0.014699056 | -0.117973602 | 0.08857549 | 7.679243035 | -0.340177268 | 0.744213596 | 0.995570006 | -5.208398039 |
| **A_51_P461504** | Eef2k | 0.032392583 | -0.195197846 | 0.259983013 | 8.58279136 | 0.340173691 | 0.74421617 | 0.995570006 | -5.208399043 |
| **A_52_P481279** | Drc1 | -0.015473558 | -0.124193393 | 0.093246276 | 7.809129404 | -0.34016572 | 0.744221905 | 0.995570006 | -5.208401283 |
| **A_55_P2408355** | BB166591 | 0.060064418 | -0.362018283 | 0.48214712 | 9.468953974 | 0.340117404 | 0.744256668 | 0.995570006 | -5.208414854 |
| **A_66_P134252** | NA | 0.019841575 | -0.119599532 | 0.159282682 | 12.87390358 | 0.340090469 | 0.744276047 | 0.995570006 | -5.208422418 |
| **A_55_P2186693** | NA | 0.021707494 | -0.130855816 | 0.174270804 | 8.703930191 | 0.340070314 | 0.744290548 | 0.995570006 | -5.208428078 |
| **A_52_P459276** | Ift81 | -0.032410167 | -0.260266653 | 0.195446319 | 9.516503901 | -0.339960924 | 0.744369257 | 0.995570006 | -5.208458793 |
| **A_55_P2143561** | Gm6813 | -0.019801725 | -0.159039095 | 0.119435646 | 14.99696076 | -0.339904052 | 0.744410179 | 0.995570006 | -5.208474758 |
| **A_55_P2034097** | Elavl1 | 0.023613204 | -0.142442633 | 0.189669041 | 13.39796024 | 0.339867711 | 0.744436328 | 0.995570006 | -5.208484958 |
| **A_55_P1984289** | Cdan1 | -0.015130831 | -0.121596798 | 0.091335136 | 7.999519946 | -0.33967306 | 0.744576396 | 0.995570006 | -5.208539575 |
| **A_51_P220407** | Rnpepl1 | 0.023966687 | -0.144698784 | 0.192632158 | 13.2932676 | 0.339618192 | 0.744615881 | 0.995570006 | -5.208554965 |

| | | | | | | | | | |
|---|---|---|---|---|---|---|---|---|---|
| **A_55_P2050044** | NA | -0.019113206 | -0.153632406 | 0.115405994 | 10.56028807 | -0.339592743 | 0.744634195 | 0.995570006 | -5.208562102 |
| **A_55_P2125947** | Agpat2 | -0.026906211 | -0.216295928 | 0.162483505 | 8.681910278 | -0.339551213 | 0.744664081 | 0.995570006 | -5.208573748 |
| **A_55_P2113683** | Raph1 | -0.025405325 | -0.204286074 | 0.153475425 | 11.88499283 | -0.339445671 | 0.744740036 | 0.995570006 | -5.20860334 |
| **A_55_P2051479** | NA | 0.053040924 | -0.320442628 | 0.426524476 | 10.20898837 | 0.339428836 | 0.744752152 | 0.995570006 | -5.208608059 |
| **A_55_P2183110** | Fmr1 | 0.022344118 | -0.135106163 | 0.179794399 | 8.598326515 | 0.339178963 | 0.74493199 | 0.995570006 | -5.208678078 |
| **A_55_P1980966** | Mettl23 | 0.020620152 | -0.124686815 | 0.165927119 | 11.31967492 | 0.339167757 | 0.744940055 | 0.995570006 | -5.208681217 |
| **A_65_P10913** | Tgfb2 | 0.03992317 | -0.241427197 | 0.321273536 | 11.45543946 | 0.33914597 | 0.744955736 | 0.995570006 | -5.20868732 |
| **A_51_P320606** | Prl5a1 | -0.015353668 | -0.123556956 | 0.09284962 | 7.644115923 | -0.339141413 | 0.744959016 | 0.995570006 | -5.208688596 |
| **A_55_P1984178** | Pde4dip | 0.029950655 | -0.181131894 | 0.241033204 | 14.02465854 | 0.33912759 | 0.744968965 | 0.995570006 | -5.208692468 |
| **A_55_P2186833** | NA | 0.025197237 | -0.152433693 | 0.202828168 | 14.87197494 | 0.339034168 | 0.745036209 | 0.995570006 | -5.20871863 |
| **A_55_P2174895** | Mrpl43 | -0.029321801 | -0.236047441 | 0.177403838 | 12.50152906 | -0.33900448 | 0.745057578 | 0.995570006 | -5.208726942 |
| **A_55_P2085400** | NA | 0.017276359 | -0.104558132 | 0.13911085 | 8.02372812 | 0.338915237 | 0.745121816 | 0.995570006 | -5.208751926 |
| **A_55_P2066505** | Mpo | -0.014472644 | -0.11653577 | 0.087590481 | 7.665327461 | -0.338912947 | 0.745123464 | 0.995570006 | -5.208752567 |
| **A_55_P2054559** | NA | 0.032802523 | -0.198591208 | 0.264196254 | 10.36372524 | 0.338816691 | 0.745192753 | 0.995570006 | -5.208779506 |
| **A_52_P455089** | NA | 0.016975565 | -0.102788764 | 0.136739893 | 9.393188658 | 0.338770725 | 0.745225842 | 0.995570006 | -5.208792368 |
| **A_55_P2101177** | Ncaph2 | -0.028070263 | -0.226132048 | 0.169991523 | 12.5228575 | -0.338730979 | 0.745254455 | 0.995570006 | -5.208803489 |
| **A_55_P2005055** | Pepd | 0.018075397 | -0.109484123 | 0.145634917 | 11.38957649 | 0.338675753 | 0.745294211 | 0.995570006 | -5.208818938 |
| **A_55_P2177173** | Gnl1 | -0.01974948 | -0.159128871 | 0.119629912 | 10.88761633 | -0.33866181 | 0.745304248 | 0.995570006 | -5.208822838 |
| **A_52_P1114640** | 2410003L1 | 0.014732505 | -0.089240374 | 0.118705383 | 7.665555873 | 0.338661362 | 0.745304571 | 0.995570006 | -5.208822963 |
| **A_55_P1965574** | A2ml1 | -0.015822946 | -0.127500016 | 0.095854125 | 7.747776112 | -0.338635481 | 0.745323202 | 0.995570006 | -5.208830202 |
| **A_55_P1965629** | Hcfc1 | 0.027713324 | -0.167947856 | 0.223374504 | 11.37362827 | 0.338526824 | 0.745401426 | 0.995570006 | -5.208860588 |
| **A_55_P2040628** | Gid8 | -0.024645042 | -0.19865571 | 0.149365626 | 12.29008056 | -0.338503236 | 0.745418408 | 0.995570006 | -5.208867183 |
| **A_52_P282762** | Myd88 | -0.021103063 | -0.170143658 | 0.127937532 | 9.232234573 | -0.338415351 | 0.745481681 | 0.995570006 | -5.189902241 |
| **A_52_P287052** | Znrd2 | -0.039702047 | -0.320168342 | 0.240764248 | 10.53551744 | -0.338330658 | 0.745542657 | 0.995570006 | -5.208915422 |
| **A_55_P2044582** | Iglon5 | 0.031451955 | -0.190748708 | 0.253652619 | 8.198883607 | 0.338307346 | 0.745559442 | 0.995570006 | -5.208921936 |
| **A_51_P412348** | Scnn1b | -0.01345875 | -0.108542596 | 0.081625096 | 7.630999755 | -0.338303989 | 0.745561859 | 0.995570006 | -5.208922874 |
| **A_55_P1953583** | Ace2 | 0.014699087 | -0.089157564 | 0.118555738 | 7.724821748 | 0.338271313 | 0.745585385 | 0.995570006 | -5.208932004 |
| **A_55_P2173837** | Mpp6 | 0.015255411 | -0.092579494 | 0.123090316 | 12.41881293 | 0.338122194 | 0.745692754 | 0.995570006 | -5.20897366 |
| **A_55_P2025911** | Sms | -0.023726888 | -0.191445881 | 0.143992105 | 9.2148812 | -0.338117513 | 0.745696125 | 0.995570006 | -5.208974967 |
| **A_51_P329949** | Fam13a | -0.029915945 | -0.241436758 | 0.181604868 | 9.335857557 | -0.338032729 | 0.745757174 | 0.995570006 | -5.208998643 |
| **A_66_P113268** | Mme | 0.021828911 | -0.132534798 | 0.176192621 | 7.771721552 | 0.337983889 | 0.745792342 | 0.995570006 | -5.209012279 |
| **A_55_P1957433** | Mprip | -0.038992721 | -0.31476004 | 0.236774597 | 9.999849012 | -0.337948001 | 0.745818185 | 0.995570006 | -5.209022297 |
| **A_55_P2140711** | Tufm | -0.040428072 | -0.326382606 | 0.245526461 | 12.92810706 | -0.337905439 | 0.745848834 | 0.995570006 | -5.209034178 |
| **A_55_P2045928** | NA | 0.053061866 | -0.322406783 | 0.428530515 | 12.76083193 | 0.337767586 | 0.745948104 | 0.995570006 | -5.209072647 |
| **A_51_P145533** | Rnf14 | 0.021765212 | -0.132268348 | 0.175798771 | 12.89652962 | 0.337719918 | 0.745982432 | 0.995570006 | -5.209085946 |
| **A_51_P167232** | Cask | 0.022550481 | -0.137073918 | 0.182174879 | 8.288602247 | 0.337649153 | 0.746033394 | 0.995570006 | -5.209105685 |
| **A_51_P292527** | P4htm | -0.02089154 | -0.168780317 | 0.126997238 | 8.984001401 | -0.337632593 | 0.746045321 | 0.995570006 | -5.209110303 |
| **A_55_P2029517** | Wnk2 | -0.023659366 | -0.191155552 | 0.14383682 | 8.52607511 | -0.337603793 | 0.746066062 | 0.995570006 | -5.209118335 |
| **A_55_P1982533** | Elovl4 | 0.023263079 | -0.141432421 | 0.187958578 | 7.855758664 | 0.337593898 | 0.746073188 | 0.995570006 | -5.209121095 |
| **A_55_P2046101** | Xlr4b | 0.038744272 | -0.235557773 | 0.313046317 | 7.915838208 | 0.337588459 | 0.746077105 | 0.995570006 | -5.209122611 |
| **A_55_P2054280** | Fam178b | 0.01720588 | -0.104609686 | 0.139021445 | 8.241605993 | 0.337585069 | 0.746079547 | 0.995570006 | -5.209123557 |
| **A_51_P327451** | Alas2 | -0.108407033 | -0.87593226 | 0.659118195 | 12.86305438 | -0.33757784 | 0.746084753 | 0.995570006 | -5.209125573 |
| **A_51_P298538** | Prmt7 | 0.024481631 | -0.14888369 | 0.197846951 | 12.48909246 | 0.337510466 | 0.746133275 | 0.995570006 | -5.209144358 |
| **A_55_P2153016** | Ccdc34 | 0.027628771 | -0.168051169 | 0.223308711 | 9.122017694 | 0.337461621 | 0.746168454 | 0.995570006 | -5.209157975 |
| **A_55_P2029549** | Dtd1 | 0.019293297 | -0.117359143 | 0.155945736 | 10.95876251 | 0.337441263 | 0.746183117 | 0.995570006 | -5.20916365 |
| **A_66_P120472** | Pglyrp2 | -0.015369627 | -0.124232715 | 0.09349346 | 7.627795279 | -0.337436318 | 0.746186678 | 0.995570006 | -5.209165029 |
| **A_66_P114768** | Bach2 | 0.030695997 | -0.186736788 | 0.248128781 | 9.337228887 | 0.337416133 | 0.746201216 | 0.995570006 | -5.209170655 |
| **A_55_P2032458** | Gm6934 | -0.020150118 | -0.162928565 | 0.122628329 | 8.437529443 | -0.337306009 | 0.746280533 | 0.995607128 | -5.209201345 |

| | | | | | | | | | |
|---|---|---|---|---|---|---|---|---|---|
| **A_51_P301311** | Ciz1 | -0.027327652 | -0.221030147 | 0.166374844 | 11.82851031 | -0.337191201 | 0.746363228 | 0.995618156 | -5.20923333 |
| **A_51_P328622** | Tlcd2 | -0.019959027 | -0.161448242 | 0.121530188 | 8.030220321 | -0.337151554 | 0.746391785 | 0.995618156 | -5.209244373 |
| **A_55_P2092968** | Traf3 | 0.02768321 | -0.168652303 | 0.224018723 | 11.09656849 | 0.336997529 | 0.746502734 | 0.995644158 | -5.209287262 |
| **A_51_P164296** | Adamdec1 | 0.01465599 | -0.089292504 | 0.118604484 | 7.812579369 | 0.336981519 | 0.746514267 | 0.995644158 | -5.209291719 |
| **A_52_P381303** | Gins2 | 0.02777989 | -0.169371798 | 0.224931579 | 9.174290586 | 0.336774469 | 0.746663423 | 0.995671558 | -5.209349341 |
| **A_55_P2107292** | Ptch1 | 0.021371965 | -0.130399074 | 0.173143005 | 8.071267023 | 0.336561691 | 0.746816717 | 0.995671558 | -5.209408523 |
| **A_55_P2038640** | Gcn1 | 0.036206915 | -0.220981547 | 0.293395377 | 10.1225915 | 0.336472192 | 0.7468812 | 0.995671558 | -5.209433405 |
| **A_55_P2062568** | Caprin2 | 0.03177998 | -0.193989578 | 0.257549537 | 9.969908325 | 0.336432054 | 0.746910119 | 0.995671558 | -5.209444562 |
| **A_55_P1956898** | NA | -0.043589401 | -0.353366389 | 0.266187586 | 13.69402874 | -0.336310855 | 0.746997446 | 0.995671558 | -5.209478243 |
| **A_55_P2037717** | Slc22a18 | 0.014810184 | -0.090475466 | 0.120095835 | 8.042993993 | 0.336202092 | 0.747075816 | 0.995671558 | -5.209508458 |
| **A_55_P1995417** | Cnppd1 | -0.022392015 | -0.181592358 | 0.136808328 | 8.725473293 | -0.336169504 | 0.747099298 | 0.995671558 | -5.20951751 |
| **A_51_P373609** | Yars2 | 0.022348748 | -0.136587103 | 0.1812846 | 10.70117018 | 0.336078295 | 0.747165022 | 0.995671558 | -5.209542839 |
| **A_55_P2216456** | Rint1 | -0.031451719 | -0.255283476 | 0.192380038 | 9.583269567 | -0.335839529 | 0.747337086 | 0.995671558 | -5.209609114 |
| **A_55_P2186668** | Cfap126 | -0.018716946 | -0.151934074 | 0.114500183 | 8.577064253 | -0.335802594 | 0.747363703 | 0.995671558 | -5.209619363 |
| **A_51_P485260** | Rgl1 | 0.02255973 | -0.138008833 | 0.183128294 | 11.48139049 | 0.335801409 | 0.747364558 | 0.995671558 | -5.209619691 |
| **A_51_P328669** | Ncbp1 | 0.017470218 | -0.10690004 | 0.141840475 | 9.007691772 | 0.335730589 | 0.747415597 | 0.995671558 | -5.209639338 |
| **A_55_P1996082** | Zfp936 | 0.023629258 | -0.144588815 | 0.191847331 | 9.876361858 | 0.335727232 | 0.747418016 | 0.995671558 | -5.20964027 |
| **A_55_P1955587** | Hspd1 | 0.016230848 | -0.099382319 | 0.131844015 | 14.84752469 | 0.335539032 | 0.747553657 | 0.995671558 | -5.209692461 |
| **A_55_P2027819** | Rpl41 | 0.017499308 | -0.10724731 | 0.142245925 | 17.66104764 | 0.335275037 | 0.747743941 | 0.995671558 | -5.209765623 |
| **A_55_P2226660** | 5033418A1 | 0.039841076 | -0.244182911 | 0.323865063 | 8.663948809 | 0.335262647 | 0.747752872 | 0.995671558 | -5.209769056 |
| **A_51_P410451** | Tube1 | 0.023154482 | -0.141957288 | 0.188266253 | 8.575528931 | 0.335170799 | 0.747819079 | 0.995671558 | -5.209794497 |
| **A_51_P208580** | Nop53 | -0.052548142 | -0.427272063 | 0.322175779 | 9.55325143 | -0.335162235 | 0.747825253 | 0.995671558 | -5.209796868 |
| **A_55_P2022845** | Csnk2a1 | 0.022802693 | -0.139864127 | 0.185469512 | 11.74421818 | 0.335039717 | 0.747913573 | 0.995671558 | -5.209830793 |
| **A_51_P114722** | Hao2 | -0.01383803 | -0.112570695 | 0.084894635 | 7.666685439 | -0.334982826 | 0.747954585 | 0.995671558 | -5.191703044 |
| **A_55_P2206605** | 5830444B0 | -0.016057668 | -0.130634432 | 0.098519097 | 7.765284308 | -0.334961613 | 0.747969878 | 0.995671558 | -5.209852414 |
| **A_51_P476018** | Sox7 | -0.022921996 | -0.186513529 | 0.140669536 | 10.58448304 | -0.3348889 | 0.748022298 | 0.995671558 | -5.209872537 |
| **A_55_P1957739** | Guca2b | 0.054215297 | -0.332758144 | 0.441188737 | 7.889710701 | 0.334849623 | 0.748050614 | 0.995671558 | -5.209883406 |
| **A_65_P18981** | Palb2 | -0.015844949 | -0.128979658 | 0.097289759 | 8.053559961 | -0.334737309 | 0.748131588 | 0.995671558 | -5.209914478 |
| **A_51_P353856** | Ubac2 | -0.023797611 | -0.193718335 | 0.146123113 | 9.007649803 | -0.334731165 | 0.748136017 | 0.995671558 | -5.209916177 |
| **A_55_P2211341** | Map3k4 | 0.021456665 | -0.13177504 | 0.174688371 | 12.51622472 | 0.334674577 | 0.748176816 | 0.995671558 | -5.209931828 |
| **A_55_P2230506** | 4931402H1 | 0.071979602 | -0.442179638 | 0.586138842 | 13.59715209 | 0.334596141 | 0.748233368 | 0.995671558 | -5.209953518 |
| **A_51_P491350** | Col4a2 | 0.024421793 | -0.150027255 | 0.19887084 | 12.98094175 | 0.334593939 | 0.748234956 | 0.995671558 | -5.209954127 |
| **A_51_P365409** | Rpusd4 | 0.019847898 | -0.121986713 | 0.161682508 | 10.5853672 | 0.334457874 | 0.748333063 | 0.995671558 | -5.209991741 |
| **A_55_P1955188** | Spata21 | -0.016769471 | -0.136719495 | 0.103180554 | 7.731050534 | -0.334139756 | 0.748562454 | 0.995671558 | -5.210079624 |
| **A_55_P2026223** | St6galnac4 | -0.026752092 | -0.218109596 | 0.164605412 | 8.727465153 | -0.33413455 | 0.748566208 | 0.995671558 | -5.210081062 |
| **A_55_P2126945** | 1700066B1 | -0.048283446 | -0.393672429 | 0.297105538 | 9.352310735 | -0.334117256 | 0.74857868 | 0.995671558 | -5.210085837 |
| **A_51_P413461** | Parg | 0.02069767 | -0.127363647 | 0.168758986 | 10.88842849 | 0.334109621 | 0.748584186 | 0.995671558 | -5.210087945 |
| **A_55_P2016426** | Nova2 | -0.022329317 | -0.182073125 | 0.137414492 | 8.909680233 | -0.334087732 | 0.748599971 | 0.995671558 | -5.210093989 |
| **A_51_P401501** | Tmem213 | -0.014336269 | -0.116915529 | 0.088242991 | 7.645674013 | -0.334030187 | 0.74864147 | 0.995671558 | -5.210109875 |
| **A_55_P1987231** | Fuom | 0.030885218 | -0.190150923 | 0.251921358 | 11.28009103 | 0.333961583 | 0.748690945 | 0.995671558 | -5.210128811 |
| **A_55_P2122380** | Zfp14 | 0.019471771 | -0.11993545 | 0.158878993 | 8.866214889 | 0.333833042 | 0.748783648 | 0.995671558 | -5.210164281 |
| **A_55_P2105366** | NA | 0.014168553 | -0.08727486 | 0.115611965 | 8.332444595 | 0.333818784 | 0.748793932 | 0.995671558 | -5.210168215 |
| **A_55_P1984745** | Pappa2 | -0.091775288 | -0.74889405 | 0.565343473 | 8.276529029 | -0.333803606 | 0.748804879 | 0.995671558 | -5.210172402 |
| **A_55_P1983903** | Thap4 | -0.035872688 | -0.292884106 | 0.221138731 | 8.357395815 | -0.333595844 | 0.748954728 | 0.995671558 | -5.2102297 |
| **A_55_P2010191** | Serpina11 | -0.049935393 | -0.407757324 | 0.307886539 | 9.035436425 | -0.33354209 | 0.7489935 | 0.995671558 | -5.210244519 |
| **A_55_P1981734** | Tut7 | 0.030969351 | -0.190951964 | 0.252890667 | 11.54338693 | 0.333535624 | 0.748998164 | 0.995671558 | -5.210246302 |
| **A_55_P1994128** | Tmem184a | -0.015427955 | -0.125985038 | 0.095129128 | 7.715227121 | -0.333526947 | 0.749004422 | 0.995671558 | -5.210248693 |
| **A_55_P2168667** | Rbpms2 | -0.027706204 | -0.226375444 | 0.170963036 | 13.80735319 | -0.333315519 | 0.749156931 | 0.995671558 | -5.210306956 |

| | | | | | | | | | |
|---|---|---|---|---|---|---|---|---|---|
| **A_55_P2020453** | NA | 0.019804609 | -0.12222041 | 0.161829628 | 8.399568319 | 0.33328099 | 0.74918184 | 0.995671558 | -5.210316467 |
| **A_55_P2114034** | NA | 0.030527787 | -0.188408588 | 0.249464162 | 8.397823582 | 0.333262567 | 0.74919513 | 0.995671558 | -5.210321542 |
| **A_55_P2004636** | Ppih | -0.023043293 | -0.188308545 | 0.142221959 | 10.72083954 | -0.333251505 | 0.749203109 | 0.995671558 | -5.210324589 |
| **A_51_P428372** | Ppbp | 0.055044424 | -0.339740204 | 0.449829052 | 8.46171841 | 0.333243917 | 0.749208583 | 0.995671558 | -5.210326679 |
| **A_55_P2039215** | Acyp2 | -0.030080064 | -0.245845708 | 0.18568558 | 9.856670332 | -0.333200461 | 0.749239932 | 0.995671558 | -5.210338647 |
| **A_55_P2088995** | Plscr1 | -0.017585143 | -0.143801569 | 0.108631283 | 9.781853412 | -0.332996108 | 0.749387355 | 0.995671558 | -5.210394907 |
| **A_51_P431894** | Radil | 0.027719693 | -0.171263421 | 0.226702807 | 8.72774923 | 0.332951767 | 0.749419345 | 0.995671558 | -5.21040711 |
| **A_55_P1955472** | Bbs1 | -0.023059521 | -0.188592233 | 0.142473191 | 8.919216628 | -0.332947364 | 0.749422522 | 0.995671558 | -5.210408322 |
| **A_55_P2027312** | Cdc34 | 0.03553871 | -0.219578222 | 0.290655641 | 14.02563895 | 0.332944238 | 0.749424776 | 0.995671558 | -5.210409182 |
| **A_55_P1973006** | Bag1 | 0.018451051 | -0.114017431 | 0.150919533 | 13.80710452 | 0.332902974 | 0.749454547 | 0.995671558 | -5.210420537 |
| **A_51_P282837** | St14 | -0.014514265 | -0.118722234 | 0.089693705 | 7.810910107 | -0.332891904 | 0.749462534 | 0.995671558 | -5.210423583 |
| **A_55_P2079116** | Rpph1 | -0.091487708 | -0.748417743 | 0.565442327 | 9.702751618 | -0.33285322 | 0.749490443 | 0.995671558 | -5.210434226 |
| **A_55_P1997106** | Large2 | 0.014115516 | -0.08724439 | 0.115475421 | 7.83656359 | 0.332843197 | 0.749497675 | 0.995671558 | -5.210436984 |
| **A_55_P1955568** | Extl2 | 0.025786564 | -0.159423138 | 0.210996266 | 10.72556212 | 0.332765947 | 0.749553411 | 0.995671558 | -5.210458233 |
| **A_52_P649285** | NA | 0.018577175 | -0.114862657 | 0.152017008 | 10.33332351 | 0.332738707 | 0.749573065 | 0.995671558 | -5.210465725 |
| **A_66_P125722** | Cyb5a | -0.019878469 | -0.16280657 | 0.123049631 | 13.10216072 | -0.332410283 | 0.749810042 | 0.995862972 | -5.210556007 |
| **A_55_P2101001** | Smad3 | 0.025983552 | -0.160917848 | 0.212884953 | 11.44878018 | 0.332273037 | 0.749909082 | 0.995862972 | -5.21059371 |
| **A_55_P2157033** | Bace2 | 0.031540001 | -0.195337977 | 0.258417979 | 11.76000057 | 0.332260331 | 0.749918251 | 0.995862972 | -5.2105972 |
| **A_51_P373461** | Brcc3 | 0.020852754 | -0.129272011 | 0.170977519 | 11.23100938 | 0.331986338 | 0.750115987 | 0.995862972 | -5.210672421 |
| **A_55_P2006519** | Clip1 | -0.024277283 | -0.199098883 | 0.150544317 | 10.20304682 | -0.331905253 | 0.750174508 | 0.995862972 | -5.210694671 |
| **A_55_P2115980** | Xlr5c | -0.014186224 | -0.116390031 | 0.088017583 | 7.644143769 | -0.331748431 | 0.750287696 | 0.995862972 | -5.210737687 |
| **A_55_P2015520** | Plekha1 | 0.018931075 | -0.117474903 | 0.155337054 | 11.82376526 | 0.331704234 | 0.750319597 | 0.995862972 | -5.210749807 |
| **A_55_P2002103** | Arhgap45 | -0.050987936 | -0.418467488 | 0.316491615 | 9.332396871 | -0.331622042 | 0.750378924 | 0.995862972 | -5.210772342 |
| **A_55_P1956863** | Egfr | -0.014265014 | -0.117078805 | 0.088548777 | 7.924574304 | -0.331611786 | 0.750386327 | 0.995862972 | -5.210775154 |
| **A_51_P208931** | N4bp3 | -0.029626164 | -0.243161723 | 0.183909395 | 11.70695163 | -0.331599865 | 0.750394932 | 0.995862972 | -5.210778421 |
| **A_66_P125560** | Macroh2a2 | 0.023952795 | -0.148694608 | 0.196600197 | 11.42680772 | 0.331592939 | 0.750399931 | 0.995862972 | -5.21078032 |
| **A_55_P2098603** | NA | 0.017172736 | -0.106635058 | 0.14098053 | 7.942776524 | 0.331513059 | 0.750457592 | 0.995862972 | -5.210802214 |
| **A_55_P2163837** | Med14 | -0.023938653 | -0.196533932 | 0.148656625 | 11.51115669 | -0.331497257 | 0.750468998 | 0.995862972 | -5.210806544 |
| **A_55_P2026014** | Mc5r | 0.015411448 | -0.095724756 | 0.126547651 | 7.900270344 | 0.331433965 | 0.750514686 | 0.995862972 | -5.210823888 |
| **A_55_P2128158** | Trappc13 | 0.021528678 | -0.133758388 | 0.176815743 | 11.17096702 | 0.33135323 | 0.750572967 | 0.995862972 | -5.210846006 |
| **A_51_P338961** | Qser1 | 0.021176473 | -0.131576854 | 0.173929799 | 10.3578785 | 0.331338637 | 0.750583502 | 0.995862972 | -5.210850003 |
| **A_51_P298355** | Dek | 0.027301877 | -0.16965042 | 0.224254175 | 11.44639474 | 0.331314616 | 0.750600843 | 0.995862972 | -5.210856583 |
| **A_55_P1995012** | Plekhm2 | 0.015951906 | -0.09914403 | 0.131047842 | 12.39118473 | 0.33125446 | 0.75064427 | 0.995862972 | -5.210873058 |
| **A_55_P2100495** | Synb | 0.014288603 | -0.088879696 | 0.117456902 | 7.665772432 | 0.331018776 | 0.750814421 | 0.996020366 | -5.210937577 |
| **A_55_P1978962** | Slc9a3r2 | 0.019267155 | -0.11989681 | 0.15843112 | 10.72389459 | 0.330902411 | 0.750898436 | 0.996050611 | -5.210969417 |
| **A_55_P2027386** | Gpr156 | 0.019675995 | -0.122476871 | 0.16182886 | 7.945025129 | 0.33081882 | 0.750958791 | 0.996050611 | -5.210992282 |
| **A_55_P2130463** | Ptgdr2 | 0.013349681 | -0.083131649 | 0.10983101 | 7.70485035 | 0.330701928 | 0.751043193 | 0.996050611 | -5.211024247 |
| **A_55_P1995151** | NA | -0.058678546 | -0.482861905 | 0.365504814 | 12.05995405 | -0.330624358 | 0.751099204 | 0.996050611 | -5.211045453 |
| **A_55_P2084278** | Ighmbp2 | 0.016508073 | -0.102837198 | 0.135853343 | 8.022864729 | 0.330598051 | 0.751118201 | 0.996050611 | -5.211052644 |
| **A_52_P641922** | Nhlrc3 | 0.01424223 | -0.088734238 | 0.117218699 | 7.917551932 | 0.33055912 | 0.751146313 | 0.996050611 | -5.211063284 |
| **A_55_P2148248** | NA | -0.016330869 | -0.134446524 | 0.101784786 | 8.356177159 | -0.330453958 | 0.751222252 | 0.996082996 | -5.21109202 |
| **A_51_P416825** | Tspan15 | 0.013604037 | -0.084936827 | 0.112144901 | 7.957119294 | 0.32995945 | 0.751579386 | 0.996172124 | -5.21122703 |
| **A_65_P16688** | Taok2 | -0.016252724 | -0.133984419 | 0.101478972 | 8.55060587 | -0.329945258 | 0.751589637 | 0.996172124 | -5.211230902 |
| **A_55_P2166399** | Stk11 | 0.040310526 | -0.251698012 | 0.332319065 | 10.03665912 | 0.329937783 | 0.751595036 | 0.996172124 | -5.211232941 |
| **A_66_P117815** | NA | 0.035628373 | -0.222525636 | 0.293782382 | 12.91323947 | 0.329857412 | 0.751653086 | 0.996172124 | -5.211254864 |
| **A_66_P100440** | Vezf1 | 0.033007481 | -0.206204246 | 0.272219208 | 11.04987184 | 0.329791187 | 0.751700921 | 0.996172124 | -5.211272925 |
| **A_66_P104390** | Arl3 | -0.022637644 | -0.186735729 | 0.14146044 | 10.79397579 | -0.329713596 | 0.751756967 | 0.996172124 | -5.211294081 |
| **A_51_P203710** | Taf8 | -0.022572044 | -0.186201505 | 0.141057416 | 10.49843452 | -0.329699684 | 0.751767016 | 0.996172124 | -5.211297873 |

| | | | | | | | | | |
|---|---|---|---|---|---|---|---|---|---|
| **A_55_P2021953** | Olfr700 | 0.017174682 | -0.107351678 | 0.141701041 | 8.004081504 | 0.329637444 | 0.751811974 | 0.996172124 | -5.211314839 |
| **A_51_P509263** | Hoxa7 | 0.016203049 | -0.101290597 | 0.133696695 | 7.899507796 | 0.329603261 | 0.751836667 | 0.996172124 | -5.211324156 |
| **A_55_P2121352** | Cdk5 | 0.033218919 | -0.207720017 | 0.274157855 | 9.696360019 | 0.329524446 | 0.751893601 | 0.996172124 | -5.211345634 |
| **A_55_P2101191** | Arrdc4 | -0.032826226 | -0.270920005 | 0.205267554 | 9.06177012 | -0.329520197 | 0.75189667 | 0.996172124 | -5.211346792 |
| **A_52_P208600** | Bms1 | 0.014245974 | -0.089087118 | 0.117579066 | 8.909224948 | 0.329504885 | 0.751907731 | 0.996172124 | -5.211350963 |
| **A_55_P1971439** | Serpinb9c | 0.016512157 | -0.103292996 | 0.13631731 | 7.679230823 | 0.329410508 | 0.75197591 | 0.996194191 | -5.211376674 |
| **A_66_P104134** | Homer1 | 0.029904317 | -0.187304568 | 0.247113201 | 8.997538573 | 0.329052682 | 0.752234426 | 0.99646839 | -5.21147409 |
| **A_55_P1996751** | NA | 0.031978246 | -0.20038394 | 0.264340433 | 10.73805683 | 0.32892609 | 0.752325892 | 0.99652128 | -5.21150853 |
| **A_55_P2152912** | NA | -0.013241182 | -0.1095053 | 0.083022936 | 7.710329096 | -0.328754303 | 0.75245002 | 0.996617423 | -5.211555245 |
| **A_51_P123920** | Mydgf | 0.032954234 | -0.206724231 | 0.272632699 | 10.99436033 | 0.328617994 | 0.752548518 | 0.996630408 | -5.186920355 |
| **A_51_P227165** | 2310030G( | 0.023818597 | -0.149444299 | 0.197081492 | 9.299072623 | 0.328563817 | 0.752587668 | 0.996630408 | -5.211607017 |
| **A_55_P1987264** | Gnb2 | -0.022248625 | -0.184160847 | 0.139663598 | 15.23355968 | -0.328422323 | 0.752689921 | 0.996630408 | -5.211645456 |
| **A_55_P1979053** | Atp6v0c | -0.027828187 | -0.230363944 | 0.17470757 | 13.01480397 | -0.328391822 | 0.752711963 | 0.996630408 | -5.211653739 |
| **A_55_P2139043** | 4833439L1 | -0.017326911 | -0.143455718 | 0.108801896 | 12.48048246 | -0.328334097 | 0.752753681 | 0.996630408 | -5.211669415 |
| **A_51_P519555** | Gnb1l | -0.02167109 | -0.179457833 | 0.136115653 | 10.26380161 | -0.328261066 | 0.752806461 | 0.996630408 | -5.211689243 |
| **A_55_P1962707** | NA | 0.044334671 | -0.278658587 | 0.367327928 | 11.58463505 | 0.328064443 | 0.752948571 | 0.996630408 | -5.211742606 |
| **A_55_P2170105** | NA | -0.028779345 | -0.238550503 | 0.180991813 | 13.24343691 | -0.327902146 | 0.753065879 | 0.996630408 | -5.21178663 |
| **A_52_P352563** | 1700073E1 | -0.022115385 | -0.18336112 | 0.139130351 | 14.87555259 | -0.327804862 | 0.753136199 | 0.996630408 | -5.211813008 |
| **A_55_P2027594** | Taf3 | 0.013795119 | -0.086799321 | 0.114389559 | 8.484721811 | 0.327763514 | 0.753166087 | 0.996630408 | -5.211824218 |
| **A_55_P2132467** | Nfx1 | 0.015029089 | -0.094636941 | 0.12469512 | 8.948600389 | 0.327544019 | 0.753324757 | 0.996630408 | -5.211883699 |
| **A_51_P108862** | Cops7a | 0.017953655 | -0.113080185 | 0.148987495 | 13.50321563 | 0.32747529 | 0.753374443 | 0.996630408 | -5.192868391 |
| **A_51_P214470** | Tdrp | -0.01417993 | -0.117702135 | 0.089342275 | 7.707350663 | -0.327378164 | 0.75344466 | 0.996630408 | -5.211928619 |
| **A_55_P1974193** | Ptges | 0.023181245 | -0.146076059 | 0.19243855 | 13.120428 | 0.327339545 | 0.753472581 | 0.996630408 | -5.211939076 |
| **A_51_P287069** | Serpinh1 | 0.022701071 | -0.143142175 | 0.188544317 | 14.37600255 | 0.327158113 | 0.753603755 | 0.996630408 | -5.211988184 |
| **A_52_P633128** | Aasdhppt | -0.034147325 | -0.283646669 | 0.215352019 | 10.18482604 | -0.327111954 | 0.75363713 | 0.996630408 | -5.212000674 |
| **A_52_P578634** | Cela1 | 0.014028481 | -0.088484794 | 0.116541756 | 7.764540282 | 0.327069219 | 0.753668029 | 0.996630408 | -5.212012236 |
| **A_55_P2145711** | Tox3 | -0.028052959 | -0.233106242 | 0.177000325 | 11.02040791 | -0.326979915 | 0.7537326 | 0.996630408 | -5.212036392 |
| **A_52_P563375** | Lgals2 | 0.037042055 | -0.233725837 | 0.307809948 | 7.880022756 | 0.32696935 | 0.75374024 | 0.996630408 | -5.212039249 |
| **A_51_P275083** | Mrps18a | 0.020633188 | -0.130194478 | 0.171460854 | 12.20329541 | 0.326959867 | 0.753747096 | 0.996630408 | -5.212041814 |
| **A_55_P2025363** | Ccnl2 | -0.023237483 | -0.193157489 | 0.146682523 | 8.798179808 | -0.326853926 | 0.753823701 | 0.996630408 | -5.21207046 |
| **A_55_P2092859** | Anxa4 | 0.02831758 | -0.178755247 | 0.235390408 | 10.26332302 | 0.326845236 | 0.753829984 | 0.996630408 | -5.21207281 |
| **A_55_P2119721** | NA | 0.023146025 | -0.146159484 | 0.192451534 | 17.05547437 | 0.326749151 | 0.753899466 | 0.996630408 | -5.212098783 |
| **A_51_P365516** | Spink1 | -0.106460696 | -0.885212189 | 0.672290796 | 8.3770399 | -0.326737922 | 0.753907586 | 0.996630408 | -5.212101818 |
| **A_55_P2082203** | Baz1a | -0.026998389 | -0.224499148 | 0.170502369 | 9.368092027 | -0.326721881 | 0.753919185 | 0.996630408 | -5.212106153 |
| **A_66_P114501** | Zcrb1 | -0.022096051 | -0.183745136 | 0.139553035 | 12.53782772 | -0.326701057 | 0.753934244 | 0.996630408 | -5.21211178 |
| **A_51_P482633** | Rps6ka4 | 0.023411978 | -0.14791194 | 0.194735896 | 9.503219084 | 0.326609822 | 0.754000221 | 0.996630408 | -5.212136432 |
| **A_52_P346113** | Foxn2 | 0.01580507 | -0.09987032 | 0.131480459 | 8.40599797 | 0.326561197 | 0.754035385 | 0.996630408 | -5.212149568 |
| **A_55_P2058681** | Tmc8 | -0.0195943 | -0.163003951 | 0.123815351 | 9.054119664 | -0.326558134 | 0.7540376 | 0.996630408 | -5.212150395 |
| **A_52_P152631** | Tmem17 | 0.01668288 | -0.105582099 | 0.138947859 | 8.884944692 | 0.326120489 | 0.754354122 | 0.996630408 | -5.212268537 |
| **A_55_P2065369** | H3f3a | -0.031356791 | -0.261230279 | 0.198516697 | 14.44126203 | -0.326025713 | 0.754422674 | 0.996630408 | -5.212294102 |
| **A_65_P03874** | Nav2 | -0.017115478 | -0.142592737 | 0.108361782 | 8.560668724 | -0.326011658 | 0.754432841 | 0.996630408 | -5.212297893 |
| **A_51_P153224** | Zdhhc12 | 0.024003742 | -0.151974253 | 0.199981738 | 9.561024196 | 0.326009098 | 0.754434692 | 0.996630408 | -5.212298583 |
| **A_55_P2097329** | Sgsm3 | -0.033509996 | -0.279202934 | 0.212182942 | 11.68877615 | -0.325979907 | 0.754455807 | 0.996630408 | -5.212306455 |
| **A_55_P2042297** | NA | 0.016636156 | -0.105341756 | 0.138614068 | 8.104406071 | 0.325972472 | 0.754461185 | 0.996630408 | -5.21230846 |
| **A_55_P2047009** | NA | -0.039830751 | -0.332070282 | 0.25240878 | 10.34209121 | -0.325753182 | 0.754619813 | 0.996630408 | -5.212367574 |
| **A_55_P2055411** | Rps7 | 0.019376185 | -0.122798214 | 0.161550584 | 15.23073056 | 0.32572868 | 0.754637537 | 0.996630408 | -5.212374177 |
| **A_55_P1967216** | Adam10 | 0.024025454 | -0.152301448 | 0.200352356 | 12.08060303 | 0.325658308 | 0.754688445 | 0.996630408 | -5.212393138 |
| **A_55_P2100222** | NA | 0.053033801 | -0.336292583 | 0.442360185 | 11.49032068 | 0.325572753 | 0.754750338 | 0.996630408 | -5.212416184 |

| | | | | | | | | | |
|---|---|---|---|---|---|---|---|---|---|
| **A_55_P2037613** | Elmsan1 | -0.03099968 | -0.25858921 | 0.19658985 | 9.161018335 | -0.325547272 | 0.754768772 | 0.996630408 | -5.212423047 |
| **A_65_P10865** | Scfd1 | 0.020682105 | -0.131169227 | 0.172533437 | 10.34072261 | 0.32552569 | 0.754784386 | 0.996630408 | -5.212428859 |
| **A_52_P268287** | Zfp507 | -0.020405184 | -0.17024888 | 0.129438511 | 9.348439353 | -0.32547016 | 0.75482456 | 0.996630408 | -5.212443812 |
| **A_51_P466824** | Tyk2 | 0.017448046 | -0.110685105 | 0.145581198 | 8.521958046 | 0.325457608 | 0.754833641 | 0.996630408 | -5.212447192 |
| **A_55_P2065754** | Krtap19-5 | -0.023557438 | -0.196557031 | 0.149442155 | 8.461175665 | -0.325455869 | 0.754834899 | 0.996630408 | -5.21244766 |
| **A_55_P2033530** | Gm6981 | -0.050710191 | -0.423304071 | 0.321883688 | 12.63436803 | -0.325288467 | 0.754956014 | 0.996630408 | -5.212492722 |
| **A_51_P288828** | Slc52a2 | 0.026965845 | -0.171168152 | 0.225099841 | 9.342375806 | 0.325285094 | 0.754958454 | 0.996630408 | -5.21249363 |
| **A_55_P2080243** | NA | 0.023839184 | -0.151325545 | 0.199003912 | 9.254472072 | 0.325277369 | 0.754964043 | 0.996630408 | -5.212495708 |
| **A_51_P121915** | Rdh16f2 | 0.013124926 | -0.083317236 | 0.109567089 | 7.663725173 | 0.325266301 | 0.754972051 | 0.996630408 | -5.202251202 |
| **A_51_P284426** | Cstad | -0.030089766 | -0.251230898 | 0.191051366 | 8.874252833 | -0.325205898 | 0.755015755 | 0.996630408 | -5.21251494 |
| **A_55_P1997628** | NA | 0.029286311 | -0.185969449 | 0.244542072 | 16.62238049 | 0.325176406 | 0.755037094 | 0.996630408 | -5.212522874 |
| **A_51_P467770** | Snx19 | 0.01631993 | -0.103693486 | 0.136333347 | 9.121842824 | 0.32501068 | 0.755157008 | 0.996633405 | -5.212567448 |
| **A_55_P1978681** | Tspan8 | -0.022281382 | -0.186384451 | 0.141821688 | 7.981322184 | -0.324514829 | 0.755515836 | 0.996633405 | -5.212700682 |
| **A_55_P2099930** | Cers5 | 0.017007599 | -0.108335942 | 0.14235114 | 11.74471105 | 0.324302411 | 0.755669574 | 0.996633405 | -5.212757698 |
| **A_55_P2148370** | NA | -0.01496752 | -0.125283887 | 0.095348847 | 8.653316866 | -0.324279159 | 0.755686403 | 0.996633405 | -5.212763937 |
| **A_55_P2040170** | Pmp22 | 0.028230617 | -0.179859349 | 0.236320583 | 11.66592175 | 0.324248789 | 0.755708385 | 0.996633405 | -5.212772085 |
| **A_55_P2109559** | Trim6 | -0.01547131 | -0.129549674 | 0.098607054 | 7.68546484 | -0.324140223 | 0.755786966 | 0.996633405 | -5.212801207 |
| **A_55_P1960765** | NA | -0.022035128 | -0.184543722 | 0.140473466 | 12.7755203 | -0.324077127 | 0.755832637 | 0.996633405 | -5.212818128 |
| **A_55_P2128373** | Cops6 | 0.021250623 | -0.135499841 | 0.178001087 | 12.96691014 | 0.324020104 | 0.755873913 | 0.996633405 | -5.212833418 |
| **A_51_P492125** | Ciapin1 | 0.027904985 | -0.177995789 | 0.233805759 | 11.58941927 | 0.323916405 | 0.755948978 | 0.996633405 | -5.212861216 |
| **A_55_P2175494** | NA | -0.017266749 | -0.144687044 | 0.110153546 | 8.674611243 | -0.323877748 | 0.755976961 | 0.996633405 | -5.212871577 |
| **A_55_P2101862** | Spryd3 | 0.020396436 | -0.130153269 | 0.17094614 | 9.511111184 | 0.323804968 | 0.756029647 | 0.996633405 | -5.212891079 |
| **A_55_P1955637** | Trp53i11 | -0.019629569 | -0.164586294 | 0.125327156 | 8.209081023 | -0.323654411 | 0.75613864 | 0.996633405 | -5.21293141 |
| **A_52_P612137** | Runx1t1 | -0.0324529 | -0.272130999 | 0.207225199 | 10.63127236 | -0.323619211 | 0.756164124 | 0.996633405 | -5.212940836 |
| **A_51_P304239** | Loxl1 | 0.036584641 | -0.233631313 | 0.306800595 | 11.37408812 | 0.323591377 | 0.756184275 | 0.996633405 | -5.21294829 |
| **A_55_P2103807** | NA | -0.015411237 | -0.129247884 | 0.098425409 | 14.59561738 | -0.323567242 | 0.756201747 | 0.996633405 | -5.212954752 |
| **A_51_P416613** | 1810058I24 | -0.021013537 | -0.17637325 | 0.134346176 | 9.307211754 | -0.323273337 | 0.75641454 | 0.996633405 | -5.21303341 |
| **A_55_P1974900** | Ndufab1 | -0.026650151 | -0.223734189 | 0.170433887 | 15.84061301 | -0.323189579 | 0.756475186 | 0.996633405 | -5.213055813 |
| **A_55_P2162815** | NA | -0.046131614 | -0.387384513 | 0.295121285 | 12.03896003 | -0.323095886 | 0.756543028 | 0.996633405 | -5.213080868 |
| **A_55_P2121537** | NA | -0.018805557 | -0.15792665 | 0.120315537 | 16.15524924 | -0.323074253 | 0.756558693 | 0.996633405 | -5.213086652 |
| **A_51_P291819** | 2310033P0 | -0.021348798 | -0.179292119 | 0.136594524 | 10.34424056 | -0.323058584 | 0.756570039 | 0.996633405 | -5.213090841 |
| **A_51_P461319** | Gatm | -0.016606845 | -0.139497457 | 0.106283766 | 8.342619294 | -0.322981442 | 0.756625899 | 0.996633405 | -5.213111462 |
| **A_51_P437144** | Dynll2 | -0.036305797 | -0.30501607 | 0.232404477 | 12.61009559 | -0.322924378 | 0.756667221 | 0.996633405 | -5.213126712 |
| **A_51_P488554** | Mrnip | 0.026664642 | -0.170698793 | 0.224028077 | 9.939879343 | 0.322907548 | 0.756679408 | 0.996633405 | -5.21313121 |
| **A_55_P2045512** | NA | 0.033321473 | -0.213399872 | 0.280042818 | 14.4304184 | 0.322794849 | 0.756761021 | 0.996633405 | -5.213161321 |
| **A_55_P1979705** | Spop | 0.029485518 | -0.188840686 | 0.247811722 | 8.696194482 | 0.322784023 | 0.756768861 | 0.996633405 | -5.213164212 |
| **A_55_P2017160** | Cdc14a | -0.02138743 | -0.179763622 | 0.136988763 | 8.530023032 | -0.322758602 | 0.756787271 | 0.996633405 | -5.213171003 |
| **A_66_P137297** | Otof | -0.076192584 | -0.640477799 | 0.48809263 | 9.6654983 | -0.322717982 | 0.756816688 | 0.996633405 | -5.213181852 |
| **A_51_P374782** | Epn3 | 0.038566871 | -0.24708236 | 0.324216102 | 10.98260765 | 0.322693694 | 0.756834278 | 0.996633405 | -5.213188338 |
| **A_55_P2154835** | NA | 0.021724477 | -0.139324392 | 0.182773347 | 11.18192186 | 0.322404275 | 0.757043889 | 0.996633405 | -5.213265595 |
| **A_55_P1997948** | Ppp4r3b | 0.035260257 | -0.226288082 | 0.296808597 | 9.955710166 | 0.322212692 | 0.757182654 | 0.996633405 | -5.213316699 |
| **A_51_P243134** | Adcy6 | 0.039218092 | -0.251692462 | 0.330128646 | 11.09098105 | 0.322207848 | 0.757186163 | 0.996633405 | -5.213317991 |
| **A_55_P2048767** | Efemp1 | 0.10013312 | -0.642663495 | 0.842929735 | 9.46543721 | 0.322193641 | 0.757196454 | 0.996633405 | -5.213321779 |
| **A_55_P2060680** | Acsf2 | 0.038177163 | -0.245072071 | 0.321426397 | 11.66628436 | 0.322139545 | 0.757235638 | 0.996633405 | -5.213336203 |
| **A_55_P2005190** | Herc4 | 0.023855316 | -0.153141101 | 0.200851733 | 10.94639301 | 0.32212901 | 0.757243269 | 0.996633405 | -5.213339012 |
| **A_52_P239403** | Ipo7 | 0.025620336 | -0.164523725 | 0.215764398 | 8.786339261 | 0.322041021 | 0.757307005 | 0.996633405 | -5.213362467 |
| **A_55_P2006467** | Serf2 | -0.024788621 | -0.20876057 | 0.159183328 | 15.45253771 | -0.322040052 | 0.757307707 | 0.996633405 | -5.213362725 |
| **A_55_P1983368** | Sh3d21 | 0.021450813 | -0.137871086 | 0.180772711 | 8.906805604 | 0.321793605 | 0.757486237 | 0.996633405 | -5.213428387 |

| | | | | | | | | | |
|---|---|---|---|---|---|---|---|---|---|
| **A_55_P1982762** | Hsd3b7 | -0.04043151 | -0.340796632 | 0.259933613 | 10.37759877 | -0.321721132 | 0.757538741 | 0.996633405 | -5.213447687 |
| **A_55_P2031668** | Gstp1 | -0.044878024 | -0.378282034 | 0.288525986 | 11.3507215 | -0.321715541 | 0.757542791 | 0.996633405 | -5.213449176 |
| **A_52_P511821** | Wdfy1 | -0.013314026 | -0.112282261 | 0.08565421 | 7.922679355 | -0.32153088 | 0.757676576 | 0.996633405 | -5.213498333 |
| **A_51_P211131** | Traf3ip2 | -0.017088619 | -0.144121344 | 0.109944106 | 8.203236542 | -0.321514447 | 0.757688482 | 0.996633405 | -5.213502706 |
| **A_55_P2213828** | 4933412E1 | -0.015413788 | -0.130047284 | 0.099219708 | 8.496480816 | -0.321371222 | 0.757792255 | 0.996633405 | -5.213540812 |
| **A_55_P1971649** | Ttc19 | -0.019066943 | -0.160875324 | 0.122741438 | 10.69938091 | -0.321357399 | 0.75780227 | 0.996633405 | -5.213544489 |
| **A_55_P2043554** | Slc12a4 | 0.024199729 | -0.155785021 | 0.204184479 | 11.87216948 | 0.321354159 | 0.757804617 | 0.996633405 | -5.213545351 |
| **A_51_P513941** | Lrpap1 | -0.013345011 | -0.112616589 | 0.085926568 | 11.04237124 | -0.32129438 | 0.757847932 | 0.996633405 | -5.21356125 |
| **A_52_P516097** | Leng9 | 0.022091757 | -0.142323631 | 0.186507144 | 8.927672891 | 0.321141864 | 0.757958444 | 0.996633405 | -5.213601801 |
| **A_52_P418644** | Plp2 | -0.038874713 | -0.328339289 | 0.250589863 | 11.11939374 | -0.320982166 | 0.758074168 | 0.996633405 | -5.213644241 |
| **A_51_P504588** | Tm2d2 | 0.026042797 | -0.167893704 | 0.219979297 | 10.52647583 | 0.320949863 | 0.758097576 | 0.996633405 | -5.213652823 |
| **A_55_P1971991** | Kdf1 | -0.013709133 | -0.115850279 | 0.088432013 | 7.707339384 | -0.320788212 | 0.758214724 | 0.996633405 | -5.213695758 |
| **A_51_P312997** | Zfp346 | 0.019651947 | -0.12678791 | 0.166091805 | 8.662776747 | 0.320741701 | 0.758248431 | 0.996633405 | -5.213708108 |
| **A_51_P483544** | Aass | -0.013767202 | -0.116358612 | 0.088824208 | 7.907192692 | -0.320733126 | 0.758254645 | 0.996633405 | -5.213710384 |
| **A_55_P2109485** | Milr1 | 0.042824414 | -0.276383865 | 0.362032693 | 15.07204913 | 0.320646433 | 0.758317475 | 0.996633405 | -5.213733398 |
| **A_55_P2008066** | Itpr1 | -0.031636853 | -0.267485294 | 0.204211589 | 11.37021733 | -0.320604338 | 0.758347984 | 0.996633405 | -5.21374457 |
| **A_55_P2207335** | Nron | 0.026067115 | -0.168265669 | 0.220399898 | 8.35220263 | 0.320594468 | 0.758355137 | 0.996633405 | -5.213747189 |
| **A_52_P267106** | Ksr2 | 0.014073291 | -0.090920741 | 0.119067322 | 8.022349365 | 0.320361394 | 0.758524067 | 0.996633405 | -5.213809021 |
| **A_52_P113537** | Xist | 0.120861727 | -0.781117238 | 1.022840692 | 7.966267166 | 0.320259129 | 0.758598193 | 0.996633405 | -5.213836138 |
| **A_52_P354298** | Ap2a2 | 0.025509304 | -0.164906681 | 0.215925288 | 9.39085736 | 0.320187468 | 0.758650137 | 0.996633405 | -5.213855134 |
| **A_51_P187171** | Smim12 | -0.025109407 | -0.212564949 | 0.162346135 | 12.1594152 | -0.320145423 | 0.758680615 | 0.996633405 | -5.213866278 |
| **A_55_P1954987** | Slc10a1 | 0.022813971 | -0.147510824 | 0.193138766 | 17.60535006 | 0.320134252 | 0.758688712 | 0.996633405 | -5.213869238 |
| **A_55_P2026753** | Zmat1 | 0.028277724 | -0.182944641 | 0.239500088 | 9.680000549 | 0.319973254 | 0.75880542 | 0.996633405 | -5.213911895 |
| **A_55_P1982986** | Cfap410 | -0.022186664 | -0.187916659 | 0.143543332 | 8.609968239 | -0.31996319 | 0.758812717 | 0.996633405 | -5.213914561 |
| **A_55_P1964138** | Pign | 0.041858401 | -0.27086335 | 0.354580153 | 10.11405453 | 0.31991432 | 0.758848144 | 0.996633405 | -5.213927505 |
| **A_55_P2114044** | Commd8 | 0.023286726 | -0.150804858 | 0.197378311 | 10.50090751 | 0.31969791 | 0.759005035 | 0.996633405 | -5.2139848 |
| **A_55_P2096917** | Mreg | 0.021288556 | -0.137910517 | 0.180487629 | 8.77739071 | 0.319605913 | 0.759071733 | 0.996633405 | -5.214009146 |
| **A_55_P2020326** | Madcam1 | 0.014704886 | -0.095336987 | 0.124746759 | 8.258702612 | 0.319383761 | 0.759232805 | 0.996633405 | -5.214067907 |
| **A_51_P520084** | Lypla2 | 0.021634297 | -0.140310098 | 0.183578692 | 12.89438143 | 0.319290503 | 0.759300426 | 0.996633405 | -5.214092562 |
| **A_55_P2061495** | Cks1b | -0.029078027 | -0.246755038 | 0.188598984 | 12.70916706 | -0.319272491 | 0.759313486 | 0.996633405 | -5.214097323 |
| **A_55_P2114482** | Rpgr | -0.027435303 | -0.23287423 | 0.178003625 | 9.224807762 | -0.319180352 | 0.759380299 | 0.996633405 | -5.214121675 |
| **A_55_P2076832** | Nrd1 | 0.018823328 | -0.122135475 | 0.15978213 | 8.764331802 | 0.319163582 | 0.759392459 | 0.996633405 | -5.214126107 |
| **A_55_P1967133** | Ppp1r3b | 0.023927179 | -0.155252116 | 0.203106474 | 12.68320086 | 0.319163195 | 0.75939274 | 0.996633405 | -5.214126209 |
| **A_55_P2004126** | NA | -0.027776601 | -0.235831358 | 0.180278156 | 10.15954182 | -0.319088081 | 0.759447209 | 0.996633405 | -5.214146055 |
| **A_55_P1967330** | Gm5069 | -0.025469701 | -0.216385223 | 0.165445821 | 16.05028335 | -0.318853906 | 0.75961703 | 0.996633405 | -5.214207898 |
| **A_55_P2071751** | NA | -0.038068613 | -0.323455282 | 0.247318057 | 9.684482783 | -0.318817754 | 0.759643248 | 0.996633405 | -5.214217442 |
| **A_55_P1975041** | Arhgef11 | 0.018755998 | -0.121851205 | 0.159363201 | 9.12388799 | 0.318817203 | 0.759643648 | 0.996633405 | -5.214217587 |
| **A_51_P434503** | Dtwd1 | 0.017582965 | -0.114264708 | 0.149430637 | 8.219507213 | 0.318734273 | 0.759703792 | 0.996633405 | -5.214239475 |
| **A_51_P189082** | Akr1c6 | -0.01322236 | -0.112400307 | 0.085955587 | 7.66214816 | -0.318641989 | 0.759770722 | 0.996633405 | -5.214263826 |
| **A_51_P393518** | Hmcn1 | -0.025676327 | -0.218272922 | 0.166920268 | 9.104398121 | -0.318634962 | 0.759775819 | 0.996633405 | -5.21426568 |
| **A_55_P1972653** | Tada3 | -0.017916727 | -0.152341662 | 0.116508209 | 10.99594788 | -0.318557597 | 0.75983193 | 0.996633405 | -5.214286088 |
| **A_51_P203192** | Mbd4 | 0.019983313 | -0.129961828 | 0.169928454 | 8.347665404 | 0.318525514 | 0.7598552 | 0.996633405 | -5.21429455 |
| **A_51_P400773** | Hdgfl2 | 0.016697186 | -0.108667893 | 0.142062265 | 11.19570573 | 0.318328726 | 0.759997937 | 0.996633405 | -5.214346434 |
| **A_52_P230904** | Wdr18 | 0.031928847 | -0.207808232 | 0.271665926 | 9.478058393 | 0.318315037 | 0.760007866 | 0.996633405 | -5.214350042 |
| **A_51_P322640** | Ccl24 | -0.015170627 | -0.129084204 | 0.09874295 | 7.836092075 | -0.318300398 | 0.760018484 | 0.996633405 | -5.214353901 |
| **A_52_P64763** | Gen1 | 0.014448601 | -0.094045355 | 0.122942557 | 8.123775542 | 0.318294682 | 0.760022631 | 0.996633405 | -5.214355407 |
| **A_52_P398211** | NA | 0.013727527 | -0.089352249 | 0.116807303 | 7.898645964 | 0.318293653 | 0.760023377 | 0.996633405 | -5.214355678 |
| **A_55_P2019203** | Spire1 | -0.018171755 | -0.154627056 | 0.118283545 | 7.993238678 | -0.318284581 | 0.760029958 | 0.996633405 | -5.214358069 |

| | | | | | | | | | |
|---|---|---|---|---|---|---|---|---|---|
| **A_52_P561736** | NA | -0.027796342 | -0.236590309 | 0.180997626 | 11.78381525 | -0.318184361 | 0.760102656 | 0.996633405 | -5.214384478 |
| **A_55_P2059337** | Fastk | 0.02546179 | -0.16581587 | 0.216739451 | 13.17717933 | 0.318151387 | 0.760126575 | 0.996633405 | -5.214393165 |
| **A_55_P2291776** | 4833447P1 | -0.016086998 | -0.136938371 | 0.104764374 | 8.417556522 | -0.318150467 | 0.760127242 | 0.996633405 | -5.214393407 |
| **A_55_P1952920** | AI854703 | 0.021666259 | -0.14111825 | 0.184450767 | 8.155747643 | 0.318111949 | 0.760155184 | 0.996633405 | -5.214403553 |
| **A_55_P1965079** | Gm10499 | 0.013031708 | -0.084886983 | 0.1109504 | 7.635753051 | 0.318086244 | 0.760173831 | 0.996633405 | -5.214410324 |
| **A_55_P1987494** | Gabrr2 | -0.02261721 | -0.192642226 | 0.147407805 | 10.93011834 | -0.317932808 | 0.760285139 | 0.996633405 | -5.214450728 |
| **A_55_P2128324** | Ubap2l | -0.01693711 | -0.1442729 | 0.110398679 | 8.721227679 | -0.317905442 | 0.760304992 | 0.996633405 | -5.214457932 |
| **A_55_P2020361** | Lzts3 | -0.064442193 | -0.549066196 | 0.420181809 | 10.98483285 | -0.317815099 | 0.760370534 | 0.996633405 | -5.214481711 |
| **A_52_P476699** | Gemin4 | -0.016470302 | -0.140358598 | 0.107417993 | 8.533574532 | -0.317746246 | 0.760420487 | 0.996633405 | -5.214499829 |
| **A_51_P206346** | Reep3 | 0.017973378 | -0.117250151 | 0.153196906 | 12.02971117 | 0.317677579 | 0.760470306 | 0.996633405 | -5.214517894 |
| **A_55_P2009351** | Prxl2c | 0.017757011 | -0.115917538 | 0.151431561 | 9.366949187 | 0.317490166 | 0.760606284 | 0.996633405 | -5.214567181 |
| **A_55_P2024461** | Tbkbp1 | 0.022150787 | -0.144604631 | 0.188906205 | 11.54527843 | 0.317481437 | 0.760612618 | 0.996633405 | -5.214569476 |
| **A_55_P2113723** | Pecr | 0.018539446 | -0.121047628 | 0.15812652 | 8.175027856 | 0.317439288 | 0.7606432 | 0.996633405 | -5.214580556 |
| **A_55_P1991069** | Milr1 | 0.030385091 | -0.198495286 | 0.259265468 | 9.389664485 | 0.317293454 | 0.760749019 | 0.996633405 | -5.214618883 |
| **A_55_P2419296** | NA | -0.021702718 | -0.185230019 | 0.141824584 | 8.18235481 | -0.317199857 | 0.760816937 | 0.996633405 | -5.214643473 |
| **A_52_P216226** | Masp1 | -0.01335356 | -0.114004877 | 0.087297756 | 7.704186912 | -0.317093066 | 0.760894431 | 0.996633405 | -5.21467152 |
| **A_52_P26299** | Slc35f1 | 0.01516732 | -0.099177127 | 0.129511767 | 8.531029148 | 0.317031862 | 0.760938846 | 0.996633405 | -5.21468759 |
| **A_52_P249856** | Cc2d2a | 0.015144908 | -0.099121215 | 0.129411031 | 8.648772135 | 0.316780382 | 0.761121353 | 0.996633405 | -5.21475359 |
| **A_51_P210956** | Vcam1 | 0.0266307 | -0.174435436 | 0.227696837 | 12.51413496 | 0.31655763 | 0.761283023 | 0.996633405 | -5.214812008 |
| **A_52_P137371** | Hmgcr | 0.022731934 | -0.148953487 | 0.194417354 | 8.312922578 | 0.3164551 | 0.761357443 | 0.996633405 | -5.214838884 |
| **A_51_P292196** | Prl7d1 | -0.041150753 | -0.352096721 | 0.269795215 | 7.813674643 | -0.316302037 | 0.761468547 | 0.996633405 | -5.21487899 |
| **A_51_P284577** | Hyal2 | 0.034989673 | -0.229410007 | 0.299389353 | 13.14515462 | 0.316291914 | 0.761475895 | 0.996633405 | -5.214881642 |
| **A_55_P2010858** | Mrps23 | -0.040641325 | -0.347920817 | 0.266638166 | 9.695549988 | -0.316113773 | 0.76160521 | 0.996633405 | -5.214928295 |
| **A_55_P1981929** | Scg5 | 0.013527355 | -0.088791664 | 0.115846374 | 7.910816908 | 0.315984405 | 0.761699124 | 0.996633405 | -5.214962158 |
| **A_55_P2077510** | Zfp583 | 0.02248493 | -0.147593127 | 0.192562986 | 8.412721009 | 0.315974754 | 0.761706131 | 0.996633405 | -5.214964684 |
| **A_55_P2029682** | Fry | 0.030640693 | -0.201130369 | 0.262411754 | 11.63291077 | 0.315971932 | 0.76170818 | 0.996633405 | -5.214965423 |
| **A_55_P2128794** | Clta | 0.017630079 | -0.115746012 | 0.151006169 | 12.63787323 | 0.315926022 | 0.761741509 | 0.996633405 | -5.214977436 |
| **A_55_P1990633** | Iigp1 | -0.012832791 | -0.109928022 | 0.084262441 | 7.736468064 | -0.315887426 | 0.761769529 | 0.996633405 | -5.214987535 |
| **A_51_P432659** | Pex7 | -0.017840119 | -0.152824666 | 0.117144429 | 10.41837046 | -0.315880502 | 0.761774557 | 0.996633405 | -5.214989347 |
| **A_55_P1986551** | Mcam | -0.024251191 | -0.207839983 | 0.1593376 | 8.881163749 | -0.315715606 | 0.761894274 | 0.996633405 | -5.215032477 |
| **A_52_P283501** | Zc3h7b | 0.019883105 | -0.130727695 | 0.170493906 | 10.30922845 | 0.31552751 | 0.762030843 | 0.996633405 | -5.21508165 |
| **A_55_P2166688** | Timm10b | -0.029294156 | -0.251217732 | 0.19262942 | 11.46314597 | -0.315490787 | 0.762057508 | 0.996633405 | -5.215091246 |
| **A_52_P670263** | Tfpi | 0.028250854 | -0.185806913 | 0.24230862 | 10.5625739 | 0.315434885 | 0.762098098 | 0.996633405 | -5.215105853 |
| **A_51_P133369** | Scly | 0.019972009 | -0.131365378 | 0.171309395 | 10.06778584 | 0.315416671 | 0.762111323 | 0.996633405 | -5.215110612 |
| **A_55_P2025153** | Hmgb1 | -0.049359113 | -0.423400654 | 0.324682429 | 14.4610706 | -0.315396332 | 0.762126092 | 0.996633405 | -5.215115926 |
| **A_55_P1975430** | St3gal3 | -0.025219859 | -0.216353345 | 0.165913626 | 11.53096745 | -0.315366103 | 0.762148042 | 0.996633405 | -5.215123822 |
| **A_51_P272817** | Map2k3 | 0.022692907 | -0.149317257 | 0.194703072 | 8.968016992 | 0.315315383 | 0.762184871 | 0.996633405 | -5.21513707 |
| **A_51_P125205** | Aqp1 | -0.049543594 | -0.425109516 | 0.326022329 | 12.54170076 | -0.31529019 | 0.762203165 | 0.996633405 | -5.21514365 |
| **A_51_P169327** | Etd | -0.01303214 | -0.111841276 | 0.085776996 | 7.705555654 | -0.315230153 | 0.762246762 | 0.996633405 | -5.215159327 |
| **A_52_P651078** | Fyttd1 | -0.018785832 | -0.161235391 | 0.123663727 | 8.291564184 | -0.315194374 | 0.762272743 | 0.996633405 | -5.215168669 |
| **A_55_P1964183** | Dysf | -0.016336102 | -0.140218751 | 0.107546547 | 11.49983844 | -0.315171614 | 0.762289271 | 0.996633405 | -5.215174611 |
| **A_51_P232207** | Hoxb6 | -0.032675664 | -0.280541034 | 0.215189705 | 8.233313327 | -0.315077728 | 0.76235745 | 0.996633405 | -5.215199118 |
| **A_55_P1968553** | Btbd9 | -0.020874579 | -0.17922434 | 0.137475183 | 10.91307992 | -0.315071728 | 0.762361808 | 0.996633405 | -5.215200684 |
| **A_51_P401958** | Taldo1 | 0.020905155 | -0.137678593 | 0.179488903 | 14.60996695 | 0.315067675 | 0.762364751 | 0.996633405 | -5.215201741 |
| **A_51_P323880** | COX2 | -0.050494001 | -0.433592325 | 0.332604324 | 16.92011151 | -0.315020387 | 0.762399092 | 0.996633405 | -5.215214082 |
| **A_55_P2067608** | Clmp | -0.024845887 | -0.213391485 | 0.16369971 | 8.610992649 | -0.314954086 | 0.762447241 | 0.996633405 | -5.215231381 |
| **A_55_P2116290** | Szrd1 | -0.026795617 | -0.230248316 | 0.176657081 | 9.905318 | -0.314781677 | 0.762572455 | 0.996633405 | -5.215276348 |
| **A_52_P331212** | Cspp1 | 0.020609635 | -0.135906871 | 0.177126141 | 8.288592631 | 0.31471634 | 0.762619909 | 0.996633405 | -5.215293384 |

| | | | | | | | | | |
|---|---|---|---|---|---|---|---|---|---|
| **A_55_P1976754** | Ppia | -0.031463239 | -0.270423173 | 0.207496695 | 17.48181833 | -0.314693283 | 0.762636655 | 0.996633405 | -5.215299394 |
| **A_52_P78922** | Ythdc2 | 0.029541925 | -0.19483851 | 0.25392236 | 8.603009878 | 0.314675503 | 0.762649569 | 0.996633405 | -5.215304029 |
| **A_55_P1975080** | Ugt2b5 | 0.013124478 | -0.086561369 | 0.112810326 | 7.659784265 | 0.314671694 | 0.762652335 | 0.996633405 | -5.215305022 |
| **A_55_P2001064** | NA | 0.016668512 | -0.109949271 | 0.143286295 | 11.77785365 | 0.314638055 | 0.762676768 | 0.996633405 | -5.21531379 |
| **A_55_P2022337** | Dhcr7 | 0.026608433 | -0.175568108 | 0.228784974 | 8.765490869 | 0.31455578 | 0.762736527 | 0.996633405 | -5.215335231 |
| **A_51_P409893** | Prkar2a | 0.029666947 | -0.196016611 | 0.255350506 | 8.738842886 | 0.314182559 | 0.76300763 | 0.996633405 | -5.215432427 |
| **A_52_P82765** | NA | 0.022930918 | -0.151567158 | 0.197428995 | 8.530115064 | 0.31407975 | 0.763082315 | 0.996633405 | -5.215459181 |
| **A_55_P2019054** | Acacb | -0.028218957 | -0.242985028 | 0.186547114 | 12.32281867 | -0.314039602 | 0.763111481 | 0.996633405 | -5.215469626 |
| **A_52_P199586** | Kpna6 | -0.026417492 | -0.227491904 | 0.174656919 | 11.47056367 | -0.314010317 | 0.763132756 | 0.996633405 | -5.215477245 |
| **A_55_P2145441** | NA | -0.024187088 | -0.208328629 | 0.159954454 | 9.504796754 | -0.313935866 | 0.763186844 | 0.996633405 | -5.21549661 |
| **A_55_P2050773** | LOC10004 | 0.013475403 | -0.089142302 | 0.116093108 | 7.803953123 | 0.313854673 | 0.763245831 | 0.996633405 | -5.215517724 |
| **A_55_P2090505** | NA | 0.070709911 | -0.467837525 | 0.609257346 | 11.84828407 | 0.313809046 | 0.763278981 | 0.996633405 | -5.215529586 |
| **A_51_P483617** | 0610040J0 | 0.017347905 | -0.11480473 | 0.14950054 | 8.196985744 | 0.313747551 | 0.763323659 | 0.996633405 | -5.215545572 |
| **A_52_P413646** | Bmp6 | 0.016314756 | -0.107991664 | 0.140621175 | 8.637335177 | 0.313686732 | 0.763367848 | 0.996633405 | -5.215561379 |
| **A_55_P2090364** | NA | 0.037749659 | -0.250017313 | 0.325516631 | 14.25969984 | 0.313531527 | 0.763480618 | 0.996633405 | -5.215601704 |
| **A_55_P2275249** | Th | -0.015544365 | -0.134112901 | 0.10302417 | 8.619907513 | -0.313337715 | 0.763621447 | 0.996633405 | -5.215652033 |
| **A_51_P101075** | Cnppd1 | -0.021420769 | -0.184817442 | 0.141975904 | 12.38351865 | -0.313329282 | 0.763627575 | 0.996633405 | -5.215654222 |
| **A_55_P1984043** | Pdcd4 | 0.048652647 | -0.322710818 | 0.420016112 | 12.7360012 | 0.313124053 | 0.763776712 | 0.996633405 | -5.215707481 |
| **A_55_P2011887** | Kif13a | 0.030564389 | -0.202737985 | 0.263866763 | 11.93353641 | 0.313116314 | 0.763782336 | 0.996633405 | -5.215709489 |
| **A_55_P2109505** | NA | -0.032101611 | -0.277158341 | 0.212955119 | 15.34039995 | -0.313090093 | 0.763801391 | 0.996633405 | -5.215716291 |
| **A_51_P404377** | Rnd2 | 0.02724725 | -0.180811575 | 0.235306075 | 8.157758182 | 0.313000957 | 0.763866168 | 0.996633405 | -5.21573941 |
| **A_55_P2179793** | NA | 0.022009809 | -0.146063264 | 0.190082881 | 8.359947965 | 0.312987693 | 0.763875808 | 0.996633405 | -5.215742849 |
| **A_55_P2009813** | NA | 0.01509982 | -0.100209835 | 0.130409475 | 7.909185038 | 0.312979021 | 0.76388211 | 0.996633405 | -5.215745098 |
| **A_55_P2037702** | Gm10638 | -0.024075054 | -0.207935299 | 0.159785191 | 9.297472825 | -0.312959807 | 0.763896074 | 0.996633405 | -5.215750081 |
| **A_55_P1976934** | Unc13d | -0.017599257 | -0.152005915 | 0.116807402 | 8.141167543 | -0.312955562 | 0.763899159 | 0.996633405 | -5.215751181 |
| **A_55_P2091676** | Fas | -0.014284156 | -0.123399249 | 0.094830937 | 8.504881447 | -0.312880785 | 0.763953504 | 0.996633405 | -5.215770569 |
| **A_55_P2173768** | Pkhd1 | 0.012465187 | -0.082766868 | 0.107697242 | 7.665594371 | 0.312841801 | 0.763981837 | 0.996633405 | -5.215780674 |
| **A_55_P1980432** | Zfp488 | 0.030015901 | -0.199304806 | 0.259336608 | 9.765087062 | 0.312836373 | 0.763985781 | 0.996633405 | -5.215782081 |
| **A_51_P497882** | Creb3l4 | -0.015008537 | -0.129709106 | 0.099692032 | 8.396668926 | -0.31273891 | 0.764056617 | 0.996633405 | -5.215807339 |
| **A_55_P2065824** | Smtn | 0.02363382 | -0.156989923 | 0.204257564 | 12.23523919 | 0.312729049 | 0.764063784 | 0.996633405 | -5.215809894 |
| **A_51_P320105** | Smyd2 | -0.020469731 | -0.176956676 | 0.136017213 | 14.63238635 | -0.312639007 | 0.764129229 | 0.996633405 | -5.215833223 |
| **A_55_P1985670** | Exosc3 | 0.022679987 | -0.150716241 | 0.196076216 | 9.066191369 | 0.312616787 | 0.764145379 | 0.996633405 | -5.215838978 |
| **A_55_P1989452** | Nenf | -0.026920751 | -0.232861812 | 0.179020311 | 13.73904418 | -0.312430447 | 0.764280824 | 0.996633405 | -5.215887231 |
| **A_55_P2000638** | Stag2 | 0.025138543 | -0.16721521 | 0.217492295 | 12.23683541 | 0.312355075 | 0.764335611 | 0.996633405 | -5.215906741 |
| **A_52_P206861** | Klf8 | 0.013843892 | -0.092097803 | 0.119785588 | 8.021251193 | 0.312320449 | 0.764360781 | 0.996633405 | -5.215915702 |
| **A_55_P1995438** | NA | 0.014563619 | -0.096913821 | 0.126041059 | 15.13546679 | 0.312242097 | 0.764417737 | 0.996633405 | -5.215935976 |
| **A_52_P176160** | Ahcy | 0.020292424 | -0.135037569 | 0.175622416 | 8.133117182 | 0.312239418 | 0.764419685 | 0.996633405 | -5.215936669 |
| **A_51_P271665** | Rev3l | 0.029463701 | -0.196092424 | 0.255019827 | 11.30053529 | 0.312206405 | 0.764443683 | 0.996633405 | -5.21594521 |
| **A_52_P506995** | Plet1 | 0.013660141 | -0.09093357 | 0.118253852 | 7.665008185 | 0.31214669 | 0.764487094 | 0.996633405 | -5.215960657 |
| **A_55_P2058122** | Kyat3 | -0.01987734 | -0.172211629 | 0.132456949 | 11.16770259 | -0.31186722 | 0.764690267 | 0.996633405 | -5.21603291 |
| **A_55_P2149931** | Arap2 | 0.01584082 | -0.105616777 | 0.137298418 | 8.014690077 | 0.311718168 | 0.764798636 | 0.996633405 | -5.21607142 |
| **A_52_P251450** | Cldn2 | -0.012568399 | -0.108953089 | 0.08381629 | 7.646097459 | -0.311659989 | 0.764840937 | 0.996633405 | -5.216086447 |
| **A_55_P2036317** | NA | -0.02295804 | -0.199123906 | 0.153207826 | 17.03554273 | -0.311474266 | 0.764975977 | 0.996633405 | -5.216134398 |
| **A_55_P2098210** | 4931406P1 | -0.022778809 | -0.197585093 | 0.152027475 | 11.50095478 | -0.31144625 | 0.764996349 | 0.996633405 | -5.216141629 |
| **A_55_P2105794** | Wdr61 | 0.014775044 | -0.098620797 | 0.128170885 | 12.26875998 | 0.311415904 | 0.765018415 | 0.996633405 | -5.216149461 |
| **A_55_P2116105** | Ap2a1 | -0.018690302 | -0.162155936 | 0.124775332 | 11.57548185 | -0.311370572 | 0.765051379 | 0.996633405 | -5.216161159 |
| **A_52_P206526** | Gtf2ird2 | -0.018242965 | -0.158275275 | 0.121789344 | 9.909804462 | -0.311369675 | 0.765052031 | 0.996633405 | -5.21616139 |
| **A_55_P2097518** | Zdhhc22 | -0.12169913 | -1.055915178 | 0.812516917 | 12.5599071 | -0.311350297 | 0.765066122 | 0.996633405 | -5.21616639 |

| | | | | | | | | | |
|---|---|---|---|---|---|---|---|---|---|
| **A_51_P151586** | Haspin | 0.01467938 | -0.098021817 | 0.127380576 | 8.715650651 | 0.311306589 | 0.765097905 | 0.996633405 | -5.216177667 |
| **A_55_P1961463** | Bnip3l | 0.023663609 | -0.158016514 | 0.205343732 | 12.80522907 | 0.31130256 | 0.765100835 | 0.996633405 | -5.216178706 |
| **A_52_P407697** | NA | 0.013559204 | -0.090553158 | 0.117671566 | 7.906958071 | 0.311272699 | 0.765122549 | 0.996633405 | -5.216186409 |
| **A_55_P2176045** | NA | 0.033328951 | -0.222626694 | 0.289284596 | 8.495967013 | 0.311218974 | 0.765161617 | 0.996633405 | -5.216200266 |
| **A_55_P1957850** | Snx9 | -0.019418652 | -0.168552908 | 0.129715603 | 10.20966733 | -0.311208042 | 0.765169567 | 0.996633405 | -5.216203086 |
| **A_66_P111666** | Nprl3 | 0.022855845 | -0.152679425 | 0.198391116 | 9.898541719 | 0.311201744 | 0.765174147 | 0.996633405 | -5.21620471 |
| **A_55_P2110200** | Ubox5 | -0.018105303 | -0.157203183 | 0.120992577 | 8.797599544 | -0.311095997 | 0.765251048 | 0.996633405 | -5.216231977 |
| **A_51_P484842** | Asgr2 | -0.013506251 | -0.117295451 | 0.090282949 | 7.931615388 | -0.311022479 | 0.765304514 | 0.996633405 | -5.216250929 |
| **A_55_P1990038** | Mark2 | -0.018099561 | -0.157210238 | 0.121011117 | 8.580812596 | -0.310968713 | 0.765343615 | 0.996633405 | -5.216264786 |
| **A_55_P2012201** | App | -0.02240349 | -0.194625432 | 0.149818451 | 13.6029568 | -0.310911174 | 0.765385462 | 0.996633405 | -5.216279613 |
| **A_55_P2126662** | Cav1 | -0.026440567 | -0.229758387 | 0.176877254 | 9.357641583 | -0.310816772 | 0.76545412 | 0.996633405 | -5.216303933 |
| **A_55_P2018211** | NA | -0.021134399 | -0.183661875 | 0.141393078 | 15.28232707 | -0.310793719 | 0.765470886 | 0.996633405 | -5.216309871 |
| **A_55_P2121541** | Gm6607 | 0.017148996 | -0.114800966 | 0.149098959 | 14.69731271 | 0.31062655 | 0.765592473 | 0.996633405 | -5.216352918 |
| **A_55_P1997604** | Pla2g4a | 0.029744798 | -0.199314412 | 0.258804009 | 8.871635314 | 0.310364758 | 0.765782897 | 0.996633405 | -5.216420285 |
| **A_51_P292550** | Chdh | 0.012644677 | -0.084756355 | 0.110045708 | 7.694896198 | 0.310279654 | 0.765844804 | 0.996633405 | -5.216442173 |
| **A_51_P345663** | Rrbp1 | -0.02598703 | -0.226274402 | 0.174300342 | 12.08949448 | -0.310107459 | 0.765970069 | 0.996633405 | -5.216486443 |
| **A_55_P2009069** | NA | 0.030003289 | -0.201244121 | 0.2612507 | 12.45013116 | 0.310099541 | 0.765975829 | 0.996633405 | -5.216488477 |
| **A_52_P299846** | Sh3rf1 | 0.018333459 | -0.122996025 | 0.159662942 | 8.231948132 | 0.310042168 | 0.766017568 | 0.996633405 | -5.216503222 |
| **A_55_P1967022** | Fgfr1 | 0.020051504 | -0.134526072 | 0.17462908 | 8.265611232 | 0.310034194 | 0.766023369 | 0.996633405 | -5.216505271 |
| **A_55_P1958652** | Arf2 | 0.015798703 | -0.106043635 | 0.137641041 | 12.08401396 | 0.309907686 | 0.766115406 | 0.996633405 | -5.216537772 |
| **A_55_P1987645** | Unc13b | -0.024157404 | -0.210480104 | 0.162165297 | 11.66626693 | -0.30988005 | 0.766135513 | 0.996633405 | -5.216544871 |
| **A_55_P1952166** | Arid3a | 0.030388339 | -0.204047106 | 0.264823783 | 8.849709017 | 0.309808137 | 0.766187833 | 0.996633405 | -5.216563339 |
| **A_51_P124126** | Cyp2d22 | -0.013441618 | -0.117146139 | 0.090262902 | 7.754611624 | -0.30978687 | 0.766203306 | 0.996633405 | -5.2165688 |
| **A_55_P2102951** | NA | 0.024033511 | -0.16139757 | 0.209464592 | 11.83559207 | 0.309773187 | 0.766213261 | 0.996633405 | -5.216572313 |
| **A_52_P380263** | Podxl | -0.022603507 | -0.197018844 | 0.151811829 | 8.523522536 | -0.309742142 | 0.766235849 | 0.996633405 | -5.216580283 |
| **A_55_P2010018** | NA | 0.01624201 | -0.109125761 | 0.141609782 | 16.28494461 | 0.309644237 | 0.766307083 | 0.996633405 | -5.216605415 |
| **A_55_P2066234** | Arfip2 | -0.020766015 | -0.18106577 | 0.139533739 | 9.278051003 | -0.309620315 | 0.766324489 | 0.996633405 | -5.216611554 |
| **A_52_P136709** | Pctp | -0.015068018 | -0.131385898 | 0.101249862 | 7.935819499 | -0.30961271 | 0.766330023 | 0.996633405 | -5.216613506 |
| **A_55_P2067151** | NA | -0.050549749 | -0.441008803 | 0.339909305 | 9.805184756 | -0.309423032 | 0.766468039 | 0.996633405 | -5.216662168 |
| **A_55_P2346654** | AU035318 | 0.031778164 | -0.213699564 | 0.277255891 | 8.285689727 | 0.309403937 | 0.766481934 | 0.996633405 | -5.216667065 |
| **A_55_P2112589** | Uxt | -0.017624462 | -0.153814746 | 0.118565821 | 11.21560996 | -0.309299265 | 0.766558101 | 0.996633405 | -5.216693905 |
| **A_55_P2010552** | Hsd17b13 | -0.016361268 | -0.142808038 | 0.110085502 | 7.704551903 | -0.309256158 | 0.76658947 | 0.996633405 | -5.216704955 |
| **A_55_P2138343** | Ndufaf5 | -0.027162956 | -0.237123351 | 0.182797438 | 8.871075283 | -0.309206623 | 0.766625517 | 0.996633405 | -5.216717652 |
| **A_55_P1989797** | NA | 0.021722987 | -0.146276979 | 0.189722954 | 14.59994435 | 0.309043408 | 0.766744294 | 0.996633405 | -5.216759474 |
| **A_55_P1986199** | Tut1 | 0.015131041 | -0.10190301 | 0.132165092 | 8.332484467 | 0.309005139 | 0.766772145 | 0.996633405 | -5.216769276 |
| **A_55_P2162592** | Lman2 | 0.017497375 | -0.11785299 | 0.15284774 | 12.37877354 | 0.308974472 | 0.766794463 | 0.996633405 | -5.216777131 |
| **A_55_P2287269** | AI415730 | 0.025475584 | -0.171616335 | 0.222567504 | 8.752896297 | 0.308933119 | 0.766824559 | 0.996633405 | -5.216787722 |
| **A_55_P2061492** | Pnldc1 | -0.013596825 | -0.118807141 | 0.091613491 | 8.11359457 | -0.308878949 | 0.766863984 | 0.996633405 | -5.216801592 |
| **A_65_P02177** | Gstm4 | -0.024027138 | -0.209948077 | 0.161893801 | 8.34481436 | -0.308875082 | 0.766866798 | 0.996633405 | -5.216802583 |
| **A_55_P2028942** | Ncam1 | -0.039743265 | -0.347425918 | 0.267939387 | 9.868702719 | -0.308723486 | 0.766977133 | 0.996633405 | -5.216841388 |
| **A_66_P106191** | Eef2 | -0.034768022 | -0.303958543 | 0.234422499 | 14.65701729 | -0.308694828 | 0.766997991 | 0.996633405 | -5.216848721 |
| **A_55_P2108933** | Dao | -0.012522298 | -0.109485785 | 0.084441188 | 7.659174545 | -0.308663274 | 0.767020958 | 0.996633405 | -5.216856795 |
| **A_51_P384441** | Tmem201 | -0.042100797 | -0.368117407 | 0.283915814 | 9.784414556 | -0.308645335 | 0.767034015 | 0.996633405 | -5.216861385 |
| **A_52_P114889** | Upk3b | 0.112461379 | -0.758408792 | 0.98333155 | 9.244441383 | 0.308644912 | 0.767034322 | 0.996633405 | -5.216861493 |
| **A_52_P334796** | Taar4 | 0.015682407 | -0.105842661 | 0.137207475 | 7.850750069 | 0.308429551 | 0.767191081 | 0.996633405 | -5.216916575 |
| **A_55_P2053616** | NA | 0.037866037 | -0.255564513 | 0.331296587 | 10.31924607 | 0.3084279 | 0.767192282 | 0.996633405 | -5.216916998 |
| **A_55_P1970650** | Dgkh | -0.014134317 | -0.123671831 | 0.095403196 | 8.535349193 | -0.30840477 | 0.767209119 | 0.996633405 | -5.216922911 |
| **A_55_P2080378** | Ttc30a2 | -0.015874782 | -0.138928354 | 0.107178789 | 8.290704555 | -0.308334906 | 0.767259975 | 0.996633405 | -5.216940771 |

| | | | | | | | | | |
|---|---|---|---|---|---|---|---|---|---|
| **A_51_P343429** | Slc25a37 | 0.11210128 | -0.757341091 | 0.981543651 | 12.45066329 | 0.308161871 | 0.767385937 | 0.996633405 | -5.216984987 |
| **A_55_P2003541** | Nrcam | 0.016039032 | -0.108360842 | 0.140438907 | 7.942727405 | 0.308153677 | 0.767391902 | 0.996633405 | -5.21698708 |
| **A_55_P2048036** | Zfx | 0.025905219 | -0.175058964 | 0.226869402 | 10.65022362 | 0.308090097 | 0.767438189 | 0.996633405 | -5.217003321 |
| **A_55_P2149219** | Aar2 | 0.025759331 | -0.1741449 | 0.225663563 | 10.34149994 | 0.307979444 | 0.767518745 | 0.996633405 | -5.217031578 |
| **A_55_P1975874** | Bcl2l15 | 0.015171396 | -0.102579596 | 0.132922388 | 7.840643676 | 0.307942828 | 0.767545403 | 0.996633405 | -5.217040926 |
| **A_52_P218976** | Cyld | 0.020058179 | -0.135660218 | 0.175776576 | 8.463529375 | 0.307865281 | 0.76760186 | 0.996633405 | -5.217060721 |
| **A_55_P2061465** | Ppp1r21 | 0.015905623 | -0.107585346 | 0.139396593 | 8.99738113 | 0.307839697 | 0.767620487 | 0.996633405 | -5.21706725 |
| **A_55_P1983588** | Pmepa1 | 0.020564172 | -0.139154456 | 0.180282799 | 10.84837974 | 0.307726422 | 0.76770296 | 0.996633405 | -5.217096154 |
| **A_51_P115085** | Nup153 | 0.025255383 | -0.171004236 | 0.221515003 | 10.28230588 | 0.307561626 | 0.76782295 | 0.996633405 | -5.217138186 |
| **A_55_P2074836** | Mapk8ip1 | -0.021355562 | -0.187384957 | 0.144673832 | 10.19954757 | -0.307422151 | 0.767924509 | 0.996633405 | -5.217173742 |
| **A_55_P2371984** | 5930436O1 | 0.033632063 | -0.22799327 | 0.295257397 | 8.548786735 | 0.307243604 | 0.768054525 | 0.996633405 | -5.217219236 |
| **A_55_P2117155** | Apoc3 | -0.012616944 | -0.110795783 | 0.085561895 | 7.663629476 | -0.307146394 | 0.768125316 | 0.996633405 | -5.217243995 |
| **A_51_P176883** | Edrf1 | -0.01492771 | -0.131166194 | 0.101310774 | 8.293478289 | -0.306939217 | 0.768276196 | 0.996633405 | -5.217296737 |
| **A_55_P1984152** | 1600029O1 | -0.014985289 | -0.131706935 | 0.101736358 | 15.06774461 | -0.30684768 | 0.768342863 | 0.996633405 | -5.217320028 |
| **A_55_P2002667** | Trim26 | 0.016997047 | -0.115398345 | 0.149392439 | 10.02426443 | 0.306838425 | 0.768349603 | 0.996633405 | -5.217322383 |
| **A_55_P2184766** | Tmc6 | 0.022272789 | -0.151314565 | 0.195860143 | 9.30377697 | 0.306666021 | 0.768475172 | 0.996633405 | -5.217366232 |
| **A_55_P2115260** | NA | 0.022496274 | -0.152902682 | 0.197895229 | 8.227412153 | 0.306543927 | 0.768564102 | 0.996633405 | -5.217397271 |
| **A_51_P249594** | Rev1 | -0.018337486 | -0.161352496 | 0.124677524 | 8.53778949 | -0.306455417 | 0.768628573 | 0.996633405 | -5.217419765 |
| **A_55_P1967191** | Gm4632 | -0.012869008 | -0.113249944 | 0.087511928 | 7.710967196 | -0.30641002 | 0.768661641 | 0.996633405 | -5.217431299 |
| **A_52_P467449** | Alox12 | -0.029990783 | -0.263982896 | 0.20400133 | 8.806684371 | -0.306334362 | 0.768716753 | 0.996633405 | -5.217450519 |
| **A_55_P2410106** | 9130230N0 | 0.01276956 | -0.086865223 | 0.112404343 | 7.882854151 | 0.306319095 | 0.768727874 | 0.996633405 | -5.217454397 |
| **A_52_P571746** | Sh2d5 | -0.01245964 | -0.109727059 | 0.084807779 | 7.692970903 | -0.306159143 | 0.768844394 | 0.996633405 | -5.217495013 |
| **A_55_P2090184** | Fgfr1op2 | 0.037788587 | -0.257665816 | 0.333242991 | 11.92026316 | 0.305688657 | 0.769187166 | 0.996633405 | -5.217614365 |
| **A_55_P2300071** | 4833412C1 | 0.04380418 | -0.298758346 | 0.386366707 | 8.918034223 | 0.305622159 | 0.769235618 | 0.996633405 | -5.217631219 |
| **A_55_P2029366** | Abhd6 | -0.017162307 | -0.151397134 | 0.11707252 | 8.252952319 | -0.305576245 | 0.769269072 | 0.996633405 | -5.217642855 |
| **A_55_P1972025** | Mycl | 0.012797022 | -0.087319652 | 0.112913695 | 7.67284302 | 0.305500288 | 0.769324418 | 0.996633405 | -5.2176621 |
| **A_55_P1985984** | Actg1 | -0.019555476 | -0.172679787 | 0.133568835 | 12.10771332 | -0.305234333 | 0.769518216 | 0.996633405 | -5.217729448 |
| **A_55_P2004208** | Defa-rs2 | 0.013546364 | -0.092566289 | 0.119659018 | 7.714893044 | 0.305115807 | 0.769604591 | 0.996633405 | -5.217759444 |
| **A_55_P1955108** | Ripor1 | 0.026764017 | -0.182907183 | 0.236435216 | 13.63311858 | 0.305085555 | 0.769626637 | 0.996633405 | -5.217767098 |
| **A_52_P613498** | 4833420G1 | 0.039839276 | -0.272279681 | 0.351958232 | 11.02222651 | 0.305070644 | 0.769637504 | 0.996633405 | -5.217770871 |
| **A_55_P2022699** | NA | -0.017766507 | -0.15709196 | 0.121558947 | 8.232495038 | -0.304775959 | 0.76985227 | 0.996633405 | -5.21784539 |
| **A_51_P295896** | 4930452B0 | -0.021870082 | -0.193376997 | 0.149636833 | 9.269160727 | -0.304773998 | 0.769853699 | 0.996633405 | -5.217845885 |
| **A_66_P133498** | Pdcd6ip | -0.020903913 | -0.1848384 | 0.143030573 | 11.10641021 | -0.304765925 | 0.769859583 | 0.996633405 | -5.217847926 |
| **A_55_P2085880** | Aco1 | -0.016361951 | -0.144744367 | 0.112020466 | 8.325858528 | -0.304606152 | 0.769976036 | 0.996633405 | -5.217888298 |
| **A_55_P2067767** | Psg25 | -0.045343793 | -0.401131503 | 0.310443918 | 7.839625714 | -0.304604301 | 0.769977385 | 0.996633405 | -5.217888766 |
| **A_55_P1981724** | Nr2c1 | -0.019470503 | -0.172285867 | 0.13334486 | 9.527093789 | -0.304522435 | 0.770037056 | 0.996633405 | -5.217909444 |
| **A_55_P2032755** | Gm10586 | -0.021084531 | -0.186725521 | 0.144556459 | 14.39897004 | -0.304232253 | 0.770248581 | 0.996633405 | -5.217982698 |
| **A_65_P03364** | Dennd4a | 0.013771208 | -0.094488028 | 0.122030444 | 7.758849581 | 0.304029843 | 0.770396138 | 0.996633405 | -5.218033754 |
| **A_55_P2028595** | AW551984 | -0.023108774 | -0.204798226 | 0.158580678 | 8.541644325 | -0.303987922 | 0.7704267 | 0.996633405 | -5.218044324 |
| **A_51_P308446** | 1700094D0 | 0.025654729 | -0.176079961 | 0.227389419 | 8.160762525 | 0.303945682 | 0.770457494 | 0.996633405 | -5.218054973 |
| **A_51_P239203** | Mapk13 | 0.015293528 | -0.105030242 | 0.135617299 | 7.931704804 | 0.303784349 | 0.770575117 | 0.996633405 | -5.218095634 |
| **A_52_P70027** | Gm20498 | -0.022214593 | -0.197004237 | 0.152575051 | 12.04833078 | -0.303760847 | 0.770592252 | 0.996633405 | -5.218101555 |
| **A_55_P2142222** | Serpina3h | 0.016557932 | -0.113725601 | 0.146841466 | 7.816740454 | 0.303756587 | 0.770595358 | 0.996633405 | -5.218102628 |
| **A_55_P2135725** | C8g | -0.015860305 | -0.140660566 | 0.108939955 | 7.886270238 | -0.303742235 | 0.770605822 | 0.996633405 | -5.218106244 |
| **A_51_P422629** | Tbpl1 | 0.015506734 | -0.106533491 | 0.13754696 | 10.46872603 | 0.303687193 | 0.770645953 | 0.996633405 | -5.21812011 |
| **A_52_P404329** | Saa4 | -0.01309322 | -0.116148014 | 0.089961575 | 7.667243404 | -0.303659857 | 0.770665885 | 0.996633405 | -5.218126995 |
| **A_55_P2050273** | Cct6b | -0.012521671 | -0.111094542 | 0.0860512 | 7.730840726 | -0.303608563 | 0.770703284 | 0.996633405 | -5.218139913 |
| **A_51_P431734** | Fam185a | -0.019949587 | -0.177070724 | 0.137171549 | 8.670896515 | -0.303464876 | 0.770808053 | 0.996633405 | -5.218176088 |

| | | | | | | | | | |
|---|---|---|---|---|---|---|---|---|---|
| **A_55_P2060444** | Apopt1 | -0.022597826 | -0.200606218 | 0.155410566 | 16.13078393 | -0.30341379 | 0.770845304 | 0.996633405 | -5.218188946 |
| **A_55_P2090600** | Mageb16-p | 0.013011054 | -0.089501379 | 0.115523487 | 7.648456723 | 0.303350744 | 0.770891277 | 0.996633405 | -5.218204811 |
| **A_55_P1972575** | Tmeff1 | 0.034029453 | -0.234167949 | 0.302226856 | 11.37326464 | 0.303256095 | 0.770960296 | 0.996633405 | -5.218228623 |
| **A_55_P2060376** | Gzmg | 0.018752316 | -0.129048465 | 0.166553097 | 7.718513918 | 0.303240578 | 0.770971611 | 0.996633405 | -5.218232526 |
| **A_66_P127484** | Abcf3 | -0.013657235 | -0.121351326 | 0.094036855 | 10.61913406 | -0.303095897 | 0.771077118 | 0.996633405 | -5.218268909 |
| **A_52_P412529** | Fbxo3 | -0.015671025 | -0.139292005 | 0.107949954 | 8.48647104 | -0.302980283 | 0.771161433 | 0.996633405 | -5.218297971 |
| **A_55_P2156504** | NA | -0.027432373 | -0.243845419 | 0.188980672 | 13.69107776 | -0.302962643 | 0.771174298 | 0.996633405 | -5.218302404 |
| **A_55_P2016049** | Gp6 | 0.018370714 | -0.126578786 | 0.163320214 | 7.903471497 | 0.302913374 | 0.77121023 | 0.996633405 | -5.218314785 |
| **A_55_P2000613** | Slc6a1 | 0.013822685 | -0.095254094 | 0.122899464 | 7.814669978 | 0.302879044 | 0.771235267 | 0.996633405 | -5.21832341 |
| **A_55_P2166414** | Uqcr10 | -0.021294739 | -0.18934339 | 0.146753912 | 16.18807894 | -0.302863143 | 0.771246864 | 0.996633405 | -5.218327405 |
| **A_51_P490700** | Ccdc40 | -0.014099899 | -0.125501782 | 0.097301984 | 7.798322053 | -0.302505038 | 0.771508053 | 0.996633405 | -5.218417319 |
| **A_55_P2079390** | 4930579G2 | 0.014940727 | -0.103105027 | 0.132986481 | 9.215369043 | 0.302503561 | 0.77150913 | 0.996633405 | -5.21841769 |
| **A_51_P353502** | Sharpin | 0.015546897 | -0.107293791 | 0.138387586 | 11.90519443 | 0.302489722 | 0.771519224 | 0.996633405 | -5.218421162 |
| **A_66_P100696** | Slc26a2 | -0.0149267 | -0.132933335 | 0.103079935 | 7.95676313 | -0.302319741 | 0.771643214 | 0.996633405 | -5.218463804 |
| **A_51_P506937** | Mrps12 | 0.013665257 | -0.094425973 | 0.121756488 | 13.13807737 | 0.302159663 | 0.771759987 | 0.996633405 | -5.21850394 |
| **A_52_P380329** | Tent4a | -0.019175753 | -0.170871633 | 0.132520127 | 10.50387766 | -0.302125766 | 0.771784715 | 0.996633405 | -5.218512436 |
| **A_66_P111449** | 1700018A0 | -0.012719989 | -0.113379461 | 0.087939482 | 8.049219057 | -0.30202385 | 0.771859064 | 0.996633405 | -5.218537976 |
| **A_55_P2160473** | LOC63486 | 0.038318077 | -0.264941052 | 0.341577206 | 13.99096213 | 0.301994443 | 0.771880518 | 0.996633405 | -5.218545343 |
| **A_55_P2176871** | NA | 0.019584785 | -0.135441237 | 0.174610806 | 15.11289989 | 0.301941861 | 0.771918878 | 0.996633405 | -5.218558515 |
| **A_55_P1969745** | Pitpnm1 | -0.02912123 | -0.259645635 | 0.201403175 | 11.76447415 | -0.301926988 | 0.771929729 | 0.996633405 | -5.218562241 |
| **A_55_P1993453** | Ftsj1 | 0.014043659 | -0.097151696 | 0.125239013 | 10.89212291 | 0.301858042 | 0.77198003 | 0.996633405 | -5.218579509 |
| **A_55_P2029945** | NA | 0.07317977 | -0.506505607 | 0.652865147 | 14.73747777 | 0.301722589 | 0.772078854 | 0.996633405 | -5.218613421 |
| **A_55_P2008244** | NA | -0.027729241 | -0.247404159 | 0.191945677 | 9.530860142 | -0.301693978 | 0.772099729 | 0.996633405 | -5.218620583 |
| **A_51_P455647** | Car2 | 0.049043368 | -0.339694387 | 0.437781123 | 9.805063029 | 0.301531505 | 0.772218274 | 0.996633405 | -5.218661238 |
| **A_55_P1975640** | Zfp217 | -0.019033077 | -0.16993029 | 0.131864137 | 10.46964468 | -0.301465007 | 0.772266796 | 0.996633405 | -5.218677871 |
| **A_52_P520397** | Timm8b | 0.013481311 | -0.093403816 | 0.120366438 | 14.38622356 | 0.301456042 | 0.772273337 | 0.996633405 | -5.218680113 |
| **A_55_P2087315** | Shroom1 | 0.016289683 | -0.1128812 | 0.145460567 | 8.319838609 | 0.301409642 | 0.772307194 | 0.996633405 | -5.218691717 |
| **A_55_P2088601** | Fbln1 | -0.022566948 | -0.201514276 | 0.15638038 | 8.394829783 | -0.301409371 | 0.772307392 | 0.996633405 | -5.218691785 |
| **A_55_P2053177** | NA | 0.016451204 | -0.114014102 | 0.146916511 | 12.35790814 | 0.301378169 | 0.772330159 | 0.996633405 | -5.218699587 |
| **A_55_P2090414** | NA | 0.03198251 | -0.221963214 | 0.285928235 | 8.600712727 | 0.301009899 | 0.772598899 | 0.996633405 | -5.218791614 |
| **A_55_P2444965** | Tmem245 | -0.018158904 | -0.162362515 | 0.126044707 | 9.087209195 | -0.300969601 | 0.772628308 | 0.996633405 | -5.218801677 |
| **A_52_P378968** | Rgs2 | -0.012762945 | -0.11412048 | 0.088594589 | 7.947801439 | -0.300956693 | 0.772637728 | 0.996633405 | -5.2188049 |
| **A_66_P136228** | Rai14 | 0.023735433 | -0.164791066 | 0.212261931 | 10.25116521 | 0.300908105 | 0.772673188 | 0.996633405 | -5.218817032 |
| **A_51_P219918** | Tmem125 | 0.01768872 | -0.122810091 | 0.158187531 | 7.774387481 | 0.300907417 | 0.77267369 | 0.996633405 | -5.218817203 |
| **A_55_P2063426** | NA | 0.023277534 | -0.161618043 | 0.20817311 | 8.647546188 | 0.300898188 | 0.772680426 | 0.996633405 | -5.218819508 |
| **A_51_P481679** | Angptl3 | -0.015631924 | -0.139846611 | 0.108582763 | 7.884037939 | -0.300779766 | 0.772766853 | 0.996633405 | -5.218849066 |
| **A_52_P183524** | Tmem86b | -0.012839159 | -0.114885525 | 0.089207207 | 7.752275837 | -0.300710201 | 0.772817625 | 0.996633405 | -5.218866425 |
| **A_52_P670874** | Ebag9 | 0.024225747 | -0.168338124 | 0.216789618 | 10.02756774 | 0.30068482 | 0.772836149 | 0.996633405 | -5.218872757 |
| **A_55_P2017870** | Pclaf | 0.025980084 | -0.180545999 | 0.232506166 | 13.53353309 | 0.30065939 | 0.77285471 | 0.996633405 | -5.218879101 |
| **A_55_P1994302** | NA | 0.013145461 | -0.091412795 | 0.117703717 | 7.890748577 | 0.300487645 | 0.772980065 | 0.996633405 | -5.218921933 |
| **A_55_P2036083** | Gm5797 | 0.045096079 | -0.313669903 | 0.40386206 | 8.97335947 | 0.300425404 | 0.773025496 | 0.996633405 | -5.21893745 |
| **A_55_P2180099** | NA | 0.02416398 | -0.16817171 | 0.21649967 | 14.01447461 | 0.300273989 | 0.773136021 | 0.996633405 | -5.218975184 |
| **A_51_P153683** | Lrrc26 | -0.012164736 | -0.109011476 | 0.084682004 | 7.649228 | -0.300211145 | 0.773181895 | 0.996633405 | -5.21899084 |
| **A_55_P2170034** | Rgl1 | 0.021923708 | -0.152623741 | 0.196471157 | 10.88285818 | 0.300199272 | 0.773190562 | 0.996633405 | -5.218993798 |
| **A_52_P59318** | Rdh10 | -0.014245488 | -0.127664058 | 0.099173082 | 8.083031276 | -0.3001942 | 0.773194264 | 0.996633405 | -5.218995061 |
| **A_55_P2026089** | Usp30 | 0.019851906 | -0.138205497 | 0.177909309 | 10.30030174 | 0.30019019 | 0.773197192 | 0.996633405 | -5.21899606 |
| **A_51_P513460** | Cfhr1 | -0.012083892 | -0.108337024 | 0.08416924 | 7.620348717 | -0.300055161 | 0.773295764 | 0.996633405 | -5.219029687 |
| **A_55_P1966664** | Cdc14b | -0.02976554 | -0.266909425 | 0.207378346 | 10.66811279 | -0.299992888 | 0.773341224 | 0.996633405 | -5.219045189 |

| | | | | | | | | | |
|---|---|---|---|---|---|---|---|---|---|
| **A_55_P2073789** | NA | -0.067344985 | -0.604083267 | 0.469393298 | 11.36544395 | -0.29988298 | 0.773421463 | 0.996633405 | -5.219072544 |
| **A_55_P2139809** | Fgf13 | 0.035809477 | -0.249688362 | 0.321307316 | 9.433105342 | 0.299781123 | 0.773495826 | 0.996633405 | -5.219097885 |
| **A_51_P231687** | Ik | 0.016381842 | -0.114236664 | 0.147000348 | 13.72033061 | 0.29975549 | 0.773514541 | 0.996633405 | -5.219104262 |
| **A_55_P2137653** | NA | -0.014526823 | -0.130360661 | 0.101307015 | 13.45493849 | -0.299739745 | 0.773526036 | 0.996633405 | -5.219108178 |
| **A_51_P303089** | Ttc28 | -0.027169626 | -0.244143155 | 0.189803902 | 9.46621716 | -0.299285753 | 0.77385752 | 0.996633405 | -5.219221014 |
| **A_52_P216672** | Klk8 | 0.020423115 | -0.142727409 | 0.18357364 | 9.895956545 | 0.299186952 | 0.773929667 | 0.996633405 | -5.219245548 |
| **A_55_P2042016** | NA | 0.012481374 | -0.087236449 | 0.112199196 | 8.192057506 | 0.2991567 | 0.773951758 | 0.996633405 | -5.219253059 |
| **A_55_P1988708** | Gstz1 | 0.025740349 | -0.179939447 | 0.231420145 | 10.98705173 | 0.299110685 | 0.77398536 | 0.996633405 | -5.219264481 |
| **A_55_P2172420** | Krit1 | 0.037517047 | -0.262292397 | 0.337326492 | 9.826994115 | 0.299083515 | 0.774005202 | 0.996633405 | -5.219271226 |
| **A_51_P204913** | Bag2 | 0.031437034 | -0.219797498 | 0.282671565 | 10.981268 | 0.299068975 | 0.774015819 | 0.996633405 | -5.219274834 |
| **A_55_P2165969** | NA | 0.014218021 | -0.099473456 | 0.127909497 | 8.59080363 | 0.29889618 | 0.774142009 | 0.996633405 | -5.219317707 |
| **A_55_P1960556** | Acox1 | 0.020780785 | -0.145391726 | 0.186953295 | 12.93304477 | 0.298890349 | 0.774146268 | 0.996633405 | -5.219319154 |
| **A_55_P1961750** | Dtd2 | 0.016906095 | -0.118300015 | 0.152112205 | 8.962098955 | 0.298851969 | 0.774174298 | 0.996633405 | -5.219328673 |
| **A_55_P1979082** | Shc1 | 0.025302817 | -0.177083982 | 0.227689616 | 11.5630455 | 0.298810481 | 0.774204597 | 0.996633405 | -5.219338962 |
| **A_55_P2149821** | Srsf11 | -0.020959328 | -0.188619909 | 0.146701253 | 12.17612955 | -0.298782751 | 0.774224849 | 0.996633405 | -5.219345838 |
| **A_55_P2117411** | NA | 0.021396997 | -0.149778672 | 0.192572667 | 11.43699767 | 0.298758268 | 0.77424273 | 0.996633405 | -5.219351909 |
| **A_51_P408471** | Cdipt | 0.024662795 | -0.17264424 | 0.22196983 | 12.20565553 | 0.298750651 | 0.774248293 | 0.996633405 | -5.219353797 |
| **A_55_P1993761** | Nsa2 | -0.017433714 | -0.15698494 | 0.122117512 | 11.26932293 | -0.298583218 | 0.77437058 | 0.996633405 | -5.219395298 |
| **A_52_P437734** | Fbxo28 | 0.019361433 | -0.135655648 | 0.174378514 | 9.201214907 | 0.298515632 | 0.774419944 | 0.996633405 | -5.219412043 |
| **A_55_P1977353** | Ogfod2 | -0.030147225 | -0.271526868 | 0.211232418 | 10.35899922 | -0.298507905 | 0.774425588 | 0.996633405 | -5.219413958 |
| **A_51_P273326** | Sec31a | -0.014744389 | -0.132830312 | 0.103341533 | 12.27286288 | -0.298426781 | 0.774484843 | 0.996633405 | -5.219434052 |
| **A_55_P2133624** | NA | -0.060146011 | -0.54191885 | 0.421626829 | 11.4333629 | -0.298382706 | 0.774517036 | 0.996633405 | -5.219444968 |
| **A_55_P1992680** | Ubtf | -0.022687667 | -0.204459532 | 0.159084197 | 14.22403009 | -0.298313093 | 0.774567884 | 0.996633405 | -5.219462204 |
| **A_52_P344424** | Lipg | 0.01210172 | -0.084924213 | 0.109127654 | 7.657574813 | 0.298104421 | 0.774720315 | 0.996633405 | -5.21951385 |
| **A_55_P2009007** | NA | 0.017638887 | -0.123846293 | 0.159124067 | 14.63813304 | 0.297967815 | 0.774820108 | 0.996633405 | -5.21954764 |
| **A_55_P1988766** | NA | -0.024744762 | -0.223274904 | 0.17378538 | 8.030124636 | -0.297896883 | 0.774871927 | 0.996633405 | -5.21956518 |
| **A_55_P2073975** | NA | 0.028206711 | -0.198119492 | 0.254532914 | 14.86411826 | 0.297869985 | 0.774891578 | 0.996633405 | -5.21957183 |
| **A_52_P90124** | Prep | 0.028788033 | -0.202268563 | 0.25984463 | 12.05302723 | 0.297784958 | 0.774953696 | 0.996633405 | -5.219592848 |
| **A_51_P228254** | Zdhhc21 | 0.019611224 | -0.137863912 | 0.17708636 | 10.95426896 | 0.297647225 | 0.775054324 | 0.996633405 | -5.219626882 |
| **A_55_P2088425** | Tcn2 | -0.021430362 | -0.193517972 | 0.150657248 | 12.97631384 | -0.29763845 | 0.775060735 | 0.996633405 | -5.21962905 |
| **A_51_P243207** | Zfp574 | -0.019593237 | -0.176929158 | 0.137742684 | 9.527214844 | -0.29763735 | 0.775061539 | 0.996633405 | -5.219629322 |
| **A_55_P1973486** | Atp6v1f | -0.018211897 | -0.16447738 | 0.128053586 | 14.0140347 | -0.297592835 | 0.775094062 | 0.996633405 | -5.219640318 |
| **A_66_P132322** | Mrpl21 | 0.01995752 | -0.140355405 | 0.180270445 | 12.90386087 | 0.297541235 | 0.775131763 | 0.996633405 | -5.219653062 |
| **A_52_P119187** | NA | 0.022972245 | -0.161566971 | 0.207511461 | 9.680445553 | 0.297525297 | 0.775143408 | 0.996633405 | -5.219656998 |
| **A_51_P458778** | Hpgd | -0.043606966 | -0.394257107 | 0.307043175 | 8.798922249 | -0.297228865 | 0.775360006 | 0.996633405 | -5.219730166 |
| **A_55_P1988010** | Gm10406 | 0.06200232 | -0.436579504 | 0.560584144 | 10.5443151 | 0.297221781 | 0.775365182 | 0.996633405 | -5.219731913 |
| **A_55_P2033084** | Zdhhc20 | -0.013916801 | -0.125844683 | 0.098011081 | 8.277615886 | -0.29717363 | 0.775400368 | 0.996633405 | -5.219743792 |
| **A_55_P2209053** | 2900001G( | -0.016095114 | -0.145603723 | 0.113413494 | 8.329893721 | -0.297032902 | 0.775503205 | 0.996633405 | -5.219778497 |
| **A_51_P148959** | Dhx36 | -0.019272012 | -0.174416194 | 0.135872169 | 10.13737506 | -0.296893507 | 0.775605073 | 0.996633405 | -5.219812857 |
| **A_55_P2069995** | NA | 0.047666969 | -0.336103273 | 0.431437212 | 11.15217043 | 0.296862523 | 0.775627716 | 0.996633405 | -5.219820492 |
| **A_52_P304323** | Isca1 | -0.027376507 | -0.247788172 | 0.193035157 | 10.20817394 | -0.296860627 | 0.775629102 | 0.996633405 | -5.21982096 |
| **A_55_P2063490** | Csprs | -0.013886745 | -0.125702186 | 0.097928696 | 7.873899625 | -0.29683002 | 0.775651469 | 0.996633405 | -5.219828501 |
| **A_55_P2023200** | Miox | 0.012792519 | -0.090217714 | 0.115802751 | 7.798533476 | 0.296814301 | 0.775662957 | 0.996633405 | -5.219832374 |
| **A_55_P2033461** | NA | 0.024296807 | -0.171365042 | 0.219958655 | 14.20251317 | 0.296791991 | 0.775679262 | 0.996633405 | -5.219837871 |
| **A_52_P302496** | Fah | 0.022773947 | -0.160823446 | 0.206371341 | 8.095915176 | 0.296470123 | 0.775914504 | 0.996633405 | -5.219917124 |
| **A_51_P233101** | Ltbp3 | 0.035373969 | -0.249816764 | 0.320564703 | 11.32927039 | 0.296454137 | 0.775926189 | 0.996633405 | -5.219921058 |
| **A_51_P223404** | Plin3 | -0.013795246 | -0.125048423 | 0.097457931 | 13.73288954 | -0.296364479 | 0.775991722 | 0.996633405 | -5.219943119 |
| **A_55_P1997585** | NA | 0.012479187 | -0.088184689 | 0.113143062 | 7.703257529 | 0.296293255 | 0.776043783 | 0.996633405 | -5.219960639 |

| | | | | | | | | | |
|---|---|---|---|---|---|---|---|---|---|
| **A_66_P130310** | A4galt | -0.020114746 | -0.182390429 | 0.142160937 | 8.024333137 | -0.296258104 | 0.776069476 | 0.996633405 | -5.219969284 |
| **A_52_P632742** | Shisa5 | -0.017989273 | -0.163137341 | 0.127158795 | 8.152965586 | -0.296218027 | 0.776098771 | 0.996633405 | -5.21997914 |
| **A_55_P2258366** | C030009H( | 0.043254663 | -0.305782962 | 0.392292289 | 11.08530534 | 0.296189612 | 0.776119541 | 0.996633405 | -5.219986126 |
| **A_55_P1966144** | Slfn14 | 0.013618301 | -0.09632994 | 0.123566541 | 7.689118994 | 0.296035486 | 0.776232207 | 0.996633405 | -5.220024013 |
| **A_55_P2032543** | Cblc | -0.013136184 | -0.119196382 | 0.092924015 | 7.916574154 | -0.296023314 | 0.776241105 | 0.996633405 | -5.220027004 |
| **A_55_P1962065** | NA | 0.02027591 | -0.143436391 | 0.183988211 | 13.53119178 | 0.296011217 | 0.776249948 | 0.996633405 | -5.220029977 |
| **A_55_P2180899** | Aldh3a2 | -0.018354713 | -0.16665637 | 0.129946944 | 8.625633832 | -0.295808558 | 0.776398101 | 0.996633405 | -5.22007976 |
| **A_55_P2443935** | Xpo7 | 0.016862435 | -0.119419898 | 0.153144768 | 8.733141358 | 0.295726233 | 0.776458287 | 0.996633405 | -5.220099974 |
| **A_52_P72654** | Slc25a17 | 0.015361846 | -0.108795494 | 0.139519186 | 11.20135503 | 0.295719622 | 0.77646312 | 0.996633405 | -5.220101597 |
| **A_55_P2009792** | Hdac6 | -0.01315999 | -0.119525715 | 0.093205735 | 11.37985278 | -0.295707939 | 0.776471661 | 0.996633405 | -5.220104465 |
| **A_55_P1952512** | Sult2a2 | -0.012844354 | -0.116709294 | 0.091020587 | 7.647362353 | -0.295564593 | 0.776576464 | 0.996633405 | -5.220139647 |
| **A_55_P2386256** | D130062J1 | 0.040787754 | -0.289053349 | 0.370628856 | 9.247120204 | 0.295552167 | 0.776585548 | 0.996633405 | -5.220142696 |
| **A_51_P503822** | Slitrk6 | 0.01421653 | -0.100752599 | 0.12918566 | 8.413732668 | 0.295543561 | 0.77659184 | 0.996633405 | -5.220144807 |
| **A_55_P2003996** | H2-T24 | -0.012608482 | -0.114588083 | 0.089371119 | 7.825861242 | -0.29550078 | 0.776623119 | 0.996633405 | -5.220155303 |
| **A_55_P2036803** | Dhx30 | 0.02303888 | -0.163355587 | 0.209433347 | 10.4142695 | 0.295418355 | 0.776683385 | 0.996633405 | -5.220175521 |
| **A_52_P352735** | Ccdc153 | 0.016617499 | -0.117828877 | 0.151063875 | 7.709222275 | 0.295410342 | 0.776689244 | 0.996633405 | -5.220177487 |
| **A_51_P155747** | Scyl3 | 0.014766739 | -0.104725334 | 0.134258813 | 8.343903023 | 0.295362015 | 0.776724579 | 0.996633405 | -5.220189338 |
| **A_52_P276343** | Pum3 | -0.012835447 | -0.116711971 | 0.091041077 | 8.738159222 | -0.29532671 | 0.776750394 | 0.996633405 | -5.220197995 |
| **A_55_P2046109** | NA | -0.013552417 | -0.123232139 | 0.096127306 | 7.774606366 | -0.295324542 | 0.776751979 | 0.996633405 | -5.220198526 |
| **A_51_P244690** | Ube2b | -0.019796972 | -0.180048943 | 0.140454999 | 13.20742538 | -0.295259936 | 0.776799218 | 0.996633405 | -5.220214365 |
| **A_55_P1991164** | Mlc1 | -0.01251506 | -0.11383955 | 0.088809431 | 7.674805081 | -0.295207667 | 0.776837438 | 0.996633405 | -5.220227176 |
| **A_52_P370443** | Riok3 | -0.019558318 | -0.177925063 | 0.138808427 | 11.47447089 | -0.295173009 | 0.776862781 | 0.996633405 | -5.22023567 |
| **A_55_P2075670** | NA | -0.017981928 | -0.163584774 | 0.127620919 | 8.255473501 | -0.295172245 | 0.776863339 | 0.996633405 | -5.220235857 |
| **A_55_P1955072** | Alad | -0.020879142 | -0.189953215 | 0.14819493 | 10.17157951 | -0.295151346 | 0.776878621 | 0.996633405 | -5.220240978 |
| **A_55_P2012369** | Eea1 | -0.032341077 | -0.294297017 | 0.229614862 | 10.21800894 | -0.295077005 | 0.776932982 | 0.996633405 | -5.220259193 |
| **A_52_P627269** | Ces2b | -0.014074553 | -0.128086705 | 0.099937599 | 7.641420993 | -0.29504795 | 0.776954229 | 0.996633405 | -5.220266311 |
| **A_55_P1985433** | Nrg1 | -0.033727281 | -0.307033487 | 0.239578926 | 8.702637383 | -0.29494495 | 0.77702955 | 0.996633405 | -5.220291537 |
| **A_55_P1983414** | NA | 0.018306189 | -0.130068736 | 0.166681114 | 8.554354023 | 0.294880846 | 0.777076428 | 0.996633405 | -5.220307233 |
| **A_55_P1957028** | Kat7 | 0.016075163 | -0.114228695 | 0.146379021 | 11.19464627 | 0.294854146 | 0.777095954 | 0.996633405 | -5.22031377 |
| **A_55_P1990573** | Snrnp70 | 0.020869197 | -0.148310137 | 0.190048531 | 12.41826032 | 0.294827205 | 0.777115656 | 0.996633405 | -5.220320365 |
| **A_55_P2070992** | Aldoc | 0.029178846 | -0.207377419 | 0.26573511 | 12.56623528 | 0.294810391 | 0.777127952 | 0.996633405 | -5.22032448 |
| **A_55_P2133337** | NA | 0.012468346 | -0.088680994 | 0.113617687 | 7.978684888 | 0.294615055 | 0.777270809 | 0.996633405 | -5.220372277 |
| **A_55_P2045367** | G630025P( | -0.029309972 | -0.267105801 | 0.208485857 | 8.490359573 | -0.294591563 | 0.77728799 | 0.996633405 | -5.220378023 |
| **A_55_P1996963** | Mtcl1 | 0.014887361 | -0.106018796 | 0.135793517 | 8.473105716 | 0.29429198 | 0.777507106 | 0.996633405 | -5.220451263 |
| **A_51_P367843** | Polr2g | 0.017812111 | -0.126864937 | 0.162489159 | 13.33588049 | 0.294255698 | 0.777533645 | 0.996633405 | -5.220460128 |
| **A_51_P434670** | Qk | -0.015620764 | -0.142502991 | 0.111261463 | 12.39114731 | -0.294246023 | 0.777540721 | 0.996633405 | -5.220462492 |
| **A_55_P1982034** | NA | -0.015090319 | -0.137726145 | 0.107545507 | 8.199814494 | -0.294096728 | 0.777649926 | 0.996633405 | -5.220498958 |
| **A_55_P2063785** | Ank1 | -0.037674168 | -0.343977646 | 0.268629309 | 10.27824384 | -0.293968555 | 0.777743685 | 0.996633405 | -5.220530251 |
| **A_52_P663585** | 10-Sep | -0.024633078 | -0.224940529 | 0.175674373 | 9.188764171 | -0.293921064 | 0.777778426 | 0.996633405 | -5.220541842 |
| **A_51_P359983** | Arhgap17 | 0.014058921 | -0.100303066 | 0.128420907 | 9.513354521 | 0.293818689 | 0.777853318 | 0.996633405 | -5.220566823 |
| **A_51_P396854** | Timm10b | 0.017331368 | -0.123665797 | 0.158328533 | 13.68225492 | 0.29378634 | 0.777876983 | 0.996633405 | -5.220574715 |
| **A_55_P2083224** | Pum2 | -0.018954639 | -0.173245692 | 0.135336414 | 10.87516984 | -0.293618832 | 0.777999529 | 0.996633405 | -5.220615566 |
| **A_55_P2009086** | Creb3l1 | -0.029114968 | -0.266122723 | 0.207892787 | 9.758856211 | -0.293604622 | 0.778009926 | 0.996633405 | -5.220619031 |
| **A_51_P181922** | Yjefn3 | 0.025149907 | -0.179596253 | 0.229896067 | 9.355952963 | 0.293582225 | 0.778026312 | 0.996633405 | -5.220624491 |
| **A_51_P507172** | 2310022A1 | -0.012547998 | -0.11470277 | 0.089606774 | 11.85657917 | -0.29357895 | 0.778028708 | 0.996633405 | -5.220625289 |
| **A_55_P2044458** | Pramef8 | 0.017986873 | -0.128486521 | 0.164460266 | 9.880557942 | 0.29349861 | 0.778087486 | 0.996633405 | -5.220644872 |
| **A_55_P2163873** | Uox | -0.011802888 | -0.107976083 | 0.084370307 | 7.631535572 | -0.293321158 | 0.77821732 | 0.996633405 | -5.220688107 |
| **A_55_P1954156** | Wdr43 | 0.02373341 | -0.169702051 | 0.217168872 | 9.49639069 | 0.293246737 | 0.778271773 | 0.996633405 | -5.220706232 |

| | | | | | | | | | |
|---|---|---|---|---|---|---|---|---|---|
| **A_55_P2165314** | Trio | -0.020716849 | -0.189646451 | 0.148212753 | 10.46245619 | -0.2931076 | 0.778373581 | 0.996633405 | -5.220740106 |
| **A_55_P2055442** | Pafah2 | 0.023653924 | -0.169378835 | 0.216686682 | 9.147517634 | 0.292874329 | 0.778544279 | 0.996633405 | -5.220796862 |
| **A_55_P2055742** | Adtrp | 0.01367199 | -0.097901502 | 0.125245482 | 7.665720716 | 0.292873334 | 0.778545007 | 0.996633405 | -5.220797104 |
| **A_51_P469578** | Psma1 | -0.013967518 | -0.128011394 | 0.100076358 | 13.46254247 | -0.292722695 | 0.778655245 | 0.996633405 | -5.220833732 |
| **A_52_P75146** | Lyrm2 | -0.021829172 | -0.200080084 | 0.156421741 | 10.97362449 | -0.292694543 | 0.778675848 | 0.996633405 | -5.220840575 |
| **A_55_P2061701** | Osgep | -0.029800733 | -0.273200616 | 0.213599149 | 10.77561332 | -0.292627891 | 0.778724626 | 0.996633405 | -5.220856774 |
| **A_52_P566840** | Adgrf1 | -0.013042724 | -0.119621611 | 0.093536163 | 7.659541914 | -0.292486789 | 0.778827894 | 0.996633405 | -5.220891056 |
| **A_52_P566605** | Hsd17b7 | 0.016455366 | -0.118021252 | 0.150931984 | 9.067483782 | 0.29246231 | 0.778845809 | 0.996633405 | -5.220897002 |
| **A_52_P244964** | Akap11 | 0.025419396 | -0.182360555 | 0.233199347 | 10.34181539 | 0.292395521 | 0.778894692 | 0.996633405 | -5.220913222 |
| **A_55_P1998656** | Entpd5 | -0.033188031 | -0.304554955 | 0.238178894 | 10.2657174 | -0.292303288 | 0.778962199 | 0.996633405 | -5.220935616 |
| **A_55_P2202539** | BC028789 | -0.012337903 | -0.113232671 | 0.088556865 | 7.788511679 | -0.292268388 | 0.778987743 | 0.996633405 | -5.220944087 |
| **A_55_P2061909** | Tceal9 | 0.025646267 | -0.184094706 | 0.235387241 | 12.41856306 | 0.292246971 | 0.779003419 | 0.996633405 | -5.220949285 |
| **A_55_P2061391** | Rhobtb3 | 0.020220612 | -0.145238745 | 0.185679968 | 9.212597757 | 0.292086906 | 0.77912058 | 0.996633405 | -5.220988124 |
| **A_55_P1957876** | NA | 0.017452514 | -0.125438027 | 0.160343054 | 10.51622427 | 0.291919859 | 0.779242858 | 0.996633405 | -5.221028636 |
| **A_55_P1976789** | Dgcr6 | -0.015269004 | -0.140310547 | 0.109772539 | 11.25188834 | -0.291853915 | 0.779291131 | 0.996633405 | -5.221044622 |
| **A_55_P2038540** | Hbb-b2 | -0.078546269 | -0.721801102 | 0.564708564 | 15.72906931 | -0.291844582 | 0.779297963 | 0.996633405 | -5.221046884 |
| **A_52_P431859** | Aopep | 0.028033751 | -0.201569122 | 0.257636625 | 9.482691987 | 0.291818639 | 0.779316954 | 0.996633405 | -5.221053172 |
| **A_66_P113490** | Sh2b2 | 0.01516502 | -0.109057591 | 0.139387632 | 8.23335739 | 0.291777283 | 0.779347229 | 0.996633405 | -5.221063194 |
| **A_51_P348592** | Slc38a7 | 0.014845687 | -0.106786828 | 0.136478203 | 8.435806401 | 0.291715661 | 0.77939234 | 0.996633405 | -5.221078126 |
| **A_52_P139588** | Edem3 | 0.02689469 | -0.193479048 | 0.247268428 | 9.720222643 | 0.291686168 | 0.779413931 | 0.996633405 | -5.221085271 |
| **A_55_P2180477** | Rpl31 | 0.014637323 | -0.105314617 | 0.134589263 | 16.153946 | 0.291651037 | 0.779439649 | 0.996633405 | -5.221093781 |
| **A_52_P661587** | Ccdc141 | 0.020127311 | -0.144835701 | 0.185090322 | 8.551800067 | 0.291613957 | 0.779466795 | 0.996633405 | -5.221102763 |
| **A_55_P2009345** | Ovch2 | -0.014381049 | -0.132251929 | 0.103489832 | 8.168549647 | -0.291603786 | 0.779474241 | 0.996633405 | -5.221105226 |
| **A_55_P2049582** | Apbb3 | 0.02281125 | -0.1641715 | 0.209794 | 9.492728317 | 0.291579289 | 0.779492176 | 0.996633405 | -5.221111159 |
| **A_51_P466371** | Pitpna | 0.022247686 | -0.160154989 | 0.204650361 | 10.63737474 | 0.29151626 | 0.77953832 | 0.996633405 | -5.221126421 |
| **A_52_P6798** | Rbm22 | 0.014604866 | -0.105178829 | 0.13438856 | 8.906212773 | 0.291413051 | 0.779613882 | 0.996633405 | -5.221151406 |
| **A_65_P10351** | Atp2b1 | 0.014842312 | -0.106921784 | 0.136606408 | 8.433693509 | 0.291334184 | 0.779671624 | 0.996633405 | -5.221170492 |
| **A_55_P1972192** | Phka1 | -0.013604047 | -0.125220549 | 0.098012455 | 10.27918316 | -0.291305613 | 0.779692544 | 0.996633405 | -5.221177406 |
| **A_55_P2006703** | Bmp1 | -0.037469165 | -0.344960901 | 0.270022571 | 11.6206106 | -0.291239112 | 0.779741234 | 0.996633405 | -5.221193494 |
| **A_55_P2044413** | Cxcl11 | -0.031630168 | -0.291252706 | 0.22799237 | 11.20949915 | -0.291184493 | 0.779781225 | 0.996633405 | -5.221206705 |
| **A_52_P33014** | Ppp1r8 | -0.025671766 | -0.236468572 | 0.185125041 | 8.584045237 | -0.291072278 | 0.77986339 | 0.996633405 | -5.221233839 |
| **A_55_P2063715** | Cfap53 | 0.012142742 | -0.087677533 | 0.111963018 | 7.67612967 | 0.290741582 | 0.780105547 | 0.996633405 | -5.221313746 |
| **A_55_P2034110** | Tgm2 | 0.029115211 | -0.210317832 | 0.268548254 | 13.71456685 | 0.290633043 | 0.780185032 | 0.996633405 | -5.221339954 |
| **A_55_P2017491** | Rbm43 | 0.01671859 | -0.120808153 | 0.154245334 | 9.98303961 | 0.290550503 | 0.780245479 | 0.996633405 | -5.221359877 |
| **A_55_P1952577** | Ces1a | -0.012455794 | -0.114928731 | 0.090017142 | 7.68295086 | -0.290516884 | 0.7802701 | 0.996633405 | -5.22136799 |
| **A_51_P141264** | Zfp24 | -0.021282843 | -0.196442013 | 0.153876326 | 8.713722477 | -0.290406216 | 0.780351151 | 0.996633405 | -5.221394692 |
| **A_55_P2025328** | Tacr2 | 0.048607715 | -0.3514752 | 0.44869063 | 12.19922834 | 0.290378414 | 0.780371512 | 0.996633405 | -5.221401398 |
| **A_55_P1980932** | Elob | -0.024344732 | -0.224882373 | 0.176192908 | 14.95248132 | -0.2901471 | 0.780540931 | 0.996633405 | -5.221457171 |
| **A_55_P1982563** | Rhbdf1 | 0.030817368 | -0.223244996 | 0.284879731 | 11.15115056 | 0.289910765 | 0.780714041 | 0.996633405 | -5.22151411 |
| **A_55_P2087281** | NA | 0.021732371 | -0.157502024 | 0.200966767 | 8.711558224 | 0.289797672 | 0.780796884 | 0.996633405 | -5.221541341 |
| **A_55_P2125841** | Cog1 | 0.022532971 | -0.163337558 | 0.2084035 | 9.313481348 | 0.289745735 | 0.780834929 | 0.996633405 | -5.221553843 |
| **A_52_P539434** | Lbh | -0.036902839 | -0.341363335 | 0.267557657 | 10.31107014 | -0.289692973 | 0.78087358 | 0.996633405 | -5.221566542 |
| **A_55_P2152035** | Tcf19 | 0.027419153 | -0.198886735 | 0.253725041 | 9.652527435 | 0.289579165 | 0.780956953 | 0.996633405 | -5.221593925 |
| **A_55_P2164348** | Hoxa10 | 0.012516316 | -0.090834739 | 0.115867371 | 7.632196518 | 0.289448128 | 0.78105295 | 0.996633405 | -5.221625441 |
| **A_51_P268529** | Csad | -0.031648522 | -0.293038304 | 0.229741259 | 9.84063636 | -0.289383632 | 0.781100202 | 0.996633405 | -5.221640948 |
| **A_55_P2035667** | Gk | 0.013318676 | -0.096712353 | 0.123349706 | 8.633291146 | 0.289304382 | 0.781158264 | 0.996633405 | -5.221659998 |
| **A_55_P2152956** | Plekhm3 | -0.015438218 | -0.142984628 | 0.112108192 | 8.369385479 | -0.289293093 | 0.781166535 | 0.996633405 | -5.221662711 |
| **A_51_P301930** | Lrrc17 | -0.031932794 | -0.295793803 | 0.231928216 | 10.68778583 | -0.289248307 | 0.781199348 | 0.996633405 | -5.221673474 |

| | | | | | | | | | |
|---|---|---|---|---|---|---|---|---|---|
| **A_52_P487436** | Nags | 0.011864842 | -0.086208675 | 0.109938359 | 7.647985673 | 0.28914744 | 0.78127325 | 0.996633405 | -5.221697708 |
| **A_55_P2094896** | Phyhd1 | 0.029921318 | -0.217428893 | 0.277271529 | 11.00383946 | 0.289119618 | 0.781293635 | 0.996633405 | -5.22170439 |
| **A_55_P2015782** | Bcas1 | 0.015923265 | -0.115721389 | 0.147567919 | 8.080587601 | 0.289093305 | 0.781312915 | 0.996633405 | -5.221710711 |
| **A_51_P156291** | F9 | -0.01545403 | -0.14324718 | 0.112339119 | 8.076137353 | -0.289030264 | 0.781359105 | 0.996633405 | -5.22172585 |
| **A_66_P120770** | Ywhaz | -0.019935461 | -0.184796752 | 0.14492583 | 8.778185713 | -0.289012562 | 0.781372076 | 0.996633405 | -5.2217301 |
| **A_51_P413111** | Ppm1l | -0.016085478 | -0.149161484 | 0.116990528 | 11.71667828 | -0.288897214 | 0.781456596 | 0.996633405 | -5.221757791 |
| **A_66_P104365** | Ccdc167 | -0.017870485 | -0.165768506 | 0.130027537 | 10.9741937 | -0.288790629 | 0.781534697 | 0.996633405 | -5.221783368 |
| **A_51_P373369** | Med4 | 0.015711238 | -0.114342721 | 0.145765197 | 10.89310467 | 0.288732697 | 0.781577148 | 0.996633405 | -5.221797266 |
| **A_55_P2168914** | NA | -0.012000485 | -0.111347515 | 0.087346546 | 7.765757431 | -0.288704149 | 0.781598068 | 0.996633405 | -5.221804114 |
| **A_55_P2094118** | Fmnl3 | -0.021728356 | -0.201615115 | 0.158158404 | 9.40306876 | -0.288693362 | 0.781605972 | 0.996633405 | -5.221806701 |
| **A_51_P156631** | Glrx5 | 0.020415375 | -0.148641528 | 0.189472278 | 13.52903259 | 0.288624762 | 0.781656243 | 0.996633405 | -5.221823153 |
| **A_55_P2069961** | NA | -0.021783964 | -0.202206859 | 0.15863893 | 16.50513236 | -0.288572142 | 0.781694804 | 0.996633405 | -5.22183577 |
| **A_55_P2211164** | 5330406M2 | -0.021503053 | -0.199655851 | 0.156649744 | 7.972198338 | -0.288480603 | 0.781761887 | 0.996633405 | -5.221857714 |
| **A_52_P148658** | Mettl22 | 0.0234185 | -0.170640532 | 0.217477533 | 10.34538066 | 0.288425961 | 0.781801932 | 0.996633405 | -5.221870809 |
| **A_52_P53470** | NA | -0.017448022 | -0.16207129 | 0.127175245 | 8.05917906 | -0.288348153 | 0.781858956 | 0.996633405 | -5.221889452 |
| **A_55_P2017710** | Igsf9 | 0.020614076 | -0.150282008 | 0.191510161 | 8.349751204 | 0.288297513 | 0.781896069 | 0.996633405 | -5.221901583 |
| **A_55_P2077546** | NA | 0.018015382 | -0.131362254 | 0.167393018 | 9.039236668 | 0.288248481 | 0.781932004 | 0.996633405 | -5.221913327 |
| **A_55_P2127692** | Timm50 | -0.048632214 | -0.451885304 | 0.354620876 | 9.216262204 | -0.288240811 | 0.781937626 | 0.996633405 | -5.221915164 |
| **A_55_P2043622** | NA | 0.031192754 | -0.227470677 | 0.289856184 | 14.17901847 | 0.288222455 | 0.781951079 | 0.996633405 | -5.22191956 |
| **A_55_P2184146** | NA | 0.013076298 | -0.095364024 | 0.121516621 | 8.763238469 | 0.28820609 | 0.781963073 | 0.996633405 | -5.221923479 |
| **A_55_P2149942** | Ninj2 | -0.011772822 | -0.109414318 | 0.085868674 | 7.693896232 | -0.288174328 | 0.781986352 | 0.996633405 | -5.221931084 |
| **A_55_P1952793** | Son | -0.027200224 | -0.252826186 | 0.198425739 | 12.23255197 | -0.288132693 | 0.782016867 | 0.996633405 | -5.221941052 |
| **A_55_P1963724** | NA | 0.014105616 | -0.102950537 | 0.13116177 | 7.934937788 | 0.288009588 | 0.782107097 | 0.996633405 | -5.221970517 |
| **A_55_P2014124** | Gjb3 | -0.020130954 | -0.187204349 | 0.146942442 | 7.789400545 | -0.287982555 | 0.782126911 | 0.996633405 | -5.221976986 |
| **A_52_P376337** | Sdhaf3 | -0.016164558 | -0.150366973 | 0.118037858 | 8.922575483 | -0.287880748 | 0.782201534 | 0.996633405 | -5.222001342 |
| **A_55_P2067645** | Gm5136 | -0.017979094 | -0.167327606 | 0.131369419 | 8.861671561 | -0.287723956 | 0.782316464 | 0.996633405 | -5.222038836 |
| **A_66_P119123** | A1cf | 0.012308049 | -0.0899795 | 0.114595598 | 7.65116698 | 0.287591189 | 0.782413789 | 0.996633405 | -5.22207057 |
| **A_55_P2046037** | Gm7083 | 0.022378556 | -0.16361757 | 0.208374681 | 13.73235334 | 0.287565834 | 0.782432375 | 0.996633405 | -5.222076629 |
| **A_55_P1986188** | Def8 | -0.019810659 | -0.184471053 | 0.144849735 | 12.31707232 | -0.287553659 | 0.782441301 | 0.996633405 | -5.222079538 |
| **A_55_P2039694** | NA | 0.03760911 | -0.275007761 | 0.350225981 | 15.25795973 | 0.287534377 | 0.782455436 | 0.996633405 | -5.222084145 |
| **A_55_P2036999** | Zic3 | -0.011952637 | -0.111364115 | 0.087458842 | 7.680268133 | -0.287366617 | 0.78257842 | 0.996633405 | -5.222124214 |
| **A_55_P2138906** | Lrfn1 | 0.015124006 | -0.110710977 | 0.140958988 | 8.524257831 | 0.287259615 | 0.782656866 | 0.996633405 | -5.22214976 |
| **A_51_P340653** | Slc16a4 | 0.012520559 | -0.091657549 | 0.116698666 | 7.793146435 | 0.28724758 | 0.78266569 | 0.996633405 | -5.222152633 |
| **A_55_P2084763** | NA | -0.045762217 | -0.426533889 | 0.335009455 | 10.13144863 | -0.28724439 | 0.782668028 | 0.996633405 | -5.222153394 |
| **A_55_P1978927** | Srgap1 | 0.013942478 | -0.102157862 | 0.130042819 | 8.010418933 | 0.287022286 | 0.782830869 | 0.996633405 | -5.222206387 |
| **A_52_P58471** | Tmem199 | -0.022075118 | -0.205923079 | 0.161772844 | 8.892041376 | -0.286981128 | 0.782861046 | 0.996633405 | -5.222216203 |
| **A_55_P1987912** | Chpf2 | 0.015881378 | -0.116414097 | 0.148176853 | 8.610361101 | 0.286914385 | 0.782909982 | 0.996633405 | -5.222232117 |
| **A_52_P253044** | Syt13 | -0.013764113 | -0.128455531 | 0.100927305 | 7.890123109 | -0.286831237 | 0.782970949 | 0.996633405 | -5.222251939 |
| **A_55_P2121877** | Gpr137b | -0.023203743 | -0.216581204 | 0.170173719 | 8.060569359 | -0.286788226 | 0.783002487 | 0.996633405 | -5.22226219 |
| **A_55_P1999849** | NA | 0.022360059 | -0.164115214 | 0.208835331 | 12.12605284 | 0.28658986 | 0.783147945 | 0.996633405 | -5.222309448 |
| **A_55_P2133220** | Arhgef39 | 0.022141349 | -0.162520483 | 0.20680318 | 10.08663603 | 0.286573525 | 0.783159924 | 0.996633405 | -5.222313338 |
| **A_51_P142744** | Sulf1 | -0.051973154 | -0.485531611 | 0.381585304 | 11.61448025 | -0.286510498 | 0.783206142 | 0.996633405 | -5.222328346 |
| **A_51_P452203** | Vash2 | -0.026212002 | -0.244903461 | 0.192479457 | 9.393680283 | -0.28646891 | 0.78323664 | 0.996633405 | -5.222338247 |
| **A_55_P2025660** | Ranbp9 | 0.022297502 | -0.163765345 | 0.208360349 | 9.691375597 | 0.28642154 | 0.783271378 | 0.996633405 | -5.222349523 |
| **A_51_P304527** | Cluap1 | -0.012419252 | -0.116153433 | 0.09131493 | 10.33733707 | -0.2861427 | 0.783475872 | 0.996633405 | -5.222415862 |
| **A_55_P1964053** | Golga4 | 0.0288126 | -0.211932764 | 0.269557964 | 12.27940818 | 0.286044531 | 0.783547871 | 0.996633405 | -5.222439203 |
| **A_52_P104761** | Rlim | 0.014542285 | -0.106968991 | 0.13605356 | 8.164661014 | 0.286038965 | 0.783551954 | 0.996633405 | -5.222440526 |
| **A_55_P2073452** | Glp2r | 0.012630492 | -0.092926878 | 0.118187862 | 7.657660872 | 0.285983409 | 0.7835927 | 0.996633405 | -5.222453731 |

| | | | | | | | | | |
|---|---|---|---|---|---|---|---|---|---|
| **A_52_P363216** | Gcnt2 | -0.019960443 | -0.186781892 | 0.146861006 | 8.528861526 | -0.285974575 | 0.78359918 | 0.996633405 | -5.22245583 |
| **A_55_P1999561** | Pram1 | -0.014140315 | -0.132332685 | 0.104052055 | 7.866676791 | -0.285942547 | 0.78362267 | 0.996633405 | -5.222463442 |
| **A_51_P497563** | Vps18 | -0.024040716 | -0.224999684 | 0.176918252 | 9.964443605 | -0.285923032 | 0.783636984 | 0.996633405 | -5.222468079 |
| **A_55_P2145942** | Atg16l1 | 0.013571553 | -0.099911916 | 0.127055023 | 8.810814894 | 0.285828862 | 0.783706055 | 0.996633405 | -5.222490452 |
| **A_55_P2197914** | C030027H1 | 0.030602878 | -0.225343719 | 0.286549475 | 11.24186275 | 0.285773558 | 0.78374662 | 0.996633405 | -5.222503588 |
| **A_52_P441036** | Col23a1 | 0.015496334 | -0.11412362 | 0.145116287 | 8.467205973 | 0.285736833 | 0.783773558 | 0.996633405 | -5.22251231 |
| **A_51_P214306** | Haus4 | -0.039187355 | -0.393682994 | 0.315308284 | 11.43657884 | -0.290710468 | 0.783786972 | 0.996633405 | -4.998805046 |
| **A_55_P2081865** | Slc2a8 | -0.017402227 | -0.162993788 | 0.128189334 | 11.19566232 | -0.285678633 | 0.783816248 | 0.996633405 | -5.222526129 |
| **A_55_P2210315** | Get4 | 0.02996203 | -0.220725898 | 0.280649958 | 9.590853556 | 0.285658362 | 0.783831118 | 0.996633405 | -5.222530942 |
| **A_51_P452820** | Rpl31 | 0.01556778 | -0.114763348 | 0.145898908 | 9.857902104 | 0.285487872 | 0.783956179 | 0.996633405 | -5.222571406 |
| **A_51_P403536** | Ltbp4 | -0.027683528 | -0.259521858 | 0.204154803 | 13.92377296 | -0.285394318 | 0.784024809 | 0.996633405 | -5.2225936 |
| **A_55_P2058953** | Rpl13a | -0.014435864 | -0.135335201 | 0.106463473 | 15.93760931 | -0.285382927 | 0.784033165 | 0.996633405 | -5.222596302 |
| **A_52_P633481** | Akap11 | -0.025448764 | -0.238594684 | 0.187697155 | 8.965399692 | -0.285363752 | 0.784047232 | 0.996633405 | -5.22260085 |
| **A_52_P312102** | NA | -0.021719289 | -0.203630807 | 0.16019223 | 10.13544241 | -0.285360942 | 0.784049293 | 0.996633405 | -5.222601516 |
| **A_51_P326994** | Calhm2 | -0.015383277 | -0.144245302 | 0.113478748 | 8.559021137 | -0.285320543 | 0.784078929 | 0.996633405 | -5.222611097 |
| **A_55_P2007196** | Aldoa | 0.024461549 | -0.180496085 | 0.229419182 | 17.95432774 | 0.285252189 | 0.784129075 | 0.996633405 | -5.222627304 |
| **A_55_P2018292** | Map2k4 | 0.020198999 | -0.149121223 | 0.189519221 | 9.391369631 | 0.28512162 | 0.784224865 | 0.996633405 | -5.222658253 |
| **A_51_P137168** | Smarcal1 | 0.019812397 | -0.146353119 | 0.185977912 | 10.24623352 | 0.284974005 | 0.784333166 | 0.996633405 | -5.222693225 |
| **A_65_P09443** | Psma4 | 0.020074078 | -0.148296282 | 0.188444438 | 12.69916747 | 0.284956847 | 0.784345754 | 0.996633405 | -5.222697289 |
| **A_55_P2136832** | Gm6981 | 0.025691454 | -0.189819893 | 0.241202801 | 15.82355366 | 0.284923109 | 0.784370507 | 0.996633405 | -5.222705279 |
| **A_51_P113403** | Slc26a11 | 0.031612664 | -0.233617039 | 0.296842367 | 9.339993859 | 0.284870886 | 0.784408824 | 0.996633405 | -5.179011268 |
| **A_55_P1983036** | Tdrd3 | -0.013702215 | -0.128690264 | 0.101285835 | 10.60596309 | -0.284804723 | 0.784457369 | 0.996633405 | -5.222733309 |
| **A_55_P2045658** | Nme6 | -0.033676519 | -0.316326443 | 0.248973405 | 9.838601418 | -0.284765555 | 0.784486107 | 0.996633405 | -5.22274258 |
| **A_55_P2121744** | Vars2 | 0.014248716 | -0.105355385 | 0.133852818 | 8.441545951 | 0.284733652 | 0.784509516 | 0.996633405 | -5.222750131 |
| **A_55_P2026400** | Capns1 | -0.025261322 | -0.237344442 | 0.186821797 | 12.66074599 | -0.284681409 | 0.78454785 | 0.996633405 | -5.222762494 |
| **A_52_P294253** | Zfp41 | -0.01777428 | -0.167008608 | 0.131460049 | 9.534796586 | -0.284663905 | 0.784560693 | 0.996633405 | -5.222766635 |
| **A_51_P142923** | Chka | 0.019677317 | -0.145539928 | 0.184894562 | 8.702299967 | 0.284655542 | 0.78456683 | 0.996633405 | -5.222768614 |
| **A_55_P2046411** | NA | 0.037112257 | -0.274561715 | 0.348786228 | 12.35282074 | 0.284594144 | 0.784611882 | 0.996633405 | -5.222783139 |
| **A_52_P128691** | Gpr4 | -0.016306876 | -0.153257547 | 0.120643795 | 8.023571053 | -0.284587447 | 0.784616796 | 0.996633405 | -5.222784723 |
| **A_51_P513254** | Rac1 | -0.013713296 | -0.128903028 | 0.101476436 | 13.02393502 | -0.28453599 | 0.784654554 | 0.996633405 | -5.222796894 |
| **A_55_P2000533** | Polk | -0.017990073 | -0.169199922 | 0.133219777 | 8.231669248 | -0.28435572 | 0.784786839 | 0.996633405 | -5.222839516 |
| **A_55_P2070825** | Nudt5 | 0.027525277 | -0.203856578 | 0.258907131 | 11.19548285 | 0.284322696 | 0.784811072 | 0.996633405 | -5.222847321 |
| **A_55_P2031157** | Efna3 | -0.017215392 | -0.161936477 | 0.127505693 | 8.770598441 | -0.284311373 | 0.784819382 | 0.996633405 | -5.222849996 |
| **A_52_P486045** | Coq4 | 0.025658548 | -0.190343382 | 0.241660478 | 10.17914701 | 0.283911889 | 0.78511256 | 0.996633405 | -5.222944339 |
| **A_55_P2071059** | Nab2 | -0.020722679 | -0.195200036 | 0.153754679 | 10.8244788 | -0.283867679 | 0.785145007 | 0.996633405 | -5.222954772 |
| **A_51_P462192** | Olr1 | -0.016460205 | -0.155084678 | 0.122164268 | 7.738891475 | -0.283794818 | 0.785198485 | 0.996633405 | -5.222971962 |
| **A_51_P236469** | Metap1d | 0.012040318 | -0.089365722 | 0.113446358 | 8.354385147 | 0.283780961 | 0.785208655 | 0.996633405 | -5.222975231 |
| **A_55_P2144095** | NA | -0.02093247 | -0.197347225 | 0.155482284 | 10.50632588 | -0.28359248 | 0.785346998 | 0.996633405 | -5.223019679 |
| **A_51_P424641** | Sirt4 | -0.02449351 | -0.230925555 | 0.181938536 | 9.090055521 | -0.283584856 | 0.785352595 | 0.996633405 | -5.223021476 |
| **A_52_P272945** | Ccnh | 0.025388279 | -0.188693207 | 0.239469766 | 8.768872374 | 0.283441402 | 0.785457895 | 0.996633405 | -5.223055285 |
| **A_55_P1985808** | Ranbp9 | -0.016930849 | -0.159733536 | 0.125871839 | 11.03626862 | -0.283368445 | 0.785511449 | 0.996633405 | -5.223072473 |
| **A_52_P338180** | Timm13 | 0.019895192 | -0.147918725 | 0.187709109 | 13.99156818 | 0.283353962 | 0.785522081 | 0.996633405 | -5.223075885 |
| **A_52_P81630** | 2810001G2 | -0.016795853 | -0.158565863 | 0.124974158 | 8.887848656 | -0.283156681 | 0.785666903 | 0.996633405 | -5.223122339 |
| **A_51_P371051** | Glipr1 | -0.018137904 | -0.171236131 | 0.134960324 | 7.949517134 | -0.283156189 | 0.785667265 | 0.996633405 | -5.223122454 |
| **A_52_P648424** | Nedd4 | 0.024825691 | -0.184810692 | 0.234462073 | 13.87164314 | 0.283037396 | 0.785754475 | 0.996633405 | -5.223150412 |
| **A_55_P2063558** | NA | 0.013458532 | -0.100205604 | 0.127122669 | 16.18811937 | 0.282998007 | 0.785783392 | 0.996633405 | -5.223159679 |
| **A_55_P2082779** | Gm10240 | 0.01311494 | -0.097660387 | 0.123890268 | 15.04283576 | 0.282964797 | 0.785807773 | 0.996633405 | -5.223167492 |
| **A_55_P2124528** | Capns1 | 0.02101396 | -0.156681813 | 0.198709734 | 14.15873725 | 0.282644113 | 0.78604322 | 0.996633405 | -5.223242887 |

| | | | | | | | | | |
|---|---|---|---|---|---|---|---|---|---|
| **A_55_P2176802** | Hipk1 | 0.014385663 | -0.107298299 | 0.136069625 | 8.334621771 | 0.282556741 | 0.786107373 | 0.996633405 | -5.223263414 |
| **A_55_P2455725** | Klhl20 | -0.026099475 | -0.246951463 | 0.194752513 | 9.250387961 | -0.28244871 | 0.786186697 | 0.996633405 | -5.223288787 |
| **A_51_P295708** | Ints1 | -0.015600198 | -0.147608929 | 0.116408533 | 10.38723713 | -0.282446752 | 0.786188134 | 0.996633405 | -5.223289247 |
| **A_55_P2022434** | Gpi1 | -0.016801874 | -0.158985123 | 0.125381375 | 16.35732859 | -0.282434937 | 0.78619681 | 0.996633405 | -5.223292021 |
| **A_55_P2138333** | Mad2l1 | -0.019983148 | -0.189103545 | 0.149137249 | 8.10830038 | -0.282408038 | 0.786216561 | 0.996633405 | -5.223298337 |
| **A_55_P2056021** | Snap91 | -0.020238723 | -0.191563614 | 0.151086167 | 10.13173245 | -0.282339592 | 0.786266822 | 0.996633405 | -5.223314405 |
| **A_51_P406466** | Ankrd35 | 0.012072665 | -0.090167737 | 0.114313068 | 7.752997062 | 0.282221272 | 0.786353707 | 0.996633405 | -5.223342173 |
| **A_52_P610403** | Sf1 | 0.013316281 | -0.099473028 | 0.12610559 | 8.337914654 | 0.28217865 | 0.786385006 | 0.996633405 | -5.223352173 |
| **A_55_P2149983** | Foxp4 | 0.026982027 | -0.201570755 | 0.255534809 | 10.81184373 | 0.282161138 | 0.786397866 | 0.996633405 | -5.223356281 |
| **A_55_P2073397** | NA | 0.018984121 | -0.14184902 | 0.179817262 | 11.64691266 | 0.282113642 | 0.786432745 | 0.996633405 | -5.223367423 |
| **A_51_P122035** | Nipa1 | -0.017166901 | -0.16262496 | 0.128291159 | 9.375332194 | -0.282074116 | 0.786461772 | 0.996633405 | -5.217272804 |
| **A_51_P513085** | Gins4 | 0.019351436 | -0.144622693 | 0.183325565 | 10.27918034 | 0.282063568 | 0.786469518 | 0.996633405 | -5.223379166 |
| **A_55_P2012331** | NA | -0.033676456 | -0.319052397 | 0.251699484 | 13.00068425 | -0.282044841 | 0.78648327 | 0.996633405 | -5.223383558 |
| **A_55_P2148182** | NA | 0.027823922 | -0.207973318 | 0.263621162 | 11.02758446 | 0.282025748 | 0.786497292 | 0.996633405 | -5.223388035 |
| **A_55_P1977498** | Htr2b | -0.016126661 | -0.152828397 | 0.120575075 | 7.713720393 | -0.281954849 | 0.786549359 | 0.996633405 | -5.223404657 |
| **A_52_P49250** | Erbb2 | 0.019225526 | -0.143760385 | 0.182211437 | 10.03988871 | 0.281927403 | 0.786569516 | 0.996633405 | -5.217307044 |
| **A_55_P2092622** | NA | -0.015222996 | -0.14434836 | 0.113902367 | 8.144871637 | -0.281771927 | 0.786683702 | 0.996633405 | -5.223447525 |
| **A_55_P2075274** | Rab11fip4 | -0.011617729 | -0.110169898 | 0.086934441 | 7.746799968 | -0.281750165 | 0.786699685 | 0.996633405 | -5.223452623 |
| **A_51_P308469** | Rngtt | 0.032125672 | -0.240499262 | 0.304750607 | 10.21725129 | 0.281640937 | 0.786779909 | 0.996633405 | -5.223478206 |
| **A_55_P2128283** | Nr2f2 | 0.030130906 | -0.225582637 | 0.285844449 | 9.698903306 | 0.281622657 | 0.786793335 | 0.996633405 | -5.223482486 |
| **A_51_P416689** | Ext1 | -0.012412652 | -0.117756791 | 0.092931488 | 9.657153327 | -0.281619892 | 0.786795366 | 0.996633405 | -5.223483134 |
| **A_51_P143915** | Rnf181 | -0.023967711 | -0.22741212 | 0.179476698 | 10.22215877 | -0.281572302 | 0.78683032 | 0.996633405 | -5.223494276 |
| **A_55_P2095251** | Ell3 | 0.01215878 | -0.091049133 | 0.115366693 | 7.755505722 | 0.281569851 | 0.78683212 | 0.996633405 | -5.22349485 |
| **A_55_P1966721** | NA | -0.037786388 | -0.358670472 | 0.283097696 | 11.65780267 | -0.281446815 | 0.78692249 | 0.996633405 | -5.223523648 |
| **A_51_P356967** | Lgi3 | 0.013150377 | -0.098540511 | 0.124841264 | 7.976838453 | 0.281403559 | 0.786954263 | 0.996633405 | -5.223533769 |
| **A_51_P477902** | Kdm5c | 0.058978699 | -0.441953063 | 0.559910462 | 11.33590885 | 0.281401075 | 0.786956088 | 0.996633405 | -5.223534351 |
| **A_55_P1958275** | Bcl11a | 0.030955837 | -0.232055821 | 0.293967495 | 10.16219943 | 0.281304503 | 0.787027024 | 0.996633405 | -5.223556943 |
| **A_52_P39083** | Ccbe1 | 0.012724662 | -0.095389069 | 0.120838392 | 7.760803801 | 0.281303102 | 0.787028053 | 0.996633405 | -5.22355727 |
| **A_55_P2089070** | 4933431K1 | 0.020546661 | -0.15403 | 0.195123322 | 9.158424168 | 0.281296423 | 0.787032959 | 0.996633405 | -5.223558832 |
| **A_55_P2089942** | Xpo1 | 0.018962768 | -0.14220411 | 0.180129646 | 10.8084604 | 0.281212788 | 0.787094394 | 0.996633405 | -5.223578391 |
| **A_51_P255682** | Mfge8 | 0.024101363 | -0.180756015 | 0.228958741 | 15.00411185 | 0.281189518 | 0.787111488 | 0.996633405 | -5.223583832 |
| **A_55_P1981756** | Vmn2r123 | 0.011932447 | -0.089502297 | 0.113367191 | 7.623263118 | 0.281158949 | 0.787133944 | 0.996633405 | -5.223590979 |
| **A_55_P2019217** | NA | -0.016671544 | -0.158492663 | 0.125149575 | 8.947174535 | -0.280959711 | 0.787280306 | 0.996633405 | -5.223637541 |
| **A_51_P183275** | Sympk | -0.02046115 | -0.194520717 | 0.153598416 | 11.08521203 | -0.280957918 | 0.787281623 | 0.996633405 | -5.223637959 |
| **A_55_P1985259** | C2cd3 | -0.016994727 | -0.161568491 | 0.127579036 | 8.482616625 | -0.280953114 | 0.787285153 | 0.996633405 | -5.223639082 |
| **A_66_P103654** | Asxl3 | 0.018722987 | -0.140567993 | 0.178013967 | 8.643480657 | 0.280926743 | 0.787304526 | 0.996633405 | -5.223645242 |
| **A_52_P146457** | NA | -0.014141892 | -0.134490844 | 0.106207059 | 14.78753622 | -0.280849946 | 0.787360945 | 0.996633405 | -5.223663179 |
| **A_52_P78023** | Limd2 | 0.016749339 | -0.1258128 | 0.159311478 | 12.2174601 | 0.280803555 | 0.787395027 | 0.996633405 | -5.223674012 |
| **A_55_P2061061** | Set | 0.020687478 | -0.155443241 | 0.196818196 | 12.39154543 | 0.280725306 | 0.787452515 | 0.996633405 | -5.22369228 |
| **A_55_P2064712** | NA | -0.018976116 | -0.180539479 | 0.142587246 | 15.19785021 | -0.280720146 | 0.787456305 | 0.996633405 | -5.223693484 |
| **A_66_P134181** | Aym1 | -0.022751669 | -0.216482308 | 0.17097897 | 8.572665236 | -0.280688149 | 0.787479814 | 0.996633405 | -5.223700953 |
| **A_51_P265008** | Ppig | 0.017745172 | -0.133416825 | 0.168907168 | 11.2598694 | 0.280573546 | 0.787564014 | 0.996633405 | -5.223727696 |
| **A_55_P2052170** | NA | -0.021594502 | -0.205573983 | 0.162384979 | 12.17744154 | -0.280532342 | 0.787594287 | 0.996633405 | -5.223737309 |
| **A_55_P2076273** | NA | 0.027243551 | -0.204966526 | 0.259453628 | 13.38516118 | 0.280408895 | 0.787684991 | 0.996633405 | -5.2237661 |
| **A_55_P1970810** | NA | 0.017565581 | -0.132186933 | 0.167318096 | 11.48447591 | 0.280348044 | 0.787729702 | 0.996633405 | -5.223780287 |
| **A_51_P438666** | Keap1 | 0.030044366 | -0.226145358 | 0.286234089 | 11.02985034 | 0.280291848 | 0.787770994 | 0.996633405 | -5.223793387 |
| **A_55_P2119478** | Ndufv2 | 0.019939452 | -0.150094707 | 0.189973611 | 14.97234744 | 0.280276171 | 0.787782513 | 0.996633405 | -5.223797041 |
| **A_52_P685938** | Efcab14 | 0.020560019 | -0.154797936 | 0.195917974 | 9.940988152 | 0.280225193 | 0.787819972 | 0.996633405 | -5.223808921 |

| | | | | | | | | | |
|---|---|---|---|---|---|---|---|---|---|
| **A_55_P2057296** | Upk3bl | 0.016600992 | -0.124991382 | 0.158193366 | 7.754016856 | 0.280222699 | 0.787821804 | 0.996633405 | -5.223809502 |
| **A_52_P598605** | Ptch1 | 0.013109717 | -0.098756194 | 0.124975629 | 7.905346414 | 0.280094573 | 0.787915954 | 0.996633405 | -5.223839352 |
| **A_66_P126189** | Ercc5 | -0.012730708 | -0.12144529 | 0.095983873 | 8.774279742 | -0.279881314 | 0.78807267 | 0.996633405 | -5.223889006 |
| **A_55_P2006008** | Serpinb1a | -0.024581373 | -0.23452615 | 0.185363404 | 9.629024058 | -0.279840263 | 0.788102838 | 0.996633405 | -5.22389856 |
| **A_51_P192344** | Ash2l | 0.013173567 | -0.099359109 | 0.125706243 | 11.08629491 | 0.279791085 | 0.788138979 | 0.996633405 | -5.223910004 |
| **A_51_P226932** | Tubgcp2 | -0.014439105 | -0.137830202 | 0.108951993 | 9.197573898 | -0.279682676 | 0.788218651 | 0.996633405 | -5.223935223 |
| **A_51_P245533** | Tex19.1 | 0.013084643 | -0.098743056 | 0.124912341 | 7.680707028 | 0.27965437 | 0.788239455 | 0.996633405 | -5.223941807 |
| **A_55_P1991552** | Atraid | 0.012843939 | -0.096963419 | 0.122651297 | 11.895145 | 0.279560577 | 0.788308388 | 0.996633405 | -5.223963616 |
| **A_66_P100853** | NA | -0.0171915 | -0.164190539 | 0.129807538 | 16.72606365 | -0.279517122 | 0.788340326 | 0.996633405 | -5.223973718 |
| **A_55_P2115981** | Xlr5c | 0.016447039 | -0.124188994 | 0.157083073 | 7.693786028 | 0.279511872 | 0.788344184 | 0.996633405 | -5.223974939 |
| **A_51_P400581** | Anp32e | 0.027880049 | -0.210545734 | 0.266305832 | 9.410620621 | 0.279479172 | 0.788368218 | 0.996633405 | -5.223982539 |
| **A_55_P2056220** | NA | -0.018760955 | -0.17924359 | 0.141721679 | 10.8248211 | -0.279406201 | 0.788421851 | 0.996633405 | -5.223999498 |
| **A_55_P1999361** | Dip2a | 0.025885453 | -0.19563918 | 0.247410086 | 10.86582266 | 0.279281955 | 0.788513174 | 0.996633405 | -5.224028362 |
| **A_55_P1976983** | Rpl36 | -0.013391283 | -0.127996004 | 0.101213437 | 16.67923212 | -0.27927292 | 0.788519814 | 0.996633405 | -5.22403046 |
| **A_51_P489313** | Sri | -0.021944345 | -0.209801769 | 0.165913079 | 12.3672177 | -0.279192269 | 0.788579096 | 0.996633405 | -5.22404919 |
| **A_52_P598300** | Gtf2f2 | -0.023510068 | -0.224797395 | 0.177777259 | 10.08203324 | -0.279155753 | 0.788605937 | 0.996633405 | -5.224057668 |
| **A_66_P107174** | NA | -0.018923514 | -0.180960508 | 0.14311348 | 8.623907752 | -0.279123719 | 0.788629484 | 0.996633405 | -5.224065105 |
| **A_55_P1957249** | Pdgfrb | -0.015714752 | -0.150290279 | 0.118860775 | 10.87634133 | -0.279094043 | 0.788651298 | 0.996633405 | -5.224071993 |
| **A_55_P1957845** | Tnfaip1 | 0.02103785 | -0.159198013 | 0.201273712 | 13.55211017 | 0.278977561 | 0.788736923 | 0.996633405 | -5.224099025 |
| **A_55_P1982171** | H2afz | -0.017795819 | -0.170325698 | 0.134734061 | 15.75062291 | -0.278850984 | 0.788829971 | 0.996633405 | -5.224128387 |
| **A_66_P120603** | Trps1 | 0.016448821 | -0.124538973 | 0.157436615 | 8.780387682 | 0.278844701 | 0.78883459 | 0.996633405 | -5.224129844 |
| **A_55_P2114009** | Ankra2 | -0.032990721 | -0.315777433 | 0.249795991 | 10.8944461 | -0.278831567 | 0.788844245 | 0.996633405 | -5.22413289 |
| **A_52_P556281** | G3bp2 | 0.017549509 | -0.132898224 | 0.167997242 | 11.81099384 | 0.278797233 | 0.788869485 | 0.996633405 | -5.224140851 |
| **A_51_P300506** | Cox6b2 | 0.017936475 | -0.135909912 | 0.171782862 | 9.358263529 | 0.278649921 | 0.788977783 | 0.996633405 | -5.224175001 |
| **A_55_P2139615** | Rtl6 | 0.014572359 | -0.11045137 | 0.139596089 | 8.727220904 | 0.278577828 | 0.789030784 | 0.996633405 | -5.224191706 |
| **A_55_P2120956** | NA | -0.01425291 | -0.136553878 | 0.108048057 | 8.677861904 | -0.278536935 | 0.789060848 | 0.996633405 | -5.224201181 |
| **A_52_P107234** | Prpf39 | 0.024945988 | -0.189133096 | 0.239025072 | 8.693808128 | 0.27850667 | 0.789083099 | 0.996633405 | -5.224208191 |
| **A_52_P267391** | Trim12a | -0.026698408 | -0.255829105 | 0.20243229 | 9.547133676 | -0.278491031 | 0.789094597 | 0.996633405 | -5.224211814 |
| **A_55_P2301063** | 9930024M1 | 0.026142983 | -0.19824298 | 0.250528947 | 8.512724333 | 0.278463702 | 0.78911469 | 0.996633405 | -5.224218144 |
| **A_51_P400217** | Vpreb1 | -0.014456105 | -0.138536828 | 0.109624618 | 8.030249856 | -0.278455703 | 0.789120571 | 0.996633405 | -5.224219996 |
| **A_51_P154973** | Stard3 | -0.020482162 | -0.196298339 | 0.155334015 | 11.24110914 | -0.27843645 | 0.789134726 | 0.996633405 | -5.224224455 |
| **A_55_P2120682** | Golga7 | 0.014719712 | -0.111694727 | 0.141134151 | 9.746911528 | 0.278299068 | 0.789235735 | 0.996633405 | -5.224256263 |
| **A_51_P256665** | Pon3 | -0.016777932 | -0.160903206 | 0.127347343 | 10.96582354 | -0.278232209 | 0.789284894 | 0.996633405 | -5.224271737 |
| **A_66_P138584** | Mnd1 | 0.016888491 | -0.128233107 | 0.162010088 | 9.151965587 | 0.278142861 | 0.78935059 | 0.996633405 | -5.224292411 |
| **A_51_P459128** | Serpinb9e | 0.01226885 | -0.093167826 | 0.117705527 | 7.698528361 | 0.278112997 | 0.789372549 | 0.996633405 | -5.224299319 |
| **A_55_P1972688** | Wnk1 | -0.028050761 | -0.269127153 | 0.21302563 | 12.33893175 | -0.278098791 | 0.789382994 | 0.996633405 | -5.224302605 |
| **A_55_P2174942** | NA | 0.023242832 | -0.176567961 | 0.223053625 | 13.59824685 | 0.278022046 | 0.789439426 | 0.996633405 | -5.224320355 |
| **A_55_P1979970** | NA | -0.018798639 | -0.180408571 | 0.142811293 | 11.00204719 | -0.278014522 | 0.789444958 | 0.996633405 | -5.224322095 |
| **A_55_P2172470** | Nrgn | 0.016857781 | -0.12820484 | 0.161920401 | 8.084897226 | 0.277749962 | 0.789639504 | 0.996633405 | -5.224383244 |
| **A_52_P375876** | Arf3 | -0.013074914 | -0.125654702 | 0.099504873 | 12.35199221 | -0.277579609 | 0.789764782 | 0.996633405 | -5.224422589 |
| **A_51_P383991** | 4-Sep | 0.02224915 | -0.169548255 | 0.214046556 | 10.7837641 | 0.277255286 | 0.790003309 | 0.996633405 | -5.22449743 |
| **A_52_P653937** | Rpl21 | -0.017713137 | -0.170440056 | 0.135013781 | 8.565855575 | -0.277197323 | 0.790045941 | 0.996633405 | -5.224510796 |
| **A_55_P2110837** | Rnf7 | 0.014706826 | -0.112141671 | 0.141555323 | 11.85537153 | 0.277103983 | 0.790114594 | 0.996633405 | -5.224532315 |
| **A_55_P1956147** | B4galnt2 | -0.011691085 | -0.112529248 | 0.089147078 | 7.701194961 | -0.277101591 | 0.790116354 | 0.996633405 | -5.224532867 |
| **A_51_P299934** | BC048403 | 0.015560634 | -0.118771286 | 0.149892554 | 8.803085951 | 0.276858072 | 0.790295478 | 0.996633405 | -5.224588975 |
| **A_55_P2119927** | NA | 0.023711382 | -0.181025271 | 0.228448034 | 8.008258987 | 0.27680275 | 0.790336172 | 0.996633405 | -5.224601715 |
| **A_55_P2100655** | Lsm6 | 0.020529352 | -0.156739862 | 0.197798567 | 13.68718753 | 0.276790425 | 0.790345239 | 0.996633405 | -5.224604553 |
| **A_55_P2147616** | NA | -0.023926567 | -0.230620437 | 0.182767303 | 11.71523726 | -0.276669908 | 0.790433894 | 0.996633405 | -5.224632297 |

| | | | | | | | | | |
|---|---|---|---|---|---|---|---|---|---|
| **A_51_P221362** | Tnks2 | 0.019799128 | -0.151279744 | 0.190878 | 12.48111713 | 0.276604227 | 0.790482211 | 0.996633405 | -5.224647412 |
| **A_51_P391716** | Ermap | 0.014063567 | -0.107489893 | 0.135617027 | 7.794248839 | 0.276526837 | 0.790539144 | 0.996633405 | -5.224665217 |
| **A_52_P350616** | Mrm1 | 0.017783286 | -0.136005277 | 0.171571849 | 11.54042456 | 0.276373948 | 0.790651622 | 0.996633405 | -5.224700378 |
| **A_51_P296487** | Lss | 0.012266961 | -0.093856206 | 0.118390128 | 8.014775081 | 0.276271397 | 0.79072707 | 0.996633405 | -5.224723952 |
| **A_51_P127841** | Pdss1 | 0.01514067 | -0.115854344 | 0.146135683 | 10.11679564 | 0.276248198 | 0.790744138 | 0.996633405 | -5.20637672 |
| **A_51_P325856** | Mcemp1 | -0.017066221 | -0.164736068 | 0.130603625 | 8.265330713 | -0.276219722 | 0.790765089 | 0.996633405 | -5.224735828 |
| **A_55_P2471798** | NA | 0.024732535 | -0.189328137 | 0.238793208 | 9.924610306 | 0.276147352 | 0.790818335 | 0.996633405 | -5.224752456 |
| **A_55_P1981959** | Trafd1 | 0.019472874 | -0.149108127 | 0.188053875 | 12.16513937 | 0.276077206 | 0.790869946 | 0.996633405 | -5.224768569 |
| **A_55_P1965303** | Dsc2 | -0.016799233 | -0.162234863 | 0.128636398 | 9.636413211 | -0.276075432 | 0.790871251 | 0.996633405 | -5.224768976 |
| **A_55_P2127973** | Gm9112 | -0.013678181 | -0.132192236 | 0.104835874 | 7.694151192 | -0.275846598 | 0.791039627 | 0.996633405 | -5.224821513 |
| **A_52_P585751** | Gns | -0.020367585 | -0.196876338 | 0.156141169 | 11.1226051 | -0.275792476 | 0.791079451 | 0.996633405 | -5.224833932 |
| **A_55_P1963687** | Tsen54 | -0.030707207 | -0.296868444 | 0.23545403 | 11.65014629 | -0.275743094 | 0.791115789 | 0.996633405 | -5.224845262 |
| **A_55_P1980539** | Cetn4 | -0.013355721 | -0.12912536 | 0.102413919 | 8.431507707 | -0.275728574 | 0.791126473 | 0.996633405 | -5.224848593 |
| **A_55_P2124712** | Ces2c | -0.012740944 | -0.123231454 | 0.097749565 | 7.744529842 | -0.275604164 | 0.791218023 | 0.996633405 | -5.224877126 |
| **A_52_P131490** | Atg4b | -0.014213349 | -0.137523496 | 0.109096798 | 8.74136338 | -0.275490566 | 0.791301619 | 0.996633405 | -5.224903168 |
| **A_55_P2143832** | Col4a4 | 0.013316473 | -0.102234215 | 0.128867161 | 8.006019828 | 0.275439237 | 0.791339393 | 0.996633405 | -5.224914932 |
| **A_51_P354744** | Cox15 | -0.026133788 | -0.252929375 | 0.2006618 | 10.29477902 | -0.275408214 | 0.791362223 | 0.996633405 | -5.224922041 |
| **A_51_P488196** | Bmper | -0.036382933 | -0.352142114 | 0.279376248 | 9.057284785 | -0.275391702 | 0.791374374 | 0.996633405 | -5.224925824 |
| **A_55_P1966593** | NA | 0.05677836 | -0.436177448 | 0.549734167 | 9.024717867 | 0.275285911 | 0.79145223 | 0.996633405 | -5.224950059 |
| **A_55_P1973334** | Mapk1 | 0.018535726 | -0.142401067 | 0.179472518 | 12.86786052 | 0.275272851 | 0.791461842 | 0.996633405 | -5.22495305 |
| **A_52_P27103** | Necap2 | 0.019215729 | -0.14770209 | 0.186133548 | 8.401083089 | 0.275146058 | 0.791555158 | 0.996633405 | -5.224982083 |
| **A_55_P2174085** | St8sia5 | -0.015615159 | -0.151296973 | 0.120066655 | 8.170666977 | -0.275064098 | 0.79161548 | 0.996633405 | -5.225000843 |
| **A_55_P2014361** | Pdpk1 | -0.015974033 | -0.154841389 | 0.122893323 | 11.87614923 | -0.27493089 | 0.791713524 | 0.996633405 | -5.225031321 |
| **A_55_P2157205** | Cdx1 | -0.012385002 | -0.120078826 | 0.095308822 | 7.695362338 | -0.274861824 | 0.79176436 | 0.996633405 | -5.225047118 |
| **A_55_P2045755** | Plcxd2 | 0.012758713 | -0.098213407 | 0.123730833 | 7.89537806 | 0.274790755 | 0.79181667 | 0.996633405 | -5.225063369 |
| **A_52_P660945** | Ctsf | 0.027439992 | -0.211276997 | 0.26615698 | 8.921261075 | 0.274732337 | 0.791859671 | 0.996633405 | -5.225076725 |
| **A_55_P1966765** | NA | -0.019979788 | -0.193853563 | 0.153893986 | 13.11593783 | -0.274641354 | 0.791926643 | 0.996633405 | -5.225097519 |
| **A_66_P121583** | Tmem267 | 0.018175601 | -0.140006598 | 0.1763578 | 8.407608255 | 0.274625146 | 0.791938574 | 0.996633405 | -5.225101223 |
| **A_55_P2018847** | Crlf2 | -0.020967074 | -0.203443725 | 0.161509576 | 11.10437533 | -0.274624745 | 0.791938868 | 0.996633405 | -5.225101314 |
| **A_55_P1963610** | NA | -0.014714531 | -0.142847963 | 0.113418902 | 15.38944411 | -0.274468866 | 0.792053614 | 0.996633405 | -5.225136923 |
| **A_55_P2032165** | NA | -0.019612102 | -0.190411669 | 0.151187465 | 14.49658315 | -0.274439431 | 0.792075283 | 0.996633405 | -5.225143645 |
| **A_55_P2117943** | NA | 0.024215328 | -0.186689521 | 0.235120176 | 9.976962383 | 0.274418206 | 0.792090908 | 0.996633405 | -5.225148492 |
| **A_65_P03606** | Rpap2 | 0.016402935 | -0.126462869 | 0.159268739 | 8.590857932 | 0.274411564 | 0.792095797 | 0.996633405 | -5.225150009 |
| **A_55_P2024993** | Hoxb4 | 0.031042374 | -0.239362219 | 0.301446967 | 8.576443153 | 0.27437845 | 0.792120174 | 0.996633405 | -5.225157569 |
| **A_55_P2172227** | Ubn2 | 0.018262747 | -0.140841827 | 0.177367322 | 8.457422895 | 0.274342173 | 0.792146881 | 0.996633405 | -5.225165851 |
| **A_55_P1984035** | Litaf | -0.017854714 | -0.173416081 | 0.137706653 | 8.295640812 | -0.274321776 | 0.792161896 | 0.996633405 | -5.225170507 |
| **A_55_P1968340** | Plekhh1 | -0.030259728 | -0.293984128 | 0.233464673 | 10.54713227 | -0.274235601 | 0.792225337 | 0.996633405 | -5.225190174 |
| **A_51_P474422** | Ift27 | -0.014022768 | -0.136277426 | 0.10823189 | 12.13318541 | -0.274143186 | 0.792293374 | 0.996633405 | -5.225211259 |
| **A_51_P447668** | Ptpn12 | 0.014841532 | -0.114562853 | 0.144245917 | 9.038158047 | 0.27411883 | 0.792311305 | 0.996633405 | -5.225216814 |
| **A_55_P2079787** | Prss50 | -0.01140835 | -0.110886038 | 0.088069338 | 7.85769969 | -0.274098266 | 0.792326445 | 0.996633405 | -5.225221505 |
| **A_55_P1963201** | Chchd7 | 0.018428113 | -0.142337729 | 0.179193954 | 13.55172673 | 0.273965712 | 0.792424037 | 0.996633405 | -5.22525173 |
| **A_51_P385351** | Slc44a3 | 0.011397889 | -0.08805729 | 0.110853067 | 7.710608083 | 0.273908898 | 0.792465867 | 0.996633405 | -5.225264681 |
| **A_55_P2021892** | Ccdc114 | -0.021866567 | -0.212686504 | 0.168953371 | 8.789104674 | -0.273883569 | 0.792484516 | 0.996633405 | -5.225270454 |
| **A_51_P249193** | Gsg1l | -0.019559548 | -0.190253468 | 0.151134373 | 8.811561509 | -0.273873424 | 0.792491985 | 0.996633405 | -5.225272766 |
| **A_51_P463187** | Tbc1d2b | 0.01643565 | -0.127001719 | 0.159873018 | 8.709950769 | 0.27386321 | 0.792499506 | 0.996633405 | -5.225275094 |
| **A_55_P2023818** | Cysltr1 | -0.018947663 | -0.184330339 | 0.146435013 | 8.230746632 | -0.273826054 | 0.792526863 | 0.996633405 | -5.22528356 |
| **A_55_P2109354** | Dot1l | 0.018087615 | -0.139818163 | 0.175993393 | 11.09017225 | 0.273774141 | 0.792565087 | 0.996633405 | -5.225295388 |
| **A_55_P2044843** | NA | 0.017295985 | -0.133712678 | 0.168304648 | 14.53255737 | 0.273749017 | 0.792583585 | 0.996633405 | -5.225301111 |

| | | | | | | | | | |
|---|---|---|---|---|---|---|---|---|---|
| **A_55_P2066878** | Kcnj13 | 0.021583135 | -0.166867958 | 0.210034228 | 8.00274354 | 0.273731638 | 0.792596382 | 0.996633405 | -5.22530507 |
| **A_55_P2148688** | NA | -0.02718475 | -0.26457708 | 0.210207579 | 9.104915178 | -0.273695591 | 0.792622924 | 0.996633405 | -5.225313281 |
| **A_55_P2394308** | Fst | 0.028720612 | -0.222177595 | 0.27961882 | 8.140636333 | 0.273593182 | 0.792698331 | 0.996633405 | -5.2253366 |
| **A_55_P1987650** | Ctnnd2 | 0.012395341 | -0.095888915 | 0.120679598 | 7.727548093 | 0.273591313 | 0.792699707 | 0.996633405 | -5.225337026 |
| **A_55_P2113886** | NA | -0.011820602 | -0.115101016 | 0.091459812 | 7.684978565 | -0.27354624 | 0.792732896 | 0.996633405 | -5.225347287 |
| **A_55_P2011300** | Fggy | -0.013015652 | -0.126742137 | 0.100710833 | 7.808358271 | -0.273535337 | 0.792740925 | 0.996633405 | -5.225349769 |
| **A_52_P486482** | R3hcc1l | 0.013093424 | -0.101327083 | 0.127513931 | 8.438911695 | 0.273500737 | 0.792766403 | 0.996633405 | -5.225357644 |
| **A_52_P606679** | Eloa | -0.018164062 | -0.176908648 | 0.140580524 | 8.671316945 | -0.273478495 | 0.792782781 | 0.996633405 | -5.225362706 |
| **A_55_P2028046** | Cyp3a59 | -0.011338115 | -0.110437174 | 0.087760945 | 7.666285254 | -0.273451588 | 0.792802595 | 0.996633405 | -5.225368829 |
| **A_55_P2080905** | NA | -0.014493686 | -0.14121918 | 0.112231809 | 16.41569077 | -0.273353084 | 0.792875132 | 0.996633405 | -5.22539124 |
| **A_51_P301994** | Trmt10c | -0.0165055 | -0.160857484 | 0.127846484 | 8.997157139 | -0.273284546 | 0.792925604 | 0.996633405 | -5.225406829 |
| **A_55_P2025837** | NA | -0.0116268 | -0.113327776 | 0.090074176 | 14.20959975 | -0.273239951 | 0.792958444 | 0.996633405 | -5.22541697 |
| **A_51_P373379** | Csnk2b | -0.015476979 | -0.15085791 | 0.119903952 | 13.8023968 | -0.27323594 | 0.792961398 | 0.996633405 | -5.225417882 |
| **A_55_P2024853** | Cep57l1 | -0.012965831 | -0.126405221 | 0.100473559 | 8.77919525 | -0.273177929 | 0.793004119 | 0.996633405 | -5.225431071 |
| **A_55_P1977934** | Itgam | -0.045542743 | -0.444041708 | 0.352956223 | 12.46424822 | -0.273149944 | 0.793024729 | 0.996633405 | -5.225437433 |
| **A_52_P206613** | Adss | -0.023602635 | -0.230140511 | 0.182935241 | 9.348789184 | -0.273130323 | 0.793039179 | 0.996633405 | -5.225441893 |
| **A_52_P344290** | F2r | 0.021636333 | -0.167716517 | 0.210989184 | 12.92759463 | 0.273099527 | 0.793061858 | 0.996633405 | -5.225448892 |
| **A_55_P1983819** | Srsf7 | 0.020585435 | -0.159591382 | 0.200762252 | 12.00874686 | 0.273067662 | 0.793085325 | 0.996633405 | -5.225456134 |
| **A_55_P1952235** | Spry1 | 0.032698759 | -0.253553947 | 0.318951466 | 11.21807053 | 0.273017702 | 0.79312212 | 0.996633405 | -5.225467486 |
| **A_55_P1968649** | Atp1b2 | -0.019315332 | -0.188421077 | 0.149790412 | 11.32462435 | -0.272993909 | 0.793139642 | 0.996633405 | -5.225472891 |
| **A_51_P403693** | Uba5 | -0.012504061 | -0.121977735 | 0.096969613 | 10.94807573 | -0.272992373 | 0.793140774 | 0.996633405 | -5.22547324 |
| **A_66_P111987** | Brwd3 | 0.016437919 | -0.127478256 | 0.160354095 | 8.567438589 | 0.272989765 | 0.793142694 | 0.996633405 | -5.225473833 |
| **A_51_P201773** | Tubgcp3 | -0.018611794 | -0.181573825 | 0.144350238 | 10.22577884 | -0.272967495 | 0.793159096 | 0.996633405 | -5.225478892 |
| **A_51_P479434** | Rfc3 | 0.016744189 | -0.129932642 | 0.163421019 | 8.629301622 | 0.272842317 | 0.79325129 | 0.996633405 | -5.225507322 |
| **A_52_P650473** | NA | 0.015189769 | -0.117872481 | 0.148252019 | 8.215578348 | 0.272838363 | 0.793254201 | 0.996633405 | -5.225508219 |
| **A_51_P357341** | Clic1 | -0.041253686 | -0.402652081 | 0.320144709 | 14.76183272 | -0.272825948 | 0.793263346 | 0.996633405 | -5.225511038 |
| **A_66_P129048** | Faap20 | -0.015178963 | -0.148190372 | 0.117832446 | 13.01739302 | -0.272748477 | 0.793320405 | 0.996633405 | -5.225528625 |
| **A_52_P312467** | Hoxb3 | 0.01408935 | -0.109468288 | 0.137646989 | 7.788490185 | 0.272540157 | 0.793473845 | 0.996633405 | -5.225575894 |
| **A_55_P1998115** | Klkb1 | 0.011266643 | -0.087662938 | 0.110196224 | 7.705450028 | 0.272193343 | 0.793729317 | 0.996633405 | -5.225654509 |
| **A_66_P106098** | Kif3a | 0.022453596 | -0.174754787 | 0.219661978 | 9.20520524 | 0.272125764 | 0.7937791 | 0.996633405 | -5.225669816 |
| **A_52_P216427** | Nkx2-1 | 0.011244944 | -0.087556558 | 0.110046445 | 7.651939769 | 0.27202127 | 0.79385608 | 0.996633405 | -5.206456528 |
| **A_55_P2009146** | Gtpbp3 | -0.020122634 | -0.196929746 | 0.156684479 | 7.966514605 | -0.272015858 | 0.793860067 | 0.996633405 | -5.225694703 |
| **A_51_P306658** | Lsm8 | -0.018592293 | -0.181973154 | 0.144788568 | 11.65744339 | -0.271982465 | 0.793884668 | 0.996633405 | -5.225702263 |
| **A_55_P2117764** | Vangl2 | 0.013952008 | -0.108738258 | 0.136642274 | 8.735981455 | 0.271791412 | 0.794025422 | 0.996633405 | -5.225745496 |
| **A_51_P475995** | Thra | 0.022629261 | -0.176480183 | 0.221738705 | 12.81035259 | 0.271636196 | 0.794139781 | 0.996633405 | -5.225780598 |
| **A_55_P2178349** | Ywhaq | 0.024912675 | -0.194297841 | 0.244123191 | 12.20860439 | 0.271623981 | 0.794148781 | 0.996633405 | -5.225783359 |
| **A_51_P519189** | Eif3i | -0.025050302 | -0.245480299 | 0.195379694 | 10.92092208 | -0.271613536 | 0.794156477 | 0.996633405 | -5.225785721 |
| **A_55_P2106763** | Pxdc1 | 0.017509275 | -0.136565259 | 0.171583808 | 8.361639518 | 0.271610417 | 0.794158776 | 0.996633405 | -5.225786426 |
| **A_55_P1971054** | Paqr9 | 0.012191677 | -0.095091333 | 0.119474688 | 7.686515976 | 0.271607423 | 0.794160981 | 0.996633405 | -5.225787103 |
| **A_55_P2130109** | Nmt2 | 0.01183969 | -0.09234889 | 0.116028269 | 8.02218516 | 0.271599735 | 0.794166646 | 0.996633405 | -5.225788841 |
| **A_55_P1989321** | Sat1 | -0.020211398 | -0.198157286 | 0.15773449 | 10.00644846 | -0.271467303 | 0.794264223 | 0.996633405 | -5.225818771 |
| **A_55_P2085616** | Nsa2 | 0.01503215 | -0.117375876 | 0.147440175 | 8.802537133 | 0.271341305 | 0.794357064 | 0.996633405 | -5.225847234 |
| **A_52_P480544** | Tnpo2 | 0.046461219 | -0.362851881 | 0.455774318 | 8.63971526 | 0.271296434 | 0.794390128 | 0.996633405 | -5.225857367 |
| **A_51_P409724** | Secisbp2 | 0.019928602 | -0.155654302 | 0.195511507 | 8.838738042 | 0.271271235 | 0.794408696 | 0.996633405 | -5.225863057 |
| **A_55_P2092329** | Adrm1 | 0.011782173 | -0.092043891 | 0.115608237 | 11.65731491 | 0.271224016 | 0.794443491 | 0.996633405 | -5.225873718 |
| **A_55_P2090943** | Larp4 | 0.016130706 | -0.126085812 | 0.158347223 | 11.98353272 | 0.271089351 | 0.794542726 | 0.996633405 | -5.225904111 |
| **A_55_P2054027** | Chst5 | 0.011781292 | -0.09220648 | 0.115769064 | 7.675038533 | 0.270781994 | 0.794769232 | 0.996633405 | -5.225973426 |
| **A_52_P35057** | Kctd17 | -0.016543819 | -0.162582728 | 0.129495089 | 10.5085683 | -0.270754902 | 0.794789199 | 0.996633405 | -5.225979533 |

| | | | | | | | | | |
|---|---|---|---|---|---|---|---|---|---|
| **A_55_P2025293** | Ss18 | 0.019209199 | -0.150425532 | 0.18884393 | 12.09803864 | 0.27064725 | 0.794868538 | 0.996633405 | -5.22600379 |
| **A_55_P2086128** | Fa2h | 0.010931474 | -0.085609154 | 0.107472103 | 7.64917669 | 0.270631132 | 0.794880418 | 0.996633405 | -5.226007421 |
| **A_51_P116601** | Cfap69 | 0.014565036 | -0.114074227 | 0.143204298 | 8.157370962 | 0.270612057 | 0.794894476 | 0.996633405 | -5.219895686 |
| **A_52_P427818** | Zscan18 | -0.0252334 | -0.248293912 | 0.197827113 | 9.101297529 | -0.270372306 | 0.795071185 | 0.996633405 | -5.2260657 |
| **A_55_P1996746** | NA | -0.011296583 | -0.11118311 | 0.088589944 | 7.631871447 | -0.270302037 | 0.795122979 | 0.996633405 | -5.226081513 |
| **A_55_P2099080** | Irak1 | 0.020496461 | -0.16077098 | 0.201763902 | 10.57520936 | 0.27025156 | 0.795160186 | 0.996633405 | -5.22609287 |
| **A_52_P452667** | Prom2 | -0.011159318 | -0.109873177 | 0.087554541 | 7.637823122 | -0.27018962 | 0.795205843 | 0.996633405 | -5.226106803 |
| **A_55_P2099961** | H2ac11 | -0.026692176 | -0.26284048 | 0.209456128 | 17.10180529 | -0.27015206 | 0.795233529 | 0.996633405 | -5.22611525 |
| **A_51_P312748** | Oxsm | -0.013731284 | -0.135278051 | 0.107815482 | 9.303857324 | -0.270008157 | 0.795339606 | 0.996633405 | -5.226147603 |
| **A_55_P2114596** | Zbtb14 | -0.017778446 | -0.175175601 | 0.139618708 | 10.00482242 | -0.26996413 | 0.795372061 | 0.996633405 | -5.226157499 |
| **A_55_P2066468** | Noa1 | -0.023102306 | -0.227657197 | 0.181452584 | 10.2246301 | -0.269932145 | 0.795395639 | 0.996633405 | -5.226164686 |
| **A_55_P1974367** | Fabp3 | 0.022898151 | -0.17985752 | 0.225653822 | 17.01194196 | 0.269920916 | 0.795403917 | 0.996633405 | -5.226167209 |
| **A_51_P300378** | Wwc2 | -0.012858998 | -0.126745294 | 0.101027297 | 12.45549285 | -0.26986391 | 0.795445941 | 0.996633405 | -5.226180017 |
| **A_66_P133043** | Mbd2 | -0.013278355 | -0.130888757 | 0.104332047 | 11.49938449 | -0.269840832 | 0.795462954 | 0.996633405 | -5.226185202 |
| **A_51_P441970** | Stox2 | 0.014871325 | -0.116946836 | 0.146689487 | 8.986721127 | 0.269639528 | 0.79561136 | 0.996633405 | -5.226230405 |
| **A_52_P507305** | Iah1 | -0.020454951 | -0.201782887 | 0.160872986 | 9.350638751 | -0.269614252 | 0.795629994 | 0.996633405 | -5.226236078 |
| **A_51_P329370** | Snrnp25 | 0.01685868 | -0.132739429 | 0.16645679 | 12.75474289 | 0.269343567 | 0.795829565 | 0.996633405 | -5.226296804 |
| **A_51_P464182** | Mat1a | 0.01141686 | -0.08990812 | 0.11274184 | 7.668458068 | 0.269301809 | 0.795860354 | 0.996633405 | -5.226306167 |
| **A_66_P108188** | Car5a | -0.011868849 | -0.117220913 | 0.093483215 | 7.655672718 | -0.269261768 | 0.795889877 | 0.996633405 | -5.226315143 |
| **A_65_P12152** | Rasa2 | 0.025730668 | -0.20269646 | 0.254157796 | 8.271007594 | 0.269223221 | 0.795918299 | 0.996633405 | -5.226323783 |
| **A_55_P2176260** | NA | 0.01147376 | -0.090432345 | 0.113379866 | 7.920831138 | 0.269100619 | 0.796008699 | 0.996633405 | -5.226351256 |
| **A_51_P353232** | Tnnc2 | -0.121107954 | -1.197214861 | 0.954998953 | 8.924440804 | -0.268983959 | 0.796094722 | 0.996633405 | -5.226377387 |
| **A_51_P490023** | Tubb2a | 0.021813969 | -0.17203188 | 0.215659818 | 13.69248308 | 0.268959764 | 0.796112564 | 0.996633405 | -5.226382805 |
| **A_55_P2176212** | NA | -0.028979691 | -0.286518086 | 0.228558705 | 12.15833218 | -0.2689434 | 0.79612463 | 0.996633405 | -5.226386469 |
| **A_51_P268145** | Naa80 | -0.012359476 | -0.122200031 | 0.097481079 | 9.179316909 | -0.268934468 | 0.796131216 | 0.996633405 | -5.226388469 |
| **A_65_P09657** | Zfp213 | -0.013484872 | -0.133343001 | 0.106373258 | 9.694335699 | -0.268898548 | 0.796157704 | 0.996633405 | -5.226396511 |
| **A_55_P2111163** | S100g | 0.12187843 | -0.9616221 | 1.205378959 | 8.192420334 | 0.268848028 | 0.796194959 | 0.996633405 | -5.22640782 |
| **A_55_P2021510** | NA | -0.04001818 | -0.395945899 | 0.315909539 | 8.99886203 | -0.268722886 | 0.796287243 | 0.996633405 | -5.226435824 |
| **A_55_P1987499** | Pttg1 | -0.012589898 | -0.124581447 | 0.099401651 | 13.27169491 | -0.268686645 | 0.796313969 | 0.996633405 | -5.226443931 |
| **A_55_P2064792** | Arpc3 | -0.024870026 | -0.246170713 | 0.196430661 | 12.14620927 | -0.268597897 | 0.796379417 | 0.996633405 | -5.226463781 |
| **A_55_P1992455** | Vdac3 | -0.01762981 | -0.174632656 | 0.139373037 | 14.99769313 | -0.268379432 | 0.796540536 | 0.996633405 | -5.226512616 |
| **A_55_P1978642** | Wasf1 | 0.024002739 | -0.189766803 | 0.237772281 | 10.16984708 | 0.268363905 | 0.796551988 | 0.996633405 | -5.226516086 |
| **A_52_P42069** | Prokr1 | 0.012720686 | -0.10062275 | 0.126064121 | 7.854037377 | 0.268239838 | 0.796643493 | 0.996633405 | -5.226543801 |
| **A_52_P582261** | Ikzf5 | 0.023394012 | -0.185207326 | 0.23199535 | 9.597878459 | 0.268038235 | 0.796792191 | 0.996633405 | -5.22658881 |
| **A_55_P2023652** | Vps35l | 0.017782881 | -0.140790806 | 0.176356569 | 11.56838553 | 0.268027985 | 0.796799752 | 0.996633405 | -5.226591097 |
| **A_55_P2076683** | Sort1 | 0.015874116 | -0.125709275 | 0.157457506 | 8.229595683 | 0.267970116 | 0.796842437 | 0.996633405 | -5.22660401 |
| **A_55_P2110671** | NA | -0.030149703 | -0.299114298 | 0.238814892 | 12.09129044 | -0.267915017 | 0.796883079 | 0.996633405 | -5.226616303 |
| **A_52_P357469** | Frk | 0.01131495 | -0.089630133 | 0.112260033 | 7.912057038 | 0.267902397 | 0.796892389 | 0.996633405 | -5.226619118 |
| **A_51_P229816** | Eif2b2 | -0.026747167 | -0.265420219 | 0.211925886 | 12.69060425 | -0.267844989 | 0.796934735 | 0.996633405 | -5.226631922 |
| **A_52_P491569** | 1700017B0 | -0.017999849 | -0.178628267 | 0.142628569 | 8.683825133 | -0.26782777 | 0.796947436 | 0.996633405 | -5.226635762 |
| **A_55_P2019776** | Gxylt1 | 0.011983743 | -0.094977102 | 0.118944588 | 8.117357182 | 0.267779163 | 0.796983292 | 0.996633405 | -5.226646601 |
| **A_51_P363791** | Ap3m1 | 0.018486718 | -0.14658161 | 0.183555045 | 10.89529249 | 0.267673383 | 0.797061323 | 0.996633405 | -5.226670181 |
| **A_55_P1991931** | Nosip | 0.015472164 | -0.122743691 | 0.153688019 | 10.44710275 | 0.267548381 | 0.797153537 | 0.996633405 | -5.226698035 |
| **A_55_P2085596** | Nudt1 | 0.025764413 | -0.204460983 | 0.255989809 | 10.14631145 | 0.267470664 | 0.79721087 | 0.996633405 | -5.226715346 |
| **A_55_P2056035** | Traf7 | 0.01672347 | -0.13274347 | 0.166190409 | 13.06543043 | 0.267417841 | 0.79724984 | 0.996633405 | -5.22672711 |
| **A_52_P684057** | Snupn | 0.016283506 | -0.12929548 | 0.161862492 | 10.26010532 | 0.267336562 | 0.797309804 | 0.996633405 | -5.226745205 |
| **A_55_P2042612** | C1s1 | -0.020757067 | -0.206360286 | 0.164846152 | 8.076857496 | -0.267294251 | 0.79734102 | 0.996633405 | -5.226754623 |
| **A_51_P121179** | Erfe | -0.011122832 | -0.110606065 | 0.088360402 | 7.782230049 | -0.267223471 | 0.79739324 | 0.996633405 | -5.220650848 |

| | | | | | | | | | |
|---|---|---|---|---|---|---|---|---|---|
| **A_55_P2032267** | Gm10767 | -0.021364118 | -0.212491336 | 0.169763099 | 11.5132042 | -0.267160083 | 0.797440007 | 0.996633405 | -5.226784478 |
| **A_66_P113662** | Tmem62 | 0.013489408 | -0.107251366 | 0.134230183 | 8.021722971 | 0.26702263 | 0.797541423 | 0.996633405 | -5.226815049 |
| **A_66_P129976** | Apba1 | -0.017460914 | -0.173778822 | 0.138856995 | 8.852158179 | -0.266973014 | 0.797578031 | 0.996633405 | -5.22682608 |
| **A_51_P382849** | Emb | 0.018449177 | -0.146731398 | 0.183629751 | 8.201302363 | 0.266948295 | 0.79759627 | 0.996633405 | -5.226831575 |
| **A_55_P1957922** | Arrdc2 | -0.016929139 | -0.168562044 | 0.134703765 | 8.832764381 | -0.266839769 | 0.797676347 | 0.996633405 | -5.226855694 |
| **A_55_P2165011** | Fut10 | -0.014265479 | -0.142170045 | 0.113639087 | 8.506478612 | -0.266568885 | 0.797876233 | 0.996633405 | -5.226915856 |
| **A_55_P2000683** | Eci2 | -0.019471598 | -0.194088387 | 0.155145191 | 13.64519726 | -0.266516886 | 0.797914605 | 0.996633405 | -5.226927398 |
| **A_52_P24843** | Pten | -0.012915903 | -0.12877681 | 0.102945003 | 9.092694869 | -0.266438507 | 0.797972446 | 0.996633405 | -5.226944791 |
| **A_66_P128701** | NA | -0.013082149 | -0.13059394 | 0.104429642 | 15.19834114 | -0.266076661 | 0.798239489 | 0.996633405 | -5.227025023 |
| **A_51_P126198** | Dapk2 | -0.026189511 | -0.261447544 | 0.209068523 | 8.366417478 | -0.266067637 | 0.79824615 | 0.996633405 | -5.227027023 |
| **A_55_P2157632** | 2510039O1 | -0.017160436 | -0.171319282 | 0.136998409 | 11.85952585 | -0.266053523 | 0.798256566 | 0.996633405 | -5.22703015 |
| **A_55_P2088495** | NA | 0.011773183 | -0.093992884 | 0.11753925 | 7.759786137 | 0.266045964 | 0.798262146 | 0.996633405 | -5.227031825 |
| **A_51_P372172** | Plod3 | -0.01890158 | -0.188710853 | 0.150907694 | 10.67197844 | -0.266039319 | 0.79826705 | 0.996633405 | -5.227033297 |
| **A_55_P1960703** | NA | -0.026202734 | -0.261611595 | 0.209206128 | 9.75510379 | -0.266031417 | 0.798272882 | 0.996633405 | -5.227035048 |
| **A_65_P02327** | Pogk | -0.018814474 | -0.187996938 | 0.15036799 | 8.96193702 | -0.265794428 | 0.798447799 | 0.996633405 | -5.227087529 |
| **A_55_P2052709** | NA | 0.02442933 | -0.195246259 | 0.244104918 | 8.587184002 | 0.265790149 | 0.798450957 | 0.996633405 | -5.227088476 |
| **A_55_P2003211** | Igsf5 | 0.011745307 | -0.093879076 | 0.11736969 | 7.708334105 | 0.265772061 | 0.798464308 | 0.996633405 | -5.22709248 |
| **A_55_P2083098** | Tatdn3 | 0.024414295 | -0.195230438 | 0.244059027 | 10.12625288 | 0.265663889 | 0.798544153 | 0.996633405 | -5.227116418 |
| **A_55_P2147160** | NA | 0.010703228 | -0.085599438 | 0.107005894 | 7.651433195 | 0.265635183 | 0.798565341 | 0.996633405 | -5.227122769 |
| **A_66_P117682** | NA | -0.029888404 | -0.299006062 | 0.239229254 | 9.676272228 | -0.265442011 | 0.798707934 | 0.996633405 | -5.227165488 |
| **A_55_P2203141** | 2900011F0 | -0.0175966 | -0.176050056 | 0.140856856 | 8.748271841 | -0.265421544 | 0.798723043 | 0.996633405 | -5.227170013 |
| **A_52_P65108** | Smad1 | -0.015695501 | -0.157071987 | 0.125680985 | 8.526093973 | -0.265342696 | 0.798781248 | 0.996633405 | -5.22718744 |
| **A_55_P1968178** | Smagp | -0.028926134 | -0.289538492 | 0.231686224 | 10.53959476 | -0.265280006 | 0.798827527 | 0.996633405 | -5.227201292 |
| **A_52_P119350** | 4732419C1 | 0.010973879 | -0.087902542 | 0.1098503 | 7.690007982 | 0.265262932 | 0.798840131 | 0.996633405 | -5.227205064 |
| **A_52_P390918** | Trip4 | 0.014056872 | -0.112630999 | 0.140744743 | 8.380176694 | 0.265193445 | 0.798891429 | 0.996633405 | -5.227220414 |
| **A_51_P381558** | Rasa4 | -0.011184487 | -0.111989096 | 0.089620123 | 7.996127077 | -0.26518246 | 0.798899539 | 0.996633405 | -5.22722284 |
| **A_55_P2023384** | NA | -0.011486409 | -0.115066134 | 0.092093317 | 7.895823701 | -0.265044408 | 0.799001457 | 0.996633405 | -5.227253322 |
| **A_51_P334318** | Chp2 | 0.011569035 | -0.09282987 | 0.11596794 | 7.767811382 | 0.264856311 | 0.799140328 | 0.996633405 | -5.22729483 |
| **A_55_P2347756** | AU040972 | -0.013755872 | -0.137904977 | 0.110393233 | 8.143193292 | -0.26482175 | 0.799165846 | 0.996633405 | -5.227302454 |
| **A_55_P2185826** | Cyp3a13 | 0.012280086 | -0.098576968 | 0.12313714 | 7.859965587 | 0.264756857 | 0.799213757 | 0.996633405 | -5.227316765 |
| **A_51_P271336** | Wipf2 | 0.017918958 | -0.143924397 | 0.179762314 | 10.03850981 | 0.264622649 | 0.799312851 | 0.996633405 | -5.227346353 |
| **A_51_P361220** | Fzd4 | 0.016708294 | -0.134245666 | 0.167662254 | 10.92276138 | 0.264543279 | 0.799371456 | 0.996633405 | -5.227363843 |
| **A_51_P520718** | Ptger3 | 0.012310169 | -0.098911661 | 0.123531999 | 7.753605265 | 0.264534986 | 0.799377579 | 0.996633405 | -5.227365671 |
| **A_55_P1981322** | Grhl2 | -0.012472352 | -0.125169384 | 0.100224679 | 8.0464482 | -0.264511782 | 0.799394713 | 0.996633405 | -5.227370783 |
| **A_55_P1989511** | Tbca | -0.023473963 | -0.235733806 | 0.188785881 | 11.97387795 | -0.264318585 | 0.799537372 | 0.996633405 | -5.227413332 |
| **A_55_P1992959** | Pfkfb4 | 0.017286076 | -0.139032554 | 0.173604706 | 8.550245787 | 0.264298562 | 0.799552158 | 0.996633405 | -5.227417741 |
| **A_55_P2002898** | Ctsj | 0.091856072 | -0.73896078 | 0.922672924 | 8.155420039 | 0.264247999 | 0.799589496 | 0.996633405 | -5.227428871 |
| **A_55_P1996008** | Gm14399 | -0.01576192 | -0.158389475 | 0.126865635 | 8.976130519 | -0.264128224 | 0.799677946 | 0.996633405 | -5.227455227 |
| **A_51_P214269** | Epha1 | -0.018697959 | -0.187925104 | 0.150529186 | 7.956667057 | -0.264078659 | 0.799714549 | 0.996633405 | -5.227466131 |
| **A_55_P2003903** | Bfsp1 | -0.011476625 | -0.115363015 | 0.092409764 | 8.266523548 | -0.264036932 | 0.799745364 | 0.996633405 | -5.227475309 |
| **A_55_P2021585** | Tff1 | 0.012444455 | -0.100246111 | 0.125135022 | 7.675082791 | 0.263935288 | 0.799820429 | 0.996633405 | -5.227497659 |
| **A_55_P2045158** | NA | 0.043060176 | -0.347132833 | 0.433253185 | 10.2329566 | 0.263757882 | 0.79995145 | 0.996633405 | -5.227536649 |
| **A_51_P151897** | R3hcc1 | -0.02899191 | -0.291897295 | 0.233913475 | 10.63490363 | -0.263564233 | 0.800094476 | 0.996633405 | -5.227579179 |
| **A_66_P103957** | Bbs7 | 0.025096845 | -0.202549524 | 0.252743213 | 9.069674663 | 0.263492059 | 0.800147785 | 0.996633405 | -5.227595022 |
| **A_55_P2038117** | Tmem104 | -0.015688307 | -0.158051479 | 0.126674866 | 8.795378062 | -0.263382879 | 0.800228429 | 0.996633405 | -5.227618982 |
| **A_51_P493886** | Gpt2 | 0.013134422 | -0.106056811 | 0.132325656 | 8.385564116 | 0.263375754 | 0.800233692 | 0.996633405 | -5.227620545 |
| **A_55_P2165324** | Acsl3 | -0.015937702 | -0.160599956 | 0.128724552 | 10.85863751 | -0.263317428 | 0.800276774 | 0.996633405 | -5.22763334 |
| **A_55_P2062029** | Tsen34 | 0.024311036 | -0.196375688 | 0.244997761 | 14.74239064 | 0.263291233 | 0.800296123 | 0.996633405 | -5.227639085 |

| | | | | | | | | | |
|---|---|---|---|---|---|---|---|---|---|
| **A_55_P2165655** | Zbtb4 | 0.018292239 | -0.147905291 | 0.184489769 | 10.64538492 | 0.263057946 | 0.80046845 | 0.996633405 | -5.227690229 |
| **A_55_P2102175** | Lztr1 | 0.019444865 | -0.157241471 | 0.196131202 | 10.7616833 | 0.263033527 | 0.800486489 | 0.996633405 | -5.22769558 |
| **A_55_P2005343** | NA | -0.017796117 | -0.179502682 | 0.143910447 | 16.5481133 | -0.263030883 | 0.800488442 | 0.996633405 | -5.227696159 |
| **A_55_P2136572** | Cdc42se2 | -0.014798842 | -0.149279254 | 0.11968157 | 11.30740837 | -0.263013374 | 0.800501376 | 0.996633405 | -5.227699996 |
| **A_51_P153973** | Zfp273 | -0.015124955 | -0.152579222 | 0.122329313 | 8.550515742 | -0.262993483 | 0.80051607 | 0.996633405 | -5.227704353 |
| **A_52_P351669** | Gcc2 | 0.017820739 | -0.144168142 | 0.17980962 | 9.16701965 | 0.262935753 | 0.800558718 | 0.996633405 | -5.227717 |
| **A_55_P2213418** | 4933417E1 | 0.011576214 | -0.093689107 | 0.116841534 | 7.694883317 | 0.26283933 | 0.80062995 | 0.996633405 | -5.227738116 |
| **A_52_P555680** | Trappc12 | 0.014461101 | -0.117051437 | 0.145973638 | 11.42930867 | 0.262810868 | 0.800650977 | 0.996633405 | -5.227744348 |
| **A_55_P2171528** | 3110082I17 | -0.015066706 | -0.152091436 | 0.121958023 | 8.223319188 | -0.262801903 | 0.8006576 | 0.996633405 | -5.227746311 |
| **A_55_P2178343** | Rap1gds1 | 0.015335501 | -0.124235595 | 0.154906596 | 13.05273296 | 0.26261021 | 0.800799222 | 0.996633405 | -5.227788263 |
| **A_55_P1959541** | NA | 0.012914748 | -0.104645375 | 0.13047487 | 10.24643334 | 0.262563911 | 0.800833428 | 0.996633405 | -5.227798391 |
| **A_52_P585448** | Ptp4a2 | 0.019378562 | -0.157118157 | 0.195875282 | 12.34057605 | 0.262418257 | 0.800941044 | 0.996633405 | -5.227830242 |
| **A_52_P326664** | Unc93a | -0.010688091 | -0.108081028 | 0.086704845 | 7.656819069 | -0.262290059 | 0.801035765 | 0.996633405 | -5.227858262 |
| **A_55_P1972034** | Muc16 | 0.116975479 | -0.949258684 | 1.183209642 | 9.360421844 | 0.262211281 | 0.801093974 | 0.996633405 | -5.227875473 |
| **A_52_P469381** | Comtd1 | 0.02119171 | -0.172098877 | 0.214482298 | 12.26375883 | 0.262038097 | 0.801221944 | 0.996633405 | -5.227913293 |
| **A_52_P152219** | Rpgrip1l | -0.012480964 | -0.126338409 | 0.10137648 | 8.668852052 | -0.261996714 | 0.801252523 | 0.996633405 | -5.227922327 |
| **A_51_P253803** | Mki67 | 0.019642776 | -0.159560787 | 0.198846339 | 13.1337091 | 0.261978323 | 0.801266113 | 0.996633405 | -5.227926341 |
| **A_52_P43848** | Sco1 | 0.014671728 | -0.119216905 | 0.148560361 | 8.614210672 | 0.261906749 | 0.801319004 | 0.996633405 | -5.227941961 |
| **A_55_P2157330** | NA | 0.018698943 | -0.151944933 | 0.189342819 | 16.38775508 | 0.261899991 | 0.801323998 | 0.996633405 | -5.227943435 |
| **A_55_P2042778** | Pde5a | 0.024233582 | -0.196942305 | 0.24540947 | 8.543603752 | 0.261871942 | 0.801344725 | 0.996633405 | -5.227949555 |
| **A_55_P1969311** | Gramd1b | -0.02533386 | -0.25658731 | 0.205919591 | 9.136400219 | -0.261831729 | 0.801374442 | 0.996633405 | -5.227958328 |
| **A_52_P444162** | Rab11a | 0.016332198 | -0.132753967 | 0.165418363 | 10.89298325 | 0.261828206 | 0.801377045 | 0.996633405 | -5.227959096 |
| **A_55_P2177755** | Cdk2ap1 | -0.022082064 | -0.223748564 | 0.179584435 | 10.78749269 | -0.261706822 | 0.801466748 | 0.996633405 | -5.227985569 |
| **A_66_P103477** | Nkapd1 | -0.013613382 | -0.137960935 | 0.110734171 | 9.051205849 | -0.261660349 | 0.801501093 | 0.996633405 | -5.227995701 |
| **A_51_P254855** | Ptgs2 | 0.01137152 | -0.0924987 | 0.115241739 | 7.688696164 | 0.261659556 | 0.801501678 | 0.996633405 | -5.203009028 |
| **A_55_P2136813** | Rab10os | 0.016392421 | -0.13335365 | 0.166138492 | 8.335632357 | 0.261635579 | 0.801519399 | 0.996633405 | -5.228001101 |
| **A_51_P133792** | Prickle3 | 0.014884351 | -0.121187413 | 0.150956114 | 9.213069045 | 0.261439387 | 0.801664394 | 0.996633405 | -5.228043851 |
| **A_55_P2051859** | Golgb1 | -0.012388955 | -0.125668134 | 0.100890223 | 10.32858686 | -0.261392866 | 0.801698777 | 0.996633405 | -5.228053984 |
| **A_55_P1998776** | NA | -0.014242911 | -0.144514736 | 0.116028915 | 15.29872065 | -0.261310811 | 0.801759423 | 0.996633405 | -5.228071852 |
| **A_55_P1991199** | Nos1ap | 0.016974384 | -0.1382927 | 0.172241467 | 8.636103338 | 0.261290579 | 0.801774377 | 0.996633405 | -5.228076256 |
| **A_55_P1978050** | Aven | -0.019951065 | -0.202509956 | 0.162607826 | 10.95610145 | -0.261199428 | 0.801841748 | 0.996633405 | -5.228096097 |
| **A_52_P148678** | Plcg1 | 0.017086457 | -0.139268991 | 0.173441904 | 10.58443451 | 0.261184936 | 0.801852459 | 0.996633405 | -5.228099251 |
| **A_55_P2153391** | NA | 0.029112953 | -0.237526885 | 0.29575279 | 10.92771358 | 0.260957843 | 0.802020316 | 0.996633405 | -5.228148649 |
| **A_66_P119074** | Tbc1d22b | -0.01441982 | -0.146525105 | 0.117685465 | 8.295964502 | -0.260884797 | 0.80207431 | 0.996633405 | -5.228164529 |
| **A_66_P139785** | Plpp3 | -0.032285662 | -0.328114867 | 0.263543544 | 11.33607987 | -0.260842153 | 0.802105833 | 0.996633405 | -5.228173798 |
| **A_55_P1967618** | Med16 | -0.017537598 | -0.178263838 | 0.143188642 | 11.77661923 | -0.260790921 | 0.802143704 | 0.996633405 | -5.228184932 |
| **A_55_P2066613** | Lcp1 | -0.014060135 | -0.142968981 | 0.114848711 | 8.608387478 | -0.260684911 | 0.80222207 | 0.996633405 | -5.228207963 |
| **A_55_P1960764** | Rps4x | 0.011661869 | -0.095263716 | 0.118587455 | 15.08557064 | 0.260672771 | 0.802231044 | 0.996633405 | -5.2282106 |
| **A_55_P2001048** | Rom1 | -0.021074542 | -0.214315626 | 0.172166541 | 9.378195308 | -0.260656057 | 0.8022434 | 0.996633405 | -5.22821423 |
| **A_51_P236267** | St8sia4 | 0.038112674 | -0.311442102 | 0.38766745 | 9.852900571 | 0.260593341 | 0.802289763 | 0.996633405 | -5.228227849 |
| **A_51_P410681** | Irx5 | -0.016034433 | -0.163162955 | 0.13109409 | 8.58580341 | -0.260474889 | 0.802377331 | 0.996633405 | -5.228253564 |
| **A_55_P2045931** | Btnl10 | -0.010459619 | -0.106442467 | 0.08552323 | 7.631904713 | -0.260454191 | 0.802392633 | 0.996633405 | -5.228258056 |
| **A_52_P420394** | Apeh | 0.017703737 | -0.144774877 | 0.180182351 | 11.75424159 | 0.260422118 | 0.802416344 | 0.996633405 | -5.228265016 |
| **A_55_P2010768** | Dpm3 | 0.013621718 | -0.111398917 | 0.138642352 | 13.15816373 | 0.260410987 | 0.802424573 | 0.996633405 | -5.228267431 |
| **A_52_P650379** | Strap | 0.017342857 | -0.141854066 | 0.176539779 | 10.90602358 | 0.260372492 | 0.802453032 | 0.996633405 | -5.228275784 |
| **A_55_P2182016** | Faf2 | -0.01519979 | -0.154741015 | 0.124341435 | 10.2346824 | -0.260341972 | 0.802475597 | 0.996633405 | -5.228282405 |
| **A_66_P122729** | Zmat2 | 0.016295201 | -0.133354068 | 0.165944469 | 11.13048958 | 0.260252107 | 0.802542036 | 0.996633405 | -5.228301896 |
| **A_55_P2105873** | Ppp2r5c | -0.018034672 | -0.183682159 | 0.147612816 | 9.694813324 | -0.260215113 | 0.802569387 | 0.996633405 | -5.228309918 |

| | | | | | | | | | |
|---|---|---|---|---|---|---|---|---|---|
| **A_51_P141104** | Med18 | -0.016705011 | -0.170157002 | 0.13674698 | 8.264002369 | -0.260185672 | 0.802591154 | 0.996633405 | -5.228316302 |
| **A_52_P408025** | Mpped2 | 0.013631051 | -0.111597389 | 0.138859492 | 11.27826973 | 0.260157002 | 0.802612351 | 0.996633405 | -5.228322517 |
| **A_52_P355014** | Rbm8a | -0.020006985 | -0.203837303 | 0.163823333 | 11.80115854 | -0.260119939 | 0.802639754 | 0.996633405 | -5.228330551 |
| **A_52_P31641** | Tmem145 | -0.010554269 | -0.107624681 | 0.086516144 | 7.654620556 | -0.259866561 | 0.802827099 | 0.996633405 | -5.228385445 |
| **A_55_P1981040** | Ddx6 | 0.017335587 | -0.142154087 | 0.17682526 | 10.71888703 | 0.259785618 | 0.802886951 | 0.996633405 | -5.22840297 |
| **A_52_P593268** | Lsm6 | -0.019161102 | -0.195498267 | 0.157176063 | 13.16823178 | -0.259708261 | 0.802944151 | 0.996633405 | -5.228419714 |
| **A_55_P2079636** | Scgb3a2 | 0.010738772 | -0.088113225 | 0.109590769 | 7.635006182 | 0.259644015 | 0.802991659 | 0.996633405 | -5.228433616 |
| **A_52_P229320** | Rptor | -0.010744099 | -0.109696207 | 0.088208009 | 11.12132035 | -0.259509989 | 0.803090768 | 0.996633405 | -5.228462608 |
| **A_51_P490348** | 1110059E2 | -0.0190398 | -0.194501028 | 0.156421428 | 10.64917809 | -0.259352445 | 0.803207274 | 0.996633405 | -5.228496668 |
| **A_55_P2050807** | Mtg2 | 0.012279743 | -0.100900372 | 0.125459858 | 10.87290036 | 0.259315388 | 0.803234679 | 0.996633405 | -5.228504676 |
| **A_55_P2126572** | Tmem25 | 0.016476533 | -0.135413091 | 0.168366156 | 8.975273841 | 0.259266766 | 0.803270638 | 0.996633405 | -5.228515183 |
| **A_51_P128987** | Akr1b8 | -0.014462724 | -0.147843778 | 0.11891833 | 11.31495566 | -0.259158281 | 0.803350868 | 0.996633405 | -5.215991981 |
| **A_55_P2040227** | Ptchd4 | 0.018470785 | -0.151928925 | 0.188870496 | 8.966171117 | 0.259075076 | 0.803412406 | 0.996633405 | -5.228556584 |
| **A_51_P280245** | Zfp207 | -0.014047715 | -0.143650864 | 0.115555433 | 11.27855584 | -0.259059355 | 0.803424033 | 0.996633405 | -5.228559978 |
| **A_52_P655265** | Bora | 0.017862364 | -0.14695584 | 0.182680567 | 8.502059186 | 0.259025718 | 0.80344891 | 0.996633405 | -5.228567239 |
| **A_55_P2037328** | Ppp1r12b | 0.019046921 | -0.156757605 | 0.194851446 | 8.616812349 | 0.258942811 | 0.80351023 | 0.996633405 | -5.228585133 |
| **A_55_P2059860** | Igsf1 | 0.010487196 | -0.086311995 | 0.107286386 | 7.736293073 | 0.258938593 | 0.803513349 | 0.996633405 | -5.228586043 |
| **A_55_P2109983** | Mpg | -0.018931472 | -0.193811408 | 0.155948465 | 8.324277068 | -0.258734012 | 0.803664667 | 0.996633405 | -5.228630174 |
| **A_55_P2087087** | Pecam1 | -0.016873641 | -0.172828632 | 0.13908135 | 8.885990847 | -0.258594132 | 0.803768133 | 0.996633405 | -5.228660327 |
| **A_51_P238722** | Cd93 | -0.015801791 | -0.161908845 | 0.130305262 | 14.40756149 | -0.258490314 | 0.803844927 | 0.996633405 | -5.228682697 |
| **A_51_P501803** | Hoxa2 | 0.011912745 | -0.098282519 | 0.122108008 | 7.808864834 | 0.258379491 | 0.803926907 | 0.996633405 | -5.228706566 |
| **A_52_P601914** | Lemd2 | -0.019441082 | -0.199344283 | 0.16046212 | 11.77533535 | -0.258279932 | 0.804000556 | 0.996633405 | -5.228728001 |
| **A_55_P1987290** | Rnu1b6 | 0.074327092 | -0.613626655 | 0.762280838 | 14.48700451 | 0.258224255 | 0.804041744 | 0.996633405 | -5.228739985 |
| **A_51_P439857** | Dnajc21 | -0.013626023 | -0.139790603 | 0.112538557 | 9.580071017 | -0.258131379 | 0.804110452 | 0.996633405 | -5.22875997 |
| **A_51_P386911** | Taco1 | 0.016655132 | -0.137568001 | 0.170878265 | 8.642412409 | 0.258111699 | 0.804125012 | 0.996633405 | -5.228764203 |
| **A_55_P2050390** | Cdh22 | -0.01197857 | -0.122899512 | 0.098942372 | 7.74405081 | -0.258107452 | 0.804128153 | 0.996633405 | -5.228765117 |
| **A_55_P1952414** | Odf2 | -0.014412522 | -0.147904017 | 0.119078973 | 9.657004018 | -0.25804504 | 0.804174327 | 0.996633405 | -5.228778541 |
| **A_55_P1979335** | NA | 0.011369536 | -0.094006095 | 0.116745166 | 8.008525603 | 0.257876449 | 0.804299056 | 0.996633405 | -5.228814788 |
| **A_51_P201971** | Kmt5a | -0.021092552 | -0.216610874 | 0.17442577 | 10.61987893 | -0.257840302 | 0.804325799 | 0.996633405 | -5.228822556 |
| **A_55_P1960157** | Bcat1 | 0.010830546 | -0.08956371 | 0.111224802 | 7.767790285 | 0.257840196 | 0.804325878 | 0.996633405 | -5.228822579 |
| **A_55_P2010121** | Rab27b | 0.012308886 | -0.101829298 | 0.12644707 | 7.999391699 | 0.257748952 | 0.804393386 | 0.996633405 | -5.228842184 |
| **A_51_P359485** | Zfpl1 | 0.015943035 | -0.131937928 | 0.163823997 | 10.0924869 | 0.257672371 | 0.804450048 | 0.996633405 | -5.228858633 |
| **A_55_P2168316** | Aldh3b2 | -0.010292628 | -0.105829266 | 0.08524401 | 7.647175721 | -0.257493012 | 0.804582758 | 0.996633405 | -5.22889714 |
| **A_52_P420665** | Uts2b | -0.074249673 | -0.763556246 | 0.6150569 | 8.206318224 | -0.25744903 | 0.804615303 | 0.996633405 | -5.228906579 |
| **A_55_P2049976** | NA | -0.011723764 | -0.12056374 | 0.097116212 | 7.765632593 | -0.257446958 | 0.804616836 | 0.996633405 | -5.228907023 |
| **A_55_P2088014** | Cdc42ep4 | -0.013807517 | -0.142021568 | 0.114406534 | 10.68220286 | -0.25738847 | 0.804660114 | 0.996633405 | -5.228919572 |
| **A_55_P1960202** | Vcan | 0.033850688 | -0.280524912 | 0.348226287 | 11.35339412 | 0.257352148 | 0.804686991 | 0.996633405 | -5.228927364 |
| **A_55_P2067895** | Krba1 | 0.021546334 | -0.178590504 | 0.221683173 | 10.93014581 | 0.257309303 | 0.804718695 | 0.996633405 | -5.228936554 |
| **A_52_P444637** | Rcbtb2 | -0.013550294 | -0.139458596 | 0.112358008 | 8.539627773 | -0.257219259 | 0.804785326 | 0.996633405 | -5.228955862 |
| **A_52_P424840** | Mrc2 | 0.020628067 | -0.171101675 | 0.212357809 | 8.211406711 | 0.257145056 | 0.804840237 | 0.996633405 | -5.228971769 |
| **A_66_P115757** | Mthfd2l | 0.024831406 | -0.206130559 | 0.255793372 | 10.19617175 | 0.256962638 | 0.804975231 | 0.996633405 | -5.229010854 |
| **A_52_P227065** | Bud31 | -0.015272933 | -0.157340888 | 0.126795022 | 10.6631359 | -0.256942207 | 0.804990351 | 0.996633405 | -5.22901523 |
| **A_55_P2154880** | NA | 0.015572853 | -0.129331917 | 0.160477624 | 10.9516882 | 0.256858916 | 0.805051992 | 0.996633405 | -5.229033065 |
| **A_55_P2184364** | B4galt3 | 0.017186604 | -0.142747179 | 0.177120387 | 11.97904837 | 0.256837834 | 0.805067594 | 0.996633405 | -5.229037579 |
| **A_55_P2133315** | Steap2 | 0.019932788 | -0.16560355 | 0.205469127 | 8.207365026 | 0.256772321 | 0.805116079 | 0.996633405 | -5.229051602 |
| **A_55_P2122648** | Dtymk | -0.025005134 | -0.257779333 | 0.207769065 | 10.79040915 | -0.256745876 | 0.805135651 | 0.996633405 | -5.229057262 |
| **A_55_P2095880** | NA | -0.021757336 | -0.224364587 | 0.180849915 | 9.22788887 | -0.25666099 | 0.805198476 | 0.996633405 | -5.229075426 |
| **A_55_P2058143** | Pclo | -0.012555845 | -0.129505411 | 0.10439372 | 8.000665659 | -0.256599894 | 0.805243694 | 0.996633405 | -5.229088495 |

| | | | | | | | | | |
|---|---|---|---|---|---|---|---|---|---|
| **A_51_P438952** | Bag4 | -0.013424413 | -0.1385135 | 0.111664674 | 8.195497379 | -0.256498607 | 0.805318661 | 0.996633405 | -5.229110156 |
| **A_51_P242859** | Akr1c12 | -0.010430997 | -0.107659281 | 0.086797288 | 7.648411363 | -0.256414346 | 0.805381028 | 0.996633405 | -5.229128169 |
| **A_65_P09032** | Rab43 | -0.017599209 | -0.181784737 | 0.14658632 | 11.23784926 | -0.256193079 | 0.805544807 | 0.996633405 | -5.229175443 |
| **A_51_P281246** | Cep57 | -0.020664908 | -0.213564699 | 0.172234884 | 9.078050119 | -0.256041782 | 0.805656803 | 0.996633405 | -5.229207744 |
| **A_66_P105422** | Lonrf3 | -0.018151571 | -0.187621686 | 0.151318544 | 8.126799644 | -0.255994259 | 0.805691982 | 0.996633405 | -5.229217887 |
| **A_52_P174661** | Gatc | -0.019261912 | -0.199132517 | 0.160608693 | 8.604501418 | -0.255945988 | 0.805727715 | 0.996633405 | -5.229228187 |
| **A_55_P2156052** | NA | 0.018461156 | -0.153943739 | 0.190866052 | 10.39429303 | 0.255928376 | 0.805740753 | 0.996633405 | -5.229231944 |
| **A_51_P169032** | 2810004N2 | -0.027405101 | -0.283391949 | 0.228581747 | 12.66196502 | -0.255872071 | 0.805782435 | 0.996633405 | -5.229243956 |
| **A_51_P395111** | Nme3 | -0.026009142 | -0.268961215 | 0.216942931 | 10.79848859 | -0.255867164 | 0.805786067 | 0.996633405 | -5.229245002 |
| **A_51_P450682** | Tepp | -0.01058226 | -0.109453196 | 0.088288676 | 7.813668593 | -0.255810829 | 0.805827771 | 0.996633405 | -5.229257017 |
| **A_55_P1986703** | Aldh3b3 | -0.021106681 | -0.218406739 | 0.176193378 | 8.321072225 | -0.255683003 | 0.805922403 | 0.996633405 | -5.229284269 |
| **A_55_P2068733** | Gm21953 | -0.018192127 | -0.188262716 | 0.151878463 | 9.076102015 | -0.255660354 | 0.805939171 | 0.996633405 | -5.229289097 |
| **A_55_P1997320** | Zim1 | 0.025725538 | -0.214812579 | 0.266263656 | 8.301679799 | 0.255616977 | 0.805971284 | 0.996633405 | -5.229298341 |
| **A_55_P2012815** | Acvr2b | 0.020496878 | -0.171218941 | 0.212212697 | 11.11459106 | 0.255528242 | 0.806036979 | 0.996633405 | -5.229317247 |
| **A_51_P327796** | Itgb5 | 0.016632826 | -0.139073797 | 0.172339449 | 12.93121087 | 0.25531015 | 0.806198451 | 0.996633405 | -5.229363685 |
| **A_55_P2078138** | Otop1 | -0.058381277 | -0.605058681 | 0.488296127 | 8.664749993 | -0.255241684 | 0.806249144 | 0.996633405 | -5.229378256 |
| **A_55_P2012799** | Rtkn | -0.023656173 | -0.245180136 | 0.197867791 | 11.14912967 | -0.255230693 | 0.806257282 | 0.996633405 | -5.229380595 |
| **A_55_P2083889** | Pea15a | 0.012244396 | -0.102479609 | 0.1269684 | 11.30370674 | 0.255089265 | 0.806362001 | 0.996633405 | -5.229410679 |
| **A_55_P1979508** | Clic3 | -0.046922271 | -0.486621814 | 0.392777271 | 8.470058647 | -0.255054004 | 0.80638811 | 0.996633405 | -5.229418178 |
| **A_51_P287100** | Cdh16 | 0.025384853 | -0.212519873 | 0.26328958 | 8.587725046 | 0.255023802 | 0.806410474 | 0.996633405 | -5.229424599 |
| **A_66_P124549** | H13 | 0.013235234 | -0.110842829 | 0.137313297 | 12.06404869 | 0.254944565 | 0.806469147 | 0.996633405 | -5.229441443 |
| **A_52_P94830** | Galnt16 | 0.044568512 | -0.373269665 | 0.462406689 | 10.28073419 | 0.254934815 | 0.806476366 | 0.996633405 | -5.229443516 |
| **A_51_P465232** | Mcfd2 | -0.017412342 | -0.180704272 | 0.145879589 | 12.45716184 | -0.254859947 | 0.806531806 | 0.996633405 | -5.229459426 |
| **A_52_P381484** | Spon2 | -0.030775672 | -0.319577469 | 0.258026124 | 11.67732375 | -0.254692872 | 0.806655529 | 0.996633405 | -5.229494914 |
| **A_51_P505322** | Mpnd | 0.016747411 | -0.140412138 | 0.173906961 | 12.78633056 | 0.254692432 | 0.806655854 | 0.996633405 | -5.229495008 |
| **A_66_P100159** | Cfl2 | -0.013251352 | -0.137631934 | 0.111129229 | 13.29207136 | -0.254634209 | 0.806698971 | 0.996633405 | -5.229507369 |
| **A_51_P123134** | Ercc6l | -0.016117927 | -0.167442804 | 0.13520695 | 8.469022841 | -0.25457045 | 0.806746189 | 0.996633405 | -5.2111012 |
| **A_55_P2008936** | Slc2a9 | -0.011387043 | -0.118351395 | 0.095577309 | 7.79902492 | -0.254437443 | 0.806844691 | 0.996633405 | -5.229549126 |
| **A_55_P1968362** | Tle1 | -0.022047685 | -0.229206703 | 0.185111334 | 9.280498913 | -0.254371397 | 0.806893605 | 0.996633405 | -5.229563135 |
| **A_51_P309534** | Pex14 | -0.012358502 | -0.128478778 | 0.103761774 | 9.812407003 | -0.254370598 | 0.806894196 | 0.996633405 | -5.229563304 |
| **A_52_P615375** | H2aw | -0.022388847 | -0.23277454 | 0.187996846 | 12.27058174 | -0.254345851 | 0.806912524 | 0.996633405 | -5.229568552 |
| **A_55_P1961736** | Rcan2 | -0.014421846 | -0.150135822 | 0.12129213 | 12.86060398 | -0.253983449 | 0.807180938 | 0.996633405 | -5.229645349 |
| **A_55_P2042356** | Rftn1 | 0.019415726 | -0.163374888 | 0.20220634 | 11.8977309 | 0.253868531 | 0.807266059 | 0.996633405 | -5.22966968 |
| **A_55_P2128085** | NA | -0.017270844 | -0.179930033 | 0.145388344 | 11.07770178 | -0.253772233 | 0.807337389 | 0.996633405 | -5.229690059 |
| **A_55_P1966585** | NA | 0.036685376 | -0.309061151 | 0.382431903 | 15.80136916 | 0.253597098 | 0.807467122 | 0.996633405 | -5.229727104 |
| **A_55_P2012296** | Gopc | 0.016056425 | -0.13551207 | 0.16762492 | 9.818758994 | 0.253191462 | 0.807767624 | 0.996633405 | -5.229812809 |
| **A_55_P2023523** | Aloxe3 | -0.012021651 | -0.125504228 | 0.101460927 | 7.751716579 | -0.25318854 | 0.807769789 | 0.996633405 | -5.229813426 |
| **A_66_P108474** | NA | 0.013728176 | -0.120049838 | 0.14750619 | 7.794070983 | 0.254914907 | 0.807811374 | 0.996633405 | -5.151246459 |
| **A_55_P2025225** | Gnaz | -0.017025745 | -0.177880171 | 0.14382868 | 8.133553755 | -0.252977707 | 0.807925992 | 0.996633405 | -5.229857919 |
| **A_55_P2423228** | AI464196 | -0.011500408 | -0.12022652 | 0.097225704 | 7.940618991 | -0.252806663 | 0.808052723 | 0.996633405 | -5.229893988 |
| **A_55_P2063256** | Lgals4 | -0.027057713 | -0.282893895 | 0.22877847 | 8.230459243 | -0.252777401 | 0.808074404 | 0.996633405 | -5.229900156 |
| **A_51_P516850** | Rbbp5 | -0.015667551 | -0.163821142 | 0.132486039 | 8.207373931 | -0.252754026 | 0.808091724 | 0.996633405 | -5.229905083 |
| **A_66_P130654** | Abcg4 | 0.01121599 | -0.094953841 | 0.117385822 | 8.66957839 | 0.252490837 | 0.808286743 | 0.996633405 | -5.229960526 |
| **A_55_P2016034** | Nlrc5 | 0.010973249 | -0.092906274 | 0.114852772 | 7.696622689 | 0.252472694 | 0.808300187 | 0.996633405 | -5.229964345 |
| **A_55_P2190994** | Cdsn | -0.010376253 | -0.108608795 | 0.087856288 | 7.97022043 | -0.252461009 | 0.808308846 | 0.996633405 | -5.229966806 |
| **A_55_P2100375** | Col11a1 | -0.025098765 | -0.262742136 | 0.212544605 | 8.413533469 | -0.252426989 | 0.808334055 | 0.996633405 | -5.229973967 |
| **A_66_P138201** | Rhox10 | -0.010624427 | -0.111222009 | 0.089973154 | 7.657803572 | -0.252421951 | 0.808337789 | 0.996633405 | -5.229975028 |
| **A_51_P315555** | Nars | 0.019520544 | -0.165335604 | 0.204376691 | 10.68531835 | 0.252387096 | 0.808363617 | 0.996633405 | -5.229982364 |

| | | | | | | | | | |
|---|---|---|---|---|---|---|---|---|---|
| **A_55_P2295951** | NA | 0.03823815 | -0.323881468 | 0.400357768 | 8.85505321 | 0.252379428 | 0.8083693 | 0.996633405 | -5.229983978 |
| **A_51_P246677** | Rec8 | 0.038438066 | -0.325660419 | 0.402536551 | 9.806386321 | 0.252320066 | 0.808413289 | 0.996633405 | -5.22999647 |
| **A_52_P18922** | Chid1 | 0.017526923 | -0.148521562 | 0.183575407 | 8.294840725 | 0.252278297 | 0.808444242 | 0.996633405 | -5.230005258 |
| **A_55_P1963817** | Rps6 | 0.013554543 | -0.114939215 | 0.142048301 | 15.57038717 | 0.252122721 | 0.808559536 | 0.996633405 | -5.230037978 |
| **A_55_P2177154** | H2bc14 | 0.037666688 | -0.319503926 | 0.394837301 | 11.58895359 | 0.252052403 | 0.808611648 | 0.996633405 | -5.23005276 |
| **A_52_P290783** | BC016579 | 0.010339563 | -0.087717482 | 0.108396609 | 7.648859074 | 0.252018557 | 0.808636732 | 0.996633405 | -5.230059874 |
| **A_52_P441070** | Apob | -0.03315788 | -0.347730314 | 0.281414553 | 7.832492026 | -0.251927299 | 0.808704365 | 0.996633405 | -5.23007905 |
| **A_55_P2174067** | Cerk | 0.01641086 | -0.139301106 | 0.172122825 | 9.207188685 | 0.251894379 | 0.808728762 | 0.996633405 | -5.230085965 |
| **A_52_P608495** | Aars2 | 0.017439457 | -0.148069108 | 0.182948023 | 10.43744283 | 0.251838211 | 0.808770392 | 0.996633405 | -5.230097763 |
| **A_55_P2039235** | Mat2b | -0.020870845 | -0.21898062 | 0.177238931 | 12.88117829 | -0.251792767 | 0.808804072 | 0.996633405 | -5.230107306 |
| **A_55_P2088203** | Hcn2 | -0.016311847 | -0.171215121 | 0.138591428 | 8.892103443 | -0.251681713 | 0.808886382 | 0.996633405 | -5.23013062 |
| **A_51_P378550** | Oplah | -0.038524019 | -0.404424506 | 0.327376467 | 10.49263551 | -0.251638878 | 0.808918131 | 0.996633405 | -5.23013961 |
| **A_51_P486681** | Ap3b2 | -0.024751711 | -0.25984431 | 0.210340889 | 7.743930175 | -0.251637518 | 0.808919139 | 0.996633405 | -5.230139896 |
| **A_55_P1962411** | Gm6813 | -0.024160342 | -0.253692856 | 0.205372173 | 15.77802958 | -0.251575297 | 0.808965257 | 0.996633405 | -5.230152952 |
| **A_55_P2151956** | NA | 0.011536798 | -0.098111951 | 0.121185546 | 7.806832772 | 0.251472637 | 0.809041351 | 0.996633405 | -5.230174486 |
| **A_66_P136372** | Mrpl9 | 0.020861142 | -0.177413432 | 0.219135716 | 10.61889822 | 0.251466528 | 0.809045879 | 0.996633405 | -5.230175767 |
| **A_52_P335508** | Tiprl | 0.01639312 | -0.139468339 | 0.17225458 | 9.802152489 | 0.25138075 | 0.809109461 | 0.996633405 | -5.230193753 |
| **A_55_P1999057** | Cps1 | -0.010450045 | -0.109810957 | 0.088910866 | 7.754737854 | -0.251369013 | 0.809118161 | 0.996633405 | -5.230196214 |
| **A_52_P740562** | Rps21 | 0.010351823 | -0.088113888 | 0.108817534 | 14.43242632 | 0.251270174 | 0.809191426 | 0.996633405 | -5.23021693 |
| **A_55_P2037525** | NA | -0.022974585 | -0.241595134 | 0.195645963 | 14.29394045 | -0.251168885 | 0.80926651 | 0.996633405 | -5.230238152 |
| **A_51_P140641** | Eva1a | 0.018664001 | -0.158971778 | 0.196299781 | 9.306892971 | 0.251121239 | 0.80930183 | 0.996633405 | -5.230248131 |
| **A_55_P2130373** | Dcaf11 | -0.024082547 | -0.253517021 | 0.205351927 | 12.51138339 | -0.250872393 | 0.809486307 | 0.996633405 | -5.230300223 |
| **A_55_P2081313** | Gnas | 0.01417337 | -0.120894464 | 0.149241205 | 15.86307898 | 0.250801618 | 0.809538776 | 0.996633405 | -5.23031503 |
| **A_51_P275123** | Psd3 | -0.023394514 | -0.246446883 | 0.199657855 | 10.6343674 | -0.250678058 | 0.809630382 | 0.996633405 | -5.230340869 |
| **A_66_P118004** | Rab3ip | -0.018733002 | -0.197397551 | 0.159931546 | 12.3496483 | -0.250598308 | 0.809689509 | 0.996633405 | -5.23035754 |
| **A_55_P2132894** | Sdcbp | -0.019104342 | -0.201331763 | 0.16312308 | 8.911936811 | -0.250569078 | 0.80971118 | 0.996633405 | -5.230363649 |
| **A_51_P436020** | Tmem167 | -0.021044991 | -0.221786324 | 0.179696341 | 9.104200209 | -0.250565388 | 0.809713916 | 0.996633405 | -5.23036442 |
| **A_52_P155554** | Cdc42ep2 | -0.018491863 | -0.194909982 | 0.157926257 | 11.04457697 | -0.250522421 | 0.809745773 | 0.996633405 | -5.230373399 |
| **A_55_P2029498** | Osbpl1a | -0.020518961 | -0.216285951 | 0.175248029 | 10.81346406 | -0.250509981 | 0.809754996 | 0.996633405 | -5.230375998 |
| **A_55_P2084238** | NA | -0.020131638 | -0.212321536 | 0.172058259 | 12.37781054 | -0.25035582 | 0.809869299 | 0.996633405 | -5.230408199 |
| **A_51_P181319** | Dcxr | -0.012880951 | -0.135865997 | 0.110104094 | 8.051477174 | -0.250325307 | 0.809891923 | 0.996633405 | -5.23041457 |
| **A_55_P2061333** | NA | -0.021052981 | -0.222091553 | 0.179985591 | 11.32070659 | -0.25028991 | 0.809918169 | 0.996633405 | -5.23042196 |
| **A_55_P2074488** | Crim1 | 0.014439117 | -0.123455975 | 0.152334209 | 8.601735568 | 0.25026549 | 0.809936276 | 0.996633405 | -5.230427057 |
| **A_55_P2097048** | NA | 0.042847067 | -0.366431338 | 0.452125473 | 11.33530833 | 0.250213884 | 0.809974541 | 0.996633405 | -5.230437828 |
| **A_66_P109220** | Etfbkmt | -0.012536822 | -0.13231303 | 0.107239387 | 8.208701203 | -0.2501647 | 0.810011012 | 0.996633405 | -5.230448092 |
| **A_51_P333518** | Sh3rf2 | -0.012934205 | -0.136522879 | 0.110654469 | 8.155737153 | -0.250132543 | 0.810034856 | 0.996633405 | -5.230454801 |
| **A_52_P672313** | Exoc5 | -0.015725652 | -0.166032556 | 0.134581251 | 10.97795574 | -0.25005692 | 0.810090932 | 0.996633405 | -5.230470576 |
| **A_51_P226142** | Dennd6a | 0.01942302 | -0.166239838 | 0.205085878 | 11.41781334 | 0.25003503 | 0.810107164 | 0.996633405 | -5.230475142 |
| **A_52_P406371** | Med14 | -0.018508116 | -0.195468514 | 0.158452283 | 10.3085459 | -0.249974238 | 0.810152244 | 0.996633405 | -5.230487818 |
| **A_55_P1978201** | Incenp | 0.013473308 | -0.115362806 | 0.142309423 | 12.23439152 | 0.249945756 | 0.810173365 | 0.996633405 | -5.230493757 |
| **A_55_P1956017** | Usp45 | -0.020512492 | -0.216735976 | 0.175710992 | 9.349522444 | -0.249848398 | 0.810245561 | 0.996633405 | -5.23051405 |
| **A_55_P2167984** | NA | -0.021556701 | -0.227814381 | 0.18470098 | 11.63081004 | -0.249793591 | 0.810286205 | 0.996633405 | -5.23052547 |
| **A_55_P2181976** | Akap11 | 0.0169348 | -0.145108239 | 0.178977838 | 9.706142538 | 0.249780667 | 0.810295789 | 0.996633405 | -5.230528163 |
| **A_52_P352525** | Qser1 | 0.012101994 | -0.103700822 | 0.127904811 | 8.453385817 | 0.249773846 | 0.810300848 | 0.996633405 | -5.230529584 |
| **A_55_P2064547** | Tuba1c | -0.024538347 | -0.259438611 | 0.210361917 | 14.53252994 | -0.249672625 | 0.810375913 | 0.996633405 | -5.230550668 |
| **A_55_P2176585** | Donson | -0.011565572 | -0.12228028 | 0.099149136 | 8.011745773 | -0.249672625 | 0.810375913 | 0.996633405 | -5.230550668 |
| **A_51_P137604** | Fcna | 0.010945899 | -0.093848134 | 0.115739932 | 7.857748129 | 0.249645652 | 0.810395917 | 0.996633405 | -5.230556285 |
| **A_55_P2154914** | Acap1 | -0.013183979 | -0.139412026 | 0.113044068 | 8.065835738 | -0.249631716 | 0.810406251 | 0.996633405 | -5.230559187 |

| | | | | | | | | | |
|---|---|---|---|---|---|---|---|---|---|
| **A_66_P124858** | Diras2 | 0.023434104 | -0.201051263 | 0.24791947 | 9.425460695 | 0.249499362 | 0.81050441 | 0.996633405 | -5.230586739 |
| **A_55_P1979090** | Ublcp1 | 0.020947123 | -0.1797185 | 0.221612745 | 11.8203966 | 0.249494243 | 0.810508206 | 0.996633405 | -5.230587804 |
| **A_55_P1962334** | Trim71 | 0.009937257 | -0.085304626 | 0.105179139 | 7.657320858 | 0.249371988 | 0.810598878 | 0.996633405 | -5.230613241 |
| **A_55_P2056280** | Wdr89 | 0.037362232 | -0.320863576 | 0.395588039 | 12.33376095 | 0.249278643 | 0.81066811 | 0.996633405 | -5.230632654 |
| **A_55_P2236382** | D8Ertd158 | 0.049258881 | -0.423096205 | 0.521613968 | 9.481827458 | 0.249244212 | 0.810693647 | 0.996633405 | -5.230639813 |
| **A_55_P2021378** | Wapl | -0.015164678 | -0.160634055 | 0.130304699 | 10.16446426 | -0.249155653 | 0.810759332 | 0.996633405 | -5.230658222 |
| **A_55_P2048334** | Ubr3 | 0.023836085 | -0.204824278 | 0.252496447 | 10.19583331 | 0.249145567 | 0.810766813 | 0.996633405 | -5.230660318 |
| **A_51_P318544** | Fdx2 | -0.017356223 | -0.183857319 | 0.149144874 | 12.18107879 | -0.249142171 | 0.810769332 | 0.996633405 | -5.230661024 |
| **A_51_P303238** | Tmed10 | -0.014571948 | -0.154391904 | 0.125248008 | 12.51659935 | -0.249090735 | 0.810807484 | 0.996633405 | -5.230671712 |
| **A_52_P510215** | Traf6 | -0.011837832 | -0.125432123 | 0.101756458 | 8.254768514 | -0.249071947 | 0.810821419 | 0.996633405 | -5.230675616 |
| **A_55_P1969982** | Styx | 0.021148033 | -0.181862618 | 0.224158683 | 9.604932906 | 0.248977605 | 0.810891397 | 0.996633405 | -5.230695213 |
| **A_55_P1972969** | NA | 0.018505094 | -0.159228829 | 0.196239018 | 14.0624119 | 0.248845682 | 0.810989253 | 0.996633405 | -5.230722605 |
| **A_55_P2020090** | Pwwp3b | 0.023374145 | -0.201173622 | 0.247921912 | 10.68708847 | 0.248791833 | 0.811029198 | 0.996633405 | -5.230733782 |
| **A_55_P2004258** | Gm7102 | -0.015150893 | -0.160707801 | 0.130406014 | 8.993385588 | -0.248779481 | 0.811038361 | 0.996633405 | -5.230736346 |
| **A_55_P1964193** | Map1b | 0.012960705 | -0.111592458 | 0.137513868 | 8.860597275 | 0.248704132 | 0.811094255 | 0.996633405 | -5.23075198 |
| **A_51_P246854** | Acta1 | 0.047153414 | -0.406057593 | 0.500364421 | 11.45608238 | 0.2486691 | 0.811120242 | 0.996633405 | -5.230759248 |
| **A_55_P2031949** | Bend6 | 0.024279317 | -0.209124812 | 0.257683446 | 8.841330339 | 0.248620572 | 0.811156241 | 0.996633405 | -5.230769313 |
| **A_51_P135268** | Corin | 0.028367617 | -0.244420026 | 0.301155261 | 10.07816061 | 0.248546299 | 0.81121134 | 0.996633405 | -5.230784715 |
| **A_55_P1967196** | Lamb3 | -0.037119046 | -0.394173438 | 0.319935346 | 9.471381542 | -0.248468628 | 0.811268961 | 0.996633405 | -5.230800817 |
| **A_51_P501312** | Natd1 | -0.019358072 | -0.205627695 | 0.166911551 | 11.9914913 | -0.248387187 | 0.811329379 | 0.996633405 | -5.230817695 |
| **A_66_P138047** | Gfm1 | 0.020996077 | -0.181063517 | 0.223055672 | 13.45778256 | 0.248352091 | 0.811355417 | 0.996633405 | -5.230824967 |
| **A_51_P410918** | Tor4a | -0.013692651 | -0.145573135 | 0.118187834 | 8.695485808 | -0.248151058 | 0.811504565 | 0.996633405 | -5.2308666 |
| **A_55_P1953226** | H2al1e | -0.009950499 | -0.105818531 | 0.085917533 | 7.769492507 | -0.248073388 | 0.811562191 | 0.996633405 | -5.230882677 |
| **A_51_P205968** | Snx15 | -0.017355892 | -0.184658343 | 0.149946558 | 9.405551516 | -0.247944096 | 0.81165812 | 0.996633405 | -5.230909427 |
| **A_55_P2151666** | NA | -0.018932593 | -0.20147641 | 0.163611225 | 8.379339223 | -0.247886054 | 0.811701186 | 0.996633405 | -5.230921432 |
| **A_51_P295237** | Lrp11 | 0.014753847 | -0.127596676 | 0.157104369 | 8.055979592 | 0.247716724 | 0.811826829 | 0.996633405 | -5.230956438 |
| **A_55_P2172314** | Scamp4 | 0.013435512 | -0.116230697 | 0.14310172 | 12.21499375 | 0.247648972 | 0.811877104 | 0.996633405 | -5.230970438 |
| **A_55_P1964787** | Slc35e4 | -0.014970115 | -0.159473841 | 0.129533611 | 9.291501831 | -0.247602611 | 0.811911505 | 0.996633405 | -5.230980015 |
| **A_52_P307893** | Pip4k2b | 0.018691124 | -0.161749748 | 0.199131997 | 10.50345954 | 0.247576629 | 0.811930785 | 0.996633405 | -5.230985382 |
| **A_51_P222359** | Emilin1 | -0.029907667 | -0.318649824 | 0.25883449 | 8.648636703 | -0.24756057 | 0.811942702 | 0.996633405 | -5.230988699 |
| **A_55_P1987146** | Adgrf4 | -0.010412583 | -0.110992993 | 0.090167827 | 7.73427349 | -0.247431055 | 0.81203881 | 0.996633405 | -5.231015441 |
| **A_55_P2028288** | Nhlrc4 | 0.01015749 | -0.088008242 | 0.108323223 | 7.689652321 | 0.247306565 | 0.812131194 | 0.996633405 | -5.231041134 |
| **A_55_P1979973** | Nuak2 | -0.013404296 | -0.143009405 | 0.116200814 | 8.150780319 | -0.247190061 | 0.812217653 | 0.996633405 | -5.231065166 |
| **A_52_P365896** | Pa2g4 | 0.018176226 | -0.157572768 | 0.19392522 | 8.710766311 | 0.247183797 | 0.812222302 | 0.996633405 | -5.231066458 |
| **A_55_P2082961** | Fnbp1l | 0.018033057 | -0.15634572 | 0.192411834 | 10.73249976 | 0.247163801 | 0.812237142 | 0.996633405 | -5.231070581 |
| **A_51_P121412** | Slc35a4 | 0.019487985 | -0.169055522 | 0.208031491 | 9.273244411 | 0.247038407 | 0.812330202 | 0.996633405 | -5.187102386 |
| **A_55_P1953361** | Luc7l2 | -0.018197479 | -0.194282269 | 0.157887312 | 14.34712596 | -0.247000886 | 0.812358049 | 0.996633405 | -5.231104166 |
| **A_55_P2023314** | Casz1 | 0.016191661 | -0.140488179 | 0.172871501 | 10.77558586 | 0.246994578 | 0.812362731 | 0.996633405 | -5.231105465 |
| **A_52_P662244** | Slc25a47 | -0.010956001 | -0.116993803 | 0.095081801 | 8.579557764 | -0.246945134 | 0.812399427 | 0.996633405 | -5.231115654 |
| **A_51_P394558** | Dchs1 | 0.021102189 | -0.183140982 | 0.22534536 | 10.2163302 | 0.246938666 | 0.812404227 | 0.996633405 | -5.231116986 |
| **A_55_P1983769** | Birc5 | -0.025449273 | -0.271818784 | 0.220920238 | 9.980713877 | -0.246886619 | 0.812442856 | 0.996633405 | -5.231127708 |
| **A_55_P2156462** | Nt5c3b | -0.024466546 | -0.261388899 | 0.212455807 | 9.2445093 | -0.24681739 | 0.812494238 | 0.996633405 | -5.231141967 |
| **A_55_P2087061** | Tmem240 | -0.014327752 | -0.153182297 | 0.124526793 | 8.93882501 | -0.246619315 | 0.812641255 | 0.996633405 | -5.23118274 |
| **A_55_P2033393** | Slc25a21 | -0.010464351 | -0.11189585 | 0.090967148 | 7.659890035 | -0.246574746 | 0.812674337 | 0.996633405 | -5.23119191 |
| **A_55_P2136776** | Sult1b1 | -0.010237748 | -0.109495129 | 0.089019633 | 7.64933566 | -0.246519203 | 0.812715564 | 0.996633405 | -5.231203336 |
| **A_51_P100298** | Stx3 | -0.010626626 | -0.113656687 | 0.092403436 | 7.82895908 | -0.246513417 | 0.812719859 | 0.996633405 | -5.200147347 |
| **A_52_P77080** | Gypa | -0.024516851 | -0.26222618 | 0.213192478 | 8.058985019 | -0.246506054 | 0.812725324 | 0.996633405 | -5.23120604 |
| **A_55_P2038111** | Tmem104 | -0.011101189 | -0.118754998 | 0.09655262 | 7.72624586 | -0.246461584 | 0.812758334 | 0.996633405 | -5.231215186 |

| | | | | | | | | | |
|---|---|---|---|---|---|---|---|---|---|
| **A_55_P2004179** | Col2a1 | 0.026416562 | -0.229853258 | 0.282686381 | 8.326873466 | 0.246370077 | 0.812826258 | 0.996633405 | -5.231234 |
| **A_51_P203501** | Vars | 0.025116573 | -0.218546443 | 0.268779589 | 9.184830285 | 0.246365504 | 0.812829653 | 0.996633405 | -5.23123494 |
| **A_55_P2176385** | St6galnac4 | 0.027879092 | -0.242761579 | 0.298519763 | 9.459302977 | 0.246203768 | 0.812949713 | 0.996633405 | -5.231268176 |
| **A_65_P01991** | Prrg4 | -0.013429635 | -0.143847467 | 0.116988198 | 7.7392711 | -0.246114022 | 0.813016335 | 0.996633405 | -5.231286609 |
| **A_55_P2182273** | NA | -0.013803137 | -0.147876143 | 0.120269868 | 7.795880472 | -0.246062595 | 0.813054513 | 0.996633405 | -5.231297169 |
| **A_55_P2033725** | Ascl2 | 0.01394046 | -0.121466635 | 0.149347555 | 7.884930001 | 0.24606215 | 0.813054843 | 0.996633405 | -5.23129726 |
| **A_55_P1983450** | S100a4 | -0.01816224 | -0.194625383 | 0.158300903 | 8.474349699 | -0.245994013 | 0.813105426 | 0.996633405 | -5.231311248 |
| **A_55_P2042923** | Sgk2 | -0.019447053 | -0.208477762 | 0.169583656 | 7.732494581 | -0.245884171 | 0.813186972 | 0.996633405 | -5.231333789 |
| **A_51_P427624** | Pgc | 0.010336179 | -0.090204206 | 0.110876565 | 7.66911325 | 0.245713274 | 0.81331385 | 0.996633405 | -5.23136884 |
| **A_52_P492901** | Stx4a | 0.021856583 | -0.190778166 | 0.234491331 | 11.37168177 | 0.245672845 | 0.813343865 | 0.996633405 | -5.231377128 |
| **A_55_P2248330** | 2700068H( | -0.020979328 | -0.225210183 | 0.183251526 | 7.976613102 | -0.245515751 | 0.813460502 | 0.996633405 | -5.231409322 |
| **A_52_P296109** | NA | -0.012726039 | -0.136616705 | 0.111164627 | 8.464449856 | -0.245506958 | 0.813467031 | 0.996633405 | -5.231411123 |
| **A_55_P1987054** | Krtap5-1 | 0.012841374 | -0.11222772 | 0.137910468 | 8.155370424 | 0.245397772 | 0.813548101 | 0.996633405 | -5.231433486 |
| **A_51_P333965** | Cisd2 | -0.01145491 | -0.123023892 | 0.100114071 | 8.43860704 | -0.245390282 | 0.813553662 | 0.996633405 | -5.23143502 |
| **A_52_P365005** | Zbtb22 | -0.01753007 | -0.18833948 | 0.15327934 | 9.068750038 | -0.245290642 | 0.813627646 | 0.996633405 | -5.231455419 |
| **A_55_P1996199** | NA | 0.011796929 | -0.103152649 | 0.126746506 | 14.66977297 | 0.24528484 | 0.813631954 | 0.996633405 | -5.231456606 |
| **A_51_P140141** | Hspbp1 | -0.040677472 | -0.437047414 | 0.35569247 | 10.25486979 | -0.245280132 | 0.81363545 | 0.996633405 | -5.23145757 |
| **A_51_P423127** | Gba | -0.014972274 | -0.160887294 | 0.130942745 | 9.046687451 | -0.245243163 | 0.813662901 | 0.996633405 | -5.231465136 |
| **A_52_P8922** | Sntb2 | 0.020657072 | -0.180712094 | 0.222026239 | 9.598826786 | 0.245179939 | 0.813709847 | 0.996633405 | -5.231478073 |
| **A_55_P2031133** | Tmem220 | 0.010541744 | -0.092254034 | 0.113337522 | 8.021237735 | 0.245101697 | 0.813767947 | 0.996633405 | -5.231494078 |
| **A_51_P505967** | Spag6l | 0.010593969 | -0.092718739 | 0.113906678 | 7.677246611 | 0.245083514 | 0.813781448 | 0.996633405 | -5.231497797 |
| **A_55_P2088715** | Ss18l2 | -0.021157884 | -0.227495065 | 0.185179297 | 10.666017 | -0.245077744 | 0.813785733 | 0.996633405 | -5.231498977 |
| **A_51_P388691** | Bbx | 0.011821839 | -0.103469704 | 0.127113382 | 8.152893153 | 0.245073713 | 0.813788726 | 0.996633405 | -5.231499801 |
| **A_51_P442152** | Itfg1 | 0.012851255 | -0.112481466 | 0.138183976 | 11.66179462 | 0.245070035 | 0.813791457 | 0.996633405 | -5.231500553 |
| **A_55_P2001481** | Pde6c | -0.009804293 | -0.105428025 | 0.08581944 | 7.642789195 | -0.245052826 | 0.813804237 | 0.996633405 | -5.231504073 |
| **A_51_P436342** | Hoxb5 | 0.011221275 | -0.098261915 | 0.120704464 | 7.78782428 | 0.244964917 | 0.813869517 | 0.996633405 | -5.231522046 |
| **A_55_P2067142** | NA | 0.012490495 | -0.109391499 | 0.13437249 | 16.88248461 | 0.244934092 | 0.813892407 | 0.996633405 | -5.231528347 |
| **A_55_P1983848** | Tc2n | -0.010058355 | -0.108242876 | 0.088126166 | 7.695082625 | -0.244846024 | 0.813957807 | 0.996633405 | -5.231546344 |
| **A_55_P2014927** | Zfp341 | 0.013130781 | -0.115086647 | 0.141348209 | 9.098804069 | 0.244766868 | 0.814016591 | 0.996633405 | -5.231562515 |
| **A_55_P2064412** | NA | 0.017423477 | -0.152860958 | 0.187707911 | 8.066953143 | 0.24455075 | 0.814177092 | 0.996633405 | -5.23160664 |
| **A_66_P134501** | Olfm1 | -0.035915141 | -0.386927795 | 0.315097512 | 9.46770283 | -0.2445479 | 0.814179208 | 0.996633405 | -5.231607221 |
| **A_52_P580707** | Zfp781 | 0.013089415 | -0.114909418 | 0.141088248 | 8.427755019 | 0.244412471 | 0.814279791 | 0.996633405 | -5.231634852 |
| **A_52_P37431** | NA | -0.022514203 | -0.242685879 | 0.197657473 | 14.3861816 | -0.244401725 | 0.814287772 | 0.996633405 | -5.231637044 |
| **A_55_P2038972** | Erich1 | 0.012590706 | -0.110572108 | 0.13575352 | 8.224944737 | 0.244331593 | 0.81433986 | 0.996633405 | -5.231651346 |
| **A_51_P174081** | Zdhhc14 | 0.010664565 | -0.093726381 | 0.115055512 | 7.993398987 | 0.2441684 | 0.814461071 | 0.996633405 | -5.231684611 |
| **A_55_P2007423** | Ppp1r12a | -0.015742198 | -0.169857464 | 0.138373069 | 8.871684624 | -0.244134316 | 0.814486387 | 0.996633405 | -5.231691555 |
| **A_55_P1953713** | Bop1 | 0.019684889 | -0.173139955 | 0.212509733 | 10.82297139 | 0.243993986 | 0.814590622 | 0.996633405 | -5.231720138 |
| **A_51_P441426** | Pf4 | 0.032219717 | -0.283463163 | 0.347902598 | 11.46717432 | 0.243938208 | 0.814632054 | 0.996633405 | -5.231731495 |
| **A_51_P470079** | Il1r2 | 0.011345978 | -0.099897023 | 0.12258898 | 7.705337486 | 0.243768949 | 0.814757783 | 0.996633405 | -5.231765941 |
| **A_55_P1956667** | Cyfip1 | 0.019760727 | -0.174065437 | 0.213586891 | 10.86768003 | 0.243668641 | 0.814832297 | 0.996633405 | -5.231786344 |
| **A_51_P192589** | Pno1 | 0.027781741 | -0.244822017 | 0.300385499 | 11.87992417 | 0.243577267 | 0.814900176 | 0.996633405 | -5.231804923 |
| **A_55_P2179974** | Spsb1 | 0.011640123 | -0.102675058 | 0.125955303 | 8.088741359 | 0.243367612 | 0.815055929 | 0.996633405 | -5.231847525 |
| **A_51_P279232** | Dennd2d | 0.010353195 | -0.091333519 | 0.112039909 | 7.860712817 | 0.243343252 | 0.815074027 | 0.996633405 | -5.231852473 |
| **A_55_P2159573** | Reep6 | -0.010486854 | -0.11351797 | 0.092544261 | 7.659940923 | -0.243268552 | 0.815129524 | 0.996633405 | -5.231867641 |
| **A_55_P1996983** | Btg1c | -0.019047112 | -0.206330101 | 0.168235878 | 8.890623368 | -0.243074792 | 0.815273481 | 0.996633405 | -5.231906966 |
| **A_52_P239292** | AI597479 | 0.01898687 | -0.16771209 | 0.205685829 | 10.49952565 | 0.243063978 | 0.815281515 | 0.996633405 | -5.23190916 |
| **A_51_P311694** | Ypel5 | 0.020339546 | -0.17967661 | 0.220355703 | 8.806419623 | 0.243044233 | 0.815296186 | 0.996633405 | -5.231913166 |
| **A_55_P1986556** | Minpp1 | -0.019684294 | -0.213281088 | 0.173912501 | 10.3284289 | -0.243013726 | 0.815318853 | 0.996633405 | -5.231919354 |

| | | | | | | | | | |
|---|---|---|---|---|---|---|---|---|---|
| **A_66_P101600** | Lamb1 | -0.012961174 | -0.140443605 | 0.114521257 | 14.36505141 | -0.242998247 | 0.815330354 | 0.996633405 | -5.231922493 |
| **A_55_P2412319** | A830052D1 | 0.01041007 | -0.092011132 | 0.112831273 | 7.675889583 | 0.242925392 | 0.815384485 | 0.996633405 | -5.231937267 |
| **A_55_P2327177** | Exoc8 | -0.01318616 | -0.142940968 | 0.116568648 | 9.39029195 | -0.242886854 | 0.81541312 | 0.996633405 | -5.23194508 |
| **A_52_P675395** | Ackr3 | 0.019333353 | -0.17095363 | 0.209620336 | 9.988861235 | 0.242832742 | 0.815453327 | 0.996633405 | -5.231956049 |
| **A_52_P601688** | Abcc1 | -0.012217866 | -0.132563206 | 0.108127475 | 8.316680426 | -0.242647155 | 0.815591228 | 0.996633405 | -5.231993649 |
| **A_52_P128964** | Nhej1 | 0.013389518 | -0.118533576 | 0.145312613 | 9.090022405 | 0.242579033 | 0.815641848 | 0.996633405 | -5.232007444 |
| **A_55_P1980583** | Leprot | -0.015442268 | -0.167604265 | 0.136719729 | 12.70292435 | -0.242557118 | 0.815658133 | 0.996633405 | -5.232011881 |
| **A_55_P2066429** | NA | -0.015444936 | -0.16763949 | 0.136749617 | 9.591083876 | -0.242547132 | 0.815665553 | 0.996633405 | -5.232013903 |
| **A_65_P05316** | Zfp182 | -0.019161125 | -0.208039825 | 0.169717574 | 8.614085333 | -0.242463943 | 0.815727371 | 0.996633405 | -5.232030741 |
| **A_55_P2115028** | Fastkd1 | -0.014407813 | -0.156441864 | 0.127626239 | 9.414859803 | -0.242445828 | 0.815740832 | 0.996633405 | -5.232034407 |
| **A_55_P2065929** | NA | -0.016255862 | -0.176517148 | 0.144005424 | 8.516144973 | -0.242432321 | 0.81575087 | 0.996633405 | -5.232037141 |
| **A_51_P196590** | Hadh | -0.024412194 | -0.265120393 | 0.216296006 | 13.4344175 | -0.242395781 | 0.815778023 | 0.996633405 | -5.232044534 |
| **A_51_P375509** | Spty2d1 | 0.011757364 | -0.10418445 | 0.127699178 | 8.051832165 | 0.242370073 | 0.815797128 | 0.996633405 | -5.232049735 |
| **A_52_P576778** | Tmem30a | -0.013886315 | -0.150829257 | 0.123056627 | 11.11154052 | -0.242357519 | 0.815806457 | 0.996633405 | -5.232052275 |
| **A_55_P2020286** | Tmem179 | 0.011954696 | -0.105971109 | 0.129880502 | 7.755092968 | 0.242291861 | 0.81585525 | 0.996633405 | -5.232065555 |
| **A_55_P2065501** | Rnf7 | 0.020411103 | -0.180959183 | 0.221781388 | 13.43028101 | 0.242259164 | 0.815879548 | 0.996633405 | -5.232072167 |
| **A_51_P166288** | Acy1 | -0.014876706 | -0.161659277 | 0.131905866 | 8.404997993 | -0.242237517 | 0.815895635 | 0.996633405 | -5.232076544 |
| **A_66_P120208** | Cnnm3 | 0.013184927 | -0.116954212 | 0.143324066 | 9.335125633 | 0.242146911 | 0.815962971 | 0.996633405 | -5.232094861 |
| **A_52_P283628** | Rps6ka3 | -0.011701578 | -0.127279042 | 0.103875886 | 8.689838859 | -0.24198052 | 0.816086631 | 0.996633405 | -5.23212848 |
| **A_55_P2080649** | Zfp691 | -0.018477851 | -0.201072734 | 0.164117033 | 9.851051753 | -0.241864415 | 0.816172923 | 0.996633405 | -5.232151926 |
| **A_55_P2127765** | Crp | 0.009744393 | -0.086601846 | 0.106090631 | 7.644551282 | 0.241729214 | 0.816273411 | 0.996633405 | -5.232179214 |
| **A_52_P101487** | Ddn | -0.01169282 | -0.127379798 | 0.103994157 | 7.792213354 | -0.24157052 | 0.816391364 | 0.996633405 | -5.232211225 |
| **A_55_P2144341** | Smim40 | -0.010775057 | -0.117416729 | 0.095866616 | 7.766738756 | -0.241491441 | 0.816450144 | 0.996633405 | -5.232227169 |
| **A_52_P67643** | Mtor | 0.011100278 | -0.098760902 | 0.120961459 | 8.058923269 | 0.241489775 | 0.816451382 | 0.996633405 | -5.232227505 |
| **A_55_P2139112** | Cdc26 | -0.013868597 | -0.151151293 | 0.1234141 | 10.73590107 | -0.241449243 | 0.81648151 | 0.996633405 | -5.232235675 |
| **A_55_P1980062** | NA | -0.01553262 | -0.169338927 | 0.138273686 | 8.154499683 | -0.241368015 | 0.816541889 | 0.996633405 | -5.232252044 |
| **A_55_P2018091** | NA | 0.025727219 | -0.229231602 | 0.280686039 | 13.78971331 | 0.241174802 | 0.816685515 | 0.996633405 | -5.232290958 |
| **A_55_P2045278** | Gm10845 | -0.020806856 | -0.227052264 | 0.185438552 | 12.74492078 | -0.241118927 | 0.816727051 | 0.996633405 | -5.232302205 |
| **A_55_P2050508** | Spta1 | 0.01471667 | -0.131239612 | 0.160672951 | 8.408336803 | 0.240988261 | 0.816824188 | 0.996633405 | -5.232328499 |
| **A_55_P2154242** | Gfpt1 | 0.019019023 | -0.169633948 | 0.207671994 | 10.73070475 | 0.240953749 | 0.816849845 | 0.996633405 | -5.232335441 |
| **A_52_P185343** | Gna13 | -0.017812569 | -0.194564976 | 0.158939837 | 9.305553522 | -0.240863153 | 0.816917196 | 0.996633405 | -5.232353661 |
| **A_55_P1968464** | Kat6a | -0.013636162 | -0.14896694 | 0.121694615 | 10.68932648 | -0.240826742 | 0.816944266 | 0.996633405 | -5.232360982 |
| **A_55_P2113066** | Zkscan5 | -0.01452067 | -0.158638893 | 0.129597554 | 10.63165841 | -0.24081131 | 0.816955739 | 0.996633405 | -5.232364084 |
| **A_66_P127262** | NA | -0.02599787 | -0.284307259 | 0.23231152 | 8.880546837 | -0.240550741 | 0.817149466 | 0.996633405 | -5.232416438 |
| **A_55_P2125200** | Mterf1b | 0.012024913 | -0.107453242 | 0.131503067 | 8.390272665 | 0.240548447 | 0.817151171 | 0.996633405 | -5.232416899 |
| **A_66_P106478** | Cct3 | 0.018891067 | -0.168823053 | 0.206605186 | 11.57417392 | 0.24052968 | 0.817165125 | 0.996633405 | -5.232420668 |
| **A_51_P129100** | Sec63 | 0.012958284 | -0.115819677 | 0.141736244 | 8.553177983 | 0.240499994 | 0.817187197 | 0.996633405 | -5.226281068 |
| **A_55_P2133652** | Cep162 | -0.012498987 | -0.136719431 | 0.111721457 | 9.043149169 | -0.24048659 | 0.817197163 | 0.996633405 | -5.232429319 |
| **A_55_P1978855** | NA | -0.010494432 | -0.11480271 | 0.093813847 | 15.12915439 | -0.240463569 | 0.817214279 | 0.996633405 | -5.232433941 |
| **A_55_P2000894** | NA | 0.014032233 | -0.125583562 | 0.153648027 | 15.3123948 | 0.240215683 | 0.817398593 | 0.996633405 | -5.232483678 |
| **A_55_P2068148** | Pias1 | -0.012758002 | -0.139712594 | 0.114196591 | 8.233394976 | -0.240183609 | 0.817422443 | 0.996633405 | -5.23249011 |
| **A_55_P1971010** | Gzme | 0.034195938 | -0.306130671 | 0.374522547 | 7.817598951 | 0.240152852 | 0.817445313 | 0.996633405 | -5.232496277 |
| **A_55_P1962214** | Kpna3 | -0.022575775 | -0.247267647 | 0.202116097 | 9.716245443 | -0.240139964 | 0.817454897 | 0.996633405 | -5.23249886 |
| **A_51_P275514** | Mprip | -0.021454036 | -0.235097747 | 0.192189675 | 11.92299819 | -0.240009272 | 0.817552079 | 0.996633405 | -5.232525055 |
| **A_55_P1972772** | Gsg1 | -0.014570675 | -0.159695292 | 0.130553942 | 7.916760035 | -0.239964899 | 0.817585075 | 0.996633405 | -5.232533945 |
| **A_55_P2124097** | Gm6981 | -0.022627877 | -0.248011657 | 0.202755904 | 16.1972105 | -0.239955261 | 0.817592242 | 0.996633405 | -5.232535876 |
| **A_55_P2083694** | Adam33 | 0.025436962 | -0.228070385 | 0.278944309 | 8.315616873 | 0.239819132 | 0.817693473 | 0.996633405 | -5.232563139 |
| **A_55_P2023727** | Limch1 | 0.02391601 | -0.214441994 | 0.262274013 | 10.3908087 | 0.239810454 | 0.817699926 | 0.996633405 | -5.232564877 |

| | | | | | | | | | |
|---|---|---|---|---|---|---|---|---|---|
| **A_66_P100268** | NA | 0.0288093 | -0.258340639 | 0.315959238 | 8.3477485 | 0.239791125 | 0.8177143 | 0.996633405 | -5.232568746 |
| **A_52_P105015** | Clcn7 | -0.020723021 | -0.227360716 | 0.185914673 | 8.826256007 | -0.239691514 | 0.817788378 | 0.996633405 | -5.232588684 |
| **A_55_P2055127** | Slc7a14 | 0.016882214 | -0.151519697 | 0.185284126 | 7.786825851 | 0.239602598 | 0.817854504 | 0.996633405 | -5.232606474 |
| **A_55_P1956557** | Rab3a | 0.01383225 | -0.124175764 | 0.151840263 | 11.24176518 | 0.239550811 | 0.817893017 | 0.996633405 | -5.232616833 |
| **A_55_P2092884** | NA | -0.019286542 | -0.21172705 | 0.173153965 | 8.387611133 | -0.239533918 | 0.817905581 | 0.996633405 | -5.232620211 |
| **A_55_P2177463** | B130024G1 | 0.016367639 | -0.147014646 | 0.179749925 | 8.017809994 | 0.239436422 | 0.817978091 | 0.996633405 | -5.232639705 |
| **A_55_P2129311** | 2210016L2 | -0.013481436 | -0.148097629 | 0.121134757 | 10.00296937 | -0.239358022 | 0.818036399 | 0.996633405 | -5.232655375 |
| **A_55_P2060878** | NA | -0.024006687 | -0.263816521 | 0.215803146 | 12.06806607 | -0.239262358 | 0.81810755 | 0.996633405 | -5.232674489 |
| **A_51_P283004** | Tinag | -0.009732167 | -0.106967353 | 0.087503019 | 7.648567718 | -0.239218761 | 0.818139976 | 0.996633405 | -5.232683198 |
| **A_55_P2051701** | Pbx1 | -0.022149522 | -0.243487583 | 0.199188538 | 11.5308421 | -0.23917589 | 0.818171862 | 0.996633405 | -5.23269176 |
| **A_51_P181565** | Hbegf | -0.02658551 | -0.292269038 | 0.239098019 | 10.11667348 | -0.23916052 | 0.818183294 | 0.996633405 | -5.232694829 |
| **A_55_P2098305** | Coq5 | -0.019455101 | -0.213880658 | 0.174970455 | 9.128307776 | -0.239160403 | 0.818183381 | 0.996633405 | -5.232694852 |
| **A_51_P189905** | Dad1 | -0.016504044 | -0.181507784 | 0.148499696 | 13.1212051 | -0.239059335 | 0.818258555 | 0.996633405 | -5.232715029 |
| **A_55_P2234704** | NA | 0.021495421 | -0.193583264 | 0.236574107 | 9.817878751 | 0.238867863 | 0.818400977 | 0.996633405 | -5.232753232 |
| **A_55_P2002482** | Sarnp | 0.014599246 | -0.131547989 | 0.16074648 | 12.49588339 | 0.238753061 | 0.818486373 | 0.996633405 | -5.232776123 |
| **A_55_P1984337** | Emd | 0.01130729 | -0.101885765 | 0.124500345 | 11.96925398 | 0.238752472 | 0.818486811 | 0.996633405 | -5.23277624 |
| **A_51_P466229** | Pdgfrl | 0.064088336 | -0.577554885 | 0.705731557 | 10.21057282 | 0.238723141 | 0.81850863 | 0.996633405 | -5.232782087 |
| **A_52_P428125** | Atr | 0.017088022 | -0.154000689 | 0.188176734 | 10.61343736 | 0.238714925 | 0.818514741 | 0.996633405 | -5.232783725 |
| **A_51_P133582** | Cbx4 | 0.011451702 | -0.103274927 | 0.126178331 | 11.11968145 | 0.238569502 | 0.818622919 | 0.996633405 | -5.232812701 |
| **A_55_P2048588** | Cdk1 | -0.021334346 | -0.235087901 | 0.192419209 | 11.48358098 | -0.238547634 | 0.818639188 | 0.996633405 | -5.232817057 |
| **A_55_P2090219** | Mrpl48 | 0.016615769 | -0.149863553 | 0.183095092 | 12.85026074 | 0.238544434 | 0.818641568 | 0.996633405 | -5.232817695 |
| **A_55_P1952475** | Rpl30 | 0.017413405 | -0.157070833 | 0.191897643 | 15.59559591 | 0.238526495 | 0.818654913 | 0.996633405 | -5.232821268 |
| **A_55_P2040295** | Ptp4a1 | 0.023249819 | -0.20977971 | 0.256279348 | 10.24062478 | 0.238461212 | 0.818703478 | 0.996633405 | -5.232834268 |
| **A_55_P2118491** | Gm10046 | 0.010823929 | -0.097706463 | 0.119354322 | 7.874619024 | 0.238365124 | 0.818774961 | 0.996633405 | -5.232853396 |
| **A_55_P1956464** | Stam | 0.014905551 | -0.134595111 | 0.164406213 | 9.3425571 | 0.238294518 | 0.818827489 | 0.996633405 | -5.232867447 |
| **A_52_P508920** | Abca6 | -0.010688595 | -0.117901362 | 0.096524172 | 7.751829491 | -0.238277613 | 0.818840065 | 0.996633405 | -5.23287081 |
| **A_55_P2161132** | NA | 0.010982927 | -0.099206891 | 0.121172744 | 8.024739932 | 0.238224127 | 0.818879857 | 0.996633405 | -5.232881451 |
| **A_66_P106585** | Sae1 | -0.019943548 | -0.220079184 | 0.180192087 | 10.67365644 | -0.238170043 | 0.818920095 | 0.996633405 | -5.232892208 |
| **A_51_P331870** | Rnf145 | -0.012176714 | -0.134382805 | 0.110029377 | 9.82921674 | -0.238147694 | 0.818936722 | 0.996633405 | -5.232896652 |
| **A_55_P2024302** | Spin1 | 0.014774137 | -0.133567942 | 0.163116216 | 8.827150316 | 0.238038329 | 0.81901809 | 0.996633405 | -5.232918395 |
| **A_66_P103229** | Foxl1 | 0.010724999 | -0.096971504 | 0.118421502 | 7.751825404 | 0.238015258 | 0.819035255 | 0.996633405 | -5.23292298 |
| **A_55_P2040988** | NA | -0.016101973 | -0.177834759 | 0.145630812 | 9.424111373 | -0.237952444 | 0.81908199 | 0.996633405 | -5.232935463 |
| **A_51_P471779** | Phf5a | -0.014044877 | -0.155129621 | 0.127039867 | 12.15826618 | -0.237928805 | 0.819099578 | 0.996633405 | -5.232940159 |
| **A_51_P297480** | Dph1 | 0.016466632 | -0.148970167 | 0.181903432 | 9.187413072 | 0.237893076 | 0.819126162 | 0.996633405 | -5.232947257 |
| **A_55_P2143946** | Slc18a2 | 0.01445409 | -0.13078216 | 0.159690341 | 8.871510346 | 0.237861893 | 0.819149364 | 0.996633405 | -5.232953452 |
| **A_51_P407323** | F5 | 0.011777032 | -0.10656813 | 0.130122193 | 7.722185619 | 0.237845243 | 0.819161752 | 0.996633405 | -5.232956758 |
| **A_55_P1986902** | Med23 | 0.018867086 | -0.170799986 | 0.208534159 | 8.971328025 | 0.237750825 | 0.819232005 | 0.996633405 | -5.232975507 |
| **A_55_P1964752** | Slc23a3 | 0.075249925 | -0.681439046 | 0.831938895 | 9.103593823 | 0.237682824 | 0.819282602 | 0.996633405 | -5.232989005 |
| **A_55_P2042396** | Alkbh4 | 0.013494886 | -0.122238315 | 0.149228087 | 9.078421194 | 0.237625071 | 0.819325576 | 0.996633405 | -5.233000466 |
| **A_52_P188261** | Camk2d | -0.028313051 | -0.313151808 | 0.256525706 | 10.19591631 | -0.237572802 | 0.81936447 | 0.996633405 | -5.233010837 |
| **A_55_P1982239** | Ctrb1 | 0.01991687 | -0.180507025 | 0.220340765 | 8.053395166 | 0.237509354 | 0.819411682 | 0.996633405 | -5.233023422 |
| **A_55_P1965539** | Ndufb4 | -0.016979758 | -0.187862942 | 0.153903425 | 16.10271394 | -0.237487805 | 0.819427717 | 0.996633405 | -5.233027696 |
| **A_55_P1957154** | Mageb16 | 0.009849681 | -0.089286141 | 0.108985504 | 7.667891996 | 0.237465597 | 0.819444242 | 0.996633405 | -5.2330321 |
| **A_55_P2167160** | Hdhd2 | 0.020963203 | -0.190081786 | 0.232008192 | 8.977381938 | 0.237406021 | 0.819488575 | 0.996633405 | -5.233043912 |
| **A_51_P500718** | Dck | 0.01339171 | -0.121488273 | 0.148271692 | 8.28878517 | 0.237299955 | 0.819567504 | 0.996633405 | -5.233064935 |
| **A_55_P2422650** | 5031425E2 | -0.019310155 | -0.213836563 | 0.175216253 | 10.163702 | -0.237255521 | 0.81960057 | 0.996633405 | -5.233073739 |
| **A_52_P656714** | Map3k5 | 0.025598223 | -0.232410126 | 0.283606572 | 8.94960854 | 0.237129284 | 0.819694513 | 0.996633405 | -5.233098743 |
| **A_66_P130582** | Ncoa1 | -0.011354766 | -0.125858849 | 0.103149318 | 10.89181035 | -0.237009818 | 0.819783421 | 0.996633405 | -5.233122394 |

| | | | | | | | | | |
|---|---|---|---|---|---|---|---|---|---|
| **A_51_P468126** | Trim39 | -0.014295872 | -0.158467044 | 0.129875299 | 8.256078131 | -0.236996192 | 0.819793562 | 0.996633405 | -5.233125091 |
| **A_55_P2047673** | Srr | -0.010394416 | -0.115240053 | 0.094451221 | 10.20598889 | -0.236951177 | 0.819827063 | 0.996633405 | -5.233133999 |
| **A_51_P358354** | Jam3 | 0.014441779 | -0.131237624 | 0.160121182 | 10.9630132 | 0.236936339 | 0.819838106 | 0.996633405 | -5.233136935 |
| **A_55_P2051476** | C8a | -0.010112461 | -0.11213116 | 0.091906238 | 7.637207881 | -0.236911545 | 0.819856558 | 0.996633405 | -5.233141841 |
| **A_55_P2078300** | Fndc7 | -0.010844863 | -0.12027275 | 0.098583024 | 7.713861829 | -0.236867342 | 0.819889457 | 0.996633405 | -5.233150585 |
| **A_55_P2036250** | Cr1l | -0.017664334 | -0.195914686 | 0.160586019 | 11.87260903 | -0.23685142 | 0.819901307 | 0.996633405 | -5.233153735 |
| **A_51_P117109** | Ilvbl | -0.022336695 | -0.247752606 | 0.203079217 | 9.828886396 | -0.236833684 | 0.819914506 | 0.996633405 | -5.233157243 |
| **A_51_P307741** | Cckar | 0.010664348 | -0.096963466 | 0.118292161 | 7.71107794 | 0.23682029 | 0.819924475 | 0.996633405 | -5.233159892 |
| **A_55_P1997335** | Vps9d1 | -0.020794418 | -0.230746436 | 0.1891576 | 11.28995482 | -0.236720485 | 0.819998756 | 0.996633405 | -5.233179626 |
| **A_52_P245365** | Rtf1 | 0.010447899 | -0.095068228 | 0.115964026 | 8.030382389 | 0.236656939 | 0.820046053 | 0.996633405 | -5.233192186 |
| **A_55_P2149382** | Gm3448 | -0.012846964 | -0.142620098 | 0.11692617 | 7.998816853 | -0.236605507 | 0.820084334 | 0.996633405 | -5.23320235 |
| **A_66_P117417** | NA | 0.018839253 | -0.17152153 | 0.209200035 | 9.471709804 | 0.236534951 | 0.820136848 | 0.996633405 | -5.233216289 |
| **A_51_P498831** | Derl2 | -0.009904712 | -0.110016046 | 0.090206622 | 11.98067574 | -0.236465462 | 0.820188571 | 0.996633405 | -5.233230014 |
| **A_51_P168524** | NA | 0.015275943 | -0.139177524 | 0.16972941 | 8.503035584 | 0.236384777 | 0.820248628 | 0.996633405 | -5.233245945 |
| **A_55_P1978740** | Rpgrip1 | 0.032247329 | -0.293855297 | 0.358349956 | 11.47750534 | 0.236346181 | 0.820277356 | 0.996633405 | -5.233253564 |
| **A_51_P149562** | Apbb2 | -0.010309175 | -0.114577099 | 0.093958748 | 11.60529034 | -0.236310126 | 0.820304194 | 0.996633405 | -5.23326068 |
| **A_55_P2067483** | NA | -0.013998633 | -0.155590618 | 0.127593352 | 10.57567236 | -0.236295842 | 0.820314827 | 0.996633405 | -5.233263499 |
| **A_55_P2083184** | Gm10272 | -0.010680981 | -0.118745899 | 0.097383937 | 8.003138112 | -0.236230266 | 0.820363639 | 0.996633405 | -5.233276438 |
| **A_55_P2024175** | Map4k3 | 0.022565828 | -0.205806735 | 0.250938391 | 10.93979948 | 0.236165515 | 0.820411839 | 0.996633405 | -5.233289211 |
| **A_55_P2041910** | NA | -0.020590334 | -0.22906348 | 0.187882812 | 8.960479683 | -0.236060001 | 0.820490384 | 0.996633405 | -5.233310017 |
| **A_52_P648573** | Gorab | 0.012824132 | -0.117091062 | 0.142739327 | 8.419997261 | 0.235926742 | 0.820589585 | 0.996633405 | -5.233336281 |
| **A_52_P633379** | Plpbp | 0.0111261 | -0.101670947 | 0.123923147 | 11.07848335 | 0.235751441 | 0.820720088 | 0.996633405 | -5.23337081 |
| **A_51_P149775** | Spcs1 | 0.014676445 | -0.134140936 | 0.163493825 | 13.90314387 | 0.235709093 | 0.820751615 | 0.996633405 | -5.233379147 |
| **A_51_P371675** | Slc7a6os | 0.011945414 | -0.109180768 | 0.133071596 | 9.06971922 | 0.235706908 | 0.820753242 | 0.996633405 | -5.233379577 |
| **A_52_P99848** | Pik3cd | -0.010932986 | -0.121804284 | 0.099938312 | 8.253274812 | -0.235683282 | 0.820770831 | 0.996633405 | -5.233384228 |
| **A_51_P345836** | Prkra | 0.011203875 | -0.102508357 | 0.124916106 | 10.3162829 | 0.235488767 | 0.820915648 | 0.996633405 | -5.233422501 |
| **A_55_P2065434** | Rprl2 | 0.040518234 | -0.370988251 | 0.452024718 | 9.265277677 | 0.235333067 | 0.821031573 | 0.996633405 | -5.233453114 |
| **A_55_P2000521** | Lonrf1 | 0.016105062 | -0.147475944 | 0.179686068 | 9.548879002 | 0.235309067 | 0.821049443 | 0.996633405 | -5.233457831 |
| **A_55_P2048096** | Mzf1 | -0.011781189 | -0.131461276 | 0.107898898 | 8.201294413 | -0.235275312 | 0.821074576 | 0.996633405 | -5.233464465 |
| **A_55_P2095529** | NA | -0.013868723 | -0.154810288 | 0.127072843 | 14.59184563 | -0.235183313 | 0.821143076 | 0.996633405 | -5.23348254 |
| **A_55_P1972381** | Gjb4 | -0.009493599 | -0.105992167 | 0.087004969 | 7.713983937 | -0.235136014 | 0.821178294 | 0.996633405 | -5.23349183 |
| **A_51_P267508** | Borcs7 | 0.016425344 | -0.150592978 | 0.183443666 | 9.696823414 | 0.235049584 | 0.82124265 | 0.996633405 | -5.233508801 |
| **A_52_P49378** | Kif1a | -0.021869836 | -0.244253618 | 0.200513947 | 10.51016316 | -0.235045281 | 0.821245854 | 0.996633405 | -5.233509645 |
| **A_51_P146149** | Napsa | -0.009918506 | -0.110817999 | 0.090980987 | 7.703297081 | -0.234945107 | 0.821320446 | 0.996633405 | -5.233529307 |
| **A_52_P401640** | Bpgm | -0.050604672 | -0.565467231 | 0.464257888 | 10.77314234 | -0.234913748 | 0.821343797 | 0.996633405 | -5.233535461 |
| **A_51_P250445** | Zfp276 | -0.015392576 | -0.172018642 | 0.141233489 | 9.043288583 | -0.234885608 | 0.821364752 | 0.996633405 | -5.233540982 |
| **A_55_P2020004** | NA | 0.063115998 | -0.579334549 | 0.705566545 | 12.14165593 | 0.234805834 | 0.821424156 | 0.996633405 | -5.23355663 |
| **A_55_P2033927** | NA | -0.009666871 | -0.108075418 | 0.088741676 | 15.96215989 | -0.234780614 | 0.821442935 | 0.996633405 | -5.233561576 |
| **A_51_P167513** | Leng1 | 0.013393139 | -0.122981502 | 0.149767779 | 10.2292449 | 0.234724207 | 0.82148494 | 0.996633405 | -5.233572636 |
| **A_51_P515623** | Qpctl | -0.011336753 | -0.12679825 | 0.104124744 | 8.54935829 | -0.234671654 | 0.821524075 | 0.996633405 | -5.233582938 |
| **A_66_P128918** | Hacl1 | 0.018806054 | -0.172774808 | 0.210386915 | 9.655290875 | 0.234614413 | 0.821566703 | 0.996633405 | -5.233594157 |
| **A_55_P2162152** | NA | -0.032833208 | -0.36731673 | 0.301650315 | 9.8203185 | -0.234610648 | 0.821569506 | 0.996633405 | -5.233594894 |
| **A_55_P2133485** | Gapvd1 | -0.014128526 | -0.158090131 | 0.12983308 | 10.06072576 | -0.234562883 | 0.821605077 | 0.996633405 | -5.233604254 |
| **A_55_P2073705** | Tfdp2 | 0.02203493 | -0.202507165 | 0.246577026 | 9.437200714 | 0.234543305 | 0.821619657 | 0.996633405 | -5.233608089 |
| **A_55_P1964682** | Rn28s1 | 0.069710228 | -0.640913524 | 0.780333981 | 16.76702238 | 0.234458466 | 0.821682839 | 0.996633405 | -5.233624707 |
| **A_55_P2121886** | Map3k9 | -0.011307003 | -0.126590046 | 0.10397604 | 7.699649816 | -0.234418133 | 0.821712876 | 0.996633405 | -5.233632605 |
| **A_51_P115178** | Scara3 | -0.023532518 | -0.263536837 | 0.216471801 | 8.779072037 | -0.234346505 | 0.82176622 | 0.996633405 | -5.233646628 |
| **A_55_P2078428** | Tstd1 | 0.013449996 | -0.123760834 | 0.150660826 | 8.229816896 | 0.234284138 | 0.821812669 | 0.996633405 | -5.233658834 |

| | | | | | | | | | |
|---|---|---|---|---|---|---|---|---|---|
| **A_55_P2126557** | NA | 0.032849442 | -0.302346592 | 0.368045476 | 9.32018336 | 0.234227704 | 0.821854699 | 0.996633405 | -5.233669877 |
| **A_51_P195958** | Phlda1 | 0.041341966 | -0.380692782 | 0.463376714 | 11.21433067 | 0.234127295 | 0.821929483 | 0.996633405 | -5.233689518 |
| **A_55_P2161400** | Sctr | -0.009527485 | -0.106796023 | 0.087741052 | 7.653213966 | -0.234107339 | 0.821944345 | 0.996633405 | -5.23369342 |
| **A_51_P517381** | Cers2 | 0.016596466 | -0.152889803 | 0.186082736 | 10.21504071 | 0.234040082 | 0.821994439 | 0.996633405 | -5.233706571 |
| **A_52_P64707** | Foxa3 | 0.010192258 | -0.093900952 | 0.114285468 | 7.638861995 | 0.234022247 | 0.822007723 | 0.996633405 | -5.233710057 |
| **A_55_P2018307** | NA | 0.037325326 | -0.344049922 | 0.418700574 | 10.95587358 | 0.233916166 | 0.822086735 | 0.996633405 | -5.23373079 |
| **A_52_P227267** | Atp1a2 | -0.046784619 | -0.525267894 | 0.431698656 | 9.081408634 | -0.233692872 | 0.822253057 | 0.996633405 | -5.233774399 |
| **A_55_P2086507** | Bglap3 | 0.00988748 | -0.091239507 | 0.111014466 | 7.712361008 | 0.233683299 | 0.822260188 | 0.996633405 | -5.233776268 |
| **A_55_P1963529** | Abcb10 | 0.013791201 | -0.127320854 | 0.154903256 | 11.2704433 | 0.233586159 | 0.822332547 | 0.996633405 | -5.233795226 |
| **A_51_P465473** | Btf3l4 | 0.018523625 | -0.171039844 | 0.208087094 | 9.159024284 | 0.233550316 | 0.822359247 | 0.996633405 | -5.23380222 |
| **A_51_P106059** | Traf4 | 0.049558618 | -0.457847289 | 0.556964525 | 10.32195577 | 0.23343867 | 0.822442415 | 0.996633405 | -5.227672011 |
| **A_51_P141071** | Cand2 | 0.012641092 | -0.116807692 | 0.142089875 | 11.29880233 | 0.233397252 | 0.822473268 | 0.996633405 | -5.233832072 |
| **A_52_P248483** | Zfx | 0.015635418 | -0.144545343 | 0.175816178 | 8.691676961 | 0.233296523 | 0.822548306 | 0.996633405 | -5.233851707 |
| **A_55_P2017418** | Cfh | 0.029204289 | -0.270097068 | 0.328505645 | 11.24893154 | 0.233209953 | 0.822612799 | 0.996633405 | -5.233868575 |
| **A_55_P2004606** | Serpina4-p | 0.01043587 | -0.096641873 | 0.117513613 | 7.663586831 | 0.232937056 | 0.822816108 | 0.996633405 | -5.23392171 |
| **A_55_P2117731** | Pak4 | 0.021752073 | -0.201496922 | 0.245001067 | 13.88756469 | 0.232873605 | 0.822863381 | 0.996633405 | -5.233934055 |
| **A_55_P2042277** | Pbrm1 | -0.015378894 | -0.17328742 | 0.142529632 | 9.381489601 | -0.232770884 | 0.822939914 | 0.996633405 | -5.233954034 |
| **A_51_P266546** | Zc2hc1a | -0.012870234 | -0.145025287 | 0.119284818 | 10.33467433 | -0.232761858 | 0.82294664 | 0.996633405 | -5.23395579 |
| **A_52_P182118** | NA | 0.017982294 | -0.166754642 | 0.202719229 | 8.800756544 | 0.232648622 | 0.823031009 | 0.996633405 | -5.233977803 |
| **A_55_P2135526** | Gzmc | 0.013864797 | -0.128672852 | 0.156402446 | 7.727011193 | 0.232483994 | 0.823153674 | 0.996633405 | -5.234009789 |
| **A_55_P2146389** | NA | -0.011148244 | -0.125883931 | 0.103587442 | 8.000549925 | -0.23222934 | 0.823343429 | 0.996633405 | -5.234059222 |
| **A_52_P632691** | Zfp395 | -0.012493726 | -0.141076868 | 0.116089416 | 9.109225452 | -0.232229333 | 0.823343434 | 0.996633405 | -5.234059223 |
| **A_55_P2157093** | Bcl2l14 | 0.010213868 | -0.094955431 | 0.115383167 | 7.639529779 | 0.232118849 | 0.823425765 | 0.996633405 | -5.234080653 |
| **A_55_P1962039** | Anxa7 | -0.014632187 | -0.165374909 | 0.136110536 | 10.13776607 | -0.231996808 | 0.82351671 | 0.996633405 | -5.234104314 |
| **A_55_P2134496** | Amfr | 0.016256777 | -0.151362692 | 0.183876247 | 11.71549291 | 0.23180304 | 0.823661114 | 0.996633405 | -5.234141856 |
| **A_51_P486810** | Gpx2 | -0.010413031 | -0.117850513 | 0.09702445 | 7.926361078 | -0.231649031 | 0.823775892 | 0.996633405 | -5.234171672 |
| **A_55_P2065829** | Cfap300 | 0.06394309 | -0.596060261 | 0.723946442 | 13.03479466 | 0.231556306 | 0.823845 | 0.996633405 | -5.234189614 |
| **A_51_P124719** | Ccdc93 | -0.010110802 | -0.114508464 | 0.09428686 | 8.427410615 | -0.231474937 | 0.823905645 | 0.996633405 | -5.234205354 |
| **A_51_P481238** | Dop1b | -0.01479896 | -0.167746799 | 0.138148879 | 9.890650153 | -0.231258113 | 0.824067253 | 0.996633405 | -5.234247267 |
| **A_55_P2051596** | NA | 0.05892629 | -0.551063966 | 0.668916546 | 9.970097697 | 0.230884796 | 0.824345524 | 0.996633405 | -5.234319342 |
| **A_66_P100050** | Ndufaf8 | -0.017522112 | -0.198971301 | 0.163927076 | 10.32622933 | -0.23080253 | 0.824406848 | 0.996633405 | -5.23433521 |
| **A_52_P26216** | Numb | -0.013506128 | -0.153434692 | 0.126422437 | 8.0497432 | -0.230692569 | 0.82448882 | 0.996633405 | -5.23435641 |
| **A_51_P356055** | Grp | -0.010634211 | -0.120816662 | 0.09954824 | 7.702474916 | -0.230675766 | 0.824501346 | 0.996633405 | -5.234359649 |
| **A_51_P116487** | Lsm4 | 0.018077075 | -0.169292224 | 0.205446375 | 13.87189707 | 0.23058915 | 0.824565918 | 0.996633405 | -5.234376341 |
| **A_55_P2104289** | NA | -0.011509748 | -0.130825883 | 0.107806387 | 8.408991145 | -0.230555655 | 0.824590888 | 0.996633405 | -5.234382794 |
| **A_55_P2289040** | NA | 0.009855745 | -0.092371611 | 0.112083101 | 7.864523033 | 0.230425985 | 0.824687559 | 0.996633405 | -5.234407767 |
| **A_66_P118579** | 3300002A1 | 0.012161415 | -0.113989662 | 0.138312493 | 7.853524363 | 0.230410516 | 0.824699092 | 0.996633405 | -5.234410745 |
| **A_51_P186899** | Egln1 | 0.014624156 | -0.13707386 | 0.166322172 | 8.4929283 | 0.230409314 | 0.824699987 | 0.996633405 | -5.234410976 |
| **A_55_P1956873** | Timm17a | 0.013139566 | -0.12327566 | 0.149554793 | 14.05571848 | 0.230211661 | 0.824847348 | 0.996633405 | -5.234449014 |
| **A_55_P2134571** | NA | 0.01784196 | -0.167408725 | 0.203092645 | 8.584753011 | 0.230192871 | 0.824861357 | 0.996633405 | -5.234452628 |
| **A_51_P467110** | Dpp4 | -0.027246163 | -0.310160708 | 0.255668382 | 8.888358639 | -0.230175559 | 0.824874265 | 0.996633405 | -5.234455958 |
| **A_55_P2095535** | NA | 0.009500687 | -0.089218935 | 0.10822031 | 7.777592383 | 0.230017389 | 0.824992194 | 0.996633405 | -5.234486369 |
| **A_51_P217336** | Scamp1 | -0.013276967 | -0.151235795 | 0.124681861 | 11.72217665 | -0.230016245 | 0.824993048 | 0.996633405 | -5.234486589 |
| **A_55_P1960698** | Pih1d2 | -0.010048208 | -0.114490604 | 0.094394188 | 7.903739007 | -0.229943383 | 0.825047375 | 0.996633405 | -5.234500591 |
| **A_51_P304683** | Clpx | -0.013011576 | -0.14825793 | 0.122234778 | 11.11019162 | -0.229939442 | 0.825050313 | 0.996633405 | -5.234501348 |
| **A_55_P1976690** | Snrnp35 | -0.018569999 | -0.211682974 | 0.174542976 | 11.26520991 | -0.229831508 | 0.825130792 | 0.996633405 | -5.234522081 |
| **A_51_P196526** | Bscl2 | -0.016842597 | -0.192093013 | 0.15840782 | 8.541421797 | -0.229699008 | 0.825229591 | 0.996633405 | -5.234547521 |
| **A_51_P122238** | Uchl5 | -0.01605614 | -0.183132318 | 0.151020039 | 11.503481 | -0.229686639 | 0.825238814 | 0.996633405 | -5.228394572 |

| | | | | | | | | | |
|---|---|---|---|---|---|---|---|---|---|
| **A_51_P446469** | Dok2 | -0.014018935 | -0.159971792 | 0.131933922 | 8.733700574 | -0.229568105 | 0.825327203 | 0.996633405 | -5.23457264 |
| **A_52_P653825** | Keg1 | -0.009234282 | -0.105394538 | 0.086925973 | 7.631808744 | -0.229517952 | 0.825364603 | 0.996633405 | -5.23458226 |
| **A_52_P221756** | Spock2 | -0.023309513 | -0.266079835 | 0.219460809 | 8.633548601 | -0.229480999 | 0.825392159 | 0.996633405 | -5.234589347 |
| **A_51_P272147** | Emc9 | -0.02360842 | -0.277489067 | 0.230272227 | 11.98302833 | -0.230996008 | 0.825453215 | 0.996633405 | -5.155887648 |
| **A_55_P2064632** | NA | 0.012988785 | -0.122339446 | 0.148317016 | 14.00604371 | 0.229397796 | 0.825454205 | 0.996633405 | -5.234605299 |
| **A_51_P482503** | Tpd52l1 | 0.018025694 | -0.169782715 | 0.205834104 | 9.33936614 | 0.229396138 | 0.825455441 | 0.996633405 | -5.234605617 |
| **A_52_P618132** | Manea | -0.01644821 | -0.187864937 | 0.154968518 | 8.894067486 | -0.229337231 | 0.825499371 | 0.996633405 | -5.234616908 |
| **A_55_P1970631** | Vps54 | 0.020528454 | -0.193412565 | 0.234469472 | 9.750516052 | 0.229335511 | 0.825500653 | 0.996633405 | -5.234617238 |
| **A_65_P10195** | Myl7 | 0.135420435 | -1.275956002 | 1.546796871 | 14.32407719 | 0.22932451 | 0.825508857 | 0.996633405 | -5.234619346 |
| **A_66_P107790** | Myl12a | -0.015870054 | -0.181349378 | 0.149609269 | 14.78223234 | -0.229215407 | 0.825590221 | 0.996633405 | -5.234640249 |
| **A_55_P2057380** | NA | 0.024399459 | -0.230040094 | 0.278839012 | 8.461959432 | 0.229194772 | 0.82560561 | 0.996633405 | -5.234644201 |
| **A_52_P391081** | Hinfp | -0.01019993 | -0.116590923 | 0.096191062 | 8.281960958 | -0.2291403 | 0.825646234 | 0.996633405 | -5.234654633 |
| **A_52_P216539** | Fbxw8 | -0.009892278 | -0.113076839 | 0.093292282 | 9.012708216 | -0.229134638 | 0.825650456 | 0.996633405 | -5.234655717 |
| **A_52_P152283** | Gabpa | 0.012530366 | -0.11820675 | 0.143267483 | 10.01172306 | 0.229073045 | 0.825696391 | 0.996633405 | -5.23466751 |
| **A_66_P108267** | Foxn4 | 0.009827352 | -0.09274484 | 0.112399544 | 7.665657303 | 0.228989735 | 0.825758524 | 0.996633405 | -5.234683455 |
| **A_51_P257743** | Dnajc22 | 0.009729931 | -0.091857607 | 0.11131747 | 7.744978316 | 0.22891722 | 0.825812607 | 0.996633405 | -5.234697329 |
| **A_55_P2071930** | NA | 0.012623545 | -0.119208109 | 0.144455199 | 8.492106853 | 0.228860459 | 0.82585494 | 0.996633405 | -5.234708186 |
| **A_55_P1963563** | Zfp689 | 0.011693443 | -0.110507098 | 0.133893983 | 8.787464815 | 0.228706443 | 0.825969813 | 0.996633405 | -5.234737633 |
| **A_51_P416546** | Ctsg | 0.01153009 | -0.108999547 | 0.132059728 | 7.859558294 | 0.228637777 | 0.826021029 | 0.996633405 | -5.234750755 |
| **A_55_P2072666** | Cyp2d37-p | -0.009809753 | -0.112374982 | 0.092755476 | 7.619033802 | -0.228595177 | 0.826052803 | 0.996633405 | -5.234758894 |
| **A_51_P487668** | H2afy3 | 0.012894305 | -0.121937246 | 0.147725857 | 10.98329108 | 0.228568064 | 0.826073027 | 0.996633405 | -5.234764074 |
| **A_51_P113773** | Slx1b | -0.01382211 | -0.158366161 | 0.130721941 | 8.464434789 | -0.228551066 | 0.826085705 | 0.996633405 | -5.23476732 |
| **A_65_P03728** | St3gal1 | 0.016201095 | -0.153238787 | 0.185640976 | 11.526193 | 0.228527175 | 0.826103526 | 0.996633405 | -5.234771883 |
| **A_55_P2111990** | Tekt1 | 0.010944051 | -0.103515987 | 0.125404088 | 7.849640517 | 0.228524806 | 0.826105292 | 0.996633405 | -5.234772336 |
| **A_55_P1971966** | Uba1 | -0.021823783 | -0.250082939 | 0.206435372 | 14.11568884 | -0.22851303 | 0.826114076 | 0.996633405 | -5.234774585 |
| **A_51_P439403** | Padi1 | -0.010892888 | -0.124840566 | 0.103054789 | 8.44001186 | -0.228479219 | 0.826139296 | 0.996633405 | -5.234781041 |
| **A_55_P2117119** | Efhd1 | 0.012518232 | -0.118496628 | 0.143533092 | 7.77215822 | 0.228366063 | 0.826223701 | 0.996633405 | -5.234802642 |
| **A_51_P508959** | Prr15 | 0.029222577 | -0.276804538 | 0.335249692 | 11.12771979 | 0.228227376 | 0.826327154 | 0.996633405 | -5.234829103 |
| **A_55_P2136101** | Meis3 | 0.01843157 | -0.17470677 | 0.211569909 | 9.136061562 | 0.228088276 | 0.826430918 | 0.996633405 | -5.234855627 |
| **A_55_P1965150** | Pnkp | -0.013857028 | -0.159126173 | 0.131412118 | 11.50360698 | -0.227984773 | 0.82650813 | 0.996633405 | -5.234875352 |
| **A_51_P415688** | Gng12 | 0.02167012 | -0.20555881 | 0.248899049 | 10.56848361 | 0.227932796 | 0.826546906 | 0.996633405 | -5.234885255 |
| **A_55_P2435563** | Etl4 | -0.020277139 | -0.232984818 | 0.192430539 | 8.479832992 | -0.227841406 | 0.826615084 | 0.996633405 | -5.234902661 |
| **A_51_P499153** | Cog3 | 0.015967499 | -0.151572695 | 0.183507693 | 9.059954406 | 0.227785992 | 0.826656425 | 0.996633405 | -5.234913211 |
| **A_55_P2167294** | NA | 0.009187035 | -0.087240465 | 0.105614536 | 7.658343308 | 0.227710792 | 0.826712528 | 0.996633405 | -5.234927525 |
| **A_52_P495869** | Mafb | 0.017035899 | -0.161817489 | 0.195889287 | 8.56705233 | 0.227654906 | 0.826754223 | 0.996633405 | -5.23493816 |
| **A_55_P2164805** | Gm6749 | -0.020996561 | -0.241448457 | 0.199455335 | 14.12521737 | -0.227637318 | 0.826767345 | 0.996633405 | -5.234941506 |
| **A_55_P2420724** | Srek1 | 0.017944998 | -0.170482745 | 0.206372741 | 8.219288849 | 0.227618575 | 0.826781328 | 0.996633405 | -5.234945072 |
| **A_55_P2177614** | Srsf3 | -0.014161487 | -0.162880501 | 0.134557526 | 11.62710084 | -0.227589127 | 0.826803299 | 0.996633405 | -5.234950674 |
| **A_51_P152612** | Nol7 | 0.015850683 | -0.150775526 | 0.182476892 | 8.561386966 | 0.22735986 | 0.826974356 | 0.996633405 | -5.234994262 |
| **A_51_P202623** | Mterf2 | -0.020609638 | -0.23756417 | 0.196344893 | 10.1945636 | -0.227044393 | 0.827209743 | 0.996633405 | -5.235054169 |
| **A_51_P226853** | Gga1 | -0.018175059 | -0.209520059 | 0.173169941 | 8.807490536 | -0.227021946 | 0.827226492 | 0.996633405 | -5.235058428 |
| **A_51_P335969** | Des | -0.038897554 | -0.448598048 | 0.37080294 | 11.53381326 | -0.226915904 | 0.827305621 | 0.996633405 | -5.235078545 |
| **A_55_P2173188** | Kank3 | -0.019586888 | -0.225937126 | 0.186763351 | 8.8070349 | -0.226866091 | 0.827342792 | 0.996633405 | -5.235087991 |
| **A_51_P279062** | Treml1 | 0.071847357 | -0.685079092 | 0.828773806 | 9.437079193 | 0.226864347 | 0.827344093 | 0.996633405 | -5.235088322 |
| **A_55_P2038101** | Dock11 | -0.017990738 | -0.207543107 | 0.171561631 | 9.550827218 | -0.226844832 | 0.827358656 | 0.996633405 | -5.235092022 |
| **A_55_P1983758** | Sdhc | 0.02629166 | -0.250780037 | 0.303363358 | 13.72544561 | 0.226795786 | 0.827395255 | 0.996633405 | -5.235101321 |
| **A_55_P1992257** | Ap1s1 | 0.011672203 | -0.111346478 | 0.134690884 | 13.49014153 | 0.226772772 | 0.827412429 | 0.996633405 | -5.235105683 |
| **A_55_P2067629** | Ush1g | 0.011610455 | -0.110757593 | 0.133978503 | 8.290601968 | 0.22677247 | 0.827412655 | 0.996633405 | -5.23510574 |

| | | | | | | | | | |
|---|---|---|---|---|---|---|---|---|---|
| **A_51_P438527** | Cyb5r1 | -0.010771604 | -0.124373399 | 0.102830192 | 10.82193237 | -0.226623165 | 0.827524074 | 0.996633405 | -5.235134031 |
| **A_55_P2207593** | 8430437O( | 0.023550681 | -0.22482956 | 0.271930921 | 8.666837606 | 0.226618601 | 0.82752748 | 0.996633405 | -5.235134896 |
| **A_55_P2181262** | NA | -0.030084542 | -0.347544759 | 0.287375676 | 13.19126448 | -0.226497402 | 0.827617928 | 0.996633405 | -5.235157847 |
| **A_55_P2153517** | Enho | 0.01371964 | -0.131117823 | 0.158557102 | 10.29596544 | 0.226397139 | 0.827692754 | 0.996633405 | -5.235176825 |
| **A_55_P2135556** | Rpgrip1 | 0.028099651 | -0.268736386 | 0.324935688 | 11.43759102 | 0.226252526 | 0.827800683 | 0.996633405 | -5.235204182 |
| **A_55_P2322709** | 4633401B( | 0.021307985 | -0.203908099 | 0.24652407 | 8.608053466 | 0.226126844 | 0.827894486 | 0.996633405 | -5.235227945 |
| **A_51_P379687** | Gm10413 | 0.010018338 | -0.095925962 | 0.115962638 | 7.681256172 | 0.226009771 | 0.827981866 | 0.996633405 | -5.235250068 |
| **A_55_P1978061** | Hnrnpa3 | 0.016101101 | -0.154203201 | 0.186405403 | 9.190646145 | 0.225963908 | 0.828016097 | 0.996633405 | -5.235258731 |
| **A_55_P2015405** | 9930111J2 | 0.010753961 | -0.103038234 | 0.124546157 | 7.941649282 | 0.225873421 | 0.828083637 | 0.996633405 | -5.235275819 |
| **A_55_P2209046** | NA | -0.010571718 | -0.122450363 | 0.101306928 | 8.132258199 | -0.225843445 | 0.828106012 | 0.996633405 | -5.235281479 |
| **A_52_P77204** | Lrrc20 | -0.017001282 | -0.196947316 | 0.162944751 | 10.69189116 | -0.225812788 | 0.828128895 | 0.996633405 | -5.235287266 |
| **A_55_P2275437** | Hoxb5os | 0.009745461 | -0.093486792 | 0.112977713 | 7.699046795 | 0.225629612 | 0.828265625 | 0.996633405 | -5.235321828 |
| **A_52_P532029** | Tbk1 | -0.010368163 | -0.120214468 | 0.099478142 | 8.335035568 | -0.225592937 | 0.828293001 | 0.996633405 | -5.235328745 |
| **A_51_P165984** | Aup1 | -0.01821763 | -0.211238272 | 0.174803011 | 9.710330034 | -0.225578271 | 0.828303949 | 0.996633405 | -5.23533151 |
| **A_51_P148122** | Klhdc3 | -0.021746127 | -0.252241521 | 0.208749266 | 12.90296555 | -0.22549079 | 0.828369251 | 0.996633405 | -5.235348003 |
| **A_55_P2063366** | Commd7 | -0.020402074 | -0.236800062 | 0.195995913 | 11.22757504 | -0.225335824 | 0.828484933 | 0.996633405 | -5.235377203 |
| **A_55_P1984344** | Rhox2h | 0.00922428 | -0.08865636 | 0.107104921 | 7.751808166 | 0.225239635 | 0.828556741 | 0.996633405 | -5.235395319 |
| **A_55_P2019257** | Trpc4ap | -0.009282588 | -0.107813842 | 0.089248666 | 13.3025031 | -0.225166712 | 0.828611181 | 0.996633405 | -5.235409047 |
| **A_55_P1958986** | Spata7 | -0.010793607 | -0.125385425 | 0.103798211 | 8.668143484 | -0.225124163 | 0.828642946 | 0.996633405 | -5.235417055 |
| **A_55_P1952071** | Npb | -0.024432983 | -0.283839903 | 0.234973936 | 10.00934728 | -0.225114813 | 0.828649926 | 0.996633405 | -5.235418815 |
| **A_55_P2177278** | DXBay18 | -0.009680662 | -0.11246347 | 0.093102145 | 7.682157337 | -0.225109453 | 0.828653927 | 0.996633405 | -5.235419823 |
| **A_55_P2051337** | Ppfia1 | -0.014070702 | -0.163592538 | 0.135451135 | 9.964006749 | -0.224915958 | 0.828798387 | 0.996633405 | -5.23545622 |
| **A_55_P2086094** | Slc35b2 | -0.017791267 | -0.207008323 | 0.171425789 | 11.67294327 | -0.224727247 | 0.82893928 | 0.996633405 | -5.235491688 |
| **A_55_P2067200** | NA | 0.015310962 | -0.147690244 | 0.178312168 | 8.670916009 | 0.224502295 | 0.829107242 | 0.996633405 | -5.235533928 |
| **A_55_P2117614** | Tnfrsf13c | 0.009245105 | -0.089217682 | 0.107707893 | 7.664976625 | 0.224413444 | 0.829173585 | 0.996633405 | -5.235550601 |
| **A_66_P121656** | Pcgf1 | -0.01236818 | -0.14409454 | 0.11935818 | 8.187319737 | -0.22441002 | 0.829176142 | 0.996633405 | -5.235551243 |
| **A_52_P655084** | Tbc1d17 | -0.014306479 | -0.166732288 | 0.138119329 | 10.27949926 | -0.224327957 | 0.829237418 | 0.996633405 | -5.235566636 |
| **A_52_P507310** | Mrps24 | -0.016686292 | -0.194472702 | 0.161100117 | 12.09605728 | -0.224321254 | 0.829242424 | 0.996633405 | -5.235567893 |
| **A_51_P511680** | Ttc38 | 0.011565664 | -0.111675808 | 0.134807137 | 8.166285239 | 0.224296678 | 0.829260775 | 0.996633405 | -5.235572502 |
| **A_55_P2049479** | NA | -0.017384973 | -0.202712794 | 0.167942848 | 9.966381596 | -0.224203576 | 0.829330296 | 0.996633405 | -5.235589956 |
| **A_55_P2160387** | Cxcl11 | -0.021801099 | -0.254276583 | 0.210674384 | 16.75784654 | -0.224135357 | 0.829381238 | 0.996633405 | -5.235602741 |
| **A_55_P2157378** | Xlr5a | -0.010084724 | -0.117650304 | 0.097480857 | 7.663202383 | -0.224078304 | 0.829423842 | 0.996633405 | -5.235613431 |
| **A_55_P2082314** | NA | -0.013538085 | -0.157984414 | 0.130908245 | 8.436991385 | -0.224006101 | 0.82947776 | 0.996633405 | -5.235626955 |
| **A_51_P154684** | Alyref | 0.02656958 | -0.266101148 | 0.319240309 | 12.414558 | 0.225513505 | 0.829513394 | 0.996633405 | -5.156888723 |
| **A_55_P1952112** | Ildr2 | -0.009216452 | -0.10758076 | 0.089147855 | 8.016499154 | -0.223941906 | 0.829525699 | 0.996633405 | -5.235638975 |
| **A_55_P1969166** | Fxr2 | 0.013154884 | -0.127291647 | 0.153601415 | 12.44131444 | 0.223864456 | 0.829583539 | 0.996633405 | -5.235653474 |
| **A_55_P2155654** | Pvt1 | 0.009614359 | -0.093131025 | 0.112359743 | 8.074780792 | 0.22364909 | 0.829744378 | 0.996633405 | -5.235693763 |
| **A_51_P240864** | Sppl3 | 0.014259292 | -0.138178731 | 0.166697315 | 13.12058902 | 0.223570138 | 0.829803343 | 0.996633405 | -5.235708523 |
| **A_66_P111401** | Fbxw8 | 0.027813446 | -0.269632351 | 0.325259243 | 9.840949024 | 0.223488972 | 0.829863962 | 0.996633405 | -5.235723692 |
| **A_55_P2006915** | Myo5c | -0.010017341 | -0.117229289 | 0.097194607 | 7.745418177 | -0.223315253 | 0.82999371 | 0.996633405 | -5.235756139 |
| **A_55_P2003927** | NA | 0.014606362 | -0.141762197 | 0.17097492 | 12.48302491 | 0.223255284 | 0.830038501 | 0.996633405 | -5.235767334 |
| **A_51_P305532** | Eif2s3x | 0.106182314 | -1.031088152 | 1.24345278 | 9.879924454 | 0.223150342 | 0.830116885 | 0.996633405 | -5.235786918 |
| **A_55_P1983152** | Gramd4 | -0.020361024 | -0.238456874 | 0.197734826 | 11.23893133 | -0.223131741 | 0.830130779 | 0.996633405 | -5.235790389 |
| **A_55_P2047648** | Coq2 | -0.014787845 | -0.173205173 | 0.143629483 | 12.67921029 | -0.223106041 | 0.830149975 | 0.996633405 | -5.235795183 |
| **A_55_P2077083** | Tmem132e | -0.013631463 | -0.159732411 | 0.132469485 | 7.925270408 | -0.22299676 | 0.830231602 | 0.996633405 | -5.235815563 |
| **A_55_P2158473** | 1700028P1 | 0.009415821 | -0.091518169 | 0.110349812 | 7.654423338 | 0.222961512 | 0.830257931 | 0.996633405 | -5.235822135 |
| **A_55_P1978306** | Zfp318 | 0.014181377 | -0.137935517 | 0.166298271 | 10.10192226 | 0.222817899 | 0.830365208 | 0.996633405 | -5.235848899 |
| **A_55_P2048378** | Zfp975 | 0.032010096 | -0.311374423 | 0.375394615 | 12.06867126 | 0.222800107 | 0.830378498 | 0.996633405 | -5.235852213 |

| | | | | | | | | | |
|---|---|---|---|---|---|---|---|---|---|
| **A_52_P350786** | Phlda2 | -0.009605541 | -0.112704696 | 0.093493614 | 7.655076064 | -0.222677258 | 0.830470268 | 0.996633405 | -5.235875092 |
| **A_55_P2150283** | E030030I0( | 0.009284935 | -0.090383155 | 0.108953025 | 12.06984041 | 0.222654701 | 0.830487119 | 0.996633405 | -5.235879292 |
| **A_52_P449871** | Id4 | -0.018263238 | -0.214322524 | 0.177796048 | 8.318742241 | -0.222638104 | 0.830499517 | 0.996633405 | -5.235882382 |
| **A_51_P407248** | Antxr1 | -0.047362748 | -0.556172958 | 0.461447462 | 10.41891741 | -0.222479605 | 0.830617923 | 0.996633405 | -5.235911877 |
| **A_55_P2068414** | 6030426L1 | 0.013166782 | -0.128446064 | 0.154779627 | 8.435121387 | 0.222221521 | 0.830810734 | 0.996633405 | -5.235959859 |
| **A_55_P2124425** | Zfp661 | 0.013393657 | -0.130660941 | 0.157448255 | 8.750050711 | 0.222218997 | 0.830812619 | 0.996633405 | -5.235960328 |
| **A_52_P510771** | Tbl3 | 0.016513772 | -0.161229727 | 0.194257272 | 8.404323453 | 0.222055583 | 0.83093471 | 0.996633405 | -5.235990682 |
| **A_55_P1975837** | NA | -0.028080079 | -0.330491039 | 0.274330881 | 15.152797 | -0.221926893 | 0.831030861 | 0.996633405 | -5.236014569 |
| **A_51_P104820** | Alkbh5 | -0.014206842 | -0.167233822 | 0.138820138 | 8.482328782 | -0.221890477 | 0.83105807 | 0.996633405 | -5.236021327 |
| **A_55_P1959164** | Gm10324 | -0.01027573 | -0.120961534 | 0.100410075 | 8.265595208 | -0.221885992 | 0.831061421 | 0.996633405 | -5.236022159 |
| **A_55_P2144975** | Ccdc90b | 0.014280436 | -0.139566573 | 0.168127444 | 9.114309552 | 0.221851073 | 0.831087511 | 0.996633405 | -5.236028637 |
| **A_55_P2127357** | NA | 0.016269139 | -0.159115722 | 0.191654 | 10.75934574 | 0.221708126 | 0.83119432 | 0.996633405 | -5.236055147 |
| **A_51_P277336** | Cavin2 | -0.019810169 | -0.233451897 | 0.193831558 | 15.02305211 | -0.221621165 | 0.831259299 | 0.996633405 | -5.236071265 |
| **A_55_P1973427** | Tex14 | 0.009439975 | -0.092414077 | 0.111294028 | 7.656990603 | 0.221514251 | 0.831339188 | 0.996633405 | -5.236091074 |
| **A_51_P499838** | Bst1 | 0.017557976 | -0.172012031 | 0.207127982 | 8.065043907 | 0.221367545 | 0.831448816 | 0.996633405 | -5.23611824 |
| **A_55_P1967776** | Slc4a4 | 0.021375822 | -0.209446085 | 0.252197728 | 8.873479059 | 0.221337463 | 0.831471295 | 0.996633405 | -5.236123808 |
| **A_51_P177583** | Puf60 | -0.00940958 | -0.111033131 | 0.092213972 | 12.02684786 | -0.221301825 | 0.831497927 | 0.996633405 | -5.236130404 |
| **A_55_P2066548** | Baiap2l2 | 0.013788018 | -0.135237689 | 0.162813725 | 9.645480349 | 0.221131076 | 0.831625527 | 0.996633405 | -5.23616199 |
| **A_55_P2014450** | B430105G( | -0.01128401 | -0.133271683 | 0.110703662 | 7.942582057 | -0.221083668 | 0.831660956 | 0.996633405 | -5.236170756 |
| **A_51_P318618** | Anks4b | -0.008962579 | -0.105898876 | 0.087973719 | 7.656891378 | -0.22098138 | 0.8317374 | 0.996633405 | -5.236189663 |
| **A_52_P451378** | Twsg1 | 0.020254048 | -0.19884457 | 0.239352666 | 11.3759865 | 0.220943549 | 0.831765672 | 0.996633405 | -5.236196653 |
| **A_55_P2044242** | Slc13a5 | 0.010825588 | -0.106323244 | 0.127974421 | 7.742984678 | 0.220862854 | 0.83182598 | 0.996633405 | -5.23621156 |
| **A_55_P1997415** | Exoc6b | 0.01550426 | -0.15231343 | 0.18332195 | 10.05085365 | 0.22081188 | 0.831864076 | 0.996633405 | -5.236220973 |
| **A_55_P2149269** | Galt | -0.011587031 | -0.137030007 | 0.113855944 | 8.639094376 | -0.220767411 | 0.831897312 | 0.996633405 | -5.236229184 |
| **A_55_P1991770** | Pdlim4 | -0.058003923 | -0.686157136 | 0.570149291 | 10.50137856 | -0.220699278 | 0.831948234 | 0.996633405 | -5.236241761 |
| **A_52_P627816** | Tgm1 | -0.009517824 | -0.112620847 | 0.093585199 | 7.68901312 | -0.220635507 | 0.831995897 | 0.996633405 | -5.236253529 |
| **A_52_P272811** | Cpsf1 | 0.012912774 | -0.126978882 | 0.15280443 | 9.634015009 | 0.220615928 | 0.83201053 | 0.996633405 | -5.236257141 |
| **A_66_P131036** | NA | -0.020903767 | -0.247367876 | 0.205560343 | 9.549053782 | -0.220614625 | 0.832011504 | 0.996633405 | -5.236257382 |
| **A_51_P230904** | Copz2 | -0.022692437 | -0.268624278 | 0.223239403 | 10.02801017 | -0.22053398 | 0.83207178 | 0.996633405 | -5.236272258 |
| **A_51_P119597** | Ap3b1 | -0.012540211 | -0.148449031 | 0.123368609 | 11.83338526 | -0.220529321 | 0.832075262 | 0.996633405 | -5.236273117 |
| **A_55_P2111419** | Eny2 | 0.020296035 | -0.199768304 | 0.240360374 | 12.88022428 | 0.220429985 | 0.832149509 | 0.996633405 | -5.236291433 |
| **A_55_P1954481** | Dock9 | -0.011661726 | -0.138164922 | 0.114841469 | 7.995913869 | -0.220328396 | 0.832225443 | 0.996633405 | -5.236310156 |
| **A_55_P2029581** | NA | 0.011552505 | -0.113800823 | 0.136905833 | 7.953281568 | 0.220267001 | 0.832271334 | 0.996633405 | -5.236321467 |
| **A_66_P126415** | NA | -0.011555127 | -0.13696328 | 0.113853026 | 7.779790984 | -0.220220672 | 0.832305964 | 0.996633405 | -5.23633 |
| **A_55_P2132039** | NA | -0.019983425 | -0.236864406 | 0.196897555 | 10.07238301 | -0.220220425 | 0.832306148 | 0.996633405 | -5.236330046 |
| **A_51_P193000** | Bbs4 | 0.023428025 | -0.230862148 | 0.277718198 | 8.92404757 | 0.220198946 | 0.832322204 | 0.996633405 | -5.236334001 |
| **A_55_P2013501** | Hmgb1 | 0.023280033 | -0.229424334 | 0.2759844 | 13.54713505 | 0.220181067 | 0.832335568 | 0.996633405 | -5.236337294 |
| **A_52_P80567** | Nbas | 0.015775661 | -0.155469623 | 0.187020944 | 8.591779998 | 0.220180103 | 0.832336289 | 0.996633405 | -5.236337471 |
| **A_55_P2059357** | Myo7a | -0.011573076 | -0.137206751 | 0.1140606 | 8.949463041 | -0.220166818 | 0.83234622 | 0.996633405 | -5.236339917 |
| **A_55_P2091476** | NA | 0.015945259 | -0.15715763 | 0.189048147 | 9.213154307 | 0.220158969 | 0.832352086 | 0.996633405 | -5.236341362 |
| **A_55_P2109128** | H2bc6 | 0.01445914 | -0.147695159 | 0.17661344 | 8.079898497 | 0.221503785 | 0.832486591 | 0.996633405 | -5.157606006 |
| **A_55_P2331974** | NA | 0.011388275 | -0.112344965 | 0.135121516 | 8.132000915 | 0.219978739 | 0.832486811 | 0.996633405 | -5.236374533 |
| **A_51_P511060** | Csgalnact2 | 0.018707641 | -0.184573008 | 0.221988289 | 9.84725939 | 0.219954126 | 0.83250521 | 0.996633405 | -5.236379061 |
| **A_55_P2052106** | BC049762 | -0.010856303 | -0.128831458 | 0.107118853 | 7.984261456 | -0.219938121 | 0.832517174 | 0.996633405 | -5.236382005 |
| **A_55_P1975832** | 1810009N( | -0.016932288 | -0.200947852 | 0.167083275 | 8.769898915 | -0.219922787 | 0.832528638 | 0.996633405 | -5.236384826 |
| **A_55_P2112030** | Tmem145 | 0.009116139 | -0.090044998 | 0.108277277 | 7.66966193 | 0.219724551 | 0.832676831 | 0.996633405 | -5.23642127 |
| **A_55_P1961541** | NA | -0.012424635 | -0.147638982 | 0.122789712 | 8.081511743 | -0.219619045 | 0.832755706 | 0.996633405 | -5.236440654 |
| **A_55_P1960097** | Epb41l3 | 0.011573363 | -0.114414598 | 0.137561324 | 12.65365532 | 0.219553137 | 0.832804979 | 0.996633405 | -5.236452758 |

| | | | | | | | | | |
|---|---|---|---|---|---|---|---|---|---|
| **A_51_P413376** | Zfp42 | -0.009208006 | -0.10948617 | 0.091070157 | 7.661002114 | -0.219466567 | 0.832869701 | 0.996633405 | -5.236468651 |
| **A_51_P220062** | Mmp15 | 0.032543282 | -0.321908729 | 0.386995294 | 10.15994948 | 0.219438609 | 0.832890603 | 0.996633405 | -5.236473782 |
| **A_52_P498985** | Zfp512 | -0.017131976 | -0.203806099 | 0.169542148 | 8.942813286 | -0.219347383 | 0.832958808 | 0.996633405 | -5.236490521 |
| **A_55_P2160383** | NA | 0.017565366 | -0.173928353 | 0.209059085 | 10.2918732 | 0.219235966 | 0.83304211 | 0.996633405 | -5.236510956 |
| **A_55_P2212161** | 2310034O( | -0.009839184 | -0.117226702 | 0.097548335 | 7.708121655 | -0.21898501 | 0.833229748 | 0.996633405 | -5.236556946 |
| **A_52_P561205** | Stard4 | 0.011482221 | -0.113844078 | 0.136808519 | 7.889757356 | 0.218974128 | 0.833237885 | 0.996633405 | -5.236558939 |
| **A_55_P1998751** | Ccdc137 | 0.009358426 | -0.092813829 | 0.111530681 | 8.462013454 | 0.21891673 | 0.833280803 | 0.996633405 | -5.23656945 |
| **A_55_P1985758** | Psmd9 | -0.015072464 | -0.179660167 | 0.14951524 | 9.187041723 | -0.218874914 | 0.83331207 | 0.996633405 | -5.236577106 |
| **A_51_P408561** | Oard1 | -0.018354104 | -0.21885958 | 0.182151372 | 10.17939893 | -0.218784263 | 0.833379854 | 0.996633405 | -5.236593698 |
| **A_52_P495031** | 9130401M( | 0.011228895 | -0.111479943 | 0.133937733 | 11.88719443 | 0.218710847 | 0.833434752 | 0.996633405 | -5.23660713 |
| **A_55_P2002389** | Nelfb | -0.012585871 | -0.150190479 | 0.125018737 | 12.584472 | -0.21860467 | 0.833514148 | 0.996633405 | -5.236626548 |
| **A_52_P256426** | NA | 0.011266918 | -0.111939791 | 0.134473628 | 8.196794838 | 0.218564652 | 0.833544074 | 0.996633405 | -5.236633865 |
| **A_51_P493987** | Moxd1 | 0.03635263 | -0.361311392 | 0.434016652 | 9.021585126 | 0.218488542 | 0.833600989 | 0.996633405 | -5.236647776 |
| **A_55_P2168587** | Cdk9 | 0.013684727 | -0.136112626 | 0.163482079 | 8.593996548 | 0.218343929 | 0.833709135 | 0.996633405 | -5.236674195 |
| **A_55_P1966445** | Krt81 | -0.010349632 | -0.12365274 | 0.102953475 | 7.812932078 | -0.218319351 | 0.833727515 | 0.996633405 | -5.236678684 |
| **A_55_P2212498** | NA | -0.029829076 | -0.356410215 | 0.296752063 | 8.859579721 | -0.218302059 | 0.833740447 | 0.996633405 | -5.236681841 |
| **A_55_P2121985** | Plcb4 | -0.015123823 | -0.180759292 | 0.150511645 | 8.848420047 | -0.218231475 | 0.833793233 | 0.996633405 | -5.236694728 |
| **A_55_P2067445** | Arf5 | 0.026754293 | -0.275741571 | 0.329250157 | 12.30599611 | 0.219705632 | 0.833820921 | 0.996633405 | -5.157923587 |
| **A_55_P2002212** | Babam2 | 0.024373383 | -0.242864111 | 0.291610877 | 10.09412053 | 0.217985471 | 0.833977215 | 0.996633405 | -5.236739608 |
| **A_55_P2117472** | Acap2 | -0.012706503 | -0.15215047 | 0.126737464 | 10.2754495 | -0.217788752 | 0.834124346 | 0.996633405 | -5.236775461 |
| **A_55_P2097972** | NA | 0.018364725 | -0.1831918 | 0.21992125 | 8.382349856 | 0.217769325 | 0.834138876 | 0.996633405 | -5.236779 |
| **A_55_P2155572** | NA | -0.016242003 | -0.194554217 | 0.162070211 | 12.47121501 | -0.217704608 | 0.834187281 | 0.996633405 | -5.236790786 |
| **A_55_P1969042** | Cops6 | -0.010668729 | -0.127886542 | 0.106549084 | 13.30970038 | -0.217534535 | 0.834314491 | 0.996633405 | -5.236821746 |
| **A_55_P2165451** | NA | -0.016792735 | -0.201306555 | 0.167721084 | 10.72576412 | -0.217521244 | 0.834324432 | 0.996633405 | -5.236824164 |
| **A_51_P251892** | Magoh | -0.015663062 | -0.187834007 | 0.156507883 | 12.65242851 | -0.217433235 | 0.834390263 | 0.996633405 | -5.236840174 |
| **A_51_P217535** | Actr3 | -0.018559952 | -0.222581113 | 0.185461209 | 13.77346439 | -0.217425644 | 0.834395941 | 0.996633405 | -5.236841555 |
| **A_51_P280158** | Sf3b2 | 0.013129943 | -0.131208875 | 0.157468761 | 12.27983658 | 0.217414661 | 0.834404157 | 0.996633405 | -5.236843553 |
| **A_55_P2155906** | Chmp3 | 0.008945968 | -0.089400638 | 0.107292573 | 10.73642177 | 0.217408782 | 0.834408554 | 0.996633405 | -5.236844622 |
| **A_51_P494929** | Diablo | -0.018775219 | -0.225243815 | 0.187693378 | 10.79415573 | -0.217340233 | 0.83445983 | 0.996633405 | -5.236857086 |
| **A_55_P2002567** | NA | 0.010202116 | -0.102076927 | 0.122481159 | 15.99276567 | 0.217170427 | 0.834586852 | 0.996633405 | -5.236887946 |
| **A_55_P1963198** | NA | -0.009585663 | -0.115101735 | 0.095930409 | 8.498596658 | -0.217126418 | 0.834619774 | 0.996633405 | -5.23689594 |
| **A_55_P1959465** | Lrrfip1 | -0.016281115 | -0.195728762 | 0.163166532 | 12.93761889 | -0.216848044 | 0.834828023 | 0.996633405 | -5.236946469 |
| **A_55_P2145376** | Fastk | -0.01419857 | -0.170696474 | 0.142299333 | 9.352451283 | -0.216842911 | 0.834831863 | 0.996633405 | -5.2369474 |
| **A_51_P346893** | Extl1 | -0.009184546 | -0.110421596 | 0.092052505 | 7.867768299 | -0.216833975 | 0.834838548 | 0.996633405 | -5.236949021 |
| **A_52_P229709** | Ube2d3 | 0.010213596 | -0.102395698 | 0.12282289 | 11.16668387 | 0.216777189 | 0.83488103 | 0.996633405 | -5.23695932 |
| **A_55_P2079516** | Mpp1 | -0.009306609 | -0.1119401 | 0.093326883 | 7.950033971 | -0.216726232 | 0.834919153 | 0.996633405 | -5.23696856 |
| **A_51_P435968** | Tnfaip8 | -0.015706553 | -0.188992844 | 0.157579738 | 10.56628587 | -0.216633595 | 0.83498846 | 0.996633405 | -5.236985352 |
| **A_51_P456857** | Ift122 | -0.013909428 | -0.167420208 | 0.139601352 | 10.70672534 | -0.216560633 | 0.835043047 | 0.996633405 | -5.236998572 |
| **A_52_P681705** | Zfp955b | 0.03363814 | -0.337775882 | 0.405052162 | 9.918653574 | 0.216462571 | 0.835116415 | 0.996633405 | -5.237016333 |
| **A_51_P244950** | Dpys | 0.009139438 | -0.091841588 | 0.110120463 | 7.660406029 | 0.216316098 | 0.835226006 | 0.996633405 | -5.237042848 |
| **A_51_P144543** | NA | 0.009408197 | -0.094551611 | 0.113368004 | 7.65715324 | 0.21629678 | 0.83524046 | 0.996633405 | -5.237046344 |
| **A_55_P2047461** | Fcho1 | -0.009567756 | -0.115316345 | 0.096180833 | 7.998136192 | -0.216244285 | 0.835279738 | 0.996633405 | -5.237055842 |
| **A_51_P181365** | Atp5md | -0.01559214 | -0.187971135 | 0.156786855 | 16.19967963 | -0.216187468 | 0.83532225 | 0.996633405 | -5.237066119 |
| **A_51_P365903** | Ankzf1 | -0.009921712 | -0.119625157 | 0.099781732 | 10.42743347 | -0.216160085 | 0.835342739 | 0.996633405 | -5.237071071 |
| **A_55_P2086153** | Bcdin3d | -0.018862898 | -0.227486064 | 0.189760268 | 10.6014351 | -0.216100126 | 0.835387604 | 0.996633405 | -5.237081912 |
| **A_51_P260513** | Polr1d | -0.011669361 | -0.140751531 | 0.117412809 | 11.12179389 | -0.216067748 | 0.835411831 | 0.996633405 | -5.237087765 |
| **A_51_P201982** | Angpt2 | 0.032969222 | -0.331732396 | 0.39767084 | 9.250492516 | 0.216062871 | 0.83541548 | 0.996633405 | -5.237088647 |
| **A_51_P411130** | Rpp25l | 0.022571603 | -0.227135359 | 0.272278564 | 12.38841309 | 0.216043337 | 0.835430097 | 0.996633405 | -5.237092178 |

| | | | | | | | | | |
|---|---|---|---|---|---|---|---|---|---|
| **A_55_P1974233** | Dtd2 | -0.01451438 | -0.175163507 | 0.146134748 | 8.480667125 | -0.215938082 | 0.835508856 | 0.996633405 | -5.237111197 |
| **A_52_P708507** | Myadml2 | -0.017920757 | -0.216339421 | 0.180497908 | 8.430858301 | -0.215865359 | 0.835563274 | 0.996633405 | -5.237124332 |
| **A_55_P2101340** | Ramp3 | 0.015184428 | -0.153059049 | 0.183427904 | 9.470219856 | 0.215709528 | 0.835679885 | 0.996633405 | -5.237152464 |
| **A_52_P468033** | Setd1b | 0.019729035 | -0.198969974 | 0.238428044 | 10.61452837 | 0.215609633 | 0.835754639 | 0.996633405 | -5.237170488 |
| **A_55_P1999992** | Galnt12 | 0.026956934 | -0.271882331 | 0.325796198 | 8.669781766 | 0.215596627 | 0.835764372 | 0.996633405 | -5.237172834 |
| **A_55_P2136318** | Slc1a5 | -0.021934419 | -0.265194254 | 0.221325416 | 8.670419231 | -0.215508724 | 0.835830155 | 0.996633405 | -5.237188686 |
| **A_55_P2047200** | NA | 0.024591938 | -0.248309582 | 0.297493458 | 9.673879463 | 0.215375318 | 0.835929993 | 0.996633405 | -5.237212732 |
| **A_52_P253317** | 4930481A1 | -0.046532875 | -0.563113517 | 0.470047767 | 10.09679787 | -0.215293486 | 0.835991235 | 0.996633405 | -5.237227474 |
| **A_52_P244132** | Isoc1 | -0.010260125 | -0.124163679 | 0.103643428 | 11.01974099 | -0.215290334 | 0.835993595 | 0.996633405 | -5.237228042 |
| **A_55_P2001861** | NA | -0.023077353 | -0.279295501 | 0.233140796 | 9.532038797 | -0.215270854 | 0.836008174 | 0.996633405 | -5.237231551 |
| **A_55_P2347384** | NA | 0.014222986 | -0.143739774 | 0.172185745 | 8.160976552 | 0.21520145 | 0.836060117 | 0.996633405 | -5.237244049 |
| **A_55_P2171658** | Stx3 | 0.013355366 | -0.134998986 | 0.161709719 | 8.839018999 | 0.215161539 | 0.836089987 | 0.996633405 | -5.237251234 |
| **A_55_P2004016** | Crispld2 | 0.04892414 | -0.49457128 | 0.59241956 | 9.548019834 | 0.215147569 | 0.836100442 | 0.996633405 | -5.237253749 |
| **A_51_P188126** | Gca | 0.019878213 | -0.201055147 | 0.240811574 | 9.021322971 | 0.215042937 | 0.836178753 | 0.996633405 | -5.237272578 |
| **A_55_P2081360** | Btbd1 | 0.012985888 | -0.131359744 | 0.15733152 | 11.91647526 | 0.215019153 | 0.836196554 | 0.996633405 | -5.237276857 |
| **A_55_P2288232** | Gucy1a2 | -0.01091833 | -0.132314129 | 0.110477469 | 8.55665296 | -0.214961995 | 0.836239335 | 0.996633405 | -5.237287139 |
| **A_52_P322927** | NA | 0.008691173 | -0.087974652 | 0.105356998 | 7.692919446 | 0.214889185 | 0.836293831 | 0.996633405 | -5.237300231 |
| **A_55_P1997485** | NA | 0.020135561 | -0.203923993 | 0.244195115 | 11.45514262 | 0.214787697 | 0.836369793 | 0.996633405 | -5.237318474 |
| **A_55_P2010699** | Map11 | 0.01239762 | -0.125578782 | 0.150374023 | 8.689945203 | 0.214754681 | 0.836394506 | 0.996633405 | -5.237324406 |
| **A_52_P606826** | Pigh | -0.012741198 | -0.15454721 | 0.129064815 | 8.098318117 | -0.214745829 | 0.836401132 | 0.996633405 | -5.237325997 |
| **A_51_P467505** | Tec | -0.017578699 | -0.213234517 | 0.17807712 | 8.837725546 | -0.214735124 | 0.836409144 | 0.996633405 | -5.23732792 |
| **A_51_P244492** | Nbl1 | -0.043533975 | -0.528118025 | 0.441050075 | 10.72826902 | -0.214717946 | 0.836422002 | 0.996633405 | -5.237331006 |
| **A_55_P1953236** | Epb42 | 0.00962288 | -0.097491307 | 0.116737068 | 7.661808999 | 0.214717391 | 0.836422417 | 0.996633405 | -5.237331106 |
| **A_52_P632191** | Nfx1 | -0.010228492 | -0.124147143 | 0.10369016 | 8.359067206 | -0.214598108 | 0.836511703 | 0.996633405 | -5.237352529 |
| **A_55_P2044119** | Klc1 | -0.02156691 | -0.261809965 | 0.218676145 | 9.569113552 | -0.214558741 | 0.836541171 | 0.996633405 | -5.237359597 |
| **A_55_P2073467** | Gtf2i | -0.014316994 | -0.173990618 | 0.145356629 | 14.4335361 | -0.21430278 | 0.836732773 | 0.996633405 | -5.23740552 |
| **A_51_P327983** | 4833427G( | 0.008849988 | -0.08985314 | 0.107553116 | 7.65257878 | 0.214299371 | 0.836735325 | 0.996633405 | -5.237406131 |
| **A_55_P2027361** | Irak1bp1 | 0.052683959 | -0.535064754 | 0.640432671 | 11.54288646 | 0.214237676 | 0.83678151 | 0.996633405 | -5.237417192 |
| **A_55_P2117928** | Mxd4 | 0.021024835 | -0.213574586 | 0.255624257 | 9.719451308 | 0.214197697 | 0.836811438 | 0.996633405 | -5.237424358 |
| **A_55_P2061737** | Tmsb4x | 0.016946579 | -0.172159898 | 0.206053055 | 15.92775124 | 0.214182883 | 0.836822528 | 0.996633405 | -5.237427013 |
| **A_55_P2046092** | NA | 0.009086366 | -0.092377384 | 0.110550117 | 7.664707313 | 0.214036809 | 0.836931882 | 0.996633405 | -5.237453182 |
| **A_55_P2048800** | Cd82 | 0.014173719 | -0.144118315 | 0.172465753 | 7.925843716 | 0.214009909 | 0.83695202 | 0.996633405 | -5.237457999 |
| **A_55_P2075140** | Sirt3 | 0.011339645 | -0.115364511 | 0.138043801 | 12.38487997 | 0.213903425 | 0.83703174 | 0.996633405 | -5.237477063 |
| **A_55_P2159801** | Rfc4 | -0.019160936 | -0.233260072 | 0.1949382 | 9.403448874 | -0.213900078 | 0.837034245 | 0.996633405 | -5.237477662 |
| **A_52_P343343** | Rpl7l1 | -0.016647029 | -0.20272028 | 0.169426221 | 8.606294891 | -0.2138267 | 0.837089181 | 0.996633405 | -5.237490792 |
| **A_52_P481423** | Cttnbp2nl | 0.014426934 | -0.146962082 | 0.175815949 | 10.65746147 | 0.213653097 | 0.837219155 | 0.996633405 | -5.237521841 |
| **A_52_P414420** | Dcun1d1 | -0.016521156 | -0.201361037 | 0.168318725 | 11.19283076 | -0.213625887 | 0.837239528 | 0.996633405 | -5.237526705 |
| **A_52_P110270** | Nelfe | 0.017397227 | -0.177267454 | 0.212061909 | 10.71376968 | 0.213600373 | 0.83725863 | 0.996633405 | -5.237531265 |
| **A_52_P590392** | Rab10 | -0.012478815 | -0.152183766 | 0.127226135 | 12.08877511 | -0.213486636 | 0.837343788 | 0.996633405 | -5.237551588 |
| **A_55_P2033650** | Cpt2 | -0.014615039 | -0.178274533 | 0.149044455 | 12.39723624 | -0.213436112 | 0.837381617 | 0.996633405 | -5.237560613 |
| **A_55_P2061591** | Psmd1 | 0.015827597 | -0.161415485 | 0.193070678 | 13.48383542 | 0.213429679 | 0.837386433 | 0.996633405 | -5.237561761 |
| **A_55_P1988259** | Gng5 | 0.013490274 | -0.137579738 | 0.164560286 | 15.67767539 | 0.213428121 | 0.8373876 | 0.996633405 | -5.23756204 |
| **A_66_P128963** | Cyp2d9 | -0.009027987 | -0.110134196 | 0.092078222 | 7.715835886 | -0.213413665 | 0.837398424 | 0.996633405 | -5.237564621 |
| **A_55_P2106351** | NA | -0.018408708 | -0.224593222 | 0.187775807 | 9.303922813 | -0.213391142 | 0.837415288 | 0.996633405 | -5.237568643 |
| **A_55_P2179869** | NA | -0.034645072 | -0.4228208 | 0.353530655 | 12.90979689 | -0.213315408 | 0.837471995 | 0.996633405 | -5.237582164 |
| **A_55_P1977066** | Klk4 | -0.008861224 | -0.1081885 | 0.090466052 | 7.708929321 | -0.213223133 | 0.837541088 | 0.996633405 | -5.237598631 |
| **A_55_P1988148** | NA | 0.014107894 | -0.144031317 | 0.172247105 | 8.228142002 | 0.213221868 | 0.837542035 | 0.996633405 | -5.237598857 |
| **A_51_P116027** | Med27 | 0.011813329 | -0.120712807 | 0.144339464 | 11.0732711 | 0.213049191 | 0.837671335 | 0.996633405 | -5.237629654 |

| | | | | | | | | | |
|---|---|---|---|---|---|---|---|---|---|
| **A_55_P2046213** | NA | 0.013550227 | -0.138484128 | 0.165584582 | 16.34459648 | 0.21301685 | 0.837695553 | 0.996633405 | -5.237635419 |
| **A_51_P296448** | Casp2 | -0.017447817 | -0.213225239 | 0.178329606 | 12.08136179 | -0.213003929 | 0.837705228 | 0.996633405 | -5.237637722 |
| **A_52_P309718** | Tasor | -0.013250307 | -0.161950962 | 0.135450347 | 8.66232267 | -0.212971851 | 0.837729249 | 0.996633405 | -5.237643439 |
| **A_55_P2180086** | Lrrc28 | 0.012878794 | -0.13165641 | 0.157413998 | 10.07831253 | 0.2129662 | 0.837733481 | 0.996633405 | -5.237644446 |
| **A_51_P256304** | Mrpl37 | -0.021217155 | -0.259692839 | 0.217258529 | 9.052506529 | -0.21264353 | 0.837975115 | 0.996633405 | -5.237701907 |
| **A_55_P2074035** | Haao | 0.009123819 | -0.093491132 | 0.11173877 | 7.90142033 | 0.21250793 | 0.838076666 | 0.996633405 | -5.237726028 |
| **A_55_P2002379** | Golga2 | 0.019738582 | -0.20237631 | 0.241853474 | 12.35108416 | 0.212396525 | 0.8381601 | 0.996633405 | -5.237745835 |
| **A_55_P2165334** | Nde1 | -0.013288168 | -0.162918409 | 0.136342073 | 11.87343795 | -0.212253506 | 0.838267214 | 0.996633405 | -5.237771247 |
| **A_55_P2182921** | NA | 0.016614403 | -0.170521992 | 0.203750798 | 8.840239725 | 0.212195225 | 0.838310864 | 0.996633405 | -5.237781597 |
| **A_55_P2101387** | Nubpl | -0.013698235 | -0.168051404 | 0.140654933 | 8.981336869 | -0.212108564 | 0.838375771 | 0.996633405 | -5.237796983 |
| **A_55_P1990190** | Arhgap28 | -0.010830224 | -0.13306481 | 0.111404362 | 8.317528637 | -0.211764157 | 0.838633737 | 0.996633405 | -5.237858068 |
| **A_55_P2131865** | Krtap10-10 | 0.015674576 | -0.161331274 | 0.192680427 | 8.966923907 | 0.211649538 | 0.838719594 | 0.996633405 | -5.237878375 |
| **A_51_P472249** | Slc7a7 | -0.024855611 | -0.305669452 | 0.255958229 | 10.02339924 | -0.211550996 | 0.83879341 | 0.996633405 | -5.237895826 |
| **A_51_P394154** | Ddx51 | 0.009690822 | -0.099810041 | 0.119191684 | 9.531706986 | 0.211520374 | 0.838816348 | 0.996633405 | -5.237901247 |
| **A_55_P2318584** | Aqp8 | 0.028003858 | -0.288430831 | 0.344438547 | 9.209307167 | 0.211515847 | 0.838819739 | 0.996633405 | -5.237902048 |
| **A_51_P453519** | Zdhhc16 | -0.010540183 | -0.129641279 | 0.108560913 | 12.23307332 | -0.21151517 | 0.838820246 | 0.996633405 | -5.237902168 |
| **A_51_P162676** | Armc7 | -0.009840337 | -0.121093571 | 0.101412897 | 8.101604622 | -0.211400721 | 0.83890598 | 0.996633405 | -5.237922422 |
| **A_55_P2004777** | Micall2 | 0.011068322 | -0.114133621 | 0.136270265 | 8.596462695 | 0.211290441 | 0.838988594 | 0.996633405 | -5.237941928 |
| **A_55_P1978176** | Abhd17c | -0.015372574 | -0.189331028 | 0.15858588 | 12.84145541 | -0.21120791 | 0.839050421 | 0.996633405 | -5.237956519 |
| **A_55_P1954266** | Zfp672 | 0.01343947 | -0.138689756 | 0.165568697 | 10.39061589 | 0.211143943 | 0.839098343 | 0.996633405 | -5.237967825 |
| **A_51_P415945** | Psma2 | 0.013841155 | -0.142835484 | 0.170517795 | 14.72485597 | 0.211143261 | 0.839098853 | 0.996633405 | -5.237967945 |
| **A_52_P603976** | Jtb | 0.011800153 | -0.121783635 | 0.145383941 | 11.91098533 | 0.21112663 | 0.839111312 | 0.996633405 | -5.237970884 |
| **A_55_P2004801** | Tacc3 | -0.012447204 | -0.153373296 | 0.128478887 | 12.23353977 | -0.211100632 | 0.839130789 | 0.996633405 | -5.237975478 |
| **A_52_P308669** | Cpn1 | -0.008584671 | -0.105805398 | 0.088636056 | 7.655566541 | -0.211044447 | 0.839172882 | 0.996633405 | -5.237985403 |
| **A_55_P2089780** | Mtfr1l | 0.018239482 | -0.188340395 | 0.224819358 | 9.343060899 | 0.211024855 | 0.83918756 | 0.996633405 | -5.237988863 |
| **A_51_P509643** | Snca | -0.083894558 | -1.034246645 | 0.86645753 | 11.54722301 | -0.210988308 | 0.83921494 | 0.996633405 | -5.237995317 |
| **A_55_P2048730** | NA | 0.016828741 | -0.173924502 | 0.207581985 | 15.53133194 | 0.210857396 | 0.839313019 | 0.996633405 | -5.238018427 |
| **A_55_P2062603** | 4932438A1 | 0.015784522 | -0.163157588 | 0.194726632 | 8.983353966 | 0.210827866 | 0.839335143 | 0.996633405 | -5.238023638 |
| **A_51_P310164** | 2810459M1 | -0.009057271 | -0.111745168 | 0.093630627 | 7.717543842 | -0.210808062 | 0.839349981 | 0.996633405 | -5.238027132 |
| **A_55_P1973447** | NA | 0.025397769 | -0.262560672 | 0.31335621 | 9.46693018 | 0.210802074 | 0.839354467 | 0.996633405 | -5.238028189 |
| **A_55_P2013236** | S100g | 0.078394934 | -0.81111077 | 0.967900637 | 8.113647359 | 0.21064366 | 0.839473157 | 0.996633405 | -5.238056127 |
| **A_55_P2122491** | Gm6222 | 0.013785764 | -0.142696373 | 0.170267901 | 16.10667778 | 0.210559675 | 0.839536083 | 0.996633405 | -5.238070931 |
| **A_55_P2349198** | NA | 0.032635259 | -0.33789769 | 0.403168207 | 9.27698944 | 0.210508365 | 0.839574529 | 0.996633405 | -5.238079972 |
| **A_52_P179238** | Macroh2a1 | -0.021414073 | -0.264599276 | 0.221771131 | 8.736486507 | -0.210460821 | 0.839610152 | 0.996633405 | -5.238088348 |
| **A_51_P346938** | Lrg1 | -0.014525236 | -0.179489734 | 0.150439262 | 8.535173178 | -0.210446566 | 0.839620833 | 0.996633405 | -5.238090859 |
| **A_55_P2042146** | Fech | 0.051636015 | -0.534843995 | 0.638116026 | 13.07751066 | 0.210430477 | 0.839632888 | 0.996633405 | -5.238093692 |
| **A_51_P118742** | Drg2 | -0.022093992 | -0.273043487 | 0.228855503 | 9.58070003 | -0.210424838 | 0.839637114 | 0.996633405 | -5.238094686 |
| **A_55_P2131027** | NA | -0.034860943 | -0.431078073 | 0.361356187 | 12.71714707 | -0.210288253 | 0.839739457 | 0.996633405 | -5.238118733 |
| **A_66_P119735** | BC049265 | -0.014658844 | -0.181324166 | 0.152006479 | 8.073422018 | -0.21021495 | 0.839794385 | 0.996633405 | -5.238131633 |
| **A_55_P2083582** | NA | 0.011440433 | -0.11864163 | 0.141522495 | 14.50303308 | 0.210200703 | 0.83980506 | 0.996633405 | -5.238134139 |
| **A_55_P1963175** | Lctl | 0.00885893 | -0.091935223 | 0.109653082 | 8.071494448 | 0.21006565 | 0.839906262 | 0.996633405 | -5.238157892 |
| **A_55_P1989901** | Ddb1 | 0.015668177 | -0.162621514 | 0.193957868 | 14.64549599 | 0.210039693 | 0.839925713 | 0.996633405 | -5.238162456 |
| **A_51_P456187** | Pdcd2 | -0.012019489 | -0.148828512 | 0.124789535 | 10.58303286 | -0.209981186 | 0.839969556 | 0.996633405 | -5.23817274 |
| **A_52_P74427** | Mms19 | 0.012550719 | -0.130386591 | 0.15548803 | 8.306607351 | 0.209861196 | 0.840059474 | 0.996633405 | -5.238193823 |
| **A_55_P2035804** | Fzd8 | 0.016365254 | -0.170052063 | 0.202782571 | 8.425167817 | 0.209819396 | 0.840090798 | 0.996633405 | -5.238201165 |
| **A_51_P361492** | Pou2f1 | -0.014361908 | -0.177985077 | 0.149261261 | 10.30234417 | -0.209785986 | 0.840115836 | 0.996633405 | -5.238207032 |
| **A_55_P2087796** | Eva1b | -0.019979325 | -0.247627882 | 0.207669232 | 11.5419202 | -0.209761143 | 0.840134453 | 0.996633405 | -5.238211394 |
| **A_55_P1954081** | Ccdc115 | 0.018222027 | -0.189429385 | 0.225873438 | 11.05611035 | 0.209735006 | 0.840154041 | 0.996633405 | -5.238215982 |

| | | | | | | | | | |
|---|---|---|---|---|---|---|---|---|---|
| **A_52_P618745** | Ermp1 | -0.013753618 | -0.170553967 | 0.143046731 | 9.130532803 | -0.209642367 | 0.840223467 | 0.996633405 | -5.238232242 |
| **A_51_P446796** | Camta1 | -0.013637543 | -0.169200148 | 0.141925061 | 11.17837626 | -0.209527039 | 0.840309898 | 0.996633405 | -5.238252473 |
| **A_55_P2113031** | Foxo4 | -0.008668725 | -0.107560635 | 0.090223186 | 7.700392132 | -0.209509426 | 0.840323098 | 0.996633405 | -5.238255562 |
| **A_55_P2132207** | 1810037I17 | 0.010354645 | -0.107810443 | 0.128519733 | 8.93654285 | 0.20943784 | 0.840376749 | 0.996633405 | -5.238268114 |
| **A_66_P134453** | I830077J02 | 0.01023109 | -0.10662779 | 0.127089969 | 7.834704792 | 0.209251853 | 0.840516143 | 0.996633405 | -5.238300704 |
| **A_51_P202074** | Ncapd2 | 0.012490767 | -0.130180305 | 0.155161838 | 12.54972993 | 0.209248475 | 0.840518675 | 0.996633405 | -5.238301296 |
| **A_55_P2000158** | Pik3r3 | 0.017511197 | -0.182740121 | 0.217762515 | 10.08201058 | 0.209001584 | 0.840703726 | 0.996633405 | -5.238344514 |
| **A_52_P479262** | NA | 0.031395228 | -0.327706166 | 0.390496622 | 11.65444087 | 0.208956404 | 0.840737591 | 0.996633405 | -5.238352418 |
| **A_52_P496726** | Rasd1 | -0.010299732 | -0.128133252 | 0.107533787 | 8.105716463 | -0.208913359 | 0.840769855 | 0.996633405 | -5.238359946 |
| **A_66_P107129** | 3300005D0 | -0.009129113 | -0.113632073 | 0.095373847 | 7.811765616 | -0.208789724 | 0.840862529 | 0.996633405 | -5.23838156 |
| **A_51_P150608** | Jagn1 | 0.018667281 | -0.195054037 | 0.232388599 | 11.61315798 | 0.208757629 | 0.840886587 | 0.996633405 | -5.238387169 |
| **A_55_P2152188** | NA | 0.048881943 | -0.510862475 | 0.608626361 | 15.23490389 | 0.208721804 | 0.840913441 | 0.996633405 | -5.238393429 |
| **A_55_P2021792** | Dyrk1b | -0.017393697 | -0.216608688 | 0.181821294 | 8.669607016 | -0.208679131 | 0.840945428 | 0.996633405 | -5.238400884 |
| **A_52_P94401** | Rtn4rl1 | 0.014523697 | -0.151964308 | 0.181011701 | 10.73918267 | 0.208498699 | 0.841080683 | 0.996633405 | -5.238432389 |
| **A_55_P2006277** | Ogdh | -0.02290599 | -0.28553711 | 0.23972513 | 9.218526482 | -0.208454857 | 0.84111355 | 0.996633405 | -5.23844004 |
| **A_55_P1960506** | Wdr83os | 0.014563337 | -0.152472696 | 0.18159937 | 12.05474163 | 0.208381839 | 0.841168287 | 0.996633405 | -5.238452779 |
| **A_51_P481832** | Rabggta | 0.021316505 | -0.223219726 | 0.265852737 | 8.976712032 | 0.208344444 | 0.841196321 | 0.996633405 | -5.238459301 |
| **A_55_P2372228** | NA | -0.008351467 | -0.104244608 | 0.087541674 | 7.692351593 | -0.208153804 | 0.84133924 | 0.996633405 | -5.238492535 |
| **A_52_P219943** | Epc1 | 0.011517993 | -0.120762812 | 0.143798797 | 8.549795926 | 0.20810816 | 0.841373459 | 0.996633405 | -5.238500488 |
| **A_55_P2158448** | Gdf3 | 0.010108623 | -0.105991091 | 0.126208337 | 8.03964893 | 0.208098996 | 0.841380329 | 0.996633405 | -5.238502084 |
| **A_55_P1967015** | Fgfr1 | -0.018166764 | -0.226930071 | 0.190596543 | 9.01199924 | -0.207985247 | 0.841465609 | 0.996633405 | -5.238521895 |
| **A_51_P245454** | Ufsp2 | -0.012769686 | -0.159571811 | 0.134032439 | 11.28452877 | -0.207901203 | 0.84152862 | 0.996633405 | -5.238536525 |
| **A_51_P254471** | Birc2 | 0.019058629 | -0.200050105 | 0.238167362 | 9.008144952 | 0.207893586 | 0.84153433 | 0.996633405 | -5.23853785 |
| **A_52_P428521** | Fgd1 | 0.012942054 | -0.135868357 | 0.161752464 | 8.681898248 | 0.207863872 | 0.841556609 | 0.996633405 | -5.238543021 |
| **A_55_P1972913** | NA | 0.026072657 | -0.273773328 | 0.325918642 | 10.79066205 | 0.207824232 | 0.841586329 | 0.996633405 | -5.238549918 |
| **A_55_P2030060** | Kdm8 | -0.013657947 | -0.170751492 | 0.143435597 | 8.601430113 | -0.207795543 | 0.841607838 | 0.996633405 | -5.238554909 |
| **A_51_P158922** | Prkaa1 | 0.014680877 | -0.154223524 | 0.183585278 | 10.20246237 | 0.207740014 | 0.841649472 | 0.996633405 | -5.238564568 |
| **A_51_P497100** | Lgals4 | 0.025807555 | -0.271124325 | 0.322739436 | 8.383281045 | 0.207729975 | 0.841656999 | 0.996633405 | -5.238566313 |
| **A_51_P248666** | Cd274 | 0.008941887 | -0.093984144 | 0.111867917 | 7.865028668 | 0.207640986 | 0.841723722 | 0.996633405 | -5.238581786 |
| **A_55_P1975660** | Hemgn | -0.0212854 | -0.266310059 | 0.22373926 | 13.58710168 | -0.207625716 | 0.841735171 | 0.996633405 | -5.23858444 |
| **A_51_P207403** | Pwp1 | 0.013531656 | -0.142296223 | 0.169359535 | 9.531913955 | 0.207546268 | 0.841794741 | 0.996633405 | -5.238598246 |
| **A_55_P1985077** | Mcrip1 | 0.023954335 | -0.251900042 | 0.299808712 | 8.72876899 | 0.207545527 | 0.841795296 | 0.996633405 | -5.238598375 |
| **A_51_P341647** | Fancc | -0.009503588 | -0.118946323 | 0.099939146 | 8.860697598 | -0.207543822 | 0.841796575 | 0.996633405 | -5.238598671 |
| **A_55_P2031192** | Gtdc1 | -0.01623524 | -0.20321112 | 0.170740639 | 10.56773618 | -0.20753066 | 0.841806444 | 0.996633405 | -5.238600958 |
| **A_55_P2043362** | NA | 0.042762245 | -0.44981532 | 0.535339809 | 9.165685287 | 0.207488963 | 0.841837708 | 0.996633405 | -5.238608202 |
| **A_55_P2047842** | Clcn3 | 0.014525002 | -0.152793949 | 0.181843952 | 11.86991773 | 0.207481887 | 0.841843014 | 0.996633405 | -5.238609431 |
| **A_55_P2013118** | NA | 0.01003111 | -0.10553253 | 0.125594751 | 8.110084288 | 0.207461213 | 0.841858516 | 0.996633405 | -5.238613022 |
| **A_55_P2074647** | Snap47 | -0.01738259 | -0.21764203 | 0.18287685 | 10.54993613 | -0.207458209 | 0.841860769 | 0.996633405 | -5.238613543 |
| **A_55_P2000548** | Cfl1 | -0.016543387 | -0.207270182 | 0.174183408 | 14.59839805 | -0.207310767 | 0.841971326 | 0.996633405 | -5.238639142 |
| **A_55_P2058359** | Macroh2a1 | -0.026971855 | -0.338014756 | 0.284071047 | 9.662360009 | -0.207252437 | 0.842015064 | 0.996633405 | -5.238649264 |
| **A_55_P2008136** | Ctr9 | -0.021947335 | -0.275095253 | 0.231200582 | 8.714807943 | -0.207212794 | 0.842044791 | 0.996633405 | -5.238656141 |
| **A_55_P2008747** | Snrpd3 | 0.012829356 | -0.135223049 | 0.160881761 | 12.47865248 | 0.207108781 | 0.842122788 | 0.996633405 | -5.238674181 |
| **A_55_P2171008** | Clk4 | -0.031324662 | -0.392815755 | 0.330166432 | 9.4634949 | -0.207108498 | 0.842123 | 0.996633405 | -5.23867423 |
| **A_51_P380807** | Ckm | -0.048255715 | -0.60522528 | 0.50871385 | 14.0701186 | -0.207074407 | 0.842148564 | 0.996633405 | -5.23868014 |
| **A_55_P2104941** | Rpl30 | 0.011575951 | -0.122038021 | 0.145189922 | 14.93550054 | 0.207068446 | 0.842153034 | 0.996633405 | -5.238681174 |
| **A_51_P328932** | Tmem165 | 0.014631555 | -0.154340001 | 0.183603112 | 11.72992837 | 0.206959812 | 0.842234499 | 0.996633405 | -5.238700001 |
| **A_66_P107038** | Prcp | -0.012664175 | -0.158956133 | 0.133627784 | 11.45666176 | -0.206902417 | 0.84227754 | 0.996633405 | -5.238709944 |
| **A_52_P315111** | Mrpl52 | -0.009905462 | -0.124340658 | 0.104529733 | 14.36806568 | -0.206882742 | 0.842292295 | 0.996633405 | -5.238713352 |

| | | | | | | | | | |
|---|---|---|---|---|---|---|---|---|---|
| **A_55_P2113439** | Caln1 | -0.010122655 | -0.127109463 | 0.106864153 | 8.030171512 | -0.206807688 | 0.84234858 | 0.996633405 | -5.23872635 |
| **A_55_P2019293** | NA | -0.008446306 | -0.106059917 | 0.089167305 | 7.628891758 | -0.206807156 | 0.842348979 | 0.996633405 | -5.238726442 |
| **A_55_P2017851** | NA | -0.021337798 | -0.268011334 | 0.225335737 | 13.68831215 | -0.206745554 | 0.842395177 | 0.996633405 | -5.238737106 |
| **A_55_P1987261** | Fam214b | 0.023791759 | -0.251251724 | 0.298835241 | 10.1448537 | 0.20674467 | 0.842395841 | 0.996633405 | -5.238737259 |
| **A_55_P2091928** | Raet1b | -0.01124393 | -0.141265401 | 0.118777542 | 8.857334686 | -0.206686535 | 0.842439439 | 0.996633405 | -5.23874732 |
| **A_55_P2142371** | Rai1 | 0.021961397 | -0.232065572 | 0.275988365 | 11.75067931 | 0.206628044 | 0.842483305 | 0.996633405 | -5.23875744 |
| **A_55_P2184180** | Ndufv3 | -0.016736097 | -0.210421049 | 0.176948855 | 16.23216806 | -0.206522535 | 0.842562434 | 0.996633405 | -5.238775688 |
| **A_55_P2393734** | 4933421H1 | -0.025617246 | -0.322186094 | 0.270951603 | 10.66532178 | -0.206450547 | 0.842616425 | 0.996633405 | -5.238788133 |
| **A_55_P2030383** | Frmd4a | 0.015151103 | -0.160261992 | 0.190564198 | 11.04907272 | 0.206438823 | 0.842625218 | 0.996633405 | -5.238790159 |
| **A_55_P2039071** | Dhdh | 0.013748869 | -0.145493118 | 0.172990856 | 10.13210876 | 0.206356678 | 0.842686827 | 0.996633405 | -5.238804354 |
| **A_55_P1958532** | Hr | -0.011787932 | -0.148410612 | 0.124834748 | 7.829151776 | -0.206216757 | 0.842791772 | 0.996633405 | -5.23882852 |
| **A_55_P1989048** | Tm9sf1 | -0.011217725 | -0.14127896 | 0.11884351 | 10.08644934 | -0.206141789 | 0.842848001 | 0.996633405 | -5.238841461 |
| **A_51_P437938** | Coa5 | 0.015820764 | -0.167640774 | 0.199282302 | 10.05401089 | 0.206106439 | 0.842874516 | 0.996633405 | -5.238847562 |
| **A_55_P2042813** | Gbp11 | -0.011294626 | -0.142297246 | 0.119707995 | 7.901731545 | -0.206063462 | 0.842906752 | 0.996633405 | -5.238854978 |
| **A_52_P189618** | Armc8 | -0.010505362 | -0.132357735 | 0.111347011 | 9.652177394 | -0.206056423 | 0.842912032 | 0.996633405 | -5.238856192 |
| **A_55_P2003586** | Unkl | 0.013560631 | -0.14374756 | 0.170868823 | 8.235872914 | 0.206033433 | 0.842929276 | 0.996633405 | -5.238860158 |
| **A_52_P486322** | Fzr1 | 0.022957985 | -0.24351596 | 0.289431931 | 11.0333063 | 0.205915082 | 0.843018049 | 0.996633405 | -5.238880568 |
| **A_66_P111781** | 4930432M1 | 0.012880912 | -0.136634335 | 0.162396159 | 9.105785265 | 0.205906603 | 0.843024409 | 0.996633405 | -5.23888203 |
| **A_51_P294643** | Cdr2 | -0.021399405 | -0.269859958 | 0.227061149 | 9.016350393 | -0.205851187 | 0.843065976 | 0.996633405 | -5.238891582 |
| **A_51_P298843** | Pcbd2 | -0.015951547 | -0.201162946 | 0.169259851 | 11.04108945 | -0.205846858 | 0.843069224 | 0.996633405 | -5.238892329 |
| **A_51_P296905** | Ppat | 0.011270879 | -0.119613606 | 0.142155365 | 9.270590298 | 0.205815827 | 0.8430925 | 0.996633405 | -5.238897676 |
| **A_55_P2063495** | Dph3 | 0.013833949 | -0.146851393 | 0.17451929 | 9.368687442 | 0.205768564 | 0.843127953 | 0.996633405 | -5.23890582 |
| **A_55_P2114562** | NA | -0.009233562 | -0.116528608 | 0.098061484 | 8.091071678 | -0.205683162 | 0.843192016 | 0.996633405 | -5.23892053 |
| **A_55_P2039637** | Ift88 | 0.013165479 | -0.13988045 | 0.166211407 | 9.769078853 | 0.205600422 | 0.843254082 | 0.996633405 | -5.238934777 |
| **A_52_P180408** | Bcas2 | 0.016702499 | -0.177464006 | 0.210869004 | 11.92474403 | 0.205596768 | 0.843256823 | 0.996633405 | -5.238935406 |
| **A_52_P417859** | Gid4 | 0.030726031 | -0.326642496 | 0.388094559 | 9.881804766 | 0.205494072 | 0.843333862 | 0.996633405 | -5.23895308 |
| **A_52_P69558** | Gm8221 | 0.009025635 | -0.096020803 | 0.114072073 | 7.645830041 | 0.20535514 | 0.843438085 | 0.996633405 | -5.238976976 |
| **A_55_P1957937** | NA | 0.008367581 | -0.089041154 | 0.105776316 | 7.830078432 | 0.205310498 | 0.843471576 | 0.996633405 | -5.238984651 |
| **A_66_P118093** | NA | 0.016247265 | -0.172895669 | 0.205390198 | 8.837307686 | 0.20530488 | 0.84347579 | 0.996633405 | -5.238985617 |
| **A_55_P1962454** | NA | 0.008915015 | -0.094907 | 0.11273703 | 7.996340544 | 0.205230427 | 0.843531646 | 0.996633405 | -5.238998414 |
| **A_55_P2131105** | NA | 0.023953086 | -0.255147328 | 0.303053501 | 11.9407027 | 0.205121003 | 0.843613738 | 0.996633405 | -5.239017212 |
| **A_55_P2061278** | Sik1 | -0.0234113 | -0.296222336 | 0.249399737 | 9.50004255 | -0.205103334 | 0.843626994 | 0.996633405 | -5.239020247 |
| **A_55_P2088905** | Hpca | -0.024762018 | -0.313330424 | 0.263806388 | 9.510211874 | -0.205090897 | 0.843636325 | 0.996633405 | -5.239022383 |
| **A_52_P309337** | Metap2 | 0.009142561 | -0.09751789 | 0.115803012 | 8.612758524 | 0.204867738 | 0.843803752 | 0.996633405 | -5.239060685 |
| **A_66_P140076** | NA | -0.034330434 | -0.43501049 | 0.366349622 | 13.23904576 | -0.204781492 | 0.843868462 | 0.996633405 | -5.239075476 |
| **A_55_P1964033** | Carm1 | 0.015516654 | -0.165838694 | 0.196872002 | 10.3832591 | 0.204492245 | 0.844085489 | 0.996633405 | -5.23912504 |
| **A_55_P2119458** | Slc25a44 | -0.008701882 | -0.110423904 | 0.093020139 | 8.014823953 | -0.20445953 | 0.844110037 | 0.996633405 | -5.239130641 |
| **A_55_P2043317** | Bag1 | -0.010711397 | -0.135973786 | 0.114550992 | 14.38504199 | -0.204378199 | 0.844171064 | 0.996633405 | -5.239144563 |
| **A_55_P2168098** | Plrg1 | -0.01085077 | -0.13778036 | 0.116078819 | 11.54932671 | -0.204318097 | 0.844216163 | 0.996633405 | -5.239154847 |
| **A_55_P2185397** | Copz2 | -0.031584829 | -0.401272992 | 0.338103334 | 9.092520724 | -0.204198299 | 0.844306057 | 0.996633405 | -5.239175337 |
| **A_66_P119376** | Kctd12 | 0.028142284 | -0.301523881 | 0.357808448 | 10.12610183 | 0.204030029 | 0.844432329 | 0.996633405 | -5.239204098 |
| **A_55_P2082478** | Kidins220 | 0.019048748 | -0.204344694 | 0.24244219 | 8.771606898 | 0.203800452 | 0.844604614 | 0.996633405 | -5.2392433 |
| **A_65_P00881** | NA | 0.008994323 | -0.09653353 | 0.114522175 | 8.734413917 | 0.203709138 | 0.844673143 | 0.996633405 | -5.239258881 |
| **A_52_P62617** | Gm9895 | -0.012103488 | -0.154114166 | 0.12990719 | 8.124351257 | -0.203703584 | 0.844677311 | 0.996633405 | -5.239259828 |
| **A_55_P2109114** | NA | -0.017237323 | -0.21949986 | 0.185025213 | 14.42874577 | -0.203687095 | 0.844689686 | 0.996633405 | -5.239262641 |
| **A_55_P2182497** | Maml3 | 0.013441712 | -0.144435036 | 0.171318461 | 9.104052102 | 0.203491162 | 0.844836734 | 0.996633405 | -5.239296045 |
| **A_55_P2120161** | Myh8 | -0.030196015 | -0.384872841 | 0.32448081 | 10.188607 | -0.20348198 | 0.844843625 | 0.996633405 | -5.23929761 |
| **A_55_P2109599** | NA | -0.015861707 | -0.202179586 | 0.170456172 | 12.24925598 | -0.203471941 | 0.84485116 | 0.996633405 | -5.239299321 |

| | | | | | | | | | |
|---|---|---|---|---|---|---|---|---|---|
| **A_55_P2043582** | 4930480E1 | 0.008604951 | -0.092481556 | 0.109691458 | 8.045548381 | 0.203453117 | 0.844865288 | 0.996633405 | -5.239302528 |
| **A_52_P195839** | Ctsc | -0.014789755 | -0.188563311 | 0.158983801 | 9.535633228 | -0.203416615 | 0.844892684 | 0.996633405 | -5.239308747 |
| **A_55_P2106911** | Wnt1 | 0.013561614 | -0.145798986 | 0.172922213 | 8.558058427 | 0.203394662 | 0.84490916 | 0.996633405 | -5.239312486 |
| **A_55_P1989846** | Tm6sf1 | 0.015502279 | -0.16668496 | 0.197689519 | 9.04856056 | 0.20336993 | 0.844927723 | 0.996633405 | -5.239316698 |
| **A_55_P2387379** | NA | -0.008715643 | -0.111157003 | 0.093725717 | 7.992858125 | -0.203344875 | 0.844946527 | 0.996633405 | -5.239320965 |
| **A_55_P2016604** | Glrx3 | 0.010705993 | -0.115131626 | 0.136543612 | 13.42543957 | 0.203341297 | 0.844949213 | 0.996633405 | -5.239321574 |
| **A_51_P176517** | Ntmt1 | -0.020957046 | -0.267427725 | 0.225513633 | 11.65031107 | -0.203223501 | 0.845037626 | 0.996633405 | -5.239341628 |
| **A_55_P2095360** | Tfpi | 0.011463726 | -0.123360903 | 0.146288355 | 9.638512816 | 0.203219643 | 0.845040521 | 0.996633405 | -5.239342284 |
| **A_52_P318955** | Ap3m1 | -0.009670637 | -0.123436493 | 0.10409522 | 9.662539364 | -0.203166576 | 0.845080352 | 0.996633405 | -5.239351314 |
| **A_51_P207591** | Anxa8 | 0.077677923 | -0.836761631 | 0.992117477 | 9.28879039 | 0.203026035 | 0.84518584 | 0.996633405 | -5.233197729 |
| **A_51_P432563** | Tlr9 | 0.01504019 | -0.16206883 | 0.192149211 | 10.1967793 | 0.202965294 | 0.845231433 | 0.996633405 | -5.239385544 |
| **A_51_P338776** | Dnajc8 | 0.014087627 | -0.151974768 | 0.180150022 | 10.85172443 | 0.202756921 | 0.845387844 | 0.996633405 | -5.239420944 |
| **A_52_P1037027** | Apela | 0.043384013 | -0.468211391 | 0.554979418 | 8.516985287 | 0.202680619 | 0.84544512 | 0.996633405 | -5.239433898 |
| **A_52_P655663** | H13 | 0.020155106 | -0.217531778 | 0.257841989 | 9.540968196 | 0.202669779 | 0.845453257 | 0.996633405 | -5.239435738 |
| **A_55_P2178023** | NA | -0.018861586 | -0.241366075 | 0.203642904 | 12.48746296 | -0.20260425 | 0.845502448 | 0.996633405 | -5.239446859 |
| **A_55_P2208682** | B230334C( | 0.019315986 | -0.208650941 | 0.247282914 | 8.422822363 | 0.20251359 | 0.845570504 | 0.996633405 | -5.239462238 |
| **A_51_P389957** | Rgs14 | -0.008652957 | -0.110775144 | 0.09346923 | 7.780202539 | -0.202513309 | 0.845570716 | 0.996633405 | -5.239462286 |
| **A_51_P502225** | Slc22a4 | 0.009764402 | -0.105517336 | 0.125046141 | 7.951172878 | 0.202439047 | 0.845626464 | 0.996633405 | -5.239474879 |
| **A_51_P160744** | Ndufb3 | 0.010114084 | -0.109301522 | 0.129529691 | 14.27720389 | 0.20242988 | 0.845633345 | 0.996633405 | -5.239476433 |
| **A_51_P293982** | Plekho2 | 0.008151628 | -0.088097882 | 0.104401139 | 8.375786599 | 0.202420724 | 0.845640219 | 0.996633405 | -5.239477985 |
| **A_55_P1972104** | Acsm3 | 0.008994783 | -0.097248784 | 0.11523835 | 7.705628164 | 0.202347182 | 0.845695428 | 0.996633405 | -5.23949045 |
| **A_51_P513568** | Stx11 | 0.011217326 | -0.121329238 | 0.14376389 | 7.97737601 | 0.202269311 | 0.845753887 | 0.996633405 | -5.239503644 |
| **A_51_P463452** | Acsl1 | -0.031781433 | -0.407668224 | 0.344105358 | 9.481570214 | -0.202081027 | 0.845895242 | 0.996633405 | -5.239535525 |
| **A_51_P492482** | Cstf2t | 0.01866868 | -0.202194986 | 0.239532345 | 11.47072616 | 0.202021903 | 0.84593963 | 0.996633405 | -5.23954553 |
| **A_52_P147666** | Slc30a7 | 0.009570993 | -0.103726417 | 0.122868403 | 8.496095809 | 0.201904566 | 0.846027725 | 0.996633405 | -5.239565377 |
| **A_51_P138060** | Rnf13 | -0.008068383 | -0.103587641 | 0.087450875 | 7.894066895 | -0.201885302 | 0.846042189 | 0.996633405 | -5.239568635 |
| **A_55_P2019615** | Neil1 | 0.01583858 | -0.171728558 | 0.203405717 | 9.041792988 | 0.201822052 | 0.846089677 | 0.996633405 | -5.239579328 |
| **A_52_P143477** | Tgoln1 | -0.014333248 | -0.184208556 | 0.15554206 | 10.56624839 | -0.201661732 | 0.846210049 | 0.996633405 | -5.239606416 |
| **A_55_P2013665** | Mybpc2 | -0.013990396 | -0.179840974 | 0.151860182 | 9.167220449 | -0.201614681 | 0.846245377 | 0.996633405 | -5.239614362 |
| **A_55_P2054817** | Rxfp1 | -0.019330743 | -0.24852069 | 0.209859204 | 8.493061079 | -0.20158681 | 0.846266304 | 0.996633405 | -5.239619068 |
| **A_66_P135987** | Ocrl | 0.017199947 | -0.186762603 | 0.221162498 | 10.85588667 | 0.2015514 | 0.846292892 | 0.996633405 | -5.239625046 |
| **A_55_P2047967** | Bfar | 0.033428152 | -0.363012507 | 0.429868812 | 13.88833785 | 0.201531675 | 0.846307702 | 0.996633405 | -5.239628376 |
| **A_55_P2064741** | Nmb | 0.016246574 | -0.176517074 | 0.209010222 | 9.646555534 | 0.201440035 | 0.846376512 | 0.996633405 | -5.23964384 |
| **A_55_P1980262** | Rtl8b | -0.018790686 | -0.241847824 | 0.204266451 | 13.66685889 | -0.201342589 | 0.846449682 | 0.996633405 | -5.239660277 |
| **A_55_P2003053** | Dct | -0.048285197 | -0.621560473 | 0.524990079 | 8.843283135 | -0.201307488 | 0.84647604 | 0.996633405 | -5.239666196 |
| **A_51_P324082** | Med1 | 0.015921347 | -0.173267089 | 0.205109783 | 10.8261875 | 0.201138104 | 0.846603233 | 0.996633405 | -5.239694744 |
| **A_51_P450518** | Hpf1 | 0.02177334 | -0.23696052 | 0.2805072 | 12.68059983 | 0.20113188 | 0.846607906 | 0.996633405 | -5.239695792 |
| **A_51_P514256** | Tubb2b | 0.020406441 | -0.222102036 | 0.262914917 | 11.08999987 | 0.201117319 | 0.846618841 | 0.996633405 | -5.239698245 |
| **A_51_P487869** | Cyp4f16 | -0.011101873 | -0.143056652 | 0.120852905 | 8.43186555 | -0.201085277 | 0.846642902 | 0.996633405 | -5.239703642 |
| **A_51_P224175** | 2210016F1 | 0.01070087 | -0.116492673 | 0.137894413 | 8.757139989 | 0.20107735 | 0.846648855 | 0.996633405 | -5.239704977 |
| **A_55_P2013730** | Islr2 | 0.008855751 | -0.096410212 | 0.114121713 | 7.797243111 | 0.201069665 | 0.846654626 | 0.996633405 | -5.239706272 |
| **A_55_P2164218** | NA | -0.015308115 | -0.197309681 | 0.166693451 | 13.21278604 | -0.201027614 | 0.846686204 | 0.996633405 | -5.239713353 |
| **A_55_P2047703** | Serac1 | 0.012117522 | -0.132005518 | 0.156240563 | 8.706663239 | 0.200950709 | 0.846743956 | 0.996633405 | -5.239726299 |
| **A_55_P1958517** | Parp6 | -0.019858066 | -0.256091356 | 0.216375224 | 10.54630206 | -0.200911589 | 0.846773334 | 0.996633405 | -5.239732883 |
| **A_55_P1962693** | Pla2g5 | 0.009139601 | -0.099701433 | 0.117980636 | 8.767297659 | 0.200698316 | 0.846933498 | 0.996633405 | -5.239768754 |
| **A_51_P263993** | Lipc | 0.008068837 | -0.088070098 | 0.104207773 | 7.649623969 | 0.200595315 | 0.847010854 | 0.996633405 | -5.239786065 |
| **A_55_P1989885** | Smarce1 | -0.020233882 | -0.261341578 | 0.220873814 | 11.86299755 | -0.200575221 | 0.847025945 | 0.996633405 | -5.239789441 |
| **A_66_P132446** | Usp15 | -0.009023514 | -0.116571512 | 0.098524485 | 7.818254196 | -0.200531442 | 0.847058824 | 0.996633405 | -5.239796795 |

| | | | | | | | | | |
|---|---|---|---|---|---|---|---|---|---|
| **A_51_P275976** | Dok1 | 0.014661985 | -0.160099591 | 0.189423561 | 8.566277247 | 0.200519193 | 0.847068024 | 0.996633405 | -5.239798853 |
| **A_55_P2113291** | Sik3 | -0.01554869 | -0.200935159 | 0.16983778 | 11.6329261 | -0.200458693 | 0.847113461 | 0.996633405 | -5.239809013 |
| **A_51_P234113** | Nod1 | 0.017363181 | -0.18969631 | 0.224422671 | 9.974252089 | 0.200421023 | 0.847141753 | 0.996633405 | -5.239815337 |
| **A_66_P118934** | Plaa | -0.019249329 | -0.248848611 | 0.210349954 | 10.43156858 | -0.200379918 | 0.847172626 | 0.996633405 | -5.239822237 |
| **A_55_P2151863** | NA | 0.010503131 | -0.114792201 | 0.135798463 | 12.1986641 | 0.200351699 | 0.84719382 | 0.996633405 | -5.239826973 |
| **A_66_P128997** | Pml | 0.011143755 | -0.121853025 | 0.144140536 | 8.801619208 | 0.200262473 | 0.847260835 | 0.996633405 | -5.239841944 |
| **A_51_P161054** | Itgb6 | 0.008852023 | -0.096831754 | 0.114535799 | 7.739301992 | 0.200190435 | 0.847314942 | 0.996633405 | -5.239854026 |
| **A_55_P2186005** | Sphk1 | 0.008558917 | -0.093641201 | 0.110759034 | 8.001831057 | 0.200159657 | 0.847338059 | 0.996633405 | -5.239859186 |
| **A_55_P2031871** | Plekhh3 | 0.010944362 | -0.119751075 | 0.141639799 | 10.7247033 | 0.200142427 | 0.847351001 | 0.996633405 | -5.239862075 |
| **A_51_P245789** | Pcolce2 | 0.018570118 | -0.203282759 | 0.240422996 | 7.964780127 | 0.200059298 | 0.847413439 | 0.996633405 | -5.239876009 |
| **A_55_P1973279** | NA | 0.016788418 | -0.183982351 | 0.217559187 | 16.08496231 | 0.199856569 | 0.847565716 | 0.996633405 | -5.239909965 |
| **A_51_P322542** | Bbx | -0.020366034 | -0.263932975 | 0.223200907 | 9.908978343 | -0.199846822 | 0.847573037 | 0.996633405 | -5.239911596 |
| **A_55_P2046550** | Adamts8 | -0.010324382 | -0.133862503 | 0.11321374 | 7.888324732 | -0.199743292 | 0.847650805 | 0.996633405 | -5.239928923 |
| **A_51_P185869** | Hbq1a | 0.016028843 | -0.175816988 | 0.207874675 | 7.967147675 | 0.199691211 | 0.847689926 | 0.996633405 | -5.239937636 |
| **A_55_P1961466** | Dct | -0.063808002 | -0.827513333 | 0.699897328 | 9.405714461 | -0.199690984 | 0.847690097 | 0.996633405 | -5.239937674 |
| **A_55_P2085984** | Igf1 | -0.017215064 | -0.223290676 | 0.188860549 | 9.371277849 | -0.199660045 | 0.847713338 | 0.996633405 | -5.239942849 |
| **A_51_P442284** | Dnajc12 | -0.014631643 | -0.190072161 | 0.160808875 | 9.531940624 | -0.199329846 | 0.847961387 | 0.996633405 | -5.239998029 |
| **A_52_P198435** | Rasgrp3 | 0.020712533 | -0.22773745 | 0.269162515 | 10.33551867 | 0.19925231 | 0.848019635 | 0.996633405 | -5.240010974 |
| **A_55_P2129309** | Foxo3 | 0.015072084 | -0.165740433 | 0.195884601 | 8.284083166 | 0.199229644 | 0.848036663 | 0.996633405 | -5.240014757 |
| **A_55_P2006754** | Mfsd12 | 0.010264142 | -0.112875405 | 0.133403688 | 11.56152894 | 0.199220591 | 0.848043465 | 0.996633405 | -5.240016268 |
| **A_55_P2102798** | Prl7a2 | 0.008059304 | -0.088762013 | 0.10488062 | 7.656389428 | 0.198946206 | 0.848249605 | 0.996633405 | -5.240062028 |
| **A_55_P2150962** | Cpne7 | -0.010295299 | -0.1339897 | 0.113399102 | 8.252315737 | -0.198928975 | 0.848262551 | 0.996633405 | -5.240064899 |
| **A_55_P2068027** | NA | -0.013673521 | -0.178012925 | 0.150665882 | 8.507060465 | -0.198860168 | 0.848314246 | 0.996633405 | -5.240076364 |
| **A_66_P119489** | Spef1l | 0.012388407 | -0.136556218 | 0.161333032 | 8.011166321 | 0.198792394 | 0.848365167 | 0.996633405 | -5.240087652 |
| **A_55_P2085060** | Vgf | -0.008818734 | -0.114846522 | 0.097209054 | 7.691246768 | -0.198790513 | 0.84836658 | 0.996633405 | -5.240087965 |
| **A_55_P2135049** | NA | 0.021464924 | -0.236613751 | 0.279543598 | 14.91120778 | 0.198786261 | 0.848369775 | 0.996633405 | -5.240088674 |
| **A_52_P514061** | Padi4 | -0.021557431 | -0.280775009 | 0.237660147 | 8.863726062 | -0.198765813 | 0.848385138 | 0.996633405 | -5.240092079 |
| **A_55_P2108808** | Tubgcp4 | 0.011171819 | -0.123168401 | 0.14551204 | 8.531424732 | 0.198759082 | 0.848390195 | 0.996633405 | -5.240093199 |
| **A_51_P408071** | Kntc1 | 0.013056267 | -0.144011389 | 0.170123923 | 9.668918895 | 0.198674163 | 0.848453999 | 0.996633405 | -5.240107336 |
| **A_51_P496054** | Zcchc14 | 0.010163188 | -0.112107659 | 0.132434035 | 9.047292838 | 0.198662625 | 0.848462668 | 0.996633405 | -5.240109256 |
| **A_55_P2055708** | Vmn1r48 | -0.009899681 | -0.129072745 | 0.109273382 | 7.790749679 | -0.198541933 | 0.848553351 | 0.996633405 | -5.240129336 |
| **A_52_P73369** | Hnrnpu | 0.01171186 | -0.129287317 | 0.152711037 | 12.69438405 | 0.198526479 | 0.848564963 | 0.996633405 | -5.240131907 |
| **A_51_P489996** | Dst | 0.014958976 | -0.165179847 | 0.195097799 | 8.284220779 | 0.198474031 | 0.848604371 | 0.996633405 | -5.240140628 |
| **A_55_P1970399** | NA | -0.024747324 | -0.322801495 | 0.273306847 | 16.44510552 | -0.198445917 | 0.848625496 | 0.996633405 | -5.240145303 |
| **A_65_P09177** | Strn3 | -0.013245166 | -0.172772611 | 0.14628228 | 10.05410152 | -0.198440864 | 0.848629292 | 0.996633405 | -5.240146142 |
| **A_55_P2091616** | Krt222 | 0.031092618 | -0.343395015 | 0.405580252 | 12.42136171 | 0.198439876 | 0.848630035 | 0.996633405 | -5.240146307 |
| **A_52_P110291** | Phf12 | 0.01231501 | -0.136079054 | 0.160709073 | 9.28346859 | 0.198347787 | 0.848699231 | 0.996633405 | -5.240161612 |
| **A_55_P2011862** | Mybpc3 | 0.014857867 | -0.164199767 | 0.193915501 | 17.25565355 | 0.198322849 | 0.84871797 | 0.996633405 | -5.240165756 |
| **A_55_P1982019** | Rbfox2 | 0.021463135 | -0.237278378 | 0.280204647 | 11.54243117 | 0.198260487 | 0.848764829 | 0.996633405 | -5.240176116 |
| **A_55_P2000487** | Srebf2 | 0.008369416 | -0.092563076 | 0.109301908 | 7.803338398 | 0.198186137 | 0.848820698 | 0.996633405 | -5.240188462 |
| **A_51_P103929** | Met | 0.008196086 | -0.090730042 | 0.107122213 | 7.786870447 | 0.198017975 | 0.848947063 | 0.996633405 | -5.214995783 |
| **A_55_P2097616** | Stxbp1 | 0.010021245 | -0.110974011 | 0.131016502 | 8.455861637 | 0.19795318 | 0.848995755 | 0.996633405 | -5.240227119 |
| **A_52_P286520** | Slc23a2 | -0.010381831 | -0.135786186 | 0.115022525 | 8.27828219 | -0.197865668 | 0.849061518 | 0.996633405 | -5.240241628 |
| **A_66_P101862** | NA | 0.008510459 | -0.094328213 | 0.111349132 | 8.008146943 | 0.19779061 | 0.849117924 | 0.996633405 | -5.240254068 |
| **A_66_P124732** | Zfp606 | -0.011625017 | -0.152119276 | 0.128869242 | 9.106113631 | -0.197762594 | 0.849138978 | 0.996633405 | -5.24025871 |
| **A_55_P2146546** | NA | -0.008244502 | -0.107926948 | 0.091437945 | 7.808290733 | -0.197676416 | 0.849203743 | 0.996633405 | -5.240272986 |
| **A_51_P293938** | Rasl11b | -0.017426108 | -0.228184215 | 0.193331998 | 11.75084402 | -0.197617435 | 0.849248068 | 0.996633405 | -5.240282752 |
| **A_55_P2057503** | Arl4c | 0.019040973 | -0.21133699 | 0.249418937 | 10.42840475 | 0.197541043 | 0.849305479 | 0.996633405 | -5.240295397 |

| | | | | | | | | | |
|---|---|---|---|---|---|---|---|---|---|
| **A_55_P2056654** | Kif22 | 0.01189485 | -0.132114379 | 0.155904079 | 11.62344837 | 0.197413918 | 0.849401021 | 0.996633405 | -5.24031643 |
| **A_66_P132777** | Cep68 | -0.011096328 | -0.145482259 | 0.123289603 | 10.66717943 | -0.197348862 | 0.849449915 | 0.996633405 | -5.240327188 |
| **A_55_P1973046** | Cttnbp2nl | 0.01191583 | -0.132397648 | 0.156229307 | 9.7963201 | 0.197345182 | 0.849452681 | 0.996633405 | -5.240327797 |
| **A_55_P1998209** | NA | -0.017306209 | -0.227137231 | 0.192524812 | 15.77512628 | -0.197124857 | 0.849618276 | 0.996633405 | -5.240364205 |
| **A_55_P2106106** | C5ar2 | -0.011860216 | -0.155737651 | 0.132017218 | 8.339465348 | -0.197019432 | 0.849697516 | 0.996633405 | -5.240381612 |
| **A_51_P380005** | Galnt3 | -0.008415941 | -0.110547812 | 0.09371593 | 7.710923006 | -0.19694752 | 0.849751569 | 0.996633405 | -5.24039348 |
| **A_52_P489119** | Nol6 | -0.013666839 | -0.179541809 | 0.152208132 | 9.431792621 | -0.196922955 | 0.849770033 | 0.996633405 | -5.240397534 |
| **A_55_P2096883** | Steap3 | 0.018591117 | -0.207279814 | 0.244462048 | 9.601072449 | 0.196722607 | 0.849920627 | 0.996633405 | -5.240430572 |
| **A_55_P2002115** | NA | -0.0183605 | -0.241699413 | 0.204978414 | 15.71814743 | -0.196484921 | 0.850099297 | 0.996633405 | -5.240469726 |
| **A_55_P2160081** | Ccdc28b | -0.010086193 | -0.132797106 | 0.112624721 | 8.360847894 | -0.196450531 | 0.850125149 | 0.996633405 | -5.240475387 |
| **A_51_P469008** | Ehd1 | 0.014263049 | -0.159284863 | 0.187810961 | 13.00696763 | 0.196427424 | 0.850142519 | 0.996633405 | -5.24047919 |
| **A_55_P2093563** | Ufm1 | 0.012224097 | -0.136531651 | 0.160979845 | 11.24633298 | 0.196404833 | 0.850159502 | 0.996633405 | -5.240482908 |
| **A_51_P301736** | Ppp2r2a | -0.008986698 | -0.118360442 | 0.100387046 | 11.58931437 | -0.196379516 | 0.850178533 | 0.996633405 | -5.240487074 |
| **A_51_P184385** | Pla2g12b | 0.008152773 | -0.091078551 | 0.107384096 | 7.701697757 | 0.196365722 | 0.850188903 | 0.996633405 | -5.240489343 |
| **A_55_P2013223** | S100z | -0.008164252 | -0.107552179 | 0.091223674 | 7.678000011 | -0.196332379 | 0.850213968 | 0.996633405 | -5.240494829 |
| **A_51_P236197** | Polr1a | 0.018392091 | -0.205620847 | 0.242405029 | 11.28868321 | 0.196230782 | 0.850290344 | 0.996633405 | -5.240511538 |
| **A_55_P2128511** | Ociad2 | -0.013438218 | -0.177132107 | 0.150255672 | 8.597549245 | -0.196208734 | 0.85030692 | 0.996633405 | -5.240515163 |
| **A_55_P1965338** | Anxa8 | 0.024423971 | -0.273193261 | 0.322041202 | 8.065574918 | 0.196140522 | 0.850358199 | 0.996633405 | -5.240526375 |
| **A_51_P343889** | Slc35f6 | -0.012440024 | -0.164045564 | 0.139165515 | 8.335245771 | -0.196117023 | 0.850375866 | 0.996633405 | -5.240530236 |
| **A_52_P629333** | Map3k21 | 0.008135937 | -0.091027829 | 0.107299704 | 7.708765622 | 0.196093732 | 0.850393375 | 0.996633405 | -5.240534063 |
| **A_51_P208240** | Tnfsf14 | -0.009651032 | -0.12728473 | 0.107982665 | 7.687545143 | -0.196088065 | 0.850397636 | 0.996633405 | -5.240534995 |
| **A_55_P2185330** | NA | 0.013719789 | -0.153579028 | 0.181018605 | 8.736003049 | 0.196003436 | 0.85046126 | 0.996633405 | -5.240548896 |
| **A_55_P2362635** | Slc9a3 | -0.008296954 | -0.109534494 | 0.092940586 | 7.749358929 | -0.195878255 | 0.850555373 | 0.996633405 | -5.240569448 |
| **A_51_P395736** | Srp72 | 0.017113005 | -0.191776316 | 0.226002325 | 10.39247047 | 0.195802919 | 0.850612012 | 0.996633405 | -5.240581811 |
| **A_55_P2419711** | 4932412D2 | -0.01914981 | -0.253003679 | 0.21470406 | 8.211423029 | -0.19571722 | 0.850676445 | 0.996633405 | -5.240595868 |
| **A_55_P2029046** | Fgd3 | -0.008066135 | -0.106589973 | 0.090457702 | 7.675562798 | -0.195674091 | 0.850708872 | 0.996633405 | -5.24060294 |
| **A_66_P135194** | Yrdc | 0.011141587 | -0.125000101 | 0.147283276 | 8.416014984 | 0.195598298 | 0.850765858 | 0.996633405 | -5.240615365 |
| **A_52_P150565** | Usp39 | 0.010594009 | -0.11901995 | 0.140207967 | 8.603662806 | 0.195351922 | 0.850951107 | 0.996633405 | -5.24065572 |
| **A_55_P1987186** | Ttll9 | 0.008076522 | -0.090992648 | 0.107145691 | 7.675614704 | 0.194847557 | 0.851330367 | 0.996633405 | -5.240738176 |
| **A_55_P2068137** | NA | 0.015955911 | -0.179907074 | 0.211818896 | 11.38384108 | 0.194705566 | 0.851437146 | 0.996633405 | -5.240761351 |
| **A_55_P1961531** | Nemp1 | 0.016688011 | -0.188227444 | 0.221603466 | 9.520897628 | 0.194643091 | 0.851484129 | 0.996633405 | -5.240771543 |
| **A_51_P473229** | Zbtb7b | -0.011034827 | -0.146571749 | 0.124502095 | 8.542810006 | -0.194588447 | 0.851525224 | 0.996633405 | -5.240780455 |
| **A_55_P1983468** | Cflar | 0.010776022 | -0.121585642 | 0.143137685 | 8.964185635 | 0.194583213 | 0.85152916 | 0.996633405 | -5.240781308 |
| **A_55_P2036495** | NA | 0.023659062 | -0.267032669 | 0.314350793 | 13.06502767 | 0.194524352 | 0.851573426 | 0.996633405 | -5.240790904 |
| **A_55_P2017826** | Myb | 0.014983984 | -0.169164364 | 0.199132333 | 8.083890976 | 0.194477162 | 0.851608915 | 0.996633405 | -5.240798596 |
| **A_51_P101635** | Stub1 | -0.018977957 | -0.25226991 | 0.214313995 | 12.77612754 | -0.194428015 | 0.851645877 | 0.996633405 | -5.240806604 |
| **A_55_P2127683** | 4933434E2 | -0.013217819 | -0.17579078 | 0.149355143 | 9.318487202 | -0.19432139 | 0.851726067 | 0.996633405 | -5.240823972 |
| **A_55_P2109717** | Kif20b | 0.019360766 | -0.218866146 | 0.257587678 | 9.856560587 | 0.194240974 | 0.851786548 | 0.996633405 | -5.240837065 |
| **A_55_P2163018** | NA | -0.011356508 | -0.151113231 | 0.128400216 | 11.06164766 | -0.194214305 | 0.851806606 | 0.996633405 | -5.240841405 |
| **A_66_P121770** | Ilf2 | 0.011687218 | -0.132204788 | 0.155579225 | 10.81464711 | 0.194125959 | 0.851873052 | 0.996633405 | -5.240855781 |
| **A_52_P676798** | Gigyf1 | -0.007939632 | -0.105698174 | 0.089818909 | 8.768809032 | -0.19411308 | 0.851882738 | 0.996633405 | -5.240857876 |
| **A_52_P633489** | C1qtnf4 | 0.01996396 | -0.225886421 | 0.265814341 | 9.674034649 | 0.194081859 | 0.851906221 | 0.996633405 | -5.240862954 |
| **A_55_P2271742** | NA | 0.018057803 | -0.204628569 | 0.240744175 | 8.477220944 | 0.19381188 | 0.852109285 | 0.996633405 | -5.240906834 |
| **A_55_P2107120** | Kansl2 | -0.01483576 | -0.197806607 | 0.168135087 | 10.11628773 | -0.193792524 | 0.852123844 | 0.996633405 | -5.240909978 |
| **A_55_P2027337** | NA | -0.03696169 | -0.493226959 | 0.41930358 | 10.91579452 | -0.193617023 | 0.852255854 | 0.996633405 | -5.240938468 |
| **A_51_P263302** | Rnf24 | 0.01179928 | -0.1338618 | 0.157460359 | 8.63150942 | 0.19360702 | 0.852263379 | 0.996633405 | -5.240940091 |
| **A_55_P2109943** | Rps15a | 0.027068527 | -0.307176269 | 0.361313323 | 15.52852829 | 0.193557096 | 0.852300932 | 0.996633405 | -5.24094819 |
| **A_55_P2111380** | Ctnnd2 | 0.00921139 | -0.104558029 | 0.12298081 | 7.699236879 | 0.193512389 | 0.852334561 | 0.996633405 | -5.240955441 |

| | | | | | | | | | |
|---|---|---|---|---|---|---|---|---|---|
| **A_66_P120277** | 9330182L0 | 0.009760132 | -0.110827891 | 0.130348154 | 7.84615722 | 0.193446391 | 0.852384207 | 0.996633405 | -5.240966142 |
| **A_55_P1978716** | Zim1 | 0.013266966 | -0.150652591 | 0.177186523 | 7.932093938 | 0.193441641 | 0.85238778 | 0.996633405 | -5.240966912 |
| **A_52_P71105** | Sertad3 | -0.019421913 | -0.259517799 | 0.220673974 | 9.274405089 | -0.193337637 | 0.852466017 | 0.996633405 | -5.240983769 |
| **A_55_P2159600** | Herc1 | -0.020600075 | -0.275347169 | 0.234147019 | 11.3061382 | -0.193271889 | 0.852515475 | 0.996633405 | -5.24099442 |
| **A_51_P159402** | Gcat | 0.015173335 | -0.172468616 | 0.202815286 | 9.925804548 | 0.193268128 | 0.852518305 | 0.996633405 | -5.240995029 |
| **A_55_P2172274** | Smc2 | 0.021182937 | -0.240908711 | 0.283274586 | 11.31690631 | 0.193171092 | 0.852591302 | 0.996633405 | -5.241010742 |
| **A_55_P2102828** | NA | -0.008190369 | -0.109538567 | 0.093157828 | 7.661072625 | -0.193150834 | 0.852606542 | 0.996633405 | -5.241014022 |
| **A_55_P1986123** | Sccpdh | -0.013911379 | -0.186103115 | 0.158280356 | 13.03393578 | -0.193093221 | 0.852649884 | 0.996633405 | -5.241023347 |
| **A_52_P277104** | Bank1 | -0.009449319 | -0.126468639 | 0.107570001 | 7.701733635 | -0.192997673 | 0.852721764 | 0.996633405 | -5.241038805 |
| **A_52_P375873** | Srp9 | 0.013287755 | -0.151291796 | 0.177867306 | 11.89518792 | 0.192967812 | 0.852744229 | 0.996633405 | -5.241043635 |
| **A_51_P480904** | Blmh | -0.0128156 | -0.171550863 | 0.145919663 | 11.09619766 | -0.192963278 | 0.85274764 | 0.996633405 | -5.241044368 |
| **A_51_P227777** | Top2b | 0.01068258 | -0.12164361 | 0.143008769 | 12.85034213 | 0.192947637 | 0.852759407 | 0.996633405 | -5.241046897 |
| **A_55_P1984098** | Mst1 | 0.008561817 | -0.097528122 | 0.114651755 | 7.699825188 | 0.192886065 | 0.852805729 | 0.996633405 | -5.241056852 |
| **A_51_P137336** | Cdh1 | 0.008950522 | -0.102088122 | 0.119989165 | 7.712787299 | 0.192656349 | 0.852978555 | 0.996633405 | -5.206984903 |
| **A_55_P2125457** | NA | 0.019062332 | -0.217489663 | 0.255614326 | 14.09223339 | 0.192601001 | 0.853020197 | 0.996633405 | -5.241102901 |
| **A_55_P2063626** | Matk | 0.011350489 | -0.129506235 | 0.152207212 | 8.427291248 | 0.192595487 | 0.853024345 | 0.996633405 | -5.241103791 |
| **A_51_P146655** | Ptgr2 | -0.014504005 | -0.194508957 | 0.165500948 | 11.76038986 | -0.192580631 | 0.853035522 | 0.996633405 | -5.241106189 |
| **A_55_P2015878** | Clps | -0.012129399 | -0.162665317 | 0.138406519 | 8.04866917 | -0.192578714 | 0.853036965 | 0.996633405 | -5.241106498 |
| **A_52_P150683** | NA | 0.008988458 | -0.102591288 | 0.120568205 | 7.757643333 | 0.192534685 | 0.853070091 | 0.996633405 | -5.241113604 |
| **A_52_P127925** | Tfec | -0.00773027 | -0.10369463 | 0.088234089 | 7.666289596 | -0.19252797 | 0.853075144 | 0.996633405 | -5.241114688 |
| **A_52_P607507** | Olfm4 | 0.008100681 | -0.092482158 | 0.108683519 | 7.658770613 | 0.192489369 | 0.853104186 | 0.996633405 | -5.241120915 |
| **A_55_P2070194** | Caprin1 | 0.015812197 | -0.180528759 | 0.212153153 | 11.85984728 | 0.192482142 | 0.853109624 | 0.996633405 | -5.241122081 |
| **A_55_P1994947** | NA | 0.022904397 | -0.261791758 | 0.307600551 | 15.80473628 | 0.192285433 | 0.853257629 | 0.996633405 | -5.241153799 |
| **A_55_P2090441** | NA | -0.01349281 | -0.181526046 | 0.154540426 | 8.108979335 | -0.191918281 | 0.853533895 | 0.996633405 | -5.241212913 |
| **A_55_P2182106** | Rpl39 | -0.014416676 | -0.19408134 | 0.165247989 | 15.81686713 | -0.191783653 | 0.853635203 | 0.996633405 | -5.241234562 |
| **A_55_P2021055** | Elp5 | 0.010271891 | -0.117748325 | 0.138292107 | 11.65702646 | 0.191770133 | 0.853645377 | 0.996633405 | -5.241236735 |
| **A_51_P502132** | Mmp23 | 0.020518712 | -0.235259694 | 0.276297117 | 9.930410125 | 0.191732327 | 0.853673826 | 0.996633405 | -5.241242811 |
| **A_52_P424767** | Rbbp4 | -0.011696068 | -0.157513213 | 0.134121077 | 8.215448256 | -0.191708089 | 0.853692066 | 0.996633405 | -5.241246706 |
| **A_52_P59264** | Sncaip | 0.020759289 | -0.238079765 | 0.279598342 | 9.942095913 | 0.191686619 | 0.853708222 | 0.996633405 | -5.241250155 |
| **A_52_P598912** | Erc1 | 0.01100761 | -0.126330539 | 0.148345759 | 8.339822483 | 0.19156271 | 0.853801468 | 0.996633405 | -5.241270057 |
| **A_51_P273679** | Dmgdh | -0.0078851 | -0.106272501 | 0.0905023 | 7.626371043 | -0.191547666 | 0.85381279 | 0.996633405 | -5.241272473 |
| **A_52_P250578** | Tcaf1 | 0.010348473 | -0.11883796 | 0.139534907 | 9.065599105 | 0.191455792 | 0.85388193 | 0.996633405 | -5.241287219 |
| **A_55_P2098697** | Tnfaip2 | -0.008950523 | -0.120728923 | 0.102827878 | 8.306965468 | -0.191381362 | 0.853937944 | 0.996633405 | -5.241299161 |
| **A_52_P354123** | Rhebl1 | -0.011082686 | -0.149556388 | 0.127391015 | 8.51221627 | -0.191287624 | 0.854008489 | 0.996633405 | -5.241314194 |
| **A_55_P2069765** | Kiss1 | 0.00918218 | -0.105549777 | 0.123914137 | 7.685947568 | 0.191280427 | 0.854013906 | 0.996633405 | -5.241315348 |
| **A_55_P2018706** | Cep19 | -0.015195627 | -0.205322852 | 0.174931598 | 10.90195543 | -0.191022029 | 0.854208379 | 0.996633405 | -5.241356749 |
| **A_51_P220343** | Ccn4 | 0.024655544 | -0.284138803 | 0.333449891 | 11.84801162 | 0.190833395 | 0.854350354 | 0.996633405 | -5.241386938 |
| **A_55_P2103756** | Hic2 | 0.011123274 | -0.128192023 | 0.150438571 | 8.4372422 | 0.19082838 | 0.854354129 | 0.996633405 | -5.24138774 |
| **A_52_P483280** | Lamp2 | 0.010078096 | -0.116171021 | 0.136327212 | 12.27836832 | 0.190791603 | 0.85438181 | 0.996633405 | -5.241393622 |
| **A_55_P2121936** | Wiz | 0.010713004 | -0.123589135 | 0.145015143 | 11.37740372 | 0.190650284 | 0.854488178 | 0.996633405 | -5.241416214 |
| **A_55_P1992031** | Dydc2 | -0.009174213 | -0.124235833 | 0.105887407 | 7.728470444 | -0.190566896 | 0.854550945 | 0.996633405 | -5.241429538 |
| **A_55_P2108012** | Fam78b | -0.012760592 | -0.172832584 | 0.147311401 | 8.526761439 | -0.190530537 | 0.854578313 | 0.996633405 | -5.241435345 |
| **A_55_P2169417** | BC021767 | -0.010370781 | -0.140492416 | 0.119750855 | 7.79396313 | -0.190489516 | 0.85460919 | 0.996633405 | -5.241441896 |
| **A_51_P403814** | Slc4a5 | -0.007707359 | -0.104430522 | 0.089015804 | 7.70323579 | -0.190451425 | 0.854637862 | 0.996633405 | -5.241447977 |
| **A_55_P2095563** | Slc8a2 | -0.015694058 | -0.212649087 | 0.181260972 | 9.017429002 | -0.190448388 | 0.854640148 | 0.996633405 | -5.241448462 |
| **A_51_P145662** | Clec4g | -0.011290519 | -0.152998837 | 0.130417799 | 8.728174871 | -0.19042663 | 0.854656526 | 0.996633405 | -5.241451935 |
| **A_65_P19784** | Asb7 | -0.014748395 | -0.199905385 | 0.170408595 | 9.934248294 | -0.190376698 | 0.854694112 | 0.996633405 | -5.241459905 |
| **A_51_P389506** | NA | -0.034239859 | -0.464618771 | 0.396139053 | 12.1120322 | -0.19014728 | 0.854866808 | 0.996633405 | -5.241496494 |

| | | | | | | | | | |
|---|---|---|---|---|---|---|---|---|---|
| **A_55_P2157384** | Xlr5b | 0.007915995 | -0.09158697 | 0.10741896 | 7.672435149 | 0.190142245 | 0.854870598 | 0.996633405 | -5.241497297 |
| **A_52_P360112** | Clip2 | -0.013184476 | -0.179028236 | 0.152659284 | 8.146909852 | -0.190008431 | 0.854971332 | 0.996633405 | -5.241518617 |
| **A_55_P1983426** | Abhd14a | -0.010451531 | -0.141950878 | 0.121047815 | 12.28469455 | -0.189961441 | 0.855006706 | 0.996633405 | -5.241526101 |
| **A_51_P472518** | Capn1 | -0.013780078 | -0.187203771 | 0.159643614 | 8.220219946 | -0.189911992 | 0.855043932 | 0.996633405 | -5.241533974 |
| **A_55_P2279997** | 4930549C1 | 0.011470394 | -0.132922136 | 0.155862924 | 7.839401792 | 0.189864051 | 0.855080023 | 0.996633405 | -5.241541605 |
| **A_66_P129444** | Pnkp | -0.015396465 | -0.209213716 | 0.178420785 | 8.773493597 | -0.189861859 | 0.855081673 | 0.996633405 | -5.241541954 |
| **A_51_P423976** | Crem | -0.009564087 | -0.130069729 | 0.110941555 | 8.687660974 | -0.189690371 | 0.855210776 | 0.996633405 | -5.241569236 |
| **A_66_P139546** | Igfbp6 | -0.118716649 | -1.614746088 | 1.377312791 | 10.51779684 | -0.189662129 | 0.855232038 | 0.996633405 | -5.241573726 |
| **A_51_P335419** | Csl | -0.02446932 | -0.332834525 | 0.283895886 | 9.969904679 | -0.189655593 | 0.855236958 | 0.996633405 | -5.241574765 |
| **A_55_P1994524** | Tpd52l2 | -0.009356739 | -0.127274572 | 0.108561093 | 12.00686074 | -0.189650582 | 0.855240731 | 0.996633405 | -5.241575562 |
| **A_55_P2032643** | Ajap1 | -0.011537873 | -0.15701113 | 0.133935384 | 9.51817854 | -0.189562191 | 0.855307278 | 0.996633405 | -5.241589611 |
| **A_55_P2020796** | Man1b1 | -0.015359042 | -0.209038912 | 0.178320828 | 11.52221876 | -0.189534716 | 0.855327963 | 0.996633405 | -5.241593977 |
| **A_55_P2061110** | Snapin | -0.0082961 | -0.112948426 | 0.096356226 | 9.274244914 | -0.189467284 | 0.855378732 | 0.996633405 | -5.241604689 |
| **A_55_P2019083** | Rps6kb1 | 0.013662452 | -0.158727902 | 0.186052807 | 12.84185283 | 0.189419564 | 0.85541466 | 0.996633405 | -5.241612268 |
| **A_55_P2142439** | Echdc2 | 0.012025792 | -0.139715361 | 0.163766945 | 9.295686111 | 0.189417217 | 0.855416427 | 0.996633405 | -5.24161264 |
| **A_52_P466147** | Rarres2 | -0.036006518 | -0.490589173 | 0.418576137 | 11.76006033 | -0.189311673 | 0.855495891 | 0.996633405 | -5.241629395 |
| **A_55_P2176240** | Pdxk | -0.013221503 | -0.180190246 | 0.15374724 | 8.418098946 | -0.189258233 | 0.855536127 | 0.996633405 | -5.241637875 |
| **A_55_P2069730** | NA | 0.00764342 | -0.088890681 | 0.104177521 | 7.834828148 | 0.18924138 | 0.855548816 | 0.996633405 | -5.241640548 |
| **A_51_P246317** | Mt2 | 0.016841673 | -0.202760268 | 0.236443614 | 13.86683087 | 0.190509312 | 0.855568988 | 0.996633405 | -5.162724916 |
| **A_51_P122582** | Daglb | 0.010101166 | -0.117498047 | 0.13770038 | 9.374489102 | 0.189205023 | 0.85557619 | 0.996633405 | -5.241646316 |
| **A_55_P2136204** | Rnf38 | 0.018838468 | -0.219188413 | 0.256865349 | 11.55727147 | 0.189159741 | 0.855610285 | 0.996633405 | -5.241653497 |
| **A_55_P1986630** | Gm3627 | -0.008525304 | -0.116248559 | 0.099197952 | 7.794271185 | -0.189151386 | 0.855616576 | 0.996633405 | -5.241654822 |
| **A_55_P1976127** | Ect2 | 0.024254073 | -0.282224159 | 0.330732304 | 10.46474325 | 0.189144698 | 0.855621611 | 0.996633405 | -5.241655882 |
| **A_55_P2002079** | NA | -0.015321488 | -0.208949371 | 0.178306395 | 15.21058842 | -0.189122051 | 0.855638663 | 0.996633405 | -5.241659473 |
| **A_55_P1978675** | NA | -0.00867057 | -0.118258932 | 0.100917791 | 7.64906652 | -0.189100366 | 0.855654991 | 0.996633405 | -5.241662911 |
| **A_66_P137462** | Nsdhl | -0.011100659 | -0.151436875 | 0.129235557 | 8.703701117 | -0.189054984 | 0.855689161 | 0.996633405 | -5.241670104 |
| **A_51_P464630** | Rad17 | 0.00894524 | -0.10417354 | 0.12206402 | 10.86217664 | 0.189002006 | 0.855729052 | 0.996633405 | -5.2416785 |
| **A_52_P670978** | Nkain1 | 0.011468412 | -0.133611294 | 0.156548117 | 8.425813237 | 0.188932094 | 0.855781693 | 0.996633405 | -5.241689575 |
| **A_51_P394676** | Mef2d | -0.014333742 | -0.19569091 | 0.167023425 | 12.96281063 | -0.18890089 | 0.85580519 | 0.996633405 | -5.241694517 |
| **A_55_P2069729** | Ormdl2 | 0.011412599 | -0.133049561 | 0.155874759 | 10.78354648 | 0.188816349 | 0.855868848 | 0.996633405 | -5.241707902 |
| **A_55_P1963339** | Washc5 | 0.009283239 | -0.108248735 | 0.126815212 | 11.93571787 | 0.188778538 | 0.85589732 | 0.996633405 | -5.241713887 |
| **A_55_P2014427** | Il17re | 0.017943801 | -0.209293678 | 0.245181279 | 7.940036484 | 0.188731161 | 0.855932995 | 0.996633405 | -5.241721384 |
| **A_55_P2133047** | Tollip | -0.009814334 | -0.134229178 | 0.11460051 | 8.428574944 | -0.188537507 | 0.85607882 | 0.996633405 | -5.241752008 |
| **A_52_P121** | Osgepl1 | 0.017618349 | -0.205743101 | 0.2409798 | 10.05563815 | 0.188523775 | 0.856089161 | 0.996633405 | -5.241754179 |
| **A_55_P2179378** | Zkscan8 | -0.015314616 | -0.209507832 | 0.1788786 | 10.11606318 | -0.188486912 | 0.856116921 | 0.996633405 | -5.241760004 |
| **A_52_P23379** | Depdc7 | -0.009290578 | -0.127108703 | 0.108527546 | 8.030578423 | -0.188468939 | 0.856130456 | 0.996633405 | -5.241762844 |
| **A_55_P1978236** | Mrps7 | -0.01322756 | -0.181027506 | 0.154572385 | 13.15994432 | -0.188407016 | 0.856177087 | 0.996633405 | -5.241772627 |
| **A_55_P2033407** | Meg3 | 0.025797188 | -0.301487428 | 0.353081804 | 8.196325965 | 0.188389159 | 0.856190535 | 0.996633405 | -5.241775447 |
| **A_55_P2042273** | Pbrm1 | -0.017601641 | -0.241010639 | 0.205807358 | 9.298660569 | -0.1883049 | 0.856253988 | 0.996633405 | -5.241788752 |
| **A_52_P476904** | Chd6 | 0.008261606 | -0.096606156 | 0.113129368 | 8.09365094 | 0.188291888 | 0.856263787 | 0.996633405 | -5.241790806 |
| **A_55_P2171912** | Gja3 | -0.013459108 | -0.184545492 | 0.157627275 | 8.544772165 | -0.188022568 | 0.856466612 | 0.996633405 | -5.241833291 |
| **A_51_P490171** | Nubp2 | 0.013943003 | -0.163360038 | 0.191246044 | 9.538052609 | 0.187953006 | 0.856519002 | 0.996633405 | -5.241844254 |
| **A_51_P434101** | Herpud1 | 0.013629708 | -0.159806081 | 0.187065497 | 11.96997362 | 0.187826545 | 0.856614245 | 0.996633405 | -5.241864175 |
| **A_55_P2094731** | Sox13 | 0.031523189 | -0.369652225 | 0.432698604 | 9.574482966 | 0.187804059 | 0.85663118 | 0.996633405 | -5.241867716 |
| **A_55_P2057370** | Slc22a19 | 0.008129869 | -0.095361107 | 0.111620846 | 7.652249405 | 0.187754443 | 0.856668549 | 0.996633405 | -5.241875527 |
| **A_55_P2167763** | NA | -0.00765873 | -0.105153835 | 0.089836375 | 7.791781573 | -0.18775136 | 0.856670871 | 0.996633405 | -5.241876012 |
| **A_66_P125173** | Aip | -0.032352823 | -0.444261629 | 0.379555984 | 10.03617838 | -0.187724186 | 0.856691338 | 0.996633405 | -5.241880289 |
| **A_55_P2013414** | Dmwd | -0.010287117 | -0.14130191 | 0.120727675 | 13.28561189 | -0.187664659 | 0.856736172 | 0.996633405 | -5.241889657 |

| | | | | | | | | | |
|---|---|---|---|---|---|---|---|---|---|
| **A_55_P1979997** | Spaca9 | 0.008717706 | -0.102309416 | 0.119744829 | 7.777671995 | 0.187664571 | 0.856736238 | 0.996633405 | -5.241889671 |
| **A_55_P2064299** | Msrb1 | -0.011218474 | -0.154116427 | 0.13167948 | 8.137860201 | -0.187636314 | 0.856757521 | 0.996633405 | -5.241894116 |
| **A_51_P172231** | Gsdmd | 0.01107755 | -0.130062862 | 0.152217961 | 8.479968393 | 0.187586448 | 0.85679508 | 0.996633405 | -5.24190196 |
| **A_55_P2091297** | NA | -0.009749692 | -0.134059226 | 0.114559841 | 7.843479673 | -0.187454386 | 0.856894549 | 0.996633405 | -5.241922723 |
| **A_51_P317433** | Mesp2 | -0.009473721 | -0.130337697 | 0.111390254 | 7.805304792 | -0.187341011 | 0.856979946 | 0.996633405 | -5.241940536 |
| **A_51_P317225** | Cmtm2a | 0.008906164 | -0.104725668 | 0.122537997 | 8.331021392 | 0.187326763 | 0.856990678 | 0.996633405 | -5.241942774 |
| **A_55_P2111018** | NA | -0.013848275 | -0.190566792 | 0.162870242 | 8.857190396 | -0.187293521 | 0.857015718 | 0.996633405 | -5.241947994 |
| **A_55_P2150343** | Piezo1 | 0.020878018 | -0.245709866 | 0.287465902 | 12.2445175 | 0.187179377 | 0.857101697 | 0.996633405 | -5.241965913 |
| **A_55_P2165046** | Klhl22 | 0.016506609 | -0.194264451 | 0.227277668 | 10.32514068 | 0.187178519 | 0.857102343 | 0.996633405 | -5.241966048 |
| **A_51_P267388** | Dxo | 0.014035388 | -0.165281767 | 0.193352542 | 9.161190228 | 0.187073258 | 0.857181633 | 0.996633405 | -5.241982563 |
| **A_51_P292943** | Ell | 0.012618693 | -0.148672054 | 0.173909439 | 10.02915382 | 0.186988131 | 0.857245758 | 0.996633405 | -5.241995912 |
| **A_55_P2177026** | Mafg | 0.015488736 | -0.182530627 | 0.213508099 | 9.083446965 | 0.186946548 | 0.857277082 | 0.996633405 | -5.242002431 |
| **A_51_P481210** | Ofd1 | -0.008559936 | -0.117999228 | 0.100879356 | 9.734676293 | -0.18694178 | 0.857280674 | 0.996633405 | -5.242003178 |
| **A_55_P2082604** | Ndor1 | -0.014535919 | -0.200454075 | 0.171382236 | 16.56470465 | -0.186865802 | 0.857337908 | 0.996633405 | -5.242015084 |
| **A_55_P2124543** | Gnptg | -0.011793048 | -0.162632026 | 0.139045931 | 9.068482877 | -0.186862264 | 0.857340574 | 0.996633405 | -5.242015639 |
| **A_51_P201884** | Dsp | 0.022665146 | -0.267299433 | 0.312629725 | 8.888887987 | 0.186819715 | 0.857372626 | 0.996633405 | -5.242022304 |
| **A_55_P1953136** | NA | -0.013197423 | -0.182227023 | 0.155832178 | 14.28171808 | -0.186610247 | 0.857530426 | 0.996633405 | -5.242055097 |
| **A_55_P2048684** | Rpl27 | -0.011703974 | -0.161616696 | 0.138208749 | 14.22733484 | -0.18659671 | 0.857540624 | 0.996633405 | -5.242057215 |
| **A_55_P2015862** | Erg | -0.010716343 | -0.147987287 | 0.126554602 | 8.637392359 | -0.18658516 | 0.857549325 | 0.996633405 | -5.242059022 |
| **A_55_P1999473** | Ppp1r26 | 0.009701757 | -0.114576744 | 0.133980257 | 8.932887089 | 0.186579318 | 0.857553726 | 0.996633405 | -5.242059936 |
| **A_51_P315890** | Kcnk6 | -0.009423222 | -0.130191925 | 0.11134548 | 9.773965267 | -0.186489407 | 0.857621462 | 0.996633405 | -5.242073999 |
| **A_51_P482043** | Epm2aip1 | -0.011753821 | -0.162520206 | 0.139012563 | 10.60942456 | -0.186330393 | 0.85774126 | 0.996633405 | -5.242098853 |
| **A_52_P575078** | Babam1 | -0.010444851 | -0.144528621 | 0.123638918 | 12.38762926 | -0.186180921 | 0.857853874 | 0.996633405 | -5.242122196 |
| **A_52_P337716** | NA | -0.012773071 | -0.176822513 | 0.151276371 | 11.34730213 | -0.186092852 | 0.857920228 | 0.996633405 | -5.242135942 |
| **A_65_P10180** | Rad23b | -0.015307135 | -0.212001301 | 0.18138703 | 10.77639396 | -0.185999413 | 0.857990629 | 0.996633405 | -5.242150518 |
| **A_55_P2064962** | Pdpn | 0.008881793 | -0.10527647 | 0.123040057 | 7.835324577 | 0.185952683 | 0.858025838 | 0.996633405 | -5.242157806 |
| **A_65_P20799** | Abi2 | 0.008739582 | -0.103592226 | 0.121071389 | 8.719857414 | 0.185950362 | 0.858027587 | 0.996633405 | -5.242158167 |
| **A_55_P2119219** | NA | -0.011582419 | -0.160572788 | 0.13740795 | 13.38645335 | -0.185801928 | 0.858139427 | 0.996633405 | -5.242181302 |
| **A_55_P1984118** | Gfra4 | -0.01687731 | -0.2339964 | 0.20024178 | 8.294296485 | -0.185786583 | 0.858150989 | 0.996633405 | -5.242183693 |
| **A_55_P2053206** | Kdm4c | -0.01132797 | -0.157119753 | 0.134463812 | 9.422044865 | -0.185706964 | 0.858210981 | 0.996633405 | -5.242196094 |
| **A_66_P105270** | Hao1 | -0.00796831 | -0.110541537 | 0.094604917 | 7.656694047 | -0.185669825 | 0.858238965 | 0.996633405 | -5.242201876 |
| **A_55_P1982916** | Tspoap1 | 0.008071733 | -0.095883211 | 0.112026678 | 7.796916422 | 0.185579835 | 0.858306774 | 0.996633405 | -5.242215884 |
| **A_52_P675530** | Fat4 | -0.016133317 | -0.22406004 | 0.191793406 | 8.661759306 | -0.185448147 | 0.858406004 | 0.996633405 | -5.242236369 |
| **A_55_P2161829** | Hmgb1-rs1 | 0.014618754 | -0.173831772 | 0.20306928 | 13.11606139 | 0.185405297 | 0.858438294 | 0.996633405 | -5.242243032 |
| **A_51_P336827** | Cyb5b | -0.012321788 | -0.171217451 | 0.146573875 | 9.253122621 | -0.185340705 | 0.858486967 | 0.996633405 | -5.242253072 |
| **A_55_P1967295** | Pemt | 0.014778565 | -0.175836951 | 0.205394082 | 8.719533803 | 0.185303304 | 0.858515151 | 0.996633405 | -5.242258884 |
| **A_55_P2066976** | Acsm1 | -0.008182442 | -0.113756823 | 0.097391939 | 7.718802068 | -0.18523947 | 0.858563254 | 0.996633405 | -5.242268801 |
| **A_55_P2036285** | Elp4 | -0.01340729 | -0.186425535 | 0.159610955 | 8.686010726 | -0.18520735 | 0.858587458 | 0.996633405 | -5.24227379 |
| **A_51_P366061** | Fscn1 | 0.016727362 | -0.199145222 | 0.232599946 | 9.600035493 | 0.185199196 | 0.858593603 | 0.996633405 | -5.242275056 |
| **A_55_P2181597** | Mfap2 | 0.023640693 | -0.28161039 | 0.328891776 | 9.668590944 | 0.185102431 | 0.858666524 | 0.996633405 | -5.24229008 |
| **A_55_P2107830** | Gemin2 | -0.010264632 | -0.142826766 | 0.122297502 | 8.114125767 | -0.185068724 | 0.858691926 | 0.996633405 | -5.242295312 |
| **A_52_P140881** | Slc26a10 | -0.0139776 | -0.194634115 | 0.166678914 | 8.937132987 | -0.184921805 | 0.858802645 | 0.996633405 | -5.242318103 |
| **A_55_P1999082** | Clec14a | -0.013451748 | -0.187386531 | 0.160483034 | 8.610626244 | -0.184842329 | 0.858862541 | 0.996633405 | -5.242330425 |
| **A_51_P337523** | Adcy2 | 0.00769864 | -0.091899404 | 0.107296685 | 7.688195381 | 0.184744849 | 0.858936006 | 0.996633405 | -5.242345531 |
| **A_51_P198835** | Dnajb9 | -0.021694212 | -0.30239098 | 0.259002555 | 10.07997739 | -0.184720718 | 0.858954192 | 0.996633405 | -5.242349269 |
| **A_55_P2000039** | Prf1 | 0.00767447 | -0.09167614 | 0.10702508 | 8.019073947 | 0.184623505 | 0.859027458 | 0.996633405 | -5.242364324 |
| **A_52_P664506** | Abhd2 | -0.00765545 | -0.106822629 | 0.09151173 | 7.778731534 | -0.184506588 | 0.859115576 | 0.996633405 | -5.24238242 |
| **A_55_P2095926** | NA | -0.020305482 | -0.283408518 | 0.242797553 | 9.000125463 | -0.184457613 | 0.859152488 | 0.996633405 | -5.242389996 |

| | | | | | | | | | |
|---|---|---|---|---|---|---|---|---|---|
| **A_52_P283524** | Rcan3 | -0.012376064 | -0.172796319 | 0.14804419 | 8.564207681 | -0.184387933 | 0.859205006 | 0.996633405 | -5.242400773 |
| **A_51_P499020** | Fbp2 | -0.007891267 | -0.110216722 | 0.094434189 | 7.670177898 | -0.184319874 | 0.859256303 | 0.996633405 | -5.242411295 |
| **A_55_P1986049** | Fip1l1 | 0.013415375 | -0.160803768 | 0.187634518 | 8.533734801 | 0.184041636 | 0.859466022 | 0.996633405 | -5.24245427 |
| **A_55_P1981381** | Lamc1 | -0.013086963 | -0.183174528 | 0.157000603 | 13.93901759 | -0.183897334 | 0.859574793 | 0.996633405 | -5.242476533 |
| **A_55_P1974827** | Pde1b | -0.009398079 | -0.131558004 | 0.112761846 | 8.788301837 | -0.183873633 | 0.859592659 | 0.996633405 | -5.242480188 |
| **A_55_P1956267** | Syt11 | 0.018538873 | -0.2224715 | 0.259549247 | 9.300882853 | 0.18384708 | 0.859612674 | 0.996633405 | -5.242484283 |
| **A_55_P1953003** | Atp6v0a4 | 0.009387069 | -0.1127406 | 0.131514738 | 7.860399612 | 0.183706732 | 0.859718469 | 0.996633405 | -5.242505913 |
| **A_55_P2055257** | Maoa | 0.01591884 | -0.197889679 | 0.229727359 | 8.376464939 | 0.18494967 | 0.859727026 | 0.996633405 | -5.163563094 |
| **A_55_P1968668** | Gpatch2l | 0.0129198 | -0.155188962 | 0.181028561 | 8.875836115 | 0.183685369 | 0.859734573 | 0.996633405 | -5.242509204 |
| **A_55_P1977776** | Rs5-8s1 | -0.071082088 | -0.996397841 | 0.854233665 | 11.13522974 | -0.183602823 | 0.859796798 | 0.996633405 | -5.242521918 |
| **A_55_P2020612** | Arl4d | -0.018918883 | -0.265203471 | 0.227365705 | 9.041735872 | -0.183597767 | 0.85980061 | 0.996633405 | -5.242522696 |
| **A_55_P2033876** | NA | -0.02083342 | -0.292148122 | 0.250481283 | 12.87424219 | -0.183525478 | 0.859855104 | 0.996633405 | -5.242533824 |
| **A_55_P2031547** | Vmn2r121 | -0.007498843 | -0.10525822 | 0.090260534 | 7.667202399 | -0.183334819 | 0.859998834 | 0.996633405 | -5.242563154 |
| **A_55_P1964483** | Cyp2c37 | 0.00755032 | -0.090928972 | 0.106029611 | 7.679562686 | 0.183243913 | 0.860067366 | 0.996633405 | -5.242577128 |
| **A_55_P2189201** | 5830426C0 | 0.009340497 | -0.11252691 | 0.131207903 | 8.464653386 | 0.183185686 | 0.860111263 | 0.996633405 | -5.242586075 |
| **A_52_P123693** | E130304I02 | -0.024654911 | -0.346354892 | 0.297045069 | 11.07125844 | -0.183173037 | 0.860120799 | 0.996633405 | -5.242588019 |
| **A_55_P2040936** | Eng | 0.020793148 | -0.250522281 | 0.292108577 | 11.63189414 | 0.183170231 | 0.860122915 | 0.996633405 | -5.24258845 |
| **A_52_P445307** | Cwc15 | 0.012538932 | -0.151134436 | 0.1762123 | 13.08084764 | 0.1831014 | 0.860174807 | 0.996633405 | -5.242599021 |
| **A_55_P2113975** | Psg20 | 0.00737005 | -0.088835619 | 0.10357572 | 7.641364032 | 0.18309603 | 0.860178855 | 0.996633405 | -5.242599846 |
| **A_55_P1960053** | Hvcn1 | -0.008356189 | -0.117484763 | 0.100772385 | 8.294335033 | -0.183011729 | 0.860242411 | 0.996633405 | -5.242612788 |
| **A_51_P341746** | Mpl | 0.012523618 | -0.151094286 | 0.176141522 | 8.046602462 | 0.182939767 | 0.860296666 | 0.996633405 | -5.242623831 |
| **A_55_P1971269** | NA | 0.017215461 | -0.207709673 | 0.242140596 | 12.28577524 | 0.182932053 | 0.860302482 | 0.996633405 | -5.242625015 |
| **A_55_P2128672** | Ikbip | 0.021010188 | -0.253613358 | 0.295633734 | 10.42103817 | 0.18285267 | 0.860362332 | 0.996633405 | -5.242637191 |
| **A_55_P2057268** | Magi1 | -0.00986885 | -0.13887367 | 0.119135969 | 10.29280906 | -0.18283939 | 0.860372344 | 0.996633405 | -5.242639228 |
| **A_55_P1966114** | Epn2 | -0.010749899 | -0.151291354 | 0.129791555 | 11.82417602 | -0.182813847 | 0.860391603 | 0.996633405 | -5.242643145 |
| **A_55_P2077353** | Ceacam12 | 0.007559264 | -0.091295733 | 0.106414261 | 7.643507472 | 0.182763742 | 0.86042938 | 0.996633405 | -5.242650826 |
| **A_55_P1972054** | NA | -0.011348782 | -0.159825304 | 0.13712774 | 8.563174562 | -0.182684036 | 0.860489476 | 0.996633405 | -5.242663041 |
| **A_55_P2057441** | NA | 0.007773022 | -0.093928603 | 0.109474646 | 15.01902316 | 0.182671636 | 0.860498825 | 0.996633405 | -5.242664941 |
| **A_55_P2092296** | Fbxo2 | 0.007898079 | -0.095499115 | 0.111295273 | 7.966165306 | 0.182566822 | 0.860577854 | 0.996633405 | -5.242680994 |
| **A_55_P2018417** | Osbpl3 | 0.015459061 | -0.186931955 | 0.217850076 | 10.35708927 | 0.182558056 | 0.860584463 | 0.996633405 | -5.242682337 |
| **A_51_P425373** | Smg9 | -0.011849531 | -0.167092364 | 0.143393301 | 9.951377853 | -0.182431045 | 0.86068023 | 0.996633405 | -5.242701777 |
| **A_55_P2077448** | Epas1 | -0.010438069 | -0.147303502 | 0.126427365 | 11.38492108 | -0.182278529 | 0.860795232 | 0.996633405 | -5.242725103 |
| **A_55_P2047639** | Cubn | -0.021492934 | -0.30354197 | 0.260556103 | 8.681878923 | -0.182129464 | 0.860907635 | 0.996633405 | -5.242747883 |
| **A_52_P642453** | Obscn | -0.021600365 | -0.305063991 | 0.26186326 | 10.13581835 | -0.182126396 | 0.860909948 | 0.996633405 | -5.242748352 |
| **A_51_P106527** | Mcrip2 | 0.023961173 | -0.290582078 | 0.338504424 | 13.45661498 | 0.182069347 | 0.860952967 | 0.996633405 | -5.242757065 |
| **A_66_P102706** | 2700012I20 | -0.013212783 | -0.186661989 | 0.160236423 | 8.504424197 | -0.182066949 | 0.860954775 | 0.996633405 | -5.242757431 |
| **A_55_P2124341** | Lysmd1 | -0.010142923 | -0.143294225 | 0.12300838 | 10.99617642 | -0.182065151 | 0.860956131 | 0.996633405 | -5.242757706 |
| **A_55_P2022574** | Gpank1 | 0.011687633 | -0.141768542 | 0.165143808 | 11.49479131 | 0.182033503 | 0.860979997 | 0.996633405 | -5.242762539 |
| **A_55_P1984023** | Arrb2 | 0.008972086 | -0.108830316 | 0.126774489 | 8.637095877 | 0.182032285 | 0.860980915 | 0.996633405 | -5.242762724 |
| **A_52_P485007** | Abca2 | -0.009112513 | -0.128824685 | 0.110599658 | 8.475256247 | -0.181931962 | 0.861056567 | 0.996633405 | -5.242778037 |
| **A_55_P2098076** | Trim25 | -0.01453122 | -0.205498789 | 0.176436349 | 12.94554977 | -0.181866032 | 0.861106285 | 0.996633405 | -5.242788096 |
| **A_51_P143901** | Tm9sf2 | -0.012789587 | -0.181048853 | 0.15546968 | 12.4161641 | -0.181671434 | 0.861253035 | 0.996633405 | -5.242817764 |
| **A_55_P1966908** | Zfp40 | 0.010516384 | -0.127974037 | 0.149006804 | 8.661841935 | 0.181491305 | 0.86138888 | 0.996633405 | -5.242845199 |
| **A_55_P2059890** | Fdx1 | 0.009819424 | -0.119539511 | 0.139178358 | 12.7608081 | 0.181425657 | 0.86143839 | 0.996633405 | -5.24285519 |
| **A_51_P170807** | Map3k6 | -0.013694377 | -0.194231701 | 0.166842947 | 9.522243205 | -0.181294408 | 0.861537376 | 0.996633405 | -5.242875156 |
| **A_52_P669005** | Lrat | -0.007272852 | -0.103174744 | 0.08862904 | 7.694402776 | -0.181253638 | 0.861568124 | 0.996633405 | -5.242881355 |
| **A_55_P2007713** | Tspan4 | 0.015371886 | -0.187337055 | 0.218080826 | 10.68854899 | 0.181243889 | 0.861575477 | 0.996633405 | -5.242882837 |
| **A_55_P2105052** | Tfb2m | 0.01530342 | -0.186508842 | 0.217115683 | 10.722474 | 0.181238344 | 0.861579659 | 0.996633405 | -5.24288368 |

| | | | | | | | | | |
|---|---|---|---|---|---|---|---|---|---|
| **A_55_P2043377** | NA | 0.009035998 | -0.110173365 | 0.128245362 | 8.107641716 | 0.181165253 | 0.861634785 | 0.996633405 | -5.242894789 |
| **A_55_P2012181** | Eps15l1 | 0.013656442 | -0.166547037 | 0.19385992 | 11.56853993 | 0.181127136 | 0.861663533 | 0.996633405 | -5.242900581 |
| **A_55_P2151026** | Ccdc50 | -0.019429423 | -0.275910244 | 0.237051398 | 8.348379958 | -0.181056509 | 0.861716802 | 0.996633405 | -5.242911309 |
| **A_51_P109050** | Paqr4 | 0.016275861 | -0.198726839 | 0.23127856 | 10.02543055 | 0.180929415 | 0.861812662 | 0.996633405 | -5.242930604 |
| **A_55_P2011410** | Pdzk1 | 0.021144355 | -0.258273241 | 0.30056195 | 7.911780811 | 0.180863036 | 0.861862729 | 0.996633405 | -5.242940676 |
| **A_55_P2078633** | C4b | 0.027240958 | -0.33288 | 0.387361915 | 8.067492349 | 0.180793622 | 0.861915086 | 0.996633405 | -5.242951205 |
| **A_55_P2079167** | Dnm1 | -0.015494381 | -0.220497957 | 0.189509196 | 9.69257188 | -0.180643324 | 0.862028453 | 0.996633405 | -5.242973988 |
| **A_51_P342567** | Akap12 | -0.01958736 | -0.278831552 | 0.239656833 | 12.99331272 | -0.180582637 | 0.862074229 | 0.996633405 | -5.242983183 |
| **A_55_P1976869** | Hsdl2 | -0.015414715 | -0.219512368 | 0.188682939 | 11.87519407 | -0.180512222 | 0.862127345 | 0.996633405 | -5.242993847 |
| **A_55_P2068356** | 1700016K1 | 0.008088852 | -0.099019286 | 0.11519699 | 7.977031547 | 0.180498483 | 0.862137708 | 0.996633405 | -5.242995927 |
| **A_55_P2125013** | H4c11 | 0.024779224 | -0.303349775 | 0.352908222 | 11.12745927 | 0.180489615 | 0.862144398 | 0.996633405 | -5.24299727 |
| **A_51_P210340** | 2610306M( | -0.014222939 | -0.202583303 | 0.174137425 | 9.456500542 | -0.180471633 | 0.862157961 | 0.996633405 | -5.242999992 |
| **A_52_P119672** | Rock1 | 0.013388981 | -0.164040953 | 0.190818916 | 9.145506208 | 0.18035566 | 0.862245444 | 0.996633405 | -5.243017543 |
| **A_55_P2004711** | Cep95 | 0.029129148 | -0.35696097 | 0.415219267 | 9.912648729 | 0.180321799 | 0.862270986 | 0.996633405 | -5.243022665 |
| **A_55_P2157307** | NA | -0.030290706 | -0.432055294 | 0.371473883 | 12.33869628 | -0.180196714 | 0.862365346 | 0.996633405 | -5.243041579 |
| **A_55_P1974976** | Sprr2a2 | 0.008553845 | -0.104901843 | 0.122009534 | 15.96121212 | 0.180195627 | 0.862366165 | 0.996633405 | -5.243041743 |
| **A_66_P105791** | Prkca | 0.014609594 | -0.179183768 | 0.208402956 | 11.02604284 | 0.180180751 | 0.862377387 | 0.996633405 | -5.243043992 |
| **A_55_P2052261** | Zfp280c | 0.01603053 | -0.196622103 | 0.228683164 | 8.945803387 | 0.180171569 | 0.862384314 | 0.996633405 | -5.24304538 |
| **A_55_P2170474** | Yipf2 | 0.017078794 | -0.209583241 | 0.243740829 | 8.996205815 | 0.180089148 | 0.86244649 | 0.996633405 | -5.243057834 |
| **A_55_P2126814** | NA | -0.033076902 | -0.472065761 | 0.405911957 | 10.95154021 | -0.180086211 | 0.862448706 | 0.996633405 | -5.243058277 |
| **A_52_P547187** | Tab2 | 0.013760294 | -0.168896309 | 0.196416898 | 11.3706595 | 0.180053462 | 0.862473412 | 0.996633405 | -5.243063224 |
| **A_55_P2144476** | NA | -0.01029196 | -0.146942119 | 0.126358198 | 8.618541155 | -0.180010191 | 0.862506055 | 0.996633405 | -5.243069759 |
| **A_55_P2070009** | Krt10 | 0.00743766 | -0.091343475 | 0.106218795 | 7.721592639 | 0.179958117 | 0.86254534 | 0.996633405 | -5.243077621 |
| **A_51_P464738** | Slc2a1 | 0.027803843 | -0.341885476 | 0.397493163 | 13.3856344 | 0.179753378 | 0.862699798 | 0.996633405 | -5.243108511 |
| **A_55_P2108983** | NA | 0.015052582 | -0.185322761 | 0.215427926 | 15.24177029 | 0.179546053 | 0.862856216 | 0.996633405 | -5.243139756 |
| **A_52_P276935** | Dnajb6 | -0.014591573 | -0.208912398 | 0.179729251 | 11.65952654 | -0.179470018 | 0.862913581 | 0.996633405 | -5.243151205 |
| **A_55_P2166572** | Strbp | -0.012560453 | -0.179902734 | 0.154781828 | 9.539786585 | -0.179394342 | 0.862970678 | 0.996633405 | -5.243162596 |
| **A_55_P2030506** | Rp9 | 0.009885242 | -0.121842603 | 0.141613087 | 11.86711747 | 0.179357213 | 0.862998692 | 0.996633405 | -5.243168184 |
| **A_55_P1977649** | Hoxd8 | -0.008847729 | -0.1267508 | 0.109055343 | 7.838308928 | -0.179355959 | 0.862999638 | 0.996633405 | -5.243168372 |
| **A_55_P2048928** | Cbfb | 0.012782125 | -0.157642285 | 0.183206534 | 12.23378156 | 0.179258756 | 0.863072978 | 0.996633405 | -5.243182994 |
| **A_51_P362661** | Spin4 | 0.00865382 | -0.106766232 | 0.124073871 | 8.314022389 | 0.17919905 | 0.863118028 | 0.996633405 | -5.243191971 |
| **A_52_P622230** | Gm4890 | 0.00943635 | -0.116423343 | 0.135296042 | 7.927895527 | 0.179195254 | 0.863120891 | 0.996633405 | -5.243192541 |
| **A_51_P412966** | Lrrn1 | -0.009990575 | -0.143384567 | 0.123403417 | 7.742484907 | -0.179004258 | 0.863265005 | 0.996633405 | -5.243221238 |
| **A_51_P336403** | Emc7 | 0.010943446 | -0.135268066 | 0.157154957 | 12.30256878 | 0.178888182 | 0.863352591 | 0.996633405 | -5.243238663 |
| **A_55_P1971174** | Cd1d2 | -0.01334144 | -0.191607086 | 0.164924205 | 8.50943538 | -0.178872756 | 0.863364231 | 0.996633405 | -5.243240978 |
| **A_55_P2112479** | Nudt16l1 | 0.017490382 | -0.2164732 | 0.251453964 | 10.46174348 | 0.17867351 | 0.86351458 | 0.996633405 | -5.243270861 |
| **A_55_P2054311** | M6pr | 0.012266154 | -0.151820107 | 0.176352415 | 9.574309663 | 0.178667403 | 0.863519188 | 0.996633405 | -5.243271776 |
| **A_52_P430247** | Nacc1 | -0.017670039 | -0.254133113 | 0.218793035 | 10.05020042 | -0.178600767 | 0.863569473 | 0.996633405 | -5.243281762 |
| **A_52_P316644** | Bet1 | 0.013852506 | -0.171649544 | 0.199354556 | 10.24528051 | 0.178479678 | 0.863660849 | 0.996633405 | -5.2432999 |
| **A_55_P2161126** | NA | 0.014658209 | -0.181667453 | 0.210983871 | 8.918811719 | 0.178448526 | 0.863684358 | 0.996633405 | -5.243304564 |
| **A_52_P469502** | Cda | -0.012876864 | -0.185394673 | 0.159640945 | 8.684817104 | -0.178396078 | 0.863723937 | 0.996633405 | -5.243312414 |
| **A_55_P2117664** | Rnf165 | 0.014466666 | -0.179368057 | 0.208301388 | 8.822297253 | 0.178379936 | 0.863736118 | 0.996633405 | -5.24331483 |
| **A_55_P2032498** | Arpin | 0.010707189 | -0.132893989 | 0.154308366 | 9.694206853 | 0.178207752 | 0.863866059 | 0.996633405 | -5.243340585 |
| **A_51_P296796** | Nt5dc2 | 0.027104345 | -0.336531378 | 0.390740067 | 9.43295912 | 0.178148226 | 0.863910982 | 0.996633405 | -5.243349484 |
| **A_51_P430033** | Ccdc13 | 0.008199791 | -0.101817784 | 0.118217366 | 7.679443663 | 0.178135243 | 0.86392078 | 0.996633405 | -5.243351424 |
| **A_51_P391934** | Cpb1 | -0.009258585 | -0.133698685 | 0.115181516 | 7.7358522 | -0.177825231 | 0.864154749 | 0.996633405 | -5.243397715 |
| **A_51_P209818** | Prtn3 | -0.007169386 | -0.103559403 | 0.089220632 | 7.69608581 | -0.177770224 | 0.864196265 | 0.996633405 | -5.24340592 |
| **A_55_P1956472** | Cybrd1 | 0.010434382 | -0.12994881 | 0.150817574 | 7.838159156 | 0.177648179 | 0.864288379 | 0.996633405 | -5.243424117 |

| | | | | | | | | | |
|---|---|---|---|---|---|---|---|---|---|
| **A_55_P2091402** | Kpna2 | 0.01295224 | -0.161543811 | 0.187448291 | 13.24927397 | 0.177406039 | 0.864471141 | 0.996633405 | -5.243460182 |
| **A_51_P293989** | Senp6 | 0.009452562 | -0.117918594 | 0.136823717 | 8.449274307 | 0.177373019 | 0.864496065 | 0.996633405 | -5.243465097 |
| **A_51_P292490** | Ttbk2 | -0.01740562 | -0.251959331 | 0.217148091 | 9.544606959 | -0.177360258 | 0.864505697 | 0.996633405 | -5.243466996 |
| **A_51_P462814** | Sars2 | 0.015756523 | -0.196578622 | 0.228091668 | 9.025780359 | 0.177356712 | 0.864508373 | 0.996633405 | -5.243467523 |
| **A_55_P2274378** | AW549542 | 0.00863687 | -0.107813928 | 0.125087668 | 7.774100909 | 0.177265028 | 0.864577577 | 0.996633405 | -5.243481162 |
| **A_55_P2076545** | F5 | -0.008691435 | -0.125990037 | 0.108607167 | 7.829624885 | -0.177095607 | 0.864705461 | 0.996633405 | -5.243506348 |
| **A_55_P1989816** | Zbtb7b | 0.015729134 | -0.19659154 | 0.228049808 | 12.06375336 | 0.17706049 | 0.864731969 | 0.996633405 | -5.243511565 |
| **A_52_P485971** | Scap | 0.010966602 | -0.137109935 | 0.159043138 | 11.72381322 | 0.17700884 | 0.864770958 | 0.996633405 | -5.243519237 |
| **A_51_P394172** | Ces1c | 0.007560317 | -0.094568441 | 0.109689074 | 7.677881876 | 0.176929835 | 0.864830596 | 0.996633405 | -5.243530968 |
| **A_55_P2109342** | Rnps1 | 0.012322433 | -0.154182979 | 0.178827845 | 10.45105944 | 0.176879393 | 0.864868673 | 0.996633405 | -5.243538455 |
| **A_52_P565575** | Arhgap20 | -0.011143832 | -0.161895983 | 0.13960832 | 8.293676198 | -0.176677056 | 0.865021416 | 0.996633405 | -5.243568467 |
| **A_55_P1965459** | Zfp521 | 0.012760858 | -0.1598895 | 0.185411216 | 8.53862559 | 0.176653203 | 0.865039424 | 0.996633405 | -5.243572002 |
| **A_55_P1981105** | Ap1b1 | -0.009574846 | -0.139148404 | 0.119998712 | 12.39063379 | -0.176613767 | 0.865069194 | 0.996633405 | -5.243577847 |
| **A_51_P402160** | Zfp750 | -0.007405657 | -0.107824526 | 0.093013211 | 7.647584176 | -0.176261486 | 0.865335147 | 0.996633405 | -5.243629999 |
| **A_55_P2111039** | NA | 0.014837105 | -0.186379094 | 0.216053305 | 10.34003584 | 0.176236302 | 0.86535416 | 0.996633405 | -5.243633723 |
| **A_55_P2028660** | Dusp12 | 0.008642927 | -0.108586225 | 0.125872079 | 8.226488763 | 0.176211544 | 0.865372852 | 0.996633405 | -5.243637384 |
| **A_55_P2147871** | NA | -0.016238114 | -0.236635149 | 0.204158921 | 8.691408956 | -0.176091733 | 0.865463309 | 0.996633405 | -5.243655093 |
| **A_51_P216075** | Ddost | -0.021909293 | -0.319332189 | 0.275513602 | 12.09415066 | -0.176060976 | 0.86548653 | 0.996633405 | -5.243659637 |
| **A_55_P2073015** | Lin37 | -0.012520349 | -0.182526979 | 0.157486282 | 9.679352338 | -0.176019059 | 0.865518177 | 0.996633405 | -5.243665828 |
| **A_51_P109171** | Os9 | -0.012444561 | -0.181475834 | 0.156586713 | 11.4217841 | -0.175963113 | 0.865560418 | 0.996633405 | -5.24367409 |
| **A_55_P1987504** | Ccnc | -0.009615293 | -0.140232276 | 0.121001689 | 8.371264164 | -0.175943015 | 0.865575592 | 0.996633405 | -5.243677057 |
| **A_51_P172573** | Sod2 | -0.014865368 | -0.21680682 | 0.187076083 | 15.57498261 | -0.175937872 | 0.865579475 | 0.996633405 | -5.243677817 |
| **A_52_P80965** | Sephs1 | 0.009510272 | -0.119706516 | 0.138727061 | 8.205536326 | 0.175907008 | 0.865602778 | 0.996633405 | -5.243682373 |
| **A_55_P1979674** | Ikzf4 | 0.010535259 | -0.132691431 | 0.15376195 | 8.575339457 | 0.175804668 | 0.865680048 | 0.996633405 | -5.243697474 |
| **A_51_P188271** | Cd248 | 0.015819851 | -0.199283862 | 0.230923563 | 11.44650061 | 0.175777632 | 0.865700462 | 0.996633405 | -5.243701462 |
| **A_55_P2093226** | Rnf121 | 0.012454777 | -0.156908728 | 0.181818281 | 8.59951232 | 0.175762106 | 0.865712185 | 0.996633405 | -5.243703752 |
| **A_51_P494863** | Vmac | 0.009481255 | -0.119491902 | 0.138454412 | 8.796693794 | 0.175701571 | 0.865757892 | 0.996633405 | -5.243712678 |
| **A_55_P2007566** | Trmt61b | 0.012537721 | -0.158016012 | 0.183091454 | 8.475994182 | 0.175697871 | 0.865760685 | 0.996633405 | -5.243713223 |
| **A_51_P477121** | Pmaip1 | 0.009669725 | -0.121881062 | 0.141220513 | 8.111906636 | 0.175683034 | 0.865771888 | 0.996633405 | -5.243715411 |
| **A_51_P146813** | Mocs3 | 0.012814262 | -0.161603494 | 0.187232018 | 9.848569833 | 0.175594958 | 0.865838391 | 0.996633405 | -5.243728391 |
| **A_52_P237669** | Nup54 | -0.01077891 | -0.157593731 | 0.136035912 | 8.626703982 | -0.175474524 | 0.865929329 | 0.996633405 | -5.24374613 |
| **A_55_P2162552** | Txndc9 | 0.010303394 | -0.130133809 | 0.150740597 | 11.74477052 | 0.175350605 | 0.866022901 | 0.996633405 | -5.24376437 |
| **A_55_P2069842** | D5Ertd579e | 0.010372636 | -0.13102059 | 0.151765863 | 8.295113009 | 0.175335428 | 0.866034361 | 0.996633405 | -5.243766603 |
| **A_51_P103054** | Adamts19 | -0.017332022 | -0.253618591 | 0.218954547 | 9.483583111 | -0.1753151 | 0.866049711 | 0.996633405 | -5.243769594 |
| **A_51_P379443** | Bclaf1 | 0.016884952 | -0.213323835 | 0.247093739 | 12.53250943 | 0.175302077 | 0.866059545 | 0.996633405 | -5.243771509 |
| **A_55_P2102350** | Stambpl1 | 0.010134671 | -0.128095333 | 0.148364676 | 8.992476222 | 0.175233234 | 0.86611153 | 0.996633405 | -5.243781634 |
| **A_51_P249537** | Wdr24 | 0.011566158 | -0.1462739 | 0.169406216 | 9.357955342 | 0.17513825 | 0.866183256 | 0.996633405 | -5.243795597 |
| **A_51_P283708** | Msh2 | -0.013161047 | -0.192816582 | 0.166494488 | 11.73914873 | -0.175089043 | 0.866220414 | 0.996633405 | -5.243802828 |
| **A_55_P2186332** | NA | 0.008495723 | -0.107515066 | 0.124506511 | 8.540796482 | 0.175029433 | 0.866265429 | 0.996633405 | -5.243811584 |
| **A_55_P2007981** | Fmn2 | 0.015465786 | -0.195732466 | 0.226664038 | 9.467135659 | 0.175021263 | 0.866271599 | 0.996633405 | -5.243812784 |
| **A_51_P483658** | Trim3 | -0.012152753 | -0.178118583 | 0.153813076 | 9.785796965 | -0.175010916 | 0.866279412 | 0.996633405 | -5.243814304 |
| **A_55_P1962134** | NA | -0.008043919 | -0.117899742 | 0.101811903 | 7.955815042 | -0.175006333 | 0.866282874 | 0.996633405 | -5.243814977 |
| **A_55_P1953169** | Saa3 | 0.009002138 | -0.113957444 | 0.131961719 | 7.723209419 | 0.174981607 | 0.866301546 | 0.996633405 | -5.243818608 |
| **A_55_P2074927** | Gm7932 | 0.007138205 | -0.0903793 | 0.104655709 | 7.885368671 | 0.174950646 | 0.866324926 | 0.996633405 | -5.243823154 |
| **A_51_P267370** | Itm2c | -0.019746696 | -0.289526486 | 0.250033093 | 10.98718761 | -0.174942033 | 0.866331431 | 0.996633405 | -5.243824418 |
| **A_55_P1994927** | Rab3il1 | 0.009099434 | -0.115297388 | 0.133496257 | 10.65881142 | 0.174829307 | 0.86641656 | 0.996633405 | -5.243840961 |
| **A_52_P495553** | Zbtb10 | 0.010961226 | -0.13889284 | 0.160815293 | 8.736932076 | 0.174823473 | 0.866420965 | 0.996633405 | -5.243841817 |
| **A_55_P2137984** | NA | -0.028785512 | -0.422678905 | 0.365107882 | 9.383617373 | -0.174664399 | 0.866541099 | 0.996633405 | -5.243865143 |

| | | | | | | | | | |
|---|---|---|---|---|---|---|---|---|---|
| **A_55_P2170670** | NA | 0.016096057 | -0.204217702 | 0.236409817 | 9.496191791 | 0.174617198 | 0.866576745 | 0.996633405 | -5.24387206 |
| **A_51_P276039** | Txndc15 | -0.00864928 | -0.127093632 | 0.109795073 | 12.59417215 | -0.17453186 | 0.866641195 | 0.996633405 | -5.243884562 |
| **A_52_P604526** | Mrpl19 | -0.018121727 | -0.26634366 | 0.230100206 | 10.75253833 | -0.174489209 | 0.866673407 | 0.996633405 | -5.243890808 |
| **A_55_P2071952** | Wdr92 | 0.033714961 | -0.428136002 | 0.495565925 | 13.39317435 | 0.174473689 | 0.866685128 | 0.996633405 | -5.24389308 |
| **A_51_P350214** | Amz2 | 0.008596511 | -0.109239954 | 0.126432977 | 11.06160832 | 0.174361929 | 0.866769535 | 0.996633405 | -5.243909438 |
| **A_55_P2091741** | Lamp3 | -0.00994908 | -0.14634443 | 0.126446269 | 12.04440213 | -0.174338181 | 0.866787471 | 0.996633405 | -5.243912913 |
| **A_66_P139876** | Psph | -0.013749626 | -0.202305926 | 0.174806674 | 8.808802386 | -0.174284584 | 0.866827951 | 0.996633405 | -5.243920753 |
| **A_51_P354165** | Apcs | -0.007147102 | -0.105204101 | 0.090909898 | 7.664137749 | -0.174204947 | 0.8668881 | 0.996633405 | -5.243932398 |
| **A_55_P2118857** | NA | 0.012846563 | -0.163445345 | 0.189138471 | 8.542109406 | 0.17416613 | 0.866917418 | 0.996633405 | -5.243938072 |
| **A_55_P1977431** | Cck | -0.01986706 | -0.292558465 | 0.252824344 | 8.032916593 | -0.174129074 | 0.866945406 | 0.996633405 | -5.243943487 |
| **A_51_P412270** | Rapgef6 | 0.010515246 | -0.133846535 | 0.154877026 | 8.494228678 | 0.174091 | 0.866974163 | 0.996633405 | -5.24394905 |
| **A_51_P354698** | Dync1h1 | -0.009625239 | -0.14178925 | 0.122538771 | 13.06166584 | -0.1740634 | 0.866995009 | 0.996633405 | -5.243953082 |
| **A_55_P2002215** | Arfip1 | 0.012752329 | -0.162396086 | 0.187900744 | 9.593464188 | 0.174017296 | 0.867029831 | 0.996633405 | -5.243959816 |
| **A_55_P2186439** | Chtop | -0.012092661 | -0.178198866 | 0.154013544 | 11.33594223 | -0.173998358 | 0.867044136 | 0.996633405 | -5.243962581 |
| **A_55_P2102060** | Gstm3 | 0.00702176 | -0.089431493 | 0.103475012 | 7.642773853 | 0.173995582 | 0.867046233 | 0.996633405 | -5.243962986 |
| **A_52_P566681** | Gpm6a | -0.037584264 | -0.553876319 | 0.47870779 | 8.838731366 | -0.17398818 | 0.867051823 | 0.996633405 | -5.243964067 |
| **A_52_P661722** | 2-Mar | -0.015835638 | -0.23375859 | 0.202087313 | 13.16160871 | -0.173676772 | 0.867287043 | 0.996633405 | -5.244009496 |
| **A_52_P217240** | Ppme1 | -0.011434039 | -0.168867632 | 0.145999554 | 9.419645934 | -0.173584686 | 0.867356602 | 0.996633405 | -5.244022914 |
| **A_55_P2311208** | C130045F1 | 0.017284984 | -0.220838554 | 0.255408522 | 8.537885002 | 0.173490538 | 0.86742772 | 0.996633405 | -5.244036625 |
| **A_51_P324640** | C330018D2 | 0.010732477 | -0.137257009 | 0.158721963 | 9.670058947 | 0.173331794 | 0.867547635 | 0.996633405 | -5.244059727 |
| **A_52_P108447** | Glis2 | -0.013151146 | -0.194517315 | 0.168215022 | 11.65242641 | -0.173307143 | 0.867566257 | 0.996633405 | -5.244063313 |
| **A_52_P252121** | Nfs1 | 0.008635436 | -0.110473747 | 0.127744618 | 8.149539438 | 0.173279877 | 0.867586854 | 0.996633405 | -5.244067278 |
| **A_55_P2010672** | Kif17 | -0.008534611 | -0.126302006 | 0.109232785 | 8.205562019 | -0.17320793 | 0.867641205 | 0.996633405 | -5.244077739 |
| **A_51_P167763** | Kcnk13 | -0.008295033 | -0.122796834 | 0.106206769 | 8.271766892 | -0.173146974 | 0.867687254 | 0.996633405 | -5.244086598 |
| **A_55_P2251082** | Qk | 0.009811758 | -0.1258631 | 0.145486616 | 10.87015933 | 0.172844903 | 0.867915458 | 0.996633405 | -5.244130455 |
| **A_55_P2037862** | NA | -0.011050083 | -0.163896809 | 0.141796643 | 7.828349598 | -0.172789971 | 0.867956959 | 0.996633405 | -5.244138422 |
| **A_52_P552092** | Adat1 | 0.009347797 | -0.119990119 | 0.138685713 | 8.328637401 | 0.172739856 | 0.867994821 | 0.996633405 | -5.244145688 |
| **A_55_P2029642** | NA | 0.008295727 | -0.106528013 | 0.123119467 | 13.68817578 | 0.172675967 | 0.868043089 | 0.996633405 | -5.244154949 |
| **A_55_P2008042** | Gm15348 | -0.013432254 | -0.199357528 | 0.17249302 | 9.543858318 | -0.172671081 | 0.868046781 | 0.996633405 | -5.244155657 |
| **A_55_P1983638** | Mapk6 | 0.015559625 | -0.199999956 | 0.231119205 | 11.16254296 | 0.172520583 | 0.868160487 | 0.996633405 | -5.244177457 |
| **A_55_P2157620** | NA | -0.010808986 | -0.160640289 | 0.139022318 | 8.281761004 | -0.172421536 | 0.86823532 | 0.996633405 | -5.244191794 |
| **A_55_P1956998** | Dis3l2 | 0.010466423 | -0.134624467 | 0.155557313 | 8.511651642 | 0.172411918 | 0.868242588 | 0.996633405 | -5.244193186 |
| **A_65_P11059** | Letm1 | -0.01061487 | -0.157852322 | 0.136622581 | 11.94863956 | -0.172308039 | 0.868321074 | 0.996633405 | -5.244208213 |
| **A_52_P426870** | Slf2 | 0.011190889 | -0.144063886 | 0.166445664 | 9.97246585 | 0.172277574 | 0.868344093 | 0.996633405 | -5.244212618 |
| **A_55_P2078088** | Pqlc1 | 0.00981603 | -0.126380218 | 0.146012278 | 10.09922442 | 0.172258184 | 0.868358743 | 0.996633405 | -5.244215422 |
| **A_52_P108502** | Slc4a7 | 0.008896861 | -0.114568568 | 0.13236229 | 8.067864249 | 0.172226727 | 0.868382512 | 0.996633405 | -5.244219969 |
| **A_51_P173601** | Bcam | 0.026144155 | -0.336757749 | 0.389046059 | 11.46441429 | 0.172184674 | 0.868414286 | 0.996633405 | -5.244226047 |
| **A_55_P1970536** | NA | -0.007065342 | -0.105218045 | 0.091087361 | 7.783168673 | -0.172044208 | 0.868520422 | 0.996633405 | -5.244246338 |
| **A_55_P2080021** | Papss2 | -0.022563501 | -0.336099861 | 0.290972859 | 9.863894765 | -0.171999706 | 0.868554048 | 0.996633405 | -5.244252764 |
| **A_51_P296100** | Rhot2 | -0.007007274 | -0.104447208 | 0.09043266 | 12.95399955 | -0.171878378 | 0.868645727 | 0.996633405 | -5.244270272 |
| **A_52_P198019** | Fuz | 0.010242659 | -0.132215708 | 0.152701026 | 8.394906347 | 0.171843803 | 0.868671853 | 0.996633405 | -5.24427526 |
| **A_55_P2039712** | Gigyf2 | -0.012054206 | -0.179758414 | 0.155650001 | 11.30724034 | -0.171792338 | 0.868710742 | 0.996633405 | -5.244282681 |
| **A_55_P2149873** | Mapk1ip1l | -0.012725801 | -0.189812531 | 0.164360929 | 8.531572028 | -0.171754543 | 0.868739302 | 0.996633405 | -5.24428813 |
| **A_55_P2028300** | Ppm1d | -0.011397264 | -0.170003455 | 0.147208928 | 9.730074126 | -0.171747169 | 0.868744874 | 0.996633405 | -5.244289193 |
| **A_51_P149960** | Letm2 | 0.007993733 | -0.103287601 | 0.119275066 | 8.29919093 | 0.171686613 | 0.868790633 | 0.996633405 | -5.244297921 |
| **A_55_P2221647** | AI605517 | 0.060846207 | -0.787068782 | 0.908761196 | 10.93108437 | 0.171510349 | 0.868923832 | 0.996633405 | -5.244323308 |
| **A_55_P2124233** | Vsig8 | -0.00875699 | -0.130868577 | 0.113354598 | 7.922382966 | -0.171398535 | 0.869008329 | 0.996633405 | -5.244339398 |
| **A_55_P2018002** | Abcg4 | 0.009383565 | -0.121465805 | 0.140232934 | 8.984031444 | 0.17139784 | 0.869008854 | 0.996633405 | -5.244339499 |

| | | | | | | | | | |
|---|---|---|---|---|---|---|---|---|---|
| **A_51_P309988** | Gprc5c | 0.008198244 | -0.106175603 | 0.122572092 | 8.698717192 | 0.171318107 | 0.869069109 | 0.996633405 | -5.244350966 |
| **A_55_P2008874** | Dnmt3b | -0.008967029 | -0.134091763 | 0.116157705 | 8.065422599 | -0.171283112 | 0.869095555 | 0.996633405 | -5.244355998 |
| **A_55_P2082837** | Spin1 | 0.008794036 | -0.113923467 | 0.131511539 | 8.883463178 | 0.17127377 | 0.869102615 | 0.996633405 | -5.244357341 |
| **A_51_P145260** | Sall4 | 0.00826707 | -0.107176972 | 0.123711111 | 8.203381973 | 0.171154847 | 0.869192489 | 0.996633405 | -5.244374431 |
| **A_55_P2179676** | Spp2 | -0.021003186 | -0.3143735 | 0.272367128 | 7.852548536 | -0.171111087 | 0.869225561 | 0.996633405 | -5.244380717 |
| **A_52_P193029** | Hadha | -0.018601307 | -0.278516948 | 0.241314334 | 11.67351197 | -0.171048855 | 0.869272592 | 0.996633405 | -5.244389653 |
| **A_66_P121474** | Chmp6 | 0.0148845 | -0.193308488 | 0.223077487 | 10.39797437 | 0.170874508 | 0.869404358 | 0.996633405 | -5.244414672 |
| **A_55_P1954774** | NA | -0.013752421 | -0.206210872 | 0.17870603 | 10.91032325 | -0.170785626 | 0.869471535 | 0.996633405 | -5.244427416 |
| **A_51_P382764** | Akr1c20 | 0.00729336 | -0.094868058 | 0.109454778 | 7.653046571 | 0.170627837 | 0.869590792 | 0.996633405 | -5.244450025 |
| **A_55_P2161773** | Mpp3 | -0.013716134 | -0.205880422 | 0.178448153 | 8.645671004 | -0.170595739 | 0.869615052 | 0.996633405 | -5.244454622 |
| **A_55_P2260052** | Gm17753 | -0.013871129 | -0.2083473 | 0.180605042 | 8.354776932 | -0.170472581 | 0.86970814 | 0.996633405 | -5.244472251 |
| **A_55_P1973643** | Raf1 | -0.012082961 | -0.181648398 | 0.157482476 | 12.49605012 | -0.170311973 | 0.869829535 | 0.996633405 | -5.244495222 |
| **A_52_P253567** | Hsd3b6 | 0.007517169 | -0.09812232 | 0.113156658 | 7.669500234 | 0.170073702 | 0.87000964 | 0.996633405 | -5.244529262 |
| **A_55_P2024909** | Dnah17 | 0.008129018 | -0.106150055 | 0.12240809 | 7.790074007 | 0.170012361 | 0.870056008 | 0.996633405 | -5.244538017 |
| **A_52_P501816** | Slc6a14 | -0.007370906 | -0.111026645 | 0.096284832 | 7.640676305 | -0.169956071 | 0.870098558 | 0.996633405 | -5.244546049 |
| **A_51_P502152** | Slc19a1 | 0.010285079 | -0.134485767 | 0.155055925 | 9.558345667 | 0.169799204 | 0.870217139 | 0.996633405 | -5.244568418 |
| **A_55_P2132358** | NA | -0.037593377 | -0.567154264 | 0.491967509 | 10.13629295 | -0.169669811 | 0.870314953 | 0.996633405 | -5.244586854 |
| **A_51_P450278** | 2010003K1 | 0.008517833 | -0.111519573 | 0.12855524 | 7.668601699 | 0.169598367 | 0.870368962 | 0.996633405 | -5.244597027 |
| **A_51_P241861** | Tcirg1 | -0.009269953 | -0.139947569 | 0.121407664 | 8.359224913 | -0.169545178 | 0.870409172 | 0.996633405 | -5.244604599 |
| **A_52_P453785** | Cthrc1 | 0.025823252 | -0.338322213 | 0.389968716 | 8.65509898 | 0.169490419 | 0.870450569 | 0.996633405 | -5.244612391 |
| **A_51_P376238** | Serping1 | -0.020851623 | -0.314945635 | 0.27324239 | 9.894827973 | -0.169458284 | 0.870474862 | 0.996633405 | -5.244616962 |
| **A_55_P1966322** | Eif3h | 0.00838757 | -0.109918669 | 0.126693809 | 15.54685359 | 0.169448467 | 0.870482284 | 0.996633405 | -5.244618359 |
| **A_55_P2231152** | Gm13372 | -0.007114139 | -0.107458861 | 0.093230582 | 7.823593754 | -0.1694482 | 0.870482486 | 0.996633405 | -5.244618397 |
| **A_51_P111488** | Gfod2 | -0.014490954 | -0.218914086 | 0.189932177 | 8.593459093 | -0.169424448 | 0.870500442 | 0.996633405 | -5.244621775 |
| **A_55_P2078830** | NA | -0.007558935 | -0.11422584 | 0.099107971 | 7.730425648 | -0.169371378 | 0.870540563 | 0.996633405 | -5.244629322 |
| **A_51_P467233** | Polr3gl | -0.015310018 | -0.231398492 | 0.200778457 | 11.41466638 | -0.169337533 | 0.87056615 | 0.996633405 | -5.244634133 |
| **A_55_P1984357** | NA | -0.007100703 | -0.107331067 | 0.093129661 | 16.08084842 | -0.169321133 | 0.870578548 | 0.996633405 | -5.244636464 |
| **A_55_P2020472** | Rufy2 | 0.012308413 | -0.161541455 | 0.18615828 | 9.180705413 | 0.169214203 | 0.87065939 | 0.996633405 | -5.244651658 |
| **A_55_P2169669** | Csf1 | 0.010548775 | -0.138529188 | 0.159626737 | 8.178757913 | 0.169121061 | 0.870729808 | 0.996633405 | -5.244664885 |
| **A_55_P2045050** | Zfp953 | -0.008074642 | -0.122238806 | 0.106089523 | 8.032911579 | -0.169045102 | 0.870787237 | 0.996633405 | -5.244675667 |
| **A_51_P403942** | A930016O | -0.018602065 | -0.281971743 | 0.244767613 | 8.394375625 | -0.168812466 | 0.870963125 | 0.996633405 | -5.244708657 |
| **A_51_P375783** | Prap1 | -0.057223516 | -0.867707199 | 0.753260167 | 8.387212061 | -0.168748297 | 0.871011643 | 0.996633405 | -5.244717749 |
| **A_55_P2049448** | Phtf1 | 0.013424706 | -0.176720836 | 0.203570247 | 10.42111082 | 0.168743772 | 0.871015064 | 0.996633405 | -5.24471839 |
| **A_52_P432580** | NA | -0.013296634 | -0.201673483 | 0.175080215 | 11.2833374 | -0.168703196 | 0.871045743 | 0.996633405 | -5.244724137 |
| **A_52_P89683** | NA | 0.033720211 | -0.44415198 | 0.511592402 | 12.36492944 | 0.16865051 | 0.87108558 | 0.996633405 | -5.244731598 |
| **A_51_P114817** | Ccna1 | -0.007169632 | -0.108934253 | 0.094594988 | 7.736705992 | -0.168387248 | 0.871284639 | 0.996633405 | -5.20507083 |
| **A_55_P1992762** | Kdelr1 | 0.01272787 | -0.16794265 | 0.193398391 | 10.08904236 | 0.168374991 | 0.871293908 | 0.996633405 | -5.244770574 |
| **A_55_P1953796** | Fam98c | -0.007932383 | -0.120565035 | 0.10470027 | 10.56578606 | -0.16832494 | 0.871331753 | 0.996633405 | -5.244777648 |
| **A_55_P2089288** | Rnf121 | 0.010638055 | -0.140433671 | 0.161709781 | 10.85827864 | 0.168301573 | 0.871349423 | 0.996633405 | -5.244780949 |
| **A_66_P103926** | Eif3m | -0.01124112 | -0.17107777 | 0.14859553 | 13.14133334 | -0.168090182 | 0.871509271 | 0.996633405 | -5.244810798 |
| **A_66_P113892** | Smim6 | -0.009270311 | -0.141127395 | 0.122586774 | 7.827463756 | -0.168035081 | 0.871550938 | 0.996633405 | -5.244818572 |
| **A_51_P124606** | Nat2 | -0.008574269 | -0.130606841 | 0.113458304 | 8.178276865 | -0.167930839 | 0.871629767 | 0.996633405 | -5.244833273 |
| **A_51_P473242** | Zfp426 | 0.007949235 | -0.105288083 | 0.121186554 | 8.581998557 | 0.167781826 | 0.871742453 | 0.996633405 | -5.244854272 |
| **A_55_P2302577** | AI426330 | -0.028602233 | -0.436489521 | 0.379285055 | 10.24443787 | -0.167598027 | 0.871881452 | 0.996633405 | -5.244880148 |
| **A_55_P2103011** | Sema4d | 0.012562774 | -0.166892918 | 0.192018466 | 9.249857925 | 0.167315985 | 0.872094755 | 0.996633405 | -5.244919799 |
| **A_51_P400659** | Bcap29 | -0.010776009 | -0.164729784 | 0.143177765 | 10.69949285 | -0.167292609 | 0.872112434 | 0.996633405 | -5.244923083 |
| **A_52_P519538** | Uba6 | 0.010255556 | -0.136311088 | 0.156822201 | 8.541995923 | 0.167237324 | 0.872154247 | 0.996633405 | -5.244930847 |
| **A_55_P2016862** | Ncoa4 | -0.01706866 | -0.261306538 | 0.227169219 | 12.40792152 | -0.167030403 | 0.872310747 | 0.996633405 | -5.244959882 |

| | | | | | | | | | |
|---|---|---|---|---|---|---|---|---|---|
| **A_55_P2083351** | Hs3st1 | 0.006953693 | -0.092550137 | 0.106457524 | 7.918764491 | 0.167026302 | 0.872313849 | 0.996633405 | -5.244960457 |
| **A_66_P127463** | NA | -0.013995712 | -0.214306406 | 0.186314983 | 8.603110529 | -0.166993659 | 0.872338538 | 0.996633405 | -5.244965034 |
| **A_52_P111866** | Naa50 | 0.01082517 | -0.144132265 | 0.165782605 | 9.14810493 | 0.166967305 | 0.872358471 | 0.996633405 | -5.244968728 |
| **A_51_P125050** | Wdr74 | -0.016970931 | -0.259955582 | 0.22601372 | 9.270944234 | -0.166930603 | 0.872386231 | 0.996633405 | -5.244973873 |
| **A_55_P1964965** | Gm1673 | 0.010678207 | -0.142253937 | 0.163610351 | 11.60845662 | 0.166881689 | 0.872423228 | 0.996633405 | -5.244980727 |
| **A_51_P152155** | Glt8d1 | -0.012089606 | -0.185246255 | 0.161067043 | 11.03154973 | -0.166871482 | 0.872430948 | 0.996633405 | -5.244982157 |
| **A_51_P494541** | Abhd17b | 0.012167646 | -0.162358035 | 0.186693328 | 10.31405303 | 0.16663122 | 0.872612679 | 0.996633405 | -5.245015795 |
| **A_55_P2098499** | Gnpda2 | -0.011564004 | -0.177463666 | 0.154335657 | 9.521729036 | -0.166598805 | 0.872637198 | 0.996633405 | -5.245020329 |
| **A_52_P259402** | Dsg3 | 0.00721752 | -0.096335192 | 0.110770232 | 7.615821598 | 0.166584909 | 0.872647708 | 0.996633405 | -5.245022273 |
| **A_55_P2157627** | NA | 0.026563593 | -0.354572764 | 0.407699949 | 12.51473344 | 0.166577203 | 0.872653537 | 0.996633405 | -5.24502335 |
| **A_52_P79947** | Ndst1 | -0.013305118 | -0.204216293 | 0.177606058 | 10.16998877 | -0.16656989 | 0.872659069 | 0.996633405 | -5.245024373 |
| **A_52_P338956** | Aspg | -0.007110633 | -0.109149403 | 0.094928136 | 7.843057247 | -0.166552901 | 0.872671919 | 0.996633405 | -5.245026749 |
| **A_55_P1971006** | Urah | -0.011756377 | -0.180684866 | 0.157172111 | 7.882643985 | -0.16633351 | 0.872837873 | 0.996633405 | -5.245057408 |
| **A_55_P1982744** | NA | 0.017219135 | -0.230296809 | 0.264735079 | 13.04194115 | 0.166271295 | 0.872884936 | 0.996633405 | -5.245066095 |
| **A_51_P215559** | Gstp3 | -0.007254779 | -0.111548605 | 0.097039047 | 7.666075534 | -0.166255006 | 0.872897258 | 0.996633405 | -5.245068369 |
| **A_52_P412585** | Ear1 | -0.007384779 | -0.113574918 | 0.09880536 | 7.691566305 | -0.166212036 | 0.872929763 | 0.996633405 | -5.245074366 |
| **A_55_P2015277** | Cyld | 0.010575914 | -0.141514054 | 0.162665881 | 10.01560588 | 0.166198258 | 0.872940185 | 0.996633405 | -5.245076289 |
| **A_55_P1961247** | NA | -0.008380417 | -0.12893072 | 0.112169885 | 15.92057124 | -0.166152348 | 0.872974915 | 0.996633405 | -5.245082695 |
| **A_55_P2139341** | Zfp975 | 0.025051968 | -0.335407315 | 0.38551125 | 12.13682349 | 0.166109613 | 0.873007243 | 0.996633405 | -5.245088656 |
| **A_55_P2106039** | Map3k6 | 0.010041303 | -0.134477954 | 0.15456056 | 9.523320764 | 0.166063234 | 0.873042328 | 0.996633405 | -5.245095123 |
| **A_55_P1973399** | NA | 0.017031899 | -0.228126407 | 0.262190205 | 12.54266683 | 0.166044922 | 0.87305618 | 0.996633405 | -5.245097676 |
| **A_55_P2175260** | Tet3 | -0.011528202 | -0.177472578 | 0.154416174 | 8.117342985 | -0.16603826 | 0.87306122 | 0.996633405 | -5.245098605 |
| **A_55_P1977289** | Zbtb9 | 0.010116817 | -0.135534552 | 0.155768187 | 9.370924776 | 0.166011613 | 0.873081378 | 0.996633405 | -5.245102319 |
| **A_52_P489202** | 4732465J0 | 0.008040975 | -0.107727844 | 0.123809793 | 7.748119141 | 0.166006936 | 0.873084916 | 0.996633405 | -5.245102971 |
| **A_52_P161297** | Tcea3 | -0.015476866 | -0.238426305 | 0.207472573 | 11.42035458 | -0.165915047 | 0.87315443 | 0.996633405 | -5.245115776 |
| **A_55_P2117830** | NA | 0.007655776 | -0.102636734 | 0.117948286 | 8.312121928 | 0.165902275 | 0.873164092 | 0.996633405 | -5.245117555 |
| **A_51_P286460** | H2ac8 | 0.007868685 | -0.105531479 | 0.121268849 | 7.868947002 | 0.165843176 | 0.873208801 | 0.996633405 | -5.245125786 |
| **A_51_P455266** | Ring1 | -0.009566323 | -0.147445827 | 0.128313181 | 8.547117353 | -0.165826706 | 0.873221261 | 0.996633405 | -5.245128079 |
| **A_55_P2113650** | Clybl | -0.020374331 | -0.314450921 | 0.273702259 | 10.06162429 | -0.165589212 | 0.873400933 | 0.996633405 | -5.245161123 |
| **A_55_P2154240** | Rps29 | -0.009864862 | -0.152357902 | 0.132628179 | 16.7694811 | -0.165465128 | 0.873494811 | 0.996633405 | -5.245178369 |
| **A_66_P103999** | Rsrc1 | -0.009531056 | -0.147212249 | 0.128150137 | 8.975591832 | -0.165453335 | 0.873503732 | 0.996633405 | -5.245180007 |
| **A_51_P232742** | Mfsd14b | 0.014960745 | -0.201168082 | 0.231089572 | 9.837116378 | 0.16544348 | 0.873511189 | 0.996633405 | -5.245181376 |
| **A_55_P2059352** | Col18a1 | -0.01059379 | -0.163641907 | 0.142454328 | 12.8674159 | -0.165436942 | 0.873516135 | 0.996633405 | -5.245182284 |
| **A_52_P520607** | Ankrd22 | -0.006691978 | -0.103377967 | 0.089994012 | 7.661895165 | -0.165424587 | 0.873525482 | 0.996633405 | -5.245184 |
| **A_55_P2077501** | NA | 0.064125524 | -0.86240644 | 0.990657489 | 11.94014712 | 0.165416815 | 0.873531363 | 0.996633405 | -5.24518508 |
| **A_66_P130646** | H4c8 | -0.007518352 | -0.116206014 | 0.10116931 | 7.964914541 | -0.165329954 | 0.87359708 | 0.996633405 | -5.245197141 |
| **A_55_P2125557** | Irx2 | -0.010950091 | -0.169328601 | 0.14742842 | 9.985176 | -0.165245861 | 0.873660704 | 0.996633405 | -5.245208812 |
| **A_55_P2108042** | Dgcr2 | 0.013498333 | -0.181793015 | 0.20878968 | 11.569354 | 0.165198561 | 0.873696492 | 0.996633405 | -5.245215374 |
| **A_55_P2085779** | Ifi27l2b | -0.009627311 | -0.14892453 | 0.129669909 | 7.721229962 | -0.165185403 | 0.873706447 | 0.996633405 | -5.245217199 |
| **A_55_P2021923** | Fkbp1a | -0.029025895 | -0.449030782 | 0.390978992 | 11.4441968 | -0.165173516 | 0.873715441 | 0.996633405 | -5.245218848 |
| **A_55_P2015994** | Fgf9 | 0.017139898 | -0.230948019 | 0.265227816 | 9.939460566 | 0.165124594 | 0.873752456 | 0.996633405 | -5.245225632 |
| **A_55_P2102961** | NA | 0.010286241 | -0.138673479 | 0.159245961 | 8.8987237 | 0.165042954 | 0.873814227 | 0.996633405 | -5.245236949 |
| **A_55_P1973254** | Cmtm7 | 0.017047922 | -0.229872659 | 0.263968503 | 12.00417901 | 0.165014952 | 0.873835415 | 0.996633405 | -5.245240829 |
| **A_55_P2091308** | Rpl38 | -0.007869482 | -0.121919261 | 0.106180297 | 16.81088426 | -0.164915255 | 0.87391085 | 0.996633405 | -5.245254639 |
| **A_55_P1966615** | Actr3b | 0.015615619 | -0.210712492 | 0.24194373 | 9.987729161 | 0.164903486 | 0.873919754 | 0.996633405 | -5.245256268 |
| **A_55_P2022971** | Tspan9 | 0.017653328 | -0.238380771 | 0.273687426 | 8.316390237 | 0.164792675 | 0.874003601 | 0.996633405 | -5.245271607 |
| **A_51_P499854** | Ghr | 0.014105867 | -0.190495523 | 0.218707256 | 11.14341258 | 0.16477842 | 0.874014387 | 0.996633405 | -5.24527358 |
| **A_55_P2173066** | NA | -0.00969199 | -0.150309347 | 0.130925367 | 9.188455128 | -0.16473397 | 0.874048021 | 0.996633405 | -5.245279729 |

| | | | | | | | | | |
|---|---|---|---|---|---|---|---|---|---|
| **A_55_P1970309** | Arhgap44 | 0.008332993 | -0.11260195 | 0.129267935 | 10.96230231 | 0.164686623 | 0.874083847 | 0.996633405 | -5.245286277 |
| **A_55_P1971815** | Clcn2 | 0.009926421 | -0.134211854 | 0.154064697 | 7.982132636 | 0.164597226 | 0.874151492 | 0.996633405 | -5.245298636 |
| **A_55_P2060333** | S100a5 | 0.028318543 | -0.383705373 | 0.440342459 | 8.329730118 | 0.164269755 | 0.874399295 | 0.996633405 | -5.245343852 |
| **A_55_P1971093** | NA | 0.029107263 | -0.394605924 | 0.452820451 | 12.57976089 | 0.164186909 | 0.874461989 | 0.996633405 | -5.245355277 |
| **A_66_P129188** | H2ac20 | 0.01308219 | -0.177480248 | 0.203644629 | 16.82670094 | 0.164078733 | 0.874543852 | 0.996633405 | -5.245370187 |
| **A_51_P329322** | Zfand1 | 0.011488323 | -0.15589131 | 0.178867955 | 11.48628221 | 0.164045056 | 0.874569338 | 0.996633405 | -5.245374826 |
| **A_55_P2069939** | Ubxn7 | -0.007721964 | -0.120245234 | 0.104801306 | 8.234933641 | -0.164019148 | 0.874588944 | 0.996633405 | -5.245378395 |
| **A_51_P425833** | Lztfl1 | 0.008303852 | -0.112798781 | 0.129406485 | 9.871689869 | 0.163883472 | 0.874691622 | 0.996633405 | -5.245397074 |
| **A_51_P308912** | Mybphl | -0.091688144 | -1.429115091 | 1.245738804 | 9.151217817 | -0.163852187 | 0.874715298 | 0.996633405 | -5.245401379 |
| **A_55_P2113703** | Spib | -0.006740819 | -0.105282995 | 0.091801357 | 7.688978313 | -0.163493176 | 0.874987007 | 0.996633405 | -5.245450723 |
| **A_51_P158400** | Rc3h2 | 0.014570215 | -0.198474881 | 0.227615311 | 10.58225224 | 0.163457008 | 0.875014381 | 0.996633405 | -5.245455688 |
| **A_51_P479894** | Cpsf3 | -0.010720786 | -0.167573588 | 0.146132017 | 11.30914417 | -0.163359151 | 0.875088446 | 0.996633405 | -5.245469117 |
| **A_55_P2062538** | 1700027A1 | 0.008402103 | -0.114640375 | 0.13144458 | 7.835431519 | 0.163208225 | 0.875202678 | 0.996633405 | -5.245489812 |
| **A_52_P153189** | Arl2bp | -0.008594426 | -0.134544555 | 0.117355703 | 10.02532062 | -0.163090016 | 0.875292151 | 0.996633405 | -5.245506007 |
| **A_55_P2090239** | Casp8ap2 | -0.011540697 | -0.180685953 | 0.157604559 | 8.295703305 | -0.163072733 | 0.875305232 | 0.996633405 | -5.245508374 |
| **A_51_P100034** | Mif4gd | -0.010208975 | -0.159946912 | 0.139528963 | 11.52153426 | -0.162951892 | 0.875396699 | 0.996633405 | -5.245524917 |
| **A_55_P2050354** | Prpmp5 | -0.010631152 | -0.16684708 | 0.145584776 | 8.417999225 | -0.162653776 | 0.875622358 | 0.996633405 | -5.245565675 |
| **A_51_P184849** | Pfkfb2 | 0.012503415 | -0.171256778 | 0.196263608 | 8.111571544 | 0.162624638 | 0.875644414 | 0.996633405 | -5.245569655 |
| **A_55_P1961188** | Gm6981 | 0.016242856 | -0.222770612 | 0.255256324 | 16.09586217 | 0.162423608 | 0.875796592 | 0.996633405 | -5.245597094 |
| **A_66_P136186** | Wee1 | -0.010947675 | -0.172077744 | 0.150182394 | 9.645710105 | -0.162388193 | 0.875823401 | 0.996633405 | -5.245601924 |
| **A_55_P1952738** | Ppp3ca | 0.018257994 | -0.250564004 | 0.287079993 | 12.19728745 | 0.162329479 | 0.875867848 | 0.996633405 | -5.24560993 |
| **A_66_P126726** | Cwc27 | 0.009873346 | -0.135500717 | 0.155247409 | 9.257828464 | 0.162325424 | 0.875870918 | 0.996633405 | -5.245610482 |
| **A_55_P2091461** | Casp4 | -0.013256941 | -0.208476076 | 0.181962194 | 8.870769478 | -0.162304322 | 0.875886892 | 0.996633405 | -5.245613359 |
| **A_55_P1958295** | NA | -0.013365805 | -0.210192749 | 0.183461139 | 12.59139582 | -0.162300439 | 0.875889831 | 0.996633405 | -5.245613888 |
| **A_55_P2085994** | Zfat | -0.011156373 | -0.175530865 | 0.15321812 | 8.699066422 | -0.162217505 | 0.875952615 | 0.996633405 | -5.24562519 |
| **A_51_P217706** | Ribc1 | -0.007521845 | -0.118347109 | 0.103303419 | 8.23562881 | -0.162216395 | 0.875953455 | 0.996633405 | -5.245625341 |
| **A_55_P1963036** | Oxr1 | -0.014533034 | -0.228894257 | 0.19982819 | 11.16723787 | -0.162038861 | 0.876087856 | 0.996633405 | -5.245649514 |
| **A_55_P2003216** | Igsf5 | -0.006865709 | -0.108171378 | 0.094439961 | 7.688025249 | -0.16197976 | 0.876132599 | 0.996633405 | -5.245657555 |
| **A_52_P517762** | Slc44a4 | -0.006755332 | -0.10643901 | 0.092928345 | 7.645523179 | -0.161968966 | 0.876140771 | 0.996633405 | -5.245659023 |
| **A_51_P285779** | Asphd2 | 0.009210318 | -0.126714759 | 0.145135396 | 8.06508255 | 0.161951207 | 0.876154216 | 0.996633405 | -5.245661439 |
| **A_55_P2002918** | Klc2 | -0.010685864 | -0.168536042 | 0.147164314 | 11.92083708 | -0.161798205 | 0.87627005 | 0.996633405 | -5.245682239 |
| **A_55_P2162234** | Tm9sf3 | 0.01402337 | -0.193247961 | 0.221294702 | 8.8199941 | 0.161704568 | 0.876340942 | 0.996633405 | -5.24569496 |
| **A_55_P2018322** | Zfp644 | -0.009752129 | -0.153992506 | 0.134488248 | 10.5729515 | -0.16159269 | 0.876425646 | 0.996633405 | -5.245710148 |
| **A_52_P475040** | Etaa1 | 0.014840526 | -0.204685728 | 0.234366781 | 8.684074153 | 0.161574184 | 0.876439658 | 0.996633405 | -5.24571266 |
| **A_51_P177552** | Ndufb9 | -0.018078418 | -0.285562746 | 0.249405911 | 16.27054632 | -0.161536699 | 0.876468039 | 0.996633405 | -5.245717746 |
| **A_51_P286496** | Il2rb | -0.007033822 | -0.111105474 | 0.097037829 | 7.693693066 | -0.161535534 | 0.876468921 | 0.996633405 | -5.245717904 |
| **A_51_P346884** | Snrnp27 | 0.007828526 | -0.108056674 | 0.123713726 | 12.15898242 | 0.161458584 | 0.876527182 | 0.996633405 | -5.245728341 |
| **A_51_P190961** | Gzmf | 0.008284201 | -0.114392531 | 0.130960932 | 7.667109256 | 0.161397776 | 0.876573222 | 0.996633405 | -5.245736585 |
| **A_55_P2049854** | Araf | -0.010328067 | -0.163295238 | 0.142639105 | 8.800679273 | -0.161372649 | 0.876592247 | 0.996633405 | -5.245739991 |
| **A_51_P219849** | Alg13 | -0.009406702 | -0.148741234 | 0.12992783 | 8.435130766 | -0.161356986 | 0.876604106 | 0.996633405 | -5.245742114 |
| **A_66_P124755** | Fut8 | 0.010723322 | -0.148201734 | 0.169648377 | 9.14707733 | 0.161267218 | 0.876672075 | 0.996633405 | -5.245754276 |
| **A_55_P2030128** | Tfpt | 0.012908959 | -0.178410052 | 0.204227971 | 8.428550508 | 0.161265787 | 0.876673159 | 0.996633405 | -5.245754469 |
| **A_51_P509669** | Uap1l1 | 0.014831784 | -0.205016259 | 0.234679826 | 8.866894112 | 0.161242643 | 0.876690682 | 0.996633405 | -5.245757604 |
| **A_55_P2164957** | Slc9a3r2 | -0.016771829 | -0.265385979 | 0.231842321 | 8.622332154 | -0.161236632 | 0.876695233 | 0.996633405 | -5.245758418 |
| **A_55_P2027127** | Triml2 | 0.007383477 | -0.102083883 | 0.116850837 | 7.661483595 | 0.161207564 | 0.876717243 | 0.996633405 | -5.245762354 |
| **A_55_P2119907** | Dnah11 | 0.006820344 | -0.094314257 | 0.107954944 | 7.678471288 | 0.161181658 | 0.876736858 | 0.996633405 | -5.245765861 |
| **A_55_P2016216** | NA | -0.008489593 | -0.134420728 | 0.117441542 | 11.69280409 | -0.161124985 | 0.87677977 | 0.996633405 | -5.245773532 |
| **A_51_P288549** | Jmjd7 | -0.01367551 | -0.21658618 | 0.18923516 | 9.505156583 | -0.16108228 | 0.876812105 | 0.996633405 | -5.24577931 |

| | | | | | | | | | |
|---|---|---|---|---|---|---|---|---|---|
| **A_51_P445473** | Fut7 | -0.006652248 | -0.105357164 | 0.092052669 | 7.734315408 | -0.161078942 | 0.876814634 | 0.996633405 | -5.245779762 |
| **A_55_P1961436** | Prkca | -0.014554692 | -0.230549289 | 0.201439904 | 10.36104585 | -0.16105317 | 0.876834148 | 0.996633405 | -5.245783248 |
| **A_51_P338461** | Numa1 | 0.014100404 | -0.195284761 | 0.223485569 | 13.15955354 | 0.160951408 | 0.876911202 | 0.996633405 | -5.245797009 |
| **A_55_P2009973** | Ttc3 | -0.009028345 | -0.14311139 | 0.125054701 | 13.6966977 | -0.160932364 | 0.876925622 | 0.996633405 | -5.245799583 |
| **A_55_P1959409** | Deup1 | -0.007254377 | -0.114999183 | 0.10049043 | 7.702460338 | -0.16092106 | 0.876934182 | 0.996633405 | -5.245801111 |
| **A_55_P2170847** | Fbxo27 | 0.010618553 | -0.147167785 | 0.168404891 | 8.929715679 | 0.16084408 | 0.876992472 | 0.996633405 | -5.245811513 |
| **A_55_P2144075** | Pofut2 | 0.019119356 | -0.265085473 | 0.303324185 | 8.715718428 | 0.160787003 | 0.877035693 | 0.996633405 | -5.245819222 |
| **A_55_P1996421** | NA | -0.011891802 | -0.18872123 | 0.164937627 | 13.9553072 | -0.160731962 | 0.877077372 | 0.996633405 | -5.245826654 |
| **A_55_P1969543** | Pin4 | -0.010566583 | -0.167734066 | 0.146600899 | 11.75550073 | -0.160687103 | 0.877111341 | 0.996633405 | -5.245832709 |
| **A_55_P2139992** | Mrps18b | 0.015633504 | -0.216964182 | 0.248231191 | 12.5160013 | 0.160642361 | 0.877145221 | 0.996633405 | -5.245838746 |
| **A_66_P137475** | Dnajb14 | -0.016405834 | -0.260554693 | 0.227743026 | 10.78388607 | -0.16060266 | 0.877175285 | 0.996633405 | -5.245844102 |
| **A_66_P116098** | Ptpn2 | 0.009077201 | -0.126101362 | 0.144255764 | 8.661643769 | 0.160491957 | 0.877259117 | 0.996633405 | -5.24585903 |
| **A_55_P2009588** | Rpl13 | -0.009708668 | -0.154364278 | 0.134946942 | 16.27007135 | -0.160410756 | 0.877320608 | 0.996633405 | -5.245869973 |
| **A_55_P1956048** | H2-T3 | -0.007037947 | -0.111902848 | 0.097826954 | 7.693026238 | -0.16040761 | 0.877322991 | 0.996633405 | -5.245870397 |
| **A_55_P2103561** | Prdx6 | -0.007739998 | -0.123233892 | 0.107753896 | 13.80118469 | -0.160173609 | 0.8775002 | 0.996633405 | -5.245901901 |
| **A_51_P128075** | Tescl | 0.00701031 | -0.097620016 | 0.111640635 | 7.828732269 | 0.160135924 | 0.87752874 | 0.996633405 | -5.225265872 |
| **A_55_P2041310** | NA | -0.008325806 | -0.132742252 | 0.11609064 | 8.490325743 | -0.159940195 | 0.877676972 | 0.996633405 | -5.245933281 |
| **A_51_P125679** | Rec114 | -0.008845319 | -0.141086094 | 0.123395457 | 7.942639873 | -0.159866413 | 0.877732851 | 0.996633405 | -5.235433941 |
| **A_55_P2321453** | D11Ertd72( | 0.017841215 | -0.249010692 | 0.284693121 | 8.509873235 | 0.15979502 | 0.877786922 | 0.996633405 | -5.245952775 |
| **A_51_P243514** | Macc1 | 0.00657428 | -0.09197855 | 0.105127111 | 7.692479973 | 0.159436685 | 0.878058322 | 0.996633405 | -5.246000817 |
| **A_55_P2132403** | Szt2 | 0.009892427 | -0.138433254 | 0.158218107 | 10.25956464 | 0.159402679 | 0.878084079 | 0.996633405 | -5.24600537 |
| **A_55_P2140147** | Ppp4r3b | 0.014185846 | -0.198741257 | 0.227112949 | 10.21697966 | 0.159233126 | 0.878212505 | 0.996633405 | -5.246028061 |
| **A_55_P1975181** | Acyp1 | 0.007913506 | -0.110932325 | 0.126759336 | 8.194403966 | 0.159145403 | 0.878278951 | 0.996633405 | -5.24603979 |
| **A_55_P2122180** | Snta1 | -0.010893211 | -0.174557242 | 0.15277082 | 12.43785085 | -0.159078618 | 0.878329538 | 0.996633405 | -5.246048716 |
| **A_55_P2003753** | Sh2d5 | 0.006868301 | -0.096330165 | 0.110066768 | 7.770591158 | 0.159068885 | 0.878336911 | 0.996633405 | -5.246050017 |
| **A_55_P2082070** | Ak6 | -0.017990155 | -0.288415115 | 0.252434805 | 10.09652827 | -0.159000052 | 0.87838905 | 0.996633405 | -5.246059212 |
| **A_55_P2065308** | Myo9b | 0.008662163 | -0.121564406 | 0.138888732 | 8.589967981 | 0.158977584 | 0.878406069 | 0.996633405 | -5.246062213 |
| **A_55_P2153721** | Ccdc127 | -0.010807639 | -0.173345401 | 0.151730123 | 8.623605154 | -0.158922609 | 0.878447712 | 0.996633405 | -5.246069553 |
| **A_55_P1985410** | Reps2 | 0.007927903 | -0.111425001 | 0.127280807 | 7.924053958 | 0.158757579 | 0.878572722 | 0.996633405 | -5.246091572 |
| **A_51_P133422** | Nck2 | 0.006973742 | -0.098035059 | 0.111982542 | 7.881272137 | 0.158726443 | 0.878596308 | 0.996633405 | -5.246095724 |
| **A_55_P2159994** | Atf1 | 0.011456052 | -0.161055467 | 0.18396757 | 10.48327325 | 0.158717931 | 0.878602756 | 0.996633405 | -5.246096859 |
| **A_66_P108247** | Ucp3 | -0.010904976 | -0.175139398 | 0.153329447 | 8.89732444 | -0.158697339 | 0.878618355 | 0.996633405 | -5.246099604 |
| **A_51_P135004** | Zfp212 | -0.013924531 | -0.223667097 | 0.195818035 | 10.73783718 | -0.158673036 | 0.878636765 | 0.996633405 | -5.246102843 |
| **A_51_P256246** | Tspan13 | -0.013811363 | -0.221877021 | 0.194254295 | 12.6204391 | -0.158651895 | 0.87865278 | 0.996633405 | -5.246105661 |
| **A_52_P218181** | Zdhhc5 | -0.010708641 | -0.17216063 | 0.150743349 | 11.72505129 | -0.158525843 | 0.878748269 | 0.996633405 | -5.246122453 |
| **A_51_P253697** | Rpain | 0.011575128 | -0.163050236 | 0.186200493 | 9.325150415 | 0.158426426 | 0.878823582 | 0.996633405 | -5.246135688 |
| **A_51_P345110** | Thap4 | -0.008190394 | -0.131791921 | 0.115411132 | 11.66404567 | -0.158376272 | 0.878861577 | 0.996633405 | -5.246142361 |
| **A_65_P16483** | Pdzk1 | -0.007047892 | -0.113536337 | 0.099440554 | 7.778729173 | -0.158185208 | 0.879006323 | 0.996633405 | -5.246167765 |
| **A_55_P2072150** | Vpreb2 | -0.007707133 | -0.124220894 | 0.108806629 | 7.718371118 | -0.158097409 | 0.87907284 | 0.996633405 | -5.246179429 |
| **A_55_P2080396** | Mrpl4 | 0.011548272 | -0.163098373 | 0.186194917 | 13.6668017 | 0.158039586 | 0.879116646 | 0.996633405 | -5.246187106 |
| **A_55_P1971360** | Ciz1 | 0.012237672 | -0.172923747 | 0.197399092 | 9.567888714 | 0.157963756 | 0.879174097 | 0.996633405 | -5.246197171 |
| **A_55_P2146708** | Zcchc9 | -0.012773783 | -0.206077705 | 0.180530139 | 13.87563116 | -0.157938501 | 0.87919323 | 0.996633405 | -5.246200522 |
| **A_51_P121455** | Slc25a39 | -0.022965123 | -0.370618711 | 0.324688465 | 9.031433658 | -0.157881484 | 0.879236428 | 0.996633405 | -5.246208086 |
| **A_55_P2041673** | Ttc38 | 0.008348279 | -0.118068192 | 0.13476475 | 17.40178291 | 0.157834675 | 0.879271893 | 0.996633405 | -5.246214293 |
| **A_66_P126459** | Gm13429 | -0.008347564 | -0.134787122 | 0.118091994 | 8.471687069 | -0.157792335 | 0.879303971 | 0.996633405 | -5.246219906 |
| **A_51_P245275** | H2afx | -0.013817885 | -0.223119136 | 0.195483367 | 13.24557558 | -0.15778978 | 0.879305907 | 0.996633405 | -5.246220245 |
| **A_51_P451338** | Jag2 | 0.01278302 | -0.180864937 | 0.206430977 | 8.788183383 | 0.157771904 | 0.879319451 | 0.996633405 | -5.246222614 |
| **A_55_P2008907** | Actg1 | 0.015261605 | -0.215939596 | 0.246462805 | 14.41060543 | 0.157768117 | 0.87932232 | 0.996633405 | -5.246223116 |

| | | | | | | | | | |
|---|---|---|---|---|---|---|---|---|---|
| **A_52_P526372** | Zeb2 | -0.00673629 | -0.108793389 | 0.095320809 | 8.000638107 | -0.157756297 | 0.879331276 | 0.996633405 | -5.246224682 |
| **A_51_P151254** | Slc35b3 | -0.012214662 | -0.197285156 | 0.172855831 | 8.97121822 | -0.157744207 | 0.879340436 | 0.996633405 | -5.246226284 |
| **A_55_P2049287** | NA | -0.011192364 | -0.180846713 | 0.158461985 | 14.46267858 | -0.157676136 | 0.87939201 | 0.996633405 | -5.246235303 |
| **A_52_P406828** | Dkc1 | 0.011109819 | -0.1573599 | 0.179579538 | 9.400501494 | 0.157613808 | 0.879439234 | 0.996633405 | -5.246243557 |
| **A_52_P306396** | Ppp3cb | -0.013540775 | -0.219014779 | 0.19193323 | 11.29000236 | -0.157505511 | 0.879521287 | 0.996633405 | -5.246257891 |
| **A_55_P2173927** | Insr | -0.014598009 | -0.236139606 | 0.206943588 | 8.44132899 | -0.157488003 | 0.879534553 | 0.996633405 | -5.246260208 |
| **A_55_P2028248** | Dnajc19 | 0.01123656 | -0.15938859 | 0.18186171 | 11.97538306 | 0.157398089 | 0.87960268 | 0.996633405 | -5.2462721 |
| **A_55_P2103101** | Snx9 | -0.010277873 | -0.166402206 | 0.145846459 | 8.833810527 | -0.157340961 | 0.879645967 | 0.996633405 | -5.246279652 |
| **A_55_P2151556** | C8g | -0.011380171 | -0.184258246 | 0.161497905 | 8.377751999 | -0.157332333 | 0.879652504 | 0.996633405 | -5.246280793 |
| **A_51_P434697** | Ube2e2 | 0.009685137 | -0.13747975 | 0.156850025 | 11.43971867 | 0.157293479 | 0.879681944 | 0.996633405 | -5.246285927 |
| **A_51_P446570** | Bbox1 | -0.006366232 | -0.103315952 | 0.090583487 | 7.66350665 | -0.15694412 | 0.879946666 | 0.996633405 | -5.24633204 |
| **A_52_P106537** | Chrac1 | -0.017981206 | -0.291985529 | 0.256023116 | 10.71348206 | -0.156844951 | 0.880021814 | 0.996633405 | -5.246345111 |
| **A_55_P2023607** | Ikzf1 | 0.006868296 | -0.097804753 | 0.111541345 | 7.674064726 | 0.15682787 | 0.880034758 | 0.996633405 | -5.246347362 |
| **A_55_P2016685** | Csnk1g1 | 0.014782118 | -0.210553026 | 0.240117262 | 9.96840359 | 0.156789462 | 0.880063862 | 0.996633405 | -5.246352422 |
| **A_51_P471520** | Stk25 | -0.015294459 | -0.248587013 | 0.217998094 | 11.81443242 | -0.1566904 | 0.88013893 | 0.996633405 | -5.246365466 |
| **A_55_P2043187** | Exosc9 | 0.008974688 | -0.127972053 | 0.14592143 | 12.21010675 | 0.15663067 | 0.880184193 | 0.996633405 | -5.246373327 |
| **A_55_P2005005** | Cdk5rap1 | -0.012630732 | -0.205548636 | 0.180287172 | 10.34108085 | -0.156482265 | 0.880296657 | 0.996633405 | -5.246392846 |
| **A_51_P327491** | Apoa4 | 0.007480243 | -0.106812798 | 0.121773283 | 7.684617254 | 0.156424591 | 0.880340364 | 0.996633405 | -5.246400427 |
| **A_51_P202050** | Dtx1 | -0.016378207 | -0.266683239 | 0.233926824 | 8.492702505 | -0.156388898 | 0.880367413 | 0.996633405 | -5.246405117 |
| **A_51_P386058** | Ppp2r5b | 0.013280971 | -0.189931195 | 0.216493137 | 11.24683752 | 0.156202959 | 0.880508327 | 0.996633405 | -5.246429532 |
| **A_55_P2053266** | St6galnac6 | 0.015837407 | -0.226555331 | 0.258230146 | 12.18597522 | 0.156161372 | 0.880539845 | 0.996633405 | -5.246434989 |
| **A_55_P2055217** | NA | -0.026306003 | -0.42921743 | 0.376605423 | 13.41623058 | -0.156046639 | 0.880626798 | 0.996633405 | -5.246450036 |
| **A_52_P308465** | Plxnb1 | 0.012495215 | -0.1789251 | 0.20391553 | 8.547965222 | 0.156014455 | 0.88065119 | 0.996633405 | -5.246454255 |
| **A_55_P1957080** | NA | -0.011343417 | -0.185170943 | 0.16248411 | 8.40996488 | -0.155967627 | 0.88068668 | 0.996633405 | -5.246460392 |
| **A_55_P2135750** | NA | -0.019638053 | -0.320711195 | 0.281435088 | 8.333306679 | -0.155896209 | 0.880740808 | 0.996633405 | -5.246469748 |
| **A_52_P527463** | Fam91a1 | 0.013034201 | -0.18681618 | 0.212884582 | 11.02724735 | 0.155879343 | 0.88075359 | 0.996633405 | -5.246471957 |
| **A_55_P1965725** | Gapdh | -0.028573326 | -0.466720901 | 0.409574248 | 14.1754549 | -0.15586533 | 0.880764211 | 0.996633405 | -5.246473792 |
| **A_52_P573552** | Trib1 | 0.012022819 | -0.1724334 | 0.196479037 | 9.504954743 | 0.155783739 | 0.88082605 | 0.996633405 | -5.246484473 |
| **A_55_P2062793** | Mfap2 | 0.028320931 | -0.406222307 | 0.462864169 | 12.88412629 | 0.155769944 | 0.880836506 | 0.996633405 | -5.246486279 |
| **A_55_P2163669** | NA | 0.011639348 | -0.167047041 | 0.190325737 | 15.35808683 | 0.15568484 | 0.880901008 | 0.996633405 | -5.246497413 |
| **A_55_P1989921** | Eml2 | 0.019858596 | -0.285382389 | 0.325099582 | 12.56799972 | 0.15549443 | 0.881045329 | 0.996633405 | -5.246522304 |
| **A_52_P235861** | Peg3 | 0.017943092 | -0.257891678 | 0.293777861 | 9.650398865 | 0.155473867 | 0.881060914 | 0.996633405 | -5.24652499 |
| **A_55_P2175195** | Stxbp4 | -0.014003764 | -0.229311874 | 0.201304346 | 12.80978715 | -0.155451005 | 0.881078243 | 0.996633405 | -5.246527976 |
| **A_52_P239787** | Ccdc25 | 0.009258692 | -0.133095423 | 0.151612806 | 8.61018726 | 0.155449283 | 0.881079548 | 0.996633405 | -5.246528201 |
| **A_55_P2151797** | Dis3l2 | -0.009344191 | -0.153013598 | 0.134325216 | 11.5843143 | -0.155448503 | 0.881080139 | 0.996633405 | -5.246528303 |
| **A_55_P2160416** | Acoxl | -0.00629777 | -0.103132773 | 0.090537233 | 7.885642949 | -0.155440273 | 0.881086377 | 0.996633405 | -5.246529378 |
| **A_55_P1964805** | Rpa1 | -0.012226126 | -0.200233808 | 0.175781557 | 13.11498282 | -0.155425544 | 0.881097541 | 0.996633405 | -5.246531301 |
| **A_51_P487628** | Pramef12 | -0.024404362 | -0.400231077 | 0.351422353 | 7.896579552 | -0.15519901 | 0.88126925 | 0.996633405 | -5.246560861 |
| **A_55_P1955656** | Ctla2a | -0.010334343 | -0.169665691 | 0.148997005 | 12.57300174 | -0.155021087 | 0.881404117 | 0.996633405 | -5.246584048 |
| **A_55_P2053933** | Foxa1 | 0.006249155 | -0.090126005 | 0.102624316 | 7.651724471 | 0.154976318 | 0.881438054 | 0.996633405 | -5.246589879 |
| **A_55_P2103837** | Gbp4 | -0.012961939 | -0.212873957 | 0.186950078 | 8.219286572 | -0.154967354 | 0.881444848 | 0.996633405 | -5.246591046 |
| **A_51_P197259** | Cct7 | 0.011998366 | -0.173106359 | 0.197103091 | 13.36679175 | 0.154922224 | 0.881479059 | 0.996633405 | -5.246596921 |
| **A_52_P477369** | Csnk1d | -0.009446264 | -0.155253109 | 0.13636058 | 8.393253537 | -0.154842909 | 0.881539183 | 0.996633405 | -5.246607242 |
| **A_51_P345316** | Cep76 | 0.009310567 | -0.134464059 | 0.153085193 | 8.465665777 | 0.154775785 | 0.881590066 | 0.996633405 | -5.246615973 |
| **A_55_P2124183** | NA | 0.016813319 | -0.242875177 | 0.276501814 | 10.85241143 | 0.154742591 | 0.881615229 | 0.996633405 | -5.246620289 |
| **A_55_P1966998** | Arhgdib | -0.01333026 | -0.219229483 | 0.192568963 | 10.14844781 | -0.154736596 | 0.881619773 | 0.996633405 | -5.246621069 |
| **A_55_P2463930** | Zmym5 | -0.015511123 | -0.255177795 | 0.224155549 | 9.406762885 | -0.154683759 | 0.881659827 | 0.996633405 | -5.246627937 |
| **A_51_P516133** | H1f2 | -0.013851012 | -0.228055418 | 0.200353393 | 11.28257024 | -0.154547595 | 0.88176305 | 0.996633405 | -5.246645626 |

| | | | | | | | | | |
|---|---|---|---|---|---|---|---|---|---|
| **A_55_P2274782** | Prl7a1 | -0.006382885 | -0.105100739 | 0.092334969 | 7.646290343 | -0.154536283 | 0.881771625 | 0.996633405 | -5.246647095 |
| **A_55_P1960416** | Nrros | -0.012610471 | -0.207740833 | 0.182519892 | 9.708794443 | -0.15445984 | 0.881829576 | 0.996633405 | -5.246657018 |
| **A_55_P2181104** | Shisa3 | 0.008558623 | -0.123889586 | 0.141006833 | 7.813568647 | 0.154442547 | 0.881842686 | 0.996633405 | -5.246659262 |
| **A_55_P2420873** | NA | 0.011920322 | -0.17256816 | 0.196408804 | 10.03849308 | 0.154428646 | 0.881853224 | 0.996633405 | -5.246661066 |
| **A_66_P112482** | Tmem184c | -0.013637413 | -0.224764061 | 0.197489235 | 10.14335941 | -0.1543825 | 0.881888208 | 0.996633405 | -5.246667053 |
| **A_55_P2132800** | Zfp367 | 0.014143687 | -0.204872781 | 0.233160154 | 8.498062922 | 0.15434586 | 0.881915985 | 0.996633405 | -5.246671805 |
| **A_55_P2274553** | NA | -0.006779792 | -0.111837763 | 0.098278178 | 7.671631541 | -0.154239824 | 0.881996373 | 0.996633405 | -5.246685551 |
| **A_55_P1953708** | Cry2 | 0.01441436 | -0.209009309 | 0.23783803 | 8.397327062 | 0.154196788 | 0.882029 | 0.996633405 | -5.246691128 |
| **A_55_P2019684** | Bspry | 0.006760893 | -0.098178573 | 0.111700358 | 7.678727746 | 0.153983547 | 0.882190668 | 0.996633405 | -5.246718737 |
| **A_51_P196844** | Osbpl3 | 0.008292708 | -0.120429988 | 0.137015403 | 9.952465705 | 0.153975055 | 0.882197106 | 0.996633405 | -5.246719835 |
| **A_51_P266958** | Nr1i2 | -0.006401174 | -0.105826228 | 0.093023881 | 7.652202007 | -0.153876711 | 0.882271668 | 0.996633405 | -5.246732555 |
| **A_55_P2054157** | Hspa4 | 0.013364282 | -0.194395646 | 0.221124211 | 13.24099515 | 0.153742162 | 0.882373681 | 0.996633405 | -5.246749943 |
| **A_55_P2068731** | Gm21953 | 0.009651061 | -0.140453994 | 0.159756116 | 9.278856679 | 0.153669921 | 0.882428454 | 0.996633405 | -5.246759274 |
| **A_55_P2433438** | Tcf12 | 0.009520323 | -0.138593922 | 0.157634568 | 9.001254667 | 0.153625743 | 0.88246195 | 0.996633405 | -5.246764977 |
| **A_55_P1965524** | NA | 0.008491559 | -0.123663241 | 0.14064636 | 15.04800354 | 0.153572565 | 0.88250227 | 0.996633405 | -5.24677184 |
| **A_55_P1980321** | Pcbp3 | 0.013993422 | -0.203948185 | 0.231935028 | 9.607130927 | 0.15345919 | 0.882588234 | 0.996633405 | -5.246786465 |
| **A_51_P141860** | Fbxw11 | -0.007892389 | -0.130824376 | 0.115039597 | 12.66602647 | -0.153444996 | 0.882598996 | 0.996633405 | -5.246788295 |
| **A_55_P2049752** | Pamr1 | -0.026704592 | -0.442663422 | 0.389254238 | 8.538982569 | -0.153442171 | 0.882601138 | 0.996633405 | -5.24678866 |
| **A_55_P2131899** | Sox4 | 0.023013239 | -0.335834982 | 0.38186146 | 8.617904784 | 0.153276678 | 0.882726622 | 0.996633405 | -5.246809985 |
| **A_52_P50417** | 1700084J1 | -0.008152828 | -0.135345901 | 0.119040245 | 8.246305315 | -0.1531983 | 0.882786053 | 0.996633405 | -5.246820077 |
| **A_55_P2001228** | Dctn6 | 0.022337344 | -0.326201874 | 0.370876561 | 13.36834994 | 0.153175396 | 0.882803421 | 0.996633405 | -5.246823026 |
| **A_52_P200207** | Rmnd5a | -0.009778724 | -0.162382472 | 0.142825025 | 8.957808635 | -0.153153211 | 0.882820243 | 0.996633405 | -5.246825881 |
| **A_55_P1958804** | Hsd3b3 | -0.006417522 | -0.10659547 | 0.093760425 | 7.685489214 | -0.153110299 | 0.882852782 | 0.996633405 | -5.246831403 |
| **A_52_P654752** | Trub1 | -0.006665361 | -0.110796704 | 0.097465982 | 8.103961611 | -0.152985876 | 0.88294713 | 0.996633405 | -5.246847404 |
| **A_51_P206655** | Lsm1 | -0.007697031 | -0.128059767 | 0.112665706 | 10.52233976 | -0.152841147 | 0.883056879 | 0.996633405 | -5.246866 |
| **A_55_P1972257** | Sars | 0.013418978 | -0.196714534 | 0.223552491 | 12.30921698 | 0.152627666 | 0.883218767 | 0.996633405 | -5.246893399 |
| **A_55_P2370384** | 1600029I1 | 0.006562596 | -0.096258691 | 0.109383883 | 7.645868472 | 0.152546323 | 0.883280454 | 0.996633405 | -5.246903829 |
| **A_55_P1974347** | Fam126b | -0.00707444 | -0.118005437 | 0.103856556 | 8.323817131 | -0.152422231 | 0.88337456 | 0.996633405 | -5.246919729 |
| **A_51_P196127** | Papss1 | -0.010363734 | -0.17294152 | 0.152214052 | 10.31654405 | -0.152357619 | 0.88342356 | 0.996633405 | -5.246928003 |
| **A_55_P2000728** | Myef2 | -0.007156738 | -0.119465617 | 0.105152141 | 9.132969385 | -0.152303593 | 0.883464532 | 0.996633405 | -5.246934919 |
| **A_55_P1971574** | Acox3 | -0.021418357 | -0.35762754 | 0.314790825 | 9.881067458 | -0.152259965 | 0.88349762 | 0.996633405 | -5.246940502 |
| **A_55_P1953093** | Pabpc4 | -0.008497248 | -0.141910485 | 0.124915988 | 12.79894252 | -0.152225891 | 0.883523461 | 0.996633405 | -5.246944861 |
| **A_66_P132232** | Ercc1 | -0.018020713 | -0.300963464 | 0.264922039 | 9.365434138 | -0.152223787 | 0.883525056 | 0.996633405 | -5.24694513 |
| **A_55_P2079471** | NA | -0.016395275 | -0.273898035 | 0.241107485 | 8.914111622 | -0.152175925 | 0.883561355 | 0.996633405 | -5.246951252 |
| **A_55_P2021629** | Abcb7 | 0.009887314 | -0.145526053 | 0.165300681 | 8.402524122 | 0.15205444 | 0.88365349 | 0.996633405 | -5.246966781 |
| **A_52_P121051** | 2610001J0 | 0.013552056 | -0.199586173 | 0.226690284 | 11.79255012 | 0.151968282 | 0.883718835 | 0.996633405 | -5.246977787 |
| **A_55_P2112701** | NA | -0.034074362 | -0.570485694 | 0.502336971 | 13.421108 | -0.151823472 | 0.883828664 | 0.996633405 | -5.246996271 |
| **A_55_P1981405** | Gsk3a | 0.008717071 | -0.128540679 | 0.14597482 | 12.26472247 | 0.151789884 | 0.883854139 | 0.996633405 | -5.247000556 |
| **A_55_P1964364** | Kctd14 | 0.009137916 | -0.134846619 | 0.15312245 | 17.44417628 | 0.151684229 | 0.883934274 | 0.996633405 | -5.247014028 |
| **A_51_P413059** | Mcpt8 | -0.007153541 | -0.119902355 | 0.105595272 | 7.725010318 | -0.15164156 | 0.883966637 | 0.996633405 | -5.247019466 |
| **A_66_P136714** | Scpep1 | -0.011617619 | -0.194901568 | 0.171666329 | 11.78515147 | -0.151496237 | 0.884076862 | 0.996633405 | -5.247037976 |
| **A_66_P111562** | Ccnd1 | -0.010812755 | -0.181561469 | 0.159935958 | 12.29940393 | -0.151351964 | 0.884186293 | 0.996633405 | -5.247056335 |
| **A_51_P321531** | Purg | 0.009728721 | -0.143973728 | 0.163431169 | 9.948186347 | 0.151280894 | 0.884240201 | 0.996633405 | -5.247065372 |
| **A_55_P1966620** | Rab5a | -0.010061973 | -0.169033976 | 0.148910029 | 9.796971609 | -0.151276562 | 0.884243486 | 0.996633405 | -5.247065923 |
| **A_55_P1990225** | Fnta | -0.010041211 | -0.168746401 | 0.14866398 | 12.34081217 | -0.151218208 | 0.88428775 | 0.996633405 | -5.24707334 |
| **A_51_P480881** | Thap11 | -0.007819313 | -0.131428425 | 0.115789799 | 11.762769 | -0.151191445 | 0.88430805 | 0.996633405 | -5.247076741 |
| **A_55_P2042698** | Hsd17b10 | -0.00952527 | -0.160201179 | 0.14115064 | 14.29664365 | -0.151092389 | 0.884383188 | 0.996633405 | -5.247089323 |
| **A_51_P498890** | Degs1 | 0.011612736 | -0.172098685 | 0.195324158 | 13.30647252 | 0.151080197 | 0.884392436 | 0.996633405 | -5.247090871 |

| | | | | | | | | | |
|---|---|---|---|---|---|---|---|---|---|
| **A_55_P2262136** | Tek | -0.009933828 | -0.16711059 | 0.147242934 | 9.188907296 | -0.151055812 | 0.884410933 | 0.996633405 | -5.247093966 |
| **A_52_P460836** | Lrrn4 | 0.02917143 | -0.432533251 | 0.490876111 | 8.546306371 | 0.151008909 | 0.884446512 | 0.996633405 | -5.24709992 |
| **A_55_P2069592** | Pitpnm3 | -0.008375193 | -0.140960714 | 0.124210329 | 7.780391599 | -0.150975974 | 0.884471495 | 0.996633405 | -5.247104099 |
| **A_55_P1981949** | Klhl5 | 0.011490908 | -0.170617693 | 0.193599508 | 9.794516848 | 0.150810997 | 0.884596642 | 0.996633405 | -5.247125019 |
| **A_55_P2069935** | Fxyd2 | 0.019936276 | -0.296054601 | 0.335927153 | 8.489323117 | 0.150792111 | 0.884610968 | 0.996633405 | -5.247127413 |
| **A_55_P2022369** | Lbh | 0.016169295 | -0.240284097 | 0.272622687 | 12.4978703 | 0.150692546 | 0.884686497 | 0.996633405 | -5.247140026 |
| **A_55_P2167000** | Dmac2 | -0.013258918 | -0.223634822 | 0.197116985 | 12.61297451 | -0.150633384 | 0.884731377 | 0.996633405 | -5.247147517 |
| **A_55_P2118674** | Tnip1 | 0.013230146 | -0.196695389 | 0.223155681 | 12.41578378 | 0.150628967 | 0.884734728 | 0.996633405 | -5.247148076 |
| **A_55_P1993879** | NA | -0.014062492 | -0.23725697 | 0.209131986 | 16.13663145 | -0.150587171 | 0.884766435 | 0.996633405 | -5.247153366 |
| **A_51_P234025** | Rbm5 | 0.014870076 | -0.221163732 | 0.250903883 | 12.67968053 | 0.150573346 | 0.884776923 | 0.996633405 | -5.247155116 |
| **A_52_P247788** | Ino80c | 0.006786239 | -0.100932735 | 0.114505212 | 8.125326292 | 0.150572642 | 0.884777457 | 0.996633405 | -5.247155205 |
| **A_51_P284244** | Tmed3 | 0.012101448 | -0.1800665 | 0.204269396 | 13.10992161 | 0.150510057 | 0.884824935 | 0.996633405 | -5.247163123 |
| **A_51_P139320** | Pcbd1 | 0.009841053 | -0.14645223 | 0.166134337 | 12.25979927 | 0.150490959 | 0.884839423 | 0.996633405 | -5.247165539 |
| **A_51_P259861** | Mettl9 | -0.00891283 | -0.150477326 | 0.132651666 | 12.46961503 | -0.150477096 | 0.88484994 | 0.996633405 | -5.247167292 |
| **A_55_P1999532** | Col9a1 | 0.029800733 | -0.443637632 | 0.503239098 | 8.305714497 | 0.150443223 | 0.884875637 | 0.996633405 | -5.247171575 |
| **A_55_P1992582** | Hmgcs2 | -0.00631233 | -0.106606943 | 0.093982283 | 7.681930537 | -0.150425418 | 0.884889145 | 0.996633405 | -5.247173826 |
| **A_55_P1992839** | Dsn1 | 0.01340592 | -0.199757045 | 0.226568886 | 9.995428141 | 0.150312126 | 0.884975092 | 0.996633405 | -5.247188143 |
| **A_51_P430950** | Rdx | 0.01020491 | -0.152098043 | 0.172507863 | 11.0713973 | 0.15027678 | 0.885001908 | 0.996633405 | -5.247192608 |
| **A_55_P2021921** | Fkbp1a | -0.01939653 | -0.327897382 | 0.289104323 | 14.78996124 | -0.15027156 | 0.885005867 | 0.996633405 | -5.247193267 |
| **A_51_P294095** | Prkar1a | -0.012770813 | -0.215995548 | 0.190453922 | 11.3793474 | -0.150193488 | 0.885065097 | 0.996633405 | -5.247203124 |
| **A_51_P380153** | Ubr4 | -0.010577882 | -0.178970904 | 0.15781514 | 12.10481071 | -0.150135625 | 0.885108996 | 0.996633405 | -5.247210426 |
| **A_55_P1966749** | Cyp2s1 | 0.056518599 | -0.843698323 | 0.95673552 | 9.926255393 | 0.150055993 | 0.885169411 | 0.996633405 | -5.247220472 |
| **A_52_P299311** | NA | 0.006717226 | -0.100309893 | 0.113744344 | 14.58107304 | 0.150004825 | 0.885208231 | 0.996633405 | -5.247226923 |
| **A_51_P355151** | Camk2n2 | -0.007017427 | -0.118845458 | 0.104810604 | 8.73785469 | -0.149981037 | 0.885226279 | 0.996633405 | -5.247229922 |
| **A_51_P223575** | Tbc1d23 | 0.012373257 | -0.184999448 | 0.209745963 | 9.716262429 | 0.14983252 | 0.885338959 | 0.996633405 | -5.247248633 |
| **A_52_P365342** | Lrrn2 | -0.021519117 | -0.365155914 | 0.322117681 | 9.311149212 | -0.149669707 | 0.885462489 | 0.996633405 | -5.247269125 |
| **A_55_P2164977** | NA | 0.006485649 | -0.097141168 | 0.110112467 | 7.760161299 | 0.14958582 | 0.885526136 | 0.996633405 | -5.247279674 |
| **A_55_P2004046** | NA | -0.010698807 | -0.181712141 | 0.160314527 | 8.416548792 | -0.149525244 | 0.885572098 | 0.996633405 | -5.247287288 |
| **A_52_P108845** | Clip3 | 0.016268337 | -0.243793451 | 0.276330125 | 9.790124932 | 0.149511897 | 0.885582225 | 0.996633405 | -5.247288965 |
| **A_55_P2123942** | Lmbrd1 | 0.010840135 | -0.162503687 | 0.184183956 | 9.260849903 | 0.149463601 | 0.88561887 | 0.996633405 | -5.247295033 |
| **A_55_P1984406** | Lrrk2 | -0.011694636 | -0.198733542 | 0.175344269 | 9.457753447 | -0.14943898 | 0.885637551 | 0.996633405 | -5.247298125 |
| **A_51_P273203** | Gpaa1 | 0.010072668 | -0.151031705 | 0.171177041 | 8.736157351 | 0.149432932 | 0.885642141 | 0.996633405 | -5.247298885 |
| **A_55_P1998539** | C1qtnf1 | -0.007277989 | -0.123705559 | 0.109149581 | 9.065348578 | -0.149404851 | 0.885663447 | 0.996633405 | -5.247302411 |
| **A_55_P2366358** | 4930444F0 | -0.006146551 | -0.104532949 | 0.092239848 | 7.666998539 | -0.149315718 | 0.885731078 | 0.996633405 | -5.2473136 |
| **A_66_P134474** | Ang3 | -0.007633273 | -0.129897111 | 0.114630566 | 7.998955375 | -0.149218229 | 0.885805051 | 0.996633405 | -5.247325831 |
| **A_55_P2098354** | Shf | 0.010818904 | -0.162475638 | 0.184113447 | 9.120847259 | 0.149213298 | 0.885808793 | 0.996633405 | -5.247326449 |
| **A_51_P163953** | Nsg2 | -0.010378063 | -0.17661833 | 0.155862203 | 8.374288213 | -0.149207006 | 0.885813567 | 0.996633405 | -5.247327238 |
| **A_51_P283754** | Cpsf7 | 0.01126623 | -0.169238057 | 0.191770518 | 11.48318611 | 0.149176437 | 0.885836764 | 0.996633405 | -5.247331071 |
| **A_55_P2020331** | Camk1g | -0.012236036 | -0.208282358 | 0.183810286 | 11.12234648 | -0.14917333 | 0.885839121 | 0.996633405 | -5.247331461 |
| **A_51_P106397** | Prkd3 | 0.008265348 | -0.124183593 | 0.140714289 | 8.352381354 | 0.149149495 | 0.885857207 | 0.996633405 | -5.213063003 |
| **A_52_P260105** | Man1a | 0.007120618 | -0.107111651 | 0.121352886 | 7.846281727 | 0.148983441 | 0.88598321 | 0.996633405 | -5.247355253 |
| **A_55_P2323913** | NA | 0.013592823 | -0.20467501 | 0.231860655 | 9.094648077 | 0.148843216 | 0.886089617 | 0.996633405 | -5.247372804 |
| **A_55_P1952434** | Tm2d1 | -0.007234448 | -0.123427391 | 0.108958494 | 11.49830749 | -0.148810918 | 0.886114126 | 0.996633405 | -5.247376844 |
| **A_55_P2046943** | Bnip3l | 0.017007854 | -0.256181584 | 0.290197292 | 11.78396077 | 0.148797199 | 0.886124537 | 0.996633405 | -5.24737856 |
| **A_55_P2122020** | Klf4 | -0.014734602 | -0.251427055 | 0.221957852 | 9.845321499 | -0.148786359 | 0.886132763 | 0.996633405 | -5.247379915 |
| **A_65_P01415** | Kremen1 | -0.011333706 | -0.193416053 | 0.170748641 | 11.16451214 | -0.148769274 | 0.886145727 | 0.996633405 | -5.247382052 |
| **A_51_P159453** | Serpina3n | 0.006447372 | -0.09714469 | 0.110039433 | 7.674491268 | 0.14875288 | 0.886158168 | 0.996633405 | -5.247384102 |
| **A_65_P15340** | Zfp251 | -0.012420101 | -0.21201272 | 0.187172519 | 9.775898261 | -0.148726984 | 0.886177819 | 0.996633405 | -5.247387339 |

| | | | | | | | | | |
|---|---|---|---|---|---|---|---|---|---|
| **A_55_P2100866** | NA | -0.009928279 | -0.169919277 | 0.150062718 | 10.96441417 | -0.148315854 | 0.886489817 | 0.996633405 | -5.247438661 |
| **A_51_P334685** | Esyt1 | -0.013733548 | -0.23506091 | 0.207593814 | 9.389849571 | -0.148305334 | 0.886497801 | 0.996633405 | -5.247439973 |
| **A_51_P274124** | Mansc1 | 0.006187771 | -0.093569433 | 0.105944974 | 9.491703669 | 0.148251495 | 0.88653866 | 0.996633405 | -5.247446683 |
| **A_55_P1966883** | Rnaseh2a | -0.015902325 | -0.272321641 | 0.240516991 | 10.16592902 | -0.148224165 | 0.886559401 | 0.996633405 | -5.247450088 |
| **A_55_P2080163** | Carhsp1 | -0.007989642 | -0.136866143 | 0.120886859 | 11.92645654 | -0.148170824 | 0.886599883 | 0.996633405 | -5.247456732 |
| **A_52_P625808** | Sf3b1 | 0.013929571 | -0.21108931 | 0.238948452 | 9.325679177 | 0.14795441 | 0.886764128 | 0.996633405 | -5.247483665 |
| **A_52_P327467** | Med7 | -0.008564122 | -0.14696623 | 0.129837985 | 10.69981992 | -0.147893571 | 0.886810302 | 0.996633405 | -5.247491229 |
| **A_55_P2087322** | Rapsn | 0.011126208 | -0.168829014 | 0.19108143 | 12.8468809 | 0.147771902 | 0.886902645 | 0.996633405 | -5.247506348 |
| **A_51_P453475** | Slc1a5 | -0.010983862 | -0.188668757 | 0.166701033 | 10.05836185 | -0.147745306 | 0.886922831 | 0.996633405 | -5.247509651 |
| **A_55_P2074193** | NA | -0.017545638 | -0.301427425 | 0.26633615 | 14.44651115 | -0.147720496 | 0.886941661 | 0.996633405 | -5.247512732 |
| **A_51_P438083** | Slc6a13 | 0.007046304 | -0.107010258 | 0.121102867 | 8.48130309 | 0.147655711 | 0.886990832 | 0.996633405 | -5.247520774 |
| **A_51_P172131** | Lurap1l | -0.010377137 | -0.178362632 | 0.157608359 | 8.488273123 | -0.147643687 | 0.886999958 | 0.996633405 | -5.247522266 |
| **A_55_P2011867** | Kif19a | 0.009004987 | -0.136929819 | 0.154939793 | 8.294568266 | 0.147480068 | 0.887124146 | 0.996633405 | -5.24754256 |
| **A_51_P341349** | Mtap | 0.010884867 | -0.165532821 | 0.187302555 | 9.247920221 | 0.147465398 | 0.88713528 | 0.996633405 | -5.247544379 |
| **A_51_P285042** | Rtf1 | 0.008796646 | -0.133868787 | 0.15146208 | 9.918874244 | 0.147369463 | 0.887208097 | 0.996633405 | -5.247556266 |
| **A_55_P2071132** | Tnfrsf23 | -0.009327613 | -0.160612381 | 0.141957154 | 8.641033265 | -0.147361643 | 0.887214032 | 0.996633405 | -5.247557235 |
| **A_55_P2064004** | Rapgef2 | -0.008834121 | -0.152120183 | 0.13445194 | 10.51141763 | -0.147356247 | 0.887218128 | 0.996633405 | -5.247557903 |
| **A_51_P337662** | Ints6l | 0.019161947 | -0.291685751 | 0.330009646 | 9.055356404 | 0.147333367 | 0.887235495 | 0.996633405 | -5.247560737 |
| **A_66_P110161** | Eppk1 | -0.027571723 | -0.475238398 | 0.420094953 | 9.868014052 | -0.147203546 | 0.887334034 | 0.996633405 | -5.247576808 |
| **A_51_P219266** | Tmprss6 | -0.007182958 | -0.12386895 | 0.109503034 | 7.763568961 | -0.147127461 | 0.887391787 | 0.996633405 | -5.247586219 |
| **A_51_P391616** | Etnppl | -0.006208827 | -0.107076733 | 0.094659079 | 7.711527252 | -0.147117957 | 0.887399001 | 0.996633405 | -5.247587395 |
| **A_51_P337543** | Glce | -0.008241825 | -0.142150012 | 0.125666362 | 9.671703257 | -0.147104312 | 0.887409358 | 0.996633405 | -5.247589082 |
| **A_52_P339054** | Nap1l4 | -0.018441577 | -0.318152333 | 0.281269179 | 10.13790034 | -0.147063488 | 0.887440347 | 0.996633405 | -5.247594129 |
| **A_55_P2033616** | Dhrs7b | 0.012030124 | -0.183548713 | 0.207608961 | 10.89853316 | 0.147013554 | 0.88747825 | 0.996633405 | -5.247600301 |
| **A_55_P2026889** | NA | -0.010194054 | -0.176063196 | 0.155675087 | 12.45896597 | -0.146889407 | 0.887572488 | 0.996633405 | -5.247615636 |
| **A_55_P2000623** | Stag1 | 0.009019439 | -0.137871882 | 0.15591076 | 10.54769992 | 0.146754869 | 0.887674615 | 0.996633405 | -5.24763224 |
| **A_55_P2013645** | Tubg2 | 0.00752424 | -0.115030695 | 0.130079175 | 8.786849282 | 0.146737444 | 0.887687843 | 0.996633405 | -5.24763439 |
| **A_51_P366770** | Rims1 | 0.011401078 | -0.174300261 | 0.197102417 | 8.221666261 | 0.146737129 | 0.887688082 | 0.996633405 | -5.247634429 |
| **A_55_P2012096** | Bmp8a | 0.008853925 | -0.135401104 | 0.153108954 | 8.211650574 | 0.146694558 | 0.887720398 | 0.996633405 | -5.247639679 |
| **A_66_P100105** | Patj | 0.008435462 | -0.129011754 | 0.145882679 | 8.521365659 | 0.146683774 | 0.887728584 | 0.996633405 | -5.247641008 |
| **A_55_P2125868** | NA | 0.008545516 | -0.130725551 | 0.147816582 | 11.54510063 | 0.146651499 | 0.887753084 | 0.996633405 | -5.247644988 |
| **A_55_P1970771** | Gm13141 | 0.007876764 | -0.120615866 | 0.136369393 | 13.03618608 | 0.146513865 | 0.887857566 | 0.996633405 | -5.247661946 |
| **A_55_P2093614** | NA | 0.006893107 | -0.105566656 | 0.11935287 | 7.848553462 | 0.146496408 | 0.887870819 | 0.996633405 | -5.247664096 |
| **A_55_P2133679** | NA | -0.025452843 | -0.440961904 | 0.390056218 | 13.27209062 | -0.146408041 | 0.887937902 | 0.996633405 | -5.247674975 |
| **A_55_P1974617** | Lysmd1 | -0.010399021 | -0.180303139 | 0.159505097 | 10.36494292 | -0.146284291 | 0.888031848 | 0.996633405 | -5.247690198 |
| **A_52_P24308** | Uchl1 | -0.01964237 | -0.340681968 | 0.301397228 | 9.303152162 | -0.146232674 | 0.888071034 | 0.996633405 | -5.247696545 |
| **A_55_P2137937** | Fxyd1 | -0.014332779 | -0.248740236 | 0.220074678 | 15.72673977 | -0.146139665 | 0.888141645 | 0.996633405 | -5.247707974 |
| **A_55_P1957880** | Jpt2 | 0.013258534 | -0.203695769 | 0.230212838 | 9.38654327 | 0.146061711 | 0.888200826 | 0.996633405 | -5.247717548 |
| **A_55_P2018249** | Lmo4 | 0.008457551 | -0.130052942 | 0.146968045 | 9.539516788 | 0.145938908 | 0.888294059 | 0.996633405 | -5.24773262 |
| **A_55_P2261772** | C230098O: | 0.008382639 | -0.129002251 | 0.145767528 | 10.15422532 | 0.145831354 | 0.888375716 | 0.996633405 | -5.24774581 |
| **A_52_P214927** | Zmynd8 | -0.009599652 | -0.166932651 | 0.147733346 | 9.676633815 | -0.145829309 | 0.888377268 | 0.996633405 | -5.24774606 |
| **A_55_P2061273** | Tbx6 | 0.008140664 | -0.125342196 | 0.141623524 | 7.913565963 | 0.145761706 | 0.888428594 | 0.996633405 | -5.247754346 |
| **A_55_P2026487** | NA | 0.006599742 | -0.101748257 | 0.114947741 | 7.650915648 | 0.145584518 | 0.888563123 | 0.996633405 | -5.247776044 |
| **A_55_P1966659** | LOC54734! | 0.007454376 | -0.114933739 | 0.129842491 | 7.745260519 | 0.145573116 | 0.888571781 | 0.996633405 | -5.247777439 |
| **A_55_P1995768** | Zfp967 | -0.008644109 | -0.150570265 | 0.133282047 | 9.479576894 | -0.145568343 | 0.888575404 | 0.996633405 | -5.247778023 |
| **A_55_P2057528** | Arl4d | 0.019427091 | -0.299827791 | 0.338681972 | 9.000538094 | 0.14543849 | 0.888673997 | 0.996633405 | -5.247793906 |
| **A_55_P2036978** | Mrps14 | 0.009532433 | -0.147291337 | 0.166356203 | 12.30834686 | 0.145278384 | 0.888795563 | 0.996633405 | -5.247813471 |
| **A_55_P2071426** | Eef2kmt | 0.008100894 | -0.125183165 | 0.141384953 | 8.577828297 | 0.145265967 | 0.888804992 | 0.996633405 | -5.247814987 |

| | | | | | | | | | |
|---|---|---|---|---|---|---|---|---|---|
| **A_66_P106072** | Orai1 | 0.010920814 | -0.168824516 | 0.190666144 | 8.729609524 | 0.145213346 | 0.888844946 | 0.996633405 | -5.247821412 |
| **A_55_P1984158** | Slc44a2 | -0.010113867 | -0.176603404 | 0.15637567 | 11.61958819 | -0.145190913 | 0.88886198 | 0.996633405 | -5.24782415 |
| **A_55_P2096081** | Myo5b | 0.042185636 | -0.652570906 | 0.736942178 | 9.540394768 | 0.145124618 | 0.888912318 | 0.996633405 | -5.24783224 |
| **A_55_P2049949** | NA | 0.01086226 | -0.168071092 | 0.189795613 | 8.691921327 | 0.145090188 | 0.888938462 | 0.996633405 | -5.24783644 |
| **A_55_P1990653** | Comt | 0.009112697 | -0.141023074 | 0.159248468 | 12.66359881 | 0.14506809 | 0.888955241 | 0.996633405 | -5.247839135 |
| **A_51_P348227** | Stoml1 | 0.007684511 | -0.118934304 | 0.134303326 | 8.864820427 | 0.145053133 | 0.888966599 | 0.996633405 | -5.247840959 |
| **A_51_P289742** | Slc27a5 | -0.006182473 | -0.108093714 | 0.095728768 | 7.691219445 | -0.144993747 | 0.889011692 | 0.996633405 | -5.247848199 |
| **A_66_P139387** | Prlr | -0.00711559 | -0.124516651 | 0.11028547 | 8.164581489 | -0.144859854 | 0.889113362 | 0.996633405 | -5.247864512 |
| **A_51_P487298** | Vasn | -0.018210479 | -0.318702875 | 0.282281916 | 11.30010013 | -0.144842838 | 0.889126283 | 0.996633405 | -5.247866584 |
| **A_55_P2158151** | NA | -0.00842381 | -0.147450282 | 0.130602663 | 15.73923659 | -0.144817208 | 0.889145745 | 0.996633405 | -5.247869705 |
| **A_52_P453374** | Lrrn4cl | 0.012706655 | -0.197019274 | 0.222432584 | 8.520908773 | 0.144806565 | 0.889153827 | 0.996633405 | -5.247871001 |
| **A_51_P184796** | Zfp101 | 0.007418792 | -0.115049642 | 0.129887226 | 8.531355286 | 0.14478319 | 0.889171577 | 0.996633405 | -5.247873846 |
| **A_55_P2148609** | NA | 0.012768009 | -0.198013435 | 0.223549453 | 8.758081364 | 0.144777124 | 0.889176183 | 0.996633405 | -5.247874584 |
| **A_55_P2102759** | NA | 0.007669353 | -0.119013034 | 0.13435174 | 8.112898264 | 0.144694361 | 0.88923903 | 0.996633405 | -5.247884655 |
| **A_55_P2186808** | NA | 0.010860867 | -0.168588589 | 0.190310323 | 12.04484373 | 0.144654342 | 0.889269419 | 0.996633405 | -5.247889522 |
| **A_52_P576049** | Inhbc | 0.006158765 | -0.095600952 | 0.107918483 | 7.682196494 | 0.144652818 | 0.889270577 | 0.996633405 | -5.247889707 |
| **A_55_P2143209** | Axin1 | -0.010013592 | -0.175511894 | 0.155484709 | 8.984124445 | -0.144612387 | 0.889301279 | 0.996633405 | -5.247894623 |
| **A_55_P2031217** | Vps52 | 0.009885722 | -0.153927548 | 0.173698993 | 11.47223241 | 0.14423427 | 0.889588422 | 0.996633405 | -5.247940533 |
| **A_55_P2079684** | NA | 0.00684586 | -0.10660138 | 0.120293101 | 14.31900922 | 0.144225892 | 0.889594784 | 0.996633405 | -5.247941549 |
| **A_55_P2056606** | NA | -0.00684456 | -0.120341973 | 0.106652852 | 8.186188621 | -0.144134762 | 0.889663992 | 0.996633405 | -5.247952595 |
| **A_55_P2057653** | NA | 0.010123076 | -0.157885542 | 0.178131694 | 15.4270915 | 0.144009144 | 0.889759392 | 0.996633405 | -5.24796781 |
| **A_51_P392214** | Pcnx4 | -0.009640735 | -0.169740148 | 0.150458678 | 9.009174152 | -0.143922778 | 0.889824983 | 0.996633405 | -5.247978263 |
| **A_51_P497768** | Gpat2 | 0.008214735 | -0.128288571 | 0.144718042 | 7.943159076 | 0.14383333 | 0.889892916 | 0.996633405 | -5.247989083 |
| **A_51_P230734** | Slc34a3 | -0.006216325 | -0.109624015 | 0.097191365 | 7.718347541 | -0.143677914 | 0.890010953 | 0.996633405 | -5.248007866 |
| **A_51_P379373** | Lactb2 | -0.008247464 | -0.145494914 | 0.128999986 | 8.619044027 | -0.143623425 | 0.890052337 | 0.996633405 | -5.248014447 |
| **A_51_P407606** | Ndc1 | -0.016279524 | -0.287380585 | 0.254821537 | 9.752048436 | -0.143522379 | 0.890129082 | 0.996633405 | -5.248026644 |
| **A_55_P2032760** | Ctif | -0.010809195 | -0.190849414 | 0.169231024 | 10.70925612 | -0.143493742 | 0.890150833 | 0.996633405 | -5.248030099 |
| **A_51_P247157** | Jpt1 | -0.011015065 | -0.194524598 | 0.172494469 | 11.34255889 | -0.14346222 | 0.890174774 | 0.996633405 | -5.248033902 |
| **A_51_P519369** | Agtpbp1 | -0.010027973 | -0.177146521 | 0.157090574 | 9.260077476 | -0.143416017 | 0.890209867 | 0.996633405 | -5.248039473 |
| **A_55_P2125588** | Pdgfa | 0.007721401 | -0.121024639 | 0.136467442 | 12.80925683 | 0.143341315 | 0.890266605 | 0.996633405 | -5.248048478 |
| **A_51_P185593** | Synm | -0.010333162 | -0.182647458 | 0.161981134 | 8.102732273 | -0.143324702 | 0.890279223 | 0.996633405 | -5.24805048 |
| **A_55_P2011322** | NA | -0.011280599 | -0.199630015 | 0.177068817 | 11.75009775 | -0.143145275 | 0.890415507 | 0.996633405 | -5.248072088 |
| **A_51_P117581** | Cables1 | 0.011322671 | -0.177748971 | 0.200394312 | 9.755938925 | 0.14313031 | 0.890426874 | 0.996633405 | -5.248073889 |
| **A_66_P110490** | Polr3h | -0.009874616 | -0.174912214 | 0.155162981 | 8.645901956 | -0.143003437 | 0.890523244 | 0.996633405 | -5.24808915 |
| **A_55_P2025008** | Hvcn1 | 0.010989752 | -0.172898729 | 0.194878234 | 8.427127344 | 0.142837589 | 0.89064922 | 0.996633405 | -5.248109078 |
| **A_55_P1960616** | Dnajc17 | 0.007859155 | -0.123779483 | 0.139497793 | 8.406736758 | 0.142692655 | 0.890759313 | 0.996633405 | -5.248126475 |
| **A_55_P2033312** | Lgi4 | 0.006572742 | -0.103545648 | 0.116691133 | 8.073313659 | 0.142657918 | 0.8907857 | 0.996633405 | -5.248130642 |
| **A_52_P637812** | Prrg4 | 0.006483575 | -0.102190091 | 0.115157241 | 7.792938288 | 0.142593381 | 0.890834724 | 0.996633405 | -5.248138381 |
| **A_55_P2042121** | Slc25a22 | 0.014768442 | -0.232885305 | 0.262422189 | 8.439099633 | 0.142527578 | 0.89088471 | 0.996633405 | -5.248146268 |
| **A_55_P2153620** | Ahnak | 0.020644169 | -0.325546702 | 0.366835041 | 9.849995697 | 0.142524962 | 0.890886697 | 0.996633405 | -5.248146582 |
| **A_55_P2067563** | NA | -0.006859032 | -0.121904797 | 0.108186732 | 8.170921705 | -0.142495568 | 0.890909027 | 0.996633405 | -5.248150104 |
| **A_55_P1954555** | NA | -0.006890631 | -0.122469891 | 0.108688629 | 7.78865199 | -0.142491256 | 0.890912302 | 0.996633405 | -5.24815062 |
| **A_51_P392244** | Tbcd | -0.005793228 | -0.103035519 | 0.091449064 | 10.41007795 | -0.14238838 | 0.890990452 | 0.996633405 | -5.248162941 |
| **A_52_P562612** | Srd5a1 | 0.006345029 | -0.1002265 | 0.112916559 | 7.70755671 | 0.142298907 | 0.891058421 | 0.996633405 | -5.248173649 |
| **A_52_P537381** | Dimt1 | 0.011726633 | -0.185271523 | 0.208724788 | 10.36526939 | 0.142272281 | 0.891078648 | 0.996633405 | -5.248176834 |
| **A_51_P451588** | Plekhb1 | 0.005797566 | -0.091623517 | 0.103218649 | 7.998048171 | 0.142233492 | 0.891108115 | 0.996633405 | -5.248181474 |
| **A_52_P244723** | Tbcel | -0.014796375 | -0.263487035 | 0.233894286 | 9.471528612 | -0.142201761 | 0.891132221 | 0.996633405 | -5.248185268 |
| **A_51_P129720** | Hectd3 | -0.006018225 | -0.107254658 | 0.095218209 | 9.06716278 | -0.142082539 | 0.891222792 | 0.996633405 | -5.248199517 |

| | | | | | | | | | |
|---|---|---|---|---|---|---|---|---|---|
| **A_55_P2101696** | Gnat2 | -0.006187307 | -0.110277528 | 0.097902913 | 7.964508832 | -0.142069512 | 0.891232688 | 0.996633405 | -5.248201073 |
| **A_55_P1953688** | Top1 | 0.007528838 | -0.119142604 | 0.134200281 | 11.81991473 | 0.14205561 | 0.89124325 | 0.996633405 | -5.248202733 |
| **A_55_P2035951** | Haus8 | 0.007728241 | -0.122341498 | 0.137797979 | 13.449553 | 0.142008225 | 0.891279248 | 0.996633405 | -5.248208392 |
| **A_55_P2047867** | NA | 0.012574569 | -0.199375532 | 0.22452467 | 10.60202133 | 0.141797516 | 0.891439327 | 0.996633405 | -5.248233533 |
| **A_55_P2096823** | NA | 0.006595618 | -0.104577915 | 0.117769152 | 8.41330654 | 0.141795761 | 0.89144066 | 0.996633405 | -5.248233742 |
| **A_55_P2089020** | Slc35g2 | -0.010670066 | -0.190833169 | 0.169493038 | 9.465943536 | -0.14155016 | 0.891627254 | 0.996633405 | -5.248262998 |
| **A_52_P425734** | Afm | -0.008598266 | -0.153874212 | 0.136677681 | 7.738788317 | -0.141457595 | 0.891697582 | 0.996633405 | -5.248274012 |
| **A_66_P133525** | NA | 0.006507912 | -0.103463409 | 0.116479234 | 7.74324224 | 0.141439718 | 0.891711164 | 0.996633405 | -5.248276138 |
| **A_51_P103865** | Ppp1r35 | 0.008627458 | -0.137160646 | 0.154415561 | 8.944380315 | 0.141439229 | 0.891711536 | 0.996633405 | -5.248276196 |
| **A_51_P349888** | Ang2 | 0.006723925 | -0.106919086 | 0.120366936 | 7.77063281 | 0.141412975 | 0.891731484 | 0.996633405 | -5.248279318 |
| **A_66_P119420** | Foxk1 | -0.006970102 | -0.124842712 | 0.110902507 | 9.812538863 | -0.141330334 | 0.891794272 | 0.996633405 | -5.248289141 |
| **A_55_P2185160** | NA | -0.010290922 | -0.184419541 | 0.163837697 | 14.50697579 | -0.141251566 | 0.89185412 | 0.996633405 | -5.248298499 |
| **A_51_P432460** | Ppp1r14a | -0.018520258 | -0.3323407 | 0.295300183 | 9.806659456 | -0.141050603 | 0.892006814 | 0.996633405 | -5.248322351 |
| **A_55_P1988882** | 9-Sep | -0.012233166 | -0.219592718 | 0.195126387 | 10.43778787 | -0.141001571 | 0.892044069 | 0.996633405 | -5.248328165 |
| **A_55_P2158866** | Nme6 | 0.00908668 | -0.145002699 | 0.16317606 | 11.79262084 | 0.140942402 | 0.892089027 | 0.996633405 | -5.248335179 |
| **A_55_P2153555** | Zkscan3 | -0.014294932 | -0.257018053 | 0.228428189 | 10.32094673 | -0.140760268 | 0.892227421 | 0.996633405 | -5.24835675 |
| **A_55_P2049612** | Ube2a | -0.010603659 | -0.190674264 | 0.169466946 | 12.00541153 | -0.140741462 | 0.892241711 | 0.996633405 | -5.248358976 |
| **A_55_P2074908** | Ube4b | 0.009782826 | -0.156390648 | 0.175956299 | 8.596963295 | 0.140705717 | 0.892268873 | 0.996633405 | -5.248363205 |
| **A_52_P598634** | 1190007I07 | 0.00863582 | -0.138163458 | 0.155435098 | 12.03651328 | 0.140601123 | 0.89234835 | 0.996633405 | -5.248375576 |
| **A_55_P1992299** | Cep295 | 0.018405859 | -0.294500436 | 0.331312155 | 8.161704982 | 0.140588865 | 0.892357665 | 0.996633405 | -5.248377025 |
| **A_55_P2021345** | Gm14717 | 0.006617932 | -0.105917556 | 0.119153419 | 8.394655067 | 0.140553583 | 0.892384475 | 0.996633405 | -5.248381195 |
| **A_55_P2040777** | Lipo3 | 0.008774512 | -0.140682383 | 0.158231407 | 9.48921872 | 0.140318895 | 0.892562813 | 0.996633405 | -5.248408909 |
| **A_55_P2093251** | NA | -0.006396442 | -0.115445759 | 0.102652875 | 9.570976104 | -0.140192457 | 0.892658896 | 0.996633405 | -5.248423821 |
| **A_55_P2119716** | Mapk1ip1 | 0.013007408 | -0.208823154 | 0.23483797 | 9.81024433 | 0.140145293 | 0.892694738 | 0.996633405 | -5.24842938 |
| **A_55_P2429432** | 2610301B2 | 0.0077878 | -0.125088529 | 0.140664128 | 8.685483597 | 0.140080052 | 0.892744317 | 0.996633405 | -5.248437067 |
| **A_55_P2056808** | Tanc1 | 0.007130842 | -0.11456985 | 0.128831535 | 8.52652748 | 0.140041554 | 0.892773573 | 0.996633405 | -5.248441601 |
| **A_55_P2187918** | Cep55 | 0.006392984 | -0.10276426 | 0.115550228 | 8.695840974 | 0.139978137 | 0.892821766 | 0.996633405 | -5.248449068 |
| **A_55_P2084723** | NA | 0.005768992 | -0.092789172 | 0.104327156 | 7.828072524 | 0.139899603 | 0.892881449 | 0.996633405 | -5.248458309 |
| **A_55_P2132507** | Bcl2l2 | 0.008479593 | -0.136393031 | 0.153352217 | 9.536256311 | 0.13989359 | 0.892886019 | 0.996633405 | -5.248459016 |
| **A_55_P2092671** | Nppc | -0.006400582 | -0.115800092 | 0.102998929 | 7.785458007 | -0.139834137 | 0.892931201 | 0.996633405 | -5.248466009 |
| **A_55_P2158873** | Ppid | -0.013171992 | -0.238572283 | 0.212228298 | 10.36017642 | -0.139670964 | 0.893055209 | 0.996633405 | -5.248485185 |
| **A_55_P2038414** | Acot8 | 0.009329086 | -0.150406708 | 0.16906488 | 10.24870703 | 0.139587319 | 0.893118779 | 0.996633405 | -5.248495006 |
| **A_52_P131372** | NA | -0.010203459 | -0.184970114 | 0.164563196 | 16.52928591 | -0.139539761 | 0.893154923 | 0.996633405 | -5.248500587 |
| **A_55_P2128969** | Mettl27 | -0.007777458 | -0.141000123 | 0.125445207 | 8.555634886 | -0.13953035 | 0.893162076 | 0.996633405 | -5.248501692 |
| **A_55_P1965635** | Setdb1 | 0.009144703 | -0.147514506 | 0.165803911 | 9.075984806 | 0.139515606 | 0.893173281 | 0.996633405 | -5.248503421 |
| **A_55_P2084656** | H2ac12 | 0.009305558 | -0.150156054 | 0.16876717 | 17.52019566 | 0.139474689 | 0.893204378 | 0.996633405 | -5.248508221 |
| **A_55_P2087785** | NA | -0.006805529 | -0.123507792 | 0.109896733 | 8.149756846 | -0.139377212 | 0.893278464 | 0.996633405 | -5.24851965 |
| **A_51_P392705** | Xpc | -0.008472027 | -0.153785695 | 0.136841642 | 9.104544079 | -0.139344548 | 0.893303289 | 0.996633405 | -5.248523478 |
| **A_55_P2115310** | Spata21 | -0.006718737 | -0.12195988 | 0.108522407 | 7.69692423 | -0.139344298 | 0.893303478 | 0.996633405 | -5.248523507 |
| **A_55_P2095039** | Ncmap | 0.007425816 | -0.119963849 | 0.134815481 | 7.82791592 | 0.139321808 | 0.893320572 | 0.996633405 | -5.248526142 |
| **A_51_P481821** | Spcs3 | 0.006516048 | -0.105284938 | 0.118317033 | 10.53568774 | 0.139298928 | 0.893337962 | 0.996633405 | -5.248528823 |
| **A_55_P2208579** | Onecut2 | 0.007074638 | -0.114439836 | 0.128589112 | 7.861291071 | 0.139150688 | 0.89345063 | 0.996633405 | -5.248546178 |
| **A_55_P2147876** | Abhd17a | 0.017390014 | -0.2815604 | 0.316340427 | 10.45330628 | 0.139030439 | 0.893542027 | 0.996633405 | -5.248560243 |
| **A_51_P154933** | Zdhhc7 | -0.007546957 | -0.137306757 | 0.122212842 | 10.6303575 | -0.139008342 | 0.893558822 | 0.996633405 | -5.248562827 |
| **A_55_P2102719** | NA | -0.010527549 | -0.19165271 | 0.170597612 | 14.24009961 | -0.138917715 | 0.893627706 | 0.996633405 | -5.248573417 |
| **A_55_P2088699** | NA | -0.009267183 | -0.168765636 | 0.150231269 | 16.43501463 | -0.138867436 | 0.893665922 | 0.996633405 | -5.24857929 |
| **A_51_P208555** | Decr1 | -0.006699478 | -0.122054929 | 0.108655973 | 9.2018528 | -0.138807203 | 0.893711705 | 0.996633405 | -5.248586323 |
| **A_51_P167971** | Bace1 | -0.013306589 | -0.242446207 | 0.215833029 | 10.97363914 | -0.138795594 | 0.893720529 | 0.996633405 | -5.248587678 |

| | | | | | | | | | |
|---|---|---|---|---|---|---|---|---|---|
| **A_51_P366121** | Klf14 | -0.008069264 | -0.147029774 | 0.130891246 | 8.078020027 | -0.138787919 | 0.893726363 | 0.996633405 | -5.248588573 |
| **A_55_P2183587** | Gli1 | -0.01419563 | -0.258814118 | 0.230422858 | 8.253508503 | -0.138699382 | 0.893793659 | 0.996633405 | -5.248598904 |
| **A_51_P392291** | Pdk3 | -0.008310728 | -0.151533416 | 0.134911959 | 11.0646823 | -0.138687207 | 0.893802914 | 0.996633405 | -5.248600324 |
| **A_55_P2144248** | Zfp790 | -0.01738363 | -0.317111061 | 0.282343801 | 8.568417563 | -0.138619113 | 0.893854673 | 0.996633405 | -5.248608263 |
| **A_55_P2113673** | Eml1 | 0.011662964 | -0.189541693 | 0.212867622 | 12.76586518 | 0.138541554 | 0.893913627 | 0.996633405 | -5.248617302 |
| **A_55_P1995622** | Pomp | 0.015235168 | -0.247607547 | 0.278077883 | 10.11127669 | 0.138535295 | 0.893918385 | 0.996633405 | -5.248618031 |
| **A_55_P1982211** | NA | -0.021301939 | -0.389181433 | 0.346577555 | 9.810187885 | -0.138395729 | 0.894024474 | 0.996633405 | -5.248634283 |
| **A_52_P337220** | Mgat4b | 0.014930913 | -0.242932738 | 0.272794564 | 9.295555233 | 0.138390211 | 0.894028669 | 0.996633405 | -5.248634925 |
| **A_51_P140211** | Ndufv3 | 0.018012482 | -0.293131913 | 0.329156877 | 10.14323475 | 0.138363238 | 0.894049172 | 0.996633405 | -5.248638064 |
| **A_55_P2038494** | Rexo2 | -0.015863844 | -0.290109687 | 0.258382 | 9.971169799 | -0.138253935 | 0.89413226 | 0.996633405 | -5.248650778 |
| **A_55_P2127971** | Gm9112 | 0.00573829 | -0.093467821 | 0.104944402 | 7.686694979 | 0.138246207 | 0.894138134 | 0.996633405 | -5.248651676 |
| **A_55_P2030717** | NA | -0.01076427 | -0.196874078 | 0.175345538 | 12.0303463 | -0.13823705 | 0.894145094 | 0.996633405 | -5.248652741 |
| **A_51_P390775** | Ube2e1 | 0.010384017 | -0.169181326 | 0.18994936 | 10.78129616 | 0.138213988 | 0.894162626 | 0.996633405 | -5.248655421 |
| **A_55_P1955542** | Olfr522 | 0.007630072 | -0.124371007 | 0.139631151 | 8.269088306 | 0.138152988 | 0.894208996 | 0.996633405 | -5.24866251 |
| **A_51_P476579** | NA | 0.007307988 | -0.119216787 | 0.133832762 | 8.915618392 | 0.138048394 | 0.894288507 | 0.996633405 | -5.248674658 |
| **A_51_P170008** | Napg | -0.006904865 | -0.126491623 | 0.112681893 | 8.844247119 | -0.138000688 | 0.894324772 | 0.996633405 | -5.248680195 |
| **A_52_P334562** | Vdr | 0.007451882 | -0.121705898 | 0.136609661 | 8.057826622 | 0.137896906 | 0.894403667 | 0.996633405 | -5.248692235 |
| **A_55_P2024808** | Abl1 | -0.009174798 | -0.168208194 | 0.149858598 | 12.00602185 | -0.137885096 | 0.894412645 | 0.996633405 | -5.248693605 |
| **A_51_P310949** | Ppox | -0.010058988 | -0.18454916 | 0.164431184 | 9.285205333 | -0.137781993 | 0.894491025 | 0.996633405 | -5.248705556 |
| **A_51_P140607** | Ints13 | -0.008853265 | -0.162449745 | 0.144743216 | 10.64820251 | -0.137762594 | 0.894505773 | 0.996633405 | -5.248707804 |
| **A_55_P1952850** | NA | 0.009169559 | -0.150017584 | 0.168356702 | 14.31648869 | 0.137673254 | 0.894573692 | 0.996633405 | -5.248718151 |
| **A_52_P120612** | Grap2 | -0.005986766 | -0.109953763 | 0.097980232 | 7.714556808 | -0.137627714 | 0.894608312 | 0.996633405 | -5.248723423 |
| **A_51_P396297** | Hsdl1 | -0.011592675 | -0.213059165 | 0.189873815 | 8.386902952 | -0.137527637 | 0.894684395 | 0.996633405 | -5.248735002 |
| **A_52_P69756** | Mterf4 | 0.011793759 | -0.193214945 | 0.216802462 | 10.29918915 | 0.137495685 | 0.894708687 | 0.996633405 | -5.248738698 |
| **A_55_P1968778** | Rasa1 | 0.008473778 | -0.138906314 | 0.15585387 | 9.874401639 | 0.137419194 | 0.89476684 | 0.996633405 | -5.24874754 |
| **A_51_P499530** | Ankrd10 | -0.014740339 | -0.271147268 | 0.241666589 | 9.622627546 | -0.137400041 | 0.894781401 | 0.996633405 | -5.248749754 |
| **A_51_P131408** | Tnfrsf12a | -0.024358639 | -0.448190685 | 0.399473407 | 12.38823072 | -0.137362555 | 0.894809901 | 0.996633405 | -5.248754085 |
| **A_55_P1988907** | Eml4 | 0.016930128 | -0.277695535 | 0.31155579 | 10.42151552 | 0.137340554 | 0.894826627 | 0.996633405 | -5.248756626 |
| **A_55_P2050513** | Pgbd1 | 0.009032219 | -0.148154554 | 0.166218993 | 8.607933406 | 0.137337017 | 0.894829316 | 0.996633405 | -5.248757035 |
| **A_55_P2043269** | Cdc42se1 | 0.008241552 | -0.135277133 | 0.151760237 | 8.691849972 | 0.137249161 | 0.894896111 | 0.996633405 | -5.248767179 |
| **A_55_P2066463** | Enah | -0.011761163 | -0.216573122 | 0.193050796 | 10.68144634 | -0.137247387 | 0.89489746 | 0.996633405 | -5.248767384 |
| **A_52_P172563** | Dffa | 0.008534588 | -0.140094527 | 0.157163704 | 8.5426142 | 0.137242248 | 0.894901367 | 0.996633405 | -5.248767977 |
| **A_55_P2168296** | Rnu7 | 0.007564544 | -0.124178225 | 0.139307313 | 8.078796726 | 0.137235067 | 0.894906826 | 0.996633405 | -5.248768806 |
| **A_55_P1975922** | Olfr629 | -0.006961175 | -0.128196225 | 0.114273875 | 8.25335062 | -0.137234559 | 0.894907213 | 0.996633405 | -5.248768864 |
| **A_55_P2162698** | Prpf40b | 0.010952094 | -0.179806145 | 0.201710334 | 9.445385277 | 0.137221763 | 0.894916942 | 0.996633405 | -5.248770341 |
| **A_51_P504134** | NA | 0.009932514 | -0.163109746 | 0.182974775 | 7.703748212 | 0.137188009 | 0.894942604 | 0.996633405 | -5.248774236 |
| **A_51_P381260** | Fxyd5 | -0.012929148 | -0.238188702 | 0.212330407 | 10.53816144 | -0.137181581 | 0.894947491 | 0.996633405 | -5.248774978 |
| **A_55_P1985633** | Lrp1 | 0.006470829 | -0.106309098 | 0.119250756 | 7.688263083 | 0.13713151 | 0.89498556 | 0.996633405 | -5.248780754 |
| **A_55_P2000628** | NA | -0.01155243 | -0.213103807 | 0.189998948 | 9.601364547 | -0.136992472 | 0.895091272 | 0.99664649 | -5.248796781 |
| **A_55_P2032363** | Tomm40l | -0.010746938 | -0.198261606 | 0.17676773 | 10.20696702 | -0.136980461 | 0.895100404 | 0.99664649 | -5.248798165 |
| **A_55_P2166069** | 2310079G1 | -0.006911231 | -0.12768286 | 0.113860397 | 7.872335591 | -0.136772765 | 0.89525832 | 0.996721468 | -5.248822074 |
| **A_55_P1987600** | Trmt112 | -0.007113116 | -0.131427596 | 0.117201363 | 12.29521201 | -0.136756297 | 0.895270842 | 0.996721468 | -5.248823968 |
| **A_55_P2131844** | NA | -0.035551521 | -0.659377485 | 0.588274444 | 9.037781702 | -0.136208393 | 0.895687454 | 0.996817786 | -5.248886861 |
| **A_55_P2074197** | Zbtb24 | -0.006318219 | -0.117185923 | 0.104549484 | 8.659990071 | -0.136206794 | 0.89568867 | 0.996817786 | -5.248887045 |
| **A_55_P1992179** | Tars2 | 0.008715138 | -0.144377512 | 0.161807789 | 12.51834385 | 0.136059561 | 0.895800628 | 0.996817786 | -5.248903902 |
| **A_55_P2071970** | Nav2 | 0.008098479 | -0.134180788 | 0.150377747 | 8.564113401 | 0.136041364 | 0.895814465 | 0.996817786 | -5.248905985 |
| **A_55_P2094574** | Rapgef6 | -0.011185584 | -0.207736721 | 0.185365553 | 10.98421348 | -0.136016701 | 0.89583322 | 0.996817786 | -5.248908806 |
| **A_52_P415155** | Wnt6 | 0.005743913 | -0.095194056 | 0.106681882 | 7.723242061 | 0.136007367 | 0.895840318 | 0.996817786 | -5.248909874 |

| | | | | | | | | | |
|---|---|---|---|---|---|---|---|---|---|
| **A_51_P428086** | Sptbn1 | -0.01165598 | -0.216528174 | 0.193216213 | 10.55453886 | -0.135979965 | 0.895861155 | 0.996817786 | -5.248913008 |
| **A_52_P301495** | Vamp2 | 0.011576587 | -0.19194286 | 0.215096035 | 10.13280516 | 0.135951426 | 0.895882857 | 0.996817786 | -5.248916272 |
| **A_55_P2000783** | Axin2 | 0.006121118 | -0.101503409 | 0.113745646 | 7.795604922 | 0.135934174 | 0.895895976 | 0.996817786 | -5.248918244 |
| **A_55_P2134566** | NA | -0.011863185 | -0.220457891 | 0.196731521 | 8.929050368 | -0.135927447 | 0.895901092 | 0.996817786 | -5.248919014 |
| **A_51_P155313** | Gsto1 | -0.008643867 | -0.160666496 | 0.143378763 | 13.00176166 | -0.135896713 | 0.895924463 | 0.996817786 | -5.248922527 |
| **A_51_P404413** | Bckdk | 0.013730401 | -0.227891496 | 0.255352299 | 11.34721075 | 0.135817608 | 0.895984619 | 0.996822808 | -5.248931566 |
| **A_55_P1957815** | Tmub1 | -0.008612732 | -0.160267745 | 0.143042281 | 11.27246386 | -0.135735455 | 0.896047093 | 0.996822808 | -5.248940948 |
| **A_51_P479352** | Klhdc9 | -0.005724795 | -0.106648193 | 0.095198604 | 7.995713401 | -0.13557424 | 0.896169692 | 0.996822808 | -5.248959342 |
| **A_55_P1986541** | NA | 0.008386068 | -0.139572956 | 0.156345091 | 15.34592405 | 0.135464671 | 0.896253018 | 0.996822808 | -5.248971831 |
| **A_55_P2100679** | Dnm2 | 0.009755659 | -0.162398509 | 0.181909828 | 8.410858132 | 0.135440395 | 0.89627148 | 0.996822808 | -5.248974597 |
| **A_55_P1994020** | Casc4 | -0.012913111 | -0.241049323 | 0.2152231 | 9.081877453 | -0.135283802 | 0.896390569 | 0.996822808 | -5.248992425 |
| **A_51_P382805** | Zpbp2 | 0.006254468 | -0.10430441 | 0.116813347 | 7.869170992 | 0.135209095 | 0.896447386 | 0.996822808 | -5.249000924 |
| **A_66_P106238** | Gm5106 | -0.04673602 | -0.873192794 | 0.779720754 | 9.774495829 | -0.135157676 | 0.896486492 | 0.996822808 | -5.24900677 |
| **A_55_P2006762** | Bzw1 | 0.011817129 | -0.197151509 | 0.220785768 | 12.19027982 | 0.135157456 | 0.896486659 | 0.996822808 | -5.249006795 |
| **A_55_P2085915** | Tnfrsf14 | 0.006282162 | -0.10481891 | 0.117383233 | 7.686207663 | 0.135145 | 0.896496132 | 0.996822808 | -5.249008211 |
| **A_51_P510939** | Chchd7 | -0.012270079 | -0.229431718 | 0.20489156 | 10.77428629 | -0.135043405 | 0.896573399 | 0.996822808 | -5.249019755 |
| **A_55_P1955242** | NA | 0.008001923 | -0.133674279 | 0.149678125 | 16.27599511 | 0.134991553 | 0.896612835 | 0.996822808 | -5.249025644 |
| **A_51_P431316** | Ube2w | -0.009490017 | -0.177536994 | 0.158556959 | 9.366647589 | -0.134972547 | 0.89662729 | 0.996822808 | -5.249027802 |
| **A_55_P2152248** | Cpe | -0.011198674 | -0.209547475 | 0.187150126 | 9.750664262 | -0.134941699 | 0.896650751 | 0.996822808 | -5.249031304 |
| **A_52_P60194** | C4bp | 0.005459727 | -0.0913807 | 0.102300153 | 7.660701972 | 0.134748314 | 0.896797834 | 0.996929002 | -5.249053238 |
| **A_55_P2069597** | Tspan17 | 0.005663069 | -0.094971058 | 0.106297195 | 8.067273028 | 0.134497958 | 0.896988253 | 0.997083355 | -5.249081587 |
| **A_51_P392967** | Zmynd10 | -0.005722246 | -0.107572829 | 0.096128337 | 7.729831762 | -0.134280251 | 0.897153846 | 0.997131211 | -5.249106196 |
| **A_51_P358700** | Olfr1221 | -0.006030196 | -0.1133882 | 0.101327809 | 7.682465129 | -0.134247468 | 0.897178782 | 0.997131211 | -5.249109899 |
| **A_55_P2066478** | Crtc3 | -0.006855961 | -0.129116147 | 0.115404225 | 10.76849646 | -0.134027044 | 0.897346448 | 0.997131211 | -5.249134769 |
| **A_51_P140942** | Rp9 | -0.008581218 | -0.161660275 | 0.144497838 | 12.90905069 | -0.133980719 | 0.897381686 | 0.997131211 | -5.249139991 |
| **A_52_P260555** | Vil1 | -0.005981302 | -0.112689856 | 0.100727252 | 7.849399991 | -0.133969398 | 0.897390297 | 0.997131211 | -5.249141267 |
| **A_55_P2021275** | Iqca | 0.005885419 | -0.099167141 | 0.110937979 | 7.639264774 | 0.13389979 | 0.897443246 | 0.997131211 | -5.249149109 |
| **A_51_P363634** | Snrpe | 0.015052388 | -0.253665092 | 0.283769868 | 12.37153001 | 0.133880901 | 0.897457615 | 0.997131211 | -5.249151236 |
| **A_55_P2048813** | NA | -0.029897905 | -0.563646053 | 0.503850242 | 11.7835428 | -0.133879325 | 0.897458813 | 0.997131211 | -5.249151414 |
| **A_55_P2326091** | 9330175M2 | 0.009034759 | -0.152315046 | 0.170384565 | 9.702995865 | 0.133831166 | 0.897495447 | 0.997131211 | -5.249156836 |
| **A_66_P110104** | Myo18b | 0.010625318 | -0.179449721 | 0.200700356 | 10.04217693 | 0.133606001 | 0.89766673 | 0.997224539 | -5.249182163 |
| **A_51_P517096** | Casp8ap2 | 0.011064273 | -0.186894093 | 0.209022638 | 9.199240306 | 0.133585137 | 0.897682602 | 0.997224539 | -5.249184507 |
| **A_55_P2005248** | Meis3 | -0.007555541 | -0.143090425 | 0.127979342 | 8.797150168 | -0.133236625 | 0.897947727 | 0.997364617 | -5.249223618 |
| **A_52_P403527** | NA | 0.01822865 | -0.308807299 | 0.3452646 | 10.44626233 | 0.133219608 | 0.897960674 | 0.997364617 | -5.249225525 |
| **A_55_P1981664** | 8-Sep | 0.010585723 | -0.179352914 | 0.20052436 | 12.61574899 | 0.133203715 | 0.897972764 | 0.997364617 | -5.249227306 |
| **A_55_P2149149** | NA | 0.008344373 | -0.141440403 | 0.158129149 | 8.329412835 | 0.133148156 | 0.898015031 | 0.997364617 | -5.24923353 |
| **A_51_P348749** | Llgl2 | 0.006969142 | -0.118267977 | 0.132206261 | 9.197274034 | 0.13300114 | 0.898126878 | 0.997431544 | -5.249249988 |
| **A_55_P2121231** | Nipbl | 0.006917209 | -0.117637968 | 0.131472387 | 8.443257101 | 0.132732806 | 0.898331026 | 0.997600964 | -5.249279979 |
| **A_51_P342535** | Rnf166 | -0.012291522 | -0.23386247 | 0.209279427 | 11.69440963 | -0.132587315 | 0.898441719 | 0.997666588 | -5.249296215 |
| **A_52_P150525** | Ndufaf6 | -0.008032827 | -0.153066698 | 0.137001043 | 11.09947748 | -0.132375653 | 0.89860276 | 0.997677483 | -5.249319804 |
| **A_51_P458218** | Wdr13 | 0.016655297 | -0.284333239 | 0.317643834 | 11.27177447 | 0.132254839 | 0.898694683 | 0.997677483 | -5.249333252 |
| **A_51_P370350** | Galnt2 | -0.005683826 | -0.108429157 | 0.097061505 | 12.26777984 | -0.132217159 | 0.898723353 | 0.997677483 | -5.249337443 |
| **A_51_P102257** | Tns1 | 0.027364126 | -0.468769053 | 0.523497306 | 9.348616039 | 0.131823381 | 0.899022978 | 0.997677483 | -5.249381177 |
| **A_52_P69789** | Exosc1 | 0.009710598 | -0.166398634 | 0.18581983 | 8.516559439 | 0.131787127 | 0.899050564 | 0.997677483 | -5.249385196 |
| **A_55_P2146105** | NA | -0.01061217 | -0.20307298 | 0.181848639 | 16.03178788 | -0.131786534 | 0.899051015 | 0.997677483 | -5.249385262 |
| **A_55_P1963508** | Slc13a5 | 0.005883871 | -0.100885844 | 0.112653586 | 7.735784927 | 0.131711657 | 0.899107991 | 0.997677483 | -5.249393561 |
| **A_51_P380514** | Tjap1 | 0.007820137 | -0.134222845 | 0.149863119 | 9.084861649 | 0.131584187 | 0.899204988 | 0.997677483 | -5.249407678 |
| **A_55_P2131303** | NA | 0.00621352 | -0.106667052 | 0.119094093 | 16.87387052 | 0.131561148 | 0.89922252 | 0.997677483 | -5.249410228 |

| | | | | | | | | | |
|---|---|---|---|---|---|---|---|---|---|
| **A_55_P2110136** | Arhgap29 | 0.007348878 | -0.126208578 | 0.140906334 | 8.962043554 | 0.131510986 | 0.899260691 | 0.997677483 | -5.249415779 |
| **A_55_P2037221** | Klhdc10 | -0.009970528 | -0.191630094 | 0.171689039 | 12.29241889 | -0.131180413 | 0.899512248 | 0.997677483 | -5.249452307 |
| **A_55_P2049856** | Araf | -0.01129467 | -0.21712571 | 0.194536371 | 10.6552265 | -0.131151059 | 0.899534586 | 0.997677483 | -5.249455546 |
| **A_52_P664617** | Zfp472 | 0.005729558 | -0.098750281 | 0.110209397 | 8.461347127 | 0.131068332 | 0.899597542 | 0.997677483 | -5.249464671 |
| **A_51_P230439** | Ppfibp2 | 0.019902946 | -0.343918813 | 0.383724704 | 9.674389319 | 0.130748832 | 0.899840689 | 0.997677483 | -5.249499858 |
| **A_51_P278653** | Rprm | -0.005973883 | -0.115240373 | 0.103292607 | 7.815102628 | -0.130670889 | 0.899900007 | 0.997677483 | -5.249508429 |
| **A_51_P206585** | Runx1 | -0.005609442 | -0.108289055 | 0.097070171 | 7.740364009 | -0.130570347 | 0.899976525 | 0.997677483 | -5.249519477 |
| **A_52_P42194** | NA | -0.019318299 | -0.37295798 | 0.334321383 | 8.835832669 | -0.130562067 | 0.899982827 | 0.997677483 | -5.249520387 |
| **A_55_P2237432** | Smarca1 | 0.008419759 | -0.145792467 | 0.162631985 | 10.46287351 | 0.130493833 | 0.900034758 | 0.997677483 | -5.24952788 |
| **A_55_P2038867** | Mcat | 0.008166898 | -0.141851985 | 0.158185781 | 11.82288453 | 0.130112893 | 0.900324689 | 0.997677483 | -5.249569641 |
| **A_65_P06572** | Smad5 | -0.005839399 | -0.113111444 | 0.101432647 | 8.071234862 | -0.130104012 | 0.900331448 | 0.997677483 | -5.249570613 |
| **A_55_P2026440** | Ceacam-ps | -0.005431547 | -0.105507551 | 0.094644456 | 7.676049657 | -0.12971875 | 0.900624686 | 0.997677483 | -5.249612722 |
| **A_52_P235102** | Zfp975 | 0.019751848 | -0.344181902 | 0.383685598 | 11.13740111 | 0.129716296 | 0.900626554 | 0.997677483 | -5.249612989 |
| **A_55_P2411907** | B230213L1 | -0.016615622 | -0.323125458 | 0.289894214 | 10.38092761 | -0.129563098 | 0.900743164 | 0.997677483 | -5.249629699 |
| **A_52_P186033** | Spn | -0.007765094 | -0.151066748 | 0.135536561 | 7.939439955 | -0.129510396 | 0.900783279 | 0.997677483 | -5.249635443 |
| **A_51_P424054** | Btbd2 | 0.009033086 | -0.157729038 | 0.17579521 | 10.61913697 | 0.129463649 | 0.900818862 | 0.997677483 | -5.249640535 |
| **A_55_P2040873** | NA | 0.006486854 | -0.113321181 | 0.126294889 | 7.695095572 | 0.129406865 | 0.900862086 | 0.997677483 | -5.249646719 |
| **A_51_P379660** | Mllt11 | 0.012542542 | -0.21925698 | 0.244342065 | 10.54648531 | 0.129324907 | 0.900924473 | 0.997677483 | -5.24965564 |
| **A_66_P105288** | Rplp0 | -0.006363175 | -0.124006283 | 0.111279932 | 15.8278142 | -0.129275594 | 0.90096201 | 0.997677483 | -5.249661004 |
| **A_51_P509452** | Tmlhe | -0.00944135 | -0.184047764 | 0.165165065 | 8.386377217 | -0.129235856 | 0.900992259 | 0.997677483 | -5.249665326 |
| **A_51_P377390** | Ndst2 | 0.006435553 | -0.112617645 | 0.125488752 | 9.550658332 | 0.12919745 | 0.901021494 | 0.997677483 | -5.249669501 |
| **A_55_P1971849** | Sh3glb2 | 0.010291981 | -0.180230135 | 0.200814097 | 13.63271352 | 0.129110846 | 0.901087419 | 0.997677483 | -5.249678912 |
| **A_55_P1962314** | NA | -0.006329536 | -0.123512618 | 0.110853547 | 15.77171295 | -0.129096977 | 0.901097977 | 0.997677483 | -5.249680419 |
| **A_55_P2098999** | NA | -0.008733363 | -0.170437411 | 0.152970686 | 14.74220826 | -0.129083204 | 0.901108461 | 0.997677483 | -5.249681915 |
| **A_52_P231635** | Oxsm | 0.007030819 | -0.123158304 | 0.137219943 | 8.239930128 | 0.12907447 | 0.901115109 | 0.997677483 | -5.249682863 |
| **A_55_P2181251** | Ccdc9b | 0.015621139 | -0.273653575 | 0.304895853 | 10.17415754 | 0.129065852 | 0.90112167 | 0.997677483 | -5.249683799 |
| **A_55_P2180096** | Gm10136 | -0.007580091 | -0.148014094 | 0.132853911 | 15.74697212 | -0.129006413 | 0.901166917 | 0.997677483 | -5.249690252 |
| **A_55_P2240823** | Epb41 | 0.007553887 | -0.132409657 | 0.14751743 | 7.845540028 | 0.12899256 | 0.901177463 | 0.997677483 | -5.249691756 |
| **A_52_P656565** | Pex6 | 0.013923034 | -0.244231308 | 0.272077377 | 9.711880528 | 0.128903168 | 0.901245512 | 0.997677483 | -5.249701454 |
| **A_66_P126836** | Zbtb44 | -0.012287761 | -0.240127395 | 0.215551872 | 10.23995638 | -0.128899914 | 0.901247989 | 0.997677483 | -5.249701807 |
| **A_55_P2154870** | Sumo2 | 0.007889755 | -0.138413501 | 0.15419301 | 13.62640278 | 0.128889824 | 0.90125567 | 0.997677483 | -5.249702901 |
| **A_55_P2148534** | Nr1d2 | 0.014097209 | -0.24755979 | 0.275754207 | 10.13205725 | 0.12876858 | 0.901347968 | 0.997677483 | -5.249716043 |
| **A_55_P1988424** | Tpi1 | 0.024586393 | -0.431893254 | 0.481066041 | 12.81677466 | 0.128730822 | 0.901376713 | 0.997677483 | -5.249720133 |
| **A_52_P391983** | Nt5c2 | 0.007673333 | -0.134863061 | 0.150209728 | 10.16944579 | 0.128667074 | 0.901425242 | 0.997677483 | -5.249727036 |
| **A_51_P510418** | Aldh1b1 | -0.026710581 | -0.523020337 | 0.469599175 | 12.25378213 | -0.128629225 | 0.901454056 | 0.997677483 | -5.249731133 |
| **A_55_P2070976** | Gm6923 | 0.00904658 | -0.159156839 | 0.177249999 | 8.051444704 | 0.128546051 | 0.901517375 | 0.997677483 | -5.249740132 |
| **A_55_P2148531** | Pithd1 | -0.008717568 | -0.171112824 | 0.153677688 | 8.3265022 | -0.128301323 | 0.901703688 | 0.997677483 | -5.249766576 |
| **A_55_P1984314** | Trim43c | -0.00702538 | -0.137942965 | 0.123892206 | 9.099114503 | -0.128256958 | 0.901737464 | 0.997677483 | -5.249771364 |
| **A_52_P381929** | Ubqln2 | 0.013490331 | -0.237911083 | 0.264891745 | 9.916773517 | 0.128251963 | 0.901741267 | 0.997677483 | -5.249771904 |
| **A_51_P119031** | Naa30 | 0.008085358 | -0.142921995 | 0.15909271 | 9.966646552 | 0.127970622 | 0.901955463 | 0.997677483 | -5.24980223 |
| **A_55_P2137496** | NA | 0.005601419 | -0.099033097 | 0.110235935 | 7.728602364 | 0.127947618 | 0.901972978 | 0.997677483 | -5.249804707 |
| **A_51_P266224** | Limd1 | 0.006436536 | -0.113808491 | 0.126681563 | 11.99906221 | 0.127936425 | 0.9019815 | 0.997677483 | -5.249805912 |
| **A_55_P2112379** | NA | 0.009252847 | -0.163610356 | 0.182116049 | 16.07395207 | 0.127932811 | 0.901984251 | 0.997677483 | -5.249806301 |
| **A_51_P138141** | Angel2 | 0.011291996 | -0.199825629 | 0.22240962 | 10.35018118 | 0.127836639 | 0.902057474 | 0.997677483 | -5.249816649 |
| **A_55_P2062632** | Smc5 | -0.007324096 | -0.144259186 | 0.129610994 | 10.45594796 | -0.127834595 | 0.902059029 | 0.997677483 | -5.249816869 |
| **A_55_P1983046** | Plch1 | -0.006291401 | -0.124027088 | 0.111444286 | 7.78540317 | -0.127716902 | 0.902148638 | 0.997677483 | -5.249829523 |
| **A_55_P1999710** | Smad2 | -0.008210042 | -0.161895483 | 0.1454754 | 9.702073657 | -0.127679675 | 0.902176983 | 0.997677483 | -5.249833523 |
| **A_51_P358872** | Disc1 | 0.007190902 | -0.127628554 | 0.142010357 | 7.912589898 | 0.127479372 | 0.902329494 | 0.997677483 | -5.249855025 |

| | | | | | | | | | |
|---|---|---|---|---|---|---|---|---|---|
| **A_55_P2096165** | NA | 0.01042785 | -0.185251288 | 0.206106988 | 11.71842468 | 0.127367723 | 0.902414505 | 0.997677483 | -5.249866996 |
| **A_55_P2039903** | Wsb2 | -0.029273962 | -0.578836264 | 0.52028834 | 11.01917694 | -0.127313279 | 0.90245596 | 0.997677483 | -5.249872829 |
| **A_51_P433789** | Agr3 | 0.005100426 | -0.09078709 | 0.100987942 | 7.647962271 | 0.127131597 | 0.9025943 | 0.997677483 | -5.249892279 |
| **A_51_P366138** | Mertk | 0.008877931 | -0.158080583 | 0.175836445 | 8.319919503 | 0.12709028 | 0.902625761 | 0.997677483 | -5.249896698 |
| **A_55_P2115235** | Bckdhb | -0.005727178 | -0.113481503 | 0.102027146 | 12.6796048 | -0.12703258 | 0.902669697 | 0.997677483 | -5.249902867 |
| **A_55_P2111331** | Klhl32 | -0.006893564 | -0.136600527 | 0.1228134 | 9.378101153 | -0.127025129 | 0.902675371 | 0.997677483 | -5.249903663 |
| **A_55_P2007210** | Abca15 | 0.005478707 | -0.09767701 | 0.108634423 | 7.665641821 | 0.126938649 | 0.902741222 | 0.997677483 | -5.249912903 |
| **A_55_P1982186** | Sgsm1 | -0.005696645 | -0.113177515 | 0.101784225 | 7.815041669 | -0.126676809 | 0.90294061 | 0.997677483 | -5.249940843 |
| **A_55_P2098335** | Map2k4 | 0.007363128 | -0.131611442 | 0.146337699 | 8.734061806 | 0.126629871 | 0.902976354 | 0.997677483 | -5.249945845 |
| **A_55_P2107502** | Fat1 | -0.012570331 | -0.249828116 | 0.224687453 | 12.29046999 | -0.126629646 | 0.902976525 | 0.997677483 | -5.249945869 |
| **A_55_P1957173** | NA | 0.011224188 | -0.200736507 | 0.223184882 | 8.234079554 | 0.126563572 | 0.90302684 | 0.997677483 | -5.249952908 |
| **A_52_P461343** | Upk3a | -0.005101146 | -0.101495721 | 0.091293429 | 7.669856858 | -0.126480699 | 0.90308995 | 0.997677483 | -5.249961731 |
| **A_51_P347862** | Actn1 | 0.007628465 | -0.136590561 | 0.151847491 | 9.101382532 | 0.126422317 | 0.90313441 | 0.997677483 | -5.249967943 |
| **A_55_P1993026** | NA | -0.008084726 | -0.161013177 | 0.144843725 | 15.46149336 | -0.126353173 | 0.903187065 | 0.997677483 | -5.249975296 |
| **A_52_P339996** | Slc6a6 | 0.013545853 | -0.242799414 | 0.269891121 | 11.15135417 | 0.126296174 | 0.903230472 | 0.997677483 | -5.249981355 |
| **A_52_P540470** | Zfp407 | 0.006873895 | -0.123750271 | 0.137498062 | 10.26430721 | 0.125773315 | 0.903628666 | 0.997677483 | -5.250036808 |
| **A_55_P2076580** | Acaa1a | -0.007533987 | -0.150715739 | 0.135647765 | 14.13497115 | -0.1257611 | 0.903637969 | 0.997677483 | -5.250038101 |
| **A_55_P2182700** | Prkab1 | -0.007126361 | -0.142565004 | 0.128312282 | 8.124241235 | -0.125757638 | 0.903640606 | 0.997677483 | -5.250038467 |
| **A_52_P298002** | Gch1 | -0.005172391 | -0.103517196 | 0.093172415 | 8.029796021 | -0.125703984 | 0.903681469 | 0.997677483 | -5.250044144 |
| **A_55_P2013716** | Napa | -0.008009883 | -0.160329337 | 0.144309571 | 12.64471972 | -0.125683983 | 0.903696702 | 0.997677483 | -5.25004626 |
| **A_51_P295816** | Cep192 | -0.012630921 | -0.252905964 | 0.227644122 | 9.21961061 | -0.125642188 | 0.903728533 | 0.997677483 | -5.250050679 |
| **A_51_P185175** | Fkbp4 | -0.017263256 | -0.345752569 | 0.311226056 | 10.14445662 | -0.125606064 | 0.903756046 | 0.997677483 | -5.250054498 |
| **A_55_P1965070** | Map7d1 | 0.010681615 | -0.192576805 | 0.213940036 | 14.15998788 | 0.125602281 | 0.903758928 | 0.997677483 | -5.250054898 |
| **A_52_P409833** | Plat | -0.005517225 | -0.110563802 | 0.099529353 | 9.230190326 | -0.125530106 | 0.903813898 | 0.997677483 | -5.250062524 |
| **A_52_P242724** | NA | -0.008819139 | -0.176938249 | 0.159299971 | 11.60524621 | -0.125377103 | 0.90393043 | 0.997677483 | -5.250078677 |
| **A_55_P2133067** | Ube2v1 | 0.011379117 | -0.205611272 | 0.228369506 | 8.629896947 | 0.12533639 | 0.903961439 | 0.997677483 | -5.250082972 |
| **A_55_P2063096** | Txnrd3 | 0.007895881 | -0.143028351 | 0.158820114 | 10.93021014 | 0.125040519 | 0.904186794 | 0.997677483 | -5.250114141 |
| **A_55_P2069337** | NA | -0.008984364 | -0.180758059 | 0.162789331 | 9.105294613 | -0.125008574 | 0.904211125 | 0.997677483 | -5.250117502 |
| **A_51_P401024** | Lrrc34 | 0.005970958 | -0.108210166 | 0.120152082 | 7.752972596 | 0.12498527 | 0.904228876 | 0.997677483 | -5.250119954 |
| **A_55_P2046793** | Eogt | 0.006953907 | -0.126035087 | 0.139942901 | 8.548274942 | 0.124974756 | 0.904236884 | 0.997677483 | -5.25012106 |
| **A_55_P2116621** | Cited2 | 0.005638112 | -0.102226231 | 0.113502455 | 8.973658977 | 0.124929481 | 0.90427137 | 0.997677483 | -5.25012582 |
| **A_55_P1965278** | Dmrtc1a | 0.005022477 | -0.091149807 | 0.101194761 | 7.685881483 | 0.124817984 | 0.904356297 | 0.997677483 | -5.250137537 |
| **A_55_P2039881** | 5-Sep | -0.006087738 | -0.122691609 | 0.110516132 | 7.882423895 | -0.124782061 | 0.90438366 | 0.997677483 | -5.25014131 |
| **A_52_P349467** | Tmlhe | -0.009837353 | -0.198363146 | 0.178688441 | 8.511479143 | -0.1247144 | 0.904435198 | 0.997677483 | -5.250148413 |
| **A_55_P2053511** | Mrfap1 | 0.006038517 | -0.109699438 | 0.121776472 | 14.76108057 | 0.124699182 | 0.90444679 | 0.997677483 | -5.25015001 |
| **A_55_P2118268** | Chpf | 0.008369466 | -0.152092188 | 0.16883112 | 12.12094755 | 0.124662442 | 0.904474775 | 0.997677483 | -5.250153865 |
| **A_55_P2175400** | Zfp467 | 0.009712041 | -0.176581278 | 0.196005359 | 9.777731852 | 0.124601233 | 0.9045214 | 0.997677483 | -5.250160285 |
| **A_55_P2138257** | Ube2d2a | -0.006316147 | -0.127506444 | 0.114874151 | 10.76246036 | -0.124564268 | 0.904549557 | 0.997677483 | -5.25016416 |
| **A_51_P288479** | Slc25a29 | -0.007253081 | -0.146477954 | 0.131971791 | 8.234032162 | -0.124513046 | 0.904588575 | 0.997677483 | -5.250169528 |
| **A_55_P2053032** | Xkr8 | 0.005952531 | -0.108360218 | 0.120265279 | 8.520277476 | 0.124456077 | 0.90463197 | 0.997677483 | -5.250175497 |
| **A_55_P2167803** | Abi3bp | 0.021483467 | -0.391255897 | 0.434222831 | 8.11036015 | 0.124404938 | 0.904670925 | 0.997677483 | -5.250180852 |
| **A_52_P209172** | Mrpl41 | -0.005891635 | -0.119131147 | 0.107347876 | 13.21453938 | -0.124350349 | 0.904712509 | 0.997677483 | -5.250186565 |
| **A_51_P198694** | Gckr | 0.005392443 | -0.098261294 | 0.109046179 | 7.705515263 | 0.12433967 | 0.904720643 | 0.997677483 | -5.250187683 |
| **A_55_P2148873** | Cat | 0.015666622 | -0.285855225 | 0.31718847 | 10.75711622 | 0.124184023 | 0.90483921 | 0.997677483 | -5.25020396 |
| **A_55_P2037524** | Rps15 | -0.006489124 | -0.131427599 | 0.118449352 | 14.97662235 | -0.124136354 | 0.904875523 | 0.997677483 | -5.25020894 |
| **A_51_P111544** | Srp14 | 0.006551052 | -0.119596309 | 0.132698414 | 12.7092279 | 0.124120075 | 0.904887924 | 0.997677483 | -5.231501984 |
| **A_55_P2083725** | NA | 0.005358688 | -0.097868673 | 0.108586049 | 7.984717808 | 0.12407171 | 0.904924768 | 0.997677483 | -5.250215692 |
| **A_66_P127224** | Krt10 | -0.008032652 | -0.162774672 | 0.146709368 | 9.449873653 | -0.124068018 | 0.90492758 | 0.997677483 | -5.250216077 |

| | | | | | | | | | |
|---|---|---|---|---|---|---|---|---|---|
| **A_55_P2035590** | NA | 0.011060813 | -0.202351628 | 0.224473254 | 13.40728752 | 0.123872945 | 0.905076187 | 0.997677483 | -5.250236429 |
| **A_55_P1979341** | Cyba | 0.015684913 | -0.287232226 | 0.318602052 | 12.98844045 | 0.123756323 | 0.905165031 | 0.997677483 | -5.250248581 |
| **A_52_P381953** | Irf2bp1 | 0.005145976 | -0.094240958 | 0.10453291 | 8.643502294 | 0.123750675 | 0.905169334 | 0.997677483 | -5.25024917 |
| **A_55_P2083954** | Scfd1 | -0.01065058 | -0.216363327 | 0.195062168 | 9.746963219 | -0.123743152 | 0.905175065 | 0.997677483 | -5.250249953 |
| **A_52_P478729** | Atg2a | 0.012465611 | -0.228527624 | 0.253458847 | 8.749812034 | 0.12362829 | 0.90526257 | 0.997677483 | -5.250261909 |
| **A_55_P2202043** | C430045I1 | -0.01263626 | -0.257099537 | 0.231827018 | 8.827356967 | -0.123541834 | 0.905328435 | 0.997677483 | -5.250270901 |
| **A_55_P2060343** | Zfp352 | 0.007666497 | -0.140860569 | 0.156193563 | 10.89052621 | 0.123367428 | 0.905461307 | 0.997677483 | -5.250289022 |
| **A_52_P288614** | Rars | -0.007165126 | -0.146010423 | 0.131680171 | 8.439684837 | -0.123339394 | 0.905482665 | 0.997677483 | -5.250291932 |
| **A_55_P2080451** | NA | 0.009710136 | -0.178472657 | 0.197892929 | 12.12091914 | 0.123325968 | 0.905492894 | 0.997677483 | -5.250293326 |
| **A_52_P561927** | Ncor1 | 0.009816527 | -0.180454643 | 0.200087696 | 9.512425983 | 0.123308778 | 0.905505991 | 0.997677483 | -5.25029511 |
| **A_55_P2021570** | C87414 | -0.006205448 | -0.126517192 | 0.114106297 | 7.933867364 | -0.123274775 | 0.905531897 | 0.997677483 | -5.250298638 |
| **A_51_P283083** | Armc1 | 0.00901588 | -0.165885642 | 0.183917402 | 12.62082076 | 0.123203679 | 0.905586063 | 0.997677483 | -5.250306012 |
| **A_52_P610808** | Hexdc | -0.007223118 | -0.147357178 | 0.132910942 | 8.56183958 | -0.123194164 | 0.905593312 | 0.997677483 | -5.250306998 |
| **A_55_P1955691** | Alg6 | 0.008262991 | -0.152052503 | 0.168578484 | 8.826180857 | 0.123188706 | 0.90559747 | 0.997677483 | -5.250307564 |
| **A_66_P120567** | Anxa1 | 0.00980212 | -0.180379732 | 0.199983973 | 9.625513455 | 0.123185643 | 0.905599804 | 0.997677483 | -5.250307882 |
| **A_51_P193302** | Mrps7 | -0.010575511 | -0.215936396 | 0.194785375 | 10.49913002 | -0.123081491 | 0.905679156 | 0.997677483 | -5.250318675 |
| **A_55_P2059765** | NA | -0.00749772 | -0.153113432 | 0.138117993 | 7.947138709 | -0.123063733 | 0.905692686 | 0.997677483 | -5.250320514 |
| **A_55_P2182358** | Arid3a | -0.009657342 | -0.197350612 | 0.178035929 | 8.634588077 | -0.122975337 | 0.905760034 | 0.997677483 | -5.250329666 |
| **A_55_P2010134** | NA | -0.010356284 | -0.211718249 | 0.191005681 | 8.853740895 | -0.122923705 | 0.905799373 | 0.997677483 | -5.250335009 |
| **A_55_P2072556** | Ncoa6 | -0.01183038 | -0.241966597 | 0.218305836 | 11.75390664 | -0.122863497 | 0.905845247 | 0.997677483 | -5.250341236 |
| **A_55_P2101591** | Vta1 | 0.016683092 | -0.307892242 | 0.341258425 | 9.120483198 | 0.122848585 | 0.905856608 | 0.997677483 | -5.250342778 |
| **A_51_P283300** | Cacnb2 | -0.007104585 | -0.145329159 | 0.131119989 | 8.498203491 | -0.122846442 | 0.905858241 | 0.997677483 | -5.250343 |
| **A_55_P2153941** | Zfp386 | 0.013697493 | -0.252819008 | 0.280213994 | 10.0652193 | 0.122836125 | 0.905866102 | 0.997677483 | -5.250344067 |
| **A_55_P2009498** | Ccdc28a | 0.014084011 | -0.260063799 | 0.28823182 | 9.484345666 | 0.122786521 | 0.905903896 | 0.997677483 | -5.250349194 |
| **A_55_P2103536** | Irf2bpl | -0.008353067 | -0.171026032 | 0.154319897 | 10.43046147 | -0.122726888 | 0.905949332 | 0.997677483 | -5.250355354 |
| **A_52_P311031** | NA | 0.015450243 | -0.285543937 | 0.316444423 | 15.43055132 | 0.122683551 | 0.905982352 | 0.997677483 | -5.25035983 |
| **A_52_P625244** | Ube3a | 0.00648092 | -0.11988841 | 0.132850251 | 9.088369813 | 0.122575627 | 0.906064584 | 0.997677483 | -5.250370969 |
| **A_52_P501049** | NA | -0.005011557 | -0.102894753 | 0.09287164 | 8.537252052 | -0.122369627 | 0.906221547 | 0.997677483 | -5.250392202 |
| **A_55_P2011877** | Eif2s3y | 0.140831008 | -2.610792962 | 2.892454979 | 9.751648394 | 0.122325877 | 0.906254884 | 0.997677483 | -5.250396707 |
| **A_55_P2121466** | Ncor1 | -0.00615865 | -0.126551961 | 0.11423466 | 8.432202046 | -0.122262231 | 0.90630338 | 0.997677483 | -5.250403258 |
| **A_55_P1991851** | Speg | 0.012057136 | -0.223845144 | 0.247959416 | 11.87993306 | 0.122157788 | 0.906382965 | 0.997677483 | -5.250414001 |
| **A_51_P212630** | Chd8 | -0.011989958 | -0.246627112 | 0.222647196 | 12.25500607 | -0.122132152 | 0.906402499 | 0.997677483 | -5.250416636 |
| **A_55_P2159189** | Decr1 | -0.011994795 | -0.246783247 | 0.222793657 | 13.53947717 | -0.122102689 | 0.90642495 | 0.997677483 | -5.250419665 |
| **A_55_P2034600** | Gm5523 | 0.00524709 | -0.097590541 | 0.10808472 | 8.47785311 | 0.121948252 | 0.906542632 | 0.997677483 | -5.250435526 |
| **A_55_P2086329** | Krt85 | -0.007201855 | -0.14838624 | 0.133982529 | 8.28509454 | -0.121917728 | 0.906565892 | 0.997677483 | -5.250438658 |
| **A_55_P2064896** | Gm8580 | 0.008821079 | -0.16414451 | 0.181786668 | 14.00739334 | 0.121890862 | 0.906586364 | 0.997677483 | -5.250441415 |
| **A_55_P2038519** | NA | 0.016052021 | -0.298802179 | 0.33090622 | 9.257940597 | 0.121851072 | 0.906616685 | 0.997677483 | -5.250445496 |
| **A_55_P1962094** | NA | -0.005564935 | -0.114861833 | 0.103731964 | 7.917777323 | -0.121691816 | 0.906738044 | 0.997677483 | -5.250461818 |
| **A_55_P1970665** | Kdm3b | 0.007288497 | -0.135936843 | 0.150513837 | 8.377034154 | 0.121626228 | 0.906788025 | 0.997677483 | -5.250468534 |
| **A_55_P2102240** | Nprl3 | -0.008937207 | -0.184656797 | 0.166782382 | 9.414260923 | -0.121560029 | 0.906838472 | 0.997677483 | -5.250475309 |
| **A_51_P406077** | Fmc1 | -0.008448901 | -0.174626346 | 0.157728545 | 14.95143059 | -0.12151706 | 0.906871217 | 0.997677483 | -5.250479705 |
| **A_55_P2003581** | Scnm1 | -0.008700393 | -0.179832683 | 0.162431897 | 11.25042826 | -0.12151112 | 0.906875743 | 0.997677483 | -5.250480312 |
| **A_55_P2030501** | Ptpa | -0.011863291 | -0.245381686 | 0.221655104 | 13.50766867 | -0.121420839 | 0.906944543 | 0.997677483 | -5.250489542 |
| **A_52_P920129** | Gnai1 | -0.010485407 | -0.21714116 | 0.196170345 | 10.07318161 | -0.121268201 | 0.907060865 | 0.997677483 | -5.250505131 |
| **A_51_P498772** | 1700019D0 | -0.004951279 | -0.10259292 | 0.092690362 | 7.781899471 | -0.121196896 | 0.907115206 | 0.997677483 | -5.250512407 |
| **A_55_P2075385** | Orai2-ps | -0.005017643 | -0.104100548 | 0.094065263 | 7.943598648 | -0.121034766 | 0.907238766 | 0.997677483 | -5.250528935 |
| **A_55_P1989698** | Mmp11 | 0.008900512 | -0.167091791 | 0.184892816 | 10.03769606 | 0.120873325 | 0.907361804 | 0.997677483 | -5.25054537 |
| **A_55_P1956223** | Chka | -0.006867316 | -0.142890686 | 0.129156053 | 8.808948649 | -0.120665373 | 0.907520292 | 0.997677483 | -5.250566509 |

| | | | | | | | | | |
|---|---|---|---|---|---|---|---|---|---|
| **A_55_P2105181** | Bhmt | 0.007109462 | -0.133732357 | 0.147951281 | 7.818309794 | 0.120646365 | 0.907534779 | 0.997677483 | -5.250568439 |
| **A_55_P2293967** | Tbc1d5 | 0.005625872 | -0.105832216 | 0.11708396 | 9.520568253 | 0.120638898 | 0.90754047 | 0.997677483 | -5.250569198 |
| **A_66_P110584** | Tpbpa | 0.005379091 | -0.101490627 | 0.112248809 | 7.647410578 | 0.120299379 | 0.907799242 | 0.997677483 | -5.250603625 |
| **A_52_P402897** | Cdh4 | -0.005358507 | -0.111945306 | 0.101228292 | 8.026056351 | -0.120157128 | 0.907907665 | 0.997677483 | -5.25061802 |
| **A_51_P253732** | Il17rd | 0.008702358 | -0.164597542 | 0.182002258 | 8.956937969 | 0.120018383 | 0.908013418 | 0.997677483 | -5.250632045 |
| **A_51_P332822** | Abce1 | -0.008325952 | -0.174153287 | 0.157501383 | 11.75506614 | -0.120001567 | 0.908026236 | 0.997677483 | -5.250633743 |
| **A_51_P213691** | Scnn1a | 0.005736399 | -0.108564889 | 0.120037686 | 8.081361614 | 0.119949193 | 0.908066156 | 0.997677483 | -5.250639033 |
| **A_52_P694988** | Zfp933 | -0.005810809 | -0.121634896 | 0.110013278 | 7.976739272 | -0.119907641 | 0.908097829 | 0.997677483 | -5.250643227 |
| **A_55_P1990319** | Ergic3 | -0.005939381 | -0.124331404 | 0.112452642 | 11.9517239 | -0.119902405 | 0.908101819 | 0.997677483 | -5.250643755 |
| **A_51_P378009** | Blzf1 | -0.007230192 | -0.151451458 | 0.136991073 | 10.28043929 | -0.1198201 | 0.908164555 | 0.997677483 | -5.250652059 |
| **A_66_P133520** | Glmn | -0.005985476 | -0.125413489 | 0.113442538 | 9.108386265 | -0.119784767 | 0.908191487 | 0.997677483 | -5.250655622 |
| **A_51_P499441** | Prkd2 | 0.006815725 | -0.12920026 | 0.142831709 | 9.400077461 | 0.119765356 | 0.908206283 | 0.997677483 | -5.250657579 |
| **A_55_P2015403** | 9930111J2 | -0.005783882 | -0.121211333 | 0.109643569 | 8.950237214 | -0.119762107 | 0.90820876 | 0.997677483 | -5.250657907 |
| **A_55_P2040416** | Pradc1 | 0.010358613 | -0.196470871 | 0.217188098 | 10.08684001 | 0.119701143 | 0.90825523 | 0.997677483 | -5.250664051 |
| **A_52_P598309** | Zfas1 | -0.017256374 | -0.36188661 | 0.327373862 | 11.72397323 | -0.119675512 | 0.908274767 | 0.997677483 | -5.250666633 |
| **A_55_P2149787** | Fcnb | 0.005005085 | -0.094960542 | 0.104970713 | 7.681188574 | 0.119665767 | 0.908282195 | 0.997677483 | -5.250667615 |
| **A_51_P279712** | Rell1 | 0.008373337 | -0.158878452 | 0.175625126 | 10.11365637 | 0.119656675 | 0.908289125 | 0.997677483 | -5.250668531 |
| **A_55_P2113523** | Nsd2 | 0.006166715 | -0.117059991 | 0.129393422 | 8.349829235 | 0.119607437 | 0.908326658 | 0.997677483 | -5.250673489 |
| **A_55_P2178803** | Zbtb5 | -0.007925186 | -0.166345666 | 0.150495294 | 11.35556523 | -0.119565878 | 0.908358337 | 0.997677483 | -5.250677672 |
| **A_52_P82510** | Nmt1 | -0.008020331 | -0.168349283 | 0.15230862 | 11.52774429 | -0.119560983 | 0.908362068 | 0.997677483 | -5.250678165 |
| **A_52_P550218** | Trmt12 | -0.005742215 | -0.12075583 | 0.1092714 | 8.359961385 | -0.119327171 | 0.908540298 | 0.997677483 | -5.250701673 |
| **A_55_P2165854** | Gemin7 | -0.008866354 | -0.186459571 | 0.168726864 | 10.48717067 | -0.119324004 | 0.908542712 | 0.997677483 | -5.250701991 |
| **A_51_P144868** | Mboat7 | -0.010659985 | -0.224253172 | 0.202933202 | 10.49463831 | -0.119282938 | 0.908574016 | 0.997677483 | -5.250706115 |
| **A_51_P265374** | Vnn3 | 0.00547189 | -0.10429203 | 0.115235809 | 7.674258918 | 0.119148028 | 0.908676859 | 0.997677483 | -5.250719653 |
| **A_51_P475938** | Mfsd6l | 0.004748621 | -0.090542717 | 0.100039959 | 7.685751596 | 0.119103139 | 0.908711078 | 0.997677483 | -5.250724154 |
| **A_52_P531175** | Fkbp3 | -0.00817565 | -0.17229121 | 0.15593991 | 14.2331397 | -0.119064333 | 0.908740661 | 0.997677483 | -5.250728044 |
| **A_55_P2096283** | NA | -0.011948365 | -0.251827961 | 0.22793123 | 13.90941744 | -0.119048608 | 0.908752648 | 0.997677483 | -5.25072962 |
| **A_51_P158120** | B3galnt2 | 0.009315247 | -0.177711356 | 0.19634185 | 9.36069163 | 0.119041977 | 0.908757703 | 0.997677483 | -5.250730285 |
| **A_51_P512820** | Dera | 0.008290386 | -0.158209999 | 0.174790772 | 8.773760356 | 0.119005945 | 0.908785171 | 0.997677483 | -5.250733895 |
| **A_55_P2173457** | 1700097N( | 0.007577269 | -0.144775328 | 0.159929867 | 9.387127373 | 0.118869929 | 0.908888861 | 0.997677483 | -5.250747512 |
| **A_55_P1992208** | Gpi1 | 0.00775001 | -0.148209544 | 0.163709563 | 16.32528847 | 0.118767987 | 0.908966575 | 0.997677483 | -5.250757709 |
| **A_55_P2048744** | Eif2ak3 | 0.008731863 | -0.166994397 | 0.184458124 | 9.185896325 | 0.118762517 | 0.908970745 | 0.997677483 | -5.250758255 |
| **A_55_P1977929** | Itgam | 0.005632804 | -0.107739247 | 0.119004855 | 7.85102964 | 0.118748401 | 0.908981507 | 0.997677483 | -5.250759667 |
| **A_51_P117236** | Zfp354a | -0.006015335 | -0.127131128 | 0.115100458 | 8.833897388 | -0.118704775 | 0.909014765 | 0.997677483 | -5.24453433 |
| **A_51_P329283** | Senp5 | 0.006471196 | -0.123941292 | 0.136883684 | 9.553477148 | 0.118597217 | 0.909096763 | 0.997677483 | -5.250774769 |
| **A_55_P2115401** | Bst1 | 0.005416426 | -0.103795247 | 0.1146281 | 7.731437228 | 0.11853672 | 0.909142885 | 0.997677483 | -5.250780807 |
| **A_66_P118863** | Ncald | -0.005333179 | -0.112962392 | 0.102296033 | 7.974007503 | -0.118430939 | 0.90922353 | 0.997677483 | -5.250791358 |
| **A_55_P2135980** | Ms4a4c | 0.009460452 | -0.181570417 | 0.200491322 | 14.16211231 | 0.118363413 | 0.90927501 | 0.997677483 | -5.250798088 |
| **A_52_P297642** | Pdia5 | -0.005709724 | -0.121231303 | 0.109811855 | 9.3477579 | -0.118130249 | 0.909452776 | 0.997677483 | -5.250821297 |
| **A_52_P630016** | Bet1l | 0.012439925 | -0.239590671 | 0.264470521 | 9.63851653 | 0.117970557 | 0.909574529 | 0.997677483 | -5.250837166 |
| **A_55_P2158483** | Nkiras2 | -0.009155511 | -0.194897895 | 0.176586873 | 8.470988175 | -0.117809606 | 0.909697244 | 0.997677483 | -5.250853139 |
| **A_55_P2120615** | Ssh3 | -0.00869344 | -0.185062871 | 0.167675992 | 8.803558869 | -0.117808725 | 0.909697916 | 0.997677483 | -5.250853226 |
| **A_51_P196509** | Plekhj1 | 0.00789537 | -0.152368315 | 0.168159056 | 8.54012442 | 0.117746094 | 0.909745669 | 0.997677483 | -5.250859436 |
| **A_51_P398406** | Kctd13 | -0.005412563 | -0.11564193 | 0.104816805 | 8.625275015 | -0.117358557 | 0.910041156 | 0.997677483 | -5.250897786 |
| **A_51_P445912** | NA | -0.010286834 | -0.219862687 | 0.199289019 | 9.148347469 | -0.117313941 | 0.910075176 | 0.997677483 | -5.250902193 |
| **A_52_P205255** | Adam11 | -0.006771185 | -0.144895795 | 0.131353425 | 8.674721078 | -0.117166313 | 0.910187744 | 0.997677483 | -5.250916763 |
| **A_55_P2175272** | NA | 0.010725271 | -0.208082493 | 0.229533034 | 16.00568485 | 0.117153337 | 0.910197638 | 0.997677483 | -5.250918043 |
| **A_51_P237040** | Nog | 0.005085511 | -0.098703374 | 0.108874397 | 8.007600412 | 0.117109712 | 0.910230903 | 0.997677483 | -5.250922345 |

| | | | | | | | | | |
|---|---|---|---|---|---|---|---|---|---|
| **A_51_P345927** | Ddx41 | -0.005464966 | -0.117065414 | 0.106135481 | 8.489814354 | -0.117039023 | 0.910284805 | 0.997677483 | -5.250929311 |
| **A_51_P410744** | Dgke | -0.009616715 | -0.206014741 | 0.186781311 | 9.056855994 | -0.117030419 | 0.910291366 | 0.997677483 | -5.250930159 |
| **A_55_P2250296** | 6330566A1 | -0.005794252 | -0.124129862 | 0.112541358 | 8.408219403 | -0.11702834 | 0.910292951 | 0.997677483 | -5.250930364 |
| **A_51_P499864** | Tubg1 | -0.007096027 | -0.152134233 | 0.13794218 | 11.35752265 | -0.116934311 | 0.910364651 | 0.997677483 | -5.250939624 |
| **A_66_P102047** | Tbc1d13 | -0.006118226 | -0.131228729 | 0.118992277 | 9.03040305 | -0.116880182 | 0.910405926 | 0.997677483 | -5.250944951 |
| **A_55_P2011126** | Mfap1a | -0.004972872 | -0.106701456 | 0.096755713 | 8.542582233 | -0.11683511 | 0.910440297 | 0.997677483 | -5.250949385 |
| **A_55_P2046241** | Arl6ip6 | -0.005804519 | -0.124659848 | 0.11305081 | 8.436370442 | -0.116723076 | 0.910525728 | 0.997677483 | -5.250960399 |
| **A_55_P2028821** | Hnrnpf | 0.008474219 | -0.165112301 | 0.182060739 | 13.71353704 | 0.116679034 | 0.910559313 | 0.997677483 | -5.250964726 |
| **A_51_P162116** | Mblac1 | 0.011448116 | -0.223183469 | 0.246079701 | 8.963520599 | 0.116615611 | 0.910607677 | 0.997677483 | -5.250970954 |
| **A_55_P2131577** | NA | -0.006079888 | -0.130768229 | 0.118608453 | 17.09591213 | -0.116541032 | 0.910664549 | 0.997677483 | -5.250978274 |
| **A_51_P439876** | Map4k4 | 0.009867538 | -0.192629887 | 0.212364963 | 12.4430018 | 0.116465801 | 0.910721919 | 0.997677483 | -5.250985652 |
| **A_51_P411909** | Sirt5 | -0.00673149 | -0.145062746 | 0.131599765 | 11.44488228 | -0.116305443 | 0.910844207 | 0.997677483 | -5.251001364 |
| **A_55_P2125623** | NA | -0.007101845 | -0.153191272 | 0.138987583 | 8.170928136 | -0.116188064 | 0.910933722 | 0.997677483 | -5.251012852 |
| **A_55_P1963109** | Prh1 | -0.007104904 | -0.153319205 | 0.139109397 | 8.303574799 | -0.116138847 | 0.910971255 | 0.997677483 | -5.251017665 |
| **A_52_P310981** | Eif4e | -0.010661462 | -0.23010993 | 0.208787006 | 11.55749498 | -0.116116339 | 0.91098842 | 0.997677483 | -5.251019865 |
| **A_55_P2114074** | Acrbp | 0.016199771 | -0.317310665 | 0.349710208 | 8.826306173 | 0.116093684 | 0.911005698 | 0.997677483 | -5.25102208 |
| **A_55_P2022139** | Pura | -0.010327645 | -0.223548759 | 0.202893469 | 11.83719406 | -0.115765785 | 0.911255767 | 0.997677483 | -5.251054082 |
| **A_51_P211903** | Golga5 | -0.007253485 | -0.157047227 | 0.142540257 | 9.781912716 | -0.115734324 | 0.911279761 | 0.997677483 | -5.251057148 |
| **A_55_P2060097** | Prdm5 | -0.008393158 | -0.181762111 | 0.164975796 | 10.38226759 | -0.115707945 | 0.911299879 | 0.997677483 | -5.251059718 |
| **A_55_P1982767** | Tmem9b | -0.008093754 | -0.175320256 | 0.159132747 | 11.91021857 | -0.115678873 | 0.911322051 | 0.997677483 | -5.25106255 |
| **A_66_P134090** | Grpel2 | 0.005616412 | -0.110557333 | 0.121790157 | 8.350307918 | 0.115547383 | 0.911422335 | 0.997677483 | -5.251075348 |
| **A_51_P191586** | Arpc1a | -0.009220882 | -0.200065155 | 0.18162339 | 11.09928081 | -0.115478859 | 0.911474597 | 0.997677483 | -5.251082012 |
| **A_55_P2037689** | NA | -0.017408849 | -0.377958301 | 0.343140604 | 12.06014848 | -0.115402267 | 0.911533014 | 0.997677483 | -5.251089456 |
| **A_55_P2062246** | Tgtp2 | -0.009154165 | -0.198798304 | 0.180489974 | 7.908875619 | -0.115368822 | 0.911558522 | 0.997677483 | -5.251092705 |
| **A_55_P2033135** | Sh3bgr | 0.008535889 | -0.168341038 | 0.185412817 | 15.22239003 | 0.115341797 | 0.911579134 | 0.997677483 | -5.251095329 |
| **A_66_P104340** | Atxn7l2 | 0.005261713 | -0.103804118 | 0.114327545 | 9.109738183 | 0.115304859 | 0.911607306 | 0.997677483 | -5.251098916 |
| **A_55_P1976395** | Nsdhl | -0.004851221 | -0.105412359 | 0.095709918 | 7.874213451 | -0.115300176 | 0.911610878 | 0.997677483 | -5.25109937 |
| **A_55_P1958457** | Nbeal2 | 0.005710352 | -0.112665848 | 0.124086551 | 8.578716437 | 0.115294234 | 0.91161541 | 0.997677483 | -5.251099947 |
| **A_55_P2277019** | Loxl3 | 0.005793152 | -0.11455804 | 0.126144343 | 8.271054343 | 0.115046558 | 0.911804317 | 0.997677483 | -5.251123963 |
| **A_55_P1998107** | NA | -0.00747986 | -0.162877337 | 0.147917617 | 14.4984612 | -0.115042583 | 0.911807349 | 0.997677483 | -5.251124348 |
| **A_55_P2139009** | NA | -0.004932353 | -0.107455119 | 0.097590413 | 15.15038698 | -0.114985466 | 0.911850914 | 0.997677483 | -5.251129878 |
| **A_51_P209807** | Ccnyl1 | -0.00579653 | -0.126402743 | 0.114809684 | 8.116138586 | -0.114870236 | 0.911938805 | 0.997677483 | -5.251141028 |
| **A_52_P343856** | Creb3l2 | 0.009998784 | -0.198318583 | 0.21831615 | 9.420871019 | 0.114717803 | 0.912055074 | 0.997677483 | -5.25115576 |
| **A_52_P4046** | Baz2a | 0.00907628 | -0.180034931 | 0.198187491 | 9.406073928 | 0.114709591 | 0.912061338 | 0.997677483 | -5.251156553 |
| **A_55_P2164047** | NA | 0.011841651 | -0.23496405 | 0.258647353 | 13.64221738 | 0.11467433 | 0.912088233 | 0.997677483 | -5.251159958 |
| **A_52_P394385** | 1600012H0 | 0.006226484 | -0.123662729 | 0.136115697 | 9.615480018 | 0.114572112 | 0.912166203 | 0.997677483 | -5.251169823 |
| **A_66_P101261** | Gm3367 | 0.007182472 | -0.142704275 | 0.157069218 | 8.189575581 | 0.114530147 | 0.912198213 | 0.997677483 | -5.25117387 |
| **A_55_P2181665** | NA | 0.00518726 | -0.103126585 | 0.113501105 | 7.763539189 | 0.114462487 | 0.912249823 | 0.997677483 | -5.251180392 |
| **A_55_P2129232** | Calm2 | 0.006992989 | -0.139058693 | 0.15304467 | 13.62927417 | 0.114436721 | 0.912269477 | 0.997677483 | -5.251182875 |
| **A_55_P2104662** | Ttc33 | -0.008901201 | -0.194979136 | 0.177176734 | 10.68287746 | -0.114330701 | 0.912350349 | 0.997677483 | -5.251193085 |
| **A_51_P314418** | Blm | 0.007993691 | -0.159214998 | 0.175202381 | 9.617787562 | 0.114260907 | 0.912403588 | 0.997677483 | -5.251199802 |
| **A_55_P2205650** | 2610507I01 | 0.00862403 | -0.17177191 | 0.189019969 | 9.454194996 | 0.114259578 | 0.912404602 | 0.997677483 | -5.251199929 |
| **A_66_P127311** | Esco1 | -0.005236889 | -0.114799158 | 0.10432538 | 8.167614122 | -0.114240873 | 0.91241887 | 0.997677483 | -5.251201729 |
| **A_52_P217675** | Slf1 | -0.007574688 | -0.166060556 | 0.15091118 | 8.664058984 | -0.114230828 | 0.912426533 | 0.997677483 | -5.251202695 |
| **A_51_P292116** | Ica1 | 0.009523118 | -0.189995423 | 0.209041659 | 9.500411924 | 0.114078828 | 0.912542482 | 0.997677483 | -5.251217304 |
| **A_55_P1999548** | Tma7 | -0.0071561 | -0.15720628 | 0.14289408 | 12.48407436 | -0.113985342 | 0.912613796 | 0.997677483 | -5.251226279 |
| **A_55_P2133806** | Tnrc6c | -0.01055778 | -0.232135521 | 0.211019961 | 10.16483896 | -0.113882136 | 0.912692525 | 0.997677483 | -5.25123618 |
| **A_51_P257419** | Lhx2 | -0.004774963 | -0.104988312 | 0.095438385 | 7.71045857 | -0.113881613 | 0.912692924 | 0.997677483 | -5.25123623 |

| | | | | | | | | | |
|---|---|---|---|---|---|---|---|---|---|
| **A_55_P2097808** | P4ha1 | 0.011786014 | -0.235808004 | 0.259380032 | 9.626470122 | 0.113772146 | 0.912776431 | 0.997677483 | -5.251246721 |
| **A_55_P2022082** | Klf13 | -0.012477969 | -0.274858064 | 0.249902127 | 10.96021764 | -0.11366379 | 0.912859092 | 0.997677483 | -5.251257095 |
| **A_55_P2087539** | Clpsl2 | 0.005385132 | -0.107876744 | 0.118647008 | 8.009145241 | 0.11363751 | 0.91287914 | 0.997677483 | -5.25125961 |
| **A_51_P433733** | Nucb1 | 0.009779693 | -0.196078015 | 0.215637401 | 11.77486209 | 0.113544791 | 0.912949873 | 0.997677483 | -5.251268478 |
| **A_51_P269203** | Trpv4 | -0.007856027 | -0.173242513 | 0.157530459 | 8.280726887 | -0.113530374 | 0.912960872 | 0.997677483 | -5.251269856 |
| **A_55_P2130600** | Musk | -0.005028528 | -0.111019515 | 0.100962459 | 7.676380462 | -0.113391651 | 0.913066703 | 0.997677483 | -5.251283108 |
| **A_55_P2032946** | Gsta1 | 0.004810822 | -0.096677282 | 0.106298927 | 7.711534686 | 0.113295672 | 0.913139925 | 0.997677483 | -5.251292268 |
| **A_51_P295449** | Mfsd5 | -0.013602136 | -0.300781239 | 0.273576966 | 10.19562131 | -0.113204431 | 0.913209533 | 0.997677483 | -5.251300968 |
| **A_55_P1983773** | Birc5 | 0.010079459 | -0.202823498 | 0.222982415 | 13.78071682 | 0.11315263 | 0.913249054 | 0.997677483 | -5.251305904 |
| **A_55_P2064451** | Pced1a | 0.008740052 | -0.175929373 | 0.193409477 | 9.118812593 | 0.113117076 | 0.913276178 | 0.997677483 | -5.251309291 |
| **A_55_P2095181** | NA | -0.008342492 | -0.184640282 | 0.167955299 | 9.247831611 | -0.113098811 | 0.913290113 | 0.997677483 | -5.251311031 |
| **A_55_P2033250** | Fdft1 | -0.012437973 | -0.275400883 | 0.250524936 | 9.513816371 | -0.113048355 | 0.913328607 | 0.997677483 | -5.251315834 |
| **A_55_P1973748** | Nrbp1 | 0.01331442 | -0.268614694 | 0.295243535 | 10.52785933 | 0.112873362 | 0.913462116 | 0.997677483 | -5.251332478 |
| **A_55_P2171484** | Lta4h | -0.011545019 | -0.25604486 | 0.232954822 | 12.86856565 | -0.112856139 | 0.913475256 | 0.997677483 | -5.251334115 |
| **A_55_P2014928** | Zfp341 | -0.004992698 | -0.110756248 | 0.100770852 | 9.085609734 | -0.1128258 | 0.913498404 | 0.997677483 | -5.251336998 |
| **A_51_P379597** | Ndufs1 | -0.009439233 | -0.209482939 | 0.190604473 | 14.99554965 | -0.112777103 | 0.913535557 | 0.997677483 | -5.251341623 |
| **A_55_P2092399** | Ccdc160 | -0.004677861 | -0.103859432 | 0.09450371 | 7.901095131 | -0.112726347 | 0.913574282 | 0.997677483 | -5.251346441 |
| **A_55_P2066304** | Limk2 | -0.004969207 | -0.11071874 | 0.100780325 | 7.986279684 | -0.112309838 | 0.913892071 | 0.997677483 | -5.251385902 |
| **A_51_P141772** | Uhrf2 | 0.008748764 | -0.177632892 | 0.19513042 | 10.96297316 | 0.112189616 | 0.913983801 | 0.997677483 | -5.251397265 |
| **A_55_P2129911** | NA | 0.004859504 | -0.098690151 | 0.108409159 | 15.98667999 | 0.112163726 | 0.914003556 | 0.997677483 | -5.25139971 |
| **A_55_P2071326** | Rpl36 | -0.007918215 | -0.176702729 | 0.160866299 | 15.07059437 | -0.112125352 | 0.914032836 | 0.997677483 | -5.251403334 |
| **A_55_P2132844** | Rps16 | 0.006095049 | -0.123990036 | 0.136180133 | 15.26646481 | 0.111984731 | 0.914140134 | 0.997677483 | -5.251416602 |
| **A_66_P128384** | Hdac1 | 0.005919625 | -0.120426021 | 0.132265272 | 11.520223 | 0.111980668 | 0.914143234 | 0.997677483 | -5.251416985 |
| **A_55_P2062936** | Abca3 | 0.006921929 | -0.140868759 | 0.154712616 | 10.9041865 | 0.111940998 | 0.914173503 | 0.997677483 | -5.251420724 |
| **A_66_P110791** | Ppp1r11 | -0.007629532 | -0.170583724 | 0.15532466 | 12.14110997 | -0.111902935 | 0.914202547 | 0.997677483 | -5.251424312 |
| **A_51_P130666** | Adk | -0.007830585 | -0.175098915 | 0.159437745 | 13.27473811 | -0.111889576 | 0.914212741 | 0.997677483 | -5.25142557 |
| **A_55_P2042590** | Polm | -0.009947251 | -0.222626651 | 0.202732149 | 9.302073158 | -0.111785835 | 0.9142919 | 0.997677483 | -5.25143534 |
| **A_51_P445487** | Mturn | -0.006870875 | -0.153806799 | 0.140065049 | 8.473038 | -0.111761748 | 0.91431028 | 0.997677483 | -5.251437607 |
| **A_55_P1969352** | Get4 | 0.007007027 | -0.142946778 | 0.156960832 | 11.65827543 | 0.111682577 | 0.914370692 | 0.997677483 | -5.251445055 |
| **A_55_P2157676** | Wdr34 | 0.013874514 | -0.283143111 | 0.31089214 | 14.91002887 | 0.111646395 | 0.914398301 | 0.997677483 | -5.251448457 |
| **A_51_P113205** | F13b | 0.004678471 | -0.095525975 | 0.104882917 | 7.656173967 | 0.111590213 | 0.914441172 | 0.997677483 | -5.219914203 |
| **A_55_P2016351** | Eif3c | -0.005360436 | -0.120173148 | 0.109452276 | 14.07945827 | -0.111588458 | 0.914442512 | 0.997677483 | -5.251453902 |
| **A_55_P2024245** | Mthfr | 0.005349157 | -0.109252952 | 0.119951266 | 8.466610312 | 0.111558293 | 0.91446553 | 0.997677483 | -5.251456736 |
| **A_55_P2051778** | Sult2b1 | 0.004665604 | -0.095374366 | 0.104705573 | 7.68329297 | 0.111466262 | 0.914535757 | 0.997677483 | -5.251465377 |
| **A_51_P138044** | Foxo1 | -0.00646172 | -0.14507671 | 0.13215327 | 10.79399717 | -0.111415868 | 0.914574212 | 0.997677483 | -5.251470106 |
| **A_51_P433824** | Tpi1 | -0.00886265 | -0.199324061 | 0.18159876 | 15.54212078 | -0.111215612 | 0.914727027 | 0.997677483 | -5.251488877 |
| **A_55_P1979551** | Pitrm1 | -0.009241786 | -0.207968178 | 0.189484606 | 12.43171684 | -0.111150008 | 0.914777091 | 0.997677483 | -5.251495019 |
| **A_55_P1956978** | Tln1 | 0.009120638 | -0.187065571 | 0.205306847 | 8.280100255 | 0.111113255 | 0.914805137 | 0.997677483 | -5.251498458 |
| **A_51_P141274** | Sec24d | -0.008057585 | -0.181559451 | 0.165444281 | 10.68876823 | -0.110996653 | 0.914894119 | 0.997677483 | -5.251509362 |
| **A_52_P596755** | Dnph1 | 0.012994092 | -0.266835032 | 0.292823216 | 9.321024425 | 0.110984453 | 0.914903429 | 0.997677483 | -5.251510502 |
| **A_52_P237652** | 2610035D1 | 0.007717856 | -0.158517945 | 0.173953657 | 8.300261997 | 0.11096378 | 0.914919205 | 0.997677483 | -5.251512434 |
| **A_51_P215540** | Rnf4 | 0.005739729 | -0.118262792 | 0.12974225 | 10.17054592 | 0.110629265 | 0.915174491 | 0.997677483 | -5.251543644 |
| **A_52_P638513** | Zbtb11os1 | 0.008942798 | -0.184545616 | 0.202431212 | 9.13711021 | 0.110465731 | 0.915299296 | 0.997677483 | -5.251558867 |
| **A_55_P1972792** | Stil | 0.006340741 | -0.130947365 | 0.143628846 | 8.429305824 | 0.110386562 | 0.915359717 | 0.997677483 | -5.251566229 |
| **A_55_P1987364** | Ap5z1 | -0.007625179 | -0.172754641 | 0.157504282 | 9.073007455 | -0.110365827 | 0.915375542 | 0.997677483 | -5.251568156 |
| **A_55_P2070838** | NA | 0.005054894 | -0.104416513 | 0.114526302 | 14.53659955 | 0.110362241 | 0.915378278 | 0.997677483 | -5.251568489 |
| **A_55_P2011425** | Atp10b | 0.004506722 | -0.093222077 | 0.10223552 | 7.690586089 | 0.110216675 | 0.915489374 | 0.997677483 | -5.251582008 |
| **A_55_P1962122** | NA | -0.006735228 | -0.153080615 | 0.139610158 | 15.21086089 | -0.109997402 | 0.915656727 | 0.997677483 | -5.251602339 |

| | | | | | | | | | |
|---|---|---|---|---|---|---|---|---|---|
| **A_55_P2147402** | NA | -0.006207461 | -0.141143587 | 0.128728664 | 8.043020556 | -0.109949919 | 0.915692968 | 0.997677483 | -5.251606737 |
| **A_55_P2053858** | Scamp3 | 0.006089561 | -0.126449134 | 0.138628256 | 13.03931502 | 0.10981267 | 0.915797722 | 0.997677483 | -5.251619436 |
| **A_55_P1963645** | NA | 0.007627039 | -0.158472172 | 0.17372625 | 10.44501082 | 0.109748231 | 0.915846905 | 0.997677483 | -5.251625394 |
| **A_52_P456158** | Fchsd1 | 0.031882239 | -0.662465486 | 0.726229963 | 9.587530315 | 0.109744035 | 0.915850107 | 0.997677483 | -5.251625781 |
| **A_55_P2119892** | NA | -0.009177306 | -0.209094322 | 0.19073971 | 10.36898286 | -0.109717167 | 0.915870615 | 0.997677483 | -5.251628264 |
| **A_66_P137152** | Zbtb8os | -0.005098063 | -0.116199237 | 0.106003112 | 11.97357667 | -0.109671969 | 0.915905112 | 0.997677483 | -5.251632439 |
| **A_55_P2185504** | Masp2 | -0.004465454 | -0.101782961 | 0.092852054 | 7.712632237 | -0.109668971 | 0.9159074 | 0.997677483 | -5.251632716 |
| **A_51_P413740** | Ftcd | -0.018638236 | -0.42496764 | 0.387691169 | 9.987394281 | -0.109631581 | 0.915935939 | 0.997677483 | -5.251636169 |
| **A_55_P2035320** | Nfil3 | 0.008589775 | -0.184828974 | 0.202008524 | 11.96582366 | 0.110319015 | 0.91596954 | 0.997677483 | -5.172436154 |
| **A_55_P2185573** | Rbms1 | 0.010063559 | -0.209671461 | 0.229798578 | 12.81226228 | 0.10946151 | 0.916065749 | 0.997677483 | -5.251651858 |
| **A_52_P138110** | 1110032A0 | -0.005898188 | -0.13487225 | 0.123075874 | 8.152878042 | -0.109301308 | 0.916188029 | 0.997677483 | -5.251666614 |
| **A_51_P369784** | Ces1e | -0.00529842 | -0.12120991 | 0.110613069 | 7.777529353 | -0.109251913 | 0.916225733 | 0.997677483 | -5.25167116 |
| **A_55_P2185332** | Il17rc | -0.004791784 | -0.109679054 | 0.100095486 | 10.47384182 | -0.109190185 | 0.916272849 | 0.997677483 | -5.251676837 |
| **A_55_P2071811** | Eml2 | 0.024676535 | -0.515526747 | 0.564879817 | 9.062034488 | 0.10917824 | 0.916281967 | 0.997677483 | -5.251677936 |
| **A_55_P2007678** | Xntrpc | -0.007518752 | -0.172164433 | 0.157126928 | 8.62256809 | -0.10914518 | 0.916307202 | 0.997677483 | -5.251680975 |
| **A_51_P219061** | Oxa1l | -0.006448966 | -0.147918882 | 0.135020951 | 10.37185501 | -0.108951974 | 0.91645468 | 0.997677483 | -5.251698717 |
| **A_52_P376360** | Pdgfc | 0.013948515 | -0.292381678 | 0.320278707 | 8.542958194 | 0.108829665 | 0.916548043 | 0.997677483 | -5.251709933 |
| **A_51_P406306** | Hecw2 | -0.010960786 | -0.252007614 | 0.230086042 | 8.849912334 | -0.108679945 | 0.916662331 | 0.997677483 | -5.251723646 |
| **A_55_P2129172** | Rab40b | -0.009748419 | -0.224235749 | 0.204738911 | 8.744913374 | -0.108627974 | 0.916702004 | 0.997677483 | -5.251728401 |
| **A_52_P543430** | NA | -0.013185327 | -0.303331229 | 0.276960574 | 12.2121575 | -0.10861345 | 0.91671309 | 0.997677483 | -5.25172973 |
| **A_51_P225179** | Smndc1 | -0.012110946 | -0.278944144 | 0.254722253 | 9.146726827 | -0.108479413 | 0.91681541 | 0.997677483 | -5.251741982 |
| **A_66_P102076** | Cep68 | 0.005247965 | -0.110421226 | 0.120917156 | 9.327187998 | 0.108438217 | 0.916846857 | 0.997677483 | -5.251745745 |
| **A_55_P1958102** | NA | -0.005951754 | -0.137171306 | 0.125267797 | 16.05902692 | -0.108406565 | 0.91687102 | 0.997677483 | -5.251748635 |
| **A_55_P2160726** | Tfcp2 | 0.004646643 | -0.09784061 | 0.107133895 | 8.01422177 | 0.108362387 | 0.916904745 | 0.997677483 | -5.251752667 |
| **A_55_P2036942** | Gm4535 | 0.010068205 | -0.212145593 | 0.232282003 | 11.49001746 | 0.108290449 | 0.916959661 | 0.997677483 | -5.25175923 |
| **A_55_P2035986** | E330020D1 | 0.015690256 | -0.330751915 | 0.362132427 | 10.29976001 | 0.108245141 | 0.916994249 | 0.997677483 | -5.251763361 |
| **A_55_P2068066** | Rapgefl1 | -0.009354666 | -0.216032552 | 0.19732322 | 8.2872283 | -0.108179111 | 0.917044657 | 0.997677483 | -5.251769379 |
| **A_55_P2020661** | Orai3 | -0.00561186 | -0.129598406 | 0.118374685 | 8.32466518 | -0.10817862 | 0.917045031 | 0.997677483 | -5.251769423 |
| **A_51_P254895** | Cyp4a10 | 0.0051175 | -0.10807722 | 0.11831222 | 7.848051924 | 0.108053988 | 0.917140176 | 0.997677483 | -5.251780771 |
| **A_55_P2102065** | Gm10639 | -0.004611682 | -0.106691014 | 0.09746765 | 7.722509069 | -0.107976861 | 0.917199057 | 0.997677483 | -5.251787787 |
| **A_55_P2111237** | Selenof | 0.007836461 | -0.16563644 | 0.181309363 | 13.4040648 | 0.107968605 | 0.917205359 | 0.997677483 | -5.251788538 |
| **A_55_P2041581** | NA | -0.012020073 | -0.278220967 | 0.25418082 | 10.44577749 | -0.107921195 | 0.917241553 | 0.997677483 | -5.251792848 |
| **A_55_P1960472** | Polg | -0.011167662 | -0.258799622 | 0.236464299 | 8.805275057 | -0.107786579 | 0.917344324 | 0.997677483 | -5.251805075 |
| **A_55_P2115392** | Mst1 | -0.00539085 | -0.12494055 | 0.114158849 | 7.697387533 | -0.10777497 | 0.917353187 | 0.997677483 | -5.251806129 |
| **A_66_P119421** | Prdx1 | -0.007183919 | -0.166529484 | 0.152161645 | 9.886595324 | -0.107753307 | 0.917369725 | 0.997677483 | -5.251808095 |
| **A_51_P145631** | Svip | 0.009557315 | -0.202681031 | 0.221795662 | 9.29720029 | 0.10762699 | 0.917466161 | 0.997677483 | -5.251819551 |
| **A_55_P2145029** | Gm826 | -0.005121865 | -0.118952813 | 0.108709083 | 8.008724524 | -0.107541699 | 0.917531277 | 0.997677483 | -5.251827279 |
| **A_55_P1995173** | Odc1 | 0.012671484 | -0.268990567 | 0.294333535 | 12.33039015 | 0.107524711 | 0.917544247 | 0.997677483 | -5.251828818 |
| **A_52_P593037** | Acsl5 | -0.007535687 | -0.17504912 | 0.159977745 | 9.373944507 | -0.107518295 | 0.917549145 | 0.997677483 | -5.251829399 |
| **A_55_P2273811** | Msh5 | -0.0063493 | -0.147745821 | 0.135047222 | 8.135030911 | -0.107323845 | 0.917697603 | 0.997677483 | -5.25184699 |
| **A_55_P2006634** | Map4k5 | 0.008393257 | -0.17855065 | 0.195337163 | 11.45206444 | 0.107307066 | 0.917710414 | 0.997677483 | -5.251848507 |
| **A_55_P2055597** | Enpp2 | 0.008169039 | -0.173996334 | 0.190334411 | 9.253248216 | 0.107180128 | 0.917807329 | 0.997677483 | -5.251859972 |
| **A_55_P2088315** | Vmn1r87 | 0.008767846 | -0.186760512 | 0.204296204 | 8.251060907 | 0.107174705 | 0.91781147 | 0.997677483 | -5.251860461 |
| **A_51_P513072** | Rdh13 | -0.00886125 | -0.206947583 | 0.189225084 | 8.97523772 | -0.106917698 | 0.918007697 | 0.997677483 | -5.251883631 |
| **A_55_P2159746** | Mfsd2b | 0.004694072 | -0.100269529 | 0.109657673 | 7.663002628 | 0.106885828 | 0.91803203 | 0.997677483 | -5.251886501 |
| **A_55_P2032039** | Bola3 | 0.008689616 | -0.185803518 | 0.203182749 | 13.02140808 | 0.106783814 | 0.918109921 | 0.997677483 | -5.251895679 |
| **A_51_P209302** | Polr1e | -0.011575705 | -0.27079494 | 0.24764353 | 10.94972513 | -0.106730699 | 0.918150476 | 0.997677483 | -5.251900455 |
| **A_52_P494982** | Dhx40 | -0.004759483 | -0.111392567 | 0.101873602 | 8.390931317 | -0.106678502 | 0.918190331 | 0.997677483 | -5.251905146 |

| | | | | | | | | | |
|---|---|---|---|---|---|---|---|---|---|
| **A_55_P2167189** | 4921504E0 | 0.015686245 | -0.336089375 | 0.367461866 | 8.373213552 | 0.106576732 | 0.918268037 | 0.997677483 | -5.251914285 |
| **A_52_P661** | Plxna4 | 0.015063428 | -0.322787336 | 0.352914193 | 10.75252721 | 0.106563397 | 0.918278219 | 0.997677483 | -5.251915482 |
| **A_55_P2115607** | Ikbip | 0.007904465 | -0.169498113 | 0.185307043 | 10.67728278 | 0.106493152 | 0.918331855 | 0.997677483 | -5.251921784 |
| **A_55_P2088038** | Vps13d | 0.005945079 | -0.127523128 | 0.139413286 | 9.813056369 | 0.106460607 | 0.918356705 | 0.997677483 | -5.251924703 |
| **A_55_P2029842** | Hnrnpa1 | 0.00873012 | -0.18739918 | 0.20485942 | 13.21556699 | 0.106386588 | 0.918413224 | 0.997677483 | -5.251931338 |
| **A_55_P1962154** | Rarb | 0.009524177 | -0.204671131 | 0.223719485 | 10.33808253 | 0.106273902 | 0.918499268 | 0.997677483 | -5.251941429 |
| **A_55_P2022895** | Slc12a8 | 0.006336503 | -0.136197731 | 0.148870737 | 8.068500159 | 0.106252606 | 0.918515529 | 0.997677483 | -5.251943335 |
| **A_51_P175205** | Rps6kb1 | -0.007582137 | -0.178246572 | 0.163082298 | 8.965342639 | -0.106183668 | 0.918568169 | 0.997677483 | -5.251949502 |
| **A_55_P2114467** | Romo1 | 0.007005828 | -0.150905687 | 0.164917342 | 14.52410988 | 0.106036354 | 0.918680658 | 0.997677483 | -5.251962667 |
| **A_51_P466964** | Pgp | 0.009019453 | -0.194416091 | 0.212454997 | 12.89925087 | 0.105965014 | 0.918735133 | 0.997677483 | -5.251969036 |
| **A_55_P2145059** | Gbp5 | 0.008283997 | -0.178732228 | 0.195300222 | 8.38946802 | 0.105869232 | 0.918808274 | 0.997677483 | -5.251977581 |
| **A_51_P316173** | Vmn1r192 | -0.006322493 | -0.149270648 | 0.136625662 | 7.88066906 | -0.105710692 | 0.918929339 | 0.997677483 | -5.251991707 |
| **A_55_P2060198** | NA | -0.017041126 | -0.402578457 | 0.368496206 | 9.456599634 | -0.10564306 | 0.918980986 | 0.997677483 | -5.251997726 |
| **A_51_P185906** | Abi3 | -0.018656817 | -0.440861183 | 0.403547549 | 8.94392953 | -0.1056146 | 0.919002719 | 0.997677483 | -5.252000258 |
| **A_55_P2000543** | Cd209f | 0.004304995 | -0.093133693 | 0.101743683 | 7.660568737 | 0.105596701 | 0.919016387 | 0.997677483 | -5.25200185 |
| **A_51_P327021** | Zbtb25 | 0.008242429 | -0.178406596 | 0.194891453 | 10.2206842 | 0.105545227 | 0.919055695 | 0.997677483 | -5.252006427 |
| **A_55_P1967325** | Tafa5 | 0.005549044 | -0.120178164 | 0.131276253 | 7.923876611 | 0.105486784 | 0.919100325 | 0.997677483 | -5.252011621 |
| **A_55_P2040026** | Itga4 | 0.011231918 | -0.243758954 | 0.266222789 | 8.424809555 | 0.10527819 | 0.919259621 | 0.997677483 | -5.252030136 |
| **A_55_P2007026** | Gm9112 | -0.005004864 | -0.118798347 | 0.10878862 | 7.83864115 | -0.105119667 | 0.919380682 | 0.997677483 | -5.252044182 |
| **A_55_P2111825** | 1700001C0 | -0.004730687 | -0.112297486 | 0.102836111 | 7.685247039 | -0.105112684 | 0.919386014 | 0.997677483 | -5.2520448 |
| **A_55_P2105638** | Ralgapa2 | 0.007157094 | -0.155635611 | 0.169949799 | 9.669624167 | 0.10507779 | 0.919412663 | 0.997677483 | -5.252047889 |
| **A_51_P141390** | Anapc1 | 0.005874983 | -0.127787043 | 0.139537009 | 11.8658554 | 0.105052815 | 0.919431736 | 0.997677483 | -5.252050099 |
| **A_55_P1993955** | Pltp | -0.004583542 | -0.108950811 | 0.099783728 | 8.056662731 | -0.104965363 | 0.919498523 | 0.997677483 | -5.252057833 |
| **A_55_P2454099** | Oga | -0.010983283 | -0.261205804 | 0.239239238 | 9.012850893 | -0.104909522 | 0.919541169 | 0.997677483 | -5.252062769 |
| **A_52_P385606** | Ckb | -0.021564321 | -0.513578834 | 0.470450193 | 11.13126729 | -0.104753127 | 0.91966061 | 0.997677483 | -5.252076578 |
| **A_51_P112734** | Slc7a8 | -0.004942686 | -0.117913518 | 0.108028146 | 8.241612697 | -0.104569687 | 0.919800709 | 0.997677483 | -5.225269826 |
| **A_51_P389988** | Slc40a1 | -0.010401636 | -0.248214362 | 0.22741109 | 9.482118034 | -0.104538366 | 0.91982463 | 0.997677483 | -5.252095507 |
| **A_51_P426458** | Ube2d2a | 0.008137051 | -0.177957494 | 0.194231597 | 10.87700066 | 0.104506318 | 0.919849106 | 0.997677483 | -5.252098329 |
| **A_65_P16542** | NA | -0.01038639 | -0.248093666 | 0.227320885 | 12.07975182 | -0.104431449 | 0.919906288 | 0.997677483 | -5.252104917 |
| **A_51_P138879** | Zgpat | -0.008642816 | -0.206480289 | 0.189194658 | 9.42275124 | -0.104413302 | 0.919920147 | 0.997677483 | -5.252106513 |
| **A_55_P2003353** | Gemin8 | -0.004844512 | -0.115826913 | 0.106137889 | 8.727499803 | -0.104328996 | 0.919984537 | 0.997677483 | -5.252113924 |
| **A_51_P250358** | Prpf39 | -0.010429983 | -0.249715208 | 0.228855243 | 8.863699914 | -0.104178196 | 0.920099713 | 0.997677483 | -5.252127166 |
| **A_55_P2128929** | Cc2d2a | -0.011742356 | -0.281185983 | 0.257701272 | 10.56771906 | -0.104158915 | 0.920114439 | 0.997677483 | -5.252128858 |
| **A_55_P2026982** | Tdrp | 0.007524688 | -0.165164301 | 0.180213677 | 8.286570566 | 0.104143693 | 0.920126065 | 0.997677483 | -5.252130193 |
| **A_55_P1991054** | Tmf1 | -0.005920216 | -0.141792889 | 0.129952456 | 9.422954314 | -0.104139285 | 0.920129432 | 0.997677483 | -5.25213058 |
| **A_55_P1974407** | Cdk14 | 0.008045685 | -0.176854176 | 0.192945546 | 9.670347044 | 0.104000539 | 0.920235404 | 0.997677483 | -5.252142742 |
| **A_51_P502082** | Rrm1 | 0.006090177 | -0.134053375 | 0.146233729 | 12.23344037 | 0.103864215 | 0.920339528 | 0.997677483 | -5.252154676 |
| **A_55_P2082050** | Ankrd12 | 0.005745583 | -0.126619628 | 0.138110793 | 8.498246918 | 0.103745519 | 0.920430189 | 0.997677483 | -5.252165055 |
| **A_55_P2008349** | Prelid1 | 0.006826882 | -0.150458626 | 0.164112391 | 13.6688364 | 0.103739189 | 0.920435024 | 0.997677483 | -5.252165608 |
| **A_55_P2161055** | Trp63 | -0.007601041 | -0.182861431 | 0.167659349 | 11.12771879 | -0.10365695 | 0.920497839 | 0.997677483 | -5.252172791 |
| **A_51_P311096** | Malsu1 | -0.007131594 | -0.171824067 | 0.15756088 | 12.85264995 | -0.103495623 | 0.920621066 | 0.997677483 | -5.252186866 |
| **A_51_P109888** | Coro1b | -0.00929772 | -0.224021527 | 0.205426088 | 9.33176942 | -0.103491669 | 0.920624086 | 0.997677483 | -5.25218721 |
| **A_51_P511734** | Snip1 | 0.004663774 | -0.103044846 | 0.112372393 | 8.404347799 | 0.103489464 | 0.92062577 | 0.997677483 | -5.252187403 |
| **A_52_P527800** | Emilin2 | -0.009685652 | -0.233571226 | 0.214199923 | 12.69459324 | -0.103397938 | 0.920695682 | 0.997677483 | -5.252195378 |
| **A_55_P1975772** | Ppp2r5a | 0.007432615 | -0.164549779 | 0.179415008 | 13.0788867 | 0.103292018 | 0.920776589 | 0.997677483 | -5.252204598 |
| **A_55_P2038002** | BC005624 | 0.006716432 | -0.148800175 | 0.162233039 | 10.37816838 | 0.103221694 | 0.920830306 | 0.997677483 | -5.252210714 |
| **A_55_P2034853** | 2610028H2 | 0.010720658 | -0.237580379 | 0.259021695 | 8.840244557 | 0.103193431 | 0.920851895 | 0.997677483 | -5.252213171 |
| **A_51_P145520** | Sigirr | -0.004652397 | -0.112466889 | 0.103162095 | 8.4549011 | -0.103135636 | 0.920896043 | 0.997677483 | -5.252218194 |

| | | | | | | | | | |
|---|---|---|---|---|---|---|---|---|---|
| **A_52_P117966** | Atpsckmt | -0.006745986 | -0.163216575 | 0.149724603 | 10.62361845 | -0.103043799 | 0.920966194 | 0.997677483 | -5.252226168 |
| **A_55_P2020786** | Prkar1a | -0.005632789 | -0.136284966 | 0.125019388 | 13.67673279 | -0.103042407 | 0.920967258 | 0.997677483 | -5.252226289 |
| **A_55_P2177968** | Snrpd2 | -0.006585491 | -0.15953722 | 0.146366238 | 14.21386209 | -0.102906527 | 0.921071054 | 0.997677483 | -5.252238075 |
| **A_55_P2082444** | Npepps | 0.005342417 | -0.118856887 | 0.12954172 | 12.36622602 | 0.102808196 | 0.921146169 | 0.997677483 | -5.252246595 |
| **A_55_P1991723** | Alkbh3 | 0.006792024 | -0.15118577 | 0.164769817 | 11.48611091 | 0.102757208 | 0.921185119 | 0.997677483 | -5.252251009 |
| **A_52_P105537** | Ccn3 | -0.009568543 | -0.232155602 | 0.213018516 | 8.409302152 | -0.10274366 | 0.921195468 | 0.997677483 | -5.252252182 |
| **A_55_P2092989** | NA | 0.007801027 | -0.173769032 | 0.189371086 | 8.983531193 | 0.102687287 | 0.921238532 | 0.997677483 | -5.252257059 |
| **A_66_P109800** | Gcsh | 0.007676214 | -0.171042597 | 0.186395024 | 12.26543076 | 0.102656372 | 0.921262148 | 0.997677483 | -5.252259733 |
| **A_52_P528634** | Ttc27 | -0.008803686 | -0.21379761 | 0.196190237 | 9.212873374 | -0.102643794 | 0.921271757 | 0.997677483 | -5.25226082 |
| **A_55_P1988168** | Proser3 | 0.013670832 | -0.304726076 | 0.33206774 | 13.70941812 | 0.102620763 | 0.921289351 | 0.997677483 | -5.252262812 |
| **A_55_P2152437** | Mitf | 0.014286147 | -0.319478949 | 0.348051242 | 11.72143817 | 0.102301812 | 0.921533007 | 0.997677483 | -5.25229034 |
| **A_55_P2095794** | Evi5 | 0.007336473 | -0.164085418 | 0.178758364 | 10.43601705 | 0.102289296 | 0.921542568 | 0.997677483 | -5.252291418 |
| **A_55_P2073845** | Mrpl47 | 0.007585764 | -0.169719771 | 0.184891298 | 9.710838686 | 0.102255374 | 0.921568483 | 0.997677483 | -5.252294341 |
| **A_52_P366803** | Cyp3a44 | 0.004081532 | -0.091363635 | 0.099526699 | 7.646376686 | 0.102206474 | 0.92160584 | 0.997677483 | -5.252298552 |
| **A_55_P2009217** | Rnu3b1 | 0.014107737 | -0.315844394 | 0.344059867 | 10.48885511 | 0.102191682 | 0.921617141 | 0.997677483 | -5.252299825 |
| **A_55_P2039586** | Grk3 | 0.012901144 | -0.288852194 | 0.314654482 | 8.446575395 | 0.102184559 | 0.921622582 | 0.997677483 | -5.252300438 |
| **A_55_P2087114** | NA | 0.011621407 | -0.260407968 | 0.283650781 | 12.86175005 | 0.102106179 | 0.921682461 | 0.997677483 | -5.252307182 |
| **A_51_P425962** | Khdrbs3 | -0.008305854 | -0.202849316 | 0.186237608 | 12.49038233 | -0.102041486 | 0.921731885 | 0.997677483 | -5.252312745 |
| **A_51_P255456** | Cyp1b1 | -0.012141884 | -0.297092059 | 0.272808291 | 7.939739347 | -0.101841853 | 0.9218844 | 0.997677483 | -5.252329888 |
| **A_51_P486155** | Ocln | 0.004507108 | -0.101427589 | 0.110441806 | 7.744753572 | 0.101687807 | 0.922002089 | 0.997677483 | -5.252343093 |
| **A_55_P1993698** | Celf1 | 0.0085499 | -0.192487429 | 0.209587229 | 10.44070656 | 0.101646742 | 0.922033463 | 0.997677483 | -5.25234661 |
| **A_55_P2062737** | Elovl2 | 0.004341164 | -0.097786484 | 0.106468811 | 7.812169804 | 0.101594913 | 0.922073061 | 0.997677483 | -5.252351047 |
| **A_55_P2000958** | Sdr39u1 | 0.006843775 | -0.15421637 | 0.167903919 | 11.04735729 | 0.101558611 | 0.922100796 | 0.997677483 | -5.252354153 |
| **A_51_P108978** | Myadm | 0.008847654 | -0.199580988 | 0.217276296 | 11.10841477 | 0.101456499 | 0.922178811 | 0.997677483 | -5.252362884 |
| **A_51_P320089** | Gclm | 0.005444306 | -0.122889907 | 0.133778519 | 8.487858035 | 0.101393282 | 0.92222711 | 0.997677483 | -5.252368286 |
| **A_52_P680038** | Cdk16 | -0.011249468 | -0.276529567 | 0.25403063 | 9.085496477 | -0.101352966 | 0.922257913 | 0.997677483 | -5.252371728 |
| **A_55_P2066274** | NA | -0.010298546 | -0.253166774 | 0.232569681 | 12.86967123 | -0.101347808 | 0.922261853 | 0.997677483 | -5.252372169 |
| **A_55_P2156625** | NA | -0.010484264 | -0.257836092 | 0.236867565 | 13.33122671 | -0.101305249 | 0.92229437 | 0.997677483 | -5.252375801 |
| **A_55_P2182690** | Orai3 | 0.005515853 | -0.124641646 | 0.135673353 | 10.83292454 | 0.101286752 | 0.922308502 | 0.997677483 | -5.25237738 |
| **A_52_P243391** | Sema4f | -0.004262224 | -0.104950998 | 0.09642655 | 7.650950705 | -0.101172937 | 0.922395462 | 0.997677483 | -5.252387085 |
| **A_55_P2101570** | Zhx3 | -0.008791265 | -0.216604423 | 0.199021893 | 10.32249034 | -0.101108453 | 0.922444731 | 0.997677483 | -5.252392579 |
| **A_55_P2111950** | Dnajc7 | 0.025752672 | -0.583429115 | 0.63493446 | 12.66115894 | 0.10103795 | 0.922498599 | 0.997677483 | -5.252398582 |
| **A_52_P177490** | Cdx2 | -0.004289821 | -0.105809644 | 0.097230002 | 7.670444932 | -0.100994442 | 0.922531841 | 0.997677483 | -5.252402285 |
| **A_66_P132295** | Tfpi | -0.007098496 | -0.175146304 | 0.160949313 | 11.17142158 | -0.100958432 | 0.922559355 | 0.997677483 | -5.252405348 |
| **A_51_P157297** | Efcab14 | -0.005081825 | -0.125520092 | 0.115356442 | 10.92797303 | -0.100847311 | 0.922644259 | 0.997677483 | -5.252414793 |
| **A_52_P121342** | Nrxn1 | -0.005728592 | -0.141541826 | 0.130084642 | 8.042038418 | -0.100812628 | 0.92267076 | 0.997677483 | -5.252417739 |
| **A_55_P2115192** | Sike1 | 0.004630583 | -0.105182153 | 0.11444332 | 9.593236937 | 0.100784124 | 0.922692539 | 0.997677483 | -5.252420159 |
| **A_55_P2031407** | NA | -0.006493097 | -0.160484253 | 0.14749806 | 17.61560445 | -0.100777884 | 0.922697307 | 0.997677483 | -5.252420689 |
| **A_51_P158584** | Poll | 0.00686399 | -0.15595895 | 0.169686931 | 9.915026351 | 0.100755831 | 0.922714157 | 0.997677483 | -5.252422561 |
| **A_52_P48546** | Rnf41 | -0.005774061 | -0.142827254 | 0.131279133 | 8.608307237 | -0.10069347 | 0.922761806 | 0.997677483 | -5.252427853 |
| **A_51_P349662** | Taf5l | 0.00482346 | -0.10968914 | 0.11933606 | 10.84101054 | 0.10067336 | 0.922777172 | 0.997677483 | -5.252429558 |
| **A_51_P196158** | Btd | -0.004996789 | -0.123657859 | 0.113664281 | 10.10388747 | -0.100644918 | 0.922798904 | 0.997677483 | -5.25243197 |
| **A_55_P2144566** | NA | 0.005919476 | -0.134665668 | 0.146504621 | 15.90513989 | 0.100635912 | 0.922805785 | 0.997677483 | -5.252432734 |
| **A_55_P2061421** | NA | -0.010963917 | -0.271563975 | 0.249636141 | 12.99952767 | -0.100554236 | 0.922868194 | 0.997677483 | -5.252439655 |
| **A_55_P2017709** | Igsf9 | -0.008316553 | -0.20634313 | 0.189710023 | 11.6954346 | -0.100375802 | 0.923004537 | 0.997677483 | -5.252454757 |
| **A_52_P94521** | 2510009E0 | -0.006369382 | -0.158088244 | 0.14534948 | 9.404972862 | -0.100338325 | 0.923033173 | 0.997677483 | -5.252457925 |
| **A_55_P1957389** | Snx25 | 0.004939257 | -0.112728024 | 0.122606538 | 10.47428669 | 0.10032635 | 0.923042324 | 0.997677483 | -5.252458937 |
| **A_55_P2126925** | Msantd3 | -0.006042897 | -0.150036395 | 0.137950601 | 11.70312462 | -0.100302427 | 0.923060604 | 0.997677483 | -5.252460959 |

| | | | | | | | | | |
|---|---|---|---|---|---|---|---|---|---|
| **A_55_P2064351** | Vipr1 | 0.007291577 | -0.166490323 | 0.181073478 | 7.835830907 | 0.10028271 | 0.92307567 | 0.997677483 | -5.252462625 |
| **A_52_P563617** | Ssbp4 | -0.007682429 | -0.19090715 | 0.175542293 | 10.85342175 | -0.100212894 | 0.923129019 | 0.997677483 | -5.252468521 |
| **A_55_P2130575** | NA | 0.008893353 | -0.20346127 | 0.221247977 | 13.86909797 | 0.100095132 | 0.923219005 | 0.997677483 | -5.252478456 |
| **A_55_P2047163** | Tmc5 | 0.004416816 | -0.101166998 | 0.110000631 | 7.782945906 | 0.099981843 | 0.923305574 | 0.997677483 | -5.252488004 |
| **A_51_P313337** | Smim10l1 | -0.006719224 | -0.167449156 | 0.154010709 | 10.59996476 | -0.09991518 | 0.923356515 | 0.997677483 | -5.252493617 |
| **A_51_P219505** | Slc41a2 | -0.00496873 | -0.123902191 | 0.113964731 | 7.983334118 | -0.099850558 | 0.923405896 | 0.997677483 | -5.252499055 |
| **A_55_P2158741** | Nos2 | 0.00712109 | -0.163735313 | 0.177977493 | 8.759232551 | 0.099614913 | 0.923585969 | 0.997677483 | -5.252518853 |
| **A_55_P2074371** | Lin28a | -0.006467375 | -0.161701658 | 0.148766908 | 8.214691365 | -0.09957482 | 0.923616608 | 0.997677483 | -5.252522217 |
| **A_55_P2164778** | Ptdss1 | 0.006565242 | -0.151163802 | 0.164294285 | 11.35350301 | 0.099482839 | 0.923686899 | 0.997677483 | -5.252529929 |
| **A_55_P2082614** | Zfp791 | 0.010004527 | -0.230504855 | 0.250513909 | 11.49683282 | 0.099419981 | 0.923734935 | 0.997677483 | -5.252535196 |
| **A_55_P2005050** | NA | -0.007143855 | -0.178958984 | 0.164671275 | 11.90847522 | -0.09937573 | 0.923768751 | 0.997677483 | -5.252538901 |
| **A_55_P2025615** | Ammecr1l | 0.005264679 | -0.121422494 | 0.131951852 | 12.86757738 | 0.099322669 | 0.923809301 | 0.997677483 | -5.252543342 |
| **A_51_P282092** | Urod | 0.010608281 | -0.244773829 | 0.265990392 | 13.55490279 | 0.099280437 | 0.923841575 | 0.997677483 | -5.252546875 |
| **A_55_P2021125** | NA | 0.009391447 | -0.216702391 | 0.235485284 | 8.489964557 | 0.099277973 | 0.923843458 | 0.997677483 | -5.252547081 |
| **A_55_P2043734** | Cdkl5 | -0.004942943 | -0.123945916 | 0.11406003 | 7.889230289 | -0.09927432 | 0.92384625 | 0.997677483 | -5.252547387 |
| **A_52_P337236** | Alyref2 | 0.00940516 | -0.217064912 | 0.235875233 | 9.53699008 | 0.09925777 | 0.923858897 | 0.997677483 | -5.252548771 |
| **A_51_P206235** | Slc39a5 | -0.004490716 | -0.112678641 | 0.103697209 | 7.697567314 | -0.099207832 | 0.923897061 | 0.997677483 | -5.252552946 |
| **A_55_P2052862** | Miga1 | 0.009120539 | -0.210946471 | 0.229187549 | 10.76973534 | 0.09905461 | 0.924014157 | 0.997677483 | -5.252565742 |
| **A_55_P2150555** | Pcgf5 | -0.009959685 | -0.250296132 | 0.230376763 | 11.45969448 | -0.099045578 | 0.924021059 | 0.997677483 | -5.252566496 |
| **A_55_P2041823** | Hyal1 | 0.011783385 | -0.273161877 | 0.296728647 | 9.658230081 | 0.098836593 | 0.924180774 | 0.997677483 | -5.252583915 |
| **A_51_P444994** | Tmem203 | -0.006156233 | -0.155148053 | 0.142835588 | 9.842227499 | -0.0987556 | 0.924242673 | 0.997677483 | -5.252590657 |
| **A_55_P2007693** | NA | -0.006907948 | -0.17438832 | 0.160572424 | 9.939699067 | -0.098581243 | 0.924375928 | 0.997677483 | -5.25260515 |
| **A_52_P325116** | Gramd1b | 0.012400865 | -0.288749518 | 0.313551248 | 10.59782624 | 0.098418717 | 0.924500144 | 0.997677483 | -5.252618637 |
| **A_51_P357085** | Dctn6 | 0.008805134 | -0.205038411 | 0.22264868 | 12.85915803 | 0.098412209 | 0.924505118 | 0.997677483 | -5.252619177 |
| **A_51_P233396** | Ppp2r5d | -0.006774817 | -0.17134384 | 0.157794205 | 11.37050757 | -0.098391739 | 0.924520763 | 0.997677483 | -5.252620874 |
| **A_51_P307964** | Krt13 | -0.004428014 | -0.112035622 | 0.103179594 | 7.737115425 | -0.098350182 | 0.924552524 | 0.997677483 | -5.252624318 |
| **A_55_P2108122** | Arcn1 | 0.00423077 | -0.098661044 | 0.107122585 | 12.09497833 | 0.098276065 | 0.924609171 | 0.997677483 | -5.252630457 |
| **A_55_P1982249** | Phactr2 | -0.012754818 | -0.323122052 | 0.297612417 | 11.20647954 | -0.098221718 | 0.924650709 | 0.997677483 | -5.252634955 |
| **A_55_P1965935** | Rwdd1 | 0.008652451 | -0.201904021 | 0.219208922 | 10.4355459 | 0.09821542 | 0.924655523 | 0.997677483 | -5.252635477 |
| **A_55_P2164998** | NA | 0.010387144 | -0.242583088 | 0.263357376 | 12.54827594 | 0.098137699 | 0.924714925 | 0.997677483 | -5.252641905 |
| **A_52_P353478** | Scn7a | 0.003925474 | -0.09172896 | 0.099579909 | 7.694220313 | 0.098083547 | 0.924756315 | 0.997677483 | -5.252646381 |
| **A_51_P241843** | Tmem202 | 0.004283271 | -0.100196766 | 0.108763308 | 7.764063316 | 0.097983152 | 0.924833048 | 0.997677483 | -5.252654673 |
| **A_52_P569038** | Cnot2 | 0.00746433 | -0.174795938 | 0.189724599 | 11.20635074 | 0.09788316 | 0.924909475 | 0.997677483 | -5.252662923 |
| **A_55_P2093439** | Mex3d | 0.00649501 | -0.15215004 | 0.16514006 | 10.69609054 | 0.09785036 | 0.924934545 | 0.997677483 | -5.252665628 |
| **A_55_P2183438** | Runx1 | 0.004102999 | -0.096140815 | 0.104346812 | 7.808440987 | 0.097825689 | 0.924953402 | 0.997677483 | -5.252667661 |
| **A_51_P220682** | Retreg3 | 0.007010883 | -0.164514865 | 0.178536631 | 10.03110409 | 0.097690543 | 0.9250567 | 0.997677483 | -5.252678792 |
| **A_55_P1958562** | Dmrtc1b | 0.004039869 | -0.094813495 | 0.102893232 | 7.643403758 | 0.097675342 | 0.925068319 | 0.997677483 | -5.252680043 |
| **A_51_P221495** | Nsun5 | 0.004733403 | -0.111093588 | 0.120560393 | 8.677131494 | 0.097672615 | 0.925070403 | 0.997677483 | -5.252680267 |
| **A_55_P2024215** | Ctnnbl1 | -0.004255117 | -0.108401277 | 0.099891042 | 12.0130072 | -0.097651164 | 0.9250868 | 0.997677483 | -5.252682032 |
| **A_55_P2082235** | NA | -0.004668479 | -0.119265406 | 0.109928449 | 9.145224641 | -0.097366943 | 0.925304048 | 0.997677483 | -5.252705382 |
| **A_51_P153557** | Hspa12a | -0.005445564 | -0.139533691 | 0.128642562 | 8.45246411 | -0.097064789 | 0.925535011 | 0.997677483 | -5.25273013 |
| **A_55_P2083629** | Tle2 | 0.008489748 | -0.201212572 | 0.218192068 | 8.622260615 | 0.096761075 | 0.925767175 | 0.997677483 | -5.252754929 |
| **A_55_P1999301** | Cyp2e1 | -0.004146107 | -0.106614424 | 0.098322209 | 7.710955408 | -0.096707479 | 0.925808145 | 0.997677483 | -5.252759297 |
| **A_55_P2096166** | NA | -0.007437689 | -0.191303991 | 0.176428613 | 10.93520304 | -0.096681866 | 0.925827725 | 0.997677483 | -5.252761384 |
| **A_55_P2012939** | Arid4b | -0.007960084 | -0.204772365 | 0.188852197 | 9.351398101 | -0.096666187 | 0.925839711 | 0.997677483 | -5.252762661 |
| **A_55_P2054062** | Chst2 | 0.005570688 | -0.132378257 | 0.143519634 | 8.08495645 | 0.096516061 | 0.925954473 | 0.997677483 | -5.252774878 |
| **A_55_P2046478** | Zfp81 | -0.003988685 | -0.102769455 | 0.094792086 | 8.362201574 | -0.09650869 | 0.925960108 | 0.997677483 | -5.252775478 |
| **A_55_P1986225** | NA | -0.006733787 | -0.173664209 | 0.160196636 | 11.87046667 | -0.096412423 | 0.9260337 | 0.997677483 | -5.252783302 |

| | | | | | | | | | |
|---|---|---|---|---|---|---|---|---|---|
| **A_51_P370341** | Zbtb48 | 0.005677222 | -0.135134266 | 0.146488711 | 8.608101895 | 0.096362254 | 0.926072052 | 0.997677483 | -5.252787376 |
| **A_55_P1952076** | Npb | -0.007569279 | -0.195327417 | 0.180188859 | 10.49253681 | -0.096352921 | 0.926079187 | 0.997677483 | -5.252788134 |
| **A_55_P2067632** | Ush1c | -0.006986266 | -0.180336786 | 0.166364255 | 8.568996374 | -0.096322794 | 0.926102218 | 0.997677483 | -5.252790579 |
| **A_52_P497388** | NA | -0.005578352 | -0.144059467 | 0.132902763 | 14.68206676 | -0.096277426 | 0.9261369 | 0.997677483 | -5.25279426 |
| **A_55_P2184877** | NA | -0.007197114 | -0.185882156 | 0.171487927 | 10.95569323 | -0.096267422 | 0.926144548 | 0.997677483 | -5.252795071 |
| **A_55_P2129975** | Rabgef1 | -0.004703822 | -0.121491089 | 0.112083446 | 9.925584043 | -0.096264128 | 0.926147066 | 0.997677483 | -5.252795339 |
| **A_51_P356525** | Trappc11 | 0.004586971 | -0.109320284 | 0.118494227 | 9.170716453 | 0.096246239 | 0.926160742 | 0.997677483 | -5.252796789 |
| **A_55_P1990336** | Add1 | 0.009292905 | -0.221523525 | 0.240109335 | 13.67229098 | 0.096226328 | 0.926175963 | 0.997677483 | -5.252798404 |
| **A_55_P2066958** | Sh3glb1 | -0.00626308 | -0.162224352 | 0.149698193 | 11.55608117 | -0.095979902 | 0.926364352 | 0.997677483 | -5.252818357 |
| **A_51_P331462** | Cgrrf1 | -0.007399003 | -0.19166695 | 0.176868943 | 9.002489915 | -0.095969351 | 0.926372418 | 0.997677483 | -5.252819211 |
| **A_51_P128575** | Scgb1a1 | 0.004069379 | -0.097563418 | 0.105702175 | 7.753290112 | 0.095698107 | 0.926579786 | 0.997677483 | -5.234095829 |
| **A_52_P173368** | NA | 0.008198622 | -0.196757306 | 0.213154549 | 8.569904415 | 0.095606956 | 0.926649473 | 0.997677483 | -5.252848459 |
| **A_55_P2122334** | Nfic | 0.00841122 | -0.201885966 | 0.218708405 | 11.77492341 | 0.095594884 | 0.926658703 | 0.997677483 | -5.252849431 |
| **A_51_P382253** | Rnf2 | 0.008767512 | -0.210518702 | 0.228053725 | 10.84395922 | 0.095559565 | 0.926685705 | 0.997677483 | -5.252852275 |
| **A_55_P2083963** | Phgr1 | 0.004392867 | -0.105546851 | 0.114332585 | 7.704923395 | 0.095499818 | 0.926731383 | 0.997677483 | -5.252857084 |
| **A_55_P2074062** | Sdhaf2 | -0.004479517 | -0.116594298 | 0.107635264 | 10.45758067 | -0.095494298 | 0.926735604 | 0.997677483 | -5.252857529 |
| **A_55_P2058384** | NA | 0.009713905 | -0.233435822 | 0.252863632 | 9.062871373 | 0.095483689 | 0.926743715 | 0.997677483 | -5.252858382 |
| **A_55_P1980035** | Tbc1d9b | 0.006065985 | -0.145900322 | 0.158032292 | 10.79153781 | 0.095403242 | 0.926805219 | 0.997677483 | -5.252864851 |
| **A_55_P2417224** | NA | -0.005189206 | -0.135233472 | 0.124855059 | 7.962966671 | -0.095371553 | 0.926829447 | 0.997677483 | -5.252867398 |
| **A_52_P273120** | Myl1 | 0.031430508 | -0.75624646 | 0.819107475 | 9.342386055 | 0.095370132 | 0.926830533 | 0.997677483 | -5.252867513 |
| **A_55_P2086143** | Cyp39a1 | -0.020953347 | -0.546190292 | 0.504283599 | 11.90268723 | -0.09534705 | 0.926848181 | 0.997677483 | -5.252869367 |
| **A_55_P2106063** | NA | -0.011885475 | -0.310289419 | 0.286518469 | 13.76133572 | -0.09519653 | 0.926963261 | 0.997677483 | -5.25288145 |
| **A_51_P155303** | Rbfa | -0.008946456 | -0.233569218 | 0.215676305 | 9.668026096 | -0.095193309 | 0.926965724 | 0.997677483 | -5.252881708 |
| **A_55_P2134356** | Msl3 | -0.00857409 | -0.223933866 | 0.206785686 | 11.68841965 | -0.09515521 | 0.926994853 | 0.997677483 | -5.252884764 |
| **A_51_P325165** | Il27 | 0.005555994 | -0.134066372 | 0.145178361 | 8.332846258 | 0.09510775 | 0.927031139 | 0.997677483 | -5.252888568 |
| **A_55_P1983358** | Gm4593 | -0.004548802 | -0.118912487 | 0.109814883 | 8.028377722 | -0.095064423 | 0.927064266 | 0.997677483 | -5.252892039 |
| **A_51_P117865** | Fam20c | 0.006688955 | -0.161486025 | 0.174863934 | 9.29147902 | 0.095061792 | 0.927066277 | 0.997677483 | -5.25289225 |
| **A_55_P1989733** | Nfkbib | -0.005167412 | -0.135355687 | 0.125020864 | 10.67284103 | -0.09486594 | 0.927216022 | 0.997677483 | -5.252907921 |
| **A_51_P416858** | Myl1 | 0.046830248 | -1.133622231 | 1.227282726 | 13.44269848 | 0.094817189 | 0.927253296 | 0.997677483 | -5.252911817 |
| **A_55_P2014209** | Scyl2 | -0.004889686 | -0.128299452 | 0.11852008 | 8.251985733 | -0.094697949 | 0.927344467 | 0.997677483 | -5.252921337 |
| **A_51_P486610** | Afg3l1 | -0.010129194 | -0.266314564 | 0.246056177 | 8.698984668 | -0.094499538 | 0.927496174 | 0.997677483 | -5.252937152 |
| **A_55_P2131820** | A530006G2 | -0.006393859 | -0.168154483 | 0.155366765 | 11.18357367 | -0.094471187 | 0.927517851 | 0.997677483 | -5.252939409 |
| **A_55_P2036337** | Ifnar1 | 0.005501151 | -0.133883375 | 0.144885678 | 10.66215923 | 0.094329636 | 0.927626085 | 0.997677483 | -5.252950669 |
| **A_55_P2060036** | Ulk2 | -0.004704577 | -0.123977314 | 0.11456816 | 9.530446552 | -0.094273261 | 0.927669192 | 0.997677483 | -5.252955148 |
| **A_55_P2149274** | Sh3bp5l | -0.007316056 | -0.192819544 | 0.178187431 | 11.99049535 | -0.094261445 | 0.927678227 | 0.997677483 | -5.252956087 |
| **A_55_P1989916** | Lin54 | 0.008878809 | -0.216287683 | 0.234045302 | 9.852472298 | 0.094245387 | 0.927690505 | 0.997677483 | -5.252957362 |
| **A_55_P1984307** | Adgrf5 | -0.007484867 | -0.197354324 | 0.18238459 | 10.06219298 | -0.09421892 | 0.927710743 | 0.997677483 | -5.252959464 |
| **A_55_P2096368** | NA | 0.004465209 | -0.10892538 | 0.117855798 | 11.16238976 | 0.094118264 | 0.92778771 | 0.997677483 | -5.25296745 |
| **A_55_P2037919** | NA | -0.009222477 | -0.243431106 | 0.224986153 | 9.376734634 | -0.094113908 | 0.92779104 | 0.997677483 | -5.252967796 |
| **A_55_P2002849** | Aifm3 | 0.007499041 | -0.183174488 | 0.19817257 | 8.357752445 | 0.093999268 | 0.9278787 | 0.997677483 | -5.252976882 |
| **A_51_P169087** | Gls2 | -0.00412989 | -0.109184294 | 0.100924515 | 7.868370128 | -0.093957905 | 0.927910329 | 0.997677483 | -5.252980157 |
| **A_52_P319606** | Gpr107 | 0.011432875 | -0.279502151 | 0.302367901 | 10.39412285 | 0.093922267 | 0.92793758 | 0.997677483 | -5.252982978 |
| **A_55_P2087162** | Specc1l | -0.007460056 | -0.197711436 | 0.182791324 | 8.478839619 | -0.093718082 | 0.928093715 | 0.997677483 | -5.25299912 |
| **A_55_P2048418** | Mdfic | 0.008808532 | -0.215833197 | 0.233450262 | 10.70621111 | 0.093717836 | 0.928093903 | 0.997677483 | -5.252999139 |
| **A_52_P364130** | Map3k14 | 0.00453294 | -0.111101888 | 0.120167767 | 8.422993118 | 0.09369155 | 0.928114003 | 0.997677483 | -5.253001215 |
| **A_55_P2127421** | NA | -0.004507155 | -0.119656003 | 0.110641694 | 7.883713925 | -0.093551771 | 0.928220891 | 0.997677483 | -5.253012242 |
| **A_51_P269020** | 1700031F0 | 0.003848557 | -0.094544734 | 0.102241849 | 7.636204735 | 0.093484929 | 0.928272005 | 0.997677483 | -5.253017509 |
| **A_55_P2042229** | NA | 0.004840303 | -0.118945183 | 0.12862579 | 12.4878608 | 0.093457033 | 0.928293338 | 0.997677483 | -5.253019706 |

| | | | | | | | | | |
|---|---|---|---|---|---|---|---|---|---|
| **A_52_P126146** | Riok1 | 0.005360984 | -0.131842905 | 0.142564873 | 9.147931049 | 0.093387176 | 0.928346758 | 0.997677483 | -5.253025205 |
| **A_51_P200144** | Pak1ip1 | 0.007194219 | -0.177073164 | 0.191461603 | 11.64928349 | 0.093313471 | 0.928403121 | 0.997677483 | -5.253031003 |
| **A_52_P460929** | BC048507 | -0.004856244 | -0.129521446 | 0.119808957 | 12.03786545 | -0.093103161 | 0.92856395 | 0.997677483 | -5.253047521 |
| **A_55_P2082180** | Tnfrsf9 | 0.005322013 | -0.131520814 | 0.14216484 | 7.825986681 | 0.092952917 | 0.928678847 | 0.997677483 | -5.253059298 |
| **A_55_P2064478** | Cdc27 | 0.006843753 | -0.169218223 | 0.18290573 | 9.494926057 | 0.092904744 | 0.928715688 | 0.997677483 | -5.253063071 |
| **A_55_P2152074** | Ctps2 | 0.003870835 | -0.09572636 | 0.103468029 | 10.66109414 | 0.092889503 | 0.928727344 | 0.997677483 | -5.253064264 |
| **A_55_P2092904** | Ddb2 | 0.006406975 | -0.158558502 | 0.171372452 | 8.251059812 | 0.092825885 | 0.928775996 | 0.997677483 | -5.253069242 |
| **A_55_P1973970** | Krt73 | -0.004532861 | -0.121268063 | 0.112202341 | 8.275978073 | -0.092806768 | 0.928790615 | 0.997677483 | -5.253070737 |
| **A_51_P312846** | Fbxo9 | -0.006892136 | -0.18443283 | 0.170648557 | 11.1950012 | -0.092782289 | 0.928809336 | 0.997677483 | -5.253072651 |
| **A_51_P208152** | 3930402G2 | -0.003728401 | -0.099875766 | 0.092418965 | 7.916468786 | -0.092681769 | 0.92888621 | 0.997677483 | -5.253080506 |
| **A_55_P2112302** | NA | -0.011035694 | -0.295774581 | 0.273703192 | 13.08536361 | -0.092632208 | 0.928924112 | 0.997677483 | -5.253084375 |
| **A_55_P2117445** | NA | -0.004707865 | -0.126197398 | 0.116781668 | 13.50149614 | -0.092617767 | 0.928935157 | 0.997677483 | -5.253085502 |
| **A_55_P2039541** | Arhgap15 | -0.004218555 | -0.113581165 | 0.105144054 | 7.783128558 | -0.092194283 | 0.929259033 | 0.997677483 | -5.253118478 |
| **A_55_P1986531** | Mpig6b | -0.005951418 | -0.160246317 | 0.148343482 | 7.8055811 | -0.092188765 | 0.929263253 | 0.997677483 | -5.253118907 |
| **A_55_P2014332** | Cntln | 0.007044405 | -0.175787708 | 0.189876517 | 9.006772798 | 0.092087556 | 0.92934066 | 0.997677483 | -5.253126765 |
| **A_55_P1956712** | Med28 | -0.004456799 | -0.120399907 | 0.11148631 | 12.48564907 | -0.091872851 | 0.929504871 | 0.997677483 | -5.253143406 |
| **A_55_P2228715** | Mm2pr | -0.003858254 | -0.104405541 | 0.096689032 | 7.929681496 | -0.091712735 | 0.929627334 | 0.997677483 | -5.253155792 |
| **A_55_P1995874** | Gm14410 | -0.011707225 | -0.316972913 | 0.293558463 | 9.106108095 | -0.09166111 | 0.929666819 | 0.997677483 | -5.253159781 |
| **A_51_P491470** | Ddx47 | -0.007392766 | -0.200225821 | 0.185440288 | 9.656834417 | -0.091629356 | 0.929691106 | 0.997677483 | -5.253162233 |
| **A_55_P2108903** | Zbtb20 | -0.011841833 | -0.320808766 | 0.297125101 | 9.000139217 | -0.091604339 | 0.929710241 | 0.997677483 | -5.253164164 |
| **A_51_P324838** | Evpl | 0.024892805 | -0.624634803 | 0.674420413 | 8.999314724 | 0.091597866 | 0.929715192 | 0.997677483 | -5.253164664 |
| **A_55_P1960655** | NA | 0.010467896 | -0.262879609 | 0.2838154 | 11.2222507 | 0.091527881 | 0.92976872 | 0.997677483 | -5.253170064 |
| **A_55_P1973357** | NA | 0.004481935 | -0.112596075 | 0.121559946 | 15.47912961 | 0.091495429 | 0.929793542 | 0.997677483 | -5.253172566 |
| **A_52_P509965** | Ager | 0.004011589 | -0.101042378 | 0.109065555 | 7.695102223 | 0.091266858 | 0.92996837 | 0.997677483 | -5.253190168 |
| **A_55_P2049122** | Tm6sf2 | -0.004851974 | -0.131935431 | 0.122231482 | 8.164919082 | -0.091251206 | 0.929980341 | 0.997677483 | -5.253191372 |
| **A_55_P2183498** | Spcs2 | 0.014431513 | -0.363826099 | 0.392689125 | 9.629059879 | 0.091187083 | 0.930029389 | 0.997677483 | -5.253196301 |
| **A_55_P2156229** | Cd3e | 0.011919841 | -0.300557885 | 0.324397567 | 9.005157888 | 0.091171801 | 0.930041078 | 0.997677483 | -5.253197475 |
| **A_51_P108853** | Lyplal1 | -0.004273235 | -0.116477094 | 0.107930624 | 7.893856125 | -0.091024455 | 0.930153782 | 0.997677483 | -5.214720618 |
| **A_51_P143142** | Mrpl12 | -0.008186377 | -0.223169148 | 0.206796395 | 14.28346217 | -0.091011693 | 0.930163544 | 0.997677483 | -5.253209765 |
| **A_55_P1985840** | Mettl16 | -0.006269874 | -0.170967173 | 0.158427426 | 10.0374328 | -0.090987437 | 0.930182097 | 0.997677483 | -5.253211625 |
| **A_51_P340355** | Tarsl2 | -0.006178914 | -0.168569505 | 0.156211677 | 10.22039296 | -0.090941144 | 0.930217507 | 0.997677483 | -5.253215174 |
| **A_52_P141628** | Slc23a1 | 0.003935279 | -0.099494473 | 0.107365031 | 7.653024635 | 0.090936695 | 0.93022091 | 0.997677483 | -5.253215515 |
| **A_51_P219444** | Plcl2 | -0.005190837 | -0.141735681 | 0.131354007 | 11.07710002 | -0.09085968 | 0.930279819 | 0.997677483 | -5.253221414 |
| **A_55_P2085574** | Nfix | -0.012396956 | -0.338545533 | 0.313751622 | 10.78412441 | -0.090846602 | 0.930289823 | 0.997677483 | -5.253222416 |
| **A_66_P118124** | Fxyd3 | -0.004465425 | -0.122093489 | 0.11316264 | 7.715643749 | -0.090732102 | 0.930377407 | 0.997677483 | -5.253231176 |
| **A_52_P529486** | Wdr77 | 0.005659849 | -0.143474256 | 0.154793955 | 9.661790405 | 0.090706207 | 0.930397214 | 0.997677483 | -5.253233156 |
| **A_55_P2087911** | Mthfsd | -0.005229464 | -0.143028092 | 0.132569163 | 8.199775827 | -0.090702956 | 0.930399701 | 0.997677483 | -5.253233404 |
| **A_52_P516021** | Ptpn1 | 0.006162046 | -0.156284097 | 0.168608189 | 9.077618708 | 0.090661872 | 0.930431127 | 0.997677483 | -5.253236544 |
| **A_55_P2034350** | Terf2 | 0.007169417 | -0.181902955 | 0.196241789 | 9.382763664 | 0.090628522 | 0.930456638 | 0.997677483 | -5.253239091 |
| **A_55_P2052719** | Nadk2 | -0.005883411 | -0.16144401 | 0.149677188 | 11.29468102 | -0.090393813 | 0.930636177 | 0.997677483 | -5.253256994 |
| **A_55_P2091055** | NA | 0.009218354 | -0.234766699 | 0.253203407 | 14.06058981 | 0.090302401 | 0.930706103 | 0.997677483 | -5.253263954 |
| **A_52_P83913** | Ssx2ip | -0.004344339 | -0.119350291 | 0.110661614 | 7.873312952 | -0.090284356 | 0.930719907 | 0.997677483 | -5.253265327 |
| **A_55_P2130716** | Ube2g2 | 0.007286049 | -0.185669366 | 0.200241463 | 8.784829486 | 0.090249383 | 0.93074666 | 0.997677483 | -5.253267988 |
| **A_51_P467346** | Muc1 | 0.003720452 | -0.094920542 | 0.102361447 | 7.643371587 | 0.090146205 | 0.930825587 | 0.997677483 | -5.25327583 |
| **A_52_P445828** | Abcf2 | -0.009897089 | -0.272339937 | 0.25254576 | 9.319937477 | -0.090132592 | 0.930836001 | 0.997677483 | -5.253276864 |
| **A_55_P2172396** | A530064D0 | 0.003931271 | -0.100447793 | 0.108310335 | 7.697692098 | 0.09001787 | 0.93092376 | 0.997677483 | -5.253285573 |
| **A_55_P2056100** | Pus1 | 0.006418582 | -0.164280746 | 0.17711791 | 9.567672598 | 0.089870348 | 0.931036613 | 0.997677483 | -5.253296755 |
| **A_55_P1972040** | Nox4 | 0.005168951 | -0.132604957 | 0.142942858 | 8.255104425 | 0.089669459 | 0.931190294 | 0.997677483 | -5.253311953 |

| | | | | | | | | | |
|---|---|---|---|---|---|---|---|---|---|
| **A_51_P132013** | Cysltr2 | -0.003748871 | -0.103723629 | 0.096225887 | 7.643647861 | -0.089622965 | 0.931225862 | 0.997677483 | -5.253315465 |
| **A_52_P342131** | Zfp532 | 0.00412097 | -0.105795311 | 0.114037251 | 8.875664193 | 0.089607952 | 0.931237347 | 0.997677483 | -5.253316599 |
| **A_55_P2009111** | NA | 0.004030048 | -0.103529511 | 0.111589606 | 7.7868576 | 0.089550975 | 0.931280935 | 0.997677483 | -5.253320901 |
| **A_55_P2056419** | Coq6 | -0.005084647 | -0.141138957 | 0.130969663 | 14.08610364 | -0.089321833 | 0.931456235 | 0.997677483 | -5.253338172 |
| **A_55_P1979863** | Syne2 | -0.010278417 | -0.285748593 | 0.265191759 | 11.30360696 | -0.089178628 | 0.931565793 | 0.997677483 | -5.253348943 |
| **A_55_P1973926** | Ipo13 | -0.004498529 | -0.125189268 | 0.116192209 | 11.49248308 | -0.08908524 | 0.93163724 | 0.997677483 | -5.253355958 |
| **A_55_P2090390** | NA | -0.005972993 | -0.16624045 | 0.154294464 | 9.153484452 | -0.08907499 | 0.931645082 | 0.997677483 | -5.253356727 |
| **A_55_P1967835** | NA | -0.005131265 | -0.142847598 | 0.132585068 | 8.470729802 | -0.089052911 | 0.931661974 | 0.997677483 | -5.253358385 |
| **A_55_P2027788** | Baiap2 | 0.004784164 | -0.123766735 | 0.133335063 | 8.232067775 | 0.088948804 | 0.931741622 | 0.997677483 | -5.253366193 |
| **A_55_P2008933** | Slc17a3 | -0.003635421 | -0.101472558 | 0.094201715 | 7.651621274 | -0.08880965 | 0.931848084 | 0.997677483 | -5.253376617 |
| **A_51_P127320** | Cabs1 | -0.003655642 | -0.102175202 | 0.094863919 | 7.637966484 | -0.088685025 | 0.931943433 | 0.997677483 | -5.246432702 |
| **A_55_P2017413** | Gm4788 | -0.008282033 | -0.231513057 | 0.214948991 | 8.257902849 | -0.088673023 | 0.931952615 | 0.997677483 | -5.253386835 |
| **A_51_P189899** | Olfr1134 | 0.003633965 | -0.094433957 | 0.101701887 | 7.722133263 | 0.088565147 | 0.93203515 | 0.997677483 | -5.253394892 |
| **A_55_P2144120** | Bcl2l2 | -0.003727323 | -0.104424814 | 0.096970167 | 7.929157033 | -0.088468278 | 0.932109265 | 0.997677483 | -5.253402118 |
| **A_55_P2043182** | Gimap6 | -0.006881409 | -0.193042565 | 0.179279747 | 11.23559829 | -0.088348147 | 0.932201178 | 0.997677483 | -5.253411069 |
| **A_55_P2001290** | Ube2e3 | -0.005053115 | -0.141995236 | 0.131889007 | 12.69835369 | -0.088192413 | 0.932320333 | 0.997677483 | -5.253422654 |
| **A_55_P2120737** | NA | 0.004241475 | -0.11071913 | 0.119202081 | 7.893148617 | 0.088181411 | 0.932328751 | 0.997677483 | -5.253423472 |
| **A_66_P101646** | NA | -0.005694823 | -0.160076097 | 0.148686451 | 17.56351108 | -0.088164695 | 0.93234154 | 0.997677483 | -5.253424714 |
| **A_52_P349844** | Obscn | -0.015220317 | -0.428034806 | 0.397594173 | 11.25297695 | -0.08812069 | 0.93237521 | 0.997677483 | -5.253427983 |
| **A_66_P123028** | NA | -0.005293379 | -0.148932505 | 0.138345748 | 8.144811363 | -0.088078388 | 0.932407577 | 0.997677483 | -5.253431124 |
| **A_51_P306017** | Dll1 | 0.004700629 | -0.122931226 | 0.132332483 | 8.047496567 | 0.088024998 | 0.932448427 | 0.997677483 | -5.253435086 |
| **A_55_P2172599** | Fam174a | -0.005206367 | -0.146574016 | 0.136161281 | 10.94637039 | -0.088022543 | 0.932450306 | 0.997677483 | -5.253435268 |
| **A_55_P2088692** | NA | 0.009997412 | -0.261661381 | 0.281656205 | 9.837373099 | 0.087957514 | 0.932500062 | 0.997677483 | -5.253440091 |
| **A_55_P1998406** | NA | 0.00632992 | -0.165781344 | 0.178441185 | 9.292007899 | 0.087901865 | 0.932542642 | 0.997677483 | -5.253444215 |
| **A_55_P2090179** | Cbx5 | -0.00626975 | -0.176791419 | 0.16425192 | 9.216231684 | -0.087877918 | 0.932560964 | 0.997677483 | -5.253445988 |
| **A_51_P394585** | Snx22 | -0.003983662 | -0.112344841 | 0.104377518 | 7.841561496 | -0.087865396 | 0.932570545 | 0.997677483 | -5.253446916 |
| **A_55_P2008106** | Otud6b | 0.006013779 | -0.15764007 | 0.169667628 | 9.297817593 | 0.087827471 | 0.932599565 | 0.997677483 | -5.253449724 |
| **A_55_P2171578** | Mkrn1 | -0.021449068 | -0.605200432 | 0.562302296 | 11.23771025 | -0.087819235 | 0.932605866 | 0.997677483 | -5.253450333 |
| **A_55_P2133385** | NA | 0.011086181 | -0.290717776 | 0.312890139 | 13.6380415 | 0.087794273 | 0.932624966 | 0.997677483 | -5.25345218 |
| **A_55_P1983708** | Insrr | -0.004765122 | -0.134506264 | 0.12497602 | 7.797984593 | -0.087781995 | 0.93263436 | 0.997677483 | -5.253453089 |
| **A_51_P237514** | Eif1ax | -0.006055157 | -0.171036721 | 0.158926407 | 13.33641963 | -0.087720103 | 0.932681718 | 0.997677483 | -5.253457666 |
| **A_55_P2025073** | Csf1r | 0.004338442 | -0.113872195 | 0.122549079 | 8.631025289 | 0.087717522 | 0.932683692 | 0.997677483 | -5.253457857 |
| **A_55_P2051176** | NA | 0.005583114 | -0.146624279 | 0.157790506 | 15.39408923 | 0.087669771 | 0.93272023 | 0.997677483 | -5.253461386 |
| **A_55_P2440441** | Wdhd1 | -0.006688629 | -0.189216331 | 0.175839073 | 8.598400761 | -0.087582523 | 0.932786989 | 0.997677483 | -5.25346783 |
| **A_55_P2031207** | Ggnbp2 | 0.007126386 | -0.18743586 | 0.201688631 | 11.48002645 | 0.08754269 | 0.932817469 | 0.997677483 | -5.253470769 |
| **A_51_P275697** | Rflnb | 0.005306855 | -0.139711529 | 0.15032524 | 8.129166012 | 0.087462789 | 0.932878606 | 0.997677483 | -5.253476662 |
| **A_51_P209122** | Agr2 | 0.00375691 | -0.09897189 | 0.106485709 | 7.682834461 | 0.087407291 | 0.932921073 | 0.997677483 | -5.253480751 |
| **A_55_P2080995** | Tgif2lx1 | 0.003825296 | -0.100774297 | 0.108424889 | 7.946893825 | 0.087406594 | 0.932921606 | 0.997677483 | -5.253480803 |
| **A_51_P399625** | Cnih1 | 0.006321658 | -0.166808709 | 0.179452025 | 13.75517673 | 0.087270383 | 0.933025834 | 0.997677483 | -5.253490829 |
| **A_55_P2066453** | NA | 0.016750738 | -0.442142064 | 0.47564354 | 9.260987717 | 0.087243248 | 0.933046597 | 0.997677483 | -5.253492825 |
| **A_55_P1983006** | Hdac5 | -0.007301048 | -0.207985942 | 0.193383846 | 13.03467058 | -0.086952014 | 0.933269452 | 0.997677483 | -5.253514204 |
| **A_55_P1969156** | Pam | -0.017951336 | -0.511462063 | 0.475559392 | 11.60940745 | -0.086937931 | 0.933280229 | 0.997677483 | -5.253515236 |
| **A_66_P115038** | Prl3d1 | 0.003570722 | -0.09467669 | 0.101818133 | 7.645135301 | 0.086864834 | 0.933336164 | 0.997677483 | -5.253520589 |
| **A_55_P2041265** | Utp6 | -0.017387676 | -0.496178618 | 0.461403267 | 14.87481976 | -0.086797004 | 0.93338807 | 0.997677483 | -5.253525553 |
| **A_55_P2115955** | Raet1e | 0.005930137 | -0.157433097 | 0.169293371 | 8.880056986 | 0.086760002 | 0.933416385 | 0.997677483 | -5.25352826 |
| **A_55_P1986647** | Podxl2 | -0.00766278 | -0.218834032 | 0.203508472 | 8.735676517 | -0.086728274 | 0.933440665 | 0.997677483 | -5.253530579 |
| **A_55_P2037146** | Igfbpl1 | -0.005758136 | -0.164860445 | 0.153344174 | 9.24130786 | -0.086499688 | 0.933615589 | 0.997677483 | -5.253547266 |
| **A_52_P616392** | Sbno2 | -0.005303942 | -0.151871965 | 0.141264081 | 8.203153851 | -0.086490553 | 0.93362258 | 0.997677483 | -5.253547932 |

| | | | | | | | | | |
|---|---|---|---|---|---|---|---|---|---|
| **A_51_P102911** | Hoxa11 | 0.003573763 | -0.095203673 | 0.102351199 | 7.669766142 | 0.086472322 | 0.933636531 | 0.997677483 | -5.228074489 |
| **A_51_P183213** | Riiad1 | -0.003572311 | -0.102391268 | 0.095246646 | 8.022344999 | -0.086400879 | 0.933691203 | 0.997677483 | -5.253554466 |
| **A_51_P417053** | Tuft1 | -0.012493817 | -0.358303289 | 0.333315655 | 9.26034368 | -0.086351001 | 0.933729374 | 0.997677483 | -5.253558097 |
| **A_51_P234833** | Strn4 | -0.004555026 | -0.130663889 | 0.121553837 | 11.78191388 | -0.08632853 | 0.93374657 | 0.997677483 | -5.253559732 |
| **A_55_P2243768** | D930030O( | 0.014094721 | -0.376273046 | 0.404462488 | 8.746893613 | 0.086296199 | 0.933771312 | 0.997677483 | -5.253562084 |
| **A_55_P1958250** | Rnf114 | -0.007066181 | -0.203191746 | 0.189059383 | 12.99963908 | -0.086111212 | 0.933912877 | 0.997677483 | -5.253575525 |
| **A_52_P235569** | Ipo5 | 0.006036621 | -0.161596699 | 0.173669941 | 9.68493155 | 0.086068194 | 0.933945799 | 0.997677483 | -5.253578646 |
| **A_55_P2054728** | Ldlrad3 | -0.00756284 | -0.217751714 | 0.202626034 | 13.15233758 | -0.085997199 | 0.934000131 | 0.997677483 | -5.253583794 |
| **A_55_P2075366** | Tead3 | 0.004190688 | -0.112404459 | 0.120785834 | 8.067710314 | 0.085904112 | 0.93407137 | 0.997677483 | -5.253590538 |
| **A_52_P130787** | Kcna6 | -0.006006095 | -0.173249118 | 0.161236928 | 9.243714645 | -0.085832812 | 0.934125936 | 0.997677483 | -5.253595698 |
| **A_55_P1974019** | Dapk1 | -0.008143772 | -0.234977391 | 0.218689848 | 9.558367455 | -0.085807905 | 0.934144997 | 0.997677483 | -5.2535975 |
| **A_55_P1981205** | Xrcc4 | 0.005484017 | -0.147356668 | 0.158324702 | 9.459702367 | 0.085756869 | 0.934184056 | 0.997677483 | -5.25360119 |
| **A_51_P273609** | Itpka | -0.009876002 | -0.285270359 | 0.265518354 | 9.47276547 | -0.085710748 | 0.934219353 | 0.997677483 | -5.253604522 |
| **A_51_P469053** | Mfsd4a | -0.005166722 | -0.149300788 | 0.138967344 | 8.943140914 | -0.085675685 | 0.934246187 | 0.997677483 | -5.253607055 |
| **A_52_P343690** | Kpna4 | -0.010529325 | -0.304271484 | 0.283212833 | 11.15105432 | -0.085672885 | 0.93424833 | 0.997677483 | -5.253607257 |
| **A_55_P2049318** | NA | -0.006752945 | -0.195239023 | 0.181733132 | 10.15385886 | -0.085629437 | 0.934281581 | 0.997677483 | -5.253610394 |
| **A_55_P2131190** | NA | -0.006461195 | -0.18684381 | 0.17392142 | 8.10709134 | -0.085610557 | 0.93429603 | 0.997677483 | -5.253611756 |
| **A_51_P388298** | Mmachc | 0.00443746 | -0.119478709 | 0.128353628 | 10.93192083 | 0.085588528 | 0.93431289 | 0.997677483 | -5.253613345 |
| **A_52_P557293** | Tmprss2 | -0.003821063 | -0.110533646 | 0.102891521 | 7.654637596 | -0.08558106 | 0.934318605 | 0.997677483 | -5.253613884 |
| **A_51_P407690** | Sppl2b | 0.007303466 | -0.196804861 | 0.211411794 | 12.17121618 | 0.08552192 | 0.934363867 | 0.997677483 | -5.253618149 |
| **A_51_P309501** | Praf2 | -0.004794602 | -0.13894934 | 0.129360136 | 8.798185718 | -0.085419242 | 0.934442449 | 0.997677483 | -5.253625546 |
| **A_52_P413502** | Rtf2 | -0.00689515 | -0.199885313 | 0.186095014 | 8.329605339 | -0.085392096 | 0.934463226 | 0.997677483 | -5.2536275 |
| **A_55_P2383897** | 8030453O2 | 0.003746101 | -0.10119972 | 0.108691923 | 7.668910877 | 0.085314632 | 0.934522512 | 0.997677483 | -5.253633073 |
| **A_52_P488437** | Plek2 | 0.003653689 | -0.098762604 | 0.106069982 | 7.801791054 | 0.085265165 | 0.934560371 | 0.997677483 | -5.253636629 |
| **A_55_P2077188** | Stxbp1 | -0.005133826 | -0.149119396 | 0.138851744 | 11.18585946 | -0.085217998 | 0.93459647 | 0.997677483 | -5.253640017 |
| **A_51_P272993** | Ntm | -0.00473541 | -0.137611053 | 0.128140233 | 8.758486486 | -0.085176809 | 0.934627994 | 0.997677483 | -5.253642975 |
| **A_52_P522977** | Ssbp2 | -0.003764648 | -0.10940238 | 0.101873083 | 7.96562037 | -0.085175452 | 0.934629033 | 0.997677483 | -5.253643073 |
| **A_51_P184024** | Tsen15 | -0.004698502 | -0.136705829 | 0.127308826 | 11.26484104 | -0.085068838 | 0.934710631 | 0.997677483 | -5.253650722 |
| **A_55_P1957038** | Gstp2 | -0.004470456 | -0.130150203 | 0.121209291 | 14.78664415 | -0.085015021 | 0.934751821 | 0.997677483 | -5.25365458 |
| **A_51_P490817** | Me2 | -0.005117229 | -0.149038975 | 0.138804516 | 9.103799401 | -0.084980167 | 0.934778497 | 0.997677483 | -5.253657076 |
| **A_51_P512172** | Tet1 | 0.007370543 | -0.200040965 | 0.214782052 | 8.659207673 | 0.084932867 | 0.934814699 | 0.997677483 | -5.253660464 |
| **A_55_P2108476** | Mavs | 0.00553349 | -0.150471113 | 0.161538093 | 10.94948451 | 0.084775595 | 0.934935071 | 0.997677483 | -5.253671712 |
| **A_51_P161463** | Ccdc103 | -0.003521283 | -0.102940955 | 0.09589839 | 7.984916879 | -0.08465209 | 0.935029601 | 0.997677483 | -5.253680531 |
| **A_55_P2017982** | Pde8b | -0.003663232 | -0.107103185 | 0.099776721 | 7.782947341 | -0.084641869 | 0.935037424 | 0.997677483 | -5.25368126 |
| **A_52_P449694** | NA | 0.004854179 | -0.132226732 | 0.141935089 | 8.265948461 | 0.084634593 | 0.935042993 | 0.997677483 | -5.253681779 |
| **A_55_P2158498** | Btk | 0.004639517 | -0.126399812 | 0.135678845 | 7.899587322 | 0.084621403 | 0.935053089 | 0.997677483 | -5.25368272 |
| **A_51_P164420** | Eef1e1 | 0.006873018 | -0.187286678 | 0.201032714 | 9.70442558 | 0.084605292 | 0.93506542 | 0.997677483 | -5.253683869 |
| **A_55_P2048221** | NA | -0.004854734 | -0.142179363 | 0.132469895 | 16.74673106 | -0.084494057 | 0.935150559 | 0.997677483 | -5.253691796 |
| **A_51_P402994** | Ddx3y | -0.081822137 | -2.397677352 | 2.234033079 | 9.404572661 | -0.084443951 | 0.935188911 | 0.997677483 | -5.253695364 |
| **A_55_P2102295** | Tmem184c | 0.005843345 | -0.159628344 | 0.171315035 | 10.24851987 | 0.084400882 | 0.935221876 | 0.997677483 | -5.253698428 |
| **A_55_P1954277** | Emid1 | -0.005565007 | -0.163381194 | 0.15225118 | 9.34097181 | -0.084279762 | 0.935314583 | 0.997677483 | -5.253707039 |
| **A_55_P2043932** | Tmem8b | -0.004432695 | -0.130163859 | 0.121298469 | 8.956147306 | -0.084262443 | 0.93532784 | 0.997677483 | -5.253708269 |
| **A_55_P2124498** | Il17rb | -0.003389344 | -0.099632588 | 0.092853899 | 7.662161679 | -0.084169469 | 0.935399004 | 0.997677483 | -5.253714869 |
| **A_55_P2172852** | Hacd4 | 0.00640299 | -0.175774471 | 0.18858045 | 8.469525724 | 0.084003481 | 0.935526058 | 0.997677483 | -5.253726633 |
| **A_55_P2166349** | Dnajc19 | 0.004770039 | -0.131018196 | 0.140558275 | 10.02746277 | 0.083959329 | 0.935559853 | 0.997677483 | -5.253729759 |
| **A_51_P386880** | Stip1 | 0.010906293 | -0.29977937 | 0.321591955 | 11.87983758 | 0.0839006 | 0.935604808 | 0.997677483 | -5.253733914 |
| **A_55_P1954221** | Emp1 | -0.00486713 | -0.143517586 | 0.133783326 | 13.65425943 | -0.08389977 | 0.935605443 | 0.997677483 | -5.253733972 |
| **A_55_P2009988** | Trib3 | -0.01686909 | -0.497474817 | 0.463736638 | 9.86575724 | -0.083890317 | 0.935612679 | 0.997677483 | -5.253734641 |

| | | | | | | | | | |
|---|---|---|---|---|---|---|---|---|---|
| **A_66_P115949** | Myo9a | 0.004205597 | -0.115661801 | 0.124072995 | 8.546998306 | 0.083856301 | 0.935638716 | 0.997677483 | -5.253737046 |
| **A_51_P108581** | Grk3 | -0.007486196 | -0.22096521 | 0.205992819 | 9.608001117 | -0.083813721 | 0.935671309 | 0.997677483 | -5.247496741 |
| **A_55_P2121349** | NA | -0.004046108 | -0.119439113 | 0.111346898 | 8.794625234 | -0.083804449 | 0.935678407 | 0.997677483 | -5.25374071 |
| **A_51_P354038** | Tmc6 | 0.005608404 | -0.154345921 | 0.165562728 | 10.78715753 | 0.083801624 | 0.935680569 | 0.997677483 | -5.253740909 |
| **A_55_P2110351** | Eppk1 | -0.018304538 | -0.540489163 | 0.503880087 | 10.34215127 | -0.083780667 | 0.935696611 | 0.997677483 | -5.253742389 |
| **A_52_P436318** | Rps25 | 0.004459838 | -0.122802183 | 0.131721858 | 8.571013332 | 0.083758593 | 0.935713508 | 0.997677483 | -5.253743948 |
| **A_52_P46085** | Mvp | -0.007564758 | -0.223477852 | 0.208348337 | 8.322855775 | -0.083738499 | 0.935728889 | 0.997677483 | -5.253745366 |
| **A_51_P126302** | Rbmx2 | -0.003697316 | -0.109237426 | 0.101842794 | 8.255345414 | -0.083729431 | 0.93573583 | 0.997677483 | -5.244736867 |
| **A_55_P2097665** | Arl4a | -0.006619022 | -0.195596106 | 0.182358062 | 8.874501189 | -0.083713174 | 0.935748274 | 0.997677483 | -5.253747154 |
| **A_55_P2400691** | Arhgap21 | -0.008284898 | -0.244884433 | 0.228314637 | 12.22071268 | -0.083691704 | 0.935764708 | 0.997677483 | -5.253748668 |
| **A_51_P219325** | Ddx28 | -0.005092003 | -0.150529654 | 0.140345647 | 8.948910896 | -0.083679859 | 0.935773775 | 0.997677483 | -5.253749504 |
| **A_55_P2042713** | Pigq | -0.006003402 | -0.177516165 | 0.165509362 | 13.57080653 | -0.083658506 | 0.93579012 | 0.997677483 | -5.25375101 |
| **A_55_P1993544** | NA | -0.005249045 | -0.15554168 | 0.14504359 | 8.691000597 | -0.083474094 | 0.935931283 | 0.997677483 | -5.253764 |
| **A_51_P354126** | Reg3g | 0.004286263 | -0.118465585 | 0.127038111 | 7.669815089 | 0.083456449 | 0.93594479 | 0.997677483 | -5.253765241 |
| **A_51_P171288** | Gli3 | -0.009992451 | -0.296201226 | 0.276216324 | 9.631044811 | -0.083444596 | 0.935953863 | 0.997677483 | -5.253766075 |
| **A_51_P258281** | Gal | -0.008648451 | -0.256862494 | 0.239565592 | 9.305016855 | -0.08327624 | 0.936082738 | 0.997677483 | -5.253777905 |
| **A_55_P2144771** | NA | 0.00385645 | -0.106846477 | 0.114559377 | 8.22706587 | 0.083260254 | 0.936094975 | 0.997677483 | -5.253779027 |
| **A_55_P1955209** | NA | 0.006395048 | -0.177287006 | 0.190077102 | 14.94162204 | 0.083212048 | 0.936131877 | 0.997677483 | -5.253782409 |
| **A_51_P145415** | Lpcat3 | 0.006336868 | -0.175760047 | 0.188433782 | 10.29837538 | 0.083172771 | 0.936161943 | 0.997677483 | -5.253785163 |
| **A_55_P2156741** | Stx4a | -0.008115173 | -0.241329471 | 0.225099125 | 13.24348747 | -0.083167132 | 0.93616626 | 0.997677483 | -5.253785559 |
| **A_55_P1970087** | Rpl35a | -0.005944154 | -0.176805848 | 0.164917539 | 15.81043455 | -0.083148515 | 0.936180512 | 0.997677483 | -5.253786864 |
| **A_55_P2024591** | Atxn7l1 | 0.004715749 | -0.130887207 | 0.140318706 | 9.381921751 | 0.083117155 | 0.936204518 | 0.997677483 | -5.253789061 |
| **A_52_P282741** | Sdc3 | 0.005142488 | -0.142904818 | 0.153189794 | 9.702009028 | 0.083019845 | 0.93627901 | 0.997677483 | -5.253795875 |
| **A_51_P207622** | Fmod | 0.016138689 | -0.448529393 | 0.480806772 | 8.780725573 | 0.083010791 | 0.93628594 | 0.997677483 | -5.253796508 |
| **A_55_P2012166** | Spata6 | -0.005366338 | -0.159914629 | 0.149181953 | 9.962726423 | -0.082989462 | 0.936302268 | 0.997677483 | -5.253798001 |
| **A_51_P178347** | Zfp326 | -0.007506713 | -0.22405395 | 0.209040524 | 9.893005574 | -0.082852633 | 0.936407014 | 0.997677483 | -5.253807565 |
| **A_55_P2029622** | Ppp1r3f | -0.00361875 | -0.108087607 | 0.100850107 | 7.955207202 | -0.082790573 | 0.936454523 | 0.997677483 | -5.253811897 |
| **A_55_P2236138** | 4930447F2 | -0.003547814 | -0.106003133 | 0.098907505 | 7.703110037 | -0.082762848 | 0.936475747 | 0.997677483 | -5.253813832 |
| **A_55_P2104835** | NA | -0.003578673 | -0.107118976 | 0.099961629 | 7.999631283 | -0.082607932 | 0.936594341 | 0.997677483 | -5.253824629 |
| **A_55_P2117487** | NA | 0.009064516 | -0.253290982 | 0.271420013 | 13.87148347 | 0.082577849 | 0.936617371 | 0.997677483 | -5.253826723 |
| **A_55_P2068096** | Rap1gap | 0.00839344 | -0.234832976 | 0.251619855 | 8.967377557 | 0.082478046 | 0.936693775 | 0.997677483 | -5.253833666 |
| **A_51_P277295** | Apmap | 0.00701777 | -0.196514068 | 0.210549608 | 9.485080618 | 0.082409241 | 0.93674645 | 0.997677483 | -5.253838448 |
| **A_55_P2068663** | Stmn1 | 0.006877146 | -0.192653175 | 0.206407466 | 15.24307248 | 0.08237747 | 0.936770772 | 0.997677483 | -5.253840655 |
| **A_66_P106363** | Itgb1bp1 | 0.007345616 | -0.205867107 | 0.220558338 | 10.55354467 | 0.082342531 | 0.93679752 | 0.997677483 | -5.25384308 |
| **A_52_P532033** | Hltf | -0.006708371 | -0.201833605 | 0.188416863 | 9.649290563 | -0.082169905 | 0.936929677 | 0.997677483 | -5.25385505 |
| **A_52_P92514** | Supt20 | 0.005977962 | -0.16807501 | 0.180030933 | 8.438686969 | 0.082088218 | 0.936992215 | 0.997677483 | -5.253860705 |
| **A_55_P2341468** | Rbms3 | -0.007828989 | -0.235987363 | 0.220329386 | 11.47304243 | -0.082012184 | 0.937050426 | 0.997677483 | -5.253865964 |
| **A_55_P2015292** | Ltc4s | -0.007953047 | -0.239779666 | 0.223873572 | 8.901574472 | -0.081993495 | 0.937064734 | 0.997677483 | -5.253867256 |
| **A_55_P2166528** | Mrps36 | -0.006210308 | -0.187492959 | 0.175072342 | 13.02648381 | -0.08187778 | 0.937153324 | 0.997677483 | -5.253875248 |
| **A_55_P1959305** | Rbm47 | 0.004433799 | -0.125018492 | 0.13388609 | 8.148687101 | 0.081860695 | 0.937166404 | 0.997677483 | -5.253876427 |
| **A_55_P2062811** | Tomm5 | 0.007307739 | -0.206061736 | 0.220677213 | 13.23215119 | 0.081857758 | 0.937168653 | 0.997677483 | -5.25387663 |
| **A_55_P2056344** | Unc93a2 | -0.042855692 | -1.295144019 | 1.209432634 | 10.87316926 | -0.081792469 | 0.937218638 | 0.997677483 | -5.253881133 |
| **A_55_P1957618** | Ak2 | 0.014065004 | -0.397256453 | 0.425386461 | 11.59674315 | 0.08172739 | 0.937268463 | 0.997677483 | -5.253885618 |
| **A_55_P2102390** | Spag9 | -0.009782896 | -0.295956741 | 0.276390949 | 12.4722794 | -0.081704623 | 0.937285893 | 0.997677483 | -5.253887187 |
| **A_52_P673499** | Shmt1 | 0.006447282 | -0.182374871 | 0.195269436 | 8.736605007 | 0.081608025 | 0.93735985 | 0.997677483 | -5.253893836 |
| **A_52_P796840** | Cfhr2 | 0.009712752 | -0.274849261 | 0.294274765 | 9.338021137 | 0.081578272 | 0.93738263 | 0.997677483 | -5.253895882 |
| **A_52_P243516** | Rmdn2 | 0.00385216 | -0.109089492 | 0.116793811 | 8.344006082 | 0.081519081 | 0.937427947 | 0.997677483 | -5.253899951 |
| **A_51_P123405** | Bub1 | -0.005487105 | -0.166505068 | 0.155530857 | 9.750897362 | -0.081447563 | 0.937482703 | 0.997677483 | -5.253904864 |

| | | | | | | | | | |
|---|---|---|---|---|---|---|---|---|---|
| **A_51_P140742** | Islr | 0.016069058 | -0.455898408 | 0.488036524 | 9.790087518 | 0.081374341 | 0.937538764 | 0.997677483 | -5.253909889 |
| **A_55_P1993512** | NA | -0.005721847 | -0.173855101 | 0.162411407 | 9.165241031 | -0.081337672 | 0.93756684 | 0.997677483 | -5.253912403 |
| **A_51_P416556** | Ssr3 | 0.006095186 | -0.173102823 | 0.185293194 | 13.17054454 | 0.081294823 | 0.937599646 | 0.997677483 | -5.253915341 |
| **A_51_P151271** | Eif4b | 0.007406454 | -0.210366496 | 0.225179403 | 9.076028802 | 0.081285954 | 0.937606437 | 0.997677483 | -5.253915948 |
| **A_55_P2068121** | Sun5 | 0.00392336 | -0.111561028 | 0.119407748 | 8.045336929 | 0.081197753 | 0.937673967 | 0.997677483 | -5.253921989 |
| **A_55_P2059372** | Ppip5k2 | 0.006257648 | -0.1781189 | 0.190634197 | 8.507989548 | 0.081117507 | 0.937735407 | 0.997677483 | -5.253927479 |
| **A_52_P158431** | Tshz2 | -0.004227112 | -0.128899885 | 0.120445661 | 8.91464662 | -0.081036604 | 0.937797351 | 0.997677483 | -5.253933008 |
| **A_55_P2016981** | Nfu1 | -0.003466922 | -0.105791128 | 0.098857283 | 12.68622525 | -0.08097946 | 0.937841104 | 0.997677483 | -5.25393691 |
| **A_66_P136095** | Ctnna3 | -0.00504468 | -0.154062132 | 0.143972772 | 8.467361128 | -0.080910635 | 0.937893801 | 0.997677483 | -5.253941607 |
| **A_55_P2138718** | H2al1a | 0.003436817 | -0.098176497 | 0.105050131 | 7.697500579 | 0.080837885 | 0.937949503 | 0.997677483 | -5.253946567 |
| **A_55_P2042520** | Cop1 | -0.003953699 | -0.120872023 | 0.112964624 | 11.8567358 | -0.08082211 | 0.937961581 | 0.997677483 | -5.253947641 |
| **A_55_P2159949** | Polr1e | 0.007490978 | -0.21419265 | 0.229174605 | 10.71062232 | 0.080763291 | 0.938006618 | 0.997677483 | -5.253951647 |
| **A_55_P2077783** | Tubb2a-ps2 | 0.01468605 | -0.42070334 | 0.450075441 | 10.11509909 | 0.080618796 | 0.938117255 | 0.997677483 | -5.253961476 |
| **A_55_P2175615** | Csde1 | -0.005738674 | -0.176036918 | 0.16455957 | 13.35210564 | -0.08053979 | 0.938177749 | 0.997677483 | -5.253966843 |
| **A_51_P245368** | Abcb1b | -0.004724731 | -0.145037421 | 0.135587959 | 8.157540587 | -0.080480244 | 0.938223343 | 0.997677483 | -5.253970884 |
| **A_55_P1972839** | NA | -0.004701808 | -0.144400439 | 0.134996824 | 16.32616902 | -0.080441815 | 0.938252768 | 0.997677483 | -5.253973491 |
| **A_55_P2156556** | NA | -0.008943387 | -0.274710165 | 0.256823392 | 10.97513962 | -0.080428587 | 0.938262897 | 0.997677483 | -5.253974388 |
| **A_51_P347177** | Snx2 | -0.008921072 | -0.274089784 | 0.256247641 | 10.88726163 | -0.080408856 | 0.938278005 | 0.997677483 | -5.253975725 |
| **A_55_P2142191** | Rpl10 | 0.004921517 | -0.141469098 | 0.151312133 | 16.49448579 | 0.080351674 | 0.93832179 | 0.997677483 | -5.2539796 |
| **A_51_P288906** | Clip1 | 0.004798713 | -0.137999304 | 0.147596731 | 10.87599839 | 0.080317793 | 0.938347732 | 0.997677483 | -5.253981895 |
| **A_55_P1983558** | Tmed1 | -0.004212167 | -0.129588296 | 0.121163963 | 12.91636409 | -0.080297091 | 0.938363584 | 0.997677483 | -5.253983296 |
| **A_55_P2203996** | 4932433N0 | 0.005926433 | -0.170537068 | 0.182389934 | 8.81168254 | 0.080268951 | 0.938385131 | 0.997677483 | -5.253985201 |
| **A_55_P2160691** | NA | -0.01008646 | -0.310438586 | 0.290265666 | 7.983336499 | -0.080263334 | 0.938389432 | 0.997677483 | -5.253985581 |
| **A_55_P2128606** | Nr1h5 | 0.003308164 | -0.095253754 | 0.101870082 | 7.661511246 | 0.080220804 | 0.938421998 | 0.997677483 | -5.253988458 |
| **A_52_P77764** | Slc7a2 | 0.003652233 | -0.105163934 | 0.1124684 | 7.77982852 | 0.080218431 | 0.938423815 | 0.997677483 | -5.253988618 |
| **A_55_P2007650** | Nt5dc1 | -0.004447612 | -0.13699947 | 0.128104246 | 8.969780133 | -0.080195535 | 0.938441347 | 0.997677483 | -5.253990166 |
| **A_55_P2136957** | Wbp1l | -0.007343075 | -0.226563073 | 0.211876923 | 13.49687806 | -0.080058408 | 0.938546348 | 0.997677483 | -5.253999429 |
| **A_55_P2111439** | Gpsm1 | -0.008419607 | -0.260034458 | 0.243195244 | 8.715580666 | -0.079976919 | 0.938608746 | 0.997677483 | -5.254004926 |
| **A_55_P2027822** | Sgf29 | 0.004114563 | -0.118910739 | 0.127139865 | 10.95008686 | 0.079935264 | 0.938640643 | 0.997677483 | -5.254007733 |
| **A_55_P2100899** | Arnt | -0.003541198 | -0.109470615 | 0.102388219 | 8.319205752 | -0.079899263 | 0.938668211 | 0.997677483 | -5.254010159 |
| **A_55_P2065325** | Tmem94 | -0.007794315 | -0.241307736 | 0.225719107 | 8.509892248 | -0.079776545 | 0.938762181 | 0.997677483 | -5.254018418 |
| **A_52_P585831** | Bap1 | 0.008405118 | -0.243440255 | 0.26025049 | 10.37246444 | 0.079766209 | 0.938770096 | 0.997677483 | -5.254019113 |
| **A_52_P58257** | Mrps6 | -0.004546415 | -0.140779379 | 0.131686548 | 12.46259596 | -0.079762 | 0.938773319 | 0.997677483 | -5.254019396 |
| **A_51_P398887** | Ctns | -0.004320107 | -0.133783416 | 0.125143201 | 9.900358651 | -0.079754825 | 0.938778813 | 0.997677483 | -5.254019879 |
| **A_51_P208870** | Zdhhc24 | 0.0038644 | -0.111972692 | 0.119701493 | 8.316286867 | 0.079734008 | 0.938794753 | 0.997677483 | -5.254021278 |
| **A_55_P2096189** | Rfx8 | 0.005059666 | -0.146635718 | 0.156755049 | 7.940430058 | 0.079718406 | 0.9388067 | 0.997677483 | -5.254022327 |
| **A_55_P2028474** | Adam22 | 0.007528217 | -0.218269198 | 0.233325631 | 8.813229328 | 0.079686046 | 0.93883148 | 0.997677483 | -5.254024501 |
| **A_52_P527822** | Slc22a3 | 0.003668568 | -0.106404907 | 0.113742043 | 7.655139881 | 0.079656835 | 0.938853849 | 0.997677483 | -5.254026463 |
| **A_66_P115453** | Coro1c | 0.005288577 | -0.153423182 | 0.164000335 | 12.40544204 | 0.079641392 | 0.938865674 | 0.997677483 | -5.2540275 |
| **A_51_P275496** | BC026762 | -0.003281107 | -0.101755756 | 0.095193543 | 8.041026007 | -0.07963519 | 0.938870424 | 0.997677483 | -5.254027916 |
| **A_51_P167876** | Tfrc | -0.003766022 | -0.116839936 | 0.109307892 | 8.044027529 | -0.079603014 | 0.938895062 | 0.997677483 | -5.254030076 |
| **A_55_P2091736** | Rassf2 | 0.008811247 | -0.255834564 | 0.273457058 | 9.758500065 | 0.079575886 | 0.938915836 | 0.997677483 | -5.254031896 |
| **A_55_P1997126** | Ctse | -0.004856831 | -0.150735527 | 0.141021866 | 7.7152677 | -0.07957382 | 0.938917418 | 0.997677483 | -5.254032035 |
| **A_52_P320193** | Clec2h | -0.004240659 | -0.131623264 | 0.123141946 | 7.659978511 | -0.079566884 | 0.93892273 | 0.997677483 | -5.2540325 |
| **A_51_P125691** | Cdca4 | -0.005393831 | -0.167584635 | 0.156796972 | 9.162181972 | -0.079484099 | 0.938986124 | 0.997677483 | -5.25403805 |
| **A_55_P2124415** | 6030468B1 | 0.003388196 | -0.098898405 | 0.105674796 | 7.681726368 | 0.079169681 | 0.939226897 | 0.997677483 | -5.254059077 |
| **A_55_P2089158** | NA | 0.004355815 | -0.127237287 | 0.135948917 | 8.794938327 | 0.079112564 | 0.939270637 | 0.997677483 | -5.254062887 |
| **A_55_P2040529** | Mtfr1 | -0.003694188 | -0.115360884 | 0.107972508 | 8.77906583 | -0.07906868 | 0.939304244 | 0.997677483 | -5.254065813 |

| | | | | | | | | | |
|---|---|---|---|---|---|---|---|---|---|
| **A_55_P2060991** | Plppr3 | -0.005390568 | -0.168368044 | 0.157586908 | 9.058127616 | -0.079052583 | 0.93931657 | 0.997677483 | -5.254066886 |
| **A_52_P117420** | Tacc1 | -0.007142179 | -0.223167386 | 0.208883028 | 10.30792103 | -0.079019711 | 0.939341744 | 0.997677483 | -5.254069076 |
| **A_55_P2047599** | Arhgdia | 0.005467189 | -0.160096381 | 0.17103076 | 13.00807409 | 0.078923888 | 0.939415125 | 0.997677483 | -5.254075456 |
| **A_55_P1975447** | NA | 0.007853631 | -0.230035314 | 0.245742577 | 10.22114103 | 0.078905157 | 0.939429469 | 0.997677483 | -5.254076702 |
| **A_52_P249733** | Tcap | -0.008704412 | -0.272750793 | 0.25534197 | 13.9024093 | -0.078789499 | 0.939518042 | 0.997677483 | -5.25408439 |
| **A_66_P130908** | Cyb5r4 | 0.004335785 | -0.127263187 | 0.135934757 | 8.994722809 | 0.078745247 | 0.939551931 | 0.997677483 | -5.254087329 |
| **A_51_P110814** | Nudt9 | 0.003736855 | -0.109721182 | 0.117194891 | 12.56902809 | 0.07871909 | 0.939571963 | 0.997677483 | -5.247844155 |
| **A_51_P306151** | Sgk3 | -0.003348748 | -0.105067186 | 0.098369691 | 7.862826159 | -0.078684991 | 0.939598076 | 0.997677483 | -5.254091327 |
| **A_52_P101333** | Sap18 | -0.003982128 | -0.1250224 | 0.117058144 | 8.638925002 | -0.078631114 | 0.939639337 | 0.997677483 | -5.2540949 |
| **A_55_P2055587** | Enpp4 | -0.005416461 | -0.170406671 | 0.159573748 | 10.23436084 | -0.078463304 | 0.939767851 | 0.997677483 | -5.254106012 |
| **A_55_P2147250** | Caprin2 | 0.003743316 | -0.110312342 | 0.117798974 | 10.01228052 | 0.07844202 | 0.939784151 | 0.997677483 | -5.25410742 |
| **A_55_P2052613** | NA | -0.005895374 | -0.185551551 | 0.173760802 | 12.37154855 | -0.078429306 | 0.939793887 | 0.997677483 | -5.25410826 |
| **A_51_P192847** | Ghdc | 0.003772736 | -0.111199924 | 0.118745397 | 8.689780578 | 0.078427973 | 0.939794908 | 0.997677483 | -5.254108349 |
| **A_55_P1992174** | Tars2 | -0.004614167 | -0.145232218 | 0.136003885 | 9.505920863 | -0.078426228 | 0.939796245 | 0.997677483 | -5.254108464 |
| **A_55_P1997097** | Bod1 | -0.008191119 | -0.257858361 | 0.241476123 | 9.33280339 | -0.078413493 | 0.939805998 | 0.997677483 | -5.254109306 |
| **A_55_P1954511** | Btbd10 | -0.004192701 | -0.132073587 | 0.123688184 | 10.18487927 | -0.078360542 | 0.93984655 | 0.997677483 | -5.254112805 |
| **A_55_P2109744** | NA | -0.004109702 | -0.129843168 | 0.121623764 | 15.10225318 | -0.078121137 | 0.940029899 | 0.997677483 | -5.254128597 |
| **A_55_P2136338** | Tomm20 | -0.006093105 | -0.192531245 | 0.180345035 | 12.54494648 | -0.078111151 | 0.940037547 | 0.997677483 | -5.254129254 |
| **A_55_P2067727** | Mxra7 | 0.009530925 | -0.282282112 | 0.301343962 | 9.949686243 | 0.078061971 | 0.940075212 | 0.997677483 | -5.254132492 |
| **A_51_P213592** | Nrde2 | 0.004614636 | -0.13674724 | 0.145976512 | 8.418838836 | 0.078021496 | 0.94010621 | 0.997677483 | -5.254135155 |
| **A_55_P1978013** | Obp1a | 0.003522321 | -0.104435304 | 0.111479945 | 8.106923181 | 0.077980263 | 0.94013779 | 0.997677483 | -5.254137866 |
| **A_51_P430552** | G6pc3 | -0.005432366 | -0.172067097 | 0.161202366 | 11.94437451 | -0.077917067 | 0.94018619 | 0.997677483 | -5.254142019 |
| **A_55_P1958697** | Rbm33 | -0.006826396 | -0.216314289 | 0.202661497 | 10.51139907 | -0.077882818 | 0.94021242 | 0.997677483 | -5.254144268 |
| **A_55_P2005549** | Tnn | -0.004076865 | -0.129196572 | 0.121042841 | 7.663599269 | -0.077877102 | 0.940216798 | 0.997677483 | -5.254144644 |
| **A_55_P2002122** | Rgs12 | 0.009767795 | -0.290008703 | 0.309544292 | 12.33159102 | 0.077876794 | 0.940217034 | 0.997677483 | -5.254144664 |
| **A_55_P2093705** | Meig1 | 0.003602245 | -0.107112444 | 0.114316933 | 7.6880627 | 0.077763735 | 0.940303624 | 0.997677483 | -5.254152081 |
| **A_55_P2040773** | NA | -0.007971725 | -0.253349395 | 0.237405946 | 9.279714328 | -0.077647306 | 0.940392796 | 0.997677483 | -5.254159709 |
| **A_55_P2065953** | Btbd19 | 0.004101551 | -0.122216806 | 0.130419909 | 8.338302543 | 0.077605199 | 0.940425045 | 0.997677483 | -5.254162464 |
| **A_55_P2080151** | Hspa2 | 0.006197721 | -0.18473353 | 0.197128971 | 10.15727837 | 0.077582565 | 0.940442381 | 0.997677483 | -5.254163945 |
| **A_51_P461452** | Ube2r2 | 0.004434249 | -0.132404313 | 0.14127281 | 13.11619413 | 0.077449872 | 0.940544011 | 0.997677483 | -5.254172617 |
| **A_66_P122372** | Ppp4r2 | 0.005643748 | -0.168561099 | 0.179848594 | 8.992546946 | 0.077431287 | 0.940558245 | 0.997677483 | -5.25417383 |
| **A_51_P126337** | Fgf12 | -0.035935813 | -1.145614606 | 1.07374298 | 8.677395116 | -0.077399707 | 0.940582432 | 0.997677483 | -5.24138677 |
| **A_55_P1979252** | Glod4 | 0.004043182 | -0.12086485 | 0.128951214 | 12.48431674 | 0.077364561 | 0.940609351 | 0.997677483 | -5.254178184 |
| **A_52_P424462** | Ero1lb | -0.003635315 | -0.116033244 | 0.108762614 | 8.010569146 | -0.077302385 | 0.940656973 | 0.997677483 | -5.254182238 |
| **A_52_P320971** | Mt3 | 0.006092456 | -0.18232623 | 0.194511142 | 9.973613331 | 0.077281859 | 0.940672694 | 0.997677483 | -5.254183575 |
| **A_51_P170562** | Cpn2 | 0.0030885 | -0.09250376 | 0.098680761 | 7.665618039 | 0.077220755 | 0.940719495 | 0.997677483 | -5.254187555 |
| **A_51_P421804** | Timm9 | -0.005962861 | -0.190532585 | 0.178606863 | 11.01678549 | -0.077215299 | 0.940723674 | 0.997677483 | -5.25418791 |
| **A_52_P174775** | Ripk1 | -0.003807219 | -0.121667447 | 0.11405301 | 8.3360187 | -0.077205756 | 0.940730983 | 0.997677483 | -5.254188532 |
| **A_55_P2151685** | Pira11 | -0.00371078 | -0.118723465 | 0.111301905 | 7.763947016 | -0.077113177 | 0.940801891 | 0.997677483 | -5.254194554 |
| **A_55_P2112549** | NA | -0.011801117 | -0.37759198 | 0.353989745 | 13.49617933 | -0.077107995 | 0.940805861 | 0.997677483 | -5.254194891 |
| **A_55_P2007326** | Cyp2a4 | -0.003363176 | -0.107629388 | 0.100903036 | 7.717143991 | -0.077093036 | 0.940817318 | 0.997677483 | -5.254195863 |
| **A_52_P542172** | Rb1cc1 | 0.006505552 | -0.19530546 | 0.208316564 | 9.943060258 | 0.0770457 | 0.940853575 | 0.997677483 | -5.254198939 |
| **A_66_P112623** | Psmc6 | -0.005503448 | -0.176228408 | 0.165221511 | 12.15193501 | -0.077045454 | 0.940853763 | 0.997677483 | -5.254198955 |
| **A_66_P111433** | NA | -0.005470863 | -0.175421874 | 0.164480148 | 10.95933758 | -0.076938058 | 0.940936021 | 0.997677483 | -5.254205926 |
| **A_51_P464146** | Sowahb | 0.003105505 | -0.093533831 | 0.099744842 | 7.686319816 | 0.076804646 | 0.941038208 | 0.997677483 | -5.254214572 |
| **A_55_P2156712** | Gpr1 | -0.003707783 | -0.119092033 | 0.111676466 | 7.762480619 | -0.07680278 | 0.941039637 | 0.997677483 | -5.254214693 |
| **A_51_P298802** | Bfsp2 | -0.003709277 | -0.119185635 | 0.111767082 | 8.251271816 | -0.076772425 | 0.941062888 | 0.997677483 | -5.254216658 |
| **A_55_P1972748** | Czib | 0.016785884 | -0.505841762 | 0.539413529 | 12.40129714 | 0.076764595 | 0.941068885 | 0.997677483 | -5.254217165 |

| | | | | | | | | | |
|---|---|---|---|---|---|---|---|---|---|
| **A_55_P2180056** | NA | 0.010735856 | -0.323655562 | 0.345127273 | 9.146652237 | 0.076734501 | 0.941091936 | 0.997677483 | -5.254219112 |
| **A_66_P133413** | NA | -0.005715096 | -0.183822387 | 0.172392195 | 15.29153425 | -0.076692162 | 0.941124366 | 0.997677483 | -5.254221851 |
| **A_55_P2021996** | Hbs1l | 0.003484548 | -0.105534575 | 0.112503672 | 7.932188484 | 0.076392886 | 0.941353602 | 0.997805175 | -5.254241164 |
| **A_52_P490863** | Nop10 | -0.004826212 | -0.15591718 | 0.146264756 | 13.05093438 | -0.076344371 | 0.941390763 | 0.997805175 | -5.254244287 |
| **A_51_P153004** | Phax | 0.004747112 | -0.143999948 | 0.153494172 | 9.779833938 | 0.076276408 | 0.941442822 | 0.997805175 | -5.25424866 |
| **A_51_P174645** | Asl | -0.007003502 | -0.226584314 | 0.212577311 | 9.022640242 | -0.076230715 | 0.941477823 | 0.997805175 | -5.254251598 |
| **A_51_P352720** | Lmod2 | -0.007859967 | -0.254399196 | 0.238679262 | 8.817471614 | -0.076198041 | 0.94150285 | 0.997805175 | -5.254253697 |
| **A_55_P1960366** | Fam47e | -0.003317748 | -0.107788378 | 0.101152881 | 7.718306381 | -0.075902894 | 0.941728933 | 0.997990076 | -5.254272622 |
| **A_51_P368823** | Grb7 | -0.003883825 | -0.126325835 | 0.118558186 | 7.781868059 | -0.075812057 | 0.941798516 | 0.998009114 | -5.254278431 |
| **A_52_P48767** | Rassf8 | 0.004625007 | -0.141505164 | 0.150755179 | 8.266175569 | 0.075645248 | 0.941926295 | 0.998037492 | -5.254289082 |
| **A_55_P2024307** | Spin1 | -0.004453509 | -0.145322645 | 0.136415626 | 12.75568652 | -0.075560642 | 0.941991107 | 0.998037492 | -5.254294475 |
| **A_55_P2042818** | Zfp639 | 0.003968315 | -0.121759416 | 0.129696047 | 10.79454145 | 0.075436967 | 0.942085846 | 0.998037492 | -5.254302347 |
| **A_52_P1026830** | Tbc1d8b | -0.009399329 | -0.307198049 | 0.28839939 | 9.239142094 | -0.075436785 | 0.942085986 | 0.998037492 | -5.254302359 |
| **A_51_P515965** | Nfe2 | -0.005036177 | -0.164600097 | 0.154527743 | 8.134671044 | -0.075435437 | 0.942087018 | 0.998037492 | -5.254302444 |
| **A_55_P2002465** | Zfp747 | -0.004535667 | -0.148361101 | 0.139289766 | 10.93631012 | -0.075372797 | 0.942135004 | 0.998037492 | -5.254306427 |
| **A_55_P2302285** | Tarbp2 | -0.004356805 | -0.142971231 | 0.134257621 | 12.49618056 | -0.075122292 | 0.942326904 | 0.998075954 | -5.254322319 |
| **A_55_P2025403** | Gphn | -0.004469681 | -0.146810711 | 0.137871348 | 10.52572507 | -0.075050847 | 0.942381636 | 0.998075954 | -5.254326842 |
| **A_55_P2095266** | NA | 0.005615336 | -0.173220096 | 0.184450769 | 13.86804856 | 0.075046654 | 0.942384848 | 0.998075954 | -5.254327108 |
| **A_51_P464822** | H1f4 | -0.016271298 | -0.554413534 | 0.521870938 | 12.93854901 | -0.075109051 | 0.942715971 | 0.998075954 | -5.175073798 |
| **A_55_P1977418** | Ugt2b34 | 0.003546571 | -0.110170871 | 0.117264013 | 7.702629714 | 0.074540223 | 0.942772817 | 0.998075954 | -5.254359044 |
| **A_52_P494622** | Nr4a2 | 0.004963326 | -0.154256248 | 0.164182899 | 8.779830684 | 0.074505003 | 0.9427998 | 0.998075954 | -5.254361257 |
| **A_55_P2014041** | Rab14 | 0.005897622 | -0.18366739 | 0.195462633 | 13.39139013 | 0.074358029 | 0.942912397 | 0.998075954 | -5.25437048 |
| **A_55_P2093874** | Pbxip1 | 0.006453375 | -0.201940493 | 0.214847243 | 13.35834712 | 0.074013525 | 0.943176331 | 0.998075954 | -5.254392029 |
| **A_55_P2111563** | NA | -0.00410769 | -0.137182341 | 0.128966961 | 7.748708701 | -0.073775384 | 0.943358781 | 0.998075954 | -5.254406866 |
| **A_52_P273929** | Mrpl17 | -0.005091093 | -0.1700691 | 0.159886915 | 10.84918893 | -0.073755447 | 0.943374056 | 0.998075954 | -5.254408106 |
| **A_52_P533304** | Eif5 | 0.003558693 | -0.111901677 | 0.119019063 | 12.88826576 | 0.073665927 | 0.943442642 | 0.998075954 | -5.25441367 |
| **A_66_P101764** | NA | -0.028048447 | -0.938428747 | 0.882331853 | 12.17814143 | -0.073636833 | 0.943464933 | 0.998075954 | -5.254415477 |
| **A_52_P684130** | 2510002D2 | 0.005667115 | -0.178507419 | 0.18984165 | 8.563311801 | 0.073543046 | 0.943536789 | 0.998075954 | -5.254421296 |
| **A_55_P1995449** | NA | -0.009265241 | -0.31047107 | 0.291940587 | 11.59060612 | -0.073519491 | 0.943554836 | 0.998075954 | -5.254422757 |
| **A_52_P84027** | Cyp7a1 | 0.002921061 | -0.092052494 | 0.097894615 | 7.64058206 | 0.073510106 | 0.943562027 | 0.998075954 | -5.254423338 |
| **A_55_P2159934** | Rnf186 | -0.003012699 | -0.101004384 | 0.094978987 | 7.665207368 | -0.073481092 | 0.943584256 | 0.998075954 | -5.254425136 |
| **A_55_P2012101** | Bmp8b | -0.003028301 | -0.101662134 | 0.095605532 | 7.711024712 | -0.073380766 | 0.943661124 | 0.998075954 | -5.254431348 |
| **A_52_P472319** | Plxnd1 | -0.007437002 | -0.249943051 | 0.235069047 | 9.590587156 | -0.073296702 | 0.943725531 | 0.998075954 | -5.254436546 |
| **A_55_P2181743** | Elmod1 | 0.002933231 | -0.092757855 | 0.098624317 | 7.747512998 | 0.073262868 | 0.943751455 | 0.998075954 | -5.254438637 |
| **A_55_P1977985** | Pigp | 0.005465308 | -0.172863016 | 0.183793632 | 12.20873754 | 0.073249294 | 0.943761855 | 0.998075954 | -5.254439475 |
| **A_52_P251366** | Neil3 | -0.005730611 | -0.192767658 | 0.181306436 | 8.881042544 | -0.073228885 | 0.943777492 | 0.998075954 | -5.254440736 |
| **A_55_P2087280** | NA | -0.004591461 | -0.154632591 | 0.145449668 | 12.35352168 | -0.073139119 | 0.94384627 | 0.998075954 | -5.254446275 |
| **A_51_P315795** | Tubb4a | -0.008124436 | -0.274065271 | 0.257816398 | 11.68757135 | -0.073015881 | 0.943940694 | 0.998075954 | -5.254453868 |
| **A_51_P146303** | Mvb12a | 0.005781534 | -0.183549959 | 0.195113027 | 11.00160281 | 0.072984286 | 0.943964903 | 0.998075954 | -5.254455813 |
| **A_55_P2146483** | Zmynd8 | -0.006519729 | -0.220273408 | 0.207233951 | 10.43151557 | -0.072899583 | 0.944029802 | 0.998075954 | -5.254461023 |
| **A_55_P2023617** | Kiss1r | -0.003270423 | -0.110512534 | 0.103971688 | 7.736201954 | -0.072886608 | 0.944039745 | 0.998075954 | -5.25446182 |
| **A_55_P1976539** | Nup62cl | -0.003066762 | -0.104095117 | 0.097961593 | 7.657777223 | -0.07255142 | 0.944296572 | 0.998075954 | -5.254482372 |
| **A_55_P2134092** | Brsk1 | 0.005112617 | -0.163345375 | 0.17357061 | 9.045383765 | 0.072537204 | 0.944307465 | 0.998075954 | -5.254483242 |
| **A_66_P114585** | NA | 0.004393573 | -0.140588593 | 0.149375739 | 8.687891659 | 0.072428984 | 0.944390387 | 0.998075954 | -5.254489856 |
| **A_55_P2009013** | Dpf2 | 0.006380427 | -0.204283317 | 0.21704417 | 11.52630277 | 0.072388414 | 0.944421473 | 0.998075954 | -5.254492333 |
| **A_51_P150014** | Rrp7a | 0.00343622 | -0.110130244 | 0.117002684 | 8.031707987 | 0.072316937 | 0.944476242 | 0.998075954 | -5.254496694 |
| **A_66_P105046** | Il18 | 0.008996369 | -0.288875026 | 0.306867763 | 8.248444706 | 0.072185102 | 0.944577261 | 0.998075954 | -5.254504725 |
| **A_55_P2133751** | Txnl4a | 0.004258775 | -0.136771669 | 0.145289218 | 12.38612408 | 0.072174024 | 0.944585749 | 0.998075954 | -5.254505399 |

| | | | | | | | | | |
|---|---|---|---|---|---|---|---|---|---|
| **A_55_P2095394** | Dhx37 | -0.004466037 | -0.152500521 | 0.143568447 | 8.520583516 | -0.07210553 | 0.944638234 | 0.998075954 | -5.254509566 |
| **A_55_P1999811** | NA | -0.003571174 | -0.122072316 | 0.114929968 | 8.57190062 | -0.072027384 | 0.944698114 | 0.998075954 | -5.254514315 |
| **A_55_P1988009** | Gm10406 | -0.014354825 | -0.491207853 | 0.462498203 | 10.58971482 | -0.071948624 | 0.944758465 | 0.998075954 | -5.254519095 |
| **A_55_P2080542** | Rbbp7 | 0.004037173 | -0.130092828 | 0.138167173 | 11.53632048 | 0.071938362 | 0.944766328 | 0.998075954 | -5.254519718 |
| **A_55_P2028898** | Prl2b1 | 0.031415656 | -1.012921076 | 1.075752387 | 8.332893295 | 0.071897652 | 0.944797523 | 0.998075954 | -5.254522187 |
| **A_51_P259009** | Cacnb3 | -0.004997386 | -0.171194485 | 0.161199714 | 8.56835778 | -0.071866842 | 0.944821132 | 0.998075954 | -5.254524054 |
| **A_55_P2087766** | Dnaja3 | -0.003851528 | -0.132028898 | 0.124325843 | 12.32221277 | -0.071817582 | 0.944858878 | 0.998075954 | -5.254527038 |
| **A_55_P2003951** | Tmem192 | 0.004034687 | -0.130664207 | 0.13873358 | 11.97895946 | 0.071590423 | 0.945032947 | 0.998075954 | -5.254540772 |
| **A_66_P131205** | NA | -0.004625858 | -0.159118811 | 0.149867094 | 16.17770127 | -0.0715637 | 0.945053425 | 0.998075954 | -5.254542385 |
| **A_51_P489522** | Ctla2b | 0.004598267 | -0.149021897 | 0.158218431 | 11.9565644 | 0.071541014 | 0.945070809 | 0.998075954 | -5.254543754 |
| **A_55_P2067712** | Hmcn2 | -0.008495348 | -0.292462401 | 0.275471704 | 9.637824892 | -0.071502689 | 0.945100177 | 0.998075954 | -5.254546065 |
| **A_55_P1992319** | Col4a3bp | 0.004623546 | -0.149984442 | 0.159231533 | 8.63719319 | 0.071474705 | 0.945121621 | 0.998075954 | -5.254547752 |
| **A_52_P419373** | Lrrc39 | -0.007387994 | -0.25451922 | 0.239743232 | 9.11791609 | -0.071450958 | 0.945139819 | 0.998075954 | -5.254549183 |
| **A_52_P40293** | Hgd | 0.002975434 | -0.096565516 | 0.102516384 | 7.639136179 | 0.071442669 | 0.94514617 | 0.998075954 | -5.254549683 |
| **A_55_P2035963** | Hipk3 | 0.004766481 | -0.154964376 | 0.164497339 | 9.164023278 | 0.07132113 | 0.945239306 | 0.998075954 | -5.254556998 |
| **A_55_P2066309** | Slc39a8 | 0.018709943 | -0.608505413 | 0.6459253 | 9.240689391 | 0.07129596 | 0.945258594 | 0.998075954 | -5.254558511 |
| **A_55_P2177347** | NA | 0.004105692 | -0.133563781 | 0.141775166 | 8.206751456 | 0.07127839 | 0.945272058 | 0.998075954 | -5.254559568 |
| **A_55_P2007485** | Prss27 | 0.003004485 | -0.097944372 | 0.103953342 | 7.691559707 | 0.071134086 | 0.945382641 | 0.998075954 | -5.254568232 |
| **A_55_P1965931** | Nat8f1 | -0.004212845 | -0.145776546 | 0.137350855 | 9.043461028 | -0.071126713 | 0.945388291 | 0.998075954 | -5.254568674 |
| **A_55_P2061159** | Ing1 | 0.003107208 | -0.101436675 | 0.107651092 | 12.031012 | 0.071036392 | 0.945457506 | 0.998075954 | -5.254574088 |
| **A_55_P2401084** | 4930558N( | 0.006433823 | -0.210147129 | 0.223014775 | 8.870368562 | 0.070999941 | 0.945485439 | 0.998075954 | -5.254576271 |
| **A_52_P253748** | Zbtb45 | 0.005212529 | -0.170504333 | 0.18092939 | 9.189203698 | 0.070899658 | 0.94556229 | 0.998075954 | -5.25458227 |
| **A_55_P2059412** | Ghr | -0.003509042 | -0.121845888 | 0.114827805 | 10.06539086 | -0.07087249 | 0.94558311 | 0.998075954 | -5.254583894 |
| **A_52_P212418** | Trmt13 | -0.004184626 | -0.14530979 | 0.136940537 | 8.367235777 | -0.070869827 | 0.94558515 | 0.998075954 | -5.254584053 |
| **A_55_P2102693** | Acvr1 | 0.003888942 | -0.12738218 | 0.135160065 | 9.392359436 | 0.070806225 | 0.945633891 | 0.998075954 | -5.254587852 |
| **A_55_P2128792** | NA | -0.005487208 | -0.190809306 | 0.179834891 | 9.282769009 | -0.070767414 | 0.945663634 | 0.998075954 | -5.254590169 |
| **A_55_P2067652** | Boc | 0.006389341 | -0.209504482 | 0.222283163 | 8.726855955 | 0.07073347 | 0.945689646 | 0.998075954 | -5.254592194 |
| **A_55_P1986276** | Tmem41b | 0.004314302 | -0.141472154 | 0.150100759 | 10.59553205 | 0.070729816 | 0.945692446 | 0.998075954 | -5.254592412 |
| **A_51_P165914** | Zfp770 | -0.004161247 | -0.144860027 | 0.136537533 | 9.965514097 | -0.070687443 | 0.945724919 | 0.998075954 | -5.254594938 |
| **A_55_P2072842** | NA | -0.013023862 | -0.4539888 | 0.427941077 | 11.56685118 | -0.07059027 | 0.945799387 | 0.998075954 | -5.254600727 |
| **A_66_P133404** | Pclaf | 0.004577529 | -0.150694949 | 0.159850007 | 8.814242519 | 0.070460501 | 0.945898837 | 0.998075954 | -5.254608444 |
| **A_55_P1971074** | Atp11a | -0.003719396 | -0.129961177 | 0.122522385 | 12.07542642 | -0.070417148 | 0.945932062 | 0.998075954 | -5.254611019 |
| **A_51_P264527** | Dipk1b | -0.006752164 | -0.236059938 | 0.222555611 | 8.557177994 | -0.070377415 | 0.945962511 | 0.998075954 | -5.254613378 |
| **A_52_P1016836** | NA | 0.002956984 | -0.09749613 | 0.103410097 | 7.633413244 | 0.070354948 | 0.94597973 | 0.998075954 | -5.254614711 |
| **A_55_P2070657** | Emc10 | -0.008531341 | -0.298609705 | 0.281547022 | 8.61378376 | -0.070292843 | 0.946027325 | 0.998075954 | -5.254618394 |
| **A_51_P202801** | Abcb9 | -0.006116094 | -0.214404476 | 0.202172288 | 9.398398902 | -0.070180787 | 0.946113202 | 0.998075954 | -5.25462503 |
| **A_51_P513661** | Slc4a8 | 0.00339498 | -0.112339955 | 0.119129914 | 7.736615233 | 0.070110299 | 0.946167222 | 0.998075954 | -5.2546292 |
| **A_55_P2130264** | NA | -0.004235823 | -0.14865335 | 0.140181705 | 16.80542189 | -0.070101449 | 0.946174005 | 0.998075954 | -5.254629723 |
| **A_52_P144263** | Rbms1 | -0.003947845 | -0.138580428 | 0.130684738 | 9.201098906 | -0.070084033 | 0.946187352 | 0.998075954 | -5.254630752 |
| **A_55_P2178632** | Tmem80 | 0.003097679 | -0.102606649 | 0.108802006 | 9.517357994 | 0.070041065 | 0.946220283 | 0.998075954 | -5.254633291 |
| **A_51_P506309** | Hrg | 0.002903274 | -0.096295793 | 0.10210234 | 7.806170079 | 0.069950287 | 0.946289855 | 0.998075954 | -5.254638649 |
| **A_55_P2135486** | NA | 0.003101353 | -0.103054144 | 0.10925685 | 7.678733753 | 0.069826103 | 0.946385029 | 0.998075954 | -5.254645967 |
| **A_55_P2024841** | Gstt1 | -0.011330812 | -0.399458672 | 0.376797049 | 9.793483454 | -0.069774276 | 0.946424749 | 0.998075954 | -5.254649018 |
| **A_51_P194628** | Tango6 | 0.003745848 | -0.124970644 | 0.132462339 | 8.39960589 | 0.069554475 | 0.946593207 | 0.998075954 | -5.25466193 |
| **A_51_P237688** | Smim14 | -0.003929125 | -0.139522834 | 0.131664584 | 10.86111916 | -0.069257282 | 0.946820984 | 0.998075954 | -5.254679324 |
| **A_55_P2091305** | NA | 0.003437684 | -0.115535347 | 0.122410716 | 8.260029588 | 0.069060001 | 0.946972188 | 0.998075954 | -5.254690829 |
| **A_55_P2055975** | Ppip5k1 | 0.003774339 | -0.127074699 | 0.134623377 | 9.072645407 | 0.068941312 | 0.947063158 | 0.998075954 | -5.254697735 |
| **A_51_P195506** | Csf1 | 0.004833763 | -0.163253739 | 0.172921266 | 9.313851801 | 0.068732017 | 0.947223575 | 0.998075954 | -5.229213026 |

| | | | | | | | | | |
|---|---|---|---|---|---|---|---|---|---|
| **A_55_P2017914** | Csf1 | -0.003953007 | -0.141626796 | 0.133720782 | 8.189451237 | -0.068625492 | 0.947305223 | 0.998075954 | -5.254716054 |
| **A_55_P2116988** | 1700024P1 | -0.006417526 | -0.230042205 | 0.217207152 | 9.512212693 | -0.068589411 | 0.947332879 | 0.998075954 | -5.254718141 |
| **A_55_P2148641** | Rab4a | -0.002895528 | -0.103807227 | 0.098016171 | 7.990049701 | -0.068579669 | 0.947340346 | 0.998075954 | -5.254718705 |
| **A_55_P2071286** | Tmc4 | 0.004917317 | -0.166522826 | 0.176357459 | 8.458495008 | 0.068552736 | 0.947360989 | 0.998075954 | -5.254720262 |
| **A_55_P2075941** | Nf1 | -0.007022186 | -0.251863715 | 0.237819344 | 9.515920979 | -0.068548248 | 0.947364429 | 0.998075954 | -5.254720522 |
| **A_51_P328489** | 1700025G( | -0.004099259 | -0.147309494 | 0.139110975 | 10.32348545 | -0.06841329 | 0.947467872 | 0.998075954 | -5.254728315 |
| **A_55_P2079324** | Sephs2 | -0.002981769 | -0.107284598 | 0.10132106 | 7.982255023 | -0.068326164 | 0.947534653 | 0.998075954 | -5.254733338 |
| **A_52_P54238** | Lmf2 | -0.00398286 | -0.143388368 | 0.135422648 | 10.02288567 | -0.068284835 | 0.947566332 | 0.998075954 | -5.254735719 |
| **A_51_P248865** | Foxf2 | -0.002898185 | -0.104440491 | 0.098644121 | 7.646405353 | -0.068216308 | 0.947618858 | 0.998075954 | -5.254739663 |
| **A_55_P2136426** | Prss57 | 0.002794907 | -0.095136359 | 0.100726174 | 7.671460008 | 0.068211121 | 0.947622833 | 0.998075954 | -5.254739961 |
| **A_55_P2125351** | Sema6d | -0.00567458 | -0.204687365 | 0.193338206 | 10.62612938 | -0.06814937 | 0.947670166 | 0.998075954 | -5.254743512 |
| **A_55_P2174339** | NA | 0.027415328 | -0.934451625 | 0.989282281 | 9.844167897 | 0.068122028 | 0.947691123 | 0.998075954 | -5.254745083 |
| **A_51_P126437** | Enc1 | 0.007036745 | -0.23999598 | 0.25406947 | 10.34525295 | 0.068081084 | 0.947722507 | 0.998075954 | -5.254747434 |
| **A_51_P496720** | Dnmt3l | 0.002885754 | -0.098504576 | 0.104276084 | 7.743384567 | 0.068025517 | 0.9477651 | 0.998075954 | -5.254750623 |
| **A_55_P2108151** | NA | 0.020817744 | -0.713673805 | 0.755309293 | 14.8148403 | 0.067741683 | 0.947982664 | 0.998075954 | -5.254766872 |
| **A_52_P655687** | Egfl6 | 0.002960313 | -0.101525014 | 0.10744564 | 7.677882838 | 0.067716025 | 0.948002332 | 0.998075954 | -5.254768337 |
| **A_55_P2414524** | NA | -0.003401356 | -0.123515761 | 0.116713049 | 12.67254111 | -0.067680901 | 0.948029255 | 0.998075954 | -5.254770342 |
| **A_55_P1994648** | Eef1d | 0.003688864 | -0.126791009 | 0.134168737 | 15.19598878 | 0.067570677 | 0.948113746 | 0.998075954 | -5.254776628 |
| **A_55_P2112504** | Kcnv2 | -0.002785004 | -0.101312183 | 0.095742175 | 7.64565786 | -0.067558333 | 0.948123208 | 0.998075954 | -5.254777332 |
| **A_55_P2034864** | Tubb2b | 0.006287821 | -0.216196824 | 0.228772466 | 13.93837725 | 0.067547497 | 0.948131515 | 0.998075954 | -5.254777949 |
| **A_55_P2056190** | Rps3a1 | -0.004678504 | -0.170282703 | 0.160925696 | 16.26236879 | -0.067521914 | 0.948151124 | 0.998075954 | -5.254779406 |
| **A_55_P2074085** | Gimap8 | 0.005774685 | -0.198659199 | 0.210208569 | 9.279985367 | 0.067512556 | 0.948158298 | 0.998075954 | -5.254779939 |
| **A_51_P508115** | Trank1 | 0.002964404 | -0.102058584 | 0.107987392 | 7.889575938 | 0.067462448 | 0.948196708 | 0.998075954 | -5.25478279 |
| **A_55_P2015984** | Odr4 | 0.007468747 | -0.257313267 | 0.272250761 | 10.92855568 | 0.067416843 | 0.948231666 | 0.998075954 | -5.254785384 |
| **A_66_P115693** | Rnf207 | -0.017674336 | -0.644574687 | 0.609226016 | 12.09546738 | -0.067383524 | 0.948257207 | 0.998075954 | -5.254787278 |
| **A_55_P2016739** | Rbm12b2 | 0.004769326 | -0.164665969 | 0.174204621 | 9.450646299 | 0.067276323 | 0.948339382 | 0.998075954 | -5.254793365 |
| **A_55_P2141429** | Park7 | -0.005708462 | -0.208514906 | 0.197097982 | 13.85712077 | -0.067273885 | 0.948341251 | 0.998075954 | -5.254793503 |
| **A_55_P2074337** | Ngf | -0.003146604 | -0.114965795 | 0.108672586 | 8.847449934 | -0.067256599 | 0.948354502 | 0.998075954 | -5.254794484 |
| **A_55_P2039566** | Arhgap23 | 0.00399844 | -0.138440915 | 0.146437795 | 12.12600808 | 0.067091836 | 0.948480803 | 0.998075954 | -5.254803817 |
| **A_55_P2158112** | Rps27 | 0.00368569 | -0.127632508 | 0.135003888 | 16.30977762 | 0.06708154 | 0.948488695 | 0.998075954 | -5.2548044 |
| **A_66_P124139** | Ikzf1 | -0.003095473 | -0.113391336 | 0.10720039 | 7.724303295 | -0.06707751 | 0.948491785 | 0.998075954 | -5.254804628 |
| **A_55_P2003123** | Jkamp | 0.003899896 | -0.135116481 | 0.142916274 | 12.97577767 | 0.067049607 | 0.948513174 | 0.998075954 | -5.254806206 |
| **A_55_P2041340** | Atp5a1 | -0.005036304 | -0.184599855 | 0.174527247 | 16.66580083 | -0.067035194 | 0.948524223 | 0.998075954 | -5.254807021 |
| **A_55_P2093940** | Zfp408 | -0.007628028 | -0.279733501 | 0.264477446 | 15.99248419 | -0.067001438 | 0.948550099 | 0.998075954 | -5.254808928 |
| **A_55_P2122666** | Pnma3 | -0.003284645 | -0.120681385 | 0.114112096 | 8.212281503 | -0.066871562 | 0.948649659 | 0.998075954 | -5.25481626 |
| **A_55_P2038737** | Slc29a1 | -0.005614451 | -0.206400334 | 0.195171432 | 11.3942559 | -0.066831811 | 0.948680131 | 0.998075954 | -5.254818501 |
| **A_55_P2118843** | Hk1 | 0.008933736 | -0.310885322 | 0.328752793 | 9.568896325 | 0.066763315 | 0.948732639 | 0.998075954 | -5.254822359 |
| **A_55_P2016652** | Cdk1 | 0.003042969 | -0.106387025 | 0.112472964 | 7.938003536 | 0.066461532 | 0.948963985 | 0.998075954 | -5.254839312 |
| **A_55_P2083382** | Snf8 | -0.002934743 | -0.108516448 | 0.102646962 | 12.79668516 | -0.066434022 | 0.948985074 | 0.998075954 | -5.254840854 |
| **A_52_P573366** | Hnrnpul1 | 0.00335562 | -0.117382346 | 0.124093586 | 9.002536839 | 0.066425996 | 0.948991226 | 0.998075954 | -5.254841303 |
| **A_52_P324767** | Tmed8 | -0.002946083 | -0.108961815 | 0.10306965 | 8.972017845 | -0.066417685 | 0.948997598 | 0.998075954 | -5.254841769 |
| **A_55_P2062508** | Ube2e1 | 0.005300861 | -0.18548274 | 0.196084462 | 11.49236727 | 0.066407097 | 0.949005715 | 0.998075954 | -5.254842362 |
| **A_51_P411019** | Tmem127 | -0.003377075 | -0.124937798 | 0.118183649 | 11.87096694 | -0.066398234 | 0.949012509 | 0.998075954 | -5.254842858 |
| **A_55_P2126087** | NA | -0.003367471 | -0.124648204 | 0.117913263 | 14.74751391 | -0.066362254 | 0.949040092 | 0.998075954 | -5.254844872 |
| **A_52_P47645** | Cul3 | 0.003815126 | -0.133645781 | 0.141276033 | 9.957845343 | 0.066334397 | 0.949061447 | 0.998075954 | -5.254846431 |
| **A_51_P155085** | Dennd2a | 0.00469428 | -0.165106885 | 0.174495445 | 11.16187636 | 0.066075047 | 0.94926027 | 0.998075954 | -5.254860911 |
| **A_51_P310576** | Smim8 | -0.004698366 | -0.175007178 | 0.165610447 | 8.650724113 | -0.065935426 | 0.949367307 | 0.998075954 | -5.254868683 |
| **A_51_P117604** | Rab5a | 0.005233197 | -0.18449752 | 0.194963913 | 11.34139329 | 0.065923242 | 0.949376648 | 0.998075954 | -5.235205273 |

| | | | | | | | | | |
|---|---|---|---|---|---|---|---|---|---|
| **A_52_P138205** | Plekha8 | 0.005728225 | -0.20211967 | 0.21357612 | 8.793468724 | 0.065869376 | 0.949417944 | 0.998075954 | -5.254872354 |
| **A_51_P195875** | Notch4 | 0.003410052 | -0.120428933 | 0.127249037 | 10.20799259 | 0.065813163 | 0.949461039 | 0.998075954 | -5.254875476 |
| **A_65_P16603** | Bad | -0.00612734 | -0.228767916 | 0.216513236 | 8.525518873 | -0.065777416 | 0.949488444 | 0.998075954 | -5.254877459 |
| **A_51_P413710** | Ap3d1 | 0.003826311 | -0.135276957 | 0.14292958 | 11.06270607 | 0.065743388 | 0.949514531 | 0.998075954 | -5.254879346 |
| **A_55_P2154425** | Cd320 | -0.002789413 | -0.104250769 | 0.098671942 | 7.748474318 | -0.065708469 | 0.949541302 | 0.998075954 | -5.254881282 |
| **A_52_P400425** | Pirt | 0.006914509 | -0.245082496 | 0.258911514 | 8.412286928 | 0.065580558 | 0.949639365 | 0.998075954 | -5.254888363 |
| **A_55_P2457912** | Psme4 | -0.003676442 | -0.137671202 | 0.130318318 | 8.738445657 | -0.065576621 | 0.949642383 | 0.998075954 | -5.254888581 |
| **A_55_P2127538** | Ppp1r21 | 0.00354175 | -0.125675616 | 0.132759116 | 8.29831297 | 0.065509785 | 0.949693624 | 0.998075954 | -5.254892275 |
| **A_55_P2040893** | Tnni2 | -0.006413175 | -0.24046121 | 0.22763486 | 10.70547352 | -0.065490337 | 0.949708533 | 0.998075954 | -5.254893349 |
| **A_55_P2028029** | Abcb11 | 0.002772556 | -0.098650563 | 0.104195674 | 7.682782618 | 0.065335988 | 0.949826867 | 0.998075954 | -5.254901864 |
| **A_55_P2066792** | Sp110 | -0.004924174 | -0.18511794 | 0.175269591 | 14.77319617 | -0.065313474 | 0.949844128 | 0.998075954 | -5.254903105 |
| **A_55_P2089913** | Cd6 | -0.004198738 | -0.157891572 | 0.149494097 | 7.841446921 | -0.065294145 | 0.949858948 | 0.998075954 | -5.254904169 |
| **A_55_P2013357** | Renbp | -0.00489861 | -0.184283305 | 0.174486084 | 9.552913498 | -0.065267451 | 0.949879413 | 0.998075954 | -5.254905639 |
| **A_55_P2138549** | Cc2d1a | -0.003224958 | -0.121407152 | 0.114957236 | 10.06709852 | -0.065220049 | 0.949915755 | 0.998075954 | -5.254908247 |
| **A_55_P2116385** | Gpr183 | -0.0030977 | -0.116689184 | 0.110493784 | 7.937295521 | -0.065178257 | 0.949947796 | 0.998075954 | -5.254910545 |
| **A_51_P455166** | Prl4a1 | 0.003083692 | -0.110196354 | 0.116363738 | 7.644078955 | 0.06506189 | 0.950037012 | 0.998075954 | -5.254916936 |
| **A_55_P1977563** | Galk2 | -0.00385087 | -0.145599589 | 0.137897848 | 8.890347512 | -0.064930517 | 0.950137735 | 0.998075954 | -5.254924137 |
| **A_51_P261999** | Ccdc166 | 0.004205134 | -0.150725992 | 0.15913626 | 8.839183353 | 0.064870954 | 0.950183401 | 0.998075954 | -5.254927397 |
| **A_55_P1993522** | NA | -0.005375197 | -0.204022468 | 0.193272073 | 12.00298223 | -0.064672689 | 0.950335411 | 0.998075954 | -5.254938227 |
| **A_55_P2010469** | NA | -0.004833594 | -0.183590859 | 0.17392367 | 11.84463959 | -0.064627245 | 0.950370254 | 0.998075954 | -5.254940705 |
| **A_55_P2073935** | Abcc5 | -0.003746005 | -0.142566911 | 0.135074902 | 8.242278241 | -0.064494486 | 0.950472042 | 0.998075954 | -5.254947934 |
| **A_55_P2161775** | Prr3 | -0.005602464 | -0.213243669 | 0.202038742 | 8.996984062 | -0.064487364 | 0.950477502 | 0.998075954 | -5.254948321 |
| **A_52_P330950** | C2cd2 | -0.004623822 | -0.176152863 | 0.166905219 | 8.677568899 | -0.064427692 | 0.950523254 | 0.998075954 | -5.254951565 |
| **A_66_P114543** | Jmjd6 | -0.005354815 | -0.204238231 | 0.1935286 | 8.779799983 | -0.064350966 | 0.950582081 | 0.998075954 | -5.254955732 |
| **A_55_P2109122** | H2bc4 | 0.008456181 | -0.30584152 | 0.322753882 | 12.85581883 | 0.064304623 | 0.950617614 | 0.998075954 | -5.254958246 |
| **A_51_P354572** | Rab35 | -0.002973018 | -0.113525642 | 0.107579606 | 12.51740671 | -0.064274337 | 0.950640835 | 0.998075954 | -5.254959888 |
| **A_52_P435561** | Prr15l | -0.002873617 | -0.10975084 | 0.104003606 | 7.686924251 | -0.064261792 | 0.950650454 | 0.998075954 | -5.254960568 |
| **A_52_P561690** | BC034090 | 0.004305064 | -0.156169108 | 0.164779236 | 8.427665575 | 0.064118533 | 0.950760295 | 0.998075954 | -5.254968323 |
| **A_55_P2053850** | NA | 0.004182298 | -0.151957155 | 0.160321752 | 10.64563636 | 0.064019379 | 0.950836321 | 0.998075954 | -5.254973681 |
| **A_52_P110812** | Ggcx | 0.00312945 | -0.114235791 | 0.12049469 | 8.591298437 | 0.063729073 | 0.951058913 | 0.998075954 | -5.25498932 |
| **A_51_P315042** | Avpr1a | -0.002808847 | -0.108338942 | 0.102721248 | 7.668653875 | -0.063615201 | 0.951146227 | 0.998075954 | -5.254995435 |
| **A_55_P1979818** | Srp54b | 0.002764345 | -0.101198646 | 0.106727337 | 8.66703856 | 0.063551038 | 0.951195425 | 0.998075954 | -5.254998876 |
| **A_55_P2042202** | Pigo | -0.003310931 | -0.127994339 | 0.121372477 | 11.14197938 | -0.063467388 | 0.951259566 | 0.998075954 | -5.255003356 |
| **A_55_P2064906** | NA | 0.003965794 | -0.145569482 | 0.153501071 | 15.31266714 | 0.063386335 | 0.951321715 | 0.998075954 | -5.255007692 |
| **A_52_P267543** | Ginm1 | -0.002773662 | -0.10761724 | 0.102069916 | 8.170294064 | -0.063229656 | 0.951441854 | 0.998075954 | -5.255016058 |
| **A_52_P161630** | St6gal1 | 0.005652249 | -0.208067738 | 0.219372235 | 9.224161036 | 0.063209922 | 0.951456986 | 0.998075954 | -5.25501711 |
| **A_52_P500244** | Atrn | -0.004427077 | -0.171940522 | 0.163086367 | 10.34588667 | -0.063165011 | 0.951491423 | 0.998075954 | -5.255019503 |
| **A_66_P122219** | Kcnd3 | 0.008350126 | -0.307646954 | 0.324347205 | 8.623233389 | 0.063156648 | 0.951497836 | 0.998075954 | -5.255019949 |
| **A_51_P471458** | Sult5a1 | 0.002889803 | -0.106474612 | 0.112254217 | 7.957729674 | 0.063154063 | 0.951499818 | 0.998075954 | -5.255020086 |
| **A_55_P2016024** | Kif3c | 0.005169113 | -0.190515341 | 0.200853567 | 8.792453189 | 0.063134803 | 0.951514587 | 0.998075954 | -5.255021112 |
| **A_51_P440327** | Efhc1 | -0.0025645 | -0.099674179 | 0.094545179 | 8.046803575 | -0.06311743 | 0.951527908 | 0.998075954 | -5.255022037 |
| **A_55_P2106801** | Bglap3 | 0.003587199 | -0.13256783 | 0.139742227 | 7.850361972 | 0.06296959 | 0.951641271 | 0.998075954 | -5.255029898 |
| **A_52_P149835** | Irak1 | -0.005220166 | -0.203750375 | 0.193310043 | 8.682892873 | -0.062844438 | 0.951737239 | 0.998075954 | -5.255036538 |
| **A_55_P2023697** | Zfp386 | -0.005205849 | -0.203201033 | 0.192789335 | 10.739674 | -0.062841435 | 0.951739541 | 0.998075954 | -5.255036697 |
| **A_55_P1992505** | Entpd4 | -0.005316537 | -0.207739049 | 0.197105975 | 12.24151122 | -0.062773908 | 0.951791322 | 0.998075954 | -5.255040274 |
| **A_55_P1997300** | Mtfr2 | 0.004569331 | -0.170002136 | 0.179140798 | 13.14976877 | 0.062558809 | 0.951956265 | 0.998075954 | -5.255051643 |
| **A_52_P58059** | Usp34 | -0.005699154 | -0.223659396 | 0.212261088 | 9.199151723 | -0.062494568 | 0.952005526 | 0.998075954 | -5.25505503 |
| **A_55_P2449750** | Fam177a2 | -0.002892302 | -0.11353582 | 0.107751217 | 8.724569299 | -0.062477948 | 0.952018271 | 0.998075954 | -5.255055906 |

| | | | | | | | | | |
|---|---|---|---|---|---|---|---|---|---|
| **A_51_P314397** | Crip2 | -0.005034502 | -0.197697707 | 0.187628704 | 17.51479878 | -0.062454945 | 0.952035911 | 0.998075954 | -5.255057118 |
| **A_55_P1959903** | Map7 | 0.00298295 | -0.11157336 | 0.11753926 | 7.894479337 | 0.062235203 | 0.952204417 | 0.998075954 | -5.255068673 |
| **A_55_P2052181** | Ccdc82 | -0.005138646 | -0.202490921 | 0.19221363 | 11.82310134 | -0.062232273 | 0.952206663 | 0.998075954 | -5.255068827 |
| **A_55_P2155848** | Gm14207 | -0.005064604 | -0.199957678 | 0.189828471 | 8.202944511 | -0.062109521 | 0.952300795 | 0.998075954 | -5.255075263 |
| **A_51_P177171** | Tie1 | 0.006122644 | -0.229492808 | 0.241738096 | 10.19547923 | 0.062107543 | 0.952302312 | 0.998075954 | -5.255075367 |
| **A_55_P2037116** | Ran | -0.003766361 | -0.149025982 | 0.141493261 | 15.2270324 | -0.061970663 | 0.952407279 | 0.998075954 | -5.255082529 |
| **A_52_P8043** | Srsf2 | -0.003568571 | -0.141306399 | 0.134169257 | 10.27893114 | -0.061922748 | 0.952444024 | 0.998075954 | -5.255085032 |
| **A_55_P2160291** | Olfm1 | -0.007719302 | -0.306533681 | 0.291095077 | 8.621325383 | -0.061742708 | 0.95258209 | 0.998075954 | -5.255094422 |
| **A_51_P186798** | NA | 0.005623558 | -0.212683762 | 0.223930877 | 8.36777576 | 0.061567566 | 0.952716402 | 0.998075954 | -5.255103529 |
| **A_51_P284665** | Plcb1 | 0.003159369 | -0.119779954 | 0.126098692 | 8.076714 | 0.061421246 | 0.952828612 | 0.998075954 | -5.255111118 |
| **A_52_P513347** | Phkb | -0.003324487 | -0.13282487 | 0.126175896 | 8.616264147 | -0.061356805 | 0.952878031 | 0.998075954 | -5.255114455 |
| **A_55_P2151091** | Phactr4 | 0.003742402 | -0.142178733 | 0.149663537 | 11.62058222 | 0.061297302 | 0.952923664 | 0.998075954 | -5.255117533 |
| **A_55_P2124871** | Npw | -0.002446851 | -0.097889367 | 0.092995665 | 7.706291879 | -0.061273804 | 0.952941685 | 0.998075954 | -5.255118747 |
| **A_55_P2119737** | NA | 0.002499224 | -0.095420056 | 0.100418505 | 7.663412727 | 0.061002292 | 0.953149908 | 0.998075954 | -5.255132748 |
| **A_51_P164014** | Cenpe | -0.00633284 | -0.254538656 | 0.241872976 | 12.10079136 | -0.060981162 | 0.953166113 | 0.998075954 | -5.255133835 |
| **A_55_P1963105** | NA | -0.003114228 | -0.125176999 | 0.118948543 | 15.3985708 | -0.06097844 | 0.9531682 | 0.998075954 | -5.255133975 |
| **A_55_P2051622** | Gm8479 | 0.004989043 | -0.190614988 | 0.200593075 | 8.565825977 | 0.060960514 | 0.953181948 | 0.998075954 | -5.255134897 |
| **A_52_P317040** | Edem2 | 0.002875985 | -0.109992583 | 0.115744553 | 7.813058618 | 0.060900762 | 0.953227772 | 0.998075954 | -5.255137968 |
| **A_51_P208919** | Acbd6 | 0.005434797 | -0.208029631 | 0.218899225 | 10.99495552 | 0.060850894 | 0.953266017 | 0.998075954 | -5.255140528 |
| **A_51_P519570** | 1600002K0 | 0.002766032 | -0.105896639 | 0.111428702 | 9.688210361 | 0.060839535 | 0.953274729 | 0.998075954 | -5.255141111 |
| **A_52_P406507** | Mus81 | -0.006534126 | -0.263318386 | 0.250250134 | 8.9604632 | -0.060817451 | 0.953291666 | 0.998075954 | -5.255142244 |
| **A_51_P327585** | Gstm4 | -0.004204514 | -0.169555669 | 0.16114664 | 8.409994602 | -0.06077399 | 0.953324997 | 0.998075954 | -5.255144473 |
| **A_55_P2115467** | Tmem63b | 0.00319926 | -0.12262559 | 0.12902411 | 9.682296489 | 0.060770417 | 0.953327737 | 0.998075954 | -5.255144656 |
| **A_55_P2341110** | Chn1os3 | 0.002481208 | -0.095170095 | 0.10013251 | 7.634859329 | 0.060728727 | 0.95335971 | 0.998075954 | -5.255146792 |
| **A_55_P1984962** | NA | -0.002825128 | -0.114065458 | 0.108415203 | 9.019730594 | -0.060699476 | 0.953382143 | 0.998075954 | -5.25514829 |
| **A_52_P570700** | Dpp8 | 0.003943868 | -0.151414203 | 0.15930194 | 9.144145361 | 0.060673317 | 0.953402205 | 0.998075954 | -5.255149629 |
| **A_55_P1992450** | Vdac2 | 0.003835622 | -0.147682227 | 0.155353472 | 15.66438466 | 0.060503596 | 0.95353237 | 0.998075954 | -5.255158302 |
| **A_55_P2006018** | Slc4a2 | 0.00311512 | -0.120012324 | 0.126242564 | 10.09713474 | 0.060468481 | 0.953559301 | 0.998075954 | -5.255160094 |
| **A_52_P64364** | Ikbkb | -0.002893149 | -0.117261508 | 0.11147521 | 8.458632412 | -0.060460823 | 0.953565174 | 0.998075954 | -5.255160484 |
| **A_55_P2079485** | Ptges3 | -0.006219632 | -0.25266359 | 0.240224326 | 12.2788926 | -0.060319209 | 0.953673784 | 0.998075954 | -5.255167697 |
| **A_55_P1998001** | DXBay18 | -0.003194011 | -0.129834728 | 0.123446706 | 7.714206228 | -0.060279856 | 0.953703965 | 0.998075954 | -5.255169699 |
| **A_55_P2143934** | NA | 0.003929932 | -0.152053962 | 0.159913825 | 8.165185985 | 0.06021634 | 0.953752679 | 0.998075954 | -5.255172927 |
| **A_51_P506297** | Cst9 | -0.025837872 | -1.051610877 | 0.999935132 | 8.895554596 | -0.060202512 | 0.953763285 | 0.998075954 | -5.255173629 |
| **A_66_P120736** | Bnip3 | -0.005596436 | -0.228528566 | 0.217335695 | 8.63194308 | -0.05999955 | 0.953918948 | 0.998075954 | -5.255183917 |
| **A_55_P1958653** | Arf2 | -0.003095424 | -0.126405039 | 0.12021419 | 12.0174126 | -0.059997396 | 0.9539206 | 0.998075954 | -5.255184026 |
| **A_52_P336768** | NA | -0.004996449 | -0.20410035 | 0.194107453 | 8.868186376 | -0.05997784 | 0.953935599 | 0.998075954 | -5.255185016 |
| **A_55_P2129291** | Dclre1c | 0.003023613 | -0.117533908 | 0.123581134 | 8.195244249 | 0.059943351 | 0.953962051 | 0.998075954 | -5.25518676 |
| **A_55_P2097259** | Zup1 | -0.004710339 | -0.192565749 | 0.183145071 | 9.460176234 | -0.059929078 | 0.953972998 | 0.998075954 | -5.255187482 |
| **A_51_P226053** | Sra1 | 0.003891768 | -0.151319192 | 0.159102728 | 12.67659303 | 0.059928535 | 0.953973414 | 0.998075954 | -5.255187509 |
| **A_55_P2383248** | 4930422N0 | 0.003172327 | -0.123508111 | 0.129852765 | 8.338391742 | 0.05985184 | 0.954032237 | 0.998075954 | -5.255191383 |
| **A_55_P1956443** | Ttc5 | 0.00357534 | -0.139356445 | 0.146507124 | 9.367845422 | 0.059785745 | 0.95408293 | 0.998075954 | -5.255194718 |
| **A_55_P2099627** | Lrrc51 | -0.002608312 | -0.107287527 | 0.102070903 | 7.836375587 | -0.059553627 | 0.95426096 | 0.998075954 | -5.255206401 |
| **A_52_P276955** | Epha3 | -0.005506173 | -0.226796719 | 0.215784374 | 8.322789755 | -0.059469747 | 0.954325296 | 0.998075954 | -5.255210611 |
| **A_66_P120558** | Pam16 | 0.003315152 | -0.130254753 | 0.136885057 | 13.15929942 | 0.059320389 | 0.954439853 | 0.998075954 | -5.255218094 |
| **A_55_P2017998** | NA | -0.004392415 | -0.181570749 | 0.172785918 | 8.623728127 | -0.059251849 | 0.954492423 | 0.998075954 | -5.255221521 |
| **A_55_P2118183** | Tnks2 | -0.003129222 | -0.129667782 | 0.123409338 | 8.733034356 | -0.059104783 | 0.954605223 | 0.998075954 | -5.255228862 |
| **A_55_P2044217** | 9930021J0 | -0.002903697 | -0.120341837 | 0.114534443 | 11.53336217 | -0.059095071 | 0.954612673 | 0.998075954 | -5.255229346 |
| **A_55_P2401403** | NA | -0.006178807 | -0.256081236 | 0.243723622 | 9.717480715 | -0.059093984 | 0.954613506 | 0.998075954 | -5.2552294 |

| | | | | | | | | | |
|---|---|---|---|---|---|---|---|---|---|
| **A_51_P184321** | Aimp1 | 0.003290735 | -0.129865741 | 0.136447211 | 12.75510913 | 0.059066302 | 0.954634738 | 0.998075954 | -5.25523078 |
| **A_55_P2081530** | Sh3bgrl3 | -0.002924305 | -0.121417469 | 0.115568859 | 12.87811702 | -0.058984587 | 0.954697415 | 0.998075954 | -5.255234848 |
| **A_51_P178872** | Ttc9c | 0.005502108 | -0.217537171 | 0.228541387 | 8.787145374 | 0.058959919 | 0.954716336 | 0.998075954 | -5.255236075 |
| **A_51_P206225** | Uroc1 | 0.002536627 | -0.100347727 | 0.105420981 | 7.684908497 | 0.058927273 | 0.954741376 | 0.998075954 | -5.255237698 |
| **A_55_P2143324** | Dyrk1b | -0.003938583 | -0.163923265 | 0.156046099 | 8.785911755 | -0.058839732 | 0.954808522 | 0.998075954 | -5.255242046 |
| **A_52_P378287** | Lca5 | 0.005708074 | -0.22659293 | 0.238009078 | 8.547135608 | 0.058728324 | 0.954893975 | 0.998075954 | -5.25524757 |
| **A_55_P2076927** | Ints10 | -0.003048861 | -0.127158781 | 0.12106106 | 9.578927757 | -0.058713804 | 0.954905112 | 0.998075954 | -5.255248289 |
| **A_55_P2125714** | Atp2b4 | -0.003495725 | -0.146046649 | 0.1390552 | 8.205814933 | -0.058610622 | 0.954984256 | 0.998075954 | -5.255253395 |
| **A_55_P1993109** | Taf6 | -0.003441904 | -0.14380399 | 0.136920181 | 12.17406181 | -0.058608165 | 0.954986141 | 0.998075954 | -5.255253516 |
| **A_55_P2452259** | Gls | -0.005282655 | -0.220779299 | 0.210213988 | 10.13976941 | -0.05858965 | 0.955000343 | 0.998075954 | -5.255254431 |
| **A_55_P2051840** | NA | 0.002746578 | -0.109328206 | 0.114821361 | 9.556119315 | 0.05857241 | 0.955013566 | 0.998075954 | -5.255255283 |
| **A_55_P2042879** | Mbtd1 | 0.002596063 | -0.103439486 | 0.108631612 | 8.089207734 | 0.058515763 | 0.955057017 | 0.998075954 | -5.25525808 |
| **A_51_P497985** | C2 | -0.011526965 | -0.482546718 | 0.459492788 | 8.793697351 | -0.058490453 | 0.955076431 | 0.998075954 | -5.255259329 |
| **A_55_P1991416** | NA | -0.002417845 | -0.101311867 | 0.096476177 | 7.821892017 | -0.058434261 | 0.955119533 | 0.998075954 | -5.2552621 |
| **A_55_P1989405** | Elf1 | 0.003857488 | -0.153995724 | 0.161710701 | 9.347448227 | 0.058406384 | 0.955140916 | 0.998075954 | -5.255263474 |
| **A_55_P2106265** | Mbnl2 | -0.003862813 | -0.162105955 | 0.154380329 | 9.068162388 | -0.058342888 | 0.95518962 | 0.998075954 | -5.2552666 |
| **A_55_P2089677** | Synrg | 0.00520528 | -0.208064172 | 0.218474732 | 9.852831244 | 0.058334378 | 0.955196148 | 0.998075954 | -5.255267019 |
| **A_51_P321331** | Sptlc2 | -0.003542269 | -0.148907623 | 0.141823084 | 8.270612517 | -0.058241127 | 0.955267676 | 0.998075954 | -5.255271604 |
| **A_51_P511176** | Triap1 | -0.00414304 | -0.174225615 | 0.165939535 | 10.0166012 | -0.058219494 | 0.95528427 | 0.998075954 | -5.255272666 |
| **A_55_P2140546** | Slc25a51 | -0.005752218 | -0.261530494 | 0.250026057 | 11.06949423 | -0.059142181 | 0.955295257 | 0.998075954 | -5.115093563 |
| **A_51_P397673** | Pcsk9 | 0.002596948 | -0.104422147 | 0.109616043 | 7.730033783 | 0.057997744 | 0.955454366 | 0.998075954 | -5.255283535 |
| **A_55_P2033105** | Cdcp1 | 0.002948379 | -0.118584339 | 0.124481097 | 7.761837245 | 0.05798281 | 0.955465821 | 0.998075954 | -5.255284266 |
| **A_55_P1956160** | Gm8909 | 0.002309162 | -0.092912176 | 0.0975305 | 7.682270897 | 0.057960115 | 0.95548323 | 0.998075954 | -5.255285375 |
| **A_52_P50287** | Bend7 | -0.002556352 | -0.108167824 | 0.10305512 | 7.870340846 | -0.057852045 | 0.955566128 | 0.998075954 | -5.255290654 |
| **A_55_P2302383** | 3300002A1 | 0.005190304 | -0.209345837 | 0.219726445 | 8.327183626 | 0.057823115 | 0.95558832 | 0.998075954 | -5.255292065 |
| **A_52_P375638** | Gtpbp1 | 0.004100841 | -0.165439804 | 0.173641486 | 9.851084587 | 0.057810704 | 0.955597839 | 0.998075954 | -5.255292671 |
| **A_55_P2177273** | Kcna6 | -0.003026018 | -0.128279762 | 0.122227726 | 8.701652246 | -0.057741744 | 0.955650738 | 0.998075954 | -5.255296031 |
| **A_51_P450924** | Pole4 | 0.00288009 | -0.116402788 | 0.122162968 | 10.42864704 | 0.057708146 | 0.95567651 | 0.998075954 | -5.255297667 |
| **A_55_P2112225** | NA | 0.002456644 | -0.099530899 | 0.104444188 | 7.849706786 | 0.05757107 | 0.955781659 | 0.998075954 | -5.255304332 |
| **A_51_P274504** | Tex264 | -0.004188579 | -0.178146082 | 0.169768924 | 11.26539214 | -0.057548325 | 0.955799106 | 0.998075954 | -5.255305436 |
| **A_55_P1990314** | Elavl4 | -0.004031868 | -0.171490308 | 0.163426573 | 8.459647678 | -0.0575451 | 0.955801581 | 0.998075954 | -5.255305593 |
| **A_52_P304947** | Cenpn | 0.004405812 | -0.178741756 | 0.187553379 | 8.704901902 | 0.057495506 | 0.955839623 | 0.998075954 | -5.255307999 |
| **A_52_P476560** | Tpcn1 | -0.003867366 | -0.164834221 | 0.157099489 | 10.87774516 | -0.057423278 | 0.955895029 | 0.998075954 | -5.2553115 |
| **A_51_P146560** | Msln | 0.026376703 | -1.072402882 | 1.125156287 | 11.54360973 | 0.057374518 | 0.955932433 | 0.998075954 | -5.25531386 |
| **A_51_P138760** | Abcd4 | -0.004613011 | -0.196930158 | 0.187704136 | 9.808726677 | -0.057329169 | 0.95596722 | 0.998075954 | -5.255316054 |
| **A_55_P2018919** | Spns1 | 0.002553821 | -0.104150736 | 0.109258378 | 13.32418567 | 0.057202714 | 0.956064224 | 0.998075954 | -5.255322163 |
| **A_55_P2249849** | Sema6a | -0.005572477 | -0.23876427 | 0.227619316 | 9.017161242 | -0.057114206 | 0.95613212 | 0.998075954 | -5.25532643 |
| **A_52_P92557** | Kdm2a | 0.003766879 | -0.153879954 | 0.161413712 | 9.830103811 | 0.057109134 | 0.956136011 | 0.998075954 | -5.255326674 |
| **A_55_P2072656** | Ckmt1 | 0.005371508 | -0.219457051 | 0.230200068 | 10.42594702 | 0.057102341 | 0.956141222 | 0.998075954 | -5.255327001 |
| **A_55_P2198568** | 4930506A1 | -0.002624771 | -0.112607669 | 0.107358126 | 8.290843267 | -0.057039469 | 0.956189452 | 0.998075954 | -5.255330028 |
| **A_55_P2053497** | Poli | -0.00407133 | -0.174742667 | 0.166600007 | 9.617912426 | -0.057014436 | 0.956208655 | 0.998075954 | -5.255331232 |
| **A_55_P2175545** | Gys2 | 0.002830525 | -0.115837249 | 0.121498299 | 7.704755806 | 0.057008989 | 0.956212833 | 0.998075954 | -5.255331494 |
| **A_55_P2123566** | NA | -0.010729994 | -0.460856075 | 0.439396087 | 9.212115917 | -0.056973701 | 0.956239903 | 0.998075954 | -5.255333191 |
| **A_55_P2419021** | 9330188P0 | -0.005150543 | -0.221468908 | 0.211167822 | 8.609684888 | -0.056907403 | 0.956290762 | 0.998075954 | -5.255336375 |
| **A_55_P1960291** | Tjp3 | -0.00319351 | -0.137488083 | 0.131101062 | 8.321873384 | -0.056835413 | 0.956345988 | 0.998075954 | -5.255339828 |
| **A_51_P343613** | Ehd2 | -0.006110737 | -0.263145532 | 0.250924058 | 9.14740066 | -0.056821256 | 0.956356848 | 0.998075954 | -5.255340507 |
| **A_51_P346165** | Agpat4 | 0.005315165 | -0.218331488 | 0.228961818 | 11.23836904 | 0.05680198 | 0.956371635 | 0.998075954 | -5.255341431 |
| **A_52_P579933** | Slc16a6 | -0.005179993 | -0.223208518 | 0.212848532 | 8.126166417 | -0.056783868 | 0.956385529 | 0.998075954 | -5.255342298 |

| | | | | | | | | | |
|---|---|---|---|---|---|---|---|---|---|
| **A_55_P2037657** | NA | -0.002349719 | -0.10163688 | 0.096937441 | 8.112979025 | -0.056562945 | 0.956555007 | 0.998075954 | -5.25535286 |
| **A_55_P2014706** | Naip5 | -0.006500128 | -0.281192933 | 0.268192677 | 13.78542976 | -0.056556658 | 0.95655983 | 0.998075954 | -5.25535316 |
| **A_65_P12359** | Pin1rt1 | -0.003702185 | -0.160356066 | 0.152951695 | 11.56616878 | -0.05648409 | 0.9566155 | 0.998075954 | -5.255356619 |
| **A_55_P2073814** | Gpatch8 | -0.003015217 | -0.130614301 | 0.124583868 | 8.498831306 | -0.0564781 | 0.956620095 | 0.998075954 | -5.255356905 |
| **A_51_P421538** | Slc9b2 | 0.002443133 | -0.100967145 | 0.10585341 | 7.729795133 | 0.056466713 | 0.95662883 | 0.998075954 | -5.255357447 |
| **A_52_P110052** | Ackr1 | -0.003135764 | -0.135890364 | 0.129618836 | 7.772804722 | -0.056455065 | 0.956637766 | 0.998075954 | -5.255358002 |
| **A_51_P365859** | Sowaha | 0.002415346 | -0.099909525 | 0.104740217 | 7.665812981 | 0.056416654 | 0.956667233 | 0.998075954 | -5.25535983 |
| **A_51_P223946** | Pds5b | 0.004986467 | -0.206854353 | 0.216827286 | 9.00409659 | 0.05625905 | 0.95678814 | 0.998075954 | -5.25536732 |
| **A_55_P2095108** | A430033K( | -0.005228444 | -0.227723933 | 0.217267045 | 8.772913342 | -0.056164305 | 0.956860824 | 0.998075954 | -5.255371813 |
| **A_52_P1130990** | Trim67 | 0.003018338 | -0.125551111 | 0.131587787 | 7.75768399 | 0.056109869 | 0.956902585 | 0.998075954 | -5.25537439 |
| **A_51_P507622** | Itpkc | -0.003580616 | -0.156111185 | 0.148949953 | 8.550410713 | -0.056106084 | 0.956905489 | 0.998075954 | -5.25537457 |
| **A_55_P2104889** | Gm11515 | 0.002277088 | -0.094947877 | 0.099502053 | 7.831478847 | 0.055977205 | 0.957004361 | 0.998075954 | -5.255380662 |
| **A_51_P236175** | Rit1 | -0.005980966 | -0.261573326 | 0.249611395 | 12.8317822 | -0.055928423 | 0.957041784 | 0.998075954 | -5.255382964 |
| **A_52_P122673** | Cep41 | 0.003509456 | -0.146581727 | 0.153600638 | 8.518202321 | 0.055884803 | 0.957075249 | 0.998075954 | -5.255385021 |
| **A_55_P1967885** | Nr3c1 | 0.002750969 | -0.11524788 | 0.120749817 | 12.11619721 | 0.055720754 | 0.957201104 | 0.998075954 | -5.255392744 |
| **A_55_P2166501** | Cd44 | 0.003201303 | -0.134248976 | 0.140651582 | 8.485130022 | 0.05566604 | 0.95724308 | 0.998075954 | -5.255395314 |
| **A_51_P215438** | Prodh | -0.003323785 | -0.146445218 | 0.139797647 | 9.163334306 | -0.055505679 | 0.957366107 | 0.998075954 | -5.255402833 |
| **A_55_P2025851** | Heatr3 | -0.006511371 | -0.287021316 | 0.273998574 | 11.46424013 | -0.055479599 | 0.957386115 | 0.998075954 | -5.255404054 |
| **A_52_P582112** | Hcfc1r1 | -0.003086296 | -0.136150746 | 0.129978155 | 13.44097157 | -0.05543507 | 0.957420278 | 0.998075954 | -5.255406137 |
| **A_55_P1964233** | Tnks | 0.003392162 | -0.143060527 | 0.149844852 | 8.65491607 | 0.055359021 | 0.957478622 | 0.998075954 | -5.255409691 |
| **A_51_P501656** | Poglut3 | 0.002345172 | -0.099145846 | 0.103836189 | 9.115006616 | 0.055227597 | 0.957579451 | 0.998075954 | -5.255415821 |
| **A_55_P2110595** | Slc35a2 | 0.002472913 | -0.104560558 | 0.109506384 | 8.47948271 | 0.055220258 | 0.957585081 | 0.998075954 | -5.255416163 |
| **A_55_P1962174** | Rab8a | -0.004002949 | -0.177442968 | 0.16943707 | 9.550153091 | -0.055161983 | 0.95762979 | 0.998075954 | -5.255418876 |
| **A_55_P1981845** | NA | 0.004888869 | -0.207043296 | 0.216821034 | 9.386370123 | 0.055134142 | 0.95765115 | 0.998075954 | -5.255420171 |
| **A_51_P242356** | Fam114a2 | -0.00332881 | -0.147724832 | 0.141067212 | 11.86842077 | -0.055098899 | 0.957678189 | 0.998075954 | -5.255421809 |
| **A_55_P2105180** | Bhmt | -0.003939612 | -0.175531195 | 0.167651971 | 7.799265162 | -0.054874001 | 0.957850735 | 0.998075954 | -5.255432241 |
| **A_55_P1987196** | Gm3448 | 0.003136294 | -0.133566556 | 0.139839143 | 8.029547258 | 0.054833795 | 0.957881582 | 0.998075954 | -5.255434101 |
| **A_52_P476075** | Clspn | -0.003689582 | -0.165074898 | 0.157695734 | 8.306455965 | -0.054641458 | 0.958029149 | 0.998075954 | -5.255442982 |
| **A_55_P2177426** | Abcb8 | 0.004560671 | -0.194994862 | 0.204116205 | 8.61816316 | 0.054622821 | 0.958043448 | 0.998075954 | -5.255443841 |
| **A_52_P546513** | NA | 0.002934045 | -0.125471364 | 0.131339453 | 7.905692882 | 0.054612559 | 0.958051321 | 0.998075954 | -5.255444314 |
| **A_55_P2170009** | Lactb2 | -0.004690512 | -0.210382697 | 0.201001673 | 9.598544672 | -0.054501895 | 0.958136227 | 0.998075954 | -5.255449407 |
| **A_52_P276607** | Ptp4a2 | -0.002930243 | -0.131539228 | 0.125678743 | 13.76620123 | -0.054455456 | 0.958171857 | 0.998075954 | -5.255451541 |
| **A_55_P2156588** | Ifnk | 0.002230853 | -0.095847023 | 0.100308728 | 7.692056816 | 0.0543637 | 0.958242256 | 0.998075954 | -5.255455753 |
| **A_51_P418116** | Tmem119 | -0.00725136 | -0.327699132 | 0.313196411 | 9.481191235 | -0.05408432 | 0.958456612 | 0.998075954 | -5.255468532 |
| **A_52_P239726** | Larp4 | 0.003329793 | -0.143833809 | 0.150493394 | 9.532530397 | 0.054078663 | 0.958460952 | 0.998075954 | -5.25546879 |
| **A_55_P1979708** | Ubap1 | 0.003183498 | -0.13758555 | 0.143952547 | 9.98257143 | 0.054051355 | 0.958481905 | 0.998075954 | -5.255470035 |
| **A_51_P336385** | Dido1 | 0.003526014 | -0.152628638 | 0.159680666 | 10.96518719 | 0.053968243 | 0.958545674 | 0.998075954 | -5.255473822 |
| **A_55_P2156973** | Aar2 | 0.00381306 | -0.165630546 | 0.173256666 | 9.953027648 | 0.053784556 | 0.958686612 | 0.998075954 | -5.25548217 |
| **A_51_P470448** | Setx | 0.00403884 | -0.17581527 | 0.183892951 | 11.65333468 | 0.053671713 | 0.958773193 | 0.998075954 | -5.255487285 |
| **A_66_P111090** | Sctr | -0.002219 | -0.101083879 | 0.09664588 | 7.678639159 | -0.053644392 | 0.958794157 | 0.998075954 | -5.255488521 |
| **A_55_P2068228** | NA | -0.006504706 | -0.296866828 | 0.283857416 | 12.48606571 | -0.053542278 | 0.958872507 | 0.998075954 | -5.255493138 |
| **A_52_P71544** | Slc25a26 | -0.002763956 | -0.126331757 | 0.120803845 | 9.975223139 | -0.053460734 | 0.958935075 | 0.998075954 | -5.255496818 |
| **A_55_P2088202** | Hcn2 | -0.010150555 | -0.46440989 | 0.444108781 | 13.4382014 | -0.053406614 | 0.958976601 | 0.998075954 | -5.255499258 |
| **A_66_P137219** | Elane | -0.003457247 | -0.158399043 | 0.151484548 | 8.667716126 | -0.05332993 | 0.95903544 | 0.998075954 | -5.25550271 |
| **A_51_P502764** | Hdac10 | -0.00359961 | -0.165126009 | 0.157926789 | 9.378767194 | -0.053262446 | 0.959087221 | 0.998075954 | -5.255505744 |
| **A_55_P2185652** | Cln3 | 0.003252661 | -0.142879339 | 0.149384662 | 8.723691813 | 0.053198901 | 0.959135979 | 0.998075954 | -5.255508598 |
| **A_55_P2003648** | Bag5 | 0.004882459 | -0.214677293 | 0.224442212 | 10.75210075 | 0.053148993 | 0.959174273 | 0.998075954 | -5.255510836 |
| **A_52_P636151** | Golph3 | 0.004243987 | -0.186619016 | 0.19510699 | 11.31450413 | 0.053144882 | 0.959177428 | 0.998075954 | -5.255511021 |

| | | | | | | | | | |
|---|---|---|---|---|---|---|---|---|---|
| **A_55_P2120714** | NA | -0.003678826 | -0.169988388 | 0.162630736 | 8.180845539 | -0.052869006 | 0.959389111 | 0.998075954 | -5.255523357 |
| **A_52_P219372** | Tmem177 | -0.004646531 | -0.214790767 | 0.205497704 | 8.464905097 | -0.05284702 | 0.959405981 | 0.998075954 | -5.255524337 |
| **A_55_P2179309** | P2ry10b | -0.002937861 | -0.135840824 | 0.129965102 | 8.121700915 | -0.052833055 | 0.959416696 | 0.998075954 | -5.255524959 |
| **A_52_P532456** | Plagl1 | 0.009226075 | -0.408334459 | 0.426786609 | 15.4245342 | 0.052808846 | 0.959435273 | 0.998075954 | -5.255526038 |
| **A_51_P306527** | Ywhah | 0.004727735 | -0.209703407 | 0.219158876 | 14.74492292 | 0.052695601 | 0.959522168 | 0.998075954 | -5.255531078 |
| **A_55_P1973573** | Spn | 0.002166158 | -0.096252772 | 0.100585087 | 7.664185354 | 0.052604222 | 0.959592286 | 0.998075954 | -5.255535136 |
| **A_51_P133052** | Glyr1 | -0.003471042 | -0.161214631 | 0.154272547 | 10.92072065 | -0.052591711 | 0.959601886 | 0.998075954 | -5.255535691 |
| **A_51_P430900** | Dusp1 | -0.002898798 | -0.134662363 | 0.128864766 | 9.647425261 | -0.052581353 | 0.959609834 | 0.998075954 | -5.255536151 |
| **A_65_P19089** | Esrrg | 0.006286995 | -0.279695075 | 0.292269064 | 10.1484788 | 0.052542827 | 0.959639397 | 0.998075954 | -5.255537859 |
| **A_66_P103418** | Tmc8 | -0.002583964 | -0.120224479 | 0.115056552 | 8.242541489 | -0.052497496 | 0.959674181 | 0.998075954 | -5.255539868 |
| **A_51_P146633** | NA | -0.00689418 | -0.320904124 | 0.307115764 | 10.78889844 | -0.052474504 | 0.959691823 | 0.998075954 | -5.255540886 |
| **A_52_P140005** | Nipal1 | -0.002109362 | -0.09842125 | 0.094202527 | 7.649435145 | -0.052345611 | 0.959790728 | 0.998075954 | -5.255546584 |
| **A_55_P1962479** | Gpbp1l1 | 0.003828542 | -0.170991104 | 0.178648188 | 9.719049278 | 0.052342239 | 0.959793316 | 0.998075954 | -5.255546733 |
| **A_51_P469568** | Cldn13 | -0.00232397 | -0.108467261 | 0.103819321 | 7.670300854 | -0.052329565 | 0.959803041 | 0.998075954 | -5.255547293 |
| **A_55_P2184905** | Rpap2 | -0.002616495 | -0.122239929 | 0.117006939 | 8.531426541 | -0.052277252 | 0.959843183 | 0.998075954 | -5.255549601 |
| **A_55_P2103452** | Npy1r | 0.014096686 | -0.630787323 | 0.658980696 | 8.551798775 | 0.052244979 | 0.959867947 | 0.998075954 | -5.255551024 |
| **A_52_P67983** | Lcmt2 | -0.003475007 | -0.162570117 | 0.155620102 | 10.12975207 | -0.052204511 | 0.959899001 | 0.998075954 | -5.255552807 |
| **A_52_P154833** | Ankrd49 | -0.00423199 | -0.198087982 | 0.189624002 | 9.676835267 | -0.052176452 | 0.959920532 | 0.998075954 | -5.255554042 |
| **A_55_P2163009** | Mapkapk5 | 0.002221458 | -0.099538389 | 0.103981306 | 11.24629811 | 0.052176012 | 0.95992087 | 0.998075954 | -5.255554062 |
| **A_55_P2243367** | NA | -0.004031423 | -0.188756226 | 0.180693379 | 8.678055092 | -0.052160568 | 0.959932721 | 0.998075954 | -5.255554741 |
| **A_51_P443702** | Mrps34 | -0.00310571 | -0.145619429 | 0.13940801 | 13.38739551 | -0.052085074 | 0.959990651 | 0.998075954 | -5.255558061 |
| **A_52_P222624** | Hspb2 | 0.004417862 | -0.205344651 | 0.214180376 | 13.98896527 | 0.052318038 | 0.960075465 | 0.998075954 | -5.176248359 |
| **A_55_P2137932** | NA | -0.004222822 | -0.198592514 | 0.190146871 | 8.037763595 | -0.051925815 | 0.96011286 | 0.998075954 | -5.255565048 |
| **A_55_P2097372** | Tctex1d4 | 0.00216713 | -0.097740285 | 0.102074544 | 7.676345271 | 0.051843737 | 0.960175844 | 0.998075954 | -5.25556864 |
| **A_55_P1961129** | Egfem1 | 0.002159371 | -0.097430448 | 0.10174919 | 7.684647368 | 0.051822874 | 0.960191853 | 0.998075954 | -5.255569553 |
| **A_66_P125035** | Duxbl2 | 0.003947403 | -0.178199583 | 0.18609439 | 8.362309937 | 0.05179629 | 0.960212253 | 0.998075954 | -5.255570714 |
| **A_55_P2062891** | Atf2 | 0.005459788 | -0.246474348 | 0.257393923 | 10.77868601 | 0.051796196 | 0.960212325 | 0.998075954 | -5.255570719 |
| **A_51_P364485** | Tnfaip2 | 0.002692718 | -0.121658524 | 0.127043961 | 8.516483541 | 0.051754716 | 0.960244155 | 0.998075954 | -5.25557253 |
| **A_55_P2143159** | Men1 | 0.004966796 | -0.224479383 | 0.234412974 | 11.20444865 | 0.051737402 | 0.960257442 | 0.998075954 | -5.255573286 |
| **A_52_P506271** | Cmtm4 | -0.002999138 | -0.141760577 | 0.135762301 | 7.893449755 | -0.051657905 | 0.960318446 | 0.998075954 | -5.255576753 |
| **A_51_P181746** | Zfyve1 | 0.003581945 | -0.162241198 | 0.169405089 | 8.589524567 | 0.051627717 | 0.960341611 | 0.998075954 | -5.255578068 |
| **A_52_P302433** | Plau | 0.002130232 | -0.096734362 | 0.100994826 | 7.85329894 | 0.051498579 | 0.960440709 | 0.998092715 | -5.255583686 |
| **A_51_P306710** | Cldn15 | 0.008856662 | -0.402394163 | 0.420107487 | 8.658105245 | 0.051472163 | 0.96046098 | 0.998092715 | -5.255584833 |
| **A_55_P2298319** | NA | 0.002264099 | -0.10313105 | 0.107659249 | 7.727281659 | 0.051343326 | 0.960559848 | 0.998119697 | -5.25559042 |
| **A_55_P2144305** | Gm3211 | 0.002720106 | -0.124030861 | 0.129471073 | 8.317854307 | 0.051291298 | 0.960599774 | 0.998119697 | -5.255592673 |
| **A_55_P2162472** | NA | -0.006876222 | -0.327635642 | 0.313883198 | 9.222583247 | -0.051236517 | 0.960641812 | 0.998119697 | -5.255595042 |
| **A_55_P1967291** | Ncaph | 0.003880449 | -0.178122627 | 0.185883525 | 11.88127344 | 0.050957998 | 0.960855548 | 0.998170723 | -5.255607048 |
| **A_55_P1963767** | Npr2 | 0.003342756 | -0.153924101 | 0.160609612 | 9.943001285 | 0.050801503 | 0.960975644 | 0.998170723 | -5.255613765 |
| **A_52_P258338** | Mob3b | 0.002211 | -0.10195801 | 0.10638001 | 7.742581796 | 0.050729359 | 0.961031008 | 0.998170723 | -5.255616855 |
| **A_51_P172842** | Med19 | 0.003608776 | -0.166535321 | 0.173752873 | 10.11713928 | 0.050693486 | 0.961058537 | 0.998170723 | -5.25561839 |
| **A_55_P2016014** | Cpsf4l | -0.002026213 | -0.097593281 | 0.093540855 | 7.635889227 | -0.050674097 | 0.961073417 | 0.998170723 | -5.255619219 |
| **A_55_P2070316** | Ankrd36 | -0.002293595 | -0.110476682 | 0.105889491 | 7.69202281 | -0.050671825 | 0.96107516 | 0.998170723 | -5.255619316 |
| **A_52_P241508** | Zbtb46 | -0.002139489 | -0.103248923 | 0.098969944 | 7.913137317 | -0.05057403 | 0.961150211 | 0.998170723 | -5.255623492 |
| **A_52_P466701** | Fuom | -0.003576806 | -0.173010641 | 0.165857028 | 11.58348437 | -0.050455021 | 0.961241541 | 0.998170723 | -5.255628564 |
| **A_55_P1980292** | Purb | 0.007037537 | -0.327193868 | 0.341268942 | 10.79956483 | 0.050324859 | 0.961341432 | 0.998170723 | -5.255634097 |
| **A_51_P281380** | Tspan5 | 0.002705342 | -0.125779583 | 0.131190267 | 9.632559371 | 0.050324462 | 0.961341737 | 0.998170723 | -5.255634114 |
| **A_55_P2017714** | Kat5 | 0.002717199 | -0.126332674 | 0.131767071 | 11.64374747 | 0.050323742 | 0.961342289 | 0.998170723 | -5.255634145 |
| **A_51_P334289** | Zfp628 | -0.002577727 | -0.1251506 | 0.119995147 | 9.249470326 | -0.050263373 | 0.961388618 | 0.998170723 | -5.255636706 |

| | | | | | | | | | |
|---|---|---|---|---|---|---|---|---|---|
| **A_55_P2126259** | Slc35e2 | -0.003791064 | -0.184176181 | 0.176594053 | 8.934676038 | -0.05023074 | 0.961413662 | 0.998170723 | -5.255638089 |
| **A_55_P1973783** | Tex9 | -0.002738242 | -0.133028375 | 0.127551891 | 8.729129221 | -0.050230726 | 0.961413673 | 0.998170723 | -5.25563809 |
| **A_55_P2000182** | Slc5a6 | 0.002683459 | -0.125435323 | 0.130802241 | 9.910169352 | 0.050060048 | 0.961544659 | 0.998206548 | -5.25564531 |
| **A_51_P341108** | Spint1 | -0.002197662 | -0.107141134 | 0.10274581 | 7.684136936 | -0.050051221 | 0.961551433 | 0.998206548 | -5.255645683 |
| **A_55_P2065586** | Gm15417 | 0.003473126 | -0.162771806 | 0.169718057 | 8.710103152 | 0.04993226 | 0.96164273 | 0.998229148 | -5.2556507 |
| **A_55_P2175190** | Cep250 | -0.00356501 | -0.174358424 | 0.167228404 | 8.575744512 | -0.049888309 | 0.961676459 | 0.998229148 | -5.255652551 |
| **A_55_P2054854** | Art4 | 0.004097624 | -0.192764081 | 0.200959328 | 8.511494123 | 0.049748495 | 0.961783761 | 0.998286644 | -5.255658427 |
| **A_51_P479132** | Zcchc3 | -0.00304439 | -0.149502668 | 0.143413888 | 10.15264582 | -0.049681584 | 0.961835112 | 0.998286644 | -5.255661233 |
| **A_55_P2153580** | Zfp655 | 0.002480654 | -0.117636414 | 0.122597723 | 8.119695572 | 0.049359491 | 0.962082309 | 0.998432246 | -5.25567469 |
| **A_55_P2043009** | Psmg2 | -0.003664797 | -0.181391048 | 0.174061454 | 11.53698366 | -0.049284176 | 0.962140112 | 0.998432246 | -5.255677823 |
| **A_51_P435333** | Dctn2 | -0.005378423 | -0.266621337 | 0.255864491 | 11.09173238 | -0.04920617 | 0.96219998 | 0.998432246 | -5.255681064 |
| **A_55_P2090394** | NA | -0.00330496 | -0.164374483 | 0.157764562 | 12.54941435 | -0.049041303 | 0.962326513 | 0.998432246 | -5.255687897 |
| **A_51_P340829** | AA986860 | 0.002106163 | -0.100567005 | 0.104779331 | 8.093891712 | 0.049027998 | 0.962336725 | 0.998432246 | -5.255688447 |
| **A_55_P1974567** | Plch2 | -0.002024976 | -0.100894197 | 0.096844245 | 7.708279824 | -0.048951713 | 0.962395273 | 0.998432246 | -5.2556916 |
| **A_55_P2077114** | NA | -0.003524811 | -0.175642182 | 0.168592561 | 14.70130149 | -0.048946341 | 0.962399396 | 0.998432246 | -5.255691822 |
| **A_55_P2029846** | Fam214a | -0.008035856 | -0.400694741 | 0.384623029 | 10.07908156 | -0.048913174 | 0.962424852 | 0.998432246 | -5.255693191 |
| **A_55_P2012849** | Dock10 | 0.002274429 | -0.1089071 | 0.113455958 | 8.006454154 | 0.048893248 | 0.962440145 | 0.998432246 | -5.255694013 |
| **A_52_P624434** | Clec1a | -0.005390561 | -0.269901008 | 0.259119885 | 8.535268076 | -0.048708001 | 0.962582322 | 0.998458633 | -5.25570164 |
| **A_51_P286792** | Zfp275 | -0.00317338 | -0.158997655 | 0.152650896 | 9.762824093 | -0.048673888 | 0.962608504 | 0.998458633 | -5.255703041 |
| **A_55_P2034550** | 1700013F0 | -0.001992689 | -0.099998464 | 0.096013085 | 7.692477249 | -0.048595609 | 0.962668584 | 0.998458633 | -5.255706253 |
| **A_51_P401683** | Tm6sf1 | -0.005193347 | -0.260874764 | 0.25048807 | 10.42006458 | -0.048546428 | 0.962706331 | 0.998458633 | -5.255708268 |
| **A_52_P58524** | Neu1 | -0.003138983 | -0.157855732 | 0.151577766 | 10.73104843 | -0.048490961 | 0.962748903 | 0.998458633 | -5.255710539 |
| **A_51_P273684** | Gbe1 | 0.004375575 | -0.211445084 | 0.220196234 | 12.366597 | 0.04845641 | 0.962775421 | 0.998458633 | -5.255711952 |
| **A_51_P475298** | Zfp110 | 0.002535213 | -0.123006209 | 0.128076634 | 9.322830461 | 0.048265467 | 0.962921974 | 0.99847104 | -5.255719742 |
| **A_51_P430423** | Ada | 0.00783448 | -0.380289994 | 0.395958955 | 8.786598993 | 0.048244555 | 0.962938024 | 0.99847104 | -5.255720594 |
| **A_55_P2061565** | Zfp229 | 0.00235733 | -0.114439617 | 0.119154278 | 8.113216367 | 0.048238977 | 0.962942306 | 0.99847104 | -5.255720821 |
| **A_55_P2114088** | Zfp385c | -0.002204125 | -0.112228771 | 0.107820522 | 7.858529107 | -0.047880133 | 0.96321773 | 0.998471826 | -5.25573537 |
| **A_51_P496400** | Med9 | -0.00343819 | -0.175965789 | 0.169089409 | 11.35178754 | -0.04762998 | 0.963409734 | 0.998471826 | -5.255745449 |
| **A_51_P201945** | Cfap97 | 0.002067757 | -0.101873416 | 0.10600893 | 9.201138652 | 0.047546768 | 0.963473604 | 0.998471826 | -5.25574879 |
| **A_55_P2021455** | NA | 0.005812679 | -0.286488473 | 0.298113831 | 14.17237765 | 0.047528589 | 0.963487557 | 0.998471826 | -5.255749519 |
| **A_55_P1972541** | Zfp960 | -0.003808228 | -0.195528318 | 0.187911862 | 10.16154986 | -0.047474941 | 0.963528735 | 0.998471826 | -5.255751669 |
| **A_55_P2043028** | Rexo5 | -0.005628085 | -0.289037676 | 0.277781505 | 8.991827853 | -0.047463008 | 0.963537894 | 0.998471826 | -5.255752147 |
| **A_52_P134556** | Cd2bp2 | -0.002135847 | -0.110167571 | 0.105895877 | 10.12887597 | -0.04725284 | 0.963699212 | 0.998471826 | -5.255760544 |
| **A_55_P2078123** | Rora | -0.002268086 | -0.116997285 | 0.112461112 | 8.491860187 | -0.047249226 | 0.963701985 | 0.998471826 | -5.255760688 |
| **A_51_P332676** | Top1mt | 0.00309968 | -0.15371169 | 0.159911051 | 9.419510723 | 0.047244198 | 0.963705844 | 0.998471826 | -5.255760888 |
| **A_51_P401263** | Eme1 | -0.002481376 | -0.128200538 | 0.123237785 | 8.726062934 | -0.047173735 | 0.96375993 | 0.998471826 | -5.255763695 |
| **A_66_P138289** | Rdh5 | -0.003852285 | -0.199130958 | 0.191426389 | 8.739373557 | -0.047149022 | 0.963778899 | 0.998471826 | -5.255764678 |
| **A_52_P358388** | Ppp4r2 | -0.004585883 | -0.237800407 | 0.228628641 | 8.988081124 | -0.046997687 | 0.96389506 | 0.998471826 | -5.255770689 |
| **A_66_P106968** | NA | -0.007621407 | -0.395552837 | 0.380310023 | 12.93930455 | -0.046955808 | 0.963927205 | 0.998471826 | -5.255772348 |
| **A_51_P325374** | Tent2 | 0.003171849 | -0.158278097 | 0.164621796 | 10.51348338 | 0.046955206 | 0.963927667 | 0.998471826 | -5.255772372 |
| **A_55_P2139256** | Rps6ka1 | -0.003696137 | -0.192563697 | 0.185171423 | 11.06734148 | -0.046773498 | 0.964067143 | 0.998471826 | -5.255779557 |
| **A_55_P2004469** | NA | -0.002156801 | -0.112698439 | 0.108384837 | 15.4222198 | -0.046632999 | 0.964174989 | 0.998471826 | -5.255785093 |
| **A_52_P574993** | Pappa2 | 0.001915579 | -0.09636579 | 0.100196948 | 7.651434388 | 0.046584134 | 0.964212497 | 0.998471826 | -5.255787014 |
| **A_55_P1977096** | Elk4 | -0.005241004 | -0.274324513 | 0.263842505 | 9.966613836 | -0.046551808 | 0.964237311 | 0.998471826 | -5.255788284 |
| **A_55_P2061084** | Wfikkn2 | 0.002185523 | -0.110102253 | 0.114473299 | 8.603291129 | 0.046519178 | 0.964262357 | 0.998471826 | -5.255789565 |
| **A_52_P79506** | Zfp667 | -0.002011829 | -0.105467086 | 0.101443428 | 7.851786963 | -0.04647802 | 0.96429395 | 0.998471826 | -5.25579118 |
| **A_55_P2104917** | Nphp1 | 0.003189073 | -0.160830825 | 0.167208972 | 8.457194743 | 0.046470473 | 0.964299743 | 0.998471826 | -5.255791476 |
| **A_52_P236705** | Ripply3 | 0.002897637 | -0.146203926 | 0.151999199 | 8.557941487 | 0.04644841 | 0.964316679 | 0.998471826 | -5.255792341 |

| | | | | | | | | | |
|---|---|---|---|---|---|---|---|---|---|
| **A_55_P2087944** | NA | 0.00375129 | -0.189278519 | 0.196781099 | 10.69950716 | 0.046447825 | 0.964317128 | 0.998471826 | -5.255792364 |
| **A_52_P284821** | Slco4c1 | 0.002045875 | -0.103348414 | 0.107440165 | 7.646514288 | 0.046395005 | 0.964357673 | 0.998471826 | -5.255794432 |
| **A_66_P114627** | S100a16 | 0.004365233 | -0.220601691 | 0.229332158 | 12.26525009 | 0.046376496 | 0.96437188 | 0.998471826 | -5.255795157 |
| **A_66_P101499** | AW822252 | 0.001919786 | -0.097166997 | 0.101006568 | 7.689083074 | 0.046306946 | 0.964425267 | 0.998471826 | -5.255797876 |
| **A_55_P2076792** | Anxa6 | -0.003255646 | -0.171731831 | 0.165220539 | 14.19427978 | -0.046185728 | 0.964518315 | 0.998471826 | -5.255802606 |
| **A_55_P2167123** | Cyp4b1-ps | -0.002726494 | -0.144022635 | 0.138569646 | 7.876502217 | -0.046119379 | 0.964569246 | 0.998471826 | -5.255805189 |
| **A_55_P1990538** | Ndfip2 | -0.002126913 | -0.112356435 | 0.108102608 | 7.872832242 | -0.046116998 | 0.964571073 | 0.998471826 | -5.255805282 |
| **A_55_P2094378** | NA | 0.002135299 | -0.108667061 | 0.112937659 | 8.321796113 | 0.046059453 | 0.964615246 | 0.998471826 | -5.255807519 |
| **A_55_P2011545** | Pyroxd1 | -0.003055023 | -0.162289215 | 0.156179169 | 8.856626526 | -0.045855062 | 0.96477214 | 0.998471826 | -5.255815444 |
| **A_55_P1994751** | NA | 0.004246731 | -0.217184139 | 0.225677602 | 8.601606174 | 0.045838011 | 0.964785229 | 0.998471826 | -5.255816104 |
| **A_51_P339962** | Cwf19l2 | 0.002573358 | -0.131900355 | 0.13704707 | 8.852332942 | 0.045737443 | 0.964862428 | 0.998471826 | -5.255819989 |
| **A_55_P1972907** | Cab39 | 0.002973985 | -0.152719733 | 0.158667703 | 12.52509594 | 0.045653788 | 0.964926644 | 0.998471826 | -5.255823214 |
| **A_55_P2144086** | NA | -0.004248487 | -0.226969348 | 0.218472374 | 17.31962151 | -0.045591358 | 0.964974567 | 0.998471826 | -5.255825617 |
| **A_51_P416822** | Hhip | 0.001830707 | -0.094150175 | 0.09781159 | 7.64544832 | 0.045587243 | 0.964977726 | 0.998471826 | -5.255825776 |
| **A_55_P2017526** | Dand5 | 0.002831076 | -0.145653268 | 0.151315421 | 8.579607503 | 0.045570102 | 0.964990884 | 0.998471826 | -5.255826435 |
| **A_66_P138001** | Atp2b1 | -0.002287326 | -0.122338332 | 0.117763681 | 9.596621001 | -0.045537724 | 0.965015739 | 0.998471826 | -5.255827679 |
| **A_55_P2037812** | Palmd | -0.00440579 | -0.235776733 | 0.226965152 | 10.39540493 | -0.045511813 | 0.965035629 | 0.998471826 | -5.255828674 |
| **A_55_P1983418** | Amy1 | 0.003348328 | -0.17254818 | 0.179244837 | 7.909992629 | 0.045496707 | 0.965047225 | 0.998471826 | -5.255829254 |
| **A_55_P2150389** | Dnajc21 | 0.003142734 | -0.162015334 | 0.168300802 | 9.230131226 | 0.045479635 | 0.96506033 | 0.998471826 | -5.25582991 |
| **A_55_P2154797** | 2810006K2 | -0.001932848 | -0.103982417 | 0.100116722 | 8.467229196 | -0.045268452 | 0.965222443 | 0.998551434 | -5.255837994 |
| **A_55_P2007646** | Cryaa | 0.002314519 | -0.119963519 | 0.124592557 | 7.800027923 | 0.045239882 | 0.965244375 | 0.998551434 | -5.255839085 |
| **A_51_P115471** | Apol8 | -0.001846612 | -0.099539277 | 0.095846053 | 7.68088023 | -0.045177571 | 0.965292208 | 0.998551434 | -5.24958854 |
| **A_51_P313581** | Fabp2 | 0.002427363 | -0.126797692 | 0.131652418 | 7.751043216 | 0.04489491 | 0.965509195 | 0.998683947 | -5.255852202 |
| **A_55_P2105200** | Abraxas1 | 0.002448463 | -0.127954428 | 0.132851353 | 9.13888347 | 0.04487613 | 0.965523611 | 0.998683947 | -5.255852913 |
| **A_55_P1976928** | Unc13d | -0.001906969 | -0.104309546 | 0.100495607 | 7.855376291 | -0.044508401 | 0.965805907 | 0.9989225 | -5.255866779 |
| **A_55_P2084652** | H2ac15 | 0.004116578 | -0.217611924 | 0.225845081 | 17.10829245 | 0.04437353 | 0.965909444 | 0.998952678 | -5.255871837 |
| **A_55_P2152225** | Ihh | 0.002035831 | -0.107712123 | 0.111783785 | 7.759711231 | 0.044335791 | 0.965938416 | 0.998952678 | -5.255873249 |
| **A_55_P2178496** | Fam53b | 0.007756321 | -0.413295347 | 0.428807988 | 9.629610478 | 0.044028055 | 0.966174662 | 0.9989817 | -5.25588472 |
| **A_52_P551707** | Kdm6a | 0.005839443 | -0.31232857 | 0.324007456 | 8.617923529 | 0.043865596 | 0.966299382 | 0.9989817 | -5.255890744 |
| **A_55_P1996314** | Amy2a5 | 0.006742633 | -0.361193686 | 0.374678953 | 7.88005579 | 0.043799184 | 0.966350367 | 0.9989817 | -5.2558932 |
| **A_55_P2131473** | Rmnd5a | -0.002343934 | -0.130795252 | 0.126107384 | 8.271939875 | -0.043613003 | 0.9664933 | 0.9989817 | -5.255900066 |
| **A_51_P350403** | Exoc4 | 0.003721566 | -0.200651673 | 0.208094805 | 8.947171756 | 0.043522199 | 0.966563011 | 0.9989817 | -5.255903404 |
| **A_51_P253883** | Fam49a | -0.004507393 | -0.252468836 | 0.24345405 | 8.889507317 | -0.043446057 | 0.966621467 | 0.9989817 | -5.255906197 |
| **A_55_P2059680** | Gm11938 | -0.002608032 | -0.146316928 | 0.141100864 | 7.964768957 | -0.043374893 | 0.966676101 | 0.9989817 | -5.255908804 |
| **A_55_P2121543** | Rps20 | -0.002058634 | -0.115817341 | 0.111700073 | 16.99414758 | -0.043251742 | 0.966770647 | 0.9989817 | -5.255913304 |
| **A_55_P2033666** | NA | -0.002132651 | -0.120342504 | 0.116077202 | 16.49449982 | -0.043119642 | 0.966872064 | 0.9989817 | -5.255918118 |
| **A_55_P2120773** | Dctn4 | 0.002664332 | -0.145041585 | 0.15037025 | 11.68086781 | 0.043112142 | 0.966877822 | 0.9989817 | -5.255918391 |
| **A_52_P476731** | Fam110c | -0.002871478 | -0.162355204 | 0.156612247 | 7.804017865 | -0.043032671 | 0.966938835 | 0.9989817 | -5.255921279 |
| **A_55_P2107302** | Aurkaip1 | 0.002750786 | -0.150502617 | 0.156004188 | 14.19899113 | 0.042899852 | 0.967040804 | 0.9989817 | -5.255926094 |
| **A_55_P2170355** | NA | -0.002370696 | -0.134474168 | 0.129732776 | 8.356384818 | -0.042891459 | 0.967047249 | 0.9989817 | -5.255926397 |
| **A_55_P1985359** | Rpl28 | 0.003199819 | -0.175471117 | 0.181870754 | 14.4243776 | 0.042803629 | 0.967114679 | 0.9989817 | -5.255929573 |
| **A_55_P2013088** | Arpc4 | 0.005992342 | -0.328892777 | 0.340877462 | 9.235098643 | 0.042767109 | 0.967142717 | 0.9989817 | -5.255930891 |
| **A_55_P2154307** | Peak1 | -0.004094817 | -0.233222946 | 0.225033312 | 12.04541373 | -0.042713506 | 0.967183871 | 0.9989817 | -5.255932824 |
| **A_55_P2026547** | Gal3st2 | -0.00197647 | -0.112860829 | 0.10890789 | 7.866239042 | -0.0426019 | 0.967269556 | 0.9989817 | -5.255936841 |
| **A_51_P520384** | Atp1b3 | 0.002209032 | -0.121924229 | 0.126342293 | 11.38045871 | 0.042532705 | 0.967322681 | 0.9989817 | -5.255939326 |
| **A_55_P2253980** | Armt1 | -0.00210464 | -0.120503558 | 0.116294279 | 8.330382853 | -0.042485342 | 0.967359044 | 0.9989817 | -5.255941025 |
| **A_51_P191649** | Ndc80 | 0.003266565 | -0.180649864 | 0.187182993 | 9.093834449 | 0.042450211 | 0.967386016 | 0.9989817 | -5.255942284 |
| **A_51_P236864** | Parp8 | 0.003709079 | -0.205486545 | 0.212904702 | 9.260111145 | 0.042376255 | 0.967442797 | 0.9989817 | -5.25594493 |

| | | | | | | | | | |
|---|---|---|---|---|---|---|---|---|---|
| **A_51_P173212** | Osbpl2 | 0.002850183 | -0.158637697 | 0.164338062 | 11.24391286 | 0.042183433 | 0.967590838 | 0.9989817 | -5.255951809 |
| **A_55_P2061437** | Lsm5 | -0.004478637 | -0.258427141 | 0.249469866 | 10.87179855 | -0.042151144 | 0.967615629 | 0.9989817 | -5.255952958 |
| **A_55_P2005440** | Dennd5a | -0.002990409 | -0.172757546 | 0.166776728 | 12.97445874 | -0.042100383 | 0.967654601 | 0.9989817 | -5.255954762 |
| **A_51_P324351** | Meltf | 0.001943749 | -0.108619294 | 0.112506792 | 7.714609102 | 0.042018379 | 0.967717562 | 0.9989817 | -5.255957672 |
| **A_52_P102783** | Dync2h1 | 0.003224062 | -0.180797534 | 0.187245657 | 9.215365333 | 0.041873924 | 0.967828471 | 0.9989817 | -5.255962785 |
| **A_65_P11425** | Bmp2k | -0.0025661 | -0.149302957 | 0.144170756 | 8.381588102 | -0.041796851 | 0.967887646 | 0.9989817 | -5.255965505 |
| **A_51_P110371** | Wwox | -0.00200744 | -0.116821374 | 0.112806495 | 8.397116197 | -0.041788522 | 0.967894041 | 0.9989817 | -5.255965799 |
| **A_55_P1994668** | Tti1 | 0.003428931 | -0.193557719 | 0.200415582 | 10.83365388 | 0.041603622 | 0.968036004 | 0.9989817 | -5.255972304 |
| **A_55_P2113356** | Miip | 0.002193039 | -0.123798394 | 0.128184472 | 8.212408087 | 0.041602022 | 0.968037233 | 0.9989817 | -5.25597236 |
| **A_51_P150521** | Mgat2 | 0.003663704 | -0.207236144 | 0.214563552 | 9.037404517 | 0.041519603 | 0.968100513 | 0.9989817 | -5.25597525 |
| **A_55_P2025523** | Arhgap6 | 0.003146474 | -0.178124351 | 0.184417299 | 9.251310458 | 0.04148636 | 0.968126037 | 0.9989817 | -5.255976415 |
| **A_55_P2046158** | NA | -0.001997452 | -0.117159226 | 0.113164322 | 13.61113384 | -0.041455017 | 0.968150102 | 0.9989817 | -5.255977511 |
| **A_55_P1953028** | Tnfsf12 | 0.004937215 | -0.280103852 | 0.289978282 | 9.007418396 | 0.041398418 | 0.968193558 | 0.9989817 | -5.25597949 |
| **A_52_P122843** | Bicral | -0.002565054 | -0.15110908 | 0.145978972 | 8.607015242 | -0.041271515 | 0.968290994 | 0.9989817 | -5.255983916 |
| **A_52_P590396** | Sort1 | 0.004071236 | -0.232400811 | 0.240543282 | 10.59504586 | 0.041148647 | 0.968385333 | 0.9989817 | -5.255988188 |
| **A_52_P370311** | Plbd2 | 0.002378743 | -0.135836622 | 0.140594107 | 11.04791179 | 0.041133935 | 0.968396629 | 0.9989817 | -5.255988699 |
| **A_55_P2061338** | Slc16a11 | -0.001697312 | -0.100483623 | 0.097088998 | 7.820738394 | -0.041065219 | 0.96844939 | 0.9989817 | -5.255991082 |
| **A_51_P290904** | Raver2 | -0.002516493 | -0.149037087 | 0.144004101 | 8.045908927 | -0.041049334 | 0.968461586 | 0.9989817 | -5.255991632 |
| **A_51_P360918** | Ehd3 | 0.002834624 | -0.162377474 | 0.168046722 | 8.625182695 | 0.041007448 | 0.968493747 | 0.9989817 | -5.255993082 |
| **A_66_P121976** | Slc4a11 | -0.001723747 | -0.102369266 | 0.098921772 | 7.74109714 | -0.04093438 | 0.968549849 | 0.9989817 | -5.255995608 |
| **A_66_P129672** | 4930453N2 | 0.002183942 | -0.1255411 | 0.129908984 | 10.5695867 | 0.040867125 | 0.968601488 | 0.9989817 | -5.255997929 |
| **A_51_P295085** | Ogn | -0.006262447 | -0.372881934 | 0.36035704 | 11.70222096 | -0.040826076 | 0.968633006 | 0.9989817 | -5.255999344 |
| **A_51_P113072** | Cdc123 | -0.003693476 | -0.219958175 | 0.212571222 | 11.56719576 | -0.040818665 | 0.968638697 | 0.9989817 | -5.255999599 |
| **A_51_P377856** | Gstt3 | 0.00281539 | -0.162348609 | 0.167979389 | 8.35386372 | 0.04074106 | 0.968698283 | 0.9989817 | -5.256002269 |
| **A_55_P2184009** | Rnd2 | 0.003314121 | -0.191283894 | 0.197912137 | 8.774325777 | 0.040704192 | 0.968726591 | 0.9989817 | -5.256003536 |
| **A_55_P1983958** | NA | -0.005805588 | -0.347892197 | 0.336281021 | 12.27965369 | -0.04056199 | 0.968835777 | 0.9989817 | -5.256008411 |
| **A_55_P2076471** | Mon1b | -0.002448721 | -0.146838047 | 0.141940605 | 10.04877225 | -0.040533431 | 0.968857705 | 0.9989817 | -5.256009389 |
| **A_51_P336952** | Cryl1 | -0.004825999 | -0.289712827 | 0.280060829 | 9.722127221 | -0.040487782 | 0.968892756 | 0.9989817 | -5.256010949 |
| **A_52_P69109** | Slc10a1 | 0.001650697 | -0.095830063 | 0.099131458 | 7.650647695 | 0.04047228 | 0.968904659 | 0.9989817 | -5.256011478 |
| **A_55_P2086983** | Atxn2 | -0.003054591 | -0.183521349 | 0.177412167 | 12.33913003 | -0.040454327 | 0.968918444 | 0.9989817 | -5.256012091 |
| **A_55_P2015675** | AU015836 | 0.001686696 | -0.098096031 | 0.101469422 | 7.685718583 | 0.040400849 | 0.968959506 | 0.9989817 | -5.256013916 |
| **A_55_P2040280** | Dhps | 0.002935703 | -0.17157343 | 0.177444837 | 8.448482566 | 0.040207138 | 0.969108244 | 0.9989817 | -5.256020503 |
| **A_55_P2305010** | A730089K1 | -0.001897951 | -0.114734581 | 0.110938679 | 7.823734494 | -0.040201671 | 0.969112442 | 0.9989817 | -5.256020689 |
| **A_51_P292008** | Gpx3 | 0.013349029 | -0.783376678 | 0.810074736 | 11.57356194 | 0.040045154 | 0.969232622 | 0.9989817 | -5.256025988 |
| **A_55_P2125786** | Lrrc8e | 0.001861346 | -0.109347009 | 0.1130697 | 7.687770513 | 0.040003567 | 0.969264555 | 0.9989817 | -5.256027392 |
| **A_55_P1973813** | NA | -0.00175611 | -0.106859652 | 0.103347432 | 7.727143925 | -0.039934056 | 0.969317929 | 0.9989817 | -5.256029736 |
| **A_55_P2106241** | 4930432K2 | 0.001873398 | -0.110296724 | 0.114043521 | 8.210497394 | 0.039917379 | 0.969330734 | 0.9989817 | -5.256030298 |
| **A_55_P2130104** | Slc52a3 | 0.001733271 | -0.102768138 | 0.10623468 | 7.794472732 | 0.039641809 | 0.969542332 | 0.9989817 | -5.256039548 |
| **A_55_P2110215** | Ckap5 | 0.003393459 | -0.201493128 | 0.208280046 | 12.84731948 | 0.039585685 | 0.969585427 | 0.9989817 | -5.256041424 |
| **A_52_P117576** | Casp3 | -0.002203052 | -0.135412746 | 0.131006643 | 8.417332399 | -0.039527373 | 0.969630203 | 0.9989817 | -5.25604337 |
| **A_55_P2077153** | Slc27a2 | 0.00343549 | -0.20432298 | 0.211193959 | 7.832067743 | 0.039522011 | 0.96963432 | 0.9989817 | -5.256043549 |
| **A_65_P01643** | Chd2 | -0.003281127 | -0.201760147 | 0.195197892 | 9.97587323 | -0.039510965 | 0.969642802 | 0.9989817 | -5.256043917 |
| **A_55_P2056973** | Treml2 | 0.002226699 | -0.13257713 | 0.137030527 | 7.741910765 | 0.039479199 | 0.969667194 | 0.9989817 | -5.256044976 |
| **A_55_P2119917** | Ikzf4 | 0.002902816 | -0.172834153 | 0.178639785 | 8.661911768 | 0.039478939 | 0.969667393 | 0.9989817 | -5.256044984 |
| **A_51_P443514** | 1700001K1 | -0.002556166 | -0.157514253 | 0.152401921 | 8.59216324 | -0.03942611 | 0.969707959 | 0.9989817 | -5.256046743 |
| **A_55_P2044724** | Zfp869 | -0.002005761 | -0.123667485 | 0.119655963 | 9.793581403 | -0.039403466 | 0.969725346 | 0.9989817 | -5.256047496 |
| **A_51_P289588** | Ube2e3 | -0.002917691 | -0.179967177 | 0.174131795 | 13.01350512 | -0.039387082 | 0.969737927 | 0.9989817 | -5.256048041 |
| **A_52_P624362** | Aamp | -0.003984722 | -0.246055516 | 0.238086073 | 10.52259117 | -0.039342751 | 0.969771968 | 0.9989817 | -5.256049513 |

| | | | | | | | | | |
|---|---|---|---|---|---|---|---|---|---|
| **A_55_P2105035** | NA | 0.002723772 | -0.162830838 | 0.168278383 | 12.1081556 | 0.039322279 | 0.969787688 | 0.9989817 | -5.256050193 |
| **A_55_P1971009** | Gzme | -0.003360819 | -0.207845224 | 0.201123585 | 7.769665942 | -0.039282047 | 0.96981858 | 0.9989817 | -5.256051527 |
| **A_52_P228079** | Atf1 | 0.001982935 | -0.118842416 | 0.122808286 | 8.62450068 | 0.039224693 | 0.969862621 | 0.9989817 | -5.256053426 |
| **A_55_P1993820** | Kdm1a | -0.00291193 | -0.180605905 | 0.174782045 | 12.13829601 | -0.039166736 | 0.969907124 | 0.9989817 | -5.256055343 |
| **A_55_P2043267** | Cdc42se1 | -0.003237834 | -0.201171874 | 0.194696207 | 9.539149359 | -0.039096978 | 0.96996069 | 0.9989817 | -5.256057647 |
| **A_55_P2013823** | Gal3st1 | -0.001788601 | -0.111252506 | 0.107675304 | 7.689773246 | -0.039052756 | 0.969994648 | 0.9989817 | -5.256059105 |
| **A_55_P2033116** | Cdcp2 | 0.00156446 | -0.09430263 | 0.097431551 | 7.668464446 | 0.039003546 | 0.970032435 | 0.9989817 | -5.256060725 |
| **A_55_P2029574** | Por | 0.002929905 | -0.176933054 | 0.182792864 | 11.22757308 | 0.038933273 | 0.970086396 | 0.9989817 | -5.256063036 |
| **A_51_P515496** | Rbm17 | -0.002475714 | -0.154907607 | 0.14995618 | 11.4818033 | -0.038818047 | 0.970174877 | 0.9989817 | -5.256066816 |
| **A_51_P383857** | Prp2 | -0.001765033 | -0.110489848 | 0.106959782 | 8.161469549 | -0.038800141 | 0.970188627 | 0.9989817 | -5.256067402 |
| **A_55_P2367117** | Pitpnb | 0.002722776 | -0.165145472 | 0.170591024 | 8.771065781 | 0.038766138 | 0.970214737 | 0.9989817 | -5.256068515 |
| **A_51_P400040** | Ubxn2a | -0.001753502 | -0.110127649 | 0.106620645 | 9.151449833 | -0.038671374 | 0.970287505 | 0.9989817 | -5.256071611 |
| **A_55_P2178207** | Abcc8 | -0.005581953 | -0.351048275 | 0.339884369 | 11.86243167 | -0.038617981 | 0.970328505 | 0.9989817 | -5.256073352 |
| **A_55_P2002893** | Pfkfb1 | -0.002594087 | -0.163355312 | 0.158167138 | 8.62515566 | -0.038566687 | 0.970367894 | 0.9989817 | -5.256075022 |
| **A_55_P1984284** | Gapdh | 0.003512626 | -0.214532606 | 0.221557857 | 16.61073595 | 0.038502984 | 0.970416811 | 0.9989817 | -5.256077093 |
| **A_51_P136888** | Rb1 | 0.00278511 | -0.170857509 | 0.17642773 | 9.746773761 | 0.038334973 | 0.970545827 | 0.9989817 | -5.25608254 |
| **A_51_P384629** | Ctsd | -0.00253986 | -0.160916894 | 0.155837175 | 13.84591903 | -0.038328929 | 0.970550469 | 0.9989817 | -5.256082736 |
| **A_51_P272563** | Naa25 | 0.002253503 | -0.138377068 | 0.142884074 | 9.563237201 | 0.038299003 | 0.970573448 | 0.9989817 | -5.256083703 |
| **A_55_P2076288** | Zfp661 | -0.002122181 | -0.135040356 | 0.130795995 | 8.576099037 | -0.038159893 | 0.970680273 | 0.9989817 | -5.25608819 |
| **A_52_P329054** | Cep70 | 0.002884744 | -0.178095038 | 0.183864525 | 9.238518206 | 0.038096605 | 0.970728871 | 0.9989817 | -5.256090227 |
| **A_55_P1957835** | Cpeb3 | -0.002899403 | -0.18510778 | 0.179308974 | 11.45292449 | -0.038032016 | 0.97077847 | 0.9989817 | -5.256092301 |
| **A_55_P2089238** | Fam76a | -0.003234903 | -0.206800116 | 0.200330311 | 10.91158834 | -0.037981036 | 0.970817619 | 0.9989817 | -5.256093936 |
| **A_55_P2078433** | Mcoln2 | -0.001929647 | -0.123461596 | 0.119602301 | 7.860947476 | -0.037948678 | 0.970842467 | 0.9989817 | -5.256094973 |
| **A_52_P237077** | Esr1 | -0.001705189 | -0.109369124 | 0.105958746 | 7.672950807 | -0.037853969 | 0.970915196 | 0.9989817 | -5.224448188 |
| **A_52_P532559** | Thrb | -0.001727097 | -0.111052554 | 0.107598359 | 7.768404083 | -0.037757625 | 0.970989181 | 0.9989817 | -5.256101075 |
| **A_55_P2030662** | Eda | 0.003122265 | -0.194518943 | 0.200763473 | 8.900198603 | 0.03775734 | 0.9709894 | 0.9989817 | -5.256101084 |
| **A_55_P2147101** | Sytl4 | 0.001833571 | -0.114581684 | 0.118248827 | 8.316861448 | 0.037644112 | 0.97107635 | 0.9989817 | -5.256104686 |
| **A_55_P2277014** | AA522020 | -0.006503695 | -0.419859913 | 0.406852522 | 12.14528294 | -0.037604934 | 0.971106436 | 0.9989817 | -5.25610593 |
| **A_51_P114616** | Batf | -0.001824657 | -0.117912712 | 0.114263398 | 8.444514157 | -0.037566681 | 0.971135811 | 0.9989817 | -5.256107143 |
| **A_51_P161342** | Abcb6 | -0.002765661 | -0.178837162 | 0.17330584 | 9.490640578 | -0.037542135 | 0.971154661 | 0.9989817 | -5.256107921 |
| **A_51_P384609** | Ddx10 | 0.002163934 | -0.136395719 | 0.140723586 | 8.996038184 | 0.03732642 | 0.971320316 | 0.9989817 | -5.256114735 |
| **A_55_P2165539** | Aebp2 | -0.003515967 | -0.229697079 | 0.222665145 | 9.286395516 | -0.037153317 | 0.971453249 | 0.9989817 | -5.256120174 |
| **A_51_P360004** | Ocstamp | 0.003130551 | -0.19906093 | 0.205322031 | 8.158518858 | 0.037005564 | 0.971566715 | 0.9989817 | -5.256124797 |
| **A_55_P1997831** | Dbndd1 | 0.001760279 | -0.112020804 | 0.115541363 | 7.833521813 | 0.036976063 | 0.971589371 | 0.9989817 | -5.256125718 |
| **A_52_P156485** | Ankrd39 | 0.002978434 | -0.190590082 | 0.196546951 | 10.813305 | 0.036775828 | 0.971743142 | 0.9989817 | -5.256131949 |
| **A_55_P2008071** | Prkcd | 0.002085262 | -0.134332731 | 0.138503255 | 9.080617963 | 0.036534074 | 0.971928799 | 0.9989817 | -5.256139426 |
| **A_51_P438479** | Zfp281 | 0.002174471 | -0.140217042 | 0.144565984 | 10.90708295 | 0.03649881 | 0.971955881 | 0.9989817 | -5.256140513 |
| **A_55_P2034784** | Snhg17 | 0.001702356 | -0.110088146 | 0.113492858 | 8.672970036 | 0.036396076 | 0.972034776 | 0.9989817 | -5.256143672 |
| **A_55_P2074972** | Zswim6 | 0.001700035 | -0.109991844 | 0.113391913 | 8.319062045 | 0.036378547 | 0.972048238 | 0.9989817 | -5.256144211 |
| **A_55_P2041592** | Rdh1 | -0.001565031 | -0.104848655 | 0.101718594 | 7.833581188 | -0.036216005 | 0.972173066 | 0.9989817 | -5.256149189 |
| **A_66_P140976** | Rpl15 | -0.003128745 | -0.209854986 | 0.203597496 | 11.77481903 | -0.03617293 | 0.972206146 | 0.9989817 | -5.256150505 |
| **A_51_P259318** | Fbrs | 0.00188333 | -0.122565908 | 0.126332569 | 8.256363946 | 0.036169576 | 0.972208722 | 0.9989817 | -5.256150607 |
| **A_55_P2085288** | Acoxl | 0.001521276 | -0.099012462 | 0.102055014 | 7.828501096 | 0.036166403 | 0.972211159 | 0.9989817 | -5.256150704 |
| **A_66_P139647** | Vwa3a | -0.001761724 | -0.118478056 | 0.114954609 | 7.83780552 | -0.036075747 | 0.97228078 | 0.9989817 | -5.256153467 |
| **A_51_P167048** | Hs6st1 | -0.002481593 | -0.167463638 | 0.162500452 | 8.601679696 | -0.035950342 | 0.972377089 | 0.9989817 | -5.256157278 |
| **A_55_P2094963** | NA | 0.003583691 | -0.235810012 | 0.242977394 | 9.436183717 | 0.035778901 | 0.972508753 | 0.9989817 | -5.256162466 |
| **A_55_P1967483** | NA | -0.001701615 | -0.115438424 | 0.112035194 | 12.67121906 | -0.03575768 | 0.972525051 | 0.9989817 | -5.256163107 |
| **A_55_P2174987** | Creb3l3 | -0.001691746 | -0.114880734 | 0.111497242 | 7.999876449 | -0.035722351 | 0.972552183 | 0.9989817 | -5.256164172 |

| | | | | | | | | | |
|---|---|---|---|---|---|---|---|---|---|
| **A_51_P480119** | Prelid2 | 0.005675212 | -0.374695594 | 0.386046019 | 12.03285635 | 0.035660223 | 0.972599897 | 0.9989817 | -5.256166043 |
| **A_55_P2173035** | Dnajc8 | 0.00422387 | -0.279245348 | 0.287693089 | 10.41884495 | 0.035613431 | 0.972635833 | 0.9989817 | -5.256167451 |
| **A_55_P2071626** | C530008M | 0.00410543 | -0.271672318 | 0.279883178 | 9.876227659 | 0.035580212 | 0.972661345 | 0.9989817 | -5.256168448 |
| **A_66_P135192** | Msl2 | 0.002762468 | -0.183576227 | 0.189101163 | 9.289750277 | 0.035432631 | 0.972774686 | 0.9989817 | -5.25617287 |
| **A_55_P2023864** | Mfap1a | -0.002108853 | -0.145156034 | 0.140938328 | 9.867848099 | -0.035235147 | 0.972926354 | 0.9989817 | -5.256178758 |
| **A_51_P210143** | Syne2 | -0.002321966 | -0.160081163 | 0.155437231 | 10.2502906 | -0.035177936 | 0.972970293 | 0.9989817 | -5.256180458 |
| **A_52_P286360** | Otc | 0.001553423 | -0.104219742 | 0.107326588 | 7.647328623 | 0.035101319 | 0.973029135 | 0.9989817 | -5.25618273 |
| **A_55_P2029294** | Cetn4 | -0.001632688 | -0.112848298 | 0.109582922 | 7.96416404 | -0.035087021 | 0.973040116 | 0.9989817 | -5.256183153 |
| **A_55_P1984523** | NA | -0.005206303 | -0.359911768 | 0.349499163 | 11.96854857 | -0.035080895 | 0.973044821 | 0.9989817 | -5.256183335 |
| **A_55_P2155764** | NA | -0.001730075 | -0.119736512 | 0.116276361 | 10.6938104 | -0.03504035 | 0.97307596 | 0.9989817 | -5.256184534 |
| **A_55_P2077526** | NA | 0.003008556 | -0.203002476 | 0.209019587 | 13.87871638 | 0.034904122 | 0.973180585 | 0.9989817 | -5.256188554 |
| **A_51_P427530** | Pgm1 | 0.001779534 | -0.120092044 | 0.123651112 | 8.365276251 | 0.034899003 | 0.973184516 | 0.9989817 | -5.256188705 |
| **A_55_P2115297** | Ap3s1 | -0.003959762 | -0.276559702 | 0.268640178 | 11.22403238 | -0.034717817 | 0.97332367 | 0.9989817 | -5.256194027 |
| **A_55_P1959496** | Sbno2 | 0.001479175 | -0.100382822 | 0.103341173 | 7.893906033 | 0.034706962 | 0.973332007 | 0.9989817 | -5.256194345 |
| **A_55_P2043322** | Rnf26 | 0.002554722 | -0.174298052 | 0.179407496 | 8.885734288 | 0.034525571 | 0.97347132 | 0.9989817 | -5.256199644 |
| **A_55_P2069917** | Uba52 | -0.002732924 | -0.192322184 | 0.186856335 | 17.12146244 | -0.034452682 | 0.973527301 | 0.9989817 | -5.256201765 |
| **A_55_P2018542** | Ptges3 | -0.003370025 | -0.237258063 | 0.230518013 | 10.91308248 | -0.034437705 | 0.973538803 | 0.9989817 | -5.2562022 |
| **A_66_P125660** | Nde1 | 0.002218311 | -0.152085952 | 0.156522575 | 8.233464012 | 0.034360041 | 0.973598452 | 0.9989817 | -5.256204455 |
| **A_55_P1963863** | Arhgef17 | 0.001474329 | -0.101440991 | 0.104389649 | 10.46190068 | 0.034239192 | 0.973691268 | 0.9989817 | -5.256207952 |
| **A_52_P558609** | Clec16a | -0.001792359 | -0.127053362 | 0.123468643 | 8.734429855 | -0.034199386 | 0.97372184 | 0.9989817 | -5.256209102 |
| **A_51_P221132** | L2hgdh | 0.001983999 | -0.13668666 | 0.140654659 | 10.13183202 | 0.034195273 | 0.973724999 | 0.9989817 | -5.25620922 |
| **A_52_P4846** | Zfp451 | -0.001744848 | -0.124183964 | 0.120694268 | 8.750421647 | -0.034060152 | 0.973828778 | 0.9989817 | -5.256213112 |
| **A_55_P2083939** | Irf3 | 0.002500569 | -0.173489746 | 0.178490884 | 12.30533399 | 0.033959342 | 0.973906204 | 0.9989817 | -5.256216005 |
| **A_51_P153995** | Gp9 | -0.01061463 | -0.759851698 | 0.738622438 | 9.068062013 | -0.033860608 | 0.973982036 | 0.9989817 | -5.25621883 |
| **A_55_P2004071** | Gm2058 | 0.002895794 | -0.201536193 | 0.207327782 | 8.768854184 | 0.033855404 | 0.973986033 | 0.9989817 | -5.256218979 |
| **A_51_P191354** | NA | 0.001617616 | -0.112582066 | 0.115817299 | 8.627025091 | 0.033854764 | 0.973986524 | 0.9989817 | -5.256218997 |
| **A_55_P2330435** | NA | 0.00168238 | -0.117269433 | 0.120634193 | 7.694057869 | 0.033803535 | 0.974025871 | 0.9989817 | -5.25622046 |
| **A_55_P2062926** | D430041D( | -0.002542822 | -0.182558427 | 0.177472784 | 8.528518082 | -0.033760966 | 0.974058566 | 0.9989817 | -5.256221673 |
| **A_52_P418734** | Rnf220 | 0.002105703 | -0.147134949 | 0.151346356 | 12.3291175 | 0.033722454 | 0.974088145 | 0.9989817 | -5.25622277 |
| **A_55_P1967708** | NA | -0.002008605 | -0.144732919 | 0.140715708 | 8.7483835 | -0.033636109 | 0.974154463 | 0.9989817 | -5.256225224 |
| **A_55_P2018904** | Kbtbd6 | -0.002018545 | -0.146212749 | 0.14217566 | 8.665457645 | -0.033457976 | 0.974291279 | 0.9989817 | -5.256230266 |
| **A_55_P1969874** | NA | 0.002223934 | -0.15682614 | 0.161274008 | 15.93059717 | 0.033419282 | 0.974320998 | 0.9989817 | -5.256231358 |
| **A_55_P2176361** | Zcchc9 | 0.001486539 | -0.104992402 | 0.107965481 | 8.574524405 | 0.033367359 | 0.974360878 | 0.9989817 | -5.256232821 |
| **A_55_P2068977** | Armcx3 | 0.00229864 | -0.162392684 | 0.166989965 | 10.91085026 | 0.033358725 | 0.974367509 | 0.9989817 | -5.256233064 |
| **A_55_P2183389** | NA | -0.00218712 | -0.159316702 | 0.154942462 | 15.20676963 | -0.033267774 | 0.974437366 | 0.9989817 | -5.256235621 |
| **A_55_P2011380** | Acvrl1 | -0.003031107 | -0.221157374 | 0.21509516 | 9.945513462 | -0.033212567 | 0.974479768 | 0.9989817 | -5.25623717 |
| **A_51_P136729** | Idh3g | 0.002410942 | -0.171605647 | 0.176427532 | 15.05744293 | 0.033113515 | 0.974555846 | 0.9989817 | -5.256239942 |
| **A_55_P1984297** | Efcab14 | 0.002979075 | -0.21349666 | 0.219454811 | 9.816895586 | 0.032891326 | 0.974726504 | 0.9989817 | -5.25624613 |
| **A_51_P418560** | Lnx2 | 0.001760349 | -0.126310385 | 0.129831084 | 9.809771094 | 0.032851716 | 0.974756927 | 0.9989817 | -5.256247228 |
| **A_55_P2000409** | Rab44 | 0.001725108 | -0.123829862 | 0.127280077 | 7.803947296 | 0.032839112 | 0.974766608 | 0.9989817 | -5.256247578 |
| **A_55_P1967224** | NA | -0.004738567 | -0.350229449 | 0.340752315 | 9.262861444 | -0.032780803 | 0.974811394 | 0.9989817 | -5.256249192 |
| **A_55_P1996344** | Dcst2 | 0.001339044 | -0.096326855 | 0.099004944 | 7.650071229 | 0.032768846 | 0.974820578 | 0.9989817 | -5.256249523 |
| **A_51_P491916** | Rassf6 | 0.001342185 | -0.096992543 | 0.099676912 | 7.687439478 | 0.032622296 | 0.97493314 | 0.9989817 | -5.256253566 |
| **A_55_P2159480** | Fam160a1 | 0.003143482 | -0.227815605 | 0.234102569 | 10.73984664 | 0.032530073 | 0.975003975 | 0.9989817 | -5.256256101 |
| **A_55_P2163774** | Crip1 | -0.00308393 | -0.230157979 | 0.223990119 | 15.7464129 | -0.032459823 | 0.975057933 | 0.9989817 | -5.256258028 |
| **A_55_P2138366** | Xpo4 | 0.002286137 | -0.166192465 | 0.170764739 | 8.496422466 | 0.032431467 | 0.975079713 | 0.9989817 | -5.256258804 |
| **A_55_P2109337** | Gba2 | -0.001858103 | -0.139215418 | 0.135499211 | 11.54972282 | -0.032331602 | 0.975156418 | 0.9989817 | -5.256261533 |
| **A_55_P1993933** | NA | -0.001423663 | -0.106900831 | 0.104053505 | 7.784242414 | -0.032259522 | 0.975211782 | 0.9989817 | -5.256263497 |

| | | | | | | | | | |
|---|---|---|---|---|---|---|---|---|---|
| **A_55_P1954910** | Fbxw2 | 0.00150465 | -0.110262351 | 0.11327165 | 8.472689159 | 0.032175921 | 0.975275996 | 0.9989817 | -5.25626577 |
| **A_55_P1988313** | NA | -0.001332035 | -0.100447549 | 0.097783479 | 7.647954196 | -0.032120572 | 0.97531851 | 0.9989817 | -5.256267272 |
| **A_55_P2046802** | Mlkl | 0.00283853 | -0.208649712 | 0.214326772 | 8.513416336 | 0.032078678 | 0.975350688 | 0.9989817 | -5.256268407 |
| **A_51_P227770** | Phykpl | -0.001506061 | -0.114039396 | 0.111027274 | 9.380374068 | -0.031986774 | 0.975421279 | 0.9989817 | -5.256270891 |
| **A_51_P111962** | Bean1 | -0.001891503 | -0.143711449 | 0.139928443 | 8.493376199 | -0.031877104 | 0.975505517 | 0.9989817 | -5.256273846 |
| **A_52_P405145** | NA | 0.001398162 | -0.103882801 | 0.106679125 | 7.728300857 | 0.03174072 | 0.975610275 | 0.9989817 | -5.256277507 |
| **A_55_P1977314** | Ulk1 | 0.003537449 | -0.263214248 | 0.270289146 | 12.4315729 | 0.031695096 | 0.975645319 | 0.9989817 | -5.256278728 |
| **A_51_P338728** | Sdhaf1 | 0.002273988 | -0.169443978 | 0.173991954 | 11.47562217 | 0.031650571 | 0.975679519 | 0.9989817 | -5.256279918 |
| **A_55_P2217309** | NA | 0.001607626 | -0.120704275 | 0.123919527 | 8.061169812 | 0.031414158 | 0.975861111 | 0.9989817 | -5.256286209 |
| **A_55_P2155593** | NA | -0.003802232 | -0.293253974 | 0.28564951 | 11.16961051 | -0.0313958 | 0.975875212 | 0.9989817 | -5.256286696 |
| **A_51_P357664** | Hras | 0.004061204 | -0.30564632 | 0.313768729 | 11.73770151 | 0.031340954 | 0.97591734 | 0.9989817 | -5.256288147 |
| **A_55_P2016708** | Msl1 | 0.001952022 | -0.147258168 | 0.151162212 | 9.882742794 | 0.031267662 | 0.975973638 | 0.9989817 | -5.256290084 |
| **A_55_P2139396** | Unc45b | 0.002633356 | -0.198976676 | 0.204243388 | 11.44831537 | 0.03121811 | 0.9760117 | 0.9989817 | -5.25629139 |
| **A_55_P1994027** | Adra2b | 0.001609553 | -0.122627244 | 0.125846351 | 8.305823194 | 0.030964513 | 0.976206494 | 0.9989817 | -5.256298044 |
| **A_55_P1984561** | Cpne2 | -0.001361799 | -0.106807929 | 0.104084332 | 8.794645895 | -0.030866785 | 0.976281562 | 0.9989817 | -5.256300593 |
| **A_55_P2033795** | Fabp6 | -0.001310499 | -0.103013619 | 0.100392622 | 7.66819895 | -0.030797212 | 0.976335003 | 0.9989817 | -5.256302404 |
| **A_51_P121390** | Anapc11 | -0.001398873 | -0.11000011 | 0.107202365 | 13.70666684 | -0.030785946 | 0.976343657 | 0.9989817 | -5.224648025 |
| **A_52_P234729** | Pkd2 | 0.003970394 | -0.304405448 | 0.312346235 | 10.4773295 | 0.03077247 | 0.976354008 | 0.9989817 | -5.256303046 |
| **A_55_P2429225** | Psrc1 | 0.001501185 | -0.115128073 | 0.118130443 | 8.63374571 | 0.030763511 | 0.97636089 | 0.9989817 | -5.256303279 |
| **A_52_P623058** | R3hdm4 | -0.001375561 | -0.108288179 | 0.105537057 | 8.866097967 | -0.030751065 | 0.976370451 | 0.9989817 | -5.256303602 |
| **A_51_P418339** | Nudcd2 | 0.002511061 | -0.192890514 | 0.197912636 | 9.894801336 | 0.030714139 | 0.976398815 | 0.9989817 | -5.25630456 |
| **A_51_P463120** | Hacd4 | 0.002421114 | -0.186861367 | 0.191703595 | 8.570483317 | 0.030571298 | 0.976508536 | 0.9989817 | -5.256308253 |
| **A_55_P1973668** | Slc39a13 | 0.001894012 | -0.146511283 | 0.150299308 | 11.72062152 | 0.030503001 | 0.976560998 | 0.9989817 | -5.256310013 |
| **A_51_P148069** | Lpgat1 | -0.002628798 | -0.209423404 | 0.204165808 | 11.47943039 | -0.030382754 | 0.976653365 | 0.9989817 | -5.256313103 |
| **A_55_P2008512** | B4galt7 | -0.001588224 | -0.126683904 | 0.123507457 | 8.38855812 | -0.030344393 | 0.976682832 | 0.9989817 | -5.256314086 |
| **A_55_P2026738** | Rgs4 | 0.002393545 | -0.186658339 | 0.19144543 | 8.14642016 | 0.030260058 | 0.976747613 | 0.9989817 | -5.256316242 |
| **A_55_P2077185** | NA | -0.002020836 | -0.162024166 | 0.157982493 | 9.880309432 | -0.030186394 | 0.976804198 | 0.9989817 | -5.256318121 |
| **A_52_P526396** | Atl1 | -0.001906618 | -0.152981482 | 0.149168246 | 8.390962456 | -0.030163421 | 0.976821845 | 0.9989817 | -5.256318706 |
| **A_55_P2301941** | A530030E2 | 0.005097218 | -0.399997006 | 0.410191442 | 13.56182844 | 0.03007366 | 0.976890795 | 0.9989817 | -5.256320987 |
| **A_55_P2054628** | Nrxn1 | 0.001296391 | -0.102451518 | 0.105044299 | 8.160501212 | 0.029865214 | 0.977050915 | 0.9989817 | -5.256326259 |
| **A_52_P552665** | Fzd7 | -0.002907959 | -0.235819807 | 0.230003889 | 8.310666515 | -0.029840479 | 0.977069915 | 0.9989817 | -5.256326882 |
| **A_55_P2161675** | Zfp133-ps | 0.002071891 | -0.164000988 | 0.168144771 | 9.510876772 | 0.029817924 | 0.977087241 | 0.9989817 | -5.25632745 |
| **A_55_P2116305** | Tnfrsf25 | 0.005055277 | -0.400749358 | 0.410859913 | 8.706250409 | 0.029773996 | 0.977120985 | 0.9989817 | -5.256328555 |
| **A_51_P227222** | Adamts2 | 0.005261832 | -0.41730418 | 0.427827843 | 11.47220367 | 0.029761274 | 0.977130757 | 0.9989817 | -5.256328874 |
| **A_51_P216605** | Hbp1 | -0.00183209 | -0.149503815 | 0.145839636 | 8.183116028 | -0.029652307 | 0.977214462 | 0.9989817 | -5.256331606 |
| **A_51_P330044** | Cyp2j9 | 0.001422554 | -0.113334075 | 0.116179182 | 7.932273175 | 0.029627838 | 0.977233257 | 0.9989817 | -5.256332218 |
| **A_55_P2038952** | Ubb | -0.004414327 | -0.36173439 | 0.352905737 | 15.84852149 | -0.029526787 | 0.977310882 | 0.9989817 | -5.25633474 |
| **A_51_P398878** | Uck2 | -0.004092413 | -0.337746148 | 0.329561322 | 11.5760051 | -0.02931518 | 0.977473432 | 0.9989817 | -5.256339994 |
| **A_55_P1974204** | Fam172a | 0.001967967 | -0.158743569 | 0.162679503 | 11.10926855 | 0.02926711 | 0.977510358 | 0.9989817 | -5.256341182 |
| **A_51_P336691** | Srsf9 | -0.001910686 | -0.158141696 | 0.154320323 | 11.5663942 | -0.029230165 | 0.977538738 | 0.9989817 | -5.256342094 |
| **A_55_P1957245** | NA | 0.002656817 | -0.214730561 | 0.220044196 | 9.00869381 | 0.029210333 | 0.977553973 | 0.9989817 | -5.256342583 |
| **A_55_P2032099** | Zfp715 | -0.001448164 | -0.119998487 | 0.11710216 | 8.14800316 | -0.029196045 | 0.977564948 | 0.9989817 | -5.256342935 |
| **A_66_P138053** | Gm9917 | -0.001442361 | -0.119631496 | 0.116746773 | 7.982312506 | -0.029167935 | 0.977586542 | 0.9989817 | -5.256343628 |
| **A_55_P2184508** | NA | 0.001650754 | -0.133681725 | 0.136983233 | 14.74678139 | 0.029153416 | 0.977597695 | 0.9989817 | -5.256343985 |
| **A_55_P2068429** | Zfp958 | 0.002783737 | -0.225883224 | 0.231450698 | 11.2216088 | 0.029096046 | 0.977641766 | 0.9989817 | -5.256345395 |
| **A_51_P489452** | Cdo1 | -0.002160737 | -0.181481323 | 0.17715985 | 8.488515281 | -0.028799229 | 0.977869775 | 0.9989817 | -5.256352646 |
| **A_55_P2006861** | Trio | -0.001801778 | -0.151592087 | 0.147988532 | 8.119355388 | -0.028749254 | 0.977908165 | 0.9989817 | -5.256353859 |
| **A_55_P1981731** | Nr2c2 | -0.002874535 | -0.242627497 | 0.236878427 | 8.822898057 | -0.02865581 | 0.977979948 | 0.9989817 | -5.256356123 |

| | | | | | | | | | |
|---|---|---|---|---|---|---|---|---|---|
| **A_55_P2015742** | Cggbp1 | 0.002188628 | -0.1816926 | 0.186069856 | 9.923243542 | 0.028447471 | 0.978139993 | 0.9989817 | -5.256361143 |
| **A_51_P384754** | Ldb1 | -0.003549842 | -0.302767359 | 0.295667676 | 10.15121236 | -0.028355095 | 0.978210956 | 0.9989817 | -5.256363357 |
| **A_55_P2039601** | Drosha | 0.001408685 | -0.117585391 | 0.120402762 | 11.29813342 | 0.028294226 | 0.978257715 | 0.9989817 | -5.256364812 |
| **A_55_P2075080** | Fat2 | -0.001393941 | -0.119179771 | 0.11639189 | 7.989715971 | -0.028285272 | 0.978264594 | 0.9989817 | -5.256365026 |
| **A_51_P311540** | Cox6a1 | 0.001309955 | -0.109776644 | 0.112396554 | 17.94230859 | 0.02818407 | 0.978342337 | 0.9989817 | -5.256367437 |
| **A_51_P421303** | Caly | -0.002017974 | -0.173539776 | 0.169503828 | 8.306721151 | -0.028119354 | 0.978392052 | 0.9989817 | -5.256368974 |
| **A_51_P394735** | Arhgef18 | -0.00139027 | -0.12022366 | 0.11744312 | 9.399204269 | -0.027962103 | 0.978512853 | 0.9989817 | -5.256372696 |
| **A_51_P154585** | Vmn1r195 | -0.001937667 | -0.168310159 | 0.164434825 | 8.083230924 | -0.027835997 | 0.978609729 | 0.9989817 | -5.256375665 |
| **A_52_P748958** | Cfl1 | 0.002979372 | -0.253506096 | 0.25946484 | 13.38140583 | 0.0277633 | 0.978665576 | 0.9989817 | -5.25637737 |
| **A_51_P439996** | Gp5 | 0.001570507 | -0.133720076 | 0.136861089 | 7.741619385 | 0.027744788 | 0.978679797 | 0.9989817 | -5.256377804 |
| **A_55_P2000304** | Terf1 | -0.002194358 | -0.19153949 | 0.187150774 | 9.201570451 | -0.027698896 | 0.978715052 | 0.9989817 | -5.256378877 |
| **A_55_P2123822** | Rbbp7 | 0.002409209 | -0.206158969 | 0.210977387 | 14.26657983 | 0.027608039 | 0.97878485 | 0.9989817 | -5.256380998 |
| **A_52_P65494** | Iqgap2 | 0.001433565 | -0.123149118 | 0.126016248 | 7.749961578 | 0.027502288 | 0.978866089 | 0.9989817 | -5.256383457 |
| **A_55_P2028171** | Dppa5a | 0.001122096 | -0.096462665 | 0.098706856 | 7.881206211 | 0.027482546 | 0.978881256 | 0.9989817 | -5.256383915 |
| **A_51_P406165** | Nxpe2 | -0.001232337 | -0.108572052 | 0.106107378 | 7.839486869 | -0.02743963 | 0.978914225 | 0.9989817 | -5.256384909 |
| **A_52_P586004** | Hk3 | 0.001230247 | -0.106073725 | 0.10853422 | 7.782243463 | 0.027402219 | 0.978942964 | 0.9989817 | -5.256385775 |
| **A_51_P282667** | Hexa | 0.002072076 | -0.178658824 | 0.182802975 | 10.95603042 | 0.027402004 | 0.978943129 | 0.9989817 | -5.25638578 |
| **A_66_P101108** | Tnrc18 | 0.002378688 | -0.205913853 | 0.210671229 | 10.54114665 | 0.027294354 | 0.979025829 | 0.9989817 | -5.256388265 |
| **A_55_P2175955** | Ano9 | -0.001227951 | -0.108889556 | 0.106433654 | 7.661948128 | -0.027260219 | 0.979052052 | 0.9989817 | -5.25638905 |
| **A_51_P131561** | Plet1 | -0.002684551 | -0.238578027 | 0.233208925 | 8.072534462 | -0.027199747 | 0.979098509 | 0.9989817 | -5.25639044 |
| **A_51_P427663** | Cnn2 | -0.003095817 | -0.275505052 | 0.269313418 | 11.86829149 | -0.02716205 | 0.979127468 | 0.9989817 | -5.256391305 |
| **A_55_P2149511** | Mff | 0.001856917 | -0.161697049 | 0.165410883 | 12.73990993 | 0.02713567 | 0.979147734 | 0.9989817 | -5.256391909 |
| **A_51_P473272** | Tspan6 | -0.001770376 | -0.158977484 | 0.155436733 | 11.8769422 | -0.026915499 | 0.979316876 | 0.9989817 | -5.256396931 |
| **A_51_P314323** | Naa38 | 0.001323537 | -0.116337032 | 0.118984106 | 13.98130017 | 0.02688527 | 0.979340099 | 0.9989817 | -5.256397617 |
| **A_52_P472799** | Ecd | 0.001641888 | -0.144378656 | 0.147662432 | 10.63953709 | 0.026874401 | 0.979348449 | 0.9989817 | -5.256397864 |
| **A_55_P2116525** | NA | 0.002024575 | -0.178182636 | 0.182231785 | 8.652682588 | 0.026851636 | 0.979365938 | 0.9989817 | -5.25639838 |
| **A_51_P156927** | Arid4a | -0.001373919 | -0.123820281 | 0.121072442 | 8.529951889 | -0.026817883 | 0.979391868 | 0.9989817 | -5.256399144 |
| **A_52_P139740** | Mdp1 | -0.002100365 | -0.190008992 | 0.185808262 | 10.80508726 | -0.026715122 | 0.979470813 | 0.9989817 | -5.256401465 |
| **A_52_P312371** | Zfp345 | 0.001260273 | -0.111711168 | 0.114231714 | 7.954704096 | 0.026662757 | 0.979511042 | 0.9989817 | -5.256402645 |
| **A_55_P2024799** | NA | -0.00235874 | -0.214427278 | 0.209709798 | 12.7933228 | -0.026583548 | 0.979571893 | 0.9989817 | -5.256404424 |
| **A_51_P270949** | H1f5 | 0.002805551 | -0.25001028 | 0.255621383 | 13.21336291 | 0.026523026 | 0.979618389 | 0.9989817 | -5.256405781 |
| **A_55_P1953241** | Mafk | -0.001418308 | -0.129752544 | 0.126915927 | 9.112180874 | -0.026414186 | 0.979702004 | 0.9989817 | -5.256408212 |
| **A_55_P2011286** | Hopx | 0.005028189 | -0.451283275 | 0.461339654 | 13.15761772 | 0.02633658 | 0.979761625 | 0.9989817 | -5.256409939 |
| **A_51_P163797** | Bdp1 | -0.001155941 | -0.106204397 | 0.103892514 | 9.15689368 | -0.026299971 | 0.97978975 | 0.9989817 | -5.256410752 |
| **A_51_P516728** | Hap1 | 0.005544633 | -0.499711996 | 0.510801263 | 10.45064786 | 0.026228288 | 0.97984482 | 0.9989817 | -5.256412341 |
| **A_51_P359262** | Sec11c | 0.001664741 | -0.150516804 | 0.153846286 | 9.524888394 | 0.026145311 | 0.979908567 | 0.9989817 | -5.256414175 |
| **A_55_P1977149** | Susd6 | 0.001773313 | -0.160476407 | 0.164023033 | 11.46615161 | 0.026122245 | 0.979926288 | 0.9989817 | -5.256414684 |
| **A_55_P2000369** | Aqp4 | -0.001951888 | -0.180616457 | 0.176712681 | 9.608128935 | -0.02611113 | 0.979934827 | 0.9989817 | -5.256414929 |
| **A_65_P16081** | NA | -0.001949089 | -0.180752844 | 0.176854666 | 8.693411249 | -0.026053391 | 0.979979185 | 0.9989817 | -5.2564162 |
| **A_51_P319460** | Osmr | 0.002256546 | -0.206091758 | 0.21060485 | 9.531876674 | 0.025885906 | 0.980107856 | 0.9989817 | -5.25641987 |
| **A_52_P48681** | Cldn1 | -0.001060772 | -0.099066699 | 0.096945155 | 7.650075662 | -0.025868953 | 0.980120881 | 0.9989817 | -5.256420241 |
| **A_55_P2117984** | Vangl2 | -0.001787246 | -0.167064086 | 0.163489594 | 8.718428453 | -0.025845293 | 0.980139058 | 0.9989817 | -5.256420757 |
| **A_51_P395842** | Timm21 | 0.002021545 | -0.184971478 | 0.189014568 | 10.70570047 | 0.025838493 | 0.980144282 | 0.9989817 | -5.256420905 |
| **A_55_P2031043** | Stat5b | -0.002031791 | -0.189996332 | 0.185932749 | 11.38010563 | -0.025835229 | 0.980146789 | 0.9989817 | -5.256420976 |
| **A_55_P2081303** | Tet2 | 0.002382813 | -0.2185338 | 0.223299426 | 10.83941641 | 0.025779276 | 0.980189775 | 0.9989817 | -5.256422195 |
| **A_51_P302377** | Fgfbp1 | -0.001213639 | -0.113762152 | 0.111334874 | 7.720375741 | -0.025772631 | 0.980194881 | 0.9989817 | -5.256422339 |
| **A_66_P133240** | Hnrnpll | 0.001798476 | -0.165014198 | 0.16861115 | 11.12178954 | 0.025768241 | 0.980198253 | 0.9989817 | -5.256422435 |
| **A_55_P2326260** | 4930525G2 | -0.00113893 | -0.106970042 | 0.104692182 | 8.128946538 | -0.025721302 | 0.980234315 | 0.9989817 | -5.256423455 |

| | | | | | | | | | |
|---|---|---|---|---|---|---|---|---|---|
| **A_55_P2062549** | Gm6524 | -0.00108713 | -0.102356367 | 0.100182107 | 9.732065501 | -0.025657415 | 0.980283397 | 0.9989817 | -5.25642484 |
| **A_52_P594927** | Atl3 | 0.00156636 | -0.144487496 | 0.147620216 | 8.171897884 | 0.025632304 | 0.980302689 | 0.9989817 | -5.256425383 |
| **A_51_P319449** | Acat1 | -0.001719556 | -0.162731436 | 0.159292323 | 13.1437191 | -0.025525108 | 0.980385043 | 0.9989817 | -5.256427697 |
| **A_52_P223704** | Faah | -0.001045839 | -0.099321388 | 0.09722971 | 7.661003837 | -0.025434818 | 0.98045441 | 0.9989817 | -5.256429639 |
| **A_55_P2186961** | NA | -0.001818786 | -0.172727263 | 0.169089691 | 9.107504522 | -0.025434726 | 0.980454481 | 0.9989817 | -5.256429641 |
| **A_55_P2102471** | Gm16378 | -0.003242853 | -0.309527665 | 0.303041959 | 12.86647507 | -0.025305269 | 0.980553939 | 0.9989817 | -5.256432413 |
| **A_52_P350950** | Dennd2c | 0.001124162 | -0.105396375 | 0.1076447 | 8.02053477 | 0.02522347 | 0.980616782 | 0.9989817 | -5.256434157 |
| **A_52_P359965** | Cpd | -0.00245968 | -0.237181322 | 0.232261961 | 10.33145391 | -0.025045786 | 0.980753292 | 0.9989817 | -5.256437926 |
| **A_51_P221256** | Slc34a2 | 0.00102549 | -0.096855347 | 0.098906327 | 7.647904707 | 0.025040498 | 0.980757355 | 0.9989817 | -5.256438037 |
| **A_55_P2008277** | Tcp10c | -0.001141532 | -0.110233632 | 0.107950568 | 7.899961565 | -0.02500943 | 0.980781223 | 0.9989817 | -5.256438694 |
| **A_51_P484764** | Fam3a | 0.00165239 | -0.156369229 | 0.159674009 | 9.725595674 | 0.024992242 | 0.980794429 | 0.9989817 | -5.256439056 |
| **A_55_P2235931** | Cacna1c | -0.003467276 | -0.335562009 | 0.328627457 | 12.42000827 | -0.024953735 | 0.980824013 | 0.9989817 | -5.256439868 |
| **A_66_P136259** | Rbm4b | 0.001171526 | -0.111403936 | 0.113746987 | 8.266580073 | 0.02487237 | 0.980886524 | 0.9989817 | -5.256441578 |
| **A_52_P613643** | Zfp3 | 0.001489775 | -0.143042854 | 0.146022405 | 8.879481135 | 0.024635651 | 0.981068391 | 0.9989817 | -5.256446524 |
| **A_55_P1958200** | P2rx1 | 0.001902991 | -0.183731106 | 0.187537089 | 8.358580143 | 0.024501247 | 0.981171652 | 0.9989817 | -5.25644931 |
| **A_55_P2210301** | Gm12992 | 0.002372496 | -0.22908865 | 0.233833642 | 8.506450946 | 0.02449833 | 0.981173893 | 0.9989817 | -5.256449371 |
| **A_52_P576100** | Sh2b1 | -0.001846474 | -0.183240713 | 0.179547766 | 11.6650578 | -0.024329255 | 0.981303792 | 0.9989817 | -5.256452854 |
| **A_66_P119884** | Wdr91 | 0.001682298 | -0.166248149 | 0.169612746 | 9.604131949 | 0.02394323 | 0.981600373 | 0.9989817 | -5.256460718 |
| **A_55_P2139402** | Amy2a5 | -0.000982612 | -0.099390088 | 0.097424865 | 7.661107503 | -0.023865084 | 0.981660413 | 0.9989817 | -5.256462294 |
| **A_55_P1957911** | Tom1 | 0.001288889 | -0.128604924 | 0.131182701 | 8.595327352 | 0.023715704 | 0.981775182 | 0.9989817 | -5.256465293 |
| **A_51_P309589** | 2700099C1 | -0.001326085 | -0.134999852 | 0.132347683 | 8.698611993 | -0.023710145 | 0.981779453 | 0.9989817 | -5.256465404 |
| **A_66_P119680** | Polr2e | 0.00118647 | -0.119003721 | 0.12137666 | 12.81368768 | 0.023593742 | 0.981868886 | 0.9989817 | -5.256467728 |
| **A_55_P2079729** | NA | -0.000998918 | -0.102485235 | 0.100487399 | 15.55125638 | -0.023525105 | 0.98192162 | 0.9989817 | -5.256469093 |
| **A_55_P2134804** | Cinp | -0.00170461 | -0.175177192 | 0.171767972 | 9.015858531 | -0.023485689 | 0.981951904 | 0.9989817 | -5.256469875 |
| **A_55_P2097072** | Gm7219 | -0.001009588 | -0.103852096 | 0.10183292 | 7.876599606 | -0.02346284 | 0.981969459 | 0.9989817 | -5.256470327 |
| **A_55_P2357860** | Atxn7l1 | -0.001468561 | -0.151629724 | 0.148692602 | 8.223308885 | -0.02337457 | 0.982037278 | 0.9989817 | -5.256472072 |
| **A_55_P2013113** | Bak1 | -0.001340952 | -0.1395365 | 0.136854596 | 12.39038595 | -0.023191468 | 0.982177957 | 0.9989817 | -5.25647567 |
| **A_55_P1978366** | NA | 0.00183877 | -0.188809596 | 0.192487137 | 11.72325413 | 0.02305173 | 0.982285321 | 0.9989817 | -5.256478397 |
| **A_55_P2372902** | 5330438D1 | 0.001873158 | -0.192569393 | 0.196315709 | 9.057940046 | 0.023024609 | 0.982306158 | 0.9989817 | -5.256478924 |
| **A_55_P2134735** | Fdxacb1 | -0.00101969 | -0.106992509 | 0.104953129 | 8.716338493 | -0.022997607 | 0.982326905 | 0.9989817 | -5.256479448 |
| **A_55_P1958306** | Thrap3 | 0.001155877 | -0.119211252 | 0.121523005 | 12.45945668 | 0.02295159 | 0.98236226 | 0.9989817 | -5.25648034 |
| **A_52_P163928** | Poldip2 | 0.003203472 | -0.330435586 | 0.336842531 | 11.43640224 | 0.022948443 | 0.982364678 | 0.9989817 | -5.256480401 |
| **A_55_P2052048** | Pih1d2 | -0.001130191 | -0.119093139 | 0.116832756 | 7.884645106 | -0.022898947 | 0.982402707 | 0.9989817 | -5.256481359 |
| **A_55_P2096144** | Gnao1 | 0.0051655 | -0.533988201 | 0.5443192 | 9.515823186 | 0.022898597 | 0.982402976 | 0.9989817 | -5.256481366 |
| **A_52_P119117** | Tmem258 | 0.001082371 | -0.112115984 | 0.114280726 | 13.23502308 | 0.022853095 | 0.982437936 | 0.9989817 | -5.256482244 |
| **A_55_P1955147** | Camk1d | -0.007938302 | -0.840984743 | 0.825108139 | 8.961378126 | -0.02277548 | 0.98249757 | 0.9989817 | -5.256483739 |
| **A_52_P444601** | Zfp277 | -0.001716479 | -0.182844356 | 0.179411397 | 10.45441784 | -0.022649696 | 0.982594212 | 0.9989817 | -5.25648615 |
| **A_55_P2164090** | Kcnj15 | 0.002479028 | -0.260139451 | 0.265097507 | 8.110078466 | 0.02256136 | 0.982662084 | 0.9989817 | -5.256487835 |
| **A_66_P132613** | C230057M | -0.002288895 | -0.247094214 | 0.242516424 | 9.058370946 | -0.022346738 | 0.982826984 | 0.9989817 | -5.256491902 |
| **A_55_P2163725** | Slc7a9 | 0.00121512 | -0.129080084 | 0.131510325 | 7.710332321 | 0.022289483 | 0.982870975 | 0.9989817 | -5.256492981 |
| **A_51_P324442** | Wdr35 | 0.001004869 | -0.10697607 | 0.108985808 | 9.36864613 | 0.022241879 | 0.982907551 | 0.9989817 | -5.256493875 |
| **A_55_P2010161** | Fastkd5 | -0.001447037 | -0.157202637 | 0.154308563 | 11.00734507 | -0.022204711 | 0.982936108 | 0.9989817 | -5.256494572 |
| **A_55_P1989568** | Tasp1 | -0.001126802 | -0.12250398 | 0.120250376 | 8.296010799 | -0.02218808 | 0.982948887 | 0.9989817 | -5.256494884 |
| **A_51_P179907** | Appl2 | -0.002461616 | -0.267800112 | 0.26287688 | 9.029522675 | -0.022173241 | 0.982960288 | 0.9989817 | -5.256495162 |
| **A_55_P2175060** | Plpp5 | -0.001122694 | -0.123097372 | 0.120851985 | 8.360622407 | -0.021998887 | 0.983094251 | 0.9989817 | -5.256498411 |
| **A_51_P457196** | Sfrp4 | 0.001060317 | -0.114138553 | 0.116259188 | 7.74761381 | 0.021998688 | 0.983094403 | 0.9989817 | -5.256498415 |
| **A_51_P382121** | Ormdl1 | 0.00170724 | -0.184070503 | 0.187484983 | 10.39252337 | 0.021963928 | 0.983121111 | 0.9989817 | -5.25649906 |
| **A_55_P2005552** | Arhgef10l | 0.001194595 | -0.129869818 | 0.132259009 | 11.76805564 | 0.021784379 | 0.983259066 | 0.9989817 | -5.256502375 |

| | | | | | | | | | |
|---|---|---|---|---|---|---|---|---|---|
| **A_51_P507778** | Sdr42e1 | -0.001150885 | -0.127721493 | 0.125419724 | 8.263132112 | -0.021732417 | 0.983298991 | 0.9989817 | -5.256503329 |
| **A_51_P349803** | Med11 | -0.001523552 | -0.169737662 | 0.166690558 | 10.49521167 | -0.021647313 | 0.98336438 | 0.9989817 | -5.256504887 |
| **A_55_P2022293** | Zfp579 | -0.006197506 | -0.691713953 | 0.679318941 | 11.69951522 | -0.021607683 | 0.98339483 | 0.9989817 | -5.25650561 |
| **A_52_P542794** | 2610507B1 | 0.001659887 | -0.182337261 | 0.185657034 | 13.08276869 | 0.021561377 | 0.983430409 | 0.9989817 | -5.256506454 |
| **A_55_P2081885** | NA | -0.003214013 | -0.363492162 | 0.357064137 | 16.3295573 | -0.02132155 | 0.98361468 | 0.9989817 | -5.256510793 |
| **A_52_P199299** | Urgcp | 0.002595652 | -0.28952535 | 0.294716655 | 10.45789556 | 0.021236984 | 0.983679656 | 0.9989817 | -5.256512312 |
| **A_55_P2016356** | Eif3b | -0.001621686 | -0.184796267 | 0.181552896 | 11.0812749 | -0.021159754 | 0.983738996 | 0.9989817 | -5.256513694 |
| **A_55_P2016959** | Nme2 | 0.000973496 | -0.109068043 | 0.111015035 | 16.60711831 | 0.021143982 | 0.983751115 | 0.9989817 | -5.256513975 |
| **A_55_P2080168** | Dgkk | 0.00223679 | -0.251159678 | 0.255633258 | 9.491706171 | 0.021097635 | 0.983786726 | 0.9989817 | -5.256514801 |
| **A_55_P1981714** | Rreb1 | 0.002155851 | -0.243259958 | 0.24757166 | 11.76814618 | 0.020995454 | 0.983865237 | 0.9989817 | -5.256516616 |
| **A_55_P1953082** | NA | 0.000880001 | -0.099661361 | 0.101421363 | 7.643840578 | 0.020919326 | 0.983923731 | 0.9989817 | -5.256517963 |
| **A_55_P2142830** | Ninj1 | -0.001398086 | -0.161499433 | 0.15870326 | 12.96684536 | -0.020871231 | 0.983960685 | 0.9989817 | -5.256518811 |
| **A_55_P1995537** | Mpo | 0.002676115 | -0.305964903 | 0.311317133 | 8.143072294 | 0.020723361 | 0.984074303 | 0.9989817 | -5.256521407 |
| **A_55_P2046178** | NA | 0.00118675 | -0.135961902 | 0.138335403 | 15.30755535 | 0.02068126 | 0.984106652 | 0.9989817 | -5.256522142 |
| **A_55_P2067351** | NA | -0.001132228 | -0.132590048 | 0.130325591 | 9.268543264 | -0.020585276 | 0.984180403 | 0.9989817 | -5.256523814 |
| **A_55_P2009477** | 9530068E0 | 0.001066612 | -0.12320362 | 0.125336843 | 13.76708411 | 0.020513908 | 0.984235239 | 0.9989817 | -5.256525051 |
| **A_51_P152404** | Ikbip | -0.002092419 | -0.246555691 | 0.242370853 | 10.15116009 | -0.020457106 | 0.984278884 | 0.9989817 | -5.256526033 |
| **A_51_P230507** | Ell2 | 0.00121796 | -0.141400519 | 0.14383644 | 7.909556712 | 0.020411105 | 0.98431423 | 0.9989817 | -5.256526827 |
| **A_55_P2030578** | Tns1 | 0.001803546 | -0.209605796 | 0.213212888 | 13.4464038 | 0.020389763 | 0.984330629 | 0.9989817 | -5.256527194 |
| **A_51_P133684** | Csrp3 | 0.002087116 | -0.243997082 | 0.248171315 | 15.04084921 | 0.020270853 | 0.984421995 | 0.9989817 | -5.256529234 |
| **A_55_P2118575** | Tgs1 | -0.001008083 | -0.120123026 | 0.11810686 | 9.975623154 | -0.020227364 | 0.984455411 | 0.9989817 | -5.256529978 |
| **A_55_P2158319** | Ntan1 | -0.000956579 | -0.114137145 | 0.112223986 | 10.39241591 | -0.020200319 | 0.984476192 | 0.9989817 | -5.256530439 |
| **A_55_P2137941** | Fxyd2 | 0.002574316 | -0.302224494 | 0.307373126 | 8.376527157 | 0.020186348 | 0.984486927 | 0.9989817 | -5.256530677 |
| **A_52_P555723** | Ywhag | -0.001408773 | -0.169247935 | 0.166430389 | 11.77054487 | -0.020061198 | 0.984583089 | 0.9989817 | -5.256532803 |
| **A_55_P2122255** | Polm | 0.0012314 | -0.145767045 | 0.148229846 | 8.286367678 | 0.020021457 | 0.984613625 | 0.9989817 | -5.256533475 |
| **A_51_P464023** | Chst12 | 0.001385654 | -0.164074597 | 0.166845904 | 8.847433188 | 0.020015674 | 0.984618069 | 0.9989817 | -5.256533573 |
| **A_55_P2357303** | 5430401H0 | -0.00083603 | -0.100884298 | 0.099212237 | 7.728455478 | -0.019972005 | 0.984651623 | 0.9989817 | -5.256534309 |
| **A_55_P2103422** | Psmc2 | 0.001162223 | -0.137996639 | 0.140321084 | 13.60536533 | 0.019961245 | 0.98465989 | 0.9989817 | -5.256534491 |
| **A_51_P427516** | Thsd1 | -0.001787086 | -0.215908839 | 0.212334667 | 9.917973138 | -0.019947743 | 0.984670265 | 0.9989817 | -5.256534718 |
| **A_55_P2118090** | NA | 0.001510853 | -0.180082237 | 0.183103943 | 10.44152459 | 0.019885289 | 0.984718253 | 0.9989817 | -5.256535768 |
| **A_55_P1985825** | NA | 0.001537606 | -0.183807782 | 0.186882994 | 7.981389857 | 0.019827709 | 0.984762497 | 0.9989817 | -5.256536733 |
| **A_55_P2110512** | Lmo7 | -0.001693808 | -0.206403591 | 0.203015975 | 12.05017083 | -0.019775827 | 0.984802361 | 0.9989817 | -5.2565376 |
| **A_55_P1957004** | Timm10 | -0.001619736 | -0.197512127 | 0.194272655 | 12.90559254 | -0.019762224 | 0.984812814 | 0.9989817 | -5.256537827 |
| **A_52_P514352** | Kcnk5 | -0.001005811 | -0.122970698 | 0.120959076 | 7.820061371 | -0.019710181 | 0.984852803 | 0.9989817 | -5.256538694 |
| **A_55_P2046253** | Tmem239 | -0.001594567 | -0.196306457 | 0.193117324 | 13.16678321 | -0.019573086 | 0.984958144 | 0.9989817 | -5.256540966 |
| **A_52_P561671** | Msx1 | -0.001100791 | -0.135885444 | 0.133683862 | 8.146673961 | -0.01951972 | 0.984999149 | 0.9989817 | -5.256541847 |
| **A_52_P66226** | Rab33a | 0.002365912 | -0.287502941 | 0.292234764 | 8.967148446 | 0.019507704 | 0.985008383 | 0.9989817 | -5.256542045 |
| **A_51_P377210** | Caap1 | 0.00102783 | -0.125177395 | 0.127233055 | 8.192762693 | 0.019464941 | 0.985041241 | 0.9989817 | -5.256542748 |
| **A_52_P869802** | Thoc3 | -0.001807677 | -0.224078361 | 0.220463007 | 9.686088573 | -0.019437829 | 0.985062074 | 0.9989817 | -5.256543193 |
| **A_66_P105843** | Cyth2 | -0.001545299 | -0.191731144 | 0.188640547 | 12.40945854 | -0.01941974 | 0.985075973 | 0.9989817 | -5.25654349 |
| **A_55_P2153381** | Ermap | 0.001917127 | -0.234066631 | 0.237900885 | 8.823623754 | 0.019416811 | 0.985078223 | 0.9989817 | -5.256543538 |
| **A_55_P2107192** | NA | 0.001683703 | -0.205621427 | 0.208988833 | 14.53565927 | 0.019411745 | 0.985082116 | 0.9989817 | -5.256543621 |
| **A_51_P281089** | S100a6 | -0.005251741 | -0.653190087 | 0.642686606 | 12.31881855 | -0.01937219 | 0.98511251 | 0.9989817 | -5.256544268 |
| **A_55_P1958902** | NA | -0.001972646 | -0.245423774 | 0.241478481 | 10.43870627 | -0.019366298 | 0.985117037 | 0.9989817 | -5.256544364 |
| **A_55_P2066523** | Akt2 | 0.00191284 | -0.234246308 | 0.238071988 | 8.87712893 | 0.019359005 | 0.985122641 | 0.9989817 | -5.256544484 |
| **A_55_P1961645** | Vcp | -0.001701358 | -0.212424563 | 0.209021848 | 14.42231168 | -0.019297118 | 0.985170195 | 0.9989817 | -5.256545493 |
| **A_55_P2018283** | Hmgb1 | -0.00090932 | -0.113908351 | 0.112089712 | 9.143091826 | -0.019233178 | 0.985219325 | 0.9989817 | -5.256546533 |
| **A_51_P177691** | Gabrr1 | -0.000881465 | -0.11072978 | 0.108966849 | 7.956216564 | -0.019178775 | 0.985261128 | 0.9989817 | -5.256547414 |

| | | | | | | | | | |
|---|---|---|---|---|---|---|---|---|---|
| **A_55_P2183808** | NA | -0.001581038 | -0.19898501 | 0.195822933 | 13.95652918 | -0.019142369 | 0.985289102 | 0.9989817 | -5.256548003 |
| **A_55_P1993728** | Tmem116 | 0.000820445 | -0.10186811 | 0.103509 | 8.200847798 | 0.019095745 | 0.985324927 | 0.9989817 | -5.256548756 |
| **A_55_P2045602** | Gpkow | -0.001067487 | -0.135888868 | 0.133753895 | 9.669420368 | -0.018923999 | 0.985456896 | 0.9989817 | -5.256551511 |
| **A_55_P2017369** | NA | 0.000902726 | -0.113361619 | 0.115167071 | 11.91331829 | 0.018882278 | 0.985488954 | 0.9989817 | -5.256552177 |
| **A_55_P2013391** | Gm4402 | -0.001589246 | -0.203383531 | 0.200205039 | 9.568967572 | -0.018823107 | 0.985534421 | 0.9989817 | -5.256553118 |
| **A_51_P425798** | Mtus2 | -0.000956854 | -0.122845243 | 0.120931536 | 10.46637149 | -0.018762567 | 0.98558094 | 0.9989817 | -5.256554078 |
| **A_55_P1986958** | Brms1l | -0.001597041 | -0.205180966 | 0.201986885 | 10.99814449 | -0.018749151 | 0.985591249 | 0.9989817 | -5.256554291 |
| **A_51_P468249** | Phex | 0.000791495 | -0.10021008 | 0.10179307 | 7.671320309 | 0.018729628 | 0.98560625 | 0.9989817 | -5.256554599 |
| **A_51_P251209** | Cacna1d | -0.001001179 | -0.128804042 | 0.126801683 | 7.956254483 | -0.018723217 | 0.985611176 | 0.9989817 | -5.256554701 |
| **A_65_P08239** | Mef2a | -0.000988413 | -0.127188563 | 0.125211738 | 9.286400199 | -0.018719215 | 0.985614252 | 0.9989817 | -5.256554764 |
| **A_55_P1998309** | Sms | -0.001810437 | -0.233657327 | 0.230036454 | 10.8881806 | -0.018663417 | 0.985657126 | 0.9989817 | -5.256555644 |
| **A_51_P371867** | 3830417A1 | -0.000880819 | -0.113717461 | 0.111955824 | 7.730634983 | -0.018657158 | 0.985661936 | 0.9989817 | -5.256555743 |
| **A_55_P2083839** | Atp13a1 | -0.00093187 | -0.120430898 | 0.118567158 | 10.4442084 | -0.018638032 | 0.985676632 | 0.9989817 | -5.256556044 |
| **A_51_P484832** | Ppm1g | -0.003158475 | -0.410832416 | 0.404515467 | 9.801302458 | -0.018517125 | 0.985769537 | 0.9989817 | -5.25655794 |
| **A_52_P573336** | Sbsn | -0.00218906 | -0.28547559 | 0.281097471 | 9.537540595 | -0.018468891 | 0.9858066 | 0.9989817 | -5.256558692 |
| **A_52_P267256** | LOC10086 | -0.003675564 | -0.4811427 | 0.473791572 | 12.95404269 | -0.018398806 | 0.985860454 | 0.9989817 | -5.256559783 |
| **A_66_P100840** | NA | 0.004744247 | -0.612600361 | 0.622088855 | 10.15861145 | 0.018367444 | 0.985884552 | 0.9989817 | -5.256560269 |
| **A_51_P467224** | Fbn1 | -0.002809823 | -0.369524712 | 0.363905065 | 11.34179178 | -0.018313004 | 0.985926384 | 0.9989817 | -5.256561112 |
| **A_51_P395164** | Pex16 | -0.001461685 | -0.19342287 | 0.190499501 | 9.146629611 | -0.018199077 | 0.986013927 | 0.9989817 | -5.256562867 |
| **A_55_P2168301** | Slc46a1 | 0.00099377 | -0.13028021 | 0.13226775 | 9.147237221 | 0.018093236 | 0.986095256 | 0.9989817 | -5.256564488 |
| **A_51_P124748** | Tgfb3 | -0.002614928 | -0.349468035 | 0.344238179 | 10.71811578 | -0.018018693 | 0.986152535 | 0.9989817 | -5.256565624 |
| **A_66_P115161** | Cep85l | 0.000948354 | -0.124848852 | 0.126745559 | 8.361937649 | 0.01801808 | 0.986153006 | 0.9989817 | -5.256565634 |
| **A_55_P2177112** | Prrg2 | 0.00095139 | -0.126025162 | 0.127927941 | 7.91065395 | 0.017907875 | 0.986237689 | 0.9989817 | -5.256567304 |
| **A_55_P1979217** | Abtb1 | 0.001867067 | -0.248294291 | 0.252028424 | 10.89100516 | 0.017838105 | 0.986291301 | 0.9989817 | -5.256568357 |
| **A_52_P91402** | Zcchc17 | 0.000840614 | -0.112216512 | 0.113897739 | 12.25536298 | 0.017770832 | 0.986342995 | 0.9989817 | -5.256569368 |
| **A_51_P134627** | NA | -0.000924241 | -0.12601964 | 0.124171158 | 8.560320798 | -0.017658471 | 0.986429334 | 0.9989817 | -5.256571048 |
| **A_52_P20727** | Nhlrc1 | 0.001457858 | -0.196788593 | 0.199704308 | 9.840241434 | 0.017575953 | 0.986492743 | 0.9989817 | -5.256572275 |
| **A_52_P637172** | Pyroxd2 | 0.001045466 | -0.141522768 | 0.1436137 | 8.135216702 | 0.01752655 | 0.986530705 | 0.9989817 | -5.256573006 |
| **A_55_P2048119** | Slc29a4 | 0.000808374 | -0.110155059 | 0.111771806 | 8.126283372 | 0.017411712 | 0.986618949 | 0.9989817 | -5.256574699 |
| **A_55_P2063871** | NA | 0.001444586 | -0.197186238 | 0.20007541 | 13.18856541 | 0.017382242 | 0.986641594 | 0.9989817 | -5.256575132 |
| **A_51_P244923** | AI837181 | -0.001727532 | -0.240381121 | 0.236926057 | 9.832149695 | -0.017300847 | 0.986704139 | 0.9989817 | -5.256576323 |
| **A_55_P2108803** | Tubgcp4 | -0.000774726 | -0.108012212 | 0.10646276 | 9.334029024 | -0.017266751 | 0.98673034 | 0.9989817 | -5.256576821 |
| **A_55_P1978191** | C2cd4c | -0.001424767 | -0.199013141 | 0.196163608 | 8.014364374 | -0.017234214 | 0.986755342 | 0.9989817 | -5.256577295 |
| **A_55_P2113466** | NA | -0.001093681 | -0.154103056 | 0.151915695 | 9.295226732 | -0.017083691 | 0.986871007 | 0.9989817 | -5.256579474 |
| **A_51_P340947** | Selenoo | -0.000970437 | -0.136739672 | 0.134798798 | 9.928180396 | -0.017083436 | 0.986871203 | 0.9989817 | -5.256579478 |
| **A_52_P506344** | B3gat3 | -0.00180529 | -0.25480725 | 0.251196671 | 8.617139969 | -0.017054232 | 0.986893644 | 0.9989817 | -5.256579899 |
| **A_52_P116628** | Fam234b | 0.000811129 | -0.113089219 | 0.114711478 | 8.093902193 | 0.017020574 | 0.986919508 | 0.9989817 | -5.256580383 |
| **A_55_P2052623** | Pgap2 | -0.000894463 | -0.126974302 | 0.125185375 | 11.08279095 | -0.016956099 | 0.986969052 | 0.9989817 | -5.256581307 |
| **A_66_P116372** | Soat2 | -0.000740958 | -0.106299864 | 0.104817948 | 7.744459563 | -0.016776748 | 0.98710687 | 0.9989817 | -5.25658386 |
| **A_55_P2093696** | NA | 0.001137705 | -0.162117541 | 0.164392952 | 8.007185536 | 0.016656042 | 0.987199624 | 0.9989817 | -5.256585563 |
| **A_51_P371152** | Sos2 | -0.001135527 | -0.164705544 | 0.162434489 | 10.3382342 | -0.016592165 | 0.987248709 | 0.9989817 | -5.256586459 |
| **A_55_P1988273** | Rnf103 | 0.000790074 | -0.113171215 | 0.114751363 | 10.77359505 | 0.016569884 | 0.987265831 | 0.9989817 | -5.256586771 |
| **A_55_P2170345** | Gm6729 | -0.000982221 | -0.142825753 | 0.140861311 | 8.362562362 | -0.016550409 | 0.987280796 | 0.9989817 | -5.256587043 |
| **A_55_P2057291** | Kansl3 | -0.000892408 | -0.129843154 | 0.128058337 | 12.47529639 | -0.016540511 | 0.987288402 | 0.9989817 | -5.256587182 |
| **A_55_P1979156** | Arap3 | 0.000874978 | -0.12685998 | 0.128609937 | 8.496410736 | 0.016371809 | 0.987418038 | 0.9989817 | -5.256589525 |
| **A_52_P161087** | 2610008E1 | -0.000889362 | -0.1313179 | 0.129539176 | 9.50343595 | -0.016297272 | 0.987475315 | 0.9989817 | -5.256590552 |
| **A_51_P375969** | Ces1d | 0.000653972 | -0.096543887 | 0.09785183 | 7.662205535 | 0.016080942 | 0.987641551 | 0.9989817 | -5.256593508 |
| **A_55_P2110601** | Slc35a2 | -0.000874829 | -0.131129786 | 0.129380128 | 9.674844177 | -0.016052322 | 0.987663544 | 0.9989817 | -5.256593896 |

| | | | | | | | | | |
|---|---|---|---|---|---|---|---|---|---|
| **A_55_P2014611** | Ccdc12 | -0.001554889 | -0.233406056 | 0.230296278 | 11.79555118 | -0.016028734 | 0.98768167 | 0.9989817 | -5.256594216 |
| **A_55_P2081656** | NA | -0.001555845 | -0.23473893 | 0.231627239 | 10.48297091 | -0.015946984 | 0.98774449 | 0.9989817 | -5.256595319 |
| **A_66_P102049** | Zfp1 | 0.00075621 | -0.112792422 | 0.114304843 | 10.99819814 | 0.015917317 | 0.987767287 | 0.9989817 | -5.256595718 |
| **A_55_P2048284** | Aasdhppt | -0.000967636 | -0.147711302 | 0.14577603 | 8.869156948 | -0.015760206 | 0.987888018 | 0.9989817 | -5.256597818 |
| **A_51_P332136** | Zmat5 | 0.00166283 | -0.250733608 | 0.254059269 | 9.129683357 | 0.01574613 | 0.987898835 | 0.9989817 | -5.256598005 |
| **A_51_P202745** | Zc3h13 | 0.001653128 | -0.249934194 | 0.253240451 | 10.95315635 | 0.015704604 | 0.987930745 | 0.9989817 | -5.256598556 |
| **A_55_P2101426** | Mrps25 | 0.00126502 | -0.191542016 | 0.194072055 | 12.40634581 | 0.015681355 | 0.98794861 | 0.9989817 | -5.256598864 |
| **A_55_P2083252** | Mtus2 | 0.001247897 | -0.189504105 | 0.1919999 | 13.78117602 | 0.015635757 | 0.98798365 | 0.9989817 | -5.256599467 |
| **A_55_P2178525** | Rpl26 | -0.0010315 | -0.160634505 | 0.158571505 | 15.67850531 | -0.015446753 | 0.988128889 | 0.9989817 | -5.256601946 |
| **A_55_P2004727** | NA | 0.001149358 | -0.17956177 | 0.181860485 | 8.062801923 | 0.015201254 | 0.988317543 | 0.9989817 | -5.256605121 |
| **A_52_P500002** | Tspyl5 | -0.000973067 | -0.154103243 | 0.152157109 | 8.506014329 | -0.015187667 | 0.988327984 | 0.9989817 | -5.256605295 |
| **A_52_P198727** | Cfap410 | 0.001549992 | -0.242795651 | 0.245895636 | 9.18273968 | 0.015161218 | 0.988348309 | 0.9989817 | -5.256605634 |
| **A_55_P2020248** | NA | 0.000997798 | -0.156506925 | 0.158502522 | 14.42102046 | 0.015141133 | 0.988363743 | 0.9989817 | -5.256605891 |
| **A_55_P2185526** | 1700016P0 | -0.001161508 | -0.185132841 | 0.182809825 | 8.654064901 | -0.015089723 | 0.988403249 | 0.9989817 | -5.256606547 |
| **A_52_P279599** | Ube2k | -0.000887271 | -0.141845959 | 0.140071416 | 10.62652447 | -0.015044363 | 0.988438106 | 0.9989817 | -5.256607123 |
| **A_55_P1971458** | Jak2 | 0.001245355 | -0.196928435 | 0.199419144 | 10.19362141 | 0.01501952 | 0.988457197 | 0.9989817 | -5.256607439 |
| **A_51_P380013** | Bcl2 | -0.001085641 | -0.174480263 | 0.172308981 | 9.393109748 | -0.014964416 | 0.988499542 | 0.9989817 | -5.256608136 |
| **A_55_P1975326** | Tmem259 | 0.000950193 | -0.151154513 | 0.153054899 | 13.38880438 | 0.014930636 | 0.9885255 | 0.9989817 | -5.256608562 |
| **A_55_P2142298** | Ywhaq | -0.001521504 | -0.24517421 | 0.242131202 | 13.80823659 | -0.014924885 | 0.98852992 | 0.9989817 | -5.256608635 |
| **A_51_P171965** | Sdhaf4 | -0.001867982 | -0.301372098 | 0.297636133 | 11.01483457 | -0.014906619 | 0.988543957 | 0.9989817 | -5.256608864 |
| **A_55_P1961135** | Atg4d | -0.001452214 | -0.235395396 | 0.232490969 | 12.20028365 | -0.014836424 | 0.988597898 | 0.9989817 | -5.256609746 |
| **A_55_P2119817** | Plekhj1 | 0.000851664 | -0.136534352 | 0.13823768 | 8.318694203 | 0.014816131 | 0.988613492 | 0.9989817 | -5.256609999 |
| **A_55_P2011237** | NA | -0.003462911 | -0.563486348 | 0.556560525 | 15.30958361 | -0.014778975 | 0.988642045 | 0.9989817 | -5.256610464 |
| **A_55_P2158384** | Aanat | 0.000794942 | -0.128106145 | 0.12969603 | 8.56102675 | 0.014739687 | 0.988672236 | 0.9989817 | -5.256610953 |
| **A_55_P2020572** | Psme3 | 0.001512734 | -0.244549714 | 0.247575183 | 10.75189833 | 0.01469354 | 0.988707698 | 0.9989817 | -5.256611526 |
| **A_55_P1986053** | Pcnx | -0.001078301 | -0.177680407 | 0.175523805 | 10.04626073 | -0.014593293 | 0.988784734 | 0.9989817 | -5.256612765 |
| **A_51_P431046** | St8sia3 | -0.000632262 | -0.104323834 | 0.103059309 | 7.656197907 | -0.014573472 | 0.988799965 | 0.9989817 | -5.256613009 |
| **A_51_P423825** | Morn1 | 0.000600863 | -0.097947859 | 0.099149585 | 8.0583334 | 0.014572479 | 0.988800728 | 0.9989817 | -5.256613021 |
| **A_55_P1995512** | Tmem139 | 0.000721721 | -0.118228447 | 0.119671889 | 7.734529205 | 0.014501517 | 0.98885526 | 0.9989817 | -5.256613892 |
| **A_55_P2071354** | Togaram2 | 0.001174284 | -0.194144869 | 0.196493436 | 8.491232954 | 0.014369357 | 0.98895682 | 0.9989817 | -5.256615502 |
| **A_55_P2160253** | 6720475J1 | 0.001278046 | -0.211505777 | 0.214061868 | 9.178498888 | 0.014355452 | 0.988967505 | 0.9989817 | -5.256615671 |
| **A_55_P2038594** | Tmem234 | 0.001055948 | -0.174895913 | 0.177007808 | 12.10186052 | 0.014343582 | 0.988976627 | 0.9989817 | -5.256615814 |
| **A_55_P2006708** | Bmp1 | -0.001215176 | -0.20371895 | 0.201288598 | 9.168414222 | -0.014342182 | 0.988977703 | 0.9989817 | -5.256615831 |
| **A_55_P1990041** | Nxt2 | -0.001305249 | -0.219611597 | 0.217001099 | 10.01672053 | -0.014290129 | 0.989017704 | 0.9989817 | -5.25661646 |
| **A_51_P195215** | Fam149a | -0.001036293 | -0.1751795 | 0.173106915 | 9.481551715 | -0.014222795 | 0.989069447 | 0.9989817 | -5.25661727 |
| **A_55_P2117574** | Ppih | 0.000697585 | -0.116877985 | 0.118273156 | 8.033647464 | 0.014180428 | 0.989102005 | 0.9989817 | -5.256617778 |
| **A_55_P2118309** | NA | -0.001275287 | -0.216703335 | 0.214152762 | 9.221041629 | -0.014148642 | 0.989126431 | 0.9989817 | -5.256618158 |
| **A_55_P2107045** | Myl4 | -0.003591781 | -0.610404747 | 0.603221184 | 17.54458743 | -0.014146994 | 0.989127698 | 0.9989817 | -5.256618178 |
| **A_52_P246753** | Chmp5 | -0.001253233 | -0.21315526 | 0.210648794 | 11.81753436 | -0.014135329 | 0.989136662 | 0.9989817 | -5.256618317 |
| **A_51_P150876** | Lypd3 | -0.000560357 | -0.097751356 | 0.096630643 | 7.666577638 | -0.01377995 | 0.989409759 | 0.9989817 | -5.256622504 |
| **A_55_P1973349** | H4c14 | 0.001442521 | -0.249418574 | 0.252303617 | 14.80485726 | 0.013743523 | 0.989437752 | 0.9989817 | -5.256622927 |
| **A_55_P2174501** | Stx17 | -0.000705573 | -0.123419471 | 0.122008325 | 8.500711715 | -0.013742241 | 0.989438737 | 0.9989817 | -5.256622942 |
| **A_51_P478098** | Epb41l5 | -0.000632453 | -0.11116983 | 0.109904924 | 7.662651995 | -0.013675026 | 0.98949039 | 0.9989817 | -5.256623719 |
| **A_55_P2092085** | Tmsb10 | 0.000740084 | -0.128978768 | 0.130458936 | 14.75940724 | 0.013636003 | 0.989520378 | 0.9989817 | -5.256624169 |
| **A_55_P2104487** | Pld1 | 0.001214679 | -0.212566469 | 0.214995828 | 9.697936148 | 0.013580049 | 0.989563377 | 0.9989817 | -5.256624812 |
| **A_55_P1957708** | NA | 0.000582982 | -0.102589719 | 0.103755682 | 8.158535369 | 0.013505149 | 0.989620936 | 0.9989817 | -5.256625668 |
| **A_55_P1965427** | Uqcrh | -0.000987546 | -0.17660119 | 0.174626097 | 16.97998479 | -0.013440283 | 0.989670783 | 0.9989817 | -5.256626405 |
| **A_66_P135941** | Fam221a | -0.00072672 | -0.129960713 | 0.128507274 | 8.025246757 | -0.013440002 | 0.989670999 | 0.9989817 | -5.256626409 |

| | | | | | | | | | |
|---|---|---|---|---|---|---|---|---|---|
| **A_55_P2052166** | Akap13 | 0.001030894 | -0.182359212 | 0.184421 | 10.59268479 | 0.013435299 | 0.989674614 | 0.9989817 | -5.256626462 |
| **A_52_P367520** | Nexn | 0.001183032 | -0.210106 | 0.212472064 | 14.34721296 | 0.013382234 | 0.989715392 | 0.9989817 | -5.256627062 |
| **A_55_P2085641** | Lmbr1l | 0.001716726 | -0.306734363 | 0.310167816 | 10.72005229 | 0.013302214 | 0.989776886 | 0.9989817 | -5.256627964 |
| **A_55_P2166011** | Tmem128 | 0.000728425 | -0.130888518 | 0.132345368 | 8.666640484 | 0.013227631 | 0.989834201 | 0.9989817 | -5.256628799 |
| **A_55_P2095814** | Kalrn | -0.001377886 | -0.253184911 | 0.250429138 | 9.594703864 | -0.013078402 | 0.98994888 | 0.9989817 | -5.256630455 |
| **A_55_P2054663** | Cox4i2 | 0.001101123 | -0.200234258 | 0.202436505 | 9.187867858 | 0.013071488 | 0.989954193 | 0.9989817 | -5.256630532 |
| **A_55_P1955325** | Zfp820 | 0.000745054 | -0.136465372 | 0.13795548 | 8.851456984 | 0.012978055 | 0.990025995 | 0.9989817 | -5.256631559 |
| **A_55_P2090349** | Mettl5 | 0.00062893 | -0.115223293 | 0.116481152 | 8.648134419 | 0.012974988 | 0.990028352 | 0.9989817 | -5.256631592 |
| **A_55_P2059958** | Snx7 | 0.00083874 | -0.155644015 | 0.157321496 | 9.097641367 | 0.01281062 | 0.990154665 | 0.9989817 | -5.256633381 |
| **A_55_P2104422** | Patj | 0.000554676 | -0.102967248 | 0.1040766 | 8.03288502 | 0.01280608 | 0.990158154 | 0.9989817 | -5.25663343 |
| **A_55_P2133910** | Ttll10 | 0.001052253 | -0.195539099 | 0.197643606 | 8.446363505 | 0.012792783 | 0.990168372 | 0.9989817 | -5.256633574 |
| **A_55_P1974352** | Fam126b | -0.00098562 | -0.1857216 | 0.183750359 | 9.614400524 | -0.012751677 | 0.990199962 | 0.9989817 | -5.256634017 |
| **A_55_P2142151** | Mapk1ip1 | -0.001147461 | -0.217337751 | 0.215042828 | 10.55049385 | -0.012685601 | 0.99025074 | 0.9989817 | -5.256634726 |
| **A_55_P2148082** | Larp7 | -0.000808533 | -0.153276316 | 0.151659249 | 10.74267448 | -0.012674447 | 0.990259312 | 0.9989817 | -5.256634845 |
| **A_52_P547662** | P2ry1 | 0.000603189 | -0.11370675 | 0.114913127 | 8.714591223 | 0.012611841 | 0.990307423 | 0.9989817 | -5.256635513 |
| **A_52_P674309** | Mocs2 | 0.000774661 | -0.146413519 | 0.147962842 | 11.58399474 | 0.012579058 | 0.990332616 | 0.9989817 | -5.256635862 |
| **A_55_P2036240** | NA | 0.002861378 | -0.542739361 | 0.548462117 | 11.73983215 | 0.012534568 | 0.990366806 | 0.9989817 | -5.256636333 |
| **A_55_P1986386** | Senp3 | 0.000896224 | -0.17099825 | 0.172790698 | 8.819298847 | 0.012461317 | 0.990423099 | 0.9989817 | -5.256637106 |
| **A_55_P2076462** | Lnx1 | -0.000697109 | -0.135719532 | 0.134325313 | 8.142562306 | -0.012339686 | 0.99051657 | 0.9989817 | -5.256638379 |
| **A_52_P217474** | Ndufa6 | 0.001051228 | -0.20395354 | 0.206055995 | 14.18043788 | 0.012255808 | 0.990581029 | 0.9989817 | -5.25663925 |
| **A_51_P383194** | Pde9a | -0.000640742 | -0.125800361 | 0.124518878 | 8.447891532 | -0.012235671 | 0.990596504 | 0.9989817 | -5.256639458 |
| **A_51_P433584** | Bcas3 | -0.000774275 | -0.155741756 | 0.154193205 | 8.423887269 | -0.011941641 | 0.990822463 | 0.9989817 | -5.256642458 |
| **A_55_P1980883** | Ephb6 | -0.003823549 | -0.778706948 | 0.77105985 | 10.14472246 | -0.011793412 | 0.990936375 | 0.9989817 | -5.256643943 |
| **A_52_P376135** | Mknk2 | -0.001144208 | -0.237448391 | 0.235159974 | 13.61148533 | -0.011572915 | 0.991105826 | 0.9989817 | -5.256646117 |
| **A_55_P2040951** | Actc1 | 0.000886204 | -0.183482968 | 0.185255375 | 18.06448585 | 0.011488262 | 0.991170881 | 0.9989817 | -5.256646941 |
| **A_55_P2028418** | Hacd2 | 0.000910556 | -0.192245547 | 0.194066658 | 9.949984532 | 0.01126697 | 0.991340943 | 0.9989817 | -5.256649066 |
| **A_55_P1977008** | Gfi1 | 0.000562251 | -0.118856825 | 0.119981327 | 7.754474105 | 0.011252923 | 0.991351739 | 0.9989817 | -5.256649199 |
| **A_52_P285024** | Sertad4 | -0.000907845 | -0.195065347 | 0.193249658 | 8.494075109 | -0.011175488 | 0.991411247 | 0.9989817 | -5.256649932 |
| **A_55_P2096942** | Nrn1l | 0.000629626 | -0.134398712 | 0.135657965 | 7.943496933 | 0.011144665 | 0.991434934 | 0.9989817 | -5.256650222 |
| **A_51_P116088** | Rexo2 | 0.000827892 | -0.178445912 | 0.180101697 | 10.25658111 | 0.011037386 | 0.991517378 | 0.9989817 | -5.250394603 |
| **A_55_P2020497** | Dcaf12l1 | -0.000828258 | -0.180899471 | 0.179242955 | 9.202141656 | -0.010993362 | 0.99155121 | 0.9989817 | -5.256651636 |
| **A_55_P2010871** | Daam2 | 0.001706415 | -0.370701533 | 0.374114362 | 9.69589274 | 0.010951529 | 0.991583359 | 0.9989817 | -5.256652023 |
| **A_55_P1961870** | NA | 0.000735247 | -0.159951695 | 0.161422188 | 14.38768865 | 0.010936081 | 0.991595231 | 0.9989817 | -5.256652166 |
| **A_55_P2006375** | Dmrtc1c2 | -0.000446704 | -0.098976534 | 0.098083125 | 7.642685638 | -0.010835813 | 0.991672287 | 0.9989817 | -5.256653087 |
| **A_55_P2053733** | NA | -0.000627045 | -0.139621243 | 0.138367153 | 15.55092723 | -0.01078229 | 0.99171342 | 0.9989817 | -5.256653576 |
| **A_55_P1964464** | NA | -0.001028634 | -0.229432154 | 0.227374887 | 15.61980808 | -0.010763837 | 0.991727601 | 0.9989817 | -5.256653743 |
| **A_55_P2328787** | NA | 0.000995765 | -0.221793253 | 0.223784783 | 8.319165116 | 0.010682483 | 0.991790121 | 0.9989817 | -5.25665448 |
| **A_55_P1961968** | Plagl1 | -0.001833309 | -0.419408449 | 0.415741831 | 15.1377055 | -0.010493255 | 0.991935544 | 0.9989817 | -5.256656171 |
| **A_51_P404825** | Ccdc58 | 0.000610562 | -0.138951435 | 0.140172558 | 12.57876322 | 0.010456144 | 0.991964064 | 0.9989817 | -5.256656499 |
| **A_51_P128775** | Zfp84 | 0.000506644 | -0.11553122 | 0.116544509 | 9.270449467 | 0.010435481 | 0.991979943 | 0.9989817 | -5.243844181 |
| **A_55_P2081183** | Slc28a1 | -0.000492104 | -0.113341778 | 0.11235757 | 8.376385401 | -0.010422341 | 0.991990042 | 0.9989817 | -5.256656797 |
| **A_55_P2121344** | Nudcd3 | -0.000532805 | -0.123048952 | 0.121983343 | 13.32000385 | -0.010394033 | 0.992011797 | 0.9989817 | -5.256657045 |
| **A_55_P2048210** | Ganab | -0.000546084 | -0.12738579 | 0.126293622 | 11.67742863 | -0.010289959 | 0.992091778 | 0.9989817 | -5.256657954 |
| **A_55_P2038412** | Acot8 | 0.00060273 | -0.139870331 | 0.141075792 | 9.172245463 | 0.010255084 | 0.99211858 | 0.9989817 | -5.256658256 |
| **A_55_P1999349** | Nudt4 | 0.001219565 | -0.284138005 | 0.286577136 | 13.47495582 | 0.010214682 | 0.992149629 | 0.9989817 | -5.256658605 |
| **A_55_P2063646** | Krcc1 | 0.001050261 | -0.244738485 | 0.246839007 | 13.12835216 | 0.010212788 | 0.992151085 | 0.9989817 | -5.256658621 |
| **A_52_P53626** | NA | -0.000554118 | -0.132168512 | 0.131060277 | 8.552410862 | -0.010062544 | 0.992266549 | 0.9989817 | -5.256659907 |
| **A_55_P2064568** | Anks1 | -0.000739848 | -0.178737957 | 0.17725826 | 10.6639389 | -0.009934278 | 0.992365122 | 0.9989817 | -5.256660989 |

| | | | | | | | | | |
|---|---|---|---|---|---|---|---|---|---|
| **A_55_P2034720** | Cfap36 | 0.001029637 | -0.24879813 | 0.250857405 | 10.80409543 | 0.009850374 | 0.992429603 | 0.9989817 | -5.25666169 |
| **A_55_P2001628** | Rps4l | -0.000560876 | -0.137996897 | 0.136875145 | 13.12635595 | -0.009753833 | 0.992503796 | 0.9989817 | -5.256662489 |
| **A_55_P2041758** | Scand1 | 0.000878537 | -0.215347472 | 0.217104546 | 14.01433573 | 0.009710939 | 0.992536761 | 0.9989817 | -5.256662841 |
| **A_52_P351925** | Ank3 | 0.000926968 | -0.227370726 | 0.229224662 | 9.031443609 | 0.009704481 | 0.992541723 | 0.9989817 | -5.256662894 |
| **A_55_P2158227** | Angpt1 | -0.003576882 | -0.890736667 | 0.883582903 | 10.24694618 | -0.009636335 | 0.992594095 | 0.9989817 | -5.25666345 |
| **A_55_P2093649** | NA | -0.000672692 | -0.167764454 | 0.166419071 | 16.87686494 | -0.009622104 | 0.992605031 | 0.9989817 | -5.256663566 |
| **A_55_P1989312** | Prkd3 | 0.000853473 | -0.213092064 | 0.214799011 | 11.63680443 | 0.009534456 | 0.99267239 | 0.9989817 | -5.256664274 |
| **A_55_P2086912** | NA | 0.001007742 | -0.251718951 | 0.253734434 | 11.8721376 | 0.009530314 | 0.992675573 | 0.9989817 | -5.256664308 |
| **A_55_P2315457** | Gripap1 | 0.000845809 | -0.212781428 | 0.214473046 | 9.864022104 | 0.009462914 | 0.992727371 | 0.9989817 | -5.256664848 |
| **A_55_P2110998** | Gpr153 | 0.000784579 | -0.201544597 | 0.203113754 | 10.49637883 | 0.009268022 | 0.992877148 | 0.9989817 | -5.256666388 |
| **A_51_P477364** | Rhob | -0.001002014 | -0.260953549 | 0.258949521 | 8.591711428 | -0.009212777 | 0.992919605 | 0.9989817 | -5.256666819 |
| **A_55_P2019557** | Mrgpra2b | -0.0004492 | -0.117465059 | 0.11656666 | 7.72981105 | -0.009174952 | 0.992948674 | 0.9989817 | -5.256667113 |
| **A_52_P590235** | Vars2 | 0.000593648 | -0.1540737 | 0.155260995 | 9.659409381 | 0.009173586 | 0.992949724 | 0.9989817 | -5.256667123 |
| **A_52_P527106** | Arhgap12 | -0.000673561 | -0.178541634 | 0.177194512 | 8.78670294 | -0.009050824 | 0.993044068 | 0.9989817 | -5.256668068 |
| **A_52_P262967** | Rab10 | 0.000760004 | -0.201246909 | 0.202766917 | 9.943273728 | 0.008992054 | 0.993089234 | 0.9989817 | -5.256668515 |
| **A_51_P256384** | Atp2b2 | -0.000482836 | -0.129559197 | 0.128593525 | 7.803371607 | -0.008940505 | 0.99312885 | 0.9989817 | -5.256668905 |
| **A_55_P2023537** | NA | 0.000488667 | -0.130604036 | 0.131581371 | 7.945938436 | 0.008909309 | 0.993152825 | 0.9989817 | -5.25666914 |
| **A_51_P501773** | Slc35f3 | 0.000363649 | -0.097314853 | 0.098042151 | 7.686190569 | 0.008897999 | 0.993161517 | 0.9989817 | -5.256669225 |
| **A_55_P2155727** | Mafa | -0.00066684 | -0.179879812 | 0.178546132 | 10.10812766 | -0.008893264 | 0.993165156 | 0.9989817 | -5.256669261 |
| **A_55_P2397599** | B930025B1 | -0.001189944 | -0.321643484 | 0.319263596 | 8.926436556 | -0.008875043 | 0.993179159 | 0.9989817 | -5.256669397 |
| **A_51_P217236** | Derl1 | -0.000419989 | -0.113729386 | 0.112889409 | 10.84475352 | -0.008858921 | 0.993191549 | 0.9989817 | -5.256669518 |
| **A_55_P1988108** | Mrc1 | -0.001126273 | -0.306577208 | 0.304324662 | 9.185072688 | -0.008812749 | 0.993227033 | 0.9989817 | -5.256669862 |
| **A_55_P1971956** | Ift46 | 0.000518835 | -0.140643443 | 0.141681114 | 11.66550633 | 0.008784561 | 0.993248697 | 0.9989817 | -5.256670072 |
| **A_52_P676359** | Fcnb | -0.000353977 | -0.096943851 | 0.096235897 | 7.659551733 | -0.008758959 | 0.993268372 | 0.9989817 | -5.256670261 |
| **A_55_P2008619** | Lancl2 | -0.000628888 | -0.186691247 | 0.185433471 | 11.39494153 | -0.008888742 | 0.993276474 | 0.9989817 | -5.116437702 |
| **A_55_P2135351** | Fam89b | -0.000835293 | -0.231148862 | 0.229478276 | 10.45000962 | -0.008668191 | 0.993338129 | 0.9989817 | -5.256670929 |
| **A_55_P2073840** | Rpl7 | 0.00036368 | -0.100927205 | 0.101654564 | 16.10578349 | 0.008581396 | 0.993404833 | 0.9989817 | -5.256671561 |
| **A_55_P2132502** | Pkn1 | 0.000431355 | -0.119762638 | 0.120625347 | 10.29853236 | 0.008577505 | 0.993407823 | 0.9989817 | -5.256671589 |
| **A_55_P2033445** | Tnfrsf1b | 0.00062775 | -0.174361251 | 0.175616751 | 8.00821239 | 0.00857403 | 0.993410494 | 0.9989817 | -5.256671614 |
| **A_66_P123055** | Gm10845 | -0.000871727 | -0.251252834 | 0.249509379 | 11.96832753 | -0.008321243 | 0.993604766 | 0.9989817 | -5.256673416 |
| **A_55_P2058137** | Pde4dip | -0.000579406 | -0.16707388 | 0.165915068 | 14.1510661 | -0.008317488 | 0.993607652 | 0.9989817 | -5.256673443 |
| **A_55_P1974421** | Itga3 | -0.000425994 | -0.123240053 | 0.122388064 | 8.251845765 | -0.008290195 | 0.993628627 | 0.9989817 | -5.256673634 |
| **A_55_P2180789** | Huwe1 | -0.000671494 | -0.200979451 | 0.199636463 | 12.36788557 | -0.008327433 | 0.993641908 | 0.9989817 | -5.177328025 |
| **A_52_P505157** | Fsd2 | -0.000722164 | -0.209926872 | 0.208482544 | 8.969932635 | -0.008250374 | 0.99365923 | 0.9989817 | -5.256673912 |
| **A_51_P186856** | Krt5 | -0.000380599 | -0.111871788 | 0.11111059 | 7.75153128 | -0.008158993 | 0.993729458 | 0.9989817 | -5.256674545 |
| **A_51_P176042** | Pklr | -0.00034928 | -0.102931824 | 0.102233264 | 7.666294131 | -0.00813784 | 0.993745715 | 0.9989817 | -5.25667469 |
| **A_55_P2129373** | Capn11 | 0.00095476 | -0.286074151 | 0.28798367 | 8.810327044 | 0.007950193 | 0.993889927 | 0.9989817 | -5.256675964 |
| **A_55_P1997509** | Zbtb32 | -0.000338253 | -0.102159964 | 0.101483457 | 7.692110247 | -0.007939825 | 0.993897894 | 0.9989817 | -5.256676034 |
| **A_51_P396273** | Nras | -0.000444822 | -0.137809818 | 0.136920175 | 9.721227026 | -0.007739608 | 0.994051767 | 0.9989817 | -5.256677359 |
| **A_52_P444253** | Gm6900 | -0.000353802 | -0.112457501 | 0.111749897 | 14.62501284 | -0.00754309 | 0.994202796 | 0.9989817 | -5.256678626 |
| **A_55_P2139089** | Slc22a28 | 0.000494951 | -0.157861146 | 0.158851048 | 8.630497382 | 0.007470276 | 0.994258756 | 0.9989817 | -5.256679087 |
| **A_51_P369762** | Nmrk2 | -0.002531505 | -0.814973855 | 0.809910845 | 11.17502039 | -0.00744724 | 0.994276459 | 0.9989817 | -5.256679232 |
| **A_51_P446417** | Ufl1 | 0.000474246 | -0.152067282 | 0.153015774 | 9.437502637 | 0.007430613 | 0.994289238 | 0.9989817 | -5.256679337 |
| **A_66_P137102** | Spryd7 | -0.000343697 | -0.111521179 | 0.110833784 | 8.480589275 | -0.007388711 | 0.994321441 | 0.9989817 | -5.256679599 |
| **A_51_P230287** | Sorcs2 | 0.000345615 | -0.112159909 | 0.112851138 | 8.959324451 | 0.007342221 | 0.99435717 | 0.9989817 | -5.256679888 |
| **A_55_P2015605** | Foxp4 | 0.00063876 | -0.209537072 | 0.210814592 | 10.74829782 | 0.007263805 | 0.994417435 | 0.9989817 | -5.256680371 |
| **A_51_P304109** | Cyp2c39 | -0.000300338 | -0.099767519 | 0.099166844 | 7.64306418 | -0.007216706 | 0.994453632 | 0.9989817 | -5.256680659 |
| **A_55_P1962149** | Dnah10 | -0.000286255 | -0.096532042 | 0.095959531 | 7.650581084 | -0.007108552 | 0.994536751 | 0.9989817 | -5.256681313 |

| | | | | | | | | | |
|---|---|---|---|---|---|---|---|---|---|
| **A_66_P106385** | Cd55b | 0.000492552 | -0.166397133 | 0.167382238 | 8.21100158 | 0.007053943 | 0.99457872 | 0.9989817 | -5.256681639 |
| **A_51_P160544** | Efemp2 | 0.000865839 | -0.295783535 | 0.297515213 | 12.20163122 | 0.006975945 | 0.994638664 | 0.9989817 | -5.256682101 |
| **A_55_P2072315** | NA | -0.001413276 | -0.493720088 | 0.490893537 | 11.16419542 | -0.006861201 | 0.994726849 | 0.9989817 | -5.256682771 |
| **A_55_P2080907** | Rpl26 | -0.000446343 | -0.156463663 | 0.155570977 | 15.26704955 | -0.006837627 | 0.994744966 | 0.9989817 | -5.256682907 |
| **A_52_P473966** | Kdelr3 | -0.000859882 | -0.301642752 | 0.299922989 | 10.06861145 | -0.006832736 | 0.994748725 | 0.9989817 | -5.256682936 |
| **A_55_P2082869** | Ezh1 | -0.000615956 | -0.217471211 | 0.2162393 | 11.2092434 | -0.006788731 | 0.994782544 | 0.9989817 | -5.256683189 |
| **A_55_P2110435** | Zscan20 | -0.000309352 | -0.109793303 | 0.109174598 | 8.316260158 | -0.00675324 | 0.99480982 | 0.9989817 | -5.256683391 |
| **A_51_P140797** | Dcaf6 | 0.000388075 | -0.138801049 | 0.139577199 | 8.922458179 | 0.006663761 | 0.994878588 | 0.9989817 | -5.256683898 |
| **A_52_P250590** | Dnajc13 | -0.000536259 | -0.195009083 | 0.193936564 | 9.355754549 | -0.0065906 | 0.994934814 | 0.9989817 | -5.256684307 |
| **A_55_P2003369** | Fkbp8 | -0.000837188 | -0.304790431 | 0.303116054 | 11.8486253 | -0.006583026 | 0.994940635 | 0.9989817 | -5.256684349 |
| **A_55_P2124582** | Xlr4a | 0.000373027 | -0.135264754 | 0.136010807 | 7.735496641 | 0.006573073 | 0.994948285 | 0.9989817 | -5.256684405 |
| **A_66_P112546** | Yeats4 | 0.000507924 | -0.186031696 | 0.187047543 | 8.899728897 | 0.006507836 | 0.994998422 | 0.9989817 | -5.256684765 |
| **A_51_P510663** | Supt3 | -0.000425541 | -0.157530382 | 0.1566793 | 9.092181579 | -0.006473823 | 0.995024562 | 0.9989817 | -5.256684951 |
| **A_55_P2404554** | Atcayos | -0.000739484 | -0.278445481 | 0.276966514 | 9.417734361 | -0.006364328 | 0.995108712 | 0.9989817 | -5.256685544 |
| **A_55_P2095663** | Pgr | -0.000284984 | -0.108450618 | 0.107880649 | 7.635700174 | -0.006297097 | 0.995160382 | 0.9989817 | -5.256685904 |
| **A_52_P162184** | Snx12 | 0.000345502 | -0.131598116 | 0.13228912 | 8.824614598 | 0.006258522 | 0.995190028 | 0.9989817 | -5.256686108 |
| **A_55_P2012592** | Ildr1 | -0.000269613 | -0.104388605 | 0.103849378 | 7.660731672 | -0.006189002 | 0.995243456 | 0.9989817 | -5.256686473 |
| **A_55_P2146177** | Cerk | -0.000453286 | -0.177692704 | 0.176786132 | 12.13557336 | -0.00611253 | 0.995302228 | 0.9989817 | -5.25668687 |
| **A_52_P498208** | H2ac15 | 0.000914391 | -0.357469876 | 0.359298659 | 13.79294226 | 0.006098069 | 0.995313342 | 0.9989817 | -5.256686945 |
| **A_55_P2014224** | Sema5a | 0.000762614 | -0.304802497 | 0.306327726 | 10.94831131 | 0.005964998 | 0.995415612 | 0.998986523 | -5.256687622 |
| **A_52_P127325** | AI182371 | 0.000235546 | -0.09730602 | 0.097777111 | 7.675091598 | 0.005771576 | 0.995564265 | 0.998986523 | -5.25668858 |
| **A_66_P106836** | Kmo | 0.00024414 | -0.101777252 | 0.102265533 | 7.718618984 | 0.00571949 | 0.995604294 | 0.998986523 | -5.256688833 |
| **A_52_P49014** | Shh | 0.000239762 | -0.100875816 | 0.10135534 | 7.641095059 | 0.005667232 | 0.995644457 | 0.998986523 | -5.218099521 |
| **A_52_P145412** | Rictor | -0.000410438 | -0.174879268 | 0.174058392 | 9.039935961 | -0.005622621 | 0.995678742 | 0.998986523 | -5.256689296 |
| **A_55_P2010271** | Samsn1 | -0.00037253 | -0.160396237 | 0.159651176 | 8.105908039 | -0.005563992 | 0.995723801 | 0.998986523 | -5.256689573 |
| **A_55_P2085932** | Lamtor2 | -0.001830685 | -0.789823345 | 0.786161975 | 11.92721379 | -0.005552653 | 0.995732515 | 0.998986523 | -5.256689626 |
| **A_55_P2452914** | N4bp2 | -0.000623496 | -0.269295574 | 0.268048582 | 10.15317556 | -0.005546515 | 0.995737233 | 0.998986523 | -5.256689655 |
| **A_51_P384318** | C1ra | -0.000481091 | -0.210046648 | 0.209084466 | 8.287910836 | -0.005486764 | 0.995783153 | 0.998986523 | -5.256689933 |
| **A_52_P535907** | Sde2 | -0.000308543 | -0.137343776 | 0.13672669 | 8.965705376 | -0.005381375 | 0.99586415 | 0.999015945 | -5.256690417 |
| **A_55_P2078735** | AI662270 | -0.000239861 | -0.108367166 | 0.107887444 | 7.766565956 | -0.005301919 | 0.995925214 | 0.999025371 | -5.256690775 |
| **A_55_P2006255** | Txnip | -0.000548826 | -0.264051002 | 0.262953351 | 8.949782741 | -0.00497805 | 0.996174122 | 0.99913985 | -5.25669218 |
| **A_55_P2108693** | NA | 0.000240174 | -0.116436471 | 0.116916818 | 7.849408816 | 0.004919837 | 0.99621886 | 0.99913985 | -5.256692423 |
| **A_55_P2060747** | Esco1 | -0.000462601 | -0.226425031 | 0.225499828 | 9.594510762 | -0.004893049 | 0.996239448 | 0.99913985 | -5.256692534 |
| **A_52_P229959** | 6-Sep | -0.00033115 | -0.163153007 | 0.162490707 | 12.58228294 | -0.004860946 | 0.996264121 | 0.99913985 | -5.256692666 |
| **A_55_P2430122** | Ilf2 | 0.000254924 | -0.126256645 | 0.126766493 | 10.47639618 | 0.004816029 | 0.996298641 | 0.99913985 | -5.25669285 |
| **A_51_P460774** | Smyd4 | -0.000385652 | -0.194434696 | 0.193663392 | 9.080240275 | -0.004749997 | 0.99634939 | 0.99913985 | -5.256693116 |
| **A_51_P122649** | Degs2 | 0.000189861 | -0.097627549 | 0.098007271 | 7.646143379 | 0.004639046 | 0.996434661 | 0.999173538 | -5.243880708 |
| **A_55_P2137701** | Rex2 | -0.000198963 | -0.105049315 | 0.104651389 | 7.902810278 | -0.004535352 | 0.996514354 | 0.99920163 | -5.256693957 |
| **A_51_P492346** | Tmem174 | -0.000270771 | -0.145281281 | 0.144739738 | 8.507373052 | -0.004462851 | 0.996570074 | 0.999205682 | -5.256694233 |
| **A_52_P659258** | Ppat | -0.000300635 | -0.163852763 | 0.163251494 | 9.179846191 | -0.00439331 | 0.996623519 | 0.999207454 | -5.256694492 |
| **A_51_P292016** | Uts2r | -0.000209671 | -0.118647242 | 0.1182279 | 8.967527841 | -0.004231146 | 0.99674815 | 0.999243097 | -5.256695083 |
| **A_55_P1989384** | Rnf181 | -0.000269635 | -0.153250916 | 0.152711647 | 10.47930249 | -0.004212563 | 0.996762432 | 0.999243097 | -5.256695149 |
| **A_51_P305843** | Chordc1 | 0.000204792 | -0.123967029 | 0.124376612 | 10.58686552 | 0.003941839 | 0.996970496 | 0.999397506 | -5.256696081 |
| **A_51_P362089** | Lrrc3b | 0.000405385 | -0.256457592 | 0.257268362 | 8.970548709 | 0.003772029 | 0.997101002 | 0.999397506 | -5.256696633 |
| **A_51_P191893** | 2410022M1 | 0.000285461 | -0.18691625 | 0.187487173 | 10.84115182 | 0.003644573 | 0.997198959 | 0.999397506 | -5.256697032 |
| **A_55_P1980981** | Apba3 | -0.000306373 | -0.213458133 | 0.212845386 | 10.1223317 | -0.003435353 | 0.997359754 | 0.999397506 | -5.256697657 |
| **A_51_P202596** | Sh3tc2 | -0.000198009 | -0.138497607 | 0.138101589 | 8.630918986 | -0.003421942 | 0.997370061 | 0.999397506 | -5.256697696 |
| **A_55_P2022812** | Cyp4f17 | 0.00016403 | -0.11596532 | 0.11629338 | 8.221194581 | 0.003375905 | 0.997405443 | 0.999397506 | -5.256697828 |

| | | | | | | | | | |
|---|---|---|---|---|---|---|---|---|---|
| **A_51_P281333** | St3gal6 | 0.000160095 | -0.114838756 | 0.115158947 | 9.394133879 | 0.003327316 | 0.997442785 | 0.999397506 | -5.256697966 |
| **A_51_P390715** | Tgfb1 | -0.000160123 | -0.117673987 | 0.117353742 | 8.589963389 | -0.003256661 | 0.997497088 | 0.999397506 | -5.256698162 |
| **A_51_P201904** | Ndufb5 | -0.000233585 | -0.178467268 | 0.178000097 | 11.9490086 | -0.003132312 | 0.997592655 | 0.999397506 | -5.256698497 |
| **A_55_P2136862** | Dpy30 | -0.000372781 | -0.289960609 | 0.289215048 | 11.0684965 | -0.003076678 | 0.997635413 | 0.999397506 | -5.256698643 |
| **A_65_P14951** | Cblb | 0.000270634 | -0.210377721 | 0.210918988 | 9.454341831 | 0.003070669 | 0.997640031 | 0.999397506 | -5.256698659 |
| **A_55_P1958210** | Ephb1 | 0.000261644 | -0.204717534 | 0.205240821 | 8.333457726 | 0.00305077 | 0.997655325 | 0.999397506 | -5.25669871 |
| **A_55_P2120277** | Gm11619 | 0.000143854 | -0.113533069 | 0.113820777 | 15.43399148 | 0.003024532 | 0.99767549 | 0.999397506 | -5.256698777 |
| **A_51_P443508** | Ppa1 | -0.000440394 | -0.366920989 | 0.366040201 | 11.51577659 | -0.002872099 | 0.997792642 | 0.999397506 | -5.256699157 |
| **A_55_P2063257** | Lgals4 | -0.000236121 | -0.199455265 | 0.198983022 | 8.116270486 | -0.002832782 | 0.997822859 | 0.999397506 | -5.256699251 |
| **A_55_P2035937** | Rab6a | -0.000168169 | -0.148642206 | 0.148305867 | 12.92762982 | -0.002707107 | 0.997919446 | 0.999397506 | -5.256699545 |
| **A_55_P2102095** | 1810010H2 | -0.000123511 | -0.109263647 | 0.109016625 | 7.870980359 | -0.002704771 | 0.997921242 | 0.999397506 | -5.256699551 |
| **A_51_P136521** | Lypd2 | 0.000188809 | -0.168628081 | 0.169005698 | 8.212510972 | 0.002673099 | 0.997945584 | 0.999397506 | -5.256699622 |
| **A_55_P2038183** | Insc | 0.000284922 | -0.26769716 | 0.268267004 | 9.068752718 | 0.002541143 | 0.998046998 | 0.999397506 | -5.256699913 |
| **A_55_P2137973** | Bnip2 | 0.000130223 | -0.126296536 | 0.126556983 | 8.230673806 | 0.002461837 | 0.998107949 | 0.999397506 | -5.25670008 |
| **A_55_P2008609** | Rxrb | -0.00014026 | -0.140189857 | 0.139909337 | 12.02561049 | -0.00239365 | 0.998160355 | 0.999397506 | -5.25670022 |
| **A_51_P401184** | Rarres1 | -0.000116419 | -0.120695969 | 0.120463131 | 7.997840175 | -0.00230759 | 0.998226496 | 0.999397506 | -5.256700391 |
| **A_55_P1967227** | Tmem158 | 0.000168863 | -0.176889625 | 0.177227351 | 10.07375525 | 0.002279437 | 0.998248133 | 0.999397506 | -5.256700445 |
| **A_51_P518488** | Atpif1 | 0.000141212 | -0.148125243 | 0.148407667 | 16.83743508 | 0.002276345 | 0.998250509 | 0.999397506 | -5.256700451 |
| **A_55_P1973259** | Gamt | 0.00018205 | -0.192002454 | 0.192366554 | 11.24658801 | 0.002264023 | 0.998259979 | 0.999397506 | -5.256700475 |
| **A_55_P2025368** | Nkain1 | -0.000108843 | -0.115036412 | 0.114818725 | 8.76195368 | -0.00226353 | 0.998260359 | 0.999397506 | -5.256700476 |
| **A_55_P1973011** | Ppm1l | 0.000132758 | -0.166295009 | 0.166560524 | 11.08805205 | 0.001906528 | 0.998534733 | 0.999554607 | -5.256701104 |
| **A_55_P2170139** | Rheb | 0.000145115 | -0.18501433 | 0.18530456 | 11.77719904 | 0.001873161 | 0.998560377 | 0.999554607 | -5.256701157 |
| **A_55_P2171977** | NA | -0.000118065 | -0.152029381 | 0.15179325 | 14.22873009 | -0.001857555 | 0.998572371 | 0.999554607 | -5.256701182 |
| **A_51_P196629** | Fam102b | 0.000144143 | -0.20133597 | 0.201624256 | 10.93522461 | 0.001709904 | 0.998685848 | 0.999580426 | -5.256701404 |
| **A_55_P2128113** | Tbc1d12 | -0.000268341 | -0.398159956 | 0.397623274 | 8.644720589 | -0.001611875 | 0.998761189 | 0.999580426 | -5.256701541 |
| **A_55_P2069755** | Tymp | -0.000166098 | -0.247799676 | 0.247467481 | 9.963553277 | -0.001603108 | 0.998767927 | 0.999580426 | -5.256701553 |
| **A_55_P2010347** | Zdhhc6 | -6.54E-05 | -0.107390422 | 0.10725965 | 8.475523579 | -0.001456103 | 0.998880908 | 0.999580426 | -5.256701743 |
| **A_55_P2032187** | NA | -0.000109377 | -0.181933063 | 0.181714309 | 10.33675797 | -0.001437757 | 0.998895008 | 0.999580426 | -5.256701766 |
| **A_55_P2008599** | Pcx | -0.000110618 | -0.192650695 | 0.192429459 | 10.51901988 | -0.001373134 | 0.998944673 | 0.999580426 | -5.256701842 |
| **A_55_P2036380** | NA | -8.33E-05 | -0.156909939 | 0.156743336 | 16.98792665 | -0.001269526 | 0.999024302 | 0.999580426 | -5.256701958 |
| **A_55_P1982727** | Dnajb5 | 0.000115148 | -0.235586943 | 0.235817239 | 11.02988742 | 0.001167619 | 0.999102623 | 0.999580426 | -5.256702063 |
| **A_55_P2090698** | NA | 7.58E-05 | -0.156505905 | 0.156657422 | 14.82781695 | 0.001156374 | 0.999111265 | 0.999580426 | -5.256702074 |
| **A_55_P1955733** | NA | 0.000217519 | -0.541370895 | 0.541805934 | 11.0701552 | 0.000959926 | 0.999262245 | 0.999580426 | -5.256702249 |
| **A_52_P637282** | Wipf1 | 6.57E-05 | -0.167346545 | 0.167477867 | 9.470963979 | 0.000937412 | 0.999279549 | 0.999580426 | -5.256702267 |
| **A_55_P2073617** | NA | 7.78E-05 | -0.211239629 | 0.211395249 | 8.53756984 | 0.000880057 | 0.999323629 | 0.999580426 | -5.256702311 |
| **A_55_P2193512** | NA | -3.63E-05 | -0.099874655 | 0.099802045 | 7.744974423 | -0.000869117 | 0.999332037 | 0.999580426 | -5.256702319 |
| **A_52_P125485** | Slc25a48 | -3.34E-05 | -0.097486533 | 0.097419661 | 7.655909423 | -0.000820026 | 0.999369767 | 0.999580426 | -5.256702354 |
| **A_52_P289685** | Ssbp1 | 6.35E-05 | -0.186261822 | 0.186388893 | 12.20715092 | 0.000814993 | 0.999373634 | 0.999580426 | -5.256702358 |
| **A_52_P336290** | Rap2a | 4.02E-05 | -0.143399603 | 0.143479924 | 10.24656014 | 0.000669175 | 0.999485703 | 0.999640807 | -5.256702449 |
| **A_55_P1993589** | NA | 3.32E-05 | -0.140388162 | 0.14045449 | 14.9859391 | 0.000564472 | 0.999566173 | 0.999669578 | -5.256702503 |
| **A_55_P2044045** | Ybx1 | -1.31E-05 | -0.182435338 | 0.182409224 | 14.61461278 | -0.000171073 | 0.999868521 | 0.999920237 | -5.256702626 |
| **A_55_P2178152** | Ccsap | -4.98E-06 | -0.182353176 | 0.182343216 | 8.701171471 | -6.53E-05 | 0.999949831 | 0.999949831 | -5.256702636 |

Rychlik_Arsenic_microarray_TableS2_Liver_100_vs_0_all_p_values

| | ID | logFC | CI.L | CI.R | AveExpr | t | P.Value | adj.P.Val | B |
|---|---|---|---|---|---|---|---|---|---|
| **A_55_P2031167** | Efna1 | -1.732199205 | -2.132527126 | -1.331871283 | 11.2628963 | -10.51393149 | 3.59E-05 | 0.479759698 | -2.785719463 |
| **A_55_P1998943** | Oas1a | -0.626870669 | -0.821736282 | -0.432005055 | 10.4380862 | -7.816744729 | 0.000200024 | 0.479759698 | -2.903156455 |
| **A_52_P319438** | Ankrd37 | -1.872557407 | -2.540827911 | -1.204286902 | 9.747137364 | -6.808728809 | 0.00043407 | 0.479759698 | -2.978891927 |
| **A_51_P245796** | Ddit4 | -0.951392027 | -1.292137437 | -0.610646618 | 11.14661019 | -6.784419324 | 0.000442751 | 0.479759698 | -2.981055998 |
| **A_51_P359570** | Ifit3 | -0.61773495 | -0.839957517 | -0.395512383 | 8.765143618 | -6.754562161 | 0.000453688 | 0.479759698 | -2.983739073 |
| **A_51_P514961** | Tiparp | -0.633734324 | -0.864733338 | -0.40273531 | 9.849038894 | -6.666229997 | 0.000487895 | 0.479759698 | -2.991842575 |
| **A_51_P333274** | Gzmb | -0.480846906 | -0.657837845 | -0.303855967 | 8.414332675 | -6.601444589 | 0.000514862 | 0.479759698 | -2.997947503 |
| **A_55_P1984830** | Fgl1 | -1.327512222 | -1.817206441 | -0.837818003 | 13.19604458 | -6.587137666 | 0.000521045 | 0.479759698 | -2.999314569 |
| **A_55_P2154107** | Gcm1 | 0.616092356 | 0.382565312 | 0.849619401 | 10.56906044 | 6.410498851 | 0.000604801 | 0.479759698 | -3.016776886 |
| **A_51_P439085** | Hilpda | -0.927625015 | -1.285822211 | -0.56942782 | 11.09583293 | -6.292648305 | 0.000669233 | 0.479759698 | -3.029056003 |
| **A_51_P387123** | Oasl2 | -0.577771864 | -0.803459119 | -0.352084608 | 8.67939803 | -6.220603813 | 0.000712468 | 0.479759698 | -3.036823781 |
| **A_55_P2270412** | C230096K16Rik | 0.790995371 | 0.478223573 | 1.103767169 | 10.48031825 | 6.14510834 | 0.000761226 | 0.479759698 | -3.045184789 |
| **A_55_P2058783** | Hilpda | -0.841319551 | -1.190956699 | -0.491682404 | 11.29553864 | -5.846912605 | 0.000994782 | 0.479759698 | -3.080573027 |
| **A_55_P2168296** | Rnu7 | 0.406194696 | 0.235597933 | 0.576791458 | 8.457030071 | 5.785576647 | 0.001052382 | 0.479759698 | -3.088350761 |
| **A_51_P345367** | Psmb8 | -0.558616758 | -0.793828371 | -0.323405145 | 9.940696438 | -5.770831445 | 0.001066789 | 0.479759698 | -3.090247333 |
| **A_52_P294510** | Fgl1 | -1.214841476 | -1.727616851 | -0.702066101 | 10.10354295 | -5.75672706 | 0.00108078 | 0.479759698 | -3.092071332 |
| **A_55_P2159934** | Rnf186 | -0.764616292 | -1.087547764 | -0.44168482 | 8.582778136 | -5.753291304 | 0.00108422 | 0.479759698 | -3.092517115 |
| **A_51_P438805** | Txnip | -0.855680164 | -1.220459967 | -0.49090036 | 11.00726954 | -5.699855795 | 0.001139349 | 0.479759698 | -3.099524982 |
| **A_51_P417016** | Srsf3 | 0.514872208 | 0.294511715 | 0.735232701 | 9.315359027 | 5.677392293 | 0.001163471 | 0.479759698 | -3.102513399 |
| **A_55_P2000062** | Irf1 | -0.535553044 | -0.771967892 | -0.299138195 | 11.55856029 | -5.504411983 | 0.001369931 | 0.479759698 | -3.126398522 |
| **A_55_P2021187** | Malat1 | 0.588469903 | 0.327593625 | 0.84934618 | 9.323895561 | 5.481164755 | 0.001400726 | 0.479759698 | -3.129730158 |
| **A_51_P162955** | Serpina7 | -0.71330225 | -1.034228754 | -0.392375746 | 9.125843525 | -5.400714225 | 0.00151352 | 0.479759698 | -3.141491578 |
| **A_51_P126177** | Map1lc3b | -0.507045801 | -0.738039692 | -0.27605191 | 12.23325956 | -5.333715973 | 0.001615351 | 0.479759698 | -3.18521784 |
| **A_51_P161890** | Fcgrt | -0.479728622 | -0.698487987 | -0.260969257 | 10.59672112 | -5.328588277 | 0.001623459 | 0.479759698 | -3.152349857 |
| **A_55_P2161923** | Rabgap1 | 0.722397885 | 0.391800932 | 1.052994839 | 8.996607056 | 5.309587813 | 0.001653907 | 0.479759698 | -3.155260957 |
| **A_55_P2050688** | Zbed4 | 0.527004172 | 0.28531092 | 0.768697424 | 11.17422712 | 5.298253294 | 0.001672379 | 0.479759698 | -3.157007754 |
| **A_66_P115389** | NA | 0.564930785 | 0.305338387 | 0.824523182 | 12.60633498 | 5.287939297 | 0.00168939 | 0.479759698 | -3.158603947 |
| **A_51_P384091** | Rad21 | 0.359736433 | 0.19382972 | 0.525643147 | 13.71227663 | 5.268702221 | 0.001721645 | 0.479759698 | -3.161598166 |
| **A_66_P108152** | Cbr1 | -0.551213092 | -0.806855736 | -0.295570448 | 11.88308564 | -5.239253502 | 0.001772382 | 0.479759698 | -3.166225293 |
| **A_51_P290207** | Insig1 | -0.65078365 | -0.955495329 | -0.346071971 | 9.695621815 | -5.189562201 | 0.00186188 | 0.479759698 | -3.174154051 |
| **A_55_P2092526** | Tgif1 | -0.708145194 | -1.047355043 | -0.368935346 | 10.52615229 | -5.072675114 | 0.002093283 | 0.479759698 | -3.193421452 |
| **A_55_P2027327** | Usp1 | 0.478359351 | 0.248989262 | 0.70772944 | 11.80180409 | 5.06758071 | 0.002104084 | 0.479759698 | -3.194281394 |
| **A_55_P2037712** | Smc4 | 0.521942948 | 0.269792772 | 0.774093124 | 13.21947651 | 5.029756354 | 0.002186277 | 0.479759698 | -3.200720196 |
| **A_55_P2004447** | Fgl1 | -1.03409989 | -1.535892258 | -0.532307523 | 10.61598923 | -5.00750775 | 0.002236315 | 0.479759698 | -3.204552441 |
| **A_55_P1973347** | A130040M12Rik | -0.730422579 | -1.085093438 | -0.37575172 | 11.57247117 | -5.004167896 | 0.002243937 | 0.479759698 | -3.205130616 |
| **A_55_P2110081** | Cisd3 | -0.492343796 | -0.734283568 | -0.250404024 | 11.6704404 | -4.944750609 | 0.00238457 | 0.479759698 | -3.215544572 |
| **A_55_P2086433** | Oasl1 | -0.513125997 | -0.765295993 | -0.260956002 | 8.614036621 | -4.944402273 | 0.002385423 | 0.479759698 | -3.215606346 |
| **A_55_P2136338** | Tomm20 | 0.336953632 | 0.169840039 | 0.504067225 | 13.07202131 | 4.899384835 | 0.002498652 | 0.479759698 | -3.223661519 |
| **A_55_P2127179** | NA | -0.447520972 | -0.669645429 | -0.225396514 | 8.314180408 | -4.895535461 | 0.002508612 | 0.479759698 | -3.22435697 |
| **A_55_P2186085** | Atp11c | 0.437861595 | 0.21952572 | 0.656197469 | 9.553127479 | 4.872983528 | 0.002567872 | 0.479759698 | -3.228452654 |
| **A_52_P621588** | Ifnlr1 | -0.431826041 | -0.647664356 | -0.215987726 | 8.865064194 | -4.861423724 | 0.002598858 | 0.479759698 | -3.23056624 |
| **A_55_P1955309** | Zfp800 | 0.828118722 | 0.41158177 | 1.244655675 | 10.02899578 | 4.830835592 | 0.002682903 | 0.479759698 | -3.236205804 |
| **A_55_P2058433** | Cyp2c68 | -2.101175405 | -2.860083293 | -1.342267516 | 13.0761655 | -8.537658694 | 0.002699231 | 0.479759698 | -4.097072581 |
| **A_52_P604629** | Csrnp1 | -0.575708649 | -0.866021724 | -0.285395574 | 9.975776255 | -4.818583039 | 0.002717425 | 0.479759698 | -3.238484047 |
| **A_66_P115053** | Anp32b | 0.483084356 | 0.239383606 | 0.726785105 | 14.17746383 | 4.816696663 | 0.002722784 | 0.479759698 | -3.238835783 |
| **A_51_P289588** | Ube2e3 | 0.397321788 | 0.196721825 | 0.597921751 | 12.90107839 | 4.81276572 | 0.00273399 | 0.479759698 | -3.239569598 |
| **A_52_P381784** | Nsd2 | 0.46922056 | 0.232108242 | 0.706332877 | 9.765000936 | 4.8084611 | 0.002746321 | 0.479759698 | -3.240374482 |
| **A_55_P2174203** | Lims2 | -0.710492314 | -1.069562123 | -0.351422506 | 9.494651599 | -4.807991414 | 0.002747671 | 0.479759698 | -3.240462388 |

| | | | | | | | | | |
|---|---|---|---|---|---|---|---|---|---|
| **A_55_P2006634** | Map4k5 | 0.325962458 | 0.160894674 | 0.491030243 | 11.63713626 | 4.798311651 | 0.002775644 | 0.479759698 | -3.242277686 |
| **A_55_P2094060** | Gzma | -0.767623134 | -1.157487553 | -0.377758715 | 11.19487773 | -4.784291609 | 0.002816732 | 0.479759698 | -3.244919324 |
| **A_52_P270164** | E2f6 | 0.437631124 | 0.214994745 | 0.660267503 | 10.87661267 | 4.776340289 | 0.002840338 | 0.479759698 | -3.246424043 |
| **A_55_P1976109** | NA | 0.385792625 | 0.188734853 | 0.582850397 | 14.72058661 | 4.757113819 | 0.002898345 | 0.479759698 | -3.25008217 |
| **A_52_P387564** | Tmpo | 0.521534251 | 0.253363187 | 0.789705315 | 13.46884708 | 4.725569006 | 0.00299643 | 0.479759698 | -3.256144888 |
| **A_55_P2028288** | Nhlrc4 | 0.460812605 | 0.223839951 | 0.697785258 | 8.938812754 | 4.725081536 | 0.002997974 | 0.479759698 | -3.256239174 |
| **A_55_P2119257** | Serpine1 | -1.994483758 | -3.033466082 | -0.955501435 | 10.51696816 | -4.664504692 | 0.003197102 | 0.479759698 | -3.268099385 |
| **A_51_P462385** | G6pc | -1.09170804 | -1.661229934 | -0.522186146 | 8.455348422 | -4.657783239 | 0.003220099 | 0.479759698 | -3.269433077 |
| **A_51_P261517** | Tyrobp | -0.330334327 | -0.502939279 | -0.157729374 | 12.45142965 | -4.6503286 | 0.003245823 | 0.479759698 | -3.270916434 |
| **A_52_P402663** | Ninl | 0.6258688 | 0.29855548 | 0.953182121 | 8.913886182 | 4.646252542 | 0.003259985 | 0.479759698 | -3.271729371 |
| **A_55_P2084496** | Ssbp3 | 0.403802428 | 0.191728777 | 0.615876079 | 10.96924365 | 4.62663674 | 0.003329122 | 0.479759698 | -3.275660108 |
| **A_55_P2133255** | Mad2l1 | 0.401022104 | 0.189381839 | 0.612662369 | 11.82132703 | 4.604189639 | 0.003410267 | 0.479759698 | -3.280196081 |
| **A_55_P2066958** | Sh3glb1 | 0.30883013 | 0.145765779 | 0.471894481 | 11.48737393 | 4.601970458 | 0.003418409 | 0.479759698 | -3.280646729 |
| **A_55_P2005025** | Mbnl1 | 0.531707541 | 0.250171826 | 0.813243256 | 12.33570526 | 4.589047072 | 0.003466262 | 0.479759698 | -3.283279022 |
| **A_55_P2015782** | Bcas1 | 0.506288511 | 0.237716714 | 0.774860309 | 9.160921571 | 4.580584038 | 0.003498006 | 0.479759698 | -3.285010181 |
| **A_55_P1957344** | Hnrnpa3 | 0.455131285 | 0.213200879 | 0.697061691 | 13.18542027 | 4.571191585 | 0.00353362 | 0.479759698 | -3.286938318 |
| **A_51_P476209** | Fscn3 | 0.404912518 | 0.18838047 | 0.621444566 | 8.658507823 | 4.5438314 | 0.003639704 | 0.479759698 | -3.292596369 |
| **A_55_P2002117** | Kbtbd4 | 0.57672236 | 0.267914668 | 0.885530053 | 10.99511212 | 4.537972599 | 0.003662883 | 0.479759698 | -3.293816021 |
| **A_55_P2023864** | Mfap1a | 0.306412501 | 0.141716626 | 0.471108377 | 9.70173886 | 4.5207131 | 0.003732133 | 0.479759698 | -3.297425649 |
| **A_55_P1979463** | Prdx5 | -0.467775833 | -0.719450392 | -0.216101273 | 13.09397201 | -4.516288245 | 0.003750123 | 0.479759698 | -3.298355075 |
| **A_52_P64687** | Camk2n1 | -0.390677233 | -0.601057565 | -0.180296901 | 9.788299427 | -4.512281193 | 0.003766498 | 0.479759698 | -3.299198162 |
| **A_65_P19395** | H2-D1 | -0.372571618 | -0.57429874 | -0.170844495 | 12.01767935 | -4.487750066 | 0.003868513 | 0.479759698 | -3.304389086 |
| **A_55_P2058240** | Ipo5 | 0.525035938 | 0.240311482 | 0.809760393 | 10.93865907 | 4.480716322 | 0.003898333 | 0.479759698 | -3.305886885 |
| **A_51_P164014** | Cenpe | 0.465579456 | 0.212653811 | 0.718505101 | 12.6190707 | 4.472847911 | 0.003931997 | 0.479759698 | -3.307567423 |
| **A_55_P1967648** | Flcn | -0.342991981 | -0.529657622 | -0.156326339 | 10.04368641 | -4.464807802 | 0.003966733 | 0.479759698 | -3.309290107 |
| **A_55_P2018941** | Fmr1 | 0.586217598 | 0.267070662 | 0.905364533 | 10.43934122 | 4.463251524 | 0.003973496 | 0.479759698 | -3.309624198 |
| **A_55_P1976664** | Gm16430 | 0.31942748 | 0.144578979 | 0.49427598 | 8.062077204 | 4.439085931 | 0.004080192 | 0.479759698 | -3.314838687 |
| **A_51_P431996** | Zfp367 | 0.450600171 | 0.203745998 | 0.697454345 | 10.17009634 | 4.43541302 | 0.004096688 | 0.479759698 | -3.315635659 |
| **A_55_P2067722** | Camsap3 | -0.31616972 | -0.48984463 | -0.142494811 | 10.73742917 | -4.423503576 | 0.004150693 | 0.479759698 | -3.318227921 |
| **A_55_P2066697** | Trim47 | -0.4815347 | -0.74642218 | -0.216647219 | 10.63973721 | -4.417223206 | 0.004179493 | 0.479759698 | -3.31959992 |
| **A_55_P2124153** | Rusc2 | -0.770674343 | -1.195877568 | -0.345471117 | 9.473618915 | -4.404103732 | 0.004240381 | 0.479759698 | -3.322477134 |
| **A_52_P70854** | Tob2 | -0.318380328 | -0.494324556 | -0.142436101 | 9.793920083 | -4.396979016 | 0.004273863 | 0.479759698 | -3.324045992 |
| **A_55_P2124085** | NA | 0.664377374 | 0.296854544 | 1.031900204 | 10.7203429 | 4.392519523 | 0.004294971 | 0.479759698 | -3.32503025 |
| **A_51_P207412** | Slf1 | 0.339364173 | 0.150986272 | 0.527742074 | 9.65384995 | 4.377430169 | 0.004367263 | 0.479759698 | -3.328373703 |
| **A_55_P2030383** | Frmd4a | 0.432148472 | 0.192075283 | 0.672221661 | 11.53501304 | 4.373936859 | 0.004384193 | 0.479759698 | -3.329150625 |
| **A_55_P1976764** | Polr3e | 0.338578135 | 0.15037995 | 0.526776319 | 9.770572686 | 4.371461593 | 0.004396233 | 0.479759698 | -3.32970179 |
| **A_52_P101238** | B230354K17Rik | 0.344568703 | 0.152742314 | 0.536395092 | 9.121138574 | 4.364662491 | 0.004429497 | 0.479759698 | -3.331218554 |
| **A_55_P2048259** | Golph3l | 0.340957677 | 0.149838954 | 0.532076399 | 11.68202893 | 4.334913424 | 0.004578394 | 0.479759698 | -3.337903876 |
| **A_55_P2062246** | Tgtp2 | -1.196586747 | -1.867669661 | -0.525503833 | 10.19106901 | -4.332625943 | 0.004590073 | 0.479759698 | -3.338421234 |
| **A_55_P1959056** | Pom121 | 0.410180573 | 0.179801212 | 0.640559935 | 12.27018116 | 4.326280829 | 0.004622643 | 0.479759698 | -3.339858791 |
| **A_51_P148612** | Cox7a1 | -0.583922487 | -0.91209081 | -0.255754164 | 10.92195587 | -4.323562405 | 0.004636676 | 0.479759698 | -3.340475799 |
| **A_51_P298355** | Dek | 0.514133649 | 0.225112646 | 0.803154653 | 12.98661092 | 4.322448356 | 0.004642441 | 0.479759698 | -3.340728853 |
| **A_55_P1952788** | NA | 0.435246602 | 0.19051646 | 0.679976745 | 9.273263894 | 4.321466148 | 0.00464753 | 0.479759698 | -3.340952052 |
| **A_51_P252677** | Card19 | -0.406911185 | -0.635763971 | -0.178058398 | 10.87093775 | -4.320426405 | 0.004652924 | 0.479759698 | -3.341188422 |
| **A_55_P2030938** | Trim59 | 0.600574997 | 0.262415341 | 0.938734654 | 12.79759946 | 4.315475385 | 0.004678706 | 0.479759698 | -3.342315314 |
| **A_55_P1957855** | NA | 0.28248484 | 0.123163723 | 0.441805956 | 9.446204218 | 4.308290643 | 0.004716405 | 0.479759698 | -3.3439546 |
| **A_55_P2014506** | Amd2 | 0.377801602 | 0.164689472 | 0.590913733 | 12.62104503 | 4.307634177 | 0.004719866 | 0.479759698 | -3.344104617 |
| **A_55_P2003823** | Asgr1 | -0.791209629 | -1.23757898 | -0.344840277 | 8.9257502 | -4.307054681 | 0.004722924 | 0.479759698 | -3.344237077 |
| **A_52_P554703** | Gprin3 | -0.363692951 | -0.569002245 | -0.158383657 | 8.149544915 | -4.304368676 | 0.004737126 | 0.479759698 | -3.344851439 |
| **A_52_P24631** | Azin1 | 0.447565197 | 0.194850501 | 0.700279894 | 10.64882952 | 4.303373085 | 0.004742403 | 0.479759698 | -3.345079326 |

| | | | | | | | | | |
|---|---|---|---|---|---|---|---|---|---|
| **A_55_P2025851** | Heatr3 | 0.347884348 | 0.151451168 | 0.544317528 | 11.52978098 | 4.303315932 | 0.004742706 | 0.479759698 | -3.345092411 |
| **A_51_P329356** | Mcts2 | 0.29401285 | 0.127715007 | 0.460310693 | 10.90963046 | 4.295986274 | 0.004781757 | 0.479759698 | -3.346772981 |
| **A_55_P2073789** | NA | -0.956405158 | -1.498420943 | -0.414389373 | 10.2718076 | -4.287589158 | 0.004826939 | 0.479759698 | -3.348704379 |
| **A_55_P2030428** | Sirt1 | 0.405933999 | 0.175505286 | 0.636362713 | 10.50359284 | 4.280574122 | 0.00486505 | 0.479759698 | -3.350322881 |
| **A_55_P2076001** | Rbm12 | 0.368160949 | 0.15899105 | 0.577330848 | 10.78851503 | 4.276827396 | 0.004885543 | 0.479759698 | -3.351189187 |
| **A_55_P2095196** | Akr1b10 | -0.34467048 | -0.540921529 | -0.148419431 | 9.70711398 | -4.267517305 | 0.004936884 | 0.479759698 | -3.35334748 |
| **A_55_P2032997** | Srp19 | 0.477291528 | 0.204799416 | 0.749783641 | 11.77944268 | 4.256111327 | 0.005000607 | 0.479759698 | -3.356002649 |
| **A_55_P2040168** | Pmp22 | -0.543128475 | -0.853534214 | -0.232722737 | 9.571117608 | -4.251635455 | 0.005025864 | 0.479759698 | -3.357047897 |
| **A_55_P2115176** | Pdhx | 0.328216085 | 0.140450546 | 0.515981624 | 9.602672822 | 4.24743911 | 0.005049673 | 0.479759698 | -3.358029572 |
| **A_52_P151320** | Tnfaip8l1 | -0.33951592 | -0.533808106 | -0.145223734 | 10.27537156 | -4.246078293 | 0.00505742 | 0.479759698 | -3.358348271 |
| **A_55_P2009857** | NA | 0.516953272 | 0.221083596 | 0.812822948 | 12.01867418 | 4.2455506 | 0.005060429 | 0.479759698 | -3.358471901 |
| **A_52_P681310** | Plaur | -0.47105233 | -0.740653729 | -0.201450931 | 9.450305345 | -4.245513285 | 0.005060641 | 0.479759698 | -3.358480645 |
| **A_55_P2054857** | Ube2v2 | 0.387851502 | 0.165684166 | 0.610018838 | 12.21808106 | 4.241978469 | 0.005080844 | 0.479759698 | -3.359309486 |
| **A_66_P139387** | Prlr | -0.966702643 | -1.521079311 | -0.412325975 | 8.533752835 | -4.237123779 | 0.005108736 | 0.479759698 | -3.360449727 |
| **A_51_P464387** | Hspb8 | -0.453044873 | -0.713144466 | -0.19294528 | 9.307271517 | -4.232380802 | 0.005136152 | 0.479759698 | -3.361565873 |
| **A_55_P2081530** | Sh3bgrl3 | -0.351306475 | -0.553282896 | -0.149330054 | 13.35572778 | -4.226381254 | 0.005171068 | 0.479759698 | -3.362980767 |
| **A_55_P1974645** | Entpd2 | -0.776085226 | -1.223288517 | -0.328881934 | 9.254510837 | -4.216844764 | 0.005227114 | 0.479759698 | -3.365236804 |
| **A_51_P485260** | Rgl1 | -0.359701937 | -0.567022867 | -0.152381007 | 10.91726646 | -4.215827343 | 0.005233133 | 0.479759698 | -3.365478003 |
| **A_51_P128876** | Ifitm3 | -1.020823885 | -1.609796883 | -0.431850888 | 11.40420059 | -4.211517678 | 0.005258716 | 0.479759698 | -3.366500785 |
| **A_55_P1980591** | Hace1 | 0.562850857 | 0.23801382 | 0.887687894 | 10.45727524 | 4.210280011 | 0.005266089 | 0.479759698 | -3.366794839 |
| **A_55_P2172022** | B4galt1 | -0.324420159 | -0.511690563 | -0.137149755 | 8.647371611 | -4.209416257 | 0.005271242 | 0.479759698 | -3.367000142 |
| **A_66_P112623** | Psmc6 | 0.318562382 | 0.134574383 | 0.502550381 | 11.79576325 | 4.207151813 | 0.005284776 | 0.479759698 | -3.367538708 |
| **A_52_P599055** | Atg14 | -0.312459495 | -0.492990488 | -0.131928501 | 9.577248071 | -4.205572679 | 0.005294237 | 0.479759698 | -3.367914572 |
| **A_51_P356762** | Mcm4 | 0.354627577 | 0.149290641 | 0.559964513 | 13.18729645 | 4.196513422 | 0.005348881 | 0.479759698 | -3.370075451 |
| **A_55_P1966833** | Xaf1 | -0.457938702 | -0.723302277 | -0.192575126 | 8.629180315 | -4.193235244 | 0.005368809 | 0.479759698 | -3.370859321 |
| **A_55_P1985890** | Tiparp | -0.425495582 | -0.672319407 | -0.178671756 | 9.602453211 | -4.18881485 | 0.005395813 | 0.479759698 | -3.371917946 |
| **A_51_P369803** | Psmb9 | -0.337371277 | -0.533896449 | -0.140846105 | 9.824698866 | -4.171316165 | 0.005504207 | 0.479759698 | -3.376127099 |
| **A_55_P2061338** | Slc16a11 | -0.420848172 | -0.66604036 | -0.175655984 | 9.427180127 | -4.170633285 | 0.005508486 | 0.479759698 | -3.376291959 |
| **A_55_P2090349** | Mettl5 | 0.426102963 | 0.177696817 | 0.67450911 | 8.756819162 | 4.168073859 | 0.005524557 | 0.479759698 | -3.376910253 |
| **A_51_P462047** | Prpf38a | 0.320365856 | 0.133137082 | 0.507594631 | 10.95053801 | 4.157735129 | 0.00559001 | 0.479759698 | -3.379414298 |
| **A_55_P2014361** | Pdpk1 | 0.432036308 | 0.179422645 | 0.684649972 | 12.00089209 | 4.155723056 | 0.005602849 | 0.479759698 | -3.37990283 |
| **A_55_P1995165** | Mdc1 | 0.511119343 | 0.2121622 | 0.810076486 | 10.43797399 | 4.15428772 | 0.005612027 | 0.479759698 | -3.38025157 |
| **A_51_P189746** | Pim3 | -0.522744667 | -0.829208926 | -0.216280408 | 9.827075758 | -4.144698722 | 0.005673778 | 0.479759698 | -3.382586541 |
| **A_55_P2066116** | Bcl3 | -0.362628112 | -0.57526895 | -0.149987273 | 10.65824884 | -4.143792317 | 0.005679654 | 0.479759698 | -3.382807719 |
| **A_55_P2109717** | Kif20b | 0.578125179 | 0.238972711 | 0.917277647 | 10.36488151 | 4.141999972 | 0.005691294 | 0.479759698 | -3.383245319 |
| **A_55_P2011341** | NA | -0.842643769 | -1.337815261 | -0.347472278 | 10.88128225 | -4.134962449 | 0.005737254 | 0.479759698 | -3.384966567 |
| **A_55_P1994042** | Zbp1 | -0.350436873 | -0.55643415 | -0.144439597 | 8.453259471 | -4.133629111 | 0.005746008 | 0.479759698 | -3.385293224 |
| **A_55_P2170076** | Taf9-ps | 0.311459733 | 0.128248802 | 0.494670663 | 12.73103325 | 4.13079568 | 0.005764661 | 0.479759698 | -3.385987971 |
| **A_55_P2066309** | Slc39a8 | 0.458982249 | 0.188787469 | 0.72917703 | 11.2515446 | 4.127642957 | 0.005785495 | 0.479759698 | -3.386761934 |
| **A_55_P2036514** | Dis3 | 0.477081329 | 0.195883963 | 0.758278695 | 10.67292645 | 4.122535097 | 0.005819428 | 0.479759698 | -3.388017941 |
| **A_55_P2115003** | Prpf4 | 0.460102659 | 0.188833159 | 0.731372159 | 8.944715048 | 4.121325963 | 0.005827493 | 0.479759698 | -3.388315639 |
| **A_55_P1999349** | Nudt4 | 0.483219107 | 0.197541326 | 0.768896888 | 13.76178671 | 4.110085189 | 0.005903069 | 0.479759698 | -3.391090116 |
| **A_51_P388737** | Slc11a2 | 0.629634782 | 0.25714286 | 1.002126705 | 9.541994405 | 4.107286906 | 0.005922052 | 0.479759698 | -3.391782739 |
| **A_55_P2076303** | 43898 | 0.423867333 | 0.173045764 | 0.674688902 | 12.86476768 | 4.106277123 | 0.005928919 | 0.479759698 | -3.392032868 |
| **A_55_P2035320** | Nfil3 | -0.642718487 | -1.02378348 | -0.261653494 | 10.76215737 | -4.098311148 | 0.005983404 | 0.479759698 | -3.394009635 |
| **A_51_P184806** | Elmod2 | 0.370526428 | 0.150735042 | 0.590317814 | 9.65086988 | 4.096299583 | 0.005997251 | 0.479759698 | -3.394509805 |
| **A_55_P2410304** | Iigp1 | -1.026324263 | -1.635265437 | -0.417383088 | 9.788146182 | -4.095363397 | 0.006003707 | 0.479759698 | -3.394742723 |
| **A_55_P2009375** | Cdk18 | -0.433641982 | -0.691037874 | -0.17624609 | 10.06393317 | -4.093670646 | 0.006015401 | 0.479759698 | -3.395164091 |
| **A_55_P2164189** | Sfpq | 0.423242604 | 0.171932129 | 0.674553079 | 10.44538615 | 4.092248289 | 0.006025246 | 0.479759698 | -3.395518372 |
| **A_55_P1994887** | Zfpm1 | 0.434913816 | 0.176662604 | 0.693165028 | 13.96914499 | 4.092079145 | 0.006026418 | 0.479759698 | -3.395560516 |

| | | | | | | | | |
|---|---|---|---|---|---|---|---|---|
| **A_55_P2400691** | Arhgap21 | 0.36256798 | 0.146751932 | 0.578384029 | 12.00028687 | 4.082149438 | 0.006095668 | 0.479759698 | -3.398039603 |
| **A_55_P2002482** | Sarnp | 0.301386747 | 0.12177404 | 0.480999454 | 12.39477159 | 4.077277607 | 0.006129968 | 0.479759698 | -3.39925953 |
| **A_51_P405565** | Msh6 | 0.400905603 | 0.161494245 | 0.640316962 | 12.14734185 | 4.06893324 | 0.006189217 | 0.479759698 | -3.401354526 |
| **A_55_P1993453** | Ftsj1 | 0.382694881 | 0.153989306 | 0.611400457 | 11.74775274 | 4.065922446 | 0.006210752 | 0.479759698 | -3.402112156 |
| **A_55_P2159994** | Atf1 | 0.48802423 | 0.19634769 | 0.77970077 | 10.79680209 | 4.06558527 | 0.006213169 | 0.479759698 | -3.40219706 |
| **A_55_P2011322** | NA | 0.30914046 | 0.12419712 | 0.494083801 | 11.76715829 | 4.061629833 | 0.006241599 | 0.479759698 | -3.40319392 |
| **A_55_P2052578** | Bclaf3 | 0.39013991 | 0.155866718 | 0.624413103 | 9.4004518 | 4.046513309 | 0.006351594 | 0.479759698 | -3.407018194 |
| **A_51_P159503** | Rnf213 | -0.477684358 | -0.764758841 | -0.190609875 | 10.46602422 | -4.043240903 | 0.006375688 | 0.479759698 | -3.407849116 |
| **A_55_P2149763** | Tapbp | -0.44473153 | -0.712039203 | -0.177423858 | 10.90134092 | -4.042682975 | 0.006379807 | 0.479759698 | -3.407990892 |
| **A_55_P1978103** | Ppip5k2 | 0.30707443 | 0.122398838 | 0.491750022 | 10.13188338 | 4.040334704 | 0.006397172 | 0.479759698 | -3.408587964 |
| **A_55_P2000833** | E2f8 | 0.812910352 | 0.321237977 | 1.304582727 | 11.97796036 | 4.017445884 | 0.006569219 | 0.479759698 | -3.414437126 |
| **A_51_P436727** | Arrb1 | 0.439888939 | 0.173739735 | 0.706038143 | 13.44061795 | 4.016068016 | 0.006579739 | 0.479759698 | -3.414790946 |
| **A_65_P12977** | Dnaja2 | 0.333669382 | 0.131536677 | 0.535802086 | 11.72821142 | 4.011095092 | 0.006617864 | 0.479759698 | -3.416069555 |
| **A_66_P128537** | Isg15 | -0.928892172 | -1.494053334 | -0.363731011 | 9.364694284 | -3.99370685 | 0.006753115 | 0.479759698 | -3.420560337 |
| **A_55_P2073253** | NA | 0.44081891 | 0.172295246 | 0.709342574 | 9.862257302 | 3.988970656 | 0.006790485 | 0.479759698 | -3.421788946 |
| **A_52_P428446** | Tsn | 0.30426201 | 0.118748021 | 0.489775998 | 11.96640621 | 3.985237926 | 0.0068201 | 0.479759698 | -3.422758887 |
| **A_52_P317923** | Zfp948 | 0.377324106 | 0.147216797 | 0.607431415 | 9.319958211 | 3.984440326 | 0.006826447 | 0.479759698 | -3.422966328 |
| **A_51_P375431** | Phip | 0.365408603 | 0.142545176 | 0.58827203 | 8.719749009 | 3.984035071 | 0.006829674 | 0.479759698 | -3.423071754 |
| **A_55_P1967216** | Adam10 | 0.447219064 | 0.174447253 | 0.719990875 | 12.86762468 | 3.983859465 | 0.006831073 | 0.479759698 | -3.423117442 |
| **A_55_P1982613** | Hnrnpa3 | 0.424213882 | 0.165430911 | 0.682996854 | 14.22377363 | 3.983202352 | 0.006836311 | 0.479759698 | -3.423288435 |
| **A_55_P1986526** | Cdh26 | 0.437277516 | 0.170523024 | 0.704032007 | 9.350559656 | 3.983167733 | 0.006836587 | 0.479759698 | -3.423297445 |
| **A_52_P309337** | Metap2 | 0.457467112 | 0.177491744 | 0.737442481 | 8.985232272 | 3.970299248 | 0.006940067 | 0.479759698 | -3.426655162 |
| **A_55_P1994939** | Hmgb2 | 0.392012545 | 0.151951947 | 0.632073143 | 14.99867231 | 3.967914182 | 0.006959437 | 0.479759698 | -3.427279383 |
| **A_55_P2043682** | Ubac1 | 0.401877054 | 0.155712898 | 0.64804121 | 13.88260126 | 3.966903026 | 0.006967667 | 0.479759698 | -3.427544203 |
| **A_51_P392546** | Haus3 | 0.435830786 | 0.168584979 | 0.703076592 | 11.19747355 | 3.962690863 | 0.007002067 | 0.479759698 | -3.428648511 |
| **A_55_P1983773** | Birc5 | 0.43825463 | 0.169515215 | 0.706994045 | 14.69840816 | 3.962582663 | 0.007002953 | 0.479759698 | -3.428676902 |
| **A_55_P2002466** | Zfp747 | -0.386318801 | -0.62332649 | -0.149311112 | 9.743250365 | -3.960651201 | 0.007018793 | 0.479759698 | -3.429183919 |
| **A_51_P329322** | Zfand1 | 0.265665008 | 0.102576585 | 0.42875343 | 11.0338559 | 3.958169721 | 0.007039201 | 0.479759698 | -3.429835891 |
| **A_55_P2025463** | Brpf3 | 0.606652945 | 0.233670619 | 0.979635272 | 11.30396596 | 3.952166619 | 0.007088846 | 0.479759698 | -3.431415788 |
| **A_55_P2009952** | Me1 | -0.453258838 | -0.732187153 | -0.174330523 | 10.23300456 | -3.948542952 | 0.007119001 | 0.479759698 | -3.432371297 |
| **A_55_P2274553** | NA | 0.630638403 | 0.242195376 | 1.019081431 | 8.450694187 | 3.944902488 | 0.00714944 | 0.479759698 | -3.433332627 |
| **A_55_P2130970** | Parp10 | -0.333192586 | -0.538654179 | -0.127730993 | 11.75227463 | -3.940468563 | 0.007186708 | 0.479759698 | -3.43450537 |
| **A_51_P230098** | Pbk | 0.326299024 | 0.124694688 | 0.527903361 | 10.5879767 | 3.932774942 | 0.007251889 | 0.479759698 | -3.436545207 |
| **A_52_P215176** | Opn3 | -0.404376933 | -0.654240898 | -0.154512969 | 8.372060783 | -3.932474698 | 0.007254446 | 0.479759698 | -3.436624938 |
| **A_51_P504442** | Sf3a1 | 0.330592655 | 0.126277917 | 0.534907394 | 10.9541896 | 3.931666692 | 0.007261332 | 0.479759698 | -3.436839556 |
| **A_55_P2105256** | Trim47 | -0.52994019 | -0.85766305 | -0.20221733 | 10.76534157 | -3.929192532 | 0.007282463 | 0.479759698 | -3.43749716 |
| **A_55_P2227154** | A530041M06Rik | 0.295601225 | 0.112575695 | 0.478626754 | 8.591847279 | 3.924440465 | 0.007323241 | 0.479759698 | -3.438762024 |
| **A_55_P1952905** | Dnm1l | 0.380256227 | 0.144703968 | 0.615808486 | 11.02631464 | 3.922583982 | 0.00733924 | 0.479759698 | -3.439256817 |
| **A_52_P463346** | Serinc3 | 0.496607998 | 0.1884805 | 0.804735496 | 12.51505976 | 3.916214229 | 0.007394432 | 0.479759698 | -3.440957281 |
| **A_55_P2061437** | Lsm5 | 0.353904318 | 0.134274553 | 0.573534084 | 11.37823081 | 3.915415472 | 0.007401385 | 0.479759698 | -3.441170821 |
| **A_52_P81783** | Gspt1 | 0.380294542 | 0.144186973 | 0.616402111 | 9.326464821 | 3.913752624 | 0.007415884 | 0.479759698 | -3.441615585 |
| **A_66_P125660** | Nde1 | 0.382070402 | 0.144661308 | 0.619479496 | 8.462882256 | 3.91047248 | 0.007444577 | 0.479759698 | -3.442493794 |
| **A_55_P2169729** | NA | 0.372791722 | 0.140942152 | 0.604641292 | 13.61515466 | 3.906997695 | 0.007475107 | 0.479759698 | -3.443425367 |
| **A_51_P502964** | Tmem63a | -0.281604164 | -0.45683806 | -0.106370267 | 10.11530437 | -3.904848087 | 0.007494063 | 0.479759698 | -3.444002312 |
| **A_52_P417654** | Tcea1 | 0.274553776 | 0.103537915 | 0.445569638 | 9.535680735 | 3.900984436 | 0.007528269 | 0.479759698 | -3.44504054 |
| **A_55_P2032318** | 4930522L14Rik | 0.406963209 | 0.153453144 | 0.660473274 | 8.639141339 | 3.900705225 | 0.007530748 | 0.479759698 | -3.445115631 |
| **A_66_P103490** | Srsf1 | 0.512113923 | 0.192855121 | 0.831372725 | 10.87917487 | 3.897686394 | 0.007557605 | 0.479759698 | -3.445928044 |
| **A_51_P355753** | Hic1 | -0.916534073 | -1.487934882 | -0.345133263 | 9.965031501 | -3.897543335 | 0.00755888 | 0.479759698 | -3.445966568 |
| **A_55_P1961049** | Cep295 | 0.468927588 | 0.176104808 | 0.761750367 | 10.43262972 | 3.891205028 | 0.007615626 | 0.479759698 | -3.447675578 |
| **A_55_P2167620** | NA | 0.40092447 | 0.150487652 | 0.651361288 | 10.10348474 | 3.889981861 | 0.007626632 | 0.479759698 | -3.448005879 |

| | | | | | | | | | |
|---|---|---|---|---|---|---|---|---|---|
| **A_55_P1955632** | Trp53i11 | -0.806373274 | -1.310082496 | -0.302664051 | 10.42540387 | -3.889908733 | 0.00762729 | 0.479759698 | -3.448025632 |
| **A_51_P326229** | Ddx25 | 0.713715317 | 0.267783866 | 1.159646768 | 9.136581889 | 3.889019338 | 0.007635304 | 0.479759698 | -3.44826591 |
| **A_55_P1996008** | Gm14399 | 0.414398344 | 0.155353713 | 0.673442974 | 8.879973348 | 3.887107984 | 0.007652558 | 0.479759698 | -3.448782568 |
| **A_55_P1993365** | NA | 0.638331911 | 0.239099055 | 1.037564766 | 8.762581699 | 3.885111345 | 0.007670629 | 0.479759698 | -3.449322698 |
| **A_55_P2430367** | Zbtb8b | -0.45558715 | -0.74107476 | -0.17009954 | 8.823664747 | -3.877639136 | 0.007738676 | 0.479759698 | -3.45134789 |
| **A_55_P2090184** | Fgfr1op2 | 0.292983041 | 0.109196644 | 0.476769438 | 12.38785679 | 3.873578017 | 0.00777594 | 0.479759698 | -3.452451101 |
| **A_66_P125831** | Pnn | 0.348496127 | 0.12987376 | 0.567118494 | 10.90185754 | 3.873348307 | 0.007778054 | 0.479759698 | -3.452513555 |
| **A_51_P481210** | Ofd1 | 0.419770016 | 0.156138605 | 0.683401426 | 10.01054828 | 3.868988592 | 0.007818292 | 0.479759698 | -3.453699973 |
| **A_55_P2055477** | Scaf11 | 0.280198716 | 0.104205362 | 0.456192071 | 10.3764153 | 3.868593171 | 0.007821953 | 0.479759698 | -3.453807681 |
| **A_55_P2043684** | Ubac1 | 0.425717668 | 0.158194027 | 0.693241309 | 14.36770496 | 3.866719755 | 0.007839324 | 0.479759698 | -3.45431821 |
| **A_55_P2077263** | Cenpk | 0.573856435 | 0.213192632 | 0.934520238 | 10.80640891 | 3.866196176 | 0.007844186 | 0.479759698 | -3.454460959 |
| **A_55_P2032543** | Cblc | -0.3667994 | -0.597400514 | -0.136198286 | 10.56104718 | -3.865008063 | 0.007855232 | 0.479759698 | -3.454784999 |
| **A_55_P2050799** | Nup35 | 0.38443414 | 0.142617447 | 0.626250833 | 11.23950427 | 3.86294798 | 0.007874426 | 0.479759698 | -3.455347217 |
| **A_55_P2115691** | NA | 0.480583315 | 0.177811279 | 0.783355352 | 11.97159868 | 3.856879905 | 0.007931266 | 0.479759698 | -3.457005936 |
| **A_55_P2058812** | Fancl | 0.401553518 | 0.148457345 | 0.654649692 | 9.958740154 | 3.855147915 | 0.007947573 | 0.479759698 | -3.457480113 |
| **A_55_P2086983** | Atxn2 | 0.292153623 | 0.107962756 | 0.476344491 | 11.5004753 | 3.854130114 | 0.007957173 | 0.479759698 | -3.457758914 |
| **A_55_P2114953** | Usp18 | -0.295582886 | -0.481935943 | -0.109229829 | 9.307839447 | -3.854126393 | 0.007957208 | 0.479759698 | -3.457759934 |
| **A_51_P351363** | Tyms-ps | 0.646699504 | 0.238940429 | 1.054458578 | 11.58607134 | 3.853737083 | 0.007960883 | 0.479759698 | -3.457866605 |
| **A_51_P351860** | C1qb | -0.380495544 | -0.620615974 | -0.140375114 | 10.68681372 | -3.850380521 | 0.007992651 | 0.479759698 | -3.458786995 |
| **A_55_P2116802** | NA | 0.492191058 | 0.181362492 | 0.803019625 | 11.32176407 | 3.847653754 | 0.008018561 | 0.479759698 | -3.459535594 |
| **A_55_P2164534** | Dtl | 0.552287396 | 0.203120661 | 0.901454132 | 10.63343924 | 3.843398881 | 0.008059178 | 0.479759698 | -3.460705335 |
| **A_55_P2124461** | NA | -0.467195584 | -0.762666364 | -0.171724804 | 13.23572501 | -3.842088635 | 0.008071732 | 0.479759698 | -3.461065944 |
| **A_51_P267494** | Cdc42ep3 | 0.277781636 | 0.102022739 | 0.453540534 | 10.77130243 | 3.840337567 | 0.008088542 | 0.479759698 | -3.461548169 |
| **A_55_P2130610** | Smc3 | 0.351655431 | 0.129139504 | 0.574171357 | 12.25570454 | 3.840072436 | 0.008091091 | 0.479759698 | -3.461621213 |
| **A_55_P1980020** | Rsl1d1 | 0.276441737 | 0.101482729 | 0.451400746 | 12.88819638 | 3.839286248 | 0.008098654 | 0.479759698 | -3.461837852 |
| **A_55_P2020911** | Rps6ka6 | 0.402931782 | 0.147877785 | 0.657985779 | 9.341106021 | 3.83868592 | 0.008104434 | 0.479759698 | -3.462003322 |
| **A_55_P2073705** | Tfdp2 | 0.721000698 | 0.264347984 | 1.177653412 | 10.79193827 | 3.836479051 | 0.008125723 | 0.479759698 | -3.462611947 |
| **A_55_P2047842** | Clcn3 | 0.525836331 | 0.192558126 | 0.859114535 | 12.96799958 | 3.833777154 | 0.008151871 | 0.479759698 | -3.463357819 |
| **A_55_P2041017** | Gm15365 | 0.288347192 | 0.10551036 | 0.471184025 | 13.48004962 | 3.832085822 | 0.008168286 | 0.479759698 | -3.463825127 |
| **A_51_P394558** | Dchs1 | -0.334539656 | -0.546834448 | -0.122244864 | 8.959148785 | -3.829053685 | 0.008197807 | 0.479759698 | -3.464663679 |
| **A_51_P155873** | Ppp1r3g | -1.129935266 | -1.847570529 | -0.412300003 | 10.45060884 | -3.825894426 | 0.008228691 | 0.479759698 | -3.465538461 |
| **A_55_P2114938** | Irf9 | -0.369147029 | -0.603628862 | -0.134665196 | 10.7924882 | -3.825369288 | 0.008233838 | 0.479759698 | -3.465683975 |
| **A_51_P341010** | Ercc8 | 0.282997354 | 0.103234671 | 0.462760036 | 9.74618901 | 3.825304607 | 0.008234472 | 0.479759698 | -3.4657019 |
| **A_51_P101573** | Klc4 | -0.324314716 | -0.530472684 | -0.118156748 | 10.45339571 | -3.822519584 | 0.008261826 | 0.479759698 | -3.466474148 |
| **A_55_P1969710** | NA | 0.43364188 | 0.1579762 | 0.70930756 | 14.13286691 | 3.822361057 | 0.008263386 | 0.479759698 | -3.466518131 |
| **A_55_P2039523** | Phf10 | 0.499649635 | 0.181929305 | 0.817369964 | 12.38168715 | 3.821235109 | 0.008274476 | 0.479759698 | -3.466830603 |
| **A_55_P1996872** | Polr2k | 0.327703799 | 0.118876753 | 0.536530845 | 12.43109257 | 3.813097591 | 0.008355116 | 0.479759698 | -3.469093067 |
| **A_51_P234544** | Azin1 | 0.43850707 | 0.159016295 | 0.717997844 | 12.55100812 | 3.812346013 | 0.008362608 | 0.479759698 | -3.469302396 |
| **A_55_P2042713** | Pigq | 0.429401965 | 0.155350834 | 0.703453095 | 14.90206591 | 3.807287024 | 0.008413231 | 0.479759698 | -3.470713043 |
| **A_51_P223404** | Plin3 | -0.342374612 | -0.561362382 | -0.123386842 | 13.24799478 | -3.798961399 | 0.008497281 | 0.479759698 | -3.473040723 |
| **A_55_P1979650** | S100a1 | -0.497841162 | -0.816310791 | -0.179371533 | 10.01399493 | -3.798446108 | 0.008502514 | 0.479759698 | -3.473185041 |
| **A_55_P2125208** | Arid5a | -0.299657645 | -0.491654484 | -0.107660806 | 9.916949212 | -3.792402916 | 0.008564147 | 0.479759698 | -3.474879758 |
| **A_55_P2018480** | Carnmt1 | 0.370890423 | 0.133199069 | 0.608581778 | 10.9380664 | 3.791538084 | 0.008573007 | 0.479759698 | -3.475122618 |
| **A_51_P336721** | Tipin | 0.392396101 | 0.140833195 | 0.643959008 | 12.98668926 | 3.790192507 | 0.008586813 | 0.479759698 | -3.475500645 |
| **A_66_P115531** | Crebzf | 0.31764536 | 0.11399544 | 0.521295281 | 10.95094023 | 3.790019648 | 0.008588589 | 0.479759698 | -3.475549223 |
| **A_55_P2010906** | Rbm45 | 0.281940911 | 0.101036542 | 0.46284528 | 9.430076007 | 3.786973206 | 0.008619945 | 0.479759698 | -3.476405895 |
| **A_55_P2123471** | Kpna2 | 0.368207109 | 0.131910991 | 0.604503227 | 14.01242962 | 3.786332687 | 0.008626554 | 0.479759698 | -3.476586143 |
| **A_55_P2028054** | Incenp | 0.377698863 | 0.135296695 | 0.620101031 | 13.61911625 | 3.786102493 | 0.00862893 | 0.479759698 | -3.476650933 |
| **A_55_P2064038** | NA | 0.566735834 | 0.202992113 | 0.930479554 | 8.6358181 | 3.785893192 | 0.008631092 | 0.479759698 | -3.476709847 |
| **A_51_P353095** | Zfyve27 | -0.244939093 | -0.402203456 | -0.087674729 | 10.2609085 | -3.784521455 | 0.008645272 | 0.479759698 | -3.477096088 |

| | | | | | | | | |
|---|---|---|---|---|---|---|---|---|
| **A_51_P210474** | Mrpl18 | 0.417528837 | 0.149372006 | 0.685685668 | 11.94265367 | 3.783387286 | 0.008657016 | 0.479759698 | -3.477415595 |
| **A_55_P2079390** | 4930579G24Rik | 0.380842537 | 0.136137401 | 0.625547672 | 9.134423626 | 3.781686769 | 0.008674658 | 0.479759698 | -3.477894916 |
| **A_55_P2018091** | NA | -0.437369434 | -0.719003917 | -0.155734952 | 12.81748676 | -3.773512482 | 0.008760018 | 0.479759698 | -3.480203492 |
| **A_51_P487690** | Ifi44 | -0.320247087 | -0.526510652 | -0.113983522 | 8.044526679 | -3.772644277 | 0.008769138 | 0.479759698 | -3.480449129 |
| **A_55_P2102936** | Pnrc1 | -0.3663481 | -0.603577963 | -0.129118237 | 10.6689089 | -3.752388312 | 0.008984944 | 0.479759698 | -3.48620403 |
| **A_51_P383218** | Tmem161a | 0.250746744 | 0.088358037 | 0.413135451 | 8.678021339 | 3.751998605 | 0.008989153 | 0.479759698 | -3.486315201 |
| **A_55_P2172420** | Krit1 | 0.334531557 | 0.117632688 | 0.551430426 | 9.717679029 | 3.747684247 | 0.009035897 | 0.479759698 | -3.487547099 |
| **A_55_P2106525** | Nmnat3 | 0.536908066 | 0.188326411 | 0.88548972 | 11.72313446 | 3.742644613 | 0.00909084 | 0.479759698 | -3.488988747 |
| **A_55_P2064771** | Ly6c1 | -0.539035311 | -0.889000698 | -0.189069924 | 12.25801263 | -3.742616364 | 0.009091149 | 0.479759698 | -3.488996836 |
| **A_55_P1998987** | NA | 0.597882325 | 0.209707138 | 0.986057511 | 9.153190298 | 3.742580354 | 0.009091543 | 0.479759698 | -3.489007147 |
| **A_55_P2001188** | Phf6 | 0.260827307 | 0.091442844 | 0.430211769 | 10.25894479 | 3.741645821 | 0.009101773 | 0.479759698 | -3.489274804 |
| **A_55_P1973809** | Hbb-bt | 0.350891526 | 0.122981293 | 0.578801758 | 17.67056951 | 3.741039081 | 0.009108422 | 0.479759698 | -3.489448631 |
| **A_55_P1972263** | Nop56 | 0.437673944 | 0.153355839 | 0.721992049 | 12.6661661 | 3.740497475 | 0.009114361 | 0.479759698 | -3.489603833 |
| **A_55_P1976127** | Ect2 | 0.649158894 | 0.227268943 | 1.071048844 | 11.60623262 | 3.738823986 | 0.00913274 | 0.479759698 | -3.490083596 |
| **A_55_P1998942** | Oas1a | -0.332139926 | -0.548385565 | -0.115894287 | 8.605107296 | -3.732131305 | 0.009206652 | 0.479759698 | -3.49200545 |
| **A_55_P2055612** | Zfp800 | 0.75280412 | 0.262065097 | 1.243543143 | 9.260826855 | 3.727473654 | 0.00925848 | 0.479759698 | -3.493345927 |
| **A_55_P2057268** | Magi1 | -0.335754365 | -0.554855994 | -0.116652736 | 9.264102578 | -3.723567659 | 0.009302193 | 0.479759698 | -3.494471976 |
| **A_55_P2058861** | Mvk | 0.328533492 | 0.114123425 | 0.542943558 | 12.91167633 | 3.72321115 | 0.009306194 | 0.479759698 | -3.49457484 |
| **A_55_P2004016** | Crispld2 | -0.677247352 | -1.119393137 | -0.235101568 | 10.29601729 | -3.721902704 | 0.009320895 | 0.479759698 | -3.494952489 |
| **A_55_P2138257** | Ube2d2a | 0.392290506 | 0.136171703 | 0.64840931 | 10.85468305 | 3.721769933 | 0.009322388 | 0.479759698 | -3.494990821 |
| **A_55_P1988228** | Aspm | 0.584533606 | 0.202553169 | 0.966514043 | 10.88554167 | 3.718361011 | 0.009360817 | 0.479759698 | -3.495975685 |
| **A_51_P490296** | Slx4 | 0.352365747 | 0.121999021 | 0.582732473 | 10.1909803 | 3.716696709 | 0.009379642 | 0.479759698 | -3.496456996 |
| **A_55_P2033255** | NA | -0.46153424 | -0.763332575 | -0.159735905 | 14.42916577 | -3.715953538 | 0.009388062 | 0.479759698 | -3.49667202 |
| **A_55_P2039566** | Arhgap23 | 0.419978449 | 0.145324087 | 0.694632812 | 12.4189025 | 3.715554994 | 0.009392581 | 0.479759698 | -3.496787358 |
| **A_55_P2078459** | Kpna2 | 0.343198866 | 0.118407387 | 0.567990345 | 12.94890829 | 3.709788828 | 0.009458227 | 0.479759698 | -3.498458108 |
| **A_55_P1978241** | Artn | 0.329642393 | 0.113672974 | 0.545611811 | 8.734035905 | 3.708804904 | 0.009469479 | 0.479759698 | -3.49874358 |
| **A_55_P2001228** | Dctn6 | 0.491407798 | 0.169164998 | 0.813650597 | 14.38697764 | 3.705458988 | 0.009507854 | 0.479759698 | -3.499715179 |
| **A_55_P2127357** | NA | -0.347369756 | -0.575171768 | -0.119567744 | 9.831928172 | -3.705251034 | 0.009510245 | 0.479759698 | -3.499775607 |
| **A_51_P164296** | Adamdec1 | -0.336085024 | -0.556574941 | -0.115595106 | 8.158844155 | -3.703766631 | 0.00952733 | 0.479759698 | -3.500207098 |
| **A_52_P681** | Cnot4 | 0.327800634 | 0.112532271 | 0.543068996 | 9.096551848 | 3.700094116 | 0.009569745 | 0.479759698 | -3.501275721 |
| **A_55_P2097612** | Zfp131 | 0.326229895 | 0.111790798 | 0.540668991 | 10.23884868 | 3.696604397 | 0.009610242 | 0.479759698 | -3.502292587 |
| **A_55_P2056344** | Unc93a2 | 0.564897927 | 0.193307331 | 0.936488523 | 15.46411586 | 3.693928249 | 0.009641426 | 0.479759698 | -3.503073333 |
| **A_55_P1995173** | Odc1 | 0.412909082 | 0.141158111 | 0.684660052 | 13.99063124 | 3.692041045 | 0.009663484 | 0.479759698 | -3.503624405 |
| **A_55_P2034705** | Nmi | -0.301280415 | -0.499749559 | -0.10281127 | 10.04903081 | -3.68859601 | 0.009703894 | 0.479759698 | -3.504631425 |
| **A_55_P1998596** | Nupl2 | 0.275393953 | 0.093970725 | 0.45681718 | 9.50180299 | 3.688456783 | 0.009705531 | 0.479759698 | -3.504672151 |
| **A_52_P627816** | Tgm1 | -0.350570591 | -0.581886625 | -0.119254558 | 8.65420182 | -3.6825863 | 0.009774836 | 0.479759698 | -3.506391396 |
| **A_55_P1972748** | Czib | 0.641749796 | 0.218014238 | 1.065485353 | 14.78291895 | 3.680052541 | 0.009804917 | 0.479759698 | -3.507134665 |
| **A_51_P304170** | Rtp4 | -0.698657193 | -1.161042707 | -0.236271679 | 10.16294389 | -3.671496379 | 0.009907255 | 0.479759698 | -3.50965006 |
| **A_55_P2028300** | Ppm1d | 0.451235848 | 0.152183001 | 0.750288695 | 10.00744089 | 3.666391536 | 0.009968874 | 0.479759698 | -3.511154845 |
| **A_55_P2147001** | Wdr55 | 0.348985938 | 0.117590234 | 0.580381643 | 11.73767453 | 3.66467803 | 0.009989652 | 0.479759698 | -3.511660621 |
| **A_55_P2027737** | Il1rl1 | 0.630608203 | 0.212032823 | 1.049183584 | 10.55362426 | 3.660742019 | 0.010037562 | 0.479759698 | -3.512823705 |
| **A_55_P2058757** | Cap1 | -0.35136083 | -0.584839357 | -0.117882303 | 11.16183549 | -3.65670216 | 0.010086999 | 0.479759698 | -3.514019344 |
| **A_55_P2032760** | Ctif | -0.335546913 | -0.558651541 | -0.112442285 | 9.504438943 | -3.654499082 | 0.010114072 | 0.479759698 | -3.514672167 |
| **A_55_P2102175** | Lztr1 | -0.428290737 | -0.713126126 | -0.143455348 | 10.69972605 | -3.653658264 | 0.010124426 | 0.479759698 | -3.514921469 |
| **A_51_P327828** | Ppib | -0.272059354 | -0.453061817 | -0.091056892 | 14.07595644 | -3.652265731 | 0.0101416 | 0.479759698 | -3.515334536 |
| **A_51_P236324** | Gtf2e1 | 0.243879332 | 0.081449833 | 0.406308831 | 9.476060135 | 3.648323 | 0.010190398 | 0.479759698 | -3.516505292 |
| **A_51_P330369** | Ssx2ip | 0.528877802 | 0.176492917 | 0.881262687 | 11.91697252 | 3.646878183 | 0.010208345 | 0.479759698 | -3.516934769 |
| **A_55_P2116165** | Pfkfb3 | -0.528979649 | -0.88187637 | -0.176082927 | 10.14077188 | -3.642290063 | 0.010265568 | 0.479759698 | -3.518300219 |
| **A_55_P2042521** | Cop1 | 0.250449959 | 0.083150387 | 0.41774953 | 11.66103703 | 3.637552947 | 0.010325021 | 0.479759698 | -3.519712588 |
| **A_52_P18068** | Txlng | 0.250134433 | 0.082937766 | 0.4173311 | 8.660239803 | 3.635206219 | 0.010354614 | 0.479759698 | -3.520413235 |

| | | | | | | | | | |
|---|---|---|---|---|---|---|---|---|---|
| **A_51_P282667** | Hexa | -0.269400471 | -0.449514438 | -0.089286504 | 10.983285 | -3.634411917 | 0.010364652 | 0.479759698 | -3.52065053 |
| **A_55_P2089565** | Tbx22 | 0.721976189 | 0.23884944 | 1.205102938 | 9.333350513 | 3.631156608 | 0.010405902 | 0.479759698 | -3.521623815 |
| **A_51_P122035** | Nipa1 | 0.465198628 | 0.153846965 | 0.776550291 | 10.97756976 | 3.630533297 | 0.010413821 | 0.479759698 | -3.544935693 |
| **A_55_P2305010** | A730089K16Rik | 0.633040528 | 0.208975474 | 1.057105583 | 10.86265605 | 3.627289498 | 0.01045514 | 0.479759698 | -3.52278163 |
| **A_55_P2095854** | Rab3c | 0.539406017 | 0.177263691 | 0.901548343 | 12.02994343 | 3.619259212 | 0.010558208 | 0.479759698 | -3.525191506 |
| **A_55_P2057946** | Hsp90aa1 | 0.298502653 | 0.09768346 | 0.499321846 | 14.08299665 | 3.611820572 | 0.010654683 | 0.479759698 | -3.52743059 |
| **A_55_P1979551** | Pitrm1 | 0.584491921 | 0.19114191 | 0.977841933 | 13.11569888 | 3.610626249 | 0.010670264 | 0.479759698 | -3.527790696 |
| **A_52_P633410** | Gabarapl1 | -0.310319827 | -0.519196802 | -0.101442851 | 13.54787209 | -3.609957944 | 0.010678993 | 0.479759698 | -3.527992274 |
| **A_51_P133562** | Serpina6 | 0.975428124 | 0.318677001 | 1.632179247 | 15.48201161 | 3.608923184 | 0.010692524 | 0.479759698 | -3.528304487 |
| **A_55_P2098539** | Itgb3bp | 0.368520825 | 0.120393559 | 0.616648091 | 10.23142804 | 3.608865803 | 0.010693275 | 0.479759698 | -3.528321804 |
| **A_52_P638459** | Ccl5 | -0.662687145 | -1.109102726 | -0.216271564 | 9.005908354 | -3.607051901 | 0.010717042 | 0.479759698 | -3.528869421 |
| **A_55_P1959305** | Rbm47 | -0.376973971 | -0.630931555 | -0.123016388 | 10.65771161 | -3.606893901 | 0.010719115 | 0.479759698 | -3.528917139 |
| **A_55_P2121252** | Amd2 | 0.443297179 | 0.144648741 | 0.741945617 | 9.981647028 | 3.606765513 | 0.010720799 | 0.479759698 | -3.528955917 |
| **A_55_P2147250** | Caprin2 | 0.341146347 | 0.111308958 | 0.570983736 | 9.428131199 | 3.606643655 | 0.010722399 | 0.479759698 | -3.528992724 |
| **A_51_P316935** | Depp1 | -0.930608641 | -1.557700922 | -0.30351636 | 11.6114289 | -3.605942645 | 0.010731604 | 0.479759698 | -3.529204496 |
| **A_52_P375876** | Arf3 | -0.473183614 | -0.782427305 | -0.163939924 | 11.84042659 | -3.892881784 | 0.01073318 | 0.479759698 | -3.714020143 |
| **A_52_P219943** | Epc1 | 0.394350453 | 0.128538884 | 0.660162021 | 8.648641698 | 3.604886985 | 0.010745484 | 0.479759698 | -3.529523517 |
| **A_52_P463977** | Tmem140 | -0.419641395 | -0.703066997 | -0.136215793 | 11.27251381 | -3.597679186 | 0.010840781 | 0.479759698 | -3.53170523 |
| **A_55_P2069989** | Klhl12 | 0.433742102 | 0.140464504 | 0.7270197 | 12.0612594 | 3.593650839 | 0.010894446 | 0.479759698 | -3.532927237 |
| **A_55_P2066230** | Hck | -0.297389601 | -0.498507837 | -0.096271365 | 11.69456929 | -3.593002467 | 0.010903111 | 0.479759698 | -3.533124102 |
| **A_66_P136834** | Usp7 | 0.384040357 | 0.124317765 | 0.643762948 | 8.931986272 | 3.59294299 | 0.010903906 | 0.479759698 | -3.533142163 |
| **A_52_P379337** | Rtn4 | -0.283287517 | -0.474962259 | -0.091612775 | 9.488297913 | -3.591250824 | 0.010926558 | 0.479759698 | -3.5336562 |
| **A_51_P126525** | D230025D16Rik | -0.294570382 | -0.493895159 | -0.095245605 | 9.835996005 | -3.590963417 | 0.010930411 | 0.479759698 | -3.615503517 |
| **A_51_P204402** | Shcbp1 | 0.50328103 | 0.162709388 | 0.843852671 | 10.50017235 | 3.590750813 | 0.010933262 | 0.479759698 | -3.533808155 |
| **A_55_P2004037** | NA | 0.694638496 | 0.22452598 | 1.164751012 | 9.607649949 | 3.590378402 | 0.010938257 | 0.479759698 | -3.533921352 |
| **A_55_P2065671** | Ccnb1 | 0.453066901 | 0.146406143 | 0.75972766 | 12.41635268 | 3.589941069 | 0.010944127 | 0.479759698 | -3.534054303 |
| **A_55_P1988083** | Prc1 | 0.344209857 | 0.111176343 | 0.57724337 | 13.03477814 | 3.589121071 | 0.010955142 | 0.479759698 | -3.534303647 |
| **A_55_P2059134** | Smarca5 | 0.425493848 | 0.137363816 | 0.71362388 | 11.62115573 | 3.588293404 | 0.010966272 | 0.479759698 | -3.534555403 |
| **A_51_P324814** | Krt18 | -0.306247488 | -0.513842847 | -0.098652129 | 14.37757309 | -3.584578406 | 0.011016384 | 0.479759698 | -3.535686419 |
| **A_55_P2016862** | Ncoa4 | 0.374147533 | 0.120476657 | 0.62781841 | 14.39999931 | 3.583896553 | 0.011025609 | 0.479759698 | -3.535894184 |
| **A_55_P2082156** | Rab11fip3 | -0.283330371 | -0.475578777 | -0.091081965 | 9.350253681 | -3.581076271 | 0.011063856 | 0.479759698 | -3.536754129 |
| **A_55_P2106150** | Cenpk | 0.66421828 | 0.213491894 | 1.114944666 | 11.64770328 | 3.580807959 | 0.011067503 | 0.479759698 | -3.53683599 |
| **A_66_P126836** | Zbtb44 | 0.32244953 | 0.103579482 | 0.541319579 | 10.10941243 | 3.579798678 | 0.01108123 | 0.479759698 | -3.537143997 |
| **A_55_P2076631** | Gypc | 0.400950443 | 0.128700403 | 0.673200482 | 14.06820394 | 3.578540483 | 0.01109837 | 0.479759698 | -3.537528135 |
| **A_55_P2010191** | Serpina11 | -0.677870789 | -1.138383189 | -0.217358389 | 12.95545598 | -3.576751599 | 0.011122789 | 0.479759698 | -3.538074621 |
| **A_51_P123405** | Bub1 | 0.443917604 | 0.142078054 | 0.745757154 | 10.06509122 | 3.573628578 | 0.011165562 | 0.479759698 | -3.539029586 |
| **A_55_P2028064** | Kdm4b | -0.246398147 | -0.414067019 | -0.078729275 | 10.06465962 | -3.570821853 | 0.011204157 | 0.479759698 | -3.539888824 |
| **A_55_P2109382** | Adora2a | -0.359964507 | -0.605063568 | -0.114865447 | 9.04904534 | -3.568627544 | 0.011234434 | 0.479759698 | -3.540561234 |
| **A_55_P1953688** | Top1 | 0.399800398 | 0.127550981 | 0.672049816 | 12.73427709 | 3.568284321 | 0.011239178 | 0.479759698 | -3.540666461 |
| **A_55_P1995924** | Zfp984 | 0.4182644 | 0.133037451 | 0.70349135 | 9.94801322 | 3.563227356 | 0.011309328 | 0.479759698 | -3.542218479 |
| **A_55_P2139748** | Phf14 | 0.266346285 | 0.084686344 | 0.448006226 | 8.532887452 | 3.562629531 | 0.011317653 | 0.479759698 | -3.542402158 |
| **A_55_P1963175** | Lctl | 0.255226228 | 0.081079885 | 0.429372572 | 8.177387495 | 3.561181681 | 0.011337843 | 0.479759698 | -3.542847179 |
| **A_55_P2124586** | Xlr4a | 0.440636993 | 0.139957917 | 0.74131607 | 8.263209511 | 3.560909503 | 0.011341643 | 0.479759698 | -3.542930865 |
| **A_55_P2105002** | Pcna | 0.401378531 | 0.126959766 | 0.675797296 | 14.06029676 | 3.554049909 | 0.011437873 | 0.479759698 | -3.545042909 |
| **A_55_P2099479** | H2afz | 0.273340822 | 0.086398442 | 0.460283202 | 14.22158568 | 3.55287492 | 0.011454447 | 0.479759698 | -3.545405249 |
| **A_55_P2062727** | Nasp | 0.455423509 | 0.143790851 | 0.767056166 | 10.40618747 | 3.551040873 | 0.011480369 | 0.479759698 | -3.545971158 |
| **A_55_P1968763** | Shc4 | 0.684964191 | 0.215864897 | 1.154063484 | 11.19785805 | 3.548021786 | 0.011523182 | 0.479759698 | -3.546903598 |
| **A_55_P1994907** | Senp2 | 0.416074855 | 0.130899679 | 0.701250032 | 11.17312534 | 3.545217959 | 0.011563099 | 0.479759698 | -3.547770536 |
| **A_55_P2000077** | Nrf1 | 0.332959467 | 0.104617612 | 0.561301322 | 10.55092241 | 3.543145863 | 0.011592697 | 0.479759698 | -3.54841183 |
| **A_55_P2105512** | Nop56 | 0.3361877 | 0.105530856 | 0.566844545 | 13.26643718 | 3.541593106 | 0.011614931 | 0.479759698 | -3.548892731 |

| | | | | | | | | | |
|---|---|---|---|---|---|---|---|---|---|
| **A_55_P2103284** | Twf2 | -0.368045765 | -0.620601545 | -0.115489985 | 9.504279389 | -3.541014473 | 0.011623229 | 0.479759698 | -3.549072013 |
| **A_51_P458384** | Slc38a2 | -0.395171978 | -0.666515511 | -0.123828444 | 11.22089006 | -3.538749761 | 0.011655767 | 0.479759698 | -3.54977409 |
| **A_55_P2101757** | Dscc1 | 0.576053051 | 0.180372395 | 0.971733707 | 9.663620171 | 3.537535862 | 0.011673249 | 0.479759698 | -3.550150662 |
| **A_52_P275069** | Epp13 | 0.811159568 | 0.253340081 | 1.368979056 | 9.911454318 | 3.533424388 | 0.011732676 | 0.479759698 | -3.551427427 |
| **A_55_P2071952** | Wdr92 | -0.384760339 | -0.64948768 | -0.120032999 | 12.60541443 | -3.531625887 | 0.011758775 | 0.479759698 | -3.551986569 |
| **A_51_P514712** | Parp14 | -0.278521489 | -0.470233045 | -0.086809932 | 9.406099689 | -3.530153611 | 0.011780188 | 0.479759698 | -3.55244458 |
| **A_55_P2294184** | Cemip2 | -0.288470945 | -0.487066601 | -0.089875288 | 9.964837286 | -3.529518971 | 0.011789432 | 0.479759698 | -3.552642091 |
| **A_55_P2023424** | Trip10 | -0.414909409 | -0.700551712 | -0.129267106 | 10.81419906 | -3.529506184 | 0.011789618 | 0.479759698 | -3.55264607 |
| **A_55_P2003824** | Asgr1 | -1.353485429 | -2.285402854 | -0.421568003 | 11.45495709 | -3.529062431 | 0.011796086 | 0.479759698 | -3.552784204 |
| **A_55_P1968438** | Arxes1 | 0.522344031 | 0.16242782 | 0.882260242 | 9.440510554 | 3.526455572 | 0.011834163 | 0.479759698 | -3.553596157 |
| **A_55_P2114908** | Nek3 | 0.303299744 | 0.094273734 | 0.512325755 | 8.769747192 | 3.525777496 | 0.011844089 | 0.479759698 | -3.55380749 |
| **A_55_P2005055** | Pepd | 0.356507256 | 0.110792463 | 0.602222049 | 12.2202523 | 3.525496268 | 0.011848209 | 0.479759698 | -3.553895155 |
| **A_55_P2108248** | Art4 | 0.529956412 | 0.164673807 | 0.895239017 | 12.92742269 | 3.525285939 | 0.011851291 | 0.479759698 | -3.553960726 |
| **A_51_P198076** | Mien1 | -0.301100481 | -0.508656684 | -0.093544278 | 12.33241701 | -3.524998375 | 0.011855506 | 0.479759698 | -3.554050384 |
| **A_51_P122845** | Naa50 | 0.503149275 | 0.156174221 | 0.850124328 | 10.07597389 | 3.523560955 | 0.011876601 | 0.479759698 | -3.598065703 |
| **A_55_P2130448** | NA | 0.39935017 | 0.123927527 | 0.674772814 | 13.24748828 | 3.523201007 | 0.01188189 | 0.479759698 | -3.554611001 |
| **A_55_P2096305** | Rc3h1 | 0.284169726 | 0.088080075 | 0.480259376 | 11.04324309 | 3.521326743 | 0.011909471 | 0.479759698 | -3.555196017 |
| **A_51_P363749** | Irf6 | -0.546139374 | -0.923060738 | -0.169218009 | 9.817338201 | -3.520755956 | 0.011917885 | 0.479759698 | -3.555374262 |
| **A_51_P115471** | Apol8 | 0.637910554 | 0.197618882 | 1.078202227 | 11.48777264 | 3.520485058 | 0.01192188 | 0.479759698 | -3.566304826 |
| **A_55_P2043466** | Eif5 | 0.383287482 | 0.118454376 | 0.648120589 | 13.25043288 | 3.516701853 | 0.011977832 | 0.479759698 | -3.556641409 |
| **A_55_P1955517** | NA | 0.442312041 | 0.136681419 | 0.747942663 | 11.10229457 | 3.516536152 | 0.011980289 | 0.479759698 | -3.556693243 |
| **A_55_P2038928** | NA | 0.299653246 | 0.092589107 | 0.506717386 | 11.75725944 | 3.516392002 | 0.011982427 | 0.479759698 | -3.556738337 |
| **A_52_P583458** | E2f3 | 0.341412637 | 0.105483636 | 0.577341637 | 9.921739404 | 3.516263829 | 0.011984329 | 0.479759698 | -3.556778436 |
| **A_52_P132165** | Hsd17b11 | -0.343366305 | -0.580653911 | -0.106078698 | 11.23831512 | -3.516137161 | 0.011986208 | 0.479759698 | -3.556818066 |
| **A_66_P113487** | Pacsin2 | 0.292077414 | 0.090202602 | 0.493952225 | 13.11943442 | 3.51559655 | 0.011994234 | 0.479759698 | -3.556987226 |
| **A_55_P1957459** | Lilrb4a | -0.311252135 | -0.526456917 | -0.096047353 | 9.546802875 | -3.514338672 | 0.012012929 | 0.479759698 | -3.55738096 |
| **A_55_P2044547** | NA | -0.327890624 | -0.554624365 | -0.101156883 | 9.572127934 | -3.513953684 | 0.012018657 | 0.479759698 | -3.557501505 |
| **A_55_P1980721** | NA | 0.347027363 | 0.107026029 | 0.587028698 | 12.96173236 | 3.51344576 | 0.01202622 | 0.479759698 | -3.557660571 |
| **A_55_P2040026** | Itga4 | 0.540712671 | 0.166614254 | 0.914811089 | 11.6866444 | 3.512075673 | 0.012046644 | 0.479759698 | -3.558089794 |
| **A_55_P1977318** | Wbp11 | 0.419046897 | 0.128845121 | 0.709248674 | 12.32374088 | 3.508696079 | 0.012097188 | 0.479759698 | -3.559149532 |
| **A_55_P2042698** | Hsd17b10 | -0.371959215 | -0.629612413 | -0.114306017 | 13.91985077 | -3.507865828 | 0.012109641 | 0.479759698 | -3.559410085 |
| **A_51_P340355** | Tarsl2 | 0.438747465 | 0.134793353 | 0.742701578 | 9.840450939 | 3.50743625 | 0.01211609 | 0.479759698 | -3.55954493 |
| **A_51_P214319** | Cox11 | 0.330410613 | 0.10134941 | 0.559471815 | 11.73608721 | 3.504980787 | 0.012153022 | 0.479759698 | -3.560316133 |
| **A_55_P1957173** | NA | 0.77867205 | 0.238642229 | 1.318701872 | 10.67226423 | 3.50364449 | 0.012173174 | 0.479759698 | -3.56073614 |
| **A_66_P114501** | Zcrb1 | 0.281218742 | 0.085907432 | 0.476530051 | 12.3295494 | 3.498646454 | 0.01224887 | 0.479759698 | -3.562308979 |
| **A_52_P105138** | Frs2 | 0.27886576 | 0.085185644 | 0.472545876 | 10.96716265 | 3.498592377 | 0.012249692 | 0.479759698 | -3.562326013 |
| **A_55_P2096630** | Ssxb9 | 0.495618729 | 0.15137511 | 0.839862348 | 8.66398289 | 3.498364033 | 0.012253163 | 0.479759698 | -3.562397945 |
| **A_55_P2027147** | Mrps18c | 0.327210592 | 0.099825976 | 0.554595208 | 12.61690493 | 3.496628267 | 0.012279583 | 0.479759698 | -3.562944946 |
| **A_55_P2062598** | E2f7 | 0.368957927 | 0.112510279 | 0.625405575 | 9.80655968 | 3.49591862 | 0.012290402 | 0.479759698 | -3.563168687 |
| **A_55_P1984123** | Kmt2c | 0.373101077 | 0.113704668 | 0.632497487 | 10.21184201 | 3.494988349 | 0.012304601 | 0.479759698 | -3.563462079 |
| **A_55_P1975090** | NA | 0.68437927 | 0.208347584 | 1.160410957 | 12.79751236 | 3.493366677 | 0.012329396 | 0.479759698 | -3.563973781 |
| **A_55_P2108067** | Dgcr8 | 0.31499826 | 0.095772447 | 0.534224073 | 9.978383312 | 3.491400374 | 0.012359534 | 0.479759698 | -3.564594656 |
| **A_55_P1962811** | Anp32b | 0.472464796 | 0.143609462 | 0.801320131 | 14.60781783 | 3.490983148 | 0.01236594 | 0.479759698 | -3.564726459 |
| **A_55_P2093665** | NA | 0.28002884 | 0.085055708 | 0.475001972 | 8.692930218 | 3.489885532 | 0.012382808 | 0.479759698 | -3.565073301 |
| **A_55_P2151506** | Nudt9 | 0.270278905 | 0.082084446 | 0.458473363 | 12.98732053 | 3.489703331 | 0.01238561 | 0.479759698 | -3.56513089 |
| **A_55_P2018949** | Fbxo7 | 0.384813331 | 0.116807034 | 0.652819628 | 13.28093953 | 3.488898213 | 0.012398003 | 0.479759698 | -3.565385415 |
| **A_55_P2031403** | NA | -0.44807816 | -0.760184919 | -0.135971402 | 8.782488667 | -3.48846036 | 0.012404748 | 0.479759698 | -3.565523868 |
| **A_55_P2110351** | Eppk1 | -0.329311999 | -0.558717161 | -0.099906837 | 8.444283432 | -3.488089005 | 0.012410472 | 0.479759698 | -3.565641313 |
| **A_51_P388847** | Stap2 | -0.658676438 | -1.117647395 | -0.199705482 | 9.219029625 | -3.487145859 | 0.012425022 | 0.479759698 | -3.565939666 |
| **A_51_P155152** | Ank | 0.458827149 | 0.138516611 | 0.779137686 | 10.18647809 | 3.480655732 | 0.012525657 | 0.479759698 | -3.567995692 |

| | | | | | | | | | |
|---|---|---|---|---|---|---|---|---|---|
| **A_51_P285042** | Rtf1 | 0.272695785 | 0.082281594 | 0.463109976 | 9.877439421 | 3.47986425 | 0.01253799 | 0.479759698 | -3.56824678 |
| **A_52_P504361** | Naglu | -0.302900582 | -0.514483555 | -0.091317609 | 9.489561346 | -3.478584701 | 0.012557958 | 0.479759698 | -3.568652863 |
| **A_55_P2088720** | NA | -0.482157539 | -0.81929307 | -0.145022008 | 12.09410648 | -3.475102549 | 0.012612475 | 0.479759698 | -3.569758988 |
| **A_55_P2182705** | Kpna2 | 0.42116217 | 0.126573237 | 0.715751103 | 13.23906977 | 3.473890431 | 0.012631512 | 0.479759698 | -3.570144372 |
| **A_55_P2098935** | Sh2d3c | -0.4735486 | -0.804963978 | -0.142133221 | 10.78040557 | -3.471963007 | 0.01266185 | 0.479759698 | -3.570757554 |
| **A_55_P1954092** | NA | -0.448692712 | -0.762790631 | -0.134594793 | 13.87832175 | -3.471100162 | 0.012675457 | 0.479759698 | -3.571032202 |
| **A_55_P2443935** | Xpo7 | 0.652209985 | 0.195549659 | 1.10887031 | 10.60004164 | 3.470382544 | 0.012686786 | 0.479759698 | -3.571260694 |
| **A_52_P20357** | Aqr | 0.295605374 | 0.08844336 | 0.502767388 | 9.843327949 | 3.467251857 | 0.012736341 | 0.479759698 | -3.572258252 |
| **A_55_P1998416** | Ifi47 | -0.876130699 | -1.490316214 | -0.261945183 | 10.899344 | -3.466191212 | 0.012753178 | 0.479759698 | -3.572596487 |
| **A_55_P1978201** | Incenp | 0.412399611 | 0.1230738 | 0.701725422 | 13.21510751 | 3.463492541 | 0.012796127 | 0.479759698 | -3.573457705 |
| **A_55_P2087944** | NA | 0.547718374 | 0.163454028 | 0.931982719 | 11.88781202 | 3.463461858 | 0.012796616 | 0.479759698 | -3.573467502 |
| **A_51_P324287** | Kif23 | 0.511681709 | 0.152644202 | 0.870719217 | 9.878774227 | 3.462926401 | 0.012805157 | 0.479759698 | -3.573638488 |
| **A_55_P1953980** | Sdhb | -0.228222577 | -0.388410513 | -0.068034642 | 12.41140932 | -3.461879848 | 0.012821869 | 0.479759698 | -3.573972785 |
| **A_55_P2169586** | Ccdc80 | -0.351987705 | -0.599117806 | -0.104857605 | 10.16725073 | -3.460868016 | 0.012838048 | 0.479759698 | -3.574296118 |
| **A_51_P390967** | Slc43a3 | 0.326018716 | 0.097104113 | 0.554933319 | 12.22512586 | 3.460606537 | 0.012842233 | 0.479759698 | -3.574379695 |
| **A_51_P155174** | Zfp672 | 0.465416865 | 0.138530152 | 0.792303577 | 10.84881449 | 3.45961712 | 0.012858082 | 0.479759698 | -3.574696019 |
| **A_55_P2154228** | Ncapd3 | 0.511269688 | 0.151994707 | 0.87054467 | 11.22251598 | 3.457850871 | 0.012886428 | 0.479759698 | -3.575261001 |
| **A_55_P2004258** | Gm7102 | 0.370570394 | 0.110057064 | 0.631083724 | 9.630757621 | 3.456399695 | 0.012909768 | 0.479759698 | -3.575725485 |
| **A_55_P2006792** | Ntrk2 | 0.675958971 | 0.200162015 | 1.151755926 | 9.949065495 | 3.452088064 | 0.01297939 | 0.479759698 | -3.577107058 |
| **A_55_P1999637** | Gm15455 | 0.382109891 | 0.113095809 | 0.651123973 | 11.40527207 | 3.451409217 | 0.01299039 | 0.479759698 | -3.577324789 |
| **A_55_P1990633** | Iigp1 | -0.681312124 | -1.161031208 | -0.201593041 | 8.510637747 | -3.450978958 | 0.012997366 | 0.479759698 | -3.577462818 |
| **A_52_P628067** | Cdca3 | 0.413788261 | 0.121999134 | 0.705577389 | 13.19138941 | 3.44581732 | 0.013081384 | 0.479759698 | -3.579120475 |
| **A_55_P2035265** | Trpm7 | 0.36763213 | 0.10832393 | 0.62694033 | 10.61398706 | 3.444929974 | 0.013095887 | 0.479759698 | -3.579405777 |
| **A_55_P2070084** | Zfp988 | 0.596496546 | 0.175207207 | 1.017785885 | 10.24164302 | 3.440413576 | 0.013169981 | 0.479759698 | -3.580859406 |
| **A_55_P2008244** | NA | 0.322923686 | 0.094725789 | 0.551121582 | 9.6495375 | 3.438519192 | 0.013201196 | 0.479759698 | -3.581469875 |
| **A_51_P331288** | Akr1b7 | -0.875276048 | -1.493914278 | -0.256637818 | 11.35549882 | -3.437886057 | 0.013211647 | 0.479759698 | -3.581674003 |
| **A_55_P2435504** | Cep295 | 0.359990474 | 0.105326016 | 0.614654932 | 9.508918039 | 3.434834847 | 0.013262139 | 0.479759698 | -3.582658432 |
| **A_51_P324491** | Cmpk1 | 0.35113087 | 0.102426328 | 0.599835413 | 10.52622473 | 3.430587375 | 0.013332779 | 0.479759698 | -3.584030736 |
| **A_52_P385615** | Rnpc3 | 0.449597679 | 0.131116409 | 0.768078949 | 9.472969898 | 3.430230886 | 0.013338726 | 0.479759698 | -3.584146015 |
| **A_66_P106661** | Slc7a1 | 0.4637058 | 0.134981665 | 0.792429934 | 11.98014782 | 3.427631507 | 0.013382182 | 0.479759698 | -3.584987057 |
| **A_55_P1988373** | Cenpu | 0.455350722 | 0.132534943 | 0.7781665 | 9.439902151 | 3.427476329 | 0.013384781 | 0.479759698 | -3.585037293 |
| **A_55_P1963718** | Lias | 0.308281587 | 0.089698318 | 0.526864856 | 11.08181357 | 3.426998062 | 0.013392795 | 0.479759698 | -3.585192139 |
| **A_55_P2185821** | Cyp3a13 | -0.788580293 | -1.348065194 | -0.229095392 | 10.06489988 | -3.424843554 | 0.013428962 | 0.479759698 | -3.585890045 |
| **A_55_P1978608** | NA | 0.447611314 | 0.130010595 | 0.765212034 | 9.569108761 | 3.424544127 | 0.013433997 | 0.479759698 | -3.585987084 |
| **A_55_P2364315** | D030002E05Rik | 0.28690146 | 0.083321983 | 0.490480937 | 8.548458259 | 3.424380014 | 0.013436758 | 0.479759698 | -3.586040274 |
| **A_51_P456208** | Tff3 | -0.86465055 | -1.478309943 | -0.250991157 | 14.91513317 | -3.423705692 | 0.013448107 | 0.479759698 | -3.586258863 |
| **A_55_P2015964** | Rif1 | 0.398885771 | 0.115687897 | 0.682083645 | 9.665826495 | 3.422486509 | 0.013468652 | 0.479759698 | -3.586654218 |
| **A_51_P424810** | Ncapg2 | 0.663697848 | 0.192240292 | 1.135155404 | 11.20667753 | 3.420668574 | 0.013499352 | 0.479759698 | -3.587244079 |
| **A_55_P2295786** | Mindy2 | 0.358806718 | 0.103871218 | 0.613742217 | 8.501540924 | 3.419900261 | 0.01351235 | 0.479759698 | -3.587493495 |
| **A_55_P2110215** | Ckap5 | 0.321119219 | 0.092892378 | 0.549346061 | 12.28800262 | 3.418871423 | 0.013529776 | 0.479759698 | -3.587827599 |
| **A_52_P272854** | NA | 0.91821632 | 0.26446672 | 1.571965921 | 10.1144151 | 3.412846537 | 0.013632323 | 0.479759698 | -3.589786756 |
| **A_55_P2119985** | Baiap2 | 0.281915741 | 0.081169124 | 0.482662358 | 8.821360722 | 3.41235557 | 0.013640717 | 0.479759698 | -3.589946607 |
| **A_55_P2042520** | Cop1 | 0.242557778 | 0.069784603 | 0.415330952 | 12.02072856 | 3.411317023 | 0.013658491 | 0.479759698 | -3.590284839 |
| **A_55_P2197338** | Tnfsf13os | -0.285667915 | -0.489239779 | -0.08209605 | 8.414826012 | -3.409784244 | 0.013684771 | 0.479759698 | -3.590784276 |
| **A_55_P2024290** | Fam149a | 0.320591127 | 0.092126295 | 0.549055959 | 8.498982858 | 3.409693405 | 0.01368633 | 0.479759698 | -3.590813884 |
| **A_55_P1953311** | Stx2 | 0.488377946 | 0.140303133 | 0.836452758 | 11.15779613 | 3.409311121 | 0.013692893 | 0.479759698 | -3.590938496 |
| **A_55_P2045476** | Hmgxb4 | 0.287518339 | 0.082580779 | 0.4924559 | 9.617695381 | 3.409001386 | 0.013698214 | 0.479759698 | -3.591039473 |
| **A_55_P2039354** | NA | 0.664079441 | 0.190497006 | 1.137661876 | 8.981696323 | 3.40727854 | 0.01372785 | 0.479759698 | -3.591601359 |
| **A_52_P656867** | Fam98b | 0.253616442 | 0.072711093 | 0.434521792 | 9.839480847 | 3.406506214 | 0.013741159 | 0.479759698 | -3.591853363 |
| **A_55_P2135880** | Fancl | 0.42589021 | 0.121931944 | 0.729848477 | 11.09957205 | 3.40460621 | 0.013773959 | 0.479759698 | -3.592473639 |

| | | | | | | | | | |
|---|---|---|---|---|---|---|---|---|---|
| **A_51_P251357** | Ctps | 0.291332116 | 0.083283304 | 0.499380929 | 11.78007409 | 3.402564071 | 0.013809308 | 0.479759698 | -3.593140816 |
| **A_55_P2128734** | Pbxip1 | -0.473534026 | -0.811879406 | -0.135188646 | 11.15443168 | -3.400745473 | 0.013840872 | 0.479759698 | -3.5937354 |
| **A_55_P2140151** | Flcn | -0.401039003 | -0.687841241 | -0.114236765 | 10.37833023 | -3.397717486 | 0.013893601 | 0.479759698 | -3.594726304 |
| **A_55_P2172274** | Smc2 | 0.589192675 | 0.167660386 | 1.010724965 | 12.29467457 | 3.396328431 | 0.013917864 | 0.479759698 | -3.595181254 |
| **A_66_P125067** | Snrpd1 | 0.325561785 | 0.092628095 | 0.558495475 | 11.11070676 | 3.396129975 | 0.013921334 | 0.479759698 | -3.595246273 |
| **A_55_P1961039** | Pnpla2 | -0.367146167 | -0.629981608 | -0.104310726 | 10.8863041 | -3.394206508 | 0.013955017 | 0.479759698 | -3.5958767 |
| **A_55_P2161077** | NA | 0.373288116 | 0.105979352 | 0.64059688 | 9.763963564 | 3.393236695 | 0.013972034 | 0.479759698 | -3.596194738 |
| **A_55_P1967633** | Lsm5 | 0.411973451 | 0.116406822 | 0.707540081 | 10.48098697 | 3.38685824 | 0.014084522 | 0.479759698 | -3.598289397 |
| **A_55_P2048518** | Zfpm1 | 0.420190872 | 0.118354696 | 0.722027048 | 13.93663182 | 3.382661288 | 0.014159079 | 0.479759698 | -3.599670438 |
| **A_55_P2126827** | NA | 0.312807332 | 0.088028054 | 0.537586609 | 13.76285886 | 3.381456791 | 0.014180556 | 0.479759698 | -3.600067194 |
| **A_51_P199567** | Med26 | 0.3140467 | 0.088245526 | 0.539847873 | 9.998958343 | 3.379490477 | 0.014215693 | 0.479759698 | -3.600715281 |
| **A_51_P494825** | Mfhas1 | 0.30763693 | 0.086301012 | 0.528972848 | 10.78767927 | 3.377300946 | 0.014254932 | 0.479759698 | -3.60143751 |
| **A_55_P2010332** | Ewsr1 | 0.265193265 | 0.074320012 | 0.456066518 | 11.78022406 | 3.37598574 | 0.014278559 | 0.479759698 | -3.601871628 |
| **A_51_P154585** | Vmn1r195 | 0.730542382 | 0.204631664 | 1.2564531 | 10.09334903 | 3.375332674 | 0.014290306 | 0.479759698 | -3.602087269 |
| **A_51_P346826** | Rragc | -0.228602286 | -0.393289929 | -0.063914643 | 10.7652332 | -3.372894402 | 0.014334261 | 0.479759698 | -3.602892857 |
| **A_52_P76034** | Rcc2 | 0.394932064 | 0.110401245 | 0.679462882 | 12.28868574 | 3.372688799 | 0.014337975 | 0.479759698 | -3.602960821 |
| **A_55_P2001871** | Uros | 0.45123322 | 0.125902441 | 0.776564 | 13.14442376 | 3.37022659 | 0.014382524 | 0.479759698 | -3.603775139 |
| **A_51_P336952** | Cryl1 | -0.346872241 | -0.596962566 | -0.096781916 | 9.803247896 | -3.370201317 | 0.014382982 | 0.479759698 | -3.603783501 |
| **A_66_P102607** | Btrc | 0.439922003 | 0.12257723 | 0.757266777 | 8.873338992 | 3.368429974 | 0.014415126 | 0.479759698 | -3.604369808 |
| **A_55_P2041050** | Zmym5 | 0.232683449 | 0.064743427 | 0.400623471 | 9.42215269 | 3.366622912 | 0.014448 | 0.479759698 | -3.604968345 |
| **A_55_P2023235** | Fen1 | 0.499962454 | 0.139083756 | 0.860841152 | 13.55189031 | 3.366350504 | 0.014452962 | 0.479759698 | -3.605058607 |
| **A_55_P2123716** | Tug1 | 0.423662014 | 0.117705749 | 0.729618279 | 9.259937604 | 3.364676887 | 0.014483493 | 0.479759698 | -3.605613367 |
| **A_52_P381665** | Aff1 | 0.441468444 | 0.122482558 | 0.76045433 | 11.71106631 | 3.362880136 | 0.014516348 | 0.479759698 | -3.606209336 |
| **A_52_P651034** | Tmx1 | 0.334849345 | 0.092661613 | 0.577037077 | 11.07388625 | 3.359545311 | 0.014577545 | 0.479759698 | -3.607316548 |
| **A_55_P2107745** | Yy1 | 0.343100154 | 0.094781791 | 0.591418516 | 10.97834657 | 3.357339544 | 0.014618178 | 0.479759698 | -3.608049666 |
| **A_55_P2113768** | NA | 0.258742069 | 0.071462006 | 0.446022131 | 12.11124205 | 3.357056796 | 0.014623396 | 0.479759698 | -3.608143686 |
| **A_52_P634822** | Nap1l4 | 0.362542064 | 0.099778562 | 0.625305566 | 8.638937298 | 3.352559933 | 0.01470665 | 0.479759698 | -3.609640342 |
| **A_51_P466148** | Bckdha | -0.521351779 | -0.899314712 | -0.143388845 | 11.97666378 | -3.351697672 | 0.014722674 | 0.479759698 | -3.609927613 |
| **A_55_P2117164** | Tmem106b | -0.290912912 | -0.501823881 | -0.080001943 | 9.080594051 | -3.351560164 | 0.014725231 | 0.479759698 | -3.609973434 |
| **A_55_P1964477** | Srsf1 | 0.480240812 | 0.131970136 | 0.828511488 | 10.45203195 | 3.3506213 | 0.014742702 | 0.479759698 | -3.610286349 |
| **A_55_P2131766** | Rsrp1 | -0.299479497 | -0.51672298 | -0.082236013 | 13.51159666 | -3.349681583 | 0.014760212 | 0.479759698 | -3.61059966 |
| **A_55_P2093291** | Coil | 0.370923386 | 0.10172587 | 0.640120902 | 9.752203172 | 3.348084086 | 0.014790031 | 0.479759698 | -3.611132538 |
| **A_51_P211341** | Eif3j1 | 0.582828457 | 0.159339514 | 1.0063174 | 11.6251554 | 3.344120066 | 0.014864309 | 0.479759698 | -3.612456209 |
| **A_52_P90363** | Ifi27l2a | -0.873531278 | -1.508466258 | -0.238596298 | 10.9854315 | -3.342969353 | 0.014885948 | 0.479759698 | -3.612840828 |
| **A_55_P2057390** | Smap1 | 0.250471262 | 0.068384959 | 0.432557565 | 11.9585374 | 3.342441493 | 0.014895885 | 0.479759698 | -3.613017319 |
| **A_55_P2131844** | NA | 0.979870251 | 0.26742446 | 1.692316042 | 12.20314248 | 3.341949911 | 0.014905146 | 0.479759698 | -3.613181712 |
| **A_55_P2182690** | Orai3 | -0.304407986 | -0.525767259 | -0.083048713 | 11.15323979 | -3.341500349 | 0.014913621 | 0.479759698 | -3.613332079 |
| **A_51_P172131** | Lurap1l | -0.52083777 | -0.89974915 | -0.141926391 | 10.27499047 | -3.340011885 | 0.014941719 | 0.479759698 | -3.613830115 |
| **A_66_P111689** | NA | 0.92593061 | 0.252119177 | 1.599742043 | 10.77912142 | 3.339052556 | 0.014959859 | 0.479759698 | -3.614151253 |
| **A_55_P2092073** | Taf11 | 0.242839517 | 0.066115268 | 0.419563766 | 9.548130575 | 3.33892301 | 0.014962311 | 0.479759698 | -3.614194627 |
| **A_55_P1954724** | A130040M12Rik | -0.620241852 | -1.071815691 | -0.168668013 | 12.16762616 | -3.337455493 | 0.014990112 | 0.479759698 | -3.614686131 |
| **A_52_P228236** | Tfrc | 0.584021346 | 0.158477071 | 1.00956562 | 12.5080273 | 3.334779767 | 0.015040947 | 0.479759698 | -3.698694971 |
| **A_66_P126348** | Smim31 | 1.129978369 | 0.306365627 | 1.95359111 | 12.3425338 | 3.333728764 | 0.015060966 | 0.479759698 | -3.615935521 |
| **A_55_P2128646** | Gmnn | 0.511965902 | 0.138786476 | 0.885145329 | 10.54110223 | 3.333546619 | 0.015064438 | 0.479759698 | -3.615996631 |
| **A_55_P2080902** | NA | 0.391079058 | 0.105929219 | 0.676228897 | 8.363551115 | 3.33253423 | 0.015083754 | 0.479759698 | -3.616336362 |
| **A_55_P2467801** | Srp54a | 0.338140512 | 0.091572527 | 0.584708498 | 9.495298492 | 3.332297051 | 0.015088283 | 0.479759698 | -3.616415973 |
| **A_55_P1991406** | Fbxo34 | 0.522963013 | 0.141586346 | 0.90433968 | 11.02922232 | 3.33196203 | 0.015094683 | 0.479759698 | -3.616528436 |
| **A_55_P1993522** | NA | 0.400909196 | 0.108473497 | 0.693344896 | 11.39510523 | 3.331185514 | 0.015109529 | 0.479759698 | -3.616789159 |
| **A_55_P2071059** | Nab2 | -0.386523818 | -0.668486049 | -0.104561588 | 10.30512086 | -3.330953058 | 0.015113976 | 0.479759698 | -3.616867223 |
| **A_55_P2035102** | Odc1 | 0.655254529 | 0.177237482 | 1.133271575 | 12.88171218 | 3.330809815 | 0.015116718 | 0.479759698 | -3.616915331 |

| | | | | | | | | | |
|---|---|---|---|---|---|---|---|---|---|
| **A_52_P27020** | Sgo2a | 0.352014835 | 0.095094926 | 0.608934744 | 8.959561345 | 3.329249899 | 0.015146604 | 0.479759698 | -3.617439394 |
| **A_55_P2398534** | Gid8 | 0.451488145 | 0.121950954 | 0.781025336 | 12.0047928 | 3.329086691 | 0.015149735 | 0.479759698 | -3.617494243 |
| **A_51_P312310** | Mtbp | 0.331536113 | 0.089324404 | 0.573747821 | 10.45562708 | 3.325974361 | 0.015209572 | 0.479759698 | -3.618540836 |
| **A_55_P2052959** | Ctbp1 | 0.269193977 | 0.072261367 | 0.466126586 | 14.13496365 | 3.321474265 | 0.015296547 | 0.479759698 | -3.620056272 |
| **A_51_P453246** | Uqcc3 | -0.300123019 | -0.519739042 | -0.080506995 | 12.56089755 | -3.320614583 | 0.015313224 | 0.479759698 | -3.620346068 |
| **A_55_P2057212** | Tpr | 0.48685339 | 0.130505189 | 0.84320159 | 11.02531234 | 3.319760991 | 0.015329803 | 0.479759698 | -3.620633904 |
| **A_55_P2068988** | NA | -0.462532636 | -0.801125985 | -0.123939287 | 10.56968963 | -3.319304902 | 0.015338669 | 0.479759698 | -3.620787738 |
| **A_51_P330133** | Pspc1 | 0.330887076 | 0.088593148 | 0.573181004 | 9.514796516 | 3.318336794 | 0.015357508 | 0.479759698 | -3.621114357 |
| **A_52_P180408** | Bcas2 | 0.266369666 | 0.0712166 | 0.461522732 | 12.51221004 | 3.316596026 | 0.015391445 | 0.479759698 | -3.621701957 |
| **A_55_P2178036** | Serpina6 | 0.962638039 | 0.256987493 | 1.668288584 | 12.93055008 | 3.314793864 | 0.015426666 | 0.479759698 | -3.622310685 |
| **A_55_P2118857** | NA | 0.497428946 | 0.132720602 | 0.86213729 | 8.957017973 | 3.314122391 | 0.015439811 | 0.479759698 | -3.622537599 |
| **A_51_P256224** | Zfp768 | -0.263076572 | -0.456230918 | -0.069922227 | 10.97975891 | -3.309488527 | 0.015530863 | 0.479759698 | -3.624105105 |
| **A_52_P30989** | Cdkn3 | 0.394501973 | 0.104742327 | 0.68426162 | 10.69659623 | 3.308220631 | 0.015555878 | 0.479759698 | -3.624534475 |
| **A_55_P2121886** | Map3k9 | -0.253153164 | -0.439098005 | -0.067208323 | 7.928537125 | -3.308128839 | 0.015557691 | 0.479759698 | -3.624565568 |
| **A_55_P2093241** | Zc3h4 | 0.335063047 | 0.088933382 | 0.581192712 | 11.86617178 | 3.307849649 | 0.015563206 | 0.479759698 | -3.624660146 |
| **A_55_P2186621** | Rbmxl1 | 0.287859119 | 0.076399632 | 0.499318606 | 12.96036427 | 3.307775357 | 0.015564674 | 0.479759698 | -3.624685315 |
| **A_51_P288447** | Nup133 | 0.43593696 | 0.115635264 | 0.756238656 | 11.69395301 | 3.307102395 | 0.015577977 | 0.479759698 | -3.624913333 |
| **A_55_P1994128** | Tmem184a | -0.518896396 | -0.900215558 | -0.137577234 | 8.665942394 | -3.306550909 | 0.015588889 | 0.479759698 | -3.625100235 |
| **A_55_P2024868** | Psg18 | -0.460625007 | -0.799127546 | -0.122122468 | 7.996643367 | -3.30650185 | 0.01558986 | 0.479759698 | -3.625116864 |
| **A_55_P2176260** | NA | 0.510621444 | 0.135337671 | 0.885905217 | 8.381150457 | 3.306149231 | 0.015596841 | 0.479759698 | -3.625236391 |
| **A_51_P267370** | Itm2c | -0.390092114 | -0.676810804 | -0.103373424 | 10.8292064 | -3.305935352 | 0.015601078 | 0.479759698 | -3.625308897 |
| **A_51_P239203** | Mapk13 | -0.47834689 | -0.83007839 | -0.126615391 | 10.05544686 | -3.304569428 | 0.015628162 | 0.479759698 | -3.625772091 |
| **A_55_P2093874** | Pbxip1 | -0.461610376 | -0.801102731 | -0.122118021 | 11.17455802 | -3.30391414 | 0.015641174 | 0.479759698 | -3.625994389 |
| **A_51_P378550** | Oplah | -0.504094336 | -0.874971317 | -0.133217354 | 9.128956305 | -3.302669584 | 0.015665919 | 0.479759698 | -3.626416737 |
| **A_52_P585233** | Ccpg1 | -0.347198851 | -0.602755113 | -0.091642588 | 9.287656417 | -3.301223587 | 0.015694723 | 0.479759698 | -3.626907694 |
| **A_55_P2006703** | Bmp1 | -0.489276393 | -0.849425224 | -0.129127562 | 11.67579053 | -3.301075377 | 0.015697678 | 0.479759698 | -3.626958031 |
| **A_51_P500344** | Mat2a | 0.341948403 | 0.090092476 | 0.593804331 | 13.26439596 | 3.299070472 | 0.015737719 | 0.479759698 | -3.627639233 |
| **A_55_P2073218** | Hps4 | -0.274154858 | -0.476169507 | -0.07214021 | 10.73145031 | -3.297587081 | 0.015767417 | 0.479759698 | -3.628143571 |
| **A_52_P222073** | Ackr2 | -0.422419831 | -0.733812986 | -0.111026676 | 8.829523473 | -3.296236972 | 0.015794499 | 0.479759698 | -3.628602839 |
| **A_55_P2047559** | Ccp110 | 0.43436298 | 0.114077814 | 0.754648147 | 10.27082166 | 3.295331939 | 0.015812682 | 0.479759698 | -3.628910836 |
| **A_55_P2103698** | Isg15 | -1.206678463 | -2.096596464 | -0.316760461 | 12.43228367 | -3.294767387 | 0.015824036 | 0.479759698 | -3.629103014 |
| **A_55_P2097072** | Gm7219 | 0.401222403 | 0.105301041 | 0.697143766 | 8.242096247 | 3.294519223 | 0.01582903 | 0.479759698 | -3.629187505 |
| **A_55_P2262287** | 5430434G16Rik | 0.561782345 | 0.147292766 | 0.976271924 | 10.40431152 | 3.293348267 | 0.015852616 | 0.479759698 | -3.629586275 |
| **A_55_P1989858** | Thrap3 | 0.341881026 | 0.089618946 | 0.594143106 | 11.27595014 | 3.293109829 | 0.015857424 | 0.479759698 | -3.629667497 |
| **A_52_P162099** | Ckap2 | 0.348774191 | 0.091260027 | 0.606288356 | 9.417472946 | 3.290988818 | 0.015900258 | 0.479759698 | -3.630390322 |
| **A_51_P502225** | Slc22a4 | 0.582240182 | 0.152198839 | 1.012281524 | 10.6522844 | 3.289842858 | 0.015923454 | 0.479759698 | -3.630781096 |
| **A_55_P2058871** | Otud5 | 0.331898108 | 0.086513091 | 0.577283125 | 12.59711157 | 3.286547586 | 0.01599036 | 0.479759698 | -3.631905725 |
| **A_55_P2160862** | Wrn | 0.29897937 | 0.077889606 | 0.520069134 | 8.578609511 | 3.285910841 | 0.016003324 | 0.479759698 | -3.632123197 |
| **A_52_P598835** | Ccdc71l | 0.537223748 | 0.139780077 | 0.934667418 | 10.66802554 | 3.284451143 | 0.016033086 | 0.479759698 | -3.632621934 |
| **A_55_P2203051** | Gt(ROSA)26Sor | 0.466429549 | 0.121313362 | 0.811545736 | 11.42933079 | 3.284005801 | 0.016042178 | 0.479759698 | -3.632774148 |
| **A_55_P1957154** | Mageb16 | 0.499212479 | 0.129815907 | 0.868609051 | 8.509538044 | 3.283792912 | 0.016046526 | 0.479759698 | -3.632846922 |
| **A_52_P477819** | Hmox2 | -0.297116436 | -0.517046391 | -0.077186482 | 11.49608919 | -3.282656814 | 0.016069753 | 0.479759698 | -3.633235378 |
| **A_51_P249286** | Rgs16 | -1.058829291 | -1.842591357 | -0.275067226 | 9.907692935 | -3.282652391 | 0.016069843 | 0.479759698 | -3.633236891 |
| **A_52_P330984** | Lin9 | 0.370778191 | 0.096235693 | 0.645320689 | 9.613777913 | 3.281616222 | 0.016091059 | 0.479759698 | -3.633591324 |
| **A_55_P2041708** | Gas2 | 0.330538577 | 0.085754417 | 0.575322736 | 8.72749625 | 3.281119376 | 0.016101243 | 0.479759698 | -3.633761325 |
| **A_51_P133137** | Kif20a | 0.4060775 | 0.105155025 | 0.706999976 | 11.72411768 | 3.278970354 | 0.016145374 | 0.479759698 | -3.634496998 |
| **A_51_P483013** | Arel1 | 0.390415511 | 0.101051235 | 0.679779787 | 11.37362868 | 3.278425647 | 0.01615658 | 0.479759698 | -3.634683561 |
| **A_55_P1974533** | Eif4e | 0.368552314 | 0.095206139 | 0.641898488 | 9.135977831 | 3.276191889 | 0.016202625 | 0.479759698 | -3.635449025 |
| **A_55_P2071846** | NA | 0.436274635 | 0.11256376 | 0.75998551 | 13.53083397 | 3.274808158 | 0.016231221 | 0.479759698 | -3.635923523 |
| **A_55_P1960416** | Nrros | -0.327800248 | -0.571196221 | -0.084404276 | 11.69719768 | -3.27249566 | 0.016279132 | 0.479759698 | -3.636717055 |

| | | | | | | | | | |
|---|---|---|---|---|---|---|---|---|---|
| **A_51_P393958** | Fbxo5 | 0.521422708 | 0.134256439 | 0.908588977 | 11.92696661 | 3.272469565 | 0.016279673 | 0.479759698 | -3.636726013 |
| **A_55_P2002465** | Zfp747 | -0.342210763 | -0.596623857 | -0.08779767 | 10.48458489 | -3.268416508 | 0.016364022 | 0.479759698 | -3.638118486 |
| **A_55_P2416494** | 8430426J06Rik | -1.084123528 | -1.890550655 | -0.277696401 | 8.980592917 | -3.266606565 | 0.016401843 | 0.479759698 | -3.638740993 |
| **A_51_P408561** | Oard1 | 0.304596029 | 0.077991621 | 0.531200436 | 11.05171592 | 3.266172159 | 0.016410934 | 0.479759698 | -3.638890464 |
| **A_52_P148212** | Mis18bp1 | 0.443794184 | 0.113435752 | 0.774152616 | 9.059668581 | 3.264219836 | 0.016451862 | 0.479759698 | -3.639562522 |
| **A_51_P223946** | Pds5b | 0.244660866 | 0.062438426 | 0.426883307 | 8.953984179 | 3.262464839 | 0.016488748 | 0.479759698 | -3.640167072 |
| **A_55_P2197998** | D4Ertd58e | 0.779534844 | 0.198903345 | 1.360166343 | 9.558354413 | 3.262256681 | 0.016493129 | 0.479759698 | -3.640238803 |
| **A_55_P2102628** | Usp4 | 0.345504226 | 0.088154859 | 0.602853592 | 11.07296885 | 3.262221519 | 0.016493869 | 0.479759698 | -3.64025092 |
| **A_55_P2002689** | Hmgb1 | 0.357432964 | 0.090949676 | 0.623916252 | 12.69752282 | 3.259175999 | 0.016558116 | 0.479759698 | -3.641301057 |
| **A_55_P1965629** | Hcfc1 | 0.325100768 | 0.08256557 | 0.567635966 | 11.71304708 | 3.257064884 | 0.01660281 | 0.479759698 | -3.642029698 |
| **A_51_P392967** | Zmynd10 | -0.490018177 | -0.855627587 | -0.124408766 | 9.422304874 | -3.256701946 | 0.016610507 | 0.479759698 | -3.642155022 |
| **A_55_P2183288** | Cryzl1 | 0.218585541 | 0.055489186 | 0.381681896 | 10.71717209 | 3.256569596 | 0.016613315 | 0.479759698 | -3.642200727 |
| **A_55_P1971963** | Tmem176b | -0.255160607 | -0.445584698 | -0.064736517 | 14.50744391 | -3.25592901 | 0.016626912 | 0.479759698 | -3.642421977 |
| **A_52_P543824** | Srsf6 | 0.38997178 | 0.098901325 | 0.681042236 | 10.09509566 | 3.255504083 | 0.016635939 | 0.479759698 | -3.64256877 |
| **A_55_P2044413** | Cxcl11 | 0.349071488 | 0.088469721 | 0.609673255 | 10.96328494 | 3.254769298 | 0.01665156 | 0.479759698 | -3.64282266 |
| **A_51_P468599** | Smchd1 | 0.328769468 | 0.088477894 | 0.569061042 | 11.50567993 | 3.480930111 | 0.016663608 | 0.479759698 | -3.796632726 |
| **A_52_P618187** | Mrs2 | 0.368136556 | 0.093201902 | 0.643071211 | 9.075735623 | 3.253588696 | 0.016676692 | 0.479759698 | -3.643230738 |
| **A_55_P2007377** | Elf2 | 0.222586711 | 0.056298724 | 0.388874697 | 10.53733632 | 3.252531764 | 0.016699227 | 0.479759698 | -3.643596222 |
| **A_55_P2133910** | Ttll10 | 0.505721008 | 0.127872513 | 0.883569503 | 9.952471939 | 3.252194323 | 0.016706428 | 0.479759698 | -3.643712938 |
| **A_51_P314111** | Psip1 | 0.456300454 | 0.115367416 | 0.797233492 | 11.03456165 | 3.252108309 | 0.016708265 | 0.479759698 | -3.643742692 |
| **A_55_P2050508** | Spta1 | 0.773135259 | 0.195381464 | 1.350889054 | 11.49771898 | 3.251590618 | 0.016719321 | 0.479759698 | -3.643921788 |
| **A_55_P2140212** | NA | -0.563535322 | -0.984983701 | -0.142086943 | 13.03181941 | -3.24907655 | 0.016773127 | 0.479759698 | -3.644792028 |
| **A_55_P2085641** | Lmbr1l | -0.291548292 | -0.50961358 | -0.073483005 | 10.48284453 | -3.248681629 | 0.016781596 | 0.479759698 | -3.644928804 |
| **A_55_P2047509** | Ufd1 | 0.273469877 | 0.068783963 | 0.47815579 | 12.44804558 | 3.246420177 | 0.016830184 | 0.479759698 | -3.645712413 |
| **A_55_P1959953** | Helz2 | -0.301313794 | -0.526846689 | -0.075780898 | 11.56149579 | -3.246327325 | 0.016832182 | 0.479759698 | -3.645744601 |
| **A_51_P347452** | Htatsf1 | 0.215443264 | 0.054119797 | 0.376766732 | 11.60422127 | 3.245028842 | 0.016860154 | 0.479759698 | -3.646194848 |
| **A_55_P2081405** | BC052688 | -0.460151986 | -0.8047163 | -0.115587672 | 13.10147361 | -3.244996205 | 0.016860857 | 0.479759698 | -3.646206168 |
| **A_51_P184100** | Capn7 | 0.261497654 | 0.065664069 | 0.45733124 | 8.886825497 | 3.244619781 | 0.016868976 | 0.479759698 | -3.646336735 |
| **A_55_P2073915** | Nos1 | 0.385130498 | 0.096661858 | 0.673599138 | 8.435501562 | 3.244087007 | 0.016880474 | 0.479759698 | -3.646521565 |
| **A_55_P2025695** | C330007P06Rik | 0.269637205 | 0.067638334 | 0.471636076 | 11.00008333 | 3.243501211 | 0.016893126 | 0.479759698 | -3.646724831 |
| **A_51_P330428** | Eif4ebp1 | -0.316975369 | -0.554451212 | -0.079499526 | 13.16525227 | -3.243316016 | 0.016897128 | 0.479759698 | -3.646789101 |
| **A_55_P2005838** | Ticrr | 0.509659763 | 0.127799954 | 0.891519572 | 9.938812599 | 3.24309437 | 0.016901919 | 0.479759698 | -3.646866027 |
| **A_55_P2048378** | Zfp975 | -0.495686761 | -0.867143851 | -0.124229671 | 10.0086885 | -3.242513974 | 0.016914472 | 0.479759698 | -3.647067494 |
| **A_55_P2039694** | NA | 0.243363919 | 0.060965755 | 0.425762083 | 15.38265052 | 3.242044087 | 0.016924642 | 0.479759698 | -3.647230633 |
| **A_51_P312121** | Xdh | -0.392608124 | -0.687000431 | -0.098215817 | 8.611705225 | -3.24052976 | 0.016957463 | 0.479759698 | -3.647756582 |
| **A_55_P2074438** | Gtf2a1 | 0.262400997 | 0.06555934 | 0.459242653 | 10.19570694 | 3.239154466 | 0.016987331 | 0.479759698 | -3.6482345 |
| **A_52_P85983** | Mrpl45 | 0.235293462 | 0.058777873 | 0.411809051 | 11.80213757 | 3.238992799 | 0.016990845 | 0.479759698 | -3.648290695 |
| **A_52_P273821** | Abhd5 | -0.874757266 | -1.53108393 | -0.218430602 | 10.06099156 | -3.238550693 | 0.017000461 | 0.479759698 | -3.648444389 |
| **A_55_P1954598** | NA | 0.487848096 | 0.121580556 | 0.854115637 | 12.66015885 | 3.236453501 | 0.017046155 | 0.479759698 | -3.649173802 |
| **A_55_P2028804** | Uchl3 | 0.340688329 | 0.084759591 | 0.596617066 | 8.116285905 | 3.234606022 | 0.01708652 | 0.479759698 | -3.649816834 |
| **A_51_P469578** | Psma1 | 0.241345776 | 0.059988107 | 0.422703444 | 13.54796627 | 3.233604995 | 0.017108434 | 0.479759698 | -3.650165435 |
| **A_55_P2139889** | Zfp783 | 0.273806013 | 0.068050457 | 0.479561568 | 8.646664438 | 3.233512914 | 0.017110452 | 0.479759698 | -3.650197508 |
| **A_55_P2094731** | Sox13 | 0.82525323 | 0.205082825 | 1.445423634 | 12.6886762 | 3.233399632 | 0.017112934 | 0.479759698 | -3.650236968 |
| **A_55_P1987264** | Gnb2 | -0.270183171 | -0.473252806 | -0.067113536 | 14.77128834 | -3.232931449 | 0.017123197 | 0.479759698 | -3.650400066 |
| **A_66_P106509** | NA | 0.73519206 | 0.182088478 | 1.288295642 | 13.32000308 | 3.229814099 | 0.017191702 | 0.479759698 | -3.651486765 |
| **A_55_P2086664** | Tspan33 | 0.482105705 | 0.119133524 | 0.845077885 | 14.84761995 | 3.227394981 | 0.01724507 | 0.479759698 | -3.652330928 |
| **A_55_P2295445** | 2600002B07Rik | 0.517686888 | 0.127895371 | 0.907478405 | 10.24761958 | 3.227141226 | 0.017250678 | 0.479759698 | -3.652419521 |
| **A_55_P2181738** | Lyz1 | -0.306590213 | -0.537438961 | -0.075741464 | 13.91952711 | -3.227111357 | 0.017251338 | 0.479759698 | -3.65242995 |
| **A_51_P481920** | Ccna2 | 0.494481062 | 0.122129889 | 0.866832234 | 13.03133632 | 3.22686002 | 0.017256896 | 0.479759698 | -3.652517708 |
| **A_51_P490257** | Fbxo4 | -0.397385557 | -0.696687322 | -0.098083792 | 9.740020491 | -3.226160651 | 0.017272369 | 0.479759698 | -3.652761946 |

| | | | | | | | | | |
|---|---|---|---|---|---|---|---|---|---|
| **A_55_P1962707** | NA | 0.469660861 | 0.115729508 | 0.823592213 | 12.92856314 | 3.224397111 | 0.017311455 | 0.479759698 | -3.653378101 |
| **A_51_P191649** | Ndc80 | 0.373414841 | 0.091903617 | 0.654926065 | 9.187430349 | 3.223139094 | 0.017339396 | 0.479759698 | -3.65381788 |
| **A_55_P1953087** | Mcm3 | 0.378732068 | 0.093129018 | 0.664335118 | 13.3755207 | 3.222199508 | 0.017360296 | 0.479759698 | -3.654146476 |
| **A_55_P2017312** | Eapp | 0.314655611 | 0.077370693 | 0.551940529 | 11.4751168 | 3.222170625 | 0.017360939 | 0.479759698 | -3.654156579 |
| **A_51_P344566** | Plk1 | 0.551527192 | 0.135580384 | 0.967474 | 12.80243862 | 3.22190197 | 0.01736692 | 0.479759698 | -3.654250556 |
| **A_51_P194853** | Sec14l4 | -0.710217707 | -1.246318887 | -0.174116527 | 10.43417408 | -3.219051478 | 0.017430522 | 0.479759698 | -3.65524825 |
| **A_55_P2184816** | Eif2s1 | 0.266845885 | 0.065339709 | 0.46835206 | 12.35431746 | 3.217772521 | 0.017459141 | 0.479759698 | -3.655696238 |
| **A_51_P228171** | Cenpp | 0.432062889 | 0.105743172 | 0.758382606 | 10.44557816 | 3.217265028 | 0.017470511 | 0.479759698 | -3.655874059 |
| **A_55_P2036240** | NA | -0.434627982 | -0.763009896 | -0.106246068 | 10.88481091 | -3.216041484 | 0.017497958 | 0.479759698 | -3.656302915 |
| **A_55_P2243566** | Redrum | 0.54973975 | 0.134009591 | 0.965469908 | 13.07449715 | 3.213133709 | 0.017563373 | 0.479759698 | -3.65732288 |
| **A_51_P296448** | Casp2 | 0.23297064 | 0.056705049 | 0.409236231 | 12.35020892 | 3.211565921 | 0.017598754 | 0.479759698 | -3.657873271 |
| **A_55_P1968129** | Eloc | 0.259361201 | 0.063095134 | 0.455627268 | 10.47625486 | 3.211019762 | 0.017611098 | 0.479759698 | -3.658065081 |
| **A_51_P237668** | Bex2 | 0.370908 | 0.090052825 | 0.651763175 | 13.68735198 | 3.208979624 | 0.017657289 | 0.479759698 | -3.658781919 |
| **A_55_P1957213** | NA | -0.608593008 | -1.069502845 | -0.14768317 | 12.14069152 | -3.208441828 | 0.017669488 | 0.479759698 | -3.658970973 |
| **A_55_P2102698** | NA | 0.408067822 | 0.098769865 | 0.71736578 | 14.02664392 | 3.205815227 | 0.017729197 | 0.479759698 | -3.659894854 |
| **A_55_P2041265** | Utp6 | 0.254170554 | 0.061474525 | 0.446866583 | 15.09251517 | 3.205056284 | 0.017746491 | 0.479759698 | -3.660161971 |
| **A_55_P2058746** | Mterf1a | 0.2671841 | 0.064472992 | 0.469895207 | 9.062057878 | 3.202699946 | 0.017800301 | 0.479759698 | -3.660991785 |
| **A_55_P2052685** | Dguok | -0.231915003 | -0.407879383 | -0.055950624 | 10.33325097 | -3.20248624 | 0.01780519 | 0.479759698 | -3.66106708 |
| **A_51_P283083** | Armc1 | 0.26825862 | 0.064641123 | 0.471876116 | 12.20567698 | 3.201266144 | 0.01783313 | 0.479759698 | -3.66149707 |
| **A_51_P492595** | Fbxo42 | 0.381300961 | 0.091806408 | 0.670795514 | 10.58144042 | 3.20044739 | 0.017851907 | 0.479759698 | -3.661785725 |
| **A_52_P424767** | Rbbp4 | 0.316208681 | 0.076034731 | 0.556382631 | 8.780115689 | 3.199124022 | 0.017882301 | 0.479759698 | -3.662252471 |
| **A_51_P372172** | Plod3 | -0.284590579 | -0.500800907 | -0.068380252 | 10.03465731 | -3.198359478 | 0.017899886 | 0.479759698 | -3.662522225 |
| **A_55_P1956752** | NA | 0.397456203 | 0.09544932 | 0.699463086 | 8.583523018 | 3.197831874 | 0.017912032 | 0.479759698 | -3.662708424 |
| **A_55_P2062444** | Memo1 | 0.374575837 | 0.089938046 | 0.659213628 | 12.49929328 | 3.197646031 | 0.017916313 | 0.479759698 | -3.662774019 |
| **A_51_P314907** | Dbf4 | 0.415556151 | 0.099710725 | 0.731401578 | 9.422362407 | 3.196968167 | 0.017931936 | 0.479759698 | -3.663013317 |
| **A_51_P317673** | Zfp764 | -0.453353721 | -0.797948086 | -0.108759357 | 10.10933623 | -3.196775982 | 0.017936368 | 0.479759698 | -3.663081172 |
| **A_51_P230439** | Ppfibp2 | -0.319068238 | -0.561636468 | -0.076500008 | 10.4521592 | -3.196191878 | 0.017949845 | 0.479759698 | -3.663287434 |
| **A_55_P2074796** | Cdkn2c | 0.43752979 | 0.104885811 | 0.770173769 | 10.01177871 | 3.196032232 | 0.01795353 | 0.479759698 | -3.663343816 |
| **A_52_P343690** | Kpna4 | 0.329303036 | 0.078906845 | 0.579699228 | 10.77835031 | 3.195591116 | 0.017963718 | 0.479759698 | -3.663499624 |
| **A_55_P2171086** | NA | 0.25091647 | 0.059981049 | 0.441851891 | 8.81047798 | 3.193198052 | 0.018019097 | 0.479759698 | -3.664345323 |
| **A_51_P362903** | Mrps22 | 0.368466897 | 0.0880231 | 0.648910694 | 11.55878744 | 3.192536183 | 0.018034446 | 0.479759698 | -3.664579357 |
| **A_55_P2087985** | NA | 0.344776944 | 0.082221214 | 0.607332675 | 11.69065837 | 3.19080235 | 0.018074722 | 0.479759698 | -3.665192703 |
| **A_55_P2062015** | Acyp1 | 0.387536463 | 0.092216477 | 0.682856449 | 8.487586078 | 3.188621144 | 0.01812553 | 0.479759698 | -3.665964863 |
| **A_55_P2219903** | AW061147 | 0.466002989 | 0.110817488 | 0.82118849 | 10.67446877 | 3.187987886 | 0.01814031 | 0.479759698 | -3.666189157 |
| **A_55_P2091230** | NA | 0.45151106 | 0.107346108 | 0.795676013 | 11.8622724 | 3.18775503 | 0.018145748 | 0.479759698 | -3.666271645 |
| **A_55_P2105152** | Trim59 | 0.597464892 | 0.142019744 | 1.05291004 | 12.80703835 | 3.187568304 | 0.01815011 | 0.479759698 | -3.666337797 |
| **A_55_P1964313** | Pla2g4b | 0.369715185 | 0.087852666 | 0.651577704 | 11.40732321 | 3.187228124 | 0.018158059 | 0.479759698 | -3.666458326 |
| **A_55_P1982034** | NA | 0.213439827 | 0.050697644 | 0.37618201 | 8.264834443 | 3.186827169 | 0.018167434 | 0.479759698 | -3.666600406 |
| **A_55_P1962419** | Ccar1 | 0.419336115 | 0.099557085 | 0.739115145 | 10.03275425 | 3.186364647 | 0.018178255 | 0.479759698 | -3.66676433 |
| **A_55_P2073377** | Mki67 | 0.509319865 | 0.120897698 | 0.897742032 | 14.64619305 | 3.186176257 | 0.018182664 | 0.479759698 | -3.666831105 |
| **A_52_P414420** | Dcun1d1 | 0.379970986 | 0.090130378 | 0.669811595 | 11.44548766 | 3.185476403 | 0.018199055 | 0.479759698 | -3.667079213 |
| **A_55_P2052016** | Crispld2 | -0.468736472 | -0.826423798 | -0.111049147 | 9.467747989 | -3.184259002 | 0.018227606 | 0.479759698 | -3.667510949 |
| **A_51_P214796** | Suz12 | 0.448790294 | 0.106298575 | 0.791282013 | 11.09231931 | 3.184025737 | 0.018233082 | 0.479759698 | -3.667593696 |
| **A_55_P1959606** | Tom1l1 | 0.413004162 | 0.097616726 | 0.728391598 | 12.33636503 | 3.181949106 | 0.018281912 | 0.479759698 | -3.668330657 |
| **A_51_P504588** | Tm2d2 | -0.208760325 | -0.368188849 | -0.049331801 | 9.996074216 | -3.181743217 | 0.018286761 | 0.479759698 | -3.668403753 |
| **A_55_P2099700** | Klc4 | -0.294731351 | -0.519876096 | -0.069586606 | 11.21292806 | -3.180883097 | 0.018307033 | 0.479759698 | -3.668709183 |
| **A_65_P05252** | Cdc25a | 0.341175286 | 0.080423823 | 0.601926748 | 10.54743794 | 3.179318229 | 0.018343978 | 0.479759698 | -3.669265118 |
| **A_55_P1979090** | Ublcp1 | 0.252607577 | 0.059359257 | 0.445855897 | 12.15360257 | 3.176243835 | 0.018416798 | 0.479759698 | -3.670358257 |
| **A_52_P329054** | Cep70 | 0.495940728 | 0.116414139 | 0.875467318 | 10.02709387 | 3.17519759 | 0.018441651 | 0.479759698 | -3.670730543 |
| **A_55_P2020072** | NA | 0.277989684 | 0.065110737 | 0.490868632 | 12.98583307 | 3.173066359 | 0.01849239 | 0.479759698 | -3.671489345 |

| | | | | | | | | | |
|---|---|---|---|---|---|---|---|---|---|
| **A_55_P2042525** | Apoo | 0.360871428 | 0.084458389 | 0.637284468 | 10.86794485 | 3.172321253 | 0.018510165 | 0.479759698 | -3.671754771 |
| **A_51_P487999** | Sgo1 | 0.476110671 | 0.111359288 | 0.840862055 | 11.45496542 | 3.171715004 | 0.018524641 | 0.479759698 | -3.671970787 |
| **A_66_P114381** | Ypel2 | -0.305704742 | -0.539920409 | -0.071489075 | 8.168739178 | -3.17153437 | 0.018528957 | 0.479759698 | -3.672035159 |
| **A_55_P2179069** | NA | 0.543675413 | 0.127054511 | 0.960296316 | 8.502059306 | 3.170894749 | 0.018544247 | 0.479759698 | -3.672263133 |
| **A_66_P119350** | 2310040G24Rik | -0.506896915 | -0.895367634 | -0.118426197 | 9.681697721 | -3.170622581 | 0.018550757 | 0.479759698 | -3.672360155 |
| **A_55_P1956842** | NA | 0.212312756 | 0.049513836 | 0.375111677 | 14.75175389 | 3.168894322 | 0.018592156 | 0.479759698 | -3.67297647 |
| **A_51_P142046** | Oxld1 | -0.33463964 | -0.591314045 | -0.077965234 | 10.73270383 | -3.167947734 | 0.018614873 | 0.479759698 | -3.673314199 |
| **A_55_P1989061** | Tsc22d3 | -0.600158628 | -1.060561081 | -0.139756175 | 10.71683061 | -3.167463467 | 0.018626506 | 0.479759698 | -3.673487024 |
| **A_55_P2114044** | Commd8 | 0.300129417 | 0.069867583 | 0.530391251 | 10.49329175 | 3.167158468 | 0.018633837 | 0.479759698 | -3.673595887 |
| **A_55_P2024155** | Zbtb16 | -0.919745129 | -1.625840833 | -0.213649425 | 9.295319296 | -3.16509767 | 0.018683454 | 0.479759698 | -3.674331767 |
| **A_51_P438821** | Pycard | -0.330918837 | -0.584992567 | -0.076845107 | 10.75443288 | -3.164790121 | 0.018690871 | 0.479759698 | -3.674441636 |
| **A_55_P2146650** | NA | 0.479712204 | 0.111217693 | 0.848206715 | 8.922665811 | 3.163245751 | 0.018728163 | 0.479759698 | -3.674993534 |
| **A_52_P278354** | Bmp7 | 0.583006401 | 0.135137379 | 1.030875424 | 9.709911585 | 3.163045853 | 0.018732996 | 0.479759698 | -3.675064993 |
| **A_55_P2169198** | NA | 0.401785998 | 0.092984621 | 0.710587374 | 8.391056283 | 3.161540565 | 0.018769433 | 0.479759698 | -3.675603264 |
| **A_55_P1964403** | Agfg2 | 0.35569859 | 0.082303388 | 0.629093792 | 8.863932546 | 3.161363568 | 0.018773723 | 0.479759698 | -3.675666575 |
| **A_51_P300936** | Rbbp8 | 0.371344518 | 0.085897008 | 0.656792027 | 11.37751248 | 3.161068796 | 0.018780869 | 0.479759698 | -3.675772023 |
| **A_55_P1960936** | NA | -0.418165973 | -0.739652775 | -0.09667917 | 13.89421048 | -3.160594131 | 0.018792382 | 0.479759698 | -3.675941848 |
| **A_66_P100232** | Paxbp1 | 0.375197741 | 0.086698381 | 0.6636971 | 9.55419374 | 3.160083443 | 0.018804778 | 0.479759698 | -3.676124594 |
| **A_51_P226053** | Sra1 | -0.26425333 | -0.467477648 | -0.061029012 | 12.84855975 | -3.159570007 | 0.018817249 | 0.479759698 | -3.676308358 |
| **A_55_P2022840** | Csnk2a1 | 0.26749629 | 0.06168346 | 0.47330912 | 11.59678964 | 3.158119123 | 0.01885254 | 0.479759698 | -3.67682783 |
| **A_51_P325485** | Utp3 | 0.294813172 | 0.067917548 | 0.521708795 | 12.13382759 | 3.157213515 | 0.018874604 | 0.479759698 | -3.677152212 |
| **A_55_P1990440** | R3hdm1 | 0.376443768 | 0.08669996 | 0.666187575 | 11.00452613 | 3.156960427 | 0.018880775 | 0.479759698 | -3.677242886 |
| **A_52_P556099** | Oip5 | 0.466841263 | 0.107510216 | 0.826172311 | 9.480435053 | 3.156877145 | 0.018882806 | 0.479759698 | -3.677272725 |
| **A_55_P2089942** | Xpo1 | 0.532829494 | 0.122393491 | 0.943265496 | 11.57597529 | 3.154467058 | 0.01894169 | 0.479759698 | -3.67813663 |
| **A_52_P206881** | Prpf18 | 0.229147737 | 0.052578458 | 0.405717017 | 10.89724004 | 3.153433063 | 0.018967013 | 0.479759698 | -3.678507502 |
| **A_51_P181451** | C1qa | -0.395072746 | -0.699503003 | -0.090642489 | 10.58917933 | -3.153351954 | 0.018969001 | 0.479759698 | -3.678536601 |
| **A_66_P115061** | Smc6 | 0.412138867 | 0.094503792 | 0.729773943 | 10.33027185 | 3.152813689 | 0.0189822 | 0.479759698 | -3.678729726 |
| **A_55_P2013336** | Melk | 0.382494751 | 0.087612116 | 0.677377386 | 9.991647067 | 3.151805945 | 0.019006938 | 0.479759698 | -3.6790914 |
| **A_55_P2359610** | AI646519 | 0.589388284 | 0.134929887 | 1.04384668 | 9.628209952 | 3.151305851 | 0.019019227 | 0.479759698 | -3.679270931 |
| **A_55_P2426549** | NA | 0.249971857 | 0.057154154 | 0.44278956 | 8.529137423 | 3.150122196 | 0.019048348 | 0.479759698 | -3.679695985 |
| **A_66_P138765** | Trappc10 | 0.457578884 | 0.104618144 | 0.810539623 | 9.990117134 | 3.15008855 | 0.019049177 | 0.479759698 | -3.67970807 |
| **A_55_P2126368** | Atad5 | 0.654568839 | 0.149092076 | 1.160045601 | 10.48072309 | 3.146569439 | 0.019136047 | 0.479759698 | -3.680972896 |
| **A_55_P2224830** | D230044B12Rik | 0.433507169 | 0.098733089 | 0.768281249 | 9.359044644 | 3.146499361 | 0.019137781 | 0.479759698 | -3.6809981 |
| **A_55_P1970676** | Sh3yl1 | 0.566633779 | 0.128956783 | 1.004310775 | 10.89700154 | 3.145805947 | 0.019154951 | 0.479759698 | -3.681247522 |
| **A_55_P2005680** | Ipo11 | 0.293457516 | 0.066553878 | 0.520361154 | 11.6731773 | 3.142584518 | 0.019234936 | 0.479759698 | -3.682407103 |
| **A_55_P2054342** | 2810408I11Rik | 0.474387186 | 0.107530694 | 0.841243678 | 10.16148891 | 3.142099471 | 0.01924701 | 0.479759698 | -3.682581818 |
| **A_55_P2000968** | Ppp2r3c | 0.327386859 | 0.074171313 | 0.580602406 | 10.21682307 | 3.141622996 | 0.01925888 | 0.479759698 | -3.682753476 |
| **A_55_P1968028** | Tdgf1 | 0.720826266 | 0.163160074 | 1.278492458 | 9.1603419 | 3.140794169 | 0.019279545 | 0.479759698 | -3.683052144 |
| **A_55_P2055087** | Prkar2b | 0.370055224 | 0.083726949 | 0.6563835 | 13.00502352 | 3.140403779 | 0.019289287 | 0.479759698 | -3.683192852 |
| **A_51_P428578** | Retreg1 | -0.268848295 | -0.47687149 | -0.0608251 | 8.300797298 | -3.140354764 | 0.019290511 | 0.479759698 | -3.683210521 |
| **A_55_P2017267** | Nanp | 0.632075323 | 0.142770294 | 1.121380352 | 10.54722879 | 3.138862885 | 0.019327792 | 0.479759698 | -3.683748433 |
| **A_65_P20433** | Parpbp | 0.454441645 | 0.102570835 | 0.806312455 | 9.620227992 | 3.138181602 | 0.019344843 | 0.479759698 | -3.683994175 |
| **A_55_P2129925** | Mybl2 | 0.495610027 | 0.111614879 | 0.879605175 | 10.43327172 | 3.136155129 | 0.019395658 | 0.479759698 | -3.68472549 |
| **A_55_P1978905** | NA | 0.251560434 | 0.056629558 | 0.446491311 | 7.95120573 | 3.135774969 | 0.019405207 | 0.479759698 | -3.684862743 |
| **A_55_P2061268** | Rnpc3 | 0.300699521 | 0.067499951 | 0.533899091 | 8.472343309 | 3.133200444 | 0.019470008 | 0.479759698 | -3.685792748 |
| **A_55_P1972277** | Samd11 | 0.503792238 | 0.113043688 | 0.894540787 | 9.265556398 | 3.1328333 | 0.019479268 | 0.479759698 | -3.685925443 |
| **A_55_P2081616** | Timeless | 0.373311855 | 0.083648904 | 0.662974805 | 11.12902234 | 3.131569255 | 0.019511187 | 0.479759698 | -3.686382438 |
| **A_66_P133328** | Tpd52 | -0.338236349 | -0.600765376 | -0.075707322 | 10.27417387 | -3.130589597 | 0.019535963 | 0.479759698 | -3.686736762 |
| **A_55_P2095488** | Syncrip | 0.357289626 | 0.079866172 | 0.634713081 | 9.514928098 | 3.129395263 | 0.019566215 | 0.479759698 | -3.687168902 |
| **A_55_P2013236** | S100g | -0.57009125 | -1.012861814 | -0.127320686 | 10.25383607 | -3.128591231 | 0.01958661 | 0.479759698 | -3.687459926 |

| | | | | | | | | | |
|---|---|---|---|---|---|---|---|---|---|
| **A_55_P1979833** | Cited1 | 1.159228786 | 0.258264037 | 2.060193535 | 11.21893437 | 3.126400035 | 0.019642308 | 0.479759698 | -3.688253473 |
| **A_51_P141836** | Ube3c | 0.265915836 | 0.059202668 | 0.472629005 | 9.743398858 | 3.125786039 | 0.019657946 | 0.479759698 | -3.688475947 |
| **A_55_P2067351** | NA | 0.433281017 | 0.096327497 | 0.770234536 | 9.433846038 | 3.124516727 | 0.019690317 | 0.479759698 | -3.688936023 |
| **A_55_P1999022** | Zfp760 | 0.279188823 | 0.061988399 | 0.496389246 | 8.998897884 | 3.12334925 | 0.019720142 | 0.479759698 | -3.689359375 |
| **A_55_P1958867** | Gm9790 | 0.231826468 | 0.051465752 | 0.412187185 | 14.63864523 | 3.123232077 | 0.019723138 | 0.479759698 | -3.689401874 |
| **A_55_P2006255** | Txnip | -0.289247921 | -0.514326218 | -0.064169624 | 8.288006921 | -3.122624868 | 0.019738672 | 0.479759698 | -3.689622141 |
| **A_55_P2012789** | Impa1 | 0.306425387 | 0.06797935 | 0.544871425 | 10.71165399 | 3.122610553 | 0.019739038 | 0.479759698 | -3.689627335 |
| **A_51_P451106** | Brca2 | 0.497970172 | 0.110460746 | 0.885479599 | 10.60797174 | 3.122512942 | 0.019741537 | 0.479759698 | -3.689662748 |
| **A_55_P2054511** | NA | 0.418978242 | 0.092740407 | 0.745216076 | 9.243244168 | 3.120616018 | 0.019790158 | 0.479759698 | -3.690351205 |
| **A_55_P2045026** | Mgrn1 | -0.25888172 | -0.460461132 | -0.057302307 | 12.39222266 | -3.120602118 | 0.019790515 | 0.479759698 | -3.690356251 |
| **A_51_P169693** | Bst2 | -0.5717486 | -1.017004127 | -0.126493073 | 10.80289431 | -3.120175193 | 0.019801476 | 0.479759698 | -3.690511262 |
| **A_51_P394802** | Fam111a | 0.423928188 | 0.093763611 | 0.754092765 | 10.7846202 | 3.119931133 | 0.019807745 | 0.479759698 | -3.690599889 |
| **A_55_P1957319** | Krr1 | 0.237304752 | 0.052474844 | 0.422134661 | 10.09674021 | 3.119732642 | 0.019812845 | 0.479759698 | -3.690671973 |
| **A_55_P1968304** | NA | 0.254270355 | 0.056188151 | 0.452352559 | 11.65825941 | 3.119129875 | 0.019828342 | 0.479759698 | -3.690890906 |
| **A_51_P359272** | Alcam | -0.267696832 | -0.47625924 | -0.059134424 | 10.30797244 | -3.118820559 | 0.019836299 | 0.479759698 | -3.691003273 |
| **A_55_P2008896** | NA | 0.372806692 | 0.082301114 | 0.66331227 | 13.4821541 | 3.118260638 | 0.019850712 | 0.479759698 | -3.691206709 |
| **A_55_P2011937** | Fignl1 | 0.500831877 | 0.110494524 | 0.891169229 | 12.65713807 | 3.117705148 | 0.019865022 | 0.479759698 | -3.691408577 |
| **A_52_P635182** | Rras | -0.457253781 | -0.813856631 | -0.100650932 | 11.20392606 | -3.11570037 | 0.019916761 | 0.479759698 | -3.692137461 |
| **A_55_P2046499** | Vmn1r70 | 0.549483923 | 0.120595541 | 0.978372305 | 9.341122272 | 3.113105917 | 0.019983936 | 0.479759698 | -3.693081521 |
| **A_51_P496751** | Nop58 | 0.412765571 | 0.090417185 | 0.735113957 | 12.40932143 | 3.111438024 | 0.02002725 | 0.479759698 | -3.693688896 |
| **A_52_P659312** | Spsb4 | 0.491010058 | 0.107379253 | 0.874640864 | 8.83574249 | 3.109997958 | 0.02006473 | 0.479759698 | -3.694213601 |
| **A_55_P1992234** | Slx | 0.239954798 | 0.052466217 | 0.427443379 | 8.009789691 | 3.109838281 | 0.02006889 | 0.479759698 | -3.694271798 |
| **A_55_P2088070** | Spata5 | 0.396285005 | 0.086432208 | 0.706137801 | 9.699220058 | 3.107673663 | 0.020125383 | 0.479759698 | -3.695061063 |
| **A_51_P333159** | Rpa1 | 0.450391151 | 0.098213668 | 0.802568634 | 11.9344125 | 3.107502257 | 0.020129864 | 0.479759698 | -3.695123587 |
| **A_55_P1969625** | Hnrnpr | 0.307950269 | 0.067132352 | 0.548768187 | 12.27346641 | 3.107241301 | 0.020136687 | 0.479759698 | -3.695218785 |
| **A_52_P535484** | Gvin1 | -0.323296991 | -0.576197369 | -0.070396613 | 8.778304885 | -3.10624261 | 0.020162826 | 0.479759698 | -3.695583194 |
| **A_55_P2028248** | Dnajc19 | 0.404267858 | 0.087994999 | 0.720540718 | 12.36490657 | 3.105921516 | 0.020171237 | 0.479759698 | -3.695700384 |
| **A_52_P123400** | Dnaaf2 | 0.426317716 | 0.092749951 | 0.759885482 | 9.381483814 | 3.105506866 | 0.020182105 | 0.479759698 | -3.695851741 |
| **A_55_P2129614** | Itprip | -0.465803446 | -0.830651141 | -0.100955751 | 11.12937519 | -3.102232046 | 0.020268163 | 0.479759698 | -3.697047918 |
| **A_55_P1964154** | Asah1 | -0.250313358 | -0.446515468 | -0.054111247 | 8.582272415 | -3.100013184 | 0.020326698 | 0.479759698 | -3.697859196 |
| **A_51_P343429** | Slc25a37 | 0.515171415 | 0.111329114 | 0.919013717 | 16.17994053 | 3.099724804 | 0.020334319 | 0.479759698 | -3.697964683 |
| **A_51_P424221** | Irf2bp2 | -0.377510797 | -0.673584152 | -0.081437441 | 10.1884294 | -3.098227044 | 0.02037395 | 0.479759698 | -3.69851273 |
| **A_55_P2066429** | NA | 0.366662358 | 0.07905252 | 0.654272195 | 9.666281683 | 3.097745818 | 0.020386702 | 0.479759698 | -3.698688879 |
| **A_55_P1986596** | Cacna1h | 0.225100669 | 0.048482698 | 0.40171864 | 8.288384758 | 3.096885038 | 0.020409532 | 0.479759698 | -3.699004036 |
| **A_55_P2118575** | Tgs1 | 0.294139622 | 0.063342886 | 0.524936359 | 10.00990961 | 3.096756482 | 0.020412944 | 0.479759698 | -3.699051113 |
| **A_55_P2175767** | Rangrf | 0.361972711 | 0.07763938 | 0.646306042 | 9.27750947 | 3.093365566 | 0.020503165 | 0.479759698 | -3.70029364 |
| **A_55_P2007650** | Nt5dc1 | 0.265253702 | 0.056780455 | 0.473726949 | 9.711242923 | 3.091678408 | 0.020548216 | 0.479759698 | -3.700912428 |
| **A_55_P2045127** | NA | 0.317801681 | 0.067935084 | 0.567668278 | 8.133794958 | 3.090517282 | 0.020579284 | 0.479759698 | -3.701338504 |
| **A_55_P2009576** | Rnu12 | 0.506569412 | 0.108225166 | 0.904913658 | 13.87854127 | 3.090036455 | 0.020592163 | 0.479759698 | -3.701514997 |
| **A_52_P199586** | Kpna6 | 0.22500682 | 0.048070381 | 0.40194326 | 10.75416651 | 3.090022112 | 0.020592548 | 0.479759698 | -3.701520262 |
| **A_55_P2026823** | Uspl1 | 0.327909616 | 0.069987657 | 0.585831575 | 10.51117812 | 3.089221311 | 0.020614019 | 0.479759698 | -3.701814274 |
| **A_51_P434171** | Myo19 | 0.289228965 | 0.061680807 | 0.516777122 | 8.922093253 | 3.088528756 | 0.020632607 | 0.479759698 | -3.702068611 |
| **A_55_P1977330** | Zbtb1 | 0.343839838 | 0.073324869 | 0.614354807 | 8.666508485 | 3.08850338 | 0.020633288 | 0.479759698 | -3.702077932 |
| **A_51_P246224** | Tram2 | -0.31957013 | -0.570998173 | -0.068142088 | 8.995837193 | -3.088415014 | 0.020635661 | 0.479759698 | -3.702110389 |
| **A_51_P210603** | Cyp4f13 | -0.334748488 | -0.598140454 | -0.071356522 | 8.961530443 | -3.088156411 | 0.020642608 | 0.479759698 | -3.702205379 |
| **A_55_P2078895** | Gin1 | 0.351544009 | 0.07491567 | 0.628172348 | 8.503057382 | 3.087921281 | 0.020648926 | 0.479759698 | -3.702291756 |
| **A_55_P2182222** | Rab3c | 0.566388658 | 0.120667392 | 1.012109924 | 11.22924809 | 3.087694906 | 0.020655012 | 0.479759698 | -3.702374923 |
| **A_55_P1965846** | Sp2 | 0.35468971 | 0.075529426 | 0.633849994 | 9.696933444 | 3.087295122 | 0.020665763 | 0.479759698 | -3.702521814 |
| **A_66_P140658** | Olfr787 | 0.588215274 | 0.125243532 | 1.051187017 | 9.319439867 | 3.087201681 | 0.020668276 | 0.479759698 | -3.702556151 |
| **A_55_P2130565** | Prdm11 | 0.262488681 | 0.055803757 | 0.469173606 | 8.386797728 | 3.085922176 | 0.020702731 | 0.479759698 | -3.703026434 |

| | | | | | | | | | |
|---|---|---|---|---|---|---|---|---|---|
| **A_51_P201773** | Tubgcp3 | 0.247509095 | 0.052555392 | 0.442462799 | 10.49037469 | 3.084912583 | 0.020729961 | 0.479759698 | -3.703397664 |
| **A_51_P201480** | Stat3 | -0.262613762 | -0.469618744 | -0.05560878 | 12.02543188 | -3.082619151 | 0.020791962 | 0.479759698 | -3.788263674 |
| **A_55_P2065364** | Tdrkh | 0.325541287 | 0.068767999 | 0.582314574 | 8.47474507 | 3.080629173 | 0.020845922 | 0.479759698 | -3.704974184 |
| **A_55_P2039611** | Eed | 0.269665225 | 0.056917124 | 0.482413327 | 12.05205778 | 3.079941316 | 0.020864609 | 0.479759698 | -3.705227578 |
| **A_55_P2186961** | NA | -0.670486871 | -1.200074241 | -0.140899502 | 12.15043205 | -3.076350741 | 0.02096245 | 0.479759698 | -3.706551295 |
| **A_55_P1964368** | Kctd14 | 0.383275195 | 0.080503599 | 0.68604679 | 9.89286859 | 3.075946464 | 0.020973498 | 0.479759698 | -3.706700444 |
| **A_55_P2035981** | Mindy1 | -0.303025135 | -0.542406444 | -0.063643827 | 12.07381006 | -3.075895711 | 0.020974885 | 0.479759698 | -3.70671917 |
| **A_66_P122927** | NA | 0.754127716 | 0.158045365 | 1.350210067 | 12.15870812 | 3.074127162 | 0.021023291 | 0.479759698 | -3.707371905 |
| **A_51_P142187** | Abhd11 | -0.207526882 | -0.371599252 | -0.043454512 | 10.20407942 | -3.073421317 | 0.021042644 | 0.479759698 | -3.707632534 |
| **A_55_P1978628** | Hnrnpa2b1 | 0.356535378 | 0.074575964 | 0.638494791 | 8.475487476 | 3.072551852 | 0.02106651 | 0.479759698 | -3.707953668 |
| **A_52_P283910** | Hnrnpdl | 0.250545227 | 0.052359765 | 0.448730689 | 10.39042796 | 3.071832493 | 0.021086277 | 0.479759698 | -3.708219438 |
| **A_55_P2022104** | Cop1 | 0.233067887 | 0.048570896 | 0.417564877 | 8.87108924 | 3.069561513 | 0.021148815 | 0.479759698 | -3.709058907 |
| **A_55_P2157695** | Tsc1 | 0.24938313 | 0.051906804 | 0.446859457 | 10.22038212 | 3.068564254 | 0.02117634 | 0.479759698 | -3.70942776 |
| **A_55_P1991688** | Rad51ap1 | 0.466519592 | 0.097031822 | 0.836007362 | 11.26404667 | 3.067983421 | 0.02119239 | 0.479759698 | -3.709642652 |
| **A_55_P2167189** | 4921504E06Rik | 0.971242041 | 0.20192715 | 1.740556932 | 11.53170685 | 3.067655054 | 0.021201469 | 0.479759698 | -3.709764158 |
| **A_52_P125350** | Anp32e | 0.416028662 | 0.086350148 | 0.745707177 | 10.52991799 | 3.066308144 | 0.021238755 | 0.479759698 | -3.710262707 |
| **A_52_P608322** | Maff | -0.727832047 | -1.304626411 | -0.151037683 | 10.66239885 | -3.066149699 | 0.021243146 | 0.479759698 | -3.71032137 |
| **A_55_P2040988** | NA | 0.666864936 | 0.138328626 | 1.195401245 | 9.721504663 | 3.06581711 | 0.021252365 | 0.479759698 | -3.71044452 |
| **A_55_P2140171** | NA | 0.363319783 | 0.075363667 | 0.651275899 | 11.18662383 | 3.065814878 | 0.021252427 | 0.479759698 | -3.710445346 |
| **A_55_P2062039** | Tsen34 | -0.397408383 | -0.712406142 | -0.082410624 | 11.90853267 | -3.06558067 | 0.021258923 | 0.479759698 | -3.710532077 |
| **A_55_P2020004** | NA | 0.600569865 | 0.124270683 | 1.076869047 | 11.97665229 | 3.0638457 | 0.021307105 | 0.479759698 | -3.711174785 |
| **A_55_P1957754** | NA | 0.240185283 | 0.049687028 | 0.430683537 | 15.18008142 | 3.063645997 | 0.021312659 | 0.479759698 | -3.711248789 |
| **A_55_P2357303** | 5430401H09Rik | 0.636420638 | 0.131564102 | 1.141277174 | 11.445465 | 3.063087905 | 0.021328187 | 0.479759698 | -3.71145563 |
| **A_52_P135873** | NA | 0.460868864 | 0.095242152 | 0.826495575 | 11.39627231 | 3.062828228 | 0.021335417 | 0.479759698 | -3.711551887 |
| **A_51_P195034** | Esco2 | 0.532947478 | 0.110115669 | 0.955779287 | 9.981660448 | 3.062668202 | 0.021339873 | 0.479759698 | -3.711611209 |
| **A_66_P102163** | NA | 0.767211881 | 0.158489472 | 1.375934291 | 9.50190489 | 3.062522012 | 0.021343945 | 0.479759698 | -3.711665405 |
| **A_55_P2000638** | Stag2 | 0.266224907 | 0.054915233 | 0.477534581 | 12.31965164 | 3.061346533 | 0.021376718 | 0.479759698 | -3.712101287 |
| **A_55_P1962982** | NA | 0.290270269 | 0.05979864 | 0.520741898 | 10.99095648 | 3.060330098 | 0.021405101 | 0.479759698 | -3.71247834 |
| **A_55_P1975322** | NA | 0.429299878 | 0.088380448 | 0.770219308 | 11.22172036 | 3.059794083 | 0.021420085 | 0.479759698 | -3.712677234 |
| **A_52_P681391** | G0s2 | -0.33505275 | -0.601175687 | -0.068929812 | 11.48449958 | -3.059243648 | 0.021435484 | 0.479759698 | -3.712881518 |
| **A_55_P2045268** | Tmem59 | -0.355572808 | -0.63801008 | -0.073135536 | 12.4807763 | -3.059072161 | 0.021440284 | 0.479759698 | -3.712945171 |
| **A_55_P2044704** | NA | 0.307123567 | 0.06310547 | 0.551141664 | 12.10793393 | 3.058259238 | 0.021463054 | 0.479759698 | -3.713246965 |
| **A_66_P139250** | Kctd3 | 0.245711799 | 0.050472476 | 0.440951122 | 10.71448413 | 3.058031173 | 0.021469446 | 0.479759698 | -3.713331649 |
| **A_51_P502054** | Gtf2b | -0.439754784 | -0.789300386 | -0.090209182 | 11.85735188 | -3.056961154 | 0.021499467 | 0.479759698 | -3.713729054 |
| **A_55_P1981301** | Nup88 | 0.328342255 | 0.067339635 | 0.589344875 | 11.70367752 | 3.056786443 | 0.021504373 | 0.479759698 | -3.713793956 |
| **A_55_P2051414** | Axl | -0.346462885 | -0.621997784 | -0.070927987 | 10.52353381 | -3.055366364 | 0.021544295 | 0.479759698 | -3.714321641 |
| **A_55_P2122648** | Dtymk | 0.453274322 | 0.092560661 | 0.813987982 | 11.49283894 | 3.053386042 | 0.0216001 | 0.479759698 | -3.715057954 |
| **A_55_P2163198** | NA | 0.294126627 | 0.060006129 | 0.528247126 | 11.35436167 | 3.052657583 | 0.021620668 | 0.479759698 | -3.715328936 |
| **A_55_P2069554** | Brix1 | 0.319249563 | 0.065120614 | 0.573378512 | 12.06857574 | 3.05252592 | 0.021624387 | 0.479759698 | -3.715377921 |
| **A_55_P2419299** | NA | 0.341366671 | 0.069501352 | 0.61323199 | 8.900237165 | 3.051058227 | 0.021665898 | 0.479759698 | -3.715924134 |
| **A_55_P2084758** | Agfg1 | 0.529451313 | 0.10771557 | 0.951187056 | 10.76564326 | 3.050484404 | 0.021682151 | 0.479759698 | -3.716137764 |
| **A_55_P2253980** | Armt1 | 0.236216858 | 0.048051051 | 0.424382666 | 8.568087306 | 3.050375956 | 0.021685224 | 0.479759698 | -3.716178143 |
| **A_51_P375187** | Rpf2 | 0.344046299 | 0.069973063 | 0.618119536 | 11.62356221 | 3.050236063 | 0.021689189 | 0.479759698 | -3.716230233 |
| **A_52_P456158** | Fchsd1 | 0.72798968 | 0.148001602 | 1.307977757 | 13.22525149 | 3.049926303 | 0.021697972 | 0.479759698 | -3.716345583 |
| **A_55_P2146535** | Cnr2 | 0.551653032 | 0.112055069 | 0.991250995 | 10.8215568 | 3.049253472 | 0.021717061 | 0.479759698 | -3.716596179 |
| **A_51_P517096** | Casp8ap2 | 0.2839361 | 0.057354098 | 0.510518102 | 9.422877297 | 3.044937563 | 0.021839942 | 0.479759698 | -3.718205066 |
| **A_55_P2015405** | 9930111J21Rik1 | 0.272148583 | 0.054965334 | 0.489331833 | 8.574444075 | 3.04482923 | 0.021843036 | 0.479759698 | -3.718245482 |
| **A_55_P1953169** | Saa3 | -0.844614832 | -1.518661943 | -0.17056772 | 11.8469413 | -3.044750011 | 0.021845299 | 0.479759698 | -3.718275038 |
| **A_66_P120745** | Armc1 | 0.249623068 | 0.050394088 | 0.448852048 | 11.63677101 | 3.044495908 | 0.021852559 | 0.479759698 | -3.718369846 |
| **A_55_P2370250** | Syn3 | 0.446820801 | 0.090174397 | 0.803467205 | 8.416872219 | 3.044238835 | 0.021859907 | 0.479759698 | -3.718465771 |

| | | | | | | | | | |
|---|---|---|---|---|---|---|---|---|---|
| **A_51_P332355** | Kif18a | 0.559236493 | 0.112727236 | 1.005745751 | 9.595834787 | 3.043324243 | 0.021886069 | 0.479759698 | -3.718807115 |
| **A_55_P2016739** | Rbm12b2 | 0.377623445 | 0.075947435 | 0.679299456 | 9.567980229 | 3.041594844 | 0.021935632 | 0.479759698 | -3.719452867 |
| **A_55_P1960738** | Gm4470 | 0.359131457 | 0.072154364 | 0.646108549 | 9.061411799 | 3.04081084 | 0.021958141 | 0.479759698 | -3.719745742 |
| **A_51_P345316** | Cep76 | 0.590081089 | 0.118394802 | 1.061767375 | 10.2361028 | 3.039776374 | 0.021987879 | 0.479759698 | -3.720132305 |
| **A_52_P176300** | Med1 | 0.265708149 | 0.053271163 | 0.478145135 | 9.172002463 | 3.039190562 | 0.022004739 | 0.479759698 | -3.720351277 |
| **A_55_P2018076** | Arap1 | -0.454203825 | -0.817400594 | -0.091007056 | 11.79174734 | -3.038729203 | 0.022018026 | 0.479759698 | -3.720523761 |
| **A_55_P2048588** | Cdk1 | 0.426499804 | 0.085400205 | 0.767599403 | 11.82198211 | 3.038231175 | 0.02203238 | 0.479759698 | -3.720709986 |
| **A_55_P1974636** | Gm16430 | 0.246272664 | 0.049293251 | 0.443252077 | 8.095424636 | 3.037935568 | 0.022040904 | 0.479759698 | -3.720820537 |
| **A_51_P175205** | Rps6kb1 | 0.217474697 | 0.043450782 | 0.391498613 | 8.757548899 | 3.036567836 | 0.022080392 | 0.479759698 | -3.721332189 |
| **A_55_P2142590** | Mark3 | 0.252840457 | 0.050412025 | 0.455268888 | 10.67301732 | 3.034996856 | 0.022125843 | 0.479759698 | -3.721920181 |
| **A_52_P30451** | Ppp1r3c | -0.990338713 | -1.783251475 | -0.197425952 | 10.66900886 | -3.0348801 | 0.022129225 | 0.479759698 | -3.721963894 |
| **A_51_P431329** | Car3 | 0.972336212 | 0.193428398 | 1.751244026 | 11.38960824 | 3.03328759 | 0.022175409 | 0.479759698 | -3.722560304 |
| **A_52_P633128** | Aasdhppt | 0.308574951 | 0.061331285 | 0.555818617 | 10.62214385 | 3.03262481 | 0.022194661 | 0.479759698 | -3.72280862 |
| **A_55_P1995127** | NA | 0.219636561 | 0.043577969 | 0.395695154 | 13.74478409 | 3.031311728 | 0.022232855 | 0.479759698 | -3.723300751 |
| **A_55_P2288117** | NA | 0.293149229 | 0.05812592 | 0.528172538 | 8.667893307 | 3.030826038 | 0.022247001 | 0.479759698 | -3.723482841 |
| **A_55_P2146136** | Gla | 0.315659987 | 0.062512071 | 0.568807902 | 15.51160886 | 3.029900504 | 0.022273984 | 0.479759698 | -3.72382992 |
| **A_55_P2063146** | NA | 0.443256618 | 0.087649693 | 0.798863542 | 12.94673237 | 3.028783337 | 0.022306601 | 0.479759698 | -3.724249014 |
| **A_51_P481788** | Mapk1 | 0.251115456 | 0.049623082 | 0.45260783 | 10.506199 | 3.028293879 | 0.022320908 | 0.479759698 | -3.818833641 |
| **A_55_P2086511** | Ppid | 0.289125514 | 0.057107909 | 0.521143118 | 12.28393715 | 3.027949775 | 0.022330972 | 0.479759698 | -3.724561824 |
| **A_55_P2457153** | G2e3 | 0.36252611 | 0.071578938 | 0.653473283 | 9.794098024 | 3.027668624 | 0.022339198 | 0.479759698 | -3.724667353 |
| **A_55_P2139430** | NA | 0.360335539 | 0.071085774 | 0.649585305 | 11.26162224 | 3.027033809 | 0.022357784 | 0.479759698 | -3.724905665 |
| **A_55_P2149001** | NA | 0.356513555 | 0.070330848 | 0.642696262 | 13.87902879 | 3.027023886 | 0.022358075 | 0.479759698 | -3.72490939 |
| **A_55_P2174401** | NA | 0.660499852 | 0.130127667 | 1.190872036 | 10.5847384 | 3.026043549 | 0.022386811 | 0.479759698 | -3.72527752 |
| **A_55_P2134790** | Qsox1 | -0.640843961 | -1.155504495 | -0.126183427 | 13.20736071 | -3.025621646 | 0.022399191 | 0.479759698 | -3.72543599 |
| **A_52_P680495** | Pdcd10 | 0.333864651 | 0.065489877 | 0.602239425 | 12.27824209 | 3.022817556 | 0.022481656 | 0.479759698 | -3.726489826 |
| **A_55_P2057912** | NA | 0.236321005 | 0.046354984 | 0.426287026 | 12.61178648 | 3.022801209 | 0.022482137 | 0.479759698 | -3.726495973 |
| **A_55_P1995195** | Fosl2 | -0.611913938 | -1.103800905 | -0.120026971 | 9.23309534 | -3.022791685 | 0.022482418 | 0.479759698 | -3.726499554 |
| **A_55_P2064538** | Zfp985 | 0.33697292 | 0.066093488 | 0.607852352 | 8.376014782 | 3.022749515 | 0.022483661 | 0.479759698 | -3.72651541 |
| **A_55_P2129319** | Mre11a | 0.353232524 | 0.069256105 | 0.637208942 | 9.91247817 | 3.022467194 | 0.022491982 | 0.479759698 | -3.726621573 |
| **A_55_P2004571** | Pitpnm2 | -0.315859381 | -0.569918043 | -0.061800718 | 10.74977267 | -3.020945994 | 0.022536879 | 0.479759698 | -3.727193781 |
| **A_55_P2022499** | Pnisr | 0.334198321 | 0.065342115 | 0.603054527 | 9.417730584 | 3.020420342 | 0.022552415 | 0.479759698 | -3.727391579 |
| **A_52_P97670** | Atad2b | 0.272678014 | 0.053224943 | 0.492131085 | 10.35559599 | 3.019198367 | 0.022588577 | 0.479759698 | -3.72785154 |
| **A_51_P133612** | Cdt1 | 0.43999963 | 0.085876883 | 0.794122377 | 13.21598091 | 3.019129039 | 0.022590631 | 0.479759698 | -3.727877642 |
| **A_51_P245090** | Aqp3 | 0.830171587 | 0.162026915 | 1.498316259 | 10.03052463 | 3.01912118 | 0.022590864 | 0.479759698 | -3.727880601 |
| **A_55_P2010788** | Slc6a9 | 0.543442896 | 0.106023984 | 0.980861807 | 13.37931935 | 3.0188362 | 0.022599307 | 0.479759698 | -3.727987901 |
| **A_51_P346938** | Lrg1 | -1.026881613 | -1.853458384 | -0.200304841 | 10.16222952 | -3.018702886 | 0.022603258 | 0.479759698 | -3.7280381 |
| **A_55_P2056192** | NA | 0.433248568 | 0.084444134 | 0.782053001 | 12.47948229 | 3.018132683 | 0.022620165 | 0.479759698 | -3.728252834 |
| **A_55_P2178203** | Derl2 | -0.3514698 | -0.634473797 | -0.068465802 | 11.54837446 | -3.017717856 | 0.022632474 | 0.479759698 | -3.728409083 |
| **A_55_P2126557** | NA | 0.825743046 | 0.160702499 | 1.490783592 | 10.66252804 | 3.017032506 | 0.022652826 | 0.479759698 | -3.728667276 |
| **A_55_P2018479** | Carnmt1 | 0.358312558 | 0.069565945 | 0.64705917 | 11.08697178 | 3.01528467 | 0.022704818 | 0.479759698 | -3.729326025 |
| **A_55_P2096043** | Acot11 | 0.70302805 | 0.136209767 | 1.269846334 | 9.761041273 | 3.013783091 | 0.022749588 | 0.479759698 | -3.729892286 |
| **A_51_P342535** | Rnf166 | -0.353826385 | -0.639182911 | -0.068469859 | 10.94142583 | -3.012906087 | 0.02277578 | 0.479759698 | -3.730223152 |
| **A_51_P253803** | Mki67 | 0.502975588 | 0.097208208 | 0.908742969 | 15.07928621 | 3.011986114 | 0.022803291 | 0.479759698 | -3.730570339 |
| **A_55_P2296088** | Sp3 | 0.522211857 | 0.100911365 | 0.943512348 | 8.911846804 | 3.011881987 | 0.022806407 | 0.479759698 | -3.730609642 |
| **A_51_P153486** | Dnajb1 | 0.289986204 | 0.05595625 | 0.524016159 | 13.58341338 | 3.010849711 | 0.022837323 | 0.479759698 | -3.73099936 |
| **A_55_P2000793** | Rbm34 | 0.395566966 | 0.076325408 | 0.714808524 | 9.584709309 | 3.01081301 | 0.022838423 | 0.479759698 | -3.731013218 |
| **A_51_P201721** | Crtap | -0.446900279 | -0.807655323 | -0.086145235 | 10.78552536 | -3.010103317 | 0.022859705 | 0.479759698 | -3.731281236 |
| **A_55_P2178862** | U2surp | 0.304636651 | 0.058679917 | 0.550593385 | 8.961539988 | 3.009585075 | 0.02287526 | 0.479759698 | -3.731476994 |
| **A_55_P1965931** | Nat8f1 | -0.634310767 | -1.146467039 | -0.122154494 | 11.94881174 | -3.009419812 | 0.022880223 | 0.479759698 | -3.731539427 |
| **A_55_P2002702** | Ddx18 | 0.311459905 | 0.059929436 | 0.562990374 | 10.69010296 | 3.008809797 | 0.022898551 | 0.479759698 | -3.73176991 |

| | | | | | | | | | |
|---|---|---|---|---|---|---|---|---|---|
| **A_55_P1956017** | Usp45 | 0.414602764 | 0.079742895 | 0.749462633 | 10.59508903 | 3.008515679 | 0.022907393 | 0.479759698 | -3.731881055 |
| **A_55_P1956862** | Egfr | -0.28530177 | -0.51582664 | -0.054776899 | 8.11932584 | -3.007252318 | 0.022945417 | 0.479759698 | -3.7323586 |
| **A_55_P2133385** | NA | 0.424639934 | 0.081280609 | 0.76799926 | 14.34384326 | 3.005074036 | 0.023011139 | 0.479759698 | -3.733182482 |
| **A_55_P2128499** | Mgl2 | 0.36115198 | 0.069034174 | 0.653269786 | 14.08480777 | 3.004105353 | 0.023040431 | 0.479759698 | -3.733549065 |
| **A_55_P2064984** | Cenpi | 0.539073049 | 0.103036421 | 0.975109677 | 9.855590039 | 3.0040547 | 0.023041963 | 0.479759698 | -3.733568237 |
| **A_51_P414889** | Ifi35 | -0.302300391 | -0.546866682 | -0.057734101 | 10.25759736 | -3.003483859 | 0.023059245 | 0.479759698 | -3.733784326 |
| **A_66_P124011** | Spint2 | -0.818184588 | -1.480131866 | -0.156237311 | 10.69337451 | -3.003385526 | 0.023062223 | 0.479759698 | -3.733821553 |
| **A_52_P578043** | NA | 0.275042907 | 0.052486298 | 0.497599516 | 8.512498037 | 3.002915716 | 0.023076459 | 0.479759698 | -3.733999436 |
| **A_55_P2056896** | Kif2a | 0.452568927 | 0.086353777 | 0.818784077 | 10.45111428 | 3.002836003 | 0.023078875 | 0.479759698 | -3.73402962 |
| **A_55_P2340206** | E230006M18Rik | 0.363483313 | 0.069333204 | 0.657633422 | 8.984900367 | 3.002608114 | 0.023085784 | 0.479759698 | -3.734115918 |
| **A_51_P202596** | Sh3tc2 | 0.245211979 | 0.04667525 | 0.443748708 | 9.478261606 | 3.001124475 | 0.023130821 | 0.479759698 | -3.734677917 |
| **A_55_P2083939** | Irf3 | -0.250162318 | -0.452802893 | -0.047521744 | 12.50034308 | -2.999705875 | 0.023173972 | 0.479759698 | -3.735215554 |
| **A_52_P609200** | Serinc3 | 0.272450502 | 0.051640491 | 0.493260513 | 15.12010737 | 2.998140918 | 0.023221677 | 0.479759698 | -3.73580897 |
| **A_51_P370678** | Gfi1b | 0.525191779 | 0.099431616 | 0.950951941 | 12.19970848 | 2.997340523 | 0.023246116 | 0.479759698 | -3.736112598 |
| **A_55_P2031436** | Ly6e | -0.529898237 | -0.959568701 | -0.100227774 | 12.24021062 | -2.99667858 | 0.023266348 | 0.479759698 | -3.736363769 |
| **A_55_P2027999** | Hk1 | 0.495433498 | 0.093580166 | 0.89728683 | 13.50254081 | 2.995718371 | 0.023295732 | 0.479759698 | -3.73672822 |
| **A_55_P2096395** | Cdc42bpa | 0.318776731 | 0.060118826 | 0.577434635 | 9.763364144 | 2.994636033 | 0.0233289 | 0.479759698 | -3.737139172 |
| **A_55_P2006940** | Sfmbt1 | 0.433361444 | 0.081668719 | 0.785054169 | 9.96397953 | 2.99412601 | 0.023344547 | 0.479759698 | -3.737332877 |
| **A_66_P120567** | Anxa1 | -0.488837941 | -0.885632831 | -0.092043051 | 11.29329585 | -2.993519039 | 0.023363184 | 0.479759698 | -3.737563447 |
| **A_55_P2041397** | Ezh2 | 0.423596446 | 0.079571029 | 0.767621862 | 11.11816183 | 2.991885484 | 0.023413421 | 0.479759698 | -3.73818423 |
| **A_51_P193813** | Fga | -0.492517463 | -0.892596608 | -0.092438319 | 13.12365845 | -2.991292696 | 0.02343168 | 0.479759698 | -3.738409588 |
| **A_66_P106654** | Camsap1 | 0.230836974 | 0.043259926 | 0.418414022 | 10.33778533 | 2.990259411 | 0.023463543 | 0.479759698 | -3.738802522 |
| **A_66_P118368** | Catip | 0.663359701 | 0.124177761 | 1.202541641 | 10.17067768 | 2.98948884 | 0.023487336 | 0.479759698 | -3.739095645 |
| **A_51_P172842** | Med19 | 0.245851318 | 0.04577417 | 0.445928467 | 10.49895232 | 2.985783003 | 0.023602124 | 0.479759698 | -3.74050644 |
| **A_52_P58059** | Usp34 | 0.235680532 | 0.043850526 | 0.427510539 | 8.967473625 | 2.985316337 | 0.023616622 | 0.479759698 | -3.740684227 |
| **A_55_P2106690** | Tal1 | 0.5637559 | 0.104885545 | 1.022626256 | 12.86739286 | 2.98527461 | 0.023617919 | 0.479759698 | -3.740700125 |
| **A_52_P244193** | Cd24a | 0.406225744 | 0.075512796 | 0.736938693 | 15.41663793 | 2.984691388 | 0.023636053 | 0.479759698 | -3.740922362 |
| **A_55_P2432388** | Prr14l | 0.303627427 | 0.05639332 | 0.550861533 | 10.28783673 | 2.984116717 | 0.023653935 | 0.479759698 | -3.741141384 |
| **A_55_P2033215** | Cabp5 | 0.552600501 | 0.102609232 | 1.00259177 | 11.79608605 | 2.983941992 | 0.023659375 | 0.479759698 | -3.741207985 |
| **A_55_P1972799** | Eif4enif1 | 0.293509819 | 0.054486426 | 0.532533213 | 10.84077578 | 2.983770501 | 0.023664715 | 0.479759698 | -3.741273357 |
| **A_55_P2083481** | Lpin1 | -0.906185648 | -1.644338747 | -0.16803255 | 9.638189304 | -2.98300459 | 0.023688583 | 0.479759698 | -3.74156537 |
| **A_55_P2074291** | Fbxo6 | -0.386642718 | -0.701603644 | -0.071681792 | 11.57454071 | -2.98288387 | 0.023692347 | 0.479759698 | -3.741611403 |
| **A_66_P128631** | Cinp | 0.415126037 | 0.076899226 | 0.753352848 | 9.014895343 | 2.982326117 | 0.023709748 | 0.479759698 | -3.741824111 |
| **A_55_P2157838** | NA | 0.436394612 | 0.080826602 | 0.791962623 | 12.3080435 | 2.982221435 | 0.023713015 | 0.479759698 | -3.741864038 |
| **A_52_P644972** | Mzt1 | 0.378047126 | 0.070012448 | 0.686081805 | 10.26523066 | 2.982150235 | 0.023715237 | 0.479759698 | -3.741891195 |
| **A_55_P2016842** | Me1 | -0.361600032 | -0.656262338 | -0.066937727 | 9.278838762 | -2.981858752 | 0.023724339 | 0.479759698 | -3.74200238 |
| **A_51_P279232** | Dennd2d | -0.200688123 | -0.364234438 | -0.037141808 | 9.15730042 | -2.981701044 | 0.023729264 | 0.479759698 | -3.742062542 |
| **A_55_P1985670** | Exosc3 | 0.353884017 | 0.065479202 | 0.642288832 | 10.12795625 | 2.981546801 | 0.023734083 | 0.479759698 | -3.742121386 |
| **A_51_P362054** | Atp5e | -0.24364984 | -0.442252723 | -0.045046957 | 15.35973979 | -2.981012314 | 0.023750789 | 0.479759698 | -3.742325316 |
| **A_55_P2065268** | NA | 0.599658593 | 0.110760092 | 1.088557094 | 9.346593808 | 2.980358754 | 0.023771234 | 0.479759698 | -3.742574729 |
| **A_55_P2095345** | Rara | -0.602031071 | -1.092913393 | -0.111148748 | 10.92789951 | -2.980057901 | 0.023780652 | 0.479759698 | -3.742689561 |
| **A_55_P1960456** | Larp1 | 0.31652869 | 0.0583841 | 0.57467328 | 13.35654172 | 2.979430375 | 0.023800308 | 0.479759698 | -3.742929117 |
| **A_66_P100780** | Katnbl1 | 0.255590541 | 0.047131767 | 0.464049315 | 10.02768352 | 2.979255757 | 0.023805781 | 0.479759698 | -3.742995787 |
| **A_51_P105709** | Trip13 | 0.386777667 | 0.07130287 | 0.702252465 | 10.50112024 | 2.979064518 | 0.023811776 | 0.479759698 | -3.743068807 |
| **A_51_P210143** | Syne2 | -0.321168148 | -0.583177833 | -0.059158463 | 8.989696618 | -2.97850485 | 0.023829332 | 0.479759698 | -3.743282531 |
| **A_55_P1963868** | Plekho2 | 0.256101708 | 0.047110458 | 0.465092959 | 12.5494494 | 2.977608251 | 0.023857484 | 0.479759698 | -3.743625008 |
| **A_55_P1970935** | Usp7 | 0.393848205 | 0.072443236 | 0.715253174 | 12.81912062 | 2.977552808 | 0.023859226 | 0.479759698 | -3.743646189 |
| **A_55_P2368625** | Sp4 | 0.286590461 | 0.052683672 | 0.52049725 | 8.968027903 | 2.977159432 | 0.02387159 | 0.479759698 | -3.743796485 |
| **A_51_P443364** | Cbll1 | 0.20595773 | 0.037841563 | 0.374073897 | 10.1794531 | 2.976814722 | 0.02388243 | 0.479759698 | -3.743928204 |
| **A_52_P476400** | NA | 0.315559792 | 0.057958313 | 0.573161272 | 9.882904073 | 2.976572727 | 0.023890044 | 0.479759698 | -3.744020684 |

| | | | | | | | | | |
|---|---|---|---|---|---|---|---|---|---|
| **A_51_P128336** | Serinc5 | 0.36318816 | 0.066652531 | 0.659723789 | 11.45636851 | 2.97603469 | 0.02390698 | 0.479759698 | -3.752660752 |
| **A_55_P1975430** | St3gal3 | -0.204541823 | -0.371566501 | -0.037517146 | 10.56389567 | -2.975669342 | 0.023918487 | 0.479759698 | -3.744365987 |
| **A_55_P1974587** | Vangl1 | 0.5541651 | 0.101437229 | 1.00689297 | 10.2286163 | 2.974302454 | 0.023961594 | 0.479759698 | -3.744888662 |
| **A_55_P2106429** | Herc2 | 0.384002265 | 0.070274145 | 0.697730384 | 10.71933883 | 2.974154521 | 0.023966265 | 0.479759698 | -3.744945244 |
| **A_55_P2068172** | Nufip2 | 0.230391105 | 0.042082749 | 0.41869946 | 9.553834935 | 2.972893179 | 0.024006126 | 0.479759698 | -3.745427806 |
| **A_55_P2017645** | Tap2 | -0.374062586 | -0.679822724 | -0.068302449 | 10.45929125 | -2.972669293 | 0.024013209 | 0.479759698 | -3.745513482 |
| **A_55_P2000668** | Stxbp2 | 0.388953305 | 0.070915478 | 0.706991132 | 10.24085042 | 2.971678882 | 0.024044568 | 0.479759698 | -3.745892571 |
| **A_51_P499776** | Denr | 0.22575228 | 0.041057853 | 0.410446706 | 8.448827189 | 2.970034838 | 0.024096721 | 0.479759698 | -3.746522132 |
| **A_51_P456957** | Pex11a | -0.537408813 | -0.977144885 | -0.097672741 | 11.06527772 | -2.969585817 | 0.024110986 | 0.479759698 | -3.773203974 |
| **A_55_P2033041** | Sirpb1b | -0.255333564 | -0.46427044 | -0.046396688 | 8.136220732 | -2.969449884 | 0.024115306 | 0.479759698 | -3.746746219 |
| **A_55_P2001238** | Luc7l3 | 0.418809196 | 0.076031777 | 0.761586614 | 12.56937761 | 2.968842744 | 0.024134613 | 0.479759698 | -3.746978852 |
| **A_51_P286373** | Ttll12 | 0.435696222 | 0.079067756 | 0.792324687 | 11.2241283 | 2.96859518 | 0.02414249 | 0.479759698 | -3.747073724 |
| **A_55_P2029417** | 2610318N02Rik | 0.432598639 | 0.078158486 | 0.787038792 | 9.990879295 | 2.965687744 | 0.024235209 | 0.479759698 | -3.748188525 |
| **A_55_P2260094** | Tcf7l2 | 0.377387924 | 0.068057198 | 0.686718649 | 8.337479675 | 2.964477289 | 0.024273924 | 0.479759698 | -3.748652983 |
| **A_55_P2166884** | NA | 0.329412666 | 0.059263674 | 0.599561659 | 13.86841989 | 2.9629214 | 0.024323785 | 0.479759698 | -3.749250274 |
| **A_55_P2102574** | Prorsd1 | 0.274689258 | 0.049387973 | 0.499990543 | 10.7007424 | 2.962519361 | 0.024336687 | 0.479759698 | -3.749404666 |
| **A_55_P1958730** | NA | 0.494875533 | 0.08886557 | 0.900885496 | 11.82615116 | 2.96170965 | 0.024362694 | 0.479759698 | -3.749715678 |
| **A_55_P2081169** | NA | 0.215296454 | 0.038558817 | 0.392034091 | 15.68850833 | 2.959995308 | 0.024417856 | 0.479759698 | -3.750374449 |
| **A_55_P2069277** | Mtf1 | 0.328402159 | 0.058722876 | 0.598081442 | 8.450920831 | 2.958977121 | 0.024450681 | 0.479759698 | -3.750765894 |
| **A_55_P2154208** | NA | 0.415088638 | 0.074204618 | 0.755972658 | 11.85590452 | 2.958812141 | 0.024456004 | 0.479759698 | -3.750829335 |
| **A_51_P476018** | Sox7 | -0.594031874 | -1.081970202 | -0.106093546 | 8.553811549 | -2.958203212 | 0.024475663 | 0.479759698 | -3.751063519 |
| **A_52_P399584** | Ckap2l | 0.358404777 | 0.063980613 | 0.652828941 | 9.090833748 | 2.957900274 | 0.02448545 | 0.479759698 | -3.751180043 |
| **A_51_P514675** | Zcchc10 | 0.260117227 | 0.046346364 | 0.473888089 | 9.696210828 | 2.9566764 | 0.02452503 | 0.479759698 | -3.751650925 |
| **A_55_P2043382** | Llgl2 | -0.371722735 | -0.677319374 | -0.066126095 | 10.32067862 | -2.955654994 | 0.024558115 | 0.479759698 | -3.752044061 |
| **A_52_P330214** | Fanca | 0.481824316 | 0.085614177 | 0.878034454 | 10.20474281 | 2.954924019 | 0.024581822 | 0.479759698 | -3.752325497 |
| **A_55_P2014284** | Serpina11 | -0.729505797 | -1.329460763 | -0.129550831 | 13.63560401 | -2.954563198 | 0.024593534 | 0.479759698 | -3.752464444 |
| **A_52_P296632** | NA | 0.715260778 | 0.126851769 | 1.303669787 | 11.71711395 | 2.953712945 | 0.024621155 | 0.479759698 | -3.752791934 |
| **A_55_P2032278** | NA | 0.279859735 | 0.049617521 | 0.510101948 | 8.641055037 | 2.953511428 | 0.024627706 | 0.479759698 | -3.752869566 |
| **A_55_P1973352** | NA | -0.662041157 | -1.206793399 | -0.117288916 | 11.17529779 | -2.953038795 | 0.024643078 | 0.479759698 | -3.753051664 |
| **A_51_P463846** | Gbp7 | -0.213798208 | -0.3897401 | -0.037856315 | 8.228401267 | -2.952690934 | 0.024654399 | 0.479759698 | -3.753185708 |
| **A_65_P06446** | Tfrc | 0.730595717 | 0.129311333 | 1.331880101 | 11.80831376 | 2.95243528 | 0.024662723 | 0.479759698 | -3.753284231 |
| **A_55_P1993518** | Tbc1d14 | 0.233145473 | 0.041203059 | 0.425087888 | 10.72231152 | 2.951475781 | 0.02469399 | 0.479759698 | -3.75365408 |
| **A_51_P474454** | Rnaseh2b | 0.367940074 | 0.06495997 | 0.670920178 | 11.80449736 | 2.950843455 | 0.024714618 | 0.479759698 | -3.753897883 |
| **A_55_P2177233** | Abhd5 | -0.819987018 | -1.495249221 | -0.144724814 | 11.53167849 | -2.95065018 | 0.024720927 | 0.479759698 | -3.753972414 |
| **A_51_P406165** | Nxpe2 | 0.702861448 | 0.124035721 | 1.281687175 | 11.92497451 | 2.950564318 | 0.024723731 | 0.479759698 | -3.754005526 |
| **A_55_P2359436** | A930026B05Rik | 0.281984409 | 0.049713731 | 0.514255087 | 9.085653353 | 2.949944864 | 0.024743965 | 0.479759698 | -3.754244441 |
| **A_51_P243914** | Skap2 | -0.330147251 | -0.602137851 | -0.058156651 | 10.95902718 | -2.949422437 | 0.024761045 | 0.479759698 | -3.754445974 |
| **A_51_P406638** | Rfc1 | 0.346299038 | 0.060723874 | 0.631874201 | 11.51497871 | 2.946551517 | 0.024855128 | 0.479759698 | -3.755554121 |
| **A_55_P2024600** | Zfp658 | 0.355575864 | 0.062241164 | 0.648910563 | 13.89094286 | 2.945452496 | 0.024891246 | 0.479759698 | -3.755978624 |
| **A_51_P209280** | Rab31 | -0.260912508 | -0.476172522 | -0.045652495 | 8.572317788 | -2.945199553 | 0.024899566 | 0.479759698 | -3.756076347 |
| **A_55_P2317665** | E2f1 | 0.462982355 | 0.080997061 | 0.844967649 | 10.64845078 | 2.945106368 | 0.024902632 | 0.479759698 | -3.756112351 |
| **A_51_P369252** | Ddias | 0.371670279 | 0.064799163 | 0.678541395 | 8.869779132 | 2.942964413 | 0.024973222 | 0.479759698 | -3.756940256 |
| **A_55_P2004511** | Cd300lf | 0.523333577 | 0.091130127 | 0.955537026 | 9.614879317 | 2.942209275 | 0.02499816 | 0.479759698 | -3.757232277 |
| **A_55_P2102322** | Yif1b | -0.275171885 | -0.502444561 | -0.047899209 | 12.4953333 | -2.941982072 | 0.025005668 | 0.479759698 | -3.757320154 |
| **A_51_P157524** | E2f4 | 0.642252815 | 0.111738941 | 1.172766689 | 9.746043226 | 2.941659893 | 0.02501632 | 0.479759698 | -3.757444777 |
| **A_55_P2018904** | Kbtbd6 | 0.374446232 | 0.065138622 | 0.683753842 | 9.235725556 | 2.941589376 | 0.025018651 | 0.479759698 | -3.757472056 |
| **A_55_P2045952** | Mysm1 | 0.227437593 | 0.039526709 | 0.415348477 | 9.0670605 | 2.940989704 | 0.025038492 | 0.479759698 | -3.75770406 |
| **A_55_P2159807** | Rfc4 | 0.489463496 | 0.085044028 | 0.893882964 | 12.27063844 | 2.940840316 | 0.025043437 | 0.479759698 | -3.757761864 |
| **A_52_P254174** | Nr3c1 | -0.256127715 | -0.467782847 | -0.044472583 | 8.862476684 | -2.940430783 | 0.025056999 | 0.479759698 | -3.757920342 |
| **A_66_P112164** | Rab29 | -0.201650404 | -0.368315734 | -0.034985074 | 10.29842865 | -2.939930189 | 0.025073587 | 0.479759698 | -3.758114089 |

| | | | | | | | | | |
|---|---|---|---|---|---|---|---|---|---|
| **A_52_P480360** | Dut | 0.38741452 | 0.067119889 | 0.70770915 | 12.04425231 | 2.939066574 | 0.025102233 | 0.479759698 | -3.758448415 |
| **A_55_P2173982** | Rrm2 | 0.554817968 | 0.095917263 | 1.013718673 | 14.4102685 | 2.93775099 | 0.025145938 | 0.479759698 | -3.758957902 |
| **A_55_P2015426** | Asnsd1 | 0.302004648 | 0.052208757 | 0.55180054 | 12.06625514 | 2.93772761 | 0.025146715 | 0.479759698 | -3.758966959 |
| **A_55_P1993404** | NA | 0.272882604 | 0.047121727 | 0.49864348 | 11.41296028 | 2.937043347 | 0.02516948 | 0.479759698 | -3.759232047 |
| **A_55_P2111478** | Polq | 0.565339703 | 0.097538903 | 1.033140504 | 9.457794261 | 2.93651156 | 0.025187188 | 0.479759698 | -3.759438109 |
| **A_55_P2171413** | Me1 | -0.321235822 | -0.587062488 | -0.055409156 | 9.247427212 | -2.93635536 | 0.025192391 | 0.479759698 | -3.759498642 |
| **A_55_P2087404** | Mapre3 | -0.363342186 | -0.664026631 | -0.06265774 | 9.323746818 | -2.936216339 | 0.025197024 | 0.479759698 | -3.75955252 |
| **A_55_P2048334** | Ubr3 | 0.276336019 | 0.047593271 | 0.505078767 | 9.34784645 | 2.935440973 | 0.025222877 | 0.479759698 | -3.759853064 |
| **A_55_P2083649** | Alas1 | -0.64329467 | -1.139938767 | -0.146650573 | 12.17690626 | -3.994203825 | 0.025234413 | 0.479759698 | -4.19639266 |
| **A_51_P234359** | Sct | -0.980514811 | -1.782115889 | -0.178913733 | 8.386744059 | -3.11199028 | 0.025275428 | 0.479759698 | -3.884664677 |
| **A_55_P1975380** | Nol11 | 0.295893877 | 0.050791683 | 0.540996072 | 10.16090517 | 2.933404594 | 0.025290913 | 0.479759698 | -3.760642779 |
| **A_55_P2067629** | Ush1g | 0.401778958 | 0.068887088 | 0.734670828 | 8.869726179 | 2.932696956 | 0.025314601 | 0.479759698 | -3.760917334 |
| **A_55_P2019054** | Acacb | -0.945535712 | -1.729203893 | -0.16186753 | 10.62174559 | -2.93176335 | 0.025345891 | 0.479759698 | -3.761279663 |
| **A_55_P2123362** | NA | 0.350848064 | 0.059993819 | 0.64170231 | 8.642213964 | 2.931074563 | 0.025369002 | 0.479759698 | -3.761547054 |
| **A_55_P2258261** | 1810008I18Rik | -0.790355716 | -1.445803633 | -0.134907798 | 9.336405624 | -2.929999857 | 0.025405107 | 0.479759698 | -3.761964387 |
| **A_55_P2117959** | Cyp4b1 | 0.294697139 | 0.05026278 | 0.539131499 | 9.591028818 | 2.929522624 | 0.025421158 | 0.479759698 | -3.762149757 |
| **A_55_P2054240** | Med8 | 0.241143232 | 0.041078829 | 0.441207636 | 13.05005551 | 2.928791415 | 0.025445771 | 0.479759698 | -3.762433837 |
| **A_55_P2404484** | 1700001L05Rik | 0.26306022 | 0.044784137 | 0.481336302 | 9.41310452 | 2.928412267 | 0.025458544 | 0.479759698 | -3.762581168 |
| **A_55_P2238079** | A830054O07Rik | 0.236752042 | 0.040251743 | 0.433252341 | 8.311030659 | 2.927613266 | 0.025485484 | 0.479759698 | -3.762891708 |
| **A_55_P2151463** | NA | 0.264727623 | 0.044873782 | 0.484581463 | 10.61382719 | 2.925825311 | 0.025545879 | 0.479759698 | -3.763586925 |
| **A_51_P493105** | Mettl15 | 0.256820416 | 0.043485753 | 0.47015508 | 9.209962408 | 2.925171322 | 0.025568008 | 0.479759698 | -3.763841325 |
| **A_55_P2065369** | H3f3a | 0.28882034 | 0.048901478 | 0.528739202 | 14.81789758 | 2.925139427 | 0.025569088 | 0.479759698 | -3.763853734 |
| **A_55_P2236607** | Ccnc | 0.276893131 | 0.046857909 | 0.506928353 | 8.624288965 | 2.924832722 | 0.025579473 | 0.479759698 | -3.763973062 |
| **A_55_P2060253** | Gm12657 | 0.272181416 | 0.04593991 | 0.498422922 | 15.03593267 | 2.923272998 | 0.02563236 | 0.479759698 | -3.764580092 |
| **A_55_P2032099** | Zfp715 | 0.372719787 | 0.062866621 | 0.682572953 | 8.625424569 | 2.922871348 | 0.025645998 | 0.479759698 | -3.764736463 |
| **A_66_P101862** | NA | 0.321213158 | 0.05416249 | 0.588263826 | 8.600855632 | 2.922690633 | 0.025652136 | 0.479759698 | -3.764806826 |
| **A_52_P502771** | Rad54b | 0.371716343 | 0.062607965 | 0.680824721 | 8.924733233 | 2.922025951 | 0.025674728 | 0.479759698 | -3.765065665 |
| **A_55_P2101806** | Esf1 | 0.451507195 | 0.076030361 | 0.826984029 | 11.01106757 | 2.921895752 | 0.025679156 | 0.479759698 | -3.765116373 |
| **A_55_P2156068** | NA | 0.278683923 | 0.046875146 | 0.5104927 | 8.768967945 | 2.921226477 | 0.02570193 | 0.479759698 | -3.765377072 |
| **A_51_P231630** | Zfp239 | 0.274221594 | 0.046099883 | 0.502343304 | 8.421890757 | 2.920910301 | 0.025712697 | 0.479759698 | -3.765500251 |
| **A_55_P2015687** | Phf11d | -0.473935426 | -0.868646154 | -0.079224698 | 8.742910494 | -2.91758442 | 0.025826244 | 0.479759698 | -3.76679679 |
| **A_55_P2081615** | Timeless | 0.363785757 | 0.060810089 | 0.666761425 | 8.641539839 | 2.917568952 | 0.025826773 | 0.479759698 | -3.766802823 |
| **A_55_P2179296** | Mad2l1 | 0.387607056 | 0.064788129 | 0.710425982 | 8.676780061 | 2.917533654 | 0.025827981 | 0.479759698 | -3.766816592 |
| **A_55_P2100526** | Ptges3 | 0.260801172 | 0.043579089 | 0.478023256 | 11.94718337 | 2.917351462 | 0.025834217 | 0.479759698 | -3.76688766 |
| **A_55_P1992475** | NA | 0.757399523 | 0.126470811 | 1.388328235 | 10.37803323 | 2.916942879 | 0.025848208 | 0.479759698 | -3.767047054 |
| **A_55_P2054913** | Olfr406 | 0.326037857 | 0.054332121 | 0.597743593 | 8.430165382 | 2.915764288 | 0.025888612 | 0.479759698 | -3.767506965 |
| **A_52_P207614** | NA | 0.248728976 | 0.041315178 | 0.456142775 | 10.40033085 | 2.913881769 | 0.025953289 | 0.479759698 | -3.768241947 |
| **A_52_P135707** | Creb3 | -0.249957589 | -0.458400965 | -0.041514213 | 10.61026805 | -2.913811264 | 0.025955714 | 0.479759698 | -3.768269483 |
| **A_55_P2091402** | Kpna2 | 0.355277928 | 0.058926962 | 0.651628893 | 13.4176733 | 2.913030752 | 0.025982584 | 0.479759698 | -3.768574361 |
| **A_55_P2010672** | Kif17 | 0.321843218 | 0.05325935 | 0.590427086 | 8.794529834 | 2.91170671 | 0.026028233 | 0.479759698 | -3.769091735 |
| **A_55_P2059055** | Zfp943 | 0.252199647 | 0.041614655 | 0.462784639 | 9.947911895 | 2.910048635 | 0.026085519 | 0.479759698 | -3.769739965 |
| **A_51_P176377** | Usp25 | 0.433301201 | 0.071285039 | 0.795317363 | 11.34917449 | 2.908339636 | 0.026144708 | 0.479759698 | -3.770408488 |
| **A_55_P2022114** | NA | 0.434981762 | 0.071538673 | 0.798424851 | 8.850292198 | 2.90815682 | 0.026151048 | 0.479759698 | -3.770480025 |
| **A_55_P1971249** | Eya3 | 0.229754512 | 0.037771912 | 0.421737112 | 11.41358186 | 2.90793951 | 0.026158586 | 0.479759698 | -3.770565065 |
| **A_66_P136099** | Coasy | -0.215696253 | -0.395936089 | -0.035456417 | 12.50756465 | -2.907870126 | 0.026160994 | 0.479759698 | -3.770592219 |
| **A_55_P2114074** | Acrbp | -0.216714799 | -0.397809359 | -0.035620239 | 8.736527714 | -2.907812191 | 0.026163004 | 0.479759698 | -3.770614892 |
| **A_51_P382928** | Cstf3 | 0.232824198 | 0.038232683 | 0.427415713 | 10.45252038 | 2.90728365 | 0.026181352 | 0.479759698 | -3.770821762 |
| **A_51_P227165** | 2310030G06Rik | -0.294146917 | -0.540161025 | -0.04813281 | 9.571448473 | -2.905276549 | 0.026251154 | 0.479759698 | -3.771607679 |
| **A_55_P2159595** | NA | 0.306157331 | 0.049839342 | 0.56247532 | 8.389595757 | 2.90234319 | 0.02635353 | 0.479759698 | -3.772757259 |
| **A_55_P2011802** | Pdk2 | -0.289670265 | -0.532186137 | -0.047154392 | 9.136481268 | -2.90233098 | 0.026353957 | 0.479759698 | -3.772762046 |

| | | | | | | | | |
|---|---|---|---|---|---|---|---|---|
| **A_55_P2137706** | Tmpo | 0.296734771 | 0.04822201 | 0.545247532 | 12.24252997 | 2.90136878 | 0.026387632 | 0.479759698 | -3.773139383 |
| **A_55_P2125766** | Ola1 | 0.568170479 | 0.092326376 | 1.044014583 | 10.31431441 | 2.901330224 | 0.026388982 | 0.479759698 | -3.773154506 |
| **A_51_P321150** | Lyz2 | -0.424679486 | -0.780497017 | -0.068861955 | 10.21989756 | -2.900127756 | 0.026431134 | 0.479759698 | -3.773626247 |
| **A_55_P2145257** | NA | 0.272143793 | 0.044080198 | 0.500207388 | 9.576270989 | 2.899516978 | 0.026452572 | 0.479759698 | -3.773865936 |
| **A_51_P195153** | Gtse1 | 0.582678372 | 0.094348876 | 1.071007869 | 9.3485175 | 2.899339884 | 0.026458791 | 0.479759698 | -3.773935443 |
| **A_51_P143190** | Lyl1 | 0.423871834 | 0.068632011 | 0.779111656 | 13.41264308 | 2.899319676 | 0.026459501 | 0.479759698 | -3.773943375 |
| **A_51_P500718** | Dck | 0.534824492 | 0.086394149 | 0.983254834 | 9.391697187 | 2.898007367 | 0.026505639 | 0.479759698 | -3.77445857 |
| **A_55_P2072906** | Gm6788 | 0.263890609 | 0.042438124 | 0.485343094 | 13.0023383 | 2.895519977 | 0.026593327 | 0.479759698 | -3.77543572 |
| **A_51_P336635** | Xpo1 | 0.273680343 | 0.043974615 | 0.503386071 | 10.88485484 | 2.895042668 | 0.026610189 | 0.479759698 | -3.775623322 |
| **A_51_P229105** | Cetn3 | 0.249221885 | 0.040033438 | 0.458410332 | 11.61031124 | 2.894887363 | 0.026615678 | 0.479759698 | -3.77568437 |
| **A_52_P449417** | Vangl1 | 0.425698805 | 0.068120472 | 0.783277139 | 11.62815869 | 2.892773459 | 0.026690512 | 0.479759698 | -3.776515628 |
| **A_52_P400224** | Ccs | -0.292286973 | -0.537895517 | -0.046678428 | 13.29811754 | -2.89167298 | 0.02672956 | 0.479759698 | -3.776948611 |
| **A_51_P360492** | Mcm6 | 0.327618899 | 0.05229399 | 0.602943807 | 14.55889641 | 2.891389716 | 0.026739621 | 0.479759698 | -3.777060087 |
| **A_55_P2411907** | B230213L16Rik | 0.408674354 | 0.065203582 | 0.752145126 | 11.02204711 | 2.891151083 | 0.026748099 | 0.479759698 | -3.777154007 |
| **A_55_P2013586** | Prss8 | -0.437807004 | -0.805777584 | -0.069836424 | 9.67179992 | -2.891031323 | 0.026752356 | 0.479759698 | -3.777201145 |
| **A_51_P393426** | Pros1 | -0.491156263 | -0.9040092 | -0.078303326 | 9.352284506 | -2.890729784 | 0.026763075 | 0.479759698 | -3.77731984 |
| **A_52_P52128** | Leo1 | 0.331507114 | 0.05282993 | 0.610184297 | 10.79293874 | 2.890511013 | 0.026770856 | 0.479759698 | -3.777405962 |
| **A_55_P2141889** | NA | 0.246902869 | 0.039188189 | 0.454617548 | 11.35885956 | 2.888298898 | 0.026849663 | 0.479759698 | -3.77827715 |
| **A_55_P2146389** | NA | 0.253295649 | 0.040200694 | 0.466390604 | 8.576805885 | 2.888269738 | 0.026850703 | 0.479759698 | -3.778288639 |
| **A_55_P1971619** | NA | 0.556939077 | 0.088183766 | 1.025694388 | 8.599968765 | 2.886986002 | 0.026896553 | 0.479759698 | -3.778794513 |
| **A_55_P2009813** | NA | 0.42387936 | 0.067088926 | 0.780669794 | 8.649279938 | 2.886770488 | 0.026904259 | 0.479759698 | -3.778879461 |
| **A_55_P1959515** | Cep135 | 0.278730619 | 0.044024026 | 0.513437213 | 8.78774817 | 2.885642852 | 0.026944615 | 0.479759698 | -3.779324036 |
| **A_55_P2099785** | Usp15 | 0.472098355 | 0.07445098 | 0.86974573 | 11.8559197 | 2.884812268 | 0.026974382 | 0.479759698 | -3.779651606 |
| **A_66_P126080** | Rpl39 | 0.227547698 | 0.035836923 | 0.419258473 | 9.502394957 | 2.88409202 | 0.027000223 | 0.479759698 | -3.779935736 |
| **A_55_P2035043** | Gm2436 | 0.254980262 | 0.040055566 | 0.469904958 | 10.37869997 | 2.882726438 | 0.027049291 | 0.479759698 | -3.780474633 |
| **A_55_P1980636** | Aurka | 0.476370251 | 0.074678972 | 0.878061529 | 12.33644165 | 2.881611431 | 0.027089426 | 0.479759698 | -3.780914832 |
| **A_51_P157270** | Plekhd1 | -0.476738676 | -0.878764782 | -0.074712571 | 8.570042836 | -2.881438275 | 0.027095664 | 0.479759698 | -3.780983208 |
| **A_55_P1999082** | Clec14a | -0.3932385 | -0.724884894 | -0.061592106 | 8.508428821 | -2.881137204 | 0.027106515 | 0.479759698 | -3.781102104 |
| **A_52_P126146** | Riok1 | 0.198496528 | 0.031000524 | 0.365992531 | 9.368739632 | 2.879596617 | 0.027162112 | 0.479759698 | -3.781710691 |
| **A_55_P2167123** | Cyp4b1-ps2 | 0.312445378 | 0.048704709 | 0.576186046 | 11.28984289 | 2.878592509 | 0.027198414 | 0.479759698 | -3.78210752 |
| **A_55_P2068177** | NA | 0.23647435 | 0.036845616 | 0.436103083 | 11.69995986 | 2.878353817 | 0.027207051 | 0.479759698 | -3.782201873 |
| **A_52_P553316** | Snrk | -0.46717147 | -0.861577745 | -0.072765195 | 10.87197347 | -2.878164991 | 0.027213886 | 0.479759698 | -3.782276519 |
| **A_51_P219028** | Wdr53 | 0.333716068 | 0.051971739 | 0.615460398 | 10.73004304 | 2.878094986 | 0.027216421 | 0.479759698 | -3.782304194 |
| **A_55_P2008870** | NA | 0.204845294 | 0.03187893 | 0.377811658 | 13.99468077 | 2.877713307 | 0.027230243 | 0.479759698 | -3.782455097 |
| **A_55_P1972913** | NA | 0.384373487 | 0.059774143 | 0.70897283 | 10.55228062 | 2.877325373 | 0.0272443 | 0.479759698 | -3.782608492 |
| **A_55_P2152035** | Tcf19 | 0.347414049 | 0.053970653 | 0.640857445 | 9.752415148 | 2.876777184 | 0.027264177 | 0.479759698 | -3.78282529 |
| **A_55_P2128821** | Selenok | -0.406436724 | -0.749772738 | -0.06310071 | 13.87549106 | -2.876449611 | 0.027276062 | 0.479759698 | -3.782954857 |
| **A_55_P2033420** | Sppl2a | -0.325189294 | -0.599961119 | -0.050417469 | 10.31481293 | -2.875724185 | 0.027302402 | 0.479759698 | -3.783241842 |
| **A_55_P2109485** | Milr1 | 0.363572725 | 0.056331041 | 0.670814409 | 15.27605264 | 2.87537404 | 0.027315126 | 0.479759698 | -3.783380387 |
| **A_55_P2142580** | NA | -0.360596968 | -0.665594156 | -0.05559978 | 9.759185229 | -2.872826674 | 0.027407882 | 0.479759698 | -3.784388823 |
| **A_55_P2003246** | Asxl2 | 0.323182763 | 0.049774557 | 0.59659097 | 10.08859768 | 2.872234097 | 0.027429508 | 0.479759698 | -3.784623533 |
| **A_51_P213030** | Macrod1 | -0.362791308 | -0.669755678 | -0.055826939 | 9.926010128 | -2.871786138 | 0.027445868 | 0.479759698 | -3.784800994 |
| **A_55_P2029746** | NA | -0.524581648 | -0.9684457 | -0.080717596 | 13.37000004 | -2.871747878 | 0.027447266 | 0.479759698 | -3.784816152 |
| **A_55_P2069012** | Pidd1 | 0.543810686 | 0.083552117 | 1.004069255 | 9.919275888 | 2.87097257 | 0.027475608 | 0.479759698 | -3.78512336 |
| **A_55_P2150108** | Pgm2l1 | 0.393621553 | 0.060465178 | 0.726777929 | 9.328837746 | 2.87087267 | 0.027479262 | 0.479759698 | -3.785162951 |
| **A_55_P1975874** | Bcl2l15 | 0.648868477 | 0.099474625 | 1.19826233 | 9.185954847 | 2.869829328 | 0.027517457 | 0.479759698 | -3.785576506 |
| **A_55_P1968649** | Atp1b2 | 0.421401446 | 0.064384244 | 0.778418648 | 10.67525239 | 2.868072125 | 0.027581914 | 0.479759698 | -3.786273348 |
| **A_55_P2095342** | Rara | -0.400323882 | -0.739658439 | -0.060989325 | 9.128972817 | -2.866596714 | 0.027636159 | 0.479759698 | -3.78685876 |
| **A_51_P178815** | H1f8 | 0.217436857 | 0.033115931 | 0.401757783 | 8.678814265 | 2.866432415 | 0.027642207 | 0.479759698 | -3.786923969 |
| **A_55_P2000728** | Myef2 | 0.267325574 | 0.04070654 | 0.493944609 | 9.196066083 | 2.866337413 | 0.027645705 | 0.479759698 | -3.786961675 |

| | | | | | | | | | |
|---|---|---|---|---|---|---|---|---|---|
| **A_55_P2049577** | Aggf1 | 0.209624929 | 0.031758088 | 0.38749177 | 12.04058714 | 2.863723781 | 0.027742116 | 0.479759698 | -3.787999516 |
| **A_55_P2009111** | NA | 0.203828536 | 0.030879556 | 0.376777515 | 8.621186784 | 2.863717474 | 0.027742349 | 0.479759698 | -3.788002021 |
| **A_55_P2061690** | Atp5d | -0.232353581 | -0.429512406 | -0.035194755 | 13.37174066 | -2.863626262 | 0.02774572 | 0.479759698 | -3.788038257 |
| **A_55_P2032294** | Zfp932 | 0.234004073 | 0.035327157 | 0.432680989 | 9.381615259 | 2.861931237 | 0.027808447 | 0.479759698 | -3.788711841 |
| **A_55_P2031496** | Rufy3 | 0.215228991 | 0.032467828 | 0.397990155 | 8.443022985 | 2.861541473 | 0.027822893 | 0.479759698 | -3.788866783 |
| **A_55_P1999601** | NA | 0.521705098 | 0.078559425 | 0.964850771 | 12.97060621 | 2.860630448 | 0.027856689 | 0.479759698 | -3.78922902 |
| **A_55_P2071799** | Slbp | 0.2820419 | 0.042397357 | 0.521686443 | 9.45351651 | 2.85975795 | 0.027889096 | 0.479759698 | -3.789576042 |
| **A_52_P378084** | NA | 0.450667335 | 0.067611403 | 0.833723267 | 9.281463101 | 2.858755923 | 0.027926365 | 0.479759698 | -3.789974709 |
| **A_51_P151586** | Haspin | 0.293961019 | 0.04408048 | 0.543841557 | 8.926815429 | 2.858515161 | 0.027935328 | 0.479759698 | -3.790070518 |
| **A_55_P2058864** | Mvk | 0.302687327 | 0.045286769 | 0.560087885 | 10.44295379 | 2.857379627 | 0.027977642 | 0.479759698 | -3.7905225 |
| **A_65_P13527** | Osbpl8 | 0.325455761 | 0.048668383 | 0.602243138 | 10.40060054 | 2.857122564 | 0.02798723 | 0.479759698 | -3.790624844 |
| **A_55_P2127688** | 4933434E20Rik | 0.20450269 | 0.03057926 | 0.37842612 | 12.51098807 | 2.857091317 | 0.027988396 | 0.479759698 | -3.790637285 |
| **A_52_P411003** | Dlgap5 | 0.410085094 | 0.061247602 | 0.758922585 | 8.860710562 | 2.856498613 | 0.028010518 | 0.479759698 | -3.790873293 |
| **A_55_P2014511** | Amd2 | 0.368304612 | 0.054998701 | 0.681610524 | 10.68832482 | 2.856417978 | 0.02801353 | 0.479759698 | -3.790905405 |
| **A_52_P662244** | Slc25a47 | -0.628219469 | -1.16263872 | -0.093800217 | 11.83791883 | -2.856356995 | 0.028015807 | 0.479759698 | -3.790929691 |
| **A_51_P239984** | Exo1 | 0.479157863 | 0.071524748 | 0.886790979 | 10.67730861 | 2.856224566 | 0.028020754 | 0.479759698 | -3.790982432 |
| **A_51_P286665** | Rbl1 | 0.373880682 | 0.055733648 | 0.692027717 | 10.06826853 | 2.855540577 | 0.028046317 | 0.479759698 | -3.791254872 |
| **A_52_P110052** | Ackr1 | 0.461964717 | 0.068817819 | 0.855111614 | 11.10757716 | 2.855203999 | 0.028058905 | 0.479759698 | -3.791388958 |
| **A_52_P69043** | Nkap | 0.253741394 | 0.037711557 | 0.469771231 | 9.202788405 | 2.854044711 | 0.028102309 | 0.479759698 | -3.79185091 |
| **A_55_P1968600** | NA | 0.289573137 | 0.042971664 | 0.536174611 | 8.718331524 | 2.853289211 | 0.028130635 | 0.479759698 | -3.792152058 |
| **A_51_P249360** | Suox | 0.265826148 | 0.039421604 | 0.492230693 | 10.66281929 | 2.852960427 | 0.028142971 | 0.479759698 | -3.792283138 |
| **A_51_P273538** | Syce2 | 0.46573857 | 0.069063985 | 0.862413154 | 11.55240201 | 2.852929359 | 0.028144137 | 0.479759698 | -3.792295525 |
| **A_55_P1957419** | Ensa | 0.229186231 | 0.033967702 | 0.424404761 | 9.117547352 | 2.852664354 | 0.028154085 | 0.479759698 | -3.792401189 |
| **A_66_P133993** | Gm5093 | 0.841191807 | 0.1246023 | 1.557781315 | 10.87845619 | 2.852382599 | 0.028164666 | 0.479759698 | -3.792513541 |
| **A_66_P131708** | Znhit1 | -0.229605859 | -0.425281858 | -0.033929861 | 13.34812189 | -2.851206006 | 0.028208897 | 0.479759698 | -3.792982835 |
| **A_51_P257258** | Cpox | 0.593539533 | 0.087668389 | 1.099410676 | 14.54771532 | 2.850971873 | 0.028217708 | 0.479759698 | -3.793076242 |
| **A_55_P2076543** | NA | 0.89077219 | 0.130891889 | 1.650652491 | 11.11367455 | 2.848424107 | 0.028313772 | 0.479759698 | -3.794093153 |
| **A_55_P1958902** | NA | 0.396838057 | 0.058243487 | 0.735432627 | 12.07004073 | 2.847846092 | 0.028335616 | 0.479759698 | -3.794323982 |
| **A_51_P363947** | Cdkn1a | -0.431215073 | -0.799151256 | -0.063278891 | 9.329826247 | -2.847768123 | 0.028338563 | 0.479759698 | -3.876719763 |
| **A_55_P2340593** | 1810019D21Rik | -0.298469566 | -0.553152207 | -0.043786925 | 9.14734938 | -2.847632222 | 0.028343702 | 0.479759698 | -3.794409401 |
| **A_55_P1987645** | Unc13b | -0.463327079 | -0.858712802 | -0.067941356 | 8.942102718 | -2.84740923 | 0.028352137 | 0.479759698 | -3.794498471 |
| **A_55_P2076927** | Ints10 | 0.268618788 | 0.039299978 | 0.497937598 | 9.530462063 | 2.846294907 | 0.028394324 | 0.479759698 | -3.794943664 |
| **A_66_P119418** | Rb1cc1 | 0.258559783 | 0.03780151 | 0.479318055 | 8.795175554 | 2.84594941 | 0.028407418 | 0.479759698 | -3.79508173 |
| **A_51_P351970** | Hells | 0.295652102 | 0.043200365 | 0.54810384 | 8.548490867 | 2.845678268 | 0.028417699 | 0.479759698 | -3.795190094 |
| **A_55_P2028394** | Atg13 | -0.269358475 | -0.499458596 | -0.039258354 | 11.79853774 | -2.844441373 | 0.028464647 | 0.479759698 | -3.795684553 |
| **A_55_P1967683** | Bex4 | 0.252863364 | 0.036849175 | 0.468877552 | 13.56434634 | 2.844374801 | 0.028467176 | 0.479759698 | -3.795711172 |
| **A_52_P195922** | Exosc6 | 0.271195455 | 0.039470767 | 0.502920143 | 11.65795903 | 2.843762279 | 0.028490458 | 0.479759698 | -3.795956114 |
| **A_66_P105689** | Trim34a | -0.25355252 | -0.470209117 | -0.036895923 | 8.52733849 | -2.843670042 | 0.028493966 | 0.479759698 | -3.795993003 |
| **A_55_P2061525** | Cdc27 | 0.291903854 | 0.042461816 | 0.541345891 | 9.073675097 | 2.843500914 | 0.028500399 | 0.479759698 | -3.796060647 |
| **A_55_P2006133** | Tmc6 | 0.669299139 | 0.096968191 | 1.241630087 | 10.90246137 | 2.841556036 | 0.028574488 | 0.479759698 | -3.796838786 |
| **A_55_P2454004** | Smim8 | 0.227905018 | 0.032808036 | 0.423002001 | 10.95693861 | 2.838484488 | 0.028691917 | 0.479759698 | -3.798068733 |
| **A_55_P1967002** | NA | 0.761603816 | 0.109205957 | 1.414001676 | 10.20675835 | 2.836610938 | 0.0287638 | 0.479759698 | -3.798819582 |
| **A_55_P2049025** | NA | 0.280905656 | 0.040248283 | 0.52156303 | 14.70112136 | 2.836249944 | 0.028777672 | 0.479759698 | -3.798964308 |
| **A_51_P145631** | Svip | 0.408486717 | 0.058457126 | 0.758516307 | 10.21158144 | 2.835674436 | 0.028799803 | 0.479759698 | -3.799195073 |
| **A_51_P503654** | Tmem205 | -0.304731728 | -0.565954386 | -0.04350907 | 11.55580306 | -2.834588486 | 0.028841611 | 0.479759698 | -3.799630632 |
| **A_55_P2079020** | Snhg7os | 0.344970482 | 0.049219677 | 0.640721286 | 10.15644516 | 2.834256748 | 0.028854396 | 0.479759698 | -3.799763718 |
| **A_66_P137297** | Otof | 1.139387227 | 0.162514815 | 2.116259639 | 12.98518073 | 2.834109925 | 0.028860057 | 0.479759698 | -3.799822626 |
| **A_55_P2153381** | Ermap | 0.543399214 | 0.07747291 | 1.009325517 | 9.912500704 | 2.8339029 | 0.02886804 | 0.479759698 | -3.799905691 |
| **A_55_P1957413** | Lsp1 | -0.385808741 | -0.716745591 | -0.054871891 | 12.56541711 | -2.832762232 | 0.028912068 | 0.479759698 | -3.800363471 |
| **A_55_P2147886** | Fam118a | 0.389660439 | 0.055417582 | 0.723903296 | 11.25509407 | 2.832744269 | 0.028912762 | 0.479759698 | -3.800370682 |

| | | | | | | | | | |
|---|---|---|---|---|---|---|---|---|---|
| **A_55_P2070174** | NA | 0.482569211 | 0.068625794 | 0.896512628 | 10.01590963 | 2.832708059 | 0.028914161 | 0.479759698 | -3.800385217 |
| **A_55_P2019203** | Spire1 | 0.507970202 | 0.072129044 | 0.94381136 | 9.501938961 | 2.83199959 | 0.028941546 | 0.479759698 | -3.800669636 |
| **A_55_P2032916** | Slc6a20a | 0.608154281 | 0.086327781 | 1.129980781 | 13.96330348 | 2.831853792 | 0.028947185 | 0.479759698 | -3.800728176 |
| **A_55_P2118609** | St6galnac1 | 0.673013936 | 0.095496901 | 1.25053097 | 10.48854085 | 2.831668743 | 0.028954344 | 0.479759698 | -3.80080248 |
| **A_55_P2081925** | Zfp53 | 0.324610416 | 0.045982635 | 0.603238196 | 8.937385252 | 2.830878458 | 0.028984938 | 0.479759698 | -3.80111986 |
| **A_55_P2152427** | Zwilch | 0.484828851 | 0.068662806 | 0.900994896 | 10.550154 | 2.830772722 | 0.028989034 | 0.479759698 | -3.80116233 |
| **A_52_P316405** | Lamtor4 | -0.252671173 | -0.469572923 | -0.035769423 | 12.12220845 | -2.830582575 | 0.028996401 | 0.479759698 | -3.801238708 |
| **A_52_P392509** | Gin1 | 0.253258816 | 0.035824975 | 0.470692657 | 8.64156096 | 2.830222784 | 0.029010347 | 0.479759698 | -3.801383242 |
| **A_55_P2013823** | Gal3st1 | -0.289892527 | -0.538832105 | -0.040952949 | 8.530905326 | -2.8296079 | 0.029034198 | 0.479759698 | -3.801630292 |
| **A_55_P2139886** | Sft2d1 | 0.425507147 | 0.060081286 | 0.790933007 | 8.366137811 | 2.829376679 | 0.029043172 | 0.479759698 | -3.801723206 |
| **A_51_P324934** | Mcm3 | 0.429524471 | 0.060640104 | 0.798408838 | 12.88340731 | 2.829312058 | 0.029045681 | 0.479759698 | -3.801749174 |
| **A_55_P2107182** | Angptl8 | -0.546817589 | -1.016612109 | -0.077023069 | 12.98528875 | -2.828249497 | 0.029086963 | 0.479759698 | -3.802176254 |
| **A_55_P2039416** | Sgk1 | 0.405083025 | 0.05705304 | 0.75311301 | 10.72262732 | 2.82820295 | 0.029088772 | 0.479759698 | -3.802194966 |
| **A_55_P2100550** | Mtif2 | 0.272037788 | 0.038307976 | 0.5057676 | 13.85273131 | 2.828123136 | 0.029091876 | 0.479759698 | -3.802227053 |
| **A_52_P313080** | Hnrnpab | 0.259744396 | 0.036573246 | 0.482915546 | 12.34021988 | 2.828077613 | 0.029093646 | 0.479759698 | -3.802245354 |
| **A_51_P382121** | Ormdl1 | 0.394456604 | 0.055510013 | 0.733403195 | 11.08456836 | 2.827816017 | 0.029103822 | 0.479759698 | -3.802350528 |
| **A_55_P1980421** | Wdr81 | 0.395757494 | 0.05561257 | 0.735902419 | 12.6020136 | 2.82714668 | 0.029129875 | 0.479759698 | -3.802619674 |
| **A_51_P485594** | Tab3 | 0.435856623 | 0.06122037 | 0.810492875 | 10.51209471 | 2.826942908 | 0.029137812 | 0.479759698 | -3.802701624 |
| **A_55_P2095301** | NA | 0.379762484 | 0.05307751 | 0.706447458 | 8.47341926 | 2.824659408 | 0.029226908 | 0.479759698 | -3.80362035 |
| **A_55_P1985495** | Gm12005 | 0.270559608 | 0.03779028 | 0.503328935 | 14.94612341 | 2.824362249 | 0.029238524 | 0.479759698 | -3.803739958 |
| **A_51_P265374** | Vnn3 | -1.188640086 | -2.211435834 | -0.165844338 | 10.55572757 | -2.823869663 | 0.02925779 | 0.479759698 | -3.803938252 |
| **A_51_P265571** | Adm | -0.308517002 | -0.574082504 | -0.0429515 | 8.414562664 | -2.822868412 | 0.029296993 | 0.479759698 | -3.804341413 |
| **A_51_P341465** | Csf2ra | -0.228758144 | -0.425821641 | -0.031694647 | 11.33223861 | -2.820678367 | 0.029382938 | 0.479759698 | -3.805223718 |
| **A_55_P2045886** | Stk3 | 0.313090026 | 0.043290808 | 0.582889245 | 9.662765772 | 2.819757337 | 0.029419162 | 0.479759698 | -3.805594965 |
| **A_55_P1953796** | Fam98c | -0.368073673 | -0.685293891 | -0.050853455 | 9.88062877 | -2.81940253 | 0.02943313 | 0.479759698 | -3.80573801 |
| **A_55_P2037862** | NA | 0.51244395 | 0.070660394 | 0.954227507 | 9.210698522 | 2.818512807 | 0.029468187 | 0.479759698 | -3.806096788 |
| **A_52_P21550** | Gcnt1 | 0.461481407 | 0.063534295 | 0.859428519 | 11.46639255 | 2.817812165 | 0.029495825 | 0.479759698 | -3.806379394 |
| **A_55_P2111163** | S100g | -0.553665617 | -1.031243281 | -0.076087954 | 10.53359291 | -2.816999289 | 0.029527925 | 0.479759698 | -3.806707352 |
| **A_51_P363791** | Ap3m1 | 0.230740295 | 0.031560082 | 0.429920508 | 10.76569339 | 2.814883589 | 0.029611649 | 0.479759698 | -3.807561353 |
| **A_55_P2023488** | NA | 0.329523383 | 0.04505414 | 0.613992626 | 11.11580715 | 2.814712936 | 0.029618413 | 0.479759698 | -3.807630263 |
| **A_52_P475870** | Tmem29 | 0.232876576 | 0.031833741 | 0.433919412 | 9.259422468 | 2.814624057 | 0.029621937 | 0.479759698 | -3.807666154 |
| **A_51_P113072** | Cdc123 | 0.192977943 | 0.026373389 | 0.359582498 | 12.12912751 | 2.814517759 | 0.029626152 | 0.479759698 | -3.807709081 |
| **A_66_P105422** | Lonrf3 | 0.334379636 | 0.045569988 | 0.623189284 | 8.964988368 | 2.81326934 | 0.0296757 | 0.479759698 | -3.808213345 |
| **A_55_P2123234** | NA | 0.483971505 | 0.065915352 | 0.902027659 | 9.785647489 | 2.812991121 | 0.029686754 | 0.479759698 | -3.808325753 |
| **A_66_P138584** | Mnd1 | 0.303806069 | 0.041362764 | 0.566249373 | 9.115087675 | 2.812834215 | 0.02969299 | 0.479759698 | -3.808389151 |
| **A_51_P288026** | Haus6 | 0.294351458 | 0.040022937 | 0.54867998 | 8.990980353 | 2.812252519 | 0.029716122 | 0.479759698 | -3.808624216 |
| **A_51_P381360** | Ndufa13 | -0.206992598 | -0.385905477 | -0.028079718 | 15.77239813 | -2.811230131 | 0.029756825 | 0.479759698 | -3.809037476 |
| **A_55_P2046943** | Bnip3l | 0.246577143 | 0.03341891 | 0.459735375 | 13.31354359 | 2.810825565 | 0.029772948 | 0.479759698 | -3.809201044 |
| **A_55_P2166555** | NA | -0.262217353 | -0.488922953 | -0.035511754 | 11.44254441 | -2.810492118 | 0.029786243 | 0.479759698 | -3.809335875 |
| **A_66_P128440** | NA | 0.530600591 | 0.071738083 | 0.989463099 | 10.20722645 | 2.809754296 | 0.029815685 | 0.479759698 | -3.80963427 |
| **A_55_P1973838** | Slc6a9 | 0.542603407 | 0.073359169 | 1.011847645 | 13.46925263 | 2.809744031 | 0.029816095 | 0.479759698 | -3.809638421 |
| **A_55_P2004168** | NA | 0.472926649 | 0.063817984 | 0.882035314 | 9.987198987 | 2.808913066 | 0.029849292 | 0.479759698 | -3.809974574 |
| **A_55_P2177198** | H2bc3 | 0.272314223 | 0.036650399 | 0.507978047 | 9.086146173 | 2.807763953 | 0.029895263 | 0.479759698 | -3.810439579 |
| **A_55_P2132099** | NA | 0.700614873 | 0.094256963 | 1.306972783 | 8.580648492 | 2.807588753 | 0.029902279 | 0.479759698 | -3.810510491 |
| **A_51_P116616** | Rbm14 | 0.296862089 | 0.039913924 | 0.553810254 | 12.13192102 | 2.807323158 | 0.029912918 | 0.479759698 | -3.834656043 |
| **A_55_P2118090** | NA | 0.392325598 | 0.052734553 | 0.731916644 | 10.95676801 | 2.807201615 | 0.029917788 | 0.479759698 | -3.810667201 |
| **A_51_P470851** | Cplx3 | 0.618305181 | 0.082955883 | 1.153654479 | 9.774860353 | 2.806395255 | 0.029950118 | 0.479759698 | -3.810993672 |
| **A_51_P115666** | Fam83h | -0.383271343 | -0.715180258 | -0.051362429 | 9.554949224 | -2.805889872 | 0.0299704 | 0.479759698 | -3.811198331 |
| **A_55_P2049313** | NA | 0.564500901 | 0.075383435 | 1.053618368 | 9.093103853 | 2.804365765 | 0.030031655 | 0.479759698 | -3.811815735 |
| **A_66_P133269** | Hesx1 | 0.45989255 | 0.0610864 | 0.858698701 | 8.618248171 | 2.802061841 | 0.030124506 | 0.479759698 | -3.812749627 |

| | | | | | | | | | |
|---|---|---|---|---|---|---|---|---|---|
| **A_52_P630632** | Rcbtb2 | -0.226709741 | -0.423328742 | -0.03009074 | 10.09837822 | -2.801740347 | 0.030137487 | 0.479759698 | -3.81288 |
| **A_55_P2036978** | Mrps14 | 0.214276947 | 0.028412299 | 0.400141596 | 12.60553683 | 2.801314342 | 0.030154697 | 0.479759698 | -3.813052776 |
| **A_51_P443475** | Bccip | 0.253134861 | 0.033480807 | 0.472788915 | 12.07554862 | 2.800244318 | 0.030197971 | 0.479759698 | -3.813486856 |
| **A_55_P2065799** | Naa50 | 0.419698928 | 0.055405366 | 0.783992489 | 10.23968213 | 2.799429613 | 0.030230964 | 0.479759698 | -3.813817463 |
| **A_55_P2046109** | NA | 0.362084934 | 0.047692199 | 0.676477669 | 8.424024298 | 2.798473133 | 0.030269747 | 0.479759698 | -3.814205714 |
| **A_51_P482029** | Mcoln1 | -0.231698321 | -0.432928242 | -0.030468399 | 11.42049891 | -2.797779713 | 0.030297897 | 0.479759698 | -3.814487261 |
| **A_55_P1971614** | NA | 0.389516097 | 0.051078221 | 0.727953973 | 10.67326651 | 2.796595397 | 0.030346041 | 0.479759698 | -3.814968274 |
| **A_55_P2038046** | NA | 0.373531946 | 0.048965085 | 0.698098808 | 10.39049412 | 2.796448103 | 0.030352034 | 0.479759698 | -3.815028111 |
| **A_55_P2107155** | Rasd2 | 0.262394399 | 0.034370177 | 0.490418621 | 8.950421797 | 2.796126193 | 0.030365137 | 0.479759698 | -3.815158894 |
| **A_55_P2098880** | Galnt10 | 0.374284791 | 0.048878404 | 0.699691178 | 11.95672629 | 2.794855099 | 0.030416934 | 0.479759698 | -3.815675439 |
| **A_55_P2051254** | Pvt1 | 0.325651728 | 0.042476634 | 0.608826822 | 8.95603047 | 2.79435464 | 0.030437353 | 0.479759698 | -3.815878874 |
| **A_52_P267391** | Trim12a | -0.251543281 | -0.470331138 | -0.032755423 | 9.584862762 | -2.793654498 | 0.030465945 | 0.479759698 | -3.816163535 |
| **A_51_P383194** | Pde9a | -0.215313179 | -0.402595116 | -0.028031243 | 8.47061441 | -2.793559407 | 0.03046983 | 0.479759698 | -3.816202202 |
| **A_55_P1963309** | G3bp1 | 0.368659344 | 0.047991373 | 0.689327316 | 12.86952296 | 2.79352686 | 0.03047116 | 0.479759698 | -3.816215437 |
| **A_51_P154842** | Oas1f | -0.319143676 | -0.596823863 | -0.041463489 | 9.429899495 | -2.792701629 | 0.030504903 | 0.479759698 | -3.816551054 |
| **A_52_P93902** | Alg5 | -0.232936572 | -0.435629854 | -0.03024329 | 10.78073784 | -2.792424975 | 0.030516224 | 0.479759698 | -3.816663588 |
| **A_52_P398211** | NA | 0.483377263 | 0.06258943 | 0.904165095 | 8.252676955 | 2.79129819 | 0.03056238 | 0.479759698 | -3.817122035 |
| **A_55_P2056729** | Igfbp2 | -1.364598585 | -2.552720377 | -0.176476793 | 10.77517251 | -2.790789909 | 0.030583225 | 0.479759698 | -3.817328891 |
| **A_52_P77227** | Rnf212 | 0.380386751 | 0.049189445 | 0.711584057 | 8.849675676 | 2.790755413 | 0.03058464 | 0.479759698 | -3.817342931 |
| **A_51_P474169** | Proser2 | -0.43168013 | -0.807593878 | -0.055766383 | 10.31850717 | -2.790339453 | 0.030601711 | 0.479759698 | -3.817512242 |
| **A_55_P2090364** | NA | 0.313525614 | 0.040473059 | 0.586578169 | 14.38560735 | 2.790037053 | 0.030614128 | 0.479759698 | -3.817635345 |
| **A_55_P2121185** | Omp | 0.372778544 | 0.048064272 | 0.697492816 | 8.500295579 | 2.789540813 | 0.030634516 | 0.479759698 | -3.817837383 |
| **A_52_P286098** | Epb41l4b | -0.497007873 | -0.930125142 | -0.063890604 | 11.262515 | -2.788309368 | 0.030685172 | 0.479759698 | -3.818338894 |
| **A_66_P116311** | Kif5b | 0.24645638 | 0.031625652 | 0.461287109 | 8.850163899 | 2.787576884 | 0.030715345 | 0.479759698 | -3.818637297 |
| **A_55_P2347976** | Mir17hg | 0.340767945 | 0.043698896 | 0.637836993 | 9.259001545 | 2.787305158 | 0.030726547 | 0.479759698 | -3.818748012 |
| **A_51_P371091** | Rcsd1 | -0.403606416 | -0.755480724 | -0.051732108 | 10.92674325 | -2.787107324 | 0.030734705 | 0.479759698 | -3.818828626 |
| **A_55_P2206461** | A930006K02Rik | 0.603475105 | 0.077211486 | 1.129738724 | 9.81086976 | 2.78637266 | 0.03076502 | 0.479759698 | -3.819128035 |
| **A_66_P131110** | Zfp473 | 0.310340866 | 0.039688204 | 0.580993528 | 8.399641045 | 2.786184379 | 0.030772795 | 0.479759698 | -3.819204779 |
| **A_55_P2076592** | NA | 0.421996714 | 0.053825376 | 0.790168052 | 8.4416574 | 2.785109551 | 0.030817217 | 0.479759698 | -3.819642976 |
| **A_51_P196972** | Slc4a1 | 0.424742274 | 0.054155118 | 0.795329431 | 16.18485527 | 2.78495584 | 0.030823575 | 0.479759698 | -3.819705656 |
| **A_55_P2114288** | Mrpl24 | -0.200389162 | -0.375240423 | -0.025537901 | 12.85108035 | -2.784765618 | 0.030831446 | 0.479759698 | -3.819783227 |
| **A_52_P638895** | Vegfa | -0.710953503 | -1.331358036 | -0.09054897 | 10.32373286 | -2.784514124 | 0.030841855 | 0.479759698 | -3.819885793 |
| **A_55_P2004833** | NA | 0.29067644 | 0.037016677 | 0.544336203 | 13.90970834 | 2.784462923 | 0.030843974 | 0.479759698 | -3.819906675 |
| **A_55_P2071191** | Prss8 | -0.554586983 | -1.038565397 | -0.07060857 | 9.823597626 | -2.784369514 | 0.030847842 | 0.479759698 | -3.819944772 |
| **A_52_P453624** | Arl6ip6 | 0.217616513 | 0.027693123 | 0.407539902 | 8.956830769 | 2.78417534 | 0.030855883 | 0.479759698 | -3.82002397 |
| **A_55_P1952312** | NA | 0.224985267 | 0.02861087 | 0.421359664 | 13.51660759 | 2.783892129 | 0.030867615 | 0.479759698 | -3.820139494 |
| **A_51_P179864** | Prkx | -0.334421119 | -0.626324905 | -0.042517334 | 9.005157382 | -2.78379441 | 0.030871664 | 0.479759698 | -3.820179356 |
| **A_66_P137495** | Btaf1 | 0.43201574 | 0.054735755 | 0.809295725 | 10.06329639 | 2.782396338 | 0.030929657 | 0.479759698 | -3.820749813 |
| **A_55_P2366273** | Amn1 | 0.224251951 | 0.028372985 | 0.420130918 | 8.489372499 | 2.78183659 | 0.030952908 | 0.479759698 | -3.82097828 |
| **A_52_P460929** | BC048507 | 0.391804633 | 0.049550547 | 0.73405872 | 12.01701985 | 2.781660443 | 0.030960229 | 0.479759698 | -3.821050185 |
| **A_51_P162471** | Dnajc9 | 0.344788885 | 0.043544926 | 0.646032844 | 13.20762009 | 2.781109571 | 0.030983136 | 0.479759698 | -3.821275083 |
| **A_65_P06453** | Nup35 | 0.309274747 | 0.039012859 | 0.579536636 | 11.07833971 | 2.78062769 | 0.031003189 | 0.479759698 | -3.821471848 |
| **A_51_P375969** | Ces1d | -0.812803352 | -1.523121777 | -0.102484926 | 11.04112982 | -2.780453232 | 0.031010452 | 0.479759698 | -3.821543092 |
| **A_55_P1971006** | Urah | -0.413930778 | -0.77570103 | -0.052160527 | 13.49987526 | -2.780212923 | 0.03102046 | 0.479759698 | -3.821641234 |
| **A_55_P2099390** | Ranbp9 | 0.362794947 | 0.045571827 | 0.680018067 | 8.811591831 | 2.778942661 | 0.031073418 | 0.479759698 | -3.822160135 |
| **A_51_P451151** | Ube2c | 0.354207959 | 0.044467008 | 0.663948909 | 13.71386086 | 2.778707785 | 0.031083221 | 0.479759698 | -3.822256105 |
| **A_51_P424854** | Ahsg | 0.379254897 | 0.04758708 | 0.710922714 | 17.46160442 | 2.778504178 | 0.031091721 | 0.479759698 | -3.822339305 |
| **A_55_P2210575** | A830073O21Rik | 0.436268825 | 0.054618903 | 0.817918746 | 12.72563078 | 2.77761591 | 0.031128835 | 0.479759698 | -3.822702341 |
| **A_52_P87210** | Dcun1d5 | 0.240609064 | 0.030119908 | 0.451098221 | 11.40991751 | 2.777572766 | 0.031130638 | 0.479759698 | -3.822719977 |
| **A_55_P1953231** | Map2k5 | -0.265131084 | -0.497081637 | -0.03318053 | 10.93496952 | -2.7774639 | 0.031135191 | 0.479759698 | -3.822764478 |

| | | | | | | | | |
|---|---|---|---|---|---|---|---|---|
| **A_52_P482925** | Serbp1 | 0.252081362 | 0.031518202 | 0.472644522 | 13.91291256 | 2.77709637 | 0.031150564 | 0.479759698 | -3.822914726 |
| **A_51_P470724** | Eif2s2 | 0.34871282 | 0.043551668 | 0.653873973 | 10.16014983 | 2.776654583 | 0.031169054 | 0.479759698 | -3.823095355 |
| **A_55_P1969078** | Adar | -0.211504645 | -0.396595748 | -0.026413541 | 10.3177475 | -2.776627072 | 0.031170206 | 0.479759698 | -3.823106604 |
| **A_55_P2103422** | Psmc2 | 0.253199783 | 0.031574832 | 0.474824733 | 13.64606433 | 2.776053722 | 0.03119422 | 0.479759698 | -3.823341064 |
| **A_51_P303180** | Fam114a1 | -0.251418963 | -0.471493951 | -0.031343974 | 9.833791342 | -2.775942921 | 0.031198864 | 0.479759698 | -3.823386379 |
| **A_55_P1960566** | Ybx3 | 0.428092304 | 0.053353762 | 0.802830846 | 14.45314031 | 2.775826047 | 0.031203762 | 0.479759698 | -3.823434179 |
| **A_52_P671812** | NA | 0.404810194 | 0.050418847 | 0.759201542 | 12.58133917 | 2.775565749 | 0.031214675 | 0.479759698 | -3.823540646 |
| **A_52_P646034** | Nfatc2ip | 0.313352204 | 0.039003959 | 0.587700449 | 9.611736784 | 2.775324414 | 0.031224796 | 0.479759698 | -3.823639363 |
| **A_55_P2107785** | Gm13105 | -0.278214671 | -0.521881786 | -0.034547556 | 9.774575096 | -2.77438224 | 0.031264343 | 0.479759698 | -3.824024834 |
| **A_52_P121** | Osgepl1 | 0.410198978 | 0.050929237 | 0.769468719 | 10.07678158 | 2.774323638 | 0.031266805 | 0.479759698 | -3.824048813 |
| **A_55_P2042918** | Trim37 | 0.265922501 | 0.032981523 | 0.498863479 | 8.552679746 | 2.773910096 | 0.031284181 | 0.479759698 | -3.824218046 |
| **A_55_P1965554** | Orc3 | 0.256090286 | 0.031721798 | 0.480458773 | 11.29197004 | 2.773412274 | 0.031305113 | 0.479759698 | -3.824421797 |
| **A_51_P462299** | Aida | 0.225740259 | 0.027936923 | 0.423543594 | 9.252603273 | 2.773055727 | 0.031320113 | 0.479759698 | -3.824567748 |
| **A_51_P146303** | Mvb12a | -0.28365799 | -0.532296252 | -0.035019728 | 11.27319233 | -2.77210866 | 0.031359996 | 0.479759698 | -3.824955506 |
| **A_51_P442964** | Knl1 | 0.402785281 | 0.049533198 | 0.756037364 | 8.946811808 | 2.770588642 | 0.03142412 | 0.479759698 | -3.825578098 |
| **A_52_P467266** | Mtf1 | 0.455967875 | 0.056025747 | 0.855910002 | 9.520073816 | 2.77025841 | 0.031438069 | 0.479759698 | -3.825713399 |
| **A_51_P178828** | Mbl2 | -1.276354846 | -2.396025838 | -0.156683854 | 12.34368058 | -2.769900661 | 0.031453189 | 0.479759698 | -3.825859992 |
| **A_55_P1983819** | Srsf7 | 0.275799996 | 0.033841886 | 0.517758106 | 12.79862086 | 2.769728885 | 0.031460451 | 0.479759698 | -3.825930385 |
| **A_51_P337089** | Gins1 | 0.489102848 | 0.059830952 | 0.918374744 | 11.16033373 | 2.768540774 | 0.031510733 | 0.479759698 | -3.82641738 |
| **A_52_P134083** | Cyhr1 | -0.286719383 | -0.538398939 | -0.035039827 | 9.966929939 | -2.768167106 | 0.031526564 | 0.479759698 | -3.826570581 |
| **A_51_P508959** | Prr15 | -0.437436435 | -0.821434014 | -0.053438856 | 9.205361574 | -2.768022754 | 0.031532682 | 0.479759698 | -3.826629769 |
| **A_66_P103398** | Ogfrl1 | 0.367337642 | 0.044692466 | 0.689982819 | 9.793487978 | 2.766453954 | 0.031599256 | 0.479759698 | -3.8272732 |
| **A_51_P136888** | Rb1 | 0.486566878 | 0.059189302 | 0.913944453 | 11.04391513 | 2.766393782 | 0.031601812 | 0.479759698 | -3.849633558 |
| **A_55_P1969982** | Styx | 0.330123868 | 0.039827107 | 0.620420629 | 8.867480381 | 2.763235656 | 0.031736297 | 0.479759698 | -3.828594183 |
| **A_51_P172251** | Ifrd2 | 0.428308651 | 0.051663032 | 0.804954271 | 13.40238721 | 2.763166883 | 0.031739232 | 0.479759698 | -3.828622426 |
| **A_55_P2065567** | Chac2 | 0.53282957 | 0.064165525 | 1.001493615 | 12.03331433 | 2.762548244 | 0.03176565 | 0.479759698 | -3.828876516 |
| **A_55_P1955691** | Alg6 | 0.258389208 | 0.030991574 | 0.485786842 | 9.783687667 | 2.761033137 | 0.03183045 | 0.479759698 | -3.829499023 |
| **A_55_P2076580** | Acaa1a | -0.255617547 | -0.480644032 | -0.030591062 | 13.84188332 | -2.760197868 | 0.031866234 | 0.479759698 | -3.829842337 |
| **A_55_P2017082** | Hnrnpa2b1 | 0.287334251 | 0.034319445 | 0.540349058 | 12.94223693 | 2.759463476 | 0.031897732 | 0.479759698 | -3.830144265 |
| **A_51_P355040** | NA | 0.22865451 | 0.027303231 | 0.43000579 | 8.538172627 | 2.759361294 | 0.031902117 | 0.479759698 | -3.830186281 |
| **A_55_P2001048** | Rom1 | 0.223222778 | 0.026580295 | 0.419865261 | 9.755435921 | 2.758318089 | 0.031946923 | 0.479759698 | -3.830615307 |
| **A_52_P335609** | Idnk | -0.241676836 | -0.454595527 | -0.028758145 | 10.22968512 | -2.758064557 | 0.031957823 | 0.479759698 | -3.830719596 |
| **A_55_P1953093** | Pabpc4 | 0.433705549 | 0.051488412 | 0.815922687 | 13.15232242 | 2.757198365 | 0.031995091 | 0.479759698 | -3.831075965 |
| **A_55_P2052172** | NA | 0.209298009 | 0.024734842 | 0.393861177 | 11.25988227 | 2.755517993 | 0.032067521 | 0.479759698 | -3.831767586 |
| **A_52_P478745** | Paxbp1 | 0.259356839 | 0.030604973 | 0.488108706 | 8.652735355 | 2.754966023 | 0.032091352 | 0.479759698 | -3.831994852 |
| **A_55_P2063336** | Tedc1 | 0.305398737 | 0.036010103 | 0.574787371 | 9.963800079 | 2.754680009 | 0.032103707 | 0.479759698 | -3.832112631 |
| **A_51_P408946** | Ccne1 | 0.54769405 | 0.064392029 | 1.030996072 | 12.66691358 | 2.75361108 | 0.032149929 | 0.479759698 | -3.832552904 |
| **A_55_P2059105** | Gm9920 | 0.365040253 | 0.042837247 | 0.687243258 | 9.5733309 | 2.752924866 | 0.032179639 | 0.479759698 | -3.832835623 |
| **A_55_P2061170** | Tbc1d2 | -0.282183206 | -0.531282676 | -0.033083736 | 8.763536731 | -2.752590133 | 0.032194142 | 0.479759698 | -3.832973556 |
| **A_55_P2058201** | Scgb2b20 | 0.863874925 | 0.101105318 | 1.626644532 | 10.62012497 | 2.751950877 | 0.032221859 | 0.479759698 | -3.833237013 |
| **A_55_P1954081** | Ccdc115 | 0.184529656 | 0.021577723 | 0.347481589 | 11.14014702 | 2.751628789 | 0.032235834 | 0.479759698 | -3.833369776 |
| **A_55_P2016652** | Cdk1 | 0.649195507 | 0.075783317 | 1.222607697 | 8.466277694 | 2.751007493 | 0.032262809 | 0.479759698 | -3.83362591 |
| **A_52_P157880** | Gm1947 | 0.520478111 | 0.060716427 | 0.980239795 | 10.16518708 | 2.75076116 | 0.03227351 | 0.479759698 | -3.833727477 |
| **A_55_P2113920** | NA | 0.383171154 | 0.044631786 | 0.721710523 | 8.712267027 | 2.750216075 | 0.032297205 | 0.479759698 | -3.833952253 |
| **A_52_P356119** | Hus1 | 0.227501382 | 0.026448293 | 0.428554472 | 8.503646472 | 2.74951743 | 0.032327601 | 0.479759698 | -3.834240409 |
| **A_55_P2214487** | E330013P08Rik | 0.338168664 | 0.039188853 | 0.637148474 | 8.9010157 | 2.748366765 | 0.032377731 | 0.479759698 | -3.834715143 |
| **A_52_P660032** | Cpsf2 | 0.341161564 | 0.039523459 | 0.64279967 | 10.05731269 | 2.748255345 | 0.032382589 | 0.479759698 | -3.834761122 |
| **A_55_P2387284** | NA | 0.478668004 | 0.055435699 | 0.901900308 | 13.13509653 | 2.748139525 | 0.03238764 | 0.479759698 | -3.834808917 |
| **A_55_P2001143** | Ehmt2 | -0.252840192 | -0.476404461 | -0.029275922 | 12.77495588 | -2.748064393 | 0.032390918 | 0.479759698 | -3.834839923 |
| **A_52_P181759** | Eif5a2 | 0.262315554 | 0.032179 | 0.492452109 | 8.843372659 | 2.899885182 | 0.032427663 | 0.479759698 | -3.942041288 |

| | | | | | | | | | |
|---|---|---|---|---|---|---|---|---|---|
| **A_51_P193146** | Ms4a6c | -0.213733832 | -0.402813208 | -0.024654456 | 8.459613397 | -2.746706787 | 0.032450195 | 0.479759698 | -3.835400316 |
| **A_55_P2135031** | NA | 0.387083837 | 0.044520048 | 0.729647627 | 8.154896011 | 2.745660103 | 0.032495975 | 0.479759698 | -3.835832533 |
| **A_55_P2013143** | Esrp2 | -0.292052081 | -0.550579989 | -0.033524173 | 9.63525761 | -2.744960267 | 0.032526624 | 0.479759698 | -3.836121604 |
| **A_55_P2092085** | Tmsb10 | -0.43472985 | -0.8196867 | -0.049772999 | 14.58165046 | -2.744041009 | 0.032566928 | 0.479759698 | -3.836501408 |
| **A_52_P631499** | NA | 0.515629436 | 0.05897431 | 0.972284562 | 11.35475513 | 2.743674295 | 0.032583022 | 0.479759698 | -3.836652952 |
| **A_55_P2186395** | NA | 0.24602471 | 0.02813623 | 0.46391319 | 9.273676842 | 2.743643297 | 0.032584382 | 0.479759698 | -3.836665762 |
| **A_55_P1970846** | Pemt | -0.936663384 | -1.766248862 | -0.107077906 | 11.22247058 | -2.743503959 | 0.0325905 | 0.479759698 | -3.836723349 |
| **A_51_P408071** | Kntc1 | 0.490744343 | 0.05605987 | 0.925428816 | 10.15698679 | 2.743243528 | 0.032601937 | 0.479759698 | -3.836830989 |
| **A_65_P08521** | Ssrp1 | 0.567996055 | 0.064864404 | 1.071127707 | 10.03533075 | 2.743133028 | 0.032606791 | 0.479759698 | -3.836876662 |
| **A_55_P2033393** | Slc25a21 | 0.508151835 | 0.057876314 | 0.958427357 | 9.169023356 | 2.742195096 | 0.032648022 | 0.479759698 | -3.837264409 |
| **A_66_P130908** | Cyb5r4 | 0.355321505 | 0.040436706 | 0.670206303 | 9.496908626 | 2.741908683 | 0.032660624 | 0.479759698 | -3.837382838 |
| **A_55_P2158653** | Cldn7 | -0.888690567 | -1.676541647 | -0.100839487 | 9.796978491 | -2.740877494 | 0.032706039 | 0.479759698 | -3.837809312 |
| **A_55_P2027361** | Irak1bp1 | 0.638043978 | 0.072218216 | 1.203869741 | 14.24540496 | 2.7400033 | 0.032744592 | 0.479759698 | -3.838170968 |
| **A_52_P529570** | Nsl1 | 0.406001188 | 0.045923268 | 0.766079107 | 10.44014729 | 2.739769345 | 0.032754918 | 0.479759698 | -3.838267773 |
| **A_55_P2051596** | NA | -0.336921421 | -0.635759087 | -0.038083755 | 10.050832 | -2.739532628 | 0.032765369 | 0.479759698 | -3.838365728 |
| **A_55_P2050773** | LOC100048452 | 0.456109421 | 0.051518614 | 0.860700229 | 8.572099298 | 2.739278703 | 0.032776584 | 0.479759698 | -3.838470812 |
| **A_51_P134142** | Cyp2c70 | -1.213300326 | -2.290079296 | -0.136521357 | 13.45083915 | -2.737946379 | 0.032835496 | 0.479759698 | -3.839022319 |
| **A_55_P1980883** | Ephb6 | 0.640163153 | 0.071690855 | 1.208635451 | 13.92347827 | 2.736305326 | 0.032908213 | 0.479759698 | -3.839701947 |
| **A_51_P389714** | Phlpp2 | 0.402085793 | 0.045027354 | 0.759144232 | 10.10473092 | 2.736293099 | 0.032908756 | 0.479759698 | -3.839707012 |
| **A_55_P2125774** | Ola1 | 0.283241344 | 0.031648791 | 0.534833898 | 11.43920274 | 2.735533596 | 0.032942469 | 0.479759698 | -3.840021676 |
| **A_55_P2053798** | NA | 0.3977218 | 0.044409863 | 0.751033738 | 14.90991033 | 2.735295646 | 0.032953039 | 0.479759698 | -3.840120275 |
| **A_55_P2020612** | Arl4d | -0.366655217 | -0.69242655 | -0.040883884 | 8.617106095 | -2.734816509 | 0.032974334 | 0.479759698 | -3.840318837 |
| **A_55_P2053516** | Cacfd1 | -0.364451609 | -0.688284459 | -0.040618759 | 11.79196084 | -2.734652571 | 0.032981623 | 0.479759698 | -3.840386782 |
| **A_55_P1982818** | Odc1 | 0.466215409 | 0.051775119 | 0.8806557 | 10.86693503 | 2.733429285 | 0.03303607 | 0.479759698 | -3.840893895 |
| **A_51_P289881** | Txndc12 | -0.250631712 | -0.473532587 | -0.027730837 | 9.940294978 | -2.732168224 | 0.033092299 | 0.479759698 | -3.841416877 |
| **A_55_P2358002** | 5-Mar | 0.325479246 | 0.035996277 | 0.614962214 | 8.562132562 | 2.732017468 | 0.033099027 | 0.479759698 | -3.841479411 |
| **A_55_P2040936** | Eng | -0.28358309 | -0.535826063 | -0.031340118 | 9.885301284 | -2.73177197 | 0.033109988 | 0.479759698 | -3.841581252 |
| **A_52_P491544** | Nedd9 | -0.710436661 | -1.342597103 | -0.078276219 | 9.068791399 | -2.730745568 | 0.033155855 | 0.479759698 | -3.842007126 |
| **A_55_P1993636** | Rab9 | -0.263424506 | -0.497893525 | -0.028955487 | 10.65583877 | -2.729944999 | 0.033191677 | 0.479759698 | -3.842339394 |
| **A_55_P2006525** | Adamtsl4 | -0.317158036 | -0.599534114 | -0.034781959 | 8.453785362 | -2.729172635 | 0.033226276 | 0.479759698 | -3.842660036 |
| **A_51_P388374** | Ppil4 | 0.23368685 | 0.025421378 | 0.441952321 | 10.50752503 | 2.726466648 | 0.033347798 | 0.479759698 | -3.843784034 |
| **A_55_P1973779** | Pde4c | 0.440555676 | 0.047850318 | 0.833261034 | 14.53234928 | 2.725945466 | 0.033371258 | 0.479759698 | -3.844000631 |
| **A_55_P1976237** | Dnajb3 | 0.420118386 | 0.045589813 | 0.794646959 | 10.04784318 | 2.725648956 | 0.033384612 | 0.479759698 | -3.844123874 |
| **A_66_P110161** | Eppk1 | -0.245874271 | -0.465089491 | -0.02665905 | 8.216008429 | -2.725370618 | 0.033397154 | 0.479759698 | -3.844239573 |
| **A_52_P620748** | Ncl | 0.36914326 | 0.03999845 | 0.698288069 | 9.227543554 | 2.725154449 | 0.033406897 | 0.479759698 | -3.844329437 |
| **A_51_P316553** | Kdr | -0.483846809 | -0.915275335 | -0.052418283 | 11.25889166 | -2.72509855 | 0.033409417 | 0.479759698 | -3.881039816 |
| **A_51_P308298** | Myl9 | -0.764227951 | -1.445677409 | -0.082778493 | 13.14777119 | -2.725037329 | 0.033412177 | 0.479759698 | -3.844378128 |
| **A_52_P73126** | Epb42 | 0.53339787 | 0.057669848 | 1.009125893 | 11.62494943 | 2.724430491 | 0.033439551 | 0.479759698 | -3.844630441 |
| **A_55_P2227321** | Ptprd | -0.190717979 | -0.360823939 | -0.02061202 | 8.202381678 | -2.724302296 | 0.033445337 | 0.479759698 | -3.844683749 |
| **A_55_P1975922** | Olfr629 | 0.499516868 | 0.053957871 | 0.945075864 | 9.027897521 | 2.724131888 | 0.033453029 | 0.479759698 | -3.844754613 |
| **A_55_P1961458** | Speer1 | 0.555653262 | 0.060015747 | 1.051290777 | 9.851690668 | 2.724099 | 0.033454514 | 0.479759698 | -3.84476829 |
| **A_52_P80593** | NA | 0.323772804 | 0.034837234 | 0.612708373 | 12.20497591 | 2.722842653 | 0.033511288 | 0.479759698 | -3.845290868 |
| **A_55_P1952877** | Snrpf | 0.287952729 | 0.030916502 | 0.544988957 | 12.93290974 | 2.72213745 | 0.033543202 | 0.479759698 | -3.845584289 |
| **A_55_P2060642** | Spidr | 0.226149131 | 0.024271457 | 0.428026806 | 10.24099012 | 2.722010646 | 0.033548943 | 0.479759698 | -3.845637057 |
| **A_51_P155142** | Cdca8 | 0.363463243 | 0.038992885 | 0.687933602 | 13.06896658 | 2.721878018 | 0.03355495 | 0.479759698 | -3.84569225 |
| **A_55_P2029420** | 2610318N02Rik | 0.382521803 | 0.041002736 | 0.72404087 | 10.86110266 | 2.721600821 | 0.033567508 | 0.479759698 | -3.845807614 |
| **A_55_P2043668** | Zfp617 | 0.203680256 | 0.021701572 | 0.38565894 | 9.343436941 | 2.719641139 | 0.03365643 | 0.479759698 | -3.846623483 |
| **A_51_P363630** | Snrpe | 0.221003389 | 0.023527149 | 0.418479629 | 14.47747214 | 2.719363551 | 0.033669046 | 0.479759698 | -3.846739091 |
| **A_55_P1957911** | Tom1 | 0.330252861 | 0.035149642 | 0.625356081 | 9.175403829 | 2.719291908 | 0.033672303 | 0.479759698 | -3.846768931 |
| **A_52_P423247** | Pde4b | -0.361864273 | -0.685217301 | -0.038511245 | 7.987356043 | -2.719267699 | 0.033673404 | 0.479759698 | -3.846779014 |

| | | | | | | | | | |
|---|---|---|---|---|---|---|---|---|---|
| **A_51_P202623** | Mterf2 | 0.314034465 | 0.033332973 | 0.594735958 | 9.022864445 | 2.718415149 | 0.033712188 | 0.479759698 | -3.847134154 |
| **A_55_P2065829** | Cfap300 | 0.614836683 | 0.06524235 | 1.164431016 | 13.0536637 | 2.718320835 | 0.033716481 | 0.479759698 | -3.847173448 |
| **A_51_P225948** | Snx33 | -0.270503012 | -0.512313851 | -0.028692173 | 8.729219403 | -2.718188245 | 0.033722518 | 0.479759698 | -3.847228691 |
| **A_55_P2173224** | Kif2a | 0.308586754 | 0.032703786 | 0.584469722 | 9.074444943 | 2.717913181 | 0.033735046 | 0.479759698 | -3.847343301 |
| **A_55_P2166123** | Atp9a | 0.448244996 | 0.04711712 | 0.849372872 | 9.425499453 | 2.715287341 | 0.033854886 | 0.479759698 | -3.848437906 |
| **A_55_P2126695** | NA | 0.566732699 | 0.059172262 | 1.074293137 | 8.353588012 | 2.713149328 | 0.033952798 | 0.479759698 | -3.84932983 |
| **A_52_P83913** | Ssx2ip | 0.427997456 | 0.044577731 | 0.811417181 | 8.820864839 | 2.712376201 | 0.033988278 | 0.479759698 | -3.849652507 |
| **A_55_P2183854** | Sctr | 0.57359624 | 0.05972677 | 1.087465709 | 9.446290259 | 2.71229341 | 0.03399208 | 0.479759698 | -3.849687066 |
| **A_66_P134542** | Anln | 0.381539244 | 0.039720085 | 0.723358403 | 11.79111772 | 2.712226795 | 0.033995139 | 0.479759698 | -3.849714873 |
| **A_52_P350111** | Cyth1 | -0.247914992 | -0.470033467 | -0.025796518 | 9.770223494 | -2.712072516 | 0.034002225 | 0.479759698 | -3.849779277 |
| **A_55_P2280821** | Dlc1 | -0.437780477 | -0.830043429 | -0.045517524 | 8.908726823 | -2.711828907 | 0.034013418 | 0.479759698 | -3.849880977 |
| **A_55_P1993293** | Ndrg3 | 0.365910107 | 0.038037378 | 0.693782835 | 10.80324528 | 2.711766551 | 0.034016283 | 0.479759698 | -3.84990701 |
| **A_51_P498831** | Derl2 | -0.312871452 | -0.593243483 | -0.032499422 | 13.66720081 | -2.711530164 | 0.034027149 | 0.479759698 | -3.850005705 |
| **A_51_P136460** | Zmat2 | 0.189961887 | 0.019727335 | 0.360196439 | 12.24508168 | 2.7114522 | 0.034030733 | 0.479759698 | -3.850038257 |
| **A_51_P262515** | Phf11a | -0.240576882 | -0.456179574 | -0.02497419 | 8.306385754 | -2.711333267 | 0.034036202 | 0.479759698 | -3.850087918 |
| **A_52_P355084** | Metrnl | -0.414684485 | -0.786479344 | -0.042889626 | 9.547531833 | -2.710176629 | 0.034089436 | 0.479759698 | -3.850570966 |
| **A_51_P127756** | Ddx20 | 0.219850829 | 0.022644828 | 0.41705683 | 10.21684308 | 2.708888758 | 0.034148813 | 0.479759698 | -3.85110903 |
| **A_52_P387009** | Egln3 | -0.684766218 | -1.299041645 | -0.070490791 | 10.16863655 | -2.708709142 | 0.034157103 | 0.479759698 | -3.85118409 |
| **A_51_P240801** | Tmem173 | -0.228725051 | -0.4339051 | -0.023545002 | 9.829247291 | -2.708705525 | 0.03415727 | 0.479759698 | -3.851185602 |
| **A_55_P2009762** | Mapkapk3 | -0.377786089 | -0.716885975 | -0.038686203 | 10.53740813 | -2.707082599 | 0.034232272 | 0.479759698 | -3.851864 |
| **A_52_P139650** | Ska1 | 0.390697278 | 0.039916655 | 0.741477902 | 11.06309677 | 2.706375036 | 0.034265027 | 0.479759698 | -3.852159877 |
| **A_55_P2051727** | Rnf8 | 0.26793552 | 0.027358753 | 0.508512288 | 11.327468 | 2.706199416 | 0.034273162 | 0.479759698 | -3.852233325 |
| **A_55_P2148906** | Cltb | 0.372592813 | 0.037978534 | 0.707207092 | 9.798261041 | 2.705659825 | 0.034298169 | 0.479759698 | -3.85245902 |
| **A_51_P519837** | Cherp | 0.223636005 | 0.022785311 | 0.424486699 | 11.11910762 | 2.705525162 | 0.034304413 | 0.479759698 | -3.852515351 |
| **A_51_P453401** | Cipc | 0.212298203 | 0.02162197 | 0.402974437 | 10.53735496 | 2.705409078 | 0.034309796 | 0.479759698 | -3.852563913 |
| **A_51_P321886** | Cmtm3 | -0.290726398 | -0.551940037 | -0.029512758 | 10.65843624 | -2.704405473 | 0.034356377 | 0.479759698 | -3.852983825 |
| **A_55_P2098578** | Tubd1 | 0.278610641 | 0.028281659 | 0.528939622 | 9.304987228 | 2.704392707 | 0.03435697 | 0.479759698 | -3.852989167 |
| **A_51_P197321** | Clta | -0.23911008 | -0.453965184 | -0.024254976 | 13.71102348 | -2.704178768 | 0.034366908 | 0.479759698 | -3.853078697 |
| **A_55_P1963454** | Cenpq | 0.310504292 | 0.031446751 | 0.589561833 | 9.144741876 | 2.703690863 | 0.034389585 | 0.479759698 | -3.853282902 |
| **A_55_P1976898** | Pnp2 | 0.394929079 | 0.039971541 | 0.749886618 | 10.48940234 | 2.703497035 | 0.034398598 | 0.479759698 | -3.853364033 |
| **A_55_P2026049** | NA | 0.282828312 | 0.028617549 | 0.537039074 | 8.247108813 | 2.703411377 | 0.034402582 | 0.479759698 | -3.85339989 |
| **A_52_P23308** | Fam241a | 0.438560626 | 0.044353573 | 0.832767679 | 11.36539567 | 2.703263851 | 0.034409445 | 0.479759698 | -3.853461646 |
| **A_55_P2092909** | Rassf1 | -0.234833476 | -0.445935529 | -0.023731423 | 11.15575346 | -2.703029207 | 0.034420363 | 0.479759698 | -3.853559876 |
| **A_55_P2013756** | NA | 0.714513041 | 0.072198807 | 1.356827275 | 10.017752 | 2.702998487 | 0.034421793 | 0.479759698 | -3.853572737 |
| **A_55_P2113208** | Snrpa1 | 0.289022866 | 0.029160836 | 0.548884897 | 12.12245755 | 2.702542702 | 0.034443012 | 0.479759698 | -3.853763568 |
| **A_51_P431046** | St8sia3 | 0.341462675 | 0.034373815 | 0.648551534 | 8.459929479 | 2.701857037 | 0.03447496 | 0.479759698 | -3.854050698 |
| **A_52_P433864** | Naa16 | 0.352315948 | 0.035112084 | 0.669519811 | 8.824856175 | 2.698839283 | 0.034615946 | 0.479759698 | -3.855315154 |
| **A_55_P2171912** | Gja3 | 0.553003317 | 0.054647496 | 1.051359138 | 9.588368269 | 2.696319733 | 0.034734126 | 0.479759698 | -3.856371779 |
| **A_55_P1979156** | Arap3 | -0.256635318 | -0.487920632 | -0.025350004 | 8.068097031 | -2.696196588 | 0.034739913 | 0.479759698 | -3.856423444 |
| **A_55_P2051949** | Elmod3 | -0.348876764 | -0.663329602 | -0.034423925 | 9.87118912 | -2.695874774 | 0.034755041 | 0.479759698 | -3.856558468 |
| **A_55_P2142172** | Olfr1229 | 0.235675176 | 0.0232511 | 0.448099253 | 9.05318285 | 2.695834897 | 0.034756916 | 0.479759698 | -3.856575201 |
| **A_55_P2024649** | Rpe | 0.474682883 | 0.046796504 | 0.902569262 | 10.37176325 | 2.695617696 | 0.034767131 | 0.479759698 | -3.856666341 |
| **A_55_P2017491** | Rbm43 | 0.219206831 | 0.021606466 | 0.416807196 | 10.8103663 | 2.695563267 | 0.034769691 | 0.479759698 | -3.856689182 |
| **A_55_P2009345** | Ovch2 | 0.304074722 | 0.029907454 | 0.578241991 | 8.655596662 | 2.694932568 | 0.034799374 | 0.479759698 | -3.856953872 |
| **A_55_P2078735** | AI662270 | 0.667595802 | 0.065626348 | 1.269565255 | 10.05397416 | 2.694773897 | 0.034806846 | 0.479759698 | -3.857020471 |
| **A_51_P483329** | Sdad1 | 0.315094223 | 0.030971913 | 0.599216534 | 11.60367139 | 2.694748838 | 0.034808026 | 0.479759698 | -3.85703099 |
| **A_55_P2018919** | Spns1 | -0.300354142 | -0.571192462 | -0.029515821 | 13.02698498 | -2.694676917 | 0.034811413 | 0.479759698 | -3.857061178 |
| **A_55_P1970905** | Lin28b | 0.656062173 | 0.064446294 | 1.247678052 | 9.89505446 | 2.694563143 | 0.034816773 | 0.479759698 | -3.857108936 |
| **A_51_P338998** | Prim2 | 0.356973947 | 0.034906896 | 0.679040997 | 10.7114986 | 2.693229817 | 0.034879645 | 0.479759698 | -3.857668741 |
| **A_55_P2101585** | Cd55 | 0.175409685 | 0.01714863 | 0.33367074 | 10.13952729 | 2.693163382 | 0.034882781 | 0.479759698 | -3.857696641 |

| | | | | | | | | | |
|---|---|---|---|---|---|---|---|---|---|
| **A_52_P485450** | Vta1 | 0.297553218 | 0.029056585 | 0.56604985 | 9.8522785 | 2.692830372 | 0.034898504 | 0.479759698 | -3.857836496 |
| **A_51_P110814** | Nudt9 | 0.265319506 | 0.025869559 | 0.504769453 | 13.19514158 | 2.692387858 | 0.034919409 | 0.479759698 | -3.879242869 |
| **A_55_P2062543** | Kifc5b | 0.502881417 | 0.049030461 | 0.956732373 | 10.71612617 | 2.692374831 | 0.034920025 | 0.479759698 | -3.858027835 |
| **A_51_P245275** | H2afx | 0.230512284 | 0.022470444 | 0.438554125 | 13.92261773 | 2.692319373 | 0.034922646 | 0.479759698 | -3.858051131 |
| **A_52_P476075** | Clspn | 0.41054769 | 0.039912437 | 0.781182942 | 8.831612887 | 2.691535285 | 0.034959724 | 0.479759698 | -3.858380539 |
| **A_51_P466162** | C1qbp | 0.245043425 | 0.023803467 | 0.466283383 | 13.61342791 | 2.6913035 | 0.034970693 | 0.479759698 | -3.858477931 |
| **A_55_P2063041** | Prelid3b | 0.209812979 | 0.020379617 | 0.399246342 | 9.767199528 | 2.691281171 | 0.034971749 | 0.479759698 | -3.858487314 |
| **A_51_P168632** | Prim1 | 0.422607967 | 0.041044648 | 0.804171286 | 12.66027439 | 2.691251292 | 0.034973164 | 0.479759698 | -3.858499869 |
| **A_55_P1968178** | Smagp | -0.302524399 | -0.575713131 | -0.029335668 | 10.8708033 | -2.690796256 | 0.03499471 | 0.479759698 | -3.85869109 |
| **A_51_P101137** | Samd14 | 0.462806932 | 0.044865796 | 0.880748067 | 11.97818546 | 2.690716399 | 0.034998492 | 0.479759698 | -3.858724652 |
| **A_51_P106859** | Lsm10 | -0.205849572 | -0.391765607 | -0.019933536 | 10.98615922 | -2.690396607 | 0.035013644 | 0.479759698 | -3.865888998 |
| **A_51_P314264** | Ssr2 | -0.233419579 | -0.444280794 | -0.022558364 | 12.27860122 | -2.689823394 | 0.035040822 | 0.479759698 | -3.859100012 |
| **A_55_P1967295** | Pemt | -0.937824187 | -1.785118683 | -0.090529692 | 12.58350266 | -2.689491855 | 0.035056551 | 0.479759698 | -3.859239396 |
| **A_55_P2018002** | Abcg4 | 0.477419628 | 0.046082102 | 0.908757154 | 11.38506149 | 2.689466975 | 0.035057732 | 0.479759698 | -3.859249856 |
| **A_55_P2010758** | Npm3 | 0.340317749 | 0.032839636 | 0.647795862 | 8.498876215 | 2.689388734 | 0.035061445 | 0.479759698 | -3.859282752 |
| **A_51_P224311** | Popdc2 | 0.45615015 | 0.04382234 | 0.868477959 | 10.35791985 | 2.688118339 | 0.035121795 | 0.479759698 | -3.859816996 |
| **A_55_P2120652** | Snrpa1 | 0.280246438 | 0.026854542 | 0.533638335 | 12.26782943 | 2.687389205 | 0.035156482 | 0.479759698 | -3.860123717 |
| **A_51_P272553** | Bhlhe40 | -0.626171161 | -1.192394412 | -0.05994791 | 9.911374819 | -2.687129223 | 0.035168859 | 0.479759698 | -3.8602331 |
| **A_55_P2122718** | Gpr19 | 0.308344342 | 0.029503364 | 0.58718532 | 9.439736711 | 2.686968503 | 0.035176513 | 0.479759698 | -3.860300723 |
| **A_55_P2068772** | Ncbp2 | 0.358924218 | 0.034337651 | 0.683510785 | 9.282457414 | 2.686924167 | 0.035178624 | 0.479759698 | -3.860319379 |
| **A_52_P578768** | Slc38a9 | 0.264782231 | 0.025281911 | 0.504282551 | 9.207183232 | 2.686370611 | 0.035205001 | 0.479759698 | -3.860552321 |
| **A_66_P132037** | Jmjd4 | 0.20793105 | 0.019844902 | 0.396017198 | 8.477089273 | 2.686245636 | 0.035210959 | 0.479759698 | -3.860604918 |
| **A_52_P126825** | Cramp1l | 0.275997788 | 0.026237476 | 0.525758101 | 10.83620219 | 2.68513028 | 0.035264178 | 0.479759698 | -3.861074412 |
| **A_55_P2053810** | NA | 0.365958821 | 0.034786882 | 0.697130761 | 8.514643688 | 2.685108734 | 0.035265207 | 0.479759698 | -3.861083483 |
| **A_55_P1982414** | NA | 0.260949771 | 0.024792855 | 0.497106687 | 8.425308873 | 2.684970014 | 0.035271832 | 0.479759698 | -3.861141887 |
| **A_55_P2459897** | A2m | -0.522172056 | -0.994778355 | -0.049565757 | 10.41332938 | -2.684709567 | 0.035284274 | 0.479759698 | -3.861251548 |
| **A_55_P1992884** | NA | 0.371433845 | 0.035177841 | 0.707689848 | 8.331999318 | 2.684074808 | 0.035314618 | 0.479759698 | -3.86151885 |
| **A_51_P480233** | Nmral1 | 0.330655129 | 0.031306184 | 0.630004073 | 10.91714142 | 2.683988936 | 0.035318725 | 0.479759698 | -3.861555015 |
| **A_55_P2119032** | Rasl2-9 | 0.651059495 | 0.061640116 | 1.240478874 | 11.58252505 | 2.683981112 | 0.0353191 | 0.479759698 | -3.861558311 |
| **A_51_P171200** | Golm1 | 0.339907696 | 0.032180421 | 0.647634971 | 12.22606268 | 2.683973325 | 0.035319472 | 0.479759698 | -3.86156159 |
| **A_55_P2038273** | NA | 0.368664288 | 0.034885791 | 0.702442784 | 9.253920468 | 2.683835574 | 0.035326062 | 0.479759698 | -3.861619607 |
| **A_55_P2161342** | Pkib | 0.578975032 | 0.054726577 | 1.103223486 | 11.11101379 | 2.683526358 | 0.035340859 | 0.479759698 | -3.861749849 |
| **A_55_P1978866** | NA | -0.423967851 | -0.807862297 | -0.040073405 | 12.26189775 | -2.683516595 | 0.035341326 | 0.479759698 | -3.861753961 |
| **A_66_P135391** | Igfbp2 | -1.475808109 | -2.812553939 | -0.139062279 | 11.41792383 | -2.682651412 | 0.035382763 | 0.479759698 | -3.862118445 |
| **A_55_P2004606** | Serpina4-ps1 | -0.873431667 | -1.664679671 | -0.082183663 | 8.64377535 | -2.682251489 | 0.035401935 | 0.479759698 | -3.862286958 |
| **A_51_P368009** | E2f2 | 0.30661266 | 0.028816098 | 0.584409222 | 14.4010497 | 2.681923621 | 0.03541766 | 0.479759698 | -3.862425125 |
| **A_55_P2025174** | Rps27a | 0.202728878 | 0.019028866 | 0.38642889 | 15.80195844 | 2.681572977 | 0.035434486 | 0.479759698 | -3.862572906 |
| **A_55_P1985945** | NA | 0.377636598 | 0.035390063 | 0.719883132 | 13.97851592 | 2.681132062 | 0.035455656 | 0.479759698 | -3.862758754 |
| **A_55_P2037116** | Ran | 0.262030905 | 0.024496962 | 0.499564849 | 15.80124622 | 2.680464299 | 0.035487744 | 0.479759698 | -3.86304027 |
| **A_55_P1990134** | Cox18 | 0.247210277 | 0.023092113 | 0.471328441 | 9.115785417 | 2.680233639 | 0.035498834 | 0.479759698 | -3.863137525 |
| **A_55_P2174905** | Cuta | -0.210893295 | -0.402134248 | -0.019652341 | 12.21623118 | -2.679569715 | 0.035530778 | 0.479759698 | -3.863417501 |
| **A_55_P2080451** | NA | 0.422004031 | 0.039249209 | 0.804758854 | 12.40704216 | 2.679039507 | 0.035556311 | 0.479759698 | -3.86364113 |
| **A_55_P2043009** | Psmg2 | 0.288996091 | 0.02677391 | 0.551218273 | 11.28805514 | 2.677970152 | 0.035607866 | 0.479759698 | -3.864092271 |
| **A_55_P2100375** | Col11a1 | 0.346956623 | 0.031998162 | 0.661915085 | 8.573832208 | 2.676733235 | 0.035667598 | 0.479759698 | -3.864614291 |
| **A_55_P2075769** | Ube2n | 0.26397539 | 0.024294704 | 0.503656077 | 9.541650467 | 2.676169334 | 0.035694864 | 0.479759698 | -3.864852342 |
| **A_51_P411770** | Nip7 | 0.177429183 | 0.016283491 | 0.338574875 | 12.1737722 | 2.67540504 | 0.035731856 | 0.479759698 | -3.865175057 |
| **A_51_P112308** | Tcim | -0.475484844 | -0.90736732 | -0.043602369 | 10.34671951 | -2.675187864 | 0.035742375 | 0.479759698 | -3.872219502 |
| **A_55_P2089655** | Eif2s2 | 0.304252468 | 0.027899376 | 0.58060556 | 13.39427808 | 2.675179762 | 0.035742767 | 0.479759698 | -3.865270192 |
| **A_55_P1970120** | Vopp1 | 0.363914199 | 0.033369574 | 0.694458824 | 13.46984521 | 2.675174346 | 0.03574303 | 0.479759698 | -3.86527248 |
| **A_55_P2040371** | Rnaset2a | -0.30189893 | -0.576211457 | -0.027586403 | 13.48341947 | -2.674232245 | 0.035788699 | 0.479759698 | -3.865670405 |

| | | | | | | | | | |
|---|---|---|---|---|---|---|---|---|---|
| **A_55_P2087225** | Hnrnpa3 | 0.247126799 | 0.022575436 | 0.471678162 | 11.02963209 | 2.674159683 | 0.035792219 | 0.479759698 | -3.865701059 |
| **A_55_P2145800** | NA | 0.250404557 | 0.022807837 | 0.478001277 | 8.329521699 | 2.673372141 | 0.035830447 | 0.479759698 | -3.866033798 |
| **A_55_P1973213** | Trmt61a | 0.378927532 | 0.034489697 | 0.723365367 | 9.968572532 | 2.673181824 | 0.035839692 | 0.479759698 | -3.86611422 |
| **A_52_P233335** | Pphln1 | 0.241880401 | 0.021979227 | 0.461781575 | 9.864412399 | 2.672737586 | 0.03586128 | 0.479759698 | -3.866301959 |
| **A_51_P492765** | Wdr75 | 0.352802903 | 0.032043507 | 0.673562298 | 10.8974194 | 2.672612234 | 0.035867375 | 0.479759698 | -3.866354939 |
| **A_55_P2076994** | Defa-rs10 | 0.259931172 | 0.023581141 | 0.496281203 | 8.589155432 | 2.672304177 | 0.035882356 | 0.479759698 | -3.866485147 |
| **A_51_P428483** | Fgb | -0.506610227 | -0.967271621 | -0.045948834 | 17.61973413 | -2.672239166 | 0.035885519 | 0.479759698 | -3.866512627 |
| **A_55_P2194064** | NA | 0.226429431 | 0.020462335 | 0.432396527 | 8.351472458 | 2.671272651 | 0.035932571 | 0.479759698 | -3.866921239 |
| **A_51_P501018** | Nek2 | 0.296872689 | 0.026808754 | 0.566936624 | 9.071345346 | 2.671079686 | 0.035941972 | 0.479759698 | -3.867002833 |
| **A_55_P2010038** | Tnfsf9 | -0.188428215 | -0.359858766 | -0.016997665 | 8.024018771 | -2.670797154 | 0.035955743 | 0.479759698 | -3.867122309 |
| **A_66_P103957** | Bbs7 | 0.449241829 | 0.040488396 | 0.857995262 | 9.598862118 | 2.670557678 | 0.035967419 | 0.479759698 | -3.867223585 |
| **A_55_P2049086** | Map3k7 | 0.238347944 | 0.021417502 | 0.455278387 | 10.89509952 | 2.669771529 | 0.036005778 | 0.479759698 | -3.867556108 |
| **A_55_P2091928** | Raet1b | 0.37094095 | 0.033277428 | 0.708604472 | 9.813149506 | 2.669339563 | 0.036026874 | 0.479759698 | -3.867738854 |
| **A_51_P203955** | Gbp2 | -0.248527957 | -0.474879164 | -0.022176751 | 9.211756891 | -2.667937341 | 0.036095443 | 0.479759698 | -3.868332242 |
| **A_66_P119376** | Kctd12 | 0.287952253 | 0.025694089 | 0.550210416 | 10.0605807 | 2.667931366 | 0.036095736 | 0.479759698 | -3.868334771 |
| **A_51_P377856** | Gstt3 | -0.242584352 | -0.463572549 | -0.021596155 | 10.6544491 | -2.667330886 | 0.036125143 | 0.479759698 | -3.86858896 |
| **A_55_P2019690** | Slc43a1 | 0.441863871 | 0.039312857 | 0.844414885 | 12.76379171 | 2.667170368 | 0.036133008 | 0.479759698 | -3.868656916 |
| **A_51_P395555** | Vkorc1 | -0.397752103 | -0.760211501 | -0.035292704 | 13.96540205 | -2.666467578 | 0.036167465 | 0.479759698 | -3.868954488 |
| **A_55_P2029106** | Bmf | -0.827437403 | -1.581531748 | -0.073343058 | 9.28810011 | -2.666199568 | 0.036180615 | 0.479759698 | -3.869067985 |
| **A_66_P122729** | Zmat2 | 0.244220816 | 0.02162403 | 0.466817602 | 11.26668464 | 2.665919164 | 0.036194378 | 0.479759698 | -3.86918674 |
| **A_55_P1955227** | Rabac1 | -0.462166612 | -0.883431425 | -0.0409018 | 12.99470938 | -2.665793915 | 0.036200527 | 0.479759698 | -3.869239788 |
| **A_55_P2409570** | Luc7l2 | 0.264412815 | 0.023238448 | 0.505587182 | 14.41311496 | 2.664001972 | 0.036288628 | 0.479759698 | -3.869998976 |
| **A_52_P650379** | Strap | 0.361615706 | 0.031688487 | 0.691542925 | 10.58398365 | 2.663252403 | 0.036325548 | 0.479759698 | -3.870316669 |
| **A_55_P2052719** | Nadk2 | 0.641238663 | 0.05606182 | 1.226415506 | 12.83792571 | 2.662660264 | 0.036354742 | 0.479759698 | -3.870567689 |
| **A_55_P1977245** | Slc22a13 | 0.204704718 | 0.017846906 | 0.39156253 | 8.274965384 | 2.661949314 | 0.036389827 | 0.479759698 | -3.870869137 |
| **A_51_P331328** | Gpihbp1 | -0.744526458 | -1.424207776 | -0.06484514 | 11.27535677 | -2.661693179 | 0.036402476 | 0.479759698 | -3.870977757 |
| **A_55_P2170139** | Rheb | 0.209229672 | 0.018210823 | 0.40024852 | 10.88970257 | 2.661523098 | 0.036410878 | 0.479759698 | -3.871049888 |
| **A_55_P2060333** | S100a5 | 0.712735965 | 0.061924974 | 1.363546957 | 10.90896533 | 2.661074198 | 0.036433062 | 0.479759698 | -3.871240283 |
| **A_55_P1956762** | Sgms1 | 0.256201954 | 0.022244234 | 0.490159673 | 9.004812699 | 2.66089812 | 0.036441768 | 0.479759698 | -3.871314972 |
| **A_51_P200667** | Clmn | -0.288947425 | -0.55287549 | -0.02501936 | 8.715377665 | -2.660213194 | 0.036475654 | 0.479759698 | -3.871605543 |
| **A_55_P2072391** | Acp1 | 0.456862327 | 0.039547801 | 0.874176853 | 9.762258579 | 2.660143311 | 0.036479113 | 0.479759698 | -3.871635193 |
| **A_55_P2077079** | Rab29 | -0.180009728 | -0.344439185 | -0.015580272 | 9.630766775 | -2.660109679 | 0.036480778 | 0.479759698 | -3.871649463 |
| **A_55_P2066707** | Trim27 | 0.284029545 | 0.02450056 | 0.543558529 | 12.13240328 | 2.659260234 | 0.036522856 | 0.479759698 | -3.872009925 |
| **A_55_P2116988** | 1700024P12Rik | 0.507737135 | 0.043745518 | 0.971728753 | 12.00957017 | 2.658961093 | 0.036537686 | 0.479759698 | -3.872136888 |
| **A_55_P2067563** | NA | -0.203226416 | -0.389005854 | -0.017446977 | 8.50894206 | -2.658065635 | 0.036582119 | 0.479759698 | -3.872517013 |
| **A_66_P131530** | Rpp30 | 0.230412174 | 0.019644777 | 0.441179571 | 8.494407225 | 2.656349273 | 0.036667444 | 0.479759698 | -3.873245909 |
| **A_66_P132613** | C230057M02Rik | 0.349340081 | 0.029772173 | 0.668907989 | 9.770834619 | 2.656246937 | 0.036672538 | 0.479759698 | -3.87328938 |
| **A_55_P2040341** | NA | 0.371110357 | 0.031548144 | 0.710672571 | 9.002186338 | 2.655625979 | 0.036703464 | 0.479759698 | -3.873553189 |
| **A_55_P2391065** | Cep128 | 0.237247432 | 0.020151785 | 0.454343078 | 8.351694323 | 2.655422274 | 0.036713615 | 0.479759698 | -3.873639742 |
| **A_55_P2142451** | Echdc2 | -0.45765403 | -0.876488232 | -0.038819827 | 12.19807474 | -2.655084458 | 0.036730456 | 0.479759698 | -3.87378329 |
| **A_51_P296036** | Nrbp2 | -0.360942164 | -0.691338927 | -0.030545401 | 9.552582509 | -2.654514017 | 0.036758913 | 0.479759698 | -3.874025722 |
| **A_55_P1961014** | Selenbp1 | -0.422500673 | -0.809275739 | -0.035725607 | 15.04378338 | -2.654312948 | 0.036768949 | 0.479759698 | -3.874111184 |
| **A_55_P2148370** | NA | 0.610661877 | 0.051579651 | 1.169744102 | 9.376778845 | 2.654045182 | 0.036782319 | 0.479759698 | -3.874225003 |
| **A_55_P1989728** | C2cd2l | 0.249671229 | 0.021071451 | 0.478271008 | 11.88094297 | 2.653847019 | 0.036792217 | 0.479759698 | -3.874309243 |
| **A_51_P138772** | Ddx27 | 0.358919201 | 0.030154452 | 0.68768395 | 10.81763998 | 2.652739698 | 0.036847576 | 0.479759698 | -3.874780061 |
| **A_55_P1984118** | Gfra4 | 0.521685971 | 0.043721113 | 0.999650828 | 10.33576189 | 2.652139601 | 0.036877614 | 0.479759698 | -3.875035281 |
| **A_55_P1963529** | Abcb10 | 0.476517895 | 0.039867056 | 0.913168735 | 13.58050592 | 2.651722689 | 0.036898498 | 0.479759698 | -3.875212621 |
| **A_55_P2069306** | Ptpn18 | -0.402303894 | -0.770988416 | -0.033619372 | 12.29138345 | -2.651444368 | 0.036912447 | 0.479759698 | -3.875331021 |
| **A_55_P2142439** | Echdc2 | -0.357304686 | -0.684752269 | -0.029857104 | 10.51884622 | -2.65142968 | 0.036913183 | 0.479759698 | -3.87533727 |
| **A_52_P73369** | Hnrnpu | 0.225853964 | 0.018772408 | 0.43293552 | 12.50702447 | 2.650144089 | 0.036977687 | 0.479759698 | -3.875884307 |

| | | | | | | | | |
|---|---|---|---|---|---|---|---|---|
| **A_55_P1956734** | NA | 0.656406631 | 0.054382738 | 1.258430525 | 10.73934034 | 2.649368818 | 0.037016644 | 0.479759698 | -3.8762143 |
| **A_55_P2024391** | NA | 0.280787393 | 0.023261606 | 0.538313179 | 9.82461872 | 2.649354488 | 0.037017365 | 0.479759698 | -3.876220401 |
| **A_52_P259402** | Dsg3 | 0.935736 | 0.077244535 | 1.794227465 | 8.39217833 | 2.648503464 | 0.037060179 | 0.479759698 | -3.87658273 |
| **A_55_P1990490** | Recql | 0.314380096 | 0.025893213 | 0.602866979 | 9.034999215 | 2.647964511 | 0.037087321 | 0.479759698 | -3.876812242 |
| **A_51_P189916** | Nol12 | 0.233533165 | 0.019191977 | 0.447874352 | 11.50921071 | 2.64743997 | 0.037113757 | 0.479759698 | -3.877035653 |
| **A_55_P2041270** | Arhgef1 | -0.238364889 | -0.457150938 | -0.01957884 | 11.56810342 | -2.647316388 | 0.037119988 | 0.479759698 | -3.877088294 |
| **A_52_P459929** | Itga1 | -0.309601784 | -0.593786931 | -0.025416638 | 8.988701783 | -2.647190947 | 0.037126314 | 0.479759698 | -3.877141729 |
| **A_55_P1983769** | Birc5 | 0.429771487 | 0.035264853 | 0.824278121 | 11.09534496 | 2.647076407 | 0.037132092 | 0.479759698 | -3.877190522 |
| **A_52_P305995** | Ammecr1 | 0.593664965 | 0.048698799 | 1.138631131 | 10.10761415 | 2.647006872 | 0.037135599 | 0.479759698 | -3.877220144 |
| **A_55_P1998811** | NA | -0.349326788 | -0.670116346 | -0.02853723 | 11.00483627 | -2.646030566 | 0.037184888 | 0.479759698 | -3.877636121 |
| **A_66_P134481** | Skp2 | 0.346602591 | 0.0283108 | 0.664894383 | 8.895333808 | 2.645998266 | 0.037186519 | 0.479759698 | -3.877649885 |
| **A_55_P2023081** | Dnaja2 | 0.251124309 | 0.020428775 | 0.481819843 | 9.785528973 | 2.645043125 | 0.037234809 | 0.479759698 | -3.878056968 |
| **A_51_P288479** | Slc25a29 | -0.200548265 | -0.384785971 | -0.016310558 | 8.290768386 | -2.644987221 | 0.037237637 | 0.479759698 | -3.878080799 |
| **A_55_P2051094** | Rorc | -0.289542408 | -0.555567899 | -0.023516918 | 8.409232051 | -2.644673837 | 0.037253497 | 0.479759698 | -3.878214392 |
| **A_55_P2178496** | Fam53b | 0.418704509 | 0.033941199 | 0.803467819 | 11.36792431 | 2.644217494 | 0.037276604 | 0.479759698 | -3.87840895 |
| **A_51_P479321** | Acss1 | 0.413178461 | 0.033472801 | 0.792884121 | 13.24823716 | 2.64407513 | 0.037283816 | 0.479759698 | -3.878469651 |
| **A_66_P133112** | Gm14635 | 0.433170829 | 0.034952401 | 0.831389258 | 9.123966673 | 2.643145299 | 0.037330955 | 0.479759698 | -3.878866179 |
| **A_51_P502082** | Rrm1 | 0.44108874 | 0.03557327 | 0.846604211 | 12.91502398 | 2.643027816 | 0.037336915 | 0.479759698 | -3.878916288 |
| **A_55_P1983868** | NA | 0.306032397 | 0.024600444 | 0.58746435 | 8.821845328 | 2.642269966 | 0.037375389 | 0.479759698 | -3.879239569 |
| **A_52_P338956** | Aspg | -0.532624766 | -1.022567424 | -0.042682107 | 8.581184394 | -2.641552769 | 0.037411837 | 0.479759698 | -3.879545578 |
| **A_55_P2063396** | Zfand2a | 0.193916998 | 0.015519569 | 0.372314426 | 9.973030763 | 2.641255887 | 0.037426936 | 0.479759698 | -3.879672269 |
| **A_66_P115996** | Kyat1 | -0.260967589 | -0.501136893 | -0.020798285 | 11.91096518 | -2.640293842 | 0.037475908 | 0.479759698 | -3.880082891 |
| **A_55_P2006579** | Rbbp6 | 0.227832529 | 0.018082435 | 0.437582623 | 9.603877849 | 2.639348609 | 0.037524091 | 0.479759698 | -3.880486454 |
| **A_55_P2070869** | Lcn2 | -0.381509482 | -0.732748302 | -0.030270662 | 13.84812673 | -2.63928334 | 0.03752742 | 0.479759698 | -3.880514325 |
| **A_55_P1965416** | Atp9a | -0.68674758 | -1.319227234 | -0.054267926 | 12.53023414 | -2.638358256 | 0.037574643 | 0.479759698 | -3.880909406 |
| **A_55_P1987494** | Gabrr2 | 0.394146836 | 0.031050433 | 0.757243239 | 11.01118333 | 2.637662877 | 0.037610182 | 0.479759698 | -3.881206459 |
| **A_55_P2171273** | Mier3 | 0.329329751 | 0.0258973 | 0.632762201 | 8.967133159 | 2.637255033 | 0.037631043 | 0.479759698 | -3.881380712 |
| **A_55_P2141699** | Rnf149 | 0.43083916 | 0.033861349 | 0.827816971 | 12.31381771 | 2.63713357 | 0.037637258 | 0.479759698 | -3.881432612 |
| **A_66_P120347** | Sstr5 | 0.315903419 | 0.024750308 | 0.60705653 | 11.14896007 | 2.636429005 | 0.03767333 | 0.479759698 | -3.881733702 |
| **A_55_P2116650** | A130040M12Rik | -0.597024548 | -1.147299206 | -0.04674989 | 12.14838161 | -2.636306286 | 0.037679617 | 0.479759698 | -3.881786152 |
| **A_55_P2092826** | Anxa1 | -0.341497053 | -0.656312989 | -0.026681117 | 13.42012421 | -2.635805994 | 0.037705257 | 0.479759698 | -3.881999995 |
| **A_52_P582261** | Ikzf5 | 0.452609866 | 0.035317025 | 0.869902707 | 10.20902555 | 2.63551973 | 0.037719937 | 0.479759698 | -3.882122369 |
| **A_55_P2083988** | Lmo2 | 0.3717774 | 0.029006003 | 0.714548797 | 13.27907763 | 2.635491379 | 0.037721391 | 0.479759698 | -3.882134489 |
| **A_55_P2171033** | NA | 0.348982607 | 0.027166554 | 0.67079866 | 8.926760509 | 2.634991799 | 0.037747026 | 0.479759698 | -3.882348081 |
| **A_55_P1957747** | Inip | 0.283759943 | 0.022085566 | 0.545434321 | 9.279999775 | 2.634954245 | 0.037748953 | 0.479759698 | -3.882364138 |
| **A_55_P1989928** | Lin52 | 0.198610757 | 0.015451172 | 0.381770343 | 9.183784646 | 2.634852475 | 0.037754178 | 0.479759698 | -3.882407654 |
| **A_51_P473888** | Il6st | -0.403369118 | -0.775359023 | -0.031379212 | 10.03512062 | -2.634842618 | 0.037754684 | 0.479759698 | -3.882411868 |
| **A_52_P600318** | Scamp2 | -0.202512831 | -0.389294271 | -0.015731392 | 10.52332796 | -2.634523131 | 0.037771091 | 0.479759698 | -3.882548486 |
| **A_55_P2001290** | Ube2e3 | 0.251982446 | 0.019524861 | 0.484440031 | 12.49181516 | 2.63396351 | 0.037799847 | 0.479759698 | -3.882787821 |
| **A_52_P655265** | Bora | 0.23848279 | 0.018475679 | 0.458489901 | 8.565688857 | 2.633925674 | 0.037801792 | 0.479759698 | -3.882804003 |
| **A_52_P150114** | NA | 0.399298279 | 0.030917292 | 0.767679267 | 8.297470105 | 2.633803801 | 0.037808058 | 0.479759698 | -3.882856131 |
| **A_55_P2013928** | Dhrs13 | 0.366150233 | 0.028347883 | 0.703952583 | 9.924373486 | 2.633782077 | 0.037809175 | 0.479759698 | -3.882865423 |
| **A_52_P220783** | Tmem229b | -0.29851069 | -0.573920067 | -0.023101314 | 10.19640461 | -2.633688187 | 0.037814003 | 0.479759698 | -3.882905584 |
| **A_55_P2058992** | Vamp4 | -0.218885739 | -0.420843796 | -0.016927682 | 8.666220451 | -2.633537296 | 0.037821764 | 0.479759698 | -3.882970128 |
| **A_55_P1971769** | NA | 0.251268689 | 0.019412685 | 0.483124693 | 12.80974715 | 2.63331745 | 0.037833074 | 0.479759698 | -3.883064173 |
| **A_55_P2202043** | C430045I18Rik | 0.338485909 | 0.026126858 | 0.65084496 | 8.820749933 | 2.63311416 | 0.037843536 | 0.479759698 | -3.883151142 |
| **A_55_P2070940** | Krtap20-2 | 0.686893289 | 0.053006129 | 1.320780448 | 11.04517191 | 2.633058496 | 0.037846401 | 0.479759698 | -3.883174957 |
| **A_55_P1996946** | Cdc20 | 0.280757677 | 0.021630053 | 0.539885301 | 12.90691282 | 2.632698445 | 0.037864939 | 0.479759698 | -3.883329005 |
| **A_55_P2036250** | Cr1l | 0.260025473 | 0.020018499 | 0.500032447 | 12.59729992 | 2.632541477 | 0.037873024 | 0.479759698 | -3.883396169 |
| **A_55_P2140195** | Gm15517 | 0.700343617 | 0.053887162 | 1.346800071 | 13.41428484 | 2.63241943 | 0.037879311 | 0.479759698 | -3.883448393 |

| | | | | | | | | | |
|---|---|---|---|---|---|---|---|---|---|
| **A_55_P1961736** | Rcan2 | -0.195502034 | -0.375964953 | -0.015039115 | 8.443025419 | -2.632367326 | 0.037881996 | 0.479759698 | -3.883470689 |
| **A_52_P350616** | Mrm1 | 0.411847688 | 0.031582528 | 0.792112848 | 11.2401171 | 2.631681239 | 0.037917364 | 0.479759698 | -3.883764308 |
| **A_51_P295312** | Gabarap | -0.365621701 | -0.703270768 | -0.027972635 | 12.12761799 | -2.631174196 | 0.037943524 | 0.479759698 | -3.883981342 |
| **A_52_P504330** | Ankib1 | 0.222521325 | 0.017008885 | 0.428033766 | 9.233783968 | 2.630974929 | 0.037953811 | 0.479759698 | -3.884066645 |
| **A_51_P263246** | Dusp8 | 0.346006322 | 0.026418683 | 0.665593962 | 9.085686198 | 2.630735886 | 0.037966154 | 0.479759698 | -3.884168982 |
| **A_55_P2007022** | Adgre5 | -0.516422369 | -0.993428624 | -0.039416114 | 12.06919029 | -2.630656606 | 0.037970249 | 0.479759698 | -3.884202925 |
| **A_55_P2152009** | Tesc | -0.387155321 | -0.744941178 | -0.029369463 | 9.816500047 | -2.629330936 | 0.038038788 | 0.479759698 | -3.884770611 |
| **A_52_P512817** | Shisa5 | -0.783002832 | -1.506659815 | -0.05934585 | 11.96675642 | -2.629140318 | 0.038048654 | 0.479759698 | -3.884852257 |
| **A_55_P2151855** | Acp1 | 0.411378499 | 0.03110551 | 0.791651489 | 13.061898 | 2.628629029 | 0.03807513 | 0.479759698 | -3.885071278 |
| **A_55_P2020128** | Dhrs3 | -0.357786386 | -0.688597907 | -0.026974864 | 13.15602823 | -2.628006123 | 0.038107413 | 0.479759698 | -3.885338159 |
| **A_55_P2133485** | Gapvd1 | 0.359285312 | 0.026981644 | 0.69158898 | 11.19641902 | 2.62716601 | 0.038150999 | 0.479759698 | -3.885698179 |
| **A_51_P513449** | Pogz | 0.217313942 | 0.016308257 | 0.418319628 | 9.592875253 | 2.627014302 | 0.038158876 | 0.479759698 | -3.885763202 |
| **A_55_P2143324** | Dyrk1b | 0.276819844 | 0.020741827 | 0.532897861 | 8.931478287 | 2.626685704 | 0.038175942 | 0.479759698 | -3.88590405 |
| **A_66_P105307** | Adnp | 0.265237352 | 0.019790785 | 0.51068392 | 11.79009436 | 2.625795566 | 0.038222213 | 0.479759698 | -3.886285665 |
| **A_52_P574190** | Psmd12 | 0.215400044 | 0.016043996 | 0.414756092 | 13.25624997 | 2.625424664 | 0.038241511 | 0.479759698 | -3.886444706 |
| **A_66_P132446** | Usp15 | 0.52920342 | 0.039414059 | 1.018992781 | 8.374856339 | 2.625406061 | 0.038242479 | 0.479759698 | -3.886452683 |
| **A_55_P2230688** | Pigm | 0.179744346 | 0.013291709 | 0.346196984 | 8.844068288 | 2.623902819 | 0.038320799 | 0.479759698 | -3.887097454 |
| **A_55_P2036813** | H2bu2 | 0.269746895 | 0.019926619 | 0.51956717 | 11.26103205 | 2.623686617 | 0.038332078 | 0.479759698 | -3.887190211 |
| **A_52_P143477** | Tgoln1 | -0.382179414 | -0.73621499 | -0.028143837 | 10.87686563 | -2.623031906 | 0.038366252 | 0.479759698 | -3.88747114 |
| **A_66_P130654** | Abcg4 | 0.483406003 | 0.035553627 | 0.931258379 | 10.80436689 | 2.622770836 | 0.038379889 | 0.479759698 | -3.887583177 |
| **A_55_P1957219** | Ptma | 0.335342058 | 0.024464673 | 0.646219443 | 14.92364573 | 2.621090911 | 0.038467759 | 0.479759698 | -3.888304325 |
| **A_52_P590392** | Rab10 | 0.2660287 | 0.019339166 | 0.512718234 | 11.60619424 | 2.620359975 | 0.038506058 | 0.479759698 | -3.888618211 |
| **A_55_P1975843** | NA | -0.308604811 | -0.59480518 | -0.022404443 | 11.96247273 | -2.620086882 | 0.038520377 | 0.479759698 | -3.888735503 |
| **A_55_P2091446** | Trim33 | 0.219396769 | 0.015919452 | 0.422874086 | 9.00416841 | 2.619976615 | 0.038526161 | 0.479759698 | -3.888782865 |
| **A_55_P2166011** | Tmem128 | 0.250106673 | 0.018037132 | 0.482176214 | 9.492004214 | 2.618727599 | 0.038591735 | 0.479759698 | -3.889319453 |
| **A_52_P555672** | Vrk1 | 0.347883637 | 0.025081626 | 0.670685647 | 9.734516766 | 2.618671127 | 0.038594703 | 0.479759698 | -3.889343719 |
| **A_55_P2105820** | Tbc1d31 | 0.249945555 | 0.017990424 | 0.481900686 | 9.77394368 | 2.618331454 | 0.038612558 | 0.479759698 | -3.889489682 |
| **A_55_P1968628** | NA | 0.446460561 | 0.032064826 | 0.860856297 | 9.686451517 | 2.617887698 | 0.038635897 | 0.479759698 | -3.889680395 |
| **A_51_P407165** | Abhd5 | -0.797360506 | -1.537621555 | -0.057099457 | 10.3722114 | -2.617297024 | 0.038666987 | 0.479759698 | -3.889934288 |
| **A_66_P107680** | Med8 | 0.243117197 | 0.017388618 | 0.468845775 | 8.683743038 | 2.617051815 | 0.038679901 | 0.479759698 | -3.890039702 |
| **A_66_P129800** | Rab8b | 0.228526989 | 0.016286411 | 0.440767568 | 11.7185375 | 2.616328467 | 0.038718023 | 0.479759698 | -3.890350707 |
| **A_55_P1979341** | Cyba | -0.321827998 | -0.620912382 | -0.022743615 | 13.65399868 | -2.614648276 | 0.038806727 | 0.479759698 | -3.891073373 |
| **A_52_P561377** | Fam160b1 | -0.189806025 | -0.366203229 | -0.013408822 | 11.51306301 | -2.614577306 | 0.038810479 | 0.479759698 | -3.891103907 |
| **A_55_P2185667** | NA | 0.274277427 | 0.019245304 | 0.529309551 | 7.969251076 | 2.613234414 | 0.038881538 | 0.479759698 | -3.891681772 |
| **A_55_P2136101** | Meis3 | 0.465133459 | 0.032608093 | 0.897658826 | 9.17097002 | 2.613058832 | 0.038890839 | 0.479759698 | -3.891757345 |
| **A_52_P219473** | Cdc6 | 0.56207539 | 0.039389464 | 1.084761315 | 9.30821179 | 2.612985221 | 0.038894739 | 0.479759698 | -3.89178903 |
| **A_55_P1999847** | Nutf2 | 0.203866068 | 0.01427651 | 0.393455626 | 13.38178481 | 2.612845454 | 0.038902145 | 0.479759698 | -3.891849191 |
| **A_55_P2023118** | Prss28 | 0.226741639 | 0.015861485 | 0.437621794 | 8.120068831 | 2.61263512 | 0.038913294 | 0.479759698 | -3.891939733 |
| **A_55_P1957954** | Rsu1 | -0.205561027 | -0.396743863 | -0.01437819 | 10.85805631 | -2.612612895 | 0.038914472 | 0.479759698 | -3.8919493 |
| **A_51_P101660** | Ncbp3 | 0.176258097 | 0.012314581 | 0.340201613 | 8.315506676 | 2.612390046 | 0.038926288 | 0.479759698 | -3.898676779 |
| **A_51_P400581** | Anp32e | 0.316133799 | 0.022085166 | 0.610182431 | 9.917729889 | 2.612371604 | 0.038927266 | 0.479759698 | -3.892053175 |
| **A_55_P2091323** | Wdr26 | 0.383476345 | 0.026765721 | 0.74018697 | 10.21353918 | 2.61219578 | 0.038936591 | 0.479759698 | -3.892128872 |
| **A_55_P1958840** | Itga6 | -0.469176045 | -0.905749169 | -0.032602922 | 10.89729392 | -2.611331597 | 0.038982461 | 0.479759698 | -3.892500982 |
| **A_51_P361220** | Fzd4 | -0.411651633 | -0.794744494 | -0.028558772 | 9.641550829 | -2.611012633 | 0.038999405 | 0.479759698 | -3.89263835 |
| **A_55_P1998072** | NA | 0.388697788 | 0.026790814 | 0.750604762 | 8.503443485 | 2.609746396 | 0.03906675 | 0.479759698 | -3.893183809 |
| **A_51_P424781** | Sf3b6 | 0.199933537 | 0.013672614 | 0.386194459 | 12.64751628 | 2.608237221 | 0.039147175 | 0.479759698 | -3.893834189 |
| **A_52_P93933** | Mcl1 | 0.257720961 | 0.01759515 | 0.497846772 | 11.23904341 | 2.607918924 | 0.03916416 | 0.479759698 | -3.893971397 |
| **A_55_P1988246** | NA | 0.378200344 | 0.025706419 | 0.730694269 | 11.81808468 | 2.607074684 | 0.039209248 | 0.479759698 | -3.894335387 |
| **A_66_P133642** | NA | 0.973650339 | 0.065795439 | 1.881505239 | 10.90423511 | 2.605972129 | 0.039268213 | 0.479759698 | -3.894810886 |
| **A_55_P2045482** | NA | -0.205879205 | -0.397870179 | -0.013888231 | 14.94513721 | -2.605642682 | 0.039285851 | 0.479759698 | -3.894952997 |

| | | | | | | | | |
|---|---|---|---|---|---|---|---|---|
| **A_51_P391716** | Ermap | 0.514871265 | 0.034729075 | 0.995013455 | 13.11533271 | 2.605625374 | 0.039286778 | 0.479759698 | -3.894960464 |
| **A_51_P151126** | Cd52 | -0.366056157 | -0.707460955 | -0.02465136 | 12.9024861 | -2.605321272 | 0.039303066 | 0.479759698 | -3.895091655 |
| **A_55_P1961302** | NA | 0.269797017 | 0.018157119 | 0.521436914 | 11.95605196 | 2.605198581 | 0.03930964 | 0.479759698 | -3.895144588 |
| **A_55_P1970389** | Srsf10 | 0.445012051 | 0.029885865 | 0.860138236 | 10.2165242 | 2.60480269 | 0.039330859 | 0.479759698 | -3.895315402 |
| **A_51_P362176** | Fgg | -0.288288986 | -0.557221773 | -0.019356199 | 17.50598144 | -2.604758635 | 0.039333221 | 0.479759698 | -3.895334412 |
| **A_55_P1977563** | Galk2 | -0.176444652 | -0.341121774 | -0.011767529 | 8.452472677 | -2.603505011 | 0.039400498 | 0.479759698 | -3.89587545 |
| **A_55_P2160322** | Anapc15 | 0.279480217 | 0.018608512 | 0.540351921 | 8.884312484 | 2.603198496 | 0.039416966 | 0.479759698 | -3.896007767 |
| **A_55_P1972541** | Zfp960 | 0.30588513 | 0.020298527 | 0.591471734 | 9.857842185 | 2.602577811 | 0.039450335 | 0.479759698 | -3.896275741 |
| **A_55_P2096834** | Fbxl19 | 0.221782298 | 0.014706181 | 0.428858416 | 11.38535694 | 2.602435979 | 0.039457965 | 0.479759698 | -3.896336983 |
| **A_51_P199135** | Cd83 | -0.646811457 | -1.250749822 | -0.042873092 | 8.78379827 | -2.602365386 | 0.039461763 | 0.479759698 | -3.896367465 |
| **A_52_P151853** | Tpd52 | -0.272912477 | -0.527761947 | -0.018063007 | 8.823092025 | -2.602093191 | 0.039476411 | 0.479759698 | -3.896485006 |
| **A_55_P1978726** | NA | 0.504307417 | 0.033348 | 0.975266833 | 10.97285864 | 2.601926712 | 0.039485372 | 0.479759698 | -3.8965569 |
| **A_55_P2149906** | NA | 0.414418728 | 0.027315986 | 0.80152147 | 14.15178819 | 2.601335184 | 0.039517232 | 0.479759698 | -3.896812383 |
| **A_55_P2169539** | NA | 0.490342011 | 0.032320234 | 0.948363788 | 8.383612009 | 2.601334292 | 0.03951728 | 0.479759698 | -3.896812768 |
| **A_55_P2042870** | Aoc2 | 0.366781875 | 0.024154047 | 0.709409702 | 14.72072662 | 2.60116812 | 0.039526235 | 0.479759698 | -3.896884547 |
| **A_65_P19780** | Klf9 | -0.838484007 | -1.621841945 | -0.05512607 | 9.492541426 | -2.600864485 | 0.039542603 | 0.479759698 | -3.897015711 |
| **A_52_P196105** | Ttyh3 | -0.320306785 | -0.619582672 | -0.021030898 | 10.97346225 | -2.600624207 | 0.039555561 | 0.479759698 | -3.897119516 |
| **A_51_P123134** | Ercc6l | 0.336177029 | 0.022003649 | 0.650350409 | 8.929480514 | 2.600050839 | 0.0395865 | 0.479759698 | -3.910386784 |
| **A_55_P2408355** | BB166591 | 0.490476472 | 0.031944857 | 0.949008087 | 11.04252293 | 2.599154424 | 0.039634923 | 0.479759698 | -3.897754649 |
| **A_51_P114910** | Cstb | -0.25184824 | -0.487320997 | -0.016375484 | 12.11201341 | -2.598851363 | 0.039651307 | 0.479759698 | -3.897885645 |
| **A_55_P1997141** | Mybl2 | 0.457659797 | 0.029746018 | 0.885573577 | 12.50621146 | 2.59878099 | 0.039655113 | 0.479759698 | -3.897916065 |
| **A_55_P2078845** | Qk | 0.230528131 | 0.014975415 | 0.446080846 | 10.82346706 | 2.598684878 | 0.039660311 | 0.479759698 | -3.897957612 |
| **A_51_P281246** | Cep57 | 0.289437518 | 0.018732306 | 0.56014273 | 9.534391707 | 2.598013462 | 0.039696645 | 0.479759698 | -3.898247883 |
| **A_51_P359636** | Lgals3bp | -0.35932366 | -0.695399426 | -0.023247894 | 12.02057617 | -2.597956093 | 0.039699752 | 0.479759698 | -3.898272688 |
| **A_55_P1983006** | Hdac5 | -0.429626965 | -0.831493791 | -0.02776014 | 11.91270323 | -2.597721357 | 0.039712464 | 0.479759698 | -3.898374186 |
| **A_51_P192800** | Grn | -0.272816287 | -0.528006881 | -0.017625693 | 11.68116144 | -2.597698957 | 0.039713677 | 0.479759698 | -3.898383872 |
| **A_55_P2064171** | NA | 0.504711506 | 0.032583167 | 0.976839845 | 11.22442576 | 2.597564409 | 0.039720966 | 0.479759698 | -3.898442053 |
| **A_55_P2056293** | Sf3b4 | 0.251370039 | 0.016219514 | 0.486520564 | 8.490283626 | 2.597471252 | 0.039726013 | 0.479759698 | -3.898482337 |
| **A_55_P1961094** | Tmem230 | 0.1803313 | 0.011629928 | 0.349032671 | 10.58019521 | 2.597381177 | 0.039730894 | 0.479759698 | -3.89852129 |
| **A_55_P2182921** | NA | 0.360361392 | 0.023111387 | 0.697611397 | 8.948150212 | 2.596387321 | 0.039784791 | 0.479759698 | -3.898951148 |
| **A_51_P214306** | Haus4 | 0.270703652 | 0.017353419 | 0.524053885 | 11.83707356 | 2.596306713 | 0.039789166 | 0.479759698 | -3.898986017 |
| **A_51_P405476** | Fcer1g | -0.37758747 | -0.731007584 | -0.024167355 | 12.69788871 | -2.596028762 | 0.039804254 | 0.479759698 | -3.899106261 |
| **A_55_P2000813** | Cops2 | 0.537181403 | 0.034311157 | 1.040051648 | 10.197957 | 2.595662473 | 0.039824148 | 0.479759698 | -3.899264736 |
| **A_55_P2035677** | Srp19 | 0.332246304 | 0.02110328 | 0.643389328 | 11.26456565 | 2.594676861 | 0.03987773 | 0.479759698 | -3.899691246 |
| **A_55_P1967592** | Ccnk | 0.257601659 | 0.016340752 | 0.498862567 | 12.25552497 | 2.594447504 | 0.03989021 | 0.479759698 | -3.899790515 |
| **A_55_P2016346** | Eif3d | 0.209816715 | 0.013249344 | 0.406384086 | 14.42888368 | 2.593652824 | 0.039933482 | 0.479759698 | -3.900134517 |
| **A_66_P116632** | Mafg | 0.325771071 | 0.020498991 | 0.631043151 | 12.66792441 | 2.593036436 | 0.03996708 | 0.479759698 | -3.900401396 |
| **A_51_P300655** | Tbccd1 | 0.431014515 | 0.027102227 | 0.834926803 | 9.964834175 | 2.592913442 | 0.039973787 | 0.479759698 | -3.900454654 |
| **A_51_P140607** | Ints13 | 0.259601549 | 0.016310178 | 0.502892921 | 10.63841386 | 2.592768622 | 0.039981687 | 0.479759698 | -3.900517367 |
| **A_55_P2097518** | Zdhhc22 | 0.383909806 | 0.02403877 | 0.743780842 | 15.9396645 | 2.592182056 | 0.040013699 | 0.479759698 | -3.900771399 |
| **A_51_P483473** | St3gal5 | 0.256747825 | 0.016037278 | 0.497458371 | 14.03325261 | 2.591760357 | 0.040036731 | 0.479759698 | -3.900954058 |
| **A_55_P2025143** | Tlk1 | 0.361357595 | 0.02250742 | 0.70020777 | 11.69012698 | 2.591269972 | 0.040063531 | 0.479759698 | -3.901166496 |
| **A_52_P460957** | Crem | -0.291062473 | -0.564017225 | -0.018107722 | 8.64899876 | -2.591067613 | 0.040074596 | 0.479759698 | -3.901254169 |
| **A_51_P275101** | Chst11 | 0.235705921 | 0.014662457 | 0.456749385 | 8.479794396 | 2.591051253 | 0.040075491 | 0.479759698 | -3.901261257 |
| **A_55_P2012815** | Acvr2b | -0.194678147 | -0.377268651 | -0.012087642 | 10.63297622 | -2.590730291 | 0.040093048 | 0.479759698 | -3.901400326 |
| **A_55_P2005135** | Tardbp | 0.304120799 | 0.018854467 | 0.589387131 | 9.565233086 | 2.590471353 | 0.040107218 | 0.479759698 | -3.90151253 |
| **A_51_P505823** | Endod1 | 0.475175246 | 0.029401079 | 0.920949414 | 11.68685087 | 2.590133206 | 0.04012573 | 0.479759698 | -3.901659071 |
| **A_55_P2019719** | Oas2 | -0.320287133 | -0.620865837 | -0.019708428 | 8.69580926 | -2.589193288 | 0.040177236 | 0.479759698 | -3.902066474 |
| **A_55_P1965757** | Eme2 | 0.2025413 | 0.012459256 | 0.392623343 | 9.530400429 | 2.589140927 | 0.040180107 | 0.479759698 | -3.902089174 |
| **A_52_P152283** | Gabpa | 0.284583683 | 0.017478416 | 0.55168895 | 9.799970566 | 2.58887292 | 0.040194807 | 0.479759698 | -3.902205363 |

| | | | | | | | | | |
|---|---|---|---|---|---|---|---|---|---|
| **A_51_P357685** | Dhx16 | 0.270730398 | 0.016597804 | 0.524862992 | 11.33913109 | 2.588569572 | 0.040211453 | 0.479759698 | -3.902336884 |
| **A_52_P613498** | 4833420G17Rik | 0.194962069 | 0.011931664 | 0.377992474 | 11.16094342 | 2.588272951 | 0.040227736 | 0.479759698 | -3.902465501 |
| **A_55_P2003673** | Mtmr12 | 0.18610639 | 0.01134906 | 0.36086372 | 10.18370517 | 2.587671093 | 0.040260797 | 0.479759698 | -3.902726505 |
| **A_55_P2163837** | Med14 | 0.229147472 | 0.013960757 | 0.444334188 | 8.869176338 | 2.587514562 | 0.0402694 | 0.479759698 | -3.902794394 |
| **A_55_P2158458** | Iqcc | 0.198152806 | 0.012014554 | 0.384291058 | 9.202304611 | 2.586710255 | 0.040313637 | 0.479759698 | -3.903143282 |
| **A_51_P169047** | Tdp2 | 0.31631022 | 0.019129215 | 0.613491225 | 9.768406272 | 2.58627896 | 0.04033738 | 0.479759698 | -3.903330401 |
| **A_51_P410949** | Polr3g | 0.370275337 | 0.022360411 | 0.718190264 | 9.058414769 | 2.58603807 | 0.040350647 | 0.479759698 | -3.903434923 |
| **A_55_P1963389** | NA | 0.251582064 | 0.01518282 | 0.487981309 | 10.3624559 | 2.585930095 | 0.040356595 | 0.479759698 | -3.903481775 |
| **A_55_P2089488** | Coq10b | 0.263543729 | 0.015898384 | 0.511189073 | 9.078769857 | 2.585864163 | 0.040360228 | 0.479759698 | -3.903510385 |
| **A_52_P671661** | Mak16 | 0.243132386 | 0.014653846 | 0.471610926 | 8.991908711 | 2.585714594 | 0.04036847 | 0.479759698 | -3.90357529 |
| **A_55_P2155921** | Hace1 | 0.32793971 | 0.019722684 | 0.636156736 | 8.79813385 | 2.585357297 | 0.040388166 | 0.479759698 | -3.903730348 |
| **A_55_P1995007** | Arhgap11a | 0.4789518 | 0.028744748 | 0.929158853 | 9.535033218 | 2.585012842 | 0.040407164 | 0.479759698 | -3.90387985 |
| **A_55_P2211341** | Map3k4 | -0.221095812 | -0.428946314 | -0.01324531 | 11.60289032 | -2.58471479 | 0.04042361 | 0.479759698 | -3.904009223 |
| **A_55_P2045931** | Btnl10 | 0.449106759 | 0.026776774 | 0.871436745 | 10.58971942 | 2.583930707 | 0.040466908 | 0.479759698 | -3.904349619 |
| **A_55_P2075200** | Tor3a | -0.317235691 | -0.615631785 | -0.018839597 | 9.2830215 | -2.58328366 | 0.040502676 | 0.479759698 | -3.904630583 |
| **A_55_P1975181** | Acyp1 | 0.460492674 | 0.027303914 | 0.893681434 | 8.427391674 | 2.583025748 | 0.040516942 | 0.479759698 | -3.90474259 |
| **A_66_P108474** | NA | 0.303525078 | 0.017966857 | 0.589083298 | 8.121767291 | 2.582754348 | 0.040531961 | 0.479759698 | -3.904860464 |
| **A_51_P486068** | Zfp280b | 0.34428601 | 0.02031582 | 0.668256199 | 9.604225483 | 2.582245409 | 0.040560139 | 0.479759698 | -3.90508153 |
| **A_51_P299062** | Kel | 0.548290519 | 0.032156408 | 1.06442463 | 13.60349511 | 2.58125769 | 0.040614885 | 0.479759698 | -3.905510658 |
| **A_55_P2026635** | Asnsd1 | 0.231162324 | 0.01347136 | 0.448853288 | 12.29588483 | 2.580238428 | 0.040671462 | 0.479759698 | -3.905953622 |
| **A_55_P1952324** | NA | 0.332768159 | 0.019387669 | 0.646148649 | 8.851590365 | 2.580197782 | 0.04067372 | 0.479759698 | -3.905971288 |
| **A_55_P2096340** | Rnf170 | 0.425846891 | 0.024735732 | 0.826958049 | 9.613800964 | 2.579716172 | 0.040700483 | 0.479759698 | -3.906180642 |
| **A_55_P1977812** | Cd244a | 0.425822618 | 0.02472844 | 0.826916796 | 9.037055865 | 2.579678336 | 0.040702587 | 0.479759698 | -3.90619709 |
| **A_55_P2133895** | Hnrnph2 | 0.224898558 | 0.013017505 | 0.436779612 | 10.13878744 | 2.57915675 | 0.040731594 | 0.479759698 | -3.906423857 |
| **A_55_P2063266** | Piga | 0.281206625 | 0.016150198 | 0.546263052 | 10.6552555 | 2.577925706 | 0.040800144 | 0.479759698 | -3.906959207 |
| **A_51_P205545** | Creld1 | -0.247697588 | -0.481214921 | -0.014180255 | 8.808364017 | -2.577423856 | 0.040828124 | 0.479759698 | -3.907177505 |
| **A_51_P492298** | Aqp2 | 0.694402267 | 0.039722897 | 1.349081636 | 10.82511954 | 2.577304091 | 0.040834805 | 0.479759698 | -3.907229606 |
| **A_51_P120461** | Riox2 | 0.24736666 | 0.014132821 | 0.480600499 | 10.09801235 | 2.577109035 | 0.040845687 | 0.479759698 | -3.947465882 |
| **A_55_P2021841** | Calm3 | 0.270690496 | 0.015459492 | 0.5259215 | 13.84744985 | 2.57704954 | 0.040849007 | 0.479759698 | -3.907340349 |
| **A_51_P165704** | Mcm7 | 0.370262669 | 0.021091549 | 0.719433788 | 13.56049595 | 2.576646268 | 0.040871518 | 0.479759698 | -3.90751581 |
| **A_55_P2137431** | Rtel1 | 0.292975576 | 0.016593934 | 0.569357218 | 10.83219483 | 2.575760114 | 0.04092103 | 0.479759698 | -3.907901442 |
| **A_55_P2039601** | Drosha | 0.335634861 | 0.018946204 | 0.652323518 | 10.95898609 | 2.575240205 | 0.040950108 | 0.479759698 | -3.908127741 |
| **A_55_P2157922** | Ndufaf1 | 0.329169131 | 0.018580155 | 0.639758106 | 11.08546859 | 2.575231367 | 0.040950602 | 0.479759698 | -3.908131588 |
| **A_51_P347240** | Lrr1 | 0.354238053 | 0.019929637 | 0.688546469 | 8.919772838 | 2.574726429 | 0.040978864 | 0.479759698 | -3.908351404 |
| **A_52_P106259** | Egfr | -0.855927369 | -1.663791586 | -0.048063153 | 9.791859537 | -2.574433812 | 0.040995252 | 0.479759698 | -3.914914407 |
| **A_52_P454994** | Pi4ka | 0.233766033 | 0.013076852 | 0.454455213 | 12.27705891 | 2.573851898 | 0.041027862 | 0.479759698 | -3.908732193 |
| **A_55_P2096367** | NA | -0.231610201 | -0.450284355 | -0.012936048 | 13.27203408 | -2.573614056 | 0.041041198 | 0.479759698 | -3.908835771 |
| **A_55_P2015989** | Rpl22 | 0.220454612 | 0.012275783 | 0.428633441 | 15.98887092 | 2.573154218 | 0.041066995 | 0.479759698 | -3.909036048 |
| **A_55_P2325729** | Arhgap1 | 0.330453609 | 0.018312234 | 0.642594984 | 10.91290845 | 2.572422791 | 0.041108064 | 0.479759698 | -3.909354667 |
| **A_65_P01319** | Pde4b | -0.635176664 | -1.235155745 | -0.035197582 | 8.483071825 | -2.57241844 | 0.041108308 | 0.479759698 | -3.909356563 |
| **A_51_P369200** | Tpx2 | 0.432282589 | 0.023926446 | 0.840638732 | 11.51411331 | 2.572242085 | 0.041118217 | 0.479759698 | -3.909433396 |
| **A_55_P2090142** | Kcna7 | 0.267752134 | 0.014745408 | 0.52075886 | 8.838924546 | 2.571485401 | 0.04116076 | 0.479759698 | -3.909763107 |
| **A_55_P2028821** | Hnrnpf | 0.260309904 | 0.014284624 | 0.506335184 | 13.99178601 | 2.570953058 | 0.041190718 | 0.479759698 | -3.90999511 |
| **A_55_P2016851** | Lce1a2 | 0.588034961 | 0.03219601 | 1.143873913 | 11.53245218 | 2.570616905 | 0.041209648 | 0.479759698 | -3.910141629 |
| **A_66_P119155** | NA | -0.377702435 | -0.734827793 | -0.020577077 | 10.93047083 | -2.569876689 | 0.041251362 | 0.479759698 | -3.910464317 |
| **A_55_P2016852** | Zyx | -0.410880821 | -0.799417512 | -0.02234413 | 12.72939904 | -2.569608863 | 0.041266467 | 0.479759698 | -3.91058109 |
| **A_55_P2085806** | Paxx | -0.294825719 | -0.573641512 | -0.016009927 | 10.63830062 | -2.569396847 | 0.041278428 | 0.479759698 | -3.910673537 |
| **A_55_P2016486** | Tmem131 | 0.212410412 | 0.011441393 | 0.41337943 | 11.40106726 | 2.568206132 | 0.04134567 | 0.479759698 | -3.911192835 |
| **A_51_P109171** | Os9 | -0.221752409 | -0.431566448 | -0.01193837 | 12.42325603 | -2.56812992 | 0.041349978 | 0.479759698 | -3.911226079 |
| **A_51_P381749** | Prelid3b | 0.214174971 | 0.011520553 | 0.416829388 | 12.64552891 | 2.568004792 | 0.041357052 | 0.479759698 | -3.911280662 |

| | | | | | | | | |
|---|---|---|---|---|---|---|---|---|
| **A_55_P2081349** | Focad | 0.271857453 | 0.014570051 | 0.529144856 | 10.31513685 | 2.567473152 | 0.041387121 | 0.479759698 | -3.911512596 |
| **A_52_P598714** | Afg3l1 | 0.220563799 | 0.011820077 | 0.429307521 | 10.03608536 | 2.567461849 | 0.041387761 | 0.479759698 | -3.911517527 |
| **A_55_P1998797** | NA | 0.27923653 | 0.014946461 | 0.543526599 | 13.10994186 | 2.567287918 | 0.041397603 | 0.479759698 | -3.911593414 |
| **A_51_P447545** | Igfbp1 | -0.829091748 | -1.613895009 | -0.044288488 | 16.56019579 | -2.566994767 | 0.041414198 | 0.479759698 | -3.911721327 |
| **A_65_P20683** | NA | 0.786903779 | 0.041767962 | 1.532039595 | 10.64208813 | 2.566075207 | 0.041466298 | 0.479759698 | -3.912122635 |
| **A_55_P2048022** | Gm6578 | 0.187382069 | 0.009908814 | 0.364855325 | 11.61976185 | 2.565537123 | 0.041496817 | 0.479759698 | -3.912357512 |
| **A_52_P276935** | Dnajb6 | 0.374505527 | 0.019799473 | 0.72921158 | 11.53963192 | 2.565504731 | 0.041498655 | 0.479759698 | -3.912371653 |
| **A_55_P2260052** | Gm17753 | 0.449777052 | 0.023680265 | 0.875873839 | 8.499163868 | 2.564910548 | 0.041532384 | 0.479759698 | -3.912631063 |
| **A_65_P03364** | Dennd4a | 0.441348669 | 0.023213149 | 0.859484189 | 8.759241999 | 2.564767188 | 0.041540526 | 0.479759698 | -3.912693659 |
| **A_51_P249051** | Pycr2 | 0.222612081 | 0.011695996 | 0.433528165 | 13.41538235 | 2.564615236 | 0.041549159 | 0.479759698 | -3.912760008 |
| **A_55_P2022699** | NA | 0.536444934 | 0.028120649 | 1.044769219 | 8.894770448 | 2.564292008 | 0.041567527 | 0.479759698 | -3.912901155 |
| **A_55_P1982494** | H2al1m | 0.354312982 | 0.018471587 | 0.690154377 | 8.69670716 | 2.563515978 | 0.041611662 | 0.479759698 | -3.913240086 |
| **A_55_P2077613** | Fam102a | -0.5489118 | -1.069315063 | -0.028508538 | 10.97122497 | -2.56298312 | 0.041641996 | 0.479759698 | -3.913472856 |
| **A_55_P2108808** | Tubgcp4 | 0.197806973 | 0.010257078 | 0.385356868 | 8.504497932 | 2.562760145 | 0.041654697 | 0.479759698 | -3.91357027 |
| **A_51_P232371** | Stab1 | -0.274751585 | -0.535256532 | -0.014246639 | 11.68636759 | -2.562756961 | 0.041654878 | 0.479759698 | -3.913571661 |
| **A_55_P1989772** | Sqor | -0.416808183 | -0.812048233 | -0.021568133 | 10.78185425 | -2.562468174 | 0.041671333 | 0.479759698 | -3.913697836 |
| **A_55_P1985259** | C2cd3 | 0.207421493 | 0.010651649 | 0.404191337 | 8.502838871 | 2.561405895 | 0.04173192 | 0.479759698 | -3.914162054 |
| **A_55_P2133427** | Vkorc1 | -0.398740696 | -0.777109452 | -0.020371941 | 12.40881233 | -2.560698728 | 0.041772304 | 0.479759698 | -3.914471166 |
| **A_55_P2169010** | Ndfip1 | -0.17645384 | -0.343896169 | -0.009011512 | 11.06217261 | -2.560643084 | 0.041775484 | 0.479759698 | -3.914495492 |
| **A_66_P108567** | Xrcc6 | 0.277376595 | 0.014001614 | 0.540751576 | 12.24345965 | 2.55904831 | 0.041866717 | 0.479759698 | -3.915192835 |
| **A_55_P1985793** | Dock6 | -0.430904769 | -0.840075838 | -0.021733699 | 11.19165126 | -2.558936858 | 0.0418731 | 0.479759698 | -3.915241582 |
| **A_51_P237575** | Akirin1 | 0.237821426 | 0.011890446 | 0.463752406 | 10.93355755 | 2.557751702 | 0.041941047 | 0.479759698 | -3.915760039 |
| **A_55_P1961567** | Ypel3 | 0.506936809 | 0.02533223 | 0.988541388 | 14.4641042 | 2.557681183 | 0.041945093 | 0.479759698 | -3.915790894 |
| **A_52_P15008** | Mapkapk2 | -0.300354519 | -0.585730154 | -0.014978884 | 10.12671274 | -2.557410646 | 0.041960622 | 0.479759698 | -3.915909271 |
| **A_55_P1986902** | Med23 | 0.235356134 | 0.011734932 | 0.458977337 | 8.848318594 | 2.557382756 | 0.041962223 | 0.479759698 | -3.915921475 |
| **A_51_P334308** | Tspan14 | -0.235025754 | -0.458458337 | -0.011593171 | 11.79643671 | -2.555948721 | 0.042044637 | 0.479759698 | -3.916549114 |
| **A_66_P112855** | AI413582 | -0.20874045 | -0.407190779 | -0.010290121 | 10.96709482 | -2.555865415 | 0.04204943 | 0.479759698 | -3.916585583 |
| **A_55_P2472435** | Gbp3 | -0.213838308 | -0.417152514 | -0.010524101 | 8.505724342 | -2.55564762 | 0.042061964 | 0.479759698 | -3.916680931 |
| **A_52_P490863** | Nop10 | -0.179638761 | -0.350439983 | -0.008837539 | 12.98101353 | -2.555596396 | 0.042064912 | 0.479759698 | -3.916703357 |
| **A_51_P469568** | Cldn13 | 0.418154108 | 0.020556277 | 0.815751939 | 11.85494007 | 2.555498031 | 0.042070574 | 0.479759698 | -3.916746423 |
| **A_55_P2076360** | NA | 0.27882485 | 0.01364131 | 0.54400839 | 8.468303109 | 2.554865854 | 0.042106983 | 0.479759698 | -3.91702323 |
| **A_55_P2067976** | Gmeb1 | 0.288560924 | 0.013977141 | 0.563144707 | 9.771420881 | 2.553558579 | 0.04218238 | 0.479759698 | -3.917595798 |
| **A_51_P303749** | Depdc1b | 0.414248463 | 0.020004519 | 0.808492407 | 11.29830803 | 2.55316607 | 0.042205045 | 0.479759698 | -3.917767754 |
| **A_55_P2140449** | B3galt4 | -0.177708672 | -0.346907243 | -0.0085101 | 8.232956421 | -2.552084894 | 0.042267544 | 0.479759698 | -3.918241513 |
| **A_55_P1998411** | Chchd3 | 0.277281873 | 0.01316635 | 0.541397396 | 11.75128871 | 2.551001661 | 0.04233026 | 0.479759698 | -3.918716322 |
| **A_66_P139703** | NA | -0.744342774 | -1.453387756 | -0.035297792 | 9.44046337 | -2.550835061 | 0.042339915 | 0.479759698 | -3.91878936 |
| **A_66_P125212** | Mapk1ip1l | 0.339239661 | 0.016061572 | 0.66241775 | 11.53311168 | 2.550632569 | 0.042351652 | 0.479759698 | -3.918878138 |
| **A_55_P2040170** | Pmp22 | -0.694251636 | -1.355778066 | -0.032725207 | 11.5217788 | -2.550074691 | 0.042384007 | 0.479759698 | -3.919122755 |
| **A_51_P405668** | 5-Mar | 0.327969569 | 0.01544864 | 0.640490498 | 11.83896749 | 2.549985035 | 0.042389209 | 0.479759698 | -3.919162071 |
| **A_55_P1998688** | Gm2012 | 0.193680243 | 0.009110808 | 0.378249678 | 8.029937824 | 2.549815322 | 0.042399058 | 0.479759698 | -3.919236496 |
| **A_51_P415905** | Pola1 | 0.546926792 | 0.025611254 | 1.06824233 | 10.84695879 | 2.549245832 | 0.042432126 | 0.479759698 | -3.919486264 |
| **A_55_P1996988** | Tent4b | 0.295413627 | 0.013770325 | 0.577056929 | 10.75430883 | 2.548673983 | 0.042465358 | 0.479759698 | -3.919737108 |
| **A_55_P2014927** | Zfp341 | 0.280671602 | 0.013033105 | 0.5483101 | 9.575394532 | 2.548197467 | 0.042493071 | 0.479759698 | -3.919946166 |
| **A_55_P2065726** | Snx29 | 0.261347527 | 0.01209754 | 0.510597514 | 8.524679516 | 2.547806491 | 0.042515823 | 0.479759698 | -3.920117717 |
| **A_55_P2151601** | Samd9l | -0.277361639 | -0.541888131 | -0.012835146 | 9.816648024 | -2.547771119 | 0.042517882 | 0.479759698 | -3.920133238 |
| **A_55_P2093974** | Mphosph10 | 0.346479306 | 0.016028222 | 0.676930389 | 8.978335054 | 2.547729462 | 0.042520307 | 0.479759698 | -3.920151518 |
| **A_55_P2327177** | Exoc8 | 0.217030026 | 0.010012761 | 0.424047292 | 8.854817907 | 2.547395892 | 0.042539731 | 0.479759698 | -3.920297899 |
| **A_55_P1956843** | NA | 0.279158174 | 0.012845257 | 0.54547109 | 13.93155084 | 2.547072501 | 0.042558571 | 0.479759698 | -3.920439827 |
| **A_55_P2006975** | NA | 0.449064722 | 0.020613862 | 0.877515583 | 9.729582913 | 2.54677811 | 0.04257573 | 0.479759698 | -3.920569038 |
| **A_52_P88695** | Dgka | -0.253647758 | -0.495659797 | -0.01163572 | 9.165762199 | -2.54669683 | 0.042580468 | 0.479759698 | -3.920604715 |

| | | | | | | | | |
|---|---|---|---|---|---|---|---|---|
| **A_55_P1996082** | Zfp936 | 0.332693201 | 0.015226123 | 0.650160278 | 9.813086554 | 2.54641052 | 0.042597165 | 0.479759698 | -3.920730394 |
| **A_52_P310981** | Eif4e | 0.184758614 | 0.008454919 | 0.361062308 | 11.52214171 | 2.546399096 | 0.042597831 | 0.479759698 | -3.920735409 |
| **A_51_P241769** | Rhd | 0.45506973 | 0.020770706 | 0.889368755 | 15.10191778 | 2.546081399 | 0.042616366 | 0.479759698 | -3.920874878 |
| **A_55_P2153621** | Ahnak | -0.275959132 | -0.53937842 | -0.012539845 | 9.83677313 | -2.545542714 | 0.042647813 | 0.479759698 | -3.92111139 |
| **A_51_P464738** | Slc2a1 | 0.515981613 | 0.023369283 | 1.008593944 | 14.29969021 | 2.545142697 | 0.042671181 | 0.479759698 | -3.921287043 |
| **A_51_P313157** | Arhgef5 | -0.32651853 | -0.638289759 | -0.014747301 | 9.298365439 | -2.544807793 | 0.042690756 | 0.479759698 | -3.92143412 |
| **A_55_P2098598** | Btg1 | -0.511417666 | -0.999784605 | -0.023050726 | 12.31903288 | -2.544559781 | 0.042705258 | 0.479759698 | -3.921543046 |
| **A_55_P1975650** | Fbxo33 | 0.269223257 | 0.012032361 | 0.526414153 | 9.410684093 | 2.543549368 | 0.042764394 | 0.479759698 | -3.921986898 |
| **A_52_P447196** | Col4a6 | 0.242778238 | 0.010836415 | 0.474720061 | 8.485855611 | 2.543395379 | 0.042773414 | 0.479759698 | -3.922054553 |
| **A_51_P367060** | Ifrd1 | 0.292532454 | 0.01299635 | 0.572068559 | 11.34825397 | 2.542841761 | 0.042805859 | 0.479759698 | -3.922297811 |
| **A_51_P335981** | Narf | 0.446200677 | 0.019800923 | 0.872600431 | 10.5072286 | 2.542707879 | 0.042813709 | 0.479759698 | -3.922356644 |
| **A_52_P267824** | Ces2g | 0.459095574 | 0.020356503 | 0.897834646 | 9.967362267 | 2.542611361 | 0.042819369 | 0.479759698 | -3.922399059 |
| **A_55_P1968365** | NA | 0.253574088 | 0.011186224 | 0.495961952 | 8.280204633 | 2.542009623 | 0.042854675 | 0.479759698 | -3.922663522 |
| **A_55_P2108404** | NA | 0.326872666 | 0.014371283 | 0.639374049 | 11.26922028 | 2.5416155 | 0.042877817 | 0.479759698 | -3.922836764 |
| **A_55_P2004797** | Tacc2 | -0.435056127 | -0.851090059 | -0.019022195 | 10.1453633 | -2.540971096 | 0.042915682 | 0.479759698 | -3.923120062 |
| **A_52_P82247** | Cycs | 0.318030452 | 0.013838162 | 0.622222741 | 12.09818433 | 2.540409297 | 0.042948722 | 0.479759698 | -3.923367087 |
| **A_51_P107782** | Kdsr | -0.294379137 | -0.576012093 | -0.01274618 | 8.883554047 | -2.539842243 | 0.042982098 | 0.479759698 | -3.942486756 |
| **A_66_P139295** | Tomm40 | 0.400112072 | 0.017248261 | 0.782975883 | 12.32598601 | 2.539338085 | 0.043011795 | 0.479759698 | -3.923838215 |
| **A_55_P2116235** | NA | 0.204047236 | 0.00879076 | 0.399303712 | 14.59099013 | 2.539267519 | 0.043015954 | 0.479759698 | -3.923869255 |
| **A_51_P152826** | Golt1b | 0.283475254 | 0.012193317 | 0.55475719 | 10.00039022 | 2.539086317 | 0.043026634 | 0.479759698 | -3.923948965 |
| **A_55_P2027822** | Sgf29 | 0.251742989 | 0.010804634 | 0.492681345 | 11.46238148 | 2.538835911 | 0.043041397 | 0.479759698 | -3.924059125 |
| **A_55_P1998354** | Wbp4 | 0.181659306 | 0.007740759 | 0.355577854 | 8.507956574 | 2.53801942 | 0.043089573 | 0.479759698 | -3.924418374 |
| **A_66_P132888** | Dhfr | 0.34790569 | 0.014815616 | 0.680995763 | 9.183842693 | 2.537949811 | 0.043093683 | 0.479759698 | -3.924449005 |
| **A_55_P2087567** | Olfr373 | 0.320454514 | 0.013612646 | 0.627296382 | 8.774477864 | 2.537668943 | 0.04311027 | 0.479759698 | -3.924572606 |
| **A_55_P2080592** | Spata31d1b | 1.023653754 | 0.04346917 | 2.003838339 | 11.06296807 | 2.537630611 | 0.043112534 | 0.479759698 | -3.924589476 |
| **A_55_P2116054** | Caprin1 | 0.228555893 | 0.009555786 | 0.447556001 | 8.425861309 | 2.5358951 | 0.043215182 | 0.479759698 | -3.925353451 |
| **A_66_P113884** | NA | 0.284262765 | 0.01179342 | 0.556732109 | 8.166424237 | 2.535044092 | 0.04326561 | 0.479759698 | -3.925728205 |
| **A_55_P2091858** | Hipk2 | 0.359975689 | 0.014929317 | 0.705022062 | 10.60577321 | 2.535005435 | 0.043267902 | 0.479759698 | -3.925745231 |
| **A_55_P2186808** | NA | 0.453540859 | 0.018793458 | 0.88828826 | 12.17876581 | 2.534910398 | 0.043273537 | 0.479759698 | -3.925787088 |
| **A_55_P2094292** | Erp27 | 0.373077891 | 0.015425517 | 0.730730265 | 9.562029765 | 2.534670968 | 0.043287739 | 0.479759698 | -3.925892545 |
| **A_55_P1956488** | Dmtn | 0.59080236 | 0.024318003 | 1.157286718 | 11.11588613 | 2.534180173 | 0.043316865 | 0.479759698 | -3.926108739 |
| **A_55_P1973941** | Slc7a5 | 0.482662446 | 0.019855688 | 0.945469205 | 10.00320254 | 2.534119008 | 0.043320497 | 0.479759698 | -3.926135684 |
| **A_51_P459661** | Lipa | -0.207046856 | -0.405584919 | -0.008508793 | 11.21707951 | -2.534008394 | 0.043327065 | 0.479759698 | -3.926184415 |
| **A_55_P1954266** | Zfp672 | 0.258224312 | 0.0104747 | 0.505973925 | 9.851815296 | 2.532604267 | 0.043410528 | 0.479759698 | -3.926803128 |
| **A_55_P2118799** | Rmi1 | 0.204664528 | 0.008274452 | 0.401054604 | 8.699469636 | 2.532247956 | 0.043431735 | 0.479759698 | -3.926960172 |
| **A_55_P2002723** | Inpp5a | -0.551372595 | -1.080483794 | -0.022261396 | 10.04751533 | -2.532103256 | 0.04344035 | 0.479759698 | -3.927023954 |
| **A_55_P1997275** | Xpnpep1 | 0.202566966 | 0.008152422 | 0.39698151 | 13.26887022 | 2.531763078 | 0.043460611 | 0.479759698 | -3.927173908 |
| **A_55_P1970536** | NA | 0.296892482 | 0.011942226 | 0.581842738 | 8.103772241 | 2.531706387 | 0.043463988 | 0.479759698 | -3.9271989 |
| **A_51_P190765** | Rtca | 0.205053878 | 0.008201681 | 0.401906076 | 10.2660718 | 2.531109356 | 0.043499574 | 0.479759698 | -3.927462118 |
| **A_55_P2052623** | Pgap2 | -0.476428734 | -0.933914583 | -0.018942884 | 10.40345001 | -2.530483268 | 0.043536925 | 0.479759698 | -3.927738196 |
| **A_55_P1964664** | Fus | 0.198040281 | 0.007857264 | 0.388223298 | 13.35367826 | 2.530259067 | 0.043550308 | 0.479759698 | -3.927837071 |
| **A_55_P2174582** | Urah | -0.392237023 | -0.768975394 | -0.015498652 | 12.91020419 | -2.529833374 | 0.043575731 | 0.479759698 | -3.928024823 |
| **A_51_P150763** | Ropn1l | -0.238597204 | -0.467853202 | -0.009341206 | 10.26550283 | -2.528877735 | 0.043632861 | 0.479759698 | -3.928446392 |
| **A_55_P2065513** | Ebf1 | -0.28372578 | -0.556591037 | -0.010860523 | 8.556399261 | -2.526584026 | 0.043770304 | 0.479759698 | -3.929458706 |
| **A_55_P2082185** | NA | 0.276392403 | 0.010563034 | 0.542221772 | 13.55354982 | 2.526424537 | 0.043779878 | 0.479759698 | -3.92952912 |
| **A_55_P2038262** | Phf11b | -0.203906855 | -0.400036064 | -0.007777645 | 8.365218604 | -2.52622912 | 0.043791611 | 0.479759698 | -3.9296154 |
| **A_55_P2083401** | Acp1 | 0.365649269 | 0.013842907 | 0.71745563 | 12.65551589 | 2.525481608 | 0.043836525 | 0.479759698 | -3.929945487 |
| **A_52_P213181** | Cul4b | 0.204606145 | 0.00769704 | 0.40151525 | 9.38038118 | 2.524852794 | 0.043874345 | 0.479759698 | -3.930223213 |
| **A_51_P173100** | Pttg1ip | -0.210959078 | -0.413983273 | -0.007934883 | 11.76086809 | -2.524838542 | 0.043875202 | 0.479759698 | -3.930229508 |
| **A_52_P480939** | Tnpo1 | 0.267670065 | 0.01003079 | 0.525309341 | 11.03752911 | 2.524474128 | 0.043897136 | 0.479759698 | -3.930390481 |

| | | | | | | | | |
|---|---|---|---|---|---|---|---|---|
| **A_51_P125135** | Cdca5 | 0.329647674 | 0.012332951 | 0.646962398 | 11.69893526 | 2.524311703 | 0.043906916 | 0.479759698 | -3.930462235 |
| **A_55_P2092329** | Adrm1 | 0.324641248 | 0.012098441 | 0.637184056 | 11.66222416 | 2.523930428 | 0.043929882 | 0.479759698 | -3.930630682 |
| **A_55_P1986158** | Klhdc4 | 0.248663715 | 0.009198746 | 0.488128685 | 11.50648718 | 2.523211277 | 0.043973234 | 0.479759698 | -3.930948452 |
| **A_51_P357573** | Cald1 | -0.22341827 | -0.438573019 | -0.008263521 | 10.03409065 | -2.523195707 | 0.043974173 | 0.479759698 | -3.930955332 |
| **A_55_P2043798** | Mrto4 | 0.320467096 | 0.011647712 | 0.62928648 | 11.63780069 | 2.52151805 | 0.044075485 | 0.479759698 | -3.931696892 |
| **A_55_P2045367** | G630025P09Rik | -0.322972122 | -0.634337935 | -0.01160631 | 8.193475275 | -2.520445441 | 0.044140387 | 0.479759698 | -3.932171193 |
| **A_52_P114282** | Armcx5 | 0.240349029 | 0.008629784 | 0.472068274 | 8.910810064 | 2.520365095 | 0.044145253 | 0.479759698 | -3.932206728 |
| **A_51_P365964** | Tor1a | -0.182944096 | -0.359342514 | -0.006545679 | 12.00028598 | -2.520036925 | 0.044165132 | 0.479759698 | -3.932351875 |
| **A_52_P150565** | Usp39 | 0.273879754 | 0.009717646 | 0.538041862 | 9.138545869 | 2.519257714 | 0.044212372 | 0.479759698 | -3.932696568 |
| **A_55_P2161773** | Mpp3 | 0.452317674 | 0.016002048 | 0.8886333 | 9.348610921 | 2.518987309 | 0.044228778 | 0.479759698 | -3.932816203 |
| **A_51_P459070** | Aplf | 0.367886113 | 0.012965299 | 0.722806927 | 10.00623559 | 2.518634308 | 0.044250205 | 0.479759698 | -3.932972394 |
| **A_55_P2182392** | Adat3 | 0.250730935 | 0.008819325 | 0.492642545 | 9.311395882 | 2.518456168 | 0.044261021 | 0.479759698 | -3.933051221 |
| **A_55_P2134735** | Fdxacb1 | 0.262621699 | 0.009225025 | 0.516018374 | 9.205992178 | 2.518331431 | 0.044268597 | 0.479759698 | -3.93310642 |
| **A_55_P2145136** | NA | 0.494292844 | 0.017360374 | 0.971225315 | 9.440186989 | 2.518318299 | 0.044269395 | 0.479759698 | -3.933112231 |
| **A_55_P1992839** | Dsn1 | 0.473512203 | 0.0165847 | 0.930439706 | 10.37515656 | 2.518065744 | 0.044284738 | 0.479759698 | -3.933223998 |
| **A_55_P2021570** | C87414 | 0.183467631 | 0.006404729 | 0.360530532 | 8.175034363 | 2.517764259 | 0.044303062 | 0.479759698 | -3.933357429 |
| **A_55_P2035613** | Tmbim1 | -0.329121941 | -0.646818145 | -0.011425738 | 11.20228496 | -2.517259568 | 0.044333753 | 0.479759698 | -3.933580821 |
| **A_55_P2105220** | Hscb | 0.287070157 | 0.009904732 | 0.564235583 | 12.78609179 | 2.516704246 | 0.04436755 | 0.479759698 | -3.93382666 |
| **A_55_P2110671** | NA | 0.35034487 | 0.012087071 | 0.688602669 | 12.56549142 | 2.516698165 | 0.04436792 | 0.479759698 | -3.933829353 |
| **A_55_P2155572** | NA | 0.274287772 | 0.009445233 | 0.539130312 | 13.49717425 | 2.516528728 | 0.044378237 | 0.479759698 | -3.93390437 |
| **A_55_P2143522** | Fstl3 | -0.497289988 | -0.977470442 | -0.017109534 | 8.708152573 | -2.516450703 | 0.044382989 | 0.479759698 | -3.933938916 |
| **A_52_P161495** | Bcl6 | -0.312870239 | -0.615064609 | -0.010675868 | 9.26125932 | -2.515712876 | 0.044427951 | 0.479759698 | -3.934265634 |
| **A_55_P1967335** | NA | 0.262344124 | 0.008912042 | 0.515776207 | 10.73378744 | 2.515318229 | 0.04445202 | 0.479759698 | -3.934440415 |
| **A_51_P286460** | H2ac8 | 0.433422035 | 0.014696612 | 0.852147458 | 8.474347929 | 2.515155532 | 0.044461946 | 0.479759698 | -3.934512476 |
| **A_52_P817257** | Smim22 | -0.470207648 | -0.924525639 | -0.015889657 | 11.52037018 | -2.514854957 | 0.044480292 | 0.479759698 | -3.934645615 |
| **A_55_P2340101** | NA | 0.504249634 | 0.016981398 | 0.99151787 | 11.20673583 | 2.514552337 | 0.044498769 | 0.479759698 | -3.93477967 |
| **A_55_P1957598** | Atad2b | 0.218527079 | 0.007355297 | 0.429698861 | 10.79542123 | 2.51450535 | 0.044501639 | 0.479759698 | -3.934800486 |
| **A_55_P1961531** | Nemp1 | 0.264319533 | 0.008761941 | 0.519877126 | 9.680420346 | 2.513180373 | 0.044582642 | 0.479759698 | -3.935387572 |
| **A_51_P324640** | C330018D20Rik | 0.35295634 | 0.011692894 | 0.694219787 | 10.33420907 | 2.513126814 | 0.04458592 | 0.479759698 | -3.935411308 |
| **A_55_P2144305** | Gm3211 | 0.276490327 | 0.009037579 | 0.543943076 | 8.60602177 | 2.511979356 | 0.044656201 | 0.479759698 | -3.935919923 |
| **A_55_P2101696** | Gnat2 | 0.341117072 | 0.011104345 | 0.671129798 | 9.058396977 | 2.5116317 | 0.044677518 | 0.479759698 | -3.936074055 |
| **A_55_P1985514** | Dlg3 | -0.39218087 | -0.771598063 | -0.012763676 | 10.40445726 | -2.511612217 | 0.044678713 | 0.479759698 | -3.936082693 |
| **A_55_P2037081** | Zfp979 | 0.187414662 | 0.006092155 | 0.36873717 | 8.591618153 | 2.511510726 | 0.044684938 | 0.479759698 | -3.936127692 |
| **A_66_P103477** | Nkapd1 | 0.168322415 | 0.005441362 | 0.331203467 | 9.06470398 | 2.511045458 | 0.044713488 | 0.479759698 | -3.936333998 |
| **A_55_P2145543** | Eef2k | 0.212005344 | 0.006844941 | 0.417165747 | 8.932921476 | 2.510940689 | 0.04471992 | 0.479759698 | -3.936380458 |
| **A_55_P2120615** | Ssh3 | -0.380259268 | -0.74832388 | -0.012194656 | 8.876318357 | -2.51037692 | 0.044754545 | 0.479759698 | -3.936630484 |
| **A_55_P1992487** | NA | 0.360471319 | 0.011533657 | 0.709408982 | 14.12371398 | 2.510186891 | 0.044766222 | 0.479759698 | -3.936714769 |
| **A_55_P1986983** | Gne | -0.395615751 | -0.778597579 | -0.012633924 | 11.57775877 | -2.510028174 | 0.044775978 | 0.479759698 | -3.93678517 |
| **A_55_P2414924** | Luc7l | 0.258557284 | 0.00824212 | 0.508872449 | 8.90133147 | 2.509879123 | 0.044785142 | 0.479759698 | -3.936851286 |
| **A_52_P330540** | Mis18a | 0.292670531 | 0.009253161 | 0.576087902 | 10.25590371 | 2.509202553 | 0.044826762 | 0.479759698 | -3.937151433 |
| **A_55_P2121760** | NA | 0.265941378 | 0.008406265 | 0.523476491 | 8.2799414 | 2.50918483 | 0.044827853 | 0.479759698 | -3.937159297 |
| **A_55_P2110245** | Fignl1 | 0.448915658 | 0.014171713 | 0.883659604 | 12.68474281 | 2.509079376 | 0.044834344 | 0.479759698 | -3.937206085 |
| **A_51_P223036** | Fbxo30 | 0.424726178 | 0.013378309 | 0.836074047 | 9.986479688 | 2.508897779 | 0.044845524 | 0.479759698 | -3.93728666 |
| **A_66_P112024** | NA | -0.329538867 | -0.648722695 | -0.010355039 | 10.93169205 | -2.508701294 | 0.044857624 | 0.479759698 | -3.937373844 |
| **A_52_P570266** | Psmb10 | -0.247943321 | -0.488181815 | -0.007704828 | 13.17535687 | -2.507800631 | 0.044913133 | 0.479759698 | -3.937773552 |
| **A_51_P302377** | Fgfbp1 | -0.297972941 | -0.586727943 | -0.00921794 | 8.287004629 | -2.507439722 | 0.044935396 | 0.479759698 | -3.93793375 |
| **A_52_P145349** | Stradb | 0.449726702 | 0.013788386 | 0.885665019 | 10.66672691 | 2.506725742 | 0.044979474 | 0.479759698 | -3.938250713 |
| **A_55_P2143204** | Pbxip1 | -0.295255765 | -0.58151815 | -0.008993381 | 9.277321066 | -2.506209032 | 0.045011401 | 0.479759698 | -3.938480141 |
| **A_51_P401504** | Col9a2 | 0.291218751 | 0.008867635 | 0.573569867 | 8.818780584 | 2.506184359 | 0.045012927 | 0.479759698 | -3.938491097 |
| **A_55_P2093689** | Pold3 | 0.332768077 | 0.009938066 | 0.655598088 | 11.68688568 | 2.50467249 | 0.045106485 | 0.479759698 | -3.939162587 |

| | | | | | | | | | |
|---|---|---|---|---|---|---|---|---|---|
| **A_55_P2145557** | NA | -0.360804756 | -0.710841095 | -0.010768417 | 13.07942294 | -2.504622682 | 0.045109571 | 0.479759698 | -3.939184714 |
| **A_55_P1965050** | Cks2 | 0.329152257 | 0.009794369 | 0.648510144 | 11.78302941 | 2.50439241 | 0.04512384 | 0.479759698 | -3.939287015 |
| **A_51_P174081** | Zdhhc14 | 0.279299402 | 0.008277986 | 0.550320817 | 8.844954463 | 2.504087992 | 0.045142711 | 0.479759698 | -3.939422266 |
| **A_55_P1977096** | Elk4 | 0.371028465 | 0.010925911 | 0.731131018 | 10.67308248 | 2.503595817 | 0.045173238 | 0.479759698 | -3.939640962 |
| **A_55_P2180884** | Psmd10 | 0.324960799 | 0.009541231 | 0.640380367 | 9.847778479 | 2.503372808 | 0.045187077 | 0.479759698 | -3.939740065 |
| **A_55_P2062678** | Hmox2 | -0.209983262 | -0.413809287 | -0.006157237 | 9.791924958 | -2.503273096 | 0.045193266 | 0.479759698 | -3.939784377 |
| **A_55_P1979432** | Prdx2 | 0.282628475 | 0.008238694 | 0.557018257 | 16.08219375 | 2.502828957 | 0.045220845 | 0.479759698 | -3.939981772 |
| **A_51_P517870** | Nfatc1 | -0.265924373 | -0.524105985 | -0.00774276 | 9.005977184 | -2.502741663 | 0.045226268 | 0.479759698 | -3.940020572 |
| **A_52_P211223** | Cdca2 | 0.35452682 | 0.010244326 | 0.698809314 | 9.412567219 | 2.502173058 | 0.045261606 | 0.479759698 | -3.940273327 |
| **A_55_P1961499** | Ly6c1 | -0.43681222 | -0.861071914 | -0.012552527 | 9.794900446 | -2.501763172 | 0.045287098 | 0.479759698 | -3.940455554 |
| **A_51_P156631** | Glrx5 | 0.341944645 | 0.009807082 | 0.674082208 | 14.87675447 | 2.501618047 | 0.045296128 | 0.479759698 | -3.940520078 |
| **A_55_P1965467** | 2300009A05Rik | 0.460484794 | 0.013205422 | 0.907764166 | 10.54915595 | 2.501610052 | 0.045296625 | 0.479759698 | -3.940523633 |
| **A_55_P2064257** | Dnaic2 | -0.368952526 | -0.727329886 | -0.010575166 | 8.369092142 | -2.501572599 | 0.045298956 | 0.479759698 | -3.940540286 |
| **A_52_P253567** | Hsd3b6 | -0.41111573 | -0.810521887 | -0.011709572 | 8.746164544 | -2.501108466 | 0.045327848 | 0.479759698 | -3.940746666 |
| **A_55_P2041310** | NA | 0.39051892 | 0.011060955 | 0.769976885 | 9.106683657 | 2.500700006 | 0.04535329 | 0.479759698 | -3.940928312 |
| **A_52_P595954** | Nelfa | 0.264689348 | 0.007487711 | 0.521890985 | 10.16905601 | 2.500609779 | 0.045358912 | 0.479759698 | -3.94096844 |
| **A_55_P2109922** | NA | 0.631858921 | 0.017873107 | 1.245844734 | 9.947622685 | 2.500604301 | 0.045359253 | 0.479759698 | -3.940970877 |
| **A_55_P2138333** | Mad2l1 | 0.262636021 | 0.00741957 | 0.517852472 | 8.434411184 | 2.500511256 | 0.045365052 | 0.479759698 | -3.941012259 |
| **A_51_P486618** | Akr1b10 | -0.255069505 | -0.502933488 | -0.007205521 | 10.31535943 | -2.500508311 | 0.045365235 | 0.479759698 | -3.941013569 |
| **A_51_P349825** | 1700021F05Rik | 0.245743926 | 0.006881741 | 0.484606111 | 11.70627585 | 2.499876649 | 0.045404621 | 0.479759698 | -3.941294532 |
| **A_51_P365019** | Gclc | 0.42462058 | 0.011835018 | 0.837406141 | 11.81106894 | 2.499537923 | 0.045425756 | 0.479759698 | -3.941445219 |
| **A_51_P202050** | Dtx1 | -0.293092637 | -0.578038213 | -0.00814706 | 7.85675681 | -2.49934482 | 0.04543781 | 0.479759698 | -3.941531129 |
| **A_51_P435968** | Tnfaip8 | 0.357652421 | 0.009841089 | 0.705463753 | 10.30707452 | 2.498622407 | 0.045482933 | 0.479759698 | -3.941852567 |
| **A_66_P100365** | NA | 0.199952824 | 0.005496785 | 0.394408864 | 8.555534147 | 2.498557198 | 0.045487008 | 0.479759698 | -3.941881586 |
| **A_51_P428505** | Krtcap2 | -0.244902318 | -0.483194315 | -0.006610321 | 13.62559537 | -2.497276487 | 0.045567127 | 0.479759698 | -3.94245161 |
| **A_55_P2128263** | Phf21a | -0.368558067 | -0.72724862 | -0.009867514 | 10.17496153 | -2.49671616 | 0.045602226 | 0.479759698 | -3.942701067 |
| **A_55_P2019814** | Rnf20 | 0.230021518 | 0.006122749 | 0.453920287 | 11.38074665 | 2.496318226 | 0.04562717 | 0.479759698 | -3.942878251 |
| **A_55_P2008576** | Ubr2 | 0.300316103 | 0.007973108 | 0.592659098 | 8.448421179 | 2.496141017 | 0.045638283 | 0.479759698 | -3.942957161 |
| **A_55_P2183055** | E4f1 | 0.301982777 | 0.008010726 | 0.595954829 | 10.33418365 | 2.496084716 | 0.045641814 | 0.479759698 | -3.942982233 |
| **A_51_P356931** | Orc1 | 0.302110304 | 0.007980316 | 0.596240293 | 8.621671992 | 2.495797939 | 0.045659806 | 0.479759698 | -3.943109943 |
| **A_55_P2015238** | Dctn4 | 0.173103155 | 0.004554404 | 0.341651907 | 11.07016245 | 2.495529076 | 0.04567668 | 0.479759698 | -3.943229686 |
| **A_51_P215097** | Klf16 | 0.391656369 | 0.01013333 | 0.773179409 | 9.266784152 | 2.494408705 | 0.045747067 | 0.479759698 | -3.943728757 |
| **A_55_P2001593** | Rnf180 | 0.49777096 | 0.012864418 | 0.982677502 | 9.568367338 | 2.49433455 | 0.04575173 | 0.479759698 | -3.943761795 |
| **A_55_P2065113** | Gm11559 | 0.394474191 | 0.010161714 | 0.778786667 | 9.639739917 | 2.494119735 | 0.04576524 | 0.479759698 | -3.943857505 |
| **A_55_P2098335** | Map2k4 | 0.304678517 | 0.007823872 | 0.601533162 | 8.640429982 | 2.493912277 | 0.045778291 | 0.479759698 | -3.943949942 |
| **A_55_P2090943** | Larp4 | 0.325005559 | 0.008303692 | 0.641707426 | 11.94560078 | 2.493580278 | 0.045799186 | 0.479759698 | -3.944097882 |
| **A_51_P239113** | Zfp35 | 0.199342021 | 0.005072975 | 0.393611067 | 10.7594741 | 2.493322395 | 0.045815423 | 0.479759698 | -3.944212805 |
| **A_55_P2020577** | Pcolce | -0.479406336 | -0.946651129 | -0.012161542 | 12.4300426 | -2.49311601 | 0.045828422 | 0.479759698 | -3.944304785 |
| **A_55_P1996299** | Pofut1 | 0.165249949 | 0.004171688 | 0.326328211 | 11.64921023 | 2.492800883 | 0.045848277 | 0.479759698 | -3.944445238 |
| **A_55_P1959541** | NA | 0.216555289 | 0.005453617 | 0.427656961 | 10.35121122 | 2.492644307 | 0.045858146 | 0.479759698 | -3.944515029 |
| **A_55_P2058851** | Fancf | 0.38280357 | 0.009630141 | 0.755976998 | 10.07916748 | 2.492576263 | 0.045862435 | 0.479759698 | -3.944545359 |
| **A_51_P208361** | Ak3 | -0.327245728 | -0.646285894 | -0.008205561 | 11.86345312 | -2.492365962 | 0.045875695 | 0.479759698 | -3.944639103 |
| **A_52_P466147** | Rarres2 | -0.363258367 | -0.717450111 | -0.009066623 | 12.73386492 | -2.492070829 | 0.04589431 | 0.479759698 | -3.944770672 |
| **A_55_P1977071** | Fam110a | 0.405424558 | 0.010110974 | 0.800738143 | 10.64597548 | 2.49201988 | 0.045897525 | 0.479759698 | -3.944793386 |
| **A_55_P2066613** | Lcp1 | -0.185699333 | -0.366784757 | -0.004613909 | 9.995001456 | -2.491781971 | 0.045912538 | 0.479759698 | -3.944899454 |
| **A_55_P2000823** | Rbm39 | 0.190309248 | 0.004696234 | 0.375922262 | 13.09466239 | 2.491349511 | 0.045939841 | 0.479759698 | -3.945092277 |
| **A_51_P114616** | Batf | -0.201020898 | -0.397115079 | -0.004926716 | 8.959802428 | -2.490919483 | 0.045967007 | 0.479759698 | -3.95128401 |
| **A_55_P2008417** | Mnd1 | 0.27256001 | 0.006642372 | 0.538477649 | 9.926334828 | 2.490566719 | 0.045989305 | 0.479759698 | -3.945441364 |
| **A_55_P2133220** | Arhgef39 | 0.29043585 | 0.007066358 | 0.573805342 | 10.65533225 | 2.490464296 | 0.045995781 | 0.479759698 | -3.945487046 |
| **A_55_P2032559** | NA | 0.415058007 | 0.010046237 | 0.820069776 | 14.16404774 | 2.490143305 | 0.046016083 | 0.479759698 | -3.945630218 |

| | | | | | | | | | |
|---|---|---|---|---|---|---|---|---|---|
| **A_51_P502798** | Nxpe4 | 0.3008251 | 0.007249434 | 0.594400765 | 8.744880277 | 2.489873051 | 0.046033184 | 0.479759698 | -3.94575077 |
| **A_55_P1987156** | Crhr2 | 0.31 | 0.007468977 | 0.612531022 | 8.430768862 | 2.489860226 | 0.046033996 | 0.479759698 | -3.945756491 |
| **A_55_P2337138** | Ago2 | 0.40697082 | 0.00978876 | 0.80415288 | 11.92028613 | 2.489756276 | 0.046040575 | 0.479759698 | -3.945802863 |
| **A_66_P106716** | Akap8 | 0.210915752 | 0.005072514 | 0.41675899 | 9.457028178 | 2.48974919 | 0.046041024 | 0.479759698 | -3.945806024 |
| **A_52_P298548** | Nbn | 0.237049761 | 0.005689486 | 0.468410036 | 8.743201915 | 2.4896249 | 0.046048892 | 0.479759698 | -3.945861471 |
| **A_51_P271865** | Clec4f | -0.509602764 | -1.007015312 | -0.012190216 | 11.24705024 | -2.489420301 | 0.046061847 | 0.479759698 | -3.945952749 |
| **A_55_P1957088** | Fam53b | 0.252841502 | 0.005956293 | 0.499726711 | 12.48601799 | 2.488493316 | 0.046120593 | 0.479759698 | -3.946366369 |
| **A_55_P1953053** | NA | 0.211675281 | 0.004967242 | 0.418383321 | 10.7311454 | 2.488261192 | 0.046135315 | 0.479759698 | -3.94646996 |
| **A_55_P2011237** | NA | 0.467710883 | 0.010959493 | 0.924462274 | 15.57896531 | 2.488174225 | 0.046140832 | 0.479759698 | -3.946508772 |
| **A_55_P2062029** | Tsen34 | -0.420851837 | -0.831858461 | -0.009845212 | 13.27505632 | -2.488075727 | 0.046147082 | 0.479759698 | -3.946552732 |
| **A_51_P478610** | Asb6 | 0.380803904 | 0.008815444 | 0.752792364 | 10.93382092 | 2.487454319 | 0.04618653 | 0.479759698 | -3.946830097 |
| **A_51_P258078** | Nop2 | 0.322982933 | 0.007455859 | 0.638510006 | 11.95515565 | 2.487288337 | 0.046197073 | 0.479759698 | -3.946904191 |
| **A_55_P2021358** | Klhl15 | 0.236692189 | 0.005463882 | 0.467920496 | 8.369102124 | 2.487288237 | 0.046197079 | 0.479759698 | -3.946904235 |
| **A_55_P2142251** | H3c15 | 0.384994465 | 0.008841877 | 0.761147053 | 8.711900375 | 2.486987607 | 0.046216181 | 0.479759698 | -3.947038444 |
| **A_55_P2122849** | NA | 0.354264307 | 0.007951881 | 0.700576733 | 11.07371464 | 2.485664517 | 0.046300347 | 0.480040109 | -3.947629239 |
| **A_52_P366653** | Hnrnph3 | 0.243152138 | 0.005456836 | 0.480847441 | 8.448652249 | 2.485654049 | 0.046301014 | 0.480040109 | -3.947633915 |
| **A_55_P2004201** | NA | 0.318771915 | 0.007072985 | 0.630470844 | 12.2200663 | 2.485008792 | 0.04634212 | 0.480040109 | -3.947922119 |
| **A_52_P683306** | Fam13b | 0.208613844 | 0.00461659 | 0.412611098 | 8.594809719 | 2.48486039 | 0.046351579 | 0.480040109 | -3.94798841 |
| **A_55_P2348815** | Casd1 | -0.226234608 | -0.44757478 | -0.004894435 | 9.111410877 | -2.483601912 | 0.046431878 | 0.480040109 | -3.948550681 |
| **A_55_P2138529** | Mrpl40 | 0.210815596 | 0.00454661 | 0.417084581 | 11.99865304 | 2.483430394 | 0.046442833 | 0.480040109 | -3.948627329 |
| **A_66_P128918** | Hacl1 | -0.594647097 | -1.176483397 | -0.012810798 | 11.4817256 | -2.483371425 | 0.0464466 | 0.480040109 | -3.948653681 |
| **A_55_P2074972** | Zswim6 | -0.196439419 | -0.38866223 | -0.004216608 | 8.096466518 | -2.483172597 | 0.046459305 | 0.480040109 | -3.948742539 |
| **A_55_P1973970** | Krt73 | 0.302147277 | 0.006428907 | 0.597865647 | 9.17374174 | 2.482696148 | 0.046489762 | 0.480040109 | -3.948955486 |
| **A_55_P2063251** | Dusp3 | -0.308965994 | -0.61139986 | -0.006532129 | 9.507940699 | -2.482352492 | 0.046511744 | 0.480040109 | -3.9491091 |
| **A_55_P2048101** | Gm6762 | -0.163696964 | -0.323937742 | -0.003456187 | 8.121242697 | -2.482280015 | 0.046516381 | 0.480040109 | -3.949141499 |
| **A_55_P1963764** | 1-Sep | 0.531090392 | 0.011125151 | 1.051055634 | 9.284463854 | 2.481860237 | 0.046543249 | 0.480061078 | -3.949329163 |
| **A_55_P2073709** | Tfdp2 | 0.460600315 | 0.009132209 | 0.912068421 | 11.1737188 | 2.479021793 | 0.046725354 | 0.480627651 | -3.950598675 |
| **A_55_P2111302** | Cp | -0.821080006 | -1.625981733 | -0.01617828 | 13.88259612 | -2.478710503 | 0.04674537 | 0.480627651 | -3.950737962 |
| **A_51_P400585** | Api5 | 0.228237491 | 0.004446925 | 0.452028058 | 10.79276864 | 2.478154612 | 0.046781138 | 0.480627651 | -3.950986727 |
| **A_55_P2004801** | Tacc3 | 0.392265909 | 0.007588922 | 0.776942895 | 12.90044819 | 2.477807423 | 0.046803492 | 0.480627651 | -3.951142115 |
| **A_55_P1964509** | Lcn12 | 0.338202167 | 0.006515778 | 0.669888555 | 9.746967257 | 2.477604178 | 0.046816583 | 0.480627651 | -3.951233086 |
| **A_55_P2003789** | Taf4 | 0.250446384 | 0.004819445 | 0.496073324 | 10.14149351 | 2.477547318 | 0.046820246 | 0.480627651 | -3.951258537 |
| **A_55_P2169529** | NA | 0.294530513 | 0.005634967 | 0.583426059 | 8.98114028 | 2.477265965 | 0.046838376 | 0.480627651 | -3.95138448 |
| **A_55_P1954820** | D1Ertd622e | 0.475754434 | 0.00908199 | 0.942426878 | 12.20142666 | 2.477158959 | 0.046845273 | 0.480627651 | -3.951432382 |
| **A_55_P2091116** | Kbtbd12 | 0.202903092 | 0.003862438 | 0.401943745 | 8.14246345 | 2.477023141 | 0.046854029 | 0.480627651 | -3.951493184 |
| **A_55_P2085412** | Ankle1 | 0.41872124 | 0.007968557 | 0.829473923 | 9.78439919 | 2.477010065 | 0.046854872 | 0.480627651 | -3.951499037 |
| **A_55_P2155654** | Pvt1 | 0.266744155 | 0.005048755 | 0.528439556 | 8.500710321 | 2.476749085 | 0.046871702 | 0.480627651 | -3.951615877 |
| **A_52_P362111** | Fam3c | -0.309957595 | -0.614277731 | -0.005637459 | 11.46498468 | -2.474883626 | 0.046992185 | 0.481205014 | -3.952451283 |
| **A_52_P527800** | Emilin2 | 0.296245923 | 0.005321565 | 0.587170281 | 11.47649445 | 2.474317842 | 0.047028791 | 0.481205014 | -3.952704742 |
| **A_52_P534583** | Ahsp | 0.307761328 | 0.005458648 | 0.610064007 | 16.86845275 | 2.473746767 | 0.04706577 | 0.481205014 | -3.952960612 |
| **A_55_P1998631** | Capg | -0.428267996 | -0.849053265 | -0.007482727 | 11.55459111 | -2.473080671 | 0.04710894 | 0.481205014 | -3.953259106 |
| **A_55_P1961210** | Trp53inp2 | -0.529987749 | -1.050775699 | -0.009199799 | 11.55786969 | -2.472794878 | 0.047127475 | 0.481205014 | -3.953387194 |
| **A_51_P419959** | Eaf1 | 0.318295319 | 0.005511945 | 0.631078693 | 10.35328629 | 2.47269062 | 0.047134239 | 0.481205014 | -3.953433924 |
| **A_55_P2106175** | Gch1 | -0.227138138 | -0.450346432 | -0.003929845 | 10.91367486 | -2.472651577 | 0.047136772 | 0.481205014 | -3.953451423 |
| **A_51_P446510** | Emp3 | -0.28095639 | -0.557123523 | -0.004789257 | 10.12055452 | -2.472009369 | 0.047178459 | 0.481205014 | -3.953739299 |
| **A_55_P2169356** | Gm1966 | -0.19593797 | -0.388565907 | -0.003310033 | 8.403532758 | -2.471624662 | 0.04720345 | 0.481205014 | -3.953911772 |
| **A_51_P234174** | Pms2 | 0.242254132 | 0.004085329 | 0.480422935 | 9.533073724 | 2.471550617 | 0.047208261 | 0.481205014 | -3.95394497 |
| **A_51_P414243** | Pomgnt2 | -0.226571676 | -0.449376709 | -0.003766644 | 10.75239096 | -2.47094917 | 0.047247364 | 0.481205014 | -3.954214654 |
| **A_55_P2156345** | Zfp51 | 0.283062486 | 0.004702676 | 0.561422296 | 9.076298254 | 2.470921645 | 0.047249154 | 0.481205014 | -3.954226997 |
| **A_55_P2160198** | NA | 0.207727104 | 0.003401746 | 0.412052463 | 10.31410843 | 2.470324962 | 0.047287982 | 0.481205014 | -3.954494591 |

| | | | | | | | | | |
|---|---|---|---|---|---|---|---|---|---|
| **A_51_P505521** | NA | 0.38739169 | 0.006307527 | 0.768475852 | 13.13946977 | 2.470088929 | 0.04730335 | 0.481205014 | -3.954600457 |
| **A_55_P1992421** | Hmgb3 | 0.324153645 | 0.005224626 | 0.643082664 | 12.99783107 | 2.469676454 | 0.04733022 | 0.481205014 | -3.954785476 |
| **A_55_P1959137** | Usp32 | 0.426716872 | 0.006841381 | 0.846592363 | 12.83150403 | 2.469462748 | 0.047344147 | 0.481205014 | -3.954881345 |
| **A_51_P481569** | Ddx42 | 0.181303011 | 0.002814659 | 0.359791364 | 11.30592144 | 2.468188507 | 0.047427281 | 0.481205014 | -3.955453086 |
| **A_51_P295816** | Cep192 | 0.301027284 | 0.004640095 | 0.597414474 | 9.513094046 | 2.467911723 | 0.047445359 | 0.481205014 | -3.955577302 |
| **A_55_P1998506** | Aplf | 0.310263816 | 0.004752045 | 0.615775586 | 8.681887252 | 2.467665965 | 0.047461416 | 0.481205014 | -3.955687603 |
| **A_55_P1970695** | NA | 0.229252088 | 0.003503448 | 0.455000729 | 8.660201745 | 2.467580593 | 0.047466996 | 0.481205014 | -3.955725922 |
| **A_52_P629895** | Adh1 | -1.175208567 | -2.332771834 | -0.0176453 | 13.79814641 | -2.466910542 | 0.047510811 | 0.481205014 | -3.956026698 |
| **A_51_P293069** | Flvcr1 | 0.286009356 | 0.004266674 | 0.567752039 | 9.297766556 | 2.466668485 | 0.04752665 | 0.481205014 | -3.956135367 |
| **A_55_P1954418** | NA | 0.182894055 | 0.002701837 | 0.363086274 | 11.72797347 | 2.466304793 | 0.047550458 | 0.481205014 | -3.956298657 |
| **A_55_P2088131** | Zfp975 | -0.33604995 | -0.667173004 | -0.004926896 | 9.788870337 | -2.46602574 | 0.047568734 | 0.481205014 | -3.956423957 |
| **A_55_P2076048** | Nuf2 | 0.29715079 | 0.004300733 | 0.590000846 | 10.12716218 | 2.465555403 | 0.047599554 | 0.481205014 | -3.956635169 |
| **A_51_P113773** | Slx1b | 0.317495364 | 0.004563604 | 0.630427124 | 9.148332913 | 2.465306577 | 0.047615868 | 0.481205014 | -3.956746919 |
| **A_52_P445474** | Lmtk2 | 0.201493064 | 0.002863302 | 0.400122825 | 8.383512491 | 2.464898083 | 0.047642662 | 0.481205014 | -3.956930394 |
| **A_52_P208613** | Fancb | 0.255001265 | 0.003575734 | 0.506426796 | 9.056006894 | 2.464428073 | 0.047673511 | 0.481205014 | -3.957141524 |
| **A_66_P121770** | Ilf2 | 0.174681729 | 0.002397644 | 0.346965814 | 10.48854864 | 2.463686879 | 0.047722202 | 0.481205014 | -3.957474526 |
| **A_55_P2102240** | Nprl3 | 0.317338107 | 0.004332433 | 0.630343782 | 9.63182047 | 2.463503621 | 0.047734249 | 0.481205014 | -3.95755687 |
| **A_55_P2067112** | Gm4924 | 0.26808906 | 0.003640068 | 0.532538051 | 10.0514214 | 2.463317346 | 0.047746497 | 0.481205014 | -3.957640574 |
| **A_55_P2169149** | Washc3 | -0.19270827 | -0.382858593 | -0.002557948 | 10.56423938 | -2.462558046 | 0.047796458 | 0.481205014 | -3.957981815 |
| **A_55_P2106666** | Adcy9 | -0.219380413 | -0.435874946 | -0.002885881 | 9.409133558 | -2.462261114 | 0.04781601 | 0.481205014 | -3.95811528 |
| **A_55_P2023607** | Ikzf1 | 0.465745028 | 0.005905969 | 0.925584087 | 8.983609822 | 2.461079022 | 0.047893933 | 0.481205014 | -3.958646714 |
| **A_55_P2000618** | Gm12690 | 0.280506156 | 0.003509867 | 0.557502444 | 8.743544166 | 2.460660146 | 0.047921577 | 0.481205014 | -3.95883507 |
| **A_51_P223709** | Pask | 0.33528859 | 0.004160972 | 0.666416208 | 9.890525342 | 2.460404761 | 0.047938439 | 0.481205014 | -3.958949919 |
| **A_55_P1979986** | Crcp | 0.259848187 | 0.003213819 | 0.516482554 | 10.83202103 | 2.46029999 | 0.047945359 | 0.481205014 | -3.958997038 |
| **A_52_P50478** | Rnf123 | 0.498621864 | 0.006035945 | 0.991207782 | 10.7139781 | 2.459645475 | 0.04798861 | 0.481205014 | -3.959291426 |
| **A_51_P144512** | Rbm19 | 0.258098762 | 0.003116088 | 0.513081436 | 10.40279402 | 2.459565761 | 0.04799388 | 0.481205014 | -3.959327283 |
| **A_55_P2052081** | Eif1ad | 0.251546882 | 0.002963229 | 0.500130536 | 12.54212474 | 2.458835989 | 0.048042158 | 0.481205014 | -3.959655589 |
| **A_55_P2143516** | NA | 0.351703122 | 0.004131313 | 0.699274932 | 10.21859313 | 2.458752803 | 0.048047664 | 0.481205014 | -3.959693016 |
| **A_55_P2033932** | Zzz3 | 0.203894258 | 0.002369376 | 0.405419141 | 9.247670321 | 2.458439405 | 0.048068415 | 0.481205014 | -3.959834028 |
| **A_55_P2003586** | Unkl | 0.418747098 | 0.004779224 | 0.832714971 | 9.076495535 | 2.457923489 | 0.048102596 | 0.481205014 | -3.96006619 |
| **A_51_P455906** | Pold4 | -0.242084809 | -0.481420422 | -0.002749197 | 9.309483282 | -2.457782241 | 0.048111958 | 0.481205014 | -3.960129758 |
| **A_55_P2069221** | Prr11 | 0.214101832 | 0.002180279 | 0.426023385 | 8.566638616 | 2.454869698 | 0.048305438 | 0.481205014 | -3.961441061 |
| **A_66_P128293** | Pheta2 | 0.496468107 | 0.005006294 | 0.98792992 | 11.32336982 | 2.454622806 | 0.048321876 | 0.481205014 | -3.961552266 |
| **A_55_P2025790** | Fanci | 0.369189613 | 0.003659257 | 0.734719968 | 9.389088838 | 2.45419583 | 0.048350319 | 0.481205014 | -3.961744602 |
| **A_55_P2055763** | Gm5878 | 0.384923399 | 0.003796907 | 0.766049892 | 10.71497364 | 2.454078008 | 0.04835817 | 0.481205014 | -3.96179768 |
| **A_51_P378667** | Wrap53 | 0.259609488 | 0.002488065 | 0.516730911 | 11.42185134 | 2.453383768 | 0.048404462 | 0.481205014 | -3.962110466 |
| **A_52_P109403** | Cebpz | 0.393480018 | 0.003757593 | 0.783202443 | 9.934966068 | 2.453298962 | 0.04841012 | 0.481205014 | -3.962148678 |
| **A_51_P155747** | Scyl3 | 0.190021529 | 0.001794785 | 0.378248272 | 8.47528795 | 2.453040216 | 0.048427387 | 0.481205014 | -3.962265273 |
| **A_55_P2169445** | Mfsd6 | -0.337366765 | -0.671565484 | -0.003168046 | 9.444592886 | -2.452904865 | 0.048436422 | 0.481205014 | -3.962326267 |
| **A_55_P2022987** | Psmd2 | 0.208593417 | 0.001854513 | 0.415332321 | 13.6699826 | 2.451667543 | 0.048519098 | 0.481205014 | -3.962883953 |
| **A_55_P2024431** | Gaa | -0.209220838 | -0.416583079 | -0.001858598 | 12.99719246 | -2.451649894 | 0.048520278 | 0.481205014 | -3.962891909 |
| **A_52_P450188** | Mtmr4 | 0.20364484 | 0.001745517 | 0.405544163 | 9.225334379 | 2.450878247 | 0.048571914 | 0.481205014 | -3.963239803 |
| **A_55_P2165334** | Nde1 | 0.201718572 | 0.00169196 | 0.401745184 | 12.12717508 | 2.450424329 | 0.048602316 | 0.481205014 | -3.963444484 |
| **A_55_P2041090** | Tyw5 | 0.188500171 | 0.001542347 | 0.375457996 | 10.22587143 | 2.449916552 | 0.048636349 | 0.481205014 | -3.96367348 |
| **A_52_P574697** | Khdrbs1 | 0.241088891 | 0.001942642 | 0.48023514 | 11.94141755 | 2.449609258 | 0.048656957 | 0.481205014 | -3.963812079 |
| **A_55_P2018666** | Thrsp | -0.370675203 | -0.738410093 | -0.002940313 | 8.130722135 | -2.449299455 | 0.048677742 | 0.481205014 | -3.96395182 |
| **A_52_P161707** | Cand1 | 0.178306441 | 0.00140099 | 0.355211891 | 8.951219746 | 2.449114026 | 0.048690187 | 0.481205014 | -3.964035467 |
| **A_66_P130582** | Ncoa1 | -0.278122897 | -0.554074637 | -0.002171156 | 9.492419642 | -2.448988779 | 0.048698595 | 0.481205014 | -3.964091968 |
| **A_52_P304947** | Cenpn | 0.254867807 | 0.001967743 | 0.50776787 | 8.838054795 | 2.448776969 | 0.048712818 | 0.481205014 | -3.964187523 |
| **A_55_P2005941** | Plcxd1 | 0.326888609 | 0.002490051 | 0.651287167 | 10.99293879 | 2.448522282 | 0.048729925 | 0.481205014 | -3.964302429 |

| | | | | | | | | | |
|---|---|---|---|---|---|---|---|---|---|
| **A_55_P2079089** | Fuca1 | -0.302317175 | -0.60233356 | -0.00230079 | 10.56712645 | -2.448505233 | 0.048731071 | 0.481205014 | -3.964310121 |
| **A_55_P2022231** | Gm826 | 0.477904505 | 0.003629153 | 0.952179857 | 8.694455409 | 2.448464202 | 0.048733827 | 0.481205014 | -3.964328634 |
| **A_55_P2020361** | Lzts3 | 0.496676671 | 0.003715619 | 0.989637723 | 11.65815653 | 2.448185619 | 0.048752549 | 0.481205014 | -3.964454331 |
| **A_55_P1991920** | Dhx32 | 0.271582441 | 0.001996875 | 0.541168007 | 9.69179615 | 2.447869385 | 0.048773809 | 0.481205014 | -3.964597029 |
| **A_66_P128702** | Nup43 | 0.275397618 | 0.002024836 | 0.5487704 | 9.977097022 | 2.447868567 | 0.048773864 | 0.481205014 | -3.964597398 |
| **A_52_P412452** | Cntn6 | 0.317703402 | 0.002335463 | 0.633071341 | 8.404933879 | 2.44786529 | 0.048774085 | 0.481205014 | -3.964598877 |
| **A_55_P2097808** | P4ha1 | -0.23240378 | -0.463103454 | -0.001704106 | 8.625454657 | -2.44781953 | 0.048777162 | 0.481205014 | -3.964619527 |
| **A_55_P2381538** | NA | 0.235599769 | 0.001721949 | 0.469477589 | 9.492357258 | 2.447761008 | 0.048781098 | 0.481205014 | -3.964645936 |
| **A_52_P6828** | Xk | 0.602424928 | 0.00432759 | 1.200522266 | 10.31299156 | 2.447452401 | 0.048801858 | 0.481205014 | -3.964785208 |
| **A_55_P2042146** | Fech | 0.276021663 | 0.001969983 | 0.550073343 | 15.67979646 | 2.447337632 | 0.048809581 | 0.481205014 | -3.964837005 |
| **A_55_P2105808** | Wdr62 | 0.619773538 | 0.004268491 | 1.235278585 | 13.62268353 | 2.446721851 | 0.04885104 | 0.481205014 | -3.965114945 |
| **A_52_P5549** | Fam133b | 0.248464962 | 0.001620277 | 0.495309647 | 9.965229405 | 2.445820394 | 0.048911799 | 0.481205014 | -3.965521912 |
| **A_55_P2061991** | Pnldc1 | 0.588755666 | 0.00379994 | 1.173711392 | 10.58760946 | 2.445655559 | 0.048922918 | 0.481205014 | -3.965596339 |
| **A_52_P131836** | Bysl | 0.379973679 | 0.002368688 | 0.757578669 | 11.6451854 | 2.44511324 | 0.048959518 | 0.481205014 | -3.965841229 |
| **A_55_P2002122** | Rgs12 | 0.262961896 | 0.001594167 | 0.524329625 | 11.69818107 | 2.444691416 | 0.048988005 | 0.481205014 | -3.966031734 |
| **A_55_P1993512** | NA | 0.466440232 | 0.002741518 | 0.930138947 | 10.49334011 | 2.444236921 | 0.049018719 | 0.481205014 | -3.966237018 |
| **A_51_P161929** | Tspo2 | 0.374753031 | 0.002182888 | 0.747323174 | 14.70979839 | 2.444107451 | 0.049027472 | 0.481205014 | -3.966295501 |
| **A_55_P1986556** | Minpp1 | 0.391829081 | 0.002202848 | 0.781455315 | 11.36001147 | 2.44360871 | 0.049061205 | 0.481205014 | -3.966520806 |
| **A_52_P276525** | Ankrd9 | 0.452423375 | 0.002466489 | 0.902380262 | 10.65968775 | 2.443190448 | 0.049089514 | 0.481205014 | -3.966709779 |
| **A_52_P372165** | Rpsa | 0.271308183 | 0.001471763 | 0.541144603 | 8.705302874 | 2.443124029 | 0.04909401 | 0.481205014 | -3.96673979 |
| **A_55_P1963900** | Mib2 | -0.444735078 | -0.887080279 | -0.002389877 | 10.24669246 | -2.442998802 | 0.04910249 | 0.481205014 | -3.966796373 |
| **A_55_P1986103** | Suco | 0.227581812 | 0.00116903 | 0.453994594 | 8.46596534 | 2.44241691 | 0.049141913 | 0.481205014 | -3.967059323 |
| **A_55_P1999102** | Pi16 | -0.315499149 | -0.629410297 | -0.001588 | 10.00571977 | -2.442162966 | 0.049159128 | 0.481205014 | -3.967174091 |
| **A_51_P256682** | Kpna1 | 0.396249636 | 0.001987706 | 0.790511565 | 9.887872188 | 2.442121248 | 0.049161956 | 0.481205014 | -3.967192945 |
| **A_51_P436878** | Sertad1 | -0.340623343 | -0.679648626 | -0.001598059 | 10.87493258 | -2.441324491 | 0.049216012 | 0.481205014 | -3.967553086 |
| **A_52_P551011** | Smim1 | 0.463458955 | 0.002149289 | 0.92476862 | 12.93314347 | 2.441191857 | 0.049225017 | 0.481205014 | -3.967613045 |
| **A_55_P2104597** | Kctd9 | 0.287998841 | 0.001324167 | 0.574673515 | 12.18876348 | 2.441094556 | 0.049231624 | 0.481205014 | -3.967657033 |
| **A_55_P1983450** | S100a4 | -0.357245932 | -0.712851657 | -0.001640207 | 8.786450086 | -2.441078449 | 0.049232718 | 0.481205014 | -3.967664315 |
| **A_55_P2183692** | Rgs19 | -0.202993887 | -0.405098255 | -0.00088952 | 9.149793424 | -2.440565407 | 0.04926757 | 0.481205014 | -3.96789627 |
| **A_52_P363216** | Gcnt2 | -0.379696552 | -0.757736548 | -0.001656555 | 8.979281711 | -2.44051843 | 0.049270763 | 0.481205014 | -3.967917511 |
| **A_55_P1992762** | Kdelr1 | -0.212149566 | -0.423414169 | -0.000884963 | 9.726584936 | -2.440049286 | 0.049302659 | 0.481205014 | -3.96812965 |
| **A_55_P2140843** | NA | 0.429523552 | 0.001773305 | 0.8572738 | 9.65073733 | 2.439944244 | 0.049309803 | 0.481205014 | -3.968177152 |
| **A_55_P2136209** | Ublcp1 | 0.20887594 | 0.000803027 | 0.416948852 | 9.778340082 | 2.439248568 | 0.049357146 | 0.481205014 | -3.968491783 |
| **A_55_P2147402** | NA | 0.183494033 | 0.000702815 | 0.366285251 | 8.219907486 | 2.439213465 | 0.049359537 | 0.481205014 | -3.968507661 |
| **A_51_P260548** | Glg1 | -0.202792205 | -0.404847816 | -0.000736594 | 12.08067207 | -2.438728938 | 0.04939254 | 0.481205014 | -3.968726832 |
| **A_51_P245789** | Pcolce2 | -0.219469501 | -0.438167159 | -0.000771844 | 10.0268889 | -2.438446519 | 0.049411788 | 0.481205014 | -3.968854596 |
| **A_55_P2151461** | NA | 0.211501873 | 0.000736509 | 0.422267238 | 12.995489 | 2.438361899 | 0.049417557 | 0.481205014 | -3.968892879 |
| **A_55_P2249055** | NA | 0.270348013 | 0.000837772 | 0.539858253 | 9.976033544 | 2.43742409 | 0.049481535 | 0.481457414 | -3.969317213 |
| **A_51_P309589** | 2700099C18Rik | 0.251952288 | 0.000639739 | 0.503264838 | 8.802752989 | 2.436056292 | 0.049575004 | 0.481457414 | -3.969936297 |
| **A_55_P2052170** | NA | 0.174328833 | 0.000437998 | 0.348219668 | 12.32021315 | 2.435991217 | 0.049579456 | 0.481457414 | -3.969965756 |
| **A_51_P509971** | Plekho1 | -0.398400084 | -0.795850762 | -0.000949406 | 11.08158396 | -2.435675166 | 0.049601081 | 0.481457414 | -3.97010884 |
| **A_52_P117420** | Tacc1 | 0.235407886 | 0.000507939 | 0.470307832 | 10.30172331 | 2.435125101 | 0.049638743 | 0.481457414 | -3.970357896 |
| **A_55_P1963489** | NA | 0.348972773 | 0.000725534 | 0.697220012 | 9.593607688 | 2.434933201 | 0.049651889 | 0.481457414 | -3.970444792 |
| **A_55_P2057380** | NA | 0.364352238 | 0.000679652 | 0.728024824 | 8.694402393 | 2.434411917 | 0.049687617 | 0.481457414 | -3.970680863 |
| **A_55_P2053838** | Tnfaip3 | -0.254376565 | -0.508279892 | -0.000473237 | 9.146352158 | -2.43439975 | 0.049688451 | 0.481457414 | -3.970686373 |
| **A_51_P125368** | Hars | 0.333612719 | 0.000612827 | 0.666612611 | 10.71375884 | 2.434342585 | 0.049692371 | 0.481457414 | -3.970712264 |
| **A_55_P2006467** | Serf2 | -0.294303455 | -0.588066629 | -0.000540282 | 14.70910665 | -2.434339796 | 0.049692562 | 0.481457414 | -3.970713527 |
| **A_55_P2059025** | U2af1 | 0.186609241 | 0.000285044 | 0.372933438 | 13.52316846 | 2.433588128 | 0.049744134 | 0.481712803 | -3.971053994 |
| **A_55_P1997766** | Acp5 | 0.313797934 | 0.000398804 | 0.627197064 | 15.22258845 | 2.432962875 | 0.049787075 | 0.481712803 | -3.971337255 |
| **A_55_P2040628** | Gid8 | 0.199528675 | 0.000201997 | 0.398855354 | 13.07668843 | 2.432333261 | 0.049830355 | 0.481712803 | -3.971622539 |

| | | | | | | | | |
|---|---|---|---|---|---|---|---|---|
| **A_55_P2040367** | Bola3 | 0.372817901 | 0.000358604 | 0.745277199 | 12.53529066 | 2.432210318 | 0.04983881 | 0.481712803 | -3.971678251 |
| **A_66_P110791** | Ppp1r11 | -0.200270162 | -0.400370969 | -0.000169355 | 12.02716231 | -2.431927353 | 0.049858278 | 0.481712803 | -3.971806485 |
| **A_51_P310949** | Ppox | 0.44132685 | 0.000239131 | 0.882414569 | 12.30992102 | 2.431188169 | 0.049909169 | 0.481712803 | -3.972141513 |
| **A_55_P2147846** | Npcd | 0.188237619 | 4.81E-05 | 0.376427133 | 8.599117967 | 2.430491975 | 0.049957151 | 0.481712803 | -3.972457118 |
| **A_55_P2110743** | Mettl7a2 | 0.324152252 | 4.85E-05 | 0.648256046 | 13.32483786 | 2.430234139 | 0.049974933 | 0.481712803 | -3.972574017 |
| **A_52_P188678** | Nectin2 | -0.474761826 | -0.949662968 | 0.000139317 | 9.453758601 | -2.429158013 | 0.050049222 | 0.481712803 | -3.973062003 |
| **A_51_P495485** | Pip4p2 | 0.194847303 | -8.84E-05 | 0.389782998 | 10.8779659 | 2.428769039 | 0.050076102 | 0.481712803 | -3.973238424 |
| **A_55_P2118717** | Ninl | 0.415068906 | -0.000199058 | 0.83033687 | 9.175055456 | 2.428706083 | 0.050080455 | 0.481712803 | -3.97326698 |
| **A_55_P2099760** | NA | 0.837049991 | -0.00045811 | 1.674558093 | 9.685906669 | 2.42854172 | 0.050091819 | 0.481712803 | -3.973341534 |
| **A_55_P2070721** | Rapgef1 | 0.218919508 | -0.000138373 | 0.437977389 | 10.39570277 | 2.428335953 | 0.050106049 | 0.481712803 | -3.973434874 |
| **A_51_P229911** | Adcy4 | -0.492015197 | -0.984350753 | 0.000320359 | 9.107009637 | -2.428289741 | 0.050109246 | 0.481712803 | -3.973455837 |
| **A_55_P1965154** | Spc25 | 0.307885762 | -0.000217189 | 0.615988713 | 13.69863829 | 2.428157966 | 0.050118362 | 0.481712803 | -3.973515617 |
| **A_55_P2151986** | Gngt2 | -0.262134264 | -0.524539206 | 0.000270678 | 11.57227188 | -2.427364363 | 0.050173301 | 0.481712803 | -3.973875674 |
| **A_55_P2009940** | Xlr | 0.220954306 | -0.000268896 | 0.442177507 | 8.045491519 | 2.426917341 | 0.050204274 | 0.481712803 | -3.974078522 |
| **A_55_P2083224** | Pum2 | 0.192781289 | -0.000242101 | 0.385804679 | 10.68281904 | 2.426823151 | 0.050210803 | 0.481712803 | -3.974121265 |
| **A_51_P284486** | Gstm2 | -1.012999255 | -2.027308264 | 0.001309754 | 10.56387043 | -2.426733202 | 0.050217039 | 0.481712803 | -3.974162086 |
| **A_52_P50417** | 1700084J12Rik | 0.228144231 | -0.000309039 | 0.456597501 | 8.597277395 | 2.426583846 | 0.050227395 | 0.481712803 | -3.974229869 |
| **A_55_P2168243** | NA | -0.454030925 | -0.908717083 | 0.000655232 | 10.35165418 | -2.426369237 | 0.050242279 | 0.481712803 | -3.97432727 |
| **A_55_P2144771** | NA | 0.435475323 | -0.000750517 | 0.871701163 | 9.346559142 | 2.425690299 | 0.050289397 | 0.481734469 | -3.974635446 |
| **A_55_P1981780** | Cul4b | 0.206577043 | -0.000362151 | 0.413516236 | 9.867598755 | 2.425618475 | 0.050294385 | 0.481734469 | -3.974668051 |
| **A_51_P311362** | 4930449E01Rik | 0.684537101 | -0.001488476 | 1.370562678 | 9.834002623 | 2.424598726 | 0.050365249 | 0.482103877 | -3.975131037 |
| **A_55_P2003492** | NA | 0.206805313 | -0.000471298 | 0.414081924 | 13.57867012 | 2.424345887 | 0.050382836 | 0.482103877 | -3.975245851 |
| **A_55_P2017729** | Gm2138 | 0.338650709 | -0.000849355 | 0.678150773 | 8.659558702 | 2.423791836 | 0.050421396 | 0.482234122 | -3.97549747 |
| **A_55_P2107901** | Dlk2 | 0.529072899 | -0.001444935 | 1.059590734 | 9.807460622 | 2.423252765 | 0.050458943 | 0.482354554 | -3.975742322 |
| **A_55_P2136413** | NA | 0.265964305 | -0.000836691 | 0.532765301 | 11.39196586 | 2.422250738 | 0.050528813 | 0.482501908 | -3.976197547 |
| **A_55_P2001718** | Pex1 | 0.270166957 | -0.000948763 | 0.541282676 | 8.940124323 | 2.421367568 | 0.05059048 | 0.482501908 | -3.976598875 |
| **A_55_P2097116** | Cnep1r1 | 0.315002114 | -0.001160922 | 0.631165149 | 8.73142619 | 2.420948572 | 0.050619763 | 0.482501908 | -3.976789306 |
| **A_51_P373890** | Guca1b | 0.277318601 | -0.001058996 | 0.555696197 | 9.006909866 | 2.420627196 | 0.050642236 | 0.482501908 | -3.976935385 |
| **A_55_P1985825** | NA | 0.51309192 | -0.002066111 | 1.028249951 | 9.57687678 | 2.420125512 | 0.050677338 | 0.482501908 | -3.977163445 |
| **A_55_P1980041** | Kpna6 | 0.355247957 | -0.001431028 | 0.711926942 | 9.431997394 | 2.420121979 | 0.050677586 | 0.482501908 | -3.977165051 |
| **A_66_P130647** | NA | -0.774544638 | -1.552575621 | 0.003486345 | 11.59326772 | -2.418982626 | 0.0507574 | 0.482501908 | -3.977683102 |
| **A_55_P1994773** | Gm16381 | 0.305142739 | -0.001391706 | 0.611677184 | 9.684007672 | 2.418838911 | 0.050767476 | 0.482501908 | -3.977748459 |
| **A_55_P2142863** | Parp9 | -0.262315274 | -0.525931849 | 0.0013013 | 9.861131174 | -2.417876177 | 0.050835034 | 0.482501908 | -3.978186342 |
| **A_51_P224517** | Jade1 | 0.297156057 | -0.001500943 | 0.595813057 | 9.385221242 | 2.417659177 | 0.050850275 | 0.482501908 | -3.978285056 |
| **A_66_P119170** | Phf21a | -0.287543504 | -0.576568331 | 0.001481322 | 8.48080624 | -2.41741716 | 0.050867278 | 0.482501908 | -3.978395156 |
| **A_55_P2027240** | Engase | 0.289045808 | -0.001495692 | 0.579587307 | 10.59568263 | 2.417361998 | 0.050871154 | 0.482501908 | -3.978420252 |
| **A_66_P111666** | Nprl3 | 0.320337543 | -0.001692452 | 0.642367537 | 10.22093564 | 2.417100478 | 0.050889535 | 0.482501908 | -3.978539235 |
| **A_55_P2132671** | NA | 0.271211806 | -0.001453325 | 0.543876937 | 8.468584398 | 2.416919448 | 0.050902264 | 0.482501908 | -3.978621603 |
| **A_55_P2129034** | Rheb | 0.210749561 | -0.001141784 | 0.422640905 | 10.42375756 | 2.416777398 | 0.050912253 | 0.482501908 | -3.978686238 |
| **A_55_P2039324** | Ccnf | 0.653971081 | -0.003587752 | 1.311529915 | 12.25229429 | 2.416613053 | 0.050923814 | 0.482501908 | -3.97876102 |
| **A_52_P192426** | Tnfrsf1a | -0.351915489 | -0.70578552 | 0.001954542 | 12.06964947 | -2.416449854 | 0.050935296 | 0.482501908 | -3.984441965 |
| **A_52_P471502** | Dcaf13 | 0.263345329 | -0.001539345 | 0.528230003 | 9.508852844 | 2.415749938 | 0.050984571 | 0.482501908 | -3.979153818 |
| **A_55_P2124831** | Brdt | 0.520236094 | -0.003187644 | 1.043659832 | 9.645291664 | 2.415072958 | 0.051032279 | 0.482501908 | -3.97946197 |
| **A_55_P2049553** | NA | 0.591206254 | -0.003726104 | 1.186138612 | 11.02679919 | 2.414652383 | 0.051061941 | 0.482501908 | -3.979653437 |
| **A_55_P2170335** | Bmi1 | 0.221325538 | -0.001398263 | 0.444049339 | 9.873168662 | 2.414616079 | 0.051064503 | 0.482501908 | -3.979669966 |
| **A_55_P1955656** | Ctla2a | -0.476262939 | -0.955562313 | 0.003036435 | 11.74513444 | -2.414477236 | 0.051074299 | 0.482501908 | -3.979733179 |
| **A_55_P1996504** | Abcg2 | 0.403574706 | -0.002580954 | 0.809730366 | 13.79809987 | 2.41443 | 0.051077633 | 0.482501908 | -3.979754686 |
| **A_55_P2379480** | NA | 0.436076966 | -0.002802516 | 0.874956449 | 9.082140007 | 2.414354613 | 0.051082953 | 0.482501908 | -3.97978901 |
| **A_55_P2005720** | NA | 0.451374682 | -0.002942065 | 0.905691428 | 10.91420005 | 2.414135477 | 0.051098422 | 0.482501908 | -3.979888788 |
| **A_66_P106760** | Adam32 | 0.366777384 | -0.002478103 | 0.736032871 | 9.301189195 | 2.413563782 | 0.051138802 | 0.482505424 | -3.980149123 |

| | | | | | | | | | |
|---|---|---|---|---|---|---|---|---|---|
| **A_55_P1964348** | Gdpd2 | 0.390744026 | -0.002721233 | 0.784209285 | 10.30824723 | 2.413065684 | 0.05117401 | 0.482505424 | -3.980375975 |
| **A_52_P235569** | Ipo5 | 0.238180883 | -0.001747327 | 0.478109094 | 9.500551159 | 2.412174798 | 0.051237046 | 0.482505424 | -3.980781792 |
| **A_52_P540470** | Zfp407 | 0.179449027 | -0.001320596 | 0.360218649 | 9.886065115 | 2.412119638 | 0.051240951 | 0.482505424 | -3.980806922 |
| **A_55_P2120906** | NA | 0.216473485 | -0.001654922 | 0.434601892 | 9.049471434 | 2.411435615 | 0.051289409 | 0.482505424 | -3.981118577 |
| **A_55_P1959628** | Pfdn4 | 0.219767387 | -0.001693564 | 0.441228339 | 9.83593511 | 2.411289043 | 0.051299799 | 0.482505424 | -3.981185366 |
| **A_55_P1955108** | Ripor1 | -0.321293101 | -0.645117657 | 0.002531456 | 12.31486186 | -2.41087564 | 0.051329115 | 0.482505424 | -3.981373755 |
| **A_51_P103397** | Vwf | -0.254187795 | -0.510385977 | 0.002010388 | 11.20477791 | -2.410803638 | 0.051334222 | 0.482505424 | -3.981406569 |
| **A_55_P2023181** | Lsm2 | 0.264749906 | -0.002256207 | 0.53175602 | 13.0818226 | 2.409338378 | 0.051438281 | 0.482505424 | -3.982074472 |
| **A_55_P2017998** | NA | 0.247374605 | -0.002255165 | 0.497004374 | 8.770098248 | 2.407919292 | 0.051539271 | 0.482505424 | -3.982721572 |
| **A_55_P2006876** | NA | -0.206932845 | -0.415766159 | 0.001900468 | 14.09944828 | -2.40775803 | 0.05155076 | 0.482505424 | -3.982795122 |
| **A_55_P2119030** | Rasl2-9 | 0.215140756 | -0.001991954 | 0.432273466 | 14.09059469 | 2.407579447 | 0.051563486 | 0.482505424 | -3.982876576 |
| **A_55_P2068027** | NA | 0.411468438 | -0.003934657 | 0.826871533 | 8.546812707 | 2.406855341 | 0.051615122 | 0.482505424 | -3.983206888 |
| **A_51_P438527** | Cyb5r1 | 0.267126705 | -0.002578363 | 0.536831773 | 10.07563443 | 2.40664143 | 0.051630387 | 0.482505424 | -3.983304479 |
| **A_52_P274028** | Ssr1 | 0.288122685 | -0.002796266 | 0.579041637 | 8.793069078 | 2.406515315 | 0.051639389 | 0.482505424 | -3.983362019 |
| **A_55_P2151609** | Sorl1 | -0.394202419 | -0.792254804 | 0.003849966 | 9.518410236 | -2.40636911 | 0.051649826 | 0.482505424 | -3.983428726 |
| **A_55_P2038587** | NA | 0.455073552 | -0.004445087 | 0.914592191 | 9.096620689 | 2.406365838 | 0.05165006 | 0.482505424 | -3.983430219 |
| **A_52_P234910** | Gm15032 | 0.189363974 | -0.001883913 | 0.380611862 | 13.01970969 | 2.405935071 | 0.051680825 | 0.482505424 | -3.983626775 |
| **A_55_P2173373** | Zfp62 | 0.277976242 | -0.002847612 | 0.558800097 | 9.067898039 | 2.405231446 | 0.051731119 | 0.482505424 | -3.983947882 |
| **A_55_P2017369** | NA | 0.197638339 | -0.002049561 | 0.397326239 | 12.15722954 | 2.40493108 | 0.051752605 | 0.482505424 | -3.984084976 |
| **A_52_P231737** | Actl6a | 0.251929351 | -0.002659081 | 0.506517783 | 10.22656181 | 2.40449175 | 0.051784047 | 0.482505424 | -3.984285515 |
| **A_55_P2133195** | Gm4951 | -0.972786684 | -1.955937514 | 0.010364145 | 10.09013665 | -2.404255711 | 0.051800948 | 0.482505424 | -3.984393268 |
| **A_51_P438200** | Sil1 | -0.260342777 | -0.523531932 | 0.002846378 | 12.81316971 | -2.403591902 | 0.05184851 | 0.482505424 | -3.984696336 |
| **A_52_P251183** | Tmem177 | 0.194139103 | -0.002127952 | 0.390406157 | 8.948405925 | 2.403525876 | 0.051853243 | 0.482505424 | -3.984726483 |
| **A_55_P1983152** | Gramd4 | 0.177201827 | -0.00196373 | 0.356367384 | 10.35240062 | 2.403238429 | 0.051873855 | 0.482505424 | -3.984857738 |
| **A_51_P400366** | Rhbg | 0.35472904 | -0.003948446 | 0.713406527 | 8.562610484 | 2.403121979 | 0.051882207 | 0.482505424 | -3.984910915 |
| **A_55_P1991718** | Hmmr | 0.447402681 | -0.005063445 | 0.899868806 | 10.87970518 | 2.402678711 | 0.051914015 | 0.482505424 | -3.985113347 |
| **A_55_P1970860** | Pemt | -0.917945996 | -1.846306598 | 0.010414606 | 13.27408063 | -2.402611877 | 0.051918812 | 0.482505424 | -3.985143871 |
| **A_55_P2440441** | Wdhd1 | 0.336440661 | -0.003907907 | 0.676789229 | 9.210864705 | 2.401970884 | 0.051964848 | 0.482505424 | -3.985436647 |
| **A_52_P179178** | Bend3 | 0.28649752 | -0.003423391 | 0.576418432 | 10.15969136 | 2.401178885 | 0.052021787 | 0.482505424 | -3.985798462 |
| **A_55_P1962781** | NA | 0.30302519 | -0.003622829 | 0.60967321 | 8.399900267 | 2.401163636 | 0.052022884 | 0.482505424 | -3.98580543 |
| **A_51_P352687** | Rnf216 | 0.258798376 | -0.003293912 | 0.520890665 | 10.47410319 | 2.39933281 | 0.05215476 | 0.482505424 | -3.986642111 |
| **A_55_P2130388** | Mical1 | -0.417326375 | -0.83997066 | 0.005317911 | 10.22880534 | -2.399297051 | 0.05215734 | 0.482505424 | -3.986658457 |
| **A_55_P2171116** | Lgals3 | -0.578246275 | -1.16388667 | 0.007394119 | 12.48502718 | -2.399192021 | 0.052164916 | 0.482505424 | -3.986706467 |
| **A_52_P406828** | Dkc1 | 0.440107109 | -0.00567766 | 0.885891879 | 10.09271804 | 2.39892321 | 0.052184312 | 0.482505424 | -3.986829351 |
| **A_55_P2131379** | Ybx1 | 0.651261853 | -0.008678737 | 1.311202444 | 11.54696324 | 2.39791613 | 0.052257045 | 0.482505424 | -3.987289802 |
| **A_55_P1981739** | Donson | 0.441982026 | -0.006009715 | 0.889973768 | 11.45794853 | 2.39727463 | 0.05230343 | 0.482505424 | -3.987583167 |
| **A_55_P2071626** | C530008M17Rik | 0.262721934 | -0.003591301 | 0.529035169 | 8.828099088 | 2.397103419 | 0.052315817 | 0.482505424 | -3.987661473 |
| **A_55_P2065149** | Kdm2b | 0.199787371 | -0.002739253 | 0.402313994 | 11.39333893 | 2.39700587 | 0.052322876 | 0.482505424 | -3.987706089 |
| **A_55_P2074864** | NA | 0.226964505 | -0.003181126 | 0.457110135 | 13.36087777 | 2.396284608 | 0.0523751 | 0.482505424 | -3.988036012 |
| **A_55_P1990870** | Mylk3 | 0.491479717 | -0.006920862 | 0.989880296 | 10.33412401 | 2.396129306 | 0.052386352 | 0.482505424 | -3.988107059 |
| **A_55_P1953608** | NA | 0.193825553 | -0.002756386 | 0.390407492 | 11.07173708 | 2.395800255 | 0.052410201 | 0.482505424 | -3.988257602 |
| **A_55_P2174022** | Brd4 | 0.276193898 | -0.003940399 | 0.556328195 | 12.25007877 | 2.395692015 | 0.052418048 | 0.482505424 | -3.988307125 |
| **A_52_P263518** | Gng2 | -0.34496882 | -0.694880541 | 0.004942902 | 10.40092242 | -2.395546146 | 0.052428626 | 0.482505424 | -3.988373867 |
| **A_51_P303620** | Whrn | -0.24480838 | -0.493205007 | 0.003588247 | 8.298343148 | -2.39476981 | 0.052484958 | 0.482505424 | -3.988729119 |
| **A_51_P302520** | Myom1 | -0.379392081 | -0.764387001 | 0.005602839 | 8.078350394 | -2.394508877 | 0.052503906 | 0.482505424 | -3.988848538 |
| **A_55_P2021413** | Celf4 | 0.408749627 | -0.006109854 | 0.823609107 | 9.900128318 | 2.39408485 | 0.052534713 | 0.482505424 | -3.989042616 |
| **A_55_P1957708** | NA | 0.432565283 | -0.006643083 | 0.871773648 | 8.95061168 | 2.393118733 | 0.052604974 | 0.482505424 | -3.989484891 |
| **A_55_P2158011** | Knstrn | 0.326795605 | -0.005082424 | 0.658673634 | 10.0278482 | 2.392659476 | 0.052638408 | 0.482505424 | -3.98969517 |
| **A_51_P292357** | Rps3a1 | 0.198688637 | -0.00315716 | 0.400534434 | 14.64317513 | 2.391864152 | 0.05269636 | 0.482505424 | -3.990059384 |
| **A_51_P468294** | Polg | -0.183745 | -0.370452164 | 0.002962164 | 9.529869054 | -2.391320222 | 0.052736032 | 0.482505424 | -3.990308517 |

| | | | | | | | | | |
|---|---|---|---|---|---|---|---|---|---|
| **A_55_P2155002** | Svbp | 0.265261368 | -0.004276483 | 0.53479922 | 12.33937829 | 2.391318548 | 0.052736155 | 0.482505424 | -3.990309283 |
| **A_52_P322927** | NA | 0.408429268 | -0.006623252 | 0.823481788 | 8.934085634 | 2.391095876 | 0.052752405 | 0.482505424 | -3.990411282 |
| **A_51_P155073** | Nudt14 | -0.432455139 | -0.871927064 | 0.007016786 | 9.007076207 | -2.391074541 | 0.052753962 | 0.482505424 | -3.990421056 |
| **A_55_P1983414** | NA | 0.283582933 | -0.004613597 | 0.571779463 | 8.73756644 | 2.390972224 | 0.052761431 | 0.482505424 | -3.990467926 |
| **A_55_P2046550** | Adamts8 | 0.335139395 | -0.005584044 | 0.675862834 | 8.733975389 | 2.390048205 | 0.052828931 | 0.482505424 | -3.990891267 |
| **A_55_P1966649** | Glmn | 0.190591618 | -0.003238267 | 0.384421503 | 9.306888483 | 2.389275601 | 0.052885439 | 0.482505424 | -3.991245313 |
| **A_55_P2076757** | Znfx1 | 0.164368856 | -0.002804314 | 0.331542025 | 11.3719006 | 2.389109987 | 0.05289756 | 0.482505424 | -3.991321215 |
| **A_52_P244964** | Akap11 | 0.26158045 | -0.004485915 | 0.527646814 | 9.81263406 | 2.388902887 | 0.052912721 | 0.482505424 | -3.991416135 |
| **A_55_P2060188** | NA | 0.286551889 | -0.004930543 | 0.578034321 | 8.582278524 | 2.388768594 | 0.052922555 | 0.482505424 | -3.991477688 |
| **A_55_P1959994** | Unc5cl | 0.351788405 | -0.006125858 | 0.709702667 | 8.482514144 | 2.388282548 | 0.052958163 | 0.482505424 | -3.991700485 |
| **A_55_P2091297** | NA | 0.414175064 | -0.007285153 | 0.835635282 | 8.99292813 | 2.387869292 | 0.052988458 | 0.482505424 | -3.991889937 |
| **A_51_P465128** | Chst15 | -0.199318032 | -0.402174567 | 0.003538503 | 8.773568452 | -2.387485688 | 0.053016595 | 0.482505424 | -3.992065814 |
| **A_51_P111757** | Asb1 | 0.331870008 | -0.006061193 | 0.669801208 | 10.22056436 | 2.386288248 | 0.053104527 | 0.482505424 | -3.992614935 |
| **A_52_P35377** | Plagl2 | 0.414304721 | -0.00757156 | 0.836181003 | 9.894646763 | 2.386261104 | 0.053106522 | 0.482505424 | -3.992627385 |
| **A_55_P1966538** | Ctu2 | 0.193251062 | -0.003535399 | 0.390037524 | 11.18794457 | 2.3862166 | 0.053109793 | 0.482505424 | -3.992647797 |
| **A_51_P429366** | Hes6 | -0.375324679 | -0.757538139 | 0.006888782 | 10.74192229 | -2.386076332 | 0.053120104 | 0.482505424 | -3.992712133 |
| **A_55_P2156425** | Upk1a | -0.193243404 | -0.390070421 | 0.003583614 | 8.282837237 | -2.385630373 | 0.053152902 | 0.482505424 | -3.992916695 |
| **A_55_P1953063** | Hspe1 | 0.470260052 | -0.008732473 | 0.949252576 | 12.07251718 | 2.385572067 | 0.053157191 | 0.482505424 | -3.992943441 |
| **A_51_P286142** | Znrd1 | -0.281412751 | -0.568059806 | 0.005234304 | 11.01352852 | -2.38550031 | 0.053162471 | 0.482505424 | -3.992976359 |
| **A_55_P2203141** | 2900011F02Rik | 0.293446975 | -0.005464456 | 0.592358405 | 9.121754368 | 2.385449914 | 0.053166179 | 0.482505424 | -3.992999478 |
| **A_55_P2361547** | 2610201A13Rik | 0.338938925 | -0.006333718 | 0.684211567 | 8.754824704 | 2.385297028 | 0.05317743 | 0.482505424 | -3.993069615 |
| **A_52_P602898** | Nkrf | 0.241877998 | -0.004536442 | 0.488292439 | 8.602309706 | 2.385137389 | 0.053189182 | 0.482505424 | -3.993142854 |
| **A_52_P305182** | Ccdc32 | 0.207854506 | -0.00390415 | 0.419613162 | 10.44899392 | 2.385071817 | 0.053194009 | 0.482505424 | -3.993172938 |
| **A_51_P117581** | Cables1 | -0.194358523 | -0.392368338 | 0.003651293 | 8.566523294 | -2.385064117 | 0.053194576 | 0.482505424 | -4.010384174 |
| **A_55_P1956502** | Hnrnph1 | 0.253330593 | -0.004824598 | 0.511485784 | 13.50162618 | 2.384459591 | 0.053239105 | 0.482505424 | -3.993453845 |
| **A_55_P1957583** | Gps2 | 0.358241112 | -0.006853753 | 0.723335978 | 10.65384203 | 2.384256021 | 0.053254108 | 0.482505424 | -3.993547258 |
| **A_51_P172231** | Gsdmd | -0.247553121 | -0.49984754 | 0.004741299 | 9.719655937 | -2.384206953 | 0.053257725 | 0.482505424 | -3.993569775 |
| **A_51_P479434** | Rfc3 | 0.198364841 | -0.003803433 | 0.400533115 | 8.883643062 | 2.384157187 | 0.053261394 | 0.482505424 | -3.993592613 |
| **A_55_P2131128** | Helq | 0.317998252 | -0.006136141 | 0.642132644 | 8.956323421 | 2.38387131 | 0.053282474 | 0.482505424 | -3.993723807 |
| **A_55_P2092869** | Anxa9 | 0.266659636 | -0.005153243 | 0.538472516 | 8.571677474 | 2.383803432 | 0.053287481 | 0.482505424 | -3.993754959 |
| **A_52_P218955** | D3Ertd751e | 0.18364922 | -0.003568962 | 0.370867401 | 8.268761729 | 2.383549927 | 0.053306183 | 0.482505424 | -3.993871307 |
| **A_55_P2097840** | NA | 0.390369083 | -0.007622227 | 0.788360392 | 9.817302018 | 2.383334578 | 0.053322075 | 0.482505424 | -3.993970149 |
| **A_52_P306357** | NA | -1.117568762 | -2.257679726 | 0.022542201 | 9.222978923 | -2.381827588 | 0.053433428 | 0.482505424 | -3.994661987 |
| **A_55_P2082276** | NA | 0.223544079 | -0.004512699 | 0.451600857 | 9.90117299 | 2.381789496 | 0.053436246 | 0.482505424 | -3.994679478 |
| **A_51_P108252** | Gpsm2 | 0.442107289 | -0.008975928 | 0.893190507 | 11.53791755 | 2.381519792 | 0.053456201 | 0.482505424 | -3.994803325 |
| **A_55_P2066081** | Prmt6 | 0.217283388 | -0.004449658 | 0.439016434 | 8.425049978 | 2.381109073 | 0.053486605 | 0.482505424 | -3.994991941 |
| **A_55_P2097309** | Nudt4 | 0.362796038 | -0.007489784 | 0.73308186 | 11.6692178 | 2.380721758 | 0.053515293 | 0.482505424 | -3.995169828 |
| **A_51_P148105** | Rad51 | 0.37623375 | -0.007924979 | 0.760392479 | 10.40307578 | 2.379743966 | 0.053587788 | 0.482505424 | -3.995618989 |
| **A_51_P297165** | Sf3a3 | 0.189685822 | -0.00399982 | 0.383371464 | 12.7797287 | 2.379691356 | 0.053591692 | 0.482505424 | -3.995643159 |
| **A_55_P1994275** | Gar1 | 0.23747019 | -0.005076116 | 0.480016496 | 12.85368847 | 2.379017431 | 0.053641721 | 0.482505424 | -3.995952804 |
| **A_55_P2158384** | Aanat | 0.347456525 | -0.007455225 | 0.702368275 | 9.459902993 | 2.378829323 | 0.053655694 | 0.482505424 | -3.996039243 |
| **A_52_P64763** | Gen1 | 0.260453307 | -0.005646919 | 0.526553532 | 8.460866974 | 2.378306498 | 0.05369455 | 0.482505424 | -3.996279511 |
| **A_55_P2084696** | H2ac1 | 0.296515467 | -0.006430973 | 0.599461906 | 9.207060772 | 2.378289333 | 0.053695826 | 0.482505424 | -3.9962874 |
| **A_55_P2007273** | Pole | 0.398763853 | -0.008696722 | 0.806224429 | 10.81433483 | 2.37800837 | 0.05371672 | 0.482505424 | -3.996416533 |
| **A_55_P2108678** | NA | 0.190505917 | -0.004168641 | 0.385180476 | 9.091871246 | 2.377839078 | 0.053729314 | 0.482505424 | -3.996494345 |
| **A_55_P1974681** | Nsun4 | -0.164286619 | -0.332171624 | 0.003598387 | 8.92432999 | -2.377789863 | 0.053732975 | 0.482505424 | -3.996516966 |
| **A_51_P108629** | Prl3b1 | -0.681228409 | -1.377384287 | 0.014927469 | 8.854609606 | -2.377767822 | 0.053734615 | 0.482505424 | -3.996527098 |
| **A_55_P2128493** | Rap1a | 0.189052781 | -0.004178229 | 0.382283791 | 9.678448561 | 2.377329812 | 0.053767216 | 0.482505424 | -3.996728441 |
| **A_55_P2069052** | Sacs | 0.283572194 | -0.006297352 | 0.573441739 | 8.665658222 | 2.377082433 | 0.053785637 | 0.482505424 | -3.996842166 |
| **A_52_P162298** | Ydjc | 0.32781714 | -0.007308868 | 0.662943148 | 9.777528441 | 2.376877027 | 0.053800938 | 0.482505424 | -3.996936599 |

| | | | | | | | | | |
|---|---|---|---|---|---|---|---|---|---|
| **A_55_P2305525** | C81189 | 0.536023942 | -0.011970441 | 1.084018324 | 9.645796925 | 2.376792513 | 0.053807235 | 0.482505424 | -3.996975456 |
| **A_55_P1953132** | Camkv | 0.255014958 | -0.005704368 | 0.515734285 | 15.16792061 | 2.37670686 | 0.053813617 | 0.482505424 | -3.997014836 |
| **A_51_P186798** | NA | 0.162741627 | -0.003643996 | 0.329127251 | 8.416134899 | 2.376654463 | 0.053817522 | 0.482505424 | -3.997038927 |
| **A_51_P453475** | Slc1a5 | 0.394558096 | -0.008996774 | 0.798112966 | 11.43899297 | 2.375699768 | 0.05388872 | 0.482505424 | -3.997477933 |
| **A_55_P2118740** | NA | 0.436435457 | -0.010043602 | 0.882914517 | 12.89712471 | 2.37521059 | 0.053925239 | 0.482505424 | -3.997702917 |
| **A_51_P208680** | Chtf18 | 0.367666749 | -0.008512501 | 0.743845999 | 10.43323665 | 2.374885677 | 0.05394951 | 0.482505424 | -3.997852367 |
| **A_55_P1995633** | NA | 0.232730263 | -0.005425371 | 0.470885898 | 12.87985066 | 2.374516482 | 0.053977102 | 0.482505424 | -3.998022201 |
| **A_55_P2230638** | Ece1 | -0.195926527 | -0.396464434 | 0.004611381 | 8.489621096 | -2.373995819 | 0.054016039 | 0.482505424 | -3.998261739 |
| **A_55_P2106039** | Map3k6 | -0.32152702 | -0.650673043 | 0.007619004 | 10.13986973 | -2.373624695 | 0.054043811 | 0.482505424 | -3.998432499 |
| **A_55_P2164139** | NA | 0.409200036 | -0.009707964 | 0.828108036 | 13.54624781 | 2.373559911 | 0.05404866 | 0.482505424 | -3.998462309 |
| **A_55_P1994074** | Mrtfa | -0.221868375 | -0.449041508 | 0.005304757 | 8.892117583 | -2.373130526 | 0.054080813 | 0.482505424 | -3.998659899 |
| **A_55_P2128324** | Ubap2l | 0.396544779 | -0.009568908 | 0.802658466 | 9.212228967 | 2.372617881 | 0.054119227 | 0.482505424 | -3.99889583 |
| **A_55_P2075941** | Nf1 | 0.511896661 | -0.01244418 | 1.036237501 | 10.05644636 | 2.372202721 | 0.054150357 | 0.482505424 | -3.99908692 |
| **A_51_P285446** | Lig1 | 0.37151351 | -0.009217291 | 0.752244311 | 13.25762089 | 2.371044955 | 0.054237269 | 0.482505424 | -3.999619922 |
| **A_55_P2185148** | Mettl7a3 | 0.277370919 | -0.006905328 | 0.561647165 | 8.72963305 | 2.37084707 | 0.054252138 | 0.482505424 | -3.999711038 |
| **A_52_P255841** | Dbr1 | 0.278327346 | -0.006992301 | 0.563646993 | 9.425453796 | 2.37032223 | 0.054291596 | 0.482505424 | -3.999952723 |
| **A_51_P117477** | Slc27a1 | 0.614081355 | -0.015446351 | 1.243609061 | 9.972075097 | 2.370250526 | 0.054296989 | 0.482505424 | -3.999985745 |
| **A_55_P1993940** | Sh3bgrl3 | -0.368272916 | -0.745908972 | 0.009363139 | 11.30374947 | -2.369624425 | 0.054344104 | 0.482505424 | -4.000274107 |
| **A_55_P1969311** | Gramd1b | -0.200963461 | -0.407040159 | 0.005113237 | 8.276650161 | -2.369580155 | 0.054347437 | 0.482505424 | -4.000294498 |
| **A_55_P2113061** | NA | 0.269835796 | -0.006885503 | 0.546557094 | 8.773063178 | 2.36940971 | 0.054360271 | 0.482505424 | -4.000373008 |
| **A_55_P2094378** | NA | 0.198027649 | -0.005144122 | 0.401199419 | 8.56934199 | 2.368348753 | 0.054440231 | 0.482505424 | -4.000861782 |
| **A_52_P235102** | Zfp975 | -0.32891462 | -0.666528722 | 0.008699483 | 9.35965768 | -2.367259061 | 0.054522484 | 0.482505424 | -4.00136393 |
| **A_55_P1992445** | Vdac1 | 0.151515485 | -0.004011013 | 0.307041983 | 14.35978476 | 2.367204718 | 0.054526589 | 0.482505424 | -4.001388976 |
| **A_52_P542290** | Pdxdc1 | -0.167629786 | -0.339701716 | 0.004442145 | 9.032196521 | -2.3671422 | 0.054531313 | 0.482505424 | -4.00141779 |
| **A_51_P426633** | Dnttip2 | 0.200065152 | -0.005322087 | 0.405452391 | 12.43014005 | 2.366906931 | 0.054549091 | 0.482505424 | -4.001526227 |
| **A_55_P2018292** | Map2k4 | 0.320477067 | -0.008542832 | 0.649496967 | 9.366462936 | 2.366780493 | 0.054558649 | 0.482505424 | -4.001584506 |
| **A_55_P2030080** | Kdm6b | -0.337959168 | -0.68501754 | 0.009099204 | 9.955805199 | -2.366164293 | 0.054605251 | 0.482505424 | -4.001868557 |
| **A_55_P1994117** | Ifnl3 | 0.239142296 | -0.006464221 | 0.484748813 | 8.500708764 | 2.365918053 | 0.054623885 | 0.482505424 | -4.001982079 |
| **A_55_P2180854** | Mrgprg | 0.297326119 | -0.008076181 | 0.602728419 | 7.918752566 | 2.365614361 | 0.054646876 | 0.482505424 | -4.002122097 |
| **A_55_P2027230** | NA | 0.295552858 | -0.008041408 | 0.599147125 | 8.193624605 | 2.365509996 | 0.054654779 | 0.482505424 | -4.002170218 |
| **A_51_P232901** | Cnp | -0.205589547 | -0.416792199 | 0.005613106 | 9.146019369 | -2.365292474 | 0.054671255 | 0.482505424 | -4.002270516 |
| **A_55_P2037408** | Lrrc57 | 0.169985725 | -0.00464364 | 0.34461509 | 9.201856604 | 2.365257167 | 0.05467393 | 0.482505424 | -4.002286797 |
| **A_55_P1973560** | NA | 0.251219302 | -0.006885352 | 0.509323956 | 12.81552205 | 2.365050174 | 0.054689614 | 0.482505424 | -4.002382247 |
| **A_51_P294778** | Zfp954 | -0.195048373 | -0.395450173 | 0.005353427 | 9.477949443 | -2.364960563 | 0.054696406 | 0.482505424 | -4.00242357 |
| **A_55_P2128042** | NA | 0.497185717 | -0.013790541 | 1.008161974 | 10.63297864 | 2.364291994 | 0.054747103 | 0.482505424 | -4.002731905 |
| **A_55_P2043182** | Gimap6 | -0.542253252 | -1.099553515 | 0.015047011 | 10.34918819 | -2.364264746 | 0.05474917 | 0.482505424 | -4.002744473 |
| **A_51_P144543** | NA | 0.232055715 | -0.006453435 | 0.470564865 | 8.504280282 | 2.364124872 | 0.054759783 | 0.482505424 | -4.002808988 |
| **A_55_P2034400** | Cdk8 | 0.486159166 | -0.013606419 | 0.985924751 | 11.75956372 | 2.36371614 | 0.054790809 | 0.482505424 | -4.002997523 |
| **A_55_P2035038** | NA | 0.51969674 | -0.01455957 | 1.05395305 | 11.67180728 | 2.363651919 | 0.054795685 | 0.482505424 | -4.003027148 |
| **A_55_P1956863** | Egfr | -0.424413671 | -0.860845008 | 0.012017665 | 8.488757137 | -2.362961404 | 0.054848147 | 0.482505424 | -4.00334571 |
| **A_51_P488175** | Phf20l1 | 0.193438518 | -0.005539409 | 0.392416446 | 8.536906715 | 2.362224896 | 0.05490416 | 0.482505424 | -4.003685552 |
| **A_55_P1992640** | Hdac3 | 0.233002994 | -0.006748463 | 0.472754451 | 9.376852306 | 2.361475451 | 0.054961218 | 0.482505424 | -4.004031428 |
| **A_51_P387235** | Nampt | -0.250321197 | -0.507904349 | 0.007261955 | 9.888551671 | -2.361366313 | 0.054969532 | 0.482505424 | -4.004081802 |
| **A_55_P2074393** | NA | 0.306026981 | -0.008906693 | 0.620960655 | 9.061884566 | 2.36115125 | 0.05498592 | 0.482505424 | -4.00418107 |
| **A_55_P2037608** | Acp5 | 0.30010554 | -0.008746527 | 0.608957606 | 15.33140082 | 2.361058186 | 0.054993013 | 0.482505424 | -4.004224028 |
| **A_55_P2034227** | NA | -0.299792442 | -0.60840502 | 0.008820136 | 13.20200481 | -2.36042522 | 0.05504128 | 0.482505424 | -4.004516229 |
| **A_55_P2033120** | Srxn1 | 0.465963855 | -0.013774977 | 0.945702686 | 10.69859239 | 2.360100762 | 0.055066039 | 0.482505424 | -4.00466603 |
| **A_55_P2168990** | NA | -0.366512886 | -0.743931892 | 0.01090612 | 14.12063101 | -2.359655875 | 0.055100006 | 0.482505424 | -4.004871451 |
| **A_55_P1999172** | Nat10 | 0.312375032 | -0.009364889 | 0.634114952 | 11.01637166 | 2.359144549 | 0.055139073 | 0.482505424 | -4.005107578 |
| **A_55_P2077133** | Oxsr1 | 0.313124619 | -0.009430071 | 0.63567931 | 9.488304548 | 2.358832172 | 0.055162954 | 0.482505424 | -4.005251846 |

| | | | | | | | | |
|---|---|---|---|---|---|---|---|---|
| **A_55_P1980746** | Dram2 | -0.253076369 | -0.51381939 | 0.007666652 | 9.778480503 | -2.358425117 | 0.055194089 | 0.482505424 | -4.005439858 |
| **A_55_P2050902** | Fetub | -1.079584391 | -2.192161371 | 0.032992589 | 13.09896353 | -2.357814943 | 0.055240794 | 0.482505424 | -4.005721723 |
| **A_55_P1962661** | Cyp2c67 | -1.05345879 | -2.139245649 | 0.032328069 | 12.74253243 | -2.357524197 | 0.055263064 | 0.482505424 | -4.005856046 |
| **A_55_P2162404** | NA | 0.460069867 | -0.014122394 | 0.934262127 | 9.241956508 | 2.357504427 | 0.055264578 | 0.482505424 | -4.00586518 |
| **A_51_P296416** | Orc6 | 0.241693539 | -0.007472826 | 0.490859904 | 12.3924462 | 2.356995822 | 0.055303558 | 0.482505424 | -4.006100178 |
| **A_55_P2171788** | NA | 0.397641254 | -0.012325268 | 0.807607777 | 11.70802427 | 2.356818998 | 0.055317116 | 0.482505424 | -4.006181885 |
| **A_52_P193236** | Tedc2 | 0.368796704 | -0.011469066 | 0.749062473 | 9.008976091 | 2.356584334 | 0.055335115 | 0.482505424 | -4.006290324 |
| **A_55_P2063376** | Add2 | 0.557679082 | -0.017376861 | 1.132735025 | 12.01633468 | 2.356445759 | 0.055345747 | 0.482505424 | -4.006354364 |
| **A_55_P2363442** | Zfp867 | -0.236373459 | -0.480143824 | 0.007396905 | 8.480080458 | -2.356139462 | 0.055369254 | 0.482505424 | -4.00649592 |
| **A_52_P212214** | Pomt2 | -0.183145702 | -0.372038205 | 0.005746801 | 8.264885677 | -2.355945279 | 0.055384163 | 0.482505424 | -4.006585668 |
| **A_51_P414396** | Mmrn2 | -0.385701833 | -0.783527997 | 0.012124331 | 11.27197882 | -2.355816991 | 0.055394014 | 0.482505424 | -4.006644963 |
| **A_66_P109296** | NA | 0.427474706 | -0.013440234 | 0.868389646 | 11.07223693 | 2.355802055 | 0.055395161 | 0.482505424 | -4.006651866 |
| **A_55_P2091974** | Gpr137 | -0.349021382 | -0.709018251 | 0.010975488 | 9.818883627 | -2.355789589 | 0.055396119 | 0.482505424 | -4.006657628 |
| **A_55_P2000107** | Atox1 | -0.255716953 | -0.519519627 | 0.008085721 | 14.75217727 | -2.355393738 | 0.055426529 | 0.482505424 | -4.006840604 |
| **A_51_P324228** | Satb1 | -0.190934358 | -0.387909549 | 0.006040834 | 8.437443191 | -2.355351575 | 0.05542977 | 0.482505424 | -4.006860094 |
| **A_55_P2032643** | Ajap1 | 0.255617108 | -0.00809116 | 0.519325377 | 8.064148156 | 2.355316962 | 0.05543243 | 0.482505424 | -4.006876095 |
| **A_55_P2164502** | NA | 0.297397634 | -0.00945169 | 0.604246959 | 9.259982234 | 2.355025028 | 0.055454871 | 0.482505424 | -4.007011051 |
| **A_52_P672313** | Exoc5 | 0.281150318 | -0.008954981 | 0.571255617 | 10.88273226 | 2.3548655 | 0.055467138 | 0.482505424 | -4.007084802 |
| **A_51_P288505** | Tradd | -0.192662162 | -0.391462998 | 0.006138674 | 11.00549262 | -2.35484004 | 0.055469096 | 0.482505424 | -4.007096573 |
| **A_55_P1955676** | Kat14 | 0.230200565 | -0.007362105 | 0.467763234 | 9.864242848 | 2.354568752 | 0.055489964 | 0.482505424 | -4.007221999 |
| **A_51_P512119** | AF067063 | 0.293108565 | -0.009457488 | 0.595674619 | 9.10337243 | 2.35391891 | 0.055539984 | 0.482505424 | -4.00752248 |
| **A_55_P2255449** | NA | 0.251069466 | -0.008272165 | 0.510411097 | 8.929372828 | 2.352365764 | 0.055659726 | 0.482505424 | -4.008240838 |
| **A_52_P431872** | Ptcd3 | 0.253196905 | -0.008406129 | 0.514799939 | 12.51656184 | 2.351791438 | 0.055704072 | 0.482505424 | -4.008506544 |
| **A_55_P1986296** | Tagln2 | -0.417175056 | -0.84825807 | 0.013907957 | 14.73599756 | -2.351476334 | 0.055728418 | 0.482505424 | -4.00865234 |
| **A_66_P101942** | Gm9706 | -0.422029473 | -0.858248549 | 0.014189603 | 9.986536495 | -2.350830501 | 0.055778352 | 0.482505424 | -4.008951197 |
| **A_52_P188261** | Camk2d | -0.210475582 | -0.428039561 | 0.007088396 | 7.992746622 | -2.350703841 | 0.05578815 | 0.482505424 | -4.009009814 |
| **A_55_P2112892** | Dab1 | 0.219395475 | -0.007423568 | 0.446214518 | 8.320529547 | 2.350343515 | 0.055816035 | 0.482505424 | -4.00917658 |
| **A_55_P1994047** | NA | -0.391485504 | -0.79621808 | 0.013247072 | 11.59666504 | -2.350340116 | 0.055816298 | 0.482505424 | -4.009178153 |
| **A_51_P414653** | Plvap | -0.557416675 | -1.133803199 | 0.018969849 | 10.78277769 | -2.349899706 | 0.0558504 | 0.482505424 | -4.009382004 |
| **A_55_P2005050** | NA | 0.292664415 | -0.009978717 | 0.595307548 | 12.32358811 | 2.349753396 | 0.055861734 | 0.482505424 | -4.009449731 |
| **A_55_P1993733** | Gm2824 | 0.228492808 | -0.007791844 | 0.46477746 | 8.294785283 | 2.349742168 | 0.055862604 | 0.482505424 | -4.009454929 |
| **A_55_P2173457** | 1700097N02Rik | 0.283073748 | -0.009658 | 0.575805496 | 10.8895376 | 2.349702925 | 0.055865644 | 0.482505424 | -4.009473095 |
| **A_66_P108672** | Mtg1 | 0.220287887 | -0.007547205 | 0.448122978 | 10.86247745 | 2.349379585 | 0.055890702 | 0.482505424 | -4.009622779 |
| **A_55_P2147310** | Mrtfa | -0.221462595 | -0.450518362 | 0.007593172 | 11.2281202 | -2.349320906 | 0.055895251 | 0.482505424 | -4.009649945 |
| **A_55_P2010768** | Dpm3 | -0.245068752 | -0.498552286 | 0.008414783 | 13.0890235 | -2.349207471 | 0.055904045 | 0.482505424 | -4.009702462 |
| **A_51_P196973** | Chaf1a | 0.390680109 | -0.01346392 | 0.794824138 | 10.74844542 | 2.348920523 | 0.055926298 | 0.482505424 | -4.009835315 |
| **A_55_P1974442** | Sumf2 | 0.198068575 | -0.006833468 | 0.402970618 | 9.516134445 | 2.348834824 | 0.055932946 | 0.482505424 | -4.009874994 |
| **A_55_P2086949** | Cntf | 0.3201706 | -0.011084203 | 0.651425402 | 8.877524486 | 2.348564302 | 0.055953936 | 0.482505424 | -4.010000254 |
| **A_52_P830312** | Iba57 | 0.514130941 | -0.018116142 | 1.046378023 | 10.49946735 | 2.347165107 | 0.056062632 | 0.482505424 | -4.010648257 |
| **A_55_P2039603** | Drosha | 0.245913541 | -0.008713853 | 0.500540934 | 11.37534743 | 2.346715851 | 0.056097579 | 0.482505424 | -4.010856367 |
| **A_55_P1994927** | Rab3il1 | 0.450390214 | -0.01597328 | 0.916753708 | 13.38830278 | 2.34664604 | 0.056103011 | 0.482505424 | -4.010888708 |
| **A_55_P1953809** | Plaat3 | 0.4193435 | -0.014873992 | 0.853560993 | 10.38589332 | 2.34663633 | 0.056103767 | 0.482505424 | -4.010893206 |
| **A_52_P455404** | Itpa | 0.229608479 | -0.008177647 | 0.467394605 | 12.67000689 | 2.346305724 | 0.056129502 | 0.482505424 | -4.011046371 |
| **A_55_P2153292** | Tubb4b | 0.287255386 | -0.010316841 | 0.584827613 | 15.46438115 | 2.345627121 | 0.056182363 | 0.482505424 | -4.011360798 |
| **A_51_P509229** | Tnrc6a | 0.190034067 | -0.006838727 | 0.386906861 | 9.324476385 | 2.345464955 | 0.056195003 | 0.482505424 | -4.011435944 |
| **A_55_P2031207** | Ggnbp2 | 0.187935106 | -0.006824954 | 0.382695166 | 11.95929341 | 2.344721155 | 0.056253016 | 0.482505424 | -4.011780654 |
| **A_55_P2181222** | Ifi208 | -0.244130449 | -0.49721984 | 0.008958943 | 8.421317919 | -2.34385746 | 0.056320459 | 0.482505424 | -4.012181008 |
| **A_55_P2017724** | NA | 0.644491708 | -0.02375227 | 1.312735685 | 12.0505797 | 2.343502762 | 0.05634818 | 0.482505424 | -4.012345448 |
| **A_51_P348749** | Llgl2 | -0.313181161 | -0.637909288 | 0.011546967 | 9.624581213 | -2.343467366 | 0.056350947 | 0.482505424 | -4.012361859 |
| **A_51_P394585** | Snx22 | 0.310316026 | -0.011480318 | 0.632112369 | 9.116607914 | 2.343183431 | 0.056373149 | 0.482505424 | -4.012493504 |

| | | | | | | | | | |
|---|---|---|---|---|---|---|---|---|---|
| **A_55_P2034003** | Gtf3c1 | -0.185488415 | -0.377859897 | 0.006883066 | 9.360689491 | -2.342929875 | 0.056392984 | 0.482505424 | -4.012611072 |
| **A_55_P2160473** | LOC634869 | 0.242154342 | -0.008995797 | 0.493304481 | 14.08299482 | 2.342836741 | 0.056400271 | 0.482505424 | -4.012654259 |
| **A_55_P2009677** | Zmiz1 | -0.35283051 | -0.718787387 | 0.013126367 | 11.08649116 | -2.342714732 | 0.056409819 | 0.482505424 | -4.012710835 |
| **A_55_P2125200** | Mterf1b | 0.182370548 | -0.006880726 | 0.371621823 | 8.549512167 | 2.341526511 | 0.056502893 | 0.482505424 | -4.013261914 |
| **A_55_P1977583** | Anxa11 | -0.237319714 | -0.483596347 | 0.00895692 | 11.12245963 | -2.341498025 | 0.056505126 | 0.482505424 | -4.013275128 |
| **A_66_P106375** | Strap | 0.217037303 | -0.008203554 | 0.442278159 | 10.1500837 | 2.341371903 | 0.056515015 | 0.482505424 | -4.013333631 |
| **A_51_P484753** | Saraf | -0.247612557 | -0.504599235 | 0.009374121 | 11.88578462 | -2.341236268 | 0.056525652 | 0.482505424 | -4.013396549 |
| **A_51_P461761** | Zcchc8 | 0.246360814 | -0.009338251 | 0.502059879 | 9.616074047 | 2.341130802 | 0.056533925 | 0.482505424 | -4.013445474 |
| **A_55_P2083764** | NA | 0.183620882 | -0.006961429 | 0.374203194 | 8.440886464 | 2.341114575 | 0.056535198 | 0.482505424 | -4.013453002 |
| **A_55_P2074055** | NA | -0.210795748 | -0.429610096 | 0.0080186 | 8.66656879 | -2.340826574 | 0.056557794 | 0.482505424 | -4.01358661 |
| **A_51_P120398** | Asb17 | 0.280210684 | -0.010676229 | 0.571097597 | 8.380754075 | 2.340688906 | 0.056568599 | 0.482505424 | -4.032110305 |
| **A_55_P2016623** | S100a11-ps | -0.666857012 | -1.359168363 | 0.025454338 | 9.981690536 | -2.340531334 | 0.056580969 | 0.482505424 | -4.013723587 |
| **A_66_P107734** | NA | 0.554570803 | -0.021243543 | 1.130385149 | 13.40428746 | 2.34022551 | 0.056604984 | 0.482505424 | -4.013865485 |
| **A_52_P56397** | Aunip | 0.448274364 | -0.017187341 | 0.91373607 | 10.12355022 | 2.340146982 | 0.056611153 | 0.482505424 | -4.013901922 |
| **A_55_P2076941** | Lrr1 | 0.317940011 | -0.012315252 | 0.648195275 | 9.078534159 | 2.339260707 | 0.056680817 | 0.482505424 | -4.014313208 |
| **A_55_P1955244** | NA | 0.455934374 | -0.01772829 | 0.929597037 | 13.19582942 | 2.33892541 | 0.056707196 | 0.482505424 | -4.01446883 |
| **A_51_P159711** | Car7 | 0.629746868 | -0.024625803 | 1.28411954 | 11.24232923 | 2.338428267 | 0.056746331 | 0.482505424 | -4.014699594 |
| **A_55_P2058962** | Mcm10 | 0.413730707 | -0.016215541 | 0.843676955 | 11.21276071 | 2.338227593 | 0.056762136 | 0.482505424 | -4.01479275 |
| **A_55_P2052759** | Scd3 | 0.18572604 | -0.007279682 | 0.378731763 | 8.47758839 | 2.338222328 | 0.056762551 | 0.482505424 | -4.014795194 |
| **A_55_P2132507** | Bcl2l2 | -0.248602789 | -0.506952092 | 0.009746513 | 8.767523197 | -2.338201275 | 0.05676421 | 0.482505424 | -4.014804968 |
| **A_51_P262340** | Rbm3 | 0.197376708 | -0.007795918 | 0.402549334 | 14.02385365 | 2.337543353 | 0.056816062 | 0.482505424 | -4.015110422 |
| **A_52_P541157** | Calu | -0.175936563 | -0.358859258 | 0.006986133 | 9.088230088 | -2.337069878 | 0.056853407 | 0.482505424 | -4.015330274 |
| **A_55_P2024075** | Prkacb | -0.277561153 | -0.566151771 | 0.011029465 | 11.59324542 | -2.33700512 | 0.056858517 | 0.482505424 | -4.015360345 |
| **A_55_P2106180** | Dnah2 | -0.23324437 | -0.475815821 | 0.00932708 | 9.193291704 | -2.336440222 | 0.056903112 | 0.482505424 | -4.015622685 |
| **A_52_P137765** | Lmna | 0.313404196 | -0.012596301 | 0.639404693 | 13.0342889 | 2.335983302 | 0.056939208 | 0.482505424 | -4.015834906 |
| **A_55_P2058384** | NA | 0.237413571 | -0.009570983 | 0.484398125 | 8.994712477 | 2.335710083 | 0.056960804 | 0.482505424 | -4.015961817 |
| **A_65_P11425** | Bmp2k | 0.502111476 | -0.020254927 | 1.024477879 | 10.28134082 | 2.33565181 | 0.056965411 | 0.482505424 | -4.015988886 |
| **A_55_P2009993** | Tomm20 | 0.265776605 | -0.010764488 | 0.542317699 | 12.2579834 | 2.335287007 | 0.056994262 | 0.482505424 | -4.016158353 |
| **A_55_P2413924** | Gm10404 | 0.265582622 | -0.010830558 | 0.541995803 | 8.738019878 | 2.334662434 | 0.057043691 | 0.482505424 | -4.016448531 |
| **A_55_P1982817** | NA | 0.501747683 | -0.020683142 | 1.024178507 | 11.90337381 | 2.333671761 | 0.057122186 | 0.482505424 | -4.016908891 |
| **A_55_P1971498** | Exoc1 | 0.181133455 | -0.007474744 | 0.369741653 | 11.06894989 | 2.333572462 | 0.05713006 | 0.482505424 | -4.01695504 |
| **A_55_P1995318** | NA | 0.357173906 | -0.014830943 | 0.729178755 | 9.41869462 | 2.332997704 | 0.057175658 | 0.482505424 | -4.017222184 |
| **A_51_P518621** | H3c1 | 0.335756796 | -0.014071628 | 0.685585221 | 9.035665474 | 2.332130812 | 0.057244504 | 0.482505424 | -4.017625182 |
| **A_55_P2031442** | Pithd1 | 0.322662211 | -0.013655211 | 0.658979632 | 12.30976958 | 2.331212853 | 0.057317499 | 0.482505424 | -4.018052012 |
| **A_55_P2056654** | Kif22 | 0.402956752 | -0.017073505 | 0.822987009 | 12.38836347 | 2.331100786 | 0.057326417 | 0.482505424 | -4.018104128 |
| **A_55_P2122666** | Pnma3 | 0.28087824 | -0.01191364 | 0.573670119 | 8.838090666 | 2.330999908 | 0.057334446 | 0.482505424 | -4.01815104 |
| **A_55_P2173005** | Dnase2a | 0.468744724 | -0.019882335 | 0.957371783 | 12.2959668 | 2.330998898 | 0.057334527 | 0.482505424 | -4.01815151 |
| **A_55_P2082070** | Ak6 | 0.205456397 | -0.00874284 | 0.419655635 | 10.05816247 | 2.330692279 | 0.057358938 | 0.482505424 | -4.01829411 |
| **A_55_P2113598** | Ikzf3 | 0.353032224 | -0.015026471 | 0.721090918 | 8.541250868 | 2.330668229 | 0.057360853 | 0.482505424 | -4.018305295 |
| **A_51_P406008** | Hgh1 | 0.232644442 | -0.009949881 | 0.475238765 | 9.05998044 | 2.330210939 | 0.057397281 | 0.482505424 | -4.018517989 |
| **A_55_P2008106** | Otud6b | 0.261642663 | -0.011212796 | 0.534498122 | 9.032825067 | 2.330017071 | 0.057412732 | 0.482505424 | -4.018608168 |
| **A_55_P2019463** | Tnks1bp1 | -0.274666401 | -0.561149447 | 0.011816646 | 9.600184674 | -2.329645279 | 0.057442375 | 0.482505424 | -4.018781121 |
| **A_55_P2095047** | Mat2a | 0.227690787 | -0.009820354 | 0.465201927 | 8.658057953 | 2.329403162 | 0.057461687 | 0.482505424 | -4.018893758 |
| **A_51_P115655** | Iqcd | 0.333691448 | -0.014418762 | 0.681801657 | 10.01384849 | 2.32922533 | 0.057475877 | 0.482505424 | -4.018976494 |
| **A_52_P342896** | Mrps33 | 0.222154228 | -0.009602032 | 0.453910488 | 9.383901696 | 2.329197409 | 0.057478105 | 0.482505424 | -4.018989485 |
| **A_55_P2117033** | Mtmr3 | 0.454455385 | -0.019647591 | 0.92855836 | 10.47366335 | 2.329173079 | 0.057480046 | 0.482505424 | -4.019000804 |
| **A_55_P1988760** | Sestd1 | -0.291183602 | -0.595039227 | 0.012672023 | 8.059386822 | -2.32853528 | 0.05753097 | 0.482505424 | -4.019297571 |
| **A_51_P254805** | Kif4 | 0.398678359 | -0.017365342 | 0.81472206 | 9.775682653 | 2.328449911 | 0.05753779 | 0.482505424 | -4.019337296 |
| **A_55_P2047778** | Krtap4-8 | 0.323047026 | -0.014116436 | 0.660210489 | 10.78607552 | 2.328136455 | 0.057562837 | 0.482505424 | -4.019483166 |
| **A_66_P108178** | Lin52 | 0.189124152 | -0.008308025 | 0.386556329 | 9.454593415 | 2.327620899 | 0.057604058 | 0.482505424 | -4.01972311 |

| | | | | | | | | | |
|---|---|---|---|---|---|---|---|---|---|
| **A_55_P2057142** | Cks1brt | 0.199379482 | -0.008772171 | 0.407531136 | 9.457053095 | 2.327468369 | 0.05761626 | 0.482505424 | -4.019794105 |
| **A_51_P357341** | Clic1 | -0.334278248 | -0.683266997 | 0.0147105 | 14.15590669 | -2.327447434 | 0.057617934 | 0.482505424 | -4.019803849 |
| **A_52_P618745** | Ermp1 | 0.308288366 | -0.013623053 | 0.630199786 | 9.657347309 | 2.327040505 | 0.0576505 | 0.482505424 | -4.019993267 |
| **A_51_P100174** | Mns1 | 0.433580929 | -0.019179749 | 0.886341608 | 12.46311707 | 2.326937182 | 0.057658772 | 0.482505424 | -4.020041364 |
| **A_51_P270213** | Ceacam11 | -0.823933565 | -1.68432973 | 0.0364626 | 8.406026487 | -2.326895706 | 0.057662093 | 0.482505424 | -4.020060672 |
| **A_51_P327564** | Glb1 | -0.261188608 | -0.533974591 | 0.011597374 | 9.988309226 | -2.326565963 | 0.057688501 | 0.482505424 | -4.02021418 |
| **A_55_P2177273** | Kcna6 | 0.192577103 | -0.008612784 | 0.393766991 | 8.862420813 | 2.325849937 | 0.057745889 | 0.482505424 | -4.02054756 |
| **A_55_P2091305** | NA | 0.240167428 | -0.010754212 | 0.491089067 | 8.527391721 | 2.325729378 | 0.057755557 | 0.482505424 | -4.020603698 |
| **A_52_P559919** | Eif2ak2 | -0.191567315 | -0.391713534 | 0.008578905 | 8.845704484 | -2.325718832 | 0.057756403 | 0.482505424 | -4.020608609 |
| **A_55_P2163873** | Uox | -0.828889369 | -1.694997203 | 0.037218465 | 11.29090118 | -2.325454207 | 0.057777631 | 0.482505424 | -4.020731836 |
| **A_55_P1970234** | Dclk2 | 0.438053667 | -0.019705028 | 0.895812363 | 9.701676121 | 2.325272774 | 0.05779219 | 0.482505424 | -4.020816329 |
| **A_55_P2052380** | Gbp6 | 0.430935983 | -0.019459311 | 0.881331278 | 8.722380418 | 2.324888363 | 0.05782305 | 0.482505424 | -4.020995358 |
| **A_55_P1959753** | NA | 0.279038055 | -0.012603421 | 0.570679531 | 8.994352353 | 2.324862853 | 0.057825098 | 0.482505424 | -4.02100724 |
| **A_55_P2286503** | A930001C03Rik | 0.368685223 | -0.016688494 | 0.754058941 | 8.812037857 | 2.324646004 | 0.057842515 | 0.482505424 | -4.02110824 |
| **A_55_P1975341** | Nasp | 0.317284706 | -0.014383345 | 0.648952756 | 12.60360197 | 2.324495385 | 0.057854615 | 0.482505424 | -4.021178396 |
| **A_55_P2027701** | NA | -0.174878088 | -0.357687646 | 0.007931469 | 13.39638362 | -2.324447213 | 0.057858485 | 0.482505424 | -4.021200834 |
| **A_66_P105736** | Net1 | 0.212741058 | -0.009682775 | 0.43516489 | 9.855158227 | 2.324091295 | 0.057887091 | 0.482505424 | -4.021366626 |
| **A_55_P1997106** | Large2 | -0.178597883 | -0.365363807 | 0.008168041 | 8.885517313 | -2.323602609 | 0.057926392 | 0.482505424 | -4.021594288 |
| **A_55_P2042818** | Zfp639 | 0.212640291 | -0.009749495 | 0.435030077 | 10.54363933 | 2.323346098 | 0.057947032 | 0.482505424 | -4.021713798 |
| **A_55_P1998912** | Txnrd2 | 0.326496983 | -0.014986492 | 0.667980457 | 13.64314884 | 2.323232475 | 0.057956177 | 0.482505424 | -4.021766738 |
| **A_55_P2088425** | Tcn2 | -0.25405113 | -0.519767072 | 0.011664812 | 12.92707095 | -2.32320059 | 0.057958743 | 0.482505424 | -4.021781595 |
| **A_55_P1995233** | Lonp2 | 0.218348408 | -0.01003072 | 0.446727536 | 11.77405691 | 2.323147627 | 0.057963007 | 0.482505424 | -4.021806272 |
| **A_55_P2053202** | Itsn2 | -0.205513875 | -0.420534826 | 0.009507076 | 10.29839035 | -2.322434952 | 0.058020408 | 0.482505424 | -4.022138368 |
| **A_55_P2157008** | Ear10 | -0.365171115 | -0.747259264 | 0.016917034 | 9.741123889 | -2.322287792 | 0.058032268 | 0.482505424 | -4.022206949 |
| **A_55_P2116664** | Hook2 | -0.33184625 | -0.679148641 | 0.015456141 | 10.83792781 | -2.321733301 | 0.058076978 | 0.482505424 | -4.022465382 |
| **A_55_P1969072** | Cbx3 | 0.263870199 | -0.012315771 | 0.54005617 | 13.54516006 | 2.32151728 | 0.058094407 | 0.482505424 | -4.022566073 |
| **A_52_P648601** | Dusp9 | 0.800755474 | -0.037428309 | 1.638939257 | 9.723296776 | 2.321367239 | 0.058106515 | 0.482505424 | -4.022636012 |
| **A_55_P2143030** | Timd2 | 0.490877223 | -0.023042543 | 1.004796989 | 8.961525801 | 2.320923087 | 0.058142373 | 0.482505424 | -4.022843063 |
| **A_55_P2212908** | Pon2 | 0.301971248 | -0.014184761 | 0.618127257 | 8.625123317 | 2.320851444 | 0.058148159 | 0.482505424 | -4.022876462 |
| **A_55_P2014665** | Racgap1 | 0.370502439 | -0.017443395 | 0.758448274 | 10.82954102 | 2.320615387 | 0.058167229 | 0.482505424 | -4.022986516 |
| **A_55_P2129706** | Srsf4 | 0.258110327 | -0.012229569 | 0.528450223 | 11.45536657 | 2.319948944 | 0.058221101 | 0.482505424 | -4.023297257 |
| **A_55_P2154977** | Rpap3 | 0.178678365 | -0.008555736 | 0.365912466 | 11.18530667 | 2.318836935 | 0.058311105 | 0.482505424 | -4.023815862 |
| **A_52_P570543** | Sdk2 | 0.239990048 | -0.011545578 | 0.491525675 | 8.683306147 | 2.318338875 | 0.058351464 | 0.482505424 | -4.024048186 |
| **A_51_P398467** | Bsdc1 | 0.368500731 | -0.017737662 | 0.754739125 | 11.91769837 | 2.318281135 | 0.058356144 | 0.482505424 | -4.024075121 |
| **A_55_P1966721** | NA | -0.33794675 | -0.692254641 | 0.016361142 | 9.627656626 | -2.317664867 | 0.058406126 | 0.482505424 | -4.024362627 |
| **A_55_P2051099** | Tnfrsf1a | -0.426331976 | -0.873347252 | 0.0206833 | 10.28581859 | -2.317441242 | 0.058424274 | 0.482505424 | -4.024466964 |
| **A_55_P2008884** | Cltc | 0.260809286 | -0.012732642 | 0.534351214 | 13.68786776 | 2.316766875 | 0.058479037 | 0.482505424 | -4.024781641 |
| **A_51_P250058** | Epas1 | -0.325840594 | -0.667597169 | 0.01591598 | 12.29089094 | -2.316709075 | 0.058483733 | 0.482505424 | -4.024808615 |
| **A_52_P8043** | Srsf2 | 0.292386023 | -0.014320212 | 0.599092259 | 10.7403656 | 2.316419391 | 0.058507275 | 0.482505424 | -4.024943806 |
| **A_51_P297963** | Recql | 0.227400731 | -0.011138682 | 0.465940145 | 8.646990862 | 2.316407166 | 0.058508269 | 0.482505424 | -4.024949511 |
| **A_55_P1981455** | NA | -0.283358613 | -0.580638686 | 0.01392146 | 8.795628264 | -2.316081342 | 0.05853476 | 0.482505424 | -4.025101581 |
| **A_65_P10357** | Srsf6 | 0.20166552 | -0.009915249 | 0.41324629 | 11.39587282 | 2.316000496 | 0.058541335 | 0.482505424 | -4.025139315 |
| **A_55_P2117830** | NA | 0.222585998 | -0.010971111 | 0.456143106 | 8.475150734 | 2.315730096 | 0.058563333 | 0.482505424 | -4.025265528 |
| **A_55_P2009477** | 9530068E07Rik | -0.287606519 | -0.589412213 | 0.014199174 | 13.24977471 | -2.315551725 | 0.058577848 | 0.482505424 | -4.02534879 |
| **A_55_P2036967** | Cox17 | 0.318090473 | -0.015758563 | 0.651939508 | 8.065286989 | 2.315174471 | 0.05860856 | 0.482505424 | -4.0255249 |
| **A_51_P457331** | Hmgn5 | 0.316492636 | -0.015680237 | 0.64866551 | 11.05607702 | 2.315168664 | 0.058609033 | 0.482505424 | -4.025527611 |
| **A_55_P2067645** | Gm5136 | 0.166721852 | -0.008263731 | 0.341707434 | 8.969878652 | 2.315119681 | 0.058613022 | 0.482505424 | -4.025550478 |
| **A_51_P242603** | E430018J23Rik | -0.169416492 | -0.347232877 | 0.008399892 | 8.31642103 | -2.315085839 | 0.058615778 | 0.482505424 | -4.025566278 |
| **A_55_P1973011** | Ppm1l | 0.269026861 | -0.013422002 | 0.551475725 | 10.13901177 | 2.314403097 | 0.058671409 | 0.482505424 | -4.025885043 |
| **A_51_P308469** | Rngtt | 0.237703908 | -0.011868157 | 0.487275974 | 9.923897748 | 2.314320689 | 0.058678128 | 0.482505424 | -4.025923522 |

| | | | | | | | | | |
|---|---|---|---|---|---|---|---|---|---|
| **A_55_P2032695** | Tut4 | -0.198043049 | -0.405987466 | 0.009901368 | 11.45676278 | -2.314171435 | 0.058690298 | 0.482505424 | -4.025993216 |
| **A_51_P338600** | Tmem14c | 0.292189412 | -0.0146618 | 0.599040625 | 13.54127632 | 2.313768055 | 0.058723203 | 0.482505424 | -4.026181585 |
| **A_52_P520466** | Kif18b | 0.327308677 | -0.016443858 | 0.671061211 | 8.974300003 | 2.31363475 | 0.058734081 | 0.482505424 | -4.02624384 |
| **A_55_P2050988** | NA | 0.203434514 | -0.010261956 | 0.417130984 | 9.450772747 | 2.313185577 | 0.058770751 | 0.482505424 | -4.026453621 |
| **A_52_P116628** | Fam234b | 0.312059445 | -0.015774244 | 0.639893134 | 9.26257764 | 2.312953706 | 0.05878969 | 0.482505424 | -4.026561924 |
| **A_51_P178872** | Ttc9c | 0.187783929 | -0.009593266 | 0.385161123 | 8.950539311 | 2.311770075 | 0.058886467 | 0.482505424 | -4.027114866 |
| **A_55_P1957922** | Arrdc2 | 0.332157358 | -0.017008499 | 0.681323216 | 10.25249353 | 2.311507446 | 0.058907962 | 0.482505424 | -4.027237576 |
| **A_55_P2160286** | Olfm1 | -0.199859446 | -0.409974167 | 0.010255274 | 8.117573648 | -2.311273761 | 0.058927096 | 0.482505424 | -4.027346769 |
| **A_55_P2151308** | G430095P16Rik | 0.362179737 | -0.018605354 | 0.742964828 | 9.84656067 | 2.31114611 | 0.05893755 | 0.482505424 | -4.027406419 |
| **A_51_P500541** | Flywch2 | -0.55817775 | -1.145185137 | 0.028829636 | 8.36331192 | -2.310532833 | 0.058987803 | 0.482505424 | -4.027693021 |
| **A_55_P1979575** | Shroom2 | -0.249348192 | -0.511611422 | 0.012915038 | 10.02978653 | -2.310212911 | 0.059014036 | 0.482505424 | -4.027842546 |
| **A_55_P2178152** | Ccsap | 0.198693238 | -0.010295163 | 0.407681639 | 8.891530653 | 2.310170821 | 0.059017488 | 0.482505424 | -4.027862219 |
| **A_51_P307243** | Ppp5c | 0.383024105 | -0.019873177 | 0.785921387 | 10.52005507 | 2.310015838 | 0.059030201 | 0.482505424 | -4.02793466 |
| **A_52_P238044** | Psmc3ip | 0.235177884 | -0.01223656 | 0.482592328 | 9.328902743 | 2.309694915 | 0.059056535 | 0.482505424 | -4.028084671 |
| **A_52_P89099** | Nolc1 | 0.234045089 | -0.01219858 | 0.480288759 | 8.61546676 | 2.309498303 | 0.059072675 | 0.482505424 | -4.028176581 |
| **A_52_P272945** | Ccnh | 0.352204186 | -0.018450133 | 0.722858506 | 9.189672984 | 2.308918677 | 0.059120282 | 0.482505424 | -4.028447562 |
| **A_55_P1990648** | Ndufaf4 | 0.306645849 | -0.016141389 | 0.629433087 | 9.685341865 | 2.308362039 | 0.059166039 | 0.482505424 | -4.028707832 |
| **A_55_P2091055** | NA | 0.27373116 | -0.014457128 | 0.561919447 | 14.67929657 | 2.307974998 | 0.059197876 | 0.482505424 | -4.028888823 |
| **A_55_P1971458** | Jak2 | 0.284656524 | -0.015115364 | 0.584428412 | 10.8731547 | 2.307349733 | 0.059249346 | 0.482505424 | -4.029181249 |
| **A_51_P204845** | Celf1 | 0.322516091 | -0.017127889 | 0.662160072 | 11.76681217 | 2.307335003 | 0.059250559 | 0.482505424 | -4.029188138 |
| **A_51_P296905** | Ppat | 0.32383662 | -0.01730923 | 0.664982471 | 9.655775896 | 2.30658283 | 0.059312539 | 0.482505424 | -4.029539976 |
| **A_55_P2462377** | Nsf | 0.231478572 | -0.012384691 | 0.475341835 | 10.63155592 | 2.306468899 | 0.059321933 | 0.482505424 | -4.029593274 |
| **A_55_P2030752** | Nedd4l | -0.727005074 | -1.492959106 | 0.038948957 | 10.58782969 | -2.306311289 | 0.05933493 | 0.482505424 | -4.029667008 |
| **A_51_P261001** | Mtf2 | 0.231463292 | -0.01241845 | 0.475345034 | 10.59066548 | 2.3061419 | 0.059348903 | 0.482505424 | -4.029746255 |
| **A_55_P1983448** | S100a4 | -0.408814464 | -0.839631919 | 0.02200299 | 9.232605654 | -2.305770891 | 0.059379519 | 0.482505424 | -4.029919841 |
| **A_51_P434697** | Ube2e2 | -0.554599809 | -1.139084479 | 0.02988486 | 10.66104612 | -2.305630882 | 0.059391076 | 0.482505424 | -4.029985351 |
| **A_55_P2035286** | Uhrf1 | 0.345737378 | -0.018740641 | 0.710215398 | 12.9578982 | 2.304932341 | 0.059448776 | 0.482505424 | -4.030312233 |
| **A_52_P106739** | Ddx46 | 0.279182869 | -0.01519125 | 0.573556987 | 10.93566474 | 2.304476747 | 0.059486439 | 0.482505424 | -4.030525457 |
| **A_52_P628590** | Pvr | -0.192553554 | -0.395586059 | 0.010478952 | 8.354895546 | -2.304459892 | 0.059487833 | 0.482505424 | -4.030533346 |
| **A_55_P2110136** | Arhgap29 | -0.195775524 | -0.4022195 | 0.010668453 | 8.398064876 | -2.304301846 | 0.059500905 | 0.482505424 | -4.030607319 |
| **A_55_P2074473** | Gtf2f1 | 0.229011539 | -0.012516225 | 0.470539304 | 13.47974495 | 2.303952357 | 0.059529821 | 0.482505424 | -4.030770907 |
| **A_55_P2002578** | Ephx1 | -0.384130601 | -0.789326288 | 0.021065086 | 11.64638497 | -2.303548079 | 0.059563288 | 0.482505424 | -4.030960157 |
| **A_55_P2030060** | Kdm8 | 0.214880729 | -0.011791253 | 0.44155271 | 8.926097524 | 2.303471355 | 0.059569642 | 0.482505424 | -4.030996076 |
| **A_66_P116372** | Soat2 | 0.539099157 | -0.029604721 | 1.107803034 | 11.35791218 | 2.30338032 | 0.059577181 | 0.482505424 | -4.031038694 |
| **A_51_P455671** | Psmd5 | 0.264686454 | -0.01456066 | 0.543933569 | 10.46646584 | 2.303171143 | 0.059594509 | 0.482505424 | -4.031136625 |
| **A_51_P295192** | Nfkbia | -0.741173375 | -1.523123013 | 0.040776263 | 11.73269974 | -2.303160565 | 0.059595386 | 0.482505424 | -4.031141577 |
| **A_55_P2127702** | Racgap1 | 0.30596347 | -0.016835787 | 0.628762727 | 11.77156406 | 2.303139479 | 0.059597133 | 0.482505424 | -4.03115145 |
| **A_55_P1997741** | Slc9a8 | 0.275356859 | -0.015209921 | 0.56592364 | 11.89743087 | 2.302677552 | 0.059635419 | 0.482505424 | -4.03136773 |
| **A_55_P2148248** | NA | 0.267040126 | -0.014755426 | 0.548835678 | 8.454467262 | 2.302637538 | 0.059638736 | 0.482505424 | -4.031386466 |
| **A_55_P2249220** | A430102J17Rik | 0.26413208 | -0.014612402 | 0.542876561 | 9.014631863 | 2.302491639 | 0.059650835 | 0.482505424 | -4.031454783 |
| **A_55_P1985463** | NA | 0.256347642 | -0.014184286 | 0.526879571 | 14.99072036 | 2.302470041 | 0.059652626 | 0.482505424 | -4.031464897 |
| **A_51_P387220** | Ddx1 | 0.274607604 | -0.015244453 | 0.564459661 | 11.53699865 | 2.30207443 | 0.059685445 | 0.482505424 | -4.031650156 |
| **A_55_P2119917** | Ikzf4 | -0.374635954 | -0.770092314 | 0.020820406 | 8.445032477 | -2.301940422 | 0.059696567 | 0.482505424 | -4.031712913 |
| **A_55_P2090798** | Mtus1 | -0.414820135 | -0.85274553 | 0.02310526 | 9.739477281 | -2.301669101 | 0.05971909 | 0.482505424 | -4.031839983 |
| **A_55_P2139814** | Fgf13 | 0.484310578 | -0.027129848 | 0.995751004 | 9.152716852 | 2.300976009 | 0.059776667 | 0.482505424 | -4.032164619 |
| **A_55_P2120773** | Dctn4 | 0.158588771 | -0.00889915 | 0.326076693 | 11.09548165 | 2.300764298 | 0.059794266 | 0.482505424 | -4.032263793 |
| **A_55_P2111770** | Klhl13 | 0.318133523 | -0.017893358 | 0.654160405 | 11.17884436 | 2.300480745 | 0.059817845 | 0.482505424 | -4.032396629 |
| **A_55_P1985143** | NA | 0.323894968 | -0.018254713 | 0.666044649 | 11.87261614 | 2.30022993 | 0.05983871 | 0.482505424 | -4.032514135 |
| **A_52_P146164** | Ube2n | 0.193974958 | -0.010981999 | 0.398931915 | 9.181250499 | 2.29967355 | 0.059885021 | 0.482505424 | -4.032774822 |
| **A_55_P1990919** | Msrb1 | -0.280666351 | -0.577246016 | 0.015913315 | 14.1771449 | -2.299493394 | 0.059900025 | 0.482505424 | -4.03285924 |

| | | | | | | | | |
|---|---|---|---|---|---|---|---|---|
| **A_55_P2075359** | Eif4a3l1 | 0.224264434 | -0.01281332 | 0.461342188 | 12.20320906 | 2.298543826 | 0.059979169 | 0.482505424 | -4.03330425 |
| **A_52_P77204** | Lrrc20 | 0.300038627 | -0.017161756 | 0.61723901 | 9.789555982 | 2.29840552 | 0.059990706 | 0.482505424 | -4.033369075 |
| **A_55_P2131238** | Ttc39a | 0.315008166 | -0.018097019 | 0.64811335 | 9.134525961 | 2.297860228 | 0.060036213 | 0.482505424 | -4.033624677 |
| **A_51_P437021** | Trak1 | -0.279690725 | -0.575456347 | 0.016074898 | 9.404168624 | -2.29780706 | 0.060040652 | 0.482505424 | -4.033649601 |
| **A_66_P123804** | Fbxo30 | 0.34016726 | -0.019610059 | 0.699944579 | 10.141433 | 2.297428051 | 0.060072306 | 0.482505424 | -4.033827281 |
| **A_55_P2061094** | Fnta | 0.198166754 | -0.011493805 | 0.407827313 | 8.411728967 | 2.296662848 | 0.060136266 | 0.482505424 | -4.034186057 |
| **A_51_P363719** | Stx7 | -0.218664969 | -0.450019895 | 0.012689957 | 10.30048631 | -2.296590962 | 0.060142278 | 0.482505424 | -4.034219765 |
| **A_55_P2065621** | Hook2 | -0.267925999 | -0.551421387 | 0.015569388 | 10.37216259 | -2.296423863 | 0.060156256 | 0.482505424 | -4.034298121 |
| **A_55_P2175190** | Cep250 | 0.294090228 | -0.017102905 | 0.605283361 | 9.182723177 | 2.296327243 | 0.06016434 | 0.482505424 | -4.03434343 |
| **A_55_P2081148** | Foxk2 | 0.206046332 | -0.011992716 | 0.424085381 | 10.42454607 | 2.296221608 | 0.060173179 | 0.482505424 | -4.034392968 |
| **A_51_P122723** | Grpel1 | -0.164037452 | -0.33764778 | 0.009572877 | 13.07891913 | -2.295887713 | 0.060201128 | 0.482505424 | -4.034549556 |
| **A_55_P1968908** | Hypk | 0.297443816 | -0.017401447 | 0.612289079 | 11.67029114 | 2.295572269 | 0.060227544 | 0.482505424 | -4.034697502 |
| **A_55_P2044359** | Rxrg | 0.30107286 | -0.017621244 | 0.619766964 | 10.1913664 | 2.295518347 | 0.060232061 | 0.482505424 | -4.034722793 |
| **A_65_P12104** | Tmtc1 | -0.432979242 | -0.89133599 | 0.025377506 | 8.035373236 | -2.295337938 | 0.060247176 | 0.482505424 | -4.034807412 |
| **A_55_P2018274** | NA | 0.284089614 | -0.016653745 | 0.584832974 | 11.93393456 | 2.295316083 | 0.060249007 | 0.482505424 | -4.034817664 |
| **A_51_P423091** | Flot1 | -0.242225974 | -0.498708028 | 0.014256081 | 10.78226972 | -2.294810959 | 0.060291349 | 0.482505424 | -4.035054609 |
| **A_52_P103391** | B4galt1 | -0.260876054 | -0.537109441 | 0.015357332 | 12.84881285 | -2.294780962 | 0.060293864 | 0.482505424 | -4.035068681 |
| **A_52_P508317** | Erlec1 | 0.230488997 | -0.01363788 | 0.474615873 | 9.147027711 | 2.294128772 | 0.060348581 | 0.482505424 | -4.035374656 |
| **A_55_P2062034** | Tsen34 | -0.432847089 | -0.891305691 | 0.025611514 | 12.9437412 | -2.294127567 | 0.060348682 | 0.482505424 | -4.035375222 |
| **A_51_P185869** | Hbq1a | 0.32750292 | -0.019448642 | 0.674454483 | 9.894186067 | 2.293662523 | 0.06038773 | 0.482505424 | -4.035593426 |
| **A_55_P1967015** | Fgfr1 | 0.218417295 | -0.013004872 | 0.449839462 | 8.836549845 | 2.293323166 | 0.060416241 | 0.482505424 | -4.035752671 |
| **A_55_P1968778** | Rasa1 | 0.256782523 | -0.015293116 | 0.528858163 | 9.461448795 | 2.29329008 | 0.060419021 | 0.482505424 | -4.035768198 |
| **A_55_P2122200** | Prkar2b | 0.282203437 | -0.016909833 | 0.581316707 | 9.498508128 | 2.292502446 | 0.06048525 | 0.482505424 | -4.036137852 |
| **A_55_P2002893** | Pfkfb1 | -0.802082759 | -1.652269947 | 0.04810443 | 12.00244743 | -2.292386349 | 0.060495019 | 0.482505424 | -4.036192345 |
| **A_55_P2158478** | Nkiras2 | -0.253591945 | -0.522404105 | 0.015220216 | 11.4992694 | -2.292290908 | 0.06050305 | 0.482505424 | -4.036237143 |
| **A_51_P492346** | Tmem174 | 0.322838533 | -0.019388196 | 0.665065263 | 9.07809249 | 2.292211191 | 0.06050976 | 0.482505424 | -4.036274562 |
| **A_55_P2254779** | NA | 0.325382613 | -0.019633088 | 0.670398315 | 9.950497726 | 2.291599253 | 0.060561289 | 0.482648684 | -4.036561825 |
| **A_52_P183524** | Tmem86b | 0.354434596 | -0.021430679 | 0.730299871 | 8.990331361 | 2.291327097 | 0.060584221 | 0.482648684 | -4.036689597 |
| **A_55_P1991199** | Nos1ap | 0.481958887 | -0.029190099 | 0.993107872 | 10.08455861 | 2.291108616 | 0.060602636 | 0.482648684 | -4.036792176 |
| **A_55_P2022251** | Cdkn2d | 0.251657949 | -0.015300799 | 0.518616696 | 13.48359565 | 2.290602263 | 0.060645339 | 0.482789848 | -4.037029932 |
| **A_55_P1968143** | Rbm25 | 0.326863318 | -0.020081774 | 0.67380841 | 10.32922466 | 2.289225771 | 0.060761581 | 0.483477094 | -4.037676404 |
| **A_55_P2059347** | Ormdl3 | 0.227192267 | -0.013983262 | 0.468367796 | 12.49795354 | 2.288987886 | 0.060781693 | 0.483477094 | -4.037788148 |
| **A_65_P18433** | Ankrd28 | 0.293723092 | -0.01812757 | 0.605573753 | 9.83648646 | 2.288624855 | 0.060812399 | 0.48352236 | -4.03795869 |
| **A_51_P476711** | Mtrex | 0.209923492 | -0.012994012 | 0.432840995 | 11.43013751 | 2.288232027 | 0.060845643 | 0.483574684 | -4.038143246 |
| **A_52_P253230** | Ankrd46 | -0.25042715 | -0.516387524 | 0.015533224 | 10.53921215 | -2.287955982 | 0.060869016 | 0.483574684 | -4.038272946 |
| **A_51_P215922** | Casp6 | -0.171573314 | -0.353852783 | 0.010706154 | 10.37287494 | -2.287152786 | 0.060937075 | 0.483916482 | -4.038650377 |
| **A_52_P271155** | Mtpap | 0.242778344 | -0.015269633 | 0.500826322 | 10.0064816 | 2.286086585 | 0.061027541 | 0.484435868 | -4.039151505 |
| **A_55_P2024808** | Abl1 | -0.250224055 | -0.516227903 | 0.015779793 | 10.70376479 | -2.285726842 | 0.061058096 | 0.48447945 | -4.039320618 |
| **A_55_P1956827** | Ogfr | -0.154767759 | -0.319352356 | 0.009816838 | 13.30123747 | -2.284938385 | 0.06112512 | 0.484600003 | -4.039691314 |
| **A_55_P2068812** | Pkm | -0.358348486 | -0.739453164 | 0.022756193 | 13.44568853 | -2.284780495 | 0.061138551 | 0.484600003 | -4.039765555 |
| **A_51_P346575** | Rhag | 0.474461795 | -0.030239261 | 0.979162852 | 10.77886484 | 2.284284659 | 0.061180749 | 0.484600003 | -4.039998718 |
| **A_55_P1981296** | Nup85 | 0.260371901 | -0.016639521 | 0.537383324 | 12.93927563 | 2.283913366 | 0.061212367 | 0.484600003 | -4.040173333 |
| **A_55_P2111416** | Rcc2 | 0.330046374 | -0.021111603 | 0.681204351 | 10.98233287 | 2.283787102 | 0.061223123 | 0.484600003 | -4.040232717 |
| **A_52_P86693** | Ifi27 | -0.675016022 | -1.393250317 | 0.043218274 | 11.90801851 | -2.28365835 | 0.061234093 | 0.484600003 | -4.040293273 |
| **A_51_P143721** | Vipas39 | -0.197117934 | -0.406890116 | 0.012654248 | 9.546966666 | -2.283291876 | 0.061265329 | 0.484600003 | -4.040465647 |
| **A_51_P436483** | Kin | 0.191522599 | -0.012303978 | 0.395349176 | 10.17743101 | 2.283191844 | 0.061273858 | 0.484600003 | -4.040512701 |
| **A_55_P1966878** | Rnaseh2a | 0.313430986 | -0.020287896 | 0.647149868 | 9.650237797 | 2.282150796 | 0.061362694 | 0.484675141 | -4.041002458 |
| **A_52_P419162** | Cyhr1 | -0.230994757 | -0.476942643 | 0.014953128 | 8.871497098 | -2.282139662 | 0.061363644 | 0.484675141 | -4.041007696 |
| **A_55_P2048607** | Hp | -0.786610566 | -1.624171285 | 0.050950153 | 15.84295732 | -2.282057923 | 0.061370625 | 0.484675141 | -4.041046155 |
| **A_66_P103271** | Irf2bp2 | -0.434165709 | -0.896613373 | 0.028281955 | 12.19587416 | -2.281267003 | 0.061438214 | 0.484675141 | -4.041418327 |

| | | | | | | | | |
|---|---|---|---|---|---|---|---|---|
| **A_55_P2018611** | Igfbp7 | 0.225437869 | -0.014690816 | 0.465566554 | 7.988407317 | 2.28121394 | 0.061442752 | 0.484675141 | -4.041443299 |
| **A_51_P423549** | Shbg | 0.709201214 | -0.046350229 | 1.464752657 | 9.881575086 | 2.280807434 | 0.061477523 | 0.484675141 | -4.041634612 |
| **A_55_P2071137** | Asap1 | 0.367453506 | -0.024030597 | 0.758937608 | 10.89539113 | 2.280717283 | 0.061485237 | 0.484675141 | -4.041677042 |
| **A_55_P2160682** | Set | 0.38961414 | -0.02548716 | 0.804715439 | 10.86727125 | 2.280677121 | 0.061488674 | 0.484675141 | -4.041695945 |
| **A_55_P2144126** | NA | -0.369679452 | -0.763583093 | 0.024224189 | 14.34093397 | -2.280439244 | 0.061509034 | 0.484675141 | -4.041807908 |
| **A_55_P1987449** | Ep400 | 0.306376751 | -0.020136827 | 0.632890328 | 10.47673292 | 2.280015239 | 0.061545343 | 0.484763625 | -4.042007492 |
| **A_52_P660385** | Trmt6 | 0.360334774 | -0.023767571 | 0.74443712 | 8.979688114 | 2.279514748 | 0.06158823 | 0.484903829 | -4.042243105 |
| **A_55_P2025611** | Psme2 | -0.182166222 | -0.376390722 | 0.012058278 | 12.04369445 | -2.279014184 | 0.061631154 | 0.484931391 | -4.04247878 |
| **A_55_P1993475** | Gm5168 | 0.16610531 | -0.011011724 | 0.343222344 | 7.937754516 | 2.278800853 | 0.061649457 | 0.484931391 | -4.042579228 |
| **A_55_P2017870** | Pclaf | 0.275695595 | -0.018303228 | 0.569694419 | 13.66569866 | 2.278596494 | 0.061666996 | 0.484931391 | -4.042675457 |
| **A_55_P2096887** | Steap3 | 0.300424675 | -0.020059831 | 0.620909181 | 9.66716913 | 2.277779873 | 0.061737131 | 0.485285484 | -4.043060033 |
| **A_55_P2086866** | NA | 0.41127816 | -0.027675502 | 0.850231822 | 13.99043956 | 2.276670398 | 0.06183255 | 0.485350587 | -4.04358264 |
| **A_55_P2060107** | Pkm | -0.411714503 | -0.851231936 | 0.02780293 | 11.9662578 | -2.276162422 | 0.061876289 | 0.485350587 | -4.043821962 |
| **A_52_P286928** | NA | 0.311776574 | -0.02112753 | 0.644680677 | 9.321097199 | 2.275660759 | 0.061919515 | 0.485350587 | -4.044058337 |
| **A_55_P2128944** | Bcl2l13 | 0.259740943 | -0.017647458 | 0.537129344 | 11.0605878 | 2.275282386 | 0.061952139 | 0.485350587 | -4.044236638 |
| **A_55_P2135039** | Hyi | -0.314834036 | -0.651108579 | 0.021440507 | 10.34930165 | -2.274944858 | 0.061981256 | 0.485350587 | -4.044395705 |
| **A_52_P469502** | Cda | -0.190363065 | -0.393697794 | 0.012971664 | 8.786392669 | -2.274858129 | 0.06198874 | 0.485350587 | -4.04443658 |
| **A_66_P140531** | Selenok | -0.409613975 | -0.84729241 | 0.02806446 | 12.71785957 | -2.274064643 | 0.062057254 | 0.485350587 | -4.044810583 |
| **A_55_P2045792** | Pes1 | 0.18572181 | -0.012736949 | 0.38418057 | 9.992206576 | 2.273923366 | 0.062069461 | 0.485350587 | -4.04487718 |
| **A_55_P1982449** | NA | 0.204523818 | -0.014031006 | 0.423078643 | 7.907594224 | 2.273875505 | 0.062073597 | 0.485350587 | -4.044899741 |
| **A_55_P2016426** | Nova2 | 0.254746856 | -0.017489382 | 0.526983093 | 8.987078959 | 2.273767668 | 0.062082917 | 0.485350587 | -4.044950577 |
| **A_66_P134878** | D11Wsu47e | 0.288039906 | -0.019783394 | 0.595863207 | 10.67951429 | 2.273706274 | 0.062088224 | 0.485350587 | -4.044979519 |
| **A_52_P354682** | Elovl7 | -0.200488728 | -0.414841871 | 0.013864415 | 8.961278466 | -2.272706189 | 0.062174734 | 0.485350587 | -4.045451036 |
| **A_55_P2004099** | NA | 0.205494112 | -0.014222082 | 0.425210307 | 8.402938117 | 2.272586928 | 0.062185058 | 0.485350587 | -4.045507273 |
| **A_55_P1973553** | Bbs12 | 0.182544681 | -0.012665272 | 0.377754635 | 8.925018894 | 2.272220189 | 0.062216819 | 0.485350587 | -4.045680213 |
| **A_55_P2121408** | Tpm2 | -0.582520996 | -1.205463747 | 0.040421755 | 10.04772503 | -2.272200421 | 0.062218531 | 0.485350587 | -4.045689536 |
| **A_55_P1970154** | Rnf10 | 0.293991734 | -0.020416023 | 0.608399492 | 12.2189359 | 2.272087522 | 0.062228312 | 0.485350587 | -4.045742778 |
| **A_55_P1959973** | Dcaf12l2 | 0.51038317 | -0.035495713 | 1.056262054 | 12.63926754 | 2.271868759 | 0.062247269 | 0.485350587 | -4.045845949 |
| **A_55_P2024853** | Cep57l1 | 0.307853298 | -0.021418384 | 0.637124981 | 8.968710822 | 2.271813195 | 0.062252085 | 0.485350587 | -4.045872154 |
| **A_52_P520408** | Serp1 | -0.213820922 | -0.442548996 | 0.014907152 | 11.76389348 | -2.271506132 | 0.062278706 | 0.485350587 | -4.046016979 |
| **A_51_P351194** | Cnfn | 0.336546485 | -0.023519483 | 0.696612452 | 8.511467931 | 2.271151851 | 0.062309435 | 0.485350587 | -4.046184087 |
| **A_51_P101196** | Psme1 | -0.228004227 | -0.471981523 | 0.015973068 | 12.93376016 | -2.270788442 | 0.062340972 | 0.485350587 | -4.046355514 |
| **A_55_P2247608** | 4931428A05Rik | 0.256606314 | -0.018019693 | 0.531232321 | 8.716426381 | 2.270433909 | 0.062371754 | 0.485350587 | -4.046522768 |
| **A_55_P2109337** | Gba2 | 0.200642954 | -0.014090775 | 0.415376683 | 11.27957842 | 2.27042331 | 0.062372675 | 0.485350587 | -4.046527768 |
| **A_55_P1979605** | Eif3m | 0.194255518 | -0.013679562 | 0.402190598 | 12.87113514 | 2.270015325 | 0.062408118 | 0.485350587 | -4.046720256 |
| **A_66_P109049** | Brd9 | 0.178803606 | -0.012597048 | 0.370204259 | 9.176546859 | 2.269948718 | 0.062413907 | 0.485350587 | -4.046751683 |
| **A_51_P181517** | Fcgr4 | -0.35845866 | -0.742184279 | 0.025266958 | 9.83945225 | -2.269872542 | 0.062420527 | 0.485350587 | -4.046787625 |
| **A_66_P121774** | Tmem59 | -0.346787503 | -0.718037762 | 0.024462755 | 14.56165534 | -2.269759611 | 0.062430344 | 0.485350587 | -4.046840911 |
| **A_55_P2062568** | Caprin2 | 0.257282825 | -0.018174333 | 0.532739983 | 9.334909447 | 2.269550877 | 0.062448492 | 0.485350587 | -4.046939405 |
| **A_55_P2010161** | Fastkd5 | 0.240338109 | -0.017086229 | 0.497762447 | 10.87054381 | 2.268591102 | 0.06253201 | 0.485433814 | -4.047392346 |
| **A_55_P1964752** | Slc23a3 | 0.582247126 | -0.041690589 | 1.206184841 | 9.971079985 | 2.267510489 | 0.062626182 | 0.485433814 | -4.047902434 |
| **A_51_P170696** | Haus1 | 0.278629176 | -0.019955991 | 0.577214343 | 12.0934059 | 2.267470003 | 0.062629713 | 0.485433814 | -4.047921547 |
| **A_55_P1982384** | NA | 0.33415384 | -0.023976603 | 0.692284283 | 12.18134373 | 2.267192546 | 0.062653917 | 0.485433814 | -4.048052539 |
| **A_55_P2080756** | Olfr704 | 0.269567975 | -0.019408698 | 0.558544647 | 9.111989635 | 2.2666721 | 0.062699345 | 0.485433814 | -4.04829827 |
| **A_52_P483301** | Mios | 0.197257898 | -0.014210941 | 0.408726738 | 8.885720679 | 2.266580816 | 0.062707316 | 0.485433814 | -4.048341374 |
| **A_55_P2074762** | Lect2 | 0.656085326 | -0.047355218 | 1.359525869 | 11.84171255 | 2.266293313 | 0.062732429 | 0.485433814 | -4.048477135 |
| **A_55_P2085225** | NA | 0.207859079 | -0.0150033 | 0.430721459 | 11.57946746 | 2.266289699 | 0.062732745 | 0.485433814 | -4.048478842 |
| **A_51_P267278** | Slc15a2 | -0.843313143 | -1.747521299 | 0.060895012 | 8.718955814 | -2.266228194 | 0.062738119 | 0.485433814 | -4.048507886 |
| **A_55_P1955975** | Afg3l2 | 0.374000427 | -0.027020335 | 0.775021189 | 11.43048987 | 2.266148833 | 0.062745053 | 0.485433814 | -4.048545364 |
| **A_55_P2174953** | Akr1b3 | -0.189503679 | -0.39270673 | 0.013699372 | 8.771129413 | -2.266055854 | 0.062753179 | 0.485433814 | -4.048589273 |

| | | | | | | | | | |
|---|---|---|---|---|---|---|---|---|---|
| **A_55_P2000039** | Prf1 | 0.222212476 | -0.016072813 | 0.460497764 | 8.442078261 | 2.26597126 | 0.062760572 | 0.485433814 | -4.048629222 |
| **A_55_P2091861** | Hipk2 | 0.355080289 | -0.025734976 | 0.735895555 | 9.53562422 | 2.265663483 | 0.06278748 | 0.485447681 | -4.048774578 |
| **A_55_P2090794** | NA | 0.251548384 | -0.01834915 | 0.521445919 | 9.246182182 | 2.264674571 | 0.062874018 | 0.485804283 | -4.049241687 |
| **A_55_P2021921** | Fkbp1a | -0.474050841 | -0.982800295 | 0.034698614 | 14.53235391 | -2.264144569 | 0.062920448 | 0.485804283 | -4.049492075 |
| **A_55_P1986185** | NA | 0.158161861 | -0.011611807 | 0.327935529 | 14.65380966 | 2.263677865 | 0.062961362 | 0.485804283 | -4.049712584 |
| **A_55_P2150294** | Synj2bp | -0.173106572 | -0.358960641 | 0.012747496 | 10.69265149 | -2.263209065 | 0.063002487 | 0.485804283 | -4.049934107 |
| **A_55_P1999309** | Zfp668 | 0.245280345 | -0.01808721 | 0.5086479 | 10.10257565 | 2.262995369 | 0.063021243 | 0.485804283 | -4.050035093 |
| **A_55_P2071389** | Erh | 0.219104526 | -0.016199808 | 0.45440886 | 14.06837038 | 2.26258348 | 0.06305741 | 0.485804283 | -4.050229753 |
| **A_55_P2045158** | NA | 0.261190399 | -0.019366486 | 0.541747283 | 10.0124512 | 2.262139938 | 0.063096381 | 0.485804283 | -4.050439392 |
| **A_51_P208801** | Ndufab1 | 0.370811833 | -0.027498174 | 0.76912184 | 10.71061343 | 2.262119562 | 0.063098172 | 0.485804283 | -4.050449023 |
| **A_55_P1968295** | Pacsin3 | 0.170388817 | -0.012637184 | 0.353414819 | 11.40859997 | 2.26209836 | 0.063100035 | 0.485804283 | -4.050459045 |
| **A_55_P2135750** | NA | 0.646607772 | -0.047978888 | 1.341194432 | 10.6983944 | 2.262026409 | 0.06310636 | 0.485804283 | -4.050493054 |
| **A_66_P121361** | C330021F23Rik | 0.348836511 | -0.025890982 | 0.723564004 | 9.106328583 | 2.261984193 | 0.063110071 | 0.485804283 | -4.050513009 |
| **A_55_P2120303** | NA | 0.228264038 | -0.016975735 | 0.473503811 | 8.440533093 | 2.261672818 | 0.06313745 | 0.485821562 | -4.050660198 |
| **A_51_P111952** | Nudt19 | 0.271484759 | -0.020258182 | 0.5632277 | 12.06086731 | 2.261144334 | 0.063183947 | 0.485985877 | -4.055684044 |
| **A_55_P2280089** | 4930570B17Rik | 0.313606171 | -0.023570996 | 0.650783338 | 10.76596894 | 2.260006204 | 0.063284202 | 0.486563383 | -4.051448193 |
| **A_55_P2171463** | Nptn | 0.182163915 | -0.013800833 | 0.378128663 | 12.79699682 | 2.258746988 | 0.063395315 | 0.486800754 | -4.052043763 |
| **A_55_P2145754** | Atp5pb | 0.192812332 | -0.014614146 | 0.40023881 | 13.06647349 | 2.258675298 | 0.063401647 | 0.486800754 | -4.052077675 |
| **A_55_P2101216** | Aldh3a1 | 0.394413749 | -0.029928979 | 0.818756477 | 9.436728764 | 2.258491556 | 0.063417879 | 0.486800754 | -4.052164595 |
| **A_55_P1985693** | Fhdc1 | 0.335311857 | -0.025536571 | 0.696160285 | 12.13194902 | 2.257913409 | 0.06346898 | 0.486800754 | -4.052438113 |
| **A_52_P238858** | Shisa9 | 0.253587519 | -0.019337168 | 0.526512207 | 8.489216317 | 2.257710445 | 0.06348693 | 0.486800754 | -4.052534142 |
| **A_55_P1958160** | Sgce | -0.179676841 | -0.373058359 | 0.013704677 | 10.60216218 | -2.2576693 | 0.06349057 | 0.486800754 | -4.05255361 |
| **A_55_P2064272** | NA | 0.33679437 | -0.025773886 | 0.699362627 | 8.261303745 | 2.257138633 | 0.063537528 | 0.486800754 | -4.052804711 |
| **A_55_P1957909** | Mrpl49 | 0.281423614 | -0.021541533 | 0.584388761 | 10.6394911 | 2.257101323 | 0.063540831 | 0.486800754 | -4.052822367 |
| **A_55_P2006277** | Ogdh | 0.249629201 | -0.019109056 | 0.518367459 | 8.873358663 | 2.257091052 | 0.063541741 | 0.486800754 | -4.052827227 |
| **A_51_P165914** | Zfp770 | 0.199760954 | -0.015352271 | 0.414874179 | 9.37449877 | 2.256455023 | 0.063598074 | 0.486951812 | -4.053128226 |
| **A_55_P1959833** | Tmem266 | 0.286092399 | -0.022066263 | 0.594251061 | 9.219609424 | 2.255875503 | 0.063649447 | 0.486951812 | -4.05340252 |
| **A_51_P345046** | Metap2 | 0.193600879 | -0.014943214 | 0.402144971 | 12.30490192 | 2.255758596 | 0.063659816 | 0.486951812 | -4.053457858 |
| **A_55_P1968553** | Btbd9 | -0.357045275 | -0.741653969 | 0.027563418 | 9.620038739 | -2.255731389 | 0.063662229 | 0.486951812 | -4.053470737 |
| **A_55_P1964183** | Dysf | -0.218306453 | -0.453633894 | 0.017020988 | 9.55764038 | -2.254120816 | 0.063805258 | 0.487264368 | -4.054233254 |
| **A_66_P109175** | Farsb | 0.242168643 | -0.01889587 | 0.503233156 | 11.55886188 | 2.253996593 | 0.063816303 | 0.487264368 | -4.054292078 |
| **A_55_P2092661** | Nup210 | 0.322899873 | -0.025215977 | 0.671015722 | 12.59850915 | 2.253861712 | 0.063828299 | 0.487264368 | -4.054355951 |
| **A_51_P455647** | Car2 | 0.384010934 | -0.030034608 | 0.798056476 | 15.35695693 | 2.253609507 | 0.063850734 | 0.487264368 | -4.054475388 |
| **A_51_P292116** | Ica1 | -0.295183285 | -0.613492696 | 0.023126126 | 9.197851741 | -2.253333488 | 0.063875298 | 0.487264368 | -4.05460611 |
| **A_51_P371152** | Sos2 | 0.279346692 | -0.021930017 | 0.580623401 | 10.49139412 | 2.252999855 | 0.063905001 | 0.487264368 | -4.05476413 |
| **A_66_P126415** | NA | 0.37811236 | -0.029700821 | 0.785925542 | 9.038149251 | 2.252904613 | 0.063913483 | 0.487264368 | -4.054809242 |
| **A_52_P607683** | Ppm1e | 0.2331655 | -0.018381478 | 0.484712478 | 8.374816411 | 2.252311097 | 0.063966367 | 0.487264368 | -4.055090385 |
| **A_52_P204331** | D630039A03Rik | -0.531109016 | -1.104303465 | 0.042085433 | 8.323394515 | -2.251463377 | 0.064041979 | 0.487264368 | -4.055492008 |
| **A_52_P155554** | Cdc42ep2 | -0.31946162 | -0.664244618 | 0.025321378 | 10.31153287 | -2.251417497 | 0.064046074 | 0.487264368 | -4.055513747 |
| **A_55_P2055107** | Agpat1 | 0.166916492 | -0.01323237 | 0.347065354 | 10.88046944 | 2.251390946 | 0.064048444 | 0.487264368 | -4.055526328 |
| **A_52_P222354** | Vps13c | 0.264589095 | -0.020986109 | 0.5501643 | 8.44280307 | 2.251306544 | 0.064055978 | 0.487264368 | -4.055566319 |
| **A_51_P230624** | Ccdc71l | 0.410596561 | -0.032740732 | 0.853933855 | 10.21778083 | 2.250423382 | 0.064134867 | 0.487264368 | -4.05598483 |
| **A_55_P2005828** | Hbb-bh1 | 0.877261037 | -0.069967655 | 1.824489728 | 11.24378786 | 2.25038687 | 0.06413813 | 0.487264368 | -4.056002133 |
| **A_55_P2183597** | Tbc1d2 | -0.410003296 | -0.852730875 | 0.032724283 | 9.206744001 | -2.25026653 | 0.064148888 | 0.487264368 | -4.056059167 |
| **A_55_P1976152** | Vrk2 | 0.31973262 | -0.025527135 | 0.664992374 | 10.18671205 | 2.250215838 | 0.06415342 | 0.487264368 | -4.056083192 |
| **A_55_P2014259** | Zfp82 | -0.187686726 | -0.390449805 | 0.015076353 | 8.796361436 | -2.249198945 | 0.064244406 | 0.487264368 | -4.0565652 |
| **A_55_P1972648** | Pheta2 | 0.453338797 | -0.036520786 | 0.94319838 | 10.44093826 | 2.248715268 | 0.06428773 | 0.487264368 | -4.056794501 |
| **A_52_P655803** | Chpt1 | -0.268988729 | -0.559657034 | 0.021679575 | 8.172818031 | -2.248638254 | 0.06429463 | 0.487264368 | -4.056831014 |
| **A_55_P2156016** | NA | 0.397965459 | -0.03207926 | 0.828010177 | 9.473145551 | 2.248614205 | 0.064296786 | 0.487264368 | -4.056842416 |
| **A_55_P2056521** | Rpl21 | 0.166360839 | -0.013454597 | 0.346176274 | 8.45183918 | 2.248057015 | 0.064346738 | 0.487264368 | -4.057106606 |

| | | | | | | | | | |
|---|---|---|---|---|---|---|---|---|---|
| **A_51_P190111** | Mcm5 | 0.372936294 | -0.030173821 | 0.776046409 | 12.6698183 | 2.247988804 | 0.064352856 | 0.487264368 | -4.05713895 |
| **A_55_P2163263** | NA | 0.197517289 | -0.016031711 | 0.411066289 | 9.366081702 | 2.247453745 | 0.064400867 | 0.487264368 | -4.057392681 |
| **A_55_P2068247** | NA | 0.490446757 | -0.039818305 | 1.020711819 | 10.95400203 | 2.247408621 | 0.064404918 | 0.487264368 | -4.05741408 |
| **A_55_P2155906** | Chmp3 | 0.191913248 | -0.015583847 | 0.399410342 | 11.2229815 | 2.247377992 | 0.064407667 | 0.487264368 | -4.057428606 |
| **A_51_P516133** | H1f2 | 0.289860683 | -0.023554014 | 0.60327538 | 12.73087842 | 2.247259073 | 0.064418344 | 0.487264368 | -4.057485004 |
| **A_51_P317695** | Trpc3 | -0.199670803 | -0.415578754 | 0.016237148 | 8.094787239 | -2.247134762 | 0.064429507 | 0.487264368 | -4.05754396 |
| **A_52_P527500** | Helb | 0.35578542 | -0.029044886 | 0.740615727 | 9.974347295 | 2.24647748 | 0.064488562 | 0.487264368 | -4.057855714 |
| **A_51_P290018** | NA | 0.258743667 | -0.021166849 | 0.538654184 | 11.98567631 | 2.246123869 | 0.064520357 | 0.487264368 | -4.058023454 |
| **A_52_P144285** | Fam72a | 0.191993918 | -0.015716553 | 0.39970439 | 8.760415178 | 2.246013019 | 0.064530327 | 0.487264368 | -4.05807604 |
| **A_55_P2088203** | Hcn2 | 0.44899653 | -0.036778416 | 0.934771476 | 10.36928458 | 2.24590334 | 0.064540193 | 0.487264368 | -4.058128071 |
| **A_52_P350950** | Dennd2c | 0.275689845 | -0.02260193 | 0.57398162 | 8.621586583 | 2.245756572 | 0.064553399 | 0.487264368 | -4.058197699 |
| **A_55_P2159890** | Zbtb7a | 0.274910457 | -0.022565895 | 0.572386809 | 10.219624 | 2.245546238 | 0.064572328 | 0.487264368 | -4.058297488 |
| **A_51_P234113** | Nod1 | -0.327467494 | -0.681864849 | 0.02692986 | 9.958601441 | -2.245230405 | 0.064600763 | 0.487264368 | -4.058447338 |
| **A_52_P235278** | Pms1 | 0.220258806 | -0.018154994 | 0.458672607 | 8.785426446 | 2.244838389 | 0.064636075 | 0.487264368 | -4.058633348 |
| **A_51_P435339** | Epor | 0.494179579 | -0.040804365 | 1.029163524 | 12.66180043 | 2.244539413 | 0.064663019 | 0.487264368 | -4.058775221 |
| **A_55_P2386256** | D130062J10Rik | 0.38153211 | -0.031508729 | 0.794572949 | 9.92252034 | 2.244508682 | 0.064665789 | 0.487264368 | -4.058789805 |
| **A_52_P425706** | Rnf168 | 0.205124317 | -0.016983703 | 0.427232338 | 10.70376509 | 2.24406843 | 0.064705488 | 0.487264368 | -4.058998737 |
| **A_51_P151820** | Urb1 | 0.280640257 | -0.023270753 | 0.584551267 | 8.845681389 | 2.243813333 | 0.064728503 | 0.487264368 | -4.059119809 |
| **A_55_P1986225** | NA | 0.192850989 | -0.016063689 | 0.401765667 | 11.93086776 | 2.243035291 | 0.064798749 | 0.487264368 | -4.059489119 |
| **A_55_P2158697** | H4c9 | 0.515647561 | -0.042998726 | 1.074293848 | 11.71379368 | 2.242844892 | 0.064815951 | 0.487264368 | -4.059579504 |
| **A_52_P113916** | NA | 0.34772592 | -0.029052339 | 0.724504179 | 9.369811862 | 2.242510158 | 0.064846205 | 0.487264368 | -4.059738417 |
| **A_55_P2025615** | Ammecr1l | -0.274725708 | -0.572573923 | 0.023122507 | 12.35134631 | -2.241235479 | 0.064961547 | 0.487264368 | -4.060343672 |
| **A_51_P320852** | Cd9 | -0.331368315 | -0.690629822 | 0.027893192 | 13.63897368 | -2.241214799 | 0.06496342 | 0.487264368 | -4.060353493 |
| **A_55_P2100209** | Kif2c | 0.237429187 | -0.020007382 | 0.494865755 | 8.585827072 | 2.241026831 | 0.064980447 | 0.487264368 | -4.06044276 |
| **A_51_P282092** | Urod | 0.306611299 | -0.025838138 | 0.639060735 | 15.83020088 | 2.241020052 | 0.064981062 | 0.487264368 | -4.06044598 |
| **A_52_P542172** | Rb1cc1 | 0.21732894 | -0.01834598 | 0.453003861 | 9.496271306 | 2.240718928 | 0.065008349 | 0.487264368 | -4.060588994 |
| **A_51_P117881** | Leap2 | 0.35624262 | -0.030148766 | 0.742634005 | 13.58705642 | 2.24027653 | 0.065048459 | 0.487264368 | -4.060799122 |
| **A_51_P302139** | Mastl | 0.192876696 | -0.016325623 | 0.402079016 | 8.366530641 | 2.240249828 | 0.065050881 | 0.487264368 | -4.060811805 |
| **A_55_P2139027** | Plec | -0.532006497 | -1.10910312 | 0.045090126 | 10.86805454 | -2.240018432 | 0.065071872 | 0.487264368 | -4.060921721 |
| **A_66_P110817** | Mterf3 | 0.162371043 | -0.013779215 | 0.338521301 | 9.067911535 | 2.239796108 | 0.065092046 | 0.487264368 | -4.061027333 |
| **A_55_P2419854** | NA | 0.233413888 | -0.019815472 | 0.486643249 | 8.348232223 | 2.23973081 | 0.065097973 | 0.487264368 | -4.061058353 |
| **A_55_P2102664** | NA | 0.505199139 | -0.042900858 | 1.053299137 | 10.00699272 | 2.239680099 | 0.065102576 | 0.487264368 | -4.061082444 |
| **A_66_P125209** | Kif2c | 0.238944647 | -0.02029952 | 0.498188814 | 8.672375353 | 2.239605378 | 0.065109359 | 0.487264368 | -4.061117941 |
| **A_51_P463428** | Pik3ip1 | -0.340878278 | -0.710731497 | 0.028974941 | 8.359181688 | -2.239510556 | 0.065117967 | 0.487264368 | -4.061162988 |
| **A_55_P2063970** | Ube2l3 | 0.229254436 | -0.019509093 | 0.478017965 | 13.09941943 | 2.23931004 | 0.065136176 | 0.487264368 | -4.061258251 |
| **A_55_P2180086** | Lrrc28 | 0.180126372 | -0.015351812 | 0.375604557 | 9.805476237 | 2.239041769 | 0.065160545 | 0.487264368 | -4.06138571 |
| **A_51_P468055** | Smyd1 | 0.424350348 | -0.036345625 | 0.885046321 | 9.9972404 | 2.238171368 | 0.065239675 | 0.487264368 | -4.061799301 |
| **A_55_P2120254** | Avpi1 | -0.35389611 | -0.738234754 | 0.030442533 | 13.63057231 | -2.237406655 | 0.065309279 | 0.487264368 | -4.062162738 |
| **A_51_P492087** | Pole3 | 0.267599994 | -0.023104856 | 0.558304844 | 10.69741595 | 2.236747757 | 0.065369313 | 0.487264368 | -4.062475934 |
| **A_52_P202029** | Trak2 | 0.47202751 | -0.041004391 | 0.985059412 | 11.21242456 | 2.235661913 | 0.065468371 | 0.487264368 | -4.062992171 |
| **A_66_P116098** | Ptpn2 | 0.188120518 | -0.016373357 | 0.392614394 | 8.698771697 | 2.235316633 | 0.065499902 | 0.487264368 | -4.063156352 |
| **A_51_P236829** | Zswim7 | 0.235730516 | -0.020529095 | 0.491990127 | 9.946062807 | 2.23521258 | 0.065509407 | 0.487264368 | -4.063205832 |
| **A_52_P472319** | Plxnd1 | -0.292545408 | -0.610568224 | 0.025477407 | 8.663531594 | -2.235209304 | 0.065509706 | 0.487264368 | -4.06320739 |
| **A_55_P1968381** | Cast | 0.2233288 | -0.019466437 | 0.466124037 | 9.638905325 | 2.235052657 | 0.065524019 | 0.487264368 | -4.063281881 |
| **A_55_P2024679** | NA | 0.378807452 | -0.033086497 | 0.7907014 | 9.919965654 | 2.234684885 | 0.065557634 | 0.487264368 | -4.063456781 |
| **A_52_P53470** | NA | 0.228400171 | -0.019989844 | 0.476790187 | 8.591425307 | 2.234320548 | 0.065590953 | 0.487264368 | -4.06363006 |
| **A_55_P2138222** | NA | 0.226622933 | -0.019839827 | 0.473085693 | 9.357195922 | 2.234270429 | 0.065595537 | 0.487264368 | -4.063653898 |
| **A_55_P2114118** | Cadm1 | -0.208454964 | -0.435159653 | 0.018249725 | 9.142173225 | -2.234266259 | 0.065595919 | 0.487264368 | -4.063655882 |
| **A_55_P2043122** | Arsg | 0.252289583 | -0.02210409 | 0.526683256 | 9.313785482 | 2.234129873 | 0.065608397 | 0.487264368 | -4.063720752 |
| **A_52_P533304** | Eif5 | 0.173021567 | -0.015160832 | 0.361203967 | 13.53852374 | 2.234109362 | 0.065610273 | 0.487264368 | -4.063730508 |

| | | | | | | | | | |
|---|---|---|---|---|---|---|---|---|---|
| **A_66_P117786** | Crxos | 0.302067343 | -0.026517684 | 0.630652369 | 8.542677137 | 2.233773812 | 0.065640984 | 0.487264368 | -4.063890116 |
| **A_55_P1984289** | Cdan1 | 0.323282963 | -0.028483529 | 0.675049455 | 8.743086143 | 2.233117317 | 0.065701112 | 0.487264368 | -4.06420242 |
| **A_55_P1996171** | Pcgf5 | 0.256607397 | -0.022620041 | 0.535834834 | 10.7639965 | 2.233028515 | 0.065709249 | 0.487264368 | -4.064244667 |
| **A_55_P2163774** | Crip1 | -0.571779346 | -1.193999299 | 0.050440606 | 13.55654173 | -2.232892008 | 0.065721761 | 0.487264368 | -4.064309613 |
| **A_51_P473953** | Arhgef26 | -0.561662448 | -1.172917078 | 0.049592181 | 8.167745772 | -2.232731074 | 0.065736514 | 0.487264368 | -4.064386182 |
| **A_55_P2012206** | Ndufb11 | -0.193285469 | -0.403656072 | 0.017085134 | 14.71824997 | -2.232530206 | 0.065754933 | 0.487264368 | -4.064481755 |
| **A_52_P430262** | Rbm15 | 0.336138015 | -0.029775502 | 0.702051531 | 9.811139528 | 2.232144817 | 0.065790286 | 0.487264368 | -4.064665134 |
| **A_51_P348665** | Ramp1 | -0.316935225 | -0.66196021 | 0.02808976 | 9.280047684 | -2.23204607 | 0.065799348 | 0.487264368 | -4.064712124 |
| **A_51_P342082** | Cnpy2 | -0.281165456 | -0.587265815 | 0.024934903 | 12.52949945 | -2.231933818 | 0.06580965 | 0.487264368 | -4.064765541 |
| **A_55_P2067682** | Atp5l | 0.326900278 | -0.029042155 | 0.68284271 | 13.58337989 | 2.231612135 | 0.065839184 | 0.487264368 | -4.064918626 |
| **A_55_P2094049** | Hectd4 | 0.435932326 | -0.03883239 | 0.910697042 | 12.68205565 | 2.23112462 | 0.065883968 | 0.487264368 | -4.065150649 |
| **A_55_P1987290** | Rnu1b6 | 0.84507536 | -0.075612878 | 1.765763597 | 16.37515121 | 2.230314117 | 0.065958492 | 0.487264368 | -4.065536447 |
| **A_66_P100268** | NA | 0.535986869 | -0.048010666 | 1.119984404 | 8.515541164 | 2.230110208 | 0.065977255 | 0.487264368 | -4.065633519 |
| **A_55_P2168321** | NA | 0.230637717 | -0.020754405 | 0.482029839 | 8.261516172 | 2.229265814 | 0.06605501 | 0.487264368 | -4.06603554 |
| **A_51_P427232** | Zbed5 | -0.218073903 | -0.455809314 | 0.019661508 | 8.119542871 | -2.228912446 | 0.066087578 | 0.487264368 | -4.066203802 |
| **A_66_P109577** | Plcb3 | -0.396410065 | -0.828568164 | 0.035748034 | 9.82888934 | -2.228872395 | 0.06609127 | 0.487264368 | -4.066222874 |
| **A_55_P2009217** | Rnu3b1 | 0.941115623 | -0.084914805 | 1.967146051 | 14.38493087 | 2.22877348 | 0.06610039 | 0.487264368 | -4.066269977 |
| **A_55_P2021345** | Gm14717 | 0.33066112 | -0.029839678 | 0.691161918 | 8.993401694 | 2.228743508 | 0.066103154 | 0.487264368 | -4.06628425 |
| **A_55_P2114863** | Mgll | 0.193363454 | -0.017467175 | 0.404194084 | 10.37297966 | 2.228557681 | 0.066120291 | 0.487264368 | -4.066372743 |
| **A_55_P2012819** | Sec61a2 | 0.214840535 | -0.019475599 | 0.449156669 | 9.471504825 | 2.22790784 | 0.066180256 | 0.487264368 | -4.066682235 |
| **A_55_P1986691** | Card6 | 0.221675153 | -0.020100999 | 0.463451305 | 8.553360625 | 2.227854091 | 0.066185218 | 0.487264368 | -4.066707835 |
| **A_51_P353173** | Fam98a | 0.236172149 | -0.021425161 | 0.493769458 | 10.96913356 | 2.227771001 | 0.06619289 | 0.487264368 | -4.066747411 |
| **A_55_P2093614** | NA | 0.377968643 | -0.034290804 | 0.79022809 | 9.212377851 | 2.227759705 | 0.066193933 | 0.487264368 | -4.066752791 |
| **A_55_P2029310** | Arf4 | -0.211536728 | -0.442335102 | 0.019261646 | 10.86291942 | -2.227082098 | 0.066256534 | 0.487264368 | -4.067075564 |
| **A_55_P2000219** | Ubap2 | 0.214232674 | -0.019530211 | 0.447995559 | 11.79229098 | 2.22686218 | 0.066276864 | 0.487264368 | -4.06718033 |
| **A_55_P2064568** | Anks1 | -0.396150619 | -0.828462514 | 0.036161276 | 10.26745265 | -2.226621214 | 0.066299147 | 0.487264368 | -4.067295129 |
| **A_66_P102260** | Irs2 | -0.448275216 | -0.937470191 | 0.040919759 | 11.86168895 | -2.226619102 | 0.066299342 | 0.487264368 | -4.067296135 |
| **A_51_P115085** | Nup153 | 0.304514853 | -0.027807863 | 0.636837569 | 10.77897752 | 2.226545838 | 0.066306119 | 0.487264368 | -4.071917635 |
| **A_55_P1979684** | Rhoc | -0.532715026 | -1.114100515 | 0.048670462 | 11.22006005 | -2.226455138 | 0.066314509 | 0.487264368 | -4.067374253 |
| **A_51_P170946** | Hnrnpm | 0.187985404 | -0.017185464 | 0.393156272 | 13.59511839 | 2.226340692 | 0.066325098 | 0.487264368 | -4.067428781 |
| **A_51_P116088** | Rexo2 | 0.275017716 | -0.025144386 | 0.575179819 | 10.73958219 | 2.226322122 | 0.066326816 | 0.487264368 | -4.067437629 |
| **A_55_P2117664** | Rnf165 | 0.212766463 | -0.019475156 | 0.445008081 | 8.501319606 | 2.226108425 | 0.066346593 | 0.487264368 | -4.067539448 |
| **A_55_P2048224** | NA | 0.152289412 | -0.013973415 | 0.31855224 | 7.971700084 | 2.225654453 | 0.066388625 | 0.487264368 | -4.067755767 |
| **A_55_P2036026** | Robo4 | -0.576680379 | -1.206274835 | 0.052914077 | 10.93444788 | -2.225653074 | 0.066388753 | 0.487264368 | -4.067756424 |
| **A_55_P2011525** | Rnf214 | 0.198831158 | -0.018264072 | 0.415926389 | 8.454408283 | 2.225447477 | 0.066407798 | 0.487264368 | -4.067854398 |
| **A_51_P342652** | Cd79b | -0.218441806 | -0.456963074 | 0.020079463 | 11.2922494 | -2.225316749 | 0.066419911 | 0.487264368 | -4.067916697 |
| **A_55_P2049976** | NA | 0.325507732 | -0.029931018 | 0.680946482 | 8.088841154 | 2.22525469 | 0.066425661 | 0.487264368 | -4.067946272 |
| **A_55_P2053374** | Sorbs1 | 0.386180644 | -0.035713547 | 0.808074835 | 11.07772417 | 2.224181099 | 0.06652523 | 0.487807834 | -4.068457969 |
| **A_55_P2049080** | Fam160a2 | -0.20764903 | -0.434620331 | 0.019322272 | 10.44912681 | -2.22301374 | 0.066633669 | 0.487807834 | -4.069014494 |
| **A_55_P2179020** | Mrpl15 | 0.218473867 | -0.020344283 | 0.457292017 | 10.30603153 | 2.222876603 | 0.06664642 | 0.487807834 | -4.069079882 |
| **A_51_P249313** | Herc3 | 0.245674546 | -0.02288075 | 0.514229842 | 9.215281397 | 2.222847301 | 0.066649145 | 0.487807834 | -4.069093854 |
| **A_55_P1990319** | Ergic3 | -0.206797984 | -0.432867024 | 0.019271056 | 12.07158555 | -2.222738637 | 0.066659251 | 0.487807834 | -4.069145667 |
| **A_51_P330213** | Asf1b | 0.328202289 | -0.030598891 | 0.687003468 | 12.03366347 | 2.222649245 | 0.066667565 | 0.487807834 | -4.069188292 |
| **A_52_P320971** | Mt3 | 0.30963963 | -0.028882739 | 0.648161999 | 9.859396838 | 2.222554185 | 0.066676408 | 0.487807834 | -4.06923362 |
| **A_55_P1969156** | Pam | -0.213104188 | -0.446232287 | 0.020023911 | 8.220990401 | -2.221163621 | 0.066805904 | 0.488164075 | -4.069896806 |
| **A_55_P2048848** | Smn1 | 0.194423142 | -0.018341503 | 0.407187786 | 10.76019914 | 2.220402375 | 0.066876905 | 0.488164075 | -4.070259942 |
| **A_55_P2016004** | Cd22 | 0.26869008 | -0.025415503 | 0.562795663 | 8.636629734 | 2.219890504 | 0.06692469 | 0.488164075 | -4.070504153 |
| **A_51_P140368** | Cmc2 | 0.238865829 | -0.022625397 | 0.500357056 | 11.85004142 | 2.219627481 | 0.066949258 | 0.488164075 | -4.07062965 |
| **A_52_P18816** | Gosr2 | 0.163806939 | -0.015525512 | 0.343139389 | 8.720549768 | 2.219507414 | 0.066960476 | 0.488164075 | -4.070686941 |
| **A_55_P2094611** | NA | 0.624770231 | -0.059344264 | 1.308884727 | 8.806402396 | 2.219089021 | 0.066999582 | 0.488164075 | -4.070886591 |

| | | | | | | | | | |
|---|---|---|---|---|---|---|---|---|---|
| **A_52_P684057** | Snupn | 0.150042787 | -0.014278637 | 0.314364211 | 10.07938777 | 2.218728298 | 0.067033317 | 0.488164075 | -4.071058736 |
| **A_55_P2051384** | Ccdc88c | 0.215044176 | -0.020529129 | 0.45061748 | 8.566772148 | 2.218118784 | 0.067090358 | 0.488164075 | -4.07134964 |
| **A_66_P119074** | Tbc1d22b | 0.262635682 | -0.025073495 | 0.55034486 | 8.687024239 | 2.21811063 | 0.067091121 | 0.488164075 | -4.071353532 |
| **A_55_P1984523** | NA | 0.297418069 | -0.028471175 | 0.623307313 | 11.65537034 | 2.217586211 | 0.06714024 | 0.488164075 | -4.071603853 |
| **A_55_P1968723** | Noct | 0.598089799 | -0.057287664 | 1.253467262 | 11.86243358 | 2.217471677 | 0.067150972 | 0.488164075 | -4.071658528 |
| **A_55_P2003927** | NA | 0.331189774 | -0.031731673 | 0.694111221 | 13.29336907 | 2.217417519 | 0.067156048 | 0.488164075 | -4.071684382 |
| **A_55_P2018249** | Lmo4 | -0.244344972 | -0.51211157 | 0.023421626 | 8.78342574 | -2.217329291 | 0.067164317 | 0.488164075 | -4.0717265 |
| **A_55_P2001920** | Flna | -0.455820475 | -0.95533576 | 0.043694811 | 13.47464822 | -2.217319291 | 0.067165254 | 0.488164075 | -4.071731274 |
| **A_52_P49406** | Pde12 | 0.260352759 | -0.024964383 | 0.5456699 | 8.931984203 | 2.217264527 | 0.067170388 | 0.488164075 | -4.071757418 |
| **A_55_P2050727** | Fcrl5 | 0.280277019 | -0.026898594 | 0.587452631 | 11.45727808 | 2.217093178 | 0.067186452 | 0.488164075 | -4.07183922 |
| **A_55_P2088725** | Usp14 | 0.269460373 | -0.025903827 | 0.564824573 | 12.51579857 | 2.216767987 | 0.06721695 | 0.488164075 | -4.071994474 |
| **A_55_P1980677** | Fam24b | 0.299067178 | -0.028751104 | 0.62688546 | 9.989355548 | 2.216760486 | 0.067217654 | 0.488164075 | -4.071998056 |
| **A_52_P73475** | Fam78a | 0.404165989 | -0.03897872 | 0.847310699 | 9.776278052 | 2.216140981 | 0.067275794 | 0.488164075 | -4.072293854 |
| **A_55_P2148082** | Larp7 | 0.307784045 | -0.029688184 | 0.645256275 | 10.23182148 | 2.216109689 | 0.067278733 | 0.488164075 | -4.072308796 |
| **A_55_P2066468** | Noa1 | 0.253160146 | -0.02442887 | 0.530749161 | 10.41854141 | 2.216033129 | 0.067285922 | 0.488164075 | -4.072345354 |
| **A_51_P396811** | Fam122a | 0.198465581 | -0.019152071 | 0.416083232 | 9.131486733 | 2.216023031 | 0.06728687 | 0.488164075 | -4.072350177 |
| **A_51_P153042** | Psg16 | -0.709171927 | -1.486850851 | 0.068506997 | 8.280330059 | -2.215819578 | 0.067305979 | 0.488164075 | -4.072447331 |
| **A_55_P2130975** | Ncoa1 | -0.28523072 | -0.5981409 | 0.02767946 | 9.884418487 | -2.214928924 | 0.067389696 | 0.488587933 | -4.072872695 |
| **A_66_P131979** | Cdc6 | 0.419010524 | -0.040767861 | 0.878788909 | 10.10099177 | 2.214417832 | 0.067437784 | 0.488753256 | -4.073116822 |
| **A_52_P576689** | Zfp944 | 0.172863385 | -0.016863759 | 0.362590529 | 8.999522176 | 2.213893538 | 0.067487152 | 0.488927719 | -4.073367283 |
| **A_55_P2022539** | NA | 0.265218591 | -0.025930928 | 0.556368109 | 8.827169802 | 2.213456929 | 0.067528291 | 0.488998858 | -4.073575877 |
| **A_55_P2044788** | NA | 0.21500229 | -0.021066182 | 0.451070763 | 9.244070115 | 2.213035016 | 0.06756807 | 0.488998858 | -4.073777469 |
| **A_51_P485161** | Cplx2 | -0.24207305 | -0.507912632 | 0.023766533 | 10.03973448 | -2.212636055 | 0.067605707 | 0.488998858 | -4.073968111 |
| **A_51_P106952** | Rbm26 | 0.222886297 | -0.021962848 | 0.467735441 | 10.12160371 | 2.21191262 | 0.06767401 | 0.488998858 | -4.074313843 |
| **A_55_P1985895** | Thumpd1 | 0.230705676 | -0.022738568 | 0.484149919 | 11.20659416 | 2.21186714 | 0.067678306 | 0.488998858 | -4.074335579 |
| **A_52_P607507** | Olfm4 | -0.285918931 | -0.600029652 | 0.028191789 | 8.850358479 | -2.21178721 | 0.067685857 | 0.488998858 | -4.074373782 |
| **A_55_P1967672** | Tas2r109 | 0.286780014 | -0.028326107 | 0.601886135 | 8.787899345 | 2.21144036 | 0.067718636 | 0.488998858 | -4.074539566 |
| **A_52_P315976** | Tpm2 | -0.496354869 | -1.041820053 | 0.049110316 | 9.887068363 | -2.211100277 | 0.067750791 | 0.488998858 | -4.074702128 |
| **A_51_P501844** | Cyp26b1 | -0.475422776 | -0.997972387 | 0.047126834 | 8.27269042 | -2.210729694 | 0.067785847 | 0.488998858 | -4.074879282 |
| **A_66_P131036** | NA | 0.28721372 | -0.028576798 | 0.603004239 | 10.23852852 | 2.209984793 | 0.06785637 | 0.488998858 | -4.075235419 |
| **A_55_P1976839** | Spata5 | 0.332057551 | -0.0330863 | 0.697201403 | 9.269038133 | 2.209696145 | 0.067883718 | 0.488998858 | -4.075373436 |
| **A_52_P672453** | Prpf3 | 0.233459364 | -0.023306606 | 0.490225334 | 10.69560893 | 2.209311853 | 0.067920145 | 0.488998858 | -4.075557199 |
| **A_55_P2083894** | NA | 0.306645585 | -0.030669609 | 0.643960779 | 8.32394348 | 2.208940418 | 0.067955372 | 0.488998858 | -4.075734829 |
| **A_55_P2078460** | NA | 0.388673266 | -0.038909157 | 0.816255689 | 10.82110008 | 2.208757384 | 0.067972738 | 0.488998858 | -4.075822366 |
| **A_55_P2058497** | Tmem50a | -0.275826147 | -0.579267875 | 0.027615581 | 11.35161611 | -2.2087335 | 0.067975005 | 0.488998858 | -4.075833788 |
| **A_55_P2034928** | BC147527 | 0.292830241 | -0.02937577 | 0.615036252 | 8.895562352 | 2.208337646 | 0.06801258 | 0.488998858 | -4.076023119 |
| **A_52_P426768** | Cited4 | 0.514164087 | -0.051755203 | 1.080083378 | 11.65877046 | 2.207651062 | 0.068077802 | 0.488998858 | -4.07635154 |
| **A_55_P2279575** | 5430400D12Rik | 0.19030764 | -0.019224241 | 0.39983952 | 8.467373586 | 2.206933777 | 0.06814601 | 0.488998858 | -4.076694697 |
| **A_55_P2040416** | Pradc1 | 0.203044104 | -0.020538159 | 0.426626367 | 9.900749469 | 2.206664078 | 0.068171674 | 0.488998858 | -4.076823738 |
| **A_55_P2069787** | NA | 0.17367842 | -0.017581294 | 0.364938133 | 13.3950889 | 2.206508203 | 0.068186511 | 0.488998858 | -4.076898322 |
| **A_66_P132855** | Baz1b | 0.240439707 | -0.024343258 | 0.505222672 | 8.873899749 | 2.206476657 | 0.068189514 | 0.488998858 | -4.076913416 |
| **A_51_P497402** | Tonsl | 0.371614343 | -0.037654381 | 0.780883066 | 9.798801431 | 2.206312873 | 0.068205109 | 0.488998858 | -4.076991787 |
| **A_51_P417854** | Rab20 | -0.167364842 | -0.351742067 | 0.017012383 | 8.065754831 | -2.20566802 | 0.068266543 | 0.488998858 | -4.077300377 |
| **A_55_P2133993** | NA | -0.318121118 | -0.668579063 | 0.032336827 | 9.615283469 | -2.205666155 | 0.068266721 | 0.488998858 | -4.07730127 |
| **A_55_P2202539** | BC028789 | 0.391641716 | -0.039836872 | 0.823120303 | 9.182786044 | 2.205529569 | 0.06827974 | 0.488998858 | -4.077366638 |
| **A_55_P2167530** | NA | 0.176642952 | -0.018065744 | 0.371351648 | 8.150656782 | 2.204419048 | 0.068385693 | 0.488998858 | -4.077898186 |
| **A_55_P2176385** | St6galnac4 | 0.315771364 | -0.0323583 | 0.663901028 | 9.251176836 | 2.204016804 | 0.068424111 | 0.488998858 | -4.078090749 |
| **A_51_P201982** | Angpt2 | -0.280714829 | -0.590198523 | 0.028768865 | 8.665064878 | -2.20399585 | 0.068426113 | 0.488998858 | -4.078100781 |
| **A_51_P122425** | Ctf1 | -0.174417999 | -0.366724834 | 0.017888836 | 8.784871194 | -2.203838514 | 0.068441147 | 0.488998858 | -4.078176106 |
| **A_55_P2065370** | NA | 0.216801566 | -0.022244307 | 0.455847439 | 13.49842343 | 2.203760289 | 0.068448623 | 0.488998858 | -4.078213557 |

| | | | | | | | | | |
|---|---|---|---|---|---|---|---|---|---|
| **A_55_P2032714** | Fhit | -0.143616658 | -0.301973151 | 0.014739834 | 8.386671223 | -2.203698275 | 0.06845455 | 0.488998858 | -4.078243248 |
| **A_55_P1971278** | Kdm2a | 0.176489161 | -0.018128515 | 0.371106836 | 9.281483656 | 2.203529888 | 0.068470647 | 0.488998858 | -4.078323869 |
| **A_55_P2319035** | AW011956 | 0.555781961 | -0.057099754 | 1.168663675 | 9.759163816 | 2.203489429 | 0.068474515 | 0.488998858 | -4.078343241 |
| **A_51_P364786** | U2af1 | 0.190371266 | -0.019565197 | 0.400307728 | 9.174322244 | 2.203417079 | 0.068481433 | 0.488998858 | -4.078377882 |
| **A_55_P1981709** | Rreb1 | 0.16981681 | -0.017453248 | 0.357086868 | 9.473897999 | 2.203411044 | 0.06848201 | 0.488998858 | -4.078380772 |
| **A_55_P2012939** | Arid4b | 0.228228028 | -0.023470866 | 0.479926922 | 9.316083643 | 2.203285925 | 0.068493975 | 0.488998858 | -4.07844068 |
| **A_55_P2339379** | Sass6 | 0.296249963 | -0.030466855 | 0.62296678 | 8.655231483 | 2.203281577 | 0.068494391 | 0.488998858 | -4.078442762 |
| **A_55_P1971124** | Mbd1 | 0.238817352 | -0.024562285 | 0.502196988 | 9.995082338 | 2.203265699 | 0.06849591 | 0.488998858 | -4.078450365 |
| **A_55_P2010008** | Zfp113 | -0.168021673 | -0.353365331 | 0.017321984 | 8.491138836 | -2.202778176 | 0.068542553 | 0.488998858 | -4.078683813 |
| **A_51_P483373** | Txndc5 | -0.224221008 | -0.471560787 | 0.023118772 | 10.1898911 | -2.202751573 | 0.068545099 | 0.488998858 | -4.078696552 |
| **A_55_P2129316** | NA | -0.242481678 | -0.51000882 | 0.025045465 | 8.5479497 | -2.202390199 | 0.068579696 | 0.488998858 | -4.078869611 |
| **A_55_P1993762** | NA | 0.14577529 | -0.015070682 | 0.306621262 | 8.221001154 | 2.202200793 | 0.068597837 | 0.488998858 | -4.078960321 |
| **A_52_P515497** | Atp1a1 | -0.226419568 | -0.476256722 | 0.023417586 | 13.05536827 | -2.202115644 | 0.068605993 | 0.488998858 | -4.079001101 |
| **A_51_P490817** | Me2 | 0.294584143 | -0.030508857 | 0.619677143 | 9.750688593 | 2.201835842 | 0.068632804 | 0.488998858 | -4.079135113 |
| **A_52_P201232** | Xpo5 | 0.290829055 | -0.030123927 | 0.611782037 | 9.429394691 | 2.201808614 | 0.068635413 | 0.488998858 | -4.079148154 |
| **A_55_P2126805** | Wdr81 | 0.336908066 | -0.034997867 | 0.708814 | 12.87809086 | 2.201210068 | 0.068692803 | 0.489084452 | -4.079434856 |
| **A_55_P1995902** | Zfp992 | 0.316459538 | -0.032882336 | 0.665801411 | 8.762073498 | 2.201155547 | 0.068698033 | 0.489084452 | -4.079460974 |
| **A_55_P2144476** | NA | 0.184548747 | -0.019235518 | 0.388333012 | 8.697115408 | 2.2005115 | 0.068759846 | 0.489098363 | -4.079769515 |
| **A_55_P2022845** | Csnk2a1 | 0.19106368 | -0.019921713 | 0.402049073 | 11.34026907 | 2.200436992 | 0.068767001 | 0.489098363 | -4.079805212 |
| **A_51_P386358** | Wwp2 | 0.167926953 | -0.017517115 | 0.353371021 | 12.23794325 | 2.200344339 | 0.068775899 | 0.489098363 | -4.079849603 |
| **A_51_P487547** | Ccdc91 | -0.216536513 | -0.455717558 | 0.022644532 | 9.021903473 | -2.199822138 | 0.068826071 | 0.489110454 | -4.080099811 |
| **A_65_P17827** | Cenpl | 0.315755127 | -0.03303404 | 0.664544294 | 10.15009636 | 2.19973625 | 0.068834327 | 0.489110454 | -4.080140966 |
| **A_52_P252121** | Nfs1 | 0.240648773 | -0.025200593 | 0.506498139 | 8.415613633 | 2.199536692 | 0.068853513 | 0.489110454 | -4.080236592 |
| **A_52_P195798** | Dmac2l | 0.258419386 | -0.027179594 | 0.544018366 | 8.786204255 | 2.198627354 | 0.068941006 | 0.48938034 | -4.080672386 |
| **A_51_P349341** | Npc1 | -0.145619952 | -0.306556535 | 0.015316631 | 8.411768275 | -2.198615557 | 0.068942142 | 0.48938034 | -4.08067804 |
| **A_51_P239673** | Hprt | 0.264442131 | -0.027901892 | 0.556786154 | 13.14359522 | 2.197959157 | 0.069005371 | 0.48951902 | -4.080992668 |
| **A_51_P129803** | Cacybp | 0.221533919 | -0.023421059 | 0.466488897 | 11.98450162 | 2.197541826 | 0.069045602 | 0.48951902 | -4.081192726 |
| **A_52_P675617** | Agbl3 | 0.298864937 | -0.031615872 | 0.629345746 | 8.938341517 | 2.19741412 | 0.069057917 | 0.48951902 | -4.081253949 |
| **A_51_P407866** | 1700020D05Rik | 0.238392705 | -0.025225025 | 0.502010436 | 8.515849236 | 2.197361618 | 0.069062981 | 0.48951902 | -4.081279119 |
| **A_55_P2114333** | NA | 0.350325489 | -0.037132375 | 0.737783354 | 15.45265359 | 2.197001967 | 0.06909768 | 0.489585433 | -4.081451549 |
| **A_55_P2004347** | NA | 0.217346347 | -0.023080793 | 0.457773487 | 12.46990609 | 2.196605381 | 0.069135963 | 0.489677184 | -4.081641701 |
| **A_66_P132493** | Siglece | 0.327746554 | -0.034888751 | 0.69038186 | 10.44323416 | 2.196095591 | 0.069185206 | 0.48978725 | -4.081886154 |
| **A_55_P2185548** | Cdk8 | 0.370938347 | -0.039519366 | 0.781396059 | 11.8900381 | 2.19591993 | 0.069202182 | 0.48978725 | -4.081970392 |
| **A_55_P2063785** | Ank1 | 0.475824982 | -0.05087346 | 1.002523425 | 10.96576391 | 2.195171195 | 0.069274589 | 0.490120254 | -4.082329485 |
| **A_55_P2102857** | Krtcap3 | -0.385199605 | -0.811682882 | 0.041283673 | 9.198979509 | -2.194658819 | 0.069324184 | 0.490291675 | -4.082575252 |
| **A_55_P2051622** | Gm8479 | 0.377529994 | -0.040610827 | 0.795670815 | 8.793160663 | 2.193876026 | 0.069400023 | 0.490648517 | -4.082950779 |
| **A_55_P2032187** | NA | 0.211610942 | -0.022818388 | 0.446040273 | 9.66407499 | 2.193357189 | 0.069450336 | 0.490824699 | -4.083199713 |
| **A_51_P457528** | Ccnb2 | 0.314682741 | -0.034041794 | 0.663407275 | 13.78081473 | 2.19267169 | 0.069516869 | 0.491115336 | -4.083528652 |
| **A_55_P2009449** | Pnma2 | 0.510951549 | -0.055545959 | 1.077449057 | 9.023089711 | 2.191618235 | 0.069619242 | 0.491658868 | -4.084034247 |
| **A_51_P120093** | Snx10 | -0.401778595 | -0.847316525 | 0.043759336 | 9.750606532 | -2.191216556 | 0.069658317 | 0.491735675 | -4.084227057 |
| **A_52_P596755** | Dnph1 | 0.365102531 | -0.039911678 | 0.77011674 | 9.559076258 | 2.190421901 | 0.069735687 | 0.491735675 | -4.084608549 |
| **A_51_P336385** | Dido1 | 0.299131736 | -0.03273277 | 0.630996243 | 10.61232648 | 2.190205546 | 0.069756767 | 0.491735675 | -4.084712426 |
| **A_55_P2073945** | Eif2b3 | 0.192556135 | -0.021082844 | 0.406195113 | 8.579921008 | 2.190080387 | 0.069768964 | 0.491735675 | -4.08477252 |
| **A_66_P100220** | NA | 0.386482847 | -0.042345724 | 0.815311419 | 8.8535741 | 2.18992731 | 0.069783886 | 0.491735675 | -4.08484602 |
| **A_52_P240453** | Polr3k | 0.228862644 | -0.025132443 | 0.482857731 | 10.65192273 | 2.189438665 | 0.069831539 | 0.491735675 | -4.08508066 |
| **A_55_P2032238** | Zbtb8os | 0.185354531 | -0.020373186 | 0.391082248 | 11.25557417 | 2.189241083 | 0.069850817 | 0.491735675 | -4.085175543 |
| **A_55_P1989981** | Fam129b | -0.825786253 | -1.742342561 | 0.090770055 | 12.08063214 | -2.189231494 | 0.069851753 | 0.491735675 | -4.085180148 |
| **A_55_P2117649** | Slfn9 | 0.228195824 | -0.02509187 | 0.481483518 | 8.34539656 | 2.18915641 | 0.06985908 | 0.491735675 | -4.085216206 |
| **A_52_P456134** | Dgat1 | -0.206744194 | -0.436277072 | 0.022788683 | 8.74409476 | -2.188626282 | 0.069910838 | 0.491920853 | -4.085470808 |
| **A_51_P363400** | Prg2 | 0.301557328 | -0.033412929 | 0.636527585 | 8.406401158 | 2.187493788 | 0.070021537 | 0.492181707 | -4.086014798 |

| | | | | | | | | | |
|---|---|---|---|---|---|---|---|---|---|
| **A_55_P2014987** | Castor2 | -0.445996157 | -0.94151701 | 0.049524697 | 9.124177595 | -2.187018059 | 0.070068093 | 0.492181707 | -4.086243351 |
| **A_55_P2048323** | Cdk10 | -0.156439024 | -0.330252437 | 0.01737439 | 10.63881236 | -2.186980938 | 0.070071727 | 0.492181707 | -4.086261186 |
| **A_55_P2120956** | NA | 0.239912164 | -0.026729469 | 0.506553798 | 8.629672296 | 2.186288629 | 0.070139537 | 0.492181707 | -4.086593833 |
| **A_55_P1972182** | Siah1b | 0.34945152 | -0.039066209 | 0.737969249 | 10.98426722 | 2.185542629 | 0.070212682 | 0.492181707 | -4.086952331 |
| **A_55_P2134004** | Gstm2 | -0.892588169 | -1.88505797 | 0.099881632 | 9.916161528 | -2.185329931 | 0.070233551 | 0.492181707 | -4.087054555 |
| **A_55_P2123331** | Cog2 | 0.399709771 | -0.04473642 | 0.844155963 | 9.338060734 | 2.185288424 | 0.070237624 | 0.492181707 | -4.087074504 |
| **A_55_P2068248** | NA | 0.173389371 | -0.019444125 | 0.366222867 | 11.48787955 | 2.184857838 | 0.070279894 | 0.492181707 | -4.087281463 |
| **A_51_P353008** | Minpp1 | 0.314225602 | -0.035257259 | 0.663708463 | 11.2774498 | 2.184735541 | 0.070291904 | 0.492181707 | -4.087340248 |
| **A_51_P519791** | Ska3 | 0.266948196 | -0.030014142 | 0.563910535 | 8.574701805 | 2.184282498 | 0.070336414 | 0.492181707 | -4.087558026 |
| **A_51_P248122** | Bbc3 | -0.32498237 | -0.686512145 | 0.036547405 | 10.37541172 | -2.184232775 | 0.070341301 | 0.492181707 | -4.087581929 |
| **A_55_P1983768** | Birc5 | 0.35315389 | -0.039734543 | 0.746042322 | 11.28696537 | 2.184127267 | 0.070351671 | 0.492181707 | -4.08763265 |
| **A_55_P2141287** | Erlin1 | 0.368904002 | -0.041525006 | 0.779333009 | 11.94007182 | 2.18402954 | 0.070361279 | 0.492181707 | -4.087679631 |
| **A_55_P2113698** | Nup205 | 0.295432854 | -0.033300784 | 0.624166492 | 11.09437612 | 2.183724434 | 0.070391282 | 0.492181707 | -4.087826315 |
| **A_55_P2045234** | NA | 0.234722515 | -0.026465031 | 0.495910061 | 9.053185652 | 2.183662289 | 0.070397394 | 0.492181707 | -4.087856193 |
| **A_55_P2083859** | NA | 0.357759153 | -0.040354005 | 0.755872311 | 13.85169671 | 2.183571466 | 0.070406329 | 0.492181707 | -4.087899859 |
| **A_51_P376238** | Serping1 | -0.479622152 | -1.013400127 | 0.054155822 | 12.37189297 | -2.18334202 | 0.070428905 | 0.492181707 | -4.088010178 |
| **A_55_P2178006** | NA | 0.39694005 | -0.044881758 | 0.838761859 | 8.825981907 | 2.183036316 | 0.070458996 | 0.492181707 | -4.08815717 |
| **A_55_P2145706** | Thoc7 | 0.205504493 | -0.023236618 | 0.434245603 | 10.43897822 | 2.183032915 | 0.070459331 | 0.492181707 | -4.088158805 |
| **A_51_P157976** | Mmab | 0.291950459 | -0.033024654 | 0.616925572 | 9.008932941 | 2.182942255 | 0.070468258 | 0.492181707 | -4.088202399 |
| **A_55_P2170034** | Rgl1 | -0.251006274 | -0.530437311 | 0.028424762 | 9.972010239 | -2.182695359 | 0.070492573 | 0.492181707 | -4.088321124 |
| **A_55_P2207335** | Nron | 0.418765527 | -0.047456037 | 0.884987091 | 9.864374924 | 2.182537703 | 0.070508104 | 0.492181707 | -4.088396938 |
| **A_55_P2027392** | Gpr146 | -0.347115319 | -0.733719759 | 0.039489121 | 11.4956554 | -2.181675388 | 0.070593116 | 0.492597234 | -4.088811659 |
| **A_55_P2089035** | Pole | 0.28454159 | -0.032409045 | 0.601492225 | 8.526080204 | 2.181410087 | 0.070619292 | 0.492602054 | -4.088939266 |
| **A_55_P2027323** | Golga7 | 0.169219111 | -0.019323624 | 0.357761845 | 12.64348567 | 2.180834939 | 0.070676073 | 0.492820279 | -4.089215933 |
| **A_55_P1957474** | Pus7 | 0.354764254 | -0.040619682 | 0.75014819 | 10.64574185 | 2.180238589 | 0.070734997 | 0.493019013 | -4.089502833 |
| **A_52_P470401** | Tmem186 | 0.178931236 | -0.020541921 | 0.378404394 | 8.521759439 | 2.1796406 | 0.070794133 | 0.493019013 | -4.089790556 |
| **A_52_P354271** | 2310057M21Rik | 0.232560329 | -0.026702512 | 0.49182317 | 8.905555143 | 2.179608773 | 0.070797282 | 0.493019013 | -4.089805871 |
| **A_51_P353386** | Micos13 | -0.288193604 | -0.609491401 | 0.033104192 | 12.38497104 | -2.179514591 | 0.0708066 | 0.493019013 | -4.08985119 |
| **A_55_P2044468** | Marf1 | 0.18591338 | -0.021455203 | 0.393281964 | 8.55594805 | 2.178466445 | 0.070910394 | 0.493199435 | -4.090355605 |
| **A_55_P2139894** | Cwc15 | 0.145326717 | -0.016773999 | 0.307427433 | 10.71191066 | 2.178430554 | 0.070913951 | 0.493199435 | -4.09037288 |
| **A_51_P141390** | Anapc1 | 0.168739126 | -0.019539286 | 0.357017538 | 12.21587679 | 2.177702034 | 0.070986189 | 0.493199435 | -4.090723543 |
| **A_55_P2002133** | Ago2 | 0.367838049 | -0.042623769 | 0.778299866 | 10.05532995 | 2.177544683 | 0.071001802 | 0.493199435 | -4.090799288 |
| **A_52_P46085** | Mvp | -0.213062843 | -0.45083665 | 0.024710965 | 8.267019545 | -2.177343221 | 0.071021796 | 0.493199435 | -4.090896272 |
| **A_51_P382369** | Zfp608 | -0.50989419 | -1.078926815 | 0.059138434 | 9.26549556 | -2.177339173 | 0.071022198 | 0.493199435 | -4.090898221 |
| **A_51_P407774** | Slc22a21 | 0.157417063 | -0.018258254 | 0.333092381 | 8.156374925 | 2.177329957 | 0.071023112 | 0.493199435 | -4.090902657 |
| **A_51_P444447** | Cebpd | -0.670971399 | -1.419959189 | 0.07801639 | 12.30762953 | -2.176769582 | 0.071078759 | 0.493199435 | -4.091172442 |
| **A_52_P232580** | Dyrk3 | 0.340969069 | -0.03966707 | 0.721605208 | 10.92287423 | 2.176647757 | 0.071090862 | 0.493199435 | -4.091231097 |
| **A_51_P209302** | Polr1e | 0.222462927 | -0.025896373 | 0.470822227 | 9.16590998 | 2.176508704 | 0.071104679 | 0.493199435 | -4.091298049 |
| **A_52_P278154** | Eif1ad | 0.25295731 | -0.029502548 | 0.535417167 | 9.949145146 | 2.176074136 | 0.071147879 | 0.493199435 | -4.091507298 |
| **A_51_P365859** | Sowaha | 0.500195132 | -0.058352635 | 1.058742899 | 10.6625226 | 2.176017229 | 0.071153539 | 0.493199435 | -4.091534701 |
| **A_55_P1964842** | Opa1 | 0.271339342 | -0.031719013 | 0.574397698 | 11.17801025 | 2.175553136 | 0.071199707 | 0.493199435 | -4.091758189 |
| **A_51_P460332** | Apoc4 | -0.278645849 | -0.589872661 | 0.032580963 | 13.77201664 | -2.17549837 | 0.071205157 | 0.493199435 | -4.091784564 |
| **A_55_P2146254** | Ifitm1 | -0.741612576 | -1.570069409 | 0.086844257 | 10.32812258 | -2.175155905 | 0.071239249 | 0.493199435 | -4.091949497 |
| **A_51_P161342** | Abcb6 | 0.413450331 | -0.048419069 | 0.875319732 | 11.16500869 | 2.175140639 | 0.071240769 | 0.493199435 | -4.09195685 |
| **A_55_P2027594** | Taf3 | 0.302724346 | -0.035532675 | 0.640981367 | 8.93830036 | 2.174621708 | 0.07129246 | 0.493333329 | -4.092206792 |
| **A_51_P474468** | Utp11 | 0.151972745 | -0.017852654 | 0.321798144 | 12.05856373 | 2.174434116 | 0.071311155 | 0.493333329 | -4.092297152 |
| **A_52_P4698** | Rfwd3 | 0.347959197 | -0.041105819 | 0.737024213 | 12.17806185 | 2.173148116 | 0.071439454 | 0.493349744 | -4.09291669 |
| **A_52_P525837** | Trmt6 | 0.293860185 | -0.034721772 | 0.622442141 | 9.06625764 | 2.173102565 | 0.071444003 | 0.493349744 | -4.092938637 |
| **A_55_P2037111** | Ran | 0.34893918 | -0.041241119 | 0.739119479 | 10.75627456 | 2.173039334 | 0.071450318 | 0.493349744 | -4.092969104 |
| **A_55_P2085181** | Chaf1b | 0.36180131 | -0.042775393 | 0.766378013 | 12.08547707 | 2.172963609 | 0.071457881 | 0.493349744 | -4.093005591 |

| | | | | | | | | |
|---|---|---|---|---|---|---|---|---|
| **A_51_P254045** | Traip | 0.447239026 | -0.052911993 | 0.947390046 | 10.70079411 | 2.172809863 | 0.07147324 | 0.493349744 | -4.093079673 |
| **A_55_P2064412** | NA | 0.335325564 | -0.039787227 | 0.710438355 | 8.444163413 | 2.17214083 | 0.071540112 | 0.493349744 | -4.093402069 |
| **A_51_P487869** | Cyp4f16 | -0.144118754 | -0.305340932 | 0.017103425 | 8.243418538 | -2.17209543 | 0.071544652 | 0.493349744 | -4.093423948 |
| **A_55_P1976934** | Unc13d | 0.330746737 | -0.039275281 | 0.700768754 | 9.54353006 | 2.171956838 | 0.071558514 | 0.493349744 | -4.09349074 |
| **A_55_P2117333** | NA | 0.285199054 | -0.033898681 | 0.604296789 | 9.79307546 | 2.171738589 | 0.071580348 | 0.493349744 | -4.093595924 |
| **A_51_P406557** | Myorg | -0.208772022 | -0.442385813 | 0.024841769 | 8.276770793 | -2.171485877 | 0.071605639 | 0.493349744 | -4.093717723 |
| **A_51_P122238** | Uchl5 | 0.279867934 | -0.033314927 | 0.593050795 | 11.89776782 | 2.171392549 | 0.071614981 | 0.493349744 | -4.098049415 |
| **A_51_P103706** | Cyp2c29 | -0.796073663 | -1.686930169 | 0.094782844 | 12.95790386 | -2.171344279 | 0.071619814 | 0.493349744 | -4.098072417 |
| **A_55_P1993813** | NA | -0.190183389 | -0.403099296 | 0.022732517 | 16.33363369 | -2.170439397 | 0.071710465 | 0.493798214 | -4.094222161 |
| **A_55_P2028710** | Rnf170 | 0.193891181 | -0.023203893 | 0.410986255 | 10.31654613 | 2.170157613 | 0.071738719 | 0.493816842 | -4.094358008 |
| **A_55_P2168736** | Relb | -0.207736284 | -0.440431629 | 0.024959062 | 8.341586377 | -2.169241225 | 0.07183068 | 0.494135151 | -4.094799849 |
| **A_51_P340947** | Selenoo | -0.273831105 | -0.580640488 | 0.032978277 | 10.55788256 | -2.168689276 | 0.071886127 | 0.494135151 | -4.095066014 |
| **A_51_P270666** | Uspl1 | 0.229426021 | -0.027673789 | 0.486525831 | 10.709856 | 2.168323644 | 0.071922881 | 0.494135151 | -4.095242348 |
| **A_55_P2036942** | Gm4535 | 0.189055452 | -0.022839855 | 0.40095076 | 9.751168294 | 2.167958957 | 0.07195956 | 0.494135151 | -4.095418239 |
| **A_52_P372032** | Ddx21 | 0.381150469 | -0.046074756 | 0.808375694 | 9.994357141 | 2.167817714 | 0.071973771 | 0.494135151 | -4.095486365 |
| **A_51_P326191** | Serpina3g | -0.214771848 | -0.455574504 | 0.026030809 | 9.314082573 | -2.167201383 | 0.072035815 | 0.494135151 | -4.095783663 |
| **A_55_P2065991** | S100a11 | -0.762611165 | -1.617733196 | 0.092510865 | 11.20252553 | -2.166996715 | 0.07205643 | 0.494135151 | -4.095882397 |
| **A_55_P2004137** | Gnai2 | -0.195000409 | -0.413656692 | 0.023655874 | 13.35104076 | -2.166989215 | 0.072057186 | 0.494135151 | -4.095886015 |
| **A_55_P2091736** | Rassf2 | -0.3827236 | -0.811879511 | 0.04643231 | 9.758050819 | -2.166972173 | 0.072058902 | 0.494135151 | -4.095894236 |
| **A_51_P214747** | Parp12 | -0.169245268 | -0.359026474 | 0.020535938 | 8.471997352 | -2.166938178 | 0.072062327 | 0.494135151 | -4.095910636 |
| **A_55_P2110032** | Akap7 | 0.219465314 | -0.026633886 | 0.465564514 | 9.785532885 | 2.166900042 | 0.07206617 | 0.494135151 | -4.095929034 |
| **A_51_P362066** | Chil1 | -1.013199874 | -2.150128812 | 0.123729065 | 10.34146367 | -2.165434217 | 0.072214013 | 0.494810862 | -4.096636293 |
| **A_52_P112958** | Ywhab | -0.181165029 | -0.384472277 | 0.022142219 | 10.89816178 | -2.165233292 | 0.072234302 | 0.494810862 | -4.096733255 |
| **A_55_P2074414** | Gtf2h1 | 0.227000163 | -0.027795487 | 0.481795812 | 10.73840101 | 2.164797858 | 0.072278293 | 0.494810862 | -4.0969434 |
| **A_66_P132846** | Exosc1 | 0.278561225 | -0.034158376 | 0.591280826 | 9.244013696 | 2.164455936 | 0.072312855 | 0.494810862 | -4.097108428 |
| **A_51_P278868** | H2-DMb1 | -0.231233027 | -0.490824766 | 0.028358712 | 8.994049284 | -2.164423231 | 0.072316162 | 0.494810862 | -4.097124214 |
| **A_66_P123014** | Twistnb | 0.223376109 | -0.027398154 | 0.474150371 | 10.27155046 | 2.164397124 | 0.072318801 | 0.494810862 | -4.097136815 |
| **A_55_P2112997** | Ddx18 | 0.244362141 | -0.030032762 | 0.518757044 | 11.04172124 | 2.163919351 | 0.072367128 | 0.494810862 | -4.097367432 |
| **A_52_P343856** | Creb3l2 | -0.222753353 | -0.472895152 | 0.027388447 | 8.31144904 | -2.16382019 | 0.072377163 | 0.494810862 | -4.0974153 |
| **A_55_P2139878** | NA | 0.281345457 | -0.034618484 | 0.597309398 | 9.051457282 | 2.163642847 | 0.072395112 | 0.494810862 | -4.097500909 |
| **A_51_P481676** | Zfp787 | -0.196841032 | -0.417961791 | 0.024279727 | 9.619343948 | -2.163063688 | 0.072453761 | 0.495036676 | -4.097780511 |
| **A_55_P1982211** | NA | 0.374936249 | -0.046413055 | 0.796285552 | 9.684635611 | 2.162212325 | 0.072540065 | 0.495189767 | -4.098191583 |
| **A_66_P134501** | Olfm1 | -0.168118348 | -0.357063429 | 0.020826733 | 8.748316221 | -2.162034968 | 0.072558057 | 0.495189767 | -4.098277227 |
| **A_51_P450518** | Hpf1 | 0.212484628 | -0.026326439 | 0.451295694 | 13.49066505 | 2.162002827 | 0.072561318 | 0.495189767 | -4.098292748 |
| **A_55_P2086475** | NA | 0.359963949 | -0.044633136 | 0.764561035 | 11.63363899 | 2.161819584 | 0.072579912 | 0.495189767 | -4.098381237 |
| **A_52_P48777** | Copg2 | -0.159318785 | -0.338411914 | 0.019774344 | 9.692281359 | -2.161579691 | 0.072604263 | 0.495189767 | -4.098497089 |
| **A_51_P219789** | H2-Q2 | -0.300014295 | -0.637387807 | 0.037359217 | 12.7852469 | -2.160797931 | 0.072683675 | 0.495224094 | -4.098874664 |
| **A_51_P488991** | Oaf | 0.293173898 | -0.036533739 | 0.622881535 | 10.72055942 | 2.160625432 | 0.072701209 | 0.495224094 | -4.098957985 |
| **A_55_P2124701** | NA | 0.287617256 | -0.035859576 | 0.611094088 | 13.03195954 | 2.160503363 | 0.07271362 | 0.495224094 | -4.09901695 |
| **A_55_P1997501** | Gm10778 | 0.237289187 | -0.029603922 | 0.504182296 | 8.71241283 | 2.160348306 | 0.072729388 | 0.495224094 | -4.09909185 |
| **A_55_P1984158** | Slc44a2 | -0.616920313 | -1.310886322 | 0.077045697 | 10.86182646 | -2.16010101 | 0.072754543 | 0.495224094 | -4.099211312 |
| **A_55_P1991802** | Kat14 | 0.197449296 | -0.024704368 | 0.419602961 | 10.48265565 | 2.159659568 | 0.072799469 | 0.495224094 | -4.099424575 |
| **A_51_P109888** | Coro1b | -0.234681941 | -0.498740062 | 0.029376181 | 9.377176009 | -2.159550333 | 0.072810591 | 0.495224094 | -4.099477351 |
| **A_55_P1984861** | NA | 0.218629441 | -0.027370926 | 0.464629807 | 15.15434793 | 2.159514274 | 0.072814262 | 0.495224094 | -4.099494772 |
| **A_52_P528963** | Eps15l1 | 0.224458358 | -0.028151751 | 0.477068467 | 9.638553435 | 2.15907756 | 0.072858744 | 0.495352327 | -4.099705775 |
| **A_51_P177171** | Tie1 | -0.49902681 | -1.060747453 | 0.062693834 | 9.039118824 | -2.15867212 | 0.072900066 | 0.495458991 | -4.099901685 |
| **A_55_P2005300** | Ranbp3 | 0.199168038 | -0.025062159 | 0.423398236 | 12.43136207 | 2.158284714 | 0.072939571 | 0.495542765 | -4.100088895 |
| **A_55_P2052097** | Ip6k2 | -0.254532482 | -0.541125227 | 0.032060264 | 8.756957115 | -2.158048536 | 0.072963667 | 0.495542765 | -4.100203034 |
| **A_55_P1973848** | Sema4b | 0.21980094 | -0.027762169 | 0.467364048 | 9.674530828 | 2.157380784 | 0.073031835 | 0.495557031 | -4.100525767 |
| **A_55_P2060444** | Apopt1 | 0.188861669 | -0.023871415 | 0.401594752 | 16.14069634 | 2.15720777 | 0.073049508 | 0.495557031 | -4.100609395 |

| | | | | | | | | | |
|---|---|---|---|---|---|---|---|---|---|
| **A_55_P2146783** | Ropn1l | -0.212550892 | -0.451969922 | 0.026868138 | 9.111422996 | -2.157185308 | 0.073051803 | 0.495557031 | -4.100620252 |
| **A_52_P194250** | Ets1 | -0.401039778 | -0.852808055 | 0.050728498 | 9.705219722 | -2.157023662 | 0.073068319 | 0.495557031 | -4.100698387 |
| **A_51_P354165** | Apcs | -0.752989678 | -1.601599158 | 0.095619802 | 11.22783181 | -2.156077328 | 0.073165088 | 0.495797863 | -4.101155869 |
| **A_55_P2113439** | Caln1 | 0.297616832 | -0.037816452 | 0.633050116 | 8.915862787 | 2.155929351 | 0.073180231 | 0.495797863 | -4.101227413 |
| **A_55_P2178044** | Inppl1 | -0.499354219 | -1.062239774 | 0.063531336 | 10.93760482 | -2.155618038 | 0.073212101 | 0.495797863 | -4.101377933 |
| **A_55_P1998384** | Wsb2 | 0.23603263 | -0.030049679 | 0.502114939 | 10.74653894 | 2.155456355 | 0.073228658 | 0.495797863 | -4.10145611 |
| **A_66_P111572** | Fam122b | 0.224391925 | -0.028571606 | 0.477355456 | 8.952435022 | 2.155422928 | 0.073232081 | 0.495797863 | -4.101472273 |
| **A_51_P279606** | Socs1 | -0.338357478 | -0.719864064 | 0.043149109 | 8.806240357 | -2.155047897 | 0.073270503 | 0.495884297 | -4.10165362 |
| **A_55_P1957080** | NA | 0.265994904 | -0.033984824 | 0.565974631 | 8.610100706 | 2.154589795 | 0.073317463 | 0.496028436 | -4.101875154 |
| **A_52_P24160** | Mrpl51 | 0.142533269 | -0.018288992 | 0.303355529 | 14.14064038 | 2.15354163 | 0.073425026 | 0.49643472 | -4.102382113 |
| **A_51_P430900** | Dusp1 | 0.164073049 | -0.021056072 | 0.349202169 | 9.611508548 | 2.153504083 | 0.073428882 | 0.49643472 | -4.102400275 |
| **A_55_P2104422** | Patj | -0.179928946 | -0.382979137 | 0.023121246 | 8.020493433 | -2.153182393 | 0.073461928 | 0.49648448 | -4.102555887 |
| **A_65_P15809** | Gm4759 | 0.237267745 | -0.03058888 | 0.50512437 | 9.157901167 | 2.152382735 | 0.07354414 | 0.496546591 | -4.10294275 |
| **A_51_P454691** | 4930430F08Rik | 0.291938847 | -0.037802163 | 0.621679856 | 9.39305515 | 2.151305632 | 0.073655025 | 0.496546591 | -4.103463934 |
| **A_52_P559545** | Cercam | 0.31347838 | -0.040600692 | 0.667557451 | 8.846213094 | 2.15124822 | 0.07366094 | 0.496546591 | -4.103491717 |
| **A_55_P2052834** | Lst1 | -0.294148693 | -0.626425098 | 0.038127712 | 12.37580523 | -2.151050515 | 0.073681313 | 0.496546591 | -4.103587395 |
| **A_55_P2075429** | Ttll4 | 0.252826085 | -0.032878671 | 0.53853084 | 8.814292151 | 2.150243281 | 0.073764558 | 0.496546591 | -4.103978087 |
| **A_51_P152612** | Nol7 | 0.222538679 | -0.028980791 | 0.47405815 | 8.619510133 | 2.149894191 | 0.073800587 | 0.496546591 | -4.104147062 |
| **A_52_P883557** | Slc30a10 | 0.432321375 | -0.056313252 | 0.920956003 | 9.22073526 | 2.149837617 | 0.073806427 | 0.496546591 | -4.104174447 |
| **A_51_P225832** | 2700097O09Rik | 0.391027175 | -0.051092955 | 0.833147305 | 11.18310562 | 2.149066431 | 0.07388609 | 0.496546591 | -4.104547781 |
| **A_51_P217382** | Kif1c | -0.194383633 | -0.414189485 | 0.025422219 | 10.44910734 | -2.148837793 | 0.073909725 | 0.496546591 | -4.104658475 |
| **A_55_P2143000** | NA | 0.194920268 | -0.025515206 | 0.415355743 | 8.646441281 | 2.148615492 | 0.073932713 | 0.496546591 | -4.104766108 |
| **A_51_P176517** | Ntmt1 | -0.184602313 | -0.393372439 | 0.024167813 | 10.88523298 | -2.148582197 | 0.073936156 | 0.496546591 | -4.104782228 |
| **A_55_P1952676** | Gtf2a2 | 0.178578117 | -0.023392944 | 0.380549179 | 12.22086704 | 2.148435307 | 0.07395135 | 0.496546591 | -4.104853351 |
| **A_55_P2040529** | Mtfr1 | 0.2607931 | -0.034191247 | 0.555777447 | 9.579440861 | 2.14822771 | 0.073972829 | 0.496546591 | -4.104953872 |
| **A_55_P2028193** | Mgat1 | -0.1769809 | -0.377202598 | 0.023240798 | 13.24743042 | -2.147822804 | 0.074014741 | 0.496546591 | -4.105149942 |
| **A_55_P2136154** | Heca | 0.219050722 | -0.028766492 | 0.466867935 | 9.752833116 | 2.147812712 | 0.074015786 | 0.496546591 | -4.105154829 |
| **A_51_P328669** | Ncbp1 | 0.259990278 | -0.034149174 | 0.55412973 | 9.147908721 | 2.147766271 | 0.074020595 | 0.496546591 | -4.105177319 |
| **A_51_P184284** | Dld | 0.469372453 | -0.061715453 | 1.000460359 | 12.03452747 | 2.147505945 | 0.074047556 | 0.496546591 | -4.105303388 |
| **A_52_P33382** | NA | 0.355082346 | -0.046731815 | 0.756896506 | 13.82875344 | 2.147271852 | 0.074071809 | 0.496546591 | -4.105416759 |
| **A_55_P2134496** | Amfr | 0.280386564 | -0.036936665 | 0.597709792 | 12.4242019 | 2.147032027 | 0.074096664 | 0.496546591 | -4.105532912 |
| **A_52_P226929** | Cflar | -0.148905701 | -0.31748069 | 0.019669287 | 8.379466154 | -2.146354118 | 0.074166969 | 0.496546591 | -4.105861266 |
| **A_51_P187665** | Rmi1 | 0.156901117 | -0.020787095 | 0.334589328 | 9.806092418 | 2.145609126 | 0.074244308 | 0.496546591 | -4.106222164 |
| **A_55_P2066827** | Hexim2 | 0.304611002 | -0.04036026 | 0.649582264 | 8.973911873 | 2.145585656 | 0.074246746 | 0.496546591 | -4.106233534 |
| **A_51_P257550** | Marcksl1 | -0.491119524 | -1.047352133 | 0.065113085 | 10.81123969 | -2.145427991 | 0.074263125 | 0.496546591 | -4.106309919 |
| **A_55_P1990874** | Banp | 0.353765426 | -0.046943442 | 0.754474295 | 10.5974753 | 2.145209054 | 0.074285875 | 0.496546591 | -4.106415993 |
| **A_51_P422124** | Fam126a | 0.387004333 | -0.051374409 | 0.825383074 | 11.04030604 | 2.145109819 | 0.074296189 | 0.496546591 | -4.106464073 |
| **A_51_P421876** | Irf7 | -0.194479308 | -0.414821273 | 0.025862656 | 7.994640965 | -2.144664555 | 0.074342485 | 0.496546591 | -4.106679819 |
| **A_55_P2147831** | Gimap4 | -0.439045929 | -0.936506399 | 0.05841454 | 9.845477045 | -2.14454206 | 0.074355227 | 0.496546591 | -4.106739175 |
| **A_55_P1969885** | NA | 0.403134642 | -0.053819435 | 0.86008872 | 9.264752672 | 2.143683928 | 0.07444455 | 0.496546591 | -4.107155033 |
| **A_55_P2056100** | Pus1 | 0.258234028 | -0.034497267 | 0.550965323 | 9.838555297 | 2.143519824 | 0.074461644 | 0.496546591 | -4.107234566 |
| **A_52_P249424** | Vegfa | -0.652569582 | -1.392407479 | 0.087268315 | 9.216363617 | -2.143253007 | 0.074489446 | 0.496546591 | -4.107363886 |
| **A_55_P2185526** | 1700016P03Rik | 0.262166109 | -0.035080686 | 0.559412903 | 9.553591667 | 2.143100596 | 0.074505332 | 0.496546591 | -4.107437759 |
| **A_55_P2131483** | Hmgb1-ps7 | 0.245883875 | -0.0329391 | 0.524706851 | 12.4186156 | 2.142815015 | 0.074535107 | 0.496546591 | -4.107576185 |
| **A_51_P281089** | S100a6 | -1.037783817 | -2.21471886 | 0.139151225 | 12.36718026 | -2.142582678 | 0.07455934 | 0.496546591 | -4.107688808 |
| **A_55_P2057821** | Golga1 | 0.15004186 | -0.020150638 | 0.320234358 | 9.552986803 | 2.142176331 | 0.074601741 | 0.496546591 | -4.107885793 |
| **A_55_P1991693** | Pprc1 | 0.2585358 | -0.034749471 | 0.551821071 | 8.582288617 | 2.14197119 | 0.074623157 | 0.496546591 | -4.107985246 |
| **A_51_P346964** | Mrap | 0.66958863 | -0.090061933 | 1.429239192 | 10.42139645 | 2.141792509 | 0.074641815 | 0.496546591 | -4.108071873 |
| **A_52_P477431** | Mnt | -0.224868226 | -0.479988129 | 0.030251676 | 11.03209439 | -2.141740964 | 0.074647199 | 0.496546591 | -4.108096864 |
| **A_55_P2129920** | Chd7 | 0.347131597 | -0.046714995 | 0.74097819 | 11.87729396 | 2.14165861 | 0.0746558 | 0.496546591 | -4.108136792 |

| | | | | | | | | | |
|---|---|---|---|---|---|---|---|---|---|
| **A_52_P32466** | Yif1a | -0.152505949 | -0.325539273 | 0.020527374 | 11.82249259 | -2.141609206 | 0.074660961 | 0.496546591 | -4.108160745 |
| **A_55_P2057441** | NA | 0.219557793 | -0.02960679 | 0.468722375 | 15.44678342 | 2.141143308 | 0.074709647 | 0.496546591 | -4.108386643 |
| **A_55_P2040743** | Zwilch | 0.329724173 | -0.044469336 | 0.703917682 | 9.501611222 | 2.141103825 | 0.074713774 | 0.496546591 | -4.108405787 |
| **A_52_P473966** | Kdelr3 | -0.34044385 | -0.726805375 | 0.045917674 | 8.470419898 | -2.141089453 | 0.074715276 | 0.496546591 | -4.108412756 |
| **A_55_P2077185** | NA | 0.213335642 | -0.028830358 | 0.455501642 | 10.07898778 | 2.140589741 | 0.074767535 | 0.496546591 | -4.108655074 |
| **A_51_P345747** | E130309D02Rik | 0.21207618 | -0.028693125 | 0.452845485 | 9.185907957 | 2.1402966 | 0.074798209 | 0.496546591 | -4.108797234 |
| **A_52_P213889** | Tmc7 | -0.257035958 | -0.548862694 | 0.034790778 | 8.340787891 | -2.140188346 | 0.074809539 | 0.496546591 | -4.108849734 |
| **A_55_P1967481** | Hbb-y | 0.887782253 | -0.120368521 | 1.895933027 | 10.76906172 | 2.139755544 | 0.074854857 | 0.496546591 | -4.109059642 |
| **A_55_P2054942** | Iffo2 | 0.196321403 | -0.026621314 | 0.41926412 | 9.103584532 | 2.139722971 | 0.074858269 | 0.496546591 | -4.10907544 |
| **A_51_P163261** | Gm6985 | 0.349984306 | -0.047518546 | 0.747487158 | 9.033878023 | 2.139397627 | 0.074892355 | 0.496546591 | -4.109233243 |
| **A_55_P2145441** | NA | 0.279971234 | -0.038060249 | 0.598002717 | 9.798814135 | 2.139077335 | 0.074925927 | 0.496546591 | -4.109388606 |
| **A_55_P2040996** | Klf7 | -0.224404549 | -0.479325206 | 0.030516108 | 10.08230615 | -2.138995233 | 0.074934535 | 0.496546591 | -4.109428433 |
| **A_51_P162098** | Ndufaf4 | 0.267830161 | -0.036464743 | 0.572125064 | 10.20685137 | 2.138690757 | 0.074966468 | 0.496546591 | -4.109576134 |
| **A_55_P2018747** | Cox8a | -0.172233872 | -0.367922683 | 0.023454939 | 15.26958924 | -2.138630515 | 0.074972787 | 0.496546591 | -4.109605359 |
| **A_55_P2156944** | Rbis | 0.309632372 | -0.042172419 | 0.661437163 | 12.38326589 | 2.138591319 | 0.074976899 | 0.496546591 | -4.109624374 |
| **A_52_P320193** | Clec2h | -0.571745408 | -1.221393318 | 0.077902502 | 8.08475628 | -2.138492979 | 0.074987217 | 0.496546591 | -4.109672082 |
| **A_55_P2008722** | Zfp984 | 0.317934971 | -0.04332114 | 0.679191083 | 9.487936653 | 2.138485385 | 0.074988014 | 0.496546591 | -4.109675766 |
| **A_51_P133582** | Cbx4 | -0.294961316 | -0.630130104 | 0.040207472 | 10.1152912 | -2.138379009 | 0.074999177 | 0.496546591 | -4.109727373 |
| **A_55_P2098057** | Sf3a2 | 0.258854235 | -0.035331894 | 0.553040363 | 12.56155453 | 2.138042194 | 0.075034533 | 0.496546591 | -4.109890783 |
| **A_66_P139493** | Tmem250-ps | 0.247882702 | -0.033845835 | 0.52961124 | 11.14945189 | 2.137955053 | 0.075043683 | 0.496546591 | -4.109933062 |
| **A_55_P1971872** | NA | 0.161910438 | -0.022126254 | 0.345947131 | 8.824498628 | 2.137733769 | 0.075066924 | 0.496546591 | -4.110040428 |
| **A_55_P2078633** | C4b | -0.577277778 | -1.23348308 | 0.078927524 | 13.45365107 | -2.13760912 | 0.075080018 | 0.496546591 | -4.11010091 |
| **A_52_P482203** | Tsen2 | 0.147215735 | -0.020128084 | 0.314559554 | 10.47275705 | 2.137606417 | 0.075080302 | 0.496546591 | -4.110102221 |
| **A_52_P670026** | Rsad2 | -0.648864569 | -1.38645283 | 0.088723693 | 11.09810267 | -2.137584308 | 0.075082625 | 0.496546591 | -4.110112949 |
| **A_55_P2102961** | NA | 0.237033011 | -0.032443191 | 0.506509214 | 9.07318444 | 2.137330111 | 0.075109337 | 0.496546591 | -4.110236293 |
| **A_55_P2125811** | Esam | -0.621669221 | -1.328446107 | 0.085107665 | 10.08195775 | -2.137274071 | 0.075115228 | 0.496546591 | -4.110263486 |
| **A_66_P114463** | NA | 0.416575003 | -0.057154889 | 0.890304894 | 12.83849433 | 2.136710116 | 0.07517453 | 0.496587847 | -4.11053716 |
| **A_55_P2154049** | Myo18a | 0.172236579 | -0.023650831 | 0.368123988 | 8.449148366 | 2.136495859 | 0.075197072 | 0.496587847 | -4.110641141 |
| **A_51_P506674** | Rpp38 | 0.234777489 | -0.032262518 | 0.501817495 | 10.13432553 | 2.136305273 | 0.07521713 | 0.496587847 | -4.110733639 |
| **A_55_P2380132** | NA | 0.196712327 | -0.027046444 | 0.420471098 | 8.31931075 | 2.13616452 | 0.075231947 | 0.496587847 | -4.110801953 |
| **A_52_P558699** | Ppp4r3a | 0.230630106 | -0.031741913 | 0.493002126 | 9.633700703 | 2.13590371 | 0.07525941 | 0.496587847 | -4.110928541 |
| **A_55_P2055134** | NA | 0.195819091 | -0.026966878 | 0.41860506 | 10.5806405 | 2.135749841 | 0.075275617 | 0.496587847 | -4.111003227 |
| **A_55_P2083967** | Trmt5 | 0.189220214 | -0.026110143 | 0.404550571 | 8.620783208 | 2.135233913 | 0.075329985 | 0.496776962 | -4.111253666 |
| **A_55_P2151091** | Phactr4 | 0.216782737 | -0.029947725 | 0.463513198 | 11.47938515 | 2.13493724 | 0.075361266 | 0.49681375 | -4.111397687 |
| **A_55_P2029592** | Dus2 | 0.292617639 | -0.04053081 | 0.625766087 | 10.14222217 | 2.134252975 | 0.075433466 | 0.497120172 | -4.111729896 |
| **A_55_P2057446** | Ccdc134 | 0.199900587 | -0.027738591 | 0.427539765 | 9.68179494 | 2.133782993 | 0.075483097 | 0.497218957 | -4.111958097 |
| **A_55_P2005873** | Zfp280d | 0.292687834 | -0.040653651 | 0.626029319 | 9.504030271 | 2.133528725 | 0.075509962 | 0.497218957 | -4.112081567 |
| **A_52_P681488** | Stx17 | 0.155446119 | -0.021603401 | 0.332495639 | 8.895411477 | 2.133380495 | 0.075525628 | 0.497218957 | -4.112153548 |
| **A_52_P232830** | Ets1 | -0.394764623 | -0.844503744 | 0.054974499 | 9.470398805 | -2.132852132 | 0.075581495 | 0.497271905 | -4.11241014 |
| **A_52_P89717** | NA | 0.238938197 | -0.033291338 | 0.511167731 | 8.563270372 | 2.132718465 | 0.075595636 | 0.497271905 | -4.112475058 |
| **A_55_P2159600** | Herc1 | 0.38660063 | -0.053987258 | 0.827188517 | 12.40946556 | 2.132127604 | 0.075658174 | 0.497271905 | -4.11276204 |
| **A_51_P505493** | Elovl5 | -0.393892425 | -0.842809282 | 0.055024433 | 11.16091767 | -2.132037814 | 0.075667682 | 0.497271905 | -4.112805653 |
| **A_55_P2005778** | Uap1 | 0.19765996 | -0.027620538 | 0.422940458 | 11.67339029 | 2.131956282 | 0.075676317 | 0.497271905 | -4.112845257 |
| **A_55_P2117574** | Ppih | 0.264908311 | -0.037064544 | 0.566881166 | 8.466528604 | 2.131625304 | 0.07571138 | 0.497271905 | -4.113006032 |
| **A_51_P348325** | Poc1b | -0.28100543 | -0.601431549 | 0.03942069 | 10.25356224 | -2.130933959 | 0.075784673 | 0.497271905 | -4.113341893 |
| **A_55_P2019293** | NA | 0.984867287 | -0.138194497 | 2.107929071 | 8.655945063 | 2.130871457 | 0.075791302 | 0.497271905 | -4.113372258 |
| **A_55_P2184128** | Ndufa3 | -0.161234205 | -0.345116347 | 0.022647938 | 14.96360591 | -2.13059456 | 0.075820681 | 0.497271905 | -4.113506791 |
| **A_55_P2082314** | NA | 0.201312711 | -0.028316169 | 0.430941591 | 9.017184444 | 2.130236781 | 0.075858657 | 0.497271905 | -4.113680631 |
| **A_55_P2150555** | Pcgf5 | 0.293583213 | -0.041356823 | 0.62852325 | 10.20202035 | 2.129841796 | 0.075900606 | 0.497271905 | -4.113872563 |
| **A_55_P1961650** | Gm2066 | 0.141075691 | -0.019886451 | 0.302037834 | 7.937556834 | 2.129666665 | 0.075919213 | 0.497271905 | -4.113957667 |

| | | | | | | | | | |
|---|---|---|---|---|---|---|---|---|---|
| **A_55_P2044202** | Vps29 | 0.180831931 | -0.025508425 | 0.387172286 | 11.10835208 | 2.129482784 | 0.075938754 | 0.497271905 | -4.114047027 |
| **A_51_P463497** | Rnh1 | -0.315136337 | -0.674739802 | 0.044467129 | 9.789314568 | -2.129402714 | 0.075947265 | 0.497271905 | -4.114085939 |
| **A_55_P2042526** | Apoo | 0.215506406 | -0.030413688 | 0.4614265 | 10.46525818 | 2.129361305 | 0.075951667 | 0.497271905 | -4.114106063 |
| **A_55_P2125963** | NA | 0.369044407 | -0.052120363 | 0.790209176 | 8.877215316 | 2.12916727 | 0.075972297 | 0.497271905 | -4.114200363 |
| **A_55_P2183672** | Pebp1 | 0.238775086 | -0.033733836 | 0.511284009 | 16.78614781 | 2.129077513 | 0.075981842 | 0.497271905 | -4.114243985 |
| **A_55_P2034953** | NA | 0.259943811 | -0.036838028 | 0.55672565 | 9.647046617 | 2.128263265 | 0.076068486 | 0.497271905 | -4.114639748 |
| **A_55_P2376423** | Igkv4-72 | 0.325322525 | -0.046196255 | 0.696841305 | 9.026148997 | 2.127730166 | 0.076125268 | 0.497271905 | -4.114898893 |
| **A_55_P2092776** | Apoo | 0.233884464 | -0.033245754 | 0.501014682 | 10.21325187 | 2.127460697 | 0.076153987 | 0.497271905 | -4.115029895 |
| **A_55_P1966438** | Gstm2 | -0.419055491 | -0.897707905 | 0.059596922 | 8.086992731 | -2.127328077 | 0.076168125 | 0.497271905 | -4.11509437 |
| **A_55_P1961320** | Tes | -0.681745294 | -1.460558143 | 0.097067554 | 9.35278889 | -2.127023217 | 0.076200634 | 0.497271905 | -4.115242589 |
| **A_55_P2207593** | 8430437O03Rik | 0.281945547 | -0.040262821 | 0.604153915 | 8.9938418 | 2.12623672 | 0.07628457 | 0.497271905 | -4.115625012 |
| **A_51_P492830** | Cenph | 0.362185446 | -0.051764536 | 0.776135427 | 10.63006068 | 2.12601496 | 0.076308253 | 0.497271905 | -4.11573285 |
| **A_52_P483129** | Utp23 | 0.198030574 | -0.028304237 | 0.424365386 | 9.369374188 | 2.126004017 | 0.076309422 | 0.497271905 | -4.115738171 |
| **A_55_P2145804** | Aen | 0.263217331 | -0.037625231 | 0.564059894 | 11.79677742 | 2.125976168 | 0.076312397 | 0.497271905 | -4.115751714 |
| **A_55_P1972842** | Klf3 | 0.27011658 | -0.038644905 | 0.578878064 | 12.48504522 | 2.125745708 | 0.076337019 | 0.497271905 | -4.115863788 |
| **A_55_P2124381** | NA | 0.207322802 | -0.029745533 | 0.444391138 | 12.68478886 | 2.124989112 | 0.076417909 | 0.497271905 | -4.11623176 |
| **A_55_P1973334** | Mapk1 | 0.199720649 | -0.02866941 | 0.428110707 | 12.6470429 | 2.124853348 | 0.076432433 | 0.497271905 | -4.116297795 |
| **A_52_P531662** | Ttc9c | 0.23637405 | -0.033935279 | 0.506683379 | 9.588174793 | 2.12481905 | 0.076436102 | 0.497271905 | -4.116314478 |
| **A_55_P1978151** | NA | 0.174293469 | -0.02503506 | 0.373621997 | 16.80788805 | 2.124686419 | 0.076450295 | 0.497271905 | -4.11637899 |
| **A_51_P416689** | Ext1 | -0.153804223 | -0.329717602 | 0.022109157 | 9.236146882 | -2.124479655 | 0.076472425 | 0.497271905 | -4.116479565 |
| **A_52_P59579** | Tmbim6 | -0.313303415 | -0.671665485 | 0.045058655 | 12.41526377 | -2.12435103 | 0.076486196 | 0.497271905 | -4.116542133 |
| **A_55_P1952738** | Ppp3ca | -0.413701357 | -0.88691302 | 0.059510305 | 11.58496706 | -2.124294358 | 0.076492264 | 0.497271905 | -4.116569702 |
| **A_55_P1987770** | Psenen | -0.207195631 | -0.444201799 | 0.029810537 | 12.94691845 | -2.124242697 | 0.076497796 | 0.497271905 | -4.116594832 |
| **A_55_P2178247** | Fam20b | 0.223363876 | -0.032156944 | 0.478884696 | 8.566544101 | 2.124074934 | 0.076515763 | 0.497271905 | -4.116676442 |
| **A_51_P221014** | Cd47 | 0.187981124 | -0.027089111 | 0.403051359 | 13.76863097 | 2.123817145 | 0.07654338 | 0.497271905 | -4.116801853 |
| **A_55_P2012037** | Lrrc8e | 0.222622109 | -0.032106014 | 0.477350232 | 8.889403672 | 2.123609145 | 0.07656567 | 0.497271905 | -4.116903045 |
| **A_55_P2153904** | Gm6132 | 0.211440728 | -0.03058179 | 0.453463246 | 13.4313984 | 2.122834122 | 0.076648784 | 0.497271905 | -4.117280132 |
| **A_51_P240723** | 1700022A21Rik | 0.198779148 | -0.028780356 | 0.426338652 | 8.276476294 | 2.122555401 | 0.076678697 | 0.497271905 | -4.117415757 |
| **A_55_P2047867** | NA | -0.305426192 | -0.655089593 | 0.04423721 | 9.818194594 | -2.122458896 | 0.076689057 | 0.497271905 | -4.117462717 |
| **A_51_P234728** | Lpar6 | -0.336800832 | -0.722422233 | 0.04882057 | 11.77550711 | -2.122243518 | 0.076712183 | 0.497271905 | -4.117567526 |
| **A_55_P1990170** | NA | 0.246163601 | -0.035699631 | 0.528026832 | 14.79749841 | 2.122113452 | 0.076726152 | 0.497271905 | -4.117630822 |
| **A_55_P2138718** | H2al1a | 0.341594006 | -0.0495499 | 0.732737912 | 8.585001188 | 2.122056105 | 0.076732312 | 0.497271905 | -4.11765873 |
| **A_52_P745** | Acox3 | -0.222181979 | -0.476598781 | 0.032234823 | 9.132439009 | -2.122004157 | 0.076737892 | 0.497271905 | -4.117684011 |
| **A_55_P2311208** | C130045F17Rik | 0.151078994 | -0.021944699 | 0.324102687 | 8.391261473 | 2.121688853 | 0.076771772 | 0.497271905 | -4.117837461 |
| **A_51_P459091** | Ybey | 0.155569644 | -0.022611955 | 0.333751243 | 8.379724022 | 2.121510547 | 0.076790938 | 0.497271905 | -4.117924243 |
| **A_51_P262670** | Rbm38 | 0.427738017 | -0.062172594 | 0.917648628 | 10.14405131 | 2.12150566 | 0.076791463 | 0.497271905 | -4.117926621 |
| **A_52_P1020860** | AW112010 | -0.559900905 | -1.201192468 | 0.081390659 | 9.607617105 | -2.12147946 | 0.07679428 | 0.497271905 | -4.117939373 |
| **A_52_P522977** | Ssbp2 | 0.429972148 | -0.062674859 | 0.922619155 | 8.881902301 | 2.120741159 | 0.076873695 | 0.497279747 | -4.118298735 |
| **A_55_P2074831** | Fam168a | -0.351789187 | -0.754894765 | 0.051316392 | 10.03863888 | -2.120541943 | 0.076895138 | 0.497279747 | -4.11839571 |
| **A_55_P2176350** | 0610010F05Rik | 0.205291329 | -0.030083107 | 0.440665765 | 9.601269722 | 2.119310078 | 0.077027866 | 0.497279747 | -4.118995443 |
| **A_55_P2102471** | Gm16378 | 0.313626124 | -0.04604598 | 0.673298228 | 13.38129914 | 2.118793657 | 0.077083578 | 0.497279747 | -4.119246904 |
| **A_51_P103222** | Slc39a4 | -0.384519886 | -0.825508352 | 0.05646858 | 13.5355358 | -2.118725841 | 0.077090897 | 0.497279747 | -4.119279927 |
| **A_55_P2050697** | Nfkbil1 | -0.191023588 | -0.410130947 | 0.02808377 | 10.68353456 | -2.118425647 | 0.077123304 | 0.497279747 | -4.119426114 |
| **A_55_P1955857** | Ap5z1 | 0.271209736 | -0.03987717 | 0.582296641 | 11.49928834 | 2.118393982 | 0.077126723 | 0.497279747 | -4.119441534 |
| **A_52_P658437** | Espl1 | 0.29767903 | -0.043894971 | 0.639253032 | 9.411947387 | 2.117613146 | 0.077211086 | 0.497279747 | -4.11982182 |
| **A_55_P2154782** | NA | 0.258305536 | -0.038111822 | 0.554722895 | 9.061030779 | 2.117450521 | 0.077228668 | 0.497279747 | -4.11990103 |
| **A_55_P2008347** | Prelid1 | -0.174092198 | -0.373893719 | 0.025709324 | 14.26119042 | -2.117208872 | 0.077254801 | 0.497279747 | -4.120018733 |
| **A_55_P2158873** | Ppid | 0.152842138 | -0.022580868 | 0.328265145 | 10.65657607 | 2.11709206 | 0.077267437 | 0.497279747 | -4.120075633 |
| **A_52_P317653** | Car1 | 0.287182183 | -0.042433468 | 0.616797835 | 9.731862636 | 2.11705849 | 0.077271069 | 0.497279747 | -4.120091985 |
| **A_55_P1969745** | Pitpnm1 | -0.202399627 | -0.43471948 | 0.029920226 | 10.61252325 | -2.116930366 | 0.077284932 | 0.497279747 | -4.120154396 |

| | | | | | | | | | |
|---|---|---|---|---|---|---|---|---|---|
| **A_55_P1956812** | Fam83g | 0.441728249 | -0.065300155 | 0.948756654 | 10.17849173 | 2.116927931 | 0.077285195 | 0.497279747 | -4.120155582 |
| **A_55_P2138422** | Obox1 | 0.324047751 | -0.047910619 | 0.696006121 | 8.952975555 | 2.116887921 | 0.077289525 | 0.497279747 | -4.120175073 |
| **A_52_P276792** | Foxp1 | -0.311202532 | -0.668422315 | 0.046017251 | 9.662876546 | -2.116853526 | 0.077293247 | 0.497279747 | -4.120191827 |
| **A_55_P2085979** | Igf1 | 0.27573548 | -0.040779951 | 0.592250912 | 9.636860501 | 2.116805489 | 0.077298445 | 0.497279747 | -4.120215227 |
| **A_52_P661071** | Snhg3 | 0.295782127 | -0.04391198 | 0.635476234 | 11.82214187 | 2.115763421 | 0.077411309 | 0.497279747 | -4.120722903 |
| **A_52_P655663** | H13 | -0.158738405 | -0.341073254 | 0.023596444 | 9.017344708 | -2.115414708 | 0.077449114 | 0.497279747 | -4.120892811 |
| **A_55_P2452914** | N4bp2 | 0.572852264 | -0.085230353 | 1.230934882 | 10.51559187 | 2.11517061 | 0.077475589 | 0.497279747 | -4.121011753 |
| **A_55_P2000658** | Gm15645 | 0.340033566 | -0.050597382 | 0.730664513 | 11.4520141 | 2.115136167 | 0.077479325 | 0.497279747 | -4.121028536 |
| **A_55_P1986306** | Ltv1 | 0.203147232 | -0.030266462 | 0.436560925 | 11.48489993 | 2.114792521 | 0.077516615 | 0.497279747 | -4.121195995 |
| **A_55_P2137840** | Etfbkmt | 0.243460804 | -0.03629975 | 0.523221358 | 9.466304564 | 2.114587989 | 0.077538817 | 0.497279747 | -4.121295669 |
| **A_52_P517001** | Rpf1 | 0.165775913 | -0.024761661 | 0.356313486 | 10.09213457 | 2.114092501 | 0.077592631 | 0.497279747 | -4.121537148 |
| **A_51_P136355** | Gng11 | -0.492456094 | -1.058545959 | 0.073633772 | 13.62359973 | -2.113806966 | 0.077623659 | 0.497279747 | -4.121676316 |
| **A_55_P2038767** | Fam49a | -0.205099694 | -0.440875602 | 0.030676214 | 8.341059705 | -2.113726419 | 0.077632414 | 0.497279747 | -4.121715575 |
| **A_55_P2079619** | Rnf43 | 0.188642701 | -0.028221512 | 0.405506914 | 10.3943811 | 2.113660853 | 0.077639542 | 0.497279747 | -4.121747533 |
| **A_55_P1982883** | Gys1 | -0.351785808 | -0.756266826 | 0.052695211 | 9.9135776 | -2.113310727 | 0.077677615 | 0.497279747 | -4.121918196 |
| **A_55_P2016316** | Xrn2 | 0.193095503 | -0.028932301 | 0.415123307 | 10.31419846 | 2.113235928 | 0.077685751 | 0.497279747 | -4.121954657 |
| **A_55_P2070352** | Csf3r | 0.156919371 | -0.023512025 | 0.337350767 | 14.37256916 | 2.113234244 | 0.077685934 | 0.497279747 | -4.121955478 |
| **A_51_P297069** | Tmod1 | 0.341235413 | -0.051130253 | 0.733601078 | 9.880057729 | 2.113227663 | 0.07768665 | 0.497279747 | -4.121958686 |
| **A_55_P2073975** | NA | 0.182124218 | -0.027304544 | 0.39155298 | 15.09945059 | 2.113073309 | 0.077703442 | 0.497279747 | -4.122033928 |
| **A_66_P125437** | Rpl30 | 0.188425022 | -0.028298179 | 0.405148223 | 10.41719087 | 2.112595532 | 0.077755444 | 0.497279747 | -4.122266841 |
| **A_55_P2037787** | NA | 0.240534437 | -0.036127878 | 0.517196752 | 8.167091312 | 2.112566772 | 0.077758575 | 0.497279747 | -4.122280861 |
| **A_55_P1967291** | Ncaph | 0.330333711 | -0.049666028 | 0.71033345 | 12.33615339 | 2.112286324 | 0.077789117 | 0.497279747 | -4.122417588 |
| **A_52_P350786** | Phlda2 | 0.359085318 | -0.054039273 | 0.772209909 | 9.218419962 | 2.112028578 | 0.077817197 | 0.497279747 | -4.122543253 |
| **A_55_P2005465** | Zfp422 | -0.280101064 | -0.602355091 | 0.042152963 | 9.579966337 | -2.112027623 | 0.077817302 | 0.497279747 | -4.122543719 |
| **A_51_P427663** | Cnn2 | -0.740177185 | -1.591784138 | 0.111429769 | 10.90220614 | -2.111930802 | 0.077827852 | 0.497279747 | -4.122590926 |
| **A_52_P486702** | G3bp1 | 0.263247678 | -0.039653546 | 0.566148901 | 9.620365474 | 2.11177046 | 0.077845329 | 0.497279747 | -4.122669106 |
| **A_55_P2101182** | Pcmt1 | 0.245567336 | -0.037002224 | 0.528136896 | 11.29834527 | 2.111681487 | 0.077855028 | 0.497279747 | -4.122712488 |
| **A_55_P2115166** | AI661453 | -0.239289814 | -0.514682651 | 0.036103024 | 10.37388042 | -2.111323394 | 0.077894077 | 0.497279747 | -4.122887099 |
| **A_55_P1964038** | Gm10032 | 0.374859543 | -0.056570197 | 0.806289283 | 8.936372846 | 2.111259815 | 0.077901012 | 0.497279747 | -4.122918102 |
| **A_55_P2025902** | NA | 0.240512723 | -0.036296763 | 0.517322208 | 9.532379307 | 2.111252981 | 0.077901758 | 0.497279747 | -4.122921434 |
| **A_55_P2081005** | Cox7c | 0.186527803 | -0.028223373 | 0.401278978 | 8.258394179 | 2.110528466 | 0.077980833 | 0.497620179 | -4.123274757 |
| **A_55_P2109932** | Tmem63b | -0.438091962 | -0.942906298 | 0.066722375 | 11.03419348 | -2.108709689 | 0.078179699 | 0.498060535 | -4.124161923 |
| **A_66_P112436** | Prkar2b | 0.309954886 | -0.047213308 | 0.667123081 | 9.518131012 | 2.108671356 | 0.078183896 | 0.498060535 | -4.124180625 |
| **A_55_P2054628** | Nrxn1 | -0.240870581 | -0.518439715 | 0.036698553 | 8.070777331 | -2.108607658 | 0.078190871 | 0.498060535 | -4.124211701 |
| **A_51_P172085** | Arhgdig | 0.454033934 | -0.069177652 | 0.97724552 | 9.983761257 | 2.108599744 | 0.078191737 | 0.498060535 | -4.124215562 |
| **A_51_P208377** | Trappc5 | -0.165478147 | -0.356210501 | 0.025254208 | 11.65349659 | -2.108140087 | 0.078242086 | 0.498060535 | -4.124439826 |
| **A_55_P2138961** | Tpk1 | 0.256615479 | -0.039233709 | 0.552464666 | 9.006022333 | 2.107636236 | 0.078297313 | 0.498060535 | -4.124685674 |
| **A_55_P2086885** | Slc45a4 | 0.292033143 | -0.044677158 | 0.628743444 | 9.391970977 | 2.107457999 | 0.078316859 | 0.498060535 | -4.124772647 |
| **A_55_P1958245** | Tcof1 | 0.282639551 | -0.043243948 | 0.60852305 | 10.62933345 | 2.107432888 | 0.078319614 | 0.498060535 | -4.124784901 |
| **A_51_P146407** | Fam104a | 0.263281637 | -0.040287257 | 0.566850531 | 9.657628122 | 2.107397647 | 0.078323479 | 0.498060535 | -4.124802098 |
| **A_55_P2002314** | Zmiz2 | 0.162515448 | -0.02488178 | 0.349912675 | 10.19893962 | 2.107243273 | 0.078340414 | 0.498060535 | -4.124877431 |
| **A_55_P2060567** | Ddx39 | 0.259423582 | -0.039718857 | 0.558566021 | 13.39152647 | 2.107242953 | 0.078340449 | 0.498060535 | -4.124877587 |
| **A_55_P2096652** | Kyat1 | -0.245139527 | -0.527833705 | 0.037554651 | 11.02638914 | -2.107073422 | 0.078359051 | 0.498060535 | -4.124960318 |
| **A_55_P2061456** | Nfyc | 0.325496668 | -0.049987657 | 0.700980992 | 9.669466821 | 2.106385833 | 0.078434542 | 0.498376483 | -4.12529589 |
| **A_51_P459350** | Dstn | -0.464953322 | -1.00153074 | 0.071624097 | 11.52480859 | -2.105523786 | 0.078529292 | 0.498486974 | -4.125716663 |
| **A_51_P403760** | Braf | 0.190798674 | -0.029412138 | 0.411009487 | 9.557887507 | 2.10532866 | 0.078550755 | 0.498486974 | -4.125811916 |
| **A_55_P2030721** | Ankle1 | 0.336127708 | -0.05185199 | 0.724107406 | 9.328728057 | 2.105127975 | 0.078572836 | 0.498486974 | -4.125909885 |
| **A_51_P466886** | Scoc | 0.217679732 | -0.033643184 | 0.469002647 | 9.169535786 | 2.104597711 | 0.078631209 | 0.498486974 | -4.126168763 |
| **A_51_P489522** | Ctla2b | -0.432186685 | -0.931260841 | 0.066887472 | 11.2448232 | -2.104211985 | 0.078673699 | 0.498486974 | -4.126357094 |
| **A_55_P2034420** | NA | 0.339261319 | -0.052515587 | 0.731038224 | 10.60161121 | 2.104159722 | 0.078679458 | 0.498486974 | -4.126382612 |

| | | | | | | | | | |
|---|---|---|---|---|---|---|---|---|---|
| **A_66_P125828** | NA | -0.543789043 | -1.171844207 | 0.084266121 | 9.553200322 | -2.103855225 | 0.078713019 | 0.498486974 | -4.126531292 |
| **A_66_P139785** | Plpp3 | -0.452866726 | -0.975915444 | 0.070181992 | 11.57156621 | -2.10383395 | 0.078715364 | 0.498486974 | -4.126541681 |
| **A_52_P376214** | Cstf3 | 0.258765848 | -0.040108394 | 0.557640089 | 9.827832974 | 2.103786476 | 0.078720598 | 0.498486974 | -4.126564863 |
| **A_55_P2035315** | Rasgef1b | -0.46361714 | -0.999106752 | 0.071872472 | 11.09779521 | -2.103737858 | 0.078725959 | 0.498486974 | -4.126588603 |
| **A_55_P2090406** | H13 | -0.185539043 | -0.399941659 | 0.028863574 | 9.914881051 | -2.102753766 | 0.078834541 | 0.498486974 | -4.127069185 |
| **A_51_P492125** | Ciapin1 | 0.218489362 | -0.034049635 | 0.471028358 | 10.94409861 | 2.10225326 | 0.078889824 | 0.498486974 | -4.127313641 |
| **A_51_P519364** | Fam83f | 0.499395343 | -0.077898532 | 1.076689219 | 11.15764666 | 2.101990397 | 0.078918875 | 0.498486974 | -4.127442037 |
| **A_55_P2017769** | B020031M17Rik | 0.230376847 | -0.035938191 | 0.496691885 | 8.451527411 | 2.101969105 | 0.078921228 | 0.498486974 | -4.127452437 |
| **A_55_P2054854** | Art4 | 0.351712151 | -0.054890316 | 0.758314618 | 9.158391977 | 2.101844355 | 0.078935019 | 0.498486974 | -4.127513374 |
| **A_55_P2087374** | NA | 0.216244381 | -0.033772025 | 0.466260788 | 15.50904426 | 2.101645741 | 0.078956981 | 0.498486974 | -4.127610393 |
| **A_55_P2016902** | Entr1 | 0.193365483 | -0.030207688 | 0.416938653 | 9.715602648 | 2.101563202 | 0.078966109 | 0.498486974 | -4.127650714 |
| **A_52_P265877** | Aldh9a1 | 0.289953918 | -0.045317839 | 0.625225675 | 11.7685361 | 2.101431315 | 0.078980697 | 0.498486974 | -4.127715141 |
| **A_52_P174661** | Gatc | 0.184551739 | -0.028857801 | 0.397961279 | 8.396359703 | 2.101297296 | 0.078995524 | 0.498486974 | -4.127780612 |
| **A_51_P466964** | Pgp | 0.308049669 | -0.048177172 | 0.664276509 | 12.88456653 | 2.101247919 | 0.079000988 | 0.498486974 | -4.127804734 |
| **A_51_P130727** | Fkbp11 | 1.040247902 | -0.162757219 | 2.243253023 | 10.37686583 | 2.101128248 | 0.079014231 | 0.498486974 | -4.127863198 |
| **A_55_P2078955** | Aqp8 | -0.621609927 | -1.340507767 | 0.097287913 | 10.67897259 | -2.101038214 | 0.079024196 | 0.498486974 | -4.127907183 |
| **A_55_P2167035** | Zfand1 | 0.182979436 | -0.02865705 | 0.394615923 | 8.845505182 | 2.100849452 | 0.079045092 | 0.498486974 | -4.127999404 |
| **A_55_P2037922** | Rpl7a | 0.251139591 | -0.039364408 | 0.54164359 | 14.61129938 | 2.100614008 | 0.079071163 | 0.498488752 | -4.128114436 |
| **A_55_P2021275** | Iqca | -0.265560244 | -0.572813825 | 0.041693336 | 8.098410897 | -2.100145075 | 0.079123116 | 0.498578611 | -4.128343559 |
| **A_55_P1953226** | H2al1e | 0.41484559 | -0.06515999 | 0.894851171 | 9.113565673 | 2.100019757 | 0.079137005 | 0.498578611 | -4.128404793 |
| **A_55_P1971991** | Kdf1 | -0.46985514 | -1.013687466 | 0.073977185 | 9.291419916 | -2.099336965 | 0.079212727 | 0.498601523 | -4.128738451 |
| **A_55_P1976007** | Psme4 | 0.197313662 | -0.031077732 | 0.425705056 | 9.560610605 | 2.099232833 | 0.079224281 | 0.498601523 | -4.128789341 |
| **A_55_P1969181** | Paox | -0.362773546 | -0.782852066 | 0.057304973 | 11.71222897 | -2.098400227 | 0.07931673 | 0.498601523 | -4.129196271 |
| **A_65_P09523** | Scrn3 | 0.277519187 | -0.043857269 | 0.598895644 | 11.1588253 | 2.098273743 | 0.079330784 | 0.498601523 | -4.129258095 |
| **A_55_P1967144** | NA | 0.167451784 | -0.026465152 | 0.36136872 | 16.63216127 | 2.098249978 | 0.079333425 | 0.498601523 | -4.129269711 |
| **A_55_P2065732** | Medag | 0.482711994 | -0.076305863 | 1.041729851 | 14.70345576 | 2.098193792 | 0.079339669 | 0.498601523 | -4.129297174 |
| **A_51_P389636** | Kcnn4 | 0.436911359 | -0.069068685 | 0.942891403 | 10.48803686 | 2.098181902 | 0.07934099 | 0.498601523 | -4.129302986 |
| **A_55_P2129309** | Foxo3 | 0.41896441 | -0.066302976 | 0.904231796 | 10.35601451 | 2.097873115 | 0.079375315 | 0.498601523 | -4.129453925 |
| **A_55_P2009604** | NA | 0.266569825 | -0.042217502 | 0.575357152 | 8.596175844 | 2.09765812 | 0.079399223 | 0.498601523 | -4.129559023 |
| **A_55_P1987984** | NA | 0.371421271 | -0.05888513 | 0.801727673 | 9.34229923 | 2.097356006 | 0.079432831 | 0.498601523 | -4.129706714 |
| **A_55_P2057528** | Arl4d | -0.35492034 | -0.766115691 | 0.056275011 | 8.471759273 | -2.097325715 | 0.079436202 | 0.498601523 | -4.129721522 |
| **A_55_P2212498** | NA | 0.288682824 | -0.045824041 | 0.623189689 | 8.832350163 | 2.097003228 | 0.079472094 | 0.498601523 | -4.129879182 |
| **A_55_P1999785** | NA | 0.201198707 | -0.031941143 | 0.434338557 | 11.17013733 | 2.096968283 | 0.079475984 | 0.498601523 | -4.129896267 |
| **A_65_P15503** | Map3k2 | 0.222244691 | -0.035367974 | 0.479857356 | 9.637720028 | 2.096270747 | 0.079553678 | 0.498704563 | -4.130237319 |
| **A_66_P110343** | Snx10 | -0.260469573 | -0.562391898 | 0.041452753 | 8.518477558 | -2.096259091 | 0.079554977 | 0.498704563 | -4.130243018 |
| **A_66_P111449** | 1700018A04Rik | 0.220030057 | -0.035054119 | 0.475114232 | 8.531344853 | 2.095953688 | 0.079589019 | 0.498704563 | -4.130392355 |
| **A_55_P2050592** | NA | 0.307293056 | -0.048966487 | 0.663552598 | 8.786951589 | 2.095894554 | 0.079595612 | 0.498704563 | -4.130421271 |
| **A_52_P475040** | Etaa1 | 0.250290145 | -0.039972592 | 0.540552883 | 8.925070416 | 2.095249049 | 0.079667618 | 0.498858976 | -4.130736942 |
| **A_51_P225506** | Pnrc2 | 0.251579948 | -0.040183892 | 0.543343788 | 12.5149964 | 2.095210908 | 0.079671874 | 0.498858976 | -4.130755595 |
| **A_55_P2000280** | Mthfd1l | 0.433164947 | -0.069324807 | 0.935654701 | 9.692380556 | 2.094639474 | 0.079735676 | 0.49909113 | -4.131035075 |
| **A_55_P1992676** | Ubtf | 0.366050528 | -0.058639082 | 0.790740138 | 9.961027671 | 2.094366054 | 0.079766223 | 0.49909113 | -4.131168811 |
| **A_55_P2119219** | NA | -0.196729857 | -0.42499434 | 0.031534626 | 13.31690608 | -2.094185382 | 0.079786414 | 0.49909113 | -4.131257186 |
| **A_51_P217498** | Slc2a4 | 0.311661975 | -0.0501609 | 0.67348485 | 10.98453936 | 2.093008476 | 0.079918067 | 0.499431186 | -4.131832931 |
| **A_55_P1967478** | Hbb-y | 0.937632061 | -0.150953618 | 2.026217741 | 11.18814538 | 2.092921896 | 0.079927761 | 0.499431186 | -4.131875291 |
| **A_55_P2026982** | Tdrp | -0.539607858 | -1.166153061 | 0.086937345 | 8.417391835 | -2.092709978 | 0.079951493 | 0.499431186 | -4.131978976 |
| **A_52_P206925** | Parp16 | 0.317354314 | -0.051169371 | 0.685878 | 9.406034751 | 2.092484213 | 0.079976784 | 0.499431186 | -4.13208944 |
| **A_55_P2168185** | NA | 0.164387381 | -0.026517606 | 0.355292367 | 16.6711876 | 2.092350286 | 0.079991791 | 0.499431186 | -4.132154972 |
| **A_55_P1990538** | Ndfip2 | 0.31743118 | -0.051211673 | 0.686074034 | 8.205381359 | 2.092314446 | 0.079995807 | 0.499431186 | -4.132172509 |
| **A_52_P219966** | Spg21 | -0.17711373 | -0.382845836 | 0.028618376 | 8.301673242 | -2.091863517 | 0.080046358 | 0.499438273 | -4.132393164 |
| **A_51_P466829** | Pheta1 | -0.485033825 | -1.048445779 | 0.078378129 | 10.63532135 | -2.091843348 | 0.08004862 | 0.499438273 | -4.132403033 |

| | | | | | | | | | |
|---|---|---|---|---|---|---|---|---|---|
| **A_55_P2121076** | Cul2 | 0.346160681 | -0.056000365 | 0.748321727 | 9.205282023 | 2.09151471 | 0.080085483 | 0.499507034 | -4.132563859 |
| **A_51_P319592** | Usp42 | 0.172204362 | -0.027962922 | 0.372371647 | 9.423975235 | 2.090423312 | 0.080208028 | 0.49973308 | -4.133098024 |
| **A_55_P1991585** | Blvrb | 0.396553896 | -0.064399784 | 0.857507575 | 14.6030317 | 2.090393872 | 0.080211336 | 0.49973308 | -4.133112434 |
| **A_55_P2017789** | Il4ra | -0.240334788 | -0.519728277 | 0.039058701 | 8.391903385 | -2.090179319 | 0.08023545 | 0.49973308 | -4.133217456 |
| **A_55_P2105271** | Pdlim7 | -0.265151791 | -0.573403334 | 0.043099753 | 11.18944316 | -2.090126124 | 0.08024143 | 0.49973308 | -4.133243495 |
| **A_51_P183971** | Barhl1 | 0.243941663 | -0.039663644 | 0.52754697 | 9.271548763 | 2.090041045 | 0.080250995 | 0.49973308 | -4.133285142 |
| **A_55_P2046101** | Xlr4b | 0.276693148 | -0.045037807 | 0.598424103 | 8.128766523 | 2.089723109 | 0.080286749 | 0.499794708 | -4.13344078 |
| **A_51_P418935** | Neurl2 | -0.174962582 | -0.378450058 | 0.028524894 | 9.002657343 | -2.089251315 | 0.080339835 | 0.499799423 | -4.133671752 |
| **A_55_P1987261** | Fam214b | 0.474531283 | -0.077505046 | 1.026567612 | 12.23647523 | 2.088720735 | 0.080399578 | 0.499799423 | -4.133931526 |
| **A_55_P2101666** | Vamp5 | 0.361595103 | -0.059140609 | 0.782330815 | 10.08419575 | 2.088316657 | 0.080445108 | 0.499799423 | -4.134129381 |
| **A_52_P214630** | Sox9 | -0.229362333 | -0.49624911 | 0.037524443 | 8.288167613 | -2.088229525 | 0.080454929 | 0.499799423 | -4.134172047 |
| **A_51_P492482** | Cstf2t | 0.193997238 | -0.031741641 | 0.419736117 | 10.77576008 | 2.088201345 | 0.080458105 | 0.499799423 | -4.134185846 |
| **A_55_P2113524** | Nsd2 | 0.189585185 | -0.031041109 | 0.410211478 | 8.298660974 | 2.087999146 | 0.080480901 | 0.499799423 | -4.134284859 |
| **A_66_P107943** | NA | -0.142565088 | -0.308484192 | 0.023354016 | 12.93592273 | -2.087853301 | 0.080497348 | 0.499799423 | -4.134356279 |
| **A_55_P2037883** | Ino80 | 0.18076434 | -0.029665635 | 0.391194315 | 8.840164751 | 2.087316693 | 0.080557891 | 0.499799423 | -4.13461907 |
| **A_55_P1965101** | Xlr3b | 0.354352725 | -0.058187499 | 0.766892948 | 8.994180079 | 2.08714521 | 0.080577248 | 0.499799423 | -4.134703054 |
| **A_55_P2123831** | Thap6 | 0.268380985 | -0.044095041 | 0.580857012 | 9.312223516 | 2.08697972 | 0.080595933 | 0.499799423 | -4.134784107 |
| **A_65_P05396** | Rad18 | 0.238243751 | -0.039200209 | 0.51568771 | 8.96149423 | 2.086553057 | 0.080644127 | 0.499799423 | -4.134993085 |
| **A_52_P77080** | Gypa | 0.291954885 | -0.048045019 | 0.631954789 | 11.84177974 | 2.086508419 | 0.080649171 | 0.499799423 | -4.13501495 |
| **A_55_P2157872** | Srsf5 | 0.175500178 | -0.028948549 | 0.379948904 | 12.14938557 | 2.08581766 | 0.080727264 | 0.499799423 | -4.135353318 |
| **A_51_P515965** | Nfe2 | 0.338324055 | -0.055806752 | 0.732454863 | 13.67115502 | 2.085814508 | 0.08072762 | 0.499799423 | -4.135354862 |
| **A_55_P2128501** | Krt8 | -0.27834979 | -0.60261709 | 0.04591751 | 10.9145207 | -2.085791686 | 0.080730202 | 0.499799423 | -4.135366042 |
| **A_55_P1982227** | NA | 0.144517015 | -0.023845637 | 0.312879667 | 12.21149513 | 2.085721961 | 0.080738089 | 0.499799423 | -4.135400199 |
| **A_52_P507382** | Unc93b1 | -0.253942942 | -0.549803876 | 0.041917992 | 10.37555414 | -2.085603334 | 0.080751509 | 0.499799423 | -4.135458313 |
| **A_51_P490509** | Bub1b | 0.28866408 | -0.0476514 | 0.62497956 | 10.18217595 | 2.085590681 | 0.080752941 | 0.499799423 | -4.135464512 |
| **A_51_P246727** | Mlxip | -0.161699279 | -0.350126621 | 0.026728064 | 8.23801848 | -2.085198233 | 0.080797357 | 0.499828444 | -4.135656778 |
| **A_55_P1980311** | Pcbp2 | 0.248558515 | -0.041186885 | 0.538303916 | 10.27258431 | 2.08446825 | 0.08088004 | 0.499828444 | -4.136014444 |
| **A_55_P2110713** | Anxa2 | -0.452685949 | -0.980434322 | 0.075062423 | 14.98413298 | -2.084266753 | 0.080902878 | 0.499828444 | -4.136113178 |
| **A_51_P369862** | Nceh1 | -0.375037954 | -0.812340096 | 0.062264188 | 9.758130664 | -2.083899664 | 0.080944501 | 0.499828444 | -4.136293062 |
| **A_51_P159814** | Smpd4 | 0.276866915 | -0.046063795 | 0.599797625 | 9.751330345 | 2.083266852 | 0.081016305 | 0.499828444 | -4.136603186 |
| **A_55_P2068152** | Anapc15 | 0.248341671 | -0.04141356 | 0.538096901 | 11.47269569 | 2.082579089 | 0.081094418 | 0.499828444 | -4.136940278 |
| **A_55_P2004777** | Micall2 | 0.375263037 | -0.062591345 | 0.813117419 | 10.53016714 | 2.082520465 | 0.08110108 | 0.499828444 | -4.136969013 |
| **A_55_P2041121** | Rn4.5s | 0.205478661 | -0.034297823 | 0.445255146 | 16.85425146 | 2.082300137 | 0.081126122 | 0.499828444 | -4.137077012 |
| **A_52_P113700** | Sfxn1 | 0.398338544 | -0.066528607 | 0.863205695 | 12.21498211 | 2.082124345 | 0.081146108 | 0.499828444 | -4.137163184 |
| **A_55_P2102002** | Noc3l | 0.181304278 | -0.030293806 | 0.392902362 | 9.081236048 | 2.08199417 | 0.08116091 | 0.499828444 | -4.137226996 |
| **A_55_P2170355** | NA | 0.189354974 | -0.031646299 | 0.410356246 | 8.458186068 | 2.081925245 | 0.081168749 | 0.499828444 | -4.137260784 |
| **A_55_P2035872** | Mical3 | 0.149361678 | -0.02498816 | 0.323711517 | 9.357393672 | 2.081616996 | 0.081203816 | 0.499828444 | -4.137411896 |
| **A_55_P1989916** | Lin54 | 0.296200693 | -0.049570524 | 0.641971909 | 10.12234464 | 2.0815192 | 0.081214945 | 0.499828444 | -4.13745984 |
| **A_55_P2078088** | Pqlc1 | 0.183826695 | -0.030807586 | 0.398460976 | 11.91958757 | 2.081098707 | 0.081262812 | 0.499828444 | -4.137665994 |
| **A_55_P2043008** | NA | 0.235722579 | -0.039576372 | 0.511021529 | 8.779123621 | 2.080557945 | 0.081324412 | 0.499828444 | -4.137931134 |
| **A_55_P1962906** | Als2cl | -0.347510951 | -0.753408777 | 0.058386875 | 9.164500701 | -2.080343063 | 0.081348903 | 0.499828444 | -4.1380365 |
| **A_52_P367760** | Calml4 | -0.298714302 | -0.64768795 | 0.050259347 | 8.335713904 | -2.079919713 | 0.081397176 | 0.499828444 | -4.138244097 |
| **A_51_P282508** | Rhoc | -0.496071588 | -1.075737909 | 0.083594733 | 12.96937689 | -2.079454753 | 0.081450228 | 0.499828444 | -4.138472117 |
| **A_51_P160344** | Cenpv | 0.220760281 | -0.037235063 | 0.478755624 | 11.41367382 | 2.079180812 | 0.0814815 | 0.499828444 | -4.138606468 |
| **A_55_P2184766** | Tmc6 | 0.332323718 | -0.056076236 | 0.720723673 | 9.994590581 | 2.079052025 | 0.081496207 | 0.499828444 | -4.138669632 |
| **A_66_P126603** | NA | 0.334144341 | -0.056430236 | 0.724718918 | 13.98355691 | 2.078802969 | 0.081524654 | 0.499828444 | -4.138791787 |
| **A_55_P2073219** | Hps4 | -0.225575837 | -0.489261476 | 0.038109802 | 11.03162477 | -2.078687904 | 0.081537801 | 0.499828444 | -4.138848225 |
| **A_51_P175555** | Pelo | 0.153935883 | -0.02602155 | 0.333893317 | 12.46982839 | 2.078515492 | 0.081557503 | 0.499828444 | -4.138932793 |
| **A_55_P1963576** | Gm11595 | 0.302488202 | -0.051140593 | 0.656116997 | 8.65573017 | 2.078471185 | 0.081562567 | 0.499828444 | -4.138954525 |
| **A_51_P223443** | Cd81 | -0.376475253 | -0.816601696 | 0.06365119 | 13.59529223 | -2.078462346 | 0.081563577 | 0.499828444 | -4.138958861 |

| | | | | | | | | | |
|---|---|---|---|---|---|---|---|---|---|
| **A_52_P238242** | Sertad2 | 0.237282118 | -0.040130674 | 0.514694911 | 10.31505214 | 2.078364499 | 0.081574762 | 0.499828444 | -4.139006857 |
| **A_52_P600625** | Rad23b | 0.272490043 | -0.046106351 | 0.591086436 | 8.815752938 | 2.078226942 | 0.081590488 | 0.499828444 | -4.139074332 |
| **A_51_P455338** | Rnase2a | -0.287812546 | -0.624351299 | 0.048726207 | 10.16399781 | -2.078058787 | 0.081609716 | 0.499828444 | -4.139156818 |
| **A_55_P1963344** | Upf3b | 0.189355503 | -0.03206909 | 0.410780097 | 9.355081531 | 2.07795082 | 0.081622064 | 0.499828444 | -4.139209781 |
| **A_55_P2186210** | NA | 0.210030458 | -0.035622819 | 0.455683735 | 13.7058759 | 2.07750896 | 0.08167262 | 0.499828444 | -4.139426546 |
| **A_55_P2159801** | Rfc4 | 0.399433354 | -0.067843455 | 0.866710164 | 9.979823303 | 2.077080308 | 0.081721696 | 0.499828444 | -4.139636848 |
| **A_66_P125110** | NA | 0.180794328 | -0.030715113 | 0.392303769 | 10.37105021 | 2.077008303 | 0.081729942 | 0.499828444 | -4.139672176 |
| **A_55_P2440032** | Usp24 | 0.147528422 | -0.025080543 | 0.320137387 | 8.302713951 | 2.076804128 | 0.081753331 | 0.499828444 | -4.139772353 |
| **A_55_P1999334** | Inpp5b | 0.15200511 | -0.025882187 | 0.329892407 | 11.72341392 | 2.076330293 | 0.081807635 | 0.499828444 | -4.140004851 |
| **A_55_P2058270** | Gm6507 | 0.239195065 | -0.04080453 | 0.519194659 | 8.189147169 | 2.075764122 | 0.081872569 | 0.499828444 | -4.140282681 |
| **A_51_P310649** | Arsa | -0.324559981 | -0.704530297 | 0.055410335 | 10.3897918 | -2.075527482 | 0.081899725 | 0.499828444 | -4.140398812 |
| **A_55_P2100088** | Ccdc92b | 0.445400815 | -0.076182059 | 0.966983689 | 11.43074745 | 2.074965469 | 0.081964256 | 0.499828444 | -4.140674639 |
| **A_55_P2153016** | Ccdc34 | 0.206951172 | -0.035400518 | 0.449302862 | 8.491198891 | 2.074937534 | 0.081967465 | 0.499828444 | -4.14068835 |
| **A_55_P2010223** | Gabarap | -0.235140213 | -0.510504633 | 0.040224206 | 13.93392349 | -2.074924378 | 0.081968976 | 0.499828444 | -4.140694807 |
| **A_51_P520857** | Selenok-ps1 | -0.32614071 | -0.708114067 | 0.055832647 | 13.36505622 | -2.074699154 | 0.081994852 | 0.499828444 | -4.140805352 |
| **A_55_P2038519** | NA | 0.390437479 | -0.066857426 | 0.847732384 | 9.350235619 | 2.074618884 | 0.082004077 | 0.499828444 | -4.140844752 |
| **A_55_P2085596** | Nudt1 | 0.175775485 | -0.030109695 | 0.381660665 | 10.36869012 | 2.074514175 | 0.082016111 | 0.499828444 | -4.140896147 |
| **A_55_P2038540** | Hbb-b2 | 0.215434036 | -0.036908908 | 0.46777698 | 16.68456485 | 2.074466093 | 0.082021638 | 0.499828444 | -4.140919748 |
| **A_55_P1957739** | Guca2b | -0.341549021 | -0.741652471 | 0.05855443 | 8.133009109 | -2.074263554 | 0.082044923 | 0.499828444 | -4.141019166 |
| **A_55_P2073482** | NA | 0.298788452 | -0.051234727 | 0.64881163 | 11.59988172 | 2.07419791 | 0.082052471 | 0.499828444 | -4.141051389 |
| **A_55_P1999364** | Krt85 | -0.473522903 | -1.028408288 | 0.081362482 | 10.04512286 | -2.073580463 | 0.082123503 | 0.499828444 | -4.141354494 |
| **A_51_P196158** | Btd | -0.320656822 | -0.696420362 | 0.055106718 | 10.75117339 | -2.073523849 | 0.08213002 | 0.499828444 | -4.141382287 |
| **A_55_P2139181** | Cald1 | -0.222569635 | -0.483406308 | 0.038267038 | 9.741150049 | -2.073387378 | 0.082145729 | 0.499828444 | -4.141449286 |
| **A_55_P2135035** | NA | 0.544889644 | -0.093685893 | 1.183465181 | 10.4766472 | 2.073382678 | 0.08214627 | 0.499828444 | -4.141451594 |
| **A_51_P219721** | Ola1 | 0.250874202 | -0.043181884 | 0.544930288 | 11.4253174 | 2.073046389 | 0.082184995 | 0.499828444 | -4.141616697 |
| **A_51_P152990** | Grem2 | -0.365786955 | -0.794575492 | 0.063001582 | 8.922474758 | -2.072851715 | 0.082207421 | 0.499828444 | -4.141712278 |
| **A_55_P1965448** | NA | 0.251988918 | -0.043406116 | 0.547383952 | 12.31536246 | 2.072819288 | 0.082211157 | 0.499828444 | -4.1417282 |
| **A_55_P1993517** | Tbc1d14 | 0.232252907 | -0.040031282 | 0.504537096 | 8.032489474 | 2.07263069 | 0.082232889 | 0.499828444 | -4.141820801 |
| **A_55_P2024342** | Tmem33 | 0.215728924 | -0.037240451 | 0.468698299 | 9.876503259 | 2.072161587 | 0.082286971 | 0.499828444 | -4.142051143 |
| **A_52_P186033** | Spn | 0.399859799 | -0.069065951 | 0.86878555 | 11.24985504 | 2.071986161 | 0.082307204 | 0.499828444 | -4.142137287 |
| **A_55_P2429225** | Psrc1 | 0.218783786 | -0.037794384 | 0.475361956 | 8.956496977 | 2.071946892 | 0.082311734 | 0.499828444 | -4.14215657 |
| **A_55_P2118734** | Rnmt | 0.272904998 | -0.047150054 | 0.592960049 | 10.10709812 | 2.071905733 | 0.082316483 | 0.499828444 | -4.142176782 |
| **A_55_P2044045** | Ybx1 | 0.238103846 | -0.041205806 | 0.517413499 | 13.80608844 | 2.071398492 | 0.082375023 | 0.499828444 | -4.142425883 |
| **A_55_P2007560** | NA | 0.235799961 | -0.040818133 | 0.512418054 | 8.610855938 | 2.071315875 | 0.082384561 | 0.499828444 | -4.142466457 |
| **A_51_P139400** | Taf1 | 0.218408721 | -0.03785731 | 0.474674753 | 8.946309517 | 2.070914276 | 0.082430945 | 0.499828444 | -4.142663697 |
| **A_66_P103418** | Tmc8 | 0.358150458 | -0.062248919 | 0.778549836 | 9.396961537 | 2.070077647 | 0.082527657 | 0.499828444 | -4.143074638 |
| **A_55_P1993544** | NA | 0.250085091 | -0.043540141 | 0.543710324 | 8.949146916 | 2.069558074 | 0.082587777 | 0.499828444 | -4.143329875 |
| **A_55_P2046852** | Gata1 | 0.397362257 | -0.069185896 | 0.86391041 | 13.33045027 | 2.069537634 | 0.082590143 | 0.499828444 | -4.143339916 |
| **A_55_P2040815** | Lrrc45 | -0.362597686 | -0.788347946 | 0.063152573 | 10.35336203 | -2.069442178 | 0.082601194 | 0.499828444 | -4.143386811 |
| **A_55_P2022693** | Tcerg1 | 0.276411227 | -0.048170449 | 0.600992903 | 9.3844214 | 2.069259083 | 0.082622393 | 0.499828444 | -4.143476763 |
| **A_55_P1966790** | Impdh2 | 0.216016076 | -0.037678254 | 0.469710405 | 12.39489765 | 2.068990519 | 0.082653499 | 0.499828444 | -4.143608709 |
| **A_51_P367310** | Chaf1b | 0.370501526 | -0.064669364 | 0.805672417 | 11.51664266 | 2.068775448 | 0.082678418 | 0.499828444 | -4.143714378 |
| **A_51_P106799** | Pparg | -0.19049407 | -0.414277916 | 0.033289776 | 8.983305867 | -2.068406611 | 0.082721171 | 0.499828444 | -4.151552166 |
| **A_55_P1953439** | Lemd3 | 0.241519282 | -0.042291456 | 0.525330019 | 9.462135397 | 2.067788785 | 0.082792834 | 0.499828444 | -4.144199197 |
| **A_55_P1973868** | Sema3b | -0.724344162 | -1.575532556 | 0.126844233 | 8.709197491 | -2.067771092 | 0.082794887 | 0.499828444 | -4.144207892 |
| **A_52_P306396** | Ppp3cb | 0.2238748 | -0.039266711 | 0.48701631 | 10.85618712 | 2.067278727 | 0.082852045 | 0.499828444 | -4.144449859 |
| **A_51_P117995** | Pfkm | 0.210562251 | -0.036943724 | 0.458068226 | 10.7089402 | 2.067178679 | 0.082863664 | 0.499828444 | -4.144499028 |
| **A_55_P2017929** | Clu | -0.298250985 | -0.648872907 | 0.052370937 | 13.76875531 | -2.066931144 | 0.082892418 | 0.499828444 | -4.144620686 |
| **A_66_P122155** | Wdr26 | 0.267802793 | -0.047032832 | 0.582638418 | 12.00481956 | 2.066875997 | 0.082898826 | 0.499828444 | -4.14464779 |
| **A_55_P2355345** | A130019P10Rik | 0.248897247 | -0.043754627 | 0.541549122 | 8.739154207 | 2.06657881 | 0.082933365 | 0.499828444 | -4.144793858 |

| | | | | | | | | | |
|---|---|---|---|---|---|---|---|---|---|
| **A_51_P507053** | Slc38a1 | 0.186066429 | -0.032719477 | 0.404852336 | 11.06301081 | 2.06648316 | 0.082944484 | 0.499828444 | -4.144840872 |
| **A_55_P2128451** | Sfpq | 0.271186601 | -0.047744398 | 0.5901176 | 11.99060135 | 2.066115919 | 0.08298719 | 0.499828444 | -4.145021386 |
| **A_55_P2150058** | Zfp593 | 0.203117344 | -0.035810257 | 0.442044944 | 9.70989076 | 2.065683955 | 0.083037451 | 0.499828444 | -4.145233727 |
| **A_55_P2128794** | Clta | -0.177357292 | -0.386024561 | 0.031309977 | 13.16839936 | -2.065275086 | 0.083085054 | 0.499828444 | -4.14543473 |
| **A_55_P2153141** | NA | 0.291843296 | -0.051590047 | 0.635276639 | 11.72376184 | 2.064859248 | 0.083133496 | 0.499828444 | -4.145639174 |
| **A_52_P586141** | Adcy7 | 0.358271343 | -0.063341526 | 0.779884212 | 11.00352645 | 2.064816215 | 0.083138511 | 0.499828444 | -4.145660332 |
| **A_55_P2102181** | NA | -0.275259512 | -0.599186668 | 0.048667643 | 13.69987976 | -2.064800837 | 0.083140303 | 0.499828444 | -4.145667893 |
| **A_55_P1988766** | NA | 0.525313228 | -0.092941722 | 1.143568178 | 9.984006996 | 2.064590498 | 0.083164819 | 0.499828444 | -4.145771311 |
| **A_55_P2055537** | Slc16a6 | 0.405310476 | -0.071741046 | 0.882361997 | 12.33232529 | 2.064456481 | 0.083180443 | 0.499828444 | -4.145837205 |
| **A_55_P2103097** | Ccr8 | 0.52026473 | -0.092090063 | 1.132619522 | 8.654156732 | 2.064450396 | 0.083181152 | 0.499828444 | -4.145840197 |
| **A_52_P354823** | Irf8 | -0.207106471 | -0.450891546 | 0.036678604 | 10.7339219 | -2.064285414 | 0.08320039 | 0.499828444 | -4.145921319 |
| **A_55_P2022364** | Nxn | -0.410056095 | -0.892755085 | 0.072642894 | 10.77380971 | -2.064191909 | 0.083211296 | 0.499828444 | -4.145967296 |
| **A_55_P2023777** | Tmx2 | 0.212930197 | -0.037775178 | 0.463635572 | 8.776376088 | 2.063748639 | 0.083263014 | 0.499828444 | -4.146185267 |
| **A_55_P2090060** | Mab21l3 | -0.265299198 | -0.577720079 | 0.047121684 | 7.865402787 | -2.063379312 | 0.08330613 | 0.499828444 | -4.14636689 |
| **A_55_P2131190** | NA | -0.272102855 | -0.592550826 | 0.048345116 | 8.700596828 | -2.063282818 | 0.083317399 | 0.499828444 | -4.146414344 |
| **A_55_P2062148** | Arl1 | -0.227417933 | -0.495259172 | 0.040423306 | 10.29668827 | -2.063148326 | 0.083333107 | 0.499828444 | -4.146480487 |
| **A_55_P2041693** | Gm5834 | -0.317929855 | -0.692393058 | 0.056533348 | 8.673472408 | -2.063029097 | 0.083347036 | 0.499828444 | -4.146539125 |
| **A_55_P2033610** | Fastkd3 | 0.170178498 | -0.0302673 | 0.370624295 | 9.826902097 | 2.062960534 | 0.083355046 | 0.499828444 | -4.146572845 |
| **A_52_P543040** | Utp14a | 0.225060152 | -0.040052541 | 0.490172844 | 9.360132408 | 2.062772228 | 0.083377051 | 0.499828444 | -4.146665459 |
| **A_66_P111840** | NA | 0.215950493 | -0.038470904 | 0.47037189 | 14.60466995 | 2.062451554 | 0.083414538 | 0.499828444 | -4.146823181 |
| **A_55_P2084686** | Pdik1l | 0.182098103 | -0.032442518 | 0.396638725 | 8.558879914 | 2.062429331 | 0.083417136 | 0.499828444 | -4.146834112 |
| **A_55_P2022404** | Gtf2h5 | 0.177782683 | -0.031689997 | 0.387255364 | 11.53093567 | 2.062268724 | 0.083435918 | 0.499828444 | -4.146913109 |
| **A_52_P652913** | Akip1 | 0.343168772 | -0.061190194 | 0.747527739 | 10.46926943 | 2.062167186 | 0.083447794 | 0.499828444 | -4.146963054 |
| **A_55_P1980302** | Psenen | -0.232497137 | -0.506453054 | 0.04145878 | 12.37718124 | -2.062149341 | 0.083449882 | 0.499828444 | -4.146971832 |
| **A_52_P68702** | Frmd4b | -0.18679814 | -0.406936676 | 0.033340396 | 8.374903162 | -2.061862319 | 0.083483464 | 0.499828444 | -4.147113017 |
| **A_55_P1981884** | Ercc6l2 | 0.165729465 | -0.029596449 | 0.361055378 | 9.485390452 | 2.061688518 | 0.083503805 | 0.499828444 | -4.147198512 |
| **A_55_P2230913** | Etv6 | -0.199013962 | -0.433573741 | 0.035545817 | 8.586083942 | -2.06164171 | 0.083509284 | 0.499828444 | -4.147221538 |
| **A_52_P267106** | Ksr2 | 0.18353371 | -0.03286747 | 0.399934891 | 8.182332584 | 2.06081691 | 0.083605892 | 0.499828444 | -4.147627308 |
| **A_52_P663757** | Tnik | -0.187882194 | -0.409415183 | 0.033650794 | 8.350002452 | -2.060774192 | 0.083610899 | 0.499828444 | -4.147648325 |
| **A_55_P2048041** | Slc44a1 | -0.175706513 | -0.382897306 | 0.031484279 | 8.783515934 | -2.060632752 | 0.083627478 | 0.499828444 | -4.147717914 |
| **A_55_P2036633** | Ssrp1 | 0.182767431 | -0.032761957 | 0.39829682 | 13.50253476 | 2.060513675 | 0.083641438 | 0.499828444 | -4.147776502 |
| **A_66_P110607** | Sart1 | 0.3130717 | -0.056154158 | 0.682297558 | 9.999954233 | 2.060321012 | 0.083664031 | 0.499828444 | -4.147871298 |
| **A_55_P2043377** | NA | 0.230611974 | -0.041370214 | 0.502594163 | 8.435693231 | 2.060272088 | 0.083669769 | 0.499828444 | -4.147895371 |
| **A_55_P2177712** | NA | -0.14864168 | -0.323952222 | 0.026668861 | 7.977250864 | -2.060230267 | 0.083674674 | 0.499828444 | -4.147915948 |
| **A_55_P2236818** | A430103D13Rik | 0.146120687 | -0.026224381 | 0.318465754 | 8.344118286 | 2.060136685 | 0.083685651 | 0.499828444 | -4.147961995 |
| **A_55_P1977593** | NA | 0.222932075 | -0.040112999 | 0.485977149 | 11.97713307 | 2.059328238 | 0.083780545 | 0.499828444 | -4.14835982 |
| **A_55_P2139753** | Jade2 | -0.25523529 | -0.556434048 | 0.045963468 | 9.450719345 | -2.05906821 | 0.08381109 | 0.499828444 | -4.148487787 |
| **A_51_P269203** | Trpv4 | 0.170537419 | -0.030742893 | 0.371817731 | 8.570228478 | 2.058740361 | 0.083849618 | 0.499828444 | -4.148649139 |
| **A_55_P1958082** | Prpsap2 | 0.171503719 | -0.030939299 | 0.373946736 | 9.164937979 | 2.058514492 | 0.083876171 | 0.499828444 | -4.148760307 |
| **A_51_P152747** | Nrip1 | 0.214522968 | -0.038707154 | 0.46775309 | 9.395990938 | 2.058456155 | 0.083883031 | 0.499828444 | -4.14878902 |
| **A_55_P1972699** | Rars2 | 0.186883321 | -0.033744209 | 0.407510851 | 10.74362254 | 2.058230595 | 0.083909559 | 0.499828444 | -4.14890004 |
| **A_52_P225260** | Ywhaz | -0.185345871 | -0.404159712 | 0.033467969 | 8.48853279 | -2.058217738 | 0.083911071 | 0.499828444 | -4.148906369 |
| **A_52_P456750** | Aph1b | -0.209235636 | -0.456254098 | 0.037782826 | 8.539253676 | -2.058208793 | 0.083912124 | 0.499828444 | -4.148910772 |
| **A_55_P2020090** | Pwwp3b | -0.631290189 | -1.376610403 | 0.114030025 | 8.74476045 | -2.058113537 | 0.08392333 | 0.499828444 | -4.148957658 |
| **A_55_P2058681** | Tmc8 | 0.297347529 | -0.053733347 | 0.648428404 | 9.915585022 | 2.05797621 | 0.083939487 | 0.499828444 | -4.149025254 |
| **A_55_P2079973** | NA | -0.165329428 | -0.360569888 | 0.029911033 | 13.59389481 | -2.05761221 | 0.083982331 | 0.499828444 | -4.149204431 |
| **A_55_P2010469** | NA | 0.247034476 | -0.04470599 | 0.538774941 | 12.05173416 | 2.057520091 | 0.083993177 | 0.499828444 | -4.149249778 |
| **A_55_P1976908** | NA | 0.335476399 | -0.060723503 | 0.731676302 | 12.80541916 | 2.057457142 | 0.084000589 | 0.499828444 | -4.149280766 |
| **A_51_P314418** | Blm | 0.371353809 | -0.067286533 | 0.809994151 | 11.07811322 | 2.057133615 | 0.084038696 | 0.499828444 | -4.149440034 |
| **A_55_P2138306** | NA | -0.251309974 | -0.548172086 | 0.045552138 | 10.4245962 | -2.057018237 | 0.08405229 | 0.499828444 | -4.149496835 |

| | | | | | | | | |
|---|---|---|---|---|---|---|---|---|
| **A_55_P2082826** | Fam160b1 | -0.156875325 | -0.342220295 | 0.028469645 | 11.26978836 | -2.056634062 | 0.08409757 | 0.499828444 | -4.149685974 |
| **A_55_P2102533** | Fuca2 | 0.209632969 | -0.038082582 | 0.457348521 | 8.407164964 | 2.056314331 | 0.084135273 | 0.499828444 | -4.149843395 |
| **A_55_P2149983** | Foxp4 | -0.155198304 | -0.338592534 | 0.028195927 | 11.63123641 | -2.0562906 | 0.084138072 | 0.499828444 | -4.149855079 |
| **A_51_P379443** | Bclaf1 | 0.234549264 | -0.042641526 | 0.511740053 | 12.15508655 | 2.056072706 | 0.084163777 | 0.499828444 | -4.149962365 |
| **A_55_P2099064** | Nsd2 | 0.191336973 | -0.034834745 | 0.417508692 | 8.33869132 | 2.055624524 | 0.084216675 | 0.499828444 | -4.150183052 |
| **A_55_P2167000** | Dmac2 | -0.179210217 | -0.39105259 | 0.032632156 | 10.49083005 | -2.055574032 | 0.084222636 | 0.499828444 | -4.150207916 |
| **A_51_P234308** | Rpa2 | 0.3372582 | -0.061431477 | 0.735947876 | 10.37488056 | 2.05546798 | 0.084235159 | 0.499828444 | -4.150260139 |
| **A_55_P2007640** | Eprs | 0.381344441 | -0.069499079 | 0.832187961 | 10.91374651 | 2.055297895 | 0.084255247 | 0.499828444 | -4.150343897 |
| **A_55_P2133751** | Txnl4a | 0.162932579 | -0.029698676 | 0.355563833 | 12.67553056 | 2.05524863 | 0.084261066 | 0.499828444 | -4.150368157 |
| **A_55_P1989469** | NA | 0.210118183 | -0.038324277 | 0.458560643 | 15.38647247 | 2.055043434 | 0.084285309 | 0.499828444 | -4.150469209 |
| **A_55_P1997335** | Vps9d1 | -0.210278598 | -0.458927038 | 0.038369842 | 10.05521008 | -2.05490866 | 0.084301235 | 0.499828444 | -4.150535582 |
| **A_55_P2152956** | Plekhm3 | 0.186570043 | -0.034092401 | 0.407232487 | 8.352415912 | 2.054455211 | 0.084354843 | 0.499828444 | -4.150758906 |
| **A_55_P1954768** | NA | 0.336105971 | -0.061437421 | 0.733649363 | 9.044977192 | 2.054352088 | 0.084367039 | 0.499828444 | -4.150809697 |
| **A_55_P2084433** | NA | 0.152257155 | -0.027848756 | 0.332363066 | 12.79955662 | 2.054153681 | 0.08439051 | 0.499828444 | -4.150907419 |
| **A_52_P214927** | Zmynd8 | 0.25415795 | -0.046497551 | 0.554813451 | 9.677619373 | 2.054081798 | 0.084399015 | 0.499828444 | -4.150942825 |
| **A_52_P572702** | Gxylt1 | 0.152363056 | -0.027882957 | 0.332609069 | 10.01335974 | 2.053984664 | 0.084410509 | 0.499828444 | -4.150990668 |
| **A_52_P200617** | Iars | 0.369917199 | -0.067747492 | 0.80758189 | 11.35637839 | 2.0537435 | 0.084439053 | 0.499828444 | -4.151109458 |
| **A_55_P2079520** | Mpp1 | 0.280539431 | -0.051438059 | 0.61251692 | 11.64931989 | 2.053375918 | 0.084482578 | 0.499828444 | -4.151290526 |
| **A_52_P658122** | Ets2 | -0.537720538 | -1.17409111 | 0.098650034 | 11.02322339 | -2.053192764 | 0.084504274 | 0.499828444 | -4.151380751 |
| **A_51_P354572** | Rab35 | 0.158436087 | -0.029073702 | 0.345945876 | 12.24145142 | 2.053115364 | 0.084513444 | 0.499828444 | -4.15141888 |
| **A_52_P86517** | Cbwd1 | 0.185128471 | -0.034006218 | 0.404263159 | 9.43592986 | 2.052793538 | 0.084551584 | 0.499828444 | -4.151577425 |
| **A_51_P133422** | Nck2 | 0.195763176 | -0.035966673 | 0.427493026 | 8.453201531 | 2.052731812 | 0.084558902 | 0.499828444 | -4.151607835 |
| **A_52_P599624** | Pfas | 0.314522192 | -0.057844898 | 0.686889282 | 8.740807802 | 2.052405605 | 0.084597582 | 0.499904209 | -4.151768549 |
| **A_55_P2061492** | Pnldc1 | 0.188158702 | -0.034683496 | 0.4110009 | 8.323479099 | 2.051682071 | 0.08468344 | 0.500258625 | -4.152125046 |
| **A_51_P401263** | Eme1 | 0.300917634 | -0.055526786 | 0.657362054 | 9.404065376 | 2.05134642 | 0.084723301 | 0.500341178 | -4.152290441 |
| **A_55_P2021216** | Crlf3 | 0.289152226 | -0.053480035 | 0.631784487 | 13.39606624 | 2.050602475 | 0.084811715 | 0.500710335 | -4.15265706 |
| **A_55_P1982030** | H3f3b | 0.246000724 | -0.045671498 | 0.537672946 | 12.03027205 | 2.049389485 | 0.084956073 | 0.501409446 | -4.153254923 |
| **A_55_P2046563** | Cym | -0.607308732 | -1.327448457 | 0.112830993 | 8.60854072 | -2.04916036 | 0.084983369 | 0.501417443 | -4.153367868 |
| **A_52_P390944** | Chst3 | -0.310458219 | -0.678705882 | 0.057789443 | 8.848036894 | -2.048548981 | 0.085056247 | 0.501493325 | -4.153669263 |
| **A_52_P572284** | Laptm4a | -0.271107149 | -0.592687137 | 0.050472838 | 11.15437856 | -2.048496118 | 0.085062552 | 0.501493325 | -4.153695325 |
| **A_55_P1980831** | Arpc2 | -0.240007341 | -0.524710992 | 0.04469631 | 14.68169102 | -2.048399583 | 0.085074065 | 0.501493325 | -4.153742917 |
| **A_55_P1971244** | Ndufa4l2 | -0.231287395 | -0.50576622 | 0.04319143 | 12.35516987 | -2.047511299 | 0.085180087 | 0.501948106 | -4.154180884 |
| **A_51_P479818** | Lonrf3 | 0.136021719 | -0.025426483 | 0.297469922 | 8.207513901 | 2.047190392 | 0.085218421 | 0.501948106 | -4.154339123 |
| **A_51_P288916** | Tmtc2 | -0.377147185 | -0.824840684 | 0.070546313 | 8.868074007 | -2.046978458 | 0.085243748 | 0.501948106 | -4.154443632 |
| **A_52_P569038** | Cnot2 | 0.153491812 | -0.028749414 | 0.335733037 | 10.94306818 | 2.046547244 | 0.085295302 | 0.501948106 | -4.154656282 |
| **A_52_P92557** | Kdm2a | 0.169526151 | -0.03176554 | 0.370817841 | 9.792882932 | 2.046416561 | 0.085310933 | 0.501948106 | -4.154720731 |
| **A_55_P2110537** | Slfn9 | 0.241564741 | -0.045299013 | 0.528428494 | 9.288229731 | 2.046166907 | 0.0853408 | 0.501948106 | -4.154843856 |
| **A_52_P653902** | Eny2 | 0.161114998 | -0.030224357 | 0.352454353 | 9.208041132 | 2.046043454 | 0.085355574 | 0.501948106 | -4.154904743 |
| **A_52_P631568** | Thoc6 | 0.224914387 | -0.042196498 | 0.492025273 | 9.831980779 | 2.046015124 | 0.085358964 | 0.501948106 | -4.154918715 |
| **A_55_P1993955** | Pltp | -0.379105536 | -0.829505186 | 0.071294114 | 9.405866193 | -2.045244675 | 0.085451225 | 0.502273624 | -4.155298729 |
| **A_66_P111301** | Mplkip | 0.187137633 | -0.035206539 | 0.409481805 | 10.32142019 | 2.045118936 | 0.085466291 | 0.502273624 | -4.155360752 |
| **A_55_P1979221** | Txndc9 | 0.21767721 | -0.041036869 | 0.476391288 | 10.25094311 | 2.044448091 | 0.085546721 | 0.502593485 | -4.155691683 |
| **A_55_P2221365** | NA | 0.151747572 | -0.028644953 | 0.332140097 | 8.54069025 | 2.044025936 | 0.085597373 | 0.502704111 | -4.155899953 |
| **A_55_P1985094** | Wrap73 | 0.193044712 | -0.03647108 | 0.422560503 | 10.7442963 | 2.04375356 | 0.085630071 | 0.502704111 | -4.156034337 |
| **A_51_P398878** | Uck2 | 0.172482578 | -0.032597651 | 0.377562808 | 11.89742359 | 2.043641106 | 0.085643574 | 0.502704111 | -4.156089821 |
| **A_51_P375783** | Prap1 | -1.373296621 | -3.006827663 | 0.260234422 | 8.471101799 | -2.042773186 | 0.085747864 | 0.503071236 | -4.156518079 |
| **A_55_P2175502** | M1ap | 0.525040465 | -0.09951936 | 1.14960029 | 9.941845523 | 2.042687448 | 0.085758173 | 0.503071236 | -4.156560388 |
| **A_55_P2076792** | Anxa6 | -0.388500437 | -0.850892057 | 0.073891183 | 12.60417959 | -2.041572213 | 0.085892385 | 0.503705676 | -4.157110776 |
| **A_66_P126552** | Prr14l | 0.236252458 | -0.045010492 | 0.517515408 | 9.145825074 | 2.041018763 | 0.085959068 | 0.503943836 | -4.15738395 |
| **A_55_P1958379** | Slit1 | 0.547271715 | -0.104502289 | 1.19904572 | 10.23011275 | 2.04027711 | 0.086048509 | 0.504257929 | -4.157750057 |

| | | | | | | | | | |
|---|---|---|---|---|---|---|---|---|---|
| **A_52_P215170** | Acot4 | 0.474309138 | -0.090625311 | 1.039243588 | 9.241141129 | 2.040077295 | 0.086072623 | 0.504257929 | -4.1578487 |
| **A_55_P2047188** | Fgf1 | -0.851838496 | -1.866757824 | 0.163080831 | 9.680587128 | -2.039430588 | 0.086150712 | 0.504257929 | -4.158167984 |
| **A_51_P162437** | Bcr | -0.25691094 | -0.563031352 | 0.049209473 | 11.23450314 | -2.039264208 | 0.086170814 | 0.504257929 | -4.158250132 |
| **A_51_P176352** | Ndrg2 | -0.464660492 | -1.01836382 | 0.089042835 | 12.00808109 | -2.039115397 | 0.086188798 | 0.504257929 | -4.158323608 |
| **A_55_P2092831** | NA | 0.275349645 | -0.052832477 | 0.603531768 | 12.46041264 | 2.038697502 | 0.086239319 | 0.504257929 | -4.158529954 |
| **A_52_P278497** | Eif4ebp3 | -0.146969004 | -0.322138584 | 0.028200575 | 10.6012105 | -2.038685591 | 0.086240759 | 0.504257929 | -4.158535835 |
| **A_55_P2023572** | NA | 0.217104308 | -0.041665167 | 0.475873783 | 8.544203549 | 2.038630818 | 0.086247383 | 0.504257929 | -4.158562882 |
| **A_55_P2063594** | Gm5801 | 0.196428021 | -0.03771969 | 0.430575731 | 11.46903468 | 2.038434277 | 0.086271157 | 0.504257929 | -4.158659935 |
| **A_66_P100105** | Patj | -0.281647356 | -0.61738213 | 0.054087418 | 8.578046911 | -2.038414699 | 0.086273525 | 0.504257929 | -4.158669602 |
| **A_55_P2156785** | Ppfia3 | 0.306514388 | -0.058950644 | 0.67197942 | 9.835124699 | 2.037925132 | 0.086332773 | 0.50444516 | -4.158911367 |
| **A_55_P2058923** | Bud23 | 0.332831343 | -0.064052255 | 0.729714941 | 11.2218769 | 2.037718813 | 0.086357754 | 0.50444516 | -4.15901326 |
| **A_55_P2107647** | NA | -0.171259362 | -0.37555648 | 0.033037757 | 11.04560413 | -2.036926079 | 0.086453807 | 0.504827813 | -4.159404791 |
| **A_51_P112355** | Igtp | -0.71386484 | -1.565577932 | 0.137848252 | 11.24748549 | -2.036600557 | 0.086493281 | 0.504827813 | -4.166865275 |
| **A_55_P2025155** | NA | 0.264665404 | -0.051174734 | 0.580505542 | 13.36998042 | 2.036165354 | 0.086546083 | 0.504827813 | -4.15978056 |
| **A_55_P1959703** | Zfp646 | 0.199560767 | -0.038589263 | 0.437710797 | 15.90886619 | 2.036140362 | 0.086549116 | 0.504827813 | -4.159792906 |
| **A_51_P294555** | Ifitm6 | -0.405092846 | -0.888528158 | 0.078342466 | 12.31933347 | -2.036101356 | 0.08655385 | 0.504827813 | -4.159812175 |
| **A_55_P2101191** | Arrdc4 | -0.171044814 | -0.375236513 | 0.033146885 | 8.777259397 | -2.035424589 | 0.086636033 | 0.504952978 | -4.160146514 |
| **A_55_P2096535** | Icmt | 0.243901176 | -0.047294486 | 0.535096837 | 8.832102894 | 2.035223847 | 0.086660425 | 0.504952978 | -4.160245692 |
| **A_51_P267346** | Topors | 0.155355868 | -0.030173409 | 0.340885144 | 10.84024508 | 2.034690692 | 0.086725242 | 0.504952978 | -4.160509117 |
| **A_55_P1976928** | Unc13d | 0.336944686 | -0.065452679 | 0.739342051 | 9.925826853 | 2.034635758 | 0.086731923 | 0.504952978 | -4.160536261 |
| **A_55_P2156052** | NA | 0.254813183 | -0.049558803 | 0.55918517 | 9.97643056 | 2.034231634 | 0.08678109 | 0.504952978 | -4.16073595 |
| **A_55_P2158990** | Jun | -0.864557462 | -1.897314331 | 0.168199408 | 10.72461136 | -2.034131194 | 0.086793314 | 0.504952978 | -4.160785582 |
| **A_51_P501069** | Spta1 | 0.494259605 | -0.096193707 | 1.084712917 | 10.36504902 | 2.034008407 | 0.086808261 | 0.504952978 | -4.160846258 |
| **A_55_P1952502** | Mapk7 | -0.363837065 | -0.798485237 | 0.070811108 | 10.52155584 | -2.034006192 | 0.08680853 | 0.504952978 | -4.160847353 |
| **A_55_P2133205** | Nmt2 | 0.313174491 | -0.060969351 | 0.687318332 | 9.05077809 | 2.033906424 | 0.086820677 | 0.504952978 | -4.160896655 |
| **A_55_P2275054** | Nrap | 0.160906265 | -0.031337832 | 0.353150362 | 8.775029596 | 2.033776053 | 0.086836551 | 0.504952978 | -4.160961081 |
| **A_52_P466701** | Fuom | -0.234401423 | -0.514485135 | 0.04568229 | 9.840381248 | -2.033553384 | 0.086863671 | 0.504958767 | -4.161071122 |
| **A_55_P1974984** | NA | -0.280675465 | -0.616302632 | 0.054951702 | 9.252411025 | -2.032031953 | 0.087049203 | 0.505776906 | -4.161823101 |
| **A_55_P2000623** | Stag1 | 0.167106735 | -0.032757882 | 0.366971352 | 10.11810951 | 2.031614136 | 0.087100224 | 0.505776906 | -4.162029642 |
| **A_55_P1992889** | Serping1 | -0.412059488 | -0.9049889 | 0.080869923 | 13.50929706 | -2.031226604 | 0.087147574 | 0.505776906 | -4.162221224 |
| **A_51_P397564** | Snrnp40 | 0.233021236 | -0.04579372 | 0.511836192 | 11.10576768 | 2.030778815 | 0.087202319 | 0.505776906 | -4.16244261 |
| **A_55_P2015217** | Alg11 | -0.253486389 | -0.556799271 | 0.049826494 | 9.108715707 | -2.030705651 | 0.087211267 | 0.505776906 | -4.162478784 |
| **A_55_P2003133** | Ngrn | 0.157094398 | -0.030882007 | 0.345070803 | 12.31055066 | 2.030675588 | 0.087214943 | 0.505776906 | -4.162493647 |
| **A_51_P350453** | Pdk4 | -0.619651164 | -1.361264943 | 0.121962616 | 9.642982365 | -2.030264719 | 0.087265212 | 0.505776906 | -4.162696797 |
| **A_55_P2136318** | Slc1a5 | 0.409501876 | -0.080609028 | 0.899612779 | 9.704778254 | 2.030227565 | 0.087269759 | 0.505776906 | -4.162715169 |
| **A_52_P609024** | Cenpw | 0.319466713 | -0.062910922 | 0.701844348 | 12.0739346 | 2.030094804 | 0.087286009 | 0.505776906 | -4.162780814 |
| **A_55_P2077535** | Zfp513 | -0.190724589 | -0.41904781 | 0.037598632 | 10.36663815 | -2.029737122 | 0.087329805 | 0.505776906 | -4.162957682 |
| **A_52_P74524** | NA | 0.187290235 | -0.036940777 | 0.411521248 | 8.367692083 | 2.029563514 | 0.08735107 | 0.505776906 | -4.163043532 |
| **A_51_P293753** | Cul7 | -0.245718779 | -0.53990869 | 0.048471132 | 12.0627093 | -2.029521993 | 0.087356157 | 0.505776906 | -4.163064064 |
| **A_55_P1953683** | Nsmaf | 0.195707757 | -0.038639545 | 0.430055059 | 11.68355153 | 2.029229978 | 0.087391939 | 0.505776906 | -4.163208473 |
| **A_55_P2075358** | NA | 0.228869262 | -0.045284063 | 0.503022586 | 12.28130872 | 2.028509944 | 0.087480233 | 0.505776906 | -4.163564576 |
| **A_55_P2155848** | Gm14207 | -0.203280188 | -0.44679722 | 0.040236844 | 8.036142711 | -2.028378048 | 0.087496417 | 0.505776906 | -4.163629812 |
| **A_55_P2050857** | D11Wsu47e | 0.243064862 | -0.04814427 | 0.534273995 | 10.02079145 | 2.028151442 | 0.087524228 | 0.505776906 | -4.163741894 |
| **A_52_P575771** | Mtfr1 | 0.217357025 | -0.043064196 | 0.477778246 | 11.44083263 | 2.028058603 | 0.087535625 | 0.505776906 | -4.163787815 |
| **A_52_P444601** | Zfp277 | -0.33595521 | -0.7385235 | 0.066613079 | 9.526880964 | -2.027799483 | 0.087567441 | 0.505776906 | -4.163915985 |
| **A_55_P1967504** | Abhd11 | -0.140676342 | -0.309253 | 0.027900315 | 10.45519988 | -2.027714554 | 0.087577872 | 0.505776906 | -4.163957996 |
| **A_55_P2354531** | 2610029K11Rik | -0.165691251 | -0.364271675 | 0.032889174 | 8.859682172 | -2.027432157 | 0.087612565 | 0.505776906 | -4.164097687 |
| **A_55_P2003981** | NA | 0.388269502 | -0.077100333 | 0.853639336 | 8.818265108 | 2.02730102 | 0.087628679 | 0.505776906 | -4.164162558 |
| **A_52_P524895** | Eif3a | 0.24512054 | -0.048679021 | 0.538920101 | 11.35371309 | 2.027270737 | 0.087632401 | 0.505776906 | -4.164177539 |
| **A_52_P454183** | Olfml2b | -0.178849832 | -0.393243367 | 0.035543702 | 8.171530058 | -2.027029379 | 0.08766207 | 0.505776906 | -4.164296939 |

| | | | | | | | | |
|---|---|---|---|---|---|---|---|---|
| **A_55_P2010134** | NA | 0.297396403 | -0.059138832 | 0.653931637 | 8.895229333 | 2.026825898 | 0.08768709 | 0.505776906 | -4.164397604 |
| **A_55_P2121222** | Clcc1 | 0.181916505 | -0.036183736 | 0.400016746 | 11.56782962 | 2.026745172 | 0.087697018 | 0.505776906 | -4.164437541 |
| **A_55_P2164615** | NA | 0.171827334 | -0.034184738 | 0.377839406 | 16.33735544 | 2.026668747 | 0.087706419 | 0.505776906 | -4.164475351 |
| **A_55_P2139112** | Cdc26 | 0.14690481 | -0.029244552 | 0.323054172 | 11.27666045 | 2.026460435 | 0.087732046 | 0.505776906 | -4.164578412 |
| **A_55_P2005555** | Celf2 | -0.178484666 | -0.3925267 | 0.035557368 | 8.228622806 | -2.026212692 | 0.087762535 | 0.505776906 | -4.164700986 |
| **A_55_P2117243** | Npm1 | 0.280607746 | -0.055903086 | 0.617118577 | 12.71558124 | 2.026206929 | 0.087763244 | 0.505776906 | -4.164703838 |
| **A_55_P2060747** | Esco1 | 0.241548557 | -0.048183341 | 0.531280455 | 9.515402446 | 2.025775552 | 0.087816357 | 0.505932152 | -4.164917278 |
| **A_51_P346165** | Agpat4 | 0.298592583 | -0.059600555 | 0.65678572 | 12.51283101 | 2.025559208 | 0.087843007 | 0.505934886 | -4.165024328 |
| **A_52_P370402** | Kcmf1 | 0.186531214 | -0.03726783 | 0.410330259 | 12.0164777 | 2.025239915 | 0.087882353 | 0.506010722 | -4.165182325 |
| **A_55_P1959218** | Usp51 | 0.156495629 | -0.031300276 | 0.344291533 | 7.953326975 | 2.024879967 | 0.08792673 | 0.506111524 | -4.165360449 |
| **A_51_P342871** | S100a10 | -0.483984694 | -1.064891043 | 0.096921655 | 11.36892594 | -2.024457638 | 0.087978826 | 0.506111524 | -4.165569456 |
| **A_55_P2032079** | Dbp | -0.302008097 | -0.664515914 | 0.060499721 | 8.718366881 | -2.024344391 | 0.087992801 | 0.506111524 | -4.165625503 |
| **A_51_P108489** | Cfap20 | -0.249354201 | -0.548710375 | 0.050001974 | 10.23885519 | -2.024005358 | 0.088034652 | 0.506111524 | -4.176336804 |
| **A_51_P423825** | Morn1 | -0.171493674 | -0.377390878 | 0.03440353 | 8.145643832 | -2.023861758 | 0.088052384 | 0.506111524 | -4.165864375 |
| **A_55_P2132942** | Adal | 0.145761613 | -0.029261121 | 0.320784348 | 9.475942729 | 2.023633642 | 0.08808056 | 0.506111524 | -4.165977284 |
| **A_51_P109939** | Naxe | -0.219158372 | -0.482314726 | 0.043997982 | 12.88174873 | -2.023612693 | 0.088083148 | 0.506111524 | -4.165987653 |
| **A_66_P136998** | Tmem43 | -0.160680409 | -0.353641133 | 0.032280315 | 9.0535373 | -2.023378805 | 0.088112047 | 0.50612712 | -4.166103423 |
| **A_55_P2067226** | Arpc5l | 0.177766015 | -0.035735989 | 0.39126802 | 10.77863215 | 2.023158786 | 0.088139242 | 0.506132917 | -4.166212332 |
| **A_51_P164420** | Eef1e1 | 0.287099676 | -0.057762899 | 0.631962251 | 10.33041912 | 2.022878619 | 0.088173882 | 0.506181459 | -4.16635102 |
| **A_52_P49601** | Fth1 | 0.179141429 | -0.036076215 | 0.394359073 | 16.49315494 | 2.022559701 | 0.088213331 | 0.506257565 | -4.166508897 |
| **A_55_P1980551** | Dennd1a | 0.251427233 | -0.050670106 | 0.553524572 | 9.072350162 | 2.022314078 | 0.088243726 | 0.506281679 | -4.166630496 |
| **A_55_P2185332** | Il17rc | -0.166044733 | -0.365606427 | 0.03351696 | 10.8948466 | -2.021767047 | 0.088311456 | 0.506346569 | -4.166901328 |
| **A_55_P1977766** | Ttc14 | 0.188655913 | -0.038116604 | 0.415428431 | 8.604425742 | 2.021450868 | 0.088350628 | 0.506346569 | -4.167057877 |
| **A_52_P410636** | Trp73 | 0.206128232 | -0.041678672 | 0.453935136 | 8.435468629 | 2.02119058 | 0.088382888 | 0.506346569 | -4.167186758 |
| **A_55_P2089358** | Hspe1 | 0.233030508 | -0.047138297 | 0.513199313 | 14.18431036 | 2.021045974 | 0.088400816 | 0.506346569 | -4.167258361 |
| **A_66_P100159** | Cfl2 | -0.196633805 | -0.433103803 | 0.039836194 | 10.53959516 | -2.020530093 | 0.088464803 | 0.506346569 | -4.16751382 |
| **A_51_P114297** | Kifc3 | -0.150127946 | -0.330671065 | 0.030415173 | 8.552345865 | -2.020522962 | 0.088465687 | 0.506346569 | -4.167517352 |
| **A_51_P314521** | Smim11 | -0.263339311 | -0.580051837 | 0.053373214 | 11.7031333 | -2.02038272 | 0.08848309 | 0.506346569 | -4.167586802 |
| **A_55_P2059686** | Rcc1 | 0.226818188 | -0.045992926 | 0.499629302 | 11.59483857 | 2.020221582 | 0.088503091 | 0.506346569 | -4.167666602 |
| **A_55_P2091846** | Tut4 | 0.316239762 | -0.064142753 | 0.696622276 | 10.03760771 | 2.02012907 | 0.088514575 | 0.506346569 | -4.167712417 |
| **A_55_P2066578** | Ifi204 | -0.291979645 | -0.643240484 | 0.059281193 | 9.601843042 | -2.019789137 | 0.088556788 | 0.506346569 | -4.16788077 |
| **A_55_P2021398** | NA | 0.212670111 | -0.043226777 | 0.468566998 | 12.0533594 | 2.019410649 | 0.088603812 | 0.506346569 | -4.168068227 |
| **A_66_P122372** | Ppp4r2 | 0.2057852 | -0.041879523 | 0.453449923 | 8.847647125 | 2.01898539 | 0.088656678 | 0.506346569 | -4.168278863 |
| **A_55_P2130666** | Armc1 | 0.189555507 | -0.038578474 | 0.417689488 | 11.00034766 | 2.018968838 | 0.088658736 | 0.506346569 | -4.168287062 |
| **A_55_P1964033** | Carm1 | 0.247355497 | -0.050348686 | 0.545059681 | 10.09734946 | 2.018923294 | 0.0886644 | 0.506346569 | -4.168309621 |
| **A_55_P2258567** | D930043N17Rik | 0.402295224 | -0.082056093 | 0.886646542 | 9.427880179 | 2.018215701 | 0.088752441 | 0.506346569 | -4.168660134 |
| **A_55_P2152771** | Lhfpl2 | -0.266643022 | -0.587702614 | 0.054416571 | 8.62034153 | -2.018030665 | 0.088775478 | 0.506346569 | -4.1687518 |
| **A_55_P2076713** | Immp1l | 0.243741404 | -0.049744928 | 0.537227736 | 10.8613993 | 2.018016052 | 0.088777298 | 0.506346569 | -4.168759039 |
| **A_55_P2101336** | Ramp2 | -0.866138926 | -1.909220582 | 0.17694273 | 11.93022016 | -2.017680692 | 0.088819067 | 0.506346569 | -4.168925181 |
| **A_55_P2033780** | Fabp3 | -0.83580667 | -1.842381558 | 0.170768218 | 8.474900966 | -2.017636521 | 0.08882457 | 0.506346569 | -4.168947065 |
| **A_52_P119672** | Rock1 | 0.178297703 | -0.036442687 | 0.393038093 | 9.053135451 | 2.017507698 | 0.088840621 | 0.506346569 | -4.169010888 |
| **A_51_P440892** | Dph2 | 0.186064963 | -0.038047545 | 0.41017747 | 9.657624351 | 2.017352049 | 0.088860019 | 0.506346569 | -4.169088004 |
| **A_55_P2129696** | Smad9 | -0.24441466 | -0.538827412 | 0.049998092 | 7.956025737 | -2.01722259 | 0.088876156 | 0.506346569 | -4.169152145 |
| **A_55_P2192729** | 5830469G19Rik | 0.231314538 | -0.047369756 | 0.509998833 | 9.435138374 | 2.016850115 | 0.088922601 | 0.506346569 | -4.169336698 |
| **A_55_P2024888** | Ctss | -0.293569692 | -0.647260281 | 0.060120896 | 12.45462666 | -2.016837479 | 0.088924177 | 0.506346569 | -4.169342959 |
| **A_55_P2080708** | Cstf2 | 0.179137196 | -0.036702281 | 0.394976673 | 8.991701875 | 2.01668506 | 0.08894319 | 0.506346569 | -4.169418482 |
| **A_55_P2033461** | NA | 0.244324796 | -0.050064515 | 0.538714107 | 14.82755791 | 2.016641485 | 0.088948627 | 0.506346569 | -4.169440073 |
| **A_51_P420655** | Reep4 | 0.219590064 | -0.045016664 | 0.484196792 | 9.682050969 | 2.016484983 | 0.088968154 | 0.506346569 | -4.169517622 |
| **A_51_P263220** | Taf5 | 0.262807667 | -0.053927328 | 0.579542661 | 8.942051428 | 2.016160818 | 0.089008617 | 0.506346569 | -4.169678256 |
| **A_51_P515452** | Fkbpl | -0.233043874 | -0.513915311 | 0.047827563 | 10.00858923 | -2.01610573 | 0.089015494 | 0.506346569 | -4.169705554 |

| | | | | | | | | |
|---|---|---|---|---|---|---|---|---|
| **A_51_P346453** | Noc4l | 0.306198231 | -0.062948528 | 0.67534499 | 10.10435377 | 2.015518581 | 0.089088835 | 0.506346569 | -4.169996527 |
| **A_52_P610403** | Sf1 | 0.267515362 | -0.055077294 | 0.590108019 | 8.947321179 | 2.015011081 | 0.089152275 | 0.506346569 | -4.17024805 |
| **A_51_P310164** | 2810459M11Rik | 0.474970266 | -0.097873203 | 1.047813736 | 9.937699605 | 2.014715121 | 0.089189293 | 0.506346569 | -4.17039474 |
| **A_55_P2180764** | Gls | 0.281872975 | -0.058085222 | 0.621831173 | 9.322359355 | 2.014703363 | 0.089190764 | 0.506346569 | -4.170400568 |
| **A_55_P2015520** | Plekha1 | -0.328939677 | -0.725669486 | 0.067790133 | 10.95048682 | -2.014673234 | 0.089194533 | 0.506346569 | -4.170415501 |
| **A_55_P1995205** | Top2a | 0.409957032 | -0.08455707 | 0.904471135 | 11.17008855 | 2.014386713 | 0.089230387 | 0.506346569 | -4.17055752 |
| **A_55_P1997951** | Htatsf1 | 0.151875129 | -0.031351377 | 0.335101635 | 9.116676634 | 2.014102414 | 0.089265978 | 0.506346569 | -4.170698444 |
| **A_51_P136699** | Ttc7b | -0.204463295 | -0.451148267 | 0.042221678 | 9.429850185 | -2.013983229 | 0.089280902 | 0.506346569 | -4.170757524 |
| **A_66_P134412** | Mettl25 | 0.182728421 | -0.03776115 | 0.403217992 | 10.01998073 | 2.013729993 | 0.089312621 | 0.506346569 | -4.170883058 |
| **A_52_P551909** | Arhgap5 | -0.18286836 | -0.403527807 | 0.037791087 | 8.546435919 | -2.013720692 | 0.089313787 | 0.506346569 | -4.170887668 |
| **A_55_P2102515** | Daam1 | 0.328191978 | -0.067848521 | 0.724232476 | 9.652895681 | 2.013592344 | 0.089329867 | 0.506346569 | -4.170951295 |
| **A_55_P2125149** | Tulp1 | 0.629614017 | -0.130220882 | 1.389448916 | 11.63352267 | 2.01343837 | 0.089349163 | 0.506346569 | -4.171027626 |
| **A_55_P1953030** | Tnfsf12 | 0.20846586 | -0.043126702 | 0.460058421 | 8.785686352 | 2.013354887 | 0.089359626 | 0.506346569 | -4.171069013 |
| **A_55_P2154387** | Bmp4 | -0.982961067 | -2.169543791 | 0.203621658 | 9.641386038 | -2.012896683 | 0.089417078 | 0.506346569 | -4.171296178 |
| **A_55_P2072041** | NA | 0.184477094 | -0.038235385 | 0.407189573 | 8.699943673 | 2.012709453 | 0.089440564 | 0.506346569 | -4.171389006 |
| **A_55_P2008619** | Lancl2 | 0.204474006 | -0.042390965 | 0.451338978 | 10.78908025 | 2.012620186 | 0.089451764 | 0.506346569 | -4.171433265 |
| **A_66_P121583** | Tmem267 | 0.208558932 | -0.043267606 | 0.460385469 | 8.515970149 | 2.012382299 | 0.089481618 | 0.506346569 | -4.171551214 |
| **A_52_P163849** | Ivns1abp | -0.140709975 | -0.310674748 | 0.029254797 | 8.194840015 | -2.011634888 | 0.089575479 | 0.506346569 | -4.171921823 |
| **A_55_P2040425** | Guf1 | 0.193401926 | -0.040228381 | 0.427032232 | 8.56601948 | 2.011475767 | 0.089595475 | 0.506346569 | -4.172000729 |
| **A_51_P142923** | Chka | -0.255169825 | -0.563436494 | 0.053096843 | 9.101333521 | -2.011342064 | 0.08961228 | 0.506346569 | -4.172067033 |
| **A_55_P2016216** | NA | 0.159422637 | -0.03318869 | 0.352033964 | 11.98594848 | 2.011181914 | 0.089632413 | 0.506346569 | -4.172146453 |
| **A_52_P511381** | Tesk1 | -0.260538844 | -0.57534109 | 0.054263402 | 10.0050574 | -2.011026757 | 0.089651923 | 0.506346569 | -4.1722234 |
| **A_52_P180842** | Exoc3 | 0.168833845 | -0.035195765 | 0.372863456 | 9.554469631 | 2.010710292 | 0.089691729 | 0.506346569 | -4.172380348 |
| **A_51_P125691** | Cdca4 | 0.1945761 | -0.040659233 | 0.429811433 | 9.670713734 | 2.009879999 | 0.089796252 | 0.506346569 | -4.172792161 |
| **A_55_P2199118** | Bend4 | -0.352733018 | -0.779181543 | 0.073715506 | 8.52229136 | -2.009845563 | 0.08980059 | 0.506346569 | -4.172809242 |
| **A_52_P375493** | Tfdp1 | 0.21990922 | -0.045985902 | 0.485804342 | 10.30179302 | 2.009630703 | 0.089827659 | 0.506346569 | -4.172915819 |
| **A_51_P439299** | Relt | 0.295407261 | -0.061778168 | 0.652592691 | 8.854445047 | 2.009604622 | 0.089830945 | 0.506346569 | -4.172928756 |
| **A_51_P485899** | Srsf1 | 0.200652255 | -0.042002807 | 0.443307317 | 11.58528915 | 2.009268054 | 0.089873366 | 0.506346569 | -4.17309571 |
| **A_51_P299805** | Slc46a3 | 0.24725365 | -0.051773265 | 0.546280565 | 10.89784605 | 2.009165077 | 0.089886349 | 0.506346569 | -4.173146794 |
| **A_51_P150014** | Rrp7a | 0.267427144 | -0.056015091 | 0.59086938 | 8.750735973 | 2.009055555 | 0.08990016 | 0.506346569 | -4.173201125 |
| **A_55_P1998406** | NA | 0.312904842 | -0.065605663 | 0.691415347 | 9.2489764 | 2.008711357 | 0.089943576 | 0.506346569 | -4.173371878 |
| **A_51_P495171** | Pbdc1 | 0.371644061 | -0.07793235 | 0.821220472 | 11.63138211 | 2.008662032 | 0.089949799 | 0.506346569 | -4.173396349 |
| **A_66_P124674** | Ubxn2a | 0.17925645 | -0.037644155 | 0.396157055 | 8.844518375 | 2.008154939 | 0.090013806 | 0.506346569 | -4.173647931 |
| **A_51_P179921** | Igdcc3 | 0.182026364 | -0.038251781 | 0.402304509 | 8.286431681 | 2.007918461 | 0.09004367 | 0.506346569 | -4.17376526 |
| **A_51_P124606** | Nat2 | 0.190724979 | -0.040105421 | 0.42155538 | 9.163064136 | 2.007695104 | 0.090071887 | 0.506346569 | -4.173876084 |
| **A_51_P519555** | Gnb1l | 0.207712854 | -0.043680561 | 0.459106268 | 10.48631542 | 2.007671551 | 0.090074863 | 0.506346569 | -4.17388777 |
| **A_51_P167763** | Kcnk13 | 0.198874712 | -0.041829276 | 0.439578701 | 9.031016819 | 2.007610542 | 0.090082572 | 0.506346569 | -4.173918042 |
| **A_51_P386612** | Pir | 0.23799742 | -0.05006527 | 0.52606011 | 9.983659991 | 2.007559499 | 0.090089022 | 0.506346569 | -4.173943369 |
| **A_52_P424826** | Txnl4b | 0.155237014 | -0.03266044 | 0.343134468 | 10.61989085 | 2.007509342 | 0.090095361 | 0.506346569 | -4.173968256 |
| **A_51_P469285** | Nrp1 | -0.413996815 | -0.915130232 | 0.087136602 | 8.375329215 | -2.007367207 | 0.090113326 | 0.506346569 | -4.174038784 |
| **A_55_P2028961** | Idi1 | 0.314562991 | -0.066231161 | 0.695357142 | 13.64989697 | 2.007245738 | 0.090128682 | 0.506346569 | -4.174099058 |
| **A_66_P134674** | Far1 | 0.208368548 | -0.043898125 | 0.460635221 | 8.778386639 | 2.00703744 | 0.090155021 | 0.506346569 | -4.17420242 |
| **A_55_P2117155** | Apoc3 | -0.747711641 | -1.653051188 | 0.157627907 | 10.98077936 | -2.006808072 | 0.090184033 | 0.506346569 | -4.174316241 |
| **A_55_P2117375** | NA | 0.174717438 | -0.036847879 | 0.386282756 | 8.527365685 | 2.006665427 | 0.090202081 | 0.506346569 | -4.174387029 |
| **A_55_P2182736** | Limd2 | -0.245443053 | -0.542693008 | 0.051806902 | 9.378424254 | -2.006375133 | 0.09023882 | 0.506346569 | -4.174531092 |
| **A_55_P1980566** | Ndufb11 | -0.233513584 | -0.516329072 | 0.049301904 | 14.892681 | -2.006282797 | 0.090250509 | 0.506346569 | -4.174576916 |
| **A_55_P2121275** | Gm4907 | -0.415374618 | -0.918502813 | 0.087753577 | 8.770372966 | -2.006062632 | 0.090278387 | 0.506346569 | -4.174686183 |
| **A_52_P337220** | Mgat4b | -0.15539927 | -0.343644265 | 0.032845725 | 8.62823621 | -2.005897445 | 0.090299309 | 0.506346569 | -4.174768166 |
| **A_55_P2039225** | Rap1gap | -0.292245608 | -0.646281316 | 0.0617901 | 9.45943119 | -2.005783779 | 0.090313708 | 0.506346569 | -4.174824581 |
| **A_51_P397548** | Mtrf1l | 0.172302004 | -0.036458905 | 0.381062914 | 9.764254178 | 2.005507718 | 0.090348689 | 0.506346569 | -4.174961599 |

| | | | | | | | | | |
|---|---|---|---|---|---|---|---|---|---|
| **A_51_P160413** | Yod1 | 0.410691725 | -0.086964646 | 0.908348096 | 9.879117872 | 2.00525484 | 0.090380745 | 0.506346569 | -4.175087115 |
| **A_55_P2144248** | Zfp790 | 0.799695551 | -0.169433205 | 1.768824307 | 9.490088696 | 2.005055455 | 0.090406027 | 0.506346569 | -4.175186084 |
| **A_55_P2077009** | Acap3 | 0.196208304 | -0.041575957 | 0.433992564 | 10.49478472 | 2.005014269 | 0.090411251 | 0.506346569 | -4.175206527 |
| **A_55_P1994302** | NA | 0.297050309 | -0.06296966 | 0.657070277 | 8.732183217 | 2.0048718 | 0.090429322 | 0.506346569 | -4.175277247 |
| **A_55_P1980080** | Rbms2 | -0.298155742 | -0.659627205 | 0.063315721 | 9.664366346 | -2.00425211 | 0.090507966 | 0.506640162 | -4.175584869 |
| **A_51_P103650** | Trim8 | -0.189775199 | -0.419876205 | 0.040325808 | 11.2419339 | -2.004029568 | 0.090536226 | 0.506651623 | -4.179066228 |
| **A_52_P279599** | Ube2k | 0.258973844 | -0.055073114 | 0.573020802 | 10.23828016 | 2.003754456 | 0.090571173 | 0.506700492 | -4.175831931 |
| **A_52_P584078** | Pop1 | 0.297111717 | -0.06328766 | 0.657511093 | 10.54909158 | 2.00317521 | 0.090644799 | 0.506858629 | -4.176119523 |
| **A_52_P1029978** | Lasp1 | -0.176268908 | -0.390090741 | 0.037552925 | 13.22017471 | -2.003119485 | 0.090651885 | 0.506858629 | -4.176147191 |
| **A_55_P2199040** | NA | 0.162076 | -0.034574283 | 0.358726282 | 8.336562477 | 2.002660458 | 0.090710277 | 0.506882868 | -4.176375115 |
| **A_55_P2170225** | Sh3bgrl2 | 0.167652287 | -0.035805022 | 0.371109597 | 8.209573156 | 2.002254943 | 0.090761894 | 0.506882868 | -4.176576481 |
| **A_55_P2100305** | Lrba | 0.254219983 | -0.054299322 | 0.562739289 | 11.38729514 | 2.002214166 | 0.090767086 | 0.506882868 | -4.17659673 |
| **A_66_P116154** | Cstf2 | 0.260281161 | -0.055597675 | 0.576159997 | 9.524103135 | 2.002190492 | 0.090770101 | 0.506882868 | -4.176608487 |
| **A_55_P2141884** | Tagln2 | -0.46831283 | -1.03675189 | 0.10012623 | 9.959621064 | -2.001867514 | 0.090811236 | 0.506882868 | -4.176768877 |
| **A_55_P2156062** | NA | -0.149115014 | -0.330115487 | 0.031885459 | 8.158629167 | -2.001819214 | 0.090817389 | 0.506882868 | -4.176792864 |
| **A_52_P595824** | Cpeb4 | 0.34497733 | -0.073803729 | 0.763758389 | 8.663522699 | 2.001643429 | 0.090839788 | 0.506882868 | -4.176880162 |
| **A_51_P340704** | Prpf31 | 0.187856619 | -0.040252756 | 0.415965993 | 12.42387357 | 2.001089699 | 0.090910379 | 0.506978541 | -4.177155171 |
| **A_51_P447976** | Tent5c | 0.226997708 | -0.048644271 | 0.502639686 | 14.42128684 | 2.001056276 | 0.090914641 | 0.506978541 | -4.177171772 |
| **A_55_P2065449** | Kdm5b | -0.385102746 | -0.852769178 | 0.082563687 | 11.16872313 | -2.000891805 | 0.09093562 | 0.506978541 | -4.177253461 |
| **A_55_P2006762** | Bzw1 | 0.183125444 | -0.039295062 | 0.40554595 | 12.12500085 | 2.000585214 | 0.09097474 | 0.507050389 | -4.177405743 |
| **A_66_P107353** | NA | 0.197944438 | -0.042581536 | 0.438470412 | 15.99104706 | 1.999698449 | 0.091087983 | 0.507069226 | -4.177846233 |
| **A_55_P2077746** | Mphosph8 | 0.194700499 | -0.041893636 | 0.431294634 | 10.52588326 | 1.999614504 | 0.09109871 | 0.507069226 | -4.177887935 |
| **A_51_P323521** | Snx21 | -0.190427383 | -0.42185177 | 0.040997004 | 10.98014243 | -1.999417398 | 0.091123903 | 0.507069226 | -4.177985854 |
| **A_55_P2087265** | NA | -0.711908137 | -1.577101345 | 0.153285071 | 9.855839481 | -1.999374021 | 0.091129449 | 0.507069226 | -4.178007403 |
| **A_55_P2040559** | NA | 0.212188758 | -0.045698579 | 0.470076096 | 15.92626886 | 1.99928884 | 0.091140339 | 0.507069226 | -4.178049721 |
| **A_51_P257885** | Mmd2 | 0.386024531 | -0.083191482 | 0.855240545 | 9.917337908 | 1.999057417 | 0.091169933 | 0.507069226 | -4.178164694 |
| **A_55_P2158448** | Gdf3 | 0.258909864 | -0.055947718 | 0.573767447 | 8.518750134 | 1.998101884 | 0.091292227 | 0.507069226 | -4.178639453 |
| **A_55_P2250296** | 6330566A10Rik | 0.20018032 | -0.043257381 | 0.443618021 | 8.488829136 | 1.998097747 | 0.091292757 | 0.507069226 | -4.178641509 |
| **A_55_P2028159** | C9orf72 | -0.157003748 | -0.34797238 | 0.033964884 | 10.00687671 | -1.997704155 | 0.091343179 | 0.507069226 | -4.178837086 |
| **A_52_P513123** | Dna2 | 0.311490609 | -0.067434956 | 0.690416173 | 9.077527791 | 1.997442286 | 0.091376742 | 0.507069226 | -4.178967215 |
| **A_51_P272817** | Map2k3 | 0.32294762 | -0.069943415 | 0.715838655 | 10.14334166 | 1.997299339 | 0.091395068 | 0.507069226 | -4.179038251 |
| **A_55_P2175692** | Reep3 | -0.323975712 | -0.718128126 | 0.070176703 | 10.75866479 | -1.997245497 | 0.091401972 | 0.507069226 | -4.179065008 |
| **A_52_P467690** | Sptbn1 | -0.35089785 | -0.777826965 | 0.076031264 | 9.67142367 | -1.997138224 | 0.091415728 | 0.507069226 | -4.179118318 |
| **A_55_P2035972** | Mark3 | 0.193227932 | -0.04187816 | 0.428334024 | 10.52603676 | 1.997051257 | 0.091426882 | 0.507069226 | -4.179161537 |
| **A_55_P2041070** | Tle3 | -0.280638266 | -0.622117628 | 0.060841096 | 11.0040332 | -1.996942756 | 0.091440799 | 0.507069226 | -4.179215459 |
| **A_52_P121342** | Nrxn1 | -0.190038295 | -0.421288937 | 0.041212347 | 7.941753066 | -1.996831264 | 0.091455103 | 0.507069226 | -4.179270867 |
| **A_55_P2092859** | Anxa4 | -0.15037077 | -0.333359922 | 0.032618383 | 11.42606907 | -1.996738843 | 0.091466961 | 0.507069226 | -4.179316799 |
| **A_55_P1952359** | Anapc5 | 0.174664489 | -0.037895958 | 0.387224935 | 14.58418309 | 1.996665671 | 0.091476351 | 0.507069226 | -4.179353165 |
| **A_51_P239924** | Polr2c | -0.192354198 | -0.426442579 | 0.041734183 | 12.47718293 | -1.996664057 | 0.091476558 | 0.507069226 | -4.179353967 |
| **A_55_P2086423** | Recql4 | 0.378626662 | -0.082408439 | 0.839661764 | 9.457068127 | 1.995539781 | 0.091620952 | 0.507530293 | -4.17991277 |
| **A_55_P2129477** | NA | 0.246818716 | -0.053731098 | 0.54736853 | 8.965645128 | 1.995468217 | 0.091630151 | 0.507530293 | -4.179948342 |
| **A_55_P2076134** | NA | 0.26809125 | -0.058372648 | 0.594555149 | 12.87643318 | 1.995403211 | 0.091638508 | 0.507530293 | -4.179980656 |
| **A_55_P2021029** | Tigd3 | 0.198476492 | -0.043271714 | 0.440224698 | 8.489613029 | 1.994936172 | 0.09169857 | 0.507607474 | -4.180212821 |
| **A_55_P2061219** | Ces3a | -0.381006546 | -0.845091454 | 0.083078362 | 8.562476488 | -1.994886452 | 0.091704967 | 0.507607474 | -4.180237538 |
| **A_55_P2013986** | Ddx31 | 0.148417203 | -0.032381796 | 0.329216201 | 9.327099686 | 1.994671629 | 0.091732608 | 0.507610563 | -4.180344333 |
| **A_55_P1976242** | Arhgef7 | -0.293502017 | -0.651075993 | 0.064071959 | 12.28024515 | -1.99447398 | 0.091758048 | 0.507610563 | -4.180442593 |
| **A_55_P2021119** | Sun2 | -0.27033237 | -0.599778714 | 0.059113975 | 12.72912071 | -1.993868659 | 0.091836003 | 0.507896452 | -4.180743542 |
| **A_55_P2141585** | NA | -0.147181997 | -0.326571566 | 0.032207572 | 11.16371891 | -1.993612248 | 0.091869044 | 0.507933855 | -4.180871031 |
| **A_55_P2078831** | NA | 0.222971377 | -0.048872794 | 0.494815547 | 9.750541531 | 1.993022861 | 0.091945039 | 0.507967689 | -4.181164095 |
| **A_52_P275627** | Kpnb1 | 0.235462716 | -0.051639725 | 0.522565157 | 10.90811621 | 1.992821748 | 0.091970984 | 0.507967689 | -4.181264101 |

| | | | | | | | | | |
|---|---|---|---|---|---|---|---|---|---|
| **A_55_P1960479** | NA | 0.216416934 | -0.047467721 | 0.480301589 | 11.76258884 | 1.992784296 | 0.091975817 | 0.507967689 | -4.181282724 |
| **A_51_P424054** | Btbd2 | -0.190095919 | -0.421890477 | 0.041698639 | 9.68164991 | -1.992749677 | 0.091980284 | 0.507967689 | -4.18129994 |
| **A_55_P1991075** | Lmf1 | -0.175067757 | -0.388585955 | 0.038450442 | 11.18026239 | -1.992298738 | 0.092038492 | 0.508143964 | -4.181524188 |
| **A_52_P282762** | Myd88 | -0.179821984 | -0.399212362 | 0.039568394 | 9.701845394 | -1.991628797 | 0.092125038 | 0.508476546 | -4.185160205 |
| **A_51_P513530** | Spag5 | 0.388505062 | -0.085603363 | 0.862613487 | 10.81408151 | 1.991141839 | 0.092187997 | 0.508546856 | -4.182099572 |
| **A_55_P2297485** | Ago2 | 0.373038086 | -0.082199606 | 0.828275777 | 11.93770414 | 1.991123283 | 0.092190397 | 0.508546856 | -4.182108802 |
| **A_51_P423008** | Klhl7 | 0.300720906 | -0.066346692 | 0.667788504 | 9.092228036 | 1.990676825 | 0.092248159 | 0.508720303 | -4.182330874 |
| **A_51_P270184** | Nhlrc2 | 0.158016159 | -0.034945829 | 0.350978148 | 10.04255116 | 1.989816031 | 0.092359629 | 0.509048892 | -4.182759081 |
| **A_55_P1975510** | H6pd | -0.314639455 | -0.698888443 | 0.069609534 | 12.93781768 | -1.989681843 | 0.092377018 | 0.509048892 | -4.182825838 |
| **A_66_P134294** | Uba1y | 0.327890829 | -0.072615064 | 0.728396722 | 9.133241812 | 1.989314956 | 0.09242458 | 0.509048892 | -4.183008368 |
| **A_51_P509999** | Prl8a9 | -0.579127764 | -1.286590055 | 0.128334527 | 8.150264494 | -1.989089291 | 0.092453846 | 0.509048892 | -4.183120643 |
| **A_51_P519251** | Nupr1 | -1.031308941 | -2.291212867 | 0.228594986 | 9.6637741 | -1.988998898 | 0.092465571 | 0.509048892 | -4.183165617 |
| **A_55_P1999633** | Cit | 0.348548374 | -0.077257869 | 0.774354616 | 8.986312522 | 1.988997448 | 0.092465759 | 0.509048892 | -4.183166339 |
| **A_55_P2060487** | Zfp593 | 0.230245631 | -0.051087832 | 0.511579094 | 9.785500687 | 1.988626376 | 0.092513909 | 0.50916895 | -4.183350969 |
| **A_55_P2157408** | NA | 0.222430968 | -0.049393891 | 0.494255828 | 8.255287325 | 1.98833368 | 0.092551907 | 0.509233081 | -4.183496609 |
| **A_55_P1992329** | Gmpr | 0.391307336 | -0.087055507 | 0.869670179 | 10.08167565 | 1.987667519 | 0.092638447 | 0.509564181 | -4.183828102 |
| **A_55_P2076176** | NA | -0.257808924 | -0.573005937 | 0.05738809 | 15.00066226 | -1.987462949 | 0.092665038 | 0.50956544 | -4.183929906 |
| **A_51_P235726** | Icam4 | 0.382803712 | -0.085266388 | 0.850873811 | 12.23850843 | 1.987231351 | 0.092695152 | 0.509586063 | -4.184045163 |
| **A_66_P111426** | Lce1a2 | 0.355019949 | -0.079137387 | 0.789177284 | 8.982661275 | 1.986958528 | 0.092730639 | 0.509636203 | -4.184180942 |
| **A_55_P2192662** | Lepr | -0.206170541 | -0.458404288 | 0.046063206 | 8.031678543 | -1.986125139 | 0.092839126 | 0.509976349 | -4.184595737 |
| **A_55_P2136752** | Ermn | 0.216713046 | -0.048424968 | 0.48185106 | 10.92666736 | 1.98607775 | 0.092845298 | 0.509976349 | -4.184619324 |
| **A_55_P2041626** | Ceacam12 | -0.534779729 | -1.189258819 | 0.11969936 | 8.120285759 | -1.985465525 | 0.092925081 | 0.510222243 | -4.184924075 |
| **A_51_P354038** | Tmc6 | 0.341075436 | -0.076385018 | 0.758535889 | 11.65621391 | 1.985264109 | 0.092951344 | 0.510222243 | -4.18502434 |
| **A_55_P2044297** | Ciao2a | 0.178078783 | -0.03995074 | 0.396108305 | 13.16262431 | 1.984632337 | 0.09303377 | 0.510222243 | -4.185338857 |
| **A_51_P296608** | Gadd45a | 0.370957044 | -0.083251319 | 0.825165406 | 11.09177269 | 1.984502659 | 0.093050698 | 0.510222243 | -4.185403419 |
| **A_51_P400752** | NA | -0.277762637 | -0.61788801 | 0.062362735 | 12.98190462 | -1.984348677 | 0.093070802 | 0.510222243 | -4.185480082 |
| **A_55_P1977144** | Cyp2c67 | -0.613104952 | -1.363892256 | 0.137682353 | 14.56711818 | -1.98427149 | 0.093080881 | 0.510222243 | -4.185518512 |
| **A_66_P136857** | Phrf1 | 0.19302148 | -0.043379604 | 0.429422565 | 10.06409078 | 1.983989485 | 0.093117716 | 0.510222243 | -4.18565892 |
| **A_51_P338317** | Rhoq | -0.398247777 | -0.886050376 | 0.089554822 | 9.597638879 | -1.98377512 | 0.093145726 | 0.510222243 | -4.185765654 |
| **A_51_P363634** | Snrpe | 0.178095536 | -0.040106511 | 0.396297583 | 12.38987669 | 1.983249721 | 0.093214412 | 0.510222243 | -4.186027268 |
| **A_51_P515623** | Qpctl | -0.132458779 | -0.29476597 | 0.029848411 | 8.834393651 | -1.983015811 | 0.093245008 | 0.510222243 | -4.186143746 |
| **A_55_P2085425** | Ophn1 | -0.235608649 | -0.524312615 | 0.053095318 | 8.888200955 | -1.982995215 | 0.093247702 | 0.510222243 | -4.186154003 |
| **A_55_P1965076** | NA | -0.197974367 | -0.440580373 | 0.044631639 | 12.65122097 | -1.982853386 | 0.093266259 | 0.510222243 | -4.18622463 |
| **A_55_P2063216** | Dusp14 | -0.412954723 | -0.919014146 | 0.0931047 | 8.33406759 | -1.982823744 | 0.093270138 | 0.510222243 | -4.186239391 |
| **A_51_P517430** | Cd1d1 | -0.492417473 | -1.095889853 | 0.111054907 | 10.06081217 | -1.982710226 | 0.093284994 | 0.510222243 | -4.186295922 |
| **A_66_P136801** | Peg13 | -0.271323474 | -0.603839838 | 0.06119289 | 9.970802513 | -1.982702416 | 0.093286016 | 0.510222243 | -4.186299811 |
| **A_55_P2016618** | Neurl1a | -0.206671128 | -0.460111343 | 0.046769086 | 9.599457087 | -1.981469868 | 0.093447473 | 0.510278114 | -4.186913663 |
| **A_55_P2056496** | Tk1 | 0.327079261 | -0.074031017 | 0.72818954 | 12.58532854 | 1.981401128 | 0.093456486 | 0.510278114 | -4.186947901 |
| **A_55_P2157102** | NA | -0.162399012 | -0.361573219 | 0.036775195 | 14.06231047 | -1.981223519 | 0.093479777 | 0.510278114 | -4.187036367 |
| **A_55_P2011943** | Gramd1a | -0.261161465 | -0.581553613 | 0.059230682 | 10.29768495 | -1.980662242 | 0.093553419 | 0.510278114 | -4.187315947 |
| **A_55_P2059606** | Stat2 | -0.181053587 | -0.403198285 | 0.041091112 | 10.30170998 | -1.980406615 | 0.093586978 | 0.510278114 | -4.187443286 |
| **A_55_P2093696** | NA | 0.287516765 | -0.065270831 | 0.640304361 | 9.041597819 | 1.980309428 | 0.09359974 | 0.510278114 | -4.1874917 |
| **A_55_P1994564** | Sap30bp | 0.153009775 | -0.034741776 | 0.340761325 | 8.738018203 | 1.980244573 | 0.093608258 | 0.510278114 | -4.187524009 |
| **A_66_P101107** | NA | 0.186581123 | -0.04238295 | 0.415545197 | 9.127961381 | 1.980083705 | 0.093629387 | 0.510278114 | -4.187604149 |
| **A_55_P2174258** | Pin4 | 0.186133833 | -0.042285796 | 0.414553462 | 12.06325588 | 1.980045126 | 0.093634455 | 0.510278114 | -4.187623367 |
| **A_55_P2028661** | Dusp12 | 0.21651843 | -0.049211849 | 0.482248709 | 11.96091761 | 1.979871546 | 0.093657262 | 0.510278114 | -4.187709842 |
| **A_55_P2149382** | Gm3448 | 0.255551615 | -0.058112479 | 0.569215709 | 8.950230439 | 1.979689193 | 0.093681226 | 0.510278114 | -4.18780069 |
| **A_55_P2061909** | Tceal9 | 0.212641081 | -0.048356705 | 0.473638868 | 12.85541354 | 1.979673353 | 0.093683308 | 0.510278114 | -4.187808582 |
| **A_55_P1964658** | NA | 0.239762092 | -0.054547377 | 0.534071561 | 12.18726527 | 1.97951808 | 0.09370372 | 0.510278114 | -4.18788594 |
| **A_52_P11174** | Hip1 | -0.266059779 | -0.592650575 | 0.060531018 | 8.964719142 | -1.979513521 | 0.093704319 | 0.510278114 | -4.187888211 |

| | | | | | | | | | |
|---|---|---|---|---|---|---|---|---|---|
| **A_51_P138044** | Foxo1 | -0.279335284 | -0.622234818 | 0.06356425 | 10.4838725 | -1.979438853 | 0.093714136 | 0.510278114 | -4.187925412 |
| **A_55_P2097578** | NA | 0.292087776 | -0.066472333 | 0.650647884 | 11.78862357 | 1.979404713 | 0.093718625 | 0.510278114 | -4.187942422 |
| **A_55_P1973254** | Cmtm7 | -0.136443074 | -0.303969782 | 0.031083634 | 11.39936537 | -1.979022038 | 0.093768957 | 0.510408381 | -4.188133085 |
| **A_51_P327751** | Ifit1 | -0.13864499 | -0.308977737 | 0.031687756 | 8.035911458 | -1.97783119 | 0.093925756 | 0.511097203 | -4.188726473 |
| **A_55_P1995874** | Gm14410 | -0.156286737 | -0.348309979 | 0.035736504 | 8.408290057 | -1.977659485 | 0.093948387 | 0.511097203 | -4.188812041 |
| **A_51_P161265** | Ccl25 | 0.42401484 | -0.097186642 | 0.945216321 | 9.551719002 | 1.976781209 | 0.094064226 | 0.511583407 | -4.189249752 |
| **A_55_P2008427** | Dnd1 | 0.174819859 | -0.040141272 | 0.38978099 | 8.369234455 | 1.976123201 | 0.094151108 | 0.511698563 | -4.189577722 |
| **A_55_P2061333** | NA | 0.272888837 | -0.062690154 | 0.608467828 | 10.9217708 | 1.975942011 | 0.094175046 | 0.511698563 | -4.189668037 |
| **A_51_P370252** | Capn10 | -0.334275778 | -0.745396723 | 0.076845168 | 11.04137302 | -1.975688597 | 0.094208536 | 0.511698563 | -4.189794357 |
| **A_55_P1965233** | Sh3tc1 | -0.310980704 | -0.693461888 | 0.07150048 | 9.366677979 | -1.975634294 | 0.094215714 | 0.511698563 | -4.189821426 |
| **A_55_P2021378** | Wapl | 0.232913211 | -0.053573269 | 0.519399692 | 11.5540647 | 1.97548247 | 0.094235785 | 0.511698563 | -4.189897109 |
| **A_55_P1957875** | NA | 0.22613632 | -0.052023492 | 0.504296133 | 9.825658385 | 1.975418538 | 0.094244238 | 0.511698563 | -4.189928979 |
| **A_52_P352362** | NA | 0.186732678 | -0.043005312 | 0.416470668 | 8.697639779 | 1.975016365 | 0.094297432 | 0.511843604 | -4.190129468 |
| **A_55_P1956882** | Cpt1b | -0.334163279 | -0.745334535 | 0.077007977 | 8.590422622 | -1.974782027 | 0.094328441 | 0.511868176 | -4.190246293 |
| **A_55_P2043277** | Siglecf | 0.280191366 | -0.064701588 | 0.62508432 | 8.473469764 | 1.974029393 | 0.094428103 | 0.512121646 | -4.190621534 |
| **A_52_P380369** | Phf11d | -0.188144364 | -0.419758909 | 0.04347018 | 8.052506194 | -1.973824681 | 0.094455229 | 0.512121646 | -4.190723604 |
| **A_52_P134666** | Srpr | -0.194137979 | -0.433151036 | 0.044875079 | 10.36142601 | -1.97365875 | 0.094477222 | 0.512121646 | -4.19080634 |
| **A_51_P508191** | Vps26c | -0.143411004 | -0.319974163 | 0.033152155 | 11.12102225 | -1.973629257 | 0.094481132 | 0.512121646 | -4.190821046 |
| **A_55_P2021892** | Ccdc114 | 0.150120679 | -0.034735668 | 0.334977025 | 8.810982946 | 1.973282854 | 0.094527063 | 0.512226968 | -4.190993775 |
| **A_51_P486155** | Ocln | -0.287561719 | -0.641836782 | 0.066713343 | 8.678851891 | -1.972303189 | 0.094657082 | 0.512527862 | -4.191482315 |
| **A_52_P84901** | Slc44a1 | -0.161291347 | -0.360012646 | 0.037429953 | 9.18822825 | -1.972194931 | 0.094671461 | 0.512527862 | -4.191536305 |
| **A_52_P78023** | Limd2 | -0.228676095 | -0.510443368 | 0.053091177 | 13.46857027 | -1.972029507 | 0.094693437 | 0.512527862 | -4.191618807 |
| **A_55_P2115582** | Slc20a1 | 0.383207742 | -0.088974643 | 0.855390128 | 12.90161588 | 1.972003504 | 0.094696892 | 0.512527862 | -4.191631775 |
| **A_55_P2167112** | Kank2 | -0.425364256 | -0.949591744 | 0.098863232 | 11.36921638 | -1.971625343 | 0.09474715 | 0.512527862 | -4.191820383 |
| **A_55_P2021665** | Zfp638 | 0.158468588 | -0.036857082 | 0.353794258 | 10.17793276 | 1.971365062 | 0.094781757 | 0.512527862 | -4.191950203 |
| **A_51_P475378** | Echdc3 | -0.474003126 | -1.058302532 | 0.11029628 | 12.25228796 | -1.9711921 | 0.094804761 | 0.512527862 | -4.192036474 |
| **A_55_P1962334** | Trim71 | 0.258720039 | -0.06022413 | 0.577664208 | 8.225930353 | 1.97105431 | 0.094823091 | 0.512527862 | -4.192105203 |
| **A_55_P1968993** | NA | 0.312434743 | -0.072728008 | 0.697597494 | 11.30261225 | 1.971052676 | 0.094823309 | 0.512527862 | -4.192106018 |
| **A_52_P251623** | Fam222a | 0.210683055 | -0.049066636 | 0.470432746 | 8.66499337 | 1.970868989 | 0.09484775 | 0.512527862 | -4.192197642 |
| **A_51_P300337** | Csrp2 | -0.715260382 | -1.59731404 | 0.166793277 | 11.70880498 | -1.970390721 | 0.094911419 | 0.512693703 | -4.192436216 |
| **A_52_P665202** | NA | 0.202135358 | -0.047155494 | 0.451426211 | 14.38614581 | 1.970240016 | 0.09493149 | 0.512693703 | -4.192511396 |
| **A_55_P2279997** | 4930549C15Rik | 0.304103939 | -0.070998944 | 0.679206822 | 8.6397234 | 1.969948319 | 0.09497035 | 0.512760308 | -4.192656913 |
| **A_55_P2061278** | Sik1 | -0.258179715 | -0.576721255 | 0.060361824 | 9.312791294 | -1.969424026 | 0.095040238 | 0.512874622 | -4.19291848 |
| **A_52_P241508** | Zbtb46 | 0.283641795 | -0.066328485 | 0.633612075 | 8.760913675 | 1.969347015 | 0.095050508 | 0.512874622 | -4.192956901 |
| **A_55_P2051825** | Ephx3 | 0.393810673 | -0.092145811 | 0.879767158 | 10.1885779 | 1.969125019 | 0.095080119 | 0.512874622 | -4.193067661 |
| **A_51_P484158** | Steap1 | 0.239722223 | -0.056114322 | 0.535558767 | 8.539315863 | 1.968972561 | 0.09510046 | 0.512874622 | -4.193143728 |
| **A_55_P1986993** | Brd8 | 0.136377373 | -0.03196032 | 0.304715065 | 8.199889254 | 1.968539521 | 0.095158259 | 0.512874622 | -4.193359796 |
| **A_55_P2003638** | Stxbp6 | 0.200240725 | -0.046951765 | 0.447433215 | 8.927557615 | 1.968340937 | 0.095184777 | 0.512874622 | -4.193458885 |
| **A_55_P2085546** | Dnaja4 | 0.2535246 | -0.059458984 | 0.566508185 | 10.83604863 | 1.968256689 | 0.095196029 | 0.512874622 | -4.193500924 |
| **A_55_P2130575** | NA | 0.363431803 | -0.0852926 | 0.812156207 | 14.49107744 | 1.968006051 | 0.095229512 | 0.512874622 | -4.193625992 |
| **A_55_P2084739** | Gcnt1 | 0.320624914 | -0.075279825 | 0.716529654 | 9.407919477 | 1.967839866 | 0.095251719 | 0.512874622 | -4.19370892 |
| **A_55_P2091206** | NA | 0.275268976 | -0.064708932 | 0.615246884 | 10.10565293 | 1.967386826 | 0.095312285 | 0.512874622 | -4.193935003 |
| **A_51_P435333** | Dctn2 | -0.289370461 | -0.646776965 | 0.068036043 | 10.67822512 | -1.967319669 | 0.095321266 | 0.512874622 | -4.193968518 |
| **A_55_P1954643** | 2310081J21Rik | 0.461710496 | -0.10858028 | 1.032001273 | 9.162488128 | 1.96723657 | 0.095332381 | 0.512874622 | -4.194009989 |
| **A_55_P2152049** | Xrcc3 | 0.290262296 | -0.068379599 | 0.648904191 | 9.159779738 | 1.966585329 | 0.09541953 | 0.512874622 | -4.194335012 |
| **A_55_P2141860** | Aen | 0.208449505 | -0.049120286 | 0.466019296 | 11.91025214 | 1.966478179 | 0.095433877 | 0.512874622 | -4.194388491 |
| **A_51_P115159** | Fam162a | 0.137839906 | -0.032507343 | 0.308187155 | 13.87543773 | 1.966178908 | 0.095473958 | 0.512874622 | -4.201046571 |
| **A_51_P357433** | Ap1g2 | 0.274427201 | -0.064731 | 0.613585403 | 9.239207039 | 1.966110954 | 0.095483062 | 0.512874622 | -4.194571782 |
| **A_52_P680710** | Kpna3 | 0.273442656 | -0.064507975 | 0.611393287 | 10.03589177 | 1.966057394 | 0.095490237 | 0.512874622 | -4.194598516 |
| **A_55_P2419483** | 4732460I02Rik | 0.240810364 | -0.056815294 | 0.538436022 | 8.310718779 | 1.966020282 | 0.09549521 | 0.512874622 | -4.19461704 |

| | | | | | | | | | |
|---|---|---|---|---|---|---|---|---|---|
| **A_55_P1958517** | Parp6 | -0.337920373 | -0.755568297 | 0.07972755 | 9.716642238 | -1.966016864 | 0.095495668 | 0.512874622 | -4.194618746 |
| **A_51_P206346** | Reep3 | -0.274642933 | -0.614261791 | 0.064975926 | 11.17091441 | -1.96498762 | 0.095633674 | 0.513146057 | -4.195132521 |
| **A_51_P453995** | Gm11961 | 0.254786537 | -0.060336345 | 0.569909419 | 9.236971437 | 1.964625268 | 0.095682308 | 0.513146057 | -4.195313416 |
| **A_51_P440047** | Lgalsl | -0.24162956 | -0.540495554 | 0.057236435 | 10.36566444 | -1.96452133 | 0.095696263 | 0.513146057 | -4.195365306 |
| **A_55_P1994229** | Hrc | -0.373500515 | -0.835474981 | 0.088473951 | 8.447269145 | -1.964519854 | 0.095696461 | 0.513146057 | -4.195366043 |
| **A_55_P1992814** | Shq1 | 0.173272421 | -0.041082443 | 0.387627285 | 8.468233128 | 1.964170974 | 0.095743317 | 0.513146057 | -4.195540223 |
| **A_55_P2095133** | Synpo2 | -0.234492911 | -0.524621037 | 0.055635215 | 7.823668949 | -1.963916751 | 0.095777475 | 0.513146057 | -4.19566715 |
| **A_51_P382688** | Zmynd19 | 0.208662261 | -0.049523277 | 0.4668478 | 10.31653561 | 1.963790639 | 0.095794424 | 0.513146057 | -4.195730117 |
| **A_52_P218181** | Zdhhc5 | 0.172730601 | -0.040996391 | 0.386457593 | 11.67106764 | 1.963781204 | 0.095795692 | 0.513146057 | -4.195734827 |
| **A_52_P514306** | Spata2 | 0.155088827 | -0.036899616 | 0.34707727 | 9.088394897 | 1.962856789 | 0.095920023 | 0.513146057 | -4.19619641 |
| **A_55_P2115607** | Ikbip | -0.25360225 | -0.567549556 | 0.060345055 | 10.00553644 | -1.96281574 | 0.095925548 | 0.513146057 | -4.196216908 |
| **A_55_P2002752** | Zswim6 | -0.224381767 | -0.502159527 | 0.053395993 | 8.277756551 | -1.962787487 | 0.095929351 | 0.513146057 | -4.196231016 |
| **A_55_P2082271** | NA | 0.237592466 | -0.056603263 | 0.531788195 | 15.59674241 | 1.962363649 | 0.095986415 | 0.513146057 | -4.19644267 |
| **A_55_P2078645** | Mettl5 | 0.191185919 | -0.045571742 | 0.427943579 | 10.27380811 | 1.962162863 | 0.09601346 | 0.513146057 | -4.196542941 |
| **A_51_P483118** | Hmga1 | 0.160107391 | -0.038185221 | 0.358400002 | 14.67094658 | 1.961950459 | 0.096042079 | 0.513146057 | -4.196649018 |
| **A_55_P2152437** | Mitf | -0.191308577 | -0.428280625 | 0.04566347 | 8.222990721 | -1.961645425 | 0.096083193 | 0.513146057 | -4.19680136 |
| **A_52_P227937** | Cystm1 | -0.268455186 | -0.601124409 | 0.064214037 | 9.418583563 | -1.960840931 | 0.096191711 | 0.513146057 | -4.197203175 |
| **A_55_P1987650** | Ctnnd2 | -0.40033033 | -0.896512065 | 0.095851404 | 7.937658765 | -1.960473205 | 0.096241355 | 0.513146057 | -4.197386855 |
| **A_51_P202074** | Ncapd2 | 0.288099643 | -0.068988555 | 0.645187841 | 13.02112569 | 1.960425816 | 0.096247754 | 0.513146057 | -4.197410526 |
| **A_51_P487360** | Hpcal1 | -0.396193983 | -0.887288163 | 0.094900196 | 10.33906991 | -1.960316874 | 0.096262467 | 0.513146057 | -4.197464945 |
| **A_55_P2049549** | Rab22a | 0.198494014 | -0.04757055 | 0.444558578 | 10.03509153 | 1.960114891 | 0.096289752 | 0.513146057 | -4.197565842 |
| **A_55_P1985015** | Pard3 | -0.185636733 | -0.415766953 | 0.044493486 | 9.117731268 | -1.960078455 | 0.096294675 | 0.513146057 | -4.197584043 |
| **A_55_P2144364** | NA | 0.164926033 | -0.039569784 | 0.36942185 | 8.190076585 | 1.959692696 | 0.096346809 | 0.513146057 | -4.19777675 |
| **A_55_P2087805** | Hsd3b1 | -0.720028194 | -1.612836466 | 0.172780079 | 13.40995442 | -1.959631832 | 0.096355038 | 0.513146057 | -4.197807156 |
| **A_55_P2083839** | Atp13a1 | 0.346632299 | -0.083185329 | 0.776449928 | 12.02353895 | 1.959602539 | 0.096358998 | 0.513146057 | -4.19782179 |
| **A_55_P2043062** | Hs3st6 | -0.221298987 | -0.495714823 | 0.053116849 | 8.553237452 | -1.959536894 | 0.096367874 | 0.513146057 | -4.197854584 |
| **A_55_P2173623** | Samd1 | 0.152239533 | -0.036549369 | 0.341028436 | 12.44965539 | 1.959449936 | 0.096379632 | 0.513146057 | -4.197898027 |
| **A_66_P108815** | Pgls | -0.242689795 | -0.543645026 | 0.058265436 | 10.68128112 | -1.95944378 | 0.096380465 | 0.513146057 | -4.197901102 |
| **A_55_P2037732** | Foxl2 | -0.416896422 | -0.933904505 | 0.100111661 | 13.81037177 | -1.959359035 | 0.096391926 | 0.513146057 | -4.197943439 |
| **A_55_P2140031** | Fam129b | -0.766350346 | -1.716874503 | 0.184173811 | 12.09242092 | -1.959058424 | 0.096432591 | 0.513146057 | -4.198093624 |
| **A_55_P2001159** | Oser1 | -0.239248157 | -0.536002711 | 0.057506396 | 11.86799361 | -1.958999833 | 0.096440519 | 0.513146057 | -4.198122897 |
| **A_55_P1978316** | Adamts14 | -0.344564253 | -0.771953041 | 0.082824535 | 9.075114875 | -1.958981271 | 0.096443031 | 0.513146057 | -4.198132171 |
| **A_55_P1961385** | NA | 0.219036976 | -0.052696124 | 0.490770076 | 8.430273593 | 1.958655612 | 0.096487109 | 0.513146057 | -4.198294877 |
| **A_52_P215106** | Pik3r6 | -0.176940271 | -0.396461321 | 0.042580779 | 10.17490846 | -1.958545682 | 0.096501992 | 0.513146057 | -4.198349803 |
| **A_55_P2020248** | NA | 0.171654742 | -0.041326164 | 0.384635649 | 14.09791086 | 1.958386129 | 0.096523599 | 0.513146057 | -4.198429523 |
| **A_55_P2169849** | Thoc6 | 0.226945782 | -0.054667795 | 0.508559359 | 9.698118961 | 1.958175965 | 0.096552066 | 0.513146057 | -4.198534533 |
| **A_55_P2133632** | Sipa1l2 | -0.658777643 | -1.476318601 | 0.158763316 | 10.03788324 | -1.95799925 | 0.096576009 | 0.513146057 | -4.198622833 |
| **A_55_P1958275** | Bcl11a | 0.346572521 | -0.083555798 | 0.776700839 | 10.75115666 | 1.957849379 | 0.096596319 | 0.513146057 | -4.198697721 |
| **A_55_P1961059** | Usp16 | 0.194660032 | -0.046944528 | 0.436264591 | 10.97583663 | 1.957739275 | 0.096611243 | 0.513146057 | -4.198752739 |
| **A_55_P2079912** | NA | 0.313670893 | -0.075710297 | 0.703052083 | 9.96615813 | 1.95741288 | 0.096655498 | 0.513146057 | -4.198915841 |
| **A_55_P2126469** | NA | 0.136840229 | -0.033089558 | 0.306770016 | 15.93955115 | 1.956714527 | 0.096750253 | 0.513146057 | -4.199264836 |
| **A_55_P2141479** | Rnu2-10 | 1.127766542 | -0.272866986 | 2.52840007 | 12.6673938 | 1.956491102 | 0.096780588 | 0.513146057 | -4.199376497 |
| **A_55_P2002103** | Arhgap45 | -0.162918797 | -0.365258876 | 0.039421283 | 11.90278294 | -1.95646673 | 0.096783898 | 0.513146057 | -4.199388678 |
| **A_52_P489295** | Adamts1 | -0.452937862 | -1.015511106 | 0.109635382 | 10.22712713 | -1.956332821 | 0.096802084 | 0.513146057 | -4.199455603 |
| **A_66_P102237** | Nemf | 0.227395244 | -0.055067221 | 0.50985771 | 10.98936976 | 1.956157508 | 0.096825899 | 0.513146057 | -4.199543223 |
| **A_55_P2097553** | C77080 | -0.294240074 | -0.659743721 | 0.071263573 | 10.80394038 | -1.956110098 | 0.09683234 | 0.513146057 | -4.199566919 |
| **A_52_P260994** | Fgd2 | -0.25029033 | -0.561212718 | 0.060632058 | 8.566360199 | -1.956028893 | 0.096843373 | 0.513146057 | -4.199607506 |
| **A_55_P2138130** | NA | 0.212835813 | -0.051573227 | 0.477244854 | 12.44195023 | 1.955922289 | 0.09685786 | 0.513146057 | -4.199660788 |
| **A_51_P317433** | Mesp2 | 0.342253276 | -0.083023871 | 0.767530423 | 8.892765736 | 1.955504216 | 0.096914693 | 0.513146057 | -4.199869753 |
| **A_55_P2121846** | NA | 0.163515063 | -0.039686874 | 0.366717 | 15.97677156 | 1.955298704 | 0.096942643 | 0.513146057 | -4.199972478 |

| | | | | | | | | |
|---|---|---|---|---|---|---|---|---|
| **A_55_P1965000** | Ddx58 | -0.264791198 | -0.593865031 | 0.064282634 | 10.53876601 | -1.955209889 | 0.096954725 | 0.513146057 | -4.200016873 |
| **A_55_P2064547** | Tuba1c | 0.220146493 | -0.053523484 | 0.49381647 | 15.51062754 | 1.954644601 | 0.097031656 | 0.513146057 | -4.200299449 |
| **A_55_P1962918** | Ifi211 | -0.243640198 | -0.546518258 | 0.059237861 | 8.832845497 | -1.954628917 | 0.097033791 | 0.513146057 | -4.20030729 |
| **A_52_P247788** | Ino80c | 0.279197391 | -0.067890826 | 0.626285608 | 8.862138206 | 1.95458551 | 0.097039701 | 0.513146057 | -4.200328989 |
| **A_55_P2080248** | Cyp2j8 | -0.267136148 | -0.59923772 | 0.064965425 | 8.547478199 | -1.954541587 | 0.097045681 | 0.513146057 | -4.200350946 |
| **A_51_P237514** | Eif1ax | 0.163278571 | -0.039775364 | 0.366332505 | 13.13145613 | 1.953893868 | 0.097133918 | 0.513146057 | -4.200674758 |
| **A_51_P513785** | Ctcf | 0.177651915 | -0.043280978 | 0.398584809 | 11.29474355 | 1.953856672 | 0.097138988 | 0.513146057 | -4.200693354 |
| **A_55_P1957468** | She | -0.413378808 | -0.92751917 | 0.100761554 | 9.43245401 | -1.953663212 | 0.097165359 | 0.513146057 | -4.200790076 |
| **A_55_P2032039** | Bola3 | 0.260196272 | -0.063449358 | 0.583841903 | 12.76696028 | 1.953504929 | 0.097186941 | 0.513146057 | -4.200869213 |
| **A_55_P2156490** | Gm3146 | 0.169764128 | -0.041406539 | 0.380934795 | 8.31936135 | 1.953419527 | 0.097198587 | 0.513146057 | -4.200911911 |
| **A_51_P168630** | Klf1 | 0.465779284 | -0.113677151 | 1.045235719 | 11.83282599 | 1.953181347 | 0.097231075 | 0.513146057 | -4.201030998 |
| **A_55_P2235059** | 6230415J03Rik | 0.403274994 | -0.098423895 | 0.904973884 | 9.687888459 | 1.953175838 | 0.097231827 | 0.513146057 | -4.201033753 |
| **A_51_P264695** | Crym | -0.214506936 | -0.481412782 | 0.05239891 | 7.818930203 | -1.95283901 | 0.09727779 | 0.513146057 | -4.201202169 |
| **A_55_P2157360** | Tagap1 | -0.133055415 | -0.298615492 | 0.032504661 | 8.393940823 | -1.952810614 | 0.097281666 | 0.513146057 | -4.201216367 |
| **A_55_P2065754** | Krtap19-5 | 0.511340403 | -0.125007362 | 1.147688168 | 9.131172438 | 1.952534766 | 0.097319325 | 0.513146057 | -4.201354299 |
| **A_55_P1961089** | Npb | 0.462791231 | -0.113165373 | 1.038747835 | 10.18100648 | 1.952443827 | 0.097331744 | 0.513146057 | -4.201399772 |
| **A_55_P2008591** | Kras | 0.233287434 | -0.057083864 | 0.523658731 | 8.531100039 | 1.952184453 | 0.097367172 | 0.513146057 | -4.201529472 |
| **A_55_P2076777** | Mdfi | -0.923420049 | -2.073178625 | 0.226338527 | 11.27600595 | -1.951532692 | 0.097456254 | 0.513146057 | -4.201855405 |
| **A_55_P2068366** | Gm9292 | -0.181352753 | -0.407156995 | 0.04445149 | 8.207560647 | -1.951530051 | 0.097456615 | 0.513146057 | -4.201856726 |
| **A_55_P2100866** | NA | 0.316619275 | -0.077623861 | 0.710862412 | 10.65295004 | 1.951445372 | 0.097468195 | 0.513146057 | -4.201899074 |
| **A_55_P2171658** | Stx3 | -0.508621185 | -1.141941554 | 0.124699184 | 8.556792222 | -1.951435397 | 0.097469559 | 0.513146057 | -4.201904063 |
| **A_55_P2047768** | Serpinb6c | -0.329439737 | -0.739699901 | 0.080820426 | 11.98192493 | -1.951191179 | 0.097502964 | 0.513146057 | -4.202026201 |
| **A_55_P2013118** | NA | 0.283610429 | -0.069599759 | 0.636820617 | 8.667673201 | 1.951066909 | 0.097519966 | 0.513146057 | -4.202088351 |
| **A_51_P228295** | Mpzl1 | -0.385246199 | -0.865078858 | 0.094586459 | 11.93507736 | -1.950885352 | 0.097544812 | 0.513146057 | -4.202179155 |
| **A_55_P1969645** | Sdcbp | 0.200495286 | -0.049249233 | 0.450239805 | 11.29629717 | 1.950704065 | 0.097569627 | 0.513146057 | -4.202269826 |
| **A_55_P2033055** | Pdp1 | 0.156991767 | -0.038577437 | 0.35256097 | 8.782510038 | 1.950561276 | 0.097589177 | 0.513146057 | -4.202341243 |
| **A_55_P2150299** | NA | 0.489650924 | -0.120359595 | 1.099661443 | 9.941723572 | 1.950439319 | 0.097605878 | 0.513146057 | -4.202402242 |
| **A_51_P417891** | Trim10 | 0.440591487 | -0.108307167 | 0.989490141 | 11.74269801 | 1.950415432 | 0.097609149 | 0.513146057 | -4.20241419 |
| **A_55_P1969820** | Slc36a1 | 0.248729811 | -0.061175535 | 0.558635158 | 8.721337633 | 1.95021261 | 0.097636931 | 0.513146057 | -4.202515638 |
| **A_66_P106133** | Magi3 | -0.184696069 | -0.414824119 | 0.045431981 | 9.084671941 | -1.950164664 | 0.097643499 | 0.513146057 | -4.20253962 |
| **A_51_P273556** | Fam83d | 0.16700875 | -0.041133888 | 0.375151389 | 8.637037496 | 1.94967113 | 0.097711139 | 0.513247152 | -4.202786489 |
| **A_55_P2116900** | NA | 0.398416529 | -0.098137852 | 0.894970911 | 9.452164395 | 1.949636822 | 0.097715843 | 0.513247152 | -4.202803651 |
| **A_51_P210560** | Pptc7 | -0.253379228 | -0.569380004 | 0.062621547 | 11.78338011 | -1.948345844 | 0.097893003 | 0.514013401 | -4.203449483 |
| **A_52_P543430** | NA | -0.252033746 | -0.566387554 | 0.062320063 | 11.29546126 | -1.948153424 | 0.097919436 | 0.514013401 | -4.203545754 |
| **A_51_P462972** | Ap5m1 | 0.181564762 | -0.04491904 | 0.408048565 | 9.564835504 | 1.947949112 | 0.09794751 | 0.514013401 | -4.203647977 |
| **A_55_P2121521** | H3c11 | 0.434529064 | -0.107576276 | 0.976634403 | 9.869555289 | 1.947683271 | 0.097984052 | 0.514013401 | -4.203780988 |
| **A_55_P1953252** | Ctnnd1 | -0.217822503 | -0.489652485 | 0.054007479 | 12.22421638 | -1.947101433 | 0.098064076 | 0.514013401 | -4.204072122 |
| **A_51_P397296** | Marveld3 | -0.207484282 | -0.466416267 | 0.051447702 | 7.77584807 | -1.94707505 | 0.098067706 | 0.514013401 | -4.204085324 |
| **A_55_P1952618** | Ear2 | -0.285927495 | -0.642755441 | 0.070900451 | 11.10887041 | -1.947064098 | 0.098069213 | 0.514013401 | -4.204090804 |
| **A_51_P124568** | Mpp1 | 0.235948063 | -0.058513001 | 0.530409126 | 13.00174933 | 1.947025901 | 0.09807447 | 0.514013401 | -4.207167981 |
| **A_51_P339962** | Cwf19l2 | 0.162676357 | -0.040366855 | 0.365719568 | 8.713114873 | 1.946790203 | 0.098106909 | 0.514044034 | -4.204227861 |
| **A_55_P2073579** | Chchd5 | -0.212459358 | -0.477715143 | 0.052796427 | 9.681005036 | -1.946230123 | 0.098184036 | 0.514308735 | -4.204508139 |
| **A_55_P2009206** | Sumo2 | 0.232100228 | -0.057708163 | 0.52190862 | 8.960577654 | 1.946022243 | 0.098212678 | 0.514319387 | -4.204612173 |
| **A_55_P1952256** | Ccnb1 | 0.259157305 | -0.064500286 | 0.582814897 | 9.198562251 | 1.945632656 | 0.098266379 | 0.514461222 | -4.204807149 |
| **A_55_P1956847** | Nolc1 | 0.304249378 | -0.075842286 | 0.684341041 | 11.66932451 | 1.945022113 | 0.098350595 | 0.514691922 | -4.205112727 |
| **A_51_P167374** | Gpatch1 | 0.143310147 | -0.035732625 | 0.322352919 | 9.077675396 | 1.944927149 | 0.0983637 | 0.514691922 | -4.205160259 |
| **A_55_P2034853** | 2610028H24Rik | 0.227902616 | -0.056945339 | 0.512750572 | 9.155927582 | 1.944103548 | 0.098477434 | 0.514977125 | -4.205572515 |
| **A_51_P192162** | NA | -0.210853576 | -0.474437895 | 0.052730743 | 8.588851457 | -1.943768727 | 0.098523708 | 0.514977125 | -4.205740123 |
| **A_52_P494930** | Fnip2 | 0.352677185 | -0.088242634 | 0.793597004 | 10.10080449 | 1.943573343 | 0.098550721 | 0.514977125 | -4.205837934 |
| **A_66_P107585** | Taf2 | -0.226383203 | -0.509447018 | 0.056680611 | 10.67783644 | -1.94331425 | 0.098586554 | 0.514977125 | -4.205967641 |

| | | | | | | | | | |
|---|---|---|---|---|---|---|---|---|---|
| **A_51_P194628** | Tango6 | 0.313826083 | -0.078576469 | 0.706228635 | 9.025313363 | 1.943302469 | 0.098588183 | 0.514977125 | -4.205973539 |
| **A_55_P2014635** | Ccdc82 | 0.243258002 | -0.060909551 | 0.547425556 | 10.54289312 | 1.943289212 | 0.098590017 | 0.514977125 | -4.205980176 |
| **A_55_P2018322** | Zfp644 | 0.18526012 | -0.0464055 | 0.416925739 | 9.903099607 | 1.943137541 | 0.098611 | 0.514977125 | -4.206056108 |
| **A_52_P169901** | Slc25a51 | 0.212806916 | -0.053346491 | 0.478960322 | 11.65197901 | 1.942839377 | 0.098652263 | 0.514977125 | -4.206205384 |
| **A_55_P2067251** | Tmpo | 0.280773829 | -0.070391932 | 0.63193959 | 9.622315101 | 1.942798001 | 0.09865799 | 0.514977125 | -4.206226099 |
| **A_51_P174854** | Maged1 | -0.271188509 | -0.610534269 | 0.068157251 | 12.05844979 | -1.941833751 | 0.098791558 | 0.515189359 | -4.206708892 |
| **A_55_P1974780** | NA | 0.347241818 | -0.087277269 | 0.781760905 | 9.362229368 | 1.941808294 | 0.098795086 | 0.515189359 | -4.20672164 |
| **A_52_P431894** | Pknox1 | 0.136140713 | -0.034251272 | 0.306532699 | 9.358416469 | 1.941431387 | 0.098847346 | 0.515189359 | -4.206910371 |
| **A_65_P16753** | Bex1 | 0.470116226 | -0.118291507 | 1.058523958 | 13.32868575 | 1.941377796 | 0.098854779 | 0.515189359 | -4.206937207 |
| **A_55_P2111855** | Gale | -0.296149364 | -0.666857589 | 0.07455886 | 12.35638883 | -1.941161961 | 0.09888472 | 0.515189359 | -4.207045288 |
| **A_55_P1971179** | Cd1d1 | -0.295075668 | -0.664466433 | 0.074315098 | 8.335269316 | -1.941022428 | 0.098904082 | 0.515189359 | -4.207115162 |
| **A_55_P2117741** | Nck2 | -0.180822226 | -0.407201559 | 0.045557106 | 11.46181786 | -1.940877949 | 0.098924133 | 0.515189359 | -4.207187514 |
| **A_55_P2176688** | NA | 0.262521942 | -0.066148926 | 0.591192811 | 14.54268883 | 1.940830396 | 0.098930734 | 0.515189359 | -4.207211328 |
| **A_55_P1959648** | Hspg2 | -0.173976884 | -0.391797872 | 0.043844104 | 7.935817396 | -1.940774211 | 0.098938533 | 0.515189359 | -4.207239464 |
| **A_51_P111932** | Tor1aip2 | -0.176333013 | -0.39720624 | 0.044540215 | 8.910038446 | -1.939874969 | 0.099063443 | 0.515651985 | -4.207689822 |
| **A_55_P2044197** | Vps29 | 0.159289411 | -0.04024794 | 0.358826761 | 11.41585101 | 1.939750595 | 0.099080732 | 0.515651985 | -4.207752114 |
| **A_55_P2015007** | Zfp46 | -0.578345862 | -1.302955108 | 0.146263384 | 10.56148682 | -1.939398032 | 0.099129757 | 0.515768256 | -4.207928701 |
| **A_51_P501107** | Cul5 | 0.204674479 | -0.051867936 | 0.461216894 | 8.933954444 | 1.938597751 | 0.099241128 | 0.516208763 | -4.208329564 |
| **A_55_P2175880** | Mgst1 | -0.434686031 | -0.979615716 | 0.110243654 | 13.64370824 | -1.938288445 | 0.099284206 | 0.516293897 | -4.208484506 |
| **A_51_P127841** | Pdss1 | 0.232963444 | -0.059133744 | 0.525060631 | 9.106137496 | 1.937954566 | 0.099330727 | 0.5163626 | -4.227411428 |
| **A_51_P151484** | Atp1b1 | -0.517838998 | -1.167183439 | 0.131505444 | 11.46285558 | -1.937772619 | 0.099356088 | 0.5163626 | -4.208742916 |
| **A_52_P489778** | Ablim1 | -0.219778682 | -0.495412061 | 0.055854697 | 8.449202982 | -1.937478736 | 0.099397065 | 0.5163626 | -4.208890147 |
| **A_51_P220993** | Pfn1 | -0.287627896 | -0.64848206 | 0.073226268 | 11.1552508 | -1.936789722 | 0.099493202 | 0.5163626 | -4.209235356 |
| **A_55_P2086094** | Slc35b2 | -0.15223155 | -0.343260099 | 0.038797 | 11.61222206 | -1.936375501 | 0.099551042 | 0.5163626 | -4.209442903 |
| **A_51_P506748** | Arhgap35 | -0.290551211 | -0.655154752 | 0.07405233 | 11.87748984 | -1.936355071 | 0.099553896 | 0.5163626 | -4.20945314 |
| **A_55_P2027107** | Rgs3 | 0.178759474 | -0.045577479 | 0.403096426 | 9.135917997 | 1.936205455 | 0.099574797 | 0.5163626 | -4.209528108 |
| **A_51_P477419** | Nfic | -0.319201606 | -0.719790924 | 0.081387713 | 10.21364119 | -1.936194097 | 0.099576384 | 0.5163626 | -4.209533799 |
| **A_55_P2174743** | Akap7 | 0.32000057 | -0.081592734 | 0.721593874 | 9.449657744 | 1.936187795 | 0.099577264 | 0.5163626 | -4.209536957 |
| **A_52_P539310** | Serp2 | -0.385963569 | -0.870437419 | 0.09851028 | 8.763470036 | -1.935794104 | 0.099632283 | 0.5163626 | -4.209734231 |
| **A_55_P2024245** | Mthfr | 0.163841137 | -0.041822721 | 0.369504995 | 8.854227889 | 1.935745082 | 0.099639136 | 0.5163626 | -4.209758796 |
| **A_51_P208121** | Klhl25 | 0.310730893 | -0.079414068 | 0.700875854 | 9.977985293 | 1.935270247 | 0.09970554 | 0.5163626 | -4.209996745 |
| **A_51_P214083** | 2310039H08Rik | -0.151247797 | -0.341157713 | 0.038662119 | 12.35997601 | -1.935194433 | 0.099716147 | 0.5163626 | -4.210034739 |
| **A_55_P2144796** | Gfer | 0.140292683 | -0.035862724 | 0.316448091 | 9.325678453 | 1.935183854 | 0.099717627 | 0.5163626 | -4.21004004 |
| **A_51_P487913** | NA | 0.280984786 | -0.071829695 | 0.633799267 | 8.623556448 | 1.93517209 | 0.099719273 | 0.5163626 | -4.210045935 |
| **A_51_P391926** | Ubxn6 | -0.285607223 | -0.644251024 | 0.073036579 | 11.79463707 | -1.935035982 | 0.099738318 | 0.5163626 | -4.210114145 |
| **A_55_P2099269** | Ift80 | 0.175605277 | -0.044955461 | 0.396166016 | 9.320307623 | 1.934606064 | 0.099798498 | 0.5163626 | -4.210329604 |
| **A_55_P2006698** | Bmp1 | -0.230346278 | -0.519677519 | 0.058984962 | 9.569338518 | -1.934501453 | 0.099813147 | 0.5163626 | -4.210382033 |
| **A_55_P2040044** | NA | 0.189835097 | -0.048648032 | 0.428318226 | 8.545977415 | 1.934202931 | 0.099854962 | 0.5163626 | -4.210531649 |
| **A_55_P2114789** | Gtf2a2 | 0.240540132 | -0.061684977 | 0.542765241 | 9.126589326 | 1.933927511 | 0.099893556 | 0.5163626 | -4.210669692 |
| **A_55_P2045549** | Dpy19l1 | 0.250403536 | -0.06421451 | 0.565021581 | 10.47316034 | 1.933926736 | 0.099893665 | 0.5163626 | -4.21067008 |
| **A_55_P1990041** | Nxt2 | -0.144935516 | -0.327062697 | 0.037191666 | 10.4836642 | -1.933673926 | 0.099929104 | 0.5163626 | -4.210796795 |
| **A_51_P166023** | Hip1 | -0.161083644 | -0.363505426 | 0.041338139 | 7.979072513 | -1.93364787 | 0.099932758 | 0.5163626 | -4.210809856 |
| **A_55_P1959164** | Gm10324 | 0.153277461 | -0.039343763 | 0.345898686 | 8.025786159 | 1.933558645 | 0.099945269 | 0.5163626 | -4.210854578 |
| **A_55_P2133057** | Pdia2 | 0.585544797 | -0.15038275 | 1.321472343 | 12.46464274 | 1.933340086 | 0.099975922 | 0.5163626 | -4.21096413 |
| **A_55_P2082203** | Baz1a | 0.363839164 | -0.093470219 | 0.821148547 | 10.9756643 | 1.933225529 | 0.099991992 | 0.5163626 | -4.211021553 |
| **A_55_P1979674** | Ikzf4 | -0.294632669 | -0.665166865 | 0.075901528 | 8.42955421 | -1.932127548 | 0.100146151 | 0.51691778 | -4.211571965 |
| **A_55_P1959800** | NA | 0.271330862 | -0.069923261 | 0.612584985 | 9.073706033 | 1.93198823 | 0.100165728 | 0.51691778 | -4.211641809 |
| **A_55_P2136289** | NA | -0.282413952 | -0.637625647 | 0.072797743 | 9.882275353 | -1.931888604 | 0.10017973 | 0.51691778 | -4.211691756 |
| **A_55_P2429480** | Atad2 | 0.335077823 | -0.086530492 | 0.756686138 | 8.996928388 | 1.931166446 | 0.100281286 | 0.516969174 | -4.212053821 |
| **A_51_P324572** | Ctnna1 | -0.344792321 | -0.778628939 | 0.089044297 | 13.02714351 | -1.931143594 | 0.100284501 | 0.516969174 | -4.212065279 |

| | | | | | | | | | |
|---|---|---|---|---|---|---|---|---|---|
| **A_51_P424338** | Nqo1 | 0.233769863 | -0.060395068 | 0.527934795 | 8.875128943 | 1.930993514 | 0.10030562 | 0.516969174 | -4.212140527 |
| **A_51_P416137** | Slc31a2 | -0.207712115 | -0.469100716 | 0.053676486 | 9.539509225 | -1.930893732 | 0.100319664 | 0.516969174 | -4.212190558 |
| **A_55_P2147101** | Sytl4 | -0.20234034 | -0.456979778 | 0.052299098 | 8.271256339 | -1.930812033 | 0.100331164 | 0.516969174 | -4.212231523 |
| **A_55_P2077068** | Malt1 | 0.267766603 | -0.069258847 | 0.604792052 | 9.456522393 | 1.930531539 | 0.100370656 | 0.516969174 | -4.212372167 |
| **A_55_P2353004** | Pitpnm2os1 | 0.318881627 | -0.082576746 | 0.720339999 | 8.840103495 | 1.930066026 | 0.100436233 | 0.516969174 | -4.212605594 |
| **A_51_P117109** | Ilvbl | -0.152646415 | -0.344827287 | 0.039534457 | 8.883695207 | -1.930010351 | 0.100444079 | 0.516969174 | -4.212633513 |
| **A_55_P1968340** | Plekhh1 | -0.378321216 | -0.854634194 | 0.097991762 | 8.584935271 | -1.929974058 | 0.100449194 | 0.516969174 | -4.212651712 |
| **A_51_P221337** | Ranbp10 | 0.344313079 | -0.089195742 | 0.777821899 | 12.47512187 | 1.929917617 | 0.100457148 | 0.516969174 | -4.212680015 |
| **A_55_P2054352** | Rbm6 | 0.163814746 | -0.042474468 | 0.370103959 | 10.2410686 | 1.929566104 | 0.100506704 | 0.517086529 | -4.21285629 |
| **A_55_P2060878** | NA | 0.335968496 | -0.08717777 | 0.759114761 | 11.43790979 | 1.929262096 | 0.100549583 | 0.517169476 | -4.213008748 |
| **A_55_P2008740** | Fcgr1 | -0.195852353 | -0.442589741 | 0.050885035 | 8.674586248 | -1.928754802 | 0.100621174 | 0.517308907 | -4.213263165 |
| **A_55_P2139473** | Ilk | -0.274445819 | -0.620250984 | 0.071359346 | 11.8251535 | -1.928449774 | 0.100664246 | 0.517308907 | -4.21341615 |
| **A_55_P2046812** | Sparc | -0.70449447 | -1.592358152 | 0.183369213 | 15.10677765 | -1.928033099 | 0.100723112 | 0.517308907 | -4.21362514 |
| **A_55_P2067752** | Psg26 | -0.252608179 | -0.570982244 | 0.065765887 | 7.773980153 | -1.927937335 | 0.100736646 | 0.517308907 | -4.213673174 |
| **A_55_P2304507** | Noxa1 | -0.381106189 | -0.861450124 | 0.099237746 | 8.579574446 | -1.927866155 | 0.100746707 | 0.517308907 | -4.213708876 |
| **A_51_P203192** | Mbd4 | 0.175426503 | -0.045691962 | 0.396544967 | 8.774456725 | 1.92776186 | 0.10076145 | 0.517308907 | -4.21376119 |
| **A_52_P549827** | Mgst1 | -0.381954222 | -0.863397625 | 0.099489181 | 13.8268944 | -1.927743573 | 0.100764035 | 0.517308907 | -4.213770362 |
| **A_51_P274124** | Mansc1 | -0.287474719 | -0.649887341 | 0.074937902 | 8.496904735 | -1.927434078 | 0.100807799 | 0.517335746 | -4.213925607 |
| **A_51_P187018** | Magohb | 0.151834364 | -0.039590177 | 0.343258905 | 10.52310911 | 1.927328079 | 0.100822792 | 0.517335746 | -4.213978778 |
| **A_55_P2409306** | Vps35l | 0.1814383 | -0.047339416 | 0.410216016 | 8.377456901 | 1.927074196 | 0.100858712 | 0.517382709 | -4.214106133 |
| **A_66_P127458** | Zfp652 | -0.236772464 | -0.535382111 | 0.061837183 | 10.98111411 | -1.926684257 | 0.100913906 | 0.517528493 | -4.214301745 |
| **A_55_P1985840** | Mettl16 | 0.179651694 | -0.047033393 | 0.40633678 | 9.997097503 | 1.925712971 | 0.101051518 | 0.518096761 | -4.214789028 |
| **A_51_P381260** | Fxyd5 | -0.295575502 | -0.66865587 | 0.077504866 | 10.21097395 | -1.925081978 | 0.101141017 | 0.518176965 | -4.215105621 |
| **A_66_P139196** | Baz1a | 0.302105126 | -0.079223637 | 0.683433889 | 9.406464202 | 1.925048692 | 0.101145741 | 0.518176965 | -4.215122322 |
| **A_55_P1978061** | Hnrnpa3 | 0.203768935 | -0.053456171 | 0.46099404 | 9.370450922 | 1.924898395 | 0.101167071 | 0.518176965 | -4.215197736 |
| **A_55_P2086659** | Rad54l | 0.345491548 | -0.090771846 | 0.781754942 | 9.069918732 | 1.924295847 | 0.101252632 | 0.518176965 | -4.215500086 |
| **A_55_P1968568** | Rapgef5 | -0.367992799 | -0.832687842 | 0.096702244 | 8.275124857 | -1.92421887 | 0.101263567 | 0.518176965 | -4.215538713 |
| **A_51_P493558** | Rnpep | -0.233909708 | -0.529307482 | 0.061488066 | 10.48197581 | -1.924084836 | 0.101282612 | 0.518176965 | -4.215605973 |
| **A_66_P106289** | Vps54 | 0.13705754 | -0.036031781 | 0.310146861 | 9.264880484 | 1.924047759 | 0.10128788 | 0.518176965 | -4.215624579 |
| **A_51_P108978** | Myadm | -0.277984614 | -0.629061712 | 0.073092485 | 9.81321703 | -1.92398396 | 0.101296947 | 0.518176965 | -4.215656595 |
| **A_51_P497692** | Mageh1 | -0.183066328 | -0.414278491 | 0.048145834 | 9.129987691 | -1.923893311 | 0.101309831 | 0.518176965 | -4.215702085 |
| **A_55_P2126950** | Zfp467 | -0.555769147 | -1.257768958 | 0.146230663 | 9.866616321 | -1.923714542 | 0.101335244 | 0.518176965 | -4.215791797 |
| **A_55_P2015675** | AU015836 | 0.254653956 | -0.067082075 | 0.576389987 | 8.504456699 | 1.923241921 | 0.10140246 | 0.518383536 | -4.216028983 |
| **A_51_P104897** | Itpr3 | -0.304672181 | -0.689668805 | 0.080324443 | 10.67466926 | -1.92291049 | 0.101449623 | 0.518487508 | -4.21619532 |
| **A_52_P546090** | Ssr4 | -0.178690354 | -0.404537076 | 0.047156367 | 14.1700166 | -1.922518414 | 0.101505443 | 0.518514227 | -4.216392103 |
| **A_55_P2049141** | Cops7b | 0.222528533 | -0.058783249 | 0.503840314 | 9.123952287 | 1.922122099 | 0.101561898 | 0.518514227 | -4.216591021 |
| **A_55_P2204491** | AI131754 | -0.130391055 | -0.295245052 | 0.034462942 | 7.902634281 | -1.921903194 | 0.101593095 | 0.518514227 | -4.216700899 |
| **A_55_P1972084** | Napb | 0.194817809 | -0.051497446 | 0.441133064 | 8.479426908 | 1.921854626 | 0.101600018 | 0.518514227 | -4.216725277 |
| **A_51_P494361** | Mbnl2 | -0.582599661 | -1.319252231 | 0.15405291 | 10.17947243 | -1.921722645 | 0.101618832 | 0.518514227 | -4.216791525 |
| **A_55_P2004119** | NA | 0.230973762 | -0.061100316 | 0.52304784 | 14.46619985 | 1.921555014 | 0.101642734 | 0.518514227 | -4.21687567 |
| **A_55_P2054300** | Alg8 | 0.495306114 | -0.13113348 | 1.121745708 | 10.50975571 | 1.921222565 | 0.101690153 | 0.518514227 | -4.217042552 |
| **A_51_P254895** | Cyp4a10 | -0.473103291 | -1.071539541 | 0.125332959 | 8.778014794 | -1.920973022 | 0.101725761 | 0.518514227 | -4.217167821 |
| **A_55_P2016034** | Nlrc5 | -0.168124609 | -0.380820366 | 0.044571147 | 7.897059335 | -1.920682826 | 0.101767186 | 0.518514227 | -4.217313502 |
| **A_51_P174434** | Cenpj | 0.150061876 | -0.039784187 | 0.339907939 | 9.400247085 | 1.920666516 | 0.101769515 | 0.518514227 | -4.21732169 |
| **A_55_P2170134** | Eif2b4 | 0.215034962 | -0.057030703 | 0.487100626 | 12.37612475 | 1.920518652 | 0.101790629 | 0.518514227 | -4.217395921 |
| **A_51_P365903** | Ankzf1 | -0.154212897 | -0.349345773 | 0.04091998 | 10.09793244 | -1.920319257 | 0.101819109 | 0.518514227 | -4.217496025 |
| **A_51_P365399** | Smg5 | 0.346574837 | -0.092057777 | 0.785207452 | 11.37089248 | 1.919903042 | 0.101878582 | 0.518514227 | -4.217704986 |
| **A_55_P1975021** | Rad18 | 0.256603267 | -0.068175294 | 0.581381827 | 9.388358739 | 1.919808972 | 0.101892028 | 0.518514227 | -4.217752216 |
| **A_52_P383653** | Cpne8 | -0.209052144 | -0.473690232 | 0.055585945 | 7.95016036 | -1.919488273 | 0.101937883 | 0.518514227 | -4.217913233 |
| **A_55_P1996998** | Polr2a | 0.35851934 | -0.095370748 | 0.812409428 | 9.732465776 | 1.919309792 | 0.101963411 | 0.518514227 | -4.218002847 |

| | | | | | | | | | |
|---|---|---|---|---|---|---|---|---|---|
| **A_52_P142191** | Aph1b | -0.166818918 | -0.378015834 | 0.044377998 | 8.478729511 | -1.919291399 | 0.101966042 | 0.518514227 | -4.218012082 |
| **A_55_P2137611** | Irgm2 | -0.533845402 | -1.209922643 | 0.142231839 | 10.98563135 | -1.918679252 | 0.10205365 | 0.518514227 | -4.218319453 |
| **A_55_P1984113** | Nup62 | 0.14878957 | -0.03967734 | 0.33725648 | 12.92158699 | 1.918317851 | 0.102105408 | 0.518514227 | -4.218500929 |
| **A_55_P2070194** | Caprin1 | 0.161546526 | -0.043146678 | 0.36623973 | 11.55715017 | 1.917685519 | 0.102196029 | 0.518514227 | -4.218818472 |
| **A_55_P2033660** | Crls1 | 0.251116299 | -0.067110698 | 0.569343296 | 10.60845317 | 1.917436856 | 0.102231687 | 0.518514227 | -4.218943351 |
| **A_51_P224593** | Arl8a | -0.203689723 | -0.461822822 | 0.054443375 | 9.719525738 | -1.917381852 | 0.102239576 | 0.518514227 | -4.218970975 |
| **A_51_P170758** | Grwd1 | 0.248897423 | -0.066536696 | 0.564331542 | 10.30762572 | 1.917321412 | 0.102248246 | 0.518514227 | -4.219001329 |
| **A_52_P55772** | Tbxa2r | -0.24319339 | -0.55140849 | 0.06502171 | 8.881745114 | -1.917260145 | 0.102257035 | 0.518514227 | -4.219032099 |
| **A_55_P1998776** | NA | 0.258779926 | -0.069198422 | 0.586758274 | 15.53743971 | 1.917205202 | 0.102264917 | 0.518514227 | -4.219059692 |
| **A_55_P2147141** | Spata2 | 0.226932338 | -0.060690405 | 0.514555082 | 11.1260162 | 1.917151145 | 0.102272673 | 0.518514227 | -4.219086841 |
| **A_55_P1955637** | Trp53i11 | -0.130530822 | -0.295987278 | 0.034925635 | 7.873573916 | -1.916957753 | 0.102300425 | 0.518514227 | -4.219183968 |
| **A_55_P2001078** | Mmel1 | 0.492055864 | -0.131712539 | 1.115824268 | 9.155807751 | 1.916788649 | 0.102324698 | 0.518514227 | -4.2192689 |
| **A_55_P2109114** | NA | -0.250957651 | -0.569094399 | 0.067179097 | 14.4125113 | -1.916769065 | 0.10232751 | 0.518514227 | -4.219278736 |
| **A_51_P200561** | Shtn1 | -0.383149639 | -0.868879079 | 0.102579801 | 9.454078056 | -1.916713415 | 0.102335499 | 0.518514227 | -4.219306686 |
| **A_55_P2013833** | Ddx3x | 0.275239271 | -0.073716493 | 0.624195034 | 12.54766725 | 1.916563496 | 0.102357025 | 0.518514227 | -4.219381984 |
| **A_55_P2091476** | NA | 0.438626097 | -0.117515619 | 0.994767813 | 10.1413364 | 1.916426571 | 0.10237669 | 0.518514227 | -4.219450757 |
| **A_55_P2111277** | NA | 0.22031011 | -0.059039682 | 0.499659903 | 10.15711457 | 1.916325433 | 0.102391217 | 0.518514227 | -4.219501556 |
| **A_55_P2007404** | Vbp1 | 0.251421011 | -0.067414636 | 0.570256658 | 9.983440204 | 1.916098748 | 0.102423785 | 0.518514227 | -4.219615415 |
| **A_55_P2103796** | Arhgap11a | 0.367231763 | -0.098525948 | 0.832989474 | 9.229946725 | 1.915858246 | 0.10245835 | 0.518514227 | -4.219736219 |
| **A_55_P1954544** | Zfp446 | 0.178191821 | -0.047809011 | 0.404192654 | 8.968779568 | 1.915847416 | 0.102459906 | 0.518514227 | -4.219741658 |
| **A_55_P2048176** | NA | 0.188796747 | -0.050659776 | 0.428253269 | 8.322552883 | 1.915803772 | 0.10246618 | 0.518514227 | -4.219763581 |
| **A_51_P224445** | Gpn3 | 0.181675244 | -0.048755598 | 0.412106086 | 9.185410876 | 1.915747792 | 0.102474228 | 0.518514227 | -4.2197917 |
| **A_51_P424232** | Mecp2 | 0.169642637 | -0.045553224 | 0.384838498 | 9.57317917 | 1.915509405 | 0.102508505 | 0.518551921 | -4.219911446 |
| **A_52_P256279** | Zbtb11 | 0.245833384 | -0.066055822 | 0.557722591 | 9.370620727 | 1.915242209 | 0.102546938 | 0.518610614 | -4.220045666 |
| **A_52_P116134** | Aldh2 | -0.295572344 | -0.670687157 | 0.079542469 | 11.81203866 | -1.91462079 | 0.102636378 | 0.518729925 | -4.220357841 |
| **A_55_P2086500** | Dennd5a | -0.282428826 | -0.640891863 | 0.076034212 | 11.55092664 | -1.91446675 | 0.10265856 | 0.518729925 | -4.220435227 |
| **A_55_P2428968** | Lrrc2 | 0.76811676 | -0.206830944 | 1.743064464 | 9.514125339 | 1.914384237 | 0.102670445 | 0.518729925 | -4.22047668 |
| **A_55_P2088440** | Arrdc2 | 0.323660066 | -0.087188026 | 0.734508157 | 10.5174705 | 1.914216402 | 0.102694622 | 0.518729925 | -4.220560999 |
| **A_52_P275408** | Ppp4c | -0.145723614 | -0.330748016 | 0.039300788 | 10.66672711 | -1.913745188 | 0.102762533 | 0.518729925 | -4.220797743 |
| **A_55_P2030209** | Cir1 | 0.180335605 | -0.048655606 | 0.409326817 | 10.36729295 | 1.913576618 | 0.102786838 | 0.518729925 | -4.220882437 |
| **A_55_P2112302** | NA | -0.186468796 | -0.423256029 | 0.050318438 | 10.52281133 | -1.913511476 | 0.102796232 | 0.518729925 | -4.220915167 |
| **A_52_P361551** | Hspa14 | 0.16669145 | -0.044988317 | 0.378371217 | 12.72645783 | 1.913450207 | 0.102805068 | 0.518729925 | -4.220945952 |
| **A_55_P2019349** | Txnrd2 | 0.298511439 | -0.080574789 | 0.677597668 | 8.946775512 | 1.913401719 | 0.102812062 | 0.518729925 | -4.220970314 |
| **A_51_P308961** | Hmgcl | -0.220555855 | -0.500768058 | 0.059656347 | 12.68877261 | -1.912558543 | 0.10293375 | 0.519208363 | -4.221393985 |
| **A_55_P1993012** | NA | 0.41920642 | -0.113482355 | 0.951895196 | 11.34564045 | 1.912218735 | 0.102982832 | 0.519232308 | -4.221564739 |
| **A_55_P2270137** | B930036G03Rik | 0.249031217 | -0.06747539 | 0.565537824 | 10.09410968 | 1.911851694 | 0.103035873 | 0.519232308 | -4.221749186 |
| **A_51_P148675** | Dag1 | -0.473368735 | -1.075039583 | 0.128302114 | 10.45992105 | -1.911717823 | 0.103055226 | 0.519232308 | -4.221816462 |
| **A_55_P2129407** | NA | 0.171401602 | -0.046489329 | 0.389292532 | 8.354733789 | 1.911432258 | 0.10309652 | 0.519232308 | -4.221959973 |
| **A_51_P175580** | Trp53inp1 | -0.346224086 | -0.786367928 | 0.093919756 | 10.02745976 | -1.911374715 | 0.103104842 | 0.519232308 | -4.221988891 |
| **A_55_P2025974** | Ruvbl2 | 0.178614375 | -0.048494159 | 0.405722909 | 11.73076399 | 1.911024006 | 0.103155583 | 0.519232308 | -4.222165148 |
| **A_51_P386539** | Rnf125 | -0.375383786 | -0.852754541 | 0.10198697 | 9.265359333 | -1.910745692 | 0.103195867 | 0.519232308 | -4.222305026 |
| **A_55_P2083184** | Gm10272 | 0.182628256 | -0.049620323 | 0.414876834 | 8.331456112 | 1.910724605 | 0.10319892 | 0.519232308 | -4.222315624 |
| **A_55_P2112230** | Pcyt2 | -0.272262859 | -0.618507288 | 0.073981569 | 14.05094547 | -1.910683688 | 0.103204844 | 0.519232308 | -4.222336189 |
| **A_55_P2103706** | NA | 0.345087284 | -0.093773658 | 0.783948226 | 11.95015393 | 1.91066793 | 0.103207126 | 0.519232308 | -4.222344109 |
| **A_55_P2172688** | Srbd1 | 0.24755974 | -0.067349585 | 0.562469065 | 9.061078976 | 1.910194921 | 0.103275634 | 0.519430616 | -4.222581851 |
| **A_51_P511375** | Ranbp2 | 0.282902771 | -0.076996852 | 0.642802393 | 10.94819635 | 1.910024767 | 0.103300289 | 0.519430616 | -4.222667377 |
| **A_51_P145260** | Sall4 | 0.218940757 | -0.059620179 | 0.497501693 | 8.782379219 | 1.909807488 | 0.103331781 | 0.519453836 | -4.222776591 |
| **A_55_P2015232** | NA | 0.3184664 | -0.086807174 | 0.723739975 | 11.15047194 | 1.909407049 | 0.103389846 | 0.519558982 | -4.222977877 |
| **A_65_P17700** | Apobec3 | 0.400441266 | -0.109231306 | 0.910113838 | 10.67513263 | 1.909109117 | 0.103433067 | 0.519558982 | -4.223127643 |
| **A_55_P2131279** | Atp6v1a | 0.145420259 | -0.039667532 | 0.33050805 | 10.34686995 | 1.909107261 | 0.103433337 | 0.519558982 | -4.223128576 |

| | | | | | | | | | |
|---|---|---|---|---|---|---|---|---|---|
| **A_52_P540219** | Timp2 | -0.270574789 | -0.62634224 | 0.085192663 | 11.09230836 | -2.076475976 | 0.103468495 | 0.519600555 | -4.284048903 |
| **A_55_P2085060** | Vgf | 0.152632759 | -0.041699383 | 0.346964901 | 7.768807487 | 1.908474254 | 0.103525228 | 0.519617071 | -4.223446795 |
| **A_55_P2178578** | Tmprss4 | -0.349102479 | -0.793580529 | 0.095375571 | 7.938260945 | -1.908472044 | 0.103525549 | 0.519617071 | -4.223447906 |
| **A_55_P1976689** | Snrnp35 | 0.253954412 | -0.069453801 | 0.577362625 | 9.744541851 | 1.9080419 | 0.103588038 | 0.519795741 | -4.223664157 |
| **A_55_P2171484** | Lta4h | 0.255556503 | -0.069929307 | 0.581042312 | 12.18737258 | 1.907822937 | 0.103619863 | 0.519820484 | -4.223774242 |
| **A_55_P2051701** | Pbx1 | -0.457739181 | -1.040982819 | 0.125504457 | 8.947277952 | -1.907002519 | 0.103739189 | 0.520284066 | -4.224186739 |
| **A_51_P481398** | Kif11 | 0.355556125 | -0.097660682 | 0.808772932 | 9.773057989 | 1.906274099 | 0.10384525 | 0.520480116 | -4.224553013 |
| **A_55_P2046378** | Eif4g1 | 0.194704934 | -0.053543848 | 0.442953715 | 13.53323481 | 1.905781121 | 0.10391709 | 0.520480116 | -4.224800915 |
| **A_55_P2139640** | Fbh1 | -0.188272367 | -0.428326434 | 0.0517817 | 9.141769109 | -1.905727069 | 0.10392497 | 0.520480116 | -4.224828097 |
| **A_55_P1988102** | Hmbs | 0.379597629 | -0.104418617 | 0.863613875 | 11.36148161 | 1.905665806 | 0.103933902 | 0.520480116 | -4.224858905 |
| **A_55_P2108883** | NA | -0.319644028 | -0.727258956 | 0.0879709 | 8.552168862 | -1.905459417 | 0.103963998 | 0.520480116 | -4.224962697 |
| **A_52_P644690** | Flt3l | 0.223262148 | -0.06144887 | 0.507973167 | 8.983489167 | 1.905434451 | 0.103967639 | 0.520480116 | -4.224975252 |
| **A_55_P1960197** | P2ry14 | -0.172774909 | -0.393103029 | 0.047553212 | 9.578564034 | -1.905434093 | 0.103967691 | 0.520480116 | -4.224975432 |
| **A_55_P1960431** | Ubn1 | 0.126209107 | -0.034769112 | 0.287187327 | 11.85469281 | 1.905051691 | 0.104023479 | 0.520480116 | -4.225167747 |
| **A_55_P2087182** | Car4 | -0.130133882 | -0.296128458 | 0.035860695 | 7.720577142 | -1.904932868 | 0.10404082 | 0.520480116 | -4.225227505 |
| **A_55_P1980446** | Sbno1 | 0.178259901 | -0.049128179 | 0.40564798 | 10.58809897 | 1.904886729 | 0.104047554 | 0.520480116 | -4.22525071 |
| **A_65_P01415** | Kremen1 | -0.309550903 | -0.704474914 | 0.085373108 | 8.877388232 | -1.904590986 | 0.104090729 | 0.520561373 | -4.225399451 |
| **A_52_P325116** | Gramd1b | -0.215865945 | -0.491377555 | 0.059645666 | 8.756175956 | -1.903826716 | 0.104202387 | 0.520757844 | -4.225783856 |
| **A_55_P2117411** | NA | 0.145671714 | -0.04025285 | 0.331596279 | 11.58507995 | 1.903801426 | 0.104206084 | 0.520757844 | -4.225796576 |
| **A_52_P51078** | Ctsh | -0.780843205 | -1.777595486 | 0.215909076 | 14.31714916 | -1.903530266 | 0.10424573 | 0.520757844 | -4.22593297 |
| **A_55_P2165544** | Aebp2 | 0.191884562 | -0.053061502 | 0.436830625 | 9.098936863 | 1.903499464 | 0.104250234 | 0.520757844 | -4.225948463 |
| **A_51_P274504** | Tex264 | -0.233319535 | -0.531193434 | 0.064554364 | 10.79035469 | -1.903276309 | 0.104282874 | 0.520757844 | -4.226060714 |
| **A_51_P467110** | Dpp4 | -0.207026668 | -0.471350298 | 0.057296961 | 9.336954692 | -1.903152076 | 0.104301049 | 0.520757844 | -4.226123206 |
| **A_55_P2047932** | Cyb5r4 | 0.309236215 | -0.085639522 | 0.704111953 | 10.21762886 | 1.902887394 | 0.104339783 | 0.520757844 | -4.226256351 |
| **A_51_P215438** | Prodh | -0.180950321 | -0.4120176 | 0.050116958 | 8.994889471 | -1.902847995 | 0.10434555 | 0.520757844 | -4.22627617 |
| **A_51_P238734** | Mfsd11 | -0.127895136 | -0.29123393 | 0.035443659 | 8.352834919 | -1.902601658 | 0.104381614 | 0.520803358 | -4.226400091 |
| **A_55_P2141013** | Siglech | 0.188853581 | -0.052361248 | 0.430068409 | 14.28934457 | 1.902411272 | 0.104409494 | 0.520808031 | -4.226495867 |
| **A_51_P499441** | Prkd2 | -0.280371767 | -0.638627691 | 0.077884156 | 9.036673075 | -1.901621543 | 0.104525225 | 0.521177689 | -4.226893174 |
| **A_55_P2067146** | NA | 0.316842798 | -0.088054309 | 0.721739906 | 8.530015217 | 1.901438814 | 0.104552021 | 0.521177689 | -4.226985108 |
| **A_55_P2065646** | Zfp189 | 0.185534745 | -0.051572873 | 0.422642362 | 9.372707043 | 1.901353784 | 0.104564493 | 0.521177689 | -4.227027889 |
| **A_55_P2007693** | NA | -0.364553311 | -0.830781243 | 0.101674621 | 9.81440929 | -1.899966515 | 0.104768174 | 0.521923509 | -4.227725917 |
| **A_55_P2019615** | Neil1 | -0.163784244 | -0.373253113 | 0.045684625 | 8.930653713 | -1.899922215 | 0.104774685 | 0.521923509 | -4.227748209 |
| **A_55_P2151056** | Rarg | -0.633369625 | -1.443465465 | 0.176726214 | 9.734227064 | -1.899783096 | 0.104795134 | 0.521923509 | -4.227818216 |
| **A_55_P2120577** | Cited2 | -0.272098362 | -0.620223046 | 0.076026321 | 10.63097105 | -1.899215733 | 0.10487857 | 0.522088557 | -4.228103731 |
| **A_55_P2016237** | Hand2 | 0.239208357 | -0.066840687 | 0.545257401 | 8.603285622 | 1.899190418 | 0.104882294 | 0.522088557 | -4.22811647 |
| **A_55_P1963634** | Yae1d1 | 0.161458972 | -0.045190107 | 0.368108051 | 9.235364427 | 1.898505667 | 0.104983085 | 0.522231076 | -4.228461083 |
| **A_55_P2360266** | 4833411C07Rik | -0.457873164 | -1.043935972 | 0.128189643 | 10.20181849 | -1.89838467 | 0.105000905 | 0.522231076 | -4.22852198 |
| **A_51_P439403** | Padi1 | 0.299960127 | -0.084058824 | 0.683979077 | 8.618100219 | 1.897990617 | 0.10505896 | 0.522231076 | -4.228720308 |
| **A_55_P2073329** | NA | 0.585791364 | -0.164165964 | 1.335748691 | 11.37000787 | 1.897971123 | 0.105061833 | 0.522231076 | -4.22873012 |
| **A_51_P137604** | Fcna | -0.248028466 | -0.565619313 | 0.06956238 | 11.10985968 | -1.897652734 | 0.105108765 | 0.522231076 | -4.228890373 |
| **A_55_P2112622** | Smad4 | -0.328290989 | -0.748671774 | 0.092089796 | 11.15881574 | -1.897576502 | 0.105120005 | 0.522231076 | -4.228928743 |
| **A_55_P1971184** | Ube2l3 | 0.191308794 | -0.053703132 | 0.436320721 | 13.10372388 | 1.897277685 | 0.105164076 | 0.522231076 | -4.229079151 |
| **A_52_P184149** | Mthfd2 | 0.438552895 | -0.123115991 | 1.000221781 | 10.54163467 | 1.897251063 | 0.105168004 | 0.522231076 | -4.229092552 |
| **A_65_P16680** | B4galt5 | -0.156849224 | -0.357737947 | 0.044039499 | 10.39106051 | -1.897186406 | 0.105177542 | 0.522231076 | -4.229125097 |
| **A_55_P2000224** | Cd63 | -0.379179138 | -0.864906607 | 0.10654833 | 14.0731133 | -1.896858609 | 0.105225915 | 0.522231076 | -4.2292901 |
| **A_55_P2107192** | NA | -0.235327414 | -0.536783798 | 0.06612897 | 11.84530814 | -1.896842298 | 0.105228322 | 0.522231076 | -4.22929831 |
| **A_52_P406371** | Med14 | 0.248242832 | -0.069818361 | 0.566304025 | 9.652856719 | 1.896484173 | 0.105281196 | 0.522231076 | -4.229478586 |
| **A_51_P332201** | Cst3 | -0.360127019 | -0.821544338 | 0.101290301 | 12.92971753 | -1.896465744 | 0.105283918 | 0.522231076 | -4.229487863 |
| **A_51_P411264** | Txndc16 | -0.225782939 | -0.515075631 | 0.063509753 | 8.829503828 | -1.896430134 | 0.105289177 | 0.522231076 | -4.229505789 |
| **A_55_P2159522** | Col14a1 | -0.148963967 | -0.339863543 | 0.041935609 | 8.264296099 | -1.896092216 | 0.105339096 | 0.522344634 | -4.2296759 |

| | | | | | | | | | |
|---|---|---|---|---|---|---|---|---|---|
| **A_55_P2143693** | Sytl1 | -0.193865185 | -0.442341493 | 0.054611123 | 8.700468233 | -1.895824043 | 0.105378728 | 0.52240714 | -4.229810905 |
| **A_55_P2025363** | Ccnl2 | -0.166404063 | -0.379713929 | 0.046905803 | 8.986391768 | -1.895554045 | 0.105418645 | 0.522471025 | -4.229946833 |
| **A_51_P389751** | Relb | -0.278031077 | -0.634475433 | 0.07841328 | 8.693455296 | -1.895329786 | 0.105451812 | 0.522501428 | -4.230059737 |
| **A_65_P12392** | Timp2 | -0.378834665 | -0.864696178 | 0.107026848 | 8.926701473 | -1.894612519 | 0.10555796 | 0.522724425 | -4.230420866 |
| **A_55_P2080804** | Slirp | 0.204801926 | -0.057891473 | 0.467495325 | 12.99557797 | 1.894384214 | 0.105591769 | 0.522724425 | -4.23053582 |
| **A_51_P350332** | Rbpms | -0.230540837 | -0.526258478 | 0.065176805 | 8.644076417 | -1.894322075 | 0.105600973 | 0.522724425 | -4.230567107 |
| **A_52_P235479** | Usp38 | 0.212595017 | -0.060109596 | 0.485299631 | 9.622778653 | 1.894278305 | 0.105607457 | 0.522724425 | -4.230589146 |
| **A_55_P1953236** | Epb42 | 0.345507332 | -0.09784062 | 0.788855284 | 9.212343272 | 1.893632726 | 0.105703131 | 0.522724425 | -4.230914218 |
| **A_55_P2043451** | NA | 0.133571809 | -0.037831352 | 0.30497497 | 11.80225771 | 1.893560437 | 0.10571385 | 0.522724425 | -4.230950619 |
| **A_55_P2088325** | Ctdsp2 | 0.243548473 | -0.069034611 | 0.556131558 | 9.962752606 | 1.893228911 | 0.10576302 | 0.522724425 | -4.231117564 |
| **A_55_P2159830** | Yaf2 | 0.167730244 | -0.047548084 | 0.383008572 | 8.646761032 | 1.893190235 | 0.105768758 | 0.522724425 | -4.23113704 |
| **A_66_P123294** | NA | 0.230789284 | -0.065458277 | 0.527036845 | 8.214098747 | 1.892971368 | 0.105801233 | 0.522724425 | -4.231247257 |
| **A_51_P418259** | Pccb | 0.131097388 | -0.03719044 | 0.299385216 | 8.934667406 | 1.892886272 | 0.105813863 | 0.522724425 | -4.231290111 |
| **A_55_P1963873** | NA | -0.204333691 | -0.466638558 | 0.057971177 | 13.16348284 | -1.892852701 | 0.105818845 | 0.522724425 | -4.231307017 |
| **A_55_P2048011** | Larp4b | 0.224537482 | -0.06370572 | 0.512780684 | 12.79116914 | 1.892835896 | 0.10582134 | 0.522724425 | -4.23131548 |
| **A_55_P2086329** | Krt85 | 0.188947382 | -0.053698768 | 0.431593533 | 8.52158548 | 1.892128656 | 0.105926368 | 0.522996088 | -4.231671656 |
| **A_55_P2010699** | Map11 | 0.23909919 | -0.067985217 | 0.546183598 | 9.409680032 | 1.891923318 | 0.10595688 | 0.522996088 | -4.231775072 |
| **A_55_P2166773** | Pitpna | -0.130509874 | -0.298154444 | 0.037134695 | 12.39696016 | -1.891633822 | 0.105999914 | 0.522996088 | -4.231920877 |
| **A_55_P2075636** | Epc2 | 0.175416229 | -0.049981789 | 0.400814247 | 9.474266737 | 1.891049367 | 0.106086845 | 0.522996088 | -4.232215253 |
| **A_51_P366770** | Rims1 | 0.326016359 | -0.092894437 | 0.744927154 | 8.602537532 | 1.891041366 | 0.106088036 | 0.522996088 | -4.232219283 |
| **A_55_P2136817** | Coro2b | -0.569474979 | -1.301333192 | 0.162383234 | 9.142825526 | -1.890735965 | 0.10613349 | 0.522996088 | -4.232373114 |
| **A_55_P2085186** | NA | -0.276493134 | -0.631856806 | 0.078870538 | 9.692734315 | -1.890577617 | 0.106157065 | 0.522996088 | -4.232452875 |
| **A_51_P352594** | St5 | -0.143076033 | -0.326974097 | 0.04082203 | 9.139042328 | -1.890483644 | 0.106171058 | 0.522996088 | -4.232500211 |
| **A_52_P110070** | 5730416F02Rik | -0.205665246 | -0.470012791 | 0.0586823 | 8.217234766 | -1.890465761 | 0.106173721 | 0.522996088 | -4.23250922 |
| **A_55_P2120348** | Pdcd7 | 0.144837208 | -0.041374607 | 0.331049023 | 9.900539524 | 1.88997518 | 0.106246804 | 0.522996088 | -4.232756342 |
| **A_55_P2402134** | NA | 0.171448318 | -0.048980065 | 0.391876701 | 9.402929909 | 1.889943857 | 0.106251472 | 0.522996088 | -4.232772121 |
| **A_55_P1988433** | Stard7 | 0.179605483 | -0.051380419 | 0.410591385 | 12.39277477 | 1.889371264 | 0.106336839 | 0.522996088 | -4.233060574 |
| **A_55_P2020532** | Selenov | 0.202699856 | -0.058014348 | 0.463414059 | 8.503301197 | 1.8891739 | 0.10636628 | 0.522996088 | -4.233160003 |
| **A_66_P104340** | Atxn7l2 | 0.166696446 | -0.047725239 | 0.381118131 | 9.222603664 | 1.88903857 | 0.106386471 | 0.522996088 | -4.233228182 |
| **A_51_P507509** | Gimap1 | -0.387630196 | -0.886277425 | 0.111017033 | 9.192692126 | -1.8888931 | 0.10640818 | 0.522996088 | -4.23330147 |
| **A_55_P2040873** | NA | 0.334386299 | -0.095790508 | 0.764563106 | 9.919947819 | 1.888794337 | 0.106422921 | 0.522996088 | -4.233351227 |
| **A_66_P122758** | Tsr1 | 0.26114067 | -0.074856267 | 0.597137608 | 9.788356851 | 1.88852346 | 0.106463362 | 0.522996088 | -4.2334877 |
| **A_51_P418908** | Larp1 | 0.30703781 | -0.088046084 | 0.702121704 | 9.748418902 | 1.888364045 | 0.106487169 | 0.522996088 | -4.233568019 |
| **A_55_P2037439** | Depdc1a | 0.250283025 | -0.071773199 | 0.57233925 | 8.764982796 | 1.888351715 | 0.106489011 | 0.522996088 | -4.23357423 |
| **A_51_P380986** | Epc1 | 0.136086503 | -0.039032167 | 0.311205172 | 9.968619984 | 1.888277385 | 0.106500113 | 0.522996088 | -4.233611681 |
| **A_55_P1987222** | NA | 0.194342876 | -0.055775672 | 0.444461424 | 8.502752284 | 1.888017064 | 0.106539006 | 0.522996088 | -4.233742842 |
| **A_55_P2077994** | Ube2v2 | 0.245590326 | -0.070493825 | 0.561674477 | 9.68851012 | 1.887955371 | 0.106548225 | 0.522996088 | -4.233773927 |
| **A_55_P2260648** | 6330418K02Rik | 0.153258555 | -0.043993921 | 0.350511032 | 9.006272578 | 1.887928107 | 0.106552299 | 0.522996088 | -4.233787664 |
| **A_52_P164161** | Cyp51 | 0.399709296 | -0.114819479 | 0.914238072 | 10.45615873 | 1.887633907 | 0.106596276 | 0.522996088 | -4.233935901 |
| **A_55_P2166282** | NA | -0.287807131 | -0.658291535 | 0.082677273 | 9.305110787 | -1.887621039 | 0.1065982 | 0.522996088 | -4.233942385 |
| **A_55_P2152926** | Mybbp1a | 0.237751944 | -0.068310697 | 0.543814584 | 13.00512699 | 1.88754339 | 0.10660981 | 0.522996088 | -4.23398151 |
| **A_55_P1992720** | NA | 0.716358174 | -0.205934018 | 1.638650366 | 9.082744784 | 1.887317113 | 0.10664365 | 0.522996088 | -4.234095528 |
| **A_52_P370443** | Riok3 | 0.250381557 | -0.071999393 | 0.572762507 | 11.69871272 | 1.887192291 | 0.106662322 | 0.522996088 | -4.234158425 |
| **A_55_P2142724** | NA | 0.487054797 | -0.140063423 | 1.114173017 | 8.913225899 | 1.887172483 | 0.106665285 | 0.522996088 | -4.234168406 |
| **A_55_P2062652** | Zfp941 | 0.137358366 | -0.039514717 | 0.31423145 | 7.981252819 | 1.887020244 | 0.106688063 | 0.522996088 | -4.234245119 |
| **A_52_P423810** | Mt1 | 0.601421107 | -0.173169822 | 1.376012037 | 10.71987228 | 1.886641782 | 0.10674471 | 0.523141101 | -4.234435832 |
| **A_51_P326393** | Gapvd1 | 0.30333345 | -0.087419088 | 0.694085989 | 11.59293834 | 1.886260568 | 0.106801799 | 0.523256831 | -4.234627939 |
| **A_52_P420394** | Apeh | 0.30236389 | -0.087168156 | 0.691895935 | 13.09015186 | 1.886122608 | 0.106822467 | 0.523256831 | -4.234697464 |
| **A_55_P2041060** | Rbp3 | 0.250970181 | -0.07244338 | 0.574383742 | 10.79219105 | 1.885589221 | 0.10690241 | 0.52349922 | -4.234966273 |
| **A_52_P526852** | Eri1 | 0.279179681 | -0.080616313 | 0.638975675 | 9.046288513 | 1.885431122 | 0.106926117 | 0.52349922 | -4.235045952 |

| | | | | | | | | | |
|---|---|---|---|---|---|---|---|---|---|
| **A_55_P1983488** | Lsr | -0.217345675 | -0.497483331 | 0.06279198 | 13.35541273 | -1.885222883 | 0.106957351 | 0.523519533 | -4.235150904 |
| **A_52_P646755** | Tex30 | 0.293526476 | -0.084893934 | 0.671946886 | 10.67391724 | 1.884759396 | 0.107026901 | 0.523685412 | -4.235384506 |
| **A_51_P208410** | Ppp1r12c | -0.155716963 | -0.356483641 | 0.045049715 | 10.73134297 | -1.88463599 | 0.107045427 | 0.523685412 | -4.235446707 |
| **A_55_P2033287** | NA | 0.1929076 | -0.055854539 | 0.44166974 | 15.65625462 | 1.884292174 | 0.107097058 | 0.523685566 | -4.235620004 |
| **A_55_P2051592** | Gfm2 | 0.31243487 | -0.090483875 | 0.715353616 | 10.19454802 | 1.884192257 | 0.107112067 | 0.523685566 | -4.235670367 |
| **A_66_P107790** | Myl12a | -0.444615003 | -1.018098276 | 0.128868271 | 13.41358228 | -1.883850984 | 0.107163347 | 0.523685566 | -4.23584239 |
| **A_51_P219385** | Six5 | -0.192776767 | -0.44143029 | 0.055876757 | 8.654191923 | -1.883836742 | 0.107165488 | 0.523685566 | -4.235849569 |
| **A_55_P2133165** | Wwc1 | -0.394560175 | -0.903532092 | 0.114411741 | 9.369799667 | -1.883660439 | 0.10719199 | 0.523685566 | -4.235938439 |
| **A_52_P593268** | Lsm6 | -0.165906439 | -0.379933354 | 0.048120477 | 13.034613 | -1.883553829 | 0.107208018 | 0.523685566 | -4.235992179 |
| **A_51_P267508** | Borcs7 | 0.249493645 | -0.072559591 | 0.57154688 | 9.224014995 | 1.882413417 | 0.107379626 | 0.524268917 | -4.23656708 |
| **A_52_P576100** | Sh2b1 | -0.18239251 | -0.417831475 | 0.053046454 | 10.93377152 | -1.882399728 | 0.107381688 | 0.524268917 | -4.236573981 |
| **A_55_P1971639** | Abitram | 0.151478245 | -0.044104413 | 0.347060902 | 9.580910172 | 1.881928462 | 0.107452684 | 0.524332052 | -4.236811574 |
| **A_66_P106585** | Sae1 | 0.176716715 | -0.05147769 | 0.404911119 | 10.04275548 | 1.881723578 | 0.107483564 | 0.524332052 | -4.236914871 |
| **A_55_P2102738** | NA | 0.236175784 | -0.068813617 | 0.541165185 | 13.83697156 | 1.881628174 | 0.107497947 | 0.524332052 | -4.236962973 |
| **A_55_P2042879** | Mbtd1 | 0.218081299 | -0.063552554 | 0.499715152 | 8.593898362 | 1.881554305 | 0.107509084 | 0.524332052 | -4.237000217 |
| **A_55_P2117420** | NA | 0.214517269 | -0.062562018 | 0.491596556 | 15.96390608 | 1.881227799 | 0.107558325 | 0.524332052 | -4.237164841 |
| **A_52_P252258** | Mapk12 | -0.302801609 | -0.693943868 | 0.08834065 | 8.666961551 | -1.881077238 | 0.107581039 | 0.524332052 | -4.237240755 |
| **A_55_P2015670** | Itga6 | -0.781068897 | -1.790022534 | 0.227884739 | 12.08972882 | -1.881054262 | 0.107584506 | 0.524332052 | -4.23725234 |
| **A_55_P1988872** | NA | 0.366008542 | -0.106935726 | 0.838952809 | 9.851085346 | 1.880461491 | 0.107673982 | 0.524473253 | -4.237551234 |
| **A_55_P2062714** | Zfp975 | 0.196147986 | -0.057364842 | 0.449660814 | 8.372033097 | 1.880040051 | 0.107737641 | 0.524473253 | -4.237763748 |
| **A_66_P108019** | Heatr1 | 0.264418589 | -0.077335078 | 0.606172257 | 11.91999432 | 1.880017923 | 0.107740984 | 0.524473253 | -4.237774907 |
| **A_55_P1960916** | Egln3 | -0.402503566 | -0.922777442 | 0.11777031 | 9.329129529 | -1.879840065 | 0.107767862 | 0.524473253 | -4.237864596 |
| **A_65_P03606** | Rpap2 | 0.138421195 | -0.040523519 | 0.317365909 | 8.228235872 | 1.879606372 | 0.107803188 | 0.524473253 | -4.237982444 |
| **A_55_P2077751** | NA | 0.282938705 | -0.082837722 | 0.648715132 | 8.997932777 | 1.879575766 | 0.107807815 | 0.524473253 | -4.237997878 |
| **A_51_P383399** | Aldh1a7 | -1.037434676 | -2.378916967 | 0.304047616 | 9.848593632 | -1.879139428 | 0.107873806 | 0.524473253 | -4.238217925 |
| **A_51_P111612** | Arrdc4 | -0.177638275 | -0.40734618 | 0.052069631 | 8.607205469 | -1.879073613 | 0.107883764 | 0.524473253 | -4.240936545 |
| **A_51_P425373** | Smg9 | 0.209723531 | -0.061482098 | 0.480929159 | 9.963278492 | 1.879021075 | 0.107891713 | 0.524473253 | -4.238277613 |
| **A_55_P2020786** | Prkar1a | -0.124266508 | -0.284981449 | 0.036448433 | 12.78663102 | -1.878802067 | 0.107924856 | 0.524473253 | -4.238388064 |
| **A_55_P2017418** | Cfh | -0.482125151 | -1.105668128 | 0.141417826 | 13.45051798 | -1.87878284 | 0.107927766 | 0.524473253 | -4.238397761 |
| **A_55_P1968053** | NA | 0.142024831 | -0.041666294 | 0.325715956 | 15.68649256 | 1.878708054 | 0.107939086 | 0.524473253 | -4.238435478 |
| **A_55_P2032081** | Dbp | -0.341853932 | -0.784183577 | 0.100475713 | 8.871090544 | -1.877922741 | 0.108058028 | 0.524919234 | -4.238831556 |
| **A_52_P42069** | Prokr1 | 0.304431006 | -0.089519821 | 0.698381833 | 8.524319789 | 1.877716642 | 0.108089265 | 0.524939048 | -4.238935509 |
| **A_55_P2069721** | Hspa8 | 0.164132892 | -0.048290544 | 0.376556328 | 14.91818687 | 1.877484589 | 0.108124447 | 0.524978002 | -4.239052554 |
| **A_55_P2044843** | NA | 0.243444294 | -0.071661435 | 0.558550023 | 15.1565579 | 1.877268916 | 0.108157155 | 0.525004932 | -4.239161341 |
| **A_55_P2014570** | Zfp961 | 0.133302567 | -0.03935351 | 0.305958645 | 10.20745645 | 1.876030231 | 0.108345198 | 0.525567516 | -4.239786185 |
| **A_52_P447424** | NA | 0.187807908 | -0.055444868 | 0.431060684 | 9.104336061 | 1.876027753 | 0.108345575 | 0.525567516 | -4.239787436 |
| **A_55_P2047310** | Pdgfb | -0.783267379 | -1.797929391 | 0.231394634 | 9.63245389 | -1.875736491 | 0.108389838 | 0.525567516 | -4.239934371 |
| **A_55_P2003643** | Tstd2 | 0.171840391 | -0.050774409 | 0.39445519 | 9.594936993 | 1.875661256 | 0.108401274 | 0.525567516 | -4.239972326 |
| **A_51_P461475** | Atn1 | -0.244039214 | -0.56024 | 0.072161572 | 8.73803134 | -1.875339326 | 0.108450224 | 0.525567516 | -4.24013474 |
| **A_51_P246816** | Napa | 0.225553218 | -0.06672611 | 0.517832546 | 10.10476476 | 1.875141805 | 0.108480269 | 0.525567516 | -4.240234392 |
| **A_55_P2134690** | Ankrd26 | 0.172906683 | -0.051204184 | 0.39701755 | 8.559236959 | 1.874701184 | 0.10854732 | 0.525567516 | -4.240456698 |
| **A_55_P2005859** | Fn3k | 0.428172549 | -0.1268441 | 0.983189197 | 10.0503736 | 1.874545552 | 0.108571013 | 0.525567516 | -4.240535221 |
| **A_51_P347961** | Npc2 | -0.760296082 | -1.745947354 | 0.225355191 | 12.56047589 | -1.874315317 | 0.108606072 | 0.525567516 | -4.240651387 |
| **A_55_P2139396** | Unc45b | -0.143094465 | -0.3286102 | 0.042421271 | 8.002267556 | -1.874240292 | 0.108617499 | 0.525567516 | -4.240689241 |
| **A_52_P123944** | Mospd2 | -0.172439738 | -0.396044852 | 0.051165375 | 8.25021486 | -1.873867217 | 0.10867434 | 0.525567516 | -4.240877485 |
| **A_55_P2026738** | Rgs4 | -0.154629196 | -0.355144627 | 0.045886236 | 7.745024142 | -1.873815752 | 0.108682183 | 0.525567516 | -4.240903453 |
| **A_51_P434426** | Rwdd4a | 0.234121592 | -0.069518232 | 0.537761416 | 9.32061165 | 1.873552759 | 0.108722273 | 0.525567516 | -4.241036157 |
| **A_55_P2019312** | Car12 | -0.123952184 | -0.284742483 | 0.036838115 | 7.900772837 | -1.873171447 | 0.108780424 | 0.525567516 | -4.24122857 |
| **A_52_P432580** | NA | -0.445510506 | -1.023558882 | 0.13253787 | 11.27776553 | -1.872737702 | 0.108846609 | 0.525567516 | -4.241447449 |
| **A_55_P1975560** | Ifi204 | -0.374718446 | -0.860920252 | 0.11148336 | 10.66575003 | -1.872715021 | 0.108850071 | 0.525567516 | -4.241458895 |

| | | | | | | | | | |
|---|---|---|---|---|---|---|---|---|---|
| **A_52_P648424** | Nedd4 | -0.418062361 | -0.960610068 | 0.124485345 | 12.27334912 | -1.872346945 | 0.108906269 | 0.525567516 | -4.241644644 |
| **A_52_P221776** | Kif12 | 0.330391453 | -0.09838959 | 0.759172496 | 8.947469959 | 1.87230422 | 0.108912794 | 0.525567516 | -4.241666206 |
| **A_55_P2078138** | Otop1 | 0.586053212 | -0.174678893 | 1.346785317 | 11.26247404 | 1.871925215 | 0.108970694 | 0.525567516 | -4.241857478 |
| **A_55_P2160323** | NA | 0.19976935 | -0.059558911 | 0.459097611 | 8.480453549 | 1.871811871 | 0.108988015 | 0.525567516 | -4.24191468 |
| **A_55_P2113165** | Camkk1 | -0.231881158 | -0.53289991 | 0.069137594 | 8.4138577 | -1.871781276 | 0.108992691 | 0.525567516 | -4.241930121 |
| **A_51_P127297** | Hsd11b1 | -0.980327412 | -2.253027487 | 0.292372663 | 12.5130407 | -1.871665633 | 0.109010367 | 0.525567516 | -4.255668553 |
| **A_66_P108792** | Rad1 | 0.199760415 | -0.059592714 | 0.459113543 | 10.47235031 | 1.871548681 | 0.109028246 | 0.525567516 | -4.24204751 |
| **A_55_P2151951** | NA | 0.168142372 | -0.050166034 | 0.386450778 | 13.87887046 | 1.8715003 | 0.109035643 | 0.525567516 | -4.242071928 |
| **A_55_P2096737** | Elmod3 | -0.271449342 | -0.623887811 | 0.080989128 | 9.858052633 | -1.871495016 | 0.109036451 | 0.525567516 | -4.242074595 |
| **A_55_P2116105** | Ap2a1 | 0.325561466 | -0.097191195 | 0.748314126 | 11.85640741 | 1.871241473 | 0.109075224 | 0.525567516 | -4.242202559 |
| **A_51_P487813** | Lxn | -0.913571043 | -2.116012667 | 0.288870581 | 10.61073771 | -1.932950312 | 0.109080035 | 0.525567516 | -4.271194188 |
| **A_51_P430552** | G6pc3 | -0.164654019 | -0.378469407 | 0.04916137 | 10.75684377 | -1.871184297 | 0.10908397 | 0.525567516 | -4.242231417 |
| **A_55_P2002597** | NA | -0.124959003 | -0.287259142 | 0.037341135 | 9.361054801 | -1.87081934 | 0.10913981 | 0.525567516 | -4.24241562 |
| **A_51_P313761** | Shmt2 | 0.28806878 | -0.086116199 | 0.662253758 | 12.4831492 | 1.870652131 | 0.109165403 | 0.525567516 | -4.242500016 |
| **A_51_P297694** | Prpf6 | -0.217028674 | -0.498954059 | 0.064896711 | 11.56237338 | -1.870536227 | 0.109183147 | 0.525567516 | -4.242558518 |
| **A_55_P2153382** | Ermap | 0.459943175 | -0.137566504 | 1.057452855 | 11.33622922 | 1.870434149 | 0.109198776 | 0.525567516 | -4.242610042 |
| **A_55_P2121945** | Cnot6l | -0.20990163 | -0.482592713 | 0.062789453 | 11.39814225 | -1.870372306 | 0.109208246 | 0.525567516 | -4.242641257 |
| **A_55_P2138097** | NA | -0.319412885 | -0.734394596 | 0.095568825 | 14.79254864 | -1.870280148 | 0.10922236 | 0.525567516 | -4.242687775 |
| **A_51_P374869** | Abca7 | 0.277069559 | -0.082902609 | 0.637041727 | 12.23865246 | 1.870264707 | 0.109224725 | 0.525567516 | -4.242695569 |
| **A_55_P2093727** | Gemin6 | 0.275693452 | -0.082582456 | 0.633969359 | 10.10992243 | 1.869786567 | 0.109297982 | 0.525751171 | -4.242936921 |
| **A_55_P2060278** | Fam45a | 0.286080097 | -0.085718771 | 0.657878964 | 10.67077191 | 1.869660576 | 0.109317293 | 0.525751171 | -4.24300052 |
| **A_55_P1959748** | Asns | 0.326618874 | -0.098005657 | 0.751243405 | 14.35438593 | 1.869043402 | 0.109411939 | 0.525800829 | -4.243312073 |
| **A_66_P113578** | Washc1 | -0.150026078 | -0.345086947 | 0.045034792 | 11.04222331 | -1.868872993 | 0.109438087 | 0.525800829 | -4.243398099 |
| **A_55_P1965070** | Map7d1 | -0.219672722 | -0.505290357 | 0.065944913 | 12.64390785 | -1.868849383 | 0.10944171 | 0.525800829 | -4.243410019 |
| **A_55_P2120433** | Wdr12 | 0.26018122 | -0.078115344 | 0.598477785 | 9.062804049 | 1.868794503 | 0.109450132 | 0.525800829 | -4.243437723 |
| **A_55_P2156731** | H2-Eb1 | -0.295502773 | -0.679743718 | 0.088738172 | 9.46537888 | -1.868706549 | 0.109463632 | 0.525800829 | -4.243482126 |
| **A_55_P2159746** | Mfsd2b | 0.324799756 | -0.09761468 | 0.747214193 | 9.078145084 | 1.868358154 | 0.10951712 | 0.525927061 | -4.243658012 |
| **A_52_P632691** | Zfp395 | 0.172120703 | -0.051828237 | 0.396069643 | 9.177355435 | 1.867528714 | 0.109644567 | 0.52631318 | -4.244076775 |
| **A_55_P2011712** | NA | 0.23759708 | -0.071560925 | 0.546755085 | 8.956472818 | 1.867427681 | 0.109660102 | 0.52631318 | -4.244127787 |
| **A_51_P382970** | Itga9 | -0.425533352 | -0.979268261 | 0.128201557 | 10.06190757 | -1.867303408 | 0.109679212 | 0.52631318 | -4.244190532 |
| **A_52_P251366** | Neil3 | 0.629032567 | -0.189746891 | 1.447812025 | 9.556201349 | 1.866763848 | 0.109762223 | 0.526505179 | -4.244462967 |
| **A_55_P2074776** | Gm8894 | -0.335016924 | -0.771108708 | 0.10107486 | 15.4662615 | -1.86668927 | 0.109773701 | 0.526505179 | -4.244500623 |
| **A_51_P177432** | NA | 0.234241182 | -0.070788182 | 0.539270545 | 8.492381045 | 1.865970574 | 0.109884379 | 0.526888429 | -4.244863532 |
| **A_52_P225898** | Kcnj8 | -0.187625613 | -0.43199313 | 0.056741904 | 8.055459426 | -1.86565715 | 0.109932681 | 0.526888429 | -4.245021804 |
| **A_55_P1982111** | Homer1 | 0.22593219 | -0.068329379 | 0.520193759 | 8.686274961 | 1.865639615 | 0.109935384 | 0.526888429 | -4.245030659 |
| **A_66_P126313** | Rhof | -0.573435522 | -1.320463382 | 0.173592337 | 9.056400346 | -1.865223948 | 0.109999476 | 0.527064915 | -4.24524057 |
| **A_55_P2091473** | Bnip3 | -0.317461762 | -0.731106984 | 0.096183459 | 14.75923912 | -1.864861574 | 0.110055381 | 0.527110207 | -4.245423574 |
| **A_51_P175771** | Flii | -0.210272084 | -0.484274071 | 0.063729902 | 10.80805787 | -1.86470913 | 0.110078907 | 0.527110207 | -4.245500563 |
| **A_55_P1954633** | NA | -0.252771382 | -0.582167025 | 0.07662426 | 10.19521639 | -1.864632471 | 0.11009074 | 0.527110207 | -4.245539278 |
| **A_52_P261322** | Tanc1 | -0.161412684 | -0.371807601 | 0.048982233 | 7.796994621 | -1.864170382 | 0.110162091 | 0.527142456 | -4.245772655 |
| **A_55_P2018847** | Crlf2 | -0.244932151 | -0.564264826 | 0.074400523 | 11.25022452 | -1.863741292 | 0.110228388 | 0.527142456 | -4.245989374 |
| **A_55_P1994020** | Casc4 | -0.180682173 | -0.41626873 | 0.054904384 | 8.690671881 | -1.863579775 | 0.110253354 | 0.527142456 | -4.246070954 |
| **A_52_P533198** | Dnajc2 | 0.350661862 | -0.106568651 | 0.807892376 | 9.698159692 | 1.863530557 | 0.110260963 | 0.527142456 | -4.246095813 |
| **A_55_P2153291** | NA | 0.161630267 | -0.049130122 | 0.372390655 | 15.33759315 | 1.863446328 | 0.110273985 | 0.527142456 | -4.246138357 |
| **A_55_P1952414** | Odf2 | 0.287272711 | -0.087351546 | 0.661896969 | 9.896326286 | 1.863295211 | 0.110297353 | 0.527142456 | -4.246214685 |
| **A_51_P468240** | Dennd4a | 0.41018032 | -0.124744501 | 0.945105141 | 9.885682882 | 1.863224811 | 0.11030824 | 0.527142456 | -4.246250244 |
| **A_55_P2090600** | Mageb16-ps1 | 0.129244241 | -0.039328164 | 0.297816645 | 7.766329894 | 1.862978769 | 0.1103463 | 0.527142456 | -4.246374522 |
| **A_51_P290576** | Plk2 | -0.488552081 | -1.125816306 | 0.148712144 | 8.969776881 | -1.862835553 | 0.11036846 | 0.527142456 | -4.246446863 |
| **A_55_P2014867** | Iws1 | 0.121163953 | -0.036882488 | 0.279210395 | 8.611297658 | 1.862824337 | 0.110370196 | 0.527142456 | -4.246452529 |
| **A_55_P2060607** | NA | 0.140557911 | -0.042885811 | 0.324001633 | 15.80872601 | 1.861811161 | 0.110527094 | 0.527231141 | -4.246964331 |

| | | | | | | | | | |
|---|---|---|---|---|---|---|---|---|---|
| **A_51_P505795** | Tapbpl | -0.216678007 | -0.499467959 | 0.066111944 | 9.401788113 | -1.861804386 | 0.110528144 | 0.527231141 | -4.246967754 |
| **A_51_P118763** | Ahctf1 | 0.358574204 | -0.109434752 | 0.82658316 | 10.35456901 | 1.861693008 | 0.110545405 | 0.527231141 | -4.260840458 |
| **A_51_P509098** | Asah1 | -0.299904777 | -0.691371229 | 0.091561676 | 9.953892997 | -1.861538497 | 0.110569356 | 0.527231141 | -4.247102075 |
| **A_55_P2089070** | 4933431K14Rik | 0.231612715 | -0.070749255 | 0.533974685 | 9.373057867 | 1.861308753 | 0.110604978 | 0.527231141 | -4.247218139 |
| **A_55_P2127765** | Crp | -0.489642625 | -1.128876879 | 0.149591629 | 10.16613994 | -1.861239957 | 0.110615647 | 0.527231141 | -4.247252895 |
| **A_51_P199168** | Cidea | -0.340504277 | -0.785066866 | 0.104058312 | 8.258584413 | -1.861113447 | 0.110635269 | 0.527231141 | -4.247316808 |
| **A_66_P117659** | Nfatc3 | 0.248038842 | -0.075858049 | 0.571935733 | 9.184354984 | 1.860784606 | 0.110686289 | 0.527231141 | -4.247482942 |
| **A_51_P147034** | Ica1l | 0.218489084 | -0.066893651 | 0.503871819 | 8.464057807 | 1.860309643 | 0.110760021 | 0.527231141 | -4.247722908 |
| **A_52_P598300** | Gtf2f2 | 0.170849516 | -0.052341479 | 0.394040511 | 9.951198674 | 1.860031395 | 0.110803238 | 0.527231141 | -4.247863493 |
| **A_55_P2150906** | Atp5c1 | 0.190749194 | -0.058472222 | 0.43997061 | 12.93601735 | 1.859775584 | 0.110842985 | 0.527231141 | -4.247992744 |
| **A_51_P332932** | NA | 0.249612222 | -0.076518324 | 0.575742769 | 8.781713847 | 1.859762806 | 0.110844971 | 0.527231141 | -4.2479992 |
| **A_55_P2172624** | Gm1123 | 0.274392908 | -0.084140046 | 0.632925863 | 8.627345397 | 1.85963192 | 0.110865314 | 0.527231141 | -4.248065333 |
| **A_55_P1957876** | NA | 0.178963337 | -0.054879306 | 0.412805979 | 10.08240443 | 1.859617166 | 0.110867607 | 0.527231141 | -4.248072788 |
| **A_51_P211934** | Larp7 | 0.2044478 | -0.062734833 | 0.471630432 | 10.65601785 | 1.859333978 | 0.110911634 | 0.527231141 | -4.248215878 |
| **A_55_P1994477** | Prrc2c | 0.126605127 | -0.038860075 | 0.292070328 | 9.199705994 | 1.85920727 | 0.110931338 | 0.527231141 | -4.248279902 |
| **A_55_P1974967** | H4c14 | 0.508185144 | -0.15603632 | 1.172406609 | 9.069776425 | 1.859055041 | 0.110955016 | 0.527231141 | -4.248356823 |
| **A_55_P2117710** | Snhg1 | 0.182781483 | -0.056144263 | 0.421707228 | 9.897183413 | 1.858884624 | 0.11098153 | 0.527231141 | -4.248442936 |
| **A_52_P150525** | Ndufaf6 | 0.287893973 | -0.088471941 | 0.664259887 | 10.91951088 | 1.858683646 | 0.111012805 | 0.527231141 | -4.248544493 |
| **A_55_P2105286** | Lrrc56 | 0.398229228 | -0.122381844 | 0.9188403 | 10.1363858 | 1.858672701 | 0.111014509 | 0.527231141 | -4.248550024 |
| **A_55_P2051313** | Gstk1 | -0.511234708 | -1.179688611 | 0.157219195 | 10.5375214 | -1.858369444 | 0.111061718 | 0.527231141 | -4.248703267 |
| **A_52_P239787** | Ccdc25 | 0.209058131 | -0.064293644 | 0.482409906 | 8.614474987 | 1.858353601 | 0.111064185 | 0.527231141 | -4.248711273 |
| **A_55_P2138366** | Xpo4 | 0.359914454 | -0.110688993 | 0.830517901 | 9.428492616 | 1.858349405 | 0.111064838 | 0.527231141 | -4.248713394 |
| **A_55_P1963080** | Rps12 | 0.2064487 | -0.063500298 | 0.476397698 | 16.69051827 | 1.858290563 | 0.111074001 | 0.527231141 | -4.248743129 |
| **A_51_P275976** | Dok1 | 0.255736636 | -0.078673559 | 0.590146831 | 9.855831876 | 1.858217851 | 0.111085325 | 0.527231141 | -4.248779873 |
| **A_55_P2154714** | Klhl9 | 0.239542997 | -0.073705502 | 0.552791496 | 12.86791824 | 1.858136736 | 0.111097958 | 0.527231141 | -4.248820864 |
| **A_55_P2088530** | Htt | 0.267035861 | -0.082231475 | 0.616303196 | 10.92542046 | 1.857782232 | 0.111153188 | 0.527297931 | -4.249000013 |
| **A_66_P115378** | Orc2 | 0.258635786 | -0.079660404 | 0.596931975 | 8.709019517 | 1.857696221 | 0.111166593 | 0.527297931 | -4.24904348 |
| **A_51_P114177** | Eci3 | -0.158840029 | -0.366648151 | 0.048968093 | 10.98288073 | -1.857293891 | 0.111229314 | 0.527465999 | -4.26531669 |
| **A_55_P2011380** | Acvrl1 | -0.767189778 | -1.771118762 | 0.236739206 | 9.588826617 | -1.856876432 | 0.111294431 | 0.527645342 | -4.249457788 |
| **A_55_P2145029** | Gm826 | 0.245559535 | -0.075808858 | 0.566927929 | 8.478875085 | 1.856679021 | 0.111325237 | 0.527661969 | -4.249557562 |
| **A_55_P2079485** | Ptges3 | 0.225195374 | -0.069648208 | 0.520038956 | 12.34860566 | 1.855884629 | 0.111449288 | 0.527811864 | -4.249959072 |
| **A_51_P284244** | Tmed3 | -0.216357186 | -0.499634515 | 0.066920144 | 13.17560798 | -1.855849238 | 0.111454817 | 0.527811864 | -4.24997696 |
| **A_55_P2081656** | NA | 0.489675689 | -0.151516965 | 1.130868343 | 12.19340188 | 1.855680457 | 0.111481193 | 0.527811864 | -4.250062272 |
| **A_51_P232142** | Thumpd3 | 0.178071576 | -0.055100568 | 0.411243719 | 11.07941389 | 1.85567162 | 0.111482574 | 0.527811864 | -4.250066738 |
| **A_55_P1962214** | Kpna3 | 0.242746458 | -0.075143888 | 0.560636804 | 9.679145776 | 1.855490567 | 0.111510874 | 0.527811864 | -4.250158254 |
| **A_55_P2010386** | Tspan32 | 0.38259159 | -0.118470306 | 0.883653485 | 10.90423737 | 1.855355906 | 0.111531928 | 0.527811864 | -4.250226321 |
| **A_52_P134556** | Cd2bp2 | 0.188527711 | -0.058413265 | 0.435468688 | 9.664256037 | 1.85509102 | 0.111573352 | 0.527811864 | -4.250360216 |
| **A_55_P2005552** | Arhgef10l | -0.21860911 | -0.504953817 | 0.067735598 | 10.91123309 | -1.855078468 | 0.111575316 | 0.527811864 | -4.250366561 |
| **A_55_P2095010** | NA | 0.205207389 | -0.063656528 | 0.474071305 | 7.983949512 | 1.854571845 | 0.111654588 | 0.527863503 | -4.250622658 |
| **A_51_P159453** | Serpina3n | -0.564588892 | -1.304327973 | 0.175150189 | 8.944472689 | -1.854543204 | 0.111659072 | 0.527863503 | -4.250637136 |
| **A_51_P230507** | Ell2 | 0.323127245 | -0.100255747 | 0.746510237 | 9.714355531 | 1.854485146 | 0.11166816 | 0.527863503 | -4.250666485 |
| **A_51_P192589** | Pno1 | 0.278044988 | -0.086416842 | 0.642506817 | 11.64583418 | 1.853728845 | 0.111786618 | 0.528170316 | -4.251048819 |
| **A_52_P380329** | Tent4a | 0.167575232 | -0.05208349 | 0.387233954 | 10.14750156 | 1.85372184 | 0.111787716 | 0.528170316 | -4.251052361 |
| **A_51_P126302** | Rbmx2 | 0.165629059 | -0.051501056 | 0.382759174 | 8.41651081 | 1.853530175 | 0.111817756 | 0.528183141 | -4.254831875 |
| **A_51_P460734** | Prkcd | -0.19101663 | -0.441457043 | 0.059423783 | 9.776780642 | -1.85331805 | 0.111851013 | 0.528211147 | -4.2512565 |
| **A_51_P349008** | Ptges2 | -0.15119731 | -0.349461787 | 0.047067168 | 10.11511122 | -1.853029538 | 0.11189626 | 0.528295754 | -4.251402365 |
| **A_51_P256202** | Ctsz | -0.151336021 | -0.349837521 | 0.047165479 | 12.76918353 | -1.852514893 | 0.111977018 | 0.528515348 | -4.251662566 |
| **A_66_P124925** | Pdcd5 | 0.20192365 | -0.062981375 | 0.466828675 | 12.96329271 | 1.852167167 | 0.112031615 | 0.528515348 | -4.25183838 |
| **A_52_P355014** | Rbm8a | 0.147555151 | -0.0460325 | 0.341142802 | 11.86236587 | 1.852080737 | 0.112045189 | 0.528515348 | -4.251882081 |
| **A_51_P461504** | Eef2k | 0.213822101 | -0.06674121 | 0.494385411 | 9.258994173 | 1.851846155 | 0.112082041 | 0.528515348 | -4.252000693 |

| | | | | | | | | | |
|---|---|---|---|---|---|---|---|---|---|
| **A_55_P1985304** | Krtap5-3 | 0.243911132 | -0.076159173 | 0.563981436 | 8.54888251 | 1.851694885 | 0.112105811 | 0.528515348 | -4.252077181 |
| **A_55_P2020306** | Aasdh | 0.127070615 | -0.039677301 | 0.293818531 | 9.434688997 | 1.851688395 | 0.11210683 | 0.528515348 | -4.252080462 |
| **A_55_P2035951** | Haus8 | 0.208578601 | -0.065164374 | 0.482321576 | 12.08416095 | 1.851441342 | 0.112145662 | 0.52853444 | -4.252205384 |
| **A_51_P516728** | Hap1 | 0.611080173 | -0.191014163 | 1.413174509 | 11.93713543 | 1.851211043 | 0.112181873 | 0.52853444 | -4.252321837 |
| **A_51_P119031** | Naa30 | 0.249471857 | -0.078002776 | 0.57694649 | 9.591436078 | 1.851088082 | 0.112201211 | 0.52853444 | -4.260451101 |
| **A_55_P2023200** | Miox | 0.47468433 | -0.14856549 | 1.09793415 | 10.28699667 | 1.850656949 | 0.112269042 | 0.52853444 | -4.25260203 |
| **A_55_P1955827** | NA | 0.273648003 | -0.085683236 | 0.632979242 | 12.45413577 | 1.850463387 | 0.112299508 | 0.52853444 | -4.252699912 |
| **A_55_P2095794** | Evi5 | 0.26308322 | -0.082446347 | 0.608612788 | 11.54223123 | 1.850082613 | 0.112359466 | 0.52853444 | -4.252892472 |
| **A_55_P1982916** | Tspoap1 | 0.267250917 | -0.083759116 | 0.618260949 | 8.606894621 | 1.850047431 | 0.112365007 | 0.52853444 | -4.252910264 |
| **A_55_P2066180** | Hid1 | -0.484301994 | -1.120544889 | 0.151940901 | 8.779753368 | -1.849594394 | 0.112436388 | 0.52853444 | -4.253139377 |
| **A_55_P1964076** | Dnajb6 | 0.138549847 | -0.043476177 | 0.32057587 | 9.465904806 | 1.849506052 | 0.112450312 | 0.52853444 | -4.253184055 |
| **A_55_P2078204** | Prr5 | 0.145438338 | -0.045638503 | 0.336515179 | 9.376675548 | 1.849498735 | 0.112451465 | 0.52853444 | -4.253187756 |
| **A_52_P429774** | Cnst | 0.145715797 | -0.045728017 | 0.33715961 | 9.356091853 | 1.849475096 | 0.112455192 | 0.52853444 | -4.253199711 |
| **A_55_P1969002** | Tbx3 | -0.935915875 | -2.16555396 | 0.29372221 | 9.756171415 | -1.849450436 | 0.112459079 | 0.52853444 | -4.253212183 |
| **A_51_P325836** | Hpx | -0.350112946 | -0.810114712 | 0.109888819 | 14.72427835 | -1.84940429 | 0.112466354 | 0.52853444 | -4.253235521 |
| **A_55_P1993609** | Slx4ip | 0.153499839 | -0.048255135 | 0.355254813 | 8.752405545 | 1.848701798 | 0.112577156 | 0.528803655 | -4.253590814 |
| **A_55_P2089677** | Synrg | 0.393819277 | -0.123845578 | 0.911484133 | 10.78884501 | 1.848551177 | 0.112600927 | 0.528803655 | -4.253666996 |
| **A_66_P127160** | Eif2b3 | 0.238948617 | -0.075148071 | 0.553045305 | 9.112186657 | 1.848520852 | 0.112605714 | 0.528803655 | -4.253682333 |
| **A_55_P2140383** | Mis18a | 0.241755339 | -0.076205832 | 0.559716509 | 9.705428797 | 1.847503096 | 0.112766473 | 0.529314703 | -4.254197123 |
| **A_52_P22781** | Zfp866 | 0.183102883 | -0.057726516 | 0.423932282 | 9.533891986 | 1.847433732 | 0.112777438 | 0.529314703 | -4.25423221 |
| **A_55_P2065465** | Mettl1 | 0.223005416 | -0.070358067 | 0.516368898 | 10.02575052 | 1.847109095 | 0.112828769 | 0.529314703 | -4.254396424 |
| **A_55_P2033520** | Med6 | 0.206379493 | -0.065117824 | 0.47787681 | 10.20335805 | 1.847073547 | 0.112834391 | 0.529314703 | -4.254414406 |
| **A_55_P2073845** | Mrpl47 | 0.215330161 | -0.067974395 | 0.498634716 | 9.040730235 | 1.846862209 | 0.112867821 | 0.529314703 | -4.254521313 |
| **A_55_P2007733** | Epn1 | 0.14756989 | -0.046591483 | 0.341731263 | 14.68126668 | 1.846792527 | 0.112878846 | 0.529314703 | -4.254556562 |
| **A_51_P215066** | Emc4 | -0.241965839 | -0.560378483 | 0.076446805 | 11.9556568 | -1.846489917 | 0.112926735 | 0.529410832 | -4.254709643 |
| **A_55_P2359807** | NA | 0.248968331 | -0.078760518 | 0.576697181 | 9.017887728 | 1.845918932 | 0.113017151 | 0.529701336 | -4.254998498 |
| **A_51_P310576** | Smim8 | 0.157705321 | -0.049908403 | 0.365319046 | 8.293822734 | 1.845752552 | 0.11304351 | 0.529701336 | -4.255082671 |
| **A_55_P2120919** | NA | 0.232280762 | -0.073555804 | 0.538117329 | 8.259118768 | 1.845470137 | 0.113088268 | 0.529782628 | -4.255225549 |
| **A_55_P1997400** | Slc7a4 | -0.25611081 | -0.593390542 | 0.081168921 | 9.26270804 | -1.845104021 | 0.113146315 | 0.529798462 | -4.255410777 |
| **A_51_P430620** | Slc25a12 | 0.222435669 | -0.070496447 | 0.515367784 | 10.49906086 | 1.845103069 | 0.113146467 | 0.529798462 | -4.255411259 |
| **A_51_P193336** | Nucb2 | -0.376619378 | -0.872708896 | 0.11947014 | 8.762334642 | -1.844700222 | 0.113210372 | 0.529907348 | -4.255615078 |
| **A_51_P344399** | Rilpl2 | -0.20548931 | -0.476260751 | 0.065282131 | 9.172626398 | -1.844036726 | 0.113315704 | 0.529907348 | -4.255950787 |
| **A_51_P131942** | Dph5 | 0.255834431 | -0.081281617 | 0.592950479 | 10.80518519 | 1.844007805 | 0.113320298 | 0.529907348 | -4.255965421 |
| **A_55_P2077618** | Csgalnact1 | -0.168600682 | -0.390789764 | 0.0535884 | 8.294466783 | -1.843825435 | 0.113349267 | 0.529907348 | -4.256057698 |
| **A_55_P1973818** | Mmel1 | 0.476625136 | -0.151493863 | 1.104744135 | 9.09205555 | 1.843818642 | 0.113350347 | 0.529907348 | -4.256061135 |
| **A_55_P1967835** | NA | 0.337354076 | -0.107279842 | 0.781987994 | 8.997975032 | 1.843599416 | 0.113385181 | 0.529907348 | -4.256172064 |
| **A_55_P1957028** | Kat7 | 0.147408959 | -0.046894146 | 0.341712065 | 11.24543111 | 1.843432868 | 0.113411652 | 0.529907348 | -4.256256338 |
| **A_55_P2214625** | 4930519A11Rik | 0.24483582 | -0.077898911 | 0.567570551 | 8.420667339 | 1.843369683 | 0.113421696 | 0.529907348 | -4.25628831 |
| **A_55_P1970788** | C5ar2 | 0.38778775 | -0.123388544 | 0.898964044 | 9.456774782 | 1.843344766 | 0.113425657 | 0.529907348 | -4.256300919 |
| **A_55_P2000289** | Mab21l2 | 0.154207775 | -0.049080509 | 0.35749606 | 8.281559912 | 1.843219729 | 0.113445537 | 0.529907348 | -4.256364189 |
| **A_51_P407683** | Ints3 | 0.136140243 | -0.043345829 | 0.315626315 | 9.789895086 | 1.843057801 | 0.113471287 | 0.529907348 | -4.256446129 |
| **A_55_P2126780** | Pgam5 | 0.169101199 | -0.053886377 | 0.392088774 | 11.67097354 | 1.842676979 | 0.113531869 | 0.529915934 | -4.256638838 |
| **A_51_P475875** | Lrwd1 | 0.26701713 | -0.085104651 | 0.61913891 | 10.81147039 | 1.842593027 | 0.113545229 | 0.529915934 | -4.256681322 |
| **A_55_P2117130** | Zfp937 | 0.15602975 | -0.049766794 | 0.361826294 | 8.569055645 | 1.842266794 | 0.113597158 | 0.529915934 | -4.256846414 |
| **A_55_P1992838** | Socs2 | -0.59833755 | -1.387603124 | 0.190928024 | 8.569383214 | -1.84207067 | 0.113628388 | 0.529915934 | -4.256945665 |
| **A_55_P2093221** | Rnf111 | 0.188396261 | -0.060151749 | 0.436944272 | 10.98038898 | 1.841811492 | 0.113669671 | 0.529915934 | -4.257076829 |
| **A_51_P217430** | Riox1 | 0.163106732 | -0.052100555 | 0.378314019 | 9.561603003 | 1.84161186 | 0.11370148 | 0.529915934 | -4.257177861 |
| **A_55_P2057459** | Mall | -0.293074636 | -0.679779126 | 0.093629855 | 7.811065544 | -1.841544457 | 0.113712222 | 0.529915934 | -4.257211973 |
| **A_55_P1973783** | Tex9 | 0.128332641 | -0.041004439 | 0.297669721 | 8.668199131 | 1.841485297 | 0.113721651 | 0.529915934 | -4.257241913 |
| **A_55_P2277825** | E430010N07Rik | 0.167855269 | -0.053652034 | 0.389362572 | 9.107088501 | 1.841323591 | 0.113747427 | 0.529915934 | -4.257323752 |

| | | | | | | | | | |
|---|---|---|---|---|---|---|---|---|---|
| **A_55_P2007196** | Aldoa | -0.252232287 | -0.58509835 | 0.080633776 | 14.98491137 | -1.841256729 | 0.113758087 | 0.529915934 | -4.257357592 |
| **A_55_P2107830** | Gemin2 | 0.137358994 | -0.043921266 | 0.318639254 | 8.449827771 | 1.841152555 | 0.113774697 | 0.529915934 | -4.257410315 |
| **A_55_P1958165** | Ms4a7 | -0.197674168 | -0.458590555 | 0.06324222 | 9.117923857 | -1.840906583 | 0.113813927 | 0.529970944 | -4.257534805 |
| **A_55_P2048794** | Cd82 | 0.172214849 | -0.055129725 | 0.399559424 | 14.22115023 | 1.840641418 | 0.113856232 | 0.530030545 | -4.257669012 |
| **A_66_P117189** | Rad54l2 | 0.155773368 | -0.049894934 | 0.36144167 | 8.988794506 | 1.840386491 | 0.113896918 | 0.530030545 | -4.25779804 |
| **A_55_P1960331** | Srbd1 | 0.203726861 | -0.065316881 | 0.472770602 | 10.06188438 | 1.839960874 | 0.113964879 | 0.530030545 | -4.258013467 |
| **A_51_P379478** | Nckap5 | -0.503451493 | -1.168315044 | 0.161412058 | 8.609948995 | -1.839959644 | 0.113965075 | 0.530030545 | -4.25801409 |
| **A_55_P2010957** | Gm4971 | 0.218515576 | -0.070063024 | 0.507094176 | 12.84192133 | 1.839930706 | 0.113969697 | 0.530030545 | -4.258028737 |
| **A_51_P183102** | Rnf139 | 0.252153758 | -0.080872918 | 0.585180434 | 10.95919046 | 1.839795751 | 0.113991255 | 0.530030545 | -4.258097047 |
| **A_52_P478629** | Arcn1 | -0.140734206 | -0.326631845 | 0.045163434 | 8.655292218 | -1.839538915 | 0.114032295 | 0.530093848 | -4.258227051 |
| **A_55_P1961270** | Cd72 | -0.301854535 | -0.700666355 | 0.096957284 | 8.608444888 | -1.839131883 | 0.114097363 | 0.530168377 | -4.258433085 |
| **A_55_P2157048** | Mllt3 | 0.235924484 | -0.075786371 | 0.54763534 | 9.297792395 | 1.839095478 | 0.114103184 | 0.530168377 | -4.258451514 |
| **A_65_P16081** | NA | 0.131148961 | -0.042171085 | 0.304469006 | 8.020338203 | 1.83865077 | 0.114174321 | 0.53021655 | -4.258676629 |
| **A_55_P2081865** | Slc2a8 | -0.140825516 | -0.326938524 | 0.045287493 | 10.4470227 | -1.838602347 | 0.114182069 | 0.53021655 | -4.258701141 |
| **A_55_P2067632** | Ush1c | -0.22205206 | -0.515541739 | 0.071437618 | 7.842458296 | -1.838421809 | 0.114210963 | 0.53021655 | -4.258792534 |
| **A_55_P2348676** | B130052P14Rik | 0.133325646 | -0.042900318 | 0.30955161 | 8.14472786 | 1.838344882 | 0.114223277 | 0.53021655 | -4.258831477 |
| **A_52_P197666** | Med12 | 0.224738846 | -0.072382806 | 0.521860498 | 11.13419775 | 1.83792182 | 0.11429102 | 0.53040363 | -4.259045648 |
| **A_55_P2430472** | Dido1 | 0.212953891 | -0.068618451 | 0.494526233 | 9.234790473 | 1.837717607 | 0.114323734 | 0.530417904 | -4.259149031 |
| **A_55_P2151897** | NA | 0.212539464 | -0.068509008 | 0.493587936 | 15.47554287 | 1.837560058 | 0.114348979 | 0.530417904 | -4.259228792 |
| **A_55_P2090239** | Casp8ap2 | 0.201111502 | -0.064870303 | 0.467093306 | 8.434337177 | 1.837249637 | 0.114398735 | 0.53043086 | -4.259385949 |
| **A_55_P1989703** | Galnt11 | 0.328411267 | -0.105990222 | 0.762812756 | 9.94737712 | 1.837003281 | 0.114438238 | 0.53043086 | -4.259510675 |
| **A_55_P1958419** | NA | 0.233715729 | -0.075443145 | 0.542874604 | 11.78234141 | 1.836916491 | 0.114452158 | 0.53043086 | -4.259554616 |
| **A_55_P1974098** | Gm12063 | 0.261353397 | -0.08437959 | 0.607086383 | 10.66538471 | 1.836836583 | 0.114464975 | 0.53043086 | -4.259595073 |
| **A_55_P2111563** | NA | 0.134760122 | -0.043526268 | 0.313046512 | 7.819583267 | 1.836649959 | 0.114494916 | 0.53043086 | -4.25968956 |
| **A_51_P449935** | Ftsj3 | 0.257201502 | -0.083098532 | 0.597501537 | 9.549982874 | 1.836515919 | 0.114516425 | 0.53043086 | -4.259757426 |
| **A_55_P2013188** | Dnajb6 | 0.217064888 | -0.070255748 | 0.504385523 | 11.65048808 | 1.835717925 | 0.114644561 | 0.530492718 | -4.26016147 |
| **A_55_P2120993** | Zfp282 | 0.344626134 | -0.111556016 | 0.800808285 | 10.21756497 | 1.835663655 | 0.11465328 | 0.530492718 | -4.26018895 |
| **A_55_P2098603** | NA | 0.234675575 | -0.075967069 | 0.545318219 | 8.549921946 | 1.835650534 | 0.114655388 | 0.530492718 | -4.260195593 |
| **A_55_P2167025** | Nectin2 | -0.467743346 | -1.087030742 | 0.151544049 | 10.24998452 | -1.835264089 | 0.114717497 | 0.530492718 | -4.260391271 |
| **A_66_P117838** | Dhx29 | 0.150560103 | -0.048781118 | 0.349901323 | 8.650213114 | 1.835253147 | 0.114719256 | 0.530492718 | -4.260396811 |
| **A_55_P2019751** | Paox | -0.297790768 | -0.692069944 | 0.096488408 | 11.68520626 | -1.835230334 | 0.114722924 | 0.530492718 | -4.260408363 |
| **A_55_P1963563** | Zfp689 | 0.144602614 | -0.046859348 | 0.336064577 | 8.880944105 | 1.835172231 | 0.114732265 | 0.530492718 | -4.260437784 |
| **A_51_P181341** | Necab2 | -0.209814499 | -0.487639407 | 0.068010408 | 7.833877358 | -1.835048334 | 0.114752187 | 0.530492718 | -4.260500522 |
| **A_52_P630867** | Abcc4 | 0.269549753 | -0.087403072 | 0.626502577 | 8.946718416 | 1.834895365 | 0.114776789 | 0.530492718 | -4.260577982 |
| **A_52_P572045** | Pfkp | 0.250224363 | -0.081228591 | 0.581677316 | 8.349327105 | 1.834386678 | 0.114858636 | 0.530744101 | -4.260835574 |
| **A_55_P1970027** | Itch | 0.154884881 | -0.050340708 | 0.36011047 | 10.55642237 | 1.83383689 | 0.114947161 | 0.531026214 | -4.261113992 |
| **A_55_P2005868** | Bbs7 | 0.221640599 | -0.072069282 | 0.51535048 | 8.712334109 | 1.833639462 | 0.114978967 | 0.531046224 | -4.261213975 |
| **A_55_P2057065** | Gtf3c5 | 0.303229719 | -0.098760944 | 0.705220381 | 10.9234712 | 1.832900909 | 0.115098024 | 0.531469115 | -4.261588011 |
| **A_55_P2131964** | Pdia2 | 0.533175648 | -0.173855308 | 1.240206604 | 12.27908735 | 1.832377986 | 0.115182395 | 0.531530286 | -4.261852856 |
| **A_51_P388801** | Tbcc | 0.22277115 | -0.072715837 | 0.518258138 | 11.59384166 | 1.831908494 | 0.115258197 | 0.531530286 | -4.26209065 |
| **A_55_P2021490** | Gm16386 | 0.17203013 | -0.056163246 | 0.400223506 | 8.44329292 | 1.831827911 | 0.115271212 | 0.531530286 | -4.262131466 |
| **A_55_P2155593** | NA | -0.152197105 | -0.354094403 | 0.049700193 | 10.58619519 | -1.831719942 | 0.115288653 | 0.531530286 | -4.262186153 |
| **A_51_P453010** | Mphosph6 | 0.13568075 | -0.04433579 | 0.31569729 | 10.00278273 | 1.831424481 | 0.115336395 | 0.531530286 | -4.262335808 |
| **A_51_P282394** | Mrps28 | 0.209790077 | -0.068565161 | 0.488145316 | 13.23713895 | 1.831338956 | 0.115350218 | 0.531530286 | -4.262379129 |
| **A_55_P2091039** | NA | 0.316933344 | -0.103584505 | 0.737451192 | 9.341917632 | 1.831330328 | 0.115351612 | 0.531530286 | -4.262383499 |
| **A_55_P1977653** | Palld | -0.625130116 | -1.454656561 | 0.204396329 | 9.80183703 | -1.831147697 | 0.115381136 | 0.531530286 | -4.262476007 |
| **A_52_P163928** | Poldip2 | -0.146660228 | -0.341276864 | 0.047956408 | 10.6737838 | -1.83111484 | 0.115386448 | 0.531530286 | -4.26249265 |
| **A_51_P241943** | Sap30l | -0.246347343 | -0.573309515 | 0.08061483 | 12.1579084 | -1.830769044 | 0.115442371 | 0.531530286 | -4.26266781 |
| **A_66_P122559** | Myct1 | -0.209611779 | -0.487888494 | 0.068664935 | 8.379186648 | -1.830298847 | 0.115518455 | 0.531530286 | -4.262905992 |
| **A_51_P118539** | 2700049A03Rik | 0.152379629 | -0.049932419 | 0.354691677 | 10.05284391 | 1.830157034 | 0.115541412 | 0.531530286 | -4.26297783 |

| | | | | | | | | | |
|---|---|---|---|---|---|---|---|---|---|
| **A_55_P2106250** | Trappc6a | -0.316306185 | -0.736284245 | 0.103671874 | 12.63871881 | -1.830055544 | 0.115557844 | 0.531530286 | -4.263029242 |
| **A_51_P402536** | Uvrag | -0.146205274 | -0.34035206 | 0.047941512 | 11.2830182 | -1.829852238 | 0.115590768 | 0.531530286 | -4.263132233 |
| **A_55_P1973858** | NA | 0.21388885 | -0.070149413 | 0.497927113 | 9.940501732 | 1.829761502 | 0.115605465 | 0.531530286 | -4.263178199 |
| **A_55_P2176917** | NA | 0.144470255 | -0.047408668 | 0.336349178 | 10.78522565 | 1.829508187 | 0.115646506 | 0.531530286 | -4.263306527 |
| **A_52_P147803** | Syne4 | -0.25540325 | -0.594631598 | 0.083825097 | 10.33863308 | -1.829437057 | 0.115658033 | 0.531530286 | -4.263342562 |
| **A_51_P247637** | Rnf144a | -0.378920717 | -0.88223151 | 0.124390076 | 8.575674515 | -1.829343644 | 0.115673172 | 0.531530286 | -4.263389885 |
| **A_55_P2081483** | Atp5g2 | -0.155823431 | -0.362813241 | 0.05116638 | 15.59581774 | -1.829224389 | 0.115692502 | 0.531530286 | -4.263450301 |
| **A_55_P2155931** | Tnfsfm13 | -0.288541133 | -0.671829434 | 0.094747168 | 8.584549494 | -1.829217544 | 0.115693612 | 0.531530286 | -4.263453769 |
| **A_55_P2092622** | NA | 0.178733846 | -0.058716549 | 0.416184241 | 8.31684139 | 1.829014271 | 0.115726569 | 0.531530286 | -4.26355675 |
| **A_55_P1959076** | Zfp930 | 0.180931896 | -0.059448605 | 0.421312396 | 8.581371211 | 1.828938435 | 0.115738866 | 0.531530286 | -4.26359517 |
| **A_55_P1998578** | Ifitm2 | -0.233496762 | -0.543720611 | 0.076727087 | 12.53304011 | -1.82889541 | 0.115745844 | 0.531530286 | -4.263616967 |
| **A_55_P2178658** | NA | -0.138020048 | -0.321417085 | 0.045376989 | 12.22826962 | -1.828660346 | 0.115783973 | 0.531530286 | -4.263736058 |
| **A_51_P130808** | Tpbpb | -0.841021313 | -1.958893463 | 0.276850837 | 8.585526139 | -1.828092026 | 0.115876208 | 0.531530286 | -4.264023995 |
| **A_55_P2006768** | NA | 0.141250699 | -0.046523324 | 0.329024721 | 13.22996145 | 1.827840452 | 0.115917061 | 0.531530286 | -4.264151458 |
| **A_51_P305320** | B4gat1 | -0.216503806 | -0.504318086 | 0.071310474 | 10.34115826 | -1.827832465 | 0.115918358 | 0.531530286 | -4.264155505 |
| **A_51_P211325** | Nudt2 | 0.126958348 | -0.041849046 | 0.295765742 | 9.405245471 | 1.827481488 | 0.115975377 | 0.531530286 | -4.264333336 |
| **A_52_P54856** | Ubald1 | -0.165673842 | -0.385965264 | 0.054617581 | 9.524223934 | -1.827424929 | 0.115984568 | 0.531530286 | -4.264361993 |
| **A_55_P1962479** | Gpbp1l1 | 0.191499719 | -0.063151272 | 0.44615071 | 9.700240718 | 1.827283614 | 0.116007535 | 0.531530286 | -4.264433596 |
| **A_55_P2059179** | Slc48a1 | 0.226101763 | -0.074615169 | 0.526818694 | 12.41285736 | 1.826960911 | 0.116059999 | 0.531530286 | -4.264597108 |
| **A_51_P167876** | Tfrc | 0.316803984 | -0.104558995 | 0.738166964 | 8.844098901 | 1.826911241 | 0.116068076 | 0.531530286 | -4.264622276 |
| **A_51_P227004** | Cks1b | 0.157718139 | -0.052081923 | 0.367518201 | 13.00176741 | 1.826666322 | 0.116107912 | 0.531530286 | -4.264746379 |
| **A_55_P2102250** | Dzip1 | 0.137569528 | -0.045431207 | 0.320570263 | 8.229867451 | 1.826638476 | 0.116112442 | 0.531530286 | -4.264760489 |
| **A_66_P128997** | Pml | 0.191740324 | -0.063327948 | 0.446808595 | 9.309167057 | 1.82658634 | 0.116120924 | 0.531530286 | -4.264786907 |
| **A_55_P2173098** | Commd1 | -0.194589367 | -0.453448189 | 0.064269455 | 12.05932342 | -1.826582631 | 0.116121528 | 0.531530286 | -4.264788787 |
| **A_51_P462814** | Sars2 | 0.217764564 | -0.071930846 | 0.507459974 | 8.98692723 | 1.826538308 | 0.116128739 | 0.531530286 | -4.264811246 |
| **A_55_P2170094** | Lrrc27 | 0.245643771 | -0.081255425 | 0.572542967 | 9.23683135 | 1.825892025 | 0.116233941 | 0.531874476 | -4.265138739 |
| **A_55_P2043627** | Fam89a | 0.191114916 | -0.063239474 | 0.445469306 | 9.202009248 | 1.825738341 | 0.116258972 | 0.531874476 | -4.265216618 |
| **A_51_P100828** | Prdx3 | 0.28885105 | -0.09564334 | 0.673345439 | 12.90301018 | 1.825438187 | 0.116307873 | 0.531972284 | -4.270298308 |
| **A_55_P2086198** | NA | 0.198066945 | -0.065668716 | 0.461802607 | 9.890502275 | 1.824846483 | 0.116404333 | 0.532287519 | -4.265668585 |
| **A_55_P2253289** | NA | 0.304827542 | -0.101160338 | 0.710815423 | 8.647506944 | 1.824417899 | 0.11647425 | 0.532392439 | -4.26588579 |
| **A_55_P2126207** | Fthl17a | -0.198611684 | -0.463152581 | 0.065929213 | 7.866565525 | -1.824295389 | 0.116494243 | 0.532392439 | -4.265947879 |
| **A_51_P507851** | Gcc1 | 0.156896603 | -0.052092873 | 0.36588608 | 8.810081975 | 1.824199414 | 0.116509908 | 0.532392439 | -4.26599652 |
| **A_55_P2074193** | NA | -0.39319572 | -0.917049288 | 0.130657847 | 14.12367208 | -1.823820384 | 0.116571794 | 0.53254933 | -4.26618862 |
| **A_51_P303725** | Tmem38a | -0.399095801 | -0.931110375 | 0.132918772 | 8.739379054 | -1.822790761 | 0.116740069 | 0.532996168 | -4.266710483 |
| **A_51_P184728** | Cnksr3 | -0.200680036 | -0.468200588 | 0.066840516 | 10.40765898 | -1.82276301 | 0.116744608 | 0.532996168 | -4.266724549 |
| **A_55_P2126951** | Zfp467 | -0.419939726 | -0.979787694 | 0.139908242 | 8.868725489 | -1.822636418 | 0.116765315 | 0.532996168 | -4.266788715 |
| **A_55_P2127804** | Map3k7 | 0.121910844 | -0.040633901 | 0.284455589 | 8.350229413 | 1.82243729 | 0.116797893 | 0.532996168 | -4.266889649 |
| **A_55_P2068096** | Rap1gap | -0.208629641 | -0.486806223 | 0.069546941 | 8.993939571 | -1.822378709 | 0.116807479 | 0.532996168 | -4.266919343 |
| **A_55_P2102454** | NA | 0.459688799 | -0.153543683 | 1.072921281 | 10.88930909 | 1.821469737 | 0.116956319 | 0.533549372 | -4.267380103 |
| **A_52_P238816** | Ppt2 | 0.20797105 | -0.069552395 | 0.485494496 | 10.39670007 | 1.820901254 | 0.1170495 | 0.533804146 | -4.267668285 |
| **A_55_P2173968** | Ctdspl | -0.461147465 | -1.076650983 | 0.154356053 | 11.72578751 | -1.82050751 | 0.117114081 | 0.533804146 | -4.267867894 |
| **A_55_P2156973** | Aar2 | 0.146076414 | -0.048905324 | 0.341058152 | 9.818566408 | 1.820410577 | 0.117129986 | 0.533804146 | -4.267917035 |
| **A_55_P1990785** | Zranb3 | 0.168759833 | -0.056516135 | 0.3940358 | 8.856455194 | 1.820276708 | 0.117151954 | 0.533804146 | -4.267984902 |
| **A_55_P2031337** | Trmt2a | 0.196235558 | -0.065749117 | 0.458220233 | 11.21850117 | 1.820057069 | 0.117188006 | 0.533804146 | -4.268096252 |
| **A_52_P101333** | Sap18 | 0.136231149 | -0.045646768 | 0.318109065 | 8.532251046 | 1.820034571 | 0.117191699 | 0.533804146 | -4.268107658 |
| **A_51_P249930** | Tceal1 | -0.32183914 | -0.751636207 | 0.107957927 | 8.218564355 | -1.819527402 | 0.117274991 | 0.533804146 | -4.268364786 |
| **A_52_P552092** | Adat1 | 0.145185289 | -0.048702322 | 0.3390729 | 8.56318682 | 1.819515433 | 0.117276957 | 0.533804146 | -4.268370854 |
| **A_55_P1959403** | Ccdc77 | 0.259031576 | -0.086929698 | 0.60499285 | 9.862067291 | 1.819317133 | 0.11730954 | 0.533804146 | -4.268471393 |
| **A_55_P2159390** | NA | 0.202049865 | -0.067815622 | 0.471915352 | 11.64021762 | 1.819258474 | 0.11731918 | 0.533804146 | -4.268501133 |
| **A_52_P204689** | Map4k2 | -0.121944837 | -0.284839656 | 0.040949981 | 8.489008627 | -1.819027802 | 0.117357096 | 0.533804146 | -4.268618086 |

| | | | | | | | | | |
|---|---|---|---|---|---|---|---|---|---|
| **A_52_P228079** | Atf1 | 0.240342233 | -0.080710138 | 0.561394605 | 9.083234535 | 1.819019684 | 0.117358431 | 0.533804146 | -4.268622202 |
| **A_55_P2133716** | Tax1bp3 | -0.628599105 | -1.468375284 | 0.211177074 | 12.0232197 | -1.81883539 | 0.117388733 | 0.533804146 | -4.268715643 |
| **A_55_P2249897** | NA | 0.154839808 | -0.052037041 | 0.361716657 | 8.647292811 | 1.81867007 | 0.117415922 | 0.533804146 | -4.268799464 |
| **A_55_P1985749** | Rad23a | 0.250425324 | -0.084281958 | 0.585132606 | 8.642300648 | 1.818010018 | 0.117524536 | 0.533804146 | -4.269134138 |
| **A_55_P2114467** | Romo1 | -0.138786812 | -0.324286399 | 0.046712775 | 13.93766565 | -1.817977243 | 0.117529932 | 0.533804146 | -4.269150757 |
| **A_55_P2269459** | Prl8a6 | -0.59904894 | -1.399734849 | 0.201636968 | 8.157587878 | -1.817955749 | 0.117533471 | 0.533804146 | -4.269161655 |
| **A_51_P277444** | Rnf19a | 0.151704652 | -0.051065696 | 0.354475 | 11.40484304 | 1.817932032 | 0.117537376 | 0.533804146 | -4.269173681 |
| **A_55_P2170309** | Ppfibp2 | -0.179767865 | -0.420049039 | 0.06051331 | 8.355665512 | -1.817923079 | 0.11753885 | 0.533804146 | -4.269178221 |
| **A_55_P2105366** | NA | 0.187660414 | -0.06320906 | 0.438529887 | 8.701527418 | 1.817640702 | 0.117585352 | 0.533804146 | -4.269321404 |
| **A_51_P381873** | Bub3 | 0.125213277 | -0.042178973 | 0.292605526 | 12.11631342 | 1.817599625 | 0.117592118 | 0.533804146 | -4.269342233 |
| **A_55_P1963434** | Gpc4 | -0.174653505 | -0.408202101 | 0.05889509 | 8.217822795 | -1.81711844 | 0.117671406 | 0.533897283 | -4.269586232 |
| **A_55_P2034670** | Fbxl12 | -0.265525481 | -0.620590679 | 0.089539716 | 11.36304816 | -1.817110293 | 0.117672749 | 0.533897283 | -4.269590363 |
| **A_51_P418725** | Plekhf1 | -0.202542752 | -0.473424807 | 0.068339302 | 8.061361013 | -1.816852465 | 0.117715256 | 0.533897283 | -4.269721106 |
| **A_55_P2004153** | NA | 0.195222898 | -0.065890007 | 0.456335803 | 10.22171144 | 1.816710003 | 0.117738749 | 0.533897283 | -4.269793348 |
| **A_55_P2030662** | Eda | 0.190538694 | -0.064342275 | 0.445419663 | 8.458067315 | 1.816473067 | 0.117777832 | 0.533897283 | -4.269913501 |
| **A_51_P416613** | 1810058I24Rik | 0.2022242 | -0.068310793 | 0.472759193 | 9.00636476 | 1.816322103 | 0.11780274 | 0.533897283 | -4.269990057 |
| **A_55_P1985288** | Bhlha9 | 0.215489969 | -0.072795056 | 0.503774994 | 7.862571398 | 1.816302428 | 0.117805987 | 0.533897283 | -4.270000035 |
| **A_52_P573366** | Hnrnpul1 | 0.138276209 | -0.046792019 | 0.323344437 | 8.229425627 | 1.815510596 | 0.117936723 | 0.534364491 | -4.270401599 |
| **A_52_P587648** | Phf3 | 0.167166798 | -0.056603704 | 0.3909373 | 10.94457755 | 1.815224639 | 0.117983972 | 0.53445329 | -4.270546624 |
| **A_55_P1952220** | Pnpo | 0.284728342 | -0.096488067 | 0.66594475 | 15.91112479 | 1.814856547 | 0.118044818 | 0.53460363 | -4.270733308 |
| **A_55_P2077228** | Cenpl | 0.287500562 | -0.097464726 | 0.67246585 | 10.38436106 | 1.814681098 | 0.118073831 | 0.534609765 | -4.270822292 |
| **A_65_P08022** | Hnrnpk | 0.189534171 | -0.064330622 | 0.443398964 | 11.01404643 | 1.814129286 | 0.118165128 | 0.534846362 | -4.271102166 |
| **A_55_P2170009** | Lactb2 | 0.260837685 | -0.088561533 | 0.610236904 | 11.08522276 | 1.813976252 | 0.118190459 | 0.534846362 | -4.271179785 |
| **A_55_P1996983** | Btg1c | -0.193619924 | -0.45300025 | 0.065760403 | 8.505712303 | -1.813828411 | 0.118214936 | 0.534846362 | -4.271254772 |
| **A_51_P108581** | Grk3 | -0.204069824 | -0.477469093 | 0.069329445 | 8.842904966 | -1.813696565 | 0.118236769 | 0.534846362 | -4.271321646 |
| **A_51_P395921** | Uchl4 | 0.371476927 | -0.126256494 | 0.869210348 | 10.5329267 | 1.8135028 | 0.118268862 | 0.534866364 | -4.271419928 |
| **A_55_P1983016** | NA | 0.13177612 | -0.044813137 | 0.308365378 | 8.268574087 | 1.813241396 | 0.118312173 | 0.534937073 | -4.271552519 |
| **A_55_P2106489** | Ankrd52 | 0.264955318 | -0.090241929 | 0.620152566 | 8.930495698 | 1.812534321 | 0.118429401 | 0.535301858 | -4.271911181 |
| **A_51_P273433** | Prkg2 | -0.266446732 | -0.623683944 | 0.09079048 | 7.837171466 | -1.812328397 | 0.118463563 | 0.535301858 | -4.272015639 |
| **A_55_P1953650** | Zdhhc18 | -0.27907925 | -0.653268846 | 0.095110346 | 11.73264217 | -1.812253837 | 0.118475935 | 0.535301858 | -4.272053461 |
| **A_55_P2139611** | Acbd4 | 0.379387618 | -0.129361651 | 0.888136886 | 9.67314138 | 1.812018152 | 0.11851505 | 0.53535345 | -4.272173018 |
| **A_55_P2146546** | NA | 0.148669071 | -0.050741726 | 0.348079869 | 7.942039859 | 1.811570111 | 0.118589445 | 0.535561818 | -4.272400304 |
| **A_55_P1970537** | Bcl7b | 0.240870498 | -0.082239827 | 0.563980823 | 10.77160792 | 1.811406672 | 0.118616594 | 0.535561818 | -4.272483217 |
| **A_52_P93467** | Sdc4 | -0.191995568 | -0.449589416 | 0.065598281 | 9.483759294 | -1.811085296 | 0.118669996 | 0.535677804 | -4.272646253 |
| **A_51_P494675** | Cotl1 | -0.199749701 | -0.467814013 | 0.068314611 | 13.6018118 | -1.81063257 | 0.118745265 | 0.535717384 | -4.272875931 |
| **A_52_P679101** | Tjp2 | -0.321121495 | -0.752162291 | 0.109919301 | 10.90823227 | -1.810231799 | 0.118811935 | 0.535717384 | -4.273079258 |
| **A_55_P2101776** | Adora1 | -0.532822937 | -1.248058851 | 0.182412976 | 10.34329596 | -1.810159268 | 0.118824005 | 0.535717384 | -4.273116056 |
| **A_55_P2008907** | Actg1 | -0.427705383 | -1.001854738 | 0.146443971 | 13.807512 | -1.810101903 | 0.118833551 | 0.535717384 | -4.27314516 |
| **A_52_P588881** | Iqgap3 | 0.240793584 | -0.082457198 | 0.564044366 | 8.913838624 | 1.810041428 | 0.118843617 | 0.535717384 | -4.273175842 |
| **A_55_P2104978** | Edc4 | 0.161906883 | -0.055451394 | 0.379265159 | 11.32759379 | 1.809973927 | 0.118854852 | 0.535717384 | -4.273210089 |
| **A_55_P2295671** | NA | 0.307630028 | -0.105521835 | 0.720781892 | 9.322016309 | 1.80926507 | 0.118972905 | 0.535717384 | -4.27356974 |
| **A_51_P328400** | Cops2 | 0.183424473 | -0.062939193 | 0.429788139 | 10.75369565 | 1.809105151 | 0.118999554 | 0.535717384 | -4.27365088 |
| **A_55_P2112617** | Smad4 | -0.177151756 | -0.415099778 | 0.060796265 | 8.96382523 | -1.809033271 | 0.119011534 | 0.535717384 | -4.273687351 |
| **A_52_P455089** | NA | 0.162290751 | -0.055759473 | 0.380340975 | 9.251647037 | 1.808507949 | 0.119099123 | 0.535717384 | -4.273953899 |
| **A_55_P1999384** | NA | -0.263011078 | -0.616392122 | 0.090369966 | 9.945216531 | -1.80848113 | 0.119103597 | 0.535717384 | -4.273967507 |
| **A_55_P2161600** | Diaph2 | -0.261138219 | -0.612061754 | 0.089785317 | 8.840609964 | -1.808177788 | 0.119154205 | 0.535717384 | -4.274121427 |
| **A_52_P359257** | NA | -0.142746306 | -0.334608602 | 0.049115991 | 7.875925733 | -1.807833496 | 0.119211671 | 0.535717384 | -4.27429613 |
| **A_51_P179131** | Vasp | -0.190423638 | -0.446372128 | 0.065524852 | 9.798328826 | -1.807804548 | 0.119216503 | 0.535717384 | -4.274310819 |
| **A_55_P2035205** | Nifk | 0.288448981 | -0.099302422 | 0.676200385 | 12.20146307 | 1.80758538 | 0.1192531 | 0.535717384 | -4.274422033 |
| **A_51_P415247** | NA | 0.151919757 | -0.052304194 | 0.356143707 | 8.717565307 | 1.807551879 | 0.119258695 | 0.535717384 | -4.274439033 |

| | | | | | | | | | |
|---|---|---|---|---|---|---|---|---|---|
| **A_55_P2124619** | Smurf2 | -0.231008494 | -0.541577785 | 0.079560798 | 9.263244506 | -1.807393125 | 0.119285212 | 0.535717384 | -4.274519592 |
| **A_52_P300730** | Hmga2 | 0.220697726 | -0.076013233 | 0.517408686 | 8.751343031 | 1.807371625 | 0.119288804 | 0.535717384 | -4.274530502 |
| **A_55_P1968789** | Rasa3 | -0.204207086 | -0.478755915 | 0.070341743 | 12.07920965 | -1.807317278 | 0.119297883 | 0.535717384 | -4.27455808 |
| **A_55_P2154132** | Tuba1b | 0.206499143 | -0.071160969 | 0.484159254 | 17.04795657 | 1.807123974 | 0.119330182 | 0.535717384 | -4.274656173 |
| **A_51_P131561** | Plet1 | -0.398693199 | -0.934794494 | 0.137408096 | 8.609344116 | -1.807070767 | 0.119339074 | 0.535717384 | -4.274683173 |
| **A_55_P2089080** | Ssbp2 | 0.121762256 | -0.041982728 | 0.285507239 | 10.45869242 | 1.806874001 | 0.119371963 | 0.535717384 | -4.274783025 |
| **A_55_P1992739** | Dync1i2 | -0.198765336 | -0.466088963 | 0.068558291 | 12.77298444 | -1.806701856 | 0.119400744 | 0.535717384 | -4.274870383 |
| **A_51_P265008** | Ppig | 0.163370977 | -0.056367362 | 0.383109316 | 11.20123624 | 1.806559452 | 0.119424557 | 0.535717384 | -4.27494265 |
| **A_51_P195875** | Notch4 | -0.379251223 | -0.889396658 | 0.130894211 | 8.617649517 | -1.80640936 | 0.119449661 | 0.535717384 | -4.275018819 |
| **A_52_P577729** | Rnf169 | -0.130224833 | -0.305405836 | 0.044956169 | 10.49976359 | -1.806300462 | 0.119467879 | 0.535717384 | -4.275074083 |
| **A_51_P460954** | Ccl6 | -0.153347838 | -0.359637387 | 0.052941711 | 10.27655242 | -1.806273956 | 0.119472313 | 0.535717384 | -4.275087534 |
| **A_52_P120612** | Grap2 | 0.363852834 | -0.125690829 | 0.853396496 | 8.858097406 | 1.80599905 | 0.119518316 | 0.535717384 | -4.275227047 |
| **A_55_P2073010** | Gm4832 | 0.174897983 | -0.060422802 | 0.410218767 | 14.27427488 | 1.805958232 | 0.119525148 | 0.535717384 | -4.275247762 |
| **A_51_P323842** | Ciao2b | -0.263778797 | -0.618708846 | 0.091151252 | 12.49477185 | -1.805844304 | 0.119544218 | 0.535717384 | -4.275305581 |
| **A_55_P2090617** | Pilra | -0.225180808 | -0.52818794 | 0.077826325 | 8.596664607 | -1.805766993 | 0.119557161 | 0.535717384 | -4.275344816 |
| **A_55_P1994817** | Taf15 | 0.327265012 | -0.113120756 | 0.76765078 | 9.722097959 | 1.805716187 | 0.119565668 | 0.535717384 | -4.275370601 |
| **A_66_P130960** | Psmd11 | 0.244750163 | -0.084662582 | 0.574162908 | 10.48693607 | 1.805368167 | 0.119623953 | 0.535854317 | -4.275547226 |
| **A_51_P413698** | Mrpl20 | 0.143001304 | -0.049499018 | 0.335501627 | 13.97608218 | 1.805060348 | 0.119675528 | 0.53596114 | -4.275703452 |
| **A_55_P2096823** | NA | 0.128964368 | -0.044690214 | 0.30261895 | 8.587598144 | 1.804540677 | 0.119762649 | 0.535994198 | -4.275967206 |
| **A_55_P2173927** | Insr | -0.126257346 | -0.296283259 | 0.043768566 | 8.022443423 | -1.804366401 | 0.119791879 | 0.535994198 | -4.27605566 |
| **A_55_P2052485** | Ushbp1 | -0.501788245 | -1.177569463 | 0.173992974 | 9.90528803 | -1.804253492 | 0.119810821 | 0.535994198 | -4.276112968 |
| **A_51_P139651** | Nos3 | -0.183704675 | -0.431134936 | 0.063725585 | 8.293416924 | -1.804058373 | 0.119843561 | 0.535994198 | -4.285679129 |
| **A_55_P2473316** | Mapre1 | 0.174973848 | -0.060747807 | 0.410695502 | 11.2686483 | 1.803669035 | 0.119908916 | 0.535994198 | -4.276409621 |
| **A_51_P236412** | Cdkl2 | -0.175153275 | -0.411141508 | 0.060834959 | 7.970036925 | -1.803479047 | 0.11994082 | 0.535994198 | -4.276506056 |
| **A_55_P2302285** | Tarbp2 | 0.158668492 | -0.055112183 | 0.372449167 | 12.21039993 | 1.80345554 | 0.119944768 | 0.535994198 | -4.276517988 |
| **A_51_P311694** | Ypel5 | 0.291650174 | -0.101308203 | 0.68460855 | 10.24521281 | 1.803428289 | 0.119949345 | 0.535994198 | -4.27653182 |
| **A_55_P2011169** | NA | 0.238805584 | -0.08296152 | 0.560572688 | 8.667678291 | 1.80337492 | 0.11995831 | 0.535994198 | -4.27655891 |
| **A_51_P513254** | Rac1 | -0.28035508 | -0.658108425 | 0.097398265 | 12.29726948 | -1.803363605 | 0.11996021 | 0.535994198 | -4.276564653 |
| **A_51_P292016** | Uts2r | 0.14987113 | -0.052105987 | 0.351848247 | 8.687967086 | 1.803013596 | 0.120019018 | 0.53607888 | -4.276742316 |
| **A_51_P152845** | Trim24 | 0.139177827 | -0.048414904 | 0.326770559 | 10.50593619 | 1.802757184 | 0.120062117 | 0.53607888 | -4.276872473 |
| **A_55_P2457154** | G2e3 | 0.206662274 | -0.071936071 | 0.485260619 | 9.855692877 | 1.802460936 | 0.120111932 | 0.53607888 | -4.277022854 |
| **A_51_P501922** | Plxna1 | 0.246295215 | -0.085735548 | 0.578325978 | 9.56464742 | 1.802440094 | 0.120115437 | 0.53607888 | -4.277033434 |
| **A_55_P2135271** | Zfp358 | -0.362387634 | -0.850958905 | 0.126183638 | 10.26106072 | -1.802306428 | 0.120137921 | 0.53607888 | -4.277101286 |
| **A_66_P119680** | Polr2e | -0.175185692 | -0.411377127 | 0.061005744 | 12.52200366 | -1.80226096 | 0.120145569 | 0.53607888 | -4.277124367 |
| **A_55_P2032600** | Rps24 | 0.154446871 | -0.053812599 | 0.362706341 | 17.14329549 | 1.802011442 | 0.120187553 | 0.536142446 | -4.277251031 |
| **A_55_P2040031** | NA | 0.226702258 | -0.079095627 | 0.532500142 | 12.76762909 | 1.801376769 | 0.120294408 | 0.53631643 | -4.277573222 |
| **A_55_P1994997** | Arhgap11a | 0.211011856 | -0.073649262 | 0.495672973 | 8.505719967 | 1.801199821 | 0.120324216 | 0.53631643 | -4.277663051 |
| **A_51_P319453** | Zfp143 | 0.191201816 | -0.066736754 | 0.449140386 | 8.723815758 | 1.801187455 | 0.1203263 | 0.53631643 | -4.277669329 |
| **A_55_P1954306** | Esrra | -0.185185651 | -0.435028592 | 0.06465729 | 11.28036274 | -1.801040333 | 0.120351089 | 0.53631643 | -4.277744019 |
| **A_51_P494125** | Alpl | 0.437869441 | -0.152975323 | 1.028714206 | 9.789523132 | 1.800754192 | 0.120399318 | 0.53631643 | -4.277889286 |
| **A_55_P1955039** | Myo1b | -0.225016495 | -0.528647286 | 0.078614295 | 8.866771644 | -1.800742999 | 0.120401204 | 0.53631643 | -4.277894968 |
| **A_55_P1970270** | B3gnt2 | -0.150095244 | -0.352642514 | 0.052452025 | 9.804806439 | -1.800626874 | 0.120420783 | 0.53631643 | -4.277953923 |
| **A_51_P350666** | Peli2 | -0.264148939 | -0.620715619 | 0.09241774 | 8.739039346 | -1.800077916 | 0.120513377 | 0.536442031 | -4.278232627 |
| **A_55_P2151466** | Casc3 | 0.174972189 | -0.061226923 | 0.411171301 | 11.29149103 | 1.800006002 | 0.120525512 | 0.536442031 | -4.278269138 |
| **A_55_P1954860** | NA | 0.348439266 | -0.122038308 | 0.818916839 | 13.86718267 | 1.799580805 | 0.120597286 | 0.536442031 | -4.278485016 |
| **A_51_P268843** | Rasip1 | -0.557764875 | -1.310912219 | 0.195382469 | 9.988600316 | -1.799510569 | 0.120609146 | 0.536442031 | -4.278520677 |
| **A_55_P2122180** | Snta1 | 0.337926473 | -0.118381729 | 0.794234675 | 10.44365131 | 1.799480435 | 0.120614235 | 0.536442031 | -4.278535976 |
| **A_55_P1973046** | Cttnbp2nl | -0.457015078 | -1.074172528 | 0.160142373 | 8.700029359 | -1.79935867 | 0.120634799 | 0.536442031 | -4.2785978 |
| **A_55_P2041624** | Ceacam12 | -0.577556704 | -1.357570406 | 0.202456998 | 8.129152518 | -1.799184026 | 0.120664301 | 0.536442031 | -4.278686472 |
| **A_51_P292008** | Gpx3 | -0.502378343 | -1.180907413 | 0.176150727 | 9.374771799 | -1.799059967 | 0.120685262 | 0.536442031 | -4.278749461 |

| | | | | | | | | | |
|---|---|---|---|---|---|---|---|---|---|
| **A_52_P590396** | Sort1 | -0.374824209 | -0.881136926 | 0.131488509 | 10.32201909 | -1.798837719 | 0.120722821 | 0.536442031 | -4.278862307 |
| **A_55_P2154430** | Cd320 | -0.264454127 | -0.621683229 | 0.092774974 | 11.01451723 | -1.798815856 | 0.120726516 | 0.536442031 | -4.278873408 |
| **A_52_P621567** | Tcf20 | 0.121508654 | -0.042644981 | 0.285662289 | 10.7762117 | 1.798621976 | 0.120759292 | 0.536464343 | -4.27897185 |
| **A_65_P05316** | Zfp182 | 0.147625818 | -0.051895521 | 0.347147157 | 8.631224904 | 1.797861182 | 0.120887989 | 0.536912671 | -4.279358157 |
| **A_55_P2187261** | NA | 0.305010679 | -0.10731986 | 0.717341217 | 9.434415516 | 1.797433089 | 0.120960465 | 0.537111149 | -4.279575537 |
| **A_55_P2050439** | Dlgap5 | 0.305587424 | -0.107669754 | 0.718844602 | 9.049976314 | 1.796793884 | 0.121068762 | 0.537468557 | -4.279900128 |
| **A_51_P354706** | Lefty1 | 0.220616684 | -0.077792687 | 0.519026055 | 8.676496351 | 1.796424973 | 0.121131307 | 0.537572169 | -4.28008747 |
| **A_55_P2136334** | NA | 0.226959937 | -0.080061263 | 0.533981136 | 12.21805807 | 1.796238607 | 0.121162916 | 0.537572169 | -4.280182112 |
| **A_55_P1981105** | Ap1b1 | 0.248903163 | -0.087848731 | 0.585655057 | 13.30349873 | 1.795988528 | 0.121205343 | 0.537572169 | -4.280309112 |
| **A_52_P530280** | Nsa2 | 0.198401357 | -0.07004235 | 0.466845064 | 11.22985643 | 1.79586878 | 0.121225664 | 0.537572169 | -4.280369925 |
| **A_51_P341130** | Iqgap1 | -0.60746641 | -1.429509829 | 0.214577008 | 10.97567016 | -1.795604567 | 0.121270513 | 0.537572169 | -4.280504106 |
| **A_55_P1980432** | Zfp488 | 0.153352185 | -0.05420336 | 0.36090773 | 9.639461034 | 1.795307382 | 0.121320978 | 0.537572169 | -4.280655035 |
| **A_55_P1962641** | Rnf26 | 0.183897369 | -0.065056658 | 0.432851397 | 13.21238939 | 1.794897073 | 0.121390686 | 0.537572169 | -4.280863419 |
| **A_55_P2052166** | Akap13 | 0.213179388 | -0.075417491 | 0.501776267 | 10.58056113 | 1.794885587 | 0.121392637 | 0.537572169 | -4.280869253 |
| **A_51_P172853** | Cd14 | -0.208746421 | -0.491366945 | 0.073874102 | 8.389694501 | -1.794727558 | 0.121419496 | 0.537572169 | -4.280949513 |
| **A_51_P407606** | Ndc1 | 0.330975547 | -0.117144468 | 0.779095562 | 9.961419412 | 1.794670629 | 0.121429173 | 0.537572169 | -4.280978426 |
| **A_51_P155085** | Dennd2a | -0.54911045 | -1.292574565 | 0.194353665 | 9.779142513 | -1.794662907 | 0.121430486 | 0.537572169 | -4.280982348 |
| **A_52_P286141** | Cisd2 | 0.158932055 | -0.056285007 | 0.374149118 | 10.32004627 | 1.794394752 | 0.121476079 | 0.537572169 | -4.281118542 |
| **A_51_P494929** | Diablo | 0.19276512 | -0.068267025 | 0.453797265 | 10.54314567 | 1.794393344 | 0.121476318 | 0.537572169 | -4.281119257 |
| **A_66_P115323** | Utp4 | 0.260576416 | -0.092288085 | 0.613440917 | 10.63380752 | 1.794363083 | 0.121481465 | 0.537572169 | -4.281134627 |
| **A_52_P315345** | Lats2 | -0.222370276 | -0.523546137 | 0.078805586 | 9.399029669 | -1.794071561 | 0.121531052 | 0.537668506 | -4.281282691 |
| **A_51_P394471** | Tcf3 | 0.144200878 | -0.051134447 | 0.339536204 | 10.87277476 | 1.793784656 | 0.121579873 | 0.537761411 | -4.281428413 |
| **A_51_P502132** | Mmp23 | -0.301691787 | -0.710527525 | 0.10714395 | 8.418929721 | -1.793072396 | 0.121701158 | 0.537959813 | -4.281790189 |
| **A_52_P37702** | NA | 0.317755245 | -0.112891962 | 0.748402453 | 8.778391433 | 1.792892631 | 0.121731787 | 0.537959813 | -4.2818815 |
| **A_55_P2099526** | H2afv | -0.327963616 | -0.772453442 | 0.116526211 | 13.62975197 | -1.792862687 | 0.12173689 | 0.537959813 | -4.28189671 |
| **A_55_P2286493** | 4933405E24Rik | 0.310357576 | -0.110287073 | 0.731002225 | 9.432255325 | 1.7927931 | 0.121748749 | 0.537959813 | -4.281932056 |
| **A_55_P2091348** | NA | 0.177839179 | -0.063207958 | 0.418886315 | 9.605941783 | 1.792704281 | 0.121763888 | 0.537959813 | -4.281977172 |
| **A_51_P148314** | Fam117a | 0.187750757 | -0.066797124 | 0.442298638 | 11.5601578 | 1.792236839 | 0.121843589 | 0.538039838 | -4.282214613 |
| **A_55_P2157966** | Map1a | 0.141369752 | -0.05031437 | 0.333053873 | 8.105250113 | 1.792064123 | 0.121873051 | 0.538039838 | -4.282302348 |
| **A_55_P2041723** | Mid1ip1 | -0.24284789 | -0.572141592 | 0.086445812 | 12.03492052 | -1.79198388 | 0.121886741 | 0.538039838 | -4.282343109 |
| **A_55_P2043322** | Rnf26 | 0.226780574 | -0.080733021 | 0.534294169 | 9.194461985 | 1.791945179 | 0.121893344 | 0.538039838 | -4.282362768 |
| **A_55_P1966573** | Gemin4 | 0.233497729 | -0.083175621 | 0.550171079 | 10.51975974 | 1.791654784 | 0.121942904 | 0.538135704 | -4.282510283 |
| **A_55_P2000943** | Nrip3 | 0.114955441 | -0.040966513 | 0.270877395 | 7.900335818 | 1.791453133 | 0.121977329 | 0.538142463 | -4.28261272 |
| **A_55_P2068154** | Anapc15 | 0.281352458 | -0.100293621 | 0.662998537 | 10.37944065 | 1.79131968 | 0.122000118 | 0.538142463 | -4.282680513 |
| **A_55_P1982049** | Icam2 | -0.711706931 | -1.677284216 | 0.253870354 | 12.18683142 | -1.791007249 | 0.122053484 | 0.538202945 | -4.282839228 |
| **A_55_P1973699** | Htra2 | 0.264019112 | -0.094196053 | 0.622234276 | 12.67494675 | 1.790913407 | 0.122069518 | 0.538202945 | -4.2828869 |
| **A_55_P1977699** | NA | 0.217686814 | -0.077789571 | 0.513163199 | 14.28670823 | 1.790162832 | 0.122197834 | 0.538558523 | -4.283268209 |
| **A_55_P1986680** | NA | 0.376064813 | -0.134444819 | 0.886574445 | 8.741678645 | 1.789954325 | 0.122233503 | 0.538558523 | -4.283374138 |
| **A_55_P1963645** | NA | 0.351780316 | -0.125763396 | 0.829324028 | 11.13719625 | 1.789952857 | 0.122233755 | 0.538558523 | -4.283374883 |
| **A_51_P205968** | Snx15 | 0.25426339 | -0.090988671 | 0.59951545 | 11.28394897 | 1.789496042 | 0.122311938 | 0.538780182 | -4.283606967 |
| **A_52_P370935** | Glcci1 | -0.191481273 | -0.451523733 | 0.068561188 | 9.284542125 | -1.789226111 | 0.122358158 | 0.538860981 | -4.283744108 |
| **A_55_P2186619** | Rbmxl1 | 0.147328797 | -0.052808289 | 0.347465883 | 12.68578875 | 1.788723691 | 0.122444235 | 0.53909906 | -4.283999373 |
| **A_55_P1987746** | NA | 0.164705481 | -0.059054104 | 0.388465065 | 15.56849573 | 1.788585037 | 0.122468 | 0.53909906 | -4.28406982 |
| **A_52_P54261** | Tmem56 | 0.495130007 | -0.177678054 | 1.167938068 | 10.30859452 | 1.788180071 | 0.122537437 | 0.539239072 | -4.28427558 |
| **A_52_P487024** | Smg7 | 0.156496149 | -0.056202308 | 0.369194605 | 9.164587722 | 1.787814707 | 0.122600116 | 0.539239072 | -4.284461222 |
| **A_52_P480544** | Tnpo2 | 0.17585858 | -0.063166993 | 0.414884153 | 9.207684168 | 1.787731868 | 0.122614331 | 0.539239072 | -4.284503313 |
| **A_55_P2051874** | Psmc2 | 0.214186806 | -0.076953733 | 0.505327345 | 13.33020133 | 1.787611841 | 0.122634932 | 0.539239072 | -4.2845643 |
| **A_55_P1982454** | Eps8 | -0.30761594 | -0.725816077 | 0.110584197 | 8.527445467 | -1.787342798 | 0.12268112 | 0.539239072 | -4.284701006 |
| **A_55_P2167645** | NA | 0.411080316 | -0.147789953 | 0.969950584 | 10.43748628 | 1.787305799 | 0.122687473 | 0.539239072 | -4.284719806 |
| **A_51_P237383** | Rnase4 | -0.348546962 | -0.822475385 | 0.125381462 | 8.612985409 | -1.787029553 | 0.122734918 | 0.539239072 | -4.284860174 |

| | | | | | | | | | |
|---|---|---|---|---|---|---|---|---|---|
| **A_55_P2055587** | Enpp4 | 0.217964444 | -0.0784116 | 0.514340488 | 9.923939072 | 1.787004912 | 0.122739151 | 0.539239072 | -4.284872695 |
| **A_51_P487298** | Vasn | -0.463410612 | -1.093554926 | 0.166733703 | 9.945756531 | -1.786936588 | 0.122750888 | 0.539239072 | -4.284907413 |
| **A_55_P2058143** | Pclo | 0.143602289 | -0.051737232 | 0.33894181 | 8.157302761 | 1.786300145 | 0.122860278 | 0.539429251 | -4.285230818 |
| **A_51_P265495** | Ly6a | -0.336354272 | -0.793933377 | 0.121224832 | 8.822371634 | -1.786133654 | 0.12288891 | 0.539429251 | -4.285315422 |
| **A_55_P2176585** | Donson | 0.201731945 | -0.072732931 | 0.476196822 | 8.480557165 | 1.785957377 | 0.122919232 | 0.539429251 | -4.285404999 |
| **A_55_P2034453** | NA | 0.164072885 | -0.059168296 | 0.387314066 | 8.431969692 | 1.785852936 | 0.1229372 | 0.539429251 | -4.285458073 |
| **A_51_P219849** | Alg13 | 0.184234891 | -0.066507804 | 0.434977586 | 8.650733578 | 1.785364035 | 0.123021347 | 0.539429251 | -4.285706521 |
| **A_51_P135654** | Eif4ebp2 | -0.189117546 | -0.446558127 | 0.068323035 | 11.64718351 | -1.784999117 | 0.123084192 | 0.539429251 | -4.285891969 |
| **A_55_P2068570** | NA | 0.135727046 | -0.049037083 | 0.320491174 | 7.951686439 | 1.784974133 | 0.123088496 | 0.539429251 | -4.285904665 |
| **A_55_P1989321** | Sat1 | -0.165477148 | -0.39074186 | 0.059787564 | 10.81212882 | -1.784958209 | 0.123091239 | 0.539429251 | -4.285912758 |
| **A_55_P1987196** | Gm3448 | 0.207140973 | -0.074844888 | 0.489126835 | 9.332890619 | 1.78493279 | 0.123095618 | 0.539429251 | -4.285925676 |
| **A_51_P507172** | 2310022A10Rik | 0.163416154 | -0.059047843 | 0.385880151 | 10.53856141 | 1.784918695 | 0.123098046 | 0.539429251 | -4.285932839 |
| **A_51_P284966** | Dolk | 0.151971183 | -0.054992954 | 0.35893532 | 10.63725396 | 1.784223831 | 0.123217812 | 0.539429251 | -4.286285974 |
| **A_55_P2099840** | NA | -0.431162968 | -1.018411305 | 0.156085368 | 9.112090207 | -1.784032851 | 0.12325075 | 0.539429251 | -4.286383035 |
| **A_52_P546513** | NA | 0.47951861 | -0.173683273 | 1.132720493 | 8.481138985 | 1.783779743 | 0.123294415 | 0.539429251 | -4.286511672 |
| **A_55_P2176838** | Mitf | -0.289578931 | -0.684044807 | 0.104886946 | 8.87928481 | -1.783777611 | 0.123294783 | 0.539429251 | -4.286512755 |
| **A_55_P2035400** | Ripk4 | -0.283124157 | -0.66881111 | 0.102562795 | 10.07784326 | -1.783713783 | 0.123305796 | 0.539429251 | -4.286545194 |
| **A_52_P449629** | Ankrd13c | 0.158186943 | -0.057323482 | 0.373697367 | 8.877322046 | 1.783551025 | 0.123333885 | 0.539429251 | -4.286627914 |
| **A_55_P2078285** | Uck1 | -0.199186476 | -0.470567668 | 0.072194716 | 12.53544557 | -1.783459662 | 0.123349656 | 0.539429251 | -4.286674349 |
| **A_66_P124420** | NA | -0.129258979 | -0.305371437 | 0.046853478 | 7.932559698 | -1.783420826 | 0.12335636 | 0.539429251 | -4.286694087 |
| **A_51_P137336** | Cdh1 | -0.260437439 | -0.615320565 | 0.094445686 | 11.07775565 | -1.783204933 | 0.123393635 | 0.539429251 | -4.286803814 |
| **A_52_P218379** | Kif26a | -0.398039663 | -0.940526311 | 0.144446985 | 8.41670473 | -1.782873316 | 0.123450912 | 0.539429251 | -4.28697236 |
| **A_51_P158210** | Mcm2 | 0.38640056 | -0.140229093 | 0.913030213 | 12.39339558 | 1.782853372 | 0.123454358 | 0.539429251 | -4.286982497 |
| **A_51_P453909** | Cyp2f2 | -0.973686963 | -2.300764307 | 0.353390381 | 10.85618414 | -1.782815123 | 0.123460966 | 0.539429251 | -4.287001937 |
| **A_66_P128079** | Krt8 | -0.180079872 | -0.425526215 | 0.065366471 | 13.38089887 | -1.782755544 | 0.12347126 | 0.539429251 | -4.287032219 |
| **A_51_P328300** | Pdia4 | -0.288062963 | -0.680726176 | 0.10460025 | 10.40244871 | -1.78258561 | 0.123500626 | 0.539429251 | -4.287118591 |
| **A_55_P2076787** | Synpo | 0.195014255 | -0.070857709 | 0.46088622 | 8.589335692 | 1.782284388 | 0.123552696 | 0.539429251 | -4.287271694 |
| **A_55_P1979027** | Tmem65 | 0.141903839 | -0.051561708 | 0.335369385 | 8.351767681 | 1.782270828 | 0.123555041 | 0.539429251 | -4.287278586 |
| **A_55_P2058657** | NA | 0.139729942 | -0.050783875 | 0.330243759 | 16.32537853 | 1.78215794 | 0.123574561 | 0.539429251 | -4.287335965 |
| **A_52_P205797** | Vgll4 | -0.242064918 | -0.572117225 | 0.087987389 | 8.020317978 | -1.782100817 | 0.12358444 | 0.539429251 | -4.287364999 |
| **A_55_P2183473** | Cldn3 | -1.529451042 | -3.615536734 | 0.556634651 | 10.04072523 | -1.781503271 | 0.123687823 | 0.539429251 | -4.287668727 |
| **A_55_P1974650** | Npdc1 | -0.326586294 | -0.772133928 | 0.11896134 | 9.848494792 | -1.781094659 | 0.123758567 | 0.539429251 | -4.287876428 |
| **A_55_P2105694** | Prr29 | 0.184284717 | -0.067130664 | 0.435700098 | 8.762029095 | 1.78106868 | 0.123763066 | 0.539429251 | -4.287889634 |
| **A_52_P255804** | NA | 0.158233608 | -0.05764304 | 0.374110256 | 11.52307363 | 1.781050578 | 0.123766202 | 0.539429251 | -4.287898835 |
| **A_55_P2147641** | Cstf1 | 0.152565185 | -0.055582209 | 0.360712578 | 11.48735128 | 1.781015302 | 0.123772311 | 0.539429251 | -4.287916767 |
| **A_52_P504302** | Pmpca | 0.237389804 | -0.086504354 | 0.561283962 | 9.877585358 | 1.780910674 | 0.123790434 | 0.539429251 | -4.287969951 |
| **A_55_P2333126** | Oga | 0.153754506 | -0.056035806 | 0.363544817 | 9.045421257 | 1.780842914 | 0.123802172 | 0.539429251 | -4.288004394 |
| **A_55_P2168058** | Glod4 | -0.139272934 | -0.329314628 | 0.05076876 | 10.89825541 | -1.780742074 | 0.123819643 | 0.539429251 | -4.288055654 |
| **A_52_P578562** | Slc41a1 | -0.151579601 | -0.358419944 | 0.055260742 | 7.849657929 | -1.780691559 | 0.123828396 | 0.539429251 | -4.288081332 |
| **A_55_P2142200** | NA | 0.126412982 | -0.046100544 | 0.298926509 | 10.88460585 | 1.780539913 | 0.123854675 | 0.539429251 | -4.288158418 |
| **A_52_P472660** | Smurf2 | -0.210107813 | -0.496881407 | 0.076665781 | 9.005836789 | -1.780271467 | 0.123901209 | 0.539510354 | -4.288294878 |
| **A_66_P135176** | Trak2 | 0.318577229 | -0.116320723 | 0.753475181 | 9.777388237 | 1.779961286 | 0.123954998 | 0.539623008 | -4.288452557 |
| **A_66_P118934** | Plaa | 0.384963723 | -0.140608691 | 0.910536137 | 10.51942547 | 1.779796844 | 0.123983523 | 0.539625653 | -4.288536151 |
| **A_55_P2026315** | Gys1 | -0.203154645 | -0.480638107 | 0.074328817 | 8.30270627 | -1.778987271 | 0.124124053 | 0.539940625 | -4.288947709 |
| **A_55_P1980651** | Ubr5 | 0.128605806 | -0.047064836 | 0.304276448 | 11.88713821 | 1.778871498 | 0.124144162 | 0.539940625 | -4.289006566 |
| **A_51_P172752** | Gpatch4 | 0.291689157 | -0.106775446 | 0.690153761 | 11.87103504 | 1.778745141 | 0.124166113 | 0.539940625 | -4.289070803 |
| **A_51_P145511** | Prl2a1 | 0.215723098 | -0.07902081 | 0.510467007 | 8.307791551 | 1.778422732 | 0.12422214 | 0.539940625 | -4.289234712 |
| **A_52_P28806** | Foxm1 | 0.242827894 | -0.088960262 | 0.574616049 | 9.629668602 | 1.778364921 | 0.124232189 | 0.539940625 | -4.289264103 |
| **A_55_P2153517** | Enho | -0.424704508 | -1.005000257 | 0.155591241 | 11.17689657 | -1.778364052 | 0.12423234 | 0.539940625 | -4.289264544 |
| **A_51_P261107** | Ogt | 0.153777247 | -0.056349539 | 0.363904034 | 9.744335361 | 1.778254237 | 0.12425143 | 0.539940625 | -4.289320374 |

| | | | | | | | | | |
|---|---|---|---|---|---|---|---|---|---|
| **A_55_P2122220** | Kcnk4 | 0.260376366 | -0.095465077 | 0.616217808 | 9.257117425 | 1.777985536 | 0.124298154 | 0.540022256 | -4.289456982 |
| **A_51_P302651** | Topbp1 | 0.266542462 | -0.097815632 | 0.630900556 | 10.46057858 | 1.777547323 | 0.12437439 | 0.540093509 | -4.289679775 |
| **A_55_P1970229** | Csnk1d | -0.216550224 | -0.512631606 | 0.079531157 | 12.78792924 | -1.777177182 | 0.124438819 | 0.540093509 | -4.289867963 |
| **A_55_P2110513** | Lmo7 | -0.89549074 | -2.120144699 | 0.32916322 | 9.513516578 | -1.776768709 | 0.124509959 | 0.540093509 | -4.290075645 |
| **A_52_P248483** | Zfx | 0.136653795 | -0.050237689 | 0.32354528 | 8.733584761 | 1.776705198 | 0.124521023 | 0.540093509 | -4.290107937 |
| **A_55_P2006861** | Trio | 0.42729447 | -0.15709886 | 1.011687799 | 8.790504505 | 1.776663625 | 0.124528266 | 0.540093509 | -4.290129075 |
| **A_52_P119187** | NA | 0.260927104 | -0.09595869 | 0.617812897 | 9.257822594 | 1.776532358 | 0.124551139 | 0.540093509 | -4.290195816 |
| **A_52_P668984** | Tmem243 | -0.506956113 | -1.200402088 | 0.186489863 | 8.794141624 | -1.776400641 | 0.124574095 | 0.540093509 | -4.290262788 |
| **A_55_P1975095** | Timmdc1 | 0.17986019 | -0.06617409 | 0.42589447 | 9.764620813 | 1.776325763 | 0.124587147 | 0.540093509 | -4.29030086 |
| **A_51_P403536** | Ltbp4 | -0.834130139 | -1.975161472 | 0.306901194 | 11.53499141 | -1.776312746 | 0.124589416 | 0.540093509 | -4.290307478 |
| **A_55_P2108122** | Arcn1 | 0.162213533 | -0.059686433 | 0.384113498 | 12.70893312 | 1.776286588 | 0.124593976 | 0.540093509 | -4.290320779 |
| **A_55_P1973573** | Spn | 0.23112733 | -0.085137188 | 0.547391849 | 8.554497005 | 1.775758983 | 0.124685982 | 0.540371155 | -4.290589046 |
| **A_52_P531175** | Fkbp3 | 0.124042754 | -0.045781522 | 0.29386703 | 12.44371486 | 1.774822059 | 0.124849532 | 0.54077777 | -4.291065456 |
| **A_55_P1954910** | Fbxw2 | 0.146417889 | -0.05404393 | 0.346879708 | 8.664113308 | 1.774784645 | 0.124856067 | 0.54077777 | -4.291084481 |
| **A_55_P2141093** | Eya2 | -0.821345428 | -1.946094912 | 0.303404055 | 8.213207546 | -1.774406953 | 0.124922059 | 0.54077777 | -4.291276539 |
| **A_55_P2175356** | Uros | 0.278692794 | -0.102963093 | 0.66034868 | 8.799641132 | 1.774340489 | 0.124933676 | 0.54077777 | -4.291310336 |
| **A_51_P385773** | Eif2ak1 | 0.369103992 | -0.136385971 | 0.874593954 | 10.34230786 | 1.774268715 | 0.124946222 | 0.54077777 | -4.291346834 |
| **A_51_P497113** | Igf2bp1 | 0.318164451 | -0.117565563 | 0.753894464 | 8.772109927 | 1.774260428 | 0.12494767 | 0.54077777 | -4.291351048 |
| **A_55_P1983708** | Insrr | 0.250775454 | -0.092809193 | 0.594360101 | 8.544283882 | 1.773513366 | 0.125078328 | 0.541108699 | -4.291730947 |
| **A_55_P2002357** | Tnrc18 | 0.174660183 | -0.064642596 | 0.413962962 | 9.128546402 | 1.773492503 | 0.125081979 | 0.541108699 | -4.291741557 |
| **A_55_P2171373** | Eloc | 0.161277087 | -0.059727371 | 0.382281544 | 13.30200244 | 1.773188171 | 0.125135244 | 0.541108699 | -4.291896321 |
| **A_51_P110395** | Setd2 | 0.138622253 | -0.051337908 | 0.328582415 | 11.20127135 | 1.773183222 | 0.125136111 | 0.541108699 | -4.305173093 |
| **A_55_P1993493** | Efr3a | -0.211256144 | -0.500799212 | 0.078286924 | 10.02578889 | -1.772880099 | 0.125189187 | 0.541217134 | -4.29205299 |
| **A_51_P265106** | Dlat | 0.137974976 | -0.051227218 | 0.327177169 | 8.987744631 | 1.771974011 | 0.125347974 | 0.541697672 | -4.292513794 |
| **A_55_P2107045** | Myl4 | -1.913882153 | -4.53841975 | 0.710655445 | 10.25057553 | -1.771926009 | 0.125356391 | 0.541697672 | -4.292538207 |
| **A_55_P2035877** | Faap20 | -0.201372842 | -0.477650253 | 0.074904569 | 10.18530715 | -1.771082168 | 0.125504456 | 0.542152785 | -4.292967376 |
| **A_55_P1982996** | Ramp1 | 0.208169394 | -0.077470983 | 0.49380977 | 8.486029819 | 1.770844673 | 0.125546159 | 0.542152785 | -4.293088167 |
| **A_52_P30877** | Gm5176 | 0.138872783 | -0.051682264 | 0.329427829 | 13.50940377 | 1.770842236 | 0.125546587 | 0.542152785 | -4.293089407 |
| **A_52_P676798** | Gigyf1 | 0.258857034 | -0.096424963 | 0.61413903 | 9.418052153 | 1.770394119 | 0.125625311 | 0.542152785 | -4.293317325 |
| **A_55_P2118309** | NA | 0.304702807 | -0.113538806 | 0.72294442 | 9.190179502 | 1.770241032 | 0.125652215 | 0.542152785 | -4.293395189 |
| **A_51_P359262** | Sec11c | 0.139714772 | -0.052061511 | 0.331491054 | 10.54752035 | 1.770233757 | 0.125653494 | 0.542152785 | -4.293398889 |
| **A_55_P1987326** | Prl3a1 | -0.204388623 | -0.484952984 | 0.076175738 | 7.89750479 | -1.770139135 | 0.125670127 | 0.542152785 | -4.293447016 |
| **A_52_P192307** | Rbak | -0.20051574 | -0.475783045 | 0.074751566 | 8.466777477 | -1.770015322 | 0.125691894 | 0.542152785 | -4.293509991 |
| **A_55_P2061045** | Rpl29 | -0.236627347 | -0.56149177 | 0.088237076 | 15.15275508 | -1.769888757 | 0.125714149 | 0.542152785 | -4.293574366 |
| **A_52_P518997** | Epha2 | -0.349763242 | -0.830023369 | 0.130496885 | 8.967133313 | -1.769623281 | 0.125760842 | 0.542233172 | -4.293709397 |
| **A_55_P1977478** | Zfp629 | -0.198402354 | -0.470914858 | 0.07411015 | 10.35117037 | -1.769064125 | 0.125859244 | 0.542325136 | -4.293993812 |
| **A_65_P05040** | Ston2 | 0.245626891 | -0.091775621 | 0.583029403 | 8.702339248 | 1.768930573 | 0.125882758 | 0.542325136 | -4.294061744 |
| **A_55_P2143015** | Sema6c | 0.210970949 | -0.078836875 | 0.500778773 | 8.822189627 | 1.768869278 | 0.125893551 | 0.542325136 | -4.294092922 |
| **A_51_P329949** | Fam13a | -0.265996589 | -0.631393376 | 0.099400197 | 9.328613733 | -1.768864368 | 0.125894416 | 0.542325136 | -4.29409542 |
| **A_55_P1982334** | Llph | 0.211831931 | -0.079225853 | 0.502889715 | 11.5736309 | 1.768460632 | 0.125965532 | 0.542325136 | -4.294300787 |
| **A_55_P1964359** | Cnot8 | -0.170591543 | -0.404987112 | 0.063804025 | 11.2813179 | -1.768443915 | 0.125968478 | 0.542325136 | -4.294309291 |
| **A_55_P2044533** | NA | -0.232070486 | -0.550949987 | 0.086809015 | 8.414475905 | -1.768383685 | 0.125979091 | 0.542325136 | -4.294339928 |
| **A_51_P369636** | Hat1 | 0.273957526 | -0.102522671 | 0.650437722 | 10.23941056 | 1.768171099 | 0.126016557 | 0.542325136 | -4.294448066 |
| **A_55_P2099232** | Bdp1 | 0.188128458 | -0.070418042 | 0.446674958 | 10.11017845 | 1.768068235 | 0.126034689 | 0.542325136 | -4.294500391 |
| **A_51_P283004** | Tinag | -0.902941862 | -2.144193206 | 0.338309483 | 9.394833443 | -1.767596956 | 0.126117798 | 0.542528118 | -4.294740126 |
| **A_52_P186937** | Cmpk2 | -0.168857072 | -0.400995656 | 0.063281513 | 9.491512513 | -1.767482452 | 0.126137998 | 0.542528118 | -4.294798374 |
| **A_52_P669964** | Cchcr1 | 0.174367473 | -0.065394818 | 0.414129764 | 8.856163039 | 1.767127084 | 0.126200711 | 0.542677094 | -4.294979151 |
| **A_55_P2120843** | NA | -0.283540283 | -0.673478037 | 0.106397472 | 8.917637295 | -1.766862163 | 0.126247482 | 0.542749749 | -4.295113919 |
| **A_52_P318073** | NA | 0.299406183 | -0.112443118 | 0.711255484 | 9.915475015 | 1.766467375 | 0.126317212 | 0.542749749 | -4.295314756 |
| **A_55_P1956557** | Rab3a | -0.23983731 | -0.569761596 | 0.090086977 | 9.162339928 | -1.766386134 | 0.126331566 | 0.542749749 | -4.295356086 |

| | | | | | | | | | |
|---|---|---|---|---|---|---|---|---|---|
| **A_55_P2059931** | Prom1 | 0.281016653 | -0.105600835 | 0.667634142 | 11.57801862 | 1.766175074 | 0.126368865 | 0.542749749 | -4.295463459 |
| **A_55_P2162782** | Ahnak | -0.196337955 | -0.466468648 | 0.073792739 | 7.942128482 | -1.766092794 | 0.126383408 | 0.542749749 | -4.295505317 |
| **A_55_P2102095** | 1810010H24Rik | -0.226415848 | -0.537931747 | 0.08510005 | 9.065094533 | -1.766077656 | 0.126386084 | 0.542749749 | -4.295513019 |
| **A_66_P107584** | Fam133b | 0.160370912 | -0.060409243 | 0.381151066 | 8.36569698 | 1.765016435 | 0.126573811 | 0.543084255 | -4.296052915 |
| **A_55_P2096762** | Arhgdig | 0.425773628 | -0.160410368 | 1.011957624 | 9.436411992 | 1.764932051 | 0.126588749 | 0.543084255 | -4.296095847 |
| **A_55_P1953998** | NA | -0.514533026 | -1.222925933 | 0.193859881 | 10.20444257 | -1.764908687 | 0.126592886 | 0.543084255 | -4.296107734 |
| **A_55_P2075048** | Atic | 0.270602509 | -0.101967599 | 0.643172617 | 10.52343808 | 1.764846755 | 0.126603852 | 0.543084255 | -4.296139243 |
| **A_55_P2008755** | NA | 0.146055225 | -0.05503647 | 0.347146921 | 13.00320878 | 1.764843306 | 0.126604462 | 0.543084255 | -4.296140998 |
| **A_55_P2172146** | Pds5a | 0.293779687 | -0.110766772 | 0.698326146 | 9.516875925 | 1.764560482 | 0.126654551 | 0.543178568 | -4.296284891 |
| **A_55_P2105843** | NA | -0.280456215 | -0.666728628 | 0.105816197 | 12.35547389 | -1.764227414 | 0.126713562 | 0.543207505 | -4.296454349 |
| **A_55_P1996653** | D030056L22Rik | 0.231928443 | -0.087518739 | 0.551375625 | 12.35924336 | 1.76416069 | 0.126725387 | 0.543207505 | -4.296488297 |
| **A_55_P2075592** | Clstn1 | -0.193871498 | -0.460920903 | 0.073177908 | 7.876463336 | -1.764028263 | 0.126748859 | 0.543207505 | -4.296555674 |
| **A_51_P370350** | Galnt2 | -0.19242653 | -0.457506622 | 0.072653562 | 11.57483895 | -1.76388808 | 0.126773711 | 0.543207505 | -4.296626999 |
| **A_55_P2301063** | 9930024M15Rik | 0.187442144 | -0.070811931 | 0.445696219 | 8.459269568 | 1.763612832 | 0.126822521 | 0.543229677 | -4.296767044 |
| **A_52_P612137** | Runx1t1 | 0.300974812 | -0.113743097 | 0.715692721 | 10.84984936 | 1.763439445 | 0.126853277 | 0.543229677 | -4.296855264 |
| **A_51_P237688** | Smim14 | -0.272184507 | -0.647249868 | 0.102880854 | 11.0303957 | -1.763354512 | 0.126868345 | 0.543229677 | -4.296898478 |
| **A_55_P1974877** | Bcl2l1 | 0.212515345 | -0.08034847 | 0.505379161 | 14.83775488 | 1.763225132 | 0.126891303 | 0.543229677 | -4.296964308 |
| **A_52_P653966** | Atf7ip | 0.1960473 | -0.074207556 | 0.466302156 | 10.49821374 | 1.762668114 | 0.126990187 | 0.543389603 | -4.297247729 |
| **A_51_P247799** | Casp8 | -0.184533732 | -0.438923281 | 0.069855817 | 9.562555679 | -1.762624036 | 0.126998015 | 0.543389603 | -4.297270157 |
| **A_51_P136792** | Calcoco1 | -0.152065054 | -0.36170647 | 0.057576362 | 8.040943857 | -1.762525974 | 0.127015432 | 0.543389603 | -4.297320053 |
| **A_55_P2055597** | Enpp2 | -0.49710761 | -1.182491185 | 0.188275966 | 12.22312127 | -1.762381429 | 0.12704111 | 0.543389603 | -4.297393602 |
| **A_51_P237040** | Nog | 0.452800097 | -0.171605831 | 1.077206026 | 8.839347461 | 1.762068076 | 0.127096793 | 0.543462586 | -4.297553048 |
| **A_55_P2195157** | Dbn1 | -0.278839017 | -0.663376317 | 0.105698283 | 8.815188367 | -1.761968986 | 0.127114406 | 0.543462586 | -4.297603469 |
| **A_51_P313483** | Dyrk1a | 0.228383076 | -0.086662276 | 0.543428427 | 9.83432963 | 1.761465049 | 0.127204017 | 0.543654597 | -4.297859897 |
| **A_55_P2282257** | NA | 0.374764226 | -0.142227149 | 0.891755602 | 10.42718324 | 1.761400109 | 0.12721557 | 0.543654597 | -4.297892942 |
| **A_55_P1972530** | Efl1 | 0.158080918 | -0.060027896 | 0.376189732 | 9.990828621 | 1.761121915 | 0.127265069 | 0.543721797 | -4.298034504 |
| **A_55_P2016119** | Sgcb | -0.312011726 | -0.742550525 | 0.118527073 | 8.676064589 | -1.760928856 | 0.127299432 | 0.543721797 | -4.298132745 |
| **A_55_P2063999** | Gpatch8 | 0.16528744 | -0.062814485 | 0.393389364 | 10.45536583 | 1.760735387 | 0.127333877 | 0.543721797 | -4.298231196 |
| **A_51_P493857** | Katna1 | 0.196358396 | -0.074658133 | 0.467374924 | 10.72815323 | 1.760503471 | 0.127375178 | 0.543721797 | -4.298349213 |
| **A_51_P501312** | Natd1 | 0.158648191 | -0.060329795 | 0.377626176 | 11.79449568 | 1.760426334 | 0.127388918 | 0.543721797 | -4.298388466 |
| **A_55_P2417212** | NA | 0.267055726 | -0.101606489 | 0.635717942 | 9.086844591 | 1.760177458 | 0.127433259 | 0.543721797 | -4.298515115 |
| **A_55_P2136204** | Rnf38 | 0.199672347 | -0.075973079 | 0.475317774 | 11.17996494 | 1.760152599 | 0.127437689 | 0.543721797 | -4.298527766 |
| **A_55_P2056260** | NA | 0.174344114 | -0.066350278 | 0.415038506 | 15.42052344 | 1.760047984 | 0.127456333 | 0.543721797 | -4.298581003 |
| **A_51_P418168** | Manf | -0.122866195 | -0.292566031 | 0.046833641 | 11.95827202 | -1.759276801 | 0.127593851 | 0.543828518 | -4.298973459 |
| **A_55_P2055299** | NA | 0.205289003 | -0.078313963 | 0.48889197 | 8.539074802 | 1.758887676 | 0.127663295 | 0.543828518 | -4.299171491 |
| **A_52_P114620** | Elmo3 | -0.188114264 | -0.448027015 | 0.071798486 | 10.40438913 | -1.758641559 | 0.127707236 | 0.543828518 | -4.299296745 |
| **A_51_P158772** | Elof1 | -0.318041353 | -0.757501744 | 0.121419038 | 13.40501295 | -1.758518914 | 0.127729139 | 0.543828518 | -4.299359162 |
| **A_51_P297336** | Chmp7 | -0.220289938 | -0.524694645 | 0.08411477 | 10.66290424 | -1.758435671 | 0.127744007 | 0.543828518 | -4.299401527 |
| **A_55_P1978696** | NA | 0.315387111 | -0.120432587 | 0.751206809 | 9.412300375 | 1.758410523 | 0.127748499 | 0.543828518 | -4.299414326 |
| **A_51_P443514** | 1700001K19Rik | 0.364896691 | -0.139340004 | 0.869133386 | 9.380129445 | 1.758404013 | 0.127749662 | 0.543828518 | -4.299417639 |
| **A_55_P1990066** | NA | 0.224587184 | -0.085776372 | 0.53495074 | 9.120612722 | 1.758318074 | 0.127765013 | 0.543828518 | -4.299461376 |
| **A_55_P2044005** | NA | 0.25813333 | -0.098618713 | 0.614885374 | 9.725884142 | 1.758169753 | 0.127791513 | 0.543828518 | -4.299536862 |
| **A_52_P323820** | Ufm1 | 0.137728699 | -0.052630821 | 0.328088219 | 8.195480322 | 1.758057334 | 0.127811602 | 0.543828518 | -4.299594077 |
| **A_55_P2179791** | NA | 0.279250403 | -0.106730785 | 0.66523159 | 8.47721393 | 1.757967568 | 0.127827645 | 0.543828518 | -4.299639762 |
| **A_51_P281024** | Rad9a | 0.258362105 | -0.098857456 | 0.615581666 | 9.439880404 | 1.757424884 | 0.127924677 | 0.543828518 | -4.299915961 |
| **A_55_P2151672** | NA | 0.201047008 | -0.076981603 | 0.479075618 | 9.133877901 | 1.757079099 | 0.12798654 | 0.543828518 | -4.300091951 |
| **A_51_P285770** | Fem1a | 0.13757713 | -0.052682054 | 0.327836314 | 11.60923849 | 1.757048728 | 0.127991975 | 0.543828518 | -4.300107409 |
| **A_51_P224175** | 2210016F16Rik | 0.129635049 | -0.049643749 | 0.308913847 | 8.253704166 | 1.757019952 | 0.127997125 | 0.543828518 | -4.300122055 |
| **A_55_P1982171** | H2afz | 0.169716413 | -0.064998314 | 0.40443114 | 15.92227768 | 1.756979493 | 0.128004366 | 0.543828518 | -4.300142648 |
| **A_51_P213359** | Has2 | -0.122363526 | -0.291591462 | 0.04686441 | 7.931367148 | -1.756965024 | 0.128006955 | 0.543828518 | -4.300150012 |

| | | | | | | | | | |
|---|---|---|---|---|---|---|---|---|---|
| **A_55_P2108803** | Tubgcp4 | 0.157374449 | -0.060287377 | 0.375036274 | 9.175374489 | 1.756851865 | 0.12802721 | 0.543828518 | -4.300207606 |
| **A_55_P2275249** | Th | 0.277512617 | -0.106376383 | 0.661401616 | 8.896850429 | 1.75654894 | 0.128081446 | 0.543828518 | -4.300361787 |
| **A_55_P2037294** | NA | -0.114109222 | -0.271971025 | 0.043752582 | 8.363584326 | -1.756413921 | 0.128105627 | 0.543828518 | -4.300430508 |
| **A_55_P2084763** | NA | 0.207474214 | -0.079585552 | 0.494533979 | 12.63813174 | 1.756204114 | 0.128143211 | 0.543828518 | -4.300537296 |
| **A_52_P496202** | Pcsk1n | 0.27893974 | -0.107052828 | 0.664932307 | 8.735792224 | 1.755960078 | 0.12818694 | 0.543828518 | -4.300661508 |
| **A_55_P2172334** | Nup155 | 0.237574313 | -0.091191108 | 0.566339734 | 10.15239994 | 1.755886898 | 0.128200057 | 0.543828518 | -4.300698756 |
| **A_55_P2162537** | Acrbp | 0.132389686 | -0.050817162 | 0.315596534 | 8.059158579 | 1.755883263 | 0.128200708 | 0.543828518 | -4.300700606 |
| **A_55_P1957439** | NA | 0.19349291 | -0.074278738 | 0.461264558 | 8.517790214 | 1.755834806 | 0.128209394 | 0.543828518 | -4.30072527 |
| **A_51_P479043** | Nif3l1 | 0.171905218 | -0.066006203 | 0.409816639 | 11.14444229 | 1.755726876 | 0.128228742 | 0.543828518 | -4.300780206 |
| **A_51_P359293** | Ubxn2b | 0.206716673 | -0.079391977 | 0.492825324 | 9.271319959 | 1.75560863 | 0.128249943 | 0.543828518 | -4.300840393 |
| **A_52_P458697** | Higd1a | 0.195645414 | -0.075159457 | 0.466450285 | 12.85783371 | 1.755482033 | 0.128272645 | 0.543828518 | -4.300904831 |
| **A_55_P1962289** | NA | -0.189703633 | -0.452310345 | 0.072903079 | 8.536793622 | -1.755306721 | 0.128304089 | 0.543828518 | -4.300994066 |
| **A_55_P2019699** | Samhd1 | -0.222178888 | -0.529761918 | 0.085404142 | 10.63297593 | -1.755187864 | 0.128325411 | 0.543828518 | -4.301054565 |
| **A_51_P226269** | Rgcc | 0.25649409 | -0.098641707 | 0.611629888 | 14.22343934 | 1.754955472 | 0.128367111 | 0.543885991 | -4.301172856 |
| **A_55_P1982762** | Hsd3b7 | -0.324165174 | -0.773100247 | 0.124769899 | 12.74440191 | -1.754551047 | 0.128439712 | 0.543899477 | -4.301378716 |
| **A_52_P276607** | Ptp4a2 | 0.116214965 | -0.044734392 | 0.277164322 | 13.11634635 | 1.754510613 | 0.128446973 | 0.543899477 | -4.301399298 |
| **A_55_P2078623** | Stxbp3 | -0.174312745 | -0.415729272 | 0.067103782 | 10.08918071 | -1.754467522 | 0.128454711 | 0.543899477 | -4.301421232 |
| **A_52_P447971** | NA | 0.184184531 | -0.070931024 | 0.439300087 | 9.784947948 | 1.754281977 | 0.128488037 | 0.543921433 | -4.30151568 |
| **A_55_P2175494** | NA | -0.167096266 | -0.398652327 | 0.064459795 | 8.720530636 | -1.753451592 | 0.128637284 | 0.544355972 | -4.30193838 |
| **A_55_P2097329** | Sgsm3 | 0.219841263 | -0.084816336 | 0.524498862 | 12.52065489 | 1.753397506 | 0.128647011 | 0.544355972 | -4.301965912 |
| **A_55_P2027157** | NA | -0.335239217 | -0.800138372 | 0.129659939 | 8.283243869 | -1.75218214 | 0.128865773 | 0.544759076 | -4.302584614 |
| **A_55_P2348799** | 1700025G04Rik | -0.167866023 | -0.400677008 | 0.064944963 | 7.907663852 | -1.752033963 | 0.128892469 | 0.544759076 | -4.302660048 |
| **A_52_P193194** | Slc35d2 | 0.303012419 | -0.117231179 | 0.723256016 | 9.061216292 | 1.752033926 | 0.128892476 | 0.544759076 | -4.302660066 |
| **A_51_P167527** | Lum | 0.392585512 | -0.151907416 | 0.93707844 | 11.5793732 | 1.751964145 | 0.12890505 | 0.544759076 | -4.302695591 |
| **A_55_P2254968** | BB212172 | 0.14492967 | -0.056103126 | 0.345962466 | 8.416411111 | 1.751755866 | 0.128942587 | 0.544759076 | -4.302801623 |
| **A_55_P2135027** | NA | 0.164956125 | -0.063890205 | 0.393802454 | 10.32111274 | 1.75149008 | 0.128990503 | 0.544759076 | -4.302936933 |
| **A_51_P469902** | Ggps1 | 0.1197598 | -0.046410541 | 0.285930142 | 8.7088251 | 1.751220132 | 0.129039188 | 0.544759076 | -4.303074363 |
| **A_55_P2005440** | Dennd5a | -0.29719354 | -0.709580605 | 0.115193525 | 11.11713125 | -1.751126498 | 0.129056079 | 0.544759076 | -4.303122032 |
| **A_55_P2136572** | Cdc42se2 | -0.15030364 | -0.358871388 | 0.058264108 | 10.49149652 | -1.751078179 | 0.129064796 | 0.544759076 | -4.303146632 |
| **A_55_P1996623** | Dnajb12 | 0.189060711 | -0.073291327 | 0.451412749 | 9.948388438 | 1.751055991 | 0.129068799 | 0.544759076 | -4.303157928 |
| **A_55_P2138906** | Lrfn1 | 0.133665101 | -0.051819466 | 0.319149668 | 8.543726785 | 1.751029409 | 0.129073595 | 0.544759076 | -4.303171461 |
| **A_51_P195958** | Phlda1 | -0.948517944 | -2.264790459 | 0.367754571 | 11.44245944 | -1.750987023 | 0.129081243 | 0.544759076 | -4.30319304 |
| **A_55_P2106394** | Knstrn | 0.435764758 | -0.169004983 | 1.040534498 | 13.67118351 | 1.75083508 | 0.129108662 | 0.544759076 | -4.303270395 |
| **A_55_P2128668** | Ccnb1 | 0.392880827 | -0.152460901 | 0.938222555 | 9.866445019 | 1.750553122 | 0.129159558 | 0.54480983 | -4.303413944 |
| **A_55_P1973178** | Haus5 | 0.281571857 | -0.109288111 | 0.672431826 | 10.6097064 | 1.750456174 | 0.129177063 | 0.54480983 | -4.303463302 |
| **A_55_P1975903** | Dnmt3b | 0.31623493 | -0.122794325 | 0.755264186 | 9.375649282 | 1.750247906 | 0.129214675 | 0.544849576 | -4.303569336 |
| **A_55_P2045876** | NA | -0.136783467 | -0.326800182 | 0.053233247 | 8.495542879 | -1.749141696 | 0.129414629 | 0.545276208 | -4.304132548 |
| **A_55_P1983999** | Desi1 | 0.232767857 | -0.090604317 | 0.556140031 | 8.930972708 | 1.749055339 | 0.129430251 | 0.545276208 | -4.304176516 |
| **A_52_P195018** | Arap3 | -0.402222109 | -0.961042013 | 0.156597794 | 11.23588698 | -1.748949472 | 0.129449405 | 0.545276208 | -4.304230419 |
| **A_52_P284889** | Prkcz | -0.380963433 | -0.91026203 | 0.148335163 | 8.435003191 | -1.748903065 | 0.129457802 | 0.545276208 | -4.304254047 |
| **A_51_P427516** | Thsd1 | -0.416405788 | -0.994963832 | 0.162152256 | 8.87260075 | -1.748851809 | 0.129467077 | 0.545276208 | -4.304280144 |
| **A_55_P2281624** | 2610034E01Rik | 0.163020362 | -0.063494543 | 0.389535266 | 8.335923425 | 1.748752137 | 0.129485115 | 0.545276208 | -4.304330893 |
| **A_55_P2109857** | Rgs2 | -0.92046564 | -2.199567363 | 0.358636084 | 11.40163081 | -1.748580722 | 0.129516143 | 0.545288069 | -4.30441817 |
| **A_55_P2018387** | Zkscan17 | -0.265537287 | -0.634607292 | 0.103532719 | 10.9149552 | -1.748235566 | 0.129578641 | 0.545380146 | -4.304593912 |
| **A_51_P215489** | Slc37a1 | -0.154682393 | -0.369686025 | 0.060321238 | 8.138325238 | -1.748148316 | 0.129594444 | 0.545380146 | -4.304638336 |
| **A_55_P2140835** | NA | 0.395163097 | -0.154239035 | 0.944565229 | 9.109148861 | 1.747709429 | 0.129673966 | 0.545596016 | -4.304861807 |
| **A_52_P118310** | Chn1 | -0.225280207 | -0.538529229 | 0.087968815 | 7.783439288 | -1.747497252 | 0.129712428 | 0.545639068 | -4.304969844 |
| **A_55_P2393483** | 6430584L05Rik | 0.212421723 | -0.083137385 | 0.507980831 | 8.60346897 | 1.746376061 | 0.12991585 | 0.545642308 | -4.305540751 |
| **A_55_P2001073** | Arid2 | 0.133241809 | -0.052153101 | 0.318636719 | 9.16301438 | 1.746328342 | 0.129924515 | 0.545642308 | -4.30556505 |
| **A_55_P2104332** | Pxdn | -0.297607544 | -0.711726116 | 0.116511027 | 8.692076037 | -1.746233912 | 0.129941663 | 0.545642308 | -4.305613136 |

| | | | | | | | | |
|---|---|---|---|---|---|---|---|---|
| **A_55_P2010931** | Cnn2 | -0.573699598 | -1.372034036 | 0.224634839 | 9.460180581 | -1.746155318 | 0.129955937 | 0.545642308 | -4.305653156 |
| **A_55_P2071802** | Slbp | 0.212177005 | -0.08309241 | 0.507446419 | 9.192571664 | 1.746075587 | 0.129970419 | 0.545642308 | -4.305693757 |
| **A_55_P2002667** | Trim26 | -0.153356434 | -0.366774896 | 0.060062027 | 10.69565448 | -1.746036049 | 0.129977602 | 0.545642308 | -4.30571389 |
| **A_66_P119174** | Cmtm6 | -0.227724649 | -0.544662643 | 0.089213345 | 12.09349517 | -1.745898233 | 0.130002639 | 0.545642308 | -4.305784069 |
| **A_52_P501049** | NA | 0.147338797 | -0.057723993 | 0.352401587 | 8.415364956 | 1.745876213 | 0.13000664 | 0.545642308 | -4.305795282 |
| **A_55_P2125743** | NA | 0.137935434 | -0.054067191 | 0.329938059 | 16.41216715 | 1.745628675 | 0.130051624 | 0.545642308 | -4.305921334 |
| **A_55_P2335903** | NA | 0.30658953 | -0.120218253 | 0.733397314 | 10.59898268 | 1.745453077 | 0.130083544 | 0.545642308 | -4.306010753 |
| **A_55_P2125714** | Atp2b4 | 0.273435441 | -0.107264674 | 0.654135556 | 8.642345926 | 1.745239305 | 0.130122414 | 0.545642308 | -4.306119612 |
| **A_55_P2079787** | Prss50 | 0.299787467 | -0.117628247 | 0.71720318 | 9.077485104 | 1.745130334 | 0.130142232 | 0.545642308 | -4.306175104 |
| **A_51_P234462** | Gpn1 | 0.140851244 | -0.055283672 | 0.33698616 | 9.731242638 | 1.744974009 | 0.130170667 | 0.545642308 | -4.306254711 |
| **A_55_P2111735** | Ppp1r2 | 0.175762509 | -0.068998225 | 0.420523243 | 10.46689955 | 1.744888522 | 0.13018622 | 0.545642308 | -4.306298244 |
| **A_52_P209484** | Tmem88 | -0.475040839 | -1.136645451 | 0.186563774 | 10.725678 | -1.744679314 | 0.130224289 | 0.545642308 | -4.306404782 |
| **A_55_P1956457** | 6030458C11Rik | 0.241667705 | -0.094916674 | 0.578252083 | 10.43985179 | 1.744648136 | 0.130229963 | 0.545642308 | -4.306420659 |
| **A_55_P2077909** | Tpgs2 | 0.303779888 | -0.119326577 | 0.726886352 | 8.677326425 | 1.744586653 | 0.130241153 | 0.545642308 | -4.30645197 |
| **A_52_P344290** | F2r | -0.313494558 | -0.750168739 | 0.123179622 | 12.10033703 | -1.744438575 | 0.130268108 | 0.545642308 | -4.306527378 |
| **A_55_P2124228** | Uaca | -0.410378131 | -0.982083348 | 0.161327087 | 10.42440676 | -1.744195824 | 0.130312309 | 0.545642308 | -4.306651 |
| **A_55_P2185273** | Umps | 0.257298729 | -0.101198133 | 0.615795592 | 12.380631 | 1.74395579 | 0.130356029 | 0.545642308 | -4.306773239 |
| **A_55_P2093143** | Casd1 | 0.143607984 | -0.056512995 | 0.343728964 | 7.991446334 | 1.743689514 | 0.130404545 | 0.545642308 | -4.306908844 |
| **A_52_P484118** | Urb2 | 0.328718681 | -0.12939771 | 0.786835073 | 9.063503559 | 1.743539311 | 0.13043192 | 0.545642308 | -4.306985338 |
| **A_55_P1966144** | Slfn14 | 0.264046403 | -0.103957335 | 0.63205014 | 8.566944338 | 1.743456897 | 0.130446943 | 0.545642308 | -4.307027309 |
| **A_51_P240453** | Nusap1 | 0.23211943 | -0.091412434 | 0.555651293 | 11.90637418 | 1.743322056 | 0.130471526 | 0.545642308 | -4.307095979 |
| **A_55_P2092364** | Ermard | 0.140073973 | -0.055163887 | 0.335311834 | 9.686884372 | 1.743317927 | 0.130472279 | 0.545642308 | -4.307098082 |
| **A_55_P2072443** | Kansl1 | 0.240343245 | -0.094674629 | 0.57536112 | 8.709227072 | 1.743199655 | 0.130493845 | 0.545642308 | -4.307158316 |
| **A_51_P505322** | Mpnd | -0.326426609 | -0.781522092 | 0.128668874 | 11.7333681 | -1.742874907 | 0.130553078 | 0.545642308 | -4.307323703 |
| **A_55_P2068199** | NA | 0.292995695 | -0.115496322 | 0.701487712 | 9.364172296 | 1.742853385 | 0.130557005 | 0.545642308 | -4.307334664 |
| **A_66_P123582** | Mog | 0.220560038 | -0.08695023 | 0.528070306 | 8.54050292 | 1.742811412 | 0.130564663 | 0.545642308 | -4.30735604 |
| **A_55_P1971956** | Ift46 | -0.330739556 | -0.79188576 | 0.130406648 | 11.26438286 | -1.742732339 | 0.130579091 | 0.545642308 | -4.307396311 |
| **A_51_P421140** | Tubb6 | 0.261040431 | -0.102935662 | 0.625016525 | 12.63659576 | 1.742681852 | 0.130588304 | 0.545642308 | -4.307422023 |
| **A_55_P1976351** | Gpcpd1 | 0.359059026 | -0.141677576 | 0.859795627 | 10.03748831 | 1.742367252 | 0.130645726 | 0.545708314 | -4.307582246 |
| **A_65_P11306** | NA | 0.147642329 | -0.058266343 | 0.353551 | 8.857191901 | 1.742285969 | 0.130660566 | 0.545708314 | -4.307623643 |
| **A_55_P2069350** | Psph | 0.433073612 | -0.17107085 | 1.037218074 | 13.19631966 | 1.741823365 | 0.130745057 | 0.545927291 | -4.307859248 |
| **A_66_P106385** | Cd55b | 0.157273167 | -0.062168004 | 0.376714339 | 9.027940949 | 1.741484886 | 0.13080691 | 0.545927291 | -4.308031638 |
| **A_51_P493037** | Dclre1a | 0.15215931 | -0.060166037 | 0.364484656 | 8.491912809 | 1.741325168 | 0.130836107 | 0.545927291 | -4.308112985 |
| **A_52_P375312** | Jaml | -0.244918482 | -0.586723552 | 0.096886588 | 8.302068385 | -1.741110154 | 0.130875422 | 0.545927291 | -4.308222496 |
| **A_55_P2156625** | NA | 0.193137502 | -0.076410459 | 0.462685464 | 14.12485285 | 1.741060042 | 0.130884587 | 0.545927291 | -4.308248019 |
| **A_51_P204913** | Bag2 | 0.269042243 | -0.106471868 | 0.644556354 | 10.5168402 | 1.740914341 | 0.130911237 | 0.545927291 | -4.308322227 |
| **A_51_P498023** | Scyl1 | -0.230609302 | -0.552542721 | 0.091324117 | 11.43045109 | -1.740579835 | 0.13097244 | 0.545927291 | -4.3084926 |
| **A_55_P1962305** | Plac8 | -0.165601037 | -0.396790146 | 0.065588073 | 12.52700776 | -1.740519401 | 0.1309835 | 0.545927291 | -4.308523381 |
| **A_51_P482503** | Tpd52l1 | -0.275604828 | -0.660376222 | 0.109166565 | 7.915950443 | -1.740472777 | 0.130992034 | 0.545927291 | -4.308547128 |
| **A_55_P1983095** | Adam19 | -0.464846631 | -1.113825087 | 0.184131825 | 9.352377447 | -1.74045419 | 0.130995436 | 0.545927291 | -4.308556595 |
| **A_55_P2151666** | NA | 0.147314035 | -0.058401566 | 0.353029637 | 8.178706895 | 1.740043418 | 0.131070644 | 0.545931708 | -4.308765817 |
| **A_51_P198434** | H2-K1 | -0.360797418 | -0.864709126 | 0.143114289 | 13.5678857 | -1.739771298 | 0.131120489 | 0.545931708 | -4.30890442 |
| **A_55_P2040583** | Map6d1 | 0.285952587 | -0.113458706 | 0.68536388 | 11.04446805 | 1.739629962 | 0.131146385 | 0.545931708 | -4.308976409 |
| **A_52_P623775** | Eri2 | 0.19098339 | -0.075790057 | 0.457756837 | 8.759939203 | 1.739547077 | 0.131161574 | 0.545931708 | -4.309018626 |
| **A_55_P1997320** | Zim1 | 0.208253065 | -0.082672015 | 0.499178145 | 8.337796321 | 1.739375822 | 0.131192963 | 0.545931708 | -4.309105856 |
| **A_51_P460890** | Rgl2 | -0.438963622 | -1.052201809 | 0.174274565 | 9.528189648 | -1.739332165 | 0.131200966 | 0.545931708 | -4.309128093 |
| **A_55_P2050044** | NA | 0.239364502 | -0.095038087 | 0.57376709 | 11.4702741 | 1.739295217 | 0.131207739 | 0.545931708 | -4.309146912 |
| **A_55_P2118843** | Hk1 | 0.265143281 | -0.10535673 | 0.635643292 | 8.596676411 | 1.738903936 | 0.13127949 | 0.545931708 | -4.309346216 |
| **A_52_P558087** | Poc5 | 0.112643564 | -0.044768418 | 0.270055545 | 9.707692985 | 1.738808625 | 0.131296974 | 0.545931708 | -4.309394764 |
| **A_65_P05238** | Knop1 | 0.184879347 | -0.073486358 | 0.443245051 | 9.646562266 | 1.738748314 | 0.131308038 | 0.545931708 | -4.309425484 |

| | | | | | | | | | |
|---|---|---|---|---|---|---|---|---|---|
| **A_52_P408757** | Fcgr3 | -0.208921967 | -0.500971539 | 0.083127604 | 8.971047982 | -1.738243935 | 0.131400605 | 0.545931708 | -4.309682401 |
| **A_66_P112356** | Agt | -0.371406207 | -0.890630066 | 0.147817652 | 10.81161132 | -1.738111794 | 0.131424866 | 0.545931708 | -4.309749711 |
| **A_55_P2131340** | Churc1 | 0.188172217 | -0.074920006 | 0.45126444 | 11.19303724 | 1.7379236 | 0.131459427 | 0.545931708 | -4.309845573 |
| **A_65_P10450** | Tmpo | 0.208078536 | -0.082846095 | 0.499003168 | 8.73208072 | 1.737920797 | 0.131459942 | 0.545931708 | -4.309847001 |
| **A_55_P1962586** | NA | 0.445266106 | -0.177352676 | 1.067884887 | 9.196749768 | 1.737723239 | 0.131496232 | 0.545931708 | -4.309947634 |
| **A_52_P412529** | Fbxo3 | 0.170285549 | -0.067852636 | 0.408423735 | 9.359471898 | 1.73752853 | 0.131532008 | 0.545931708 | -4.310046817 |
| **A_55_P2038737** | Slc29a1 | 0.269928869 | -0.107568614 | 0.647426352 | 12.03698257 | 1.7374746 | 0.131541919 | 0.545931708 | -4.310074288 |
| **A_55_P1964332** | NA | 0.21425724 | -0.085398364 | 0.513912844 | 10.73517387 | 1.737385892 | 0.131558223 | 0.545931708 | -4.310119475 |
| **A_55_P2070823** | Ino80d | 0.169577965 | -0.067590412 | 0.406746342 | 10.19901366 | 1.737384037 | 0.131558564 | 0.545931708 | -4.31012042 |
| **A_55_P2032818** | Trim2 | -0.16073369 | -0.385536768 | 0.064069388 | 10.65013653 | -1.737352128 | 0.131564429 | 0.545931708 | -4.310136674 |
| **A_52_P591153** | Pld4 | -0.159416186 | -0.382394185 | 0.063561813 | 8.475149999 | -1.737215072 | 0.131589623 | 0.545931708 | -4.31020649 |
| **A_55_P2008181** | Il3ra | -0.170313633 | -0.408592047 | 0.067964781 | 10.05659633 | -1.736792364 | 0.131667358 | 0.546033129 | -4.310421818 |
| **A_52_P651948** | Fam229b | -0.198854874 | -0.477066895 | 0.079357148 | 7.788145221 | -1.736774912 | 0.131670568 | 0.546033129 | -4.310430708 |
| **A_55_P2135331** | Evl | -0.558327664 | -1.339859352 | 0.223204024 | 11.6716061 | -1.735904161 | 0.131830841 | 0.546553661 | -4.310874282 |
| **A_55_P2100628** | Gm12216 | 0.329708365 | -0.131839795 | 0.791256524 | 10.91517406 | 1.735785798 | 0.131852642 | 0.546553661 | -4.310934579 |
| **A_55_P2068260** | H3c8 | 0.324643438 | -0.129935583 | 0.779222459 | 9.459651983 | 1.735323423 | 0.131937839 | 0.546646849 | -4.311170126 |
| **A_55_P2045802** | Nsmf | -0.314292131 | -0.754376905 | 0.125792643 | 11.37721476 | -1.735323123 | 0.131937894 | 0.546646849 | -4.311170278 |
| **A_55_P1991960** | Nwd1 | 0.274212691 | -0.109777699 | 0.65820308 | 8.509366827 | 1.735203379 | 0.131959967 | 0.546646849 | -4.311231281 |
| **A_51_P429295** | Pus10 | 0.196794295 | -0.078889907 | 0.472478498 | 9.636524676 | 1.734537977 | 0.132082687 | 0.547037979 | -4.311570264 |
| **A_55_P2382070** | 4930593A02Rik | 0.153588409 | -0.061677097 | 0.368853915 | 8.404617878 | 1.733673 | 0.13224238 | 0.547225988 | -4.312010932 |
| **A_66_P100789** | Kansl2 | 0.1222004 | -0.0490726 | 0.2934734 | 9.960623281 | 1.733671903 | 0.132242582 | 0.547225988 | -4.312011491 |
| **A_55_P1967266** | Plxnb2 | -0.286656781 | -0.688443091 | 0.115129529 | 12.16042181 | -1.733605492 | 0.132254851 | 0.547225988 | -4.312045325 |
| **A_52_P327467** | Med7 | 0.138701482 | -0.055717599 | 0.333120562 | 11.11281553 | 1.73350622 | 0.132273192 | 0.547225988 | -4.312095901 |
| **A_55_P2138792** | Fdx1 | -0.618452732 | -1.485400444 | 0.248494981 | 12.82925017 | -1.733392033 | 0.132294292 | 0.547225988 | -4.312154076 |
| **A_51_P169516** | Ppp1r3d | -0.205599144 | -0.493811538 | 0.082613249 | 8.958109618 | -1.73337225 | 0.132297948 | 0.547225988 | -4.312164154 |
| **A_55_P1958457** | Nbeal2 | -0.178595539 | -0.428996989 | 0.07180591 | 8.710554128 | -1.733073407 | 0.132353187 | 0.547337345 | -4.312316407 |
| **A_55_P1980035** | Tbc1d9b | -0.130238201 | -0.312923808 | 0.052447405 | 10.33032109 | -1.73227663 | 0.132500574 | 0.547547187 | -4.312722351 |
| **A_55_P1969522** | NA | 0.162682112 | -0.065515094 | 0.390879319 | 9.343215207 | 1.732258369 | 0.132503954 | 0.547547187 | -4.312731655 |
| **A_66_P124551** | Akap13 | 0.17898476 | -0.072088952 | 0.430058473 | 11.67419063 | 1.732199859 | 0.132514784 | 0.547547187 | -4.312761465 |
| **A_51_P107362** | Socs2 | -0.681571164 | -1.637663256 | 0.274520927 | 9.247477813 | -1.732186587 | 0.13251724 | 0.547547187 | -4.312768227 |
| **A_55_P1957228** | Sgk3 | 0.119900085 | -0.048341067 | 0.288141237 | 11.83465419 | 1.731691189 | 0.13260897 | 0.547630122 | -4.31302063 |
| **A_51_P185292** | 4930581F22Rik | 0.324479391 | -0.13082294 | 0.779781722 | 8.683592427 | 1.731691135 | 0.13260898 | 0.547630122 | -4.313020658 |
| **A_66_P107174** | NA | 0.17143279 | -0.069127919 | 0.4119935 | 8.682399109 | 1.731619181 | 0.132622309 | 0.547630122 | -4.313057318 |
| **A_55_P2053077** | Chuk | 0.168422624 | -0.068004954 | 0.404850202 | 11.47885766 | 1.730953832 | 0.132745616 | 0.548022216 | -4.313396319 |
| **A_55_P1997415** | Exoc6b | -0.30030487 | -0.721916195 | 0.121306456 | 9.349070924 | -1.730745835 | 0.132784186 | 0.54806439 | -4.313502297 |
| **A_66_P120901** | Atxn10 | -0.15039432 | -0.361580126 | 0.060791486 | 11.7858969 | -1.73041351 | 0.132845834 | 0.548124916 | -4.313671623 |
| **A_51_P181565** | Hbegf | -0.244356614 | -0.587535706 | 0.098822478 | 8.656678558 | -1.730160793 | 0.132892733 | 0.548124916 | -4.313800389 |
| **A_55_P1975120** | NA | 0.193694334 | -0.078366479 | 0.465755148 | 8.6275878 | 1.729952242 | 0.132931448 | 0.548124916 | -4.313906652 |
| **A_55_P2069515** | Dennd4a | 0.374840606 | -0.151721639 | 0.901402851 | 11.41328809 | 1.729737113 | 0.132971395 | 0.548124916 | -4.314016268 |
| **A_55_P2166069** | 2310079G19Rik | 0.245894667 | -0.099539984 | 0.591329318 | 8.628180082 | 1.729682529 | 0.132981533 | 0.548124916 | -4.31404408 |
| **A_55_P2108042** | Dgcr2 | -0.173095738 | -0.41627403 | 0.070082553 | 10.87012688 | -1.729596359 | 0.132997538 | 0.548124916 | -4.314087987 |
| **A_51_P212068** | Aoc1 | -0.274776998 | -0.660826839 | 0.111272843 | 7.813295192 | -1.729498483 | 0.13301572 | 0.548124916 | -4.314137858 |
| **A_55_P2117146** | Pa2g4 | 0.211068766 | -0.085482976 | 0.507620508 | 11.56788807 | 1.729444702 | 0.133025712 | 0.548124916 | -4.314165262 |
| **A_51_P372702** | Il16 | -0.205818548 | -0.495040089 | 0.083402993 | 9.523006854 | -1.729167496 | 0.133077223 | 0.548220301 | -4.31430651 |
| **A_55_P2160043** | A330069E16Rik | 0.140792877 | -0.057109449 | 0.338695204 | 8.292098341 | 1.728673497 | 0.133169069 | 0.548246779 | -4.314558227 |
| **A_55_P2012599** | NA | 0.291376237 | -0.118206936 | 0.70095941 | 9.41344871 | 1.728602802 | 0.133182218 | 0.548246779 | -4.31459425 |
| **A_52_P406036** | Cdkn2aip | 0.232681127 | -0.094415861 | 0.559778115 | 8.407173976 | 1.728493707 | 0.133202511 | 0.548246779 | -4.31464984 |
| **A_55_P2118784** | Rcor2 | -0.165911391 | -0.399153005 | 0.067330224 | 9.09245618 | -1.728436199 | 0.13321321 | 0.548246779 | -4.314679144 |
| **A_55_P2097729** | Gtpbp4 | 0.258979409 | -0.105119109 | 0.623077927 | 9.138265277 | 1.728341321 | 0.133230862 | 0.548246779 | -4.31472749 |
| **A_52_P434052** | Wdr37 | 0.149387955 | -0.060664625 | 0.359440535 | 8.568720439 | 1.728107488 | 0.133274378 | 0.548246779 | -4.314846642 |

| | | | | | | | | | |
|---|---|---|---|---|---|---|---|---|---|
| **A_51_P484526** | Wif1 | -0.322923568 | -0.776993494 | 0.131146358 | 8.415126482 | -1.728065472 | 0.133282199 | 0.548246779 | -4.314868052 |
| **A_55_P2010586** | Gm6252 | 0.287353116 | -0.116749247 | 0.691455479 | 10.87964502 | 1.727856654 | 0.133321073 | 0.548290004 | -4.314974459 |
| **A_55_P2144566** | NA | -0.16388581 | -0.394398427 | 0.066626807 | 15.12045564 | -1.727546874 | 0.133378763 | 0.548410575 | -4.315132313 |
| **A_51_P209930** | Rtn2 | -0.250192331 | -0.60216184 | 0.101777178 | 8.424168212 | -1.727237825 | 0.133436341 | 0.548530633 | -4.315289797 |
| **A_55_P2067332** | NA | 0.155484555 | -0.063280315 | 0.374249425 | 8.444917775 | 1.727002082 | 0.133480278 | 0.548594576 | -4.315409927 |
| **A_55_P2060454** | Nsd3 | 0.186856114 | -0.076078208 | 0.449790436 | 10.58809636 | 1.726804702 | 0.133517076 | 0.548629157 | -4.315510509 |
| **A_52_P682382** | Scd1 | -0.402840406 | -0.969912667 | 0.164231854 | 11.59895751 | -1.726147131 | 0.133639738 | 0.54901647 | -4.315845601 |
| **A_55_P2002093** | Dph7 | 0.153404329 | -0.062642639 | 0.369451296 | 8.747454829 | 1.725331807 | 0.133791979 | 0.54940884 | -4.316261093 |
| **A_55_P2470474** | 9530082P21Rik | -0.167317436 | -0.402976753 | 0.06834188 | 7.972642597 | -1.725201301 | 0.133816363 | 0.54940884 | -4.316327601 |
| **A_55_P2027212** | NA | 0.212324059 | -0.086753834 | 0.511401952 | 8.637582505 | 1.725035691 | 0.133847312 | 0.54940884 | -4.316411997 |
| **A_55_P2157992** | NA | -0.158972088 | -0.382945204 | 0.065001027 | 15.07419213 | -1.724678612 | 0.133914067 | 0.54940884 | -4.316593971 |
| **A_55_P2083121** | Vill | -0.138910245 | -0.334625661 | 0.056805172 | 10.98335134 | -1.724616068 | 0.133925763 | 0.54940884 | -4.316625845 |
| **A_52_P334796** | Taar4 | 0.283907447 | -0.116127401 | 0.683942296 | 8.742191226 | 1.724495829 | 0.133948251 | 0.54940884 | -4.316687122 |
| **A_55_P1964068** | Slc25a38 | 0.260802567 | -0.106685671 | 0.628290806 | 12.88926598 | 1.724453973 | 0.133956079 | 0.54940884 | -4.316708452 |
| **A_51_P491350** | Col4a2 | -0.834240099 | -2.009764031 | 0.341283834 | 10.3784693 | -1.724418902 | 0.13396264 | 0.54940884 | -4.316726326 |
| **A_55_P2010116** | Rab27b | -0.213962201 | -0.515546683 | 0.087622282 | 9.501392212 | -1.723896758 | 0.134060345 | 0.549590638 | -4.316992425 |
| **A_55_P1991783** | NA | 0.447420189 | -0.183281372 | 1.07812175 | 11.9346083 | 1.723752307 | 0.134087388 | 0.549590638 | -4.317066043 |
| **A_55_P2106106** | C5ar2 | 0.550362202 | -0.225502847 | 1.326227251 | 10.76789998 | 1.723636176 | 0.134109133 | 0.549590638 | -4.317125227 |
| **A_55_P2158107** | Ddx31 | 0.178349945 | -0.073090204 | 0.429790093 | 9.736672686 | 1.723540701 | 0.134127012 | 0.549590638 | -4.317173885 |
| **A_55_P2255628** | 9430013L14Rik | 0.151520791 | -0.062140574 | 0.365182156 | 8.862386029 | 1.72317513 | 0.134195493 | 0.549590638 | -4.317360195 |
| **A_55_P2020395** | 2-Mar | 0.230750461 | -0.094642945 | 0.556143867 | 8.705244202 | 1.723125933 | 0.134204712 | 0.549590638 | -4.317385268 |
| **A_51_P255682** | Mfge8 | -0.250981075 | -0.604904151 | 0.102942002 | 14.18267539 | -1.723119047 | 0.134206002 | 0.549590638 | -4.317388778 |
| **A_55_P2038597** | NA | 0.310123555 | -0.127271541 | 0.74751865 | 12.06335283 | 1.722836379 | 0.134258981 | 0.549593761 | -4.317532839 |
| **A_51_P375987** | Fign | 0.264881897 | -0.108741671 | 0.638505464 | 9.528746462 | 1.722666483 | 0.134290833 | 0.549593761 | -4.317619427 |
| **A_51_P269687** | Pole2 | 0.259363231 | -0.106477495 | 0.625203958 | 9.469320873 | 1.722659909 | 0.134292065 | 0.549593761 | -4.317622777 |
| **A_55_P2149873** | Mapk1ip1l | 0.443563291 | -0.182178477 | 1.06930506 | 9.18005791 | 1.722438171 | 0.134333648 | 0.549647563 | -4.317735787 |
| **A_52_P421897** | NA | 0.269685629 | -0.110834407 | 0.650205665 | 8.501930022 | 1.722120215 | 0.134393297 | 0.549775246 | -4.317897836 |
| **A_66_P118093** | NA | -0.342138028 | -0.825091858 | 0.140815802 | 8.723011776 | -1.721388599 | 0.134530646 | 0.550122297 | -4.318270717 |
| **A_55_P1993054** | Psip1 | 0.298278227 | -0.122790735 | 0.719347189 | 9.794530024 | 1.721279959 | 0.134551053 | 0.550122297 | -4.318326088 |
| **A_55_P2133568** | Snhg5 | 0.12631097 | -0.052004691 | 0.30462663 | 8.36823029 | 1.721213611 | 0.134563517 | 0.550122297 | -4.318359904 |
| **A_52_P585751** | Gns | -0.220180915 | -0.531085717 | 0.090723887 | 10.92559218 | -1.720819943 | 0.134637495 | 0.550308342 | -4.318560548 |
| **A_55_P2024858** | Fars2 | -0.134000871 | -0.323255264 | 0.055253523 | 11.5975789 | -1.720461027 | 0.134704977 | 0.550345511 | -4.318743482 |
| **A_55_P1959379** | BC080695 | 0.157525689 | -0.065001043 | 0.380052422 | 8.247318763 | 1.720094816 | 0.134773865 | 0.550345511 | -4.318930136 |
| **A_55_P2086682** | Nfib | -0.563506609 | -1.359538017 | 0.232524799 | 11.52587439 | -1.720093282 | 0.134774153 | 0.550345511 | -4.318930917 |
| **A_55_P2033500** | Ltbp4 | -0.703408727 | -1.697087082 | 0.290269629 | 9.459424301 | -1.720065998 | 0.134779287 | 0.550345511 | -4.318944824 |
| **A_66_P123186** | Mapk1ip1l | 0.198615973 | -0.081969524 | 0.479201469 | 9.297583714 | 1.720014636 | 0.134788952 | 0.550345511 | -4.318971003 |
| **A_52_P167278** | Mthfd1l | 0.347287629 | -0.143383718 | 0.837958975 | 11.4860416 | 1.719815284 | 0.13482647 | 0.550382437 | -4.319072612 |
| **A_55_P2127243** | Tpsb2 | 0.124987214 | -0.051629443 | 0.30160387 | 8.274577065 | 1.71955914 | 0.134874691 | 0.550429595 | -4.319203168 |
| **A_55_P2451183** | Pdpk1 | 0.221077187 | -0.091365681 | 0.533520055 | 10.29972005 | 1.71931916 | 0.134919885 | 0.550429595 | -4.319325486 |
| **A_55_P2034027** | Nos3 | -0.288499366 | -0.696240705 | 0.119241974 | 10.08683717 | -1.719266919 | 0.134929725 | 0.550429595 | -4.319352114 |
| **A_55_P2013011** | NA | 0.216398615 | -0.089489357 | 0.522286587 | 8.366748228 | 1.718997578 | 0.134980469 | 0.550429595 | -4.319489399 |
| **A_55_P1999246** | Gm10318 | 0.233357665 | -0.096511824 | 0.563227154 | 11.75712586 | 1.718949478 | 0.134989533 | 0.550429595 | -4.319513916 |
| **A_55_P2108837** | Tuba1c | 0.273698738 | -0.113219155 | 0.660616632 | 15.7785335 | 1.718846798 | 0.135008884 | 0.550429595 | -4.319566253 |
| **A_55_P2002703** | NA | 0.20350761 | -0.084230548 | 0.491245768 | 10.01714466 | 1.718566668 | 0.13506169 | 0.550528767 | -4.319709039 |
| **A_52_P370203** | Jarid2 | -0.144735623 | -0.349395199 | 0.059923954 | 9.975793601 | -1.718409051 | 0.135091411 | 0.550533816 | -4.319789379 |
| **A_55_P2159094** | P3h1 | 0.126175654 | -0.052274627 | 0.304625936 | 9.439323198 | 1.718072623 | 0.135154871 | 0.550676327 | -4.319960863 |
| **A_55_P2173035** | Dnajc8 | 0.205616038 | -0.085302726 | 0.496534803 | 10.71765316 | 1.717388071 | 0.135284085 | 0.55103489 | -4.320309798 |
| **A_55_P1978466** | Pnma5 | 0.144179722 | -0.059824942 | 0.348184387 | 7.719168078 | 1.717304376 | 0.135299891 | 0.55103489 | -4.320352461 |
| **A_51_P133792** | Prickle3 | -0.124091275 | -0.299689036 | 0.051506486 | 8.317474369 | -1.717139041 | 0.13533112 | 0.551045972 | -4.320436738 |
| **A_51_P506683** | Fbxw7 | 0.143643468 | -0.05966867 | 0.346955606 | 9.137410173 | 1.716744895 | 0.135405597 | 0.551195778 | -4.320637649 |

| | | | | | | | | |
|---|---|---|---|---|---|---|---|---|
| **A_55_P1979645** | S100a1 | -0.369364972 | -0.892193198 | 0.153463254 | 9.58270669 | -1.716642543 | 0.135424944 | 0.551195778 | -4.320689822 |
| **A_55_P2152946** | NA | 0.323267668 | -0.134394131 | 0.780929467 | 9.14985265 | 1.716330009 | 0.135484036 | 0.551320198 | -4.320849135 |
| **A_66_P130310** | A4galt | 0.197896168 | -0.082336105 | 0.478128441 | 8.213582706 | 1.715941288 | 0.135557568 | 0.55141532 | -4.321047286 |
| **A_55_P2114680** | Car4 | 0.244367446 | -0.101697881 | 0.590432773 | 10.55802745 | 1.715807057 | 0.135582969 | 0.55141532 | -4.321115711 |
| **A_55_P1968355** | Tle1 | -0.198116161 | -0.478690568 | 0.082458246 | 11.31536183 | -1.715754076 | 0.135592996 | 0.55141532 | -4.321142718 |
| **A_55_P1955717** | Hccs | 0.144407324 | -0.060297335 | 0.349111982 | 16.02317593 | 1.714133656 | 0.135900015 | 0.55254762 | -4.321968755 |
| **A_52_P469381** | Comtd1 | 0.222743213 | -0.093042093 | 0.538528519 | 11.34016454 | 1.713940542 | 0.135936648 | 0.5525732 | -4.3220672 |
| **A_52_P107234** | Prpf39 | 0.181060384 | -0.075664637 | 0.437785406 | 8.905447699 | 1.71371433 | 0.135979573 | 0.5525732 | -4.322182519 |
| **A_55_P2253885** | AI503301 | 0.176125073 | -0.073611779 | 0.425861926 | 8.628011545 | 1.713648487 | 0.13599207 | 0.5525732 | -4.322216085 |
| **A_55_P2027771** | Zfp260 | -0.356030649 | -0.861035843 | 0.148974545 | 11.05474018 | -1.713068501 | 0.136102194 | 0.552574322 | -4.322511755 |
| **A_55_P1965478** | Kcnd1 | 0.260161592 | -0.108919575 | 0.629242759 | 9.621237451 | 1.712791444 | 0.136154831 | 0.552574322 | -4.322652997 |
| **A_51_P134301** | Pfdn5 | -0.298439099 | -0.721846317 | 0.124968118 | 12.37207642 | -1.712697456 | 0.136172691 | 0.552574322 | -4.322700912 |
| **A_51_P229602** | Plxna2 | -0.22900508 | -0.553930245 | 0.095920085 | 10.07473182 | -1.71255669 | 0.136199445 | 0.552574322 | -4.322772674 |
| **A_55_P1996202** | NA | 0.177300689 | -0.074276883 | 0.42887826 | 11.39430842 | 1.712464951 | 0.136216884 | 0.552574322 | -4.322819443 |
| **A_51_P371750** | Marco | 0.32429858 | -0.135885163 | 0.784482323 | 13.87669656 | 1.71236745 | 0.136235421 | 0.552574322 | -4.322869149 |
| **A_52_P529486** | Wdr77 | 0.221483682 | -0.092849306 | 0.53581667 | 9.49498501 | 1.712123009 | 0.136281904 | 0.552574322 | -4.322993766 |
| **A_65_P03022** | Cdc14b | 0.334177035 | -0.140110013 | 0.808464084 | 9.671921865 | 1.712058207 | 0.136294229 | 0.552574322 | -4.323026802 |
| **A_51_P464630** | Rad17 | 0.199654062 | -0.083723174 | 0.483031298 | 10.59078489 | 1.711970905 | 0.136310836 | 0.552574322 | -4.323071309 |
| **A_55_P2115467** | Tmem63b | -0.211500217 | -0.511697714 | 0.088697281 | 8.796144871 | -1.711933687 | 0.136317916 | 0.552574322 | -4.323090283 |
| **A_55_P2068554** | NA | 0.137383352 | -0.057615051 | 0.332381756 | 16.55486968 | 1.711930947 | 0.136318437 | 0.552574322 | -4.32309168 |
| **A_51_P136781** | Pnliprp1 | 0.510866808 | -0.214282191 | 1.236015806 | 10.3658616 | 1.711841788 | 0.1363354 | 0.552574322 | -4.323137134 |
| **A_51_P332822** | Abce1 | 0.259554371 | -0.108963511 | 0.628072253 | 11.7676636 | 1.711405679 | 0.1364184 | 0.55279481 | -4.323359467 |
| **A_52_P595642** | Smim7 | -0.153377014 | -0.371186271 | 0.064432242 | 10.10268382 | -1.711067472 | 0.136482801 | 0.552809026 | -4.323531891 |
| **A_55_P1994917** | Wdr5 | 0.173179965 | -0.072752951 | 0.41911288 | 11.51114961 | 1.71105582 | 0.13648502 | 0.552809026 | -4.323537831 |
| **A_52_P629333** | Map3k21 | -0.134960935 | -0.3266485 | 0.05672663 | 7.864280406 | -1.710792462 | 0.136535189 | 0.552809026 | -4.323672097 |
| **A_51_P500550** | Mfsd1 | -0.172569478 | -0.417674007 | 0.072535051 | 8.778273958 | -1.710786592 | 0.136536308 | 0.552809026 | -4.323675089 |
| **A_55_P2424988** | C78228 | 0.16699086 | -0.070215282 | 0.404197003 | 8.027319177 | 1.71060588 | 0.136570744 | 0.55283265 | -4.323767221 |
| **A_55_P2011286** | Hopx | -1.511216312 | -3.659315818 | 0.636883193 | 9.281201107 | -1.709446159 | 0.136791937 | 0.553612093 | -4.324358484 |
| **A_51_P188772** | Arhgef28 | -0.544894342 | -1.319703388 | 0.229914705 | 8.224430512 | -1.708837652 | 0.136908136 | 0.553966374 | -4.324668727 |
| **A_51_P181319** | Dcxr | -0.501838325 | -1.215548547 | 0.211871897 | 9.730303291 | -1.708539789 | 0.13696505 | 0.554080673 | -4.324820591 |
| **A_51_P159565** | Arhgef9 | 0.149762705 | -0.063250424 | 0.362775834 | 8.353771061 | 1.708364319 | 0.136998589 | 0.554100381 | -4.324910055 |
| **A_55_P2056674** | Lrrk1 | -0.208466355 | -0.505018709 | 0.088086 | 10.57955279 | -1.708117666 | 0.137045746 | 0.554175152 | -4.325035812 |
| **A_55_P2133943** | NA | 0.30062267 | -0.127111086 | 0.728356426 | 8.728857779 | 1.707777908 | 0.13711073 | 0.554321962 | -4.32520904 |
| **A_51_P279127** | Fam173a | -0.185547236 | -0.449576923 | 0.078482451 | 12.82745294 | -1.707595167 | 0.137145694 | 0.554326424 | -4.325302213 |
| **A_55_P2179413** | Lgals3bp | -0.279428008 | -0.677104314 | 0.118248298 | 10.90376904 | -1.70735334 | 0.137191977 | 0.554326424 | -4.325425511 |
| **A_51_P163797** | Bdp1 | 0.162832556 | -0.068923953 | 0.394589066 | 9.159576276 | 1.707231787 | 0.137215246 | 0.554326424 | -4.325487487 |
| **A_65_P15518** | NA | 0.14722142 | -0.062343115 | 0.356785956 | 8.394169942 | 1.707011329 | 0.137257459 | 0.554326424 | -4.325599891 |
| **A_51_P187901** | Nop56 | 0.292038856 | -0.123684141 | 0.707761853 | 10.91499069 | 1.706945985 | 0.137269973 | 0.554326424 | -4.325633208 |
| **A_51_P475076** | Vbp1 | 0.145091536 | -0.061491874 | 0.351674947 | 10.49056369 | 1.706592471 | 0.137337696 | 0.554326424 | -4.325813455 |
| **A_52_P249588** | Cnot11 | 0.161729055 | -0.06854351 | 0.392001619 | 8.837207976 | 1.706589379 | 0.137338288 | 0.554326424 | -4.325815031 |
| **A_52_P304128** | Mmp14 | -0.181896405 | -0.440919121 | 0.077126312 | 8.754783003 | -1.706355235 | 0.137383161 | 0.554326424 | -4.325934415 |
| **A_52_P220229** | Plekhb2 | -0.141700178 | -0.343576914 | 0.060176558 | 8.308149116 | -1.705561209 | 0.137535438 | 0.554326424 | -4.326339274 |
| **A_51_P519369** | Agtpbp1 | 0.159434513 | -0.067734726 | 0.386603752 | 8.966045158 | 1.705359738 | 0.137574102 | 0.554326424 | -4.326442001 |
| **A_52_P99807** | Dpy19l3 | 0.17452993 | -0.074176013 | 0.423235873 | 8.555223589 | 1.705167082 | 0.137611084 | 0.554326424 | -4.326540234 |
| **A_55_P2087589** | Rps11 | -0.258297077 | -0.626371733 | 0.10977758 | 15.20343491 | -1.705166393 | 0.137611216 | 0.554326424 | -4.326540586 |
| **A_55_P2007413** | Agps | 0.163341836 | -0.069438359 | 0.396122032 | 11.34643402 | 1.705040088 | 0.137635467 | 0.554326424 | -4.326604987 |
| **A_55_P2145480** | NA | 0.193138349 | -0.082112523 | 0.468389221 | 11.359064 | 1.704994569 | 0.137644207 | 0.554326424 | -4.326628197 |
| **A_52_P528600** | Hhex | 0.251582504 | -0.106988811 | 0.610153818 | 11.09950627 | 1.704857485 | 0.137670534 | 0.554326424 | -4.326698095 |
| **A_55_P2131438** | H2ac19 | 0.304755016 | -0.129617656 | 0.739127688 | 17.10310558 | 1.704792623 | 0.137682992 | 0.554326424 | -4.326731168 |
| **A_66_P104713** | Lrp5 | 0.243436508 | -0.103609034 | 0.59048205 | 10.10404749 | 1.704442788 | 0.137750204 | 0.554326424 | -4.326909547 |

| | | | | | | | | | |
|---|---|---|---|---|---|---|---|---|---|
| **A_55_P2072908** | NA | 0.170829114 | -0.072712527 | 0.414370756 | 13.68169362 | 1.704401272 | 0.137758182 | 0.554326424 | -4.326930716 |
| **A_55_P1966987** | Limk1 | -0.272355588 | -0.6606801 | 0.115968924 | 9.545778954 | -1.704216143 | 0.137793765 | 0.554326424 | -4.327025114 |
| **A_55_P2059890** | Fdx1 | -0.575271213 | -1.395594606 | 0.245052181 | 12.85551581 | -1.704004488 | 0.137834457 | 0.554326424 | -4.327133037 |
| **A_55_P2112701** | NA | -0.253194609 | -0.614283278 | 0.10789406 | 11.60910056 | -1.703820277 | 0.137869882 | 0.554326424 | -4.327226966 |
| **A_52_P604195** | Mbp | 0.17560041 | -0.07483244 | 0.42603326 | 10.97215421 | 1.703795311 | 0.137874684 | 0.554326424 | -4.327239697 |
| **A_51_P448489** | Mea1 | -0.201765766 | -0.489521325 | 0.085989793 | 10.78649855 | -1.703754232 | 0.137882586 | 0.554326424 | -4.327260643 |
| **A_55_P1959152** | Gm10324 | 0.139549842 | -0.059497936 | 0.338597621 | 8.170313094 | 1.703551256 | 0.137921633 | 0.554326424 | -4.327364142 |
| **A_55_P2078257** | NA | 0.22196728 | -0.094643368 | 0.538577929 | 8.532677384 | 1.703517631 | 0.137928103 | 0.554326424 | -4.327381288 |
| **A_51_P215995** | Zswim6 | -0.225826106 | -0.548007595 | 0.096355382 | 9.051565894 | -1.70316511 | 0.137995947 | 0.554326424 | -4.327561042 |
| **A_55_P1967553** | Tmem254a | -0.167355596 | -0.406130475 | 0.071419283 | 11.40407851 | -1.703079002 | 0.138012524 | 0.554326424 | -4.327604949 |
| **A_51_P259902** | Arl6ip5 | -0.277961614 | -0.674553581 | 0.118630353 | 10.70820845 | -1.703037065 | 0.138020599 | 0.554326424 | -4.327626334 |
| **A_55_P2133115** | NA | -0.201277835 | -0.488504884 | 0.085949215 | 14.71206058 | -1.702761427 | 0.138073678 | 0.554326424 | -4.327766885 |
| **A_55_P2082708** | Pwp2 | 0.192018579 | -0.08201445 | 0.466051608 | 10.72114392 | 1.702642737 | 0.138096541 | 0.554326424 | -4.327827408 |
| **A_55_P2262136** | Tek | -0.134056843 | -0.32537443 | 0.057260743 | 8.123967181 | -1.70261825 | 0.138101258 | 0.554326424 | -4.327839894 |
| **A_51_P265545** | Maco1 | 0.127421303 | -0.054446787 | 0.309289392 | 10.51796842 | 1.70242789 | 0.138137934 | 0.554326424 | -4.327936962 |
| **A_55_P2143279** | Nkd2 | 0.265287395 | -0.113384666 | 0.643959456 | 9.5221012 | 1.702301731 | 0.138162246 | 0.554326424 | -4.328001293 |
| **A_52_P422859** | Hspg2 | -0.258491657 | -0.627490379 | 0.110507065 | 11.76142387 | -1.702177546 | 0.138186181 | 0.554326424 | -4.328064617 |
| **A_55_P1968153** | Pcgf2 | -0.140855501 | -0.341932398 | 0.060221396 | 8.46577312 | -1.702138235 | 0.138193759 | 0.554326424 | -4.328084663 |
| **A_55_P2083794** | Tmem159 | -0.583427871 | -1.416297177 | 0.249441435 | 8.327377959 | -1.702133047 | 0.138194759 | 0.554326424 | -4.328087309 |
| **A_55_P2050628** | Cyp4a31 | -0.487158927 | -1.182687397 | 0.208369543 | 8.875239697 | -1.70191922 | 0.138235985 | 0.554326424 | -4.328196344 |
| **A_55_P2083624** | Wbp2 | 0.201479343 | -0.086179883 | 0.48913857 | 13.2890193 | 1.701905365 | 0.138238656 | 0.554326424 | -4.328203409 |
| **A_55_P2002884** | Zcchc24 | -0.19394816 | -0.470906684 | 0.083010364 | 12.21244615 | -1.701586833 | 0.138300091 | 0.554326424 | -4.328365837 |
| **A_55_P1964805** | Rpa1 | 0.18496978 | -0.079169608 | 0.449109167 | 13.04663228 | 1.701573846 | 0.138302597 | 0.554326424 | -4.32837246 |
| **A_51_P398723** | Flt1 | -0.24592244 | -0.597115917 | 0.105271037 | 11.41618772 | -1.701511575 | 0.13831461 | 0.554326424 | -4.328404214 |
| **A_55_P2014066** | NA | 0.167477068 | -0.071707608 | 0.406661745 | 8.264541342 | 1.701395132 | 0.138337077 | 0.554326424 | -4.328463591 |
| **A_55_P2358679** | NA | 0.163190692 | -0.069884079 | 0.396265463 | 11.3034493 | 1.701309422 | 0.138353617 | 0.554326424 | -4.328507297 |
| **A_51_P189269** | Mbnl2 | -0.266551818 | -0.647352758 | 0.114249122 | 8.327561081 | -1.700853179 | 0.138441691 | 0.554326424 | -4.328739951 |
| **A_55_P2021721** | NA | 0.371728885 | -0.15933611 | 0.90279388 | 10.56662046 | 1.700833581 | 0.138445476 | 0.554326424 | -4.328749945 |
| **A_52_P239536** | Ppp1r9a | -0.210077455 | -0.510230158 | 0.090075249 | 9.43686195 | -1.700671276 | 0.138476821 | 0.554326424 | -4.32883271 |
| **A_52_P582858** | Dgkd | -0.172198863 | -0.418232471 | 0.073834746 | 10.40380181 | -1.700666008 | 0.138477839 | 0.554326424 | -4.328835396 |
| **A_52_P512575** | Hopx | -1.506492047 | -3.659000994 | 0.646016899 | 9.617412406 | -1.700611326 | 0.138488401 | 0.554326424 | -4.328863281 |
| **A_52_P608495** | Aars2 | 0.165353075 | -0.07093885 | 0.401645 | 9.293787516 | 1.70038233 | 0.138532642 | 0.554334579 | -4.328980055 |
| **A_51_P474701** | Fbp1 | -0.868272066 | -2.109102582 | 0.372558451 | 15.61125718 | -1.700303905 | 0.138547796 | 0.554334579 | -4.329020047 |
| **A_55_P1973208** | Tax1bp3 | -0.380517696 | -0.924367631 | 0.16333224 | 8.411780374 | -1.700117609 | 0.138583801 | 0.554363885 | -4.329115048 |
| **A_52_P500488** | Lars | 0.232762406 | -0.099958632 | 0.565483445 | 12.262832 | 1.699870216 | 0.138631628 | 0.55441646 | -4.329241204 |
| **A_55_P1966383** | NA | 0.286670293 | -0.123137378 | 0.696477964 | 11.0234317 | 1.699752919 | 0.13865431 | 0.55441646 | -4.329301019 |
| **A_55_P2045085** | NA | 0.186188331 | -0.080029538 | 0.4524062 | 14.49682884 | 1.699411078 | 0.138720434 | 0.554566135 | -4.329475341 |
| **A_55_P1981992** | NA | 0.329858154 | -0.141847297 | 0.801563605 | 9.665770234 | 1.699180512 | 0.13876505 | 0.554592334 | -4.329592918 |
| **A_55_P1963807** | Actg2 | -0.613696807 | -1.491370125 | 0.263976511 | 9.299994384 | -1.699042165 | 0.138791827 | 0.554592334 | -4.329663469 |
| **A_55_P2038002** | BC005624 | 0.159367879 | -0.068570916 | 0.387306675 | 10.8234577 | 1.69889185 | 0.138820927 | 0.554592334 | -4.329740123 |
| **A_55_P2015984** | Odr4 | 0.213619324 | -0.091932931 | 0.519171579 | 10.87369564 | 1.698784277 | 0.138841756 | 0.554592334 | -4.329794981 |
| **A_55_P2045622** | Setdb2 | 0.212804155 | -0.091611768 | 0.517220078 | 9.405042776 | 1.698618801 | 0.138873802 | 0.554605729 | -4.329879366 |
| **A_55_P2069579** | NA | 0.186946599 | -0.080626465 | 0.454519663 | 8.223224711 | 1.697689904 | 0.139053827 | 0.555173809 | -4.330353069 |
| **A_55_P2071581** | NA | 0.165741276 | -0.07149519 | 0.402977742 | 8.473103037 | 1.697588488 | 0.139073495 | 0.555173809 | -4.330404788 |
| **A_55_P2121082** | Cul2 | 0.184431146 | -0.079599479 | 0.44846177 | 10.44141574 | 1.697317741 | 0.139126016 | 0.555245446 | -4.330542861 |
| **A_55_P2069856** | Tnpo3 | 0.153559689 | -0.066313725 | 0.373433104 | 11.65634044 | 1.697022859 | 0.139183241 | 0.555245446 | -4.330693242 |
| **A_55_P2180944** | Tmco4 | -0.190598206 | -0.463522554 | 0.082326142 | 10.77668459 | -1.696913548 | 0.13920446 | 0.555245446 | -4.330748988 |
| **A_55_P1968483** | Gtpbp6 | -0.147260084 | -0.358140055 | 0.063619888 | 10.02177354 | -1.696808758 | 0.139224804 | 0.555245446 | -4.330802428 |
| **A_51_P205385** | Uox | -0.64668039 | -1.572853058 | 0.279492278 | 14.19774652 | -1.696605694 | 0.139264236 | 0.555245446 | -4.330905986 |
| **A_51_P443359** | Trappc2l | -0.266214018 | -0.647517388 | 0.115089353 | 11.42796483 | -1.696459377 | 0.139292654 | 0.555245446 | -4.330980604 |

| | | | | | | | | | |
|---|---|---|---|---|---|---|---|---|---|
| **A_55_P2093794** | Usp3 | -0.188113005 | -0.45756305 | 0.081337039 | 10.09060312 | -1.696382373 | 0.139307613 | 0.555245446 | -4.331019875 |
| **A_51_P116665** | 2610005L07Rik | 0.160877228 | -0.069570262 | 0.391324717 | 9.38718954 | 1.696312185 | 0.139321249 | 0.555245446 | -4.331055669 |
| **A_51_P514412** | Gsk3b | -0.145725846 | -0.354520492 | 0.063068801 | 8.378575092 | -1.695900682 | 0.13940122 | 0.555399333 | -4.331265528 |
| **A_52_P318955** | Ap3m1 | 0.228725192 | -0.099006166 | 0.55645655 | 9.612638208 | 1.695817814 | 0.13941733 | 0.555399333 | -4.331307789 |
| **A_55_P1974900** | Ndufab1 | 0.172815072 | -0.074832038 | 0.420462181 | 14.27653714 | 1.695631756 | 0.139453506 | 0.555428977 | -4.331402676 |
| **A_66_P124952** | Hspe1 | 0.169031606 | -0.073217347 | 0.41128056 | 14.55969544 | 1.695466441 | 0.139485657 | 0.555442577 | -4.331486984 |
| **A_55_P2080703** | Cstf2 | 0.246761854 | -0.106934164 | 0.600457872 | 9.096807367 | 1.695239419 | 0.139529821 | 0.555503999 | -4.331602763 |
| **A_52_P677891** | Fam174b | -0.591438758 | -1.439292021 | 0.256414505 | 8.461561373 | -1.695010038 | 0.139574458 | 0.555539254 | -4.331719745 |
| **A_55_P1953583** | Ace2 | -0.419080173 | -1.019889403 | 0.181729058 | 8.136337298 | -1.694898544 | 0.139596159 | 0.555539254 | -4.331776606 |
| **A_55_P2139407** | Myl12b | -0.355270802 | -0.864779355 | 0.15423775 | 12.65719582 | -1.694303575 | 0.139712019 | 0.555885881 | -4.332080037 |
| **A_55_P1980956** | Osbp | 0.190762918 | -0.082972556 | 0.464498391 | 8.234573734 | 1.693347394 | 0.139898412 | 0.556162181 | -4.332567691 |
| **A_51_P466875** | Sbds | 0.128579634 | -0.055943751 | 0.313103018 | 11.62842805 | 1.693183241 | 0.139930435 | 0.556162181 | -4.33265141 |
| **A_55_P1991911** | NA | 0.141862235 | -0.061745166 | 0.345469637 | 14.06264915 | 1.692997929 | 0.139966595 | 0.556162181 | -4.33274592 |
| **A_55_P2078433** | Mcoln2 | 0.510692347 | -0.222314685 | 1.243699378 | 10.34201542 | 1.692912056 | 0.139983354 | 0.556162181 | -4.332789716 |
| **A_52_P228318** | Smu1 | 0.109297575 | -0.047609609 | 0.266204759 | 13.20527138 | 1.692586555 | 0.140046898 | 0.556162181 | -4.332955725 |
| **A_55_P2109658** | NA | 0.156355904 | -0.068116816 | 0.380828624 | 10.99842378 | 1.69252037 | 0.140059822 | 0.556162181 | -4.33298948 |
| **A_55_P2055819** | Adam15 | -0.478837486 | -1.166306407 | 0.208631434 | 11.53908463 | -1.692459411 | 0.140071726 | 0.556162181 | -4.33302057 |
| **A_55_P2009146** | Gtpbp3 | 0.31341859 | -0.136586264 | 0.763423445 | 8.733600217 | 1.692352171 | 0.140092671 | 0.556162181 | -4.333075263 |
| **A_55_P1974367** | Fabp3 | -0.791715496 | -1.928462401 | 0.345031409 | 8.479542399 | -1.69234364 | 0.140094337 | 0.556162181 | -4.333079614 |
| **A_52_P740562** | Rps21 | 0.13955557 | -0.060884412 | 0.339995552 | 14.95800983 | 1.691788266 | 0.140202855 | 0.556162181 | -4.333362864 |
| **A_55_P1999240** | NA | 0.250703203 | -0.109414923 | 0.610821328 | 9.94115245 | 1.691601611 | 0.140239346 | 0.556162181 | -4.333458061 |
| **A_55_P2091472** | Bnip3 | -0.274643796 | -0.669194914 | 0.119907322 | 14.00628484 | -1.691413152 | 0.140276197 | 0.556162181 | -4.333554179 |
| **A_51_P321651** | Fam192a | 0.132679418 | -0.05794228 | 0.323301117 | 10.4618479 | 1.691275715 | 0.140303078 | 0.556162181 | -4.333624275 |
| **A_55_P2043028** | Rexo5 | 0.194316487 | -0.084861104 | 0.473494078 | 9.597448057 | 1.691267426 | 0.1403047 | 0.556162181 | -4.333628502 |
| **A_55_P2266977** | Ahcyl2 | 0.147450785 | -0.064395679 | 0.35929725 | 9.884215929 | 1.691254861 | 0.140307157 | 0.556162181 | -4.333634911 |
| **A_52_P530291** | Pim1 | 0.32210505 | -0.14067862 | 0.78488872 | 9.751707975 | 1.69123009 | 0.140312003 | 0.556162181 | -4.333647545 |
| **A_55_P2021555** | Tbrg1 | -0.173378845 | -0.42248437 | 0.07572668 | 13.74903478 | -1.691203755 | 0.140317155 | 0.556162181 | -4.333660976 |
| **A_55_P2125006** | Gpatch1 | 0.136874616 | -0.059783603 | 0.333532836 | 9.665721447 | 1.691196227 | 0.140318627 | 0.556162181 | -4.333664815 |
| **A_55_P2169415** | Cgn | 0.151217839 | -0.066054643 | 0.368490321 | 11.4011045 | 1.691147512 | 0.140328157 | 0.556162181 | -4.333689661 |
| **A_55_P2016967** | Gm8213 | 0.154899413 | -0.067685735 | 0.377484562 | 15.30140674 | 1.690973406 | 0.140362223 | 0.556183151 | -4.333778459 |
| **A_51_P421846** | Ech1 | -0.303888821 | -0.740672041 | 0.1328944 | 12.08005017 | -1.690565364 | 0.140442092 | 0.55623584 | -4.333986572 |
| **A_55_P2077343** | Ahdc1 | -0.18314678 | -0.446427006 | 0.080133445 | 8.450501636 | -1.690301729 | 0.140493719 | 0.55623584 | -4.334121033 |
| **A_55_P1953382** | Prl8a8 | -0.434721141 | -1.059770629 | 0.190328347 | 8.074703085 | -1.689972144 | 0.140558286 | 0.55623584 | -4.334289131 |
| **A_55_P2038458** | Lancl2 | 0.181108983 | -0.079296566 | 0.441514532 | 8.832144508 | 1.689946466 | 0.140563317 | 0.55623584 | -4.334302228 |
| **A_55_P2075731** | NA | 0.39993478 | -0.175124429 | 0.974993989 | 11.65592222 | 1.689895307 | 0.140573343 | 0.55623584 | -4.33432832 |
| **A_55_P2248320** | C430003N24Rik | 0.127971711 | -0.056040967 | 0.311984388 | 8.42286043 | 1.689854919 | 0.140581257 | 0.55623584 | -4.334348919 |
| **A_51_P167105** | B3galt4 | -0.187316865 | -0.456675261 | 0.082041532 | 8.396775077 | -1.689777608 | 0.14059641 | 0.55623584 | -4.334388351 |
| **A_66_P125401** | NA | 0.193765374 | -0.084873715 | 0.472404463 | 9.062896252 | 1.689730013 | 0.140605738 | 0.55623584 | -4.334412625 |
| **A_52_P150683** | NA | 0.239877527 | -0.105123522 | 0.584878577 | 8.550808234 | 1.689477203 | 0.1406553 | 0.556261471 | -4.334541567 |
| **A_52_P158211** | NA | 0.248477116 | -0.108907806 | 0.605862039 | 9.643677171 | 1.689403387 | 0.140669775 | 0.556261471 | -4.334579216 |
| **A_51_P195240** | Gpatch11 | 0.156677786 | -0.068750314 | 0.382105885 | 10.16854155 | 1.688816895 | 0.140784829 | 0.556335188 | -4.334878349 |
| **A_55_P2063126** | Patj | -0.361562415 | -0.881796021 | 0.158671191 | 8.784724305 | -1.688760508 | 0.140795895 | 0.556335188 | -4.334907109 |
| **A_51_P465772** | Ruvbl1 | 0.11828158 | -0.051918394 | 0.288481555 | 12.71081129 | 1.688654562 | 0.140816691 | 0.556335188 | -4.334961146 |
| **A_66_P119247** | Gatad1 | 0.144193292 | -0.063306032 | 0.351692616 | 10.93508573 | 1.688540805 | 0.140839022 | 0.556335188 | -4.335019167 |
| **A_55_P1995012** | Plekhm2 | -0.185501833 | -0.452491902 | 0.081488235 | 11.02711694 | -1.688248172 | 0.140896484 | 0.556335188 | -4.335168422 |
| **A_51_P189905** | Dad1 | -0.148447394 | -0.362115091 | 0.065220304 | 12.94737035 | -1.688172792 | 0.14091129 | 0.556335188 | -4.335206869 |
| **A_66_P120603** | Trps1 | -0.249577015 | -0.608839333 | 0.109685303 | 8.051708388 | -1.688014246 | 0.140942435 | 0.556335188 | -4.335287734 |
| **A_66_P110742** | Gm9949 | 0.154609114 | -0.067955435 | 0.377173663 | 8.413512631 | 1.687960549 | 0.140952985 | 0.556335188 | -4.335315122 |
| **A_55_P2322039** | NA | 0.168615104 | -0.07413609 | 0.411366297 | 8.325326564 | 1.68778953 | 0.14098659 | 0.556335188 | -4.33540235 |
| **A_55_P2002642** | Deup1 | -0.822660138 | -2.007086458 | 0.361766182 | 8.469384113 | -1.687701333 | 0.141003924 | 0.556335188 | -4.335447335 |

| | | | | | | | | | |
|---|---|---|---|---|---|---|---|---|---|
| **A_55_P2054739** | Polr3c | 0.168068025 | -0.073960265 | 0.410096315 | 9.792458135 | 1.68733826 | 0.141075301 | 0.556335188 | -4.33563252 |
| **A_52_P564951** | C2cd3 | 0.214167581 | -0.09427992 | 0.522615081 | 8.735411051 | 1.687157646 | 0.141110822 | 0.556335188 | -4.335724642 |
| **A_55_P2159740** | Mfsd2b | 0.141611517 | -0.062346412 | 0.345569446 | 15.59476309 | 1.687101339 | 0.141121897 | 0.556335188 | -4.335753361 |
| **A_55_P1996941** | Ube2c | 0.245198967 | -0.107991618 | 0.598389552 | 13.11253006 | 1.686913084 | 0.141158932 | 0.556335188 | -4.335849381 |
| **A_55_P1958103** | NA | -0.168205804 | -0.410502135 | 0.074090526 | 15.59489453 | -1.686853357 | 0.141170684 | 0.556335188 | -4.335879845 |
| **A_51_P412225** | Tbc1d16 | -0.572180533 | -1.396552832 | 0.252191765 | 9.787292535 | -1.686525367 | 0.141235237 | 0.556335188 | -4.336047138 |
| **A_51_P396917** | Zfyve21 | -0.195566785 | -0.477368089 | 0.086234518 | 11.00416464 | -1.686301742 | 0.141279265 | 0.556335188 | -4.336161199 |
| **A_55_P1998380** | Traf6 | 0.118355047 | -0.052196452 | 0.288906547 | 8.640902397 | 1.686220756 | 0.141295213 | 0.556335188 | -4.336202506 |
| **A_55_P2119817** | Plekhj1 | 0.196972521 | -0.0868728 | 0.480817841 | 8.709959966 | 1.686192267 | 0.141300824 | 0.556335188 | -4.336217037 |
| **A_55_P2080850** | Ptpn2 | 0.135874094 | -0.059937065 | 0.331685253 | 10.35527998 | 1.686096438 | 0.141319698 | 0.556335188 | -4.336265916 |
| **A_51_P134007** | Ncl | 0.174400859 | -0.076967466 | 0.425769185 | 13.29889853 | 1.685859034 | 0.141366467 | 0.556335188 | -4.336387006 |
| **A_55_P2121142** | Tbc1d22b | 0.18407769 | -0.081239591 | 0.449394972 | 9.554464241 | 1.685849519 | 0.141368342 | 0.556335188 | -4.336391859 |
| **A_55_P2068137** | NA | -0.270343036 | -0.66000942 | 0.119323348 | 10.8288131 | -1.685797611 | 0.14137857 | 0.556335188 | -4.336418335 |
| **A_55_P2085445** | Acsm1 | -0.330131068 | -0.805975475 | 0.145713338 | 8.972455854 | -1.685794441 | 0.141379194 | 0.556335188 | -4.336419952 |
| **A_55_P2149896** | Lrrc75a | -0.503040289 | -1.228201005 | 0.222120426 | 9.885769303 | -1.685589006 | 0.141419681 | 0.55634665 | -4.336524736 |
| **A_55_P2186648** | Bard1 | 0.321348992 | -0.141958056 | 0.784656039 | 10.46022502 | 1.685354341 | 0.141465941 | 0.55634665 | -4.33664443 |
| **A_55_P2205156** | Aamdc | -0.12565523 | -0.306820864 | 0.055510405 | 8.560782761 | -1.685341586 | 0.141468456 | 0.55634665 | -4.336650936 |
| **A_52_P1172945** | Ankrd34b | -0.189664327 | -0.463195426 | 0.083866772 | 7.762896796 | -1.684853453 | 0.141564732 | 0.556566686 | -4.336899915 |
| **A_66_P122173** | Mrtfb | -0.196793168 | -0.480620116 | 0.08703378 | 10.55431981 | -1.684765957 | 0.141581996 | 0.556566686 | -4.336944544 |
| **A_52_P343343** | Rpl7l1 | 0.180012585 | -0.079666578 | 0.439691748 | 9.029522643 | 1.684414433 | 0.141651375 | 0.556687662 | -4.337123845 |
| **A_55_P2011126** | Mfap1a | 0.126333641 | -0.055955063 | 0.308622345 | 8.541755737 | 1.684001386 | 0.141732939 | 0.556687662 | -4.337334527 |
| **A_55_P2068673** | Stmn2 | -0.24663927 | -0.602541818 | 0.109263278 | 7.82663586 | -1.683892327 | 0.141754482 | 0.556687662 | -4.337390155 |
| **A_55_P2273811** | Msh5 | 0.208697481 | -0.092455913 | 0.509850874 | 8.560995913 | 1.6838858 | 0.141755772 | 0.556687662 | -4.337393485 |
| **A_66_P113749** | Il2rg | -0.119306671 | -0.291468833 | 0.05285549 | 8.229143999 | -1.683876408 | 0.141757627 | 0.556687662 | -4.337398275 |
| **A_55_P2088825** | Foxp1 | -0.271779268 | -0.663995701 | 0.120437164 | 11.36479567 | -1.683734958 | 0.141785574 | 0.556687662 | -4.337470424 |
| **A_55_P1974593** | NA | 0.198400477 | -0.087970891 | 0.484771844 | 15.91054682 | 1.683434822 | 0.141844891 | 0.556807454 | -4.337623516 |
| **A_52_P180565** | 2310061I04Rik | 0.219785733 | -0.097596624 | 0.537168089 | 9.914090431 | 1.682673694 | 0.141995423 | 0.557180308 | -4.338011749 |
| **A_51_P154753** | Klc3 | -0.679558811 | -1.660884293 | 0.30176667 | 8.563575971 | -1.682663061 | 0.141997527 | 0.557180308 | -4.338017173 |
| **A_52_P144925** | Taf1d | 0.15940345 | -0.070860766 | 0.389667666 | 9.720002348 | 1.682110241 | 0.142106958 | 0.55736827 | -4.338299155 |
| **A_55_P1995572** | Meis1 | -0.451537632 | -1.103814425 | 0.20073916 | 10.19525099 | -1.682074447 | 0.142114047 | 0.55736827 | -4.338317413 |
| **A_55_P2061386** | Chd3 | -0.614049305 | -1.501181107 | 0.273082497 | 11.24758039 | -1.681892697 | 0.142150044 | 0.55736827 | -4.33841012 |
| **A_55_P1960491** | Snta1 | 0.211360841 | -0.094007084 | 0.516728767 | 11.05454381 | 1.681838537 | 0.142160772 | 0.55736827 | -4.338437746 |
| **A_51_P234692** | Neat1 | -0.169357152 | -0.414070329 | 0.075356025 | 10.13592321 | -1.681625852 | 0.142202911 | 0.557420413 | -4.338546233 |
| **A_55_P2158281** | NA | 0.151566076 | -0.067483243 | 0.370615395 | 16.26420676 | 1.681292552 | 0.14226897 | 0.557534935 | -4.338716244 |
| **A_66_P115116** | Wwtr1 | -0.80998992 | -1.980860532 | 0.360880692 | 9.132065854 | -1.680946524 | 0.142337583 | 0.557534935 | -4.338892748 |
| **A_51_P284608** | Cd74 | -0.368819118 | -0.902068323 | 0.164430088 | 8.715822221 | -1.680607884 | 0.142404762 | 0.557534935 | -4.339065485 |
| **A_55_P2029191** | Pde4a | -0.153157128 | -0.374610675 | 0.068296418 | 8.945676276 | -1.680497086 | 0.142426749 | 0.557534935 | -4.339122001 |
| **A_55_P1964174** | Nme1 | 0.269220413 | -0.120097668 | 0.658538495 | 11.96932791 | 1.680299126 | 0.14246604 | 0.557534935 | -4.339222979 |
| **A_51_P239654** | Nr4a1 | -0.439901069 | -1.07604143 | 0.196239292 | 8.985244671 | -1.680293919 | 0.142467073 | 0.557534935 | -4.339225635 |
| **A_52_P503730** | Fchsd2 | -0.215753927 | -0.527769112 | 0.096261259 | 8.997378189 | -1.680220064 | 0.142481735 | 0.557534935 | -4.339263307 |
| **A_55_P1978465** | H2-Q8 | 0.424741328 | -0.189547847 | 1.039030502 | 10.16344074 | 1.680098901 | 0.142505791 | 0.557534935 | -4.339325112 |
| **A_52_P604526** | Mrpl19 | 0.165320526 | -0.073797084 | 0.404438136 | 10.07677917 | 1.679957932 | 0.142533785 | 0.557534935 | -4.339397019 |
| **A_52_P412585** | Ear1 | -0.206026381 | -0.504063745 | 0.092010984 | 9.32285586 | -1.679713867 | 0.142582263 | 0.557534935 | -4.339521514 |
| **A_55_P2124415** | 6030468B19Rik | 0.357345407 | -0.159639041 | 0.874329855 | 10.62703622 | 1.679553781 | 0.142614069 | 0.557534935 | -4.339603173 |
| **A_55_P2160032** | Atp5pb | 0.140179774 | -0.062627191 | 0.34298674 | 13.8785541 | 1.679521926 | 0.142620399 | 0.557534935 | -4.339619422 |
| **A_55_P2182497** | Maml3 | -0.275831031 | -0.674926659 | 0.123264596 | 9.145285181 | -1.679381416 | 0.142648323 | 0.557534935 | -4.339691096 |
| **A_55_P2060021** | Otub2 | 0.272809672 | -0.121932448 | 0.667551792 | 11.38855163 | 1.679304622 | 0.142663587 | 0.557534935 | -4.339730268 |
| **A_52_P579933** | Slc16a6 | 0.371830563 | -0.166201525 | 0.909862652 | 9.307879055 | 1.679268322 | 0.142670802 | 0.557534935 | -4.339748784 |
| **A_55_P1996223** | Sf3b4 | 0.254890844 | -0.114003723 | 0.623785411 | 11.52860307 | 1.678939958 | 0.142736089 | 0.557534935 | -4.339916281 |
| **A_55_P1970710** | Mllt6 | -0.640255382 | -1.566951291 | 0.286440527 | 10.3737812 | -1.678800853 | 0.142763755 | 0.557534935 | -4.339987239 |

| | | | | | | | | |
|---|---|---|---|---|---|---|---|---|
| **A_52_P195839** | Ctsc | -0.494540886 | -1.210380548 | 0.221298776 | 10.71762986 | -1.67868664 | 0.142786474 | 0.557534935 | -4.340045498 |
| **A_51_P134062** | Stxbp2 | -0.206413484 | -0.505204735 | 0.092377767 | 10.97513288 | -1.678623799 | 0.142798976 | 0.557534935 | -4.340077553 |
| **A_55_P2038414** | Acot8 | 0.180339764 | -0.080726414 | 0.441405942 | 11.16747821 | 1.67851055 | 0.142821509 | 0.557534935 | -4.340135321 |
| **A_51_P428086** | Sptbn1 | -0.423542975 | -1.036744522 | 0.189658572 | 8.082371483 | -1.678330279 | 0.142857384 | 0.557534935 | -4.340227277 |
| **A_51_P517120** | Zfp930 | 0.19784579 | -0.088618652 | 0.484310232 | 10.61496494 | 1.678182856 | 0.142886729 | 0.557534935 | -4.340302478 |
| **A_51_P132170** | Ccdc141 | -0.134384725 | -0.329006727 | 0.060237278 | 7.825286508 | -1.677803739 | 0.142962218 | 0.557534935 | -4.340495866 |
| **A_55_P2135025** | NA | 0.11211463 | -0.050333739 | 0.274563 | 11.17872641 | 1.676988647 | 0.143124649 | 0.557534935 | -4.340911646 |
| **A_55_P2023114** | Hsd3b2 | -0.325795108 | -0.79786287 | 0.146272655 | 8.537237656 | -1.676962704 | 0.143129822 | 0.557534935 | -4.34092488 |
| **A_55_P2041470** | NA | 0.144065762 | -0.064699704 | 0.352831228 | 8.714521371 | 1.676815619 | 0.143159153 | 0.557534935 | -4.340999909 |
| **A_51_P367125** | Pold1 | 0.198168781 | -0.089032968 | 0.48537053 | 11.30935683 | 1.676607278 | 0.143200709 | 0.557534935 | -4.341106185 |
| **A_55_P2122555** | Csnk1e | -0.401172214 | -0.982592732 | 0.180248304 | 8.69489165 | -1.676577681 | 0.143206614 | 0.557534935 | -4.341121282 |
| **A_51_P146149** | Napsa | -1.084903373 | -2.657315666 | 0.48750892 | 9.420856272 | -1.676516446 | 0.143218831 | 0.557534935 | -4.341152519 |
| **A_52_P321140** | Defb1 | -0.481976577 | -1.180550811 | 0.216597657 | 8.852721669 | -1.676472982 | 0.143227503 | 0.557534935 | -4.34117469 |
| **A_55_P2042356** | Rftn1 | -0.174183069 | -0.426649955 | 0.078283816 | 9.901233307 | -1.676427225 | 0.143236633 | 0.557534935 | -4.341198031 |
| **A_51_P448618** | Slc16a10 | 0.357831313 | -0.160915394 | 0.876578021 | 10.65931711 | 1.67612413 | 0.143297124 | 0.557534935 | -4.341352642 |
| **A_52_P123354** | Tusc2 | 0.194097523 | -0.08729504 | 0.475490085 | 9.714995206 | 1.67606388 | 0.143309151 | 0.557534935 | -4.341383376 |
| **A_55_P2130905** | NA | 0.32140191 | -0.144559042 | 0.787362863 | 13.25832398 | 1.676031275 | 0.143315661 | 0.557534935 | -4.341400008 |
| **A_55_P2086912** | NA | 0.184396172 | -0.082993637 | 0.45178598 | 12.22299859 | 1.67567673 | 0.14338646 | 0.557534935 | -4.341580864 |
| **A_51_P148196** | Sdf2 | -0.229163653 | -0.561478435 | 0.103151128 | 13.30127377 | -1.675634396 | 0.143394916 | 0.557534935 | -4.34160246 |
| **A_51_P378825** | Actr8 | -0.111508336 | -0.273210469 | 0.050193798 | 10.51954565 | -1.67561706 | 0.143398379 | 0.557534935 | -4.341611303 |
| **A_55_P2051894** | Zfp598 | 0.234883502 | -0.105748349 | 0.575515353 | 9.608244071 | 1.675523208 | 0.143417128 | 0.557534935 | -4.341659178 |
| **A_55_P1992834** | Socs2 | -0.604720175 | -1.48171172 | 0.27227137 | 8.953621928 | -1.675491544 | 0.143423454 | 0.557534935 | -4.34167533 |
| **A_51_P109258** | Cys1 | 0.179037661 | -0.08064031 | 0.438715633 | 8.384707939 | 1.675299564 | 0.143461814 | 0.557534935 | -4.341773261 |
| **A_55_P2158404** | Cmpk2 | -0.185525712 | -0.454627538 | 0.083576114 | 8.651332309 | -1.67521538 | 0.143478638 | 0.557534935 | -4.341816204 |
| **A_55_P2054420** | Tcf4 | -0.484089732 | -1.186272095 | 0.218092632 | 10.23120775 | -1.675170987 | 0.143487511 | 0.557534935 | -4.341838849 |
| **A_55_P2036180** | Mcf2l | -0.525650891 | -1.288174324 | 0.236872542 | 8.693502321 | -1.675048552 | 0.143511985 | 0.557534935 | -4.341901305 |
| **A_51_P202430** | Pdap1 | 0.1363697 | -0.061468891 | 0.334208292 | 13.64546492 | 1.674904561 | 0.143540772 | 0.557534935 | -4.341974757 |
| **A_52_P466887** | Hnrnpd | 0.167332285 | -0.075455098 | 0.410119667 | 12.58001251 | 1.674699216 | 0.143581835 | 0.557534935 | -4.342079505 |
| **A_55_P1975695** | Snx18 | -0.217097479 | -0.532100456 | 0.097905497 | 10.83541454 | -1.674647141 | 0.143592251 | 0.557534935 | -4.342106069 |
| **A_55_P2084656** | H2ac12 | 0.147933682 | -0.066734748 | 0.362602113 | 17.883921 | 1.674488143 | 0.143624056 | 0.557534935 | -4.342187177 |
| **A_52_P256817** | Zik1 | 0.135905801 | -0.061310471 | 0.333122073 | 8.393760062 | 1.674474114 | 0.143626863 | 0.557534935 | -4.342194333 |
| **A_52_P138110** | 1110032A03Rik | 0.134718406 | -0.060791689 | 0.330228501 | 8.234869687 | 1.674329536 | 0.14365579 | 0.557534935 | -4.342268084 |
| **A_55_P2045414** | Hipk4 | 0.312860215 | -0.141203343 | 0.766923773 | 9.386131657 | 1.674236787 | 0.14367435 | 0.557534935 | -4.342315397 |
| **A_51_P165082** | Tmem191c | 0.136856341 | -0.061791508 | 0.33550419 | 8.332116701 | 1.674033896 | 0.143714959 | 0.557580578 | -4.342418895 |
| **A_55_P1977875** | Arglu1 | 0.185654713 | -0.083906413 | 0.455215839 | 11.04656538 | 1.673523846 | 0.143817095 | 0.557864865 | -4.342679079 |
| **A_52_P325443** | Arhgap29 | -0.25816355 | -0.633077572 | 0.116750472 | 9.382147974 | -1.673194508 | 0.143883081 | 0.55791188 | -4.34284708 |
| **A_55_P2068121** | Sun5 | 0.160563047 | -0.07261484 | 0.393740934 | 8.300481206 | 1.673175236 | 0.143886943 | 0.55791188 | -4.342856911 |
| **A_51_P391825** | Exosc10 | 0.191619031 | -0.086723359 | 0.469961422 | 10.14900919 | 1.672794056 | 0.143963354 | 0.557919515 | -4.343051358 |
| **A_66_P118256** | Strn3 | 0.144423376 | -0.065373016 | 0.354219768 | 10.91231095 | 1.672717758 | 0.143978654 | 0.557919515 | -4.343090279 |
| **A_55_P2034415** | Lrrc8d | 0.311782366 | -0.141144325 | 0.764709058 | 10.27753333 | 1.672656732 | 0.143990891 | 0.557919515 | -4.343121409 |
| **A_51_P409724** | Secisbp2 | 0.118694785 | -0.053740228 | 0.291129799 | 8.669750456 | 1.672589528 | 0.14400437 | 0.557919515 | -4.343155691 |
| **A_51_P261999** | Ccdc166 | -0.153743278 | -0.37712499 | 0.069638434 | 8.781092312 | -1.672367469 | 0.144048914 | 0.557930501 | -4.343268968 |
| **A_51_P278034** | Ufsp1 | 0.281719137 | -0.127624991 | 0.691063266 | 9.823153957 | 1.672287616 | 0.144064935 | 0.557930501 | -4.343309703 |
| **A_51_P394172** | Ces1c | -0.701508392 | -1.721203184 | 0.3181864 | 10.90113023 | -1.671651948 | 0.144192534 | 0.558093668 | -4.34363397 |
| **A_55_P1989312** | Prkd3 | -0.222896738 | -0.546897973 | 0.101104496 | 11.35612696 | -1.67163031 | 0.144196879 | 0.558093668 | -4.343645008 |
| **A_55_P1959763** | Dach1 | -0.590878634 | -1.449776859 | 0.268019591 | 9.036910475 | -1.671628513 | 0.14419724 | 0.558093668 | -4.343645925 |
| **A_55_P2034784** | Snhg17 | 0.293007351 | -0.132962596 | 0.718977298 | 8.844313172 | 1.671409036 | 0.144241323 | 0.558093668 | -4.343757885 |
| **A_51_P232399** | Acy3 | 0.191149968 | -0.086755016 | 0.469054952 | 10.38499964 | 1.671325671 | 0.14425807 | 0.558093668 | -4.343800411 |
| **A_51_P429209** | Smpd1 | -0.184882828 | -0.453711484 | 0.083945829 | 12.11492383 | -1.671106782 | 0.144302053 | 0.558093668 | -4.343912071 |
| **A_55_P2136765** | Uqcr10 | -0.146759856 | -0.360167356 | 0.066647645 | 13.31311674 | -1.671016681 | 0.144320161 | 0.558093668 | -4.343958034 |

| | | | | | | | | |
|---|---|---|---|---|---|---|---|---|
| **A_55_P2070936** | Tra2b | 0.176910036 | -0.080388363 | 0.434208434 | 8.241342043 | 1.670700397 | 0.144383744 | 0.558093668 | -4.344119378 |
| **A_55_P2176963** | Hsph1 | 0.234947926 | -0.106797151 | 0.576693003 | 10.72285507 | 1.670523299 | 0.144419357 | 0.558093668 | -4.34420972 |
| **A_55_P2077666** | Porcn | -0.259324219 | -0.63655108 | 0.117902642 | 8.515744454 | -1.670412215 | 0.1444417 | 0.558093668 | -4.344266386 |
| **A_55_P2143306** | Rest | -0.159649339 | -0.391905936 | 0.072607257 | 9.446747826 | -1.670252984 | 0.144473733 | 0.558093668 | -4.344347614 |
| **A_51_P106428** | Nsmce4a | 0.136258335 | -0.061981679 | 0.334498349 | 12.67543031 | 1.670147962 | 0.144494864 | 0.558093668 | -4.347604403 |
| **A_55_P1970597** | Actg1 | -0.263332015 | -0.646457893 | 0.119793862 | 16.3163933 | -1.670111113 | 0.144502279 | 0.558093668 | -4.344419986 |
| **A_52_P585448** | Ptp4a2 | 0.165715041 | -0.075423056 | 0.406853139 | 11.42288793 | 1.669857031 | 0.144553417 | 0.558093668 | -4.3445496 |
| **A_55_P2128853** | Ear12 | -0.285678127 | -0.701388286 | 0.130032033 | 9.629662596 | -1.669819544 | 0.144560964 | 0.558093668 | -4.344568723 |
| **A_55_P1963031** | Ampd2 | 0.385191383 | -0.17537143 | 0.945754195 | 9.80524725 | 1.669688546 | 0.144587337 | 0.558093668 | -4.344635548 |
| **A_55_P2161545** | Ddx21 | 0.417386158 | -0.19005518 | 1.024827497 | 10.5328336 | 1.669617116 | 0.14460172 | 0.558093668 | -4.344671987 |
| **A_55_P2102060** | Gstm3 | 0.558828905 | -0.25452123 | 1.37217904 | 9.945373093 | 1.669492635 | 0.144626788 | 0.558093668 | -4.344735488 |
| **A_55_P2046593** | Gm15226 | -0.128330243 | -0.31516945 | 0.058508964 | 9.612861026 | -1.668953316 | 0.144735445 | 0.558318995 | -4.345010609 |
| **A_51_P150802** | Mpv17 | -0.128120454 | -0.314687814 | 0.058446906 | 10.32181571 | -1.668652836 | 0.144796017 | 0.558318995 | -4.345163892 |
| **A_55_P2027812** | Arxes2 | 0.160527042 | -0.073232443 | 0.394286528 | 10.22140773 | 1.668638078 | 0.144798992 | 0.558318995 | -4.34517142 |
| **A_51_P429472** | Mindy3 | 0.194961607 | -0.089046205 | 0.478969418 | 9.739582262 | 1.668022863 | 0.144923088 | 0.558318995 | -4.345485259 |
| **A_55_P1979082** | Shc1 | 0.114043109 | -0.052095794 | 0.280182011 | 11.79729788 | 1.667941827 | 0.144939441 | 0.558318995 | -4.345526598 |
| **A_55_P2077628** | NA | -0.523006225 | -1.284948751 | 0.238936302 | 9.256743431 | -1.667891645 | 0.144949569 | 0.558318995 | -4.345552197 |
| **A_51_P291501** | Ccno | -0.665756637 | -1.635706011 | 0.304192736 | 8.460979584 | -1.667821726 | 0.144963682 | 0.558318995 | -4.345587865 |
| **A_52_P27103** | Necap2 | 0.173970666 | -0.079511134 | 0.427452466 | 9.47926974 | 1.667678899 | 0.144992514 | 0.558318995 | -4.345660725 |
| **A_51_P499061** | Ube2o | 0.236409911 | -0.108057128 | 0.580876951 | 14.40870043 | 1.667635749 | 0.145001225 | 0.558318995 | -4.345682737 |
| **A_55_P2043833** | Srgap2 | 0.305280974 | -0.139589431 | 0.75015138 | 9.340189412 | 1.667436918 | 0.145041374 | 0.558318995 | -4.345784167 |
| **A_52_P268231** | NA | 0.19620014 | -0.089717759 | 0.482118039 | 8.689088616 | 1.667405228 | 0.145047774 | 0.558318995 | -4.345800333 |
| **A_51_P215559** | Gstp3 | 0.293208125 | -0.134080788 | 0.720497038 | 10.82817424 | 1.66739143 | 0.145050561 | 0.558318995 | -4.345807372 |
| **A_55_P2082604** | Ndor1 | 0.150420597 | -0.068792243 | 0.369633436 | 16.36192094 | 1.667341301 | 0.145060686 | 0.558318995 | -4.345832944 |
| **A_55_P2102828** | NA | 0.352371633 | -0.161238554 | 0.86598182 | 9.520964068 | 1.66705719 | 0.145118081 | 0.558347923 | -4.345977877 |
| **A_66_P121196** | Katnb1 | 0.198490871 | -0.090883313 | 0.487865054 | 10.61405559 | 1.666724973 | 0.145185221 | 0.558347923 | -4.346147351 |
| **A_55_P2000818** | Cab39l | 0.189470271 | -0.086758868 | 0.46569941 | 9.664610474 | 1.666689789 | 0.145192334 | 0.558347923 | -4.3461653 |
| **A_52_P227065** | Bud31 | 0.159309823 | -0.072959409 | 0.391579056 | 11.26721841 | 1.666610293 | 0.145208405 | 0.558347923 | -4.346205853 |
| **A_55_P1960708** | Setd1a | 0.20994612 | -0.096186751 | 0.516078991 | 10.19236257 | 1.666406988 | 0.145249514 | 0.558347923 | -4.346309565 |
| **A_51_P424641** | Sirt4 | -0.188966268 | -0.464522068 | 0.086589533 | 8.459905519 | -1.666318121 | 0.145267486 | 0.558347923 | -4.346354899 |
| **A_55_P1975670** | Prps1l3 | 0.142547487 | -0.065320974 | 0.350415949 | 11.30902861 | 1.666303679 | 0.145270407 | 0.558347923 | -4.346362267 |
| **A_51_P427934** | Megf8 | -0.341373313 | -0.839302012 | 0.156555386 | 8.768404186 | -1.665887223 | 0.145354662 | 0.55856069 | -4.346574714 |
| **A_55_P2043723** | Rabac1 | -0.509515706 | -1.252877586 | 0.233846175 | 11.4449873 | -1.66548405 | 0.145436275 | 0.558653258 | -4.346780385 |
| **A_55_P2178084** | Dnmt3a | -0.29255206 | -0.719390592 | 0.134286471 | 10.46265358 | -1.665416 | 0.145450054 | 0.558653258 | -4.346815099 |
| **A_51_P292757** | Acbd4 | 0.342867463 | -0.157454103 | 0.843189029 | 11.87880451 | 1.665176371 | 0.145498586 | 0.558653258 | -4.346937342 |
| **A_55_P2102897** | Gm12657 | 0.150614878 | -0.06918022 | 0.370409976 | 14.32912926 | 1.665072169 | 0.145519695 | 0.558653258 | -4.346990499 |
| **A_55_P1989048** | Tm9sf1 | 0.115886801 | -0.053230756 | 0.285004358 | 10.21072915 | 1.665054556 | 0.145523263 | 0.558653258 | -4.346999483 |
| **A_55_P2240468** | 1110019D14Rik | 0.126194243 | -0.057997859 | 0.310386345 | 8.30012358 | 1.664760367 | 0.145582876 | 0.558771132 | -4.347149559 |
| **A_51_P161021** | Ifit2 | -0.151283611 | -0.372125018 | 0.069557795 | 8.233356576 | -1.664541271 | 0.145627288 | 0.558830624 | -4.347261327 |
| **A_55_P2347384** | NA | 0.321339635 | -0.147827838 | 0.790507107 | 10.26341347 | 1.664253924 | 0.145685555 | 0.558837843 | -4.347407912 |
| **A_51_P322994** | Mrpl22 | 0.181493895 | -0.083494876 | 0.446482666 | 10.40800379 | 1.664246831 | 0.145686993 | 0.558837843 | -4.347411531 |
| **A_51_P380178** | Id3 | -0.626762287 | -1.542077023 | 0.28855245 | 10.39879226 | -1.663855439 | 0.145766394 | 0.558841836 | -4.347611192 |
| **A_55_P1955577** | Ccm2 | 0.108252908 | -0.049840815 | 0.266346631 | 12.46502182 | 1.663826875 | 0.14577219 | 0.558841836 | -4.347625764 |
| **A_51_P400555** | Cts3 | -0.320359774 | -0.788239329 | 0.14751978 | 7.857777808 | -1.663746291 | 0.145788544 | 0.558841836 | -4.347666872 |
| **A_55_P1958919** | Olfr1347 | 0.131947538 | -0.060768685 | 0.324663762 | 8.129602536 | 1.663666246 | 0.14580479 | 0.558841836 | -4.347707706 |
| **A_55_P2069875** | Trim44 | -0.258773759 | -0.636781231 | 0.119233714 | 12.30036928 | -1.663424284 | 0.145853909 | 0.558841836 | -4.347831139 |
| **A_51_P126437** | Enc1 | -0.216571341 | -0.532946015 | 0.099803332 | 9.550802748 | -1.66334549 | 0.145869908 | 0.558841836 | -4.347871334 |
| **A_55_P2120189** | Gas2l3 | 0.220619155 | -0.101700237 | 0.542938546 | 8.922375659 | 1.663182746 | 0.145902958 | 0.558841836 | -4.347954355 |
| **A_51_P479052** | Tars | 0.24801761 | -0.114363513 | 0.610398733 | 12.47643523 | 1.66303022 | 0.14593394 | 0.558841836 | -4.348032164 |
| **A_55_P2028595** | AW551984 | 0.205454475 | -0.094826839 | 0.505735789 | 8.921506366 | 1.662533812 | 0.146034815 | 0.558841836 | -4.348285398 |

| | | | | | | | | | |
|---|---|---|---|---|---|---|---|---|---|
| **A_55_P2077555** | C330021F23Rik | 0.209335918 | -0.096632055 | 0.515303891 | 8.677043323 | 1.662459106 | 0.146050002 | 0.558841836 | -4.348323508 |
| **A_55_P2046241** | Arl6ip6 | 0.153750412 | -0.070977348 | 0.378478172 | 8.45464268 | 1.662427654 | 0.146056396 | 0.558841836 | -4.348339552 |
| **A_52_P103550** | Tfdp1 | 0.204517179 | -0.094451257 | 0.503485615 | 10.00768456 | 1.662216708 | 0.146099289 | 0.558841836 | -4.348447163 |
| **A_51_P450573** | Tgfbr2 | -0.450618696 | -1.109522541 | 0.208285149 | 11.09826034 | -1.661767856 | 0.146190597 | 0.558841836 | -4.348676137 |
| **A_55_P1992319** | Col4a3bp | 0.186773939 | -0.086361503 | 0.459909382 | 8.755192966 | 1.661580584 | 0.146228709 | 0.558841836 | -4.348771671 |
| **A_55_P2098995** | Cfhr2 | -0.30879335 | -0.760542741 | 0.14295604 | 11.03672774 | -1.660938505 | 0.146359452 | 0.558841836 | -4.349099216 |
| **A_55_P2046245** | Hes1 | -0.133179582 | -0.328037455 | 0.06167829 | 8.618456854 | -1.660744723 | 0.146398933 | 0.558841836 | -4.349198071 |
| **A_55_P2229645** | NA | 0.249655619 | -0.115642202 | 0.614953441 | 8.84910723 | 1.660647486 | 0.146418748 | 0.558841836 | -4.349247675 |
| **A_55_P2062603** | 4932438A13Rik | 0.214431802 | -0.099382589 | 0.528246194 | 9.230740238 | 1.66034955 | 0.146479477 | 0.558841836 | -4.349399662 |
| **A_52_P657123** | Gm10270 | 0.192582817 | -0.089271386 | 0.474437021 | 10.22096392 | 1.660260398 | 0.146497653 | 0.558841836 | -4.349445141 |
| **A_55_P2034275** | Arfgap2 | 0.184535224 | -0.085541408 | 0.454611856 | 12.50678619 | 1.660257518 | 0.14649824 | 0.558841836 | -4.34944661 |
| **A_55_P2106911** | Wnt1 | 0.22893202 | -0.106177277 | 0.564041317 | 8.904954387 | 1.659981517 | 0.146554527 | 0.558841836 | -4.349587407 |
| **A_52_P67212** | 2900052N01Rik | 0.227585982 | -0.10557932 | 0.560751283 | 8.627943739 | 1.659850345 | 0.146581284 | 0.558841836 | -4.349654322 |
| **A_66_P109315** | Snapc2 | -0.165558167 | -0.407982956 | 0.076866621 | 11.14412046 | -1.659421734 | 0.146668749 | 0.558841836 | -4.34987297 |
| **A_52_P5652** | Sae1 | 0.282641261 | -0.131261307 | 0.696543828 | 9.444219571 | 1.659283635 | 0.146696941 | 0.558841836 | -4.349943419 |
| **A_55_P2032912** | NA | -0.454751172 | -1.120723518 | 0.211221174 | 12.35644315 | -1.65920795 | 0.146712394 | 0.558841836 | -4.349982028 |
| **A_51_P254646** | Jdp2 | -0.252781435 | -0.62299368 | 0.117430809 | 9.4179289 | -1.659119186 | 0.146730519 | 0.558841836 | -4.35002731 |
| **A_55_P2108773** | Tedc1 | 0.233962276 | -0.10870923 | 0.576633783 | 9.310097969 | 1.659017755 | 0.146751233 | 0.558841836 | -4.350079053 |
| **A_55_P1981929** | Scg5 | -0.124224042 | -0.306180005 | 0.057731921 | 7.833627578 | -1.65890896 | 0.146773455 | 0.558841836 | -4.350134552 |
| **A_52_P481423** | Cttnbp2nl | -0.53380844 | -1.315839137 | 0.248222257 | 9.302893793 | -1.658612082 | 0.146834108 | 0.558841836 | -4.350285999 |
| **A_55_P2079430** | NA | -0.27959891 | -0.689237114 | 0.130039294 | 9.244884299 | -1.65851044 | 0.146854879 | 0.558841836 | -4.35033785 |
| **A_55_P1993019** | NA | 0.150899329 | -0.070188609 | 0.371987266 | 10.61304358 | 1.658461707 | 0.146864839 | 0.558841836 | -4.35036271 |
| **A_66_P123824** | Sorl1 | -0.180361771 | -0.444617345 | 0.083893804 | 8.468060706 | -1.65845435 | 0.146866343 | 0.558841836 | -4.350366463 |
| **A_51_P426754** | Anxa5 | -0.355879474 | -0.877326417 | 0.165567469 | 12.16594057 | -1.658349266 | 0.146887823 | 0.558841836 | -4.35042007 |
| **A_55_P2110037** | Akap7 | 0.175626953 | -0.081714448 | 0.432968354 | 9.575341039 | 1.658306084 | 0.14689665 | 0.558841836 | -4.350442098 |
| **A_55_P2068977** | Armcx3 | -0.191070513 | -0.471042036 | 0.088901009 | 9.801359118 | -1.658299616 | 0.146897972 | 0.558841836 | -4.350445398 |
| **A_55_P1967708** | NA | 0.215316414 | -0.100239067 | 0.530871896 | 8.864548059 | 1.658000277 | 0.146959178 | 0.558841836 | -4.3505981 |
| **A_55_P2005972** | NA | -0.387418703 | -0.955201234 | 0.180363828 | 8.301105205 | -1.657989383 | 0.146961406 | 0.558841836 | -4.350603657 |
| **A_66_P136186** | Wee1 | 0.167263033 | -0.077875154 | 0.41240122 | 9.266773666 | 1.65795289 | 0.14696887 | 0.558841836 | -4.350622273 |
| **A_55_P2188464** | Uhrf1bp1l | -0.156693727 | -0.386342953 | 0.0729555 | 11.07436104 | -1.657943826 | 0.146970723 | 0.558841836 | -4.350626897 |
| **A_52_P148678** | Plcg1 | -0.254732735 | -0.628073909 | 0.11860844 | 9.605668414 | -1.657914226 | 0.146976778 | 0.558841836 | -4.350641997 |
| **A_51_P180724** | Mlh1 | 0.186948007 | -0.087061497 | 0.460957511 | 10.50525486 | 1.657823925 | 0.146995248 | 0.558841836 | -4.350688062 |
| **A_51_P275454** | Trim30a | -0.214645185 | -0.529327362 | 0.100036991 | 8.88144543 | -1.657418549 | 0.147078194 | 0.558841836 | -4.350894857 |
| **A_51_P438952** | Bag4 | 0.125382782 | -0.058443707 | 0.309209271 | 8.396023113 | 1.657345293 | 0.147093188 | 0.558841836 | -4.350932226 |
| **A_55_P2092399** | Ccdc160 | -0.137932084 | -0.340171625 | 0.064307457 | 7.868557284 | -1.657228585 | 0.147117079 | 0.558841836 | -4.350991763 |
| **A_55_P2172934** | Rgs7bp | 0.374885658 | -0.174810838 | 0.924582153 | 12.0427146 | 1.657139413 | 0.147135336 | 0.558841836 | -4.351037252 |
| **A_55_P2037797** | Tex44 | 0.224591775 | -0.104753008 | 0.553936559 | 8.759237648 | 1.657014266 | 0.147160961 | 0.558841836 | -4.351101093 |
| **A_66_P106113** | Rhoj | -0.396904629 | -0.978936602 | 0.185127343 | 9.723846746 | -1.657000012 | 0.14716388 | 0.558841836 | -4.351108365 |
| **A_55_P2079167** | Dnm1 | -0.370274448 | -0.913255749 | 0.172706853 | 8.606117682 | -1.656998283 | 0.147164234 | 0.558841836 | -4.351109247 |
| **A_55_P2100864** | Tma7 | 0.165977702 | -0.077459857 | 0.409415261 | 14.23601876 | 1.656705645 | 0.147224174 | 0.558841836 | -4.351258529 |
| **A_55_P1984406** | Lrrk2 | -0.359905402 | -0.887825072 | 0.168014269 | 9.759047792 | -1.656546799 | 0.147256719 | 0.558841836 | -4.351339561 |
| **A_55_P2041910** | NA | 0.614511072 | -0.286944554 | 1.515966697 | 10.15410355 | 1.656412686 | 0.147284203 | 0.558841836 | -4.351407976 |
| **A_52_P552589** | Map4k1 | 0.287614135 | -0.134313658 | 0.709541929 | 11.57243074 | 1.656362085 | 0.147294573 | 0.558841836 | -4.351433789 |
| **A_52_P72237** | Actg1 | -0.331960655 | -0.818967494 | 0.155046183 | 11.298024 | -1.656283756 | 0.147310628 | 0.558841836 | -4.351473747 |
| **A_55_P2001342** | Cacul1 | 0.145644242 | -0.068032523 | 0.359321007 | 8.79668727 | 1.656224515 | 0.147322772 | 0.558841836 | -4.351503967 |
| **A_55_P2101944** | NA | 0.276104577 | -0.128978695 | 0.681187848 | 11.88227067 | 1.656198877 | 0.147328028 | 0.558841836 | -4.351517046 |
| **A_51_P343323** | Cox6c | 0.126664309 | -0.059171516 | 0.312500135 | 14.88271478 | 1.656181801 | 0.147331529 | 0.558841836 | -4.351525757 |
| **A_55_P2116853** | Brpf1 | 0.125119512 | -0.05845229 | 0.308691314 | 11.37888932 | 1.656159876 | 0.147336023 | 0.558841836 | -4.351536942 |
| **A_55_P2082961** | Fnbp1l | -0.315692066 | -0.778963187 | 0.147579056 | 10.02979738 | -1.655814293 | 0.147406889 | 0.559000278 | -4.351713233 |
| **A_51_P424532** | Vnn1 | -0.616776991 | -1.521960191 | 0.28840621 | 8.569589498 | -1.655674148 | 0.147435637 | 0.559000278 | -4.351784725 |

| | | | | | | | | |
|---|---|---|---|---|---|---|---|---|
| **A_55_P2126870** | Nsf | 0.121023633 | -0.056636041 | 0.298683307 | 8.188371531 | 1.655253498 | 0.147521956 | 0.559033678 | -4.351999309 |
| **A_51_P301215** | Knop1 | 0.15485039 | -0.072518838 | 0.382219619 | 11.03349252 | 1.654869708 | 0.147600753 | 0.559033678 | -4.35219509 |
| **A_51_P513163** | Tra2a | 0.12491106 | -0.05852592 | 0.30834804 | 12.67714086 | 1.654615888 | 0.147652888 | 0.559033678 | -4.352324571 |
| **A_55_P1955998** | Nlrp1a | 0.281249471 | -0.131785855 | 0.694284798 | 13.02286136 | 1.654579752 | 0.147660312 | 0.559033678 | -4.352343004 |
| **A_55_P2048348** | Aig1 | -0.25170529 | -0.621376522 | 0.117965943 | 9.724305047 | -1.654473733 | 0.147682095 | 0.559033678 | -4.352397087 |
| **A_52_P18765** | Hsbp1l1 | -0.278549499 | -0.687656413 | 0.130557416 | 8.560724179 | -1.654431346 | 0.147690804 | 0.559033678 | -4.35241871 |
| **A_52_P212686** | Lrrk1 | -0.152662697 | -0.376886829 | 0.071561434 | 8.262617069 | -1.654374281 | 0.147702531 | 0.559033678 | -4.35244782 |
| **A_51_P249594** | Rev1 | 0.212854063 | -0.099792265 | 0.525500391 | 8.682401746 | 1.654290597 | 0.147719729 | 0.559033678 | -4.352490509 |
| **A_55_P2099358** | Cars | 0.219052758 | -0.1027143 | 0.540819817 | 12.32603186 | 1.65420883 | 0.147736535 | 0.559033678 | -4.352532221 |
| **A_55_P2120682** | Golga7 | 0.206661356 | -0.096948877 | 0.510271589 | 10.27495742 | 1.653964022 | 0.147786863 | 0.559033678 | -4.352657103 |
| **A_55_P1978618** | Grcc10 | -0.379157339 | -0.936189507 | 0.17787483 | 12.65985623 | -1.653950011 | 0.147789744 | 0.559033678 | -4.35266425 |
| **A_51_P384609** | Ddx10 | 0.190358323 | -0.08936303 | 0.470079676 | 9.232086808 | 1.653596098 | 0.147862532 | 0.559033678 | -4.352844789 |
| **A_52_P683441** | Capn5 | 0.315367069 | -0.148063056 | 0.778797194 | 10.1090302 | 1.653542149 | 0.147873631 | 0.559033678 | -4.35287231 |
| **A_55_P2139673** | Rpl11 | 0.193940816 | -0.091086564 | 0.478968197 | 15.73076897 | 1.653353909 | 0.147912362 | 0.559033678 | -4.352968335 |
| **A_55_P2049266** | Mettl6 | 0.159105548 | -0.07472844 | 0.392939535 | 10.02551036 | 1.653335063 | 0.14791624 | 0.559033678 | -4.352977949 |
| **A_55_P2150831** | Cdh4 | -0.225224687 | -0.556240528 | 0.105791154 | 8.020403704 | -1.653295196 | 0.147924444 | 0.559033678 | -4.352998286 |
| **A_55_P2007326** | Cyp2a4 | -0.812705729 | -2.007209416 | 0.381797957 | 11.24546267 | -1.653213777 | 0.147941201 | 0.559033678 | -4.353039819 |
| **A_51_P222359** | Emilin1 | -0.187130942 | -0.462212848 | 0.087950964 | 8.361545616 | -1.652976838 | 0.147989976 | 0.559033678 | -4.353160687 |
| **A_51_P192344** | Ash2l | 0.116828773 | -0.054937544 | 0.288595091 | 10.744504 | 1.652703705 | 0.14804622 | 0.559033678 | -4.353300018 |
| **A_51_P312497** | Atp7a | -0.257077638 | -0.635067064 | 0.120911789 | 11.62908368 | -1.652600341 | 0.14806751 | 0.559033678 | -4.353352746 |
| **A_55_P2047698** | Pop7 | 0.121671651 | -0.057235409 | 0.30057871 | 12.41283428 | 1.652513862 | 0.148085325 | 0.559033678 | -4.35339686 |
| **A_55_P1983558** | Tmed1 | -0.214047589 | -0.528788681 | 0.100693502 | 12.48481912 | -1.65249473 | 0.148089266 | 0.559033678 | -4.35340662 |
| **A_55_P2026785** | Dcaf1 | 0.160472923 | -0.075504728 | 0.396450575 | 9.776593296 | 1.652395785 | 0.148109652 | 0.559033678 | -4.353457094 |
| **A_52_P117999** | Selenot | -0.160607602 | -0.396865878 | 0.075650673 | 8.697389446 | -1.651818242 | 0.148228697 | 0.55921184 | -4.353751709 |
| **A_52_P233441** | Gata2 | -0.187506645 | -0.463362755 | 0.088349464 | 8.359475465 | -1.651647052 | 0.148264001 | 0.55921184 | -4.353839036 |
| **A_55_P2008071** | Prkcd | -0.125332763 | -0.309742665 | 0.059077138 | 9.035886101 | -1.651442922 | 0.148306109 | 0.55921184 | -4.353943166 |
| **A_55_P2032598** | NA | 0.149443607 | -0.070443701 | 0.369330914 | 15.81796455 | 1.65143075 | 0.14830862 | 0.55921184 | -4.353949375 |
| **A_51_P495077** | Ift22 | -0.307665795 | -0.760385168 | 0.145053579 | 8.361254374 | -1.651327924 | 0.148329835 | 0.55921184 | -4.354001829 |
| **A_55_P2023797** | Cep112 | -0.216349798 | -0.534718571 | 0.102018975 | 8.102201522 | -1.651236268 | 0.148348748 | 0.55921184 | -4.354048584 |
| **A_55_P1989071** | Oas1d | 0.201319083 | -0.094953982 | 0.497592148 | 8.066021128 | 1.651109827 | 0.148374843 | 0.55921184 | -4.354113083 |
| **A_55_P2315457** | Gripap1 | 0.189937425 | -0.08959676 | 0.46947161 | 10.73629249 | 1.651044613 | 0.148388304 | 0.55921184 | -4.35414635 |
| **A_55_P2031989** | Rin2 | -0.616517008 | -1.523984348 | 0.290950333 | 9.147507946 | -1.650810594 | 0.148436617 | 0.55926275 | -4.354265727 |
| **A_55_P1962234** | Bad | -0.354208945 | -0.875613536 | 0.167195647 | 11.80831235 | -1.6506989 | 0.148459681 | 0.55926275 | -4.354322703 |
| **A_55_P1967168** | Lrp4 | -0.203527603 | -0.503170201 | 0.096114995 | 8.399751115 | -1.65045221 | 0.148510633 | 0.559345679 | -4.354448543 |
| **A_55_P2092611** | Slc45a3 | 0.242611666 | -0.114678558 | 0.599901889 | 9.602377259 | 1.649961214 | 0.148612095 | 0.559358756 | -4.354699006 |
| **A_51_P327874** | Pth1r | -0.534393106 | -1.321399681 | 0.25261347 | 9.247156671 | -1.649930591 | 0.148618426 | 0.559358756 | -4.354714627 |
| **A_52_P35064** | Tm7sf2 | -0.208555396 | -0.515764567 | 0.098653774 | 13.5366508 | -1.649568843 | 0.148693225 | 0.559358756 | -4.354899159 |
| **A_51_P331870** | Rnf145 | 0.121362059 | -0.057420605 | 0.300144722 | 10.42539565 | 1.649455947 | 0.148716576 | 0.559358756 | -4.354956749 |
| **A_55_P2096967** | Gjb1 | -0.404663049 | -1.000790387 | 0.191464289 | 14.41265128 | -1.64944447 | 0.14871895 | 0.559358756 | -4.354962603 |
| **A_52_P40954** | Dubr | -0.194150888 | -0.4801673 | 0.091865524 | 8.273255749 | -1.649421363 | 0.14872373 | 0.559358756 | -4.35497439 |
| **A_55_P2053032** | Xkr8 | 0.190046524 | -0.089960458 | 0.470053506 | 8.787605487 | 1.649203542 | 0.148768795 | 0.559358756 | -4.355085503 |
| **A_66_P131604** | Otud4 | 0.216014091 | -0.102256724 | 0.534284906 | 11.47496606 | 1.649181501 | 0.148773356 | 0.559358756 | -4.355096746 |
| **A_55_P2131579** | NA | -0.260727968 | -0.644925932 | 0.123469997 | 14.64959307 | -1.648981369 | 0.148814774 | 0.559358756 | -4.355198835 |
| **A_52_P372151** | Ctnnal1 | 0.1309929 | -0.062057288 | 0.324043087 | 9.996057099 | 1.648772429 | 0.148858027 | 0.559358756 | -4.355305417 |
| **A_55_P2179726** | Mettl18 | 0.126328622 | -0.059848606 | 0.31250585 | 9.002903684 | 1.648763591 | 0.148859857 | 0.559358756 | -4.355309925 |
| **A_55_P2117928** | Mxd4 | -0.456039698 | -1.128132703 | 0.216053308 | 9.391444376 | -1.648756278 | 0.148861371 | 0.559358756 | -4.355313656 |
| **A_55_P2134012** | Oxa1l | -0.131693925 | -0.325798985 | 0.062411135 | 12.1332036 | -1.648587768 | 0.148896264 | 0.559381125 | -4.355399614 |
| **A_55_P2122619** | Asb11 | -0.11084689 | -0.274269586 | 0.052575805 | 7.792397878 | -1.648140887 | 0.148988837 | 0.559608154 | -4.355627571 |
| **A_55_P2113523** | Nsd2 | 0.157503099 | -0.074722834 | 0.389729032 | 8.321214114 | 1.648016574 | 0.149014598 | 0.559608154 | -4.355690983 |
| **A_55_P2005229** | NA | 0.212599926 | -0.100927647 | 0.5261275 | 10.07564514 | 1.647671227 | 0.149086187 | 0.559764172 | -4.355867146 |

| | | | | | | | | | |
|---|---|---|---|---|---|---|---|---|---|
| **A_52_P338110** | Chmp1b | 0.114018943 | -0.054141978 | 0.282179863 | 9.280449707 | 1.647536798 | 0.149114063 | 0.559764172 | -4.355935718 |
| **A_55_P2003813** | Scn3b | 0.363476291 | -0.172995227 | 0.89994781 | 8.857050291 | 1.646313756 | 0.149367905 | 0.560491757 | -4.356559592 |
| **A_55_P2131865** | Krtap10-10 | 0.316291478 | -0.150593014 | 0.783175969 | 9.473541367 | 1.646119013 | 0.149408362 | 0.560491757 | -4.35665893 |
| **A_51_P353502** | Sharpin | -0.137061184 | -0.339383901 | 0.065261533 | 12.07784721 | -1.646087891 | 0.149414828 | 0.560491757 | -4.356674805 |
| **A_51_P208152** | 3930402G23Rik | -0.472127109 | -1.169075453 | 0.224821235 | 9.588752384 | -1.646044366 | 0.149423872 | 0.560491757 | -4.356697007 |
| **A_55_P2047852** | Clcn3 | 0.304651661 | -0.145122066 | 0.754425388 | 11.82579605 | 1.645859111 | 0.149462371 | 0.560527392 | -4.356791505 |
| **A_55_P2160686** | Tsc22d1 | -0.392865637 | -0.972980672 | 0.187249398 | 10.89682552 | -1.645557685 | 0.149525034 | 0.560653613 | -4.356945261 |
| **A_55_P1998081** | Trmt1 | -0.163649121 | -0.405432065 | 0.078133822 | 12.66593826 | -1.644641353 | 0.14971568 | 0.561177829 | -4.357412674 |
| **A_55_P2101570** | Zhx3 | -0.301460498 | -0.746861883 | 0.143940886 | 10.3045886 | -1.644606641 | 0.149722907 | 0.561177829 | -4.35743038 |
| **A_55_P2042958** | Rad50 | 0.147349226 | -0.07037584 | 0.365074291 | 9.634088249 | 1.644457361 | 0.149753988 | 0.561185506 | -4.357506526 |
| **A_55_P1957937** | NA | 0.276323942 | -0.132027478 | 0.684675361 | 8.910645887 | 1.644249183 | 0.149797344 | 0.561197144 | -4.357612715 |
| **A_55_P2091681** | Nudt1 | 0.177398388 | -0.084774524 | 0.4395713 | 10.67002295 | 1.644163643 | 0.149815162 | 0.561197144 | -4.357656348 |
| **A_55_P1956728** | NA | 0.191235735 | -0.091494629 | 0.473966098 | 14.14012717 | 1.643538139 | 0.149945516 | 0.561475841 | -4.357975409 |
| **A_55_P1955502** | Tasor2 | 0.188957611 | -0.090406432 | 0.468321655 | 9.194700834 | 1.643527864 | 0.149947658 | 0.561475841 | -4.35798065 |
| **A_51_P342716** | Ppp1cb | 0.129981974 | -0.062225484 | 0.322189431 | 13.30092659 | 1.643221395 | 0.150011567 | 0.561518247 | -4.358136974 |
| **A_55_P2171185** | NA | -0.298801622 | -0.74065385 | 0.143050607 | 12.91368703 | -1.643194942 | 0.150017084 | 0.561518247 | -4.358150468 |
| **A_51_P282523** | Adgra2 | -0.396588126 | -0.983244651 | 0.1900684 | 9.78111992 | -1.642627125 | 0.150135565 | 0.561626099 | -4.3584401 |
| **A_55_P2012369** | Eea1 | 0.252190373 | -0.120934162 | 0.625314908 | 9.529822497 | 1.642320395 | 0.150199605 | 0.561626099 | -4.358596556 |
| **A_55_P1986863** | Ccdc126 | -0.185620205 | -0.46025622 | 0.08901581 | 9.078063147 | -1.642294158 | 0.150205084 | 0.561626099 | -4.35860994 |
| **A_55_P2062926** | D430041D05Rik | 0.158855925 | -0.076216918 | 0.393928769 | 9.071989092 | 1.642041566 | 0.150257841 | 0.561626099 | -4.358738781 |
| **A_51_P415809** | Tusc1 | 0.183982778 | -0.088305195 | 0.456270751 | 9.972746433 | 1.641844048 | 0.150299108 | 0.561626099 | -4.35883953 |
| **A_51_P204053** | Hba-x | 0.505030886 | -0.242440382 | 1.252502154 | 9.151846232 | 1.641748487 | 0.150319078 | 0.561626099 | -4.358888273 |
| **A_51_P195573** | Dnmt1 | 0.275803339 | -0.132418215 | 0.684024893 | 10.27772518 | 1.641673457 | 0.150334759 | 0.561626099 | -4.358926544 |
| **A_55_P2091247** | NA | -0.161059933 | -0.399462574 | 0.077342708 | 13.81685172 | -1.641570889 | 0.150356197 | 0.561626099 | -4.358978861 |
| **A_51_P193794** | Lrp1 | -0.165537992 | -0.410592725 | 0.079516741 | 11.13629429 | -1.641412651 | 0.150389277 | 0.561626099 | -4.359059574 |
| **A_52_P362981** | NA | -0.243439407 | -0.603844228 | 0.116965413 | 8.857188984 | -1.641283032 | 0.15041638 | 0.561626099 | -4.359125689 |
| **A_55_P2159234** | Tatdn2 | 0.168693988 | -0.081070108 | 0.418458083 | 8.304820715 | 1.64116704 | 0.150440637 | 0.561626099 | -4.359184853 |
| **A_55_P2039210** | Cpt1b | -0.257949137 | -0.639949703 | 0.124051429 | 8.636831214 | -1.640790985 | 0.150519306 | 0.561626099 | -4.359376667 |
| **A_55_P2179974** | Spsb1 | 0.141806641 | -0.068201078 | 0.35181436 | 8.192441491 | 1.64075789 | 0.150526231 | 0.561626099 | -4.359393548 |
| **A_55_P2109354** | Dot1l | 0.170263102 | -0.08188714 | 0.422413344 | 10.47338439 | 1.640757285 | 0.150526358 | 0.561626099 | -4.359393857 |
| **A_55_P2065971** | Setd3 | 0.152664498 | -0.073432075 | 0.378761071 | 11.42020715 | 1.640692771 | 0.150539859 | 0.561626099 | -4.359426763 |
| **A_52_P671769** | Arhgap18 | -0.247211962 | -0.613335796 | 0.118911872 | 8.721016468 | -1.640682966 | 0.150541911 | 0.561626099 | -4.359431765 |
| **A_55_P2014645** | Knop1 | 0.202473864 | -0.097425104 | 0.502372831 | 10.05187998 | 1.640503602 | 0.150579452 | 0.561626099 | -4.359523252 |
| **A_55_P2169621** | Prorsd1 | 0.296191492 | -0.14252389 | 0.734906874 | 15.4476691 | 1.640487427 | 0.150582838 | 0.561626099 | -4.359531502 |
| **A_51_P386899** | Flvcr2 | 0.203161515 | -0.097773703 | 0.504096734 | 10.18122989 | 1.640407009 | 0.150599673 | 0.561626099 | -4.359572521 |
| **A_55_P2077522** | NA | 0.301322955 | -0.145050514 | 0.747696425 | 9.994806263 | 1.640276388 | 0.150627021 | 0.561626099 | -4.359639146 |
| **A_51_P385952** | Man2a2 | -0.172061266 | -0.427060153 | 0.08293762 | 8.542990297 | -1.639562662 | 0.15077654 | 0.562075168 | -4.360003192 |
| **A_55_P2132467** | Nfx1 | 0.145044354 | -0.069971122 | 0.36005983 | 9.092493322 | 1.639133394 | 0.150866536 | 0.562098163 | -4.360222144 |
| **A_55_P2142728** | NA | 0.24306616 | -0.11729504 | 0.603427361 | 11.997648 | 1.638964944 | 0.150901865 | 0.562098163 | -4.360308063 |
| **A_66_P129469** | NA | 0.178911874 | -0.086343971 | 0.444167718 | 7.911410452 | 1.638918627 | 0.150911581 | 0.562098163 | -4.360331687 |
| **A_51_P413111** | Ppm1l | 0.15494766 | -0.074793644 | 0.384688963 | 10.2428124 | 1.638811978 | 0.150933954 | 0.562098163 | -4.360386084 |
| **A_51_P224227** | Rlf | 0.139847535 | -0.067551774 | 0.347246843 | 10.239239 | 1.638440593 | 0.151011889 | 0.562098163 | -4.360575511 |
| **A_51_P352283** | Cyb561d1 | 0.111892944 | -0.054053855 | 0.277839743 | 9.107894379 | 1.638388947 | 0.151022731 | 0.562098163 | -4.360601854 |
| **A_52_P156190** | Ednra | -0.794451297 | -1.972708188 | 0.383805593 | 10.21428635 | -1.638364312 | 0.151027902 | 0.562098163 | -4.360614419 |
| **A_51_P102789** | C1qc | -0.285609096 | -0.709229704 | 0.138011513 | 12.12995804 | -1.63824233 | 0.151053511 | 0.562098163 | -4.363528927 |
| **A_66_P124052** | Bok | -0.481736342 | -1.19629692 | 0.232824236 | 9.501680797 | -1.63814955 | 0.151072992 | 0.562098163 | -4.360723959 |
| **A_51_P498212** | Ube2j1 | 0.149816238 | -0.072417872 | 0.372050348 | 8.409403329 | 1.638065854 | 0.151090568 | 0.562098163 | -4.360766648 |
| **A_55_P1976207** | Arf5 | 0.165444913 | -0.079984228 | 0.410874054 | 13.47737823 | 1.637987111 | 0.151107105 | 0.562098163 | -4.360806811 |
| **A_66_P139310** | Cog1 | 0.148587399 | -0.071850217 | 0.369025016 | 10.78504705 | 1.637870135 | 0.151131675 | 0.562098163 | -4.360866475 |
| **A_55_P1969356** | Ccm2l | -0.286778791 | -0.71236227 | 0.138804687 | 8.32658687 | -1.637364834 | 0.151237855 | 0.56216274 | -4.361124202 |

| | | | | | | | | | |
|---|---|---|---|---|---|---|---|---|---|
| **A_51_P182131** | 5330417C22Rik | -0.458465299 | -1.138846607 | 0.22191601 | 8.13760559 | -1.637334016 | 0.151244333 | 0.56216274 | -4.361139921 |
| **A_55_P2083844** | Dis3l | 0.191549231 | -0.0927461 | 0.475844561 | 9.987578262 | 1.637170364 | 0.151278738 | 0.56216274 | -4.361223391 |
| **A_52_P317040** | Edem2 | -0.138817921 | -0.344850115 | 0.067214272 | 8.136070139 | -1.637169478 | 0.151278924 | 0.56216274 | -4.361223843 |
| **A_55_P2032845** | Rbm10 | 0.143543663 | -0.069512044 | 0.356599369 | 11.49279414 | 1.637095603 | 0.151294458 | 0.56216274 | -4.361261522 |
| **A_55_P2097269** | NA | 0.196549882 | -0.095324669 | 0.488424433 | 12.7527519 | 1.636288001 | 0.151464369 | 0.562541389 | -4.361673433 |
| **A_51_P393078** | Tigd2 | -0.162292515 | -0.403306694 | 0.078721664 | 9.115548517 | -1.636210165 | 0.151480754 | 0.562541389 | -4.361713132 |
| **A_55_P2037657** | NA | 0.182756471 | -0.088684956 | 0.454197898 | 8.347395846 | 1.635986903 | 0.151527763 | 0.562541389 | -4.361827005 |
| **A_55_P2106265** | Mbnl2 | -0.200196777 | -0.497545997 | 0.097152443 | 8.359775014 | -1.635962961 | 0.151532805 | 0.562541389 | -4.361839216 |
| **A_55_P2160716** | NA | 0.137508604 | -0.066736208 | 0.341753416 | 15.5829774 | 1.635919869 | 0.151541881 | 0.562541389 | -4.361861195 |
| **A_55_P2158072** | NA | 0.105785097 | -0.051379606 | 0.262949799 | 7.911754758 | 1.635507959 | 0.151628655 | 0.562744353 | -4.362071283 |
| **A_55_P2063558** | NA | 0.122550201 | -0.059547933 | 0.304648335 | 16.40092476 | 1.635278475 | 0.151677019 | 0.562744353 | -4.362188327 |
| **A_52_P257547** | Dele1 | 0.126183985 | -0.061320264 | 0.313688234 | 10.35006888 | 1.635220468 | 0.151689246 | 0.562744353 | -4.362217912 |
| **A_55_P2063711** | Snu13 | 0.145804687 | -0.070870062 | 0.362479435 | 13.1320634 | 1.635107733 | 0.151713013 | 0.562744353 | -4.362275411 |
| **A_55_P1999301** | Cyp2e1 | -0.671648706 | -1.669876247 | 0.326578834 | 9.149296 | -1.63491743 | 0.15175314 | 0.562785196 | -4.362372471 |
| **A_55_P2035717** | Pgap2 | -0.427346742 | -1.062591301 | 0.207897817 | 10.53237084 | -1.634641923 | 0.151811251 | 0.562799957 | -4.362512987 |
| **A_51_P411297** | Nup50 | 0.277901756 | -0.135259642 | 0.691063155 | 10.16306708 | 1.634386409 | 0.151865164 | 0.562799957 | -4.362643306 |
| **A_55_P1972230** | NA | 0.165114524 | -0.080378402 | 0.41060745 | 11.7426441 | 1.634291354 | 0.151885225 | 0.562799957 | -4.362691786 |
| **A_52_P227391** | Kif15 | 0.257517449 | -0.125366165 | 0.640401062 | 8.847433851 | 1.634267221 | 0.151890318 | 0.562799957 | -4.362704094 |
| **A_55_P2056808** | Tanc1 | -0.127049188 | -0.315969525 | 0.06187115 | 7.952302497 | -1.634091493 | 0.151927413 | 0.562799957 | -4.36279372 |
| **A_51_P420400** | Lef1 | -0.22407787 | -0.557327523 | 0.109171784 | 8.732783478 | -1.633851005 | 0.151978192 | 0.562799957 | -4.362916374 |
| **A_55_P2147842** | Zglp1 | 0.171917281 | -0.083763583 | 0.427598145 | 9.430000411 | 1.633821088 | 0.15198451 | 0.562799957 | -4.362931632 |
| **A_51_P230734** | Slc34a3 | -0.184135703 | -0.457992594 | 0.089721189 | 7.770791093 | -1.633794834 | 0.151990055 | 0.562799957 | -4.362945022 |
| **A_55_P2079064** | NA | 0.110081547 | -0.053656821 | 0.273819915 | 8.572972182 | 1.633605759 | 0.152029993 | 0.562840017 | -4.363041454 |
| **A_55_P2035539** | D3Ertd751e | 0.128792655 | -0.062864565 | 0.320449875 | 8.507317669 | 1.632860569 | 0.152187494 | 0.562997669 | -4.363421513 |
| **A_55_P2095394** | Dhx37 | 0.167355776 | -0.081687884 | 0.416399436 | 8.507189929 | 1.632857953 | 0.152188047 | 0.562997669 | -4.363422847 |
| **A_55_P2062255** | Syne3 | 0.301259392 | -0.147073847 | 0.749592632 | 8.88026514 | 1.632761856 | 0.152208369 | 0.562997669 | -4.363471857 |
| **A_55_P2038303** | NA | 0.319526061 | -0.156043251 | 0.795095373 | 10.15842409 | 1.632584437 | 0.152245895 | 0.562997669 | -4.363562343 |
| **A_55_P2074843** | Ccdc122 | 0.202236963 | -0.098812422 | 0.503286349 | 8.244277451 | 1.632322546 | 0.152301305 | 0.562997669 | -4.363695909 |
| **A_66_P134785** | 2610044O15Rik8 | 0.168451796 | -0.082344014 | 0.419247607 | 8.470874781 | 1.632069159 | 0.152354934 | 0.562997669 | -4.363825139 |
| **A_51_P244213** | Hapln3 | 0.160557893 | -0.07849157 | 0.399607356 | 8.289758218 | 1.632026016 | 0.152364067 | 0.562997669 | -4.363847142 |
| **A_55_P2055881** | NA | -0.240550438 | -0.598698254 | 0.117597377 | 16.18971568 | -1.632025855 | 0.152364101 | 0.562997669 | -4.363847224 |
| **A_55_P2053258** | Poc1a | 0.233801648 | -0.114299688 | 0.581902985 | 11.92945219 | 1.632018462 | 0.152365666 | 0.562997669 | -4.363850994 |
| **A_55_P2032458** | Gm6934 | 0.19736625 | -0.09650011 | 0.49123261 | 8.676768219 | 1.63194758 | 0.152380672 | 0.562997669 | -4.363887144 |
| **A_52_P253044** | Syt13 | -0.131999558 | -0.32855943 | 0.064560314 | 7.957664197 | -1.631776987 | 0.152416794 | 0.562997669 | -4.363974148 |
| **A_51_P173212** | Osbpl2 | -0.114977921 | -0.286203023 | 0.056247181 | 10.30885189 | -1.631662027 | 0.15244114 | 0.562997669 | -4.364032778 |
| **A_52_P345247** | Tfb1m | 0.241418522 | -0.1181124 | 0.600949445 | 10.56566026 | 1.631614394 | 0.152451229 | 0.562997669 | -4.364057071 |
| **A_51_P411917** | Gata6 | -0.358157603 | -0.891678437 | 0.175363231 | 9.497683067 | -1.631195371 | 0.152540007 | 0.563181884 | -4.364270772 |
| **A_55_P2095153** | Pdf | 0.130057996 | -0.06369062 | 0.323806612 | 10.34210989 | 1.631103941 | 0.152559385 | 0.563181884 | -4.364317401 |
| **A_55_P1987904** | Chpf2 | -0.142134078 | -0.353897697 | 0.069629542 | 8.183812885 | -1.630910217 | 0.152600451 | 0.563203382 | -4.3644162 |
| **A_66_P113059** | Itgb1 | -0.256537311 | -0.638774204 | 0.125699583 | 10.55530638 | -1.630801583 | 0.152623484 | 0.563203382 | -4.364471603 |
| **A_51_P429682** | Ebp | -0.312626439 | -0.77870184 | 0.153448961 | 14.81573987 | -1.629869045 | 0.15282134 | 0.563591855 | -4.364947189 |
| **A_55_P2031871** | Plekhh3 | -0.376186487 | -0.937031147 | 0.184658172 | 9.495780574 | -1.62983557 | 0.152828447 | 0.563591855 | -4.364964261 |
| **A_55_P2021149** | NA | 0.126901079 | -0.062301314 | 0.316103473 | 8.005432456 | 1.629753338 | 0.152845906 | 0.563591855 | -4.365006198 |
| **A_55_P1952076** | Npb | 0.413440337 | -0.203092405 | 1.02997308 | 10.48689114 | 1.629445689 | 0.152911244 | 0.563591855 | -4.365163094 |
| **A_55_P1980863** | NA | 0.138828645 | -0.06820408 | 0.345861369 | 8.235830198 | 1.629383357 | 0.152924485 | 0.563591855 | -4.365194882 |
| **A_55_P1978962** | Slc9a3r2 | -0.523206706 | -1.30353661 | 0.257123199 | 9.617375896 | -1.6292144 | 0.152960382 | 0.563591855 | -4.365281047 |
| **A_55_P2136353** | NA | 0.179007721 | -0.087989534 | 0.446004976 | 9.213054562 | 1.629101545 | 0.152984364 | 0.563591855 | -4.365338601 |
| **A_55_P2155062** | Naa40 | 0.171265394 | -0.084202538 | 0.426733325 | 9.888339198 | 1.628982483 | 0.153009668 | 0.563591855 | -4.36539932 |
| **A_51_P101545** | Hgfac | -0.360147211 | -0.897375793 | 0.17708137 | 13.73008652 | -1.628936427 | 0.153019458 | 0.563591855 | -4.365422807 |
| **A_55_P2049429** | Phtf2 | 0.167963396 | -0.082586831 | 0.418513624 | 9.935556742 | 1.6289323 | 0.153020335 | 0.563591855 | -4.365424912 |

| | | | | | | | | | |
|---|---|---|---|---|---|---|---|---|---|
| **A_55_P2010347** | Zdhhc6 | 0.162623857 | -0.080028132 | 0.405275846 | 8.874867326 | 1.628484352 | 0.153115581 | 0.563835219 | -4.365653355 |
| **A_52_P15490** | Nectin3 | -0.138546326 | -0.345312955 | 0.068220304 | 8.291813959 | -1.628162519 | 0.153184046 | 0.563873153 | -4.365817481 |
| **A_51_P284946** | Rnd3 | 0.174026215 | -0.08569184 | 0.43374427 | 10.33966348 | 1.628154901 | 0.153185667 | 0.563873153 | -4.365821366 |
| **A_51_P432851** | Dhrs11 | 0.343089991 | -0.168981125 | 0.855161107 | 10.39717485 | 1.628024582 | 0.153213399 | 0.563873153 | -4.365887825 |
| **A_51_P194498** | Wfdc1 | -0.276734521 | -0.689877052 | 0.13640801 | 8.522041252 | -1.627596029 | 0.15330463 | 0.563995755 | -4.366106373 |
| **A_51_P225592** | Tpm4 | -0.519878595 | -1.296075839 | 0.256318648 | 13.07675477 | -1.627470142 | 0.153331439 | 0.563995755 | -4.366170571 |
| **A_55_P1979863** | Syne2 | -0.573572024 | -1.429942411 | 0.282798362 | 10.21827467 | -1.627456948 | 0.153334249 | 0.563995755 | -4.366177299 |
| **A_55_P1980336** | NA | 0.132595017 | -0.065417047 | 0.330607081 | 7.982938271 | 1.627116845 | 0.1534067 | 0.564154891 | -4.366350739 |
| **A_51_P465292** | Hnmt | -0.121591241 | -0.303198008 | 0.060015526 | 7.92966772 | -1.626872254 | 0.153458825 | 0.564239229 | -4.36647547 |
| **A_52_P476560** | Tpcn1 | 0.192925659 | -0.095275369 | 0.481126687 | 11.81075265 | 1.626588344 | 0.15351935 | 0.564354416 | -4.366620252 |
| **A_51_P246677** | Rec8 | -0.344446364 | -0.859094946 | 0.170202217 | 9.725908271 | -1.626275105 | 0.153586154 | 0.564427215 | -4.36677999 |
| **A_52_P633752** | Apc | 0.233472775 | -0.115401524 | 0.582347074 | 8.608642377 | 1.626112012 | 0.153620948 | 0.564427215 | -4.366863159 |
| **A_55_P2062777** | Mfap5 | -0.480513615 | -1.198548756 | 0.237521526 | 8.248374517 | -1.626084788 | 0.153626757 | 0.564427215 | -4.366877042 |
| **A_55_P2091364** | Cep164 | -0.214994868 | -0.536309396 | 0.10631966 | 10.24390396 | -1.625851665 | 0.153676505 | 0.564502692 | -4.366995923 |
| **A_66_P105711** | Ap4s1 | 0.109974248 | -0.054412326 | 0.274360821 | 8.589468036 | 1.625578121 | 0.153734899 | 0.564609892 | -4.367135416 |
| **A_51_P419389** | Bmpr2 | -0.234086358 | -0.58408293 | 0.115910214 | 8.604757333 | -1.625157676 | 0.153824693 | 0.56483235 | -4.367349819 |
| **A_51_P322265** | Grap | -0.239260459 | -0.597065015 | 0.118544098 | 8.853839716 | -1.624831213 | 0.15389445 | 0.564917629 | -4.367516295 |
| **A_51_P510891** | Afp | 0.128748935 | -0.063796616 | 0.321294486 | 17.71827988 | 1.624775439 | 0.15390637 | 0.564917629 | -4.367544736 |
| **A_66_P112399** | BC029722 | -0.217072938 | -0.54179194 | 0.107646065 | 10.35224301 | -1.624355818 | 0.153996084 | 0.565072541 | -4.367758716 |
| **A_52_P544555** | Hsbp1 | -0.266388845 | -0.664891994 | 0.132114304 | 11.70920761 | -1.624304571 | 0.154007044 | 0.565072541 | -4.367784848 |
| **A_55_P2086510** | Ppid | 0.214184433 | -0.106257305 | 0.53462617 | 12.12533651 | 1.624134581 | 0.154043403 | 0.56509868 | -4.367871532 |
| **A_51_P342271** | Setd6 | 0.313866917 | -0.155791635 | 0.783525469 | 10.34144071 | 1.623852193 | 0.154103823 | 0.565213053 | -4.368015529 |
| **A_51_P490795** | Mxd1 | 0.279322749 | -0.13875384 | 0.697399338 | 11.55428789 | 1.6234303 | 0.154194132 | 0.565361956 | -4.368230663 |
| **A_51_P278427** | 3830406C13Rik | 0.118588486 | -0.058967063 | 0.296144036 | 9.598345126 | 1.622898892 | 0.154307956 | 0.565361956 | -4.368501638 |
| **A_52_P454295** | Ttn | -0.132168289 | -0.330057288 | 0.06572071 | 7.698531301 | -1.622888956 | 0.154310085 | 0.565361956 | -4.368506705 |
| **A_55_P2096368** | NA | -0.124929555 | -0.311995352 | 0.062136242 | 10.88398223 | -1.622758872 | 0.15433796 | 0.565361956 | -4.368573037 |
| **A_51_P239036** | Ubn2 | 0.191784352 | -0.095405184 | 0.478973889 | 10.96552087 | 1.622660805 | 0.154358978 | 0.565361956 | -4.368623042 |
| **A_52_P489119** | Nol6 | 0.225219932 | -0.112044506 | 0.562484371 | 9.327755326 | 1.622629854 | 0.154365612 | 0.565361956 | -4.368638825 |
| **A_52_P125485** | Slc25a48 | 0.135730903 | -0.067525444 | 0.338987251 | 8.018154443 | 1.622623689 | 0.154366934 | 0.565361956 | -4.368641969 |
| **A_55_P1990196** | NA | -0.314499895 | -0.785477459 | 0.156477669 | 12.24977473 | -1.622570122 | 0.154378416 | 0.565361956 | -4.368669283 |
| **A_55_P2042923** | Sgk2 | 0.488691148 | -0.243430947 | 1.220813243 | 10.48995158 | 1.621937622 | 0.154514056 | 0.565657412 | -4.368991801 |
| **A_55_P1968448** | NA | 0.192383932 | -0.095834858 | 0.480602722 | 15.08694695 | 1.621920996 | 0.154517623 | 0.565657412 | -4.369000279 |
| **A_65_P15245** | Nrp2 | -0.304593649 | -0.760959531 | 0.151772234 | 8.069103593 | -1.621775976 | 0.154548739 | 0.565664188 | -4.369074225 |
| **A_51_P411757** | Ankrd13b | -0.420794344 | -1.051402352 | 0.209813665 | 9.658777789 | -1.621412811 | 0.154626688 | 0.56576654 | -4.369259403 |
| **A_66_P139618** | Stfa2 | -0.377968481 | -0.944434558 | 0.188497596 | 15.43010132 | -1.621305541 | 0.154649719 | 0.56576654 | -4.369314101 |
| **A_55_P1986566** | Gm10318 | 0.288966291 | -0.144145957 | 0.722078538 | 8.938185086 | 1.621175035 | 0.154677744 | 0.56576654 | -4.369380646 |
| **A_55_P2024337** | Tmem53 | -0.18214343 | -0.455158614 | 0.090871754 | 10.54067707 | -1.621100346 | 0.154693785 | 0.56576654 | -4.369418729 |
| **A_55_P2058794** | Cmip | -0.161051949 | -0.40248865 | 0.080384752 | 13.17131888 | -1.62086142 | 0.154745109 | 0.565847182 | -4.369540556 |
| **A_52_P198473** | NA | 0.200893166 | -0.100342737 | 0.502129068 | 8.303041631 | 1.620472331 | 0.154828724 | 0.566045851 | -4.369738949 |
| **A_52_P346581** | Colgalt1 | -0.190566125 | -0.476383998 | 0.095251748 | 10.66294849 | -1.62009137 | 0.154910635 | 0.566149904 | -4.369933197 |
| **A_51_P400040** | Ubxn2a | 0.184227453 | -0.092087523 | 0.460542429 | 9.148486853 | 1.620067512 | 0.154915766 | 0.566149904 | -4.369945361 |
| **A_55_P1982019** | Rbfox2 | 0.123880694 | -0.06199709 | 0.309758478 | 10.37123804 | 1.619419378 | 0.155055221 | 0.566253113 | -4.370275832 |
| **A_66_P117417** | NA | -0.306161705 | -0.76561201 | 0.1532886 | 8.681933206 | -1.619181422 | 0.15510645 | 0.566253113 | -4.37039716 |
| **A_55_P2055232** | Sdf4 | -0.237325488 | -0.593481108 | 0.118830132 | 10.00769494 | -1.619152559 | 0.155112665 | 0.566253113 | -4.370411877 |
| **A_55_P2018191** | Ccm2l | -0.229372027 | -0.573664486 | 0.114920431 | 8.27089963 | -1.618810944 | 0.155186242 | 0.566253113 | -4.370586056 |
| **A_51_P312846** | Fbxo9 | 0.200206671 | -0.10030877 | 0.500722113 | 11.90277736 | 1.61880651 | 0.155187197 | 0.566253113 | -4.370588317 |
| **A_52_P331762** | Lmo1 | -0.271232844 | -0.678389256 | 0.135923568 | 8.230848632 | -1.618691881 | 0.155211893 | 0.566253113 | -4.370646763 |
| **A_55_P1984386** | Depdc5 | 0.213551826 | -0.107025564 | 0.534129217 | 8.735431228 | 1.618652376 | 0.155220405 | 0.566253113 | -4.370666905 |
| **A_55_P1956160** | Gm8909 | 0.174000985 | -0.08722436 | 0.435226331 | 8.429926707 | 1.618525646 | 0.155247715 | 0.566253113 | -4.37073152 |
| **A_52_P288251** | Tmem204 | -0.658878924 | -1.648112854 | 0.330355007 | 8.671055348 | -1.618414647 | 0.155271638 | 0.566253113 | -4.370788115 |

| | | | | | | | | | |
|---|---|---|---|---|---|---|---|---|---|
| **A_55_P2040121** | Cdv3 | 0.176803248 | -0.088669059 | 0.442275555 | 10.3127236 | 1.618282002 | 0.155300231 | 0.566253113 | -4.370855746 |
| **A_55_P2187266** | Pop1 | 0.228993428 | -0.114854586 | 0.572841441 | 10.31872701 | 1.618227908 | 0.155311893 | 0.566253113 | -4.370883327 |
| **A_66_P106191** | Eef2 | -0.1359037 | -0.339982925 | 0.068175525 | 14.53475429 | -1.618138435 | 0.155331184 | 0.566253113 | -4.370928945 |
| **A_51_P161636** | Kng2 | -0.406666112 | -1.01735623 | 0.204024006 | 14.98087507 | -1.618081082 | 0.155343551 | 0.566253113 | -4.370958187 |
| **A_55_P2404157** | Rbm27 | 0.121565339 | -0.060994759 | 0.304125437 | 8.356698483 | 1.618031955 | 0.155354144 | 0.566253113 | -4.370983235 |
| **A_55_P2086954** | NA | 0.179488495 | -0.090094923 | 0.449071913 | 11.87565912 | 1.617806701 | 0.155402728 | 0.566292899 | -4.371098083 |
| **A_51_P479583** | Obsl1 | -0.272162042 | -0.68096896 | 0.136644876 | 8.984696854 | -1.617679597 | 0.155430148 | 0.566292899 | -4.371162888 |
| **A_55_P2094901** | Phyhd1 | 0.215831207 | -0.108383937 | 0.540046351 | 16.35954356 | 1.617573906 | 0.155452953 | 0.566292899 | -4.371216775 |
| **A_52_P199928** | NA | 0.19976398 | -0.10035032 | 0.49987828 | 12.56206445 | 1.617386006 | 0.155493503 | 0.566333883 | -4.371312576 |
| **A_55_P2198568** | 4930506A18Rik | 0.137080366 | -0.068890549 | 0.34305128 | 8.450692163 | 1.617158341 | 0.155542648 | 0.56640615 | -4.371428651 |
| **A_55_P2152976** | NA | 0.209969656 | -0.105550228 | 0.525489539 | 15.01458338 | 1.617011068 | 0.155574447 | 0.566415236 | -4.371503738 |
| **A_52_P22617** | NA | 0.246943473 | -0.124184291 | 0.618071238 | 8.486614442 | 1.616803705 | 0.155619232 | 0.566471587 | -4.371609461 |
| **A_55_P2060072** | Clk2 | -0.159358729 | -0.398885412 | 0.080167954 | 10.02579718 | -1.616609575 | 0.155661169 | 0.566517555 | -4.371708437 |
| **A_55_P1986551** | Mcam | -0.179412671 | -0.449125912 | 0.090300571 | 7.895670686 | -1.616344876 | 0.155718369 | 0.566534636 | -4.371843391 |
| **A_55_P1956973** | Tln1 | -0.142195914 | -0.355964496 | 0.071572668 | 13.77890989 | -1.616316589 | 0.155724482 | 0.566534636 | -4.371857813 |
| **A_55_P2069337** | NA | 0.227293231 | -0.114448347 | 0.569034808 | 9.418415302 | 1.616113548 | 0.155768373 | 0.566587672 | -4.371961331 |
| **A_55_P2086148** | NA | 0.206215946 | -0.103888936 | 0.516320829 | 11.80281781 | 1.61583433 | 0.155828751 | 0.566676589 | -4.372103685 |
| **A_66_P128839** | Gm13262 | 0.166984315 | -0.084140859 | 0.418109489 | 8.138658066 | 1.615729365 | 0.155851454 | 0.566676589 | -4.3721572 |
| **A_52_P447944** | Epcam | -0.558426385 | -1.398412334 | 0.281559564 | 8.130033783 | -1.615388911 | 0.155925113 | 0.566837785 | -4.372330773 |
| **A_55_P2125623** | NA | 0.211044067 | -0.106504737 | 0.528592872 | 8.346535838 | 1.614900819 | 0.156030772 | 0.567115229 | -4.372579614 |
| **A_55_P2088385** | Tnnt1 | -0.191774324 | -0.480377511 | 0.096828863 | 10.13338421 | -1.614628174 | 0.156089822 | 0.56717242 | -4.372718613 |
| **A_55_P1969276** | Hhip | -0.930268983 | -2.330349037 | 0.469811072 | 9.929237857 | -1.614503018 | 0.156116935 | 0.56717242 | -4.372782419 |
| **A_55_P2172924** | NA | 0.157480601 | -0.079556231 | 0.394517432 | 8.370487439 | 1.614337808 | 0.156152733 | 0.56717242 | -4.372866645 |
| **A_51_P494006** | Scaf8 | 0.110719103 | -0.055938513 | 0.277376719 | 10.70594829 | 1.614286375 | 0.156163879 | 0.56717242 | -4.372892866 |
| **A_55_P2039712** | Gigyf2 | 0.193777277 | -0.097927445 | 0.485481998 | 11.57725344 | 1.614145126 | 0.156194493 | 0.567177036 | -4.372964877 |
| **A_55_P2008790** | Zfp30 | -0.154603032 | -0.38736697 | 0.078160906 | 8.254035041 | -1.613932989 | 0.156240482 | 0.567237469 | -4.373073026 |
| **A_55_P2013534** | Slc25a19 | -0.161686029 | -0.405205608 | 0.08183355 | 10.20069569 | -1.613324761 | 0.156372411 | 0.567609828 | -4.373383102 |
| **A_55_P1964985** | Gm10240 | 0.151958077 | -0.076968866 | 0.38088502 | 15.95790751 | 1.61290976 | 0.156462488 | 0.56780314 | -4.373594667 |
| **A_51_P456857** | Ift122 | -0.261375014 | -0.655193553 | 0.132443524 | 8.953824161 | -1.612690779 | 0.156510039 | 0.56780314 | -4.373706302 |
| **A_55_P2050877** | Plekhg2 | -0.418399844 | -1.04881734 | 0.212017651 | 10.96609991 | -1.612673487 | 0.156513794 | 0.56780314 | -4.373715117 |
| **A_55_P2068451** | NA | 0.236141182 | -0.119753715 | 0.592036079 | 14.64085138 | 1.612252879 | 0.156605167 | 0.568028013 | -4.373929537 |
| **A_55_P1994355** | Ptges3 | 0.194768091 | -0.098807711 | 0.488343894 | 13.61972076 | 1.612058287 | 0.156647458 | 0.568074805 | -4.374028737 |
| **A_51_P194479** | 1110059G10Rik | 0.178729955 | -0.090748006 | 0.448207917 | 9.93064652 | 1.611600087 | 0.156747081 | 0.568251865 | -4.374262317 |
| **A_55_P2029846** | Fam214a | -0.246444657 | -0.618034796 | 0.125145483 | 9.332240268 | -1.611530073 | 0.156762308 | 0.568251865 | -4.374298008 |
| **A_52_P254149** | Irgq | -0.137509881 | -0.344860888 | 0.069841126 | 8.790439512 | -1.61142815 | 0.156784479 | 0.568251865 | -4.374349966 |
| **A_51_P153063** | Fabp12 | -0.505757815 | -1.268548292 | 0.257032663 | 9.782024031 | -1.611092694 | 0.156857469 | 0.568334754 | -4.374520971 |
| **A_51_P215475** | Ptprb | -0.496905389 | -1.246363089 | 0.252552311 | 10.19849955 | -1.611052786 | 0.156866155 | 0.568334754 | -4.374541315 |
| **A_51_P214470** | Tdrp | -0.151935466 | -0.381117018 | 0.077246086 | 7.749352764 | -1.610878169 | 0.156904164 | 0.568349171 | -4.374630328 |
| **A_55_P1972025** | Mycl | -0.124484444 | -0.312291704 | 0.063322816 | 8.053930129 | -1.610593333 | 0.156966183 | 0.568349171 | -4.374775527 |
| **A_55_P2114995** | NA | -0.277953048 | -0.697326667 | 0.141420572 | 13.85250346 | -1.610473271 | 0.156992331 | 0.568349171 | -4.374836729 |
| **A_52_P488437** | Plek2 | 0.300519379 | -0.15293066 | 0.753969418 | 9.702806249 | 1.610372065 | 0.157014377 | 0.568349171 | -4.37488832 |
| **A_51_P505868** | Lhfp | -0.305410096 | -0.766243328 | 0.155423135 | 8.103545013 | -1.610359316 | 0.157017154 | 0.568349171 | -4.381405972 |
| **A_52_P179068** | Gucy1b1 | -0.465775616 | -1.16887855 | 0.237327317 | 9.575616022 | -1.609685486 | 0.157164009 | 0.568774224 | -4.375238304 |
| **A_55_P2075474** | Gid8 | 0.224113805 | -0.114226648 | 0.562454258 | 10.14412729 | 1.609525535 | 0.157198888 | 0.568793954 | -4.375319839 |
| **A_55_P1958230** | Dhodh | 0.120017351 | -0.061201849 | 0.30123655 | 10.58449999 | 1.609248142 | 0.157259394 | 0.568906387 | -4.375461237 |
| **A_52_P183152** | Txnl1 | 0.180566233 | -0.092149152 | 0.453281619 | 8.580327455 | 1.608829747 | 0.157350697 | 0.569106866 | -4.375674509 |
| **A_52_P355075** | Rab40c | -0.213743368 | -0.536588462 | 0.109101727 | 10.60742663 | -1.608724385 | 0.157373698 | 0.569106866 | -4.375728215 |
| **A_55_P2020661** | Orai3 | -0.136499115 | -0.342696989 | 0.06969876 | 8.55318822 | -1.608528792 | 0.157416404 | 0.56915482 | -4.375827915 |
| **A_55_P2165869** | Cebpb | -0.368458113 | -0.925160194 | 0.188243969 | 12.15548334 | -1.608231139 | 0.157481415 | 0.569235022 | -4.375979636 |
| **A_55_P1980262** | Rtl8b | -0.840479258 | -2.110414536 | 0.42945602 | 10.9641992 | -1.608157577 | 0.157497486 | 0.569235022 | -4.376017133 |

| | | | | | | | | | |
|---|---|---|---|---|---|---|---|---|---|
| **A_55_P2030486** | Srsf2 | 0.265701326 | -0.135848625 | 0.667251276 | 9.989684653 | 1.607819656 | 0.15757133 | 0.569395445 | -4.376189378 |
| **A_51_P454008** | Lbp | -0.445573282 | -1.119237455 | 0.228090891 | 11.7415954 | -1.607159129 | 0.157715767 | 0.569721298 | -4.376526057 |
| **A_55_P2159480** | Fam160a1 | -0.240498089 | -0.604113037 | 0.12311686 | 8.220027317 | -1.607137701 | 0.157720454 | 0.569721298 | -4.376536979 |
| **A_55_P1968703** | Gfra2 | -0.523013082 | -1.313961341 | 0.267935178 | 10.62665362 | -1.606747623 | 0.157805814 | 0.569731139 | -4.376735804 |
| **A_55_P2030284** | Trem1 | 0.113867172 | -0.05834141 | 0.286075754 | 8.936828362 | 1.606670923 | 0.157822603 | 0.569731139 | -4.376774898 |
| **A_52_P312371** | Zfp345 | 0.136468385 | -0.069928979 | 0.342865749 | 7.97833099 | 1.606612323 | 0.157835431 | 0.569731139 | -4.376804766 |
| **A_55_P2372228** | NA | 0.359623748 | -0.184313665 | 0.90356116 | 9.896103216 | 1.606506993 | 0.157858492 | 0.569731139 | -4.376858453 |
| **A_52_P35057** | Kctd17 | -0.251709311 | -0.632437246 | 0.129018625 | 9.591084716 | -1.606451896 | 0.157870556 | 0.569731139 | -4.376886536 |
| **A_66_P104221** | NA | 0.155398922 | -0.079692812 | 0.390490656 | 8.314540655 | 1.606178582 | 0.157930414 | 0.569762179 | -4.377025842 |
| **A_51_P314763** | Rabggtb | 0.204556572 | -0.104908969 | 0.514022114 | 10.18138283 | 1.606143442 | 0.157938111 | 0.569762179 | -4.377043753 |
| **A_55_P2039454** | Gm5088 | 0.431448308 | -0.221537721 | 1.084434337 | 8.940804458 | 1.605491719 | 0.158080937 | 0.569832196 | -4.377375927 |
| **A_55_P2033864** | Plekha8 | 0.158777581 | -0.08157055 | 0.399125712 | 9.51296529 | 1.605209132 | 0.158142904 | 0.569832196 | -4.377519956 |
| **A_55_P2060349** | Lzts2 | -0.511894282 | -1.286833814 | 0.26304525 | 10.2945207 | -1.605076186 | 0.158172065 | 0.569832196 | -4.377587715 |
| **A_55_P2151026** | Ccdc50 | 0.134679483 | -0.069211618 | 0.338570584 | 8.284890823 | 1.605041841 | 0.1581796 | 0.569832196 | -4.37760522 |
| **A_55_P1968231** | Rbpms | -0.162602163 | -0.408811055 | 0.08360673 | 8.255431854 | -1.604744041 | 0.158244942 | 0.569832196 | -4.377757001 |
| **A_55_P1975305** | NA | 0.162425609 | -0.083516499 | 0.408367718 | 11.55657028 | 1.604740458 | 0.158245728 | 0.569832196 | -4.377758827 |
| **A_55_P1962314** | NA | 0.15330616 | -0.078830109 | 0.385442429 | 16.12801573 | 1.604721958 | 0.158249789 | 0.569832196 | -4.377768256 |
| **A_51_P203710** | Taf8 | 0.133329135 | -0.068563558 | 0.335221828 | 10.57780439 | 1.604677084 | 0.158259637 | 0.569832196 | -4.377791127 |
| **A_55_P2001218** | Cd80 | 0.193286306 | -0.099420896 | 0.485993507 | 9.146090667 | 1.604541176 | 0.158289469 | 0.569832196 | -4.377860395 |
| **A_51_P320539** | Otulin | 0.143556413 | -0.07387597 | 0.360988796 | 10.8235816 | 1.604285143 | 0.158345683 | 0.569832196 | -4.377990885 |
| **A_55_P1958532** | Hr | -0.190906211 | -0.480056909 | 0.098244487 | 7.80654531 | -1.604275692 | 0.158347758 | 0.569832196 | -4.377995702 |
| **A_55_P1967613** | NA | 0.275233895 | -0.141656881 | 0.69212467 | 8.943880583 | 1.604215908 | 0.158360887 | 0.569832196 | -4.378026171 |
| **A_55_P2086811** | Sp140 | -0.166665237 | -0.419116393 | 0.08578592 | 9.356828243 | -1.604171692 | 0.158370598 | 0.569832196 | -4.378048707 |
| **A_55_P2055324** | Spint1 | -0.400532406 | -1.007231171 | 0.20616636 | 7.964923589 | -1.604160199 | 0.158373122 | 0.569832196 | -4.378054564 |
| **A_52_P238468** | Gipr | -0.137774464 | -0.346481448 | 0.070932519 | 8.218210413 | -1.604039057 | 0.15839973 | 0.569832196 | -4.378116306 |
| **A_51_P383629** | Vps4a | 0.124700032 | -0.064241352 | 0.313641415 | 8.45445768 | 1.603698276 | 0.158474605 | 0.569995465 | -4.378289986 |
| **A_55_P2095384** | Glrx2 | 0.149908835 | -0.077285068 | 0.377102737 | 11.53784118 | 1.603296137 | 0.158563003 | 0.570140957 | -4.378494936 |
| **A_51_P261351** | Mrpl49 | 0.158148838 | -0.081540687 | 0.397838363 | 10.7492871 | 1.603245905 | 0.158574049 | 0.570140957 | -4.378520537 |
| **A_52_P364130** | Map3k14 | -0.181946426 | -0.45773966 | 0.093846808 | 8.302283833 | -1.603035388 | 0.158620346 | 0.570201353 | -4.378627825 |
| **A_55_P1964638** | Cxadr | 0.220214829 | -0.113647487 | 0.554077146 | 12.68923388 | 1.602737315 | 0.158685922 | 0.570331011 | -4.378779734 |
| **A_55_P2279977** | NA | 0.425959261 | -0.220027451 | 1.071945973 | 9.327829082 | 1.602240367 | 0.158795306 | 0.570506172 | -4.379032995 |
| **A_51_P138923** | Peg12 | 0.180240797 | -0.093158778 | 0.453640373 | 9.052034799 | 1.601911257 | 0.158867787 | 0.570506172 | -4.379200718 |
| **A_55_P2074836** | Mapk8ip1 | -0.316785662 | -0.797320242 | 0.163748918 | 9.993613459 | -1.601858168 | 0.158879482 | 0.570506172 | -4.379227773 |
| **A_55_P1966337** | Polr2h | 0.167083692 | -0.0864265 | 0.420593884 | 11.00209225 | 1.601481137 | 0.158962561 | 0.570506172 | -4.379419915 |
| **A_51_P141772** | Uhrf2 | 0.197137591 | -0.101996492 | 0.496271675 | 11.39332529 | 1.601351736 | 0.158991084 | 0.570506172 | -4.37948586 |
| **A_55_P2118674** | Tnip1 | -0.306940235 | -0.772695219 | 0.158814749 | 11.21051483 | -1.601325056 | 0.158996965 | 0.570506172 | -4.379499456 |
| **A_55_P2066727** | Serpinb6a | -0.221785549 | -0.558326674 | 0.114755576 | 12.14655325 | -1.601320605 | 0.158997946 | 0.570506172 | -4.379501724 |
| **A_52_P509744** | Nup160 | 0.275991285 | -0.142839535 | 0.694822106 | 9.729019507 | 1.601179146 | 0.159029134 | 0.570506172 | -4.379573814 |
| **A_52_P493091** | Fmr1 | 0.111272882 | -0.057590272 | 0.280136036 | 8.396781161 | 1.601170685 | 0.159031 | 0.570506172 | -4.379578126 |
| **A_51_P251717** | Hikeshi | 0.129349944 | -0.066949189 | 0.325649078 | 9.921334154 | 1.601146438 | 0.159036346 | 0.570506172 | -4.379590482 |
| **A_52_P301374** | Aaas | 0.195978872 | -0.101465478 | 0.493423221 | 9.464661729 | 1.600982996 | 0.159072389 | 0.570506172 | -4.379673773 |
| **A_52_P579640** | Rrs1 | 0.268076602 | -0.138812898 | 0.674966102 | 10.58334118 | 1.600905203 | 0.159089547 | 0.570506172 | -4.379713417 |
| **A_55_P2003338** | Clint1 | 0.274706789 | -0.1422801 | 0.691693679 | 11.10897465 | 1.600774588 | 0.15911836 | 0.570506172 | -4.379779979 |
| **A_55_P2042184** | NA | 0.144079841 | -0.074665809 | 0.362825491 | 8.234518669 | 1.600467963 | 0.159186018 | 0.570603159 | -4.379936235 |
| **A_55_P1978226** | Prkn | 0.537199455 | -0.278468642 | 1.352867553 | 9.889914415 | 1.600314264 | 0.159219942 | 0.570603159 | -4.380014559 |
| **A_55_P2107357** | Gm15421 | 0.119067274 | -0.061748387 | 0.299882935 | 14.94507647 | 1.600072108 | 0.159273405 | 0.570603159 | -4.38013796 |
| **A_55_P2012389** | Sfxn3 | -0.386653153 | -0.973854185 | 0.20054788 | 9.25997578 | -1.599992454 | 0.159290995 | 0.570603159 | -4.380178551 |
| **A_55_P2034042** | Gga3 | 0.175722972 | -0.091251784 | 0.442697727 | 10.94006474 | 1.599342694 | 0.159434547 | 0.570603159 | -4.380509658 |
| **A_51_P221632** | 1700037H04Rik | 0.217960356 | -0.113228176 | 0.549148888 | 13.41177868 | 1.599136028 | 0.159480231 | 0.570603159 | -4.38061497 |
| **A_55_P1970636** | Unc45a | -0.253896932 | -0.639710221 | 0.131916356 | 10.61323019 | -1.599055216 | 0.159498098 | 0.570603159 | -4.380656149 |

| | | | | | | | | | |
|---|---|---|---|---|---|---|---|---|---|
| **A_51_P370163** | Ctr9 | 0.163033467 | -0.084709806 | 0.410776741 | 9.427726593 | 1.599035414 | 0.159502477 | 0.570603159 | -4.38066624 |
| **A_55_P2134800** | Cinp | 0.241601914 | -0.125546933 | 0.608750761 | 9.781922412 | 1.59897396 | 0.159516066 | 0.570603159 | -4.380697555 |
| **A_55_P2174198** | H4c9 | 0.325856959 | -0.169366368 | 0.821080286 | 11.64726396 | 1.598855062 | 0.15954236 | 0.570603159 | -4.380758142 |
| **A_55_P2091120** | Kbtbd12 | 0.134398536 | -0.069860552 | 0.338657623 | 7.912000874 | 1.598808093 | 0.159552748 | 0.570603159 | -4.380782076 |
| **A_55_P2174501** | Stx17 | 0.136527436 | -0.070977216 | 0.344032088 | 8.550648174 | 1.598730596 | 0.15956989 | 0.570603159 | -4.380821566 |
| **A_55_P2141943** | NA | 0.347195681 | -0.180509452 | 0.874900814 | 10.26842678 | 1.59869709 | 0.159577302 | 0.570603159 | -4.380838639 |
| **A_55_P1963124** | NA | 0.205270413 | -0.106730506 | 0.517271333 | 14.90638503 | 1.59865103 | 0.159587491 | 0.570603159 | -4.38086211 |
| **A_52_P14453** | Atxn1 | -0.150556534 | -0.379433062 | 0.078319995 | 8.321620911 | -1.59838553 | 0.159646237 | 0.570603159 | -4.380997399 |
| **A_51_P124254** | Col4a1 | -0.851403339 | -2.145926669 | 0.443119991 | 10.80592566 | -1.598117313 | 0.159705605 | 0.570603159 | -4.385969181 |
| **A_51_P134812** | Chac1 | 0.526953823 | -0.274384886 | 1.328292531 | 8.478396744 | 1.597863318 | 0.159761844 | 0.570603159 | -4.381263495 |
| **A_51_P149325** | Psmd8 | -0.153790728 | -0.387692752 | 0.080111295 | 11.55695752 | -1.597641616 | 0.159810948 | 0.570603159 | -4.381376463 |
| **A_52_P1197913** | Gadd45b | -0.692182905 | -1.745008414 | 0.360642603 | 9.666653821 | -1.597524987 | 0.159836786 | 0.570603159 | -4.381435891 |
| **A_55_P2255399** | F630048H11Rik | 0.106788875 | -0.055640026 | 0.269217776 | 8.166339971 | 1.597518491 | 0.159838225 | 0.570603159 | -4.381439201 |
| **A_55_P2129685** | Pkig | 0.113822805 | -0.059325904 | 0.286971513 | 8.431480463 | 1.597324726 | 0.15988116 | 0.570603159 | -4.381537932 |
| **A_55_P2007919** | Akr1c19 | 0.486091445 | -0.253473764 | 1.225656654 | 11.70345038 | 1.597072729 | 0.159937015 | 0.570603159 | -4.381666335 |
| **A_66_P128963** | Cyp2d9 | -0.315448983 | -0.795407109 | 0.164509143 | 8.513081968 | -1.597014913 | 0.159949832 | 0.570603159 | -4.381695794 |
| **A_55_P2136657** | Thbs3 | -0.544078545 | -1.371916246 | 0.283759156 | 8.570327393 | -1.59698041 | 0.159957482 | 0.570603159 | -4.381713374 |
| **A_55_P2172121** | Mxd4 | -0.253100523 | -0.638264298 | 0.132063252 | 8.275079721 | -1.596727469 | 0.160013571 | 0.570603159 | -4.381842256 |
| **A_55_P1956048** | H2-T3 | 0.299029493 | -0.156029906 | 0.754088891 | 8.317318313 | 1.596721321 | 0.160014935 | 0.570603159 | -4.381845389 |
| **A_55_P2005585** | Trps1 | -0.287573165 | -0.7252506 | 0.150104271 | 8.229562821 | -1.596531123 | 0.160057124 | 0.570603159 | -4.3819423 |
| **A_55_P2168899** | NA | 0.143658192 | -0.075001675 | 0.362318058 | 8.36163875 | 1.596410245 | 0.160083942 | 0.570603159 | -4.38200389 |
| **A_55_P2001948** | Cab39l | 0.150229517 | -0.078453282 | 0.378912316 | 11.47585994 | 1.596264889 | 0.160116196 | 0.570603159 | -4.382077952 |
| **A_55_P2240823** | Epb41 | 0.284463588 | -0.148645136 | 0.717572313 | 8.643351247 | 1.595926701 | 0.160191264 | 0.570603159 | -4.382250265 |
| **A_55_P1989748** | NA | 0.130510885 | -0.068202094 | 0.329223865 | 8.400039198 | 1.595892709 | 0.160198811 | 0.570603159 | -4.382267585 |
| **A_52_P536735** | Hpcal1 | -0.259974533 | -0.655873226 | 0.13592416 | 8.579298847 | -1.595621676 | 0.160258999 | 0.570603159 | -4.382405679 |
| **A_55_P2128153** | Col6a1 | -0.976352415 | -2.463201713 | 0.510496882 | 11.12614142 | -1.595595644 | 0.160264781 | 0.570603159 | -4.382418943 |
| **A_55_P1962304** | Plac8 | -0.192156332 | -0.484789308 | 0.100476644 | 9.033171602 | -1.595565455 | 0.160271486 | 0.570603159 | -4.382434324 |
| **A_55_P2125456** | NA | 0.285955076 | -0.149583768 | 0.72149392 | 14.51530486 | 1.595343124 | 0.160320879 | 0.570603159 | -4.382547603 |
| **A_55_P2053988** | Epas1 | -0.144245611 | -0.363972506 | 0.075481284 | 8.464401838 | -1.595153859 | 0.160362936 | 0.570603159 | -4.382644034 |
| **A_51_P413866** | Cfb | -0.33456047 | -0.84420714 | 0.1750862 | 14.91525116 | -1.595102604 | 0.160374328 | 0.570603159 | -4.382670149 |
| **A_55_P2018706** | Cep19 | -0.262164952 | -0.661564332 | 0.137234428 | 10.0669498 | -1.594962345 | 0.160405505 | 0.570603159 | -4.38274161 |
| **A_55_P2012096** | Bmp8a | 0.149915764 | -0.078476881 | 0.37830841 | 8.296508205 | 1.594954792 | 0.160407184 | 0.570603159 | -4.382745459 |
| **A_66_P121369** | Gars | 0.136130275 | -0.071270707 | 0.343531258 | 14.32783553 | 1.59487666 | 0.160424553 | 0.570603159 | -4.382785266 |
| **A_52_P42834** | Usp25 | 0.252618409 | -0.132277378 | 0.637514196 | 9.642861764 | 1.594795595 | 0.160442577 | 0.570603159 | -4.382826568 |
| **A_51_P278266** | Gltpd2 | 0.397089274 | -0.208078934 | 1.002257482 | 12.44778475 | 1.594392493 | 0.16053223 | 0.570603159 | -4.383031945 |
| **A_51_P481482** | Dram1 | -0.574065558 | -1.448949881 | 0.300818764 | 8.178589964 | -1.594388108 | 0.160533206 | 0.570603159 | -4.383034179 |
| **A_55_P2027436** | Bri3 | -0.494574533 | -1.248431289 | 0.259282223 | 13.57251226 | -1.594138708 | 0.160588698 | 0.570603159 | -4.383161244 |
| **A_55_P1961690** | Tctex1d2 | -0.273768461 | -0.691081712 | 0.14354479 | 9.499990595 | -1.594059134 | 0.160606408 | 0.570603159 | -4.383201786 |
| **A_55_P2000390** | Arpc5 | -0.146725006 | -0.370393501 | 0.076943489 | 13.68069271 | -1.593978681 | 0.160624315 | 0.570603159 | -4.383242774 |
| **A_66_P129508** | Hyls1 | 0.113952894 | -0.059781905 | 0.287687693 | 8.246728469 | 1.593755627 | 0.160673971 | 0.570603159 | -4.383356415 |
| **A_55_P2008609** | Rxrb | -0.190941161 | -0.48206468 | 0.100182359 | 11.93145569 | -1.593695896 | 0.160687271 | 0.570603159 | -4.383386847 |
| **A_55_P1970910** | Stard13 | -0.394385025 | -0.995714425 | 0.206944375 | 8.772654007 | -1.59364347 | 0.160698945 | 0.570603159 | -4.383413556 |
| **A_55_P1994898** | NA | -0.151195832 | -0.381746607 | 0.079354944 | 13.42988148 | -1.593515971 | 0.160727339 | 0.570603159 | -4.383478513 |
| **A_55_P2055742** | Adtrp | -0.252159634 | -0.636671884 | 0.132352615 | 8.205283378 | -1.593487185 | 0.160733751 | 0.570603159 | -4.383493178 |
| **A_55_P1960472** | Polg | -0.144047929 | -0.363709515 | 0.075613656 | 8.983752755 | -1.593441393 | 0.16074395 | 0.570603159 | -4.383516508 |
| **A_55_P2431118** | Itsn1 | 0.263644261 | -0.138401643 | 0.665690164 | 11.0826592 | 1.593403877 | 0.160752307 | 0.570603159 | -4.383535621 |
| **A_55_P2225087** | 5430439M09Rik | 0.149637216 | -0.07856904 | 0.377843472 | 9.081130456 | 1.593291585 | 0.160777323 | 0.570603159 | -4.38359283 |
| **A_55_P2102569** | Cox7a2l | -0.380379287 | -0.960485428 | 0.199726854 | 11.36057825 | -1.593281768 | 0.16077951 | 0.570603159 | -4.383597831 |
| **A_51_P193446** | Prpf40a | 0.182920207 | -0.096064797 | 0.461905211 | 9.380809826 | 1.593176944 | 0.160802865 | 0.570603159 | -4.383651235 |
| **A_55_P2130178** | Fn1 | -0.28239807 | -0.713134644 | 0.148338504 | 16.28221753 | -1.59306378 | 0.160828083 | 0.570603159 | -4.383708888 |

| | | | | | | | | | |
|---|---|---|---|---|---|---|---|---|---|
| **A_51_P472481** | Dync2li1 | -0.302799372 | -0.764749176 | 0.159150433 | 8.090757564 | -1.592734438 | 0.160901494 | 0.57074531 | -4.383876673 |
| **A_55_P2037613** | Elmsan1 | 0.174527788 | -0.091792233 | 0.440847809 | 9.72258112 | 1.592369889 | 0.16098279 | 0.57074531 | -4.384062393 |
| **A_52_P327402** | Cds1 | -0.187387529 | -0.473334098 | 0.09855904 | 7.775501184 | -1.592351656 | 0.160986857 | 0.57074531 | -4.384071682 |
| **A_66_P126504** | Trabd2b | -0.381739676 | -0.964297331 | 0.200817979 | 8.178657913 | -1.592251167 | 0.161009274 | 0.57074531 | -4.384122875 |
| **A_55_P2005719** | NA | -0.173814907 | -0.439077713 | 0.091447899 | 13.00778447 | -1.592186177 | 0.161023773 | 0.57074531 | -4.384155984 |
| **A_55_P2060680** | Acsf2 | 0.16952125 | -0.089292609 | 0.42833511 | 11.24082163 | 1.591548241 | 0.161166164 | 0.57074531 | -4.384480972 |
| **A_55_P2021854** | NA | 0.134878168 | -0.071086138 | 0.340842473 | 8.504182438 | 1.591229732 | 0.161237301 | 0.57074531 | -4.384643229 |
| **A_52_P920129** | Gnai1 | -0.321258043 | -0.811857941 | 0.169341855 | 8.417013523 | -1.591144948 | 0.161256242 | 0.57074531 | -4.384686419 |
| **A_66_P117477** | Dlk1 | 0.1883197 | -0.099337926 | 0.475977326 | 15.97002439 | 1.590754102 | 0.161343586 | 0.57074531 | -4.384885523 |
| **A_55_P2004046** | NA | 0.285157191 | -0.150439314 | 0.720753695 | 9.300688066 | 1.590681134 | 0.161359898 | 0.57074531 | -4.384922694 |
| **A_55_P2145879** | Tpt1 | -0.19853698 | -0.501820171 | 0.104746212 | 17.29916083 | -1.590655964 | 0.161365525 | 0.57074531 | -4.384935516 |
| **A_55_P1994032** | Xbp1 | -0.246391412 | -0.622783671 | 0.130000846 | 14.05516639 | -1.590625988 | 0.161372226 | 0.57074531 | -4.384950786 |
| **A_55_P2161702** | Nrip2 | 0.212426062 | -0.112088625 | 0.536940749 | 8.538748037 | 1.590584073 | 0.161381597 | 0.57074531 | -4.384972138 |
| **A_55_P2011678** | Pdzk1ip1 | -0.252918834 | -0.639295069 | 0.133457401 | 12.27252673 | -1.590574272 | 0.161383789 | 0.57074531 | -4.384977131 |
| **A_55_P2083654** | Prmt2 | -0.576226581 | -1.456575439 | 0.304122277 | 10.02100698 | -1.590456047 | 0.161410224 | 0.57074531 | -4.385037355 |
| **A_55_P2048563** | Cdk2ap1 | 0.191315898 | -0.100998286 | 0.483630081 | 13.22426057 | 1.590319415 | 0.16144078 | 0.57074531 | -4.385106957 |
| **A_51_P502872** | 2200002D01Rik | -0.56727143 | -1.43402697 | 0.299484111 | 10.23952988 | -1.590294195 | 0.161446421 | 0.57074531 | -4.385119804 |
| **A_55_P2090070** | Myh14 | -1.419462916 | -3.588609302 | 0.74968347 | 9.604697721 | -1.590077815 | 0.161494824 | 0.57074531 | -4.385230028 |
| **A_55_P1997604** | Pla2g4a | 0.147850928 | -0.078093603 | 0.373795458 | 9.712954184 | 1.590030333 | 0.161505448 | 0.57074531 | -4.385254216 |
| **A_55_P2197638** | 1110046J04Rik | -0.19590135 | -0.495290183 | 0.103487483 | 7.991751363 | -1.589955688 | 0.16152215 | 0.57074531 | -4.38529224 |
| **A_51_P208919** | Acbd6 | 0.155876103 | -0.082367453 | 0.394119659 | 10.89308704 | 1.589796606 | 0.161557751 | 0.57074531 | -4.385373275 |
| **A_51_P506787** | Nap1l1 | 0.148756022 | -0.078610788 | 0.376122832 | 10.4991533 | 1.589756746 | 0.161566672 | 0.57074531 | -4.38539358 |
| **A_51_P387845** | Fam120a | 0.207815453 | -0.109828186 | 0.525459091 | 10.24653868 | 1.58972083 | 0.161574711 | 0.57074531 | -4.385411875 |
| **A_55_P2069979** | Large1 | -0.19610518 | -0.495855024 | 0.103644663 | 10.03933723 | -1.589693102 | 0.161580918 | 0.57074531 | -4.385426 |
| **A_55_P2140107** | Cyp21a1 | 0.125925997 | -0.066578584 | 0.318430577 | 8.176115787 | 1.589488971 | 0.161626617 | 0.57074531 | -4.385529982 |
| **A_55_P2154173** | NA | -0.293811495 | -0.743013101 | 0.15539011 | 14.78132929 | -1.589317527 | 0.161665008 | 0.57074531 | -4.385617313 |
| **A_55_P1952547** | Sh3rf1 | -0.147779597 | -0.37373529 | 0.078176096 | 8.04228209 | -1.589184716 | 0.161694754 | 0.57074531 | -4.385684964 |
| **A_52_P451834** | Stk35 | 0.228376649 | -0.12084381 | 0.577597108 | 9.429138487 | 1.589041381 | 0.161726863 | 0.57074531 | -4.385757976 |
| **A_52_P223571** | Mlec | 0.157842482 | -0.083526387 | 0.39921135 | 8.419852108 | 1.589007092 | 0.161734545 | 0.57074531 | -4.385775443 |
| **A_55_P2106844** | NA | 0.151832881 | -0.080358932 | 0.384024694 | 8.31890321 | 1.58892032 | 0.161753987 | 0.57074531 | -4.385819642 |
| **A_51_P288103** | Psmd6 | 0.121834159 | -0.064530916 | 0.308199235 | 12.07499737 | 1.588501873 | 0.161847773 | 0.570873968 | -4.386032787 |
| **A_51_P388298** | Mmachc | 0.215717368 | -0.114258895 | 0.545693631 | 9.467727689 | 1.588494088 | 0.161849519 | 0.570873968 | -4.386036753 |
| **A_55_P1974317** | Surf6 | 0.180523679 | -0.095664142 | 0.456711501 | 9.457809326 | 1.588227976 | 0.16190919 | 0.570980247 | -4.386172301 |
| **A_55_P2152467** | NA | 0.1628002 | -0.086298257 | 0.411898656 | 8.9326481 | 1.588060657 | 0.161946719 | 0.571008415 | -4.386257526 |
| **A_52_P429265** | Ppp6r2 | 0.196528549 | -0.104342294 | 0.497399393 | 8.526669576 | 1.587189324 | 0.16214229 | 0.571355923 | -4.386701338 |
| **A_52_P244064** | Ube2g2 | 0.185468491 | -0.09849084 | 0.469427821 | 11.0795035 | 1.587074021 | 0.162168186 | 0.571355923 | -4.386760067 |
| **A_52_P183038** | Impdh1 | -0.149217217 | -0.377675167 | 0.079240733 | 9.677449353 | -1.587069152 | 0.162169279 | 0.571355923 | -4.386762546 |
| **A_55_P1993933** | NA | -0.173258758 | -0.43853435 | 0.092016834 | 7.863426596 | -1.587015224 | 0.162181393 | 0.571355923 | -4.386790014 |
| **A_52_P293120** | Prrg3 | -0.358724683 | -0.908037321 | 0.190587956 | 8.172894615 | -1.586809741 | 0.162227557 | 0.571355923 | -4.386894673 |
| **A_55_P2097448** | Rbpms | -0.135630823 | -0.343328709 | 0.072067063 | 9.323110701 | -1.586753663 | 0.162240157 | 0.571355923 | -4.386923236 |
| **A_55_P2130930** | Nhp2 | 0.150439962 | -0.079945501 | 0.380825425 | 13.89151716 | 1.586687253 | 0.162255081 | 0.571355923 | -4.38695706 |
| **A_55_P1968464** | Kat6a | 0.171385871 | -0.091096027 | 0.433867769 | 10.29495447 | 1.58656857 | 0.162281754 | 0.571355923 | -4.387017509 |
| **A_55_P1982156** | Nelfcd | 0.209415524 | -0.11134961 | 0.530180659 | 13.65599758 | 1.586371525 | 0.162326047 | 0.571407789 | -4.387117868 |
| **A_55_P1977468** | Dab2ip | -0.455202349 | -1.152770224 | 0.242365525 | 11.62440684 | -1.585627657 | 0.162493364 | 0.571892614 | -4.38749673 |
| **A_55_P2018994** | Elk3 | -0.506219856 | -1.297570874 | 0.285131162 | 10.56298452 | -1.627467695 | 0.162551022 | 0.57199139 | -4.394840096 |
| **A_55_P2041457** | NA | 0.187540318 | -0.099994479 | 0.475075114 | 10.98170868 | 1.58484731 | 0.162669061 | 0.572302564 | -4.387894159 |
| **A_52_P49321** | Adamts9 | -0.533170752 | -1.350723986 | 0.284382482 | 9.010018573 | -1.584650403 | 0.162713424 | 0.572317069 | -4.387994441 |
| **A_55_P2044439** | Mfng | -0.501835906 | -1.271519197 | 0.267847385 | 9.819871505 | -1.584283364 | 0.162796147 | 0.572317069 | -4.388181366 |
| **A_55_P2010539** | NA | 0.186765577 | -0.099690734 | 0.473221888 | 8.667235128 | 1.584242383 | 0.162805386 | 0.572317069 | -4.388202237 |
| **A_51_P312748** | Oxsm | 0.212744484 | -0.113569279 | 0.539058248 | 9.179863857 | 1.584185762 | 0.162818151 | 0.572317069 | -4.388231072 |

| | | | | | | | | |
|---|---|---|---|---|---|---|---|---|
| **A_51_P337662** | Ints6l | 0.289530252 | -0.154650739 | 0.733711243 | 10.7700472 | 1.583861376 | 0.162891304 | 0.572317069 | -4.388396272 |
| **A_55_P2067747** | Psg21 | -0.240353267 | -0.609109214 | 0.12840268 | 7.867382176 | -1.583777559 | 0.162910211 | 0.572317069 | -4.388438958 |
| **A_55_P1980951** | 1110012L19Rik | -0.249157743 | -0.631428679 | 0.133113194 | 10.87152061 | -1.58374879 | 0.1629167 | 0.572317069 | -4.388453608 |
| **A_52_P167189** | Isca1 | 0.199938578 | -0.106820422 | 0.506697577 | 12.30609158 | 1.583734855 | 0.162919844 | 0.572317069 | -4.388460705 |
| **A_51_P513085** | Gins4 | 0.243606922 | -0.13021102 | 0.617424863 | 10.8869154 | 1.583480325 | 0.162977275 | 0.572317069 | -4.388590327 |
| **A_51_P215887** | Aldh18a1 | 0.230752753 | -0.123378718 | 0.584884225 | 8.826383011 | 1.583308551 | 0.163016043 | 0.572317069 | -4.388677804 |
| **A_52_P141628** | Slc23a1 | 0.284972768 | -0.152385155 | 0.722330691 | 8.64147002 | 1.583250201 | 0.163029215 | 0.572317069 | -4.388707519 |
| **A_55_P2396375** | 1810073O08Rik | 0.25275 | -0.135162559 | 0.640662558 | 8.79985488 | 1.583217247 | 0.163036654 | 0.572317069 | -4.388724301 |
| **A_52_P509185** | Zfp318 | -0.134362893 | -0.340591109 | 0.071865324 | 8.760268942 | -1.583122234 | 0.163058104 | 0.572317069 | -4.388772686 |
| **A_55_P2140546** | Slc25a51 | 0.188523608 | -0.100938969 | 0.477986184 | 10.92606233 | 1.582546601 | 0.163188118 | 0.572506664 | -4.389065823 |
| **A_55_P1977917** | Cln3 | -0.167127054 | -0.423744483 | 0.089490375 | 10.04735265 | -1.582500289 | 0.163198582 | 0.572506664 | -4.389089407 |
| **A_55_P2023783** | Leng8 | 0.224688353 | -0.12031464 | 0.569691345 | 9.124038994 | 1.582489684 | 0.163200979 | 0.572506664 | -4.389094808 |
| **A_51_P290819** | Hint3 | -0.108672636 | -0.275558095 | 0.058212823 | 11.42419469 | -1.582285668 | 0.163247085 | 0.572511678 | -4.3891987 |
| **A_55_P2002963** | Coro1a | -0.23027404 | -0.583945994 | 0.123397914 | 12.89709967 | -1.582076746 | 0.163294313 | 0.572511678 | -4.389305088 |
| **A_55_P2141301** | Synj2bp | 0.160458686 | -0.086003056 | 0.406920428 | 10.68642838 | 1.581965133 | 0.163319549 | 0.572511678 | -4.389361925 |
| **A_55_P2043367** | Apol9a | -0.119928074 | -0.304136192 | 0.064280043 | 8.126294851 | -1.581959222 | 0.163320885 | 0.572511678 | -4.389364934 |
| **A_55_P2179106** | Ppig | 0.161985269 | -0.086941731 | 0.410912269 | 9.125574183 | 1.581199634 | 0.163492729 | 0.572958689 | -4.389751728 |
| **A_55_P2085536** | Smarcal1 | 0.104787728 | -0.056256121 | 0.265831576 | 10.31596145 | 1.581064076 | 0.163523415 | 0.572958689 | -4.389820755 |
| **A_55_P2240116** | 2310005E17Rik | 0.369722273 | -0.198510421 | 0.937954968 | 8.338753731 | 1.581002605 | 0.163537332 | 0.572958689 | -4.389852056 |
| **A_52_P659762** | Eid2 | -0.122000856 | -0.309522118 | 0.065520407 | 8.094394067 | -1.580867781 | 0.163567859 | 0.572961788 | -4.389920708 |
| **A_55_P1963201** | Chchd7 | 0.115497949 | -0.062062414 | 0.293058313 | 12.96819029 | 1.58056164 | 0.163637197 | 0.5730448 | -4.390076594 |
| **A_52_P473953** | Ctdspl | -0.32328684 | -0.820309673 | 0.173735992 | 9.37346765 | -1.580501368 | 0.163650851 | 0.5730448 | -4.390107283 |
| **A_55_P2141938** | 1810058I24Rik | 0.215018412 | -0.115637829 | 0.545674653 | 9.523631439 | 1.580091057 | 0.163743833 | 0.573071002 | -4.390316207 |
| **A_52_P121960** | NA | 0.1729677 | -0.093041999 | 0.438977399 | 15.07423678 | 1.579976868 | 0.163769718 | 0.573071002 | -4.39037435 |
| **A_55_P1960735** | Gdf15 | 0.368159846 | -0.198069333 | 0.934389025 | 9.579372269 | 1.579891864 | 0.16378899 | 0.573071002 | -4.390417632 |
| **A_51_P259555** | Gpatch3 | 0.163359483 | -0.087904702 | 0.414623668 | 8.594946621 | 1.579781235 | 0.163814076 | 0.573071002 | -4.390473962 |
| **A_55_P2096422** | Inhbb | -0.248691159 | -0.631210115 | 0.133827797 | 8.500953519 | -1.579758038 | 0.163819336 | 0.573071002 | -4.390485773 |
| **A_52_P403527** | NA | 0.258130034 | -0.13892584 | 0.655185908 | 11.16335748 | 1.579683573 | 0.163836223 | 0.573071002 | -4.390523688 |
| **A_55_P2156229** | Cd3e | 0.369230549 | -0.198772498 | 0.937233596 | 9.93936841 | 1.579538258 | 0.163869183 | 0.573082583 | -4.390597678 |
| **A_55_P2051429** | NA | 0.211584341 | -0.113932293 | 0.537100975 | 8.464863075 | 1.579405066 | 0.163899399 | 0.573084565 | -4.390665495 |
| **A_55_P2093301** | NA | 0.108840531 | -0.058643937 | 0.276324998 | 7.907429321 | 1.579062435 | 0.163977151 | 0.573252732 | -4.390839949 |
| **A_65_P01834** | Lima1 | -0.159934375 | -0.406126296 | 0.086257546 | 8.509851593 | -1.578524069 | 0.164099392 | 0.573422125 | -4.391114057 |
| **A_52_P657435** | Psmc5 | 0.164754531 | -0.088879789 | 0.418388852 | 12.43023207 | 1.578383518 | 0.164131319 | 0.573422125 | -4.391185617 |
| **A_55_P2062797** | 4921524J17Rik | 0.178611059 | -0.096360666 | 0.453582785 | 8.758968714 | 1.57835066 | 0.164138784 | 0.573422125 | -4.391202347 |
| **A_55_P2000047** | 9930104L06Rik | 0.122267742 | -0.065966327 | 0.310501812 | 8.376612751 | 1.578326509 | 0.164144271 | 0.573422125 | -4.391214643 |
| **A_51_P252410** | Cope | -0.22119266 | -0.561796045 | 0.119410725 | 12.1200668 | -1.577992521 | 0.164220169 | 0.573583601 | -4.391384688 |
| **A_55_P2096204** | Kdm1b | 0.187210299 | -0.101141431 | 0.475562028 | 9.92060495 | 1.577576271 | 0.164314807 | 0.573766625 | -4.391596611 |
| **A_51_P187716** | Zc3h11a | -0.18471768 | -0.469253415 | 0.099818054 | 12.3213354 | -1.577447227 | 0.164344156 | 0.573766625 | -4.391662309 |
| **A_55_P1979708** | Ubap1 | 0.279580923 | -0.15120834 | 0.710370187 | 10.92007561 | 1.576978794 | 0.164450738 | 0.573766625 | -4.391900793 |
| **A_55_P2017112** | Mpnd | -0.197982599 | -0.503103784 | 0.107138586 | 10.35899026 | -1.576659266 | 0.164523477 | 0.573766625 | -4.392063466 |
| **A_55_P2053459** | Timd2 | 0.478780269 | -0.25913306 | 1.216693598 | 12.34506808 | 1.576572978 | 0.164543125 | 0.573766625 | -4.392107395 |
| **A_51_P196925** | Cx3cl1 | -0.28650265 | -0.72808873 | 0.155083429 | 8.322875593 | -1.576509017 | 0.164557691 | 0.573766625 | -4.392139957 |
| **A_55_P2146560** | H2-Ab1 | -0.205029553 | -0.521048651 | 0.110989545 | 10.40242396 | -1.576472226 | 0.16456607 | 0.573766625 | -4.392158687 |
| **A_52_P425860** | Iqsec1 | -0.18898164 | -0.480276211 | 0.10231293 | 9.770973607 | -1.576414472 | 0.164579224 | 0.573766625 | -4.392188089 |
| **A_55_P2017764** | Cep89 | 0.192449878 | -0.104227388 | 0.489127143 | 10.54228985 | 1.576219012 | 0.164623749 | 0.573766625 | -4.392287595 |
| **A_52_P219266** | Frmd8 | -0.263560139 | -0.669866438 | 0.14274616 | 11.03955417 | -1.576192881 | 0.164629702 | 0.573766625 | -4.392300898 |
| **A_51_P298666** | Scaf4 | 0.263887233 | -0.142930578 | 0.670705043 | 9.8545939 | 1.576164746 | 0.164636112 | 0.573766625 | -4.392315222 |
| **A_52_P289893** | Prdm4 | 0.120470183 | -0.065253116 | 0.306193483 | 9.023889224 | 1.576145729 | 0.164640445 | 0.573766625 | -4.392324903 |
| **A_51_P343252** | Cd151 | -0.36411637 | -0.925527138 | 0.197294399 | 11.73997981 | -1.575950798 | 0.164684865 | 0.573766625 | -4.392424139 |
| **A_51_P319460** | Osmr | -0.317432024 | -0.80688688 | 0.172022832 | 8.788330934 | -1.575873256 | 0.164702538 | 0.573766625 | -4.392463614 |

| | | | | | | | | | |
|---|---|---|---|---|---|---|---|---|---|
| **A_55_P1961391** | NA | 0.118025853 | -0.06397947 | 0.300031177 | 8.035147138 | 1.57570984 | 0.164739789 | 0.573766625 | -4.392546805 |
| **A_52_P420504** | Acta2 | -0.919652746 | -2.337861488 | 0.498555996 | 10.73095562 | -1.575675938 | 0.164747518 | 0.573766625 | -4.398140926 |
| **A_55_P2098305** | Coq5 | 0.164914463 | -0.089486491 | 0.419315416 | 8.267801774 | 1.575154648 | 0.164866405 | 0.573945804 | -4.392829435 |
| **A_51_P498403** | Cnpy3 | -0.116250396 | -0.295605617 | 0.063104825 | 9.934357106 | -1.574938521 | 0.164915719 | 0.573945804 | -4.392939456 |
| **A_55_P2184526** | NA | 0.162213031 | -0.088117291 | 0.412543354 | 8.233691684 | 1.574546426 | 0.16500522 | 0.573945804 | -4.393139052 |
| **A_51_P201971** | Kmt5a | 0.238418445 | -0.129525035 | 0.606361924 | 10.8433696 | 1.574497332 | 0.165016429 | 0.573945804 | -4.393164043 |
| **A_52_P470316** | Laptm4b | -0.233793322 | -0.594600377 | 0.127013733 | 9.179534832 | -1.574491319 | 0.165017802 | 0.573945804 | -4.393167103 |
| **A_55_P2024307** | Spin1 | -0.242729238 | -0.617334846 | 0.13187637 | 11.68761673 | -1.574457736 | 0.165025471 | 0.573945804 | -4.393184199 |
| **A_55_P2326727** | 9430085M18Rik | 0.195292997 | -0.106110314 | 0.496696309 | 10.47994757 | 1.574424503 | 0.16503306 | 0.573945804 | -4.393201116 |
| **A_55_P2117487** | NA | 0.276532516 | -0.150255019 | 0.703320051 | 15.04238258 | 1.574409376 | 0.165036514 | 0.573945804 | -4.393208816 |
| **A_55_P1999902** | Pip5k1a | 0.391022753 | -0.212582532 | 0.994628039 | 15.97675746 | 1.574099512 | 0.16510729 | 0.574077492 | -4.393366548 |
| **A_55_P2020602** | Pacsin1 | 0.358760156 | -0.195165882 | 0.912686194 | 12.42446997 | 1.573749529 | 0.165187263 | 0.574077492 | -4.3935447 |
| **A_52_P78922** | Ythdc2 | 0.18111808 | -0.098531454 | 0.460767615 | 8.539080896 | 1.573732432 | 0.165191171 | 0.574077492 | -4.393553403 |
| **A_52_P361081** | Arhgef16 | -0.11539471 | -0.293567189 | 0.062777769 | 8.957665328 | -1.573723636 | 0.165193182 | 0.574077492 | -4.39355788 |
| **A_52_P229959** | 6-Sep | -0.116000463 | -0.295165564 | 0.063164638 | 8.399639748 | -1.573220119 | 0.165308307 | 0.574201716 | -4.393814179 |
| **A_55_P2012349** | Fermt2 | -0.244646647 | -0.622526612 | 0.133233318 | 10.7821356 | -1.573144406 | 0.165325625 | 0.574201716 | -4.393852718 |
| **A_55_P2023942** | Ccdc77 | 0.242059212 | -0.131906494 | 0.616024918 | 9.58832622 | 1.572798284 | 0.165404815 | 0.574201716 | -4.394028896 |
| **A_55_P1978013** | Obp1a | 0.186591814 | -0.101716164 | 0.474899792 | 8.369545126 | 1.57260306 | 0.165449496 | 0.574201716 | -4.394128264 |
| **A_55_P2179869** | NA | -0.162719587 | -0.414143501 | 0.088704327 | 12.89730433 | -1.572593363 | 0.165451716 | 0.574201716 | -4.3941332 |
| **A_55_P2066976** | Acsm1 | -0.447232808 | -1.138269529 | 0.243803913 | 11.95151592 | -1.572590753 | 0.165452313 | 0.574201716 | -4.394134528 |
| **A_55_P2021055** | Elp5 | 0.139860778 | -0.076245034 | 0.35596659 | 11.14134874 | 1.572579758 | 0.16545483 | 0.574201716 | -4.394140125 |
| **A_55_P2162234** | Tm9sf3 | -0.128085006 | -0.32600195 | 0.069831939 | 8.556181566 | -1.572528422 | 0.165466582 | 0.574201716 | -4.394166255 |
| **A_55_P2060404** | Pus3 | 0.17438453 | -0.095170418 | 0.443939479 | 9.581642247 | 1.57196849 | 0.16559481 | 0.574523196 | -4.394451252 |
| **A_55_P2175206** | Rbbp6 | -0.143462325 | -0.365246274 | 0.078321623 | 8.167148555 | -1.57177705 | 0.165638673 | 0.574523196 | -4.394548691 |
| **A_51_P391805** | Dkkl1 | 0.208801445 | -0.114001483 | 0.531604373 | 8.426760324 | 1.571734638 | 0.165648392 | 0.574523196 | -4.394570277 |
| **A_52_P93910** | Nrp2 | -0.589880051 | -1.501996109 | 0.322236008 | 10.07279851 | -1.571436354 | 0.16571676 | 0.574606618 | -4.394722095 |
| **A_55_P1972192** | Phka1 | -0.17587118 | -0.447827654 | 0.096085294 | 8.589095002 | -1.571370028 | 0.165731966 | 0.574606618 | -4.394755852 |
| **A_51_P481159** | Cbr3 | -0.54674492 | -1.392344889 | 0.298855049 | 8.788623542 | -1.571096956 | 0.165794585 | 0.574606618 | -4.394894835 |
| **A_52_P408530** | St3gal2 | 0.15729447 | -0.085982083 | 0.400571023 | 10.88769099 | 1.571073086 | 0.165800059 | 0.574606618 | -4.394906983 |
| **A_55_P1980341** | Romo1 | -0.141014069 | -0.363555051 | 0.081526912 | 13.99552198 | -1.612110526 | 0.165843222 | 0.574606618 | -4.401188433 |
| **A_51_P177518** | Emc6 | 0.141145253 | -0.077185162 | 0.359475667 | 12.34508645 | 1.570851843 | 0.165850811 | 0.574606618 | -4.395019586 |
| **A_55_P2108255** | Lgmn | 0.143691046 | -0.078666704 | 0.366048796 | 13.46580638 | 1.570220433 | 0.165995732 | 0.574887876 | -4.395340936 |
| **A_51_P169920** | Noc2l | 0.237823263 | -0.130230738 | 0.605877264 | 11.16568436 | 1.570095177 | 0.166024495 | 0.574887876 | -4.395404683 |
| **A_55_P2031966** | Fbxo39 | 0.322232496 | -0.176557016 | 0.821022008 | 8.379657901 | 1.569767059 | 0.166099864 | 0.574887876 | -4.395571671 |
| **A_55_P1973462** | Pex11b | -0.138608566 | -0.35316678 | 0.075949649 | 9.933822002 | -1.569741401 | 0.166105759 | 0.574887876 | -4.395584729 |
| **A_55_P1954925** | Endov | 0.143889168 | -0.078843965 | 0.366622301 | 15.9514258 | 1.569735439 | 0.166107129 | 0.574887876 | -4.395587763 |
| **A_55_P2172299** | Timm17b | -0.125528251 | -0.319850081 | 0.068793579 | 13.31457283 | -1.569650906 | 0.166126552 | 0.574887876 | -4.395630783 |
| **A_55_P2083464** | Lpin3 | -0.125614813 | -0.320077995 | 0.068848369 | 8.316378243 | -1.56959157 | 0.166140187 | 0.574887876 | -4.39566098 |
| **A_55_P1966089** | NA | 0.158437815 | -0.086933201 | 0.40380883 | 8.341661234 | 1.56898493 | 0.16627965 | 0.575051951 | -4.395969705 |
| **A_51_P512629** | Ttll5 | 0.134128967 | -0.073607812 | 0.341865745 | 8.69636217 | 1.568889572 | 0.166301582 | 0.575051951 | -4.396018232 |
| **A_51_P478895** | Ppp6r1 | 0.239930473 | -0.131688555 | 0.611549502 | 9.509122206 | 1.568811112 | 0.166319629 | 0.575051951 | -4.39605816 |
| **A_51_P336161** | Man2b1 | -0.223597026 | -0.569924111 | 0.122730058 | 11.94347757 | -1.568782569 | 0.166326196 | 0.575051951 | -4.396072685 |
| **A_55_P2036894** | Med29 | 0.16618946 | -0.09124562 | 0.423624539 | 10.75498264 | 1.5686243 | 0.166362608 | 0.575051951 | -4.396153227 |
| **A_51_P156263** | Zfp511 | 0.166384235 | -0.09135506 | 0.424123529 | 10.18340632 | 1.568609088 | 0.166366108 | 0.575051951 | -4.396160968 |
| **A_51_P109541** | E130308A19Rik | 0.113251441 | -0.062199833 | 0.288702716 | 9.643676613 | 1.568448996 | 0.166402949 | 0.575076453 | -4.396242437 |
| **A_55_P2031382** | Crebzf | 0.130578096 | -0.071734755 | 0.332890947 | 8.63433476 | 1.568303278 | 0.166436489 | 0.57508954 | -4.396316591 |
| **A_55_P2093595** | Mmd | -0.1905302 | -0.485869298 | 0.104808898 | 10.57409163 | -1.567566842 | 0.166606089 | 0.575572671 | -4.396691342 |
| **A_51_P458866** | F11r | -0.180751224 | -0.461023923 | 0.099521475 | 11.81190593 | -1.567052839 | 0.16672456 | 0.575872542 | -4.396952895 |
| **A_55_P2063366** | Commd7 | -0.198760075 | -0.507041125 | 0.109520975 | 11.11765113 | -1.566626653 | 0.166822851 | 0.575872542 | -4.397169757 |
| **A_55_P2049186** | Plvap | -0.221651664 | -0.565445977 | 0.12214265 | 8.328414409 | -1.566590523 | 0.166831186 | 0.575872542 | -4.397188141 |

| | | | | | | | | | |
|---|---|---|---|---|---|---|---|---|---|
| **A_55_P1958623** | Ncln | -0.143984036 | -0.367334249 | 0.079366177 | 12.23942698 | -1.566430612 | 0.166868082 | 0.575872542 | -4.397269509 |
| **A_52_P502577** | S1pr3 | -0.847647318 | -2.162539813 | 0.467245177 | 8.692494757 | -1.566419694 | 0.166870601 | 0.575872542 | -4.397275064 |
| **A_52_P299358** | Lclat1 | -0.259913421 | -0.663099028 | 0.143272186 | 8.889036365 | -1.566415151 | 0.166871649 | 0.575872542 | -4.397277376 |
| **A_51_P429276** | Tmod3 | -0.322322442 | -0.822436781 | 0.177791898 | 9.482625979 | -1.566045681 | 0.166956927 | 0.575970905 | -4.397465371 |
| **A_55_P2177614** | Srsf3 | 0.22258138 | -0.122777783 | 0.567940543 | 11.98614198 | 1.56603346 | 0.166959749 | 0.575970905 | -4.39747159 |
| **A_55_P2118268** | Chpf | -0.248994065 | -0.635426237 | 0.137438108 | 10.38942699 | -1.565665232 | 0.167044782 | 0.576092681 | -4.397658949 |
| **A_51_P184969** | Rilpl1 | -0.411868343 | -1.051094524 | 0.227357837 | 8.477208129 | -1.565622479 | 0.167054657 | 0.576092681 | -4.397680702 |
| **A_51_P319449** | Acat1 | -0.349997561 | -0.893386421 | 0.193391299 | 12.64967142 | -1.565083367 | 0.167179233 | 0.576318796 | -4.397955003 |
| **A_55_P2136533** | Kmt2d | 0.157852561 | -0.087221923 | 0.402927045 | 8.649416665 | 1.56508066 | 0.167179859 | 0.576318796 | -4.39795638 |
| **A_55_P2061428** | Fads6 | 0.491533513 | -0.271842202 | 1.254909227 | 9.550237411 | 1.564580751 | 0.167295454 | 0.576540737 | -4.398210727 |
| **A_55_P2091676** | Fas | -0.267006908 | -0.681691438 | 0.147677622 | 10.37780648 | -1.564544257 | 0.167303896 | 0.576540737 | -4.398229295 |
| **A_55_P2146525** | Camta2 | -0.121260512 | -0.309622658 | 0.067101635 | 10.77927232 | -1.564260054 | 0.167369649 | 0.576586824 | -4.398373889 |
| **A_51_P114693** | Parm1 | -0.340184248 | -0.868668889 | 0.188300393 | 8.851188756 | -1.56410181 | 0.167406271 | 0.576586824 | -4.403720767 |
| **A_55_P2049256** | Mapkapk5 | 0.138569589 | -0.076704167 | 0.353843346 | 10.66126965 | 1.564083844 | 0.16741043 | 0.576586824 | -4.398463539 |
| **A_51_P514922** | Ccar2 | -0.116638912 | -0.297855522 | 0.064577698 | 9.248190241 | -1.563970828 | 0.16743659 | 0.576586824 | -4.398521038 |
| **A_52_P267256** | LOC100861738 | -0.200845091 | -0.513030347 | 0.111340165 | 12.74365759 | -1.563262905 | 0.167600544 | 0.576960638 | -4.398881195 |
| **A_52_P65719** | Eme2 | 0.115358248 | -0.063951942 | 0.294668439 | 8.629905903 | 1.563244355 | 0.167604842 | 0.576960638 | -4.398890632 |
| **A_55_P2032608** | Dipk2a | 0.130608468 | -0.072486734 | 0.33370367 | 10.32578356 | 1.562625336 | 0.167748333 | 0.577351767 | -4.399205547 |
| **A_52_P176619** | Atat1 | -0.198431532 | -0.507095806 | 0.110232742 | 8.49692886 | -1.562095243 | 0.167871302 | 0.577491015 | -4.399475213 |
| **A_51_P255387** | Med31 | -0.214304106 | -0.547669504 | 0.119061292 | 10.62503124 | -1.562043636 | 0.167883278 | 0.577491015 | -4.399501466 |
| **A_51_P148959** | Dhx36 | 0.223181642 | -0.12399738 | 0.570360663 | 10.21117757 | 1.562025724 | 0.167887435 | 0.577491015 | -4.399510578 |
| **A_51_P477941** | Fbxw4 | 0.143073513 | -0.079502953 | 0.36564998 | 9.948189468 | 1.561935831 | 0.167908299 | 0.577491015 | -4.399556307 |
| **A_55_P2086203** | Kcnq1 | -0.386642386 | -0.988319759 | 0.215034987 | 8.288323648 | -1.561453201 | 0.168020355 | 0.577720325 | -4.399801819 |
| **A_52_P253748** | Zbtb45 | 0.166054179 | -0.09236293 | 0.424471288 | 9.202538856 | 1.561391226 | 0.168034749 | 0.577720325 | -4.399833345 |
| **A_52_P490272** | Nup98 | 0.140815963 | -0.078361854 | 0.359993779 | 8.717250687 | 1.561127888 | 0.168095925 | 0.577817816 | -4.399967301 |
| **A_55_P2142758** | Slc9a7 | -0.142599479 | -0.364625562 | 0.079426604 | 7.875178584 | -1.560619867 | 0.168214002 | 0.577817816 | -4.400225717 |
| **A_55_P2031481** | Fam131a | -0.241207792 | -0.616775514 | 0.134359931 | 9.118488246 | -1.560580806 | 0.168223085 | 0.577817816 | -4.400245586 |
| **A_55_P2165969** | NA | 0.120867766 | -0.067358415 | 0.309093947 | 8.41635515 | 1.560319922 | 0.168283754 | 0.577817816 | -4.400378287 |
| **A_52_P119350** | 4732419C18Rik | -0.119109343 | -0.304612733 | 0.066394046 | 7.998632421 | -1.560188849 | 0.168314244 | 0.577817816 | -4.400444958 |
| **A_55_P2447879** | Asxl2 | 0.24819753 | -0.138369164 | 0.634764223 | 9.235532162 | 1.560113559 | 0.168331759 | 0.577817816 | -4.400483254 |
| **A_52_P486322** | Fzr1 | 0.217562131 | -0.121310632 | 0.556434893 | 12.32863909 | 1.560018789 | 0.16835381 | 0.577817816 | -4.400531459 |
| **A_55_P2029151** | Snap47 | -0.114774315 | -0.293568473 | 0.064019843 | 8.376998531 | -1.559820327 | 0.168399995 | 0.577817816 | -4.400632405 |
| **A_66_P100853** | NA | 0.218983527 | -0.122151178 | 0.560118233 | 17.18817733 | 1.559799337 | 0.16840488 | 0.577817816 | -4.400643081 |
| **A_51_P142153** | Filip1l | -0.251308957 | -0.642810812 | 0.140192897 | 9.685968758 | -1.559758401 | 0.168414408 | 0.577817816 | -4.400663903 |
| **A_55_P1995210** | Gm11849 | 0.125905339 | -0.070244804 | 0.322055483 | 16.37970947 | 1.559691505 | 0.16842998 | 0.577817816 | -4.400697929 |
| **A_51_P436719** | Eftud2 | 0.214044088 | -0.119425818 | 0.547513994 | 9.964813099 | 1.55965944 | 0.168437444 | 0.577817816 | -4.400714238 |
| **A_55_P2148935** | NA | 0.289547972 | -0.161570925 | 0.740666869 | 11.55938062 | 1.559598096 | 0.168451725 | 0.577817816 | -4.40074544 |
| **A_52_P154833** | Ankrd49 | 0.212553837 | -0.118640958 | 0.543748633 | 9.953812674 | 1.559439875 | 0.168488565 | 0.577841637 | -4.400825916 |
| **A_55_P2016287** | Tpst1 | 0.237810741 | -0.132813152 | 0.608434634 | 8.402120182 | 1.559126102 | 0.168561645 | 0.577989717 | -4.400985509 |
| **A_55_P2031357** | Zfpm2 | -0.119425798 | -0.305584749 | 0.066733154 | 8.013418803 | -1.558825187 | 0.168631758 | 0.578126744 | -4.401138559 |
| **A_52_P423364** | Fbxl7 | -0.344908485 | -0.882617005 | 0.192800035 | 8.91064356 | -1.558619656 | 0.168679663 | 0.578126744 | -4.401243094 |
| **A_51_P497100** | Lgals4 | -1.024893861 | -2.622743413 | 0.572955691 | 9.265476233 | -1.558569581 | 0.168691336 | 0.578126744 | -4.401268563 |
| **A_55_P2053320** | Zfr2 | -0.23311767 | -0.596633486 | 0.130398145 | 8.046789418 | -1.5582426 | 0.168767579 | 0.578285507 | -4.401434865 |
| **A_55_P2053958** | Rabep2 | -0.305526177 | -0.782061253 | 0.171008899 | 10.10821497 | -1.557889827 | 0.168849873 | 0.578400555 | -4.401614281 |
| **A_55_P2016708** | Msl1 | 0.10631715 | -0.059540687 | 0.272174988 | 9.474094792 | 1.557580555 | 0.16892205 | 0.578400555 | -4.40177157 |
| **A_55_P2067453** | Tspan17 | -0.445824388 | -1.14133116 | 0.249682384 | 8.990813574 | -1.557563095 | 0.168926125 | 0.578400555 | -4.40178045 |
| **A_55_P1967069** | Lpar6 | -0.244517248 | -0.625986455 | 0.136951958 | 11.41759857 | -1.557518461 | 0.168936545 | 0.578400555 | -4.40180315 |
| **A_52_P686785** | Lyve1 | -0.220701181 | -0.565028627 | 0.123626265 | 10.87535427 | -1.557457503 | 0.168950775 | 0.578400555 | -4.401834151 |
| **A_55_P2111907** | Noxo1 | -0.33922056 | -0.868541471 | 0.190100352 | 8.062575779 | -1.557206845 | 0.169009304 | 0.578498465 | -4.401961628 |
| **A_55_P2014928** | Zfp341 | 0.181591545 | -0.101859823 | 0.465042914 | 9.465502142 | 1.556683261 | 0.169131621 | 0.578814086 | -4.402227899 |

| | | | | | | | | | |
|---|---|---|---|---|---|---|---|---|---|
| **A_66_P133520** | Glmn | 0.131394396 | -0.073719616 | 0.336508407 | 8.945915747 | 1.556555831 | 0.169161404 | 0.578814086 | -4.402292703 |
| **A_51_P279100** | Ptgs1 | -0.269134618 | -0.689348359 | 0.151079123 | 11.47081041 | -1.556261245 | 0.169230272 | 0.578849562 | -4.402442511 |
| **A_55_P2072193** | Firre | 0.275984974 | -0.154954658 | 0.706924605 | 8.364114134 | 1.556152626 | 0.169255671 | 0.578849562 | -4.402497747 |
| **A_55_P2000553** | Slc6a8 | -0.479046845 | -1.227135621 | 0.269041931 | 9.21384096 | -1.555994417 | 0.169292673 | 0.578849562 | -4.402578201 |
| **A_55_P2130219** | Tpr | 0.197191377 | -0.1107748 | 0.505157554 | 9.01112012 | 1.555851297 | 0.169326153 | 0.578849562 | -4.40265098 |
| **A_51_P521176** | Rassf7 | -0.24488314 | -0.627361685 | 0.137595405 | 11.85897066 | -1.555732755 | 0.169353888 | 0.578849562 | -4.402711261 |
| **A_55_P2064811** | Nxt1 | 0.149011055 | -0.083729926 | 0.381752037 | 12.45748101 | 1.555710624 | 0.169359066 | 0.578849562 | -4.402722516 |
| **A_55_P1968399** | Rdh7 | -0.638165139 | -1.634979062 | 0.358648783 | 13.24856783 | -1.555615173 | 0.169381402 | 0.578849562 | -4.402771054 |
| **A_55_P2007510** | Pef1 | -0.125102486 | -0.320566705 | 0.070361733 | 11.84962439 | -1.555184292 | 0.169482265 | 0.578875746 | -4.40299016 |
| **A_51_P134262** | 1700052K11Rik | 0.122535261 | -0.068946382 | 0.314016905 | 10.11507278 | 1.55495249 | 0.16953655 | 0.578875746 | -4.40310803 |
| **A_52_P164136** | Arrdc3 | -0.172971156 | -0.443303279 | 0.097360968 | 9.281516083 | -1.554745183 | 0.169585113 | 0.578875746 | -4.403213444 |
| **A_55_P2024461** | Tbkbp1 | -0.352017123 | -0.902270591 | 0.198236345 | 10.88885787 | -1.554476604 | 0.169648047 | 0.578875746 | -4.403350012 |
| **A_55_P2134251** | Hmgb1 | 0.166113008 | -0.093554917 | 0.425780932 | 12.98675951 | 1.554420532 | 0.169661189 | 0.578875746 | -4.403378523 |
| **A_55_P2157676** | Wdr34 | 0.158231365 | -0.089125258 | 0.405587988 | 15.09384235 | 1.554362177 | 0.169674867 | 0.578875746 | -4.403408195 |
| **A_51_P463765** | Timp3 | -1.050835434 | -2.6936576 | 0.591986732 | 10.48307232 | -1.554273147 | 0.169695737 | 0.578875746 | -4.403453465 |
| **A_66_P117289** | Cdk13 | 0.174421375 | -0.098260805 | 0.447103555 | 8.715139279 | 1.554268827 | 0.16969675 | 0.578875746 | -4.403455661 |
| **A_55_P2144095** | NA | -0.187438052 | -0.480506005 | 0.1056299 | 9.863209434 | -1.554077317 | 0.169741651 | 0.578875746 | -4.403553038 |
| **A_55_P2000127** | Tgfb1i1 | -0.730752144 | -1.873331868 | 0.41182758 | 10.15443802 | -1.554056393 | 0.169746558 | 0.578875746 | -4.403563677 |
| **A_55_P2110027** | NA | -0.174110882 | -0.446354957 | 0.098133193 | 13.02779505 | -1.553998761 | 0.169760073 | 0.578875746 | -4.403592981 |
| **A_51_P453088** | Coa6 | 0.247927404 | -0.13976434 | 0.635619147 | 11.38982887 | 1.55389321 | 0.169784827 | 0.578875746 | -4.40364665 |
| **A_55_P2061737** | Tmsb4x | -0.379635526 | -0.973324537 | 0.214053485 | 15.72873327 | -1.55378536 | 0.169810125 | 0.578875746 | -4.403701487 |
| **A_55_P2013078** | Thoc5 | 0.187189491 | -0.105608665 | 0.479987648 | 9.874126019 | 1.55344655 | 0.169889619 | 0.578875746 | -4.403873755 |
| **A_55_P2113359** | Zfp523 | -0.173909067 | -0.445936233 | 0.098118099 | 10.44077302 | -1.553435186 | 0.169892286 | 0.578875746 | -4.403879533 |
| **A_55_P2008691** | NA | 0.159885321 | -0.090213833 | 0.409984476 | 15.22081131 | 1.553386617 | 0.169903685 | 0.578875746 | -4.403904228 |
| **A_51_P191909** | Rem2 | -0.158548187 | -0.406579319 | 0.089482945 | 7.831358751 | -1.553238951 | 0.169938345 | 0.578875746 | -4.403979308 |
| **A_55_P2077879** | Kat6b | -0.13119418 | -0.33643313 | 0.07404477 | 7.990432074 | -1.553237881 | 0.169938596 | 0.578875746 | -4.403979852 |
| **A_52_P62617** | Gm9895 | 0.334400015 | -0.188760211 | 0.857560241 | 9.628123652 | 1.553154856 | 0.169958087 | 0.578875746 | -4.404022065 |
| **A_55_P2171493** | BC030867 | 0.281566746 | -0.159026535 | 0.722160027 | 10.42838354 | 1.552839896 | 0.170032045 | 0.579025615 | -4.4041822 |
| **A_51_P231738** | Dnajc4 | -0.174109848 | -0.446624352 | 0.098404655 | 10.53469624 | -1.552447441 | 0.170124242 | 0.579050033 | -4.404381732 |
| **A_55_P1976958** | NA | 0.182718311 | -0.103293631 | 0.468730254 | 10.3313392 | 1.552319434 | 0.170154324 | 0.579050033 | -4.404446813 |
| **A_52_P529360** | Rab11fip5 | -0.388156435 | -0.995810501 | 0.219497631 | 8.998730599 | -1.552149578 | 0.170194249 | 0.579050033 | -4.404533169 |
| **A_55_P1966664** | Cdc14b | 0.265244587 | -0.150026089 | 0.680515263 | 10.81295429 | 1.552024078 | 0.170223753 | 0.579050033 | -4.404596974 |
| **A_55_P1956064** | NA | 0.194076695 | -0.109798373 | 0.497951763 | 12.15038759 | 1.551892048 | 0.170254798 | 0.579050033 | -4.404664097 |
| **A_55_P2052281** | Rnf208 | -0.183683657 | -0.471304715 | 0.1039374 | 7.920178648 | -1.551790286 | 0.170278729 | 0.579050033 | -4.404715832 |
| **A_55_P1957967** | Dhx32 | 0.142821496 | -0.080829837 | 0.366472829 | 9.753618863 | 1.551691124 | 0.170302052 | 0.579050033 | -4.404766246 |
| **A_55_P1999838** | Cox7c | 0.105012685 | -0.059455927 | 0.269481297 | 14.48600262 | 1.55146479 | 0.170355297 | 0.579050033 | -4.404881311 |
| **A_51_P350817** | Cnn1 | -0.673950166 | -1.729489706 | 0.381589374 | 8.506261698 | -1.551445297 | 0.170359883 | 0.579050033 | -4.404891221 |
| **A_51_P217336** | Scamp1 | -0.225571587 | -0.578870958 | 0.127727785 | 11.32211007 | -1.551403325 | 0.170369759 | 0.579050033 | -4.404912558 |
| **A_52_P252322** | Pdcd11 | 0.142085438 | -0.080476994 | 0.36464787 | 8.428801236 | 1.551246806 | 0.170406593 | 0.579050033 | -4.404992129 |
| **A_55_P2164998** | NA | 0.177514242 | -0.100545491 | 0.455573976 | 12.96756324 | 1.551237486 | 0.170408786 | 0.579050033 | -4.404996867 |
| **A_52_P646783** | Fbxl22 | -0.121113661 | -0.3108374 | 0.068610078 | 7.822083589 | -1.551153033 | 0.170428663 | 0.579050033 | -4.405039801 |
| **A_52_P323830** | Zfp830 | 0.111280416 | -0.063082093 | 0.285642925 | 8.974615034 | 1.550775099 | 0.170517643 | 0.579054832 | -4.40523193 |
| **A_55_P1975837** | NA | -0.188007071 | -0.482596242 | 0.1065821 | 12.41795411 | -1.550745728 | 0.17052456 | 0.579054832 | -4.405246861 |
| **A_55_P1990081** | Snrpg | 0.15926691 | -0.090321452 | 0.408855273 | 14.93755045 | 1.550545133 | 0.170571807 | 0.579054832 | -4.405348835 |
| **A_55_P1979728** | Atf4 | 0.23282354 | -0.173966811 | 0.639613891 | 15.26078934 | 1.76490687 | 0.170585158 | 0.579054832 | -4.44531988 |
| **A_51_P216593** | Wfs1 | -0.138826296 | -0.35640171 | 0.078749118 | 9.238273519 | -1.550404813 | 0.170604864 | 0.579054832 | -4.405420167 |
| **A_55_P2084122** | Dus4l | 0.186698528 | -0.105919772 | 0.479316827 | 8.779532194 | 1.550324464 | 0.170623796 | 0.579054832 | -4.405461012 |
| **A_51_P295610** | Cyc1 | -0.151139789 | -0.388036158 | 0.085756579 | 12.98975908 | -1.550256631 | 0.170639781 | 0.579054832 | -4.405495494 |
| **A_55_P2408330** | Rbm33 | 0.152762497 | -0.086708306 | 0.392233299 | 8.651119956 | 1.550055921 | 0.170687085 | 0.579079779 | -4.405597523 |
| **A_55_P2113683** | Raph1 | -0.275628492 | -0.707749577 | 0.156492592 | 11.54348802 | -1.549893444 | 0.170725387 | 0.579079779 | -4.405680116 |

| | | | | | | | | |
|---|---|---|---|---|---|---|---|---|
| **A_51_P361492** | Pou2f1 | 0.112275295 | -0.063751737 | 0.288302327 | 9.905131543 | 1.549844148 | 0.17073701 | 0.579079779 | -4.405705175 |
| **A_55_P1985984** | Actg1 | -0.302696328 | -0.777448637 | 0.172055982 | 11.8966259 | -1.549256243 | 0.170875678 | 0.579357481 | -4.406004019 |
| **A_52_P159256** | Lpp | -0.145879864 | -0.37468238 | 0.082922653 | 8.521704552 | -1.549236571 | 0.17088032 | 0.579357481 | -4.406014018 |
| **A_55_P1952071** | Npb | 0.401170911 | -0.228111153 | 1.030452975 | 10.14308489 | 1.549056542 | 0.170922806 | 0.579357481 | -4.406105528 |
| **A_55_P1997585** | NA | -0.776676258 | -1.99509798 | 0.441745464 | 10.92299079 | -1.548907866 | 0.1709579 | 0.579357481 | -4.406181101 |
| **A_55_P1954718** | Cyb561 | -1.001867937 | -2.573608912 | 0.569873038 | 8.977169674 | -1.548861881 | 0.170968755 | 0.579357481 | -4.406204475 |
| **A_52_P86158** | Mtx2 | 0.157394624 | -0.089600218 | 0.404389466 | 11.35511198 | 1.548407258 | 0.171076114 | 0.579474123 | -4.406435555 |
| **A_55_P2065824** | Smtn | -0.312277931 | -0.802342776 | 0.177786914 | 10.30200203 | -1.548356398 | 0.171088129 | 0.579474123 | -4.406461406 |
| **A_51_P237754** | H2-T23 | -0.285792733 | -0.734325981 | 0.162740515 | 12.95928302 | -1.548245152 | 0.171114411 | 0.579474123 | -4.40651795 |
| **A_52_P621418** | NA | -0.128280582 | -0.329624577 | 0.073063414 | 12.62204445 | -1.548122874 | 0.171143304 | 0.579474123 | -4.406580102 |
| **A_52_P97699** | D430019H16Rik | -0.361803945 | -0.929718114 | 0.206110224 | 8.627459854 | -1.548010076 | 0.171169961 | 0.579474123 | -4.406637434 |
| **A_51_P507290** | Klf5 | -0.630844164 | -1.621099151 | 0.359410822 | 8.781396322 | -1.547954678 | 0.171183054 | 0.579474123 | -4.40666559 |
| **A_51_P151254** | Slc35b3 | -0.129753224 | -0.333494762 | 0.073988315 | 9.554439425 | -1.547468312 | 0.171298046 | 0.579570591 | -4.406912791 |
| **A_55_P2112479** | Nudt16l1 | -0.140837214 | -0.361983362 | 0.080308933 | 9.903586177 | -1.547466435 | 0.17129849 | 0.579570591 | -4.406913745 |
| **A_55_P1990609** | Ube2j1 | 0.152675247 | -0.087061253 | 0.392411747 | 11.5565733 | 1.547453688 | 0.171301505 | 0.579570591 | -4.406920223 |
| **A_51_P103509** | Cyb561d2 | -0.140874609 | -0.362134053 | 0.080384836 | 10.38579184 | -1.547084711 | 0.171388793 | 0.579645536 | -4.407107754 |
| **A_66_P107818** | Gdf3 | 0.303698876 | -0.173321535 | 0.780719286 | 8.571290215 | 1.546996785 | 0.1714096 | 0.579645536 | -4.407152441 |
| **A_55_P1955791** | Fopnl | 0.177445769 | -0.101271705 | 0.456163242 | 12.36858165 | 1.546979791 | 0.171413621 | 0.579645536 | -4.407161079 |
| **A_55_P1975685** | Nup37 | 0.117550216 | -0.067120691 | 0.302221123 | 11.6220671 | 1.546707307 | 0.171478118 | 0.579762206 | -4.407299563 |
| **A_51_P222092** | Igbp1 | -0.170773959 | -0.439215112 | 0.097667194 | 9.692679963 | -1.545808677 | 0.171690983 | 0.580256714 | -4.407756256 |
| **A_55_P2160387** | Cxcl11 | 0.133123054 | -0.076144599 | 0.342390707 | 16.59422317 | 1.545732565 | 0.171709024 | 0.580256714 | -4.407794936 |
| **A_51_P163578** | Ugt2b35 | 0.330889925 | -0.189272283 | 0.851052133 | 9.609787733 | 1.545709716 | 0.17171444 | 0.580256714 | -4.407806548 |
| **A_51_P358872** | Disc1 | 0.233906014 | -0.133987298 | 0.601799326 | 8.494756943 | 1.54490822 | 0.17190453 | 0.580267241 | -4.408213851 |
| **A_52_P571746** | Sh2d5 | 0.116281433 | -0.066609893 | 0.29917276 | 8.082023872 | 1.544900292 | 0.171906411 | 0.580267241 | -4.40821788 |
| **A_55_P1964965** | Gm1673 | -0.459021627 | -1.181002098 | 0.262958844 | 9.657525345 | -1.544866255 | 0.171914488 | 0.580267241 | -4.408235176 |
| **A_66_P101600** | Lamb1 | -0.699344664 | -1.799529875 | 0.400840547 | 11.40957312 | -1.544573758 | 0.171983912 | 0.580267241 | -4.408383811 |
| **A_55_P2100050** | NA | 0.232353416 | -0.133178816 | 0.597885647 | 10.70923613 | 1.544566361 | 0.171985668 | 0.580267241 | -4.40838757 |
| **A_51_P450924** | Pole4 | 0.154408216 | -0.08852713 | 0.397343563 | 11.05209888 | 1.544411001 | 0.172022554 | 0.580267241 | -4.408466516 |
| **A_55_P2029021** | NA | 0.27310741 | -0.156595504 | 0.702810324 | 15.33603707 | 1.544359393 | 0.172034808 | 0.580267241 | -4.40849274 |
| **A_52_P340073** | Efnb2 | -0.492616285 | -1.267692645 | 0.282460075 | 8.532829902 | -1.544356154 | 0.172035577 | 0.580267241 | -4.408494386 |
| **A_55_P2020962** | NA | -0.154055249 | -0.396454406 | 0.088343907 | 16.82358224 | -1.544289026 | 0.172051518 | 0.580267241 | -4.408528497 |
| **A_55_P2245920** | Lrrc8a | -0.15228518 | -0.391906677 | 0.087336317 | 11.90828053 | -1.544240905 | 0.172062947 | 0.580267241 | -4.40855295 |
| **A_55_P2150044** | Cdkl2 | -0.163003609 | -0.419495487 | 0.093488269 | 8.253548626 | -1.544211534 | 0.172069922 | 0.580267241 | -4.408567874 |
| **A_55_P2016604** | Glrx3 | 0.144020033 | -0.082606582 | 0.370646648 | 12.88499494 | 1.54417026 | 0.172079725 | 0.580267241 | -4.408588847 |
| **A_55_P2084408** | Uba1 | 0.151146131 | -0.086714385 | 0.389006647 | 8.357074927 | 1.544037586 | 0.172111241 | 0.580267241 | -4.408656264 |
| **A_52_P207335** | Tmem246 | -0.257398593 | -0.662499286 | 0.147702099 | 10.03659863 | -1.543925614 | 0.172137843 | 0.580267241 | -4.40871316 |
| **A_52_P99531** | Ippk | 0.231917015 | -0.133330158 | 0.597164187 | 9.710632314 | 1.542868594 | 0.172389156 | 0.580877162 | -4.409250246 |
| **A_51_P289239** | F8 | -0.180065277 | -0.463650936 | 0.103520383 | 8.448790295 | -1.542868445 | 0.172389192 | 0.580877162 | -4.409250322 |
| **A_55_P2061591** | Psmd1 | 0.165264833 | -0.095054063 | 0.425583729 | 13.32567027 | 1.542616398 | 0.172449168 | 0.580877162 | -4.409378384 |
| **A_55_P2436715** | Paics | 0.111385306 | -0.064064691 | 0.286835302 | 11.89252349 | 1.542615622 | 0.172449353 | 0.580877162 | -4.409378778 |
| **A_51_P101228** | Gmeb2 | 0.127601454 | -0.073422224 | 0.328625132 | 8.456216103 | 1.542380759 | 0.172505258 | 0.580877162 | -4.41098504 |
| **A_55_P2027136** | H3c7 | 0.236707584 | -0.136223947 | 0.609639114 | 9.968993426 | 1.542290762 | 0.172526685 | 0.580877162 | -4.409543833 |
| **A_55_P2002113** | Rpl36 | -0.154700385 | -0.398440322 | 0.089039552 | 16.60685515 | -1.542225534 | 0.172542216 | 0.580877162 | -4.409576974 |
| **A_51_P435922** | Rsph9 | -0.548500848 | -1.412752464 | 0.315750768 | 8.85668548 | -1.542127538 | 0.172565553 | 0.580877162 | -4.409626762 |
| **A_52_P552832** | Ndufa4 | 0.142794362 | -0.082216137 | 0.36780486 | 13.61418858 | 1.542025183 | 0.17258993 | 0.580877162 | -4.409678766 |
| **A_55_P2001970** | Zswim9 | 0.114128175 | -0.065734042 | 0.293990391 | 9.158706928 | 1.541828677 | 0.172636739 | 0.580877162 | -4.409778603 |
| **A_52_P244723** | Tbcel | 0.309902842 | -0.178510523 | 0.798316208 | 11.50626225 | 1.541775744 | 0.172649351 | 0.580877162 | -4.409805496 |
| **A_51_P506309** | Hrg | -0.919919231 | -2.37011509 | 0.530276629 | 10.64320724 | -1.541367601 | 0.172746619 | 0.580973277 | -4.410012852 |
| **A_51_P477364** | Rhob | -0.114905929 | -0.296062825 | 0.066250966 | 7.961123392 | -1.541241726 | 0.172776627 | 0.580973277 | -4.410076802 |
| **A_52_P648573** | Gorab | 0.283356924 | -0.163470361 | 0.730184208 | 8.515054067 | 1.540910211 | 0.172855684 | 0.580973277 | -4.410245222 |

| | | | | | | | | | |
|---|---|---|---|---|---|---|---|---|---|
| **A_55_P2064598** | Cnot6 | -0.172144054 | -0.45838761 | 0.114099503 | 12.63403087 | -1.641958219 | 0.172892659 | 0.580973277 | -4.418692976 |
| **A_55_P2051778** | Sult2b1 | -0.153601524 | -0.395869741 | 0.088666693 | 7.794996561 | -1.540572958 | 0.172936145 | 0.580973277 | -4.410416553 |
| **A_51_P392291** | Pdk3 | 0.154127792 | -0.088978005 | 0.397233588 | 9.811851984 | 1.540525287 | 0.172947521 | 0.580973277 | -4.410440771 |
| **A_55_P2038952** | Ubb | 0.12563223 | -0.072535368 | 0.323799827 | 15.67875749 | 1.540464212 | 0.172962096 | 0.580973277 | -4.410471797 |
| **A_52_P100199** | Abhd10 | 0.151540716 | -0.087500978 | 0.390582409 | 8.457545896 | 1.540418997 | 0.172972888 | 0.580973277 | -4.410494767 |
| **A_55_P2127959** | Zfp36l3 | 0.318806032 | -0.184183451 | 0.821795515 | 9.000646095 | 1.540106714 | 0.173047437 | 0.580973277 | -4.410653408 |
| **A_66_P119518** | Tuba8 | -0.265682468 | -0.684866144 | 0.153501208 | 10.6315497 | -1.540074477 | 0.173055134 | 0.580973277 | -4.410669784 |
| **A_51_P512992** | Entpd3 | -0.232894686 | -0.600410166 | 0.134620794 | 7.773971086 | -1.539809987 | 0.1731183 | 0.580973277 | -4.410804142 |
| **A_55_P1971849** | Sh3glb2 | -0.210912825 | -0.543745773 | 0.121920123 | 12.11184603 | -1.539784224 | 0.173124454 | 0.580973277 | -4.410817229 |
| **A_51_P254656** | Hdc | -0.39326122 | -1.013857628 | 0.227335189 | 10.04476376 | -1.53976716 | 0.17312853 | 0.580973277 | -4.410825898 |
| **A_51_P369971** | Fra10ac1 | 0.161453843 | -0.093356146 | 0.416263832 | 10.94614266 | 1.539625609 | 0.173162346 | 0.580973277 | -4.410897803 |
| **A_55_P1952901** | Gm833 | 0.130479602 | -0.075465039 | 0.336424242 | 8.022225827 | 1.539484488 | 0.173196066 | 0.580973277 | -4.410969489 |
| **A_55_P2124871** | Npw | -0.189603178 | -0.488916702 | 0.109710345 | 7.968803378 | -1.539226254 | 0.173257784 | 0.580973277 | -4.411100664 |
| **A_55_P1996456** | Srrm1 | 0.234405502 | -0.135642403 | 0.604453407 | 9.449112465 | 1.539192861 | 0.173265767 | 0.580973277 | -4.411117627 |
| **A_66_P123735** | NA | 0.197840767 | -0.114488199 | 0.510169734 | 9.143623978 | 1.539170436 | 0.173271128 | 0.580973277 | -4.411129018 |
| **A_55_P2216996** | Tmem87a | 0.170001327 | -0.098377848 | 0.438380503 | 8.608758561 | 1.539170342 | 0.17327115 | 0.580973277 | -4.411129066 |
| **A_55_P2063163** | Agk | 0.114083701 | -0.066022821 | 0.294190224 | 9.578019516 | 1.539137257 | 0.17327906 | 0.580973277 | -4.411145871 |
| **A_51_P349691** | Ano10 | -0.134347649 | -0.346464327 | 0.07776903 | 9.287911293 | -1.538999367 | 0.173312028 | 0.580983037 | -4.411215914 |
| **A_55_P2121856** | Ier5l | -0.608435747 | -1.569258259 | 0.352386766 | 10.07283615 | -1.538702787 | 0.173382958 | 0.581023479 | -4.411366562 |
| **A_55_P2019014** | Tmem120b | 0.180512084 | -0.104562654 | 0.465586821 | 10.34186534 | 1.538617738 | 0.173403303 | 0.581023479 | -4.411409762 |
| **A_55_P2067772** | Cplane1 | -0.104065827 | -0.268436522 | 0.060304868 | 8.213047637 | -1.538391727 | 0.17345738 | 0.581023479 | -4.411524562 |
| **A_52_P295067** | Snn | -0.367353562 | -0.947585584 | 0.21287846 | 9.133285264 | -1.53838753 | 0.173458384 | 0.581023479 | -4.411526694 |
| **A_51_P464981** | Map1lc3a | -0.453706753 | -1.17038484 | 0.262971335 | 9.146023897 | -1.538276149 | 0.17348504 | 0.581023479 | -4.411583269 |
| **A_51_P310821** | Hoxa5 | -0.679631835 | -1.753249288 | 0.393985618 | 8.127868356 | -1.538180635 | 0.173507901 | 0.581023479 | -4.411631782 |
| **A_55_P1985298** | Krtap5-2 | 0.163668932 | -0.094898298 | 0.422236161 | 8.447942794 | 1.538069479 | 0.17353451 | 0.581023479 | -4.411688242 |
| **A_55_P1992450** | Vdac2 | 0.123044724 | -0.071361945 | 0.317451394 | 13.74748269 | 1.537924541 | 0.173569211 | 0.581039018 | -4.411761859 |
| **A_52_P587831** | Ddx56 | 0.156258224 | -0.090659097 | 0.403175545 | 10.54186766 | 1.537710273 | 0.173620524 | 0.581071274 | -4.411870689 |
| **A_55_P2150283** | E030030I06Rik | 0.124650546 | -0.072384171 | 0.321685262 | 11.73164602 | 1.537215019 | 0.173739181 | 0.581071274 | -4.412122229 |
| **A_51_P145772** | Rnf113a2 | 0.143850552 | -0.083586579 | 0.371287683 | 9.392044206 | 1.536856625 | 0.173825095 | 0.581071274 | -4.412304252 |
| **A_52_P684378** | Gpx1 | 0.204513604 | -0.118847968 | 0.527875176 | 17.15729485 | 1.536798698 | 0.173838985 | 0.581071274 | -4.412333672 |
| **A_55_P2056876** | Akap13 | 0.167730524 | -0.097485336 | 0.432946385 | 9.070092379 | 1.536723745 | 0.17385696 | 0.581071274 | -4.412371738 |
| **A_55_P1987944** | Rai1 | -0.203208341 | -0.524532588 | 0.118115907 | 10.01527926 | -1.536672145 | 0.173869335 | 0.581071274 | -4.412397945 |
| **A_55_P2177488** | H2bc21 | 0.120497762 | -0.070041975 | 0.311037499 | 8.441283207 | 1.536655834 | 0.173873246 | 0.581071274 | -4.412406228 |
| **A_51_P434503** | Dtwd1 | 0.193993673 | -0.112790366 | 0.500777712 | 8.777779933 | 1.536519208 | 0.173906018 | 0.581071274 | -4.412475617 |
| **A_52_P638605** | Ap1m2 | -0.260358153 | -0.672100673 | 0.151384366 | 7.797763594 | -1.536486163 | 0.173913945 | 0.581071274 | -4.412492399 |
| **A_51_P485756** | Nts | -0.110267185 | -0.284657923 | 0.064123553 | 7.775494657 | -1.536406232 | 0.17393312 | 0.581071274 | -4.412532993 |
| **A_51_P330615** | Zfp119b | 0.187554481 | -0.109114722 | 0.484223685 | 9.700822182 | 1.536166073 | 0.173990747 | 0.581071274 | -4.41265496 |
| **A_55_P2166128** | NA | 0.155344809 | -0.090395759 | 0.401085377 | 8.45020997 | 1.536041946 | 0.174020539 | 0.581071274 | -4.412717998 |
| **A_55_P2070661** | Taf7l | 0.224746121 | -0.130839841 | 0.580332082 | 10.23095654 | 1.535786291 | 0.174081913 | 0.581071274 | -4.412847831 |
| **A_55_P2064043** | Cd44 | -0.130173303 | -0.33613145 | 0.075784843 | 10.22354133 | -1.535769863 | 0.174085858 | 0.581071274 | -4.412856174 |
| **A_55_P1982737** | Dnajc5 | 0.154143032 | -0.089766459 | 0.398052524 | 14.5577776 | 1.535601 | 0.174126408 | 0.581071274 | -4.412941928 |
| **A_55_P2052490** | Ushbp1 | -0.140006993 | -0.361587088 | 0.081573101 | 8.131866425 | -1.535331549 | 0.174191132 | 0.581071274 | -4.413078763 |
| **A_55_P2140941** | Tinagl1 | -0.127195887 | -0.328522728 | 0.074130954 | 7.98583013 | -1.535163293 | 0.17423156 | 0.581071274 | -4.413164207 |
| **A_55_P2032445** | Slc25a37 | 0.20336863 | -0.118527331 | 0.525264591 | 13.16747963 | 1.535152853 | 0.174234069 | 0.581071274 | -4.413169508 |
| **A_52_P589550** | Nr2c1 | -0.113247325 | -0.292506804 | 0.066012154 | 8.337179021 | -1.535072923 | 0.174253277 | 0.581071274 | -4.413210098 |
| **A_55_P2007964** | Cx3cr1 | -0.194172207 | -0.501562551 | 0.113218137 | 9.900321268 | -1.53489982 | 0.174294884 | 0.581071274 | -4.413298001 |
| **A_51_P113003** | Cenpq | 0.223854983 | -0.130574784 | 0.578284751 | 9.851689805 | 1.534686829 | 0.17434609 | 0.581071274 | -4.418031761 |
| **A_55_P2015605** | Foxp4 | -0.150994775 | -0.390087233 | 0.088097684 | 11.52064636 | -1.534543589 | 0.174380535 | 0.581071274 | -4.413478895 |
| **A_55_P2266178** | Il2rb | -0.182236884 | -0.470809455 | 0.106335688 | 8.268444277 | -1.534491263 | 0.174393119 | 0.581071274 | -4.413505466 |
| **A_55_P2093994** | Maged2 | -0.642257612 | -1.65932132 | 0.374806096 | 12.19945031 | -1.534420144 | 0.174410225 | 0.581071274 | -4.413541579 |

| | | | | | | | | | |
|---|---|---|---|---|---|---|---|---|---|
| **A_55_P2045188** | NA | -0.166363731 | -0.429816264 | 0.097088803 | 7.878475063 | -1.534403074 | 0.174414331 | 0.581071274 | -4.413550248 |
| **A_51_P117236** | Zfp354a | -0.125674886 | -0.324705162 | 0.073355389 | 8.354056324 | -1.534307986 | 0.174437205 | 0.581071274 | -4.413598532 |
| **A_52_P335508** | Tiprl | 0.137516747 | -0.080276987 | 0.355310481 | 9.419846939 | 1.53424034 | 0.174453479 | 0.581071274 | -4.413632881 |
| **A_55_P2145496** | Wdr45 | -0.630112531 | -1.62806543 | 0.367840368 | 10.61293609 | -1.534232794 | 0.174455294 | 0.581071274 | -4.413636713 |
| **A_55_P1953459** | Yap1 | -0.299588053 | -0.774084858 | 0.174908753 | 11.58432107 | -1.534173179 | 0.174469637 | 0.581071274 | -4.413666984 |
| **A_55_P2043987** | Luzp1 | -0.168014762 | -0.434133545 | 0.098104021 | 10.56705626 | -1.534105056 | 0.174486029 | 0.581071274 | -4.413701576 |
| **A_51_P464308** | Gnb4 | -0.407245767 | -1.052326063 | 0.23783453 | 8.367479938 | -1.534002229 | 0.174510774 | 0.581071274 | -4.413753788 |
| **A_52_P552665** | Fzd7 | -0.179142703 | -0.463003297 | 0.104717892 | 8.423620384 | -1.533476775 | 0.174637273 | 0.581288072 | -4.414020595 |
| **A_51_P206268** | Seh1l | 0.21311442 | -0.124579585 | 0.550808426 | 10.54246619 | 1.533460787 | 0.174641124 | 0.581288072 | -4.414028713 |
| **A_51_P154973** | Stard3 | -0.309184456 | -0.79918835 | 0.180819438 | 11.07904373 | -1.533208821 | 0.174701814 | 0.581288072 | -4.414156648 |
| **A_52_P582150** | Gosr2 | 0.09943319 | -0.058152548 | 0.257018928 | 11.25364366 | 1.533195913 | 0.174704924 | 0.581288072 | -4.414163202 |
| **A_51_P487501** | NA | 0.127936213 | -0.07487365 | 0.330746077 | 11.37710575 | 1.532807469 | 0.174798528 | 0.581288072 | -4.414360428 |
| **A_66_P137943** | Tctn2 | -0.51708841 | -1.336887704 | 0.302710885 | 9.044110361 | -1.532641045 | 0.174838646 | 0.581288072 | -4.414444926 |
| **A_52_P164709** | Poc1a | 0.236628738 | -0.138560552 | 0.611818027 | 10.77071094 | 1.532499155 | 0.174872856 | 0.581288072 | -4.414516966 |
| **A_55_P2159666** | NA | 0.222205042 | -0.130136219 | 0.574546303 | 8.823439059 | 1.53240512 | 0.174895532 | 0.581288072 | -4.414564709 |
| **A_55_P1991896** | Ceacam5 | -0.237316859 | -0.613664457 | 0.139030739 | 7.82839383 | -1.532225312 | 0.1749389 | 0.581288072 | -4.414655999 |
| **A_55_P1958246** | Tcof1 | 0.167580083 | -0.098189124 | 0.43334929 | 11.27872597 | 1.532148742 | 0.17495737 | 0.581288072 | -4.414694874 |
| **A_51_P427674** | Cpt1a | -0.397142185 | -1.027001919 | 0.232717548 | 10.10383077 | -1.532093835 | 0.174970616 | 0.581288072 | -4.41472275 |
| **A_55_P2143885** | Etl4 | -0.194992159 | -0.50424977 | 0.114265453 | 9.11966786 | -1.532074692 | 0.174975235 | 0.581288072 | -4.414732469 |
| **A_51_P511734** | Snip1 | 0.129972426 | -0.076168463 | 0.336113314 | 8.612199685 | 1.532040579 | 0.174983465 | 0.581288072 | -4.414749788 |
| **A_55_P2003931** | NA | 0.288101204 | -0.168930123 | 0.745132531 | 8.453232066 | 1.53173026 | 0.175058351 | 0.581288072 | -4.414907335 |
| **A_51_P396854** | Timm10b | -0.117462858 | -0.303804093 | 0.068878377 | 13.3244648 | -1.531703772 | 0.175064745 | 0.581288072 | -4.414920783 |
| **A_55_P2088615** | Fbln5 | -0.897054979 | -2.320206247 | 0.526096289 | 10.34596512 | -1.531620554 | 0.175084832 | 0.581288072 | -4.414963031 |
| **A_55_P2010003** | Zfp113 | -0.134737106 | -0.348509536 | 0.079035324 | 8.635197401 | -1.531506031 | 0.175112481 | 0.581288072 | -4.415021172 |
| **A_55_P2025078** | Csf1r | -0.219899889 | -0.56881028 | 0.129010501 | 13.78227703 | -1.53141994 | 0.175133267 | 0.581288072 | -4.415064878 |
| **A_66_P123787** | Lars2 | 0.191419291 | -0.112360224 | 0.495198807 | 10.43720675 | 1.531124157 | 0.175204702 | 0.581288072 | -4.415215038 |
| **A_55_P1975732** | Sipa1l3 | -0.374672451 | -0.969282159 | 0.219937257 | 9.388729425 | -1.531097878 | 0.17521105 | 0.581288072 | -4.415228379 |
| **A_55_P2171348** | Grm2 | 0.227075921 | -0.13332771 | 0.587479552 | 9.155646103 | 1.530964481 | 0.175243277 | 0.581288072 | -4.4152961 |
| **A_55_P2078238** | NA | 0.160316724 | -0.094146361 | 0.41477981 | 8.468910174 | 1.530866188 | 0.175267027 | 0.581288072 | -4.415345998 |
| **A_55_P2059860** | Igsf1 | 0.222460011 | -0.130664464 | 0.575584485 | 8.459002007 | 1.530760771 | 0.175292501 | 0.581288072 | -4.415399513 |
| **A_51_P513392** | Azi2 | 0.196378618 | -0.11534959 | 0.508106827 | 9.567463588 | 1.530739487 | 0.175297645 | 0.581288072 | -4.415410318 |
| **A_55_P2107686** | Ttll1 | -0.102714591 | -0.265786414 | 0.060357231 | 7.869870695 | -1.530510832 | 0.175352913 | 0.581297789 | -4.415526393 |
| **A_52_P539434** | Lbh | -0.26456251 | -0.684643157 | 0.155518137 | 8.01050576 | -1.530307889 | 0.175401981 | 0.581297789 | -4.415629415 |
| **A_52_P635338** | Fes | -0.143083671 | -0.370277175 | 0.084109833 | 10.76242201 | -1.530302732 | 0.175403228 | 0.581297789 | -4.415632032 |
| **A_51_P267796** | Ddr2 | -0.329589266 | -0.852948148 | 0.193769617 | 8.46344009 | -1.530229776 | 0.17542087 | 0.581297789 | -4.415669067 |
| **A_55_P2073489** | NA | 0.195767498 | -0.115317529 | 0.506852525 | 9.155567897 | 1.529130921 | 0.1756868 | 0.581907393 | -4.416226853 |
| **A_55_P2045837** | NA | 0.108902745 | -0.064155322 | 0.281960811 | 8.552832411 | 1.529079856 | 0.175699168 | 0.581907393 | -4.416252773 |
| **A_55_P2001628** | Rps4l | -0.210897871 | -0.546039623 | 0.124243882 | 12.46118394 | -1.529068169 | 0.175701998 | 0.581907393 | -4.416258706 |
| **A_51_P307325** | Rbbp4 | 0.098328342 | -0.057942373 | 0.254599058 | 13.61156445 | 1.528918392 | 0.175738278 | 0.581907393 | -4.41633473 |
| **A_52_P304323** | Isca1 | 0.233564441 | -0.137650663 | 0.604779545 | 11.39637509 | 1.528847876 | 0.17575536 | 0.581907393 | -4.416370522 |
| **A_51_P250358** | Prpf39 | 0.134339942 | -0.079234693 | 0.347914577 | 8.733617138 | 1.528405789 | 0.175862494 | 0.581984652 | -4.41659491 |
| **A_55_P2065966** | Ctdp1 | 0.2050321 | -0.120934406 | 0.530998605 | 9.526077574 | 1.528382553 | 0.175868126 | 0.581984652 | -4.416606703 |
| **A_52_P600531** | Bahd1 | 0.241725412 | -0.142609201 | 0.626060026 | 9.134815182 | 1.528255612 | 0.175898901 | 0.581984652 | -4.416671133 |
| **A_55_P2431862** | Srpk2 | -0.158354933 | -0.410133674 | 0.093423808 | 9.911946245 | -1.528254655 | 0.175899133 | 0.581984652 | -4.416671618 |
| **A_51_P462516** | Senp7 | 0.172502272 | -0.101834889 | 0.446839434 | 8.754004713 | 1.527894504 | 0.175986472 | 0.582173973 | -4.416854411 |
| **A_51_P275949** | Loxl2 | -0.216720524 | -0.561467798 | 0.12802675 | 8.569248886 | -1.527504122 | 0.176081189 | 0.58228661 | -4.417052541 |
| **A_51_P109144** | Grtp1 | 0.230390242 | -0.136108236 | 0.596888719 | 9.105092834 | 1.52747846 | 0.176087417 | 0.58228661 | -4.417065565 |
| **A_55_P2186615** | Acp1 | 0.235580729 | -0.139250437 | 0.610411895 | 11.67637785 | 1.527169552 | 0.176162402 | 0.58228661 | -4.417222341 |
| **A_55_P2025273** | Trabd | -0.16427999 | -0.425727831 | 0.097167852 | 10.09517288 | -1.526802264 | 0.176251597 | 0.58228661 | -4.41740874 |
| **A_55_P1974134** | Mrpl58 | -0.185761843 | -0.481402044 | 0.109878358 | 12.76829463 | -1.526779117 | 0.17625722 | 0.58228661 | -4.417420486 |

| | | | | | | | | | |
|---|---|---|---|---|---|---|---|---|---|
| **A_52_P8903** | Tmem251 | 0.176254814 | -0.104288215 | 0.456797843 | 9.170507624 | 1.526598022 | 0.176301215 | 0.58228661 | -4.41751239 |
| **A_55_P2042487** | Dpysl3 | -0.476143337 | -1.234104916 | 0.281818242 | 9.368742238 | -1.526418809 | 0.176344764 | 0.58228661 | -4.417603338 |
| **A_55_P2078266** | H3f3a | 0.131694527 | -0.077954987 | 0.341344041 | 14.05876152 | 1.526360279 | 0.176358988 | 0.58228661 | -4.41763304 |
| **A_52_P255224** | NA | 0.124494365 | -0.073702156 | 0.322690887 | 7.975550173 | 1.526289288 | 0.176376243 | 0.58228661 | -4.417669066 |
| **A_55_P2154240** | Rps29 | -0.180882777 | -0.468867072 | 0.107101517 | 16.55003342 | -1.526200541 | 0.176397816 | 0.58228661 | -4.417714103 |
| **A_55_P2026669** | Atp5l | 0.141682192 | -0.083893728 | 0.367258112 | 14.73815044 | 1.526179867 | 0.176402842 | 0.58228661 | -4.417724594 |
| **A_65_P09032** | Rab43 | 0.157524787 | -0.09327999 | 0.408329565 | 13.25864134 | 1.526146712 | 0.176410902 | 0.58228661 | -4.417741419 |
| **A_55_P2127020** | Nup54 | 0.164790776 | -0.097583504 | 0.427165057 | 10.64195679 | 1.526141591 | 0.176412147 | 0.58228661 | -4.417744018 |
| **A_55_P1961140** | Mettl3 | 0.146532473 | -0.086822455 | 0.3798874 | 10.32024035 | 1.525808716 | 0.17649309 | 0.582454318 | -4.417912939 |
| **A_51_P394190** | Lmo4 | -0.27397447 | -0.710358995 | 0.162410056 | 11.92957791 | -1.525541202 | 0.176558165 | 0.582469846 | -4.418048688 |
| **A_55_P2058494** | NA | -0.178144038 | -0.461895406 | 0.105607329 | 13.69161965 | -1.52551513 | 0.176564509 | 0.582469846 | -4.418061919 |
| **A_55_P2050236** | Hikeshi | 0.116108158 | -0.068861103 | 0.30107742 | 11.20500831 | 1.52526872 | 0.176624472 | 0.582469846 | -4.418186956 |
| **A_52_P407692** | H2afj | 0.163912599 | -0.097217177 | 0.425042375 | 17.59585389 | 1.525243313 | 0.176630656 | 0.582469846 | -4.418199848 |
| **A_52_P600535** | Zfp39 | 0.211754731 | -0.125613501 | 0.549122963 | 8.681033168 | 1.525148481 | 0.176653739 | 0.582469846 | -4.418247969 |
| **A_52_P83623** | Rapgef6 | 0.16164737 | -0.09591068 | 0.419205419 | 8.980288621 | 1.525024087 | 0.176684022 | 0.582469846 | -4.41831109 |
| **A_51_P468464** | Gdpd5 | -0.395410539 | -1.025505334 | 0.234684256 | 8.47063268 | -1.524844427 | 0.176727767 | 0.582469846 | -4.418402252 |
| **A_55_P2133315** | Steap2 | 0.25446467 | -0.151042122 | 0.659971463 | 9.060064275 | 1.524798829 | 0.176738872 | 0.582469846 | -4.41842539 |
| **A_55_P2064686** | Lrrc61 | -0.120494779 | -0.312556459 | 0.0715669 | 8.858168601 | -1.524441273 | 0.176825968 | 0.582657542 | -4.418606815 |
| **A_52_P796840** | Cfhr2 | -0.421466095 | -1.093396676 | 0.250464487 | 12.48096414 | -1.52412794 | 0.176902326 | 0.582751531 | -4.418765798 |
| **A_51_P120615** | Rabl3 | 0.149004135 | -0.088583595 | 0.386591864 | 8.572068999 | 1.523903623 | 0.17695701 | 0.582751531 | -4.42043334 |
| **A_55_P2037186** | Alkbh2 | -0.132835547 | -0.344648515 | 0.078977421 | 9.660444949 | -1.523859586 | 0.176967747 | 0.582751531 | -4.418901955 |
| **A_55_P2062707** | Mthfsl | -0.158530024 | -0.411340248 | 0.094280199 | 9.987714488 | -1.523702158 | 0.177006136 | 0.582751531 | -4.41898183 |
| **A_55_P2074521** | Psg28 | -0.290202221 | -0.753000983 | 0.172596541 | 7.877149414 | -1.523672864 | 0.17701328 | 0.582751531 | -4.418996693 |
| **A_55_P1952235** | Spry1 | -0.383585373 | -0.995375665 | 0.228204918 | 10.76774263 | -1.523500659 | 0.177055283 | 0.582751531 | -4.419084063 |
| **A_55_P1961863** | Pcdhga9 | -0.667590103 | -1.732376302 | 0.397196095 | 11.22183477 | -1.523458631 | 0.177065536 | 0.582751531 | -4.419105387 |
| **A_52_P382876** | Bloc1s3 | 0.28119647 | -0.167349439 | 0.72974238 | 8.678028285 | 1.523302495 | 0.17710363 | 0.582777674 | -4.419184603 |
| **A_51_P488196** | Bmper | -0.609385953 | -1.581686471 | 0.362914565 | 8.374270941 | -1.52291306 | 0.177198677 | 0.582987012 | -4.41938218 |
| **A_55_P2162698** | Prpf40b | -0.168173525 | -0.436522201 | 0.10017515 | 8.542038934 | -1.522794718 | 0.177227569 | 0.582987012 | -4.419442219 |
| **A_55_P2210301** | Gm12992 | 0.487531516 | -0.290661003 | 1.265724035 | 9.471287937 | 1.522295043 | 0.17734961 | 0.583289197 | -4.419695715 |
| **A_55_P2250249** | Taok1 | -0.240952134 | -0.625596839 | 0.143692571 | 12.3224419 | -1.522138629 | 0.177387829 | 0.583315642 | -4.419775065 |
| **A_52_P131490** | Atg4b | 0.187534157 | -0.111864736 | 0.486933051 | 9.269431591 | 1.521995539 | 0.177422798 | 0.583322074 | -4.419847654 |
| **A_55_P1973965** | Ppp1r13b | -0.384356036 | -0.998026804 | 0.229314733 | 10.52153251 | -1.521883673 | 0.177450142 | 0.583322074 | -4.419904404 |
| **A_55_P2010641** | Slc38a5 | 0.226478355 | -0.135174211 | 0.58813092 | 13.18465652 | 1.521662509 | 0.177504213 | 0.5834006 | -4.420016597 |
| **A_51_P294643** | Cdr2 | 0.312779754 | -0.186735171 | 0.812294679 | 11.44146705 | 1.521504895 | 0.177542756 | 0.583422378 | -4.420096552 |
| **A_55_P1979893** | Tef | -0.441278671 | -1.146070819 | 0.263513477 | 12.1257962 | -1.521370774 | 0.177575561 | 0.583422378 | -4.420164588 |
| **A_55_P1958280** | Arrdc1 | -0.491346227 | -1.276158683 | 0.29346623 | 10.31794386 | -1.521265186 | 0.17760139 | 0.583422378 | -4.42021815 |
| **A_55_P1969776** | Pdhb | 0.130018592 | -0.077723399 | 0.337760584 | 12.89166342 | 1.520772878 | 0.177721868 | 0.58361234 | -4.420467877 |
| **A_51_P256066** | Tiam2 | -0.154957323 | -0.402561106 | 0.092646459 | 8.380231722 | -1.52068065 | 0.177744447 | 0.58361234 | -4.420514659 |
| **A_52_P539632** | Plxna3 | 0.116601597 | -0.069726951 | 0.302930145 | 8.283667641 | 1.520576547 | 0.177769935 | 0.58361234 | -4.420567464 |
| **A_66_P126138** | Ncoa1 | -0.143275512 | -0.372264489 | 0.085713465 | 8.74253371 | -1.520339509 | 0.177827985 | 0.58361234 | -4.420687698 |
| **A_52_P47255** | Gmps | 0.169466718 | -0.101389231 | 0.440322666 | 9.275078395 | 1.520299766 | 0.17783772 | 0.58361234 | -4.420707856 |
| **A_55_P2004152** | NA | 0.145993286 | -0.087373674 | 0.379360246 | 12.5379675 | 1.520115907 | 0.177882761 | 0.58361234 | -4.420801114 |
| **A_51_P135340** | Panx1 | -0.171199077 | -0.444873542 | 0.102475388 | 9.985825659 | -1.520023596 | 0.177905379 | 0.58361234 | -4.420847936 |
| **A_52_P322535** | Mplkip | 0.125303199 | -0.075008339 | 0.325614738 | 8.421701976 | 1.519985282 | 0.177914768 | 0.58361234 | -4.420867369 |
| **A_51_P335419** | Csl | 0.210738315 | -0.126165702 | 0.547642332 | 9.150498607 | 1.519919206 | 0.17793096 | 0.58361234 | -4.420900884 |
| **A_55_P2113774** | Yif1b | -0.231346852 | -0.601288213 | 0.138594508 | 9.79765665 | -1.51954615 | 0.178022405 | 0.583721694 | -4.421090099 |
| **A_55_P1966204** | Cxcl12 | 0.289946793 | -0.17370322 | 0.753596806 | 10.21071815 | 1.519536799 | 0.178024698 | 0.583721694 | -4.421094842 |
| **A_52_P359965** | Cpd | -0.141447854 | -0.367681945 | 0.084786238 | 9.851163103 | -1.51922291 | 0.178101675 | 0.58378546 | -4.421254043 |
| **A_55_P2015941** | NA | 0.354258009 | -0.21239744 | 0.920913457 | 11.39126322 | 1.519091023 | 0.178134028 | 0.58378546 | -4.421320933 |
| **A_55_P2067151** | NA | 0.236100595 | -0.141556213 | 0.613757402 | 8.95388745 | 1.519088067 | 0.178134753 | 0.58378546 | -4.421322432 |

| | | | | | | | | | |
|---|---|---|---|---|---|---|---|---|---|
| **A_52_P574759** | Rbpjl | -0.860262337 | -2.23660223 | 0.516077556 | 8.269710205 | -1.518757378 | 0.178215898 | 0.583863427 | -4.421490148 |
| **A_55_P2081981** | NA | 0.125903412 | -0.075554068 | 0.327360892 | 8.343573483 | 1.518578657 | 0.178259767 | 0.583863427 | -4.421580788 |
| **A_55_P2003483** | Gldc | 0.555806017 | -0.333554632 | 1.445166666 | 12.09188418 | 1.518547997 | 0.178267294 | 0.583863427 | -4.421596337 |
| **A_65_P09177** | Strn3 | 0.177660695 | -0.106629223 | 0.461950612 | 9.64105216 | 1.518494027 | 0.178280544 | 0.583863427 | -4.421623708 |
| **A_55_P1958394** | Slit2 | -0.714447197 | -1.857798093 | 0.428903698 | 8.369493603 | -1.518356629 | 0.17831428 | 0.583863427 | -4.421693389 |
| **A_66_P128593** | Lims1 | -0.146179673 | -0.380131312 | 0.087771965 | 10.33672818 | -1.518252777 | 0.178339783 | 0.583863427 | -4.421746057 |
| **A_55_P2117525** | Rbm3 | 0.160556726 | -0.096427887 | 0.41754134 | 12.68770492 | 1.518114652 | 0.178373709 | 0.5838756 | -4.421816105 |
| **A_55_P2027788** | Baiap2 | -0.123091459 | -0.320190161 | 0.074007242 | 8.460957481 | -1.517495274 | 0.17852591 | 0.584003593 | -4.422130205 |
| **A_52_P550049** | Nfya | 0.109319538 | -0.065729383 | 0.284368459 | 9.014486252 | 1.517474974 | 0.178530901 | 0.584003593 | -4.422140499 |
| **A_55_P1967025** | NA | 0.1362469 | -0.081921247 | 0.354415048 | 8.00771256 | 1.517464277 | 0.178533531 | 0.584003593 | -4.422145924 |
| **A_55_P2094641** | NA | -0.334511445 | -0.870254387 | 0.201231496 | 8.150781063 | -1.517182111 | 0.178602911 | 0.584003593 | -4.42228901 |
| **A_55_P2176391** | NA | 0.165919899 | -0.099850366 | 0.431690165 | 11.77344823 | 1.516964 | 0.178656559 | 0.584003593 | -4.422399612 |
| **A_52_P275354** | Osbpl7 | -0.122340278 | -0.318311825 | 0.073631269 | 9.676866971 | -1.51690936 | 0.178670001 | 0.584003593 | -4.422427319 |
| **A_55_P2028511** | NA | -0.222876628 | -0.579919084 | 0.134165828 | 12.47235055 | -1.516798383 | 0.178697305 | 0.584003593 | -4.422483593 |
| **A_55_P1997145** | Nlrx1 | 0.139265573 | -0.08384308 | 0.362374227 | 8.490742911 | 1.516737925 | 0.178712181 | 0.584003593 | -4.42251425 |
| **A_55_P2029775** | Gm38396 | 0.207868559 | -0.125153627 | 0.540890744 | 8.594942282 | 1.516696998 | 0.178722252 | 0.584003593 | -4.422535004 |
| **A_55_P2098697** | Tnfaip2 | 0.293673825 | -0.176918527 | 0.764266177 | 10.63613692 | 1.516364345 | 0.17880413 | 0.584003593 | -4.422703681 |
| **A_55_P2071097** | Adrm1 | 0.127569991 | -0.076860505 | 0.332000487 | 13.42602748 | 1.516303133 | 0.1788192 | 0.584003593 | -4.422734719 |
| **A_55_P2105794** | Wdr61 | 0.105931407 | -0.063831417 | 0.275694232 | 12.75670836 | 1.516230881 | 0.17883699 | 0.584003593 | -4.422771355 |
| **A_66_P102076** | Cep68 | -0.116746963 | -0.303879784 | 0.070385858 | 9.347563084 | -1.515928846 | 0.178911375 | 0.584003593 | -4.4229245 |
| **A_55_P2110887** | NA | 0.136369048 | -0.082244032 | 0.354982128 | 11.39760915 | 1.515733514 | 0.178959496 | 0.584003593 | -4.423023541 |
| **A_55_P2132237** | Fastkd3 | 0.141398971 | -0.085283078 | 0.36808102 | 9.101438372 | 1.515696715 | 0.178968563 | 0.584003593 | -4.423042199 |
| **A_55_P2115846** | Gm10836 | -0.166203538 | -0.432662823 | 0.100255747 | 7.917134401 | -1.515627912 | 0.178985517 | 0.584003593 | -4.423077084 |
| **A_55_P2144746** | Apip | 0.223344781 | -0.134729951 | 0.581419513 | 10.57206474 | 1.515602533 | 0.178991771 | 0.584003593 | -4.423089952 |
| **A_55_P2064478** | Cdc27 | 0.16115302 | -0.097217684 | 0.419523725 | 9.220304294 | 1.515578269 | 0.17899775 | 0.584003593 | -4.423102255 |
| **A_52_P318673** | Saa1 | -0.423786685 | -1.103235906 | 0.255662537 | 11.18856093 | -1.515561244 | 0.179001946 | 0.584003593 | -4.423110887 |
| **A_55_P1984314** | Trim43c | 0.181928231 | -0.109765705 | 0.473622167 | 9.342344496 | 1.515499805 | 0.179017088 | 0.584003593 | -4.423142037 |
| **A_55_P2008452** | Wtap | 0.154141284 | -0.093072649 | 0.401355217 | 10.95209774 | 1.515057856 | 0.179126042 | 0.584211421 | -4.423366111 |
| **A_51_P467224** | Fbn1 | -0.633695943 | -1.650067651 | 0.382675765 | 9.181152962 | -1.514996216 | 0.179141243 | 0.584211421 | -4.423397363 |
| **A_55_P1959738** | Zfp511 | 0.197785814 | -0.119539379 | 0.515111007 | 9.610069953 | 1.514515685 | 0.179259789 | 0.584446494 | -4.423640987 |
| **A_51_P303160** | Arg1 | -0.454097636 | -1.18267376 | 0.274478487 | 12.65390632 | -1.514458913 | 0.179273799 | 0.584446494 | -4.423669769 |
| **A_51_P469053** | Mfsd4a | -0.185719757 | -0.483725603 | 0.11228609 | 8.318002061 | -1.514315996 | 0.179309073 | 0.584462026 | -4.423742224 |
| **A_55_P1987761** | Prkcz | -0.274030759 | -0.713775655 | 0.165714137 | 8.217432546 | -1.514194608 | 0.179339038 | 0.584462026 | -4.423803764 |
| **A_52_P400999** | Arhgap31 | -0.4994878 | -1.301262676 | 0.302287076 | 9.192027276 | -1.513755141 | 0.179447562 | 0.584594367 | -4.424026554 |
| **A_55_P2180415** | Cd72 | -0.220918416 | -0.575556575 | 0.133719742 | 8.420424713 | -1.513664575 | 0.179469935 | 0.584594367 | -4.424072466 |
| **A_52_P626756** | Trnt1 | 0.261039468 | -0.158064025 | 0.68014296 | 9.955202886 | 1.513450022 | 0.179522946 | 0.584594367 | -4.424181231 |
| **A_66_P114627** | S100a16 | -0.502491168 | -1.309358555 | 0.304376218 | 11.61692534 | -1.513245741 | 0.179573432 | 0.584594367 | -4.424284786 |
| **A_52_P423128** | Arglu1 | 0.129999404 | -0.078757212 | 0.338756021 | 11.15879839 | 1.513158084 | 0.1795951 | 0.584594367 | -4.424329222 |
| **A_66_P134428** | Sfrp1 | 0.123665925 | -0.074932543 | 0.322264393 | 7.979762946 | 1.51306416 | 0.17961832 | 0.584594367 | -4.424376833 |
| **A_52_P464437** | Trp53bp1 | 0.186023175 | -0.11271743 | 0.48476378 | 9.635791313 | 1.513059428 | 0.17961949 | 0.584594367 | -4.424379232 |
| **A_65_P03719** | Fuca2 | 0.154177006 | -0.093422209 | 0.401776221 | 8.636857981 | 1.513050885 | 0.179621602 | 0.584594367 | -4.424383562 |
| **A_55_P2143934** | NA | 0.225692297 | -0.136838076 | 0.58822267 | 8.652134625 | 1.512709476 | 0.17970603 | 0.584770681 | -4.424556625 |
| **A_51_P518163** | Rrp9 | 0.201436643 | -0.122171419 | 0.525044705 | 9.424967781 | 1.5125242 | 0.179751862 | 0.584821367 | -4.42465054 |
| **A_51_P235878** | Guk1 | -0.19387073 | -0.505357628 | 0.117616167 | 11.15784111 | -1.512361638 | 0.179792085 | 0.584853789 | -4.424732941 |
| **A_52_P222096** | 4931428F04Rik | -0.131177516 | -0.34197982 | 0.079624787 | 8.032179342 | -1.512053788 | 0.179868279 | 0.584934496 | -4.424888983 |
| **A_51_P456721** | Azgp1 | -0.775405782 | -2.021551814 | 0.47074025 | 13.05581913 | -1.511970387 | 0.179888927 | 0.584934496 | -4.424931257 |
| **A_55_P2021923** | Fkbp1a | 0.529924462 | -0.321753932 | 1.381602855 | 12.82624171 | 1.511894637 | 0.179907682 | 0.584934496 | -4.424969651 |
| **A_52_P67643** | Mtor | 0.185155872 | -0.11250531 | 0.482817054 | 8.689781018 | 1.511466328 | 0.180013762 | 0.585180963 | -4.425186741 |
| **A_51_P137007** | Zw10 | 0.176370207 | -0.107232925 | 0.459973339 | 9.788648408 | 1.511114563 | 0.180100928 | 0.585197979 | -4.425365029 |
| **A_51_P317443** | Cd3eap | 0.176699962 | -0.107452068 | 0.460851993 | 9.341698342 | 1.511015369 | 0.180125515 | 0.585197979 | -4.425415303 |

| | | | | | | | | | |
|---|---|---|---|---|---|---|---|---|---|
| **A_55_P1996751** | NA | 0.160263543 | -0.097466803 | 0.417993889 | 11.01041151 | 1.510957932 | 0.180139753 | 0.585197979 | -4.425444413 |
| **A_51_P324704** | Mrpl16 | 0.165330983 | -0.100586878 | 0.431248843 | 11.93361105 | 1.510740696 | 0.180193614 | 0.585197979 | -4.425554511 |
| **A_55_P2114776** | Greb1 | 0.145286309 | -0.08841776 | 0.378990377 | 8.741561492 | 1.510572611 | 0.180235299 | 0.585197979 | -4.425639698 |
| **A_55_P2081437** | NA | 0.17875967 | -0.108813645 | 0.466332985 | 8.994433534 | 1.510442336 | 0.180267613 | 0.585197979 | -4.425705722 |
| **A_51_P108334** | Slc25a4 | -0.326476563 | -0.851712694 | 0.198759568 | 13.16226783 | -1.510360453 | 0.180287926 | 0.585197979 | -4.42574722 |
| **A_52_P459276** | Ift81 | -0.226275615 | -0.590314368 | 0.137763137 | 8.06280902 | -1.510335138 | 0.180294207 | 0.585197979 | -4.425760049 |
| **A_51_P280401** | Tcta | 0.262857788 | -0.160049633 | 0.685765209 | 8.195884736 | 1.510284382 | 0.1803068 | 0.585197979 | -4.425785772 |
| **A_55_P2123982** | Rhoh | 0.141404196 | -0.086107687 | 0.368916079 | 8.401342013 | 1.510224117 | 0.180321753 | 0.585197979 | -4.425816314 |
| **A_55_P2036952** | NA | 0.179364946 | -0.109287916 | 0.468017807 | 8.532577731 | 1.509888551 | 0.180405036 | 0.585216466 | -4.425986372 |
| **A_55_P1962731** | Clcn4 | -0.184820176 | -0.482275501 | 0.112635149 | 11.67156691 | -1.509770103 | 0.180434442 | 0.585216466 | -4.426046397 |
| **A_55_P2008016** | Armc3 | -0.224707634 | -0.586362564 | 0.136947296 | 8.261707671 | -1.509755519 | 0.180438063 | 0.585216466 | -4.426053788 |
| **A_55_P1974362** | Hp1bp3 | -0.126125135 | -0.329130239 | 0.07687997 | 12.89076162 | -1.509655571 | 0.18046288 | 0.585216466 | -4.426104438 |
| **A_55_P2074337** | Ngf | 0.390742758 | -0.238205231 | 1.019690748 | 9.216437519 | 1.509591333 | 0.180478833 | 0.585216466 | -4.426136992 |
| **A_55_P2154307** | Peak1 | -0.447945417 | -1.1693975 | 0.273506667 | 10.83170009 | -1.508692719 | 0.180702119 | 0.58552908 | -4.426592354 |
| **A_55_P2412319** | A830052D11Rik | 0.142121307 | -0.086792297 | 0.37103491 | 8.539950503 | 1.508588453 | 0.180728044 | 0.58552908 | -4.426645187 |
| **A_51_P418116** | Tmem119 | -0.434866224 | -1.135357709 | 0.265625261 | 8.887345228 | -1.508467668 | 0.18075808 | 0.58552908 | -4.42670639 |
| **A_55_P2140592** | Dmkn | -0.622132371 | -1.624303041 | 0.380038299 | 8.351465695 | -1.508427009 | 0.180768192 | 0.58552908 | -4.426726993 |
| **A_51_P391380** | Mindy2 | 0.19525404 | -0.119282765 | 0.509790845 | 10.58784929 | 1.508383405 | 0.180779037 | 0.58552908 | -4.426749087 |
| **A_55_P2060608** | NA | 0.208979443 | -0.127696095 | 0.545654981 | 14.74068069 | 1.508256457 | 0.180810614 | 0.58552908 | -4.426813412 |
| **A_52_P84275** | Derl2 | -0.251943885 | -0.657841204 | 0.153953434 | 10.57923874 | -1.5082413 | 0.180814385 | 0.58552908 | -4.426821092 |
| **A_55_P2048249** | NA | -0.13923559 | -0.363554725 | 0.085083545 | 12.89190877 | -1.50822844 | 0.180817584 | 0.58552908 | -4.426827608 |
| **A_51_P296100** | Rhot2 | 0.169775745 | -0.103804271 | 0.443355761 | 11.28129857 | 1.507906672 | 0.180897647 | 0.58569022 | -4.426990645 |
| **A_51_P131164** | Enkur | -0.295124217 | -0.770818368 | 0.180569935 | 7.859021166 | -1.507510079 | 0.180996373 | 0.585760201 | -4.427191587 |
| **A_55_P2086692** | Nfib | -0.472851551 | -1.235080927 | 0.289377825 | 9.900556221 | -1.507378528 | 0.181029132 | 0.585760201 | -4.427258239 |
| **A_51_P436342** | Hoxb5 | -0.478264887 | -1.249253306 | 0.292723531 | 8.193187976 | -1.507314345 | 0.181045117 | 0.585760201 | -4.427290758 |
| **A_55_P1994492** | Usp31 | 0.159305487 | -0.097530933 | 0.416141907 | 8.855810186 | 1.507152908 | 0.181085329 | 0.585760201 | -4.42737255 |
| **A_52_P141161** | Hnrnpf | 0.296851098 | -0.181840006 | 0.775542201 | 9.959973342 | 1.50683775 | 0.181163855 | 0.585760201 | -4.427532222 |
| **A_51_P427964** | Lmod3 | 0.103336404 | -0.063307708 | 0.269980516 | 8.129412518 | 1.506768594 | 0.18118109 | 0.585760201 | -4.427567258 |
| **A_51_P424959** | Bcl6b | -0.364081831 | -0.951249195 | 0.223085534 | 8.38614121 | -1.506677441 | 0.18120381 | 0.585760201 | -4.427613439 |
| **A_51_P193302** | Mrps7 | 0.200420232 | -0.122816212 | 0.523656677 | 10.41670584 | 1.506622433 | 0.181217522 | 0.585760201 | -4.427641307 |
| **A_55_P2055213** | Abcf1 | 0.133833516 | -0.082096136 | 0.349763167 | 8.323860932 | 1.506037521 | 0.181363385 | 0.585760201 | -4.427937629 |
| **A_66_P101108** | Tnrc18 | -0.25984539 | -0.679132568 | 0.159441788 | 9.340414749 | -1.505867027 | 0.181405922 | 0.585760201 | -4.428024 |
| **A_55_P2084626** | Rab5b | -0.138075811 | -0.360878316 | 0.084726695 | 11.35343656 | -1.50584656 | 0.181411029 | 0.585760201 | -4.428034368 |
| **A_55_P2218334** | 9430011C21Rik | 0.151804169 | -0.093156844 | 0.396765181 | 8.047108004 | 1.505809104 | 0.181420376 | 0.585760201 | -4.428053343 |
| **A_55_P2125311** | Rab3b | 0.120940291 | -0.074221106 | 0.316101689 | 8.447328822 | 1.505775685 | 0.181428716 | 0.585760201 | -4.428070273 |
| **A_52_P180826** | Ndufaf7 | 0.124025105 | -0.076117674 | 0.324167885 | 10.65814965 | 1.505749986 | 0.181435129 | 0.585760201 | -4.428083291 |
| **A_55_P2099952** | Car1 | 0.216163778 | -0.132666239 | 0.564993795 | 8.980933554 | 1.505747888 | 0.181435653 | 0.585760201 | -4.428084354 |
| **A_55_P1964433** | Zmynd12 | -0.310593469 | -0.811815626 | 0.190628689 | 8.063947221 | -1.505723562 | 0.181441723 | 0.585760201 | -4.428096677 |
| **A_55_P1982951** | Dcaf10 | 0.111829294 | -0.068645205 | 0.292303794 | 8.298577904 | 1.505646183 | 0.181461036 | 0.585760201 | -4.428135876 |
| **A_55_P1992079** | Ptprf | -0.345936661 | -0.904228319 | 0.212354997 | 11.13789097 | -1.505631317 | 0.181464746 | 0.585760201 | -4.428143406 |
| **A_52_P94521** | 2510009E07Rik | -0.107324682 | -0.280548017 | 0.065898653 | 8.249275572 | -1.505484906 | 0.181501294 | 0.585780349 | -4.428217575 |
| **A_55_P2009427** | Nrsn2 | 0.228094858 | -0.140142187 | 0.596331903 | 9.319723251 | 1.505120282 | 0.181592342 | 0.585950686 | -4.428402279 |
| **A_55_P1988498** | Naa20 | 0.151193011 | -0.092907972 | 0.395293994 | 11.09715874 | 1.505030765 | 0.181614701 | 0.585950686 | -4.428447624 |
| **A_55_P2061645** | NA | 0.307890018 | -0.189271278 | 0.805051314 | 9.326611337 | 1.50480937 | 0.181670011 | 0.586031316 | -4.42855977 |
| **A_51_P451588** | Plekhb1 | -0.71572528 | -1.871691568 | 0.440241008 | 8.422257737 | -1.504472928 | 0.181754093 | 0.586204718 | -4.428730188 |
| **A_55_P2025626** | NA | -0.098765038 | -0.258296521 | 0.060766444 | 8.100089806 | -1.504319286 | 0.181792503 | 0.586226859 | -4.42880801 |
| **A_55_P2167840** | Gm14151 | 0.248604022 | -0.152987867 | 0.65019591 | 8.625355839 | 1.504202849 | 0.181821616 | 0.586226859 | -4.428866987 |
| **A_55_P2001773** | Copg2 | -0.126240226 | -0.330219442 | 0.07773899 | 9.732349995 | -1.503817154 | 0.181918085 | 0.586374893 | -4.429062342 |
| **A_52_P306351** | Asph | 0.265205671 | -0.163405011 | 0.693816354 | 11.22874128 | 1.503498521 | 0.181997817 | 0.586374893 | -4.429223724 |
| **A_52_P190647** | Mxd3 | 0.308696911 | -0.190202606 | 0.807596428 | 9.281236738 | 1.50349639 | 0.18199835 | 0.586374893 | -4.429224804 |

| | | | | | | | | | |
|---|---|---|---|---|---|---|---|---|---|
| **A_52_P463143** | Cdc37 | -0.157858362 | -0.412997637 | 0.097280912 | 14.31797047 | -1.503396261 | 0.182023412 | 0.586374893 | -4.429275516 |
| **A_55_P2133266** | Rtl8c | -0.766617677 | -2.005743185 | 0.472507831 | 11.31203917 | -1.503303682 | 0.182046587 | 0.586374893 | -4.429322405 |
| **A_55_P2087281** | NA | 0.271197501 | -0.167181296 | 0.709576298 | 9.288751502 | 1.503208875 | 0.182070323 | 0.586374893 | -4.429370421 |
| **A_52_P171033** | Mcmbp | 0.138102159 | -0.085139561 | 0.36134388 | 8.876293495 | 1.503170683 | 0.182079885 | 0.586374893 | -4.429389764 |
| **A_55_P2184567** | NA | 0.132605891 | -0.081829993 | 0.347041775 | 8.162482138 | 1.502617851 | 0.182218356 | 0.586643348 | -4.429669744 |
| **A_55_P1955612** | Nup93 | 0.153247244 | -0.094575892 | 0.40107038 | 11.56366808 | 1.502567579 | 0.182230952 | 0.586643348 | -4.429695204 |
| **A_55_P2182006** | Cacng6 | 0.109553913 | -0.067638269 | 0.286746095 | 7.93468631 | 1.502334108 | 0.182289464 | 0.586643348 | -4.42981344 |
| **A_55_P2153923** | Slc17a5 | -0.107070994 | -0.280256525 | 0.066114538 | 8.039666193 | -1.502254164 | 0.182309503 | 0.586643348 | -4.429853925 |
| **A_55_P2262753** | 9530053J19Rik | 0.2186991 | -0.135048177 | 0.572446377 | 8.874917247 | 1.502232246 | 0.182314998 | 0.586643348 | -4.429865025 |
| **A_51_P237752** | Cavin1 | -0.639572142 | -1.674298023 | 0.395153739 | 13.16140795 | -1.501922129 | 0.182392755 | 0.586795867 | -4.430022071 |
| **A_55_P2060996** | NA | -0.098792346 | -0.258650949 | 0.061066258 | 8.076741786 | -1.501656049 | 0.182459496 | 0.586867177 | -4.430156812 |
| **A_55_P2047155** | Meis2 | -0.225783484 | -0.591165361 | 0.139598393 | 9.035625359 | -1.501510442 | 0.182496028 | 0.586867177 | -4.430230545 |
| **A_55_P2027969** | Myl6 | -0.253899565 | -0.664819603 | 0.157020473 | 15.49830868 | -1.501370319 | 0.18253119 | 0.586867177 | -4.430301501 |
| **A_55_P1988728** | Fbxl19 | 0.18452551 | -0.114121208 | 0.483172228 | 9.385466804 | 1.501349685 | 0.182536369 | 0.586867177 | -4.430311949 |
| **A_55_P2096127** | NA | 0.277051141 | -0.171437896 | 0.725540177 | 11.67475007 | 1.501036664 | 0.182614943 | 0.586949905 | -4.430470452 |
| **A_55_P1957584** | Gps2 | 0.312320209 | -0.193298361 | 0.817938779 | 10.63558604 | 1.500929385 | 0.18264188 | 0.586949905 | -4.430524774 |
| **A_55_P2106358** | NA | 0.110335353 | -0.068293763 | 0.288964469 | 8.041420836 | 1.500878818 | 0.182654578 | 0.586949905 | -4.430550379 |
| **A_52_P346458** | Tollip | -0.127993204 | -0.335237347 | 0.079250939 | 11.11831893 | -1.5006791 | 0.182704737 | 0.586949905 | -4.430651505 |
| **A_52_P178904** | Seh1l | 0.19799948 | -0.122644419 | 0.51864338 | 8.805909648 | 1.500459432 | 0.182759922 | 0.586949905 | -4.430762731 |
| **A_51_P441942** | Tial1 | 0.166628158 | -0.103222285 | 0.436478602 | 10.00771984 | 1.500404807 | 0.182773647 | 0.586949905 | -4.430790389 |
| **A_55_P1971199** | Slc4a1ap | 0.125698072 | -0.077894069 | 0.329290213 | 10.65510226 | 1.50020565 | 0.182823696 | 0.586949905 | -4.430891227 |
| **A_51_P511482** | Ift57 | -0.384532009 | -1.007375557 | 0.23831154 | 9.218176545 | -1.500157009 | 0.182835922 | 0.586949905 | -4.430915855 |
| **A_55_P2023441** | Elp4 | 0.1412508 | -0.087576081 | 0.370077682 | 9.198882478 | 1.49991644 | 0.182896399 | 0.586949905 | -4.431037657 |
| **A_51_P121275** | Cenpo | 0.177107004 | -0.109812918 | 0.464026927 | 9.412786228 | 1.499885897 | 0.182904078 | 0.586949905 | -4.432358195 |
| **A_55_P2129217** | NA | 0.166626936 | -0.103319541 | 0.436573414 | 15.59041119 | 1.499860037 | 0.182910581 | 0.586949905 | -4.431066214 |
| **A_52_P38157** | Lypla1 | -0.121743368 | -0.318983867 | 0.075497131 | 8.297055918 | -1.499796748 | 0.182926495 | 0.586949905 | -4.431098257 |
| **A_55_P1954258** | Zfp652 | 0.111677208 | -0.06930355 | 0.292657965 | 8.70898698 | 1.499392498 | 0.183028177 | 0.58701535 | -4.431302925 |
| **A_55_P2054362** | NA | -0.206856277 | -0.542097956 | 0.128385402 | 9.729925347 | -1.499318453 | 0.183046808 | 0.58701535 | -4.431340412 |
| **A_66_P126425** | Eif1 | -0.139007018 | -0.364289888 | 0.086275853 | 14.33395175 | -1.499311056 | 0.183048669 | 0.58701535 | -4.431344157 |
| **A_52_P102559** | Dars2 | 0.257546157 | -0.159869946 | 0.67496226 | 9.905096106 | 1.499232763 | 0.18306837 | 0.58701535 | -4.431383794 |
| **A_52_P191537** | NA | 0.183062198 | -0.113676464 | 0.47980086 | 9.286117519 | 1.499021034 | 0.183121659 | 0.587066639 | -4.431490986 |
| **A_66_P113490** | Sh2b2 | -0.147526875 | -0.386685212 | 0.091631463 | 8.811286141 | -1.498886697 | 0.183155477 | 0.587066639 | -4.431558995 |
| **A_55_P2085994** | Zfat | 0.110775485 | -0.068814661 | 0.290365632 | 8.496809104 | 1.498802281 | 0.183176731 | 0.587066639 | -4.43160173 |
| **A_66_P133498** | Pdcd6ip | 0.214683771 | -0.133390221 | 0.562757764 | 11.51293689 | 1.49868662 | 0.183205855 | 0.587066639 | -4.431660284 |
| **A_66_P111781** | 4930432M17Rik | 0.365779761 | -0.227374734 | 0.958934255 | 9.677429892 | 1.498425086 | 0.183271727 | 0.587158416 | -4.431792682 |
| **A_55_P1998023** | NA | 0.138701763 | -0.086233321 | 0.363636848 | 8.777007525 | 1.498331711 | 0.18329525 | 0.587158416 | -4.431839951 |
| **A_55_P1992455** | Vdac3 | 0.113738544 | -0.070847078 | 0.298324165 | 13.72076207 | 1.497245389 | 0.183569126 | 0.587562697 | -4.432389848 |
| **A_55_P1953419** | Rab13 | -0.202516733 | -0.531186159 | 0.126152693 | 8.003151354 | -1.497217156 | 0.183576249 | 0.587562697 | -4.432404138 |
| **A_52_P668715** | Nutf2 | 0.116069832 | -0.072310987 | 0.304450652 | 8.406857822 | 1.497151897 | 0.183592715 | 0.587562697 | -4.432437171 |
| **A_55_P1955072** | Alad | 0.237358178 | -0.147886755 | 0.622603111 | 12.97545356 | 1.497098769 | 0.183606121 | 0.587562697 | -4.432464063 |
| **A_55_P2107288** | Tgfbr2 | -0.132472551 | -0.347484256 | 0.082539154 | 8.254341792 | -1.497086814 | 0.183609137 | 0.587562697 | -4.432470114 |
| **A_55_P2063257** | Lgals4 | -0.833208958 | -2.185584048 | 0.519166132 | 8.792234565 | -1.497062586 | 0.183615251 | 0.587562697 | -4.432482377 |
| **A_55_P2064328** | Lama2 | -0.422731204 | -1.108908554 | 0.263446146 | 8.401218514 | -1.496963177 | 0.183640338 | 0.587562697 | -4.432532694 |
| **A_51_P407248** | Antxr1 | -0.565142043 | -1.482539691 | 0.352255606 | 8.965005978 | -1.496866894 | 0.183664639 | 0.587562697 | -4.432581428 |
| **A_51_P217706** | Ribc1 | 0.119012398 | -0.0742311 | 0.312255897 | 8.455411323 | 1.496478582 | 0.183762678 | 0.587779049 | -4.432777971 |
| **A_52_P193925** | Sulf2 | -0.300166251 | -0.787604613 | 0.187272111 | 8.271304905 | -1.496322976 | 0.183801978 | 0.587784385 | -4.432856728 |
| **A_55_P1962699** | NA | 0.171207237 | -0.106848462 | 0.449262936 | 11.55242586 | 1.496144382 | 0.183847093 | 0.587784385 | -4.432947118 |
| **A_55_P2094721** | Sox11 | -0.209974587 | -0.550999544 | 0.13105037 | 8.121031841 | -1.496110813 | 0.183855574 | 0.587784385 | -4.432964108 |
| **A_51_P417321** | Zfp236 | 0.117431234 | -0.073310557 | 0.308173024 | 8.792935894 | 1.495963361 | 0.183892832 | 0.587806276 | -4.433038736 |
| **A_51_P405397** | Ecm1 | -0.385472592 | -1.011651243 | 0.240706058 | 11.89759808 | -1.495816907 | 0.183929845 | 0.587827376 | -4.433112857 |

| | | | | | | | | | |
|---|---|---|---|---|---|---|---|---|---|
| **A_66_P135534** | Zfp950 | 0.261930149 | -0.163641017 | 0.687501316 | 8.619920291 | 1.495534665 | 0.184001194 | 0.587958187 | -4.433255699 |
| **A_52_P358349** | Mtdh | -0.161586753 | -0.424178521 | 0.101005015 | 11.53149677 | -1.495229431 | 0.184078384 | 0.588074714 | -4.433410172 |
| **A_51_P100573** | Map3k3 | -0.130904382 | -0.343646093 | 0.081837329 | 8.113066308 | -1.495149868 | 0.18409851 | 0.588074714 | -4.433450436 |
| **A_65_P15675** | Ipo9 | 0.132749961 | -0.083174697 | 0.348674618 | 8.827912684 | 1.493878751 | 0.184420317 | 0.589005339 | -4.434093669 |
| **A_55_P2168426** | Pced1b | 0.152815753 | -0.095867975 | 0.40149948 | 9.190975825 | 1.493151741 | 0.184604608 | 0.589212093 | -4.434461526 |
| **A_51_P246387** | Nktr | 0.102589854 | -0.064361042 | 0.269540749 | 9.750473724 | 1.493134208 | 0.184609055 | 0.589212093 | -4.434470397 |
| **A_52_P799815** | Tmem171 | -0.667449928 | -1.753741232 | 0.418841376 | 8.151873932 | -1.492985455 | 0.184646785 | 0.589212093 | -4.434545661 |
| **A_55_P1996421** | NA | -0.277421831 | -0.728990826 | 0.174147164 | 13.9212052 | -1.492793405 | 0.184695507 | 0.589212093 | -4.43464283 |
| **A_55_P2138739** | NA | 0.436328194 | -0.273913238 | 1.146569626 | 9.240113698 | 1.492761625 | 0.18470357 | 0.589212093 | -4.434658909 |
| **A_51_P102503** | Metap1 | 0.241898729 | -0.151949386 | 0.635746844 | 10.81498675 | 1.492409498 | 0.184792937 | 0.589212093 | -4.434837064 |
| **A_55_P1989594** | Mynn | 0.15306912 | -0.096157019 | 0.402295259 | 9.1579366 | 1.492372323 | 0.184802374 | 0.589212093 | -4.434855872 |
| **A_55_P2037141** | Jsrp1 | 0.101376685 | -0.063695773 | 0.266449143 | 8.6515233 | 1.492267416 | 0.184829008 | 0.589212093 | -4.434908947 |
| **A_55_P2005005** | Cdk5rap1 | 0.44463128 | -0.279419478 | 1.168682038 | 10.17664891 | 1.492155858 | 0.184857334 | 0.589212093 | -4.434965386 |
| **A_55_P2004636** | Ppih | 0.235590284 | -0.148078826 | 0.619259395 | 11.53917266 | 1.492051213 | 0.184883908 | 0.589212093 | -4.435018328 |
| **A_55_P2042723** | Gm2769 | 0.102361888 | -0.064351301 | 0.269075076 | 8.148022674 | 1.491940552 | 0.184912014 | 0.589212093 | -4.435074313 |
| **A_55_P2140328** | NA | 0.101552275 | -0.063859833 | 0.266964384 | 12.77824702 | 1.49178264 | 0.184952128 | 0.589212093 | -4.435154201 |
| **A_55_P2085485** | Pidd1 | 0.173979026 | -0.109409666 | 0.457367718 | 8.475288594 | 1.491755222 | 0.184959094 | 0.589212093 | -4.435168071 |
| **A_55_P1999710** | Smad2 | 0.149841942 | -0.094232606 | 0.39391649 | 9.431363014 | 1.491743272 | 0.18496213 | 0.589212093 | -4.435174117 |
| **A_52_P556281** | G3bp2 | 0.16906603 | -0.106327348 | 0.444459408 | 11.01370889 | 1.491715665 | 0.184969144 | 0.589212093 | -4.435188083 |
| **A_51_P341108** | Spint1 | -0.749737931 | -1.971214455 | 0.471738594 | 8.215477154 | -1.491446048 | 0.185037659 | 0.589212093 | -4.435324479 |
| **A_52_P403443** | Tns3 | -0.586253903 | -1.541432686 | 0.368924881 | 10.35568343 | -1.491366106 | 0.185057978 | 0.589212093 | -4.43536492 |
| **A_66_P116360** | NA | 0.153900343 | -0.09685975 | 0.404660436 | 8.619557033 | 1.491297727 | 0.18507536 | 0.589212093 | -4.435399512 |
| **A_55_P2120466** | Snx11 | 0.137109947 | -0.086319131 | 0.360539026 | 8.972133895 | 1.491119535 | 0.185120663 | 0.589212093 | -4.435489653 |
| **A_55_P2047461** | Fcho1 | 0.194404603 | -0.122407141 | 0.511216347 | 10.54413587 | 1.491037134 | 0.185141616 | 0.589212093 | -4.435531337 |
| **A_55_P2122588** | Ubr2 | 0.159500506 | -0.100429878 | 0.419430889 | 9.016961283 | 1.491036264 | 0.185141837 | 0.589212093 | -4.435531777 |
| **A_55_P2139009** | NA | 0.119429994 | -0.075206492 | 0.314066481 | 15.49860007 | 1.490981806 | 0.185155686 | 0.589212093 | -4.435559325 |
| **A_55_P2002215** | Arfip1 | 0.142528616 | -0.089776315 | 0.374833547 | 9.02174646 | 1.490825553 | 0.185195427 | 0.589241548 | -4.435638366 |
| **A_52_P625808** | Sf3b1 | 0.168674707 | -0.106302392 | 0.443651806 | 8.878629561 | 1.490515947 | 0.185274194 | 0.589395145 | -4.435794978 |
| **A_55_P2011551** | Anapc5 | 0.207944648 | -0.13112496 | 0.547014257 | 10.789181 | 1.490191468 | 0.185356779 | 0.58950004 | -4.435959108 |
| **A_55_P2075794** | Tti2 | -0.105564522 | -0.277703196 | 0.066574153 | 11.19229041 | -1.490125062 | 0.185373685 | 0.58950004 | -4.435992696 |
| **A_52_P68028** | Ace | -0.338965209 | -0.891734856 | 0.213804438 | 7.96620646 | -1.490026959 | 0.185398662 | 0.58950004 | -4.436042318 |
| **A_55_P2011659** | Anapc5 | 0.16473827 | -0.103944737 | 0.433421276 | 11.61515899 | 1.489832659 | 0.185448141 | 0.589560382 | -4.436140596 |
| **A_55_P2147971** | Ranbp1 | 0.202478689 | -0.127952349 | 0.532909727 | 12.29624843 | 1.488955348 | 0.185671703 | 0.590107166 | -4.436584318 |
| **A_55_P1952420** | Odf2 | 0.200012713 | -0.126435377 | 0.526460803 | 8.698478954 | 1.488766741 | 0.185719798 | 0.590107166 | -4.436679705 |
| **A_55_P1997872** | Apip | 0.2329771 | -0.147340762 | 0.613294962 | 11.07706569 | 1.488502953 | 0.185787083 | 0.590107166 | -4.436813112 |
| **A_66_P136899** | Nadk | 0.275389815 | -0.174177541 | 0.72495717 | 10.81841892 | 1.488457005 | 0.185798806 | 0.590107166 | -4.43683635 |
| **A_55_P2113071** | Ago4 | -0.326728986 | -0.860107504 | 0.206649532 | 8.394097579 | -1.488453712 | 0.185799646 | 0.590107166 | -4.436838015 |
| **A_51_P328489** | 1700025G04Rik | -0.252135717 | -0.663783435 | 0.159512 | 8.830388563 | -1.488304686 | 0.18583767 | 0.590107166 | -4.436913381 |
| **A_51_P508444** | St13 | -0.139536952 | -0.367377104 | 0.088303201 | 11.75434993 | -1.488134407 | 0.185881127 | 0.590107166 | -4.436999493 |
| **A_55_P1972817** | Trmt11 | 0.139419907 | -0.08823576 | 0.367075574 | 9.690160212 | 1.488091077 | 0.185892187 | 0.590107166 | -4.437021406 |
| **A_55_P2162855** | Nipa2 | 0.129400545 | -0.081896238 | 0.340697327 | 10.02890134 | 1.488080445 | 0.185894901 | 0.590107166 | -4.437026782 |
| **A_55_P2184339** | Mdga1 | -0.182393477 | -0.480256402 | 0.115469449 | 8.133228153 | -1.487907866 | 0.185938957 | 0.590150098 | -4.437114056 |
| **A_55_P2028721** | Rnf130 | -0.159796768 | -0.420813703 | 0.101220168 | 11.66005147 | -1.487587408 | 0.186020789 | 0.590305857 | -4.437276108 |
| **A_52_P558368** | Zdhhc1 | -0.383026647 | -1.008720742 | 0.242667448 | 8.244410247 | -1.487476527 | 0.186049112 | 0.590305857 | -4.437332178 |
| **A_51_P467837** | Ap3m2 | -0.103669592 | -0.273052385 | 0.0657132 | 7.896889887 | -1.48718601 | 0.186123338 | 0.590379113 | -4.437479083 |
| **A_52_P82765** | NA | 0.405297167 | -0.256980309 | 1.067574644 | 9.331234749 | 1.487019871 | 0.186165798 | 0.590379113 | -4.437563092 |
| **A_55_P2078271** | Rbis | 0.316314853 | -0.200641548 | 0.833271253 | 11.97965507 | 1.48678735 | 0.186225239 | 0.590379113 | -4.437680665 |
| **A_55_P1999878** | Sec24b | 0.111865453 | -0.070969777 | 0.294700683 | 11.08210886 | 1.486686138 | 0.186251117 | 0.590379113 | -4.437731842 |
| **A_52_P602669** | Serpinb6d | 0.133573604 | -0.084767173 | 0.351914381 | 8.018655148 | 1.486513921 | 0.186295159 | 0.590379113 | -4.43781892 |
| **A_52_P487615** | Otulinl | -0.114482256 | -0.301619543 | 0.072655031 | 8.089813261 | -1.486486743 | 0.18630211 | 0.590379113 | -4.437832661 |

| | | | | | | | | | |
|---|---|---|---|---|---|---|---|---|---|
| **A_55_P1977418** | Ugt2b34 | 0.423847825 | -0.269049619 | 1.116745268 | 13.24001345 | 1.486360614 | 0.186334373 | 0.590379113 | -4.437896435 |
| **A_55_P2055419** | Arid4a | 0.130659336 | -0.08294408 | 0.344262752 | 8.769500103 | 1.486330679 | 0.186342031 | 0.590379113 | -4.43791157 |
| **A_55_P2076871** | Lef1 | -0.214300104 | -0.564653746 | 0.136053538 | 9.618640263 | -1.486274183 | 0.186356484 | 0.590379113 | -4.437940135 |
| **A_55_P2139763** | Trim56 | 0.172491625 | -0.109526114 | 0.454509364 | 11.44600613 | 1.486191508 | 0.186377637 | 0.590379113 | -4.437981937 |
| **A_51_P157982** | Tcaf1 | -0.142371072 | -0.375172538 | 0.090430394 | 8.137383743 | -1.48600145 | 0.186426273 | 0.590436412 | -4.438078031 |
| **A_55_P2009064** | Psmc1 | 0.125363747 | -0.07968184 | 0.330409335 | 12.00840833 | 1.485609702 | 0.186526557 | 0.590440577 | -4.438276093 |
| **A_51_P200477** | Hspa4 | 0.150003277 | -0.095368548 | 0.395375102 | 10.7659855 | 1.485454123 | 0.186566398 | 0.590440577 | -4.438354749 |
| **A_51_P234140** | Cdkl1 | 0.133011743 | -0.084566796 | 0.350590282 | 8.343854278 | 1.485446853 | 0.18656826 | 0.590440577 | -4.438358425 |
| **A_55_P2149293** | Agfg2 | 0.199403794 | -0.126789709 | 0.525597298 | 8.458676799 | 1.485392751 | 0.186582117 | 0.590440577 | -4.438385777 |
| **A_55_P2182412** | Adat1 | 0.187017158 | -0.118940086 | 0.492974402 | 8.528153752 | 1.485264845 | 0.18661488 | 0.590440577 | -4.438450441 |
| **A_55_P2117169** | Zmym1 | 0.211815052 | -0.134735936 | 0.558366039 | 10.08449863 | 1.48515871 | 0.18664207 | 0.590440577 | -4.438504098 |
| **A_55_P2047489** | Calm1 | -0.364379651 | -0.960608547 | 0.231849246 | 14.35879448 | -1.484992581 | 0.186684637 | 0.590440577 | -4.438588084 |
| **A_51_P490700** | Ccdc40 | -0.22035162 | -0.580926224 | 0.140222984 | 7.849659297 | -1.484924258 | 0.186702146 | 0.590440577 | -4.438622625 |
| **A_55_P1955479** | NA | 0.139727976 | -0.088949875 | 0.368405827 | 10.36845566 | 1.484712808 | 0.186756344 | 0.590440577 | -4.43872952 |
| **A_52_P382914** | Trip6 | -0.340818816 | -0.898649004 | 0.217011373 | 11.05000106 | -1.484583874 | 0.186789399 | 0.590440577 | -4.438794699 |
| **A_52_P55190** | BC005537 | 0.208398242 | -0.132698409 | 0.549494894 | 9.654738969 | 1.484566939 | 0.186793741 | 0.590440577 | -4.43880326 |
| **A_52_P236705** | Ripply3 | -0.371476932 | -0.979493931 | 0.236540067 | 8.440354637 | -1.484565341 | 0.18679415 | 0.590440577 | -4.438804068 |
| **A_52_P418644** | Plp2 | -0.560338095 | -1.47758907 | 0.35691288 | 9.719853366 | -1.484380212 | 0.186841623 | 0.590483338 | -4.438897654 |
| **A_55_P2095880** | NA | -0.27037631 | -0.713024253 | 0.172271633 | 8.147611451 | -1.484203241 | 0.186887013 | 0.590483338 | -4.438987113 |
| **A_51_P498720** | Cacna2d1 | -0.265755133 | -0.700880095 | 0.169369829 | 7.993957987 | -1.484057926 | 0.186924292 | 0.590483338 | -4.43906057 |
| **A_55_P2072032** | NA | 0.117730389 | -0.07504771 | 0.310508488 | 15.9076436 | 1.483932259 | 0.186956536 | 0.590483338 | -4.439124093 |
| **A_55_P1969831** | Ptpmt1 | -0.113788926 | -0.300114963 | 0.072537111 | 12.54267315 | -1.483917108 | 0.186960424 | 0.590483338 | -4.439131752 |
| **A_55_P1987504** | Ccnc | 0.112108311 | -0.071522079 | 0.295738701 | 8.338834142 | 1.483462056 | 0.187077227 | 0.590677629 | -4.439361768 |
| **A_52_P302041** | NA | 0.144033716 | -0.091895861 | 0.379963292 | 10.76070117 | 1.483422853 | 0.187087293 | 0.590677629 | -4.439381584 |
| **A_51_P360004** | Ocstamp | 0.27958593 | -0.178412036 | 0.737583896 | 11.19445918 | 1.483320337 | 0.187113618 | 0.590677629 | -4.439433401 |
| **A_51_P191469** | Rnf31 | -0.141108469 | -0.372321523 | 0.090104585 | 9.919438098 | -1.48294116 | 0.187211015 | 0.590732208 | -4.439625054 |
| **A_66_P121965** | 1110032F04Rik | 0.111889407 | -0.071452354 | 0.295231168 | 8.044502234 | 1.482896237 | 0.187222558 | 0.590732208 | -4.439647759 |
| **A_55_P1981381** | Lamc1 | -0.558003547 | -1.472437597 | 0.356430502 | 11.77359563 | -1.48274941 | 0.187260287 | 0.590732208 | -4.439721969 |
| **A_51_P289414** | Spg11 | 0.118283505 | -0.075562496 | 0.312129506 | 10.07353599 | 1.482690582 | 0.187275406 | 0.590732208 | -4.439751702 |
| **A_55_P2107502** | Fat1 | -0.341025823 | -0.899919759 | 0.217868114 | 10.32443642 | -1.482658243 | 0.187283717 | 0.590732208 | -4.439768047 |
| **A_55_P1995045** | Zc3hav1 | 0.221371007 | -0.141456453 | 0.584198467 | 9.282744074 | 1.482530992 | 0.187316426 | 0.590738978 | -4.439832361 |
| **A_55_P2007535** | Edf1 | -0.104955963 | -0.277008874 | 0.067096948 | 14.38072547 | -1.482273288 | 0.187382682 | 0.590829477 | -4.439962606 |
| **A_52_P175157** | NA | -0.23403225 | -0.617772502 | 0.149708002 | 9.097971412 | -1.481909017 | 0.187476374 | 0.590829477 | -4.440146704 |
| **A_55_P2021054** | Elp5 | 0.218855949 | -0.14001164 | 0.577723538 | 11.83738309 | 1.481860455 | 0.187488868 | 0.590829477 | -4.440171246 |
| **A_52_P650387** | Ccnjl | -0.152742264 | -0.403225918 | 0.097741389 | 8.742909274 | -1.481709361 | 0.187527745 | 0.590829477 | -4.440247604 |
| **A_55_P1978191** | C2cd4c | 0.265714031 | -0.170053496 | 0.701481557 | 9.393720646 | 1.481640405 | 0.18754549 | 0.590829477 | -4.440282452 |
| **A_55_P1954086** | Postn | -0.389094513 | -1.027242742 | 0.249053716 | 10.59257061 | -1.481551412 | 0.187568394 | 0.590829477 | -4.440327426 |
| **A_52_P472324** | Slpi | -0.324778737 | -0.857493154 | 0.207935681 | 11.6617025 | -1.481413597 | 0.187603868 | 0.590829477 | -4.440397071 |
| **A_52_P78123** | Dcaf12 | 0.268611879 | -0.172061658 | 0.709285417 | 11.6394261 | 1.481124045 | 0.18767842 | 0.590829477 | -4.440543393 |
| **A_55_P1967659** | Lactb | 0.241558881 | -0.154768814 | 0.637886576 | 9.685400301 | 1.480988809 | 0.187713249 | 0.590829477 | -4.440611732 |
| **A_51_P359237** | Tmem9 | -0.151512584 | -0.400102249 | 0.097077081 | 12.90775545 | -1.480978747 | 0.18771584 | 0.590829477 | -4.440616817 |
| **A_55_P2139809** | Fgf13 | 0.16377037 | -0.10494032 | 0.432481061 | 7.81561154 | 1.480926741 | 0.187729236 | 0.590829477 | -4.440643097 |
| **A_55_P2176908** | Tdpoz8 | 0.448163828 | -0.287208875 | 1.183536531 | 11.17069913 | 1.480854826 | 0.187747761 | 0.590829477 | -4.440679437 |
| **A_55_P2077816** | Hmgxb3 | 0.103276084 | -0.066188789 | 0.272740956 | 11.36934085 | 1.480823376 | 0.187755863 | 0.590829477 | -4.440695329 |
| **A_55_P1990314** | Elavl4 | 0.439664131 | -0.282003125 | 1.161331387 | 9.476981958 | 1.480359602 | 0.187875375 | 0.590829477 | -4.440929677 |
| **A_51_P174961** | F10 | -0.240797499 | -0.636046301 | 0.154451303 | 14.37683408 | -1.480350652 | 0.187877682 | 0.590829477 | -4.440934199 |
| **A_55_P2331974** | NA | 0.152995377 | -0.09815984 | 0.404150595 | 8.4348505 | 1.480196227 | 0.187917493 | 0.590829477 | -4.441012228 |
| **A_52_P364232** | Pik3c2a | -0.108417393 | -0.286394025 | 0.069559239 | 8.165217974 | -1.480195788 | 0.187917606 | 0.590829477 | -4.44101245 |
| **A_52_P654752** | Trub1 | 0.158714284 | -0.101844437 | 0.419273005 | 8.460415171 | 1.480108629 | 0.187940079 | 0.590829477 | -4.44105649 |
| **A_55_P1960989** | Nlgn2 | -0.267807339 | -0.707558333 | 0.171943654 | 9.771132059 | -1.479785729 | 0.188023357 | 0.590829477 | -4.441219641 |

| | | | | | | | | | |
|---|---|---|---|---|---|---|---|---|---|
| **A_55_P2004732** | Ttc39a | 0.193775287 | -0.124432931 | 0.511983506 | 8.456091266 | 1.479688121 | 0.188048537 | 0.590829477 | -4.441268958 |
| **A_52_P122843** | Bicral | 0.168976997 | -0.108518794 | 0.446472789 | 8.567991027 | 1.479634251 | 0.188062435 | 0.590829477 | -4.441296176 |
| **A_51_P371119** | Cops4 | 0.142218612 | -0.091339322 | 0.375776545 | 12.1660038 | 1.479602306 | 0.188070677 | 0.590829477 | -4.441312316 |
| **A_55_P2019217** | NA | 0.133522071 | -0.085764001 | 0.352808144 | 8.535352605 | 1.479534855 | 0.188088082 | 0.590829477 | -4.441346396 |
| **A_55_P2129000** | Cyyr1 | -0.329029703 | -0.869405289 | 0.211345883 | 8.13716422 | -1.47952591 | 0.18809039 | 0.590829477 | -4.441350915 |
| **A_55_P2307621** | 5730526G10Rik | 0.132604278 | -0.0851868 | 0.350395356 | 8.513086529 | 1.47945118 | 0.188109674 | 0.590829477 | -4.441388672 |
| **A_55_P1993503** | Adora2b | -0.204248887 | -0.539758086 | 0.131260311 | 8.092146381 | -1.479239369 | 0.188164344 | 0.590829477 | -4.441495687 |
| **A_55_P2183551** | NA | -0.201775936 | -0.533235471 | 0.129683599 | 14.80640639 | -1.479183461 | 0.188178776 | 0.590829477 | -4.441523933 |
| **A_55_P2185826** | Cyp3a13 | -0.214254359 | -0.566232819 | 0.1377241 | 8.657532222 | -1.479097386 | 0.188200998 | 0.590829477 | -4.44156742 |
| **A_51_P394394** | Tspan2 | -0.53840104 | -1.423218063 | 0.346415984 | 9.383065814 | -1.478548615 | 0.188342731 | 0.591111443 | -4.441844663 |
| **A_55_P1967022** | Fgfr1 | -0.11161267 | -0.295047871 | 0.071822531 | 7.832195649 | -1.478475068 | 0.188361734 | 0.591111443 | -4.441881818 |
| **A_55_P1991552** | Atraid | -0.144821508 | -0.382874711 | 0.093231694 | 11.80236334 | -1.478230733 | 0.188424877 | 0.591111443 | -4.442005252 |
| **A_51_P227770** | Phykpl | 0.203283254 | -0.130883073 | 0.537449581 | 9.95811332 | 1.478162255 | 0.188442577 | 0.591111443 | -4.442039845 |
| **A_51_P364485** | Tnfaip2 | 0.299223895 | -0.192668259 | 0.791116048 | 11.32975476 | 1.478119564 | 0.188453613 | 0.591111443 | -4.442061411 |
| **A_55_P2081313** | Gnas | -0.34180266 | -0.903801663 | 0.220196343 | 14.33411744 | -1.47782525 | 0.188529708 | 0.591111443 | -4.442210087 |
| **A_55_P2097259** | Zup1 | 0.117832686 | -0.075931385 | 0.311596757 | 9.96080567 | 1.477664079 | 0.188571391 | 0.591111443 | -4.442291502 |
| **A_51_P433870** | Vegfc | -0.188074515 | -0.497344581 | 0.12119555 | 9.417340479 | -1.477662508 | 0.188571798 | 0.591111443 | -4.442292296 |
| **A_51_P170911** | Ptpn9 | -0.12174216 | -0.32195634 | 0.078472021 | 8.242281399 | -1.477506354 | 0.188612192 | 0.591111443 | -4.442371175 |
| **A_51_P320089** | Gclm | 0.297249596 | -0.19164101 | 0.786140203 | 10.56923251 | 1.477381886 | 0.188644395 | 0.591111443 | -4.442434048 |
| **A_55_P2046348** | Itpripl2 | -0.29697489 | -0.785422206 | 0.191472426 | 10.61788651 | -1.477356106 | 0.188651065 | 0.591111443 | -4.44244707 |
| **A_55_P1960291** | Tjp3 | -0.912923953 | -2.414475493 | 0.588627586 | 8.863299599 | -1.477330104 | 0.188657794 | 0.591111443 | -4.442460204 |
| **A_55_P2137932** | NA | 0.262330329 | -0.169360636 | 0.694021293 | 8.942047973 | 1.476585958 | 0.188850442 | 0.591300859 | -4.442836074 |
| **A_51_P201137** | Lbr | 0.262690507 | -0.169599138 | 0.694980152 | 11.66467712 | 1.476565563 | 0.188855725 | 0.591300859 | -4.442846375 |
| **A_55_P1957835** | Cpeb3 | 0.314990552 | -0.203383387 | 0.83336449 | 10.01255909 | 1.476513959 | 0.188869091 | 0.591300859 | -4.442872439 |
| **A_55_P2063490** | Csprs | 0.125131164 | -0.080810119 | 0.331072447 | 8.075668572 | 1.476404156 | 0.188897536 | 0.591300859 | -4.442927899 |
| **A_55_P2025073** | Csf1r | 0.18989282 | -0.122637811 | 0.502423451 | 9.287800591 | 1.476383368 | 0.188902921 | 0.591300859 | -4.442938398 |
| **A_55_P2122334** | Nfic | -0.240097793 | -0.635328466 | 0.15513288 | 10.38813175 | -1.476116775 | 0.188972 | 0.591300859 | -4.443073044 |
| **A_55_P2084512** | NA | 0.101808941 | -0.06578626 | 0.269404141 | 7.97718436 | 1.476071961 | 0.188983615 | 0.591300859 | -4.443095678 |
| **A_55_P2013088** | Arpc4 | -0.165651071 | -0.438380244 | 0.107078103 | 9.247999649 | -1.475862302 | 0.189037961 | 0.591300859 | -4.443201566 |
| **A_51_P344008** | Slc25a20 | 0.219785598 | -0.142111291 | 0.581682487 | 8.926939447 | 1.475698278 | 0.189080488 | 0.591300859 | -4.443284404 |
| **A_55_P1965635** | Setdb1 | 0.157793358 | -0.102058569 | 0.417645285 | 9.462454067 | 1.475522937 | 0.189125958 | 0.591300859 | -4.443372955 |
| **A_66_P103301** | Dicer1 | 0.126506348 | -0.081878213 | 0.334890909 | 9.703648005 | 1.475128892 | 0.189228182 | 0.591300859 | -4.443571952 |
| **A_55_P2046153** | NA | 0.119115527 | -0.07710165 | 0.315332704 | 14.9628062 | 1.475076495 | 0.189241779 | 0.591300859 | -4.443598413 |
| **A_52_P243516** | Rmdn2 | -0.210201148 | -0.556471473 | 0.136069177 | 8.339036537 | -1.475037284 | 0.189251955 | 0.591300859 | -4.443618214 |
| **A_55_P2140751** | NA | -0.268194137 | -0.709999179 | 0.173610906 | 11.45681091 | -1.475033214 | 0.189253011 | 0.591300859 | -4.44362027 |
| **A_55_P2186352** | Ttc12 | -0.428312296 | -1.133956465 | 0.277331872 | 8.2757808 | -1.474884375 | 0.189291641 | 0.591300859 | -4.443695432 |
| **A_51_P148037** | Sh3pxd2b | -0.276839581 | -0.732955046 | 0.179275884 | 8.209347563 | -1.474811703 | 0.189310505 | 0.591300859 | -4.44373213 |
| **A_52_P436238** | Odc1 | 0.239190157 | -0.15494319 | 0.633323505 | 10.80166622 | 1.474630836 | 0.189357462 | 0.591300859 | -4.443823464 |
| **A_55_P1986711** | NA | 0.175795431 | -0.11389836 | 0.465489222 | 9.134123564 | 1.474523119 | 0.189385433 | 0.591300859 | -4.443877857 |
| **A_51_P435068** | Acadsb | -0.320948964 | -0.849876141 | 0.207978213 | 9.505640917 | -1.474427032 | 0.189410387 | 0.591300859 | -4.443926378 |
| **A_55_P2163812** | 4-Sep | -0.584032974 | -1.546534692 | 0.378468745 | 8.645864295 | -1.474412632 | 0.189414127 | 0.591300859 | -4.443933649 |
| **A_55_P2076489** | Rac1 | -0.1082712 | -0.286715859 | 0.070173459 | 12.66107328 | -1.47432281 | 0.189437457 | 0.591300859 | -4.443979005 |
| **A_55_P1970670** | Scp2 | -0.17179205 | -0.454928724 | 0.111344623 | 14.31886539 | -1.474314464 | 0.189439625 | 0.591300859 | -4.44398322 |
| **A_55_P2078830** | NA | 0.112556223 | -0.072965693 | 0.298078139 | 7.861977276 | 1.474203644 | 0.189468413 | 0.591300859 | -4.444039178 |
| **A_51_P140731** | Cops5 | 0.107552634 | -0.069725968 | 0.284831236 | 13.11147716 | 1.474171198 | 0.189476842 | 0.591300859 | -4.444055562 |
| **A_55_P1967315** | Ppp1r9b | 0.129068366 | -0.08368822 | 0.341824951 | 11.47096385 | 1.47407638 | 0.189501477 | 0.591300859 | -4.44410344 |
| **A_55_P2175813** | Ythdf2 | 0.122701854 | -0.079566574 | 0.324970283 | 9.732884111 | 1.474029637 | 0.189513623 | 0.591300859 | -4.444127042 |
| **A_55_P1966659** | LOC547349 | 0.108187071 | -0.070169123 | 0.286543265 | 8.045928954 | 1.473907935 | 0.189545249 | 0.591304088 | -4.444188493 |
| **A_55_P1993976** | Aimp2 | 0.149014815 | -0.096712713 | 0.394742343 | 12.90314554 | 1.47352947 | 0.189643632 | 0.591510654 | -4.444379588 |
| **A_55_P2181084** | Rpl17 | 0.211967022 | -0.137616189 | 0.561550233 | 15.07523935 | 1.473333014 | 0.189694719 | 0.591510654 | -4.444478779 |

| | | | | | | | | | |
|---|---|---|---|---|---|---|---|---|---|
| **A_52_P64356** | Sparcl1 | -1.419861757 | -3.761706652 | 0.921983137 | 11.66115337 | -1.473231932 | 0.18972101 | 0.591510654 | -4.444529814 |
| **A_55_P2121618** | Tox4 | -1.066220189 | -2.82487078 | 0.692430402 | 9.347983782 | -1.47316207 | 0.189739183 | 0.591510654 | -4.444565087 |
| **A_55_P2136786** | Zkscan1 | 0.19833366 | -0.128824495 | 0.525491815 | 14.52485889 | 1.473064843 | 0.189764477 | 0.591510654 | -4.444614176 |
| **A_51_P333622** | 2010300C02Rik | -0.744059039 | -1.971835541 | 0.483717462 | 8.797845577 | -1.472554133 | 0.189897389 | 0.591829511 | -4.444872017 |
| **A_55_P2067142** | NA | 0.119155808 | -0.077507176 | 0.315818793 | 16.97250794 | 1.472230401 | 0.189981685 | 0.591894181 | -4.44503545 |
| **A_55_P1982563** | Rhbdf1 | -0.440917622 | -1.168673816 | 0.286838573 | 9.861257256 | -1.472159054 | 0.190000267 | 0.591894181 | -4.445071469 |
| **A_51_P349961** | Gc | -0.478900253 | -1.269368722 | 0.311568215 | 14.75863381 | -1.472121667 | 0.190010005 | 0.591894181 | -4.445090343 |
| **A_55_P2134236** | Foxa2 | -0.40473867 | -1.072899809 | 0.26342247 | 11.10452146 | -1.471894477 | 0.190069192 | 0.591983146 | -4.445205033 |
| **A_51_P224771** | Tmem184b | -0.205519146 | -0.54482985 | 0.133791558 | 10.94528959 | -1.471763116 | 0.190103421 | 0.591994365 | -4.445271346 |
| **A_55_P2070766** | NA | 0.3522495 | -0.229443606 | 0.933942606 | 11.92991056 | 1.471430173 | 0.190190203 | 0.592146479 | -4.445439414 |
| **A_55_P1963483** | Vps16 | 0.178487399 | -0.116278684 | 0.473253483 | 12.02041868 | 1.471340668 | 0.190213539 | 0.592146479 | -4.445484595 |
| **A_51_P324871** | Cyb561a3 | -0.126024918 | -0.334181845 | 0.08213201 | 11.04849435 | -1.47112218 | 0.190270514 | 0.592201279 | -4.445594882 |
| **A_55_P1974671** | Sft2d1 | -0.095840103 | -0.254171787 | 0.062491582 | 9.62281242 | -1.470830502 | 0.190346599 | 0.592201279 | -4.44574211 |
| **A_51_P446825** | Trmt9b | -0.112763755 | -0.299055222 | 0.073527712 | 7.877414025 | -1.470820774 | 0.190349138 | 0.592201279 | -4.445747021 |
| **A_51_P467410** | Npm1 | 0.177516004 | -0.115774722 | 0.470806731 | 15.32009482 | 1.470694174 | 0.190382171 | 0.592201279 | -4.445810922 |
| **A_55_P2004452** | Tceal8 | 0.247194235 | -0.161220686 | 0.655609157 | 10.23198125 | 1.470685893 | 0.190384332 | 0.592201279 | -4.445815101 |
| **A_55_P2170670** | NA | 0.120165887 | -0.078398879 | 0.318730653 | 9.944055611 | 1.470490414 | 0.190435348 | 0.592264658 | -4.445913767 |
| **A_55_P1986703** | Aldh3b3 | 0.397707116 | -0.259590556 | 1.055004788 | 8.967386643 | 1.470227212 | 0.19050406 | 0.592383039 | -4.446046612 |
| **A_52_P100252** | Fasn | -0.203717762 | -0.540460656 | 0.133025133 | 9.390513856 | -1.469987509 | 0.190566656 | 0.592397129 | -4.446167593 |
| **A_51_P213691** | Scnn1a | -0.888544635 | -2.357310085 | 0.580220815 | 8.991837712 | -1.469975139 | 0.190569887 | 0.592397129 | -4.446173836 |
| **A_55_P1995512** | Tmem139 | -0.572097655 | -1.51785296 | 0.37365765 | 8.115056124 | -1.46985526 | 0.1906012 | 0.592399196 | -4.446234339 |
| **A_51_P384718** | Efna4 | -0.148077974 | -0.392952456 | 0.096796509 | 7.910017329 | -1.469366451 | 0.190728929 | 0.592447387 | -4.446481032 |
| **A_55_P2069842** | D5Ertd579e | 0.228537421 | -0.149395315 | 0.606470157 | 8.621641054 | 1.469352514 | 0.190732572 | 0.592447387 | -4.446488065 |
| **A_55_P2154982** | Cd8b1 | 0.20558774 | -0.134474974 | 0.545650454 | 8.511885621 | 1.468998608 | 0.190825101 | 0.592447387 | -4.446666666 |
| **A_55_P2121608** | NA | -0.549121905 | -1.457428754 | 0.359184944 | 10.53699018 | -1.46899179 | 0.190826884 | 0.592447387 | -4.446670107 |
| **A_55_P1999849** | NA | 0.182080394 | -0.119103539 | 0.483264326 | 11.7530872 | 1.468975569 | 0.190831126 | 0.592447387 | -4.446678293 |
| **A_51_P398235** | Dnajc11 | 0.195507814 | -0.128011276 | 0.519026905 | 9.151686327 | 1.468410216 | 0.190979028 | 0.592447387 | -4.452667175 |
| **A_52_P605812** | Ptrh1 | 0.153838921 | -0.100735986 | 0.408413829 | 9.011868215 | 1.468364313 | 0.190991041 | 0.592447387 | -4.446986749 |
| **A_55_P2059019** | Mapk8ip3 | 0.173796557 | -0.11384082 | 0.461433935 | 11.57213369 | 1.468179105 | 0.191039519 | 0.592447387 | -4.447080205 |
| **A_51_P227222** | Adamts2 | -0.434908609 | -1.154734748 | 0.28491753 | 10.1630315 | -1.468093043 | 0.19106205 | 0.592447387 | -4.447123632 |
| **A_55_P2125229** | Arhgef2 | -0.315488064 | -0.837675669 | 0.206699541 | 11.17967912 | -1.468045658 | 0.191074456 | 0.592447387 | -4.447147542 |
| **A_55_P2008066** | Itpr1 | -0.168876345 | -0.448409406 | 0.110656716 | 10.02057135 | -1.467975578 | 0.191092805 | 0.592447387 | -4.447182903 |
| **A_55_P2067895** | Krba1 | -0.273715356 | -0.726788578 | 0.179357865 | 9.114885596 | -1.46795911 | 0.191097117 | 0.592447387 | -4.447191213 |
| **A_51_P187842** | Eif4e3 | -0.136455189 | -0.362330433 | 0.089420055 | 8.068595159 | -1.467927513 | 0.191105391 | 0.592447387 | -4.447207156 |
| **A_55_P2087087** | Pecam1 | -0.175359472 | -0.46564655 | 0.114927606 | 8.222436896 | -1.467860263 | 0.191123002 | 0.592447387 | -4.447241089 |
| **A_51_P414448** | Naa15 | 0.220029593 | -0.144223745 | 0.584282932 | 10.03590335 | 1.467779252 | 0.191144219 | 0.592447387 | -4.447281965 |
| **A_55_P1974421** | Itga3 | -0.205105801 | -0.544714018 | 0.134502415 | 8.04513601 | -1.467516337 | 0.191213091 | 0.592447387 | -4.447414623 |
| **A_52_P413584** | NA | 0.143746633 | -0.094272264 | 0.381765531 | 8.498211716 | 1.467470674 | 0.191225055 | 0.592447387 | -4.447437663 |
| **A_55_P2145720** | Thoc2 | 0.166286396 | -0.109090401 | 0.441663194 | 8.893041841 | 1.46727854 | 0.191275403 | 0.592447387 | -4.447534604 |
| **A_55_P2132502** | Pkn1 | -0.183506263 | -0.487410054 | 0.120397528 | 10.57391183 | -1.467229204 | 0.191288333 | 0.592447387 | -4.447559496 |
| **A_51_P314277** | Parp1 | 0.163604997 | -0.107357666 | 0.43456766 | 9.64487652 | 1.46713576 | 0.191312826 | 0.592447387 | -4.447606642 |
| **A_55_P2313033** | Ptprb | -0.159218544 | -0.422934112 | 0.104497025 | 8.393583155 | -1.467037 | 0.191338715 | 0.592447387 | -4.44765647 |
| **A_66_P116195** | Psmd13 | 0.229306108 | -0.1505566 | 0.609168817 | 12.55009115 | 1.466804225 | 0.191399748 | 0.592447387 | -4.447773911 |
| **A_55_P2106079** | NA | 0.136558547 | -0.089709287 | 0.362826381 | 8.134667919 | 1.466490511 | 0.191482031 | 0.592447387 | -4.447932181 |
| **A_52_P384901** | Wdr43 | 0.258439314 | -0.169813017 | 0.686691644 | 9.815719796 | 1.466364821 | 0.191515007 | 0.592447387 | -4.447995591 |
| **A_51_P222475** | Ehbp1l1 | 0.23287624 | -0.1530242 | 0.61877668 | 12.26485937 | 1.466334645 | 0.191522925 | 0.592447387 | -4.448010815 |
| **A_55_P1991618** | Ppp1r16a | -0.249771073 | -0.663687697 | 0.164145552 | 10.59981052 | -1.466264967 | 0.191541208 | 0.592447387 | -4.448045966 |
| **A_55_P1981099** | NA | -0.149074428 | -0.396135503 | 0.097986647 | 14.65377049 | -1.466162185 | 0.191568182 | 0.592447387 | -4.448097818 |
| **A_51_P225771** | NA | 0.259284436 | -0.170478626 | 0.689047499 | 12.46137916 | 1.465988461 | 0.19161378 | 0.592447387 | -4.448185457 |
| **A_52_P599264** | Mdfic | -0.391802728 | -1.041257377 | 0.257651921 | 8.816686292 | -1.46589146 | 0.191639245 | 0.592447387 | -4.44823439 |

| | | | | | | | | | |
|---|---|---|---|---|---|---|---|---|---|
| **A_55_P2156186** | BC028528 | -0.461404193 | -1.226235867 | 0.303427481 | 8.514669757 | -1.46588149 | 0.191641863 | 0.592447387 | -4.44823942 |
| **A_66_P115949** | Myo9a | 0.115203407 | -0.075783195 | 0.306190009 | 8.403337601 | 1.465701762 | 0.191689054 | 0.592447387 | -4.448330084 |
| **A_55_P2015585** | Zcchc24 | -0.194642222 | -0.517354385 | 0.128069941 | 12.01392218 | -1.465564406 | 0.191725127 | 0.592447387 | -4.448399373 |
| **A_51_P293781** | Fkbp1b | -0.151739866 | -0.403383607 | 0.099903875 | 9.826483769 | -1.465199467 | 0.191820999 | 0.592447387 | -4.448583459 |
| **A_55_P2010048** | NA | -0.171133892 | -0.454959804 | 0.11269202 | 8.653625703 | -1.465099684 | 0.19184722 | 0.592447387 | -4.448633792 |
| **A_55_P2064567** | Anks1 | -0.125105306 | -0.332629076 | 0.082418464 | 8.5668776 | -1.464842963 | 0.191914698 | 0.592447387 | -4.448763284 |
| **A_51_P272147** | Emc9 | 0.184477441 | -0.121544142 | 0.490499024 | 11.65479597 | 1.464786731 | 0.191929481 | 0.592447387 | -4.448791647 |
| **A_55_P2008258** | Larp1b | 0.318067028 | -0.209617364 | 0.84575142 | 11.66253699 | 1.464628875 | 0.191970986 | 0.592447387 | -4.448871268 |
| **A_55_P2259500** | D130012P04Rik | 0.111326735 | -0.073369198 | 0.296022668 | 8.234396891 | 1.464621239 | 0.191972994 | 0.592447387 | -4.448875119 |
| **A_66_P116678** | Rps8 | 0.309603373 | -0.204062351 | 0.823269098 | 10.08699572 | 1.464563766 | 0.191988108 | 0.592447387 | -4.448904107 |
| **A_55_P2080119** | NA | -0.194859158 | -0.51819195 | 0.128473634 | 14.57211181 | -1.464381582 | 0.192036024 | 0.592447387 | -4.448995997 |
| **A_51_P324442** | Wdr35 | -0.242855586 | -0.645839811 | 0.160128638 | 8.155822959 | -1.464344436 | 0.192045795 | 0.592447387 | -4.449014732 |
| **A_55_P1961009** | Apbb1ip | 0.344117526 | -0.226975083 | 0.915210134 | 10.22079859 | 1.464142817 | 0.192098838 | 0.592447387 | -4.449116421 |
| **A_55_P2168914** | NA | 0.422753689 | -0.278860492 | 1.124367869 | 9.277511612 | 1.464105043 | 0.192108777 | 0.592447387 | -4.449135473 |
| **A_55_P2065519** | NA | 0.116199272 | -0.076650382 | 0.309048926 | 9.883676631 | 1.464089857 | 0.192112773 | 0.592447387 | -4.449143132 |
| **A_55_P2012498** | Cdca7 | 0.152622292 | -0.100678703 | 0.405923287 | 11.74153945 | 1.46407817 | 0.192115848 | 0.592447387 | -4.449149026 |
| **A_66_P125035** | Duxbl2 | -0.194130538 | -0.51633177 | 0.128070693 | 7.820798687 | -1.46402958 | 0.192128635 | 0.592447387 | -4.449173533 |
| **A_55_P2068391** | Gm9222 | -0.195952378 | -0.52122953 | 0.129324774 | 9.03966707 | -1.463794692 | 0.192190456 | 0.592447387 | -4.449291997 |
| **A_55_P2280868** | 0610005C13Rik | 0.296987608 | -0.196008986 | 0.789984201 | 9.537645682 | 1.463786032 | 0.192192736 | 0.592447387 | -4.449296364 |
| **A_55_P2050353** | Prpmp5 | 0.162684239 | -0.107377666 | 0.432746144 | 8.552321379 | 1.463744726 | 0.192203609 | 0.592447387 | -4.449317196 |
| **A_51_P107234** | Abcf1 | 0.183255858 | -0.120973379 | 0.487485094 | 12.20524843 | 1.463659674 | 0.192226001 | 0.592447387 | -4.449360091 |
| **A_51_P128463** | Grrp1 | -0.487240286 | -1.296157925 | 0.321677352 | 9.113229618 | -1.463598897 | 0.192242003 | 0.592447387 | -4.450849296 |
| **A_55_P1966585** | NA | 0.127196664 | -0.08400994 | 0.338403269 | 15.72782633 | 1.463360798 | 0.192304703 | 0.592447387 | -4.449510819 |
| **A_51_P214825** | Orc5 | 0.142459489 | -0.094093714 | 0.379012693 | 8.931041892 | 1.463341664 | 0.192309743 | 0.592447387 | -4.449520469 |
| **A_65_P11059** | Letm1 | 0.098379359 | -0.064986295 | 0.261745014 | 10.78175931 | 1.463276583 | 0.192326885 | 0.592447387 | -4.449553289 |
| **A_51_P240693** | Tecpr1 | -0.546604477 | -1.454400675 | 0.361191721 | 9.880385367 | -1.463079799 | 0.192378725 | 0.592447387 | -4.449652527 |
| **A_55_P1998957** | Oas1c | -0.139355732 | -0.370801624 | 0.092090159 | 8.013596203 | -1.463047923 | 0.192387124 | 0.592447387 | -4.449668602 |
| **A_52_P138806** | Dlgap3 | 0.179417181 | -0.118568539 | 0.4774029 | 8.412644698 | 1.463025058 | 0.192393149 | 0.592447387 | -4.449680132 |
| **A_55_P2027496** | Adck5 | -0.237053884 | -0.630767072 | 0.156659303 | 11.69298568 | -1.463020134 | 0.192394446 | 0.592447387 | -4.449682615 |
| **A_51_P251205** | Psd | 0.33436384 | -0.221085434 | 0.889813113 | 9.452717557 | 1.462709526 | 0.192476305 | 0.592605049 | -4.449839248 |
| **A_55_P2253348** | Galnt7 | 0.238319582 | -0.157638568 | 0.634277733 | 10.91319762 | 1.462492444 | 0.192533535 | 0.592685882 | -4.449948715 |
| **A_55_P2149363** | 6430548M08Rik | -0.146235982 | -0.389231382 | 0.096759418 | 8.516721712 | -1.462309774 | 0.192581704 | 0.592685882 | -4.450040826 |
| **A_52_P427759** | NA | 0.19188998 | -0.126977749 | 0.510757709 | 13.28016873 | 1.462261073 | 0.192594548 | 0.592685882 | -4.450065384 |
| **A_55_P2167416** | NA | 0.281169867 | -0.186306841 | 0.748646575 | 8.59859749 | 1.461477007 | 0.192801444 | 0.592917678 | -4.450460724 |
| **A_52_P161806** | Ovca2 | 0.125655826 | -0.083369545 | 0.334681198 | 9.516169584 | 1.460719461 | 0.193001536 | 0.592917678 | -4.450842656 |
| **A_52_P28792** | Isg20l2 | 0.172662697 | -0.114563696 | 0.459889091 | 9.926702642 | 1.460687678 | 0.193009935 | 0.592917678 | -4.45085868 |
| **A_51_P473242** | Zfp426 | 0.099839166 | -0.06624709 | 0.265925422 | 8.608816136 | 1.460664375 | 0.193016093 | 0.592917678 | -4.450870428 |
| **A_66_P129263** | Lekr1 | -0.097586401 | -0.259958227 | 0.064785426 | 7.946088575 | -1.46036633 | 0.193094874 | 0.592917678 | -4.451020683 |
| **A_55_P1999923** | Nyap1 | -0.155335924 | -0.413801998 | 0.10313015 | 8.399852535 | -1.46033182 | 0.193103998 | 0.592917678 | -4.45103808 |
| **A_52_P612382** | Cdc25b | 0.185498137 | -0.123172443 | 0.494168717 | 13.25362948 | 1.460250968 | 0.193125375 | 0.592917678 | -4.45107884 |
| **A_55_P1981210** | Pex26 | 0.179778617 | -0.119416074 | 0.478973307 | 8.134487686 | 1.460048695 | 0.193178866 | 0.592917678 | -4.451180808 |
| **A_65_P03031** | Ncapd3 | 0.225422992 | -0.149739069 | 0.600585053 | 8.996679211 | 1.460032373 | 0.193183183 | 0.592917678 | -4.451189036 |
| **A_52_P458000** | AI467606 | -0.132452925 | -0.352895832 | 0.087989982 | 9.319046916 | -1.459985734 | 0.193195519 | 0.592917678 | -4.451212547 |
| **A_55_P2184741** | Katnal1 | 0.160686393 | -0.106767903 | 0.428140688 | 10.24950444 | 1.459865055 | 0.193227441 | 0.592917678 | -4.451273381 |
| **A_55_P2035812** | Rnf208 | -0.130532319 | -0.347814127 | 0.08674949 | 7.878481346 | -1.45974795 | 0.193258423 | 0.592917678 | -4.451332412 |
| **A_51_P202033** | Wls | -0.656573101 | -1.749531389 | 0.436385186 | 9.185756953 | -1.459696907 | 0.193271928 | 0.592917678 | -4.451358142 |
| **A_55_P2123491** | B3gnt5 | -0.134938774 | -0.359569482 | 0.089691934 | 7.944159238 | -1.459657032 | 0.193282479 | 0.592917678 | -4.451378243 |
| **A_52_P189132** | Thap1 | 0.153258355 | -0.101880027 | 0.408396736 | 9.84148848 | 1.45959226 | 0.193299619 | 0.592917678 | -4.451410893 |
| **A_51_P406020** | Abhd4 | -0.304905434 | -0.81257799 | 0.202767123 | 12.96668157 | -1.459367484 | 0.193359111 | 0.592917678 | -4.451524196 |
| **A_55_P1963737** | Aktip | -0.157226459 | -0.419013031 | 0.104560114 | 8.06054955 | -1.459356694 | 0.193361967 | 0.592917678 | -4.451529635 |

| | | | | | | | | | |
|---|---|---|---|---|---|---|---|---|---|
| **A_55_P2278604** | A330009N23Rik | -0.272000262 | -0.724921463 | 0.18092094 | 8.347919249 | -1.459250532 | 0.193390072 | 0.592917678 | -4.451583147 |
| **A_52_P672803** | Ctsa | 0.26327167 | -0.175142321 | 0.701685661 | 8.52303777 | 1.459159986 | 0.193414045 | 0.592917678 | -4.451628787 |
| **A_52_P53626** | NA | 0.119005763 | -0.079222634 | 0.317234161 | 8.656484356 | 1.458764929 | 0.193518674 | 0.592917678 | -4.451827911 |
| **A_55_P1969543** | Pin4 | 0.163861965 | -0.109093429 | 0.436817359 | 11.96684524 | 1.458712371 | 0.193532597 | 0.592917678 | -4.451854402 |
| **A_55_P1989846** | Tm6sf1 | -0.146275207 | -0.389943703 | 0.097393288 | 8.521236971 | -1.45866153 | 0.193546067 | 0.592917678 | -4.451880026 |
| **A_52_P493810** | Srpk1 | 0.168060163 | -0.11189946 | 0.448019785 | 9.081131474 | 1.458654947 | 0.193547811 | 0.592917678 | -4.451883345 |
| **A_55_P2013029** | NA | 0.169956204 | -0.113162822 | 0.45307523 | 8.636482071 | 1.458650198 | 0.193549069 | 0.592917678 | -4.451885738 |
| **A_55_P2168692** | Cep131 | -0.135388454 | -0.360931531 | 0.090154624 | 9.182030795 | -1.458596996 | 0.193563166 | 0.592917678 | -4.451912553 |
| **A_55_P1993128** | Gas8 | -0.289985962 | -0.773082419 | 0.193110496 | 9.206515929 | -1.458566754 | 0.193571179 | 0.592917678 | -4.451927796 |
| **A_55_P2175942** | NA | -0.141347166 | -0.376837193 | 0.09414286 | 13.12530574 | -1.458470927 | 0.193596572 | 0.592917678 | -4.451976093 |
| **A_51_P345836** | Prkra | -0.272189094 | -0.725702188 | 0.181323999 | 9.545386553 | -1.458357768 | 0.193626562 | 0.592917678 | -4.452033126 |
| **A_52_P576049** | Inhbc | 0.219202409 | -0.146031107 | 0.584435925 | 8.487924001 | 1.458336976 | 0.193632073 | 0.592917678 | -4.452043605 |
| **A_55_P2140955** | NA | 0.110767011 | -0.073804367 | 0.295338389 | 16.06753899 | 1.458240891 | 0.193657542 | 0.592917678 | -4.452092032 |
| **A_55_P2022263** | NA | 0.166211073 | -0.110759299 | 0.443181445 | 17.27837739 | 1.458175603 | 0.19367485 | 0.592917678 | -4.452124936 |
| **A_51_P161830** | Enpep | -0.382822348 | -1.020759229 | 0.255114533 | 9.055300822 | -1.458151876 | 0.19368114 | 0.592917678 | -4.452136894 |
| **A_55_P2148116** | Tsr2 | 0.14098823 | -0.093973069 | 0.375949529 | 8.176629788 | 1.458040922 | 0.193710558 | 0.592917678 | -4.452192813 |
| **A_51_P156857** | Smim3 | 0.160683222 | -0.107101966 | 0.42846841 | 11.16809017 | 1.458032383 | 0.193712822 | 0.592917678 | -4.452197117 |
| **A_55_P2191002** | 2610027K06Rik | 0.156140505 | -0.104130902 | 0.416411913 | 8.470951004 | 1.457713949 | 0.193797273 | 0.592923962 | -4.452357599 |
| **A_51_P409429** | Aars | 0.298568431 | -0.19914739 | 0.796284252 | 12.34227818 | 1.457624399 | 0.193821029 | 0.592923962 | -4.452402728 |
| **A_52_P451378** | Twsg1 | -0.282174605 | -0.752574692 | 0.188225481 | 10.05075151 | -1.457584437 | 0.193831631 | 0.592923962 | -4.452422867 |
| **A_51_P406365** | Maf1 | -0.247250069 | -0.659435433 | 0.164935294 | 11.62653446 | -1.457562026 | 0.193837576 | 0.592923962 | -4.452434161 |
| **A_51_P280532** | Supt16 | 0.143418495 | -0.095741247 | 0.382578236 | 8.458779004 | 1.457136624 | 0.193950471 | 0.593071042 | -4.452648538 |
| **A_55_P2091196** | Hrct1 | -0.282594681 | -0.753840824 | 0.188651463 | 8.049357854 | -1.457133567 | 0.193951282 | 0.593071042 | -4.452650078 |
| **A_51_P308948** | Tamm41 | 0.120960858 | -0.080763506 | 0.322685222 | 10.96444287 | 1.457034023 | 0.193977709 | 0.593071042 | -4.45270024 |
| **A_52_P435561** | Prr15l | -0.364909247 | -0.973530883 | 0.24371239 | 8.161833054 | -1.456869563 | 0.194021375 | 0.593089412 | -4.452783114 |
| **A_55_P2050354** | Prpmp5 | 0.211083488 | -0.140998167 | 0.563165143 | 8.781159439 | 1.456780283 | 0.194045085 | 0.593089412 | -4.452828103 |
| **A_55_P2088716** | Ss18l2 | 0.123989744 | -0.082861639 | 0.330841127 | 12.04085288 | 1.456500112 | 0.194119504 | 0.59313604 | -4.452969279 |
| **A_66_P130241** | Slc25a1 | -0.164630351 | -0.439283797 | 0.110023094 | 11.12642973 | -1.456491795 | 0.194121713 | 0.59313604 | -4.452973469 |
| **A_51_P275514** | Mprip | -0.414431499 | -1.10591426 | 0.277051261 | 10.63363857 | -1.45631254 | 0.194169341 | 0.593137843 | -4.453063793 |
| **A_55_P2087911** | Mthfsd | 0.336413493 | -0.224923426 | 0.897750412 | 9.182865613 | 1.45624011 | 0.194188589 | 0.593137843 | -4.453100288 |
| **A_55_P2055217** | NA | -0.349619122 | -0.933074864 | 0.23383662 | 12.63738415 | -1.456030421 | 0.194244322 | 0.593137843 | -4.453205942 |
| **A_51_P347862** | Actn1 | -0.124177358 | -0.331408955 | 0.083054238 | 8.565664298 | -1.456027688 | 0.194245049 | 0.593137843 | -4.453207319 |
| **A_55_P1966332** | Plac9a | -1.216817638 | -3.24827631 | 0.814641033 | 11.51144803 | -1.455461407 | 0.194395634 | 0.593503904 | -4.453492632 |
| **A_55_P1988689** | Rbpjl | -0.345449353 | -0.922223432 | 0.231324725 | 7.959929234 | -1.455331197 | 0.194430275 | 0.593515916 | -4.453558234 |
| **A_55_P1954277** | Emid1 | -0.668594187 | -1.785187723 | 0.447999349 | 9.643695003 | -1.454958735 | 0.194529395 | 0.593724724 | -4.45374588 |
| **A_66_P106611** | Gna14 | -0.158073892 | -0.422129184 | 0.105981399 | 7.914688136 | -1.454616337 | 0.194620555 | 0.593821118 | -4.453918372 |
| **A_55_P1967736** | Nkd1 | -0.887501302 | -2.370039253 | 0.595036649 | 9.069331696 | -1.454609329 | 0.194622421 | 0.593821118 | -4.453921903 |
| **A_51_P470448** | Setx | 0.131959744 | -0.08859839 | 0.352517879 | 11.01690732 | 1.453789658 | 0.194840813 | 0.593861612 | -4.454334803 |
| **A_55_P2048409** | Apool | 0.115631006 | -0.077637332 | 0.308899345 | 10.55305874 | 1.45377361 | 0.194845091 | 0.593861612 | -4.454342886 |
| **A_55_P2122546** | Csnk1e | -0.305121848 | -0.815192053 | 0.204948358 | 9.583474416 | -1.453538498 | 0.194907777 | 0.593861612 | -4.454461313 |
| **A_55_P2063256** | Lgals4 | -0.921273008 | -2.461409624 | 0.618863608 | 9.028449731 | -1.453490809 | 0.194920494 | 0.593861612 | -4.454485334 |
| **A_55_P2151006** | C1qtnf7 | -0.633984446 | -1.693873087 | 0.425904194 | 8.532107623 | -1.453454882 | 0.194930075 | 0.593861612 | -4.45450343 |
| **A_51_P212420** | Lama4 | -1.070775409 | -2.861058994 | 0.719508175 | 8.86085318 | -1.453314974 | 0.19496739 | 0.593861612 | -4.4545739 |
| **A_51_P394154** | Ddx51 | 0.181896658 | -0.122233429 | 0.486026745 | 9.92366332 | 1.453277409 | 0.19497741 | 0.593861612 | -4.454592821 |
| **A_51_P141535** | Oat | -0.239347738 | -0.639537585 | 0.160842109 | 10.50988612 | -1.453270475 | 0.19497926 | 0.593861612 | -4.454596313 |
| **A_55_P2369474** | Nr1h4 | -0.439430152 | -1.174206257 | 0.295345952 | 10.57617483 | -1.453175336 | 0.195004639 | 0.593861612 | -4.454644232 |
| **A_55_P2023042** | Ift43 | -0.552105569 | -1.475292852 | 0.371081714 | 9.572190775 | -1.453166922 | 0.195006884 | 0.593861612 | -4.45464847 |
| **A_55_P1969746** | Pitpnm1 | -0.119333818 | -0.318884403 | 0.080216767 | 9.958507274 | -1.453094035 | 0.19502633 | 0.593861612 | -4.454685181 |
| **A_55_P1990331** | Add1 | 0.267403845 | -0.179753124 | 0.714560814 | 8.811251595 | 1.453084377 | 0.195028907 | 0.593861612 | -4.454690046 |
| **A_55_P2059958** | Snx7 | 0.167306709 | -0.112470706 | 0.447084123 | 8.986991742 | 1.453061154 | 0.195035103 | 0.593861612 | -4.454701743 |

| | | | | | | | | | |
|---|---|---|---|---|---|---|---|---|---|
| **A_51_P334685** | Esyt1 | -0.124754477 | -0.3333931 | 0.083884146 | 9.062886336 | -1.452929767 | 0.195070163 | 0.593874812 | -4.454767917 |
| **A_51_P273508** | Ube2g1 | 0.116150392 | -0.078149034 | 0.310449819 | 8.587823974 | 1.452554217 | 0.195170407 | 0.594086426 | -4.454957061 |
| **A_55_P2178553** | Apoa2 | 0.158789137 | -0.106864226 | 0.424442499 | 17.68716924 | 1.452408089 | 0.195209426 | 0.594111634 | -4.455030656 |
| **A_52_P541833** | Vps37b | -0.113632084 | -0.303762463 | 0.076498295 | 10.38567124 | -1.452220785 | 0.195259449 | 0.594113676 | -4.455124985 |
| **A_55_P2031021** | Bex1 | 0.34679383 | -0.233501675 | 0.927089336 | 14.31393672 | 1.45212949 | 0.195283836 | 0.594113676 | -4.455170962 |
| **A_66_P118696** | Nup107 | 0.171047762 | -0.115208401 | 0.457303925 | 12.13953132 | 1.451930207 | 0.195337077 | 0.594113676 | -4.455271321 |
| **A_55_P1954161** | Plagl2 | 0.266262464 | -0.179409886 | 0.711934815 | 8.620475601 | 1.451701899 | 0.19539809 | 0.594113676 | -4.455386294 |
| **A_55_P2115477** | Parva | -0.256669924 | -0.686302687 | 0.172962838 | 8.23250483 | -1.451646195 | 0.195412979 | 0.594113676 | -4.455414345 |
| **A_55_P1992019** | Ptpru | -0.237997795 | -0.636379746 | 0.160384156 | 9.42499634 | -1.451631784 | 0.195416831 | 0.594113676 | -4.455421602 |
| **A_55_P2121827** | Dgcr2 | 0.365061347 | -0.246032767 | 0.976155461 | 9.738589051 | 1.45157988 | 0.195430705 | 0.594113676 | -4.45544774 |
| **A_55_P2089761** | NA | 0.155132693 | -0.104584103 | 0.41484949 | 10.45367895 | 1.45139788 | 0.195479363 | 0.594113676 | -4.455539388 |
| **A_55_P2014427** | Il17re | -0.44053129 | -1.178064716 | 0.297002137 | 7.985435191 | -1.451370331 | 0.195486729 | 0.594113676 | -4.455553261 |
| **A_51_P235687** | Alox5ap | -0.257572249 | -0.688876475 | 0.173731976 | 11.44266324 | -1.451104025 | 0.195557948 | 0.59418627 | -4.455687359 |
| **A_55_P2137309** | Kctd11 | -0.275433315 | -0.736662672 | 0.185796042 | 9.402122763 | -1.451051131 | 0.195572096 | 0.59418627 | -4.455713993 |
| **A_51_P432460** | Ppp1r14a | -1.365981786 | -3.65390344 | 0.921939868 | 9.27250458 | -1.450731192 | 0.195657696 | 0.594352915 | -4.455875091 |
| **A_51_P158922** | Prkaa1 | 0.208545528 | -0.140807165 | 0.557898222 | 10.06867281 | 1.450507487 | 0.195717568 | 0.59444137 | -4.455987728 |
| **A_55_P2064741** | Nmb | -0.236726083 | -0.633354684 | 0.159902518 | 8.935822786 | -1.450258012 | 0.195784358 | 0.594468096 | -4.456113337 |
| **A_51_P112223** | Gsta4 | 0.110200883 | -0.0744396 | 0.294841367 | 9.542028731 | 1.450244865 | 0.195787878 | 0.594468096 | -4.459087086 |
| **A_55_P1968664** | NA | 0.121609608 | -0.082163691 | 0.325382906 | 13.07433986 | 1.450119528 | 0.195821442 | 0.594476622 | -4.456183061 |
| **A_55_P2061341** | Ift74 | -0.400675693 | -1.072153917 | 0.270802532 | 8.977722011 | -1.449920706 | 0.195874695 | 0.594519858 | -4.456283162 |
| **A_55_P1972448** | NA | 0.169980073 | -0.114900026 | 0.454860171 | 9.270971557 | 1.449836698 | 0.1958972 | 0.594519858 | -4.456325456 |
| **A_51_P116601** | Cfap69 | -0.351157274 | -0.939833851 | 0.237519302 | 8.127202948 | -1.449466234 | 0.195996472 | 0.594677221 | -4.458363865 |
| **A_51_P401730** | Ints14 | 0.113966941 | -0.07709305 | 0.305026932 | 10.25323645 | 1.449413584 | 0.196010584 | 0.594677221 | -4.45653847 |
| **A_66_P119123** | A1cf | 0.261788317 | -0.177133223 | 0.700709857 | 9.129464893 | 1.449260835 | 0.196051532 | 0.594708106 | -4.456615368 |
| **A_51_P416191** | Hectd1 | 0.221417067 | -0.149888191 | 0.592722324 | 10.9608079 | 1.448982642 | 0.196126128 | 0.594757516 | -4.456755413 |
| **A_51_P419971** | Tmem106b | -0.241822813 | -0.647350852 | 0.163705226 | 11.04377472 | -1.448970588 | 0.196129361 | 0.594757516 | -4.456761481 |
| **A_55_P2181655** | Tns3 | -0.150101618 | -0.401847991 | 0.101644755 | 7.953786041 | -1.44878967 | 0.196177888 | 0.594806837 | -4.456852555 |
| **A_51_P114049** | Tmem109 | -0.377347289 | -1.010271639 | 0.25557706 | 11.8934338 | -1.448680519 | 0.19620717 | 0.594806837 | -4.4569075 |
| **A_55_P2066399** | Fcho2 | 0.120534469 | -0.081738355 | 0.322807292 | 8.28544858 | 1.44796115 | 0.196400259 | 0.595122264 | -4.457269599 |
| **A_51_P346523** | Zcchc7 | 0.099420539 | -0.06742122 | 0.266262299 | 9.146293199 | 1.447953255 | 0.196402379 | 0.595122264 | -4.457273573 |
| **A_51_P392005** | Car8 | 0.161276266 | -0.109369013 | 0.431921546 | 8.663575283 | 1.44794876 | 0.196403587 | 0.595122264 | -4.457275836 |
| **A_55_P2063465** | Thbs3 | -0.46273211 | -1.239405732 | 0.313941513 | 8.542500857 | -1.447685652 | 0.196474254 | 0.595203954 | -4.457408265 |
| **A_55_P2077213** | Cenpb | 0.175090747 | -0.118857191 | 0.469038685 | 10.99237641 | 1.447357999 | 0.196562289 | 0.595203954 | -4.457573174 |
| **A_51_P464182** | Mat1a | -0.548744032 | -1.470037892 | 0.372549829 | 11.50645403 | -1.447287536 | 0.196581226 | 0.595203954 | -4.457608638 |
| **A_52_P559817** | Fbln7 | -0.127041466 | -0.340347195 | 0.086264263 | 7.976511224 | -1.447192047 | 0.196606891 | 0.595203954 | -4.457656696 |
| **A_55_P2008898** | NA | 0.136444116 | -0.092665721 | 0.365553952 | 13.9854153 | 1.447085739 | 0.196635468 | 0.595203954 | -4.457710199 |
| **A_55_P2117943** | NA | 0.30681905 | -0.208399928 | 0.822038029 | 10.21135103 | 1.447017083 | 0.196653926 | 0.595203954 | -4.457744752 |
| **A_52_P79947** | Ndst1 | -0.187413905 | -0.502167404 | 0.127339593 | 10.23236402 | -1.446819766 | 0.196706982 | 0.595203954 | -4.457844054 |
| **A_55_P2136732** | Cd99 | -0.549674899 | -1.472843572 | 0.373493773 | 12.42140582 | -1.446798455 | 0.196712713 | 0.595203954 | -4.457854779 |
| **A_55_P1998751** | Ccdc137 | 0.140760869 | -0.095664658 | 0.377186396 | 8.770029466 | 1.446674286 | 0.196746109 | 0.595203954 | -4.457917268 |
| **A_51_P479408** | Hyal1 | -0.116538533 | -0.312287615 | 0.079210549 | 7.962872571 | -1.44661512 | 0.196762023 | 0.595203954 | -4.457947043 |
| **A_55_P2122643** | Dtymk | 0.208438262 | -0.141680774 | 0.558557298 | 12.74142576 | 1.44658817 | 0.196769272 | 0.595203954 | -4.457960605 |
| **A_55_P2083963** | Phgr1 | -0.369436331 | -0.990053993 | 0.251181331 | 7.896608442 | -1.44643413 | 0.196810713 | 0.595236156 | -4.458038124 |
| **A_55_P2076057** | Hmga1 | 0.34394179 | -0.234066742 | 0.921950322 | 8.769153372 | 1.445885449 | 0.196958388 | 0.595400407 | -4.458314229 |
| **A_55_P2021996** | Hbs1l | 0.17503982 | -0.11912552 | 0.469205161 | 8.647816117 | 1.445867671 | 0.196963174 | 0.595400407 | -4.458323175 |
| **A_55_P2026223** | St6galnac4 | -0.120024123 | -0.321767068 | 0.081718823 | 7.997137086 | -1.445617418 | 0.197030563 | 0.595400407 | -4.458449099 |
| **A_55_P1993880** | Rps24 | 0.158080176 | -0.107631262 | 0.423791614 | 15.22629807 | 1.445607358 | 0.197033273 | 0.595400407 | -4.458454161 |
| **A_51_P452820** | Rpl31 | 0.119516556 | -0.081375386 | 0.320408499 | 10.06149582 | 1.44560201 | 0.197034713 | 0.595400407 | -4.458456852 |
| **A_55_P1977289** | Zbtb9 | 0.099879348 | -0.068020557 | 0.267779253 | 9.445531587 | 1.445467859 | 0.197070848 | 0.595400407 | -4.458524353 |
| **A_55_P1993698** | Celf1 | 0.309278043 | -0.210646186 | 0.829202272 | 10.55935836 | 1.44541388 | 0.197085389 | 0.595400407 | -4.458551513 |

| | | | | | | | | | |
|---|---|---|---|---|---|---|---|---|---|
| **A_55_P2106096** | Fam171a1 | -0.413899157 | -1.109761755 | 0.28196344 | 8.58355746 | -1.445287469 | 0.197119446 | 0.595400407 | -4.458615118 |
| **A_55_P1980826** | Kcnrg | -0.24796012 | -0.664864276 | 0.168944036 | 7.825137454 | -1.445202829 | 0.197142253 | 0.595400407 | -4.458657706 |
| **A_51_P421094** | NA | 0.125354784 | -0.085468469 | 0.336178037 | 10.51345091 | 1.444792878 | 0.19725275 | 0.595641057 | -4.458863967 |
| **A_52_P634447** | Hs1bp3 | -0.12215812 | -0.32764717 | 0.083330931 | 9.192575176 | -1.444497659 | 0.197332358 | 0.595788369 | -4.459012496 |
| **A_55_P1981217** | Aatf | 0.200035561 | -0.136525198 | 0.536596321 | 9.273052717 | 1.444198599 | 0.197413031 | 0.595926412 | -4.459162951 |
| **A_51_P438701** | Atpaf2 | 0.158163272 | -0.107965744 | 0.424292288 | 9.508689386 | 1.444097784 | 0.197440234 | 0.595926412 | -4.45921367 |
| **A_55_P2131954** | Gm2590 | 0.115830533 | -0.079090778 | 0.310751844 | 8.390551135 | 1.44393259 | 0.197484814 | 0.595926412 | -4.459296774 |
| **A_52_P139588** | Edem3 | 0.174028893 | -0.118857613 | 0.4669154 | 9.72074164 | 1.443793836 | 0.197522267 | 0.595926412 | -4.459366576 |
| **A_55_P2085574** | Nfix | -0.551476031 | -1.479668032 | 0.37671597 | 8.987043117 | -1.443683554 | 0.197552039 | 0.595926412 | -4.459422053 |
| **A_55_P1985554** | B4galt4 | -0.229893032 | -0.616839679 | 0.157053614 | 8.106941556 | -1.443636689 | 0.197564692 | 0.595926412 | -4.459445629 |
| **A_52_P552879** | Cracr2b | -0.314993581 | -0.845307662 | 0.215320501 | 8.563923663 | -1.443283788 | 0.197659996 | 0.595926412 | -4.45962315 |
| **A_55_P2025760** | Adam9 | -0.176756594 | -0.474348635 | 0.120835447 | 10.76001725 | -1.443236493 | 0.197672771 | 0.595926412 | -4.45964694 |
| **A_55_P1991351** | S100a13 | -0.270033304 | -0.724675251 | 0.184608644 | 10.88483445 | -1.443214939 | 0.197678594 | 0.595926412 | -4.459657783 |
| **A_51_P380807** | Ckm | -0.235545984 | -0.632131234 | 0.161039266 | 7.775643604 | -1.443186094 | 0.197686386 | 0.595926412 | -4.459672292 |
| **A_55_P2279685** | 2900076A07Rik | 0.131355661 | -0.089880866 | 0.352592188 | 8.388197682 | 1.442697077 | 0.197818535 | 0.596168935 | -4.459918267 |
| **A_55_P1971076** | Atp11a | -0.694950141 | -1.865453891 | 0.475553608 | 9.086039046 | -1.442660123 | 0.197828524 | 0.596168935 | -4.459936854 |
| **A_55_P2129469** | NA | -0.227943615 | -0.61198709 | 0.15609986 | 9.057617793 | -1.442215738 | 0.197948688 | 0.596251045 | -4.460160364 |
| **A_51_P307220** | Ankrd54 | 0.191649389 | -0.131249054 | 0.514547832 | 8.797905562 | 1.442197297 | 0.197953676 | 0.596251045 | -4.460169639 |
| **A_55_P2076190** | NA | 0.193592135 | -0.132600121 | 0.51978439 | 15.02916691 | 1.442106225 | 0.197978311 | 0.596251045 | -4.460215444 |
| **A_55_P1989865** | Thrap3 | 0.248129427 | -0.169986324 | 0.666245177 | 9.599977545 | 1.441998915 | 0.198007342 | 0.596251045 | -4.460269414 |
| **A_52_P135822** | Myo18a | -0.222156338 | -0.596525162 | 0.152212485 | 11.63791276 | -1.44192351 | 0.198027744 | 0.596251045 | -4.460307337 |
| **A_52_P429534** | Elavl1 | 0.108559751 | -0.074386873 | 0.291506375 | 8.374188187 | 1.441875051 | 0.198040856 | 0.596251045 | -4.460331709 |
| **A_55_P2067200** | NA | 0.184182372 | -0.126325064 | 0.494689808 | 8.787579566 | 1.441316124 | 0.198192152 | 0.596369013 | -4.460612798 |
| **A_55_P2019083** | Rps6kb1 | 0.115856627 | -0.079471852 | 0.311185107 | 12.59843186 | 1.44124728 | 0.198210795 | 0.596369013 | -4.460647418 |
| **A_66_P109941** | Mamdc4 | -0.748095016 | -2.009499114 | 0.513309082 | 8.475637315 | -1.441072109 | 0.198258237 | 0.596369013 | -4.460735508 |
| **A_52_P595225** | Vps35 | 0.151215393 | -0.10376677 | 0.406197556 | 8.875467091 | 1.441017948 | 0.198272909 | 0.596369013 | -4.460762744 |
| **A_55_P1955279** | Specc1 | 0.274108126 | -0.188104015 | 0.736320266 | 12.46731454 | 1.440999235 | 0.198277978 | 0.596369013 | -4.460772154 |
| **A_51_P482633** | Rps6ka4 | -0.326817256 | -0.877957494 | 0.224322982 | 9.113044428 | -1.440874146 | 0.198311866 | 0.596369013 | -4.460835057 |
| **A_66_P127956** | Zfp947 | 0.187200195 | -0.128496376 | 0.502896766 | 11.18905739 | 1.440852819 | 0.198317644 | 0.596369013 | -4.460845781 |
| **A_52_P546635** | Zeb1 | -0.182603175 | -0.490554554 | 0.125348204 | 9.487111797 | -1.440818779 | 0.198326867 | 0.596369013 | -4.460862899 |
| **A_52_P263095** | Ahnak | -0.598910933 | -1.609102052 | 0.411280185 | 11.39794375 | -1.440594939 | 0.198387526 | 0.596458623 | -4.460975456 |
| **A_51_P165683** | Hirip3 | 0.18825821 | -0.129576685 | 0.506093104 | 11.04177966 | 1.439247676 | 0.198752982 | 0.597179403 | -4.461652851 |
| **A_51_P439876** | Map4k4 | -0.230542145 | -0.619783229 | 0.15869894 | 10.90849905 | -1.439179103 | 0.198771599 | 0.597179403 | -4.461687326 |
| **A_55_P1959828** | Slc35g1 | 0.284765671 | -0.196039488 | 0.765570829 | 10.23019472 | 1.439135556 | 0.198783423 | 0.597179403 | -4.461709218 |
| **A_51_P298190** | Sbf2 | -0.170857378 | -0.459345635 | 0.11763088 | 9.037106724 | -1.439092749 | 0.198795047 | 0.597179403 | -4.461730739 |
| **A_51_P325318** | Serinc2 | -0.225607339 | -0.60654383 | 0.155329153 | 7.876831828 | -1.439076344 | 0.198799501 | 0.597179403 | -4.461738987 |
| **A_52_P172563** | Dffa | 0.115897816 | -0.079807333 | 0.311602965 | 8.341298595 | 1.438984745 | 0.198824376 | 0.597179403 | -4.461785036 |
| **A_55_P2105371** | Dlk1 | 0.213729979 | -0.14721412 | 0.574674078 | 12.89723913 | 1.438827356 | 0.198867123 | 0.597179403 | -4.461864159 |
| **A_55_P2090557** | Lrrc75a | -0.444489127 | -1.195190178 | 0.306211924 | 9.944300203 | -1.4387234 | 0.198895363 | 0.597179403 | -4.46191642 |
| **A_52_P504068** | Cdk8 | 0.279259773 | -0.192591387 | 0.751110932 | 9.211433866 | 1.438091576 | 0.199067075 | 0.597179403 | -4.46223403 |
| **A_55_P2160416** | Acoxl | -0.26241554 | -0.705805918 | 0.180974838 | 7.813079576 | -1.438091352 | 0.199067136 | 0.597179403 | -4.462234143 |
| **A_51_P114878** | Elp1 | 0.134518431 | -0.092783986 | 0.361820848 | 10.27494771 | 1.438006763 | 0.199090136 | 0.597179403 | -4.467014141 |
| **A_52_P398989** | Cytip | -0.16393549 | -0.440948011 | 0.113077031 | 8.913099037 | -1.437992998 | 0.199093878 | 0.597179403 | -4.462283582 |
| **A_52_P464629** | Nme7 | -0.099373174 | -0.267293506 | 0.068547158 | 8.315402199 | -1.437967491 | 0.199100814 | 0.597179403 | -4.462296404 |
| **A_66_P110104** | Myo18b | 0.171908513 | -0.11858532 | 0.462402345 | 8.88371703 | 1.43794957 | 0.199105687 | 0.597179403 | -4.462305411 |
| **A_55_P1977776** | Rs5-8s1 | 1.256506793 | -0.867113442 | 3.380127027 | 14.61280917 | 1.437709608 | 0.19917095 | 0.597179403 | -4.462426028 |
| **A_55_P2171091** | Sap30bp | 0.141886616 | -0.097919318 | 0.381692549 | 10.12615578 | 1.437688156 | 0.199176785 | 0.597179403 | -4.462436811 |
| **A_51_P495269** | Lor | 0.16240375 | -0.11211022 | 0.43691772 | 8.159583893 | 1.437522962 | 0.199221725 | 0.597179403 | -4.462519843 |
| **A_51_P140141** | Hspbp1 | 0.162020488 | -0.1118471 | 0.435888075 | 10.08290788 | 1.437515341 | 0.199223799 | 0.597179403 | -4.462523674 |
| **A_51_P216702** | Eogt | -0.192907824 | -0.518995725 | 0.133180076 | 8.388029866 | -1.437468534 | 0.199236534 | 0.597179403 | -4.4625472 |

| | | | | | | | | | |
|---|---|---|---|---|---|---|---|---|---|
| **A_51_P365008** | Tns1 | -0.706991729 | -1.902104173 | 0.488120715 | 11.47660227 | -1.437436781 | 0.199245174 | 0.597179403 | -4.462563159 |
| **A_52_P387724** | Tcf7l1 | -0.097528335 | -0.262466258 | 0.067409589 | 8.929866986 | -1.436790588 | 0.199421077 | 0.597613952 | -4.462887935 |
| **A_55_P2113650** | Clybl | 0.207185493 | -0.143331027 | 0.557702013 | 8.994078598 | 1.436263225 | 0.199564737 | 0.597873426 | -4.463152966 |
| **A_55_P2183208** | Prl2c1 | -0.623604281 | -1.678707383 | 0.431498821 | 8.561746253 | -1.43614198 | 0.19959778 | 0.597873426 | -4.463213896 |
| **A_55_P2154817** | Ccdc28a | 0.185444927 | -0.128353178 | 0.499243032 | 9.563328532 | 1.435978142 | 0.199642437 | 0.597873426 | -4.463296229 |
| **A_55_P2039637** | Ift88 | -0.296533313 | -0.798317216 | 0.205250591 | 8.978899523 | -1.435952098 | 0.199649537 | 0.597873426 | -4.463309317 |
| **A_51_P161362** | Gnl2 | 0.118112178 | -0.081759885 | 0.317984242 | 11.07322931 | 1.435905211 | 0.199662319 | 0.597873426 | -4.463332878 |
| **A_55_P2111399** | Pcp2 | 0.13414577 | -0.092916861 | 0.3612084 | 8.270568856 | 1.435537382 | 0.199762622 | 0.59808112 | -4.463517713 |
| **A_51_P520139** | Ccpg1os | -0.373072328 | -1.004643015 | 0.258498359 | 8.714014467 | -1.43533826 | 0.199816939 | 0.598151094 | -4.463617768 |
| **A_65_P18167** | Hnrnpul2 | 0.13864731 | -0.09608933 | 0.37338395 | 10.128009 | 1.435204385 | 0.199853466 | 0.598167797 | -4.463685036 |
| **A_55_P1970655** | Kdm3b | 0.120433811 | -0.083518327 | 0.32438595 | 8.497721326 | 1.43483961 | 0.199953022 | 0.598373118 | -4.463868319 |
| **A_51_P283266** | Zfp874b | 0.139952374 | -0.097076589 | 0.376981338 | 8.710910866 | 1.434703117 | 0.199990287 | 0.598378314 | -4.463936898 |
| **A_55_P2222870** | Mylk | -0.463151593 | -1.247616578 | 0.321313392 | 8.275768774 | -1.434606479 | 0.200016674 | 0.598378314 | -4.463985451 |
| **A_55_P2102126** | Lamp2 | -0.248882209 | -0.670535221 | 0.172770804 | 11.10046203 | -1.434240011 | 0.200116767 | 0.59851695 | -4.464169568 |
| **A_51_P247359** | Ptprcap | 0.254696433 | -0.176847589 | 0.686240455 | 11.00481666 | 1.434104988 | 0.200153658 | 0.59851695 | -4.464237403 |
| **A_55_P2061319** | Mthfs | -0.124113884 | -0.334436391 | 0.086208622 | 8.922016561 | -1.43389651 | 0.20021063 | 0.59851695 | -4.464342138 |
| **A_55_P2095529** | NA | 0.144773084 | -0.100565795 | 0.390111964 | 15.08303371 | 1.433853029 | 0.200222514 | 0.59851695 | -4.464363982 |
| **A_55_P1968085** | NA | 0.163300914 | -0.113454585 | 0.440056414 | 8.273296383 | 1.433756983 | 0.200248768 | 0.59851695 | -4.464412232 |
| **A_51_P489903** | Ogg1 | 0.097245247 | -0.067568602 | 0.262059095 | 9.324910624 | 1.433698629 | 0.20026472 | 0.59851695 | -4.464441547 |
| **A_55_P2257341** | 2410004I01Rik | -0.259682721 | -0.69981685 | 0.180451408 | 7.815242268 | -1.433643585 | 0.200279768 | 0.59851695 | -4.464469199 |
| **A_55_P1995996** | Zfp984 | 0.185153551 | -0.12871318 | 0.499020281 | 8.657915807 | 1.433408416 | 0.200344072 | 0.598616567 | -4.464587335 |
| **A_55_P2099141** | NA | -0.17650009 | -0.475768021 | 0.122767842 | 14.6278578 | -1.433071757 | 0.20043616 | 0.598799156 | -4.464756448 |
| **A_55_P2075268** | Tmem216 | -0.170300509 | -0.459143342 | 0.118542324 | 10.15449238 | -1.432641535 | 0.200553897 | 0.59894539 | -4.464972548 |
| **A_55_P2002859** | Aifm2 | -0.134863221 | -0.363610053 | 0.093883612 | 7.949623508 | -1.43258905 | 0.200568265 | 0.59894539 | -4.464998911 |
| **A_52_P515769** | Pcdh12 | -0.580063278 | -1.563966849 | 0.403840293 | 9.949145421 | -1.432537584 | 0.200582355 | 0.59894539 | -4.465024761 |
| **A_51_P348227** | Stoml1 | -0.177519162 | -0.478647791 | 0.123609467 | 8.3096843 | -1.432439806 | 0.200609125 | 0.59894539 | -4.465073872 |
| **A_55_P2089158** | NA | 0.12745946 | -0.08876913 | 0.343688051 | 8.903276314 | 1.432326896 | 0.200640044 | 0.59894539 | -4.465130583 |
| **A_51_P233160** | Lysmd2 | -0.187481197 | -0.505561722 | 0.130599329 | 9.194778863 | -1.432200516 | 0.200674655 | 0.598956209 | -4.465194059 |
| **A_51_P510849** | Tmem50b | -0.52299599 | -1.410666034 | 0.364674055 | 10.64192375 | -1.431627339 | 0.200831699 | 0.599253932 | -4.465481926 |
| **A_52_P409833** | Plat | -0.122907897 | -0.331519378 | 0.085703585 | 7.731295817 | -1.431610151 | 0.20083641 | 0.599253932 | -4.465490559 |
| **A_52_P26626** | Fam92b | -0.553933593 | -1.494367532 | 0.386500346 | 8.051604611 | -1.431240439 | 0.200937769 | 0.599306752 | -4.465676227 |
| **A_55_P2153743** | Slc26a6 | 0.244820406 | -0.170830533 | 0.660471344 | 9.282071377 | 1.431205633 | 0.200947314 | 0.599306752 | -4.465693706 |
| **A_51_P414126** | Rab19 | -0.403760905 | -1.089303198 | 0.281781387 | 8.321675838 | -1.431110613 | 0.200973373 | 0.599306752 | -4.465741423 |
| **A_52_P3412** | Med17 | 0.15629901 | -0.109097458 | 0.421695477 | 9.465743308 | 1.431015301 | 0.200999515 | 0.599306752 | -4.465789286 |
| **A_55_P2015178** | Nap1l1 | 0.114573903 | -0.079977904 | 0.309125711 | 13.51419995 | 1.43098021 | 0.20100914 | 0.599306752 | -4.465806907 |
| **A_55_P2073467** | Gtf2i | -0.220887239 | -0.59601465 | 0.154240172 | 13.33021558 | -1.430787102 | 0.201062118 | 0.599372251 | -4.465903878 |
| **A_51_P295582** | Cutc | 0.130693028 | -0.091300912 | 0.352686969 | 8.473109022 | 1.43052183 | 0.201134914 | 0.5994968 | -4.466037082 |
| **A_55_P1989251** | Chfr | 0.144492277 | -0.100982841 | 0.389967394 | 10.98501794 | 1.430277632 | 0.201201948 | 0.599573233 | -4.4661597 |
| **A_51_P207591** | Anxa8 | -0.915388531 | -2.470616548 | 0.639839486 | 8.618305112 | -1.430192791 | 0.201225242 | 0.599573233 | -4.46681669 |
| **A_55_P1988009** | Gm10406 | -0.326088744 | -0.88014746 | 0.227969971 | 8.009371793 | -1.430089463 | 0.201253616 | 0.599573233 | -4.466254181 |
| **A_55_P2159665** | Zfp534 | 0.232100684 | -0.162436543 | 0.62663791 | 10.28569724 | 1.429458731 | 0.201426892 | 0.599938431 | -4.466570858 |
| **A_55_P2077188** | Stxbp1 | -0.215602377 | -0.582150819 | 0.150946066 | 9.250974453 | -1.429240634 | 0.201486839 | 0.599938431 | -4.466680354 |
| **A_55_P1993805** | Zfp947 | 0.130863392 | -0.091652146 | 0.353378929 | 10.44061191 | 1.429028923 | 0.201545047 | 0.599938431 | -4.466786639 |
| **A_51_P511707** | Ntpcr | -0.161893493 | -0.437207153 | 0.113420166 | 9.967395819 | -1.428844028 | 0.201595895 | 0.599938431 | -4.46687946 |
| **A_55_P1985519** | NA | 0.131872785 | -0.092390154 | 0.356135724 | 16.45825691 | 1.428830978 | 0.201599484 | 0.599938431 | -4.466886011 |
| **A_51_P505617** | Il18r1 | -0.550314051 | -1.486207042 | 0.38557894 | 8.196975807 | -1.428787349 | 0.201611485 | 0.599938431 | -4.466907913 |
| **A_51_P217535** | Actr3 | -0.197957554 | -0.534620214 | 0.138705105 | 13.1044414 | -1.428763409 | 0.20161807 | 0.599938431 | -4.466919932 |
| **A_55_P2067700** | Tas2r113 | 0.394038022 | -0.276163382 | 1.064239425 | 8.566581565 | 1.428617568 | 0.20165819 | 0.599938431 | -4.466993144 |
| **A_52_P623751** | Fem1b | 0.114220397 | -0.080055051 | 0.308495845 | 12.23645616 | 1.428594372 | 0.201664571 | 0.599938431 | -4.467004788 |
| **A_55_P2082979** | Irak2 | 0.130949829 | -0.091792886 | 0.353692544 | 12.49035776 | 1.428514378 | 0.201686581 | 0.599938431 | -4.467044944 |

| | | | | | | | | | |
|---|---|---|---|---|---|---|---|---|---|
| **A_55_P2069832** | Prpsap1 | -0.117798039 | -0.31822924 | 0.082633161 | 12.77801428 | -1.428091136 | 0.201803068 | 0.600131836 | -4.4672574 |
| **A_55_P2012201** | App | -0.65602152 | -1.772312654 | 0.460269615 | 12.01219877 | -1.427985506 | 0.20183215 | 0.600131836 | -4.467310421 |
| **A_52_P162509** | Ahcyl2 | 0.116233658 | -0.081557028 | 0.314024343 | 8.29976807 | 1.427937695 | 0.201845315 | 0.600131836 | -4.46733442 |
| **A_55_P1956819** | Gm4832 | 0.179178425 | -0.125778597 | 0.484135447 | 15.70407243 | 1.427677995 | 0.201916835 | 0.600131836 | -4.467464773 |
| **A_55_P2009285** | Fbxo25 | -0.264390575 | -0.714382991 | 0.185601841 | 9.635337395 | -1.427657279 | 0.201922541 | 0.600131836 | -4.46747517 |
| **A_55_P2060938** | NA | 0.272363555 | -0.191216955 | 0.735944066 | 11.65141678 | 1.427601561 | 0.201937889 | 0.600131836 | -4.467503137 |
| **A_55_P1963206** | NA | 0.167020109 | -0.117323742 | 0.451363961 | 8.819423976 | 1.427276488 | 0.202027455 | 0.600269765 | -4.467666294 |
| **A_66_P119643** | Atrx | -0.105279394 | -0.284521231 | 0.073962443 | 8.176745622 | -1.427207701 | 0.202046412 | 0.600269765 | -4.467700818 |
| **A_51_P491976** | Fbxw17 | -0.241664456 | -0.653170893 | 0.169841982 | 9.494521721 | -1.426984756 | 0.202107864 | 0.600325039 | -4.46781271 |
| **A_52_P117576** | Casp3 | 0.182209178 | -0.128085953 | 0.49250431 | 8.559449382 | 1.426850517 | 0.202144874 | 0.600325039 | -4.467880081 |
| **A_55_P2029902** | Gab3 | 0.336389359 | -0.236498307 | 0.909277025 | 11.42694852 | 1.426776561 | 0.202165267 | 0.600325039 | -4.467917197 |
| **A_55_P2064492** | Pcmtd2 | 0.118707163 | -0.083469313 | 0.320883639 | 10.31563251 | 1.426689594 | 0.202189249 | 0.600325039 | -4.467960841 |
| **A_55_P2117764** | Vangl2 | -0.104351185 | -0.282093553 | 0.073391184 | 8.411051716 | -1.426558575 | 0.202225385 | 0.600340111 | -4.468026593 |
| **A_66_P137265** | Gm9933 | 0.22289962 | -0.156814208 | 0.602613448 | 9.045707051 | 1.426382835 | 0.202273864 | 0.600391818 | -4.468114787 |
| **A_51_P333099** | Mrps15 | 0.11052704 | -0.077802889 | 0.298856969 | 11.24648063 | 1.426042232 | 0.202367852 | 0.600578567 | -4.468285708 |
| **A_51_P412835** | Daxx | -0.132357247 | -0.357942089 | 0.093227594 | 11.36445586 | -1.425676536 | 0.202468808 | 0.600713312 | -4.468469212 |
| **A_55_P2051864** | Golgb1 | -0.311917648 | -0.843580081 | 0.219744784 | 13.64102349 | -1.425565457 | 0.202499483 | 0.600713312 | -4.468524949 |
| **A_55_P2024993** | Hoxb4 | -0.458766831 | -1.240801876 | 0.323268213 | 8.85697461 | -1.425440143 | 0.202534093 | 0.600713312 | -4.468587827 |
| **A_55_P1973698** | Htra2 | 0.226754045 | -0.159798641 | 0.613306731 | 10.88542515 | 1.425376313 | 0.202551724 | 0.600713312 | -4.468619854 |
| **A_51_P334942** | Aldh1a1 | -0.677647249 | -1.832897301 | 0.477602802 | 8.950885397 | -1.425315055 | 0.202568646 | 0.600713312 | -4.468650591 |
| **A_52_P447284** | Clic6 | -0.77281473 | -2.090421981 | 0.54479252 | 8.78324085 | -1.425189469 | 0.202603343 | 0.600724039 | -4.468713603 |
| **A_66_P118849** | NA | -0.215464693 | -0.582936318 | 0.152006932 | 8.187692565 | -1.424739595 | 0.202727676 | 0.600920348 | -4.468939318 |
| **A_55_P2083128** | Sec16a | -0.866148768 | -2.343442835 | 0.611145298 | 10.38507112 | -1.424651788 | 0.202751952 | 0.600920348 | -4.468983371 |
| **A_51_P367298** | NA | 0.109378864 | -0.077184851 | 0.295942579 | 8.31035563 | 1.424588439 | 0.202769468 | 0.600920348 | -4.469015153 |
| **A_55_P1960765** | NA | 0.176751741 | -0.124746247 | 0.47824973 | 13.29978861 | 1.424500057 | 0.202793907 | 0.600920348 | -4.469059494 |
| **A_51_P108020** | Zmynd15 | -0.118663175 | -0.321104143 | 0.083777793 | 8.170689211 | -1.424297613 | 0.202849897 | 0.600977013 | -4.469161057 |
| **A_55_P2159949** | Polr1e | 0.136022935 | -0.096050022 | 0.368095893 | 8.907552948 | 1.424199388 | 0.202877068 | 0.600977013 | -4.469210335 |
| **A_55_P2177885** | Dact3 | -0.376557757 | -1.019071537 | 0.265956024 | 8.483026102 | -1.424073289 | 0.202911954 | 0.600977013 | -4.469273594 |
| **A_55_P1953846** | Abca8b | -0.363089982 | -0.982663945 | 0.256483981 | 12.46976374 | -1.423981335 | 0.202937398 | 0.600977013 | -4.469319724 |
| **A_51_P214343** | Sephs1 | 0.114309389 | -0.080787724 | 0.309406503 | 9.173344296 | 1.423686114 | 0.203019104 | 0.60112688 | -4.469467819 |
| **A_55_P2154797** | 2810006K23Rik | 0.155901579 | -0.110226082 | 0.422029239 | 8.517358765 | 1.423454817 | 0.20308314 | 0.601224386 | -4.469583843 |
| **A_52_P18081** | Pacc1 | -0.149619058 | -0.405051963 | 0.105813847 | 10.59381955 | -1.423289556 | 0.203128905 | 0.60126778 | -4.46966674 |
| **A_52_P480351** | Loxl2 | -0.213577948 | -0.578293272 | 0.151137375 | 10.62031198 | -1.422936726 | 0.203226643 | 0.601464981 | -4.469843716 |
| **A_55_P2056220** | NA | -0.257113362 | -0.696361975 | 0.182135251 | 10.21690297 | -1.422320393 | 0.203397478 | 0.601613079 | -4.47015284 |
| **A_51_P348183** | Tmem141 | -0.16479788 | -0.446339211 | 0.11674345 | 11.31197926 | -1.422304724 | 0.203401823 | 0.601613079 | -4.470160699 |
| **A_55_P2056035** | Traf7 | -0.160579388 | -0.434976324 | 0.113817548 | 12.25825106 | -1.421980794 | 0.203491664 | 0.601613079 | -4.470323155 |
| **A_51_P384629** | Ctsd | -0.104013573 | -0.281771148 | 0.073744002 | 14.16716676 | -1.421821528 | 0.20353585 | 0.601613079 | -4.470403027 |
| **A_55_P1975537** | Nutf2 | 0.151844628 | -0.107669397 | 0.411358653 | 11.393996 | 1.421745257 | 0.203557013 | 0.601613079 | -4.470441277 |
| **A_51_P176365** | Gimap5 | -0.258974804 | -0.701620874 | 0.183671267 | 8.728323721 | -1.421621844 | 0.203591261 | 0.601613079 | -4.470503166 |
| **A_52_P99411** | Itm2b | -0.170502145 | -0.461956409 | 0.120952119 | 13.88145559 | -1.421486116 | 0.203628932 | 0.601613079 | -4.47057123 |
| **A_55_P2143802** | NA | 0.22230354 | -0.157705272 | 0.602312352 | 8.475852544 | 1.421464112 | 0.20363504 | 0.601613079 | -4.470582264 |
| **A_55_P2048202** | Ss18 | -0.177609413 | -0.481218405 | 0.12599958 | 11.53149092 | -1.421459652 | 0.203636278 | 0.601613079 | -4.470584501 |
| **A_55_P1988531** | Myh11 | -0.215047242 | -0.582666773 | 0.152572289 | 7.845485137 | -1.421407129 | 0.203650858 | 0.601613079 | -4.470610839 |
| **A_55_P2143561** | Gm6813 | 0.111303559 | -0.078971016 | 0.301578135 | 15.37362145 | 1.421384186 | 0.203657227 | 0.601613079 | -4.470622344 |
| **A_55_P2044109** | NA | -0.090766948 | -0.245957606 | 0.064423709 | 16.33900256 | -1.421167776 | 0.203717313 | 0.601613079 | -4.470730863 |
| **A_55_P1962359** | Trim11 | -0.11511215 | -0.311936797 | 0.081712497 | 11.21170287 | -1.421100761 | 0.203735922 | 0.601613079 | -4.470764468 |
| **A_55_P2037368** | Pdxdc1 | -0.112097647 | -0.303769998 | 0.079574705 | 9.397192645 | -1.421085518 | 0.203740156 | 0.601613079 | -4.470772111 |
| **A_51_P433127** | Rdm1 | -0.368807692 | -0.999427057 | 0.261811672 | 11.25575972 | -1.421071263 | 0.203744114 | 0.601613079 | -4.470779259 |
| **A_51_P176883** | Edrf1 | 0.120400785 | -0.085499399 | 0.326300969 | 8.388173886 | 1.420874676 | 0.203798717 | 0.601613079 | -4.470877834 |
| **A_51_P357422** | Coprs | -0.217613708 | -0.589766658 | 0.154539242 | 8.572553901 | -1.420849153 | 0.203805807 | 0.601613079 | -4.470890631 |

| | | | | | | | | | |
|---|---|---|---|---|---|---|---|---|---|
| **A_55_P1963777** | Gtf3c6 | 0.132913206 | -0.094430995 | 0.360257407 | 11.6790105 | 1.420585711 | 0.203879001 | 0.601616109 | -4.471022724 |
| **A_51_P258409** | Hey1 | -0.789260663 | -2.139314446 | 0.56079312 | 8.865277764 | -1.420537088 | 0.203892514 | 0.601616109 | -4.471047104 |
| **A_51_P255657** | Misp3 | -0.506625731 | -1.373243679 | 0.359992217 | 7.958908512 | -1.420504955 | 0.203901444 | 0.601616109 | -4.471063215 |
| **A_55_P2028532** | Ate1 | 0.126259616 | -0.089734916 | 0.342254148 | 8.66231431 | 1.420381143 | 0.203935855 | 0.601616109 | -4.471125293 |
| **A_55_P1956517** | Polr2l | 0.148001286 | -0.10524723 | 0.401249802 | 14.55026769 | 1.420043897 | 0.204029614 | 0.601616109 | -4.471294379 |
| **A_55_P2087671** | Abhd12 | -0.094745677 | -0.256871601 | 0.067380247 | 8.278845512 | -1.420005827 | 0.204040201 | 0.601616109 | -4.471313466 |
| **A_55_P2123933** | D2hgdh | 0.107020176 | -0.076110809 | 0.290151162 | 8.551093183 | 1.419995667 | 0.204043026 | 0.601616109 | -4.47131856 |
| **A_55_P2036220** | Chadl | -0.171806302 | -0.465807666 | 0.122195062 | 9.258294162 | -1.419949614 | 0.204055834 | 0.601616109 | -4.471341649 |
| **A_55_P2035986** | E330020D12Rik | 0.252811365 | -0.179902109 | 0.685524839 | 8.981729512 | 1.419643713 | 0.204140923 | 0.601688812 | -4.471495009 |
| **A_55_P2062133** | Etv3 | -0.109430643 | -0.296733666 | 0.07787238 | 8.26876042 | -1.419637142 | 0.204142751 | 0.601688812 | -4.471498303 |
| **A_52_P551743** | Aff4 | 0.213714207 | -0.152136974 | 0.579565389 | 9.22763113 | 1.419423924 | 0.204202079 | 0.601716937 | -4.471605194 |
| **A_55_P2145005** | NA | 0.185476112 | -0.132052643 | 0.503004866 | 8.872803442 | 1.419345457 | 0.204223916 | 0.601716937 | -4.471644531 |
| **A_55_P1972039** | Nox4 | -0.290796184 | -0.788656731 | 0.207064362 | 7.819678382 | -1.419267249 | 0.204245684 | 0.601716937 | -4.471683736 |
| **A_55_P2001347** | Rpl7a | 0.136740313 | -0.097408006 | 0.370888632 | 15.74088108 | 1.419020648 | 0.204314333 | 0.601781398 | -4.471807355 |
| **A_51_P434670** | Qk | -0.344973791 | -0.935776216 | 0.245828635 | 10.59749825 | -1.418819081 | 0.204370462 | 0.601781398 | -4.471908396 |
| **A_52_P424784** | Clstn2 | -0.470412933 | -1.276060187 | 0.33523432 | 8.182714184 | -1.418788017 | 0.204379114 | 0.601781398 | -4.471923967 |
| **A_51_P283876** | Bphl | -0.214634604 | -0.582238148 | 0.15296894 | 12.62563704 | -1.418741394 | 0.204392099 | 0.601781398 | -4.471947337 |
| **A_55_P2325378** | NA | -0.114540791 | -0.31076574 | 0.081684158 | 8.468848862 | -1.418368706 | 0.204495926 | 0.601995392 | -4.472134146 |
| **A_55_P2143150** | Qk | -0.434572136 | -1.179244783 | 0.310100511 | 8.771618589 | -1.418011208 | 0.204595565 | 0.602056104 | -4.472313331 |
| **A_55_P1958804** | Hsd3b3 | -0.529747963 | -1.437523944 | 0.378028019 | 12.14075625 | -1.417992051 | 0.204600906 | 0.602056104 | -4.472322932 |
| **A_55_P2151442** | C330007P06Rik | 0.184130604 | -0.131413701 | 0.499674909 | 10.734457 | 1.417910503 | 0.204623641 | 0.602056104 | -4.472363804 |
| **A_55_P2069176** | NA | 0.236675735 | -0.168933026 | 0.642284497 | 11.93712935 | 1.417847743 | 0.20464114 | 0.602056104 | -4.47239526 |
| **A_52_P408025** | Mpped2 | -0.331142554 | -0.89893463 | 0.236649521 | 8.463388368 | -1.417127273 | 0.204842121 | 0.602555678 | -4.472756334 |
| **A_66_P116126** | Sdhb | -0.146459731 | -0.397679629 | 0.104760166 | 14.32031258 | -1.416600492 | 0.204989185 | 0.602815933 | -4.473020313 |
| **A_55_P2014282** | NA | 0.108129323 | -0.077363185 | 0.29362183 | 17.18829493 | 1.416446908 | 0.20503208 | 0.602815933 | -4.473097273 |
| **A_51_P306066** | Eif4g2 | 0.793600725 | -0.567818165 | 2.155019615 | 11.41354475 | 1.416424638 | 0.2050383 | 0.602815933 | -4.473108432 |
| **A_55_P2129786** | Slc38a10 | 0.335444939 | -0.240167827 | 0.911057706 | 9.27298413 | 1.416035091 | 0.205147138 | 0.602815933 | -4.473303622 |
| **A_55_P2024595** | Atxn7l1 | 0.237501332 | -0.170061184 | 0.645063848 | 9.649370564 | 1.415973103 | 0.205164462 | 0.602815933 | -4.473334681 |
| **A_55_P1996469** | Zfp592 | 0.101051608 | -0.072401895 | 0.274505112 | 11.33008813 | 1.415609089 | 0.205266221 | 0.602815933 | -4.473517065 |
| **A_55_P2059154** | Dtx3l | -0.223695079 | -0.607715176 | 0.160325019 | 9.893378764 | -1.415421098 | 0.205318791 | 0.602815933 | -4.473611251 |
| **A_51_P324410** | Vti1a | 0.116910068 | -0.083791297 | 0.317611434 | 9.763010359 | 1.415418205 | 0.2053196 | 0.602815933 | -4.473612701 |
| **A_55_P1977747** | Picalm | 0.131942841 | -0.094577048 | 0.358462729 | 13.90520688 | 1.415346187 | 0.205339742 | 0.602815933 | -4.473648782 |
| **A_52_P598634** | 1190007I07Rik | 0.131158842 | -0.094092955 | 0.356410638 | 12.05006426 | 1.414856837 | 0.205476656 | 0.602815933 | -4.473893938 |
| **A_51_P238722** | Cd93 | -0.575502972 | -1.563956909 | 0.412950966 | 12.21631818 | -1.414732477 | 0.205511463 | 0.602815933 | -4.473956236 |
| **A_55_P2414479** | NA | 0.10210227 | -0.073267165 | 0.277471705 | 8.104834747 | 1.414701073 | 0.205520254 | 0.602815933 | -4.473971968 |
| **A_55_P2020472** | Rufy2 | -0.116355986 | -0.316237328 | 0.083525356 | 8.383510743 | -1.41448929 | 0.205579545 | 0.602815933 | -4.47407806 |
| **A_52_P597775** | Gprc5a | -1.397199183 | -3.797681546 | 1.00328318 | 10.28668147 | -1.414304726 | 0.205631229 | 0.602815933 | -4.474170513 |
| **A_51_P174016** | Slc39a7 | -0.095378326 | -0.259257054 | 0.068500401 | 13.47964395 | -1.414198273 | 0.205661044 | 0.602815933 | -4.474223837 |
| **A_51_P297865** | Kctd6 | 0.132500408 | -0.095164742 | 0.360165559 | 8.610411809 | 1.41417726 | 0.20566693 | 0.602815933 | -4.474234362 |
| **A_55_P1971074** | Atp11a | -0.758078727 | -2.060635564 | 0.544478111 | 9.677030458 | -1.414167381 | 0.205669697 | 0.602815933 | -4.474239311 |
| **A_52_P639774** | Gart | 0.231755609 | -0.166501465 | 0.630012684 | 9.012917847 | 1.414001743 | 0.205716099 | 0.602815933 | -4.47432228 |
| **A_55_P2185712** | Copa | -0.143259908 | -0.389472411 | 0.102952595 | 11.59491977 | -1.41383183 | 0.205763708 | 0.602815933 | -4.474407387 |
| **A_55_P2041823** | Hyal1 | 0.17738806 | -0.127484743 | 0.482260864 | 9.376561075 | 1.413802972 | 0.205771795 | 0.602815933 | -4.474421842 |
| **A_55_P2121484** | Smo | 0.232367269 | -0.167010139 | 0.631744677 | 9.148701703 | 1.413756613 | 0.205784787 | 0.602815933 | -4.474445062 |
| **A_51_P286106** | NA | -0.146406471 | -0.398063547 | 0.105250606 | 8.892821028 | -1.413625314 | 0.205821587 | 0.602815933 | -4.474510826 |
| **A_65_P13209** | Ctsd | -0.225616223 | -0.613440081 | 0.162207636 | 9.361640265 | -1.413575436 | 0.205835568 | 0.602815933 | -4.474535808 |
| **A_55_P1979014** | NA | 0.140775319 | -0.101243351 | 0.382793989 | 8.321604335 | 1.413386176 | 0.205888628 | 0.602815933 | -4.4746306 |
| **A_55_P2129354** | Pilra | -0.217032574 | -0.590184966 | 0.156119818 | 9.881068965 | -1.413259391 | 0.205924179 | 0.602815933 | -4.4746941 |
| **A_55_P2435563** | Etl4 | -0.100577219 | -0.273507658 | 0.07235322 | 8.246720245 | -1.413225182 | 0.205933772 | 0.602815933 | -4.474711233 |
| **A_55_P1979998** | Spaca9 | -0.33653648 | -0.915196112 | 0.242123151 | 9.234544461 | -1.413162653 | 0.205951308 | 0.602815933 | -4.474742549 |

| | | | | | | | | |
|---|---|---|---|---|---|---|---|---|
| **A_55_P2125841** | Cog1 | 0.134854012 | -0.09703141 | 0.366739434 | 9.439879593 | 1.413102338 | 0.205968225 | 0.602815933 | -4.474772757 |
| **A_52_P516097** | Leng9 | -0.198865247 | -0.540835225 | 0.143104731 | 8.528542715 | -1.413038849 | 0.205986033 | 0.602815933 | -4.474804554 |
| **A_51_P464234** | Fkbp2 | -0.119348692 | -0.324586173 | 0.085888789 | 12.78968709 | -1.413006558 | 0.205995091 | 0.602815933 | -4.474820726 |
| **A_55_P2136020** | Uba2 | 0.136892188 | -0.098543641 | 0.372328016 | 13.03102727 | 1.412828018 | 0.206045179 | 0.602815933 | -4.474910141 |
| **A_51_P416822** | Hhip | -0.251001057 | -0.682719329 | 0.180717214 | 7.864489239 | -1.41272721 | 0.206073466 | 0.602815933 | -4.474960625 |
| **A_55_P2027022** | Prdm1 | -0.151833548 | -0.412997425 | 0.109330329 | 8.097269248 | -1.412660566 | 0.206092167 | 0.602815933 | -4.474994 |
| **A_51_P520936** | Bcar3 | -0.268470225 | -0.730315496 | 0.193375046 | 10.16635048 | -1.412481652 | 0.206142382 | 0.602815933 | -4.475083598 |
| **A_55_P2111355** | Egfl7 | -0.602277409 | -1.638422846 | 0.433868028 | 13.65153809 | -1.412404342 | 0.206164084 | 0.602815933 | -4.475122313 |
| **A_52_P358860** | Gss | 0.338114569 | -0.257265856 | 0.933494993 | 10.67669137 | 1.444813193 | 0.206177542 | 0.602815933 | -4.470627934 |
| **A_51_P247157** | Jpt1 | 0.162976307 | -0.117475061 | 0.443427674 | 10.54602177 | 1.412050078 | 0.206263557 | 0.602815933 | -4.475299714 |
| **A_55_P2090179** | Cbx5 | 0.112750253 | -0.081285579 | 0.306786086 | 8.697802709 | 1.411948243 | 0.206292159 | 0.602815933 | -4.475350706 |
| **A_51_P340699** | Rasl11a | -0.234500217 | -0.638090471 | 0.169090037 | 7.945034849 | -1.411840926 | 0.206322304 | 0.602815933 | -4.475404443 |
| **A_55_P1972040** | Nox4 | -0.417941331 | -1.137270858 | 0.301388196 | 7.887338208 | -1.411791695 | 0.206336135 | 0.602815933 | -4.475429095 |
| **A_55_P1999548** | Tma7 | 0.182649871 | -0.131869245 | 0.497168987 | 11.96696377 | 1.411092594 | 0.206532624 | 0.602815933 | -4.475779132 |
| **A_55_P2110512** | Lmo7 | -0.606231187 | -1.650290023 | 0.437827649 | 8.700897909 | -1.410900835 | 0.206586549 | 0.602815933 | -4.475875138 |
| **A_51_P470715** | Cish | -0.317535819 | -0.86442536 | 0.229353723 | 11.76447756 | -1.410835218 | 0.206605005 | 0.602815933 | -4.47590799 |
| **A_55_P2113723** | Pecr | 0.204857759 | -0.148004844 | 0.557720363 | 8.867706419 | 1.410684754 | 0.20664733 | 0.602815933 | -4.475983319 |
| **A_52_P10041** | Akr1b3 | -0.185284614 | -0.504436154 | 0.133866925 | 11.53462355 | -1.410670561 | 0.206651323 | 0.602815933 | -4.475990424 |
| **A_55_P2166384** | Cnot1 | 0.147352766 | -0.106478578 | 0.401184109 | 11.44616222 | 1.410575158 | 0.206678164 | 0.602815933 | -4.476038186 |
| **A_65_P19319** | Soga1 | -0.363786165 | -0.990449075 | 0.262876745 | 9.206097755 | -1.410572381 | 0.206678945 | 0.602815933 | -4.476039576 |
| **A_55_P2025730** | Hif3a | 0.189347404 | -0.136839731 | 0.515534538 | 8.996545953 | 1.410508524 | 0.206696913 | 0.602815933 | -4.476071544 |
| **A_52_P494622** | Nr4a2 | -0.167332793 | -0.455599335 | 0.120933749 | 7.990274101 | -1.410489994 | 0.206702127 | 0.602815933 | -4.476080821 |
| **A_66_P135842** | Fip1l1 | 0.122019398 | -0.088185568 | 0.332224363 | 11.26991185 | 1.410487025 | 0.206702962 | 0.602815933 | -4.476082307 |
| **A_66_P104309** | Myl2 | -0.457277983 | -1.245041985 | 0.33048602 | 8.01748517 | -1.410481351 | 0.206704559 | 0.602815933 | -4.476085148 |
| **A_55_P2019113** | Apol7b | 0.137367738 | -0.099289768 | 0.374025244 | 8.334740774 | 1.410417381 | 0.20672256 | 0.602815933 | -4.476117172 |
| **A_55_P1984932** | Zc3h18 | 0.155603213 | -0.112485345 | 0.423691771 | 11.97516647 | 1.410338853 | 0.206744661 | 0.602815933 | -4.476156485 |
| **A_55_P2104327** | Pxdn | -0.700467518 | -1.907501365 | 0.506566328 | 11.2154089 | -1.410105939 | 0.206810222 | 0.602815933 | -4.476273081 |
| **A_55_P1970901** | Gramd2 | -0.194467496 | -0.529607026 | 0.140672034 | 7.734269291 | -1.409952736 | 0.206853357 | 0.602815933 | -4.476349773 |
| **A_55_P2057519** | Rpl17 | 0.305749005 | -0.221180512 | 0.832678522 | 14.14703454 | 1.409924035 | 0.206861439 | 0.602815933 | -4.47636414 |
| **A_55_P2137521** | Nmral1 | 0.189819038 | -0.137316844 | 0.51695492 | 8.980083608 | 1.409920983 | 0.206862298 | 0.602815933 | -4.476365668 |
| **A_52_P39247** | Hbp1 | -0.195381938 | -0.532130185 | 0.141366309 | 11.63361734 | -1.409815429 | 0.206892023 | 0.602815933 | -4.476418505 |
| **A_55_P2000643** | NA | -0.203196917 | -0.553459535 | 0.147065701 | 8.270016468 | -1.40963448 | 0.206942989 | 0.602815933 | -4.476509081 |
| **A_55_P2049647** | H2-K1 | -0.278879073 | -0.759646348 | 0.201888202 | 12.24044085 | -1.409497198 | 0.206981663 | 0.602815933 | -4.476577797 |
| **A_55_P2147871** | NA | 0.196399217 | -0.142229327 | 0.535027761 | 9.035678585 | 1.409286782 | 0.207040953 | 0.602815933 | -4.476683118 |
| **A_52_P421344** | Paqr5 | -0.332040701 | -0.904634208 | 0.240552806 | 8.10502047 | -1.409055476 | 0.207106147 | 0.602815933 | -4.47679889 |
| **A_55_P1966209** | Fez2 | 0.14860556 | -0.107676687 | 0.404887807 | 10.71957668 | 1.408963443 | 0.207132092 | 0.602815933 | -4.476844953 |
| **A_55_P2052261** | Zfp280c | 0.121287332 | -0.087887317 | 0.33046198 | 9.153328318 | 1.40893054 | 0.207141369 | 0.602815933 | -4.476861421 |
| **A_55_P2239317** | Afap1l1 | -0.537428244 | -1.464595159 | 0.38973867 | 9.150144792 | -1.408463997 | 0.207272942 | 0.602815933 | -4.477094916 |
| **A_52_P26953** | Ergic2 | 0.101987259 | -0.073964165 | 0.277938684 | 9.394776258 | 1.408433428 | 0.207281565 | 0.602815933 | -4.477110215 |
| **A_55_P2216456** | Rint1 | 0.196476759 | -0.142494794 | 0.535448311 | 9.650175566 | 1.408416556 | 0.207286325 | 0.602815933 | -4.477118658 |
| **A_51_P451346** | Klf6 | -0.284090087 | -0.7742213 | 0.206041125 | 11.62750395 | -1.408402896 | 0.207290179 | 0.602815933 | -4.477125495 |
| **A_52_P161087** | 2610008E11Rik | -0.17917005 | -0.48828632 | 0.12994622 | 8.717560132 | -1.408402345 | 0.207290334 | 0.602815933 | -4.477125771 |
| **A_55_P1986113** | Ppil1 | 0.229792469 | -0.166681728 | 0.626266666 | 10.94776216 | 1.408328772 | 0.207311092 | 0.602815933 | -4.477162591 |
| **A_51_P374818** | Shd | -0.134036657 | -0.365303746 | 0.097230431 | 7.756401799 | -1.408292753 | 0.207321255 | 0.602815933 | -4.477180616 |
| **A_55_P2081223** | Sptbn1 | -0.424060866 | -1.155849429 | 0.307727697 | 13.01086983 | -1.408074934 | 0.207382722 | 0.602815933 | -4.477289621 |
| **A_52_P9703** | Cyp2a4 | -0.618564202 | -1.686013918 | 0.448885514 | 9.755142591 | -1.408058004 | 0.207387501 | 0.602815933 | -4.477298093 |
| **A_55_P2269497** | Pdp1 | 0.14775851 | -0.107248136 | 0.402765155 | 8.768158217 | 1.407940145 | 0.207420768 | 0.602815933 | -4.477357073 |
| **A_55_P2007756** | Krt16 | 0.247228403 | -0.179479029 | 0.673935834 | 8.823584768 | 1.407833661 | 0.207450828 | 0.602815933 | -4.477410359 |
| **A_52_P368306** | Tmem100 | -0.205081656 | -0.559053168 | 0.148889856 | 7.770074604 | -1.407802375 | 0.207459661 | 0.602815933 | -4.477426015 |
| **A_55_P1998892** | Smox | 0.226457495 | -0.164418308 | 0.617333298 | 12.38095241 | 1.407768042 | 0.207469355 | 0.602815933 | -4.477443195 |

| | | | | | | | | | |
|---|---|---|---|---|---|---|---|---|---|
| **A_52_P174915** | Gja1 | -0.333199632 | -0.908320938 | 0.241921674 | 8.065041885 | -1.407758781 | 0.207471969 | 0.602815933 | -4.47744783 |
| **A_55_P2006930** | 2610005L07Rik | 0.131132924 | -0.095210132 | 0.357475979 | 11.33565536 | 1.407757208 | 0.207472413 | 0.602815933 | -4.477448617 |
| **A_55_P1992074** | Ptpra | 0.115803921 | -0.084088103 | 0.315695946 | 11.97974008 | 1.407702844 | 0.207487763 | 0.602815933 | -4.47747582 |
| **A_51_P418317** | Mis12 | 0.122939399 | -0.08927686 | 0.335155657 | 8.527413794 | 1.407653033 | 0.207501829 | 0.602815933 | -4.477500746 |
| **A_55_P2078428** | Tstd1 | -0.538928994 | -1.469258733 | 0.391400745 | 9.913940755 | -1.407595384 | 0.207518108 | 0.602815933 | -4.477529593 |
| **A_55_P2045481** | NA | 0.136822161 | -0.099386217 | 0.37303054 | 14.08826629 | 1.407486819 | 0.207548769 | 0.602815933 | -4.477583918 |
| **A_55_P2111005** | Pafah1b1 | 0.146375152 | -0.106326395 | 0.399076698 | 10.49261229 | 1.407481347 | 0.207550315 | 0.602815933 | -4.477586656 |
| **A_55_P1954365** | Ttc28 | -0.356727229 | -0.972651716 | 0.259197257 | 9.106209194 | -1.407317148 | 0.207596696 | 0.602860058 | -4.477668817 |
| **A_51_P221495** | Nsun5 | 0.116200984 | -0.084515058 | 0.316917026 | 8.518144158 | 1.406730518 | 0.207762479 | 0.603162926 | -4.477962335 |
| **A_51_P172688** | Stard3nl | -0.255743009 | -0.697493515 | 0.186007497 | 9.522420187 | -1.406727261 | 0.2077634 | 0.603162926 | -4.477963964 |
| **A_51_P167971** | Bace1 | -0.148765165 | -0.405791659 | 0.108261329 | 10.64696362 | -1.406392507 | 0.207858057 | 0.603347106 | -4.478131445 |
| **A_55_P1956989** | BC004004 | -0.187749785 | -0.512234799 | 0.13673523 | 13.9367174 | -1.405943899 | 0.207984969 | 0.603390546 | -4.478355873 |
| **A_55_P2419296** | NA | 0.128801694 | -0.093809167 | 0.351412555 | 8.416940797 | 1.405912894 | 0.207993743 | 0.603390546 | -4.478371383 |
| **A_55_P1962876** | Adam11 | 0.326110992 | -0.237521039 | 0.889743024 | 9.253373853 | 1.405895241 | 0.207998739 | 0.603390546 | -4.478380215 |
| **A_55_P1984640** | Ube2f | 0.201564898 | -0.146832363 | 0.549962159 | 11.50968186 | 1.40579942 | 0.208025857 | 0.603390546 | -4.478428149 |
| **A_55_P2112991** | Slc4a3 | -0.198586692 | -0.541862899 | 0.144689515 | 8.011358778 | -1.405690235 | 0.208056762 | 0.603390546 | -4.478482768 |
| **A_51_P341736** | Mmp2 | -0.679922618 | -1.85524481 | 0.495399574 | 8.825746771 | -1.405677654 | 0.208060323 | 0.603390546 | -4.478489061 |
| **A_55_P1961034** | Gm13272 | -0.146615064 | -0.40008668 | 0.106856553 | 8.044398391 | -1.405505173 | 0.208109153 | 0.603441618 | -4.478575342 |
| **A_55_P1999541** | Ccdc42 | 0.115577254 | -0.084283306 | 0.315437813 | 8.241390028 | 1.405168679 | 0.208204446 | 0.603526843 | -4.478743659 |
| **A_51_P270519** | Cenps | 0.17366823 | -0.126654094 | 0.473990554 | 8.517948418 | 1.405128203 | 0.208215911 | 0.603526843 | -4.478763905 |
| **A_51_P184910** | Phpt1 | -0.290724355 | -0.793545122 | 0.212096412 | 12.47008384 | -1.404919365 | 0.208275076 | 0.603526843 | -4.478868363 |
| **A_55_P1963878** | NA | -0.095152276 | -0.259724896 | 0.069420344 | 11.25741385 | -1.404897979 | 0.208281135 | 0.603526843 | -4.478879059 |
| **A_52_P295104** | Smim5 | 0.22277735 | -0.162544995 | 0.608099695 | 9.88025954 | 1.404850233 | 0.208294664 | 0.603526843 | -4.478902941 |
| **A_55_P2039180** | Zswim8 | -0.214234933 | -0.58485467 | 0.156384803 | 11.18293861 | -1.404575003 | 0.208372668 | 0.603662365 | -4.4790406 |
| **A_52_P1026777** | Ecscr | -0.472260516 | -1.289407217 | 0.344886186 | 9.878031302 | -1.404315839 | 0.208446142 | 0.603784728 | -4.479170218 |
| **A_55_P2062632** | Smc5 | 0.274392017 | -0.200499057 | 0.749283092 | 10.75535187 | 1.403979138 | 0.208541634 | 0.603865378 | -4.479338606 |
| **A_55_P1980621** | NA | 0.144408685 | -0.10556512 | 0.394382489 | 14.0054911 | 1.403724895 | 0.208613766 | 0.603865378 | -4.479465751 |
| **A_52_P560628** | 1300002E11Rik | 0.152014634 | -0.11117475 | 0.415204018 | 9.282515349 | 1.403460586 | 0.208688778 | 0.603865378 | -4.479597923 |
| **A_51_P511236** | Pik3r1 | 0.11212974 | -0.082027891 | 0.306287371 | 9.037911202 | 1.403296814 | 0.20873527 | 0.603865378 | -4.479679818 |
| **A_55_P2162160** | Anp32a | 0.221360284 | -0.161938247 | 0.604658816 | 12.43395305 | 1.40328453 | 0.208738758 | 0.603865378 | -4.47968596 |
| **A_51_P386270** | Cyp2d12 | -0.489595137 | -1.337378561 | 0.358188286 | 12.18484307 | -1.403251012 | 0.208748274 | 0.603865378 | -4.47970272 |
| **A_55_P2154709** | Pter | 0.349392316 | -0.255658985 | 0.954443617 | 11.58441657 | 1.403150771 | 0.208776737 | 0.603865378 | -4.479752845 |
| **A_51_P467448** | Pif1 | 0.218403708 | -0.159812716 | 0.596620131 | 10.30170888 | 1.403145839 | 0.208778137 | 0.603865378 | -4.479755311 |
| **A_52_P131559** | Srprb | -0.098138134 | -0.268113607 | 0.071837338 | 8.357021347 | -1.402925895 | 0.208840601 | 0.603865378 | -4.479865288 |
| **A_55_P2341890** | NA | -0.090655572 | -0.247682452 | 0.066371308 | 8.018601045 | -1.402825622 | 0.208869085 | 0.603865378 | -4.479915425 |
| **A_51_P288138** | Fpr2 | -0.338893383 | -0.925914244 | 0.248127479 | 9.909792365 | -1.402790262 | 0.20887913 | 0.603865378 | -4.479933106 |
| **A_66_P117543** | NA | 0.289067149 | -0.211658924 | 0.789793223 | 9.440875666 | 1.402754668 | 0.208889242 | 0.603865378 | -4.479950903 |
| **A_55_P2011116** | L3mbtl3 | -0.15779619 | -0.431147618 | 0.115555238 | 8.725491377 | -1.402679194 | 0.208910685 | 0.603865378 | -4.47998864 |
| **A_52_P444162** | Rab11a | -0.10190423 | -0.278458608 | 0.074650149 | 10.85015191 | -1.402480744 | 0.208967076 | 0.603865378 | -4.480087861 |
| **A_55_P2177463** | B130024G19Rik | -0.335935855 | -0.917970687 | 0.246098977 | 8.295964862 | -1.402460288 | 0.20897289 | 0.603865378 | -4.480098089 |
| **A_55_P2124319** | NA | 0.113327786 | -0.083021834 | 0.309677407 | 7.8419139 | 1.402456916 | 0.208973848 | 0.603865378 | -4.480099775 |
| **A_66_P106861** | B3glct | 0.115280129 | -0.084534904 | 0.315095161 | 9.232139795 | 1.401875623 | 0.209139113 | 0.604140097 | -4.480390394 |
| **A_51_P116906** | Rapgef3 | -0.507319515 | -1.386764538 | 0.372125508 | 9.583394893 | -1.40170319 | 0.209188159 | 0.604140097 | -4.480476596 |
| **A_51_P181400** | Efcab2 | 0.11735039 | -0.086105429 | 0.320806209 | 8.384217844 | 1.401514552 | 0.209241827 | 0.604140097 | -4.480570898 |
| **A_55_P2055975** | Ppip5k1 | 0.111580998 | -0.081883351 | 0.305045347 | 8.327591782 | 1.40143347 | 0.209264899 | 0.604140097 | -4.480611431 |
| **A_51_P180001** | Ago2 | 0.189775749 | -0.139295992 | 0.518847489 | 11.81171988 | 1.401307074 | 0.20930087 | 0.604140097 | -4.480674614 |
| **A_51_P453351** | Lpgat1 | -0.184617424 | -0.504751931 | 0.135517083 | 11.38605508 | -1.401275039 | 0.209309987 | 0.604140097 | -4.480690628 |
| **A_55_P2028837** | Tspan2 | -0.359027294 | -0.981651081 | 0.263596493 | 8.323983045 | -1.401151016 | 0.209345289 | 0.604140097 | -4.480752624 |
| **A_55_P2066878** | Kcnj13 | -0.2488657 | -0.680453429 | 0.182722028 | 7.890491077 | -1.401132304 | 0.209350616 | 0.604140097 | -4.480761977 |
| **A_55_P2048498** | NA | 0.152019164 | -0.111637958 | 0.415676285 | 8.457711103 | 1.401012536 | 0.209384713 | 0.604140097 | -4.480821845 |

| | | | | | | | | |
|---|---|---|---|---|---|---|---|---|
| **A_51_P502456** | H2-K2 | -0.257804059 | -0.704953826 | 0.189345709 | 12.93656136 | -1.400941272 | 0.209405003 | 0.604140097 | -4.480857467 |
| **A_55_P1960311** | Dnase1l1 | 0.245611744 | -0.180399243 | 0.671622732 | 10.34875292 | 1.400914137 | 0.20941273 | 0.604140097 | -4.48087103 |
| **A_66_P109395** | Rpia | 0.157527564 | -0.11576795 | 0.430823077 | 10.43776292 | 1.400577814 | 0.209508516 | 0.604326236 | -4.481039136 |
| **A_55_P2225082** | Mettl4 | 0.174583938 | -0.128350315 | 0.477518192 | 8.79670298 | 1.40035805 | 0.209571128 | 0.60441664 | -4.481148977 |
| **A_51_P338397** | Kctd10 | -0.260470434 | -0.712501158 | 0.191560289 | 8.306753032 | -1.400147114 | 0.20963124 | 0.604499811 | -4.481254402 |
| **A_51_P185846** | Dnal4 | -0.32145642 | -0.879381912 | 0.236469071 | 9.91990917 | -1.400003395 | 0.209672205 | 0.604515372 | -4.481326229 |
| **A_51_P233027** | Rmdn3 | 0.179665779 | -0.13218643 | 0.491517989 | 9.300995346 | 1.399908753 | 0.209699186 | 0.604515372 | -4.481373529 |
| **A_55_P1956418** | Efr3b | -0.245013565 | -0.670390699 | 0.180363569 | 8.105986583 | -1.399584672 | 0.2097916 | 0.60463326 | -4.481535489 |
| **A_51_P361448** | Scara5 | -0.188931106 | -0.516971945 | 0.139109734 | 7.782120233 | -1.399454361 | 0.20982877 | 0.60463326 | -4.48160061 |
| **A_66_P133404** | Pclaf | 0.195518871 | -0.1439866 | 0.535024342 | 9.131678426 | 1.399345941 | 0.209859699 | 0.60463326 | -4.48165479 |
| **A_51_P455946** | Rac3 | -0.456174363 | -1.248301623 | 0.335952898 | 8.536552718 | -1.399326644 | 0.209865205 | 0.60463326 | -4.481664433 |
| **A_55_P2090519** | Ppdpf | -0.158421525 | -0.433572605 | 0.116729554 | 13.23462415 | -1.399027202 | 0.209950653 | 0.60475676 | -4.481814066 |
| **A_55_P2108850** | 2-Mar | 0.168315016 | -0.12403584 | 0.460665873 | 9.726244727 | 1.398948354 | 0.209973158 | 0.60475676 | -4.481853465 |
| **A_55_P2046393** | Rcn3 | -0.454186645 | -1.243164138 | 0.334790847 | 11.83212772 | -1.398791342 | 0.21001798 | 0.60475676 | -4.481931921 |
| **A_55_P2029643** | NA | -0.264391114 | -0.723688194 | 0.194905965 | 11.72612422 | -1.398737957 | 0.210033221 | 0.60475676 | -4.481958596 |
| **A_55_P2042156** | Aatk | -0.152592934 | -0.417776803 | 0.112590936 | 8.573838063 | -1.398203897 | 0.210185752 | 0.605105809 | -4.482225438 |
| **A_55_P2131105** | NA | 0.195879582 | -0.144637547 | 0.536396712 | 12.57681642 | 1.397762531 | 0.210311885 | 0.605378766 | -4.482445947 |
| **A_52_P28057** | Gm14124 | 0.110013227 | -0.08125189 | 0.301278344 | 8.043251266 | 1.397630346 | 0.210349673 | 0.605397385 | -4.482511984 |
| **A_51_P215077** | Mgst3 | 0.163616288 | -0.120899605 | 0.448132181 | 12.84009409 | 1.397343545 | 0.210431685 | 0.605543254 | -4.48265526 |
| **A_51_P211903** | Golga5 | 0.205591265 | -0.151951687 | 0.563134218 | 10.17643385 | 1.397203378 | 0.210471777 | 0.605568469 | -4.48272528 |
| **A_55_P2018181** | A730008H23Rik | 0.112153759 | -0.082966327 | 0.307273845 | 8.418410256 | 1.396673985 | 0.210623261 | 0.605863454 | -4.482989722 |
| **A_55_P2041581** | NA | 0.140685544 | -0.104093739 | 0.385464828 | 9.58066338 | 1.396554874 | 0.210657358 | 0.605863454 | -4.483049216 |
| **A_51_P466824** | Tyk2 | -0.11575051 | -0.317154596 | 0.085653575 | 8.477065024 | -1.396489992 | 0.210675933 | 0.605863454 | -4.483081624 |
| **A_55_P2163928** | Rtl8c | -0.137730792 | -0.37739446 | 0.101932876 | 7.856490099 | -1.396407048 | 0.210699681 | 0.605863454 | -4.483123053 |
| **A_65_P11853** | Arhgap28 | -0.281731706 | -0.772018036 | 0.208554625 | 8.107682558 | -1.396269105 | 0.210739183 | 0.605878146 | -4.483191951 |
| **A_55_P2054708** | NA | -0.211596902 | -0.579894359 | 0.156700555 | 7.886195768 | -1.396026856 | 0.210808569 | 0.605878146 | -4.483312943 |
| **A_66_P126726** | Cwc27 | 0.100808095 | -0.074661918 | 0.276278107 | 8.779722575 | 1.395968726 | 0.210825222 | 0.605878146 | -4.483341975 |
| **A_51_P279308** | Acbd5 | 0.247030895 | -0.182964841 | 0.677026631 | 9.955434184 | 1.395951444 | 0.210830173 | 0.605878146 | -4.483350606 |
| **A_51_P173086** | Eif3l | 0.12590002 | -0.093267888 | 0.345067928 | 14.34720936 | 1.395828385 | 0.210865432 | 0.60588939 | -4.483412066 |
| **A_55_P2137973** | Bnip2 | 0.136290101 | -0.101016189 | 0.37359639 | 8.439059072 | 1.395527029 | 0.210951798 | 0.605968086 | -4.483562567 |
| **A_52_P452019** | Usp31 | 0.12699578 | -0.094129435 | 0.348120995 | 8.293109133 | 1.395514042 | 0.21095552 | 0.605968086 | -4.483569053 |
| **A_55_P2045050** | Zfp953 | 0.112056449 | -0.083148591 | 0.307261489 | 8.454155276 | 1.394854856 | 0.211144553 | 0.606272175 | -4.483898232 |
| **A_55_P2056926** | Mtss2 | -0.516794896 | -1.417144342 | 0.383554551 | 9.05585763 | -1.394730515 | 0.211180227 | 0.606272175 | -4.48396032 |
| **A_52_P33014** | Ppp1r8 | 0.17506099 | -0.129931 | 0.48005298 | 8.847890168 | 1.394710711 | 0.211185909 | 0.606272175 | -4.483970208 |
| **A_55_P2096257** | Myl6 | -0.300324024 | -0.823562812 | 0.222914763 | 14.0560221 | -1.39467602 | 0.211195863 | 0.606272175 | -4.483987531 |
| **A_51_P277006** | Chst8 | -0.252697908 | -0.692984797 | 0.187588982 | 8.073608644 | -1.394598138 | 0.211218212 | 0.606272175 | -4.484026419 |
| **A_55_P2094412** | Ssbp1 | 0.123691425 | -0.09185954 | 0.339242391 | 9.944618336 | 1.394353241 | 0.211288502 | 0.606383883 | -4.484148699 |
| **A_55_P2127421** | NA | 0.179216034 | -0.133153277 | 0.491585346 | 8.436157202 | 1.394092825 | 0.211363269 | 0.606397351 | -4.484278722 |
| **A_55_P2032034** | Tbp | 0.106910393 | -0.079475506 | 0.293296292 | 9.3707571 | 1.393766636 | 0.211456954 | 0.606397351 | -4.484441577 |
| **A_55_P2101340** | Ramp3 | -0.097909515 | -0.268605611 | 0.072786581 | 7.91437982 | -1.39374878 | 0.211462083 | 0.606397351 | -4.484450492 |
| **A_55_P2070179** | Ing4 | -0.252183037 | -0.691894089 | 0.187528014 | 10.59170042 | -1.393579275 | 0.211510782 | 0.606397351 | -4.484535116 |
| **A_55_P2024749** | Hspb11 | -0.205973752 | -0.56511813 | 0.153170626 | 10.06560944 | -1.393561041 | 0.211516022 | 0.606397351 | -4.484544219 |
| **A_51_P369628** | Ptov1 | -0.469771442 | -1.288910149 | 0.349367265 | 10.66141419 | -1.393517261 | 0.211528602 | 0.606397351 | -4.484566076 |
| **A_55_P1990336** | Add1 | 0.137755562 | -0.102487169 | 0.377998293 | 13.86346899 | 1.393291785 | 0.211593403 | 0.606397351 | -4.484678638 |
| **A_51_P427476** | Mxra8 | -0.439259552 | -1.20539442 | 0.326875316 | 8.292817949 | -1.393154157 | 0.211632965 | 0.606397351 | -4.484747342 |
| **A_55_P1998299** | Itgb4 | -0.123465098 | -0.338839437 | 0.091909241 | 8.109103738 | -1.392943294 | 0.211693593 | 0.606397351 | -4.484852603 |
| **A_55_P2075829** | NA | -0.146121229 | -0.401042968 | 0.10880051 | 8.797512447 | -1.392802806 | 0.211733995 | 0.606397351 | -4.484922731 |
| **A_51_P293339** | Tob1 | -0.126721651 | -0.347804922 | 0.09436162 | 9.597591421 | -1.392765916 | 0.211744605 | 0.606397351 | -4.484941146 |
| **A_55_P2073617** | NA | 0.408606293 | -0.304275409 | 1.121487996 | 9.310668867 | 1.392742321 | 0.211751392 | 0.606397351 | -4.484952923 |
| **A_55_P2023692** | Neurl1a | -0.18242246 | -0.500716395 | 0.135871474 | 9.558294243 | -1.392621625 | 0.21178611 | 0.606397351 | -4.48501317 |

| | | | | | | | | | |
|---|---|---|---|---|---|---|---|---|---|
| **A_55_P2139937** | Lrrc56 | 0.337738746 | -0.251768802 | 0.927246293 | 10.01373722 | 1.392113693 | 0.211932274 | 0.606397351 | -4.485266695 |
| **A_52_P239052** | Zfp148 | -0.147367276 | -0.404602894 | 0.109868342 | 10.55553828 | -1.392044572 | 0.211952172 | 0.606397351 | -4.485301194 |
| **A_55_P2081050** | Kcnj13 | -0.379194641 | -1.041118978 | 0.282729696 | 7.850332669 | -1.391992934 | 0.211967037 | 0.606397351 | -4.485326966 |
| **A_52_P200265** | Mocs1 | 0.203915049 | -0.152040777 | 0.559870875 | 10.80803728 | 1.39199079 | 0.211967655 | 0.606397351 | -4.485328036 |
| **A_51_P314323** | Naa38 | -0.147466068 | -0.404884254 | 0.109952117 | 13.60032055 | -1.391989839 | 0.211967929 | 0.606397351 | -4.485328511 |
| **A_66_P111089** | Ube2f | 0.17698253 | -0.131964084 | 0.485929143 | 9.692532688 | 1.391970874 | 0.211973389 | 0.606397351 | -4.485337977 |
| **A_51_P395473** | Tns2 | -0.333073049 | -0.914527757 | 0.248381659 | 8.82821375 | -1.391896013 | 0.211994942 | 0.606397351 | -4.48537534 |
| **A_51_P113182** | Lifr | -0.097101829 | -0.266616268 | 0.07241261 | 7.881829279 | -1.391886759 | 0.211997607 | 0.606397351 | -4.485379958 |
| **A_65_P07627** | Dach1 | -0.253340452 | -0.6956306 | 0.188949695 | 7.992285359 | -1.391811644 | 0.212019236 | 0.606397351 | -4.485417447 |
| **A_55_P2021099** | Tmem181a | -0.194355276 | -0.533690837 | 0.144980286 | 9.599524345 | -1.391714486 | 0.212047215 | 0.606397351 | -4.485465937 |
| **A_51_P256093** | Map2k6 | 0.149149705 | -0.111295185 | 0.409594594 | 9.229899511 | 1.391520938 | 0.212102963 | 0.606397351 | -4.48556253 |
| **A_55_P2065874** | D430042O09Rik | 0.171696319 | -0.12818 | 0.471572639 | 9.461276324 | 1.391239829 | 0.212183954 | 0.606397351 | -4.485702817 |
| **A_55_P2175550** | Atf2 | -0.117781539 | -0.323501153 | 0.087938075 | 8.513917821 | -1.39118444 | 0.212199916 | 0.606397351 | -4.485730458 |
| **A_51_P163444** | Carhsp1 | 0.11663154 | -0.087082714 | 0.320345794 | 12.44661133 | 1.391162234 | 0.212206315 | 0.606397351 | -4.485741539 |
| **A_55_P1991792** | Slc25a46 | 0.138167781 | -0.103194361 | 0.379529923 | 11.60323371 | 1.390979793 | 0.212258899 | 0.606397351 | -4.485832581 |
| **A_52_P262219** | Fos | -0.12160757 | -0.334111294 | 0.090896154 | 7.945001209 | -1.390520046 | 0.21239146 | 0.606397351 | -4.486395742 |
| **A_51_P151902** | Spon1 | -0.236566644 | -0.649963609 | 0.176830321 | 7.799114697 | -1.390494943 | 0.212398701 | 0.606397351 | -4.486074518 |
| **A_55_P2022181** | Arvcf | -0.093383667 | -0.256584189 | 0.069816855 | 8.323324434 | -1.390376985 | 0.212432725 | 0.606397351 | -4.486133375 |
| **A_51_P368496** | Tmem98 | -0.37225509 | -1.022821235 | 0.278311055 | 9.63112532 | -1.390376359 | 0.212432906 | 0.606397351 | -4.486133688 |
| **A_51_P278887** | Hint2 | -0.115065573 | -0.316172437 | 0.086041291 | 12.37864414 | -1.390278157 | 0.212461236 | 0.606397351 | -4.486182686 |
| **A_55_P2153620** | Ahnak | 0.269317295 | -0.201394448 | 0.740029039 | 9.128107222 | 1.390248387 | 0.212469825 | 0.606397351 | -4.48619754 |
| **A_55_P2073814** | Gpatch8 | 0.192840856 | -0.144225313 | 0.529907026 | 8.945990586 | 1.39016732 | 0.212493215 | 0.606397351 | -4.486237988 |
| **A_55_P2056325** | Anxa3 | -0.479250523 | -1.316936443 | 0.358435396 | 8.824735849 | -1.390159299 | 0.21249553 | 0.606397351 | -4.48624199 |
| **A_51_P473272** | Tspan6 | -0.352049506 | -0.96740567 | 0.263306658 | 9.830092274 | -1.390145869 | 0.212499405 | 0.606397351 | -4.486248691 |
| **A_55_P2090055** | Phrf1 | 0.114975951 | -0.086041749 | 0.31599365 | 10.4532178 | 1.389811498 | 0.212595908 | 0.606397351 | -4.486415518 |
| **A_65_P17602** | Pgm5 | -0.278632125 | -0.765874387 | 0.208610136 | 7.842004592 | -1.389534795 | 0.212675797 | 0.606397351 | -4.486553565 |
| **A_51_P370510** | Clba1 | -0.246586815 | -0.677795184 | 0.184621554 | 7.858838409 | -1.389523381 | 0.212679093 | 0.606397351 | -4.486559259 |
| **A_52_P503208** | NA | 0.180245351 | -0.1349516 | 0.495442302 | 12.20792725 | 1.389521442 | 0.212679653 | 0.606397351 | -4.486560227 |
| **A_55_P2042062** | Mier1 | 0.161130338 | -0.120662376 | 0.442923053 | 8.432685352 | 1.389411048 | 0.212711534 | 0.606397351 | -4.4866153 |
| **A_55_P2144666** | Prpsap1 | -0.144312429 | -0.39669604 | 0.108071183 | 12.53056103 | -1.389395136 | 0.21271613 | 0.606397351 | -4.486623238 |
| **A_55_P2040490** | NA | 0.140194966 | -0.104989795 | 0.385379727 | 8.345089108 | 1.389383495 | 0.212719492 | 0.606397351 | -4.486629046 |
| **A_55_P1984622** | Smarcc1 | 0.195173245 | -0.146192662 | 0.536539152 | 8.394122314 | 1.389259348 | 0.212755351 | 0.606397351 | -4.486690979 |
| **A_55_P2182867** | Sgcb | -0.220432176 | -0.605985763 | 0.165121411 | 8.486323798 | -1.389227683 | 0.212764498 | 0.606397351 | -4.486706776 |
| **A_55_P2151156** | Farsa | 0.180039057 | -0.134888048 | 0.494966163 | 11.53760927 | 1.389120364 | 0.212795502 | 0.606397351 | -4.486760313 |
| **A_55_P2158473** | 1700028P14Rik | -0.295413271 | -0.812213797 | 0.221387256 | 7.774006631 | -1.388961611 | 0.212841373 | 0.606397351 | -4.486839506 |
| **A_55_P1971854** | Vcpkmt | 0.119757046 | -0.089766402 | 0.329280495 | 10.82502723 | 1.388838128 | 0.212877058 | 0.606397351 | -4.486901103 |
| **A_51_P380005** | Galnt3 | -0.176651905 | -0.48574105 | 0.13243724 | 8.379082189 | -1.388729819 | 0.212908363 | 0.606397351 | -4.48695513 |
| **A_51_P407999** | 1500011B03Rik | -0.11882262 | -0.326729447 | 0.089084208 | 8.733557148 | -1.388716389 | 0.212912246 | 0.606397351 | -4.48696183 |
| **A_52_P622230** | Gm4890 | 0.135503711 | -0.101599295 | 0.372606717 | 8.213063897 | 1.388664452 | 0.212927259 | 0.606397351 | -4.486987736 |
| **A_52_P508991** | Fmo1 | -0.368740125 | -1.014003908 | 0.276523658 | 10.65053667 | -1.388565268 | 0.212955933 | 0.606397351 | -4.487037211 |
| **A_55_P1984770** | Fam220a | 0.185103009 | -0.138904676 | 0.509110694 | 13.61958357 | 1.38816585 | 0.213071438 | 0.606636886 | -4.487236435 |
| **A_51_P145662** | Clec4g | -0.23282877 | -0.640423054 | 0.174765513 | 11.09042965 | -1.388007297 | 0.213117304 | 0.606678111 | -4.487315516 |
| **A_55_P2037454** | Etv5 | -0.911098688 | -2.506504441 | 0.684307065 | 10.88566308 | -1.387642064 | 0.213222994 | 0.606878998 | -4.487497672 |
| **A_51_P334289** | Zfp628 | 0.116357498 | -0.087430454 | 0.32014545 | 9.16090996 | 1.387391591 | 0.213295502 | 0.606878998 | -4.487622587 |
| **A_55_P1988048** | Emcn | -0.499856803 | -1.375379821 | 0.375666214 | 8.177560483 | -1.387270747 | 0.213330493 | 0.606878998 | -4.487682851 |
| **A_55_P2336173** | Mirg | 0.252547499 | -0.189854103 | 0.694949102 | 9.550826243 | 1.387105746 | 0.213378277 | 0.606878998 | -4.487765134 |
| **A_66_P115034** | Bend5 | -0.215919601 | -0.594200861 | 0.16236166 | 7.976974647 | -1.386948804 | 0.213423737 | 0.606878998 | -4.487843397 |
| **A_55_P1979997** | Spaca9 | -0.312217794 | -0.859217565 | 0.234781976 | 8.514186037 | -1.386927299 | 0.213429967 | 0.606878998 | -4.487854121 |
| **A_55_P2049572** | Pld2 | -0.267674966 | -0.736640406 | 0.201290474 | 10.10632668 | -1.386915834 | 0.213433289 | 0.606878998 | -4.487859838 |
| **A_52_P376337** | Sdhaf3 | 0.140326776 | -0.105528685 | 0.386182237 | 8.619362655 | 1.386895941 | 0.213439052 | 0.606878998 | -4.487869758 |

| | | | | | | | | |
|---|---|---|---|---|---|---|---|---|
| **A_55_P2080562** | Arhgap26 | 0.463090454 | -0.348479845 | 1.274660753 | 10.31371045 | 1.386509575 | 0.213551011 | 0.60707131 | -4.488062416 |
| **A_51_P332141** | Clcn2 | 0.237388059 | -0.178681374 | 0.653457492 | 9.890539588 | 1.38636073 | 0.213594156 | 0.60707131 | -4.488136633 |
| **A_51_P208031** | Cnot6l | -0.142576277 | -0.39247992 | 0.107327366 | 8.600862465 | -1.386302069 | 0.213611163 | 0.60707131 | -4.488165882 |
| **A_55_P1989452** | Nenf | -0.25985495 | -0.715396647 | 0.195686747 | 12.74485075 | -1.386072823 | 0.213677635 | 0.60707131 | -4.488280184 |
| **A_66_P113245** | Mfng | -0.473207156 | -1.30279015 | 0.356375837 | 10.47643627 | -1.386036453 | 0.213688182 | 0.60707131 | -4.488298317 |
| **A_51_P475228** | Armc6 | 0.192706564 | -0.145141078 | 0.530554206 | 9.624988084 | 1.385985876 | 0.213702851 | 0.60707131 | -4.488323534 |
| **A_51_P268559** | Idh3a | 0.168264339 | -0.126749277 | 0.463277956 | 10.96584921 | 1.385904206 | 0.213726538 | 0.60707131 | -4.488364253 |
| **A_52_P381430** | Tbc1d4 | -0.097339897 | -0.26802675 | 0.073346956 | 7.768903401 | -1.385715257 | 0.213781351 | 0.607090304 | -4.488458456 |
| **A_55_P2137591** | Rnf220 | 0.117009151 | -0.088203255 | 0.322221556 | 9.040572723 | 1.38547727 | 0.213850408 | 0.607090304 | -4.488577103 |
| **A_55_P1998461** | Rpl17 | 0.215403111 | -0.162428496 | 0.593234717 | 11.18495026 | 1.385277802 | 0.213908303 | 0.607090304 | -4.488676543 |
| **A_51_P212754** | Tgfbi | -0.770501675 | -2.122392061 | 0.581388711 | 11.71424045 | -1.384890055 | 0.214020886 | 0.607090304 | -4.488869836 |
| **A_55_P1996817** | NA | -0.198455949 | -0.546662038 | 0.14975014 | 12.94826787 | -1.384876192 | 0.214024912 | 0.607090304 | -4.488876746 |
| **A_52_P535255** | Syf2 | 0.184860651 | -0.139497913 | 0.509219215 | 10.68055579 | 1.384848603 | 0.214032925 | 0.607090304 | -4.488890499 |
| **A_66_P129048** | Faap20 | -0.167603164 | -0.46168844 | 0.126482111 | 11.9770868 | -1.384816159 | 0.214042348 | 0.607090304 | -4.488906671 |
| **A_55_P2061061** | Set | 0.11099518 | -0.083779678 | 0.305770038 | 12.4935413 | 1.384695915 | 0.214077275 | 0.607090304 | -4.488966609 |
| **A_55_P1985808** | Ranbp9 | 0.161081131 | -0.121594461 | 0.443756724 | 10.90553907 | 1.384648532 | 0.21409104 | 0.607090304 | -4.488990228 |
| **A_55_P1997614** | Gm9693 | 0.094144357 | -0.071071036 | 0.25935975 | 7.948158571 | 1.384608444 | 0.214102686 | 0.607090304 | -4.489010211 |
| **A_51_P155755** | Pld6 | -0.124157672 | -0.342084701 | 0.093769356 | 8.703334318 | -1.3843492 | 0.214178014 | 0.607090304 | -4.48913943 |
| **A_55_P2126890** | NA | 0.190877846 | -0.144163347 | 0.525919039 | 8.647983718 | 1.384332799 | 0.21418278 | 0.607090304 | -4.489147605 |
| **A_55_P2103536** | Irf2bpl | -0.294102229 | -0.810346028 | 0.222141571 | 8.901393633 | -1.384288643 | 0.214195613 | 0.607090304 | -4.489169613 |
| **A_51_P143805** | Tmem42 | -0.255520475 | -0.704048954 | 0.193008003 | 10.66072819 | -1.384263834 | 0.214202824 | 0.607090304 | -4.489181979 |
| **A_55_P2118675** | Tnip1 | -0.242305537 | -0.667671263 | 0.183060188 | 10.63383283 | -1.384152799 | 0.214235098 | 0.607090304 | -4.489237322 |
| **A_51_P125679** | Rec114 | 0.135534867 | -0.102396062 | 0.373465796 | 7.818127118 | 1.384150527 | 0.214235758 | 0.607090304 | -4.489341212 |
| **A_55_P1984009** | Rsph3b | -0.275082451 | -0.758233514 | 0.208068612 | 11.60275601 | -1.383448939 | 0.214439787 | 0.607552527 | -4.489588117 |
| **A_55_P1972927** | Elk3 | -0.321744045 | -0.886881713 | 0.243393622 | 9.900024437 | -1.3833735 | 0.214461736 | 0.607552527 | -4.489625712 |
| **A_55_P2084666** | H2ac10 | 0.157272263 | -0.119001546 | 0.433546073 | 17.35484635 | 1.383233856 | 0.214502371 | 0.607578593 | -4.489695302 |
| **A_55_P2184146** | NA | 0.250115441 | -0.1893487 | 0.689579581 | 9.028414606 | 1.382930163 | 0.214590765 | 0.607592277 | -4.48984664 |
| **A_52_P16752** | Aox3 | -0.393869071 | -1.085925146 | 0.298187004 | 9.011629224 | -1.382909572 | 0.21459676 | 0.607592277 | -4.489856901 |
| **A_51_P407690** | Sppl2b | 0.316228255 | -0.239479917 | 0.871936427 | 13.10074123 | 1.382729017 | 0.21464933 | 0.607592277 | -4.489946871 |
| **A_52_P306537** | Emc1 | 0.173749872 | -0.13159695 | 0.479096694 | 8.692621657 | 1.382656434 | 0.214670467 | 0.607592277 | -4.489983038 |
| **A_52_P577662** | Ednrb | -0.345712115 | -0.953331969 | 0.261907738 | 8.639768433 | -1.382502206 | 0.214715385 | 0.607592277 | -4.490059887 |
| **A_55_P2000853** | Mvb12b | -0.236747112 | -0.652852051 | 0.179357827 | 11.84211837 | -1.382499582 | 0.214716149 | 0.607592277 | -4.490061194 |
| **A_55_P2023762** | Epha4 | -0.22077036 | -0.608815824 | 0.167275104 | 7.821564177 | -1.382424252 | 0.214738092 | 0.607592277 | -4.490098729 |
| **A_55_P2057291** | Kansl3 | 0.09055859 | -0.06863007 | 0.249747251 | 11.22836298 | 1.38229493 | 0.214775767 | 0.607592277 | -4.490163165 |
| **A_55_P2112240** | Cdk5rap2 | 0.177496277 | -0.134603283 | 0.489595836 | 10.49474346 | 1.381908477 | 0.214888386 | 0.607592277 | -4.49035571 |
| **A_55_P1990735** | Ift122 | -0.178058496 | -0.491156397 | 0.135039404 | 8.54907479 | -1.381865375 | 0.21490095 | 0.607592277 | -4.490377184 |
| **A_55_P2077866** | Nktr | 0.166033697 | -0.125926593 | 0.457993987 | 10.94969573 | 1.381833258 | 0.214910312 | 0.607592277 | -4.490393185 |
| **A_55_P2172989** | Ptpn23 | -0.095482692 | -0.263390116 | 0.072424732 | 10.53391832 | -1.381776958 | 0.214926725 | 0.607592277 | -4.490421234 |
| **A_55_P2162472** | NA | 0.530239607 | -0.402304876 | 1.462784091 | 10.36641515 | 1.381611046 | 0.214975098 | 0.607592277 | -4.490503892 |
| **A_52_P570240** | Kbtbd11 | -0.413118119 | -1.139679069 | 0.313442831 | 8.402427757 | -1.381609719 | 0.214975485 | 0.607592277 | -4.490504553 |
| **A_55_P2145656** | NA | -0.299430297 | -0.826049912 | 0.227189319 | 8.071171269 | -1.381598643 | 0.214978715 | 0.607592277 | -4.490510071 |
| **A_51_P459944** | Tcf21 | -1.12960726 | -3.117073061 | 0.857858542 | 8.764839082 | -1.381055079 | 0.215137269 | 0.607911598 | -4.490780857 |
| **A_55_P1983358** | Gm4593 | 0.106301141 | -0.08076984 | 0.293372121 | 7.909520626 | 1.38074885 | 0.215226641 | 0.607911598 | -4.490933398 |
| **A_55_P2068289** | Slc17a2 | -0.173091998 | -0.477739534 | 0.131555538 | 8.799796809 | -1.380582964 | 0.215275068 | 0.607911598 | -4.491016027 |
| **A_55_P1974892** | Zbtb18 | 0.220435719 | -0.167542549 | 0.608413987 | 9.917461732 | 1.380567855 | 0.215279479 | 0.607911598 | -4.491023553 |
| **A_55_P2116149** | NA | 0.211209106 | -0.160536546 | 0.582954758 | 12.06184857 | 1.380542976 | 0.215286743 | 0.607911598 | -4.491035945 |
| **A_55_P2136880** | Ppp1r15a | 0.215794319 | -0.16403747 | 0.595626107 | 14.41991071 | 1.38048562 | 0.21530349 | 0.607911598 | -4.491064514 |
| **A_51_P333949** | Ints7 | 0.167235186 | -0.127143136 | 0.461613508 | 8.992434364 | 1.380400225 | 0.215328426 | 0.607911598 | -4.491107048 |
| **A_51_P383857** | Prp2 | 0.155457334 | -0.118213354 | 0.429128023 | 8.653523087 | 1.380276585 | 0.215364535 | 0.607911598 | -4.491168631 |
| **A_55_P2081482** | Atp5g2 | -0.122272933 | -0.337530489 | 0.092984624 | 15.30529065 | -1.380241597 | 0.215374754 | 0.607911598 | -4.491186057 |

| | | | | | | | | | |
|---|---|---|---|---|---|---|---|---|---|
| **A_55_P2078048** | BC004004 | -0.192366719 | -0.531086279 | 0.146352841 | 13.39513691 | -1.379980182 | 0.215451121 | 0.608038103 | -4.491316257 |
| **A_55_P2087414** | Sema3e | -0.171072462 | -0.472320501 | 0.130175578 | 7.748655577 | -1.379872833 | 0.215482488 | 0.608038103 | -4.491369721 |
| **A_55_P1962411** | Gm6813 | 0.1347995 | -0.102615003 | 0.372214004 | 16.05623129 | 1.379635071 | 0.215551975 | 0.608110238 | -4.491488133 |
| **A_52_P678056** | Eva1b | -0.532804458 | -1.471391303 | 0.405782388 | 11.99918031 | -1.379356659 | 0.215633369 | 0.608110238 | -4.491626783 |
| **A_66_P103198** | NA | 0.132386175 | -0.100850863 | 0.365623213 | 8.200444128 | 1.379203356 | 0.215678199 | 0.608110238 | -4.491703125 |
| **A_51_P328652** | Acot7 | -0.137725569 | -0.380389341 | 0.104938203 | 10.45963037 | -1.379090673 | 0.215711156 | 0.608110238 | -4.491759238 |
| **A_51_P246924** | Tppp3 | -0.697820174 | -1.927434434 | 0.531794087 | 8.119624781 | -1.378979526 | 0.215743668 | 0.608110238 | -4.491814585 |
| **A_55_P2141038** | Gnpnat1 | 0.243668191 | -0.185730047 | 0.67306643 | 9.118770987 | 1.378865071 | 0.215777152 | 0.608110238 | -4.491871578 |
| **A_51_P332602** | Surf1 | -0.137031089 | -0.378515505 | 0.104453327 | 11.00294358 | -1.378837829 | 0.215785122 | 0.608110238 | -4.491885143 |
| **A_55_P2007713** | Tspan4 | -0.111542774 | -0.308129272 | 0.085043723 | 11.6182713 | -1.378703716 | 0.215824365 | 0.608110238 | -4.491951923 |
| **A_52_P15212** | Ctbp2 | -0.376498876 | -1.040157593 | 0.287159841 | 9.77648002 | -1.378485081 | 0.215888353 | 0.608110238 | -4.492060786 |
| **A_55_P1958374** | NA | 0.258720766 | -0.197343075 | 0.714784606 | 12.17941055 | 1.378443078 | 0.215900648 | 0.608110238 | -4.4920817 |
| **A_52_P462125** | Smarca5-ps | 0.248732197 | -0.189739223 | 0.687203617 | 10.91834063 | 1.378395684 | 0.215914522 | 0.608110238 | -4.492105298 |
| **A_51_P295237** | Lrp11 | -0.098586689 | -0.272378628 | 0.075205249 | 7.928185287 | -1.378389147 | 0.215916436 | 0.608110238 | -4.492108552 |
| **A_51_P268234** | Il34 | -0.319646425 | -0.883177565 | 0.243884716 | 7.919479332 | -1.378272593 | 0.215950559 | 0.608110238 | -4.492166585 |
| **A_55_P2055404** | Slc15a4 | -0.159574761 | -0.440911443 | 0.12176192 | 11.49448378 | -1.378227885 | 0.21596365 | 0.608110238 | -4.492188844 |
| **A_55_P1990866** | Mylk3 | 0.176339056 | -0.134566318 | 0.487244429 | 8.490270279 | 1.378172158 | 0.215979967 | 0.608110238 | -4.49221659 |
| **A_51_P410451** | Tube1 | 0.141060097 | -0.10769865 | 0.389818844 | 8.554342988 | 1.37787242 | 0.216067754 | 0.608133012 | -4.49236582 |
| **A_55_P2042272** | Pbrm1 | 0.102159213 | -0.078007864 | 0.28232629 | 7.864387817 | 1.37779719 | 0.216089792 | 0.608133012 | -4.492403274 |
| **A_55_P2033521** | Med6 | 0.114672637 | -0.087573307 | 0.316918582 | 9.87749607 | 1.377726996 | 0.216110357 | 0.608133012 | -4.49243822 |
| **A_55_P2056973** | Treml2 | 0.243742991 | -0.186193904 | 0.673679887 | 8.831636617 | 1.377560272 | 0.216159209 | 0.608133012 | -4.492521221 |
| **A_55_P2158227** | Angpt1 | -0.37388636 | -1.033420533 | 0.285647814 | 8.703275575 | -1.377480651 | 0.216182542 | 0.608133012 | -4.492560858 |
| **A_55_P1971962** | Tmem176b | -0.189859811 | -0.524790634 | 0.145071012 | 9.772868562 | -1.377403293 | 0.216205215 | 0.608133012 | -4.492599368 |
| **A_52_P108845** | Clip3 | -0.283668477 | -0.784105541 | 0.216768588 | 8.295497136 | -1.377351536 | 0.216220385 | 0.608133012 | -4.492625133 |
| **A_51_P394515** | Tkt | 0.11765278 | -0.089915707 | 0.325221267 | 14.21610354 | 1.377285462 | 0.216239753 | 0.608133012 | -4.492658025 |
| **A_52_P392674** | Acot9 | -0.218953926 | -0.605359709 | 0.167451856 | 8.761155663 | -1.376868009 | 0.216362158 | 0.608316268 | -4.492865826 |
| **A_66_P102733** | Ccdc28b | -0.37538514 | -1.03791772 | 0.287147441 | 11.58843959 | -1.376743471 | 0.216398686 | 0.608316268 | -4.492927817 |
| **A_55_P2082570** | NA | 0.161921357 | -0.123860721 | 0.447703434 | 8.524107863 | 1.376741275 | 0.21639933 | 0.608316268 | -4.49292891 |
| **A_55_P1987459** | Pigt | 0.224443838 | -0.171730435 | 0.620618112 | 8.130533825 | 1.376589986 | 0.216443713 | 0.608352557 | -4.493004213 |
| **A_55_P2061979** | NA | -0.159548995 | -0.441281102 | 0.122183113 | 15.68453519 | -1.376071239 | 0.216595958 | 0.608691955 | -4.493262402 |
| **A_51_P351238** | Plgrkt | -0.096730699 | -0.26756049 | 0.074099093 | 10.88714828 | -1.375890597 | 0.216648996 | 0.608752499 | -4.493352305 |
| **A_55_P2041075** | Ccnd3 | 0.166974494 | -0.127961932 | 0.46191092 | 14.482354 | 1.375640367 | 0.216722485 | 0.608870482 | -4.493476835 |
| **A_52_P336290** | Rap2a | -0.552569225 | -1.528834687 | 0.423696237 | 9.585025358 | -1.375314295 | 0.216818282 | 0.609004664 | -4.493639101 |
| **A_55_P2047345** | Sat2 | 0.197759759 | -0.151656123 | 0.547175641 | 9.944503058 | 1.375239925 | 0.216840137 | 0.609004664 | -4.493676109 |
| **A_55_P1966327** | NA | -0.730447661 | -2.021227024 | 0.560331702 | 8.47241445 | -1.375055661 | 0.216894294 | 0.609004664 | -4.4937678 |
| **A_55_P2067652** | Boc | -0.410496505 | -1.135898919 | 0.314905908 | 8.042041503 | -1.375034697 | 0.216900456 | 0.609004664 | -4.493778232 |
| **A_55_P2014326** | NA | 0.19433497 | -0.149118214 | 0.537788154 | 9.028172231 | 1.374885715 | 0.216944253 | 0.609004664 | -4.493852364 |
| **A_55_P2107499** | Galnt18 | -0.603208545 | -1.66937738 | 0.462960291 | 8.642632766 | -1.374753044 | 0.216983262 | 0.609004664 | -4.493918378 |
| **A_55_P2108988** | Kcnip3 | -0.150754527 | -0.41725438 | 0.115745327 | 8.45146699 | -1.374537448 | 0.217046667 | 0.609004664 | -4.49402565 |
| **A_66_P104118** | NA | 0.126048711 | -0.096777231 | 0.348874653 | 8.311989575 | 1.374535143 | 0.217047345 | 0.609004664 | -4.494026797 |
| **A_51_P392861** | Med25 | -0.283087537 | -0.783531019 | 0.217355945 | 9.62760238 | -1.374513157 | 0.217053812 | 0.609004664 | -4.494037736 |
| **A_51_P478311** | Nmd3 | 0.198309989 | -0.1523399 | 0.548959878 | 8.843219084 | 1.374213069 | 0.217142095 | 0.609076605 | -4.494187041 |
| **A_66_P114054** | Sco2 | -0.222235355 | -0.615190222 | 0.170719512 | 10.00298247 | -1.374211781 | 0.217142474 | 0.609076605 | -4.494187682 |
| **A_66_P135165** | Nfib | -0.157674938 | -0.436526195 | 0.121176319 | 8.059316099 | -1.373957349 | 0.217217351 | 0.609198228 | -4.494314264 |
| **A_55_P2085940** | NA | 0.114917464 | -0.08838322 | 0.318218147 | 11.32004482 | 1.373505437 | 0.217350403 | 0.609482945 | -4.49453908 |
| **A_55_P2047639** | Cubn | -0.365976207 | -1.013584551 | 0.281632136 | 8.236952927 | -1.373167784 | 0.217449862 | 0.609668204 | -4.494707043 |
| **A_51_P123047** | Smc1a | 0.097708629 | -0.075203508 | 0.270620765 | 13.32167516 | 1.37306352 | 0.217480583 | 0.609668204 | -4.494758906 |
| **A_51_P160824** | Cspg4 | -0.542374355 | -1.50255307 | 0.417804361 | 8.178647972 | -1.372556595 | 0.21763 | 0.609668204 | -4.495011047 |
| **A_51_P382805** | Zpbp2 | 0.14364021 | -0.110651181 | 0.397931602 | 8.343831211 | 1.37254807 | 0.217632513 | 0.609668204 | -4.495015287 |
| **A_55_P2000090** | Arf1 | -0.160291263 | -0.444065545 | 0.12348302 | 14.00131759 | -1.37252418 | 0.217639557 | 0.609668204 | -4.495027169 |

| | | | | | | | | | |
|---|---|---|---|---|---|---|---|---|---|
| **A_55_P2182358** | Arid3a | 0.272763584 | -0.210188579 | 0.755715746 | 11.18511003 | 1.372351817 | 0.217690384 | 0.609668204 | -4.495112895 |
| **A_55_P2208260** | D230019N24Rik | 0.177582231 | -0.136930647 | 0.492095109 | 9.244054028 | 1.371968894 | 0.217803341 | 0.609668204 | -4.495303335 |
| **A_55_P2074811** | Zfp609 | 0.179463651 | -0.138396516 | 0.497323819 | 9.049491286 | 1.371903553 | 0.217822621 | 0.609668204 | -4.49533583 |
| **A_55_P2246650** | 9330159M07Rik | 0.140593918 | -0.108442925 | 0.389630762 | 8.56942915 | 1.371785224 | 0.21785754 | 0.609668204 | -4.495394675 |
| **A_52_P394385** | 1600012H06Rik | -0.102869435 | -0.285084202 | 0.079345333 | 9.468161359 | -1.371784751 | 0.217857679 | 0.609668204 | -4.495394911 |
| **A_51_P122649** | Degs2 | -0.154820017 | -0.429085233 | 0.119445198 | 7.73249526 | -1.371638196 | 0.217900934 | 0.609668204 | -4.495514831 |
| **A_51_P387302** | Zmym2 | -0.146392633 | -0.405772208 | 0.112986941 | 9.218870579 | -1.371407873 | 0.217968929 | 0.609668204 | -4.495582324 |
| **A_55_P1970920** | Timm17b | -0.097517383 | -0.270313589 | 0.075278824 | 13.46700369 | -1.371295411 | 0.218002137 | 0.609668204 | -4.495638247 |
| **A_55_P2047837** | NA | 0.166757507 | -0.128746239 | 0.462261253 | 8.133392877 | 1.371215115 | 0.218025849 | 0.609668204 | -4.495678174 |
| **A_66_P122877** | NA | 0.118804957 | -0.09177553 | 0.329385445 | 16.33938873 | 1.370880579 | 0.218124666 | 0.609668204 | -4.495844515 |
| **A_55_P2171303** | Bin3 | 0.128496525 | -0.099283325 | 0.356276376 | 8.461096358 | 1.370753201 | 0.218162302 | 0.609668204 | -4.495907848 |
| **A_55_P2088360** | NA | 0.123632622 | -0.095546562 | 0.342811805 | 14.67439339 | 1.370619679 | 0.218201761 | 0.609668204 | -4.495974235 |
| **A_51_P221949** | Csnk1g3 | -0.171883032 | -0.476617439 | 0.132851374 | 10.29365914 | -1.370549432 | 0.218222522 | 0.609668204 | -4.496009161 |
| **A_66_P129672** | 4930453N24Rik | 0.104416968 | -0.080740937 | 0.289574873 | 10.09924377 | 1.370288493 | 0.218299659 | 0.609668204 | -4.496138891 |
| **A_52_P343627** | Rbp7 | -0.225166011 | -0.624445578 | 0.174113557 | 7.753856692 | -1.370278791 | 0.218302528 | 0.609668204 | -4.496143715 |
| **A_52_P676271** | Tuba4a | 0.190242402 | -0.14711516 | 0.527599965 | 15.00672786 | 1.37025079 | 0.218310807 | 0.609668204 | -4.496157636 |
| **A_55_P2095814** | Kalrn | -0.11518144 | -0.31943849 | 0.089075609 | 7.734578595 | -1.370214756 | 0.218321461 | 0.609668204 | -4.496175551 |
| **A_51_P375201** | Plk3 | -0.174299276 | -0.483448734 | 0.134850182 | 9.05769869 | -1.369967559 | 0.218394565 | 0.609668204 | -4.496298442 |
| **A_55_P2043362** | NA | -0.116118164 | -0.322093273 | 0.089856945 | 7.904540494 | -1.369836101 | 0.21843345 | 0.609668204 | -4.496363792 |
| **A_52_P103977** | Ints8 | 0.129520793 | -0.100234867 | 0.359276453 | 8.511722873 | 1.369797806 | 0.218444779 | 0.609668204 | -4.496382829 |
| **A_51_P117794** | Bik | -0.244358377 | -0.677834697 | 0.189117944 | 8.056288122 | -1.369761772 | 0.21845544 | 0.609668204 | -4.496400742 |
| **A_55_P1981830** | NA | 0.297358064 | -0.230190271 | 0.8249064 | 10.15318083 | 1.369621779 | 0.21849686 | 0.609668204 | -4.496470333 |
| **A_51_P483658** | Trim3 | -0.354774747 | -0.984336678 | 0.274787184 | 8.820753117 | -1.369296283 | 0.218593194 | 0.609668204 | -4.496632131 |
| **A_55_P2008006** | Ap1g1 | 0.15495283 | -0.120017394 | 0.429923054 | 9.543246929 | 1.36929504 | 0.218593562 | 0.609668204 | -4.496632749 |
| **A_55_P2120526** | NA | 0.327429672 | -0.25367429 | 0.908533634 | 9.966208435 | 1.369138509 | 0.218639902 | 0.609668204 | -4.496710554 |
| **A_52_P216539** | Fbxw8 | -0.094348631 | -0.261803898 | 0.073106636 | 8.434433148 | -1.369052114 | 0.218665483 | 0.609668204 | -4.496753496 |
| **A_52_P144297** | Tspyl3 | 0.115191911 | -0.089299009 | 0.319682831 | 9.030016315 | 1.368772095 | 0.218748413 | 0.609668204 | -4.496892673 |
| **A_55_P1978511** | H2-Q7 | -0.235093422 | -0.652525954 | 0.18233911 | 13.3214906 | -1.368476596 | 0.218835958 | 0.609668204 | -4.497039537 |
| **A_65_P18981** | Palb2 | 0.176029414 | -0.136559711 | 0.48861854 | 8.48013912 | 1.368341719 | 0.218875927 | 0.609668204 | -4.497106569 |
| **A_51_P503937** | Calm1 | -0.306811979 | -0.851652074 | 0.238028116 | 9.08382712 | -1.368316111 | 0.218883517 | 0.609668204 | -4.497119295 |
| **A_66_P112546** | Yeats4 | 0.149623118 | -0.116101176 | 0.415347413 | 9.108878969 | 1.368203282 | 0.218916958 | 0.609668204 | -4.497175367 |
| **A_55_P1967885** | Nr3c1 | -0.257307119 | -0.714394646 | 0.199780408 | 12.03140141 | -1.36784101 | 0.219024364 | 0.609668204 | -4.497355397 |
| **A_55_P2060663** | NA | 0.129300578 | -0.100399049 | 0.359000205 | 15.78968023 | 1.367802415 | 0.21903581 | 0.609668204 | -4.497374576 |
| **A_55_P2047200** | NA | 0.189456395 | -0.147137985 | 0.526050775 | 8.66991589 | 1.367683466 | 0.219071088 | 0.609668204 | -4.497433684 |
| **A_55_P2150343** | Piezo1 | 0.167053117 | -0.129746407 | 0.463852641 | 12.18593841 | 1.367648752 | 0.219081384 | 0.609668204 | -4.497450934 |
| **A_51_P395164** | Pex16 | 0.182415337 | -0.141681772 | 0.506512446 | 10.02159864 | 1.367632401 | 0.219086234 | 0.609668204 | -4.497459059 |
| **A_55_P2162678** | Dnajc10 | -0.165359137 | -0.459161214 | 0.128442939 | 11.00625575 | -1.367591918 | 0.219098243 | 0.609668204 | -4.497479175 |
| **A_55_P1973447** | NA | -0.179639674 | -0.498826225 | 0.139546877 | 7.894076109 | -1.367542597 | 0.219112873 | 0.609668204 | -4.497503683 |
| **A_52_P104824** | Diaph3 | 0.275228984 | -0.213862781 | 0.764320749 | 10.64230882 | 1.367373018 | 0.219163185 | 0.609668204 | -4.497587945 |
| **A_51_P181851** | Eml3 | -0.125716076 | -0.349121533 | 0.097689381 | 10.44449119 | -1.367351679 | 0.219169516 | 0.609668204 | -4.497598548 |
| **A_55_P2049318** | NA | -0.1412845 | -0.392377591 | 0.10980859 | 9.20781413 | -1.367234307 | 0.219204345 | 0.609668204 | -4.497656867 |
| **A_51_P471458** | Sult5a1 | -0.407049774 | -1.130579526 | 0.316479979 | 8.029218919 | -1.367018248 | 0.219268473 | 0.609668204 | -4.497764219 |
| **A_52_P220810** | Trib2 | -0.24257773 | -0.673783875 | 0.188628416 | 8.314240556 | -1.366939125 | 0.219291961 | 0.609668204 | -4.497803531 |
| **A_51_P304510** | BC017158 | 0.161537653 | -0.125618182 | 0.448693488 | 11.10604631 | 1.366908088 | 0.219301175 | 0.609668204 | -4.497818952 |
| **A_55_P2125561** | Mrps36 | 0.133141076 | -0.10355082 | 0.369832972 | 12.00736671 | 1.366821697 | 0.219326824 | 0.609668204 | -4.497861874 |
| **A_66_P140599** | NA | 0.109483016 | -0.085151884 | 0.304117916 | 8.32594994 | 1.366813395 | 0.219329289 | 0.609668204 | -4.497865998 |
| **A_51_P121818** | Prkag1 | 0.163299197 | -0.127011467 | 0.453609861 | 11.62789603 | 1.366797732 | 0.21933394 | 0.609668204 | -4.498076424 |
| **A_52_P649224** | Sash1 | -0.292864384 | -0.813519288 | 0.22779052 | 10.42351178 | -1.36678368 | 0.219338112 | 0.609668204 | -4.497880761 |
| **A_51_P130110** | Idh3b | -0.101143708 | -0.280959855 | 0.078672439 | 10.68753125 | -1.366763498 | 0.219344105 | 0.609668204 | -4.497890788 |
| **A_55_P2007663** | Alkbh8 | 0.226237127 | -0.176006058 | 0.628480311 | 9.616405771 | 1.366653352 | 0.219376813 | 0.609668204 | -4.497945511 |

| | | | | | | | | | |
|---|---|---|---|---|---|---|---|---|---|
| **A_55_P2033075** | NA | 0.109639314 | -0.085339962 | 0.304618589 | 16.09016369 | 1.36634712 | 0.219467773 | 0.609668204 | -4.498097648 |
| **A_55_P2020592** | Trim3 | -0.184292332 | -0.512051554 | 0.14346689 | 7.904273223 | -1.366266859 | 0.219491619 | 0.609668204 | -4.49813752 |
| **A_55_P2016968** | NA | 0.110290562 | -0.085899019 | 0.306480143 | 15.98396069 | 1.365983956 | 0.219575688 | 0.609668204 | -4.498278057 |
| **A_55_P2169669** | Csf1 | 0.127116049 | -0.099012406 | 0.353244504 | 8.334383538 | 1.365929734 | 0.219591804 | 0.609668204 | -4.498304992 |
| **A_66_P114769** | NA | 0.147533826 | -0.114926395 | 0.409994047 | 10.88139811 | 1.365876092 | 0.219607749 | 0.609668204 | -4.498331638 |
| **A_55_P1986541** | NA | -0.11367797 | -0.315928506 | 0.088572565 | 15.49921816 | -1.365745628 | 0.219646533 | 0.609668204 | -4.498396444 |
| **A_52_P563450** | Brix1 | 0.185413638 | -0.144467794 | 0.515295069 | 9.001672394 | 1.365736742 | 0.219649175 | 0.609668204 | -4.498400858 |
| **A_55_P2099105** | Rhobtb1 | 0.19750863 | -0.153903967 | 0.548921227 | 9.201632883 | 1.365689405 | 0.219663249 | 0.609668204 | -4.498424372 |
| **A_55_P2066657** | Vps13a | 0.123231578 | -0.096035919 | 0.342499075 | 8.451431769 | 1.36562337 | 0.219682884 | 0.609668204 | -4.498457173 |
| **A_55_P2155312** | Dock3 | 0.203224854 | -0.158382847 | 0.564832554 | 10.30747948 | 1.365596322 | 0.219690927 | 0.609668204 | -4.498470608 |
| **A_55_P1986421** | Tcp11 | -0.260788952 | -0.724847692 | 0.203269787 | 7.823003505 | -1.365524268 | 0.219712353 | 0.609668204 | -4.498506398 |
| **A_55_P2181251** | Ccdc9b | -0.223814389 | -0.622107248 | 0.17447847 | 8.86265363 | -1.365427587 | 0.219741106 | 0.609668204 | -4.49855442 |
| **A_55_P2024591** | Atxn7l1 | 0.100819391 | -0.078598479 | 0.280237262 | 9.245586636 | 1.36540523 | 0.219747756 | 0.609668204 | -4.498565524 |
| **A_51_P188271** | Cd248 | -0.69035444 | -1.918946306 | 0.538237426 | 10.17358219 | -1.365361572 | 0.219760741 | 0.609668204 | -4.498587209 |
| **A_52_P207635** | Zfc3h1 | 0.158776095 | -0.123809348 | 0.441361537 | 9.513031772 | 1.365269911 | 0.219788007 | 0.609668204 | -4.498632736 |
| **A_55_P2116993** | Hnrnpdl | 0.13865735 | -0.108141531 | 0.385456231 | 12.27472819 | 1.365157937 | 0.219821318 | 0.609668204 | -4.498688351 |
| **A_55_P2293013** | Ces2a | -0.424608125 | -1.180388585 | 0.331172335 | 12.83951016 | -1.365135717 | 0.219827929 | 0.609668204 | -4.498699388 |
| **A_55_P2151071** | Cox7a2l | -0.306870173 | -0.853096979 | 0.239356633 | 14.4364259 | -1.365101228 | 0.219838191 | 0.609668204 | -4.498716517 |
| **A_52_P232314** | NA | 0.165983616 | -0.129479053 | 0.461446284 | 9.320545723 | 1.36504131 | 0.219856019 | 0.609668204 | -4.498746277 |
| **A_55_P2012669** | NA | 0.138417799 | -0.107982484 | 0.384818082 | 17.12409965 | 1.365004006 | 0.21986712 | 0.609668204 | -4.498764804 |
| **A_52_P366525** | Coq10b | 0.130343703 | -0.101689108 | 0.362376514 | 8.490888121 | 1.364972313 | 0.219876551 | 0.609668204 | -4.498780544 |
| **A_55_P2256686** | NA | 0.510186604 | -0.398049658 | 1.418422867 | 8.679631219 | 1.364939502 | 0.219886315 | 0.609668204 | -4.49879684 |
| **A_51_P279552** | Cav2 | -0.689926386 | -1.918253237 | 0.538400465 | 8.428315192 | -1.364809378 | 0.219925042 | 0.609668204 | -4.498920865 |
| **A_55_P2006668** | Pi4kb | 0.121843178 | -0.095104417 | 0.338790774 | 8.973507355 | 1.364676039 | 0.219964733 | 0.609668204 | -4.498927686 |
| **A_55_P2127825** | Bcas3 | -0.179364527 | -0.498734377 | 0.140005323 | 8.612685236 | -1.364664305 | 0.219968226 | 0.609668204 | -4.498933514 |
| **A_52_P645632** | Senp8 | 0.097580771 | -0.076208414 | 0.271369956 | 7.989798868 | 1.364346526 | 0.220062846 | 0.609668204 | -4.499091327 |
| **A_55_P2040011** | Itga7 | -0.643082086 | -1.788465358 | 0.502301186 | 8.201944451 | -1.364265086 | 0.220087101 | 0.609668204 | -4.499131769 |
| **A_55_P2423513** | B230369F24Rik | 0.169358956 | -0.132285175 | 0.471003087 | 10.62982829 | 1.3642579 | 0.220089241 | 0.609668204 | -4.499135338 |
| **A_55_P2033586** | Mmab | 0.172831929 | -0.135001191 | 0.480665049 | 8.65635793 | 1.364243279 | 0.220093596 | 0.609668204 | -4.499142599 |
| **A_55_P1977787** | NA | 0.214421155 | -0.167554799 | 0.596397109 | 10.41662652 | 1.364001337 | 0.220165668 | 0.609668204 | -4.499262741 |
| **A_55_P2148609** | NA | 0.127501057 | -0.099640761 | 0.354642875 | 8.812156009 | 1.363954477 | 0.22017963 | 0.609668204 | -4.499286009 |
| **A_52_P89567** | Rhob | -0.18816369 | -0.523389377 | 0.147061997 | 8.105036178 | -1.36389746 | 0.220196619 | 0.609668204 | -4.499314322 |
| **A_55_P2087114** | NA | 0.154695164 | -0.120936099 | 0.430326426 | 11.97081648 | 1.363739598 | 0.220243662 | 0.609668204 | -4.499392707 |
| **A_52_P547447** | Sirt2 | -0.140437179 | -0.390669686 | 0.109795328 | 10.10320265 | -1.363708533 | 0.220252921 | 0.609668204 | -4.499408133 |
| **A_55_P1957249** | Pdgfrb | -0.422067521 | -1.17416115 | 0.330026109 | 9.437135584 | -1.363619529 | 0.220279449 | 0.609668204 | -4.499452326 |
| **A_55_P2027213** | Krtap20-2 | 0.173003126 | -0.135281407 | 0.48128766 | 8.777685831 | 1.36359501 | 0.220286758 | 0.609668204 | -4.4994645 |
| **A_51_P445841** | Deptor | -0.254584073 | -0.708378969 | 0.199210823 | 9.492842539 | -1.363185041 | 0.220408993 | 0.609919173 | -4.499668052 |
| **A_55_P2008687** | Nlk | -0.130672971 | -0.363629179 | 0.102283236 | 9.624898287 | -1.362996273 | 0.220465296 | 0.609987648 | -4.49976177 |
| **A_55_P2023484** | Osbpl5 | -0.860947305 | -2.396116004 | 0.674221395 | 10.16916799 | -1.362710659 | 0.22055051 | 0.610118378 | -4.499903565 |
| **A_51_P440790** | Dpp7 | -0.154670944 | -0.430484208 | 0.12114232 | 10.01354444 | -1.36262633 | 0.220575675 | 0.610118378 | -4.499945428 |
| **A_52_P605834** | Tm9sf3 | -0.138067077 | -0.384330512 | 0.108196358 | 10.54737168 | -1.362301975 | 0.220672493 | 0.610298844 | -4.500106443 |
| **A_51_P415688** | Gng12 | -0.13490847 | -0.375595812 | 0.105778872 | 10.72449863 | -1.361975058 | 0.220770115 | 0.610377865 | -4.50026872 |
| **A_52_P362033** | Polr2e | -0.187271841 | -0.521388201 | 0.146844519 | 10.00873221 | -1.36193985 | 0.220780631 | 0.610377865 | -4.500286196 |
| **A_55_P2052924** | Selenow | -0.187019396 | -0.520697806 | 0.146659014 | 13.19540868 | -1.361889062 | 0.220795801 | 0.610377865 | -4.500311405 |
| **A_55_P2046867** | Gata5 | 0.204606482 | -0.160541948 | 0.569754913 | 10.15845694 | 1.361548573 | 0.220897527 | 0.610481472 | -4.500480405 |
| **A_55_P2181665** | NA | 0.148495754 | -0.11652111 | 0.413512618 | 8.254492992 | 1.361519024 | 0.220906357 | 0.610481472 | -4.500495071 |
| **A_55_P1998188** | Cdk4 | -0.160035075 | -0.445661106 | 0.125590957 | 12.50698505 | -1.361446502 | 0.22092803 | 0.610481472 | -4.500531065 |
| **A_55_P2149515** | Rsrc1 | 0.12573813 | -0.098742415 | 0.350218674 | 8.93212924 | 1.361041847 | 0.221048997 | 0.610701535 | -4.500731894 |
| **A_55_P1978136** | Tnip2 | -0.109317163 | -0.304506486 | 0.085872159 | 9.010817031 | -1.360866388 | 0.221101467 | 0.610701535 | -4.50081897 |
| **A_55_P2289765** | Fam160a2 | 0.173169821 | -0.136037846 | 0.482377487 | 8.876013802 | 1.360833973 | 0.221111161 | 0.610701535 | -4.500835056 |

| | | | | | | | | | |
|---|---|---|---|---|---|---|---|---|---|
| **A_52_P80806** | Syncrip | 0.236544518 | -0.185847163 | 0.658936198 | 9.950606267 | 1.36075745 | 0.221134049 | 0.610701535 | -4.500873031 |
| **A_51_P112445** | Wdr3 | 0.173588346 | -0.136418063 | 0.483594754 | 9.857229941 | 1.360608193 | 0.221178698 | 0.61073758 | -4.500887417 |
| **A_55_P2016252** | Ctnnal1 | 0.121009972 | -0.095188626 | 0.33720857 | 8.472912599 | 1.360039358 | 0.221348935 | 0.610986098 | -4.50122936 |
| **A_66_P123432** | NA | 0.364567406 | -0.286874982 | 1.016009793 | 8.801262091 | 1.359831238 | 0.221411249 | 0.610986098 | -4.501332623 |
| **A_52_P486253** | Polr2b | 0.164824265 | -0.129717279 | 0.459365808 | 11.08180172 | 1.359745961 | 0.221436787 | 0.610986098 | -4.501374934 |
| **A_55_P2093650** | NA | -0.147513369 | -0.411158735 | 0.116131997 | 12.74811633 | -1.359547636 | 0.221496189 | 0.610986098 | -4.501473331 |
| **A_55_P2151868** | Mest | -0.289478437 | -0.806852982 | 0.227896108 | 8.547189133 | -1.359547392 | 0.221496262 | 0.610986098 | -4.501473452 |
| **A_51_P198775** | BC055324 | 0.217721434 | -0.171414763 | 0.606857632 | 10.78319594 | 1.35951106 | 0.221507146 | 0.610986098 | -4.501491478 |
| **A_51_P139069** | Wdr45 | -0.565777532 | -1.57701812 | 0.445463056 | 10.43411257 | -1.359484916 | 0.221514978 | 0.610986098 | -4.501504449 |
| **A_52_P515094** | Tmem176a | -0.162166365 | -0.452037792 | 0.127705063 | 10.87260103 | -1.359372754 | 0.221548581 | 0.610986098 | -4.501560095 |
| **A_55_P2175400** | Zfp467 | -0.311003629 | -0.866950085 | 0.244942826 | 9.154793696 | -1.359301138 | 0.221570039 | 0.610986098 | -4.501595624 |
| **A_52_P78373** | Sorbs3 | -0.134854557 | -0.375939763 | 0.106230649 | 7.864677457 | -1.359184002 | 0.221605141 | 0.610986098 | -4.501653736 |
| **A_55_P2173737** | Hspa4l | 0.199777016 | -0.157427147 | 0.55698118 | 10.03391776 | 1.35897729 | 0.221667098 | 0.610986098 | -4.501756283 |
| **A_51_P216922** | Ddx59 | 0.125284418 | -0.098731513 | 0.349300349 | 8.605850868 | 1.358943322 | 0.22167728 | 0.610986098 | -4.501773134 |
| **A_55_P2069949** | Dynlrb1 | -0.176691377 | -0.492636014 | 0.13925326 | 12.85738751 | -1.35890018 | 0.221690214 | 0.610986098 | -4.501794535 |
| **A_51_P479352** | Klhdc9 | -0.301324384 | -0.840189752 | 0.237540984 | 8.051269756 | -1.358742603 | 0.221737458 | 0.610986098 | -4.501872703 |
| **A_55_P1962284** | Klhl24 | -0.096584787 | -0.269323244 | 0.076153671 | 7.828542694 | -1.358635247 | 0.221769651 | 0.610986098 | -4.501925957 |
| **A_66_P107255** | Nom1 | 0.181640832 | -0.143221508 | 0.506503172 | 9.314840744 | 1.358617808 | 0.221774881 | 0.610986098 | -4.501934608 |
| **A_55_P1974695** | NA | 0.151842947 | -0.119750488 | 0.423436382 | 8.903965977 | 1.358496567 | 0.221811243 | 0.610986098 | -4.501994748 |
| **A_51_P455807** | Ehd4 | -0.266502195 | -0.743211476 | 0.210207086 | 11.66091115 | -1.358408444 | 0.221837676 | 0.610986098 | -4.502038459 |
| **A_51_P148355** | Zfp710 | -0.139287652 | -0.388531381 | 0.109956077 | 11.6303247 | -1.357911808 | 0.221986697 | 0.61112263 | -4.50228479 |
| **A_52_P123693** | E130304I02Rik | 0.341497419 | -0.269620063 | 0.952614901 | 11.93416719 | 1.357831585 | 0.222010777 | 0.61112263 | -4.502324578 |
| **A_51_P437786** | Rbm28 | 0.116980615 | -0.092366168 | 0.326327398 | 11.61965544 | 1.35778435 | 0.222024956 | 0.61112263 | -4.502348005 |
| **A_52_P300416** | Dcp1a | 0.112504919 | -0.088839699 | 0.313849538 | 8.829100257 | 1.357733945 | 0.222040088 | 0.61112263 | -4.502373003 |
| **A_55_P2020453** | NA | 0.158859376 | -0.125498058 | 0.44321681 | 8.66689734 | 1.357473795 | 0.222118202 | 0.61112263 | -4.502502023 |
| **A_51_P416869** | Nedd8 | -0.150999439 | -0.421295558 | 0.11929668 | 12.99330255 | -1.357433967 | 0.222130163 | 0.61112263 | -4.502521775 |
| **A_55_P2129658** | Smad6 | -0.121766249 | -0.339735561 | 0.096203064 | 9.253896658 | -1.35742162 | 0.222133871 | 0.61112263 | -4.502527898 |
| **A_55_P2021266** | Hpse | -0.161636214 | -0.450979775 | 0.127707346 | 8.905301021 | -1.357400603 | 0.222140183 | 0.61112263 | -4.502538321 |
| **A_55_P2058646** | Nhlh1 | 0.291556751 | -0.230414408 | 0.81352791 | 8.755649309 | 1.357249793 | 0.222185481 | 0.611142028 | -4.50261311 |
| **A_51_P519108** | Dzank1 | -0.178051305 | -0.496834326 | 0.140731717 | 7.775688343 | -1.357166611 | 0.22221047 | 0.611142028 | -4.502654361 |
| **A_55_P2110101** | Tm2d3 | -0.134321565 | -0.374859821 | 0.106216691 | 11.11699755 | -1.356890415 | 0.222293459 | 0.611161993 | -4.502791322 |
| **A_55_P2028600** | Myh9 | -0.176264705 | -0.491919405 | 0.139389996 | 13.56142051 | -1.356863894 | 0.222301429 | 0.611161993 | -4.502804473 |
| **A_55_P2024575** | Sec24c | -0.115617512 | -0.322700139 | 0.091465116 | 13.54824738 | -1.356635383 | 0.222370114 | 0.611161993 | -4.502917783 |
| **A_55_P2009201** | NA | 0.121370303 | -0.096022568 | 0.338763174 | 11.64216528 | 1.356595363 | 0.222382145 | 0.611161993 | -4.502937627 |
| **A_55_P2009351** | Prxl2c | -0.096138822 | -0.268341718 | 0.076064073 | 9.421946164 | -1.356567902 | 0.222390401 | 0.611161993 | -4.502951243 |
| **A_51_P123077** | Nubp1 | 0.15121268 | -0.119703442 | 0.422128803 | 11.58696545 | 1.35624 | 0.222489001 | 0.611161993 | -4.503024502 |
| **A_55_P2092296** | Fbxo2 | 0.093439397 | -0.073969821 | 0.260848615 | 8.103336687 | 1.356231562 | 0.222491539 | 0.611161993 | -4.503118009 |
| **A_55_P2053551** | Mast4 | -0.220412633 | -0.615319515 | 0.17449425 | 8.122130033 | -1.356203836 | 0.222499879 | 0.611161993 | -4.503131756 |
| **A_51_P416858** | Myl1 | -0.891096941 | -2.487658804 | 0.705464921 | 8.179617953 | -1.356195788 | 0.222502299 | 0.611161993 | -4.503135746 |
| **A_55_P2133405** | Dmkn | -0.608100819 | -1.69793293 | 0.481731292 | 8.490928913 | -1.355811076 | 0.222618039 | 0.61139302 | -4.503326481 |
| **A_51_P472726** | Pdlim2 | -1.088146812 | -3.039230029 | 0.862936406 | 10.44025335 | -1.355173466 | 0.222809982 | 0.611735466 | -4.503642567 |
| **A_55_P2118391** | Lrp6 | -0.100495282 | -0.280700709 | 0.079710144 | 11.77716117 | -1.355067719 | 0.222841829 | 0.611735466 | -4.503694985 |
| **A_55_P2122987** | Dlx3 | 0.279348295 | -0.221625543 | 0.780322134 | 10.6742874 | 1.354921603 | 0.222885842 | 0.611735466 | -4.503767413 |
| **A_55_P1959119** | NA | 0.138082805 | -0.109595072 | 0.385760682 | 13.39762962 | 1.354676429 | 0.22295971 | 0.611735466 | -4.503888938 |
| **A_51_P474422** | Ift27 | -0.240798113 | -0.672726497 | 0.191130271 | 11.77742391 | -1.354641959 | 0.222970097 | 0.611735466 | -4.503906023 |
| **A_52_P298088** | Itgb1bp2 | -0.123298205 | -0.34448123 | 0.097884819 | 7.731315023 | -1.354528515 | 0.223004285 | 0.611735466 | -4.503962251 |
| **A_55_P2386592** | Cep63 | 0.130672708 | -0.103753557 | 0.365098974 | 10.77204738 | 1.354446366 | 0.223029045 | 0.611735466 | -4.504002967 |
| **A_66_P132062** | Gm5608 | 0.090211259 | -0.071628373 | 0.25205089 | 7.836707275 | 1.354437756 | 0.22303164 | 0.611735466 | -4.504007234 |
| **A_55_P2042615** | NA | 0.216669336 | -0.172071549 | 0.605410221 | 8.818876398 | 1.354317287 | 0.223067955 | 0.611735466 | -4.504066942 |
| **A_51_P244492** | Nbl1 | -0.793966902 | -2.218809474 | 0.63087567 | 9.488632601 | -1.354000126 | 0.223163586 | 0.611735466 | -4.504224128 |

| | | | | | | | | | |
|---|---|---|---|---|---|---|---|---|---|
| A_55_P2032453 | Mrpl23 | -0.14667965 | -0.409913581 | 0.116554282 | 13.72951592 | -1.353976677 | 0.223170658 | 0.611735466 | -4.504235749 |
| A_55_P2043582 | 4930480E11Rik | 0.101092975 | -0.080357359 | 0.282543308 | 8.040010383 | 1.353774703 | 0.223231578 | 0.611735466 | -4.504335842 |
| A_55_P2203996 | 4932433N03Rik | 0.161406849 | -0.128321966 | 0.451135663 | 9.572082638 | 1.353672038 | 0.22326255 | 0.611735466 | -4.504386719 |
| A_55_P1978972 | Rp2 | -0.098212114 | -0.274506417 | 0.078082189 | 8.349128981 | -1.353661162 | 0.223265831 | 0.611735466 | -4.504392108 |
| A_55_P2143261 | Rpain | 0.252931359 | -0.201138855 | 0.707001574 | 12.34045724 | 1.353514313 | 0.22331014 | 0.611735466 | -4.504464879 |
| A_55_P2275279 | Mid2 | -0.178489074 | -0.498920677 | 0.14194253 | 7.799344238 | -1.353503807 | 0.22331331 | 0.611735466 | -4.504470085 |
| A_51_P490023 | Tubb2a | -0.128250719 | -0.35853991 | 0.102038471 | 13.44784799 | -1.353223233 | 0.223397992 | 0.611735466 | -4.504609116 |
| A_51_P140211 | Ndufv3 | -0.109811409 | -0.307002857 | 0.087380039 | 9.168295596 | -1.353139518 | 0.223423263 | 0.611735466 | -4.504650597 |
| A_65_P11455 | Pphln1 | 0.096420441 | -0.076730861 | 0.269571743 | 8.426610459 | 1.353089558 | 0.223438347 | 0.611735466 | -4.504675353 |
| A_55_P2010014 | Map7d1 | -0.151332524 | -0.423102229 | 0.120437182 | 11.97915837 | -1.353051791 | 0.223449749 | 0.611735466 | -4.504694066 |
| A_52_P465121 | NA | -0.197975273 | -0.553525985 | 0.157575439 | 11.67960225 | -1.352983771 | 0.223470287 | 0.611735466 | -4.504727769 |
| A_52_P644534 | Dhrsx | 0.156434323 | -0.124527341 | 0.437395987 | 9.291855389 | 1.352907705 | 0.223493256 | 0.611735466 | -4.504765459 |
| A_55_P2152387 | Wwox | -0.160910434 | -0.44992679 | 0.128105923 | 8.897271344 | -1.352835443 | 0.223515079 | 0.611735466 | -4.504801262 |
| A_51_P345649 | Pdgfra | -0.518884854 | -1.451114404 | 0.413344695 | 8.849539377 | -1.352481453 | 0.223622009 | 0.611735466 | -4.504976648 |
| A_55_P1970033 | Per1 | -0.232202595 | -0.659698061 | 0.19529287 | 11.85911263 | -1.381904156 | 0.223643618 | 0.611735466 | -4.496714821 |
| A_51_P318830 | Syt10 | -0.291547805 | -0.815418965 | 0.232323354 | 7.921764485 | -1.352285762 | 0.223681141 | 0.611735466 | -4.505073599 |
| A_55_P2067712 | Hmcn2 | 0.462318756 | -0.368434579 | 1.293072091 | 10.5311983 | 1.352236358 | 0.223696072 | 0.611735466 | -4.505098074 |
| A_55_P2143790 | Gm14410 | -0.115394631 | -0.322773846 | 0.091984585 | 9.309683623 | -1.352083657 | 0.223742227 | 0.611735466 | -4.505173723 |
| A_52_P596592 | Rassf9 | -0.200704753 | -0.561433446 | 0.16002394 | 7.781504934 | -1.35194853 | 0.223783076 | 0.611735466 | -4.505240663 |
| A_55_P2167769 | Arid1a | 0.160585037 | -0.128043261 | 0.449213335 | 11.02458979 | 1.351914907 | 0.223793242 | 0.611735466 | -4.50525732 |
| A_55_P2002210 | Babam2 | 0.130828052 | -0.104351115 | 0.366007219 | 11.85634753 | 1.351715262 | 0.223853611 | 0.611735466 | -4.505356218 |
| A_55_P2042590 | Polm | -0.138696803 | -0.388025573 | 0.110631967 | 9.088779856 | -1.351690451 | 0.223861114 | 0.611735466 | -4.505368508 |
| A_52_P575668 | NA | -0.136877743 | -0.382938717 | 0.109183231 | 8.562606482 | -1.351678128 | 0.223864841 | 0.611735466 | -4.505374612 |
| A_51_P335770 | Afap1 | -0.382029951 | -1.068828715 | 0.304768813 | 9.213619617 | -1.351609069 | 0.223885727 | 0.611735466 | -4.505408821 |
| A_52_P201551 | Nav1 | -0.148817552 | -0.416363367 | 0.118728264 | 8.42574534 | -1.351571982 | 0.223896945 | 0.611735466 | -4.505427192 |
| A_51_P298538 | Prmt7 | 0.105642409 | -0.084296746 | 0.295581563 | 12.09351396 | 1.351471782 | 0.223927254 | 0.611735466 | -4.505476825 |
| A_55_P2085915 | Tnfrsf14 | 0.202423791 | -0.161567878 | 0.566415461 | 8.842046031 | 1.351304737 | 0.223977791 | 0.611735466 | -4.505559566 |
| A_55_P2092864 | Emp2 | -1.130457679 | -3.16322844 | 0.902313082 | 9.072793116 | -1.351291646 | 0.223981752 | 0.611735466 | -4.505566051 |
| A_55_P2148708 | NA | -0.338625664 | -0.947564841 | 0.270313512 | 11.04282196 | -1.351229578 | 0.224000533 | 0.611735466 | -4.505596794 |
| A_55_P2170109 | Bivm | 0.133543896 | -0.106629382 | 0.373717175 | 9.930676135 | 1.35108461 | 0.224044403 | 0.611735466 | -4.505668597 |
| A_55_P2094362 | Erp29 | -0.127830389 | -0.357737614 | 0.102076837 | 9.369222752 | -1.351029021 | 0.224061228 | 0.611735466 | -4.50569613 |
| A_55_P2153067 | 2610028H24Rik | 0.132319418 | -0.105667729 | 0.370306565 | 17.02207924 | 1.350993531 | 0.22407197 | 0.611735466 | -4.505713708 |
| A_55_P2143233 | Rasgrp2 | -0.193527896 | -0.541676906 | 0.154621114 | 10.96671229 | -1.350708397 | 0.224158289 | 0.611804469 | -4.505854927 |
| A_55_P2003123 | Jkamp | -0.117918697 | -0.330063934 | 0.09422654 | 12.04513039 | -1.350618129 | 0.224185623 | 0.611804469 | -4.505899632 |
| A_55_P2093822 | Ube2d2a | 0.128321254 | -0.102543021 | 0.359185529 | 12.21434542 | 1.35059473 | 0.224192708 | 0.611804469 | -4.505911221 |
| A_52_P268279 | Fbxl17 | -0.147804331 | -0.413763548 | 0.118154886 | 9.499269905 | -1.350377843 | 0.224258397 | 0.611804469 | -4.506018631 |
| A_55_P2012296 | Gopc | 0.135501459 | -0.108358674 | 0.379361591 | 9.511239097 | 1.350163469 | 0.22432334 | 0.611804469 | -4.506124792 |
| A_51_P439824 | Gipc1 | -0.245517188 | -0.687398108 | 0.196363732 | 10.00041726 | -1.350081046 | 0.224348315 | 0.611804469 | -4.506165608 |
| A_51_P446315 | Cdc37l1 | 0.158296238 | -0.126651676 | 0.443244152 | 9.892270986 | 1.349858669 | 0.224415707 | 0.611804469 | -4.506275725 |
| A_52_P198009 | Mrpl44 | 0.089532758 | -0.071652822 | 0.250718339 | 10.9543398 | 1.349705342 | 0.224462185 | 0.611804469 | -4.506351647 |
| A_51_P275435 | Mab21l4 | -0.538531647 | -1.5080614 | 0.430998107 | 8.010507851 | -1.34968766 | 0.224467545 | 0.611804469 | -4.506360403 |
| A_55_P1974019 | Dapk1 | -0.305616023 | -0.855879388 | 0.244647341 | 10.74067097 | -1.349549163 | 0.224509536 | 0.611804469 | -4.50642898 |
| A_51_P421958 | Sgms1 | -0.109342348 | -0.306217995 | 0.087533298 | 10.99858106 | -1.349520821 | 0.22451813 | 0.611804469 | -4.506443013 |
| A_55_P1978176 | Abhd17c | -0.238423946 | -0.667754795 | 0.190906903 | 12.46593618 | -1.349400806 | 0.224554523 | 0.611804469 | -4.506502436 |
| A_51_P502614 | Dusp6 | -0.219218066 | -0.613975687 | 0.175539555 | 12.61300018 | -1.349363653 | 0.224565791 | 0.611804469 | -4.506520831 |
| A_52_P200207 | Rmnd5a | 0.157479387 | -0.126105646 | 0.441064421 | 9.080410939 | 1.349346849 | 0.224570887 | 0.611804469 | -4.506529151 |
| A_52_P86073 | Mrpl1 | 0.208869311 | -0.167276227 | 0.585014849 | 9.735776323 | 1.349279457 | 0.224591327 | 0.611804469 | -4.506562518 |
| A_51_P368823 | Grb7 | -0.248445244 | -0.69589727 | 0.199006782 | 11.87898407 | -1.349172242 | 0.224623848 | 0.611804469 | -4.506615601 |
| A_52_P513347 | Phkb | 0.088591595 | -0.070988712 | 0.248171901 | 8.101388242 | 1.348951742 | 0.224690746 | 0.611804469 | -4.506724769 |
| A_55_P1968362 | Tle1 | 0.126835501 | -0.101653052 | 0.355324054 | 8.508336921 | 1.348837307 | 0.224725472 | 0.611804469 | -4.506781423 |

| | | | | | | | | |
|---|---|---|---|---|---|---|---|---|
| **A_55_P2052145** | Ly6g6d | -0.402384594 | -1.127292173 | 0.322522984 | 8.228634437 | -1.348782413 | 0.224742131 | 0.611804469 | -4.506808599 |
| **A_55_P1953157** | Mettl18 | 0.101351732 | -0.081277511 | 0.283980976 | 9.442482973 | 1.348478559 | 0.224834366 | 0.611804469 | -4.506959022 |
| **A_51_P140901** | Pmm2 | 0.140597052 | -0.112752182 | 0.393946285 | 9.167396582 | 1.348465401 | 0.22483836 | 0.611804469 | -4.506965535 |
| **A_55_P1974640** | NA | 0.13892905 | -0.111421126 | 0.389279226 | 8.022350551 | 1.348429836 | 0.224849158 | 0.611804469 | -4.506983141 |
| **A_55_P1970954** | Pknox2 | -0.268772012 | -0.75310477 | 0.215560745 | 7.897519336 | -1.348414422 | 0.224853839 | 0.611804469 | -4.506990771 |
| **A_52_P39157** | Bmi1 | 0.123981618 | -0.099463604 | 0.347426841 | 9.011408654 | 1.348246856 | 0.224904721 | 0.611804469 | -4.507073719 |
| **A_55_P2057777** | Fgfr1 | -0.444040255 | -1.244372238 | 0.356291728 | 9.594917102 | -1.348141134 | 0.22493683 | 0.611804469 | -4.507126052 |
| **A_51_P246773** | Sesn3 | -0.118618747 | -0.332420437 | 0.095182942 | 8.716282591 | -1.348110185 | 0.224946231 | 0.611804469 | -4.507141372 |
| **A_51_P326030** | Zfp959 | 0.127085575 | -0.101980102 | 0.356151252 | 8.455345762 | 1.348091675 | 0.224951853 | 0.611804469 | -4.507150534 |
| **A_51_P164207** | Mta2 | 0.189680896 | -0.152238095 | 0.531599888 | 12.05295065 | 1.34798034 | 0.224985673 | 0.611810364 | -4.507205644 |
| **A_55_P2153862** | Irak1 | 0.115587456 | -0.092841387 | 0.324016299 | 9.107542046 | 1.347522654 | 0.225124751 | 0.611950675 | -4.50743218 |
| **A_55_P2130373** | Dcaf11 | -0.104229378 | -0.292189906 | 0.083731151 | 12.47300852 | -1.347431434 | 0.225152479 | 0.611950675 | -4.507477327 |
| **A_51_P472323** | Fbh1 | -0.141062673 | -0.395448057 | 0.113322711 | 10.9585982 | -1.347420477 | 0.22515581 | 0.611950675 | -4.50748275 |
| **A_55_P2036673** | Ddx52 | 0.15782013 | -0.126811648 | 0.442451908 | 10.50155685 | 1.347293455 | 0.225194427 | 0.611950675 | -4.507545616 |
| **A_55_P1960703** | NA | 0.229991912 | -0.184814105 | 0.644797929 | 9.281942733 | 1.347257794 | 0.22520527 | 0.611950675 | -4.507563265 |
| **A_51_P409010** | Comp | -0.437052657 | -1.225388017 | 0.351282702 | 8.945362473 | -1.347118956 | 0.225247487 | 0.611950675 | -4.507631976 |
| **A_55_P2095251** | Ell3 | 0.201209553 | -0.161733029 | 0.564152134 | 9.247492314 | 1.347081466 | 0.225258888 | 0.611950675 | -4.50765053 |
| **A_55_P1970017** | Zer1 | -0.137718879 | -0.386166595 | 0.110728836 | 11.50616771 | -1.346919564 | 0.22530813 | 0.611998434 | -4.507730652 |
| **A_51_P161086** | Ppic | -0.832336979 | -2.334241373 | 0.669567414 | 11.51270229 | -1.346604593 | 0.225403956 | 0.612172695 | -4.507886518 |
| **A_55_P2168761** | Sf1 | 0.140837024 | -0.113377139 | 0.395051187 | 12.58251421 | 1.346171172 | 0.225535878 | 0.612286928 | -4.508100983 |
| **A_51_P454993** | Tmcc2 | 0.326955006 | -0.263251439 | 0.91716145 | 11.37404712 | 1.346068721 | 0.225567071 | 0.612286928 | -4.508151675 |
| **A_55_P2066548** | Baiap2l2 | 0.279553873 | -0.225105005 | 0.784212751 | 10.35720786 | 1.346017744 | 0.225582594 | 0.612286928 | -4.508176898 |
| **A_55_P1987534** | Tnrc6b | 0.20276527 | -0.163284989 | 0.568815528 | 9.297468597 | 1.345972048 | 0.225596509 | 0.612286928 | -4.508199507 |
| **A_55_P2083629** | Tle2 | -0.233333942 | -0.654596631 | 0.187928747 | 7.994239891 | -1.345885491 | 0.22562287 | 0.612286928 | -4.508242333 |
| **A_51_P504314** | Prpf38b | 0.098056018 | -0.078989927 | 0.275101964 | 9.495286135 | 1.345771905 | 0.225657466 | 0.612286928 | -4.508298531 |
| **A_52_P53906** | Ccnd2 | -0.293051169 | -0.822185687 | 0.236083348 | 9.647040992 | -1.345738119 | 0.225667757 | 0.612286928 | -4.508315247 |
| **A_55_P2129856** | Mtfmt | 0.097334875 | -0.078460826 | 0.273130576 | 10.12431157 | 1.345375189 | 0.225778334 | 0.612500972 | -4.508494801 |
| **A_55_P2137461** | NA | 0.10467168 | -0.084397123 | 0.293740484 | 15.96339658 | 1.345217505 | 0.225826393 | 0.61252943 | -4.508572809 |
| **A_51_P129317** | Mob4 | 0.116333175 | -0.093813052 | 0.326479402 | 10.78524581 | 1.345132829 | 0.225852204 | 0.61252943 | -4.508182995 |
| **A_55_P1962092** | Mpc1 | -0.257535044 | -0.722801692 | 0.207731604 | 13.37655894 | -1.344985495 | 0.22589712 | 0.612530278 | -4.508687581 |
| **A_55_P1957159** | NA | -0.225798142 | -0.633747107 | 0.182150824 | 15.49273654 | -1.344923916 | 0.225915896 | 0.612530278 | -4.508718043 |
| **A_55_P1983426** | Abhd14a | -0.112961817 | -0.317124588 | 0.091200954 | 11.58579708 | -1.34443035 | 0.226066436 | 0.612761848 | -4.508962183 |
| **A_55_P2096917** | Mreg | 0.35095759 | -0.283387556 | 0.985302736 | 9.532962629 | 1.344349554 | 0.226091087 | 0.612761848 | -4.509002145 |
| **A_55_P2180909** | Ccdc166 | -0.102288927 | -0.287198646 | 0.082620793 | 8.532107211 | -1.344163415 | 0.226147889 | 0.612761848 | -4.50909421 |
| **A_55_P2053157** | Zc3h3 | -0.101116773 | -0.283911961 | 0.081678415 | 10.42741503 | -1.34413111 | 0.226157749 | 0.612761848 | -4.509110188 |
| **A_55_P2008042** | Gm15348 | 0.124115325 | -0.100256929 | 0.34848758 | 10.01203561 | 1.344124346 | 0.226159813 | 0.612761848 | -4.509113533 |
| **A_51_P294807** | Stom | 0.226408584 | -0.182944292 | 0.635761461 | 11.68358854 | 1.343934899 | 0.22621764 | 0.612832623 | -4.509207229 |
| **A_55_P2057681** | Cdyl2 | -0.099619321 | -0.279781663 | 0.080543021 | 8.011568566 | -1.343577577 | 0.226326746 | 0.612934419 | -4.509383943 |
| **A_51_P111544** | Srp14 | -0.174504195 | -0.49010665 | 0.14109826 | 12.42250618 | -1.343534082 | 0.226340031 | 0.612934419 | -4.509277775 |
| **A_55_P2006250** | Myo5a | 0.108883598 | -0.08804459 | 0.305811786 | 8.784354436 | 1.343500301 | 0.226350348 | 0.612934419 | -4.509422158 |
| **A_51_P159610** | Ccdc120 | -0.118815621 | -0.333728127 | 0.096096885 | 7.908011769 | -1.343368133 | 0.226390721 | 0.612957871 | -4.509487517 |
| **A_55_P2038077** | Rpsa | 0.136022491 | -0.11004177 | 0.382086751 | 16.51655058 | 1.343214501 | 0.226437657 | 0.612999087 | -4.509563488 |
| **A_52_P225570** | Parva | -0.300481011 | -0.844148199 | 0.243186176 | 9.068326531 | -1.342972435 | 0.22651163 | 0.613113471 | -4.509683185 |
| **A_55_P2088571** | Repin1 | -0.218803533 | -0.614796689 | 0.177189624 | 10.61602437 | -1.342609879 | 0.226622463 | 0.613195378 | -4.509862451 |
| **A_55_P2007678** | Xntrpc | 0.3486271 | -0.282323971 | 0.979578172 | 9.273081443 | 1.342606207 | 0.226623586 | 0.613195378 | -4.509864267 |
| **A_51_P212023** | Sart3 | 0.093489296 | -0.075714727 | 0.262693319 | 11.84298108 | 1.342562134 | 0.226637062 | 0.613195378 | -4.509886058 |
| **A_55_P2038489** | NA | 0.097155961 | -0.078728607 | 0.27304053 | 8.069273711 | 1.342223707 | 0.226740569 | 0.613389567 | -4.510053379 |
| **A_55_P1984601** | Vapb | 0.111662509 | -0.090527849 | 0.313852867 | 8.81951774 | 1.341930829 | 0.226830179 | 0.613392034 | -4.510198171 |
| **A_55_P2253284** | NA | -0.137126104 | -0.385438472 | 0.111186264 | 7.767301001 | -1.341853103 | 0.226853966 | 0.613392034 | -4.510236595 |
| **A_51_P424878** | Prxl2b | -0.191681342 | -0.538794747 | 0.155432062 | 10.74078845 | -1.341811921 | 0.22686657 | 0.613392034 | -4.510256953 |

| | | | | | | | | | |
|---|---|---|---|---|---|---|---|---|---|
| **A_55_P2035827** | Vma21 | 0.119680022 | -0.097057369 | 0.336417414 | 10.49627363 | 1.341748158 | 0.226886086 | 0.613392034 | -4.510288474 |
| **A_55_P2105873** | Ppp2r5c | 0.146281771 | -0.118639766 | 0.411203308 | 9.937979338 | 1.341702203 | 0.226900152 | 0.613392034 | -4.510311191 |
| **A_55_P2110355** | Zfp637 | -0.165674684 | -0.465749978 | 0.13440061 | 9.413341935 | -1.341556905 | 0.226944632 | 0.613426485 | -4.510383016 |
| **A_55_P2038697** | Zfp235 | -0.091562149 | -0.257433519 | 0.074309222 | 8.449287493 | -1.341305584 | 0.227021587 | 0.613426779 | -4.510507246 |
| **A_55_P2083879** | Vsig2 | -1.297674398 | -3.64855205 | 1.053203255 | 8.72500265 | -1.341278298 | 0.227029944 | 0.613426779 | -4.510520734 |
| **A_52_P161630** | St6gal1 | -0.397993818 | -1.119040663 | 0.323053028 | 11.64357018 | -1.341207687 | 0.22705157 | 0.613426779 | -4.510555636 |
| **A_51_P246317** | Mt2 | 0.297845837 | -0.241831225 | 0.8375229 | 17.41534587 | 1.34103701 | 0.227103851 | 0.613426779 | -4.510639998 |
| **A_51_P185175** | Fkbp4 | 0.143428925 | -0.116459818 | 0.403317669 | 9.953737963 | 1.341011383 | 0.227111702 | 0.613426779 | -4.510652664 |
| **A_55_P2036764** | Prr15l | -0.221922814 | -0.624064387 | 0.180218759 | 7.982728128 | -1.340930184 | 0.227136579 | 0.613426779 | -4.510692798 |
| **A_55_P2000930** | Ccnyl1 | 0.145488626 | -0.118174455 | 0.409151706 | 8.987750689 | 1.340796629 | 0.227177502 | 0.613426779 | -4.510758807 |
| **A_66_P117251** | Snx6 | -0.139431481 | -0.392157663 | 0.113294701 | 11.53875468 | -1.340583262 | 0.227242894 | 0.613426779 | -4.51086426 |
| **A_55_P1958537** | Map6 | -0.698850859 | -1.965668563 | 0.567966844 | 8.218163768 | -1.340459105 | 0.227280953 | 0.613426779 | -4.51092562 |
| **A_55_P2038298** | Rpl31 | -0.210996581 | -0.59347728 | 0.171484117 | 15.93012277 | -1.340445262 | 0.227285196 | 0.613426779 | -4.510932461 |
| **A_55_P2131353** | 1110059G10Rik | 0.147096655 | -0.119556075 | 0.413749384 | 9.37383048 | 1.340417074 | 0.227293838 | 0.613426779 | -4.510946392 |
| **A_52_P3845** | Rnf217 | -0.146659553 | -0.412574794 | 0.119255687 | 8.389012229 | -1.340140455 | 0.227378657 | 0.613570022 | -4.511083094 |
| **A_52_P636830** | G3bp2 | 0.22786433 | -0.18533969 | 0.641068351 | 10.40916647 | 1.33996976 | 0.227431011 | 0.61362511 | -4.511167445 |
| **A_55_P2122709** | Arhgef3 | -0.312650551 | -0.87969031 | 0.254389208 | 10.39682964 | -1.339765765 | 0.227493593 | 0.61362511 | -4.511268248 |
| **A_66_P105032** | Gm13889 | -0.657730936 | -1.850629021 | 0.535167148 | 10.66497477 | -1.339763424 | 0.227494311 | 0.61362511 | -4.511269405 |
| **A_51_P267454** | Slc25a11 | -0.142497294 | -0.40096974 | 0.115975153 | 13.32440221 | -1.339601275 | 0.227544066 | 0.613673679 | -4.511349527 |
| **A_55_P1974407** | Cdk14 | -0.439027085 | -1.235436142 | 0.357381973 | 8.21198631 | -1.339486412 | 0.227579318 | 0.613683124 | -4.511406282 |
| **A_55_P2104583** | C87436 | 0.137067849 | -0.111626798 | 0.385762495 | 9.249249154 | 1.339221303 | 0.227660698 | 0.613715317 | -4.51153727 |
| **A_51_P430357** | Lrp10 | -0.174265154 | -0.490457538 | 0.14192723 | 9.901537289 | -1.339190433 | 0.227670176 | 0.613715317 | -4.511552522 |
| **A_51_P371331** | Mpp5 | -0.111552105 | -0.313963927 | 0.090859716 | 8.81445858 | -1.339137237 | 0.227686509 | 0.613715317 | -4.511578805 |
| **A_55_P2159070** | 2810025M15Rik | -0.382919919 | -1.077910784 | 0.312070946 | 10.32892508 | -1.33878874 | 0.227793538 | 0.613835383 | -4.51175098 |
| **A_51_P381321** | Ddx17 | -0.229793886 | -0.646873943 | 0.18728617 | 13.76397177 | -1.338758483 | 0.227802833 | 0.613835383 | -4.511765928 |
| **A_66_P124497** | Imp3 | -0.122600873 | -0.345136384 | 0.099934638 | 13.36414905 | -1.338682012 | 0.227826325 | 0.613835383 | -4.511803707 |
| **A_55_P2000369** | Aqp4 | -0.133798903 | -0.376691161 | 0.109093355 | 8.280056477 | -1.338511387 | 0.22787875 | 0.61389106 | -4.511887997 |
| **A_55_P1964225** | Elac2 | 0.200158689 | -0.163263427 | 0.563580806 | 10.15747344 | 1.338277832 | 0.227950527 | 0.61392441 | -4.512003371 |
| **A_51_P377760** | Rnasel | 0.110823545 | -0.090397453 | 0.312044542 | 9.851819821 | 1.338264409 | 0.227954653 | 0.61392441 | -4.512010002 |
| **A_51_P292566** | Xpot | 0.169951198 | -0.138663631 | 0.478566028 | 10.75886186 | 1.338106341 | 0.228003243 | 0.613969726 | -4.512088082 |
| **A_51_P372895** | Zfp433 | 0.146979991 | -0.119951745 | 0.413911727 | 8.995917239 | 1.337954037 | 0.228050071 | 0.614010284 | -4.512163314 |
| **A_55_P2047967** | Bfar | 0.10630451 | -0.086821056 | 0.299430076 | 13.59372968 | 1.337504061 | 0.228188471 | 0.614037221 | -4.512385566 |
| **A_51_P509997** | Cox6a2 | -1.025518337 | -2.888639773 | 0.837603099 | 8.820113552 | -1.337474334 | 0.228197617 | 0.614037221 | -4.512400248 |
| **A_51_P327983** | 4833427G06Rik | -0.355254477 | -1.000709763 | 0.29020081 | 7.849498296 | -1.337385427 | 0.228224972 | 0.614037221 | -4.512444158 |
| **A_55_P1993599** | Oaz2 | -0.645984012 | -1.819794146 | 0.527826123 | 13.20712655 | -1.337233055 | 0.228271861 | 0.614037221 | -4.512519411 |
| **A_55_P1974855** | Zfp868 | 0.090511537 | -0.073962634 | 0.254985707 | 8.890176978 | 1.337178618 | 0.228288615 | 0.614037221 | -4.512546295 |
| **A_55_P2075000** | NA | 0.108255397 | -0.088465735 | 0.304976529 | 12.38148813 | 1.337154936 | 0.228295904 | 0.614037221 | -4.512557991 |
| **A_55_P2234361** | Rnf150 | -0.398780612 | -1.123519385 | 0.325958162 | 9.119293555 | -1.3370133 | 0.228339501 | 0.614037221 | -4.512627938 |
| **A_51_P146813** | Mocs3 | 0.123959032 | -0.101330317 | 0.34924838 | 9.845073643 | 1.336967054 | 0.228353738 | 0.614037221 | -4.512650777 |
| **A_55_P2008599** | Pcx | -0.280358131 | -0.789929428 | 0.229213166 | 14.20559186 | -1.336876804 | 0.228381524 | 0.614037221 | -4.512695345 |
| **A_51_P340402** | Eefsec | 0.190317736 | -0.155645799 | 0.536281272 | 9.788280753 | 1.336694391 | 0.228437692 | 0.614037221 | -4.512785424 |
| **A_55_P1955242** | NA | 0.12611474 | -0.103141629 | 0.355371109 | 16.36774321 | 1.336680547 | 0.228441956 | 0.614037221 | -4.512792261 |
| **A_55_P2045911** | Spata32 | 0.120754629 | -0.098770406 | 0.340279664 | 8.120070642 | 1.336604513 | 0.228465372 | 0.614037221 | -4.512829807 |
| **A_51_P281806** | Cactin | 0.157348993 | -0.128753495 | 0.443451482 | 12.30073243 | 1.336366323 | 0.228538743 | 0.614037221 | -4.512947423 |
| **A_52_P69109** | Slc10a1 | -0.456938816 | -1.287814192 | 0.37393656 | 9.514835851 | -1.336304261 | 0.228557864 | 0.614037221 | -4.512978067 |
| **A_55_P2186446** | NA | -0.175734835 | -0.495300633 | 0.143830964 | 14.48233296 | -1.33622857 | 0.228581185 | 0.614037221 | -4.51301544 |
| **A_52_P229320** | Rptor | 0.137419959 | -0.112482112 | 0.387322031 | 10.74639362 | 1.336174406 | 0.228597875 | 0.614037221 | -4.513042185 |
| **A_55_P2124791** | Col18a1 | -0.233987619 | -0.659503029 | 0.191527791 | 13.81299936 | -1.336167101 | 0.228600126 | 0.614037221 | -4.513045792 |
| **A_55_P1974233** | Dtd2 | -0.085083202 | -0.239876698 | 0.069710293 | 8.524782226 | -1.335593541 | 0.228776931 | 0.614377488 | -4.513328972 |
| **A_55_P2041082** | NA | 0.168940676 | -0.138437553 | 0.476318905 | 8.614459405 | 1.335501292 | 0.228805379 | 0.614377488 | -4.513374514 |

| | | | | | | | | | |
|---|---|---|---|---|---|---|---|---|---|
| **A_55_P2113857** | Serpina1e | 0.09488715 | -0.077768469 | 0.26754277 | 17.71901292 | 1.335395395 | 0.228838039 | 0.614377488 | -4.513426794 |
| **A_51_P466229** | Pdgfrl | -0.241095008 | -0.679805785 | 0.19761577 | 8.304150184 | -1.335343826 | 0.228853946 | 0.614377488 | -4.513452252 |
| **A_55_P2042778** | Pde5a | -0.12989252 | -0.366287949 | 0.106502908 | 7.920598413 | -1.335144464 | 0.228915448 | 0.614457254 | -4.513550668 |
| **A_52_P194805** | Zfp706 | -0.109660775 | -0.309259624 | 0.089938073 | 12.63423573 | -1.334985256 | 0.228964573 | 0.61450378 | -4.513629259 |
| **A_55_P2043486** | Msx2 | 0.090051815 | -0.073883301 | 0.253986932 | 7.891441383 | 1.334761484 | 0.229033636 | 0.614566202 | -4.513739717 |
| **A_52_P656545** | Ogt | 0.138103568 | -0.113343003 | 0.38955014 | 8.501805632 | 1.33457311 | 0.229091789 | 0.614566202 | -4.513832698 |
| **A_51_P490348** | 1110059E24Rik | -0.211718453 | -0.597280706 | 0.173843801 | 10.28476233 | -1.33428127 | 0.229181908 | 0.614566202 | -4.513976741 |
| **A_55_P2186928** | NA | 0.275117258 | -0.225966482 | 0.776200999 | 9.795527917 | 1.334107154 | 0.22923569 | 0.614566202 | -4.514062675 |
| **A_51_P233815** | Tmem126a | 0.121257903 | -0.099599636 | 0.342115441 | 10.21996508 | 1.334077358 | 0.229244895 | 0.614566202 | -4.514077381 |
| **A_55_P2061495** | Cks1b | 0.096869703 | -0.079570448 | 0.273309854 | 12.8398656 | 1.334055007 | 0.2292518 | 0.614566202 | -4.514088412 |
| **A_65_P07569** | Rock1 | 0.166958535 | -0.137145854 | 0.471062924 | 9.088828208 | 1.334040841 | 0.229256176 | 0.614566202 | -4.514095403 |
| **A_55_P1996953** | NA | 0.226276328 | -0.185885464 | 0.63843812 | 9.328661496 | 1.333996168 | 0.229269978 | 0.614566202 | -4.51411745 |
| **A_55_P2279140** | F830014O18Rik | -0.196942554 | -0.555708748 | 0.161823639 | 8.66610112 | -1.333863051 | 0.229311109 | 0.614566202 | -4.514183146 |
| **A_55_P2076533** | Pmvk | -0.158841688 | -0.448211899 | 0.130528523 | 13.75734212 | -1.333809673 | 0.229327603 | 0.614566202 | -4.514209488 |
| **A_55_P2033135** | Sh3bgr | -0.429569027 | -1.212156325 | 0.353018271 | 7.970593124 | -1.333777401 | 0.229337577 | 0.614566202 | -4.514225415 |
| **A_55_P1963533** | Pcdhac2 | -0.168781962 | -0.476424715 | 0.138860791 | 7.789237332 | -1.333099395 | 0.229547192 | 0.614959132 | -4.514559986 |
| **A_52_P260555** | Vil1 | -0.371508178 | -1.048665713 | 0.305649356 | 8.56851572 | -1.333097312 | 0.229547837 | 0.614959132 | -4.514561014 |
| **A_66_P111714** | Utp20 | 0.159700443 | -0.13143554 | 0.450836425 | 8.518026825 | 1.33288728 | 0.229612806 | 0.61504794 | -4.514664647 |
| **A_55_P1983036** | Tdrd3 | 0.101828172 | -0.083826491 | 0.287482835 | 9.686022078 | 1.332739518 | 0.229658524 | 0.615058409 | -4.514737553 |
| **A_52_P32903** | Zzef1 | 0.091661554 | -0.075466024 | 0.258789132 | 9.792472124 | 1.332668964 | 0.229680356 | 0.615058409 | -4.514772363 |
| **A_51_P451052** | Pgm5 | -0.49212275 | -1.389552878 | 0.405307377 | 8.1915603 | -1.332465541 | 0.229743313 | 0.615066588 | -4.514872726 |
| **A_55_P1997509** | Zbtb32 | -0.238090328 | -0.67227513 | 0.196094474 | 7.862250704 | -1.332448168 | 0.229748691 | 0.615066588 | -4.514881297 |
| **A_55_P2003414** | Kng1 | -0.257032423 | -0.725794594 | 0.211729747 | 17.35309932 | -1.332350665 | 0.229778873 | 0.615066588 | -4.514929401 |
| **A_51_P356642** | Krt19 | -1.246021197 | -3.518701851 | 1.026659457 | 9.334290148 | -1.332202378 | 0.229824782 | 0.615093575 | -4.515002556 |
| **A_55_P1981734** | Tut7 | 0.164746972 | -0.135763571 | 0.465257515 | 12.54294252 | 1.332112542 | 0.229852599 | 0.615093575 | -4.515046875 |
| **A_55_P2150737** | NA | 0.122412021 | -0.100894417 | 0.34571846 | 9.364861559 | 1.332005484 | 0.229885753 | 0.615097137 | -4.515099688 |
| **A_55_P2111331** | Klhl32 | -0.107703612 | -0.304202867 | 0.088795644 | 8.117299659 | -1.331841509 | 0.229936541 | 0.615147874 | -4.515180577 |
| **A_55_P1956223** | Chka | -0.146351755 | -0.413408887 | 0.120705377 | 9.672674034 | -1.331609676 | 0.230008363 | 0.615254865 | -4.515294936 |
| **A_52_P89477** | Bcl9 | -0.137317956 | -0.387921571 | 0.113285658 | 8.235720837 | -1.331444874 | 0.230059431 | 0.615306316 | -4.515376226 |
| **A_55_P2071691** | Kank3 | -0.471412306 | -1.331806885 | 0.388982273 | 10.42820488 | -1.331332209 | 0.230094349 | 0.615314566 | -4.515431797 |
| **A_52_P96748** | Ttc30b | -0.248988024 | -0.703520139 | 0.20554409 | 8.140173858 | -1.331058291 | 0.230179264 | 0.615405735 | -4.5155669 |
| **A_51_P496054** | Zcchc14 | -0.143359137 | -0.405080476 | 0.118362203 | 8.272186895 | -1.330973571 | 0.230205533 | 0.615405735 | -4.515608684 |
| **A_55_P2174288** | Snap29 | 0.135085341 | -0.111545026 | 0.381715707 | 10.27497204 | 1.330898279 | 0.230228881 | 0.615405735 | -4.515645818 |
| **A_51_P143142** | Mrpl12 | 0.111650771 | -0.092207517 | 0.31550906 | 13.64018112 | 1.330811493 | 0.230255796 | 0.615405735 | -4.51568862 |
| **A_55_P1965313** | Mctp2 | -0.139674666 | -0.394723727 | 0.115374394 | 8.290640244 | -1.330690644 | 0.230293279 | 0.61542082 | -4.515748219 |
| **A_52_P13730** | Cbx2 | -0.13084099 | -0.36985503 | 0.10817305 | 8.514405043 | -1.330159123 | 0.230458204 | 0.615776419 | -4.516010334 |
| **A_51_P415607** | Fkbp7 | -0.429036624 | -1.212949024 | 0.354875776 | 9.194550615 | -1.329872549 | 0.230547168 | 0.615816933 | -4.516151642 |
| **A_52_P497424** | Aanat | 0.248878229 | -0.205861564 | 0.703618023 | 8.818636366 | 1.329863717 | 0.23054991 | 0.615816933 | -4.516155997 |
| **A_51_P116581** | Ccdc22 | 0.110088298 | -0.091078649 | 0.311255245 | 10.97295974 | 1.32974303 | 0.230587385 | 0.615816933 | -4.516215504 |
| **A_52_P654965** | Eif3j1 | 0.251797598 | -0.208341131 | 0.711936327 | 12.25161134 | 1.329676469 | 0.230608056 | 0.615816933 | -4.516248323 |
| **A_55_P2128672** | Ikbip | -0.103080771 | -0.291463353 | 0.085301811 | 9.904295301 | -1.329597235 | 0.230632665 | 0.615816933 | -4.516287389 |
| **A_55_P2164805** | Gm6749 | 0.120668813 | -0.099903522 | 0.341241148 | 14.70175021 | 1.329312809 | 0.230721022 | 0.615837886 | -4.516427622 |
| **A_66_P107129** | 3300005D01Rik | 0.275109921 | -0.227779405 | 0.777999247 | 8.442836927 | 1.329281693 | 0.23073069 | 0.615837886 | -4.516442962 |
| **A_55_P2012430** | NA | 0.145402758 | -0.120390826 | 0.411196343 | 8.309718727 | 1.329264298 | 0.230736094 | 0.615837886 | -4.516451538 |
| **A_51_P333965** | Cisd2 | 0.118527537 | -0.09817271 | 0.335227784 | 8.47667425 | 1.32905527 | 0.230801052 | 0.615876976 | -4.516554589 |
| **A_55_P2053083** | Gm6098 | 0.155111037 | -0.128482906 | 0.43870498 | 10.5824034 | 1.329012111 | 0.230814466 | 0.615876976 | -4.516575867 |
| **A_55_P1957845** | Tnfaip1 | -0.256103631 | -0.724448379 | 0.212241117 | 12.17169628 | -1.328719383 | 0.230905466 | 0.615880995 | -4.516720172 |
| **A_55_P2077671** | NA | 0.120079205 | -0.099514583 | 0.339672993 | 13.61922398 | 1.328712261 | 0.230907681 | 0.615880995 | -4.516723683 |
| **A_55_P1957245** | NA | -0.501944677 | -1.419880204 | 0.41599085 | 8.319989923 | -1.32869978 | 0.230911561 | 0.615880995 | -4.516729836 |
| **A_55_P2162920** | NA | 0.142671972 | -0.118283854 | 0.403627799 | 16.08577656 | 1.328479498 | 0.230980063 | 0.615948432 | -4.516838422 |

| | | | | | | | | | |
|---|---|---|---|---|---|---|---|---|---|
| **A_55_P1979814** | Abcc8 | -0.190978084 | -0.540349355 | 0.158393188 | 7.86774055 | -1.328249098 | 0.23105173 | 0.615948432 | -4.51695199 |
| **A_55_P2157902** | Igsf10 | -0.13843416 | -0.391756352 | 0.114888033 | 7.925260037 | -1.327862845 | 0.231171921 | 0.615948432 | -4.517142367 |
| **A_55_P2085779** | Ifi27l2b | -0.197630096 | -0.55927777 | 0.164017579 | 7.735709325 | -1.327854817 | 0.23117442 | 0.615948432 | -4.517146324 |
| **A_55_P1996963** | Mtcl1 | -0.449661967 | -1.272557765 | 0.37323383 | 7.945227756 | -1.327775042 | 0.231199251 | 0.615948432 | -4.517185641 |
| **A_55_P2248651** | Aak1 | -0.113779817 | -0.322011402 | 0.094451768 | 9.947031641 | -1.327705688 | 0.23122084 | 0.615948432 | -4.517219822 |
| **A_55_P2159264** | Lifr | -0.295798709 | -0.837184876 | 0.245587458 | 11.22834344 | -1.327615484 | 0.231248922 | 0.615948432 | -4.517264278 |
| **A_52_P276955** | Epha3 | -0.283660157 | -0.802888659 | 0.235568345 | 7.946907822 | -1.327464769 | 0.23129585 | 0.615948432 | -4.517338554 |
| **A_55_P2156462** | Nt5c3b | -0.173789883 | -0.491937566 | 0.144357799 | 8.194826472 | -1.327330018 | 0.231337814 | 0.615948432 | -4.517404961 |
| **A_51_P269792** | Rad51b | 0.164216007 | -0.136410484 | 0.464842498 | 8.509138783 | 1.327307136 | 0.23134494 | 0.615948432 | -4.517416238 |
| **A_51_P254583** | Foxred1 | 0.153030028 | -0.127147803 | 0.433207859 | 10.20213859 | 1.327168539 | 0.23138811 | 0.615948432 | -4.517484537 |
| **A_51_P161691** | Coq7 | 0.123159292 | -0.102330605 | 0.348649188 | 10.58532911 | 1.327160002 | 0.231390769 | 0.615948432 | -4.517488744 |
| **A_55_P1958255** | Rnf114 | -0.186551599 | -0.528105911 | 0.155002713 | 8.812620189 | -1.327157274 | 0.231391619 | 0.615948432 | -4.517490088 |
| **A_52_P361993** | Zc2hc1a | -0.225719161 | -0.63900331 | 0.187564989 | 8.496274452 | -1.327097609 | 0.231410206 | 0.615948432 | -4.51751949 |
| **A_51_P509489** | Kras | 0.156545524 | -0.130087251 | 0.443178298 | 9.839261905 | 1.327082721 | 0.231414844 | 0.615948432 | -4.517526826 |
| **A_55_P1965338** | Anxa8 | -0.251704396 | -0.712629723 | 0.209220931 | 7.925644675 | -1.326915957 | 0.231466802 | 0.616001902 | -4.517609003 |
| **A_55_P2060592** | Hoxa1 | -0.154781855 | -0.438278317 | 0.128714607 | 8.391681137 | -1.326647656 | 0.231550417 | 0.616139595 | -4.517741206 |
| **A_55_P2182806** | NA | -0.117621359 | -0.333075372 | 0.097832653 | 9.850929627 | -1.326523037 | 0.231589264 | 0.616158139 | -4.517802608 |
| **A_55_P1981879** | Ercc6l2 | 0.132771423 | -0.110455819 | 0.375998665 | 10.45362376 | 1.326403267 | 0.231626604 | 0.616172671 | -4.51786162 |
| **A_51_P450527** | Tagln | -0.582748259 | -1.650415227 | 0.484918708 | 11.85986576 | -1.326259072 | 0.231671566 | 0.616177586 | -4.517932664 |
| **A_55_P2130535** | Dnmt3b | 0.180199354 | -0.149973281 | 0.510371989 | 10.28275188 | 1.326158227 | 0.231703016 | 0.616177586 | -4.517982348 |
| **A_51_P443618** | Mettl26 | 0.152984186 | -0.127337379 | 0.433305751 | 12.00185451 | 1.32609067 | 0.231724087 | 0.616177586 | -4.518015631 |
| **A_66_P105075** | Mmp11 | -0.486248809 | -1.377446234 | 0.404948616 | 8.196277665 | -1.325768869 | 0.231824478 | 0.616296074 | -4.518174167 |
| **A_51_P164030** | Tcp1 | 0.107277574 | -0.089352305 | 0.303907452 | 12.34192733 | 1.325691954 | 0.231848479 | 0.616296074 | -4.518212057 |
| **A_51_P163378** | Zwint | 0.115227517 | -0.096021309 | 0.326476344 | 13.28638498 | 1.325394267 | 0.23194139 | 0.616296074 | -4.518358699 |
| **A_51_P392687** | Vim | -0.455900431 | -1.291759382 | 0.379958521 | 13.1059644 | -1.325318296 | 0.231965107 | 0.616296074 | -4.518396122 |
| **A_55_P2114569** | Zfp24 | 0.130351495 | -0.108640363 | 0.369343352 | 10.73473891 | 1.325305801 | 0.231969008 | 0.616296074 | -4.518402277 |
| **A_55_P2037585** | Trf | 0.123875214 | -0.103253052 | 0.351003479 | 17.70083076 | 1.32524575 | 0.231987756 | 0.616296074 | -4.518431856 |
| **A_55_P2056473** | Spc24 | 0.194001088 | -0.161736937 | 0.549739113 | 10.7048724 | 1.325125661 | 0.232025253 | 0.616296074 | -4.518491009 |
| **A_51_P238786** | Mcl1 | -0.096838074 | -0.274431024 | 0.080754876 | 13.25925698 | -1.324962572 | 0.232076186 | 0.616296074 | -4.518571338 |
| **A_51_P118765** | Eef1a1 | 0.099656036 | -0.083105853 | 0.282417925 | 16.15381336 | 1.324955094 | 0.232078521 | 0.616296074 | -4.518575021 |
| **A_55_P2072611** | 1-Mar | 0.302848483 | -0.252565547 | 0.858262512 | 9.584146464 | 1.324926374 | 0.232087492 | 0.616296074 | -4.518589167 |
| **A_51_P226932** | Tubgcp2 | 0.153753485 | -0.128318861 | 0.435825831 | 9.052577698 | 1.324486836 | 0.232224814 | 0.616381949 | -4.518805646 |
| **A_55_P1952793** | Son | 0.097622068 | -0.081482078 | 0.276726215 | 11.915162 | 1.324419463 | 0.232245869 | 0.616381949 | -4.518838826 |
| **A_55_P1979242** | Kcne2 | -0.329336545 | -0.93358775 | 0.274914659 | 7.823278746 | -1.32435858 | 0.232264898 | 0.616381949 | -4.51886881 |
| **A_55_P2081164** | Cd300lf | -0.277045637 | -0.785358879 | 0.231267605 | 10.2310789 | -1.324350929 | 0.232267289 | 0.616381949 | -4.518872578 |
| **A_52_P149577** | Rab25 | -0.758828413 | -2.151139458 | 0.633482632 | 8.119173851 | -1.324312581 | 0.232279276 | 0.616381949 | -4.518891463 |
| **A_55_P1996862** | Psme2 | -0.134625143 | -0.381686315 | 0.112436028 | 14.56510145 | -1.324051482 | 0.2323609 | 0.61642329 | -4.519020043 |
| **A_55_P2318584** | Aqp8 | -0.463297931 | -1.313612466 | 0.387016604 | 15.62199669 | -1.323926718 | 0.232399913 | 0.61642329 | -4.519081481 |
| **A_55_P1990067** | NA | -0.13779995 | -0.390726288 | 0.115126389 | 12.51829964 | -1.323848204 | 0.232424467 | 0.61642329 | -4.519120144 |
| **A_51_P269320** | Adsl | 0.131442529 | -0.109853887 | 0.372738945 | 9.145791944 | 1.323634946 | 0.232491171 | 0.61642329 | -4.519225153 |
| **A_52_P445360** | Krt20 | 0.388220662 | -0.324591073 | 1.101032398 | 12.87571739 | 1.323387396 | 0.232568623 | 0.61642329 | -4.519347043 |
| **A_52_P389347** | Snap29 | 0.086351426 | -0.07220509 | 0.244907942 | 10.90071882 | 1.323331377 | 0.232586153 | 0.61642329 | -4.519374624 |
| **A_55_P2010298** | Serpina1d | -0.433553292 | -1.229651839 | 0.362545255 | 17.03478789 | -1.323301626 | 0.232595463 | 0.61642329 | -4.519389273 |
| **A_51_P377452** | Ncf4 | -0.214880021 | -0.609454048 | 0.179694005 | 12.86323715 | -1.323276907 | 0.232603199 | 0.61642329 | -4.519401443 |
| **A_55_P2089913** | Cd6 | 0.098317259 | -0.082231401 | 0.278865918 | 7.796184688 | 1.323179245 | 0.232633765 | 0.61642329 | -4.519449527 |
| **A_55_P2126281** | NA | 0.101039907 | -0.084522726 | 0.286602541 | 8.374292339 | 1.323078466 | 0.232665311 | 0.61642329 | -4.519499145 |
| **A_55_P2108708** | Kcne3 | -0.19867509 | -0.563554474 | 0.166204294 | 8.110178321 | -1.323053122 | 0.232673244 | 0.61642329 | -4.519511623 |
| **A_55_P2095311** | NA | -0.205892951 | -0.58405336 | 0.172267458 | 8.399786872 | -1.322965772 | 0.23270059 | 0.61642329 | -4.519554627 |
| **A_55_P2177911** | Lepr | -0.126615118 | -0.359204286 | 0.105974051 | 7.972161217 | -1.322754555 | 0.232766725 | 0.61642329 | -4.519658611 |
| **A_55_P1963631** | Yae1d1 | 0.110686331 | -0.09265762 | 0.314030282 | 9.886638797 | 1.322653006 | 0.232798528 | 0.61642329 | -4.519708603 |

| | | | | | | | | | |
|---|---|---|---|---|---|---|---|---|---|
| **A_52_P259335** | Fkbp15 | 0.145801056 | -0.122062202 | 0.413664315 | 10.99984448 | 1.32260668 | 0.232813037 | 0.61642329 | -4.519731409 |
| **A_55_P2013296** | Prkag2 | -0.152368644 | -0.43229821 | 0.127560921 | 10.32064101 | -1.32260458 | 0.232813695 | 0.61642329 | -4.519732443 |
| **A_52_P655084** | Tbc1d17 | -0.125795991 | -0.356950134 | 0.105358152 | 10.05744679 | -1.32235575 | 0.232891643 | 0.61642329 | -4.519854934 |
| **A_51_P183995** | Paf1 | -0.133390768 | -0.378533176 | 0.11175164 | 11.54886436 | -1.322179793 | 0.232946778 | 0.61642329 | -4.519941548 |
| **A_55_P2136743** | Slc36a4 | -0.099742467 | -0.283056087 | 0.083571153 | 8.085080106 | -1.322112954 | 0.232967724 | 0.61642329 | -4.519974448 |
| **A_55_P2063715** | Cfap53 | -0.194364177 | -0.551621453 | 0.162893099 | 7.747419469 | -1.321960048 | 0.233015649 | 0.61642329 | -4.520049711 |
| **A_55_P2029549** | Dtd1 | 0.125507962 | -0.105187191 | 0.356203116 | 10.89264558 | 1.321952943 | 0.233017875 | 0.61642329 | -4.520053208 |
| **A_55_P1986486** | Gtf2ird1 | -0.173685293 | -0.492973555 | 0.145602969 | 10.66765782 | -1.321792494 | 0.233068174 | 0.61642329 | -4.520132181 |
| **A_55_P2141709** | Gabarapl2 | -0.135324495 | -0.384117485 | 0.113468494 | 13.45577476 | -1.321665233 | 0.233108076 | 0.61642329 | -4.520194817 |
| **A_55_P2112737** | Actb | -0.183057806 | -0.519629977 | 0.153514366 | 16.78866781 | -1.321579328 | 0.233135014 | 0.61642329 | -4.520237097 |
| **A_55_P2064766** | NA | 0.096802757 | -0.081186334 | 0.274791848 | 11.07075792 | 1.321531538 | 0.233150001 | 0.61642329 | -4.520260617 |
| **A_55_P2102390** | Spag9 | -0.174496937 | -0.495399321 | 0.146405447 | 11.66093889 | -1.321289714 | 0.233225852 | 0.61642329 | -4.520379631 |
| **A_51_P315646** | Nfs1 | 0.185525639 | -0.155684206 | 0.526735485 | 11.28173056 | 1.321190896 | 0.233256853 | 0.61642329 | -4.520428262 |
| **A_55_P2022074** | Klf10 | -0.160981144 | -0.45712183 | 0.135159542 | 11.09428582 | -1.320870134 | 0.233357509 | 0.61642329 | -4.520586111 |
| **A_51_P359046** | Slurp1 | -0.29963269 | -0.850847569 | 0.251582188 | 7.842910346 | -1.320843766 | 0.233365785 | 0.61642329 | -4.520599086 |
| **A_55_P2049867** | Ccrl2 | 0.104706693 | -0.087945889 | 0.297359275 | 8.629412659 | 1.320634982 | 0.233431325 | 0.61642329 | -4.520701824 |
| **A_51_P153013** | Map2k3os | 0.106785829 | -0.089694258 | 0.303265915 | 8.357050361 | 1.320621218 | 0.233435647 | 0.61642329 | -4.520708596 |
| **A_55_P2103756** | Hic2 | 0.15478262 | -0.130009323 | 0.439574564 | 8.469070959 | 1.320619434 | 0.233436207 | 0.61642329 | -4.520709474 |
| **A_55_P1989921** | Eml2 | -0.247901294 | -0.704070162 | 0.208267573 | 11.04891461 | -1.320493722 | 0.233475678 | 0.61642329 | -4.520771331 |
| **A_55_P2023727** | Limch1 | -1.25255405 | -3.557643421 | 1.052535322 | 8.778914317 | -1.320358593 | 0.233518112 | 0.61642329 | -4.52083782 |
| **A_51_P139678** | Sprr1a | 0.374934275 | -0.315076525 | 1.064945074 | 8.649425915 | 1.320329858 | 0.233527137 | 0.61642329 | -4.520851958 |
| **A_55_P1981670** | 8-Sep | 0.149771734 | -0.125887966 | 0.425431434 | 8.148058143 | 1.320200121 | 0.233567886 | 0.61642329 | -4.520915792 |
| **A_55_P2015753** | Enho | -0.358093584 | -1.017181022 | 0.300993854 | 10.60576963 | -1.320190778 | 0.233570821 | 0.61642329 | -4.520920388 |
| **A_55_P2107203** | Gpr137b | -0.128926463 | -0.366257692 | 0.108404765 | 8.172458769 | -1.319989182 | 0.233634154 | 0.61642329 | -4.521019574 |
| **A_51_P299287** | Impa2 | -0.117795323 | -0.334652831 | 0.099062184 | 10.32806597 | -1.319887073 | 0.233666238 | 0.61642329 | -4.52106981 |
| **A_55_P2075303** | Zfp609 | -0.168952785 | -0.480029358 | 0.142123788 | 8.49137173 | -1.319718298 | 0.233719278 | 0.61642329 | -4.521152842 |
| **A_52_P235241** | Fntb | 0.09921742 | -0.083474094 | 0.281908934 | 8.089346737 | 1.319631708 | 0.233746494 | 0.61642329 | -4.52119544 |
| **A_51_P337675** | Cd53 | -0.158468764 | -0.450270518 | 0.133332991 | 8.410518811 | -1.31958983 | 0.233759658 | 0.61642329 | -4.521216042 |
| **A_52_P158710** | Dcbld1 | -0.097237911 | -0.276295282 | 0.081819461 | 8.792040164 | -1.319552284 | 0.23377146 | 0.61642329 | -4.521234512 |
| **A_55_P2136340** | NA | 0.131177608 | -0.110382521 | 0.372737738 | 14.09604196 | 1.319525062 | 0.233780018 | 0.61642329 | -4.521247904 |
| **A_55_P2001903** | Smim4 | -0.486312967 | -1.381850553 | 0.409224619 | 12.28331337 | -1.319517701 | 0.233782332 | 0.61642329 | -4.521251525 |
| **A_51_P481238** | Dop1b | 0.165869882 | -0.139707116 | 0.47144688 | 11.29194592 | 1.31895526 | 0.233959207 | 0.61642329 | -4.521528192 |
| **A_55_P2041634** | Vps51 | 0.147482665 | -0.124226746 | 0.419192077 | 10.51536257 | 1.318923133 | 0.233969314 | 0.61642329 | -4.521543995 |
| **A_55_P2330435** | NA | -0.209796907 | -0.59631822 | 0.176724405 | 7.763949596 | -1.31889076 | 0.233979498 | 0.61642329 | -4.521559918 |
| **A_51_P114843** | H1f10 | -0.169487687 | -0.481745568 | 0.142770194 | 8.000587646 | -1.31888805 | 0.233980351 | 0.61642329 | -4.521561251 |
| **A_55_P2184364** | B4galt3 | -0.204040038 | -0.579976328 | 0.171896251 | 11.68950302 | -1.318816387 | 0.234002898 | 0.61642329 | -4.521596499 |
| **A_55_P1981836** | Ccm2l | -0.364633657 | -1.036481493 | 0.307214179 | 9.51480093 | -1.318769881 | 0.23401753 | 0.61642329 | -4.521619373 |
| **A_55_P1964902** | NA | 0.088054923 | -0.07419416 | 0.250304006 | 7.929208576 | 1.318726029 | 0.234031329 | 0.61642329 | -4.521640942 |
| **A_55_P2126269** | Nmb | -0.195550853 | -0.555892531 | 0.164790826 | 8.825204937 | -1.318646558 | 0.234056337 | 0.61642329 | -4.52168003 |
| **A_55_P2163344** | NA | 0.194077349 | -0.163595452 | 0.551750151 | 9.040003024 | 1.318475684 | 0.234110117 | 0.61642329 | -4.52176407 |
| **A_51_P367162** | Psmb1 | -0.17880577 | -0.508335067 | 0.150723527 | 12.51057434 | -1.318471316 | 0.234111491 | 0.61642329 | -4.521766218 |
| **A_55_P2041758** | Scand1 | -0.137678316 | -0.391413728 | 0.116057097 | 13.91959579 | -1.3184621 | 0.234114392 | 0.61642329 | -4.521770751 |
| **A_52_P37498** | Dpcd | -0.318604809 | -0.905811088 | 0.26860147 | 9.525795351 | -1.318392807 | 0.234136204 | 0.61642329 | -4.52180483 |
| **A_55_P2110507** | NA | 0.170097217 | -0.143423753 | 0.483618187 | 8.392243765 | 1.318298637 | 0.23416585 | 0.61642329 | -4.521851143 |
| **A_51_P371174** | Bag3 | -0.179262172 | -0.50968429 | 0.151159946 | 10.2059184 | -1.318265032 | 0.23417643 | 0.61642329 | -4.52186767 |
| **A_55_P2173188** | Kank3 | -0.275123069 | -0.782338808 | 0.232092669 | 8.364010498 | -1.31800627 | 0.234257912 | 0.616553811 | -4.521994924 |
| **A_55_P2015485** | Cfap45 | -0.67476913 | -1.9189718 | 0.56943354 | 8.236244597 | -1.317793211 | 0.234325022 | 0.616626047 | -4.522099697 |
| **A_55_P2064451** | Pced1a | -0.093462549 | -0.265811137 | 0.078886038 | 9.365473745 | -1.3176895 | 0.234357695 | 0.616626047 | -4.522150696 |
| **A_55_P1954486** | Mapt | -0.606785444 | -1.725921383 | 0.512350494 | 8.155628484 | -1.317454124 | 0.234431863 | 0.616626047 | -4.522266434 |
| **A_55_P2139043** | 4833439L19Rik | 0.127219398 | -0.107426127 | 0.361864924 | 11.81357725 | 1.317419989 | 0.23444262 | 0.616626047 | -4.522283218 |

| | | | | | | | | | |
|---|---|---|---|---|---|---|---|---|---|
| **A_55_P2003129** | Nemp1 | 0.166683541 | -0.140751954 | 0.474119037 | 8.631217962 | 1.317412863 | 0.234444866 | 0.616626047 | -4.522286722 |
| **A_55_P2162344** | Lrsam1 | -0.090963575 | -0.258765977 | 0.076838827 | 9.245167008 | -1.317202472 | 0.234511182 | 0.61671655 | -4.522390169 |
| **A_55_P2079757** | NA | 0.211186158 | -0.178468853 | 0.60084117 | 9.542962964 | 1.316947228 | 0.234591658 | 0.616792427 | -4.522515662 |
| **A_55_P2149511** | Mff | 0.109081792 | -0.092194683 | 0.310358267 | 12.38411808 | 1.316868576 | 0.234616461 | 0.616792427 | -4.522554331 |
| **A_55_P2126662** | Cav1 | -0.348673071 | -0.992079101 | 0.294732958 | 7.901262039 | -1.316789849 | 0.234641291 | 0.616792427 | -4.522593035 |
| **A_52_P544043** | Pcsk5 | -0.358363768 | -1.019764147 | 0.30303661 | 8.352257511 | -1.316566635 | 0.234711701 | 0.616792427 | -4.522702771 |
| **A_55_P2103880** | Gtf2ird1 | -0.10444334 | -0.297223151 | 0.088336471 | 9.662471761 | -1.316443999 | 0.234750394 | 0.616792427 | -4.522763059 |
| **A_51_P418339** | Nudcd2 | 0.227114957 | -0.192094183 | 0.646324096 | 10.52308579 | 1.316431246 | 0.234754418 | 0.616792427 | -4.522769328 |
| **A_51_P430259** | Cfap126 | -0.627814214 | -1.78699038 | 0.531361953 | 9.959876068 | -1.316027274 | 0.234881915 | 0.616792427 | -4.522967906 |
| **A_55_P1969530** | Zfp146 | 0.09068979 | -0.076788172 | 0.258167752 | 9.2202112 | 1.315781929 | 0.234959378 | 0.616792427 | -4.5230885 |
| **A_51_P227345** | Dpep1 | -0.859459154 | -2.446796268 | 0.727877961 | 8.189344241 | -1.315646636 | 0.235002104 | 0.616792427 | -4.523154997 |
| **A_55_P1957762** | Tsr1 | 0.206180884 | -0.17461513 | 0.586976898 | 9.786330164 | 1.315646433 | 0.235002168 | 0.616792427 | -4.523155097 |
| **A_55_P1974554** | Eif4b | -0.160832225 | -0.45794131 | 0.13627686 | 11.75984867 | -1.315346969 | 0.235096765 | 0.616792427 | -4.523302278 |
| **A_55_P2033312** | Lgi4 | 0.09257751 | -0.078457765 | 0.263612785 | 8.051506063 | 1.315233895 | 0.235132492 | 0.616792427 | -4.52335785 |
| **A_52_P426870** | Slf2 | 0.155298778 | -0.131664289 | 0.442261845 | 9.270599311 | 1.314998393 | 0.235206918 | 0.616792427 | -4.523473584 |
| **A_55_P2051521** | NA | 0.088477661 | -0.075020643 | 0.251975964 | 8.313209271 | 1.314932838 | 0.235227639 | 0.616792427 | -4.523505799 |
| **A_52_P17112** | Znrf1 | -0.106674793 | -0.303813636 | 0.090464049 | 9.71505076 | -1.314839668 | 0.235257091 | 0.616792427 | -4.523551584 |
| **A_51_P161554** | Arid3b | 0.202630424 | -0.171864494 | 0.577125342 | 8.791326764 | 1.314746168 | 0.235286651 | 0.616792427 | -4.52359753 |
| **A_55_P1989813** | Gcm1 | 0.228358717 | -0.193724531 | 0.650441964 | 10.40129932 | 1.314627361 | 0.235324217 | 0.616792427 | -4.52365591 |
| **A_55_P2038412** | Acot8 | 0.146389037 | -0.124227938 | 0.417006011 | 9.994737843 | 1.314427715 | 0.235387355 | 0.616792427 | -4.523754011 |
| **A_55_P2158102** | Camp | -0.215055954 | -0.612620808 | 0.182508901 | 14.81622394 | -1.314397349 | 0.23539696 | 0.616792427 | -4.523768932 |
| **A_52_P471395** | Ifngr2 | -0.187507872 | -0.534203634 | 0.159187891 | 11.11721318 | -1.314177899 | 0.235466381 | 0.616792427 | -4.523876757 |
| **A_55_P1973926** | Ipo13 | 0.152499287 | -0.129481431 | 0.434480005 | 10.74957487 | 1.314109609 | 0.235487988 | 0.616792427 | -4.52391031 |
| **A_55_P2075894** | NA | 0.114610315 | -0.097341432 | 0.326562062 | 12.34644646 | 1.313922941 | 0.235547058 | 0.616792427 | -4.524002022 |
| **A_52_P339996** | Slc6a6 | -0.144802711 | -0.412591997 | 0.122986575 | 10.65530622 | -1.31391323 | 0.235550131 | 0.616792427 | -4.524006793 |
| **A_55_P2004711** | Cep95 | 0.132391049 | -0.112473942 | 0.377256039 | 10.23945802 | 1.313757219 | 0.235599511 | 0.616792427 | -4.52408344 |
| **A_66_P126332** | Zfp703 | -0.293823242 | -0.837296579 | 0.249650094 | 11.47385906 | -1.313684554 | 0.235622513 | 0.616792427 | -4.524119138 |
| **A_51_P207403** | Pwp1 | 0.155760307 | -0.132346762 | 0.443867376 | 9.640208529 | 1.313669359 | 0.235627323 | 0.616792427 | -4.524126603 |
| **A_55_P1973563** | Inava | -0.539333367 | -1.53706021 | 0.458393476 | 8.374293311 | -1.313496204 | 0.235682146 | 0.616792427 | -4.524211668 |
| **A_51_P319732** | Mrps31 | 0.135467799 | -0.115142454 | 0.386078053 | 11.83120383 | 1.313470821 | 0.235690183 | 0.616792427 | -4.524224137 |
| **A_55_P2164221** | Tpt1 | -0.117836105 | -0.335851433 | 0.100179223 | 17.33952046 | -1.31333204 | 0.235734132 | 0.616792427 | -4.524292313 |
| **A_55_P1960216** | Cd151 | -0.251480989 | -0.716803717 | 0.213841739 | 8.6355258 | -1.313209705 | 0.235772879 | 0.616792427 | -4.524352407 |
| **A_51_P189814** | Cldn5 | -0.764460478 | -2.178964994 | 0.650044039 | 10.74161944 | -1.313209113 | 0.235773066 | 0.616792427 | -4.524352697 |
| **A_52_P594917** | Pip4k2c | 0.186220161 | -0.158379531 | 0.530819854 | 11.19616406 | 1.313091535 | 0.235810311 | 0.616792427 | -4.524410453 |
| **A_55_P2020200** | Snx6 | -0.120560489 | -0.343658076 | 0.102537097 | 8.512115104 | -1.313086447 | 0.235811923 | 0.616792427 | -4.524412953 |
| **A_55_P2142830** | Ninj1 | -0.125562826 | -0.357920081 | 0.106794429 | 13.4165154 | -1.313070468 | 0.235816986 | 0.616792427 | -4.524420802 |
| **A_55_P2069530** | NA | -0.267621234 | -0.76288308 | 0.227640613 | 11.25229707 | -1.313012574 | 0.235835327 | 0.616792427 | -4.524449239 |
| **A_51_P368591** | Tle6 | -0.375863045 | -1.071456349 | 0.319730259 | 9.354462524 | -1.312977925 | 0.235846305 | 0.616792427 | -4.524466259 |
| **A_51_P377948** | Gimap9 | 0.111812145 | -0.095115679 | 0.318739969 | 8.531005116 | 1.312965392 | 0.235850276 | 0.616792427 | -4.524472415 |
| **A_55_P2072373** | Msn | -0.247289209 | -0.704959088 | 0.210380669 | 12.87221161 | -1.31291323 | 0.235866803 | 0.616792427 | -4.524498036 |
| **A_55_P1985079** | Mcrip1 | -0.267412213 | -0.762325369 | 0.227500943 | 12.75517125 | -1.312911429 | 0.235867374 | 0.616792427 | -4.524498921 |
| **A_55_P2046158** | NA | -0.451302765 | -1.286562936 | 0.383957407 | 11.69235676 | -1.312893234 | 0.235873139 | 0.616792427 | -4.524507858 |
| **A_55_P2172239** | NA | 0.152016395 | -0.129336293 | 0.433369083 | 16.48180501 | 1.312872493 | 0.235879712 | 0.616792427 | -4.524518045 |
| **A_52_P311031** | NA | 0.264415707 | -0.224966828 | 0.753798241 | 16.13458931 | 1.31287075 | 0.235880264 | 0.616792427 | -4.524518901 |
| **A_51_P128065** | Prcc | 0.12799905 | -0.108929064 | 0.364927164 | 10.42471767 | 1.31272374 | 0.235926852 | 0.616816035 | -4.524591109 |
| **A_66_P121745** | Arhgef28 | -0.157311001 | -0.448514347 | 0.133892344 | 7.783416789 | -1.312640878 | 0.235953115 | 0.616816035 | -4.524631807 |
| **A_52_P578634** | Cela1 | -0.11619737 | -0.331401578 | 0.099006838 | 8.927994457 | -1.311984576 | 0.236161221 | 0.616899416 | -4.524954128 |
| **A_52_P159429** | AU021092 | -0.522158656 | -1.489398623 | 0.445081311 | 7.989032413 | -1.311751102 | 0.236235292 | 0.616899416 | -4.52506878 |
| **A_51_P271503** | Il1r1 | -0.243682239 | -0.695102935 | 0.207738458 | 10.25925037 | -1.311673059 | 0.236260056 | 0.616899416 | -4.525107102 |
| **A_51_P374782** | Epn3 | 0.373089772 | -0.318062711 | 1.064242254 | 11.99323023 | 1.311664184 | 0.236262872 | 0.616899416 | -4.52511146 |

| | | | | | | | | | |
|---|---|---|---|---|---|---|---|---|---|
| **A_55_P2127952** | NA | 0.221649455 | -0.188971096 | 0.632270007 | 8.51959294 | 1.311623459 | 0.236275796 | 0.616899416 | -4.525131458 |
| **A_55_P2098802** | Uqcrq | 0.178988058 | -0.152612617 | 0.510588733 | 14.40037951 | 1.311571103 | 0.236292412 | 0.616899416 | -4.525157166 |
| **A_51_P406583** | Myo1e | -0.282874316 | -0.806978167 | 0.241229536 | 9.032362873 | -1.311473 | 0.236323548 | 0.616899416 | -4.525205338 |
| **A_51_P109369** | Fbxo32 | -0.091727847 | -0.261687054 | 0.078231359 | 7.883636762 | -1.311413635 | 0.236342392 | 0.616899416 | -4.525234487 |
| **A_51_P412914** | Efs | -0.566421092 | -1.615951055 | 0.483108871 | 8.481572851 | -1.311377609 | 0.236353828 | 0.616899416 | -4.525252176 |
| **A_52_P617327** | Rcan1 | -0.2097783 | -0.59848436 | 0.178927761 | 8.725097584 | -1.311361524 | 0.236358934 | 0.616899416 | -4.525260074 |
| **A_55_P2133170** | Luc7l2 | 0.230774624 | -0.196866234 | 0.658415482 | 9.545175864 | 1.311269771 | 0.236388062 | 0.616899416 | -4.525305125 |
| **A_55_P2169839** | Rufy3 | 0.12948048 | -0.110475609 | 0.369436569 | 9.983519411 | 1.311160069 | 0.236422893 | 0.616899416 | -4.525358987 |
| **A_65_P11603** | Ldb3 | -0.209972709 | -0.59914837 | 0.179202951 | 7.774147579 | -1.31099299 | 0.23647595 | 0.616899416 | -4.525441019 |
| **A_66_P123527** | Ndufb4 | 0.113640447 | -0.097004978 | 0.324285873 | 14.11552041 | 1.310883483 | 0.23651073 | 0.616899416 | -4.525494782 |
| **A_55_P1960074** | Pafah1b3 | -0.251010195 | -0.716288975 | 0.214268584 | 10.98967542 | -1.310875073 | 0.236513401 | 0.616899416 | -4.525498911 |
| **A_55_P2142191** | Rpl10 | 0.087514084 | -0.074704597 | 0.249732766 | 16.54508532 | 1.310871963 | 0.236514389 | 0.616899416 | -4.525500438 |
| **A_55_P2060445** | Apopt1 | 0.134700721 | -0.115021816 | 0.384423258 | 14.05355569 | 1.310676071 | 0.236576618 | 0.616899416 | -4.525596609 |
| **A_55_P2049612** | Ube2a | 0.129220536 | -0.110349401 | 0.368790473 | 11.42408857 | 1.310636956 | 0.236589046 | 0.616899416 | -4.525615812 |
| **A_51_P440327** | Efhc1 | -0.349730735 | -0.998121837 | 0.298660367 | 7.916859141 | -1.310629513 | 0.23659141 | 0.616899416 | -4.525619465 |
| **A_51_P413710** | Ap3d1 | -0.106334657 | -0.303518634 | 0.090849321 | 10.72522099 | -1.310347242 | 0.23668111 | 0.617050064 | -4.525758034 |
| **A_55_P2044684** | Rsph1 | -0.535098217 | -1.527459062 | 0.457262628 | 8.486033011 | -1.310228593 | 0.236718823 | 0.617065156 | -4.525816277 |
| **A_51_P474459** | Socs3 | 0.138281599 | -0.118212907 | 0.394776106 | 10.78864518 | 1.309994632 | 0.236793204 | 0.617175815 | -4.52593112 |
| **A_55_P2091461** | Casp4 | -0.155254772 | -0.443297745 | 0.1327882 | 9.225045485 | -1.309697093 | 0.236887828 | 0.617225099 | -4.526077161 |
| **A_55_P2471798** | NA | 0.153284571 | -0.131187605 | 0.437756748 | 9.918859609 | 1.309308046 | 0.237011604 | 0.617225099 | -4.526268102 |
| **A_55_P2165655** | Zbtb4 | -0.203791205 | -0.582004283 | 0.174421872 | 9.670809347 | -1.309278648 | 0.237020959 | 0.617225099 | -4.52628253 |
| **A_51_P353232** | Tnnc2 | -0.225108904 | -0.642886945 | 0.192669136 | 8.144078536 | -1.309273128 | 0.237022716 | 0.617225099 | -4.526285239 |
| **A_52_P359621** | Luzp1 | 0.120086713 | -0.102824589 | 0.342998015 | 9.824662975 | 1.309019323 | 0.237103499 | 0.617225099 | -4.526409794 |
| **A_55_P2034864** | Tubb2b | -0.459763225 | -1.313238792 | 0.393712341 | 9.395933664 | -1.308959855 | 0.23712243 | 0.617225099 | -4.526438976 |
| **A_55_P1981241** | Grasp | -0.521422571 | -1.489405287 | 0.446560145 | 8.971965698 | -1.30889682 | 0.237142498 | 0.617225099 | -4.526469909 |
| **A_51_P519592** | Srsf10 | 0.135885895 | -0.116378574 | 0.388150363 | 8.380637849 | 1.308884979 | 0.237146269 | 0.617225099 | -4.52647572 |
| **A_52_P329917** | Prrg2 | -0.174280765 | -0.497840362 | 0.149278831 | 10.61916495 | -1.308815297 | 0.237168455 | 0.617225099 | -4.526509914 |
| **A_55_P2001168** | NA | 0.207163635 | -0.177445641 | 0.591772911 | 10.50659177 | 1.308811062 | 0.237169804 | 0.617225099 | -4.526511992 |
| **A_55_P2003043** | P4ha1 | -0.093471326 | -0.267039752 | 0.0800971 | 7.922115894 | -1.308551647 | 0.237252418 | 0.617225099 | -4.526639286 |
| **A_55_P2062054** | Dbndd2 | -0.307545276 | -0.878631916 | 0.263541365 | 11.60530628 | -1.308549779 | 0.237253013 | 0.617225099 | -4.526640203 |
| **A_52_P509710** | Zwint | 0.181821485 | -0.15591844 | 0.51956141 | 11.55616307 | 1.308115187 | 0.237391472 | 0.617225099 | -4.526853438 |
| **A_52_P117313** | Azin1 | 0.090678663 | -0.077768631 | 0.259125957 | 7.873925129 | 1.308049745 | 0.237412327 | 0.617225099 | -4.526885546 |
| **A_51_P240614** | Tm4sf1 | -0.681963516 | -1.948986122 | 0.58505909 | 10.50054719 | -1.307856113 | 0.237474045 | 0.617225099 | -4.526980543 |
| **A_55_P1985464** | Gm12271 | 0.100403847 | -0.086161162 | 0.286968855 | 14.69048774 | 1.307685621 | 0.2375284 | 0.617225099 | -4.527064185 |
| **A_55_P1995135** | Casz1 | -0.139458936 | -0.398594428 | 0.119676557 | 8.239533767 | -1.307683473 | 0.237529085 | 0.617225099 | -4.527065239 |
| **A_55_P2093232** | Myh7 | -1.158667278 | -3.311716107 | 0.994381551 | 8.444426273 | -1.307639564 | 0.237543085 | 0.617225099 | -4.52708678 |
| **A_52_P424462** | Ero1lb | -0.18592377 | -0.531419788 | 0.159572248 | 8.352819781 | -1.307600447 | 0.237555558 | 0.617225099 | -4.527105969 |
| **A_55_P2146185** | Rs1 | 0.195706055 | -0.167997835 | 0.559409944 | 13.5433119 | 1.307493399 | 0.237589695 | 0.617225099 | -4.527158483 |
| **A_51_P448391** | Nkiras1 | 0.143397717 | -0.123104397 | 0.409899831 | 8.649522178 | 1.307449029 | 0.237603845 | 0.617225099 | -4.527180249 |
| **A_51_P245405** | Ppp3cc | -0.096545301 | -0.275991365 | 0.082900763 | 8.336784082 | -1.307315451 | 0.237646451 | 0.617225099 | -4.527245775 |
| **A_52_P97572** | Aplnr | -0.439427195 | -1.256219688 | 0.377365299 | 8.034730088 | -1.30724919 | 0.237667588 | 0.617225099 | -4.527278278 |
| **A_55_P2087539** | Clpsl2 | 0.222532922 | -0.1911272 | 0.636193044 | 8.432979157 | 1.307175213 | 0.237691188 | 0.617225099 | -4.527314566 |
| **A_52_P559498** | Clk3 | 0.163885409 | -0.140762602 | 0.46853342 | 10.552556 | 1.307149106 | 0.237699517 | 0.617225099 | -4.527327372 |
| **A_51_P227108** | Bri3bp | 0.10925084 | -0.0938384 | 0.31234008 | 11.98898077 | 1.307136849 | 0.237703428 | 0.617225099 | -4.527333384 |
| **A_55_P1956443** | Ttc5 | -0.136693439 | -0.390796806 | 0.117409928 | 9.197114512 | -1.307134986 | 0.237704022 | 0.617225099 | -4.527334298 |
| **A_51_P491595** | AW549877 | -0.1837394 | -0.525299662 | 0.157820863 | 9.16317891 | -1.307128079 | 0.237706226 | 0.617225099 | -4.527337686 |
| **A_51_P358243** | Tstd3 | 0.207978032 | -0.178684318 | 0.594640383 | 9.712155362 | 1.306979474 | 0.237753643 | 0.617259258 | -4.527410578 |
| **A_52_P226127** | Dido1 | 0.151995523 | -0.130615554 | 0.434606601 | 9.242469876 | 1.306847178 | 0.237795863 | 0.617259258 | -4.527475467 |
| **A_55_P2050652** | Ikbkg | 0.195799387 | -0.168274751 | 0.559873524 | 9.140806352 | 1.306786642 | 0.237815184 | 0.617259258 | -4.527505159 |
| **A_55_P2092953** | Traf5 | -0.16979254 | -0.485566763 | 0.145981683 | 8.961199958 | -1.306547245 | 0.237891606 | 0.617374712 | -4.527622574 |

| | | | | | | | | | |
|---|---|---|---|---|---|---|---|---|---|
| **A_51_P267740** | Zc3h15 | 0.15131863 | -0.130164962 | 0.432802222 | 9.945253814 | 1.306238575 | 0.237990174 | 0.617418986 | -4.527773954 |
| **A_55_P2145401** | H3c3 | 0.157789868 | -0.135736006 | 0.451315743 | 11.53263087 | 1.306218747 | 0.237996507 | 0.617418986 | -4.527783678 |
| **A_55_P2008933** | Slc17a3 | 0.362078912 | -0.311485018 | 1.035642842 | 9.716776163 | 1.306193741 | 0.238004494 | 0.617418986 | -4.527795941 |
| **A_55_P2004588** | Nabp2 | 0.151931413 | -0.130737598 | 0.434600425 | 11.03439959 | 1.306028239 | 0.238057362 | 0.617473261 | -4.527877103 |
| **A_55_P1959838** | NA | 0.130392129 | -0.112278541 | 0.373062799 | 15.31493223 | 1.305621447 | 0.238187351 | 0.617605379 | -4.528076578 |
| **A_55_P2081373** | Glis1 | 0.103663938 | -0.089265213 | 0.296593089 | 8.185685862 | 1.30560871 | 0.238191422 | 0.617605379 | -4.528082823 |
| **A_55_P2125786** | Lrrc8e | -0.307668024 | -0.880287037 | 0.264950988 | 8.135927576 | -1.305568876 | 0.238204154 | 0.617605379 | -4.528102355 |
| **A_52_P312102** | NA | -0.597180062 | -1.708941517 | 0.514581392 | 8.324010834 | -1.305199432 | 0.238322271 | 0.617828752 | -4.528283497 |
| **A_51_P464238** | Atf4 | 0.215435078 | -0.185748647 | 0.616618803 | 9.451935713 | 1.304837115 | 0.238438159 | 0.618026793 | -4.528461129 |
| **A_66_P136372** | Mrpl9 | 0.117984977 | -0.101739803 | 0.337709756 | 10.38482162 | 1.304760686 | 0.238462612 | 0.618026793 | -4.528498598 |
| **A_55_P1974283** | Rbmx | 0.153488964 | -0.132386541 | 0.43936447 | 10.31077105 | 1.304618098 | 0.238508237 | 0.618062168 | -4.528568498 |
| **A_51_P105927** | Rasl12 | -1.063486642 | -3.044533823 | 0.91756054 | 8.323411036 | -1.304428891 | 0.238568791 | 0.618117086 | -4.52770228 |
| **A_55_P1976356** | Fgf18 | -0.546714786 | -1.565349654 | 0.471920083 | 8.146644824 | -1.304143767 | 0.238660068 | 0.618117086 | -4.528801012 |
| **A_52_P413502** | Rtf2 | -0.119003534 | -0.34073501 | 0.102727941 | 8.488346332 | -1.304114433 | 0.238669461 | 0.618117086 | -4.52881539 |
| **A_55_P2054261** | C2cd4b | -0.301224652 | -0.862556841 | 0.260107536 | 7.79221903 | -1.303928428 | 0.238729026 | 0.618117086 | -4.52890656 |
| **A_55_P2064896** | Gm8580 | -0.16688311 | -0.47790021 | 0.14413399 | 13.75466829 | -1.303800988 | 0.238769844 | 0.618117086 | -4.528969023 |
| **A_52_P129443** | Wdpcp | -0.240268271 | -0.688066873 | 0.20753033 | 8.186951468 | -1.303757679 | 0.238783717 | 0.618117086 | -4.528990249 |
| **A_55_P2073507** | 2610528J11Rik | -0.087657449 | -0.251044424 | 0.075729526 | 8.05382671 | -1.303630721 | 0.238824389 | 0.618117086 | -4.529052473 |
| **A_55_P2172999** | Ptpn13 | -0.402168558 | -1.15178305 | 0.347445933 | 8.325916065 | -1.303626949 | 0.238825597 | 0.618117086 | -4.529054321 |
| **A_51_P143031** | Eps8l3 | -0.539975999 | -1.546476045 | 0.466524048 | 7.958619271 | -1.303598482 | 0.238834718 | 0.618117086 | -4.529068273 |
| **A_55_P1956130** | LOC68395 | -0.185646139 | -0.531699208 | 0.160406929 | 11.94778018 | -1.303546135 | 0.23885149 | 0.618117086 | -4.529093928 |
| **A_55_P2145606** | Kmo | 0.35623421 | -0.307883329 | 1.02035175 | 9.157992188 | 1.303388435 | 0.238902024 | 0.618117086 | -4.529171214 |
| **A_51_P177691** | Gabrr1 | 0.281136816 | -0.243055698 | 0.80532933 | 8.991672177 | 1.303197076 | 0.238963356 | 0.618117086 | -4.529264992 |
| **A_55_P2017982** | Pde8b | -0.363603124 | -1.041614789 | 0.314408541 | 8.125824886 | -1.303087651 | 0.238998434 | 0.618117086 | -4.529318615 |
| **A_51_P153683** | Lrrc26 | -0.338851942 | -0.970766991 | 0.293063107 | 7.811807048 | -1.30297016 | 0.239036103 | 0.618117086 | -4.52937619 |
| **A_55_P2114838** | NA | 0.134848634 | -0.116631456 | 0.386328724 | 15.1857946 | 1.302945153 | 0.239044121 | 0.618117086 | -4.529388444 |
| **A_51_P406306** | Hecw2 | -0.121254448 | -0.347383996 | 0.104875099 | 7.879807008 | -1.302937413 | 0.239046603 | 0.618117086 | -4.529392236 |
| **A_66_P111534** | 5430431A17Rik | 0.175802155 | -0.152102823 | 0.503707134 | 8.379650652 | 1.302744876 | 0.239108346 | 0.618117086 | -4.529486581 |
| **A_55_P2070079** | Lgals9 | -0.237877936 | -0.681571124 | 0.205815252 | 10.65928737 | -1.302730524 | 0.239112949 | 0.618117086 | -4.529493613 |
| **A_55_P1988202** | Ifi203 | -0.193734769 | -0.555144531 | 0.167674992 | 9.373159895 | -1.302539434 | 0.239174243 | 0.618117086 | -4.529587245 |
| **A_52_P245766** | NA | 0.439049008 | -0.380103916 | 1.258201932 | 10.85528081 | 1.302360464 | 0.239231663 | 0.618117086 | -4.529674934 |
| **A_51_P229925** | Chst7 | -0.159675492 | -0.457642269 | 0.138291285 | 8.836836207 | -1.302127793 | 0.239306329 | 0.618117086 | -4.529788928 |
| **A_51_P122582** | Daglb | 0.169640513 | -0.146930153 | 0.486211179 | 9.5785137 | 1.302093278 | 0.239317407 | 0.618117086 | -4.529805837 |
| **A_51_P519570** | 1600002K03Rik | 0.112352914 | -0.097323171 | 0.322029 | 10.73335476 | 1.302022923 | 0.239339989 | 0.618117086 | -4.529840306 |
| **A_52_P342131** | Zfp532 | -0.141805656 | -0.406461441 | 0.122850129 | 8.049560414 | -1.301953134 | 0.239362392 | 0.618117086 | -4.529874496 |
| **A_51_P343309** | Cmas | 0.223024232 | -0.193289001 | 0.639337465 | 12.27617514 | 1.301712351 | 0.239439699 | 0.618117086 | -4.529992453 |
| **A_51_P443443** | 5830417I10Rik | 0.119172395 | -0.103296112 | 0.341640903 | 10.41874421 | 1.301638295 | 0.239463481 | 0.618117086 | -4.530028731 |
| **A_52_P589699** | Polr3a | 0.110339372 | -0.095642605 | 0.316321348 | 8.616745757 | 1.301620782 | 0.239469105 | 0.618117086 | -4.53003731 |
| **A_55_P2079636** | Scgb3a2 | -0.193548392 | -0.554868207 | 0.167771422 | 7.736903638 | -1.301610308 | 0.239472468 | 0.618117086 | -4.53004244 |
| **A_66_P109562** | Gm12271 | 0.143000474 | -0.123965883 | 0.409966831 | 11.83462014 | 1.301559815 | 0.239488685 | 0.618117086 | -4.530067175 |
| **A_52_P198435** | Rasgrp3 | -0.288801459 | -0.827962562 | 0.250359643 | 9.175426815 | -1.301559482 | 0.239488792 | 0.618117086 | -4.530067338 |
| **A_55_P2176953** | Usp51 | 0.084316746 | -0.073117777 | 0.24175127 | 7.843428497 | 1.301358804 | 0.239553251 | 0.618200907 | -4.530165639 |
| **A_55_P2159895** | Zfp52 | -0.084633052 | -0.242727118 | 0.073461013 | 7.906108004 | -1.300791307 | 0.239735619 | 0.618425927 | -4.530443598 |
| **A_55_P2052799** | Cask | -0.191539963 | -0.549337097 | 0.166257171 | 9.641413188 | -1.30078563 | 0.239737444 | 0.618425927 | -4.530446379 |
| **A_52_P672740** | Rala | -0.140739982 | -0.403657686 | 0.122177721 | 8.137439945 | -1.300711112 | 0.2397614 | 0.618425927 | -4.530482875 |
| **A_55_P2174601** | Plekha7 | 0.133438311 | -0.115843259 | 0.382719882 | 9.018428383 | 1.30068926 | 0.239768426 | 0.618425927 | -4.530493577 |
| **A_55_P1954755** | NA | 0.18357553 | -0.159525206 | 0.526676266 | 16.16614627 | 1.300098719 | 0.239958352 | 0.618501666 | -4.530782775 |
| **A_51_P150087** | Abhd10 | 0.119388379 | -0.103765435 | 0.342542193 | 8.710438794 | 1.299992755 | 0.239992446 | 0.618501666 | -4.530834663 |
| **A_55_P2145691** | Hlcs | -0.125540458 | -0.360198515 | 0.109117599 | 8.269409852 | -1.299964307 | 0.2400016 | 0.618501666 | -4.530848593 |
| **A_52_P579531** | Pdlim3 | -0.567935735 | -1.629628596 | 0.493757126 | 8.620422275 | -1.299820816 | 0.240047776 | 0.618501666 | -4.530918854 |

| | | | | | | | | | |
|---|---|---|---|---|---|---|---|---|---|
| **A_55_P2035932** | Fgf7 | -0.326578643 | -0.937165957 | 0.284008671 | 7.88706627 | -1.299640369 | 0.240105856 | 0.618501666 | -4.531007208 |
| **A_51_P220062** | Mmp15 | -0.19858799 | -0.569886022 | 0.172710041 | 7.911140587 | -1.299611431 | 0.240115171 | 0.618501666 | -4.531021377 |
| **A_51_P198387** | Uckl1 | 0.122542023 | -0.106585436 | 0.351669482 | 9.664219205 | 1.299544321 | 0.240136776 | 0.618501666 | -4.531054235 |
| **A_55_P2052634** | Rpl6 | -0.107374679 | -0.308167952 | 0.093418595 | 15.68354689 | -1.299379189 | 0.240189944 | 0.618501666 | -4.531135085 |
| **A_51_P320980** | Sfxn4 | -0.101001671 | -0.289895744 | 0.087892403 | 7.910355028 | -1.299252062 | 0.240230882 | 0.618501666 | -4.531197324 |
| **A_55_P2011672** | NA | 0.163138283 | -0.141975756 | 0.468252321 | 13.94929068 | 1.299202612 | 0.240246808 | 0.618501666 | -4.531221534 |
| **A_55_P2113459** | Washc1 | -0.133289633 | -0.382584105 | 0.116004839 | 11.3191916 | -1.299172783 | 0.240256415 | 0.618501666 | -4.531236137 |
| **A_52_P160518** | Sfmbt1 | 0.204278859 | -0.177798235 | 0.586355952 | 9.318537229 | 1.299138971 | 0.240267305 | 0.618501666 | -4.531252691 |
| **A_52_P452689** | Atf3 | -0.109297706 | -0.313725205 | 0.095129793 | 8.102188702 | -1.299136904 | 0.240267971 | 0.618501666 | -4.531253703 |
| **A_52_P266530** | Ide | 0.348312179 | -0.303262094 | 0.999886453 | 10.22519028 | 1.298936502 | 0.240332527 | 0.618501666 | -4.531351811 |
| **A_55_P1967196** | Lamb3 | -0.928408921 | -2.665262441 | 0.808444599 | 8.748481107 | -1.298850904 | 0.240360106 | 0.618501666 | -4.531393714 |
| **A_51_P240019** | Gyg | -0.95120642 | -2.730788336 | 0.828375496 | 12.12443035 | -1.298793116 | 0.240378726 | 0.618501666 | -4.531422003 |
| **A_66_P135185** | Etv1 | -0.734267015 | -2.108232694 | 0.639698665 | 8.831254426 | -1.298557914 | 0.240454524 | 0.618501666 | -4.531537137 |
| **A_55_P2025090** | Gins3 | 0.2434238 | -0.212076948 | 0.698924548 | 13.05256866 | 1.298545382 | 0.240458564 | 0.618501666 | -4.531543271 |
| **A_55_P2162269** | Bri3 | -0.513447103 | -1.47424221 | 0.447348004 | 11.14750356 | -1.298518418 | 0.240467255 | 0.618501666 | -4.53155647 |
| **A_55_P1972792** | Stil | 0.142651285 | -0.124293518 | 0.409596087 | 8.852654376 | 1.298486407 | 0.240477573 | 0.618501666 | -4.531572139 |
| **A_51_P246066** | Slamf9 | 0.191013326 | -0.166445724 | 0.548472377 | 8.767684609 | 1.298436033 | 0.240493811 | 0.618501666 | -4.531596796 |
| **A_52_P126266** | Prkab2 | -0.098645198 | -0.283252642 | 0.085962246 | 8.209114486 | -1.298404253 | 0.240504056 | 0.618501666 | -4.531612352 |
| **A_51_P399985** | Myo9b | -0.092387785 | -0.265304029 | 0.080528459 | 12.08243785 | -1.298260819 | 0.2405503 | 0.618501666 | -4.531682559 |
| **A_51_P301998** | Fmo2 | -1.087638215 | -3.123375515 | 0.948099085 | 8.277505717 | -1.298212879 | 0.240565757 | 0.618501666 | -4.531706023 |
| **A_51_P323610** | Selenos | -0.161841514 | -0.464813759 | 0.14113073 | 12.24311336 | -1.297986806 | 0.240638664 | 0.618564695 | -4.531816673 |
| **A_52_P69998** | Kdelr2 | -0.170146059 | -0.488676544 | 0.148384425 | 10.79466055 | -1.297938402 | 0.240654276 | 0.618564695 | -4.531840363 |
| **A_55_P2007249** | Spdef | -0.251020769 | -0.721002096 | 0.218960557 | 7.962358652 | -1.297813367 | 0.240694609 | 0.618586106 | -4.531901557 |
| **A_51_P479769** | Ampd2 | 0.274386027 | -0.239428644 | 0.788200698 | 12.36505825 | 1.297593556 | 0.24076553 | 0.618686111 | -4.532009131 |
| **A_55_P2179582** | Nmur2 | -0.115185278 | -0.330901482 | 0.100530925 | 7.817092639 | -1.297470214 | 0.240805333 | 0.618694993 | -4.532069491 |
| **A_51_P264495** | Pgam2 | -0.599869584 | -1.723711397 | 0.523972229 | 8.385054613 | -1.296984675 | 0.240962077 | 0.618694993 | -4.532307083 |
| **A_55_P2077546** | NA | 0.175357101 | -0.153173492 | 0.503887695 | 9.052162633 | 1.296972382 | 0.240966047 | 0.618694993 | -4.532313098 |
| **A_51_P214107** | Tbc1d20 | -0.130954965 | -0.376334795 | 0.114424865 | 8.787545305 | -1.296779978 | 0.241028185 | 0.618694993 | -4.53240724 |
| **A_52_P602996** | Cdip1 | -0.122064129 | -0.350791437 | 0.10666318 | 8.212004306 | -1.29674095 | 0.241040791 | 0.618694993 | -4.532426336 |
| **A_51_P379976** | Arl4a | 0.168824866 | -0.147530435 | 0.485180167 | 12.38543315 | 1.296714855 | 0.24104922 | 0.618694993 | -4.532439103 |
| **A_55_P1973448** | Ybx2 | 0.10839964 | -0.094758721 | 0.311558001 | 8.172487142 | 1.29651137 | 0.241114958 | 0.618694993 | -4.532538661 |
| **A_55_P2150048** | Sypl | -0.199389668 | -0.573080673 | 0.174301338 | 10.16882235 | -1.296502009 | 0.241117983 | 0.618694993 | -4.532543241 |
| **A_55_P2125588** | Pdgfa | -0.639713776 | -1.838705878 | 0.559278326 | 10.0893598 | -1.296440441 | 0.241137876 | 0.618694993 | -4.532573362 |
| **A_52_P179238** | Macroh2a1 | -0.196079405 | -0.563589535 | 0.171430725 | 8.555777684 | -1.296420395 | 0.241144354 | 0.618694993 | -4.53258317 |
| **A_55_P2035484** | Gucd1 | 0.192271454 | -0.16812322 | 0.552666129 | 13.1663769 | 1.296342129 | 0.241169645 | 0.618694993 | -4.53262146 |
| **A_55_P2079561** | Lilra6 | -0.292864159 | -0.841830861 | 0.256102543 | 10.02408928 | -1.296293704 | 0.241185295 | 0.618694993 | -4.532645151 |
| **A_55_P1969845** | Lrpprc | 0.245190865 | -0.214432539 | 0.70481427 | 13.28733784 | 1.296239764 | 0.241202728 | 0.618694993 | -4.53267154 |
| **A_55_P2028773** | Slc22a17 | -0.325266542 | -0.935017018 | 0.284483934 | 8.787841469 | -1.296195274 | 0.241217107 | 0.618694993 | -4.532693305 |
| **A_55_P2082869** | Ezh1 | -0.102875932 | -0.295763085 | 0.090011222 | 11.60408676 | -1.295966177 | 0.241291166 | 0.618802832 | -4.532805378 |
| **A_51_P382925** | Fadd | 0.104429737 | -0.091388828 | 0.300248302 | 9.026261578 | 1.29584635 | 0.241329909 | 0.618820087 | -4.532863995 |
| **A_52_P370022** | NA | -0.196437585 | -0.564811947 | 0.171936777 | 14.41296903 | -1.295741531 | 0.241363805 | 0.618824909 | -4.532915268 |
| **A_55_P2120277** | Gm11619 | -0.198515615 | -0.570821418 | 0.173790189 | 15.61135347 | -1.295621229 | 0.241402713 | 0.618842577 | -4.532974114 |
| **A_51_P248705** | Cse1l | 0.157277849 | -0.137717739 | 0.452273438 | 9.918638114 | 1.295493473 | 0.241444037 | 0.618866436 | -4.533036604 |
| **A_55_P2079388** | Snora74a | 0.286508382 | -0.250945874 | 0.823962639 | 8.904636676 | 1.295325797 | 0.241498283 | 0.618923405 | -4.533118617 |
| **A_52_P547234** | Tomm70a | 0.236975832 | -0.207703774 | 0.681655439 | 10.78283676 | 1.294911338 | 0.241632415 | 0.61907924 | -4.53332132 |
| **A_66_P139546** | Igfbp6 | -0.951793113 | -2.737873306 | 0.83428708 | 9.808407559 | -1.294865897 | 0.241647126 | 0.61907924 | -4.533343544 |
| **A_55_P2145034** | Swt1 | 0.139437737 | -0.122235515 | 0.401110989 | 10.32885787 | 1.294804448 | 0.241667019 | 0.61907924 | -4.533373595 |
| **A_66_P138137** | Pnp | -0.128653121 | -0.370099247 | 0.112793006 | 13.24132556 | -1.294742107 | 0.241687203 | 0.61907924 | -4.533404082 |
| **A_55_P1972892** | NA | -0.205268094 | -0.590569358 | 0.180033171 | 12.80340928 | -1.294506404 | 0.241763528 | 0.619138877 | -4.533519346 |
| **A_55_P2007981** | Fmn2 | 0.178281559 | -0.156373061 | 0.51293618 | 8.647060962 | 1.294472378 | 0.241774548 | 0.619138877 | -4.533535985 |

| | | | | | | | | |
|---|---|---|---|---|---|---|---|---|
| **A_55_P2156002** | Sumo2 | 0.124109247 | -0.108922252 | 0.357140747 | 14.50283658 | 1.294114495 | 0.241890484 | 0.61935371 | -4.533710982 |
| **A_55_P2087796** | Eva1b | -0.490352039 | -1.411269782 | 0.430565703 | 11.70897815 | -1.293809497 | 0.241989326 | 0.619485069 | -4.533860108 |
| **A_51_P420547** | Clic5 | -1.097956968 | -3.160089178 | 0.964175242 | 8.732865262 | -1.293754884 | 0.242007028 | 0.619485069 | -4.533886809 |
| **A_55_P2072985** | Sptan1 | -0.250566383 | -0.721210386 | 0.220077621 | 11.62510035 | -1.293640079 | 0.242044245 | 0.619485069 | -4.533942938 |
| **A_55_P2308835** | E130110O22Rik | 0.15030437 | -0.132032755 | 0.432641496 | 8.47459973 | 1.293560689 | 0.242069984 | 0.619485069 | -4.533981751 |
| **A_55_P2345116** | NA | 0.110084601 | -0.09672947 | 0.316898673 | 8.359607826 | 1.293390536 | 0.242125158 | 0.619544239 | -4.534064935 |
| **A_55_P2030046** | Barx2 | 0.469500336 | -0.412645982 | 1.351646654 | 9.625042933 | 1.29323804 | 0.242174616 | 0.619578073 | -4.534139485 |
| **A_66_P103092** | NA | 0.288844248 | -0.253947882 | 0.831636377 | 12.22624055 | 1.293044198 | 0.242237496 | 0.619578073 | -4.534234242 |
| **A_55_P1955276** | Specc1 | 0.194999329 | -0.171463524 | 0.561462182 | 8.857731713 | 1.292963746 | 0.242263599 | 0.619578073 | -4.534273569 |
| **A_51_P468249** | Phex | -0.4483487 | -1.291146036 | 0.394448637 | 7.84846847 | -1.292635114 | 0.242370246 | 0.619578073 | -4.534434203 |
| **A_55_P1962039** | Anxa7 | 0.190001825 | -0.167199513 | 0.547203162 | 10.98797671 | 1.292492062 | 0.242416682 | 0.619578073 | -4.534504122 |
| **A_51_P409893** | Prkar2a | 0.095497306 | -0.084045099 | 0.275039711 | 8.231717821 | 1.292430717 | 0.242436598 | 0.619578073 | -4.534534105 |
| **A_55_P1976709** | NA | 0.110092142 | -0.096906145 | 0.317090429 | 11.81016075 | 1.292328015 | 0.242469944 | 0.619578073 | -4.5345843 |
| **A_51_P461108** | Osbpl10 | -0.194588854 | -0.560465273 | 0.171287564 | 8.045146927 | -1.292310077 | 0.242475768 | 0.619578073 | -4.534593067 |
| **A_51_P151038** | Pmpcb | -0.103681443 | -0.298632372 | 0.091269485 | 12.22970504 | -1.29228682 | 0.24248332 | 0.619578073 | -4.534604434 |
| **A_52_P481686** | Wtip | -0.421443037 | -1.213959545 | 0.371073471 | 11.00366796 | -1.292152448 | 0.242526956 | 0.619578073 | -4.534670105 |
| **A_55_P2014640** | Knop1 | 0.241628631 | -0.212840262 | 0.696097525 | 11.63240622 | 1.291895602 | 0.242610384 | 0.619578073 | -4.534795626 |
| **A_52_P49042** | Arfgef3 | -0.130929163 | -0.377210612 | 0.115352286 | 7.714965209 | -1.291777991 | 0.242648595 | 0.619578073 | -4.5348531 |
| **A_55_P2084980** | Dcaf12l2 | 0.399285019 | -0.351810309 | 1.150380347 | 11.69897214 | 1.291728211 | 0.242664769 | 0.619578073 | -4.534877426 |
| **A_66_P137660** | Fam166b | -0.290021944 | -0.835599059 | 0.255555171 | 7.994423775 | -1.291688829 | 0.242677566 | 0.619578073 | -4.53489667 |
| **A_51_P307316** | Syde1 | -0.330961887 | -0.95356051 | 0.291636736 | 8.895745114 | -1.291674295 | 0.242682289 | 0.619578073 | -4.534903772 |
| **A_55_P2144716** | Hecw2 | -0.208825704 | -0.601711554 | 0.184060146 | 8.585907341 | -1.291518866 | 0.242732801 | 0.619578073 | -4.534979723 |
| **A_55_P2140953** | NA | 0.123220466 | -0.108619288 | 0.35506022 | 15.21452577 | 1.291451579 | 0.242754671 | 0.619578073 | -4.535012602 |
| **A_55_P1954393** | Susd4 | -0.676922521 | -1.950590495 | 0.596745453 | 9.294974424 | -1.291415288 | 0.242766467 | 0.619578073 | -4.535030334 |
| **A_51_P214197** | Stk17b | 0.230497813 | -0.203205717 | 0.664201343 | 10.1843225 | 1.291388877 | 0.242775052 | 0.619578073 | -4.53504324 |
| **A_52_P639551** | Tinf2 | 0.149501647 | -0.131802862 | 0.430806156 | 9.401736629 | 1.291375294 | 0.242779467 | 0.619578073 | -4.535049876 |
| **A_55_P2145307** | Ints11 | 0.154620305 | -0.136386714 | 0.445627323 | 10.07042081 | 1.291059477 | 0.242882148 | 0.619667316 | -4.535204185 |
| **A_51_P440210** | Prkg1 | -0.166499176 | -0.479922882 | 0.146924531 | 8.029726727 | -1.290813309 | 0.242962211 | 0.619667316 | -4.535324455 |
| **A_51_P336599** | Kcne3 | -0.323461008 | -0.93240485 | 0.285482835 | 8.275927859 | -1.290707641 | 0.242996585 | 0.619667316 | -4.535376079 |
| **A_55_P2111018** | NA | 0.153869131 | -0.135808926 | 0.443547187 | 9.162524813 | 1.290681517 | 0.243005084 | 0.619667316 | -4.535388842 |
| **A_55_P1996613** | NA | 0.242528415 | -0.214073882 | 0.699130712 | 14.74970234 | 1.290647741 | 0.243016073 | 0.619667316 | -4.535405342 |
| **A_52_P891775** | Cdr2l | -0.226614663 | -0.653308024 | 0.200078698 | 9.453782917 | -1.290491983 | 0.243066752 | 0.619667316 | -4.535481433 |
| **A_51_P171999** | Apoe | -0.270130031 | -0.778788943 | 0.238528881 | 16.81568275 | -1.29041499 | 0.243091808 | 0.619667316 | -4.53469809 |
| **A_55_P2011352** | NA | -0.108578951 | -0.313052101 | 0.095894199 | 11.42116559 | -1.290305484 | 0.243127447 | 0.619667316 | -4.535572538 |
| **A_55_P1988148** | NA | 0.184232891 | -0.162739219 | 0.531205001 | 8.514965685 | 1.29019629 | 0.243162989 | 0.619667316 | -4.535625878 |
| **A_52_P222593** | Cebpzos | 0.193813109 | -0.171223829 | 0.558850047 | 11.75547224 | 1.290118265 | 0.243188389 | 0.619667316 | -4.535663991 |
| **A_51_P316949** | Ccdc6 | 0.144395085 | -0.127574455 | 0.416364624 | 9.286854504 | 1.290076111 | 0.243202113 | 0.619667316 | -4.535684582 |
| **A_55_P1964757** | Stam2 | 0.106353995 | -0.093975293 | 0.306683283 | 8.209676459 | 1.290008436 | 0.243224147 | 0.619667316 | -4.535717638 |
| **A_55_P1970159** | Rnf11 | 0.180081569 | -0.159127302 | 0.51929044 | 11.33434 | 1.289986763 | 0.243231203 | 0.619667316 | -4.535728224 |
| **A_55_P2038947** | Lig3 | 0.147335659 | -0.130250672 | 0.424921991 | 9.324810935 | 1.289712717 | 0.243320447 | 0.619796876 | -4.535862077 |
| **A_55_P2040115** | Cdv3 | 0.140876064 | -0.124577096 | 0.406329224 | 13.13902457 | 1.289533111 | 0.243378952 | 0.619796876 | -4.535949798 |
| **A_55_P2177910** | Lepr | 0.533405412 | -0.471763087 | 1.538573911 | 12.25034302 | 1.289441778 | 0.243408708 | 0.619796876 | -4.535994404 |
| **A_51_P118168** | Zbtb43 | -0.114296109 | -0.32968125 | 0.101089032 | 9.45226816 | -1.289433343 | 0.243411456 | 0.619796876 | -4.535578622 |
| **A_55_P2046453** | Aarsd1 | 0.135934961 | -0.120268474 | 0.392138397 | 12.63059806 | 1.289227047 | 0.243478678 | 0.619796876 | -4.536099272 |
| **A_52_P234354** | Hic2 | 0.245281758 | -0.217024899 | 0.707588416 | 10.08076595 | 1.289194049 | 0.243489432 | 0.619796876 | -4.536115386 |
| **A_55_P2173358** | NA | 0.098043195 | -0.086791158 | 0.282877547 | 8.011084056 | 1.28889623 | 0.24358651 | 0.619796876 | -4.536260822 |
| **A_55_P1982578** | Kdm3a | -0.144275787 | -0.416280444 | 0.127728869 | 10.86923247 | -1.288843847 | 0.243603588 | 0.619796876 | -4.536286401 |
| **A_55_P2043207** | Magef1 | 0.310448253 | -0.274844086 | 0.895740593 | 10.29493512 | 1.288841672 | 0.243604298 | 0.619796876 | -4.536287463 |
| **A_55_P2164111** | Nploc4 | 0.167703462 | -0.148485356 | 0.48389228 | 10.7750237 | 1.28877977 | 0.243624481 | 0.619796876 | -4.53631769 |
| **A_55_P2052696** | Synm | -0.339801947 | -0.980569862 | 0.300965969 | 7.880650025 | -1.288570823 | 0.243692619 | 0.619796876 | -4.536419716 |

| | | | | | | | | | |
|---|---|---|---|---|---|---|---|---|---|
| **A_55_P1958137** | NA | -0.141132798 | -0.407284562 | 0.125018967 | 9.415491107 | -1.288492187 | 0.243718267 | 0.619796876 | -4.536458112 |
| **A_55_P2071354** | Togaram2 | -0.234272993 | -0.676077957 | 0.20753197 | 7.804958398 | -1.288471524 | 0.243725007 | 0.619796876 | -4.536468201 |
| **A_55_P1967928** | Srfbp1 | 0.148345079 | -0.131416247 | 0.428106406 | 9.240241485 | 1.288453222 | 0.243730977 | 0.619796876 | -4.536477137 |
| **A_51_P161946** | Epop | -0.153111959 | -0.441919084 | 0.135695166 | 8.71126687 | -1.288203275 | 0.243812519 | 0.619922677 | -4.536599172 |
| **A_66_P106072** | Orai1 | -0.091215837 | -0.263314462 | 0.080882789 | 8.610690202 | -1.287881878 | 0.243917407 | 0.620072376 | -4.536756081 |
| **A_55_P2102210** | NA | 0.152584401 | -0.135325537 | 0.440494339 | 10.93525967 | 1.28776516 | 0.243955507 | 0.620072376 | -4.536813061 |
| **A_55_P1975110** | Pnpt1 | 0.208327828 | -0.184775148 | 0.601430804 | 10.83242239 | 1.287728012 | 0.243967635 | 0.620072376 | -4.536831196 |
| **A_52_P274496** | Tspan18 | -0.458979798 | -1.325244064 | 0.407284467 | 10.50838699 | -1.287438108 | 0.244062296 | 0.620164719 | -4.536972713 |
| **A_55_P2101246** | Aldh6a1 | -0.183487797 | -0.529820611 | 0.162845018 | 8.273181392 | -1.287350282 | 0.24409098 | 0.620164719 | -4.537015583 |
| **A_55_P2153021** | Ttn | -0.328842124 | -0.949586903 | 0.291902655 | 7.818780229 | -1.287234166 | 0.244128908 | 0.620164719 | -4.537072261 |
| **A_66_P124139** | Ikzf1 | 0.301282572 | -0.267567543 | 0.870132687 | 9.661983408 | 1.286943108 | 0.244224001 | 0.620164719 | -4.537214324 |
| **A_66_P135694** | NA | 0.111697864 | -0.099226198 | 0.322621927 | 16.28025671 | 1.286772978 | 0.2442796 | 0.620164719 | -4.537297358 |
| **A_51_P468762** | Alkbh6 | -0.230507805 | -0.665800807 | 0.204785196 | 10.24179742 | -1.286729152 | 0.244293925 | 0.620164719 | -4.537318747 |
| **A_51_P317214** | Hpdl | 0.106259399 | -0.094456009 | 0.306974807 | 7.880421117 | 1.28638164 | 0.244407534 | 0.620164719 | -4.537488341 |
| **A_55_P2086631** | NA | -0.169124754 | -0.488601056 | 0.150351547 | 14.64325792 | -1.286327988 | 0.244425078 | 0.620164719 | -4.537514523 |
| **A_52_P468068** | Tchh | -0.480333487 | -1.38772207 | 0.427055095 | 8.013347019 | -1.286271787 | 0.244443457 | 0.620164719 | -4.537541948 |
| **A_66_P121053** | Sftpd | -1.266752993 | -3.659837063 | 1.126331077 | 8.384045157 | -1.286225669 | 0.244458539 | 0.620164719 | -4.537564454 |
| **A_51_P430327** | Gmpr2 | -0.088176108 | -0.25476634 | 0.078414124 | 9.208563885 | -1.286129155 | 0.244490106 | 0.620164719 | -4.53761155 |
| **A_51_P152155** | Glt8d1 | -0.167917222 | -0.485208519 | 0.149374076 | 10.42427547 | -1.285938706 | 0.244552406 | 0.620164719 | -4.537704481 |
| **A_55_P1981704** | Ccdc113 | -0.569099993 | -1.644476918 | 0.506276932 | 7.924557482 | -1.285911427 | 0.24456133 | 0.620164719 | -4.537717791 |
| **A_51_P433789** | Agr3 | -0.210236599 | -0.607510059 | 0.18703686 | 7.746725004 | -1.285884497 | 0.244570141 | 0.620164719 | -4.537730932 |
| **A_52_P177490** | Cdx2 | -0.168557389 | -0.48710433 | 0.149989553 | 7.713589025 | -1.285752994 | 0.24461317 | 0.620164719 | -4.537795096 |
| **A_51_P131800** | Cyba | -0.281303624 | -0.812941162 | 0.250333913 | 10.2167402 | -1.285709579 | 0.244627377 | 0.620164719 | -4.537816279 |
| **A_51_P284823** | Polr1b | 0.172132243 | -0.153183781 | 0.497448267 | 9.42127525 | 1.28570094 | 0.244630204 | 0.620164719 | -4.537820495 |
| **A_51_P248644** | Tdp1 | 0.117372611 | -0.104454619 | 0.33919984 | 10.41545276 | 1.285686543 | 0.244634915 | 0.620164719 | -4.537827519 |
| **A_55_P1989698** | Mmp11 | -0.653611768 | -1.889001848 | 0.581778311 | 8.697311583 | -1.285579513 | 0.244669944 | 0.620164719 | -4.53787974 |
| **A_66_P124788** | Tbcb | -0.221815037 | -0.641068359 | 0.197438285 | 10.27153244 | -1.285575717 | 0.244671186 | 0.620164719 | -4.537881592 |
| **A_55_P2077053** | Itih4 | -0.282006337 | -0.815053218 | 0.251040544 | 11.77470844 | -1.285513524 | 0.244691543 | 0.620164719 | -4.537911936 |
| **A_52_P505501** | Nsmce2 | 0.126351059 | -0.112498637 | 0.365200755 | 8.659330206 | 1.285397296 | 0.24472959 | 0.620164719 | -4.537968642 |
| **A_55_P2159026** | Rpl22l1 | 0.08409911 | -0.074884278 | 0.243082499 | 15.12976066 | 1.28535426 | 0.244743678 | 0.620164719 | -4.537989638 |
| **A_55_P2031112** | Dhps | 0.162022747 | -0.144291807 | 0.4683373 | 10.7047633 | 1.285261648 | 0.244774 | 0.620164719 | -4.53803482 |
| **A_51_P225793** | Prr5l | -0.221233754 | -0.639639628 | 0.19717212 | 10.24314703 | -1.284803777 | 0.244923957 | 0.620325075 | -4.538258185 |
| **A_51_P317317** | Gatd1 | -0.093541638 | -0.270485465 | 0.083402189 | 8.239531646 | -1.284555121 | 0.245005427 | 0.620325075 | -4.538379476 |
| **A_51_P499891** | Krit1 | 0.125753465 | -0.112133612 | 0.363640542 | 8.581442213 | 1.28449465 | 0.245025244 | 0.620325075 | -4.538408972 |
| **A_51_P151835** | Tmtc3 | 0.103999883 | -0.092742808 | 0.300742575 | 8.855075527 | 1.284450677 | 0.245039655 | 0.620325075 | -4.53843042 |
| **A_55_P1961135** | Atg4d | 0.139952368 | -0.124846123 | 0.404750859 | 13.22405192 | 1.284245153 | 0.24510702 | 0.620325075 | -4.538530664 |
| **A_51_P331752** | Ccl11 | -0.498133883 | -1.440779261 | 0.444511495 | 7.95822196 | -1.284047027 | 0.245171976 | 0.620325075 | -4.538627294 |
| **A_51_P407193** | Clp1 | 0.182720544 | -0.163054331 | 0.528495418 | 8.897109498 | 1.284035811 | 0.245175654 | 0.620325075 | -4.538632764 |
| **A_66_P134383** | NA | 0.194532989 | -0.173603106 | 0.562669085 | 10.81839547 | 1.284008941 | 0.245184464 | 0.620325075 | -4.538645869 |
| **A_52_P173442** | Wscd2 | -0.547105958 | -1.582485465 | 0.488273549 | 8.13738058 | -1.283970567 | 0.245197048 | 0.620325075 | -4.538664584 |
| **A_51_P309534** | Pex14 | 0.170096416 | -0.151823249 | 0.49201608 | 10.29677297 | 1.283898952 | 0.245220532 | 0.620325075 | -4.53869951 |
| **A_51_P413785** | Commd3 | 0.09350425 | -0.083482932 | 0.270491432 | 11.52567212 | 1.283727147 | 0.245276881 | 0.620325075 | -4.538783296 |
| **A_55_P2004230** | Emc6 | -0.112168359 | -0.324512566 | 0.100175848 | 11.78784479 | -1.283551027 | 0.245334656 | 0.620325075 | -4.538869182 |
| **A_52_P471143** | Pusl1 | 0.133304887 | -0.119064473 | 0.385674246 | 9.18720047 | 1.283490427 | 0.245354539 | 0.620325075 | -4.538898733 |
| **A_55_P2051716** | Tbx1 | -0.634204742 | -1.835054268 | 0.566644784 | 7.980280731 | -1.283287852 | 0.245421012 | 0.620325075 | -4.538997514 |
| **A_55_P2172440** | Scarf2 | -0.632773528 | -1.83101949 | 0.565472433 | 11.29769377 | -1.2831739 | 0.245458412 | 0.620325075 | -4.539053078 |
| **A_55_P2022369** | Lbh | -0.525808353 | -1.521515206 | 0.469898499 | 9.03568686 | -1.283155159 | 0.245464563 | 0.620325075 | -4.539062216 |
| **A_55_P2153541** | Sik3 | -0.17231126 | -0.498625854 | 0.154003335 | 11.65885732 | -1.283099538 | 0.245482821 | 0.620325075 | -4.539089336 |
| **A_51_P281380** | Tspan5 | -0.277985938 | -0.804458467 | 0.248486591 | 8.981782023 | -1.283010769 | 0.245511961 | 0.620325075 | -4.539132618 |
| **A_51_P193000** | Bbs4 | -0.232447503 | -0.672762177 | 0.207867172 | 8.521312336 | -1.282758539 | 0.245594778 | 0.620325075 | -4.539255596 |

| | | | | | | | | | |
|---|---|---|---|---|---|---|---|---|---|
| **A_55_P2015430** | Gpn2 | 0.107152431 | -0.095830683 | 0.310135544 | 10.01497269 | 1.282700623 | 0.245613797 | 0.620325075 | -4.539283833 |
| **A_55_P2137527** | Fam183b | -1.151596486 | -3.333658254 | 1.030465282 | 8.280576899 | -1.282379243 | 0.245719361 | 0.620325075 | -4.539440511 |
| **A_55_P2155616** | Gm9457 | 0.120920717 | -0.108217067 | 0.350058501 | 15.37456094 | 1.282292772 | 0.245747771 | 0.620325075 | -4.539482665 |
| **A_55_P2040773** | NA | 0.304164787 | -0.272250881 | 0.880580455 | 9.655061203 | 1.282201695 | 0.245777698 | 0.620325075 | -4.539527063 |
| **A_55_P2136957** | Wbp1l | -0.212894487 | -0.616383133 | 0.190594158 | 13.32993583 | -1.282083429 | 0.245816563 | 0.620325075 | -4.539584714 |
| **A_51_P112627** | St6galnac2 | -0.860358278 | -2.491397436 | 0.770680879 | 8.528812238 | -1.281734704 | 0.245931194 | 0.620325075 | -4.539297225 |
| **A_55_P1955417** | Atad3a | 0.162167612 | -0.145281945 | 0.469617168 | 13.34113787 | 1.281661798 | 0.245955165 | 0.620325075 | -4.539790231 |
| **A_51_P354744** | Cox15 | 0.21215768 | -0.190101515 | 0.614416875 | 9.521854399 | 1.281551214 | 0.245991529 | 0.620325075 | -4.53984413 |
| **A_51_P238448** | Ccnd3 | 0.272392321 | -0.244154485 | 0.788939127 | 12.55053086 | 1.281351758 | 0.246057129 | 0.620325075 | -4.539941341 |
| **A_55_P1978311** | NA | 0.118447801 | -0.106170167 | 0.343065769 | 14.40500399 | 1.281343876 | 0.246059721 | 0.620325075 | -4.539945182 |
| **A_51_P248865** | Foxf2 | -0.621718325 | -1.800713626 | 0.557276976 | 7.943413111 | -1.281341178 | 0.246060609 | 0.620325075 | -4.539946497 |
| **A_55_P1963300** | Vsig10 | -0.234716264 | -0.679885543 | 0.210453015 | 8.524822345 | -1.281153557 | 0.246122331 | 0.620325075 | -4.540037935 |
| **A_55_P2098354** | Shf | 0.101919311 | -0.091397013 | 0.295235635 | 9.187071194 | 1.281064928 | 0.246151492 | 0.620325075 | -4.540081127 |
| **A_55_P2027117** | Rgs3 | -0.26661684 | -0.772362822 | 0.239129142 | 8.892297698 | -1.280968129 | 0.246183344 | 0.620325075 | -4.5401283 |
| **A_55_P2066384** | Akt1 | -0.290366439 | -0.841193592 | 0.260460713 | 12.2486665 | -1.280897174 | 0.246206695 | 0.620325075 | -4.540162877 |
| **A_52_P222967** | Adck2 | -0.084238535 | -0.244116161 | 0.07563909 | 8.603043772 | -1.280283966 | 0.246408579 | 0.620325075 | -4.540461675 |
| **A_51_P200517** | Rasgef1a | -0.608418077 | -1.763202724 | 0.54636657 | 8.054609167 | -1.280219084 | 0.246429948 | 0.620325075 | -4.540493287 |
| **A_55_P2139256** | Rps6ka1 | 0.129513307 | -0.116308501 | 0.375335114 | 13.48166131 | 1.280198085 | 0.246436865 | 0.620325075 | -4.540503519 |
| **A_55_P2001891** | Prps1 | 0.216171056 | -0.194132944 | 0.626475056 | 11.61233681 | 1.280191625 | 0.246438992 | 0.620325075 | -4.540506666 |
| **A_51_P491227** | Suclg1 | -0.194541927 | -0.563811326 | 0.174727473 | 12.9983968 | -1.280127069 | 0.246460257 | 0.620325075 | -4.540538119 |
| **A_55_P2052420** | Setd5 | -0.212742769 | -0.616631484 | 0.191145946 | 11.90764896 | -1.279900704 | 0.246534832 | 0.620325075 | -4.540648403 |
| **A_66_P134704** | Gm4952 | -0.52402881 | -1.51891879 | 0.47086117 | 11.74424786 | -1.279862448 | 0.246547438 | 0.620325075 | -4.540667041 |
| **A_52_P1042732** | Akap5 | -1.550423804 | -4.494338293 | 1.393490686 | 8.689729951 | -1.279700753 | 0.246600722 | 0.620325075 | -4.540745813 |
| **A_55_P2152468** | NA | 0.154042386 | -0.13845668 | 0.446541451 | 10.08860132 | 1.27967281 | 0.246609932 | 0.620325075 | -4.540759426 |
| **A_51_P431785** | Myom2 | -0.512495704 | -1.485643594 | 0.460652185 | 8.060527273 | -1.279659937 | 0.246614174 | 0.620325075 | -4.540765697 |
| **A_55_P2022585** | Pira6 | -0.205433412 | -0.595521675 | 0.184654852 | 8.795263064 | -1.279650538 | 0.246617272 | 0.620325075 | -4.540770275 |
| **A_52_P563617** | Ssbp4 | -0.200043094 | -0.580008982 | 0.179922794 | 10.4110812 | -1.279269787 | 0.246742792 | 0.620325075 | -4.540955749 |
| **A_52_P216427** | Nkx2-1 | -1.150825032 | -3.33687733 | 1.035227266 | 8.205581978 | -1.279180826 | 0.246772127 | 0.620325075 | -4.540529173 |
| **A_55_P2009121** | Agrn | -0.623739148 | -1.808584356 | 0.56110606 | 11.87360375 | -1.279159131 | 0.246779281 | 0.620325075 | -4.541009649 |
| **A_52_P428745** | Camk2d | -0.187562779 | -0.543890802 | 0.168765244 | 9.749579652 | -1.279027465 | 0.246822706 | 0.620325075 | -4.541073781 |
| **A_55_P2175615** | Csde1 | 0.120537423 | -0.108487686 | 0.349562532 | 12.39859051 | 1.278857021 | 0.246878929 | 0.620325075 | -4.541156797 |
| **A_55_P2029121** | Pex11g | 0.145854783 | -0.131281987 | 0.422991553 | 9.023437374 | 1.278820866 | 0.246890856 | 0.620325075 | -4.541174406 |
| **A_55_P1984284** | Gapdh | -0.144605599 | -0.419369041 | 0.130157843 | 15.05995457 | -1.278819793 | 0.246891211 | 0.620325075 | -4.541174929 |
| **A_55_P2090194** | NA | -0.432235161 | -1.253627395 | 0.389157072 | 12.8053992 | -1.278652964 | 0.246946255 | 0.620325075 | -4.54125618 |
| **A_55_P2045086** | NA | 0.152958944 | -0.137730641 | 0.44364853 | 12.46983918 | 1.278582022 | 0.246969665 | 0.620325075 | -4.54129073 |
| **A_51_P453167** | Naxd | -0.355217533 | -1.030323456 | 0.31988839 | 11.94763648 | -1.278514521 | 0.246991941 | 0.620325075 | -4.541323604 |
| **A_55_P2110567** | Proca1 | -0.305922921 | -0.887348202 | 0.27550236 | 8.114967204 | -1.278501655 | 0.246996188 | 0.620325075 | -4.54132987 |
| **A_55_P2141851** | Pls3 | -0.706979106 | -2.051018462 | 0.637060249 | 9.904378711 | -1.278138105 | 0.247116197 | 0.620325075 | -4.541506912 |
| **A_66_P129976** | Apba1 | -0.097717948 | -0.283523333 | 0.088087437 | 7.938006437 | -1.277906947 | 0.24719253 | 0.620325075 | -4.541619472 |
| **A_55_P2059765** | NA | -0.831049248 | -2.411306839 | 0.749208343 | 10.40636465 | -1.277856436 | 0.247209213 | 0.620325075 | -4.541644067 |
| **A_55_P2069765** | Kiss1 | -0.418750143 | -1.215029027 | 0.377528741 | 8.479259486 | -1.277829641 | 0.247218063 | 0.620325075 | -4.541657114 |
| **A_52_P133436** | Plekhm1 | 0.105647547 | -0.095255711 | 0.306550804 | 9.433339534 | 1.277778649 | 0.247234906 | 0.620325075 | -4.541681943 |
| **A_55_P2121613** | Sox4 | -0.389851354 | -1.131235318 | 0.35153261 | 9.062208804 | -1.277729872 | 0.247251018 | 0.620325075 | -4.541705693 |
| **A_52_P647607** | Ubr4 | 0.118019886 | -0.106468564 | 0.342508337 | 8.366636433 | 1.277451376 | 0.24734303 | 0.620325075 | -4.541841289 |
| **A_55_P1988183** | Cdkn2b | -0.841948357 | -2.443443061 | 0.759546347 | 8.113545936 | -1.277447721 | 0.247344237 | 0.620325075 | -4.541843068 |
| **A_51_P158018** | Bend5 | -0.365174825 | -1.059857649 | 0.329507999 | 8.3272004 | -1.277313368 | 0.247388637 | 0.620325075 | -4.541908479 |
| **A_55_P1999760** | Lrrc51 | -0.62003569 | -1.799554828 | 0.559483448 | 8.78603994 | -1.277305804 | 0.247391137 | 0.620325075 | -4.541912162 |
| **A_55_P2152547** | Myl3 | -0.506709268 | -1.470647501 | 0.457228965 | 8.391627153 | -1.27729976 | 0.247393135 | 0.620325075 | -4.541915104 |
| **A_52_P58257** | Mrps6 | -0.103811519 | -0.301317999 | 0.093694961 | 11.64902017 | -1.277166115 | 0.247437308 | 0.620325075 | -4.541980168 |
| **A_51_P437978** | Agtr2 | -0.153707707 | -0.446225944 | 0.13881053 | 7.728336408 | -1.276808854 | 0.247555426 | 0.620325075 | -4.542154086 |

| | | | | | | | | | |
|---|---|---|---|---|---|---|---|---|---|
| **A_55_P2072841** | NA | 0.208783664 | -0.188553583 | 0.606120912 | 10.34602812 | 1.276792803 | 0.247560734 | 0.620325075 | -4.542161899 |
| **A_51_P272106** | Cirbp | -0.149309416 | -0.433467624 | 0.134848791 | 10.44595868 | -1.276762687 | 0.247570694 | 0.620325075 | -4.542176559 |
| **A_55_P1960999** | Pigr | 0.189220968 | -0.170906489 | 0.549348425 | 12.54869303 | 1.276721625 | 0.247584273 | 0.620325075 | -4.542196547 |
| **A_51_P175758** | Atg101 | -0.124440823 | -0.361304139 | 0.112422493 | 11.26861685 | -1.276580654 | 0.2476309 | 0.620325075 | -4.542265167 |
| **A_52_P572447** | Agpat5 | 0.174767592 | -0.157892126 | 0.50742731 | 11.02309253 | 1.276567774 | 0.24763516 | 0.620325075 | -4.542271436 |
| **A_51_P400659** | Bcap29 | 0.122112803 | -0.11032799 | 0.354553597 | 10.09256673 | 1.276532984 | 0.247646668 | 0.620325075 | -4.54228837 |
| **A_55_P2134566** | NA | 0.141084199 | -0.127474676 | 0.409643074 | 9.444916398 | 1.276503636 | 0.247656377 | 0.620325075 | -4.542302655 |
| **A_51_P427768** | Diaph1 | 0.116891097 | -0.105619547 | 0.339401741 | 8.674231801 | 1.27647947 | 0.247664371 | 0.620325075 | -4.542314418 |
| **A_55_P2074536** | Ppm1b | 0.203940594 | -0.184283912 | 0.592165099 | 12.89320218 | 1.27645034 | 0.247674008 | 0.620325075 | -4.542328597 |
| **A_55_P2071384** | Crh | -0.762629307 | -2.214425301 | 0.689166687 | 7.996964272 | -1.276412609 | 0.247686491 | 0.620325075 | -4.542346962 |
| **A_52_P177293** | Snx24 | 0.117709488 | -0.106377566 | 0.341796543 | 8.513102158 | 1.276373835 | 0.24769932 | 0.620325075 | -4.542365835 |
| **A_52_P458870** | Slc25a16 | 0.14486674 | -0.13092305 | 0.420656529 | 8.592619924 | 1.276361488 | 0.247703405 | 0.620325075 | -4.542371844 |
| **A_55_P1998962** | Oas1b | -0.157583802 | -0.457590885 | 0.142423281 | 13.79278147 | -1.276330817 | 0.247713553 | 0.620325075 | -4.542386773 |
| **A_55_P2009498** | Ccdc28a | -0.099878816 | -0.290028855 | 0.090271222 | 9.870708132 | -1.276321717 | 0.247716564 | 0.620325075 | -4.542391202 |
| **A_52_P229709** | Ube2d3 | 0.116749258 | -0.105529175 | 0.33902769 | 11.22774159 | 1.276262447 | 0.247736176 | 0.620325075 | -4.542420049 |
| **A_51_P412270** | Rapgef6 | 0.132233781 | -0.11954089 | 0.384008451 | 8.490068141 | 1.276184801 | 0.24776187 | 0.620325075 | -4.542457841 |
| **A_55_P2106255** | Mbnl3 | 0.138700117 | -0.125414499 | 0.402814734 | 10.50389196 | 1.276049673 | 0.247806592 | 0.620325075 | -4.542523607 |
| **A_55_P1955382** | Rgma | 0.245421722 | -0.22199593 | 0.712839373 | 9.278073919 | 1.275824912 | 0.247880995 | 0.620325075 | -4.542632992 |
| **A_52_P284821** | Slco4c1 | -0.188452778 | -0.547370874 | 0.170465319 | 8.247553868 | -1.275822849 | 0.247881678 | 0.620325075 | -4.542633996 |
| **A_55_P1972097** | NA | -0.216336777 | -0.628390092 | 0.195716537 | 16.24849546 | -1.275734009 | 0.247911092 | 0.620325075 | -4.542677231 |
| **A_55_P2084797** | Aff3 | 0.231495994 | -0.209522106 | 0.672514093 | 8.769519891 | 1.275470021 | 0.247998514 | 0.620325075 | -4.542805695 |
| **A_51_P256304** | Mrpl37 | 0.132852739 | -0.12024862 | 0.385954099 | 8.812635871 | 1.275437626 | 0.248009244 | 0.620325075 | -4.542821459 |
| **A_51_P520639** | Mslnl | -0.342471408 | -0.994945131 | 0.310002315 | 7.82529795 | -1.275394331 | 0.248023585 | 0.620325075 | -4.542842527 |
| **A_51_P499153** | Cog3 | 0.120690528 | -0.109249773 | 0.350630828 | 9.17997567 | 1.275384925 | 0.2480267 | 0.620325075 | -4.542847103 |
| **A_66_P102090** | Pkmyt1 | 0.159586014 | -0.144462288 | 0.463634316 | 11.34411181 | 1.275367759 | 0.248032387 | 0.620325075 | -4.542855456 |
| **A_51_P472274** | Sox18 | -0.362649591 | -1.053626235 | 0.328327054 | 9.134270549 | -1.275284298 | 0.248060035 | 0.620325075 | -4.542896068 |
| **A_51_P260300** | Ikbkg | 0.175131963 | -0.158590213 | 0.508854139 | 8.713615914 | 1.275156641 | 0.248102328 | 0.620325075 | -4.542958183 |
| **A_51_P426195** | Nppb | -0.555690009 | -1.614714786 | 0.503334769 | 7.918761362 | -1.274998448 | 0.248154748 | 0.620325075 | -4.543035154 |
| **A_55_P2047626** | Cutal | 0.144547855 | -0.130929609 | 0.420025319 | 8.943157995 | 1.274995832 | 0.248155615 | 0.620325075 | -4.543036427 |
| **A_51_P518298** | Rnf187 | 0.121122765 | -0.10972784 | 0.351973369 | 13.1310349 | 1.274905365 | 0.248185597 | 0.620325075 | -4.543080443 |
| **A_55_P2230968** | NA | 0.12543421 | -0.113638062 | 0.364506481 | 8.667478873 | 1.274881971 | 0.24819335 | 0.620325075 | -4.543091825 |
| **A_55_P1953377** | NA | -0.644389707 | -1.872782013 | 0.5840026 | 9.737798059 | -1.274660993 | 0.2482666 | 0.620325075 | -4.543199336 |
| **A_51_P272563** | Naa25 | 0.19721433 | -0.178779528 | 0.573208187 | 9.995526397 | 1.27450313 | 0.248318941 | 0.620325075 | -4.543276136 |
| **A_55_P2101426** | Mrps25 | 0.130058905 | -0.117910332 | 0.378028143 | 11.54061172 | 1.274457849 | 0.248333956 | 0.620325075 | -4.543298165 |
| **A_55_P1962951** | Stx5a | -0.182514416 | -0.530555629 | 0.165526797 | 12.40967294 | -1.274235468 | 0.248407708 | 0.620325075 | -4.543406346 |
| **A_66_P139333** | Atf6 | -0.091675697 | -0.266504005 | 0.083152611 | 8.392843709 | -1.274164958 | 0.248431097 | 0.620325075 | -4.543440645 |
| **A_55_P2006226** | NA | 0.083118519 | -0.07541687 | 0.241653909 | 16.4822681 | 1.273956984 | 0.248500094 | 0.620325075 | -4.54354181 |
| **A_55_P2058122** | Kyat3 | -0.283434982 | -0.824080793 | 0.257210828 | 11.56935484 | -1.273866152 | 0.248530233 | 0.620325075 | -4.543585992 |
| **A_66_P124858** | Diras2 | -0.145446388 | -0.422890316 | 0.131997541 | 8.467747266 | -1.273828329 | 0.248542785 | 0.620325075 | -4.543604389 |
| **A_51_P245631** | Rftn2 | -0.252296118 | -0.733596294 | 0.229004058 | 8.278627225 | -1.273731053 | 0.248575068 | 0.620325075 | -4.543651703 |
| **A_55_P2112445** | Tbx4 | -1.264029143 | -3.67549098 | 1.147432695 | 8.319241669 | -1.27367869 | 0.248592447 | 0.620325075 | -4.543677172 |
| **A_55_P2102679** | NA | 0.171829615 | -0.155997115 | 0.499656346 | 8.426181318 | 1.273611127 | 0.248614872 | 0.620325075 | -4.543710033 |
| **A_51_P146560** | Msln | -0.996573122 | -2.898173574 | 0.905027329 | 10.08879199 | -1.273424165 | 0.248676938 | 0.620325075 | -4.543800963 |
| **A_52_P656565** | Pex6 | -0.138323115 | -0.40226374 | 0.12561751 | 10.28244383 | -1.273420123 | 0.24867828 | 0.620325075 | -4.543802929 |
| **A_55_P2028922** | Stoml2 | 0.120905947 | -0.109801307 | 0.3516132 | 12.30629216 | 1.273413948 | 0.24868033 | 0.620325075 | -4.543805933 |
| **A_51_P477682** | Prss12 | -0.584717304 | -1.700454152 | 0.531019544 | 9.082423836 | -1.27340737 | 0.248682514 | 0.620325075 | -4.543809131 |
| **A_52_P355057** | Wdr4 | 0.18439617 | -0.167487601 | 0.536279941 | 11.14176945 | 1.273314979 | 0.248713191 | 0.620325075 | -4.543854065 |
| **A_55_P2037338** | Plcd3 | -0.186662327 | -0.542907558 | 0.169582903 | 8.222219171 | -1.273182927 | 0.248757042 | 0.620325075 | -4.543918285 |
| **A_55_P2079568** | NA | 0.121071826 | -0.110001555 | 0.352145208 | 8.589705905 | 1.273140587 | 0.248771103 | 0.620325075 | -4.543938876 |
| **A_55_P1989276** | Romo1 | -0.113233288 | -0.329380258 | 0.102913681 | 14.02103175 | -1.272940655 | 0.248837512 | 0.620325075 | -4.544036102 |

| | | | | | | | | | |
|---|---|---|---|---|---|---|---|---|---|
| **A_55_P2007601** | Sftpd | -1.972244183 | -5.737176066 | 1.792687701 | 8.718003693 | -1.27287791 | 0.248858356 | 0.620325075 | -4.544066614 |
| **A_51_P472217** | Sapcd2 | 0.212409246 | -0.193088632 | 0.617907123 | 11.13958433 | 1.272823018 | 0.248876593 | 0.620325075 | -4.544093306 |
| **A_55_P2179796** | NA | -0.138946288 | -0.404246167 | 0.126353591 | 13.5806222 | -1.272603422 | 0.248949561 | 0.620325075 | -4.544200086 |
| **A_55_P2186693** | NA | 0.172547935 | -0.156944515 | 0.502040385 | 9.19197165 | 1.272469811 | 0.248993967 | 0.620325075 | -4.544265052 |
| **A_55_P1954511** | Btbd10 | 0.135254426 | -0.123042485 | 0.393551336 | 9.388278182 | 1.27237598 | 0.249025156 | 0.620325075 | -4.544310674 |
| **A_51_P322972** | Hkdc1 | 0.324585286 | -0.295325066 | 0.944495639 | 11.80158253 | 1.272281258 | 0.249056645 | 0.620325075 | -4.544356728 |
| **A_51_P120254** | Fbxl8 | -0.100364998 | -0.292049424 | 0.091319429 | 8.206050523 | -1.272268097 | 0.24906102 | 0.620325075 | -4.544363127 |
| **A_51_P436020** | Tmem167 | 0.200779389 | -0.18270706 | 0.584265838 | 9.334997075 | 1.272190928 | 0.249086677 | 0.620325075 | -4.544400646 |
| **A_55_P1978424** | Bcl2a1d | -0.484161556 | -1.408954599 | 0.440631487 | 9.118213397 | -1.272122509 | 0.249109426 | 0.620325075 | -4.54443391 |
| **A_55_P1960411** | Lrrc23 | -1.069508804 | -3.112410665 | 0.973393057 | 8.2524763 | -1.272096474 | 0.249118083 | 0.620325075 | -4.544446568 |
| **A_55_P2143431** | Nfib | -0.392104286 | -1.141159375 | 0.356950803 | 10.98980641 | -1.271952869 | 0.249165839 | 0.620325075 | -4.544516383 |
| **A_55_P2169963** | Zfp982 | 0.111357745 | -0.101393251 | 0.32410874 | 8.09740458 | 1.271838637 | 0.249203833 | 0.620325075 | -4.544571917 |
| **A_51_P424561** | Zfp93 | -0.149740679 | -0.43585489 | 0.136373531 | 8.204111974 | -1.271696744 | 0.249251034 | 0.620325075 | -4.544640896 |
| **A_66_P129419** | Cox7a2l | -0.28579769 | -0.831900314 | 0.260304934 | 14.22472389 | -1.271650131 | 0.249266542 | 0.620325075 | -4.544663555 |
| **A_52_P120925** | Metap1 | 0.122825145 | -0.111871647 | 0.357521936 | 8.447817498 | 1.271637485 | 0.249270749 | 0.620325075 | -4.544669702 |
| **A_51_P248666** | Cd274 | -0.26325727 | -0.766295211 | 0.239780671 | 8.668378379 | -1.271636017 | 0.249271237 | 0.620325075 | -4.544670416 |
| **A_51_P269084** | Chchd10 | -0.184269146 | -0.536382202 | 0.167843911 | 15.64921997 | -1.27160926 | 0.24928014 | 0.620325075 | -4.544683423 |
| **A_66_P132832** | Myzap | -0.194057058 | -0.564884215 | 0.176770099 | 7.882861067 | -1.271572424 | 0.249292396 | 0.620325075 | -4.544701329 |
| **A_51_P422369** | Odf3b | -0.515630206 | -1.501142028 | 0.469881616 | 8.376468427 | -1.271334117 | 0.249371698 | 0.620325075 | -4.544817168 |
| **A_66_P124776** | NA | 0.188863017 | -0.172115132 | 0.549841165 | 8.837017659 | 1.271303371 | 0.249381931 | 0.620325075 | -4.544832112 |
| **A_51_P513541** | Rfk | 0.170006142 | -0.154937523 | 0.494949808 | 10.64751813 | 1.271275644 | 0.24939116 | 0.620325075 | -4.544845589 |
| **A_52_P365768** | Nsun2 | 0.182173207 | -0.166057761 | 0.530404174 | 12.11756112 | 1.271160248 | 0.249429571 | 0.620325075 | -4.544901679 |
| **A_51_P495379** | Fhod1 | -0.637514548 | -1.856181905 | 0.581152809 | 9.91386849 | -1.271124561 | 0.249441451 | 0.620325075 | -4.544919024 |
| **A_66_P102395** | Elac2 | 0.139171093 | -0.126868714 | 0.4052109 | 8.71303603 | 1.271117221 | 0.249443895 | 0.620325075 | -4.544922592 |
| **A_55_P2065434** | Rprl2 | 0.671142034 | -0.611909205 | 1.954193273 | 11.43610308 | 1.27102364 | 0.24947505 | 0.620325075 | -4.544968076 |
| **A_55_P2218223** | Pacs2 | 0.105140306 | -0.09586843 | 0.306149043 | 8.715667278 | 1.270976419 | 0.249490772 | 0.620325075 | -4.544991027 |
| **A_51_P189985** | Ppan | 0.195174068 | -0.177998763 | 0.5683469 | 11.91419902 | 1.270852905 | 0.2495319 | 0.620347537 | -4.545051057 |
| **A_51_P227077** | Mdh1b | -0.222250235 | -0.647296993 | 0.202796524 | 8.248942751 | -1.270541071 | 0.249635762 | 0.620409161 | -4.545202606 |
| **A_51_P110341** | Scgb3a1 | -1.11410489 | -3.244817519 | 1.016607739 | 8.281283945 | -1.270528435 | 0.249639972 | 0.620409161 | -4.545208746 |
| **A_55_P2084646** | Csrnp3 | 0.274592841 | -0.250601885 | 0.799787567 | 8.619916993 | 1.270433811 | 0.249671496 | 0.620409161 | -4.54525473 |
| **A_51_P314397** | Crip2 | -0.695705102 | -2.026708115 | 0.635297911 | 13.15957975 | -1.270074915 | 0.249791095 | 0.620409161 | -4.545429129 |
| **A_55_P2059986** | Chst14 | -0.117991927 | -0.343730892 | 0.107747038 | 9.168112199 | -1.270073787 | 0.249791471 | 0.620409161 | -4.545429677 |
| **A_55_P2089780** | Mtfr1l | 0.154971091 | -0.141531365 | 0.451473548 | 8.874434723 | 1.27000545 | 0.24981425 | 0.620409161 | -4.545462883 |
| **A_66_P139876** | Psph | 0.403050943 | -0.36809724 | 1.174199126 | 11.20378519 | 1.27000459 | 0.249814536 | 0.620409161 | -4.5454633 |
| **A_55_P2002782** | Vps26a | 0.10061525 | -0.091915475 | 0.293145975 | 10.8656733 | 1.269834007 | 0.249871404 | 0.620409161 | -4.545546185 |
| **A_51_P400375** | Erg28 | 0.165538298 | -0.151252423 | 0.482329019 | 14.1963405 | 1.269723689 | 0.249908187 | 0.620409161 | -4.545599785 |
| **A_55_P2040245** | Piezo2 | -0.341791761 | -0.995928619 | 0.312345096 | 7.994372131 | -1.269627028 | 0.24994042 | 0.620409161 | -4.545646748 |
| **A_55_P2097206** | Camk4 | -0.208442624 | -0.607383412 | 0.190498164 | 7.801659164 | -1.269583528 | 0.249954927 | 0.620409161 | -4.545667883 |
| **A_55_P2158946** | Lama2 | -0.450556398 | -1.312955349 | 0.411842553 | 8.49769067 | -1.269474936 | 0.249991145 | 0.620409161 | -4.545720641 |
| **A_55_P2001063** | Fcf1 | 0.107640751 | -0.098431072 | 0.313712573 | 12.407887 | 1.269232824 | 0.250071912 | 0.620409161 | -4.545838262 |
| **A_55_P2026275** | Ppp1r1b | -0.124077842 | -0.361634077 | 0.113478392 | 8.481990597 | -1.269144256 | 0.250101463 | 0.620409161 | -4.545881288 |
| **A_52_P276343** | Pum3 | 0.106554828 | -0.097454791 | 0.310564446 | 8.570644962 | 1.26912873 | 0.250106644 | 0.620409161 | -4.54588883 |
| **A_51_P272123** | Ndufa10 | -0.089392202 | -0.260565669 | 0.081781266 | 11.40489936 | -1.268955448 | 0.250164471 | 0.620409161 | -4.545973006 |
| **A_55_P1981829** | Rhox8 | 0.158115689 | -0.14468585 | 0.460917228 | 12.03196882 | 1.268820175 | 0.250209621 | 0.620409161 | -4.546038715 |
| **A_51_P195506** | Csf1 | 0.146755472 | -0.134307458 | 0.427818401 | 9.380096599 | 1.268743771 | 0.250235126 | 0.620409161 | -4.544151208 |
| **A_55_P2041320** | NA | -0.109607631 | -0.319545806 | 0.100330544 | 12.08844162 | -1.26862295 | 0.250275463 | 0.620409161 | -4.546134513 |
| **A_55_P2081178** | Msl3l2 | -0.298642093 | -0.870737353 | 0.273453167 | 8.319599001 | -1.268428116 | 0.250340521 | 0.620409161 | -4.546229145 |
| **A_52_P322181** | Adrb1 | -0.157810538 | -0.460139718 | 0.144518641 | 8.167337055 | -1.268350034 | 0.250366598 | 0.620409161 | -4.546267068 |
| **A_55_P1984496** | Usp48 | 0.14999728 | -0.137454096 | 0.437448655 | 9.797420998 | 1.267950155 | 0.250500183 | 0.620409161 | -4.546461272 |
| **A_66_P120414** | Adcy5 | -0.087410141 | -0.254924118 | 0.080103836 | 7.987482371 | -1.267926151 | 0.250508203 | 0.620409161 | -4.546472928 |

| | | | | | | | | | |
|---|---|---|---|---|---|---|---|---|---|
| **A_55_P2122605** | Cbr2 | -2.423846299 | -7.069397688 | 2.22170509 | 9.000114901 | -1.26780073 | 0.250550116 | 0.620409161 | -4.546533835 |
| **A_55_P2078008** | Ddx46 | 0.141829501 | -0.130028992 | 0.413687994 | 9.833299976 | 1.267671888 | 0.250593178 | 0.620409161 | -4.546596401 |
| **A_55_P1974567** | Plch2 | -0.433785892 | -1.265373257 | 0.397801474 | 7.951801001 | -1.26750806 | 0.250647943 | 0.620409161 | -4.546675953 |
| **A_55_P2049582** | Apbb3 | -0.089647249 | -0.261534817 | 0.082240319 | 9.938306684 | -1.267289074 | 0.250721162 | 0.620409161 | -4.546782282 |
| **A_51_P357085** | Dctn6 | 0.105834114 | -0.097101773 | 0.308770001 | 12.28384062 | 1.267214152 | 0.250746217 | 0.620409161 | -4.54681866 |
| **A_51_P196895** | Ist1 | -0.114272739 | -0.333402222 | 0.104856743 | 9.500564223 | -1.267141207 | 0.250770613 | 0.620409161 | -4.546854077 |
| **A_55_P2093231** | NA | -1.254411903 | -3.659945481 | 1.151121674 | 8.368829794 | -1.267103038 | 0.250783379 | 0.620409161 | -4.546872608 |
| **A_66_P115443** | Snx4 | -0.117130683 | -0.34175346 | 0.107492095 | 8.479962104 | -1.267068429 | 0.250794955 | 0.620409161 | -4.546889411 |
| **A_55_P1959550** | Carmil2 | 0.137387148 | -0.126088696 | 0.400862993 | 8.693691156 | 1.267034653 | 0.250806253 | 0.620409161 | -4.54690581 |
| **A_55_P1962747** | H2-Ab1 | -0.202724358 | -0.591503931 | 0.186055215 | 10.58180796 | -1.267026457 | 0.250808994 | 0.620409161 | -4.546909789 |
| **A_51_P205215** | Klhl30 | -0.112234316 | -0.327496399 | 0.103027767 | 7.766205928 | -1.266897022 | 0.250852294 | 0.620409161 | -4.546972629 |
| **A_55_P1984481** | Mfn2 | 0.092678273 | -0.08508525 | 0.270441797 | 8.53546491 | 1.26683039 | 0.250874587 | 0.620409161 | -4.547004978 |
| **A_51_P160439** | Crybg3 | -0.239436691 | -0.698736429 | 0.219863047 | 9.094874249 | -1.26671144 | 0.250914387 | 0.620409161 | -4.547062725 |
| **A_51_P513032** | Trps1 | -0.316548238 | -0.923782877 | 0.290686401 | 8.342242767 | -1.266678945 | 0.250925261 | 0.620409161 | -4.5470785 |
| **A_52_P365896** | Pa2g4 | 0.140922153 | -0.129409637 | 0.411253943 | 8.716261902 | 1.266675404 | 0.250926446 | 0.620409161 | -4.547080219 |
| **A_55_P1964193** | Map1b | -0.171846795 | -0.501515751 | 0.157822162 | 7.778103568 | -1.266620672 | 0.250944762 | 0.620409161 | -4.547106789 |
| **A_55_P2067181** | Usp21 | -0.086051292 | -0.251145861 | 0.079043278 | 9.684965804 | -1.266507583 | 0.250982611 | 0.620409161 | -4.547161688 |
| **A_55_P2074647** | Snap47 | -0.159903719 | -0.466752805 | 0.146945367 | 9.7540962 | -1.2662426 | 0.251071316 | 0.620409161 | -4.547290316 |
| **A_55_P2007457** | Muc5b | -0.405930354 | -1.185012164 | 0.373151456 | 7.944861901 | -1.266052315 | 0.251135032 | 0.620409161 | -4.547382678 |
| **A_55_P2039436** | Ilkap | -0.094525173 | -0.275942634 | 0.086892288 | 11.90575306 | -1.266052123 | 0.251135096 | 0.620409161 | -4.547382771 |
| **A_55_P2186668** | Cfap126 | -0.446569076 | -1.303696154 | 0.410558002 | 8.883873096 | -1.265979344 | 0.25115947 | 0.620409161 | -4.547418096 |
| **A_51_P167843** | Timm17a | 0.090990608 | -0.083653302 | 0.265634518 | 11.65769542 | 1.265978447 | 0.25115977 | 0.620409161 | -4.547418531 |
| **A_52_P371718** | Tgs1 | 0.146773558 | -0.134944439 | 0.428491555 | 9.894771445 | 1.265949608 | 0.251169429 | 0.620409161 | -4.547432529 |
| **A_55_P2172807** | NA | -0.232211779 | -0.677950543 | 0.213526984 | 7.788414045 | -1.265863948 | 0.25119812 | 0.620409161 | -4.547474105 |
| **A_55_P2088601** | Fbln1 | -0.152683815 | -0.445770815 | 0.140403185 | 7.768944473 | -1.265842394 | 0.251205339 | 0.620409161 | -4.547484566 |
| **A_52_P552547** | Lppos | -0.100363079 | -0.293033316 | 0.092307157 | 8.815579254 | -1.265734261 | 0.251241562 | 0.620409161 | -4.547537047 |
| **A_55_P2120807** | Dld | 0.289104429 | -0.265913228 | 0.844122087 | 10.68287488 | 1.265701032 | 0.251252695 | 0.620409161 | -4.547553175 |
| **A_52_P216672** | Klk8 | -0.232710795 | -0.679542415 | 0.214120826 | 8.998875609 | -1.265481556 | 0.251326233 | 0.620409161 | -4.547659689 |
| **A_55_P2088038** | Vps13d | 0.149869444 | -0.137909343 | 0.43764823 | 9.401375657 | 1.265428197 | 0.251344114 | 0.620409161 | -4.547685584 |
| **A_52_P522864** | Zfp442 | 0.173889626 | -0.160038914 | 0.507818166 | 11.07083828 | 1.265328599 | 0.251377494 | 0.620409161 | -4.547733918 |
| **A_55_P2176792** | Sh3gl3 | -0.097583624 | -0.285003158 | 0.089835909 | 7.777069458 | -1.265159498 | 0.251434176 | 0.620409161 | -4.547815977 |
| **A_55_P2084723** | NA | 0.119026072 | -0.109615398 | 0.347667541 | 8.383798188 | 1.264941052 | 0.251507416 | 0.620409161 | -4.547921977 |
| **A_66_P125226** | NA | 0.095058711 | -0.087544092 | 0.277661513 | 12.58698142 | 1.264933427 | 0.251509973 | 0.620409161 | -4.547925677 |
| **A_55_P1992910** | Gyg | -0.974346715 | -2.846254122 | 0.897560692 | 12.06577772 | -1.264772317 | 0.251564001 | 0.620409161 | -4.54800385 |
| **A_55_P1959985** | Alas1 | 0.175909389 | -0.162060386 | 0.513879164 | 9.283109246 | 1.264719886 | 0.251581586 | 0.620409161 | -4.548029289 |
| **A_55_P2093705** | Meig1 | -0.632325085 | -1.847217034 | 0.582566864 | 7.967067034 | -1.264695422 | 0.251589792 | 0.620409161 | -4.54804116 |
| **A_55_P1990810** | 4932412D23Rik | 0.102660433 | -0.094591659 | 0.299912526 | 8.061208846 | 1.264633446 | 0.25161058 | 0.620409161 | -4.548071229 |
| **A_55_P2068356** | 1700016K19Rik | -0.743801534 | -2.172995902 | 0.685392834 | 8.40622748 | -1.264587727 | 0.251625916 | 0.620409161 | -4.548093412 |
| **A_55_P2042655** | NA | -0.313922017 | -0.91715074 | 0.289306706 | 10.71295477 | -1.264511991 | 0.251651323 | 0.620409161 | -4.548130157 |
| **A_55_P2167984** | NA | -0.170531228 | -0.498229559 | 0.157167102 | 10.13072558 | -1.264482665 | 0.251661162 | 0.620409161 | -4.548144385 |
| **A_55_P2081378** | NA | 0.103597234 | -0.09549904 | 0.302693508 | 15.78848283 | 1.264352635 | 0.25170479 | 0.620409161 | -4.54820747 |
| **A_55_P2094445** | Ms4a10 | -0.499213713 | -1.458626372 | 0.460198946 | 7.932807648 | -1.264341087 | 0.251708665 | 0.620409161 | -4.548213073 |
| **A_55_P2076846** | Tead2 | -0.398572351 | -1.164603565 | 0.367458862 | 9.80442807 | -1.264281816 | 0.251728554 | 0.620409161 | -4.548241828 |
| **A_66_P132249** | Akr1c13 | 0.243252915 | -0.224316336 | 0.710822166 | 13.09186099 | 1.264140368 | 0.251776024 | 0.620409161 | -4.548310448 |
| **A_65_P20104** | Mical2 | -0.089792561 | -0.26239037 | 0.082805249 | 7.749712197 | -1.264119897 | 0.251782895 | 0.620409161 | -4.548320379 |
| **A_65_P12530** | NA | 0.117282469 | -0.108160246 | 0.342725183 | 8.16289033 | 1.264096074 | 0.251790891 | 0.620409161 | -4.548331936 |
| **A_66_P101703** | Krtap5-5 | 0.166919894 | -0.153968454 | 0.487808243 | 8.598504141 | 1.263971675 | 0.251832649 | 0.620409161 | -4.548392283 |
| **A_55_P2123421** | Dnal4 | -0.170537468 | -0.498382552 | 0.157307617 | 8.166795649 | -1.263962888 | 0.251835598 | 0.620409161 | -4.548396546 |
| **A_55_P2066414** | Dffa | 0.129600659 | -0.119572884 | 0.378774201 | 8.673455911 | 1.263829451 | 0.251880397 | 0.620440447 | -4.548461275 |
| **A_52_P609972** | Eln | -1.059949934 | -3.098046496 | 0.978146628 | 8.610204852 | -1.263699416 | 0.25192406 | 0.620468928 | -4.548524351 |

| | | | | | | | | | |
|---|---|---|---|---|---|---|---|---|---|
| **A_55_P2068414** | 6030426L16Rik | -0.107317656 | -0.3136916 | 0.099056288 | 7.774419536 | -1.263570572 | 0.251967329 | 0.62048072 | -4.548586847 |
| **A_55_P2046037** | Gm7083 | 0.108769452 | -0.100420308 | 0.317959212 | 13.8661077 | 1.263425697 | 0.25201599 | 0.62048072 | -4.548657116 |
| **A_55_P1989568** | Tasp1 | 0.119889509 | -0.110705654 | 0.350484673 | 8.47370389 | 1.263322343 | 0.25205071 | 0.62048072 | -4.548707245 |
| **A_55_P2046661** | Tnfrsf14 | 0.177304916 | -0.163728144 | 0.518337976 | 8.504900801 | 1.263302871 | 0.252057251 | 0.62048072 | -4.548716689 |
| **A_55_P2005345** | Epb41 | 0.269623658 | -0.249153279 | 0.788400595 | 8.534264028 | 1.262875463 | 0.252200879 | 0.620684349 | -4.548923974 |
| **A_55_P2064333** | Lama3 | -0.768609171 | -2.247484683 | 0.710266342 | 8.099896322 | -1.262865599 | 0.252204195 | 0.620684349 | -4.548928757 |
| **A_51_P230977** | Cdc42 | -0.115211843 | -0.336917373 | 0.106493686 | 13.76388722 | -1.262710496 | 0.252256334 | 0.620733632 | -4.549003973 |
| **A_55_P2121372** | Trp53i13 | -0.307024764 | -0.897886358 | 0.283836831 | 10.51904801 | -1.262614674 | 0.25228855 | 0.620733883 | -4.549050439 |
| **A_55_P2047361** | Ppih | 0.209844368 | -0.194081251 | 0.613769987 | 10.24090862 | 1.262348033 | 0.252378217 | 0.62084781 | -4.549179733 |
| **A_51_P388478** | Efnb1 | -0.388628169 | -1.13672827 | 0.359471932 | 10.9840471 | -1.262285961 | 0.252399094 | 0.62084781 | -4.54920983 |
| **A_55_P1956873** | Timm17a | 0.091025953 | -0.084228071 | 0.266279977 | 12.4823857 | 1.262061225 | 0.252474696 | 0.620920273 | -4.549318795 |
| **A_66_P139946** | Sfta3-ps | -1.44917375 | -4.239498597 | 1.341151097 | 8.359892616 | -1.261969566 | 0.252505536 | 0.620920273 | -4.549363235 |
| **A_55_P2005248** | Meis3 | 0.162292274 | -0.15022687 | 0.474811418 | 9.245210695 | 1.261840345 | 0.252549019 | 0.620920273 | -4.549425884 |
| **A_55_P2035657** | Psmb6 | -0.113293298 | -0.331485971 | 0.104899374 | 14.64389919 | -1.261674276 | 0.252604912 | 0.620920273 | -4.549506394 |
| **A_55_P1970596** | NA | -0.137152488 | -0.401314275 | 0.127009299 | 15.37974497 | -1.261586074 | 0.252634603 | 0.620920273 | -4.549549153 |
| **A_52_P40367** | Mrpl42 | 0.101625213 | -0.094117369 | 0.297367795 | 10.75125871 | 1.26153512 | 0.252651756 | 0.620920273 | -4.549573855 |
| **A_52_P587948** | Zfp11 | 0.088041299 | -0.081549388 | 0.257631987 | 8.215605314 | 1.261443005 | 0.252682768 | 0.620920273 | -4.549618509 |
| **A_55_P2372902** | 5330438D12Rik | 0.138260433 | -0.128067248 | 0.404588114 | 8.936435264 | 1.261434762 | 0.252685544 | 0.620920273 | -4.549622504 |
| **A_51_P287100** | Cdh16 | -1.353245791 | -3.96029332 | 1.253801738 | 8.468415302 | -1.261278304 | 0.252738226 | 0.620970787 | -4.549698347 |
| **A_51_P202745** | Zc3h13 | 0.168071121 | -0.155766174 | 0.491908415 | 10.56478944 | 1.2610997 | 0.252798378 | 0.621039635 | -4.549784921 |
| **A_51_P511680** | Ttc38 | 0.140813556 | -0.130538566 | 0.412165677 | 8.47017618 | 1.260940033 | 0.252852162 | 0.621092826 | -4.549862311 |
| **A_55_P1983848** | Tc2n | -0.166974479 | -0.488774624 | 0.154825666 | 7.750167531 | -1.260802472 | 0.252898509 | 0.621099325 | -4.549928985 |
| **A_51_P394728** | Kti12 | 0.182793237 | -0.169510529 | 0.535097002 | 10.39766325 | 1.260741437 | 0.252919074 | 0.621099325 | -4.549958566 |
| **A_55_P2077448** | Epas1 | 0.12956643 | -0.120191095 | 0.379323955 | 11.94927846 | 1.260541356 | 0.252986502 | 0.621158355 | -4.550055536 |
| **A_51_P437938** | Coa5 | 0.156581308 | -0.145266031 | 0.458428647 | 9.765056148 | 1.260479402 | 0.253007384 | 0.621158355 | -4.550085561 |
| **A_51_P315666** | Nid2 | -0.162020894 | -0.474437627 | 0.150395839 | 9.552293659 | -1.260143277 | 0.253120703 | 0.621170937 | -4.550248449 |
| **A_55_P1978740** | Rpgrip1 | 0.241899913 | -0.224550553 | 0.708350379 | 11.7819007 | 1.260124252 | 0.253127118 | 0.621170937 | -4.550257668 |
| **A_52_P59264** | Sncaip | -0.337909553 | -0.989573035 | 0.313753929 | 8.262618343 | -1.259970202 | 0.253179069 | 0.621170937 | -4.550332316 |
| **A_55_P2119430** | Usp10 | 0.17581506 | -0.163256008 | 0.514886128 | 11.90301844 | 1.259936126 | 0.253190562 | 0.621170937 | -4.550348828 |
| **A_55_P1999289** | Ovol2 | 0.10453627 | -0.097080413 | 0.306152952 | 11.20748517 | 1.259864165 | 0.253214834 | 0.621170937 | -4.550383696 |
| **A_55_P1963374** | Klhdc7a | 0.210577483 | -0.195621993 | 0.61677696 | 10.8287417 | 1.259667025 | 0.253281339 | 0.621170937 | -4.550479217 |
| **A_55_P1953043** | E130311K13Rik | 0.154437983 | -0.143508453 | 0.452384419 | 9.697047753 | 1.259502737 | 0.253336772 | 0.621170937 | -4.550558816 |
| **A_52_P297075** | Heatr5a | 0.1415867 | -0.131577675 | 0.414751075 | 10.39228264 | 1.259451908 | 0.253353925 | 0.621170937 | -4.550583442 |
| **A_51_P487048** | Rhox6 | -0.097613368 | -0.285954924 | 0.090728189 | 7.733841618 | -1.259349659 | 0.253388433 | 0.621170937 | -4.55063298 |
| **A_55_P2211164** | 5330406M23Rik | 0.107179175 | -0.099629128 | 0.313987479 | 8.136238222 | 1.259289632 | 0.253408694 | 0.621170937 | -4.550662061 |
| **A_51_P466673** | Srsf7 | 0.127980985 | -0.118981768 | 0.374943738 | 9.38080042 | 1.259207145 | 0.253436537 | 0.621170937 | -4.550702023 |
| **A_55_P1972481** | NA | -0.166914608 | -0.489046284 | 0.155217068 | 12.6879232 | -1.259053267 | 0.253488486 | 0.621170937 | -4.550776568 |
| **A_51_P337944** | Bmp2k | 0.188916276 | -0.175678979 | 0.553511531 | 11.46680138 | 1.25904587 | 0.253490983 | 0.621170937 | -4.550780152 |
| **A_51_P480241** | Elf3 | -0.250078365 | -0.732717063 | 0.232560333 | 7.940824166 | -1.259033164 | 0.253495273 | 0.621170937 | -4.550786307 |
| **A_55_P2279807** | NA | -0.177799467 | -0.520956091 | 0.165357156 | 9.496369422 | -1.258987037 | 0.253510848 | 0.621170937 | -4.550808652 |
| **A_51_P334104** | Dcn | -0.211245023 | -0.618967458 | 0.196477411 | 13.58863865 | -1.258940099 | 0.253526697 | 0.621170937 | -4.55083139 |
| **A_52_P342159** | Nfatc4 | -0.411056601 | -1.204657172 | 0.38254397 | 8.706729995 | -1.258585849 | 0.253646342 | 0.621180573 | -4.551002988 |
| **A_55_P2110200** | Ubox5 | 0.098830745 | -0.091979355 | 0.289640845 | 8.834441341 | 1.258559922 | 0.2536551 | 0.621180573 | -4.551015546 |
| **A_55_P1993549** | Npnt | -0.254502965 | -0.745877016 | 0.236871085 | 7.847908366 | -1.258530713 | 0.253664968 | 0.621180573 | -4.551029694 |
| **A_52_P540434** | Ppp1cc | 0.124488046 | -0.115900821 | 0.364876912 | 10.98806534 | 1.258335613 | 0.253730886 | 0.621180573 | -4.551124191 |
| **A_51_P518488** | Atpif1 | 0.126577292 | -0.117849002 | 0.371003586 | 16.64595813 | 1.258319906 | 0.253736194 | 0.621180573 | -4.551131798 |
| **A_55_P2101357** | Aftph | 0.140308774 | -0.130639584 | 0.411257132 | 11.45403828 | 1.258292173 | 0.253745565 | 0.621180573 | -4.55114523 |
| **A_51_P354126** | Reg3g | -0.963585283 | -2.82439554 | 0.897224974 | 8.128227747 | -1.258262507 | 0.25375559 | 0.621180573 | -4.551159598 |
| **A_55_P1990086** | Mks1 | -0.163100676 | -0.478228265 | 0.152026912 | 8.578654962 | -1.257628946 | 0.253969774 | 0.621461116 | -4.551466422 |
| **A_55_P2067445** | Arf5 | 0.282925169 | -0.449677524 | 1.015527861 | 13.79251406 | 1.537716516 | 0.253973269 | 0.621461116 | -4.523196522 |

| | | | | | | | | | |
|---|---|---|---|---|---|---|---|---|---|
| **A_55_P2289800** | Fryl | 0.339486235 | -0.316443341 | 0.995415811 | 12.25658535 | 1.257616264 | 0.253974063 | 0.621461116 | -4.551472563 |
| **A_55_P2038347** | Acot3 | 0.208575346 | -0.194466105 | 0.611616797 | 8.992011727 | 1.25746657 | 0.254024692 | 0.621461116 | -4.551545049 |
| **A_55_P1963863** | Arhgef17 | -0.442047478 | -1.29633218 | 0.412237224 | 8.95702128 | -1.257330575 | 0.254070696 | 0.621461116 | -4.551610899 |
| **A_55_P2022082** | Klf13 | -0.118965382 | -0.348917667 | 0.110986903 | 10.68680463 | -1.257089105 | 0.254152398 | 0.621461116 | -4.551727815 |
| **A_55_P2000683** | Eci2 | -0.119069844 | -0.349225043 | 0.111085355 | 12.80549606 | -1.25708367 | 0.254154237 | 0.621461116 | -4.551730446 |
| **A_55_P2068882** | Ccng2 | -0.083034883 | -0.243544093 | 0.077474327 | 8.348977299 | -1.257024699 | 0.254174193 | 0.621461116 | -4.551758998 |
| **A_55_P1982559** | 1700007K13Rik | -0.176323456 | -0.517188702 | 0.164541791 | 7.732860688 | -1.256928443 | 0.254206771 | 0.621461116 | -4.551805601 |
| **A_51_P367780** | Adamtsl2 | -0.455760154 | -1.33693093 | 0.425410622 | 7.985526229 | -1.256780566 | 0.254256826 | 0.621461116 | -4.551877193 |
| **A_55_P2024327** | Kmt5c | 0.109533728 | -0.102245462 | 0.321312917 | 13.14941236 | 1.256746759 | 0.254268271 | 0.621461116 | -4.55189356 |
| **A_55_P2146050** | Dennd5b | 0.154983024 | -0.14468305 | 0.454649099 | 11.65203718 | 1.256694578 | 0.254285936 | 0.621461116 | -4.551918822 |
| **A_51_P329370** | Snrnp25 | 0.220739758 | -0.206078606 | 0.647558122 | 13.84112316 | 1.256668283 | 0.254294839 | 0.621461116 | -4.551931551 |
| **A_55_P2087825** | Ncoa2 | 0.147677893 | -0.137946187 | 0.433301973 | 10.3805334 | 1.256330368 | 0.254409267 | 0.621461116 | -4.552095132 |
| **A_55_P1986386** | Senp3 | 0.103565687 | -0.096752323 | 0.303883697 | 8.922990962 | 1.2562587 | 0.254433542 | 0.621461116 | -4.552129823 |
| **A_51_P282837** | St14 | -0.4170274 | -1.223797652 | 0.389742851 | 8.122402729 | -1.256023901 | 0.254513085 | 0.621461116 | -4.552243474 |
| **A_55_P1981845** | NA | 0.197413898 | -0.184504751 | 0.579332546 | 9.746919005 | 1.256001178 | 0.254520784 | 0.621461116 | -4.552254473 |
| **A_52_P657360** | Tnni1 | -0.532180956 | -1.561939549 | 0.497577636 | 8.012803914 | -1.255761298 | 0.254602074 | 0.621461116 | -4.552370575 |
| **A_55_P2182931** | Sim2 | 0.137912007 | -0.128946389 | 0.404770403 | 8.42576211 | 1.255753495 | 0.254604718 | 0.621461116 | -4.552374352 |
| **A_51_P303906** | Actr2 | 0.126188031 | -0.117985831 | 0.370361893 | 11.87234928 | 1.255747091 | 0.254606889 | 0.621461116 | -4.552377451 |
| **A_51_P127464** | Ppp1r37 | 0.110065269 | -0.10291135 | 0.323041888 | 9.205782017 | 1.255745298 | 0.254607497 | 0.621461116 | -4.552378319 |
| **A_52_P24439** | Sftpa1 | -1.846592335 | -5.42026173 | 1.72707706 | 8.552038168 | -1.255566862 | 0.25466798 | 0.621461116 | -4.552464677 |
| **A_66_P118124** | Fxyd3 | -0.585697144 | -1.719208048 | 0.547813761 | 8.109456393 | -1.255540113 | 0.254677048 | 0.621461116 | -4.552477622 |
| **A_55_P1978506** | H2-Q6 | -0.158667211 | -0.46575 | 0.148415578 | 11.48901401 | -1.255494751 | 0.254692427 | 0.621461116 | -4.552499575 |
| **A_55_P2126391** | Srl | -0.163137851 | -0.478880024 | 0.152604322 | 7.889405757 | -1.255467092 | 0.254701804 | 0.621461116 | -4.552512961 |
| **A_51_P301636** | Kazn | -0.232611874 | -0.682844243 | 0.217620494 | 8.295766932 | -1.255389104 | 0.254728246 | 0.621461116 | -4.552550702 |
| **A_51_P494863** | Vmac | -0.167784573 | -0.492590332 | 0.157021185 | 8.344558107 | -1.255195857 | 0.254793777 | 0.621461116 | -4.552644219 |
| **A_51_P414837** | Faf1 | 0.16856028 | -0.157774394 | 0.494894954 | 10.08893871 | 1.255090988 | 0.254829345 | 0.621461116 | -4.552694965 |
| **A_55_P2126489** | Atxn7l3b | -0.207815525 | -0.610245346 | 0.194614295 | 12.70290126 | -1.254789927 | 0.254931479 | 0.621461116 | -4.55284064 |
| **A_55_P2066007** | Crocc | -0.188881032 | -0.554645328 | 0.176883264 | 8.830786642 | -1.254787622 | 0.254932261 | 0.621461116 | -4.552841755 |
| **A_55_P2089020** | Slc35g2 | -0.172275092 | -0.505889024 | 0.161338839 | 8.192459113 | -1.254762416 | 0.254940814 | 0.621461116 | -4.552853951 |
| **A_55_P2083929** | Robo2 | -0.161483317 | -0.474201874 | 0.15123524 | 8.005217366 | -1.254749982 | 0.254945033 | 0.621461116 | -4.552859968 |
| **A_55_P2185652** | Cln3 | 0.126129103 | -0.118142628 | 0.370400833 | 9.235290728 | 1.254657784 | 0.254976319 | 0.621461116 | -4.552904577 |
| **A_55_P2059864** | Igsf1 | 0.247418117 | -0.231755533 | 0.726591766 | 12.4305552 | 1.254647594 | 0.254979777 | 0.621461116 | -4.552909507 |
| **A_55_P2064074** | Vps37d | -0.452295136 | -1.328285623 | 0.423695352 | 8.4739106 | -1.25460125 | 0.254995505 | 0.621461116 | -4.55293193 |
| **A_55_P1995698** | Thumpd1 | 0.146088014 | -0.136888498 | 0.429064527 | 10.42686394 | 1.254432752 | 0.255052695 | 0.621522131 | -4.553013451 |
| **A_55_P2086825** | Rgl3 | -0.503365753 | -1.478488792 | 0.471757286 | 8.261946893 | -1.254317368 | 0.255091864 | 0.621539222 | -4.553069273 |
| **A_66_P136813** | Smim41 | -1.189595786 | -3.494346317 | 1.115154745 | 8.949098338 | -1.254176568 | 0.255139669 | 0.621544037 | -4.553137389 |
| **A_55_P2078675** | NA | 0.095791307 | -0.089805053 | 0.281387667 | 8.012455038 | 1.254122132 | 0.255158153 | 0.621544037 | -4.553163723 |
| **A_55_P2017075** | Zfp54 | 0.112100262 | -0.105122809 | 0.329323333 | 8.204033881 | 1.253960532 | 0.255213032 | 0.621555506 | -4.553241898 |
| **A_51_P346104** | Sgta | -0.106163886 | -0.311901984 | 0.099574211 | 11.34125509 | -1.253849114 | 0.255250876 | 0.621555506 | -4.553295794 |
| **A_55_P2021953** | Olfr700 | 0.171596333 | -0.16095494 | 0.504147606 | 8.774999648 | 1.253812451 | 0.25526333 | 0.621555506 | -4.553313528 |
| **A_52_P136782** | Rgs5 | -0.158527013 | -0.465770441 | 0.148716414 | 7.725229093 | -1.253729562 | 0.255291488 | 0.621555506 | -4.553353623 |
| **A_55_P2175260** | Tet3 | 0.110892322 | -0.104075283 | 0.325859926 | 8.052793021 | 1.253463374 | 0.255381932 | 0.621590444 | -4.553482374 |
| **A_52_P242251** | Utp25 | 0.102205958 | -0.095927625 | 0.300339541 | 8.490243387 | 1.253433539 | 0.255392071 | 0.621590444 | -4.553496805 |
| **A_51_P280245** | Zfp207 | 0.108452523 | -0.101795529 | 0.318700574 | 11.45122687 | 1.253403401 | 0.255402313 | 0.621590444 | -4.553511381 |
| **A_55_P2063316** | Mgll | 0.146550488 | -0.137605804 | 0.430706779 | 11.48120508 | 1.253179207 | 0.255478516 | 0.621697624 | -4.553619812 |
| **A_55_P2124528** | Capns1 | -0.122836812 | -0.367367947 | 0.121694323 | 13.51114204 | -1.278017318 | 0.255531297 | 0.621747789 | -4.53947556 |
| **A_55_P2005213** | Ces2c | -0.392650286 | -1.154171732 | 0.368871161 | 11.18179557 | -1.252872764 | 0.255582706 | 0.621794605 | -4.55376801 |
| **A_55_P2030702** | NA | -0.141577367 | -0.416205863 | 0.133051129 | 9.491256509 | -1.252654843 | 0.255656822 | 0.621799372 | -4.553873391 |
| **A_66_P118592** | Atrx | 0.117747408 | -0.1106761 | 0.346170915 | 8.655426666 | 1.252546182 | 0.255693785 | 0.621799372 | -4.553925934 |
| **A_55_P2046338** | Mapk14 | 0.084984815 | -0.079884119 | 0.249853749 | 11.68950028 | 1.252522945 | 0.25570169 | 0.621799372 | -4.55393717 |

| | | | | | | | | | |
|---|---|---|---|---|---|---|---|---|---|
| **A_55_P2113837** | Tbc1d1 | -0.235004164 | -0.690951825 | 0.220943498 | 10.73979547 | -1.252402004 | 0.255742837 | 0.621799372 | -4.553995648 |
| **A_55_P1964138** | Pign | 0.204908213 | -0.19265073 | 0.602467156 | 8.916484567 | 1.252394142 | 0.255745512 | 0.621799372 | -4.55399945 |
| **A_52_P90289** | Oaz2 | -0.32015355 | -0.941470881 | 0.30116378 | 8.594361889 | -1.252068367 | 0.255856378 | 0.621990684 | -4.554156962 |
| **A_55_P1980051** | Il17d | 0.312261756 | -0.293835915 | 0.918359427 | 9.732396214 | 1.251870401 | 0.255923769 | 0.621994569 | -4.554252671 |
| **A_51_P181722** | Rbks | 0.211873074 | -0.199398428 | 0.623144575 | 11.04442816 | 1.251786718 | 0.255952261 | 0.621994569 | -4.554293127 |
| **A_55_P1977761** | Ttc14 | 0.118764177 | -0.111812461 | 0.349340816 | 8.330844344 | 1.251564828 | 0.256027822 | 0.621994569 | -4.554400393 |
| **A_55_P2096266** | Sftpb | -2.255811848 | -6.636028938 | 2.124405242 | 8.820713488 | -1.251383508 | 0.256089582 | 0.621994569 | -4.554488042 |
| **A_51_P157042** | Ccn2 | -0.665509089 | -1.957796917 | 0.626778739 | 9.849585612 | -1.251347488 | 0.256101853 | 0.621994569 | -4.554505453 |
| **A_51_P125446** | Lzic | 0.099800008 | -0.093995617 | 0.293595632 | 9.216957767 | 1.251324059 | 0.256109834 | 0.621994569 | -4.554516778 |
| **A_52_P287992** | Tbcel | 0.165442738 | -0.15585727 | 0.486742745 | 9.215283342 | 1.25118106 | 0.256158554 | 0.621994569 | -4.554585899 |
| **A_55_P2108424** | NA | -0.122713692 | -0.361047622 | 0.115620238 | 10.24051307 | -1.25109514 | 0.256187831 | 0.621994569 | -4.554627428 |
| **A_55_P1995768** | Zfp967 | 0.116985399 | -0.110223304 | 0.344194102 | 9.056914976 | 1.251093846 | 0.256188272 | 0.621994569 | -4.554628054 |
| **A_51_P324651** | Adgrl1 | -0.228699332 | -0.672921316 | 0.215522652 | 8.287364088 | -1.25097329 | 0.256229356 | 0.621994569 | -4.554686322 |
| **A_51_P257951** | Retnla | -1.963764626 | -5.778167689 | 1.850638437 | 8.607952323 | -1.250967535 | 0.256231317 | 0.621994569 | -4.554689104 |
| **A_55_P2050448** | Slc25a3 | 0.227851301 | -0.214738816 | 0.670441418 | 11.08466205 | 1.250929947 | 0.256244128 | 0.621994569 | -4.554707271 |
| **A_52_P277139** | Ankrd17 | 0.16671681 | -0.157170924 | 0.490604545 | 10.98863239 | 1.250742997 | 0.256307853 | 0.622071131 | -4.554797624 |
| **A_55_P2064776** | 4930412O13Rik | 0.1618085 | -0.152592311 | 0.476209312 | 7.747561306 | 1.250549431 | 0.256373847 | 0.622151901 | -4.554891171 |
| **A_55_P1996212** | Zbtb2 | 0.099742251 | -0.094075583 | 0.293560085 | 9.365075339 | 1.250456587 | 0.256405507 | 0.622151901 | -4.554936039 |
| **A_52_P1016836** | NA | -0.192878178 | -0.567714321 | 0.181957965 | 7.76161006 | -1.250330496 | 0.256448509 | 0.622158812 | -4.554996971 |
| **A_55_P2083009** | Catsperd | 0.16558086 | -0.156229753 | 0.487391473 | 8.95261382 | 1.250238765 | 0.256479797 | 0.622158812 | -4.555041298 |
| **A_55_P2091756** | Nxnl1 | 0.215658104 | -0.203543345 | 0.634859552 | 8.75708805 | 1.250046581 | 0.256545359 | 0.622158812 | -4.555134163 |
| **A_55_P1967149** | Irx3 | -0.990288636 | -2.915312746 | 0.934735475 | 8.197622302 | -1.24999654 | 0.256562432 | 0.622158812 | -4.555158342 |
| **A_55_P2099448** | Slc12a2 | -0.890772081 | -2.622648836 | 0.841104674 | 9.811931407 | -1.24977779 | 0.256637078 | 0.622158812 | -4.555264036 |
| **A_51_P180974** | Cavin3 | -0.627499314 | -1.847555569 | 0.59255694 | 9.333754849 | -1.249731133 | 0.256653002 | 0.622158812 | -4.555286579 |
| **A_51_P352385** | Ism1 | -0.382782349 | -1.127161546 | 0.361596847 | 7.909156338 | -1.249513249 | 0.256727376 | 0.622158812 | -4.555391847 |
| **A_51_P521052** | Ly6k | 0.174171402 | -0.164592789 | 0.512935593 | 7.969457078 | 1.24928792 | 0.256804311 | 0.622158812 | -4.555500704 |
| **A_55_P2183237** | Ccdc191 | -0.089679507 | -0.264112154 | 0.084753139 | 8.746071956 | -1.24924791 | 0.256817974 | 0.622158812 | -4.555520032 |
| **A_51_P184321** | Aimp1 | 0.131029624 | -0.123872547 | 0.385931794 | 12.53915492 | 1.249048057 | 0.25688623 | 0.622158812 | -4.555616575 |
| **A_52_P432685** | Ctdsp2 | 0.160916573 | -0.152127969 | 0.473961116 | 9.102936257 | 1.24904426 | 0.256887527 | 0.622158812 | -4.555618409 |
| **A_55_P2013601** | Ldb2 | -0.285959129 | -0.842288956 | 0.270370699 | 9.54484493 | -1.248978059 | 0.256910141 | 0.622158812 | -4.555650388 |
| **A_55_P2346639** | Sec14l3 | -1.621831763 | -4.777168372 | 1.533504846 | 8.44584464 | -1.248944944 | 0.256921453 | 0.622158812 | -4.555666384 |
| **A_55_P2073308** | Bin1 | -0.092208366 | -0.271604883 | 0.08718815 | 11.4015072 | -1.248934063 | 0.25692517 | 0.622158812 | -4.55567164 |
| **A_55_P2081318** | Gnas | -0.148915788 | -0.438643401 | 0.140811825 | 7.885338817 | -1.248918343 | 0.25693054 | 0.622158812 | -4.555679233 |
| **A_51_P329928** | Phlda3 | -0.483871863 | -1.425324993 | 0.457581268 | 10.02990627 | -1.248863156 | 0.256949394 | 0.622158812 | -4.55570589 |
| **A_65_P14951** | Cblb | -0.182970651 | -0.5389756 | 0.173034297 | 9.06988598 | -1.248845142 | 0.256955548 | 0.622158812 | -4.555714591 |
| **A_51_P397925** | Smap2 | 0.189800312 | -0.179573104 | 0.559173728 | 10.14277487 | 1.248574541 | 0.257048015 | 0.622189187 | -4.555845294 |
| **A_51_P296249** | Foxs1 | -0.187850854 | -0.553432784 | 0.177731077 | 8.167054734 | -1.248566389 | 0.257050801 | 0.622189187 | -4.555849232 |
| **A_51_P516637** | Bmp5 | -0.092188143 | -0.271603958 | 0.087227672 | 8.069566557 | -1.248525832 | 0.257064662 | 0.622189187 | -4.55586882 |
| **A_55_P2067483** | NA | -0.127404882 | -0.375395288 | 0.120585524 | 10.9908914 | -1.248344292 | 0.257126715 | 0.62226146 | -4.555956498 |
| **A_51_P518600** | Atp6v1c2 | -0.533347034 | -1.571740658 | 0.505046589 | 7.92141373 | -1.248047346 | 0.257228244 | 0.622372374 | -4.556099904 |
| **A_55_P1970105** | Ltbp2 | -1.097744528 | -3.235271359 | 1.039782302 | 9.243980088 | -1.247880204 | 0.257285407 | 0.622372374 | -4.556180617 |
| **A_51_P176505** | Rps19bp1 | 0.104020161 | -0.098537696 | 0.306578018 | 11.61309797 | 1.247819067 | 0.257306319 | 0.622372374 | -4.55621014 |
| **A_51_P305547** | Snai2 | -0.491879419 | -1.449734007 | 0.465975169 | 9.261894814 | -1.247792172 | 0.257315519 | 0.622372374 | -4.556223127 |
| **A_51_P193686** | 1700012B09Rik | -0.536538962 | -1.581407512 | 0.508329587 | 7.92298383 | -1.247736262 | 0.257334644 | 0.622372374 | -4.556250125 |
| **A_52_P315111** | Mrpl52 | 0.123178345 | -0.116719526 | 0.363076216 | 14.3387523 | 1.247645368 | 0.25736574 | 0.622372374 | -4.556294014 |
| **A_52_P284778** | Rae1 | 0.135026357 | -0.128035187 | 0.3980879 | 11.07799957 | 1.247223758 | 0.25751002 | 0.622534077 | -4.55649758 |
| **A_51_P326994** | Calhm2 | -0.109187063 | -0.32191469 | 0.103540564 | 8.451254349 | -1.247183846 | 0.257523682 | 0.622534077 | -4.556516849 |
| **A_66_P129188** | H2ac20 | 0.149556991 | -0.141826589 | 0.440940572 | 17.38281639 | 1.247167639 | 0.25752923 | 0.622534077 | -4.556524674 |
| **A_51_P260169** | Gstm5 | 0.20701243 | -0.196384228 | 0.610409087 | 12.43209647 | 1.246945053 | 0.257605435 | 0.622578935 | -4.556632133 |
| **A_55_P2019684** | Bspry | -0.900351777 | -2.654940041 | 0.854236488 | 8.180784048 | -1.246867183 | 0.257632099 | 0.622578935 | -4.556669725 |

| | | | | | | | | | |
|---|---|---|---|---|---|---|---|---|---|
| **A_55_P2144781** | Erich5 | 0.141959309 | -0.134696248 | 0.418614865 | 9.337432868 | 1.246831218 | 0.257644415 | 0.622578935 | -4.556687087 |
| **A_52_P327664** | Gbp5 | -0.100772234 | -0.297175956 | 0.095631489 | 7.858387422 | -1.246735594 | 0.257677163 | 0.622580237 | -4.556733249 |
| **A_52_P139097** | NA | 0.092513843 | -0.087834972 | 0.272862658 | 11.03540145 | 1.246455041 | 0.257773265 | 0.622734589 | -4.556868674 |
| **A_55_P2167294** | NA | 0.236457185 | -0.224578857 | 0.697493227 | 8.780825528 | 1.24623753 | 0.257847794 | 0.622836792 | -4.556973662 |
| **A_55_P2122329** | Nfic | -0.103703065 | -0.30594389 | 0.09853776 | 8.269322845 | -1.245965315 | 0.257941093 | 0.622853773 | -4.557105044 |
| **A_55_P2052495** | Tmx2 | 0.1039499 | -0.098775043 | 0.306674842 | 11.85086809 | 1.245948459 | 0.257946871 | 0.622853773 | -4.557113179 |
| **A_55_P2046793** | Eogt | -0.104932299 | -0.30958768 | 0.099723081 | 7.987706281 | -1.24585991 | 0.257977227 | 0.622853773 | -4.557155914 |
| **A_55_P2100968** | Dnah7b | -0.255762398 | -0.75468235 | 0.243157555 | 7.855520611 | -1.245629862 | 0.258056106 | 0.622853773 | -4.557266933 |
| **A_51_P327491** | Apoa4 | -0.436489666 | -1.287963372 | 0.41498404 | 7.888365879 | -1.245620978 | 0.258059153 | 0.622853773 | -4.55727122 |
| **A_55_P1962483** | NA | 0.165177384 | -0.157042309 | 0.487397077 | 10.01766639 | 1.245608874 | 0.258063304 | 0.622853773 | -4.557277061 |
| **A_55_P1970650** | Dgkh | -0.112612215 | -0.332299221 | 0.10707479 | 8.144423169 | -1.245559051 | 0.258080391 | 0.622853773 | -4.557301105 |
| **A_55_P1996578** | Fgfr3 | -0.612963546 | -1.80893415 | 0.583007059 | 10.17563456 | -1.245366934 | 0.258146285 | 0.622922515 | -4.557393811 |
| **A_51_P232708** | Npff | 0.125451461 | -0.119335947 | 0.370238869 | 8.697770603 | 1.245288101 | 0.258173329 | 0.622922515 | -4.55743185 |
| **A_51_P386304** | Ccnl2 | -0.088505284 | -0.261260761 | 0.084250194 | 11.18792401 | -1.244860142 | 0.258320182 | 0.623124939 | -4.557638339 |
| **A_55_P2036863** | Rabl2 | -0.330138436 | -0.974545452 | 0.314268579 | 9.923665514 | -1.24485572 | 0.2583217 | 0.623124939 | -4.557640473 |
| **A_55_P2100155** | Mapt | -0.346723719 | -1.023587541 | 0.330140103 | 8.017388276 | -1.244702148 | 0.258374416 | 0.623135389 | -4.557714564 |
| **A_55_P1954758** | Prickle1 | -0.496494838 | -1.465773928 | 0.472784252 | 10.49852583 | -1.244655271 | 0.258390509 | 0.623135389 | -4.557737179 |
| **A_51_P383991** | 4-Sep | -0.673336793 | -1.988367778 | 0.641694192 | 10.29993114 | -1.244169496 | 0.258557329 | 0.623459907 | -4.557971519 |
| **A_55_P2164090** | Kcnj15 | -0.23781644 | -0.702407952 | 0.226775072 | 7.899392638 | -1.243809277 | 0.258681092 | 0.623637223 | -4.558145268 |
| **A_55_P2093535** | Rps8 | -0.148732845 | -0.439302872 | 0.141837182 | 16.42536861 | -1.243767664 | 0.258695393 | 0.623637223 | -4.558165338 |
| **A_55_P1968493** | Telo2 | 0.116999711 | -0.111637711 | 0.345637132 | 9.775545565 | 1.243428055 | 0.258812128 | 0.623759562 | -4.558329127 |
| **A_55_P2183338** | Marveld3 | -0.18155223 | -0.536407938 | 0.173303478 | 7.990819812 | -1.243177043 | 0.258898439 | 0.623759562 | -4.558450177 |
| **A_52_P570700** | Dpp8 | 0.092147565 | -0.087962439 | 0.27225757 | 8.952103797 | 1.243166267 | 0.258902145 | 0.623759562 | -4.558455373 |
| **A_51_P269134** | Sftpb | -0.720007654 | -2.127412272 | 0.687396964 | 8.031246922 | -1.243086445 | 0.258929597 | 0.623759562 | -4.558493865 |
| **A_55_P2101001** | Smad3 | -0.204457808 | -0.60412659 | 0.195210973 | 10.44187291 | -1.243044464 | 0.258944036 | 0.623759562 | -4.558514108 |
| **A_52_P406507** | Mus81 | -0.123569808 | -0.365128017 | 0.117988401 | 8.438992516 | -1.243007534 | 0.258956739 | 0.623759562 | -4.558531917 |
| **A_55_P2011862** | Mybpc3 | -0.787181044 | -2.326159259 | 0.751797172 | 8.221670951 | -1.242868966 | 0.259004406 | 0.623759562 | -4.558598733 |
| **A_55_P1979067** | A430073D23Rik | 0.133504121 | -0.127540838 | 0.394549081 | 12.63635831 | 1.242689275 | 0.25906623 | 0.623759562 | -4.558685375 |
| **A_51_P450123** | Mrpl36 | 0.083105159 | -0.07939944 | 0.245609758 | 12.00691109 | 1.242640536 | 0.259083001 | 0.623759562 | -4.558708874 |
| **A_52_P486482** | R3hcc1l | 0.094795453 | -0.090571295 | 0.280162201 | 8.604482792 | 1.242621505 | 0.25908955 | 0.623759562 | -4.55871805 |
| **A_55_P2004213** | Gprasp2 | -0.244201918 | -0.721753949 | 0.233350114 | 8.540351352 | -1.242543388 | 0.259116433 | 0.623759562 | -4.558755714 |
| **A_55_P1981494** | H2-Bl | -0.253448756 | -0.749103206 | 0.242205695 | 10.37664779 | -1.242494119 | 0.259133389 | 0.623759562 | -4.558779468 |
| **A_52_P312467** | Hoxb3 | -0.18533178 | -0.547824595 | 0.177161035 | 7.896541194 | -1.24232059 | 0.259193119 | 0.623772484 | -4.558863129 |
| **A_51_P239286** | Bcl2l12 | -0.118434147 | -0.350102682 | 0.113234389 | 9.309868977 | -1.242204423 | 0.259233112 | 0.623772484 | -4.558919133 |
| **A_55_P2071486** | Suv39h1 | 0.25530617 | -0.244250214 | 0.754862553 | 9.636578377 | 1.241823821 | 0.259364177 | 0.623772484 | -4.559102607 |
| **A_55_P2029842** | Hnrnpa1 | 0.126037663 | -0.120589251 | 0.372664577 | 12.54523959 | 1.241775428 | 0.259380846 | 0.623772484 | -4.559125934 |
| **A_55_P1999078** | NA | 0.164424398 | -0.157317776 | 0.486166571 | 9.328436997 | 1.24177084 | 0.259382426 | 0.623772484 | -4.559128146 |
| **A_55_P2139089** | Slc22a28 | 0.099723054 | -0.095416902 | 0.294863011 | 8.557210375 | 1.241745392 | 0.259391192 | 0.623772484 | -4.559140412 |
| **A_51_P158678** | Fam181b | -0.30934396 | -0.914720636 | 0.296032716 | 7.849009726 | -1.241649865 | 0.2594241 | 0.623772484 | -4.559186458 |
| **A_55_P2045203** | Chp1 | -0.192624873 | -0.569628797 | 0.184379051 | 12.98125413 | -1.241508462 | 0.259472819 | 0.623772484 | -4.559254614 |
| **A_51_P295215** | Alkbh5 | 0.172321737 | -0.164969492 | 0.509612966 | 11.4059917 | 1.241418477 | 0.259503826 | 0.623772484 | -4.559297985 |
| **A_51_P308446** | 1700094D03Rik | -0.398130582 | -1.177458742 | 0.381197577 | 8.334322605 | -1.241333166 | 0.259533226 | 0.623772484 | -4.559339102 |
| **A_55_P2009861** | NA | -0.237362861 | -0.702040397 | 0.227314674 | 11.65876534 | -1.241207183 | 0.259576647 | 0.623772484 | -4.559399821 |
| **A_55_P2054157** | Hspa4 | 0.132023052 | -0.126452229 | 0.390498333 | 12.92905722 | 1.241120474 | 0.259606535 | 0.623772484 | -4.559441609 |
| **A_55_P2017944** | Etfb | -0.137893247 | -0.407863403 | 0.13207691 | 15.44986185 | -1.241110437 | 0.259609995 | 0.623772484 | -4.559446446 |
| **A_51_P227962** | Dynlrb2 | -1.43179209 | -4.23527557 | 1.37169139 | 8.516397961 | -1.240981041 | 0.259654605 | 0.623772484 | -4.559508805 |
| **A_51_P308048** | Cmtm8 | -0.116793339 | -0.34548803 | 0.111901353 | 12.90530952 | -1.240923985 | 0.259674277 | 0.623772484 | -4.559536301 |
| **A_51_P330044** | Cyp2j9 | -0.105870084 | -0.313183204 | 0.101443035 | 7.90838866 | -1.240879647 | 0.259689565 | 0.623772484 | -4.559557668 |
| **A_52_P588483** | Fbln1 | -0.419590113 | -1.241277314 | 0.402097089 | 8.280608942 | -1.240800364 | 0.259716904 | 0.623772484 | -4.559595874 |
| **A_52_P181468** | Kmt5a | 0.232038564 | -0.222367611 | 0.686444738 | 9.88641893 | 1.240792425 | 0.259719641 | 0.623772484 | -4.559599699 |

| | | | | | | | | | |
|---|---|---|---|---|---|---|---|---|---|
| **A_55_P2125557** | Irx2 | -0.696969333 | -2.0621236 | 0.668184933 | 7.998751653 | -1.240552442 | 0.259802411 | 0.623892955 | -4.55971534 |
| **A_55_P2007470** | Pdgfa | -0.67193546 | -1.988154121 | 0.6442832 | 10.56969408 | -1.240459834 | 0.259834357 | 0.623892955 | -4.559759963 |
| **A_51_P419286** | Batf3 | -0.146615589 | -0.433851544 | 0.140620366 | 8.608776375 | -1.240293696 | 0.259891677 | 0.623943486 | -4.559840013 |
| **A_51_P460774** | Smyd4 | 0.147309311 | -0.141322678 | 0.435941301 | 8.513277503 | 1.240134886 | 0.259946479 | 0.623943486 | -4.559916528 |
| **A_55_P1984152** | 1600029O15Rik | -0.173419544 | -0.513242122 | 0.166403034 | 15.48933225 | -1.24002088 | 0.259985827 | 0.623943486 | -4.559971454 |
| **A_66_P104442** | Cog6 | -0.086706751 | -0.256624982 | 0.083211479 | 8.02304744 | -1.239927028 | 0.260018222 | 0.623943486 | -4.560016669 |
| **A_51_P403636** | Smad7 | -0.179606121 | -0.531591521 | 0.17237928 | 9.521013977 | -1.239880047 | 0.26003444 | 0.623943486 | -4.560039303 |
| **A_55_P1966114** | Epn2 | -0.38701621 | -1.145589742 | 0.371557321 | 10.48843002 | -1.239694458 | 0.260098514 | 0.623943486 | -4.560128709 |
| **A_52_P108089** | Mosmo | -0.127668686 | -0.377908236 | 0.122570864 | 9.996594379 | -1.239685802 | 0.260101503 | 0.623943486 | -4.560132879 |
| **A_51_P362209** | Commd10 | -0.121863923 | -0.360732506 | 0.11700466 | 9.770839692 | -1.239650644 | 0.260113643 | 0.623943486 | -4.560149815 |
| **A_51_P149562** | Apbb2 | -0.457082626 | -1.353145153 | 0.438979901 | 9.94861073 | -1.239480182 | 0.26017251 | 0.624007253 | -4.560231929 |
| **A_52_P523712** | Tbx5 | -0.868619295 | -2.57167001 | 0.834431419 | 8.094002565 | -1.23932463 | 0.260226238 | 0.624013015 | -4.560306856 |
| **A_51_P157462** | Rgn | -0.295112498 | -0.873780813 | 0.283555818 | 14.10800781 | -1.239199094 | 0.260269606 | 0.624013015 | -4.560367323 |
| **A_51_P401501** | Tmem213 | -1.559411252 | -4.617326657 | 1.498504153 | 8.408821344 | -1.239134321 | 0.260291985 | 0.624013015 | -4.560398521 |
| **A_55_P2157914** | NA | 0.121391687 | -0.116657108 | 0.359440482 | 8.261211811 | 1.239099408 | 0.260304048 | 0.624013015 | -4.560415337 |
| **A_52_P1004509** | BC065397 | 0.082081113 | -0.078925885 | 0.243088111 | 8.069166844 | 1.238744311 | 0.260426767 | 0.624131086 | -4.560586358 |
| **A_51_P337523** | Adcy2 | -0.177706222 | -0.526305995 | 0.17089355 | 7.847407436 | -1.238678856 | 0.260449394 | 0.624131086 | -4.560617881 |
| **A_51_P379710** | Rpn1 | -0.111337515 | -0.329744626 | 0.107069597 | 13.74501523 | -1.238676609 | 0.26045017 | 0.624131086 | -4.560618962 |
| **A_66_P104980** | Tceal5 | 0.331489428 | -0.318902177 | 0.981881033 | 12.59545223 | 1.238448478 | 0.260529043 | 0.624183091 | -4.560728823 |
| **A_51_P491635** | Ly6g6e | -0.206938466 | -0.612985353 | 0.199108422 | 8.001337121 | -1.238363742 | 0.260558345 | 0.624183091 | -4.560769627 |
| **A_55_P2019397** | Mtch1 | -0.295703268 | -0.875935215 | 0.284528679 | 14.21427776 | -1.238333656 | 0.26056875 | 0.624183091 | -4.560784115 |
| **A_55_P2133067** | Ube2v1 | 0.174732955 | -0.168264997 | 0.517730906 | 8.729187432 | 1.237845618 | 0.260737575 | 0.624203289 | -4.561019104 |
| **A_51_P278653** | Rprm | -0.542647851 | -1.6078569 | 0.522561198 | 7.956528305 | -1.237845463 | 0.260737628 | 0.624203289 | -4.561019179 |
| **A_55_P2065631** | Hook3 | -0.104803069 | -0.310537951 | 0.100931813 | 10.32050726 | -1.237796522 | 0.260754563 | 0.624203289 | -4.561042742 |
| **A_52_P93837** | Mme | -0.102843815 | -0.304754445 | 0.099066815 | 7.709497005 | -1.237662358 | 0.260800993 | 0.624203289 | -4.561107334 |
| **A_55_P1987201** | Capn2 | -0.185108331 | -0.548677393 | 0.178460731 | 12.03975086 | -1.237149642 | 0.260978495 | 0.624203289 | -4.561354156 |
| **A_55_P2144386** | Adgre1 | 0.181363152 | -0.174891243 | 0.537617546 | 10.96062168 | 1.23700659 | 0.261028038 | 0.624203289 | -4.561423015 |
| **A_55_P1988010** | Gm10406 | -0.25371408 | -0.75209531 | 0.24466715 | 7.960435488 | -1.236989694 | 0.26103389 | 0.624203289 | -4.561431147 |
| **A_52_P154101** | Calca | -0.304005526 | -0.901191346 | 0.293180294 | 7.799393392 | -1.236958646 | 0.261044644 | 0.624203289 | -4.561446092 |
| **A_51_P149714** | Ms4a6d | -0.08522006 | -0.252635936 | 0.082195817 | 8.646670273 | -1.23688232 | 0.261071083 | 0.624203289 | -4.56148283 |
| **A_55_P2007738** | Mir22hg | 0.143013351 | -0.137957105 | 0.423983807 | 8.06190984 | 1.236798975 | 0.261099956 | 0.624203289 | -4.561522945 |
| **A_55_P1986276** | Tmem41b | 0.10017627 | -0.096661471 | 0.297014012 | 10.65678 | 1.236629702 | 0.261158605 | 0.624203289 | -4.561604416 |
| **A_55_P2054027** | Chst5 | -0.460661434 | -1.365905078 | 0.44458221 | 7.874389381 | -1.236515487 | 0.261198184 | 0.624203289 | -4.561659386 |
| **A_55_P2032678** | Lox | -0.29337927 | -0.869927749 | 0.283169209 | 7.909006659 | -1.236450635 | 0.261220659 | 0.624203289 | -4.561690597 |
| **A_55_P2086978** | Col23a1 | -0.168513556 | -0.499682881 | 0.162655769 | 7.817646464 | -1.236425434 | 0.261229393 | 0.624203289 | -4.561702725 |
| **A_55_P1960202** | Vcan | -0.37583847 | -1.1144671 | 0.362790159 | 8.393973806 | -1.236397971 | 0.261238912 | 0.624203289 | -4.561715942 |
| **A_52_P642488** | Kcnk1 | -0.29982518 | -0.88910786 | 0.2894575 | 7.845661501 | -1.236310664 | 0.261269174 | 0.624203289 | -4.561757958 |
| **A_51_P341918** | Tsc22d1 | -0.313106798 | -0.928533923 | 0.302320328 | 11.95272172 | -1.236229353 | 0.261297361 | 0.624203289 | -4.561797087 |
| **A_55_P2088014** | Cdc42ep4 | -0.319554452 | -0.947759707 | 0.308650803 | 11.25378684 | -1.236022842 | 0.26136896 | 0.624203289 | -4.561896464 |
| **A_51_P417053** | Tuft1 | 0.168846215 | -0.163101784 | 0.500794215 | 11.01718172 | 1.235960135 | 0.261390704 | 0.624203289 | -4.561926638 |
| **A_52_P519538** | Uba6 | 0.108741089 | -0.105061122 | 0.3225433 | 8.22781292 | 1.235846911 | 0.26142997 | 0.624203289 | -4.561981119 |
| **A_51_P356967** | Lgi3 | -0.924732345 | -2.743178671 | 0.89371398 | 8.107785566 | -1.235659325 | 0.261495036 | 0.624203289 | -4.562071378 |
| **A_55_P1967568** | Chtop | 0.081629965 | -0.078920604 | 0.242180534 | 12.05316565 | 1.235437982 | 0.261571829 | 0.624203289 | -4.562177873 |
| **A_51_P288558** | Cfap52 | -0.484997724 | -1.438939674 | 0.468944227 | 7.911362859 | -1.235381069 | 0.261591578 | 0.624203289 | -4.562205255 |
| **A_51_P419117** | Rab15 | -1.16371337 | -3.45269013 | 1.12526339 | 8.714483462 | -1.235343771 | 0.26160452 | 0.624203289 | -4.562223199 |
| **A_55_P2026945** | Zranb2 | 0.185585703 | -0.179454763 | 0.550626169 | 10.82791288 | 1.235340542 | 0.261605641 | 0.624203289 | -4.562224752 |
| **A_55_P2169276** | 4732491K20Rik | 0.091176723 | -0.088166394 | 0.27051984 | 8.135201197 | 1.235328486 | 0.261609825 | 0.624203289 | -4.562230552 |
| **A_51_P516362** | Vps11 | -0.112997779 | -0.335332041 | 0.109336484 | 10.03238635 | -1.234942396 | 0.261743838 | 0.624203289 | -4.562416289 |
| **A_55_P1982451** | Hmgn5 | 0.158696807 | -0.153560483 | 0.470954097 | 11.24799708 | 1.234919906 | 0.261751646 | 0.624203289 | -4.562427108 |
| **A_55_P2089219** | NA | -1.337970595 | -3.970976474 | 1.295035284 | 8.325224503 | -1.234746856 | 0.261811734 | 0.624203289 | -4.562510349 |

| | | | | | | | | | |
|---|---|---|---|---|---|---|---|---|---|
| **A_55_P2040951** | Actc1 | -1.75763156 | -5.216746309 | 1.701483189 | 8.766385194 | -1.234656258 | 0.261843197 | 0.624203289 | -4.562553928 |
| **A_51_P208580** | Nop53 | -0.236891937 | -0.703146051 | 0.229362176 | 9.446263775 | -1.234555994 | 0.26187802 | 0.624203289 | -4.562602154 |
| **A_51_P353252** | Mal2 | -0.451550859 | -1.340315746 | 0.437214028 | 11.63476422 | -1.234533767 | 0.26188574 | 0.624203289 | -4.562612845 |
| **A_51_P516452** | Eif3f | -0.30151068 | -0.894983879 | 0.291962519 | 12.2308422 | -1.234482046 | 0.261903706 | 0.624203289 | -4.562637722 |
| **A_52_P232637** | Dhh | -0.153474597 | -0.455588712 | 0.148639518 | 7.810233144 | -1.234379425 | 0.261939355 | 0.624203289 | -4.562687079 |
| **A_55_P1966644** | Trpc6 | -0.087129951 | -0.258663853 | 0.08440395 | 8.41323568 | -1.234243063 | 0.261986732 | 0.624203289 | -4.562752662 |
| **A_51_P184353** | Nr2f6 | -0.105739626 | -0.313912148 | 0.102432896 | 12.64999337 | -1.234234141 | 0.261989832 | 0.624203289 | -4.562756953 |
| **A_51_P160544** | Efemp2 | -0.480082071 | -1.425301962 | 0.465137821 | 11.03376759 | -1.234143964 | 0.262021167 | 0.624203289 | -4.562800322 |
| **A_51_P128575** | Scgb1a1 | -1.491879493 | -4.429472437 | 1.445713452 | 8.565515571 | -1.23402886 | 0.262061168 | 0.624203289 | -4.562168006 |
| **A_51_P437717** | Glmp | -0.110956619 | -0.329439378 | 0.107526139 | 10.27829739 | -1.234011578 | 0.262067175 | 0.624203289 | -4.562863989 |
| **A_52_P42269** | Fgf10 | -0.301210798 | -0.89440016 | 0.291978564 | 7.794615678 | -1.233844336 | 0.262125306 | 0.624203289 | -4.562255861 |
| **A_55_P1962149** | Dnah10 | -0.302040398 | -0.896875725 | 0.292794929 | 7.793848407 | -1.233819044 | 0.262134098 | 0.624203289 | -4.562956577 |
| **A_55_P2147291** | NA | 0.096979786 | -0.094015219 | 0.287974791 | 16.03113471 | 1.233793283 | 0.262143053 | 0.624203289 | -4.562968965 |
| **A_51_P224468** | Maip1 | 0.13064391 | -0.126650557 | 0.387938377 | 10.72043753 | 1.233791893 | 0.262143537 | 0.624203289 | -4.562969633 |
| **A_51_P335969** | Des | -0.138969315 | -0.412709123 | 0.134770492 | 7.843253247 | -1.233570995 | 0.26222034 | 0.624203289 | -4.561736668 |
| **A_51_P486091** | Ift52 | 0.08316283 | -0.080689397 | 0.247015057 | 9.979964699 | 1.233275489 | 0.262323114 | 0.624203289 | -4.56321794 |
| **A_55_P2021530** | Clic4 | 0.103052165 | -0.099987366 | 0.306091696 | 8.184182932 | 1.233274378 | 0.2623235 | 0.624203289 | -4.563218474 |
| **A_55_P1960023** | NA | 0.091375891 | -0.088660197 | 0.271411979 | 8.372194319 | 1.233261707 | 0.262327908 | 0.624203289 | -4.563224566 |
| **A_55_P2066478** | Crtc3 | -0.136929556 | -0.406737142 | 0.13287803 | 9.830692829 | -1.233179322 | 0.262356567 | 0.624203289 | -4.563264176 |
| **A_51_P511511** | Stk33 | -0.390524362 | -1.160146475 | 0.379097751 | 7.935505194 | -1.232973615 | 0.262428138 | 0.624203289 | -4.563363074 |
| **A_55_P2004727** | NA | 0.265135233 | -0.257447978 | 0.787718445 | 8.571689032 | 1.23280725 | 0.262486034 | 0.624203289 | -4.563443053 |
| **A_55_P2007525** | Smu1 | 0.196160033 | -0.190493505 | 0.582813571 | 9.822683108 | 1.232740676 | 0.262509204 | 0.624203289 | -4.563475056 |
| **A_51_P159612** | Hebp2 | -0.09490699 | -0.281983014 | 0.092169035 | 8.302686795 | -1.232716633 | 0.262517573 | 0.624203289 | -4.563486614 |
| **A_55_P2006261** | Krt15 | -0.700622435 | -2.081814838 | 0.680569967 | 8.131356643 | -1.23257413 | 0.262567178 | 0.624203289 | -4.563555116 |
| **A_65_P01643** | Chd2 | 0.181797686 | -0.176620454 | 0.540215825 | 10.21414035 | 1.23248476 | 0.262598292 | 0.624203289 | -4.563598076 |
| **A_52_P442710** | Ndnf | -0.440902654 | -1.31019562 | 0.428390312 | 7.822778789 | -1.232422836 | 0.262619853 | 0.624203289 | -4.563627842 |
| **A_52_P5891** | Rflnb | -0.132579074 | -0.394017334 | 0.128859186 | 8.016631577 | -1.232222196 | 0.262689722 | 0.624203289 | -4.563724281 |
| **A_51_P209183** | Cxcl14 | -0.467695669 | -1.389997664 | 0.454606325 | 7.931084133 | -1.232177828 | 0.262705174 | 0.624203289 | -4.563745606 |
| **A_55_P2454099** | Oga | 0.23054036 | -0.224123363 | 0.685204083 | 8.928552846 | 1.23208268 | 0.262738315 | 0.624203289 | -4.563791337 |
| **A_55_P2027609** | Scgb3a2 | -1.725065871 | -5.127429736 | 1.677297994 | 8.501303667 | -1.231992645 | 0.262769678 | 0.624203289 | -4.56383461 |
| **A_52_P250578** | Tcaf1 | -0.127118463 | -0.377855014 | 0.123618088 | 7.902332867 | -1.231896367 | 0.262803219 | 0.624203289 | -4.563880881 |
| **A_55_P2162965** | Ptms | -0.170225071 | -0.505988688 | 0.165538547 | 13.1671435 | -1.231893252 | 0.262804305 | 0.624203289 | -4.563882379 |
| **A_55_P2159432** | Fndc4 | -0.164318795 | -0.488436868 | 0.159799277 | 10.36813307 | -1.231876537 | 0.262810128 | 0.624203289 | -4.563890412 |
| **A_55_P2171206** | Matn4 | -0.572980479 | -1.703248711 | 0.557287754 | 8.135387956 | -1.23180367 | 0.262835517 | 0.624203289 | -4.563925431 |
| **A_55_P2083814** | Zzz3 | 0.138382044 | -0.13463932 | 0.411403408 | 10.90867447 | 1.231590409 | 0.262909834 | 0.624203289 | -4.564027917 |
| **A_55_P2235970** | C130068B02Rik | 0.083404468 | -0.081152105 | 0.247961041 | 8.111924311 | 1.231564813 | 0.262918755 | 0.624203289 | -4.564040217 |
| **A_55_P2072801** | Lypd5 | 0.107487692 | -0.104585403 | 0.319560786 | 8.269858605 | 1.231562201 | 0.262919665 | 0.624203289 | -4.564041473 |
| **A_52_P24951** | Ado | 0.131537613 | -0.127995661 | 0.391070888 | 9.268827564 | 1.23151612 | 0.262935726 | 0.624203289 | -4.564063616 |
| **A_55_P2044167** | NA | 0.230049782 | -0.223880659 | 0.683980224 | 9.202496836 | 1.231446949 | 0.262959837 | 0.624203289 | -4.564096855 |
| **A_55_P2034300** | Tmem40 | 0.243203256 | -0.236711641 | 0.723118153 | 12.35739191 | 1.231369362 | 0.262986883 | 0.624203289 | -4.564134138 |
| **A_55_P2180551** | Sinhcaf | -0.160216819 | -0.476379539 | 0.155945902 | 8.357903722 | -1.231347502 | 0.262994504 | 0.624203289 | -4.564144642 |
| **A_66_P127411** | Acat2 | 0.296681925 | -0.28880876 | 0.882172611 | 10.36829575 | 1.231272805 | 0.263020546 | 0.624203289 | -4.564180535 |
| **A_55_P1956978** | Tln1 | 0.33049376 | -0.321731165 | 0.982718684 | 8.80559709 | 1.231257989 | 0.263025712 | 0.624203289 | -4.564187654 |
| **A_55_P2069251** | Prr18 | -0.688333799 | -2.046952901 | 0.670285304 | 8.04632805 | -1.231075156 | 0.263089464 | 0.624203289 | -4.564275504 |
| **A_55_P2096485** | Bcs1l | 0.123455038 | -0.120229088 | 0.367139164 | 11.05819883 | 1.23101903 | 0.263109038 | 0.624203289 | -4.564302471 |
| **A_55_P2140780** | Clip4 | -0.130936601 | -0.389391588 | 0.127518387 | 7.750856987 | -1.231003632 | 0.263114408 | 0.624203289 | -4.564309869 |
| **A_55_P2179299** | Ceacam13 | -0.170459698 | -0.506966153 | 0.166046757 | 7.870450778 | -1.230868066 | 0.263161691 | 0.624203289 | -4.564375003 |
| **A_55_P1955325** | Zfp820 | 0.100344712 | -0.097747086 | 0.298436511 | 8.407491148 | 1.230867165 | 0.263162005 | 0.624203289 | -4.564375435 |
| **A_51_P480982** | Glo1 | 0.166094864 | -0.161817087 | 0.494006816 | 12.84965192 | 1.230784866 | 0.263190714 | 0.624203289 | -4.564414975 |
| **A_52_P384314** | Tmem198b | -0.163432329 | -0.486119444 | 0.159254785 | 9.005774274 | -1.230664112 | 0.263232841 | 0.624203289 | -4.564472988 |

| | | | | | | | | | |
|---|---|---|---|---|---|---|---|---|---|
| **A_55_P2109585** | Plekha7 | 0.086783411 | -0.084594554 | 0.258161376 | 8.06428192 | 1.230452698 | 0.263306611 | 0.624203289 | -4.564574552 |
| **A_51_P130994** | NA | 0.086901631 | -0.084716658 | 0.25851992 | 11.25486789 | 1.230403474 | 0.26332379 | 0.624203289 | -4.564598198 |
| **A_55_P2022604** | Exoc3l4 | 0.134828039 | -0.131449138 | 0.401105217 | 10.61952276 | 1.230352239 | 0.263341671 | 0.624203289 | -4.56462281 |
| **A_51_P135389** | Zfp474 | -0.188083946 | -0.559580396 | 0.183412504 | 7.717104088 | -1.230212823 | 0.263390334 | 0.624203289 | -4.56468978 |
| **A_55_P2146171** | Cox20 | 0.18737454 | -0.208855444 | 0.583604524 | 12.59972592 | 1.291127226 | 0.263459022 | 0.624203289 | -4.541052548 |
| **A_55_P1970309** | Arhgap44 | -0.474322487 | -1.41138575 | 0.462740776 | 9.220166024 | -1.229951513 | 0.263481564 | 0.624203289 | -4.564815295 |
| **A_51_P296528** | Cers4 | -0.444700692 | -1.323383723 | 0.433982339 | 9.16380045 | -1.229755447 | 0.263550034 | 0.624203289 | -4.564909465 |
| **A_55_P1987121** | Rtraf | -0.214569677 | -0.702754444 | 0.27361509 | 13.61925182 | -1.355344419 | 0.263582745 | 0.624203289 | -4.532575247 |
| **A_51_P256246** | Tspan13 | -0.400226606 | -1.191207727 | 0.390754515 | 11.68276716 | -1.229484412 | 0.26364471 | 0.624203289 | -4.565039633 |
| **A_51_P455866** | Elf5 | -0.897960776 | -2.672810991 | 0.876889439 | 8.118498487 | -1.229359349 | 0.263688406 | 0.624203289 | -4.565099692 |
| **A_55_P2031999** | Dglucy | -0.121172835 | -0.36069347 | 0.118347799 | 8.624007262 | -1.229265024 | 0.263721366 | 0.624203289 | -4.565144988 |
| **A_55_P2452259** | Gls | 0.168432084 | -0.164522601 | 0.501386768 | 9.439235344 | 1.229200933 | 0.263743764 | 0.624203289 | -4.565175765 |
| **A_52_P550147** | NA | -0.453011245 | -1.348532424 | 0.442509933 | 8.182769377 | -1.229182336 | 0.263750263 | 0.624203289 | -4.565184695 |
| **A_55_P2011545** | Pyroxd1 | 0.112499543 | -0.109892389 | 0.334891476 | 8.545441252 | 1.229178399 | 0.263751639 | 0.624203289 | -4.565186586 |
| **A_51_P268734** | Kptn | -0.180747553 | -0.538171757 | 0.17667665 | 10.39394367 | -1.228772995 | 0.263893356 | 0.624203289 | -4.565381247 |
| **A_55_P2143517** | Fstl1 | -0.851338652 | -2.534877123 | 0.832199819 | 11.89252263 | -1.228747071 | 0.263902421 | 0.624203289 | -4.565393695 |
| **A_51_P195100** | Vps4b | -0.142576493 | -0.424535027 | 0.139382041 | 10.73514895 | -1.228700043 | 0.263918865 | 0.624203289 | -4.565416274 |
| **A_51_P483280** | Prnp | -0.141827076 | -0.422361012 | 0.138706859 | 10.68420516 | -1.228448445 | 0.264006857 | 0.624203289 | -4.565537068 |
| **A_55_P2084965** | Klk13 | -0.244921315 | -0.729459606 | 0.239616977 | 7.833841557 | -1.228235562 | 0.264081329 | 0.624203289 | -4.565639268 |
| **A_55_P2076462** | Lnx1 | -0.180134635 | -0.536536981 | 0.176267711 | 7.810380098 | -1.228117326 | 0.264122699 | 0.624203289 | -4.565696027 |
| **A_55_P2035946** | Penk | -0.604957962 | -1.801946805 | 0.59203088 | 8.35076723 | -1.228056318 | 0.264144047 | 0.624203289 | -4.565725313 |
| **A_51_P276943** | Gpc1 | -0.181560733 | -0.540818714 | 0.177697249 | 10.86195408 | -1.228000913 | 0.264163436 | 0.624203289 | -4.565751909 |
| **A_55_P2135601** | Polr2i | 0.15865433 | -0.155280157 | 0.472588817 | 10.24148023 | 1.227993563 | 0.264166008 | 0.624203289 | -4.565755437 |
| **A_55_P2066559** | Endou | -0.080524949 | -0.239876223 | 0.078826324 | 7.976134418 | -1.227886177 | 0.264203592 | 0.624203289 | -4.565806984 |
| **A_51_P346132** | Rhox9 | -0.098063277 | -0.292131006 | 0.096004452 | 7.729376168 | -1.227824425 | 0.264225206 | 0.624203289 | -4.565836625 |
| **A_55_P2083028** | NA | 0.209772863 | -0.205384377 | 0.624930102 | 11.48523639 | 1.227778088 | 0.264241426 | 0.624203289 | -4.565858866 |
| **A_51_P336509** | Zfpm2 | -0.10079665 | -0.300303087 | 0.098709788 | 8.253762961 | -1.227643791 | 0.264288441 | 0.624203289 | -4.565923326 |
| **A_55_P1989765** | NA | 0.188633455 | -0.184731392 | 0.561998303 | 8.694325052 | 1.227632799 | 0.264292289 | 0.624203289 | -4.565928602 |
| **A_51_P182471** | Celf2 | -0.201205821 | -0.599507902 | 0.197096259 | 10.04571373 | -1.227470763 | 0.264349024 | 0.624203289 | -4.566006372 |
| **A_51_P209122** | Agr2 | -0.174569966 | -0.520173427 | 0.171033495 | 7.753764098 | -1.227367538 | 0.264385173 | 0.624203289 | -4.566055913 |
| **A_55_P2091506** | Srp54c | -0.133261374 | -0.397116251 | 0.130593502 | 11.64316182 | -1.227219794 | 0.26443692 | 0.624203289 | -4.566126818 |
| **A_51_P125183** | Coq5 | 0.14508055 | -0.142182203 | 0.432343302 | 8.688651077 | 1.227193551 | 0.264446112 | 0.624203289 | -4.560823511 |
| **A_52_P479269** | Sdc1 | -0.307408641 | -0.916085876 | 0.301268593 | 9.790543848 | -1.227191114 | 0.264446966 | 0.624203289 | -4.566140582 |
| **A_55_P2025820** | NA | -0.093403934 | -0.278363436 | 0.091555568 | 7.880806395 | -1.227076698 | 0.264487047 | 0.624203289 | -4.566195489 |
| **A_55_P2023732** | Limch1 | -1.38389071 | -4.12471707 | 1.35693565 | 9.087185353 | -1.226883881 | 0.264554605 | 0.624203289 | -4.566288017 |
| **A_55_P1991134** | Fkbp1a | -0.230946175 | -0.688355683 | 0.226463333 | 13.0503709 | -1.226842393 | 0.264569143 | 0.624203289 | -4.566307925 |
| **A_55_P1982404** | Gpm6b | -0.437563866 | -1.304243506 | 0.429115774 | 9.6019017 | -1.226778188 | 0.264591643 | 0.624203289 | -4.566338733 |
| **A_55_P2182483** | Cd47 | 0.12098092 | -0.118653434 | 0.360615273 | 13.74393041 | 1.226735669 | 0.264606544 | 0.624203289 | -4.566359135 |
| **A_55_P2109454** | Gtf3c5 | 0.222865038 | -0.218582216 | 0.664312293 | 10.98788503 | 1.226722449 | 0.264611177 | 0.624203289 | -4.566365479 |
| **A_55_P2148243** | Wdr12 | 0.20383649 | -0.199933072 | 0.607606051 | 11.32433454 | 1.226680733 | 0.264625798 | 0.624203289 | -4.566385495 |
| **A_52_P8324** | Tmem178 | -0.391223084 | -1.166216592 | 0.383770424 | 8.019360863 | -1.226618745 | 0.264647525 | 0.624203289 | -4.566415238 |
| **A_51_P335460** | Scin | -0.098237634 | -0.292863541 | 0.096388274 | 7.725726141 | -1.226479891 | 0.2646962 | 0.624203289 | -4.566481862 |
| **A_66_P117682** | NA | 0.122914034 | -0.120630662 | 0.36645873 | 9.42163489 | 1.22632614 | 0.264750105 | 0.624203289 | -4.566555629 |
| **A_55_P2127934** | Palm3 | -0.152787174 | -0.455561688 | 0.149987339 | 7.808425257 | -1.226170244 | 0.264804772 | 0.624203289 | -4.566630422 |
| **A_55_P2056212** | Gpr3 | 0.117652559 | -0.115502161 | 0.350807279 | 8.738361735 | 1.226140827 | 0.264815089 | 0.624203289 | -4.566644535 |
| **A_55_P2011867** | Kif19a | 0.102277085 | -0.100409794 | 0.304963965 | 8.456495433 | 1.226128241 | 0.264819503 | 0.624203289 | -4.566650573 |
| **A_52_P449871** | Id4 | -0.105736766 | -0.315282941 | 0.103809409 | 7.8662087 | -1.226110113 | 0.264825861 | 0.624203289 | -4.56665927 |
| **A_52_P399934** | Dusp2 | -0.14508698 | -0.432652413 | 0.142478452 | 10.07485879 | -1.225956191 | 0.264879849 | 0.624203289 | -4.56673311 |
| **A_55_P1966749** | Cyp2s1 | -0.663759626 | -1.979354749 | 0.651835498 | 9.046063529 | -1.2259472 | 0.264883003 | 0.624203289 | -4.566737424 |
| **A_55_P1961152** | Pou3f1 | 0.097601533 | -0.095871288 | 0.291074355 | 8.095106454 | 1.225800695 | 0.2649344 | 0.624203289 | -4.566807703 |

| | | | | | | | | | |
|---|---|---|---|---|---|---|---|---|---|
| **A_55_P1967776** | Slc4a4 | -0.105745396 | -0.315383942 | 0.103893151 | 8.203114305 | -1.225669886 | 0.264980297 | 0.624203289 | -4.56687045 |
| **A_55_P2102540** | Olfml3 | -0.243546336 | -0.726384941 | 0.239292268 | 7.771533428 | -1.225639654 | 0.264990905 | 0.624203289 | -4.566884952 |
| **A_52_P601914** | Lemd2 | -0.118733555 | -0.35415098 | 0.11668387 | 11.66395214 | -1.225513371 | 0.265035223 | 0.624203289 | -4.566945525 |
| **A_51_P262721** | 0610009L18Rik | -0.292158647 | -0.871434163 | 0.287116869 | 9.478140454 | -1.225509722 | 0.265036503 | 0.624203289 | -4.566947275 |
| **A_51_P417074** | Arhgap8 | -0.764088257 | -2.279254482 | 0.751077969 | 8.535980216 | -1.225367714 | 0.265086347 | 0.624203289 | -4.567015388 |
| **A_51_P473240** | Zfp426 | -0.087688816 | -0.261587164 | 0.086209533 | 10.2298584 | -1.225270379 | 0.265120515 | 0.624203289 | -4.567062072 |
| **A_55_P2178632** | Tmem80 | 0.176773216 | -0.173877146 | 0.527423579 | 9.758826756 | 1.22496974 | 0.265226075 | 0.624203289 | -4.567206257 |
| **A_55_P2001155** | Rpl31 | -0.135398146 | -0.403981513 | 0.133185221 | 15.00491922 | -1.224945576 | 0.265234561 | 0.624203289 | -4.567217845 |
| **A_51_P328001** | Sftpc | -3.38223246 | -10.09174939 | 3.327284465 | 9.374112671 | -1.224885207 | 0.265255763 | 0.624203289 | -4.567246796 |
| **A_52_P222350** | Flnb | -0.108193103 | -0.322842536 | 0.106456329 | 9.873538388 | -1.224765721 | 0.265297731 | 0.624203289 | -4.567304095 |
| **A_55_P2154536** | Hagh | -0.156172078 | -0.466011766 | 0.15366761 | 12.44789652 | -1.224755876 | 0.265301189 | 0.624203289 | -4.567308816 |
| **A_55_P2131920** | Prrx1 | -0.171961513 | -0.51313278 | 0.169209755 | 7.826165009 | -1.224734629 | 0.265308652 | 0.624203289 | -4.567319005 |
| **A_52_P350664** | Pygb | -0.139392544 | -0.415950185 | 0.137165098 | 9.325405441 | -1.224720733 | 0.265313534 | 0.624203289 | -4.567325668 |
| **A_55_P2017640** | Phf7 | 0.104287215 | -0.102647243 | 0.311221673 | 9.935768553 | 1.224563878 | 0.265368639 | 0.624203289 | -4.567400884 |
| **A_66_P108345** | Sgcb | -0.237346982 | -0.708331152 | 0.233637189 | 8.640641017 | -1.224505078 | 0.265389298 | 0.624203289 | -4.567429079 |
| **A_55_P2155499** | Tmem9 | -0.155829444 | -0.465096272 | 0.153437383 | 9.955477591 | -1.224332484 | 0.265449948 | 0.624203289 | -4.567511835 |
| **A_55_P2092046** | NA | 0.250936066 | -0.247099463 | 0.748971596 | 9.680326831 | 1.224294643 | 0.265463247 | 0.624203289 | -4.56752998 |
| **A_55_P1997977** | Rab2b | -0.135223908 | -0.403606418 | 0.133158602 | 9.151062649 | -1.224284811 | 0.265466702 | 0.624203289 | -4.567534693 |
| **A_55_P2002115** | NA | -0.203930075 | -0.608696427 | 0.200836276 | 15.71600281 | -1.224221682 | 0.26548889 | 0.624203289 | -4.567564962 |
| **A_51_P225179** | Smndc1 | 0.144516912 | -0.142328874 | 0.431362697 | 8.852272871 | 1.224202853 | 0.265495508 | 0.624203289 | -4.567573989 |
| **A_52_P608132** | Snx32 | 0.111977151 | -0.110301367 | 0.334255669 | 8.287920713 | 1.22409496 | 0.265533434 | 0.624203289 | -4.567625718 |
| **A_55_P2000027** | Spink2 | -0.144963389 | -0.432725434 | 0.142798656 | 7.759870918 | -1.224074952 | 0.265540467 | 0.624203289 | -4.567635311 |
| **A_55_P2147186** | A430005L14Rik | 0.100368036 | -0.098878408 | 0.299614481 | 12.64812045 | 1.224018653 | 0.265560259 | 0.624203289 | -4.567662302 |
| **A_66_P136228** | Rai14 | -0.319793952 | -0.95466562 | 0.315077717 | 10.71021546 | -1.223960742 | 0.265580619 | 0.624203289 | -4.567690067 |
| **A_55_P2179030** | Ostf1 | -0.100745224 | -0.300751871 | 0.099261422 | 11.80826265 | -1.223948741 | 0.265584839 | 0.624203289 | -4.56769582 |
| **A_55_P2177639** | NA | 0.184527215 | -0.18181568 | 0.550870109 | 15.96619243 | 1.223927921 | 0.265592158 | 0.624203289 | -4.567705801 |
| **A_55_P2134092** | Brsk1 | -0.098112791 | -0.292924145 | 0.096698562 | 7.889323927 | -1.223755223 | 0.265652885 | 0.624203289 | -4.567788593 |
| **A_55_P2169714** | Gcn1 | 0.107325914 | -0.105782593 | 0.320434421 | 12.88197543 | 1.223733916 | 0.265660378 | 0.624203289 | -4.567798807 |
| **A_55_P2039379** | Slc27a6 | -0.409036405 | -1.221431716 | 0.403358905 | 7.880232902 | -1.223426108 | 0.265768646 | 0.624203289 | -4.567946359 |
| **A_55_P2098226** | C1rl | -2.795439036 | -8.34753941 | 2.756661337 | 8.620693785 | -1.223420929 | 0.265770468 | 0.624203289 | -4.567948842 |
| **A_55_P2101392** | Tmem237 | 0.101034823 | -0.099643016 | 0.301712662 | 8.21523628 | 1.223361635 | 0.265791328 | 0.624203289 | -4.567977263 |
| **A_55_P1997931** | Map9 | 0.228845227 | -0.225705887 | 0.68339634 | 9.63345776 | 1.223326325 | 0.265803752 | 0.624203289 | -4.567994188 |
| **A_55_P2026794** | Unc50 | 0.096631305 | -0.095324845 | 0.288587454 | 11.70823209 | 1.22320431 | 0.265846684 | 0.624203289 | -4.568052672 |
| **A_55_P1954271** | Col26a1 | -0.245522923 | -0.733260307 | 0.242214461 | 7.890256636 | -1.223176674 | 0.26585641 | 0.624203289 | -4.568065918 |
| **A_55_P1983177** | Acsl4 | 0.226558997 | -0.223534797 | 0.676652792 | 12.29560694 | 1.223098625 | 0.265883876 | 0.624203289 | -4.568103327 |
| **A_55_P1990121** | Aqp5 | -1.098119544 | -3.279810734 | 1.083571647 | 8.234416045 | -1.223036821 | 0.265905628 | 0.624203289 | -4.568132949 |
| **A_55_P2004876** | Srpx2 | -0.196562867 | -0.587143226 | 0.194017492 | 7.835136452 | -1.222853038 | 0.265970318 | 0.624260633 | -4.56822103 |
| **A_55_P2066453** | NA | -0.17707683 | -0.528963363 | 0.174809703 | 7.840462237 | -1.222762977 | 0.266002024 | 0.624260633 | -4.568264191 |
| **A_55_P2124941** | Brinp1 | -0.491079271 | -1.467021266 | 0.484862723 | 8.010427194 | -1.222674306 | 0.266033244 | 0.624260633 | -4.568306686 |
| **A_55_P2065866** | Cygb | -0.485039815 | -1.449037563 | 0.478957933 | 10.08068319 | -1.222600472 | 0.266059242 | 0.624260633 | -4.568342068 |
| **A_55_P2111523** | Higd1b | -1.45615819 | -4.350613713 | 1.438297333 | 8.817604413 | -1.222432438 | 0.266118418 | 0.624318466 | -4.56842259 |
| **A_55_P2048406** | NA | -0.163508279 | -0.488544553 | 0.161527995 | 12.00909758 | -1.222337414 | 0.266151888 | 0.624318466 | -4.568468125 |
| **A_55_P2081183** | Slc28a1 | 0.181455506 | -0.179290414 | 0.542201426 | 8.837295612 | 1.22222711 | 0.266190744 | 0.624318466 | -4.568520979 |
| **A_52_P66226** | Rab33a | 0.097493462 | -0.096372844 | 0.291359768 | 8.164842267 | 1.221958188 | 0.266285495 | 0.624318466 | -4.568649829 |
| **A_51_P121179** | Erfe | 0.195336412 | -0.193152413 | 0.583825237 | 8.694245822 | 1.221765518 | 0.266353397 | 0.624318466 | -4.567996042 |
| **A_55_P2079103** | Pan2 | 0.084280524 | -0.083363422 | 0.251924471 | 10.98210412 | 1.221581767 | 0.266418171 | 0.624318466 | -4.568830169 |
| **A_51_P433026** | Plpp6 | -0.134682423 | -0.402592298 | 0.133227452 | 9.042541151 | -1.221533518 | 0.266435181 | 0.624318466 | -4.568853283 |
| **A_55_P1953713** | Bop1 | 0.150475582 | -0.14886069 | 0.449811854 | 11.00211941 | 1.221489883 | 0.266450566 | 0.624318466 | -4.568874187 |
| **A_51_P134452** | Lpar3 | -0.111169923 | -0.33231907 | 0.109979224 | 7.687434202 | -1.221476811 | 0.266455175 | 0.624318466 | -4.568880449 |
| **A_55_P2283207** | Maz | 0.139619722 | -0.138136947 | 0.417376391 | 8.169114283 | 1.221421226 | 0.266474774 | 0.624318466 | -4.568907076 |

| | | | | | | | | | |
|---|---|---|---|---|---|---|---|---|---|
| **A_55_P2110177** | NA | 0.169219843 | -0.167431002 | 0.505870687 | 13.78848583 | 1.221391148 | 0.26648538 | 0.624318466 | -4.568921485 |
| **A_51_P467629** | Efcab10 | -0.450459593 | -1.346682833 | 0.445763648 | 7.88780477 | -1.221301322 | 0.266517056 | 0.624318466 | -4.568964514 |
| **A_55_P2134246** | Serpinb9 | -0.437512527 | -1.308003726 | 0.432978671 | 8.204225488 | -1.221263275 | 0.266530473 | 0.624318466 | -4.568982739 |
| **A_55_P2173952** | Myh6 | -1.855285502 | -5.546678639 | 1.836107636 | 8.771460637 | -1.221247364 | 0.266536085 | 0.624318466 | -4.56899036 |
| **A_55_P2085288** | Acoxl | -0.207643447 | -0.620840347 | 0.205553453 | 7.817135759 | -1.221080692 | 0.266594872 | 0.624380502 | -4.569070196 |
| **A_55_P2144120** | Bcl2l2 | 0.094281974 | -0.093358586 | 0.281922534 | 7.85281409 | 1.220914172 | 0.266653617 | 0.624394928 | -4.569149955 |
| **A_55_P2058601** | Gm7097 | -0.12570396 | -0.375887731 | 0.124479811 | 7.931082077 | -1.220880098 | 0.266665639 | 0.624394928 | -4.569166275 |
| **A_51_P289889** | Npnt | -0.258916259 | -0.774282359 | 0.256449841 | 7.888213239 | -1.220749806 | 0.266711613 | 0.624415627 | -4.569228678 |
| **A_52_P550218** | Trmt12 | 0.138245721 | -0.136980117 | 0.413471559 | 8.690665546 | 1.220522201 | 0.26679194 | 0.624415627 | -4.569337682 |
| **A_51_P487278** | Yipf3 | -0.166542678 | -0.498126088 | 0.165040732 | 11.06068908 | -1.220438609 | 0.266821447 | 0.624415627 | -4.569377714 |
| **A_52_P465886** | Gramd3 | -0.24134902 | -0.721871842 | 0.239173801 | 9.090806723 | -1.22043516 | 0.266822664 | 0.624415627 | -4.569379366 |
| **A_51_P153423** | Fndc1 | -0.65463803 | -1.958090046 | 0.648813986 | 8.196777907 | -1.220363956 | 0.266847801 | 0.624415627 | -4.569413464 |
| **A_55_P2175284** | Ei24 | -0.128015198 | -0.382921312 | 0.126890917 | 9.044147889 | -1.220293976 | 0.266872507 | 0.624415627 | -4.569446976 |
| **A_55_P2012899** | Vsig4 | -0.381189519 | -1.140270309 | 0.377891272 | 10.4377859 | -1.220214379 | 0.266900611 | 0.624415627 | -4.569485091 |
| **A_55_P2135098** | Rplp0 | 0.205833685 | -0.204095271 | 0.615762642 | 17.31862122 | 1.22008768 | 0.266945351 | 0.624444717 | -4.569545761 |
| **A_66_P111011** | Gata3 | 0.090284611 | -0.089541808 | 0.27011103 | 8.181291253 | 1.219953912 | 0.266992594 | 0.624479653 | -4.569609812 |
| **A_55_P2104119** | NA | -0.142482447 | -0.426312076 | 0.141347181 | 8.591508232 | -1.219794937 | 0.26704875 | 0.624535424 | -4.56968593 |
| **A_51_P447988** | Atp6ap1 | 0.119353437 | -0.118467959 | 0.357174833 | 9.767100585 | 1.219458977 | 0.267167455 | 0.624737446 | -4.569846776 |
| **A_51_P170807** | Map3k6 | -0.221916922 | -0.664218686 | 0.220384843 | 10.15209795 | -1.219143806 | 0.267278856 | 0.624747089 | -4.569997654 |
| **A_52_P613241** | Icam1 | -0.152564282 | -0.456664169 | 0.151535605 | 7.911667537 | -1.219045177 | 0.267313726 | 0.624747089 | -4.570044866 |
| **A_51_P254986** | Serf1 | -0.316826757 | -0.948350838 | 0.314697323 | 11.99807607 | -1.219032057 | 0.267318365 | 0.624747089 | -4.570051147 |
| **A_55_P2108644** | NA | 0.157647375 | -0.156589603 | 0.471884353 | 13.12856398 | 1.219025085 | 0.26732083 | 0.624747089 | -4.570054484 |
| **A_55_P1979508** | Clic3 | -1.33926718 | -4.008891952 | 1.330357593 | 8.670252159 | -1.218990136 | 0.267333187 | 0.624747089 | -4.570071213 |
| **A_55_P1971889** | F3 | -0.612569963 | -1.833776037 | 0.60863611 | 8.550844701 | -1.218849073 | 0.267383069 | 0.624749358 | -4.570138734 |
| **A_51_P291839** | Cox14 | -0.122128795 | -0.365610974 | 0.121353384 | 13.76031442 | -1.218804588 | 0.267398802 | 0.624749358 | -4.570160027 |
| **A_55_P2009752** | Hlf | 0.249373352 | -0.247937231 | 0.746683935 | 9.486493854 | 1.218443881 | 0.267526397 | 0.624942829 | -4.570332666 |
| **A_52_P1092823** | Irx1 | -1.403708709 | -4.203267008 | 1.395849591 | 8.338767434 | -1.218346072 | 0.267561005 | 0.624942829 | -4.570379475 |
| **A_55_P2012047** | Gigyf1 | -0.100135028 | -0.299853435 | 0.09958338 | 9.675979971 | -1.218291223 | 0.267580414 | 0.624942829 | -4.570405724 |
| **A_51_P425824** | C1qtnf12 | 0.197568508 | -0.196507996 | 0.591645013 | 13.02632489 | 1.218204972 | 0.267610937 | 0.624942829 | -4.570447 |
| **A_51_P143468** | Klhl26 | 0.110066643 | -0.109517283 | 0.329650569 | 8.998155317 | 1.217974973 | 0.267692345 | 0.62499569 | -4.570557062 |
| **A_66_P125634** | Wdr63 | 0.171964199 | -0.17111059 | 0.515038989 | 9.580571917 | 1.217958316 | 0.267698242 | 0.62499569 | -4.570565032 |
| **A_52_P131372** | NA | -0.158665002 | -0.475302738 | 0.157972734 | 16.54166796 | -1.217591645 | 0.267828073 | 0.62518258 | -4.570740479 |
| **A_55_P2100065** | Fsd1l | 0.133101863 | -0.132553751 | 0.398757477 | 8.133850565 | 1.217442123 | 0.267881031 | 0.62518258 | -4.570812017 |
| **A_55_P1981731** | Nr2c2 | -0.124540476 | -0.373111635 | 0.124030684 | 8.37720465 | -1.217427116 | 0.267886346 | 0.62518258 | -4.570819197 |
| **A_55_P2232988** | Scube2 | -0.652242271 | -1.954474735 | 0.649990193 | 8.224327638 | -1.217036526 | 0.268024732 | 0.62518258 | -4.571006056 |
| **A_55_P1979595** | Selenon | -0.114462595 | -0.342993038 | 0.114067847 | 10.73867152 | -1.217034015 | 0.268025622 | 0.62518258 | -4.571007257 |
| **A_66_P121117** | Serpina1d | -0.358121084 | -1.073166751 | 0.356924582 | 15.02735298 | -1.216968398 | 0.268048876 | 0.62518258 | -4.571038646 |
| **A_51_P461040** | Crct1 | 0.169778597 | -0.169223603 | 0.508780797 | 7.967559556 | 1.216924439 | 0.268064456 | 0.62518258 | -4.571059674 |
| **A_51_P221256** | Slc34a2 | -0.530611482 | -1.590106121 | 0.528883158 | 7.925947098 | -1.21691731 | 0.268066983 | 0.62518258 | -4.571063084 |
| **A_52_P410685** | Krt7 | -0.898276311 | -2.691913047 | 0.895360426 | 8.672903395 | -1.216910519 | 0.268069389 | 0.62518258 | -4.571066333 |
| **A_55_P2170813** | Pkig | 0.119602692 | -0.119242577 | 0.358447962 | 12.87332296 | 1.216767216 | 0.268120184 | 0.625225605 | -4.57113488 |
| **A_55_P2005570** | Celf2 | -0.287267111 | -0.861111039 | 0.286576816 | 10.93276363 | -1.216396905 | 0.268251482 | 0.625390854 | -4.571312001 |
| **A_51_P328089** | Crb3 | -0.164606014 | -0.493425757 | 0.16421373 | 8.995680195 | -1.216384845 | 0.268255759 | 0.625390854 | -4.571317769 |
| **A_55_P2016166** | Wfdc2 | -1.325837903 | -3.974978958 | 1.323303152 | 8.656033587 | -1.216097894 | 0.268357541 | 0.62555269 | -4.571455004 |
| **A_55_P1979616** | Tmem189 | 0.145299429 | -0.145061826 | 0.435660684 | 8.765092411 | 1.215929605 | 0.268417248 | 0.625616422 | -4.571535482 |
| **A_55_P2003903** | Bfsp1 | -0.139708466 | -0.418956035 | 0.139539103 | 7.89390346 | -1.215672275 | 0.268508569 | 0.625655287 | -4.571658532 |
| **A_55_P2136763** | NA | -0.08926628 | -0.2677247 | 0.089192141 | 13.31693555 | -1.215440155 | 0.268590966 | 0.625655287 | -4.57176952 |
| **A_55_P2033922** | Tsga8 | 0.105504448 | -0.105422968 | 0.316431864 | 8.067404095 | 1.215404742 | 0.268603538 | 0.625655287 | -4.571786451 |
| **A_52_P344376** | Eif4a2 | -0.12130307 | -0.363827804 | 0.121221663 | 9.074294529 | -1.215343232 | 0.268625378 | 0.625655287 | -4.57181586 |
| **A_55_P1960556** | Acox1 | -0.392640976 | -1.177659957 | 0.392378006 | 14.07744167 | -1.215342405 | 0.268625671 | 0.625655287 | -4.571816256 |

| | | | | | | | | | |
|---|---|---|---|---|---|---|---|---|---|
| **A_51_P377210** | Caap1 | 0.106381927 | -0.106319824 | 0.319083678 | 8.410061945 | 1.215290149 | 0.268644226 | 0.625655287 | -4.57184124 |
| **A_52_P461343** | Upk3a | -0.437770659 | -1.313332056 | 0.437790737 | 7.898261301 | -1.214907559 | 0.268780107 | 0.625655287 | -4.572024146 |
| **A_55_P1977910** | NA | -0.373483093 | -1.120482171 | 0.373515984 | 8.343275184 | -1.214881926 | 0.268789213 | 0.625655287 | -4.5720364 |
| **A_55_P2041961** | Bmp2 | 0.196431902 | -0.196476353 | 0.589340158 | 9.350400502 | 1.214797972 | 0.268819039 | 0.625655287 | -4.572076533 |
| **A_55_P2069885** | Slc50a1 | -0.103601402 | -0.310841341 | 0.103638536 | 9.021100747 | -1.214717728 | 0.26884755 | 0.625655287 | -4.572114892 |
| **A_55_P1958697** | Rbm33 | 0.185685435 | -0.18576197 | 0.55713284 | 10.82658559 | 1.21468509 | 0.268859147 | 0.625655287 | -4.572130493 |
| **A_51_P118603** | Wdr82 | 0.099407243 | -0.09945921 | 0.298273695 | 10.30409886 | 1.214617939 | 0.268883009 | 0.625655287 | -4.566476231 |
| **A_52_P252737** | Rangap1 | 0.187539013 | -0.187650608 | 0.562728633 | 11.4630574 | 1.214574053 | 0.268898604 | 0.625655287 | -4.572183569 |
| **A_51_P233947** | Tbc1d10b | 0.107462735 | -0.107540233 | 0.322465703 | 12.16675459 | 1.214497493 | 0.268925813 | 0.625655287 | -4.572220164 |
| **A_66_P118165** | NA | 0.466801434 | -0.467218471 | 1.400821339 | 10.83540031 | 1.214392954 | 0.268962968 | 0.625655287 | -4.57227013 |
| **A_55_P2096883** | Steap3 | 0.27698905 | -0.277261454 | 0.831239555 | 10.46428601 | 1.214338301 | 0.268982395 | 0.625655287 | -4.572296253 |
| **A_55_P2058152** | Ppp4r3b | 0.122496126 | -0.122634473 | 0.367626726 | 9.577081747 | 1.214249735 | 0.26901388 | 0.625655287 | -4.572338583 |
| **A_55_P2135632** | Rps18 | -0.141418352 | -0.424450034 | 0.141613329 | 15.95871031 | -1.214098465 | 0.269067661 | 0.625655287 | -4.57241088 |
| **A_52_P413395** | Sln | -1.240141256 | -3.722144364 | 1.241861853 | 8.363897728 | -1.214093191 | 0.269069537 | 0.625655287 | -4.5724134 |
| **A_55_P2082478** | Kidins220 | 0.37981526 | -0.380362933 | 1.139993453 | 9.595983885 | 1.214060115 | 0.269081298 | 0.625655287 | -4.572429208 |
| **A_55_P2003513** | Hsph1 | 0.171785625 | -0.172073114 | 0.515644363 | 9.811254403 | 1.213919652 | 0.269131248 | 0.625696162 | -4.572496336 |
| **A_55_P2090209** | Tpm3 | -0.153531579 | -0.460907027 | 0.153843868 | 14.61998947 | -1.213701061 | 0.269208997 | 0.625760974 | -4.572600795 |
| **A_51_P314679** | Rrn3 | 0.107029671 | -0.107270701 | 0.321330044 | 10.7834809 | 1.213568945 | 0.269255998 | 0.625760974 | -4.572663927 |
| **A_55_P1968235** | Rab33b | -0.155207353 | -0.466004037 | 0.155589331 | 9.630890999 | -1.213442226 | 0.269301085 | 0.625760974 | -4.572724477 |
| **A_55_P1965827** | Ubb | -0.10252799 | -0.307845814 | 0.102789834 | 16.10536001 | -1.213386002 | 0.269321092 | 0.625760974 | -4.572751342 |
| **A_55_P2206605** | 5830444B04Rik | -0.08254375 | -0.2478566 | 0.0827691 | 7.793042389 | -1.213279248 | 0.269359084 | 0.625760974 | -4.572802349 |
| **A_66_P119735** | BC049265 | 0.253328843 | -0.254032787 | 0.760690473 | 8.556972629 | 1.213249744 | 0.269369584 | 0.625760974 | -4.572816446 |
| **A_66_P100050** | Ndufaf8 | 0.114994852 | -0.115323099 | 0.345312804 | 9.856205577 | 1.213203904 | 0.269385899 | 0.625760974 | -4.572838348 |
| **A_55_P1981959** | Trafd1 | -0.226812658 | -0.681121254 | 0.227495938 | 12.84014208 | -1.213108155 | 0.269419981 | 0.625760974 | -4.572884094 |
| **A_55_P2093764** | Eif2s3x | 0.235648708 | -0.23639196 | 0.707689376 | 9.608568882 | 1.213022443 | 0.269450493 | 0.625760974 | -4.572925044 |
| **A_51_P268193** | Slc7a10 | -0.275654623 | -0.827912019 | 0.276602773 | 7.876818825 | -1.212849541 | 0.269512053 | 0.625762988 | -4.573007647 |
| **A_55_P1973289** | Mapk9 | 0.086349863 | -0.086673012 | 0.259372737 | 10.32731829 | 1.212666325 | 0.269577298 | 0.625762988 | -4.573095171 |
| **A_51_P142091** | Asb8 | 0.122763264 | -0.123226866 | 0.368753395 | 10.95419706 | 1.212645707 | 0.269584641 | 0.625762988 | -4.573105021 |
| **A_52_P670812** | Snord123 | -0.536972346 | -1.613024702 | 0.539080011 | 10.04972532 | -1.212555724 | 0.26961669 | 0.625762988 | -4.573148004 |
| **A_55_P2036854** | Pnkd | -0.116425135 | -0.349771479 | 0.116921209 | 11.09762795 | -1.212352572 | 0.26968906 | 0.625762988 | -4.573245044 |
| **A_66_P124549** | H13 | -0.14332003 | -0.430637294 | 0.143997234 | 12.8136901 | -1.212071826 | 0.269789098 | 0.625762988 | -4.573379136 |
| **A_55_P2049597** | Tpp2 | 0.137034678 | -0.137703729 | 0.411773086 | 10.03496328 | 1.211976774 | 0.269822975 | 0.625762988 | -4.573424533 |
| **A_52_P667477** | Fyco1 | -0.105249196 | -0.316276019 | 0.105777628 | 8.372446783 | -1.211893099 | 0.269852801 | 0.625762988 | -4.573464495 |
| **A_66_P137605** | Cgas | -0.126987314 | -0.381636071 | 0.127661442 | 9.256092256 | -1.211719138 | 0.269914817 | 0.625762988 | -4.573547573 |
| **A_66_P137654** | Smpd2 | -0.096815842 | -0.290961759 | 0.097330074 | 8.110447595 | -1.211717433 | 0.269915425 | 0.625762988 | -4.573548388 |
| **A_52_P116372** | Ppp1r15b | 0.114481581 | -0.115105239 | 0.3440684 | 8.909491202 | 1.211635124 | 0.269944772 | 0.625762988 | -4.573587694 |
| **A_51_P386983** | Tnni3 | -0.829583605 | -2.493329679 | 0.83416247 | 8.140218217 | -1.211591745 | 0.26996024 | 0.625762988 | -4.573608409 |
| **A_51_P155843** | Igsf10 | -0.209942263 | -0.630992062 | 0.211107536 | 8.390777711 | -1.211573034 | 0.269966912 | 0.625762988 | -4.573617344 |
| **A_51_P293938** | Rasl11b | -0.837849529 | -2.518255127 | 0.842556069 | 9.135505926 | -1.211532584 | 0.269981336 | 0.625762988 | -4.57363666 |
| **A_55_P2003096** | Pla2g6 | -0.201804371 | -0.606568052 | 0.20295931 | 9.85455752 | -1.211468764 | 0.270004096 | 0.625762988 | -4.573667135 |
| **A_55_P2166399** | Stk11 | 0.146972604 | -0.147824837 | 0.441770046 | 10.49984854 | 1.21142315 | 0.270020364 | 0.625762988 | -4.573688917 |
| **A_55_P2071236** | Aqp5 | -0.350874629 | -1.054724789 | 0.352975532 | 7.895301717 | -1.211308991 | 0.270061081 | 0.625762988 | -4.573743427 |
| **A_55_P2340448** | B230114P17Rik | -0.234003183 | -0.70347985 | 0.235473484 | 8.207497835 | -1.211130501 | 0.270124754 | 0.625762988 | -4.573828652 |
| **A_52_P24783** | Pds5b | 0.111717638 | -0.112448668 | 0.335883944 | 9.345514627 | 1.21097339 | 0.270180812 | 0.625762988 | -4.573903665 |
| **A_55_P2048418** | Mdfic | -0.225638864 | -0.678399351 | 0.227121622 | 10.04546738 | -1.210956594 | 0.270186805 | 0.625762988 | -4.573911684 |
| **A_52_P257812** | Lpl | -0.290431041 | -0.873207407 | 0.292345325 | 11.13392956 | -1.210944641 | 0.270191071 | 0.625762988 | -4.573917391 |
| **A_66_P115467** | Sfta2 | -2.032793584 | -6.112021082 | 2.046433914 | 9.199102349 | -1.210872856 | 0.270216688 | 0.625762988 | -4.573951663 |
| **A_55_P2065731** | Medag | 0.122500576 | -0.123323312 | 0.368324464 | 16.45966758 | 1.210869209 | 0.270217989 | 0.625762988 | -4.573953404 |
| **A_55_P2020552** | Iscu | 0.082090104 | -0.082651227 | 0.246831435 | 12.01759506 | 1.210797252 | 0.27024367 | 0.625762988 | -4.573987757 |
| **A_65_P10913** | Tgfb2 | -0.848263412 | -2.55088838 | 0.854361555 | 8.839686483 | -1.210583992 | 0.270319793 | 0.625762988 | -4.574089566 |

| | | | | | | | | | |
|---|---|---|---|---|---|---|---|---|---|
| **A_55_P2059680** | Gm11938 | 0.220095598 | -0.221681483 | 0.661872679 | 8.898382465 | 1.210574061 | 0.270323338 | 0.625762988 | -4.574094307 |
| **A_55_P2033510** | Ppm1m | -0.174006629 | -0.523275402 | 0.175262144 | 10.31446536 | -1.210568099 | 0.270325467 | 0.625762988 | -4.574097153 |
| **A_51_P324838** | Evpl | -0.661240851 | -1.988895387 | 0.666413686 | 8.514996981 | -1.210201763 | 0.270456275 | 0.625807474 | -4.574272022 |
| **A_55_P1998442** | Sema3f | -0.395115199 | -1.188453483 | 0.398223085 | 9.752846997 | -1.210175937 | 0.270465499 | 0.625807474 | -4.57428435 |
| **A_55_P1969523** | Sumo2 | 0.097334308 | -0.098101109 | 0.292769725 | 11.00615994 | 1.21016856 | 0.270468134 | 0.625807474 | -4.574287871 |
| **A_55_P1978795** | Pja2 | -0.116153986 | -0.349380289 | 0.117072317 | 10.90986524 | -1.2101516 | 0.270474191 | 0.625807474 | -4.574295966 |
| **A_51_P371745** | Havcr1 | 0.302550976 | -0.30513624 | 0.910238193 | 9.211708641 | 1.209766759 | 0.270611674 | 0.626050633 | -4.574479643 |
| **A_51_P518893** | Ccin | 0.290278953 | -0.29288395 | 0.873441856 | 9.932265576 | 1.209508286 | 0.270704045 | 0.626076763 | -4.574602994 |
| **A_55_P2070838** | NA | -0.180727262 | -0.543807586 | 0.182353063 | 14.54092173 | -1.209495183 | 0.270708729 | 0.626076763 | -4.574609247 |
| **A_55_P1982733** | Dnajc5 | 0.156550361 | -0.157966971 | 0.471067694 | 9.097241302 | 1.209463259 | 0.27072014 | 0.626076763 | -4.574624481 |
| **A_52_P509965** | Ager | -1.187833918 | -3.575247697 | 1.19957986 | 8.3113034 | -1.208958005 | 0.270900793 | 0.626166439 | -4.57486557 |
| **A_55_P2102424** | Msto1 | 0.17077838 | -0.172477375 | 0.514034136 | 9.95672582 | 1.208921918 | 0.2709137 | 0.626166439 | -4.574882788 |
| **A_51_P203501** | Vars | 0.190891336 | -0.192819525 | 0.574602197 | 10.12433996 | 1.208830237 | 0.270946492 | 0.626166439 | -4.57492653 |
| **A_52_P655687** | Egfl6 | -0.293156573 | -0.882459215 | 0.296146068 | 7.789126128 | -1.208772129 | 0.270967279 | 0.626166439 | -4.574954253 |
| **A_55_P1978471** | NA | -0.139516126 | -0.419975406 | 0.140943153 | 11.71029883 | -1.208753612 | 0.270973903 | 0.626166439 | -4.574963087 |
| **A_55_P2069470** | 1110017D15Rik | -0.923713919 | -2.780613146 | 0.933185308 | 8.402697161 | -1.208738462 | 0.270979322 | 0.626166439 | -4.574970315 |
| **A_55_P1989578** | NA | 0.091060258 | -0.091996646 | 0.274117162 | 8.75661688 | 1.208720683 | 0.270985683 | 0.626166439 | -4.574978797 |
| **A_55_P2134597** | NA | 0.183546369 | -0.185511557 | 0.552604294 | 8.900174565 | 1.208466036 | 0.271076795 | 0.626302101 | -4.575100281 |
| **A_55_P1985273** | Lmcd1 | 0.131411088 | -0.132869024 | 0.3956912 | 9.571429195 | 1.208233068 | 0.271160174 | 0.626335064 | -4.575211414 |
| **A_55_P2016089** | Uggt1 | -0.096068409 | -0.289307536 | 0.097170719 | 12.01727905 | -1.20800496 | 0.271241835 | 0.626335064 | -4.57532022 |
| **A_51_P249215** | Ptger2 | -0.21744513 | -0.654857644 | 0.219967384 | 8.020533694 | -1.207929732 | 0.271268771 | 0.626335064 | -4.575356101 |
| **A_52_P820196** | Cxcl15 | -1.312057509 | -3.951605015 | 1.327489996 | 8.350973726 | -1.207832128 | 0.271303722 | 0.626335064 | -4.575402654 |
| **A_55_P2098120** | NA | -0.239009746 | -0.719852548 | 0.241833056 | 7.923577123 | -1.207801821 | 0.271314575 | 0.626335064 | -4.575417109 |
| **A_55_P2032960** | Brap | -0.124160671 | -0.37395747 | 0.125636128 | 11.53674394 | -1.207759251 | 0.271329821 | 0.626335064 | -4.575437412 |
| **A_52_P94845** | Nicn1 | -0.122947442 | -0.370358849 | 0.124463964 | 8.358881628 | -1.207488408 | 0.271426835 | 0.626335064 | -4.575566581 |
| **A_55_P2151952** | NA | 0.112673568 | -0.114081506 | 0.339428642 | 13.62710492 | 1.207391802 | 0.271461446 | 0.626335064 | -4.575612651 |
| **A_51_P404193** | Sp5 | -0.494397792 | -1.489416118 | 0.500620534 | 7.914597115 | -1.207337339 | 0.27148096 | 0.626335064 | -4.575638623 |
| **A_51_P339200** | Abhd16a | -0.100087127 | -0.301524975 | 0.10135072 | 8.492927353 | -1.207314289 | 0.271489219 | 0.626335064 | -4.575649615 |
| **A_55_P2050426** | Ankrd50 | -0.167858396 | -0.505728227 | 0.170011435 | 8.856092986 | -1.207193373 | 0.271532549 | 0.626335064 | -4.575707275 |
| **A_55_P1996907** | BC067074 | -0.293092612 | -0.883057084 | 0.29687186 | 7.918425099 | -1.207152675 | 0.271547134 | 0.626335064 | -4.575726682 |
| **A_55_P2025829** | NA | -0.19707089 | -0.593782926 | 0.199641145 | 14.05200613 | -1.207063985 | 0.271578921 | 0.626335064 | -4.575768972 |
| **A_55_P2124233** | Vsig8 | -0.159048913 | -0.479224707 | 0.161126882 | 7.84424949 | -1.20705038 | 0.271583797 | 0.626335064 | -4.575775459 |
| **A_51_P227445** | NA | -0.19551766 | -0.589150514 | 0.198115195 | 8.285453881 | -1.20691821 | 0.271631174 | 0.626335064 | -4.57583848 |
| **A_52_P306305** | NA | -0.645603241 | -1.945405979 | 0.654199497 | 10.10937061 | -1.206900433 | 0.271637547 | 0.626335064 | -4.575846956 |
| **A_52_P624434** | Clec1a | -0.498492737 | -1.502125846 | 0.505140371 | 7.955177667 | -1.20688821 | 0.271641929 | 0.626335064 | -4.575852784 |
| **A_52_P573336** | Sbsn | -0.127742728 | -0.385176851 | 0.129691396 | 8.958937579 | -1.205738868 | 0.272054228 | 0.62708027 | -4.576400687 |
| **A_55_P2079860** | Asap3 | -0.100161743 | -0.30201328 | 0.101689794 | 8.600368681 | -1.205738146 | 0.272054487 | 0.62708027 | -4.576401031 |
| **A_51_P492410** | Pmvk | -0.101848626 | -0.307103441 | 0.103406189 | 12.41255053 | -1.205715957 | 0.272062453 | 0.62708027 | -4.576411607 |
| **A_55_P2121662** | Mapk3 | -0.252289409 | -0.76097499 | 0.256396172 | 8.967527002 | -1.205126901 | 0.272273977 | 0.627390576 | -4.576692333 |
| **A_55_P2200628** | 4733401D01Rik | 0.178040214 | -0.180950119 | 0.537030548 | 9.474554301 | 1.205087392 | 0.272288169 | 0.627390576 | -4.57671116 |
| **A_51_P156438** | Slc25a33 | -0.285119759 | -0.860030347 | 0.28979083 | 10.94752963 | -1.205064231 | 0.272296489 | 0.627390576 | -4.576722196 |
| **A_55_P2164724** | Gnaq | -0.167237832 | -0.504477028 | 0.170001364 | 9.870000707 | -1.20497954 | 0.272326915 | 0.627390576 | -4.576762552 |
| **A_55_P2144556** | Flrt3 | -0.4539721 | -1.369734338 | 0.461790139 | 8.291853676 | -1.204563281 | 0.272476499 | 0.627619979 | -4.576960886 |
| **A_51_P218975** | Kcns3 | -0.292105295 | -0.881398711 | 0.29718812 | 7.861330531 | -1.204456254 | 0.272514971 | 0.627619979 | -4.577011876 |
| **A_51_P149960** | Letm2 | 0.100113575 | -0.101912355 | 0.302139505 | 8.049549692 | 1.204117986 | 0.272636596 | 0.627619979 | -4.577173024 |
| **A_55_P1971759** | Zfp688 | 0.182357143 | -0.185634873 | 0.550349159 | 9.685825592 | 1.204113907 | 0.272638063 | 0.627619979 | -4.577174967 |
| **A_55_P1989885** | Smarce1 | 0.181792672 | -0.185076718 | 0.548662062 | 11.68977468 | 1.204059878 | 0.272657493 | 0.627619979 | -4.577200704 |
| **A_55_P2075385** | Orai2-ps | 0.081374553 | -0.082861108 | 0.245610214 | 8.387794683 | 1.203938606 | 0.272701111 | 0.627619979 | -4.577258471 |
| **A_51_P493720** | Tfam | 0.138682348 | -0.141221397 | 0.418586093 | 11.06075169 | 1.20391456 | 0.272709761 | 0.627619979 | -4.577269925 |
| **A_55_P2166528** | Mrps36 | 0.093844369 | -0.095569594 | 0.283258332 | 11.77373227 | 1.203869519 | 0.272725962 | 0.627619979 | -4.577291379 |

| | | | | | | | | | |
|---|---|---|---|---|---|---|---|---|---|
| **A_51_P193716** | Supv3l1 | 0.115844447 | -0.117993846 | 0.34968274 | 9.110407724 | 1.203767953 | 0.2727625 | 0.627619979 | -4.577339757 |
| **A_55_P2170345** | Gm6729 | 0.200574548 | -0.204306539 | 0.605455635 | 8.437866469 | 1.203736755 | 0.272773724 | 0.627619979 | -4.577354617 |
| **A_55_P1994027** | Adra2b | 0.284357572 | -0.28970321 | 0.858418354 | 9.914502625 | 1.203621973 | 0.272815023 | 0.627619979 | -4.577409287 |
| **A_55_P1992959** | Pfkfb4 | -0.10871026 | -0.328174989 | 0.110754468 | 9.642802618 | -1.203618883 | 0.272816134 | 0.627619979 | -4.577410758 |
| **A_52_P129697** | Stradb | 0.137066178 | -0.139689507 | 0.413821863 | 8.434866647 | 1.203419216 | 0.272887988 | 0.62766984 | -4.577505853 |
| **A_55_P2060284** | Itpripl2 | -0.10441105 | -0.315250854 | 0.106428754 | 7.941721681 | -1.203308676 | 0.272927774 | 0.62766984 | -4.577558498 |
| **A_55_P2022094** | Mta3 | 0.147622787 | -0.150513188 | 0.445758762 | 10.79924297 | 1.203156731 | 0.272982472 | 0.62766984 | -4.577630858 |
| **A_55_P2058928** | Ralgds | -0.391514093 | -1.18223024 | 0.399202054 | 9.159544935 | -1.203122865 | 0.272994664 | 0.62766984 | -4.577646985 |
| **A_51_P230287** | Sorcs2 | -0.182448905 | -0.550964299 | 0.186066489 | 8.079789073 | -1.203008832 | 0.273035722 | 0.62766984 | -4.577701287 |
| **A_52_P203560** | Fzd10 | -0.247317911 | -0.746858009 | 0.252222187 | 7.768314237 | -1.203007691 | 0.273036132 | 0.62766984 | -4.577701831 |
| **A_55_P1960366** | Fam47e | 0.137062001 | -0.139893678 | 0.414017679 | 8.460805899 | 1.202513557 | 0.273214107 | 0.62766984 | -4.577937112 |
| **A_51_P111462** | Arl15 | -0.137906648 | -0.41660793 | 0.140794633 | 9.51327647 | -1.202345903 | 0.273274514 | 0.62766984 | -4.573721529 |
| **A_52_P141488** | Grk5 | -0.185227605 | -0.559592444 | 0.189137234 | 8.526049513 | -1.202247405 | 0.273310009 | 0.62766984 | -4.578063824 |
| **A_55_P2089530** | Fkbp10 | -0.549979014 | -1.661564362 | 0.561606335 | 11.10978853 | -1.202227044 | 0.273317347 | 0.62766984 | -4.578073517 |
| **A_55_P2016064** | NA | 0.135830641 | -0.138720714 | 0.410381996 | 8.307498339 | 1.202146368 | 0.273346423 | 0.62766984 | -4.578111923 |
| **A_55_P2166262** | Rnf215 | -0.30006482 | -0.90660226 | 0.30647262 | 10.17323133 | -1.202100162 | 0.273363077 | 0.62766984 | -4.57813392 |
| **A_51_P163173** | Rbm12b1 | 0.108649281 | -0.110969997 | 0.32826856 | 8.397202912 | 1.202097202 | 0.273364144 | 0.62766984 | -4.578135328 |
| **A_66_P127175** | Ankrd16 | -0.16219252 | -0.490044179 | 0.16565914 | 10.54860715 | -1.202089 | 0.273367101 | 0.62766984 | -4.578139233 |
| **A_52_P621617** | Dnajb4 | 0.134994921 | -0.137895891 | 0.407885732 | 8.924230285 | 1.202020027 | 0.273391963 | 0.62766984 | -4.578172067 |
| **A_55_P1973770** | Unc5b | -0.435321596 | -1.315358793 | 0.4447156 | 9.185610885 | -1.201966526 | 0.273411249 | 0.62766984 | -4.578197535 |
| **A_51_P430563** | Mtpn | 0.138235789 | -0.141223465 | 0.417695042 | 9.332819418 | 1.201946644 | 0.273418417 | 0.62766984 | -4.578206999 |
| **A_55_P2116604** | NA | -0.14796291 | -0.447089333 | 0.151163513 | 13.49747081 | -1.201935812 | 0.273422322 | 0.62766984 | -4.578212155 |
| **A_66_P106098** | Kif3a | -0.109877713 | -0.332041945 | 0.112286519 | 8.549576522 | -1.201762539 | 0.273484795 | 0.627698728 | -4.578294634 |
| **A_51_P182572** | Phactr1 | -0.458366026 | -1.385229986 | 0.468497933 | 8.194833705 | -1.201654493 | 0.273523756 | 0.627698728 | -4.578346062 |
| **A_51_P103594** | Fam162b | -0.839675082 | -2.53761935 | 0.858269186 | 8.099329894 | -1.201630722 | 0.273532329 | 0.627698728 | -4.576662844 |
| **A_55_P1978047** | Aven | 0.118950004 | -0.121631687 | 0.359531696 | 8.383836719 | 1.201392946 | 0.273618091 | 0.627775137 | -4.578470546 |
| **A_51_P288876** | Tmem45a | -0.384719722 | -1.162891452 | 0.393452008 | 8.129586381 | -1.201301972 | 0.27365091 | 0.627775137 | -4.578513843 |
| **A_55_P2370384** | 1600029I14Rik | -0.45651134 | -1.379989651 | 0.466966972 | 7.887345451 | -1.201179907 | 0.27369495 | 0.627775137 | -4.578571934 |
| **A_51_P143915** | Rnf181 | -0.119053972 | -0.359888978 | 0.121781033 | 10.79176923 | -1.201178273 | 0.273695539 | 0.627775137 | -4.578572711 |
| **A_55_P2018457** | Hdac7 | -0.291649612 | -0.881736059 | 0.298436835 | 11.39256815 | -1.200961131 | 0.273773898 | 0.627880361 | -4.578676045 |
| **A_66_P135106** | Slco3a1 | -0.30377004 | -0.91845448 | 0.3109144 | 8.277198141 | -1.200814457 | 0.273826839 | 0.62792727 | -4.57874584 |
| **A_51_P275350** | Pten | -0.196211217 | -0.593352552 | 0.200930118 | 10.55746938 | -1.200499352 | 0.273940602 | 0.628095405 | -4.578895772 |
| **A_55_P2003541** | Nrcam | -0.710681195 | -2.1492381 | 0.727875709 | 8.065301114 | -1.200413767 | 0.273971508 | 0.628095405 | -4.578936492 |
| **A_51_P205106** | Mlf1 | -0.547272842 | -1.655249189 | 0.560703505 | 8.125205731 | -1.200208221 | 0.274045746 | 0.628095405 | -4.579034282 |
| **A_55_P2055291** | Gpn2 | 0.090810043 | -0.093041689 | 0.274661775 | 9.306151368 | 1.200188177 | 0.274052986 | 0.628095405 | -4.579043818 |
| **A_55_P1953728** | Nes | -0.339750113 | -1.027630339 | 0.348130113 | 9.686160446 | -1.200134646 | 0.274072323 | 0.628095405 | -4.579069284 |
| **A_55_P2112125** | Tnk2 | 0.151964263 | -0.155750861 | 0.459679386 | 11.9391682 | 1.199984995 | 0.274126389 | 0.628095405 | -4.579140476 |
| **A_51_P108266** | Actn2 | -0.239401074 | -0.724169923 | 0.245367775 | 7.766306594 | -1.199981577 | 0.274127624 | 0.628095405 | -4.579142102 |
| **A_55_P2060198** | NA | 0.27466332 | -0.281616752 | 0.830943392 | 10.21218543 | 1.199748876 | 0.274211711 | 0.628110855 | -4.579252795 |
| **A_55_P1970414** | Mccc1 | -0.082609999 | -0.249931769 | 0.08471177 | 8.676486602 | -1.199674309 | 0.274238661 | 0.628110855 | -4.579288264 |
| **A_55_P2064595** | NA | 0.103509512 | -0.106164966 | 0.31318399 | 8.650293863 | 1.199548685 | 0.27428407 | 0.628110855 | -4.579348017 |
| **A_55_P2079344** | Zfp362 | -0.154585796 | -0.467729183 | 0.158557591 | 9.193449769 | -1.199525627 | 0.274292405 | 0.628110855 | -4.579358984 |
| **A_65_P11092** | Tspan15 | -0.261611798 | -0.791578762 | 0.268355167 | 8.380718783 | -1.199476422 | 0.274310192 | 0.628110855 | -4.579382387 |
| **A_51_P433430** | Phc1 | -0.117761096 | -0.356329263 | 0.12080707 | 8.222277338 | -1.199423455 | 0.274329341 | 0.628110855 | -4.579407579 |
| **A_55_P2012734** | Fndc10 | -0.169882774 | -0.514088403 | 0.174322856 | 8.360423566 | -1.199263354 | 0.274387229 | 0.628153175 | -4.579483724 |
| **A_55_P2042121** | Slc25a22 | 0.321389846 | -0.329828169 | 0.97260786 | 9.277077791 | 1.199192584 | 0.274412821 | 0.628153175 | -4.579517381 |
| **A_55_P2121349** | NA | 0.133394455 | -0.136938618 | 0.403727529 | 8.558042549 | 1.199007184 | 0.274479874 | 0.628232266 | -4.579605551 |
| **A_55_P1999992** | Galnt12 | -0.197493552 | -0.597810386 | 0.202823283 | 8.331166243 | -1.198760034 | 0.274569282 | 0.62834918 | -4.579723079 |
| **A_55_P2147126** | Zfp712 | -0.101663256 | -0.307751458 | 0.104424946 | 8.244633232 | -1.198654644 | 0.274607416 | 0.62834918 | -4.579773192 |
| **A_51_P428134** | Lrig3 | -0.172645925 | -0.522644711 | 0.177352862 | 9.406736168 | -1.198596435 | 0.274628479 | 0.62834918 | -4.57980087 |

| | | | | | | | | | |
|---|---|---|---|---|---|---|---|---|---|
| **A_66_P138046** | Gm12298 | 0.085257506 | -0.087618217 | 0.25813323 | 7.980484132 | 1.198344822 | 0.274719544 | 0.628466947 | -4.579920503 |
| **A_55_P1981322** | Grhl2 | -0.534768839 | -1.619181058 | 0.54964338 | 8.053047182 | -1.198270533 | 0.274746436 | 0.628466947 | -4.579955822 |
| **A_55_P2019964** | Lancl1 | -0.231682621 | -0.701525728 | 0.238160486 | 11.05028426 | -1.198184747 | 0.274777493 | 0.628466947 | -4.579996608 |
| **A_51_P307741** | Cckar | -0.37743634 | -1.14317632 | 0.38830364 | 7.897465558 | -1.197693187 | 0.274955509 | 0.628652137 | -4.580230285 |
| **A_55_P1993463** | NA | 0.324519982 | -0.3338912 | 0.982931164 | 9.642451216 | 1.197643147 | 0.274973636 | 0.628652137 | -4.580254071 |
| **A_51_P495232** | Gpx8 | -0.684538356 | -2.073500812 | 0.704424101 | 9.51556392 | -1.197541216 | 0.275010564 | 0.628652137 | -4.580302521 |
| **A_55_P1994258** | Gapdh | -0.167557205 | -0.507572268 | 0.172457858 | 11.73395005 | -1.1974245 | 0.275052853 | 0.628652137 | -4.580357996 |
| **A_55_P2041592** | Rdh1 | 0.268986415 | -0.27688276 | 0.81485559 | 8.645036688 | 1.197360606 | 0.275076006 | 0.628652137 | -4.580388365 |
| **A_55_P2035495** | Zbtb20 | -0.194879908 | -0.59038002 | 0.200620203 | 9.223614264 | -1.197301828 | 0.275097307 | 0.628652137 | -4.580416301 |
| **A_52_P350554** | Kcnb1 | -0.083491756 | -0.252959709 | 0.085976198 | 7.848059231 | -1.197124164 | 0.2751617 | 0.628652137 | -4.580500738 |
| **A_52_P159885** | Ddx49 | 0.146878483 | -0.151267364 | 0.445024331 | 9.284415487 | 1.197050861 | 0.275188271 | 0.628652137 | -4.580535575 |
| **A_55_P2113160** | Mal | -1.264276948 | -3.830773736 | 1.302219841 | 8.806982518 | -1.196973907 | 0.275216169 | 0.628652137 | -4.580572146 |
| **A_66_P138929** | Lox | -0.459466257 | -1.392256645 | 0.473324131 | 8.219244667 | -1.196885896 | 0.275248078 | 0.628652137 | -4.58061397 |
| **A_55_P2121936** | Wiz | 0.091876477 | -0.094658867 | 0.278411822 | 10.79218079 | 1.196813258 | 0.275274416 | 0.628652137 | -4.580648488 |
| **A_55_P2093251** | NA | 0.148053607 | -0.15254552 | 0.448652734 | 9.563379042 | 1.196780399 | 0.275286331 | 0.628652137 | -4.580664102 |
| **A_51_P295085** | Ogn | -0.946210486 | -2.867388585 | 0.974967613 | 8.86456196 | -1.196749676 | 0.275297472 | 0.628652137 | -4.580678702 |
| **A_51_P238383** | B4galt4 | -0.394738496 | -1.196290599 | 0.406813608 | 9.255044654 | -1.196632827 | 0.275339847 | 0.628652137 | -4.580734226 |
| **A_55_P2262560** | 5730471H19Rik | -0.116826233 | -0.354056785 | 0.120404319 | 8.319231284 | -1.196610868 | 0.275347811 | 0.628652137 | -4.58074466 |
| **A_55_P2136678** | NA | 0.086368521 | -0.089029738 | 0.26176678 | 8.043818941 | 1.196501897 | 0.275387336 | 0.628652137 | -4.580796438 |
| **A_52_P5901** | Saal1 | 0.110383867 | -0.11380434 | 0.334572073 | 9.125173717 | 1.196398968 | 0.275424674 | 0.628652137 | -4.580845344 |
| **A_55_P1990970** | Casp9 | 0.08575712 | -0.088422193 | 0.259936433 | 10.92234414 | 1.196346001 | 0.27544389 | 0.628652137 | -4.58087051 |
| **A_51_P419439** | Gnmt | -0.356223073 | -1.079908202 | 0.367462057 | 12.73839151 | -1.196067213 | 0.275545049 | 0.62871769 | -4.581002962 |
| **A_52_P251450** | Cldn2 | -0.1376349 | -0.417328301 | 0.142058501 | 7.997848045 | -1.195720133 | 0.275671032 | 0.62871769 | -4.581167842 |
| **A_55_P2020331** | Camk1g | -0.854889322 | -2.592296042 | 0.882517399 | 8.85439367 | -1.195615631 | 0.275708974 | 0.62871769 | -4.581217482 |
| **A_55_P1983754** | Pcp4l1 | -0.168347516 | -0.510498005 | 0.173802972 | 9.178678552 | -1.195563749 | 0.275727813 | 0.62871769 | -4.581242126 |
| **A_55_P2115028** | Fastkd1 | 0.090810004 | -0.093754238 | 0.275374246 | 8.636423848 | 1.19555434 | 0.275731229 | 0.62871769 | -4.581246595 |
| **A_55_P2017684** | Macf1 | -0.548336988 | -1.662826946 | 0.566152969 | 10.93033131 | -1.195513741 | 0.275745972 | 0.62871769 | -4.58126588 |
| **A_51_P397920** | Ilrun | 0.121179667 | -0.12512068 | 0.367480014 | 11.31803752 | 1.195495428 | 0.275752622 | 0.62871769 | -4.581274578 |
| **A_52_P281702** | Igfbp5 | -0.426801856 | -1.294317391 | 0.440713679 | 9.275847916 | -1.195452234 | 0.275768308 | 0.62871769 | -4.581295094 |
| **A_52_P82660** | Mcm9 | 0.089233448 | -0.092152568 | 0.270619464 | 8.170231662 | 1.195382958 | 0.275793467 | 0.62871769 | -4.581327999 |
| **A_55_P2044947** | NA | 0.100708274 | -0.104004856 | 0.305421404 | 8.372014499 | 1.195370798 | 0.275797884 | 0.62871769 | -4.581333775 |
| **A_66_P139568** | Helz | 0.08931463 | -0.09227658 | 0.27090584 | 10.16260073 | 1.195118507 | 0.275889528 | 0.628852439 | -4.581453599 |
| **A_55_P2045818** | NA | -0.196593038 | -0.596340158 | 0.203154081 | 14.62757908 | -1.194994705 | 0.275934508 | 0.628861206 | -4.581512394 |
| **A_66_P105764** | Tor1b | 0.108097539 | -0.111717274 | 0.327912351 | 9.418287967 | 1.194928833 | 0.275958443 | 0.628861206 | -4.581543677 |
| **A_52_P537566** | Cenpt | 0.144111411 | -0.149007944 | 0.437230765 | 9.049158438 | 1.194640028 | 0.276063405 | 0.628910709 | -4.581680822 |
| **A_55_P2132800** | Zfp367 | 0.349405565 | -0.361312876 | 1.060124006 | 9.409825287 | 1.194580503 | 0.276085043 | 0.628910709 | -4.581709087 |
| **A_55_P2146374** | Nmnat1 | -0.102931537 | -0.312331431 | 0.106468358 | 8.916577373 | -1.194414831 | 0.276145273 | 0.628910709 | -4.581787752 |
| **A_51_P337918** | Aldh4a1 | 0.264485298 | -0.273699119 | 0.802669714 | 12.58547868 | 1.19413549 | 0.276246854 | 0.628910709 | -4.58192038 |
| **A_55_P2065529** | Mok | -0.511230204 | -1.551617371 | 0.529156963 | 8.2451705 | -1.194001044 | 0.276295755 | 0.628910709 | -4.581984208 |
| **A_55_P2325698** | 2010002M09Rik | -0.189171995 | -0.574149996 | 0.195806007 | 8.825772242 | -1.193999428 | 0.276296343 | 0.628910709 | -4.581984976 |
| **A_52_P249856** | Cc2d2a | -0.104075248 | -0.315878012 | 0.107727515 | 8.134388644 | -1.193985416 | 0.27630144 | 0.628910709 | -4.581991628 |
| **A_55_P2212161** | 2310034O05Rik | -0.139133027 | -0.422285581 | 0.144019528 | 8.073035577 | -1.193968688 | 0.276307525 | 0.628910709 | -4.581999569 |
| **A_55_P1972426** | Pdzd11 | 0.146409225 | -0.151558216 | 0.444376667 | 9.742668791 | 1.193940873 | 0.276317644 | 0.628910709 | -4.582012774 |
| **A_66_P111660** | Mt1 | 0.162920721 | -0.168665902 | 0.494507345 | 17.20592583 | 1.193885041 | 0.276337955 | 0.628910709 | -4.582039279 |
| **A_55_P2069730** | NA | 0.119184018 | -0.123420374 | 0.36178841 | 8.299971703 | 1.193720227 | 0.276397921 | 0.628910709 | -4.582117518 |
| **A_55_P1996046** | Zfp931 | -0.126015418 | -0.382526438 | 0.130495602 | 8.930279667 | -1.193715536 | 0.276399628 | 0.628910709 | -4.582119745 |
| **A_55_P2006281** | Ogdh | 0.185505838 | -0.192137446 | 0.563149122 | 12.68951912 | 1.193600533 | 0.276441477 | 0.628910709 | -4.582174335 |
| **A_51_P201904** | Ndufb5 | 0.088013736 | -0.091173675 | 0.267201147 | 10.23113719 | 1.193510244 | 0.276474336 | 0.628910709 | -4.582217193 |
| **A_51_P446583** | Zfp445 | -0.093379343 | -0.28353946 | 0.096780775 | 9.42919824 | -1.193203628 | 0.276585951 | 0.628910709 | -4.582362725 |
| **A_51_P204740** | Cd34 | -0.88663217 | -2.692255328 | 0.918990989 | 11.59015712 | -1.193162396 | 0.276600963 | 0.628910709 | -4.582382294 |

| | | | | | | | | | |
|---|---|---|---|---|---|---|---|---|---|
| **A_55_P2122419** | Arl1 | -0.094523428 | -0.287030285 | 0.097983429 | 10.03901907 | -1.193098906 | 0.276624081 | 0.628910709 | -4.582412427 |
| **A_51_P311990** | P4hb | -0.166247945 | -0.504834321 | 0.172338431 | 14.00712201 | -1.193081184 | 0.276630534 | 0.628910709 | -4.582420837 |
| **A_55_P1956799** | Smpx | -0.293367425 | -0.8908819 | 0.30414705 | 7.802290712 | -1.193017043 | 0.276653891 | 0.628910709 | -4.582451278 |
| **A_51_P404077** | Fzd2 | -0.828329813 | -2.515818073 | 0.859158448 | 8.556793183 | -1.192739828 | 0.276754857 | 0.628910709 | -4.582582834 |
| **A_52_P560728** | Serhl | -0.229626136 | -0.697432404 | 0.238180133 | 9.47398821 | -1.192719912 | 0.276762112 | 0.628910709 | -4.582592285 |
| **A_55_P2090025** | Mest | -0.267802276 | -0.813406322 | 0.277801769 | 11.46059017 | -1.192668838 | 0.276780718 | 0.628910709 | -4.582616521 |
| **A_55_P2150442** | NA | 0.17905982 | -0.185784964 | 0.543904605 | 13.42512917 | 1.192540648 | 0.276827421 | 0.628910709 | -4.582677349 |
| **A_51_P240136** | Gng8 | -0.208385326 | -0.632989817 | 0.216219166 | 7.772878054 | -1.192520183 | 0.276834877 | 0.628910709 | -4.58268706 |
| **A_55_P2087923** | Otud3 | 0.094905082 | -0.098480923 | 0.288291086 | 8.773136596 | 1.192470423 | 0.276853009 | 0.628910709 | -4.582710671 |
| **A_55_P1965427** | Uqcrh | -0.085574796 | -0.259956028 | 0.088806435 | 15.4504244 | -1.192420193 | 0.276871312 | 0.628910709 | -4.582734504 |
| **A_55_P2007871** | Ptpdc1 | -0.281984809 | -0.856632966 | 0.292663347 | 8.616470549 | -1.192358589 | 0.276893762 | 0.628910709 | -4.582763734 |
| **A_55_P2010773** | Tbx2 | -0.987591796 | -3.000190053 | 1.02500646 | 8.924056379 | -1.192349491 | 0.276897077 | 0.628910709 | -4.582768051 |
| **A_55_P2114596** | Zbtb14 | 0.122757787 | -0.127423788 | 0.372939362 | 9.578514882 | 1.192276322 | 0.276923744 | 0.628910709 | -4.582802768 |
| **A_52_P82209** | Wnk1 | 0.121631961 | -0.12628013 | 0.369544051 | 10.24421527 | 1.192156275 | 0.276967499 | 0.628936183 | -4.582859724 |
| **A_55_P2040723** | Dnali1 | -0.410637618 | -1.247695717 | 0.426420481 | 7.888958355 | -1.192027619 | 0.2770144 | 0.628955622 | -4.582920763 |
| **A_51_P416439** | Zdhhc6 | 0.148614682 | -0.15434533 | 0.451574694 | 10.9544413 | 1.191954277 | 0.277041139 | 0.628955622 | -4.582955558 |
| **A_55_P2080794** | Taf9 | 0.1527722 | -0.181322365 | 0.486866764 | 12.36427155 | 1.248477492 | 0.277236609 | 0.629038741 | -4.551880999 |
| **A_66_P123028** | NA | 0.212399607 | -0.220792228 | 0.645591443 | 8.674835667 | 1.191397366 | 0.27724425 | 0.629038741 | -4.583219737 |
| **A_55_P2026109** | Rpap1 | 0.118489947 | -0.123174467 | 0.360154361 | 10.83317349 | 1.191384622 | 0.277248899 | 0.629038741 | -4.583225781 |
| **A_55_P2050513** | Pgbd1 | 0.07817997 | -0.08127388 | 0.23763382 | 8.213062312 | 1.191361828 | 0.277257215 | 0.629038741 | -4.583236593 |
| **A_51_P176474** | Kndc1 | -0.187049993 | -0.568590565 | 0.194490579 | 7.727096299 | -1.191242444 | 0.277300774 | 0.629038741 | -4.583293218 |
| **A_66_P128909** | Numbl | -0.143035766 | -0.4348174 | 0.148745867 | 8.613655884 | -1.191159405 | 0.277331075 | 0.629038741 | -4.583332602 |
| **A_55_P2168504** | Mcmbp | 0.103991083 | -0.108152147 | 0.316134312 | 9.711388212 | 1.191105178 | 0.277350865 | 0.629038741 | -4.583358321 |
| **A_51_P423976** | Crem | -0.09976812 | -0.303315101 | 0.103778861 | 8.2529597 | -1.190996026 | 0.277390701 | 0.629038741 | -4.583410087 |
| **A_55_P2184877** | NA | -0.167599446 | -0.50954381 | 0.174344919 | 9.765065042 | -1.190968615 | 0.277400706 | 0.629038741 | -4.583423087 |
| **A_52_P479262** | NA | -0.68151084 | -2.071969624 | 0.708947944 | 10.4611166 | -1.190961815 | 0.277403189 | 0.629038741 | -4.583426312 |
| **A_51_P430082** | Tst | -0.173081655 | -0.526307248 | 0.180143938 | 13.2809888 | -1.190644375 | 0.277519076 | 0.62917352 | -4.583576849 |
| **A_55_P2146249** | NA | 0.125353264 | -0.130477362 | 0.38118389 | 13.46912697 | 1.190601163 | 0.277534854 | 0.62917352 | -4.58359734 |
| **A_55_P2078300** | Fndc7 | -0.187578942 | -0.570426925 | 0.195269042 | 7.80772394 | -1.190531542 | 0.277560278 | 0.62917352 | -4.583630353 |
| **A_55_P2082625** | NA | 0.149615805 | -0.155790814 | 0.455022424 | 9.53884087 | 1.190370668 | 0.277619031 | 0.629174896 | -4.583706634 |
| **A_51_P451338** | Jag2 | -0.218220934 | -0.663707378 | 0.22726551 | 7.833975434 | -1.190268953 | 0.277656184 | 0.629174896 | -4.583754861 |
| **A_55_P2122604** | Cbr2 | -1.320778018 | -4.017270419 | 1.375714382 | 8.399392901 | -1.190183214 | 0.277687505 | 0.629174896 | -4.583795513 |
| **A_51_P437240** | Emp2 | -1.187530814 | -3.612057712 | 1.236996085 | 9.951647161 | -1.19014827 | 0.277700271 | 0.629174896 | -4.58381208 |
| **A_51_P199624** | Sfswap | 0.187574278 | -0.195479012 | 0.570627568 | 9.008570623 | 1.189863865 | 0.277804192 | 0.629174896 | -4.583946915 |
| **A_55_P2091201** | Thoc2 | 0.087508795 | -0.091197957 | 0.266215547 | 8.543107724 | 1.189854704 | 0.277807539 | 0.629174896 | -4.583951257 |
| **A_55_P1974827** | Pde1b | -0.097985718 | -0.298093029 | 0.102121593 | 8.476985604 | -1.189824788 | 0.277818473 | 0.629174896 | -4.58396544 |
| **A_51_P133684** | Csrp3 | -0.486002747 | -1.478592934 | 0.50658744 | 7.94506831 | -1.18973965 | 0.27784959 | 0.629174896 | -4.5840058 |
| **A_55_P2076560** | Nr5a1 | 0.135547909 | -0.141291761 | 0.41238758 | 8.432752789 | 1.189728016 | 0.277853842 | 0.629174896 | -4.584011315 |
| **A_52_P121491** | Msrb3 | -0.105488202 | -0.320999399 | 0.110022996 | 8.02603082 | -1.189370707 | 0.277984468 | 0.62933641 | -4.584180685 |
| **A_51_P228276** | Nipsnap2 | -0.142750105 | -0.434390851 | 0.14889064 | 9.015894084 | -1.189354792 | 0.277990288 | 0.62933641 | -4.584188229 |
| **A_55_P2065819** | Fbxo22 | 0.140249252 | -0.146336583 | 0.426835088 | 11.18588655 | 1.189129142 | 0.27807281 | 0.629434162 | -4.58429518 |
| **A_55_P2087186** | Ier3ip1 | -0.109345402 | -0.332817954 | 0.11412715 | 13.25172499 | -1.18893887 | 0.278142411 | 0.629434162 | -4.584385356 |
| **A_55_P2422243** | 2900072G11Rik | 0.333556312 | -0.348165984 | 1.015278607 | 10.1380917 | 1.188898705 | 0.278157105 | 0.629434162 | -4.58440439 |
| **A_66_P120572** | Gm14488 | 0.108985248 | -0.113774431 | 0.331744928 | 8.273357607 | 1.18881513 | 0.278187683 | 0.629434162 | -4.584443997 |
| **A_51_P104527** | Atrip | 0.096091585 | -0.100318061 | 0.292501231 | 9.865541357 | 1.188791611 | 0.278196288 | 0.629434162 | -4.584455143 |
| **A_55_P1998656** | Entpd5 | 0.328120664 | -0.342762483 | 0.999003811 | 9.63960836 | 1.188419827 | 0.278332351 | 0.629668306 | -4.584631317 |
| **A_55_P1971709** | Dnajc8 | 0.189241095 | -0.197726025 | 0.576208216 | 10.02267181 | 1.188295838 | 0.27837774 | 0.629678772 | -4.584690066 |
| **A_51_P338461** | Numa1 | 0.092488571 | -0.096652735 | 0.281629878 | 13.14023171 | 1.188187217 | 0.278417508 | 0.629678772 | -4.584741531 |
| **A_51_P280893** | Pla2g1b | -0.372449679 | -1.134148156 | 0.389248798 | 7.842012329 | -1.188140244 | 0.278434708 | 0.629678772 | -4.584763786 |
| **A_55_P2102759** | NA | 0.08833658 | -0.092334557 | 0.269007716 | 8.183387278 | 1.188050752 | 0.278467478 | 0.629679209 | -4.584806185 |

| | | | | | | | | | |
|---|---|---|---|---|---|---|---|---|---|
| **A_55_P1994586** | Itsn1 | 0.240914353 | -0.251867669 | 0.733696376 | 8.727883665 | 1.187930434 | 0.278511541 | 0.62970518 | -4.584863187 |
| **A_55_P2109486** | Milr1 | 0.125822455 | -0.13157199 | 0.3832169 | 16.96766114 | 1.187796861 | 0.278560466 | 0.629742134 | -4.584926466 |
| **A_51_P333923** | Tspan1 | -1.13165851 | -3.44812016 | 1.184803139 | 8.314168327 | -1.187062179 | 0.278829694 | 0.630265242 | -4.585274462 |
| **A_51_P146044** | Ccdc92 | -0.277856724 | -0.846654585 | 0.290941136 | 8.043552416 | -1.186987503 | 0.278857072 | 0.630265242 | -4.585309829 |
| **A_55_P2062549** | Gm6524 | 0.11729808 | -0.122885075 | 0.357481235 | 10.0745327 | 1.186674331 | 0.278971913 | 0.630353161 | -4.585458138 |
| **A_52_P352735** | Ccdc153 | -1.025644807 | -3.125880502 | 1.074590887 | 8.27468792 | -1.186621299 | 0.278991364 | 0.630353161 | -4.58548325 |
| **A_52_P480301** | Sppl3 | 0.094840091 | -0.099367164 | 0.289047345 | 8.686043772 | 1.186614639 | 0.278993807 | 0.630353161 | -4.585486404 |
| **A_51_P220723** | B230118H07Rik | -0.182382272 | -0.555883947 | 0.191119402 | 11.85944624 | -1.18651507 | 0.27903033 | 0.630361998 | -4.585533553 |
| **A_55_P1961943** | Cd68 | -0.207031152 | -0.631090311 | 0.217028007 | 10.88329784 | -1.186294289 | 0.27911133 | 0.630471298 | -4.585638092 |
| **A_52_P353478** | Scn7a | -0.347957814 | -1.061050782 | 0.365135154 | 7.775937976 | -1.185669447 | 0.279340679 | 0.630797412 | -4.585933911 |
| **A_52_P582309** | Nub1 | -0.105939525 | -0.323055053 | 0.111176003 | 10.22152062 | -1.185633125 | 0.279354016 | 0.630797412 | -4.585951105 |
| **A_52_P337427** | Izumo4 | -0.087624123 | -0.267219871 | 0.091971625 | 8.372613083 | -1.185525291 | 0.279393615 | 0.630797412 | -4.58600215 |
| **A_55_P2021540** | Myocd | -0.177114769 | -0.54016051 | 0.185930973 | 7.754714175 | -1.185431924 | 0.279427905 | 0.630797412 | -4.586046344 |
| **A_52_P487436** | Nags | 0.213139745 | -0.223768496 | 0.650047987 | 9.432477656 | 1.185379452 | 0.279447177 | 0.630797412 | -4.586071181 |
| **A_55_P1962139** | Mff | 0.099403777 | -0.104368559 | 0.303176114 | 11.67331461 | 1.185334299 | 0.279463762 | 0.630797412 | -4.586092553 |
| **A_55_P2091671** | Psmg1 | 0.119540328 | -0.125540599 | 0.364621254 | 11.28939708 | 1.18519038 | 0.279516631 | 0.630797412 | -4.586160672 |
| **A_52_P491861** | Stag1 | 0.167349152 | -0.175749282 | 0.510447586 | 9.692893493 | 1.185189975 | 0.27951678 | 0.630797412 | -4.586160863 |
| **A_55_P2054842** | Clpb | -0.149735149 | -0.456790036 | 0.157319738 | 10.613232 | -1.184925193 | 0.279614069 | 0.630943304 | -4.586286178 |
| **A_51_P128147** | Chmp1a | -0.11729218 | -0.357843328 | 0.123258968 | 10.91164271 | -1.184799367 | 0.279660312 | 0.63096058 | -4.579487005 |
| **A_55_P1964213** | Tprkb | 0.35407817 | -0.372135058 | 1.080291398 | 10.10108111 | 1.184726725 | 0.279687012 | 0.63096058 | -4.586380101 |
| **A_55_P2071271** | Abracl | 0.090634409 | -0.095296312 | 0.276565131 | 11.32212803 | 1.18447294 | 0.279780309 | 0.630997542 | -4.586500192 |
| **A_51_P115005** | Edn1 | -0.143900936 | -0.439113403 | 0.151311532 | 7.857116443 | -1.184437399 | 0.279793376 | 0.630997542 | -4.583826005 |
| **A_55_P2142808** | NA | 0.309801866 | -0.325792168 | 0.9453959 | 15.37420241 | 1.184370023 | 0.279818151 | 0.630997542 | -4.586548889 |
| **A_52_P647906** | NA | -0.121776335 | -0.371789479 | 0.128236809 | 11.15441943 | -1.183540833 | 0.280123199 | 0.630997542 | -4.586941173 |
| **A_51_P236013** | Rnf40 | 0.131965316 | -0.138997336 | 0.402927967 | 10.17219548 | 1.183405426 | 0.280173041 | 0.630997542 | -4.587005223 |
| **A_52_P425839** | Retnlg | -0.428206576 | -1.307481166 | 0.451068014 | 11.73536024 | -1.183346685 | 0.280194665 | 0.630997542 | -4.587033007 |
| **A_51_P329441** | Bsg | -0.212056505 | -0.647496354 | 0.223383345 | 12.12640864 | -1.183332021 | 0.280200063 | 0.630997542 | -4.587039943 |
| **A_55_P2036495** | NA | 0.089120792 | -0.093893402 | 0.272134986 | 13.19854937 | 1.183252557 | 0.280229319 | 0.630997542 | -4.587077528 |
| **A_52_P509906** | Ift172 | -0.163472093 | -0.499178336 | 0.17223415 | 8.370096862 | -1.183225156 | 0.280239408 | 0.630997542 | -4.587090488 |
| **A_66_P105244** | Tor1aip1 | 0.112430851 | -0.118482471 | 0.343344172 | 8.300377408 | 1.183095212 | 0.280287255 | 0.630997542 | -4.587151946 |
| **A_51_P498388** | Sbk1 | -0.100637488 | -0.307343578 | 0.106068603 | 8.540034522 | -1.183013503 | 0.280317345 | 0.630997542 | -4.58719059 |
| **A_51_P425048** | H2-Q2 | -0.201444229 | -0.615213894 | 0.212325436 | 11.01074856 | -1.182985363 | 0.280327708 | 0.630997542 | -4.587203898 |
| **A_55_P1964559** | Smarca5-ps | 0.149048941 | -0.157110784 | 0.455208666 | 10.27843236 | 1.182943559 | 0.280343105 | 0.630997542 | -4.587223668 |
| **A_51_P316801** | Wdr60 | -0.136193506 | -0.415953004 | 0.143565992 | 8.828989563 | -1.182918294 | 0.28035241 | 0.630997542 | -4.587235617 |
| **A_55_P2034550** | 1700013F07Rik | -0.210119588 | -0.641744729 | 0.221505553 | 7.831029976 | -1.182886284 | 0.2803642 | 0.630997542 | -4.587250755 |
| **A_55_P2060036** | Ulk2 | -0.108242557 | -0.330594359 | 0.114109245 | 9.101077642 | -1.1828797 | 0.280366625 | 0.630997542 | -4.587253869 |
| **A_55_P2013840** | Krt42 | -0.757257839 | -2.312891512 | 0.798375833 | 8.373179011 | -1.182822649 | 0.280387639 | 0.630997542 | -4.587280849 |
| **A_51_P253883** | Fam49a | -0.114164042 | -0.348700337 | 0.120372254 | 8.170810151 | -1.182775892 | 0.280404863 | 0.630997542 | -4.58730296 |
| **A_55_P2104259** | Plin5 | -0.521541066 | -1.592996611 | 0.549914479 | 10.97276491 | -1.182762489 | 0.2804098 | 0.630997542 | -4.587309298 |
| **A_51_P149699** | P3h3 | -0.299330851 | -0.914345762 | 0.31568406 | 8.81894853 | -1.182630362 | 0.280458477 | 0.630997542 | -4.587371779 |
| **A_55_P2084631** | Hist1h2an | 0.130234924 | -0.137350993 | 0.397820841 | 17.54454906 | 1.182625933 | 0.280460109 | 0.630997542 | -4.587373873 |
| **A_52_P510771** | Tbl3 | 0.128406176 | -0.135425967 | 0.392238319 | 8.289150272 | 1.182609589 | 0.28046613 | 0.630997542 | -4.587381602 |
| **A_52_P502267** | Epc2 | 0.084284385 | -0.088894206 | 0.257462976 | 9.56979434 | 1.182595192 | 0.280471435 | 0.630997542 | -4.58738841 |
| **A_51_P269173** | Alg9 | 0.180660058 | -0.190554174 | 0.55187429 | 9.267543224 | 1.182553277 | 0.280486879 | 0.630997542 | -4.58740823 |
| **A_51_P253732** | Il17rd | -0.17787242 | -0.54345803 | 0.18771319 | 8.060078266 | -1.182231998 | 0.280605281 | 0.631190442 | -4.587560141 |
| **A_52_P160970** | Akap10 | 0.128341697 | -0.135515325 | 0.392198719 | 9.321552401 | 1.181904294 | 0.280726094 | 0.631227512 | -4.587715072 |
| **A_52_P335259** | Cdkal1 | -0.123051043 | -0.376041103 | 0.129939016 | 9.510704865 | -1.181857271 | 0.280743433 | 0.631227512 | -4.587737303 |
| **A_51_P353882** | Zfp580 | -0.31869404 | -0.974050245 | 0.336662164 | 8.998832552 | -1.181625122 | 0.28082905 | 0.631227512 | -4.587847045 |
| **A_51_P136521** | Lypd2 | -0.763411465 | -2.333295322 | 0.806472391 | 8.574648269 | -1.181610506 | 0.280834441 | 0.631227512 | -4.587853954 |
| **A_66_P128942** | Mipep | 0.083029135 | -0.087722463 | 0.253780733 | 7.968781032 | 1.181541354 | 0.280859949 | 0.631227512 | -4.587886641 |

| | | | | | | | | | |
|---|---|---|---|---|---|---|---|---|---|
| **A_66_P111773** | Gatad2a | 0.164798654 | -0.174127676 | 0.503724983 | 9.645810933 | 1.181494054 | 0.280877398 | 0.631227512 | -4.587909 |
| **A_51_P142896** | Cd59a | 0.20305666 | -0.214555368 | 0.620668688 | 11.17175076 | 1.18148287 | 0.280881524 | 0.631227512 | -4.587914286 |
| **A_52_P140356** | Calm3 | 0.141613189 | -0.149633468 | 0.432859846 | 10.8230728 | 1.181478824 | 0.280883017 | 0.631227512 | -4.587916199 |
| **A_55_P1988699** | Cacna2d1 | -0.232692141 | -0.711331123 | 0.245946842 | 8.144444417 | -1.181290843 | 0.280952372 | 0.631258914 | -4.58800505 |
| **A_51_P383032** | Clec4d | -0.169409696 | -0.517920134 | 0.179100742 | 8.814497775 | -1.181151652 | 0.281003736 | 0.631258914 | -4.588070837 |
| **A_55_P2157388** | Skint9 | 0.119369481 | -0.126199899 | 0.364938862 | 8.455687372 | 1.18114246 | 0.281007128 | 0.631258914 | -4.588075182 |
| **A_51_P510418** | Aldh1b1 | 0.446135658 | -0.471758781 | 1.364030098 | 11.52814096 | 1.181020365 | 0.28105219 | 0.631258914 | -4.588132885 |
| **A_55_P2003158** | Wwp1 | 0.152427982 | -0.161188274 | 0.466044237 | 8.448328473 | 1.180998438 | 0.281060283 | 0.631258914 | -4.588143248 |
| **A_55_P2171977** | NA | -0.191253312 | -0.584789074 | 0.202282449 | 14.19620708 | -1.180885937 | 0.281101811 | 0.631278831 | -4.588196415 |
| **A_55_P2143041** | Phgdh | 0.228097749 | -0.241352661 | 0.69754816 | 12.34695615 | 1.180631771 | 0.281195651 | 0.631348316 | -4.588316523 |
| **A_52_P390918** | Trip4 | 0.082162296 | -0.08694137 | 0.251265962 | 8.653100907 | 1.180599879 | 0.281207427 | 0.631348316 | -4.588331593 |
| **A_55_P2038422** | Adamts17 | -0.931202732 | -2.847894184 | 0.98548872 | 8.372068488 | -1.180525098 | 0.281235043 | 0.631348316 | -4.588366929 |
| **A_55_P2099560** | Apobec1 | -0.238725766 | -0.730157332 | 0.2527058 | 9.930782986 | -1.180373459 | 0.281291049 | 0.631348316 | -4.58843858 |
| **A_55_P2172619** | NA | -0.138045398 | -0.422271767 | 0.14618097 | 16.3759466 | -1.180159636 | 0.281370037 | 0.631348316 | -4.588539606 |
| **A_55_P2042486** | Dpysl3 | -0.104564292 | -0.319863157 | 0.110734573 | 7.73092278 | -1.18011641 | 0.281386008 | 0.631348316 | -4.588560029 |
| **A_55_P2099540** | H2afj | 0.162735071 | -0.172347385 | 0.497817527 | 9.578743607 | 1.180083278 | 0.281398249 | 0.631348316 | -4.588575682 |
| **A_51_P201785** | Morn5 | -0.685744156 | -2.097824707 | 0.726336395 | 8.081480304 | -1.180010394 | 0.28142518 | 0.631348316 | -4.588610115 |
| **A_52_P367520** | Nexn | -0.778180417 | -2.380610865 | 0.824250032 | 8.747234506 | -1.180006223 | 0.281426721 | 0.631348316 | -4.588612085 |
| **A_52_P129428** | Fzd2 | -0.236822628 | -0.724577509 | 0.250932252 | 7.842469305 | -1.17979014 | 0.281506578 | 0.631352006 | -4.588714166 |
| **A_52_P120803** | Ankrd1 | -0.652596744 | -1.996675855 | 0.691482366 | 8.045291931 | -1.17978606 | 0.281508085 | 0.631352006 | -4.588716094 |
| **A_51_P487668** | H2afy3 | -0.21170879 | -0.647759517 | 0.224341937 | 9.690489947 | -1.179736629 | 0.281526356 | 0.631352006 | -4.588739444 |
| **A_55_P2235911** | 0610009E02Rik | -0.126881822 | -0.388243026 | 0.134479382 | 8.437912673 | -1.179618225 | 0.281570125 | 0.631364584 | -4.588795376 |
| **A_52_P335064** | Mustn1 | -0.100438702 | -0.307374992 | 0.106497589 | 7.980464678 | -1.17936333 | 0.281664367 | 0.631364584 | -4.588915775 |
| **A_55_P2151546** | Adarb1 | -0.60180416 | -1.841719226 | 0.638110907 | 9.545318863 | -1.179360117 | 0.281665556 | 0.631364584 | -4.588917292 |
| **A_55_P1999800** | Gm14066 | -0.190628268 | -0.583421629 | 0.202165092 | 7.767326948 | -1.179251273 | 0.281705807 | 0.631364584 | -4.588968701 |
| **A_55_P2117689** | Rmnd1 | 0.091312234 | -0.096869576 | 0.279494044 | 10.59856899 | 1.17905623 | 0.281777947 | 0.631364584 | -4.589060817 |
| **A_55_P2057792** | NA | 0.104308502 | -0.110666144 | 0.319283148 | 12.2143255 | 1.179005022 | 0.281796891 | 0.631364584 | -4.589085002 |
| **A_51_P511250** | Tent5b | -0.347878884 | -1.064891965 | 0.369134197 | 7.92775619 | -1.178919573 | 0.281828502 | 0.631364584 | -4.589125355 |
| **A_55_P2012984** | Ldha | 0.137136966 | -0.145523859 | 0.419797791 | 16.69156199 | 1.178886796 | 0.281840629 | 0.631364584 | -4.589140834 |
| **A_55_P2366358** | 4930444F02Rik | -0.131815961 | -0.403527835 | 0.139895914 | 7.816521629 | -1.178806629 | 0.28187029 | 0.631364584 | -4.589178692 |
| **A_52_P568805** | Ubxn1 | -0.085286551 | -0.261090206 | 0.090517104 | 13.41921765 | -1.178788365 | 0.281877049 | 0.631364584 | -4.589187317 |
| **A_55_P1989663** | Slco3a1 | -0.206512352 | -0.632216165 | 0.219191462 | 8.832599049 | -1.178749932 | 0.28189127 | 0.631364584 | -4.589205466 |
| **A_51_P327904** | Yipf1 | -0.105698423 | -0.323621237 | 0.112224391 | 8.561958509 | -1.178552676 | 0.28196427 | 0.631371587 | -4.589298611 |
| **A_55_P2132403** | Szt2 | 0.106614828 | -0.113202956 | 0.326432612 | 10.185122 | 1.17852276 | 0.281975343 | 0.631371587 | -4.589312737 |
| **A_55_P2039699** | Camp | -0.17852546 | -0.546622519 | 0.189571599 | 15.89428459 | -1.178476704 | 0.28199239 | 0.631371587 | -4.589334484 |
| **A_51_P233101** | Ltbp3 | -0.884520423 | -2.708635855 | 0.939595008 | 9.117163044 | -1.178253495 | 0.282075021 | 0.631430859 | -4.589439873 |
| **A_51_P512591** | Cdc42se1 | -0.130794084 | -0.400531809 | 0.13894364 | 10.29445808 | -1.17822871 | 0.282084198 | 0.631430859 | -4.589451575 |
| **A_55_P2209767** | 2610002J23Rik | 0.107342087 | -0.114066684 | 0.328750859 | 9.45295434 | 1.178035571 | 0.282155716 | 0.631517814 | -4.58954276 |
| **A_52_P589169** | Aopep | -0.335926273 | -1.02897163 | 0.357119083 | 8.674024174 | -1.1777836 | 0.282249042 | 0.631537083 | -4.58966171 |
| **A_55_P1968035** | 1810009A15Rik | 0.119054132 | -0.126595315 | 0.364703578 | 12.49853706 | 1.177638162 | 0.282302923 | 0.631537083 | -4.589730363 |
| **A_55_P2005190** | Herc4 | 0.132491815 | -0.140891662 | 0.405875292 | 11.33449637 | 1.17760591 | 0.282314872 | 0.631537083 | -4.589745587 |
| **A_55_P1987439** | Pigt | -0.14516263 | -0.444713731 | 0.154388471 | 11.35210675 | -1.177516757 | 0.282347906 | 0.631537083 | -4.58978767 |
| **A_51_P302167** | Cobl | 0.090466704 | -0.09622222 | 0.277155629 | 7.928006985 | 1.177479637 | 0.282361661 | 0.631537083 | -4.58980519 |
| **A_52_P561736** | NA | 0.103845596 | -0.110463818 | 0.318155009 | 11.63653452 | 1.17741624 | 0.282385154 | 0.631537083 | -4.589835114 |
| **A_52_P148658** | Mettl22 | 0.107897006 | -0.114777453 | 0.330571465 | 8.810727121 | 1.17739497 | 0.282393037 | 0.631537083 | -4.589845153 |
| **A_55_P2132024** | Tekt1 | -0.785328864 | -2.406680263 | 0.836022534 | 8.103195868 | -1.176948876 | 0.2825584 | 0.631768977 | -4.59005569 |
| **A_55_P1999818** | NA | 0.418687075 | -0.445721021 | 1.283095172 | 14.06211502 | 1.176938901 | 0.282562099 | 0.631768977 | -4.590060397 |
| **A_51_P440743** | Celsr1 | -0.757302472 | -2.321432578 | 0.806827633 | 8.664757154 | -1.176466834 | 0.282737181 | 0.632087321 | -4.590283154 |
| **A_55_P1989405** | Elf1 | 0.11217061 | -0.119528789 | 0.343870009 | 10.08903343 | 1.176352181 | 0.282779719 | 0.632109307 | -4.59033725 |
| **A_52_P496260** | Exosc8 | 0.093231585 | -0.09937512 | 0.28583829 | 11.27771927 | 1.176182882 | 0.28284254 | 0.632124123 | -4.590417126 |

| | | | | | | | | |
|---|---|---|---|---|---|---|---|---|
| **A_55_P2061465** | Ppp1r21 | 0.090323649 | -0.096279504 | 0.276926802 | 9.275056669 | 1.176158054 | 0.282851753 | 0.632124123 | -4.590428839 |
| **A_55_P1995152** | NA | -0.165918321 | -0.508740344 | 0.176903702 | 15.125048 | -1.176004058 | 0.282908908 | 0.632178761 | -4.59050149 |
| **A_55_P2088028** | Mtss1 | -0.097729071 | -0.299694166 | 0.104236024 | 8.354247353 | -1.175792382 | 0.282987486 | 0.632216681 | -4.590601346 |
| **A_51_P457492** | Tmem223 | -0.220398716 | -0.675880211 | 0.235082779 | 12.88567556 | -1.175767664 | 0.282996663 | 0.632216681 | -4.590613006 |
| **A_51_P293607** | Mrps14 | 0.146466568 | -0.156243859 | 0.449176995 | 8.621692221 | 1.17569403 | 0.283024002 | 0.632216681 | -4.59064774 |
| **A_55_P2106975** | Clcn5 | 0.15413646 | -0.164468728 | 0.472741648 | 8.628588837 | 1.175535438 | 0.283082893 | 0.632275161 | -4.590722546 |
| **A_51_P426270** | Mgp | -1.836754561 | -5.634313364 | 1.960804243 | 9.52119771 | -1.175248778 | 0.283189367 | 0.632382148 | -4.59085775 |
| **A_52_P161400** | Cnpy4 | 0.091466952 | -0.097647367 | 0.280581272 | 9.093014213 | 1.175230312 | 0.283196227 | 0.632382148 | -4.590866459 |
| **A_55_P1991164** | Mlc1 | -0.742277236 | -2.277245124 | 0.792690652 | 8.083857335 | -1.175032927 | 0.283269563 | 0.63247284 | -4.590959548 |
| **A_55_P1958295** | NA | -0.188669216 | -0.578887205 | 0.201548773 | 13.02729606 | -1.174835191 | 0.283343045 | 0.632563839 | -4.591052795 |
| **A_65_P15412** | Usp13 | -0.101109311 | -0.310303817 | 0.108085194 | 7.702654799 | -1.174421699 | 0.283496758 | 0.632802315 | -4.591247766 |
| **A_55_P1953341** | Wfdc2 | -1.623720717 | -4.983604183 | 1.736162749 | 9.531859152 | -1.174276328 | 0.283550816 | 0.632802315 | -4.591316305 |
| **A_51_P141926** | Fxyd4 | -0.410514527 | -1.260002447 | 0.438973393 | 7.899338061 | -1.174233624 | 0.283566698 | 0.632802315 | -4.591336439 |
| **A_52_P10732** | Map6 | 0.090422118 | -0.096706857 | 0.277551094 | 8.5945961 | 1.174131736 | 0.283604593 | 0.632802315 | -4.591384473 |
| **A_55_P2009732** | Prr9 | 0.102002171 | -0.109121424 | 0.313125766 | 8.037403347 | 1.173966848 | 0.283665929 | 0.632802315 | -4.591462205 |
| **A_55_P2109875** | Rpl23a | -0.173831877 | -0.533641581 | 0.185977827 | 14.98630687 | -1.173923339 | 0.283682116 | 0.632802315 | -4.591482716 |
| **A_52_P302544** | Col8a2 | 0.091386222 | -0.09779455 | 0.280566993 | 8.16862896 | 1.173780574 | 0.283735234 | 0.632802315 | -4.591550014 |
| **A_66_P122362** | Aldh16a1 | 0.111466477 | -0.11929786 | 0.342230814 | 12.1396984 | 1.173704509 | 0.283763538 | 0.632802315 | -4.591585869 |
| **A_51_P103780** | Sft2d2 | -0.109491339 | -0.336186104 | 0.117203427 | 9.800343153 | -1.17360368 | 0.283801062 | 0.632802315 | -4.588790522 |
| **A_55_P2090324** | Zfp956 | 0.148948833 | -0.159445241 | 0.457342908 | 9.667481955 | 1.173584244 | 0.283808295 | 0.632802315 | -4.591642556 |
| **A_55_P2026530** | Erc2 | -0.165692619 | -0.508775442 | 0.177390204 | 8.495118017 | -1.173511574 | 0.283835343 | 0.632802315 | -4.591676809 |
| **A_51_P114854** | Clns1a | 0.084799143 | -0.090798969 | 0.260397256 | 9.89400309 | 1.173423578 | 0.283868097 | 0.632802315 | -4.586931345 |
| **A_52_P51548** | Pard3 | -0.093292377 | -0.286501113 | 0.099916359 | 9.354619381 | -1.173282482 | 0.283920624 | 0.632802315 | -4.591784783 |
| **A_52_P501816** | Slc6a14 | -0.174462564 | -0.535779375 | 0.186854248 | 7.718520259 | -1.173268117 | 0.283925972 | 0.632802315 | -4.591791553 |
| **A_51_P118650** | Wdr6 | -0.229449343 | -0.704661589 | 0.245762902 | 8.977135626 | -1.173227907 | 0.283940943 | 0.632802315 | -4.584565779 |
| **A_55_P2032388** | Trim24 | 0.111558366 | -0.119531918 | 0.342648651 | 9.471616789 | 1.17301522 | 0.284020141 | 0.63286418 | -4.591910735 |
| **A_55_P1971025** | Paqr6 | 0.226411902 | -0.242617032 | 0.695440836 | 8.886103814 | 1.172958935 | 0.284041103 | 0.63286418 | -4.591937259 |
| **A_51_P461429** | Cyp7b1 | -0.158844021 | -0.487920564 | 0.170232522 | 7.912213867 | -1.172889602 | 0.284066926 | 0.63286418 | -4.591969932 |
| **A_51_P211786** | Chst13 | 0.151556174 | -0.162474453 | 0.4655868 | 8.480580324 | 1.172694305 | 0.284139676 | 0.632879342 | -4.592061957 |
| **A_51_P105068** | Lypd6b | -0.197110872 | -0.605538075 | 0.21131633 | 7.750094121 | -1.172678895 | 0.284145417 | 0.632879342 | -4.590161987 |
| **A_55_P2076989** | NA | 0.174296138 | -0.18688075 | 0.535473025 | 10.47694739 | 1.172602999 | 0.284173693 | 0.632879342 | -4.592104979 |
| **A_55_P2072284** | Dnaaf3 | -0.30851253 | -0.947981514 | 0.330956453 | 8.07487679 | -1.172293919 | 0.284288872 | 0.632879342 | -4.592250603 |
| **A_51_P247184** | Npr3 | -0.227916686 | -0.700357902 | 0.24452453 | 8.316589938 | -1.172226494 | 0.284314003 | 0.632879342 | -4.592282368 |
| **A_51_P279639** | Col6a2 | -0.69848119 | -2.146624693 | 0.749662313 | 9.094552835 | -1.171996471 | 0.284399753 | 0.632879342 | -4.592390731 |
| **A_55_P1971513** | NA | 0.096386868 | -0.103476291 | 0.296250027 | 16.12166963 | 1.171839976 | 0.284458106 | 0.632879342 | -4.592464449 |
| **A_52_P580582** | Nppa | -1.83468764 | -5.639181684 | 1.969806404 | 8.748490717 | -1.171786299 | 0.284478123 | 0.632879342 | -4.591508898 |
| **A_51_P129012** | B2m | -0.240028496 | -0.737786403 | 0.257729411 | 14.03443952 | -1.171730746 | 0.28449884 | 0.632879342 | -4.5925159 |
| **A_55_P2159299** | Ing1 | 0.113196457 | -0.12156503 | 0.347957944 | 10.99835428 | 1.17162646 | 0.284537736 | 0.632879342 | -4.592565021 |
| **A_51_P113722** | Fbxl14 | 0.15605385 | -0.167598329 | 0.47970603 | 10.58712592 | 1.171599403 | 0.284547828 | 0.632879342 | -4.591596063 |
| **A_55_P2006018** | Slc4a2 | 0.218295833 | -0.23444862 | 0.671040285 | 10.74731673 | 1.17158957 | 0.284551496 | 0.632879342 | -4.592582396 |
| **A_55_P1984307** | Adgrf5 | -0.211580301 | -0.650457701 | 0.227297098 | 8.828142676 | -1.171426929 | 0.284612167 | 0.632879342 | -4.592658999 |
| **A_55_P2006392** | NA | 0.09370603 | -0.100671565 | 0.288083624 | 16.60689341 | 1.171398121 | 0.284622915 | 0.632879342 | -4.592672566 |
| **A_55_P2024525** | Prdm8 | -0.192054686 | -0.590493437 | 0.206384064 | 7.747223667 | -1.171241708 | 0.284681276 | 0.632879342 | -4.592746231 |
| **A_51_P185713** | Ptdss2 | 0.21320977 | -0.229123712 | 0.655543252 | 8.867597343 | 1.171225385 | 0.284687367 | 0.632879342 | -4.592753918 |
| **A_55_P2036883** | H2bc18 | 0.208865235 | -0.22447396 | 0.64220443 | 13.21007367 | 1.171173873 | 0.284706589 | 0.632879342 | -4.592778177 |
| **A_51_P234847** | 1700001L19Rik | -0.300695008 | -0.924633198 | 0.323243181 | 7.793566097 | -1.17102951 | 0.284760467 | 0.632879342 | -4.59284616 |
| **A_55_P2274378** | AW549542 | -0.16456454 | -0.506045726 | 0.176916647 | 7.800445355 | -1.1709886 | 0.284775736 | 0.632879342 | -4.592865425 |
| **A_65_P20371** | Bclaf1 | 0.174011068 | -0.187139203 | 0.535161338 | 10.87511178 | 1.170771432 | 0.284856805 | 0.632879342 | -4.592967685 |
| **A_51_P291078** | Sel1l3 | -0.130169837 | -0.400333523 | 0.139993849 | 7.748989107 | -1.170756497 | 0.284862382 | 0.632879342 | -4.592974718 |
| **A_55_P2095039** | Ncmap | -0.224989321 | -0.691960723 | 0.241982082 | 7.796479761 | -1.170724774 | 0.284874226 | 0.632879342 | -4.592989655 |

| | | | | | | | | |
|---|---|---|---|---|---|---|---|---|
| **A_55_P1998947** | Trim16 | -0.294256188 | -0.90508486 | 0.316572484 | 7.955282523 | -1.17054841 | 0.28494008 | 0.632879342 | -4.593072694 |
| **A_51_P336874** | Atp6v0e | -0.09387665 | -0.288752178 | 0.100998878 | 13.57544578 | -1.170532473 | 0.284946032 | 0.632879342 | -4.593080197 |
| **A_55_P1978050** | Aven | 0.147367273 | -0.158579799 | 0.453314345 | 10.99803583 | 1.170409758 | 0.284991862 | 0.632879342 | -4.593137973 |
| **A_55_P2113673** | Eml1 | -0.568389583 | -1.74848976 | 0.611710594 | 10.35919944 | -1.17033562 | 0.285019554 | 0.632879342 | -4.593172877 |
| **A_55_P2175752** | Mfap1b | 0.211228141 | -0.227337537 | 0.649793819 | 9.333504673 | 1.170308409 | 0.285029718 | 0.632879342 | -4.593185688 |
| **A_55_P2162204** | Kctd15 | -0.515291953 | -1.585224238 | 0.554640333 | 9.139320885 | -1.17025433 | 0.285049919 | 0.632879342 | -4.593211147 |
| **A_51_P295896** | 4930452B06Rik | -0.11974254 | -0.368376097 | 0.128891017 | 8.502637538 | -1.170231842 | 0.28505832 | 0.632879342 | -4.593221734 |
| **A_55_P2091486** | Tmem219 | -0.09650015 | -0.296899121 | 0.10389882 | 11.30426491 | -1.17008037 | 0.28511491 | 0.632879342 | -4.59329304 |
| **A_55_P1981306** | Hdgfl3 | -0.191267572 | -0.588494796 | 0.205959653 | 8.222701231 | -1.1699991 | 0.285145277 | 0.632879342 | -4.593331297 |
| **A_55_P2164469** | Prcd | -0.783425314 | -2.41051307 | 0.843662442 | 8.148030005 | -1.169956763 | 0.285161097 | 0.632879342 | -4.593351226 |
| **A_55_P1990190** | Arhgap28 | 0.080498824 | -0.086714285 | 0.247711933 | 8.099458471 | 1.169775179 | 0.285228959 | 0.632879342 | -4.593436699 |
| **A_52_P707503** | Tmem212 | -0.44448561 | -1.367807979 | 0.478836759 | 7.895384765 | -1.169735142 | 0.285243924 | 0.632879342 | -4.593455544 |
| **A_55_P2117681** | Rnf115 | 0.143391082 | -0.154497247 | 0.44127941 | 10.00553969 | 1.169639003 | 0.28527986 | 0.632879342 | -4.593500794 |
| **A_55_P2067757** | Psg27 | -0.138641611 | -0.426685967 | 0.149402745 | 7.787322848 | -1.169546285 | 0.285314522 | 0.632879342 | -4.593544433 |
| **A_52_P423814** | Cox8b | -0.524776567 | -1.615143644 | 0.56559051 | 7.936439045 | -1.169458712 | 0.285347263 | 0.632879342 | -4.593585649 |
| **A_51_P461578** | Fdxr | 0.20349753 | -0.219338091 | 0.626333151 | 10.84811559 | 1.169420666 | 0.285361488 | 0.632879342 | -4.593603555 |
| **A_55_P2094352** | Zfp523 | 0.0765646 | -0.08253328 | 0.23566248 | 8.001464972 | 1.169356176 | 0.285385603 | 0.632879342 | -4.593633906 |
| **A_51_P143296** | Myh8 | -0.582562781 | -1.793215488 | 0.628089927 | 8.019628109 | -1.16924722 | 0.285426348 | 0.632879342 | -4.593685181 |
| **A_51_P317640** | Tgfb2 | -0.684649629 | -2.10749286 | 0.738193602 | 8.47437646 | -1.169215365 | 0.285438261 | 0.632879342 | -4.593700172 |
| **A_52_P39418** | Ccpg1 | -0.090992794 | -0.280099207 | 0.098113619 | 7.922617974 | -1.169186874 | 0.285448916 | 0.632879342 | -4.59371358 |
| **A_55_P2167107** | Cdk14 | -0.641112991 | -1.973772468 | 0.691546486 | 9.125871 | -1.168957102 | 0.285534863 | 0.63299729 | -4.593821704 |
| **A_52_P353417** | Gm2a | -0.210074786 | -0.646817691 | 0.22666812 | 9.361252829 | -1.168775931 | 0.285602647 | 0.633074949 | -4.593906951 |
| **A_51_P131408** | Tnfrsf12a | -0.194728239 | -0.599626903 | 0.210170424 | 10.41464258 | -1.168599732 | 0.285668583 | 0.633148496 | -4.593989854 |
| **A_55_P2183735** | Pisd | -0.085146922 | -0.262213103 | 0.091919259 | 10.30522873 | -1.168467185 | 0.285718192 | 0.633185843 | -4.594052214 |
| **A_51_P349888** | Ang2 | 0.081004778 | -0.087483979 | 0.249493536 | 7.945009264 | 1.168215326 | 0.285812477 | 0.633322179 | -4.5941707 |
| **A_51_P342773** | Vipr2 | -0.20071866 | -0.618302371 | 0.216865052 | 7.899295757 | -1.167958432 | 0.285908674 | 0.633356574 | -4.594291543 |
| **A_55_P1959425** | Slc16a3 | 0.253579104 | -0.273998619 | 0.781156828 | 12.20819936 | 1.167912222 | 0.285925981 | 0.633356574 | -4.594313279 |
| **A_55_P2148688** | NA | 0.125619273 | -0.135751385 | 0.386989931 | 9.523431646 | 1.167838084 | 0.285953749 | 0.633356574 | -4.594348151 |
| **A_55_P2021659** | Abcb8 | 0.174931267 | -0.189120589 | 0.538983124 | 11.09401068 | 1.167581974 | 0.286049693 | 0.633356574 | -4.594468608 |
| **A_52_P584293** | Atrnl1 | -0.169277794 | -0.521598542 | 0.183042954 | 8.535464337 | -1.16746793 | 0.286092425 | 0.633356574 | -4.594522244 |
| **A_55_P2129373** | Capn11 | 0.332184408 | -0.359348343 | 1.023717158 | 9.515676784 | 1.167211827 | 0.286188406 | 0.633356574 | -4.594642681 |
| **A_55_P1992582** | Hmgcs2 | -0.253942069 | -0.782616927 | 0.274732789 | 8.701654269 | -1.167156748 | 0.286209052 | 0.633356574 | -4.594668582 |
| **A_51_P386389** | Actr6 | 0.116278403 | -0.12583506 | 0.358391865 | 10.36939504 | 1.166979716 | 0.286275419 | 0.633356574 | -4.594751827 |
| **A_55_P2111429** | Btf3l4 | -0.098917219 | -0.30488306 | 0.107048622 | 10.15788792 | -1.166970529 | 0.286278864 | 0.633356574 | -4.594756147 |
| **A_52_P410765** | Sema7a | -0.683917435 | -2.108042434 | 0.740207564 | 8.354391115 | -1.166913742 | 0.286300155 | 0.633356574 | -4.594782848 |
| **A_55_P2052939** | Dyrk2 | -0.092507228 | -0.285138192 | 0.100123735 | 8.141199766 | -1.166897638 | 0.286306194 | 0.633356574 | -4.59479042 |
| **A_55_P2023314** | Casz1 | -0.176421657 | -0.543812208 | 0.190968894 | 7.964827372 | -1.166828699 | 0.286332043 | 0.633356574 | -4.594822834 |
| **A_55_P2037962** | Trpm6 | -0.437764837 | -1.349463847 | 0.473934174 | 7.905918195 | -1.166735949 | 0.286366825 | 0.633356574 | -4.594866443 |
| **A_55_P2054559** | NA | 0.37822447 | -0.409480072 | 1.165929012 | 9.686172299 | 1.166727575 | 0.286369966 | 0.633356574 | -4.59487038 |
| **A_51_P268697** | Slc1a3 | -0.190748454 | -0.588032298 | 0.20653539 | 7.807300723 | -1.166657325 | 0.286396313 | 0.633356574 | -4.594903409 |
| **A_51_P309184** | Pxmp2 | 0.219138259 | -0.237284842 | 0.675561359 | 10.92965636 | 1.166631715 | 0.286405918 | 0.633356574 | -4.59491545 |
| **A_55_P1982249** | Phactr2 | -0.146841128 | -0.452736161 | 0.159053904 | 9.091714735 | -1.166429455 | 0.286481788 | 0.633356574 | -4.595010538 |
| **A_52_P338180** | Timm13 | -0.116301287 | -0.35858339 | 0.125980815 | 13.63062495 | -1.166396957 | 0.28649398 | 0.633356574 | -4.595025816 |
| **A_51_P108901** | Ccdc86 | 0.136583995 | -0.147955435 | 0.421123425 | 10.61525383 | 1.166381284 | 0.28649986 | 0.633356574 | -4.595033184 |
| **A_51_P192964** | Clec12b | -0.144759808 | -0.4463473 | 0.156827684 | 7.730227846 | -1.166320375 | 0.286522713 | 0.633356574 | -4.595061817 |
| **A_51_P303089** | Ttc28 | -0.172127555 | -0.530752515 | 0.186497405 | 8.241865862 | -1.166253809 | 0.286547689 | 0.633356574 | -4.595093109 |
| **A_51_P202801** | Abcb9 | 0.160478089 | -0.173876307 | 0.494832486 | 10.2527191 | 1.166250642 | 0.286548877 | 0.633356574 | -4.595094598 |
| **A_55_P2082361** | Ppt1 | -0.077633468 | -0.239409996 | 0.08414306 | 8.138027747 | -1.166048638 | 0.286624683 | 0.633407046 | -4.595189553 |
| **A_52_P378968** | Rgs2 | -0.183041004 | -0.564482013 | 0.198400006 | 7.839859765 | -1.166015152 | 0.286637251 | 0.633407046 | -4.595205293 |
| **A_55_P2025225** | Gnaz | 0.250916776 | -0.272011517 | 0.773845069 | 8.565128827 | 1.165925357 | 0.286670955 | 0.63340911 | -4.595247499 |

| | | | | | | | | | |
|---|---|---|---|---|---|---|---|---|---|
| **A_52_P15197** | Dcun1d2 | 0.104026808 | -0.11282323 | 0.320876847 | 10.20864006 | 1.165652124 | 0.286773532 | 0.633415877 | -4.595375919 |
| **A_55_P1980321** | Pcbp3 | -0.180994952 | -0.558296716 | 0.196306812 | 9.175199851 | -1.165630267 | 0.286781739 | 0.633415877 | -4.595386192 |
| **A_55_P1971159** | NA | 0.129232207 | -0.140178974 | 0.398643388 | 9.573480254 | 1.165569931 | 0.286804395 | 0.633415877 | -4.595414548 |
| **A_55_P2067308** | NA | 0.106795533 | -0.115843114 | 0.329434179 | 13.07259824 | 1.165562918 | 0.286807029 | 0.633415877 | -4.595417844 |
| **A_55_P2001558** | Zfp128 | -0.091341095 | -0.28179472 | 0.09911253 | 8.318803777 | -1.165360144 | 0.286883182 | 0.633415877 | -4.595513136 |
| **A_66_P130887** | Pcdh18 | -0.467769296 | -1.443118532 | 0.50757994 | 9.039185963 | -1.165345632 | 0.286888633 | 0.633415877 | -4.595519956 |
| **A_55_P1952696** | NA | 0.114942287 | -0.124732821 | 0.354617395 | 13.11289375 | 1.165306292 | 0.286903409 | 0.633415877 | -4.595538443 |
| **A_55_P2168628** | Sac3d1 | 0.18720464 | -0.203209806 | 0.577619086 | 11.77249551 | 1.165128751 | 0.286970104 | 0.63347469 | -4.59562187 |
| **A_51_P408649** | Shisa2 | -0.25191965 | -0.777327558 | 0.273488258 | 8.020428172 | -1.165060903 | 0.286995595 | 0.63347469 | -4.59565375 |
| **A_51_P394814** | Svep1 | -0.48909543 | -1.50946078 | 0.53126992 | 9.030865672 | -1.164718816 | 0.28712415 | 0.633686081 | -4.595814478 |
| **A_55_P1953578** | Top3a | 0.151323216 | -0.164460046 | 0.467106477 | 9.544616949 | 1.164393159 | 0.287246575 | 0.633880019 | -4.595967467 |
| **A_55_P2147566** | Tac2 | 0.139760061 | -0.151913734 | 0.431433857 | 9.928112801 | 1.164310619 | 0.287277612 | 0.633880019 | -4.596006241 |
| **A_55_P2042217** | Fam161a | -0.149092555 | -0.460324809 | 0.162139699 | 7.889844759 | -1.164004205 | 0.287392855 | 0.634061921 | -4.596150169 |
| **A_55_P1973886** | Socs4 | 0.124052883 | -0.13497782 | 0.383083587 | 8.768696141 | 1.16369403 | 0.287509552 | 0.63424699 | -4.596295847 |
| **A_51_P216075** | Ddost | -0.094177133 | -0.290852649 | 0.102498382 | 12.9536298 | -1.16353207 | 0.287570502 | 0.634274481 | -4.596371907 |
| **A_52_P613808** | Zbtb7a | 0.229225236 | -0.249508637 | 0.707959108 | 8.696545136 | 1.163460009 | 0.287597624 | 0.634274481 | -4.596405747 |
| **A_51_P225224** | Htra1 | -1.094645796 | -3.380918269 | 1.191626677 | 8.792024451 | -1.163399346 | 0.287620458 | 0.634274481 | -4.596434234 |
| **A_52_P418515** | Cfap299 | -0.378712393 | -1.169771713 | 0.412346926 | 7.880792475 | -1.163278376 | 0.287665996 | 0.634286883 | -4.596491038 |
| **A_55_P2159555** | NA | -0.203168613 | -0.627592264 | 0.221255039 | 9.031930549 | -1.163162057 | 0.287709789 | 0.634286883 | -4.596545656 |
| **A_51_P485240** | Adipor1 | 0.121531392 | -0.132372168 | 0.375434951 | 11.97774903 | 1.163062011 | 0.28774746 | 0.634286883 | -4.596592631 |
| **A_55_P2073809** | AU015836 | 0.239242339 | -0.260622593 | 0.739107271 | 8.560712723 | 1.162970125 | 0.287782062 | 0.634286883 | -4.596635773 |
| **A_52_P336958** | Jcad | -0.341200435 | -1.05414365 | 0.37174278 | 10.90000661 | -1.162887828 | 0.287813056 | 0.634286883 | -4.596674412 |
| **A_66_P140858** | NA | -0.211761257 | -0.654260888 | 0.230738374 | 11.93700845 | -1.162831486 | 0.287834277 | 0.634286883 | -4.596700864 |
| **A_55_P2203478** | Scn5a | 0.371939716 | -0.405309498 | 1.149188929 | 8.957789584 | 1.162774375 | 0.287855789 | 0.634286883 | -4.596727677 |
| **A_55_P2035122** | NA | -0.162046286 | -0.500720628 | 0.176628055 | 10.43762123 | -1.162625855 | 0.287911737 | 0.634337852 | -4.596797401 |
| **A_52_P462350** | Dr1 | 0.116789947 | -0.12732098 | 0.360900874 | 9.879319375 | 1.162522672 | 0.287950613 | 0.634351196 | -4.596845839 |
| **A_55_P2069083** | Galnt12 | -0.150008315 | -0.463600203 | 0.163583573 | 8.517964594 | -1.162341388 | 0.288018925 | 0.634429378 | -4.596930937 |
| **A_55_P2037982** | Agap1 | -0.230766752 | -0.71324529 | 0.251711787 | 9.386877813 | -1.162193457 | 0.288074678 | 0.634479884 | -4.597000374 |
| **A_52_P45834** | Pafah1b2 | 0.120739438 | -0.131717636 | 0.373196511 | 9.266441403 | 1.16210346 | 0.288108602 | 0.634482302 | -4.597042616 |
| **A_66_P124445** | Rmc1 | 0.138877923 | -0.151589024 | 0.42934487 | 9.379184898 | 1.161768725 | 0.288234806 | 0.63462362 | -4.597199716 |
| **A_51_P496804** | Adat2 | 0.097198019 | -0.106095963 | 0.300492001 | 9.150250785 | 1.161759094 | 0.288238438 | 0.63462362 | -4.597204236 |
| **A_55_P2125457** | NA | 0.111503037 | -0.121779729 | 0.344785802 | 14.49231396 | 1.16141446 | 0.288368425 | 0.634821779 | -4.597365962 |
| **A_55_P2118630** | Lasp1 | 0.100034628 | -0.10926661 | 0.309335866 | 8.056378628 | 1.161346337 | 0.288394125 | 0.634821779 | -4.597397927 |
| **A_55_P1971174** | Cd1d2 | -0.133702428 | -0.413510213 | 0.146105356 | 8.13658686 | -1.161081464 | 0.288494069 | 0.63492399 | -4.597522206 |
| **A_55_P2138878** | Lrfn4 | -0.392023821 | -1.212461045 | 0.428413403 | 9.134323558 | -1.161048308 | 0.288506582 | 0.63492399 | -4.597537762 |
| **A_55_P1980548** | NA | -0.182403843 | -0.564171485 | 0.199363799 | 11.63055616 | -1.16096214 | 0.288539103 | 0.63492399 | -4.597578189 |
| **A_52_P81468** | Psma4 | -0.087401859 | -0.270372968 | 0.095569251 | 11.61591654 | -1.160703613 | 0.288636694 | 0.634984041 | -4.597699473 |
| **A_51_P116027** | Med27 | -0.11896244 | -0.368015111 | 0.130090231 | 10.89698412 | -1.160651533 | 0.288656357 | 0.634984041 | -4.597723904 |
| **A_51_P356055** | Grp | -0.302779959 | -0.936674453 | 0.331114534 | 7.801077123 | -1.160628782 | 0.288664948 | 0.634984041 | -4.597734577 |
| **A_55_P2086835** | NA | -0.54156391 | -1.675591484 | 0.592463663 | 9.629222293 | -1.160404195 | 0.288749756 | 0.635051339 | -4.597839926 |
| **A_55_P2010018** | NA | -0.169330772 | -0.52394756 | 0.185286016 | 16.38769087 | -1.16027193 | 0.288799712 | 0.635051339 | -4.597901964 |
| **A_51_P508115** | Trank1 | 0.084080686 | -0.09201208 | 0.260173452 | 7.86405405 | 1.160213519 | 0.288821776 | 0.635051339 | -4.597929361 |
| **A_55_P1983268** | 4930444P10Rik | -0.25155129 | -0.778455018 | 0.275352438 | 7.89099314 | -1.160054697 | 0.288881776 | 0.635051339 | -4.598003851 |
| **A_51_P255304** | Cav3 | -0.327674833 | -1.014052755 | 0.358703088 | 7.826725521 | -1.160013306 | 0.288897414 | 0.635051339 | -4.598023262 |
| **A_55_P2109874** | Rpl23a | -0.09998001 | -0.309417384 | 0.109457364 | 16.05767588 | -1.159957774 | 0.288918397 | 0.635051339 | -4.598049306 |
| **A_52_P497188** | Prrg1 | -0.14913331 | -0.461548713 | 0.163282093 | 8.552425632 | -1.159912979 | 0.288935323 | 0.635051339 | -4.598070314 |
| **A_52_P609120** | Pdxp | 0.176796227 | -0.193588968 | 0.547181421 | 8.862518759 | 1.159851964 | 0.28895838 | 0.635051339 | -4.598098928 |
| **A_55_P2077313** | Ddr1 | -0.362392941 | -1.121839081 | 0.397053199 | 8.656431887 | -1.159487148 | 0.289096271 | 0.635221226 | -4.598270001 |
| **A_52_P281145** | Kank4 | -0.161804187 | -0.500924384 | 0.177316009 | 7.766178675 | -1.159362611 | 0.289143356 | 0.635221226 | -4.598328395 |
| **A_55_P1976789** | Dgcr6 | -0.198017699 | -0.613051991 | 0.217016594 | 10.8084625 | -1.159319701 | 0.289159581 | 0.635221226 | -4.598348514 |

| | | | | | | | | | |
|---|---|---|---|---|---|---|---|---|---|
| **A_55_P1954331** | Cavin4 | -0.252478434 | -0.781668576 | 0.276711708 | 7.779329665 | -1.159299721 | 0.289167136 | 0.635221226 | -4.598357882 |
| **A_55_P2007566** | Trmt61b | 0.114061983 | -0.125050463 | 0.353174429 | 8.388087814 | 1.159102717 | 0.289241637 | 0.635312682 | -4.598450247 |
| **A_52_P517063** | Immp2l | -0.116937868 | -0.362120541 | 0.128244805 | 8.962483975 | -1.158907001 | 0.289315667 | 0.63534413 | -4.598542001 |
| **A_55_P2047748** | NA | 0.15707886 | -0.17227163 | 0.486429349 | 11.17234942 | 1.158891068 | 0.289321695 | 0.63534413 | -4.59854947 |
| **A_55_P2112185** | Nhsl1 | -0.107422148 | -0.332689338 | 0.117845041 | 8.575358409 | -1.158721544 | 0.289385832 | 0.635412785 | -4.598628939 |
| **A_55_P2133096** | Tgds | -0.078815163 | -0.244174439 | 0.086544113 | 8.308379712 | -1.15814891 | 0.289602569 | 0.635816453 | -4.598897339 |
| **A_55_P2117472** | Acap2 | -0.136194375 | -0.422024935 | 0.149636186 | 10.31586561 | -1.157800407 | 0.289734542 | 0.635840793 | -4.599060658 |
| **A_51_P110841** | Mrps35 | 0.088459532 | -0.097191702 | 0.274110766 | 10.53083888 | 1.157790513 | 0.289738289 | 0.635840793 | -4.593899955 |
| **A_51_P203182** | Apobr | -0.17343591 | -0.537432125 | 0.190560305 | 8.83830725 | -1.157778137 | 0.289742977 | 0.635840793 | -4.599071094 |
| **A_51_P127915** | Rnasek | -0.149082875 | -0.462014366 | 0.163848616 | 10.63342639 | -1.157608428 | 0.289807263 | 0.635840793 | -4.591237106 |
| **A_55_P1960813** | NA | -0.227251214 | -0.704265447 | 0.249763019 | 14.40156886 | -1.157598788 | 0.289810915 | 0.635840793 | -4.599155133 |
| **A_51_P410681** | Irx5 | -0.250676657 | -0.776862468 | 0.275509155 | 7.780482971 | -1.157598485 | 0.28981103 | 0.635840793 | -4.599155275 |
| **A_55_P2088625** | Fbln7 | -0.108753067 | -0.337096563 | 0.11959043 | 7.980291166 | -1.157273624 | 0.289934123 | 0.635987929 | -4.599307484 |
| **A_51_P241667** | Prkdc | 0.094344694 | -0.103760283 | 0.29244967 | 8.49573027 | 1.157191626 | 0.2899652 | 0.635987929 | -4.597296806 |
| **A_55_P2000603** | Slc50a1 | -0.08497065 | -0.26341105 | 0.093469749 | 10.37637242 | -1.157068165 | 0.290011996 | 0.635987929 | -4.599403739 |
| **A_55_P1973349** | H4c14 | -0.149826984 | -0.464473241 | 0.164819273 | 15.63127861 | -1.157046085 | 0.290020366 | 0.635987929 | -4.599414082 |
| **A_55_P2360661** | Cbx5 | 0.171882238 | -0.189122275 | 0.532886751 | 10.17478441 | 1.156915282 | 0.290069953 | 0.635987929 | -4.599475358 |
| **A_51_P349015** | Pacrg | -0.16675999 | -0.517017372 | 0.183497391 | 7.783775222 | -1.156878509 | 0.290083895 | 0.635987929 | -4.599492584 |
| **A_55_P1993026** | NA | -0.133130649 | -0.412830059 | 0.146568761 | 15.63735675 | -1.15656405 | 0.290203141 | 0.635987929 | -4.599639879 |
| **A_55_P2343151** | Akap1 | 0.15524858 | -0.170929305 | 0.481426465 | 9.822375623 | 1.156528428 | 0.290216652 | 0.635987929 | -4.599656563 |
| **A_55_P1966102** | Nme5 | -0.638375692 | -1.979806563 | 0.70305518 | 8.163385661 | -1.156355135 | 0.290282387 | 0.635987929 | -4.599737726 |
| **A_55_P2034740** | Exosc2 | 0.092405192 | -0.101808332 | 0.286618716 | 17.62848628 | 1.156112493 | 0.290374448 | 0.635987929 | -4.59985136 |
| **A_52_P378287** | Lca5 | -0.135572386 | -0.420519003 | 0.149374231 | 7.965721867 | -1.156088081 | 0.290383712 | 0.635987929 | -4.599862793 |
| **A_55_P2132844** | Rps16 | -0.115500233 | -0.358302307 | 0.12730184 | 15.51321363 | -1.155882422 | 0.290461763 | 0.635987929 | -4.599959098 |
| **A_51_P477676** | Mcph1 | 0.149817058 | -0.165126833 | 0.46476095 | 8.811188346 | 1.15587605 | 0.290464182 | 0.635987929 | -4.599962082 |
| **A_55_P2177755** | Cdk2ap1 | 0.112867554 | -0.124403274 | 0.350138382 | 10.44102342 | 1.155867241 | 0.290467525 | 0.635987929 | -4.599966207 |
| **A_55_P2164977** | NA | 0.183989826 | -0.202829741 | 0.570809394 | 8.603731067 | 1.155762407 | 0.290507319 | 0.635987929 | -4.600015295 |
| **A_51_P214251** | Rnls | -0.090114374 | -0.279576582 | 0.099347834 | 7.795546068 | -1.155725415 | 0.290521362 | 0.635987929 | -4.600032616 |
| **A_51_P211305** | Alpk3 | -0.081660273 | -0.253354258 | 0.090033713 | 7.724382884 | -1.155683554 | 0.290537254 | 0.635987929 | -4.600052216 |
| **A_55_P2043932** | Tmem8b | -0.139018889 | -0.431343086 | 0.153305307 | 7.925285988 | -1.155559304 | 0.290584428 | 0.635987929 | -4.600110391 |
| **A_55_P2024625** | Ccl21a | -1.325126232 | -4.11168313 | 1.461430665 | 8.940585025 | -1.155506853 | 0.290604344 | 0.635987929 | -4.600134949 |
| **A_52_P358093** | Fbxo10 | 0.101622112 | -0.112081836 | 0.31532606 | 9.551818292 | 1.155470496 | 0.29061815 | 0.635987929 | -4.600151971 |
| **A_51_P233153** | Cadps2 | -0.169845231 | -0.527039104 | 0.187348643 | 7.741933292 | -1.15540048 | 0.290644738 | 0.635987929 | -4.600184751 |
| **A_51_P239750** | Inhba | -0.330604159 | -1.025979148 | 0.364770831 | 10.10360429 | -1.155240577 | 0.290705469 | 0.635987929 | -4.60025961 |
| **A_51_P183213** | Riiad1 | -0.343213312 | -1.065148925 | 0.3787223 | 7.883256414 | -1.155177837 | 0.2907293 | 0.635987929 | -4.600288982 |
| **A_55_P2080333** | NA | 0.091593605 | -0.101071941 | 0.284259152 | 8.30358178 | 1.155165698 | 0.290733911 | 0.635987929 | -4.600294664 |
| **A_52_P554267** | Ilf3 | 0.102365442 | -0.112961428 | 0.317692311 | 9.787972779 | 1.155149854 | 0.29073993 | 0.635987929 | -4.600302081 |
| **A_52_P254795** | Ndufc1 | 0.090971337 | -0.10040529 | 0.282347964 | 14.38594958 | 1.155044914 | 0.290779795 | 0.635987929 | -4.600351205 |
| **A_55_P2122425** | NA | -0.107380937 | -0.333282776 | 0.118520902 | 12.5659871 | -1.155022943 | 0.290788143 | 0.635987929 | -4.60036149 |
| **A_66_P112305** | Myo1f | -0.232790015 | -0.722532809 | 0.256952778 | 10.66943466 | -1.154993351 | 0.290799385 | 0.635987929 | -4.600375342 |
| **A_55_P2020458** | NA | 0.168577881 | -0.186179145 | 0.523334907 | 11.48751634 | 1.15465642 | 0.290927421 | 0.636099904 | -4.60053305 |
| **A_55_P2149500** | Kifc2 | -0.147264292 | -0.457208175 | 0.162679591 | 8.113178825 | -1.154509667 | 0.290983202 | 0.636099904 | -4.600601734 |
| **A_55_P1975989** | Wdr63 | -0.364606765 | -1.131998542 | 0.402785012 | 7.921452248 | -1.154491582 | 0.290990077 | 0.636099904 | -4.600610198 |
| **A_52_P239726** | Larp4 | 0.233276834 | -0.257757603 | 0.724311271 | 9.813600538 | 1.154364202 | 0.291038503 | 0.636099904 | -4.600669812 |
| **A_55_P2083725** | NA | 0.147144452 | -0.162605671 | 0.456894575 | 8.230856728 | 1.154291752 | 0.29106605 | 0.636099904 | -4.600703717 |
| **A_55_P1985559** | Gatad1 | 0.09477672 | -0.104739014 | 0.294292454 | 8.518268057 | 1.154270809 | 0.291074013 | 0.636099904 | -4.600713518 |
| **A_55_P2145696** | P3h4 | -0.502143061 | -1.559227805 | 0.554941683 | 9.695366004 | -1.154252569 | 0.291080949 | 0.636099904 | -4.600722054 |
| **A_51_P377526** | Sts | -0.094710419 | -0.294124716 | 0.104703879 | 9.066575287 | -1.154050074 | 0.291157955 | 0.636131431 | -4.600816811 |
| **A_55_P2106241** | 4930432K21Rik | 0.113571571 | -0.12555695 | 0.352700092 | 8.378449507 | 1.15404155 | 0.291161197 | 0.636131431 | -4.6008208 |
| **A_51_P391764** | Cbx6 | -0.388202292 | -1.205732584 | 0.429328001 | 10.9180978 | -1.153818319 | 0.291246109 | 0.636163288 | -4.600925251 |

| | | | | | | | | |
|---|---|---|---|---|---|---|---|---|
| **A_55_P2064321** | Lama5 | -0.850803513 | -2.642643825 | 0.941036799 | 10.22319201 | -1.153753843 | 0.291270638 | 0.636163288 | -4.600955419 |
| **A_52_P253004** | Ralb | -0.226859401 | -0.704662298 | 0.250943496 | 12.32249718 | -1.153695481 | 0.291292843 | 0.636163288 | -4.600982725 |
| **A_52_P631547** | Cytl1 | -0.229420165 | -0.712632503 | 0.253792173 | 7.791986514 | -1.153657151 | 0.291307427 | 0.636163288 | -4.601000658 |
| **A_51_P175699** | Mtfp1 | 0.101508828 | -0.112343263 | 0.315360919 | 11.13382651 | 1.15338288 | 0.291411801 | 0.636319329 | -4.601128972 |
| **A_51_P335081** | Gnb1 | -0.131059856 | -0.407226525 | 0.145106812 | 11.07113673 | -1.15313888 | 0.291504682 | 0.636450243 | -4.601243113 |
| **A_55_P2021821** | Prmt1 | 0.137039649 | -0.151764109 | 0.425843407 | 12.43092975 | 1.152992778 | 0.29156031 | 0.636499799 | -4.601311453 |
| **A_65_P11773** | Adi1 | -0.138200346 | -0.42951734 | 0.153116648 | 10.69479075 | -1.15272709 | 0.291661491 | 0.636586727 | -4.60143572 |
| **A_55_P2209053** | 2900001G08Rik | -0.394382747 | -1.225723468 | 0.436957974 | 8.288769351 | -1.15271526 | 0.291665997 | 0.636586727 | -4.601441252 |
| **A_51_P373609** | Yars2 | 0.216678106 | -0.240182287 | 0.6735385 | 10.35481338 | 1.152430412 | 0.29177451 | 0.636751665 | -4.601574466 |
| **A_52_P106537** | Chrac1 | -0.152807112 | -0.475066705 | 0.169452481 | 10.237986 | -1.152181513 | 0.291869355 | 0.636883479 | -4.601690856 |
| **A_55_P1978615** | Rpl17 | -0.196323359 | -0.610385353 | 0.217738636 | 15.62138936 | -1.152098985 | 0.291900809 | 0.636883479 | -4.601729445 |
| **A_55_P2037812** | Palmd | -0.278643279 | -0.866389672 | 0.309103114 | 9.780801736 | -1.151971644 | 0.291949349 | 0.636909012 | -4.601788987 |
| **A_51_P292030** | Nus1 | 0.104424944 | -0.115854715 | 0.324704602 | 10.22555853 | 1.151895402 | 0.291978413 | 0.636909012 | -4.601824634 |
| **A_52_P533146** | Ddit3 | 0.137601041 | -0.152699863 | 0.427901944 | 9.323084323 | 1.151745491 | 0.292035569 | 0.636961805 | -4.601894722 |
| **A_51_P398406** | Kctd13 | -0.080103623 | -0.249113825 | 0.08890658 | 8.209335177 | -1.151655068 | 0.292070049 | 0.636965133 | -4.601936995 |
| **A_52_P413492** | Ldlrap1 | 0.160565317 | -0.178383682 | 0.499514317 | 9.884734916 | 1.151066925 | 0.292294401 | 0.637348739 | -4.602211922 |
| **A_55_P2161347** | Acmsd | 0.19265406 | -0.214049595 | 0.599357716 | 8.607677145 | 1.151021086 | 0.292311893 | 0.637348739 | -4.602233347 |
| **A_51_P467346** | Muc1 | -0.276076969 | -0.858937508 | 0.30678357 | 7.774581108 | -1.15092948 | 0.292346851 | 0.637353066 | -4.602276161 |
| **A_55_P1985159** | Mef2a | 0.321539585 | -0.357450146 | 1.000529317 | 11.5734568 | 1.150679643 | 0.292442212 | 0.637489062 | -4.602392923 |
| **A_52_P157274** | Abi1 | -0.085172766 | -0.265087348 | 0.094741817 | 8.200743178 | -1.150317095 | 0.29258064 | 0.637718899 | -4.602562339 |
| **A_52_P398925** | Stfa2l1 | -0.299433495 | -0.932061974 | 0.333194983 | 14.96534234 | -1.150097953 | 0.29266434 | 0.63781965 | -4.602664731 |
| **A_55_P2026420** | Pou6f1 | -0.142245301 | -0.442813348 | 0.158322745 | 7.930559652 | -1.149948288 | 0.292721515 | 0.63781965 | -4.602734656 |
| **A_52_P920158** | Rbm24 | -0.189909551 | -0.591216899 | 0.211397797 | 7.760258688 | -1.149880962 | 0.292747238 | 0.63781965 | -4.602766109 |
| **A_55_P2093704** | Meig1 | -0.265110122 | -0.825342363 | 0.29512212 | 7.78501199 | -1.149850555 | 0.292758856 | 0.63781965 | -4.602780315 |
| **A_51_P127681** | Clic4 | 0.116297174 | -0.129536194 | 0.362130542 | 12.92732754 | 1.149506732 | 0.292890253 | 0.637967642 | -4.596668428 |
| **A_51_P223498** | Slc39a10 | -0.090677244 | -0.28236748 | 0.101012991 | 7.926638561 | -1.149427311 | 0.292920612 | 0.637967642 | -4.602978028 |
| **A_55_P2107347** | Ro60 | -0.143647591 | -0.447329349 | 0.160034167 | 8.292657651 | -1.149377871 | 0.292939513 | 0.637967642 | -4.603001122 |
| **A_52_P18807** | Eif3c | 0.134706081 | -0.150085162 | 0.419497324 | 13.84543308 | 1.1493274 | 0.292958808 | 0.637967642 | -4.603024696 |
| **A_66_P110251** | Tmem91 | 0.136408005 | -0.152030538 | 0.424846548 | 9.292792832 | 1.149131565 | 0.293033686 | 0.638051427 | -4.603116163 |
| **A_51_P271984** | Tmem45b | -0.101568451 | -0.316364688 | 0.113227786 | 7.7056238 | -1.148987619 | 0.293088735 | 0.638051427 | -4.603183391 |
| **A_55_P2009008** | NA | -0.183929368 | -0.572908593 | 0.205049858 | 13.91940258 | -1.148967805 | 0.293096313 | 0.638051427 | -4.603192644 |
| **A_52_P494115** | Gatad2b | 0.149659785 | -0.166913231 | 0.466232801 | 8.653306071 | 1.148720606 | 0.29319087 | 0.638185398 | -4.603308085 |
| **A_55_P2008136** | Ctr9 | 0.139857366 | -0.156039257 | 0.435753989 | 8.919640477 | 1.148493456 | 0.293277781 | 0.638302695 | -4.603414153 |
| **A_55_P1953489** | Lrrc8b | 0.198763103 | -0.221885702 | 0.619411909 | 9.465936715 | 1.148151764 | 0.293408559 | 0.638368836 | -4.603573689 |
| **A_55_P2201822** | Naf1 | 0.204601309 | -0.228476596 | 0.637679213 | 10.51875245 | 1.147956863 | 0.293483177 | 0.638368836 | -4.603664679 |
| **A_51_P275591** | Zfp292 | -0.113909057 | -0.355058804 | 0.12724069 | 9.334081807 | -1.147769384 | 0.293554968 | 0.638368836 | -4.603752197 |
| **A_66_P126492** | Brk1 | -0.156722902 | -0.488517654 | 0.17507185 | 12.04688439 | -1.147746936 | 0.293563565 | 0.638368836 | -4.603762676 |
| **A_51_P430423** | Ada | 0.116632241 | -0.130304807 | 0.36356929 | 9.394970366 | 1.147666113 | 0.29359452 | 0.638368836 | -4.603800403 |
| **A_55_P2153600** | Mfsd4b3-ps | 0.133808851 | -0.149522725 | 0.417140426 | 8.312699996 | 1.147553794 | 0.293637542 | 0.638368836 | -4.60385283 |
| **A_55_P2127683** | 4933434E20Rik | 0.08842983 | -0.098852655 | 0.275712315 | 8.999634036 | 1.147320667 | 0.293726855 | 0.638368836 | -4.60396164 |
| **A_51_P123676** | Synpo | -0.124659583 | -0.388672543 | 0.139353376 | 8.111811568 | -1.147317492 | 0.293728071 | 0.638368836 | -4.600608732 |
| **A_51_P279062** | Treml1 | -0.197530154 | -0.615880953 | 0.220820644 | 12.20371361 | -1.147297349 | 0.293735789 | 0.638368836 | -4.603972523 |
| **A_55_P2151398** | Tmem116 | -0.163471068 | -0.509704882 | 0.182762746 | 7.84195261 | -1.147240867 | 0.293757432 | 0.638368836 | -4.603998884 |
| **A_55_P2152305** | Atp5h | -0.079582913 | -0.248155106 | 0.08898928 | 15.40114856 | -1.147141746 | 0.293795416 | 0.638368836 | -4.604045143 |
| **A_52_P49014** | Shh | -0.244636978 | -0.762833269 | 0.273559312 | 7.774074557 | -1.147125655 | 0.293801583 | 0.638368836 | -4.599527714 |
| **A_52_P78770** | Chpt1 | -0.101080185 | -0.315221378 | 0.113061009 | 7.786493185 | -1.146961911 | 0.293864342 | 0.638368836 | -4.604129066 |
| **A_51_P171288** | Gli3 | -0.243915461 | -0.760701908 | 0.272870985 | 8.336322195 | -1.146862635 | 0.293902397 | 0.638368836 | -4.604175393 |
| **A_55_P2094262** | Tph1 | -0.111090862 | -0.34647154 | 0.124289816 | 7.691448181 | -1.14680801 | 0.293923339 | 0.638368836 | -4.604200882 |
| **A_52_P548202** | NA | 0.357039998 | -0.399469396 | 1.113549392 | 10.01957466 | 1.146794854 | 0.293928382 | 0.638368836 | -4.604207021 |
| **A_55_P2060604** | Gm5797 | -0.18024423 | -0.562166109 | 0.20167765 | 7.953037258 | -1.146753357 | 0.293944292 | 0.638368836 | -4.604226384 |

| | | | | | | | | | |
|---|---|---|---|---|---|---|---|---|---|
| **A_65_P08768** | Fbln1 | -0.298249681 | -0.930249669 | 0.333750308 | 8.381265009 | -1.146690213 | 0.293968502 | 0.638368836 | -4.604255848 |
| **A_52_P334670** | Clock | 0.110187658 | -0.123309356 | 0.343684671 | 8.662384312 | 1.146660386 | 0.293979938 | 0.638368836 | -4.604269765 |
| **A_55_P2020497** | Dcaf12l1 | 0.118394656 | -0.132497508 | 0.369286819 | 8.263642972 | 1.146642912 | 0.293986639 | 0.638368836 | -4.604277919 |
| **A_55_P2151061** | NA | -0.081770046 | -0.255056314 | 0.091516221 | 8.345423266 | -1.146603563 | 0.294001727 | 0.638368836 | -4.604296279 |
| **A_51_P343833** | Traf1 | -0.427346816 | -1.333253421 | 0.478559789 | 8.220081843 | -1.14625234 | 0.294136433 | 0.638428539 | -4.604460144 |
| **A_55_P1968977** | Stk38l | -0.180762382 | -0.563955227 | 0.202430463 | 8.99554118 | -1.146235496 | 0.294142894 | 0.638428539 | -4.604468002 |
| **A_51_P505229** | Gtpbp8 | 0.222954132 | -0.249805853 | 0.695714117 | 9.703038032 | 1.145929775 | 0.294260193 | 0.638428539 | -4.604610618 |
| **A_51_P386911** | Taco1 | 0.129517252 | -0.145122718 | 0.404157222 | 8.585491403 | 1.145900914 | 0.294271268 | 0.638428539 | -4.604624081 |
| **A_51_P324351** | Meltf | -0.539005932 | -1.682031945 | 0.604020081 | 8.020474263 | -1.145831137 | 0.294298047 | 0.638428539 | -4.604656629 |
| **A_52_P229728** | Cdc42 | -0.126609989 | -0.395113504 | 0.141893525 | 11.46903312 | -1.145779865 | 0.294317725 | 0.638428539 | -4.604680544 |
| **A_51_P505967** | Spag6l | -0.221511013 | -0.691317345 | 0.248295319 | 7.779739931 | -1.145670278 | 0.294359787 | 0.638428539 | -4.604731659 |
| **A_52_P305279** | Spata13 | 0.138858229 | -0.15565264 | 0.433369098 | 9.376614373 | 1.145654022 | 0.294366027 | 0.638428539 | -4.604739241 |
| **A_55_P2014124** | Gjb3 | -0.098199888 | -0.306495814 | 0.110096038 | 7.963825457 | -1.1455483 | 0.294406612 | 0.638428539 | -4.60478855 |
| **A_51_P101006** | Ak8 | -0.149494446 | -0.466708334 | 0.167719442 | 8.03754399 | -1.145133328 | 0.294565958 | 0.638428539 | -4.603878372 |
| **A_55_P2151822** | Plxnd1 | -0.256808902 | -0.80180014 | 0.288182337 | 12.66376387 | -1.144995399 | 0.294618937 | 0.638428539 | -4.605046392 |
| **A_55_P2081055** | 1700056E22Rik | 0.100497818 | -0.112783933 | 0.313779569 | 8.544068987 | 1.144948951 | 0.29463678 | 0.638428539 | -4.605068051 |
| **A_55_P2045859** | NA | 0.112686709 | -0.126467177 | 0.351840595 | 10.92805527 | 1.1449287 | 0.29464456 | 0.638428539 | -4.605077494 |
| **A_52_P375638** | Gtpbp1 | 0.209989258 | -0.235706401 | 0.655684917 | 9.872909913 | 1.144832275 | 0.294681605 | 0.638428539 | -4.605122454 |
| **A_51_P227777** | Top2b | 0.116728611 | -0.131028507 | 0.364485729 | 12.34475344 | 1.144812506 | 0.2946892 | 0.638428539 | -4.605131672 |
| **A_55_P1991124** | Tom1l2 | -0.156085691 | -0.487421139 | 0.175249757 | 10.06838333 | -1.144664933 | 0.294745905 | 0.638428539 | -4.605200477 |
| **A_55_P1979335** | NA | -0.092577369 | -0.289106109 | 0.103951371 | 8.271533493 | -1.144621647 | 0.294762539 | 0.638428539 | -4.605220658 |
| **A_55_P2329313** | C530014P21Rik | 0.117236379 | -0.131667588 | 0.366140345 | 8.154607661 | 1.144494648 | 0.294811347 | 0.638428539 | -4.605279867 |
| **A_55_P1964628** | NA | -0.372855671 | -1.16450382 | 0.418792478 | 12.16654316 | -1.144436607 | 0.294833656 | 0.638428539 | -4.605306926 |
| **A_51_P317272** | Bod1 | -0.272788675 | -0.852030851 | 0.306453502 | 11.16877951 | -1.1443249 | 0.294876596 | 0.638428539 | -4.605359001 |
| **A_55_P1953003** | Atp6v0a4 | -0.581831364 | -1.81733918 | 0.653676452 | 8.243883745 | -1.144286621 | 0.294891311 | 0.638428539 | -4.605376846 |
| **A_55_P1970715** | NA | 0.076283648 | -0.085710113 | 0.23827741 | 8.014434651 | 1.144237967 | 0.294910016 | 0.638428539 | -4.605399526 |
| **A_55_P2101177** | Ncaph2 | 0.095307675 | -0.107122181 | 0.297737532 | 11.96764293 | 1.14402759 | 0.294990906 | 0.638428539 | -4.605497591 |
| **A_52_P207303** | Bmp3 | -0.154099201 | -0.481440506 | 0.173242105 | 7.76543133 | -1.143886054 | 0.295045337 | 0.638428539 | -4.605563561 |
| **A_55_P2111148** | Gemin5 | 0.175633488 | -0.197453505 | 0.548720481 | 9.542784569 | 1.143879846 | 0.295047725 | 0.638428539 | -4.605566455 |
| **A_55_P1953241** | Mafk | 0.21449232 | -0.241144881 | 0.670129522 | 9.989839379 | 1.143867606 | 0.295052433 | 0.638428539 | -4.60557216 |
| **A_66_P124755** | Fut8 | -0.129398637 | -0.404279385 | 0.145482112 | 9.011572049 | -1.143848653 | 0.295059722 | 0.638428539 | -4.605580994 |
| **A_55_P2035963** | Hipk3 | 0.100936485 | -0.113488098 | 0.315361067 | 8.255132058 | 1.143817643 | 0.29507165 | 0.638428539 | -4.605595447 |
| **A_52_P606679** | Eloa | 0.118286493 | -0.133004371 | 0.369577356 | 8.552605203 | 1.143777751 | 0.295086994 | 0.638428539 | -4.605614039 |
| **A_55_P2448065** | Sfxn2 | 0.146553657 | -0.164795572 | 0.457902886 | 11.38958213 | 1.143752493 | 0.29509671 | 0.638428539 | -4.605625812 |
| **A_52_P198727** | Cfap410 | -0.107969375 | -0.337356221 | 0.121417471 | 8.349669371 | -1.14370828 | 0.295113717 | 0.638428539 | -4.605646417 |
| **A_51_P100298** | Stx3 | -0.086725442 | -0.2709834 | 0.097532515 | 7.776308613 | -1.143677191 | 0.295125676 | 0.638428539 | -4.603488927 |
| **A_51_P320614** | Txnrd1 | 0.153676034 | -0.172828926 | 0.480180994 | 11.94500735 | 1.143666892 | 0.295129638 | 0.638428539 | -4.605665707 |
| **A_55_P2079746** | Pls3 | -0.636628608 | -1.989419509 | 0.716162293 | 9.972189187 | -1.143506575 | 0.295191317 | 0.638428539 | -4.60574042 |
| **A_55_P2039038** | Zfp664 | -0.195233949 | -0.610144531 | 0.219676632 | 10.17709903 | -1.143362695 | 0.295246681 | 0.638428539 | -4.605807469 |
| **A_55_P2096867** | Gap43 | -0.091852645 | -0.287058002 | 0.103352712 | 8.389523369 | -1.143360346 | 0.295247586 | 0.638428539 | -4.605808564 |
| **A_52_P680870** | Ctxn1 | -0.129352903 | -0.404266466 | 0.145560659 | 7.894037327 | -1.1433079 | 0.295267769 | 0.638428539 | -4.605833003 |
| **A_66_P101620** | Cfap97 | 0.126409056 | -0.14226243 | 0.395080542 | 9.023093871 | 1.143246284 | 0.295291483 | 0.638428539 | -4.605861716 |
| **A_55_P2170847** | Fbxo27 | -0.159281138 | -0.497876706 | 0.179314429 | 8.029462711 | -1.143052744 | 0.295365979 | 0.638428539 | -4.605951897 |
| **A_55_P2073084** | Gtf3c2 | 0.156569817 | -0.176279314 | 0.489418947 | 10.56534046 | 1.142993617 | 0.295388741 | 0.638428539 | -4.605979446 |
| **A_51_P199899** | Fto | -0.141639357 | -0.442884873 | 0.159606159 | 9.431163977 | -1.142474575 | 0.29558862 | 0.638428539 | -4.606221258 |
| **A_51_P202014** | Taf12 | 0.134675359 | -0.151801175 | 0.421151892 | 10.79889616 | 1.142305525 | 0.295653744 | 0.638428539 | -4.606300004 |
| **A_55_P2100241** | Rn18s | 0.14598355 | -0.164551317 | 0.456518418 | 17.53514384 | 1.142290961 | 0.295659355 | 0.638428539 | -4.606306788 |
| **A_55_P1966760** | NA | -0.105702831 | -0.330560368 | 0.119154706 | 17.30522791 | -1.142253134 | 0.295673929 | 0.638428539 | -4.606324407 |
| **A_55_P2142218** | NA | -0.216292315 | -0.67645101 | 0.243866381 | 14.21370277 | -1.142132905 | 0.295720256 | 0.638428539 | -4.606380408 |
| **A_55_P2124736** | Col14a1 | -0.14145745 | -0.442422152 | 0.159507252 | 12.2608544 | -1.142071915 | 0.295743759 | 0.638428539 | -4.606408815 |

| | | | | | | | | | |
|---|---|---|---|---|---|---|---|---|---|
| **A_55_P1963579** | NA | 0.187052083 | -0.210929584 | 0.58503375 | 10.16482567 | 1.142043564 | 0.295754684 | 0.638428539 | -4.606422019 |
| **A_55_P2069592** | Pitpnm3 | -0.26613672 | -0.832399975 | 0.300126535 | 7.772645559 | -1.142009216 | 0.295767922 | 0.638428539 | -4.606438017 |
| **A_55_P2053923** | Rab17 | -0.172406909 | -0.539254059 | 0.194440241 | 9.919411956 | -1.141964767 | 0.295785053 | 0.638428539 | -4.606458719 |
| **A_51_P248638** | Myoz2 | -0.140611529 | -0.439804942 | 0.158581884 | 7.745732979 | -1.141963156 | 0.295785674 | 0.638428539 | -4.606459469 |
| **A_51_P105017** | Rad52 | 0.088302868 | -0.099610108 | 0.276215844 | 8.50091119 | 1.141829418 | 0.295837222 | 0.638428539 | -4.60189222 |
| **A_52_P648304** | Pigs | 0.093131162 | -0.105080529 | 0.291342854 | 11.71934718 | 1.141691965 | 0.295890211 | 0.638428539 | -4.606585767 |
| **A_52_P11441** | Rab6b | -0.409063521 | -1.279715306 | 0.461588264 | 8.569407363 | -1.141640708 | 0.295909972 | 0.638428539 | -4.606609636 |
| **A_55_P1984655** | Smtnl2 | -0.734213002 | -2.296946942 | 0.828520939 | 10.09720219 | -1.141616443 | 0.295919328 | 0.638428539 | -4.606620936 |
| **A_55_P2178343** | Rap1gds1 | -0.127795812 | -0.399806911 | 0.144215287 | 11.84639339 | -1.141597965 | 0.295926452 | 0.638428539 | -4.60662954 |
| **A_51_P103850** | Fbn2 | -0.44051894 | -1.37819716 | 0.497159279 | 8.343383894 | -1.141547394 | 0.295945952 | 0.638428539 | -4.60665309 |
| **A_55_P1953414** | Rab12 | -0.410495036 | -1.284286855 | 0.463296783 | 10.61328258 | -1.141518947 | 0.295956921 | 0.638428539 | -4.606666336 |
| **A_51_P378051** | Aoc3 | -0.366953295 | -1.148157383 | 0.414250793 | 8.040130193 | -1.141377937 | 0.296011299 | 0.638428539 | -4.606731997 |
| **A_55_P2316300** | Neo1 | -0.358012982 | -1.120221625 | 0.404195661 | 8.837632316 | -1.141321753 | 0.296032968 | 0.638428539 | -4.606758157 |
| **A_52_P669922** | Hamp | 0.317511547 | -0.358519614 | 0.993542707 | 9.856716758 | 1.141237407 | 0.2960655 | 0.638428539 | -4.60679743 |
| **A_51_P304239** | Loxl1 | -0.617393228 | -1.932077597 | 0.697291141 | 8.811331085 | -1.141099595 | 0.296118662 | 0.638428539 | -4.606861594 |
| **A_55_P2064025** | A430073D23Rik | 0.17824041 | -0.201376311 | 0.557857132 | 12.02051373 | 1.14089067 | 0.29619927 | 0.638428539 | -4.606958861 |
| **A_55_P2001438** | Coq2 | -0.090798 | -0.284233211 | 0.102637211 | 10.84538737 | -1.140575238 | 0.296321007 | 0.638428539 | -4.607105699 |
| **A_55_P2024366** | Raver2 | -0.699714343 | -2.190814051 | 0.791385366 | 8.574883069 | -1.140242646 | 0.296449412 | 0.638428539 | -4.607260504 |
| **A_55_P2123502** | Jam2 | -0.219778219 | -0.688159895 | 0.248603456 | 9.021195277 | -1.140165627 | 0.296479153 | 0.638428539 | -4.60729635 |
| **A_55_P2040295** | Ptp4a1 | -0.147614689 | -0.46221283 | 0.166983452 | 9.981474896 | -1.140135881 | 0.296490641 | 0.638428539 | -4.607310194 |
| **A_51_P111164** | Rnd1 | -0.116618535 | -0.365166294 | 0.131929224 | 8.76810564 | -1.140094682 | 0.296506551 | 0.638428539 | -4.607329368 |
| **A_55_P2169396** | NA | -0.12944952 | -0.405344091 | 0.146445052 | 15.58541498 | -1.140093517 | 0.296507001 | 0.638428539 | -4.60732991 |
| **A_55_P2087162** | Specc1l | -0.182708077 | -0.5721225 | 0.206706347 | 8.183403369 | -1.140063132 | 0.296518737 | 0.638428539 | -4.607344051 |
| **A_52_P515826** | Med13 | 0.226618063 | -0.256471397 | 0.709707523 | 8.53770827 | 1.139856423 | 0.296598581 | 0.638428539 | -4.607440247 |
| **A_51_P159352** | Sub1 | 0.091239132 | -0.10326187 | 0.285740134 | 8.297644512 | 1.139836324 | 0.296606345 | 0.638428539 | -4.6074496 |
| **A_55_P2105436** | Foxp1 | -0.0993052 | -0.311002762 | 0.112392362 | 8.243672313 | -1.139828006 | 0.296609558 | 0.638428539 | -4.607453471 |
| **A_55_P2060922** | Unc5a | -0.123932492 | -0.388134659 | 0.140269675 | 8.938848827 | -1.139808777 | 0.296616987 | 0.638428539 | -4.607462419 |
| **A_52_P664404** | Zfp286 | -0.194513388 | -0.60919281 | 0.220166033 | 8.108873097 | -1.139777826 | 0.296628944 | 0.638428539 | -4.607476821 |
| **A_52_P413855** | Slc35a3 | 0.138120189 | -0.15637629 | 0.432616669 | 8.518487301 | 1.139620482 | 0.296689738 | 0.638428539 | -4.607550037 |
| **A_52_P525317** | Gja5 | -0.297692726 | -0.932501509 | 0.337116057 | 8.611540231 | -1.139484667 | 0.296742222 | 0.638428539 | -4.607613231 |
| **A_55_P2064906** | NA | 0.134400806 | -0.152227094 | 0.421028706 | 15.6121923 | 1.139374776 | 0.296784693 | 0.638428539 | -4.60766436 |
| **A_55_P2020743** | Abcc10 | -0.092943772 | -0.29116344 | 0.105275896 | 8.856535953 | -1.139348899 | 0.296794695 | 0.638428539 | -4.6076764 |
| **A_66_P133413** | NA | -0.180760088 | -0.56628195 | 0.204761773 | 15.52078347 | -1.139296395 | 0.296814989 | 0.638428539 | -4.607700827 |
| **A_51_P327021** | Zbtb25 | 0.078370261 | -0.088799813 | 0.245540336 | 9.51605792 | 1.139136972 | 0.296876619 | 0.638428539 | -4.607774996 |
| **A_55_P1957227** | Las1l | 0.121719136 | -0.137932292 | 0.381370564 | 9.823272168 | 1.13907241 | 0.29690158 | 0.638428539 | -4.607805031 |
| **A_51_P173601** | Bcam | -0.681330682 | -2.134780745 | 0.77211938 | 8.629008786 | -1.139045364 | 0.296912037 | 0.638428539 | -4.607817612 |
| **A_55_P2048656** | Atp5l | 0.102530014 | -0.11620221 | 0.321262238 | 14.85285014 | 1.138994006 | 0.296931895 | 0.638428539 | -4.607841504 |
| **A_66_P102581** | Ube2z | 0.110429228 | -0.125157996 | 0.346016451 | 10.76021472 | 1.138978405 | 0.296937928 | 0.638428539 | -4.607848762 |
| **A_52_P20966** | Hnrnpab | 0.085917304 | -0.097393919 | 0.269228527 | 13.62264226 | 1.138871631 | 0.296979217 | 0.638428539 | -4.60789843 |
| **A_55_P2074908** | Ube4b | 0.096636915 | -0.109583806 | 0.302857635 | 8.467178928 | 1.138659687 | 0.29706119 | 0.638428539 | -4.607997016 |
| **A_55_P2323913** | NA | 0.118551071 | -0.134437146 | 0.371539288 | 9.340225224 | 1.138645083 | 0.297066839 | 0.638428539 | -4.608003809 |
| **A_55_P2137286** | Gm10845 | 0.22868119 | -0.259328003 | 0.716690384 | 11.41800106 | 1.138637884 | 0.297069624 | 0.638428539 | -4.608007157 |
| **A_55_P1952638** | Fgd6 | 0.178611336 | -0.202557012 | 0.559779683 | 9.244111469 | 1.138611008 | 0.297080021 | 0.638428539 | -4.608019658 |
| **A_55_P2109613** | Ahr | -0.256277823 | -0.803204237 | 0.290648592 | 10.69973101 | -1.138584627 | 0.297090226 | 0.638428539 | -4.608031928 |
| **A_55_P2079041** | NA | 0.090083131 | -0.102172332 | 0.282338594 | 16.31585029 | 1.138539156 | 0.297107816 | 0.638428539 | -4.608053077 |
| **A_51_P280437** | Slc12a6 | -0.098798697 | -0.309669383 | 0.112071989 | 9.043329644 | -1.138461099 | 0.297138015 | 0.638428539 | -4.608089381 |
| **A_52_P385206** | Harbi1 | -0.114258233 | -0.358136175 | 0.129619709 | 8.765630286 | -1.138408612 | 0.297158323 | 0.638428539 | -4.608113792 |
| **A_55_P2023617** | Kiss1r | -0.080685406 | -0.252905394 | 0.091534582 | 7.782893979 | -1.138399312 | 0.297161921 | 0.638428539 | -4.608118118 |
| **A_55_P2111825** | 1700001C02Rik | -0.549520434 | -1.722464899 | 0.623424031 | 7.972401177 | -1.138386102 | 0.297167032 | 0.638428539 | -4.608124261 |
| **A_55_P2341468** | Rbms3 | -0.542788057 | -1.701700085 | 0.616123971 | 9.70744237 | -1.138054347 | 0.297295421 | 0.638613285 | -4.608278543 |

| | | | | | | | | | |
|---|---|---|---|---|---|---|---|---|---|
| **A_55_P1955921** | NA | -0.182618269 | -0.572553394 | 0.207316856 | 14.43647607 | -1.137981108 | 0.297323771 | 0.638613285 | -4.6083126 |
| **A_55_P2036285** | Elp4 | 0.108262949 | -0.122967368 | 0.339493266 | 8.826297289 | 1.13767514 | 0.29744223 | 0.638613285 | -4.608454867 |
| **A_55_P1956718** | Rab38 | -0.377159651 | -1.18275032 | 0.428431018 | 8.943037282 | -1.137611535 | 0.297466861 | 0.638613285 | -4.60848444 |
| **A_51_P437608** | Tulp3 | -0.203175724 | -0.637171741 | 0.230820294 | 8.385720989 | -1.137546764 | 0.297491945 | 0.638613285 | -4.608514553 |
| **A_52_P519882** | Etf1 | 0.133945142 | -0.152180766 | 0.420071049 | 10.24625837 | 1.137504104 | 0.297508467 | 0.638613285 | -4.608534387 |
| **A_52_P155124** | Zfp229 | 0.091094668 | -0.103508168 | 0.285697504 | 8.031678058 | 1.13743603 | 0.297534833 | 0.638613285 | -4.608566035 |
| **A_66_P120125** | Nrep | -0.661886876 | -2.075885951 | 0.752112198 | 10.64572521 | -1.137412074 | 0.297544112 | 0.638613285 | -4.608577172 |
| **A_55_P2248330** | 2700068H02Rik | 0.19486639 | -0.221435585 | 0.611168364 | 8.352856338 | 1.137395898 | 0.297550377 | 0.638613285 | -4.608584692 |
| **A_51_P397375** | Gatb | 0.145022523 | -0.164839787 | 0.454884834 | 9.751259959 | 1.137234148 | 0.297613037 | 0.63867685 | -4.608659886 |
| **A_55_P2053177** | NA | 0.113473679 | -0.129023674 | 0.355971031 | 11.12768731 | 1.137028427 | 0.297692745 | 0.638776983 | -4.608755515 |
| **A_55_P2142371** | Rai1 | -0.139798326 | -0.43862334 | 0.159026688 | 10.45952323 | -1.136758503 | 0.297797357 | 0.638930526 | -4.608880975 |
| **A_66_P108267** | Foxn4 | -0.323682624 | -1.015811631 | 0.368446383 | 7.827038367 | -1.136358917 | 0.297952277 | 0.63919196 | -4.609066679 |
| **A_51_P117739** | Vegfd | -0.883918499 | -2.774235099 | 1.006398101 | 8.93868932 | -1.136215904 | 0.29800774 | 0.639229906 | -4.603605908 |
| **A_55_P2034245** | Meox1 | -0.269212636 | -0.844978245 | 0.306552974 | 8.047789215 | -1.136142766 | 0.298036107 | 0.639229906 | -4.60916712 |
| **A_52_P67200** | Stt3b | 0.166940739 | -0.190245999 | 0.524127478 | 11.98589266 | 1.135664879 | 0.298221517 | 0.639492007 | -4.609389154 |
| **A_66_P116373** | Pwwp2b | -0.339415997 | -1.065733264 | 0.38690127 | 9.282130166 | -1.1355052 | 0.298283491 | 0.639492007 | -4.609463334 |
| **A_66_P111798** | Znrf2 | 0.128288083 | -0.146238296 | 0.402814462 | 9.417469733 | 1.135495512 | 0.298287251 | 0.639492007 | -4.609467834 |
| **A_55_P2043782** | Trpm1 | 0.140342244 | -0.159999524 | 0.440684011 | 8.368847772 | 1.135418254 | 0.298317241 | 0.639492007 | -4.609503723 |
| **A_55_P2065751** | Krtap19-5 | 0.365680982 | -0.416952582 | 1.148314545 | 9.987656499 | 1.13534302 | 0.298346446 | 0.639492007 | -4.609538671 |
| **A_55_P2044582** | Iglon5 | -0.153335142 | -0.481512046 | 0.174841762 | 7.776727907 | -1.135316307 | 0.298356817 | 0.639492007 | -4.609551079 |
| **A_55_P2066463** | Enah | -0.260469618 | -0.818039032 | 0.297099796 | 8.093179802 | -1.135118809 | 0.298433499 | 0.639585443 | -4.609642814 |
| **A_51_P515605** | Col3a1 | -0.448758309 | -1.409494349 | 0.51197773 | 12.45239405 | -1.134988889 | 0.298483953 | 0.639622652 | -4.609703157 |
| **A_51_P181365** | Atp5md | 0.106913312 | -0.122042678 | 0.335869302 | 15.15729834 | 1.134652727 | 0.298614531 | 0.639807121 | -4.609859275 |
| **A_51_P437144** | Dynll2 | -0.123011849 | -0.386465911 | 0.140442213 | 10.88517023 | -1.13455417 | 0.298652823 | 0.639807121 | -4.609905042 |
| **A_55_P1980416** | Zkscan1 | -0.129967661 | -0.408329391 | 0.148394069 | 10.77918247 | -1.134511664 | 0.298669339 | 0.639807121 | -4.609924781 |
| **A_52_P243249** | Mapk3 | -0.293776479 | -0.923037202 | 0.335484244 | 10.78375126 | -1.134408797 | 0.298709312 | 0.639821841 | -4.609972547 |
| **A_55_P2038757** | Fblim1 | -0.537687838 | -1.689672575 | 0.614296899 | 8.844389164 | -1.134140024 | 0.298813776 | 0.639969352 | -4.610097341 |
| **A_55_P2142156** | Gm16441 | 0.170577749 | -0.194906829 | 0.536062328 | 9.536862637 | 1.134061252 | 0.298844398 | 0.639969352 | -4.610133913 |
| **A_66_P121446** | Gm7969 | 0.257433465 | -0.294238518 | 0.809105449 | 10.33702362 | 1.133880439 | 0.298914698 | 0.639979027 | -4.610217856 |
| **A_55_P2088640** | Trip4 | 0.139471419 | -0.159432794 | 0.438375632 | 9.563101031 | 1.133799791 | 0.298946058 | 0.639979027 | -4.610255296 |
| **A_55_P2050226** | Ackr4 | 0.088610891 | -0.101293967 | 0.278515749 | 7.900439693 | 1.133794167 | 0.298948245 | 0.639979027 | -4.610257907 |
| **A_51_P386526** | Rcc1l | 0.126965762 | -0.145192858 | 0.399124382 | 8.820146707 | 1.13356837 | 0.299036062 | 0.640013541 | -4.610362722 |
| **A_55_P2067727** | Mxra7 | -0.239745456 | -0.753783685 | 0.274292773 | 8.392526906 | -1.133282427 | 0.299147303 | 0.640013541 | -4.610495443 |
| **A_66_P105843** | Cyth2 | -0.21924417 | -0.68932832 | 0.25083998 | 11.57895986 | -1.133275852 | 0.299149861 | 0.640013541 | -4.610498495 |
| **A_55_P2178803** | Zbtb5 | -0.160273309 | -0.503932186 | 0.183385569 | 10.44500961 | -1.133226766 | 0.29916896 | 0.640013541 | -4.610521276 |
| **A_55_P2161132** | NA | 0.091277562 | -0.104458465 | 0.287013589 | 7.933971725 | 1.133121425 | 0.299209952 | 0.640013541 | -4.610570165 |
| **A_51_P163778** | 2810025M15Rik | -0.256999957 | -0.808155307 | 0.294155393 | 10.75550569 | -1.133032095 | 0.299244718 | 0.640013541 | -4.610611623 |
| **A_52_P319123** | Vps13b | 0.17736464 | -0.203007399 | 0.557736679 | 10.19853852 | 1.133030618 | 0.299245292 | 0.640013541 | -4.610612308 |
| **A_51_P196629** | Fam102b | -0.219298554 | -0.689614717 | 0.25101761 | 9.981254698 | -1.132997763 | 0.29925808 | 0.640013541 | -4.610627555 |
| **A_66_P138567** | NA | 0.117337416 | -0.134311395 | 0.368986226 | 10.14283628 | 1.132986736 | 0.299262372 | 0.640013541 | -4.610632672 |
| **A_55_P2113912** | NA | 0.156156547 | -0.1788339 | 0.491146995 | 14.05839219 | 1.132689734 | 0.299377988 | 0.640073254 | -4.610770494 |
| **A_55_P2005833** | Adamts8 | -0.329191057 | -1.035387588 | 0.377005473 | 7.926773316 | -1.132675843 | 0.299383397 | 0.640073254 | -4.610776939 |
| **A_51_P487918** | Rinl | 0.126262426 | -0.144605652 | 0.397130504 | 9.494010621 | 1.132659815 | 0.299389637 | 0.640073254 | -4.610784376 |
| **A_55_P2008407** | Ppp1r14c | -0.099489654 | -0.313007833 | 0.114028525 | 10.52708908 | -1.132208088 | 0.299565562 | 0.64031619 | -4.610993963 |
| **A_55_P2444515** | NA | -0.23678069 | -0.74502454 | 0.271463161 | 8.298486174 | -1.132028439 | 0.29963555 | 0.64031619 | -4.611077303 |
| **A_55_P2174935** | Mn1 | 0.115456748 | -0.13237453 | 0.363288025 | 8.74556723 | 1.131999911 | 0.299646666 | 0.64031619 | -4.611090537 |
| **A_55_P1995417** | Cnppd1 | 0.161424116 | -0.185091784 | 0.507940017 | 9.216681387 | 1.131953115 | 0.2996649 | 0.64031619 | -4.611112245 |
| **A_55_P2072095** | Mucl3 | -0.119503048 | -0.376032762 | 0.137026665 | 7.681766099 | -1.131942837 | 0.299668904 | 0.64031619 | -4.611117012 |
| **A_51_P255565** | Smarcad1 | 0.108785326 | -0.124782864 | 0.342353515 | 8.976171007 | 1.131722139 | 0.299754912 | 0.640353052 | -4.611219384 |
| **A_51_P116755** | Cap2 | -0.101089165 | -0.318144854 | 0.115966524 | 7.768220802 | -1.131661722 | 0.29977846 | 0.640353052 | -4.606417875 |

| | | | | | | | | | |
|---|---|---|---|---|---|---|---|---|---|
| **A_51_P160536** | Cfap161 | -0.237774049 | -0.748323709 | 0.27277561 | 7.773859128 | -1.131643549 | 0.299785544 | 0.640353052 | -4.611255836 |
| **A_55_P2002903** | Smoc2 | -0.115113333 | -0.362373921 | 0.132147256 | 11.43019731 | -1.131237826 | 0.299943724 | 0.640586954 | -4.611444002 |
| **A_51_P475938** | Mfsd6l | -0.116368975 | -0.366336628 | 0.133598678 | 7.728919402 | -1.131192681 | 0.299961329 | 0.640586954 | -4.611464938 |
| **A_55_P2170105** | NA | -0.23806747 | -0.749493238 | 0.273358298 | 12.092666 | -1.131099055 | 0.299997843 | 0.640594156 | -4.611508354 |
| **A_51_P351697** | Mrpl35 | 0.161037653 | -0.184966957 | 0.507042263 | 8.833111354 | 1.130911801 | 0.300070882 | 0.640619196 | -4.611595183 |
| **A_55_P2055854** | Ppara | -0.311564166 | -0.981098069 | 0.357969737 | 12.11952309 | -1.130727927 | 0.300142618 | 0.640619196 | -4.611680439 |
| **A_51_P486512** | Letmd1 | 0.124608571 | -0.143193776 | 0.392410919 | 11.26091241 | 1.130620163 | 0.300184667 | 0.640619196 | -4.611730402 |
| **A_55_P2084344** | NA | 0.144368532 | -0.165904542 | 0.454641607 | 13.89115424 | 1.130606927 | 0.300189832 | 0.640619196 | -4.611736539 |
| **A_55_P2062851** | 7-Mar | 0.132498734 | -0.152295366 | 0.417292835 | 11.28215628 | 1.130482726 | 0.300238302 | 0.640619196 | -4.611794119 |
| **A_55_P1962224** | Afap1l2 | -0.370488945 | -1.166896686 | 0.425918796 | 8.551882509 | -1.130376108 | 0.300279915 | 0.640619196 | -4.611843546 |
| **A_55_P2243768** | D930030O05Rik | 0.092708712 | -0.106603066 | 0.292020489 | 8.274564533 | 1.13024026 | 0.300332943 | 0.640619196 | -4.611906521 |
| **A_55_P2090330** | Kcnmb4 | -0.244583747 | -0.770414962 | 0.281247467 | 7.911955735 | -1.130223727 | 0.300339398 | 0.640619196 | -4.611914184 |
| **A_55_P2009086** | Creb3l1 | -0.249889804 | -0.787146432 | 0.287366824 | 8.53463342 | -1.130186054 | 0.300354105 | 0.640619196 | -4.611931648 |
| **A_55_P2112085** | Vwa8 | 0.154232098 | -0.177362974 | 0.48582717 | 9.315320986 | 1.130185908 | 0.300354162 | 0.640619196 | -4.611931715 |
| **A_55_P2108933** | Dao | -0.196103834 | -0.61774116 | 0.225533492 | 7.755526177 | -1.130134735 | 0.300374141 | 0.640619196 | -4.611955436 |
| **A_55_P1988368** | Upp1 | -0.143224414 | -0.451269346 | 0.164820519 | 8.517499493 | -1.129759944 | 0.300520499 | 0.640860627 | -4.612129151 |
| **A_52_P49250** | Erbb2 | -0.215451597 | -0.678978651 | 0.248075456 | 8.452131741 | -1.129426102 | 0.300650917 | 0.641049735 | -4.612283864 |
| **A_55_P2215880** | Ephb2 | 0.210686985 | -0.242636363 | 0.664010333 | 8.780662588 | 1.129309055 | 0.300696653 | 0.641049735 | -4.612338102 |
| **A_55_P2158866** | Nme6 | -0.170532976 | -0.537485125 | 0.196419172 | 12.92742744 | -1.129229269 | 0.300727833 | 0.641049735 | -4.612375072 |
| **A_55_P2096081** | Myo5b | -0.129690289 | -0.40876617 | 0.149385592 | 10.53279427 | -1.129193432 | 0.300741838 | 0.641049735 | -4.612391677 |
| **A_55_P2048210** | Ganab | -0.080972359 | -0.255229022 | 0.093284305 | 12.50452387 | -1.129095263 | 0.300780208 | 0.641060826 | -4.612437163 |
| **A_52_P432919** | Rab3d | -0.088059415 | -0.27767552 | 0.101556689 | 9.989379362 | -1.128453756 | 0.30103104 | 0.641524694 | -4.612734358 |
| **A_55_P2009567** | Ptbp3 | 0.111581252 | -0.128727713 | 0.351890218 | 10.72890482 | 1.128247675 | 0.301111656 | 0.641579593 | -4.612829814 |
| **A_52_P360330** | Map1b | -0.079080722 | -0.249424314 | 0.091262869 | 7.721347781 | -1.128049135 | 0.301189338 | 0.641579593 | -4.612921769 |
| **A_51_P341379** | Acad9 | 0.099514641 | -0.11486298 | 0.313892261 | 10.49914909 | 1.127952267 | 0.301227246 | 0.641579593 | -4.612966631 |
| **A_51_P184062** | Hbs1l | 0.147280003 | -0.170020468 | 0.464580474 | 11.39297632 | 1.127862756 | 0.301262278 | 0.641579593 | -4.613008085 |
| **A_55_P1977633** | Irf2bpl | -0.38339913 | -1.209443185 | 0.442644926 | 10.58990406 | -1.127797433 | 0.301287846 | 0.641579593 | -4.613038336 |
| **A_66_P104479** | NA | -0.102132588 | -0.32219671 | 0.117931534 | 16.79082104 | -1.127712208 | 0.301321206 | 0.641579593 | -4.613077802 |
| **A_52_P46419** | Grk6 | 0.132955698 | -0.153546791 | 0.419458187 | 10.05786341 | 1.127617333 | 0.301358348 | 0.641579593 | -4.613121735 |
| **A_55_P2110362** | Zfp637 | -0.142481105 | -0.449550311 | 0.164588101 | 8.54557754 | -1.127467929 | 0.301416844 | 0.641579593 | -4.613190916 |
| **A_55_P1967855** | Kctd20 | 0.13995649 | -0.16172518 | 0.441638161 | 11.52585511 | 1.127268334 | 0.301495005 | 0.641579593 | -4.61328333 |
| **A_55_P2165314** | Trio | -0.105780192 | -0.333801954 | 0.122241571 | 9.286060165 | -1.12722663 | 0.301511339 | 0.641579593 | -4.613302638 |
| **A_55_P2130695** | Armc4 | -0.212555527 | -0.670810105 | 0.245699051 | 7.757540659 | -1.127064521 | 0.301574836 | 0.641579593 | -4.613377689 |
| **A_55_P2098734** | Ttc21b | -0.22299803 | -0.703776925 | 0.257780866 | 8.858391674 | -1.127038675 | 0.301584961 | 0.641579593 | -4.613389654 |
| **A_55_P2170732** | Fndc10 | -0.133684328 | -0.421918047 | 0.154549391 | 8.220821281 | -1.126986985 | 0.301605211 | 0.641579593 | -4.613413584 |
| **A_55_P2064150** | NA | -0.092832463 | -0.293007376 | 0.10734245 | 15.5268541 | -1.126868952 | 0.301651455 | 0.641579593 | -4.613468225 |
| **A_55_P2134571** | NA | 0.092484863 | -0.10695136 | 0.291921086 | 8.51532858 | 1.1268077 | 0.301675455 | 0.641579593 | -4.613496578 |
| **A_55_P2045512** | NA | 0.171985826 | -0.198949593 | 0.542921244 | 14.66490225 | 1.12662022 | 0.301748925 | 0.641579593 | -4.61358336 |
| **A_66_P132232** | Ercc1 | -0.159378916 | -0.503145034 | 0.184387203 | 8.616664064 | -1.126551334 | 0.301775924 | 0.641579593 | -4.613615245 |
| **A_52_P84096** | Pgd | -0.116015079 | -0.366249475 | 0.134219318 | 12.0862836 | -1.126550391 | 0.301776294 | 0.641579593 | -4.613615681 |
| **A_55_P2048928** | Cbfb | -0.212253114 | -0.670113081 | 0.245606854 | 11.15703794 | -1.12643098 | 0.3018231 | 0.641579593 | -4.613670949 |
| **A_55_P2121096** | Fbn2 | -0.623487342 | -1.968461061 | 0.721486378 | 9.23431511 | -1.126411385 | 0.301830781 | 0.641579593 | -4.613680018 |
| **A_51_P393518** | Hmcn1 | -0.107724371 | -0.340114425 | 0.124665682 | 8.005900049 | -1.126366228 | 0.301848484 | 0.641579593 | -4.613700918 |
| **A_55_P2016069** | Adamtsl1 | -0.139492969 | -0.440418249 | 0.16143231 | 7.767649623 | -1.126359004 | 0.301851316 | 0.641579593 | -4.613704261 |
| **A_51_P509263** | Hoxa7 | -0.291576974 | -0.920661813 | 0.337507866 | 8.387510612 | -1.126230266 | 0.301901789 | 0.641579593 | -4.613763842 |
| **A_66_P115406** | Pabpc1 | 0.181611095 | -0.210256625 | 0.573478814 | 15.811606 | 1.12612364 | 0.301943598 | 0.641579593 | -4.613813187 |
| **A_55_P1952915** | Sh3bgr | -0.363136158 | -1.146802989 | 0.420530673 | 7.998815244 | -1.125955478 | 0.302009546 | 0.641579593 | -4.613891005 |
| **A_66_P118579** | 3300002A11Rik | -0.501244617 | -1.583013128 | 0.580523894 | 8.342313301 | -1.125896775 | 0.30203257 | 0.641579593 | -4.613918168 |
| **A_52_P65494** | Iqgap2 | 0.220597368 | -0.255585066 | 0.696779802 | 8.791700168 | 1.125667546 | 0.302122491 | 0.641579593 | -4.614024234 |
| **A_55_P2072035** | Thy1 | -0.174081491 | -0.549861131 | 0.20169815 | 8.561483449 | -1.125647834 | 0.302130225 | 0.641579593 | -4.614033354 |

| | | | | | | | | | |
|---|---|---|---|---|---|---|---|---|---|
| **A_55_P2034372** | Eif4g3 | -0.120965288 | -0.38209519 | 0.140164615 | 10.49259258 | -1.12560845 | 0.302145677 | 0.641579593 | -4.614051577 |
| **A_55_P2181356** | Cdh11 | -0.264875312 | -0.836682608 | 0.306931984 | 7.987488946 | -1.125576397 | 0.302158253 | 0.641579593 | -4.614066407 |
| **A_52_P87503** | Hbq1b | 0.165346706 | -0.191615909 | 0.522309321 | 10.83751958 | 1.125527216 | 0.302177551 | 0.641579593 | -4.614089161 |
| **A_52_P327615** | Rab18 | 0.074997022 | -0.086925514 | 0.236919558 | 11.55829081 | 1.125433688 | 0.302214251 | 0.641579593 | -4.614132431 |
| **A_51_P443702** | Mrps34 | -0.080213984 | -0.253401223 | 0.092973256 | 13.03702318 | -1.125427142 | 0.30221682 | 0.641579593 | -4.61413546 |
| **A_55_P2105035** | NA | 0.148722313 | -0.172383882 | 0.469828507 | 12.45879052 | 1.125409652 | 0.302223684 | 0.641579593 | -4.614143551 |
| **A_52_P214891** | Stx6 | -0.327615511 | -1.03501425 | 0.379783228 | 9.288482576 | -1.125338982 | 0.302251419 | 0.641579593 | -4.614176245 |
| **A_52_P142208** | Akirin1 | -0.162913978 | -0.514702282 | 0.188874326 | 9.80691618 | -1.125278813 | 0.302275033 | 0.641579593 | -4.61420408 |
| **A_51_P212679** | Cwf19l1 | 0.179778251 | -0.2084612 | 0.568017702 | 9.05220044 | 1.125176558 | 0.30231517 | 0.641579593 | -4.614251384 |
| **A_55_P2046145** | Gm10693 | -0.221982229 | -0.701388305 | 0.257423847 | 9.313456916 | -1.125117457 | 0.30233837 | 0.641579593 | -4.614278723 |
| **A_55_P2169332** | NA | -0.127830574 | -0.403929621 | 0.148268474 | 15.59258399 | -1.125001286 | 0.302383977 | 0.641579593 | -4.61433246 |
| **A_51_P439626** | Map2k2 | 0.121550715 | -0.140984888 | 0.384086319 | 11.31494182 | 1.124999942 | 0.302384505 | 0.641579593 | -4.614333082 |
| **A_51_P319070** | Retsat | -0.169254298 | -0.534900584 | 0.196391987 | 9.778222886 | -1.124764834 | 0.302476822 | 0.641705027 | -4.614441827 |
| **A_52_P639402** | Kcnk3 | -0.267708885 | -0.846164973 | 0.310747203 | 7.965896194 | -1.124541734 | 0.302564447 | 0.641820477 | -4.614545009 |
| **A_55_P2074062** | Sdhaf2 | 0.099999761 | -0.11612032 | 0.316119841 | 10.0299254 | 1.124312476 | 0.302654512 | 0.64194108 | -4.614651029 |
| **A_55_P2022149** | NA | 0.114455068 | -0.132946154 | 0.361856289 | 8.59825628 | 1.12412958 | 0.302726379 | 0.641972646 | -4.614735601 |
| **A_52_P539161** | Rdh11 | -0.25326932 | -0.800738535 | 0.294199895 | 11.99249119 | -1.124102904 | 0.302736862 | 0.641972646 | -4.614747936 |
| **A_55_P2015337** | NA | -0.127107844 | -0.401887423 | 0.147671735 | 14.70090115 | -1.124012361 | 0.302772447 | 0.641972646 | -4.614789801 |
| **A_55_P1958480** | NA | -0.405623771 | -1.282553502 | 0.47130596 | 8.603432134 | -1.123936546 | 0.302802246 | 0.641972646 | -4.614824855 |
| **A_55_P2143499** | Pgbd5 | -0.206639185 | -0.653567238 | 0.240288869 | 7.946044251 | -1.12346165 | 0.302988958 | 0.642204334 | -4.615044403 |
| **A_55_P2091368** | Cep164 | 0.079354234 | -0.092305897 | 0.251014365 | 9.173361477 | 1.123269204 | 0.303064649 | 0.642204334 | -4.61513336 |
| **A_51_P209518** | Galnt1 | 0.1365569 | -0.158854478 | 0.431968279 | 9.490097003 | 1.123232393 | 0.303079128 | 0.642204334 | -4.615150375 |
| **A_55_P2013371** | Ttc29 | -0.305150452 | -0.965327012 | 0.355026107 | 7.820461412 | -1.123148308 | 0.303112206 | 0.642204334 | -4.61518924 |
| **A_55_P2129271** | Tead2 | -0.424579283 | -1.343145614 | 0.493987048 | 10.86502088 | -1.123133718 | 0.303117946 | 0.642204334 | -4.615195983 |
| **A_51_P456187** | Pdcd2 | 0.088602384 | -0.103094825 | 0.280299592 | 10.83826937 | 1.12308546 | 0.303136931 | 0.642204334 | -4.615218288 |
| **A_51_P181538** | Brwd1 | -0.122995783 | -0.38910974 | 0.143118174 | 8.533371361 | -1.123067235 | 0.303144102 | 0.642204334 | -4.615226712 |
| **A_55_P2091284** | Ltbp3 | -0.635340086 | -2.010278899 | 0.739598727 | 8.79896262 | -1.12280949 | 0.303245521 | 0.642241789 | -4.615345832 |
| **A_55_P2034081** | Hrh1 | 0.102746406 | -0.119614027 | 0.325106839 | 8.142874119 | 1.122773926 | 0.303259517 | 0.642241789 | -4.615362267 |
| **A_55_P2015022** | NA | 0.146752251 | -0.170878005 | 0.464382507 | 14.34450662 | 1.122654432 | 0.303306548 | 0.642241789 | -4.615417487 |
| **A_55_P2092884** | NA | 0.3584111 | -0.417376375 | 1.134198575 | 9.183460594 | 1.122591829 | 0.303331191 | 0.642241789 | -4.615446416 |
| **A_55_P1996329** | Kat5 | -0.120017602 | -0.379812723 | 0.139777519 | 8.998242142 | -1.122527899 | 0.303356357 | 0.642241789 | -4.615475958 |
| **A_55_P2101591** | Vta1 | 0.169038125 | -0.196872833 | 0.534949083 | 9.641686401 | 1.122515741 | 0.303361143 | 0.642241789 | -4.615481576 |
| **A_51_P288257** | Cdk5rap3 | -0.087877876 | -0.278119011 | 0.10236326 | 11.61812122 | -1.122427525 | 0.303395873 | 0.64224497 | -4.615522338 |
| **A_55_P2096189** | Rfx8 | 0.075013735 | -0.087404972 | 0.237432441 | 7.716702176 | 1.12224565 | 0.303467485 | 0.642326218 | -4.615606373 |
| **A_55_P2069485** | Ptpn13 | -0.471828177 | -1.493617058 | 0.549960703 | 8.727373894 | -1.122033671 | 0.303550969 | 0.642432571 | -4.615704308 |
| **A_55_P2165074** | Rnf31 | -0.084188978 | -0.266541497 | 0.098163542 | 9.726650792 | -1.121828985 | 0.303631598 | 0.642449832 | -4.615798867 |
| **A_55_P1955188** | Spata21 | -0.076002559 | -0.240641762 | 0.088636645 | 7.698782966 | -1.121703686 | 0.303680965 | 0.642449832 | -4.615856747 |
| **A_55_P2302577** | AI426330 | -0.263515818 | -0.834382903 | 0.307351267 | 8.959856549 | -1.121643581 | 0.303704648 | 0.642449832 | -4.615884511 |
| **A_55_P2014357** | Pdpk1 | 0.120920145 | -0.141066138 | 0.382906429 | 10.15581709 | 1.121510373 | 0.303757141 | 0.642449832 | -4.61594604 |
| **A_55_P2161126** | NA | 0.1497162 | -0.174665839 | 0.474098238 | 9.053219272 | 1.121489431 | 0.303765394 | 0.642449832 | -4.615955713 |
| **A_55_P2087322** | Rapsn | 0.267335413 | -0.311947939 | 0.846618764 | 12.77741917 | 1.121369225 | 0.303812771 | 0.642449832 | -4.616011232 |
| **A_55_P1957815** | Tmub1 | -0.102208485 | -0.32368743 | 0.11927046 | 10.45218077 | -1.121340981 | 0.303823904 | 0.642449832 | -4.616024277 |
| **A_55_P2015495** | Abat | 0.336926466 | -0.393242805 | 1.067095737 | 12.73159356 | 1.121230142 | 0.303867596 | 0.642449832 | -4.616075467 |
| **A_55_P2058591** | NA | 0.150055154 | -0.17516883 | 0.475279138 | 8.604435503 | 1.121118555 | 0.303911588 | 0.642449832 | -4.616127001 |
| **A_51_P201945** | Cfap97 | 0.135070142 | -0.157710823 | 0.427851107 | 8.985615292 | 1.120984759 | 0.303964344 | 0.642449832 | -4.616188788 |
| **A_51_P256665** | Pon3 | -0.151047934 | -0.478518815 | 0.176422947 | 11.96544603 | -1.12079269 | 0.304040088 | 0.642449832 | -4.616277479 |
| **A_55_P2007768** | Gm8709 | -0.134792848 | -0.427034527 | 0.157448832 | 15.09162845 | -1.120747768 | 0.304057806 | 0.642449832 | -4.616298221 |
| **A_52_P302496** | Fah | -0.324676411 | -1.028633085 | 0.379280262 | 12.45528217 | -1.120696449 | 0.304078048 | 0.642449832 | -4.616321917 |
| **A_55_P2074154** | Dhx57 | 0.169632378 | -0.198177959 | 0.537442716 | 8.978036664 | 1.120644873 | 0.304098393 | 0.642449832 | -4.616345731 |
| **A_55_P2102135** | Rdh12 | 0.198689091 | -0.232147369 | 0.62952555 | 10.35339746 | 1.120584892 | 0.304122054 | 0.642449832 | -4.616373425 |

| | | | | | | | | | |
|---|---|---|---|---|---|---|---|---|---|
| **A_55_P2026119** | Dlec1 | 0.139973582 | -0.163567256 | 0.443514421 | 9.282458553 | 1.120500714 | 0.304155263 | 0.642449832 | -4.61641229 |
| **A_55_P1996821** | Myo15b | -0.226127006 | -0.71653891 | 0.264284897 | 7.9032441 | -1.120403919 | 0.304193454 | 0.642449832 | -4.616456979 |
| **A_55_P2181597** | Mfap2 | -0.473975045 | -1.50195045 | 0.55400036 | 8.487094031 | -1.120355734 | 0.304212466 | 0.642449832 | -4.616479224 |
| **A_51_P503822** | Slitrk6 | -0.138307741 | -0.438288371 | 0.161672889 | 7.80636822 | -1.120305491 | 0.304232292 | 0.642449832 | -4.616502419 |
| **A_55_P2048684** | Rpl27 | -0.160790889 | -0.509538826 | 0.187957049 | 14.3339317 | -1.120296493 | 0.304235843 | 0.642449832 | -4.616506574 |
| **A_52_P217710** | Fzd6 | -0.139858599 | -0.443219971 | 0.163502774 | 7.92397246 | -1.120242591 | 0.304257114 | 0.642449832 | -4.616531457 |
| **A_51_P487678** | Dtnbp1 | -0.097909032 | -0.31032911 | 0.114511045 | 9.355645001 | -1.119980305 | 0.304360638 | 0.642587217 | -4.616652533 |
| **A_55_P2433218** | Bloc1s6 | 0.085992625 | -0.100600734 | 0.272585983 | 10.61620964 | 1.11981998 | 0.304423932 | 0.642587217 | -4.616726536 |
| **A_55_P2138070** | NA | 0.146991976 | -0.171970526 | 0.465954478 | 14.16803767 | 1.119791553 | 0.304435155 | 0.642587217 | -4.616739656 |
| **A_52_P221756** | Spock2 | -0.249164295 | -0.78985818 | 0.29152959 | 7.820690605 | -1.119740895 | 0.304455157 | 0.642587217 | -4.616763038 |
| **A_55_P1990285** | Rplp2 | 0.131446798 | -0.15382934 | 0.416722935 | 17.17141812 | 1.119612538 | 0.304505843 | 0.642624024 | -4.616822278 |
| **A_66_P139647** | Vwa3a | -0.15067363 | -0.477746224 | 0.176398965 | 7.746865698 | -1.119376754 | 0.304598968 | 0.642651326 | -4.616931092 |
| **A_55_P2060483** | Rnf141 | 0.155950296 | -0.182581554 | 0.494482146 | 9.143947874 | 1.119360194 | 0.304605509 | 0.642651326 | -4.616938735 |
| **A_55_P1971840** | Muc13 | -0.51896961 | -1.64564229 | 0.60770307 | 9.060573431 | -1.119250643 | 0.304648786 | 0.642651326 | -4.616989288 |
| **A_55_P2109983** | Mpg | 0.110366861 | -0.129239013 | 0.349972735 | 8.720660526 | 1.119243085 | 0.304651772 | 0.642651326 | -4.616992775 |
| **A_52_P456977** | Deaf1 | 0.118307697 | -0.138570681 | 0.375186074 | 8.395255107 | 1.11909934 | 0.304708565 | 0.642700988 | -4.617059105 |
| **A_52_P456271** | Tmem167 | 0.108638779 | -0.127294515 | 0.344572074 | 12.09051301 | 1.11886795 | 0.304800005 | 0.642806603 | -4.617165868 |
| **A_55_P2129975** | Rabgef1 | 0.099748735 | -0.116890168 | 0.316387639 | 10.35534379 | 1.118804331 | 0.30482515 | 0.642806603 | -4.61719522 |
| **A_55_P1957433** | Mprip | -0.245681012 | -0.779352168 | 0.287990145 | 9.062361516 | -1.118616058 | 0.304899573 | 0.642893405 | -4.617282079 |
| **A_51_P165934** | Phb | -0.102238739 | -0.32442346 | 0.119945983 | 11.54472 | -1.11810987 | 0.305099741 | 0.642958934 | -4.617515573 |
| **A_55_P1966928** | Lnx1 | -0.146991923 | -0.466446279 | 0.172462433 | 7.763508126 | -1.118067045 | 0.305116681 | 0.642958934 | -4.617535325 |
| **A_52_P660922** | Dock1 | -0.218009555 | -0.691812938 | 0.255793828 | 10.13435485 | -1.118048285 | 0.305124102 | 0.642958934 | -4.617543978 |
| **A_52_P537381** | Dimt1 | 0.116977034 | -0.137260964 | 0.371215032 | 10.32319579 | 1.11800394 | 0.305141644 | 0.642958934 | -4.61756443 |
| **A_55_P1990964** | Osbpl11 | -0.134803508 | -0.427788908 | 0.158181893 | 10.37582651 | -1.117991244 | 0.305146666 | 0.642958934 | -4.617570285 |
| **A_55_P1993840** | Tmem132c | -0.355734637 | -1.128911937 | 0.417442664 | 7.885950426 | -1.11797025 | 0.305154972 | 0.642958934 | -4.617579968 |
| **A_55_P2068877** | Ibtk | 0.197043494 | -0.231232188 | 0.625319176 | 11.1765716 | 1.117948698 | 0.305163498 | 0.642958934 | -4.617589908 |
| **A_55_P2012592** | Ildr1 | -0.100515942 | -0.319033454 | 0.118001571 | 7.75004032 | -1.11771708 | 0.305255142 | 0.642997819 | -4.617696724 |
| **A_55_P2095732** | Fbrsl1 | -0.091752236 | -0.291222997 | 0.107718526 | 8.669720097 | -1.117688029 | 0.305266638 | 0.642997819 | -4.617710121 |
| **A_55_P1979305** | Gle1 | 0.109628029 | -0.128713123 | 0.34796918 | 9.325136924 | 1.117649839 | 0.305281752 | 0.642997819 | -4.617727733 |
| **A_55_P2124582** | Xlr4a | 0.077336434 | -0.090824526 | 0.245497394 | 7.794150382 | 1.117486163 | 0.305346532 | 0.643064187 | -4.617803207 |
| **A_55_P1967693** | Wipi2 | 0.133765964 | -0.157164898 | 0.424696825 | 8.784531028 | 1.117220816 | 0.305451575 | 0.643142256 | -4.617925554 |
| **A_55_P1964364** | Kctd14 | 0.116127134 | -0.136443191 | 0.368697459 | 17.19094259 | 1.117209379 | 0.305456104 | 0.643142256 | -4.617930828 |
| **A_55_P2189201** | 5830426C09Rik | 0.077733627 | -0.09135927 | 0.246826525 | 8.377372717 | 1.117034935 | 0.305525179 | 0.643142256 | -4.618011253 |
| **A_51_P381218** | Gulp1 | -0.582050077 | -1.848209173 | 0.684109019 | 8.644655816 | -1.117005369 | 0.305536887 | 0.643142256 | -4.618024883 |
| **A_55_P2038187** | BC051019 | 0.145789279 | -0.171405897 | 0.462984454 | 10.89966405 | 1.11681748 | 0.305611303 | 0.643142256 | -4.618111499 |
| **A_52_P565575** | Arhgap20 | -0.2679705 | -0.851053516 | 0.315112516 | 7.92939088 | -1.116708404 | 0.305654511 | 0.643142256 | -4.61816178 |
| **A_55_P2162552** | Txndc9 | 0.10523553 | -0.123753327 | 0.334224386 | 11.57035694 | 1.116686415 | 0.305663222 | 0.643142256 | -4.618171916 |
| **A_51_P349192** | NA | -0.289493325 | -0.919492819 | 0.340506169 | 11.37486482 | -1.116558657 | 0.305713838 | 0.643142256 | -4.618230805 |
| **A_55_P1961760** | NA | 0.304923641 | -0.358715438 | 0.96856272 | 11.88921535 | 1.116457855 | 0.305753779 | 0.643142256 | -4.618277266 |
| **A_55_P2038007** | Csrp1 | -0.560888976 | -1.781657071 | 0.659879119 | 14.07653502 | -1.116418239 | 0.305769478 | 0.643142256 | -4.618295526 |
| **A_51_P330630** | Usp3 | -0.108499662 | -0.344650168 | 0.127650844 | 8.663799302 | -1.11640737 | 0.305773785 | 0.643142256 | -4.618300535 |
| **A_55_P1961873** | Akt3 | -0.155655794 | -0.49444902 | 0.183137431 | 8.469914902 | -1.116384413 | 0.305782882 | 0.643142256 | -4.618311116 |
| **A_55_P2125823** | NA | 0.15410286 | -0.18137088 | 0.489576601 | 14.45757257 | 1.116182884 | 0.305862754 | 0.643240255 | -4.618403995 |
| **A_51_P394339** | Eif2b1 | 0.14169453 | -0.166792191 | 0.450181251 | 10.88426421 | 1.116091501 | 0.305898978 | 0.643246448 | -4.618446109 |
| **A_52_P391983** | Nt5c2 | 0.121879697 | -0.14355185 | 0.387311243 | 9.072691892 | 1.115737465 | 0.306039349 | 0.643471618 | -4.61860925 |
| **A_55_P2169659** | Vwf | 0.143938449 | -0.169568222 | 0.457445119 | 8.597083138 | 1.115612112 | 0.306089063 | 0.643503612 | -4.618667007 |
| **A_51_P157214** | Mfsd14a | 0.149485939 | -0.176127114 | 0.475098993 | 10.70952312 | 1.115531214 | 0.30612115 | 0.643503612 | -4.618704279 |
| **A_55_P2375905** | 1110035E04Rik | 0.079471953 | -0.093655797 | 0.252599702 | 8.08272958 | 1.115399354 | 0.306173457 | 0.643543578 | -4.61876503 |
| **A_55_P1954476** | NA | 0.099733147 | -0.117673129 | 0.317139423 | 8.258910937 | 1.114681092 | 0.306458509 | 0.643847944 | -4.619095884 |
| **A_55_P2130725** | Rhox8 | 0.109063828 | -0.128684442 | 0.346812097 | 8.29061746 | 1.114670632 | 0.306462661 | 0.643847944 | -4.619100702 |

| | | | | | | | | | |
|---|---|---|---|---|---|---|---|---|---|
| **A_51_P273755** | Exoc7 | -0.115949749 | -0.368721571 | 0.136822073 | 9.358592938 | -1.114613617 | 0.306485298 | 0.643847944 | -4.61912696 |
| **A_55_P2155041** | Ssbp2 | 0.094178946 | -0.111150795 | 0.299508686 | 11.51610476 | 1.114513042 | 0.306525233 | 0.643847944 | -4.619173279 |
| **A_52_P424840** | Mrc2 | -0.236021113 | -0.7506044 | 0.278562175 | 7.928580181 | -1.114495619 | 0.306532152 | 0.643847944 | -4.619181302 |
| **A_66_P137048** | Men1 | -0.081832533 | -0.260247876 | 0.09658281 | 11.2997772 | -1.114492071 | 0.30653356 | 0.643847944 | -4.619182936 |
| **A_55_P2159216** | NA | 0.09117214 | -0.107613904 | 0.289958184 | 11.21387031 | 1.114447068 | 0.306551432 | 0.643847944 | -4.619203661 |
| **A_55_P2087315** | Shroom1 | 0.16466846 | -0.194414161 | 0.523751081 | 8.810565571 | 1.114292549 | 0.306612799 | 0.643893362 | -4.619274818 |
| **A_55_P1990525** | Ube2v1 | 0.107521781 | -0.126975439 | 0.342019001 | 13.88833542 | 1.114145574 | 0.306671179 | 0.643893362 | -4.619342495 |
| **A_55_P2026475** | NA | 0.107194335 | -0.12660118 | 0.34098985 | 11.82715334 | 1.114086339 | 0.306694711 | 0.643893362 | -4.61936977 |
| **A_66_P111437** | Ube2j2 | 0.090599513 | -0.107007148 | 0.288206174 | 13.77234323 | 1.114057153 | 0.306706306 | 0.643893362 | -4.619383208 |
| **A_55_P2027102** | Rgs3 | -0.301517722 | -0.95925841 | 0.356222967 | 10.26391358 | -1.113887482 | 0.30677372 | 0.643945501 | -4.619461329 |
| **A_55_P2009792** | Hdac6 | 0.120094467 | -0.141897794 | 0.382086728 | 10.6587942 | 1.113826955 | 0.306797771 | 0.643945501 | -4.619489195 |
| **A_55_P2028798** | Nqo2 | 0.144029204 | -0.170224197 | 0.458282604 | 11.28997312 | 1.113662927 | 0.306862959 | 0.644012393 | -4.619564711 |
| **A_52_P556290** | Arih1 | 0.117159411 | -0.138592096 | 0.372910917 | 9.02307967 | 1.113120464 | 0.307078626 | 0.644388342 | -4.619814412 |
| **A_52_P208727** | Zmym4 | -0.088221696 | -0.280838478 | 0.104395085 | 8.697845846 | -1.112921342 | 0.307157822 | 0.644388342 | -4.619906055 |
| **A_51_P271425** | Lhfpl4 | 0.429476436 | -0.508220749 | 1.367173622 | 10.49080336 | 1.112909674 | 0.307162463 | 0.644388342 | -4.619911424 |
| **A_55_P2111790** | Gem | -0.224925551 | -0.716042709 | 0.266191608 | 7.983366315 | -1.112850625 | 0.307185952 | 0.644388342 | -4.619938599 |
| **A_51_P114817** | Ccna1 | -0.168477657 | -0.536406816 | 0.199451501 | 7.77191316 | -1.11265698 | 0.307262992 | 0.644388342 | -4.620027712 |
| **A_55_P2148519** | Teddm2 | 0.102161733 | -0.120944992 | 0.325268458 | 8.702899356 | 1.112650532 | 0.307265558 | 0.644388342 | -4.620030679 |
| **A_55_P2111292** | 3110009E18Rik | 0.125863175 | -0.149055159 | 0.400781509 | 9.434421889 | 1.112444028 | 0.307347732 | 0.644388342 | -4.6201257 |
| **A_55_P2092286** | NA | -0.150658947 | -0.479775656 | 0.178457761 | 7.765465387 | -1.112316006 | 0.307398685 | 0.644388342 | -4.620184605 |
| **A_55_P2277019** | Loxl3 | -0.093518476 | -0.297825706 | 0.110788753 | 8.075830963 | -1.112235816 | 0.307430605 | 0.644388342 | -4.620221499 |
| **A_55_P2250424** | 9530029O12Rik | 0.259679767 | -0.307725747 | 0.827085281 | 10.70596827 | 1.112058796 | 0.307501076 | 0.644388342 | -4.62030294 |
| **A_55_P1978306** | Zfp318 | -0.123755231 | -0.394169627 | 0.146659164 | 9.399956749 | -1.112031136 | 0.307512089 | 0.644388342 | -4.620315665 |
| **A_55_P2065953** | Btbd19 | 0.149369153 | -0.17702281 | 0.475761116 | 8.785943407 | 1.111999648 | 0.307524626 | 0.644388342 | -4.62033015 |
| **A_55_P2067221** | Iqsec3 | -0.4662357 | -1.485081131 | 0.55260973 | 10.77680524 | -1.111937589 | 0.307549337 | 0.644388342 | -4.620358699 |
| **A_52_P354123** | Rhebl1 | 0.088686922 | -0.105145419 | 0.282519263 | 8.519190883 | 1.111774038 | 0.307614467 | 0.644388342 | -4.620433934 |
| **A_52_P94401** | Rtn4rl1 | -0.113309475 | -0.360956587 | 0.134337636 | 8.172639509 | -1.11177307 | 0.307614853 | 0.644388342 | -4.62043438 |
| **A_51_P370050** | Nagpa | 0.202096852 | -0.239623712 | 0.643817415 | 9.250013019 | 1.111719233 | 0.307636295 | 0.644388342 | -4.620459144 |
| **A_55_P1952828** | NA | 0.085193591 | -0.10101655 | 0.271403733 | 8.406511425 | 1.1116979 | 0.307644792 | 0.644388342 | -4.620468957 |
| **A_55_P2035667** | Gk | -0.224664966 | -0.715754233 | 0.266424301 | 9.0977348 | -1.111624478 | 0.307674036 | 0.644388342 | -4.620502728 |
| **A_51_P291417** | Thbd | -0.765353019 | -2.438323724 | 0.907617686 | 9.53817514 | -1.111620769 | 0.307675514 | 0.644388342 | -4.620504434 |
| **A_51_P410918** | Tor4a | -0.076495339 | -0.243719713 | 0.090729035 | 9.22070934 | -1.111523333 | 0.307714327 | 0.644399808 | -4.62054925 |
| **A_55_P2051914** | Zfp568 | -0.139595817 | -0.444818052 | 0.165626417 | 10.4310663 | -1.111320759 | 0.307795035 | 0.644444936 | -4.620642418 |
| **A_51_P234025** | Rbm5 | 0.110429469 | -0.131025487 | 0.351884426 | 12.46660386 | 1.111301879 | 0.307802558 | 0.644444936 | -4.620651102 |
| **A_55_P2143219** | Rasgrp2 | -0.160567453 | -0.511762779 | 0.190627873 | 11.71312159 | -1.11094352 | 0.307945379 | 0.644638046 | -4.620815898 |
| **A_51_P414637** | Col5a1 | -0.151720996 | -0.483579305 | 0.180137314 | 10.87689691 | -1.110903095 | 0.307961494 | 0.644638046 | -4.620834486 |
| **A_51_P304125** | Slc29a2 | 0.090307168 | -0.107238767 | 0.287853102 | 8.088323501 | 1.110803693 | 0.308001121 | 0.644651182 | -4.620880192 |
| **A_55_P2090817** | Macrod2 | 0.075508594 | -0.089693823 | 0.24071101 | 8.916438224 | 1.110614081 | 0.308076723 | 0.644717579 | -4.620967372 |
| **A_55_P2167299** | NA | -0.125435576 | -0.399892643 | 0.149021492 | 15.39253883 | -1.11052796 | 0.308111066 | 0.644717579 | -4.621006967 |
| **A_52_P588539** | Snapin | -0.093910004 | -0.299415236 | 0.111595228 | 8.139587934 | -1.11038137 | 0.30816953 | 0.644717579 | -4.621074359 |
| **A_66_P106836** | Kmo | 0.429707471 | -0.510673754 | 1.370088696 | 11.00072874 | 1.110330179 | 0.308189949 | 0.644717579 | -4.621097892 |
| **A_55_P2053511** | Mrfap1 | 0.091967866 | -0.109301017 | 0.293236749 | 14.22381591 | 1.110305937 | 0.308199619 | 0.644717579 | -4.621109036 |
| **A_55_P1999294** | Gpbp1 | 0.079996269 | -0.095098272 | 0.255090811 | 11.69254598 | 1.110146554 | 0.308263201 | 0.644729122 | -4.621182302 |
| **A_55_P2004920** | NA | 0.149055531 | -0.177201177 | 0.475312239 | 9.154690434 | 1.11012488 | 0.308271848 | 0.644729122 | -4.621192265 |
| **A_51_P134627** | NA | 0.172417611 | -0.205164889 | 0.550000111 | 9.08735313 | 1.109565526 | 0.30849508 | 0.644835469 | -4.62144935 |
| **A_55_P2022885** | Zfp866 | 0.07684696 | -0.091444494 | 0.245138414 | 8.579369741 | 1.109552404 | 0.308500318 | 0.644835469 | -4.621455381 |
| **A_55_P2055875** | Josd2 | 0.228940026 | -0.272484182 | 0.730364234 | 10.11673486 | 1.109429269 | 0.308549479 | 0.644835469 | -4.621511966 |
| **A_55_P2171578** | Mkrn1 | 0.139329245 | -0.165831629 | 0.44449012 | 13.88559913 | 1.109421614 | 0.308552535 | 0.644835469 | -4.621515484 |
| **A_55_P2197847** | Lhx1os | 0.251856818 | -0.299785709 | 0.803499346 | 8.957937165 | 1.109377011 | 0.308570345 | 0.644835469 | -4.62153598 |
| **A_55_P2067842** | 5-Mar | 0.129810649 | -0.154524378 | 0.414145677 | 11.51290578 | 1.109336101 | 0.30858668 | 0.644835469 | -4.621554779 |

| | | | | | | | | | |
|---|---|---|---|---|---|---|---|---|---|
| A_55_P2071496 | Fcf1 | 0.100679214 | -0.119852853 | 0.321211282 | 12.11350328 | 1.109305733 | 0.308598806 | 0.644835469 | -4.621568733 |
| A_55_P2157307 | NA | -0.263283573 | -0.840001969 | 0.313434822 | 9.681240275 | -1.10928502 | 0.308607078 | 0.644835469 | -4.621578251 |
| A_51_P416585 | Hdlbp | -0.149752831 | -0.477797256 | 0.178291593 | 12.13991357 | -1.109240124 | 0.308625006 | 0.644835469 | -4.62159888 |
| A_55_P1978875 | Kmt2b | -0.076848839 | -0.245210493 | 0.091512816 | 10.26053934 | -1.109116872 | 0.30867423 | 0.644835469 | -4.621655512 |
| A_51_P479914 | Pik3cb | 0.198532441 | -0.236430739 | 0.63349562 | 9.406953692 | 1.10907822 | 0.308689668 | 0.644835469 | -4.621673271 |
| A_55_P2018017 | Tnfsf10 | -0.281689404 | -0.898913802 | 0.335534994 | 8.238579776 | -1.108946552 | 0.308742262 | 0.644875642 | -4.621733765 |
| A_51_P338262 | Tnnt2 | -1.020688455 | -3.257428159 | 1.216051249 | 8.856634885 | -1.108819729 | 0.308792929 | 0.644911779 | -4.621792031 |
| A_55_P2046092 | NA | 0.071256298 | -0.084922437 | 0.227435032 | 7.709128024 | 1.108624683 | 0.308870863 | 0.645004853 | -4.621881633 |
| A_55_P2017774 | NA | -0.169175302 | -0.540022815 | 0.201672211 | 17.08786562 | -1.108472132 | 0.30893183 | 0.645007655 | -4.621951708 |
| A_55_P2079516 | Mpp1 | 0.134239542 | -0.160030362 | 0.428509446 | 8.810318102 | 1.10845433 | 0.308938945 | 0.645007655 | -4.621959885 |
| A_55_P2013716 | Napa | 0.133148546 | -0.158753559 | 0.425050651 | 13.10157521 | 1.108363946 | 0.308975072 | 0.64501341 | -4.622001401 |
| A_55_P2133840 | Sipa1 | 0.122903994 | -0.146615223 | 0.392423211 | 12.30910784 | 1.108050232 | 0.309100492 | 0.645153481 | -4.622145485 |
| A_55_P2132207 | 1810037I17Rik | 0.086253633 | -0.10290001 | 0.275407277 | 8.663407777 | 1.108015602 | 0.30911434 | 0.645153481 | -4.622161389 |
| A_55_P2126644 | NA | 0.090444589 | -0.107918308 | 0.288807487 | 10.44093784 | 1.10791218 | 0.309155698 | 0.645153481 | -4.622208885 |
| A_51_P363642 | 1700088E04Rik | -0.335765376 | -1.072198553 | 0.400667802 | 7.965681593 | -1.107862219 | 0.309175679 | 0.645153481 | -4.622231828 |
| A_51_P238523 | Shisa4 | -0.118228592 | -0.377667239 | 0.141210055 | 8.026303126 | -1.107314627 | 0.309394748 | 0.645465841 | -4.622483263 |
| A_55_P2052062 | Cd200 | -0.725853096 | -2.318703153 | 0.866996961 | 8.251009813 | -1.107278907 | 0.309409042 | 0.645465841 | -4.622499662 |
| A_55_P2165324 | Acsl3 | 0.195044306 | -0.233047626 | 0.623136237 | 10.0715195 | 1.107081064 | 0.309488226 | 0.645465841 | -4.622590488 |
| A_51_P171616 | Wnt10a | -0.083457191 | -0.266644772 | 0.09973039 | 7.67078041 | -1.107008424 | 0.309517303 | 0.645465841 | -4.622623834 |
| A_51_P228254 | Zdhhc21 | -0.098630355 | -0.315136415 | 0.117875706 | 10.15980004 | -1.106939096 | 0.309545056 | 0.645465841 | -4.622655658 |
| A_51_P479786 | Smtn | -0.16579296 | -0.529731795 | 0.198145875 | 8.637661478 | -1.106931824 | 0.309547968 | 0.645465841 | -4.622658997 |
| A_51_P191782 | Olfml3 | -0.769551293 | -2.458926918 | 0.919824332 | 8.627613472 | -1.106864701 | 0.309574841 | 0.645465841 | -4.622689808 |
| A_55_P2247633 | Gpatch2 | 0.147150886 | -0.175906531 | 0.470208303 | 9.103720161 | 1.106792875 | 0.309603598 | 0.645465841 | -4.622722777 |
| A_55_P2048164 | Msantd3 | -0.153406338 | -0.490214032 | 0.183401357 | 8.214306059 | -1.106737145 | 0.309625913 | 0.645465841 | -4.622748357 |
| A_55_P1953503 | Lrrc4b | -0.332711048 | -1.063364406 | 0.39794231 | 7.924625895 | -1.10646843 | 0.309733529 | 0.645564584 | -4.622871689 |
| A_55_P1975535 | Nutf2 | 0.129752494 | -0.155195952 | 0.414700939 | 12.52693212 | 1.106452082 | 0.309740077 | 0.645564584 | -4.622879191 |
| A_52_P196568 | Meg3 | 0.118718642 | -0.142078793 | 0.379516078 | 8.305154745 | 1.106111211 | 0.309876637 | 0.645735245 | -4.623035619 |
| A_52_P444457 | Dnlz | 0.082132822 | -0.098299553 | 0.262565197 | 9.214397542 | 1.10607727 | 0.309890237 | 0.645735245 | -4.623051193 |
| A_55_P2123484 | Gm11627 | -0.427035914 | -1.365250617 | 0.511178789 | 8.054560398 | -1.105975115 | 0.309931173 | 0.645735245 | -4.623098067 |
| A_55_P1963578 | NA | 0.135090328 | -0.161737554 | 0.431918209 | 15.73496718 | 1.105866626 | 0.309974653 | 0.645735245 | -4.623147845 |
| A_55_P2010962 | Surf6 | 0.150955385 | -0.180742806 | 0.482653575 | 10.76831033 | 1.105830836 | 0.309988998 | 0.645735245 | -4.623164266 |
| A_51_P372456 | Wdr86 | -0.566085329 | -1.810080451 | 0.677909794 | 9.318602105 | -1.105723171 | 0.310032154 | 0.64575555 | -4.623213663 |
| A_55_P2077083 | Tmem132e | -0.13990705 | -0.447426832 | 0.167612733 | 9.033060362 | -1.105477044 | 0.310130829 | 0.645891477 | -4.623326579 |
| A_55_P2045946 | Tepsin | 0.101494497 | -0.121675828 | 0.324664823 | 9.618712704 | 1.105068602 | 0.310294636 | 0.646072612 | -4.623513932 |
| A_51_P158120 | B3galnt2 | 0.171042339 | -0.205078287 | 0.547162966 | 10.09370284 | 1.104993354 | 0.310324823 | 0.646072612 | -4.623548445 |
| A_65_P19138 | 7-Mar | 0.125028906 | -0.149920477 | 0.399978289 | 11.10039106 | 1.104945534 | 0.310344007 | 0.646072612 | -4.623570377 |
| A_55_P2031157 | Efna3 | 0.231653696 | -0.277790507 | 0.741097899 | 9.00213465 | 1.104907185 | 0.310359393 | 0.646072612 | -4.623587966 |
| A_55_P2002517 | Fhl1 | -0.277215612 | -0.886906977 | 0.332475752 | 7.928997315 | -1.104818228 | 0.310395085 | 0.646072612 | -4.623628763 |
| A_52_P266320 | Akap17b | 0.08754693 | -0.105008635 | 0.280102494 | 8.236168498 | 1.104760241 | 0.310418353 | 0.646072612 | -4.623655357 |
| A_55_P2185397 | Copz2 | -0.123214802 | -0.394265389 | 0.147835784 | 8.497000777 | -1.104576303 | 0.31049217 | 0.646088815 | -4.623739707 |
| A_55_P1964053 | Golga4 | 0.090034698 | -0.108025944 | 0.288095339 | 11.436239 | 1.10457426 | 0.31049299 | 0.646088815 | -4.623740644 |
| A_55_P2026487 | NA | 0.172839897 | -0.207408031 | 0.553087825 | 8.714930597 | 1.104486294 | 0.310528297 | 0.646092729 | -4.623780981 |
| A_55_P1955722 | Med15 | 0.100228308 | -0.120305601 | 0.320762217 | 10.23315102 | 1.104328328 | 0.310591709 | 0.646155111 | -4.623853413 |
| A_51_P124345 | Cpne5 | 0.074937258 | -0.089968131 | 0.239842646 | 8.463477466 | 1.104195924 | 0.310644867 | 0.646173284 | -4.623914121 |
| A_55_P1990573 | Snrnp70 | 0.105482866 | -0.126652307 | 0.33761804 | 12.68554495 | 1.104140045 | 0.310667304 | 0.646173284 | -4.62393974 |
| A_51_P271439 | Garem1 | 0.139880915 | -0.167992217 | 0.447754047 | 9.145885051 | 1.104002008 | 0.310722736 | 0.646194114 | -4.624003025 |
| A_51_P100289 | Srf | -0.217095466 | -0.694976657 | 0.260785725 | 9.766238576 | -1.103860024 | 0.310779761 | 0.646194114 | -4.624068116 |
| A_55_P2134082 | Clip4 | -0.123580716 | -0.395645656 | 0.148484223 | 7.7385005 | -1.103726117 | 0.31083355 | 0.646194114 | -4.624129499 |
| A_55_P2116794 | NA | 0.129426585 | -0.155510916 | 0.414364085 | 11.44929871 | 1.103715318 | 0.310837888 | 0.646194114 | -4.62413445 |
| A_51_P234044 | 1190005I06Rik | 0.076525514 | -0.091959754 | 0.245010783 | 8.657480513 | 1.1036402 | 0.310868066 | 0.646194114 | -4.624168882 |

| | | | | | | | | | |
|---|---|---|---|---|---|---|---|---|---|
| **A_55_P1958250** | Rnf114 | -0.181570097 | -0.581351483 | 0.218211288 | 13.10802817 | -1.103582861 | 0.310891103 | 0.646194114 | -4.624195165 |
| **A_55_P2033282** | Cept1 | -0.194502269 | -0.622788507 | 0.23378397 | 12.41484147 | -1.103503564 | 0.310922965 | 0.646194114 | -4.624231512 |
| **A_55_P2106901** | Nkain4 | -0.685504706 | -2.195092187 | 0.824082774 | 8.754636155 | -1.103406008 | 0.310962166 | 0.646194114 | -4.624276225 |
| **A_55_P1959228** | Slc4a4 | -0.085897536 | -0.275064048 | 0.103268976 | 8.295219692 | -1.103366106 | 0.310978201 | 0.646194114 | -4.624294513 |
| **A_52_P91274** | 1700018G05Rik | 0.071772669 | -0.086308275 | 0.229853613 | 7.683621045 | 1.103221627 | 0.311036268 | 0.6462453 | -4.624360729 |
| **A_51_P394735** | Arhgef18 | -0.077944669 | -0.249689738 | 0.0938004 | 9.44376657 | -1.102770976 | 0.311217444 | 0.646521776 | -4.624567239 |
| **A_55_P2145711** | Tox3 | -0.277372639 | -0.888567761 | 0.333822483 | 7.936951685 | -1.102724257 | 0.311236232 | 0.646521776 | -4.624588645 |
| **A_52_P580533** | Spata33 | -0.173844479 | -0.556951538 | 0.20926258 | 7.808716387 | -1.102615105 | 0.31128013 | 0.646540959 | -4.624638657 |
| **A_55_P2151685** | Pira11 | -0.190342263 | -0.609836549 | 0.229152022 | 8.960926493 | -1.102534961 | 0.311312365 | 0.646540959 | -4.624675376 |
| **A_66_P120249** | Nr1h3 | -0.095352292 | -0.305609041 | 0.114904458 | 11.35971083 | -1.101956314 | 0.311545187 | 0.646765115 | -4.624940452 |
| **A_55_P1983588** | Pmepa1 | -0.47567764 | -1.524601954 | 0.573246674 | 8.383264107 | -1.101924333 | 0.311558059 | 0.646765115 | -4.624955101 |
| **A_55_P2137927** | Cd82 | 0.162277104 | -0.195566022 | 0.520120229 | 10.30397546 | 1.101914145 | 0.31156216 | 0.646765115 | -4.624959767 |
| **A_52_P434820** | Gnpda2 | -0.137735801 | -0.441470731 | 0.16599913 | 8.158007028 | -1.101882501 | 0.311574897 | 0.646765115 | -4.624974261 |
| **A_55_P2107972** | Smad5 | -0.273784622 | -0.877551652 | 0.329982409 | 11.05220132 | -1.101850937 | 0.311587602 | 0.646765115 | -4.624988718 |
| **A_52_P603654** | Stx8 | -0.100437425 | -0.321971478 | 0.121096628 | 8.977155077 | -1.101636374 | 0.311673979 | 0.646781769 | -4.625086987 |
| **A_52_P462472** | Nexmif | -0.192655393 | -0.617696857 | 0.23238607 | 7.780743021 | -1.101369543 | 0.311781425 | 0.646781769 | -4.625209181 |
| **A_55_P2072228** | Cdk9 | 0.084471918 | -0.101903896 | 0.270847731 | 12.07856498 | 1.101300888 | 0.311809076 | 0.646781769 | -4.625240619 |
| **A_51_P294515** | Gde1 | -0.303183474 | -0.972179909 | 0.365812961 | 9.466488455 | -1.101196722 | 0.311851032 | 0.646781769 | -4.625288316 |
| **A_55_P2005470** | Mfap4 | -1.693926118 | -5.431726237 | 2.043874001 | 10.83883582 | -1.101188278 | 0.311854434 | 0.646781769 | -4.625292183 |
| **A_52_P527463** | Fam91a1 | -0.13547383 | -0.434426294 | 0.163478633 | 11.63293671 | -1.101124596 | 0.311880086 | 0.646781769 | -4.625321342 |
| **A_52_P75384** | B230219D22Rik | -0.164996511 | -0.529103484 | 0.199110461 | 11.10706898 | -1.10110556 | 0.311887755 | 0.646781769 | -4.625330058 |
| **A_51_P363103** | Blvra | 0.144656024 | -0.174565942 | 0.46387799 | 10.59795178 | 1.101100462 | 0.311889809 | 0.646781769 | -4.625332392 |
| **A_55_P2023537** | NA | 0.150971823 | -0.182205788 | 0.484149435 | 8.613626774 | 1.10104046 | 0.311913981 | 0.646781769 | -4.625359865 |
| **A_55_P2417569** | Prrx2 | -0.313233795 | -1.004530431 | 0.378062841 | 7.898721795 | -1.101000099 | 0.311930242 | 0.646781769 | -4.625378344 |
| **A_55_P2122898** | Rnf25 | 0.23538298 | -0.284328709 | 0.755094669 | 10.71522464 | 1.10051448 | 0.312125945 | 0.647118137 | -4.625600659 |
| **A_55_P2156435** | NA | 0.187789392 | -0.226892246 | 0.602471029 | 10.97791931 | 1.100371767 | 0.312183477 | 0.647168 | -4.625665984 |
| **A_55_P2156464** | Nt5c3b | -0.228320555 | -0.73258273 | 0.27594162 | 8.628406764 | -1.100200423 | 0.312252563 | 0.647192174 | -4.625744408 |
| **A_55_P2009973** | Ttc3 | -0.196244705 | -0.62968294 | 0.237193529 | 11.88993493 | -1.100155106 | 0.312270837 | 0.647192174 | -4.625765149 |
| **A_55_P1992049** | Gucy1a1 | -0.470617291 | -1.51027035 | 0.569035769 | 10.85269667 | -1.099923884 | 0.312364091 | 0.647192174 | -4.625870968 |
| **A_51_P386280** | Olfr187 | 0.303013701 | -0.366462726 | 0.972490127 | 8.782659822 | 1.099791009 | 0.31241769 | 0.647192174 | -4.625931774 |
| **A_52_P494514** | Insr | -0.174577693 | -0.560295994 | 0.211140607 | 9.16326343 | -1.099769562 | 0.312426342 | 0.647192174 | -4.625941588 |
| **A_55_P1993423** | NA | 0.096843127 | -0.117131306 | 0.31081756 | 13.49348156 | 1.099740219 | 0.31243818 | 0.647192174 | -4.625955015 |
| **A_52_P145767** | Vcpip1 | 0.121397195 | -0.146829521 | 0.389623912 | 9.555838603 | 1.099739463 | 0.312438485 | 0.647192174 | -4.625955361 |
| **A_55_P1979620** | Eif3h | -0.309635534 | -0.993855516 | 0.374584448 | 13.07363977 | -1.099608862 | 0.312491178 | 0.647192174 | -4.626015121 |
| **A_52_P27122** | Ago4 | -0.109275119 | -0.350760597 | 0.132210358 | 7.905662413 | -1.099546145 | 0.312516484 | 0.647192174 | -4.626043818 |
| **A_51_P196207** | Capsl | -0.240555991 | -0.772173288 | 0.291061307 | 8.307820703 | -1.099512731 | 0.312529968 | 0.647192174 | -4.626059107 |
| **A_55_P2061804** | NA | 0.095626726 | -0.115743192 | 0.306996644 | 9.213300733 | 1.099307773 | 0.312612685 | 0.647253889 | -4.62615288 |
| **A_52_P262967** | Rab10 | 0.170007868 | -0.205805721 | 0.545821457 | 10.46739016 | 1.099207622 | 0.31265311 | 0.647253889 | -4.626198699 |
| **A_51_P506045** | Cpq | -0.198822672 | -0.638340332 | 0.240694989 | 10.97642031 | -1.099189987 | 0.312660229 | 0.647253889 | -4.626206766 |
| **A_55_P2176492** | Tollip | 0.086485121 | -0.104713821 | 0.277684063 | 8.29571363 | 1.099104797 | 0.312694619 | 0.647255761 | -4.626245739 |
| **A_55_P2070316** | Ankrd36 | 0.141422245 | -0.171255261 | 0.454099752 | 8.4311996 | 1.099016665 | 0.3127302 | 0.647260097 | -4.626286055 |
| **A_55_P2111380** | Ctnnd2 | -0.215217587 | -0.691091132 | 0.260655958 | 7.823977855 | -1.098928371 | 0.312765851 | 0.647264575 | -4.626326444 |
| **A_51_P138895** | Ccdc102a | -0.313326177 | -1.006258361 | 0.379606006 | 9.469550799 | -1.098725329 | 0.312847845 | 0.647357674 | -4.626419317 |
| **A_52_P1077588** | Heatr6 | 0.135537442 | -0.164228766 | 0.435303649 | 8.768249452 | 1.098651112 | 0.31287782 | 0.647357674 | -4.626453262 |
| **A_55_P2076906** | NA | -0.74711142 | -2.399867205 | 0.905644366 | 9.294202494 | -1.098398365 | 0.31297992 | 0.647499612 | -4.626568855 |
| **A_55_P1970002** | Sh3kbp1 | 0.072797567 | -0.08828007 | 0.233875204 | 7.96094684 | 1.098157937 | 0.313077068 | 0.647631277 | -4.626678802 |
| **A_55_P2128373** | Cops6 | -0.075077411 | -0.241245484 | 0.091090663 | 12.76048884 | -1.097854765 | 0.313199606 | 0.647761606 | -4.626817425 |
| **A_55_P1961451** | Adamts7 | -0.082529684 | -0.265194881 | 0.100135513 | 7.894560809 | -1.097836235 | 0.313207096 | 0.647761606 | -4.626825897 |
| **A_55_P2015936** | NA | 0.08056825 | -0.097793047 | 0.258929547 | 16.56816012 | 1.097606063 | 0.313300157 | 0.647871896 | -4.626931128 |
| **A_52_P111031** | Pcdh17 | -0.510575372 | -1.640952267 | 0.619801523 | 9.457790646 | -1.097538541 | 0.31332746 | 0.647871896 | -4.626961996 |

| | | | | | | | | | |
|---|---|---|---|---|---|---|---|---|---|
| **A_65_P09285** | 2610524H06Rik | 0.125135455 | -0.152024991 | 0.4022959 | 8.81987787 | 1.097064886 | 0.313519048 | 0.648192272 | -4.627178502 |
| **A_51_P265806** | Clca3a2 | -0.163063983 | -0.524256453 | 0.198128488 | 9.483766336 | -1.096989691 | 0.313549472 | 0.648192272 | -4.62721287 |
| **A_55_P2073977** | NA | -0.09149655 | -0.294196097 | 0.111202998 | 14.54451678 | -1.096819408 | 0.313618378 | 0.648265387 | -4.627290692 |
| **A_52_P686136** | Prrc2b | -0.169554547 | -0.545233177 | 0.206124082 | 11.6022771 | -1.096670448 | 0.313678667 | 0.648300518 | -4.627358765 |
| **A_55_P2135633** | NA | -0.10919848 | -0.351182354 | 0.132785395 | 16.41098268 | -1.096511913 | 0.313742841 | 0.648300518 | -4.627431208 |
| **A_52_P404363** | Cdc42bpb | -0.082195827 | -0.264344762 | 0.099953109 | 8.197305466 | -1.096494151 | 0.313750031 | 0.648300518 | -4.627439324 |
| **A_55_P1957004** | Timm10 | 0.119436918 | -0.145279617 | 0.384153453 | 11.81855407 | 1.096328511 | 0.313817094 | 0.648300518 | -4.627515008 |
| **A_55_P2296118** | 9630010G10Rik | -0.22357148 | -0.719093858 | 0.271950899 | 8.94108132 | -1.096317426 | 0.313821583 | 0.648300518 | -4.627520073 |
| **A_51_P278489** | Itfg2 | 0.075764048 | -0.092171668 | 0.243699763 | 8.701243989 | 1.096234055 | 0.313855342 | 0.648300518 | -4.627558164 |
| **A_55_P2142028** | Clip4 | -0.232164637 | -0.746789142 | 0.282459868 | 7.90433304 | -1.096197473 | 0.313870156 | 0.648300518 | -4.627574877 |
| **A_55_P1986123** | Sccpdh | -0.187016098 | -0.601634717 | 0.22760252 | 10.3485713 | -1.096007136 | 0.313947244 | 0.648390456 | -4.627661834 |
| **A_55_P2032165** | NA | -0.151965204 | -0.488926106 | 0.184995698 | 15.02729301 | -1.095841729 | 0.314014247 | 0.648401349 | -4.627737395 |
| **A_55_P2149440** | Ehmt1 | -0.135914989 | -0.437290497 | 0.165460519 | 10.0389883 | -1.095828493 | 0.314019609 | 0.648401349 | -4.627743441 |
| **A_55_P2007656** | Cryab | -0.480564817 | -1.546305323 | 0.585175688 | 8.340506463 | -1.095679886 | 0.314079818 | 0.648456399 | -4.627811322 |
| **A_52_P260818** | Card10 | -0.099213446 | -0.319274712 | 0.12084782 | 8.669917173 | -1.095494284 | 0.314155029 | 0.648487579 | -4.627896096 |
| **A_55_P1970399** | NA | -0.100317778 | -0.322855907 | 0.122220351 | 14.7388262 | -1.095359457 | 0.314209674 | 0.648487579 | -4.627957674 |
| **A_55_P2170681** | Cdca8 | 0.191014257 | -0.232731003 | 0.614759517 | 9.371344091 | 1.095327824 | 0.314222496 | 0.648487579 | -4.62797212 |
| **A_51_P160482** | Vsig2 | -0.264580055 | -0.851596509 | 0.322436399 | 7.901773983 | -1.095191381 | 0.314277806 | 0.648487579 | -4.628034432 |
| **A_51_P277336** | Cavin2 | -0.322552145 | -1.038267595 | 0.393163305 | 13.17692691 | -1.095072143 | 0.314326148 | 0.648487579 | -4.628088883 |
| **A_55_P2175341** | NA | -0.215061628 | -0.692264969 | 0.262141714 | 8.333879264 | -1.095071917 | 0.31432624 | 0.648487579 | -4.628088986 |
| **A_55_P1980601** | Oscp1 | -0.313449279 | -1.009056686 | 0.382158129 | 8.538913061 | -1.094929775 | 0.314383876 | 0.648487579 | -4.628153892 |
| **A_55_P2030282** | Gm5796 | -0.305075364 | -0.982110333 | 0.371959606 | 8.239023562 | -1.094912024 | 0.314391073 | 0.648487579 | -4.628161997 |
| **A_51_P341349** | Mtap | 0.085215071 | -0.103899958 | 0.274330099 | 8.28982191 | 1.09489773 | 0.31439687 | 0.648487579 | -4.628168524 |
| **A_55_P2097972** | NA | 0.330015172 | -0.402548549 | 1.062578892 | 9.103097446 | 1.094640943 | 0.314501018 | 0.648633181 | -4.628285769 |
| **A_52_P217437** | Htra3 | 0.078703073 | -0.096030347 | 0.253436493 | 7.951525803 | 1.094457506 | 0.314575434 | 0.64871744 | -4.628369515 |
| **A_55_P2025248** | Mxd1 | 0.185815858 | -0.226888249 | 0.598519965 | 11.49755281 | 1.094024815 | 0.314751024 | 0.649010299 | -4.628567027 |
| **A_66_P117022** | Atp6v1e1 | -0.118606951 | -0.382060073 | 0.144846172 | 12.49789192 | -1.093931124 | 0.314789055 | 0.649019482 | -4.62860979 |
| **A_55_P1965219** | Syt6 | 0.203756853 | -0.248960345 | 0.65647405 | 9.177973514 | 1.093624977 | 0.314913354 | 0.649206507 | -4.628749509 |
| **A_55_P2419711** | 4932412D23Rik | 0.265437467 | -0.324390925 | 0.85526586 | 8.937208868 | 1.093502398 | 0.314963133 | 0.649239885 | -4.628805446 |
| **A_55_P2035344** | Tcf25 | -0.120805524 | -0.389303229 | 0.147692181 | 10.92103926 | -1.093274968 | 0.31505551 | 0.649361053 | -4.628909223 |
| **A_55_P2014249** | Sema3a | -0.291147476 | -0.938360893 | 0.356065941 | 8.334035648 | -1.093071842 | 0.315138034 | 0.64946189 | -4.629001901 |
| **A_51_P351896** | Gask1b | -0.220714622 | -0.711443827 | 0.270014584 | 7.87953895 | -1.092879774 | 0.315216082 | 0.649553482 | -4.629089525 |
| **A_52_P591166** | Dpysl2 | -0.528526206 | -1.703804785 | 0.646752373 | 10.91111026 | -1.092720005 | 0.315281018 | 0.649618035 | -4.629162409 |
| **A_55_P1993473** | Gm5935 | 0.210919061 | -0.258135653 | 0.679973776 | 8.745146808 | 1.09263602 | 0.315315156 | 0.649619128 | -4.629200719 |
| **A_52_P295313** | Wdr46 | 0.126908111 | -0.155361519 | 0.409177741 | 11.63551074 | 1.092467217 | 0.315383781 | 0.649691261 | -4.629277715 |
| **A_51_P383001** | Meox2 | -0.935403506 | -3.016277375 | 1.145470364 | 8.338399145 | -1.092286147 | 0.315457407 | 0.649704768 | -4.6293603 |
| **A_55_P2043287** | Tle5 | -0.180280102 | -0.581326584 | 0.22076638 | 10.19481734 | -1.092285762 | 0.315457564 | 0.649704768 | -4.629360475 |
| **A_51_P161054** | Itgb6 | -0.358734162 | -1.156947228 | 0.439478904 | 7.883472377 | -1.092036346 | 0.315559003 | 0.649805675 | -4.629474221 |
| **A_55_P2176212** | NA | 0.181796967 | -0.222737653 | 0.586331588 | 12.69862485 | 1.091978602 | 0.315582492 | 0.649805675 | -4.629500553 |
| **A_55_P1999532** | Col9a1 | -0.092217141 | -0.29744303 | 0.113008747 | 7.730362358 | -1.0918493 | 0.315635094 | 0.649805675 | -4.629559515 |
| **A_52_P309737** | Cdk12 | 0.151062061 | -0.185125545 | 0.487249668 | 8.623186375 | 1.091834708 | 0.315641031 | 0.649805675 | -4.629566169 |
| **A_51_P320105** | Smyd2 | -0.087166378 | -0.281188716 | 0.106855961 | 11.75018324 | -1.091642543 | 0.315719221 | 0.649855918 | -4.629653789 |
| **A_55_P2141395** | Ly6g6c | 0.118258275 | -0.144984117 | 0.381500667 | 8.493439374 | 1.091588373 | 0.315741265 | 0.649855918 | -4.629678487 |
| **A_55_P1982668** | Nadk | 0.141394034 | -0.173373184 | 0.456161251 | 11.32708528 | 1.09150261 | 0.315776169 | 0.649855918 | -4.629717588 |
| **A_51_P396708** | Med21 | 0.079944847 | -0.098035523 | 0.257925217 | 10.52939029 | 1.091444256 | 0.315799919 | 0.649855918 | -4.629744192 |
| **A_55_P2213348** | 5330421C15Rik | 0.170185338 | -0.208768382 | 0.549139057 | 8.473433064 | 1.091237182 | 0.315884211 | 0.649960179 | -4.629838593 |
| **A_55_P1995442** | NA | 0.101588341 | -0.124665417 | 0.327842099 | 8.627333617 | 1.091016344 | 0.315974127 | 0.650075987 | -4.629939259 |
| **A_55_P2002577** | Ephx1 | 0.11213104 | -0.13768064 | 0.36194272 | 8.849956225 | 1.090677364 | 0.316112186 | 0.650167703 | -4.630093759 |
| **A_55_P2159852** | NA | 0.103059236 | -0.126548903 | 0.332667375 | 8.973107165 | 1.090643535 | 0.316125966 | 0.650167703 | -4.630109177 |
| **A_55_P2159865** | Zfp59 | -0.087776054 | -0.283338807 | 0.107786699 | 8.27049947 | -1.090619101 | 0.31613592 | 0.650167703 | -4.630120312 |

| | | | | | | | | | |
|---|---|---|---|---|---|---|---|---|---|
| **A_51_P244923** | AI837181 | -0.098378235 | -0.317595095 | 0.120838624 | 9.396587378 | -1.090456296 | 0.316202248 | 0.650167703 | -4.630194505 |
| **A_55_P2037513** | Pld3 | 0.109462093 | -0.134454463 | 0.353378648 | 11.57342353 | 1.090449751 | 0.316204915 | 0.650167703 | -4.630197487 |
| **A_55_P1972772** | Gsg1 | -0.201058309 | -0.649096274 | 0.246979657 | 7.958308691 | -1.090411434 | 0.316220527 | 0.650167703 | -4.630214948 |
| **A_52_P649332** | Sdf4 | -0.116656551 | -0.376660183 | 0.143347082 | 11.35504691 | -1.090216884 | 0.316299808 | 0.650185757 | -4.630303598 |
| **A_51_P336622** | Dexi | -0.300122242 | -0.969063397 | 0.368818913 | 11.34254893 | -1.090168064 | 0.316319705 | 0.650185757 | -4.630325843 |
| **A_51_P214423** | Tead1 | -0.260825748 | -0.84219288 | 0.320541384 | 9.518608985 | -1.090142261 | 0.316330222 | 0.650185757 | -4.630337599 |
| **A_55_P1956097** | Cyb5d2 | 0.148686911 | -0.182826084 | 0.480199906 | 9.515511847 | 1.089821501 | 0.31646098 | 0.650385358 | -4.630483738 |
| **A_55_P1959393** | Hhat | -0.103393847 | -0.333959758 | 0.127172064 | 8.305323894 | -1.089639371 | 0.316535246 | 0.65042844 | -4.630566707 |
| **A_51_P517012** | Tysnd1 | 0.101933957 | -0.125383593 | 0.329251506 | 11.35435111 | 1.089605045 | 0.316549244 | 0.65042844 | -4.630582344 |
| **A_52_P281033** | Socs5 | -0.110707729 | -0.357660267 | 0.136244809 | 8.123672672 | -1.089300331 | 0.316673531 | 0.650572697 | -4.630721138 |
| **A_55_P2072150** | Vpreb2 | 0.159128922 | -0.195878845 | 0.51413669 | 9.829972986 | 1.089166953 | 0.316727946 | 0.650572697 | -4.630781885 |
| **A_55_P1953758** | Serinc2 | -0.278111567 | -0.89868577 | 0.342462636 | 8.046416592 | -1.088951463 | 0.316815877 | 0.650572697 | -4.630880022 |
| **A_55_P2176240** | Pdxk | 0.134168892 | -0.165247532 | 0.433585315 | 8.891501714 | 1.088828306 | 0.31686614 | 0.650572697 | -4.630936104 |
| **A_55_P1954221** | Emp1 | -0.984480264 | -3.181813732 | 1.212853204 | 10.42283547 | -1.08866493 | 0.316932828 | 0.650572697 | -4.631010497 |
| **A_51_P421303** | Caly | 0.121828023 | -0.150092306 | 0.393748351 | 8.560457657 | 1.088651082 | 0.316938481 | 0.650572697 | -4.631016802 |
| **A_55_P1954156** | Wdr43 | 0.187649395 | -0.231210933 | 0.606509724 | 10.30136843 | 1.08858195 | 0.316966704 | 0.650572697 | -4.631048279 |
| **A_55_P2071028** | NA | 0.188631433 | -0.23242662 | 0.609689485 | 11.7699614 | 1.088567276 | 0.316972695 | 0.650572697 | -4.63105496 |
| **A_55_P2012181** | Eps15l1 | 0.121945402 | -0.150259749 | 0.394150554 | 10.8212571 | 1.088559771 | 0.316975759 | 0.650572697 | -4.631058377 |
| **A_51_P314339** | Chtop | 0.082972883 | -0.102239515 | 0.268185282 | 10.69566176 | 1.088552341 | 0.316978792 | 0.650572697 | -4.631061761 |
| **A_52_P424272** | Gatc | 0.102543701 | -0.126369012 | 0.331456413 | 8.51494806 | 1.088484539 | 0.317006475 | 0.650572697 | -4.631092631 |
| **A_51_P426276** | Pdk2 | -0.083734569 | -0.270665944 | 0.103196806 | 8.125556058 | -1.088443222 | 0.317023345 | 0.650572697 | -4.631111442 |
| **A_55_P2097508** | Mcc | 0.171317441 | -0.211214946 | 0.553849827 | 11.35227781 | 1.088219632 | 0.317114651 | 0.650690987 | -4.631213233 |
| **A_52_P2732** | Mfn1 | 0.114470648 | -0.141158349 | 0.370099645 | 10.32970871 | 1.088096003 | 0.317165147 | 0.65072552 | -4.631269512 |
| **A_51_P177984** | Fam234a | -0.102152264 | -0.330295537 | 0.125991008 | 11.07769707 | -1.087986533 | 0.317209864 | 0.650734385 | -4.631319343 |
| **A_55_P2076524** | 1700028I16Rik | 0.37454727 | -0.462004119 | 1.211098659 | 11.01746588 | 1.0879206 | 0.3172368 | 0.650734385 | -4.631349354 |
| **A_52_P876147** | Crocc2 | -0.195433901 | -0.631975478 | 0.241107676 | 7.753288812 | -1.087821096 | 0.317277454 | 0.650748717 | -4.631394645 |
| **A_55_P2247235** | Rab26os | 0.216987491 | -0.267821566 | 0.701796548 | 8.678973575 | 1.087544816 | 0.317390355 | 0.650787219 | -4.631520387 |
| **A_55_P2079981** | NA | -0.090444916 | -0.292547887 | 0.111658055 | 8.339074662 | -1.087413331 | 0.317444098 | 0.650787219 | -4.631580224 |
| **A_55_P2388197** | 1010001B22Rik | 0.19748582 | -0.243819257 | 0.638790897 | 10.19189197 | 1.087377101 | 0.317458907 | 0.650787219 | -4.631596711 |
| **A_51_P345792** | Mfsd13a | 0.223123522 | -0.27547697 | 0.721724014 | 10.50606428 | 1.087366234 | 0.31746335 | 0.650787219 | -4.631601656 |
| **A_55_P1994194** | Heph | -0.233007361 | -0.753742646 | 0.287727923 | 9.102752362 | -1.087266043 | 0.317504308 | 0.650787219 | -4.631647248 |
| **A_55_P1972720** | Pmm1 | 0.16933302 | -0.209103145 | 0.547769186 | 11.16599488 | 1.087256996 | 0.317508007 | 0.650787219 | -4.631651364 |
| **A_55_P2031192** | Gtdc1 | 0.102897249 | -0.127122549 | 0.332917047 | 9.936064698 | 1.086980458 | 0.317621081 | 0.650787219 | -4.631777192 |
| **A_52_P537852** | Fam120b | 0.094346166 | -0.116587416 | 0.305279748 | 8.324422357 | 1.08683025 | 0.317682514 | 0.650787219 | -4.631845531 |
| **A_55_P1955015** | Myo1g | -0.141650376 | -0.458343977 | 0.175043226 | 11.63420438 | -1.086830032 | 0.317682603 | 0.650787219 | -4.63184563 |
| **A_55_P1999057** | Cps1 | -0.496401566 | -1.606263913 | 0.61346078 | 10.08655755 | -1.086793956 | 0.317697359 | 0.650787219 | -4.631862043 |
| **A_55_P2173073** | Zfp931 | -0.120580965 | -0.390210481 | 0.14904855 | 8.445457586 | -1.086662083 | 0.317751303 | 0.650787219 | -4.631922035 |
| **A_51_P142813** | Aspn | -0.239556025 | -0.775282268 | 0.296170218 | 7.876083862 | -1.086544119 | 0.317799564 | 0.650787219 | -4.631975697 |
| **A_55_P2012146** | NA | -0.165928623 | -0.537011943 | 0.205154698 | 7.849239597 | -1.086508339 | 0.317814204 | 0.650787219 | -4.631991974 |
| **A_55_P2117984** | Vangl2 | 0.152048194 | -0.188035839 | 0.492132227 | 8.580566212 | 1.086371121 | 0.317870352 | 0.650787219 | -4.63205439 |
| **A_55_P2414524** | NA | -0.103481219 | -0.334957778 | 0.12799534 | 7.69097317 | -1.086269799 | 0.317911816 | 0.650787219 | -4.632100475 |
| **A_55_P2231747** | NA | -0.247501449 | -0.801160286 | 0.306157387 | 7.97656775 | -1.086222263 | 0.317931271 | 0.650787219 | -4.632122096 |
| **A_55_P2001233** | Ifi208 | 0.082077903 | -0.101539997 | 0.265695802 | 10.49564223 | 1.086161549 | 0.317956121 | 0.650787219 | -4.632149709 |
| **A_55_P2079009** | Slco2b1 | -0.122058909 | -0.395123104 | 0.151005287 | 7.969199511 | -1.086145262 | 0.317962788 | 0.650787219 | -4.632157117 |
| **A_52_P527625** | Colec12 | -0.220121981 | -0.712568951 | 0.272324988 | 9.000245034 | -1.086143314 | 0.317963585 | 0.650787219 | -4.632158003 |
| **A_66_P133383** | NA | -0.155980527 | -0.504937852 | 0.192976799 | 7.764945438 | -1.086128606 | 0.317969605 | 0.650787219 | -4.632164692 |
| **A_55_P1959683** | Xlr3b | 0.224891231 | -0.278409014 | 0.728191476 | 8.581116334 | 1.085746826 | 0.318125907 | 0.650921621 | -4.632338312 |
| **A_52_P82510** | Nmt1 | 0.119960802 | -0.148525692 | 0.388447296 | 11.46192105 | 1.085675673 | 0.318155045 | 0.650921621 | -4.632370666 |
| **A_55_P2019890** | Rasef | -0.110758288 | -0.358658081 | 0.137141505 | 7.75543948 | -1.085633561 | 0.318172291 | 0.650921621 | -4.632389815 |
| **A_51_P171965** | Sdhaf4 | 0.112505016 | -0.139313068 | 0.364323099 | 10.21858103 | 1.085595813 | 0.31818775 | 0.650921621 | -4.632406979 |

| | | | | | | | | |
|---|---|---|---|---|---|---|---|---|
| **A_51_P124126** | Cyp2d22 | -0.259435573 | -0.840180365 | 0.321309218 | 9.540647795 | -1.085493909 | 0.318229487 | 0.650921621 | -4.632453312 |
| **A_52_P239707** | Nfyb | 0.219918779 | -0.272376615 | 0.712214173 | 9.7989257 | 1.085474767 | 0.318237328 | 0.650921621 | -4.632462016 |
| **A_52_P51029** | Ralgapa1 | 0.130807103 | -0.162052223 | 0.42366643 | 9.309089274 | 1.085314132 | 0.31830313 | 0.650987325 | -4.632535048 |
| **A_55_P1972575** | Tmeff1 | -0.597302002 | -1.93481007 | 0.740206067 | 8.48105173 | -1.085127447 | 0.318379618 | 0.651021736 | -4.632619918 |
| **A_55_P2170834** | NA | 0.442717576 | -0.548704363 | 1.434139516 | 11.22233717 | 1.085054189 | 0.318409638 | 0.651021736 | -4.632653221 |
| **A_52_P360650** | Abcb7 | 0.212885174 | -0.263880217 | 0.689650564 | 9.016518609 | 1.084985375 | 0.318437838 | 0.651021736 | -4.632684502 |
| **A_51_P285271** | Ccdc61 | -0.137281403 | -0.444745848 | 0.170183041 | 9.601865398 | -1.084925704 | 0.318462293 | 0.651021736 | -4.632711625 |
| **A_52_P285024** | Sertad4 | -0.094498764 | -0.306156809 | 0.117159281 | 7.746372431 | -1.084862099 | 0.318488361 | 0.651021736 | -4.632740537 |
| **A_55_P2115141** | Lcor | 0.1201624 | -0.149112458 | 0.389437257 | 8.280006534 | 1.084316277 | 0.318712143 | 0.651410278 | -4.632988603 |
| **A_66_P139116** | Rpl8 | 0.108046481 | -0.134121227 | 0.350214189 | 16.65007883 | 1.084120573 | 0.318792411 | 0.651470001 | -4.633077532 |
| **A_51_P460048** | Cnrip1 | -0.193291502 | -0.626537282 | 0.239954278 | 9.031466747 | -1.084080689 | 0.318808771 | 0.651470001 | -4.633095655 |
| **A_51_P349073** | Endog | -0.193108263 | -0.625998436 | 0.239781911 | 10.68417403 | -1.083942683 | 0.318865387 | 0.651516814 | -4.63315836 |
| **A_55_P2065834** | Dhx8 | 0.181937498 | -0.225942925 | 0.589817921 | 10.8079308 | 1.083858394 | 0.31889997 | 0.651518604 | -4.633196655 |
| **A_52_P576442** | Dap | -0.107575805 | -0.34879639 | 0.133644779 | 11.84969832 | -1.083636014 | 0.318991224 | 0.651571627 | -4.633297684 |
| **A_55_P2119864** | Cd2bp2 | 0.101329567 | -0.125886616 | 0.32854575 | 9.153397181 | 1.083627742 | 0.318994619 | 0.651571627 | -4.633301442 |
| **A_55_P2172375** | NA | 0.096108074 | -0.119429847 | 0.311645995 | 8.411958089 | 1.083476196 | 0.31905682 | 0.651571627 | -4.633370284 |
| **A_55_P2273929** | Gm20558 | -0.172231576 | -0.558492201 | 0.214029049 | 7.761543002 | -1.083466594 | 0.319060761 | 0.651571627 | -4.633374646 |
| **A_51_P286946** | Lhpp | -0.14434181 | -0.468159297 | 0.179475676 | 10.21522025 | -1.083116169 | 0.319204629 | 0.65163206 | -4.633533813 |
| **A_55_P2079158** | Saxo2 | -0.11174695 | -0.362449743 | 0.138955842 | 7.73492343 | -1.08307791 | 0.319220339 | 0.65163206 | -4.633551189 |
| **A_55_P1952743** | NA | 0.226853557 | -0.282121918 | 0.735829032 | 12.38155718 | 1.083008653 | 0.319248781 | 0.65163206 | -4.633582643 |
| **A_55_P1959869** | Ppp1r2 | 0.16058405 | -0.199716619 | 0.52088472 | 11.40717522 | 1.082980228 | 0.319260454 | 0.65163206 | -4.633595552 |
| **A_51_P342707** | Pold2 | 0.126367287 | -0.1571803 | 0.409914873 | 9.210445575 | 1.08290883 | 0.319289777 | 0.65163206 | -4.633627977 |
| **A_51_P407879** | Fam193b | -0.083724135 | -0.271589126 | 0.104140855 | 9.778427539 | -1.082899135 | 0.319293759 | 0.65163206 | -4.633632379 |
| **A_55_P1965539** | Ndufb4 | 0.11454945 | -0.142516272 | 0.371615171 | 14.94308571 | 1.08275956 | 0.31935109 | 0.65163206 | -4.633695763 |
| **A_51_P209327** | Apln | -0.327395814 | -1.062134722 | 0.407343093 | 7.933898865 | -1.082737737 | 0.319360055 | 0.65163206 | -4.633705672 |
| **A_55_P2052479** | NA | -0.122026179 | -0.395938478 | 0.151886119 | 15.14871695 | -1.082491939 | 0.31946104 | 0.651656745 | -4.633817282 |
| **A_55_P2068330** | Cryzl1 | 0.084788913 | -0.105550534 | 0.27512836 | 9.345081398 | 1.08241413 | 0.319493013 | 0.651656745 | -4.63385261 |
| **A_55_P1952744** | Timm8a1 | 0.228549468 | -0.284545368 | 0.741644303 | 12.35295591 | 1.082345111 | 0.319521377 | 0.651656745 | -4.633883946 |
| **A_55_P1958349** | Nipsnap3b | -0.135111557 | -0.43844045 | 0.168217335 | 8.828019422 | -1.082335514 | 0.319525321 | 0.651656745 | -4.633888303 |
| **A_55_P2022211** | Plxdc2 | -0.881311228 | -2.85994593 | 1.097323474 | 9.729015765 | -1.082298037 | 0.319540723 | 0.651656745 | -4.633905318 |
| **A_51_P108659** | Pon1 | -0.535154267 | -1.736797173 | 0.666488639 | 10.65946747 | -1.082148233 | 0.319602295 | 0.651713551 | -4.633973327 |
| **A_55_P2010871** | Daam2 | -0.287795639 | -0.934230042 | 0.358638763 | 8.419672459 | -1.081789936 | 0.3197496 | 0.651737133 | -4.634135971 |
| **A_55_P2026125** | Srrm2 | 0.153663354 | -0.191533242 | 0.498859949 | 8.549175892 | 1.081650595 | 0.319806902 | 0.651737133 | -4.634199216 |
| **A_66_P105965** | Drp2 | 0.075236215 | -0.093780934 | 0.244253364 | 7.784195545 | 1.081631578 | 0.319814723 | 0.651737133 | -4.634207847 |
| **A_51_P516850** | Rbbp5 | 0.1324582 | -0.165120391 | 0.430036791 | 8.665071344 | 1.081584253 | 0.319834187 | 0.651737133 | -4.634229325 |
| **A_55_P1962937** | Trem2 | 0.099281947 | -0.123770207 | 0.322334101 | 9.354274952 | 1.081551122 | 0.319847814 | 0.651737133 | -4.634244362 |
| **A_51_P453856** | Csnk1a1 | 0.098061456 | -0.122249305 | 0.318372218 | 12.03137948 | 1.081548045 | 0.31984908 | 0.651737133 | -4.634245759 |
| **A_52_P305685** | NA | -0.076767722 | -0.249274036 | 0.095738592 | 8.44364142 | -1.081326505 | 0.319940212 | 0.651737133 | -4.634346298 |
| **A_51_P175567** | Dact1 | -0.122922936 | -0.399154757 | 0.153308885 | 7.880710529 | -1.081290548 | 0.319955005 | 0.651737133 | -4.634362615 |
| **A_55_P2185595** | Samd4b | -0.11238285 | -0.364929445 | 0.140163746 | 10.89377742 | -1.081288818 | 0.319955717 | 0.651737133 | -4.634363401 |
| **A_51_P290904** | Raver2 | 0.173596677 | -0.216558927 | 0.563752281 | 8.263057164 | 1.081151976 | 0.320012021 | 0.651737133 | -4.634425496 |
| **A_55_P1959624** | NA | 0.272973023 | -0.340578858 | 0.886524905 | 9.651102832 | 1.081064551 | 0.320047996 | 0.651737133 | -4.634465166 |
| **A_51_P191520** | Stard10 | -0.113705168 | -0.369291806 | 0.14188147 | 11.48095411 | -1.080998888 | 0.320075019 | 0.651737133 | -4.63449496 |
| **A_55_P1961395** | Pdpn | -0.893858895 | -2.903278778 | 1.115560989 | 9.334864371 | -1.080889903 | 0.320119874 | 0.651737133 | -4.634544409 |
| **A_51_P414548** | Casp7 | 0.071208473 | -0.088881893 | 0.231298839 | 9.531863999 | 1.080810769 | 0.320152447 | 0.651737133 | -4.634580312 |
| **A_51_P504065** | Spa17 | -0.25851292 | -0.839756709 | 0.322730868 | 8.592346552 | -1.080704893 | 0.320196031 | 0.651737133 | -4.634628346 |
| **A_55_P2001998** | Zfp503 | -0.382045812 | -1.241056948 | 0.476965324 | 8.662405307 | -1.080686778 | 0.320203489 | 0.651737133 | -4.634636564 |
| **A_55_P2130090** | Cnn3 | -0.109202091 | -0.354752988 | 0.136348806 | 12.67726862 | -1.080619049 | 0.320231373 | 0.651737133 | -4.63466729 |
| **A_55_P2019512** | Mocs2 | 0.113685185 | -0.142004643 | 0.369375013 | 12.14205859 | 1.080372726 | 0.3203328 | 0.651737133 | -4.634779028 |
| **A_51_P417469** | Rnaseh2c | 0.106727883 | -0.133327356 | 0.346783123 | 14.30980685 | 1.08031373 | 0.320357097 | 0.651737133 | -4.634805788 |

| | | | | | | | | | |
|---|---|---|---|---|---|---|---|---|---|
| **A_55_P2149931** | Arap2 | -0.128992307 | -0.419131259 | 0.161146644 | 8.476396157 | -1.080291509 | 0.320366249 | 0.651737133 | -4.634815867 |
| **A_55_P1975743** | Dap3 | 0.090633375 | -0.113262554 | 0.294529304 | 13.29198383 | 1.08009707 | 0.320446338 | 0.651737133 | -4.634904057 |
| **A_51_P349546** | Cd109 | -0.17556689 | -0.570545794 | 0.219412015 | 7.746788393 | -1.080069999 | 0.32045749 | 0.651737133 | -4.634916335 |
| **A_51_P342567** | Akap12 | -0.320404749 | -1.041241697 | 0.400432199 | 11.36335753 | -1.080053066 | 0.320464465 | 0.651737133 | -4.634924015 |
| **A_52_P380301** | Unc5c | -0.296310652 | -0.962951918 | 0.370330614 | 7.830570878 | -1.08003607 | 0.320471467 | 0.651737133 | -4.634931723 |
| **A_52_P597634** | Fzd1 | -0.17069552 | -0.554747597 | 0.213356557 | 10.09463329 | -1.079978711 | 0.320495098 | 0.651737133 | -4.634957737 |
| **A_55_P2184508** | NA | -0.107286814 | -0.348686849 | 0.13411322 | 14.22485277 | -1.079921557 | 0.320518645 | 0.651737133 | -4.634983657 |
| **A_55_P1957373** | Pdcl | -0.12666287 | -0.411665955 | 0.158340216 | 9.429096338 | -1.079898533 | 0.320528132 | 0.651737133 | -4.634994098 |
| **A_55_P2050997** | Serinc4 | -0.074822099 | -0.243189979 | 0.093545781 | 8.138436036 | -1.079826128 | 0.320557966 | 0.651737133 | -4.635026933 |
| **A_51_P331021** | Ttc32 | 0.138396736 | -0.173101175 | 0.449894647 | 9.981073246 | 1.079577682 | 0.320660353 | 0.651876732 | -4.635139593 |
| **A_65_P10195** | Myl7 | -1.141782071 | -3.711876832 | 1.42831269 | 8.617575396 | -1.07948664 | 0.32069788 | 0.651884459 | -4.635180874 |
| **A_55_P1969131** | Cidec | -0.147242306 | -0.478722796 | 0.184238183 | 7.789936925 | -1.079338898 | 0.320758785 | 0.651938418 | -4.635247859 |
| **A_55_P2100149** | Mapt | -0.510829578 | -1.660930509 | 0.639271354 | 8.417659137 | -1.079253012 | 0.320794195 | 0.651938418 | -4.635286798 |
| **A_55_P1961129** | Egfem1 | -0.143911523 | -0.467965892 | 0.180142847 | 7.757779449 | -1.079097972 | 0.320858124 | 0.651938418 | -4.635357085 |
| **A_55_P2168316** | Aldh3b2 | -0.144394454 | -0.469559885 | 0.180770977 | 7.713972953 | -1.079019599 | 0.320890445 | 0.651938418 | -4.635392613 |
| **A_51_P301809** | Slit3 | -0.143933949 | -0.468064267 | 0.18019637 | 7.779993935 | -1.079013238 | 0.320893068 | 0.651938418 | -4.635395497 |
| **A_52_P612803** | Ccng1 | 0.135790431 | -0.1700373 | 0.441618163 | 9.446983624 | 1.07888584 | 0.320945612 | 0.651967626 | -4.635453246 |
| **A_55_P1964459** | NA | -0.082510165 | -0.268352107 | 0.103331778 | 8.087961036 | -1.078814825 | 0.320974904 | 0.651967626 | -4.635485436 |
| **A_52_P264229** | Ing5 | 0.109806865 | -0.137542125 | 0.357155856 | 9.347404513 | 1.078704625 | 0.321020364 | 0.651987939 | -4.635535385 |
| **A_55_P1967443** | NA | 0.243773854 | -0.30539538 | 0.792943089 | 11.74209626 | 1.078609185 | 0.32105974 | 0.651987939 | -4.635578642 |
| **A_52_P182118** | NA | -0.080995177 | -0.263470421 | 0.101480068 | 8.716593455 | -1.078545301 | 0.321086098 | 0.651987939 | -4.635607596 |
| **A_55_P2003517** | Zfp536 | -0.190726044 | -0.620523794 | 0.239071706 | 7.740375699 | -1.0782738 | 0.32119814 | 0.652049668 | -4.635730636 |
| **A_51_P380432** | Scx | -0.784131332 | -2.551189262 | 0.982926597 | 8.458065703 | -1.078254327 | 0.321206177 | 0.652049668 | -4.635739461 |
| **A_55_P2075294** | NA | 0.149627176 | -0.187632509 | 0.486886862 | 9.313079268 | 1.078026008 | 0.321300425 | 0.652049668 | -4.635842918 |
| **A_55_P1983135** | NA | 0.202192077 | -0.253562138 | 0.657946291 | 9.13769537 | 1.077994706 | 0.321313348 | 0.652049668 | -4.635857101 |
| **A_55_P1967539** | Hunk | -0.383085089 | -1.246623203 | 0.480453024 | 9.873402205 | -1.077945805 | 0.321333538 | 0.652049668 | -4.635879258 |
| **A_51_P206824** | Hjv | -0.195762979 | -0.637087973 | 0.245562015 | 8.562190602 | -1.077842316 | 0.321376269 | 0.652049668 | -4.635926147 |
| **A_52_P91402** | Zcchc17 | 0.115294053 | -0.144632608 | 0.375220713 | 12.06329969 | 1.077802698 | 0.321392628 | 0.652049668 | -4.635944096 |
| **A_55_P2061021** | Smg6 | -0.095623756 | -0.311211566 | 0.119964055 | 11.13019692 | -1.077766756 | 0.32140747 | 0.652049668 | -4.63596038 |
| **A_52_P429749** | Tmem115 | -0.116941306 | -0.380597923 | 0.146715311 | 9.005547375 | -1.077736157 | 0.321420106 | 0.652049668 | -4.635974242 |
| **A_55_P2011747** | Lsg1 | 0.131962182 | -0.16561642 | 0.429540783 | 11.18370573 | 1.077533988 | 0.321503605 | 0.652110779 | -4.63606583 |
| **A_55_P2062936** | Abca3 | -0.307146022 | -0.999791281 | 0.385499236 | 12.39103945 | -1.077499851 | 0.321517705 | 0.652110779 | -4.636081294 |
| **A_55_P2041828** | Tubb3 | 0.12319257 | -0.154679811 | 0.401064951 | 7.84655721 | 1.077264434 | 0.321614961 | 0.652239596 | -4.63618793 |
| **A_55_P2096777** | Tprn | 0.109899951 | -0.138011 | 0.357810901 | 8.706576262 | 1.077171803 | 0.321653236 | 0.652248782 | -4.636229886 |
| **A_55_P2090633** | Ptp4a1 | -0.099868507 | -0.325174784 | 0.12543777 | 9.204666731 | -1.077056422 | 0.321700916 | 0.652264104 | -4.636282143 |
| **A_66_P135700** | Cyp2d11 | -0.338438091 | -1.102117354 | 0.425241173 | 14.47014918 | -1.076840615 | 0.321790111 | 0.652264104 | -4.636379878 |
| **A_55_P2039289** | Hspb6 | -0.111396692 | -0.362778194 | 0.139984811 | 8.921721794 | -1.076768057 | 0.321820104 | 0.652264104 | -4.636412735 |
| **A_51_P359603** | Itgb7 | 0.171868067 | -0.215987044 | 0.559723177 | 8.888025645 | 1.076735078 | 0.321833738 | 0.652264104 | -4.636427669 |
| **A_55_P2111439** | Gpsm1 | -0.082856588 | -0.269845164 | 0.104131988 | 7.770363042 | -1.076701112 | 0.32184778 | 0.652264104 | -4.63644305 |
| **A_55_P1958314** | Mospd1 | 0.114753226 | -0.14423261 | 0.373739062 | 11.19906982 | 1.076643888 | 0.321871438 | 0.652264104 | -4.636468962 |
| **A_55_P1975456** | Nfyb | 0.215981203 | -0.271505482 | 0.703467888 | 9.984439749 | 1.076555406 | 0.321908022 | 0.652264104 | -4.636509027 |
| **A_51_P407675** | Zrsr1 | 0.138054782 | -0.173561712 | 0.449671276 | 9.920971879 | 1.076500429 | 0.321930755 | 0.652264104 | -4.63653392 |
| **A_55_P1990046** | NA | 0.096225163 | -0.12102143 | 0.313471757 | 8.264024232 | 1.076264141 | 0.322028473 | 0.652393708 | -4.636640901 |
| **A_55_P1956672** | Rab34 | -0.26866075 | -0.875264339 | 0.337942839 | 11.42515098 | -1.076173854 | 0.322065819 | 0.652400986 | -4.636681776 |
| **A_51_P370286** | Med24 | 0.135571904 | -0.170644183 | 0.441787992 | 10.66266925 | 1.07578351 | 0.322227317 | 0.652605382 | -4.636858473 |
| **A_51_P399853** | Zfp704 | -0.16889123 | -0.550371573 | 0.212589114 | 8.742339367 | -1.075766763 | 0.322234247 | 0.652605382 | -4.636866053 |
| **A_55_P2149719** | NA | 0.104384142 | -0.131438471 | 0.340206754 | 15.01694832 | 1.075554119 | 0.322322255 | 0.652608233 | -4.636962297 |
| **A_55_P1982628** | Tma16 | 0.173098083 | -0.217981321 | 0.564177486 | 9.832586946 | 1.075500217 | 0.322344566 | 0.652608233 | -4.636986692 |
| **A_51_P292754** | Yaf2 | 0.095521185 | -0.120313043 | 0.311355412 | 10.72947725 | 1.075381528 | 0.322393699 | 0.652608233 | -4.637040405 |
| **A_51_P246854** | Acta1 | -0.191258022 | -0.623429346 | 0.240913303 | 8.006698254 | -1.075342724 | 0.322409765 | 0.652608233 | -4.637057966 |

| | | | | | | | | | |
|---|---|---|---|---|---|---|---|---|---|
| **A_52_P32369** | Hmgn3 | -0.229527172 | -0.74820936 | 0.289155017 | 8.379414902 | -1.075266114 | 0.322441483 | 0.652608233 | -4.637092634 |
| **A_51_P246705** | Nop14 | 0.127026788 | -0.160041685 | 0.414095261 | 8.474071463 | 1.075209287 | 0.322465013 | 0.652608233 | -4.637118349 |
| **A_55_P2003753** | Sh2d5 | 0.103720008 | -0.130689893 | 0.338129908 | 8.205682661 | 1.07515178 | 0.322488825 | 0.652608233 | -4.637144371 |
| **A_55_P1999715** | Tcrb-J | -0.095438413 | -0.3111399 | 0.120263074 | 8.255008573 | -1.075110884 | 0.322505761 | 0.652608233 | -4.637162876 |
| **A_51_P135732** | Ccdc136 | -0.131384238 | -0.42837054 | 0.165602064 | 7.748850828 | -1.074954388 | 0.322570572 | 0.652621585 | -4.637233686 |
| **A_52_P467726** | Nsg1 | -0.363609251 | -1.185543659 | 0.458325157 | 8.693307036 | -1.0749319 | 0.322579886 | 0.652621585 | -4.63724386 |
| **A_51_P296057** | Upf1 | 0.143152396 | -0.180517228 | 0.466822019 | 9.797110516 | 1.074681733 | 0.322683517 | 0.652754152 | -4.637357041 |
| **A_52_P625277** | Accs | -0.109513976 | -0.35714302 | 0.138115068 | 8.554088919 | -1.074610684 | 0.322712953 | 0.652754152 | -4.637389183 |
| **A_55_P2054062** | Chst2 | -0.196170233 | -0.639802891 | 0.247462424 | 7.906690363 | -1.074466277 | 0.32277279 | 0.652779246 | -4.637454508 |
| **A_55_P1963980** | Carmil1 | -0.072352029 | -0.235981202 | 0.091277144 | 8.095228456 | -1.07441774 | 0.322792904 | 0.652779246 | -4.637476463 |
| **A_55_P2052690** | Synm | -0.116079727 | -0.378637586 | 0.146478131 | 7.752475661 | -1.074272719 | 0.322853007 | 0.652808766 | -4.63754206 |
| **A_51_P257419** | Lhx2 | 0.20001828 | -0.252420506 | 0.652457067 | 9.292555157 | 1.074219543 | 0.322875048 | 0.652808766 | -4.637566112 |
| **A_52_P257728** | Cnnm4 | 0.075454191 | -0.095281761 | 0.246190143 | 8.875276541 | 1.073844946 | 0.323030349 | 0.652913803 | -4.637735526 |
| **A_55_P2114269** | Fam177a2 | 0.096196157 | -0.121483333 | 0.313875648 | 16.36213064 | 1.073800006 | 0.323048984 | 0.652913803 | -4.637755848 |
| **A_55_P2133337** | NA | -0.445343 | -1.453115608 | 0.562429609 | 8.320849578 | -1.073779897 | 0.323057323 | 0.652913803 | -4.637764941 |
| **A_55_P1964108** | B230118H07Rik | -0.150360554 | -0.490617116 | 0.189896008 | 10.21216678 | -1.073768343 | 0.323062114 | 0.652913803 | -4.637770166 |
| **A_66_P106536** | Atp5mpl | 0.088915271 | -0.112328529 | 0.290159072 | 14.90372943 | 1.073586489 | 0.323137536 | 0.65295765 | -4.637852398 |
| **A_55_P2068822** | Cdk20 | -0.122449912 | -0.399685923 | 0.154786098 | 9.468091657 | -1.073228081 | 0.323286223 | 0.65295765 | -4.638014443 |
| **A_55_P2010312** | Serpina1a | -0.322870248 | -1.053957234 | 0.408216738 | 17.19475734 | -1.073104864 | 0.323337353 | 0.65295765 | -4.638070145 |
| **A_55_P2006915** | Myo5c | -0.207129698 | -0.676172358 | 0.261912962 | 8.062868497 | -1.073033343 | 0.323367034 | 0.65295765 | -4.638102477 |
| **A_51_P478688** | Samd10 | 0.078795665 | -0.099637928 | 0.257229258 | 10.30760373 | 1.07302266 | 0.323371468 | 0.65295765 | -4.638107306 |
| **A_55_P2092310** | Phgdh | 0.218834437 | -0.276745838 | 0.714414712 | 12.55747469 | 1.07296324 | 0.323396129 | 0.65295765 | -4.638134166 |
| **A_51_P246339** | Rfc5 | 0.143600343 | -0.181613913 | 0.468814598 | 11.98728515 | 1.072924323 | 0.323412282 | 0.65295765 | -4.638151757 |
| **A_55_P1955587** | Hspd1 | 0.250829141 | -0.317282426 | 0.818940707 | 14.46448741 | 1.072821696 | 0.323454881 | 0.65295765 | -4.638198146 |
| **A_55_P2123841** | Zfhx4 | 0.159735509 | -0.202092023 | 0.521563041 | 8.533938728 | 1.072711778 | 0.323500512 | 0.65295765 | -4.638247827 |
| **A_51_P503542** | Zfp414 | 0.117105128 | -0.148159838 | 0.382370094 | 11.08595126 | 1.072702288 | 0.323504452 | 0.65295765 | -4.638252117 |
| **A_55_P1962901** | Osr1 | -0.16248685 | -0.530571335 | 0.205597635 | 7.850031408 | -1.072639775 | 0.323530405 | 0.65295765 | -4.63828037 |
| **A_55_P2034067** | Myom3 | -0.139497424 | -0.455575679 | 0.176580832 | 7.768745781 | -1.072394939 | 0.323632071 | 0.65295765 | -4.63839102 |
| **A_66_P120277** | 9330182L06Rik | -0.120705366 | -0.394213606 | 0.152802874 | 7.746530086 | -1.072356904 | 0.323647867 | 0.65295765 | -4.638408208 |
| **A_55_P2018307** | NA | 0.376783853 | -0.476977839 | 1.230545544 | 12.31219379 | 1.072355558 | 0.323648426 | 0.65295765 | -4.638408816 |
| **A_51_P499020** | Fbp2 | -0.109603382 | -0.35798267 | 0.138775907 | 7.703205661 | -1.072239405 | 0.323696669 | 0.65295765 | -4.638461304 |
| **A_55_P2022812** | Cyp4f17 | -0.113367115 | -0.37028373 | 0.1435495 | 9.173052811 | -1.07220565 | 0.32371069 | 0.65295765 | -4.638476558 |
| **A_55_P2183378** | Snx11 | 0.122745375 | -0.155428605 | 0.400919355 | 11.08001566 | 1.072190174 | 0.323717119 | 0.65295765 | -4.63848355 |
| **A_51_P424256** | Gm6211 | 0.098491585 | -0.124779166 | 0.321762336 | 12.75749104 | 1.071890647 | 0.323841557 | 0.65295765 | -4.638618886 |
| **A_55_P2094242** | NA | 0.084329076 | -0.106872131 | 0.275530284 | 8.14130029 | 1.071691785 | 0.323924196 | 0.65295765 | -4.638708728 |
| **A_55_P2313861** | Zbed6 | -0.078359415 | -0.256043083 | 0.099324254 | 8.315906523 | -1.071585578 | 0.323968338 | 0.65295765 | -4.638756707 |
| **A_55_P2085335** | Mia | -1.006789583 | -3.290198122 | 1.276618957 | 9.241607606 | -1.07136704 | 0.324059184 | 0.65295765 | -4.638855423 |
| **A_55_P2180445** | NA | 0.181200659 | -0.229775407 | 0.592176726 | 11.20707038 | 1.071337807 | 0.324071337 | 0.65295765 | -4.638868627 |
| **A_55_P1991985** | Ptprs | -0.392457375 | -1.282650531 | 0.497735781 | 11.66759374 | -1.071251475 | 0.324107232 | 0.65295765 | -4.638907621 |
| **A_51_P116651** | Dpt | -0.209521192 | -0.684775007 | 0.265732624 | 7.809949094 | -1.071236921 | 0.324113284 | 0.65295765 | -4.637478883 |
| **A_52_P155708** | Creg1 | 0.136166809 | -0.17270395 | 0.445037568 | 8.953910629 | 1.071217485 | 0.324121365 | 0.65295765 | -4.638922973 |
| **A_55_P2117726** | Stk33 | -0.140425533 | -0.45900492 | 0.178153855 | 7.713557597 | -1.071054563 | 0.324189115 | 0.65295765 | -4.638996554 |
| **A_55_P2014555** | Lrp8 | 0.076416874 | -0.096960144 | 0.249793892 | 7.81636265 | 1.070978939 | 0.324220567 | 0.65295765 | -4.639030707 |
| **A_55_P1976754** | Ppia | 0.090505235 | -0.114852533 | 0.295863003 | 17.30328628 | 1.070892202 | 0.324256644 | 0.65295765 | -4.639069877 |
| **A_55_P2143025** | Sema3c | -0.452698659 | -1.47991867 | 0.574521352 | 8.359458464 | -1.070850701 | 0.324273907 | 0.65295765 | -4.639088618 |
| **A_55_P2023449** | NA | -0.165687746 | -0.541678057 | 0.210302566 | 7.936167063 | -1.070771797 | 0.32430673 | 0.65295765 | -4.639124249 |
| **A_51_P466221** | Amhr2 | -0.318429244 | -1.041046554 | 0.404188067 | 7.905988604 | -1.070749236 | 0.324316115 | 0.65295765 | -4.639134437 |
| **A_55_P1962035** | Coa5 | 0.140804368 | -0.178726999 | 0.460335735 | 9.61080875 | 1.070744418 | 0.32431812 | 0.65295765 | -4.639136612 |
| **A_55_P1992989** | Mrps26 | 0.150852374 | -0.191489479 | 0.493194228 | 12.91446012 | 1.070718588 | 0.324328865 | 0.65295765 | -4.639148276 |
| **A_55_P1965674** | Alg2 | 0.092899582 | -0.117934432 | 0.303733597 | 11.03294538 | 1.070671573 | 0.324348426 | 0.65295765 | -4.639169505 |

| | | | | | | | | | |
|---|---|---|---|---|---|---|---|---|---|
| **A_51_P395736** | Srp72 | 0.124073347 | -0.157557482 | 0.405704175 | 11.59824679 | 1.070487234 | 0.324425126 | 0.65295765 | -4.639252737 |
| **A_55_P1976012** | Cnih4 | -0.073345342 | -0.239840863 | 0.09315018 | 9.125798909 | -1.070417422 | 0.324454178 | 0.65295765 | -4.639284257 |
| **A_52_P360112** | Clip2 | -0.090551135 | -0.296116391 | 0.115014122 | 7.929755618 | -1.070353839 | 0.32448064 | 0.65295765 | -4.639312963 |
| **A_52_P670874** | Ebag9 | 0.100550875 | -0.127734407 | 0.328836156 | 10.04121616 | 1.070264525 | 0.324517813 | 0.65295765 | -4.639353285 |
| **A_52_P402897** | Cdh4 | -0.109257558 | -0.357332639 | 0.138817522 | 7.918593285 | -1.070166964 | 0.324558422 | 0.65295765 | -4.639397328 |
| **A_52_P532227** | S1pr1 | -0.169077173 | -0.55301439 | 0.214860043 | 9.235904067 | -1.070059572 | 0.324603129 | 0.65295765 | -4.639445807 |
| **A_55_P2038514** | NA | -0.173499664 | -0.567483816 | 0.220484489 | 7.813662931 | -1.070047541 | 0.324608137 | 0.65295765 | -4.639451238 |
| **A_55_P2002319** | Ap5s1 | -0.118147286 | -0.386437322 | 0.15014275 | 11.33249279 | -1.070045872 | 0.324608832 | 0.65295765 | -4.639451992 |
| **A_52_P656714** | Map3k5 | -0.098638046 | -0.322630156 | 0.125354065 | 8.345458496 | -1.07002747 | 0.324616494 | 0.65295765 | -4.639460298 |
| **A_55_P1990864** | NA | 0.128361742 | -0.163148535 | 0.419872019 | 8.198039641 | 1.069953541 | 0.324647273 | 0.65295765 | -4.639493669 |
| **A_55_P2115255** | Fer | 0.085568625 | -0.108761371 | 0.27989862 | 8.563720517 | 1.069936245 | 0.324654475 | 0.65295765 | -4.639501476 |
| **A_55_P2154027** | Ptgis | -0.33538188 | -1.097064317 | 0.426300558 | 8.869727803 | -1.069913929 | 0.324663767 | 0.65295765 | -4.639511549 |
| **A_66_P117389** | Rcn2 | -0.187354966 | -0.61286237 | 0.238152439 | 9.141237963 | -1.069895288 | 0.324671528 | 0.65295765 | -4.639519963 |
| **A_55_P2161640** | Slc38a1 | 0.086932026 | -0.110527728 | 0.28439178 | 8.411854606 | 1.069755179 | 0.324729872 | 0.653007043 | -4.639583203 |
| **A_51_P121228** | Ascc2 | 0.161584039 | -0.205497643 | 0.528665721 | 11.78621819 | 1.069593944 | 0.324797024 | 0.653056704 | -4.639655972 |
| **A_52_P485376** | Ube2k | 0.079779745 | -0.101479501 | 0.261038992 | 12.47826437 | 1.06948738 | 0.324841412 | 0.653056704 | -4.639704064 |
| **A_51_P293269** | Pde7a | -0.382150874 | -1.250440886 | 0.486139137 | 8.494998258 | -1.069432164 | 0.324864414 | 0.653056704 | -4.639728982 |
| **A_52_P316933** | Sh3bgrl2 | 0.193743273 | -0.246507555 | 0.633994101 | 9.843067719 | 1.069324803 | 0.324909142 | 0.653056704 | -4.639777429 |
| **A_55_P1955733** | NA | 0.341413784 | -0.434466099 | 1.117293667 | 12.59089858 | 1.069226584 | 0.324950065 | 0.653056704 | -4.639821749 |
| **A_55_P2104487** | Pld1 | -0.133715952 | -0.43759706 | 0.170165155 | 10.31792694 | -1.069209256 | 0.324957286 | 0.653056704 | -4.639829569 |
| **A_55_P2186929** | NA | 0.20252421 | -0.2578156 | 0.662864019 | 12.34929443 | 1.069009589 | 0.325040493 | 0.65308337 | -4.639919659 |
| **A_52_P476029** | Nol4l | 0.074604921 | -0.094990698 | 0.244200541 | 9.890106308 | 1.068897436 | 0.325087238 | 0.65308337 | -4.639970259 |
| **A_55_P1988795** | Acss2 | -0.220153028 | -0.720631099 | 0.280325043 | 8.951595501 | -1.068864861 | 0.325100816 | 0.65308337 | -4.639984955 |
| **A_51_P149422** | Gng13 | -0.147618084 | -0.483204769 | 0.1879686 | 7.79662096 | -1.068853131 | 0.325105706 | 0.65308337 | -4.639990247 |
| **A_55_P2008692** | Eef1a1 | 0.075706192 | -0.096455779 | 0.247868162 | 15.76115884 | 1.06850698 | 0.325250022 | 0.65330538 | -4.6401464 |
| **A_55_P1957098** | Mtfr1l | -0.097563897 | -0.319494899 | 0.124367105 | 11.35187207 | -1.068204377 | 0.325376225 | 0.653424072 | -4.640282886 |
| **A_55_P1981205** | Xrcc4 | 0.113295141 | -0.144420398 | 0.37101068 | 9.080970008 | 1.068203181 | 0.325376724 | 0.653424072 | -4.640283425 |
| **A_55_P2062793** | Mfap2 | -0.726839481 | -2.380494352 | 0.92681539 | 10.89632859 | -1.068013702 | 0.325455769 | 0.653453749 | -4.640368878 |
| **A_55_P2208071** | Nbeal1 | -0.120107304 | -0.39336918 | 0.153154572 | 8.872436094 | -1.068005679 | 0.325459116 | 0.653453749 | -4.640372496 |
| **A_55_P1965368** | Ctc1 | 0.096498144 | -0.123101604 | 0.316097892 | 11.47957207 | 1.067751798 | 0.325565052 | 0.653463124 | -4.640486981 |
| **A_55_P1978642** | Wasf1 | 0.075572841 | -0.096416614 | 0.247562297 | 8.50436978 | 1.067694775 | 0.32558885 | 0.653463124 | -4.640512693 |
| **A_51_P150678** | Tnfaip8l2 | -0.135227387 | -0.442979649 | 0.172524876 | 8.986438326 | -1.067693475 | 0.325589392 | 0.653463124 | -4.640513279 |
| **A_51_P297586** | Birc6 | 0.149585742 | -0.190850807 | 0.490022292 | 9.699456653 | 1.06767042 | 0.325599014 | 0.653463124 | -4.640523674 |
| **A_55_P2119716** | Mapk1ip1 | 0.115519168 | -0.147425825 | 0.37846416 | 9.273540065 | 1.06751094 | 0.32566558 | 0.653528862 | -4.64059558 |
| **A_55_P2099627** | Lrrc51 | -0.194345651 | -0.636769062 | 0.24807776 | 7.843188168 | -1.067382103 | 0.325719364 | 0.653529105 | -4.640653666 |
| **A_55_P2077055** | Itih4 | -0.280868625 | -0.920279627 | 0.358542377 | 15.82621138 | -1.067348667 | 0.325733323 | 0.653529105 | -4.640668739 |
| **A_51_P419226** | S100a14 | -0.916696631 | -3.004027417 | 1.170634155 | 8.477186689 | -1.067130532 | 0.325824405 | 0.653617026 | -4.640767075 |
| **A_55_P2161775** | Prr3 | 0.142231722 | -0.181654076 | 0.466117521 | 9.205131028 | 1.067057329 | 0.325854975 | 0.653617026 | -4.640800072 |
| **A_51_P390538** | Mpeg1 | -0.179995419 | -0.589897291 | 0.229906454 | 11.59051428 | -1.067000784 | 0.32587859 | 0.653617026 | -4.64082556 |
| **A_55_P1997097** | Bod1 | -0.109321539 | -0.358355782 | 0.139712703 | 8.20185736 | -1.066669458 | 0.326016993 | 0.653706294 | -4.640974893 |
| **A_55_P2053456** | Gm6548 | -0.12028865 | -0.394327261 | 0.153749961 | 16.01777693 | -1.066586498 | 0.326051655 | 0.653706294 | -4.64101228 |
| **A_55_P2163138** | Tm4sf5 | 0.137089961 | -0.175237925 | 0.449417847 | 8.576013973 | 1.066542288 | 0.326070128 | 0.653706294 | -4.641032204 |
| **A_55_P2169391** | Atp6v1a | -0.137250835 | -0.449951838 | 0.175450168 | 11.28233145 | -1.066519767 | 0.326079538 | 0.653706294 | -4.641042353 |
| **A_55_P2062891** | Atf2 | 0.099592906 | -0.127317818 | 0.326503629 | 10.68208056 | 1.066489472 | 0.326092198 | 0.653706294 | -4.641056005 |
| **A_51_P193490** | Fbxo21 | -0.194429192 | -0.637492898 | 0.248634513 | 11.41473291 | -1.066297732 | 0.326172329 | 0.653771597 | -4.641142406 |
| **A_51_P477833** | Klhl21 | -0.107541202 | -0.35261629 | 0.137533886 | 9.310669982 | -1.066249665 | 0.32619242 | 0.653771597 | -4.641164065 |
| **A_51_P236469** | Metap1d | 0.082871901 | -0.106068967 | 0.27181277 | 8.538008584 | 1.065772684 | 0.326391838 | 0.653792444 | -4.641378961 |
| **A_51_P348022** | Nudt21 | 0.093985027 | -0.120298543 | 0.308268597 | 11.96027748 | 1.065744222 | 0.32640374 | 0.653792444 | -4.641391783 |
| **A_55_P2055463** | Snapin | -0.162908292 | -0.534390935 | 0.208574351 | 10.48840487 | -1.065584398 | 0.326470584 | 0.653792444 | -4.641463777 |
| **A_51_P431903** | Pdcd6 | -0.08777785 | -0.287947727 | 0.112392026 | 12.65112825 | -1.065539144 | 0.326489513 | 0.653792444 | -4.641484161 |

| | | | | | | | | | |
|---|---|---|---|---|---|---|---|---|---|
| **A_55_P2423646** | Nf2 | 0.075955244 | -0.097293862 | 0.24920435 | 8.549498961 | 1.065295153 | 0.326591585 | 0.653792444 | -4.641594056 |
| **A_55_P1981366** | Lamc2 | -1.053112619 | -3.455205456 | 1.348980218 | 9.187567342 | -1.065290901 | 0.326593364 | 0.653792444 | -4.641595971 |
| **A_66_P139267** | Rbpj | -0.155838167 | -0.51130392 | 0.199627586 | 9.68561965 | -1.065268916 | 0.326602563 | 0.653792444 | -4.641605872 |
| **A_55_P2166639** | Cdk4 | -0.139236166 | -0.456856155 | 0.178383823 | 11.32894404 | -1.065190831 | 0.326635236 | 0.653792444 | -4.641641039 |
| **A_55_P2024080** | Rpl7a | 0.109715043 | -0.140563794 | 0.35999388 | 15.46333714 | 1.065185485 | 0.326637473 | 0.653792444 | -4.641643447 |
| **A_55_P2085460** | 5530601H04Rik | 0.133704806 | -0.17130177 | 0.438711382 | 8.620589624 | 1.065175096 | 0.32664182 | 0.653792444 | -4.641648125 |
| **A_55_P1992849** | Adrb3 | -0.217828679 | -0.71480723 | 0.279149871 | 7.878217851 | -1.065026963 | 0.326703811 | 0.653792444 | -4.641714835 |
| **A_55_P2009360** | Psg-ps1 | -0.183721822 | -0.602903452 | 0.235459807 | 7.825616558 | -1.064980589 | 0.32672322 | 0.653792444 | -4.641735718 |
| **A_55_P1993109** | Taf6 | 0.097481452 | -0.124936677 | 0.319899581 | 11.8932719 | 1.064964171 | 0.326730091 | 0.653792444 | -4.641743111 |
| **A_52_P507305** | Iah1 | -0.097800886 | -0.320954858 | 0.125353086 | 10.29305834 | -1.064930722 | 0.326744091 | 0.653792444 | -4.641758173 |
| **A_55_P2271742** | NA | 0.11754886 | -0.1506758 | 0.385773521 | 8.217154331 | 1.064885486 | 0.326763025 | 0.653792444 | -4.641778543 |
| **A_55_P2004637** | NA | 0.163628583 | -0.209745732 | 0.537002898 | 11.2092449 | 1.064873256 | 0.326768145 | 0.653792444 | -4.64178405 |
| **A_51_P111488** | Gfod2 | -0.203059075 | -0.666427436 | 0.260309286 | 8.269723009 | -1.064827397 | 0.326787341 | 0.653792444 | -4.6418047 |
| **A_55_P2080603** | Hspa8 | 0.122780478 | -0.157412333 | 0.40297329 | 15.84590232 | 1.064769301 | 0.326811661 | 0.653792444 | -4.641830859 |
| **A_52_P412321** | Pcgf6 | 0.189451908 | -0.242937159 | 0.621840975 | 9.099061953 | 1.064651495 | 0.326860981 | 0.65380235 | -4.641883901 |
| **A_52_P127024** | Tnks | 0.068594153 | -0.087969754 | 0.22515806 | 8.297677844 | 1.064580813 | 0.326890575 | 0.65380235 | -4.641915724 |
| **A_55_P2118931** | Foxj3 | -0.113262049 | -0.371807192 | 0.145283094 | 10.82442955 | -1.064464592 | 0.326939241 | 0.65380235 | -4.641968048 |
| **A_51_P482711** | Dhcr24 | 0.239292659 | -0.30701314 | 0.785598459 | 14.42078023 | 1.064331105 | 0.326995144 | 0.65380235 | -4.642028142 |
| **A_55_P2000022** | Ccdc151 | -0.293851798 | -0.964731086 | 0.37702749 | 7.955626287 | -1.064307585 | 0.327004994 | 0.65380235 | -4.64203873 |
| **A_52_P649296** | Nras | 0.149120815 | -0.191341065 | 0.489582695 | 10.06297763 | 1.064272803 | 0.327019562 | 0.65380235 | -4.642054388 |
| **A_51_P284577** | Hyal2 | 0.108173035 | -0.138832723 | 0.355178793 | 13.21917611 | 1.064131078 | 0.327078927 | 0.653853406 | -4.642118184 |
| **A_51_P103718** | Ip6k1 | 0.116009412 | -0.148940618 | 0.380959442 | 11.00597924 | 1.063928499 | 0.327163798 | 0.653955435 | -4.636480289 |
| **A_55_P1971328** | Lage3 | 0.161575558 | -0.207469566 | 0.530620683 | 11.74624904 | 1.063847512 | 0.327197733 | 0.653955638 | -4.642245816 |
| **A_55_P1991516** | E4f1 | 0.087227236 | -0.112058571 | 0.286513043 | 9.494098781 | 1.063552495 | 0.327321373 | 0.654002658 | -4.642378583 |
| **A_55_P2056739** | Col16a1 | -0.453472332 | -1.489530122 | 0.582585459 | 10.3361749 | -1.06353063 | 0.327330538 | 0.654002658 | -4.642388422 |
| **A_55_P1959338** | Spindoc | -0.216459318 | -0.711036954 | 0.278118317 | 9.266661593 | -1.063469408 | 0.327356201 | 0.654002658 | -4.642415971 |
| **A_55_P2072213** | NA | 0.102462622 | -0.131655415 | 0.336580659 | 11.5851297 | 1.063441928 | 0.327367721 | 0.654002658 | -4.642428337 |
| **A_55_P2383902** | 8030453O22Rik | -0.076220337 | -0.250385947 | 0.097945272 | 7.805709701 | -1.063387746 | 0.327390435 | 0.654002658 | -4.642452717 |
| **A_55_P1966863** | Mad2l2 | -0.114431723 | -0.375936707 | 0.147073261 | 10.68939148 | -1.063284923 | 0.327433544 | 0.654010677 | -4.642498983 |
| **A_55_P2107528** | NA | -0.289163029 | -0.950014918 | 0.371688861 | 10.66436036 | -1.063216768 | 0.327462121 | 0.654010677 | -4.642529648 |
| **A_51_P296878** | Triqk | -0.22334012 | -0.734034374 | 0.287354135 | 8.06086576 | -1.062646856 | 0.32770116 | 0.65435561 | -4.642786032 |
| **A_51_P401958** | Taldo1 | 0.113081479 | -0.145493827 | 0.371656785 | 15.46048608 | 1.062643578 | 0.327702535 | 0.65435561 | -4.642787506 |
| **A_55_P2160291** | Olfm1 | -0.084051883 | -0.276269897 | 0.108166132 | 8.071517421 | -1.062518614 | 0.327754968 | 0.654392705 | -4.642843714 |
| **A_52_P317393** | Adgrg1 | 0.240712545 | -0.309952048 | 0.791377139 | 11.47717327 | 1.062171785 | 0.327900528 | 0.654533134 | -4.642999696 |
| **A_55_P2107120** | Kansl2 | 0.101091404 | -0.130206456 | 0.332389265 | 9.537037897 | 1.062003148 | 0.327971322 | 0.654533134 | -4.643075529 |
| **A_55_P2105242** | Arhgap39 | -0.095691218 | -0.314660298 | 0.123277862 | 10.33174453 | -1.061872753 | 0.328026071 | 0.654533134 | -4.643134161 |
| **A_55_P2335529** | 4921531C22Rik | 0.090423762 | -0.11652457 | 0.297372094 | 8.209033552 | 1.061704925 | 0.328096547 | 0.654533134 | -4.643209619 |
| **A_52_P407854** | Dda1 | 0.118701754 | -0.152987422 | 0.390390929 | 8.461850045 | 1.061617302 | 0.328133348 | 0.654533134 | -4.643249013 |
| **A_55_P2030204** | Supt6 | 0.069152354 | -0.089144126 | 0.227448834 | 12.36755095 | 1.06149732 | 0.328183744 | 0.654533134 | -4.643302953 |
| **A_51_P254471** | Birc2 | 0.078855615 | -0.101666113 | 0.259377343 | 9.046290647 | 1.061417708 | 0.328217188 | 0.654533134 | -4.643338742 |
| **A_51_P413916** | Pgrmc2 | 0.127066663 | -0.163834876 | 0.417968202 | 9.55347266 | 1.061374853 | 0.328235191 | 0.654533134 | -4.643358006 |
| **A_55_P2010622** | Sec16b | 0.27153859 | -0.350139591 | 0.893216771 | 10.59920614 | 1.061326779 | 0.328255388 | 0.654533134 | -4.643379617 |
| **A_55_P2068134** | NA | 0.117780683 | -0.151888187 | 0.387449552 | 8.39954888 | 1.061271352 | 0.328278676 | 0.654533134 | -4.643404532 |
| **A_51_P259214** | Slc39a6 | -0.100715894 | -0.331332641 | 0.129900852 | 9.286455476 | -1.06118319 | 0.32831572 | 0.654533134 | -4.64344416 |
| **A_51_P167426** | Herc1 | 0.169000486 | -0.217977925 | 0.555978898 | 8.463749471 | 1.061168639 | 0.328321834 | 0.654533134 | -4.6434507 |
| **A_55_P1993789** | Hmgn2 | -0.100134653 | -0.329435116 | 0.129165809 | 13.96026325 | -1.061115499 | 0.328344164 | 0.654533134 | -4.643474585 |
| **A_55_P2074246** | Zfp184 | -0.111002626 | -0.365198055 | 0.143192802 | 8.062789856 | -1.061081414 | 0.328358488 | 0.654533134 | -4.643489905 |
| **A_55_P2095271** | Pkn3 | -0.13898541 | -0.457288259 | 0.179317438 | 8.821197077 | -1.060991434 | 0.328396303 | 0.654533134 | -4.643530347 |
| **A_55_P1999829** | Thoc7 | 0.117384485 | -0.15146081 | 0.386229779 | 12.95763939 | 1.060941524 | 0.32841728 | 0.654533134 | -4.643552779 |
| **A_52_P682745** | Dock4 | -0.241950696 | -0.796097856 | 0.312196464 | 8.519544229 | -1.060925659 | 0.328423948 | 0.654533134 | -4.643559909 |

| | | | | | | | | | |
|---|---|---|---|---|---|---|---|---|---|
| **A_55_P2118906** | Atp5j2 | -0.104533751 | -0.34399036 | 0.134922857 | 15.27061132 | -1.060749652 | 0.328497932 | 0.654533134 | -4.643639008 |
| **A_55_P2211319** | Fgf18 | -0.582633257 | -1.91735957 | 0.752093056 | 8.391060038 | -1.060684536 | 0.328525307 | 0.654533134 | -4.643668269 |
| **A_55_P2079471** | NA | 0.101867147 | -0.131513655 | 0.335247949 | 9.182767053 | 1.060601415 | 0.328560254 | 0.654533134 | -4.643705621 |
| **A_51_P359549** | Fem1c | 0.085757461 | -0.110731064 | 0.282245986 | 9.150853076 | 1.060517673 | 0.328595465 | 0.654533134 | -4.64374325 |
| **A_55_P2068496** | NA | 0.316833206 | -0.409131193 | 1.042797605 | 12.39926415 | 1.060470415 | 0.328615337 | 0.654533134 | -4.643764485 |
| **A_51_P330090** | Trappc9 | -0.078492385 | -0.258376421 | 0.101391652 | 10.95995211 | -1.060273941 | 0.328697965 | 0.654533134 | -4.643852762 |
| **A_55_P2069939** | Ubxn7 | -0.08106157 | -0.266844291 | 0.104721151 | 8.332651289 | -1.060212406 | 0.328723847 | 0.654533134 | -4.643880408 |
| **A_55_P1965655** | Mtmr14 | 0.085043741 | -0.109867921 | 0.279955402 | 8.723810048 | 1.060199803 | 0.328729148 | 0.654533134 | -4.64388607 |
| **A_51_P169745** | Tuba1a | -0.523729764 | -1.724136841 | 0.676677313 | 12.59055704 | -1.06013677 | 0.328755662 | 0.654533134 | -4.643914389 |
| **A_51_P442206** | Ankrd42 | -0.120499949 | -0.396701828 | 0.155701929 | 7.822537866 | -1.060091676 | 0.328774632 | 0.654533134 | -4.643934647 |
| **A_55_P2022293** | Zfp579 | -0.346489605 | -1.140748584 | 0.447769374 | 10.79877588 | -1.060013182 | 0.328807654 | 0.654533134 | -4.64396991 |
| **A_55_P2148288** | Tmem79 | -0.135791155 | -0.447074197 | 0.175491887 | 8.191680083 | -1.059983754 | 0.328820035 | 0.654533134 | -4.643983129 |
| **A_55_P2159850** | NA | 0.154307086 | -0.199438025 | 0.508052198 | 13.27756793 | 1.059933485 | 0.328841185 | 0.654533134 | -4.644005711 |
| **A_55_P2096283** | NA | -0.212265279 | -0.698955076 | 0.274424517 | 13.9727867 | -1.059765821 | 0.328911735 | 0.654606149 | -4.644081024 |
| **A_66_P109368** | Prex2 | -0.118787125 | -0.39118321 | 0.153608961 | 7.967976653 | -1.059623782 | 0.328971512 | 0.654657711 | -4.644144822 |
| **A_55_P2086087** | NA | -0.184249645 | -0.606924431 | 0.238425141 | 14.56649759 | -1.0592135 | 0.329144229 | 0.654933992 | -4.644329079 |
| **A_55_P2016681** | Alkbh7 | -0.168225583 | -0.554257793 | 0.217806626 | 10.46337345 | -1.058892053 | 0.3292796 | 0.655078976 | -4.644473414 |
| **A_55_P2120055** | Fgfr4 | -0.17870149 | -0.58877792 | 0.231374941 | 10.70260295 | -1.058879531 | 0.329284874 | 0.655078976 | -4.644479036 |
| **A_55_P2100108** | Zfp148 | -0.073899303 | -0.243541839 | 0.095743233 | 8.248820422 | -1.058494917 | 0.329446909 | 0.65523139 | -4.644651703 |
| **A_65_P10673** | Tmem44 | -0.224088049 | -0.738515189 | 0.290339091 | 8.231386765 | -1.058468681 | 0.329457965 | 0.65523139 | -4.64466348 |
| **A_65_P11718** | Nol8 | 0.126257938 | -0.163589157 | 0.416105034 | 8.437824752 | 1.058456295 | 0.329463184 | 0.65523139 | -4.64466904 |
| **A_51_P287198** | Krt23 | -0.285419124 | -0.940776253 | 0.369938006 | 9.076123023 | -1.058249884 | 0.329550174 | 0.65528336 | -4.644761691 |
| **A_55_P2149575** | Sdr39u1 | -0.069739507 | -0.229873554 | 0.090394541 | 7.999800801 | -1.058225883 | 0.329560289 | 0.65528336 | -4.644772463 |
| **A_51_P138141** | Angel2 | 0.079789477 | -0.103447873 | 0.263026827 | 9.898138978 | 1.058070979 | 0.329625586 | 0.65528336 | -4.644841987 |
| **A_52_P134023** | Rtp3 | 0.247723063 | -0.321205195 | 0.816651321 | 9.789287945 | 1.05801573 | 0.329648877 | 0.65528336 | -4.644866782 |
| **A_51_P423880** | Smarcd3 | -0.328958233 | -1.08452972 | 0.426613255 | 8.221807185 | -1.057909185 | 0.329693798 | 0.65528336 | -4.644914598 |
| **A_55_P2038121** | Rbx1 | -0.123925174 | -0.408601348 | 0.160750999 | 13.74599386 | -1.057770882 | 0.329752115 | 0.65528336 | -4.644976661 |
| **A_51_P173678** | Slc10a6 | -0.16131829 | -0.531907412 | 0.209270832 | 7.74454518 | -1.057728316 | 0.329770065 | 0.65528336 | -4.644995761 |
| **A_55_P2176325** | Prex1 | 0.175434984 | -0.227602508 | 0.578472475 | 9.617764908 | 1.057679149 | 0.3297908 | 0.65528336 | -4.645017824 |
| **A_51_P114826** | Cdh13 | -0.094167538 | -0.310506129 | 0.122171053 | 7.897025499 | -1.057670544 | 0.329794429 | 0.65528336 | -4.645021685 |
| **A_52_P727235** | Trabd2b | -0.151545453 | -0.499775342 | 0.196684435 | 7.795754624 | -1.057450523 | 0.329887231 | 0.655342393 | -4.645120406 |
| **A_52_P662013** | Plg | -0.235058575 | -0.775195446 | 0.305078296 | 14.90081173 | -1.057439339 | 0.329891949 | 0.655342393 | -4.645125424 |
| **A_51_P191354** | NA | -0.209216709 | -0.690071437 | 0.271638018 | 8.82283065 | -1.05722072 | 0.329984182 | 0.655448202 | -4.645223505 |
| **A_52_P14666** | NA | 0.111203628 | -0.144402804 | 0.36681006 | 8.315019948 | 1.057134793 | 0.330020439 | 0.655448202 | -4.645262052 |
| **A_51_P242076** | Ubr7 | 0.087203245 | -0.113264375 | 0.287670865 | 11.17859262 | 1.05699176 | 0.330080799 | 0.655448202 | -4.645326213 |
| **A_55_P2155546** | Nlrc3 | 0.200011703 | -0.259786264 | 0.659809671 | 9.768186965 | 1.056991634 | 0.330080853 | 0.655448202 | -4.64532627 |
| **A_65_P16796** | Psd4 | 0.095797682 | -0.124459028 | 0.316054393 | 8.17740961 | 1.056839512 | 0.330145059 | 0.655508355 | -4.645394504 |
| **A_52_P60194** | C4bp | -0.185405532 | -0.611761318 | 0.240950254 | 10.29864319 | -1.056656224 | 0.330222432 | 0.655550433 | -4.64547671 |
| **A_55_P2093196** | NA | 0.085117865 | -0.110634176 | 0.280869907 | 15.99752338 | 1.056568389 | 0.330259517 | 0.655550433 | -4.645516103 |
| **A_55_P2275437** | Hoxb5os | -0.464501908 | -1.532772652 | 0.603768835 | 8.067123923 | -1.056548303 | 0.330267997 | 0.655550433 | -4.64552511 |
| **A_55_P1959263** | Dnpep | -0.119401466 | -0.39408989 | 0.155286959 | 12.95303811 | -1.056215383 | 0.330408591 | 0.655699024 | -4.645674401 |
| **A_51_P507899** | Ttc8 | -0.159451268 | -0.526277797 | 0.20737526 | 8.650162435 | -1.05621038 | 0.330410704 | 0.655699024 | -4.645676645 |
| **A_55_P2069425** | Ttll3 | -0.074026901 | -0.244354189 | 0.096300386 | 8.317194981 | -1.056059844 | 0.330474292 | 0.655757888 | -4.645744141 |
| **A_55_P2036377** | NA | 0.087496787 | -0.113845605 | 0.288839179 | 16.41030525 | 1.055942019 | 0.330524069 | 0.655789339 | -4.645796967 |
| **A_55_P2048478** | Olfml1 | -0.188707193 | -0.623078785 | 0.245664398 | 8.28038465 | -1.055626371 | 0.330657452 | 0.65586877 | -4.645938471 |
| **A_51_P209319** | Slc35b1 | 0.103731395 | -0.135049832 | 0.342512622 | 12.47285614 | 1.055585041 | 0.330674919 | 0.65586877 | -4.645956998 |
| **A_52_P441036** | Col23a1 | -0.185838372 | -0.613623219 | 0.241946474 | 7.767288071 | -1.05558494 | 0.330674962 | 0.65586877 | -4.645957043 |
| **A_52_P630571** | Epg5 | 0.153832164 | -0.200304685 | 0.507969013 | 9.313989446 | 1.055502386 | 0.330709855 | 0.65586877 | -4.645994047 |
| **A_55_P1984541** | NA | -0.127345243 | -0.42052229 | 0.165831805 | 8.038464252 | -1.055445827 | 0.330733763 | 0.65586877 | -4.646019398 |
| **A_52_P127643** | NA | -0.099033129 | -0.327092955 | 0.129026697 | 9.349474368 | -1.055151695 | 0.330858116 | 0.655991706 | -4.646151225 |

| | | | | | | | | | |
|---|---|---|---|---|---|---|---|---|---|
| **A_66_P119884** | Wdr91 | 0.140108755 | -0.182563736 | 0.462781245 | 9.827989242 | 1.055082746 | 0.330887272 | 0.655991706 | -4.646182125 |
| **A_55_P2068233** | Phactr1 | -0.266015396 | -0.878709791 | 0.346678999 | 7.968784457 | -1.05498444 | 0.330928845 | 0.655991706 | -4.646226179 |
| **A_51_P208870** | Zdhhc24 | 0.139209936 | -0.18142435 | 0.459844221 | 8.733375546 | 1.054978142 | 0.330931508 | 0.655991706 | -4.646229001 |
| **A_51_P379775** | Nrm | 0.099411366 | -0.129634219 | 0.328456951 | 8.864605481 | 1.054623166 | 0.331081662 | 0.656186294 | -4.646388058 |
| **A_55_P1959496** | Sbno2 | 0.101102144 | -0.131847328 | 0.334051616 | 8.229760547 | 1.054585567 | 0.33109757 | 0.656186294 | -4.646404904 |
| **A_52_P641216** | Galk2 | -0.121480259 | -0.401481997 | 0.158521479 | 10.40361115 | -1.054212523 | 0.331255432 | 0.656343987 | -4.646572024 |
| **A_51_P206585** | Runx1 | -0.075517921 | -0.249584839 | 0.098548998 | 8.303167867 | -1.054185336 | 0.33126694 | 0.656343987 | -4.646584203 |
| **A_51_P435817** | Zbtb8a | -0.138086832 | -0.456382441 | 0.180208777 | 8.247041404 | -1.054155812 | 0.331279436 | 0.656343987 | -4.646597427 |
| **A_51_P373901** | Rsph1 | -0.144240175 | -0.476745006 | 0.188264655 | 7.805128225 | -1.054074899 | 0.331313687 | 0.656343987 | -4.646633671 |
| **A_52_P109304** | Tbl2 | 0.137069678 | -0.178964222 | 0.453103578 | 9.125237364 | 1.053879391 | 0.331396457 | 0.656343987 | -4.646721238 |
| **A_52_P330289** | Inpp4b | -0.127526161 | -0.421563647 | 0.166511324 | 7.932166351 | -1.053852369 | 0.331407899 | 0.656343987 | -4.646733341 |
| **A_55_P2030524** | Vldlr | -0.379246647 | -1.253699007 | 0.495205712 | 10.43283109 | -1.053825699 | 0.331419191 | 0.656343987 | -4.646745285 |
| **A_52_P555235** | Rgs19 | -0.101027934 | -0.333989717 | 0.13193385 | 9.429038694 | -1.053755798 | 0.33144879 | 0.656343987 | -4.64677659 |
| **A_55_P2097022** | Eid1 | 0.084301438 | -0.110118736 | 0.278721612 | 8.250729791 | 1.053602625 | 0.331513657 | 0.656379217 | -4.646845186 |
| **A_55_P1959174** | Pmepa1 | -0.389264905 | -1.28704894 | 0.50851913 | 8.198526657 | -1.05355342 | 0.331534497 | 0.656379217 | -4.64686722 |
| **A_52_P365011** | Fes | -0.138163549 | -0.456879392 | 0.180552293 | 9.785569439 | -1.053350774 | 0.331620336 | 0.656477153 | -4.646957961 |
| **A_52_P436318** | Rps25 | 0.086387011 | -0.112904743 | 0.285678765 | 8.884419135 | 1.053276291 | 0.331651891 | 0.656477153 | -4.646991311 |
| **A_55_P2293381** | Cldn18 | -0.225680772 | -0.746387595 | 0.295026051 | 7.799076177 | -1.05313605 | 0.331711311 | 0.656527537 | -4.6470541 |
| **A_52_P234729** | Pkd2 | -0.245860221 | -0.813170858 | 0.321450417 | 9.346895128 | -1.053053726 | 0.331746196 | 0.656529356 | -4.647090957 |
| **A_55_P2097279** | Pirb | -0.178783565 | -0.591386932 | 0.233819802 | 11.37880575 | -1.05287791 | 0.331820707 | 0.656609587 | -4.647169665 |
| **A_55_P2086075** | Gm2058 | 0.149753503 | -0.1959296 | 0.495436607 | 9.236132161 | 1.052645234 | 0.331919337 | 0.656698621 | -4.647273817 |
| **A_51_P315634** | Nedd1 | 0.115482004 | -0.151115817 | 0.382079825 | 9.223178868 | 1.052545563 | 0.331961594 | 0.656698621 | -4.647318429 |
| **A_55_P2119155** | Set | 0.1056667 | -0.13830521 | 0.349638609 | 11.08499978 | 1.052401617 | 0.33202263 | 0.656698621 | -4.647382853 |
| **A_51_P512820** | Dera | 0.187139767 | -0.245011174 | 0.619290708 | 9.512628273 | 1.052237586 | 0.332092194 | 0.656698621 | -4.647456262 |
| **A_55_P2154235** | 1700008J07Rik | 0.213734706 | -0.279900873 | 0.707370286 | 8.947159794 | 1.052087312 | 0.332155934 | 0.656698621 | -4.647523509 |
| **A_55_P2076832** | Nrd1 | 0.095768366 | -0.125416228 | 0.31695296 | 8.865827707 | 1.05208394 | 0.332157364 | 0.656698621 | -4.647525018 |
| **A_55_P1968943** | NA | 0.112040233 | -0.146726274 | 0.370806741 | 16.12462263 | 1.052080884 | 0.332158661 | 0.656698621 | -4.647526385 |
| **A_55_P2000158** | Pik3r3 | -0.299634264 | -0.991682718 | 0.39241419 | 8.375771098 | -1.052054314 | 0.332169932 | 0.656698621 | -4.647538275 |
| **A_52_P659258** | Ppat | 0.140914473 | -0.184561284 | 0.46639023 | 9.10561677 | 1.052010671 | 0.332188446 | 0.656698621 | -4.647557803 |
| **A_55_P2177941** | Tmem208 | -0.098485643 | -0.325970513 | 0.128999226 | 11.78410676 | -1.051970591 | 0.332205449 | 0.656698621 | -4.647575737 |
| **A_51_P155196** | Abtb2 | -0.117613063 | -0.389306725 | 0.1540806 | 9.80945587 | -1.05186315 | 0.332251032 | 0.656721567 | -4.64762381 |
| **A_55_P2114218** | Zcchc7 | 0.115912985 | -0.151963065 | 0.383789035 | 10.14900458 | 1.051432491 | 0.332433797 | 0.65701563 | -4.647816478 |
| **A_55_P1961715** | Nme5 | -0.221212131 | -0.732566089 | 0.290141828 | 7.793208954 | -1.051164064 | 0.332547754 | 0.657106488 | -4.647936546 |
| **A_52_P457411** | Ubl5 | -0.12838001 | -0.425143245 | 0.168383225 | 13.11363462 | -1.051164049 | 0.332547761 | 0.657106488 | -4.647936553 |
| **A_51_P511949** | Setd7 | -0.148878393 | -0.493080023 | 0.195323237 | 9.427350258 | -1.050998121 | 0.332618219 | 0.657152564 | -4.648010764 |
| **A_55_P2007001** | Ctnnbip1 | -0.10486046 | -0.3473055 | 0.137584579 | 9.674430911 | -1.050949011 | 0.332639075 | 0.657152564 | -4.648032728 |
| **A_55_P2129348** | Pilra | -0.12627322 | -0.418298816 | 0.165752375 | 9.230516116 | -1.050687407 | 0.332750191 | 0.6573049 | -4.648149715 |
| **A_66_P100601** | NA | -0.141670041 | -0.46932975 | 0.185989668 | 13.5556993 | -1.050601862 | 0.332786533 | 0.657309513 | -4.648187967 |
| **A_51_P513062** | B630019K06Rik | -0.152620289 | -0.505648291 | 0.200407713 | 8.061697103 | -1.050476415 | 0.332839832 | 0.657347615 | -4.648244059 |
| **A_55_P2127659** | Mtmr2 | -0.076027894 | -0.251924485 | 0.099868696 | 8.289462562 | -1.050264609 | 0.332929837 | 0.657383301 | -4.648338756 |
| **A_55_P2014807** | Rpl4 | 0.084479782 | -0.110972534 | 0.279932097 | 15.49086089 | 1.050255952 | 0.332933517 | 0.657383301 | -4.648342626 |
| **A_51_P337771** | Pcdhb17 | -0.124688451 | -0.413184564 | 0.163807663 | 7.827093532 | -1.050193801 | 0.332959932 | 0.657383301 | -4.648370411 |
| **A_55_P1988678** | Znhit6 | 0.143322363 | -0.188318948 | 0.474963674 | 11.24137658 | 1.050094845 | 0.333001992 | 0.657399194 | -4.648414649 |
| **A_52_P108346** | Myc | 0.24850331 | -0.326590167 | 0.823596787 | 10.29778162 | 1.049970083 | 0.333055028 | 0.657436748 | -4.64847042 |
| **A_52_P18596** | NA | 0.146760198 | -0.193013844 | 0.486534241 | 12.06281813 | 1.049545525 | 0.333235556 | 0.65767214 | -4.648660179 |
| **A_55_P2139992** | Mrps18b | 0.154418926 | -0.203091756 | 0.511929608 | 12.7281549 | 1.049529607 | 0.333242327 | 0.65767214 | -4.648667293 |
| **A_55_P2107292** | Ptch1 | -0.123731807 | -0.410227137 | 0.162763522 | 8.086465217 | -1.049414351 | 0.33329135 | 0.657701736 | -4.6487188 |
| **A_55_P2033665** | NA | 0.353507284 | -0.465187481 | 1.172202048 | 9.882571137 | 1.049203045 | 0.333381242 | 0.657705551 | -4.648813223 |
| **A_55_P1996674** | Itih3 | -0.250225259 | -0.829846158 | 0.32939564 | 15.87496958 | -1.04898747 | 0.333472971 | 0.657705551 | -4.648909544 |
| **A_52_P660242** | Isy1 | 0.085772036 | -0.112916037 | 0.28446011 | 10.85917314 | 1.048955613 | 0.333486528 | 0.657705551 | -4.648923777 |

| | | | | | | | | | |
|---|---|---|---|---|---|---|---|---|---|
| **A_55_P2044953** | Lst1 | 0.135578091 | -0.178499518 | 0.4496557 | 11.73402147 | 1.048903968 | 0.333508507 | 0.657705551 | -4.648946851 |
| **A_52_P222624** | Hspb2 | -0.497711251 | -1.650713513 | 0.65529101 | 8.322892439 | -1.048891313 | 0.333513893 | 0.657705551 | -4.648952505 |
| **A_55_P2120545** | Wdtc1 | 0.11248882 | -0.148107592 | 0.373085231 | 8.468834015 | 1.048875925 | 0.333520442 | 0.657705551 | -4.64895938 |
| **A_55_P2030269** | Dnah6 | -0.329044238 | -1.091341031 | 0.433252555 | 8.24580822 | -1.04885001 | 0.333531471 | 0.657705551 | -4.648970957 |
| **A_55_P1953884** | Dpm1 | 0.106364259 | -0.14009463 | 0.352823148 | 11.01661725 | 1.048659318 | 0.333612641 | 0.657785272 | -4.649056145 |
| **A_55_P2022509** | Mgat5b | 0.105410917 | -0.138853912 | 0.349675746 | 9.129243258 | 1.048595143 | 0.333639961 | 0.657785272 | -4.649084812 |
| **A_55_P2204804** | Dclk1 | -0.121717677 | -0.403795806 | 0.160360452 | 7.79384974 | -1.048497574 | 0.333681501 | 0.657800074 | -4.649128395 |
| **A_51_P312485** | Fpr1 | -0.243301952 | -0.807473841 | 0.320869938 | 9.999803368 | -1.047893963 | 0.333938581 | 0.658239733 | -4.649397972 |
| **A_51_P479808** | Mapk6 | 0.113800946 | -0.150119803 | 0.377721694 | 8.507336362 | 1.04774483 | 0.334002121 | 0.658297849 | -4.649464563 |
| **A_55_P2015934** | NA | 0.08368657 | -0.110428125 | 0.277801264 | 16.70724705 | 1.047563949 | 0.334079202 | 0.658304903 | -4.649545324 |
| **A_55_P2082688** | Ak1 | -0.657889267 | -2.183954294 | 0.86817576 | 9.225653209 | -1.047521512 | 0.334097288 | 0.658304903 | -4.64956427 |
| **A_55_P2028444** | Ube2j2 | 0.089192391 | -0.117706586 | 0.296091369 | 12.59595528 | 1.047496675 | 0.334107874 | 0.658304903 | -4.649575359 |
| **A_55_P2088375** | Tnnt3 | -0.104921563 | -0.348369722 | 0.138526595 | 7.7998906 | -1.047228488 | 0.334222194 | 0.6584193 | -4.649695083 |
| **A_55_P1962299** | Hba-a2 | 0.104925993 | -0.138538914 | 0.3483909 | 17.66137452 | 1.047200653 | 0.334234062 | 0.6584193 | -4.649707509 |
| **A_55_P2062480** | NA | 0.11108706 | -0.146699043 | 0.368873164 | 13.68611054 | 1.047097596 | 0.334278001 | 0.658427334 | -4.64975351 |
| **A_55_P2088416** | Lamtor5 | -0.136175206 | -0.452200295 | 0.179849884 | 12.45208229 | -1.047031304 | 0.334306268 | 0.658427334 | -4.6497831 |
| **A_55_P1990944** | Casp1 | -0.160338833 | -0.53248322 | 0.211805554 | 9.178900639 | -1.046912617 | 0.334356881 | 0.658459924 | -4.649836074 |
| **A_55_P2073248** | Slc25a34 | -0.09139465 | -0.303583697 | 0.120794396 | 7.737274929 | -1.046600656 | 0.334489944 | 0.658523041 | -4.649975297 |
| **A_55_P2347529** | 3010003L21Rik | -0.132133325 | -0.438911844 | 0.174645194 | 8.424438695 | -1.046575601 | 0.334500633 | 0.658523041 | -4.649986477 |
| **A_55_P2052744** | NA | -0.087860364 | -0.291878729 | 0.116158001 | 12.0245643 | -1.046422149 | 0.334566103 | 0.658523041 | -4.650054952 |
| **A_55_P2143832** | Col4a4 | -0.243073463 | -0.807574046 | 0.32142712 | 7.933904074 | -1.046300283 | 0.334618105 | 0.658523041 | -4.650109328 |
| **A_51_P405912** | Lmcd1 | -0.239587439 | -0.796023529 | 0.31684865 | 8.072656295 | -1.046241505 | 0.334643188 | 0.658523041 | -4.650135553 |
| **A_66_P131398** | Ror2 | -0.393001099 | -1.305743877 | 0.519741679 | 8.250122708 | -1.046233323 | 0.33464668 | 0.658523041 | -4.650139203 |
| **A_55_P2111990** | Tekt1 | -0.609142443 | -2.024014701 | 0.805729814 | 8.120150466 | -1.046127982 | 0.334691639 | 0.658523041 | -4.650186202 |
| **A_55_P2006708** | Bmp1 | 0.110378238 | -0.14601663 | 0.366773106 | 9.166970966 | 1.04606174 | 0.334719913 | 0.658523041 | -4.650215755 |
| **A_55_P2073557** | Slc27a5 | -0.301867977 | -1.003094918 | 0.399358964 | 13.01492871 | -1.04602398 | 0.334736031 | 0.658523041 | -4.6502326 |
| **A_51_P209372** | Msmo1 | 0.219457252 | -0.29037824 | 0.729292745 | 14.14719219 | 1.04593106 | 0.334775696 | 0.658523041 | -4.650274052 |
| **A_55_P2105200** | Abraxas1 | 0.107809214 | -0.142651877 | 0.358270306 | 8.493665721 | 1.045920805 | 0.334780074 | 0.658523041 | -4.650278628 |
| **A_51_P419925** | Arhgef40 | -0.11598574 | -0.385467886 | 0.153496405 | 8.302873319 | -1.045822043 | 0.334822238 | 0.658523041 | -4.650322684 |
| **A_51_P316951** | Wipf3 | -0.107522441 | -0.357346261 | 0.142301378 | 8.123374575 | -1.045799577 | 0.33483183 | 0.658523041 | -4.650332705 |
| **A_55_P2080907** | Rpl26 | -0.139338625 | -0.463146886 | 0.184469636 | 15.28107024 | -1.045602914 | 0.334915806 | 0.658621184 | -4.650420426 |
| **A_55_P1993758** | Nsa2 | 0.170227343 | -0.225435599 | 0.565890285 | 11.0776891 | 1.045411164 | 0.3349977 | 0.658674858 | -4.650505947 |
| **A_55_P2036083** | Gm5797 | -0.091507534 | -0.304208326 | 0.121193258 | 7.75912016 | -1.045372169 | 0.335014356 | 0.658674858 | -4.650523337 |
| **A_51_P317181** | Pcdhb21 | -0.148869115 | -0.494925596 | 0.197187366 | 7.904294509 | -1.045299656 | 0.335045331 | 0.658674858 | -4.650555676 |
| **A_51_P196127** | Papss1 | -0.134676588 | -0.447767095 | 0.178413918 | 10.09856164 | -1.04521443 | 0.335081739 | 0.658679441 | -4.650593682 |
| **A_51_P218953** | Zfp536 | -0.221739002 | -0.737280732 | 0.293802727 | 7.773522228 | -1.045108678 | 0.335126921 | 0.658701267 | -4.650640839 |
| **A_55_P2006677** | Gm7969 | 0.21952756 | -0.290996577 | 0.730051698 | 9.951276867 | 1.044854842 | 0.33523539 | 0.658833759 | -4.65075402 |
| **A_52_P467389** | Slc15a3 | -0.102257971 | -0.340079282 | 0.135563339 | 8.691022785 | -1.04479141 | 0.3352625 | 0.658833759 | -4.650782301 |
| **A_66_P128586** | Cwc22 | 0.120514801 | -0.159849176 | 0.400878778 | 9.530130591 | 1.044482974 | 0.335394347 | 0.659025855 | -4.650919804 |
| **A_55_P1997948** | Ppp4r3b | 0.119609317 | -0.158687414 | 0.397906048 | 10.74583219 | 1.044335626 | 0.335457349 | 0.659038612 | -4.650985485 |
| **A_55_P2069217** | Prr13 | 0.139935613 | -0.185706207 | 0.465577432 | 9.011457782 | 1.04417014 | 0.335528118 | 0.659038612 | -4.651059245 |
| **A_55_P2014332** | Cntln | 0.098303426 | -0.130465667 | 0.327072519 | 8.832080928 | 1.044129804 | 0.335545369 | 0.659038612 | -4.651077223 |
| **A_66_P129852** | Fxn | 0.149477413 | -0.198389206 | 0.497344031 | 10.68048858 | 1.0441094 | 0.335554096 | 0.659038612 | -4.651086317 |
| **A_66_P132322** | Mrpl21 | 0.084552601 | -0.112227376 | 0.281332578 | 12.31048127 | 1.044069132 | 0.335571319 | 0.659038612 | -4.651104264 |
| **A_51_P209736** | Atoh8 | -0.15389708 | -0.512226691 | 0.204432531 | 7.808033117 | -1.043592311 | 0.335775315 | 0.659166367 | -4.651316746 |
| **A_55_P2258366** | C030009H01Rik | 0.422018508 | -0.560606068 | 1.404643084 | 12.42980089 | 1.043583167 | 0.335779229 | 0.659166367 | -4.651320821 |
| **A_55_P2106063** | NA | 0.11557896 | -0.153573374 | 0.384731294 | 12.03304667 | 1.043431208 | 0.335844262 | 0.659166367 | -4.651388526 |
| **A_55_P2160726** | Tfcp2 | 0.079448421 | -0.105608369 | 0.264505211 | 8.148309435 | 1.04319005 | 0.335947491 | 0.659166367 | -4.651495963 |
| **A_52_P155805** | 2010204K13Rik | 0.0852365 | -0.1133034 | 0.283776399 | 8.319647225 | 1.043184193 | 0.335949999 | 0.659166367 | -4.651498573 |
| **A_55_P2083098** | Tatdn3 | 0.104271039 | -0.138623102 | 0.347165181 | 9.501092404 | 1.04310938 | 0.335982028 | 0.659166367 | -4.651531899 |

| | | | | | | | | | |
|---|---|---|---|---|---|---|---|---|---|
| **A_52_P59206** | Cst6 | -0.114893304 | -0.382533645 | 0.152747038 | 7.726943304 | -1.043100926 | 0.335985648 | 0.659166367 | -4.651535665 |
| **A_55_P2030225** | NA | 0.099768754 | -0.132639602 | 0.33217711 | 16.1540334 | 1.043100132 | 0.335985988 | 0.659166367 | -4.651536019 |
| **A_55_P2157205** | Cdx1 | -0.148995601 | -0.496088139 | 0.198096937 | 7.82614673 | -1.043064967 | 0.336001044 | 0.659166367 | -4.651551684 |
| **A_51_P139184** | Fank1 | -0.159062583 | -0.529629783 | 0.211504618 | 7.757828325 | -1.042999841 | 0.33602893 | 0.659166367 | -4.651580693 |
| **A_52_P109010** | Hipk3 | 0.099334061 | -0.132102041 | 0.330770162 | 11.61251088 | 1.042918264 | 0.336063862 | 0.659166367 | -4.65161703 |
| **A_66_P124239** | Gm6557 | 0.104108175 | -0.138469798 | 0.346686148 | 8.599687756 | 1.042837545 | 0.336098429 | 0.659166367 | -4.651652983 |
| **A_55_P1954965** | Npm1 | 0.102696787 | -0.136602065 | 0.34199564 | 14.37033879 | 1.042796179 | 0.336116146 | 0.659166367 | -4.651671407 |
| **A_66_P134181** | Aym1 | 0.244178635 | -0.324916797 | 0.813274068 | 8.915804736 | 1.042571266 | 0.336212483 | 0.659166367 | -4.651771575 |
| **A_55_P2154252** | Gfpt2 | -0.078936808 | -0.262913379 | 0.105039763 | 7.892537252 | -1.04255801 | 0.336218162 | 0.659166367 | -4.651777479 |
| **A_55_P2024763** | Lama3 | -0.423790246 | -1.411556958 | 0.563976466 | 8.26640422 | -1.042508871 | 0.336239213 | 0.659166367 | -4.651799361 |
| **A_55_P2184414** | Zfp330 | 0.075433651 | -0.100399686 | 0.251266987 | 11.13691304 | 1.042430475 | 0.3362728 | 0.659166367 | -4.651834273 |
| **A_55_P1981599** | Ccdc125 | 0.126993492 | -0.169037986 | 0.42302497 | 9.351398393 | 1.04238166 | 0.336293715 | 0.659166367 | -4.65185601 |
| **A_55_P1964573** | Arhgef6 | -0.129318422 | -0.430814304 | 0.17217746 | 10.14303492 | -1.042226718 | 0.336360108 | 0.659166367 | -4.651925002 |
| **A_55_P2108143** | NA | 0.309030447 | -0.41146165 | 1.029522544 | 12.3864814 | 1.042210005 | 0.33636727 | 0.659166367 | -4.651932444 |
| **A_52_P657317** | Tmem160 | -0.213615127 | -0.711658105 | 0.284427851 | 10.94562887 | -1.042193527 | 0.336374331 | 0.659166367 | -4.65193978 |
| **A_55_P2010611** | Slc30a5 | 0.090316267 | -0.120261613 | 0.300894146 | 10.43429799 | 1.042164844 | 0.336386624 | 0.659166367 | -4.651952552 |
| **A_55_P2002898** | Ctsj | -0.158532956 | -0.528199343 | 0.211133431 | 7.838668859 | -1.042060137 | 0.336431499 | 0.659187476 | -4.651999171 |
| **A_55_P2046709** | NA | 0.090950763 | -0.121180408 | 0.303081933 | 17.77104422 | 1.041801664 | 0.336542298 | 0.659337733 | -4.652114241 |
| **A_55_P1986441** | Gcdh | 0.28015199 | -0.373381127 | 0.933685108 | 10.95818576 | 1.041619978 | 0.336620198 | 0.659343622 | -4.652195117 |
| **A_51_P259029** | Dusp26 | -0.169436277 | -0.564713825 | 0.225841272 | 7.774690199 | -1.041567549 | 0.33664268 | 0.659343622 | -4.652218455 |
| **A_52_P49378** | Kif1a | -0.157082227 | -0.523543106 | 0.209378652 | 7.840505158 | -1.041555986 | 0.336647639 | 0.659343622 | -4.652223601 |
| **A_52_P667913** | Pcdhga9 | -0.096741829 | -0.322493526 | 0.129009868 | 8.3807083 | -1.041277438 | 0.336767106 | 0.659510778 | -4.652347576 |
| **A_51_P246471** | Pbx2 | -0.110747833 | -0.369213001 | 0.147717336 | 10.49551413 | -1.041157425 | 0.336818589 | 0.659544778 | -4.652400986 |
| **A_55_P2104289** | NA | 0.122177034 | -0.163078715 | 0.407432783 | 8.597467152 | 1.040730689 | 0.337001702 | 0.659836496 | -4.65259087 |
| **A_55_P1983968** | Slc8b1 | 0.135832322 | -0.181376845 | 0.453041489 | 8.496811632 | 1.040496406 | 0.337102267 | 0.659966547 | -4.652695101 |
| **A_52_P439159** | Thap12 | 0.088055561 | -0.117634632 | 0.293745754 | 9.552311626 | 1.04022286 | 0.337219716 | 0.660129622 | -4.652816784 |
| **A_55_P2394308** | Fst | -0.26985602 | -0.900302895 | 0.360590854 | 10.39537751 | -1.040080143 | 0.337281006 | 0.660182448 | -4.652880263 |
| **A_55_P1955715** | Gm14326 | 0.117230277 | -0.156695104 | 0.391155658 | 10.08544134 | 1.039897911 | 0.337359279 | 0.660182448 | -4.652961311 |
| **A_51_P364421** | Vamp7 | 0.160784058 | -0.214970029 | 0.536538146 | 9.578287483 | 1.039734518 | 0.337429472 | 0.660182448 | -4.653033974 |
| **A_52_P370473** | Msl1 | -0.129514613 | -0.432198395 | 0.173169168 | 11.31360696 | -1.039711413 | 0.337439399 | 0.660182448 | -4.653044248 |
| **A_51_P110931** | Znrd1as | 0.085278314 | -0.114024611 | 0.28458124 | 9.460228619 | 1.039700186 | 0.337444223 | 0.660182448 | -4.649855782 |
| **A_66_P108247** | Ucp3 | 0.189823914 | -0.253837267 | 0.633485095 | 11.42902838 | 1.039639284 | 0.33747039 | 0.660182448 | -4.653076323 |
| **A_66_P122219** | Kcnd3 | 0.071872665 | -0.096115708 | 0.239861038 | 8.086190788 | 1.039603451 | 0.337485787 | 0.660182448 | -4.653092256 |
| **A_55_P2055123** | Frrs1 | 0.211711987 | -0.283161666 | 0.70658564 | 10.02631491 | 1.039523482 | 0.33752015 | 0.660182856 | -4.653127815 |
| **A_55_P2051334** | H2-K2 | -0.16831447 | -0.561797492 | 0.225168552 | 13.64998637 | -1.039390266 | 0.337577401 | 0.660228026 | -4.653187047 |
| **A_52_P194316** | Atp13a3 | 0.16916123 | -0.226447037 | 0.564769497 | 9.321704922 | 1.03900746 | 0.337741958 | 0.660463772 | -4.653357231 |
| **A_52_P130727** | Rtl6 | -0.091999548 | -0.307165655 | 0.123166558 | 8.066820116 | -1.038950899 | 0.337766278 | 0.660463772 | -4.653382374 |
| **A_55_P2180056** | NA | 0.373224378 | -0.499752579 | 1.246201335 | 10.21597245 | 1.038844181 | 0.337812167 | 0.660480743 | -4.65342981 |
| **A_55_P1978451** | Ttc1 | 0.111547147 | -0.149381348 | 0.372475643 | 10.87109361 | 1.038771791 | 0.337843298 | 0.660480743 | -4.653461986 |
| **A_55_P2157627** | NA | -0.263489124 | -0.879903368 | 0.352925121 | 9.933214619 | -1.038659544 | 0.337891574 | 0.660482717 | -4.653511876 |
| **A_55_P2146749** | Rps13 | 0.0684663 | -0.091713343 | 0.228645944 | 16.40691358 | 1.038610546 | 0.337912649 | 0.660482717 | -4.653533653 |
| **A_55_P1994388** | Lsm14a | 0.219536907 | -0.294134171 | 0.733207985 | 12.03500515 | 1.038497886 | 0.33796111 | 0.660510647 | -4.653583721 |
| **A_55_P2101441** | Gpbar1 | 0.072888155 | -0.097677587 | 0.243453897 | 7.937328692 | 1.038360929 | 0.33802003 | 0.66055871 | -4.653644584 |
| **A_52_P535012** | NA | 0.105841586 | -0.141857419 | 0.35354059 | 15.23795456 | 1.038281858 | 0.338054051 | 0.66055871 | -4.653679721 |
| **A_51_P369371** | Ubtd1 | -0.158241289 | -0.528625239 | 0.212142661 | 8.931420623 | -1.038127848 | 0.338120324 | 0.660619221 | -4.653748155 |
| **A_55_P2004501** | Immp1l | 0.146627046 | -0.196597676 | 0.489851767 | 12.24692241 | 1.03805105 | 0.338153375 | 0.660619221 | -4.653782277 |
| **A_55_P1984144** | NA | 0.112983444 | -0.151508749 | 0.377475637 | 15.60654163 | 1.037970809 | 0.33818791 | 0.66061992 | -4.653817928 |
| **A_55_P2166421** | Rpusd3 | 0.156087964 | -0.20934373 | 0.521519658 | 10.0368728 | 1.037878209 | 0.338227768 | 0.660631016 | -4.653859069 |
| **A_51_P219444** | Plcl2 | -0.104994202 | -0.350923907 | 0.140935504 | 10.12659504 | -1.037379152 | 0.338442645 | 0.660939844 | -4.654080756 |
| **A_51_P183051** | Upb1 | 0.256026313 | -0.34373515 | 0.855787776 | 13.57685197 | 1.03726383 | 0.338492314 | 0.660939844 | -4.654131975 |

| | | | | | | | | | |
|---|---|---|---|---|---|---|---|---|---|
| **A_55_P2036007** | Rai2 | -0.069890515 | -0.233618145 | 0.093837116 | 7.948997202 | -1.037240462 | 0.33850238 | 0.660939844 | -4.654142353 |
| **A_55_P2055638** | Gsn | -0.146055007 | -0.488223408 | 0.196113394 | 12.70331856 | -1.037193385 | 0.338522658 | 0.660939844 | -4.654163261 |
| **A_66_P138053** | Gm9917 | -0.145794261 | -0.487382787 | 0.195794265 | 7.887104072 | -1.037099305 | 0.338563186 | 0.660952208 | -4.654205042 |
| **A_55_P2020463** | Rufy2 | -0.112813503 | -0.377179965 | 0.151552959 | 8.790229711 | -1.036902486 | 0.338647984 | 0.661050987 | -4.654292443 |
| **A_51_P325374** | Tent2 | 0.092241232 | -0.123946668 | 0.308429133 | 11.38956963 | 1.036756818 | 0.338710754 | 0.661106753 | -4.654357124 |
| **A_51_P220162** | Notch3 | -0.544872907 | -1.822220465 | 0.732474651 | 9.039118094 | -1.036500034 | 0.33882143 | 0.661256 | -4.654471132 |
| **A_55_P2097828** | Gmfg | -0.109592999 | -0.366564018 | 0.14737802 | 12.78853811 | -1.036291304 | 0.338911416 | 0.661333338 | -4.654563793 |
| **A_55_P1966124** | Erh | 0.114467211 | -0.153943598 | 0.382878021 | 14.67881136 | 1.036249394 | 0.338929486 | 0.661333338 | -4.654582397 |
| **A_55_P2082529** | Sav1 | 0.097496614 | -0.131163943 | 0.326157172 | 11.10942323 | 1.036051793 | 0.339014696 | 0.661371869 | -4.654670107 |
| **A_51_P231320** | Mmp8 | -0.361338495 | -1.208860623 | 0.486183632 | 10.70660435 | -1.035968082 | 0.339050799 | 0.661371869 | -4.654707261 |
| **A_55_P2090390** | NA | 0.116195195 | -0.156342161 | 0.388732551 | 9.267236603 | 1.035965567 | 0.339051883 | 0.661371869 | -4.654708377 |
| **A_55_P2178808** | Zbtb5 | 0.079127373 | -0.106486356 | 0.264741102 | 8.886058795 | 1.035857088 | 0.339098673 | 0.661396392 | -4.654756522 |
| **A_51_P139108** | Cpxm1 | -0.263546928 | -0.881876021 | 0.354782166 | 8.699940759 | -1.035670165 | 0.33917931 | 0.661430669 | -4.654839476 |
| **A_55_P2012630** | NA | -0.093521996 | -0.312952903 | 0.12590891 | 15.08151252 | -1.035616976 | 0.339202258 | 0.661430669 | -4.654863079 |
| **A_55_P2008197** | Esr2 | 0.123135108 | -0.165794985 | 0.412065201 | 10.17197057 | 1.035552944 | 0.339229885 | 0.661430669 | -4.654891493 |
| **A_55_P2152592** | Agpat3 | -0.179459961 | -0.600575279 | 0.241655356 | 12.33482727 | -1.035499086 | 0.339253125 | 0.661430669 | -4.654915391 |
| **A_55_P1955457** | Grb10 | -0.180865856 | -0.60533703 | 0.243605319 | 12.13065789 | -1.035360453 | 0.33931295 | 0.661443061 | -4.654976903 |
| **A_51_P342669** | Pgam1 | -0.135566606 | -0.453746466 | 0.182613253 | 14.86364979 | -1.035292883 | 0.339342112 | 0.661443061 | -4.655006883 |
| **A_55_P2130716** | Ube2g2 | 0.186957087 | -0.251857807 | 0.625771981 | 8.984566344 | 1.035246477 | 0.339362142 | 0.661443061 | -4.655027472 |
| **A_55_P2034858** | Pias4 | 0.137032255 | -0.18462992 | 0.458694431 | 8.639273257 | 1.035156469 | 0.339400993 | 0.661452086 | -4.655067405 |
| **A_55_P2183438** | Runx1 | 0.102063387 | -0.137611203 | 0.341737977 | 8.721491407 | 1.034739843 | 0.339580871 | 0.661735927 | -4.655252219 |
| **A_51_P508580** | Tcf15 | -0.192556145 | -0.644844601 | 0.259732311 | 7.845403327 | -1.034487073 | 0.339690042 | 0.661881938 | -4.655364327 |
| **A_51_P165087** | Snai1 | -0.607182901 | -2.033574356 | 0.819208555 | 8.654064744 | -1.034341603 | 0.339752883 | 0.661920191 | -4.655428839 |
| **A_55_P2332731** | D130062J21Rik | -0.168803512 | -0.56540306 | 0.227796036 | 8.655598493 | -1.034218856 | 0.339805916 | 0.661920191 | -4.655483271 |
| **A_55_P2162811** | NA | 0.280955447 | -0.379151805 | 0.941062699 | 15.27342505 | 1.034203828 | 0.339812409 | 0.661920191 | -4.655489935 |
| **A_51_P435198** | Cryz | 0.157364614 | -0.21242134 | 0.527150569 | 12.30625241 | 1.034046001 | 0.339880608 | 0.661986324 | -4.655559916 |
| **A_55_P2146483** | Zmynd8 | 0.110993216 | -0.149866886 | 0.371853319 | 10.21122413 | 1.033884358 | 0.339950468 | 0.662055677 | -4.655631584 |
| **A_51_P123920** | Mydgf | 0.100861469 | -0.136236506 | 0.337959444 | 11.77762169 | 1.033666958 | 0.340044444 | 0.662171978 | -4.654129483 |
| **A_55_P2067505** | Slc16a3 | 0.217211059 | -0.293473406 | 0.727895524 | 12.11551752 | 1.033504747 | 0.340114576 | 0.662195774 | -4.655799868 |
| **A_55_P2051124** | Scaf1 | 0.15172965 | -0.205010077 | 0.508469376 | 13.24909788 | 1.033480218 | 0.340125182 | 0.662195774 | -4.655810741 |
| **A_55_P2127425** | Sys1 | -0.143776238 | -0.48193333 | 0.194380854 | 10.85604067 | -1.033122463 | 0.340279904 | 0.662335177 | -4.655969304 |
| **A_55_P2067153** | NA | -0.116013363 | -0.388881549 | 0.156854823 | 13.76435431 | -1.033090342 | 0.340293799 | 0.662335177 | -4.65598354 |
| **A_55_P2126925** | Msantd3 | -0.239341146 | -0.802288621 | 0.323606329 | 9.371840122 | -1.03307697 | 0.340299583 | 0.662335177 | -4.655989466 |
| **A_55_P2092103** | NA | 0.074877676 | -0.101273947 | 0.251029299 | 8.246981868 | 1.032877691 | 0.340385795 | 0.662336068 | -4.656077774 |
| **A_51_P249118** | Abcb4 | 0.210881522 | -0.285234916 | 0.706997961 | 12.52837092 | 1.032852012 | 0.340396906 | 0.662336068 | -4.656089153 |
| **A_55_P2062418** | Atp6v0a1 | 0.115574376 | -0.156339912 | 0.387488663 | 11.6269661 | 1.032791648 | 0.340423025 | 0.662336068 | -4.656115901 |
| **A_55_P2024302** | Spin1 | -0.078377673 | -0.262784298 | 0.106028953 | 8.347128724 | -1.032759106 | 0.340437107 | 0.662336068 | -4.65613032 |
| **A_55_P2071751** | NA | 0.1409399 | -0.190783168 | 0.472662969 | 9.591529134 | 1.0323845 | 0.340599237 | 0.662528805 | -4.656296289 |
| **A_51_P369784** | Ces1e | 0.364358703 | -0.493224358 | 1.221941764 | 9.816932996 | 1.032371822 | 0.340604725 | 0.662528805 | -4.656301905 |
| **A_51_P214985** | Zfp521 | -0.21092433 | -0.707416419 | 0.28556776 | 8.141275516 | -1.032280048 | 0.340644455 | 0.662539412 | -4.65634256 |
| **A_55_P2003548** | NA | 0.082481509 | -0.111691598 | 0.276654615 | 17.08351207 | 1.032168751 | 0.340692642 | 0.662566463 | -4.656391861 |
| **A_55_P2133365** | 1110017D15Rik | -0.304539613 | -1.021649952 | 0.412570726 | 7.865085966 | -1.031908041 | 0.34080554 | 0.662719343 | -4.656507336 |
| **A_52_P627269** | Ces2b | -0.223124859 | -0.748669978 | 0.302420259 | 8.681538719 | -1.031623299 | 0.340928878 | 0.662782225 | -4.656633438 |
| **A_66_P138047** | Gfm1 | 0.176034241 | -0.238612802 | 0.590681283 | 11.79208103 | 1.031577279 | 0.340948816 | 0.662782225 | -4.656653817 |
| **A_55_P1993719** | Tmem216 | 0.112373343 | -0.152342302 | 0.377088987 | 8.624094044 | 1.031494412 | 0.340984719 | 0.662782225 | -4.656690511 |
| **A_55_P2219765** | Wnt5a | -0.381995979 | -1.281964196 | 0.517972238 | 8.083226319 | -1.031370744 | 0.341038305 | 0.662782225 | -4.656745269 |
| **A_66_P130601** | Foxp2 | -0.138093059 | -0.463441912 | 0.187255794 | 7.802163926 | -1.031349257 | 0.341047616 | 0.662782225 | -4.656754783 |
| **A_55_P2114994** | NA | 0.159539348 | -0.216356026 | 0.535434722 | 14.83782853 | 1.031297634 | 0.341069987 | 0.662782225 | -4.65677764 |
| **A_55_P2133624** | NA | 0.37560363 | -0.509792028 | 1.260999288 | 12.88213639 | 1.030802783 | 0.341284494 | 0.662782225 | -4.656996709 |
| **A_55_P2075631** | Slc20a2 | 0.254815189 | -0.34586258 | 0.855492957 | 11.08657417 | 1.030782274 | 0.341293386 | 0.662782225 | -4.657005786 |

| | | | | | | | | | |
|---|---|---|---|---|---|---|---|---|---|
| **A_52_P504787** | Epha7 | -0.106852168 | -0.358746595 | 0.14504226 | 7.968023118 | -1.030737237 | 0.341312915 | 0.662782225 | -4.657025721 |
| **A_55_P2017038** | Cbx3 | 0.191665341 | -0.260227952 | 0.643558635 | 11.50563356 | 1.030601761 | 0.341371663 | 0.662782225 | -4.657085684 |
| **A_51_P489229** | Eif3g | 0.11055088 | -0.150110043 | 0.371211802 | 14.44935585 | 1.03055094 | 0.341393703 | 0.662782225 | -4.657108177 |
| **A_55_P2095181** | NA | 0.372167437 | -0.505349757 | 1.24968463 | 11.10830586 | 1.030542544 | 0.341397344 | 0.662782225 | -4.657111892 |
| **A_55_P2072706** | Chsy1 | 0.099918337 | -0.135692735 | 0.335529409 | 11.26331297 | 1.030463687 | 0.341431546 | 0.662782225 | -4.657146792 |
| **A_55_P2108151** | NA | 0.148187233 | -0.201255334 | 0.497629799 | 16.90543966 | 1.030429231 | 0.341446492 | 0.662782225 | -4.657162041 |
| **A_51_P322989** | Ptrhd1 | -0.112656271 | -0.378318557 | 0.153006015 | 11.21222316 | -1.030406657 | 0.341456283 | 0.662782225 | -4.657172031 |
| **A_55_P2181976** | Akap11 | 0.115705704 | -0.157160364 | 0.388571772 | 9.199606687 | 1.03035866 | 0.341477103 | 0.662782225 | -4.657193272 |
| **A_52_P354298** | Ap2a2 | 0.156483661 | -0.212574678 | 0.525542 | 9.701500153 | 1.030284498 | 0.341509274 | 0.662782225 | -4.65722609 |
| **A_55_P2145059** | Gbp5 | 0.092091696 | -0.125117956 | 0.309301348 | 8.500601978 | 1.030207102 | 0.341542851 | 0.662782225 | -4.657260339 |
| **A_55_P2074736** | Prkar1b | -0.291378493 | -0.978692751 | 0.395935765 | 8.152161166 | -1.030114093 | 0.341583205 | 0.662782225 | -4.657301494 |
| **A_55_P1960941** | Wac | 0.115767278 | -0.157314323 | 0.38884888 | 11.81018144 | 1.030093319 | 0.341592219 | 0.662782225 | -4.657310687 |
| **A_51_P148060** | Ptgfrn | -0.149308343 | -0.50151344 | 0.202896754 | 7.966907037 | -1.030081596 | 0.341597306 | 0.662782225 | -4.657315874 |
| **A_55_P1964568** | Megf10 | -0.130492075 | -0.438336686 | 0.177352535 | 7.737957782 | -1.029996558 | 0.341634206 | 0.662782225 | -4.6573535 |
| **A_55_P2073841** | NA | 0.102041758 | -0.138757338 | 0.342840854 | 14.42546239 | 1.029689468 | 0.341767486 | 0.662782225 | -4.657489361 |
| **A_55_P2040048** | Swi5 | -0.125117534 | -0.420380072 | 0.170145004 | 13.48377445 | -1.029658042 | 0.341781128 | 0.662782225 | -4.657503263 |
| **A_55_P2106434** | Zfp114 | -0.068733703 | -0.230953465 | 0.09348606 | 7.908996767 | -1.029554086 | 0.341826257 | 0.662782225 | -4.657549249 |
| **A_55_P1964282** | Sbk3 | -0.200508247 | -0.673735553 | 0.272719059 | 7.779599435 | -1.02954571 | 0.341829893 | 0.662782225 | -4.657552954 |
| **A_55_P2059412** | Ghr | -0.184301296 | -0.619309678 | 0.250707087 | 10.21725019 | -1.029470607 | 0.3418625 | 0.662782225 | -4.657586176 |
| **A_55_P1989215** | Entpd8 | -0.228548443 | -0.768010108 | 0.310913222 | 10.22060601 | -1.029439593 | 0.341875966 | 0.662782225 | -4.657599894 |
| **A_55_P2030756** | Nedd4l | -0.231000969 | -0.776273593 | 0.314271655 | 8.024356283 | -1.029397945 | 0.341894049 | 0.662782225 | -4.657618315 |
| **A_55_P2071326** | Rpl36 | -0.126437878 | -0.424904961 | 0.172029205 | 15.26673837 | -1.029352077 | 0.341913966 | 0.662782225 | -4.657638603 |
| **A_51_P196395** | Cfap298 | -0.15647935 | -0.525870902 | 0.212912202 | 11.21431762 | -1.02932676 | 0.341924959 | 0.662782225 | -4.657649801 |
| **A_55_P2005549** | Tnn | -0.09549976 | -0.321019487 | 0.130019968 | 7.690263765 | -1.028965775 | 0.342081743 | 0.662782225 | -4.657809448 |
| **A_51_P343517** | Ly6d | -0.16258177 | -0.546519252 | 0.221355712 | 9.020047258 | -1.028950601 | 0.342088334 | 0.662782225 | -4.657816158 |
| **A_55_P2066096** | Zfand3 | -0.099925117 | -0.335899923 | 0.13604969 | 11.61482781 | -1.028945126 | 0.342090713 | 0.662782225 | -4.657818579 |
| **A_55_P2130890** | Parva | -0.145965823 | -0.490671355 | 0.198739709 | 9.479682262 | -1.028930677 | 0.342096989 | 0.662782225 | -4.657824968 |
| **A_55_P2083023** | Sac3d1 | 0.114253924 | -0.155567931 | 0.384075778 | 11.04022305 | 1.02890953 | 0.342106176 | 0.662782225 | -4.65783432 |
| **A_55_P2110935** | Mlip | -0.071047289 | -0.238842787 | 0.096748208 | 7.695132572 | -1.028846061 | 0.34213375 | 0.662782225 | -4.657862385 |
| **A_55_P2363030** | Cmtm4 | -0.157532895 | -0.529591173 | 0.214525382 | 8.993803415 | -1.028829653 | 0.342140878 | 0.662782225 | -4.657869641 |
| **A_55_P2033927** | NA | 0.06762042 | -0.092102284 | 0.227343124 | 16.24177616 | 1.028713411 | 0.342191384 | 0.662813634 | -4.657921039 |
| **A_51_P290601** | Znhit2 | 0.077099202 | -0.10503656 | 0.259234963 | 11.16525768 | 1.028579449 | 0.342249596 | 0.662859964 | -4.657980269 |
| **A_51_P284665** | Plcb1 | -0.095853546 | -0.322352449 | 0.130645357 | 8.565725537 | -1.02831287 | 0.342365459 | 0.663017931 | -4.658098121 |
| **A_55_P2094118** | Fmnl3 | -0.195711956 | -0.658236009 | 0.266812098 | 8.888775943 | -1.028173066 | 0.342426235 | 0.663056107 | -4.65815992 |
| **A_55_P2043612** | Prdm6 | -0.376470395 | -1.266233838 | 0.513293047 | 7.990437492 | -1.02810971 | 0.34245378 | 0.663056107 | -4.658187925 |
| **A_55_P1972907** | Cab39 | 0.10653832 | -0.145288739 | 0.358365378 | 12.18174775 | 1.027984675 | 0.342508146 | 0.663092739 | -4.65824319 |
| **A_55_P1958464** | NA | 0.225045194 | -0.306956365 | 0.757046753 | 12.43519885 | 1.027874347 | 0.342556124 | 0.663092739 | -4.658291952 |
| **A_55_P1970775** | Hdac1 | 0.092542206 | -0.126234937 | 0.311319349 | 13.4788204 | 1.027829525 | 0.342575616 | 0.663092739 | -4.658311761 |
| **A_51_P464822** | H1f4 | 0.189666649 | -0.258862041 | 0.638195339 | 13.94223677 | 1.027504972 | 0.342716789 | 0.663129476 | -4.658455184 |
| **A_55_P2040888** | Dhx15 | 0.141161512 | -0.192663235 | 0.474986259 | 9.71785957 | 1.027497945 | 0.342719847 | 0.663129476 | -4.65845829 |
| **A_52_P220573** | Serpini1 | 0.07002099 | -0.095570563 | 0.235612543 | 7.805976285 | 1.027479713 | 0.342727778 | 0.663129476 | -4.658466345 |
| **A_55_P2070686** | Wdfy1 | 0.140057415 | -0.191165193 | 0.471280023 | 8.974800872 | 1.027470409 | 0.342731826 | 0.663129476 | -4.658470457 |
| **A_51_P295708** | Ints1 | 0.13624408 | -0.186048634 | 0.458536793 | 9.937393049 | 1.027188958 | 0.342854293 | 0.663179869 | -4.65859481 |
| **A_52_P497388** | NA | 0.14287817 | -0.19517876 | 0.480935101 | 15.15199917 | 1.026973472 | 0.34294808 | 0.663179869 | -4.658690006 |
| **A_52_P654618** | Znhit3 | 0.1359328 | -0.185701053 | 0.457566654 | 9.366120467 | 1.026941485 | 0.342962004 | 0.663179869 | -4.658704136 |
| **A_55_P1970164** | Rnf11 | 0.115393153 | -0.157675245 | 0.388461551 | 10.85035642 | 1.026814012 | 0.343017496 | 0.663179869 | -4.658760444 |
| **A_55_P2076984** | Farp1 | -0.089089943 | -0.299929769 | 0.121749884 | 8.310763192 | -1.026737015 | 0.343051018 | 0.663179869 | -4.658794454 |
| **A_55_P2145109** | Rpn2 | 0.132919039 | -0.181661234 | 0.447499311 | 14.37903394 | 1.02668897 | 0.343071937 | 0.663179869 | -4.658815675 |
| **A_55_P2071532** | NA | -0.108967843 | -0.366867574 | 0.148931889 | 14.74277095 | -1.026669483 | 0.343080422 | 0.663179869 | -4.658824282 |
| **A_55_P2046411** | NA | -0.144799708 | -0.487525902 | 0.197926486 | 15.10472418 | -1.026605479 | 0.343108292 | 0.663179869 | -4.65885255 |

| | | | | | | | | | |
|---|---|---|---|---|---|---|---|---|---|
| **A_51_P441426** | Pf4 | -0.159362649 | -0.53656444 | 0.217839142 | 14.45740603 | -1.026587529 | 0.343116108 | 0.663179869 | -4.658860478 |
| **A_55_P2244142** | Klf13 | -0.199804607 | -0.67275699 | 0.273147777 | 9.789180057 | -1.026529106 | 0.343141549 | 0.663179869 | -4.658886281 |
| **A_52_P304720** | Crlf1 | -0.130233259 | -0.438515631 | 0.178049114 | 7.728572815 | -1.026493972 | 0.34315685 | 0.663179869 | -4.658901797 |
| **A_55_P2072945** | NA | -0.109375654 | -0.368299896 | 0.149548589 | 16.33804512 | -1.026434251 | 0.343182859 | 0.663179869 | -4.658928172 |
| **A_55_P2037398** | NA | 0.136884957 | -0.187177128 | 0.460947043 | 15.44130453 | 1.026385934 | 0.343203903 | 0.663179869 | -4.658949509 |
| **A_51_P450278** | 2010003K11Rik | -0.139223465 | -0.468898344 | 0.190451414 | 7.841934937 | -1.026147456 | 0.343307783 | 0.663278071 | -4.659054817 |
| **A_65_P16952** | Ywhaq | -0.15543756 | -0.52355145 | 0.21267633 | 11.86487158 | -1.026022662 | 0.343362153 | 0.663278071 | -4.659109919 |
| **A_55_P2120714** | NA | 0.146351142 | -0.200269323 | 0.492971606 | 8.480843924 | 1.02594742 | 0.343394938 | 0.663278071 | -4.659143139 |
| **A_51_P173285** | Nkx2-5 | -0.109029059 | -0.367344157 | 0.14928604 | 7.772156667 | -1.025594445 | 0.34354877 | 0.663278071 | -4.659298966 |
| **A_55_P2037912** | Cyp2d10 | -0.281000025 | -0.946775545 | 0.384775495 | 13.24044477 | -1.025561538 | 0.343563114 | 0.663278071 | -4.659313491 |
| **A_55_P2117630** | Snrnp25 | 0.202365713 | -0.277141343 | 0.681872769 | 12.5091896 | 1.025475098 | 0.343600796 | 0.663278071 | -4.659351647 |
| **A_55_P2357860** | Atxn7l1 | 0.085658487 | -0.117312378 | 0.288629352 | 8.579567084 | 1.02546274 | 0.343606184 | 0.663278071 | -4.659357101 |
| **A_52_P15300** | Hoxb2 | -0.308624504 | -1.040013654 | 0.422764647 | 8.292687027 | -1.025333341 | 0.3436626 | 0.663278071 | -4.659414215 |
| **A_51_P462746** | Tex10 | 0.08775832 | -0.120233356 | 0.295749996 | 9.615295389 | 1.02523998 | 0.343703308 | 0.663278071 | -4.659455421 |
| **A_51_P354077** | Svil | -0.157912608 | -0.532174847 | 0.216349631 | 11.14868375 | -1.025236323 | 0.343704903 | 0.663278071 | -4.659457035 |
| **A_51_P368903** | Anapc13 | -0.132003421 | -0.444881636 | 0.180874795 | 14.32124405 | -1.025163295 | 0.343736748 | 0.663278071 | -4.659489264 |
| **A_55_P2151011** | NA | -0.104597113 | -0.352532385 | 0.143338158 | 7.844879482 | -1.025096081 | 0.34376606 | 0.663278071 | -4.659518927 |
| **A_51_P306527** | Ywhah | 0.096253646 | -0.131909646 | 0.324416938 | 13.91617385 | 1.025072552 | 0.343776322 | 0.663278071 | -4.659529311 |
| **A_51_P235801** | Zfp36l1 | -0.244186807 | -0.823072805 | 0.334699192 | 12.0660624 | -1.024972795 | 0.343819831 | 0.663278071 | -4.659573332 |
| **A_55_P2325803** | Tnfrsf11a | -0.099031328 | -0.333804451 | 0.135741796 | 9.125202884 | -1.024961168 | 0.343824902 | 0.663278071 | -4.659578463 |
| **A_55_P1958140** | B9d1 | -0.626833474 | -2.112900765 | 0.859233816 | 9.120347478 | -1.024936347 | 0.343835729 | 0.663278071 | -4.659589416 |
| **A_51_P199608** | Ttll1 | -0.087976721 | -0.296562686 | 0.120609245 | 8.474370514 | -1.024863143 | 0.34386766 | 0.663278071 | -4.659621718 |
| **A_55_P2024260** | Vps33b | 0.08701649 | -0.119330803 | 0.293363784 | 8.907762457 | 1.02467461 | 0.343949911 | 0.663278071 | -4.659704905 |
| **A_51_P327295** | Akr1a1 | -0.105658361 | -0.356245005 | 0.144928283 | 13.26989577 | -1.024540517 | 0.34400842 | 0.663278071 | -4.659764066 |
| **A_55_P2094686** | Smim10l1 | 0.131495182 | -0.180390224 | 0.443380588 | 9.028329803 | 1.024467004 | 0.3440405 | 0.663278071 | -4.659796498 |
| **A_55_P2025712** | Immt | 0.108542054 | -0.148924407 | 0.366008515 | 10.19645033 | 1.024378752 | 0.344079014 | 0.663278071 | -4.65983543 |
| **A_55_P2045658** | Nme6 | -0.147703437 | -0.498075926 | 0.202669052 | 10.66288316 | -1.024339196 | 0.344096278 | 0.663278071 | -4.659852879 |
| **A_51_P167048** | Hs6st1 | 0.116976736 | -0.160513487 | 0.394466959 | 9.230287462 | 1.024318464 | 0.344105327 | 0.663278071 | -4.659862025 |
| **A_55_P1966528** | Pmf1 | 0.150448062 | -0.206453394 | 0.507349518 | 10.7890794 | 1.024286542 | 0.34411926 | 0.663278071 | -4.659876106 |
| **A_65_P19112** | Zfp397 | 0.069369602 | -0.095205622 | 0.233944826 | 8.25847856 | 1.024207464 | 0.344153777 | 0.663278071 | -4.659910988 |
| **A_55_P2153797** | Slc38a6 | 0.142304969 | -0.195338986 | 0.479948924 | 9.788193387 | 1.024104502 | 0.344198723 | 0.663278071 | -4.659956403 |
| **A_55_P1989539** | Cklf | 0.092748251 | -0.127317101 | 0.312813603 | 8.319646571 | 1.024087926 | 0.344205959 | 0.663278071 | -4.659963715 |
| **A_55_P2094520** | Fam171a1 | -0.430206633 | -1.45098712 | 0.590573854 | 9.826506377 | -1.024065962 | 0.344215548 | 0.663278071 | -4.659973402 |
| **A_51_P372473** | Auh | -0.074317114 | -0.250720615 | 0.102086386 | 11.31679617 | -1.023681434 | 0.344383456 | 0.663535469 | -4.660142987 |
| **A_51_P379597** | Ndufs1 | 0.140224831 | -0.192694709 | 0.473144371 | 12.97949571 | 1.023455182 | 0.344482282 | 0.663659726 | -4.660242753 |
| **A_55_P2127204** | Amigo1 | -0.088937478 | -0.300124676 | 0.122249719 | 7.996996249 | -1.023293966 | 0.344552714 | 0.66366493 | -4.660313834 |
| **A_51_P299934** | BC048403 | 0.143365074 | -0.19706432 | 0.483794469 | 9.199656393 | 1.023291816 | 0.344553654 | 0.66366493 | -4.660314782 |
| **A_55_P2125982** | Lrrc4b | -0.325823489 | -1.099682519 | 0.448035541 | 7.960600392 | -1.02306617 | 0.344652254 | 0.663788703 | -4.66041426 |
| **A_55_P2120354** | Cib2 | -0.245207621 | -0.82768537 | 0.337270127 | 9.692648798 | -1.022910918 | 0.344720107 | 0.66383824 | -4.660482697 |
| **A_55_P2174987** | Creb3l3 | -0.224385824 | -0.757434153 | 0.308662505 | 12.92314953 | -1.022850163 | 0.344746663 | 0.66383824 | -4.660509477 |
| **A_55_P1973387** | Srrm1 | 0.126820305 | -0.174572337 | 0.428212947 | 12.95133615 | 1.022443545 | 0.344924439 | 0.663972261 | -4.660688686 |
| **A_52_P94830** | Galnt16 | -0.469532155 | -1.585511746 | 0.646447436 | 8.302809993 | -1.022332756 | 0.344972889 | 0.663972261 | -4.660737508 |
| **A_55_P2048119** | Slc29a4 | -0.117196442 | -0.395751239 | 0.161358355 | 7.836174717 | -1.022320277 | 0.344978347 | 0.663972261 | -4.660743007 |
| **A_51_P254234** | Chchd4 | 0.107848954 | -0.148496182 | 0.36419409 | 9.850120227 | 1.022289844 | 0.344991657 | 0.663972261 | -4.660756417 |
| **A_55_P2052881** | NA | 0.22516671 | -0.310041663 | 0.760375084 | 9.481867353 | 1.022267308 | 0.345001514 | 0.663972261 | -4.660766348 |
| **A_51_P126835** | Arl14ep | 0.068791908 | -0.094730145 | 0.232313961 | 8.497382573 | 1.022219623 | 0.34502237 | 0.663972261 | -4.66078736 |
| **A_55_P1988754** | H4f16 | -0.21552575 | -0.727955736 | 0.296904236 | 14.64275575 | -1.021992758 | 0.345121611 | 0.664085904 | -4.660887317 |
| **A_51_P258698** | Phf8 | 0.118277457 | -0.162954752 | 0.399509666 | 10.44737565 | 1.021927558 | 0.345150137 | 0.664085904 | -4.660916043 |
| **A_51_P387188** | Vrk3 | -0.103979238 | -0.351234984 | 0.143276508 | 11.11919811 | -1.021841241 | 0.345187904 | 0.664092465 | -4.66095407 |
| **A_52_P527106** | Arhgap12 | -0.08113035 | -0.274127004 | 0.111866303 | 8.933307308 | -1.021449175 | 0.345359492 | 0.664217482 | -4.661126773 |

| | | | | | | | | | |
|---|---|---|---|---|---|---|---|---|---|
| **A_55_P1985498** | Gm12005 | 0.100166253 | -0.138155348 | 0.338487854 | 14.59957445 | 1.021271494 | 0.345437276 | 0.664217482 | -4.661205029 |
| **A_51_P461724** | Atp6ap2 | 0.117998928 | -0.162761432 | 0.398759288 | 10.6974078 | 1.021234458 | 0.345453492 | 0.664217482 | -4.66122134 |
| **A_52_P243391** | Sema4f | -0.205415137 | -0.694247821 | 0.283417546 | 7.764870397 | -1.021069722 | 0.345525624 | 0.664217482 | -4.661293887 |
| **A_55_P2083889** | Pea15a | -0.150718866 | -0.509403874 | 0.207966143 | 9.747829044 | -1.021027832 | 0.345543968 | 0.664217482 | -4.661312333 |
| **A_55_P2115136** | Vsir | -0.165459427 | -0.559229841 | 0.228310987 | 9.114439266 | -1.021013825 | 0.345550103 | 0.664217482 | -4.661318501 |
| **A_55_P1992004** | NA | -0.09338426 | -0.315637314 | 0.128868793 | 10.7954327 | -1.020960961 | 0.345573254 | 0.664217482 | -4.661341779 |
| **A_52_P110291** | Phf12 | 0.077812845 | -0.107386948 | 0.263012639 | 8.82086103 | 1.020925352 | 0.345588849 | 0.664217482 | -4.661357459 |
| **A_55_P2124751** | Col16a1 | -0.271407289 | -0.917375435 | 0.374560857 | 8.690851584 | -1.020924424 | 0.345589256 | 0.664217482 | -4.661357868 |
| **A_55_P2017914** | Csf1 | 0.122102836 | -0.168515094 | 0.412720766 | 8.299267028 | 1.020907828 | 0.345596524 | 0.664217482 | -4.661365175 |
| **A_55_P2020143** | Slc39a3 | -0.092666022 | -0.313260704 | 0.12792866 | 9.066792766 | -1.020724805 | 0.345676692 | 0.664305506 | -4.661445761 |
| **A_51_P372463** | Fxr1 | 0.097990127 | -0.135306272 | 0.331286525 | 11.5214872 | 1.020604487 | 0.345729401 | 0.664330886 | -4.661498732 |
| **A_55_P1991728** | Zfp956 | 0.096091543 | -0.132699638 | 0.324882724 | 9.426976177 | 1.020537757 | 0.345758638 | 0.664330886 | -4.66152811 |
| **A_52_P511821** | Wdfy1 | 0.092238251 | -0.127422607 | 0.311899109 | 7.991733041 | 1.020332153 | 0.345848731 | 0.66443794 | -4.661618619 |
| **A_52_P171791** | Smg1 | 0.144193345 | -0.199315289 | 0.487701979 | 10.11035765 | 1.019977868 | 0.346004018 | 0.664627421 | -4.661774555 |
| **A_51_P209782** | Cyp2c23 | 0.317830775 | -0.439375407 | 1.075036957 | 11.9231366 | 1.019917363 | 0.346030544 | 0.664627421 | -4.661801183 |
| **A_55_P2080243** | NA | 0.16330995 | -0.225783989 | 0.552403889 | 9.732975912 | 1.019861903 | 0.346054859 | 0.664627421 | -4.66182559 |
| **A_51_P335480** | 1810055G02Rik | -0.252182098 | -0.853111623 | 0.348747427 | 9.818534739 | -1.019703476 | 0.346124326 | 0.664627421 | -4.661895307 |
| **A_55_P1962084** | H2ac18 | -0.284218026 | -0.961487184 | 0.393051131 | 12.69588933 | -1.019702561 | 0.346124727 | 0.664627421 | -4.66189571 |
| **A_51_P283473** | Fibin | -0.893869007 | -3.024142281 | 1.236404268 | 8.737809521 | -1.019581037 | 0.346178021 | 0.664627421 | -4.661949183 |
| **A_55_P2115713** | Nid1 | -0.086721294 | -0.29340057 | 0.119957983 | 8.020625511 | -1.019558162 | 0.346188053 | 0.664627421 | -4.661959248 |
| **A_55_P2023703** | Zfp386 | -0.101006571 | -0.341762423 | 0.13974928 | 8.015656501 | -1.019426613 | 0.346245751 | 0.664672175 | -4.662017128 |
| **A_51_P185757** | Casz1 | -0.214771009 | -0.726765456 | 0.297223438 | 8.066786956 | -1.019280219 | 0.34630997 | 0.664729434 | -4.662081535 |
| **A_55_P2079054** | Ndufv1 | 0.097902615 | -0.135537314 | 0.331342543 | 14.09072892 | 1.019066064 | 0.34640393 | 0.664745745 | -4.662175744 |
| **A_55_P1953356** | Triobp | 0.154604887 | -0.214044582 | 0.523254356 | 9.829120919 | 1.019043664 | 0.346413759 | 0.664745745 | -4.662185598 |
| **A_55_P2114402** | Notum | 0.124499899 | -0.172377677 | 0.421377475 | 10.49831016 | 1.019001428 | 0.346432293 | 0.664745745 | -4.662204176 |
| **A_55_P2113733** | NA | -0.067312436 | -0.22784268 | 0.093217808 | 10.37444581 | -1.018876698 | 0.346487031 | 0.664745745 | -4.66225904 |
| **A_55_P2035018** | Clec4n | 0.206418371 | -0.285862771 | 0.698699514 | 11.37682895 | 1.01886897 | 0.346490423 | 0.664745745 | -4.662262439 |
| **A_52_P359502** | Pdxk | -0.168411224 | -0.570099449 | 0.233277001 | 11.69780171 | -1.018744134 | 0.346545215 | 0.664781536 | -4.662317344 |
| **A_51_P314186** | Syne1 | -0.106427321 | -0.360305316 | 0.147450674 | 8.282033757 | -1.018617795 | 0.346600674 | 0.664781536 | -4.662372908 |
| **A_65_P03442** | Pgk1 | -0.388033434 | -1.313776861 | 0.537709992 | 14.04639748 | -1.018501563 | 0.346651702 | 0.664781536 | -4.662424023 |
| **A_55_P1978366** | NA | 0.140154517 | -0.194235878 | 0.474544912 | 11.43882912 | 1.018442451 | 0.346677656 | 0.664781536 | -4.662450017 |
| **A_51_P356467** | Cldn4 | -0.097525624 | -0.330210805 | 0.135159556 | 7.787017982 | -1.018434736 | 0.346681043 | 0.664781536 | -4.662453409 |
| **A_66_P120472** | Pglyrp2 | -0.189908627 | -0.643072034 | 0.263254779 | 8.658640321 | -1.018293685 | 0.34674298 | 0.664834347 | -4.662515432 |
| **A_51_P278550** | Tecr | -0.12000483 | -0.406454434 | 0.166444773 | 11.68553334 | -1.017966978 | 0.346886474 | 0.665019298 | -4.662659074 |
| **A_52_P637730** | Ubl4a | 0.117068695 | -0.16238601 | 0.396523401 | 12.83459216 | 1.017917406 | 0.346908251 | 0.665019298 | -4.662680867 |
| **A_55_P2099961** | H2ac11 | 0.11175243 | -0.155084708 | 0.378589568 | 17.35095781 | 1.017639351 | 0.34703042 | 0.665187523 | -4.662803094 |
| **A_51_P378381** | Crnde | -0.222034592 | -0.75235576 | 0.308286577 | 8.020455581 | -1.017337062 | 0.347163275 | 0.665360517 | -4.662935954 |
| **A_52_P123864** | Rab21 | -0.118968186 | -0.403159479 | 0.165223107 | 8.460354918 | -1.017192763 | 0.347226709 | 0.665360517 | -4.662999367 |
| **A_55_P2006008** | Serpinb1a | -0.175153753 | -0.593587623 | 0.243280117 | 11.49780966 | -1.017128457 | 0.347254981 | 0.665360517 | -4.663027626 |
| **A_52_P168567** | Cebpa | -0.261960145 | -0.887823021 | 0.363902731 | 10.90786417 | -1.017042776 | 0.347292653 | 0.665360517 | -4.663065275 |
| **A_55_P2036907** | Mycbpap | -0.123633054 | -0.419011429 | 0.171745321 | 7.732085123 | -1.017042474 | 0.347292786 | 0.665360517 | -4.663065408 |
| **A_51_P507023** | Nt5dc3 | -0.199557798 | -0.676415537 | 0.277299942 | 8.914750505 | -1.016864431 | 0.347371077 | 0.665401064 | -4.663143637 |
| **A_55_P2082241** | NA | -0.137854196 | -0.467275366 | 0.191566975 | 13.39493166 | -1.016837776 | 0.3473828 | 0.665401064 | -4.663155348 |
| **A_55_P1981544** | Lactb | 0.153458047 | -0.213315072 | 0.520231167 | 10.02997534 | 1.016659114 | 0.34746138 | 0.665485634 | -4.663233841 |
| **A_55_P2080378** | Ttc30a2 | -0.076719227 | -0.260102456 | 0.106664002 | 7.826428751 | -1.016547768 | 0.347510361 | 0.665492378 | -4.663282755 |
| **A_51_P112237** | Angel1 | -0.10042671 | -0.340490886 | 0.139637467 | 8.293634678 | -1.016494577 | 0.347533761 | 0.665492378 | -4.663306121 |
| **A_55_P1981011** | Slc35c2 | 0.145319047 | -0.202103093 | 0.492741186 | 10.33646934 | 1.016361577 | 0.347592277 | 0.665538496 | -4.663364543 |
| **A_51_P201338** | Mtss1 | -0.098062449 | -0.332527276 | 0.136402378 | 9.094398763 | -1.016267931 | 0.347633483 | 0.665551465 | -4.663405675 |
| **A_55_P2119100** | Yipf1 | -0.140460746 | -0.476382384 | 0.195460893 | 11.95859055 | -1.01601514 | 0.347744736 | 0.665647339 | -4.663516698 |
| **A_66_P112005** | Olfr1252 | 0.078716842 | -0.109586156 | 0.26701984 | 8.182119247 | 1.015765873 | 0.347854466 | 0.665647339 | -4.66362616 |

| | | | | | | | | | |
|---|---|---|---|---|---|---|---|---|---|
| **A_52_P680038** | Cdk16 | 0.094720839 | -0.131874084 | 0.321315762 | 8.48690982 | 1.015730631 | 0.347869982 | 0.665647339 | -4.663641634 |
| **A_66_P133204** | NA | -0.106698144 | -0.361950019 | 0.148553732 | 8.45483307 | -1.015713234 | 0.347877641 | 0.665647339 | -4.663649273 |
| **A_55_P2179604** | Vash1 | -0.116890502 | -0.396538556 | 0.162757552 | 14.28895368 | -1.015665288 | 0.347898752 | 0.665647339 | -4.663670325 |
| **A_52_P219904** | Afmid | 0.163330828 | -0.227460367 | 0.554122023 | 8.885601445 | 1.01556233 | 0.347944088 | 0.665647339 | -4.66371553 |
| **A_51_P249127** | Pkd1 | -0.207039627 | -0.702464826 | 0.288385571 | 9.679278482 | -1.01545007 | 0.347993525 | 0.665647339 | -4.663764817 |
| **A_55_P2139390** | NA | -0.130849456 | -0.444006126 | 0.182307214 | 16.37814287 | -1.015297798 | 0.348060592 | 0.665647339 | -4.663831666 |
| **A_51_P141104** | Med18 | 0.109548244 | -0.152654135 | 0.371750623 | 8.410730538 | 1.015200873 | 0.348103287 | 0.665647339 | -4.663874214 |
| **A_55_P2119478** | Ndufv2 | 0.09843381 | -0.137170053 | 0.334037672 | 13.74811002 | 1.015184731 | 0.348110397 | 0.665647339 | -4.663881299 |
| **A_51_P374101** | Pard6b | -0.071851737 | -0.243834031 | 0.100130556 | 7.745767146 | -1.015165216 | 0.348118994 | 0.665647339 | -4.663889866 |
| **A_52_P676956** | Tirap | -0.216763309 | -0.735616823 | 0.302090204 | 9.473243582 | -1.01513593 | 0.348131896 | 0.665647339 | -4.663902721 |
| **A_51_P358354** | Jam3 | -0.339562474 | -1.152454278 | 0.473329331 | 8.82938426 | -1.01500956 | 0.348187571 | 0.665647339 | -4.663958189 |
| **A_55_P2094986** | Atp1a3 | 0.100158901 | -0.139638701 | 0.339956503 | 8.391942408 | 1.014910867 | 0.348231057 | 0.665647339 | -4.664001506 |
| **A_52_P119117** | Tmem258 | -0.102624004 | -0.348328769 | 0.143080762 | 13.19368723 | -1.014889043 | 0.348240675 | 0.665647339 | -4.664011085 |
| **A_55_P2112599** | Lgals8 | 0.108547745 | -0.151394949 | 0.368490439 | 9.779357634 | 1.014673641 | 0.348335603 | 0.665647339 | -4.664105618 |
| **A_66_P100249** | Snhg12 | 0.106325538 | -0.14830226 | 0.360953336 | 11.20996129 | 1.01464697 | 0.348347358 | 0.665647339 | -4.664117322 |
| **A_55_P2083609** | Dkk3 | -0.653797228 | -2.219554665 | 0.911960209 | 9.318010429 | -1.014616173 | 0.348360933 | 0.665647339 | -4.664130836 |
| **A_55_P2068459** | Hspa1a | -0.06891683 | -0.233970021 | 0.096136361 | 7.715447429 | -1.014575934 | 0.34837867 | 0.665647339 | -4.664148494 |
| **A_52_P432969** | Pcdh19 | 0.120937266 | -0.168709398 | 0.41058393 | 8.488276813 | 1.014553149 | 0.348388713 | 0.665647339 | -4.664158492 |
| **A_52_P279687** | Lrp8 | 0.071009787 | -0.099066636 | 0.241086209 | 7.710039191 | 1.014512225 | 0.348406753 | 0.665647339 | -4.66417645 |
| **A_55_P1991600** | Aplp2 | -0.095452486 | -0.324189491 | 0.133284518 | 13.14485589 | -1.013990775 | 0.348636681 | 0.665871106 | -4.664405231 |
| **A_55_P2121543** | Rps20 | 0.078438485 | -0.109539313 | 0.266416284 | 16.99748103 | 1.013924994 | 0.348665695 | 0.665871106 | -4.664434087 |
| **A_55_P2139615** | Rtl6 | 0.131275823 | -0.183345764 | 0.445897409 | 8.715077799 | 1.013863342 | 0.34869289 | 0.665871106 | -4.664461131 |
| **A_52_P470956** | NA | -0.168450976 | -0.57219122 | 0.235289269 | 16.11048236 | -1.013805582 | 0.348718369 | 0.665871106 | -4.664486467 |
| **A_51_P409039** | Echs1 | -0.132490464 | -0.450068574 | 0.185087646 | 13.29394946 | -1.013718216 | 0.348756911 | 0.665871106 | -4.664524788 |
| **A_55_P2091153** | BC051142 | 0.119356536 | -0.166749736 | 0.405462807 | 8.798999704 | 1.013682658 | 0.348772599 | 0.665871106 | -4.664540384 |
| **A_51_P178347** | Zfp326 | 0.123204323 | -0.172152686 | 0.418561331 | 9.316799881 | 1.013588919 | 0.348813958 | 0.665871106 | -4.664581497 |
| **A_51_P150521** | Mgat2 | -0.071879046 | -0.244220835 | 0.100462743 | 9.073749458 | -1.013432659 | 0.348882911 | 0.665871106 | -4.664650027 |
| **A_55_P2433438** | Tcf12 | 0.082154468 | -0.114825855 | 0.279134792 | 9.010366892 | 1.013424808 | 0.348886375 | 0.665871106 | -4.66465347 |
| **A_55_P2105833** | Chchd5 | 0.072626199 | -0.101509913 | 0.24676231 | 9.750632725 | 1.013415775 | 0.348890362 | 0.665871106 | -4.664657431 |
| **A_55_P2135759** | NA | 0.068653317 | -0.09596949 | 0.233276124 | 16.36245726 | 1.013338897 | 0.34892429 | 0.665871106 | -4.664691144 |
| **A_55_P2042071** | Slc25a3 | 0.09106481 | -0.127304531 | 0.30943415 | 13.69893955 | 1.013309495 | 0.348937267 | 0.665871106 | -4.664704038 |
| **A_55_P2134877** | Eml1 | -0.110046279 | -0.373957902 | 0.153865344 | 7.893007882 | -1.013211319 | 0.3489806 | 0.665888057 | -4.664747088 |
| **A_55_P1988358** | Lrrc47 | 0.151279834 | -0.211647287 | 0.514206955 | 11.48317445 | 1.012849235 | 0.349140454 | 0.666100853 | -4.664905843 |
| **A_55_P2055042** | Uqcc2 | 0.154854568 | -0.216665605 | 0.526374741 | 13.72230132 | 1.012802604 | 0.349161045 | 0.666100853 | -4.664926286 |
| **A_55_P2052769** | Mast2 | 0.110053735 | -0.154039031 | 0.374146501 | 10.96114672 | 1.012584957 | 0.349257165 | 0.666211679 | -4.665021695 |
| **A_55_P1999408** | NA | 0.135799166 | -0.190096687 | 0.461695019 | 12.01051545 | 1.012514983 | 0.349288073 | 0.666211679 | -4.665052367 |
| **A_51_P212560** | Cog7 | 0.096812169 | -0.135547334 | 0.329171673 | 9.496822551 | 1.012401316 | 0.349338284 | 0.666241706 | -4.665102188 |
| **A_66_P139664** | Ppp2r5c | 0.102937608 | -0.144155822 | 0.350031039 | 8.541773727 | 1.012269296 | 0.34939661 | 0.666274406 | -4.66516005 |
| **A_55_P2059640** | Abhd14b | -0.207612315 | -0.706025537 | 0.290800908 | 12.32839084 | -1.012154346 | 0.3494474 | 0.666274406 | -4.665210427 |
| **A_52_P50329** | Stimate | -0.181644039 | -0.617726585 | 0.254438507 | 8.827665243 | -1.012128455 | 0.34945884 | 0.666274406 | -4.665221773 |
| **A_66_P127262** | NA | 0.377633334 | -0.529091093 | 1.284357761 | 9.768981631 | 1.011994602 | 0.349517992 | 0.666301111 | -4.665280429 |
| **A_55_P2059650** | Atpaf2 | 0.152674184 | -0.213937328 | 0.519285696 | 10.51981803 | 1.011911887 | 0.349554549 | 0.666301111 | -4.665316674 |
| **A_52_P84803** | NA | -0.080976132 | -0.275430904 | 0.113478639 | 16.23175798 | -1.011862763 | 0.349576262 | 0.666301111 | -4.665338199 |
| **A_51_P106259** | Camk1 | -0.09893019 | -0.3365694 | 0.138709021 | 8.012530253 | -1.011565316 | 0.349707754 | 0.666406841 | -4.658833582 |
| **A_51_P113906** | Gkn2 | -0.080969245 | -0.275471768 | 0.113533278 | 7.693409282 | -1.011528304 | 0.349724119 | 0.666406841 | -4.658849167 |
| **A_55_P2181888** | Tmem54 | 0.160778197 | -0.225449158 | 0.547005553 | 8.488330059 | 1.011503323 | 0.349735164 | 0.666406841 | -4.665495678 |
| **A_52_P299311** | NA | 0.136644871 | -0.191668739 | 0.464958482 | 15.39989508 | 1.011317768 | 0.349817218 | 0.666464797 | -4.665576962 |
| **A_55_P2179335** | Usp33 | 0.087260266 | -0.122406165 | 0.296926697 | 10.96893136 | 1.0112786 | 0.34983454 | 0.666464797 | -4.665594119 |
| **A_55_P1999641** | Frem1 | -0.650937621 | -2.215267697 | 0.913392456 | 9.054573114 | -1.01110013 | 0.349913479 | 0.666540649 | -4.66567229 |
| **A_55_P1955427** | Cux1 | -0.163601422 | -0.556793787 | 0.229590944 | 8.291453819 | -1.011032661 | 0.349943324 | 0.666540649 | -4.665701839 |

| | | | | | | | | | |
|---|---|---|---|---|---|---|---|---|---|
| **A_51_P518246** | Fibp | -0.127297663 | -0.433266127 | 0.178670801 | 9.726288989 | -1.010943662 | 0.349982696 | 0.666549959 | -4.665740817 |
| **A_51_P232742** | Mfsd14b | 0.146624324 | -0.205957585 | 0.499206233 | 10.26373525 | 1.01048341 | 0.350186365 | 0.666872143 | -4.665942357 |
| **A_55_P2056186** | Siva1 | 0.161568576 | -0.227008241 | 0.550145394 | 12.69452262 | 1.010329886 | 0.350254324 | 0.666915282 | -4.666009573 |
| **A_55_P1957733** | Cysrt1 | 0.086855159 | -0.122053376 | 0.295763694 | 7.827408852 | 1.010235497 | 0.35029611 | 0.666915282 | -4.666050895 |
| **A_52_P645159** | Cpsf6 | 0.102159659 | -0.143578928 | 0.347898246 | 8.903673953 | 1.010157902 | 0.350330465 | 0.666915282 | -4.666084863 |
| **A_55_P1971574** | Acox3 | 0.342217923 | -0.481097915 | 1.165533761 | 10.99314147 | 1.009995572 | 0.350402345 | 0.666915282 | -4.666155921 |
| **A_51_P250807** | Spata2l | -0.117714991 | -0.400942947 | 0.165512964 | 8.412050767 | -1.009901102 | 0.350444182 | 0.666915282 | -4.666197271 |
| **A_55_P2080151** | Hspa2 | -0.168861282 | -0.575202426 | 0.237479861 | 8.474921468 | -1.009770022 | 0.350502238 | 0.666915282 | -4.666254642 |
| **A_55_P2252184** | NA | 0.085771353 | -0.120626144 | 0.29216885 | 8.179639408 | 1.009766657 | 0.350503729 | 0.666915282 | -4.666256115 |
| **A_55_P2033650** | Cpt2 | 0.152877848 | -0.215003769 | 0.520759465 | 11.98260676 | 1.009763487 | 0.350505133 | 0.666915282 | -4.666257502 |
| **A_55_P2025233** | Llph | 0.139915009 | -0.196835953 | 0.47666597 | 9.922614254 | 1.009575141 | 0.350588567 | 0.666915282 | -4.66633993 |
| **A_55_P1998539** | C1qtnf1 | -0.11078867 | -0.377451887 | 0.155874548 | 8.568249871 | -1.00952115 | 0.350612487 | 0.666915282 | -4.666363557 |
| **A_55_P2040227** | Ptchd4 | 0.142152435 | -0.200026415 | 0.484331286 | 9.745783441 | 1.009448879 | 0.350644507 | 0.666915282 | -4.666395182 |
| **A_55_P1961761** | Dcc | 0.22342772 | -0.314424112 | 0.761279551 | 11.97846084 | 1.009386729 | 0.350672046 | 0.666915282 | -4.666422378 |
| **A_55_P2097372** | Tctex1d4 | -0.347509805 | -1.184149629 | 0.489130019 | 7.917159501 | -1.00928012 | 0.350719288 | 0.666915282 | -4.666469025 |
| **A_66_P139799** | Cenpb | 0.111631462 | -0.157126429 | 0.380389353 | 8.576747829 | 1.009272816 | 0.350722524 | 0.666915282 | -4.666472221 |
| **A_51_P517381** | Cers2 | 0.08280661 | -0.116559813 | 0.282173033 | 10.92316354 | 1.009244008 | 0.350735291 | 0.666915282 | -4.666484826 |
| **A_51_P509211** | Rnf34 | -0.069479542 | -0.236781073 | 0.097821989 | 10.31252077 | -1.00911397 | 0.350792924 | 0.666915282 | -4.66654172 |
| **A_55_P2019483** | Ndrg4 | -0.290775084 | -0.990943852 | 0.409393684 | 8.27725204 | -1.009107988 | 0.350795575 | 0.666915282 | -4.666544337 |
| **A_51_P328932** | Tmem165 | -0.132668538 | -0.452176201 | 0.186839124 | 11.42137624 | -1.00895049 | 0.350865389 | 0.666982406 | -4.66661324 |
| **A_51_P346923** | P3h1 | -0.092575838 | -0.315578102 | 0.130426427 | 8.285788371 | -1.008722171 | 0.350966615 | 0.667109225 | -4.666713115 |
| **A_55_P2015292** | Ltc4s | 0.252162542 | -0.355415581 | 0.859740664 | 9.71786227 | 1.008466868 | 0.351079832 | 0.667258808 | -4.666824779 |
| **A_51_P412640** | Ccdc51 | 0.101203592 | -0.142687216 | 0.3450944 | 9.624625546 | 1.008285875 | 0.351160113 | 0.667315308 | -4.666903933 |
| **A_51_P401024** | Lrrc34 | -0.218417751 | -0.744840102 | 0.3080046 | 7.794143905 | -1.008177032 | 0.351208399 | 0.667315308 | -4.666951529 |
| **A_55_P2077497** | NA | 0.13427257 | -0.189349616 | 0.457894755 | 11.7604426 | 1.008166363 | 0.351213132 | 0.667315308 | -4.666956194 |
| **A_55_P1988952** | NA | -0.182039928 | -0.620869538 | 0.256789683 | 14.83120274 | -1.007984651 | 0.351293756 | 0.667372589 | -4.667035648 |
| **A_55_P2128326** | Ubap2l | 0.123515114 | -0.17424561 | 0.421275839 | 9.336840834 | 1.007942787 | 0.351312333 | 0.667372589 | -4.667053952 |
| **A_55_P2008567** | Rps6 | -0.099082521 | -0.338024778 | 0.139859735 | 11.81612137 | -1.007597957 | 0.351465379 | 0.667560583 | -4.667204705 |
| **A_66_P117695** | Zpr1 | 0.111794147 | -0.15781184 | 0.381400133 | 12.20199034 | 1.007564189 | 0.351480369 | 0.667560583 | -4.667219466 |
| **A_51_P110759** | Slc1a1 | -0.097683924 | -0.333291644 | 0.137923795 | 7.695545281 | -1.007434388 | 0.351537994 | 0.66760443 | -4.658874563 |
| **A_55_P1971579** | Acox3 | -0.089698997 | -0.306183385 | 0.12678539 | 8.334034777 | -1.006802294 | 0.35181872 | 0.668047573 | -4.667552443 |
| **A_55_P2012780** | Rnf167 | 0.083332624 | -0.117810389 | 0.284475637 | 11.38787546 | 1.006684299 | 0.351871144 | 0.668047573 | -4.667603999 |
| **A_52_P79639** | Vat1 | -0.135638013 | -0.463035302 | 0.191759277 | 8.725213052 | -1.006675566 | 0.351875024 | 0.668047573 | -4.667607814 |
| **A_55_P2273705** | NA | 0.118854149 | -0.168114185 | 0.405822483 | 9.034112516 | 1.006383617 | 0.352004761 | 0.668190057 | -4.667735361 |
| **A_55_P2022805** | Rubcn | -0.159291964 | -0.543908124 | 0.225324196 | 11.15966278 | -1.006351104 | 0.352019212 | 0.668190057 | -4.667749564 |
| **A_55_P2142151** | Mapk1ip1 | -0.131195564 | -0.448015763 | 0.185624635 | 9.732502826 | -1.006211963 | 0.352081059 | 0.66824183 | -4.667810343 |
| **A_52_P407871** | Lsm12 | 0.106085636 | -0.15012665 | 0.362297923 | 10.96142391 | 1.006096929 | 0.352132198 | 0.668273269 | -4.667860589 |
| **A_55_P2160451** | Stk26 | 0.115760073 | -0.163872918 | 0.395393064 | 8.459342922 | 1.00589714 | 0.352221028 | 0.668319838 | -4.667947846 |
| **A_51_P439857** | Dnajc21 | 0.088767675 | -0.125672415 | 0.303207766 | 9.323091711 | 1.005847301 | 0.352243191 | 0.668319838 | -4.667969612 |
| **A_55_P2074771** | Exosc7 | 0.082130466 | -0.116288443 | 0.280549376 | 12.167732 | 1.005783296 | 0.352271653 | 0.668319838 | -4.667997563 |
| **A_55_P2125049** | Gm11127 | -0.154092896 | -0.526388203 | 0.218202411 | 12.48615739 | -1.00572268 | 0.352298611 | 0.668319838 | -4.668024034 |
| **A_55_P2060854** | NA | -0.119829266 | -0.409362186 | 0.169703653 | 12.26644118 | -1.005652968 | 0.352329616 | 0.668319838 | -4.668054475 |
| **A_51_P451516** | Arhgef25 | -0.40637284 | -1.388626865 | 0.575881184 | 10.8928252 | -1.005273066 | 0.352498619 | 0.668519 | -4.668220348 |
| **A_51_P412338** | Vps41 | 0.077106112 | -0.109271165 | 0.263483388 | 9.729227303 | 1.005261457 | 0.352503785 | 0.668519 | -4.668225416 |
| **A_52_P442676** | Cux1 | -0.280094802 | -0.957255734 | 0.39706613 | 10.89602876 | -1.00507008 | 0.352588946 | 0.668537728 | -4.668308962 |
| **A_55_P2077872** | Deaf1 | 0.119687284 | -0.169670846 | 0.409045415 | 9.783536466 | 1.005068154 | 0.352589803 | 0.668537728 | -4.668309802 |
| **A_55_P2175451** | NA | 0.085752256 | -0.121576742 | 0.293081254 | 12.8021266 | 1.005006097 | 0.352617422 | 0.668537728 | -4.668336891 |
| **A_55_P2143953** | Pank2 | 0.111870433 | -0.158643711 | 0.382384577 | 9.79889403 | 1.00486688 | 0.352679387 | 0.66858963 | -4.668397659 |
| **A_55_P2118491** | Gm10046 | -0.146829297 | -0.501959879 | 0.208301284 | 7.75218639 | -1.004633918 | 0.352783096 | 0.66866763 | -4.668499334 |
| **A_55_P2054883** | Trappc1 | 0.099935332 | -0.141778132 | 0.341648796 | 9.314677732 | 1.004619041 | 0.35278972 | 0.66866763 | -4.668505827 |

| | | | | | | | | | |
|---|---|---|---|---|---|---|---|---|---|
| **A_55_P2162369** | Cacfd1 | -0.099107839 | -0.338892448 | 0.140676771 | 8.384270179 | -1.00431486 | 0.352925173 | 0.668814813 | -4.668638564 |
| **A_55_P1967677** | Cped1 | -0.107180953 | -0.366504521 | 0.152142615 | 8.236732484 | -1.004289255 | 0.352936577 | 0.668814813 | -4.668649737 |
| **A_51_P301930** | Lrrc17 | -0.654116382 | -2.236971459 | 0.928738696 | 8.904976689 | -1.004146459 | 0.35300018 | 0.668869766 | -4.668712041 |
| **A_66_P133450** | Dnajb9 | -0.075562539 | -0.2584915 | 0.107366423 | 8.071804686 | -1.003707708 | 0.353195664 | 0.669174572 | -4.668903444 |
| **A_51_P310398** | Tk2 | -0.090194186 | -0.308581644 | 0.128193272 | 8.575068035 | -1.003538498 | 0.353271077 | 0.669251853 | -4.668977249 |
| **A_55_P2108655** | Mtch2 | -0.070006185 | -0.239533244 | 0.099520873 | 13.38460551 | -1.003414967 | 0.353326141 | 0.669290571 | -4.669031125 |
| **A_55_P2115442** | Clec2d | -0.141169156 | -0.483083935 | 0.200745624 | 9.449272794 | -1.003240678 | 0.353403842 | 0.669326789 | -4.669107133 |
| **A_51_P260035** | Sar1b | -0.149049288 | -0.510058537 | 0.211959961 | 11.37905289 | -1.003216733 | 0.353414517 | 0.669326789 | -4.669117575 |
| **A_55_P2004126** | NA | 0.123719657 | -0.175962847 | 0.423402161 | 10.77180073 | 1.003137595 | 0.353449804 | 0.669328035 | -4.669152084 |
| **A_51_P381558** | Rasa4 | -0.10796476 | -0.369558553 | 0.153629032 | 8.594309148 | -1.002854159 | 0.353576206 | 0.669344624 | -4.669275668 |
| **A_55_P2055284** | Chrm2 | -0.066426539 | -0.227382541 | 0.094529464 | 7.707269455 | -1.002807641 | 0.353596954 | 0.669344624 | -4.669295948 |
| **A_55_P2095603** | Ccdc65 | -0.132580966 | -0.453842374 | 0.188680442 | 7.717692173 | -1.002780337 | 0.353609133 | 0.669344624 | -4.669307852 |
| **A_55_P2014269** | Zfp771 | -0.114830754 | -0.393092507 | 0.163430999 | 12.30543693 | -1.002738958 | 0.353627591 | 0.669344624 | -4.669325892 |
| **A_51_P312336** | Slc14a1 | 0.217926799 | -0.310226582 | 0.746080181 | 10.24005085 | 1.002614001 | 0.353683335 | 0.669344624 | -4.669380366 |
| **A_52_P176737** | Lcorl | -0.088133959 | -0.301744514 | 0.125476596 | 8.322855669 | -1.002544732 | 0.353714239 | 0.669344624 | -4.669410561 |
| **A_51_P425737** | Tbc1d7 | 0.132295532 | -0.188420605 | 0.453011668 | 8.63641909 | 1.002322673 | 0.353813325 | 0.669344624 | -4.669507353 |
| **A_55_P2146780** | Sdhc | 0.106444655 | -0.151622511 | 0.364511821 | 12.91202967 | 1.002245918 | 0.353847579 | 0.669344624 | -4.669540806 |
| **A_55_P2106426** | NA | -0.111907853 | -0.383274792 | 0.159459086 | 14.92344225 | -1.002044058 | 0.353937678 | 0.669344624 | -4.669628779 |
| **A_55_P1969660** | Pex26 | -0.124796012 | -0.427424684 | 0.17783266 | 11.20532839 | -1.002014081 | 0.353951059 | 0.669344624 | -4.669641842 |
| **A_55_P2100899** | Arnt | 0.121380307 | -0.172981574 | 0.415742187 | 8.635070452 | 1.001958767 | 0.353975752 | 0.669344624 | -4.669665946 |
| **A_51_P260871** | Sdhd | 0.121102941 | -0.172604754 | 0.414810636 | 14.33802287 | 1.001895795 | 0.354003865 | 0.669344624 | -4.669693387 |
| **A_55_P1953972** | Pdhb | 0.075648125 | -0.107843683 | 0.259139934 | 13.09208872 | 1.001762289 | 0.354063473 | 0.669344624 | -4.66975156 |
| **A_55_P2099019** | NA | 0.100947124 | -0.143957223 | 0.345851471 | 11.13611582 | 1.001568473 | 0.354150023 | 0.669344624 | -4.669836005 |
| **A_55_P1999202** | Vmn1r217 | 0.281616877 | -0.401636634 | 0.964870388 | 9.273371215 | 1.00152085 | 0.354171292 | 0.669344624 | -4.669856752 |
| **A_52_P182520** | Timm9 | 0.083602024 | -0.119237197 | 0.286441244 | 8.703040993 | 1.001493296 | 0.354183598 | 0.669344624 | -4.669868756 |
| **A_55_P2013038** | Serpinb6c | -0.121535165 | -0.416476552 | 0.173406223 | 10.92898763 | -1.001265897 | 0.354285174 | 0.669344624 | -4.669967817 |
| **A_51_P221248** | Tspan32 | 0.210093556 | -0.299762361 | 0.719949473 | 8.831467532 | 1.001263667 | 0.35428617 | 0.669344624 | -4.669968788 |
| **A_51_P318551** | Nle1 | 0.130620625 | -0.186397998 | 0.447639248 | 8.70054223 | 1.001175404 | 0.354325602 | 0.669344624 | -4.670007234 |
| **A_55_P1994000** | Snx16 | 0.085506924 | -0.122025096 | 0.293038944 | 9.15778315 | 1.001150482 | 0.354336737 | 0.669344624 | -4.670018089 |
| **A_52_P128068** | Kazn | -0.07847922 | -0.268972365 | 0.112013925 | 7.768839608 | -1.001056323 | 0.354378808 | 0.669344624 | -4.670059101 |
| **A_55_P2096055** | NA | 0.099906533 | -0.142607306 | 0.342420372 | 13.32527137 | 1.001014923 | 0.354397307 | 0.669344624 | -4.670077132 |
| **A_51_P142515** | Antxr2 | -0.116206662 | -0.398310765 | 0.16589744 | 10.32853676 | -1.000932553 | 0.354434115 | 0.669344624 | -4.670113006 |
| **A_55_P1953402** | Mlana | 0.068231563 | -0.097425607 | 0.233888734 | 7.807655419 | 1.000825291 | 0.354482052 | 0.669344624 | -4.670159719 |
| **A_55_P2017851** | NA | -0.145716566 | -0.499499085 | 0.208065954 | 14.10372974 | -1.000819471 | 0.354484653 | 0.669344624 | -4.670162253 |
| **A_66_P112853** | Eif1a | -0.141090798 | -0.483645264 | 0.201463669 | 11.66874791 | -1.0008114 | 0.35448826 | 0.669344624 | -4.670165768 |
| **A_51_P196509** | Plekhj1 | 0.174684063 | -0.249439345 | 0.598807471 | 9.026425783 | 1.000792935 | 0.354496513 | 0.669344624 | -4.670173809 |
| **A_55_P1994694** | Serinc2 | -0.091688211 | -0.314302405 | 0.130925983 | 7.764209045 | -1.000792024 | 0.35449692 | 0.669344624 | -4.670174206 |
| **A_55_P2113291** | Sik3 | -0.135352161 | -0.463991276 | 0.193286953 | 11.18679313 | -1.000758144 | 0.354512063 | 0.669344624 | -4.67018896 |
| **A_55_P2037525** | NA | -0.161607428 | -0.554005116 | 0.230790259 | 14.64169092 | -1.00073265 | 0.354523458 | 0.669344624 | -4.670200062 |
| **A_55_P2108903** | Zbtb20 | -0.078956401 | -0.27069453 | 0.112781728 | 8.13141656 | -1.000603569 | 0.354581157 | 0.669344624 | -4.67025627 |
| **A_51_P490678** | Trim68 | -0.083426101 | -0.286028148 | 0.119175946 | 8.039053158 | -1.000555782 | 0.35460252 | 0.669344624 | -4.670277078 |
| **A_55_P1969441** | Eef1akmt2 | 0.087439785 | -0.124932466 | 0.299812036 | 9.63602192 | 1.000447953 | 0.354650728 | 0.669344624 | -4.670324027 |
| **A_55_P1965571** | Rpl8 | -0.114290429 | -0.391877467 | 0.163296608 | 15.19965303 | -1.000446506 | 0.354651375 | 0.669344624 | -4.670324657 |
| **A_66_P110088** | NA | 0.18519619 | -0.264651029 | 0.635043408 | 9.408534254 | 1.0003459 | 0.354696358 | 0.669344624 | -4.67036846 |
| **A_51_P341177** | Ddah2 | -0.481418356 | -1.650859769 | 0.688023057 | 11.65911263 | -1.000293313 | 0.354719873 | 0.669344624 | -4.670391354 |
| **A_51_P312437** | Dhrs7 | -0.194726601 | -0.667769462 | 0.278316259 | 14.16008261 | -1.000248667 | 0.354739838 | 0.669344624 | -4.670410791 |
| **A_51_P418737** | Rbm33 | 0.085574156 | -0.122352662 | 0.293500974 | 8.380401165 | 1.000035244 | 0.354835288 | 0.669367882 | -4.670503698 |
| **A_55_P2339601** | NA | 0.29500501 | -0.421837327 | 1.011847346 | 9.115907394 | 0.999974519 | 0.354862451 | 0.669367882 | -4.670530131 |
| **A_55_P2101503** | NA | 0.079564905 | -0.113775351 | 0.272905161 | 8.355129663 | 0.999959588 | 0.35486913 | 0.669367882 | -4.67053663 |
| **A_51_P277718** | Cenpc1 | 0.153230259 | -0.219171759 | 0.525632277 | 9.776596315 | 0.99980591 | 0.354937878 | 0.669367882 | -4.670603519 |

| | | | | | | | | | |
|---|---|---|---|---|---|---|---|---|---|
| A_55_P2135311 | Nol3 | -0.065150736 | -0.223509543 | 0.093208071 | 7.722501132 | -0.999678365 | 0.354994944 | 0.669367882 | -4.67065903 |
| A_51_P499698 | Asprv1 | -0.188307097 | -0.646025816 | 0.269411622 | 8.913943245 | -0.999657443 | 0.355004305 | 0.669367882 | -4.670668135 |
| A_55_P2181191 | Btg1 | 0.069451005 | -0.099385829 | 0.23828784 | 7.964167995 | 0.999526991 | 0.355062681 | 0.669367882 | -4.670724906 |
| A_51_P320304 | Plod1 | 0.102927035 | -0.147291811 | 0.353145882 | 8.20584462 | 0.999522641 | 0.355064627 | 0.669367882 | -4.670726799 |
| A_51_P446469 | Dok2 | -0.140848417 | -0.483266032 | 0.201569198 | 10.92729115 | -0.9994914 | 0.355078608 | 0.669367882 | -4.670740394 |
| A_52_P88818 | Clybl | 0.103457787 | -0.148070361 | 0.354985935 | 8.42463625 | 0.999447028 | 0.355098466 | 0.669367882 | -4.670759703 |
| A_51_P345110 | Thap4 | -0.103226073 | -0.354304573 | 0.147852427 | 10.15835893 | -0.998994435 | 0.35530107 | 0.66966553 | -4.670956623 |
| A_66_P130914 | NA | 0.132545759 | -0.189881749 | 0.454973266 | 15.51559907 | 0.998888329 | 0.355348581 | 0.66966553 | -4.671002782 |
| A_51_P345663 | Rrbp1 | -0.155852961 | -0.534986925 | 0.223281003 | 12.76700149 | -0.998862147 | 0.355360305 | 0.66966553 | -4.671014171 |
| A_55_P1987914 | Map4k4 | -0.215980883 | -0.741461361 | 0.309499596 | 9.329978865 | -0.998715784 | 0.355425853 | 0.669723759 | -4.671077837 |
| A_55_P2168301 | Slc46a1 | -0.174399962 | -0.59879762 | 0.249997696 | 10.53395925 | -0.998519604 | 0.355513726 | 0.669754507 | -4.671163165 |
| A_55_P2015403 | 9930111J21Rik1 | 0.167315865 | -0.239845709 | 0.574477438 | 8.86170531 | 0.998512548 | 0.355516887 | 0.669754507 | -4.671166234 |
| A_51_P320401 | Rab24 | 0.080628315 | -0.115592751 | 0.276849381 | 11.01508444 | 0.998447288 | 0.355546122 | 0.669754507 | -4.671194616 |
| A_52_P282035 | Rnf5 | -0.105545748 | -0.362473393 | 0.151381897 | 11.32571244 | -0.998189721 | 0.355661526 | 0.669895758 | -4.671306625 |
| A_51_P436201 | Gart | 0.130710456 | -0.18749563 | 0.448916543 | 9.409286049 | 0.99812524 | 0.355690422 | 0.669895758 | -4.671334663 |
| A_52_P610987 | Slc28a3 | 0.076249408 | -0.109396636 | 0.261895451 | 7.976999416 | 0.998007867 | 0.355743024 | 0.669929551 | -4.671385698 |
| A_55_P2123220 | Slc35e1 | 0.189139061 | -0.271417626 | 0.649695747 | 10.62053079 | 0.997886909 | 0.35579724 | 0.669966376 | -4.671438288 |
| A_55_P2042396 | Alkbh4 | 0.099498457 | -0.142872978 | 0.341869891 | 9.450142807 | 0.997511933 | 0.355965353 | 0.670152682 | -4.671601297 |
| A_51_P412966 | Lrrn1 | -0.110257361 | -0.378850576 | 0.158335853 | 7.763099673 | -0.997460591 | 0.355988376 | 0.670152682 | -4.671623614 |
| A_51_P124535 | Mest | -0.221546527 | -0.761260749 | 0.318167695 | 13.33449586 | -0.99743424 | 0.356000194 | 0.670152682 | -4.669882912 |
| A_55_P1990755 | Ppp3r1 | -0.118494777 | -0.407205967 | 0.170216412 | 11.08851388 | -0.997283842 | 0.356067644 | 0.670175108 | -4.671700436 |
| A_55_P2002849 | Aifm3 | -0.200660105 | -0.68958128 | 0.28826107 | 10.30883998 | -0.99725306 | 0.356081451 | 0.670175108 | -4.671713815 |
| A_52_P98625 | Sowahc | -0.093082878 | -0.319914575 | 0.133748818 | 8.465355942 | -0.997124188 | 0.356139257 | 0.670218645 | -4.671769821 |
| A_55_P2016356 | Eif3b | 0.113499604 | -0.163131789 | 0.390130997 | 11.31552972 | 0.996956183 | 0.356214629 | 0.670284295 | -4.671842829 |
| A_51_P368713 | Tmbim4 | -0.141314126 | -0.48575979 | 0.203131539 | 10.97375343 | -0.996891842 | 0.356243497 | 0.670284295 | -4.671870787 |
| A_51_P125050 | Wdr74 | 0.082236496 | -0.118233792 | 0.282706784 | 9.647279279 | 0.996776454 | 0.356295274 | 0.670316464 | -4.670166013 |
| A_52_P522255 | Pitx1 | 0.12480917 | -0.179502071 | 0.429120412 | 7.750344364 | 0.996578902 | 0.356383933 | 0.670392767 | -4.672006754 |
| A_51_P343613 | Ehd2 | -0.181894908 | -0.625425857 | 0.261636041 | 7.945580289 | -0.996505731 | 0.356416776 | 0.670392767 | -4.672038542 |
| A_55_P2082658 | NA | -0.080531258 | -0.27691258 | 0.115850063 | 8.069486184 | -0.996431609 | 0.356450048 | 0.670392767 | -4.672070742 |
| A_55_P2118684 | Idh2 | -0.176439712 | -0.606743499 | 0.253864075 | 14.86926827 | -0.996332645 | 0.356494474 | 0.670392767 | -4.672113731 |
| A_52_P577388 | Epdr1 | 0.07681494 | -0.110535302 | 0.264165183 | 10.12282844 | 0.996264439 | 0.356525096 | 0.670392767 | -4.672143358 |
| A_55_P1971594 | Pqbp1 | -0.075763942 | -0.260558598 | 0.109030715 | 10.02577859 | -0.996222489 | 0.356543931 | 0.670392767 | -4.672161579 |
| A_52_P181709 | Crkl | 0.080610479 | -0.116053264 | 0.277274222 | 8.59121523 | 0.995979481 | 0.356653053 | 0.670532716 | -4.672267123 |
| A_51_P362959 | Fbxo36 | -0.176618045 | -0.607567133 | 0.254331044 | 8.450293786 | -0.995846256 | 0.356712888 | 0.670533234 | -4.67232498 |
| A_55_P2345853 | NA | 0.152535857 | -0.21966415 | 0.524735864 | 8.794119484 | 0.995815218 | 0.356726829 | 0.670533234 | -4.672338458 |
| A_51_P512157 | Slc30a6 | 0.110378172 | -0.159020395 | 0.379776739 | 8.862623942 | 0.995568405 | 0.356837705 | 0.670533234 | -4.672445631 |
| A_55_P1981674 | NA | 0.067915553 | -0.097862149 | 0.233693256 | 8.314809958 | 0.995465736 | 0.356883835 | 0.670533234 | -4.672490207 |
| A_55_P2027278 | Gas2l1 | -0.154212181 | -0.530659417 | 0.222235054 | 11.26484552 | -0.995400276 | 0.356913249 | 0.670533234 | -4.672518628 |
| A_55_P2003211 | Igsf5 | -0.09201115 | -0.316631283 | 0.132608983 | 9.706270452 | -0.995348044 | 0.356936721 | 0.670533234 | -4.672541304 |
| A_55_P2132014 | Tekt2 | -0.26175077 | -0.900820079 | 0.377318539 | 8.019892855 | -0.995229398 | 0.356990042 | 0.670533234 | -4.672592811 |
| A_51_P248044 | Lamb2 | -0.319759832 | -1.100482875 | 0.46096321 | 9.134262179 | -0.995199385 | 0.357003531 | 0.670533234 | -4.67260584 |
| A_51_P520718 | Ptger3 | -0.0744974 | -0.256417619 | 0.107422819 | 8.216009752 | -0.995046401 | 0.357072295 | 0.670533234 | -4.672672247 |
| A_55_P2008639 | NA | -0.087216333 | -0.300207112 | 0.125774446 | 12.18897059 | -0.994993421 | 0.357096111 | 0.670533234 | -4.672695244 |
| A_51_P183853 | Polr2m | 0.077020577 | -0.111095239 | 0.265136394 | 13.38591753 | 0.994866133 | 0.357153336 | 0.670533234 | -4.672750491 |
| A_55_P2039556 | Pak6 | 0.071949099 | -0.103815399 | 0.247713596 | 7.865310121 | 0.994666266 | 0.357243205 | 0.670533234 | -4.672837232 |
| A_55_P1968618 | NA | -0.138913099 | -0.478313689 | 0.20048749 | 8.053479399 | -0.994520633 | 0.357308699 | 0.670533234 | -4.67290043 |
| A_51_P521374 | Nudt13 | -0.178273889 | -0.613881421 | 0.257333642 | 9.758003534 | -0.994433047 | 0.357348093 | 0.670533234 | -4.672938435 |
| A_55_P1967500 | Nell1 | -0.155579942 | -0.535760874 | 0.224600991 | 7.754374208 | -0.994366447 | 0.35737805 | 0.670533234 | -4.672967334 |
| A_51_P312997 | Zfp346 | 0.073489597 | -0.106095222 | 0.253074416 | 8.195620758 | 0.994350358 | 0.357385287 | 0.670533234 | -4.672974314 |
| A_55_P1969273 | Zfp758 | 0.12829057 | -0.185221792 | 0.441802932 | 8.747982858 | 0.994313312 | 0.357401952 | 0.670533234 | -4.672990388 |

| | | | | | | | | | |
|---|---|---|---|---|---|---|---|---|---|
| **A_51_P326685** | Lrtm1 | -0.102766218 | -0.353912899 | 0.148380464 | 7.93349436 | -0.99427408 | 0.357419601 | 0.670533234 | -4.67300741 |
| **A_51_P349662** | Taf5l | 0.126811494 | -0.183132267 | 0.436755255 | 11.11432108 | 0.994166009 | 0.357468221 | 0.670533234 | -4.673054298 |
| **A_55_P2013138** | NA | 0.088490696 | -0.127814696 | 0.304796089 | 11.96256831 | 0.994061958 | 0.357515037 | 0.670533234 | -4.673099438 |
| **A_55_P2101920** | Eva1c | -0.221377498 | -0.762517778 | 0.319762783 | 8.829080865 | -0.994046729 | 0.357521889 | 0.670533234 | -4.673106045 |
| **A_51_P422685** | Zmat4 | -0.103797791 | -0.357524992 | 0.14992941 | 7.7247872 | -0.994040941 | 0.357524494 | 0.670533234 | -4.673108556 |
| **A_52_P328078** | Atp5b | 0.108119509 | -0.156194421 | 0.372433439 | 16.19349426 | 0.993956097 | 0.357562673 | 0.670533234 | -4.673145362 |
| **A_52_P209311** | Snx12 | -0.097724213 | -0.344017947 | 0.14856952 | 11.32241335 | -1.009464817 | 0.357566664 | 0.670533234 | -4.645604615 |
| **A_55_P1974288** | Rela | -0.093337292 | -0.321542111 | 0.134867527 | 12.97700609 | -0.993833368 | 0.357617905 | 0.670533234 | -4.673198598 |
| **A_55_P2108483** | NA | -0.16475061 | -0.567568674 | 0.238067455 | 11.72800349 | -0.993805238 | 0.357630565 | 0.670533234 | -4.6732108 |
| **A_51_P140871** | Ankhd1 | 0.083824095 | -0.121152955 | 0.288801144 | 11.37525227 | 0.993680628 | 0.357686653 | 0.670533234 | -4.673264849 |
| **A_55_P1954896** | Rpl17 | -0.13151732 | -0.453130206 | 0.190095565 | 15.42761398 | -0.993648314 | 0.357701198 | 0.670533234 | -4.673278864 |
| **A_55_P1982902** | Tceal3 | 0.172622981 | -0.249539134 | 0.594785097 | 8.710363746 | 0.993579322 | 0.357732256 | 0.670533234 | -4.673308787 |
| **A_52_P438082** | Cebpg | -0.100915556 | -0.34773217 | 0.145901057 | 9.356184233 | -0.993497819 | 0.357768948 | 0.670533234 | -4.673344133 |
| **A_55_P2104662** | Ttc33 | 0.085651788 | -0.123836017 | 0.295139594 | 10.24730708 | 0.993484 | 0.35777517 | 0.670533234 | -4.673350127 |
| **A_51_P166288** | Acy1 | 0.174814121 | -0.252753963 | 0.602382204 | 8.840464169 | 0.993469227 | 0.357781821 | 0.670533234 | -4.673356533 |
| **A_55_P2107362** | Gjc1 | -0.447400316 | -1.541778183 | 0.646977551 | 8.458952488 | -0.993372595 | 0.357825329 | 0.670533234 | -4.673398439 |
| **A_51_P271370** | Wsb1 | -0.078151841 | -0.269332014 | 0.113028332 | 8.223163162 | -0.993297979 | 0.357858927 | 0.670533234 | -4.673430795 |
| **A_55_P1982096** | Drap1 | -0.186583799 | -0.643047978 | 0.269880379 | 13.60577689 | -0.993231352 | 0.357888931 | 0.670533234 | -4.673459686 |
| **A_51_P398191** | Auts2 | 0.07801927 | -0.112855058 | 0.268893598 | 8.42323288 | 0.993201915 | 0.357902187 | 0.670533234 | -4.67347245 |
| **A_66_P125173** | Aip | -0.181598679 | -0.626008312 | 0.262810953 | 9.842740145 | -0.992915776 | 0.358031067 | 0.670701479 | -4.673596512 |
| **A_55_P2039284** | Hspb1 | -0.764389873 | -2.635137241 | 1.106357496 | 10.74469071 | -0.992848469 | 0.358061388 | 0.670701479 | -4.673625691 |
| **A_55_P1983683** | Rtl5 | -0.122942099 | -0.423873055 | 0.177988856 | 8.068302842 | -0.99269755 | 0.358129382 | 0.67076384 | -4.673691114 |
| **A_55_P2005691** | NA | -0.138167262 | -0.476557182 | 0.200222658 | 15.56432395 | -0.992135346 | 0.358382766 | 0.671173383 | -4.673934779 |
| **A_55_P2028898** | Prl2b1 | -0.139290419 | -0.480551027 | 0.20197019 | 7.878500705 | -0.991786682 | 0.358539979 | 0.671361364 | -4.674085855 |
| **A_52_P603976** | Jtb | -0.095667869 | -0.330071321 | 0.138735582 | 11.52619179 | -0.991711363 | 0.358573948 | 0.671361364 | -4.674118486 |
| **A_51_P264527** | Dipk1b | -0.089012672 | -0.307130942 | 0.129105598 | 7.837247964 | -0.991614758 | 0.35861752 | 0.671361364 | -4.674160338 |
| **A_52_P585652** | Fndc3b | -0.224890359 | -0.775971397 | 0.326190679 | 10.20001418 | -0.99160466 | 0.358622075 | 0.671361364 | -4.674164712 |
| **A_55_P2039444** | Cmss1 | 0.088775426 | -0.128784343 | 0.306335195 | 10.58732907 | 0.99151061 | 0.358664499 | 0.67137576 | -4.674205454 |
| **A_55_P2011385** | NA | 0.142888504 | -0.207403517 | 0.493180525 | 9.810945908 | 0.991174757 | 0.358816028 | 0.671481435 | -4.674350927 |
| **A_51_P184796** | Zfp101 | 0.081029379 | -0.117618194 | 0.279676953 | 8.814057612 | 0.991156962 | 0.358824059 | 0.671481435 | -4.674358634 |
| **A_51_P417251** | 6330403K07Rik | -0.57616747 | -1.988877124 | 0.836542185 | 8.853149955 | -0.991012221 | 0.358889379 | 0.671481435 | -4.674421318 |
| **A_55_P2070799** | Tyw1 | 0.087608237 | -0.127221317 | 0.302437791 | 9.489578211 | 0.990909751 | 0.358935629 | 0.671481435 | -4.674465692 |
| **A_55_P1998107** | NA | -0.090385411 | -0.312028893 | 0.13125807 | 13.14966535 | -0.990892554 | 0.358943391 | 0.671481435 | -4.674473139 |
| **A_51_P359333** | Fh1 | 0.093986281 | -0.13650475 | 0.324477312 | 12.91436271 | 0.990817396 | 0.358977317 | 0.671481435 | -4.674505684 |
| **A_52_P189005** | Gk5 | -0.063470194 | -0.219128662 | 0.092188274 | 7.757372249 | -0.990786912 | 0.358991079 | 0.671481435 | -4.674518884 |
| **A_66_P120484** | Mbtd1 | -0.11534061 | -0.398248714 | 0.167567494 | 10.35995483 | -0.990649544 | 0.359053094 | 0.671481435 | -4.674578363 |
| **A_51_P408471** | Cdipt | -0.139549392 | -0.481853629 | 0.202754845 | 11.81097411 | -0.990601229 | 0.359074909 | 0.671481435 | -4.674599282 |
| **A_55_P1986998** | Enoph1 | 0.121857223 | -0.177056271 | 0.420770718 | 10.85287078 | 0.990578609 | 0.359085122 | 0.671481435 | -4.674609075 |
| **A_55_P2076635** | Vps25 | -0.099453218 | -0.343451019 | 0.144544584 | 9.51792881 | -0.990412502 | 0.359160129 | 0.671481435 | -4.674680988 |
| **A_51_P130332** | Hspa12b | -0.297354506 | -1.026889135 | 0.432180124 | 8.798355509 | -0.990402667 | 0.359164571 | 0.671481435 | -4.674685246 |
| **A_51_P109335** | Copa | -0.096309378 | -0.332600678 | 0.139981922 | 11.89336607 | -0.990384959 | 0.359172567 | 0.671481435 | -4.674692912 |
| **A_52_P484081** | Ube2v1 | 0.167986381 | -0.244268014 | 0.580240776 | 9.639646607 | 0.990129429 | 0.359287982 | 0.671632243 | -4.674803524 |
| **A_52_P58471** | Tmem199 | 0.092864604 | -0.13506948 | 0.320798687 | 8.818492085 | 0.989974769 | 0.35935785 | 0.671641103 | -4.674870463 |
| **A_55_P2075274** | Rab11fip4 | 0.090457876 | -0.131581824 | 0.312497575 | 8.027653146 | 0.989917364 | 0.359383786 | 0.671641103 | -4.674895308 |
| **A_66_P128384** | Hdac1 | 0.089875353 | -0.130740975 | 0.31049168 | 11.97396657 | 0.989888197 | 0.359396964 | 0.671641103 | -4.674907931 |
| **A_55_P2073099** | Trex1 | -0.148178368 | -0.511950989 | 0.215594252 | 10.62234101 | -0.989778438 | 0.35944656 | 0.671668847 | -4.674955431 |
| **A_55_P1976539** | Nup62cl | -0.06797226 | -0.234864957 | 0.098920438 | 7.718142492 | -0.989640737 | 0.359508788 | 0.67172019 | -4.675015019 |
| **A_51_P446558** | Timm29 | 0.081393117 | -0.118483078 | 0.281269312 | 8.226573405 | 0.989486325 | 0.359578579 | 0.671785651 | -4.675081833 |
| **A_55_P2034230** | Polr2c | 0.114120972 | -0.166213602 | 0.394455546 | 10.82319241 | 0.989172395 | 0.3597205 | 0.671985846 | -4.675217652 |
| **A_55_P2145389** | Flrt3 | -0.194905889 | -0.673754076 | 0.283942298 | 7.855925541 | -0.989031909 | 0.359784026 | 0.672039566 | -4.675278424 |

| | | | | | | | | |
|---|---|---|---|---|---|---|---|---|
| **A_55_P1963639** | Sycn | -0.173899037 | -0.60121911 | 0.253421036 | 8.494994209 | -0.988842381 | 0.359869741 | 0.672134721 | -4.675360403 |
| **A_52_P217875** | H3f3b | 0.074365943 | -0.108388922 | 0.257120808 | 8.142439888 | 0.988754174 | 0.359909639 | 0.672144291 | -4.675398553 |
| **A_55_P1996583** | Ppdpf | -0.160286257 | -0.554282741 | 0.233710228 | 11.51160599 | -0.988523799 | 0.360013859 | 0.672273971 | -4.675498183 |
| **A_51_P391934** | Cpb1 | -0.200618537 | -0.693801798 | 0.292564725 | 7.929354483 | -0.988430002 | 0.360056298 | 0.672288272 | -4.675538743 |
| **A_55_P2014224** | Sema5a | -0.389605259 | -1.347477172 | 0.568266653 | 9.073077743 | -0.988326775 | 0.360103009 | 0.672310545 | -4.675583378 |
| **A_65_P04980** | Ilf3 | 0.115059231 | -0.167879365 | 0.397997826 | 9.487158434 | 0.988126307 | 0.360193736 | 0.672395386 | -4.675670054 |
| **A_52_P273120** | Myl1 | -0.076557241 | -0.26482701 | 0.111712527 | 7.675352544 | -0.988072644 | 0.360218026 | 0.672395386 | -4.675693254 |
| **A_55_P2089710** | Ednrb | -0.144732232 | -0.50072112 | 0.211256656 | 8.050932425 | -0.987897775 | 0.360297187 | 0.672478208 | -4.67576885 |
| **A_55_P2060097** | Prdm5 | -0.208061503 | -0.719865472 | 0.303742465 | 8.777987499 | -0.987805119 | 0.360339136 | 0.6724881 | -4.675808903 |
| **A_55_P1992555** | Gys2 | 0.247523445 | -0.361396554 | 0.856443443 | 13.63310783 | 0.987732381 | 0.36037207 | 0.6724881 | -4.675840343 |
| **A_51_P110035** | Pik3r4 | -0.07053368 | -0.244098522 | 0.103031163 | 9.340928641 | -0.987456495 | 0.360497008 | 0.672584108 | -4.658453226 |
| **A_51_P170641** | Dcaf4 | -0.22573581 | -0.78122702 | 0.329755399 | 10.4471406 | -0.98743032 | 0.360508863 | 0.672584108 | -4.675970895 |
| **A_51_P389864** | B3gnt9 | -0.232992079 | -0.806432829 | 0.340448671 | 8.390406825 | -0.987269668 | 0.360581634 | 0.672584108 | -4.67604032 |
| **A_51_P155303** | Rbfa | -0.104429618 | -0.361457528 | 0.152598292 | 9.671970595 | -0.98724875 | 0.36059111 | 0.672584108 | -4.676049359 |
| **A_55_P1969650** | Rasgrp1 | 0.112791372 | -0.164847833 | 0.390430576 | 8.512568553 | 0.98713892 | 0.360640867 | 0.672584108 | -4.676096817 |
| **A_51_P273609** | Itpka | 0.113801157 | -0.166345534 | 0.393947848 | 10.60642154 | 0.987061855 | 0.360675784 | 0.672584108 | -4.676130116 |
| **A_55_P1956284** | NA | 0.067174927 | -0.098193151 | 0.232543006 | 8.11591964 | 0.987049001 | 0.360681608 | 0.672584108 | -4.676135669 |
| **A_51_P352738** | Mpv17l | -0.149387576 | -0.517159569 | 0.218384417 | 8.595311438 | -0.987004236 | 0.360701892 | 0.672584108 | -4.676155011 |
| **A_55_P2049979** | Dbi | -0.174206609 | -0.60313626 | 0.254723043 | 15.72579231 | -0.986874087 | 0.360760869 | 0.672629193 | -4.67621124 |
| **A_66_P134739** | Clpp | 0.085234016 | -0.124676115 | 0.295144147 | 10.03894322 | 0.986649138 | 0.360862823 | 0.672695536 | -4.676308417 |
| **A_51_P329283** | Senp5 | 0.084967276 | -0.124287453 | 0.294222004 | 9.370567938 | 0.986642005 | 0.360866057 | 0.672695536 | -4.676311499 |
| **A_51_P144868** | Mboat7 | -0.086336588 | -0.299022858 | 0.126349682 | 10.46247105 | -0.986367189 | 0.360990644 | 0.672822219 | -4.6764302 |
| **A_55_P2013357** | Renbp | -0.111087349 | -0.38475885 | 0.162584151 | 8.998635285 | -0.986320867 | 0.361011648 | 0.672822219 | -4.676450206 |
| **A_55_P2058628** | Hnrnpa3 | 0.139478418 | -0.204157065 | 0.483113902 | 10.25351743 | 0.986261775 | 0.361038443 | 0.672822219 | -4.676475727 |
| **A_51_P400543** | Aif1 | -0.124426238 | -0.431007747 | 0.18215527 | 11.37197708 | -0.986164134 | 0.361082721 | 0.672839864 | -4.676517894 |
| **A_55_P2021351** | NA | -0.101939956 | -0.353170759 | 0.149290847 | 16.34189239 | -0.985949664 | 0.361179994 | 0.672956246 | -4.676610506 |
| **A_55_P2047325** | Mbtps1 | -0.158568177 | -0.549501331 | 0.232364976 | 12.47943026 | -0.98559098 | 0.361342721 | 0.67319455 | -4.676765367 |
| **A_52_P377299** | Zbtb11 | -0.078447448 | -0.271873522 | 0.114978626 | 8.511690265 | -0.985478129 | 0.361393931 | 0.673225066 | -4.676814083 |
| **A_55_P2114187** | NA | -0.349122418 | -1.21007775 | 0.511832915 | 9.209365513 | -0.985326822 | 0.3614626 | 0.673236053 | -4.676879395 |
| **A_55_P2074281** | Nr2f1 | -0.251210451 | -0.870810605 | 0.368389704 | 9.552847328 | -0.985165908 | 0.361535642 | 0.673236053 | -4.676948849 |
| **A_55_P2040838** | Gm14548 | -0.206642375 | -0.716318019 | 0.303033268 | 9.477035546 | -0.985164365 | 0.361536342 | 0.673236053 | -4.676949514 |
| **A_66_P125962** | Hipk1 | 0.164798937 | -0.241713723 | 0.571311598 | 9.002287056 | 0.985061897 | 0.36158286 | 0.673236053 | -4.676993738 |
| **A_51_P483351** | Lix1l | -0.10492359 | -0.363752719 | 0.153905539 | 8.330353983 | -0.985015763 | 0.361603805 | 0.673236053 | -4.677013647 |
| **A_55_P2049856** | Araf | 0.09967838 | -0.146242414 | 0.345599174 | 9.664285435 | 0.984892679 | 0.361659691 | 0.673236053 | -4.677066763 |
| **A_55_P2043554** | Slc12a4 | -0.080756133 | -0.280017248 | 0.118504983 | 10.73228929 | -0.984773027 | 0.361714025 | 0.673236053 | -4.677118394 |
| **A_55_P2171406** | Tm7sf2 | -0.164271553 | -0.569664622 | 0.241121516 | 10.79944678 | -0.984621313 | 0.361782927 | 0.673236053 | -4.677183855 |
| **A_55_P2098801** | Gm15393 | -0.085150775 | -0.295325681 | 0.125024132 | 15.99036852 | -0.984443801 | 0.361863559 | 0.673236053 | -4.67726044 |
| **A_55_P1981415** | Gsk3b | -0.095332086 | -0.330718146 | 0.140053974 | 11.32015038 | -0.984105242 | 0.362017384 | 0.673236053 | -4.677406484 |
| **A_55_P2064578** | Prkci | -0.226533938 | -0.785892669 | 0.332824793 | 11.38929411 | -0.984070113 | 0.362033347 | 0.673236053 | -4.677421636 |
| **A_51_P480328** | Adgrl4 | -0.104234841 | -0.361616185 | 0.153146502 | 8.024035006 | -0.984054238 | 0.362040562 | 0.673236053 | -4.677428483 |
| **A_55_P1966274** | Pdzd4 | -0.184750875 | -0.640946768 | 0.271445018 | 8.166439534 | -0.98405262 | 0.362041297 | 0.673236053 | -4.677429181 |
| **A_55_P2071466** | Ncf1 | -0.175646448 | -0.609370554 | 0.258077658 | 10.0060848 | -0.984031496 | 0.362050897 | 0.673236053 | -4.677438292 |
| **A_52_P706060** | Mex3a | -0.224962184 | -0.78046877 | 0.330544402 | 8.833065068 | -0.984019027 | 0.362056564 | 0.673236053 | -4.67744367 |
| **A_51_P106144** | Zfp64 | -0.090156408 | -0.312798464 | 0.132485649 | 9.875564697 | -0.983948988 | 0.362088395 | 0.673236053 | -4.664777931 |
| **A_55_P1960097** | Epb41l3 | -0.078036257 | -0.270772849 | 0.114700336 | 9.972620183 | -0.983819533 | 0.362147235 | 0.673236053 | -4.677529708 |
| **A_52_P43848** | Sco1 | 0.091144679 | -0.133973626 | 0.316262984 | 8.291377659 | 0.983792932 | 0.362159327 | 0.673236053 | -4.67754118 |
| **A_55_P2174967** | NA | 0.149102158 | -0.219228834 | 0.51743315 | 9.676045652 | 0.983623407 | 0.362236394 | 0.673236053 | -4.677614283 |
| **A_55_P2036615** | Lrif1 | 0.122726808 | -0.180452756 | 0.425906373 | 9.294451019 | 0.983609477 | 0.362242727 | 0.673236053 | -4.67762029 |
| **A_51_P163694** | Rufy1 | -0.064344871 | -0.223303852 | 0.09461411 | 9.232644844 | -0.983585354 | 0.362253695 | 0.673236053 | -4.677630692 |
| **A_55_P2133027** | Kirrel3 | 0.147474028 | -0.216860322 | 0.511808378 | 8.475717537 | 0.983554915 | 0.362267535 | 0.673236053 | -4.677643817 |

| | | | | | | | | | |
|---|---|---|---|---|---|---|---|---|---|
| **A_52_P127465** | Ypel1 | -0.152593467 | -0.529580506 | 0.224393572 | 7.828735734 | -0.983541548 | 0.362273612 | 0.673236053 | -4.67764958 |
| **A_55_P2095365** | Meaf6 | 0.084896669 | -0.124853728 | 0.294647066 | 8.304480055 | 0.983492486 | 0.36229592 | 0.673236053 | -4.677670735 |
| **A_52_P602091** | Csf1r | -0.15062817 | -0.522803062 | 0.221546722 | 8.960488459 | -0.98342743 | 0.362325502 | 0.673236053 | -4.677698784 |
| **A_55_P1997696** | Plbd1 | -0.480292823 | -1.875390459 | 0.914804813 | 12.73227844 | -1.06161499 | 0.362327625 | 0.673236053 | -4.600841193 |
| **A_51_P242414** | Micall1 | -0.201540379 | -0.699526528 | 0.29644577 | 9.110839593 | -0.983395003 | 0.362340248 | 0.673236053 | -4.677712766 |
| **A_55_P2136906** | Vpreb3 | -0.177948945 | -0.617746714 | 0.261848824 | 12.12863112 | -0.983163133 | 0.362445702 | 0.673364965 | -4.677812729 |
| **A_51_P234627** | Nubpl | 0.084279932 | -0.124032128 | 0.292591992 | 9.707858828 | 0.983089261 | 0.362479304 | 0.673364965 | -4.677844573 |
| **A_66_P104134** | Homer1 | 0.072143524 | -0.106200192 | 0.250487241 | 8.007979607 | 0.982930319 | 0.362551609 | 0.673404371 | -4.677913086 |
| **A_51_P134551** | Coq9 | -0.085132188 | -0.295604029 | 0.125339653 | 10.0494986 | -0.982840368 | 0.362592535 | 0.673404371 | -4.677951856 |
| **A_55_P1992179** | Tars2 | 0.10745962 | -0.158224173 | 0.373143413 | 12.71866538 | 0.982796103 | 0.362612675 | 0.673404371 | -4.677970934 |
| **A_51_P253207** | NA | -0.330090581 | -1.146300785 | 0.486119623 | 8.015190584 | -0.982684943 | 0.362663258 | 0.673404371 | -4.678018842 |
| **A_52_P404329** | Saa4 | 0.283005389 | -0.416857085 | 0.982867863 | 10.84187704 | 0.982573816 | 0.36271383 | 0.673404371 | -4.678066733 |
| **A_55_P2125241** | NA | 0.080250618 | -0.118209724 | 0.278710959 | 8.283021433 | 0.982557189 | 0.362721398 | 0.673404371 | -4.678073898 |
| **A_51_P509643** | Snca | 0.16231136 | -0.239116565 | 0.563739284 | 14.50511321 | 0.982481824 | 0.362755699 | 0.673404371 | -4.678106375 |
| **A_65_P02321** | Usp36 | 0.073503893 | -0.108295261 | 0.255303048 | 8.663123631 | 0.98243013 | 0.362779229 | 0.673404371 | -4.678128651 |
| **A_55_P1981724** | Nr2c1 | -0.106719409 | -0.370700738 | 0.15726192 | 9.447831361 | -0.982320908 | 0.362828947 | 0.673431987 | -4.678175713 |
| **A_55_P2059090** | Tagap | -0.066013505 | -0.229355112 | 0.097328102 | 9.426504558 | -0.982017345 | 0.362967158 | 0.673484309 | -4.678306499 |
| **A_51_P146655** | Ptgr2 | 0.155458561 | -0.229213823 | 0.540130944 | 10.24037121 | 0.981989453 | 0.362979859 | 0.673484309 | -4.678318515 |
| **A_55_P2327518** | 1500032P08Rik | -0.086919109 | -0.301997501 | 0.128159284 | 8.250577343 | -0.981977804 | 0.362985164 | 0.673484309 | -4.678323533 |
| **A_55_P2183894** | Gm5617 | -0.084332961 | -0.29303723 | 0.124371308 | 11.28465615 | -0.981859184 | 0.363039184 | 0.673484309 | -4.678374631 |
| **A_51_P243122** | Sec14l3 | -0.21071228 | -0.732205441 | 0.310780881 | 7.889800379 | -0.981803143 | 0.363064708 | 0.673484309 | -4.678398771 |
| **A_52_P177341** | Cox19 | -0.089964222 | -0.312617988 | 0.132689544 | 11.58866959 | -0.981799876 | 0.363066196 | 0.673484309 | -4.678400178 |
| **A_52_P351882** | Impad1 | -0.103549006 | -0.359866299 | 0.152768286 | 11.77346503 | -0.981637676 | 0.363140078 | 0.673520772 | -4.678470042 |
| **A_55_P1953143** | Wt1 | 0.100610807 | -0.148442105 | 0.349663719 | 8.62341505 | 0.981603727 | 0.363155543 | 0.673520772 | -4.678484663 |
| **A_55_P2030899** | NA | 0.103812041 | -0.153243224 | 0.360867305 | 11.61088517 | 0.981305909 | 0.363291234 | 0.673677716 | -4.678612921 |
| **A_55_P2143209** | Axin1 | 0.079692876 | -0.1176703 | 0.277056052 | 9.150120787 | 0.981152608 | 0.363361096 | 0.673677716 | -4.678678932 |
| **A_55_P1991151** | Rpl13 | 0.130023952 | -0.191987459 | 0.452035364 | 15.27359669 | 0.981149734 | 0.363362406 | 0.673677716 | -4.678680169 |
| **A_55_P2136813** | Rab10os | 0.209788249 | -0.309811737 | 0.729388235 | 10.27929314 | 0.981059204 | 0.363403667 | 0.673677716 | -4.678719148 |
| **A_55_P2184385** | NA | 0.131520729 | -0.194249053 | 0.457290512 | 8.467676981 | 0.980994561 | 0.363433132 | 0.673677716 | -4.678746981 |
| **A_51_P411355** | Taf1c | 0.098146681 | -0.144985721 | 0.341279083 | 8.546237267 | 0.980880195 | 0.363485266 | 0.673677716 | -4.678796218 |
| **A_55_P2045136** | Ly6a2 | -0.163870419 | -0.56983162 | 0.242090782 | 8.477253672 | -0.980842383 | 0.363502504 | 0.673677716 | -4.678812496 |
| **A_55_P1973829** | NA | 0.083174438 | -0.122894037 | 0.289242912 | 8.544739359 | 0.980757202 | 0.363541339 | 0.673677716 | -4.678849166 |
| **A_55_P2068723** | Ccl27a | -0.07676169 | -0.266964645 | 0.113441266 | 9.956052841 | -0.980641917 | 0.363593904 | 0.673677716 | -4.678898792 |
| **A_55_P2139587** | NA | -0.15345675 | -0.533715043 | 0.226801542 | 8.541628873 | -0.980596846 | 0.363614456 | 0.673677716 | -4.678918192 |
| **A_51_P308681** | Mcur1 | 0.105553717 | -0.15600853 | 0.367115963 | 9.371240533 | 0.980576902 | 0.363623551 | 0.673677716 | -4.678926777 |
| **A_52_P65108** | Smad1 | -0.11405872 | -0.396722072 | 0.168604633 | 8.446088895 | -0.980487757 | 0.363664203 | 0.673688459 | -4.678965147 |
| **A_55_P2024061** | Gtpbp10 | 0.104911035 | -0.15514408 | 0.36496615 | 9.381155944 | 0.980254759 | 0.363770474 | 0.673798729 | -4.679065424 |
| **A_65_P19149** | Nln | 0.22252245 | -0.32909802 | 0.774142919 | 9.978016382 | 0.9802044 | 0.363793447 | 0.673798729 | -4.679087096 |
| **A_52_P263201** | Tmem33 | 0.077090256 | -0.114041568 | 0.268222081 | 8.481568075 | 0.980053252 | 0.363862403 | 0.673861874 | -4.679152138 |
| **A_66_P111747** | NA | 0.134811307 | -0.199522308 | 0.469144921 | 14.01845755 | 0.979782017 | 0.363986169 | 0.673950735 | -4.679268841 |
| **A_55_P2093679** | Zfp599 | 0.090214539 | -0.133527056 | 0.313956134 | 8.248114198 | 0.979744862 | 0.364003126 | 0.673950735 | -4.679284826 |
| **A_55_P2031242** | Sun1 | 0.074288129 | -0.109970591 | 0.258546848 | 8.254665611 | 0.979658153 | 0.364042701 | 0.673950735 | -4.67932213 |
| **A_52_P1130990** | Trim67 | -0.149352121 | -0.519823484 | 0.221119243 | 7.71544868 | -0.979580067 | 0.364078342 | 0.673950735 | -4.679355721 |
| **A_51_P365656** | Tmem101 | 0.124189946 | -0.183885276 | 0.432265169 | 9.599496383 | 0.979518985 | 0.364106225 | 0.673950735 | -4.679381997 |
| **A_51_P209527** | Bcl10 | -0.101600908 | -0.35366797 | 0.150466154 | 10.14178376 | -0.979410323 | 0.36415583 | 0.673950735 | -4.679428738 |
| **A_51_P333034** | Ier3ip1 | -0.141151744 | -0.49134416 | 0.209040672 | 11.48648056 | -0.979405864 | 0.364157866 | 0.673950735 | -4.679430656 |
| **A_52_P33453** | NA | -0.076880838 | -0.267632861 | 0.113871185 | 12.27153902 | -0.979336963 | 0.364189323 | 0.673950735 | -4.679460292 |
| **A_52_P72497** | Acbd3 | 0.105363325 | -0.156088711 | 0.366815361 | 10.5404969 | 0.979220793 | 0.364242365 | 0.673984365 | -4.679510257 |
| **A_55_P2082614** | Zfp791 | 0.249454659 | -0.369656405 | 0.868565723 | 13.30316854 | 0.979053093 | 0.364318947 | 0.674061542 | -4.679582379 |
| **A_55_P2065327** | Rpl27a | -0.124662433 | -0.434104859 | 0.184779993 | 15.55290144 | -0.978901354 | 0.364388251 | 0.67412524 | -4.679647631 |

| | | | | | | | | |
|---|---|---|---|---|---|---|---|---|
| **A_55_P2059342** | Lamp1 | -0.076820094 | -0.267524752 | 0.113884564 | 14.70617863 | -0.978806227 | 0.364431704 | 0.674135343 | -4.679688535 |
| **A_55_P1981155** | Phldb1 | -0.339595751 | -1.182775011 | 0.50358351 | 10.16592411 | -0.978645764 | 0.36450501 | 0.674135343 | -4.679757528 |
| **A_55_P1978236** | Mrps7 | 0.091355861 | -0.135497621 | 0.318209343 | 13.29366253 | 0.978530019 | 0.364557895 | 0.674135343 | -4.67980729 |
| **A_55_P2026753** | Zmat1 | 0.123154227 | -0.18266193 | 0.428970384 | 8.682799156 | 0.97852536 | 0.364560024 | 0.674135343 | -4.679809293 |
| **A_55_P2022595** | Gm10653 | 0.122704626 | -0.182007083 | 0.427416335 | 15.55117549 | 0.978486828 | 0.364577631 | 0.674135343 | -4.679825858 |
| **A_52_P187132** | Ybx1 | -0.075589755 | -0.263320406 | 0.112140896 | 12.57268026 | -0.978387605 | 0.364622973 | 0.674135343 | -4.679868513 |
| **A_52_P559808** | Arhgef12 | 0.107390914 | -0.15932826 | 0.374110089 | 11.31329348 | 0.97835505 | 0.364637851 | 0.674135343 | -4.679882507 |
| **A_55_P2122618** | Asb11 | -0.066649498 | -0.232203013 | 0.098904016 | 7.787908899 | -0.978231561 | 0.36469429 | 0.674175204 | -4.679935589 |
| **A_51_P383140** | Slk | 0.094657381 | -0.14049437 | 0.329809132 | 8.941811032 | 0.978113958 | 0.364748046 | 0.674210096 | -4.679986137 |
| **A_51_P210928** | Tsta3 | -0.10770281 | -0.37532023 | 0.15991461 | 9.670046757 | -0.977903144 | 0.364844423 | 0.674323757 | -4.680076741 |
| **A_55_P1987364** | Ap5z1 | 0.084247562 | -0.125150411 | 0.293645534 | 9.079932068 | 0.977615451 | 0.364975978 | 0.674455507 | -4.680200367 |
| **A_55_P2141084** | Tenm4 | -0.223818191 | -0.78021259 | 0.332576209 | 11.89219631 | -0.977452855 | 0.365050347 | 0.674455507 | -4.680270227 |
| **A_52_P623058** | R3hdm4 | 0.081581366 | -0.121229747 | 0.284392479 | 8.698085139 | 0.977422681 | 0.365064149 | 0.674455507 | -4.680283191 |
| **A_55_P2024654** | Trrap | 0.087728575 | -0.130384423 | 0.305841572 | 9.767025949 | 0.977333346 | 0.365105015 | 0.674455507 | -4.680321571 |
| **A_55_P2114562** | NA | 0.07882517 | -0.117177554 | 0.274827895 | 8.143367614 | 0.977205716 | 0.365163406 | 0.674455507 | -4.6803764 |
| **A_55_P2003951** | Tmem192 | -0.098698974 | -0.344127702 | 0.146729753 | 11.45020583 | -0.977170693 | 0.36517943 | 0.674455507 | -4.680391445 |
| **A_52_P280899** | Wdr33 | 0.083065265 | -0.123490983 | 0.289621513 | 9.671541249 | 0.977156912 | 0.365185736 | 0.674455507 | -4.680397364 |
| **A_55_P2139665** | NA | 0.083054663 | -0.123500619 | 0.289609944 | 16.56966935 | 0.977036759 | 0.365240715 | 0.674455507 | -4.680448976 |
| **A_55_P2070277** | Mfn1 | 0.137310004 | -0.204202196 | 0.478822205 | 10.04722449 | 0.976965318 | 0.365273408 | 0.674455507 | -4.680479661 |
| **A_55_P2053266** | St6galnac6 | 0.151898432 | -0.225918424 | 0.529715287 | 11.28975335 | 0.976911336 | 0.365298112 | 0.674455507 | -4.680502846 |
| **A_51_P423518** | Amph | -0.218949383 | -0.763543753 | 0.325644987 | 7.916888772 | -0.97690823 | 0.365299534 | 0.674455507 | -4.680504181 |
| **A_51_P142989** | Zfp429 | -0.086373374 | -0.301231051 | 0.128484303 | 8.779862263 | -0.976814728 | 0.365342329 | 0.674470092 | -4.680544339 |
| **A_55_P2076174** | NA | -0.15148756 | -0.52841081 | 0.225435689 | 14.42342763 | -0.97657867 | 0.365450386 | 0.674605148 | -4.680645712 |
| **A_52_P66313** | Psmf1 | 0.115153078 | -0.171427469 | 0.401733624 | 9.285390353 | 0.976364619 | 0.365548391 | 0.674644253 | -4.680737623 |
| **A_51_P201661** | Xrcc5 | 0.114756468 | -0.170844547 | 0.400357482 | 10.22055651 | 0.976338949 | 0.365560145 | 0.674644253 | -4.680748644 |
| **A_55_P2012439** | Tnfrsf19 | -0.283098836 | -0.987688602 | 0.42149093 | 9.574124405 | -0.976303715 | 0.36557628 | 0.674644253 | -4.680763772 |
| **A_51_P386058** | Ppp2r5b | -0.245820344 | -0.857704752 | 0.366064064 | 9.841745029 | -0.976183864 | 0.365631167 | 0.674681129 | -4.680815227 |
| **A_55_P1982084** | NA | -0.136136619 | -0.475103437 | 0.2028302 | 9.798276977 | -0.97589021 | 0.365765676 | 0.674806839 | -4.680941285 |
| **A_66_P133270** | Mfsd1 | -0.093061901 | -0.324778684 | 0.138654881 | 8.492539694 | -0.975882701 | 0.365769116 | 0.674806839 | -4.680944509 |
| **A_51_P164835** | Ppl | -0.315896581 | -1.102685633 | 0.470892472 | 8.395108014 | -0.975595539 | 0.36590069 | 0.67494573 | -4.681067758 |
| **A_55_P2076683** | Sort1 | 0.117516714 | -0.175189706 | 0.410223134 | 8.466540929 | 0.975552351 | 0.365920482 | 0.67494573 | -4.681086292 |
| **A_55_P2057786** | Rft1 | -0.105359315 | -0.367801355 | 0.157082724 | 8.944561169 | -0.975489781 | 0.365949157 | 0.67494573 | -4.681113144 |
| **A_55_P2157023** | Sox17 | -0.442232713 | -1.544170019 | 0.659704594 | 8.824489375 | -0.975162894 | 0.366098993 | 0.675154045 | -4.681253409 |
| **A_55_P2174632** | Rpl7 | 0.086518198 | -0.129080193 | 0.302116589 | 16.14896081 | 0.975090982 | 0.366131962 | 0.675154045 | -4.681284262 |
| **A_55_P2041011** | Slc2a13 | -0.157345398 | -0.549649664 | 0.234958869 | 8.287412543 | -0.97457261 | 0.366369686 | 0.67548732 | -4.681506626 |
| **A_52_P299846** | Sh3rf1 | -0.112951811 | -0.394592263 | 0.168688641 | 8.014620214 | -0.974498906 | 0.366403496 | 0.67548732 | -4.681538236 |
| **A_52_P420712** | Pja2 | -0.101901033 | -0.355994827 | 0.152192761 | 8.629549248 | -0.974468302 | 0.366417536 | 0.67548732 | -4.681551362 |
| **A_51_P490171** | Nubp2 | 0.125277216 | -0.187185122 | 0.437739554 | 10.42363368 | 0.974221266 | 0.366530879 | 0.67563183 | -4.681657301 |
| **A_55_P2018989** | Elk3 | -0.068696924 | -0.240085097 | 0.102691248 | 7.975535648 | -0.973956664 | 0.366652313 | 0.675791224 | -4.681770756 |
| **A_55_P1996952** | NA | 0.217406322 | -0.325068412 | 0.759881055 | 9.912189806 | 0.973813614 | 0.366717977 | 0.675799588 | -4.681832084 |
| **A_55_P2187043** | Tpm1 | -0.754605826 | -2.637543961 | 1.128332309 | 9.664203625 | -0.973794442 | 0.366726777 | 0.675799588 | -4.681840303 |
| **A_55_P1957593** | Ino80b | 0.121691249 | -0.182016155 | 0.425398652 | 9.76300884 | 0.97361478 | 0.366809261 | 0.67588715 | -4.68191732 |
| **A_52_P421713** | 9930012K11Rik | -0.159500959 | -0.557699067 | 0.238697149 | 7.92095614 | -0.973301283 | 0.366953223 | 0.6760074 | -4.682051689 |
| **A_55_P1967643** | Plb1 | -0.611416918 | -2.137983418 | 0.915149582 | 8.2253504 | -0.973206303 | 0.366996849 | 0.6760074 | -4.682092394 |
| **A_52_P658974** | NA | 0.097262356 | -0.145611909 | 0.34013662 | 13.61300937 | 0.973075359 | 0.367056999 | 0.6760074 | -4.682148507 |
| **A_55_P2150908** | Atp5c1 | -0.065712041 | -0.229801965 | 0.098377884 | 14.66078415 | -0.973074807 | 0.367057252 | 0.6760074 | -4.682148744 |
| **A_55_P2166506** | NA | 0.080428172 | -0.12045202 | 0.281308364 | 14.90297467 | 0.972868793 | 0.367151902 | 0.6760074 | -4.682237018 |
| **A_55_P1988033** | Atp6v1g1 | -0.114827234 | -0.401624817 | 0.171970349 | 13.51169377 | -0.972865059 | 0.367153618 | 0.6760074 | -4.682238618 |
| **A_55_P2108389** | D830030K20Rik | 0.098581398 | -0.147640926 | 0.344803722 | 9.822470172 | 0.97286087 | 0.367155543 | 0.6760074 | -4.682240413 |
| **A_51_P390775** | Ube2e1 | -0.125594209 | -0.43930872 | 0.188120302 | 11.05726861 | -0.972787981 | 0.367189035 | 0.6760074 | -4.682271642 |

| | | | | | | | | | |
|---|---|---|---|---|---|---|---|---|---|
| **A_52_P379277** | Enpp3 | 0.275499304 | -0.41265486 | 0.963653469 | 9.214116949 | 0.972787436 | 0.367189285 | 0.6760074 | -4.682271876 |
| **A_55_P1992403** | Simc1 | 0.079428835 | -0.119095393 | 0.277953063 | 8.751612039 | 0.97218265 | 0.367467277 | 0.676454762 | -4.682530943 |
| **A_51_P230987** | Pi4k2b | -0.166607831 | -0.583153719 | 0.249938056 | 9.583601578 | -0.971886948 | 0.367603258 | 0.676640641 | -4.682657577 |
| **A_55_P2025660** | Ranbp9 | 0.107907817 | -0.161938495 | 0.377754129 | 9.652498165 | 0.971671824 | 0.367702208 | 0.67675833 | -4.682749688 |
| **A_55_P1973995** | NA | 0.226088664 | -0.339448931 | 0.79162626 | 12.40130759 | 0.971405362 | 0.367824802 | 0.676816133 | -4.682863765 |
| **A_52_P385606** | Ckb | -0.15407862 | -0.539516182 | 0.231358943 | 8.780362351 | -0.971340579 | 0.367854612 | 0.676816133 | -4.682891497 |
| **A_55_P2137653** | NA | -0.074920792 | -0.262342873 | 0.11250129 | 13.26357094 | -0.971325501 | 0.367861551 | 0.676816133 | -4.682897952 |
| **A_55_P1977538** | Apba3 | -0.114865448 | -0.402221 | 0.172490103 | 9.909275835 | -0.971299151 | 0.367873676 | 0.676816133 | -4.682909231 |
| **A_55_P2011235** | Rpl13 | 0.105766588 | -0.158856631 | 0.370389807 | 16.35031929 | 0.971188955 | 0.36792439 | 0.676845011 | -4.6829564 |
| **A_55_P2153570** | Zfp655 | 0.082011453 | -0.123194003 | 0.287216909 | 8.098808717 | 0.971110815 | 0.367960354 | 0.676846753 | -4.682989845 |
| **A_55_P2016838** | NA | 0.1117144 | -0.16793515 | 0.391363949 | 14.90833731 | 0.970684785 | 0.368156485 | 0.677143087 | -4.683172166 |
| **A_51_P424272** | Mt4 | 0.157384354 | -0.236674869 | 0.551443576 | 8.312084743 | 0.970472531 | 0.36825423 | 0.677258422 | -4.683262983 |
| **A_51_P372141** | Pnkd | -0.068132619 | -0.238755142 | 0.102489905 | 7.8160201 | -0.97029079 | 0.36833794 | 0.677283558 | -4.683340735 |
| **A_55_P1976494** | Bbof1 | -0.134999264 | -0.473073999 | 0.203075471 | 7.957776731 | -0.97029071 | 0.368337977 | 0.677283558 | -4.683340769 |
| **A_55_P2057913** | Hmgb1-ps7 | 0.107619242 | -0.161909985 | 0.377148469 | 13.13552472 | 0.970213364 | 0.368373607 | 0.677284643 | -4.683373857 |
| **A_55_P1984344** | Rhox2h | 0.108279363 | -0.162941292 | 0.379500018 | 7.795037236 | 0.970076807 | 0.36843652 | 0.677335885 | -4.68343227 |
| **A_52_P407697** | NA | 0.06874622 | -0.103467169 | 0.240959609 | 7.954106464 | 0.969985176 | 0.36847874 | 0.677349079 | -4.683471463 |
| **A_55_P2000798** | Ccdc144b | 0.10905605 | -0.164191565 | 0.382303664 | 8.389764153 | 0.969787475 | 0.368569845 | 0.677452124 | -4.683556018 |
| **A_51_P228865** | Zfp72 | 0.075398797 | -0.113566185 | 0.264363778 | 8.255011001 | 0.969541211 | 0.368683354 | 0.677596325 | -4.683661329 |
| **A_52_P75777** | NA | -0.098664825 | -0.345969832 | 0.148640181 | 7.69234839 | -0.969421463 | 0.368738558 | 0.677598176 | -4.683712531 |
| **A_55_P2126814** | NA | 0.192751697 | -0.290407689 | 0.675911083 | 10.000735 | 0.969373134 | 0.36876084 | 0.677598176 | -4.683733195 |
| **A_51_P109840** | Vtn | -0.248416769 | -0.871148502 | 0.374314964 | 15.48042178 | -0.969310911 | 0.36878953 | 0.677598176 | -4.683759799 |
| **A_52_P604613** | S1pr2 | -0.290343095 | -1.018362713 | 0.437676524 | 8.533820513 | -0.969062099 | 0.368904267 | 0.677635062 | -4.683866168 |
| **A_55_P2052475** | NA | 0.109566651 | -0.16516881 | 0.384302112 | 15.48192833 | 0.969051498 | 0.368909156 | 0.677635062 | -4.6838707 |
| **A_52_P495869** | Mafb | 0.188808245 | -0.284702373 | 0.662318863 | 9.163471951 | 0.968889886 | 0.368983697 | 0.677635062 | -4.683939782 |
| **A_55_P2044932** | Gpr84 | 0.106104781 | -0.159996875 | 0.372206437 | 8.483322744 | 0.968881284 | 0.368987665 | 0.677635062 | -4.683943458 |
| **A_55_P1960683** | F2 | -0.200908141 | -0.704797546 | 0.302981265 | 15.69250644 | -0.968825355 | 0.369013464 | 0.677635062 | -4.683967364 |
| **A_66_P114768** | Bach2 | -0.191221507 | -0.670823231 | 0.288380216 | 9.160956188 | -0.968811291 | 0.369019952 | 0.677635062 | -4.683973375 |
| **A_55_P2105180** | Bhmt | 0.343308717 | -0.51807447 | 1.204691905 | 14.22644933 | 0.96843757 | 0.369192384 | 0.677887299 | -4.68413309 |
| **A_55_P2060962** | Tm2d3 | -0.069260073 | -0.243065305 | 0.104545159 | 11.17237331 | -0.968285185 | 0.369262712 | 0.677952028 | -4.684198203 |
| **A_51_P208931** | N4bp3 | -0.377565009 | -1.325303755 | 0.570173737 | 10.2875437 | -0.968024372 | 0.369383104 | 0.678046586 | -4.684309633 |
| **A_55_P2229137** | Tmem116 | -0.075316739 | -0.264375313 | 0.113741835 | 7.708287191 | -0.968006603 | 0.369391307 | 0.678046586 | -4.684317224 |
| **A_55_P2010552** | Hsd17b13 | -0.370903179 | -1.302005269 | 0.560198911 | 12.8019729 | -0.967935556 | 0.369424109 | 0.678046586 | -4.684347575 |
| **A_55_P2088202** | Hcn2 | 0.266954933 | -0.403251642 | 0.937161507 | 14.10257208 | 0.967859808 | 0.369459083 | 0.678046586 | -4.684379933 |
| **A_55_P2014706** | Naip5 | 0.095933381 | -0.144929653 | 0.336796416 | 14.14290229 | 0.967793696 | 0.369489611 | 0.678046586 | -4.684408173 |
| **A_51_P270899** | Zfp61 | -0.190855132 | -0.670246554 | 0.288536289 | 8.648396287 | -0.967379264 | 0.369681021 | 0.678187927 | -4.684585175 |
| **A_52_P113672** | Chml | -0.15605645 | -0.548078075 | 0.235965174 | 9.625871691 | -0.967285972 | 0.369724119 | 0.678187927 | -4.684625013 |
| **A_55_P2069768** | Kiss1 | -0.155308884 | -0.545453654 | 0.234835887 | 7.924981044 | -0.967283316 | 0.369725346 | 0.678187927 | -4.684626147 |
| **A_55_P2082841** | NA | 0.181956388 | -0.275148821 | 0.639061598 | 11.76649655 | 0.967240172 | 0.369745279 | 0.678187927 | -4.68464457 |
| **A_55_P1976057** | Smurf2 | 0.198160182 | -0.299674334 | 0.695994699 | 8.732055933 | 0.967196192 | 0.369765599 | 0.678187927 | -4.68466335 |
| **A_55_P2129911** | NA | 0.087369909 | -0.132133729 | 0.306873547 | 16.22559387 | 0.967171189 | 0.369777152 | 0.678187927 | -4.684674026 |
| **A_55_P1973583** | Hpn | -0.222451141 | -0.78139048 | 0.336488198 | 13.61036385 | -0.967059397 | 0.369828807 | 0.678218313 | -4.684721758 |
| **A_55_P2031125** | Ube2l6 | 0.110008861 | -0.166471863 | 0.386489585 | 13.55799531 | 0.966820832 | 0.369939061 | 0.678352169 | -4.684823607 |
| **A_55_P2042426** | Rpl6 | 0.078674254 | -0.119069067 | 0.276417574 | 16.37861537 | 0.966749593 | 0.369971989 | 0.678352169 | -4.684854018 |
| **A_55_P1982568** | Blcap | 0.300732004 | -0.455392579 | 1.056856586 | 11.58676717 | 0.966427944 | 0.370120691 | 0.678542597 | -4.684991309 |
| **A_55_P1999842** | NA | 0.121126293 | -0.18343646 | 0.425689045 | 9.866018638 | 0.966373085 | 0.370146057 | 0.678542597 | -4.685014723 |
| **A_55_P2118173** | Igf2 | -0.123900208 | -0.435462852 | 0.187662435 | 11.43938662 | -0.966295252 | 0.370182049 | 0.678544223 | -4.685047939 |
| **A_52_P337236** | Alyref2 | 0.111226808 | -0.168506922 | 0.390960538 | 9.476480761 | 0.966157271 | 0.370245862 | 0.678575202 | -4.685106821 |
| **A_55_P2136556** | Pkn2 | 0.106223546 | -0.160940988 | 0.373388081 | 9.35824196 | 0.966106891 | 0.370269163 | 0.678575202 | -4.685128319 |
| **A_55_P1983988** | Enox2 | -0.078380557 | -0.275595099 | 0.118833986 | 8.586142251 | -0.965723044 | 0.370446736 | 0.678812442 | -4.685292091 |

| | | | | | | | | | |
|---|---|---|---|---|---|---|---|---|---|
| **A_55_P2119737** | NA | 0.19837735 | -0.300787704 | 0.697542403 | 8.88365259 | 0.965675248 | 0.370468853 | 0.678812442 | -4.685312481 |
| **A_51_P519756** | Rusc1 | -0.154230245 | -0.542350005 | 0.233889515 | 9.698251238 | -0.965577161 | 0.370514242 | 0.678831258 | -4.685354323 |
| **A_51_P123795** | Zfp637 | -0.186522334 | -0.655986722 | 0.282942054 | 11.62902698 | -0.965409074 | 0.370592033 | 0.678909431 | -4.677489957 |
| **A_55_P2403453** | BC037032 | 0.130463528 | -0.198020453 | 0.458947509 | 10.27687265 | 0.965068439 | 0.37074972 | 0.679131277 | -4.685571296 |
| **A_55_P2056774** | Hsd17b14 | 0.070398338 | -0.106865529 | 0.247662205 | 8.113199427 | 0.964995694 | 0.370783402 | 0.679131277 | -4.685602316 |
| **A_51_P306658** | Lsm8 | 0.07001941 | -0.106310157 | 0.246348977 | 11.41683355 | 0.964887084 | 0.370833694 | 0.679159036 | -4.685648628 |
| **A_55_P1976033** | Cnih2 | 0.087025962 | -0.132173019 | 0.306224944 | 8.279752903 | 0.964702692 | 0.370919089 | 0.679251073 | -4.685727247 |
| **A_51_P139030** | Slc38a3 | -0.2326894 | -0.818925822 | 0.353547022 | 11.79124228 | -0.964466154 | 0.371028657 | 0.679291474 | -4.685828086 |
| **A_55_P2182458** | Kcnh4 | 0.103521387 | -0.157297351 | 0.364340125 | 8.37003295 | 0.964438374 | 0.371041527 | 0.679291474 | -4.685839928 |
| **A_55_P2057102** | Sft2d3 | -0.085512754 | -0.300961771 | 0.129936262 | 8.66844693 | -0.964427463 | 0.371046582 | 0.679291474 | -4.685844579 |
| **A_55_P2083149** | NA | 0.252498762 | -0.383885182 | 0.888882707 | 9.85398171 | 0.964102543 | 0.371197136 | 0.679461321 | -4.685983069 |
| **A_55_P2048867** | NA | -0.314776069 | -1.108159426 | 0.478607289 | 8.387695733 | -0.964054997 | 0.371219171 | 0.679461321 | -4.686003333 |
| **A_55_P2008349** | Prelid1 | -0.076176085 | -0.268198138 | 0.115845969 | 13.57383194 | -0.963941607 | 0.371271725 | 0.679461321 | -4.686051655 |
| **A_55_P2148531** | Pithd1 | 0.159469095 | -0.242539862 | 0.561478052 | 8.898826366 | 0.963882263 | 0.371299232 | 0.679461321 | -4.686076943 |
| **A_51_P309501** | Praf2 | -0.093204657 | -0.32817457 | 0.141765256 | 8.035019129 | -0.96384799 | 0.371315119 | 0.679461321 | -4.686091548 |
| **A_52_P258557** | Slc26a9 | -0.165959394 | -0.584504707 | 0.252585919 | 7.787673694 | -0.963479652 | 0.371485891 | 0.679709466 | -4.686248486 |
| **A_51_P357207** | Zfp518b | 0.062259323 | -0.094779204 | 0.219297849 | 8.032639355 | 0.963343933 | 0.37154883 | 0.679709625 | -4.686306303 |
| **A_51_P487105** | Bud31 | 0.099352983 | -0.151274726 | 0.349980692 | 10.80926721 | 0.963241127 | 0.371596511 | 0.679709625 | -4.686350096 |
| **A_66_P118195** | Ciao1 | -0.090055023 | -0.317234382 | 0.137124336 | 10.46497939 | -0.963212831 | 0.371609636 | 0.679709625 | -4.686362149 |
| **A_51_P237695** | Gnai3 | -0.095813109 | -0.337527434 | 0.145901217 | 12.06445421 | -0.963176172 | 0.37162664 | 0.679709625 | -4.686377764 |
| **A_55_P2103596** | Slc30a4 | -0.15769126 | -0.555580848 | 0.240198327 | 10.053843 | -0.963004328 | 0.371706356 | 0.67972694 | -4.686450956 |
| **A_55_P1995092** | H2-Q10 | -0.188803226 | -0.665195205 | 0.287588753 | 11.3024059 | -0.963004151 | 0.371706439 | 0.67972694 | -4.686451031 |
| **A_51_P517094** | Sh3gl1 | -0.117945172 | -0.415617597 | 0.179727253 | 10.50309295 | -0.962774885 | 0.371812813 | 0.679857144 | -4.686548668 |
| **A_51_P421734** | Slc12a7 | 0.15726572 | -0.239793074 | 0.554324515 | 9.874081556 | 0.962415121 | 0.371979784 | 0.68002687 | -4.686701853 |
| **A_52_P152631** | Tmem17 | -0.133901238 | -0.472001077 | 0.2041986 | 8.248586409 | -0.962327329 | 0.372020538 | 0.68002687 | -4.686739229 |
| **A_51_P433824** | Tpi1 | 0.098498764 | -0.150217871 | 0.3472154 | 14.32267189 | 0.962297012 | 0.372034613 | 0.68002687 | -4.686752135 |
| **A_51_P160673** | Kcne1l | -0.27097892 | -0.955238649 | 0.41328081 | 8.102596726 | -0.962271702 | 0.372046363 | 0.68002687 | -4.68676291 |
| **A_52_P230167** | Setbp1 | -0.246951625 | -0.87069965 | 0.376796399 | 7.929367017 | -0.962023973 | 0.372161387 | 0.680172792 | -4.686868362 |
| **A_55_P2041843** | Nde1 | 0.125520291 | -0.191567341 | 0.442607922 | 8.589935156 | 0.961873197 | 0.372231408 | 0.680236446 | -4.686932535 |
| **A_55_P2039439** | Ppp1ca | -0.116958391 | -0.412474312 | 0.178557531 | 14.77740699 | -0.961686873 | 0.372317952 | 0.680254024 | -4.687011831 |
| **A_51_P249867** | Snrnp48 | 0.078668607 | -0.12011552 | 0.277452734 | 9.499867423 | 0.961618797 | 0.372349576 | 0.680254024 | -4.6870408 |
| **A_51_P251209** | Cacna1d | -0.12007003 | -0.423477146 | 0.183337085 | 7.731252633 | -0.96159467 | 0.372360784 | 0.680254024 | -4.687051067 |
| **A_55_P2114148** | NA | 0.128992988 | -0.196977045 | 0.45496302 | 9.750255767 | 0.961549431 | 0.372381801 | 0.680254024 | -4.687070317 |
| **A_55_P1962174** | Rab8a | 0.08648242 | -0.132093873 | 0.305058714 | 10.01034427 | 0.961408523 | 0.372447269 | 0.680295223 | -4.687130274 |
| **A_55_P2167323** | Cort | 0.06402228 | -0.097807638 | 0.225852199 | 7.935683099 | 0.961292409 | 0.372501224 | 0.680295223 | -4.687179676 |
| **A_51_P305437** | Rcn1 | -0.209402189 | -0.738748779 | 0.319944401 | 11.22622948 | -0.961223293 | 0.372533344 | 0.680295223 | -4.687209081 |
| **A_55_P1982986** | Cfap410 | -0.084893137 | -0.299499674 | 0.129713401 | 8.022297951 | -0.961197919 | 0.372545136 | 0.680295223 | -4.687219875 |
| **A_55_P2073699** | Bnip3l | 0.088606786 | -0.13549874 | 0.312712311 | 12.62986076 | 0.960721713 | 0.3727665 | 0.680542021 | -4.687422436 |
| **A_55_P2017183** | NA | -0.076044078 | -0.268382765 | 0.116294609 | 10.35344714 | -0.960687057 | 0.372782614 | 0.680542021 | -4.687437175 |
| **A_55_P1983149** | Rps29 | -0.086233698 | -0.304346657 | 0.13187926 | 17.106873 | -0.960679963 | 0.372785913 | 0.680542021 | -4.687440192 |
| **A_55_P2061620** | Chst1 | -0.150120227 | -0.529938517 | 0.229698063 | 8.375577373 | -0.960387563 | 0.372921891 | 0.680725965 | -4.687564535 |
| **A_51_P253527** | Rffl | 0.107005166 | -0.163755729 | 0.377766062 | 8.596739414 | 0.960289086 | 0.372967696 | 0.680745288 | -4.687606407 |
| **A_55_P1984243** | Naip2 | 0.138734894 | -0.212370115 | 0.489839903 | 9.334083769 | 0.960134046 | 0.373039819 | 0.680762539 | -4.687672325 |
| **A_55_P2155833** | NA | 0.069442829 | -0.106311909 | 0.245197568 | 16.04531275 | 0.960071445 | 0.373068943 | 0.680762539 | -4.687698939 |
| **A_55_P2017845** | Ehbp1 | 0.069307543 | -0.106131307 | 0.244746394 | 8.041896033 | 0.959926368 | 0.373136446 | 0.680762539 | -4.687760613 |
| **A_51_P440399** | Bcap31 | -0.086354014 | -0.304945375 | 0.132237347 | 12.26600284 | -0.959914885 | 0.373141789 | 0.680762539 | -4.687765494 |
| **A_55_P2010966** | Zfp688 | 0.139195775 | -0.213165081 | 0.491556631 | 9.434018328 | 0.959890263 | 0.373153246 | 0.680762539 | -4.68777596 |
| **A_55_P2173066** | NA | -0.128040249 | -0.452195239 | 0.196114741 | 8.714457034 | -0.959791697 | 0.373199114 | 0.680781963 | -4.687817857 |
| **A_51_P123805** | Sbno1 | 0.10638368 | -0.162986717 | 0.375754078 | 8.789128971 | 0.959639979 | 0.373269726 | 0.68081128 | -4.687882343 |
| **A_52_P481182** | Stard5 | -0.183268031 | -0.647331129 | 0.280795067 | 8.677502195 | -0.959605809 | 0.37328563 | 0.68081128 | -4.687896866 |

| | | | | | | | | | |
|---|---|---|---|---|---|---|---|---|---|
| **A_55_P2185870** | Bcl7a | 0.079360358 | -0.121685873 | 0.28040659 | 9.052304686 | 0.959159587 | 0.373493375 | 0.681125903 | -4.688086487 |
| **A_55_P2141786** | Pard6g | -0.404722245 | -1.430149747 | 0.620705256 | 9.336228052 | -0.95903687 | 0.373550523 | 0.681158875 | -4.688138626 |
| **A_55_P1955726** | Ptpn5 | -0.176057057 | -0.622229782 | 0.270115668 | 7.755823108 | -0.95881233 | 0.373655107 | 0.681158875 | -4.688234016 |
| **A_51_P260658** | Amdhd2 | -0.104645919 | -0.369845183 | 0.160553344 | 11.44686667 | -0.958811362 | 0.373655558 | 0.681158875 | -4.688234427 |
| **A_55_P1966159** | Dlg5 | -0.233645477 | -0.825807377 | 0.358516422 | 9.598707905 | -0.958738366 | 0.373689562 | 0.681158875 | -4.688265435 |
| **A_55_P2161690** | Tomm40 | 0.121216949 | -0.186014155 | 0.428448052 | 8.906387609 | 0.958696974 | 0.373708845 | 0.681158875 | -4.688283017 |
| **A_51_P350214** | Amz2 | -0.123625787 | -0.436972082 | 0.189720509 | 10.12036398 | -0.958666812 | 0.373722897 | 0.681158875 | -4.688295829 |
| **A_55_P1989573** | Tmem175 | 0.087944801 | -0.135013334 | 0.310902935 | 9.617071505 | 0.95845127 | 0.373823326 | 0.681251599 | -4.688387376 |
| **A_55_P2079142** | Fam124b | 0.126991804 | -0.19498997 | 0.448973578 | 8.60010973 | 0.958357602 | 0.373866975 | 0.681251599 | -4.688427156 |
| **A_51_P289697** | Nek4 | -0.093206966 | -0.329535495 | 0.143121563 | 9.023997672 | -0.958330715 | 0.373879505 | 0.681251599 | -4.688438574 |
| **A_51_P173709** | Gprc5b | -0.117016069 | -0.413758467 | 0.179726329 | 8.120758603 | -0.95818439 | 0.373947703 | 0.681282892 | -4.688500711 |
| **A_55_P2349198** | NA | -0.108412041 | -0.383347372 | 0.166523289 | 8.31816665 | -0.95814262 | 0.373967173 | 0.681282892 | -4.688518448 |
| **A_55_P1954006** | Ldhb | -0.509776644 | -1.802707624 | 0.783154336 | 9.863053305 | -0.958049131 | 0.374010752 | 0.681298071 | -4.688558144 |
| **A_55_P1957003** | Timm10 | 0.113474468 | -0.174365809 | 0.401314745 | 12.28030744 | 0.957921187 | 0.374070399 | 0.681342512 | -4.688612466 |
| **A_51_P411345** | Mogat2 | 0.210379828 | -0.323430601 | 0.744190257 | 11.45448314 | 0.957635484 | 0.374203619 | 0.681520941 | -4.688733754 |
| **A_55_P1960698** | Pih1d2 | -0.157224959 | -0.556257096 | 0.241807177 | 7.954307835 | -0.957407457 | 0.374309971 | 0.681603073 | -4.688830541 |
| **A_55_P2040962** | Gm1604b | -0.101832894 | -0.360287031 | 0.156621242 | 9.593086551 | -0.957387583 | 0.374319242 | 0.681603073 | -4.688838977 |
| **A_52_P366105** | Pip4p1 | -0.089181952 | -0.315632347 | 0.137268442 | 8.705223692 | -0.956945232 | 0.374525627 | 0.681914642 | -4.689026694 |
| **A_51_P223458** | Polr3d | 0.119026689 | -0.183276955 | 0.421330333 | 9.012438081 | 0.956718471 | 0.374631459 | 0.681965003 | -4.689122902 |
| **A_66_P103792** | AI606181 | 0.189294806 | -0.29154532 | 0.670134933 | 10.04249477 | 0.956579753 | 0.374696213 | 0.681965003 | -4.68918175 |
| **A_51_P481832** | Rabggta | -0.087135562 | -0.308504198 | 0.134233075 | 8.771388991 | -0.956450579 | 0.374756519 | 0.681965003 | -4.689236544 |
| **A_52_P549166** | Arsj | -0.069338611 | -0.245497961 | 0.106820739 | 7.666057428 | -0.95642876 | 0.374766706 | 0.681965003 | -4.689245799 |
| **A_55_P2181419** | Tcf4 | -0.074131777 | -0.262470521 | 0.114206967 | 7.776874317 | -0.956418413 | 0.374771537 | 0.681965003 | -4.689250188 |
| **A_55_P2020256** | Sik2 | 0.113944305 | -0.175548668 | 0.403437278 | 9.077002539 | 0.956396077 | 0.374781966 | 0.681965003 | -4.689259662 |
| **A_51_P124748** | Tgfb3 | -0.363848997 | -1.288337433 | 0.560639439 | 9.061197955 | -0.956319228 | 0.374817849 | 0.681965003 | -4.685382411 |
| **A_52_P547187** | Tab2 | 0.121925529 | -0.187882135 | 0.431733192 | 11.25719107 | 0.956281337 | 0.374835542 | 0.681965003 | -4.689308328 |
| **A_51_P364210** | Ms4a3 | 0.097275903 | -0.149953836 | 0.344505642 | 11.50967488 | 0.956065722 | 0.374936237 | 0.682061053 | -4.68939977 |
| **A_55_P2326582** | 5730408K05Rik | 0.066873715 | -0.103096545 | 0.236843974 | 7.855335338 | 0.956017181 | 0.374958909 | 0.682061053 | -4.689420355 |
| **A_52_P437795** | Cdh11 | -0.61489009 | -2.178250915 | 0.948470735 | 8.425292846 | -0.955699718 | 0.375107214 | 0.682266617 | -4.689554965 |
| **A_52_P88808** | Vps37a | 0.125162827 | -0.193101702 | 0.443427355 | 9.178859099 | 0.955587179 | 0.375159798 | 0.682298055 | -4.689602676 |
| **A_52_P344978** | NA | 0.093904014 | -0.144911099 | 0.332719128 | 8.761080637 | 0.955444664 | 0.375226397 | 0.682346942 | -4.689663092 |
| **A_55_P2029642** | NA | -0.18635291 | -0.660315286 | 0.287609466 | 13.15942938 | -0.95537858 | 0.375257281 | 0.682346942 | -4.689691105 |
| **A_55_P2055869** | Dvl2 | -0.076807652 | -0.272214364 | 0.11859906 | 8.989098596 | -0.955098587 | 0.37538816 | 0.682520717 | -4.68980978 |
| **A_55_P1976869** | Hsdl2 | 0.132177799 | -0.204160208 | 0.468515806 | 10.42080613 | 0.954917291 | 0.375472923 | 0.682540462 | -4.689886612 |
| **A_55_P2065050** | NA | 0.125565769 | -0.193951082 | 0.445082621 | 8.680428597 | 0.954906135 | 0.375478139 | 0.682540462 | -4.689891339 |
| **A_66_P122099** | Zc3h8 | 0.093071388 | -0.143791581 | 0.329934357 | 8.382051343 | 0.954777578 | 0.375538255 | 0.682540462 | -4.689945815 |
| **A_51_P327632** | Igsf3 | -0.275657657 | -0.977216418 | 0.425901104 | 8.96291514 | -0.954748967 | 0.375551634 | 0.682540462 | -4.689957938 |
| **A_55_P2001064** | NA | 0.078758831 | -0.121722725 | 0.279240388 | 12.04280013 | 0.954570543 | 0.375635083 | 0.682540462 | -4.690033536 |
| **A_51_P257065** | Prpf40b | -0.088883883 | -0.315141186 | 0.13737342 | 8.434269705 | -0.954560818 | 0.375639631 | 0.682540462 | -4.690037656 |
| **A_55_P2026748** | Zmat1 | -0.1353058 | -0.479736926 | 0.209125325 | 7.964217119 | -0.954546771 | 0.375646202 | 0.682540462 | -4.690043607 |
| **A_55_P2030373** | Atp1a2 | -0.077787438 | -0.275857168 | 0.120282292 | 7.73109554 | -0.954277198 | 0.37577231 | 0.682646187 | -4.690157807 |
| **A_52_P592951** | Ostc | -0.109716733 | -0.389107685 | 0.169674219 | 12.71621492 | -0.954209464 | 0.375804003 | 0.682646187 | -4.690186499 |
| **A_66_P121059** | Mirg | 0.142565357 | -0.220481202 | 0.505611915 | 10.63722731 | 0.954190021 | 0.3758131 | 0.682646187 | -4.690194734 |
| **A_51_P158552** | Zscan12 | 0.066533219 | -0.102907788 | 0.235974225 | 10.02124305 | 0.954120441 | 0.375845658 | 0.682646187 | -4.690224206 |
| **A_55_P2100928** | Ptgds | -0.369735448 | -1.31145819 | 0.571987295 | 8.308653275 | -0.954006251 | 0.375899095 | 0.682650941 | -4.69027257 |
| **A_52_P303928** | Rab2a | -0.163915096 | -0.581428284 | 0.253598091 | 10.48561767 | -0.953963911 | 0.37591891 | 0.682650941 | -4.690290501 |
| **A_55_P2101526** | L1td1 | 0.070208425 | -0.108671755 | 0.249088606 | 8.109636418 | 0.95369652 | 0.376044069 | 0.682799859 | -4.690403735 |
| **A_55_P2069597** | Tspan17 | -0.110420374 | -0.391850393 | 0.171009644 | 7.952224819 | -0.953371105 | 0.37619643 | 0.682799859 | -4.690541515 |
| **A_52_P633489** | C1qtnf4 | 0.110264066 | -0.17076929 | 0.391297422 | 8.406980052 | 0.953365263 | 0.376199166 | 0.682799859 | -4.690543989 |
| **A_55_P2130256** | Polr2l | 0.234771129 | -0.363619994 | 0.833162252 | 9.529607384 | 0.953328847 | 0.376216219 | 0.682799859 | -4.690559405 |

| | | | | | | | | | |
|---|---|---|---|---|---|---|---|---|---|
| **A_55_P2119721** | NA | 0.094827962 | -0.146884822 | 0.336540746 | 17.10963613 | 0.95327891 | 0.376239605 | 0.682799859 | -4.690580545 |
| **A_51_P498631** | Gsdme | 0.167195051 | -0.258979179 | 0.593369281 | 9.515242228 | 0.953277673 | 0.376240184 | 0.682799859 | -4.690581069 |
| **A_55_P1953693** | Pex2 | 0.099078764 | -0.153474007 | 0.351631534 | 10.54126012 | 0.953260574 | 0.376248192 | 0.682799859 | -4.690588307 |
| **A_55_P2089462** | Ncoa5 | 0.155615142 | -0.241128996 | 0.552359281 | 12.97943264 | 0.953069397 | 0.376337734 | 0.68285869 | -4.690669233 |
| **A_52_P443167** | Naa35 | 0.087448379 | -0.135516912 | 0.31041367 | 11.06769409 | 0.953010514 | 0.376365317 | 0.68285869 | -4.690694156 |
| **A_55_P1989511** | Tbca | 0.094643975 | -0.146679276 | 0.335967226 | 11.64560124 | 0.952965092 | 0.376386595 | 0.68285869 | -4.690713381 |
| **A_51_P275697** | Rflnb | -0.071705748 | -0.254613822 | 0.111202326 | 8.025692457 | -0.952586192 | 0.376564128 | 0.683116661 | -4.69087373 |
| **A_55_P2038525** | C3 | -0.250978209 | -0.891329312 | 0.389372894 | 16.45043928 | -0.952359774 | 0.376670247 | 0.683202306 | -4.690969531 |
| **A_55_P2012974** | Sdha | 0.092015554 | -0.142805524 | 0.326836631 | 12.84333118 | 0.95215435 | 0.376766546 | 0.683202306 | -4.691056438 |
| **A_51_P156800** | Mob2 | 0.153226669 | -0.237876338 | 0.544329676 | 10.68349716 | 0.951976867 | 0.376849763 | 0.683202306 | -4.691131514 |
| **A_66_P121495** | Psat1 | -0.151397949 | -0.537839993 | 0.235044094 | 9.665209681 | -0.951960244 | 0.376857558 | 0.683202306 | -4.691138546 |
| **A_55_P2074488** | Crim1 | -0.093909996 | -0.333628986 | 0.145808994 | 8.223005392 | -0.951902727 | 0.376884529 | 0.683202306 | -4.691162873 |
| **A_55_P1996816** | NA | -0.188315241 | -0.669027502 | 0.29239702 | 13.98093783 | -0.951882757 | 0.376893894 | 0.683202306 | -4.69117132 |
| **A_51_P195215** | Fam149a | 0.094182238 | -0.146242606 | 0.334607082 | 9.477262776 | 0.951859508 | 0.376904797 | 0.683202306 | -4.691181153 |
| **A_55_P1960451** | Thsd7a | -0.066532504 | -0.236374761 | 0.103309754 | 7.808791618 | -0.951856109 | 0.376906391 | 0.683202306 | -4.691182591 |
| **A_55_P2018482** | Adh5 | 0.197334517 | -0.306491378 | 0.701160412 | 8.86699754 | 0.951712472 | 0.376973756 | 0.683202306 | -4.69124334 |
| **A_55_P2081168** | Rpl7 | 0.068697424 | -0.106704207 | 0.244099054 | 16.26146208 | 0.951677966 | 0.376989941 | 0.683202306 | -4.691257933 |
| **A_51_P394676** | Mef2d | -0.092070518 | -0.327165941 | 0.143024905 | 11.15350522 | -0.951611323 | 0.3770212 | 0.683202306 | -4.691286115 |
| **A_55_P2032362** | NA | 0.086304413 | -0.134074733 | 0.306683559 | 8.135726273 | 0.951580858 | 0.377035491 | 0.683202306 | -4.691298998 |
| **A_51_P438967** | Gpnmb | 0.062621828 | -0.09732564 | 0.222569296 | 7.733451559 | 0.951330806 | 0.377152802 | 0.683350816 | -4.691404732 |
| **A_55_P2039903** | Wsb2 | -0.362707998 | -1.289480572 | 0.564064576 | 11.13668342 | -0.950970725 | 0.377321782 | 0.683567274 | -4.69155696 |
| **A_55_P2022658** | Cdk11b | 0.085893459 | -0.13360442 | 0.305391338 | 11.88267134 | 0.950852065 | 0.37737748 | 0.683567274 | -4.691607118 |
| **A_55_P2053497** | Poli | 0.104593017 | -0.16269149 | 0.371877524 | 10.139214 | 0.950850183 | 0.377378363 | 0.683567274 | -4.691607914 |
| **A_55_P2006906** | Armcx3 | 0.185065428 | -0.28792718 | 0.658058035 | 9.101105473 | 0.950723285 | 0.377437935 | 0.683585248 | -4.691661549 |
| **A_52_P83557** | Arfgef2 | 0.072048255 | -0.11210403 | 0.25620054 | 8.22894396 | 0.950669463 | 0.377463204 | 0.683585248 | -4.691684297 |
| **A_55_P2380806** | Gm2115 | -0.09709905 | -0.345297442 | 0.151099342 | 7.747319765 | -0.950603056 | 0.377494384 | 0.683585248 | -4.691712362 |
| **A_55_P2137701** | Rex2 | 0.061488012 | -0.095739368 | 0.218715393 | 7.962477633 | 0.950266598 | 0.377652386 | 0.683684293 | -4.691854539 |
| **A_55_P2003266** | Fam107a | 0.161548736 | -0.251601442 | 0.574698914 | 9.339619153 | 0.950120763 | 0.377720888 | 0.683684293 | -4.691916155 |
| **A_51_P376789** | Pgghg | -0.08932447 | -0.317773308 | 0.139124368 | 9.8336788 | -0.950089865 | 0.377735402 | 0.683684293 | -4.69192921 |
| **A_55_P1982593** | Rad51d | 0.062512749 | -0.09737368 | 0.222399178 | 8.339561571 | 0.950036268 | 0.377760581 | 0.683684293 | -4.691951853 |
| **A_66_P108468** | 2610528A11Rik | -0.063801362 | -0.226991026 | 0.099388302 | 7.779546232 | -0.949993185 | 0.377780821 | 0.683684293 | -4.691970053 |
| **A_55_P2325663** | Cers3 | -0.281553613 | -1.001713047 | 0.438605821 | 7.887473666 | -0.949982574 | 0.377785806 | 0.683684293 | -4.691974536 |
| **A_51_P176042** | Pklr | 0.252423468 | -0.393326109 | 0.898173044 | 9.688193255 | 0.949836354 | 0.377854507 | 0.683684293 | -4.692036304 |
| **A_52_P516409** | Col4a6 | -0.289394316 | -1.029729859 | 0.450941227 | 8.053947224 | -0.949827163 | 0.377858826 | 0.683684293 | -4.692040186 |
| **A_51_P116932** | Lad1 | -0.068156801 | -0.242520535 | 0.106206933 | 7.700140073 | -0.949808879 | 0.377867417 | 0.683684293 | -4.69204791 |
| **A_55_P2074381** | NA | -0.179475653 | -0.638676918 | 0.279725613 | 10.41359137 | -0.949698286 | 0.377919387 | 0.683714323 | -4.692094623 |
| **A_55_P2084278** | Ighmbp2 | 0.072954607 | -0.11372577 | 0.259634984 | 7.883681039 | 0.949592425 | 0.377969138 | 0.683740333 | -4.692139334 |
| **A_55_P2187038** | Tpm1 | -0.76850228 | -2.735304996 | 1.198300436 | 9.985058176 | -0.949440055 | 0.378040755 | 0.683796829 | -4.692203684 |
| **A_55_P2097048** | NA | -0.183562261 | -0.653379112 | 0.286254589 | 11.92019609 | -0.949375455 | 0.378071122 | 0.683796829 | -4.692230964 |
| **A_51_P172573** | Sod2 | -0.110231663 | -0.392415231 | 0.171951904 | 13.77287419 | -0.949200221 | 0.378153504 | 0.683881837 | -4.69230496 |
| **A_52_P460836** | Lrrn4 | 0.093725395 | -0.146234311 | 0.333685101 | 7.938596962 | 0.949078525 | 0.378210725 | 0.68392133 | -4.692356343 |
| **A_55_P2143195** | Dyrk1a | 0.083173706 | -0.129806128 | 0.29615354 | 9.067691451 | 0.948922532 | 0.378284082 | 0.683989992 | -4.692422201 |
| **A_66_P140067** | Wdyhv1 | 0.094443615 | -0.147437705 | 0.336324934 | 9.406801067 | 0.94875365 | 0.378363512 | 0.684069621 | -4.692493494 |
| **A_66_P123989** | Ikzf5 | 0.12707966 | -0.198466282 | 0.452625603 | 8.985698084 | 0.948520993 | 0.378472958 | 0.684203499 | -4.692591696 |
| **A_55_P2235931** | Cacna1c | -0.349153473 | -1.243705851 | 0.545398904 | 8.13716406 | -0.948404883 | 0.378527588 | 0.684212553 | -4.692640699 |
| **A_52_P601385** | Cd99l2 | -0.123465621 | -0.439807043 | 0.1928758 | 10.51684659 | -0.948359882 | 0.378548763 | 0.684212553 | -4.692659691 |
| **A_55_P2009910** | Stxbp3 | -0.072974528 | -0.259991114 | 0.114042058 | 10.33333211 | -0.948144122 | 0.378650299 | 0.684332082 | -4.692750738 |
| **A_55_P2181371** | NA | 0.109316641 | -0.17088571 | 0.389518992 | 8.595883761 | 0.947976767 | 0.37872907 | 0.684410451 | -4.692821351 |
| **A_55_P2115057** | Tuba1a | -0.452030304 | -1.611052667 | 0.706992059 | 11.81249247 | -0.947673909 | 0.378871651 | 0.684562407 | -4.692949118 |
| **A_55_P2009266** | Gm16060 | -0.144992627 | -0.516769347 | 0.226784093 | 14.62037212 | -0.947647706 | 0.378883989 | 0.684562407 | -4.692960171 |

| | | | | | | | | | |
|---|---|---|---|---|---|---|---|---|---|
| A_55_P2209308 | 9530083O12Rik | 0.333672208 | -0.522008306 | 1.189352723 | 12.30340653 | 0.947526974 | 0.378940842 | 0.684601134 | -4.693011098 |
| A_55_P1962384 | Churc1 | 0.1009688 | -0.158003132 | 0.359940731 | 11.54566219 | 0.947365767 | 0.379016764 | 0.684669001 | -4.69307909 |
| A_52_P515282 | 1110051M20Rik | -0.15653837 | -0.558150582 | 0.245073843 | 8.102508036 | -0.947102722 | 0.379140673 | 0.684669001 | -4.693190019 |
| A_55_P2002819 | Rabgap1l | 0.160557978 | -0.251376795 | 0.572492751 | 9.954364404 | 0.947079914 | 0.379151418 | 0.684669001 | -4.693199637 |
| A_51_P248304 | Vps26b | 0.068874969 | -0.10785023 | 0.245600168 | 9.020088405 | 0.946991594 | 0.379193029 | 0.684669001 | -4.693236877 |
| A_52_P418489 | Tinagl1 | -0.151443038 | -0.540053341 | 0.237167265 | 9.017498532 | -0.94693069 | 0.379221727 | 0.684669001 | -4.693262557 |
| A_55_P2072288 | Tmem167b | 0.067270087 | -0.105371974 | 0.239912148 | 9.435415191 | 0.946800694 | 0.379282984 | 0.684669001 | -4.693317365 |
| A_55_P2176802 | Hipk1 | 0.165329084 | -0.258983599 | 0.589641767 | 9.374245306 | 0.946774244 | 0.379295448 | 0.684669001 | -4.693328516 |
| A_51_P118885 | Amacr | 0.234610432 | -0.367553004 | 0.836773869 | 11.90853424 | 0.946708175 | 0.379326585 | 0.684669001 | -4.691395456 |
| A_51_P293982 | Plekho2 | 0.106207701 | -0.166423972 | 0.378839374 | 9.092107481 | 0.946592131 | 0.379381279 | 0.684669001 | -4.693405289 |
| A_51_P501656 | Poglut3 | -0.188832462 | -0.673582347 | 0.295917422 | 8.551803563 | -0.946546886 | 0.379402606 | 0.684669001 | -4.693424361 |
| A_51_P181891 | Amer2 | 0.120434253 | -0.188736213 | 0.429604719 | 8.449362155 | 0.946531808 | 0.379409713 | 0.684669001 | -4.693430717 |
| A_55_P2011031 | Cdk5rap1 | 0.145200697 | -0.227563402 | 0.517964797 | 8.000569286 | 0.946493882 | 0.379427591 | 0.684669001 | -4.693446703 |
| A_66_P127567 | Nxf1 | 0.074535505 | -0.116819652 | 0.265890661 | 12.68111944 | 0.94646861 | 0.379439504 | 0.684669001 | -4.693457356 |
| A_51_P390285 | Larp6 | -0.070346617 | -0.250973354 | 0.110280121 | 7.831789416 | -0.946333833 | 0.379503042 | 0.684669001 | -4.693514163 |
| A_55_P1985077 | Mcrip1 | -0.151442757 | -0.540303271 | 0.237417756 | 8.313064472 | -0.946319637 | 0.379509735 | 0.684669001 | -4.693520146 |
| A_66_P104473 | Polg2 | 0.08133667 | -0.127564446 | 0.290237786 | 9.426387868 | 0.946082079 | 0.379621749 | 0.684770577 | -4.693620261 |
| A_55_P2024215 | Ctnnbl1 | -0.077359827 | -0.276053758 | 0.121334104 | 12.02125998 | -0.946049967 | 0.379636893 | 0.684770577 | -4.693633793 |
| A_55_P2137910 | NA | -0.186698799 | -0.666304697 | 0.292907099 | 13.02667109 | -0.945889049 | 0.379712787 | 0.684843562 | -4.6937016 |
| A_51_P473919 | Mycbp2 | 0.098785975 | -0.155089059 | 0.35266101 | 9.37555565 | 0.945493363 | 0.379899453 | 0.684987065 | -4.693868301 |
| A_55_P2149951 | Prx | -0.715640043 | -2.555106211 | 1.123826124 | 8.905545943 | -0.945335611 | 0.379973893 | 0.684987065 | -4.69393475 |
| A_52_P14456 | Srrm4 | -0.200256825 | -0.71499572 | 0.31448207 | 8.47243875 | -0.945330192 | 0.379976451 | 0.684987065 | -4.693937033 |
| A_52_P677956 | Pex11g | -0.116903453 | -0.417408203 | 0.183601297 | 11.8783434 | -0.945277208 | 0.380001455 | 0.684987065 | -4.693959349 |
| A_55_P1968841 | NA | 0.206969727 | -0.32508998 | 0.739029433 | 13.02128681 | 0.94521291 | 0.380031801 | 0.684987065 | -4.69398643 |
| A_55_P1973643 | Raf1 | -0.076026254 | -0.271468453 | 0.119415945 | 11.86463428 | -0.945210295 | 0.380033036 | 0.684987065 | -4.693987531 |
| A_52_P279152 | Vamp8 | -0.138235551 | -0.493620495 | 0.217149393 | 12.61377121 | -0.945156905 | 0.380058234 | 0.684987065 | -4.694010017 |
| A_52_P574945 | 1500026H17Rik | -0.088826142 | -0.317195188 | 0.139542903 | 8.431229276 | -0.945119566 | 0.380075859 | 0.684987065 | -4.694025742 |
| A_51_P115626 | Shank3 | -0.100725194 | -0.359720701 | 0.158270313 | 8.104047918 | -0.944994041 | 0.380135112 | 0.685007713 | -4.69211077 |
| A_55_P2074206 | Dlgap4 | -0.157808033 | -0.563602104 | 0.247986038 | 10.6012685 | -0.944945146 | 0.380158194 | 0.685007713 | -4.694099194 |
| A_51_P399889 | Ap4b1 | 0.108980298 | -0.171334102 | 0.389294698 | 11.30003666 | 0.944682285 | 0.380282304 | 0.685053469 | -4.694209874 |
| A_51_P137611 | Card14 | 0.06702895 | -0.105386152 | 0.239444051 | 7.917268068 | 0.944648631 | 0.380298197 | 0.685053469 | -4.694224044 |
| A_65_P09090 | Arih2 | 0.093851963 | -0.147561557 | 0.335265483 | 11.30131084 | 0.944637021 | 0.380303679 | 0.685053469 | -4.694228931 |
| A_66_P100932 | Tmem256 | 0.176543956 | -0.277606017 | 0.630693929 | 15.1811459 | 0.944575661 | 0.380332656 | 0.685053469 | -4.694254764 |
| A_55_P1983262 | Gm4026 | 0.165060294 | -0.259575352 | 0.589695939 | 8.856156015 | 0.944516078 | 0.380360796 | 0.685053469 | -4.694279848 |
| A_55_P2028243 | Dnajc19 | 0.285526863 | -0.449098811 | 1.020152538 | 12.08251235 | 0.944417576 | 0.380407319 | 0.685073425 | -4.694321314 |
| A_52_P273169 | Sh3d19 | -0.096675582 | -0.345476893 | 0.152125729 | 8.425740184 | -0.944163749 | 0.380527224 | 0.685225519 | -4.694428155 |
| A_55_P2186110 | R3hdm2 | -0.116974449 | -0.418087059 | 0.184138161 | 11.49758362 | -0.943941876 | 0.380632059 | 0.685299269 | -4.694521531 |
| A_55_P2133680 | NA | -0.144401883 | -0.516123345 | 0.227319579 | 11.44922563 | -0.943926998 | 0.380639089 | 0.685299269 | -4.694527792 |
| A_52_P505827 | Cyb5r4 | 0.134246607 | -0.211407358 | 0.479900572 | 9.192810661 | 0.943723924 | 0.380735062 | 0.6853657 | -4.694613244 |
| A_52_P15388 | Ltf | -0.268374594 | -0.95941483 | 0.422665642 | 13.48278948 | -0.943672401 | 0.380759414 | 0.6853657 | -4.694634923 |
| A_51_P236864 | Parp8 | -0.094721149 | -0.338632062 | 0.149189765 | 8.748826093 | -0.943623856 | 0.380782361 | 0.6853657 | -4.694655348 |
| A_66_P133043 | Mbd2 | 0.118041198 | -0.185957071 | 0.422039466 | 11.89861182 | 0.943508216 | 0.380837026 | 0.685400268 | -4.694704001 |
| A_51_P485479 | Prrc2a | 0.113377574 | -0.178689938 | 0.405445086 | 9.432586445 | 0.943250615 | 0.38095882 | 0.685546683 | -4.694812366 |
| A_55_P1992031 | Dydc2 | -0.166732561 | -0.596275124 | 0.262810003 | 7.771448439 | -0.943186127 | 0.380989315 | 0.685546683 | -4.694839492 |
| A_51_P375862 | Srpk3 | -0.067253358 | -0.240541736 | 0.106035021 | 7.668249136 | -0.943034809 | 0.381060877 | 0.685611625 | -4.694903137 |
| A_51_P401184 | Rarres1 | 0.174696489 | -0.275524722 | 0.6249177 | 8.950053077 | 0.942847412 | 0.381149515 | 0.685654301 | -4.694981947 |
| A_51_P123297 | Ppp1r7 | 0.076341173 | -0.120424567 | 0.273106913 | 8.333862565 | 0.942741304 | 0.381199711 | 0.685654301 | -4.695026567 |
| A_55_P2149122 | Farp2 | -0.126259756 | -0.451690985 | 0.199171474 | 9.04714543 | -0.942733427 | 0.381203438 | 0.685654301 | -4.69502988 |
| A_55_P2048660 | NA | 0.297779501 | -0.469779069 | 1.065338071 | 11.90391423 | 0.942684708 | 0.381226487 | 0.685654301 | -4.695050365 |
| A_52_P84027 | Cyp7a1 | -0.45643522 | -1.633084842 | 0.720214403 | 9.206389118 | -0.942573395 | 0.381279154 | 0.685685222 | -4.695097169 |

| | | | | | | | | | |
|---|---|---|---|---|---|---|---|---|---|
| **A_55_P1985359** | Rpl28 | -0.166826259 | -0.597021227 | 0.26336871 | 14.78398292 | -0.942284988 | 0.381415637 | 0.68579829 | -4.69521842 |
| **A_51_P494686** | Sumf1 | -0.060215747 | -0.215496396 | 0.095064902 | 9.746973327 | -0.942271226 | 0.381422151 | 0.68579829 | -4.695224205 |
| **A_51_P168792** | Xylb | 0.187603163 | -0.296211862 | 0.671418188 | 9.963820262 | 0.942201939 | 0.381454946 | 0.68579829 | -4.695253331 |
| **A_55_P2176361** | Zcchc9 | 0.066104944 | -0.104386024 | 0.236595912 | 8.42544779 | 0.942140672 | 0.381483947 | 0.68579829 | -4.695279084 |
| **A_55_P2137206** | Tgfbr1 | 0.1140344 | -0.180127738 | 0.408196538 | 10.10013298 | 0.941959644 | 0.381569646 | 0.68588856 | -4.695355172 |
| **A_55_P2045278** | Gm10845 | 0.069046443 | -0.109109039 | 0.247201926 | 12.65688404 | 0.941727623 | 0.381679507 | 0.685978156 | -4.695452681 |
| **A_55_P2425235** | NA | 0.125762093 | -0.198740604 | 0.45026479 | 9.276667007 | 0.941704476 | 0.381690469 | 0.685978156 | -4.695462408 |
| **A_55_P2109752** | Nans | 0.089198453 | -0.141009298 | 0.319406203 | 12.85393428 | 0.941500531 | 0.381787059 | 0.686087956 | -4.695548104 |
| **A_55_P1987988** | Atg2b | 0.080689719 | -0.127588448 | 0.288967886 | 8.221732684 | 0.941364133 | 0.381851668 | 0.686103863 | -4.695605411 |
| **A_51_P212038** | Atp6v0e2 | -0.130596093 | -0.467712578 | 0.206520391 | 8.235463429 | -0.941311544 | 0.381876581 | 0.686103863 | -4.695627504 |
| **A_55_P2007802** | Smarca2 | -0.247088189 | -0.884956348 | 0.39077997 | 8.795934766 | -0.941248403 | 0.381906494 | 0.686103863 | -4.69565403 |
| **A_52_P578790** | Dok3 | -0.141680904 | -0.507464541 | 0.224102732 | 9.090905224 | -0.941174683 | 0.381941422 | 0.686103863 | -4.695684999 |
| **A_51_P388696** | Ndufs4 | -0.062487267 | -0.223834682 | 0.098860148 | 13.08564911 | -0.94105002 | 0.38200049 | 0.686103863 | -4.695737365 |
| **A_55_P2123381** | Fga | -0.1596935 | -0.572043405 | 0.252656404 | 13.57719863 | -0.941032301 | 0.382008887 | 0.686103863 | -4.695744807 |
| **A_55_P2031979** | Sspn | -0.303542347 | -1.087500808 | 0.480416114 | 8.673194314 | -0.940826248 | 0.382106538 | 0.686215485 | -4.695831351 |
| **A_55_P2133356** | NA | -0.121972692 | -0.437080512 | 0.193135127 | 15.53792447 | -0.940560246 | 0.382232628 | 0.686293628 | -4.695943058 |
| **A_55_P2027386** | Gpr156 | -0.067608235 | -0.242276357 | 0.107059886 | 7.723730579 | -0.940522391 | 0.382250574 | 0.686293628 | -4.695958953 |
| **A_51_P262878** | Tmed9 | 0.103024711 | -0.163146629 | 0.369196051 | 14.28062754 | 0.940509748 | 0.382256568 | 0.686293628 | -4.695964262 |
| **A_51_P301736** | Ppp2r2a | -0.098309621 | -0.35232455 | 0.155705309 | 10.71168863 | -0.94041591 | 0.382301059 | 0.686309758 | -4.696003662 |
| **A_52_P356780** | Pdzd8 | 0.099733882 | -0.157999602 | 0.357467365 | 8.748245486 | 0.940275387 | 0.382367692 | 0.686324332 | -4.696062661 |
| **A_55_P2018557** | NA | 0.070493843 | -0.111682292 | 0.252669978 | 11.71751719 | 0.940249026 | 0.382380192 | 0.686324332 | -4.696073728 |
| **A_52_P445828** | Abcf2 | 0.118915553 | -0.188420814 | 0.42625192 | 9.184323394 | 0.940173262 | 0.382416122 | 0.686325091 | -4.696105535 |
| **A_52_P428125** | Atr | 0.098935737 | -0.156808619 | 0.354680094 | 10.53122638 | 0.940005348 | 0.382495763 | 0.686404289 | -4.696176022 |
| **A_55_P2136832** | Gm6981 | -0.098970414 | -0.354865291 | 0.156924464 | 14.54768372 | -0.939781696 | 0.382601859 | 0.686530944 | -4.696269895 |
| **A_55_P1968245** | Ftl1 | 0.144020817 | -0.228552918 | 0.516594551 | 16.69585762 | 0.939282484 | 0.382838758 | 0.686747926 | -4.696479378 |
| **A_55_P1968450** | Gm5766 | 0.091565765 | -0.145370063 | 0.328501593 | 15.47613872 | 0.939043217 | 0.382952341 | 0.686747926 | -4.696579758 |
| **A_51_P423743** | Cldn6 | -0.126900518 | -0.455286153 | 0.201485118 | 7.798806629 | -0.938993165 | 0.382976104 | 0.686747926 | -4.696600754 |
| **A_55_P2010271** | Samsn1 | 0.199850552 | -0.317313728 | 0.717014832 | 9.930392753 | 0.938987953 | 0.382978579 | 0.686747926 | -4.69660294 |
| **A_55_P1973521** | Rnf7 | 0.080952196 | -0.128548006 | 0.290452397 | 9.736468572 | 0.938917376 | 0.383012089 | 0.686747926 | -4.696632545 |
| **A_55_P1961235** | Rpl23 | -0.115744087 | -0.415286078 | 0.183797904 | 15.87005124 | -0.938910705 | 0.383015257 | 0.686747926 | -4.696635343 |
| **A_51_P174645** | Asl | -0.160560168 | -0.576095499 | 0.254975163 | 10.78500568 | -0.938886396 | 0.383026799 | 0.686747926 | -4.69664554 |
| **A_51_P106488** | Speg | -0.145719211 | -0.52289771 | 0.231459288 | 10.2147255 | -0.938756747 | 0.383088365 | 0.686747926 | -4.696699919 |
| **A_51_P485421** | NA | -0.157716141 | -0.565971205 | 0.250538922 | 8.592829491 | -0.938702019 | 0.383114356 | 0.686747926 | -4.696722872 |
| **A_52_P325527** | Edil3 | -0.154537595 | -0.554581431 | 0.24550624 | 7.774670938 | -0.938663124 | 0.383132829 | 0.686747926 | -4.696739185 |
| **A_55_P2011220** | Armcx1 | -0.12823197 | -0.460201798 | 0.203737859 | 9.03521784 | -0.938600729 | 0.383162463 | 0.686747926 | -4.696765352 |
| **A_55_P1952720** | Kctd2 | -0.131683514 | -0.472606695 | 0.209239667 | 11.18069944 | -0.938551406 | 0.383185891 | 0.686747926 | -4.696786037 |
| **A_55_P2063151** | NA | -0.073777992 | -0.264795976 | 0.117239992 | 10.21205174 | -0.938503207 | 0.383208786 | 0.686747926 | -4.696806249 |
| **A_55_P2090505** | NA | -0.085892626 | -0.308301684 | 0.136516431 | 11.51818891 | -0.938396979 | 0.383259247 | 0.686747926 | -4.696850794 |
| **A_55_P2076782** | 1700028P14Rik | -0.159264513 | -0.571683817 | 0.253154791 | 7.754727454 | -0.938346465 | 0.383283245 | 0.686747926 | -4.696871975 |
| **A_55_P1999361** | Dip2a | -0.181909661 | -0.652995272 | 0.289175951 | 9.387578308 | -0.938294375 | 0.383307993 | 0.686747926 | -4.696893817 |
| **A_51_P165451** | Pbld2 | 0.16710636 | -0.265731312 | 0.599944033 | 8.809884214 | 0.938104277 | 0.383398318 | 0.686747926 | -4.696973518 |
| **A_51_P412348** | Scnn1b | -0.079644054 | -0.285942974 | 0.126654865 | 7.685371895 | -0.938079395 | 0.383410142 | 0.686747926 | -4.69698395 |
| **A_51_P262701** | Cpb2 | -0.2499778 | -0.897509167 | 0.397553568 | 14.03229613 | -0.938045316 | 0.383426336 | 0.686747926 | -4.696998236 |
| **A_51_P425962** | Khdrbs3 | 0.098170046 | -0.156138162 | 0.352478254 | 9.948579905 | 0.937997769 | 0.383448932 | 0.686747926 | -4.697018169 |
| **A_52_P265002** | Pik3ca | -0.064079502 | -0.230084043 | 0.101925038 | 8.357130636 | -0.937955753 | 0.3834689 | 0.686747926 | -4.697035782 |
| **A_51_P511060** | Csgalnact2 | 0.126854997 | -0.201903585 | 0.455613579 | 10.06866668 | 0.93759152 | 0.383642036 | 0.686802374 | -4.69718845 |
| **A_55_P2312783** | D030029J20Rik | -0.123077713 | -0.442081371 | 0.195925945 | 10.22377291 | -0.937490644 | 0.383689997 | 0.686802374 | -4.697230726 |
| **A_55_P2187206** | NA | -0.070373249 | -0.252773013 | 0.112026514 | 8.28969976 | -0.93748974 | 0.383690427 | 0.686802374 | -4.697231105 |
| **A_52_P337259** | Heyl | -0.208091112 | -0.747453379 | 0.331271154 | 8.728239928 | -0.937467369 | 0.383701063 | 0.686802374 | -4.697240479 |
| **A_55_P2032936** | Cisd1 | -0.104442165 | -0.375167412 | 0.166283083 | 12.40892424 | -0.937411535 | 0.383727612 | 0.686802374 | -4.697263877 |

| | | | | | | | | |
|---|---|---|---|---|---|---|---|---|
| **A_51_P114634** | Amz1 | -0.10732404 | -0.385526288 | 0.170878208 | 8.841193138 | -0.937388384 | 0.383738621 | 0.686802374 | -4.697273578 |
| **A_55_P1987839** | Erbin | -0.092184794 | -0.331161944 | 0.146792356 | 10.39723707 | -0.937316152 | 0.383772969 | 0.686802374 | -4.697303846 |
| **A_55_P1993589** | NA | 0.092069736 | -0.146614813 | 0.330754285 | 15.33076538 | 0.937293881 | 0.38378356 | 0.686802374 | -4.697313178 |
| **A_52_P605517** | Phactr1 | -0.159797331 | -0.574236926 | 0.254642264 | 7.87699316 | -0.936896183 | 0.383972725 | 0.687000252 | -4.697479799 |
| **A_55_P2209298** | NA | -0.094573611 | -0.339863372 | 0.150716151 | 7.895055446 | -0.93685793 | 0.383990924 | 0.687000252 | -4.697495823 |
| **A_51_P461319** | Gatm | 0.118124363 | -0.188267067 | 0.424515794 | 9.627223603 | 0.936798216 | 0.384019335 | 0.687000252 | -4.697520837 |
| **A_66_P131645** | Ahnak2 | 0.080975485 | -0.129067038 | 0.291018008 | 7.859626548 | 0.93676255 | 0.384036304 | 0.687000252 | -4.697535777 |
| **A_55_P2078325** | Smco3 | -0.099620421 | -0.358121219 | 0.158880377 | 7.768114117 | -0.936417828 | 0.38420035 | 0.687127043 | -4.697680155 |
| **A_66_P119164** | Zfp354c | -0.206465742 | -0.742229913 | 0.329298429 | 8.323525291 | -0.936391631 | 0.384212819 | 0.687127043 | -4.697691126 |
| **A_55_P2096281** | NA | -0.066440345 | -0.238848788 | 0.105968098 | 16.31373605 | -0.936389511 | 0.384213828 | 0.687127043 | -4.697692014 |
| **A_55_P2039071** | Dhdh | -0.137292084 | -0.493588176 | 0.219004008 | 10.27420737 | -0.936305616 | 0.384253761 | 0.687134882 | -4.697727145 |
| **A_51_P306067** | Lpp | -0.183073846 | -0.658305887 | 0.292158194 | 10.05252271 | -0.936060204 | 0.384370593 | 0.687280221 | -4.697829903 |
| **A_55_P2175245** | Tusc3 | -0.085294193 | -0.306770587 | 0.1361822 | 10.16975332 | -0.935783132 | 0.384502529 | 0.687402282 | -4.697945897 |
| **A_55_P1971298** | Lonp1 | 0.106464709 | -0.170013954 | 0.382943371 | 13.54490851 | 0.935679768 | 0.384551758 | 0.687402282 | -4.697989164 |
| **A_55_P2049289** | NA | 0.066054833 | -0.10549466 | 0.237604326 | 16.32240309 | 0.935617529 | 0.384581403 | 0.687402282 | -4.698015215 |
| **A_55_P1999439** | Ino80e | 0.079089038 | -0.126348118 | 0.284526193 | 9.892040075 | 0.935449803 | 0.384661301 | 0.687402282 | -4.698085414 |
| **A_55_P2048759** | Mgarp | 0.283952639 | -0.453640208 | 1.021545485 | 8.877236693 | 0.935432387 | 0.384669598 | 0.687402282 | -4.698092703 |
| **A_51_P173384** | Tada1 | 0.078827228 | -0.125958802 | 0.283613258 | 9.649985348 | 0.935317627 | 0.384724273 | 0.687402282 | -4.698140729 |
| **A_55_P2103963** | Bhmt | 0.352092447 | -0.562682627 | 1.266867521 | 11.52824266 | 0.935245389 | 0.384758692 | 0.687402282 | -4.698170959 |
| **A_55_P2038687** | Tmco1 | 0.109772839 | -0.175435669 | 0.394981347 | 12.38750361 | 0.935223925 | 0.38476892 | 0.687402282 | -4.69817994 |
| **A_55_P2033666** | NA | -0.080106175 | -0.288240289 | 0.128027939 | 16.53694754 | -0.935203052 | 0.384778866 | 0.687402282 | -4.698188675 |
| **A_55_P2009915** | Stxbp3 | 0.085818404 | -0.137165201 | 0.308802009 | 8.264274825 | 0.935170264 | 0.384794489 | 0.687402282 | -4.698202394 |
| **A_51_P155763** | NA | 0.095933218 | -0.153362197 | 0.345228633 | 8.735914391 | 0.935056622 | 0.384848645 | 0.687435493 | -4.698249944 |
| **A_55_P1972187** | Slc25a14 | -0.087138707 | -0.313715666 | 0.139438252 | 9.58966945 | -0.934498387 | 0.385114757 | 0.687847268 | -4.69848347 |
| **A_55_P2055523** | NA | 0.07705758 | -0.123367699 | 0.277482859 | 13.21500955 | 0.934213329 | 0.385250698 | 0.688026495 | -4.698602685 |
| **A_55_P2011887** | Kif13a | -0.157573237 | -0.567501301 | 0.252354827 | 9.608197905 | -0.934023909 | 0.385341052 | 0.688059835 | -4.69868189 |
| **A_55_P1971522** | Kif16b | 0.066923989 | -0.107195766 | 0.241043744 | 8.684987107 | 0.933935661 | 0.385383151 | 0.688059835 | -4.698718788 |
| **A_55_P2056315** | Tbca | -0.09337325 | -0.336312663 | 0.149566162 | 13.14585023 | -0.933915727 | 0.385392662 | 0.688059835 | -4.698727122 |
| **A_52_P541270** | Crebl2 | -0.15838271 | -0.570482013 | 0.253716594 | 9.611828736 | -0.933875705 | 0.385411756 | 0.688059835 | -4.698743855 |
| **A_55_P2065537** | NA | 0.080061559 | -0.128270356 | 0.288393473 | 8.517410403 | 0.933794743 | 0.385450385 | 0.688065247 | -4.698777702 |
| **A_55_P2098911** | Lrrcc1 | 0.103969549 | -0.166598721 | 0.374537819 | 9.093090148 | 0.933711021 | 0.385490335 | 0.688073015 | -4.698812702 |
| **A_55_P2075013** | Tpt1 | -0.139649539 | -0.503227654 | 0.223928576 | 16.35528 | -0.933307943 | 0.385682714 | 0.688184116 | -4.698981181 |
| **A_55_P2126790** | 1500009L16Rik | -0.112665724 | -0.40600409 | 0.180672641 | 7.776931845 | -0.933267482 | 0.385702029 | 0.688184116 | -4.69899809 |
| **A_51_P144319** | Efna5 | 0.122958421 | -0.197238928 | 0.44315577 | 8.94105336 | 0.933090428 | 0.385786559 | 0.688184116 | -4.699072079 |
| **A_55_P2127131** | Trmt112 | -0.101085455 | -0.364336481 | 0.162165571 | 11.93529857 | -0.933043277 | 0.385809073 | 0.688184116 | -4.699091782 |
| **A_55_P2005743** | Ubxn11 | -0.103729545 | -0.373870161 | 0.16641107 | 8.034789197 | -0.933030368 | 0.385815237 | 0.688184116 | -4.699097176 |
| **A_52_P473975** | Cpsf4 | 0.087709413 | -0.140718531 | 0.316137356 | 10.50437726 | 0.932996817 | 0.385831258 | 0.688184116 | -4.699111195 |
| **A_51_P202498** | Sar1a | 0.088064096 | -0.141287631 | 0.317415823 | 10.74565763 | 0.932996588 | 0.385831367 | 0.688184116 | -4.69911129 |
| **A_55_P2142393** | Cacna1a | -0.142656371 | -0.51421151 | 0.228898768 | 8.115318755 | -0.932934361 | 0.385861082 | 0.688184116 | -4.699137291 |
| **A_55_P2037478** | Kifap3 | -0.219394318 | -0.790832302 | 0.352043667 | 9.390778688 | -0.932909379 | 0.385873012 | 0.688184116 | -4.699147729 |
| **A_55_P2000958** | Sdr39u1 | -0.090062314 | -0.324766451 | 0.144641823 | 9.665562877 | -0.932407047 | 0.386112957 | 0.688517309 | -4.699357576 |
| **A_55_P2030578** | Tns1 | -0.478142344 | -1.724241252 | 0.767956564 | 11.05922011 | -0.932369117 | 0.386131079 | 0.688517309 | -4.699373418 |
| **A_55_P2032498** | Arpin | -0.101859872 | -0.367392117 | 0.163672373 | 8.377808536 | -0.932114035 | 0.38625297 | 0.688671125 | -4.699479948 |
| **A_55_P2132681** | Fzr1 | 0.177173692 | -0.284730331 | 0.639077715 | 10.92679056 | 0.932031692 | 0.386292324 | 0.688677766 | -4.699514333 |
| **A_55_P1999182** | Cdk12 | 0.160706194 | -0.258391297 | 0.579803686 | 9.097381654 | 0.931752882 | 0.386425598 | 0.688822082 | -4.699630746 |
| **A_55_P2023384** | NA | -0.072214777 | -0.26057248 | 0.116142927 | 8.414646093 | -0.931592268 | 0.386502389 | 0.688822082 | -4.699697798 |
| **A_55_P1976984** | NA | -0.128590434 | -0.464021412 | 0.206840543 | 16.01936474 | -0.931512492 | 0.386540535 | 0.688822082 | -4.6997311 |
| **A_52_P599238** | Rspry1 | 0.074977762 | -0.120633609 | 0.270589132 | 9.255412052 | 0.93136854 | 0.386609375 | 0.688822082 | -4.699791187 |
| **A_52_P981697** | Zfp874a | 0.100967568 | -0.162452656 | 0.364387792 | 8.751120237 | 0.931356542 | 0.386615113 | 0.688822082 | -4.699796195 |
| **A_52_P487362** | Ppp4r4 | 0.076022021 | -0.122319416 | 0.274363457 | 8.252343164 | 0.931341904 | 0.386622114 | 0.688822082 | -4.699802305 |

| | | | | | | | | | |
|---|---|---|---|---|---|---|---|---|---|
| **A_55_P2186005** | Sphk1 | 0.147123637 | -0.236722434 | 0.530969709 | 9.735787943 | 0.931340614 | 0.386622731 | 0.688822082 | -4.699802843 |
| **A_55_P2072315** | NA | 0.350482132 | -0.564024434 | 1.264988699 | 12.48737614 | 0.931241332 | 0.386670216 | 0.688836161 | -4.69984428 |
| **A_55_P1964093** | Rangrf | 0.095239093 | -0.153284199 | 0.343762384 | 9.83875212 | 0.931175074 | 0.386701908 | 0.688836161 | -4.699871933 |
| **A_52_P428496** | Rhoa | -0.110460032 | -0.398754714 | 0.177834651 | 10.2543439 | -0.931004369 | 0.386783569 | 0.688886153 | -4.69994317 |
| **A_55_P2152245** | Rpl19 | -0.148965503 | -0.537772812 | 0.239841806 | 15.61169899 | -0.930967407 | 0.386801253 | 0.688886153 | -4.699958594 |
| **A_55_P2182586** | Esrp1 | -0.15386751 | -0.555669633 | 0.247934612 | 8.201593027 | -0.930503239 | 0.387023373 | 0.689218241 | -4.700152252 |
| **A_55_P2176871** | NA | 0.104080494 | -0.167765849 | 0.375926836 | 15.54632172 | 0.930312888 | 0.387114491 | 0.689307441 | -4.700231652 |
| **A_55_P2152872** | Fgfrl1 | -0.095630493 | -0.345431664 | 0.154170677 | 10.97540595 | -0.930218806 | 0.387159532 | 0.689307441 | -4.700270892 |
| **A_52_P594049** | Actr5 | 0.07772173 | -0.125328115 | 0.280771576 | 9.884507978 | 0.930085739 | 0.387223245 | 0.689307441 | -4.700326388 |
| **A_55_P2175579** | Olfr1466 | 0.331588641 | -0.53476296 | 1.197940242 | 9.451436185 | 0.930012213 | 0.387258453 | 0.689307441 | -4.700357051 |
| **A_55_P2025765** | Adam8 | -0.184049277 | -0.664943651 | 0.296845096 | 9.485985906 | -0.929967153 | 0.387280031 | 0.689307441 | -4.700375841 |
| **A_51_P484832** | Ppm1g | 0.179159819 | -0.288966884 | 0.647286523 | 9.975911577 | 0.929951694 | 0.387287434 | 0.689307441 | -4.700382287 |
| **A_51_P184385** | Pla2g12b | 0.221690991 | -0.35782562 | 0.801207602 | 11.08735045 | 0.929534139 | 0.387487437 | 0.689593569 | -4.700556383 |
| **A_51_P130544** | Gas5 | 0.129739052 | -0.209432862 | 0.468910967 | 8.887373856 | 0.929467113 | 0.387519548 | 0.689593569 | -4.700584325 |
| **A_55_P2019214** | NA | 0.109216476 | -0.176359747 | 0.394792698 | 8.822822001 | 0.929285801 | 0.387606424 | 0.689610533 | -4.700659903 |
| **A_55_P2014690** | NA | 0.136585135 | -0.220639046 | 0.493809315 | 9.326539384 | 0.929064308 | 0.387712573 | 0.689610533 | -4.700752217 |
| **A_51_P270891** | 1700056E22Rik | 0.102305175 | -0.165269198 | 0.369879549 | 9.165297786 | 0.929043984 | 0.387722314 | 0.689610533 | -4.700760687 |
| **A_55_P1968188** | Btf3 | 0.085499458 | -0.138134295 | 0.309133212 | 14.3835192 | 0.928986063 | 0.387750077 | 0.689610533 | -4.700784826 |
| **A_51_P470311** | Cby1 | -0.148561721 | -0.537173588 | 0.240050146 | 8.362301324 | -0.928910886 | 0.387786113 | 0.689610533 | -4.700816153 |
| **A_55_P1977741** | NA | 0.270228167 | -0.436733022 | 0.977189356 | 9.739221898 | 0.9287915 | 0.387843345 | 0.689610533 | -4.700865901 |
| **A_51_P149373** | Fbxl20 | -0.108584604 | -0.392668954 | 0.175499747 | 9.427013991 | -0.928761341 | 0.387857804 | 0.689610533 | -4.700878467 |
| **A_55_P2231152** | Gm13372 | 0.080057577 | -0.129397463 | 0.289512617 | 8.126200414 | 0.928741425 | 0.387867352 | 0.689610533 | -4.700886765 |
| **A_55_P2122997** | Tet1 | -0.102317643 | -0.370022305 | 0.165387019 | 8.301770873 | -0.928704993 | 0.38788482 | 0.689610533 | -4.700901945 |
| **A_66_P113268** | Mme | -0.072170723 | -0.260999194 | 0.116657748 | 7.656570224 | -0.928702831 | 0.387885856 | 0.689610533 | -4.700902846 |
| **A_55_P1959006** | Imp4 | 0.128456645 | -0.207675316 | 0.464588606 | 11.58185544 | 0.928602728 | 0.387933854 | 0.689632435 | -4.700944553 |
| **A_55_P2370096** | NA | 0.097518116 | -0.157698219 | 0.352734452 | 8.426239781 | 0.928453213 | 0.388005552 | 0.68969646 | -4.701006841 |
| **A_55_P1993807** | Nudt10 | -0.0889092 | -0.321626687 | 0.143808287 | 7.746462814 | -0.928326768 | 0.388066195 | 0.689740826 | -4.701059514 |
| **A_51_P340816** | Ipp | 0.139349126 | -0.225467514 | 0.504165766 | 9.521288843 | 0.928138522 | 0.388156491 | 0.689837883 | -4.701137923 |
| **A_51_P375406** | Rprd1a | -0.095132217 | -0.344222949 | 0.153958515 | 9.884226105 | -0.928011242 | 0.388217552 | 0.68988297 | -4.701190932 |
| **A_52_P494686** | Kansl1l | -0.066961688 | -0.242305803 | 0.108382428 | 8.197190862 | -0.927936772 | 0.388253282 | 0.689883038 | -4.701221945 |
| **A_55_P2058117** | Slc35d1 | 0.278078003 | -0.450385045 | 1.006541051 | 10.39735774 | 0.927560613 | 0.388433797 | 0.69014035 | -4.701378574 |
| **A_55_P2064243** | Rhot2 | -0.06724693 | -0.243431933 | 0.108938074 | 8.249916903 | -0.927441898 | 0.388490781 | 0.690178153 | -4.701427997 |
| **A_55_P2061381** | Ghitm | 0.097305167 | -0.157681073 | 0.352291408 | 13.40744037 | 0.927261756 | 0.388577262 | 0.690181312 | -4.701502986 |
| **A_55_P1961255** | Ppp1cc | 0.083688462 | -0.135618023 | 0.302994947 | 13.78739115 | 0.92725098 | 0.388582435 | 0.690181312 | -4.701507471 |
| **A_55_P2128181** | NA | 0.132050185 | -0.214002102 | 0.478102471 | 10.04446603 | 0.927215063 | 0.38859968 | 0.690181312 | -4.701522422 |
| **A_55_P2002150** | Pgs1 | -0.137133299 | -0.496542842 | 0.222276243 | 11.96607275 | -0.927121196 | 0.388644751 | 0.690197941 | -4.701561492 |
| **A_55_P2047571** | Rps9 | -0.099910074 | -0.361787793 | 0.161967646 | 16.12229388 | -0.927030276 | 0.38868841 | 0.690204346 | -4.701599333 |
| **A_55_P2153431** | Ralgds | -0.164379175 | -0.595269422 | 0.266511072 | 8.330658538 | -0.926964967 | 0.388719774 | 0.690204346 | -4.701626513 |
| **A_55_P2156504** | NA | 0.119733795 | -0.194190534 | 0.433658124 | 12.32885335 | 0.926776392 | 0.388810344 | 0.690204907 | -4.701704987 |
| **A_55_P2003793** | Tspan3 | -0.351054686 | -1.271514016 | 0.569404645 | 11.27356016 | -0.926730291 | 0.388832488 | 0.690204907 | -4.701724171 |
| **A_51_P377390** | Ndst2 | 0.08342267 | -0.135319292 | 0.302164632 | 10.02730944 | 0.926691484 | 0.38885113 | 0.690204907 | -4.701740318 |
| **A_55_P2020050** | 6720475J19Rik | -0.117755239 | -0.426599237 | 0.191088758 | 8.382236947 | -0.926454857 | 0.388964811 | 0.690204907 | -4.701838769 |
| **A_52_P93066** | Klhl42 | -0.094029782 | -0.340663838 | 0.152604274 | 8.400834468 | -0.926393659 | 0.388994216 | 0.690204907 | -4.701864229 |
| **A_51_P309158** | Snx20 | -0.1506926 | -0.545955495 | 0.244570296 | 10.38673132 | -0.926379779 | 0.389000886 | 0.690204907 | -4.701870003 |
| **A_55_P2074080** | NA | 0.083528369 | -0.135643731 | 0.302700468 | 14.49183366 | 0.926044634 | 0.389161952 | 0.690204907 | -4.702009411 |
| **A_52_P652212** | Psmd14 | 0.104218902 | -0.169263326 | 0.377701131 | 9.967769346 | 0.925977799 | 0.389194079 | 0.690204907 | -4.702037208 |
| **A_55_P1969615** | Pnck | -0.144683504 | -0.524400952 | 0.235033945 | 8.62480864 | -0.925852177 | 0.389254468 | 0.690204907 | -4.702089451 |
| **A_51_P177552** | Ndufb9 | -0.087300135 | -0.316418006 | 0.141817737 | 14.69340866 | -0.925846812 | 0.389257047 | 0.690204907 | -4.702091682 |
| **A_55_P2154595** | Mrpl39 | 0.064874844 | -0.105388515 | 0.235138203 | 11.43077393 | 0.925845075 | 0.389257882 | 0.690204907 | -4.702092405 |
| **A_55_P2018549** | Gcsh | 0.143933996 | -0.233876021 | 0.521744013 | 13.16856626 | 0.925706053 | 0.389324723 | 0.690204907 | -4.702150216 |

| | | | | | | | | | |
|---|---|---|---|---|---|---|---|---|---|
| **A_51_P292073** | Haghl | -0.232338042 | -0.842211105 | 0.37753502 | 9.971155777 | -0.925686784 | 0.389333987 | 0.690204907 | -4.702158228 |
| **A_55_P2261772** | C230098O21Rik | 0.066574778 | -0.108185523 | 0.241335079 | 9.494608663 | 0.925657093 | 0.389348264 | 0.690204907 | -4.702170574 |
| **A_66_P117578** | Mtmr7 | 0.126682413 | -0.205862816 | 0.459227641 | 8.651465672 | 0.92565424 | 0.389349636 | 0.690204907 | -4.70217176 |
| **A_55_P2021455** | NA | -0.143693854 | -0.520980969 | 0.23359326 | 11.22093577 | -0.925442438 | 0.389451489 | 0.690204907 | -4.702259822 |
| **A_55_P2058290** | Cluh | 0.147984092 | -0.240576105 | 0.536544289 | 8.998118789 | 0.925422195 | 0.389461225 | 0.690204907 | -4.702268238 |
| **A_55_P2045602** | Gpkow | -0.064863744 | -0.235176432 | 0.105448944 | 9.274728018 | -0.925418544 | 0.389462981 | 0.690204907 | -4.702269756 |
| **A_52_P75465** | Gdpd1 | -0.222040049 | -0.805083881 | 0.361003783 | 9.116385906 | -0.925365487 | 0.389488499 | 0.690204907 | -4.702291813 |
| **A_55_P2125972** | Gorasp1 | 0.087962461 | -0.143052138 | 0.31897706 | 9.896242485 | 0.925211743 | 0.389562451 | 0.690204907 | -4.702355725 |
| **A_51_P306151** | Sgk3 | 0.110544047 | -0.179789024 | 0.400877119 | 8.330846175 | 0.925171066 | 0.389582019 | 0.690204907 | -4.702372634 |
| **A_52_P106620** | Tnfrsf11b | -0.15386454 | -0.557983629 | 0.250254549 | 8.614388659 | -0.925150455 | 0.389591934 | 0.690204907 | -4.702381201 |
| **A_51_P222386** | Tbc1d10a | -0.120540912 | -0.437139963 | 0.196058138 | 9.159436553 | -0.92514127 | 0.389596353 | 0.690204907 | -4.702385019 |
| **A_51_P496381** | Ptk7 | 0.343980956 | -0.559570932 | 1.247532845 | 10.94646239 | 0.925048474 | 0.389640997 | 0.690204907 | -4.70242359 |
| **A_51_P487818** | Fabp1 | -0.15631027 | -0.56690533 | 0.25428479 | 15.42257425 | -0.925032481 | 0.389648691 | 0.690204907 | -4.702430237 |
| **A_51_P113403** | Slc26a11 | -0.090925855 | -0.329795519 | 0.147943809 | 8.899731504 | -0.924931525 | 0.389697266 | 0.690204907 | -4.688310659 |
| **A_51_P359137** | Doc2g | -0.065205672 | -0.236515493 | 0.10610415 | 7.820765879 | -0.924881942 | 0.389721124 | 0.690204907 | -4.702492803 |
| **A_55_P2005433** | NA | 0.089120016 | -0.145033156 | 0.323273188 | 8.321682462 | 0.924822527 | 0.389749715 | 0.690204907 | -4.702517495 |
| **A_51_P456465** | Cldn10 | -0.624532134 | -2.265515374 | 1.016451105 | 8.512490477 | -0.924770214 | 0.38977489 | 0.690204907 | -4.702539234 |
| **A_52_P4846** | Zfp451 | 0.069832357 | -0.113661745 | 0.253326459 | 8.703340214 | 0.924736037 | 0.389791338 | 0.690204907 | -4.702553437 |
| **A_52_P463183** | Asap1 | 0.103936683 | -0.169281124 | 0.377154489 | 9.042375616 | 0.924364033 | 0.389970399 | 0.690458719 | -4.702708004 |
| **A_55_P2074718** | Fh1 | 0.095379806 | -0.155381879 | 0.34614149 | 13.02675447 | 0.924226557 | 0.390036588 | 0.69050769 | -4.702765115 |
| **A_51_P363338** | Prkd1 | -0.073036156 | -0.265068674 | 0.118996362 | 7.729211676 | -0.924158194 | 0.390069505 | 0.69050769 | -4.702793513 |
| **A_55_P2022158** | Gc | -0.285052464 | -1.034736734 | 0.464631806 | 16.18294514 | -0.923909836 | 0.390189109 | 0.690656162 | -4.702896669 |
| **A_55_P1975947** | Fam234b | 0.266319962 | -0.434321358 | 0.966961282 | 12.47327415 | 0.923615394 | 0.390330942 | 0.690802051 | -4.703018944 |
| **A_55_P2052555** | Dpysl5 | 0.236782929 | -0.386226007 | 0.859791865 | 12.89912359 | 0.923505109 | 0.390384076 | 0.690802051 | -4.703064737 |
| **A_52_P420308** | NA | 0.105683253 | -0.172395174 | 0.38376168 | 10.24582833 | 0.923468451 | 0.390401739 | 0.690802051 | -4.703079957 |
| **A_51_P375390** | Gemin7 | -0.113927055 | -0.413705523 | 0.185851414 | 11.53943991 | -0.923441995 | 0.390414486 | 0.690802051 | -4.703090942 |
| **A_51_P186053** | Rrp15 | 0.127867536 | -0.208658974 | 0.464394046 | 10.55591651 | 0.923260391 | 0.390501999 | 0.690805146 | -4.703166337 |
| **A_55_P2277171** | Gm10134 | 0.069525709 | -0.113455183 | 0.2525066 | 8.475560162 | 0.923257565 | 0.390503361 | 0.690805146 | -4.703167511 |
| **A_55_P1962756** | Ttll2 | -0.064998263 | -0.236073104 | 0.106076578 | 7.751625278 | -0.923206377 | 0.390528031 | 0.690805146 | -4.70318876 |
| **A_51_P167292** | Chil3 | -0.395250583 | -1.435642566 | 0.6451414 | 12.56206729 | -0.923121172 | 0.390569097 | 0.690805146 | -4.70322413 |
| **A_55_P2056995** | Gcgr | -0.191290074 | -0.69487756 | 0.312297411 | 11.31104003 | -0.922997864 | 0.390628534 | 0.690805146 | -4.703275313 |
| **A_55_P2236382** | D8Ertd158e | -0.138355434 | -0.502589701 | 0.225878832 | 9.099126169 | -0.922993431 | 0.390630671 | 0.690805146 | -4.703277153 |
| **A_55_P2073642** | 1600014C10Rik | -0.122397833 | -0.444708031 | 0.199912365 | 8.639109612 | -0.922747488 | 0.390749242 | 0.690914145 | -4.703379226 |
| **A_55_P1993121** | NA | -0.072177323 | -0.262248077 | 0.117893431 | 15.63944858 | -0.922717301 | 0.390763797 | 0.690914145 | -4.703391753 |
| **A_52_P88091** | Dsg2 | 0.075343747 | -0.123110775 | 0.273798269 | 8.090275016 | 0.922506436 | 0.390865481 | 0.691030722 | -4.703479253 |
| **A_51_P422223** | 1700006J14Rik | 0.100295033 | -0.163921092 | 0.364511159 | 8.376712982 | 0.922366021 | 0.390933204 | 0.691070456 | -4.703537512 |
| **A_55_P2069974** | Kctd1 | -0.075361907 | -0.273906201 | 0.123182386 | 7.793121537 | -0.922311583 | 0.390959462 | 0.691070456 | -4.703560097 |
| **A_55_P2066440** | Myh3 | -0.078637891 | -0.285855216 | 0.128579434 | 7.703781046 | -0.922123275 | 0.391050302 | 0.69116782 | -4.703638216 |
| **A_55_P2056140** | Flad1 | 0.099785329 | -0.163184983 | 0.362755641 | 10.1660977 | 0.922025984 | 0.391097242 | 0.691187582 | -4.703678573 |
| **A_55_P1976734** | Cacna2d2 | -0.146198774 | -0.531581399 | 0.239183851 | 8.368579201 | -0.921795935 | 0.39120825 | 0.691257143 | -4.703773988 |
| **A_55_P2133679** | NA | -0.128366693 | -0.466746682 | 0.210013296 | 11.94421434 | -0.921787616 | 0.391212265 | 0.691257143 | -4.703777438 |
| **A_55_P1968928** | Def6 | 0.173473913 | -0.283843112 | 0.630790939 | 11.21732553 | 0.921722087 | 0.39124389 | 0.691257143 | -4.703804614 |
| **A_51_P295625** | Arf4 | -0.135181602 | -0.491615424 | 0.221252221 | 11.17594409 | -0.921556291 | 0.391323914 | 0.691335338 | -4.703873367 |
| **A_52_P566348** | Nras | 0.088742175 | -0.145316231 | 0.322800582 | 10.08761552 | 0.921274427 | 0.391459988 | 0.691433579 | -4.703990234 |
| **A_55_P1981405** | Gsk3a | 0.10398156 | -0.170288827 | 0.378251948 | 12.38556702 | 0.921214146 | 0.391489095 | 0.691433579 | -4.704015225 |
| **A_52_P19016** | Clasrp | -0.096800796 | -0.352160756 | 0.158559164 | 11.5171931 | -0.921105373 | 0.391541619 | 0.691433579 | -4.704060316 |
| **A_55_P2173313** | Plcb1 | 0.106831682 | -0.175006439 | 0.388669804 | 8.761801676 | 0.921050666 | 0.391568039 | 0.691433579 | -4.704082994 |
| **A_55_P2161465** | Gm10516 | 0.065326603 | -0.107023307 | 0.237676513 | 8.21514907 | 0.921005456 | 0.391589873 | 0.691433579 | -4.704101734 |
| **A_55_P1993064** | Mbd6 | 0.083651947 | -0.137047487 | 0.30435138 | 10.98780111 | 0.920996593 | 0.391594153 | 0.691433579 | -4.704105408 |
| **A_55_P2167999** | Ldlr | 0.122632628 | -0.200999804 | 0.446265061 | 11.67941594 | 0.920740375 | 0.39171791 | 0.691554928 | -4.704211602 |

| | | | | | | | | | |
|---|---|---|---|---|---|---|---|---|---|
| **A_55_P2185347** | NA | 0.102221977 | -0.167576298 | 0.372020251 | 7.899753988 | 0.920636727 | 0.391767983 | 0.691554928 | -4.704254556 |
| **A_55_P1967720** | Polr2i | -0.153120912 | -0.557260672 | 0.251018848 | 12.57539337 | -0.920632108 | 0.391770214 | 0.691554928 | -4.70425647 |
| **A_66_P115453** | Coro1c | -0.089213631 | -0.324701448 | 0.146274187 | 12.20229386 | -0.920546982 | 0.391811342 | 0.691564371 | -4.704291745 |
| **A_55_P2011912** | Ecm2 | -0.077306917 | -0.281441828 | 0.126827994 | 7.782131175 | -0.920204304 | 0.391976937 | 0.691740083 | -4.704433726 |
| **A_51_P491470** | Ddx47 | -0.076253585 | -0.277609593 | 0.125102424 | 9.754766701 | -0.920192861 | 0.391982468 | 0.691740083 | -4.704438467 |
| **A_55_P2057132** | Scn8a | 0.322481824 | -0.52918109 | 1.174144737 | 12.49773599 | 0.920069628 | 0.392042033 | 0.691782039 | -4.704489518 |
| **A_55_P2022139** | Pura | 0.148138905 | -0.243131636 | 0.539409447 | 10.79342011 | 0.919973185 | 0.392088654 | 0.691801149 | -4.704529467 |
| **A_55_P2161219** | Thsd7a | -0.060373085 | -0.219892089 | 0.099145919 | 8.116791219 | -0.919632118 | 0.392253559 | 0.69189176 | -4.704670726 |
| **A_55_P2026492** | NA | 0.120828727 | -0.198439117 | 0.440096572 | 8.459928725 | 0.919598406 | 0.392269862 | 0.69189176 | -4.704684687 |
| **A_55_P2106058** | Gm11110 | -0.066471407 | -0.24211115 | 0.109168335 | 7.729879419 | -0.919592182 | 0.392272872 | 0.69189176 | -4.704687264 |
| **A_51_P358906** | U2af2 | 0.132938496 | -0.218337729 | 0.484214722 | 8.871243959 | 0.919570844 | 0.392283191 | 0.69189176 | -4.704696101 |
| **A_55_P1962454** | NA | 0.060912001 | -0.100083898 | 0.2219079 | 7.920110309 | 0.919329603 | 0.39239987 | 0.692034406 | -4.704795991 |
| **A_55_P2009518** | NA | 0.094786181 | -0.155852779 | 0.34542514 | 16.01010834 | 0.918924084 | 0.392596063 | 0.692317243 | -4.704963867 |
| **A_55_P2089840** | Eif2d | 0.080578188 | -0.13256578 | 0.293722156 | 9.593070267 | 0.918602537 | 0.392751682 | 0.692388777 | -4.705096948 |
| **A_52_P821** | Fli1 | 0.081616138 | -0.134278416 | 0.297510692 | 10.60694587 | 0.918581176 | 0.392762023 | 0.692388777 | -4.705105788 |
| **A_51_P462448** | Gpx4 | -0.147611598 | -0.538100783 | 0.242877587 | 14.89214649 | -0.918532784 | 0.392785448 | 0.692388777 | -4.705125813 |
| **A_52_P257625** | Esm1 | -0.302454518 | -1.102607689 | 0.497698653 | 8.570649509 | -0.918480911 | 0.392810559 | 0.692388777 | -4.705147278 |
| **A_55_P1959238** | Nagk | -0.092071838 | -0.335662238 | 0.151518561 | 10.48003504 | -0.918437983 | 0.392831341 | 0.692388777 | -4.705165041 |
| **A_55_P2005930** | NA | -0.197044235 | -0.718439074 | 0.324350604 | 13.19064183 | -0.918290716 | 0.39290264 | 0.692388777 | -4.705225975 |
| **A_52_P308669** | Cpn1 | -0.185979534 | -0.678144022 | 0.306184954 | 11.37064618 | -0.918201653 | 0.392945765 | 0.692388777 | -4.705262823 |
| **A_55_P2006093** | Nebl | -0.070227111 | -0.25607195 | 0.115617729 | 7.679672262 | -0.918200411 | 0.392946367 | 0.692388777 | -4.705263337 |
| **A_55_P2213828** | 4933412E12Rik | -0.094114832 | -0.3432168 | 0.154987136 | 8.245277597 | -0.918045282 | 0.39302149 | 0.692388777 | -4.705327513 |
| **A_55_P1984745** | Pappa2 | -0.077027141 | -0.280930808 | 0.126876526 | 7.837920197 | -0.917913867 | 0.393085138 | 0.692388777 | -4.705381874 |
| **A_55_P1991851** | Speg | -0.080038748 | -0.291922478 | 0.131844983 | 8.119663378 | -0.917879905 | 0.393101588 | 0.692388777 | -4.705395921 |
| **A_55_P1955542** | Olfr522 | 0.111610188 | -0.183873697 | 0.407094073 | 8.497277086 | 0.917810933 | 0.393134997 | 0.692388777 | -4.705424449 |
| **A_52_P641892** | Hacd3 | -0.152403122 | -0.55593091 | 0.251124665 | 14.29097525 | -0.917706076 | 0.393185793 | 0.692388777 | -4.705467818 |
| **A_52_P405206** | Tm7sf3 | 0.126736449 | -0.208837867 | 0.462310765 | 11.63361926 | 0.917690025 | 0.393193569 | 0.692388777 | -4.705474456 |
| **A_66_P119191** | Inpp5e | -0.060905092 | -0.222181064 | 0.10037088 | 8.593472138 | -0.917628994 | 0.393223137 | 0.692388777 | -4.705499696 |
| **A_51_P367563** | Rsph3a | -0.068735216 | -0.250753332 | 0.113282901 | 8.218700456 | -0.917588311 | 0.393242848 | 0.692388777 | -4.70551652 |
| **A_55_P1963547** | Prdm16 | -0.159127464 | -0.580516407 | 0.262261479 | 8.256390204 | -0.917582653 | 0.393245589 | 0.692388777 | -4.70551886 |
| **A_51_P487389** | Drc3 | -0.126803968 | -0.462671916 | 0.209063979 | 7.772065133 | -0.917376211 | 0.393345622 | 0.692453757 | -4.705604227 |
| **A_51_P211131** | Traf3ip2 | -0.145191352 | -0.529796074 | 0.239413371 | 7.917634799 | -0.91729563 | 0.393384674 | 0.692453757 | -4.705637545 |
| **A_55_P2037156** | Pja1 | -0.111858404 | -0.408204442 | 0.184487635 | 11.31699875 | -0.917175996 | 0.393442656 | 0.692453757 | -4.705687007 |
| **A_52_P311245** | Scarb2 | -0.090554416 | -0.330477456 | 0.149368625 | 9.228804131 | -0.917108809 | 0.393475222 | 0.692453757 | -4.705714783 |
| **A_52_P497534** | Wipi1 | 0.100348016 | -0.165553447 | 0.366249479 | 10.07102371 | 0.917004048 | 0.393526005 | 0.692453757 | -4.705758091 |
| **A_55_P2021565** | Ntf3 | -0.073293601 | -0.267511647 | 0.120924446 | 7.959236871 | -0.916979581 | 0.393537866 | 0.692453757 | -4.705768205 |
| **A_51_P486239** | Clec3b | -0.316999536 | -1.15710857 | 0.523109499 | 8.158876348 | -0.916866617 | 0.393592632 | 0.692453757 | -4.7058149 |
| **A_55_P2079729** | NA | -0.089265478 | -0.32583605 | 0.147305094 | 15.88604466 | -0.91686629 | 0.393592791 | 0.692453757 | -4.705815035 |
| **A_55_P1980521** | Ccdc85a | -0.145342174 | -0.530537209 | 0.239852862 | 7.784960711 | -0.916841279 | 0.393604917 | 0.692453757 | -4.705825373 |
| **A_55_P2095727** | Pml | -0.146640468 | -0.535349648 | 0.242068712 | 11.08285594 | -0.916668334 | 0.393688775 | 0.692538254 | -4.705896853 |
| **A_52_P330950** | C2cd2 | -0.064416337 | -0.235188866 | 0.106356192 | 8.330838572 | -0.916560639 | 0.393741002 | 0.692567097 | -4.70594136 |
| **A_55_P1966169** | Ppdpf | -0.10578952 | -0.386292726 | 0.174713685 | 13.16107376 | -0.916406178 | 0.393815917 | 0.692598511 | -4.706005188 |
| **A_55_P1985433** | Nrg1 | -0.061220271 | -0.223560219 | 0.101119676 | 8.074693144 | -0.916332387 | 0.39385171 | 0.692598511 | -4.706035678 |
| **A_51_P234714** | Ceacam14 | -0.065812788 | -0.240347886 | 0.10872231 | 7.684678548 | -0.916243072 | 0.393895037 | 0.692598511 | -4.706072582 |
| **A_52_P351021** | NA | 0.067284162 | -0.111192044 | 0.245760368 | 10.18947128 | 0.916042687 | 0.393992256 | 0.692598511 | -4.706155368 |
| **A_55_P2043187** | Exosc9 | 0.213597217 | -0.353007503 | 0.780201936 | 13.35733506 | 0.916006574 | 0.394009779 | 0.692598511 | -4.706170286 |
| **A_55_P2178317** | Gm9199 | 0.171136861 | -0.28287695 | 0.625150671 | 9.383528335 | 0.915920303 | 0.394051641 | 0.692598511 | -4.706205924 |
| **A_55_P2112549** | NA | 0.105366907 | -0.174174879 | 0.384908693 | 13.80461052 | 0.915884445 | 0.394069042 | 0.692598511 | -4.706220735 |
| **A_51_P337269** | Aldob | -0.2331439 | -0.851722801 | 0.385435001 | 15.85139284 | -0.915824258 | 0.39409825 | 0.692598511 | -4.706245596 |
| **A_52_P249733** | Tcap | -0.248458525 | -0.907673012 | 0.410755962 | 7.901976915 | -0.915820475 | 0.394100087 | 0.692598511 | -4.706247158 |

| | | | | | | | | | |
|---|---|---|---|---|---|---|---|---|---|
| **A_55_P2087726** | Cops9 | -0.150599077 | -0.550186543 | 0.248988388 | 14.63992059 | -0.915785248 | 0.394117183 | 0.692598511 | -4.706261709 |
| **A_55_P1979252** | Glod4 | -0.076942278 | -0.281126225 | 0.127241669 | 12.14568184 | -0.915643957 | 0.39418576 | 0.69265605 | -4.706320064 |
| **A_55_P1968488** | Cblif | -0.24637117 | -0.900300117 | 0.407557777 | 7.864787794 | -0.915466617 | 0.394271846 | 0.692744342 | -4.7063933 |
| **A_51_P164203** | Nme4 | 0.206675421 | -0.341953488 | 0.755304331 | 11.51647118 | 0.915362955 | 0.394322173 | 0.692769795 | -4.706436105 |
| **A_55_P1977929** | Itgam | -0.121869619 | -0.445412021 | 0.201672783 | 8.442824121 | -0.915266224 | 0.39436914 | 0.69278934 | -4.706476046 |
| **A_52_P89425** | Pcnt | 0.090042104 | -0.149023332 | 0.329107541 | 10.90244047 | 0.915191618 | 0.394405367 | 0.692790017 | -4.706506849 |
| **A_66_P138001** | Atp2b1 | -0.15358806 | -0.561405793 | 0.254229673 | 9.145273547 | -0.915112607 | 0.394443736 | 0.692794455 | -4.706539469 |
| **A_55_P1973526** | Fam216b | -0.133744773 | -0.488921845 | 0.221432299 | 7.882299153 | -0.914987338 | 0.394504574 | 0.692820229 | -4.706591183 |
| **A_55_P2068104** | NA | 0.202663366 | -0.335597696 | 0.740924429 | 10.1168555 | 0.914882829 | 0.394555336 | 0.692820229 | -4.706634324 |
| **A_55_P2074081** | NA | 0.098814705 | -0.163637156 | 0.361266566 | 14.41168267 | 0.914860995 | 0.394565941 | 0.692820229 | -4.706643337 |
| **A_55_P2026818** | Slc4a7 | 0.098401834 | -0.16300416 | 0.359807829 | 9.233998453 | 0.914683491 | 0.39465217 | 0.692908693 | -4.706716601 |
| **A_51_P470079** | Il1r2 | -0.094455897 | -0.345428942 | 0.156517148 | 8.265160584 | -0.914503109 | 0.394739812 | 0.692967618 | -4.706791044 |
| **A_66_P127255** | Ing3 | -0.178568586 | -0.653060594 | 0.295923422 | 9.828134783 | -0.914448698 | 0.394766252 | 0.692967618 | -4.706813498 |
| **A_51_P354698** | Dync1h1 | -0.088439686 | -0.323459879 | 0.146580507 | 11.80920204 | -0.914376808 | 0.394801186 | 0.692967618 | -4.706843163 |
| **A_55_P2184246** | Rps26 | -0.131064545 | -0.47939513 | 0.21726604 | 16.42002568 | -0.914274915 | 0.394850706 | 0.692967618 | -4.706885206 |
| **A_55_P2050083** | Gak | 0.093023064 | -0.154223228 | 0.340269356 | 10.71422988 | 0.914205948 | 0.394884226 | 0.692967618 | -4.706913662 |
| **A_51_P190437** | Ccdc181 | -0.230065336 | -0.84161835 | 0.381487678 | 9.462823508 | -0.914113803 | 0.394929014 | 0.692967618 | -4.706951679 |
| **A_55_P2167050** | Paip2 | -0.122702886 | -0.44889225 | 0.203486477 | 11.9233455 | -0.914046253 | 0.39496185 | 0.692967618 | -4.706979546 |
| **A_66_P114333** | Tlr12 | -0.177690708 | -0.650069272 | 0.294687857 | 9.861865464 | -0.914024261 | 0.394972541 | 0.692967618 | -4.706988619 |
| **A_55_P1996852** | NA | 0.070905713 | -0.117671698 | 0.259483124 | 10.06719937 | 0.913639247 | 0.395159741 | 0.693136505 | -4.70714743 |
| **A_51_P221031** | Slc16a12 | 0.171722524 | -0.284993277 | 0.628438326 | 9.473341668 | 0.913617512 | 0.39517031 | 0.693136505 | -4.707156394 |
| **A_51_P228193** | Ociad1 | 0.065230987 | -0.108260695 | 0.238722669 | 11.01268483 | 0.913605027 | 0.395176382 | 0.693136505 | -4.707161543 |
| **A_55_P2032363** | Tomm40l | 0.089769952 | -0.149019244 | 0.328559147 | 8.892301043 | 0.913480976 | 0.395236714 | 0.693179425 | -4.707212702 |
| **A_55_P2280581** | A130088B03Rik | -0.076988459 | -0.281796491 | 0.127819572 | 7.805962385 | -0.91340174 | 0.395275253 | 0.69318281 | -4.707245377 |
| **A_51_P193563** | Cdk7 | -0.091579287 | -0.335221725 | 0.152063151 | 8.801490671 | -0.913329549 | 0.395310369 | 0.69318281 | -4.707275144 |
| **A_51_P192089** | Pinx1 | 0.119464881 | -0.198456915 | 0.437386677 | 9.681943652 | 0.913068036 | 0.395437596 | 0.693343005 | -4.707382967 |
| **A_51_P185108** | Dpy30 | 0.134726186 | -0.223922697 | 0.49337507 | 11.1848809 | 0.912779172 | 0.395578165 | 0.693526563 | -4.707502044 |
| **A_55_P2162592** | Lman2 | -0.066965465 | -0.24525615 | 0.11132522 | 13.72240715 | -0.912652453 | 0.395639842 | 0.693551018 | -4.707554273 |
| **A_55_P1986813** | Ibsp | -0.087510771 | -0.320524179 | 0.145502637 | 7.704624531 | -0.912564956 | 0.395682433 | 0.693551018 | -4.707590333 |
| **A_52_P482897** | Areg | -0.153880542 | -0.563631566 | 0.255870482 | 7.784489011 | -0.912529365 | 0.395699758 | 0.693551018 | -4.707605001 |
| **A_55_P2093770** | NA | 0.127336331 | -0.21180297 | 0.466475631 | 13.13934239 | 0.912341437 | 0.395791251 | 0.693646832 | -4.707682443 |
| **A_55_P1976112** | C130074G19Rik | 0.15821587 | -0.263199138 | 0.579630877 | 11.01829345 | 0.912269667 | 0.395826197 | 0.693646832 | -4.707712016 |
| **A_55_P2057405** | Cpt1a | -0.070386334 | -0.258032677 | 0.11726001 | 7.950646962 | -0.911447013 | 0.396226922 | 0.694286118 | -4.708050884 |
| **A_51_P513568** | Stx11 | 0.110669597 | -0.184424165 | 0.405763359 | 8.550573155 | 0.911279265 | 0.396308671 | 0.694366417 | -4.708119959 |
| **A_55_P1974976** | Sprr2a2 | 0.069349769 | -0.115611167 | 0.254310705 | 16.16837865 | 0.911062547 | 0.396414305 | 0.694487881 | -4.708209187 |
| **A_55_P2009994** | NA | 0.106541503 | -0.177636866 | 0.390719872 | 11.31242608 | 0.910984507 | 0.396452349 | 0.694487881 | -4.708241315 |
| **A_55_P2069917** | Uba52 | 0.068150125 | -0.113640529 | 0.249940778 | 16.89491999 | 0.91091592 | 0.396485787 | 0.694487881 | -4.70826955 |
| **A_51_P243304** | Galk1 | 0.103881048 | -0.173256849 | 0.381018945 | 13.25307234 | 0.910801203 | 0.396541719 | 0.694522914 | -4.708316771 |
| **A_51_P464911** | Msrb2 | -0.123278786 | -0.452243419 | 0.205685847 | 10.37388304 | -0.910588852 | 0.39664527 | 0.694641336 | -4.708404173 |
| **A_65_P15812** | Pum1 | 0.118068847 | -0.197081257 | 0.433218951 | 10.59853759 | 0.910334615 | 0.396769273 | 0.69479555 | -4.708508797 |
| **A_55_P2105181** | Bhmt | 0.314984194 | -0.526100989 | 1.156069377 | 13.95232598 | 0.909980254 | 0.396942161 | 0.695035332 | -4.708654594 |
| **A_55_P1955034** | Myo1c | -0.188097499 | -0.690472341 | 0.314277342 | 10.98418706 | -0.909784071 | 0.3970379 | 0.695072292 | -4.708735295 |
| **A_55_P2061874** | Prelid1 | -0.073406344 | -0.269464871 | 0.122652183 | 13.82547929 | -0.909768821 | 0.397045344 | 0.695072292 | -4.708741568 |
| **A_55_P2150389** | Dnajc21 | 0.084930615 | -0.141967246 | 0.311828477 | 9.100445869 | 0.909530061 | 0.397161886 | 0.695072292 | -4.708839768 |
| **A_55_P1985001** | NA | -0.170719285 | -0.626850653 | 0.285412084 | 8.97643096 | -0.909443727 | 0.397204034 | 0.695072292 | -4.708875272 |
| **A_55_P2048779** | Lratd2 | -0.105490819 | -0.387343537 | 0.1763619 | 8.220877986 | -0.909443291 | 0.397204247 | 0.695072292 | -4.708875452 |
| **A_65_P13270** | Csnk1g1 | 0.062704568 | -0.104836031 | 0.230245166 | 8.43389065 | 0.909415403 | 0.397217862 | 0.695072292 | -4.70888692 |
| **A_55_P2004752** | Prl2c5 | -0.064628586 | -0.237328742 | 0.10807157 | 7.720967919 | -0.909316592 | 0.397266106 | 0.695072292 | -4.708927552 |
| **A_55_P1978216** | Cd300c2 | -0.122734041 | -0.450706045 | 0.205237963 | 10.7991238 | -0.909308918 | 0.397269853 | 0.695072292 | -4.708930707 |
| **A_55_P2044592** | NA | 0.083193952 | -0.139126891 | 0.305514795 | 13.12608943 | 0.909273985 | 0.39728691 | 0.695072292 | -4.708945071 |

| | | | | | | | | | |
|---|---|---|---|---|---|---|---|---|---|
| **A_51_P408363** | Cfp | -0.101766756 | -0.373763405 | 0.170229893 | 12.81545863 | -0.909129117 | 0.397357652 | 0.695096811 | -4.709004636 |
| **A_52_P324754** | Nupr1l | -0.088069946 | -0.323467735 | 0.147327844 | 7.957352686 | -0.909093468 | 0.397375062 | 0.695096811 | -4.709019292 |
| **A_55_P2158351** | Rpl18a | -0.11498613 | -0.422373022 | 0.192400762 | 15.8282108 | -0.908956925 | 0.39744175 | 0.695096811 | -4.709075427 |
| **A_52_P558859** | Rras2 | -0.112384032 | -0.412816904 | 0.18804884 | 8.768634619 | -0.908950741 | 0.39744477 | 0.695096811 | -4.709077969 |
| **A_52_P108502** | Slc4a7 | -0.114629072 | -0.421088985 | 0.191830842 | 7.995277449 | -0.908875276 | 0.397481631 | 0.695098385 | -4.709108991 |
| **A_52_P337126** | Bves | -0.164530675 | -0.604515115 | 0.275453765 | 7.811802341 | -0.908641881 | 0.39759565 | 0.695182521 | -4.709204925 |
| **A_55_P2117119** | Efhd1 | -0.063485367 | -0.233268863 | 0.106298129 | 7.725358775 | -0.908576187 | 0.397627747 | 0.695182521 | -4.709231925 |
| **A_55_P2096187** | Hax1 | -0.087139888 | -0.320189572 | 0.145909795 | 13.0884516 | -0.908555939 | 0.39763764 | 0.695182521 | -4.709240246 |
| **A_52_P112178** | Srgap2 | -0.064787515 | -0.238088982 | 0.108513951 | 8.252300084 | -0.90838986 | 0.397718795 | 0.695184818 | -4.709308497 |
| **A_51_P415395** | C2cd4b | -0.205974034 | -0.756967315 | 0.345019247 | 7.867585828 | -0.908341928 | 0.39774222 | 0.695184818 | -4.709328194 |
| **A_55_P2130104** | Slc52a3 | -0.184336185 | -0.677452026 | 0.308779656 | 7.955311563 | -0.90833245 | 0.397746852 | 0.695184818 | -4.709332088 |
| **A_55_P2179221** | Man1a2 | -0.120371608 | -0.442414847 | 0.201671632 | 10.83787563 | -0.908224185 | 0.397799765 | 0.695201574 | -4.709376574 |
| **A_55_P2058942** | Aldh3b1 | -0.109817115 | -0.403641707 | 0.184007478 | 10.25721272 | -0.908165657 | 0.397828373 | 0.695201574 | -4.709400622 |
| **A_55_P1960281** | Med7 | 0.076298797 | -0.127866593 | 0.280464187 | 10.00539981 | 0.908068805 | 0.397875715 | 0.695221451 | -4.709440414 |
| **A_52_P58145** | Aldh1a2 | -0.211658771 | -0.778213759 | 0.354896218 | 8.314212182 | -0.907773271 | 0.398020203 | 0.69527829 | -4.709561818 |
| **A_66_P126459** | Gm13429 | -0.059708048 | -0.219531031 | 0.100114936 | 7.727332091 | -0.907772088 | 0.398020781 | 0.69527829 | -4.709562304 |
| **A_55_P2078516** | NA | 0.092631497 | -0.155331578 | 0.340594573 | 9.429743157 | 0.907726173 | 0.398043233 | 0.69527829 | -4.709581164 |
| **A_66_P134040** | Vps25 | -0.109072019 | -0.401061674 | 0.182917635 | 9.949796756 | -0.907672291 | 0.398069582 | 0.69527829 | -4.709603295 |
| **A_52_P598309** | Zfas1 | -0.276016039 | -1.014988096 | 0.462956019 | 12.35072512 | -0.907589558 | 0.39811004 | 0.69527829 | -4.709637274 |
| **A_52_P472302** | Fxyd6 | -0.285014083 | -1.04827812 | 0.478249955 | 9.555090932 | -0.907349717 | 0.398227349 | 0.69527829 | -4.709735769 |
| **A_51_P459320** | Gkap1 | 0.09420646 | -0.15809941 | 0.346512331 | 10.75535952 | 0.907269935 | 0.398266377 | 0.69527829 | -4.709768529 |
| **A_52_P326664** | Unc93a | -0.262599979 | -0.965972858 | 0.440772901 | 8.809725615 | -0.907177472 | 0.398311612 | 0.69527829 | -4.709806494 |
| **A_51_P441387** | Kansl1l | -0.12450339 | -0.458032465 | 0.209025684 | 8.412034663 | -0.907048833 | 0.398374551 | 0.69527829 | -4.709859309 |
| **A_52_P150236** | Apom | 0.161396801 | -0.270972235 | 0.593765836 | 15.89952715 | 0.907033917 | 0.398381849 | 0.69527829 | -4.709865432 |
| **A_55_P1978023** | NA | 0.11764799 | -0.197546815 | 0.432842794 | 9.659127945 | 0.906961077 | 0.398417492 | 0.69527829 | -4.709895336 |
| **A_55_P2158376** | NA | -0.173489821 | -0.638293509 | 0.291313867 | 13.81687877 | -0.906958931 | 0.398418542 | 0.69527829 | -4.709896216 |
| **A_55_P2104689** | NA | -0.078849374 | -0.290099395 | 0.132400647 | 7.721840239 | -0.906952787 | 0.398421549 | 0.69527829 | -4.709898739 |
| **A_52_P675530** | Fat4 | -0.122651748 | -0.451263 | 0.205959504 | 7.870081512 | -0.906931533 | 0.398431949 | 0.69527829 | -4.709907464 |
| **A_51_P305843** | Chordc1 | 0.058261374 | -0.097847008 | 0.214369756 | 10.38522929 | 0.906854662 | 0.398469568 | 0.69527829 | -4.70993902 |
| **A_55_P2174576** | Nova1 | 0.105632947 | -0.17741421 | 0.388680103 | 8.113429882 | 0.90682563 | 0.398483776 | 0.69527829 | -4.709950937 |
| **A_55_P2018507** | Gorasp2 | 0.105854986 | -0.177816527 | 0.389526499 | 9.096733459 | 0.906731667 | 0.398529763 | 0.695295766 | -4.709989506 |
| **A_55_P2105321** | Acin1 | 0.120464719 | -0.202433241 | 0.443362679 | 12.28032716 | 0.906520773 | 0.398632994 | 0.695327882 | -4.710076062 |
| **A_55_P2065059** | Wnt2 | -0.118226314 | -0.435145422 | 0.198692795 | 7.907118763 | -0.906460558 | 0.398662473 | 0.695327882 | -4.710100773 |
| **A_55_P2095578** | NA | 0.077421331 | -0.130136624 | 0.284979287 | 8.53667625 | 0.906367743 | 0.398707914 | 0.695327882 | -4.710138861 |
| **A_65_P16542** | NA | -0.127218851 | -0.468288786 | 0.213851084 | 10.7623024 | -0.906340149 | 0.398721425 | 0.695327882 | -4.710150184 |
| **A_55_P1961620** | Vax2 | 0.394570158 | -0.663293997 | 1.452434313 | 11.26426972 | 0.906311568 | 0.398735418 | 0.695327882 | -4.710161912 |
| **A_55_P2018211** | NA | -0.112296064 | -0.413387347 | 0.188795219 | 15.35228215 | -0.906253174 | 0.398764011 | 0.695327882 | -4.710185873 |
| **A_51_P112677** | Mrpl50 | 0.167682218 | -0.28199755 | 0.617361987 | 12.04690011 | 0.90608064 | 0.398848501 | 0.695400157 | -4.708134187 |
| **A_55_P2056493** | NA | 0.162162546 | -0.272764913 | 0.597090006 | 9.871091182 | 0.905976464 | 0.398899523 | 0.695400157 | -4.710299401 |
| **A_51_P346871** | Ift20 | -0.131125503 | -0.482821168 | 0.220570161 | 11.81058282 | -0.90594815 | 0.398913391 | 0.695400157 | -4.710311016 |
| **A_55_P1975011** | Ufc1 | -0.097056818 | -0.357493065 | 0.163379429 | 13.0673512 | -0.905540356 | 0.399113167 | 0.69568567 | -4.710478282 |
| **A_52_P498355** | Lyrm4 | 0.102187065 | -0.172089531 | 0.376463662 | 10.35256347 | 0.905295509 | 0.399233153 | 0.695787411 | -4.710578688 |
| **A_55_P2052240** | Crlf1 | -1.06266042 | -3.915090978 | 1.789770138 | 9.336451666 | -0.905237661 | 0.399261504 | 0.695787411 | -4.710602408 |
| **A_55_P1987245** | Ywhaz | -0.136316749 | -0.502237833 | 0.229604336 | 9.298394931 | -0.905200892 | 0.399279526 | 0.695787411 | -4.710617484 |
| **A_55_P1999154** | Gm4987 | -0.099286426 | -0.365835054 | 0.167262202 | 14.06219613 | -0.905100104 | 0.399328928 | 0.695810769 | -4.710658807 |
| **A_55_P1995055** | Prr7 | -0.062525698 | -0.230403452 | 0.105352057 | 8.048856131 | -0.905000006 | 0.399377997 | 0.695833542 | -4.710699844 |
| **A_55_P2172832** | Kcne1l | 0.186176657 | -0.313763429 | 0.686116742 | 8.851859566 | 0.904878889 | 0.399437377 | 0.695874273 | -4.710749496 |
| **A_52_P573255** | Cdc42ep1 | -0.316076492 | -1.16506464 | 0.532911657 | 11.83744033 | -0.904635773 | 0.399556587 | 0.69601922 | -4.710849146 |
| **A_55_P2018283** | Hmgb1 | 0.085198983 | -0.14370053 | 0.314098497 | 9.29999329 | 0.904425363 | 0.399659781 | 0.696078484 | -4.710935376 |
| **A_55_P2124543** | Gnptg | 0.061873409 | -0.104364781 | 0.228111599 | 9.707078305 | 0.904391418 | 0.399676432 | 0.696078484 | -4.710949287 |

| | | | | | | | | | |
|---|---|---|---|---|---|---|---|---|---|
| **A_55_P1980214** | NA | 0.099246401 | -0.167416912 | 0.365909715 | 10.20321597 | 0.904346135 | 0.399698644 | 0.696078484 | -4.710967843 |
| **A_55_P2122688** | Zmym3 | -0.119136784 | -0.439294853 | 0.201021285 | 10.39686263 | -0.904200221 | 0.399770224 | 0.69614042 | -4.711027631 |
| **A_55_P1972019** | 4632415L05Rik | -0.077436853 | -0.285679301 | 0.130805594 | 8.072301358 | -0.903569635 | 0.400079678 | 0.696563929 | -4.711285942 |
| **A_66_P102628** | 1700016J18Rik | -0.13770812 | -0.508064851 | 0.232648612 | 7.749361391 | -0.903488221 | 0.400119644 | 0.696563929 | -4.711319283 |
| **A_55_P2090535** | Thrap3 | 0.103716857 | -0.175223852 | 0.382657566 | 8.409223862 | 0.903484353 | 0.400121543 | 0.696563929 | -4.711320868 |
| **A_55_P2153631** | NA | 0.138036834 | -0.233311103 | 0.50938477 | 8.667425663 | 0.903227523 | 0.400247641 | 0.696684705 | -4.711426035 |
| **A_66_P132510** | Scamp3 | -0.101851626 | -0.37587205 | 0.172168798 | 12.36263843 | -0.903167336 | 0.400277197 | 0.696684705 | -4.711450678 |
| **A_55_P2175545** | Gys2 | 0.278738489 | -0.471213233 | 1.02869021 | 11.37167789 | 0.903122836 | 0.40029905 | 0.696684705 | -4.711468898 |
| **A_55_P2079599** | NA | 0.099572707 | -0.168392059 | 0.367537473 | 12.20181741 | 0.902912798 | 0.400402206 | 0.696762039 | -4.711554884 |
| **A_52_P365005** | Zbtb22 | 0.095117637 | -0.160867404 | 0.351102679 | 8.821254179 | 0.902879214 | 0.400418703 | 0.696762039 | -4.711568632 |
| **A_52_P610808** | Hexdc | 0.061388826 | -0.103844975 | 0.226622627 | 8.300654058 | 0.902762734 | 0.40047592 | 0.696762039 | -4.711616311 |
| **A_51_P267239** | Litaf | -0.142751612 | -0.526991192 | 0.241487967 | 11.99913687 | -0.902738808 | 0.400487674 | 0.696762039 | -4.711626104 |
| **A_51_P510106** | B3gntl1 | -0.08046382 | -0.297108563 | 0.136180923 | 7.911026422 | -0.902476036 | 0.40061678 | 0.696798849 | -4.711733648 |
| **A_65_P01418** | Wnk2 | -0.084672413 | -0.312665682 | 0.143320856 | 8.353963772 | -0.902408338 | 0.400650047 | 0.696798849 | -4.711761352 |
| **A_55_P2027930** | Nfx1 | 0.073542698 | -0.124491408 | 0.271576804 | 8.293971384 | 0.902366069 | 0.400670818 | 0.696798849 | -4.711778648 |
| **A_65_P17218** | Mndal | -0.0941247 | -0.347623536 | 0.159374135 | 8.908178871 | -0.902216628 | 0.400744263 | 0.696798849 | -4.711839796 |
| **A_55_P1990457** | Rpl15 | -0.119153112 | -0.44006558 | 0.201759356 | 15.73393838 | -0.902198266 | 0.400753288 | 0.696798849 | -4.711847309 |
| **A_55_P2028259** | Rhbdf2 | -0.062743548 | -0.231733241 | 0.106246146 | 9.437392153 | -0.902177607 | 0.400763443 | 0.696798849 | -4.711855761 |
| **A_52_P507214** | Mmp9 | -0.18771713 | -0.693335124 | 0.317900864 | 8.439510393 | -0.902120546 | 0.40079149 | 0.696798849 | -4.707526378 |
| **A_55_P2125912** | Cct6a | 0.109545976 | -0.185539296 | 0.404631247 | 10.6754549 | 0.902053059 | 0.400824663 | 0.696798849 | -4.711906717 |
| **A_55_P2170256** | Ptgr2 | 0.094008897 | -0.159250035 | 0.347267828 | 8.85907953 | 0.9019602 | 0.400870313 | 0.696798849 | -4.711944705 |
| **A_55_P2086258** | Pan3 | 0.090244215 | -0.1528861 | 0.33337453 | 10.49397294 | 0.901910505 | 0.400894744 | 0.696798849 | -4.711965034 |
| **A_55_P2182452** | Tnfsf15 | -0.178818847 | -0.660614331 | 0.302976636 | 7.924259369 | -0.90184885 | 0.400925057 | 0.696798849 | -4.711990255 |
| **A_55_P2040110** | Cdv3 | 0.100335835 | -0.170011152 | 0.370682821 | 8.950852028 | 0.90181556 | 0.400941425 | 0.696798849 | -4.712003872 |
| **A_55_P2353079** | NA | -0.075530996 | -0.27911881 | 0.128056818 | 7.699224616 | -0.901481088 | 0.401105904 | 0.697022028 | -4.712140668 |
| **A_55_P2177539** | Sod1 | -0.078408681 | -0.289785344 | 0.132967983 | 15.64999308 | -0.901343429 | 0.401173614 | 0.6970663 | -4.71219696 |
| **A_55_P2110925** | Flrt3 | -0.208573891 | -0.770892062 | 0.35374428 | 7.98425673 | -0.901282659 | 0.401203507 | 0.6970663 | -4.712221808 |
| **A_55_P2046802** | Mlkl | -0.105010282 | -0.388230221 | 0.178209656 | 8.915568704 | -0.900930295 | 0.401376872 | 0.697304832 | -4.712365867 |
| **A_52_P4046** | Baz2a | 0.131198611 | -0.222804838 | 0.48520206 | 9.490572119 | 0.900543991 | 0.401567001 | 0.697545705 | -4.71252376 |
| **A_55_P2013218** | Paip1 | 0.104287551 | -0.177116925 | 0.385692027 | 8.10659483 | 0.900501948 | 0.401587697 | 0.697545705 | -4.712540941 |
| **A_55_P1960326** | NA | 0.074282818 | -0.126262521 | 0.274828158 | 10.60284199 | 0.900034149 | 0.401818035 | 0.697848769 | -4.71273208 |
| **A_55_P2077558** | Sod3 | -0.223661284 | -0.827514011 | 0.380191443 | 11.37704294 | -0.900000955 | 0.401834383 | 0.697848769 | -4.71274564 |
| **A_55_P2317341** | NA | 0.073576146 | -0.125109436 | 0.272261727 | 8.323685419 | 0.899816329 | 0.401925321 | 0.697884271 | -4.712821057 |
| **A_55_P1956572** | Dcaf8 | 0.072683515 | -0.123592361 | 0.268959391 | 12.2013838 | 0.899812845 | 0.401927037 | 0.697884271 | -4.71282248 |
| **A_51_P494541** | Abhd17b | -0.14414042 | -0.533443716 | 0.245162877 | 10.26927238 | -0.899665135 | 0.401999803 | 0.697939881 | -4.712882811 |
| **A_55_P2119463** | Ggh | 0.102776015 | -0.174829841 | 0.380381871 | 11.10992592 | 0.899593568 | 0.402035062 | 0.697939881 | -4.712912039 |
| **A_55_P2054887** | Trappc1 | 0.10246509 | -0.17432112 | 0.379251301 | 12.45236283 | 0.899527957 | 0.40206739 | 0.697939881 | -4.712938834 |
| **A_66_P109002** | Kcng2 | 0.1059965 | -0.180376594 | 0.392369593 | 7.756895613 | 0.899378499 | 0.402141036 | 0.697958553 | -4.712999866 |
| **A_51_P227392** | Rhou | -0.078452189 | -0.29041267 | 0.133508291 | 10.98666875 | -0.899359569 | 0.402150365 | 0.697958553 | -4.713007595 |
| **A_55_P2091212** | NA | 0.120119645 | -0.204523257 | 0.444762548 | 16.83044352 | 0.899065469 | 0.402295317 | 0.698018407 | -4.713127672 |
| **A_51_P207622** | Fmod | -0.254899496 | -0.943812321 | 0.43401333 | 8.35047949 | -0.899058384 | 0.40229881 | 0.698018407 | -4.713130565 |
| **A_55_P2043269** | Cdc42se1 | 0.076106019 | -0.129594851 | 0.281806888 | 9.097257167 | 0.899013193 | 0.402321086 | 0.698018407 | -4.713149013 |
| **A_55_P2170210** | Ap1s3 | 0.063487102 | -0.108131154 | 0.235105357 | 9.323436308 | 0.89888722 | 0.40238319 | 0.698018407 | -4.713200436 |
| **A_55_P2104761** | NA | 0.095958263 | -0.163444054 | 0.35536058 | 15.32169815 | 0.898859303 | 0.402396953 | 0.698018407 | -4.713211832 |
| **A_51_P512561** | Zfp68 | -0.168297273 | -0.623257082 | 0.286662536 | 10.38218632 | -0.898850027 | 0.402401527 | 0.698018407 | -4.713215618 |
| **A_51_P479865** | Glb1l | 0.065167186 | -0.111036288 | 0.241370661 | 8.207530706 | 0.898664722 | 0.402492896 | 0.69805499 | -4.713291251 |
| **A_55_P2174506** | Stx17 | 0.066291652 | -0.112956835 | 0.245540139 | 8.394650541 | 0.898641626 | 0.402504285 | 0.69805499 | -4.713300677 |
| **A_55_P2094469** | Kdr | 0.310291476 | -0.528767803 | 1.149350756 | 12.47502068 | 0.898587535 | 0.402530959 | 0.69805499 | -4.713322752 |
| **A_55_P1991901** | NA | -0.066502345 | -0.246437631 | 0.113432942 | 7.703375004 | -0.898056804 | 0.402792754 | 0.698367614 | -4.713539304 |
| **A_66_P120208** | Cnnm3 | -0.086950161 | -0.322226123 | 0.148325801 | 8.913713909 | -0.89799935 | 0.402821103 | 0.698367614 | -4.713562742 |

| | | | | | | | | | |
|---|---|---|---|---|---|---|---|---|---|
| **A_55_P2088490** | Acad10 | 0.171064395 | -0.291845538 | 0.633974329 | 12.23788604 | 0.897937928 | 0.40285141 | 0.698367614 | -4.713587798 |
| **A_55_P2146520** | Carns1 | -0.060646527 | -0.224761069 | 0.103468015 | 8.074578868 | -0.897929123 | 0.402855755 | 0.698367614 | -4.713591389 |
| **A_51_P264053** | Pigf | 0.125134406 | -0.213543355 | 0.463812168 | 10.13854843 | 0.897786862 | 0.402925959 | 0.698383465 | -4.713649416 |
| **A_51_P106527** | Mcrip2 | 0.112389517 | -0.191817143 | 0.416596176 | 11.41087792 | 0.897718708 | 0.402959595 | 0.698383465 | -4.713677214 |
| **A_55_P2367117** | Pitpnb | 0.06693939 | -0.114259224 | 0.248138004 | 8.62203109 | 0.897656264 | 0.402990415 | 0.698383465 | -4.713702681 |
| **A_51_P472621** | Zfp719 | -0.060794951 | -0.22539048 | 0.103800578 | 8.114092389 | -0.8974963 | 0.403069375 | 0.698383465 | -4.713767916 |
| **A_55_P2018417** | Osbpl3 | 0.154486507 | -0.263795289 | 0.572768304 | 8.188469145 | 0.89743867 | 0.403097825 | 0.698383465 | -4.713791416 |
| **A_55_P2067767** | Psg25 | -0.059452607 | -0.220430333 | 0.101525118 | 7.698178568 | -0.897404637 | 0.403114626 | 0.698383465 | -4.713805293 |
| **A_51_P469661** | Stk38 | 0.081533302 | -0.139233131 | 0.302299734 | 10.33129157 | 0.89739817 | 0.403117819 | 0.698383465 | -4.71380793 |
| **A_55_P2093940** | Zfp408 | 0.065262768 | -0.111481017 | 0.242006552 | 15.94775015 | 0.897231531 | 0.403200094 | 0.69840113 | -4.713875875 |
| **A_55_P2041045** | Csnk1g2 | -0.083523599 | -0.309721225 | 0.142674028 | 12.52485091 | -0.897231154 | 0.40320028 | 0.69840113 | -4.713876028 |
| **A_55_P2073294** | Hadhb | -0.187103793 | -0.693968445 | 0.319760858 | 13.20211354 | -0.896961448 | 0.403333469 | 0.698569232 | -4.713985979 |
| **A_52_P506529** | Rabgap1l | 0.074773677 | -0.127817178 | 0.277364531 | 9.931895177 | 0.896834048 | 0.403396395 | 0.698615618 | -4.714037909 |
| **A_55_P1976224** | Ckb | -0.107749355 | -0.399756779 | 0.184258068 | 9.880760135 | -0.896610821 | 0.403506669 | 0.698743989 | -4.714128887 |
| **A_52_P171724** | 2310009B15Rik | -0.176429719 | -0.654793187 | 0.301933749 | 10.08116431 | -0.896183464 | 0.403717847 | 0.699047054 | -4.714303021 |
| **A_55_P1998389** | NA | -0.099509392 | -0.369422171 | 0.170403387 | 15.02171362 | -0.895826311 | 0.403894398 | 0.699290113 | -4.714448508 |
| **A_55_P2012221** | Armcx2 | -0.138408668 | -0.513923082 | 0.237105747 | 11.33130512 | -0.895611919 | 0.404000406 | 0.699344981 | -4.714535823 |
| **A_55_P1969352** | Get4 | 0.111916249 | -0.191757966 | 0.415590464 | 11.01567655 | 0.89550583 | 0.40405287 | 0.699344981 | -4.714579024 |
| **A_55_P2219657** | Tmf1 | 0.068466663 | -0.117323643 | 0.254256969 | 8.270034223 | 0.895445793 | 0.404082563 | 0.699344981 | -4.714603471 |
| **A_55_P2144055** | NA | 0.088290713 | -0.151300359 | 0.327881785 | 15.6364393 | 0.895421629 | 0.404094514 | 0.699344981 | -4.714613311 |
| **A_55_P2048096** | Mzf1 | -0.07086342 | -0.263173866 | 0.121447026 | 8.044463819 | -0.89536976 | 0.404120168 | 0.699344981 | -4.71463443 |
| **A_55_P2137376** | Dip2c | -0.084862865 | -0.315177227 | 0.145451496 | 9.058091458 | -0.895323247 | 0.404143175 | 0.699344981 | -4.714653369 |
| **A_52_P561650** | Vwa1 | -0.131874169 | -0.48995706 | 0.226208722 | 7.839156251 | -0.894868775 | 0.404368021 | 0.699644108 | -4.714838378 |
| **A_55_P2002449** | NA | 0.265458283 | -0.455425582 | 0.986342147 | 10.4029789 | 0.894775667 | 0.404414098 | 0.699644108 | -4.714876273 |
| **A_55_P2156288** | NA | 0.098737392 | -0.16943838 | 0.366913165 | 8.528997562 | 0.8946338 | 0.40448431 | 0.699644108 | -4.714934009 |
| **A_55_P2101431** | Mrps25 | -0.084402953 | -0.313652977 | 0.14484707 | 8.320556204 | -0.89460525 | 0.404498442 | 0.699644108 | -4.714945628 |
| **A_52_P88773** | Tor2a | -0.080450288 | -0.298971672 | 0.138071097 | 10.55202246 | -0.894575185 | 0.404513323 | 0.699644108 | -4.714957862 |
| **A_55_P2155942** | Ezh1 | -0.069215285 | -0.257228278 | 0.118797709 | 9.658012685 | -0.894534994 | 0.404533217 | 0.699644108 | -4.714974217 |
| **A_55_P2134042** | NA | 0.070671969 | -0.121319364 | 0.262663303 | 16.23523039 | 0.894434945 | 0.404582743 | 0.699657723 | -4.715014928 |
| **A_55_P2049854** | Araf | 0.065492164 | -0.11243978 | 0.243424108 | 8.000820214 | 0.89437285 | 0.404613484 | 0.699657723 | -4.715040193 |
| **A_55_P1984357** | NA | -0.093988311 | -0.349399607 | 0.161422985 | 16.17518823 | -0.89416349 | 0.404717142 | 0.699713326 | -4.71512537 |
| **A_66_P138673** | Whamm | 0.136247099 | -0.234038889 | 0.506533087 | 8.583834168 | 0.894073403 | 0.404761752 | 0.699713326 | -4.715162017 |
| **A_55_P1999823** | Zmynd19 | 0.106133313 | -0.18232171 | 0.394588336 | 8.611640315 | 0.8940397 | 0.404778442 | 0.699713326 | -4.715175727 |
| **A_55_P2099750** | NA | 0.141739618 | -0.243498642 | 0.526977879 | 12.32342498 | 0.894015473 | 0.40479044 | 0.699713326 | -4.715185582 |
| **A_51_P376057** | Bcorl1 | 0.103795761 | -0.178346121 | 0.385937643 | 9.102585815 | 0.893912985 | 0.404841198 | 0.69971623 | -4.71522727 |
| **A_55_P1964852** | Kbtbd2 | -0.070379096 | -0.261696512 | 0.12093832 | 11.91590554 | -0.893865896 | 0.404864521 | 0.69971623 | -4.715246422 |
| **A_55_P2129291** | Dclre1c | 0.110810687 | -0.190549365 | 0.41217074 | 8.759669978 | 0.893468312 | 0.405061481 | 0.69994582 | -4.715408107 |
| **A_55_P2036693** | Cd40 | -0.330619964 | -1.229830239 | 0.56859031 | 8.537191825 | -0.893410399 | 0.405090176 | 0.69994582 | -4.715431655 |
| **A_55_P2131677** | NA | 0.061162399 | -0.105198309 | 0.227523107 | 7.761320529 | 0.893340334 | 0.405124895 | 0.69994582 | -4.715460142 |
| **A_55_P2110595** | Slc35a2 | 0.06461721 | -0.111147852 | 0.240382271 | 8.109040521 | 0.893303098 | 0.405143348 | 0.69994582 | -4.715475281 |
| **A_55_P2053398** | NA | 0.081896274 | -0.140893022 | 0.304685569 | 17.16514391 | 0.893208834 | 0.405190063 | 0.69994582 | -4.715513604 |
| **A_55_P2023953** | Arpc3 | 0.081998821 | -0.141094878 | 0.305092521 | 13.63137678 | 0.893106997 | 0.405240536 | 0.69994582 | -4.715555002 |
| **A_51_P150608** | Jagn1 | -0.06831404 | -0.254179969 | 0.117551888 | 11.43706315 | -0.89308619 | 0.405250849 | 0.69994582 | -4.715563461 |
| **A_55_P2029726** | NA | -0.121693615 | -0.452855309 | 0.209468078 | 15.14225341 | -0.892916581 | 0.405334924 | 0.699988997 | -4.715632402 |
| **A_55_P1980981** | Apba3 | -0.095839245 | -0.356664102 | 0.164985611 | 9.901067263 | -0.892848139 | 0.405368854 | 0.699988997 | -4.71566022 |
| **A_55_P1977792** | Folr1 | -0.261385534 | -0.972766896 | 0.449995828 | 8.089743601 | -0.892816597 | 0.405384491 | 0.699988997 | -4.715673039 |
| **A_51_P133097** | Gm4013 | 0.11407954 | -0.196530735 | 0.424689814 | 9.522149562 | 0.892431995 | 0.405575203 | 0.700255748 | -4.715829327 |
| **A_51_P410650** | Dpp9 | 0.17545673 | -0.302395546 | 0.653309007 | 9.610588959 | 0.892194541 | 0.405692983 | 0.70039654 | -4.715925798 |
| **A_66_P126984** | Gm10523 | 0.062769897 | -0.108197372 | 0.233737165 | 8.339026545 | 0.892116619 | 0.405731638 | 0.700400718 | -4.715957452 |
| **A_51_P430033** | Ccdc13 | -0.115349318 | -0.429603928 | 0.198905292 | 7.736318996 | -0.891900817 | 0.405838708 | 0.700522985 | -4.716045106 |

| | | | | | | | | |
|---|---|---|---|---|---|---|---|---|
| **A_55_P1987186** | Ttll9 | -0.103464531 | -0.385394935 | 0.178465873 | 7.831723934 | -0.891728747 | 0.405924095 | 0.700584627 | -4.716114988 |
| **A_55_P1956647** | Sfmbt2 | 0.079111501 | -0.13647604 | 0.294699042 | 7.989422599 | 0.891659731 | 0.405958348 | 0.700584627 | -4.716143015 |
| **A_55_P2157200** | NA | -0.161245295 | -0.600681091 | 0.278190502 | 10.60039787 | -0.891609748 | 0.405983155 | 0.700584627 | -4.716163312 |
| **A_66_P108770** | Oxct1 | -0.257367331 | -0.958886639 | 0.444151977 | 11.09591564 | -0.891449978 | 0.40606246 | 0.700658925 | -4.716228185 |
| **A_55_P1966189** | Il4ra | 0.162559856 | -0.280582368 | 0.605702079 | 8.645648181 | 0.891360453 | 0.406106902 | 0.700673062 | -4.716264533 |
| **A_52_P154880** | Ing3 | 0.070563491 | -0.121828551 | 0.262955532 | 8.577213866 | 0.891201979 | 0.406185581 | 0.700709266 | -4.716328868 |
| **A_66_P138780** | Elob | -0.111338543 | -0.414936926 | 0.192259841 | 13.13661696 | -0.891105793 | 0.406233341 | 0.700709266 | -4.716367913 |
| **A_55_P2261882** | D230040J21Rik | 0.057790449 | -0.099793953 | 0.215374851 | 8.132059204 | 0.891099147 | 0.406236641 | 0.700709266 | -4.716370611 |
| **A_55_P2088296** | Fam160b2 | 0.209686315 | -0.362177372 | 0.781550002 | 14.61984481 | 0.890965232 | 0.406303142 | 0.700761438 | -4.716424966 |
| **A_55_P2059382** | Arl6 | -0.214666755 | -0.800261731 | 0.370928221 | 8.383852375 | -0.890739351 | 0.406415332 | 0.700883081 | -4.716516638 |
| **A_51_P493987** | Moxd1 | -0.082752797 | -0.308540064 | 0.143034471 | 7.71150023 | -0.89056664 | 0.406501129 | 0.700883081 | -4.716586722 |
| **A_51_P232207** | Hoxb6 | -0.137160407 | -0.511407768 | 0.237086954 | 7.790949491 | -0.890539541 | 0.406514592 | 0.700883081 | -4.716597717 |
| **A_55_P2173183** | Cdk5r1 | 0.07411253 | -0.128120667 | 0.276345727 | 8.297119433 | 0.890476329 | 0.406545997 | 0.700883081 | -4.716623365 |
| **A_55_P2040763** | Sec62 | 0.103384311 | -0.178746754 | 0.385515376 | 12.56428944 | 0.890403623 | 0.406582122 | 0.700883081 | -4.716652863 |
| **A_55_P1980426** | Nnat | -0.39112515 | -1.458568076 | 0.676317777 | 8.113454295 | -0.890336684 | 0.406615384 | 0.700883081 | -4.716680021 |
| **A_51_P361426** | Arl8b | -0.080036079 | -0.29848918 | 0.138417022 | 10.85299277 | -0.890247534 | 0.406659685 | 0.700883081 | -4.716716187 |
| **A_55_P2016249** | Ppid | 0.082833088 | -0.143267923 | 0.308934099 | 9.690919548 | 0.890193745 | 0.406686416 | 0.700883081 | -4.716738006 |
| **A_52_P532456** | Plagl1 | -0.207828793 | -0.775135239 | 0.359477652 | 13.2633715 | -0.890166378 | 0.406700017 | 0.700883081 | -4.716749108 |
| **A_51_P389885** | Spic | 0.131738775 | -0.228013411 | 0.491490961 | 10.54758254 | 0.889801981 | 0.406881148 | 0.700998328 | -4.716896902 |
| **A_55_P1955237** | NA | 0.06297859 | -0.109006285 | 0.234963465 | 16.81638043 | 0.889786613 | 0.406888788 | 0.700998328 | -4.716903135 |
| **A_51_P192751** | Mrpl14 | -0.162216318 | -0.605256212 | 0.280823577 | 14.0359384 | -0.88968218 | 0.406940711 | 0.700998328 | -4.716945484 |
| **A_55_P1960837** | NA | 0.08261669 | -0.143026654 | 0.308260033 | 13.16023926 | 0.889668988 | 0.40694727 | 0.700998328 | -4.716950833 |
| **A_51_P144926** | Cops8 | 0.069568389 | -0.120445796 | 0.259582573 | 9.475780645 | 0.889629368 | 0.40696697 | 0.700998328 | -4.716966898 |
| **A_51_P378856** | Pfkp | -0.059188814 | -0.220859363 | 0.102481736 | 8.2570116 | -0.889594131 | 0.406984491 | 0.700998328 | -4.716981186 |
| **A_55_P2058157** | Ppp4r3b | 0.109838593 | -0.190217303 | 0.409894489 | 9.794419259 | 0.889479588 | 0.407041451 | 0.701018216 | -4.717027629 |
| **A_55_P1965728** | Abca17 | 0.302822036 | -0.524474694 | 1.130118766 | 9.052290355 | 0.88942505 | 0.407068573 | 0.701018216 | -4.71704974 |
| **A_55_P2037677** | Cttn | -0.143377614 | -0.535202582 | 0.248447354 | 11.5189878 | -0.889144672 | 0.40720803 | 0.701126298 | -4.7171634 |
| **A_51_P315904** | Gadd45g | -0.518050455 | -1.933844164 | 0.897743253 | 11.6398654 | -0.889109541 | 0.407225506 | 0.701126298 | -4.717177639 |
| **A_55_P2102719** | NA | -0.17854669 | -0.666517793 | 0.309424413 | 14.32983165 | -0.889080096 | 0.407240155 | 0.701126298 | -4.717189574 |
| **A_55_P2429432** | 2610301B20Rik | 0.073918345 | -0.128133373 | 0.275970063 | 8.484704746 | 0.888940873 | 0.40730942 | 0.701139596 | -4.717246002 |
| **A_66_P123526** | 4930504D19Rik | 0.14773165 | -0.256094714 | 0.551558014 | 8.491316962 | 0.888918752 | 0.407320426 | 0.701139596 | -4.717254966 |
| **A_51_P366138** | Mertk | -0.078098124 | -0.291681682 | 0.135485435 | 9.045192193 | -0.888497011 | 0.407530307 | 0.701385387 | -4.717425861 |
| **A_52_P641849** | Khnyn | -0.070149394 | -0.261997065 | 0.121698276 | 8.198328617 | -0.888485995 | 0.40753579 | 0.701385387 | -4.717430324 |
| **A_55_P2182810** | Arl16 | 0.068561405 | -0.118989561 | 0.25611237 | 8.570897531 | 0.888267131 | 0.407644741 | 0.701510434 | -4.717518988 |
| **A_51_P487308** | Pstk | 0.062927179 | -0.109232778 | 0.235087135 | 10.40620172 | 0.888156108 | 0.407700017 | 0.701526192 | -4.71756396 |
| **A_55_P1995438** | NA | -0.091802102 | -0.342974841 | 0.159370637 | 15.36965836 | -0.888102949 | 0.407726485 | 0.701526192 | -4.717585491 |
| **A_55_P2152364** | NA | -0.057963053 | -0.216601416 | 0.10067531 | 7.802484594 | -0.887822655 | 0.407866069 | 0.701654739 | -4.717699008 |
| **A_55_P2023136** | Rnf214 | 0.076512085 | -0.132896553 | 0.285920723 | 8.578619502 | 0.887807138 | 0.407873798 | 0.701654739 | -4.717705291 |
| **A_52_P514061** | Padi4 | -0.175147484 | -0.654628457 | 0.304333488 | 9.062933733 | -0.887596775 | 0.407978582 | 0.701726666 | -4.71779047 |
| **A_51_P380153** | Ubr4 | 0.105712378 | -0.183694755 | 0.395119511 | 11.6559715 | 0.887564247 | 0.407994786 | 0.701726666 | -4.71780364 |
| **A_51_P392928** | Mthfd1 | 0.163158464 | -0.283553427 | 0.609870355 | 12.72145757 | 0.88749371 | 0.408029927 | 0.701726666 | -4.717832198 |
| **A_55_P2109599** | NA | -0.16396667 | -0.612949198 | 0.285015858 | 11.8513952 | -0.887379363 | 0.408086898 | 0.701726666 | -4.71787849 |
| **A_51_P288618** | Dusp11 | 0.079925355 | -0.138935702 | 0.298786413 | 8.245722315 | 0.887358826 | 0.408097131 | 0.701726666 | -4.717886803 |
| **A_55_P2093483** | NA | -0.091141581 | -0.340772016 | 0.158488855 | 15.83062745 | -0.887160531 | 0.408195944 | 0.70183414 | -4.71796707 |
| **A_51_P184949** | F12 | -0.15038729 | -0.562330036 | 0.261555457 | 13.30987216 | -0.887069121 | 0.4082415 | 0.701850037 | -4.718004067 |
| **A_55_P2024021** | Atp5l | 0.073613557 | -0.128135836 | 0.275362949 | 14.82817951 | 0.886602095 | 0.408474315 | 0.702084686 | -4.718193053 |
| **A_51_P300506** | Cox6b2 | 0.158645385 | -0.276154271 | 0.593445042 | 11.15513347 | 0.886587165 | 0.408481759 | 0.702084686 | -4.718199093 |
| **A_52_P685938** | Efcab14 | 0.082754413 | -0.144053402 | 0.309562227 | 9.298522382 | 0.886576743 | 0.408486956 | 0.702084686 | -4.71820331 |
| **A_51_P439836** | NA | 0.121847304 | -0.212314113 | 0.45600872 | 12.42425574 | 0.886018539 | 0.408765361 | 0.702408869 | -4.718429103 |
| **A_51_P108226** | Wfdc21 | -0.181507717 | -0.679299925 | 0.316284491 | 12.38958218 | -0.885992791 | 0.408778206 | 0.702408869 | -4.700864005 |

| | | | | | | | | | |
|---|---|---|---|---|---|---|---|---|---|
| **A_52_P432780** | Mrpl50 | 0.104812991 | -0.182732789 | 0.392358771 | 9.988178867 | 0.885709505 | 0.408919552 | 0.702408869 | -4.718554067 |
| **A_52_P377723** | Peg10 | 0.092203288 | -0.160776341 | 0.345182916 | 7.977374485 | 0.885613125 | 0.40896765 | 0.702408869 | -4.718593034 |
| **A_55_P2056342** | Mgst3 | 0.117201776 | -0.204369555 | 0.438773107 | 13.51150884 | 0.885604997 | 0.408971706 | 0.702408869 | -4.71859632 |
| **A_51_P348617** | Borcs8 | -0.119702711 | -0.448142806 | 0.208737383 | 12.3016373 | -0.885586542 | 0.408980916 | 0.702408869 | -4.718603781 |
| **A_51_P259296** | Lpl | -0.301934257 | -1.130383068 | 0.526514555 | 12.85775213 | -0.885584282 | 0.408982045 | 0.702408869 | -4.718604695 |
| **A_55_P1994132** | Tmem184a | 0.144782281 | -0.252491869 | 0.54205643 | 8.861662929 | 0.88554023 | 0.40900403 | 0.702408869 | -4.718622505 |
| **A_55_P2037221** | Klhdc10 | -0.080833807 | -0.302665768 | 0.140998154 | 11.44199202 | -0.885425661 | 0.409061214 | 0.702408869 | -4.71866882 |
| **A_52_P583155** | Ykt6 | 0.098991209 | -0.17267754 | 0.370659958 | 9.509276894 | 0.885401259 | 0.409073394 | 0.702408869 | -4.718678684 |
| **A_55_P2033680** | NA | 0.131949901 | -0.230194348 | 0.49409415 | 9.900926196 | 0.885341182 | 0.409103383 | 0.702408869 | -4.718702969 |
| **A_51_P221362** | Tnks2 | -0.140231728 | -0.525113135 | 0.244649678 | 11.86197353 | -0.885324625 | 0.409111648 | 0.702408869 | -4.718709661 |
| **A_52_P89064** | Dusp18 | -0.084904632 | -0.317954894 | 0.148145629 | 7.740852893 | -0.885248053 | 0.409149874 | 0.702412107 | -4.718740612 |
| **A_51_P476579** | NA | 0.096399584 | -0.168277069 | 0.361076237 | 9.152097635 | 0.884998871 | 0.409274287 | 0.702519104 | -4.718841319 |
| **A_55_P2082778** | NA | -0.108555317 | -0.406614147 | 0.189503513 | 14.38479417 | -0.884977638 | 0.409284889 | 0.702519104 | -4.718849899 |
| **A_66_P112816** | B4galt6 | 0.137149767 | -0.239605385 | 0.51390492 | 8.919882472 | 0.88454323 | 0.409501855 | 0.702829102 | -4.719025418 |
| **A_51_P452153** | Mzb1 | -0.090272618 | -0.338313724 | 0.157768487 | 9.577510489 | -0.884332466 | 0.409607152 | 0.702935058 | -4.710407842 |
| **A_55_P2373121** | Dcaf17 | 0.148681858 | -0.259876662 | 0.557240378 | 9.421156277 | 0.884274084 | 0.409636323 | 0.702935058 | -4.719134137 |
| **A_55_P1999473** | Ppp1r26 | -0.153042551 | -0.573673756 | 0.267588654 | 7.789133786 | -0.884084744 | 0.409730939 | 0.703034949 | -4.719210606 |
| **A_55_P2113356** | Miip | 0.102544947 | -0.179327521 | 0.384417415 | 8.636120733 | 0.883984794 | 0.409780892 | 0.703034949 | -4.719250968 |
| **A_55_P1991069** | Milr1 | 0.122542929 | -0.2143328 | 0.459418658 | 9.630038443 | 0.883897127 | 0.40982471 | 0.703034949 | -4.719286368 |
| **A_51_P198675** | Ttc36 | -0.166854828 | -0.625561755 | 0.291852098 | 10.77328487 | -0.883866492 | 0.409840023 | 0.703034949 | -4.719298738 |
| **A_51_P276039** | Txndc15 | -0.085589661 | -0.320958497 | 0.149779174 | 12.31024513 | -0.883599659 | 0.409973418 | 0.703201367 | -4.719406469 |
| **A_51_P210395** | Glrx | -0.130424905 | -0.489176065 | 0.228326255 | 9.919383999 | -0.88338578 | 0.410080363 | 0.703322391 | -4.719492804 |
| **A_55_P2083806** | Sp8 | 0.062839564 | -0.110030202 | 0.235709331 | 7.942672916 | 0.883277786 | 0.410134371 | 0.70335261 | -4.719536393 |
| **A_51_P113274** | Hps6 | -0.127056767 | -0.476673015 | 0.22255948 | 9.764975822 | -0.883058312 | 0.410244147 | 0.703449629 | -4.719624966 |
| **A_55_P2075627** | Apc2 | 0.061549063 | -0.107820165 | 0.230918291 | 8.249385035 | 0.883019162 | 0.410263731 | 0.703449629 | -4.719640764 |
| **A_51_P132718** | 2310010J17Rik | 0.132067373 | -0.231413189 | 0.495547935 | 11.76012221 | 0.882871582 | 0.410337562 | 0.703513814 | -4.719700313 |
| **A_55_P2178378** | Gas5 | 0.135375557 | -0.237260066 | 0.50801118 | 9.339675323 | 0.882752745 | 0.410397021 | 0.70355335 | -4.71974826 |
| **A_52_P387954** | Dctn5 | 0.06783627 | -0.118938608 | 0.254611149 | 11.39262973 | 0.882524332 | 0.410511323 | 0.703684834 | -4.719840405 |
| **A_52_P484838** | Rfxank | -0.11472762 | -0.430634513 | 0.201179274 | 9.174697159 | -0.882453986 | 0.41054653 | 0.703684834 | -4.71986878 |
| **A_55_P2169069** | Gm10405 | 0.209151157 | -0.366876889 | 0.785179203 | 13.64060991 | 0.882266587 | 0.410640332 | 0.703783204 | -4.719944363 |
| **A_51_P122246** | Creld2 | -0.113890834 | -0.427642803 | 0.199861135 | 12.66898437 | -0.882034356 | 0.410756596 | 0.70392005 | -4.717822376 |
| **A_55_P2186287** | Sub1 | -0.1122466 | -0.421669485 | 0.197176285 | 14.38520997 | -0.881462728 | 0.41104288 | 0.704348212 | -4.720268465 |
| **A_55_P2020443** | Hnrnpa3 | 0.083093157 | -0.145991219 | 0.312177532 | 9.845750492 | 0.881359272 | 0.411094709 | 0.70437458 | -4.720310162 |
| **A_51_P112557** | Polr1c | -0.169392451 | -0.636473964 | 0.297689062 | 11.36646142 | -0.881220443 | 0.411164267 | 0.704431318 | -4.715847675 |
| **A_55_P2293351** | Slc2a4rg-ps | 0.119248919 | -0.209595227 | 0.448093066 | 8.932661002 | 0.881145293 | 0.411201924 | 0.704433394 | -4.720396396 |
| **A_51_P282616** | Maats1 | -0.114188885 | -0.429135675 | 0.200757905 | 7.841483271 | -0.880987681 | 0.411280908 | 0.704506263 | -4.720459904 |
| **A_55_P2031466** | Lrrc32 | -0.170672248 | -0.641633969 | 0.300289473 | 10.94227949 | -0.880563112 | 0.411493731 | 0.704732217 | -4.720630945 |
| **A_55_P2027337** | NA | 0.33069541 | -0.581986289 | 1.243377109 | 12.15500334 | 0.880424286 | 0.411563337 | 0.704732217 | -4.72068686 |
| **A_51_P364984** | Gabpa | -0.058576274 | -0.220245296 | 0.103092749 | 8.36481299 | -0.880396118 | 0.411577461 | 0.704732217 | -4.720698205 |
| **A_52_P334562** | Vdr | -0.073531743 | -0.276480955 | 0.129417469 | 7.7736525 | -0.880381042 | 0.411585021 | 0.704732217 | -4.720704277 |
| **A_52_P589119** | Rhno1 | 0.096667167 | -0.170161559 | 0.363495893 | 8.324460849 | 0.880297763 | 0.411626783 | 0.704732217 | -4.720737815 |
| **A_55_P2437253** | Nae1 | 0.085749591 | -0.150946106 | 0.322445288 | 8.740745521 | 0.880288205 | 0.411631576 | 0.704732217 | -4.720741665 |
| **A_55_P2114133** | Garnl3 | -0.105778088 | -0.397931905 | 0.186375729 | 8.12008998 | -0.879766329 | 0.411893353 | 0.705003102 | -4.720951789 |
| **A_52_P412046** | NA | -0.106027338 | -0.398885668 | 0.186830992 | 14.54591737 | -0.879717973 | 0.411917615 | 0.705003102 | -4.720971255 |
| **A_55_P2044458** | Pramef8 | -0.101781444 | -0.382920251 | 0.179357363 | 9.214783007 | -0.879692723 | 0.411930284 | 0.705003102 | -4.720981419 |
| **A_52_P290725** | Arfgef2 | 0.120116511 | -0.211690732 | 0.451923755 | 8.833571692 | 0.879630009 | 0.411961753 | 0.705003102 | -4.721006663 |
| **A_55_P2183032** | Zfp787 | 0.119934613 | -0.211417519 | 0.451286745 | 9.547226401 | 0.87950428 | 0.412024846 | 0.705003102 | -4.721057268 |
| **A_55_P2092750** | Car9 | 0.064699302 | -0.114064494 | 0.243463099 | 8.586085027 | 0.879433928 | 0.412060153 | 0.705003102 | -4.721085583 |
| **A_51_P265348** | Ldb3 | 0.121874299 | -0.214884577 | 0.458633175 | 8.20264762 | 0.879379361 | 0.412087539 | 0.705003102 | -4.721107543 |
| **A_55_P2088315** | Vmn1r87 | 0.181522132 | -0.320080369 | 0.683124633 | 8.849892544 | 0.87933241 | 0.412111105 | 0.705003102 | -4.721126438 |

| | | | | | | | | | |
|---|---|---|---|---|---|---|---|---|---|
| **A_55_P1999561** | Pram1 | -0.204583174 | -0.769919468 | 0.360753121 | 10.16775692 | -0.879318545 | 0.412118064 | 0.705003102 | -4.721132017 |
| **A_55_P2132894** | Sdcbp | 0.107843891 | -0.19022631 | 0.405914092 | 9.294620786 | 0.879144326 | 0.412205516 | 0.705063731 | -4.721202122 |
| **A_66_P120507** | Plcd3 | -0.103723967 | -0.390420697 | 0.182972763 | 8.147847186 | -0.87910261 | 0.412226459 | 0.705063731 | -4.721218907 |
| **A_55_P2021515** | Rpl3 | -0.059156566 | -0.222684406 | 0.104371274 | 16.37986289 | -0.879011273 | 0.412272314 | 0.705072904 | -4.721255656 |
| **A_55_P2060498** | Zfp553 | -0.079430124 | -0.299025036 | 0.140164789 | 9.343160688 | -0.878913536 | 0.412321387 | 0.705072904 | -4.721294977 |
| **A_55_P2001193** | Stx18 | -0.077662898 | -0.292381751 | 0.137055955 | 10.38329689 | -0.87887397 | 0.412341254 | 0.705072904 | -4.721310894 |
| **A_55_P1965114** | Spsb3 | 0.076832743 | -0.135621935 | 0.289287421 | 12.91517407 | 0.878745734 | 0.41240565 | 0.705120638 | -4.72136248 |
| **A_51_P351394** | Vps45 | 0.098862032 | -0.174579819 | 0.372303882 | 9.273618202 | 0.878512078 | 0.412523003 | 0.705258899 | -4.72145646 |
| **A_55_P2194958** | Nfia | -0.071158054 | -0.268014909 | 0.1256988 | 8.14633642 | -0.878327968 | 0.412615488 | 0.705354627 | -4.7215305 |
| **A_55_P2186332** | NA | 0.069296654 | -0.122436783 | 0.261030092 | 8.024043714 | 0.878208424 | 0.412675548 | 0.705394912 | -4.721578569 |
| **A_55_P2152962** | Mtr | 0.098993557 | -0.174972515 | 0.37295963 | 9.331002742 | 0.877997615 | 0.412781476 | 0.705513587 | -4.721663326 |
| **A_55_P2071406** | NA | -0.062525882 | -0.235605528 | 0.110553765 | 7.802279677 | -0.877802906 | 0.412879333 | 0.705618446 | -4.721741599 |
| **A_51_P210082** | Ercc4 | -0.062232246 | -0.234547653 | 0.110083161 | 9.480845099 | -0.877555423 | 0.413003737 | 0.705662725 | -4.72184107 |
| **A_55_P1997926** | NA | 0.187305831 | -0.331365812 | 0.705977474 | 11.23203125 | 0.877489608 | 0.413036826 | 0.705662725 | -4.72186752 |
| **A_51_P236267** | St8sia4 | -0.162890556 | -0.613963639 | 0.288182526 | 9.921139188 | -0.877469813 | 0.413046778 | 0.705662725 | -4.721875475 |
| **A_55_P1957880** | Jpt2 | 0.146652815 | -0.259459195 | 0.552764825 | 9.605432518 | 0.87746087 | 0.413051274 | 0.705662725 | -4.721879069 |
| **A_55_P1962065** | NA | 0.11363125 | -0.201071858 | 0.428334358 | 14.26557872 | 0.877364265 | 0.413099847 | 0.705683335 | -4.72191789 |
| **A_55_P2186558** | Tmem242 | -0.17494454 | -0.659578384 | 0.309689305 | 9.780381795 | -0.877141869 | 0.413211684 | 0.705758856 | -4.722007249 |
| **A_51_P349803** | Med11 | -0.105080919 | -0.396218454 | 0.186056616 | 10.71879783 | -0.877018696 | 0.413273634 | 0.705758856 | -4.722056734 |
| **A_55_P2151956** | NA | -0.139521348 | -0.526083614 | 0.247040917 | 8.614090681 | -0.877009698 | 0.413278159 | 0.705758856 | -4.722060349 |
| **A_55_P2047598** | Arhgdia | -0.091756642 | -0.345987387 | 0.162474102 | 14.20354881 | -0.876985944 | 0.413290107 | 0.705758856 | -4.722069891 |
| **A_55_P2049949** | NA | 0.168556181 | -0.298609042 | 0.635721405 | 9.923274184 | 0.87671284 | 0.413427494 | 0.705892078 | -4.722179592 |
| **A_55_P2042753** | Cltrn | 0.137934344 | -0.244396553 | 0.520265241 | 8.717478774 | 0.876629755 | 0.413469297 | 0.705892078 | -4.722212961 |
| **A_55_P2032966** | Hmgcs1 | 0.228871807 | -0.405577935 | 0.863321548 | 15.24300546 | 0.876553164 | 0.413507836 | 0.705892078 | -4.72224372 |
| **A_55_P1959989** | D430042O09Rik | -0.138497763 | -0.522429448 | 0.245433922 | 9.388912227 | -0.876540513 | 0.413514201 | 0.705892078 | -4.722248801 |
| **A_51_P192149** | Slc9a6 | -0.06626984 | -0.250001701 | 0.117462022 | 8.480432797 | -0.876424751 | 0.413572456 | 0.705929176 | -4.722295288 |
| **A_55_P2135860** | Prss36 | -0.089690555 | -0.338447628 | 0.159066518 | 8.86329092 | -0.87610158 | 0.413735115 | 0.706144462 | -4.722425042 |
| **A_52_P67995** | Eif4a3 | -0.078383203 | -0.295826949 | 0.139060542 | 10.15036489 | -0.875909579 | 0.413831776 | 0.706186637 | -4.722502117 |
| **A_51_P173564** | Ahsa1 | -0.073618494 | -0.27785933 | 0.130622343 | 13.69513975 | -0.875845565 | 0.413864007 | 0.706186637 | -4.722527812 |
| **A_55_P2000613** | Slc6a1 | 0.068904533 | -0.122260646 | 0.260069712 | 8.000548821 | 0.875834794 | 0.413869431 | 0.706186637 | -4.722532135 |
| **A_52_P422494** | Cd300lf | -0.180328791 | -0.680893869 | 0.320236287 | 10.02705596 | -0.875362046 | 0.414107519 | 0.706472022 | -4.722721851 |
| **A_55_P1984023** | Arrb2 | -0.132537168 | -0.500441916 | 0.235367581 | 10.28285999 | -0.87535755 | 0.414109783 | 0.706472022 | -4.722723655 |
| **A_55_P1958306** | Thrap3 | 0.087871845 | -0.156094897 | 0.331838587 | 12.35869141 | 0.875189923 | 0.41419423 | 0.706553725 | -4.722790908 |
| **A_55_P2064712** | NA | 0.06881476 | -0.122343346 | 0.259972865 | 15.53753342 | 0.874726065 | 0.414427976 | 0.706890077 | -4.722976968 |
| **A_55_P2176145** | Rgs17 | -0.089523094 | -0.338341908 | 0.15929572 | 7.792198534 | -0.874248828 | 0.414668567 | 0.707238042 | -4.723168326 |
| **A_55_P1992470** | NA | -0.070508669 | -0.266500814 | 0.125483476 | 14.57995981 | -0.874152167 | 0.41471731 | 0.707239296 | -4.723207076 |
| **A_55_P2032258** | Pdlim1 | -0.064269864 | -0.242940115 | 0.114400388 | 12.41761193 | -0.874054111 | 0.41476676 | 0.707239296 | -4.723246383 |
| **A_55_P2044380** | NA | 0.07607662 | -0.135422326 | 0.287575565 | 8.055656068 | 0.874029701 | 0.414779071 | 0.707239296 | -4.723256167 |
| **A_52_P664617** | Zfp472 | 0.079790395 | -0.142102033 | 0.301682824 | 8.792735679 | 0.873758316 | 0.414915958 | 0.707341255 | -4.723364936 |
| **A_51_P286563** | Gna13 | -0.078649957 | -0.297372549 | 0.140072634 | 10.47242533 | -0.873751706 | 0.414919292 | 0.707341255 | -4.723367585 |
| **A_55_P2018825** | NA | 0.078056625 | -0.139040634 | 0.295153883 | 9.15342575 | 0.873652285 | 0.414969449 | 0.707341255 | -4.723407426 |
| **A_51_P324303** | Mylip | 0.065035163 | -0.115854646 | 0.245924971 | 10.00637153 | 0.873609447 | 0.414991062 | 0.707341255 | -4.723424592 |
| **A_55_P2039196** | Gfra4 | 0.304781308 | -0.54300134 | 1.152563957 | 14.21271574 | 0.873548444 | 0.415021841 | 0.707341255 | -4.723449036 |
| **A_55_P2075237** | Ulk3 | 0.092658981 | -0.165120703 | 0.350438665 | 9.644952232 | 0.873417766 | 0.41508778 | 0.707388601 | -4.723501394 |
| **A_66_P117730** | Hapln1 | -0.17340808 | -0.655873297 | 0.309057138 | 7.822598649 | -0.873346347 | 0.41512382 | 0.707388601 | -4.723530007 |
| **A_55_P2158498** | Btk | 0.165264969 | -0.294613031 | 0.625142969 | 10.22029838 | 0.873215351 | 0.415189932 | 0.707388601 | -4.723582485 |
| **A_51_P466910** | Cacna1g | -0.293504487 | -1.110260251 | 0.523251278 | 7.989048188 | -0.873183912 | 0.4152058 | 0.707388601 | -4.723595079 |
| **A_55_P1964525** | NA | 0.083182089 | -0.148315556 | 0.314679734 | 16.87175892 | 0.873104918 | 0.415245672 | 0.707388601 | -4.723626722 |
| **A_55_P2079967** | Map3k5 | 0.111903593 | -0.19957057 | 0.423377755 | 9.411465889 | 0.872981804 | 0.415307818 | 0.707388601 | -4.723676033 |
| **A_51_P184024** | Tsen15 | 0.09458818 | -0.168699436 | 0.357875796 | 10.90378171 | 0.872950519 | 0.415323612 | 0.707388601 | -4.723688563 |

| | | | | | | | | | |
|---|---|---|---|---|---|---|---|---|---|
| **A_55_P2098300** | Chmp6 | -0.082163369 | -0.310879297 | 0.146552559 | 8.159661011 | -0.872901055 | 0.415348584 | 0.707388601 | -4.723708373 |
| **A_52_P495553** | Zbtb10 | -0.06841933 | -0.258889497 | 0.122050836 | 8.023951168 | -0.872840815 | 0.415378997 | 0.707388601 | -4.723732498 |
| **A_55_P2079579** | Pira7 | -0.209274317 | -0.792047975 | 0.37349934 | 9.655156029 | -0.872567856 | 0.415516826 | 0.707560984 | -4.7238418 |
| **A_66_P121787** | Samd9l | -0.074681673 | -0.282687035 | 0.133323689 | 9.592497797 | -0.872414141 | 0.415594459 | 0.707615524 | -4.723903343 |
| **A_55_P2039586** | Grk3 | -0.074812882 | -0.283253379 | 0.133627615 | 8.086862254 | -0.872122465 | 0.415741797 | 0.707615524 | -4.724020101 |
| **A_55_P2316335** | Gfm2 | 0.068685453 | -0.12269774 | 0.260068646 | 8.283878673 | 0.872055567 | 0.415775596 | 0.707615524 | -4.724046877 |
| **A_55_P1960053** | Hvcn1 | 0.142178484 | -0.253992762 | 0.53834973 | 9.431100588 | 0.872035401 | 0.415785785 | 0.707615524 | -4.724054948 |
| **A_55_P2383248** | 4930422N03Rik | 0.064233296 | -0.114755944 | 0.243222536 | 8.015459054 | 0.871999974 | 0.415803684 | 0.707615524 | -4.724069127 |
| **A_55_P2140390** | NA | 0.079578481 | -0.142189457 | 0.30134642 | 13.51877415 | 0.8719269 | 0.415840608 | 0.707615524 | -4.724098372 |
| **A_51_P409988** | Arl13b | -0.085702361 | -0.324550333 | 0.153145611 | 8.48146839 | -0.871875388 | 0.415866637 | 0.707615524 | -4.724118986 |
| **A_51_P475291** | Moap1 | 0.067602791 | -0.120806983 | 0.256012564 | 8.409359553 | 0.871855246 | 0.415876815 | 0.707615524 | -4.724127047 |
| **A_55_P2183750** | Ncoa3 | 0.077572323 | -0.138629752 | 0.293774399 | 11.00134048 | 0.871826625 | 0.415891278 | 0.707615524 | -4.7241385 |
| **A_52_P485891** | Dnah5 | -0.081100812 | -0.307149292 | 0.144947668 | 7.681683296 | -0.871779794 | 0.415914945 | 0.707615524 | -4.72415724 |
| **A_65_P12359** | Pin1rt1 | 0.144922174 | -0.259124451 | 0.548968798 | 11.31683256 | 0.871538437 | 0.416036932 | 0.707735809 | -4.724253811 |
| **A_51_P243207** | Zfp574 | 0.095712289 | -0.171149245 | 0.362573822 | 9.801148232 | 0.871495027 | 0.416058875 | 0.707735809 | -4.724271178 |
| **A_55_P2080308** | Gm12538 | 0.080060308 | -0.143236188 | 0.303356805 | 8.345233952 | 0.871201348 | 0.416207349 | 0.707879796 | -4.724388657 |
| **A_55_P1983368** | Sh3d21 | -0.174333668 | -0.660601915 | 0.311934579 | 9.510104014 | -0.871141182 | 0.416237771 | 0.707879796 | -4.724412721 |
| **A_51_P157255** | Sdc2 | -0.270844314 | -1.026371535 | 0.484682907 | 10.94024138 | -0.871069475 | 0.416274031 | 0.707879796 | -4.7244414 |
| **A_51_P450021** | Higd2a | -0.10486849 | -0.397424161 | 0.18768718 | 12.67500845 | -0.87100307 | 0.416307613 | 0.707879796 | -4.724467958 |
| **A_52_P399990** | B4galt2 | -0.166711669 | -0.631828715 | 0.298405378 | 8.520555884 | -0.870937381 | 0.416340834 | 0.707879796 | -4.724494227 |
| **A_55_P2143139** | Myadml2 | -0.05898541 | -0.223560071 | 0.105589251 | 7.96521064 | -0.870893045 | 0.416363257 | 0.707879796 | -4.724511957 |
| **A_51_P427828** | Mlip | -0.065514518 | -0.248340516 | 0.11731148 | 7.673361282 | -0.87072855 | 0.416446461 | 0.707947814 | -4.724577732 |
| **A_52_P441974** | Evc2 | -0.149205659 | -0.565610084 | 0.267198766 | 9.18880369 | -0.870669134 | 0.416476517 | 0.707947814 | -4.724601488 |
| **A_51_P420100** | Emc2 | 0.078835831 | -0.141231296 | 0.298902959 | 10.22197029 | 0.870465704 | 0.416579436 | 0.708033284 | -4.724682817 |
| **A_55_P1955931** | Slc25a42 | 0.173404895 | -0.31069681 | 0.657506599 | 10.57977319 | 0.870378049 | 0.416623788 | 0.708033284 | -4.724717856 |
| **A_55_P2062310** | NA | -0.129655352 | -0.491630229 | 0.232319525 | 14.76796637 | -0.870352553 | 0.416636689 | 0.708033284 | -4.724728047 |
| **A_51_P115178** | Scara3 | -0.099585835 | -0.377677786 | 0.178506115 | 7.928646883 | -0.870146425 | 0.416741003 | 0.70811169 | -4.720205375 |
| **A_51_P158876** | Dctpp1 | -0.08744936 | -0.331658756 | 0.156760036 | 13.84567869 | -0.870116603 | 0.416756096 | 0.70811169 | -4.724822353 |
| **A_55_P2084611** | B4galt6 | 0.108205026 | -0.194005919 | 0.410415971 | 8.266690572 | 0.870002367 | 0.416813917 | 0.708147684 | -4.724868006 |
| **A_55_P2349148** | 5830408B19Rik | -0.092548302 | -0.351062579 | 0.165965975 | 8.546419089 | -0.869895556 | 0.416867985 | 0.708177297 | -4.724910687 |
| **A_55_P2047788** | Katnal1 | 0.099852012 | -0.179138958 | 0.378842982 | 8.62959245 | 0.869660736 | 0.416986869 | 0.708264955 | -4.725004509 |
| **A_55_P2112120** | Tnk2 | 0.087749199 | -0.157429313 | 0.33292771 | 8.3507267 | 0.869648885 | 0.41699287 | 0.708264955 | -4.725009243 |
| **A_66_P134474** | Ang3 | 0.092590658 | -0.166177506 | 0.351358823 | 8.365872425 | 0.869439797 | 0.417098749 | 0.708321651 | -4.725092769 |
| **A_55_P2066996** | Cav2 | -0.109584841 | -0.415869285 | 0.196699604 | 7.76221214 | -0.869378167 | 0.417129962 | 0.708321651 | -4.725117386 |
| **A_55_P2109589** | Slc1a4 | 0.267068872 | -0.479390237 | 1.013527981 | 12.0927177 | 0.86936157 | 0.417138367 | 0.708321651 | -4.725124015 |
| **A_55_P1987775** | Pla2g4d | -0.092387661 | -0.350646199 | 0.165870877 | 7.723292482 | -0.869245542 | 0.417197135 | 0.708321651 | -4.725170358 |
| **A_51_P176071** | Cmtr2 | 0.0784003 | -0.140767641 | 0.29756824 | 9.293696633 | 0.86920834 | 0.417215979 | 0.708321651 | -4.725185215 |
| **A_52_P418665** | Ttpal | 0.076490616 | -0.137354473 | 0.290335705 | 10.47153934 | 0.869144663 | 0.417248235 | 0.708321651 | -4.725210646 |
| **A_55_P1952166** | Arid3a | 0.180919871 | -0.324918134 | 0.686757876 | 11.30910237 | 0.869076492 | 0.417282769 | 0.708321651 | -4.725237869 |
| **A_55_P2158547** | Cyp2c54 | -0.38614761 | -1.465933857 | 0.693638637 | 11.06392401 | -0.868957924 | 0.417342838 | 0.708361409 | -4.725285215 |
| **A_55_P1962279** | Plac1 | -0.218817217 | -0.830789038 | 0.393154603 | 8.038594178 | -0.868826893 | 0.41740923 | 0.70840397 | -4.725337533 |
| **A_52_P114494** | Stat6 | -0.064989989 | -0.246762393 | 0.116782414 | 9.105350811 | -0.868763775 | 0.417441213 | 0.70840397 | -4.725362733 |
| **A_52_P346113** | Foxn2 | 0.107246322 | -0.192758202 | 0.407250845 | 8.701082977 | 0.868635933 | 0.417506 | 0.708451714 | -4.72541377 |
| **A_66_P102706** | 2700012I20Rik | 0.061840099 | -0.111166495 | 0.234846692 | 8.477516488 | 0.868541767 | 0.417553725 | 0.708470501 | -4.72545136 |
| **A_55_P2031668** | Gstp1 | -0.138692802 | -0.526835275 | 0.249449671 | 12.07352839 | -0.868252303 | 0.417700457 | 0.708624523 | -4.725566893 |
| **A_55_P2007966** | Tmem215 | 0.20122614 | -0.361950633 | 0.764402913 | 12.15837021 | 0.868206136 | 0.417723863 | 0.708624523 | -4.725585317 |
| **A_52_P742248** | Fam117b | -0.198204528 | -0.752967004 | 0.356557947 | 10.41656699 | -0.868139834 | 0.417757479 | 0.708624523 | -4.725611775 |
| **A_55_P2054491** | Slc31a2 | -0.062100415 | -0.235929011 | 0.111728182 | 8.280847941 | -0.868073435 | 0.417791146 | 0.708624523 | -4.725638271 |
| **A_55_P1997866** | Mtr | 0.082881862 | -0.149282635 | 0.31504636 | 10.1587319 | 0.867454857 | 0.418104887 | 0.709073774 | -4.725885044 |
| **A_51_P133369** | Scly | 0.186949747 | -0.336753616 | 0.710653109 | 11.63248863 | 0.867406571 | 0.418129385 | 0.709073774 | -4.725904302 |

| | | | | | | | | | |
|---|---|---|---|---|---|---|---|---|---|
| **A_55_P2004208** | Defa-rs2 | 0.076260577 | -0.137438078 | 0.289959232 | 8.032455896 | 0.867124565 | 0.418272482 | 0.709254215 | -4.726016761 |
| **A_55_P2144512** | NA | 0.152424868 | -0.274744027 | 0.579593762 | 8.719173014 | 0.867040521 | 0.418315134 | 0.709264318 | -4.726050272 |
| **A_51_P466378** | Itgal | 0.140581645 | -0.2535076 | 0.534670889 | 8.817968511 | 0.866796655 | 0.418438916 | 0.709411965 | -4.726147496 |
| **A_55_P1975285** | Lrig2 | 0.107694938 | -0.194232233 | 0.409622109 | 9.182618497 | 0.866714937 | 0.418480401 | 0.709419739 | -4.726180071 |
| **A_52_P482983** | Cd164 | -0.122017053 | -0.464166254 | 0.220132147 | 13.5600258 | -0.866539158 | 0.418569647 | 0.709419739 | -4.726250135 |
| **A_55_P2022569** | Pak3 | -0.079645701 | -0.302990735 | 0.143699333 | 7.710015456 | -0.866501319 | 0.41858886 | 0.709419739 | -4.726265216 |
| **A_55_P2141955** | NA | -0.103044484 | -0.39202372 | 0.185934751 | 16.14354416 | -0.866445601 | 0.418617153 | 0.709419739 | -4.726287421 |
| **A_55_P2070825** | Nudt5 | 0.100283515 | -0.180959127 | 0.381526156 | 11.96142622 | 0.866426183 | 0.418627013 | 0.709419739 | -4.72629516 |
| **A_55_P2114034** | NA | 0.171760496 | -0.31002259 | 0.653543582 | 8.791480832 | 0.866273294 | 0.418704657 | 0.709467611 | -4.726356087 |
| **A_51_P322285** | Tecpr2 | 0.076685473 | -0.138426741 | 0.291797687 | 8.542931934 | 0.866226009 | 0.418728672 | 0.709467611 | -4.726374929 |
| **A_55_P1970785** | NA | 0.124881351 | -0.225481364 | 0.475244067 | 11.65366334 | 0.866089744 | 0.418797885 | 0.709468546 | -4.726429223 |
| **A_51_P297480** | Dph1 | 0.095046086 | -0.171633306 | 0.361725478 | 9.392107072 | 0.866020097 | 0.418833264 | 0.709468546 | -4.726456972 |
| **A_55_P2061724** | NA | -0.077415983 | -0.294649328 | 0.139817362 | 7.91493323 | -0.865939063 | 0.41887443 | 0.709468546 | -4.726489255 |
| **A_52_P2947** | Ndufaf2 | 0.082829728 | -0.149595724 | 0.315255179 | 11.29696449 | 0.865935888 | 0.418876043 | 0.709468546 | -4.72649052 |
| **A_55_P1977431** | Cck | -0.20183624 | -0.768331368 | 0.364658889 | 7.944562563 | -0.865737354 | 0.418976914 | 0.709536301 | -4.726569605 |
| **A_55_P2004071** | Gm2058 | -0.066918438 | -0.254770773 | 0.120933897 | 8.758041801 | -0.865590312 | 0.419051633 | 0.709536301 | -4.726628171 |
| **A_55_P2178064** | NA | 0.082373991 | -0.148882301 | 0.313630282 | 8.052321419 | 0.865525241 | 0.419084703 | 0.709536301 | -4.726654087 |
| **A_51_P404463** | Ecrg4 | -0.186969569 | -0.711878319 | 0.337939182 | 8.050495131 | -0.865506437 | 0.419094259 | 0.709536301 | -4.726661575 |
| **A_52_P266132** | Fgl2 | -0.071453744 | -0.272059384 | 0.129151896 | 8.075856771 | -0.865495951 | 0.419099589 | 0.709536301 | -4.726665751 |
| **A_51_P365578** | Ppp2r1a | -0.110762379 | -0.421825154 | 0.200300396 | 9.195939535 | -0.865221739 | 0.419238969 | 0.709637518 | -4.726774943 |
| **A_52_P134924** | Wdr20 | 0.100368326 | -0.181516076 | 0.382252728 | 9.084484684 | 0.865184689 | 0.419257804 | 0.709637518 | -4.726789695 |
| **A_51_P158388** | Cdk19 | 0.060991101 | -0.110306941 | 0.232289143 | 9.482774934 | 0.865161653 | 0.419269515 | 0.709637518 | -4.726798867 |
| **A_55_P2121231** | Nipbl | 0.072714004 | -0.131573104 | 0.277001112 | 8.427127188 | 0.864888832 | 0.419408229 | 0.709810145 | -4.726907476 |
| **A_52_P582112** | Hcfc1r1 | -0.172556693 | -0.657443305 | 0.312329919 | 10.89429759 | -0.864718607 | 0.419494797 | 0.709881386 | -4.726975232 |
| **A_55_P1970578** | Aamdc | -0.097942022 | -0.373192549 | 0.177308504 | 9.76010219 | -0.864617653 | 0.419546143 | 0.709881386 | -4.72701541 |
| **A_51_P186487** | NA | -0.056076772 | -0.213676796 | 0.101523252 | 7.687124591 | -0.864589421 | 0.419560503 | 0.709881386 | -4.727026646 |
| **A_55_P1967761** | Foxc1 | -0.066160592 | -0.25214542 | 0.119824235 | 7.723850242 | -0.864380696 | 0.419666679 | 0.709998884 | -4.727109705 |
| **A_52_P408657** | Mycbp | 0.089179793 | -0.161633099 | 0.339992685 | 8.337222041 | 0.863972249 | 0.41987451 | 0.710288326 | -4.727272202 |
| **A_55_P2007303** | Snrpg | 0.113595266 | -0.205936711 | 0.433127244 | 14.57944214 | 0.863831616 | 0.419946086 | 0.710291065 | -4.727328139 |
| **A_55_P2320085** | 4833415N18Rik | 0.162364142 | -0.29438269 | 0.619110974 | 8.832760078 | 0.863769304 | 0.419977803 | 0.710291065 | -4.727352922 |
| **A_51_P141554** | Med22 | 0.075165207 | -0.136286281 | 0.286616695 | 10.46190162 | 0.863752471 | 0.419986372 | 0.710291065 | -4.727359617 |
| **A_52_P394395** | Abcd3 | -0.109544107 | -0.417767563 | 0.198679348 | 12.32748907 | -0.863587853 | 0.420070173 | 0.710370636 | -4.727425084 |
| **A_51_P513661** | Slc4a8 | -0.062150615 | -0.237052112 | 0.112750882 | 7.720472381 | -0.863445818 | 0.420142487 | 0.71043077 | -4.727481562 |
| **A_55_P1961099** | Atg4c | 0.096463176 | -0.17505413 | 0.367980482 | 8.870888999 | 0.863271153 | 0.420231428 | 0.71047603 | -4.727551007 |
| **A_55_P2109943** | Rps15a | 0.114236861 | -0.207316774 | 0.435790496 | 15.81657994 | 0.863248886 | 0.420242767 | 0.71047603 | -4.72755986 |
| **A_55_P2144931** | NA | -0.1256116 | -0.479219849 | 0.227996648 | 14.76859667 | -0.8631585 | 0.420288799 | 0.710491708 | -4.727595792 |
| **A_51_P496720** | Dnmt3l | -0.059472394 | -0.226922494 | 0.107977705 | 7.771601063 | -0.863004782 | 0.420367092 | 0.710561917 | -4.727656896 |
| **A_55_P2008991** | Syp | 0.109427525 | -0.198719998 | 0.417575048 | 10.25114745 | 0.862881356 | 0.420429964 | 0.710606049 | -4.727705953 |
| **A_55_P1958678** | Brd2 | -0.069074124 | -0.26362305 | 0.125474802 | 12.37121133 | -0.862719743 | 0.4205123 | 0.710663155 | -4.727770181 |
| **A_55_P1997791** | Fbl | 0.113858283 | -0.206867665 | 0.434584231 | 12.48984515 | 0.862608478 | 0.420568992 | 0.710663155 | -4.727814395 |
| **A_55_P2092984** | NA | -0.147385049 | -0.562556896 | 0.267786798 | 14.21914727 | -0.862598549 | 0.420574051 | 0.710663155 | -4.727818341 |
| **A_51_P169880** | Zbtb3 | -0.066304187 | -0.253105466 | 0.120497093 | 8.216726206 | -0.862470591 | 0.420639256 | 0.710711205 | -4.727869183 |
| **A_51_P465232** | Mcfd2 | -0.19339364 | -0.738311213 | 0.351523933 | 13.81710059 | -0.862371832 | 0.420689587 | 0.710734116 | -4.72790842 |
| **A_55_P1976504** | NA | 0.409185327 | -0.743858002 | 1.562228656 | 12.77453615 | 0.862298466 | 0.420726979 | 0.710735167 | -4.727937567 |
| **A_55_P1988260** | NA | 0.2603327 | -0.473376076 | 0.994041475 | 14.25370452 | 0.86216065 | 0.420797227 | 0.710781239 | -4.727992314 |
| **A_51_P178735** | Hibch | 0.151581256 | -0.275657561 | 0.578820074 | 11.21426417 | 0.86210068 | 0.420827798 | 0.710781239 | -4.728016135 |
| **A_51_P495780** | Plin4 | -0.076927089 | -0.2937841 | 0.139929922 | 8.09611058 | -0.861963785 | 0.420897588 | 0.710837001 | -4.728070507 |
| **A_55_P2014978** | Dvl3 | 0.294289135 | -0.535398387 | 1.123976656 | 13.37352467 | 0.861872172 | 0.420944298 | 0.710853777 | -4.728106891 |
| **A_51_P156434** | Slc25a33 | -0.174517878 | -0.666637944 | 0.317602188 | 8.623438597 | -0.861691957 | 0.421036194 | 0.71094685 | -4.728178456 |
| **A_55_P2017600** | Crmp1 | -0.082941112 | -0.316910308 | 0.151028083 | 7.805447814 | -0.861379163 | 0.42119573 | 0.711154111 | -4.728302645 |

| | | | | | | | | | |
|---|---|---|---|---|---|---|---|---|---|
| **A_51_P165435** | Cox4i1 | -0.087307802 | -0.333678782 | 0.159063178 | 15.39894878 | -0.861086323 | 0.421345129 | 0.711303295 | -4.728418884 |
| **A_55_P1968774** | Ptk2b | 0.15667544 | -0.285454407 | 0.598805287 | 8.770208612 | 0.861061712 | 0.421357687 | 0.711303295 | -4.728428652 |
| **A_55_P1995447** | NA | -0.160219428 | -0.612465683 | 0.292026826 | 13.72630824 | -0.860841881 | 0.421469867 | 0.711430535 | -4.728515892 |
| **A_55_P2039250** | Mboat2 | -0.222793257 | -0.851761147 | 0.406174633 | 8.762658384 | -0.860709819 | 0.421537269 | 0.711482176 | -4.728568294 |
| **A_55_P2178207** | Abcc8 | -0.109961913 | -0.420456404 | 0.200532578 | 8.233357482 | -0.860541019 | 0.421623433 | 0.711502236 | -4.728635266 |
| **A_55_P1971828** | Csnk2a1 | 0.151291769 | -0.275949697 | 0.578533236 | 8.428515795 | 0.86044892 | 0.421670451 | 0.711502236 | -4.728671803 |
| **A_55_P2159475** | Sbf1 | 0.112516001 | -0.20525461 | 0.430286612 | 11.81564692 | 0.860367007 | 0.421712272 | 0.711502236 | -4.728704296 |
| **A_51_P221144** | Vwa8 | 0.180092509 | -0.328550686 | 0.688735703 | 10.1951426 | 0.860331052 | 0.42173063 | 0.711502236 | -4.728718559 |
| **A_51_P436817** | Cbarp | -0.185442773 | -0.709220843 | 0.338335297 | 8.425754393 | -0.860291814 | 0.421750664 | 0.711502236 | -4.728734122 |
| **A_55_P2061076** | Gm3219 | 0.232267933 | -0.423795431 | 0.888331298 | 9.449769134 | 0.860253916 | 0.421770015 | 0.711502236 | -4.728749154 |
| **A_52_P586004** | Hk3 | -0.16028894 | -0.613095 | 0.29251712 | 8.908583323 | -0.860150639 | 0.421822753 | 0.711529103 | -4.728790116 |
| **A_52_P467046** | Wdr48 | 0.082696151 | -0.150984997 | 0.3163773 | 9.340821057 | 0.859893782 | 0.421953936 | 0.711688275 | -4.728891977 |
| **A_55_P2033958** | Sltm | 0.106979138 | -0.195415868 | 0.409374144 | 11.15971627 | 0.859622292 | 0.422092626 | 0.711833374 | -4.728999619 |
| **A_55_P2078994** | Lrrc31 | 0.073677231 | -0.134612059 | 0.281966521 | 16.11642451 | 0.859507253 | 0.422151404 | 0.711833374 | -4.729045223 |
| **A_55_P2072661** | Prex1 | 0.119403616 | -0.218159178 | 0.456966411 | 10.8425669 | 0.859500425 | 0.422154892 | 0.711833374 | -4.72904793 |
| **A_51_P438083** | Slc6a13 | 0.174592768 | -0.319030206 | 0.668215742 | 11.99832983 | 0.859437055 | 0.422187273 | 0.711833374 | -4.729073049 |
| **A_55_P2053943** | Olfr1368 | 0.110542528 | -0.202067795 | 0.423152851 | 8.534321816 | 0.859229671 | 0.422293254 | 0.711949962 | -4.729155247 |
| **A_55_P2057803** | Stum | 0.196220493 | -0.35880966 | 0.751250646 | 8.558877123 | 0.859035227 | 0.422392641 | 0.712055412 | -4.729232303 |
| **A_55_P2157770** | Neu2 | -0.119823007 | -0.458849884 | 0.21920387 | 8.325716186 | -0.858794539 | 0.422515688 | 0.712200726 | -4.72932767 |
| **A_55_P2083426** | NA | 0.088570962 | -0.162094368 | 0.339236291 | 13.68469566 | 0.858579037 | 0.422625881 | 0.712324351 | -4.729413042 |
| **A_66_P114451** | Adgrg6 | -0.332298703 | -1.273059539 | 0.608462133 | 8.628589639 | -0.85828714 | 0.422775172 | 0.712513846 | -4.729528656 |
| **A_55_P2132651** | Ccn4 | -0.084805567 | -0.324952386 | 0.155341251 | 7.745630687 | -0.8580858 | 0.42287817 | 0.712625296 | -4.729608386 |
| **A_51_P315042** | Avpr1a | 0.141623835 | -0.259557253 | 0.542804924 | 8.458678289 | 0.857786265 | 0.423031435 | 0.712814249 | -4.729726979 |
| **A_55_P2025293** | Ss18 | -0.119539722 | -0.458234085 | 0.21915464 | 11.00868969 | -0.857605315 | 0.423124043 | 0.712814249 | -4.729798608 |
| **A_55_P2168664** | Atp5k | 0.080655585 | -0.147874086 | 0.309185257 | 14.89604046 | 0.857580784 | 0.423136599 | 0.712814249 | -4.729808318 |
| **A_55_P1978441** | Kif3a | -0.138094926 | -0.529393491 | 0.253203639 | 9.48004598 | -0.857536583 | 0.423159223 | 0.712814249 | -4.729825813 |
| **A_51_P431870** | Map1s | 0.06100206 | -0.111863354 | 0.233867474 | 10.07714804 | 0.857471278 | 0.423192651 | 0.712814249 | -4.72985166 |
| **A_51_P359983** | Arhgap17 | 0.087466995 | -0.160404467 | 0.335338458 | 9.896584687 | 0.85743433 | 0.423211564 | 0.712814249 | -4.729866283 |
| **A_51_P434758** | Fam83e | -0.127733101 | -0.48981845 | 0.234352247 | 7.938650741 | -0.857187234 | 0.423338068 | 0.712965192 | -4.729964066 |
| **A_55_P2032849** | NA | 0.209311791 | -0.384198783 | 0.802822366 | 9.277139471 | 0.856936068 | 0.423466683 | 0.713119666 | -4.730063441 |
| **A_51_P155345** | Chmp2a | -0.120198903 | -0.461093556 | 0.22069575 | 11.75193919 | -0.856768527 | 0.423552493 | 0.713175707 | -4.730129718 |
| **A_55_P2063426** | NA | 0.104178461 | -0.191295126 | 0.399652049 | 8.786576255 | 0.856727017 | 0.423573755 | 0.713175707 | -4.730146138 |
| **A_51_P350048** | Gstt2 | -0.091276023 | -0.350185556 | 0.16763351 | 10.78139049 | -0.856627196 | 0.423624889 | 0.713187238 | -4.73018562 |
| **A_52_P506995** | Plet1 | -0.056918495 | -0.218381838 | 0.104544849 | 7.672307222 | -0.856569592 | 0.423654399 | 0.713187238 | -4.730208404 |
| **A_52_P134141** | Pyurf | 0.130566485 | -0.239942234 | 0.501075204 | 9.507924367 | 0.856281319 | 0.423802099 | 0.713347037 | -4.730322403 |
| **A_55_P2001250** | Cpz | 0.099162554 | -0.182323109 | 0.380648217 | 8.816240315 | 0.856001673 | 0.423945416 | 0.713347037 | -4.730432966 |
| **A_51_P299216** | Wdr90 | 0.062581737 | -0.11506983 | 0.240233305 | 9.147884517 | 0.85597634 | 0.423958401 | 0.713347037 | -4.73044298 |
| **A_55_P2148478** | Klhl25 | -0.089278998 | -0.342766928 | 0.164208932 | 12.35537118 | -0.855805771 | 0.424045836 | 0.713347037 | -4.730510405 |
| **A_55_P2182433** | Atg10 | -0.075321666 | -0.289182654 | 0.138539323 | 8.059889907 | -0.855798526 | 0.42404955 | 0.713347037 | -4.730513269 |
| **A_55_P2066484** | Crtc3 | -0.074765012 | -0.28705607 | 0.137526047 | 8.601961886 | -0.855755881 | 0.424071413 | 0.713347037 | -4.730530124 |
| **A_55_P2119927** | NA | -0.078087151 | -0.299817969 | 0.143643667 | 8.578893592 | -0.855729903 | 0.424084731 | 0.713347037 | -4.730540392 |
| **A_51_P451458** | Mamdc2 | -0.068784594 | -0.264104608 | 0.12653542 | 7.857080231 | -0.855711995 | 0.424093912 | 0.713347037 | -4.73054747 |
| **A_51_P242201** | Naaa | -0.073706023 | -0.283008929 | 0.135596884 | 9.17650346 | -0.855679065 | 0.424110795 | 0.713347037 | -4.730560485 |
| **A_55_P2023912** | NA | 0.167468297 | -0.308099451 | 0.643036045 | 12.56901262 | 0.855664274 | 0.424118379 | 0.713347037 | -4.730566331 |
| **A_55_P2061421** | NA | 0.185630667 | -0.341606765 | 0.712868099 | 14.44914791 | 0.855513128 | 0.424195878 | 0.713415308 | -4.730626063 |
| **A_51_P339503** | Cct4 | -0.086060576 | -0.330521118 | 0.158399965 | 11.58844092 | -0.855418561 | 0.424244373 | 0.713434791 | -4.730663432 |
| **A_51_P260504** | Arhgef4 | -0.059950841 | -0.230295636 | 0.110393954 | 7.891076701 | -0.855164377 | 0.424374739 | 0.713487459 | -4.730763862 |
| **A_55_P2002379** | Golga2 | -0.068231143 | -0.262117749 | 0.125655462 | 11.64265859 | -0.855102215 | 0.424406625 | 0.713487459 | -4.730788419 |
| **A_52_P258338** | Mob3b | -0.111745564 | -0.429287061 | 0.205795934 | 8.338239138 | -0.85509229 | 0.424411717 | 0.713487459 | -4.73079234 |
| **A_52_P366932** | Gna11 | -0.061429827 | -0.236002838 | 0.113143184 | 10.35903228 | -0.855037929 | 0.424439603 | 0.713487459 | -4.730813814 |

| | | | | | | | | | |
|---|---|---|---|---|---|---|---|---|---|
| **A_51_P226142** | Dennd6a | -0.083055002 | -0.319094156 | 0.152984153 | 11.11699457 | -0.85499767 | 0.424460256 | 0.713487459 | -4.730829718 |
| **A_52_P491569** | 1700017B05Rik | 0.152568867 | -0.281137351 | 0.586275086 | 9.663291573 | 0.854778249 | 0.424572833 | 0.713530624 | -4.730916385 |
| **A_55_P2094706** | Sox17 | -0.2535973 | -0.97455221 | 0.46735761 | 8.401882179 | -0.854711821 | 0.42460692 | 0.713530624 | -4.730942619 |
| **A_51_P361951** | Uqcrfs1 | 0.078600834 | -0.144860512 | 0.302062181 | 13.82667914 | 0.854688645 | 0.424618812 | 0.713530624 | -4.730951772 |
| **A_55_P2039274** | Hspb3 | -0.075850863 | -0.291501337 | 0.139799612 | 7.701522834 | -0.854659836 | 0.424633596 | 0.713530624 | -4.730963149 |
| **A_55_P2119907** | Dnah11 | -0.139913412 | -0.537856471 | 0.258029647 | 7.901707072 | -0.854322024 | 0.424806977 | 0.713759914 | -4.731096536 |
| **A_55_P1954835** | NA | -0.062488394 | -0.240249072 | 0.115272283 | 7.848859215 | -0.854175 | 0.424882453 | 0.713777428 | -4.731154579 |
| **A_51_P239766** | Plcd1 | -0.186859111 | -0.718427725 | 0.344709502 | 8.986674444 | -0.854157853 | 0.424891256 | 0.713777428 | -4.731161348 |
| **A_55_P2136902** | Cisd3 | -0.114578422 | -0.440626679 | 0.211469834 | 12.58676408 | -0.853894359 | 0.425026549 | 0.713942658 | -4.731265352 |
| **A_55_P2017362** | Prr36 | -0.062312562 | -0.239769956 | 0.115144832 | 8.677889537 | -0.853227214 | 0.425369242 | 0.714356272 | -4.731528587 |
| **A_55_P2040893** | Tnni2 | -0.063327548 | -0.243678296 | 0.117023201 | 9.115426345 | -0.853213878 | 0.425376094 | 0.714356272 | -4.731533847 |
| **A_55_P2040971** | Tom1 | -0.083577308 | -0.321620977 | 0.15446636 | 11.41761246 | -0.85312945 | 0.425419477 | 0.714356272 | -4.73156715 |
| **A_51_P497768** | Gpat2 | 0.116298455 | -0.214967885 | 0.447564796 | 8.447680197 | 0.853060487 | 0.425454916 | 0.714356272 | -4.73159435 |
| **A_55_P2047275** | Tspan7 | -0.077191113 | -0.29706521 | 0.142682983 | 12.0395291 | -0.85305381 | 0.425458347 | 0.714356272 | -4.731596984 |
| **A_51_P338776** | Dnajc8 | 0.113099119 | -0.209083293 | 0.43528153 | 11.13591011 | 0.852983405 | 0.42549453 | 0.714356272 | -4.731624752 |
| **A_55_P2046184** | Ccdc90b | 0.081179874 | -0.150176002 | 0.312535749 | 9.967739702 | 0.852611186 | 0.425685857 | 0.714615419 | -4.731771529 |
| **A_51_P384946** | Ndufa2 | -0.072836734 | -0.28043767 | 0.134764203 | 14.91035839 | -0.852519541 | 0.425732974 | 0.714632449 | -4.731807661 |
| **A_55_P2020153** | Cycs | 0.085325715 | -0.157949243 | 0.328600673 | 13.78177794 | 0.852247467 | 0.425872876 | 0.714805212 | -4.731914913 |
| **A_51_P286496** | Il2rb | -0.069477421 | -0.267604114 | 0.128649273 | 7.996391075 | -0.852086892 | 0.425955461 | 0.714823425 | -4.731978201 |
| **A_51_P115326** | Surf4 | 0.091270766 | -0.169016435 | 0.351557966 | 13.73897495 | 0.852044092 | 0.425977475 | 0.714823425 | -4.731995069 |
| **A_55_P2037425** | Gapdh | -0.098078018 | -0.377826933 | 0.181670897 | 15.19386047 | -0.851895761 | 0.426053776 | 0.714823425 | -4.732053523 |
| **A_52_P677147** | Prmt5 | 0.089765728 | -0.166288092 | 0.345819549 | 10.62806948 | 0.851848746 | 0.426077962 | 0.714823425 | -4.732072049 |
| **A_52_P367791** | Mri1 | -0.069591134 | -0.268118732 | 0.128936463 | 11.14104517 | -0.851757994 | 0.426124651 | 0.714823425 | -4.732107807 |
| **A_52_P237077** | Esr1 | -0.085072449 | -0.327808918 | 0.157664019 | 7.920940884 | -0.851602832 | 0.426204487 | 0.714823425 | -4.720611056 |
| **A_55_P2083411** | Cyp4f18 | -0.080455367 | -0.310020518 | 0.149109784 | 9.492928343 | -0.851593324 | 0.426209379 | 0.714823425 | -4.732172684 |
| **A_55_P2087013** | Hipk1 | 0.095616232 | -0.177208936 | 0.368441401 | 8.388286286 | 0.851589668 | 0.42621126 | 0.714823425 | -4.732174124 |
| **A_52_P144818** | Ppp1r1a | -0.054333352 | -0.20937442 | 0.100707717 | 7.699254034 | -0.851535843 | 0.426238957 | 0.714823425 | -4.732195328 |
| **A_51_P115817** | Ralgps2 | -0.080327164 | -0.309566447 | 0.148912118 | 9.154761292 | -0.851444969 | 0.426285723 | 0.714823425 | -4.725296052 |
| **A_55_P1974243** | Dbh | -0.05628748 | -0.21692417 | 0.10434921 | 7.663773254 | -0.85143255 | 0.426292114 | 0.714823425 | -4.732236018 |
| **A_55_P2165839** | Cebpe | -0.260605887 | -1.004399159 | 0.483187384 | 12.31998452 | -0.851363774 | 0.42632751 | 0.714823425 | -4.732263108 |
| **A_55_P2046262** | Phgdh | 0.161897741 | -0.300242388 | 0.624037871 | 13.11903461 | 0.851236617 | 0.426392958 | 0.714871149 | -4.732313191 |
| **A_55_P2044207** | Bphl | -0.108213178 | -0.417170937 | 0.200744581 | 8.816589912 | -0.851067946 | 0.426479784 | 0.7149074 | -4.732379617 |
| **A_55_P2057984** | Tbc1d19 | -0.115014787 | -0.443398284 | 0.213368711 | 8.231667484 | -0.851050914 | 0.426488552 | 0.7149074 | -4.732386324 |
| **A_51_P504134** | NA | -0.077443729 | -0.298614931 | 0.143727474 | 7.676874649 | -0.850826221 | 0.426604239 | 0.714998658 | -4.732474797 |
| **A_55_P2185330** | NA | 0.242726864 | -0.450495865 | 0.935949593 | 10.21897399 | 0.850801486 | 0.426616976 | 0.714998658 | -4.732484536 |
| **A_55_P1967538** | Hunk | -0.301544367 | -1.162864193 | 0.559775459 | 9.873701742 | -0.85068733 | 0.426675761 | 0.715013015 | -4.732529478 |
| **A_55_P1965459** | Zfp521 | 0.075190895 | -0.139606918 | 0.289988708 | 7.811839604 | 0.850586702 | 0.426727584 | 0.715013015 | -4.732569091 |
| **A_55_P2003410** | Kng1 | -0.206554601 | -0.796631116 | 0.383521914 | 13.31316096 | -0.850569357 | 0.426736517 | 0.715013015 | -4.732575919 |
| **A_51_P306710** | Cldn15 | 0.115452048 | -0.214471174 | 0.44537527 | 8.990353156 | 0.85029954 | 0.426875499 | 0.715100543 | -4.732682117 |
| **A_52_P445307** | Cwc15 | 0.070078736 | -0.130185047 | 0.27034252 | 12.70561912 | 0.850289927 | 0.426880451 | 0.715100543 | -4.732685901 |
| **A_55_P2057370** | Slc22a19 | -0.129466047 | -0.499457056 | 0.240524963 | 7.732840489 | -0.850252476 | 0.426899745 | 0.715100543 | -4.732700639 |
| **A_55_P2143095** | NA | 0.144390895 | -0.268544829 | 0.557326619 | 8.959177464 | 0.849650937 | 0.427209726 | 0.715557782 | -4.732937311 |
| **A_55_P2113081** | Tmem132a | -0.398361066 | -1.537842669 | 0.741120537 | 10.67701658 | -0.849479214 | 0.427298248 | 0.715586996 | -4.733004854 |
| **A_51_P493016** | Cmya5 | -0.085196625 | -0.328911006 | 0.158517757 | 7.711212373 | -0.849423789 | 0.427326822 | 0.715586996 | -4.733026652 |
| **A_55_P1967983** | Use1 | -0.222225275 | -0.857941854 | 0.413491304 | 12.35796176 | -0.849401657 | 0.427338232 | 0.715586996 | -4.733035356 |
| **A_55_P1969987** | Fam204a | 0.113739004 | -0.2116698 | 0.439147809 | 8.895788898 | 0.849304278 | 0.42738844 | 0.715609075 | -4.733073651 |
| **A_55_P2123727** | Grb2 | 0.076128523 | -0.141751413 | 0.294008458 | 8.352982949 | 0.84901107 | 0.42753964 | 0.715800235 | -4.73318894 |
| **A_55_P2119441** | Vma21 | -0.078209983 | -0.302068504 | 0.145648538 | 10.44424624 | -0.848929746 | 0.427581584 | 0.715808457 | -4.733220911 |
| **A_55_P1984297** | Efcab14 | 0.070747603 | -0.131808505 | 0.273303711 | 9.843089526 | 0.848690964 | 0.427704756 | 0.715882519 | -4.733314774 |
| **A_55_P2108845** | NA | 0.119245564 | -0.222165788 | 0.460656916 | 8.152111632 | 0.848686831 | 0.427706889 | 0.715882519 | -4.733316398 |

| | | | | | | | | | |
|---|---|---|---|---|---|---|---|---|---|
| **A_51_P421664** | Tmem123 | -0.141399864 | -0.546268862 | 0.263469133 | 10.32289119 | -0.848628591 | 0.427736935 | 0.715882519 | -4.733339289 |
| **A_55_P2046728** | C920009B18Rik | 0.075111081 | -0.13998685 | 0.290209013 | 11.03767516 | 0.848498288 | 0.427804164 | 0.715910318 | -4.733390499 |
| **A_55_P2009022** | NA | 0.065729038 | -0.12251133 | 0.253969407 | 8.653046947 | 0.848452829 | 0.427827621 | 0.715910318 | -4.733408364 |
| **A_55_P2075996** | Cpne1 | -0.099415279 | -0.384278938 | 0.18544838 | 11.24024577 | -0.848006687 | 0.428057874 | 0.716233609 | -4.733583656 |
| **A_55_P2034998** | Tjp1 | -0.263461119 | -1.018505105 | 0.491582867 | 10.77408261 | -0.847866484 | 0.428130251 | 0.716292706 | -4.733638729 |
| **A_52_P286360** | Otc | 0.334129998 | -0.623543391 | 1.291803386 | 11.66379229 | 0.847776233 | 0.428176847 | 0.716308661 | -4.733674178 |
| **A_55_P2168667** | Rbpms2 | 0.195276205 | -0.364518725 | 0.755071134 | 11.52555431 | 0.847624603 | 0.428255139 | 0.716309451 | -4.733733729 |
| **A_55_P1957918** | Asap2 | -0.07403073 | -0.286273837 | 0.138212376 | 8.976515523 | -0.847542782 | 0.428297391 | 0.716309451 | -4.733765861 |
| **A_52_P456640** | Fgr | -0.121650608 | -0.470432425 | 0.227131209 | 8.71704457 | -0.847507669 | 0.428315525 | 0.716309451 | -4.733779649 |
| **A_55_P1990038** | Mark2 | 0.076373696 | -0.142600687 | 0.295348079 | 8.480483785 | 0.847488248 | 0.428325554 | 0.716309451 | -4.733787275 |
| **A_55_P2047225** | Uxs1 | 0.06112428 | -0.11419617 | 0.23644473 | 8.207750366 | 0.847157902 | 0.428496182 | 0.716386496 | -4.733916977 |
| **A_55_P2457912** | Psme4 | 0.07645665 | -0.142868752 | 0.295782052 | 8.685360516 | 0.847050927 | 0.428551446 | 0.716386496 | -4.73395897 |
| **A_55_P2099080** | Irak1 | 0.072603475 | -0.135684231 | 0.280891181 | 10.08570675 | 0.84698742 | 0.428584257 | 0.716386496 | -4.733983898 |
| **A_55_P2142668** | NA | 0.307631687 | -0.574960091 | 1.190223465 | 8.827750333 | 0.846943382 | 0.428607011 | 0.716386496 | -4.734001184 |
| **A_55_P2009042** | NA | 0.093026663 | -0.173911546 | 0.359964871 | 10.99114861 | 0.846798127 | 0.428682067 | 0.716386496 | -4.734058194 |
| **A_66_P128431** | Fam122b | 0.069432262 | -0.129813196 | 0.26867772 | 7.948032439 | 0.846751691 | 0.428706063 | 0.716386496 | -4.734076417 |
| **A_55_P1970583** | Ttc38 | 0.157814172 | -0.295055866 | 0.61068421 | 10.19045722 | 0.846750772 | 0.428706538 | 0.716386496 | -4.734076778 |
| **A_55_P1960748** | Arl13b | -0.077925899 | -0.301556637 | 0.145704838 | 8.5637918 | -0.846707713 | 0.42872879 | 0.716386496 | -4.734093676 |
| **A_52_P147666** | Slc30a7 | -0.08534997 | -0.330290666 | 0.159590727 | 8.570134218 | -0.8466923 | 0.428736755 | 0.716386496 | -4.734099724 |
| **A_51_P156158** | Phlpp1 | -0.075230233 | -0.291148754 | 0.140688288 | 8.156482918 | -0.846614492 | 0.428776968 | 0.716386496 | -4.734130257 |
| **A_55_P2134804** | Cinp | 0.10745697 | -0.200957261 | 0.4158712 | 9.276799701 | 0.846609952 | 0.428779315 | 0.716386496 | -4.734132039 |
| **A_52_P432153** | Yipf6 | -0.067559361 | -0.261516595 | 0.126397873 | 8.715528014 | -0.846374831 | 0.428900847 | 0.71641873 | -4.734224291 |
| **A_51_P166740** | Kcnj4 | 0.054621785 | -0.102194116 | 0.211437687 | 7.74855337 | 0.846367505 | 0.428904634 | 0.71641873 | -4.734227165 |
| **A_55_P2147191** | NA | -0.088359626 | -0.342082126 | 0.165362874 | 16.66551815 | -0.846209849 | 0.428986139 | 0.71641873 | -4.734289013 |
| **A_66_P121636** | Ablim3 | -0.672122277 | -2.602130838 | 1.257886285 | 8.625350848 | -0.846198484 | 0.428992015 | 0.71641873 | -4.734293472 |
| **A_55_P2006615** | Rbbp9 | -0.172281801 | -0.667061801 | 0.322498198 | 9.962373839 | -0.846078106 | 0.429054257 | 0.71641873 | -4.73434069 |
| **A_55_P2187225** | NA | -0.096458506 | -0.373503901 | 0.180586888 | 9.374904737 | -0.846004722 | 0.429092203 | 0.71641873 | -4.734369473 |
| **A_51_P275642** | NA | 0.071927277 | -0.134672276 | 0.278526829 | 8.010135803 | 0.845955328 | 0.429117746 | 0.71641873 | -4.734388845 |
| **A_55_P2073840** | Rpl7 | 0.061262299 | -0.114708729 | 0.237233326 | 16.11643275 | 0.845931717 | 0.429129956 | 0.71641873 | -4.734398105 |
| **A_55_P2163961** | Kcnj5 | -0.118910946 | -0.460474821 | 0.222652928 | 9.004711831 | -0.845927402 | 0.429132188 | 0.71641873 | -4.734399797 |
| **A_52_P636151** | Golph3 | 0.082992201 | -0.155481 | 0.321465402 | 11.16517124 | 0.84563099 | 0.429285497 | 0.716560828 | -4.734516032 |
| **A_51_P297105** | Ucp2 | 0.142744852 | -0.267434895 | 0.5529246 | 11.73765156 | 0.845608677 | 0.429297039 | 0.716560828 | -4.73452478 |
| **A_55_P2014041** | Rab14 | -0.091567104 | -0.354705663 | 0.171571455 | 12.75595965 | -0.845547823 | 0.429328519 | 0.716560828 | -4.73454864 |
| **A_51_P504354** | Nrarp | -0.073299987 | -0.28398432 | 0.137384346 | 7.904941529 | -0.845385602 | 0.429412446 | 0.716584144 | -4.734612237 |
| **A_55_P2042277** | Pbrm1 | 0.096439323 | -0.180756483 | 0.373635128 | 9.053635506 | 0.845377505 | 0.429416635 | 0.716584144 | -4.734615411 |
| **A_51_P104933** | Trub2 | 0.102539471 | -0.192243071 | 0.397322013 | 12.37604438 | 0.845225327 | 0.429495377 | 0.716653673 | -4.708842476 |
| **A_55_P2019068** | Flii | 0.058159675 | -0.109095083 | 0.225414432 | 8.288189194 | 0.844941574 | 0.429642228 | 0.716836826 | -4.734786271 |
| **A_55_P2090345** | BC085271 | 0.081909358 | -0.153710431 | 0.317529147 | 11.62367078 | 0.844704774 | 0.429764808 | 0.716979456 | -4.734879058 |
| **A_55_P2090414** | NA | -0.067776763 | -0.262882527 | 0.127329 | 8.70692997 | -0.844100029 | 0.430077971 | 0.717439984 | -4.73511594 |
| **A_55_P1977968** | Epb41 | 0.161894319 | -0.319648466 | 0.643437105 | 15.34863418 | 0.855340618 | 0.430149681 | 0.717497685 | -4.701356063 |
| **A_55_P2186480** | NA | 0.076170503 | -0.143182173 | 0.295523178 | 9.303895359 | 0.84377582 | 0.430245929 | 0.717568336 | -4.735242887 |
| **A_52_P665817** | Abhd13 | 0.069945577 | -0.131515201 | 0.271406355 | 8.202259749 | 0.843631795 | 0.430320557 | 0.717568336 | -4.735299271 |
| **A_51_P186856** | Krt5 | 0.061348348 | -0.115351813 | 0.238048508 | 7.822735948 | 0.843624369 | 0.430324405 | 0.717568336 | -4.735302178 |
| **A_55_P2443568** | Ube4a | -0.067152988 | -0.260601353 | 0.126295376 | 9.573096251 | -0.843496859 | 0.430390484 | 0.717568336 | -4.735352091 |
| **A_52_P325237** | Rgs6 | -0.056323214 | -0.218574873 | 0.105928446 | 7.659094414 | -0.843492971 | 0.430392499 | 0.717568336 | -4.735353612 |
| **A_55_P1980125** | Srsf3 | 0.079716284 | -0.14994146 | 0.309374028 | 12.06323857 | 0.843430186 | 0.430425039 | 0.717568336 | -4.735378187 |
| **A_52_P652193** | Sar1a | -0.080972167 | -0.314262371 | 0.152318037 | 12.71155428 | -0.843378352 | 0.430451904 | 0.717568336 | -4.735398474 |
| **A_55_P2047703** | Serac1 | 0.055382605 | -0.104248685 | 0.215013894 | 8.141617192 | 0.843021293 | 0.430637 | 0.717754669 | -4.735538201 |
| **A_55_P2116391** | Tesk2 | 0.154584218 | -0.290984032 | 0.600152469 | 9.250523328 | 0.843012678 | 0.430641467 | 0.717754669 | -4.735541572 |
| **A_55_P2112360** | Arhgap17 | 0.123085779 | -0.231719736 | 0.477891293 | 11.65820592 | 0.842947847 | 0.430675081 | 0.717754669 | -4.735566937 |

| | | | | | | | | | |
|---|---|---|---|---|---|---|---|---|---|
| **A_51_P162116** | Mblac1 | -0.083643188 | -0.324834918 | 0.157548543 | 8.629264782 | -0.842658003 | 0.430825386 | 0.717886729 | -4.735680325 |
| **A_55_P2157478** | NA | 0.081812975 | -0.154102929 | 0.317728878 | 8.400050767 | 0.842651801 | 0.430828602 | 0.717886729 | -4.73568275 |
| **A_55_P1985788** | Usp2 | -0.088274563 | -0.342849617 | 0.166300492 | 8.424900742 | -0.842564038 | 0.430874122 | 0.717900689 | -4.735717078 |
| **A_51_P392705** | Xpc | -0.06860952 | -0.26653117 | 0.129312129 | 9.233964062 | -0.842314489 | 0.431003572 | 0.71803975 | -4.735814674 |
| **A_55_P2168549** | Ap5b1 | 0.0859631 | -0.162035438 | 0.333961638 | 9.806316472 | 0.842259924 | 0.431031881 | 0.71803975 | -4.735836011 |
| **A_51_P179907** | Appl2 | -0.100578065 | -0.390808969 | 0.189652839 | 8.925355479 | -0.842059556 | 0.431135844 | 0.718096479 | -4.735914354 |
| **A_51_P454666** | Arl6ip4 | 0.10748779 | -0.20268513 | 0.417660709 | 10.51552625 | 0.842051092 | 0.431140237 | 0.718096479 | -4.735917663 |
| **A_66_P125734** | Tnip2 | -0.059309008 | -0.230507553 | 0.111889538 | 8.723570335 | -0.841790031 | 0.43127572 | 0.718198634 | -4.736019718 |
| **A_55_P2086687** | Nfib | -0.083987372 | -0.326435296 | 0.158460552 | 8.115859424 | -0.841741448 | 0.431300937 | 0.718198634 | -4.736038708 |
| **A_51_P122085** | Usp20 | -0.061885522 | -0.240536568 | 0.116765525 | 9.753251985 | -0.841718131 | 0.43131304 | 0.718198634 | -4.726698364 |
| **A_52_P661** | Plxna4 | -0.080175842 | -0.31168658 | 0.151334896 | 8.442382652 | -0.841502832 | 0.431424805 | 0.718312573 | -4.736131966 |
| **A_52_P493620** | Fgfr1op2 | 0.106970276 | -0.201933214 | 0.415873766 | 10.34934421 | 0.841440653 | 0.431457086 | 0.718312573 | -4.736156264 |
| **A_55_P2184925** | Tmem185b | 0.085448878 | -0.161345619 | 0.332243376 | 9.824475637 | 0.841306186 | 0.431526905 | 0.718312573 | -4.736208807 |
| **A_55_P2174067** | Cerk | -0.058011928 | -0.225582069 | 0.109558213 | 8.83617335 | -0.84120889 | 0.431577428 | 0.718312573 | -4.736246822 |
| **A_55_P2183389** | NA | -0.087292234 | -0.339455439 | 0.164870971 | 15.60321835 | -0.841157038 | 0.431604355 | 0.718312573 | -4.73626708 |
| **A_55_P2034350** | Terf2 | 0.080814575 | -0.152636504 | 0.314265654 | 9.018439897 | 0.841156875 | 0.43160444 | 0.718312573 | -4.736267144 |
| **A_51_P468329** | Psmb7 | 0.132750813 | -0.250916593 | 0.516418218 | 11.16325641 | 0.840747282 | 0.431817189 | 0.718604773 | -4.736427138 |
| **A_52_P248604** | Cdh5 | -0.206991753 | -0.805426056 | 0.391442549 | 11.27144678 | -0.840465234 | 0.431963733 | 0.718786759 | -4.73653728 |
| **A_51_P448032** | Cox6b1 | -0.074298427 | -0.289124658 | 0.140527805 | 12.18250071 | -0.840379593 | 0.432008236 | 0.718798933 | -4.736570718 |
| **A_55_P1953028** | Tnfsf12 | 0.240531188 | -0.455104194 | 0.936166569 | 9.791192143 | 0.840181127 | 0.432111383 | 0.71884661 | -4.7366482 |
| **A_52_P456279** | Cct8 | 0.095980346 | -0.181613373 | 0.373574064 | 12.84147589 | 0.840148128 | 0.432128535 | 0.71884661 | -4.736661082 |
| **A_51_P237834** | Tmcc1 | -0.137594177 | -0.535624688 | 0.260436333 | 8.824715169 | -0.839976009 | 0.432218006 | 0.71884661 | -4.736728266 |
| **A_55_P2166414** | Uqcr10 | -0.06964694 | -0.271122823 | 0.131828943 | 14.92461564 | -0.839966876 | 0.432222754 | 0.71884661 | -4.736731831 |
| **A_51_P193185** | Mb | -0.075137132 | -0.292510266 | 0.142236001 | 7.711155725 | -0.839908428 | 0.43225314 | 0.71884661 | -4.736754643 |
| **A_51_P183894** | Fbxo15 | -0.06107502 | -0.23777103 | 0.11562099 | 7.681772983 | -0.839885463 | 0.432265079 | 0.71884661 | -4.736763605 |
| **A_55_P2115318** | Fmr1nb | 0.097126268 | -0.183890215 | 0.37814275 | 8.048523879 | 0.83982364 | 0.432297222 | 0.71884661 | -4.736787733 |
| **A_55_P2109544** | Trnt1 | 0.089492548 | -0.169492786 | 0.348477882 | 8.695746728 | 0.839643427 | 0.432390927 | 0.718940579 | -4.736858058 |
| **A_65_P16059** | Tgfbr3 | -0.116483136 | -0.453696709 | 0.220730437 | 10.41908429 | -0.83934633 | 0.43254544 | 0.719040654 | -4.736973972 |
| **A_55_P2131481** | NA | 0.085209018 | -0.161474652 | 0.331892689 | 14.29037446 | 0.839321505 | 0.432558353 | 0.719040654 | -4.736983657 |
| **A_52_P500244** | Atrn | -0.1293043 | -0.503649394 | 0.245040794 | 10.20527149 | -0.839313118 | 0.432562716 | 0.719040654 | -4.736986928 |
| **A_51_P242356** | Fam114a2 | -0.077030665 | -0.300063139 | 0.14600181 | 11.42898681 | -0.839225618 | 0.432608231 | 0.719054476 | -4.73702106 |
| **A_55_P2131393** | Eef1akmt2 | 0.075107851 | -0.142418264 | 0.292633967 | 9.56203968 | 0.838990652 | 0.432730473 | 0.719195814 | -4.737112705 |
| **A_65_P16483** | Pdzk1 | 0.25802526 | -0.489455178 | 1.005505699 | 10.54325226 | 0.838775202 | 0.432842584 | 0.719244664 | -4.737196723 |
| **A_51_P295420** | Zc3h7a | -0.05794038 | -0.225801727 | 0.109920968 | 10.59841507 | -0.838713862 | 0.432874507 | 0.719244664 | -4.73722064 |
| **A_55_P2144511** | NA | 0.14389559 | -0.273020212 | 0.560811392 | 8.661113619 | 0.838653023 | 0.43290617 | 0.719244664 | -4.737244361 |
| **A_55_P2026014** | Mc5r | 0.105522986 | -0.200215752 | 0.411261724 | 8.415837733 | 0.838648146 | 0.432908708 | 0.719244664 | -4.737246262 |
| **A_51_P332799** | Clptm1 | 0.090429919 | -0.171612856 | 0.352472694 | 11.79400034 | 0.838538757 | 0.432965644 | 0.719277433 | -4.73728891 |
| **A_52_P557129** | Slc12a5 | -0.071034631 | -0.276925787 | 0.134856525 | 8.301701303 | -0.838331195 | 0.433073693 | 0.719344041 | -4.737369822 |
| **A_55_P2004096** | Asb8 | 0.063315503 | -0.120204779 | 0.246835785 | 8.160187616 | 0.838318754 | 0.43308017 | 0.719344041 | -4.737374671 |
| **A_55_P1994668** | Tti1 | 0.092443408 | -0.175627775 | 0.360514591 | 10.22776338 | 0.837932445 | 0.433281321 | 0.719616314 | -4.737525224 |
| **A_55_P2047009** | NA | 0.295869957 | -0.562663576 | 1.15440349 | 11.32143246 | 0.837388119 | 0.433564867 | 0.719969488 | -4.737737279 |
| **A_55_P2132631** | NA | 0.081351266 | -0.154752608 | 0.31745514 | 12.14737609 | 0.837229251 | 0.43364765 | 0.719969488 | -4.737799152 |
| **A_51_P167513** | Leng1 | 0.063473932 | -0.120749033 | 0.247696897 | 10.07916538 | 0.837210802 | 0.433657264 | 0.719969488 | -4.737806337 |
| **A_55_P1962134** | NA | 0.130685443 | -0.248631067 | 0.510001952 | 8.441942695 | 0.837160361 | 0.43368355 | 0.719969488 | -4.73782598 |
| **A_55_P1965303** | Dsc2 | 0.247876124 | -0.471617009 | 0.967369257 | 9.532076404 | 0.837126774 | 0.433701053 | 0.719969488 | -4.737839059 |
| **A_52_P257523** | 1110065P20Rik | -0.169567571 | -0.661778311 | 0.322643168 | 11.41414343 | -0.837095301 | 0.433717456 | 0.719969488 | -4.737851315 |
| **A_52_P365342** | Lrrn2 | -0.056742907 | -0.22146787 | 0.107982056 | 7.710904616 | -0.837019069 | 0.433757187 | 0.719973608 | -4.737880998 |
| **A_55_P2015177** | NA | -0.105068296 | -0.410183805 | 0.200047213 | 11.56332699 | -0.836740122 | 0.433902593 | 0.720153119 | -4.737989599 |
| **A_51_P123879** | Tecrl | -0.087683076 | -0.34239096 | 0.167024808 | 7.696294722 | -0.836481959 | 0.434037197 | 0.720314672 | -4.733226826 |
| **A_55_P1965945** | NA | 0.056470632 | -0.107593652 | 0.220534916 | 16.25419941 | 0.836357184 | 0.434102265 | 0.720328106 | -4.738138646 |

| | | | | | | | | | |
|---|---|---|---|---|---|---|---|---|---|
| **A_51_P431554** | Ten1 | -0.148512994 | -0.580023124 | 0.282997136 | 9.125187789 | -0.836289506 | 0.43413756 | 0.720328106 | -4.738164983 |
| **A_55_P2023747** | Mfsd10 | -0.061747152 | -0.241180841 | 0.117686538 | 10.726967 | -0.836172983 | 0.434198335 | 0.720328106 | -4.738210325 |
| **A_51_P329966** | D6Wsu163e | 0.075462759 | -0.143854426 | 0.294779943 | 9.368852377 | 0.836071086 | 0.434251485 | 0.720328106 | -4.738249971 |
| **A_55_P2147220** | Ak2 | 0.107152479 | -0.20430191 | 0.418606868 | 11.84008243 | 0.83597051 | 0.434303952 | 0.720328106 | -4.7382891 |
| **A_55_P1962466** | NA | 0.117424639 | -0.223891781 | 0.45874106 | 16.95963783 | 0.835959507 | 0.434309692 | 0.720328106 | -4.738293381 |
| **A_51_P416059** | Slc49a4 | 0.065122038 | -0.124177402 | 0.254421478 | 7.935129951 | 0.835914471 | 0.434333187 | 0.720328106 | -4.738310901 |
| **A_52_P288614** | Rars | 0.082692956 | -0.157688 | 0.323073912 | 8.443773248 | 0.835894845 | 0.434343426 | 0.720328106 | -4.738318536 |
| **A_51_P256945** | Opa3 | 0.073019303 | -0.139314264 | 0.28535287 | 8.493087635 | 0.835607284 | 0.43449347 | 0.720480635 | -4.738430387 |
| **A_55_P2019650** | NA | 0.119811466 | -0.228603964 | 0.468226896 | 9.785656466 | 0.835572605 | 0.434511568 | 0.720480635 | -4.738443875 |
| **A_55_P2045036** | NA | 0.084277256 | -0.160826081 | 0.329380593 | 8.999461259 | 0.835495956 | 0.434551569 | 0.720480635 | -4.738473683 |
| **A_55_P1965079** | Gm10499 | -0.102979297 | -0.40249635 | 0.196537756 | 7.696521069 | -0.835432868 | 0.434584496 | 0.720480635 | -4.738498217 |
| **A_52_P140789** | Hspa13 | -0.081542736 | -0.318770925 | 0.155685452 | 10.54398162 | -0.835222485 | 0.434694311 | 0.720600886 | -4.73858002 |
| **A_55_P2060303** | Mras | 0.115621088 | -0.220805835 | 0.452048011 | 8.756718408 | 0.835082719 | 0.434767276 | 0.720660035 | -4.738634358 |
| **A_55_P2000783** | Axin2 | -0.092760364 | -0.362723601 | 0.177202873 | 7.757875143 | -0.834912582 | 0.434856109 | 0.720745475 | -4.738700495 |
| **A_55_P2137203** | B4galt3 | -0.122955193 | -0.480863652 | 0.234953266 | 11.76523394 | -0.83475322 | 0.434939328 | 0.720788166 | -4.738762435 |
| **A_55_P2049071** | NA | -0.082541203 | -0.322818608 | 0.157736203 | 13.66614005 | -0.834720438 | 0.434956447 | 0.720788166 | -4.738775175 |
| **A_51_P300378** | Wwc2 | -0.284089841 | -1.111439135 | 0.543259453 | 10.793835 | -0.834353308 | 0.435148213 | 0.721044132 | -4.738917834 |
| **A_51_P243514** | Macc1 | -0.107825866 | -0.421936426 | 0.206284694 | 7.759719673 | -0.834110533 | 0.435275056 | 0.721192487 | -4.739012148 |
| **A_51_P396983** | Ssu2 | -0.126775576 | -0.496210655 | 0.242659504 | 8.023303466 | -0.833836013 | 0.435418518 | 0.721336662 | -4.739118771 |
| **A_52_P27271** | Dhdds | 0.072827003 | -0.139414815 | 0.28506882 | 9.618274233 | 0.833766936 | 0.435454622 | 0.721336662 | -4.739145597 |
| **A_52_P445622** | 2610002M06Rik | 0.096942751 | -0.185592865 | 0.379478368 | 8.790226017 | 0.833729807 | 0.435474029 | 0.721336662 | -4.739160015 |
| **A_55_P2129232** | Calm2 | -0.085482799 | -0.334715735 | 0.163750137 | 13.68648216 | -0.833405748 | 0.43564344 | 0.721555445 | -4.739285838 |
| **A_55_P2031949** | Bend6 | 0.055560327 | -0.106499187 | 0.217619841 | 7.759593749 | 0.83305457 | 0.435827081 | 0.721754737 | -4.739422152 |
| **A_55_P2099930** | Cers5 | -0.185719844 | -0.727445462 | 0.356005775 | 10.54445453 | -0.833032844 | 0.435838444 | 0.721754737 | -4.739430584 |
| **A_55_P2072556** | Ncoa6 | 0.096313296 | -0.184665547 | 0.37729214 | 11.07343763 | 0.832905664 | 0.435904965 | 0.721756013 | -4.73947994 |
| **A_51_P109421** | Eif2d | 0.068832486 | -0.131979546 | 0.269644519 | 9.534194912 | 0.832888591 | 0.435913896 | 0.721756013 | -4.737086432 |
| **A_55_P2135261** | Ccdc174 | 0.073910221 | -0.141781984 | 0.289602426 | 9.925105674 | 0.832632271 | 0.436047988 | 0.721893854 | -4.73958602 |
| **A_55_P1968606** | Synrg | 0.09459447 | -0.181485002 | 0.370673941 | 16.56114632 | 0.832558617 | 0.436086525 | 0.721893854 | -4.739614595 |
| **A_52_P401311** | NA | -0.11835409 | -0.463795372 | 0.227087191 | 11.04770262 | -0.832515301 | 0.43610919 | 0.721893854 | -4.739631399 |
| **A_55_P2029574** | Por | -0.19823114 | -0.776866668 | 0.380404388 | 12.83590121 | -0.83243431 | 0.43615157 | 0.721902184 | -4.739662817 |
| **A_51_P200776** | Grsf1 | 0.083949117 | -0.161131047 | 0.32902928 | 10.31226951 | 0.832321588 | 0.43621056 | 0.721938 | -4.739706541 |
| **A_55_P2069460** | NA | -0.098710922 | -0.386921991 | 0.189500147 | 15.84840413 | -0.832219222 | 0.436264135 | 0.721938363 | -4.739746244 |
| **A_51_P258620** | Rab4b | -0.108102714 | -0.423750921 | 0.207545493 | 8.417576996 | -0.832178442 | 0.436285479 | 0.721938363 | -4.739762059 |
| **A_66_P116998** | Tro | -0.121368951 | -0.475804964 | 0.233067061 | 9.166164339 | -0.832056748 | 0.436349178 | 0.72198196 | -4.739809253 |
| **A_55_P2108012** | Fam78b | -0.052852697 | -0.207249904 | 0.10154451 | 7.681273444 | -0.831784655 | 0.436491625 | 0.722118307 | -4.739914755 |
| **A_52_P637812** | Prrg4 | -0.054579777 | -0.214027632 | 0.104868077 | 7.840105956 | -0.831756626 | 0.436506301 | 0.722118307 | -4.739925622 |
| **A_55_P2082837** | Spin1 | -0.055097227 | -0.216099957 | 0.105905503 | 8.432740015 | -0.83153339 | 0.436623199 | 0.722249877 | -4.740012161 |
| **A_55_P2103101** | Snx9 | 0.110840483 | -0.213239373 | 0.434920339 | 8.885698076 | 0.831054604 | 0.436873992 | 0.722595374 | -4.740197711 |
| **A_55_P1978860** | Trp53bp2 | 0.111055501 | -0.213677526 | 0.435788528 | 9.158662533 | 0.830991927 | 0.436906831 | 0.722595374 | -4.740221996 |
| **A_66_P133202** | Nfyc | 0.109240704 | -0.210287242 | 0.428768651 | 12.27157606 | 0.830727969 | 0.437045147 | 0.722719475 | -4.740324254 |
| **A_66_P140976** | Rpl15 | 0.088126466 | -0.169653807 | 0.345906738 | 11.24902665 | 0.830691685 | 0.437064162 | 0.722719475 | -4.740338309 |
| **A_55_P2097178** | Idua | 0.084126987 | -0.161971227 | 0.3302252 | 9.040241349 | 0.830634681 | 0.437094038 | 0.722719475 | -4.740360388 |
| **A_52_P650453** | Spink4 | 0.06408216 | -0.123405092 | 0.251569412 | 7.925551967 | 0.830517121 | 0.437155656 | 0.722759531 | -4.740405921 |
| **A_51_P189082** | Akr1c6 | -0.511108858 | -2.006675873 | 0.984458157 | 10.82535429 | -0.83040646 | 0.437213663 | 0.722793611 | -4.740448777 |
| **A_55_P2052535** | Gm14393 | -0.068933081 | -0.270717314 | 0.132851152 | 8.514181178 | -0.830087076 | 0.437381112 | 0.723008548 | -4.740572445 |
| **A_55_P2178608** | NA | 0.070563434 | -0.1360365 | 0.277163368 | 8.919990884 | 0.829913284 | 0.437472249 | 0.723008548 | -4.740639725 |
| **A_55_P1975210** | Fubp3 | -0.106131879 | -0.416877852 | 0.204614095 | 8.967662491 | -0.829895731 | 0.437481454 | 0.723008548 | -4.740646519 |
| **A_55_P2073526** | H2-Ke6 | -0.085497713 | -0.335835251 | 0.164839825 | 12.01817572 | -0.829873145 | 0.437493299 | 0.723008548 | -4.740655262 |
| **A_66_P129893** | 1700047M11Rik | -0.106490875 | -0.418396265 | 0.205414514 | 8.68437921 | -0.829607571 | 0.437632597 | 0.723126981 | -4.740758051 |
| **A_51_P181002** | Vps50 | 0.059066503 | -0.113938602 | 0.232071608 | 8.753785488 | 0.829593867 | 0.437639786 | 0.723126981 | -4.740763354 |

| | | | | | | | | | |
|---|---|---|---|---|---|---|---|---|---|
| **A_52_P5213** | Chd1l | 0.113968769 | -0.219914777 | 0.447852315 | 9.35406133 | 0.829419096 | 0.437731475 | 0.723190724 | -4.740830985 |
| **A_55_P2092968** | Traf3 | -0.096888781 | -0.380748863 | 0.186971301 | 10.12378376 | -0.829377703 | 0.437753193 | 0.723190724 | -4.740847002 |
| **A_52_P310530** | Rbfox2 | 0.153111551 | -0.295567787 | 0.601790889 | 10.30633162 | 0.829191945 | 0.437850665 | 0.723252683 | -4.740918871 |
| **A_55_P2040863** | Cnn3 | -0.133797804 | -0.525893378 | 0.25829777 | 11.4322763 | -0.829163611 | 0.437865534 | 0.723252683 | -4.740929832 |
| **A_55_P2000533** | Polk | 0.053921295 | -0.104132352 | 0.211974943 | 8.188149723 | 0.828970321 | 0.437966977 | 0.723358429 | -4.741004602 |
| **A_55_P1983162** | Pstpip2 | 0.156028899 | -0.301471041 | 0.613528839 | 9.737978575 | 0.828699719 | 0.438109024 | 0.723379876 | -4.741109259 |
| **A_55_P1988259** | Gng5 | -0.071446916 | -0.280941311 | 0.138047479 | 15.22351855 | -0.828694139 | 0.438111953 | 0.723379876 | -4.741111417 |
| **A_51_P165293** | Ckmt2 | -0.056028426 | -0.220315328 | 0.108258475 | 7.674125106 | -0.828683468 | 0.438117555 | 0.723379876 | -4.741115543 |
| **A_55_P2145521** | Stk38l | 0.070197716 | -0.135642191 | 0.276037622 | 9.33107478 | 0.828660412 | 0.43812966 | 0.723379876 | -4.741124459 |
| **A_55_P2145227** | Slmap | 0.069033062 | -0.133417869 | 0.271483993 | 11.1577794 | 0.828553484 | 0.438185801 | 0.723381327 | -4.741165806 |
| **A_65_P02543** | Srp72 | 0.094601011 | -0.182903683 | 0.372105704 | 9.548634881 | 0.828339997 | 0.438297904 | 0.723381327 | -4.741248346 |
| **A_51_P423578** | Slfn2 | -0.107222911 | -0.421763159 | 0.207317336 | 8.865226872 | -0.828313156 | 0.438312 | 0.723381327 | -4.741258722 |
| **A_52_P517762** | Slc44a4 | -0.088746624 | -0.349086482 | 0.171593234 | 8.102418187 | -0.828312786 | 0.438312194 | 0.723381327 | -4.741258865 |
| **A_55_P1983314** | Crat | 0.05534481 | -0.107023615 | 0.217713236 | 7.767671005 | 0.82824441 | 0.438348104 | 0.723381327 | -4.741285298 |
| **A_51_P325856** | Mcemp1 | -0.146191012 | -0.575087332 | 0.282705308 | 10.58185548 | -0.828231114 | 0.438355087 | 0.723381327 | -4.741290438 |
| **A_55_P2012399** | NA | -0.072564835 | -0.285496326 | 0.140366657 | 11.12867712 | -0.828074676 | 0.438437255 | 0.72344911 | -4.741350906 |
| **A_51_P403814** | Slc4a5 | -0.23702191 | -0.932604539 | 0.45856072 | 7.966157572 | -0.827985925 | 0.438483875 | 0.72344911 | -4.741385208 |
| **A_55_P1973339** | Mapk1 | 0.132920729 | -0.257186012 | 0.52302747 | 10.5454734 | 0.827927767 | 0.438514427 | 0.72344911 | -4.741407684 |
| **A_51_P346445** | Hspb7 | -0.062615724 | -0.246398812 | 0.121167365 | 7.692792242 | -0.827867907 | 0.438545875 | 0.72344911 | -4.739012296 |
| **A_55_P2326091** | 9330175M20Rik | 0.156962747 | -0.303806098 | 0.617731591 | 11.17987928 | 0.827745204 | 0.438610343 | 0.723493712 | -4.741478233 |
| **A_55_P2014337** | Bach1 | -0.124741681 | -0.490990215 | 0.241506853 | 10.20587564 | -0.827596958 | 0.43868824 | 0.723510418 | -4.741535512 |
| **A_51_P170562** | Cpn2 | -0.240665824 | -0.947285634 | 0.465953986 | 10.68975442 | -0.827583459 | 0.438695333 | 0.723510418 | -4.741540727 |
| **A_51_P498890** | Degs1 | -0.067221793 | -0.26464303 | 0.130199444 | 12.59205451 | -0.827369317 | 0.438807875 | 0.723615465 | -4.741623453 |
| **A_55_P2060672** | Hoxa3 | -0.176749786 | -0.695870868 | 0.342371296 | 8.365483761 | -0.827319802 | 0.438833901 | 0.723615465 | -4.74164258 |
| **A_55_P1994648** | Eef1d | 0.062395905 | -0.120889842 | 0.245681651 | 15.48418146 | 0.827200107 | 0.438896818 | 0.723657477 | -4.741688812 |
| **A_55_P1961463** | Bnip3l | 0.086818721 | -0.168281061 | 0.341918504 | 13.75241297 | 0.826963777 | 0.439021063 | 0.723800591 | -4.74178008 |
| **A_52_P112651** | Lman2 | -0.099533394 | -0.392037034 | 0.192970245 | 9.601286189 | -0.826838574 | 0.439086896 | 0.723847387 | -4.741828424 |
| **A_51_P487628** | Pramef12 | -0.057694612 | -0.227263357 | 0.111874134 | 7.708858267 | -0.826747014 | 0.439135044 | 0.723865023 | -4.741863775 |
| **A_55_P2039771** | Itgav | -0.05555766 | -0.218921445 | 0.107806125 | 7.890546621 | -0.826363921 | 0.439336537 | 0.724061105 | -4.742011656 |
| **A_51_P371255** | Rwdd2b | 0.069082594 | -0.134057631 | 0.272222818 | 8.825354788 | 0.826334519 | 0.439352005 | 0.724061105 | -4.742023004 |
| **A_51_P223524** | Ric1 | -0.07179792 | -0.28293837 | 0.13934253 | 9.00873834 | -0.826273087 | 0.439384323 | 0.724061105 | -4.742046713 |
| **A_51_P440865** | Fam110b | -0.083022055 | -0.327180954 | 0.161136844 | 7.808069376 | -0.826235995 | 0.439403837 | 0.724061105 | -4.742061028 |
| **A_52_P178998** | Fam168b | -0.073724137 | -0.29056279 | 0.143114516 | 12.01750894 | -0.826144822 | 0.439451806 | 0.724078421 | -4.742096212 |
| **A_52_P251615** | Arhgef40 | -0.202784856 | -0.799357895 | 0.393788183 | 9.959798464 | -0.825952526 | 0.439552992 | 0.724183411 | -4.74217041 |
| **A_52_P376135** | Mknk2 | 0.063452149 | -0.123293324 | 0.250197622 | 13.43106066 | 0.825618553 | 0.439728767 | 0.724411263 | -4.742299247 |
| **A_52_P491073** | Nek8 | 0.137709739 | -0.267621529 | 0.543041008 | 9.79032856 | 0.825539271 | 0.439770502 | 0.724418275 | -4.742329827 |
| **A_51_P505696** | Inpp5k | -0.086596985 | -0.341539947 | 0.168345978 | 9.550525655 | -0.825359075 | 0.43986537 | 0.724452885 | -4.742399321 |
| **A_51_P352303** | Homer2 | -0.088020779 | -0.347162921 | 0.171121364 | 8.071822561 | -0.825335168 | 0.439877957 | 0.724452885 | -4.742408541 |
| **A_55_P2008021** | Srpx2 | -0.254847976 | -1.005213854 | 0.495517901 | 9.475995948 | -0.825260963 | 0.439917029 | 0.724452885 | -4.742437155 |
| **A_52_P460537** | Lpar2 | -0.064778588 | -0.255521214 | 0.125964037 | 7.76631102 | -0.825214617 | 0.439941433 | 0.724452885 | -4.742455026 |
| **A_51_P450296** | Thap3 | -0.133745798 | -0.527697553 | 0.260205957 | 8.489988067 | -0.824936072 | 0.440088126 | 0.72463271 | -4.742562416 |
| **A_55_P2184541** | Cdadc1 | 0.058553915 | -0.113981311 | 0.23108914 | 10.58655797 | 0.824634213 | 0.440247136 | 0.724832785 | -4.742678767 |
| **A_52_P408690** | Sft2d2 | -0.076309587 | -0.301239856 | 0.148620683 | 8.919858917 | -0.824355209 | 0.440394145 | 0.725013067 | -4.742786282 |
| **A_55_P1985758** | Psmd9 | 0.091969651 | -0.179170309 | 0.36310961 | 10.11471226 | 0.824203015 | 0.440474351 | 0.725062008 | -4.74284492 |
| **A_51_P160113** | Copb1 | -0.135992223 | -0.536939811 | 0.264955364 | 11.46193743 | -0.824156444 | 0.440498896 | 0.725062008 | -4.742862862 |
| **A_51_P259975** | Aspa | 0.084930916 | -0.165497488 | 0.33535932 | 8.423783527 | 0.824072477 | 0.440543154 | 0.725067163 | -4.742895209 |
| **A_55_P1955548** | Ezr | -0.137320576 | -0.542308834 | 0.267667683 | 10.52740373 | -0.823903545 | 0.440632203 | 0.725067163 | -4.742960279 |
| **A_66_P112065** | Sertad2 | 0.113279563 | -0.220814198 | 0.447373323 | 9.482510061 | 0.823884606 | 0.440642188 | 0.725067163 | -4.742967574 |
| **A_55_P2147876** | Abhd17a | -0.164802525 | -0.650937087 | 0.321332036 | 10.14726312 | -0.823740755 | 0.440718028 | 0.725067163 | -4.743022976 |
| **A_55_P2052210** | Gdi2 | -0.06521728 | -0.25761451 | 0.12717995 | 13.61634155 | -0.823658259 | 0.440761525 | 0.725067163 | -4.743054744 |

| | | | | | | | | | |
|---|---|---|---|---|---|---|---|---|---|
| **A_55_P1993744** | Myo10 | -0.058982355 | -0.232988907 | 0.115024198 | 8.10974941 | -0.823644295 | 0.440768888 | 0.725067163 | -4.743060122 |
| **A_55_P2118520** | Col1a1 | -0.340036908 | -1.343239672 | 0.663165856 | 12.46842807 | -0.823607944 | 0.440788055 | 0.725067163 | -4.743074119 |
| **A_55_P2042086** | Nrtn | -0.140152477 | -0.553654376 | 0.273349422 | 10.2177043 | -0.823581266 | 0.440802123 | 0.725067163 | -4.743084392 |
| **A_55_P1976682** | 4932431P20Rik | 0.098561322 | -0.192275981 | 0.389398625 | 12.45172668 | 0.823454486 | 0.44086898 | 0.72510096 | -4.743133207 |
| **A_55_P2121466** | Ncor1 | 0.064138503 | -0.125135569 | 0.253412575 | 8.106283902 | 0.823400036 | 0.440897697 | 0.72510096 | -4.743154171 |
| **A_55_P2027906** | Nsa2 | -0.153599156 | -0.607060831 | 0.29986252 | 12.68508763 | -0.823059872 | 0.441077126 | 0.725308053 | -4.743285115 |
| **A_52_P283501** | Zc3h7b | -0.095946857 | -0.379238336 | 0.187344622 | 9.322852525 | -0.822963228 | 0.441128113 | 0.725308053 | -4.743322311 |
| **A_51_P371675** | Slc7a6os | 0.069533158 | -0.135773387 | 0.274839703 | 9.419280251 | 0.822947914 | 0.441136193 | 0.725308053 | -4.743328204 |
| **A_51_P462692** | Ptbp1 | 0.054559435 | -0.106560915 | 0.215679784 | 12.82907901 | 0.822815865 | 0.441205867 | 0.725360908 | -4.743379021 |
| **A_52_P319606** | Gpr107 | -0.060258745 | -0.238255648 | 0.117738159 | 9.524570387 | -0.822604008 | 0.441317667 | 0.725483006 | -4.743460538 |
| **A_55_P1993629** | NA | -0.061083634 | -0.241558081 | 0.119390814 | 15.19004408 | -0.822417478 | 0.441416118 | 0.725583141 | -4.743532299 |
| **A_66_P138289** | Rdh5 | 0.229375032 | -0.448449669 | 0.907199733 | 11.13143249 | 0.822265257 | 0.441496473 | 0.725653515 | -4.743590852 |
| **A_55_P2190994** | Cdsn | 0.053349521 | -0.104330499 | 0.211029542 | 7.76653274 | 0.822123476 | 0.441571326 | 0.725714834 | -4.743645382 |
| **A_55_P1960924** | Egln2 | -0.078901843 | -0.312169932 | 0.154366247 | 13.08086177 | -0.821892473 | 0.441693304 | 0.725714864 | -4.743734214 |
| **A_51_P182303** | Col1a2 | -0.382264034 | -1.512467809 | 0.747939741 | 13.59676103 | -0.821844918 | 0.441718417 | 0.725714864 | -4.743752498 |
| **A_51_P327585** | Gstm4 | 0.105397551 | -0.206274612 | 0.417069714 | 8.719496653 | 0.821704553 | 0.44179255 | 0.725714864 | -4.743806465 |
| **A_55_P2138343** | Ndufaf5 | -0.105417799 | -0.417161225 | 0.206325627 | 8.79279753 | -0.821674539 | 0.441808403 | 0.725714864 | -4.743818003 |
| **A_55_P2373852** | 2310058N22Rik | 0.097069876 | -0.190020871 | 0.384160623 | 8.558183283 | 0.821577368 | 0.441859729 | 0.725714864 | -4.743855358 |
| **A_55_P1988113** | Cyth2 | -0.100489113 | -0.397701935 | 0.196723709 | 10.08629179 | -0.82155125 | 0.441873526 | 0.725714864 | -4.743865398 |
| **A_52_P220485** | Uhrf2 | 0.071618072 | -0.140242178 | 0.283478321 | 9.359667686 | 0.821403091 | 0.441951794 | 0.725714864 | -4.743922347 |
| **A_51_P515496** | Rbm17 | 0.058248424 | -0.114069302 | 0.23056615 | 11.47541613 | 0.821367309 | 0.441970698 | 0.725714864 | -4.7439361 |
| **A_66_P102954** | Gm2155 | 0.058931118 | -0.115408588 | 0.233270824 | 7.909123127 | 0.821356232 | 0.441976551 | 0.725714864 | -4.743940357 |
| **A_55_P2393734** | 4933421H12Rik | 0.162367675 | -0.318049598 | 0.642784949 | 9.299499372 | 0.821228755 | 0.442043905 | 0.725714864 | -4.743989348 |
| **A_51_P286321** | Trir | -0.060042375 | -0.237701999 | 0.117617249 | 13.01216272 | -0.821206375 | 0.44205573 | 0.725714864 | -4.743997949 |
| **A_55_P1980316** | Pcbp2 | 0.076573716 | -0.150002197 | 0.303149629 | 13.86060482 | 0.821200441 | 0.442058866 | 0.725714864 | -4.744000229 |
| **A_55_P2168189** | NA | 0.159744297 | -0.312927779 | 0.632416373 | 12.93706739 | 0.821199365 | 0.442059434 | 0.725714864 | -4.744000643 |
| **A_55_P2058933** | Kif1b | -0.124868093 | -0.494471419 | 0.244735233 | 10.80069596 | -0.820916147 | 0.442209105 | 0.725758431 | -4.744109466 |
| **A_55_P1967227** | Tmem158 | 0.074779436 | -0.146578076 | 0.296136949 | 10.43570093 | 0.820863812 | 0.442236766 | 0.725758431 | -4.744129572 |
| **A_55_P2182077** | NA | -0.119679622 | -0.473959742 | 0.234600497 | 15.07763427 | -0.820836417 | 0.442251247 | 0.725758431 | -4.744140097 |
| **A_55_P1982683** | Cib1 | -0.064446374 | -0.255223918 | 0.12633117 | 13.19115598 | -0.820832274 | 0.442253436 | 0.725758431 | -4.744141689 |
| **A_55_P2176731** | Prdx6 | -0.097383944 | -0.385694228 | 0.190926339 | 12.11832313 | -0.820749103 | 0.442297399 | 0.725758431 | -4.744173639 |
| **A_55_P1969874** | NA | 0.087228269 | -0.171032315 | 0.345488853 | 16.02959583 | 0.820695996 | 0.442325473 | 0.725758431 | -4.744194039 |
| **A_55_P1998159** | Deaf1 | 0.074353204 | -0.145845773 | 0.294552181 | 8.407525935 | 0.820479205 | 0.442440085 | 0.725758431 | -4.744277305 |
| **A_51_P369897** | Zfp410 | 0.093592854 | -0.183590623 | 0.370776332 | 9.661133701 | 0.820462136 | 0.442449111 | 0.725758431 | -4.744283861 |
| **A_55_P2063495** | Dph3 | 0.091590133 | -0.179668899 | 0.362849165 | 8.964719721 | 0.820441596 | 0.442459971 | 0.725758431 | -4.744291749 |
| **A_51_P481221** | Bace2 | 0.06467874 | -0.126894133 | 0.256251612 | 8.10382211 | 0.820371808 | 0.442496872 | 0.725758431 | -4.744318549 |
| **A_52_P549977** | Fam32a | 0.113036502 | -0.221778582 | 0.447851586 | 9.831561068 | 0.820345656 | 0.442510701 | 0.725758431 | -4.744328592 |
| **A_55_P2177513** | Chrna2 | 0.090193571 | -0.176976474 | 0.357363617 | 8.234227139 | 0.820296784 | 0.442536545 | 0.725758431 | -4.744347359 |
| **A_51_P327796** | Itgb5 | -0.090381556 | -0.358154196 | 0.177391085 | 12.17932268 | -0.820156631 | 0.442610664 | 0.725818403 | -4.744401173 |
| **A_51_P304527** | Cluap1 | -0.093444714 | -0.370336793 | 0.183447364 | 10.09009247 | -0.820025577 | 0.442679979 | 0.725823673 | -4.744451488 |
| **A_55_P2003115** | Rnf7 | 0.096556204 | -0.189576324 | 0.382688733 | 11.96718106 | 0.819966562 | 0.442711195 | 0.725823673 | -4.744474144 |
| **A_51_P125842** | Tmem51 | 0.087005405 | -0.170833611 | 0.34484442 | 10.99754882 | 0.81993757 | 0.442726531 | 0.725823673 | -4.744485273 |
| **A_55_P1998471** | S100a9 | 0.092976884 | -0.182657375 | 0.368611143 | 17.23180378 | 0.819643467 | 0.442882123 | 0.726017179 | -4.744598158 |
| **A_55_P2183498** | Spcs2 | 0.314166252 | -0.617311625 | 1.24564413 | 10.74570265 | 0.819540038 | 0.442936851 | 0.726041022 | -4.74463785 |
| **A_52_P504421** | Rnf138 | 0.099225552 | -0.194993999 | 0.393445102 | 9.316713361 | 0.819474008 | 0.442971792 | 0.726041022 | -4.744663188 |
| **A_55_P2110998** | Gpr153 | -0.207712706 | -0.823672932 | 0.408247519 | 8.600986064 | -0.819395519 | 0.443013329 | 0.726047535 | -4.744693305 |
| **A_55_P2003494** | NA | 0.119875478 | -0.235777739 | 0.475528695 | 14.2821547 | 0.819005467 | 0.443219787 | 0.726191674 | -4.744842943 |
| **A_55_P2080048** | NA | -0.114214875 | -0.45307747 | 0.22464772 | 15.30737661 | -0.818996839 | 0.443224355 | 0.726191674 | -4.744846252 |
| **A_52_P516021** | Ptpn1 | -0.064786539 | -0.257009708 | 0.12743663 | 9.046275706 | -0.818959146 | 0.44324431 | 0.726191674 | -4.74486071 |
| **A_66_P101755** | Sp1 | 0.072138245 | -0.141906204 | 0.286182693 | 10.95855841 | 0.81892625 | 0.443261726 | 0.726191674 | -4.744873327 |

| | | | | | | | | | |
|---|---|---|---|---|---|---|---|---|---|
| **A_51_P183630** | Bcor | -0.071959153 | -0.285502581 | 0.141584276 | 10.01996148 | -0.818809773 | 0.443323396 | 0.726191674 | -4.744917999 |
| **A_55_P2126552** | Zdhhc13 | 0.06954549 | -0.136840408 | 0.275931388 | 9.375168897 | 0.818789266 | 0.443334254 | 0.726191674 | -4.744925864 |
| **A_55_P2090914** | Tpst2 | -0.069720688 | -0.276642493 | 0.137201118 | 11.37321339 | -0.818726018 | 0.443367745 | 0.726191674 | -4.744950119 |
| **A_55_P2042273** | Pbrm1 | 0.111542082 | -0.21952614 | 0.442610304 | 9.0378305 | 0.818661634 | 0.443401838 | 0.726191674 | -4.744974808 |
| **A_55_P1989328** | Rps12 | 0.084638712 | -0.166638929 | 0.335916353 | 17.17890191 | 0.818461754 | 0.443507695 | 0.726303503 | -4.745051448 |
| **A_55_P2185722** | Fermt2 | -0.094070716 | -0.373403371 | 0.185261939 | 9.215103239 | -0.818306364 | 0.443590002 | 0.726343495 | -4.745111019 |
| **A_52_P655948** | Mosmo | 0.091151395 | -0.179541455 | 0.361844245 | 9.488771589 | 0.818219308 | 0.443636119 | 0.726343495 | -4.74514439 |
| **A_55_P2001683** | Atp5g1 | 0.095304559 | -0.187727669 | 0.378336788 | 15.06378556 | 0.818202827 | 0.44364485 | 0.726343495 | -4.745150708 |
| **A_55_P2134581** | Papola | 0.150041158 | -0.295629496 | 0.595711812 | 9.669947069 | 0.818049452 | 0.443726108 | 0.726370372 | -4.745209494 |
| **A_55_P1989816** | Zbtb7b | 0.220335174 | -0.434146969 | 0.874817318 | 13.23841354 | 0.81802998 | 0.443736425 | 0.726370372 | -4.745216957 |
| **A_55_P1993851** | Gm10021 | -0.066695496 | -0.264840127 | 0.131449135 | 7.783224093 | -0.817894684 | 0.443808114 | 0.726391449 | -4.745268806 |
| **A_51_P503131** | Wnt11 | -0.09747421 | -0.387069777 | 0.192121356 | 7.844900367 | -0.817863836 | 0.443824461 | 0.726391449 | -4.745280627 |
| **A_55_P2160094** | Plscr2 | 0.059397935 | -0.117108859 | 0.235904728 | 8.139194034 | 0.817698324 | 0.443912175 | 0.726473494 | -4.745344047 |
| **A_55_P2184009** | Rnd2 | 0.073352039 | -0.144666375 | 0.291370453 | 8.523859734 | 0.817527188 | 0.444002882 | 0.726502426 | -4.745409612 |
| **A_51_P282268** | Snapc1 | 0.056092315 | -0.110646872 | 0.222831501 | 8.561851218 | 0.817426797 | 0.444056099 | 0.726502426 | -4.745448069 |
| **A_52_P504743** | Cdyl | 0.102532906 | -0.202276603 | 0.407342414 | 8.889554081 | 0.817368587 | 0.444086958 | 0.726502426 | -4.745470366 |
| **A_51_P203474** | Tsg101 | 0.06564262 | -0.129518986 | 0.260804226 | 10.21642202 | 0.817287227 | 0.444130092 | 0.726502426 | -4.745501529 |
| **A_55_P2008422** | Mpc1 | -0.153100034 | -0.608310984 | 0.302110916 | 14.20101488 | -0.817232777 | 0.444158961 | 0.726502426 | -4.745522384 |
| **A_51_P275496** | BC026762 | 0.115480473 | -0.227900034 | 0.458860979 | 9.05364504 | 0.817176945 | 0.444188564 | 0.726502426 | -4.745543767 |
| **A_51_P447258** | Alox12b | 0.06559078 | -0.129444997 | 0.260626558 | 7.949199228 | 0.817168659 | 0.444192957 | 0.726502426 | -4.74554694 |
| **A_55_P2051991** | Brsk1 | -0.065442876 | -0.260071094 | 0.129185342 | 7.925566793 | -0.817033306 | 0.44426473 | 0.726558335 | -4.745598773 |
| **A_51_P190254** | NA | -0.073365916 | -0.291589081 | 0.144857249 | 7.787865746 | -0.816914648 | 0.444327657 | 0.72659977 | -4.745644208 |
| **A_52_P632742** | Shisa5 | 0.055249524 | -0.109102991 | 0.219602038 | 8.156623478 | 0.816836949 | 0.444368866 | 0.726605686 | -4.745673958 |
| **A_55_P1984976** | Wnt5b | -0.216064388 | -0.859147523 | 0.427018748 | 8.520489312 | -0.816392976 | 0.444604387 | 0.726876918 | -4.745843906 |
| **A_51_P262171** | Irgm1 | -0.12359439 | -0.491459219 | 0.24427044 | 10.08088695 | -0.816382483 | 0.444609954 | 0.726876918 | -4.745847922 |
| **A_51_P510663** | Supt3 | -0.059312271 | -0.235967859 | 0.117343317 | 8.493876324 | -0.815831299 | 0.444902476 | 0.727254897 | -4.746058819 |
| **A_55_P2087872** | NA | -0.105044331 | -0.417946504 | 0.207857843 | 14.14279757 | -0.815731487 | 0.444955463 | 0.727254897 | -4.746096999 |
| **A_52_P168271** | Spg20 | -0.056779649 | -0.22592939 | 0.112370092 | 7.962439739 | -0.815651343 | 0.444998012 | 0.727254897 | -4.746127653 |
| **A_55_P2077964** | Vps8 | -0.058549179 | -0.232976817 | 0.11587846 | 8.055549503 | -0.815621556 | 0.445013826 | 0.727254897 | -4.746139046 |
| **A_55_P2038299** | Rpl31 | -0.092480704 | -0.368005771 | 0.183044363 | 15.18542113 | -0.815592453 | 0.445029279 | 0.727254897 | -4.746150177 |
| **A_55_P2102155** | Rgs20 | -0.084414883 | -0.335963955 | 0.16713419 | 11.02199516 | -0.81541649 | 0.445122713 | 0.727346092 | -4.74621747 |
| **A_55_P2006236** | Ugdh | -0.132062252 | -0.525671881 | 0.261547376 | 11.26069751 | -0.815260075 | 0.445205779 | 0.727350152 | -4.746277279 |
| **A_52_P8685** | Tmem183a | 0.124461998 | -0.246495622 | 0.495419619 | 9.711697554 | 0.815259111 | 0.445206291 | 0.727350152 | -4.746277647 |
| **A_52_P299363** | NA | 0.068022194 | -0.134758977 | 0.270803365 | 8.641121153 | 0.81509119 | 0.445295481 | 0.727350152 | -4.746341847 |
| **A_55_P2001553** | Igdcc4 | 0.211381367 | -0.418807825 | 0.841570558 | 10.76123565 | 0.815040031 | 0.445322656 | 0.727350152 | -4.746361404 |
| **A_55_P2144766** | Pigc | 0.058667608 | -0.116246529 | 0.233581744 | 8.34892497 | 0.814998214 | 0.44534487 | 0.727350152 | -4.74637739 |
| **A_55_P2028660** | Dusp12 | 0.085070207 | -0.168583688 | 0.338724101 | 8.611843554 | 0.814927816 | 0.445382267 | 0.727350152 | -4.746404299 |
| **A_52_P366572** | Slain2 | -0.084837536 | -0.337805313 | 0.168130241 | 8.455536074 | -0.814903213 | 0.445395338 | 0.727350152 | -4.746413704 |
| **A_55_P1961461** | Bnip3l | 0.086356822 | -0.171159527 | 0.34387317 | 13.53351132 | 0.814845051 | 0.445426238 | 0.727350152 | -4.746435934 |
| **A_55_P2012406** | NA | -0.104202147 | -0.415225645 | 0.206821352 | 15.10893176 | -0.81407919 | 0.445833265 | 0.727953301 | -4.746728559 |
| **A_55_P2043837** | Cmtr1 | 0.059769647 | -0.118649875 | 0.238189168 | 12.61127458 | 0.813994568 | 0.445878254 | 0.727965265 | -4.74676088 |
| **A_55_P2169135** | NA | -0.089849487 | -0.358095794 | 0.17839682 | 13.52916925 | -0.813888736 | 0.445934525 | 0.727972981 | -4.746801299 |
| **A_55_P2082444** | Npepps | -0.086511034 | -0.344813809 | 0.171791742 | 10.86443897 | -0.813814865 | 0.445973805 | 0.727972981 | -4.74682951 |
| **A_55_P2150962** | Cpne7 | 0.128761809 | -0.255712107 | 0.513235724 | 8.803137013 | 0.813773189 | 0.445995967 | 0.727972981 | -4.746845424 |
| **A_55_P2092983** | NA | -0.093366629 | -0.372220801 | 0.185487542 | 14.66878129 | -0.813575241 | 0.44610124 | 0.728037552 | -4.746921006 |
| **A_52_P209514** | Cacnb2 | 0.066314132 | -0.131747889 | 0.264376153 | 7.776667164 | 0.813557158 | 0.446110857 | 0.728037552 | -4.74692791 |
| **A_52_P536494** | Mycn | 0.074587236 | -0.148209323 | 0.297383794 | 8.661463983 | 0.813465658 | 0.446159526 | 0.728055507 | -4.746962843 |
| **A_52_P453785** | Cthrc1 | -0.137032936 | -0.546462148 | 0.272396275 | 7.942507133 | -0.813259843 | 0.446269011 | 0.728146532 | -4.747041407 |
| **A_55_P2012955** | NA | 0.087807564 | -0.17455841 | 0.350173538 | 16.22538611 | 0.813219172 | 0.44629065 | 0.728146532 | -4.747056931 |
| **A_52_P79187** | Ankrd12 | -0.073218745 | -0.292026504 | 0.145589015 | 8.749080392 | -0.813097733 | 0.446355262 | 0.728183573 | -4.74710328 |

| | | | | | | | | | |
|---|---|---|---|---|---|---|---|---|---|
| **A_55_P2123002** | Tet1 | -0.062316498 | -0.248582835 | 0.123949838 | 7.798462634 | -0.812927579 | 0.446445804 | 0.728183573 | -4.747168212 |
| **A_55_P1991440** | Zscan21 | -0.104848917 | -0.418285509 | 0.208587676 | 11.45799199 | -0.812825723 | 0.44650001 | 0.728183573 | -4.747207077 |
| **A_55_P2056542** | Nin | 0.08096603 | -0.16109076 | 0.323022819 | 9.122463635 | 0.812772056 | 0.446528572 | 0.728183573 | -4.747227553 |
| **A_55_P1952269** | Xylt2 | -0.10656617 | -0.425189117 | 0.212056776 | 9.989005665 | -0.812691091 | 0.446571666 | 0.728183573 | -4.747258443 |
| **A_52_P575078** | Babam1 | 0.060708301 | -0.120813339 | 0.242229941 | 12.27809063 | 0.81264873 | 0.446594214 | 0.728183573 | -4.747274603 |
| **A_51_P485458** | Txlna | 0.096233355 | -0.191533958 | 0.384000668 | 8.454233036 | 0.812582292 | 0.446629579 | 0.728183573 | -4.747299948 |
| **A_55_P2104889** | Gm11515 | -0.149768221 | -0.597662961 | 0.298126519 | 8.273991195 | -0.812506602 | 0.446669871 | 0.728183573 | -4.74732882 |
| **A_55_P2013043** | Serpinb6b | -0.515989363 | -2.059162665 | 1.02718394 | 9.846032041 | -0.812473559 | 0.446687462 | 0.728183573 | -4.747341424 |
| **A_51_P453519** | Zdhhc16 | 0.099599677 | -0.198287148 | 0.397486502 | 12.12017818 | 0.812437243 | 0.446706796 | 0.728183573 | -4.747355276 |
| **A_55_P2110978** | Atp6ap1 | -0.077812135 | -0.310547157 | 0.154922888 | 8.627779692 | -0.812397871 | 0.446727757 | 0.728183573 | -4.747370293 |
| **A_66_P114784** | Pla2g7 | -0.147412783 | -0.588404087 | 0.29357852 | 8.378787149 | -0.812247363 | 0.446807892 | 0.728252783 | -4.747427695 |
| **A_55_P1962831** | Ssr1 | 0.113357389 | -0.225810965 | 0.452525743 | 13.1649848 | 0.81211531 | 0.44687821 | 0.728276124 | -4.747478051 |
| **A_55_P2058107** | Pigyl | -0.156590631 | -0.625134981 | 0.311953719 | 11.84863964 | -0.812078959 | 0.446897569 | 0.728276124 | -4.747491912 |
| **A_55_P2005754** | Manbal | -0.127391923 | -0.508603359 | 0.253819513 | 10.22755905 | -0.812005857 | 0.4469365 | 0.728278167 | -4.747519786 |
| **A_55_P2015878** | Clps | 0.334161936 | -0.666078858 | 1.334402731 | 8.912880821 | 0.811774874 | 0.447059529 | 0.728415132 | -4.747607846 |
| **A_52_P630016** | Bet1l | -0.108334243 | -0.432640235 | 0.215971748 | 10.12525149 | -0.811697055 | 0.447100983 | 0.728415132 | -4.74763751 |
| **A_55_P2092989** | NA | 0.097294933 | -0.193991429 | 0.388581296 | 8.956885373 | 0.811620975 | 0.447141514 | 0.728415132 | -4.747666509 |
| **A_55_P2104542** | NA | 0.063987845 | -0.12759531 | 0.255570999 | 8.155614159 | 0.811565076 | 0.447171294 | 0.728415132 | -4.747687815 |
| **A_51_P405227** | Nradd | -0.102569323 | -0.409710159 | 0.204571514 | 7.937908825 | -0.811452522 | 0.447231264 | 0.728451428 | -4.747730711 |
| **A_55_P2171746** | NA | -0.171339285 | -0.684506308 | 0.341827738 | 11.16189087 | -0.811299858 | 0.447312613 | 0.728490984 | -4.747788887 |
| **A_51_P240594** | Zfp7 | -0.059047041 | -0.235951654 | 0.117857573 | 9.098254822 | -0.811039802 | 0.447451212 | 0.728490984 | -4.747887969 |
| **A_55_P2098210** | 4931406P16Rik | -0.142871847 | -0.570934443 | 0.285190749 | 10.94335708 | -0.811003199 | 0.447470722 | 0.728490984 | -4.747901914 |
| **A_55_P2160563** | Mrps17 | 0.060233585 | -0.120235924 | 0.240703094 | 11.74704599 | 0.81099479 | 0.447475204 | 0.728490984 | -4.747905117 |
| **A_55_P1969882** | Morf4l2 | 0.090952978 | -0.18156099 | 0.363466947 | 14.22964334 | 0.810982245 | 0.447481891 | 0.728490984 | -4.747909896 |
| **A_55_P2134852** | Cnot2 | 0.07054707 | -0.140827791 | 0.28192193 | 10.52603345 | 0.810977554 | 0.447484392 | 0.728490984 | -4.747911683 |
| **A_55_P1958597** | Slc27a3 | -0.266246409 | -1.064044987 | 0.531552168 | 9.232867615 | -0.810911932 | 0.447519372 | 0.728490984 | -4.74793668 |
| **A_55_P1971815** | Clcn2 | 0.089823673 | -0.179412693 | 0.359060039 | 8.81071077 | 0.810662866 | 0.447652157 | 0.728645772 | -4.748031544 |
| **A_52_P684138** | Dpf3 | 0.076484283 | -0.152842518 | 0.305811084 | 8.079401759 | 0.810402134 | 0.447791191 | 0.728791645 | -4.748130829 |
| **A_55_P1984070** | NA | -0.102463252 | -0.409708384 | 0.204781879 | 13.33538104 | -0.810338207 | 0.447825285 | 0.728791645 | -4.748155169 |
| **A_55_P1992650** | Bloc1s1 | -0.111810499 | -0.447130835 | 0.223509837 | 13.62377636 | -0.810225452 | 0.447885424 | 0.728791645 | -4.748198096 |
| **A_55_P1961870** | NA | -0.077974497 | -0.311838774 | 0.15588978 | 14.73790388 | -0.810162028 | 0.447919255 | 0.728791645 | -4.748222241 |
| **A_55_P2034260** | Ctdsp1 | 0.06679212 | -0.133538637 | 0.267122876 | 11.96698627 | 0.810141323 | 0.447930299 | 0.728791645 | -4.748230122 |
| **A_66_P106363** | Itgb1bp1 | -0.132482434 | -0.529899172 | 0.264934304 | 10.18435679 | -0.810019238 | 0.447995425 | 0.728831721 | -4.748276594 |
| **A_55_P2066553** | Oacyl | 0.174968392 | -0.34996656 | 0.699903343 | 10.26302465 | 0.809911002 | 0.44805317 | 0.728831721 | -4.748317789 |
| **A_51_P182296** | Washc3 | -0.065033533 | -0.260151919 | 0.130084853 | 11.05956324 | -0.809883112 | 0.44806805 | 0.728831721 | -4.748328403 |
| **A_55_P2000548** | Cfl1 | -0.069189204 | -0.276831842 | 0.138453434 | 14.3172258 | -0.809664292 | 0.44818481 | 0.728877368 | -4.748411674 |
| **A_51_P431894** | Radil | -0.113401393 | -0.453729289 | 0.226926503 | 7.732289692 | -0.809662509 | 0.448185762 | 0.728877368 | -4.748412353 |
| **A_51_P254912** | Rfc2 | 0.111364074 | -0.222867803 | 0.445595951 | 11.10690605 | 0.809618515 | 0.44820924 | 0.728877368 | -4.748429092 |
| **A_55_P2091145** | H4c12 | -0.117806821 | -0.471420769 | 0.235807126 | 16.11532782 | -0.80951377 | 0.44826514 | 0.728897787 | -4.748468945 |
| **A_66_P116888** | Arid1a | 0.09660827 | -0.193396737 | 0.386613276 | 8.723953785 | 0.809453672 | 0.448297216 | 0.728897787 | -4.74849181 |
| **A_55_P2078403** | Rgs19 | -0.072749328 | -0.291208329 | 0.145709673 | 10.80325905 | -0.809174582 | 0.448446194 | 0.729028279 | -4.748597973 |
| **A_51_P501248** | Sphk1 | 0.13186647 | -0.264121586 | 0.527854526 | 12.22213179 | 0.809162008 | 0.448452907 | 0.729028279 | -4.748602756 |
| **A_52_P227445** | Parp2 | 0.08456512 | -0.169408034 | 0.338538274 | 8.769136845 | 0.809071025 | 0.448501482 | 0.72904593 | -4.748637359 |
| **A_51_P466288** | Frmd5 | -0.069096239 | -0.27672132 | 0.138528841 | 7.702167334 | -0.808644779 | 0.4487291 | 0.72935459 | -4.748799436 |
| **A_55_P1965725** | Gapdh | -0.11528913 | -0.461984771 | 0.231406511 | 12.48659011 | -0.808021969 | 0.449061833 | 0.729761724 | -4.749036146 |
| **A_55_P2017636** | Thbs1 | -0.156614017 | -0.62761015 | 0.314382116 | 11.88611797 | -0.80797231 | 0.449088371 | 0.729761724 | -4.749055015 |
| **A_55_P2061565** | Zfp229 | -0.063191461 | -0.253233469 | 0.126850547 | 7.97553555 | -0.807963928 | 0.44909285 | 0.729761724 | -4.749058199 |
| **A_51_P249608** | Trdn | -0.074746384 | -0.299644931 | 0.150152164 | 7.690183989 | -0.807582174 | 0.449296897 | 0.729988813 | -4.749203221 |
| **A_66_P130882** | Eif3e | -0.098743273 | -0.395851914 | 0.198365369 | 10.79321687 | -0.807561157 | 0.449308133 | 0.729988813 | -4.749211204 |
| **A_55_P2062733** | NA | -0.089862915 | -0.360320114 | 0.180594283 | 14.5586101 | -0.807356132 | 0.449417748 | 0.730105535 | -4.749289068 |

| | | | | | | | | | |
|---|---|---|---|---|---|---|---|---|---|
| **A_51_P323878** | Coro7 | 0.09927899 | -0.199584544 | 0.398142523 | 11.4699631 | 0.80717483 | 0.449514695 | 0.730201659 | -4.749357911 |
| **A_55_P1998097** | Tmem214 | 0.099919639 | -0.200908615 | 0.400747893 | 9.300750571 | 0.807077842 | 0.449566564 | 0.730224548 | -4.749394735 |
| **A_55_P2064292** | Col7a1 | 0.098677056 | -0.19847756 | 0.395831671 | 8.235772835 | 0.806894755 | 0.449664489 | 0.730322234 | -4.749464239 |
| **A_55_P2075298** | Atp8a1 | 0.245399383 | -0.493735366 | 0.984534131 | 11.070467 | 0.806738967 | 0.449747825 | 0.730396212 | -4.749523371 |
| **A_55_P2009582** | Nkg7 | 0.111957319 | -0.225284534 | 0.449199173 | 10.58113145 | 0.806666855 | 0.449786404 | 0.730397497 | -4.749550739 |
| **A_55_P2066409** | Gabpb1 | 0.065070674 | -0.130971699 | 0.261113047 | 10.36726853 | 0.80652632 | 0.449861595 | 0.73045823 | -4.749604072 |
| **A_51_P410927** | NA | -0.149427153 | -0.599774968 | 0.300920661 | 8.732684537 | -0.806240578 | 0.450014504 | 0.73061766 | -4.749712489 |
| **A_55_P1984721** | Ube2i | 0.094743007 | -0.190830205 | 0.38031622 | 12.41559942 | 0.806144488 | 0.450065933 | 0.73061766 | -4.749748942 |
| **A_51_P399175** | Ppm1j | -0.055883134 | -0.224342889 | 0.112576621 | 7.793964602 | -0.806060755 | 0.450110752 | 0.73061766 | -4.749780704 |
| **A_52_P652316** | Mapre2 | 0.060592673 | -0.122064088 | 0.243249435 | 9.760874463 | 0.806060331 | 0.450110978 | 0.73061766 | -4.749780865 |
| **A_51_P446417** | Ufl1 | -0.072382233 | -0.290620938 | 0.145856471 | 9.596292902 | -0.80590415 | 0.450194584 | 0.730692007 | -4.749840103 |
| **A_51_P305532** | Eif2s3x | 0.328252787 | -0.661837208 | 1.318342783 | 10.04218457 | 0.805595328 | 0.450359932 | 0.730853855 | -4.749957212 |
| **A_55_P2078474** | Fam50a | 0.069129955 | -0.139387581 | 0.277647492 | 12.44554939 | 0.805576668 | 0.450369925 | 0.730853855 | -4.749964287 |
| **A_55_P2207255** | Srpx | -0.22384787 | -0.899212598 | 0.451516859 | 8.346144321 | -0.805374322 | 0.450478289 | 0.730968339 | -4.750041001 |
| **A_55_P2027012** | Serpinb8 | 0.230534533 | -0.465194366 | 0.926263433 | 13.11434354 | 0.805154337 | 0.450596122 | 0.730973445 | -4.750124387 |
| **A_52_P81630** | 2810001G20Rik | 0.057629815 | -0.116291109 | 0.231550738 | 8.675700242 | 0.805153303 | 0.450596675 | 0.730973445 | -4.750124779 |
| **A_52_P242973** | Sec11a | -0.074788896 | -0.300506117 | 0.150928324 | 12.86702853 | -0.80511074 | 0.450619477 | 0.730973445 | -4.750140911 |
| **A_55_P2006476** | NA | -0.079548304 | -0.319637058 | 0.160540449 | 12.11135136 | -0.805086045 | 0.450632706 | 0.730973445 | -4.75015027 |
| **A_66_P109220** | Etfbkmt | 0.067015576 | -0.13527574 | 0.269306893 | 8.478031806 | 0.804973726 | 0.45069288 | 0.73097751 | -4.750192836 |
| **A_55_P2295951** | NA | 0.065200445 | -0.131619966 | 0.262020855 | 8.701941918 | 0.804940191 | 0.450710847 | 0.73097751 | -4.750205544 |
| **A_52_P87964** | Pla2g12a | 0.145363809 | -0.2935595 | 0.584287118 | 11.46161765 | 0.804731197 | 0.450822833 | 0.731045753 | -4.750284734 |
| **A_55_P2069415** | Fxr1 | 0.070205756 | -0.141782033 | 0.282193545 | 10.59615942 | 0.804720496 | 0.450828567 | 0.731045753 | -4.750288788 |
| **A_55_P2016478** | Ddx39b | 0.126205398 | -0.254938805 | 0.507349601 | 9.634665529 | 0.804584757 | 0.450901312 | 0.731102378 | -4.750340213 |
| **A_51_P172502** | Cxcl12 | -0.109692734 | -0.441042872 | 0.221657404 | 11.97774766 | -0.804403393 | 0.45099852 | 0.73110539 | -4.750408913 |
| **A_52_P293556** | Wdr41 | -0.054634127 | -0.219676726 | 0.110408472 | 8.921003255 | -0.804361258 | 0.451021106 | 0.73110539 | -4.750424872 |
| **A_55_P2061809** | Ndufc2 | -0.081890874 | -0.329296329 | 0.165514582 | 14.04069226 | -0.804283985 | 0.45106253 | 0.73110539 | -4.750454138 |
| **A_52_P575178** | Cdon | -0.122399453 | -0.492198678 | 0.247399772 | 8.446580932 | -0.804260367 | 0.451075191 | 0.73110539 | -4.750463083 |
| **A_55_P2062642** | Magt1 | -0.117266564 | -0.471572104 | 0.237038977 | 10.92155297 | -0.804228471 | 0.45109229 | 0.73110539 | -4.750475162 |
| **A_66_P124555** | Rps15 | -0.17153578 | -0.689904875 | 0.346833315 | 14.0793119 | -0.804079165 | 0.451172339 | 0.731173819 | -4.750531702 |
| **A_55_P2141311** | AU022252 | -0.072152252 | -0.290250724 | 0.14594622 | 9.542262722 | -0.803860075 | 0.451289821 | 0.731302896 | -4.750614654 |
| **A_51_P196862** | Amdhd1 | 0.246382782 | -0.498505952 | 0.991271517 | 9.981244998 | 0.80371512 | 0.451367561 | 0.731367557 | -4.750669529 |
| **A_51_P268529** | Csad | -0.241484175 | -0.971785078 | 0.488816728 | 12.28988139 | -0.803470669 | 0.451498683 | 0.731460093 | -4.750762053 |
| **A_55_P2048036** | Zfx | 0.095724304 | -0.193788905 | 0.385237513 | 10.22370851 | 0.803409611 | 0.451531439 | 0.731460093 | -4.75078516 |
| **A_55_P2017055** | Kctd12b | -0.208967919 | -0.8409905 | 0.423054662 | 8.600539277 | -0.803397011 | 0.451538198 | 0.731460093 | -4.750789928 |
| **A_52_P522372** | Aard | -0.070775553 | -0.284873345 | 0.143322239 | 8.009868361 | -0.803256546 | 0.45161356 | 0.731520867 | -4.750843081 |
| **A_55_P2077114** | NA | -0.097689096 | -0.393238492 | 0.1978603 | 14.91367923 | -0.803154698 | 0.451668209 | 0.731544942 | -4.750881617 |
| **A_55_P2090415** | Nat9 | -0.059963376 | -0.241396527 | 0.121469775 | 8.075363449 | -0.803068553 | 0.451714435 | 0.731544942 | -4.750914209 |
| **A_51_P205099** | Dazap1 | 0.05680108 | -0.115089553 | 0.228691712 | 11.77376196 | 0.802948275 | 0.451778984 | 0.731544942 | -4.750959711 |
| **A_55_P2093226** | Rnf121 | 0.071710595 | -0.145299422 | 0.288720612 | 8.760267393 | 0.802946733 | 0.451779812 | 0.731544942 | -4.750960294 |
| **A_52_P675395** | Ackr3 | -0.216923237 | -0.873599682 | 0.439753207 | 8.447849185 | -0.802671473 | 0.451927558 | 0.731590868 | -4.751064407 |
| **A_55_P2196027** | NA | -0.063736046 | -0.256691534 | 0.129219442 | 8.053663521 | -0.802622206 | 0.451954005 | 0.731590868 | -4.751083039 |
| **A_55_P2279035** | 9030419F21Rik | -0.066299058 | -0.267050706 | 0.13445259 | 8.599292949 | -0.802474846 | 0.452033118 | 0.731590868 | -4.751138763 |
| **A_55_P2156409** | Rpl26 | -0.1378552 | -0.55529793 | 0.279587531 | 13.83750245 | -0.802434214 | 0.452054934 | 0.731590868 | -4.751154127 |
| **A_55_P2061701** | Osgep | -0.108918539 | -0.438745628 | 0.22090855 | 10.46222536 | -0.802414328 | 0.452065611 | 0.731590868 | -4.751161646 |
| **A_52_P428521** | Fgd1 | -0.066347134 | -0.267266121 | 0.134571852 | 8.09922587 | -0.802387915 | 0.452079793 | 0.731590868 | -4.751171632 |
| **A_55_P2149288** | Cmya5 | 0.072388303 | -0.146825996 | 0.291602603 | 8.074995387 | 0.802384826 | 0.452081452 | 0.731590868 | -4.7511728 |
| **A_51_P392214** | Pcnx4 | -0.079672581 | -0.32096247 | 0.161617308 | 8.371388747 | -0.802329854 | 0.452110969 | 0.731590868 | -4.751193584 |
| **A_55_P2017677** | Cap2 | -0.05678262 | -0.228773323 | 0.115208083 | 7.740714475 | -0.802220294 | 0.452169802 | 0.731624821 | -4.751235005 |
| **A_55_P2106608** | NA | 0.135238889 | -0.274502225 | 0.544980003 | 8.455852637 | 0.80200161 | 0.45228725 | 0.731745666 | -4.751317668 |
| **A_55_P2128626** | Gm4349 | 0.06123926 | -0.124315076 | 0.246793596 | 8.378327442 | 0.801940259 | 0.452320204 | 0.731745666 | -4.751340856 |

| | | | | | | | | | |
|---|---|---|---|---|---|---|---|---|---|
| **A_51_P331207** | Tmem218 | -0.106305992 | -0.428459471 | 0.215847487 | 9.832619913 | -0.801822259 | 0.45238359 | 0.731760219 | -4.751385451 |
| **A_51_P282179** | Mtor | 0.085479037 | -0.173594703 | 0.344552776 | 8.603369818 | 0.801713902 | 0.452441802 | 0.731760219 | -4.751426398 |
| **A_51_P343900** | Xrcc1 | 0.100128625 | -0.203346435 | 0.403603684 | 11.36551946 | 0.801712092 | 0.452442774 | 0.731760219 | -4.751427082 |
| **A_51_P308912** | Mybphl | -0.073632601 | -0.296848125 | 0.149582922 | 7.831752626 | -0.801546899 | 0.452531529 | 0.731842531 | -4.751489499 |
| **A_55_P2110601** | Slc35a2 | -0.094876334 | -0.382539738 | 0.19278707 | 9.774370844 | -0.801413156 | 0.452603397 | 0.731877496 | -4.751540026 |
| **A_51_P329158** | Zc3h10 | 0.075829841 | -0.154110525 | 0.305770207 | 9.047717915 | 0.801323935 | 0.452651344 | 0.731877496 | -4.75157373 |
| **A_55_P2019909** | Tmem35b | -0.054180708 | -0.218487773 | 0.110126357 | 9.286708772 | -0.801256613 | 0.452687526 | 0.731877496 | -4.75159916 |
| **A_55_P2015742** | Cggbp1 | 0.086697891 | -0.176230401 | 0.349626183 | 9.821260948 | 0.801224833 | 0.452704606 | 0.731877496 | -4.751611163 |
| **A_51_P427812** | Tmem108 | 0.076911928 | -0.156467991 | 0.310291847 | 8.261136996 | 0.800780336 | 0.452943555 | 0.731906256 | -4.75177902 |
| **A_55_P2010076** | Miga2 | -0.074757573 | -0.301619891 | 0.152104746 | 10.4826568 | -0.800711407 | 0.452980617 | 0.731906256 | -4.751805044 |
| **A_52_P430169** | Dip2b | -0.117150381 | -0.472667603 | 0.23836684 | 9.188291199 | -0.800693407 | 0.452990295 | 0.731906256 | -4.75181184 |
| **A_55_P1986088** | Prr14 | 0.054089106 | -0.110056535 | 0.218234748 | 11.46149747 | 0.800688587 | 0.452992887 | 0.731906256 | -4.751813659 |
| **A_51_P506937** | Mrps12 | -0.052627367 | -0.212338199 | 0.107083464 | 12.94257243 | -0.800682732 | 0.452996036 | 0.731906256 | -4.75181587 |
| **A_55_P2012523** | Twf1 | 0.107158841 | -0.218046551 | 0.432364234 | 11.09148247 | 0.800669823 | 0.453002977 | 0.731906256 | -4.751820743 |
| **A_55_P2000182** | Slc5a6 | 0.126136555 | -0.256719938 | 0.508993048 | 9.422471624 | 0.800549404 | 0.453067732 | 0.731906256 | -4.751866202 |
| **A_55_P2157330** | NA | 0.054942142 | -0.111822404 | 0.221706688 | 16.60218265 | 0.800543708 | 0.453070795 | 0.731906256 | -4.751868352 |
| **A_55_P1984098** | Mst1 | -0.167045193 | -0.674191882 | 0.340101496 | 12.71697401 | -0.800356687 | 0.453171378 | 0.731906256 | -4.751938943 |
| **A_51_P517695** | Ly6f | -0.065414306 | -0.264016737 | 0.133188125 | 7.729646048 | -0.800334189 | 0.45318348 | 0.731906256 | -4.751947434 |
| **A_51_P244504** | LTO1 | 0.081954449 | -0.166898153 | 0.330807051 | 9.845048582 | 0.800227623 | 0.453240801 | 0.731906256 | -4.751987651 |
| **A_55_P2003053** | Dct | -0.125478388 | -0.506506059 | 0.255549282 | 8.173397404 | -0.800194578 | 0.453258577 | 0.731906256 | -4.752000121 |
| **A_55_P2028310** | Trappc1 | 0.086336944 | -0.175867872 | 0.348541759 | 8.338730132 | 0.800090651 | 0.453314486 | 0.731906256 | -4.752039338 |
| **A_55_P2130792** | Tuft1 | 0.105447987 | -0.214827334 | 0.425723307 | 9.349186755 | 0.800014775 | 0.453355307 | 0.731906256 | -4.752067967 |
| **A_55_P1985351** | Slc35f2 | -0.066972645 | -0.270394534 | 0.136449244 | 7.950071805 | -0.799987049 | 0.453370224 | 0.731906256 | -4.752078428 |
| **A_55_P2022678** | C1qtnf1 | -0.069437091 | -0.280346609 | 0.141472427 | 8.093779421 | -0.799978896 | 0.45337461 | 0.731906256 | -4.752081504 |
| **A_55_P1973357** | NA | 0.067285466 | -0.137109506 | 0.271680438 | 15.75767243 | 0.799897329 | 0.453418497 | 0.731906256 | -4.752112278 |
| **A_51_P262101** | Ptcd2 | 0.075209131 | -0.153262779 | 0.303681042 | 9.380087398 | 0.799872835 | 0.453431677 | 0.731906256 | -4.752121518 |
| **A_55_P1955562** | Extl3 | 0.07969118 | -0.162402114 | 0.321784474 | 11.56659451 | 0.79985394 | 0.453441844 | 0.731906256 | -4.752128647 |
| **A_55_P2040748** | Scrib | 0.061658778 | -0.125744382 | 0.249061937 | 11.83388357 | 0.799468197 | 0.453649442 | 0.732180199 | -4.752274143 |
| **A_51_P272250** | Snf8 | 0.081484305 | -0.16619886 | 0.32916747 | 11.32588031 | 0.7993936 | 0.453689597 | 0.73218387 | -4.752302274 |
| **A_55_P2287910** | Rragd | 0.05169695 | -0.105476711 | 0.20887061 | 7.707375832 | 0.799223676 | 0.453781073 | 0.732270359 | -4.752366347 |
| **A_55_P1995774** | NA | 0.092859137 | -0.189550364 | 0.375268638 | 9.227307295 | 0.798966425 | 0.453919586 | 0.73243273 | -4.75246333 |
| **A_55_P1968863** | Synpo2l | 0.190493791 | -0.388987914 | 0.769975497 | 14.06565369 | 0.79877467 | 0.454022852 | 0.732538207 | -4.752535606 |
| **A_55_P1974028** | Stab2 | -0.13487816 | -0.545214045 | 0.275457726 | 12.29736085 | -0.79870301 | 0.454061448 | 0.732539331 | -4.752562613 |
| **A_55_P1974193** | Ptges | -0.074678943 | -0.301968295 | 0.15261041 | 13.54862122 | -0.798366413 | 0.454242769 | 0.73265747 | -4.752689446 |
| **A_52_P381929** | Ubqln2 | 0.119043366 | -0.243292209 | 0.481378942 | 9.651141441 | 0.798320739 | 0.454267377 | 0.73265747 | -4.752706654 |
| **A_55_P2026089** | Usp30 | -0.052942418 | -0.214122332 | 0.108237497 | 10.04342968 | -0.798134413 | 0.454367774 | 0.73265747 | -4.752776843 |
| **A_55_P2046703** | Atp5g3 | 0.086578282 | -0.17700581 | 0.350162374 | 15.55657753 | 0.798128752 | 0.454370825 | 0.73265747 | -4.752778976 |
| **A_55_P2165234** | Slurp2 | 0.081075433 | -0.1657585 | 0.327909366 | 8.455607351 | 0.798118916 | 0.454376125 | 0.73265747 | -4.752782681 |
| **A_55_P1984366** | Cdadc1 | 0.061810961 | -0.126389103 | 0.250011026 | 9.16995197 | 0.798047829 | 0.454414433 | 0.73265747 | -4.752809456 |
| **A_51_P309854** | Kcnn2 | -0.051978294 | -0.210243461 | 0.106286872 | 7.746658387 | -0.798031207 | 0.454423391 | 0.73265747 | -4.750289712 |
| **A_55_P2100715** | Auh | 0.124786275 | -0.255208117 | 0.504780667 | 9.074762376 | 0.797944752 | 0.454469985 | 0.73265747 | -4.752848278 |
| **A_55_P2020341** | Camk2g | -0.152438212 | -0.616643519 | 0.311767095 | 10.86231097 | -0.797933931 | 0.454475817 | 0.73265747 | -4.752852353 |
| **A_55_P2069755** | Tymp | -0.138447468 | -0.560132001 | 0.283237064 | 9.848296201 | -0.797775209 | 0.454561367 | 0.732734273 | -4.752912125 |
| **A_55_P1980539** | Cetn4 | 0.086375579 | -0.176776697 | 0.349527854 | 8.886283779 | 0.797566728 | 0.454673755 | 0.732854321 | -4.752990622 |
| **A_66_P139393** | Psmd10 | 0.107491421 | -0.220171631 | 0.435154474 | 10.13630179 | 0.797130675 | 0.454908886 | 0.733172171 | -4.753154757 |
| **A_51_P205278** | Tmem161b | 0.083503128 | -0.171168211 | 0.338174466 | 8.558987002 | 0.796720258 | 0.455130271 | 0.733457745 | -4.753309184 |
| **A_55_P2046838** | Amfr | 0.138175654 | -0.283301491 | 0.5596528 | 11.37393704 | 0.796600709 | 0.455194772 | 0.733457745 | -4.753354157 |
| **A_55_P2156583** | Cdk5r1 | 0.056168784 | -0.115164885 | 0.227502452 | 8.892890922 | 0.796591183 | 0.455199912 | 0.733457745 | -4.75335774 |
| **A_55_P2039105** | Rbck1 | 0.082233901 | -0.168702279 | 0.333170081 | 11.60136159 | 0.796289153 | 0.455362899 | 0.73359532 | -4.753471335 |
| **A_66_P102299** | Ascc1 | -0.095094577 | -0.385305598 | 0.195116445 | 8.519042192 | -0.796205248 | 0.455408184 | 0.73359532 | -4.753502887 |

| | | | | | | | | | |
|---|---|---|---|---|---|---|---|---|---|
| **A_55_P2083153** | 1700001J03Rik | 0.052292698 | -0.107318351 | 0.211903748 | 7.760419326 | 0.796088389 | 0.455471261 | 0.73359532 | -4.753546827 |
| **A_55_P2069714** | Sqstm1 | -0.095101551 | -0.385385225 | 0.195182124 | 9.560969091 | -0.796064349 | 0.455484238 | 0.73359532 | -4.753555866 |
| **A_51_P161354** | Sesn2 | 0.116729811 | -0.239576999 | 0.473036621 | 9.776026132 | 0.796050921 | 0.455491486 | 0.73359532 | -4.753560914 |
| **A_55_P2144075** | Pofut2 | -0.081747638 | -0.331287142 | 0.167791867 | 8.081690102 | -0.796011043 | 0.455513013 | 0.73359532 | -4.753575907 |
| **A_51_P278994** | Swap70 | -0.078947377 | -0.320033263 | 0.16213851 | 9.438163854 | -0.795699537 | 0.455681196 | 0.733805035 | -4.753693005 |
| **A_52_P30550** | Acp2 | -0.076272715 | -0.309216276 | 0.156670846 | 8.745769979 | -0.79561266 | 0.455728109 | 0.733819445 | -4.753725657 |
| **A_51_P315682** | Igf2bp2 | -0.099955872 | -0.405310099 | 0.205398354 | 9.517846109 | -0.795403628 | 0.455840998 | 0.73394008 | -4.75380421 |
| **A_51_P200610** | Zbtb12 | 0.063594146 | -0.130729698 | 0.257917989 | 9.986577556 | 0.795196088 | 0.455953101 | 0.734059429 | -4.753882188 |
| **A_55_P1991906** | Acadvl | -0.100629608 | -0.408155205 | 0.206895988 | 14.06172158 | -0.795110889 | 0.455999128 | 0.734072386 | -4.753914195 |
| **A_55_P2049287** | NA | -0.116642851 | -0.473172993 | 0.239887291 | 14.57525766 | -0.794959607 | 0.456080861 | 0.734117495 | -4.753971021 |
| **A_51_P245546** | Synrg | -0.069153946 | -0.280554624 | 0.142246733 | 8.85068409 | -0.794865736 | 0.456131582 | 0.734117495 | -4.754006279 |
| **A_55_P1984690** | Ptprr | 0.066371362 | -0.136535805 | 0.269278529 | 7.781188743 | 0.794815874 | 0.456158526 | 0.734117495 | -4.754025005 |
| **A_66_P101835** | Cd38 | -0.115524923 | -0.468722757 | 0.237672911 | 9.605634301 | -0.794768864 | 0.456183929 | 0.734117495 | -4.75404266 |
| **A_55_P2081805** | Slc46a1 | -0.113617227 | -0.461009395 | 0.23377494 | 10.02378235 | -0.794707576 | 0.45621705 | 0.734117495 | -4.754065676 |
| **A_55_P2160750** | Col25a1 | -0.104197596 | -0.42283579 | 0.214440598 | 7.97299593 | -0.794589928 | 0.456280632 | 0.734117669 | -4.754109853 |
| **A_51_P140237** | Fhl2 | -0.207151184 | -0.840651085 | 0.426348718 | 9.55972008 | -0.794555168 | 0.456299419 | 0.734117669 | -4.754122905 |
| **A_55_P1974830** | Pde1c | 0.053389781 | -0.109896354 | 0.216675916 | 7.744016535 | 0.794496558 | 0.456331098 | 0.734117669 | -4.75414491 |
| **A_55_P2087865** | Tinf2 | 0.117106363 | -0.241119889 | 0.475332614 | 9.155955725 | 0.794339707 | 0.456415884 | 0.734160357 | -4.754203797 |
| **A_51_P144160** | Colec10 | 0.233405041 | -0.480606223 | 0.947416304 | 12.07567401 | 0.79430694 | 0.456433598 | 0.734160357 | -4.754216097 |
| **A_52_P138598** | Pum2 | 0.059964039 | -0.123522467 | 0.243450545 | 8.498822319 | 0.794090381 | 0.456550681 | 0.734279263 | -4.754297383 |
| **A_55_P2052330** | NA | -0.055033882 | -0.223447266 | 0.113379501 | 12.17144797 | -0.794029686 | 0.456583499 | 0.734279263 | -4.754320162 |
| **A_52_P203691** | Arl5c | -0.082091665 | -0.333346808 | 0.169163477 | 9.137096656 | -0.793902729 | 0.456652152 | 0.734328573 | -4.754367806 |
| **A_55_P2092730** | Wdfy1 | -0.094477473 | -0.383716313 | 0.194761367 | 8.757170421 | -0.7936972 | 0.456763309 | 0.734420544 | -4.754444924 |
| **A_52_P370311** | Plbd2 | -0.066649189 | -0.270717035 | 0.137418657 | 11.09572938 | -0.793603326 | 0.456814085 | 0.734420544 | -4.754480143 |
| **A_55_P2031471** | Rufy4 | -0.05946594 | -0.241565934 | 0.122634053 | 7.785080878 | -0.793490168 | 0.456875298 | 0.734420544 | -4.754522592 |
| **A_52_P45738** | Trim35 | 0.067232856 | -0.138655324 | 0.273121035 | 7.921526101 | 0.793475156 | 0.456883419 | 0.734420544 | -4.754528223 |
| **A_55_P2175275** | Hmbox1 | 0.246330729 | -0.508100831 | 1.000762289 | 10.35486997 | 0.793381251 | 0.456934221 | 0.734420544 | -4.754563446 |
| **A_51_P296796** | Nt5dc2 | -0.114273234 | -0.464257814 | 0.235711346 | 9.194231578 | -0.793375523 | 0.45693732 | 0.734420544 | -4.754565595 |
| **A_55_P1952112** | Ildr2 | 0.078575125 | -0.162171935 | 0.319322184 | 8.276183627 | 0.793062249 | 0.457106831 | 0.734590842 | -4.75468308 |
| **A_55_P2059372** | Ppip5k2 | 0.064523666 | -0.133176733 | 0.262224064 | 8.058757686 | 0.793039237 | 0.457119285 | 0.734590842 | -4.754691708 |
| **A_55_P2166985** | Dedd2 | 0.100949834 | -0.208467536 | 0.410367204 | 13.03865003 | 0.792764343 | 0.457268068 | 0.734768848 | -4.75479477 |
| **A_55_P1960576** | Kdm5b | -0.052203428 | -0.212242657 | 0.1078358 | 8.00138304 | -0.792603093 | 0.457355358 | 0.73484544 | -4.754855213 |
| **A_55_P2088272** | Golga3 | -0.078366351 | -0.318633227 | 0.161900526 | 11.04546722 | -0.79253584 | 0.457391768 | 0.73484544 | -4.75488042 |
| **A_55_P2016193** | Prxl2c | -0.086235223 | -0.350723839 | 0.178253392 | 10.0793922 | -0.792247538 | 0.457547875 | 0.735035145 | -4.754988459 |
| **A_55_P2087766** | Dnaja3 | 0.13481326 | -0.27893963 | 0.54856615 | 12.18481362 | 0.791725729 | 0.457830514 | 0.735287204 | -4.755183932 |
| **A_55_P2383877** | 2810040C05Rik | 0.05949954 | -0.123113813 | 0.242112892 | 8.47605219 | 0.791706598 | 0.457840878 | 0.735287204 | -4.755191097 |
| **A_55_P2173768** | Pkhd1 | 0.216795191 | -0.448588071 | 0.882178454 | 9.768043996 | 0.791700578 | 0.45784414 | 0.735287204 | -4.755193352 |
| **A_55_P2146177** | Cerk | -0.082205661 | -0.334517084 | 0.170105762 | 11.24308707 | -0.791676953 | 0.45785694 | 0.735287204 | -4.755202199 |
| **A_51_P422893** | Tmem14a | 0.087923752 | -0.181972235 | 0.35781974 | 8.999477992 | 0.791576651 | 0.457911285 | 0.735292887 | -4.755239761 |
| **A_51_P395743** | Spopl | 0.073679986 | -0.152542009 | 0.299901981 | 8.639110619 | 0.791403371 | 0.458005181 | 0.735292887 | -4.755304644 |
| **A_55_P2079855** | Prl6a1 | 0.051658685 | -0.106955796 | 0.210273165 | 7.717760883 | 0.791377503 | 0.4580192 | 0.735292887 | -4.755314329 |
| **A_52_P694988** | Zfp933 | 0.063340024 | -0.131184601 | 0.25786465 | 7.917035368 | 0.791200999 | 0.45811486 | 0.735292887 | -4.755380407 |
| **A_52_P400677** | AW209491 | -0.053192045 | -0.216566967 | 0.110182877 | 9.804926503 | -0.791123859 | 0.458156671 | 0.735292887 | -4.755409283 |
| **A_55_P2019019** | Rpl36a | -0.068278963 | -0.277997643 | 0.141439717 | 15.41741443 | -0.791102922 | 0.45816802 | 0.735292887 | -4.75541712 |
| **A_52_P605556** | Nxnl2 | -0.070702923 | -0.287868752 | 0.146462907 | 7.760489913 | -0.791095777 | 0.458171893 | 0.735292887 | -4.755419794 |
| **A_51_P349495** | Mboat1 | -0.186664305 | -0.760033459 | 0.38670485 | 8.85840696 | -0.791061303 | 0.45819058 | 0.735292887 | -4.755432698 |
| **A_51_P230904** | Copz2 | -0.101880648 | -0.414832246 | 0.211070949 | 9.249987499 | -0.791038674 | 0.458202846 | 0.735292887 | -4.755441167 |
| **A_55_P1992388** | NA | -0.066489222 | -0.270831931 | 0.137853487 | 13.82427285 | -0.790633643 | 0.458422443 | 0.735584211 | -4.755592737 |
| **A_55_P1969914** | Tcf25 | -0.065709082 | -0.267733466 | 0.136315301 | 12.01801557 | -0.790323327 | 0.458590738 | 0.735764947 | -4.755708825 |
| **A_66_P120736** | Bnip3 | -0.121235556 | -0.493995414 | 0.251524302 | 8.721513966 | -0.790285585 | 0.45861121 | 0.735764947 | -4.755722942 |

| | | | | | | | | | |
|---|---|---|---|---|---|---|---|---|---|
| **A_52_P31641** | Tmem145 | 0.065402134 | -0.13571042 | 0.266514687 | 7.829731796 | 0.790198001 | 0.45865872 | 0.735780097 | -4.7557557 |
| **A_55_P2002160** | Sumo3 | -0.072166271 | -0.294135691 | 0.149803149 | 9.805890489 | -0.789994931 | 0.458768887 | 0.735786973 | -4.755831642 |
| **A_66_P113892** | Smim6 | -0.383324825 | -1.56249162 | 0.795841969 | 8.651353301 | -0.789905058 | 0.45881765 | 0.735786973 | -4.755865247 |
| **A_51_P487933** | Psmd7 | 0.057754192 | -0.119906723 | 0.235415107 | 12.29317146 | 0.789904896 | 0.458817737 | 0.735786973 | -4.755865308 |
| **A_55_P2036460** | Synb | -0.081855229 | -0.333658741 | 0.169948283 | 7.819468957 | -0.789892214 | 0.458824619 | 0.735786973 | -4.755870049 |
| **A_55_P2019577** | 1500011B03Rik | -0.074109383 | -0.302103787 | 0.15388502 | 8.406802695 | -0.78982741 | 0.458859782 | 0.735786973 | -4.755894279 |
| **A_51_P194004** | Glyctk | 0.202120937 | -0.419741516 | 0.823983391 | 9.899516145 | 0.789769135 | 0.458891405 | 0.735786973 | -4.755916066 |
| **A_52_P321318** | NA | -0.170420021 | -0.694875854 | 0.354035812 | 15.29638602 | -0.789577717 | 0.458995287 | 0.735892493 | -4.755987622 |
| **A_55_P2040276** | Prcd | -0.509987635 | -2.079938818 | 1.059963549 | 8.284613883 | -0.789326505 | 0.459131644 | 0.736050058 | -4.756081513 |
| **A_51_P420236** | Srrt | 0.10074228 | -0.209469308 | 0.410953867 | 8.745774459 | 0.789108911 | 0.459249777 | 0.736178383 | -4.756162821 |
| **A_65_P20699** | Tlk2 | 0.062145147 | -0.129244531 | 0.253534825 | 9.186996132 | 0.788990719 | 0.459313953 | 0.736220201 | -4.756206979 |
| **A_51_P338443** | Angptl4 | -0.175447115 | -0.716091017 | 0.365196788 | 12.73956933 | -0.788529799 | 0.459564284 | 0.736560369 | -4.756379139 |
| **A_55_P2151138** | Dynlt1c | -0.166119452 | -0.678124859 | 0.345885955 | 13.19629094 | -0.78836826 | 0.459652041 | 0.736619942 | -4.75643946 |
| **A_51_P205390** | Cabcoco1 | -0.069542396 | -0.283922729 | 0.144837937 | 8.129239378 | -0.788220808 | 0.459732154 | 0.736619942 | -4.756494512 |
| **A_51_P349023** | Lsm3 | 0.10110989 | -0.210622255 | 0.412842035 | 11.30520924 | 0.788125246 | 0.459784081 | 0.736619942 | -4.756530186 |
| **A_55_P2062911** | Epha5 | 0.181940944 | -0.379006423 | 0.742888312 | 13.96496185 | 0.788118495 | 0.459787749 | 0.736619942 | -4.756532707 |
| **A_51_P255456** | Cyp1b1 | -0.080113039 | -0.327114312 | 0.166888233 | 7.801345977 | -0.788110669 | 0.459792002 | 0.736619942 | -4.756535628 |
| **A_51_P167360** | Ptpn7 | 0.118804487 | -0.247672363 | 0.485281337 | 9.034435479 | 0.787715674 | 0.46000668 | 0.736902794 | -4.756683051 |
| **A_52_P597461** | Skil | -0.10420269 | -0.425700532 | 0.217295152 | 8.157577973 | -0.787560735 | 0.460090907 | 0.736946901 | -4.756740864 |
| **A_51_P266774** | Mfn2 | 0.062286788 | -0.129907291 | 0.254480868 | 10.13588221 | 0.787479257 | 0.460135205 | 0.736946901 | -4.756771263 |
| **A_52_P31490** | Vamp3 | -0.093127103 | -0.380492017 | 0.19423781 | 8.490557172 | -0.787454633 | 0.460148593 | 0.736946901 | -4.75678045 |
| **A_55_P2094925** | Srgn | -0.104968879 | -0.428944566 | 0.219006807 | 11.70965162 | -0.787283828 | 0.460241466 | 0.737006545 | -4.756844167 |
| **A_55_P2158522** | Pbx1 | 0.272997061 | -0.569717284 | 1.115711407 | 14.40133861 | 0.787155935 | 0.460311016 | 0.737006545 | -4.75689187 |
| **A_66_P130777** | Ephb4 | -0.071155723 | -0.290812671 | 0.148501225 | 8.279271169 | -0.787132922 | 0.460323531 | 0.737006545 | -4.756900453 |
| **A_55_P2134032** | NA | 0.252591967 | -0.527183708 | 1.032367641 | 9.828730086 | 0.787105669 | 0.460338353 | 0.737006545 | -4.756910617 |
| **A_55_P2051047** | Rps10 | 0.106557183 | -0.222498569 | 0.435612935 | 17.37578206 | 0.786858126 | 0.460472995 | 0.737137067 | -4.757002929 |
| **A_55_P2151899** | NA | -0.137058124 | -0.560333781 | 0.286217534 | 12.85861651 | -0.786800592 | 0.460504293 | 0.737137067 | -4.757024381 |
| **A_55_P2081123** | Srcin1 | -0.067015006 | -0.273991489 | 0.139961478 | 7.740496086 | -0.786745458 | 0.460534286 | 0.737137067 | -4.757044937 |
| **A_51_P105380** | Csta2 | -0.23303775 | -0.952993806 | 0.486918307 | 13.76084267 | -0.786508604 | 0.460663153 | 0.737282279 | -4.757133234 |
| **A_52_P453884** | Foxo1 | -0.093295767 | -0.381720016 | 0.195128483 | 10.37068886 | -0.785983367 | 0.460949012 | 0.737634966 | -4.757328969 |
| **A_55_P2064233** | Ipo8 | 0.09071663 | -0.189779194 | 0.371212454 | 9.686569723 | 0.785857308 | 0.461017638 | 0.737634966 | -4.757375932 |
| **A_66_P105791** | Prkca | -0.154529901 | -0.632342815 | 0.323283012 | 10.10056912 | -0.785846699 | 0.461023414 | 0.737634966 | -4.757379885 |
| **A_55_P1984168** | Casp4 | -0.085633354 | -0.350423156 | 0.179156447 | 8.972271104 | -0.785823279 | 0.461036165 | 0.737634966 | -4.757388609 |
| **A_55_P2088495** | NA | 0.070103515 | -0.146912009 | 0.287119038 | 8.20815578 | 0.784932262 | 0.461521449 | 0.738287467 | -4.757720393 |
| **A_55_P2299524** | Scube1 | -0.061304712 | -0.25111174 | 0.128502317 | 7.739919573 | -0.784810406 | 0.461587844 | 0.738287467 | -4.757765746 |
| **A_55_P1973813** | NA | 0.150637502 | -0.315757627 | 0.617032632 | 8.378197453 | 0.784805953 | 0.461590271 | 0.738287467 | -4.757767404 |
| **A_55_P2176729** | Prdx6 | -0.094735019 | -0.388086052 | 0.198616015 | 13.1517787 | -0.784704442 | 0.461645586 | 0.738287467 | -4.757805181 |
| **A_55_P2131153** | Smarcc1 | -0.080269981 | -0.328857463 | 0.168317502 | 11.32942829 | -0.784615876 | 0.461693852 | 0.738287467 | -4.757838139 |
| **A_55_P2009013** | Dpf2 | 0.07719692 | -0.161877396 | 0.316271236 | 11.17198098 | 0.784603501 | 0.461700596 | 0.738287467 | -4.757842743 |
| **A_55_P2425661** | Ncbp3 | 0.246660912 | -0.517264495 | 1.010586319 | 11.64427622 | 0.784571571 | 0.461717998 | 0.738287467 | -4.757854624 |
| **A_51_P279465** | Farsa | 0.089752894 | -0.188246682 | 0.367752469 | 9.037455525 | 0.784490186 | 0.461762354 | 0.738287467 | -4.757884906 |
| **A_55_P1977850** | Sall1 | 0.144151673 | -0.302368577 | 0.590671924 | 8.454248695 | 0.784443587 | 0.461787753 | 0.738287467 | -4.757902243 |
| **A_51_P282227** | As3mt | 0.086816138 | -0.182245349 | 0.355877625 | 10.19987877 | 0.784028975 | 0.462013782 | 0.738521527 | -4.758056469 |
| **A_51_P165504** | Twist2 | -0.099451493 | -0.407682584 | 0.208779599 | 7.823945027 | -0.784003591 | 0.462027623 | 0.738521527 | -4.758065909 |
| **A_55_P2083841** | Atp13a1 | -0.067111947 | -0.275123028 | 0.140899133 | 12.24375252 | -0.783964793 | 0.462048778 | 0.738521527 | -4.758080338 |
| **A_55_P2109505** | NA | -0.087703668 | -0.359576373 | 0.184169037 | 14.6795768 | -0.783854286 | 0.462109038 | 0.738556771 | -4.758121431 |
| **A_51_P337856** | Ubl7 | 0.083439937 | -0.175268618 | 0.342148493 | 9.533813832 | 0.783693721 | 0.462196605 | 0.738587481 | -4.758181133 |
| **A_55_P2115068** | Psmb5 | -0.099085068 | -0.406307719 | 0.208137584 | 12.80801041 | -0.783678923 | 0.462204675 | 0.738587481 | -4.758186634 |
| **A_55_P2029612** | Dhx35 | -0.076958376 | -0.315631236 | 0.161714483 | 9.311906826 | -0.783494675 | 0.462305173 | 0.738674049 | -4.758255129 |
| **A_52_P525070** | Vamp4 | -0.060679637 | -0.24888442 | 0.127525147 | 8.168712879 | -0.78342153 | 0.462345074 | 0.738674049 | -4.758282318 |

| | | | | | | | | | |
|---|---|---|---|---|---|---|---|---|---|
| **A_55_P1987302** | Eny2 | -0.068889589 | -0.28259034 | 0.144811162 | 12.78491157 | -0.783304709 | 0.462408806 | 0.738674049 | -4.758325738 |
| **A_55_P2032668** | NA | 0.085694458 | -0.180138081 | 0.351526996 | 12.58081178 | 0.783299382 | 0.462411713 | 0.738674049 | -4.758327718 |
| **A_66_P101407** | Acat3 | 0.167203598 | -0.351567888 | 0.685975083 | 10.00728742 | 0.783163989 | 0.462485584 | 0.738678592 | -4.758378034 |
| **A_55_P1955289** | Slc18b1 | -0.123722592 | -0.507593307 | 0.260148123 | 8.457504586 | -0.783154085 | 0.462490989 | 0.738678592 | -4.758381715 |
| **A_66_P114585** | NA | -0.051891538 | -0.21291426 | 0.109131183 | 7.972679089 | -0.783055548 | 0.462544757 | 0.738685592 | -4.75841833 |
| **A_55_P1999213** | Tdrd7 | -0.074390718 | -0.305256963 | 0.156475526 | 10.83349242 | -0.782963473 | 0.462595003 | 0.738685592 | -4.758452541 |
| **A_55_P2014214** | NA | -0.162170448 | -0.665482955 | 0.341142059 | 14.37214517 | -0.782919631 | 0.46261893 | 0.738685592 | -4.75846883 |
| **A_55_P2144636** | Ccdc163 | -0.085158074 | -0.349472963 | 0.179156815 | 9.84442991 | -0.782865931 | 0.462648237 | 0.738685592 | -4.75848878 |
| **A_55_P2009507** | Ocrl | -0.066572047 | -0.273228 | 0.140083906 | 10.43606223 | -0.782757401 | 0.462707473 | 0.738719149 | -4.758529098 |
| **A_66_P127036** | NA | -0.094184694 | -0.386606487 | 0.198237098 | 15.2013785 | -0.782625128 | 0.462779674 | 0.738741472 | -4.758578231 |
| **A_55_P1988789** | Acss2 | -0.21176593 | -0.869292308 | 0.445760448 | 12.27629129 | -0.782575233 | 0.462806912 | 0.738741472 | -4.758596762 |
| **A_55_P1972997** | A530016L24Rik | 0.051958167 | -0.109381291 | 0.213297625 | 7.73901373 | 0.782521744 | 0.462836113 | 0.738741472 | -4.758616628 |
| **A_55_P1967880** | Chtf8 | -0.151884775 | -0.623613133 | 0.319843582 | 13.38914242 | -0.782357856 | 0.46292559 | 0.738823279 | -4.75867749 |
| **A_55_P2096297** | NA | 0.070514007 | -0.148543484 | 0.289571497 | 8.866682718 | 0.782168771 | 0.46302884 | 0.738899202 | -4.758747698 |
| **A_51_P488673** | Rack1 | 0.075438891 | -0.158929553 | 0.309807336 | 16.88898928 | 0.782130725 | 0.463049616 | 0.738899202 | -4.758761823 |
| **A_55_P1993864** | Sh2b3 | -0.076113762 | -0.312609563 | 0.160382038 | 9.343420427 | -0.782029156 | 0.463105086 | 0.738926714 | -4.758799529 |
| **A_52_P357133** | Selenom | -0.203761004 | -0.837059123 | 0.429537115 | 9.942301352 | -0.781800714 | 0.463229862 | 0.739064796 | -4.758884323 |
| **A_51_P281700** | Khsrp | 0.20504041 | -0.43233598 | 0.8424168 | 12.61944421 | 0.78167582 | 0.46329809 | 0.739098161 | -4.758930674 |
| **A_52_P331901** | Trp53rkb | 0.071271837 | -0.150294675 | 0.29283835 | 8.417765799 | 0.781622446 | 0.46332725 | 0.739098161 | -4.758950481 |
| **A_66_P128608** | Fut11 | -0.059702431 | -0.245371198 | 0.125966335 | 8.329440075 | -0.781333338 | 0.463485219 | 0.73928914 | -4.759057749 |
| **A_55_P2012849** | Dock10 | -0.082030413 | -0.337202154 | 0.173141328 | 8.493733754 | -0.781133946 | 0.463594189 | 0.739360552 | -4.759131714 |
| **A_55_P2117400** | Cacng7 | -0.160780649 | -0.660934905 | 0.339373608 | 8.19417661 | -0.781111436 | 0.463606492 | 0.739360552 | -4.759140063 |
| **A_51_P287069** | Serpinh1 | -0.387371066 | -1.592718561 | 0.817976428 | 11.90562853 | -0.780904811 | 0.463719436 | 0.739475699 | -4.759216694 |
| **A_51_P455266** | Ring1 | 0.069562245 | -0.146912052 | 0.286036542 | 8.332110241 | 0.780819118 | 0.463766283 | 0.739475699 | -4.759248471 |
| **A_55_P2029750** | Ndufs5 | -0.077673131 | -0.319403503 | 0.164057241 | 14.3174663 | -0.780769397 | 0.463793466 | 0.739475699 | -4.759266908 |
| **A_51_P161874** | Med30 | 0.076904051 | -0.162522176 | 0.316330278 | 10.87020766 | 0.780478033 | 0.46395278 | 0.739620235 | -4.759374928 |
| **A_55_P2106569** | NA | -0.119623491 | -0.492055487 | 0.252808504 | 17.02481165 | -0.780463647 | 0.463960647 | 0.739620235 | -4.759380261 |
| **A_55_P2124115** | Invs | -0.080118671 | -0.329591038 | 0.169353697 | 9.379171295 | -0.780359057 | 0.464017846 | 0.739650416 | -4.759419029 |
| **A_51_P489452** | Cdo1 | -0.173575166 | -0.714306257 | 0.367155925 | 11.08696721 | -0.779990722 | 0.464219322 | 0.739766057 | -4.75955553 |
| **A_51_P302358** | Ltb | -0.112485175 | -0.462911111 | 0.237940761 | 10.03986603 | -0.77997779 | 0.464226396 | 0.739766057 | -4.759560321 |
| **A_55_P2068607** | Rnf19b | 0.074361575 | -0.157312709 | 0.306035858 | 10.21952807 | 0.779926972 | 0.464254199 | 0.739766057 | -4.75957915 |
| **A_55_P2094574** | Rapgef6 | 0.094797306 | -0.200552553 | 0.390147164 | 10.74738225 | 0.779906277 | 0.464265521 | 0.739766057 | -4.759586817 |
| **A_66_P122087** | Zfp622 | -0.076489879 | -0.314816373 | 0.161836616 | 13.17982526 | -0.779856749 | 0.464292619 | 0.739766057 | -4.759605167 |
| **A_55_P1955209** | NA | -0.110653903 | -0.455450513 | 0.234142708 | 14.72760018 | -0.779806655 | 0.464320027 | 0.739766057 | -4.759623725 |
| **A_55_P2054714** | NA | 0.059742121 | -0.12650233 | 0.245986571 | 9.131815721 | 0.779436037 | 0.464522844 | 0.740028192 | -4.759761001 |
| **A_51_P109050** | Paqr4 | -0.189697651 | -0.781201525 | 0.401806223 | 9.676506766 | -0.779269268 | 0.464614127 | 0.74005621 | -4.759822756 |
| **A_55_P2041673** | Ttc38 | 0.0968064 | -0.205051565 | 0.398664365 | 17.2119016 | 0.77926401 | 0.464617006 | 0.74005621 | -4.759824703 |
| **A_55_P1969058** | Epha7 | -0.06799133 | -0.280033959 | 0.144051298 | 7.797120793 | -0.779136497 | 0.46468681 | 0.740106406 | -4.759871915 |
| **A_55_P1977589** | Nfu1 | -0.056490741 | -0.232732238 | 0.119750756 | 11.81229404 | -0.778847244 | 0.464845183 | 0.74021057 | -4.759978991 |
| **A_51_P256957** | Ankmy2 | -0.070344807 | -0.289814487 | 0.149124872 | 8.615782545 | -0.778826469 | 0.464856559 | 0.74021057 | -4.759986681 |
| **A_66_P111216** | 4632427E13Rik | 0.065177477 | -0.138176245 | 0.268531199 | 8.233893768 | 0.778804779 | 0.464868437 | 0.74021057 | -4.759994708 |
| **A_51_P398887** | Ctns | -0.132011147 | -0.543921605 | 0.279899311 | 9.805144941 | -0.778737296 | 0.464905392 | 0.74021057 | -4.760019684 |
| **A_55_P1995892** | NA | -0.086937207 | -0.358254469 | 0.184380055 | 8.772515224 | -0.778594708 | 0.464983484 | 0.740273927 | -4.760072452 |
| **A_66_P137134** | NA | 0.230927345 | -0.489884388 | 0.951739079 | 9.714466216 | 0.778460722 | 0.465056872 | 0.740309931 | -4.76012203 |
| **A_52_P23379** | Depdc7 | 0.137712466 | -0.292192421 | 0.567617354 | 8.915182004 | 0.778366367 | 0.465108558 | 0.740309931 | -4.760156939 |
| **A_55_P2063166** | Ogfod1 | 0.203423583 | -0.431663929 | 0.838511094 | 14.26653341 | 0.778306961 | 0.465141102 | 0.740309931 | -4.760178917 |
| **A_55_P2170509** | Yipf4 | 0.152400745 | -0.415094059 | 0.719895549 | 12.29368361 | 0.828115186 | 0.465206039 | 0.740309931 | -4.654330548 |
| **A_55_P2353462** | 9530036O11Rik | -0.267421047 | -1.102518532 | 0.567676437 | 8.116664891 | -0.778111079 | 0.465248421 | 0.740309931 | -4.760251376 |
| **A_55_P1978089** | Dcaf17 | 0.116650821 | -0.247641653 | 0.480943294 | 10.16965094 | 0.778073796 | 0.46526885 | 0.740309931 | -4.760265166 |
| **A_55_P2163659** | Rspo3 | 0.162666088 | -0.345418412 | 0.670750588 | 9.068322393 | 0.777936707 | 0.465343971 | 0.740309931 | -4.760315867 |

| | | | | | | | | | |
|---|---|---|---|---|---|---|---|---|---|
| **A_55_P2041893** | NA | -0.079731727 | -0.328783988 | 0.169320534 | 15.36565303 | -0.777900179 | 0.465363988 | 0.740309931 | -4.760329375 |
| **A_55_P2403325** | NA | -0.057830039 | -0.238495752 | 0.122835674 | 7.867521492 | -0.777787459 | 0.465425764 | 0.740309931 | -4.760371058 |
| **A_51_P125648** | Vwce | 0.23952833 | -0.508846375 | 0.987903036 | 11.10644088 | 0.777715896 | 0.465464986 | 0.740309931 | -4.760397519 |
| **A_51_P174681** | Bzw2 | -0.05711776 | -0.235599994 | 0.121364474 | 10.71709474 | -0.777605567 | 0.46552546 | 0.740309931 | -4.76043831 |
| **A_52_P565847** | AU018091 | 0.055481184 | -0.117888168 | 0.228850536 | 7.694599503 | 0.777600594 | 0.465528186 | 0.740309931 | -4.760440148 |
| **A_55_P2017714** | Kat5 | -0.072771744 | -0.300196541 | 0.154653052 | 11.45614428 | -0.777513896 | 0.465575712 | 0.740309931 | -4.7604722 |
| **A_52_P214752** | NA | 0.128004623 | -0.272034419 | 0.528043664 | 9.50274097 | 0.777510861 | 0.465577375 | 0.740309931 | -4.760473322 |
| **A_55_P2172747** | Mfn2 | 0.076834331 | -0.163290854 | 0.316959517 | 8.939129263 | 0.777500706 | 0.465582942 | 0.740309931 | -4.760477076 |
| **A_55_P2301312** | Wdr78 | -0.27653667 | -1.140877624 | 0.587804285 | 8.505183238 | -0.777411259 | 0.465631978 | 0.740309931 | -4.76051014 |
| **A_55_P2058718** | Ephb4 | -0.073629777 | -0.303793024 | 0.15653347 | 8.917003712 | -0.777321529 | 0.465681174 | 0.740309931 | -4.760543306 |
| **A_55_P2000628** | NA | 0.081506743 | -0.173346225 | 0.336359711 | 8.786387589 | 0.777118116 | 0.46579271 | 0.740309931 | -4.760618482 |
| **A_55_P1965150** | Pnkp | -0.090939922 | -0.375296932 | 0.193417088 | 11.88092558 | -0.777094484 | 0.465805669 | 0.740309931 | -4.760627215 |
| **A_55_P2075230** | BC051226 | -0.143724887 | -0.593188971 | 0.305739196 | 8.779778236 | -0.776998486 | 0.465858314 | 0.740309931 | -4.760662687 |
| **A_55_P2021065** | E130308A19Rik | 0.073824959 | -0.157045203 | 0.30469512 | 8.750442685 | 0.776995663 | 0.465859863 | 0.740309931 | -4.76066373 |
| **A_51_P291460** | Gm6498 | -0.093713869 | -0.386909277 | 0.19948154 | 14.76594553 | -0.776658127 | 0.466045002 | 0.740309931 | -4.760788429 |
| **A_52_P585124** | Cxcr4 | -0.122420833 | -0.505460659 | 0.260618994 | 8.91064446 | -0.776594992 | 0.466079637 | 0.740309931 | -4.760811748 |
| **A_52_P569549** | Lig4 | 0.090546648 | -0.19277066 | 0.373863957 | 8.720234099 | 0.776573309 | 0.466091532 | 0.740309931 | -4.760819757 |
| **A_55_P2096486** | Bcs1l | 0.073042724 | -0.155525006 | 0.301610455 | 9.859485219 | 0.776506749 | 0.466128049 | 0.740309931 | -4.76084434 |
| **A_55_P1978321** | Fbxo8 | 0.125000185 | -0.266166054 | 0.516166424 | 10.65298777 | 0.776483946 | 0.46614056 | 0.740309931 | -4.760852762 |
| **A_55_P2115192** | Sike1 | -0.080083452 | -0.330695426 | 0.170528522 | 8.967857006 | -0.77646906 | 0.466148727 | 0.740309931 | -4.76085826 |
| **A_55_P2002562** | Sec11a | -0.066227879 | -0.273489272 | 0.141033513 | 10.99959946 | -0.776435931 | 0.466166904 | 0.740309931 | -4.760870494 |
| **A_51_P508485** | Tmtc4 | -0.096005111 | -0.396470484 | 0.204460262 | 8.817860464 | -0.776395687 | 0.466188986 | 0.740309931 | -4.760885356 |
| **A_52_P625431** | Hnrnpa0 | -0.132022581 | -0.545214201 | 0.281169038 | 8.59499822 | -0.77638995 | 0.466192133 | 0.740309931 | -4.760887475 |
| **A_55_P1998309** | Sms | 0.104634973 | -0.222842518 | 0.432112464 | 10.79766244 | 0.776387618 | 0.466193413 | 0.740309931 | -4.760888336 |
| **A_55_P2139913** | Ssb | 0.088553692 | -0.188630684 | 0.365738069 | 11.8632694 | 0.776284859 | 0.466249799 | 0.740338648 | -4.760926282 |
| **A_55_P2078710** | Morc4 | 0.20255623 | -0.431574395 | 0.836686855 | 12.94530962 | 0.776157872 | 0.466319486 | 0.740370209 | -4.760973169 |
| **A_52_P487599** | Thnsl2 | -0.181992196 | -0.751780079 | 0.387795687 | 10.66214763 | -0.776109046 | 0.466346283 | 0.740370209 | -4.760991195 |
| **A_55_P2181743** | Elmod1 | -0.066325207 | -0.274018158 | 0.141367745 | 7.688086336 | -0.775961266 | 0.466427393 | 0.740380353 | -4.76104575 |
| **A_51_P226429** | Rhobtb2 | -0.063455667 | -0.262169914 | 0.135258579 | 8.564492185 | -0.775933677 | 0.466442537 | 0.740380353 | -4.761055934 |
| **A_55_P2003719** | NA | 0.076148191 | -0.162327282 | 0.314623664 | 16.06050264 | 0.775888045 | 0.466467584 | 0.740380353 | -4.761072778 |
| **A_55_P2032267** | Gm10767 | 0.063893582 | -0.136235455 | 0.264022619 | 11.72735579 | 0.775765246 | 0.466534995 | 0.740426548 | -4.761118102 |
| **A_55_P2030761** | NA | 0.189930705 | -0.405068183 | 0.784929593 | 14.13954339 | 0.775643603 | 0.466601778 | 0.740471738 | -4.761162994 |
| **A_55_P2070769** | Dus1l | 0.078780245 | -0.190092784 | 0.347653275 | 12.6546544 | 0.799974023 | 0.4667411 | 0.740550406 | -4.697334923 |
| **A_51_P461031** | Ostm1 | 0.096973774 | -0.206919357 | 0.400866905 | 9.883998004 | 0.775383587 | 0.466744551 | 0.740550406 | -4.761258936 |
| **A_55_P1973033** | Ltbp1 | -0.10820018 | -0.447307053 | 0.230906693 | 9.036125236 | -0.775308563 | 0.466785752 | 0.740550406 | -4.761286615 |
| **A_55_P2112797** | NA | 0.144654984 | -0.308723846 | 0.598033814 | 9.649505213 | 0.775274241 | 0.466804602 | 0.740550406 | -4.761299276 |
| **A_55_P1978676** | Mospd3 | 0.076140304 | -0.162533222 | 0.31481383 | 10.72917915 | 0.775163912 | 0.466865197 | 0.740585753 | -4.761339975 |
| **A_51_P489138** | Sptb | -0.058723046 | -0.242857899 | 0.125411808 | 7.752846282 | -0.774918018 | 0.467000268 | 0.740739223 | -4.761430665 |
| **A_55_P2055168** | Gm2016 | -0.085928302 | -0.355501382 | 0.183644778 | 9.248801756 | -0.774538302 | 0.467208902 | 0.741009343 | -4.761570672 |
| **A_55_P1966322** | Eif3h | -0.197061961 | -0.815386331 | 0.421262409 | 15.07970863 | -0.774407635 | 0.467280711 | 0.741030397 | -4.761618839 |
| **A_66_P135130** | Tmem161a | 0.222107207 | -0.474897392 | 0.919111805 | 10.3414062 | 0.77430167 | 0.467338952 | 0.741030397 | -4.761657896 |
| **A_55_P1966620** | Rab5a | -0.074873365 | -0.309858764 | 0.160112033 | 9.741902975 | -0.774229414 | 0.467378668 | 0.741030397 | -4.761684526 |
| **A_52_P35048** | Serpinf1 | -0.158823827 | -0.657317185 | 0.339669531 | 10.54905937 | -0.774175581 | 0.467408259 | 0.741030397 | -4.761704365 |
| **A_66_P135192** | Msl2 | 0.052579894 | -0.112452481 | 0.217612269 | 9.326755458 | 0.774165381 | 0.467413865 | 0.741030397 | -4.761708124 |
| **A_55_P1971674** | Golga7b | 0.237419016 | -0.508010206 | 0.982848238 | 10.88116974 | 0.773913237 | 0.467552483 | 0.741189367 | -4.761801033 |
| **A_55_P1971649** | Ttc19 | 0.115171089 | -0.246494476 | 0.476836654 | 9.256071094 | 0.773783564 | 0.467623784 | 0.741241604 | -4.761848806 |
| **A_55_P1969166** | Fxr2 | 0.054075514 | -0.115767388 | 0.223918415 | 11.73262206 | 0.77363559 | 0.467705155 | 0.741309795 | -4.761903314 |
| **A_55_P2161170** | Srp68 | 0.088151999 | -0.188745577 | 0.365049575 | 11.78631807 | 0.773563911 | 0.467744575 | 0.741311487 | -4.761929714 |
| **A_52_P48767** | Rassf8 | -0.056641732 | -0.234602002 | 0.121318538 | 7.917360659 | -0.77338663 | 0.467842082 | 0.741405231 | -4.761995003 |
| **A_51_P501773** | Slc35f3 | -0.050142347 | -0.207697447 | 0.107412753 | 7.706410793 | -0.773313121 | 0.467882517 | 0.741408524 | -4.762022072 |

| | | | | | | | | | |
|---|---|---|---|---|---|---|---|---|---|
| **A_52_P446457** | Ly6g6f | -0.126378489 | -0.523623181 | 0.270866202 | 8.553133147 | -0.773033378 | 0.468036417 | 0.741591598 | -4.762125067 |
| **A_55_P2056419** | Coq6 | 0.066696002 | -0.143154659 | 0.276546664 | 12.18688379 | 0.772276198 | 0.468453155 | 0.742159434 | -4.762403706 |
| **A_55_P2055442** | Pafah2 | -0.086029464 | -0.356722186 | 0.184663258 | 9.553552579 | -0.772242724 | 0.468471584 | 0.742159434 | -4.76241602 |
| **A_52_P105840** | Fnbp1 | -0.051864376 | -0.215098999 | 0.111370247 | 8.113906943 | -0.772040473 | 0.468582947 | 0.742275018 | -4.762490412 |
| **A_55_P2050268** | Nfkbid | -0.086387573 | -0.358431839 | 0.185656693 | 8.636186331 | -0.771604736 | 0.468822932 | 0.742571418 | -4.762650636 |
| **A_52_P253317** | 4930481A15Rik | 0.150195172 | -0.322813125 | 0.623203469 | 9.566091103 | 0.77156124 | 0.468846892 | 0.742571418 | -4.762666626 |
| **A_51_P205820** | Klf11 | -0.06494888 | -0.26954616 | 0.139648399 | 8.266673938 | -0.771356249 | 0.468959826 | 0.742585788 | -4.762741977 |
| **A_55_P2052185** | Lamtor1 | 0.093214101 | -0.200443959 | 0.386872162 | 12.33024659 | 0.771299199 | 0.468991259 | 0.742585788 | -4.762762945 |
| **A_55_P2080895** | Rab7 | -0.092378411 | -0.383409872 | 0.198653051 | 9.832837655 | -0.771282954 | 0.46900021 | 0.742585788 | -4.762768915 |
| **A_55_P1992655** | Hdac6 | 0.080300533 | -0.172686068 | 0.333287135 | 10.72669272 | 0.771265843 | 0.469009639 | 0.742585788 | -4.762775204 |
| **A_55_P2117711** | Rnf185 | -0.0789612 | -0.327767767 | 0.169845367 | 10.63455492 | -0.771143299 | 0.469077164 | 0.742631871 | -4.762820238 |
| **A_55_P2157423** | NA | 0.098594561 | -0.212193728 | 0.40938285 | 8.482654432 | 0.770852883 | 0.469237221 | 0.74266895 | -4.762926943 |
| **A_55_P2024358** | Rpl34 | -0.080923425 | -0.336012111 | 0.174165262 | 16.05744408 | -0.770843555 | 0.469242362 | 0.74266895 | -4.76293037 |
| **A_55_P2083093** | Haus7 | 0.114299703 | -0.2460016 | 0.474601006 | 9.12989061 | 0.770836831 | 0.469246068 | 0.74266895 | -4.76293284 |
| **A_51_P417747** | Kap | -0.049543426 | -0.205719734 | 0.106632882 | 7.6992856 | -0.770821942 | 0.469254275 | 0.74266895 | -4.76293831 |
| **A_51_P425632** | Mad2l1bp | 0.07367883 | -0.158601043 | 0.305958703 | 12.79275231 | 0.770751416 | 0.469293151 | 0.742669668 | -4.762964218 |
| **A_52_P484956** | Nnat | -0.426671805 | -1.772009435 | 0.918665824 | 8.597210038 | -0.770629881 | 0.469360149 | 0.742707716 | -4.763008859 |
| **A_55_P1992299** | Cep295 | 0.118385812 | -0.25492593 | 0.491697554 | 8.465427164 | 0.770568402 | 0.469394043 | 0.742707716 | -4.76303144 |
| **A_55_P2177105** | NA | -0.091435956 | -0.379861479 | 0.196989567 | 12.071729 | -0.770311727 | 0.469535568 | 0.742834085 | -4.763125698 |
| **A_55_P2038831** | NA | -0.094711038 | -0.393478172 | 0.204056095 | 15.3690627 | -0.770284158 | 0.469550771 | 0.742834085 | -4.763135821 |
| **A_52_P175242** | Irs1 | 0.117860053 | -0.254024039 | 0.489744145 | 10.44646448 | 0.770091304 | 0.469657129 | 0.742941537 | -4.763206625 |
| **A_55_P2081488** | Pglyrp1 | -0.199922326 | -0.830945348 | 0.431100695 | 11.39030892 | -0.769837889 | 0.46979691 | 0.742995773 | -4.763299645 |
| **A_52_P161297** | Tcea3 | -0.145633546 | -0.605347508 | 0.314080415 | 10.9073886 | -0.769762802 | 0.469838334 | 0.742995773 | -4.763327202 |
| **A_55_P2104572** | Tmem268 | -0.126904469 | -0.527498221 | 0.273689283 | 10.33884737 | -0.769761053 | 0.469839298 | 0.742995773 | -4.763327844 |
| **A_55_P2102335** | Zfp423 | 0.16175206 | -0.348850611 | 0.672354731 | 8.123067775 | 0.769750406 | 0.469845172 | 0.742995773 | -4.763331751 |
| **A_55_P2028214** | NA | 0.120588697 | -0.260197084 | 0.501374477 | 8.575259648 | 0.769500787 | 0.469982898 | 0.743040517 | -4.763423348 |
| **A_52_P356204** | Nostrin | -0.050737477 | -0.210956929 | 0.109481975 | 8.689815164 | -0.769479072 | 0.469994881 | 0.743040517 | -4.763431315 |
| **A_66_P101261** | Gm3367 | 0.05127939 | -0.110665189 | 0.21322397 | 7.669063001 | 0.769413186 | 0.470031239 | 0.743040517 | -4.763455488 |
| **A_51_P220260** | Phf1 | -0.075579648 | -0.314317856 | 0.16315856 | 8.658502856 | -0.769247556 | 0.470122646 | 0.743040517 | -4.763516248 |
| **A_55_P1991530** | Cep126 | -0.099069669 | -0.412014056 | 0.213874718 | 7.748551232 | -0.769230924 | 0.470131826 | 0.743040517 | -4.763522349 |
| **A_55_P2026400** | Capns1 | -0.115397286 | -0.479940223 | 0.24914565 | 11.92178807 | -0.769183743 | 0.470157867 | 0.743040517 | -4.763539655 |
| **A_52_P112110** | Tmem82 | 0.103085932 | -0.222601243 | 0.428773106 | 8.586002726 | 0.769098443 | 0.470204949 | 0.743040517 | -4.763570941 |
| **A_51_P162671** | Folr2 | -0.10942422 | -0.455145952 | 0.236297513 | 9.276021922 | -0.769077254 | 0.470216645 | 0.743040517 | -4.763578712 |
| **A_51_P218518** | Fiz1 | 0.072777717 | -0.157162274 | 0.302717708 | 8.576341322 | 0.769072186 | 0.470219443 | 0.743040517 | -4.763580571 |
| **A_51_P279183** | Tkfc | -0.259885481 | -1.081121485 | 0.561350524 | 11.99845759 | -0.768948448 | 0.470287749 | 0.743085783 | -4.76362595 |
| **A_52_P417825** | Osgin2 | 0.062150727 | -0.134262289 | 0.258563743 | 8.532558427 | 0.768881015 | 0.470324977 | 0.743085783 | -4.763650677 |
| **A_55_P2151797** | Dis3l2 | -0.058091648 | -0.241700437 | 0.125517142 | 11.36611249 | -0.76878237 | 0.470379438 | 0.743111088 | -4.763686847 |
| **A_55_P2169888** | Prag1 | -0.147933942 | -0.615724913 | 0.319857029 | 8.922762138 | -0.768420929 | 0.470579029 | 0.743365646 | -4.763819348 |
| **A_51_P371001** | Tm4sf4 | -0.223941604 | -0.932246976 | 0.484363769 | 12.22403774 | -0.768240922 | 0.470678452 | 0.743383346 | -4.76388532 |
| **A_51_P216605** | Hbp1 | -0.065349474 | -0.272048152 | 0.141349205 | 8.26861201 | -0.768223493 | 0.470688079 | 0.743383346 | -4.763891706 |
| **A_55_P1956171** | NA | -0.122260383 | -0.508982757 | 0.264461992 | 14.57310186 | -0.768191753 | 0.470705612 | 0.743383346 | -4.763903338 |
| **A_51_P110301** | C3 | -0.186926185 | -0.778322178 | 0.404469808 | 17.49700191 | -0.768024287 | 0.470798125 | 0.743468706 | -4.750496966 |
| **A_52_P584188** | Frat1 | 0.123254343 | -0.266794581 | 0.513303267 | 9.881582533 | 0.767832224 | 0.470904241 | 0.743575532 | -4.764035064 |
| **A_51_P499854** | Ghr | -0.176431399 | -0.734854354 | 0.381991557 | 11.00789314 | -0.767707535 | 0.470973142 | 0.743580904 | -4.764080737 |
| **A_55_P2148873** | Cat | 0.109526829 | -0.237145847 | 0.456199506 | 11.53915245 | 0.767686832 | 0.470984583 | 0.743580904 | -4.76408832 |
| **A_55_P2073452** | Glp2r | 0.176515171 | -0.382300659 | 0.735331001 | 8.627930236 | 0.767532062 | 0.471070117 | 0.743592685 | -4.764145004 |
| **A_51_P116487** | Lsm4 | 0.129722538 | -0.280981277 | 0.540426353 | 13.65860716 | 0.767484986 | 0.471096136 | 0.743592685 | -4.764162244 |
| **A_55_P2090782** | NA | 0.19309906 | -0.41827782 | 0.804475941 | 13.95895662 | 0.767457505 | 0.471111326 | 0.743592685 | -4.764172307 |
| **A_55_P1987409** | Nbeal1 | 0.087623615 | -0.189826842 | 0.365074073 | 8.2789948 | 0.767394906 | 0.471145926 | 0.743592685 | -4.76419523 |
| **A_55_P1959818** | Tmem40 | 0.174383013 | -0.377926813 | 0.72669284 | 12.12785634 | 0.767192938 | 0.471257573 | 0.743690241 | -4.764269177 |

| | | | | | | | | | |
|---|---|---|---|---|---|---|---|---|---|
| **A_51_P501897** | Pitx2 | 0.05384047 | -0.1167053 | 0.224386241 | 7.722269848 | 0.767098405 | 0.471309836 | 0.743690241 | -4.764303784 |
| **A_55_P2454784** | Ifnar1 | 0.060678008 | -0.131550883 | 0.252906899 | 8.341929732 | 0.767000843 | 0.471363779 | 0.743690241 | -4.764339496 |
| **A_55_P2067608** | Clmp | -0.065480796 | -0.27293969 | 0.141978098 | 7.942710619 | -0.766946519 | 0.471393817 | 0.743690241 | -4.76435938 |
| **A_55_P2024689** | Rsbn1 | 0.081830995 | -0.177433108 | 0.341095097 | 8.947262431 | 0.766935129 | 0.471400115 | 0.743690241 | -4.764363549 |
| **A_55_P2360501** | NA | 0.057732672 | -0.125204188 | 0.240669531 | 8.228507998 | 0.766838 | 0.471453825 | 0.743714273 | -4.764399097 |
| **A_51_P160713** | Alb | -0.092882768 | -0.387250156 | 0.201484619 | 17.99496801 | -0.766705621 | 0.471527035 | 0.74376906 | -4.764447542 |
| **A_55_P2185203** | Snrnp40 | 0.091822352 | -0.199231851 | 0.382876555 | 9.091283984 | 0.766580427 | 0.471596278 | 0.743817581 | -4.764493351 |
| **A_55_P2148171** | Ncmap | -0.062442528 | -0.260400988 | 0.135515932 | 7.695720627 | -0.766460184 | 0.47166279 | 0.74383495 | -4.764537344 |
| **A_55_P2049752** | Pamr1 | 0.073406334 | -0.159321921 | 0.306134589 | 10.16808827 | 0.76642138 | 0.471684256 | 0.74383495 | -4.76455154 |
| **A_52_P411601** | Ep300 | 0.095496853 | -0.20729705 | 0.398290757 | 8.97862368 | 0.766346406 | 0.471725732 | 0.743839669 | -4.764578967 |
| **A_51_P473229** | Zbtb7b | -0.057248669 | -0.23881108 | 0.124313742 | 8.720150414 | -0.76616559 | 0.47182577 | 0.743922552 | -4.764645104 |
| **A_55_P2057321** | Myh7b | 0.104375635 | -0.226680649 | 0.435431919 | 8.010661317 | 0.766091219 | 0.471866921 | 0.743922552 | -4.764672304 |
| **A_55_P2053491** | Pdia6 | -0.083160296 | -0.346957613 | 0.180637022 | 14.70608435 | -0.766000123 | 0.47191733 | 0.743922552 | -4.764705618 |
| **A_55_P1971897** | Mdk | -0.395569683 | -1.650422035 | 0.859282669 | 10.02526293 | -0.765973173 | 0.471932244 | 0.743922552 | -4.764715473 |
| **A_55_P2184897** | Lbr | 0.176561586 | -0.383667658 | 0.736790829 | 8.833060239 | 0.765796953 | 0.472029769 | 0.743943672 | -4.764779906 |
| **A_55_P2091494** | Tango2 | 0.103745689 | -0.225441122 | 0.4329325 | 11.59160495 | 0.765791993 | 0.472032514 | 0.743943672 | -4.76478172 |
| **A_55_P1993879** | NA | 0.05358247 | -0.116460121 | 0.22362506 | 16.42062454 | 0.765681588 | 0.472093623 | 0.743943672 | -4.764822083 |
| **A_55_P1964302** | Timm8a1 | 0.112700389 | -0.24496824 | 0.470369018 | 9.709748295 | 0.765645536 | 0.472113579 | 0.743943672 | -4.764835262 |
| **A_66_P109595** | Ube4b | -0.100037555 | -0.417556698 | 0.217481589 | 10.62302291 | -0.765554901 | 0.47216375 | 0.743943672 | -4.764868393 |
| **A_55_P1975080** | Ugt2b5 | 0.241643847 | -0.525356798 | 1.008644492 | 11.15342749 | 0.765531738 | 0.472176573 | 0.743943672 | -4.76487686 |
| **A_51_P362738** | Dusp18 | -0.051294102 | -0.214129307 | 0.111541103 | 7.684545421 | -0.76542442 | 0.472235985 | 0.743965334 | -4.764916084 |
| **A_52_P175679** | Gm13023 | 0.241468536 | -0.525139764 | 1.008076835 | 9.678156544 | 0.765367859 | 0.4722673 | 0.743965334 | -4.764936756 |
| **A_66_P140335** | Mbnl3 | -0.059681642 | -0.249177768 | 0.129814483 | 8.272087765 | -0.765285737 | 0.472312769 | 0.743976328 | -4.764966767 |
| **A_55_P2055063** | Mcts1 | 0.088822651 | -0.193287575 | 0.370932877 | 11.25614541 | 0.765046956 | 0.472444995 | 0.744112522 | -4.765054016 |
| **A_66_P116807** | Selenot | 0.094564064 | -0.205803644 | 0.394931772 | 12.16254737 | 0.764990563 | 0.472476226 | 0.744112522 | -4.765074619 |
| **A_66_P110490** | Polr3h | 0.098146249 | -0.213724786 | 0.410017283 | 9.18478702 | 0.764683735 | 0.472646179 | 0.744319537 | -4.765186696 |
| **A_55_P2025687** | Muc4 | -0.163228993 | -0.681988417 | 0.35553043 | 8.285604603 | -0.764565141 | 0.472711879 | 0.744362356 | -4.765230007 |
| **A_55_P2042481** | Gm10653 | 0.09197405 | -0.200401475 | 0.384349575 | 15.85584667 | 0.764376781 | 0.472816243 | 0.744376486 | -4.765298786 |
| **A_52_P504478** | Adprh | -0.094545583 | -0.39513473 | 0.206043563 | 11.44410649 | -0.764277614 | 0.472871194 | 0.744376486 | -4.765334992 |
| **A_55_P1996716** | Cachd1 | -0.129651761 | -0.541883277 | 0.282579756 | 8.826224414 | -0.76422355 | 0.472901155 | 0.744376486 | -4.765354729 |
| **A_55_P2144285** | Nnt | 0.183910721 | -0.400845727 | 0.768667168 | 11.50543334 | 0.764214399 | 0.472906226 | 0.744376486 | -4.76535807 |
| **A_55_P1983284** | Flot2 | 0.17259493 | -0.37619151 | 0.721381371 | 12.16743767 | 0.764201441 | 0.472913407 | 0.744376486 | -4.7653628 |
| **A_51_P473259** | Dpyd | -0.184393284 | -0.770918576 | 0.402132008 | 9.613975322 | -0.763908854 | 0.473075574 | 0.744571107 | -4.765469595 |
| **A_51_P463087** | Cenpm | 0.102298604 | -0.22316854 | 0.427765748 | 8.469057811 | 0.763740365 | 0.473168977 | 0.744600786 | -4.765531081 |
| **A_51_P396570** | Plod2 | -0.086160555 | -0.36028696 | 0.18796585 | 8.417182654 | -0.763731679 | 0.473173792 | 0.744600786 | -4.76553425 |
| **A_55_P2031133** | Tmem220 | 0.097564868 | -0.212871779 | 0.408001516 | 8.951996085 | 0.763666374 | 0.473209998 | 0.744600786 | -4.765558079 |
| **A_52_P37431** | NA | 0.070081849 | -0.152975043 | 0.293138741 | 14.93639251 | 0.763436803 | 0.47333729 | 0.744682389 | -4.765641833 |
| **A_55_P1961983** | Uqcr11 | -0.071466025 | -0.298940383 | 0.156008333 | 15.27036736 | -0.763396856 | 0.473359442 | 0.744682389 | -4.765656404 |
| **A_52_P678816** | Ctsm | -0.06666096 | -0.278849952 | 0.145528031 | 7.762385307 | -0.763364404 | 0.473377439 | 0.744682389 | -4.765668242 |
| **A_66_P124806** | Tlr4 | -0.082675712 | -0.345871962 | 0.180520537 | 8.353133624 | -0.763275702 | 0.473426631 | 0.744687871 | -4.765700596 |
| **A_55_P2175272** | NA | -0.076296716 | -0.319203563 | 0.166610131 | 14.55398822 | -0.763219184 | 0.473457977 | 0.744687871 | -4.765721209 |
| **A_51_P155458** | Dok7 | -0.077696345 | -0.325104991 | 0.169712301 | 7.77259147 | -0.76307795 | 0.473536314 | 0.744750481 | -4.765772715 |
| **A_51_P480881** | Thap11 | 0.06880713 | -0.150414276 | 0.288028536 | 11.40853718 | 0.762664748 | 0.473765551 | 0.74505039 | -4.765923364 |
| **A_55_P2139402** | Amy2a5 | -0.219059855 | -0.917135701 | 0.479015991 | 9.431974991 | -0.76250619 | 0.473853537 | 0.745064293 | -4.765981156 |
| **A_52_P614472** | Rpusd1 | -0.080004799 | -0.334966535 | 0.174956937 | 8.436397282 | -0.762472563 | 0.473872199 | 0.745064293 | -4.765993412 |
| **A_55_P1993390** | NA | 0.069435512 | -0.151853022 | 0.290724045 | 15.06913317 | 0.762440431 | 0.473890031 | 0.745064293 | -4.766005122 |
| **A_52_P598912** | Erc1 | 0.064411724 | -0.141064105 | 0.269887554 | 8.127447495 | 0.761705994 | 0.474297749 | 0.745623445 | -4.766272685 |
| **A_51_P421538** | Slc9b2 | -0.053508902 | -0.224214438 | 0.117196634 | 7.770512698 | -0.761660838 | 0.474322825 | 0.745623445 | -4.766289129 |
| **A_55_P2124026** | Synj2 | 0.267148756 | -0.585310208 | 1.11960772 | 10.77957266 | 0.761487648 | 0.474419009 | 0.745713997 | -4.766352194 |
| **A_55_P2044369** | Hs2st1 | 0.314830408 | -0.689974743 | 1.319635558 | 11.5634401 | 0.761338879 | 0.474501641 | 0.745779162 | -4.766406357 |

| | | | | | | | | | |
|---|---|---|---|---|---|---|---|---|---|
| **A_51_P405167** | Maf | -0.078768239 | -0.330184452 | 0.172647974 | 10.19698539 | -0.761274084 | 0.474537634 | 0.745779162 | -4.766429945 |
| **A_51_P267544** | Frg1 | -0.113786709 | -0.477167042 | 0.249593623 | 10.58602692 | -0.760874989 | 0.474759367 | 0.74601467 | -4.766575199 |
| **A_66_P100565** | Ccdc107 | -0.122831769 | -0.515114293 | 0.269450755 | 13.00273008 | -0.760842799 | 0.474777255 | 0.74601467 | -4.766586913 |
| **A_51_P130079** | Lrmp | -0.146389291 | -0.613968475 | 0.321189894 | 9.802944577 | -0.760741881 | 0.474833337 | 0.74601467 | -4.766623633 |
| **A_51_P427624** | Pgc | -0.068962849 | -0.289240189 | 0.151314491 | 7.713991519 | -0.760726528 | 0.474841869 | 0.74601467 | -4.766629219 |
| **A_55_P2135556** | Rpgrip1 | 0.154723601 | -0.33965221 | 0.649099412 | 11.74118837 | 0.760470797 | 0.474984005 | 0.746117617 | -4.766722251 |
| **A_51_P168524** | NA | 0.069047032 | -0.151573674 | 0.289667738 | 8.949480845 | 0.760469733 | 0.474984597 | 0.746117617 | -4.766722638 |
| **A_51_P204080** | Hk2 | 0.071965406 | -0.158062091 | 0.301992903 | 9.572967112 | 0.760198858 | 0.475135183 | 0.746236183 | -4.766821155 |
| **A_55_P2041988** | NA | -0.070564243 | -0.296125949 | 0.154997463 | 7.785681194 | -0.760155606 | 0.475159231 | 0.746236183 | -4.766836883 |
| **A_55_P1966444** | NA | -0.063582423 | -0.266834453 | 0.139669606 | 15.10577334 | -0.760125628 | 0.475175898 | 0.746236183 | -4.766847784 |
| **A_55_P2171158** | Hmgn1 | -0.105937232 | -0.444651682 | 0.232777219 | 13.91885485 | -0.759972863 | 0.475260844 | 0.746308949 | -4.76690333 |
| **A_51_P266958** | Nr1i2 | 0.164296724 | -0.361147591 | 0.689741038 | 9.243817691 | 0.759775692 | 0.475370497 | 0.746325062 | -4.766975009 |
| **A_55_P2143436** | NA | 0.059523097 | -0.130864637 | 0.24991083 | 7.869526344 | 0.759678336 | 0.475424646 | 0.746325062 | -4.767010396 |
| **A_51_P509669** | Uap1l1 | -0.114080633 | -0.479033192 | 0.250871927 | 9.298561828 | -0.759554072 | 0.475493767 | 0.746325062 | -4.767055559 |
| **A_55_P2109263** | Ppp1r14b | -0.060252304 | -0.253012561 | 0.132507953 | 12.68472122 | -0.759520237 | 0.475512589 | 0.746325062 | -4.767067855 |
| **A_55_P2091147** | H4c12 | -0.098497964 | -0.413624253 | 0.216628326 | 13.64283614 | -0.759496549 | 0.475525767 | 0.746325062 | -4.767076464 |
| **A_55_P2105542** | Pde2a | -0.147363199 | -0.618841615 | 0.324115216 | 12.21529661 | -0.759469637 | 0.475540738 | 0.746325062 | -4.767086244 |
| **A_55_P2183668** | NA | 0.380991737 | -0.83804467 | 1.600028144 | 14.04205043 | 0.759420068 | 0.475568314 | 0.746325062 | -4.767104256 |
| **A_55_P2039541** | Arhgap15 | 0.139081031 | -0.305947903 | 0.584109964 | 9.330119048 | 0.759386447 | 0.475587019 | 0.746325062 | -4.767116473 |
| **A_52_P282741** | Sdc3 | -0.116904833 | -0.491002747 | 0.257193082 | 9.960596932 | -0.759329665 | 0.47561861 | 0.746325062 | -4.767137105 |
| **A_55_P1987146** | Adgrf4 | 0.081347816 | -0.17902544 | 0.341721072 | 7.766057807 | 0.759158946 | 0.475713601 | 0.746413523 | -4.76719913 |
| **A_51_P472901** | Slc3a2 | 0.106763341 | -0.235019671 | 0.448546353 | 12.21827305 | 0.759022888 | 0.475789314 | 0.746419641 | -4.767248555 |
| **A_55_P2164841** | NA | -0.130060107 | -0.546478651 | 0.286358438 | 8.879820416 | -0.758922159 | 0.475845374 | 0.746419641 | -4.767285141 |
| **A_52_P232648** | Slc38a1 | 0.054914984 | -0.120909827 | 0.230739794 | 7.786632745 | 0.758916316 | 0.475848626 | 0.746419641 | -4.767287264 |
| **A_55_P2022773** | Glcci1 | 0.081752796 | -0.180029199 | 0.343534792 | 9.12283122 | 0.75883269 | 0.47589517 | 0.746419641 | -4.767317635 |
| **A_55_P2159573** | Reep6 | 0.098892092 | -0.217783484 | 0.415567668 | 8.552960497 | 0.758804999 | 0.475910583 | 0.746419641 | -4.767327692 |
| **A_55_P2131288** | Ank2 | 0.204947638 | -0.45174648 | 0.861641756 | 9.886467089 | 0.758338282 | 0.476170411 | 0.746766562 | -4.767497147 |
| **A_55_P2168663** | Atp5k | 0.063641886 | -0.140351392 | 0.267635164 | 14.8663351 | 0.758071858 | 0.476318778 | 0.746904943 | -4.767593845 |
| **A_51_P517051** | Castor1 | 0.080353326 | -0.177216104 | 0.337922755 | 7.745811255 | 0.758041057 | 0.476335932 | 0.746904943 | -4.767605023 |
| **A_55_P1955528** | NA | 0.06535311 | -0.144167523 | 0.274873742 | 9.286583668 | 0.757918747 | 0.476404057 | 0.746905693 | -4.767649405 |
| **A_51_P252784** | Fam216a | -0.123922566 | -0.521224657 | 0.273379525 | 10.41444349 | -0.757901445 | 0.476413694 | 0.746905693 | -4.767655683 |
| **A_55_P2072656** | Ckmt1 | 0.187901523 | -0.414773788 | 0.790576834 | 10.90225115 | 0.757582771 | 0.476591223 | 0.746961621 | -4.767771294 |
| **A_51_P456870** | Foxj1 | -0.330943646 | -1.392581545 | 0.730694253 | 8.391849906 | -0.757461952 | 0.476658542 | 0.746961621 | -4.767815116 |
| **A_52_P86928** | Cog2 | 0.058283641 | -0.128690208 | 0.24525749 | 8.843078274 | 0.757441322 | 0.476670037 | 0.746961621 | -4.767822598 |
| **A_52_P604515** | Zfp319 | 0.051674133 | -0.114098599 | 0.217446864 | 8.175881537 | 0.757431375 | 0.47667558 | 0.746961621 | -4.767826206 |
| **A_51_P270286** | Dync1li1 | -0.050072154 | -0.2107114 | 0.110567092 | 8.48635765 | -0.757404366 | 0.47669063 | 0.746961621 | -4.767836001 |
| **A_55_P1988970** | Mkks | -0.076418265 | -0.321598426 | 0.168761896 | 9.889432702 | -0.75734722 | 0.476722475 | 0.746961621 | -4.767856725 |
| **A_52_P122696** | Arl2 | -0.124902748 | -0.525683069 | 0.275877572 | 8.777060072 | -0.757266587 | 0.47676741 | 0.746961621 | -4.767885965 |
| **A_55_P1965194** | Fbxl5 | -0.072506745 | -0.305167394 | 0.160153905 | 8.748915743 | -0.757248915 | 0.476777259 | 0.746961621 | -4.767892374 |
| **A_52_P603421** | Ccdc189 | -0.108779676 | -0.457855614 | 0.240296263 | 8.001515512 | -0.75720075 | 0.476804102 | 0.746961621 | -4.767909838 |
| **A_55_P2092501** | Med1 | 0.077037774 | -0.170196407 | 0.324271955 | 11.56148563 | 0.757143852 | 0.476835814 | 0.746961621 | -4.767930469 |
| **A_55_P2003216** | Igsf5 | -0.062544983 | -0.263406102 | 0.138316136 | 8.422909488 | -0.756623435 | 0.477125932 | 0.747355522 | -4.768119111 |
| **A_55_P2163098** | Akr1c18 | -0.113007209 | -0.475961946 | 0.249947529 | 8.20929638 | -0.756548662 | 0.477167626 | 0.747360266 | -4.768146207 |
| **A_51_P265406** | Ric8a | -0.089449173 | -0.376769636 | 0.19787129 | 11.53241074 | -0.756472181 | 0.477210275 | 0.747366505 | -4.76817392 |
| **A_55_P2054456** | Adprm | -0.075747946 | -0.319095518 | 0.167599625 | 10.68819794 | -0.756357356 | 0.477274311 | 0.747406235 | -4.768215524 |
| **A_55_P2152248** | Cpe | -0.052148074 | -0.219721201 | 0.115425053 | 7.812644129 | -0.756165896 | 0.477381098 | 0.747426864 | -4.768284882 |
| **A_55_P2091616** | Krt222 | 0.16354224 | -0.362036713 | 0.689121193 | 10.80501506 | 0.75609291 | 0.47742181 | 0.747426864 | -4.768311319 |
| **A_51_P193173** | Slc25a25 | -0.061497624 | -0.259133768 | 0.13613852 | 9.73430403 | -0.756092892 | 0.47742182 | 0.747426864 | -4.768311326 |
| **A_55_P2046374** | NA | 0.08598422 | -0.190358313 | 0.362326754 | 8.461008976 | 0.756056432 | 0.477442159 | 0.747426864 | -4.768324531 |
| **A_55_P2230506** | 4931402H11Rik | 0.218501046 | -0.483919182 | 0.920921275 | 14.49744814 | 0.755857105 | 0.477553361 | 0.747540405 | -4.768396719 |

| | | | | | | | | | |
|---|---|---|---|---|---|---|---|---|---|
| **A_55_P1995497** | Atad3a | 0.118610449 | -0.262758557 | 0.499979455 | 8.993237625 | 0.755719701 | 0.477630027 | 0.74759987 | -4.768446472 |
| **A_55_P1970866** | Rplp0 | 0.115440305 | -0.255918512 | 0.486799122 | 16.49904619 | 0.755347706 | 0.47783763 | 0.747864255 | -4.768581137 |
| **A_55_P2097244** | Bptf | -0.069643131 | -0.293701145 | 0.154414883 | 11.63796945 | -0.755267842 | 0.477882209 | 0.747872796 | -4.768610042 |
| **A_55_P2109122** | H2bc4 | 0.176713924 | -0.391867127 | 0.745294975 | 15.06969018 | 0.755199298 | 0.47792047 | 0.747872796 | -4.768634848 |
| **A_55_P2109033** | Hmgcs2 | -0.261255297 | -1.102102567 | 0.579591974 | 12.49830582 | -0.754972572 | 0.478047047 | 0.74793558 | -4.768716888 |
| **A_55_P2100120** | Nme1 | 0.082979957 | -0.184120795 | 0.350080709 | 12.86322364 | 0.754885848 | 0.478095469 | 0.74793558 | -4.768748264 |
| **A_55_P1959409** | Deup1 | -0.068820151 | -0.290349503 | 0.152709202 | 7.748489052 | -0.754861944 | 0.478108816 | 0.74793558 | -4.768756912 |
| **A_55_P2322709** | 4633401B06Rik | -0.050342141 | -0.212401345 | 0.111717062 | 8.105648204 | -0.754816132 | 0.478134398 | 0.74793558 | -4.768773485 |
| **A_52_P57308** | Cand1 | 0.054877766 | -0.121795446 | 0.231550979 | 12.18509314 | 0.75476006 | 0.478165709 | 0.74793558 | -4.768793769 |
| **A_66_P102719** | Spop | 0.061409873 | -0.136305462 | 0.259125208 | 11.66088933 | 0.754711615 | 0.478192762 | 0.74793558 | -4.768811292 |
| **A_52_P418734** | Rnf220 | -0.070057795 | -0.29567777 | 0.155562181 | 12.20079962 | -0.754504967 | 0.478308174 | 0.747990251 | -4.768886033 |
| **A_66_P130911** | Proser2 | -0.080379996 | -0.339243495 | 0.178483504 | 8.197051768 | -0.754501921 | 0.478309875 | 0.747990251 | -4.768887134 |
| **A_55_P2103115** | Tor1aip2 | 0.089945906 | -0.199747901 | 0.379639712 | 11.42273081 | 0.754441167 | 0.478343809 | 0.747990251 | -4.768909105 |
| **A_52_P594756** | Asb4 | 0.126108 | -0.28013037 | 0.532346369 | 9.199878701 | 0.75430135 | 0.478421911 | 0.747995801 | -4.768959662 |
| **A_52_P243054** | Nufip2 | 0.059593778 | -0.132396964 | 0.251584521 | 8.630481131 | 0.754230034 | 0.478461752 | 0.747995801 | -4.768985447 |
| **A_55_P2137941** | Fxyd2 | -0.125759861 | -0.530916695 | 0.279396973 | 8.434396135 | -0.754226988 | 0.478463453 | 0.747995801 | -4.768986548 |
| **A_55_P2117445** | NA | 0.071712044 | -0.159345688 | 0.302769776 | 13.06505419 | 0.754144873 | 0.478509329 | 0.747999037 | -4.769016235 |
| **A_52_P558609** | Clec16a | 0.08877349 | -0.197279388 | 0.374826367 | 8.337148946 | 0.754084752 | 0.47854292 | 0.747999037 | -4.769037969 |
| **A_66_P107703** | Smox | 0.140315443 | -0.31195053 | 0.592581416 | 10.05471033 | 0.753867024 | 0.478664581 | 0.748084999 | -4.769116668 |
| **A_55_P2034893** | Socs7 | -0.078880686 | -0.333160804 | 0.175399432 | 10.43553729 | -0.75377454 | 0.478716266 | 0.748084999 | -4.769150092 |
| **A_55_P2006662** | Zfp967 | 0.065631161 | -0.145939447 | 0.277201769 | 8.186611204 | 0.753768427 | 0.478719682 | 0.748084999 | -4.769152301 |
| **A_52_P399095** | Mettl24 | -0.06134649 | -0.259120417 | 0.136427436 | 7.771973205 | -0.753709302 | 0.478752726 | 0.748084999 | -4.769173667 |
| **A_52_P493519** | Igll1 | -0.089921499 | -0.379909968 | 0.200066971 | 9.713615826 | -0.753470053 | 0.478886454 | 0.748229664 | -4.769260112 |
| **A_55_P2176345** | Psma5 | 0.070021579 | -0.15581095 | 0.295854108 | 13.11607016 | 0.753405165 | 0.478922728 | 0.748229664 | -4.769283554 |
| **A_55_P2179027** | Gem | -0.067641914 | -0.285827433 | 0.150543604 | 7.80473817 | -0.753309002 | 0.478976488 | 0.748247837 | -4.769318291 |
| **A_52_P59228** | Zswim3 | 0.056290933 | -0.125312292 | 0.237894159 | 8.175990532 | 0.753178787 | 0.479049291 | 0.748247837 | -4.769365324 |
| **A_51_P401683** | Tm6sf1 | 0.107488863 | -0.23930281 | 0.454280536 | 9.458208302 | 0.753143961 | 0.479068764 | 0.748247837 | -4.769377902 |
| **A_55_P1980501** | Znhit1 | -0.08628864 | -0.364702813 | 0.192125533 | 10.72987973 | -0.753087556 | 0.479100304 | 0.748247837 | -4.769398273 |
| **A_51_P386810** | Gmppb | -0.082475637 | -0.348609688 | 0.183658413 | 10.79525056 | -0.75302332 | 0.479136224 | 0.748247837 | -4.769421471 |
| **A_55_P2126850** | Nudt11 | -0.106558332 | -0.450427733 | 0.237311069 | 7.800897625 | -0.752968953 | 0.479166627 | 0.748247837 | -4.769441103 |
| **A_55_P2057916** | Pex5 | 0.121232898 | -0.270052998 | 0.512518794 | 10.37944217 | 0.752851783 | 0.479232155 | 0.74828971 | -4.76948341 |
| **A_55_P2033876** | NA | -0.127230356 | -0.537962318 | 0.283501606 | 13.21443915 | -0.752688763 | 0.479323334 | 0.748371626 | -4.769542265 |
| **A_51_P172054** | Gas6 | -0.314887343 | -1.331666074 | 0.701891387 | 11.01343718 | -0.75250942 | 0.479423657 | 0.748467803 | -4.769607002 |
| **A_51_P154469** | Ccndbp1 | 0.089159166 | -0.198805816 | 0.377124149 | 12.31124059 | 0.752331956 | 0.479522944 | 0.748562347 | -4.769671049 |
| **A_55_P2048855** | Sprr2a2 | -0.108404072 | -0.458619535 | 0.241811391 | 8.017997629 | -0.75213096 | 0.479635413 | 0.748677453 | -4.769743576 |
| **A_55_P2032853** | Entpd1 | 0.054174242 | -0.120891619 | 0.229240103 | 8.049668858 | 0.751925079 | 0.479750634 | 0.748771409 | -4.76981785 |
| **A_55_P1992216** | Chchd2 | -0.058893879 | -0.249252831 | 0.131465073 | 16.7424291 | -0.751761436 | 0.47984223 | 0.748771409 | -4.769876876 |
| **A_55_P2401084** | 4930558N01Rik | 0.082759363 | -0.184739143 | 0.350257869 | 8.861018231 | 0.751759575 | 0.479843272 | 0.748771409 | -4.769877547 |
| **A_66_P132295** | Tfpi | -0.071354417 | -0.301993602 | 0.159284768 | 10.24373319 | -0.751745707 | 0.479851035 | 0.748771409 | -4.769882549 |
| **A_55_P2112510** | NA | -0.141463989 | -0.598760218 | 0.315832241 | 15.02212052 | -0.751677357 | 0.479889297 | 0.748771409 | -4.7699072 |
| **A_66_P114782** | Nr2f1 | -0.165582544 | -0.70094394 | 0.369778852 | 8.727364316 | -0.751537555 | 0.479967563 | 0.74883308 | -4.769957614 |
| **A_55_P2005343** | NA | 0.066012054 | -0.147461715 | 0.279485824 | 16.76755277 | 0.751383959 | 0.480053562 | 0.748906804 | -4.770012995 |
| **A_52_P101399** | Tmem130 | 0.057817122 | -0.129174817 | 0.24480906 | 7.80242714 | 0.751305854 | 0.480097298 | 0.748914588 | -4.770041154 |
| **A_55_P1966883** | Rnaseh2a | 0.148093708 | -0.331086775 | 0.62727419 | 10.94831324 | 0.75096669 | 0.480287246 | 0.749099205 | -4.770163405 |
| **A_51_P413059** | Mcpt8 | 0.235279006 | -0.526013924 | 0.996571937 | 8.914575511 | 0.750956135 | 0.480293158 | 0.749099205 | -4.770167208 |
| **A_51_P449911** | NA | 0.059586698 | -0.133250604 | 0.252424 | 11.83155633 | 0.750829729 | 0.480363965 | 0.749149192 | -4.77021276 |
| **A_55_P2360427** | BC005561 | 0.054791544 | -0.122544749 | 0.232127837 | 8.394856163 | 0.7507565 | 0.480404988 | 0.749152724 | -4.770239146 |
| **A_55_P2071858** | Mgmt | -0.073474773 | -0.311409032 | 0.164459486 | 10.80829156 | -0.750350997 | 0.480632196 | 0.749379002 | -4.770385225 |
| **A_52_P380263** | Podxl | -0.077914298 | -0.330236696 | 0.174408099 | 8.043912653 | -0.750316591 | 0.480651477 | 0.749379002 | -4.770397617 |
| **A_55_P2000439** | Ptprz1 | -0.073129404 | -0.309964517 | 0.163705708 | 7.768167335 | -0.750289961 | 0.480666401 | 0.749379002 | -4.770407208 |

| | | | | | | | | | |
|---|---|---|---|---|---|---|---|---|---|
| **A_52_P384574** | Stard4 | 0.078652075 | -0.17619047 | 0.33349462 | 11.64151701 | 0.749931233 | 0.480867471 | 0.749609164 | -4.770536379 |
| **A_52_P367060** | Atxn2l | -0.071315721 | -0.302408415 | 0.159776973 | 12.31731637 | -0.749863566 | 0.480905406 | 0.749609164 | -4.77056074 |
| **A_55_P2109128** | H2bc6 | 0.05845894 | -0.130996509 | 0.247914389 | 8.178097194 | 0.749768212 | 0.480958865 | 0.749609164 | -4.770595065 |
| **A_55_P2133806** | Tnrc6c | 0.058800692 | -0.131766984 | 0.249368368 | 9.477727177 | 0.749749854 | 0.480969158 | 0.749609164 | -4.770601673 |
| **A_52_P31543** | Btg2 | 0.081956859 | -0.183773906 | 0.347687624 | 11.39602498 | 0.749422379 | 0.481152787 | 0.749834896 | -4.77071953 |
| **A_55_P2155504** | Lck | 0.085775736 | -0.192381257 | 0.363932729 | 9.692581478 | 0.749303325 | 0.481219557 | 0.749846038 | -4.770762368 |
| **A_55_P2055708** | Vmn1r48 | 0.177893469 | -0.399011261 | 0.7547982 | 8.894524811 | 0.749271292 | 0.481237524 | 0.749846038 | -4.770773893 |
| **A_52_P589568** | Foxo6 | -0.131561759 | -0.558271108 | 0.295147591 | 8.14840283 | -0.749170556 | 0.481294027 | 0.749849715 | -4.770810134 |
| **A_55_P2168383** | Myo18b | -0.094756297 | -0.402128983 | 0.212616388 | 7.729758549 | -0.749076202 | 0.481346955 | 0.749849715 | -4.770844076 |
| **A_55_P2118810** | NA | -0.13612903 | -0.577717334 | 0.305459274 | 13.09068423 | -0.749059605 | 0.481356266 | 0.749849715 | -4.770850046 |
| **A_55_P2050298** | NA | -0.128348206 | -0.544735324 | 0.288038912 | 14.69638411 | -0.748989458 | 0.481395618 | 0.749850584 | -4.770875278 |
| **A_55_P1974347** | Fam126b | 0.124304973 | -0.279169232 | 0.527779177 | 8.997939445 | 0.748610506 | 0.481608246 | 0.750109551 | -4.771011555 |
| **A_51_P471779** | Phf5a | 0.07250657 | -0.162855746 | 0.307868887 | 12.50290884 | 0.748554837 | 0.481639487 | 0.750109551 | -4.77103157 |
| **A_55_P2115127** | Mphosph10 | 0.125145106 | -0.281134424 | 0.531424637 | 8.881871449 | 0.748466074 | 0.481689303 | 0.750126695 | -4.771063482 |
| **A_55_P1994608** | Itsn1 | 0.099006372 | -0.222466993 | 0.420479738 | 9.633746725 | 0.748344102 | 0.481757762 | 0.750172866 | -4.771107328 |
| **A_55_P2010292** | Serpina1e | -0.209088127 | -0.888149141 | 0.469972887 | 17.50893626 | -0.748175984 | 0.481852133 | 0.750225961 | -4.771167753 |
| **A_55_P2180301** | Clcn5 | 0.109455502 | -0.246040702 | 0.464951706 | 8.651471645 | 0.748145072 | 0.481869486 | 0.750225961 | -4.771178863 |
| **A_55_P2118183** | Tnks2 | -0.059267342 | -0.251844524 | 0.133309839 | 8.297779439 | -0.747814382 | 0.482055156 | 0.750454584 | -4.771297687 |
| **A_51_P128667** | Lynx1 | -0.096584631 | -0.410447301 | 0.217278039 | 8.123657749 | -0.74774161 | 0.482096021 | 0.750457759 | -4.771323831 |
| **A_55_P1971054** | Paqr9 | -0.131288819 | -0.558022405 | 0.295444767 | 8.605204527 | -0.747573858 | 0.482190232 | 0.750494621 | -4.771384089 |
| **A_51_P244856** | Atp2a1 | -0.048224926 | -0.204975095 | 0.108525244 | 7.777157661 | -0.747561174 | 0.482197356 | 0.750494621 | -4.771388645 |
| **A_51_P423509** | Faap100 | -0.055400747 | -0.235497775 | 0.124696282 | 7.965151806 | -0.747467405 | 0.482250023 | 0.750516159 | -4.771422322 |
| **A_51_P214487** | 2-Sep | -0.083727502 | -0.355968394 | 0.18851339 | 13.22030001 | -0.747305132 | 0.482341177 | 0.750597585 | -4.771480596 |
| **A_55_P1970771** | Gm13141 | -0.109419831 | -0.465276779 | 0.246437117 | 13.21510119 | -0.747143079 | 0.482432218 | 0.750645032 | -4.771538782 |
| **A_55_P1955869** | NA | 0.063987179 | -0.144126633 | 0.272100991 | 12.11210702 | 0.747093998 | 0.482459794 | 0.750645032 | -4.771556403 |
| **A_51_P187171** | Smim12 | -0.0795057 | -0.338123678 | 0.179112277 | 11.67852526 | -0.747003687 | 0.482510538 | 0.750645032 | -4.771588824 |
| **A_51_P443293** | Decr2 | -0.125652014 | -0.534391826 | 0.283087797 | 10.42047924 | -0.746974376 | 0.482527007 | 0.750645032 | -4.771599345 |
| **A_55_P2085974** | Igf1 | -0.224998003 | -0.956998931 | 0.507002926 | 9.570617876 | -0.746878951 | 0.48258063 | 0.750666338 | -4.771633598 |
| **A_51_P188126** | Gca | 0.110496994 | -0.249054495 | 0.470048482 | 8.47402637 | 0.746745406 | 0.48265568 | 0.750666338 | -4.771681528 |
| **A_55_P2115458** | Pot1a | 0.276681629 | -0.623629335 | 1.176992594 | 12.60397603 | 0.74674268 | 0.482657212 | 0.750666338 | -4.771682506 |
| **A_51_P288341** | Hpd | -0.272850146 | -1.160811763 | 0.615111471 | 11.43161613 | -0.746643324 | 0.482713054 | 0.750692784 | -4.771718162 |
| **A_55_P2110875** | 2700046G09Rik | -0.057184006 | -0.243375454 | 0.129007443 | 9.196984437 | -0.74627352 | 0.482920937 | 0.750948128 | -4.77185084 |
| **A_55_P2135023** | NA | 0.143620982 | -0.324059291 | 0.611301255 | 11.77355202 | 0.746194475 | 0.482965379 | 0.750948128 | -4.771879193 |
| **A_55_P2097126** | Cnep1r1 | 0.049625354 | -0.111983144 | 0.211233852 | 9.822628124 | 0.746143932 | 0.482993798 | 0.750948128 | -4.771897322 |
| **A_55_P2068701** | NA | 0.08211641 | -0.185328391 | 0.349561211 | 8.454421843 | 0.746068984 | 0.483035941 | 0.750953246 | -4.771924202 |
| **A_51_P224983** | Cln8 | 0.125708304 | -0.283752616 | 0.535169224 | 11.40432725 | 0.745992907 | 0.483078722 | 0.750959354 | -4.771951486 |
| **A_55_P2097656** | Rps11 | -0.073250482 | -0.311915299 | 0.165414336 | 15.40185753 | -0.745770621 | 0.483203735 | 0.751093285 | -4.772031192 |
| **A_52_P520495** | Vcam1 | 0.117930701 | -0.266406669 | 0.502268072 | 9.401876948 | 0.745585505 | 0.483307862 | 0.75119473 | -4.772097556 |
| **A_51_P250123** | Zfp763 | -0.066487634 | -0.283251136 | 0.150275868 | 8.137524334 | -0.745311648 | 0.483461932 | 0.751324967 | -4.772195711 |
| **A_51_P502150** | Slc9a3r1 | 0.190350786 | -0.430243372 | 0.810944943 | 10.50349523 | 0.745298384 | 0.483469395 | 0.751324967 | -4.772200465 |
| **A_51_P467751** | Hgsnat | -0.154046989 | -0.656369983 | 0.348276005 | 11.40379593 | -0.745166538 | 0.483543583 | 0.751379847 | -4.77224771 |
| **A_55_P1960764** | Rps4x | -0.07136251 | -0.304100054 | 0.161375035 | 15.27986591 | -0.745052463 | 0.483607779 | 0.751419192 | -4.772288583 |
| **A_55_P2095325** | AI182371 | 0.138072346 | -0.312275966 | 0.588420658 | 8.540894078 | 0.744974409 | 0.483651707 | 0.751427043 | -4.772316547 |
| **A_52_P72654** | Slc25a17 | -0.076098871 | -0.324351684 | 0.172153942 | 11.59082259 | -0.744847262 | 0.483723269 | 0.751448333 | -4.772362095 |
| **A_52_P431859** | Aopep | -0.130503332 | -0.556261099 | 0.295254436 | 9.021860217 | -0.744804356 | 0.48374742 | 0.751448333 | -4.772377463 |
| **A_55_P2027077** | Shc2 | -0.052589704 | -0.224174014 | 0.118994606 | 7.798343289 | -0.744742852 | 0.48378204 | 0.751448333 | -4.772399493 |
| **A_51_P493117** | Slc16a9 | -0.153877955 | -0.656011385 | 0.348255475 | 8.463882301 | -0.744629879 | 0.483845637 | 0.751477787 | -4.772439954 |
| **A_51_P309920** | Itga8 | -0.286980209 | -1.223525952 | 0.649565534 | 9.574463492 | -0.744571043 | 0.483878759 | 0.751477787 | -4.772461024 |
| **A_55_P2054663** | Cox4i2 | -0.240402869 | -1.025092625 | 0.544286888 | 8.673028659 | -0.744431689 | 0.483957218 | 0.751539253 | -4.772510924 |
| **A_51_P409452** | Cldn11 | -0.050748041 | -0.2164543 | 0.114958218 | 7.726431452 | -0.744155267 | 0.484112875 | 0.751674854 | -4.772609884 |

| | | | | | | | | | |
|---|---|---|---|---|---|---|---|---|---|
| **A_55_P2112295** | Hmg20b | -0.116849639 | -0.498408832 | 0.264709554 | 12.30948888 | -0.744129707 | 0.484127269 | 0.751674854 | -4.772619033 |
| **A_52_P28651** | Nectin1 | 0.067665556 | -0.153306507 | 0.288637619 | 10.70194504 | 0.744069451 | 0.484161205 | 0.751674854 | -4.772640601 |
| **A_55_P2020378** | Ehd4 | -0.053757782 | -0.229451503 | 0.12193594 | 8.8226442 | -0.743478277 | 0.484494239 | 0.75208867 | -4.772852134 |
| **A_55_P1962094** | NA | 0.052967302 | -0.120147619 | 0.226082222 | 7.978202263 | 0.743458171 | 0.484505568 | 0.75208867 | -4.772859326 |
| **A_55_P1962946** | Syk | -0.164424023 | -0.701942546 | 0.373094501 | 10.40605282 | -0.743284409 | 0.484603486 | 0.752180261 | -4.772921476 |
| **A_55_P1954774** | NA | 0.112915733 | -0.256305174 | 0.48213664 | 11.55871664 | 0.743107016 | 0.484703465 | 0.752201354 | -4.772984914 |
| **A_66_P119421** | Prdx1 | 0.146409317 | -0.332338143 | 0.625156777 | 10.60146916 | 0.743096851 | 0.484709195 | 0.752201354 | -4.772988549 |
| **A_66_P106231** | Nccrp1 | 0.074645651 | -0.169467564 | 0.318758866 | 8.081501128 | 0.743012997 | 0.48475646 | 0.752201354 | -4.773018532 |
| **A_55_P2134356** | Msl3 | 0.067567498 | -0.153406664 | 0.288541659 | 10.94163946 | 0.742984119 | 0.484772739 | 0.752201354 | -4.773028857 |
| **A_55_P1985070** | Robo1 | 0.10703162 | -0.243107535 | 0.457170775 | 10.58508451 | 0.742770434 | 0.484893202 | 0.752327877 | -4.773105248 |
| **A_52_P489729** | Sec61a1 | -0.107200017 | -0.457945684 | 0.243545649 | 14.13527393 | -0.742652644 | 0.484959614 | 0.752351891 | -4.773147351 |
| **A_55_P2259125** | D7Ertd143e | -0.047661873 | -0.203626204 | 0.108302457 | 7.999946809 | -0.742555659 | 0.4850143 | 0.752351891 | -4.773182013 |
| **A_52_P69756** | Mterf4 | -0.055587726 | -0.237492729 | 0.126317278 | 10.26485192 | -0.742535887 | 0.485025449 | 0.752351891 | -4.773189079 |
| **A_55_P2098471** | Rps19 | -0.144786094 | -0.618664891 | 0.329092702 | 15.25580364 | -0.742408208 | 0.485097451 | 0.752360246 | -4.773234706 |
| **A_51_P118223** | Gm1943 | 0.048641604 | -0.11056459 | 0.207847797 | 8.105826333 | 0.74238829 | 0.485108684 | 0.752360246 | -4.770523822 |
| **A_55_P2121985** | Plcb4 | -0.049409174 | -0.211216473 | 0.112398124 | 7.842900022 | -0.741980821 | 0.485338518 | 0.752656307 | -4.773387391 |
| **A_55_P2166049** | Vmn1r65 | -0.063110159 | -0.269864127 | 0.143643809 | 7.888154723 | -0.741700568 | 0.485496638 | 0.752841116 | -4.773487476 |
| **A_52_P595717** | Smug1 | -0.052819192 | -0.225910667 | 0.120272282 | 8.441234933 | -0.7414797 | 0.485621277 | 0.752859301 | -4.773566333 |
| **A_55_P2081630** | Gpa33 | 0.078789152 | -0.17940897 | 0.336987274 | 8.94290264 | 0.741475041 | 0.485623907 | 0.752859301 | -4.773567997 |
| **A_51_P376347** | Hebp1 | 0.147711277 | -0.336389151 | 0.631811705 | 13.35496853 | 0.741415013 | 0.485657786 | 0.752859301 | -4.773589426 |
| **A_51_P151897** | R3hcc1 | -0.06583134 | -0.281606585 | 0.149943905 | 9.415531773 | -0.74133459 | 0.485703178 | 0.752859301 | -4.773618133 |
| **A_55_P2007699** | Nanog | 0.159119252 | -0.362482407 | 0.680720912 | 9.450341479 | 0.741253836 | 0.485748759 | 0.752859301 | -4.773646957 |
| **A_55_P1984416** | NA | 0.162539369 | -0.370281141 | 0.695359879 | 11.86804514 | 0.741243375 | 0.485754664 | 0.752859301 | -4.77365069 |
| **A_55_P2131785** | Kiz | -0.070362244 | -0.301061649 | 0.160337161 | 10.2392873 | -0.741099286 | 0.485836003 | 0.752859301 | -4.773702113 |
| **A_55_P2002472** | Zfp777 | -0.09648373 | -0.412847335 | 0.219879875 | 9.983852417 | -0.741055542 | 0.485860698 | 0.752859301 | -4.773717723 |
| **A_51_P441782** | Micu2 | -0.070221085 | -0.30049059 | 0.16004842 | 10.96406671 | -0.740993331 | 0.485895821 | 0.752859301 | -4.773739922 |
| **A_55_P2022201** | NA | 0.090724831 | -0.206796632 | 0.388246294 | 8.836458991 | 0.740953676 | 0.48591821 | 0.752859301 | -4.773754072 |
| **A_55_P2006554** | Eif4e2 | -0.096641881 | -0.41358899 | 0.220305227 | 11.71329992 | -0.740903712 | 0.48594642 | 0.752859301 | -4.773771899 |
| **A_55_P2131672** | Mical2 | -0.062004893 | -0.265370659 | 0.141360872 | 7.753086325 | -0.740851747 | 0.485975762 | 0.752859301 | -4.773790439 |
| **A_55_P2134427** | NA | 0.084359265 | -0.192414513 | 0.361133044 | 10.01768594 | 0.740612496 | 0.486110867 | 0.752953055 | -4.773875786 |
| **A_55_P1986371** | NA | -0.110932324 | -0.474892375 | 0.253027726 | 15.45574814 | -0.740606612 | 0.48611419 | 0.752953055 | -4.773877884 |
| **A_52_P227267** | Atp1a2 | -0.068629428 | -0.293840279 | 0.156581424 | 7.77118406 | -0.740464522 | 0.486194441 | 0.753017015 | -4.773928562 |
| **A_51_P391616** | Etnppl | -0.329971388 | -1.412929292 | 0.752986516 | 8.977680856 | -0.74036844 | 0.486248712 | 0.75304073 | -4.773962826 |
| **A_66_P140711** | Rps10 | 0.05660365 | -0.129210091 | 0.242417392 | 16.26708992 | 0.740201225 | 0.486343172 | 0.753095831 | -4.774022449 |
| **A_55_P2005982** | NA | 0.076628686 | -0.174941519 | 0.328198891 | 10.56667573 | 0.740142537 | 0.486376327 | 0.753095831 | -4.774043372 |
| **A_55_P2095894** | 6430548M08Rik | 0.144905614 | -0.330875006 | 0.620686234 | 10.49355201 | 0.740051007 | 0.486428041 | 0.753095831 | -4.774076003 |
| **A_55_P2104769** | Utf1 | 0.071781452 | -0.163918341 | 0.307481244 | 8.029785503 | 0.740007678 | 0.486452522 | 0.753095831 | -4.774091448 |
| **A_51_P329413** | Pomgnt1 | -0.119625126 | -0.512462194 | 0.273211941 | 9.41494576 | -0.739934263 | 0.486494004 | 0.753095831 | -4.774117617 |
| **A_55_P2151082** | NA | 0.102442 | -0.234049136 | 0.438933136 | 12.53918905 | 0.739754489 | 0.486595594 | 0.753095831 | -4.77418169 |
| **A_55_P2032946** | Gsta1 | 0.182446905 | -0.416842889 | 0.781736699 | 8.550599195 | 0.73974631 | 0.486600216 | 0.753095831 | -4.774184604 |
| **A_52_P354373** | Lockd | 0.136788942 | -0.312616187 | 0.58619407 | 12.71200651 | 0.739598727 | 0.486683626 | 0.753095831 | -4.774237195 |
| **A_55_P2030568** | Ifit1bl2 | 0.088262758 | -0.201739146 | 0.378264662 | 9.026718024 | 0.739536876 | 0.486718586 | 0.753095831 | -4.774259233 |
| **A_55_P2180096** | Gm10136 | 0.05323252 | -0.121684347 | 0.228149388 | 15.88400628 | 0.739483569 | 0.486748718 | 0.753095831 | -4.774278225 |
| **A_55_P2103412** | Golt1a | 0.272877079 | -0.623829771 | 1.169583928 | 10.85576433 | 0.739434584 | 0.486776407 | 0.753095831 | -4.774295677 |
| **A_55_P2171008** | Clk4 | -0.081964655 | -0.3513286 | 0.18739929 | 10.06134075 | -0.739384498 | 0.48680472 | 0.753095831 | -4.77431352 |
| **A_55_P2170514** | Sfn | -0.055652778 | -0.238565544 | 0.127259988 | 8.097998479 | -0.739309042 | 0.486847377 | 0.753095831 | -4.774340399 |
| **A_55_P1967325** | Tafa5 | -0.064928508 | -0.278329144 | 0.148472127 | 7.74495045 | -0.739303745 | 0.486850371 | 0.753095831 | -4.774342286 |
| **A_52_P274238** | Maea | 0.074403441 | -0.170149208 | 0.318956091 | 11.2008546 | 0.739271289 | 0.486868721 | 0.753095831 | -4.774353847 |
| **A_66_P116804** | Smchd1 | 0.098543298 | -0.225416077 | 0.422502673 | 10.7143457 | 0.739128128 | 0.486949662 | 0.753138162 | -4.774404838 |
| **A_51_P276939** | Gmip | -0.080837332 | -0.346604139 | 0.184929475 | 8.510987123 | -0.739085055 | 0.486974016 | 0.753138162 | -4.774420177 |

| | | | | | | | | | |
|---|---|---|---|---|---|---|---|---|---|
| **A_66_P138829** | Ndufs8 | -0.100050542 | -0.429095495 | 0.228994411 | 11.41340703 | -0.7388349 | 0.487115477 | 0.753290481 | -4.774509255 |
| **A_51_P412508** | Irx4 | 0.070993352 | -0.162508224 | 0.304494927 | 7.919134537 | 0.738773064 | 0.487150448 | 0.753290481 | -4.77453127 |
| **A_55_P2034320** | NA | -0.29327415 | -1.258275417 | 0.671727117 | 10.49529134 | -0.738463595 | 0.487325498 | 0.753500884 | -4.77464143 |
| **A_55_P2371984** | 5930436O19Rik | 0.066179982 | -0.151603237 | 0.283963201 | 8.32269823 | 0.738389348 | 0.487367501 | 0.753505554 | -4.774667854 |
| **A_52_P516034** | Ptp4a1 | -0.137322443 | -0.589349001 | 0.314704116 | 9.111441346 | -0.73817742 | 0.487487409 | 0.753599767 | -4.774743267 |
| **A_55_P2111950** | Dnajc7 | 0.086247298 | -0.197673367 | 0.370167964 | 11.78063161 | 0.738128008 | 0.487515369 | 0.753599767 | -4.774760848 |
| **A_51_P211822** | Neurl4 | -0.074773156 | -0.320942345 | 0.171396032 | 11.05707831 | -0.738066014 | 0.48755045 | 0.753599767 | -4.774782904 |
| **A_55_P2020326** | Madcam1 | 0.080895114 | -0.18545051 | 0.347240738 | 8.190817864 | 0.738006038 | 0.487584391 | 0.753599767 | -4.774804241 |
| **A_66_P105288** | Rplp0 | 0.060724503 | -0.139290613 | 0.260739619 | 16.10094208 | 0.737707737 | 0.487753224 | 0.753794834 | -4.774910343 |
| **A_55_P1961188** | Gm6981 | -0.095104381 | -0.408386902 | 0.218178139 | 14.52127339 | -0.737645248 | 0.487788597 | 0.753794834 | -4.774932566 |
| **A_51_P374137** | Bco1 | 0.107526532 | -0.246722017 | 0.461775081 | 8.558996014 | 0.737548779 | 0.487843208 | 0.753818959 | -4.77496687 |
| **A_55_P2104653** | Vma21 | -0.058534237 | -0.251396688 | 0.134328214 | 8.193142539 | -0.737471881 | 0.487886742 | 0.753825967 | -4.774994212 |
| **A_55_P2135701** | BC024978 | 0.069228435 | -0.166780911 | 0.305237782 | 8.149861858 | 0.746272916 | 0.487977557 | 0.753906018 | -4.737431084 |
| **A_52_P37757** | Eps8l1 | -0.076944058 | -0.330644492 | 0.176756377 | 7.81305573 | -0.736948372 | 0.488183191 | 0.754137513 | -4.775180296 |
| **A_55_P2176045** | NA | -0.069585718 | -0.299036411 | 0.159864975 | 8.105915128 | -0.736909114 | 0.488205427 | 0.754137513 | -4.775194247 |
| **A_55_P2138095** | NA | -0.078521035 | -0.337821243 | 0.180779173 | 16.14052006 | -0.735811105 | 0.488827614 | 0.755038274 | -4.775584203 |
| **A_51_P334789** | Nudt16 | -0.056256735 | -0.242070931 | 0.129557462 | 8.426501923 | -0.735662837 | 0.488911672 | 0.755107767 | -4.775636826 |
| **A_55_P2154835** | NA | 0.090089572 | -0.207534909 | 0.387714054 | 10.95924212 | 0.735510813 | 0.488997869 | 0.755146118 | -4.775690775 |
| **A_51_P310896** | Pfkl | 0.130503765 | -0.300701806 | 0.561709337 | 10.27259388 | 0.735397024 | 0.489062393 | 0.755146118 | -4.77573115 |
| **A_51_P105408** | Alg14 | -0.086249899 | -0.37124377 | 0.198743973 | 10.30507979 | -0.735370598 | 0.489077379 | 0.755146118 | -4.753627798 |
| **A_52_P111866** | Naa50 | 0.073311839 | -0.168992555 | 0.315616233 | 9.026958376 | 0.735183945 | 0.489183236 | 0.755146118 | -4.775806742 |
| **A_52_P479500** | Npepps | -0.060549374 | -0.260679353 | 0.139580605 | 10.55305337 | -0.735158015 | 0.489197943 | 0.755146118 | -4.775815939 |
| **A_55_P2109222** | Rps3 | -0.061151439 | -0.263273334 | 0.140970457 | 15.698728 | -0.73515092 | 0.489201967 | 0.755146118 | -4.775818456 |
| **A_55_P2174895** | Mrpl43 | -0.085457837 | -0.367928917 | 0.197013242 | 12.17139799 | -0.735124837 | 0.489216761 | 0.755146118 | -4.775827708 |
| **A_51_P333071** | Anapc16 | -0.070897868 | -0.305260787 | 0.163465051 | 10.84494501 | -0.735067916 | 0.489249047 | 0.755146118 | -4.775847897 |
| **A_51_P353856** | Ubac2 | 0.06317861 | -0.145794611 | 0.272151831 | 9.028995595 | 0.734619781 | 0.489503286 | 0.755340084 | -4.776006805 |
| **A_55_P2005475** | Sult1a1 | -0.266531127 | -1.148136609 | 0.615074355 | 13.00688303 | -0.734610011 | 0.48950883 | 0.755340084 | -4.776010269 |
| **A_55_P1954886** | NA | -0.055406129 | -0.238703641 | 0.127891383 | 16.41885824 | -0.734487531 | 0.489578333 | 0.755340084 | -4.776053687 |
| **A_55_P1958808** | Stoml2 | 0.07655189 | -0.176702492 | 0.329806272 | 12.64735292 | 0.734483656 | 0.489580532 | 0.755340084 | -4.776055061 |
| **A_51_P216219** | Ptk2 | -0.128474621 | -0.553512312 | 0.296563071 | 9.32049553 | -0.734468356 | 0.489589214 | 0.755340084 | -4.776060484 |
| **A_51_P308254** | Surf2 | 0.071139692 | -0.164225854 | 0.306505238 | 8.969309198 | 0.734433167 | 0.489609184 | 0.755340084 | -4.776072957 |
| **A_55_P1981274** | NA | -0.068143122 | -0.293633871 | 0.157347627 | 16.89203099 | -0.734305001 | 0.489681923 | 0.755340383 | -4.776118383 |
| **A_51_P337195** | Pipox | 0.186311572 | -0.430262248 | 0.802885391 | 12.21379848 | 0.734239828 | 0.489718914 | 0.755340383 | -4.77614148 |
| **A_51_P499864** | Tubg1 | 0.084113058 | -0.194253265 | 0.362479381 | 10.97282972 | 0.734226265 | 0.489726612 | 0.755340383 | -4.776146286 |
| **A_55_P2277014** | AA522020 | 0.147519269 | -0.340751778 | 0.635790316 | 10.62919667 | 0.734126612 | 0.489783178 | 0.755367353 | -4.776181599 |
| **A_51_P282144** | Adam23 | -0.081404607 | -0.350995405 | 0.18818619 | 8.582124114 | -0.733714515 | 0.490017141 | 0.755654737 | -4.77632759 |
| **A_66_P126842** | Frg2f1 | -0.062888598 | -0.271181875 | 0.14540468 | 8.338966168 | -0.733634671 | 0.49006248 | 0.755654737 | -4.776355869 |
| **A_55_P2122429** | Cym | -0.092708419 | -0.399785832 | 0.214368993 | 8.633014176 | -0.733591842 | 0.490086801 | 0.755654737 | -4.776371037 |
| **A_55_P2001510** | NA | 0.094163758 | -0.217851527 | 0.406179044 | 16.76407418 | 0.733315901 | 0.490243521 | 0.755819047 | -4.776468746 |
| **A_51_P259009** | Cacnb3 | -0.103648592 | -0.447157954 | 0.239860769 | 7.895772753 | -0.733175629 | 0.4903232 | 0.755819047 | -4.776518405 |
| **A_55_P2136862** | Dpy30 | 0.152057133 | -0.351932212 | 0.656046478 | 10.83407649 | 0.733109135 | 0.490360974 | 0.755819047 | -4.776541943 |
| **A_55_P2032128** | Dnase1l1 | 0.083846629 | -0.194074985 | 0.361768242 | 8.653914613 | 0.73307173 | 0.490382225 | 0.755819047 | -4.776555183 |
| **A_51_P438692** | Vti1b | 0.058948614 | -0.136448101 | 0.254345329 | 12.12925515 | 0.733060015 | 0.49038888 | 0.755819047 | -4.776559329 |
| **A_55_P2069955** | NA | 0.068600756 | -0.158841658 | 0.29604317 | 17.58159796 | 0.732893103 | 0.490483712 | 0.755904934 | -4.776618403 |
| **A_52_P205789** | Unc119b | 0.090959395 | -0.210793934 | 0.392712725 | 8.641093124 | 0.73245118 | 0.490734854 | 0.75607644 | -4.77677476 |
| **A_55_P2084251** | Kcna4 | 0.241218749 | -0.559018462 | 1.04145596 | 13.1475419 | 0.732445824 | 0.490737898 | 0.75607644 | -4.776776655 |
| **A_52_P510877** | Bcl2l1 | 0.11384114 | -0.263832011 | 0.49151429 | 8.559002464 | 0.732430317 | 0.490746713 | 0.75607644 | -4.77678214 |
| **A_52_P573552** | Trib1 | -0.152514805 | -0.658497895 | 0.353468284 | 9.862195774 | -0.732418308 | 0.490753539 | 0.75607644 | -4.776786388 |
| **A_66_P122086** | 9030619P08Rik | -0.057881363 | -0.249925646 | 0.134162921 | 7.89753891 | -0.732353146 | 0.490790578 | 0.75607644 | -4.776809436 |
| **A_55_P2016675** | Ogfr | -0.085450619 | -0.369033869 | 0.198132632 | 10.83702371 | -0.732179935 | 0.490889044 | 0.756121508 | -4.776870695 |

| | | | | | | | | | |
|---|---|---|---|---|---|---|---|---|---|
| **A_55_P2125820** | Gm14288 | 0.093758281 | -0.21744847 | 0.404965031 | 9.984745711 | 0.732055175 | 0.490959975 | 0.756121508 | -4.776914811 |
| **A_51_P393761** | Ndufs2 | -0.084649651 | -0.365637484 | 0.196338182 | 10.49209739 | -0.73201646 | 0.490981988 | 0.756121508 | -4.7769285 |
| **A_55_P2081555** | Phf23 | -0.08253851 | -0.356558465 | 0.191481445 | 9.741543885 | -0.731909903 | 0.491042577 | 0.756121508 | -4.776966174 |
| **A_51_P283344** | ccdc198 | 0.245623617 | -0.569846459 | 1.061093694 | 8.892745674 | 0.731889106 | 0.491054403 | 0.756121508 | -4.776973526 |
| **A_66_P134453** | I830077J02Rik | 0.105990595 | -0.245898156 | 0.457879346 | 8.797230871 | 0.731888858 | 0.491054544 | 0.756121508 | -4.776973614 |
| **A_51_P472932** | Atp8b1 | -0.08055642 | -0.348073948 | 0.186961108 | 8.090292959 | -0.731696709 | 0.491163816 | 0.756127055 | -4.777041536 |
| **A_55_P2140147** | Ppp4r3b | 0.060795387 | -0.141100351 | 0.262691125 | 10.67952489 | 0.731689236 | 0.491168066 | 0.756127055 | -4.777044178 |
| **A_55_P2044252** | Zfp512b | -0.067261648 | -0.29063522 | 0.156111924 | 10.14808393 | -0.73167616 | 0.491175503 | 0.756127055 | -4.7770488 |
| **A_51_P499838** | Bst1 | -0.092815593 | -0.401113638 | 0.215482453 | 8.75394935 | -0.731532051 | 0.491257468 | 0.756148387 | -4.777099731 |
| **A_55_P2035804** | Fzd8 | -0.055652222 | -0.240512208 | 0.129207764 | 7.87471757 | -0.731514238 | 0.4912676 | 0.756148387 | -4.777106026 |
| **A_52_P46310** | Dcaf7 | 0.115434879 | -0.268086528 | 0.498956286 | 8.909635704 | 0.731359035 | 0.491355886 | 0.756175353 | -4.777160869 |
| **A_55_P2133225** | Mmgt2 | -0.074006282 | -0.319910009 | 0.171897444 | 9.591780934 | -0.731285002 | 0.491398003 | 0.756175353 | -4.777187026 |
| **A_65_P02323** | Sec22b | 0.061162604 | -0.142067092 | 0.2643923 | 8.10452935 | 0.731277126 | 0.491402484 | 0.756175353 | -4.777189809 |
| **A_51_P483261** | Pomt1 | 0.084712115 | -0.196797157 | 0.366221386 | 8.963122023 | 0.731199706 | 0.49144653 | 0.756182932 | -4.777217161 |
| **A_51_P116813** | Cyp17a1 | 0.294883311 | -0.685261962 | 1.275028585 | 12.4496953 | 0.731043018 | 0.491535683 | 0.756259908 | -4.77727251 |
| **A_55_P2053718** | Pink1 | -0.086746414 | -0.375120018 | 0.20162719 | 11.99236344 | -0.730935769 | 0.491596712 | 0.756293605 | -4.77731039 |
| **A_51_P248444** | Poglut1 | -0.052983756 | -0.229191127 | 0.123223615 | 8.261016805 | -0.730637332 | 0.491766563 | 0.7564503 | -4.777415776 |
| **A_52_P165654** | Star | 0.048892657 | -0.113726838 | 0.211512152 | 7.813988063 | 0.730557191 | 0.49181218 | 0.7564503 | -4.77744407 |
| **A_55_P2142527** | NA | 0.065279519 | -0.151845503 | 0.282404541 | 8.968565814 | 0.73055053 | 0.491815972 | 0.7564503 | -4.777446422 |
| **A_55_P2122314** | Emc1 | 0.08906054 | -0.207320618 | 0.385441699 | 8.771195308 | 0.730159808 | 0.492038419 | 0.756732225 | -4.777584334 |
| **A_52_P536481** | Mogs | 0.094712502 | -0.22054001 | 0.409965015 | 13.7293673 | 0.730015269 | 0.492120726 | 0.756747632 | -4.777635338 |
| **A_51_P147714** | Sugp2 | 0.059789672 | -0.13922436 | 0.258803704 | 8.29720023 | 0.73000471 | 0.492126739 | 0.756747632 | -4.777639063 |
| **A_55_P1953400** | Rbfox1 | -0.056413113 | -0.244242278 | 0.131416052 | 7.785193993 | -0.729793895 | 0.492246804 | 0.756818651 | -4.777713439 |
| **A_51_P416919** | Ube2j2 | 0.067114741 | -0.156365978 | 0.29059546 | 9.804972808 | 0.729728058 | 0.492284304 | 0.756818651 | -4.777736663 |
| **A_51_P267194** | Faim | -0.059108193 | -0.255932276 | 0.13771589 | 9.555665792 | -0.729713928 | 0.492292352 | 0.756818651 | -4.777741647 |
| **A_51_P321341** | Sult1a1 | -0.259027201 | -1.121637788 | 0.603583386 | 13.32289065 | -0.72964864 | 0.492329542 | 0.756818651 | -4.777764676 |
| **A_51_P425080** | Hdgf | 0.059740808 | -0.139260328 | 0.258741944 | 13.92642389 | 0.72945537 | 0.492439644 | 0.756927704 | -4.777832838 |
| **A_55_P1964464** | NA | -0.079492186 | -0.34438647 | 0.185402099 | 15.76523783 | -0.729180491 | 0.492596266 | 0.75707182 | -4.777929757 |
| **A_55_P2039598** | Rpl7a | -0.124769948 | -0.540569788 | 0.291029891 | 13.48976447 | -0.729136547 | 0.492621308 | 0.75707182 | -4.777945249 |
| **A_55_P2058737** | Mex3d | -0.121158984 | -0.524953934 | 0.282635965 | 9.218551371 | -0.729084611 | 0.492650905 | 0.75707182 | -4.777963557 |
| **A_65_P08971** | F3 | -0.057687667 | -0.249967473 | 0.134592138 | 7.721651526 | -0.729008332 | 0.492694377 | 0.757078433 | -4.777990444 |
| **A_51_P158400** | Rc3h2 | 0.075008325 | -0.175229512 | 0.325246162 | 9.901439136 | 0.728349254 | 0.493070097 | 0.757595542 | -4.778222675 |
| **A_55_P2029687** | Hmox1 | 0.126979606 | -0.296693921 | 0.550653133 | 8.80138222 | 0.728258961 | 0.493121586 | 0.757614429 | -4.778254478 |
| **A_51_P141610** | Cables2 | 0.057207151 | -0.133724828 | 0.24813913 | 8.89134902 | 0.72803932 | 0.493246848 | 0.75772142 | -4.778331827 |
| **A_55_P2040357** | Bola1 | 0.051490562 | -0.120371418 | 0.223352542 | 8.930282798 | 0.727999381 | 0.493269628 | 0.75772142 | -4.778345891 |
| **A_55_P1973930** | Krt79 | 0.051432485 | -0.12025562 | 0.223120591 | 8.183392863 | 0.727914704 | 0.493317927 | 0.757735394 | -4.778375705 |
| **A_51_P401527** | Mrm3 | 0.092675795 | -0.216718842 | 0.402070433 | 8.677952201 | 0.727841356 | 0.493359766 | 0.757739446 | -4.778401528 |
| **A_52_P200458** | Fut4 | -0.060938051 | -0.264418884 | 0.142542783 | 7.967430964 | -0.72769307 | 0.49344436 | 0.75779128 | -4.778453728 |
| **A_55_P2071536** | NA | -0.050532509 | -0.219278961 | 0.118213943 | 7.928359814 | -0.727644753 | 0.493471925 | 0.75779128 | -4.778470735 |
| **A_55_P2083406** | Myh3 | -0.058456316 | -0.253849272 | 0.136936641 | 7.673642885 | -0.726951982 | 0.493867278 | 0.758281024 | -4.778714488 |
| **A_55_P2059957** | Snx7 | -0.07506191 | -0.325961063 | 0.175837243 | 8.73626423 | -0.72694843 | 0.493869305 | 0.758281024 | -4.778715738 |
| **A_55_P2179453** | Lig3 | 0.060847889 | -0.142564851 | 0.264260629 | 8.225610305 | 0.726859639 | 0.493919993 | 0.758298613 | -4.778746966 |
| **A_51_P245156** | Gdf2 | 0.144665099 | -0.339039457 | 0.628369656 | 10.50663096 | 0.726719444 | 0.494000031 | 0.758311999 | -4.778796268 |
| **A_55_P2181508** | Gm10653 | 0.068996716 | -0.161705788 | 0.299699221 | 15.91967779 | 0.726706931 | 0.494007175 | 0.758311999 | -4.778800668 |
| **A_55_P2008895** | Tmsb15b1 | -0.04911554 | -0.213423336 | 0.115192257 | 8.432355505 | -0.726346651 | 0.494212904 | 0.758567556 | -4.778927332 |
| **A_51_P146760** | Pcdhb20 | -0.081253273 | -0.353130243 | 0.190623697 | 7.845615531 | -0.726192284 | 0.494301069 | 0.758608304 | -4.778981588 |
| **A_66_P106814** | NA | 0.144862768 | -0.339874052 | 0.629599588 | 12.74639378 | 0.726162736 | 0.494317946 | 0.758608304 | -4.778991972 |
| **A_51_P385351** | Slc44a3 | -0.06473755 | -0.281421371 | 0.151946272 | 9.060902697 | -0.725960448 | 0.4944335 | 0.758666252 | -4.779063056 |
| **A_55_P2106961** | 2900097C17Rik | 0.094061937 | -0.220774425 | 0.408898298 | 8.344859327 | 0.725959211 | 0.494434206 | 0.758666252 | -4.779063491 |
| **A_55_P1952950** | Cic | -0.072477436 | -0.315121016 | 0.170166144 | 13.31779637 | -0.725800401 | 0.494524937 | 0.758745237 | -4.779119286 |

| | | | | | | | | | |
|---|---|---|---|---|---|---|---|---|---|
| **A_55_P1971194** | Timm23 | 0.055401993 | -0.130118239 | 0.240922225 | 12.99400307 | 0.725633458 | 0.494620325 | 0.758831357 | -4.779177929 |
| **A_51_P285413** | Rbbp6 | -0.052615125 | -0.22882663 | 0.12359638 | 7.932291218 | -0.725536945 | 0.494675477 | 0.758855738 | -4.779211826 |
| **A_51_P333518** | Sh3rf2 | -0.062161125 | -0.270419598 | 0.146097348 | 7.731565121 | -0.725269433 | 0.494828367 | 0.759030038 | -4.779305766 |
| **A_52_P397204** | Csk | 0.063936162 | -0.150355915 | 0.278228239 | 9.233448227 | 0.724976008 | 0.494996103 | 0.759227081 | -4.779408774 |
| **A_66_P121459** | Cenpa | 0.116935989 | -0.275191451 | 0.509063429 | 11.35061144 | 0.724609709 | 0.495205551 | 0.759445306 | -4.779537321 |
| **A_55_P1954986** | Elk1 | 0.070700257 | -0.166409124 | 0.307809637 | 17.42166248 | 0.724528453 | 0.495252021 | 0.759445306 | -4.77956583 |
| **A_55_P1999958** | NA | 0.100726096 | -0.237085209 | 0.438537401 | 8.807757634 | 0.724521056 | 0.495256251 | 0.759445306 | -4.779568425 |
| **A_55_P1968969** | NA | 0.068580441 | -0.161475604 | 0.298636486 | 15.71159088 | 0.724352254 | 0.495352798 | 0.759483688 | -4.779627642 |
| **A_51_P392303** | Lrrn3 | 0.074212143 | -0.174739935 | 0.323164222 | 8.713051469 | 0.724339897 | 0.495359866 | 0.759483688 | -4.779631976 |
| **A_51_P460710** | Tdrd3 | 0.069346623 | -0.163399825 | 0.302093071 | 9.496047298 | 0.723978124 | 0.495566828 | 0.759740737 | -4.77975885 |
| **A_66_P128434** | Kit | -0.059822415 | -0.260629008 | 0.140984178 | 10.00988729 | -0.723884307 | 0.495620508 | 0.759762773 | -4.779791745 |
| **A_51_P443754** | Hivep2 | 0.078266772 | -0.184521907 | 0.34105545 | 8.150679941 | 0.723692309 | 0.495730377 | 0.759855008 | -4.779859053 |
| **A_55_P1973399** | NA | 0.180209462 | -0.424904469 | 0.785323393 | 14.29676009 | 0.723641771 | 0.4957593 | 0.759855008 | -4.779876767 |
| **A_51_P119401** | Tug1 | 0.064306773 | -0.151649595 | 0.28026314 | 8.746270083 | 0.7235589 | 0.495806729 | 0.759867449 | -4.777125408 |
| **A_66_P114102** | Zfp36l2 | -0.055189697 | -0.240556208 | 0.130176814 | 12.23352815 | -0.72345234 | 0.49586772 | 0.759900671 | -4.779943159 |
| **A_51_P339444** | Rab9 | -0.06864513 | -0.299264423 | 0.161974162 | 12.1113506 | -0.723264728 | 0.495975115 | 0.75994769 | -4.780008899 |
| **A_51_P206405** | Ptprz1 | -0.075774262 | -0.33034707 | 0.178798545 | 7.740106537 | -0.723257413 | 0.495979303 | 0.75994769 | -4.780011462 |
| **A_55_P2085400** | NA | -0.09905281 | -0.431862494 | 0.233756875 | 9.083001741 | -0.723192699 | 0.496016351 | 0.75994769 | -4.780034136 |
| **A_55_P1990127** | Dolpp1 | 0.049512106 | -0.116895573 | 0.215919785 | 10.99217409 | 0.722971577 | 0.496142955 | 0.760068504 | -4.780111596 |
| **A_55_P2030638** | Dusp23 | -0.103619584 | -0.451910853 | 0.244671684 | 10.07118191 | -0.722907028 | 0.496179917 | 0.760068504 | -4.780134205 |
| **A_51_P481644** | Mbip | -0.339065336 | -1.478840074 | 0.800709402 | 9.45534429 | -0.722848951 | 0.496213175 | 0.760068504 | -4.780154545 |
| **A_66_P140076** | NA | -0.102702653 | -0.447974193 | 0.242568887 | 11.57983383 | -0.722776575 | 0.496254622 | 0.760071759 | -4.780179892 |
| **A_51_P278018** | Vps36 | -0.149651275 | -0.652810272 | 0.353507723 | 11.84875639 | -0.722700517 | 0.496298181 | 0.760078246 | -4.780206526 |
| **A_55_P2040280** | Dhps | 0.063053036 | -0.148972975 | 0.275079047 | 8.287243164 | 0.722603481 | 0.496353758 | 0.760103137 | -4.780240503 |
| **A_55_P2020572** | Psme3 | 0.244335264 | -0.577493236 | 1.066163764 | 12.51406827 | 0.722417309 | 0.496460399 | 0.760144639 | -4.780305681 |
| **A_52_P106463** | Jak1 | 0.075558002 | -0.178625068 | 0.329741072 | 12.1315363 | 0.722299035 | 0.496528155 | 0.760144639 | -4.780347082 |
| **A_51_P492446** | Ldah | 0.058812179 | -0.139077402 | 0.25670176 | 10.83994786 | 0.722150191 | 0.496613433 | 0.760144639 | -4.780399176 |
| **A_55_P1964078** | NA | -0.081717322 | -0.356694733 | 0.193260089 | 12.31344079 | -0.722104909 | 0.496639379 | 0.760144639 | -4.780415023 |
| **A_55_P2127303** | Spsb2 | -0.112155977 | -0.489563926 | 0.265251972 | 10.97746287 | -0.722095384 | 0.496644837 | 0.760144639 | -4.780418356 |
| **A_66_P129029** | Taf10 | 0.079512625 | -0.188061264 | 0.347086514 | 12.28844079 | 0.72206376 | 0.496662957 | 0.760144639 | -4.780429422 |
| **A_55_P1966583** | Bpifc | 0.160874502 | -0.380607271 | 0.702356275 | 13.47782077 | 0.721915824 | 0.49674773 | 0.760144639 | -4.780481185 |
| **A_66_P125862** | A930005H10Rik | -0.054859461 | -0.239549517 | 0.129830595 | 8.733686378 | -0.72175734 | 0.496838558 | 0.760144639 | -4.780536631 |
| **A_55_P2113984** | Mospd2 | -0.052740107 | -0.230304649 | 0.124824436 | 8.304095686 | -0.721718682 | 0.496860715 | 0.760144639 | -4.780550154 |
| **A_55_P1990608** | Ube2j1 | 0.075060131 | -0.17765654 | 0.327776803 | 9.251045207 | 0.721703175 | 0.496869603 | 0.760144639 | -4.780555578 |
| **A_51_P168695** | Mast4 | -0.086373914 | -0.377189196 | 0.204441368 | 8.216086536 | -0.721686471 | 0.496879177 | 0.760144639 | -4.780561421 |
| **A_52_P495104** | NA | 0.063137652 | -0.149494875 | 0.275770179 | 10.40527314 | 0.721509269 | 0.496980752 | 0.760144639 | -4.7806234 |
| **A_55_P1989673** | Slco2a1 | -0.16244923 | -0.709585481 | 0.384687021 | 11.89677913 | -0.721448535 | 0.497015568 | 0.760144639 | -4.780644639 |
| **A_55_P2016685** | Csnk1g1 | 0.057345595 | -0.135797934 | 0.250489124 | 9.468739341 | 0.721444772 | 0.497017725 | 0.760144639 | -4.780645955 |
| **A_51_P143712** | Tdh | 0.0537042 | -0.127222411 | 0.234630811 | 7.821089933 | 0.721255256 | 0.497126379 | 0.760144639 | -4.780712223 |
| **A_55_P1966377** | 4930502E18Rik | 0.045712163 | -0.10829629 | 0.199720616 | 7.66707038 | 0.721224386 | 0.497144079 | 0.760144639 | -4.780723017 |
| **A_55_P2107750** | Hectd1 | 0.099905404 | -0.236686114 | 0.436496923 | 9.315620645 | 0.721222062 | 0.497145411 | 0.760144639 | -4.780723829 |
| **A_55_P2064876** | Fam89b | -0.113894164 | -0.49762906 | 0.269840733 | 12.08116899 | -0.721196091 | 0.497160303 | 0.760144639 | -4.780732909 |
| **A_66_P126563** | Ptges3l | -0.153434963 | -0.670499615 | 0.363629688 | 8.199984757 | -0.721045505 | 0.497246653 | 0.760144639 | -4.780785552 |
| **A_52_P193233** | Trnau1ap | -0.059342112 | -0.259336149 | 0.140651925 | 9.899086693 | -0.720989836 | 0.497278577 | 0.760144639 | -4.78080501 |
| **A_55_P2124712** | Ces2c | -0.162365516 | -0.709598504 | 0.384867473 | 8.752561061 | -0.720949285 | 0.497301833 | 0.760144639 | -4.780819184 |
| **A_51_P120295** | Cyp2d26 | -0.252514777 | -1.103589323 | 0.598559769 | 16.24747192 | -0.720945418 | 0.497304051 | 0.760144639 | -4.780820536 |
| **A_51_P368285** | Rflna | -0.181133816 | -0.791658977 | 0.429391345 | 7.863406577 | -0.720906861 | 0.497326163 | 0.760144639 | -4.780834012 |
| **A_51_P121915** | Rdh16f2 | -0.150853072 | -0.659373 | 0.357666856 | 8.483100074 | -0.720824222 | 0.49737356 | 0.760144639 | -4.780862894 |
| **A_66_P128694** | NA | 0.123666833 | -0.29323555 | 0.540569217 | 9.128000761 | 0.720778878 | 0.497399568 | 0.760144639 | -4.78087874 |
| **A_55_P1954400** | NA | 0.093598738 | -0.221941759 | 0.409139234 | 12.57963214 | 0.72077228 | 0.497403352 | 0.760144639 | -4.780881046 |

| | | | | | | | | | |
|---|---|---|---|---|---|---|---|---|---|
| **A_51_P483557** | Ube2b | -0.062165676 | -0.27178689 | 0.147455538 | 10.85662945 | -0.720607233 | 0.497498025 | 0.760192178 | -4.780938717 |
| **A_51_P470989** | Paip1 | -0.064434685 | -0.281717827 | 0.152848457 | 8.678507496 | -0.720571147 | 0.497518726 | 0.760192178 | -4.780951326 |
| **A_51_P211506** | Muc20 | 0.063330453 | -0.150264943 | 0.276925849 | 8.510372077 | 0.720450083 | 0.49758818 | 0.760192178 | -4.78099362 |
| **A_55_P2116744** | Xirp1 | -0.049354856 | -0.215821487 | 0.117111775 | 7.668444958 | -0.720420207 | 0.49760532 | 0.760192178 | -4.781004057 |
| **A_51_P137778** | NA | 0.056883349 | -0.135000022 | 0.24876672 | 7.723595065 | 0.720329183 | 0.497657546 | 0.760192178 | -4.781035853 |
| **A_52_P393306** | Snap23 | -0.100233352 | -0.438369743 | 0.23790304 | 9.63106613 | -0.720283604 | 0.497683699 | 0.760192178 | -4.781051773 |
| **A_66_P138319** | Acox2 | 0.144218136 | -0.342358152 | 0.630794425 | 13.4195173 | 0.720198358 | 0.497732614 | 0.760192178 | -4.781081546 |
| **A_51_P244690** | Ube2b | -0.060181009 | -0.263233277 | 0.142871259 | 13.34693464 | -0.720169642 | 0.497749092 | 0.760192178 | -4.781091575 |
| **A_55_P2015877** | Clps | 0.160132615 | -0.380272206 | 0.700537436 | 8.020564587 | 0.72001869 | 0.49783572 | 0.76026441 | -4.781144288 |
| **A_51_P519189** | Eif3i | -0.11165721 | -0.488552419 | 0.265237999 | 10.99767028 | -0.719862156 | 0.497925563 | 0.760303357 | -4.781198942 |
| **A_52_P628117** | Abcc5 | 0.05353139 | -0.127186307 | 0.234249087 | 7.965575849 | 0.719765497 | 0.497981045 | 0.760303357 | -4.781232686 |
| **A_55_P1975793** | Srd5a3 | -0.066480148 | -0.290926395 | 0.157966099 | 9.4226547 | -0.719718753 | 0.498007878 | 0.760303357 | -4.781249003 |
| **A_55_P2094896** | Phyhd1 | 0.081692819 | -0.194120658 | 0.357506295 | 12.02907793 | 0.719700139 | 0.498018563 | 0.760303357 | -4.781255501 |
| **A_51_P414115** | Naprt | 0.177483819 | -0.421996686 | 0.776964325 | 12.14893499 | 0.719394131 | 0.498194249 | 0.760511501 | -4.781362299 |
| **A_52_P172014** | Ramp1 | -0.049989043 | -0.218864242 | 0.118886156 | 7.778752414 | -0.719270322 | 0.498265342 | 0.760559961 | -4.781405499 |
| **A_55_P2142222** | Serpina3h | 0.051052621 | -0.121516522 | 0.223621765 | 7.976974129 | 0.718849691 | 0.498506925 | 0.760824079 | -4.781552226 |
| **A_52_P18922** | Chid1 | -0.058546135 | -0.256464646 | 0.139372376 | 8.196427569 | -0.718778376 | 0.498547892 | 0.760824079 | -4.781577096 |
| **A_52_P83959** | Taf7 | 0.046666328 | -0.111095099 | 0.204427756 | 8.558469207 | 0.718763464 | 0.498556459 | 0.760824079 | -4.781582296 |
| **A_65_P00881** | NA | -0.06864333 | -0.300856104 | 0.163569444 | 8.347074122 | -0.718282733 | 0.498832674 | 0.761140123 | -4.781749892 |
| **A_55_P2156556** | NA | 0.212078907 | -0.505476631 | 0.929634445 | 12.73845947 | 0.718166504 | 0.498899472 | 0.761140123 | -4.7817904 |
| **A_55_P1967133** | Ppp1r3b | -0.137754381 | -0.603851957 | 0.328343195 | 13.40940231 | -0.718144378 | 0.498912188 | 0.761140123 | -4.78179811 |
| **A_55_P2049261** | NA | -0.065472044 | -0.287004146 | 0.156060058 | 7.866321406 | -0.718128924 | 0.49892107 | 0.761140123 | -4.781803496 |
| **A_55_P1956807** | Gm14391 | -0.096215116 | -0.42189518 | 0.229464949 | 9.357741532 | -0.717852672 | 0.499079863 | 0.761250816 | -4.781899751 |
| **A_55_P2089238** | Fam76a | 0.062939681 | -0.150108327 | 0.275987689 | 10.58567032 | 0.717844286 | 0.499084684 | 0.761250816 | -4.781902672 |
| **A_55_P2048744** | Eif2ak3 | 0.092343649 | -0.220256002 | 0.404943299 | 9.926304891 | 0.717797152 | 0.49911178 | 0.761250816 | -4.781919092 |
| **A_51_P295442** | Nek7 | -0.060716756 | -0.266343249 | 0.144909736 | 10.63572376 | -0.717484767 | 0.49929139 | 0.761464674 | -4.782027896 |
| **A_55_P1958354** | Rps29 | -0.06590692 | -0.289149464 | 0.157335625 | 16.98619124 | -0.717360136 | 0.499363061 | 0.761513893 | -4.782071295 |
| **A_55_P2026214** | NA | -0.080844073 | -0.354870989 | 0.193182844 | 11.81079601 | -0.716866273 | 0.49964713 | 0.761886981 | -4.782243212 |
| **A_55_P1993348** | NA | 0.06062797 | -0.144940621 | 0.266196561 | 15.83101125 | 0.716637379 | 0.499778826 | 0.761910471 | -4.78232286 |
| **A_55_P2061110** | Snapin | 0.054233395 | -0.129653651 | 0.238120442 | 8.913714293 | 0.716636375 | 0.499779404 | 0.761910471 | -4.782323209 |
| **A_51_P286814** | Ncor2 | -0.183522138 | -0.805785543 | 0.438741266 | 11.36499103 | -0.716633968 | 0.499780789 | 0.761910471 | -4.782324047 |
| **A_52_P452667** | Prom2 | -0.05817919 | -0.255475587 | 0.139117206 | 7.682508816 | -0.716525592 | 0.499843152 | 0.761945211 | -4.782361752 |
| **A_55_P2131522** | Arfrp1 | -0.052298138 | -0.229684201 | 0.125087926 | 10.91663369 | -0.716390661 | 0.499920803 | 0.761945211 | -4.782408689 |
| **A_52_P102002** | Srrd | 0.063233999 | -0.151251596 | 0.277719594 | 8.580509347 | 0.716367225 | 0.499934291 | 0.761945211 | -4.782416841 |
| **A_55_P2069022** | Vstm4 | -0.060713691 | -0.266664036 | 0.145236653 | 7.901651908 | -0.716320374 | 0.499961256 | 0.761945211 | -4.782433137 |
| **A_52_P63709** | Fam49b | 0.050450316 | -0.120715525 | 0.221616158 | 7.798252307 | 0.716192856 | 0.500034652 | 0.761996988 | -4.782477486 |
| **A_66_P125496** | NA | -0.086779244 | -0.381307443 | 0.207748954 | 12.9124654 | -0.715932655 | 0.50018444 | 0.76209234 | -4.782567963 |
| **A_51_P430950** | Rdx | 0.094517033 | -0.226311132 | 0.415345198 | 10.98668691 | 0.715847946 | 0.50023321 | 0.76209234 | -4.782597412 |
| **A_52_P590235** | Vars2 | -0.055316646 | -0.243089777 | 0.132456485 | 8.762887407 | -0.715822888 | 0.500247638 | 0.76209234 | -4.782606123 |
| **A_55_P2059432** | Crabp2 | 0.054082204 | -0.129503845 | 0.237668252 | 8.50436456 | 0.715810217 | 0.500254934 | 0.76209234 | -4.782610528 |
| **A_66_P112886** | Ubl3 | -0.115125301 | -0.506145741 | 0.275895138 | 10.30315477 | -0.715409182 | 0.500485874 | 0.762132818 | -4.782749907 |
| **A_55_P2077048** | Itih5 | 0.117407997 | -0.281383585 | 0.516199579 | 9.181344519 | 0.715376855 | 0.500504493 | 0.762132818 | -4.78276114 |
| **A_52_P245415** | AW554918 | -0.058129904 | -0.255578279 | 0.13931847 | 8.464445476 | -0.715367547 | 0.500509855 | 0.762132818 | -4.782764374 |
| **A_55_P1972490** | Gm4055 | 0.12063913 | -0.289137175 | 0.530415435 | 11.66503757 | 0.715359821 | 0.500514305 | 0.762132818 | -4.782767058 |
| **A_51_P228658** | Isyna1 | -0.051332805 | -0.225705209 | 0.123039599 | 11.32225582 | -0.715320104 | 0.500537181 | 0.762132818 | -4.782780858 |
| **A_55_P2057503** | Arl4c | 0.088372415 | -0.211833727 | 0.388578558 | 9.868664274 | 0.715287012 | 0.500556242 | 0.762132818 | -4.782792355 |
| **A_55_P2138416** | NA | -0.074990847 | -0.329764331 | 0.179782637 | 13.09819507 | -0.715216001 | 0.500597146 | 0.762132818 | -4.782817026 |
| **A_51_P502348** | Tfip11 | 0.097491769 | -0.233735969 | 0.428719506 | 9.770093381 | 0.715194952 | 0.500609271 | 0.762132818 | -4.782824338 |
| **A_51_P276479** | 4930486L24Rik | -0.047639044 | -0.209532317 | 0.114254229 | 7.853665436 | -0.715018743 | 0.500710783 | 0.762132818 | -4.782885547 |
| **A_55_P2132549** | Cd48 | -0.096228905 | -0.423248169 | 0.230790358 | 8.954109298 | -0.715015404 | 0.500712706 | 0.762132818 | -4.782886706 |

| | | | | | | | | | |
|---|---|---|---|---|---|---|---|---|---|
| **A_55_P2066301** | Gpr137b-ps | -0.064732245 | -0.284717102 | 0.155252611 | 8.082506933 | -0.715008289 | 0.500716805 | 0.762132818 | -4.782889178 |
| **A_55_P2006280** | Ogdh | 0.136129319 | -0.326532533 | 0.59879117 | 12.73675772 | 0.714942589 | 0.500754658 | 0.762132818 | -4.782911996 |
| **A_55_P2106351** | NA | 0.113036909 | -0.271222831 | 0.497296648 | 9.502481772 | 0.714790179 | 0.500842475 | 0.762206456 | -4.782964924 |
| **A_51_P482425** | Lins1 | 0.054418827 | -0.130695235 | 0.239532889 | 8.533773129 | 0.714320237 | 0.501113316 | 0.762558596 | -4.783128068 |
| **A_55_P1959061** | NA | -0.160252201 | -0.705441889 | 0.384937486 | 12.61841628 | -0.714232423 | 0.501163937 | 0.762575592 | -4.783158544 |
| **A_51_P475613** | Fahd2a | 0.067478932 | -0.162140142 | 0.297098006 | 10.19221927 | 0.714074337 | 0.501255075 | 0.76265423 | -4.783213401 |
| **A_55_P2045114** | Tmem242 | -0.151546224 | -0.667631966 | 0.364539518 | 10.40434104 | -0.713520489 | 0.501574459 | 0.762897008 | -4.783405518 |
| **A_51_P153004** | Phax | -0.059514848 | -0.262222185 | 0.143192489 | 9.404334547 | -0.713409765 | 0.501638326 | 0.762897008 | -4.783443913 |
| **A_55_P1967867** | Xpa | -0.050657649 | -0.223201873 | 0.121886575 | 9.277645431 | -0.713391273 | 0.501648992 | 0.762897008 | -4.783450324 |
| **A_55_P2150757** | Gzmm | 0.050058934 | -0.120448123 | 0.220565991 | 8.017581358 | 0.713382462 | 0.501654075 | 0.762897008 | -4.783453379 |
| **A_55_P2115225** | Fap | -0.081748813 | -0.360214328 | 0.196716702 | 7.993134447 | -0.713334492 | 0.501681747 | 0.762897008 | -4.783470011 |
| **A_51_P148069** | Lpgat1 | -0.09951588 | -0.438527558 | 0.239495799 | 11.26884741 | -0.713281427 | 0.501712359 | 0.762897008 | -4.783488408 |
| **A_52_P297009** | Itpk1 | -0.054286574 | -0.239239493 | 0.130666346 | 8.209063399 | -0.713205084 | 0.501756402 | 0.762897008 | -4.783514874 |
| **A_51_P121390** | Anapc11 | -0.12767934 | -0.562818849 | 0.307460169 | 12.50823509 | -0.712976641 | 0.501888208 | 0.762897008 | -4.772243678 |
| **A_55_P2125491** | Tnfrsf4 | -0.202040249 | -0.89065613 | 0.486575632 | 8.744343901 | -0.712925338 | 0.501917812 | 0.762897008 | -4.783611835 |
| **A_55_P2077664** | Gm5458 | -0.068835861 | -0.303468048 | 0.165796327 | 7.829209984 | -0.712870013 | 0.501949738 | 0.762897008 | -4.783631008 |
| **A_52_P474089** | Capn6 | -0.10681615 | -0.470938677 | 0.257306376 | 9.53748101 | -0.712807998 | 0.501985526 | 0.762897008 | -4.783652497 |
| **A_52_P160955** | Sox11 | -0.099387781 | -0.438194887 | 0.239419325 | 7.960630171 | -0.712793407 | 0.501993947 | 0.762897008 | -4.783657553 |
| **A_55_P2281095** | 2610002M06Rik | 0.098509747 | -0.237311172 | 0.434330665 | 9.426075938 | 0.712778591 | 0.502002497 | 0.762897008 | -4.783662687 |
| **A_51_P286792** | Zfp275 | -0.062421387 | -0.275244925 | 0.15040215 | 8.932621657 | -0.712683898 | 0.502057148 | 0.762897008 | -4.783695496 |
| **A_52_P80944** | Zfp36 | -0.06065041 | -0.26744161 | 0.14614079 | 8.095688475 | -0.712664092 | 0.502068579 | 0.762897008 | -4.783702358 |
| **A_51_P304397** | Cpm | 0.135034623 | -0.325388933 | 0.595458178 | 12.09069222 | 0.712640974 | 0.502081922 | 0.762897008 | -4.783710367 |
| **A_55_P2020831** | NA | 0.095837042 | -0.230945639 | 0.422619722 | 12.45075667 | 0.712619263 | 0.502094453 | 0.762897008 | -4.783717889 |
| **A_55_P2141429** | Park7 | -0.092430826 | -0.407622553 | 0.2227609 | 13.32195939 | -0.712566196 | 0.502125083 | 0.762897008 | -4.783736273 |
| **A_51_P510764** | Agmo | 0.198291683 | -0.477969515 | 0.87455288 | 9.021866565 | 0.712480887 | 0.502174326 | 0.762911856 | -4.783765824 |
| **A_51_P472329** | Meak7 | -0.051669644 | -0.227936349 | 0.124597061 | 8.575348556 | -0.712276102 | 0.502292546 | 0.763031486 | -4.783836751 |
| **A_51_P506312** | Cdkn2aipnl | 0.055394583 | -0.133736093 | 0.244525258 | 11.36098884 | 0.711686142 | 0.502633228 | 0.763470098 | -4.784040996 |
| **A_55_P2148348** | NA | 0.052764018 | -0.12739712 | 0.232925156 | 13.45180268 | 0.711639316 | 0.502660276 | 0.763470098 | -4.784057202 |
| **A_55_P2180292** | Dmrtc1c2 | -0.051491064 | -0.227324524 | 0.124342395 | 7.741894886 | -0.711563296 | 0.502704187 | 0.7634768 | -4.78408351 |
| **A_55_P2115507** | Egfl8 | 0.064555552 | -0.156076164 | 0.285187268 | 9.084037532 | 0.710966022 | 0.503049281 | 0.763918036 | -4.784290127 |
| **A_51_P291682** | Tmed4 | -0.126779499 | -0.560110677 | 0.30655168 | 10.04793352 | -0.710906165 | 0.503083874 | 0.763918036 | -4.784310826 |
| **A_55_P2096530** | Icmt | 0.062061112 | -0.150078389 | 0.274200612 | 9.05960971 | 0.710855285 | 0.50311328 | 0.763918036 | -4.78432842 |
| **A_52_P137378** | Prkaa2 | -0.058160955 | -0.257004664 | 0.140682754 | 8.044308379 | -0.710727084 | 0.50318738 | 0.763970533 | -4.784372747 |
| **A_55_P2095266** | NA | 0.127972882 | -0.309680363 | 0.565626128 | 14.73918127 | 0.710511297 | 0.503312119 | 0.764099902 | -4.784447343 |
| **A_55_P2021465** | Phldb2 | -0.169679515 | -0.750072265 | 0.410713235 | 10.33352931 | -0.710379833 | 0.503388124 | 0.76415527 | -4.784492781 |
| **A_52_P324767** | Tmed8 | -0.062477164 | -0.276216598 | 0.151262269 | 8.314558919 | -0.710264068 | 0.503455059 | 0.764196862 | -4.784532788 |
| **A_55_P2037702** | Gm10638 | -0.107375705 | -0.474826717 | 0.260075307 | 8.354394195 | -0.71005137 | 0.503578056 | 0.764253309 | -4.784606279 |
| **A_51_P195374** | Baiap2l1 | -0.124546017 | -0.550781198 | 0.301689165 | 9.080468829 | -0.71000881 | 0.50360267 | 0.764253309 | -4.784620983 |
| **A_52_P430247** | Nacc1 | 0.068362377 | -0.165627759 | 0.302352512 | 9.43518785 | 0.70990918 | 0.503660292 | 0.764253309 | -4.7846554 |
| **A_55_P2037538** | NA | -0.124061619 | -0.54870825 | 0.300585013 | 9.665959859 | -0.70989309 | 0.503669598 | 0.764253309 | -4.784660958 |
| **A_55_P2030501** | Ptpa | -0.091638678 | -0.405321383 | 0.222044027 | 13.90413649 | -0.709857915 | 0.503689943 | 0.764253309 | -4.784673108 |
| **A_51_P197259** | Cct7 | -0.081058781 | -0.358625648 | 0.196508085 | 13.21315395 | -0.709603318 | 0.503837217 | 0.764318711 | -4.784761036 |
| **A_55_P2063505** | NA | -0.501309331 | -2.218202865 | 1.215584204 | 11.05779728 | -0.709488911 | 0.503903406 | 0.764318711 | -4.78480054 |
| **A_51_P291388** | Pafah1b1 | 0.091278305 | -0.221349151 | 0.403905761 | 11.00743484 | 0.709453015 | 0.503924175 | 0.764318711 | -4.784812934 |
| **A_52_P255849** | NA | -0.050981587 | -0.225596815 | 0.123633641 | 8.069311252 | -0.709437961 | 0.503932885 | 0.764318711 | -4.784818131 |
| **A_51_P375146** | Cd36 | -0.154806361 | -0.685036403 | 0.37542368 | 10.29881161 | -0.709426916 | 0.503939275 | 0.764318711 | -4.784821944 |
| **A_55_P1974189** | Ptges | -0.056071837 | -0.24813901 | 0.135995335 | 7.798354556 | -0.709373292 | 0.503970303 | 0.764318711 | -4.784840458 |
| **A_55_P1976108** | Rps27a | 0.07818955 | -0.18977103 | 0.346150129 | 17.20695874 | 0.709024095 | 0.504172381 | 0.764565193 | -4.784960988 |
| **A_51_P300709** | Srm | 0.109107141 | -0.264880196 | 0.483094477 | 12.37367192 | 0.708891007 | 0.504249412 | 0.764609712 | -4.785006913 |
| **A_55_P2136501** | Midn | -0.079352311 | -0.351369644 | 0.192665022 | 11.93480779 | -0.70883669 | 0.504280853 | 0.764609712 | -4.785025655 |

| | | | | | | | | | |
|---|---|---|---|---|---|---|---|---|---|
| **A_51_P386562** | Ebf2 | -0.047649176 | -0.211020023 | 0.11572167 | 7.673647922 | -0.708702603 | 0.504358474 | 0.764637477 | -4.785071915 |
| **A_51_P119429** | Nckap1l | 0.133024898 | -0.323144876 | 0.589194672 | 9.974019402 | 0.708581189 | 0.504428765 | 0.764637477 | -4.785113798 |
| **A_51_P355413** | Nisch | -0.089969142 | -0.398507398 | 0.218569114 | 14.45022698 | -0.708545505 | 0.504449425 | 0.764637477 | -4.785126106 |
| **A_66_P111562** | Ccnd1 | 0.271227444 | -0.658932252 | 1.201387141 | 10.97695995 | 0.708531728 | 0.504457401 | 0.764637477 | -4.785130858 |
| **A_55_P1957424** | Sulf1 | -0.056017848 | -0.248156223 | 0.136120526 | 7.724996677 | -0.708427641 | 0.504517669 | 0.764668864 | -4.785166757 |
| **A_55_P2123566** | NA | -0.053549639 | -0.237271323 | 0.130172045 | 8.04834908 | -0.708238152 | 0.504627398 | 0.764775205 | -4.785232101 |
| **A_51_P160907** | Morc3 | 0.112965637 | -0.274650691 | 0.500581965 | 9.69551717 | 0.708153622 | 0.504676352 | 0.764789432 | -4.785261246 |
| **A_66_P135194** | Yrdc | 0.054638073 | -0.132889834 | 0.242165981 | 8.728856054 | 0.707966417 | 0.504784781 | 0.764865466 | -4.785325783 |
| **A_55_P2083063** | Mob1b | 0.056458599 | -0.137327559 | 0.250244758 | 8.121227338 | 0.707930357 | 0.504805668 | 0.764865466 | -4.785338212 |
| **A_51_P110371** | Wwox | -0.054557477 | -0.241872968 | 0.132758015 | 8.002864608 | -0.707723746 | 0.504925357 | 0.764936218 | -4.785409421 |
| **A_55_P2042685** | NA | -0.063520669 | -0.281636412 | 0.154595074 | 10.45185538 | -0.707638148 | 0.50497495 | 0.764936218 | -4.785438918 |
| **A_51_P289742** | Slc27a5 | -0.222018853 | -0.984588441 | 0.540550735 | 11.49939421 | -0.707446434 | 0.505086033 | 0.764936218 | -4.785504972 |
| **A_55_P1952434** | Tm2d1 | 0.06042093 | -0.147115676 | 0.267957536 | 10.8340041 | 0.707417639 | 0.505102719 | 0.764936218 | -4.785514892 |
| **A_52_P501875** | Ccdc173 | -0.075991932 | -0.337014077 | 0.185030214 | 7.821050743 | -0.70741346 | 0.505105141 | 0.764936218 | -4.785516332 |
| **A_66_P106065** | Ttc3 | -0.101496212 | -0.450125887 | 0.247133463 | 8.684595 | -0.707405891 | 0.505109527 | 0.764936218 | -4.78551894 |
| **A_55_P1961645** | Vcp | 0.061631251 | -0.150076394 | 0.273338897 | 13.5030273 | 0.707371623 | 0.505129385 | 0.764936218 | -4.785530744 |
| **A_55_P1989102** | Hmgn1 | -0.110553723 | -0.490473927 | 0.269366481 | 10.7946972 | -0.707072869 | 0.505302532 | 0.765138476 | -4.785633643 |
| **A_52_P559975** | Cxcr2 | -0.177543528 | -0.78774103 | 0.432653973 | 9.092373699 | -0.706997064 | 0.505346472 | 0.76514507 | -4.785659747 |
| **A_55_P2030775** | Fnbp1 | 0.198999777 | -0.485123516 | 0.883123071 | 11.40074866 | 0.706807911 | 0.505456126 | 0.765230116 | -4.785724873 |
| **A_55_P2011151** | Tmprss3 | 0.216323832 | -0.527402895 | 0.96005056 | 12.38224445 | 0.70676359 | 0.505481821 | 0.765230116 | -4.785740131 |
| **A_55_P1975475** | Mdm2 | -0.052300969 | -0.232269097 | 0.127667159 | 11.71681755 | -0.706150586 | 0.505837303 | 0.765708296 | -4.78595109 |
| **A_52_P124173** | Zdhhc8 | -0.051664596 | -0.229514822 | 0.12618563 | 7.750436291 | -0.705865255 | 0.506002824 | 0.765821124 | -4.786049235 |
| **A_66_P107379** | Mdk | -0.449864674 | -1.99849202 | 1.098762671 | 9.346802421 | -0.705859325 | 0.506006264 | 0.765821124 | -4.786051275 |
| **A_55_P2048769** | Zfp454 | -0.070496667 | -0.313191673 | 0.17219834 | 8.046028698 | -0.70581508 | 0.506031934 | 0.765821124 | -4.786066491 |
| **A_55_P2128606** | Nr1h5 | 0.138933995 | -0.339411283 | 0.617279274 | 8.83192396 | 0.70574892 | 0.50607032 | 0.765821124 | -4.786089243 |
| **A_52_P539414** | Gtf2h3 | -0.068666036 | -0.305110109 | 0.167778036 | 9.79014621 | -0.705662008 | 0.506120749 | 0.765837479 | -4.786119128 |
| **A_55_P1957267** | Rab9 | -0.066250999 | -0.294414344 | 0.161912346 | 11.67174131 | -0.705553162 | 0.50618391 | 0.765873095 | -4.786156551 |
| **A_55_P2070331** | NA | -0.057822247 | -0.257018321 | 0.141373828 | 7.967472418 | -0.705338153 | 0.50630869 | 0.765914834 | -4.786230463 |
| **A_52_P19532** | Fgd3 | 0.064320951 | -0.157264014 | 0.285905916 | 7.984129314 | 0.705334874 | 0.506310593 | 0.765914834 | -4.78623159 |
| **A_52_P16877** | Tmcc3 | 0.061581074 | -0.150575296 | 0.273737445 | 7.949503932 | 0.705300796 | 0.506330372 | 0.765914834 | -4.786243303 |
| **A_55_P2115257** | Fer | -0.05535925 | -0.24612115 | 0.135402649 | 9.392512976 | -0.705150391 | 0.506417673 | 0.765986948 | -4.786294994 |
| **A_55_P2173333** | Plcb4 | 0.097157427 | -0.237699292 | 0.432014147 | 9.277396198 | 0.705017955 | 0.506494553 | 0.766043288 | -4.786340502 |
| **A_52_P650855** | Myo1d | -0.140466033 | -0.624725447 | 0.343793381 | 12.12831985 | -0.704817103 | 0.506611164 | 0.766159705 | -4.786409507 |
| **A_55_P1960083** | NA | 0.076612545 | -0.187576271 | 0.340801361 | 14.66272353 | 0.7046422 | 0.506712724 | 0.766253343 | -4.786469585 |
| **A_55_P2009895** | Gm5113 | -0.046782942 | -0.208152178 | 0.114586294 | 7.841014453 | -0.704449683 | 0.506824527 | 0.766362456 | -4.7865357 |
| **A_55_P2015032** | Kazald1 | -0.095758471 | -0.426172017 | 0.234655074 | 7.891079065 | -0.704210589 | 0.506963402 | 0.766395067 | -4.786617791 |
| **A_51_P495825** | Apof | -0.221876315 | -0.987539071 | 0.54378644 | 14.20950257 | -0.704136102 | 0.507006672 | 0.766395067 | -4.786643361 |
| **A_52_P297332** | Aida | 0.076234374 | -0.186864522 | 0.33933327 | 8.491051653 | 0.704068646 | 0.50704586 | 0.766395067 | -4.786666515 |
| **A_51_P348397** | Hexim1 | 0.083663214 | -0.205085398 | 0.372411826 | 9.206076542 | 0.704040803 | 0.507062035 | 0.766395067 | -4.786676072 |
| **A_55_P2125322** | NA | -0.095417104 | -0.424744091 | 0.233909883 | 15.74790778 | -0.704015304 | 0.507076849 | 0.766395067 | -4.786684824 |
| **A_55_P2105888** | G6pc3 | -0.072303498 | -0.321882923 | 0.177275927 | 9.797558008 | -0.703936877 | 0.507122415 | 0.766395067 | -4.786711741 |
| **A_52_P73559** | NA | 0.050368688 | -0.123496011 | 0.224233387 | 8.666001001 | 0.703934762 | 0.507123643 | 0.766395067 | -4.786712467 |
| **A_55_P2130855** | Nap1l1 | 0.103227014 | -0.25314377 | 0.459597798 | 11.09775047 | 0.703840836 | 0.507178218 | 0.766417619 | -4.7867447 |
| **A_55_P1966908** | Zfp40 | -0.071753048 | -0.319500689 | 0.175994594 | 8.113721801 | -0.703742876 | 0.50723514 | 0.766432512 | -4.786778314 |
| **A_51_P461877** | Thbs2 | -0.052508372 | -0.233822633 | 0.12880589 | 8.002237841 | -0.703687401 | 0.507267377 | 0.766432512 | -4.786797348 |
| **A_55_P2038747** | Ano1 | -0.644613716 | -2.871002212 | 1.581774779 | 9.586615828 | -0.70352864 | 0.507359642 | 0.766499395 | -4.786851814 |
| **A_55_P1963687** | Tsen54 | -0.094803757 | -0.422265232 | 0.232657718 | 11.31291964 | -0.703474764 | 0.507390954 | 0.766499395 | -4.786870295 |
| **A_55_P2121603** | 2810047C21Rik1 | 0.050487945 | -0.123950962 | 0.224926852 | 8.085050221 | 0.703278798 | 0.507504862 | 0.766611556 | -4.786937508 |
| **A_55_P2011087** | Gnptg | -0.057894251 | -0.258013986 | 0.142225484 | 10.9112497 | -0.702956921 | 0.507691991 | 0.766792291 | -4.787047875 |
| **A_51_P166244** | Osbpl1a | 0.097991253 | -0.240766848 | 0.436749353 | 11.94646109 | 0.702879392 | 0.507737072 | 0.766792291 | -4.787074453 |

| | | | | | | | | | |
|---|---|---|---|---|---|---|---|---|---|
| **A_66_P104190** | Gsk3b | 0.074972704 | -0.184213813 | 0.334159221 | 8.856937404 | 0.702868299 | 0.507743522 | 0.766792291 | -4.787078256 |
| **A_55_P2171196** | Gm3411 | -0.09717954 | -0.433237551 | 0.238878472 | 7.873141723 | -0.702657641 | 0.507866025 | 0.766863519 | -4.787150459 |
| **A_55_P2067952** | NA | 0.235374889 | -0.578586502 | 1.049336281 | 13.14603017 | 0.702650747 | 0.507870035 | 0.766863519 | -4.787152822 |
| **A_55_P1958200** | P2rx1 | 0.145384478 | -0.357601581 | 0.648370536 | 10.08418451 | 0.702336569 | 0.508052775 | 0.767079526 | -4.787260476 |
| **A_52_P535907** | Sde2 | 0.098113764 | -0.241388358 | 0.437615887 | 9.450339338 | 0.70221586 | 0.508122997 | 0.767101771 | -4.787301827 |
| **A_52_P488427** | Sec14l2 | -0.103250903 | -0.460549905 | 0.2540481 | 10.56370901 | -0.702174804 | 0.508146882 | 0.767101771 | -4.78731589 |
| **A_55_P2168823** | NA | 0.130561807 | -0.321365009 | 0.582488624 | 13.51301959 | 0.701990493 | 0.508254119 | 0.767203738 | -4.787379016 |
| **A_55_P1976376** | Gimap4 | -0.061496763 | -0.274401875 | 0.151408348 | 8.371540987 | -0.701858169 | 0.508331118 | 0.767260048 | -4.787424329 |
| **A_55_P2185359** | NA | -0.076498333 | -0.341447727 | 0.188451062 | 14.10349292 | -0.701571967 | 0.508497684 | 0.767324526 | -4.787522313 |
| **A_55_P2123673** | Trnp1 | -0.124807059 | -0.557112274 | 0.307498156 | 8.113567172 | -0.701506764 | 0.508535636 | 0.767324526 | -4.787544631 |
| **A_51_P511315** | Pstpip1 | -0.100918844 | -0.450517537 | 0.248679848 | 8.719352288 | -0.701432135 | 0.508579077 | 0.767324526 | -4.787570174 |
| **A_55_P2168048** | Gramd4 | 0.048191951 | -0.118754059 | 0.21513796 | 8.184708362 | 0.701425664 | 0.508582844 | 0.767324526 | -4.787572389 |
| **A_55_P1962122** | NA | 0.063893537 | -0.157462368 | 0.285249442 | 15.5129562 | 0.701372944 | 0.508613534 | 0.767324526 | -4.787590432 |
| **A_55_P2121081** | Cul2 | 0.064725594 | -0.159528982 | 0.288980171 | 9.200797917 | 0.701322738 | 0.508642761 | 0.767324526 | -4.787607613 |
| **A_55_P2083382** | Snf8 | 0.063298266 | -0.156021077 | 0.28261761 | 13.09145988 | 0.701290677 | 0.508661427 | 0.767324526 | -4.787618585 |
| **A_51_P328883** | Zfp810 | -0.061221199 | -0.273382121 | 0.150939723 | 8.334243854 | -0.701164024 | 0.508735164 | 0.767324526 | -4.787661922 |
| **A_55_P1968982** | Pbx3 | -0.095782127 | -0.427733089 | 0.236168835 | 9.261020926 | -0.701122225 | 0.508759501 | 0.767324526 | -4.787676223 |
| **A_55_P2039463** | Timmdc1 | 0.086672984 | -0.213716857 | 0.387062824 | 12.16698982 | 0.70110279 | 0.508770817 | 0.767324526 | -4.787682872 |
| **A_55_P2114009** | Ankra2 | 0.058091178 | -0.14326121 | 0.259443566 | 10.15993204 | 0.701029973 | 0.508813216 | 0.7673286 | -4.787707784 |
| **A_51_P113195** | Upk1b | -0.091715736 | -0.409656349 | 0.226224877 | 7.942599427 | -0.70094031 | 0.508865428 | 0.767347469 | -4.787738456 |
| **A_55_P2054350** | Fbxo44 | -0.055847695 | -0.249497252 | 0.137801862 | 7.880938569 | -0.700764246 | 0.508967962 | 0.767396927 | -4.787798676 |
| **A_51_P365694** | Rala | -0.113104608 | -0.505353382 | 0.279144166 | 11.28604047 | -0.700651239 | 0.509033781 | 0.767396927 | -4.787837322 |
| **A_51_P268343** | Aagab | 0.046740869 | -0.115363131 | 0.208844868 | 11.61199784 | 0.700625982 | 0.509048492 | 0.767396927 | -4.787845959 |
| **A_55_P2062469** | Col12a1 | 0.08780394 | -0.216719022 | 0.392326903 | 8.576469591 | 0.700611318 | 0.509057034 | 0.767396927 | -4.787850973 |
| **A_65_P09443** | Psma4 | 0.062717331 | -0.154897193 | 0.280331855 | 12.64339834 | 0.700297987 | 0.509239562 | 0.767570475 | -4.787958096 |
| **A_52_P52618** | Csf2rb | -0.0948763 | -0.42413846 | 0.234385859 | 10.73715792 | -0.700162923 | 0.509318256 | 0.767570475 | -4.788004261 |
| **A_51_P466371** | Pitpna | 0.069990153 | -0.172924363 | 0.312904669 | 10.0370218 | 0.700110619 | 0.509348732 | 0.767570475 | -4.788022136 |
| **A_55_P1980817** | Gm6548 | 0.065644231 | -0.162197249 | 0.293485711 | 15.52934241 | 0.70007886 | 0.509367238 | 0.767570475 | -4.78803299 |
| **A_55_P1960167** | Bcat2 | 0.072531553 | -0.179216951 | 0.324280056 | 9.753878078 | 0.700072899 | 0.509370712 | 0.767570475 | -4.788035027 |
| **A_51_P149775** | Spcs1 | 0.056138489 | -0.13878897 | 0.251065948 | 14.06808496 | 0.699795081 | 0.509532616 | 0.767750648 | -4.788129954 |
| **A_55_P1972839** | NA | -0.073681358 | -0.329545505 | 0.182182788 | 16.3526346 | -0.699731425 | 0.509569717 | 0.767750648 | -4.7881517 |
| **A_55_P2039190** | Gfra4 | 0.049796331 | -0.12318345 | 0.222776113 | 8.17964657 | 0.699495929 | 0.509706991 | 0.767877198 | -4.788232138 |
| **A_51_P171772** | Bcl11b | -0.053020857 | -0.237213641 | 0.131171928 | 7.656952162 | -0.699451033 | 0.509733164 | 0.767877198 | -4.78824747 |
| **A_51_P357877** | Fbxo46 | 0.048626287 | -0.120360295 | 0.217612869 | 9.072519814 | 0.699201058 | 0.509878908 | 0.767947433 | -4.788332826 |
| **A_51_P503866** | Edem1 | -0.156734284 | -0.70151414 | 0.388045573 | 10.28811442 | -0.699078832 | 0.509950181 | 0.767947433 | -4.788374553 |
| **A_55_P1964347** | NA | 0.062578267 | -0.154935967 | 0.280092502 | 8.138949731 | 0.699067384 | 0.509956857 | 0.767947433 | -4.788378461 |
| **A_55_P2012689** | Stk16 | -0.061215898 | -0.273998619 | 0.151566824 | 10.30256806 | -0.699054528 | 0.509964354 | 0.767947433 | -4.788382849 |
| **A_55_P2120931** | Dleu2 | 0.154219221 | -0.381898344 | 0.690336785 | 9.279776806 | 0.698975023 | 0.510010719 | 0.767947433 | -4.788409987 |
| **A_55_P2178525** | Rpl26 | -0.078023273 | -0.349263211 | 0.193216664 | 15.65506179 | -0.698962248 | 0.51001817 | 0.767947433 | -4.788414348 |
| **A_55_P2131303** | NA | -0.079947435 | -0.358044082 | 0.198149212 | 16.79640268 | -0.698541111 | 0.510263815 | 0.76825746 | -4.788558057 |
| **A_51_P221132** | L2hgdh | 0.126807116 | -0.31439173 | 0.568005961 | 9.892821548 | 0.69838105 | 0.510357197 | 0.768338208 | -4.788612659 |
| **A_55_P2074005** | Tctn1 | 0.053171076 | -0.131855843 | 0.238197995 | 8.370957214 | 0.698270546 | 0.510421673 | 0.76837543 | -4.788650349 |
| **A_51_P211968** | Lcmt1 | 0.051667647 | -0.128163269 | 0.231498563 | 9.998508122 | 0.698131953 | 0.510502547 | 0.768437328 | -4.788697614 |
| **A_51_P221651** | Coq8a | -0.152456104 | -0.683295042 | 0.378382834 | 9.624529419 | -0.697855063 | 0.510664146 | 0.768620719 | -4.788792021 |
| **A_55_P2110718** | Npm1 | 0.109651987 | -0.272186115 | 0.491490089 | 11.83459154 | 0.697783076 | 0.510706164 | 0.76862411 | -4.788816561 |
| **A_55_P1967231** | Stbd1 | -0.104602444 | -0.469019965 | 0.259815077 | 13.26180733 | -0.697470386 | 0.510888708 | 0.768838978 | -4.78892313 |
| **A_55_P2068566** | NA | -0.078960655 | -0.354214803 | 0.196293493 | 14.28951846 | -0.697043786 | 0.511137819 | 0.769009974 | -4.789068463 |
| **A_55_P2036638** | Ints11 | 0.078035393 | -0.19403232 | 0.350103106 | 10.54830935 | 0.696943874 | 0.511196173 | 0.769009974 | -4.78910249 |
| **A_55_P2049072** | NA | -0.080198769 | -0.359832476 | 0.199434938 | 13.23177313 | -0.696885409 | 0.511230322 | 0.769009974 | -4.7891224 |
| **A_55_P2229351** | NA | 0.059351042 | -0.147659215 | 0.266361298 | 8.355614509 | 0.696658067 | 0.511363126 | 0.769009974 | -4.789199808 |

| | | | | | | | | | |
|---|---|---|---|---|---|---|---|---|---|
| **A_55_P2053933** | Foxa1 | -0.247875095 | -1.112445826 | 0.616695636 | 10.08784032 | -0.696651464 | 0.511366984 | 0.769009974 | -4.789202057 |
| **A_55_P1997126** | Ctse | 0.112140154 | -0.279059697 | 0.503340005 | 9.841001159 | 0.696539349 | 0.511432485 | 0.769009974 | -4.789240223 |
| **A_55_P2128224** | NA | -0.071610963 | -0.321431201 | 0.178209275 | 16.33754384 | -0.696522397 | 0.51144239 | 0.769009974 | -4.789245994 |
| **A_55_P1983858** | Scel | -0.061018844 | -0.273897794 | 0.151860106 | 7.842226718 | -0.696489289 | 0.511461733 | 0.769009974 | -4.789257263 |
| **A_55_P2011991** | St7 | -0.082454383 | -0.370131946 | 0.20522318 | 10.25441346 | -0.696451608 | 0.511483751 | 0.769009974 | -4.789270089 |
| **A_52_P298002** | Gch1 | -0.064688518 | -0.290386261 | 0.161009225 | 8.271799159 | -0.696439149 | 0.51149103 | 0.769009974 | -4.78927433 |
| **A_51_P256384** | Atp2b2 | 0.155433985 | -0.386894738 | 0.697762708 | 9.492949686 | 0.69641251 | 0.511506595 | 0.769009974 | -4.789283397 |
| **A_51_P429335** | Prss16 | -0.061019431 | -0.273950659 | 0.151911796 | 7.872972489 | -0.696324999 | 0.511557731 | 0.769009974 | -4.78931318 |
| **A_55_P2044385** | Fgfbp3 | 0.107595055 | -0.267866 | 0.48305611 | 8.793127417 | 0.696322785 | 0.511559025 | 0.769009974 | -4.789313934 |
| **A_55_P2155826** | Rplp1 | -0.054496415 | -0.244666312 | 0.135673481 | 16.66089133 | -0.69632078 | 0.511560197 | 0.769009974 | -4.789314616 |
| **A_55_P2126627** | Phf20l1 | 0.056244514 | -0.140068436 | 0.252557464 | 8.360951545 | 0.696168567 | 0.511649148 | 0.769009974 | -4.789366413 |
| **A_55_P1989653** | Slco4a1 | -0.121697675 | -0.54649103 | 0.303095681 | 8.490629058 | -0.696125838 | 0.51167412 | 0.769009974 | -4.789380952 |
| **A_55_P2029763** | Peg10 | 0.0518129 | -0.129045301 | 0.232671101 | 7.796594483 | 0.696118031 | 0.511678683 | 0.769009974 | -4.789383608 |
| **A_55_P1997894** | NA | -0.074526578 | -0.334703286 | 0.185650131 | 15.35999566 | -0.696026784 | 0.511732014 | 0.76903033 | -4.789414653 |
| **A_55_P2136662** | 8030462N17Rik | 0.072426606 | -0.180517587 | 0.3253708 | 8.35208271 | 0.69575544 | 0.511890626 | 0.76911376 | -4.789506953 |
| **A_51_P181746** | Zfyve1 | -0.071041578 | -0.319155085 | 0.177071928 | 8.710254363 | -0.695737456 | 0.51190114 | 0.76911376 | -4.789513069 |
| **A_51_P385635** | Mrpl53 | -0.123873632 | -0.556572373 | 0.308825108 | 13.35144618 | -0.695627 | 0.511965717 | 0.76911376 | -4.789550633 |
| **A_51_P456914** | Bmt2 | -0.086688829 | -0.389502003 | 0.216124346 | 8.47274914 | -0.695619196 | 0.511970279 | 0.76911376 | -4.789553287 |
| **A_52_P596008** | Zeb2 | -0.118227484 | -0.531295153 | 0.294840185 | 10.24324652 | -0.695473253 | 0.512055611 | 0.76911376 | -4.789602912 |
| **A_51_P475249** | Rnf44 | 0.116601095 | -0.290797473 | 0.523999664 | 10.44135063 | 0.695450656 | 0.512068825 | 0.76911376 | -4.789610595 |
| **A_51_P348804** | Amotl2 | -0.122254124 | -0.549457503 | 0.304949255 | 9.282183291 | -0.695363719 | 0.512119662 | 0.76911376 | -4.789640151 |
| **A_66_P127463** | NA | 0.061557076 | -0.15357395 | 0.276688102 | 7.995593504 | 0.695277413 | 0.512170133 | 0.76911376 | -4.78966949 |
| **A_55_P2152567** | Plekha5 | -0.050233541 | -0.225829088 | 0.125362005 | 7.971339576 | -0.695125929 | 0.512258727 | 0.76911376 | -4.789720979 |
| **A_55_P2096144** | Gnao1 | -0.05537585 | -0.248955784 | 0.138204084 | 7.738909041 | -0.69509355 | 0.512277666 | 0.76911376 | -4.789731983 |
| **A_55_P2183777** | NA | 0.088554307 | -0.221013492 | 0.398122107 | 9.380219103 | 0.695083691 | 0.512283432 | 0.76911376 | -4.789735334 |
| **A_55_P2051322** | Efhd2 | -0.057318136 | -0.257693468 | 0.143057196 | 11.34656058 | -0.695073919 | 0.512289148 | 0.76911376 | -4.789738655 |
| **A_52_P649285** | NA | 0.075872168 | -0.189375405 | 0.34111974 | 9.859269386 | 0.695047143 | 0.512304809 | 0.76911376 | -4.789747755 |
| **A_55_P1977533** | Eno3 | -0.194233326 | -0.873442304 | 0.484975652 | 9.308871259 | -0.694869931 | 0.51240847 | 0.76920964 | -4.789807972 |
| **A_51_P141264** | Zfp24 | -0.047354473 | -0.213050293 | 0.118341348 | 8.309691738 | -0.694436661 | 0.51266197 | 0.769317884 | -4.789955149 |
| **A_55_P2115995** | Sms | 0.056098584 | -0.14021866 | 0.252415828 | 9.548920327 | 0.694347119 | 0.51271437 | 0.769317884 | -4.789985557 |
| **A_51_P176387** | Hook3 | -0.063549613 | -0.285946235 | 0.158847009 | 8.230806335 | -0.69433317 | 0.512722534 | 0.769317884 | -4.789990294 |
| **A_52_P250555** | Dynll1 | 0.058467856 | -0.146183479 | 0.26311919 | 8.399541007 | 0.694201861 | 0.512799383 | 0.769317884 | -4.790034879 |
| **A_51_P425696** | Nt5c2 | -0.091503964 | -0.411804696 | 0.228796768 | 9.193578007 | -0.694168925 | 0.51281866 | 0.769317884 | -4.790046061 |
| **A_55_P2146620** | Atf2 | -0.055093607 | -0.247948882 | 0.137761668 | 8.29648811 | -0.694149276 | 0.512830161 | 0.769317884 | -4.790052732 |
| **A_55_P2152566** | Sp110 | -0.099155364 | -0.446259952 | 0.247949224 | 10.14657348 | -0.694127173 | 0.512843098 | 0.769317884 | -4.790060235 |
| **A_52_P91618** | Lysmd3 | 0.10659849 | -0.266631462 | 0.479828442 | 9.416101231 | 0.693997257 | 0.512919144 | 0.769317884 | -4.790104338 |
| **A_52_P430934** | Dcun1d1 | 0.057981748 | -0.145030541 | 0.260994038 | 9.590869299 | 0.69398833 | 0.512924369 | 0.769317884 | -4.790107368 |
| **A_55_P2234744** | Spice1 | -0.067244977 | -0.302696767 | 0.168206813 | 8.923386608 | -0.693970551 | 0.512934776 | 0.769317884 | -4.790113402 |
| **A_66_P126189** | Ercc5 | 0.045452061 | -0.113698122 | 0.204602244 | 8.588872916 | 0.693952319 | 0.51294545 | 0.769317884 | -4.790119591 |
| **A_51_P236197** | Polr1a | 0.120525679 | -0.301507665 | 0.542559022 | 10.97350673 | 0.693930553 | 0.512958191 | 0.769317884 | -4.790126979 |
| **A_55_P2207055** | NA | 0.062291984 | -0.155871902 | 0.28045587 | 8.001639825 | 0.693797114 | 0.513036311 | 0.769375347 | -4.790172266 |
| **A_51_P497317** | Tcp11l2 | 0.104852472 | -0.26242275 | 0.472127694 | 11.15271898 | 0.693697666 | 0.513094536 | 0.769402971 | -4.790206014 |
| **A_51_P271725** | Fundc1 | -0.090252168 | -0.406442484 | 0.225938148 | 10.22885639 | -0.693573142 | 0.513167449 | 0.769428299 | -4.790248265 |
| **A_55_P2090698** | NA | -0.09875302 | -0.444753019 | 0.247246979 | 15.18984577 | -0.693517584 | 0.513199982 | 0.769428299 | -4.790267114 |
| **A_55_P1973279** | NA | 0.066918467 | -0.167569689 | 0.301406623 | 16.73893362 | 0.693439 | 0.513246001 | 0.769428299 | -4.790293773 |
| **A_52_P503334** | Erlin2 | -0.063587512 | -0.286417239 | 0.159242214 | 9.725820838 | -0.693396902 | 0.513270655 | 0.769428299 | -4.790308054 |
| **A_55_P1997837** | Osbp2 | -0.044007471 | -0.198247751 | 0.110232809 | 7.743707859 | -0.693284994 | 0.513336195 | 0.769466871 | -4.790346012 |
| **A_55_P1978052** | Gatb | 0.065745249 | -0.164732601 | 0.296223098 | 8.322151785 | 0.69313586 | 0.513423545 | 0.769492996 | -4.790396589 |
| **A_55_P2219059** | Lmnb2 | 0.110611239 | -0.277175208 | 0.498397686 | 9.612828858 | 0.693090299 | 0.513450233 | 0.769492996 | -4.790412039 |
| **A_55_P2022434** | Gpi1 | -0.114101484 | -0.514179476 | 0.285976509 | 14.8586698 | -0.692994549 | 0.513506323 | 0.769492996 | -4.790444506 |

| | | | | | | | | | |
|---|---|---|---|---|---|---|---|---|---|
| **A_66_P132777** | Cep68 | 0.06123664 | -0.153494764 | 0.275968044 | 9.488270517 | 0.692945338 | 0.513535152 | 0.769492996 | -4.790461191 |
| **A_55_P2084238** | NA | 0.060522404 | -0.151713634 | 0.272758442 | 13.03031744 | 0.692915428 | 0.513552674 | 0.769492996 | -4.790471332 |
| **A_55_P1990261** | Chchd6 | -0.157561673 | -0.710143208 | 0.395019862 | 10.70614682 | -0.692847102 | 0.513592704 | 0.769493324 | -4.790494495 |
| **A_51_P336391** | Tmem18 | 0.133545877 | -0.334919528 | 0.602011282 | 10.37557949 | 0.692685584 | 0.513687339 | 0.769575459 | -4.790549245 |
| **A_51_P316173** | Vmn1r192 | 0.185932703 | -0.466442766 | 0.838308172 | 9.090286916 | 0.692534398 | 0.513775931 | 0.769648529 | -4.790600484 |
| **A_55_P2013948** | Gsn | -0.103458222 | -0.466559402 | 0.259642957 | 13.36941674 | -0.692341781 | 0.513888816 | 0.769757976 | -4.790665752 |
| **A_51_P149349** | Nfxl1 | 0.081909351 | -0.205669875 | 0.369488577 | 8.518099591 | 0.692084564 | 0.514039584 | 0.769924148 | -4.790752887 |
| **A_55_P1963767** | Npr2 | -0.113357792 | -0.511425058 | 0.284709474 | 10.93026775 | -0.691955398 | 0.514115307 | 0.769977899 | -4.790796634 |
| **A_52_P451888** | Tlk2 | -0.065230605 | -0.294369445 | 0.163908236 | 8.449991839 | -0.691728839 | 0.514248142 | 0.770117172 | -4.790873352 |
| **A_52_P154501** | Ube2q2 | 0.063049301 | -0.15847479 | 0.284573393 | 8.701953343 | 0.691580125 | 0.514335348 | 0.770188096 | -4.790923699 |
| **A_52_P601980** | Kdm5b | -0.064995489 | -0.293423432 | 0.163432454 | 8.15979904 | -0.691380581 | 0.514452376 | 0.770263758 | -4.790991242 |
| **A_51_P348456** | Rest | 0.074627253 | -0.187660254 | 0.336914761 | 8.716766928 | 0.691358075 | 0.514465576 | 0.770263758 | -4.790998858 |
| **A_55_P2000117** | Ceacam13 | -0.088095361 | -0.39779475 | 0.221604028 | 7.872545917 | -0.691187508 | 0.514565624 | 0.770353881 | -4.791056579 |
| **A_55_P2068265** | Mif | -0.09947062 | -0.449267476 | 0.250326235 | 15.49965865 | -0.69097465 | 0.514690498 | 0.770437595 | -4.791128596 |
| **A_55_P2011387** | Tifa | 0.120662196 | -0.303717249 | 0.545041641 | 9.630329605 | 0.690875949 | 0.514748408 | 0.770437595 | -4.791161983 |
| **A_51_P210877** | Pcnp | 0.054764421 | -0.137865051 | 0.247393893 | 11.77086402 | 0.690810542 | 0.514786785 | 0.770437595 | -4.791184106 |
| **A_55_P2038972** | Erich1 | 0.051855977 | -0.130543324 | 0.234255279 | 8.224327149 | 0.690810359 | 0.514786892 | 0.770437595 | -4.791184168 |
| **A_55_P2136962** | NA | -0.096489174 | -0.435910643 | 0.242932294 | 10.4213147 | -0.690752508 | 0.514820838 | 0.770437595 | -4.791203735 |
| **A_55_P2054280** | Fam178b | -0.101836862 | -0.460109217 | 0.256435492 | 8.248048119 | -0.690676854 | 0.514865233 | 0.770444383 | -4.79122932 |
| **A_55_P2077608** | Hdac1 | 0.101580947 | -0.255918233 | 0.459080126 | 9.787500386 | 0.69043118 | 0.515009414 | 0.7705566 | -4.79131239 |
| **A_55_P1957927** | Nono | 0.057671643 | -0.145300489 | 0.260643774 | 13.39350273 | 0.69041322 | 0.515019956 | 0.7705566 | -4.791318462 |
| **A_55_P1979053** | Atp6v0c | -0.055221662 | -0.249697733 | 0.13925441 | 12.93655943 | -0.689964088 | 0.515283614 | 0.770891406 | -4.791470264 |
| **A_51_P279712** | Rell1 | 0.07299024 | -0.184091652 | 0.330072131 | 11.11208658 | 0.689884668 | 0.515330246 | 0.770901503 | -4.7914971 |
| **A_55_P2033445** | Tnfrsf1b | 0.102171738 | -0.257770917 | 0.462114393 | 8.705939746 | 0.68973244 | 0.515419635 | 0.770945662 | -4.791548529 |
| **A_55_P2009267** | Gm16060 | -0.101287592 | -0.458133009 | 0.255557824 | 14.86823045 | -0.689698552 | 0.515439536 | 0.770945662 | -4.791559977 |
| **A_55_P2169514** | Fgf23 | 0.18532712 | -0.467720297 | 0.838374537 | 10.60959468 | 0.689568558 | 0.515515879 | 0.770954186 | -4.791603886 |
| **A_55_P2108451** | Galns | 0.118963457 | -0.300248106 | 0.53817502 | 11.76584991 | 0.689546426 | 0.515528877 | 0.770954186 | -4.791611362 |
| **A_55_P2410007** | NA | 0.071094115 | -0.179454468 | 0.321642698 | 9.029846041 | 0.689485106 | 0.515564893 | 0.770954186 | -4.791632071 |
| **A_55_P1997291** | Fam53a | 0.045469134 | -0.1148038 | 0.205742067 | 11.550597 | 0.689349845 | 0.515644342 | 0.771013342 | -4.791677748 |
| **A_51_P140803** | Slco1b2 | -0.337440349 | -1.527453609 | 0.852572911 | 10.42730926 | -0.689014562 | 0.515841314 | 0.771248202 | -4.791790943 |
| **A_55_P1989798** | NA | -0.094598554 | -0.42824533 | 0.239048223 | 14.99459357 | -0.688938972 | 0.515885728 | 0.771254949 | -4.791816456 |
| **A_55_P2089288** | Rnf121 | 0.068571822 | -0.173334629 | 0.310478274 | 11.11176401 | 0.688781428 | 0.515978304 | 0.771298706 | -4.791869625 |
| **A_55_P2127973** | Gm9112 | -0.050263754 | -0.227590557 | 0.127063049 | 7.707057777 | -0.688753351 | 0.515994804 | 0.771298706 | -4.791879099 |
| **A_55_P2105052** | Tfb2m | 0.089726935 | -0.226974553 | 0.406428422 | 9.904687053 | 0.688423865 | 0.516188456 | 0.771440725 | -4.791990261 |
| **A_55_P2112693** | Sipa1l1 | 0.064694014 | -0.163675757 | 0.293063784 | 8.535988761 | 0.688348973 | 0.516232479 | 0.771440725 | -4.792015523 |
| **A_51_P257675** | Tspyl4 | -0.106307492 | -0.481620297 | 0.269005314 | 7.945281055 | -0.688261819 | 0.516283713 | 0.771440725 | -4.792044917 |
| **A_51_P253547** | Ctsl | -0.122755349 | -0.556152236 | 0.310641538 | 14.41176554 | -0.688236699 | 0.516298482 | 0.771440725 | -4.792053389 |
| **A_51_P102257** | Tns1 | -0.116217616 | -0.52653742 | 0.294102189 | 8.385609386 | -0.688228528 | 0.516303285 | 0.771440725 | -4.792056145 |
| **A_55_P2044364** | Mms22l | 0.16492211 | -0.417391987 | 0.747236208 | 14.7595692 | 0.688184312 | 0.51632928 | 0.771440725 | -4.792071056 |
| **A_52_P316644** | Bet1 | -0.125810571 | -0.570112617 | 0.318491475 | 11.08196429 | -0.688053186 | 0.516406375 | 0.771496276 | -4.792115272 |
| **A_55_P2068733** | Gm21953 | -0.093343385 | -0.423100662 | 0.236413893 | 8.35184389 | -0.68781611 | 0.516545781 | 0.771617781 | -4.792195197 |
| **A_52_P498193** | Aldh1l2 | -0.094575804 | -0.42870487 | 0.239553262 | 8.130743787 | -0.687779101 | 0.516567545 | 0.771617781 | -4.792207672 |
| **A_52_P367745** | Acads | -0.111965341 | -0.507645507 | 0.283714825 | 8.68685515 | -0.687578858 | 0.516685316 | 0.77173406 | -4.79227516 |
| **A_51_P458218** | Wdr13 | -0.064856332 | -0.294141312 | 0.164428648 | 9.854713856 | -0.687321558 | 0.516836669 | 0.771868 | -4.792361856 |
| **A_55_P1965402** | Uqcrb | -0.059138622 | -0.268232575 | 0.149955332 | 13.1911796 | -0.687247095 | 0.516880476 | 0.771868 | -4.792386942 |
| **A_51_P493700** | Cep350 | 0.094865924 | -0.240559435 | 0.430291284 | 9.637511725 | 0.687222766 | 0.51689479 | 0.771868 | -4.792395137 |
| **A_52_P665386** | Ube2m | 0.114564086 | -0.304804044 | 0.533932216 | 10.4008493 | 0.695017016 | 0.516991677 | 0.771952839 | -4.75322406 |
| **A_51_P455997** | Meg3 | -0.188770828 | -0.856449291 | 0.478907635 | 12.03799058 | -0.686990454 | 0.517031479 | 0.771952839 | -4.792473383 |
| **A_55_P2056241** | Pdlim5 | -0.082614207 | -0.37496507 | 0.209736657 | 8.666655037 | -0.686647027 | 0.517233589 | 0.77213622 | -4.792589016 |
| **A_51_P265219** | Ppm1k | -0.166944236 | -0.757718716 | 0.423830244 | 9.585997801 | -0.686645996 | 0.517234196 | 0.77213622 | -4.792589363 |

| | | | | | | | | | |
|---|---|---|---|---|---|---|---|---|---|
| **A_55_P2159575** | Reep6 | 0.153967703 | -0.391005755 | 0.698941161 | 13.94148235 | 0.686495143 | 0.517322991 | 0.77216308 | -4.792640142 |
| **A_51_P438666** | Keap1 | -0.071889854 | -0.326351946 | 0.182572239 | 10.88567235 | -0.686479693 | 0.517332086 | 0.77216308 | -4.792645342 |
| **A_55_P2170737** | Igf2bp2 | -0.057838862 | -0.262586585 | 0.146908862 | 9.016883017 | -0.686410383 | 0.517372887 | 0.772164353 | -4.792668669 |
| **A_66_P120260** | Taf10 | 0.053332761 | -0.135494055 | 0.242159578 | 13.65337044 | 0.686299348 | 0.517438255 | 0.772202287 | -4.792706035 |
| **A_52_P111963** | 1810030O07Rik | -0.053213378 | -0.241644728 | 0.135217971 | 9.334460044 | -0.686200234 | 0.517496609 | 0.77222975 | -4.792739385 |
| **A_55_P2143042** | Gm8096 | 0.153485332 | -0.390149692 | 0.697120356 | 10.31592368 | 0.686029259 | 0.517597282 | 0.772320354 | -4.792796907 |
| **A_55_P2180099** | NA | -0.105649122 | -0.479894522 | 0.268596278 | 13.98498859 | -0.685950235 | 0.517643817 | 0.772330169 | -4.79282349 |
| **A_51_P159896** | Dnajc1 | -0.05698252 | -0.258921415 | 0.144956374 | 10.60051133 | -0.685653769 | 0.517818422 | 0.77253105 | -4.792923195 |
| **A_55_P1997661** | Smim15 | 0.072640129 | -0.184824944 | 0.330105202 | 11.35218522 | 0.685553694 | 0.517877371 | 0.772545544 | -4.792956845 |
| **A_51_P173961** | Pdrg1 | -0.135314021 | -0.614956382 | 0.344328341 | 11.82640158 | -0.685501574 | 0.517908074 | 0.772545544 | -4.792974368 |
| **A_55_P2010341** | Rpl11 | 0.071216096 | -0.181323329 | 0.323755522 | 15.99971186 | 0.685223368 | 0.518071977 | 0.772675047 | -4.793067886 |
| **A_55_P2088905** | Hpca | 0.275625838 | -0.701777981 | 1.253029658 | 11.588701 | 0.685218508 | 0.518074841 | 0.772675047 | -4.793069519 |
| **A_52_P653825** | Keg1 | -0.198028713 | -0.90050573 | 0.504448304 | 9.005792665 | -0.684982118 | 0.518214136 | 0.772778952 | -4.793148957 |
| **A_55_P2148268** | Xxylt1 | 0.045637344 | -0.116258401 | 0.20753309 | 8.301472144 | 0.684964583 | 0.51822447 | 0.772778952 | -4.793154848 |
| **A_55_P2011132** | Armc10 | -0.105341883 | -0.479191413 | 0.268507646 | 10.35020202 | -0.684679664 | 0.518392396 | 0.772969731 | -4.793250564 |
| **A_55_P2361647** | 4831440E17Rik | -0.071963026 | -0.327404877 | 0.183478825 | 8.256067222 | -0.684542717 | 0.518473123 | 0.773030469 | -4.793296559 |
| **A_52_P611798** | Ddx5 | 0.08829535 | -0.225175165 | 0.401765866 | 12.77945918 | 0.684422574 | 0.518543951 | 0.773076438 | -4.793336903 |
| **A_55_P2067533** | Foxa2 | -0.152879051 | -0.695803676 | 0.390045575 | 9.618199995 | -0.68421348 | 0.518667233 | 0.77308932 | -4.793407106 |
| **A_51_P102507** | Vps33a | 0.049278568 | -0.125729331 | 0.224286468 | 8.714140508 | 0.684200863 | 0.518674672 | 0.77308932 | -4.790504815 |
| **A_52_P121051** | 2610001J05Rik | -0.084680974 | -0.385450222 | 0.216088274 | 10.88046467 | -0.684125224 | 0.518719275 | 0.77308932 | -4.793436732 |
| **A_51_P284686** | Aak1 | 0.049474504 | -0.126255723 | 0.225204731 | 7.844296592 | 0.68409776 | 0.51873547 | 0.77308932 | -4.793445951 |
| **A_51_P380309** | Ncam1 | -0.073972518 | -0.336757779 | 0.188812744 | 9.423692665 | -0.683994462 | 0.518796387 | 0.77308932 | -4.793480622 |
| **A_55_P1998209** | NA | 0.084907456 | -0.216789057 | 0.386603969 | 15.94770108 | 0.683846655 | 0.51888356 | 0.77308932 | -4.793530225 |
| **A_55_P1991500** | Stn1 | -0.167424107 | -0.76241382 | 0.427565606 | 9.965654728 | -0.683741157 | 0.518945785 | 0.77308932 | -4.793565624 |
| **A_52_P213696** | Fndc5 | -0.251263903 | -1.144368525 | 0.641840719 | 8.65786372 | -0.683614009 | 0.519020788 | 0.77308932 | -4.793608282 |
| **A_52_P343641** | Siah1a | 0.045669738 | -0.11666419 | 0.208003667 | 8.150693385 | 0.683600564 | 0.519028719 | 0.77308932 | -4.793612793 |
| **A_51_P350073** | Ndel1 | -0.047255487 | -0.215249896 | 0.120738923 | 11.86003803 | -0.683503274 | 0.519086114 | 0.77308932 | -4.793645429 |
| **A_52_P229972** | Slc22a1 | -0.11701927 | -0.5330393 | 0.299000759 | 8.087527997 | -0.683480826 | 0.519099358 | 0.77308932 | -4.793652959 |
| **A_55_P2024909** | Dnah17 | -0.147581174 | -0.672296413 | 0.377134065 | 8.9238122 | -0.683424387 | 0.519132655 | 0.77308932 | -4.793671889 |
| **A_55_P2106414** | Haus7 | 0.097061415 | -0.248091838 | 0.442214667 | 9.107359537 | 0.683310093 | 0.519200091 | 0.77308932 | -4.793710221 |
| **A_55_P2025153** | Hmgb1 | 0.070323568 | -0.179778475 | 0.32042561 | 14.69980379 | 0.683229869 | 0.519247427 | 0.77308932 | -4.793737123 |
| **A_65_P16688** | Taok2 | -0.046129455 | -0.210217925 | 0.117959014 | 8.359927797 | -0.683098689 | 0.519324838 | 0.77308932 | -4.793781108 |
| **A_55_P2007500** | Tmem151b | -0.053545984 | -0.244030043 | 0.136938076 | 7.961761508 | -0.683048356 | 0.519354541 | 0.77308932 | -4.793797983 |
| **A_55_P2082180** | Tnfrsf9 | -0.045680558 | -0.208252484 | 0.116891368 | 7.68628665 | -0.682761526 | 0.519523833 | 0.77308932 | -4.793894129 |
| **A_52_P614960** | Skiv2l | 0.087008598 | -0.222689217 | 0.396706413 | 9.370390053 | 0.682664345 | 0.519581199 | 0.77308932 | -4.793926697 |
| **A_52_P203981** | Srek1ip1 | 0.049442873 | -0.126568333 | 0.225454078 | 10.04333884 | 0.682569008 | 0.51963748 | 0.77308932 | -4.793958644 |
| **A_55_P2117843** | Coasy | -0.059566841 | -0.271632717 | 0.152499035 | 12.22830938 | -0.682522493 | 0.519664942 | 0.77308932 | -4.79397423 |
| **A_55_P2038111** | Tmem104 | 0.047538739 | -0.121737085 | 0.216814563 | 7.836178059 | 0.682395117 | 0.519740146 | 0.77308932 | -4.794016905 |
| **A_51_P367843** | Polr2g | -0.057707827 | -0.26319691 | 0.147781257 | 12.87046412 | -0.682384499 | 0.519746416 | 0.77308932 | -4.794020462 |
| **A_51_P365912** | Ciao3 | -0.060800277 | -0.277309034 | 0.15570848 | 11.03400736 | -0.682359561 | 0.51976114 | 0.77308932 | -4.794028816 |
| **A_51_P135004** | Zfp212 | -0.099920603 | -0.455754308 | 0.255913103 | 10.67718781 | -0.68232479 | 0.519781671 | 0.77308932 | -4.794040463 |
| **A_55_P2100222** | NA | 0.160316773 | -0.410745498 | 0.731379043 | 10.11748919 | 0.682148114 | 0.519886 | 0.77308932 | -4.79409964 |
| **A_55_P2154870** | Sumo2 | 0.070889106 | -0.181655064 | 0.323433276 | 13.22758578 | 0.682064334 | 0.519935477 | 0.77308932 | -4.794127698 |
| **A_51_P467505** | Tec | -0.090738197 | -0.414013124 | 0.23253673 | 8.484197672 | -0.682026599 | 0.519957763 | 0.77308932 | -4.794140334 |
| **A_51_P129720** | Hectd3 | 0.078710218 | -0.201712468 | 0.359132904 | 9.577084533 | 0.682026358 | 0.519957905 | 0.77308932 | -4.794140415 |
| **A_51_P159415** | Sin3a | 0.098860751 | -0.253372003 | 0.451093504 | 9.589394675 | 0.681988977 | 0.519979983 | 0.77308932 | -4.794152932 |
| **A_55_P2022971** | Tspan9 | -0.048024013 | -0.219153893 | 0.123105867 | 7.844159979 | -0.68189231 | 0.520037078 | 0.77308932 | -4.7941853 |
| **A_66_P122110** | Mitd1 | -0.05698473 | -0.260051249 | 0.146081789 | 9.231037373 | -0.681872787 | 0.52004861 | 0.77308932 | -4.794191836 |
| **A_52_P464003** | Atg3 | 0.058640529 | -0.150338466 | 0.267619523 | 10.07905227 | 0.681833648 | 0.520071728 | 0.77308932 | -4.79420494 |
| **A_55_P1954987** | Slc10a1 | 0.074893763 | -0.192008104 | 0.341795629 | 17.20721693 | 0.681831764 | 0.520072841 | 0.77308932 | -4.79420557 |

| | | | | | | | | |
|---|---|---|---|---|---|---|---|---|
| **A_51_P272806** | Ctsr | -0.046849099 | -0.213812214 | 0.120114015 | 7.689762529 | -0.681810834 | 0.520085204 | 0.77308932 | -4.794212578 |
| **A_65_P10180** | Rad23b | 0.093762339 | -0.240433036 | 0.427957714 | 10.8200857 | 0.681728085 | 0.520134085 | 0.77308932 | -4.794240279 |
| **A_51_P513941** | Lrpap1 | 0.072824257 | -0.186778972 | 0.332427486 | 11.00905725 | 0.681630727 | 0.520191599 | 0.77308932 | -4.794272868 |
| **A_51_P516870** | Itm2a | -0.092822433 | -0.423795743 | 0.238150878 | 12.14156209 | -0.681464381 | 0.520289878 | 0.77308932 | -4.794328541 |
| **A_52_P547491** | Lurap1 | -0.087037798 | -0.39740258 | 0.223326985 | 7.9247324 | -0.681425917 | 0.520312605 | 0.77308932 | -4.794341412 |
| **A_51_P422751** | Clca1 | -0.060716258 | -0.27722474 | 0.155792224 | 7.707401087 | -0.681417486 | 0.520317586 | 0.77308932 | -4.794344233 |
| **A_51_P337543** | Glce | 0.055418093 | -0.142208642 | 0.253044827 | 9.933951646 | 0.681379508 | 0.520340026 | 0.77308932 | -4.794356942 |
| **A_55_P2069246** | Ddx39b | -0.055043046 | -0.251344407 | 0.141258315 | 13.6953109 | -0.681337567 | 0.520364809 | 0.77308932 | -4.794370976 |
| **A_51_P258766** | Smo | -0.128798468 | -0.58816091 | 0.330563975 | 10.70250258 | -0.681300019 | 0.520386996 | 0.77308932 | -4.794383539 |
| **A_51_P477440** | Sec23ip | 0.07288456 | -0.187067329 | 0.332836449 | 10.07282219 | 0.68128017 | 0.520398725 | 0.77308932 | -4.79439018 |
| **A_55_P2097017** | Eid1 | 0.121761739 | -0.312535873 | 0.556059351 | 12.44389788 | 0.681250118 | 0.520416484 | 0.77308932 | -4.794400234 |
| **A_52_P240542** | Id2 | 0.080304468 | -0.206160791 | 0.366769726 | 12.05069813 | 0.681162823 | 0.520468071 | 0.77308932 | -4.794429439 |
| **A_55_P2087384** | Gps1 | -0.07187244 | -0.328282494 | 0.184537613 | 12.94267431 | -0.681099451 | 0.520505524 | 0.77308932 | -4.794450638 |
| **A_55_P2033917** | Fut10 | -0.113460882 | -0.518297155 | 0.29137539 | 8.396114376 | -0.681004414 | 0.520561693 | 0.77308932 | -4.794482427 |
| **A_51_P144783** | Cct5 | 0.049075658 | -0.126041725 | 0.22419304 | 14.64070986 | 0.680957584 | 0.520589372 | 0.77308932 | -4.79449809 |
| **A_55_P1979217** | Abtb1 | 0.086623973 | -0.222496196 | 0.395744141 | 12.28379707 | 0.680916637 | 0.520613575 | 0.77308932 | -4.794511785 |
| **A_55_P1974114** | NA | -0.07186858 | -0.328336279 | 0.184599119 | 15.12756084 | -0.680909788 | 0.520617623 | 0.77308932 | -4.794514075 |
| **A_55_P2001068** | NA | -0.104919473 | -0.479345042 | 0.269506096 | 11.79425761 | -0.680885039 | 0.520632252 | 0.77308932 | -4.794522352 |
| **A_55_P2121913** | Adi1 | -0.07226306 | -0.330147956 | 0.185621836 | 9.977962309 | -0.680884786 | 0.520632401 | 0.77308932 | -4.794522436 |
| **A_55_P1969367** | Col4a5 | -0.319802456 | -1.46169562 | 0.822090708 | 8.603766226 | -0.680517834 | 0.520849335 | 0.77327456 | -4.794645128 |
| **A_55_P1953387** | Fabp5 | 0.093419311 | -0.24014609 | 0.426984713 | 17.50498417 | 0.680516802 | 0.520849945 | 0.77327456 | -4.794645473 |
| **A_55_P2034555** | Pdcd5 | 0.070972437 | -0.182460724 | 0.324405598 | 14.06078984 | 0.68047076 | 0.520877168 | 0.77327456 | -4.794660864 |
| **A_55_P2103105** | Hacd1 | 0.076271309 | -0.196154724 | 0.348697343 | 12.48110536 | 0.680292658 | 0.520982483 | 0.773371508 | -4.794720391 |
| **A_55_P1960496** | Ppp1r2 | 0.05072087 | -0.130521976 | 0.231963716 | 8.607810506 | 0.680000151 | 0.521155476 | 0.773568898 | -4.79481813 |
| **A_55_P2033845** | NA | -0.084645935 | -0.38719655 | 0.217904679 | 15.66586855 | -0.679815807 | 0.52126452 | 0.773671343 | -4.79487971 |
| **A_51_P301994** | Trmt10c | 0.121539425 | -0.313180065 | 0.556258916 | 8.870367882 | 0.679346365 | 0.521542271 | 0.773931967 | -4.795036467 |
| **A_52_P641477** | Wars2 | -0.048361433 | -0.221364079 | 0.124641214 | 8.159011978 | -0.67924993 | 0.52159934 | 0.773931967 | -4.795068658 |
| **A_55_P2013730** | Islr2 | 0.045086956 | -0.116205492 | 0.206379403 | 7.755968167 | 0.679235024 | 0.521608161 | 0.773931967 | -4.795073634 |
| **A_51_P490388** | Nudt8 | -0.05775393 | -0.264369398 | 0.148861539 | 9.383668361 | -0.679206602 | 0.521624983 | 0.773931967 | -4.795083121 |
| **A_55_P2040161** | Upf3a | -0.072009416 | -0.329633904 | 0.185615073 | 9.517273494 | -0.679180694 | 0.521640316 | 0.773931967 | -4.795091768 |
| **A_55_P1961127** | H19 | -0.074192757 | -0.339716432 | 0.191330918 | 17.77143966 | -0.678955717 | 0.521773477 | 0.774070115 | -4.795166849 |
| **A_55_P2180477** | Rpl31 | -0.053185176 | -0.243551446 | 0.137181093 | 16.0102937 | -0.678865586 | 0.52182683 | 0.774089854 | -4.795196922 |
| **A_55_P2041539** | Pigx | -0.14843831 | -0.679922239 | 0.383045618 | 11.35818335 | -0.678639376 | 0.521960751 | 0.774188661 | -4.795272386 |
| **A_55_P2019833** | NA | 0.276156844 | -0.712655256 | 1.264968944 | 11.56014851 | 0.67861777 | 0.521973544 | 0.774188661 | -4.795279593 |
| **A_55_P1993761** | Nsa2 | -0.125489143 | -0.574888532 | 0.323910245 | 10.89834501 | -0.678510958 | 0.522036788 | 0.774223055 | -4.795315218 |
| **A_51_P462918** | Ehhadh | -0.27385785 | -1.254770536 | 0.707054836 | 10.11960572 | -0.678387804 | 0.522109714 | 0.774223831 | -4.795356288 |
| **A_55_P2208682** | B230334C09Rik | 0.048773423 | -0.125939949 | 0.223486794 | 7.954093818 | 0.678328831 | 0.522144638 | 0.774223831 | -4.795375953 |
| **A_65_P04472** | Atp6v1c1 | 0.092340805 | -0.23844773 | 0.423129339 | 9.674210578 | 0.678307151 | 0.522157477 | 0.774223831 | -4.795383182 |
| **A_55_P1994603** | Itsn1 | 0.151675175 | -0.391862402 | 0.695212752 | 15.40599994 | 0.67805999 | 0.522303863 | 0.774381481 | -4.795465581 |
| **A_55_P2397240** | B430316J06Rik | -0.050730768 | -0.232597907 | 0.131136372 | 7.979007358 | -0.677798162 | 0.522458965 | 0.774552027 | -4.795552844 |
| **A_51_P406077** | Fmc1 | -0.062608308 | -0.287085721 | 0.161869106 | 13.82243346 | -0.677707832 | 0.522512482 | 0.774569294 | -4.795582944 |
| **A_51_P273326** | Sec31a | -0.176371008 | -0.935071749 | 0.582329734 | 11.78939995 | -0.716840007 | 0.522561652 | 0.774569294 | -4.678185967 |
| **A_51_P214127** | Cpa3 | 0.078443632 | -0.202865044 | 0.359752308 | 8.276010927 | 0.677575599 | 0.522590831 | 0.774569294 | -4.795627 |
| **A_52_P214612** | NA | -0.056223312 | -0.257880036 | 0.145433412 | 8.83175401 | -0.67746507 | 0.522656325 | 0.774606971 | -4.79566382 |
| **A_55_P2152074** | Ctps2 | -0.092373774 | -0.42373294 | 0.238985392 | 11.16972212 | -0.677380811 | 0.522706257 | 0.774621579 | -4.795691886 |
| **A_52_P634122** | Rexo1 | 0.083153929 | -0.215183211 | 0.381491068 | 9.052677663 | 0.677265013 | 0.522774885 | 0.774663887 | -4.795730453 |
| **A_55_P2100963** | Hmces | -0.058641292 | -0.269099323 | 0.151816738 | 11.11314903 | -0.677050742 | 0.522901886 | 0.774792684 | -4.795801802 |
| **A_55_P1974783** | Tsnax | 0.053251026 | -0.137939742 | 0.244441794 | 12.06414723 | 0.676774916 | 0.523065402 | 0.77497556 | -4.795893622 |
| **A_55_P2087352** | NA | -0.078415872 | -0.360053688 | 0.203221943 | 13.64454904 | -0.67654424 | 0.523202178 | 0.775073252 | -4.79597039 |
| **A_51_P377376** | Gnl3 | 0.143733164 | -0.372552309 | 0.660018637 | 10.17811043 | 0.676472692 | 0.523244605 | 0.775073252 | -4.795994197 |

| | | | | | | | | |
|---|---|---|---|---|---|---|---|---|
| **A_55_P2010778** | Il11ra1 | -0.050388127 | -0.231396683 | 0.130620429 | 13.19493186 | -0.676413554 | 0.523279675 | 0.775073252 | -4.796013872 |
| **A_55_P2066799** | Sctr | 0.047262649 | -0.122523401 | 0.217048699 | 7.858473086 | 0.67639322 | 0.523291734 | 0.775073252 | -4.796020637 |
| **A_55_P2060376** | Gzmg | -0.043754029 | -0.200991005 | 0.113482947 | 7.652724106 | -0.676155454 | 0.523432753 | 0.775205705 | -4.796099729 |
| **A_55_P2056014** | NA | 0.164727934 | -0.427342227 | 0.756798096 | 9.505914549 | 0.676047587 | 0.523496737 | 0.775205705 | -4.796135603 |
| **A_51_P122660** | Liph | -0.0533448 | -0.245080571 | 0.138390972 | 7.746089321 | -0.676039596 | 0.523501478 | 0.775205705 | -4.781521759 |
| **A_55_P1973486** | Atp6v1f | -0.083145969 | -0.382162814 | 0.215870876 | 14.05457745 | -0.675660814 | 0.523726203 | 0.775425234 | -4.796264198 |
| **A_55_P2143311** | Pdxk | 0.098608303 | -0.256018873 | 0.453235479 | 8.941152362 | 0.675654479 | 0.523729962 | 0.775425234 | -4.796266304 |
| **A_52_P570652** | Flywch1 | -0.071992665 | -0.330950771 | 0.186965442 | 8.159563583 | -0.675525781 | 0.523806331 | 0.775478904 | -4.796309081 |
| **A_55_P1952202** | NA | 0.073878705 | -0.191920542 | 0.339677953 | 8.86188772 | 0.67538081 | 0.523892364 | 0.775546872 | -4.796357258 |
| **A_55_P2006911** | Pik3c3 | -0.051298596 | -0.23588678 | 0.133289588 | 8.205951224 | -0.675281372 | 0.523951382 | 0.775570067 | -4.7963903 |
| **A_55_P2078765** | Kmt2e | -0.061772166 | -0.284067953 | 0.160523621 | 10.33275117 | -0.675219203 | 0.523988282 | 0.775570067 | -4.796410955 |
| **A_51_P130905** | Gcm1 | 0.051992481 | -0.135160993 | 0.239145954 | 7.658069717 | 0.675034291 | 0.524098045 | 0.775673133 | -4.796472383 |
| **A_51_P265576** | Rbm18 | 0.052867539 | -0.13748849 | 0.243223568 | 8.107997811 | 0.674847504 | 0.524208936 | 0.775691684 | -4.79653442 |
| **A_55_P2096827** | Art3 | -0.0574919 | -0.264500222 | 0.149516423 | 7.757404291 | -0.674841905 | 0.524212261 | 0.775691684 | -4.79653628 |
| **A_55_P1957618** | Ak2 | 0.072558037 | -0.188710403 | 0.333826477 | 10.58657375 | 0.674810389 | 0.524230972 | 0.775691684 | -4.796546746 |
| **A_55_P2151209** | Krtap19-4 | 0.249654197 | -0.649747483 | 1.149055877 | 12.55554722 | 0.674478898 | 0.524427814 | 0.775923547 | -4.796656805 |
| **A_55_P2097657** | Rps11 | -0.102219038 | -0.470585541 | 0.266147466 | 15.00353119 | -0.674271566 | 0.524550953 | 0.776046336 | -4.796725621 |
| **A_51_P283754** | Cpsf7 | -0.069182863 | -0.318601447 | 0.180235721 | 11.24408217 | -0.673989158 | 0.524718712 | 0.776102573 | -4.796819327 |
| **A_55_P2078780** | Etv4 | -0.059332154 | -0.273268205 | 0.154603897 | 8.414942972 | -0.673890494 | 0.524777329 | 0.776102573 | -4.796852058 |
| **A_55_P2387379** | NA | 0.073923937 | -0.192631496 | 0.340479369 | 8.119825583 | 0.673877162 | 0.52478525 | 0.776102573 | -4.796856481 |
| **A_55_P2060722** | Uso1 | 0.098380927 | -0.25637877 | 0.453140624 | 11.46495246 | 0.673844709 | 0.524804532 | 0.776102573 | -4.796867246 |
| **A_55_P2064557** | Furin | -0.155181518 | -0.714790592 | 0.404427557 | 11.63783306 | -0.673811527 | 0.524824248 | 0.776102573 | -4.796878252 |
| **A_52_P485007** | Abca2 | -0.046388527 | -0.213675237 | 0.120898183 | 8.008946894 | -0.673802049 | 0.524829879 | 0.776102573 | -4.796881396 |
| **A_51_P292550** | Chdh | 0.189682018 | -0.494579742 | 0.873943777 | 9.433921637 | 0.673576738 | 0.524963763 | 0.77619475 | -4.796956119 |
| **A_52_P138205** | Plekha8 | 0.05831342 | -0.152051893 | 0.268678734 | 8.535882957 | 0.673561992 | 0.524972527 | 0.77619475 | -4.796961008 |
| **A_55_P2121344** | Nudcd3 | -0.078082688 | -0.359864423 | 0.203699047 | 12.38968468 | -0.673325564 | 0.525113042 | 0.776318795 | -4.797039395 |
| **A_55_P2117614** | Tnfrsf13c | 0.120161892 | -0.313507782 | 0.553831567 | 8.890929715 | 0.673272529 | 0.525144565 | 0.776318795 | -4.797056976 |
| **A_66_P127484** | Abcf3 | 0.064621311 | -0.168626186 | 0.297868807 | 10.54914021 | 0.673196673 | 0.525189656 | 0.776318795 | -4.79708212 |
| **A_52_P342620** | Csrnp2 | -0.050224148 | -0.231518205 | 0.131069909 | 7.795238592 | -0.673150543 | 0.525217077 | 0.776318795 | -4.797097409 |
| **A_66_P103229** | Foxl1 | -0.08420827 | -0.388365747 | 0.219949206 | 7.756252724 | -0.672727901 | 0.525468357 | 0.776630591 | -4.797237452 |
| **A_51_P484919** | Polrmt | 0.049157661 | -0.128421785 | 0.226737107 | 9.888435595 | 0.67263847 | 0.525521538 | 0.776630591 | -4.797267076 |
| **A_55_P2052563** | Id1 | -0.166329111 | -0.767281676 | 0.434623455 | 11.45590587 | -0.672529378 | 0.525586415 | 0.776630591 | -4.797303209 |
| **A_51_P413376** | Zfp42 | 0.058237336 | -0.152192127 | 0.268666799 | 7.677244119 | 0.672478093 | 0.525616916 | 0.776630591 | -4.797320194 |
| **A_51_P220407** | Rnpepl1 | -0.0888325 | -0.409866185 | 0.232201185 | 12.63655833 | -0.672364027 | 0.525684758 | 0.776630591 | -4.797357967 |
| **A_55_P2181262** | NA | -0.101380144 | -0.467761935 | 0.265001648 | 13.51516253 | -0.672360529 | 0.525686839 | 0.776630591 | -4.797359125 |
| **A_55_P2045163** | Ehmt1 | 0.182168892 | -0.476262757 | 0.840600541 | 13.08399902 | 0.672274608 | 0.525737946 | 0.776630591 | -4.797387574 |
| **A_55_P1982239** | Ctrb1 | 0.493335837 | -1.289999805 | 2.276671479 | 9.082048409 | 0.672191109 | 0.525787616 | 0.776630591 | -4.797415219 |
| **A_55_P1978716** | Zim1 | 0.047542503 | -0.12431744 | 0.219402445 | 7.86309922 | 0.67218771 | 0.525789638 | 0.776630591 | -4.797416344 |
| **A_51_P366024** | Ercc3 | 0.048839501 | -0.127764418 | 0.22544342 | 8.977399917 | 0.67197648 | 0.525915302 | 0.776756848 | -4.797486265 |
| **A_55_P2108690** | Gm8709 | -0.104278071 | -0.481658419 | 0.273102278 | 12.06597231 | -0.671424052 | 0.526244041 | 0.777182998 | -4.797669047 |
| **A_52_P58524** | Neu1 | -0.05599498 | -0.258667666 | 0.146677706 | 12.07284564 | -0.671331556 | 0.526299097 | 0.777193327 | -4.79769964 |
| **A_52_P492901** | Stx4a | -0.06969597 | -0.321997523 | 0.182605582 | 10.66457569 | -0.671229346 | 0.526359939 | 0.777193327 | -4.797733441 |
| **A_55_P1979705** | Spop | 0.054772032 | -0.143510223 | 0.253054288 | 8.118788065 | 0.671209654 | 0.526371661 | 0.777193327 | -4.797739953 |
| **A_55_P2071426** | Eef2kmt | -0.053802208 | -0.248641265 | 0.141036849 | 8.569695123 | -0.670976438 | 0.526510505 | 0.777338952 | -4.797817062 |
| **A_66_P105564** | Gjc2 | -0.168839467 | -0.780626609 | 0.442947675 | 8.625843565 | -0.670589604 | 0.526740857 | 0.777619646 | -4.797944916 |
| **A_51_P157083** | Gas1 | -0.076302715 | -0.352848718 | 0.200243287 | 8.064408916 | -0.670433639 | 0.526833749 | 0.777697383 | -4.797996447 |
| **A_55_P2031547** | Vmn2r121 | 0.045496161 | -0.119426461 | 0.210418783 | 7.697074924 | 0.6703131 | 0.526905549 | 0.777700914 | -4.798036268 |
| **A_52_P6451** | Sec24a | 0.158232113 | -0.415371847 | 0.731836073 | 9.943603757 | 0.670294529 | 0.526916611 | 0.777700914 | -4.798042402 |
| **A_52_P676076** | Rbx1 | 0.075527141 | -0.198336568 | 0.349390851 | 11.13769019 | 0.67011872 | 0.527021345 | 0.777720651 | -4.79810047 |
| **A_51_P352720** | Lmod2 | -0.047766045 | -0.22097437 | 0.12544228 | 7.684787189 | -0.670090883 | 0.52703793 | 0.777720651 | -4.798109663 |

| | | | | | | | | | |
|---|---|---|---|---|---|---|---|---|---|
| **A_52_P352563** | 1700073E17Rik | -0.097502721 | -0.45107654 | 0.256071098 | 14.82551782 | -0.670069464 | 0.527050691 | 0.777720651 | -4.798116737 |
| **A_55_P2132845** | NA | -0.059080499 | -0.27337144 | 0.155210442 | 16.05583273 | -0.669920912 | 0.527139201 | 0.77779188 | -4.79816579 |
| **A_66_P100440** | Vezf1 | -0.112614744 | -0.521222698 | 0.29599321 | 9.982138415 | -0.669686626 | 0.527278813 | 0.777938491 | -4.798243135 |
| **A_55_P1986188** | Def8 | 0.064063037 | -0.168429199 | 0.296555273 | 11.32098929 | 0.669548835 | 0.527360934 | 0.778000266 | -4.798288614 |
| **A_55_P2119877** | Pnisr | 0.067560341 | -0.17771878 | 0.312839463 | 9.142925427 | 0.669290163 | 0.52751512 | 0.778168339 | -4.798373971 |
| **A_51_P373461** | Brcc3 | 0.058351001 | -0.15359297 | 0.270294972 | 11.09541887 | 0.668975835 | 0.52770252 | 0.778322876 | -4.798477658 |
| **A_55_P2017709** | Igsf9 | 0.185768756 | -0.489023853 | 0.860561365 | 13.4777088 | 0.668937502 | 0.527725376 | 0.778322876 | -4.7984903 |
| **A_55_P2052425** | Setd4 | -0.054293909 | -0.25152048 | 0.142932661 | 9.95631071 | -0.668911836 | 0.52774068 | 0.778322876 | -4.798498765 |
| **A_55_P2004427** | Fbxl15 | 0.085463409 | -0.225035335 | 0.395962153 | 10.5972314 | 0.668811227 | 0.527800675 | 0.778351968 | -4.798531942 |
| **A_51_P384441** | Tmem201 | 0.046718248 | -0.123080002 | 0.216516499 | 7.918727384 | 0.668554055 | 0.527954048 | 0.778518752 | -4.79861673 |
| **A_65_P19623** | Ssu72 | -0.075676137 | -0.351003311 | 0.199651036 | 11.90831115 | -0.667871743 | 0.528361109 | 0.778927494 | -4.798841559 |
| **A_55_P1984361** | NA | 0.120325075 | -0.317466389 | 0.55811654 | 14.51743499 | 0.667839406 | 0.528380405 | 0.778927494 | -4.79885221 |
| **A_55_P2124183** | NA | 0.067851693 | -0.179025831 | 0.314729218 | 9.310384846 | 0.667824464 | 0.528389322 | 0.778927494 | -4.798857131 |
| **A_51_P200083** | Grina | 0.11614891 | -0.306460723 | 0.538758544 | 11.87586625 | 0.667819255 | 0.528392431 | 0.778927494 | -4.798858847 |
| **A_52_P276794** | Tox4 | 0.043936719 | -0.115947296 | 0.203820734 | 9.170870217 | 0.667737496 | 0.528441223 | 0.778940013 | -4.798885774 |
| **A_55_P2041894** | NA | 0.064189283 | -0.169420166 | 0.297798732 | 16.63096614 | 0.667659923 | 0.528487519 | 0.778948853 | -4.798911319 |
| **A_55_P2116621** | Cited2 | -0.089065636 | -0.413277529 | 0.235146257 | 9.395473676 | -0.667520215 | 0.528570905 | 0.778958766 | -4.798957321 |
| **A_52_P655285** | Zfp462 | -0.098098107 | -0.45519304 | 0.258996826 | 7.997125197 | -0.667513615 | 0.528574845 | 0.778958766 | -4.798959494 |
| **A_66_P139167** | Safb | 0.068974376 | -0.182275789 | 0.320224542 | 10.87453636 | 0.667059562 | 0.52884591 | 0.77919625 | -4.799108945 |
| **A_52_P55717** | Adnp | 0.058732447 | -0.155230708 | 0.272695602 | 8.483010105 | 0.666994562 | 0.528884721 | 0.77919625 | -4.799130333 |
| **A_55_P1981550** | Park7 | -0.091583737 | -0.425227201 | 0.242059726 | 12.80014037 | -0.666989396 | 0.528887806 | 0.77919625 | -4.799132032 |
| **A_55_P2035937** | Rab6a | -0.065356161 | -0.30348535 | 0.172773028 | 12.39029186 | -0.666894433 | 0.528944512 | 0.77919625 | -4.799163277 |
| **A_55_P2160383** | NA | 0.090198048 | -0.238448208 | 0.418844305 | 10.87217607 | 0.666886061 | 0.528949511 | 0.77919625 | -4.799166031 |
| **A_52_P373893** | NA | -0.057069818 | -0.265024622 | 0.150884987 | 9.841389223 | -0.666838578 | 0.528977867 | 0.77919625 | -4.799181652 |
| **A_55_P1955346** | NA | -0.066563466 | -0.309140022 | 0.17601309 | 14.57792625 | -0.666761154 | 0.529024104 | 0.779204977 | -4.799207121 |
| **A_51_P388691** | Bbx | -0.044194285 | -0.205288667 | 0.116900097 | 7.932785955 | -0.666605518 | 0.529117058 | 0.779282507 | -4.799258312 |
| **A_55_P2021476** | NA | 0.071049267 | -0.187978487 | 0.330077021 | 13.62595877 | 0.666494376 | 0.529183444 | 0.779320898 | -4.799294862 |
| **A_55_P2130965** | NA | -0.107993316 | -0.501789431 | 0.285802799 | 12.87271274 | -0.666359567 | 0.529263973 | 0.779380111 | -4.799339189 |
| **A_52_P661587** | Ccdc141 | 0.086186733 | -0.228168273 | 0.400541738 | 7.767307948 | 0.666197848 | 0.529360588 | 0.779463001 | -4.799392355 |
| **A_52_P11833** | NA | 0.067481055 | -0.178730765 | 0.313692875 | 10.12487458 | 0.665972281 | 0.529495367 | 0.779500811 | -4.799466494 |
| **A_55_P1978351** | Apobec2 | -0.045779125 | -0.212828485 | 0.121270236 | 7.668583691 | -0.665895154 | 0.529541456 | 0.779500811 | -4.79949184 |
| **A_55_P2022444** | Pdss2 | 0.081415965 | -0.215679458 | 0.378511389 | 8.541820682 | 0.665881278 | 0.529549748 | 0.779500811 | -4.7994964 |
| **A_55_P2049365** | NA | 0.091578719 | -0.24261381 | 0.425771249 | 12.70661929 | 0.665857074 | 0.529564212 | 0.779500811 | -4.799504353 |
| **A_55_P1981949** | Klhl5 | -0.105602374 | -0.490993503 | 0.279788755 | 9.615047629 | -0.665817425 | 0.529587907 | 0.779500811 | -4.799517381 |
| **A_55_P2065325** | Tmem94 | 0.079450965 | -0.210546534 | 0.369448464 | 8.064206396 | 0.665714649 | 0.529649331 | 0.77953186 | -4.799551148 |
| **A_55_P1970665** | Kdm3b | 0.072815259 | -0.193009055 | 0.338639573 | 8.42578364 | 0.665596282 | 0.529720078 | 0.779576624 | -4.799590033 |
| **A_55_P2082653** | Zfp711 | -0.10010843 | -0.465713078 | 0.265496218 | 8.039754373 | -0.665337699 | 0.529874652 | 0.779744739 | -4.799674961 |
| **A_55_P2052754** | Pcm1 | -0.046788311 | -0.217698373 | 0.124121751 | 8.56839 | -0.665201049 | 0.52995635 | 0.779805594 | -4.799719832 |
| **A_51_P279571** | Syap1 | -0.076706084 | -0.35703176 | 0.203619592 | 10.92496798 | -0.66489049 | 0.53014205 | 0.779929 | -4.799821779 |
| **A_55_P2171623** | Ptbp1 | 0.116383977 | -0.308974811 | 0.541742766 | 9.501312683 | 0.664845868 | 0.530168736 | 0.779929 | -4.799836424 |
| **A_55_P2081510** | Atp5j2 | -0.058858457 | -0.273986962 | 0.156270047 | 14.29837679 | -0.664804738 | 0.530193334 | 0.779929 | -4.799849923 |
| **A_51_P459340** | Pacrgl | 0.051014155 | -0.135447197 | 0.237475507 | 9.002312528 | 0.664790888 | 0.530201617 | 0.779929 | -4.799854468 |
| **A_52_P965502** | Ssr1 | 0.068972492 | -0.183261289 | 0.321206272 | 8.67381063 | 0.66444013 | 0.53041142 | 0.780178246 | -4.799969553 |
| **A_55_P2066543** | Orai2 | -0.060645887 | -0.282462611 | 0.161170838 | 9.174471116 | -0.664339767 | 0.530471461 | 0.780207189 | -4.800002474 |
| **A_52_P8227** | Lypla1 | 0.109838173 | -0.291945693 | 0.511622039 | 9.933282562 | 0.664269014 | 0.530513791 | 0.780210079 | -4.800025679 |
| **A_51_P479902** | Mars | 0.113014702 | -0.300535875 | 0.526565279 | 10.53624883 | 0.664032756 | 0.530655154 | 0.780348039 | -4.800103153 |
| **A_55_P2024175** | Map4k3 | -0.159904411 | -0.745085665 | 0.425276843 | 10.11704581 | -0.663977294 | 0.530688343 | 0.780348039 | -4.800121338 |
| **A_51_P348119** | Ube2q1 | -0.06786072 | -0.316324022 | 0.180602582 | 10.09033681 | -0.66365046 | 0.530883948 | 0.780576285 | -4.80022847 |
| **A_51_P225083** | Snx17 | -0.138643894 | -0.646476833 | 0.369189045 | 12.30214652 | -0.663381062 | 0.531045215 | 0.780716712 | -4.800316744 |
| **A_51_P255441** | Mrpl43 | -0.072687304 | -0.338953056 | 0.193578448 | 12.96040716 | -0.663325117 | 0.531078708 | 0.780716712 | -4.800335073 |

| | | | | | | | | | |
|---|---|---|---|---|---|---|---|---|---|
| **A_51_P156274** | Ltbr | -0.089504401 | -0.417392088 | 0.238383286 | 11.85617929 | -0.663288505 | 0.531100628 | 0.780716712 | -4.800347066 |
| **A_55_P2100884** | Fjx1 | -0.092338892 | -0.430779648 | 0.246101865 | 8.29647804 | -0.662956737 | 0.531299285 | 0.780919764 | -4.800455726 |
| **A_55_P2005285** | Klhl22 | -0.074981467 | -0.349817779 | 0.199854845 | 11.15950243 | -0.662922879 | 0.531319562 | 0.780919764 | -4.800466813 |
| **A_55_P2145833** | Dmpk | -0.060382295 | -0.281848476 | 0.161083885 | 10.23628628 | -0.662499251 | 0.531573299 | 0.781233295 | -4.80060549 |
| **A_55_P2019118** | NA | 0.04851483 | -0.129507499 | 0.226537159 | 7.918770989 | 0.662190919 | 0.531758027 | 0.781438854 | -4.80070638 |
| **A_55_P2007646** | Cryaa | -0.047534122 | -0.221973626 | 0.126905382 | 7.852762484 | -0.662130844 | 0.531794024 | 0.781438854 | -4.800726033 |
| **A_55_P2028894** | Gata4 | 0.108452904 | -0.289594038 | 0.506499845 | 9.473866293 | 0.66204892 | 0.531843115 | 0.781451583 | -4.800752832 |
| **A_55_P2067463** | 1700110I01Rik | 0.206723104 | -0.552174144 | 0.965620353 | 13.2825786 | 0.661895195 | 0.531935239 | 0.781527534 | -4.80080311 |
| **A_55_P2002079** | NA | -0.076238583 | -0.356195952 | 0.203718787 | 15.37639061 | -0.661707564 | 0.532047697 | 0.781633346 | -4.800864465 |
| **A_51_P158638** | Ggnbp1 | -0.055476823 | -0.259238335 | 0.14828469 | 8.568793865 | -0.66156514 | 0.53213307 | 0.781636146 | -4.800911029 |
| **A_55_P2141724** | Gm13275 | -0.052315051 | -0.244475192 | 0.139845091 | 7.983330694 | -0.661525408 | 0.532156887 | 0.781636146 | -4.800924017 |
| **A_55_P1973854** | Sema6b | -0.092400546 | -0.431834887 | 0.247033795 | 8.99049785 | -0.661457504 | 0.532197595 | 0.781636146 | -4.800946213 |
| **A_55_P2119145** | NA | 0.042637303 | -0.113996698 | 0.199271305 | 7.705866266 | 0.661434549 | 0.532211357 | 0.781636146 | -4.800953716 |
| **A_55_P2041131** | NA | -0.127719091 | -0.597026258 | 0.341588075 | 12.99358983 | -0.661274572 | 0.53230727 | 0.781717613 | -4.801006 |
| **A_52_P651078** | Fyttd1 | 0.051719677 | -0.138349425 | 0.241788779 | 7.915578514 | 0.661191814 | 0.532356892 | 0.781731092 | -4.801033044 |
| **A_52_P289692** | NA | -0.079000768 | -0.369374163 | 0.211372626 | 10.12352812 | -0.661085578 | 0.532420595 | 0.781765244 | -4.801067755 |
| **A_51_P509760** | Actr1a | -0.065872062 | -0.308048068 | 0.176303945 | 12.00340306 | -0.660926754 | 0.532515841 | 0.781845704 | -4.801119641 |
| **A_51_P452798** | Zfp524 | 0.072297979 | -0.193696217 | 0.338292175 | 10.55533009 | 0.660445804 | 0.532804331 | 0.78212709 | -4.8012767 |
| **A_52_P168097** | Atp2a2 | 0.079083948 | -0.211890932 | 0.370058828 | 8.059733555 | 0.660413636 | 0.53282363 | 0.78212709 | -4.801287201 |
| **A_52_P210338** | Ndufb6 | -0.072930532 | -0.341283662 | 0.195422598 | 12.77556345 | -0.660367828 | 0.532851113 | 0.78212709 | -4.801302155 |
| **A_51_P247614** | Smim10l2a | -0.045175997 | -0.211411973 | 0.121059979 | 8.432800967 | -0.660337433 | 0.532869349 | 0.78212709 | -4.801312077 |
| **A_66_P137383** | Ddx55 | 0.061028373 | -0.163578759 | 0.285635505 | 8.468257248 | 0.660224199 | 0.532937291 | 0.782167418 | -4.801349037 |
| **A_52_P496655** | Leprotl1 | -0.048127725 | -0.225335652 | 0.129080201 | 8.127491899 | -0.659926211 | 0.533116112 | 0.782222319 | -4.801446276 |
| **A_51_P105124** | P2ry6 | -0.075434412 | -0.353205674 | 0.202336851 | 9.447253847 | -0.659880635 | 0.533143466 | 0.782222319 | -4.79847967 |
| **A_55_P1972688** | Wnk1 | 0.052492658 | -0.140831939 | 0.245817256 | 11.40129111 | 0.659773158 | 0.533207974 | 0.782222319 | -4.801496207 |
| **A_52_P538470** | Spats2l | -0.070264706 | -0.329066158 | 0.188536747 | 8.37886372 | -0.659710979 | 0.533245296 | 0.782222319 | -4.801516489 |
| **A_55_P2012729** | Kcnh2 | -0.063671261 | -0.298196956 | 0.170854435 | 7.746735776 | -0.659684384 | 0.53326126 | 0.782222319 | -4.801525163 |
| **A_55_P2068734** | Ccl27a | -0.054521432 | -0.255345563 | 0.1463027 | 9.872996001 | -0.659681859 | 0.533262775 | 0.782222319 | -4.801525987 |
| **A_55_P2181246** | NA | -0.091302219 | -0.427703763 | 0.245099325 | 8.796170386 | -0.659487462 | 0.533379474 | 0.782222319 | -4.801589385 |
| **A_51_P428372** | Ppbp | -0.179671315 | -0.841670765 | 0.482328135 | 12.35610895 | -0.659484066 | 0.533381512 | 0.782222319 | -4.801590492 |
| **A_52_P99888** | Cxcl16 | -0.055001298 | -0.257669801 | 0.147667204 | 8.340842565 | -0.659431777 | 0.533412905 | 0.782222319 | -4.801607543 |
| **A_55_P2015410** | 0610010K14Rik | -0.061311325 | -0.287238927 | 0.164616277 | 13.36784977 | -0.65940859 | 0.533426825 | 0.782222319 | -4.801615103 |
| **A_55_P2068148** | Pias1 | -0.04612643 | -0.216113578 | 0.123860717 | 8.232413475 | -0.659351426 | 0.533461146 | 0.782222319 | -4.801633741 |
| **A_51_P140819** | Wdr36 | 0.078986718 | -0.212124976 | 0.370098411 | 9.573970123 | 0.659291694 | 0.533497011 | 0.782222319 | -4.801653215 |
| **A_55_P2182106** | Rpl39 | -0.059189513 | -0.277339152 | 0.158960127 | 16.00230035 | -0.659285395 | 0.533500793 | 0.782222319 | -4.801655269 |
| **A_55_P2336943** | Dph6 | 0.056673973 | -0.152255473 | 0.265603419 | 8.969824483 | 0.659124103 | 0.533597644 | 0.782281501 | -4.801707846 |
| **A_55_P2062622** | Nono | 0.066013318 | -0.17738168 | 0.309408316 | 12.97258908 | 0.659026838 | 0.533656054 | 0.782281501 | -4.801739547 |
| **A_55_P2027312** | Cdc34 | 0.076451132 | -0.205445274 | 0.358347537 | 14.47373515 | 0.658988095 | 0.533679321 | 0.782281501 | -4.801752174 |
| **A_55_P2083351** | Hs3st1 | 0.042067027 | -0.113055001 | 0.197189056 | 7.773042006 | 0.658948594 | 0.533703045 | 0.782281501 | -4.801765047 |
| **A_55_P1967332** | NA | -0.082872813 | -0.388514057 | 0.222768431 | 14.01366385 | -0.658845085 | 0.533765212 | 0.782313299 | -4.801798775 |
| **A_55_P2102715** | NA | -0.071494248 | -0.335230225 | 0.19224173 | 15.97959068 | -0.658695825 | 0.533854867 | 0.782385375 | -4.801847405 |
| **A_55_P2023391** | Grhl3 | -0.043837732 | -0.205607953 | 0.117932489 | 7.844231201 | -0.658464987 | 0.53399354 | 0.782529274 | -4.801922596 |
| **A_51_P462428** | NA | -0.049635452 | -0.232860441 | 0.133589536 | 7.689761636 | -0.658249396 | 0.534123074 | 0.782607513 | -4.801992802 |
| **A_52_P175086** | Usp48 | 0.08621803 | -0.23210867 | 0.404544729 | 9.757167912 | 0.658124739 | 0.534197982 | 0.782607513 | -4.802033387 |
| **A_52_P9437** | Psmb3 | -0.079001033 | -0.370690693 | 0.212688627 | 14.11964346 | -0.658104596 | 0.534210086 | 0.782607513 | -4.802039945 |
| **A_55_P2001583** | Tspan18 | -0.062030261 | -0.291082013 | 0.167021491 | 7.79148541 | -0.658041342 | 0.534248099 | 0.782607513 | -4.802060536 |
| **A_55_P1987600** | Trmt112 | -0.080883228 | -0.379552045 | 0.217785589 | 12.80330726 | -0.658039222 | 0.534249373 | 0.782607513 | -4.802061226 |
| **A_52_P197095** | Cdk17 | 0.066140751 | -0.178198759 | 0.31048026 | 8.349160563 | 0.657746599 | 0.53442525 | 0.78275532 | -4.802156463 |
| **A_55_P2143657** | Gm5784 | -0.053910455 | -0.253089806 | 0.145268896 | 7.871924964 | -0.657675814 | 0.534467799 | 0.78275532 | -4.802179496 |
| **A_55_P2167451** | Cnnm2 | 0.046154682 | -0.124386894 | 0.216696258 | 9.339529105 | 0.657610408 | 0.534507117 | 0.78275532 | -4.802200776 |

| | | | | | | | | | |
|---|---|---|---|---|---|---|---|---|---|
| **A_52_P674357** | Thtpa | -0.047260674 | -0.221924272 | 0.127402924 | 9.560525711 | -0.657477204 | 0.534587198 | 0.78275532 | -4.80224411 |
| **A_55_P2050807** | Mtg2 | -0.071592959 | -0.336185191 | 0.192999272 | 10.44166858 | -0.657470715 | 0.534591099 | 0.78275532 | -4.802246221 |
| **A_55_P2128283** | Nr2f2 | -0.142325943 | -0.668334988 | 0.383683102 | 9.749939702 | -0.657467133 | 0.534593252 | 0.78275532 | -4.802247386 |
| **A_55_P1965892** | Gnpda1 | 0.103762784 | -0.279773573 | 0.487299141 | 11.45098128 | 0.657382692 | 0.534644022 | 0.78277036 | -4.802274853 |
| **A_51_P219266** | Tmprss6 | -0.141236825 | -0.663514654 | 0.381041004 | 12.05954506 | -0.65709709 | 0.534815758 | 0.782950303 | -4.802367731 |
| **A_51_P102438** | Ugt2b36 | 0.236146148 | -0.63716676 | 1.109459057 | 11.06468892 | 0.657043579 | 0.534847939 | 0.782950303 | -4.802385129 |
| **A_66_P132663** | Uqcrb | 0.051118821 | -0.137958163 | 0.240195806 | 14.33159837 | 0.656939465 | 0.534910555 | 0.782970079 | -4.802418977 |
| **A_52_P47645** | Cul3 | 0.050552174 | -0.136444011 | 0.237548359 | 9.526571081 | 0.656886414 | 0.534942463 | 0.782970079 | -4.802436223 |
| **A_52_P349467** | Tmlhe | 0.090847129 | -0.245257833 | 0.426952091 | 8.505442462 | 0.656779321 | 0.535006878 | 0.783005069 | -4.802471032 |
| **A_55_P2177026** | Mafg | 0.125425396 | -0.338725304 | 0.589576096 | 9.913255757 | 0.656613276 | 0.535106762 | 0.78309196 | -4.802524995 |
| **A_52_P484405** | Twf1 | 0.063718903 | -0.172141471 | 0.299579278 | 11.18334322 | 0.656442207 | 0.53520968 | 0.783183277 | -4.802580579 |
| **A_55_P1979281** | Rab1a | -0.051431127 | -0.241951848 | 0.139089594 | 12.89814389 | -0.655944375 | 0.535509258 | 0.783562335 | -4.802742269 |
| **A_66_P108276** | Gsg1l | -0.045491366 | -0.2140383 | 0.123055569 | 7.695000129 | -0.655830039 | 0.535578076 | 0.783585198 | -4.80277939 |
| **A_55_P1989076** | Slc22a23 | 0.137582294 | -0.372343073 | 0.647507662 | 11.34222022 | 0.655600263 | 0.535716394 | 0.783585198 | -4.802853975 |
| **A_66_P133244** | Vcl | -0.051154631 | -0.240769887 | 0.138460626 | 7.913341372 | -0.655533459 | 0.535756612 | 0.783585198 | -4.802875655 |
| **A_55_P2115704** | Aip | -0.145946381 | -0.686940123 | 0.395047362 | 8.991574316 | -0.65551748 | 0.535766232 | 0.783585198 | -4.802880841 |
| **A_51_P425833** | Lztfl1 | 0.053701093 | -0.145362839 | 0.252765025 | 9.796595841 | 0.655501572 | 0.535775809 | 0.783585198 | -4.802886003 |
| **A_55_P1986647** | Podxl2 | -0.066244285 | -0.311842436 | 0.179353866 | 7.718398723 | -0.655400113 | 0.535836896 | 0.783585198 | -4.802918926 |
| **A_65_P20799** | Abi2 | -0.050296517 | -0.236790073 | 0.136197039 | 7.936210024 | -0.655325808 | 0.535881636 | 0.783585198 | -4.802943035 |
| **A_52_P156485** | Ankrd39 | 0.080318711 | -0.21749555 | 0.378132972 | 9.820937126 | 0.655321518 | 0.53588422 | 0.783585198 | -4.802944427 |
| **A_55_P2131088** | Vwa5a | 0.051708597 | -0.140027241 | 0.243444435 | 9.411154016 | 0.655303744 | 0.535894922 | 0.783585198 | -4.802950194 |
| **A_51_P513224** | Cep164 | 0.054163126 | -0.146692119 | 0.255018371 | 9.004164197 | 0.655245031 | 0.535930277 | 0.783585198 | -4.802969242 |
| **A_66_P132969** | NA | 0.077944703 | -0.211171608 | 0.367061014 | 14.85910571 | 0.655084316 | 0.53602706 | 0.783667426 | -4.803021375 |
| **A_55_P2131820** | A530006G24Rik | 0.19528709 | -0.529244673 | 0.919818853 | 13.10667001 | 0.654936648 | 0.536115996 | 0.78373817 | -4.803069266 |
| **A_52_P437230** | Rbx1 | -0.078130253 | -0.368039777 | 0.211779272 | 14.09266994 | -0.65484714 | 0.536169908 | 0.783757707 | -4.803098292 |
| **A_55_P2162152** | NA | 0.075721927 | -0.205579733 | 0.357023587 | 9.710161501 | 0.654082531 | 0.536630589 | 0.784371797 | -4.803346105 |
| **A_55_P2200618** | NA | 0.062701636 | -0.170294323 | 0.295697596 | 9.420289097 | 0.653903518 | 0.536738481 | 0.784470177 | -4.80340409 |
| **A_52_P51429** | Dennd1c | -0.098045997 | -0.462481089 | 0.266389094 | 8.606110735 | -0.65372165 | 0.536848108 | 0.784536327 | -4.803462987 |
| **A_52_P90684** | Klhl31 | -0.044211793 | -0.208565865 | 0.120142278 | 7.687070601 | -0.653643356 | 0.536895307 | 0.784536327 | -4.803488338 |
| **A_51_P259318** | Fbrs | 0.054530791 | -0.14821088 | 0.257272462 | 8.248889033 | 0.653554739 | 0.536948732 | 0.784536327 | -4.803517029 |
| **A_55_P2077783** | Tubb2a-ps2 | -0.107118463 | -0.505403227 | 0.291166302 | 9.223475353 | -0.653512392 | 0.536974263 | 0.784536327 | -4.803530738 |
| **A_55_P2062836** | Ttf2 | 0.063184931 | -0.171781979 | 0.298151841 | 10.55638943 | 0.653416355 | 0.537032167 | 0.784536327 | -4.803561826 |
| **A_65_P01641** | Rnf6 | -0.047104896 | -0.222284425 | 0.128074633 | 12.5877335 | -0.653380068 | 0.537054046 | 0.784536327 | -4.803573572 |
| **A_55_P2029187** | Pde4a | -0.050840732 | -0.239920182 | 0.138238718 | 8.261434581 | -0.653357158 | 0.53706786 | 0.784536327 | -4.803580987 |
| **A_55_P2324976** | 5033406O09Rik | 0.062160877 | -0.169044036 | 0.29336579 | 8.471641358 | 0.653285869 | 0.537110847 | 0.78453983 | -4.80360406 |
| **A_52_P560146** | NA | 0.05826907 | -0.15863771 | 0.275175849 | 8.012841966 | 0.652751902 | 0.537432895 | 0.78490818 | -4.803776815 |
| **A_55_P1962801** | Max | -0.061247588 | -0.289248482 | 0.166753305 | 11.64945262 | -0.652733092 | 0.537444242 | 0.78490818 | -4.803782898 |
| **A_52_P619248** | Prl2c5 | -0.044966102 | -0.212376309 | 0.122444104 | 7.709958309 | -0.652659256 | 0.537488785 | 0.784913925 | -4.803806777 |
| **A_55_P2152246** | Rpl19 | 0.057602306 | -0.156888763 | 0.272093375 | 16.80212706 | 0.652550078 | 0.537554652 | 0.784950809 | -4.803842082 |
| **A_55_P1965592** | NA | -0.060722191 | -0.286958198 | 0.165513816 | 8.717975595 | -0.652182127 | 0.537776674 | 0.785143104 | -4.80396103 |
| **A_55_P2064996** | Capzb | -0.073961224 | -0.349543191 | 0.201620742 | 10.78196054 | -0.652133462 | 0.537806043 | 0.785143104 | -4.803976758 |
| **A_51_P426739** | Gpt | 0.075324555 | -0.205360991 | 0.356010101 | 9.005616327 | 0.652078251 | 0.537839364 | 0.785143104 | -4.8039946 |
| **A_52_P440709** | Ryr2 | -0.052941033 | -0.250222512 | 0.144340447 | 7.693034319 | -0.652062583 | 0.53784882 | 0.785143104 | -4.803999664 |
| **A_55_P2112105** | NA | 0.093887647 | -0.256025862 | 0.443801156 | 11.6944571 | 0.651974985 | 0.53790169 | 0.785160985 | -4.804027969 |
| **A_52_P597726** | Ccdc84 | 0.066504291 | -0.181436383 | 0.314444965 | 9.318551339 | 0.651756061 | 0.538033836 | 0.785289048 | -4.804098698 |
| **A_51_P469252** | Zfp647 | -0.079160544 | -0.374313781 | 0.215992693 | 8.222525047 | -0.65169503 | 0.538070679 | 0.785289048 | -4.804118412 |
| **A_55_P1973451** | Stra6 | 0.064027184 | -0.174758686 | 0.302813054 | 7.713110237 | 0.651536822 | 0.538166193 | 0.785328748 | -4.804169509 |
| **A_55_P2098450** | Snrnp200 | 0.048047972 | -0.131150203 | 0.227246148 | 12.0746218 | 0.651515378 | 0.53817914 | 0.785328748 | -4.804176434 |
| **A_52_P128691** | Gpr4 | -0.045680629 | -0.216088801 | 0.124727543 | 7.738326974 | -0.651365641 | 0.538269551 | 0.785401385 | -4.804224785 |
| **A_55_P1976490** | Lman2l | -0.057067704 | -0.270042076 | 0.155906667 | 10.25695077 | -0.651097827 | 0.538431281 | 0.785578066 | -4.804311241 |

| | | | | | | | | | |
|---|---|---|---|---|---|---|---|---|---|
| **A_55_P1962026** | Pigg | 0.041158278 | -0.11249188 | 0.194808435 | 8.224585481 | 0.650889659 | 0.538557012 | 0.785702203 | -4.804378423 |
| **A_51_P302458** | Plscr3 | -0.103616063 | -0.490703586 | 0.28347146 | 10.9125774 | -0.650430805 | 0.538834221 | 0.786002579 | -4.804526447 |
| **A_55_P2078123** | Rora | -0.152967429 | -0.724435695 | 0.418500837 | 9.117137995 | -0.650414235 | 0.538844233 | 0.786002579 | -4.80453179 |
| **A_55_P2135220** | NA | -0.05293897 | -0.25089419 | 0.145016249 | 7.851168878 | -0.649817977 | 0.53920459 | 0.786425567 | -4.804724011 |
| **A_55_P1976212** | Selplg | -0.071790306 | -0.340244026 | 0.196663414 | 8.379638489 | -0.649799789 | 0.539215585 | 0.786425567 | -4.804729872 |
| **A_55_P2008747** | Snrpd3 | 0.055840538 | -0.15302454 | 0.264705616 | 12.6813747 | 0.649631316 | 0.539317433 | 0.786514763 | -4.804784157 |
| **A_52_P368262** | Bcl7c | 0.047249227 | -0.129500447 | 0.2239989 | 8.183154143 | 0.649560003 | 0.539360547 | 0.786518297 | -4.804807132 |
| **A_52_P69020** | Slc24a5 | 0.094387525 | -0.258801095 | 0.447576145 | 8.651498298 | 0.649368305 | 0.539476457 | 0.786627975 | -4.804868882 |
| **A_51_P133052** | Glyr1 | -0.059956586 | -0.284393252 | 0.164480081 | 10.95478722 | -0.649121917 | 0.539625456 | 0.786785882 | -4.804948226 |
| **A_55_P2183172** | Rdx | 0.066190388 | -0.181661729 | 0.314042505 | 12.09197663 | 0.648911518 | 0.539752713 | 0.786819764 | -4.805015962 |
| **A_55_P2028029** | Abcb11 | -0.192077974 | -0.911330711 | 0.527174764 | 11.83130145 | -0.648902177 | 0.539758363 | 0.786819764 | -4.805018969 |
| **A_55_P2088525** | Myh10 | -0.116585031 | -0.55318911 | 0.320019047 | 9.502283799 | -0.648840865 | 0.539795451 | 0.786819764 | -4.805038704 |
| **A_51_P367285** | Rnf135 | -0.054945692 | -0.260722497 | 0.150831114 | 8.5664349 | -0.648814298 | 0.539811521 | 0.786819764 | -4.805047255 |
| **A_51_P304109** | Cyp2c39 | 0.304474575 | -0.836181189 | 1.445130339 | 10.30995164 | 0.648604 | 0.539938744 | 0.786869421 | -4.805114933 |
| **A_55_P2005783** | Ifih1 | -0.081437419 | -0.386534386 | 0.223659548 | 9.870022098 | -0.648588581 | 0.539948073 | 0.786869421 | -4.805119894 |
| **A_55_P1974845** | Pde1a | -0.051060925 | -0.242377138 | 0.140255288 | 8.321636146 | -0.6485151 | 0.539992532 | 0.786869421 | -4.805143538 |
| **A_55_P2026437** | Gm4802 | -0.04739581 | -0.224986681 | 0.130195061 | 15.95923221 | -0.64848883 | 0.540008426 | 0.786869421 | -4.805151989 |
| **A_51_P310548** | Osgep | -0.073734655 | -0.350155263 | 0.202685954 | 11.60648961 | -0.648163276 | 0.540205429 | 0.787059593 | -4.805256709 |
| **A_55_P2027412** | NA | 0.145914171 | -0.401127466 | 0.692955807 | 9.611144814 | 0.648127244 | 0.540227235 | 0.787059593 | -4.805268296 |
| **A_55_P1968133** | Ttc4 | 0.063483076 | -0.174544007 | 0.301510159 | 11.38251812 | 0.648059341 | 0.540268332 | 0.787059593 | -4.805290132 |
| **A_52_P670978** | Nkain1 | 0.044192977 | -0.121521106 | 0.209907059 | 7.730504437 | 0.648003015 | 0.540302424 | 0.787059593 | -4.805308244 |
| **A_51_P187082** | G6pdx | 0.078539626 | -0.215997948 | 0.373077201 | 11.67496951 | 0.647934812 | 0.540343706 | 0.787059593 | -4.805330173 |
| **A_55_P2156895** | Tmem39b | 0.091129306 | -0.25065627 | 0.432914883 | 8.870664019 | 0.647869482 | 0.540383251 | 0.787059593 | -4.805351176 |
| **A_55_P2121195** | Lcorl | 0.048390131 | -0.133134199 | 0.229914461 | 8.613766858 | 0.647746605 | 0.540457635 | 0.787108621 | -4.805390676 |
| **A_55_P2075140** | Sirt3 | -0.056334112 | -0.267691944 | 0.155023721 | 11.62252558 | -0.647643918 | 0.540519801 | 0.787139851 | -4.805423681 |
| **A_55_P2099378** | Tfe3 | -0.046654853 | -0.221716027 | 0.128406322 | 11.49927378 | -0.647575146 | 0.540561438 | 0.78714118 | -4.805445783 |
| **A_55_P2152225** | Ihh | -0.174443385 | -0.829101184 | 0.480214414 | 10.46542619 | -0.647475512 | 0.540621763 | 0.787149178 | -4.8054778 |
| **A_55_P2111459** | Trpv2 | 0.092921927 | -0.255822706 | 0.441666561 | 9.634512101 | 0.647431559 | 0.540648377 | 0.787149178 | -4.805491923 |
| **A_52_P10017** | Tmem64 | 0.121391015 | -0.334305144 | 0.577087173 | 9.332376934 | 0.647283243 | 0.540738189 | 0.787155645 | -4.805539574 |
| **A_66_P126300** | Mrpl47 | 0.055317813 | -0.152359184 | 0.262994809 | 10.93004638 | 0.647231722 | 0.540769389 | 0.787155645 | -4.805556124 |
| **A_52_P615312** | Mrps27 | 0.056343837 | -0.155188137 | 0.267875812 | 9.075804526 | 0.64722247 | 0.540774992 | 0.787155645 | -4.805559096 |
| **A_55_P1964309** | Mlf2 | -0.057629145 | -0.274043178 | 0.158784888 | 13.18592888 | -0.647053138 | 0.540877546 | 0.787245639 | -4.805613484 |
| **A_55_P2163143** | Cdk4 | -0.04922305 | -0.2341469 | 0.1357008 | 13.43091265 | -0.646783279 | 0.541041007 | 0.787363864 | -4.805700137 |
| **A_55_P2048730** | NA | -0.069411656 | -0.330227484 | 0.191404172 | 15.8755278 | -0.64666842 | 0.54111059 | 0.787363864 | -4.80573701 |
| **A_55_P2132711** | Tmem248 | 0.074119226 | -0.20440805 | 0.352646503 | 10.10721625 | 0.646615832 | 0.54114245 | 0.787363864 | -4.80575389 |
| **A_55_P2002155** | Sumo1 | -0.076746228 | -0.365151195 | 0.211658739 | 13.06102842 | -0.646602671 | 0.541150424 | 0.787363864 | -4.805758115 |
| **A_52_P391718** | Rpl22 | -0.093066229 | -0.442810928 | 0.25667847 | 11.12501896 | -0.646582827 | 0.541162447 | 0.787363864 | -4.805764484 |
| **A_55_P1970867** | Gm5779 | -0.130019688 | -0.618687454 | 0.358648079 | 13.80037639 | -0.646515013 | 0.541203535 | 0.787364377 | -4.805786249 |
| **A_51_P473219** | Siae | -0.045925457 | -0.218554151 | 0.126703237 | 9.277326017 | -0.646433259 | 0.541253071 | 0.787377181 | -4.805812486 |
| **A_55_P2144975** | Ccdc90b | -0.068464911 | -0.325875033 | 0.188945212 | 8.548529915 | -0.646287288 | 0.541341525 | 0.787446593 | -4.805859325 |
| **A_51_P137388** | Zadh2 | -0.080213888 | -0.381933114 | 0.221505339 | 10.83968245 | -0.645995909 | 0.541518118 | 0.7876031 | -4.805952798 |
| **A_55_P1953899** | Mccc1 | -0.0626062 | -0.298110788 | 0.172898387 | 10.83461238 | -0.645953364 | 0.541543906 | 0.7876031 | -4.805966443 |
| **A_52_P24308** | Uchl1 | -0.057254833 | -0.272644385 | 0.158134718 | 7.767213957 | -0.645908073 | 0.541571359 | 0.7876031 | -4.805980969 |
| **A_55_P1990324** | Ptprc | -0.094873865 | -0.451980702 | 0.262232972 | 8.366503628 | -0.645552576 | 0.541786874 | 0.787857244 | -4.806094951 |
| **A_55_P1970062** | Igsf9b | 0.07767702 | -0.214762337 | 0.370116378 | 8.899699307 | 0.645416294 | 0.541869508 | 0.787918131 | -4.806138634 |
| **A_52_P286520** | Slc23a2 | -0.052427488 | -0.249860817 | 0.14500584 | 8.019097344 | -0.645240729 | 0.541975971 | 0.788013657 | -4.806194897 |
| **A_55_P2042016** | NA | -0.082637792 | -0.393990097 | 0.228714513 | 7.82927521 | -0.644925885 | 0.542166928 | 0.788232008 | -4.806295764 |
| **A_52_P331212** | Cspp1 | 0.047812063 | -0.132354608 | 0.227978734 | 8.457355236 | 0.644831458 | 0.542224208 | 0.788255995 | -4.806326008 |
| **A_55_P2119962** | Shld1 | 0.078040165 | -0.216094915 | 0.372175245 | 10.46420334 | 0.644695361 | 0.54230677 | 0.788268987 | -4.806369592 |
| **A_55_P1961168** | NA | -0.070920181 | -0.338225312 | 0.19638495 | 11.71872068 | -0.644682277 | 0.542314708 | 0.788268987 | -4.806373782 |

| | | | | | | | | | |
|---|---|---|---|---|---|---|---|---|---|
| **A_55_P2091751** | NA | -0.065074424 | -0.31043429 | 0.180285441 | 9.054635846 | -0.644451145 | 0.542454943 | 0.788413534 | -4.806447782 |
| **A_51_P212419** | Lrrc41 | -0.054890207 | -0.261992996 | 0.152212582 | 9.247969009 | -0.644009258 | 0.542723114 | 0.78874399 | -4.806589199 |
| **A_55_P2055632** | Rtf2 | -0.064344479 | -0.307171004 | 0.178482045 | 8.582848202 | -0.643870244 | 0.542807495 | 0.788807313 | -4.806633671 |
| **A_55_P2156394** | Btbd16 | 0.087900759 | -0.244226421 | 0.42002794 | 9.644401094 | 0.643089467 | 0.543281578 | 0.789370094 | -4.806883308 |
| **A_51_P517495** | Cox8a | -0.070399222 | -0.336407331 | 0.195608888 | 15.87069342 | -0.643066923 | 0.543295271 | 0.789370094 | -4.806890512 |
| **A_66_P119017** | Smarca1 | 0.079371309 | -0.220560436 | 0.379303054 | 9.195336319 | 0.643019729 | 0.543323935 | 0.789370094 | -4.806905594 |
| **A_55_P1992505** | Entpd4 | -0.058230559 | -0.278293983 | 0.161832864 | 11.32120593 | -0.642963449 | 0.54335812 | 0.789370094 | -4.806923577 |
| **A_55_P2085295** | Inf2 | 0.069044675 | -0.191993163 | 0.330082512 | 11.06931348 | 0.642702388 | 0.543516706 | 0.789487552 | -4.807006978 |
| **A_55_P1994751** | NA | 0.044302859 | -0.123195064 | 0.211800781 | 8.514372148 | 0.642695879 | 0.543520661 | 0.789487552 | -4.807009057 |
| **A_52_P550185** | Urm1 | -0.094569177 | -0.452150771 | 0.263012417 | 11.2542748 | -0.642625039 | 0.543563699 | 0.789490738 | -4.807031683 |
| **A_52_P90124** | Prep | -0.048098756 | -0.229994969 | 0.133797456 | 11.3796329 | -0.642529957 | 0.543621469 | 0.789515319 | -4.807062049 |
| **A_55_P2160077** | NA | -0.070507196 | -0.337297763 | 0.19628337 | 16.92520385 | -0.642164312 | 0.543843663 | 0.789724846 | -4.807178791 |
| **A_55_P2047882** | Zc3h7b | -0.081458822 | -0.38972072 | 0.226803077 | 9.652472207 | -0.642098215 | 0.543883834 | 0.789724846 | -4.807199888 |
| **A_52_P303891** | Nr1d2 | -0.049982968 | -0.239134012 | 0.139168077 | 8.05659043 | -0.64209085 | 0.54388831 | 0.789724846 | -4.807202239 |
| **A_51_P232858** | Rpl14 | 0.079075075 | -0.220257678 | 0.378407827 | 10.91381692 | 0.641901751 | 0.54400325 | 0.789832405 | -4.807262587 |
| **A_52_P52263** | Ilrun | 0.048409419 | -0.134886709 | 0.231705546 | 8.305758837 | 0.641740974 | 0.544100986 | 0.789885263 | -4.807313885 |
| **A_66_P124732** | Zfp606 | -0.054427818 | -0.260522628 | 0.151666992 | 8.166895162 | -0.641707417 | 0.544121386 | 0.789885263 | -4.80732459 |
| **A_55_P2315921** | 5330431K02Rik | -0.045620194 | -0.218473572 | 0.127233183 | 7.736635272 | -0.641301787 | 0.544368024 | 0.790176736 | -4.807453961 |
| **A_55_P2088912** | Gapdh | 0.166999224 | -0.4658758 | 0.799874247 | 14.5812749 | 0.641179584 | 0.544442341 | 0.790176736 | -4.807492923 |
| **A_55_P2038247** | Tyro3 | -0.25554419 | -1.22398318 | 0.712894801 | 9.330838907 | -0.641175521 | 0.544444812 | 0.790176736 | -4.807494218 |
| **A_55_P2279957** | 5730414N17Rik | -0.04453119 | -0.213326058 | 0.124263678 | 8.124709146 | -0.641044602 | 0.544524437 | 0.7902094 | -4.807535952 |
| **A_51_P145533** | Rnf14 | 0.055498024 | -0.154879778 | 0.265875827 | 12.17224999 | 0.641004083 | 0.544549082 | 0.7902094 | -4.807548868 |
| **A_55_P2090394** | NA | 0.089125652 | -0.24883589 | 0.427087194 | 12.4558269 | 0.640794277 | 0.544676705 | 0.790335263 | -4.807615732 |
| **A_51_P290626** | Gulo | 0.18705308 | -0.522475687 | 0.896581847 | 11.04152591 | 0.640586888 | 0.544802877 | 0.790459 | -4.807681808 |
| **A_51_P402435** | Trap1 | 0.094541549 | -0.264125063 | 0.453208161 | 10.97368859 | 0.640493831 | 0.544859496 | 0.790481814 | -4.807711451 |
| **A_51_P392867** | Pex12 | 0.069100133 | -0.193198388 | 0.331398655 | 8.980809841 | 0.640127127 | 0.545082651 | 0.790697243 | -4.807828231 |
| **A_55_P2101862** | Spryd3 | 0.060657226 | -0.169596929 | 0.290911382 | 9.87080784 | 0.640115379 | 0.545089801 | 0.790697243 | -4.807831971 |
| **A_55_P2081105** | NA | -0.105227676 | -0.504957832 | 0.29450248 | 9.434184526 | -0.639655672 | 0.545369635 | 0.791033785 | -4.80797829 |
| **A_51_P321331** | Sptlc2 | -0.042600772 | -0.204457132 | 0.119255588 | 8.288277096 | -0.639544673 | 0.545437215 | 0.791033785 | -4.808013607 |
| **A_55_P2103352** | Ndufb5 | -0.043192447 | -0.207299908 | 0.120915014 | 12.98991573 | -0.639532577 | 0.54544458 | 0.791033785 | -4.808017455 |
| **A_55_P2175314** | NA | 0.072315508 | -0.202542327 | 0.347173343 | 15.52476567 | 0.639302656 | 0.545584584 | 0.791110136 | -4.808090593 |
| **A_55_P1992571** | NA | 0.071839732 | -0.201231132 | 0.344910596 | 12.85113277 | 0.639252635 | 0.545615045 | 0.791110136 | -4.808106501 |
| **A_51_P348548** | Ecsit | 0.080310862 | -0.224969681 | 0.385591405 | 12.33669923 | 0.63923177 | 0.545627752 | 0.791110136 | -4.808113137 |
| **A_51_P284466** | Tsc22d4 | -0.052084098 | -0.250084927 | 0.145916732 | 10.25674477 | -0.639177274 | 0.545660941 | 0.791110136 | -4.808130468 |
| **A_51_P382152** | Procr | -0.105625282 | -0.507270615 | 0.296020051 | 8.529271055 | -0.639011019 | 0.5457622 | 0.791197597 | -4.808183331 |
| **A_51_P520927** | Ccdc97 | 0.058656359 | -0.164444593 | 0.281757311 | 10.39024596 | 0.638847011 | 0.545862101 | 0.791283078 | -4.80823547 |
| **A_65_P02177** | Gstm4 | -0.088620746 | -0.425768749 | 0.248527257 | 9.156227868 | -0.63870159 | 0.545950691 | 0.791350415 | -4.80828169 |
| **A_55_P2108689** | NA | -0.068968236 | -0.33139206 | 0.193455587 | 14.3371638 | -0.638600201 | 0.546012462 | 0.791350415 | -4.808313911 |
| **A_55_P2098140** | Tmed2 | 0.055913561 | -0.156847625 | 0.268674747 | 13.85928816 | 0.638569156 | 0.546031377 | 0.791350415 | -4.808323776 |
| **A_66_P101764** | NA | -0.264296514 | -1.270113431 | 0.741520402 | 12.68661075 | -0.638492336 | 0.546078183 | 0.791358914 | -4.808348185 |
| **A_55_P2046807** | Sparc | -0.23966045 | -1.152374114 | 0.673053214 | 11.88089479 | -0.638035741 | 0.546356436 | 0.791608544 | -4.808493215 |
| **A_55_P2131473** | Rmnd5a | 0.047741295 | -0.134081162 | 0.229563752 | 8.297843245 | 0.638013491 | 0.546369998 | 0.791608544 | -4.80850028 |
| **A_55_P1957018** | Caskin2 | 0.082430134 | -0.231534702 | 0.39639497 | 9.635621487 | 0.637952266 | 0.546407316 | 0.791608544 | -4.80851972 |
| **A_55_P2115955** | Raet1e | 0.082973188 | -0.233084218 | 0.399030595 | 9.352352255 | 0.637903515 | 0.546437032 | 0.791608544 | -4.808535199 |
| **A_55_P2107477** | Rab5c | -0.082766859 | -0.398065636 | 0.232531918 | 13.00093687 | -0.637848263 | 0.546470712 | 0.791608544 | -4.80855274 |
| **A_55_P2062508** | Ube2e1 | -0.086298559 | -0.415072805 | 0.242475686 | 11.85697485 | -0.637806506 | 0.546496167 | 0.791608544 | -4.808565996 |
| **A_51_P295967** | Proc | -0.136991094 | -0.658985923 | 0.385003735 | 14.62201996 | -0.637689584 | 0.546567447 | 0.791652467 | -4.80860311 |
| **A_52_P283055** | Slc5a3 | 0.054917849 | -0.154440996 | 0.264276694 | 8.093881672 | 0.637390214 | 0.546749977 | 0.791738239 | -4.808698113 |
| **A_55_P1980267** | Sidt2 | -0.045317777 | -0.218083768 | 0.127448214 | 11.12049856 | -0.637372806 | 0.546760592 | 0.791738239 | -4.808703636 |
| **A_55_P2033160** | Pomgnt1 | -0.164783107 | -0.793067087 | 0.463500873 | 10.87717717 | -0.637294089 | 0.546808594 | 0.791738239 | -4.80872861 |

| | | | | | | | | | |
|---|---|---|---|---|---|---|---|---|---|
| **A_55_P2105362** | Tmed5 | 0.098300603 | -0.276502872 | 0.473104079 | 9.071766909 | 0.637288032 | 0.546812288 | 0.791738239 | -4.808730532 |
| **A_55_P2057469** | Caskin1 | 0.044907586 | -0.126325851 | 0.216141023 | 7.713825379 | 0.637256576 | 0.546831471 | 0.791738239 | -4.808740511 |
| **A_55_P2157378** | Xlr5a | 0.046398151 | -0.130582899 | 0.223379201 | 7.849238798 | 0.637025905 | 0.546972154 | 0.791781985 | -4.808813675 |
| **A_55_P2130576** | NA | -0.095231811 | -0.458492565 | 0.268028943 | 14.7148009 | -0.637010738 | 0.546981404 | 0.791781985 | -4.808818485 |
| **A_55_P2169879** | NA | -0.079058879 | -0.380630612 | 0.222512854 | 14.14145566 | -0.63700554 | 0.546984575 | 0.791781985 | -4.808820134 |
| **A_51_P106059** | Traf4 | 0.117015042 | -0.32946111 | 0.563491194 | 11.33166539 | 0.636834545 | 0.547088878 | 0.791792037 | -4.808874355 |
| **A_55_P2049147** | NA | -0.096629036 | -0.465330062 | 0.27207199 | 15.35774473 | -0.636819701 | 0.547097933 | 0.791792037 | -4.808879061 |
| **A_51_P150394** | Vps28 | -0.062006029 | -0.298636526 | 0.174624467 | 13.91859032 | -0.636716925 | 0.547160632 | 0.791792037 | -4.808911645 |
| **A_55_P2107207** | Gpr137b-ps | -0.058198674 | -0.280319868 | 0.163922519 | 8.793595048 | -0.636658119 | 0.547196507 | 0.791792037 | -4.808930286 |
| **A_55_P1971439** | Serpinb9c | -0.041716367 | -0.200948486 | 0.117515751 | 7.65387725 | -0.636588806 | 0.547238796 | 0.791792037 | -4.808952256 |
| **A_55_P2002712** | Mllt10 | 0.081484511 | -0.229559139 | 0.39252816 | 9.886424893 | 0.636556433 | 0.547258547 | 0.791792037 | -4.808962517 |
| **A_52_P476699** | Gemin4 | 0.064846393 | -0.182698604 | 0.31239139 | 8.509151163 | 0.636524114 | 0.547278267 | 0.791792037 | -4.80897276 |
| **A_55_P2046253** | Tmem239 | 0.156480103 | -0.440971069 | 0.753931275 | 14.85564227 | 0.63641425 | 0.547345302 | 0.791829754 | -4.809007578 |
| **A_66_P108059** | Ttc39c | -0.11348376 | -0.546918654 | 0.319951134 | 9.244949486 | -0.636199078 | 0.547476609 | 0.791960438 | -4.809075754 |
| **A_65_P16603** | Bad | -0.100583272 | -0.484890944 | 0.283724399 | 8.284768242 | -0.63596014 | 0.547622441 | 0.792112112 | -4.809151438 |
| **A_52_P606826** | Pigh | -0.050454509 | -0.243271421 | 0.142362402 | 8.030390075 | -0.635825664 | 0.547704527 | 0.792165092 | -4.809194024 |
| **A_55_P2054728** | Ldlrad3 | 0.160357962 | -0.452523542 | 0.773239466 | 14.79925934 | 0.63576586 | 0.547741035 | 0.792165092 | -4.809212961 |
| **A_55_P2137476** | Mpv17 | -0.060456282 | -0.291559097 | 0.170646532 | 10.69774946 | -0.635651961 | 0.547810569 | 0.792182211 | -4.809249021 |
| **A_51_P391727** | Slc3a1 | -0.243279399 | -1.173307976 | 0.686749178 | 9.797120159 | -0.635612207 | 0.547834839 | 0.792182211 | -4.809261606 |
| **A_51_P465350** | Ly86 | -0.064260818 | -0.30998207 | 0.181460434 | 11.52406495 | -0.635457806 | 0.547929111 | 0.79225926 | -4.809310479 |
| **A_55_P2031407** | NA | -0.076422454 | -0.368920711 | 0.216075803 | 17.32918908 | -0.63486427 | 0.548291598 | 0.792724085 | -4.809498264 |
| **A_55_P2035554** | Strip1 | -0.069290954 | -0.334566165 | 0.195984256 | 11.39940412 | -0.634692058 | 0.5483968 | 0.792749932 | -4.809552723 |
| **A_51_P155522** | Psmb4 | -0.051912783 | -0.250720167 | 0.146894601 | 14.37314145 | -0.634490305 | 0.548520064 | 0.792749932 | -4.809616508 |
| **A_55_P2029581** | NA | -0.074102572 | -0.357948416 | 0.209743272 | 7.806715471 | -0.634357281 | 0.548601346 | 0.792749932 | -4.809658554 |
| **A_55_P2056871** | Akap13 | 0.055249847 | -0.156387483 | 0.266887177 | 11.38239156 | 0.63433985 | 0.548611997 | 0.792749932 | -4.809664063 |
| **A_51_P112762** | Slc5a3 | -0.042035607 | -0.203063153 | 0.118991939 | 7.852618357 | -0.634308223 | 0.548631324 | 0.792749932 | -4.80055057 |
| **A_51_P137094** | Sall2 | -0.097381202 | -0.470582032 | 0.275819628 | 8.676620459 | -0.634038633 | 0.548796082 | 0.792749932 | -4.809759245 |
| **A_52_P417859** | Gid4 | 0.057628407 | -0.163231435 | 0.278488249 | 9.19041284 | 0.634020133 | 0.548807389 | 0.792749932 | -4.80976509 |
| **A_55_P2140745** | NA | -0.076007031 | -0.367334037 | 0.215319975 | 8.607930789 | -0.633951758 | 0.548849181 | 0.792749932 | -4.80978669 |
| **A_55_P2266880** | Kirrel | -0.130658629 | -0.631473656 | 0.370156399 | 8.648931233 | -0.633933836 | 0.548860136 | 0.792749932 | -4.809792351 |
| **A_55_P1953851** | Stk39 | -0.076543096 | -0.36997769 | 0.216891498 | 8.757963828 | -0.633837454 | 0.548919051 | 0.792749932 | -4.809822796 |
| **A_51_P202633** | Ebi3 | -0.116205514 | -0.561738959 | 0.329327931 | 10.94972365 | -0.633766989 | 0.548962126 | 0.792749932 | -4.809845051 |
| **A_55_P1975087** | NA | -0.067155283 | -0.324629884 | 0.190319318 | 14.46061692 | -0.633766061 | 0.548962693 | 0.792749932 | -4.809845344 |
| **A_55_P2137586** | Nphp4 | 0.125447886 | -0.355536431 | 0.606432203 | 8.248141382 | 0.633746566 | 0.54897461 | 0.792749932 | -4.8098515 |
| **A_51_P449048** | Cndp2 | -0.048655644 | -0.235213936 | 0.137902648 | 9.284608303 | -0.633726489 | 0.548986884 | 0.792749932 | -4.809857841 |
| **A_52_P179729** | Txnl4a | -0.09630994 | -0.465608313 | 0.272988434 | 10.94717398 | -0.633690074 | 0.549009146 | 0.792749932 | -4.809869341 |
| **A_55_P2155901** | Snhg10 | 0.081038786 | -0.229741352 | 0.391818924 | 9.065610354 | 0.633611223 | 0.549057352 | 0.792749932 | -4.80989424 |
| **A_55_P1981681** | Syt8 | -0.070289509 | -0.339963501 | 0.199384484 | 7.758874755 | -0.633336666 | 0.549225227 | 0.792749932 | -4.809980918 |
| **A_55_P2016079** | Erich4 | 0.126607305 | -0.359142296 | 0.612356905 | 8.307927507 | 0.633329183 | 0.549229803 | 0.792749932 | -4.80998328 |
| **A_55_P2052749** | Pcm1 | 0.071347191 | -0.202410542 | 0.345104924 | 9.823905808 | 0.633276938 | 0.549261751 | 0.792749932 | -4.80999977 |
| **A_55_P1998848** | NA | -0.104685227 | -0.506422818 | 0.297052363 | 14.94667963 | -0.63317844 | 0.549321986 | 0.792749932 | -4.810030856 |
| **A_55_P2083396** | Radil | -0.065402632 | -0.316408178 | 0.185602914 | 7.810015302 | -0.633133214 | 0.549349645 | 0.792749932 | -4.810045128 |
| **A_51_P272283** | Cmbl | -0.142216163 | -0.688119792 | 0.403687466 | 11.13518404 | -0.633018153 | 0.549420018 | 0.792749932 | -4.810081435 |
| **A_52_P367294** | Fsd1l | 0.052955326 | -0.150331366 | 0.256242017 | 8.266352968 | 0.632971103 | 0.549448795 | 0.792749932 | -4.810096279 |
| **A_55_P2103501** | Ankrd29 | -0.046753722 | -0.226240833 | 0.132733389 | 7.73185653 | -0.632945197 | 0.549464641 | 0.792749932 | -4.810104452 |
| **A_55_P2181029** | NA | 0.061076517 | -0.173399545 | 0.295552579 | 8.160313012 | 0.632934749 | 0.549471031 | 0.792749932 | -4.810107749 |
| **A_55_P1956179** | Git2 | 0.090753671 | -0.257660977 | 0.43916832 | 10.8484286 | 0.632923157 | 0.549478122 | 0.792749932 | -4.810111406 |
| **A_55_P2076805** | Ddx5 | 0.110006091 | -0.312350108 | 0.532362289 | 9.405956571 | 0.632879529 | 0.549504808 | 0.792749932 | -4.810125169 |
| **A_55_P2183587** | Gli1 | -0.267920759 | -1.296578615 | 0.760737097 | 8.558870895 | -0.632875971 | 0.549506984 | 0.792749932 | -4.810126291 |
| **A_55_P2095535** | NA | -0.041799342 | -0.202299509 | 0.118700824 | 7.699321831 | -0.632815561 | 0.549543937 | 0.792749932 | -4.810145347 |

| | | | | | | | | | |
|---|---|---|---|---|---|---|---|---|---|
| **A_66_P123448** | Pigx | -0.105256266 | -0.509423891 | 0.298911358 | 11.18359587 | -0.632804601 | 0.549550642 | 0.792749932 | -4.810148805 |
| **A_51_P408082** | Apoa1 | -0.152548153 | -0.738376007 | 0.433279702 | 14.72992513 | -0.632732474 | 0.549594764 | 0.792749932 | -4.794890982 |
| **A_55_P1965620** | Bud31 | 0.085188255 | -0.242019978 | 0.412396487 | 13.10235872 | 0.632613838 | 0.549667343 | 0.792749932 | -4.810208969 |
| **A_51_P143470** | Timm44 | 0.080460256 | -0.228587957 | 0.389508469 | 11.59799293 | 0.632613366 | 0.549667632 | 0.792749932 | -4.810209118 |
| **A_55_P2084741** | Zkscan6 | -0.043748833 | -0.211833578 | 0.124335911 | 9.347729782 | -0.632442964 | 0.54977189 | 0.792749932 | -4.810262849 |
| **A_51_P384673** | Lrrc59 | 0.082458551 | -0.234352381 | 0.399269484 | 10.84784027 | 0.632439127 | 0.549774238 | 0.792749932 | -4.810264059 |
| **A_55_P2109377** | Camk2b | 0.128665232 | -0.365710757 | 0.623041221 | 10.20629575 | 0.63239296 | 0.549802486 | 0.792749932 | -4.810278614 |
| **A_51_P322273** | Ptp4a3 | 0.113916778 | -0.323818288 | 0.551651844 | 11.1877399 | 0.63235294 | 0.549826974 | 0.792749932 | -4.810291231 |
| **A_55_P2110290** | Tmem254a | -0.086200921 | -0.417752579 | 0.245350736 | 10.57929135 | -0.631748027 | 0.550197204 | 0.793201265 | -4.810481854 |
| **A_52_P536293** | NA | -0.078990665 | -0.3828293 | 0.224847971 | 14.53918447 | -0.631707394 | 0.550222079 | 0.793201265 | -4.810494653 |
| **A_66_P134728** | Psmg1 | 0.058319269 | -0.16606148 | 0.282700019 | 12.80459792 | 0.631552808 | 0.550316719 | 0.793244336 | -4.81054334 |
| **A_55_P2062070** | Pdzd2 | -0.049940677 | -0.242093775 | 0.142212421 | 7.701510979 | -0.631524528 | 0.550334034 | 0.793244336 | -4.810552247 |
| **A_55_P2163404** | NA | -0.129267891 | -0.626795454 | 0.368259673 | 14.5283479 | -0.631330405 | 0.550452896 | 0.793321833 | -4.810613371 |
| **A_55_P2028847** | NA | -0.066956783 | -0.324672054 | 0.190758487 | 11.94364163 | -0.631302658 | 0.550469886 | 0.793321833 | -4.810622106 |
| **A_55_P2005315** | NA | 0.210649989 | -0.60048083 | 1.021780807 | 12.39567189 | 0.631035405 | 0.550633555 | 0.793498544 | -4.810706229 |
| **A_55_P2128225** | Rpl18 | -0.074754497 | -0.362638831 | 0.213129838 | 15.46361329 | -0.630960946 | 0.550679159 | 0.793505105 | -4.810729661 |
| **A_55_P2023707** | Camta1 | -0.042516583 | -0.206310139 | 0.121276972 | 7.688832672 | -0.630731811 | 0.550819515 | 0.793559527 | -4.810801755 |
| **A_55_P2115392** | Mst1 | -0.138032135 | -0.669834867 | 0.393770597 | 12.3520818 | -0.630685479 | 0.550847899 | 0.793559527 | -4.81081633 |
| **A_51_P472518** | Capn1 | 0.103364975 | -0.294882599 | 0.501612549 | 8.786077833 | 0.630671862 | 0.55085624 | 0.793559527 | -4.810820614 |
| **A_51_P386880** | Stip1 | 0.087106412 | -0.24852249 | 0.422735314 | 11.61554394 | 0.630629034 | 0.550882478 | 0.793559527 | -4.810834085 |
| **A_55_P2065303** | NA | 0.082390595 | -0.235100483 | 0.399881673 | 15.91379278 | 0.63056419 | 0.550922205 | 0.793559527 | -4.810854481 |
| **A_55_P1966109** | Epn2 | -0.131215787 | -0.636978638 | 0.374547064 | 8.637288939 | -0.630408924 | 0.551017336 | 0.793585272 | -4.810903311 |
| **A_55_P2096933** | Cntln | -0.110108712 | -0.534521026 | 0.314303603 | 9.818783889 | -0.630401 | 0.551022191 | 0.793585272 | -4.810905803 |
| **A_55_P2155764** | NA | 0.052411472 | -0.149716258 | 0.254539202 | 10.45050578 | 0.630062521 | 0.551229613 | 0.793785233 | -4.811012216 |
| **A_55_P2108864** | Fer | 0.045182904 | -0.129073572 | 0.219439379 | 8.170635712 | 0.630040406 | 0.551243167 | 0.793785233 | -4.811019167 |
| **A_55_P1964408** | Cryab | -0.396548921 | -1.926887401 | 1.133789559 | 10.72356261 | -0.629640222 | 0.551488468 | 0.794074418 | -4.811144917 |
| **A_55_P2177386** | Cradd | -0.054740236 | -0.266011192 | 0.156530719 | 9.542527003 | -0.629578748 | 0.551526155 | 0.794074418 | -4.811164227 |
| **A_55_P2106424** | NA | 0.069866395 | -0.199855888 | 0.339588678 | 16.63447201 | 0.629411533 | 0.551628678 | 0.794110421 | -4.811216748 |
| **A_55_P2064792** | Arpc3 | -0.050974956 | -0.24776838 | 0.145818469 | 12.35483198 | -0.629403947 | 0.551633329 | 0.794110421 | -4.81121913 |
| **A_55_P2138816** | Mplkip | 0.063780996 | -0.182631297 | 0.310193289 | 9.528747249 | 0.6289442 | 0.55191527 | 0.794440671 | -4.811363472 |
| **A_55_P2023446** | NA | -0.052769977 | -0.256657844 | 0.15111789 | 11.48735023 | -0.628895824 | 0.551944941 | 0.794440671 | -4.811378654 |
| **A_55_P1994539** | Cldn12 | 0.122280914 | -0.35032032 | 0.594882148 | 11.23722062 | 0.628705145 | 0.552061906 | 0.794549857 | -4.811438491 |
| **A_51_P213153** | Tmem208 | 0.058683461 | -0.168311942 | 0.285678864 | 13.45568246 | 0.628176736 | 0.552386116 | 0.794814111 | -4.811604231 |
| **A_52_P96782** | Wasl | -0.084504589 | -0.41138202 | 0.242372843 | 9.582836926 | -0.628171958 | 0.552389048 | 0.794814111 | -4.81160573 |
| **A_51_P342387** | Apoh | -0.217926754 | -1.060909065 | 0.625055557 | 16.26731154 | -0.628167232 | 0.552391948 | 0.794814111 | -4.811607211 |
| **A_55_P2029061** | Tnfrsf9 | 0.18368523 | -0.526877587 | 0.894248047 | 12.85498802 | 0.628137828 | 0.552409993 | 0.794814111 | -4.811616431 |
| **A_55_P2181288** | Ppp1r2 | 0.067860088 | -0.194695053 | 0.330415228 | 9.943520821 | 0.628025213 | 0.552479106 | 0.794844597 | -4.811651737 |
| **A_55_P1987730** | Prxl2a | -0.106660196 | -0.519372254 | 0.306051862 | 14.05403138 | -0.627969295 | 0.552513426 | 0.794844597 | -4.811669267 |
| **A_51_P219061** | Oxa1l | 0.053039414 | -0.152225401 | 0.258304228 | 9.083163899 | 0.627866619 | 0.552576446 | 0.794864545 | -4.81170145 |
| **A_55_P1958833** | Gm9079 | 0.095727545 | -0.274773986 | 0.466229076 | 10.09967973 | 0.627812708 | 0.552609538 | 0.794864545 | -4.811718347 |
| **A_52_P123485** | Tcf25 | -0.056201548 | -0.273775334 | 0.161372239 | 13.05613127 | -0.627660638 | 0.552702887 | 0.794939661 | -4.811766001 |
| **A_51_P484111** | Matn2 | 0.1063473 | -0.305417034 | 0.518111633 | 10.65131486 | 0.627568202 | 0.552759634 | 0.794962125 | -4.811794964 |
| **A_51_P172323** | Pak2 | -0.050882095 | -0.247999525 | 0.146235334 | 10.63088593 | -0.627224698 | 0.552970545 | 0.7950721 | -4.811902561 |
| **A_55_P2098100** | Arhgef9 | 0.04444141 | -0.12772762 | 0.21661044 | 7.859512823 | 0.627214345 | 0.552976903 | 0.7950721 | -4.811905803 |
| **A_51_P259879** | Fkrp | 0.060772671 | -0.174683162 | 0.296228504 | 9.38809964 | 0.627165352 | 0.553006989 | 0.7950721 | -4.811921146 |
| **A_55_P2121961** | Psmc3 | -0.060396734 | -0.294438082 | 0.173644614 | 13.1003155 | -0.627052716 | 0.553076161 | 0.7950721 | -4.811956414 |
| **A_55_P1993728** | Tmem116 | -0.052844679 | -0.257647209 | 0.151957851 | 8.394410539 | -0.626973426 | 0.553124858 | 0.7950721 | -4.811981238 |
| **A_66_P122415** | Snhg6 | -0.074489933 | -0.36318154 | 0.214201673 | 12.98431257 | -0.626969792 | 0.553127089 | 0.7950721 | -4.811982375 |
| **A_52_P222230** | Gm40812 | -0.182557999 | -0.890126888 | 0.525010891 | 10.45679153 | -0.626924621 | 0.553154833 | 0.7950721 | -4.811996517 |
| **A_52_P468041** | Alkbh5 | 0.097549179 | -0.280547743 | 0.475646101 | 10.009751 | 0.626907789 | 0.553165172 | 0.7950721 | -4.812001786 |

| | | | | | | | | | |
|---|---|---|---|---|---|---|---|---|---|
| **A_55_P2116385** | Gpr183 | -0.043729889 | -0.213273096 | 0.125813318 | 7.845525354 | -0.626730989 | 0.55327377 | 0.795169061 | -4.812057124 |
| **A_51_P411271** | Nfe2l2 | -0.097794497 | -0.477041195 | 0.281452201 | 11.79389537 | -0.626578945 | 0.553367172 | 0.795244168 | -4.812104703 |
| **A_51_P114062** | Ncs1 | 0.045686029 | -0.131517151 | 0.222889209 | 7.880003211 | 0.626462514 | 0.553438703 | 0.795287837 | -4.799038681 |
| **A_55_P2145531** | Mpst | 0.049649038 | -0.143002247 | 0.242300322 | 13.28754513 | 0.626213051 | 0.553591985 | 0.795448965 | -4.812219165 |
| **A_55_P2094964** | NA | -0.072146036 | -0.352175082 | 0.20788301 | 10.33282335 | -0.626026307 | 0.553706746 | 0.795554723 | -4.812277563 |
| **A_55_P2042973** | 1010001N08Rik | -0.04532756 | -0.221295801 | 0.13064068 | 8.034574441 | -0.625909068 | 0.553778801 | 0.795599111 | -4.812314218 |
| **A_51_P491095** | Gzf1 | 0.083200039 | -0.239943282 | 0.406343361 | 9.686690936 | 0.625621313 | 0.55395568 | 0.795794079 | -4.812404162 |
| **A_52_P420466** | H2ac4 | -0.073656016 | -0.359888754 | 0.212576722 | 16.26512249 | -0.6252765 | 0.554167677 | 0.796039464 | -4.812511896 |
| **A_51_P300207** | Rsl24d1 | 0.043861157 | -0.126650685 | 0.214372999 | 9.29097607 | 0.625041315 | 0.554312302 | 0.796188042 | -4.81258535 |
| **A_55_P2162935** | Ntn1 | -0.053238618 | -0.260239059 | 0.153761823 | 7.883084862 | -0.624940532 | 0.554374285 | 0.796217904 | -4.81261682 |
| **A_55_P2054409** | Pira2 | -0.086038651 | -0.420610574 | 0.248533273 | 8.555610343 | -0.624866563 | 0.554419779 | 0.796224081 | -4.812639914 |
| **A_51_P439452** | Insig2 | -0.059974169 | -0.293233441 | 0.173285103 | 10.01780327 | -0.624753232 | 0.554489487 | 0.79626503 | -4.812675294 |
| **A_55_P1957613** | Ak2 | 0.089993568 | -0.260072772 | 0.440059908 | 14.01696845 | 0.624660874 | 0.5545463 | 0.796268555 | -4.812704123 |
| **A_65_P08239** | Mef2a | 0.043234322 | -0.124967078 | 0.211435721 | 8.105635249 | 0.624571601 | 0.554601218 | 0.796268555 | -4.812731985 |
| **A_55_P2006713** | NA | 0.068137985 | -0.197032845 | 0.333308816 | 16.73901667 | 0.624376759 | 0.554721089 | 0.796268555 | -4.812792784 |
| **A_51_P483311** | Mpv17l2 | 0.114786477 | -0.331945394 | 0.561518347 | 9.369679483 | 0.624348363 | 0.554738561 | 0.796268555 | -4.812801644 |
| **A_55_P2015734** | Syvn1 | -0.052108645 | -0.254917822 | 0.150700532 | 12.89096129 | -0.624317294 | 0.554757677 | 0.796268555 | -4.812811337 |
| **A_55_P1975041** | Arhgef11 | -0.055970778 | -0.273853476 | 0.16191192 | 8.866016695 | -0.624197157 | 0.5548316 | 0.796268555 | -4.812848814 |
| **A_55_P2036337** | Ifnar1 | 0.045409439 | -0.13136587 | 0.222184747 | 10.59905328 | 0.624176937 | 0.554844042 | 0.796268555 | -4.812855121 |
| **A_55_P1996088** | NA | 0.096042445 | -0.277886531 | 0.46997142 | 11.75035281 | 0.624104445 | 0.554888652 | 0.796268555 | -4.812877732 |
| **A_52_P108447** | Glis2 | -0.173321498 | -0.848126953 | 0.501483957 | 9.988954694 | -0.624104103 | 0.554888862 | 0.796268555 | -4.812877839 |
| **A_51_P111259** | Cdk2ap2 | -0.079684723 | -0.389957195 | 0.230587749 | 12.02537955 | -0.624043712 | 0.554926026 | 0.796268555 | -4.812896674 |
| **A_55_P2096226** | Loxl1 | -0.122858525 | -0.601262783 | 0.355545733 | 8.004866384 | -0.62401273 | 0.554945093 | 0.796268555 | -4.812906336 |
| **A_55_P2124346** | Ivns1abp | -0.059911749 | -0.293257191 | 0.173433693 | 11.54351916 | -0.623872532 | 0.555031378 | 0.796317042 | -4.812950053 |
| **A_55_P2143075** | Ubc | 0.067319339 | -0.194897749 | 0.329536427 | 15.48545899 | 0.623823948 | 0.555061281 | 0.796317042 | -4.812965201 |
| **A_52_P430274** | Dnaaf5 | 0.059445529 | -0.17214017 | 0.291031229 | 9.695180947 | 0.623721406 | 0.555124398 | 0.796318879 | -4.812997169 |
| **A_52_P32081** | Crbn | 0.048869003 | -0.141534375 | 0.23927238 | 8.868127639 | 0.623651566 | 0.555167389 | 0.796318879 | -4.81301894 |
| **A_55_P2209046** | NA | -0.097685246 | -0.478305049 | 0.282934556 | 8.646662821 | -0.623621079 | 0.555186156 | 0.796318879 | -4.813028443 |
| **A_51_P415220** | Zmat3 | 0.061160001 | -0.1771993 | 0.299519303 | 8.689807543 | 0.623474322 | 0.555276502 | 0.796389368 | -4.813074182 |
| **A_55_P2053929** | NA | -0.053234417 | -0.260794521 | 0.154325687 | 11.57967853 | -0.623206263 | 0.555441547 | 0.796410832 | -4.813157704 |
| **A_66_P113008** | Pla2g2e | 0.042942372 | -0.124493781 | 0.210378524 | 7.760462657 | 0.623189287 | 0.555452 | 0.796410832 | -4.813162993 |
| **A_51_P464029** | Fads3 | -0.239931585 | -1.175557731 | 0.695694562 | 10.60366317 | -0.623115079 | 0.555497696 | 0.796410832 | -4.813186109 |
| **A_52_P173368** | NA | 0.054006172 | -0.156607102 | 0.264619447 | 8.486052874 | 0.623075744 | 0.555521919 | 0.796410832 | -4.813198361 |
| **A_52_P197627** | Umps | 0.103939007 | -0.301420595 | 0.50929861 | 9.513744959 | 0.623047689 | 0.555539196 | 0.796410832 | -4.813207099 |
| **A_51_P277795** | Resf1 | 0.061904142 | -0.179549575 | 0.303357859 | 10.73288805 | 0.622972682 | 0.555585388 | 0.796410832 | -4.81323046 |
| **A_52_P177847** | Tril | -0.081688326 | -0.400323588 | 0.236946936 | 7.809308407 | -0.622944491 | 0.55560275 | 0.796410832 | -4.813239239 |
| **A_66_P100561** | Bnip2 | -0.072723841 | -0.356405597 | 0.210957916 | 12.58143072 | -0.622914711 | 0.555621091 | 0.796410832 | -4.813248513 |
| **A_55_P2166538** | Mrps16 | 0.092276078 | -0.267894434 | 0.45244659 | 13.03721774 | 0.622535556 | 0.555854636 | 0.796686508 | -4.813366555 |
| **A_55_P1955970** | Cbfa2t2 | -0.068632253 | -0.336555094 | 0.199290588 | 10.13470091 | -0.622446037 | 0.555909785 | 0.796706475 | -4.813394416 |
| **A_52_P588559** | Ube2d3 | 0.074033353 | -0.215079362 | 0.363146068 | 11.95727414 | 0.62221921 | 0.556049539 | 0.796847683 | -4.813464998 |
| **A_51_P159402** | Gcat | 0.070825447 | -0.20584715 | 0.347498043 | 13.35292676 | 0.622022889 | 0.556170515 | 0.796961961 | -4.81352607 |
| **A_51_P389664** | Scrt2 | 0.053998789 | -0.156999602 | 0.264997181 | 7.957450734 | 0.62185348 | 0.55627492 | 0.797049294 | -4.813578757 |
| **A_55_P2168267** | NA | -0.097124965 | -0.476776784 | 0.282526855 | 15.15272806 | -0.621625151 | 0.556415657 | 0.797049294 | -4.813649751 |
| **A_51_P361580** | Rab11b | -0.04206176 | -0.206479238 | 0.122355717 | 11.65856747 | -0.621616669 | 0.556420886 | 0.797049294 | -4.813652388 |
| **A_51_P138879** | Zgpat | 0.051677448 | -0.150328932 | 0.253683827 | 9.084915574 | 0.621611673 | 0.556423965 | 0.797049294 | -4.813653941 |
| **A_55_P2178510** | Mc3r | 0.12601719 | -0.366599724 | 0.618634104 | 8.045218826 | 0.621589487 | 0.556437641 | 0.797049294 | -4.813660838 |
| **A_52_P122673** | Cep41 | 0.04841647 | -0.140940932 | 0.237773871 | 8.233295354 | 0.621289513 | 0.556622576 | 0.797255114 | -4.813754071 |
| **A_55_P1986049** | Fip1l1 | 0.061969046 | -0.180554136 | 0.304492228 | 8.421002041 | 0.620875813 | 0.556877684 | 0.797561407 | -4.81388259 |
| **A_51_P144770** | 2410002F23Rik | 0.053520239 | -0.156044641 | 0.263085118 | 11.56555081 | 0.620558502 | 0.557073402 | 0.797782603 | -4.813981118 |
| **A_52_P664699** | Cul1 | -0.051122584 | -0.251357732 | 0.149112563 | 8.756872829 | -0.620376984 | 0.557185382 | 0.797883853 | -4.814037462 |

| | | | | | | | | | |
|---|---|---|---|---|---|---|---|---|---|
| **A_55_P1961894** | Fbxw11 | -0.077774275 | -0.382585633 | 0.227037082 | 11.76628828 | -0.619994756 | 0.557421227 | 0.798151626 | -4.814156063 |
| **A_55_P1972157** | Espnl | 0.084052728 | -0.245394063 | 0.413499519 | 8.675903179 | 0.619940091 | 0.557454961 | 0.798151626 | -4.814173021 |
| **A_55_P2147591** | Med27 | -0.044539662 | -0.219172775 | 0.13009345 | 8.254737466 | -0.619731421 | 0.557583746 | 0.798276886 | -4.814237739 |
| **A_55_P2266295** | Fzd3 | -0.051222762 | -0.252175121 | 0.149729596 | 8.129764435 | -0.619374149 | 0.557804287 | 0.798533481 | -4.814348504 |
| **A_51_P363556** | Abi2 | -0.073722717 | -0.363008219 | 0.215562786 | 8.942197248 | -0.619238358 | 0.557888124 | 0.798544856 | -4.81439059 |
| **A_52_P126055** | Pon2 | 0.05309227 | -0.155243714 | 0.261428255 | 8.922805605 | 0.619227449 | 0.557894859 | 0.798544856 | -4.81439397 |
| **A_52_P13897** | Hook1 | -0.068121433 | -0.335652149 | 0.199409283 | 8.482975217 | -0.618718803 | 0.558208965 | 0.798935289 | -4.814551545 |
| **A_51_P428364** | Psma6 | -0.061537937 | -0.303336541 | 0.180260667 | 11.58178569 | -0.618404058 | 0.558403384 | 0.799154377 | -4.814648998 |
| **A_51_P277745** | Abcc2 | -0.175999209 | -0.867709767 | 0.515711348 | 10.45841973 | -0.618257654 | 0.558493833 | 0.799224646 | -4.814694314 |
| **A_66_P134809** | Gas5 | 0.073593187 | -0.215731208 | 0.362917581 | 13.20305393 | 0.618067267 | 0.558611468 | 0.799333807 | -4.814753231 |
| **A_55_P2034517** | NA | -0.045474179 | -0.224362032 | 0.133413673 | 7.693544935 | -0.617685219 | 0.55884757 | 0.799453426 | -4.814871415 |
| **A_51_P236175** | Rit1 | -0.085856998 | -0.423624797 | 0.2519108 | 12.84732343 | -0.617647442 | 0.558870919 | 0.799453426 | -4.814883098 |
| **A_52_P581440** | Wasf2 | -0.10143362 | -0.500506908 | 0.297639668 | 12.3332035 | -0.617607349 | 0.558895701 | 0.799453426 | -4.814895496 |
| **A_55_P1962274** | Ccnt1 | 0.063351962 | -0.185945065 | 0.312648988 | 8.990528571 | 0.617484639 | 0.558971551 | 0.799453426 | -4.814933439 |
| **A_55_P2053850** | NA | 0.054252292 | -0.159245297 | 0.267749882 | 10.99786309 | 0.617459257 | 0.558987241 | 0.799453426 | -4.814941286 |
| **A_52_P628162** | Zfp592 | 0.061845076 | -0.181534249 | 0.305224402 | 8.357893747 | 0.61745404 | 0.558990467 | 0.799453426 | -4.814942899 |
| **A_51_P434101** | Herpud1 | 0.102036039 | -0.299511297 | 0.503583376 | 13.43255055 | 0.617447494 | 0.558994513 | 0.799453426 | -4.814944923 |
| **A_55_P2035159** | Pcdh1 | 0.055086123 | -0.161714802 | 0.271887048 | 9.851917622 | 0.617396647 | 0.559025946 | 0.799453426 | -4.814960643 |
| **A_52_P576778** | Tmem30a | -0.090183875 | -0.445189228 | 0.264821479 | 11.14781682 | -0.617272854 | 0.559102477 | 0.79950372 | -4.81499891 |
| **A_55_P1981714** | Rreb1 | -0.1048277 | -0.517635783 | 0.307980384 | 11.22933802 | -0.61703678 | 0.55924844 | 0.799653284 | -4.815071867 |
| **A_55_P2111444** | Trpv6 | -0.079494919 | -0.392592104 | 0.233602267 | 7.890032979 | -0.616940663 | 0.559307875 | 0.799678162 | -4.815101565 |
| **A_66_P134252** | NA | 0.068855939 | -0.202383651 | 0.340095529 | 13.27021321 | 0.616838562 | 0.559371014 | 0.799678162 | -4.815133108 |
| **A_51_P338485** | Aldh6a1 | -0.158215957 | -0.781526855 | 0.465094941 | 12.31836571 | -0.616777827 | 0.559408575 | 0.799678162 | -4.815151869 |
| **A_55_P2161675** | Zfp133-ps | 0.19039888 | -0.559808318 | 0.940606079 | 10.99183281 | 0.616689213 | 0.55946338 | 0.799678162 | -4.81517924 |
| **A_66_P119801** | Optn | 0.063345211 | -0.186252865 | 0.312943286 | 8.257924408 | 0.616674147 | 0.559472699 | 0.799678162 | -4.815183893 |
| **A_52_P295533** | Slc25a28 | 0.073320134 | -0.215653183 | 0.36229345 | 9.763565618 | 0.616522166 | 0.559566703 | 0.799753386 | -4.815230827 |
| **A_55_P2051840** | NA | 0.056290422 | -0.165639252 | 0.278220097 | 9.415340926 | 0.616314409 | 0.559695221 | 0.799868912 | -4.815294971 |
| **A_51_P162162** | Inmt | -0.113991434 | -0.563453496 | 0.335470627 | 7.902989423 | -0.616257713 | 0.559730297 | 0.799868912 | -4.815312472 |
| **A_55_P2165451** | NA | -0.070685097 | -0.349441741 | 0.208071547 | 10.90791034 | -0.616149104 | 0.559797492 | 0.799905797 | -4.815345995 |
| **A_55_P2002389** | Nelfb | -0.066475785 | -0.328674293 | 0.195722723 | 12.31430651 | -0.616050687 | 0.559858386 | 0.799933673 | -4.815376367 |
| **A_55_P2095213** | Dusp22 | 0.049474529 | -0.145733067 | 0.244682124 | 9.00021008 | 0.615840353 | 0.55998854 | 0.800060498 | -4.815441266 |
| **A_51_P340653** | Slc16a4 | 0.039527882 | -0.116492622 | 0.195548386 | 7.702969128 | 0.615609137 | 0.560131637 | 0.800204137 | -4.815512586 |
| **A_55_P2024640** | Eral1 | 0.072817422 | -0.214630581 | 0.360265424 | 10.07758763 | 0.615544126 | 0.560171875 | 0.800204137 | -4.815532635 |
| **A_55_P2088218** | Uqcrb | 0.049736923 | -0.146654264 | 0.24612811 | 12.96660811 | 0.615375364 | 0.560276339 | 0.800294218 | -4.815584673 |
| **A_55_P2010843** | Fahd1 | 0.112320603 | -0.331246513 | 0.555887718 | 10.23584283 | 0.615294839 | 0.560326188 | 0.800306281 | -4.815609498 |
| **A_66_P126320** | Mrpl46 | 0.060772098 | -0.179316556 | 0.300860751 | 10.61415328 | 0.615057588 | 0.560473075 | 0.800456928 | -4.815682627 |
| **A_52_P296109** | NA | 0.040842319 | -0.12055791 | 0.202242548 | 8.426335584 | 0.614878676 | 0.560583858 | 0.800555995 | -4.815737758 |
| **A_52_P96552** | Vkorc1l1 | 0.053628968 | -0.158374998 | 0.265632934 | 9.751842488 | 0.614665226 | 0.560716045 | 0.800685612 | -4.815803514 |
| **A_52_P642453** | Obscn | -0.041509559 | -0.205685072 | 0.122665953 | 7.702559858 | -0.614359997 | 0.560905102 | 0.800861294 | -4.815897512 |
| **A_55_P2005903** | Lhx6 | 0.170864181 | -0.505071577 | 0.84679994 | 10.69241422 | 0.61422685 | 0.560987585 | 0.800861294 | -4.815938504 |
| **A_55_P2141088** | NA | -0.150069671 | -0.743788564 | 0.443649222 | 10.89940964 | -0.614179405 | 0.561016978 | 0.800861294 | -4.815953109 |
| **A_52_P441044** | Rfxap | 0.049689748 | -0.146916326 | 0.246295823 | 8.195833998 | 0.614119735 | 0.561053947 | 0.800861294 | -4.815971476 |
| **A_66_P103654** | Asxl3 | -0.054437882 | -0.269839866 | 0.160964101 | 7.93177319 | -0.614093802 | 0.561070014 | 0.800861294 | -4.815979458 |
| **A_51_P465148** | Ctsb | 0.141972749 | -0.419816761 | 0.70376226 | 14.78129704 | 0.614065299 | 0.561087674 | 0.800861294 | -4.815988231 |
| **A_51_P204247** | C8a | 0.187749664 | -0.555329117 | 0.930828444 | 10.55519433 | 0.613942215 | 0.561163937 | 0.800911005 | -4.816026109 |
| **A_55_P2025837** | NA | -0.070523005 | -0.349776317 | 0.208730308 | 14.35747782 | -0.613642828 | 0.561349466 | 0.801092969 | -4.816118219 |
| **A_52_P37681** | Fkbp9 | -0.127446228 | -0.632134134 | 0.377241678 | 11.02461686 | -0.613602723 | 0.561374322 | 0.801092969 | -4.816130555 |
| **A_55_P2140516** | Pex13 | 0.0820313 | -0.242924108 | 0.406986708 | 11.09296456 | 0.61339328 | 0.561504139 | 0.801189879 | -4.816194968 |
| **A_52_P555680** | Trappc12 | 0.05427745 | -0.160746873 | 0.269301772 | 11.04625454 | 0.613359411 | 0.561525132 | 0.801189879 | -4.816205382 |
| **A_55_P2022297** | Ptrh2 | 0.045474888 | -0.134730749 | 0.225680525 | 9.525548087 | 0.613177847 | 0.561637687 | 0.801234971 | -4.816261203 |

| | | | | | | | | | |
|---|---|---|---|---|---|---|---|---|---|
| **A_55_P1986743** | NA | -0.069037904 | -0.342644416 | 0.204568609 | 10.58824748 | -0.613118407 | 0.561674537 | 0.801234971 | -4.816279474 |
| **A_55_P2129578** | Pkp3 | -0.060319502 | -0.299377939 | 0.178738934 | 11.27660865 | -0.613107832 | 0.561681094 | 0.801234971 | -4.816282725 |
| **A_55_P1992084** | Ptprg | -0.092286269 | -0.458168582 | 0.273596044 | 8.535390199 | -0.612884816 | 0.56181937 | 0.801349767 | -4.816351265 |
| **A_55_P2018631** | NA | -0.067397681 | -0.334635541 | 0.199840178 | 12.90632574 | -0.612816089 | 0.561861986 | 0.801349767 | -4.816372383 |
| **A_55_P1996553** | Ttc17 | 0.046736588 | -0.138589864 | 0.232063039 | 10.18348622 | 0.612777455 | 0.561885943 | 0.801349767 | -4.816384253 |
| **A_55_P2100008** | NA | -0.10516286 | -0.522513255 | 0.312187534 | 11.00833266 | -0.612272496 | 0.562199131 | 0.801559726 | -4.816539346 |
| **A_55_P2047753** | Tec | -0.077681683 | -0.386028332 | 0.230664966 | 8.14267015 | -0.612156664 | 0.562270987 | 0.801559726 | -4.816574908 |
| **A_55_P1962315** | NA | 0.049855281 | -0.148051162 | 0.247761723 | 16.29355135 | 0.612116972 | 0.562295612 | 0.801559726 | -4.816587092 |
| **A_51_P453043** | Aacs | 0.069539204 | -0.206506189 | 0.345584597 | 11.66956849 | 0.612114122 | 0.562297379 | 0.801559726 | -4.816587967 |
| **A_55_P2076772** | Hspa5 | -0.090078344 | -0.447657758 | 0.26750107 | 15.24580114 | -0.612112252 | 0.56229854 | 0.801559726 | -4.816588541 |
| **A_55_P2157610** | Rpusd3 | 0.09121294 | -0.27093659 | 0.453362469 | 10.80169518 | 0.612000416 | 0.562367925 | 0.801559726 | -4.816622869 |
| **A_55_P2002602** | NA | 0.051035767 | -0.151602422 | 0.253673956 | 15.28848441 | 0.611979028 | 0.562381195 | 0.801559726 | -4.816629433 |
| **A_55_P2149219** | Aar2 | 0.046715959 | -0.138781813 | 0.232213732 | 10.13903471 | 0.611941298 | 0.562404605 | 0.801559726 | -4.816641012 |
| **A_55_P1961466** | Dct | -0.125690215 | -0.62487997 | 0.373499541 | 8.452122909 | -0.611813411 | 0.562483958 | 0.801559726 | -4.816680257 |
| **A_51_P481821** | Spcs3 | 0.079983583 | -0.237701503 | 0.397668669 | 11.20413956 | 0.611768649 | 0.562511734 | 0.801559726 | -4.816693992 |
| **A_55_P2002918** | Klc2 | -0.127369529 | -0.633274781 | 0.378535723 | 9.739642125 | -0.61175784 | 0.562518441 | 0.801559726 | -4.816697308 |
| **A_55_P2127439** | Kri1 | 0.060200426 | -0.178920366 | 0.299321218 | 11.83690295 | 0.611737936 | 0.562530793 | 0.801559726 | -4.816703415 |
| **A_66_P129111** | NA | 0.044385566 | -0.131977706 | 0.220748837 | 7.799649518 | 0.61152864 | 0.562660681 | 0.801685706 | -4.816767621 |
| **A_55_P1960371** | NA | 0.077270295 | -0.229869139 | 0.38440973 | 15.48724495 | 0.611308141 | 0.56279754 | 0.801781699 | -4.816835244 |
| **A_55_P2067518** | Slc13a3 | -0.212489118 | -1.057135884 | 0.632157648 | 12.78927213 | -0.611286435 | 0.562811014 | 0.801781699 | -4.8168419 |
| **A_51_P106397** | Prkd3 | -0.080625655 | -0.401161482 | 0.239910172 | 8.434611931 | -0.611195103 | 0.562867709 | 0.801803372 | -4.811592397 |
| **A_51_P285487** | Rnf113a1 | -0.045003497 | -0.224028005 | 0.134021012 | 8.394969422 | -0.61082522 | 0.563097352 | 0.802071387 | -4.816983277 |
| **A_55_P1966615** | Actr3b | 0.043803647 | -0.130513362 | 0.218120656 | 8.077388882 | 0.610595633 | 0.563239921 | 0.802215343 | -4.81705362 |
| **A_55_P2080264** | NA | 0.040491578 | -0.120710635 | 0.201693792 | 7.862787327 | 0.610347113 | 0.56339427 | 0.802325721 | -4.817129739 |
| **A_55_P2058903** | Sgpp1 | 0.092677614 | -0.276318404 | 0.461673632 | 12.10232803 | 0.610290142 | 0.563429657 | 0.802325721 | -4.817147185 |
| **A_55_P2042953** | Acd | -0.067622467 | -0.336870118 | 0.201625184 | 10.785482 | -0.610270362 | 0.563441944 | 0.802325721 | -4.817153242 |
| **A_51_P501803** | Hoxa2 | -0.067802088 | -0.337815568 | 0.202211393 | 7.93400377 | -0.610155892 | 0.563513051 | 0.802367865 | -4.81718829 |
| **A_55_P2041355** | Gm4750 | -0.125249299 | -0.624274546 | 0.373775948 | 11.83926265 | -0.609868181 | 0.563691796 | 0.802544073 | -4.817276359 |
| **A_55_P2001714** | Blvra | 0.06614941 | -0.197426271 | 0.329725091 | 9.459216003 | 0.609823039 | 0.563719844 | 0.802544073 | -4.817290173 |
| **A_55_P2124273** | Myo1f | -0.10843739 | -0.540590737 | 0.323715958 | 9.406395552 | -0.609711466 | 0.563789173 | 0.802583659 | -4.817324315 |
| **A_51_P220896** | Chmp2b | -0.059559301 | -0.296989332 | 0.17787073 | 9.746624849 | -0.609532874 | 0.563900155 | 0.802682531 | -4.817378953 |
| **A_55_P2146384** | Filip1 | -0.055712667 | -0.277980272 | 0.166554939 | 7.809437092 | -0.609061242 | 0.564193305 | 0.802988435 | -4.81752318 |
| **A_55_P1962966** | NA | -0.079474748 | -0.396546052 | 0.237596555 | 11.63310902 | -0.609053457 | 0.564198144 | 0.802988435 | -4.81752556 |
| **A_55_P2066782** | Fam104a | 0.069344465 | -0.207369029 | 0.346057959 | 12.51538665 | 0.608926191 | 0.564277265 | 0.803041912 | -4.817564462 |
| **A_52_P400857** | Zfp595 | 0.057907973 | -0.173342304 | 0.289158249 | 8.30533186 | 0.608470168 | 0.564560826 | 0.803386306 | -4.817703802 |
| **A_55_P1956464** | Stam | 0.05172428 | -0.154880055 | 0.258328615 | 9.195202114 | 0.608328567 | 0.564648892 | 0.803398303 | -4.817747052 |
| **A_51_P154933** | Zdhhc7 | 0.06743578 | -0.201945657 | 0.336817217 | 9.799525973 | 0.608283323 | 0.564677033 | 0.803398303 | -4.817760869 |
| **A_55_P2091691** | Nudt3 | -0.054616943 | -0.272801539 | 0.163567653 | 12.31032197 | -0.608256126 | 0.564693949 | 0.803398303 | -4.817769174 |
| **A_52_P109319** | E2f5 | 0.052414007 | -0.157095389 | 0.261923403 | 8.303298204 | 0.607892869 | 0.564919921 | 0.803533499 | -4.817880075 |
| **A_55_P2039215** | Acyp2 | 0.099031109 | -0.296857479 | 0.494919697 | 8.380720653 | 0.607829604 | 0.564959282 | 0.803533499 | -4.817899385 |
| **A_55_P2182975** | P2rx2 | -0.055834714 | -0.279048938 | 0.167379511 | 7.870419927 | -0.607806884 | 0.564973418 | 0.803533499 | -4.817906318 |
| **A_55_P2036380** | NA | -0.063768599 | -0.318718223 | 0.191181025 | 17.03969735 | -0.607765004 | 0.564999476 | 0.803533499 | -4.817919099 |
| **A_55_P2056606** | NA | -0.075907199 | -0.379398119 | 0.227583721 | 9.069033573 | -0.60774368 | 0.565012743 | 0.803533499 | -4.817925606 |
| **A_51_P472829** | Aif1l | -0.197113596 | -0.985310459 | 0.591083267 | 8.625256971 | -0.607666182 | 0.565060964 | 0.803533499 | -4.817949255 |
| **A_55_P1973548** | NA | -0.071636484 | -0.358103234 | 0.214830266 | 15.21870526 | -0.607635629 | 0.565079976 | 0.803533499 | -4.817958577 |
| **A_55_P1976694** | 11-Sep | 0.060200672 | -0.180610857 | 0.3010122 | 8.353994583 | 0.607445407 | 0.56519835 | 0.803642703 | -4.818016609 |
| **A_52_P114722** | Ptpn6 | -0.087119025 | -0.435714837 | 0.261476787 | 10.74370375 | -0.607259096 | 0.565314304 | 0.803748451 | -4.818073433 |
| **A_55_P2156293** | Pcyox1 | -0.089570147 | -0.448107447 | 0.268967153 | 12.49656186 | -0.607032764 | 0.565455186 | 0.803859625 | -4.818142445 |
| **A_51_P284426** | Cstad | -0.072751409 | -0.364005765 | 0.218502947 | 7.961000343 | -0.606948956 | 0.565507358 | 0.803859625 | -4.818167993 |
| **A_55_P2062437** | Begain | -0.039623589 | -0.198257565 | 0.119010388 | 8.31766106 | -0.606933046 | 0.565517263 | 0.803859625 | -4.818172843 |

| | | | | | | | | | |
|---|---|---|---|---|---|---|---|---|---|
| **A_55_P2039125** | Fam219a | 0.041916746 | -0.125930354 | 0.209763845 | 7.877674561 | 0.606815837 | 0.565590234 | 0.80387013 | -4.818208569 |
| **A_55_P2067016** | Phykpl | 0.080847714 | -0.242905624 | 0.404601052 | 9.201280118 | 0.606787573 | 0.565607831 | 0.80387013 | -4.818217183 |
| **A_55_P2057686** | Cdc42ep5 | -0.08562184 | -0.428545324 | 0.257301644 | 9.320368551 | -0.606695145 | 0.565665379 | 0.80389281 | -4.81824535 |
| **A_65_P18858** | Btf3 | -0.080882174 | -0.404913022 | 0.243148673 | 12.52041935 | -0.606526318 | 0.565770504 | 0.803983095 | -4.818296789 |
| **A_55_P2112967** | Diaph1 | 0.072779212 | -0.218908148 | 0.364466572 | 8.865674764 | 0.606279562 | 0.565924174 | 0.804075514 | -4.818371952 |
| **A_55_P2032768** | NA | 0.074020979 | -0.222668296 | 0.370710254 | 14.17191527 | 0.606228243 | 0.565956138 | 0.804075514 | -4.818387581 |
| **A_55_P2161829** | Hmgb1-rs17 | -0.076225786 | -0.381755718 | 0.229304147 | 13.19599749 | -0.606221499 | 0.565960338 | 0.804075514 | -4.818389634 |
| **A_55_P1979669** | Zfp946 | 0.061476896 | -0.185015102 | 0.307968893 | 8.958341688 | 0.606027445 | 0.56608121 | 0.80418813 | -4.818448721 |
| **A_51_P506417** | Krt14 | 0.045295282 | -0.136366971 | 0.226957534 | 7.834270776 | 0.605858854 | 0.566186235 | 0.804221438 | -4.818500043 |
| **A_55_P2025855** | Hnrnpul1 | 0.08843646 | -0.266250285 | 0.443123204 | 9.685105662 | 0.60585623 | 0.566187869 | 0.804221438 | -4.818500841 |
| **A_51_P386670** | Dse | 0.044036732 | -0.132616538 | 0.220690002 | 8.212650619 | 0.605726526 | 0.566268677 | 0.804255656 | -4.818540317 |
| **A_51_P126337** | Fgf12 | -0.047133959 | -0.236225477 | 0.141957558 | 7.699409433 | -0.605682555 | 0.566296073 | 0.804255656 | -4.812145832 |
| **A_55_P1966958** | Mef2d | 0.046350506 | -0.139618023 | 0.232319035 | 7.962151779 | 0.605617214 | 0.566336786 | 0.804255656 | -4.81857358 |
| **A_55_P2159869** | Zfp59 | -0.048835598 | -0.244807129 | 0.147135932 | 8.660458207 | -0.605517526 | 0.566398902 | 0.804284777 | -4.818603911 |
| **A_51_P480499** | Emc8 | 0.111548126 | -0.336146821 | 0.559243074 | 10.31995758 | 0.605429079 | 0.566454018 | 0.804303954 | -4.818630818 |
| **A_52_P641367** | Desi1 | 0.075966294 | -0.228994212 | 0.380926801 | 9.073041055 | 0.605285862 | 0.566543271 | 0.804337919 | -4.818674381 |
| **A_55_P2099890** | Atxn7 | 0.158328846 | -0.477299575 | 0.793957267 | 14.03398099 | 0.605257149 | 0.566561165 | 0.804337919 | -4.818683113 |
| **A_51_P376711** | Gm15498 | 0.040736635 | -0.122838901 | 0.204312171 | 7.694704866 | 0.605131818 | 0.56663928 | 0.804389735 | -4.818721226 |
| **A_51_P368543** | Abhd8 | -0.105393987 | -0.52866598 | 0.317878007 | 8.610997278 | -0.605033593 | 0.566700504 | 0.80441757 | -4.818751092 |
| **A_51_P443508** | Ppa1 | 0.141391373 | -0.426655313 | 0.70943806 | 12.15512696 | 0.604814328 | 0.566837189 | 0.804515045 | -4.818817746 |
| **A_55_P2144850** | Psmb5 | -0.050562092 | -0.253724386 | 0.152600201 | 15.27032091 | -0.604735019 | 0.566886633 | 0.804515045 | -4.81884185 |
| **A_66_P111430** | 2410006H16Rik | -0.173555054 | -0.870926029 | 0.523815921 | 12.72632859 | -0.604723138 | 0.566894041 | 0.804515045 | -4.818845461 |
| **A_55_P2064962** | Pdpn | -0.062185757 | -0.31211386 | 0.187742347 | 7.736688524 | -0.604587299 | 0.566978735 | 0.804576166 | -4.81888674 |
| **A_51_P126626** | Zfp503 | -0.235301735 | -1.181145897 | 0.710542427 | 8.951113369 | -0.604489458 | 0.567039742 | 0.804577518 | -4.802761556 |
| **A_51_P471520** | Stk25 | -0.072921523 | -0.366062759 | 0.220219713 | 11.59104744 | -0.604452259 | 0.567062938 | 0.804577518 | -4.818927767 |
| **A_55_P2116924** | Dbi | -0.107877726 | -0.541603225 | 0.325847774 | 16.30636658 | -0.604365987 | 0.567116736 | 0.804594788 | -4.818953975 |
| **A_55_P2166995** | NA | 0.048108275 | -0.145382592 | 0.241599142 | 15.61249081 | 0.604146835 | 0.567253411 | 0.804729628 | -4.819020534 |
| **A_51_P270206** | Cep83 | 0.044466435 | -0.134451569 | 0.223384438 | 10.31602939 | 0.60389503 | 0.567410474 | 0.804816108 | -4.819096985 |
| **A_55_P2116496** | Tuba1b | 0.050471135 | -0.152625273 | 0.253567543 | 14.24607153 | 0.603842977 | 0.567442945 | 0.804816108 | -4.819112785 |
| **A_52_P69194** | Rimkla | -0.04108535 | -0.206413507 | 0.124242807 | 7.665472502 | -0.603842049 | 0.567443525 | 0.804816108 | -4.819113067 |
| **A_52_P574668** | Nt5e | 0.162134948 | -0.490406606 | 0.814676501 | 10.68182006 | 0.603742366 | 0.567505711 | 0.804816108 | -4.819143323 |
| **A_55_P2214841** | Irs3 | 0.041499792 | -0.125531131 | 0.208530715 | 7.804602432 | 0.60371536 | 0.56752256 | 0.804816108 | -4.819151519 |
| **A_55_P1955891** | NA | 0.140702311 | -0.425738828 | 0.70714345 | 13.73213961 | 0.603572762 | 0.567611527 | 0.804830063 | -4.819194791 |
| **A_55_P1960655** | NA | 0.163987526 | -0.496202798 | 0.82417785 | 12.05134933 | 0.603566111 | 0.567615677 | 0.804830063 | -4.819196809 |
| **A_55_P2052936** | NA | -0.066681309 | -0.335221775 | 0.201859156 | 16.09298555 | -0.603361467 | 0.567743371 | 0.804911553 | -4.819258894 |
| **A_55_P1955437** | Cmtm5 | 0.055438291 | -0.167846406 | 0.278722988 | 8.092487417 | 0.603301029 | 0.567781086 | 0.804911553 | -4.819277227 |
| **A_52_P117352** | Gja4 | -0.273238877 | -1.373848313 | 0.827370559 | 9.014010797 | -0.603243219 | 0.567817163 | 0.804911553 | -4.819294761 |
| **A_55_P2016959** | Nme2 | -0.067054267 | -0.337177515 | 0.20306898 | 16.01210854 | -0.603180994 | 0.567855997 | 0.804911553 | -4.819313632 |
| **A_55_P2105858** | Atf5 | -0.178768669 | -0.898973799 | 0.541436462 | 15.46612364 | -0.603140351 | 0.567881362 | 0.804911553 | -4.819325957 |
| **A_55_P2133047** | Tollip | 0.072994388 | -0.221207529 | 0.367196305 | 8.559107645 | 0.602874846 | 0.568047082 | 0.80505681 | -4.819406456 |
| **A_52_P99848** | Pik3cd | 0.105730199 | -0.32046397 | 0.531924367 | 9.77009449 | 0.602802068 | 0.568092513 | 0.80505681 | -4.819428516 |
| **A_52_P185485** | Arpc4 | -0.091145036 | -0.458623365 | 0.276333294 | 10.38278848 | -0.602676801 | 0.568170715 | 0.80505681 | -4.819466482 |
| **A_55_P1993640** | Gm6994 | 0.156542984 | -0.474632137 | 0.787718104 | 14.1646517 | 0.602652448 | 0.568185918 | 0.80505681 | -4.819473863 |
| **A_55_P2051859** | Golgb1 | 0.057471056 | -0.174253774 | 0.289195887 | 10.36633448 | 0.602642554 | 0.568192095 | 0.80505681 | -4.819476861 |
| **A_66_P140408** | D8Ertd738e | -0.084163685 | -0.423640266 | 0.255312895 | 10.30817734 | -0.602418242 | 0.568332148 | 0.805196224 | -4.819544826 |
| **A_55_P2089362** | Wfdc13 | 0.078306802 | -0.237621466 | 0.39423507 | 8.626977137 | 0.602274102 | 0.568422155 | 0.805207242 | -4.819588488 |
| **A_52_P587738** | P2ry2 | -0.102321996 | -0.515242957 | 0.310598965 | 9.057295556 | -0.602123064 | 0.568516478 | 0.805207242 | -4.81963423 |
| **A_55_P2185543** | Tle2 | 0.099478236 | -0.301977134 | 0.500933606 | 10.46800166 | 0.602107437 | 0.568526238 | 0.805207242 | -4.819638962 |
| **A_52_P375873** | Srp9 | 0.052683447 | -0.159957523 | 0.265324417 | 11.9046135 | 0.602019316 | 0.568581275 | 0.805207242 | -4.819665645 |
| **A_51_P272586** | Dym | 0.059504148 | -0.180670063 | 0.299678358 | 9.781847875 | 0.602010484 | 0.568586791 | 0.805207242 | -4.819668319 |

| | | | | | | | | | |
|---|---|---|---|---|---|---|---|---|---|
| **A_55_P1979178** | NA | -0.051744079 | -0.260598346 | 0.157110189 | 13.49213102 | -0.60200555 | 0.568589873 | 0.805207242 | -4.819669813 |
| **A_55_P2115990** | 2010001K21Rik | 0.063819362 | -0.193947291 | 0.321586015 | 8.352420393 | 0.601601505 | 0.568842266 | 0.805436841 | -4.819792113 |
| **A_51_P284793** | Ints12 | -0.062510317 | -0.314991247 | 0.189970614 | 8.774553477 | -0.601597894 | 0.568844521 | 0.805436841 | -4.819793206 |
| **A_55_P2037930** | NA | 0.063099049 | -0.191781818 | 0.317979916 | 8.125377948 | 0.601545891 | 0.568877011 | 0.805436841 | -4.819808942 |
| **A_52_P389230** | Smarcc2 | 0.050221948 | -0.152682938 | 0.253126835 | 8.289281292 | 0.601428827 | 0.568950152 | 0.805481395 | -4.81984436 |
| **A_51_P221428** | Cbx6 | -0.234508271 | -1.182155582 | 0.713139041 | 9.986145521 | -0.601304727 | 0.569027696 | 0.805532176 | -4.819881902 |
| **A_52_P321627** | NA | 0.084160347 | -0.256066822 | 0.424387516 | 14.60874655 | 0.601065381 | 0.569177268 | 0.805614264 | -4.819954287 |
| **A_55_P2060111** | NA | -0.130279067 | -0.657073222 | 0.396515089 | 14.00223926 | -0.600920306 | 0.56926794 | 0.805614264 | -4.819998151 |
| **A_55_P2452384** | Mga | -0.045008547 | -0.227029701 | 0.137012607 | 8.674398113 | -0.600836515 | 0.569320313 | 0.805614264 | -4.820023481 |
| **A_55_P2183433** | Rab30 | 0.093819049 | -0.285603393 | 0.473241491 | 8.5174158 | 0.600829432 | 0.56932474 | 0.805614264 | -4.820025622 |
| **A_55_P2006422** | Gm6900 | 0.066378926 | -0.20207698 | 0.334834833 | 15.76113504 | 0.600814562 | 0.569334035 | 0.805614264 | -4.820030117 |
| **A_51_P141818** | Mrpl32 | 0.046834785 | -0.142579731 | 0.2362493 | 11.56054766 | 0.600811807 | 0.569335757 | 0.805614264 | -4.82003095 |
| **A_55_P2103626** | Slc27a4 | -0.049996092 | -0.25224052 | 0.152248336 | 8.20613937 | -0.600679321 | 0.569418575 | 0.805672471 | -4.820070994 |
| **A_55_P2131173** | Sphk2 | 0.077928352 | -0.237376 | 0.393232703 | 10.97819538 | 0.600549371 | 0.569499813 | 0.805728436 | -4.820110263 |
| **A_55_P2116111** | Prag1 | -0.18776858 | -0.947898016 | 0.572360857 | 11.27618376 | -0.60023119 | 0.569698756 | 0.805915603 | -4.820206386 |
| **A_55_P2070005** | NA | -0.112566158 | -0.568315015 | 0.343182699 | 12.87172803 | -0.600157784 | 0.569744659 | 0.805915603 | -4.820228556 |
| **A_55_P2093439** | Mex3d | -0.099156777 | -0.500655286 | 0.302341733 | 9.442411789 | -0.600097273 | 0.5697825 | 0.805915603 | -4.820246829 |
| **A_51_P318381** | Pgf | 0.039980623 | -0.121913116 | 0.201874363 | 8.001215838 | 0.600071074 | 0.569798884 | 0.805915603 | -4.820254741 |
| **A_51_P366227** | Zfp799 | -0.067903614 | -0.342969942 | 0.207162715 | 8.074873633 | -0.59984445 | 0.56994062 | 0.806020504 | -4.820323163 |
| **A_55_P2091308** | Rpl38 | 0.042790666 | -0.130554573 | 0.216135905 | 16.8491802 | 0.59981914 | 0.569956451 | 0.806020504 | -4.820330803 |
| **A_55_P1990495** | Stat5a | 0.099962003 | -0.305074369 | 0.504998376 | 9.744906024 | 0.599686284 | 0.570039554 | 0.806028709 | -4.820370903 |
| **A_52_P491244** | Zfp287 | -0.055262338 | -0.279204796 | 0.16868012 | 8.001370125 | -0.599619848 | 0.570081113 | 0.806028709 | -4.820390953 |
| **A_55_P2038484** | 4933427D06Rik | 0.164403648 | -0.50182893 | 0.830636225 | 13.71904674 | 0.59960987 | 0.570087355 | 0.806028709 | -4.820393964 |
| **A_55_P2103856** | Hgs | -0.055002606 | -0.277945314 | 0.167940102 | 11.91724239 | -0.599477905 | 0.570169912 | 0.806086471 | -4.820433784 |
| **A_52_P302433** | Plau | -0.039569763 | -0.199978381 | 0.120838855 | 7.806048926 | -0.599403037 | 0.570216752 | 0.806093733 | -4.820456372 |
| **A_55_P1958220** | Crebl2 | -0.052522038 | -0.265490319 | 0.160446242 | 8.049258567 | -0.599252476 | 0.570310956 | 0.806155118 | -4.820501789 |
| **A_52_P427240** | Ltn1 | 0.051899652 | -0.158587148 | 0.262386452 | 8.303513815 | 0.599132352 | 0.570386123 | 0.806155118 | -4.820538019 |
| **A_66_P131389** | Ugt2b38 | 0.24335483 | -0.743926991 | 1.230636651 | 11.63045404 | 0.598938208 | 0.570507619 | 0.806155118 | -4.82059656 |
| **A_55_P1991770** | Pdlim4 | -0.051210363 | -0.258977766 | 0.15655704 | 9.74300555 | -0.598912852 | 0.570523489 | 0.806155118 | -4.820604204 |
| **A_51_P173906** | Mto1 | 0.075431568 | -0.230629478 | 0.381492614 | 8.659105475 | 0.598864085 | 0.57055401 | 0.806155118 | -4.820618906 |
| **A_52_P576720** | Calm4 | -0.039768789 | -0.201137166 | 0.121599588 | 7.695624992 | -0.598834929 | 0.570572258 | 0.806155118 | -4.820627695 |
| **A_55_P2184106** | Gpn2 | 0.053483602 | -0.163540245 | 0.270507449 | 8.67328653 | 0.598820113 | 0.570581532 | 0.806155118 | -4.820632161 |
| **A_51_P130427** | Psmd3 | 0.055481527 | -0.169657496 | 0.280620551 | 14.06459291 | 0.598798657 | 0.570594961 | 0.806155118 | -4.820638629 |
| **A_55_P2112922** | Impdh2 | 0.049904916 | -0.152695993 | 0.252505825 | 13.14072273 | 0.59852891 | 0.570763812 | 0.806155118 | -4.820719925 |
| **A_55_P1972634** | Cmc4 | 0.07206188 | -0.220520317 | 0.364644077 | 9.619165211 | 0.598467923 | 0.570801991 | 0.806155118 | -4.820738301 |
| **A_55_P2085781** | Smim26 | -0.049892252 | -0.252495866 | 0.152711362 | 11.7083044 | -0.598369026 | 0.570863906 | 0.806155118 | -4.820768096 |
| **A_55_P2152292** | Map2k2 | 0.09615884 | -0.294409018 | 0.486726698 | 9.956235737 | 0.598240628 | 0.570944297 | 0.806155118 | -4.820806773 |
| **A_55_P2060183** | Lamtor3 | -0.100011103 | -0.506238441 | 0.306216234 | 12.12595459 | -0.598221835 | 0.570956064 | 0.806155118 | -4.820812433 |
| **A_52_P546660** | 1700029I15Rik | -0.067532072 | -0.341836802 | 0.206772659 | 8.447073525 | -0.598218674 | 0.570958043 | 0.806155118 | -4.820813385 |
| **A_55_P2186672** | Rps14 | 0.060143144 | -0.184170883 | 0.304457171 | 17.55324216 | 0.598164888 | 0.570991722 | 0.806155118 | -4.820829584 |
| **A_55_P2171993** | Mrgbp | 0.04126965 | -0.126402545 | 0.208941845 | 8.242512122 | 0.59807125 | 0.571050357 | 0.806155118 | -4.820857784 |
| **A_55_P2228327** | 4921529L05Rik | 0.061104951 | -0.187209213 | 0.309419116 | 8.728045039 | 0.597940677 | 0.571132125 | 0.806155118 | -4.8208971 |
| **A_55_P2092958** | Traf2 | 0.041956705 | -0.128550233 | 0.212463643 | 8.036273711 | 0.597919213 | 0.571145567 | 0.806155118 | -4.820903562 |
| **A_55_P2014611** | Ccdc12 | -0.07605107 | -0.385122194 | 0.233020055 | 11.38603765 | -0.597902105 | 0.571156282 | 0.806155118 | -4.820908712 |
| **A_55_P2326517** | Ube2i | 0.072058994 | -0.220792989 | 0.364910976 | 9.481642843 | 0.597892648 | 0.571162204 | 0.806155118 | -4.820911559 |
| **A_51_P451075** | Atp2a2 | -0.142006406 | -0.719142867 | 0.435130055 | 9.017695324 | -0.597878056 | 0.571171343 | 0.806155118 | -4.820915952 |
| **A_66_P107973** | Prpf38b | 0.100326771 | -0.307479264 | 0.508132807 | 8.626188845 | 0.597786876 | 0.571228449 | 0.806155118 | -4.8209434 |
| **A_55_P2187121** | Gpr137b-ps | -0.087773513 | -0.444572537 | 0.26902551 | 9.049417487 | -0.597754719 | 0.57124859 | 0.806155118 | -4.82095308 |
| **A_51_P450100** | Rab5if | -0.067150518 | -0.340154193 | 0.205853157 | 13.26236738 | -0.59767359 | 0.571299405 | 0.806155118 | -4.820977498 |
| **A_55_P1990879** | Banp | 0.076577791 | -0.234784126 | 0.387939708 | 8.425720271 | 0.597613683 | 0.571336929 | 0.806155118 | -4.820995527 |

| | | | | | | | | | |
|---|---|---|---|---|---|---|---|---|---|
| **A_51_P250445** | Zfp276 | -0.053996547 | -0.273548595 | 0.165555502 | 9.230775579 | -0.597601504 | 0.571344558 | 0.806155118 | -4.820999192 |
| **A_55_P2013760** | Slc39a14 | -0.185934564 | -0.942259319 | 0.570390191 | 11.44630002 | -0.59735844 | 0.571496825 | 0.806311104 | -4.821072326 |
| **A_66_P113505** | Fgd4 | 0.055971509 | -0.171752368 | 0.283695385 | 9.902748933 | 0.597230025 | 0.571577279 | 0.806338061 | -4.821110953 |
| **A_52_P239023** | Zfp955b | -0.044798022 | -0.227072823 | 0.137476779 | 8.058124167 | -0.597193946 | 0.571599885 | 0.806338061 | -4.821121805 |
| **A_55_P2052897** | Arhgap30 | 0.082194587 | -0.25227669 | 0.416665865 | 8.946048755 | 0.597128198 | 0.571641081 | 0.806338061 | -4.821141578 |
| **A_52_P451073** | Tnfrsf21 | 0.072278337 | -0.221916628 | 0.366473301 | 8.773573119 | 0.596974946 | 0.571737112 | 0.80641467 | -4.821187662 |
| **A_52_P474765** | Denr | 0.046951008 | -0.144178485 | 0.238080502 | 10.11975441 | 0.596898382 | 0.571785093 | 0.806423499 | -4.821210681 |
| **A_51_P302856** | Ugt2b37 | 0.185657263 | -0.570443204 | 0.94175773 | 10.48446847 | 0.596644479 | 0.571944223 | 0.806479271 | -4.821287001 |
| **A_55_P2124016** | Nipsnap1 | 0.150216056 | -0.461553174 | 0.761985285 | 12.43144833 | 0.596639379 | 0.57194742 | 0.806479271 | -4.821288533 |
| **A_55_P2009091** | Mad1l1 | 0.083554739 | -0.256732085 | 0.423841562 | 10.10905351 | 0.596635569 | 0.571949808 | 0.806479271 | -4.821289678 |
| **A_55_P1984886** | Hcst | -0.113968485 | -0.578262416 | 0.350325446 | 10.26717802 | -0.596451255 | 0.572065342 | 0.806577215 | -4.821345063 |
| **A_55_P1989544** | Cklf | -0.065532476 | -0.332530615 | 0.201465663 | 9.493010733 | -0.596391617 | 0.572102728 | 0.806577215 | -4.82136298 |
| **A_52_P376360** | Pdgfc | -0.063022775 | -0.319846779 | 0.193801228 | 8.129416785 | -0.596272941 | 0.572177128 | 0.806623273 | -4.821398631 |
| **A_52_P80444** | Dpm2 | -0.061509761 | -0.312218116 | 0.189198593 | 11.38345567 | -0.596153949 | 0.572251732 | 0.806669612 | -4.82143437 |
| **A_55_P2094626** | Ralgps1 | 0.191108211 | -0.588023833 | 0.970240255 | 12.8121217 | 0.596007151 | 0.572343777 | 0.806740529 | -4.821478453 |
| **A_51_P507622** | Itpkc | -0.050316339 | -0.255489921 | 0.154857244 | 8.750642714 | -0.595896424 | 0.572413212 | 0.806747307 | -4.821511698 |
| **A_55_P1955483** | Grb14 | 0.068527351 | -0.210918884 | 0.347973586 | 9.921211639 | 0.595866365 | 0.572432061 | 0.806747307 | -4.821520722 |
| **A_52_P89267** | Castor2 | 0.081755965 | -0.25174334 | 0.41525527 | 7.930042093 | 0.59567271 | 0.572553512 | 0.806767794 | -4.821578851 |
| **A_51_P468073** | Ggt1 | -0.200395348 | -1.017893348 | 0.617102651 | 9.549348313 | -0.595640372 | 0.572573794 | 0.806767794 | -4.821588556 |
| **A_51_P342481** | Gpd2 | 0.038677268 | -0.119109181 | 0.196463718 | 8.564811179 | 0.595620009 | 0.572586566 | 0.806767794 | -4.821594667 |
| **A_52_P137371** | Hmgcr | 0.117941009 | -0.363259075 | 0.599141093 | 10.24343548 | 0.595555629 | 0.572626947 | 0.806767794 | -4.821613987 |
| **A_52_P102248** | Mex3b | -0.128708655 | -0.653880654 | 0.396463344 | 8.789245037 | -0.595510439 | 0.572655292 | 0.806767794 | -4.821627547 |
| **A_51_P261059** | Cpne2 | -0.056047549 | -0.284857986 | 0.172762889 | 8.508900332 | -0.595201449 | 0.572849127 | 0.806851684 | -4.821720241 |
| **A_55_P2034175** | Usp28 | 0.059293488 | -0.182769631 | 0.301356607 | 10.04572744 | 0.595198141 | 0.572851203 | 0.806851684 | -4.821721233 |
| **A_55_P2004469** | NA | -0.082208665 | -0.417843137 | 0.253425808 | 15.87810021 | -0.595160668 | 0.572874713 | 0.806851684 | -4.821732472 |
| **A_55_P2093173** | NA | 0.08515421 | -0.262561667 | 0.432870086 | 11.3705852 | 0.595065526 | 0.572934406 | 0.806851684 | -4.821761004 |
| **A_52_P282987** | Yeats2 | -0.095300607 | -0.484467299 | 0.293866086 | 9.469687754 | -0.595035932 | 0.572952975 | 0.806851684 | -4.821769878 |
| **A_51_P248857** | Inka1 | 0.061272611 | -0.188995598 | 0.31154082 | 8.51827599 | 0.594899892 | 0.573038337 | 0.806851684 | -4.821810667 |
| **A_51_P313581** | Fabp2 | -0.5867024 | -2.983199448 | 1.809794648 | 10.57750292 | -0.59487286 | 0.5730553 | 0.806851684 | -4.821818771 |
| **A_55_P2245422** | NA | 0.054046897 | -0.166745927 | 0.274839721 | 8.830220181 | 0.594797314 | 0.573102707 | 0.806851684 | -4.821841418 |
| **A_55_P2074035** | Haao | -0.148916411 | -0.757288152 | 0.459455331 | 12.63384647 | -0.594780492 | 0.573113264 | 0.806851684 | -4.82184646 |
| **A_55_P1978576** | Dtx2 | -0.097859351 | -0.497711217 | 0.301992516 | 10.57188906 | -0.594684187 | 0.573173702 | 0.806851684 | -4.821875326 |
| **A_52_P229044** | Slc20a2 | 0.079821683 | -0.246328809 | 0.405972175 | 8.485029124 | 0.594683694 | 0.573174012 | 0.806851684 | -4.821875473 |
| **A_51_P125205** | Aqp1 | -0.105564049 | -0.536950633 | 0.325822535 | 13.36445693 | -0.594610526 | 0.573219933 | 0.806857565 | -4.821897402 |
| **A_51_P104891** | Selenoi | 0.110323597 | -0.340607726 | 0.56125492 | 9.370428306 | 0.594485406 | 0.573298464 | 0.806876443 | -4.821934894 |
| **A_55_P1974204** | Fam172a | -0.051208042 | -0.260523612 | 0.158107529 | 10.79122885 | -0.594456142 | 0.573316833 | 0.806876443 | -4.821943662 |
| **A_55_P1998651** | Entpd5 | 0.166560487 | -0.514397708 | 0.847518681 | 10.98344585 | 0.59433967 | 0.573389944 | 0.806920584 | -4.821978556 |
| **A_55_P2032727** | NA | -0.113424136 | -0.577350146 | 0.350501874 | 15.27644078 | -0.594073181 | 0.573557243 | 0.807097259 | -4.822058372 |
| **A_52_P464129** | Gon7 | 0.061418375 | -0.18991245 | 0.312749199 | 9.542333904 | 0.593793929 | 0.573732586 | 0.807285226 | -4.822141979 |
| **A_55_P2156101** | Skp1a | -0.056282129 | -0.286664198 | 0.174099939 | 13.41797503 | -0.593615233 | 0.573844806 | 0.807384355 | -4.822195462 |
| **A_51_P511386** | Zfp267 | 0.045006436 | -0.139276622 | 0.229289493 | 8.086038484 | 0.593433964 | 0.573958656 | 0.807437232 | -4.822249702 |
| **A_52_P207964** | Mrps18b | 0.080398424 | -0.248806868 | 0.409603717 | 9.194252019 | 0.593422374 | 0.573965936 | 0.807437232 | -4.822253169 |
| **A_55_P2068892** | Il6ra | -0.141464094 | -0.721048741 | 0.438120552 | 11.52123043 | -0.593078992 | 0.574181642 | 0.807495995 | -4.822355878 |
| **A_55_P2103206** | NA | 0.059889706 | -0.185502142 | 0.305281555 | 8.113812463 | 0.593028055 | 0.574213644 | 0.807495995 | -4.822371109 |
| **A_55_P2012166** | Spata6 | -0.146496407 | -0.746776541 | 0.453783727 | 9.093745632 | -0.593002045 | 0.574229985 | 0.807495995 | -4.822378886 |
| **A_55_P1997747** | Cox16 | 0.042377218 | -0.131267902 | 0.216022337 | 12.24410297 | 0.592997753 | 0.574232682 | 0.807495995 | -4.822380169 |
| **A_55_P1954555** | NA | -0.047552525 | -0.242424955 | 0.147319905 | 7.723985728 | -0.59293402 | 0.574272725 | 0.807495995 | -4.822399225 |
| **A_55_P2033616** | Dhrs7b | 0.071355804 | -0.221083724 | 0.363795332 | 11.14467268 | 0.592893131 | 0.574298416 | 0.807495995 | -4.822411449 |
| **A_55_P2124090** | Ambra1 | -0.060087063 | -0.306344731 | 0.186170604 | 10.35947423 | -0.592890385 | 0.574300142 | 0.807495995 | -4.82241227 |
| **A_55_P1954302** | Esrrb | 0.046885176 | -0.145339603 | 0.239109955 | 8.150977939 | 0.592665119 | 0.574441693 | 0.807575112 | -4.822479604 |

| | | | | | | | | | |
|---|---|---|---|---|---|---|---|---|---|
| **A_51_P278540** | Myg1 | 0.058507476 | -0.181406384 | 0.298421335 | 9.996826594 | 0.592569387 | 0.574501855 | 0.807575112 | -4.822508213 |
| **A_55_P1970755** | Hdac9 | 0.070043096 | -0.217209945 | 0.357296138 | 8.563177488 | 0.592493909 | 0.574549292 | 0.807575112 | -4.822530766 |
| **A_55_P2081585** | Dennd5b | 0.133576033 | -0.414270332 | 0.681422397 | 10.15869199 | 0.592451694 | 0.574575824 | 0.807575112 | -4.822543379 |
| **A_52_P429450** | Ngp | -0.177024294 | -0.903140049 | 0.54909146 | 12.68408464 | -0.592393387 | 0.574612471 | 0.807575112 | -4.822560799 |
| **A_55_P2183110** | Fmr1 | 0.051344148 | -0.159266243 | 0.261954538 | 8.46409861 | 0.592371758 | 0.574626066 | 0.807575112 | -4.82256726 |
| **A_65_P15215** | Mrpl15 | 0.047346967 | -0.14691326 | 0.241607193 | 11.5162749 | 0.592231441 | 0.574714264 | 0.807575112 | -4.822609174 |
| **A_51_P130757** | Acer3 | 0.058685519 | -0.182113577 | 0.299484615 | 9.007054269 | 0.592187573 | 0.574741841 | 0.807575112 | -4.822622276 |
| **A_55_P2052290** | Psat1 | 0.134632653 | -0.417812136 | 0.687077442 | 14.76216037 | 0.592167697 | 0.574754335 | 0.807575112 | -4.822628212 |
| **A_55_P2105457** | Zfp7 | -0.038167351 | -0.194789685 | 0.118454983 | 8.160822797 | -0.592136072 | 0.574774216 | 0.807575112 | -4.822637657 |
| **A_55_P2038207** | Zfp687 | 0.049496447 | -0.153718565 | 0.252711459 | 9.79696712 | 0.591836071 | 0.574962826 | 0.807750246 | -4.82272723 |
| **A_51_P332652** | Pqlc3 | 0.110663586 | -0.343706144 | 0.565033316 | 8.919501593 | 0.591804872 | 0.574982443 | 0.807750246 | -4.822736543 |
| **A_55_P2389528** | D130061D10Rik | -0.047836613 | -0.244281226 | 0.148607999 | 7.985384378 | -0.59170262 | 0.575046739 | 0.807767862 | -4.822767063 |
| **A_55_P2150976** | NA | 0.056120661 | -0.174362723 | 0.286604045 | 17.09693382 | 0.59165201 | 0.575078563 | 0.807767862 | -4.822782167 |
| **A_66_P116412** | Flt3 | 0.056218154 | -0.174692462 | 0.287128769 | 8.159773138 | 0.591583249 | 0.575121804 | 0.807769899 | -4.822802687 |
| **A_55_P2405784** | BC023202 | -0.088622275 | -0.452766261 | 0.27552171 | 7.975485515 | -0.591361359 | 0.575261352 | 0.80781937 | -4.82286889 |
| **A_55_P2035593** | Hsf1 | 0.0483504 | -0.150346328 | 0.247047127 | 10.04768023 | 0.591279118 | 0.57531308 | 0.80781937 | -4.822893422 |
| **A_51_P224064** | Klhl38 | -0.041158489 | -0.210323062 | 0.128006084 | 7.758342581 | -0.591198325 | 0.575363899 | 0.80781937 | -4.82291752 |
| **A_55_P1961423** | Gsta3 | -0.184323951 | -0.941936722 | 0.573288821 | 14.51289375 | -0.591177194 | 0.575377191 | 0.80781937 | -4.822923822 |
| **A_51_P143232** | Wdr59 | 0.040862878 | -0.127127981 | 0.208853738 | 8.480606137 | 0.591053089 | 0.57545526 | 0.80781937 | -4.822960831 |
| **A_51_P186547** | Pah | 0.186442206 | -0.5801026 | 0.952987012 | 12.14411418 | 0.591003261 | 0.575486606 | 0.80781937 | -4.822975688 |
| **A_52_P613643** | Zfp3 | 0.044237862 | -0.137647063 | 0.226122787 | 7.912370812 | 0.590990656 | 0.575494536 | 0.80781937 | -4.822979446 |
| **A_55_P2028365** | Lrrc36 | -0.089771298 | -0.458922204 | 0.279379608 | 7.969843626 | -0.590903761 | 0.575549204 | 0.80781937 | -4.823005353 |
| **A_55_P2117425** | Syt5 | -0.086864722 | -0.444091426 | 0.270361982 | 8.326890505 | -0.590857437 | 0.575578349 | 0.80781937 | -4.823019163 |
| **A_55_P2076288** | Zfp661 | 0.04260333 | -0.132606641 | 0.217813301 | 8.574636797 | 0.590837317 | 0.575591008 | 0.80781937 | -4.823025161 |
| **A_55_P2128229** | NA | 0.153499571 | -0.477824742 | 0.784823884 | 10.3450123 | 0.590796401 | 0.575616751 | 0.80781937 | -4.823037357 |
| **A_52_P347591** | Ndufa7 | -0.085079231 | -0.435060144 | 0.264901683 | 14.96483134 | -0.590693759 | 0.575681334 | 0.80785135 | -4.82306795 |
| **A_55_P2008874** | Dnmt3b | -0.068424307 | -0.349943169 | 0.213094555 | 8.416965435 | -0.590590014 | 0.575746615 | 0.807884307 | -4.823098867 |
| **A_51_P136729** | Idh3g | -0.045563293 | -0.233119937 | 0.14199335 | 13.24201644 | -0.590290569 | 0.575935066 | 0.808051119 | -4.823188079 |
| **A_55_P2007423** | Ppp1r12a | 0.066159707 | -0.206199886 | 0.338519299 | 8.705424931 | 0.590247401 | 0.575962235 | 0.808051119 | -4.823200936 |
| **A_55_P2128113** | Tbc1d12 | -0.090066989 | -0.460874288 | 0.280740309 | 7.925345488 | -0.590201843 | 0.575990911 | 0.808051119 | -4.823214505 |
| **A_55_P2133637** | Etfa | -0.137359405 | -0.703034306 | 0.428315496 | 9.848609361 | -0.590030798 | 0.576098578 | 0.808109068 | -4.82326544 |
| **A_51_P220806** | Gdf9 | -0.05659378 | -0.289681491 | 0.176493931 | 8.049147935 | -0.58997351 | 0.576134641 | 0.808109068 | -4.823282497 |
| **A_52_P1055590** | Bnip1 | -0.048437247 | -0.247943735 | 0.151069241 | 9.901333512 | -0.589936972 | 0.576157643 | 0.808109068 | -4.823293375 |
| **A_55_P2066269** | Aak1 | 0.054025123 | -0.168566033 | 0.27661628 | 8.165030129 | 0.589754209 | 0.576272706 | 0.808211807 | -4.823347779 |
| **A_51_P442053** | Ptdss1 | 0.044484314 | -0.138858141 | 0.227826769 | 8.60503826 | 0.589558685 | 0.576395818 | 0.808289006 | -4.823405966 |
| **A_66_P129747** | Dnajc24 | 0.064123598 | -0.200173407 | 0.328420603 | 10.03705796 | 0.589533964 | 0.576411385 | 0.808289006 | -4.823413322 |
| **A_55_P2114898** | Gdap2 | 0.081473345 | -0.254384534 | 0.417331224 | 9.738617377 | 0.589444873 | 0.576467487 | 0.808309035 | -4.823439828 |
| **A_55_P2115235** | Bckdhb | 0.186382817 | -0.582206592 | 0.954972225 | 12.2919972 | 0.589243315 | 0.576594424 | 0.808413905 | -4.823499784 |
| **A_55_P2189571** | NA | 0.070504383 | -0.220260202 | 0.361268968 | 8.89676985 | 0.589193299 | 0.576625926 | 0.808413905 | -4.823514659 |
| **A_55_P1967105** | Adgrb2 | -0.065752298 | -0.336998914 | 0.205494318 | 8.076724118 | -0.589019665 | 0.576735293 | 0.808508593 | -4.823566291 |
| **A_55_P2102182** | NA | 0.103676436 | -0.32410487 | 0.531457742 | 9.047224714 | 0.588899853 | 0.576810768 | 0.808548711 | -4.823601912 |
| **A_55_P2057523** | Ppm1b | 0.088569569 | -0.276916387 | 0.454055524 | 10.10067716 | 0.588839622 | 0.576848712 | 0.808548711 | -4.823619816 |
| **A_52_P399054** | Tmed2 | 0.07106317 | -0.222214083 | 0.364340422 | 8.897099475 | 0.588775032 | 0.576889403 | 0.808548711 | -4.823639014 |
| **A_55_P2013780** | NA | -0.045746724 | -0.234651043 | 0.143157596 | 13.36113381 | -0.588438793 | 0.577101261 | 0.808715736 | -4.823738928 |
| **A_52_P494372** | Zxdc | 0.039750175 | -0.124395054 | 0.203895404 | 8.102956819 | 0.588428861 | 0.577107519 | 0.808715736 | -4.823741879 |
| **A_51_P310254** | Fam118a | -0.045241758 | -0.232077439 | 0.141593923 | 8.403352675 | -0.588386693 | 0.577134092 | 0.808715736 | -4.823754405 |
| **A_55_P2167160** | Hdhd2 | 0.049782496 | -0.155846024 | 0.255411017 | 8.398570199 | 0.588269741 | 0.577207794 | 0.808760381 | -4.823789144 |
| **A_55_P2058613** | NA | -0.049727273 | -0.255163283 | 0.155708736 | 8.112088834 | -0.588167827 | 0.577272024 | 0.808771477 | -4.823819411 |
| **A_66_P120558** | Pam16 | 0.041821796 | -0.130967435 | 0.214611027 | 12.7715891 | 0.588124395 | 0.577299398 | 0.808771477 | -4.823832309 |
| **A_51_P451428** | Marveld2 | 0.111103671 | -0.348005072 | 0.570212414 | 9.229173621 | 0.588025329 | 0.577361839 | 0.808800333 | -4.823861724 |

| | | | | | | | | | |
|---|---|---|---|---|---|---|---|---|---|
| A_52_P352525 | Qser1 | 0.039766743 | -0.124644513 | 0.204178 | 7.950884109 | 0.58772162 | 0.577553291 | 0.808929767 | -4.823951877 |
| A_52_P520397 | Timm8b | -0.06203246 | -0.318597823 | 0.194532903 | 13.96695238 | -0.587494989 | 0.577696179 | 0.808929767 | -4.824019125 |
| A_55_P2059318 | Dgkz | -0.055933274 | -0.287301355 | 0.175434806 | 11.21205629 | -0.587421706 | 0.577742388 | 0.808929767 | -4.824040866 |
| A_55_P2038183 | Insc | 0.105027139 | -0.329470382 | 0.539524659 | 9.523105764 | 0.587350605 | 0.577787222 | 0.808929767 | -4.824061957 |
| A_55_P1996973 | Gvin1 | -0.104645879 | -0.537614617 | 0.328322858 | 9.450661863 | -0.587284829 | 0.577828702 | 0.808929767 | -4.824081467 |
| A_51_P518592 | Atxn7 | 0.05260559 | -0.165074011 | 0.270285192 | 8.950652838 | 0.587215289 | 0.577872556 | 0.808929767 | -4.824102091 |
| A_55_P2173702 | Ebpl | -0.065859986 | -0.348280315 | 0.216560343 | 12.09612261 | -0.59329139 | 0.577950551 | 0.808929767 | -4.782069788 |
| A_51_P163252 | Schip1 | -0.056673025 | -0.291283188 | 0.177937137 | 7.802355988 | -0.586965757 | 0.578029935 | 0.808929767 | -4.824176079 |
| A_55_P1999666 | Cntrl | 0.079898466 | -0.250862171 | 0.410659102 | 9.921614499 | 0.58695906 | 0.578034159 | 0.808929767 | -4.824178065 |
| A_55_P2153887 | Tatdn3 | 0.060086235 | -0.188679256 | 0.308851726 | 9.177166013 | 0.586905319 | 0.578068057 | 0.808929767 | -4.824193996 |
| A_55_P2174816 | Zfp266 | -0.063179911 | -0.324787948 | 0.198428126 | 8.764956091 | -0.58682839 | 0.578116583 | 0.808929767 | -4.824216799 |
| A_65_P03874 | Nav2 | -0.040638777 | -0.208916511 | 0.127638958 | 7.862073866 | -0.58680953 | 0.57812848 | 0.808929767 | -4.824222389 |
| A_55_P1960178 | NA | -0.097076674 | -0.499053736 | 0.304900387 | 17.02786602 | -0.586809055 | 0.57812878 | 0.808929767 | -4.82422253 |
| A_55_P2025523 | Arhgap6 | -0.082976916 | -0.426570004 | 0.260616173 | 8.690206789 | -0.586808042 | 0.578129419 | 0.808929767 | -4.82422283 |
| A_55_P1982201 | Sgsm2 | 0.061285029 | -0.192493087 | 0.315063146 | 9.181695374 | 0.586790964 | 0.578140192 | 0.808929767 | -4.824227892 |
| A_55_P2002212 | Babam2 | 0.065333414 | -0.205223056 | 0.335889884 | 9.788741715 | 0.58676016 | 0.578159624 | 0.808929767 | -4.824237021 |
| A_55_P2012960 | Gpx7 | -0.092673646 | -0.476483182 | 0.291135889 | 8.987979937 | -0.586710256 | 0.578191106 | 0.808929767 | -4.824251811 |
| A_55_P1973159 | Serpinf1 | -0.157710894 | -0.810922698 | 0.495500911 | 14.6139839 | -0.586665916 | 0.578219078 | 0.808929767 | -4.824264951 |
| A_51_P432520 | Wrnip1 | 0.051665769 | -0.162342625 | 0.265674163 | 10.24049776 | 0.586617858 | 0.578249397 | 0.808929767 | -4.824279191 |
| A_55_P2081388 | Mdm4 | 0.063890268 | -0.200888356 | 0.328668892 | 8.577098927 | 0.586320364 | 0.578437101 | 0.809050621 | -4.824367323 |
| A_55_P1962665 | Exoc3l2 | 0.045423726 | -0.142839307 | 0.233686758 | 8.111336587 | 0.586274346 | 0.578466139 | 0.809050621 | -4.824380952 |
| A_55_P1961853 | Zfand3 | 0.066596618 | -0.209439037 | 0.342632273 | 9.123921324 | 0.586232892 | 0.578492298 | 0.809050621 | -4.824393229 |
| A_55_P1997486 | Snu13 | 0.045633932 | -0.143519277 | 0.234787142 | 12.40628617 | 0.586215594 | 0.578503214 | 0.809050621 | -4.824398352 |
| A_55_P2037544 | NA | -0.075808165 | -0.390107202 | 0.238490871 | 15.84452386 | -0.586078953 | 0.578589447 | 0.8090768 | -4.824438813 |
| A_51_P365409 | Rpusd4 | -0.039678935 | -0.204194379 | 0.12483651 | 10.21313199 | -0.586052494 | 0.578606145 | 0.8090768 | -4.824446647 |
| A_52_P451574 | Xpr1 | 0.062286029 | -0.19599106 | 0.320563118 | 10.76118268 | 0.585986957 | 0.578647508 | 0.8090768 | -4.82446605 |
| A_55_P2060415 | NA | 0.041633456 | -0.13104426 | 0.214311172 | 16.33091705 | 0.585853945 | 0.578731462 | 0.809135655 | -4.824505423 |
| A_55_P1957957 | Dhx34 | -0.058159407 | -0.299437109 | 0.183118296 | 9.256512256 | -0.58571449 | 0.57881949 | 0.809173631 | -4.824546696 |
| A_51_P182257 | Catsperz | -0.057900964 | -0.298168941 | 0.182367012 | 7.918560704 | -0.585562281 | 0.578915578 | 0.809173631 | -4.824591734 |
| A_55_P2177953 | Aqp11 | -0.172250221 | -0.887076362 | 0.542575919 | 10.51490438 | -0.585521101 | 0.578941576 | 0.809173631 | -4.824603917 |
| A_55_P2161958 | Gm10324 | 0.061959934 | -0.195225323 | 0.319145191 | 8.659182498 | 0.585393731 | 0.579021993 | 0.809173631 | -4.824641596 |
| A_51_P142164 | Hspa13 | -0.040762185 | -0.209966327 | 0.128441957 | 7.827219853 | -0.585368912 | 0.579037663 | 0.809173631 | -4.824648937 |
| A_55_P2051721 | Psg22 | -0.037287006 | -0.192069388 | 0.117495377 | 7.721076039 | -0.585354784 | 0.579046584 | 0.809173631 | -4.824653116 |
| A_51_P497061 | Ppp2r2d | -0.047390289 | -0.244123465 | 0.149342886 | 11.7473158 | -0.585322136 | 0.579067198 | 0.809173631 | -4.824662772 |
| A_55_P2097736 | Patz1 | -0.043883334 | -0.22607174 | 0.138305071 | 8.255856816 | -0.585277828 | 0.579095175 | 0.809173631 | -4.824675876 |
| A_55_P2155574 | NA | -0.097984305 | -0.504836178 | 0.308867568 | 14.29831511 | -0.585198746 | 0.579145112 | 0.809173631 | -4.824699263 |
| A_55_P1977934 | Itgam | 0.111905233 | -0.3529488 | 0.576759266 | 11.18281129 | 0.584947626 | 0.5793037 | 0.809173631 | -4.824773508 |
| A_55_P2038987 | Gas7 | -0.047166363 | -0.24310834 | 0.148775614 | 8.293063989 | -0.584908717 | 0.579328274 | 0.809173631 | -4.82478501 |
| A_51_P416509 | H1f1 | 0.205220889 | -0.647326728 | 1.057768506 | 11.14847183 | 0.584906043 | 0.579329962 | 0.809173631 | -4.8247858 |
| A_55_P2133524 | Nbr1 | 0.068291731 | -0.215425475 | 0.352008937 | 8.529761019 | 0.584878472 | 0.579347376 | 0.809173631 | -4.82479395 |
| A_51_P429212 | Npl | -0.085239582 | -0.439374707 | 0.268895542 | 8.495408122 | -0.584864819 | 0.579355999 | 0.809173631 | -4.824797985 |
| A_51_P453111 | Hexb | 0.084632097 | -0.267045518 | 0.436309711 | 11.27825951 | 0.584754489 | 0.579425687 | 0.809173631 | -4.824830592 |
| A_55_P2316612 | 6720407P12Rik | -0.066348198 | -0.342051514 | 0.209355118 | 8.151243 | -0.584750138 | 0.579428435 | 0.809173631 | -4.824831878 |
| A_55_P2103952 | NA | -0.040195852 | -0.207365077 | 0.126973374 | 7.6942054 | -0.584262611 | 0.57973643 | 0.809294829 | -4.824975902 |
| A_55_P2048454 | Rps2 | -0.046397844 | -0.239361168 | 0.146565481 | 15.43948855 | -0.584260082 | 0.579738028 | 0.809294829 | -4.824976648 |
| A_51_P162676 | Armc7 | -0.053306355 | -0.275021177 | 0.168408467 | 8.395762348 | -0.584207934 | 0.579770979 | 0.809294829 | -4.824992048 |
| A_51_P251736 | Reep5 | -0.045996025 | -0.237307179 | 0.145315129 | 8.884893266 | -0.584202213 | 0.579774593 | 0.809294829 | -4.824993737 |
| A_51_P281593 | Plpp5 | 0.106020796 | -0.33496443 | 0.547006023 | 11.13976924 | 0.584184743 | 0.579785633 | 0.809294829 | -4.824998896 |
| A_55_P2011410 | Pdzk1 | 0.157591965 | -0.497931234 | 0.813115164 | 12.54882003 | 0.584156474 | 0.579803495 | 0.809294829 | -4.825007243 |
| A_51_P275123 | Psd3 | 0.081883212 | -0.258724541 | 0.422490965 | 10.33450378 | 0.584148857 | 0.579808308 | 0.809294829 | -4.825009492 |

| | | | | | | | | | |
|---|---|---|---|---|---|---|---|---|---|
| **A_51_P241861** | Tcirg1 | -0.07598325 | -0.392129007 | 0.240162507 | 9.543970409 | -0.584001145 | 0.579901652 | 0.80932742 | -4.825053102 |
| **A_55_P2038594** | Tmem234 | 0.104358203 | -0.32987703 | 0.538593435 | 12.21325631 | 0.583962181 | 0.579926275 | 0.80932742 | -4.825064604 |
| **A_66_P101519** | Abcc9 | 0.099216219 | -0.313684757 | 0.512117194 | 9.976757371 | 0.583875095 | 0.579981312 | 0.80932742 | -4.825090308 |
| **A_51_P184849** | Pfkfb2 | 0.05034474 | -0.159181454 | 0.259870933 | 8.050478643 | 0.583846884 | 0.579999142 | 0.80932742 | -4.825098635 |
| **A_55_P1976395** | Nsdhl | 0.10993405 | -0.347777008 | 0.567645109 | 8.736240472 | 0.583611731 | 0.580147774 | 0.809476383 | -4.825168025 |
| **A_52_P521507** | Trio | -0.083320159 | -0.430284431 | 0.263644112 | 10.17366936 | -0.583510297 | 0.580211894 | 0.809507413 | -4.82519795 |
| **A_55_P1984841** | NA | -0.097020476 | -0.501101303 | 0.307060352 | 10.51509858 | -0.583416011 | 0.5802715 | 0.809532142 | -4.825225762 |
| **A_51_P169032** | 2810004N23Rik | 0.054744489 | -0.173402506 | 0.282891484 | 12.23411471 | 0.583054084 | 0.580500334 | 0.809733386 | -4.825332486 |
| **A_55_P1989797** | NA | -0.07548446 | -0.390101343 | 0.239132423 | 14.85563901 | -0.582986795 | 0.580542885 | 0.809733386 | -4.825352322 |
| **A_55_P2035087** | Arhgef15 | 0.166673848 | -0.528046244 | 0.86139394 | 12.62007038 | 0.582962733 | 0.580558101 | 0.809733386 | -4.825359415 |
| **A_51_P446106** | Lrrc38 | 0.044602139 | -0.141318582 | 0.230522861 | 8.035415693 | 0.582922853 | 0.58058332 | 0.809733386 | -4.825371169 |
| **A_55_P2035590** | NA | -0.113868204 | -0.588674946 | 0.360938538 | 14.0921166 | -0.582731887 | 0.580704093 | 0.809843393 | -4.825427448 |
| **A_52_P558939** | Rab3gap1 | 0.062712417 | -0.198948878 | 0.324373712 | 8.622580188 | 0.582367646 | 0.580934492 | 0.810101637 | -4.825534748 |
| **A_51_P129149** | Gatad2a | 0.0520607 | -0.165182264 | 0.269303664 | 9.707271972 | 0.582300916 | 0.580976708 | 0.810101637 | -4.8255544 |
| **A_55_P2104791** | NA | -0.055736617 | -0.288342913 | 0.17686968 | 14.70337647 | -0.582240385 | 0.581015003 | 0.810101637 | -4.825572224 |
| **A_51_P472888** | Enkd1 | -0.108696079 | -0.562432393 | 0.345040234 | 9.463604024 | -0.582094546 | 0.581107276 | 0.810117033 | -4.825615163 |
| **A_51_P208697** | Ttl | -0.079352355 | -0.410636647 | 0.251931936 | 8.63010501 | -0.58202571 | 0.581150832 | 0.810117033 | -4.825635427 |
| **A_51_P311904** | Ptk2b | -0.139970425 | -0.724327653 | 0.444386803 | 9.679521942 | -0.582024209 | 0.581151781 | 0.810117033 | -4.825635869 |
| **A_51_P281582** | Lrrc42 | -0.040299066 | -0.208626499 | 0.128028366 | 8.385847879 | -0.581732425 | 0.581336427 | 0.810315988 | -4.825721741 |
| **A_55_P2179483** | NA | 0.093328283 | -0.296669166 | 0.483325732 | 14.83815017 | 0.581479889 | 0.581496264 | 0.810478383 | -4.825796034 |
| **A_55_P1974228** | NA | 0.058079815 | -0.184649113 | 0.300808743 | 15.10250868 | 0.58141586 | 0.581536794 | 0.810478383 | -4.825814866 |
| **A_52_P609868** | Timd4 | 0.129080186 | -0.410578599 | 0.668738972 | 9.502525442 | 0.58119721 | 0.58167521 | 0.810564598 | -4.825879162 |
| **A_55_P2169829** | Armh4 | -0.043347506 | -0.224578479 | 0.137883466 | 7.780451378 | -0.581185655 | 0.581682526 | 0.810564598 | -4.825882559 |
| **A_51_P262533** | Cxxc1 | -0.040296513 | -0.208836084 | 0.128243059 | 9.449778871 | -0.580963392 | 0.58182325 | 0.810702249 | -4.825947895 |
| **A_55_P2144391** | NA | -0.077804321 | -0.403296584 | 0.247687942 | 10.97357946 | -0.580826251 | 0.58191009 | 0.810764804 | -4.825988199 |
| **A_51_P130095** | Fcgr2b | -0.08263802 | -0.428566917 | 0.263290878 | 8.560183146 | -0.58046528 | 0.582138699 | 0.810830045 | -4.826094244 |
| **A_51_P498257** | NA | 0.0823924 | -0.262527102 | 0.427311903 | 10.08313332 | 0.580433665 | 0.582158724 | 0.810830045 | -4.826103529 |
| **A_51_P257762** | Kat2b | 0.05103341 | -0.162612791 | 0.264679611 | 8.31242002 | 0.580420311 | 0.582167183 | 0.810830045 | -4.826107451 |
| **A_52_P632191** | Nfx1 | 0.045048746 | -0.143545555 | 0.233643047 | 8.448737452 | 0.580413264 | 0.582171646 | 0.810830045 | -4.826109521 |
| **A_51_P520849** | Sfrp2 | -0.146080807 | -0.757679778 | 0.465518163 | 8.223353384 | -0.580376213 | 0.582195115 | 0.810830045 | -4.826120402 |
| **A_55_P2122491** | Gm6222 | 0.054370434 | -0.17334276 | 0.282083629 | 15.63909919 | 0.580173379 | 0.582323604 | 0.810830045 | -4.826179958 |
| **A_51_P401343** | Cldn14 | -0.102283264 | -0.530701957 | 0.326135429 | 8.451201785 | -0.580122024 | 0.582356138 | 0.810830045 | -4.826195033 |
| **A_55_P2198946** | Gm10768 | -0.184946173 | -0.959623421 | 0.589731076 | 9.920821038 | -0.580106506 | 0.58236597 | 0.810830045 | -4.826199589 |
| **A_55_P2131607** | Trnau1ap | -0.050928677 | -0.264257627 | 0.162400274 | 9.971104215 | -0.580090542 | 0.582376083 | 0.810830045 | -4.826204275 |
| **A_55_P2415472** | 4931408D14Rik | 0.379043366 | -1.208861442 | 1.966948173 | 10.27534557 | 0.580026219 | 0.582416836 | 0.810830045 | -4.826223156 |
| **A_51_P267700** | Gkn3 | -0.069354398 | -0.359898029 | 0.221189234 | 8.518702833 | -0.580023824 | 0.582418353 | 0.810830045 | -4.826223859 |
| **A_51_P501815** | Smdt1 | -0.058753403 | -0.305022121 | 0.187515315 | 14.63225311 | -0.579704892 | 0.582620439 | 0.810970094 | -4.826317448 |
| **A_55_P2114799** | NA | -0.04414023 | -0.229162035 | 0.140881575 | 8.229541517 | -0.579688744 | 0.582630671 | 0.810970094 | -4.826322185 |
| **A_52_P351816** | Itgb1 | -0.080608991 | -0.418510035 | 0.257292053 | 8.967470959 | -0.579665082 | 0.582645667 | 0.810970094 | -4.826329127 |
| **A_55_P2003005** | Capza1 | 0.13058962 | -0.416985444 | 0.678164683 | 13.99325576 | 0.579492985 | 0.582754733 | 0.810970094 | -4.826379607 |
| **A_51_P140742** | Islr | -0.186397721 | -0.968000699 | 0.595205256 | 8.944038703 | -0.579478841 | 0.582763697 | 0.810970094 | -4.826383755 |
| **A_52_P264902** | Dnajc3 | -0.049905492 | -0.259173199 | 0.159362215 | 10.20965038 | -0.579467813 | 0.582770687 | 0.810970094 | -4.826386989 |
| **A_55_P2059010** | Rbp1 | -0.077504144 | -0.402661827 | 0.247653539 | 15.10015511 | -0.579180715 | 0.582952664 | 0.811012387 | -4.826471171 |
| **A_51_P371867** | 3830417A13Rik | -0.036911129 | -0.191770099 | 0.11794784 | 7.679230134 | -0.579167466 | 0.582961063 | 0.811012387 | -4.826475055 |
| **A_52_P76402** | Ago1 | -0.047229105 | -0.245394142 | 0.150935931 | 8.784953925 | -0.579116417 | 0.582993424 | 0.811012387 | -4.826490019 |
| **A_55_P2084706** | Acaca | 0.095132305 | -0.304032738 | 0.494297348 | 11.91934489 | 0.579106859 | 0.582999483 | 0.811012387 | -4.826492821 |
| **A_52_P8922** | Sntb2 | -0.082012715 | -0.426140026 | 0.262114596 | 8.571211698 | -0.579088897 | 0.58301087 | 0.811012387 | -4.826498086 |
| **A_55_P1957389** | Snx25 | -0.086067462 | -0.447498537 | 0.275363613 | 10.67884528 | -0.578624334 | 0.583305423 | 0.811256436 | -4.82663421 |
| **A_52_P175028** | NA | -0.109284144 | -0.568216435 | 0.349648146 | 8.978540118 | -0.578617721 | 0.583309616 | 0.811256436 | -4.826636147 |
| **A_52_P190973** | Vcl | -0.066982354 | -0.348272779 | 0.214308071 | 10.02836031 | -0.578613611 | 0.583312222 | 0.811256436 | -4.826637351 |

| | | | | | | | | | |
|---|---|---|---|---|---|---|---|---|---|
| **A_52_P466853** | Bmyc | -0.058034219 | -0.30182902 | 0.185760581 | 9.063753106 | -0.578419461 | 0.583435349 | 0.811369297 | -4.826694212 |
| **A_55_P2063381** | Tmem74b | -0.050068341 | -0.260434249 | 0.160297568 | 7.728655654 | -0.578323748 | 0.583496053 | 0.811395339 | -4.826722238 |
| **A_55_P2101473** | NA | -0.085083522 | -0.442645818 | 0.272478774 | 15.18447453 | -0.578198461 | 0.58357552 | 0.811447467 | -4.826758917 |
| **A_55_P1985764** | NA | 0.10732592 | -0.343808304 | 0.558460144 | 14.4661737 | 0.578072132 | 0.583655655 | 0.811500515 | -4.826795895 |
| **A_51_P102860** | Herpud2 | 0.048087419 | -0.154162172 | 0.25033701 | 9.197739222 | 0.577732775 | 0.583870953 | 0.811736255 | -4.80104272 |
| **A_55_P2113539** | NA | 0.041656894 | -0.133565004 | 0.216878792 | 8.053819553 | 0.577672499 | 0.583909198 | 0.811736255 | -4.826912826 |
| **A_55_P2129691** | AB124611 | 0.081223543 | -0.260490738 | 0.422937824 | 12.36387962 | 0.577566494 | 0.583976463 | 0.811771381 | -4.826943831 |
| **A_55_P1975660** | Hemgn | 0.126855732 | -0.406985671 | 0.660697135 | 15.25893624 | 0.577405654 | 0.584078531 | 0.811854878 | -4.826990866 |
| **A_55_P2117028** | Mtmr3 | 0.086342864 | -0.27713065 | 0.449816378 | 12.01731705 | 0.577214019 | 0.584200156 | 0.811942004 | -4.827046892 |
| **A_55_P2168124** | NA | -0.098568067 | -0.513538486 | 0.316402351 | 14.07977056 | -0.577168063 | 0.584229324 | 0.811942004 | -4.827060325 |
| **A_51_P243930** | Qrsl1 | 0.060407752 | -0.193948987 | 0.314764491 | 8.403773896 | 0.577075468 | 0.584288099 | 0.811942004 | -4.827087389 |
| **A_52_P460703** | Rassf3 | -0.090572113 | -0.471962852 | 0.290818626 | 8.997456444 | -0.577042164 | 0.584309239 | 0.811942004 | -4.827097122 |
| **A_51_P353592** | Commd4 | -0.069070562 | -0.36004481 | 0.221903687 | 10.74705434 | -0.576795179 | 0.584466031 | 0.812101496 | -4.827169288 |
| **A_55_P2158701** | H4c4 | -0.090151325 | -0.470312572 | 0.290009921 | 14.88278322 | -0.576218849 | 0.584831995 | 0.812551584 | -4.827337585 |
| **A_55_P2062262** | Aspscr1 | 0.05748272 | -0.184960036 | 0.299925476 | 9.126776371 | 0.576117789 | 0.584896181 | 0.812582353 | -4.827367081 |
| **A_55_P1994524** | Tpd52l2 | 0.062515484 | -0.201256845 | 0.326287813 | 12.15889053 | 0.57589267 | 0.585039175 | 0.812722597 | -4.82743277 |
| **A_55_P1955821** | Ccl21b | -0.26771151 | -1.397777535 | 0.862354515 | 11.84414215 | -0.575633969 | 0.585203525 | 0.812835776 | -4.827508233 |
| **A_51_P484764** | Fam3a | 0.042470897 | -0.136808205 | 0.22175 | 9.757944633 | 0.575632038 | 0.585204752 | 0.812835776 | -4.827508796 |
| **A_55_P1974617** | Lysmd1 | -0.053167078 | -0.277687127 | 0.171352971 | 10.65931774 | -0.575401322 | 0.585351347 | 0.812980972 | -4.827576071 |
| **A_55_P2134616** | Med12l | 0.038836 | -0.125201406 | 0.202873406 | 8.006543684 | 0.575274062 | 0.585432216 | 0.813034869 | -4.827613169 |
| **A_51_P440460** | Hip1r | 0.066391535 | -0.2141664 | 0.34694947 | 9.242498938 | 0.575007275 | 0.58560177 | 0.81315989 | -4.827690919 |
| **A_51_P311905** | Elp6 | 0.038164842 | -0.123116513 | 0.199446197 | 8.490187495 | 0.57499292 | 0.585610895 | 0.81315989 | -4.827695102 |
| **A_55_P2040993** | NA | -0.145827197 | -0.76214377 | 0.470489376 | 8.842921994 | -0.574933839 | 0.585648447 | 0.81315989 | -4.827712315 |
| **A_51_P140942** | Rp9 | -0.077656942 | -0.405985694 | 0.25067181 | 12.81040522 | -0.574717681 | 0.585785852 | 0.813292253 | -4.827775281 |
| **A_55_P2038494** | Rexo2 | 0.084039098 | -0.271318604 | 0.439396799 | 10.08588201 | 0.574643948 | 0.585832726 | 0.813298913 | -4.827796755 |
| **A_55_P2106645** | Adcy5 | -0.04998696 | -0.2613828 | 0.161408881 | 7.862983434 | -0.574570699 | 0.585879295 | 0.813305149 | -4.827818085 |
| **A_55_P1957633** | Gpsm3 | -0.075101811 | -0.392810586 | 0.242606965 | 11.9728756 | -0.574386714 | 0.585996274 | 0.813336385 | -4.827871652 |
| **A_55_P1953508** | Mob3c | 0.071877029 | -0.232199447 | 0.375953505 | 8.467329384 | 0.574368328 | 0.586007965 | 0.813336385 | -4.827877004 |
| **A_55_P2094963** | NA | 0.07790381 | -0.25168713 | 0.40749475 | 9.495390728 | 0.574336768 | 0.586028032 | 0.813336385 | -4.827886191 |
| **A_55_P2121352** | Cdk5 | -0.075590697 | -0.395500443 | 0.24431905 | 8.965698101 | -0.574148276 | 0.586147895 | 0.813371207 | -4.827941051 |
| **A_55_P2075366** | Tead3 | -0.055602724 | -0.290964725 | 0.179759278 | 7.835097079 | -0.57404099 | 0.586216125 | 0.813371207 | -4.827972269 |
| **A_55_P1953866** | Sema4c | 0.129701467 | -0.419376756 | 0.67877969 | 15.23696608 | 0.573976166 | 0.586257353 | 0.813371207 | -4.827991129 |
| **A_55_P2043662** | Zfp607a | 0.039628412 | -0.128165868 | 0.207422693 | 8.282329591 | 0.573868923 | 0.586325563 | 0.813371207 | -4.828022328 |
| **A_51_P184197** | Smarcd2 | -0.053035269 | -0.277610474 | 0.171539937 | 10.0696918 | -0.573833839 | 0.586347879 | 0.813371207 | -4.828032533 |
| **A_52_P507498** | Plxnc1 | -0.110619166 | -0.579030822 | 0.35779249 | 9.48295756 | -0.573833471 | 0.586348113 | 0.813371207 | -4.82803264 |
| **A_52_P56394** | Psmg4 | 0.052851416 | -0.171015941 | 0.276718773 | 10.78962994 | 0.573652702 | 0.586463101 | 0.813371207 | -4.828085213 |
| **A_52_P512749** | Ears2 | 0.095909383 | -0.310529095 | 0.502347861 | 9.260363379 | 0.573389149 | 0.586630772 | 0.813371207 | -4.828161838 |
| **A_55_P2025702** | C330007P06Rik | 0.052296813 | -0.169330888 | 0.273924514 | 8.484202344 | 0.573369213 | 0.586643456 | 0.813371207 | -4.828167633 |
| **A_66_P113662** | Tmem62 | 0.05253961 | -0.1701756 | 0.27525482 | 8.474173363 | 0.573218448 | 0.586739386 | 0.813371207 | -4.828211451 |
| **A_51_P432199** | Sap30 | -0.041188272 | -0.215793197 | 0.133416653 | 9.183361522 | -0.573192194 | 0.586756092 | 0.813371207 | -4.828219081 |
| **A_52_P674759** | Btbd3 | -0.18487928 | -0.968804595 | 0.599046035 | 9.111004072 | -0.573055575 | 0.58684303 | 0.813371207 | -4.828258778 |
| **A_52_P477369** | Csnk1d | 0.054319822 | -0.176022771 | 0.284662414 | 8.357317459 | 0.573016693 | 0.586867774 | 0.813371207 | -4.828270075 |
| **A_52_P515247** | Arhgdib | -0.123763065 | -0.64859034 | 0.401064209 | 13.20841525 | -0.573004259 | 0.586875687 | 0.813371207 | -4.828273687 |
| **A_52_P118560** | Kynu | 0.13069471 | -0.423528257 | 0.684917677 | 9.201313822 | 0.573002714 | 0.58687667 | 0.813371207 | -4.828274136 |
| **A_55_P1972719** | Pmm1 | 0.057822686 | -0.187382049 | 0.303027422 | 8.398421537 | 0.572997334 | 0.586880094 | 0.813371207 | -4.828275699 |
| **A_51_P241995** | Col5a3 | -0.051652016 | -0.270714163 | 0.167410132 | 8.18210979 | -0.572932056 | 0.586921639 | 0.813371207 | -4.828294662 |
| **A_55_P2032723** | Dnajc10 | -0.040429762 | -0.211900881 | 0.131041357 | 8.536968229 | -0.572919219 | 0.586929808 | 0.813371207 | -4.828298391 |
| **A_66_P133240** | Hnrnpll | -0.044178929 | -0.231557235 | 0.143199377 | 11.24103391 | -0.572900316 | 0.586941839 | 0.813371207 | -4.828303882 |
| **A_52_P357484** | Nfrkb | -0.041041118 | -0.215135334 | 0.133053099 | 8.332010704 | -0.572819805 | 0.586993081 | 0.813371207 | -4.828327266 |
| **A_55_P2038573** | NA | -0.054725058 | -0.286870122 | 0.177420007 | 15.49980635 | -0.572809168 | 0.586999851 | 0.813371207 | -4.828330356 |

| | | | | | | | | | |
|---|---|---|---|---|---|---|---|---|---|
| **A_51_P352968** | Marcks | -0.06673109 | -0.349834002 | 0.216371822 | 13.33600056 | -0.572752604 | 0.587035854 | 0.813371207 | -4.828346783 |
| **A_65_P10351** | Atp2b1 | -0.042883807 | -0.224819346 | 0.139051731 | 8.21680229 | -0.572741939 | 0.587042642 | 0.813371207 | -4.828349881 |
| **A_55_P2002557** | Srebf1 | -0.089869374 | -0.471163606 | 0.291424857 | 9.117000331 | -0.572709876 | 0.587063051 | 0.813371207 | -4.828359192 |
| **A_55_P2049479** | NA | 0.116613403 | -0.378249522 | 0.611476328 | 10.57370663 | 0.572593931 | 0.587136855 | 0.813415157 | -4.828392858 |
| **A_55_P2302383** | 3300002A11Rik | -0.042236455 | -0.221522106 | 0.137049196 | 8.156374109 | -0.572433597 | 0.587238923 | 0.813437647 | -4.828439404 |
| **A_55_P2167763** | NA | -0.076996578 | -0.403841141 | 0.249847986 | 8.466334417 | -0.572418083 | 0.5872488 | 0.813437647 | -4.828443907 |
| **A_51_P242859** | Akr1c12 | 0.142254574 | -0.461655786 | 0.746164933 | 9.799206809 | 0.572370113 | 0.58727934 | 0.813437647 | -4.828457831 |
| **A_55_P2052472** | NA | -0.054941789 | -0.288391716 | 0.178508137 | 14.22973711 | -0.571863329 | 0.587602039 | 0.81375492 | -4.828604867 |
| **A_55_P2014315** | NA | -0.038097217 | -0.199987034 | 0.123792601 | 7.703017249 | -0.571816792 | 0.587631678 | 0.81375492 | -4.828618364 |
| **A_51_P235705** | Tsfm | 0.077415011 | -0.251581378 | 0.4064114 | 9.541330274 | 0.571764568 | 0.587664939 | 0.81375492 | -4.828633509 |
| **A_52_P222709** | Pigu | -0.041822128 | -0.219562557 | 0.135918301 | 10.09960396 | -0.571745939 | 0.587676803 | 0.81375492 | -4.828638911 |
| **A_66_P116635** | Igf2os | 0.129663931 | -0.421594578 | 0.68092244 | 10.18884458 | 0.571540576 | 0.587807609 | 0.813805352 | -4.828698453 |
| **A_55_P2157705** | Uqcr10 | -0.045076354 | -0.236753565 | 0.146600856 | 14.25774848 | -0.571427965 | 0.587879344 | 0.813805352 | -4.828731094 |
| **A_52_P520607** | Ankrd22 | -0.107987671 | -0.567254739 | 0.351279398 | 8.434232073 | -0.571336614 | 0.587937539 | 0.813805352 | -4.82875757 |
| **A_55_P2148896** | NA | 0.086164137 | -0.280293543 | 0.452621817 | 9.098166693 | 0.571328521 | 0.587942695 | 0.813805352 | -4.828759915 |
| **A_52_P167382** | Tmem8 | -0.042859778 | -0.225158852 | 0.139439296 | 8.054264526 | -0.571279508 | 0.587973921 | 0.813805352 | -4.828774119 |
| **A_55_P2181542** | Sult1d1 | -0.145937483 | -0.766694996 | 0.474820029 | 9.029259905 | -0.571252426 | 0.587991175 | 0.813805352 | -4.828781966 |
| **A_52_P206613** | Adss | 0.061896414 | -0.201397384 | 0.325190211 | 9.786895574 | 0.571226106 | 0.588007944 | 0.813805352 | -4.828789593 |
| **A_66_P128446** | Ttpa | 0.17831279 | -0.581038046 | 0.937663626 | 9.645124252 | 0.570588756 | 0.588414091 | 0.814309155 | -4.828974181 |
| **A_51_P371485** | Abraxas2 | 0.074752721 | -0.243682973 | 0.393188415 | 9.352035915 | 0.570411735 | 0.588526926 | 0.814406999 | -4.829025418 |
| **A_52_P566840** | Adgrf1 | -0.08687286 | -0.457019789 | 0.28327407 | 8.488830747 | -0.570286584 | 0.588606706 | 0.814459089 | -4.829061634 |
| **A_51_P322473** | Perm1 | -0.038586687 | -0.203026507 | 0.125853133 | 7.719852031 | -0.570182246 | 0.588673223 | 0.814492823 | -4.829091822 |
| **A_55_P2180846** | Wdsub1 | -0.047832102 | -0.251842431 | 0.156178227 | 8.109154043 | -0.569705615 | 0.588977139 | 0.814552298 | -4.829229666 |
| **A_52_P537466** | Zfp330 | 0.053304881 | -0.174092046 | 0.280701807 | 9.869412661 | 0.569594222 | 0.589048179 | 0.814552298 | -4.829261867 |
| **A_51_P416825** | Tspan15 | -0.048031156 | -0.252941688 | 0.156879375 | 8.090938424 | -0.569563237 | 0.589067941 | 0.814552298 | -4.829270824 |
| **A_55_P2127884** | Btbd1 | -0.041909689 | -0.220712732 | 0.136893354 | 7.948338425 | -0.569538021 | 0.589084023 | 0.814552298 | -4.829278112 |
| **A_55_P1960157** | Bcat1 | 0.037568227 | -0.122727309 | 0.197863763 | 7.696888697 | 0.569485228 | 0.589117695 | 0.814552298 | -4.82929337 |
| **A_51_P108459** | Gpr65 | 0.087112886 | -0.284590203 | 0.458815974 | 8.941098393 | 0.569468124 | 0.589128604 | 0.814552298 | -4.812640104 |
| **A_51_P136303** | Cyp4f15 | 0.109758414 | -0.358580939 | 0.578097766 | 9.152066639 | 0.569456243 | 0.589136182 | 0.814552298 | -4.829301747 |
| **A_55_P2210213** | 4732423E21Rik | -0.072094525 | -0.379750928 | 0.235561879 | 8.688241998 | -0.569402687 | 0.589170342 | 0.814552298 | -4.829317224 |
| **A_55_P1984675** | Abhd6 | 0.065054923 | -0.212562689 | 0.342672534 | 8.60123438 | 0.569398529 | 0.589172994 | 0.814552298 | -4.829318425 |
| **A_55_P2099418** | Neil2 | 0.03650441 | -0.119275952 | 0.192284771 | 7.917195528 | 0.569397832 | 0.589173439 | 0.814552298 | -4.829318627 |
| **A_55_P2024515** | Crtc2 | -0.042288436 | -0.222776673 | 0.138199801 | 9.743755539 | -0.569319304 | 0.58922353 | 0.814552298 | -4.829341318 |
| **A_55_P2031043** | Stat5b | 0.075660803 | -0.247290189 | 0.398611795 | 11.48426951 | 0.56926897 | 0.589255637 | 0.814552298 | -4.82935586 |
| **A_66_P104390** | Arl3 | -0.087931153 | -0.463296369 | 0.287434063 | 9.734321817 | -0.569209228 | 0.589293747 | 0.814552298 | -4.82937312 |
| **A_66_P118233** | Hnrnpa0 | -0.062332748 | -0.328431242 | 0.203765745 | 12.6575385 | -0.569189724 | 0.58930619 | 0.814552298 | -4.829378754 |
| **A_55_P2004806** | Hexim2 | 0.148977188 | -0.487094651 | 0.785049028 | 11.34722382 | 0.569110756 | 0.589356567 | 0.814563681 | -4.829401565 |
| **A_51_P190845** | Atp6v1b2 | 0.066167526 | -0.216462123 | 0.348797175 | 8.662267066 | 0.568866513 | 0.589512398 | 0.814676471 | -4.829472101 |
| **A_55_P2003803** | Scml2 | 0.044683577 | -0.146212106 | 0.23557926 | 8.277947682 | 0.568767816 | 0.589575375 | 0.814676471 | -4.829500597 |
| **A_55_P1954356** | Ttc23 | -0.061992214 | -0.326842101 | 0.202857672 | 9.692428367 | -0.568748871 | 0.589587464 | 0.814676471 | -4.829506066 |
| **A_52_P869802** | Thoc3 | 0.04600888 | -0.150577718 | 0.242595478 | 9.587130552 | 0.568683913 | 0.589628915 | 0.814676471 | -4.829524818 |
| **A_55_P2167447** | Cnnm3 | 0.049835268 | -0.163128121 | 0.262798657 | 12.74997273 | 0.568610712 | 0.589675628 | 0.814676471 | -4.829545948 |
| **A_55_P2147276** | Orai1 | -0.043810737 | -0.231037179 | 0.143415704 | 8.947448596 | -0.56858653 | 0.58969106 | 0.814676471 | -4.829552927 |
| **A_52_P369123** | Fam189a2 | -0.047152848 | -0.248686404 | 0.154380708 | 7.7105228 | -0.568517381 | 0.589735191 | 0.81467921 | -4.829572884 |
| **A_51_P168203** | Aig1 | -0.073895472 | -0.389845539 | 0.242054595 | 8.737260893 | -0.568306425 | 0.589869833 | 0.814806974 | -4.829633755 |
| **A_55_P2131348** | Loxl2 | -0.050956876 | -0.268885282 | 0.16697153 | 9.188792811 | -0.568161946 | 0.589962056 | 0.814876131 | -4.829675432 |
| **A_55_P2036459** | Med24 | 0.061639298 | -0.202139294 | 0.32541789 | 8.809280467 | 0.567807763 | 0.590188172 | 0.815128131 | -4.829777566 |
| **A_51_P293688** | Rab32 | -0.081353268 | -0.429574076 | 0.26686754 | 10.61320045 | -0.567679841 | 0.590269852 | 0.815128131 | -4.82981444 |
| **A_51_P382764** | Akr1c20 | -0.24368926 | -1.286843749 | 0.79946523 | 9.298896098 | -0.567637327 | 0.590297 | 0.815128131 | -4.829826694 |
| **A_51_P104392** | Rpp25 | -0.040499267 | -0.213887078 | 0.132888544 | 7.809882677 | -0.567560014 | 0.590346369 | 0.815128131 | -4.829848975 |

| | | | | | | | | | |
|---|---|---|---|---|---|---|---|---|---|
| **A_51_P111962** | Bean1 | -0.036308374 | -0.19177435 | 0.119157603 | 7.676570217 | -0.567485317 | 0.590394071 | 0.815128131 | -4.8298705 |
| **A_51_P401964** | Aph1a | -0.045284282 | -0.239185347 | 0.148616782 | 11.08733138 | -0.567479902 | 0.590397529 | 0.815128131 | -4.82987206 |
| **A_66_P101631** | Zfp27 | 0.038005965 | -0.124790384 | 0.200802314 | 7.976421485 | 0.56727062 | 0.590531189 | 0.815147724 | -4.829932354 |
| **A_55_P1990678** | NA | 0.065776635 | -0.215995555 | 0.347548824 | 8.463437432 | 0.567226761 | 0.590559202 | 0.815147724 | -4.829944988 |
| **A_55_P2151437** | NA | 0.094831095 | -0.311406966 | 0.501069156 | 9.163796838 | 0.567222361 | 0.590562013 | 0.815147724 | -4.829946255 |
| **A_55_P1956667** | Cyfip1 | -0.074592786 | -0.394150073 | 0.244964502 | 10.48280919 | -0.567193559 | 0.590580409 | 0.815147724 | -4.829954551 |
| **A_55_P2047163** | Tmc5 | -0.042404443 | -0.224150721 | 0.139341835 | 7.696209757 | -0.566929458 | 0.590749113 | 0.815277676 | -4.830030603 |
| **A_55_P1968718** | Plekha6 | -0.045456933 | -0.240291516 | 0.149377651 | 8.201403398 | -0.566914111 | 0.590758918 | 0.815277676 | -4.830035022 |
| **A_52_P482124** | Fam32a | -0.052032514 | -0.275128506 | 0.171063478 | 10.09073896 | -0.566716983 | 0.59088486 | 0.815361087 | -4.830091767 |
| **A_51_P327121** | Was | 0.09946904 | -0.327039286 | 0.525977365 | 10.74548101 | 0.566687457 | 0.590903725 | 0.815361087 | -4.830100265 |
| **A_55_P2106459** | Zfhx3 | 0.076934458 | -0.253038512 | 0.406907429 | 9.409612108 | 0.566533667 | 0.591001992 | 0.815438468 | -4.830144521 |
| **A_51_P321531** | Purg | -0.107852081 | -0.570515401 | 0.354811238 | 9.066438629 | -0.566430525 | 0.591067902 | 0.815445667 | -4.830174197 |
| **A_55_P2099650** | Slc24a3 | -0.062245125 | -0.329300144 | 0.204809894 | 9.418959115 | -0.566353759 | 0.591116959 | 0.815445667 | -4.830196281 |
| **A_55_P2031217** | Vps52 | 0.046844397 | -0.15414907 | 0.247837864 | 11.01615046 | 0.566316089 | 0.591141033 | 0.815445667 | -4.830207116 |
| **A_55_P2156712** | Gpr1 | -0.036353957 | -0.19235155 | 0.119643635 | 7.687046685 | -0.566261439 | 0.59117596 | 0.815445667 | -4.830222836 |
| **A_55_P2091105** | NA | -0.060962982 | -0.322643177 | 0.200717214 | 16.11916106 | -0.566080941 | 0.591291324 | 0.815546596 | -4.830274743 |
| **A_55_P1963150** | Creg1 | 0.083631854 | -0.275671578 | 0.442935286 | 12.77236946 | 0.565579352 | 0.591611978 | 0.815915667 | -4.830418918 |
| **A_51_P144249** | Ubr1 | 0.046414703 | -0.153032128 | 0.245861534 | 8.245004493 | 0.565472677 | 0.591680187 | 0.815915667 | -4.830449566 |
| **A_51_P268903** | Mon1a | -0.050445383 | -0.267227345 | 0.16633658 | 9.496489837 | -0.565433411 | 0.591705294 | 0.815915667 | -4.830460846 |
| **A_55_P2028362** | NA | -0.100393282 | -0.531923434 | 0.33113687 | 8.736861644 | -0.56529702 | 0.591792512 | 0.815915667 | -4.830500022 |
| **A_55_P2279690** | 2900076A07Rik | 0.037112662 | -0.122431725 | 0.19665705 | 7.824475772 | 0.56522813 | 0.591836567 | 0.815915667 | -4.830519807 |
| **A_55_P1968668** | Gpatch2l | 0.041478886 | -0.136845689 | 0.219803461 | 8.590315863 | 0.565195993 | 0.59185712 | 0.815915667 | -4.830529035 |
| **A_51_P439612** | Dnajb2 | 0.070688161 | -0.233234371 | 0.374610693 | 10.77330576 | 0.565154217 | 0.591883837 | 0.815915667 | -4.830541032 |
| **A_66_P135987** | Ocrl | -0.064950759 | -0.344215296 | 0.214313778 | 10.0796122 | -0.565134253 | 0.591896605 | 0.815915667 | -4.830546764 |
| **A_55_P2001861** | NA | -0.145070721 | -0.768906225 | 0.478764783 | 9.725189283 | -0.565057797 | 0.591945505 | 0.815924885 | -4.830568716 |
| **A_66_P123938** | Smim13 | -0.059122769 | -0.313428639 | 0.195183102 | 10.76453496 | -0.564912997 | 0.592038121 | 0.815994355 | -4.830610284 |
| **A_55_P2014304** | Kank1 | 0.098138897 | -0.324169727 | 0.520447522 | 11.03676718 | 0.564669606 | 0.592193817 | 0.816150751 | -4.830680134 |
| **A_51_P286034** | Ccdc117 | -0.046346171 | -0.245852608 | 0.153160265 | 8.940619541 | -0.56446906 | 0.592322122 | 0.816237949 | -4.830737669 |
| **A_52_P144263** | Rbms1 | 0.060196546 | -0.198945556 | 0.319338648 | 9.09505818 | 0.564438705 | 0.592341544 | 0.816237949 | -4.830746376 |
| **A_55_P2037584** | Trf | 0.07320191 | -0.241986533 | 0.388390354 | 17.32954122 | 0.564332833 | 0.592409287 | 0.816255626 | -4.830776741 |
| **A_55_P2007088** | Rnf128 | -0.122764497 | -0.670602201 | 0.425073207 | 14.12205574 | -0.570116008 | 0.592444905 | 0.816255626 | -4.788140377 |
| **A_52_P569348** | Dbt | 0.147182631 | -0.486673594 | 0.781038856 | 10.03732853 | 0.564220668 | 0.592481061 | 0.816255626 | -4.830808906 |
| **A_51_P419300** | Pdcd2l | -0.060930711 | -0.32337293 | 0.201511508 | 11.27761007 | -0.564138494 | 0.592533647 | 0.816269893 | -4.830832467 |
| **A_55_P2007384** | Sgca | -0.036350962 | -0.192967441 | 0.120265518 | 7.6797831 | -0.56397732 | 0.592636797 | 0.816353809 | -4.830878672 |
| **A_51_P506755** | Tep1 | 0.042954813 | -0.142318847 | 0.228228473 | 9.780020651 | 0.563353946 | 0.593035845 | 0.816845282 | -4.831057269 |
| **A_52_P60353** | Greb1l | 0.138935731 | -0.460480492 | 0.738351953 | 9.114600259 | 0.563207781 | 0.593129434 | 0.81686381 | -4.831099121 |
| **A_55_P2058791** | 4833420G17Rik | 0.03883816 | -0.1287278 | 0.206404121 | 8.603631786 | 0.563191433 | 0.593139902 | 0.81686381 | -4.831103802 |
| **A_52_P217474** | Ndufa6 | -0.041046776 | -0.218159504 | 0.136065952 | 13.55191065 | -0.563134936 | 0.593176079 | 0.81686381 | -4.831119976 |
| **A_52_P258218** | Proser1 | 0.036676306 | -0.121636293 | 0.194988904 | 8.227578588 | 0.562928575 | 0.593308232 | 0.816896901 | -4.831179042 |
| **A_51_P185247** | Gdf10 | 0.11796622 | -0.391256391 | 0.627188832 | 9.784109838 | 0.562902495 | 0.593324935 | 0.816896901 | -4.831186506 |
| **A_52_P476431** | Evc | 0.054550324 | -0.180942285 | 0.290042932 | 8.052768618 | 0.562863702 | 0.59334978 | 0.816896901 | -4.831197607 |
| **A_55_P2467415** | B3galt6 | 0.04831304 | -0.160264556 | 0.256890635 | 8.480480983 | 0.562833442 | 0.59336916 | 0.816896901 | -4.831206265 |
| **A_51_P468418** | Ice2 | 0.053387896 | -0.177305616 | 0.284081407 | 8.643527912 | 0.562329169 | 0.593692185 | 0.817181659 | -4.831350505 |
| **A_51_P336691** | Srsf9 | -0.038090682 | -0.202703692 | 0.126522328 | 12.03736501 | -0.562260773 | 0.593736005 | 0.817181659 | -4.831370059 |
| **A_51_P368483** | 1700123O20Rik | 0.06767233 | -0.224810875 | 0.360155535 | 10.65816802 | 0.562203292 | 0.593772834 | 0.817181659 | -4.831386492 |
| **A_55_P2004781** | Ass1 | -0.137365312 | -0.731098994 | 0.45636837 | 14.94708297 | -0.562171182 | 0.593793408 | 0.817181659 | -4.831395671 |
| **A_52_P629037** | Gm6987 | 0.10119434 | -0.33629003 | 0.538678711 | 12.27443505 | 0.562052483 | 0.593869466 | 0.817181659 | -4.831429599 |
| **A_55_P1961888** | Twistnb | 0.071112642 | -0.236351763 | 0.378577047 | 10.52608365 | 0.561998504 | 0.593904056 | 0.817181659 | -4.831445026 |
| **A_52_P613977** | NA | -0.069841022 | -0.371836819 | 0.232154775 | 11.8284026 | -0.56194379 | 0.593939117 | 0.817181659 | -4.831460661 |
| **A_55_P1994957** | NA | 0.054236452 | -0.180291867 | 0.28876477 | 15.09395777 | 0.561926055 | 0.593950483 | 0.817181659 | -4.831465729 |

| | | | | | | | | |
|---|---|---|---|---|---|---|---|---|
| **A_51_P296429** | Tmem170 | 0.048419304 | -0.160976004 | 0.257814612 | 8.094823318 | 0.561868631 | 0.593987282 | 0.817181659 | -4.831482138 |
| **A_55_P1971448** | Snrpd1 | 0.06208524 | -0.20641868 | 0.330589161 | 13.82297153 | 0.561850698 | 0.593998775 | 0.817181659 | -4.831487261 |
| **A_55_P2077526** | NA | -0.05465973 | -0.291092334 | 0.181772874 | 14.13828387 | -0.56175029 | 0.594063125 | 0.817202108 | -4.831515947 |
| **A_55_P1988273** | Rnf103 | -0.10947253 | -0.583045979 | 0.364100919 | 10.72859393 | -0.561695571 | 0.594098196 | 0.817202108 | -4.831531579 |
| **A_55_P2069729** | Ormdl2 | -0.059506675 | -0.316990546 | 0.197977196 | 12.10253332 | -0.561563462 | 0.594182871 | 0.81725073 | -4.831569312 |
| **A_55_P1991239** | NA | 0.073891849 | -0.245885628 | 0.393669327 | 8.696938734 | 0.561476847 | 0.594238391 | 0.81725073 | -4.831594047 |
| **A_55_P2048284** | Aasdhppt | 0.072766048 | -0.242158609 | 0.387690704 | 8.638317106 | 0.561442534 | 0.594260386 | 0.81725073 | -4.831603845 |
| **A_51_P362089** | Lrrc3b | 0.072611742 | -0.241738552 | 0.386962036 | 7.938656442 | 0.561275614 | 0.594367392 | 0.81730821 | -4.831651501 |
| **A_52_P558577** | Zfp263 | -0.035972096 | -0.191710647 | 0.119766454 | 10.12367406 | -0.561245418 | 0.594386751 | 0.81730821 | -4.831660121 |
| **A_51_P291819** | 2310033P09Rik | 0.049946676 | -0.16635941 | 0.266252761 | 10.83197894 | 0.561075157 | 0.594495912 | 0.817400163 | -4.831708717 |
| **A_51_P507778** | Sdr42e1 | -0.158190423 | -0.843473758 | 0.527092912 | 10.75599284 | -0.56091003 | 0.594601793 | 0.817442398 | -4.831755835 |
| **A_66_P106593** | Vhl | -0.047138532 | -0.251348723 | 0.157071659 | 10.52955399 | -0.560895341 | 0.594611212 | 0.817442398 | -4.831760026 |
| **A_55_P2088741** | Mtif3 | 0.060404629 | -0.201358367 | 0.322167626 | 8.706579718 | 0.560718852 | 0.594724391 | 0.817539845 | -4.831810372 |
| **A_51_P362661** | Spin4 | -0.049769797 | -0.265495403 | 0.16595581 | 8.055918167 | -0.560592594 | 0.594805365 | 0.81759301 | -4.831846381 |
| **A_55_P2081571** | Raver1 | 0.082821456 | -0.276519527 | 0.44216244 | 14.10744309 | 0.560040327 | 0.595159631 | 0.81784631 | -4.832003807 |
| **A_55_P1984561** | Cpne2 | -0.039917435 | -0.213135026 | 0.133300156 | 9.104612193 | -0.559955895 | 0.595213804 | 0.81784631 | -4.832027863 |
| **A_55_P2037777** | NA | -0.04054607 | -0.216497467 | 0.135405326 | 8.545709492 | -0.559937097 | 0.595225865 | 0.81784631 | -4.832033218 |
| **A_55_P2111985** | Prss35 | -0.046676129 | -0.249236993 | 0.155884735 | 7.774697569 | -0.559915485 | 0.595239731 | 0.81784631 | -4.832039375 |
| **A_55_P2110915** | Flrt1 | -0.057730148 | -0.30828921 | 0.192828914 | 8.040643036 | -0.559855241 | 0.595278387 | 0.81784631 | -4.832056537 |
| **A_52_P681705** | Zfp955b | 0.088791298 | -0.296580261 | 0.474162858 | 9.638174016 | 0.559852902 | 0.595279887 | 0.81784631 | -4.832057203 |
| **A_51_P215038** | Tmem59l | -0.062424004 | -0.333360843 | 0.208512834 | 8.065000859 | -0.559843647 | 0.595285826 | 0.81784631 | -4.832059839 |
| **A_55_P1984730** | Sgpl1 | 0.052670757 | -0.175975453 | 0.281316967 | 10.83073758 | 0.559743092 | 0.59535035 | 0.817863302 | -4.83208848 |
| **A_51_P273679** | Dmgdh | 0.151822194 | -0.507329562 | 0.81097395 | 11.13366567 | 0.559671303 | 0.595396419 | 0.817863302 | -4.832108925 |
| **A_52_P66130** | Myo5b | 0.056201917 | -0.187826632 | 0.300230465 | 8.665369022 | 0.559620582 | 0.595428969 | 0.817863302 | -4.832123368 |
| **A_51_P340668** | Bcl9l | -0.04052978 | -0.216528816 | 0.135469256 | 7.99124616 | -0.559560628 | 0.595467445 | 0.817863302 | -4.832140439 |
| **A_55_P2052831** | 0610009B22Rik | 0.055899558 | -0.187064984 | 0.2988641 | 9.838909782 | 0.559047442 | 0.595796848 | 0.818257587 | -4.8322865 |
| **A_51_P507942** | Atp13a2 | -0.045490147 | -0.243322323 | 0.152342028 | 11.84541965 | -0.558732077 | 0.595999325 | 0.818461741 | -4.832376201 |
| **A_55_P2125947** | Agpat2 | -0.071411373 | -0.381999126 | 0.23917638 | 9.968846847 | -0.558684016 | 0.596030186 | 0.818461741 | -4.832389867 |
| **A_52_P639229** | Fgd4 | 0.058180863 | -0.194901276 | 0.311263003 | 8.235163686 | 0.558601184 | 0.596083376 | 0.818476634 | -4.832413419 |
| **A_55_P2017826** | Myb | 0.087996365 | -0.294872366 | 0.470865097 | 11.27665848 | 0.558467653 | 0.596169127 | 0.818536231 | -4.832451379 |
| **A_55_P1956593** | Plekha4 | -0.054836156 | -0.29349488 | 0.183822569 | 8.464978909 | -0.558306743 | 0.59627247 | 0.818619972 | -4.832497113 |
| **A_55_P2427150** | C430002N11Rik | 0.038205589 | -0.128129983 | 0.20454116 | 7.728300626 | 0.558116615 | 0.596394592 | 0.818729479 | -4.832551136 |
| **A_55_P1956898** | NA | 0.071570302 | -0.240180606 | 0.383321211 | 11.86165971 | 0.55783828 | 0.596573396 | 0.818888882 | -4.832630194 |
| **A_52_P438036** | Lpar4 | -0.129767696 | -0.695199182 | 0.43566379 | 8.582952224 | -0.557660384 | 0.596687693 | 0.818888882 | -4.832680706 |
| **A_55_P2095118** | Dancr | -0.135727291 | -0.727210471 | 0.455755889 | 9.85766536 | -0.557581007 | 0.596738696 | 0.818888882 | -4.83270324 |
| **A_55_P1965391** | Klhl18 | -0.043992205 | -0.235747143 | 0.147762734 | 9.348673273 | -0.557458263 | 0.59681757 | 0.818888882 | -4.83273808 |
| **A_51_P390377** | Rcor3 | -0.07510417 | -0.402565531 | 0.252357192 | 9.051491432 | -0.557297603 | 0.596920817 | 0.818888882 | -4.832783672 |
| **A_55_P2109559** | Trim6 | -0.036256557 | -0.194346987 | 0.121833874 | 7.651586828 | -0.557268073 | 0.596939795 | 0.818888882 | -4.832792051 |
| **A_55_P2017526** | Dand5 | 0.047653824 | -0.160158357 | 0.255466006 | 8.268480597 | 0.557198511 | 0.596984502 | 0.818888882 | -4.832811786 |
| **A_51_P137029** | Plce1 | -0.039410223 | -0.211274261 | 0.132453816 | 7.852832332 | -0.557194812 | 0.59698688 | 0.818888882 | -4.832812836 |
| **A_52_P216226** | Masp1 | 0.090574716 | -0.304415807 | 0.485565239 | 8.733511795 | 0.557190231 | 0.596989824 | 0.818888882 | -4.832814136 |
| **A_52_P315030** | NA | 0.052084666 | -0.175073098 | 0.27924243 | 15.45675128 | 0.557141474 | 0.597021161 | 0.818888882 | -4.832827967 |
| **A_55_P2028571** | Sgsh | 0.051040286 | -0.171566829 | 0.2736474 | 9.211012792 | 0.557130899 | 0.597027959 | 0.818888882 | -4.832830967 |
| **A_52_P322141** | Ccdc88b | 0.05588016 | -0.187845036 | 0.299605357 | 11.40621588 | 0.557109291 | 0.597041847 | 0.818888882 | -4.832837096 |
| **A_51_P354272** | Cluh | 0.089760646 | -0.301758091 | 0.481279384 | 13.06363582 | 0.557078772 | 0.597061463 | 0.818888882 | -4.832845753 |
| **A_55_P2060793** | NA | -0.068149909 | -0.365497898 | 0.22919808 | 13.40067238 | -0.556908009 | 0.597171229 | 0.818894668 | -4.832894184 |
| **A_55_P2158319** | Ntan1 | -0.071340334 | -0.382609413 | 0.239928744 | 10.02562122 | -0.556906579 | 0.597172148 | 0.818894668 | -4.832894589 |
| **A_55_P1976983** | Rpl36 | -0.04545967 | -0.243833514 | 0.152914173 | 16.58470851 | -0.556833129 | 0.597219365 | 0.818894668 | -4.832915417 |
| **A_52_P474636** | Stk4 | 0.040612967 | -0.136653053 | 0.217878987 | 8.15298833 | 0.556701527 | 0.59730397 | 0.818894668 | -4.832952728 |
| **A_55_P2035892** | Prkcsh | -0.063672167 | -0.341595282 | 0.214250947 | 14.35091888 | -0.556683249 | 0.59731572 | 0.818894668 | -4.83295791 |

| | | | | | | | | | |
|---|---|---|---|---|---|---|---|---|---|
| **A_52_P237159** | Spty2d1 | 0.092215676 | -0.310310548 | 0.494741899 | 9.556611138 | 0.556664803 | 0.59732758 | 0.818894668 | -4.832963138 |
| **A_51_P170959** | NA | -0.149763751 | -0.803553643 | 0.504026141 | 13.39819152 | -0.556610887 | 0.597362244 | 0.818894668 | -4.832978421 |
| **A_55_P2068066** | Rapgefl1 | -0.052891464 | -0.28388387 | 0.178100942 | 8.100775281 | -0.556379439 | 0.597511063 | 0.819030476 | -4.833044013 |
| **A_52_P336825** | Matr3 | 0.062992988 | -0.212169497 | 0.338155473 | 9.976623443 | 0.556270686 | 0.597580998 | 0.819030476 | -4.833074825 |
| **A_55_P2156126** | Sema6c | -0.046708079 | -0.250739967 | 0.157323809 | 8.869328853 | -0.556259126 | 0.597588432 | 0.819030476 | -4.8330781 |
| **A_55_P2436438** | Dusp22 | -0.042900593 | -0.230511238 | 0.144710051 | 8.144533419 | -0.555634255 | 0.597990355 | 0.819303197 | -4.833255037 |
| **A_55_P2183808** | NA | 0.074170109 | -0.250233982 | 0.3985742 | 13.8691284 | 0.555553368 | 0.598042393 | 0.819303197 | -4.833277929 |
| **A_52_P526372** | Zeb2 | -0.073307298 | -0.393973786 | 0.247359191 | 8.015764441 | -0.555490741 | 0.598082686 | 0.819303197 | -4.83329565 |
| **A_55_P2134487** | NA | 0.062273125 | -0.210129025 | 0.334675274 | 11.97670744 | 0.555486255 | 0.598085572 | 0.819303197 | -4.83329692 |
| **A_55_P1952512** | Sult2a2 | -0.163623482 | -0.879420684 | 0.552173721 | 8.566789432 | -0.555442134 | 0.598113959 | 0.819303197 | -4.833309404 |
| **A_51_P173022** | Nsmce1 | 0.078558213 | -0.265124499 | 0.422240926 | 11.6401291 | 0.555414353 | 0.598131834 | 0.819303197 | -4.833317264 |
| **A_55_P2087785** | NA | 0.044821822 | -0.151278431 | 0.240922074 | 8.018801553 | 0.555385502 | 0.598150398 | 0.819303197 | -4.833325426 |
| **A_55_P2003581** | Scnm1 | -0.056060414 | -0.301368747 | 0.189247919 | 10.91585574 | -0.555299378 | 0.598205814 | 0.819303197 | -4.83334979 |
| **A_55_P2123942** | Lmbrd1 | 0.071155744 | -0.240213661 | 0.382525148 | 9.195451475 | 0.55528663 | 0.598214017 | 0.819303197 | -4.833353397 |
| **A_52_P592909** | Dgat2 | 0.101924339 | -0.344138184 | 0.547986862 | 14.16245668 | 0.555220324 | 0.598256683 | 0.819303197 | -4.833372152 |
| **A_51_P244531** | NA | -0.070563139 | -0.379406958 | 0.23828068 | 8.29594018 | -0.555165111 | 0.598292214 | 0.819303197 | -4.833387768 |
| **A_51_P168395** | Ttc30a1 | 0.06256361 | -0.211352914 | 0.336480134 | 8.652897712 | 0.554992044 | 0.598403591 | 0.819303197 | -4.833436707 |
| **A_55_P1998761** | Coq8b | 0.087069493 | -0.294153203 | 0.468292189 | 11.15922097 | 0.554971214 | 0.598416997 | 0.819303197 | -4.833442597 |
| **A_55_P2167005** | NA | -0.057305509 | -0.3082242 | 0.193613182 | 14.76926745 | -0.554940664 | 0.598436659 | 0.819303197 | -4.833451234 |
| **A_55_P2116290** | Szrd1 | 0.158596943 | -0.535889573 | 0.853083458 | 10.97582017 | 0.554899306 | 0.598463278 | 0.819303197 | -4.833462927 |
| **A_55_P2044119** | Klc1 | -0.078414961 | -0.421791628 | 0.264961705 | 8.564566191 | -0.554895678 | 0.598465613 | 0.819303197 | -4.833463952 |
| **A_52_P289213** | Notch2 | -0.049447037 | -0.266017471 | 0.167123397 | 10.72785628 | -0.554784469 | 0.598537192 | 0.819343158 | -4.833495389 |
| **A_51_P242733** | Tex261 | 0.046002445 | -0.155524953 | 0.247529843 | 11.40322314 | 0.554664033 | 0.598614716 | 0.81939125 | -4.833529427 |
| **A_51_P441687** | Lrrc10 | -0.039542462 | -0.212846579 | 0.133761655 | 7.6731232 | -0.554418887 | 0.598772532 | 0.819474081 | -4.833598692 |
| **A_55_P2004741** | Cyb5rl | 0.045089328 | -0.152528899 | 0.242707555 | 8.52516209 | 0.554408595 | 0.598779158 | 0.819474081 | -4.8336016 |
| **A_55_P2161055** | Trp63 | 0.087692928 | -0.296674084 | 0.47205994 | 11.28976053 | 0.554372468 | 0.598802417 | 0.819474081 | -4.833611805 |
| **A_55_P1955472** | Bbs1 | -0.066773412 | -0.359590075 | 0.226043251 | 8.366940213 | -0.554103595 | 0.598975541 | 0.819637934 | -4.833687738 |
| **A_52_P381484** | Spon2 | -0.055951458 | -0.301359051 | 0.189456135 | 11.73990788 | -0.553995963 | 0.599044851 | 0.819637934 | -4.833718126 |
| **A_55_P2015299** | NA | -0.087559241 | -0.471605969 | 0.296487486 | 15.78704353 | -0.55398896 | 0.599049361 | 0.819637934 | -4.833720103 |
| **A_51_P393768** | Osbp | 0.076116388 | -0.257821659 | 0.410054434 | 9.164075741 | 0.553854204 | 0.599136145 | 0.819662229 | -4.833758142 |
| **A_51_P321743** | NA | 0.041468399 | -0.140469937 | 0.223406735 | 7.730401505 | 0.553829697 | 0.599151929 | 0.819662229 | -4.833765059 |
| **A_51_P415225** | Zfp105 | -0.088352396 | -0.476101638 | 0.299396846 | 9.018814165 | -0.553669453 | 0.599255138 | 0.819694474 | -4.83381028 |
| **A_55_P1975396** | Ighmbp2 | -0.047527381 | -0.256131718 | 0.161076955 | 8.813286284 | -0.553609766 | 0.599293583 | 0.819694474 | -4.833827121 |
| **A_55_P2042312** | NA | 0.056638464 | -0.191962063 | 0.305238992 | 16.36438474 | 0.553595579 | 0.599302722 | 0.819694474 | -4.833831124 |
| **A_51_P325165** | Il27 | 0.091321138 | -0.309658025 | 0.4923003 | 9.193644344 | 0.553391771 | 0.59943401 | 0.819816033 | -4.833888616 |
| **A_55_P1966372** | Srek1 | 0.139410368 | -0.472951389 | 0.751772125 | 14.70221658 | 0.553184754 | 0.599567384 | 0.819940424 | -4.833946995 |
| **A_52_P633481** | Akap11 | 0.07653369 | -0.259760396 | 0.412827776 | 8.823868745 | 0.552989153 | 0.599693417 | 0.820054761 | -4.834002138 |
| **A_55_P2146002** | Pank4 | 0.057415703 | -0.194907667 | 0.309739074 | 11.02969491 | 0.552912491 | 0.599742817 | 0.820064298 | -4.834023745 |
| **A_51_P327451** | Alas2 | 0.083598914 | -0.283995272 | 0.451193099 | 15.51778541 | 0.552605482 | 0.599940675 | 0.820276812 | -4.834110251 |
| **A_51_P489337** | Becn1 | 0.051943243 | -0.176559431 | 0.280445917 | 9.214508415 | 0.552358398 | 0.600099939 | 0.820436534 | -4.834179843 |
| **A_52_P627925** | Usp36 | 0.059348359 | -0.201947664 | 0.320644383 | 8.920828671 | 0.551898363 | 0.600396531 | 0.820662848 | -4.83430934 |
| **A_55_P2014531** | Hes7 | 0.04321352 | -0.147058053 | 0.233485093 | 8.633212362 | 0.551860009 | 0.600421262 | 0.820662848 | -4.834320133 |
| **A_52_P665675** | Abca1 | 0.124925994 | -0.425131135 | 0.674983123 | 12.44387026 | 0.551859095 | 0.600421852 | 0.820662848 | -4.83432039 |
| **A_51_P283292** | Gm14326 | -0.053904935 | -0.291311909 | 0.183502039 | 8.590146272 | -0.551719385 | 0.600511944 | 0.820662848 | -4.834359697 |
| **A_55_P1997751** | Prpf8 | 0.04914171 | -0.167329528 | 0.265612949 | 14.35824789 | 0.55161143 | 0.600581564 | 0.820662848 | -4.834390064 |
| **A_52_P162184** | Snx12 | 0.048200471 | -0.164139836 | 0.260540778 | 8.853922538 | 0.551571773 | 0.60060714 | 0.820662848 | -4.834401218 |
| **A_66_P106090** | Rpa3 | 0.08126095 | -0.276726062 | 0.439247961 | 12.42634778 | 0.551566413 | 0.600610597 | 0.820662848 | -4.834402726 |
| **A_55_P2124425** | Zfp661 | 0.04671108 | -0.159070287 | 0.252492448 | 8.781667611 | 0.551565446 | 0.60061122 | 0.820662848 | -4.834402998 |
| **A_55_P2053444** | Map4 | -0.049572193 | -0.26798026 | 0.168835874 | 10.64151947 | -0.551509051 | 0.600647593 | 0.820662848 | -4.834418858 |
| **A_52_P316437** | 1810013L24Rik | 0.040689594 | -0.138717403 | 0.22009659 | 8.386989284 | 0.551095884 | 0.600914102 | 0.820968948 | -4.834535014 |

| | | | | | | | | | |
|---|---|---|---|---|---|---|---|---|---|
| **A_52_P629451** | Gon4l | 0.050269263 | -0.171413113 | 0.271951639 | 8.530691011 | 0.551003731 | 0.600973554 | 0.820992143 | -4.834560911 |
| **A_55_P2098499** | Gnpda2 | -0.086210268 | -0.466487959 | 0.294067422 | 8.753640857 | -0.550860128 | 0.601066205 | 0.821060684 | -4.83460126 |
| **A_55_P2006340** | Uba52 | -0.041215666 | -0.22314268 | 0.140711348 | 17.09487423 | -0.550488587 | 0.601305956 | 0.821235404 | -4.834705611 |
| **A_55_P2163009** | Mapkapk5 | 0.05338278 | -0.18226063 | 0.28902619 | 11.32314751 | 0.550464195 | 0.601321698 | 0.821235404 | -4.83471246 |
| **A_55_P1971419** | Serpinb9g | -0.035058171 | -0.189841412 | 0.11972507 | 7.647540611 | -0.550362088 | 0.601387597 | 0.821235404 | -4.834741126 |
| **A_51_P218653** | Jph2 | -0.034816178 | -0.188566141 | 0.118933786 | 7.679960595 | -0.550236326 | 0.601468769 | 0.821235404 | -4.834776428 |
| **A_55_P1970810** | NA | -0.084438102 | -0.457412784 | 0.28853658 | 13.31939184 | -0.550100827 | 0.601556231 | 0.821235404 | -4.834814454 |
| **A_51_P151271** | Eif4b | -0.056929541 | -0.308439132 | 0.19458005 | 9.137430636 | -0.550004599 | 0.60161835 | 0.821235404 | -4.834841455 |
| **A_52_P641922** | Nhlrc3 | 0.070159991 | -0.239806256 | 0.380126237 | 8.270499353 | 0.549994449 | 0.601624902 | 0.821235404 | -4.834844302 |
| **A_52_P229210** | NA | -0.077616052 | -0.420541169 | 0.265309066 | 12.25122985 | -0.54996549 | 0.601643597 | 0.821235404 | -4.834852427 |
| **A_55_P2017967** | Usf2 | 0.08776882 | -0.300057301 | 0.475594941 | 13.77702139 | 0.549903385 | 0.601683691 | 0.821235404 | -4.83486985 |
| **A_55_P2128032** | Hsf2 | 0.101054148 | -0.345487899 | 0.547596195 | 9.836822655 | 0.549888926 | 0.601693025 | 0.821235404 | -4.834873906 |
| **A_55_P2329923** | Camkk2 | -0.078527039 | -0.425529767 | 0.26847569 | 11.3702494 | -0.54988202 | 0.601697484 | 0.821235404 | -4.834875843 |
| **A_52_P523445** | Cdhr5 | -0.05869978 | -0.318101992 | 0.200702432 | 7.791032213 | -0.549852227 | 0.601716719 | 0.821235404 | -4.834884201 |
| **A_51_P366161** | Mpg | 0.111134507 | -0.380024932 | 0.602293945 | 11.13653059 | 0.549806185 | 0.601746444 | 0.821235404 | -4.834897115 |
| **A_55_P2031939** | Aqp4 | -0.086940133 | -0.471375766 | 0.2974955 | 8.319468461 | -0.549515384 | 0.601934211 | 0.821433661 | -4.834978661 |
| **A_55_P1966660** | LOC547349 | -0.099965343 | -0.542106265 | 0.342175579 | 11.70900613 | -0.549378851 | 0.60202238 | 0.821483981 | -4.835016935 |
| **A_66_P102047** | Tbc1d13 | 0.054679644 | -0.187188233 | 0.296547522 | 9.220591667 | 0.54932666 | 0.602056086 | 0.821483981 | -4.835031563 |
| **A_51_P483159** | Gchfr | -0.125151578 | -0.678824924 | 0.428521767 | 13.16364292 | -0.549244737 | 0.602108995 | 0.821498183 | -4.835054522 |
| **A_66_P111433** | NA | 0.084111428 | -0.288118845 | 0.4563417 | 10.85516418 | 0.549068467 | 0.602222845 | 0.821565663 | -4.835103913 |
| **A_55_P2059904** | Chsy1 | 0.049687065 | -0.170212958 | 0.269587087 | 10.94489595 | 0.549036548 | 0.602243462 | 0.821565663 | -4.835112855 |
| **A_55_P2015862** | Erg | -0.071996743 | -0.390758065 | 0.246764579 | 8.498337826 | -0.548820621 | 0.602382946 | 0.821572383 | -4.835173336 |
| **A_55_P2003473** | Zc4h2 | -0.041424445 | -0.224834608 | 0.141985719 | 8.033451086 | -0.548803011 | 0.602394322 | 0.821572383 | -4.835178267 |
| **A_55_P2049999** | Slc9a1 | -0.049097771 | -0.266512073 | 0.16831653 | 10.97245026 | -0.548727668 | 0.602442997 | 0.821572383 | -4.835199366 |
| **A_55_P2079684** | NA | 0.05728589 | -0.196401165 | 0.310972945 | 14.26685945 | 0.548696951 | 0.602462842 | 0.821572383 | -4.835207967 |
| **A_55_P1991475** | Sesn1 | -0.049468837 | -0.268586635 | 0.169648961 | 10.8370747 | -0.548576541 | 0.602540638 | 0.821572383 | -4.835241678 |
| **A_55_P2098683** | St7l | 0.054906666 | -0.18830058 | 0.298113911 | 10.53333259 | 0.548569616 | 0.602545112 | 0.821572383 | -4.835243616 |
| **A_55_P1999269** | Grk2 | -0.115009996 | -0.624558084 | 0.394538092 | 11.52342653 | -0.548445656 | 0.602625207 | 0.821572383 | -4.835278315 |
| **A_55_P2161535** | Ddx23 | 0.113132345 | -0.388206775 | 0.614471465 | 8.61552092 | 0.548325427 | 0.602702898 | 0.821572383 | -4.835311962 |
| **A_52_P152952** | Leng1 | -0.050978204 | -0.2768867 | 0.174930291 | 8.436520547 | -0.548321356 | 0.602705529 | 0.821572383 | -4.835313101 |
| **A_52_P593037** | Acsl5 | -0.113375713 | -0.615803068 | 0.389051643 | 12.05241902 | -0.548314767 | 0.602709787 | 0.821572383 | -4.835314945 |
| **A_51_P520106** | Adprhl2 | -0.052921772 | -0.287461025 | 0.181617481 | 9.465379297 | -0.548279524 | 0.602732561 | 0.821572383 | -4.835324807 |
| **A_51_P378558** | Rpap2 | -0.035771815 | -0.194317297 | 0.122773666 | 8.880332554 | -0.548239473 | 0.602758444 | 0.821572383 | -4.835336013 |
| **A_52_P541175** | Marveld1 | -0.055199476 | -0.299930746 | 0.189531793 | 8.867395449 | -0.54806073 | 0.602873963 | 0.821636444 | -4.835386018 |
| **A_55_P2081116** | Fam89a | 0.034862803 | -0.119711405 | 0.189437011 | 7.778776883 | 0.548035206 | 0.602890459 | 0.821636444 | -4.835393158 |
| **A_55_P1992680** | Ubtf | 0.124069045 | -0.42611099 | 0.674249081 | 15.56646728 | 0.547951099 | 0.602944821 | 0.821652598 | -4.835416681 |
| **A_51_P192130** | Stk10 | 0.104518906 | -0.359154476 | 0.568192288 | 10.06973779 | 0.54772918 | 0.60308827 | 0.821770849 | -4.835478735 |
| **A_55_P2094966** | H3c14 | 0.09933061 | -0.341367721 | 0.540028942 | 13.23982708 | 0.547677484 | 0.603121689 | 0.821770849 | -4.835493187 |
| **A_52_P78257** | Plekha2 | -0.050622916 | -0.275244454 | 0.173998623 | 9.267493269 | -0.547619552 | 0.603159141 | 0.821770849 | -4.835509381 |
| **A_55_P2135631** | Rps18 | -0.113840213 | -0.619238331 | 0.391557905 | 15.03825729 | -0.547324979 | 0.603349596 | 0.821942964 | -4.835591703 |
| **A_55_P2093418** | Zmpste24 | 0.094687212 | -0.325705252 | 0.515079676 | 11.33361701 | 0.547292626 | 0.603370517 | 0.821942964 | -4.835600743 |
| **A_55_P2099338** | Akt1s1 | -0.042197784 | -0.229621639 | 0.145226071 | 8.197585944 | -0.547076382 | 0.603510352 | 0.82199974 | -4.835661147 |
| **A_51_P511015** | Fzd9 | -0.03801417 | -0.206869816 | 0.130841476 | 7.933880612 | -0.547032482 | 0.603538743 | 0.82199974 | -4.835673407 |
| **A_51_P229893** | Cts6 | -0.0390311 | -0.212404369 | 0.134342168 | 7.702047619 | -0.547030887 | 0.603539775 | 0.82199974 | -4.835673853 |
| **A_51_P511176** | Triap1 | 0.073284267 | -0.252368037 | 0.398936571 | 10.30091716 | 0.546814197 | 0.603679922 | 0.822132686 | -4.835734357 |
| **A_51_P423290** | Mmrn1 | -0.095700935 | -0.521073331 | 0.32967146 | 9.380027263 | -0.546676077 | 0.603769263 | 0.822196426 | -4.835772913 |
| **A_55_P1958887** | DXBay18 | -0.177474789 | -0.966728559 | 0.61177898 | 9.719528285 | -0.546390568 | 0.603953962 | 0.822286817 | -4.835852584 |
| **A_55_P1982291** | Clca3a1 | -0.087188038 | -0.47493171 | 0.300555635 | 9.964031594 | -0.546380728 | 0.603960329 | 0.822286817 | -4.835855329 |
| **A_52_P326657** | Fam167b | -0.063072764 | -0.343578431 | 0.217432903 | 9.904503445 | -0.54636568 | 0.603970064 | 0.822286817 | -4.835859527 |
| **A_66_P119999** | Atg7 | -0.053377066 | -0.290786726 | 0.184032595 | 9.323954641 | -0.546310437 | 0.604005806 | 0.822286817 | -4.835874938 |

| | | | | | | | | |
|---|---|---|---|---|---|---|---|---|
| **A_51_P423127** | Gba | -0.059544287 | -0.324438206 | 0.205349632 | 9.333653105 | -0.546199501 | 0.604077585 | 0.822326617 | -4.835905881 |
| **A_51_P394115** | Aadac | 0.151756655 | -0.523526788 | 0.827040099 | 15.09911087 | 0.546065619 | 0.604164216 | 0.822379397 | -4.835943218 |
| **A_55_P2108486** | NA | -0.053184066 | -0.289866295 | 0.183498164 | 10.84548857 | -0.546008081 | 0.604201449 | 0.822379397 | -4.835959261 |
| **A_55_P1958921** | Ankrd29 | -0.050266975 | -0.274000373 | 0.173466422 | 8.012976669 | -0.545927691 | 0.604253473 | 0.822392295 | -4.835981674 |
| **A_55_P2129172** | Rab40b | -0.046548044 | -0.253775805 | 0.160679717 | 7.961134451 | -0.545803967 | 0.604333544 | 0.822443362 | -4.836016163 |
| **A_51_P175232** | Fbxl4 | -0.076879822 | -0.419229208 | 0.265469564 | 9.158455044 | -0.54566488 | 0.604423565 | 0.822474276 | -4.836054927 |
| **A_51_P365980** | 2310022B05Rik | -0.105904119 | -0.577523801 | 0.365715564 | 9.98504057 | -0.545637384 | 0.604441362 | 0.822474276 | -4.836062589 |
| **A_55_P2201676** | Cyp2r1 | 0.098665645 | -0.340778438 | 0.538109728 | 9.532377415 | 0.545563777 | 0.604489006 | 0.822481205 | -4.836083099 |
| **A_51_P263965** | Hmox1 | 0.151086357 | -0.522087561 | 0.824260275 | 12.5197945 | 0.545357335 | 0.604622644 | 0.82260513 | -4.836140609 |
| **A_52_P308465** | Plxnb1 | -0.035936518 | -0.196241271 | 0.124368236 | 8.45131761 | -0.544719317 | 0.605035761 | 0.82310925 | -4.83631823 |
| **A_52_P494982** | Dhx40 | 0.041388412 | -0.143268198 | 0.226045021 | 8.254326141 | 0.5446244 | 0.605097233 | 0.823134944 | -4.83634464 |
| **A_55_P2412369** | Ubxn7 | -0.040199547 | -0.219589583 | 0.139190488 | 8.57226718 | -0.544510223 | 0.605171184 | 0.823177609 | -4.836376402 |
| **A_55_P2046084** | Dcun1d4 | -0.07709231 | -0.421188394 | 0.267003774 | 8.678056587 | -0.54439549 | 0.6052455 | 0.823181959 | -4.836408314 |
| **A_51_P494481** | Snx30 | -0.042537397 | -0.2324328 | 0.147358006 | 8.30311438 | -0.544301643 | 0.605306291 | 0.823181959 | -4.836434412 |
| **A_52_P432570** | Chdh | 0.093958431 | -0.325514289 | 0.513431152 | 8.491506361 | 0.544271036 | 0.605326118 | 0.823181959 | -4.836442923 |
| **A_55_P2145982** | NA | 0.036720709 | -0.127225716 | 0.200667133 | 8.095205332 | 0.5442423 | 0.605344734 | 0.823181959 | -4.836450913 |
| **A_52_P229316** | Galnt4 | -0.070618222 | -0.386160303 | 0.244923858 | 8.987445695 | -0.543804359 | 0.605628474 | 0.823509868 | -4.836572638 |
| **A_55_P2045258** | Ehd1 | 0.065277817 | -0.226646151 | 0.357201784 | 9.99472126 | 0.54334923 | 0.605923431 | 0.823826796 | -4.836699052 |
| **A_52_P357469** | Frk | 0.04295753 | -0.149162322 | 0.235077382 | 7.919489193 | 0.543313187 | 0.605946792 | 0.823826796 | -4.836709059 |
| **A_51_P205740** | Ube3b | -0.043228345 | -0.236587888 | 0.150131198 | 11.57021823 | -0.543233054 | 0.605998734 | 0.823839467 | -4.836731306 |
| **A_55_P1975832** | 1810009N02Rik | -0.064207579 | -0.351463912 | 0.223048755 | 9.49576895 | -0.543125094 | 0.606068717 | 0.82387666 | -4.836761273 |
| **A_55_P2002864** | Aifm1 | 0.108047479 | -0.375612797 | 0.591707754 | 12.33344195 | 0.542821959 | 0.606265242 | 0.824043344 | -4.83684539 |
| **A_55_P2109554** | Serinc3 | 0.069116617 | -0.240284779 | 0.378518013 | 12.3870191 | 0.542804441 | 0.606276599 | 0.824043344 | -4.836850249 |
| **A_55_P1963002** | Ppfia4 | -0.048665729 | -0.266551358 | 0.169219901 | 8.6113605 | -0.542722508 | 0.606329724 | 0.824057603 | -4.836872977 |
| **A_51_P429246** | Fbxl6 | -0.038575061 | -0.211317043 | 0.134166921 | 8.988490683 | -0.542615147 | 0.60639934 | 0.824094273 | -4.836902755 |
| **A_55_P2084571** | NA | -0.049881466 | -0.273348691 | 0.173585758 | 16.01104507 | -0.542386118 | 0.606547863 | 0.824238164 | -4.83696626 |
| **A_55_P1987979** | Plrg1 | 0.038944743 | -0.135592134 | 0.213481621 | 11.84541147 | 0.542181669 | 0.606680464 | 0.824310011 | -4.83702293 |
| **A_55_P1983209** | Rcbtb1 | 0.095736625 | -0.333350378 | 0.524823628 | 10.86136811 | 0.542145607 | 0.606703855 | 0.824310011 | -4.837032924 |
| **A_52_P555723** | Ywhag | 0.056510544 | -0.196812536 | 0.309833624 | 10.90025503 | 0.542048214 | 0.606767029 | 0.824310011 | -4.837059912 |
| **A_66_P106606** | 2900097C17Rik | -0.080172862 | -0.439576309 | 0.279230586 | 10.17995581 | -0.542036254 | 0.606774787 | 0.824310011 | -4.837063226 |
| **A_55_P2143968** | Gps1 | -0.056600189 | -0.310358998 | 0.197158621 | 12.48104617 | -0.541975857 | 0.606813966 | 0.824310011 | -4.83707996 |
| **A_55_P2097474** | Creb5 | 0.104984102 | -0.365912136 | 0.575880339 | 15.45176098 | 0.541728276 | 0.606974582 | 0.824470253 | -4.837148538 |
| **A_51_P504815** | Stfa3 | -0.136233894 | -0.747394412 | 0.474926623 | 15.09129142 | -0.541642919 | 0.607029963 | 0.824487538 | -4.837172176 |
| **A_51_P189361** | Osgin1 | -0.16448079 | -0.902468734 | 0.573507154 | 12.27625063 | -0.541563151 | 0.60708172 | 0.824499899 | -4.837194263 |
| **A_55_P2086746** | Fam3a | -0.038655211 | -0.212126208 | 0.134815786 | 8.151457459 | -0.541457485 | 0.607150284 | 0.824512067 | -4.837223516 |
| **A_55_P1974005** | Tnfrsf23 | 0.045755933 | -0.15961936 | 0.251131226 | 7.725386368 | 0.54135532 | 0.607216581 | 0.824512067 | -4.837251795 |
| **A_52_P16563** | Lsm14b | -0.054853582 | -0.301090687 | 0.191383523 | 9.8617377 | -0.541295837 | 0.607255183 | 0.824512067 | -4.837268258 |
| **A_51_P117130** | Naga | -0.048859109 | -0.268200747 | 0.170482529 | 8.203111018 | -0.541262139 | 0.607277051 | 0.824512067 | -4.833588261 |
| **A_51_P320444** | Rock2 | -0.041399086 | -0.227264922 | 0.14446675 | 8.369777611 | -0.541220672 | 0.607303963 | 0.824512067 | -4.837289058 |
| **A_51_P413461** | Parg | -0.074606379 | -0.409665209 | 0.260452452 | 10.31940024 | -0.541050848 | 0.607414183 | 0.824603789 | -4.837336045 |
| **A_52_P303176** | 1810037I17Rik | 0.046543962 | -0.162546527 | 0.25563445 | 9.455902771 | 0.540894115 | 0.607515916 | 0.824641988 | -4.837379399 |
| **A_55_P2181114** | NA | -0.088921699 | -0.488407853 | 0.310564455 | 14.03527134 | -0.540865413 | 0.607534547 | 0.824641988 | -4.837387337 |
| **A_52_P88829** | Ctsq | -0.042146525 | -0.231511646 | 0.147218597 | 7.73174881 | -0.540810319 | 0.607570311 | 0.824641988 | -4.837402573 |
| **A_55_P1970642** | NA | -0.091308189 | -0.501666652 | 0.319050273 | 9.097906981 | -0.540666581 | 0.607663623 | 0.824710726 | -4.837442317 |
| **A_55_P2065334** | Gm14326 | 0.043056144 | -0.15051571 | 0.236627998 | 8.758741912 | 0.540475624 | 0.6077876 | 0.824821072 | -4.837495104 |
| **A_55_P1971966** | Uba1 | 0.049272338 | -0.172345444 | 0.27089012 | 13.68399984 | 0.540233804 | 0.607944621 | 0.824899171 | -4.837561928 |
| **A_55_P2152188** | NA | 0.203866118 | -0.713142053 | 1.120874289 | 16.05201434 | 0.540200568 | 0.607966204 | 0.824899171 | -4.83757111 |
| **A_55_P1977473** | Dab2 | 0.073473399 | -0.257035989 | 0.403982788 | 9.375590419 | 0.540168833 | 0.607986813 | 0.824899171 | -4.837579877 |
| **A_55_P2131577** | NA | -0.04435489 | -0.243895423 | 0.155185644 | 16.95101879 | -0.54012411 | 0.608015856 | 0.824899171 | -4.837592232 |
| **A_51_P394014** | Qprt | 0.105468053 | -0.36927887 | 0.580214977 | 12.35443248 | 0.539811286 | 0.608219028 | 0.825073524 | -4.837678623 |

| | | | | | | | | | |
|---|---|---|---|---|---|---|---|---|---|
| **A_55_P1996818** | NA | -0.081513113 | -0.44844206 | 0.285415834 | 14.48457364 | -0.539794796 | 0.60822974 | 0.825073524 | -4.837683176 |
| **A_55_P2310898** | Kantr | -0.04970591 | -0.273506707 | 0.174094886 | 8.588181064 | -0.539671634 | 0.608309742 | 0.825124141 | -4.837717176 |
| **A_66_P137556** | Tle1 | -0.065937988 | -0.362868229 | 0.230992253 | 8.684431765 | -0.539590691 | 0.608362323 | 0.825131008 | -4.837739518 |
| **A_55_P2012799** | Rtkn | -0.048729485 | -0.268204437 | 0.170745468 | 11.57741474 | -0.539498253 | 0.608422375 | 0.825131008 | -4.837765029 |
| **A_55_P1987043** | Tpp1 | 0.083621207 | -0.293166734 | 0.460409148 | 9.92093199 | 0.539265486 | 0.608573605 | 0.825131008 | -4.837829251 |
| **A_51_P306047** | Sec13 | -0.041774592 | -0.230007991 | 0.146458806 | 10.30892125 | -0.539260644 | 0.608576752 | 0.825131008 | -4.837830587 |
| **A_55_P2185573** | Rbms1 | -0.055997713 | -0.308329871 | 0.196334445 | 12.4363606 | -0.539238482 | 0.608591152 | 0.825131008 | -4.837836701 |
| **A_55_P2041584** | NA | 0.076921188 | -0.269793313 | 0.423635688 | 8.736817341 | 0.539084896 | 0.608690951 | 0.825131008 | -4.837879061 |
| **A_52_P323852** | Tnfrsf22 | 0.036636113 | -0.128501459 | 0.201773685 | 7.871973369 | 0.539071885 | 0.608699406 | 0.825131008 | -4.837882649 |
| **A_55_P2085664** | Meaf6 | 0.05519077 | -0.193595716 | 0.303977257 | 10.63182415 | 0.539042314 | 0.608718621 | 0.825131008 | -4.837890804 |
| **A_55_P2003658** | Gnpda1 | 0.14193265 | -0.49798037 | 0.781845669 | 9.773502916 | 0.538945132 | 0.608781776 | 0.825131008 | -4.8379176 |
| **A_51_P165475** | C1galt1 | -0.055433506 | -0.305448621 | 0.194581608 | 8.975478738 | -0.538752471 | 0.608906989 | 0.825131008 | -4.837970712 |
| **A_55_P2152304** | Atp5h | -0.035347398 | -0.194790044 | 0.124095248 | 15.33290812 | -0.538686567 | 0.608949824 | 0.825131008 | -4.837988876 |
| **A_51_P450812** | Cog4 | -0.06016445 | -0.331563478 | 0.211234578 | 10.81215593 | -0.538660157 | 0.60896699 | 0.825131008 | -4.837996155 |
| **A_51_P406429** | Pdk1 | 0.082164871 | -0.288484251 | 0.452813993 | 10.11803492 | 0.538649662 | 0.608973812 | 0.825131008 | -4.837999047 |
| **A_55_P2078680** | Tpm1 | 0.100143017 | -0.351661423 | 0.551947457 | 9.844552164 | 0.5385839 | 0.609016558 | 0.825131008 | -4.838017169 |
| **A_55_P2164957** | Slc9a3r2 | -0.087062322 | -0.479857691 | 0.305733047 | 8.293743375 | -0.538576096 | 0.60902163 | 0.825131008 | -4.838019319 |
| **A_55_P1997544** | NA | -0.046246197 | -0.25489472 | 0.162402326 | 7.68192651 | -0.538572163 | 0.609024187 | 0.825131008 | -4.838020403 |
| **A_51_P231687** | Ik | -0.042909094 | -0.236538663 | 0.150720474 | 13.11377651 | -0.53846919 | 0.609091124 | 0.825131008 | -4.838048775 |
| **A_66_P138976** | Lpin2 | -0.103510341 | -0.570740139 | 0.363719456 | 9.869443853 | -0.538314896 | 0.609191429 | 0.825131008 | -4.838091279 |
| **A_52_P6798** | Rbm22 | 0.040526185 | -0.142430304 | 0.223482675 | 9.005956358 | 0.538233961 | 0.609244048 | 0.825131008 | -4.83811357 |
| **A_55_P2121364** | Tdg | 0.037262956 | -0.130967535 | 0.205493447 | 8.118253731 | 0.538214979 | 0.609256389 | 0.825131008 | -4.838118797 |
| **A_51_P191893** | 2410022M11Rik | -0.087715267 | -0.483755939 | 0.308325405 | 10.27774617 | -0.538168891 | 0.609286354 | 0.825131008 | -4.838131489 |
| **A_55_P2073940** | Abcc5 | 0.040012297 | -0.140648323 | 0.220672918 | 8.125778065 | 0.538162187 | 0.609290713 | 0.825131008 | -4.838133335 |
| **A_55_P1981775** | NA | 0.077988285 | -0.27414473 | 0.4301213 | 13.4689631 | 0.538153062 | 0.609296646 | 0.825131008 | -4.838135848 |
| **A_55_P1991778** | Fam219a | 0.152812484 | -0.537703328 | 0.843328295 | 14.44887421 | 0.537735112 | 0.609568424 | 0.825370641 | -4.838250899 |
| **A_55_P1985623** | Abcc3 | 0.095181415 | -0.334988219 | 0.525351049 | 10.6593896 | 0.537644982 | 0.609627042 | 0.825370641 | -4.838275699 |
| **A_55_P2026445** | Tfam | 0.062222994 | -0.219003487 | 0.343449474 | 10.25239405 | 0.537623048 | 0.609641307 | 0.825370641 | -4.838281734 |
| **A_55_P2009875** | NA | -0.067246069 | -0.371177809 | 0.236685671 | 11.21006301 | -0.53761829 | 0.609644402 | 0.825370641 | -4.838283043 |
| **A_51_P428754** | Grik5 | -0.068704292 | -0.379308946 | 0.241900361 | 8.416720337 | -0.53747603 | 0.609736931 | 0.825386884 | -4.838322179 |
| **A_52_P351925** | Ank3 | -0.046966565 | -0.259321945 | 0.165388815 | 8.339330281 | -0.537413677 | 0.609777488 | 0.825386884 | -4.838339329 |
| **A_51_P334281** | Gucy1b2 | 0.047770391 | -0.168227728 | 0.263768509 | 7.913275822 | 0.537393009 | 0.609790933 | 0.825386884 | -4.838345014 |
| **A_55_P2034600** | Gm5523 | 0.040548558 | -0.142829539 | 0.223926656 | 8.386977097 | 0.537292949 | 0.609856021 | 0.825386884 | -4.838372531 |
| **A_55_P2026889** | NA | 0.071605284 | -0.252246926 | 0.395457495 | 11.82719808 | 0.537256152 | 0.609879959 | 0.825386884 | -4.838382649 |
| **A_52_P140005** | Nipal1 | -0.167400745 | -0.924581827 | 0.589780337 | 10.03652473 | -0.537205959 | 0.609912612 | 0.825386884 | -4.83839645 |
| **A_55_P2102624** | Eaf2 | 0.036034289 | -0.12704055 | 0.199109129 | 7.869584695 | 0.536923224 | 0.610096561 | 0.825578019 | -4.83847417 |
| **A_55_P2029945** | NA | 0.138527418 | -0.488741609 | 0.765796445 | 15.30260911 | 0.536617814 | 0.610295298 | 0.82578209 | -4.838558083 |
| **A_55_P2058359** | Macroh2a1 | -0.075727303 | -0.418696864 | 0.267242258 | 9.188414127 | -0.536512815 | 0.610363631 | 0.82578209 | -4.838586923 |
| **A_55_P2079659** | NA | -0.118685559 | -0.656231733 | 0.418860615 | 8.720453815 | -0.536494524 | 0.610375535 | 0.82578209 | -4.838591946 |
| **A_51_P518051** | Dusp4 | -0.037715199 | -0.208561909 | 0.133131511 | 7.756307575 | -0.536405192 | 0.610433677 | 0.825802949 | -4.838616478 |
| **A_55_P2003443** | Lsm7 | 0.070220965 | -0.247999334 | 0.388441264 | 14.38634472 | 0.536194188 | 0.610571019 | 0.825930942 | -4.838674409 |
| **A_52_P59457** | Nprl2 | -0.039707538 | -0.219904085 | 0.140489009 | 9.488248815 | -0.535438617 | 0.611062965 | 0.826500647 | -4.838881688 |
| **A_51_P252199** | Mmgt1 | 0.061625826 | -0.218049786 | 0.341301438 | 9.266792064 | 0.535416 | 0.611077694 | 0.826500647 | -4.838887889 |
| **A_55_P2119892** | NA | -0.035200158 | -0.194988312 | 0.124587997 | 7.732867271 | -0.535282708 | 0.611164503 | 0.826560221 | -4.838924428 |
| **A_55_P2397599** | B930025B16Rik | -0.041559436 | -0.230300398 | 0.147181526 | 8.092562441 | -0.535040512 | 0.611322257 | 0.826698569 | -4.8389908 |
| **A_55_P1983758** | Sdhc | -0.054450561 | -0.301758315 | 0.192857194 | 12.48419088 | -0.534992644 | 0.611353438 | 0.826698569 | -4.839003915 |
| **A_52_P459564** | Itga2b | -0.119427295 | -0.661916258 | 0.423061667 | 8.672906686 | -0.534928676 | 0.611395108 | 0.826698569 | -4.839021439 |
| **A_51_P309740** | Usp40 | 0.037921631 | -0.134457817 | 0.210301079 | 8.724916887 | 0.534545538 | 0.611644725 | 0.826905759 | -4.839126364 |
| **A_55_P2025468** | Tst | -0.063114601 | -0.350104994 | 0.223875791 | 12.43771339 | -0.534374435 | 0.611756219 | 0.826905759 | -4.839173201 |
| **A_55_P2091350** | NA | 0.080056959 | -0.283983141 | 0.444097058 | 14.48218936 | 0.534358906 | 0.611766338 | 0.826905759 | -4.839177451 |

| | | | | | | | | | |
|---|---|---|---|---|---|---|---|---|---|
| **A_66_P102510** | Fam114a2 | -0.048862054 | -0.271078871 | 0.173354762 | 8.608588041 | -0.53429116 | 0.611810486 | 0.826905759 | -4.839195992 |
| **A_55_P2424459** | Atg9a | -0.057352041 | -0.318238478 | 0.203534395 | 8.839174823 | -0.534171323 | 0.611888584 | 0.826905759 | -4.839228783 |
| **A_51_P337125** | Inpp5d | 0.097742702 | -0.346892054 | 0.542377458 | 10.68889553 | 0.534151094 | 0.611901768 | 0.826905759 | -4.839234318 |
| **A_52_P500002** | Tspyl5 | -0.034703798 | -0.192577574 | 0.123169979 | 7.710134405 | -0.534133964 | 0.611912933 | 0.826905759 | -4.839239005 |
| **A_55_P2080774** | Qdpr | -0.109896985 | -0.609846079 | 0.390052109 | 11.36858642 | -0.534125338 | 0.611918554 | 0.826905759 | -4.839241364 |
| **A_55_P2320263** | NA | 0.119159887 | -0.422951546 | 0.66127132 | 10.39925234 | 0.534102616 | 0.611933363 | 0.826905759 | -4.839247581 |
| **A_55_P2168431** | Amer1 | -0.056416844 | -0.313190276 | 0.200356589 | 9.433644398 | -0.533877831 | 0.612079878 | 0.827007789 | -4.839309066 |
| **A_55_P2424767** | Hecw1 | 0.128916846 | -0.457854835 | 0.715688527 | 13.7410014 | 0.533855491 | 0.61209444 | 0.827007789 | -4.839315175 |
| **A_51_P389988** | Slc40a1 | 0.126039242 | -0.447782738 | 0.699861222 | 13.10339407 | 0.533717928 | 0.612184114 | 0.827071136 | -4.83935279 |
| **A_55_P2062495** | Smyd1 | -0.033829373 | -0.187875185 | 0.120216439 | 7.685012148 | -0.533614035 | 0.612251844 | 0.827104829 | -4.839381192 |
| **A_51_P221677** | Trim62 | -0.03819197 | -0.212205043 | 0.135821102 | 7.8259205 | -0.533302206 | 0.612455158 | 0.827265482 | -4.839466412 |
| **A_55_P2025103** | Snx3 | 0.039287754 | -0.139718628 | 0.218294135 | 11.53444993 | 0.533300356 | 0.612456364 | 0.827265482 | -4.839466917 |
| **A_55_P1953485** | Copz1 | 0.049714712 | -0.176887078 | 0.276316503 | 14.06513378 | 0.533095211 | 0.61259014 | 0.827342017 | -4.839522958 |
| **A_52_P69558** | Gm8221 | -0.110887995 | -0.616332579 | 0.394556589 | 8.394121809 | -0.53308219 | 0.612598632 | 0.827342017 | -4.839526515 |
| **A_55_P2017804** | Cox5b | 0.050603813 | -0.180096004 | 0.281303629 | 14.74744998 | 0.532990144 | 0.612658661 | 0.82736528 | -4.839551653 |
| **A_55_P2126470** | Rpl7 | 0.065094705 | -0.23170671 | 0.36189612 | 15.0711264 | 0.532921066 | 0.612703713 | 0.827368315 | -4.839570516 |
| **A_51_P502152** | Slc19a1 | 0.059535056 | -0.212007644 | 0.331077756 | 10.19299075 | 0.532743086 | 0.6128198 | 0.827439591 | -4.839619107 |
| **A_55_P2086153** | Bcdin3d | -0.048383317 | -0.269121201 | 0.172354567 | 10.12552081 | -0.532600972 | 0.612912501 | 0.827439591 | -4.839657895 |
| **A_55_P2103857** | Hgs | -0.057986994 | -0.32254289 | 0.206568902 | 11.73360539 | -0.532594087 | 0.612916993 | 0.827439591 | -4.839659775 |
| **A_51_P210956** | Vcam1 | -0.112009813 | -0.623133551 | 0.399113926 | 13.37275767 | -0.532492149 | 0.612983493 | 0.827439591 | -4.839687592 |
| **A_55_P1998001** | DXBay18 | -0.139229482 | -0.774692798 | 0.496233834 | 9.15989979 | -0.532382671 | 0.613054915 | 0.827439591 | -4.839717462 |
| **A_51_P186899** | Egln1 | 0.036743409 | -0.130969147 | 0.204455964 | 7.853746393 | 0.532349753 | 0.613076392 | 0.827439591 | -4.839726442 |
| **A_51_P380013** | Bcl2 | -0.08476209 | -0.47165584 | 0.30213166 | 8.402807565 | -0.532344939 | 0.613079533 | 0.827439591 | -4.839727755 |
| **A_55_P2169721** | Rps17 | 0.04837476 | -0.172473305 | 0.269222825 | 15.51621201 | 0.532241105 | 0.61314728 | 0.827439591 | -4.839756078 |
| **A_55_P2002632** | Rap1b | 0.041990426 | -0.149720593 | 0.233701445 | 13.30098093 | 0.532214119 | 0.613164888 | 0.827439591 | -4.839763439 |
| **A_55_P1973683** | Morn4 | -0.12342523 | -0.686966033 | 0.440115574 | 8.654340546 | -0.532183942 | 0.613184577 | 0.827439591 | -4.839771669 |
| **A_55_P1974788** | Cfl1 | -0.095419833 | -0.531367282 | 0.340527617 | 11.0457858 | -0.531848206 | 0.613403665 | 0.827619854 | -4.839863208 |
| **A_52_P485766** | Zdhhc17 | -0.060744638 | -0.338293365 | 0.21680409 | 8.459954962 | -0.53180436 | 0.613432281 | 0.827619854 | -4.839875159 |
| **A_52_P563375** | Lgals2 | 0.21603236 | -0.771085476 | 1.203150196 | 10.33104882 | 0.531781225 | 0.61344738 | 0.827619854 | -4.839881465 |
| **A_55_P1982747** | Slc8a3 | 0.130414156 | -0.465599105 | 0.726427416 | 10.52924688 | 0.531682053 | 0.613512106 | 0.827619854 | -4.839908492 |
| **A_66_P118513** | 11-Sep | -0.054728813 | -0.304862642 | 0.195405016 | 11.86243053 | -0.531651189 | 0.613532251 | 0.827619854 | -4.839916902 |
| **A_55_P1970929** | Hnf1a | 0.147009511 | -0.525248448 | 0.819267469 | 10.58583575 | 0.531364662 | 0.613719283 | 0.827814377 | -4.83999496 |
| **A_55_P2044091** | Tdg | -0.086952288 | -0.484631848 | 0.310727272 | 8.598892059 | -0.531289134 | 0.613768589 | 0.827823115 | -4.840015529 |
| **A_55_P2031032** | NA | 0.058473209 | -0.208994291 | 0.325940709 | 10.93209524 | 0.531213495 | 0.613817971 | 0.827831954 | -4.840036127 |
| **A_55_P2426272** | Scaper | -0.037019074 | -0.206394423 | 0.132356275 | 8.023168424 | -0.53107828 | 0.613906252 | 0.82789325 | -4.840072941 |
| **A_51_P449824** | NA | -0.045079079 | -0.251402473 | 0.161244315 | 8.504815199 | -0.530896364 | 0.614025035 | 0.827966699 | -4.840122458 |
| **A_55_P1968276** | Tomm22 | 0.036980158 | -0.132285538 | 0.206245855 | 12.62066312 | 0.530863665 | 0.614046388 | 0.827966699 | -4.840131357 |
| **A_51_P320481** | Mcee | -0.081394297 | -0.454141175 | 0.291352581 | 10.97863926 | -0.530594998 | 0.614221842 | 0.828145507 | -4.840204456 |
| **A_51_P372312** | Mrpl54 | -0.12759181 | -0.712215612 | 0.457031991 | 12.21562779 | -0.530309608 | 0.614408248 | 0.828246832 | -4.84028207 |
| **A_52_P25812** | Prdm2 | 0.043576487 | -0.156095754 | 0.243248728 | 8.169362111 | 0.530295224 | 0.614417644 | 0.828246832 | -4.840285981 |
| **A_55_P1998115** | Klkb1 | -0.160234953 | -0.894559524 | 0.574089618 | 12.41448378 | -0.53021546 | 0.614469749 | 0.828246832 | -4.840307666 |
| **A_52_P273594** | Zc3h14 | -0.051694563 | -0.288604047 | 0.185214922 | 9.814533265 | -0.530207183 | 0.614475156 | 0.828246832 | -4.840309917 |
| **A_55_P2062737** | Elovl2 | -0.195420924 | -1.091178123 | 0.700336276 | 12.18384941 | -0.530107493 | 0.614540281 | 0.828246832 | -4.840337015 |
| **A_51_P150120** | Mgst2 | 0.110045756 | -0.39439465 | 0.614486163 | 12.35390113 | 0.530086351 | 0.614554093 | 0.828246832 | -4.840342761 |
| **A_52_P297457** | 0610030E20Rik | 0.054887094 | -0.196781759 | 0.306555947 | 9.994980656 | 0.529936652 | 0.614651896 | 0.828320888 | -4.840383444 |
| **A_55_P2025601** | Lsm6 | -0.039960914 | -0.223242656 | 0.143320829 | 14.17302821 | -0.529784678 | 0.614751194 | 0.828396949 | -4.840424734 |
| **A_55_P1963184** | NA | -0.049223937 | -0.275065763 | 0.17661789 | 8.224479565 | -0.529608753 | 0.614866153 | 0.8284941 | -4.84047252 |
| **A_55_P2029533** | Zfp57 | 0.03840272 | -0.137852165 | 0.214657605 | 8.019304888 | 0.529424474 | 0.614986584 | 0.828598611 | -4.840522559 |
| **A_55_P2031631** | Igf1 | -0.039810099 | -0.222551639 | 0.142931441 | 10.23898755 | -0.529345426 | 0.615038247 | 0.82861046 | -4.840544019 |
| **A_55_P2121537** | NA | 0.051366188 | -0.184502427 | 0.287234804 | 16.40437597 | 0.529164098 | 0.615156767 | 0.828712374 | -4.840593237 |

| | | | | | | | | |
|---|---|---|---|---|---|---|---|---|
| **A_55_P2046877** | Foxq1 | -0.183001855 | -1.023636445 | 0.657632735 | 10.35595294 | -0.528970467 | 0.615283343 | 0.828801942 | -4.840645777 |
| **A_55_P2070406** | NA | 0.041881255 | -0.150525368 | 0.234287879 | 8.057812661 | 0.528911318 | 0.615322011 | 0.828801942 | -4.840661823 |
| **A_51_P514256** | Tubb2b | -0.060225447 | -0.336931009 | 0.216480115 | 8.251935212 | -0.528865616 | 0.61535189 | 0.828801942 | -4.840674221 |
| **A_55_P2060343** | Zfp352 | 0.135819258 | -0.488451196 | 0.760089712 | 9.92509492 | 0.528654291 | 0.615490056 | 0.828930274 | -4.840731533 |
| **A_52_P658034** | Tspyl1 | 0.046877692 | -0.168618165 | 0.262373549 | 11.3250804 | 0.528579707 | 0.615538824 | 0.828938197 | -4.840751756 |
| **A_51_P200144** | Pak1ip1 | 0.082840113 | -0.298229027 | 0.463909253 | 12.15691375 | 0.528226386 | 0.615769879 | 0.829191583 | -4.840847523 |
| **A_55_P2000909** | Atp5l | -0.040814668 | -0.228852956 | 0.147223621 | 15.30344335 | -0.527415834 | 0.61630012 | 0.829829628 | -4.841067011 |
| **A_51_P378789** | Cxcl13 | -0.040526454 | -0.227252836 | 0.146199928 | 7.675476786 | -0.527370837 | 0.616329564 | 0.829829628 | -4.841079187 |
| **A_66_P123683** | Nabp1 | -0.037104448 | -0.208096856 | 0.13388796 | 7.992563777 | -0.52726912 | 0.616396124 | 0.829833256 | -4.841106708 |
| **A_55_P2119011** | Osbpl9 | 0.06109136 | -0.220469017 | 0.342651738 | 8.640118881 | 0.527219478 | 0.616428609 | 0.829833256 | -4.841120138 |
| **A_51_P419017** | Pde10a | -0.038160459 | -0.214075231 | 0.137754314 | 7.910786095 | -0.527101757 | 0.616505649 | 0.829833256 | -4.841151981 |
| **A_65_P09657** | Zfp213 | 0.038786783 | -0.140019074 | 0.217592641 | 9.118976921 | 0.527090527 | 0.616512999 | 0.829833256 | -4.841155018 |
| **A_51_P490305** | Ifi30 | 0.042308313 | -0.152785348 | 0.237401975 | 11.13433115 | 0.526945551 | 0.616607883 | 0.829833256 | -4.841194224 |
| **A_55_P2005920** | NA | -0.052196078 | -0.292917693 | 0.188525538 | 14.50028328 | -0.52687303 | 0.61665535 | 0.829833256 | -4.841213833 |
| **A_51_P479894** | Cpsf3 | -0.05700308 | -0.31990432 | 0.20589816 | 11.24075763 | -0.526852296 | 0.616668921 | 0.829833256 | -4.841219438 |
| **A_55_P2040777** | Lipo3 | -0.083796281 | -0.470301501 | 0.302708939 | 8.904751375 | -0.526808252 | 0.61669775 | 0.829833256 | -4.841231346 |
| **A_51_P496400** | Med9 | 0.06877783 | -0.248517399 | 0.386073059 | 10.85327027 | 0.526705818 | 0.616764802 | 0.829833256 | -4.841259035 |
| **A_55_P2050932** | Tlr7 | 0.069021181 | -0.249401762 | 0.387444124 | 8.796917421 | 0.526697461 | 0.616770272 | 0.829833256 | -4.841261294 |
| **A_55_P2127692** | Timm50 | 0.070448506 | -0.25459155 | 0.395488563 | 9.055548298 | 0.526645157 | 0.616804511 | 0.829833256 | -4.84127543 |
| **A_55_P1996698** | Gm16381 | -0.079783741 | -0.448005954 | 0.288438473 | 12.87376397 | -0.526486936 | 0.616908091 | 0.829914845 | -4.841318186 |
| **A_55_P2147487** | Cyth3 | 0.09108417 | -0.329667909 | 0.51183625 | 9.523023128 | 0.526017054 | 0.617215758 | 0.830172505 | -4.841445097 |
| **A_55_P2099695** | Msantd4 | 0.052727592 | -0.190844963 | 0.296300146 | 10.55458045 | 0.526008513 | 0.617221352 | 0.830172505 | -4.841447403 |
| **A_55_P2362601** | D4Ertd681e | 0.070341888 | -0.254606911 | 0.395290688 | 8.614433605 | 0.525995797 | 0.617229679 | 0.830172505 | -4.841450836 |
| **A_55_P1962404** | Ugt1a6b | -0.119561944 | -0.672010722 | 0.432886834 | 12.46865126 | -0.525876956 | 0.617307507 | 0.830172505 | -4.841482917 |
| **A_66_P126094** | Pycrl | 0.060135869 | -0.217733893 | 0.338005631 | 9.619343426 | 0.525866481 | 0.617314368 | 0.830172505 | -4.841485744 |
| **A_55_P1989419** | AI987944 | 0.044847057 | -0.162457596 | 0.252151709 | 8.263613894 | 0.525663818 | 0.617447104 | 0.830245489 | -4.841540438 |
| **A_55_P2001608** | Klhl4 | -0.042332414 | -0.238039932 | 0.153375104 | 7.940277194 | -0.525591967 | 0.617494168 | 0.830245489 | -4.841559824 |
| **A_55_P2124498** | Il17rb | 0.097823845 | -0.354515404 | 0.550163094 | 9.029558672 | 0.525489019 | 0.617561604 | 0.830245489 | -4.841587597 |
| **A_55_P2126259** | Slc35e2 | 0.053550008 | -0.194078362 | 0.301178377 | 8.281080488 | 0.525463225 | 0.6175785 | 0.830245489 | -4.841594554 |
| **A_51_P185906** | Abi3 | 0.10025869 | -0.363368692 | 0.563886073 | 9.497481632 | 0.525455737 | 0.617583405 | 0.830245489 | -4.841596574 |
| **A_55_P1988774** | Slc17a5 | 0.123017813 | -0.446537202 | 0.692572828 | 10.77541423 | 0.524826203 | 0.617995877 | 0.830718904 | -4.841766293 |
| **A_55_P1961913** | Pygm | -0.093542852 | -0.526742294 | 0.339656591 | 8.727776016 | -0.524693767 | 0.618082669 | 0.830718904 | -4.841801975 |
| **A_55_P2045066** | Eif5a | 0.042939094 | -0.155925504 | 0.241803691 | 13.39596987 | 0.524660763 | 0.618104299 | 0.830718904 | -4.841810866 |
| **A_55_P2089253** | Atg4c | 0.063200318 | -0.229503151 | 0.355903786 | 9.349364464 | 0.524655924 | 0.61810747 | 0.830718904 | -4.841812169 |
| **A_55_P2006504** | Eif4a2 | -0.10953557 | -0.616973296 | 0.397902156 | 12.39315279 | -0.524512217 | 0.618201657 | 0.8307701 | -4.841850876 |
| **A_55_P2125013** | H4c11 | 0.083275603 | -0.302562879 | 0.469114086 | 12.12126075 | 0.524439549 | 0.618249288 | 0.8307701 | -4.841870445 |
| **A_55_P2232754** | Smarcal1 | 0.03791081 | -0.137758568 | 0.213580187 | 8.527485335 | 0.524384914 | 0.618285101 | 0.8307701 | -4.841885157 |
| **A_51_P205779** | Cd5l | 0.06199988 | -0.225319399 | 0.349319159 | 13.73329522 | 0.52433551 | 0.618317485 | 0.8307701 | -4.841898459 |
| **A_55_P2420724** | Srek1 | 0.036754697 | -0.133622516 | 0.207131909 | 8.14426755 | 0.524184924 | 0.618416199 | 0.830825518 | -4.841938997 |
| **A_51_P175424** | Car14 | 0.121111304 | -0.440349393 | 0.682572002 | 10.37912683 | 0.524141454 | 0.618444697 | 0.830825518 | -4.841950697 |
| **A_55_P2088995** | Plscr1 | -0.063431571 | -0.357572468 | 0.230709325 | 10.37580331 | -0.524002365 | 0.618535885 | 0.830847676 | -4.841988128 |
| **A_52_P1037027** | Apela | -0.034319229 | -0.193467425 | 0.124828967 | 7.749315949 | -0.523985168 | 0.61854716 | 0.830847676 | -4.841992756 |
| **A_55_P2064351** | Vipr1 | -0.142846318 | -0.805796795 | 0.520104158 | 8.985215627 | -0.52356566 | 0.618822242 | 0.831107594 | -4.842105598 |
| **A_52_P651248** | Zdhhc2 | -0.037328895 | -0.210574713 | 0.135916923 | 8.266919059 | -0.523558923 | 0.618826659 | 0.831107594 | -4.842107409 |
| **A_52_P635271** | Btbd6 | -0.060633738 | -0.342151483 | 0.220884006 | 8.431218496 | -0.523349436 | 0.618964051 | 0.831162568 | -4.842163728 |
| **A_55_P2082045** | Vps13a | 0.061128438 | -0.222703095 | 0.344959971 | 8.462784164 | 0.523318207 | 0.618984533 | 0.831162568 | -4.842172122 |
| **A_51_P220682** | Retreg3 | -0.04606275 | -0.259971138 | 0.167845638 | 10.35316302 | -0.523245176 | 0.619032436 | 0.831162568 | -4.84219175 |
| **A_55_P2074371** | Lin28a | 0.102261304 | -0.372634511 | 0.577157119 | 8.870954057 | 0.523234261 | 0.619039595 | 0.831162568 | -4.842194684 |
| **A_52_P634090** | Jag1 | -0.07490005 | -0.422806293 | 0.273006193 | 8.438165548 | -0.523122111 | 0.61911316 | 0.831203603 | -4.84222482 |
| **A_51_P128775** | Zfp84 | 0.046069012 | -0.167959565 | 0.260097588 | 8.678397651 | 0.523022438 | 0.619178544 | 0.831208986 | -4.811673983 |

| | | | | | | | | | |
|---|---|---|---|---|---|---|---|---|---|
| **A_51_P414706** | Ddt | -0.08678286 | -0.489989806 | 0.316424086 | 14.66796436 | -0.522984891 | 0.619203176 | 0.831208986 | -4.842261686 |
| **A_55_P2048671** | Rpl27a | 0.057763442 | -0.210746334 | 0.326273217 | 16.4265366 | 0.522728463 | 0.619371412 | 0.831377085 | -4.842330557 |
| **A_55_P2140201** | Dupd1 | -0.035903252 | -0.202856177 | 0.131049673 | 7.703395019 | -0.522544091 | 0.619492389 | 0.831481729 | -4.842380056 |
| **A_55_P1997628** | NA | -0.07184438 | -0.406045214 | 0.262356454 | 14.85765712 | -0.522358256 | 0.619614339 | 0.831531089 | -4.842429933 |
| **A_55_P2176894** | NA | 0.044530808 | -0.162625612 | 0.251687227 | 7.999680734 | 0.522330474 | 0.619632571 | 0.831531089 | -4.842437388 |
| **A_52_P468033** | Setd1b | 0.052104131 | -0.190301376 | 0.294509638 | 10.47754071 | 0.522291386 | 0.619658224 | 0.831531089 | -4.842447877 |
| **A_55_P2052775** | Morc2a | -0.042915287 | -0.242620888 | 0.156790315 | 10.10245447 | -0.522161634 | 0.619743382 | 0.831587632 | -4.842482689 |
| **A_55_P2425190** | Nhsl2 | -0.056793147 | -0.321191289 | 0.207604994 | 8.196528303 | -0.521940176 | 0.619888743 | 0.831683212 | -4.842542087 |
| **A_55_P2190152** | 4921509J17Rik | 0.033523354 | -0.122549849 | 0.189596557 | 7.993755874 | 0.521918039 | 0.619903274 | 0.831683212 | -4.842548023 |
| **A_66_P104527** | Tbl1x | -0.079071088 | -0.447264685 | 0.289122509 | 10.51638043 | -0.521824749 | 0.619964515 | 0.831683212 | -4.842573038 |
| **A_55_P2159485** | NA | -0.064587407 | -0.365357412 | 0.236182598 | 9.105371008 | -0.521790917 | 0.619986725 | 0.831683212 | -4.842582108 |
| **A_51_P292196** | Prl7d1 | -0.033971411 | -0.192217194 | 0.124274373 | 7.675593638 | -0.521632477 | 0.620090742 | 0.831716496 | -4.84262458 |
| **A_51_P209818** | Prtn3 | -0.083991868 | -0.475450876 | 0.307467141 | 10.11709362 | -0.521355712 | 0.620272464 | 0.831716496 | -4.842698744 |
| **A_55_P2427377** | NA | 0.147923751 | -0.541559888 | 0.837407391 | 9.053552188 | 0.521311296 | 0.62030163 | 0.831716496 | -4.842710643 |
| **A_55_P2169059** | Strada | 0.040607526 | -0.148674438 | 0.229889491 | 10.15476215 | 0.521291314 | 0.620314752 | 0.831716496 | -4.842715996 |
| **A_66_P102202** | Gm648 | -0.033334443 | -0.188733993 | 0.122065106 | 7.669796364 | -0.521226682 | 0.620357195 | 0.831716496 | -4.842733308 |
| **A_55_P2097869** | Frem2 | -0.127991689 | -0.724738662 | 0.468755283 | 8.892851207 | -0.521164393 | 0.620398101 | 0.831716496 | -4.842749991 |
| **A_52_P124472** | Kcnd2 | -0.035125283 | -0.198906772 | 0.128656206 | 7.716595876 | -0.521120562 | 0.620426885 | 0.831716496 | -4.842761729 |
| **A_55_P2088715** | Ss18l2 | 0.067538434 | -0.247421713 | 0.382498582 | 9.762928229 | 0.521049007 | 0.620473879 | 0.831716496 | -4.84278089 |
| **A_55_P1972257** | Sars | 0.070119359 | -0.256883201 | 0.39712192 | 12.44237379 | 0.52103869 | 0.620480655 | 0.831716496 | -4.842783653 |
| **A_52_P48689** | Bzw1 | 0.055262244 | -0.202462725 | 0.312987213 | 9.718331323 | 0.521020978 | 0.620492288 | 0.831716496 | -4.842788396 |
| **A_55_P1974542** | Gxylt2 | 0.169024104 | -0.619313794 | 0.957362003 | 12.77884738 | 0.520978051 | 0.620520481 | 0.831716496 | -4.842799889 |
| **A_52_P186672** | NA | 0.062613392 | -0.229425303 | 0.354652087 | 8.724222224 | 0.52096677 | 0.620527891 | 0.831716496 | -4.842802909 |
| **A_55_P2082050** | Ankrd12 | -0.046580206 | -0.263936413 | 0.170776 | 8.199703883 | -0.52072994 | 0.62068345 | 0.831865866 | -4.842866304 |
| **A_52_P279557** | NA | 0.033125141 | -0.121480297 | 0.187730578 | 7.677556299 | 0.520614373 | 0.620759366 | 0.831865866 | -4.842897229 |
| **A_55_P2108599** | Tlcd1 | 0.065893462 | -0.241660189 | 0.373447113 | 9.311002484 | 0.520600555 | 0.620768444 | 0.831865866 | -4.842900926 |
| **A_55_P1976310** | Tbx20 | -0.034314685 | -0.194514321 | 0.125884951 | 7.702073409 | -0.520477162 | 0.620849508 | 0.831916821 | -4.842933939 |
| **A_55_P2172599** | Fam174a | -0.086220519 | -0.488824381 | 0.316383344 | 10.1412331 | -0.520374352 | 0.620917054 | 0.831949656 | -4.842961439 |
| **A_55_P2162988** | Tmem179 | -0.037303742 | -0.211548097 | 0.136940614 | 7.720564459 | -0.520207807 | 0.621026483 | 0.8320386 | -4.843005978 |
| **A_55_P2032558** | NA | -0.06536137 | -0.370845114 | 0.240122373 | 14.88170146 | -0.519895709 | 0.621231575 | 0.832217634 | -4.843089408 |
| **A_55_P2038452** | NA | -0.057806451 | -0.327991855 | 0.212378952 | 9.317050728 | -0.519873422 | 0.621246222 | 0.832217634 | -4.843095364 |
| **A_51_P235117** | Zc3h11a | -0.060217201 | -0.341714379 | 0.221279977 | 13.04014238 | -0.519792141 | 0.621299643 | 0.832221654 | -4.843117085 |
| **A_51_P169327** | Etd | 0.034722935 | -0.127630518 | 0.197076388 | 7.711744665 | 0.519682495 | 0.621371709 | 0.832221654 | -4.84314638 |
| **A_55_P2101739** | NA | -0.054827097 | -0.311202188 | 0.201547995 | 13.60405316 | -0.519640044 | 0.621399612 | 0.832221654 | -4.843157721 |
| **A_55_P2158350** | Gm15427 | 0.036761665 | -0.135168228 | 0.208691559 | 16.17580081 | 0.519549547 | 0.621459097 | 0.832221654 | -4.843181894 |
| **A_55_P2017710** | Igsf9 | -0.075740706 | -0.430056563 | 0.27857515 | 8.599964263 | -0.519423927 | 0.621541674 | 0.832221654 | -4.843215443 |
| **A_55_P1985768** | Ttyh2 | 0.102957746 | -0.378711251 | 0.584626743 | 10.65923518 | 0.519389924 | 0.621564027 | 0.832221654 | -4.843224523 |
| **A_52_P339742** | Cyb5r3 | -0.080088132 | -0.454779282 | 0.294603018 | 8.51473599 | -0.519371264 | 0.621576294 | 0.832221654 | -4.843229505 |
| **A_51_P156243** | Eps15 | 0.055831476 | -0.205388558 | 0.317051511 | 9.124554874 | 0.519344836 | 0.621593668 | 0.832221654 | -4.843236562 |
| **A_55_P2099379** | Tfe3 | 0.041965764 | -0.15442743 | 0.238358958 | 9.035137047 | 0.519220574 | 0.621675363 | 0.83224576 | -4.843269737 |
| **A_55_P2129286** | Zc3hc1 | -0.071319292 | -0.405104318 | 0.262465734 | 9.558704786 | -0.519186466 | 0.621697787 | 0.83224576 | -4.843278842 |
| **A_66_P137475** | Dnajb14 | -0.066976315 | -0.380733947 | 0.246781318 | 10.93248849 | -0.5186927 | 0.622022471 | 0.832566163 | -4.84341059 |
| **A_55_P2058165** | Morn4 | -0.087987842 | -0.500189391 | 0.324213707 | 8.354402244 | -0.5186761 | 0.622033388 | 0.832566163 | -4.843415017 |
| **A_66_P101538** | Lrrk2 | -0.085149452 | -0.48409239 | 0.313793486 | 8.620444251 | -0.518625978 | 0.622066352 | 0.832566163 | -4.843428384 |
| **A_52_P679152** | Clk1 | -0.034350473 | -0.19534297 | 0.126642023 | 10.14348871 | -0.518454057 | 0.622179427 | 0.832649606 | -4.843474226 |
| **A_52_P95740** | Smim19 | -0.041000429 | -0.233195233 | 0.151194375 | 9.230568061 | -0.518358171 | 0.622242498 | 0.832649606 | -4.843499788 |
| **A_55_P1956482** | NA | 0.18334038 | -0.707713123 | 1.074393884 | 12.11670479 | 0.523475924 | 0.622266202 | 0.832649606 | -4.799755116 |
| **A_52_P39083** | Ccbe1 | -0.045047152 | -0.256247733 | 0.166153429 | 8.256270411 | -0.518269221 | 0.622301009 | 0.832649606 | -4.843523497 |
| **A_52_P66371** | Nlrx1 | -0.049000377 | -0.278833219 | 0.180832465 | 8.449861162 | -0.518048622 | 0.622446134 | 0.832760618 | -4.84358228 |
| **A_55_P2141878** | Ldha | -0.122960942 | -0.69981351 | 0.453891626 | 13.38080174 | -0.517947262 | 0.622512821 | 0.832760618 | -4.843609283 |

| | | | | | | | | | |
|---|---|---|---|---|---|---|---|---|---|
| **A_55_P2184113** | Aurkaip1 | 0.055846407 | -0.206171694 | 0.317864509 | 14.60064715 | 0.517901457 | 0.622542958 | 0.832760618 | -4.843621484 |
| **A_55_P2127497** | Trim35 | -0.143487119 | -0.816721039 | 0.529746801 | 11.29717956 | -0.517881163 | 0.62255631 | 0.832760618 | -4.843626889 |
| **A_52_P185343** | Gna13 | 0.041531701 | -0.153490528 | 0.236553929 | 9.343927122 | 0.517462389 | 0.622831883 | 0.833071585 | -4.843738389 |
| **A_55_P1994204** | Ythdc1 | -0.045102823 | -0.256924397 | 0.166718752 | 11.73676527 | -0.517388438 | 0.622880554 | 0.833079036 | -4.843758071 |
| **A_55_P1966432** | Gstm1 | -0.08184832 | -0.466320155 | 0.302623516 | 11.68950765 | -0.517283262 | 0.622949778 | 0.833113973 | -4.843786059 |
| **A_51_P484054** | F7 | 0.107375646 | -0.397248487 | 0.61199978 | 9.27781371 | 0.517036215 | 0.623112394 | 0.833273797 | -4.843851779 |
| **A_55_P2152836** | NA | 0.123283705 | -0.456419725 | 0.702987136 | 11.01022142 | 0.516752988 | 0.623298854 | 0.833458494 | -4.843927091 |
| **A_55_P2033010** | Mrrf | 0.036315344 | -0.13448698 | 0.207117667 | 7.88806149 | 0.516629943 | 0.623379869 | 0.833458494 | -4.843959798 |
| **A_55_P2176292** | NA | -0.054312033 | -0.309803607 | 0.201179541 | 13.9096635 | -0.516538464 | 0.623440105 | 0.833458494 | -4.84398411 |
| **A_51_P340829** | AA986860 | -0.143385469 | -0.817905191 | 0.531134253 | 9.479617069 | -0.516527773 | 0.623447144 | 0.833458494 | -4.843986951 |
| **A_55_P1972018** | H4c3 | 0.112811909 | -0.417912023 | 0.643535841 | 11.62826627 | 0.516498977 | 0.623466106 | 0.833458494 | -4.843994603 |
| **A_51_P416546** | Ctsg | 0.067853605 | -0.251420161 | 0.387127372 | 12.88407713 | 0.516407906 | 0.623526077 | 0.83348102 | -4.844018802 |
| **A_55_P2115981** | Xlr5c | -0.034849571 | -0.19887129 | 0.129172148 | 7.685991642 | -0.516272826 | 0.623615034 | 0.833542285 | -4.844054686 |
| **A_55_P1983903** | Thap4 | 0.039063156 | -0.144877085 | 0.223003397 | 7.976371238 | 0.516028594 | 0.623775891 | 0.83369964 | -4.844119548 |
| **A_55_P2060707** | Il15ra | 0.151359895 | -0.561670151 | 0.864389941 | 13.13566924 | 0.515805746 | 0.623922684 | 0.833800823 | -4.844178706 |
| **A_51_P140797** | Dcaf6 | 0.05087365 | -0.188793946 | 0.290541246 | 8.918347222 | 0.515782692 | 0.62393787 | 0.833800823 | -4.844184825 |
| **A_52_P35961** | NA | -0.043514079 | -0.248541546 | 0.161513389 | 8.182706947 | -0.515704516 | 0.623989371 | 0.833811998 | -4.844205572 |
| **A_51_P503261** | Lrrc8c | 0.034230293 | -0.127077202 | 0.195537787 | 8.699535736 | 0.515631282 | 0.624037618 | 0.833818825 | -4.844225005 |
| **A_55_P2065716** | Glud1 | 0.093654476 | -0.34778796 | 0.535096912 | 14.94481738 | 0.515510657 | 0.624117091 | 0.833867371 | -4.844257009 |
| **A_51_P199866** | Pin1 | 0.084934158 | -0.315572252 | 0.485440568 | 11.58399957 | 0.515295207 | 0.624259052 | 0.833947526 | -4.844314153 |
| **A_55_P2075884** | Senp8 | 0.035253023 | -0.131007305 | 0.20151335 | 8.014086762 | 0.515217871 | 0.624310014 | 0.833947526 | -4.844334661 |
| **A_55_P2131979** | 1-Mar | 0.134638845 | -0.500392269 | 0.76966996 | 10.03339499 | 0.515179488 | 0.624335308 | 0.833947526 | -4.844344838 |
| **A_55_P2106803** | Bglap2 | 0.038292798 | -0.14232481 | 0.218910405 | 7.767018863 | 0.515157703 | 0.624349664 | 0.833947526 | -4.844350613 |
| **A_52_P69789** | Exosc1 | 0.062369814 | -0.231982106 | 0.356721733 | 8.73528295 | 0.514861909 | 0.624544608 | 0.834104095 | -4.844429016 |
| **A_51_P141071** | Cand2 | 0.050438327 | -0.187621614 | 0.288498268 | 8.375546665 | 0.514822523 | 0.624570568 | 0.834104095 | -4.844439453 |
| **A_51_P245525** | NA | 0.075945547 | -0.28257448 | 0.434465574 | 15.66121323 | 0.514721235 | 0.624637332 | 0.834104095 | -4.84446629 |
| **A_55_P2008538** | Chmp4b | -0.325664864 | -1.863156846 | 1.211827118 | 12.32808879 | -0.514684672 | 0.624661433 | 0.834104095 | -4.844475976 |
| **A_51_P129866** | Atxn1l | -0.069217813 | -0.396021528 | 0.257585902 | 9.981791835 | -0.51465249 | 0.624682647 | 0.834104095 | -4.844484501 |
| **A_55_P1958434** | Got1 | -0.144619599 | -0.827538307 | 0.538299108 | 11.93060708 | -0.514566291 | 0.62473947 | 0.834122347 | -4.844507333 |
| **A_51_P180452** | Crk | -0.060100384 | -0.343982154 | 0.223781386 | 10.53567131 | -0.514426023 | 0.624831941 | 0.834188189 | -4.84454448 |
| **A_55_P1999197** | Clasp1 | 0.035308851 | -0.131593008 | 0.20221071 | 10.1375531 | 0.514050281 | 0.625079686 | 0.834369656 | -4.844643944 |
| **A_55_P2072453** | 1-Mar | -0.095313157 | -0.545859109 | 0.355232795 | 13.62171356 | -0.514040044 | 0.625086436 | 0.834369656 | -4.844646653 |
| **A_52_P10254** | Psmc4 | 0.057405038 | -0.213957724 | 0.328767801 | 12.09038949 | 0.514023469 | 0.625097366 | 0.834369656 | -4.844651039 |
| **A_51_P271665** | Rev3l | -0.0970542 | -0.555950513 | 0.361842113 | 10.42282859 | -0.513905131 | 0.625175404 | 0.834416198 | -4.84468235 |
| **A_51_P308814** | Toe1 | 0.051459587 | -0.192019811 | 0.294938986 | 8.950025488 | 0.51355536 | 0.625406091 | 0.83466646 | -4.844774858 |
| **A_51_P341025** | Eif3k | -0.073810506 | -0.423111932 | 0.275490919 | 11.43733727 | -0.513453378 | 0.625473361 | 0.834684318 | -4.844801821 |
| **A_51_P376857** | Det1 | 0.047073013 | -0.175717048 | 0.269863073 | 9.369022604 | 0.513404145 | 0.625505838 | 0.834684318 | -4.844814835 |
| **A_55_P2112111** | Ube2b | 0.063032292 | -0.235590208 | 0.361654792 | 13.46403171 | 0.512889445 | 0.625845415 | 0.834980623 | -4.84495083 |
| **A_55_P1977985** | Pigp | 0.045286903 | -0.169267655 | 0.25984146 | 11.51234062 | 0.512882719 | 0.625849854 | 0.834980623 | -4.844952606 |
| **A_55_P2011146** | Reep1 | -0.056610779 | -0.324820211 | 0.211598652 | 9.264336099 | -0.512871159 | 0.625857482 | 0.834980623 | -4.844955659 |
| **A_55_P2119420** | Ndufb2 | 0.071384117 | -0.266936201 | 0.409704435 | 13.80780071 | 0.512692187 | 0.625975584 | 0.835008237 | -4.845002918 |
| **A_51_P385697** | Mgat3 | 0.081890362 | -0.306247106 | 0.47002783 | 8.272933315 | 0.512661153 | 0.625996065 | 0.835008237 | -4.845011111 |
| **A_55_P2122380** | Zfp14 | -0.035593287 | -0.204301381 | 0.133114806 | 8.327576872 | -0.512643403 | 0.626007779 | 0.835008237 | -4.845015797 |
| **A_55_P2228715** | Mm2pr | 0.037966318 | -0.14202519 | 0.217957825 | 7.942176485 | 0.512542229 | 0.626074551 | 0.835039676 | -4.845042504 |
| **A_51_P232439** | Ice1 | 0.046677551 | -0.174660698 | 0.2680158 | 9.572844839 | 0.512430278 | 0.626148439 | 0.835078429 | -4.845072051 |
| **A_55_P2107854** | Hnrnpk | -0.051833572 | -0.297651145 | 0.193984002 | 11.06538918 | -0.512367292 | 0.626190012 | 0.835078429 | -4.845088672 |
| **A_55_P2019557** | Mrgpra2b | 0.068286223 | -0.255880553 | 0.392452999 | 8.561671809 | 0.511855978 | 0.626527556 | 0.835270413 | -4.845223533 |
| **A_55_P2040775** | NA | 0.053279425 | -0.199667353 | 0.306226204 | 13.3894082 | 0.511815658 | 0.626554177 | 0.835270413 | -4.845234163 |
| **A_55_P2056280** | Wdr89 | 0.129522154 | -0.485472285 | 0.744516593 | 14.05274687 | 0.511747889 | 0.626598924 | 0.835270413 | -4.845252027 |
| **A_51_P286488** | Ier3 | -0.098901335 | -0.568507261 | 0.370704592 | 9.192534469 | -0.511742837 | 0.626602259 | 0.835270413 | -4.845253359 |

| | | | | | | | | | |
|---|---|---|---|---|---|---|---|---|---|
| **A_55_P1977353** | Ogfod2 | -0.034815709 | -0.200157516 | 0.130526098 | 10.3759568 | -0.511653268 | 0.626661402 | 0.835270413 | -4.845276966 |
| **A_66_P136364** | NA | -0.05669613 | -0.325975385 | 0.212583125 | 13.70413664 | -0.511603733 | 0.626694111 | 0.835270413 | -4.845290021 |
| **A_55_P2017850** | Rpsa | -0.102591548 | -0.589920298 | 0.384737203 | 13.37144742 | -0.511531916 | 0.626741536 | 0.835270413 | -4.845308945 |
| **A_52_P675887** | NA | 0.05289351 | -0.198360638 | 0.304147657 | 15.85616614 | 0.511531444 | 0.626741848 | 0.835270413 | -4.84530907 |
| **A_51_P401876** | Kif1bp | 0.048714463 | -0.182708988 | 0.280137915 | 10.75497731 | 0.511485994 | 0.626771862 | 0.835270413 | -4.845321045 |
| **A_55_P1982186** | Sgsm1 | -0.080214472 | -0.461336957 | 0.300908014 | 8.980996617 | -0.511412505 | 0.626820393 | 0.835270413 | -4.845340406 |
| **A_52_P700056** | Wfdc17 | -0.080195886 | -0.461293871 | 0.300902099 | 10.06696424 | -0.511326882 | 0.626876942 | 0.835270413 | -4.845362961 |
| **A_55_P2090995** | Ebna1bp2 | 0.046685273 | -0.175179411 | 0.268549958 | 9.700683351 | 0.511298966 | 0.626895379 | 0.835270413 | -4.845370314 |
| **A_55_P2011852** | Cdk2 | 0.057207054 | -0.214661097 | 0.329075205 | 8.98258847 | 0.511298409 | 0.626895747 | 0.835270413 | -4.845370461 |
| **A_52_P110088** | Ndufaf3 | -0.070311012 | -0.404498478 | 0.263876453 | 9.659059711 | -0.511230062 | 0.626940888 | 0.835272982 | -4.845388461 |
| **A_55_P2101910** | Rpl13a | -0.126014806 | -0.725202449 | 0.473172837 | 13.36759209 | -0.511024728 | 0.627076515 | 0.835296275 | -4.845442528 |
| **A_52_P440729** | Atf7ip | 0.041018282 | -0.154026815 | 0.23606338 | 9.844779289 | 0.511005555 | 0.627089179 | 0.835296275 | -4.845447575 |
| **A_51_P257156** | Ddrgk1 | -0.054271181 | -0.31235961 | 0.203817247 | 10.60716948 | -0.510956503 | 0.627121582 | 0.835296275 | -4.845460488 |
| **A_55_P2244112** | Amotl1 | -0.039386281 | -0.226723952 | 0.147951389 | 7.890600545 | -0.510861359 | 0.627184434 | 0.835296275 | -4.845485531 |
| **A_52_P312398** | Ssr3 | -0.06915211 | -0.398113766 | 0.259809546 | 10.35013801 | -0.510791131 | 0.627230829 | 0.835296275 | -4.845504014 |
| **A_55_P1980237** | Acad11 | -0.148974434 | -0.857700015 | 0.559751148 | 12.09811096 | -0.510759935 | 0.627251439 | 0.835296275 | -4.845512223 |
| **A_55_P2015994** | Fgf9 | -0.067899929 | -0.390933625 | 0.255133767 | 7.872690446 | -0.510745654 | 0.627260874 | 0.835296275 | -4.845515981 |
| **A_55_P2073734** | Bfar | -0.036087444 | -0.207811146 | 0.135636258 | 11.01292826 | -0.51063322 | 0.627335157 | 0.835337645 | -4.845545564 |
| **A_52_P58181** | Suclg2 | -0.112131325 | -0.646128595 | 0.421865945 | 10.33535763 | -0.510236012 | 0.627597625 | 0.835559852 | -4.845650029 |
| **A_51_P346884** | Snrnp27 | 0.048108146 | -0.181053385 | 0.277269678 | 12.11046724 | 0.510105604 | 0.627683808 | 0.835559852 | -4.845684311 |
| **A_55_P2227967** | NA | -0.056446465 | -0.32533651 | 0.212443579 | 8.351058038 | -0.51008813 | 0.627695357 | 0.835559852 | -4.845688904 |
| **A_55_P1964787** | Slc35e4 | -0.040795817 | -0.235150365 | 0.153558731 | 8.688220151 | -0.510039857 | 0.627727262 | 0.835559852 | -4.845701591 |
| **A_55_P2253746** | Nip7 | 0.071982755 | -0.270959682 | 0.414925192 | 9.262437976 | 0.510023776 | 0.62773789 | 0.835559852 | -4.845705818 |
| **A_55_P2349265** | Ptk2 | 0.069539299 | -0.261785979 | 0.400864577 | 8.626757586 | 0.509986788 | 0.627762337 | 0.835559852 | -4.845715538 |
| **A_55_P2026863** | Trim32 | -0.065178851 | -0.375767412 | 0.245409711 | 10.33050444 | -0.509922798 | 0.627804632 | 0.835559852 | -4.845732354 |
| **A_55_P2061135** | Rpl17 | -0.079167814 | -0.456505832 | 0.298170205 | 14.74651649 | -0.509801697 | 0.627884679 | 0.835608852 | -4.845764172 |
| **A_51_P438479** | Zfp281 | -0.042275894 | -0.243909929 | 0.159358141 | 10.50463889 | -0.509462408 | 0.628108977 | 0.835814035 | -4.845853282 |
| **A_65_P02327** | Pogk | -0.051870824 | -0.29927971 | 0.195538061 | 8.267416587 | -0.509437661 | 0.628125339 | 0.835814035 | -4.845859779 |
| **A_52_P415155** | Wnt6 | -0.033743143 | -0.19476825 | 0.127281965 | 7.725804657 | -0.509184433 | 0.628292773 | 0.835954221 | -4.845926249 |
| **A_55_P2208579** | Onecut2 | -0.156391078 | -0.902835203 | 0.590053047 | 11.0187563 | -0.509093859 | 0.628352667 | 0.835954221 | -4.845950017 |
| **A_51_P319379** | Tmx4 | 0.085917314 | -0.324178574 | 0.496013202 | 9.522241857 | 0.509071127 | 0.628367699 | 0.835954221 | -4.845955982 |
| **A_55_P2115281** | Chchd2-ps | 0.045445663 | -0.17150292 | 0.262394246 | 15.55073427 | 0.509001212 | 0.628413934 | 0.835954221 | -4.845974325 |
| **A_55_P2179216** | Man1a2 | 0.055721939 | -0.210344737 | 0.321788614 | 9.074540168 | 0.508884151 | 0.628491352 | 0.835954221 | -4.846005034 |
| **A_55_P1982727** | Dnajb5 | -0.06427583 | -0.371200972 | 0.242649311 | 8.651785994 | -0.508860126 | 0.628507241 | 0.835954221 | -4.846011335 |
| **A_51_P108935** | Iars2 | 0.058849913 | -0.222187664 | 0.33988749 | 9.560363963 | 0.508820526 | 0.628533431 | 0.835954221 | -4.846021721 |
| **A_55_P1994175** | Ppil3 | 0.044519102 | -0.168181921 | 0.257220125 | 9.523704067 | 0.508580853 | 0.628691959 | 0.836057398 | -4.846084567 |
| **A_55_P2157551** | NA | 0.058609296 | -0.221415739 | 0.33863433 | 8.875233123 | 0.508572453 | 0.628697515 | 0.836057398 | -4.84608677 |
| **A_51_P165060** | Slc22a5 | 0.042886384 | -0.162078783 | 0.247851552 | 8.153470552 | 0.508419922 | 0.628798416 | 0.836115519 | -4.846126752 |
| **A_55_P1971010** | Gzme | -0.034936594 | -0.201975239 | 0.132102051 | 7.643816067 | -0.508214195 | 0.62893452 | 0.836115519 | -4.846180661 |
| **A_55_P2161818** | NA | -0.058072936 | -0.335768717 | 0.219622845 | 16.22366498 | -0.508145038 | 0.628980276 | 0.836115519 | -4.846198779 |
| **A_55_P2071300** | Rnf126 | 0.061521046 | -0.232707292 | 0.355749385 | 10.80303635 | 0.508068654 | 0.629030815 | 0.836115519 | -4.846218787 |
| **A_52_P283524** | Rcan3 | -0.037009542 | -0.214012005 | 0.139992921 | 8.174866729 | -0.508063021 | 0.629034542 | 0.836115519 | -4.846220263 |
| **A_55_P2132512** | Ngef | 0.087385909 | -0.330578263 | 0.50535008 | 11.31989035 | 0.508025534 | 0.629059347 | 0.836115519 | -4.846230081 |
| **A_51_P478172** | Tcrb-J | -0.047111956 | -0.272473417 | 0.178249505 | 8.066756409 | -0.507966035 | 0.629098717 | 0.836115519 | -4.846245664 |
| **A_51_P207153** | 1110008P14Rik | 0.083204047 | -0.314806381 | 0.481214475 | 10.51651173 | 0.507964298 | 0.629099867 | 0.836115519 | -4.846246119 |
| **A_51_P292877** | Snrnp200 | 0.049923048 | -0.188907963 | 0.288754059 | 10.35494076 | 0.507917952 | 0.629130535 | 0.836115519 | -4.846258256 |
| **A_55_P2241553** | 9330133O14Rik | 0.080632964 | -0.305223667 | 0.466489596 | 9.035837727 | 0.507773282 | 0.62922627 | 0.836161713 | -4.846296134 |
| **A_55_P2011440** | Ralgapb | 0.057973358 | -0.219501188 | 0.335447905 | 10.20967177 | 0.50767818 | 0.629289209 | 0.836161713 | -4.84632103 |
| **A_55_P2021049** | Ugp2 | -0.110309183 | -0.638284884 | 0.417666519 | 12.48707134 | -0.507669321 | 0.629295071 | 0.836161713 | -4.846323349 |
| **A_51_P453948** | Drg1 | 0.040988255 | -0.15525999 | 0.2372365 | 10.76407503 | 0.507500925 | 0.629406525 | 0.836200805 | -4.84636742 |

| | | | | | | | | | |
|---|---|---|---|---|---|---|---|---|---|
| **A_55_P2066533** | NA | 0.065325808 | -0.247452732 | 0.378104348 | 15.42562869 | 0.50749414 | 0.629411015 | 0.836200805 | -4.846369195 |
| **A_52_P618132** | Manea | 0.057700465 | -0.21867038 | 0.334071311 | 8.544327832 | 0.507306328 | 0.629535332 | 0.836272958 | -4.846418332 |
| **A_55_P2124096** | NA | -0.067495511 | -0.390798088 | 0.255807067 | 14.23995662 | -0.507281366 | 0.629551855 | 0.836272958 | -4.846424862 |
| **A_55_P2093649** | NA | -0.059876537 | -0.346754607 | 0.227001533 | 15.3160613 | -0.507157105 | 0.629634114 | 0.836324752 | -4.846457362 |
| **A_55_P2135967** | NA | 0.049505439 | -0.187769184 | 0.286780061 | 12.79903442 | 0.50697298 | 0.629756013 | 0.836389024 | -4.846505506 |
| **A_55_P2088630** | Ap3s2 | -0.043071831 | -0.249518826 | 0.163375165 | 10.23655278 | -0.506953299 | 0.629769044 | 0.836389024 | -4.846510652 |
| **A_55_P1994947** | NA | -0.081472234 | -0.472133209 | 0.309188742 | 15.34829034 | -0.506748861 | 0.629904407 | 0.836432435 | -4.846564087 |
| **A_55_P2115320** | Metap1d | 0.038088138 | -0.144558842 | 0.220735117 | 8.336149428 | 0.506711119 | 0.629929399 | 0.836432435 | -4.846573951 |
| **A_65_P12993** | Kif1b | -0.050881547 | -0.294930933 | 0.193167839 | 8.21368933 | -0.506600693 | 0.630002523 | 0.836432435 | -4.846602804 |
| **A_52_P149391** | Usp40 | -0.047392896 | -0.274717236 | 0.179931443 | 8.841971171 | -0.506582874 | 0.630014323 | 0.836432435 | -4.846607459 |
| **A_55_P2025665** | Mrpl40 | 0.069171127 | -0.262688681 | 0.401030936 | 11.60883934 | 0.506469602 | 0.630089337 | 0.836432435 | -4.84663705 |
| **A_55_P2166543** | Pam16 | -0.03857038 | -0.223620827 | 0.146480066 | 13.251293 | -0.506462124 | 0.630094289 | 0.836432435 | -4.846639003 |
| **A_55_P2082186** | NA | 0.065518225 | -0.248886016 | 0.379922465 | 13.38922643 | 0.506357115 | 0.630163836 | 0.836432435 | -4.846666429 |
| **A_55_P1959313** | Rbm42 | -0.044223122 | -0.256439055 | 0.16799281 | 12.5478801 | -0.506354422 | 0.630165619 | 0.836432435 | -4.846667132 |
| **A_55_P2016462** | Cxcl10 | -0.202792676 | -1.176019265 | 0.770433913 | 10.97369689 | -0.506315811 | 0.630191192 | 0.836432435 | -4.846677216 |
| **A_55_P2053733** | NA | 0.050679423 | -0.192578661 | 0.293937508 | 15.74614658 | 0.506229641 | 0.630248266 | 0.836435584 | -4.846699716 |
| **A_52_P220879** | Tgm2 | 0.050907907 | -0.193470094 | 0.295285907 | 13.54723919 | 0.506181562 | 0.630280112 | 0.836435584 | -4.846712269 |
| **A_55_P2038640** | Gcn1 | 0.055386705 | -0.210575056 | 0.321348466 | 9.860943203 | 0.506022141 | 0.630385713 | 0.836518292 | -4.846753884 |
| **A_55_P2171715** | Mrps33 | 0.084178011 | -0.320093184 | 0.488449207 | 12.58869109 | 0.505951691 | 0.630432382 | 0.836522793 | -4.846772271 |
| **A_55_P2016049** | Gp6 | -0.097901736 | -0.568756448 | 0.372952976 | 8.867895968 | -0.505227126 | 0.630912476 | 0.836981642 | -4.846961244 |
| **A_55_P2007630** | Sez6l2 | 0.037550259 | -0.143047271 | 0.218147789 | 7.875567484 | 0.505224404 | 0.630914281 | 0.836981642 | -4.846961954 |
| **A_51_P253537** | Atf6b | 0.040848471 | -0.155643515 | 0.237340457 | 8.517918144 | 0.505142781 | 0.630968376 | 0.836981642 | -4.846983227 |
| **A_66_P106583** | NA | 0.04799401 | -0.182885971 | 0.278873991 | 13.68622161 | 0.50510765 | 0.63099166 | 0.836981642 | -4.846992382 |
| **A_55_P2129234** | Calm2 | -0.067044155 | -0.389569691 | 0.255481381 | 14.15982913 | -0.505103069 | 0.630994696 | 0.836981642 | -4.846993576 |
| **A_51_P283499** | Drd4 | -0.032141035 | -0.186841275 | 0.122559204 | 7.644165011 | -0.504838035 | 0.631170368 | 0.837121593 | -4.847062624 |
| **A_55_P2124097** | Gm6981 | -0.054658261 | -0.317763689 | 0.208447166 | 14.6779277 | -0.504788199 | 0.631203404 | 0.837121593 | -4.847075605 |
| **A_55_P2066017** | NA | 0.050461686 | -0.192483395 | 0.293406768 | 7.848002338 | 0.504704107 | 0.631259149 | 0.837121593 | -4.847097504 |
| **A_55_P2049687** | Efna2 | -0.056157568 | -0.326536714 | 0.214221578 | 7.915478078 | -0.504682549 | 0.631273441 | 0.837121593 | -4.847103118 |
| **A_52_P654437** | NA | -0.076516786 | -0.444978672 | 0.291945101 | 12.01855845 | -0.504600102 | 0.631328099 | 0.837136642 | -4.847124586 |
| **A_51_P211491** | Gusb | 0.049567474 | -0.189175817 | 0.288310765 | 11.00912401 | 0.504485633 | 0.63140399 | 0.837179841 | -4.815594668 |
| **A_55_P2107398** | NA | 0.054718251 | -0.20887851 | 0.318315013 | 10.88961201 | 0.504400293 | 0.631460573 | 0.837197008 | -4.847176599 |
| **A_55_P2249849** | Sema6a | -0.044580081 | -0.259365372 | 0.170205211 | 8.663794046 | -0.504335456 | 0.631503563 | 0.837197008 | -4.847193473 |
| **A_52_P134639** | Tgoln2 | -0.033579753 | -0.195407965 | 0.128248458 | 8.130401826 | -0.504204195 | 0.631590601 | 0.837211985 | -4.847227629 |
| **A_55_P2070576** | Ccl27a | -0.082969207 | -0.482829071 | 0.316890657 | 8.59655325 | -0.504187777 | 0.631601488 | 0.837211985 | -4.8472319 |
| **A_55_P2064632** | NA | -0.067577144 | -0.393313502 | 0.258159213 | 14.31069267 | -0.504100105 | 0.631659627 | 0.837231635 | -4.847254709 |
| **A_55_P2170515** | NA | -0.051305529 | -0.298668512 | 0.196057455 | 7.825637502 | -0.503979238 | 0.631739783 | 0.837280462 | -4.847286147 |
| **A_55_P2111039** | NA | 0.079321057 | -0.303391772 | 0.462033887 | 10.27324206 | 0.503615007 | 0.631981364 | 0.837543215 | -4.847380846 |
| **A_55_P2008918** | Cbll1 | 0.048431582 | -0.185287669 | 0.282150833 | 9.023678778 | 0.503520732 | 0.632043902 | 0.837568667 | -4.847405347 |
| **A_51_P358112** | Fads1 | -0.115818883 | -0.675120345 | 0.443482578 | 10.64152165 | -0.503172164 | 0.632275154 | 0.837727419 | -4.847495902 |
| **A_51_P391367** | Tfap4 | 0.054045365 | -0.207012333 | 0.315103063 | 8.406739147 | 0.503043033 | 0.632360835 | 0.837727419 | -4.847529436 |
| **A_55_P2103807** | NA | -0.08892777 | -0.518521238 | 0.340665697 | 14.86664283 | -0.502994139 | 0.632393279 | 0.837727419 | -4.847542131 |
| **A_55_P2035424** | Hpgd | -0.129247303 | -0.753798573 | 0.495303966 | 9.02039333 | -0.502847834 | 0.632490366 | 0.837727419 | -4.847580111 |
| **A_55_P2350022** | Akap6 | -0.032647564 | -0.190437561 | 0.125142434 | 7.680554312 | -0.502752798 | 0.632553435 | 0.837727419 | -4.847604777 |
| **A_55_P2080758** | Slc8a1 | 0.032729883 | -0.125470134 | 0.190929899 | 7.789792445 | 0.502714154 | 0.632579082 | 0.837727419 | -4.847614806 |
| **A_55_P2093013** | NA | 0.060336837 | -0.231320448 | 0.351994122 | 13.98382935 | 0.502681499 | 0.632600754 | 0.837727419 | -4.84762328 |
| **A_51_P299632** | Spef1 | -0.130493555 | -0.761285296 | 0.500298185 | 10.88793232 | -0.502673806 | 0.63260586 | 0.837727419 | -4.847625276 |
| **A_55_P1981994** | Krt17 | 0.033625592 | -0.128919354 | 0.196170538 | 7.775521627 | 0.502666174 | 0.632610925 | 0.837727419 | -4.847627257 |
| **A_51_P393862** | Spryd4 | 0.106904802 | -0.409885176 | 0.623694781 | 10.77393766 | 0.502650733 | 0.632621173 | 0.837727419 | -4.847631263 |
| **A_55_P2084338** | 2310009B15Rik | -0.076317387 | -0.445291757 | 0.292656982 | 9.441461763 | -0.502586113 | 0.632664062 | 0.837727419 | -4.84764803 |
| **A_51_P268094** | Serpine2 | -0.157247193 | -0.917695678 | 0.603201291 | 8.128685971 | -0.502453982 | 0.632751762 | 0.837727419 | -4.847682307 |

| | | | | | | | | | |
|---|---|---|---|---|---|---|---|---|---|
| **A_52_P525183** | Acot2 | -0.035188497 | -0.205382747 | 0.135005753 | 7.911305422 | -0.502387728 | 0.63279574 | 0.837727419 | -4.847699491 |
| **A_55_P2144456** | Kmt5b | 0.040261215 | -0.154484025 | 0.235006455 | 8.305798001 | 0.502346308 | 0.632823235 | 0.837727419 | -4.847710234 |
| **A_51_P384148** | Ryk | -0.080296901 | -0.468730429 | 0.308136627 | 10.12156788 | -0.502302414 | 0.632852373 | 0.837727419 | -4.847721617 |
| **A_52_P237669** | Nup54 | 0.055878333 | -0.214435071 | 0.326191737 | 8.521835138 | 0.502295223 | 0.632857146 | 0.837727419 | -4.847723481 |
| **A_51_P404236** | Atp6v0b | 0.099558068 | -0.382161177 | 0.581277313 | 11.00656068 | 0.502187215 | 0.632928847 | 0.837728181 | -4.847751487 |
| **A_51_P485862** | Eef1a2 | -0.034467316 | -0.201265981 | 0.13233135 | 7.678462064 | -0.502109086 | 0.632980715 | 0.837728181 | -4.847771742 |
| **A_52_P272811** | Cpsf1 | 0.075041673 | -0.288117297 | 0.438200644 | 9.775524419 | 0.5020985 | 0.632987743 | 0.837728181 | -4.847774486 |
| **A_55_P2157068** | Cdhr2 | 0.131992384 | -0.506976884 | 0.770961652 | 11.9066288 | 0.501940329 | 0.633092757 | 0.837789871 | -4.847815482 |
| **A_55_P2161695** | Poglut2 | -0.066298101 | -0.387271507 | 0.254675304 | 9.057465553 | -0.501897727 | 0.633121043 | 0.837789871 | -4.847826522 |
| **A_55_P2087280** | NA | -0.071033313 | -0.415091067 | 0.273024441 | 12.61249763 | -0.501665122 | 0.633275497 | 0.837896962 | -4.847886786 |
| **A_52_P48546** | Rnf41 | -0.038146132 | -0.222932234 | 0.146639969 | 8.453775461 | -0.501607937 | 0.633313472 | 0.837896962 | -4.847901598 |
| **A_51_P262208** | Itgb2 | -0.115777299 | -0.676652692 | 0.445098094 | 9.72237657 | -0.501580007 | 0.63333202 | 0.837896962 | -4.847908832 |
| **A_51_P356283** | Fbxo31 | 0.068214084 | -0.262353846 | 0.398782013 | 11.54302813 | 0.501414075 | 0.633442219 | 0.837985398 | -4.8479518 |
| **A_55_P2160081** | Ccdc28b | -0.033649681 | -0.196855776 | 0.129556415 | 7.923349148 | -0.500988505 | 0.633724896 | 0.838230131 | -4.848061945 |
| **A_52_P296913** | Cnnm1 | 0.042283455 | -0.162847516 | 0.247414427 | 7.935130427 | 0.500867003 | 0.633805614 | 0.838230131 | -4.848093377 |
| **A_55_P2079251** | Ap2m1 | 0.050264213 | -0.193600971 | 0.294129398 | 12.02938104 | 0.500832238 | 0.63382871 | 0.838230131 | -4.84810237 |
| **A_55_P2161585** | Coq10a | -0.033376836 | -0.195341664 | 0.128587993 | 8.508932974 | -0.500734638 | 0.633893554 | 0.838230131 | -4.848127612 |
| **A_52_P146457** | NA | 0.048405787 | -0.186489576 | 0.283301151 | 15.25689453 | 0.500732789 | 0.633894783 | 0.838230131 | -4.84812809 |
| **A_55_P2127265** | NA | -0.061256305 | -0.358550777 | 0.236038168 | 14.94004376 | -0.500664903 | 0.633939888 | 0.838230131 | -4.848145644 |
| **A_55_P2116059** | Crybg1 | -0.123827249 | -0.724894559 | 0.47724006 | 9.681024259 | -0.500583242 | 0.633994147 | 0.838230131 | -4.848166758 |
| **A_51_P220343** | Ccn4 | -0.142495763 | -0.834208902 | 0.549217376 | 8.54828325 | -0.500563427 | 0.634007313 | 0.838230131 | -4.848171881 |
| **A_55_P2070131** | NA | -0.088496681 | -0.518161451 | 0.341168089 | 14.90200991 | -0.500472741 | 0.634067573 | 0.838230131 | -4.848195325 |
| **A_55_P2014319** | Armcx6 | -0.054234117 | -0.317573386 | 0.209105153 | 9.387295788 | -0.50042631 | 0.634098427 | 0.838230131 | -4.848207326 |
| **A_52_P451644** | Ndn | -0.063982849 | -0.374687246 | 0.246721548 | 8.082298863 | -0.500379335 | 0.634129643 | 0.838230131 | -4.848219467 |
| **A_51_P315555** | Nars | 0.108427651 | -0.437548582 | 0.654403884 | 11.51413711 | 0.505252742 | 0.634147613 | 0.838230131 | -4.804067724 |
| **A_55_P2371801** | A430046D13Rik | 0.035503986 | -0.136972007 | 0.207979978 | 7.909510608 | 0.500186131 | 0.634258042 | 0.838297167 | -4.848269391 |
| **A_51_P224564** | Ppm1f | -0.114289136 | -0.669543274 | 0.440965001 | 11.02154306 | -0.500145467 | 0.634285068 | 0.838297167 | -4.848279896 |
| **A_55_P2143494** | Doc2b | -0.032390275 | -0.189836425 | 0.125055874 | 7.730246095 | -0.499880029 | 0.634461497 | 0.838473013 | -4.848348453 |
| **A_55_P2108476** | Mavs | 0.072474869 | -0.279953251 | 0.424902989 | 11.24694428 | 0.499689331 | 0.634588265 | 0.838583208 | -4.848397687 |
| **A_55_P2109782** | Psg17 | 0.035625145 | -0.137653151 | 0.20890344 | 7.729306792 | 0.499569203 | 0.634668128 | 0.838631409 | -4.848428692 |
| **A_51_P288549** | Jmjd7 | -0.069394731 | -0.406977668 | 0.268188206 | 8.951087764 | -0.499492762 | 0.63471895 | 0.838641233 | -4.848448418 |
| **A_55_P2077143** | Mcidas | 0.141921698 | -0.548645826 | 0.832489221 | 10.70530094 | 0.499373896 | 0.634797983 | 0.838688326 | -4.848479087 |
| **A_55_P2160712** | NA | 0.056455382 | -0.218342515 | 0.331253279 | 8.448376013 | 0.49920064 | 0.634913188 | 0.838696087 | -4.848523779 |
| **A_52_P101487** | Ddn | 0.040131776 | -0.155241214 | 0.235504767 | 7.925353321 | 0.499122385 | 0.634965227 | 0.838696087 | -4.84854396 |
| **A_55_P2121141** | Tbc1d22b | 0.035021464 | -0.135489638 | 0.205532567 | 9.212095463 | 0.499073865 | 0.634997493 | 0.838696087 | -4.848556471 |
| **A_55_P2077548** | Git1 | 0.094695452 | -0.366392163 | 0.555783066 | 10.3142458 | 0.499032525 | 0.635024985 | 0.838696087 | -4.84856713 |
| **A_55_P2114088** | Zfp385c | 0.03236885 | -0.125252102 | 0.189989803 | 7.67577651 | 0.498995369 | 0.635049696 | 0.838696087 | -4.84857671 |
| **A_55_P1960375** | NA | -0.083359994 | -0.489334667 | 0.32261468 | 14.76971212 | -0.498932644 | 0.635091412 | 0.838696087 | -4.848592881 |
| **A_51_P282424** | Eapp | 0.042206346 | -0.163354406 | 0.247767097 | 10.04220536 | 0.498908318 | 0.635107591 | 0.838696087 | -4.848599152 |
| **A_52_P42380** | Tmem106c | -0.045244791 | -0.26569236 | 0.175202778 | 8.300856623 | -0.498708141 | 0.635240732 | 0.83876826 | -4.848650743 |
| **A_55_P1974954** | NA | -0.054042473 | -0.317361846 | 0.2092769 | 15.70723987 | -0.498695662 | 0.635249033 | 0.83876826 | -4.848653959 |
| **A_52_P72587** | Prkcq | -0.077059454 | -0.452732007 | 0.298613099 | 8.539307062 | -0.498424807 | 0.635429209 | 0.838932739 | -4.848723736 |
| **A_55_P2107367** | NA | 0.074399632 | -0.288340153 | 0.437139416 | 10.10428444 | 0.498377909 | 0.635460409 | 0.838932739 | -4.848735814 |
| **A_55_P2165719** | Pcyt1a | 0.057850449 | -0.224326461 | 0.340027359 | 12.4376661 | 0.49815954 | 0.635605694 | 0.839067236 | -4.848792041 |
| **A_51_P144349** | Dtx4 | -0.103206371 | -0.606716432 | 0.40030369 | 10.20296281 | -0.498059861 | 0.635672018 | 0.839097483 | -4.848817699 |
| **A_51_P420415** | Srd5a1 | -0.069215598 | -0.406985095 | 0.268553898 | 9.859765139 | -0.49792822 | 0.635759615 | 0.839155805 | -4.848851578 |
| **A_55_P2093543** | Snapc4 | 0.050689133 | -0.196728675 | 0.298106941 | 10.26870443 | 0.497813991 | 0.635835631 | 0.839198833 | -4.84888097 |
| **A_55_P2079761** | Ggps1 | 0.03476806 | -0.135016375 | 0.204552496 | 8.62635986 | 0.497583277 | 0.635989178 | 0.839232861 | -4.848940315 |
| **A_52_P494126** | Usp46 | 0.035214091 | -0.136758978 | 0.207187159 | 9.10569943 | 0.497552861 | 0.636009422 | 0.839232861 | -4.848948137 |
| **A_51_P463440** | Elovl6 | -0.141348281 | -0.831690864 | 0.548994303 | 12.09819829 | -0.497518296 | 0.636032429 | 0.839232861 | -4.848957026 |

| | | | | | | | | | |
|---|---|---|---|---|---|---|---|---|---|
| **A_51_P396752** | Arl2bp | -0.096934421 | -0.570364275 | 0.376495432 | 11.58854056 | -0.497514304 | 0.636035086 | 0.839232861 | -4.848958052 |
| **A_51_P120066** | 9330151L19Rik | 0.034018113 | -0.132278774 | 0.200315001 | 8.257721787 | 0.497060547 | 0.636337144 | 0.839574106 | -4.842324124 |
| **A_55_P2164047** | NA | -0.086320999 | -0.508427193 | 0.335785194 | 13.85583659 | -0.496910214 | 0.636437235 | 0.839647227 | -4.849113305 |
| **A_55_P2141654** | NA | -0.065294649 | -0.384654315 | 0.254065017 | 15.87959116 | -0.496799003 | 0.636511284 | 0.839647227 | -4.849141869 |
| **A_52_P660945** | Ctsf | 0.085532997 | -0.332869088 | 0.503935082 | 9.464086008 | 0.49673303 | 0.636555214 | 0.839647227 | -4.84915881 |
| **A_55_P2163128** | NA | 0.040976912 | -0.159476734 | 0.241430557 | 8.761400176 | 0.496716346 | 0.636566323 | 0.839647227 | -4.849163094 |
| **A_55_P2163725** | Slc7a9 | 0.129392115 | -0.50389292 | 0.76267715 | 8.481847678 | 0.496468588 | 0.636731316 | 0.839718375 | -4.849226699 |
| **A_55_P2095933** | Zranb1 | -0.054947856 | -0.323916335 | 0.214020622 | 9.76075387 | -0.496400895 | 0.636776399 | 0.839718375 | -4.849244073 |
| **A_51_P384618** | Dhdds | 0.048981364 | -0.190786748 | 0.288749477 | 9.937865579 | 0.496389564 | 0.636783945 | 0.839718375 | -4.84924698 |
| **A_52_P157170** | Rnf157 | -0.035428274 | -0.208858105 | 0.138001557 | 7.98475392 | -0.496374412 | 0.636794037 | 0.839718375 | -4.849250869 |
| **A_51_P126563** | Otc | 0.161963365 | -0.631240778 | 0.955167508 | 9.441088622 | 0.496152296 | 0.63694198 | 0.839807542 | -4.83218833 |
| **A_55_P1964228** | Elac1 | -0.039489874 | -0.23289259 | 0.153912842 | 8.14351251 | -0.496142429 | 0.636948553 | 0.839807542 | -4.849310388 |
| **A_55_P2054743** | Sh3bgrl2 | -0.038091699 | -0.224693476 | 0.148510079 | 7.998644202 | -0.496018363 | 0.637031196 | 0.839859218 | -4.84934221 |
| **A_51_P147373** | Dpp3 | -0.039703921 | -0.234312955 | 0.154905114 | 11.8813983 | -0.495739569 | 0.63721693 | 0.84002892 | -4.849413691 |
| **A_55_P2048751** | Prox1 | 0.05150476 | -0.200969026 | 0.303978546 | 8.21352403 | 0.495694686 | 0.637246834 | 0.84002892 | -4.849425196 |
| **A_52_P23225** | Gpc3 | 0.099351497 | -0.387979223 | 0.586682216 | 13.19484894 | 0.495374691 | 0.637460056 | 0.840252689 | -4.849507191 |
| **A_55_P2086143** | Cyp39a1 | -0.148492498 | -0.877186203 | 0.580201207 | 12.43815934 | -0.495156727 | 0.637605314 | 0.840386847 | -4.849563014 |
| **A_55_P2298029** | Nsd3 | 0.058719392 | -0.229535161 | 0.346973946 | 9.125308973 | 0.494981042 | 0.637722408 | 0.840455722 | -4.849607994 |
| **A_55_P2044463** | NA | 0.032818129 | -0.12829746 | 0.193933718 | 8.356290024 | 0.494947849 | 0.637744533 | 0.840455722 | -4.849616491 |
| **A_55_P2079116** | Rpph1 | 0.225270207 | -0.881439775 | 1.33198019 | 11.36024672 | 0.494598871 | 0.637977166 | 0.840688652 | -4.849705791 |
| **A_52_P313856** | Slc25a26 | 0.032370918 | -0.126676291 | 0.191418127 | 8.062370287 | 0.494552216 | 0.63800827 | 0.840688652 | -4.849717726 |
| **A_55_P2113066** | Zkscan5 | 0.048525101 | -0.189925776 | 0.286975978 | 10.09665384 | 0.494482256 | 0.638054913 | 0.840692802 | -4.84973562 |
| **A_55_P1972322** | Btg3 | -0.06830287 | -0.403992273 | 0.267386533 | 12.35122646 | -0.494406885 | 0.638105165 | 0.840701707 | -4.849754895 |
| **A_55_P2102065** | Gm10639 | 0.137847615 | -0.539806473 | 0.815501702 | 8.719578169 | 0.49428153 | 0.638188748 | 0.84075452 | -4.849786948 |
| **A_55_P2029235** | Btn1a1 | 0.142033699 | -0.556781031 | 0.84084843 | 14.11956231 | 0.493869876 | 0.638463267 | 0.840985061 | -4.849892155 |
| **A_52_P313102** | Yipf7 | -0.031957525 | -0.189193805 | 0.125278756 | 7.629670884 | -0.493859668 | 0.638470075 | 0.840985061 | -4.849894763 |
| **A_55_P2175405** | Zfp467 | -0.059563689 | -0.352648369 | 0.233520992 | 8.56331896 | -0.493823389 | 0.638494271 | 0.840985061 | -4.849904031 |
| **A_52_P273929** | Mrpl17 | 0.047365083 | -0.185792772 | 0.280522937 | 11.76506309 | 0.493618511 | 0.638630925 | 0.841107737 | -4.849956359 |
| **A_55_P2156340** | NA | 0.058458419 | -0.229447184 | 0.346364022 | 16.41672427 | 0.493378405 | 0.638791095 | 0.841188048 | -4.85001766 |
| **A_55_P1962380** | NA | -0.089772918 | -0.531904499 | 0.352358662 | 13.30650376 | -0.493374838 | 0.638793475 | 0.841188048 | -4.85001857 |
| **A_55_P1955287** | NA | 0.04495724 | -0.176495021 | 0.266409501 | 16.62748283 | 0.493290455 | 0.63884977 | 0.841188048 | -4.850040108 |
| **A_55_P2097820** | NA | -0.043415137 | -0.257300349 | 0.170470075 | 12.60043194 | -0.493223327 | 0.638894556 | 0.841188048 | -4.850057238 |
| **A_55_P2103452** | Npy1r | -0.033816435 | -0.200470591 | 0.132837721 | 7.687530441 | -0.493054427 | 0.639007248 | 0.841188048 | -4.850100332 |
| **A_55_P2086770** | Uevld | 0.036853344 | -0.144799154 | 0.218505842 | 8.385763016 | 0.492967986 | 0.639064927 | 0.841188048 | -4.850122382 |
| **A_51_P245989** | Ccr2 | -0.050099216 | -0.297089585 | 0.196891153 | 8.500284785 | -0.492871948 | 0.639129012 | 0.841188048 | -4.850146876 |
| **A_51_P314161** | Qtrt1 | 0.037881442 | -0.148876768 | 0.224639652 | 8.495692345 | 0.492867279 | 0.639132128 | 0.841188048 | -4.850148067 |
| **A_51_P503303** | C1galt1c1 | -0.058959706 | -0.349650762 | 0.231731351 | 10.03701951 | -0.492840996 | 0.639149667 | 0.841188048 | -4.850154769 |
| **A_51_P384640** | Mrpl10 | 0.041392495 | -0.162694478 | 0.245479468 | 10.0110378 | 0.492821347 | 0.639162779 | 0.841188048 | -4.85015978 |
| **A_51_P189208** | 4933417G07Rik | 0.044471715 | -0.174829673 | 0.263773104 | 9.264083065 | 0.492748926 | 0.639211109 | 0.841188048 | -4.850178246 |
| **A_55_P1968433** | Gpat3 | 0.159190576 | -0.625889206 | 0.944270357 | 10.31574033 | 0.49270475 | 0.639240591 | 0.841188048 | -4.850189509 |
| **A_55_P2074406** | Ldoc1 | 0.032585992 | -0.128126607 | 0.193298591 | 7.691094096 | 0.492679179 | 0.639257656 | 0.841188048 | -4.850196028 |
| **A_55_P1966982** | Gsr | 0.081319553 | -0.319873187 | 0.482512292 | 12.90205519 | 0.492521399 | 0.63936296 | 0.841198409 | -4.850236246 |
| **A_51_P329332** | Slc19a2 | 0.140464842 | -0.552559221 | 0.833488906 | 11.71775546 | 0.492495777 | 0.639380061 | 0.841198409 | -4.850242776 |
| **A_51_P185135** | Gpr108 | -0.053044836 | -0.314828082 | 0.208738409 | 10.16275384 | -0.492361919 | 0.639469408 | 0.841198409 | -4.850276885 |
| **A_55_P2088103** | Ash2l | 0.040772967 | -0.160448818 | 0.241994753 | 11.13309496 | 0.492357445 | 0.639472394 | 0.841198409 | -4.850278025 |
| **A_52_P127682** | Dagla | -0.058208593 | -0.345520393 | 0.229103206 | 8.130695649 | -0.492285257 | 0.639520581 | 0.841198409 | -4.850296416 |
| **A_55_P2172852** | Hacd4 | 0.088482846 | -0.348267709 | 0.525233401 | 8.86492966 | 0.492276165 | 0.63952665 | 0.841198409 | -4.850298733 |
| **A_55_P1976574** | Tmod4 | -0.037989722 | -0.225568543 | 0.149589098 | 8.497658131 | -0.492113761 | 0.639635065 | 0.841204401 | -4.850340099 |
| **A_55_P2177659** | Mast1 | 0.035196975 | -0.138604857 | 0.208998807 | 7.827923113 | 0.492078264 | 0.639658762 | 0.841204401 | -4.850349139 |
| **A_51_P462102** | Adgra3 | -0.109184231 | -0.648338311 | 0.42996985 | 9.646038689 | -0.492073764 | 0.639661767 | 0.841204401 | -4.850350285 |

| | | | | | | | | | |
|---|---|---|---|---|---|---|---|---|---|
| **A_55_P2114809** | Gtf2e2 | 0.035747048 | -0.140824951 | 0.212319048 | 11.33042048 | 0.491928001 | 0.639759082 | 0.841275142 | -4.850387399 |
| **A_65_P11193** | Flad1 | 0.055311111 | -0.218046993 | 0.328669215 | 9.962559542 | 0.491658576 | 0.63993898 | 0.841454459 | -4.850455976 |
| **A_65_P20174** | Jade1 | 0.044188444 | -0.17441115 | 0.262788038 | 8.200418593 | 0.491182114 | 0.640257182 | 0.841815596 | -4.850577168 |
| **A_51_P476261** | Zfp120 | 0.060708407 | -0.239732241 | 0.361149055 | 8.817445047 | 0.490990781 | 0.640384986 | 0.841926364 | -4.850625805 |
| **A_55_P1972841** | Rps28 | -0.056432125 | -0.335797942 | 0.222933692 | 17.07379915 | -0.490835907 | 0.640488447 | 0.842005114 | -4.850665163 |
| **A_52_P339054** | Nap1l4 | 0.062756853 | -0.247967336 | 0.373481043 | 9.327898565 | 0.490760143 | 0.640539063 | 0.842014387 | -4.850684413 |
| **A_66_P128701** | NA | -0.058913604 | -0.350770673 | 0.232943465 | 15.48455277 | -0.490488201 | 0.640720757 | 0.842170329 | -4.850753484 |
| **A_55_P2045693** | NA | 0.033423493 | -0.132175543 | 0.199022528 | 8.060345292 | 0.490430212 | 0.640759505 | 0.842170329 | -4.850768208 |
| **A_52_P657338** | Mrps17 | -0.035171312 | -0.209445405 | 0.139102782 | 12.22312372 | -0.490386967 | 0.640788402 | 0.842170329 | -4.850779188 |
| **A_51_P222657** | Tspo | -0.096220528 | -0.573215769 | 0.380774712 | 11.99023058 | -0.490158887 | 0.64094082 | 0.842266877 | -4.850837082 |
| **A_51_P258435** | Zdhhc3 | -0.073015894 | -0.435004099 | 0.288972311 | 9.497714963 | -0.490124232 | 0.640963981 | 0.842266877 | -4.850845877 |
| **A_51_P317191** | Eepd1 | -0.049282211 | -0.293628152 | 0.19506373 | 8.260299195 | -0.490081426 | 0.640992589 | 0.842266877 | -4.850856739 |
| **A_55_P2132781** | Slc16a2 | -0.081959476 | -0.48858793 | 0.324668978 | 10.46193259 | -0.489761447 | 0.641206461 | 0.842490632 | -4.850937908 |
| **A_52_P56751** | Lcp1 | -0.083717082 | -0.499386621 | 0.331952457 | 11.40284583 | -0.489383218 | 0.641459315 | 0.842762608 | -4.851033794 |
| **A_55_P2180491** | Rrbp1 | -0.09782632 | -0.583612003 | 0.387959364 | 12.67347862 | -0.489321381 | 0.64150066 | 0.842762608 | -4.851049464 |
| **A_66_P124017** | Map7d3 | 0.041673606 | -0.165411133 | 0.248758346 | 8.270341182 | 0.488985723 | 0.641725106 | 0.842936578 | -4.851134493 |
| **A_55_P2025403** | Gphn | 0.069847642 | -0.277295112 | 0.416990396 | 10.25373505 | 0.488907649 | 0.641777318 | 0.842936578 | -4.851154263 |
| **A_55_P2051849** | Snap25 | 0.133738545 | -0.530955812 | 0.798432902 | 10.22451014 | 0.488897471 | 0.641784125 | 0.842936578 | -4.85115684 |
| **A_55_P1980292** | Purb | 0.103705469 | -0.411880117 | 0.619291054 | 10.14478777 | 0.488746973 | 0.641884777 | 0.842936578 | -4.851194941 |
| **A_55_P2114697** | Gcm1 | 0.202277921 | -1.015189994 | 1.419745835 | 11.65694319 | 0.512337924 | 0.641905093 | 0.842936578 | -4.71677598 |
| **A_66_P106968** | NA | -0.083275914 | -0.497382136 | 0.330830309 | 11.32455357 | -0.488642052 | 0.641954953 | 0.842936578 | -4.851221497 |
| **A_51_P311540** | Cox6a1 | 0.043228982 | -0.171747792 | 0.258205755 | 17.3957218 | 0.488614842 | 0.641973153 | 0.842936578 | -4.851228384 |
| **A_55_P2016387** | Eif3k | -0.073286261 | -0.437747044 | 0.291174523 | 14.79574678 | -0.48860167 | 0.641981963 | 0.842936578 | -4.851231717 |
| **A_55_P2025368** | Nkain1 | 0.052953488 | -0.210481837 | 0.316388813 | 8.871852972 | 0.488431596 | 0.642095727 | 0.843028685 | -4.85127475 |
| **A_55_P2145149** | Cpn2 | 0.072203303 | -0.287078376 | 0.431484982 | 9.240633832 | 0.488320756 | 0.642169874 | 0.843054815 | -4.851302789 |
| **A_51_P219918** | Tmem125 | -0.043854724 | -0.26210965 | 0.174400202 | 9.397559021 | -0.488242429 | 0.642222275 | 0.843054815 | -4.851322599 |
| **A_55_P2097665** | Arl4a | 0.069851047 | -0.277828845 | 0.41753094 | 10.11187721 | 0.488176126 | 0.642266633 | 0.843054815 | -4.851339366 |
| **A_55_P2172196** | Thnsl1 | 0.032300709 | -0.128485911 | 0.193087329 | 7.801082515 | 0.488141061 | 0.642290093 | 0.843054815 | -4.851348233 |
| **A_55_P2175469** | Chd2 | 0.066332463 | -0.263986993 | 0.396651919 | 10.39065681 | 0.487949817 | 0.64241805 | 0.843133048 | -4.851396582 |
| **A_55_P2129523** | NA | 0.031467298 | -0.125260966 | 0.188195561 | 7.74409588 | 0.487860119 | 0.64247807 | 0.843133048 | -4.851419253 |
| **A_51_P475501** | Ndufa8 | 0.037917532 | -0.150938667 | 0.226773732 | 12.74271222 | 0.487856404 | 0.642480556 | 0.843133048 | -4.851420191 |
| **A_55_P2128695** | Nckap1 | -0.074028455 | -0.442990818 | 0.294933909 | 9.43938214 | -0.487528271 | 0.642700146 | 0.843363959 | -4.851503094 |
| **A_51_P503192** | Rbm15b | -0.054112003 | -0.323853479 | 0.215629472 | 9.000940373 | -0.487448876 | 0.642753283 | 0.843376431 | -4.851523145 |
| **A_55_P2049592** | NA | 0.17120982 | -0.682764315 | 1.025183955 | 9.798455521 | 0.487154976 | 0.642950006 | 0.843536405 | -4.851597346 |
| **A_55_P1999841** | Cox7c | -0.039084959 | -0.234043525 | 0.155873608 | 13.63482302 | -0.487136336 | 0.642962484 | 0.843536405 | -4.851602051 |
| **A_55_P2106033** | Myo18b | -0.032964948 | -0.197475856 | 0.13154596 | 7.68387254 | -0.486901245 | 0.643119866 | 0.843685618 | -4.851661374 |
| **A_55_P1970474** | Tmem67 | -0.039422829 | -0.236202593 | 0.157356935 | 8.062379401 | -0.486799963 | 0.643187676 | 0.843686133 | -4.851686924 |
| **A_55_P2126087** | NA | 0.040480843 | -0.161616413 | 0.2425781 | 14.46190945 | 0.486712299 | 0.643246372 | 0.843686133 | -4.851709034 |
| **A_51_P262230** | Ggact | -0.069007733 | -0.413528233 | 0.275512767 | 10.60249548 | -0.48670508 | 0.643251205 | 0.843686133 | -4.851710855 |
| **A_55_P2087061** | Tmem240 | 0.158289036 | -0.632226504 | 0.948804577 | 9.899497333 | 0.48654567 | 0.643357946 | 0.843768879 | -4.851751051 |
| **A_55_P2004333** | Rpl24 | -0.047285367 | -0.283573866 | 0.189003133 | 15.49743127 | -0.486258679 | 0.643550138 | 0.843945482 | -4.851823389 |
| **A_55_P1956948** | Rpl27 | 0.042183635 | -0.168630413 | 0.252997682 | 16.20967073 | 0.486214201 | 0.643579927 | 0.843945482 | -4.851834597 |
| **A_55_P1955568** | Extl2 | 0.05854186 | -0.234171179 | 0.351254899 | 9.456778702 | 0.485967958 | 0.643744859 | 0.844104496 | -4.851896629 |
| **A_51_P304859** | Zfand6 | 0.071293033 | -0.285399577 | 0.427985642 | 10.06985166 | 0.485664284 | 0.643948288 | 0.844313964 | -4.85197309 |
| **A_66_P115757** | Mthfd2l | 0.048088242 | -0.192564011 | 0.288740495 | 8.844914601 | 0.485547989 | 0.644026202 | 0.844358846 | -4.85200236 |
| **A_55_P2033530** | Gm6981 | -0.071013383 | -0.42644408 | 0.284417315 | 11.04998963 | -0.485476771 | 0.644073919 | 0.844364133 | -4.852020282 |
| **A_55_P1975370** | Apoc1 | -0.079252638 | -0.476076037 | 0.31757076 | 16.60249029 | -0.485288104 | 0.644200335 | 0.844472586 | -4.852067749 |
| **A_55_P2050390** | Cdh22 | -0.040593983 | -0.243959373 | 0.162771407 | 7.729088826 | -0.485029118 | 0.64437389 | 0.844561986 | -4.85213288 |
| **A_55_P2087528** | Tmem131l | 0.049739218 | -0.199478875 | 0.29895731 | 11.91057042 | 0.484956261 | 0.644422718 | 0.844561986 | -4.852151196 |
| **A_55_P2185900** | Nrg4 | -0.121368219 | -0.729598894 | 0.486862457 | 8.443116337 | -0.484863897 | 0.644484623 | 0.844561986 | -4.852174414 |

| | | | | | | | | | |
|---|---|---|---|---|---|---|---|---|---|
| **A_55_P2012478** | Zfp618 | -0.065071804 | -0.391191723 | 0.261048115 | 11.36087684 | -0.484840297 | 0.644500441 | 0.844561986 | -4.852180346 |
| **A_55_P2070992** | Aldoc | -0.105409171 | -0.633707975 | 0.422889633 | 13.32848594 | -0.484821598 | 0.644512974 | 0.844561986 | -4.852185046 |
| **A_55_P2379599** | Asph | 0.067630311 | -0.271368226 | 0.406628848 | 9.293449624 | 0.484759971 | 0.64455428 | 0.844561986 | -4.852200533 |
| **A_52_P284348** | Smim20 | -0.061036386 | -0.367001651 | 0.24492888 | 10.70433789 | -0.484729967 | 0.644574391 | 0.844561986 | -4.852208073 |
| **A_51_P235088** | Dusp27 | -0.032997057 | -0.198502384 | 0.13250827 | 7.672051193 | -0.484447172 | 0.64476396 | 0.844712605 | -4.852279118 |
| **A_51_P404084** | Psmb2 | -0.063910416 | -0.38448236 | 0.256661528 | 10.41478463 | -0.484428096 | 0.644776749 | 0.844712605 | -4.852283909 |
| **A_51_P462271** | Acan | -0.033222816 | -0.199905226 | 0.133459593 | 7.75213121 | -0.484317165 | 0.644851119 | 0.844752781 | -4.852311766 |
| **A_55_P2018345** | Amz2 | 0.032377441 | -0.130144513 | 0.194899395 | 9.354599982 | 0.484076134 | 0.645012726 | 0.844879482 | -4.852372275 |
| **A_55_P1982533** | Elovl4 | -0.031085609 | -0.187133913 | 0.124962695 | 7.657265473 | -0.48404253 | 0.645035259 | 0.844879482 | -4.852380709 |
| **A_52_P364140** | Itga5 | -0.048299521 | -0.290836583 | 0.19423754 | 8.452154213 | -0.4838914 | 0.645136601 | 0.844954964 | -4.852418633 |
| **A_51_P479132** | Zcchc3 | 0.041494926 | -0.166998413 | 0.249988266 | 9.180794526 | 0.483599677 | 0.645332244 | 0.845142208 | -4.852491807 |
| **A_52_P393589** | Clmp | -0.057442018 | -0.346093258 | 0.231209223 | 7.973903337 | -0.483547839 | 0.645367012 | 0.845142208 | -4.852504806 |
| **A_55_P2118103** | Zfp68 | -0.039527517 | -0.238311064 | 0.159256029 | 8.403593067 | -0.483172593 | 0.645618722 | 0.845336007 | -4.852598864 |
| **A_55_P2037919** | NA | -0.096447278 | -0.58150358 | 0.388609024 | 9.664693298 | -0.48314892 | 0.645634604 | 0.845336007 | -4.852604796 |
| **A_55_P1975445** | NA | -0.106760651 | -0.643704578 | 0.430183275 | 12.70613261 | -0.483131629 | 0.645646204 | 0.845336007 | -4.852609128 |
| **A_55_P2060863** | NA | 0.043657258 | -0.175972438 | 0.263286954 | 11.96268546 | 0.483001621 | 0.645733425 | 0.845392941 | -4.852641698 |
| **A_55_P2132358** | NA | 0.103784601 | -0.418510443 | 0.626079645 | 10.1316579 | 0.482836624 | 0.645844129 | 0.845480607 | -4.852683022 |
| **A_55_P2177112** | Prrg2 | 0.034015708 | -0.137216641 | 0.205248058 | 8.182033148 | 0.482699546 | 0.645936108 | 0.845513677 | -4.852717344 |
| **A_52_P311104** | Brd7 | 0.059530986 | -0.240162442 | 0.359224414 | 12.81916419 | 0.482668597 | 0.645956877 | 0.845513677 | -4.852725092 |
| **A_52_P654043** | Msto1 | -0.063239641 | -0.381661113 | 0.255181832 | 10.66006494 | -0.48258102 | 0.646015645 | 0.845516574 | -4.852747014 |
| **A_55_P2024012** | Rpl5 | -0.068865364 | -0.415646357 | 0.277915628 | 14.86760365 | -0.482534927 | 0.646046577 | 0.845516574 | -4.852758551 |
| **A_55_P1982744** | NA | 0.089035584 | -0.359455271 | 0.537526438 | 13.04487707 | 0.482384348 | 0.646147633 | 0.845591577 | -4.852796232 |
| **A_52_P504268** | B3galnt1 | 0.050867646 | -0.205432104 | 0.307167396 | 9.512758815 | 0.482254897 | 0.646234515 | 0.845592621 | -4.852828618 |
| **A_55_P2023076** | Arpc1b | -0.05407546 | -0.326539162 | 0.218388243 | 13.00474308 | -0.482252797 | 0.646235925 | 0.845592621 | -4.852829143 |
| **A_52_P584335** | Folr1 | 0.032754131 | -0.132351978 | 0.19786024 | 7.710506879 | 0.482043384 | 0.646376488 | 0.84571705 | -4.852881517 |
| **A_51_P232411** | Rag1 | -0.109974365 | -0.664473211 | 0.444524481 | 8.575500321 | -0.481918953 | 0.646460018 | 0.84571705 | -4.852912628 |
| **A_51_P345699** | Tnnt3 | -0.036777632 | -0.222214459 | 0.148659194 | 7.752837488 | -0.481915582 | 0.64646228 | 0.84571705 | -4.85291347 |
| **A_55_P1956627** | Gpr160 | 0.055250056 | -0.223457612 | 0.333957724 | 8.468849297 | 0.481689292 | 0.6466142 | 0.845858546 | -4.852970029 |
| **A_51_P141521** | Sap130 | -0.041645138 | -0.251931034 | 0.168640758 | 10.87611232 | -0.481212997 | 0.646934022 | 0.846171761 | -4.853088997 |
| **A_51_P343356** | Isoc2b | -0.058360658 | -0.353057617 | 0.236336301 | 9.19429803 | -0.481202322 | 0.646941191 | 0.846171761 | -4.853091662 |
| **A_51_P403693** | Uba5 | 0.039024887 | -0.158067377 | 0.236117151 | 11.31789922 | 0.481122055 | 0.646995097 | 0.846185008 | -4.8531117 |
| **A_55_P2029558** | Adrb2 | 0.06724432 | -0.272787549 | 0.407276189 | 8.852537507 | 0.480528527 | 0.647393772 | 0.846602054 | -4.853259779 |
| **A_51_P466759** | Mau2 | -0.041329721 | -0.250350294 | 0.167690852 | 8.977623021 | -0.480459329 | 0.647440261 | 0.846602054 | -4.853277032 |
| **A_55_P2122789** | Homer1 | 0.038628763 | -0.156748534 | 0.234006061 | 8.046303783 | 0.480418692 | 0.647467563 | 0.846602054 | -4.853287163 |
| **A_55_P1960049** | 2810408A11Rik | 0.049586954 | -0.201244971 | 0.300418878 | 9.412228369 | 0.480361076 | 0.647506273 | 0.846602054 | -4.853301527 |
| **A_55_P2162815** | NA | 0.100276057 | -0.407004891 | 0.607557004 | 10.61810135 | 0.480321342 | 0.64753297 | 0.846602054 | -4.853311431 |
| **A_55_P2250246** | Taok3 | 0.071513515 | -0.290366177 | 0.433393207 | 8.840358951 | 0.480183355 | 0.647625685 | 0.846666004 | -4.853345821 |
| **A_51_P483544** | Aass | -0.116414601 | -0.762918144 | 0.530088943 | 12.27088673 | -0.491634485 | 0.647689865 | 0.846675671 | -4.735970852 |
| **A_52_P551707** | Kdm6a | 0.138824098 | -0.563874093 | 0.841522289 | 8.997751356 | 0.480041976 | 0.647720686 | 0.846675671 | -4.853381047 |
| **A_51_P185939** | Gper1 | -0.054999248 | -0.333445041 | 0.223446544 | 7.945331346 | -0.479953633 | 0.647780053 | 0.846696013 | -4.853403053 |
| **A_55_P2009329** | Gm8709 | 0.082600697 | -0.335994307 | 0.501195701 | 9.96507992 | 0.479482608 | 0.648096631 | 0.847021346 | -4.853520328 |
| **A_52_P290783** | BC016579 | -0.031326089 | -0.190105431 | 0.127453253 | 7.746719093 | -0.479397067 | 0.648154133 | 0.847021346 | -4.853541615 |
| **A_51_P506297** | Cst9 | -0.037869542 | -0.229861286 | 0.154122201 | 7.677483459 | -0.479281529 | 0.648231802 | 0.847021346 | -4.853570361 |
| **A_55_P2089233** | Pou2af1 | -0.130069292 | -0.78952347 | 0.529384886 | 9.904231919 | -0.47926238 | 0.648244675 | 0.847021346 | -4.853575125 |
| **A_52_P501721** | Ccdc127 | 0.039733089 | -0.161728094 | 0.241194272 | 8.353861036 | 0.479230155 | 0.648266339 | 0.847021346 | -4.853583141 |
| **A_55_P2066299** | Gpr137b | -0.044534055 | -0.270381781 | 0.18131367 | 8.557117673 | -0.479137005 | 0.648328964 | 0.847021346 | -4.85360631 |
| **A_55_P2063795** | Bag6 | 0.045963892 | -0.187139911 | 0.279067696 | 12.62487952 | 0.479126981 | 0.648335703 | 0.847021346 | -4.853608804 |
| **A_51_P457196** | Sfrp4 | -0.036026743 | -0.218771047 | 0.146717561 | 7.969858487 | -0.4790318 | 0.648399697 | 0.847037041 | -4.853632473 |
| **A_55_P2031100** | Ahcy | -0.060805172 | -0.369271284 | 0.24766094 | 8.699619361 | -0.478978756 | 0.648435362 | 0.847037041 | -4.853645663 |
| **A_65_P06872** | Fam210a | 0.046157909 | -0.188091385 | 0.280407203 | 8.385499169 | 0.478796562 | 0.64855787 | 0.847125904 | -4.853690955 |

| | | | | | | | | |
|---|---|---|---|---|---|---|---|---|
| **A_51_P465273** | Mettl17 | 0.048144649 | -0.196212436 | 0.292501733 | 8.438731554 | 0.478747231 | 0.648591042 | 0.847125904 | -4.853703216 |
| **A_55_P2025328** | Tacr2 | 0.107872156 | -0.439753986 | 0.655498297 | 10.56069805 | 0.478639323 | 0.648663608 | 0.847163437 | -4.853730032 |
| **A_51_P336827** | Cyb5b | 0.048236746 | -0.19679672 | 0.293270211 | 9.741373061 | 0.478338995 | 0.648865592 | 0.84732575 | -4.853804636 |
| **A_55_P2043337** | Colq | -0.048297264 | -0.293682478 | 0.19708795 | 8.139609877 | -0.478252584 | 0.648923714 | 0.84732575 | -4.853826094 |
| **A_55_P2087385** | Gps1 | -0.053539995 | -0.325593026 | 0.218513037 | 12.71450138 | -0.478198207 | 0.64896029 | 0.84732575 | -4.853839595 |
| **A_55_P1956063** | Pbld1 | 0.035415635 | -0.144543648 | 0.215374917 | 16.63761924 | 0.478193827 | 0.648963237 | 0.84732575 | -4.853840682 |
| **A_55_P1963062** | Rps16 | -0.057219989 | -0.348047436 | 0.233607457 | 17.18743021 | -0.478074491 | 0.649043512 | 0.847373322 | -4.853870307 |
| **A_55_P2021350** | NA | -0.044819219 | -0.272696813 | 0.183058374 | 15.92602185 | -0.477909708 | 0.649154365 | 0.847460808 | -4.853911203 |
| **A_52_P36928** | Morn2 | -0.066160709 | -0.402660248 | 0.27033883 | 8.61472836 | -0.477747986 | 0.64926317 | 0.847471984 | -4.853951327 |
| **A_66_P132890** | NA | -0.03373345 | -0.205314842 | 0.137847942 | 8.197412226 | -0.477720374 | 0.649281748 | 0.847471984 | -4.853958176 |
| **A_55_P1953919** | Me3 | -0.036432889 | -0.221751999 | 0.148886222 | 7.738164498 | -0.477701481 | 0.64929446 | 0.847471984 | -4.853962863 |
| **A_55_P1967342** | NA | -0.064148063 | -0.390596217 | 0.262300091 | 14.94090259 | -0.477477068 | 0.649445461 | 0.847514719 | -4.854018516 |
| **A_55_P1976864** | Fbxl20 | 0.098958111 | -0.404643784 | 0.602560006 | 9.60091272 | 0.477471253 | 0.649449374 | 0.847514719 | -4.854019958 |
| **A_52_P498985** | Zfp512 | 0.031589594 | -0.12922955 | 0.192408738 | 8.474617783 | 0.477297855 | 0.649566061 | 0.847514719 | -4.854062943 |
| **A_51_P456366** | Pard6a | -0.048056465 | -0.292723679 | 0.196610748 | 8.397190536 | -0.477264616 | 0.649588431 | 0.847514719 | -4.854071181 |
| **A_55_P2114993** | Actg1 | -0.047792891 | -0.291129864 | 0.195544083 | 15.26700741 | -0.477241701 | 0.649603852 | 0.847514719 | -4.85407686 |
| **A_51_P153995** | Gp9 | -0.11147634 | -0.679134415 | 0.456181735 | 11.37680358 | -0.477176526 | 0.649647716 | 0.847514719 | -4.854093012 |
| **A_51_P103209** | Ttc7 | 0.067382583 | -0.275764799 | 0.410529965 | 10.47936619 | 0.477144756 | 0.649669098 | 0.847514719 | -4.854100884 |
| **A_55_P2132517** | Gmppa | -0.058853047 | -0.358733253 | 0.241027159 | 9.564382943 | -0.476874765 | 0.649850824 | 0.847514719 | -4.854167768 |
| **A_52_P417620** | Pacs1 | 0.034304363 | -0.140497827 | 0.209106552 | 8.313053845 | 0.476854273 | 0.649864618 | 0.847514719 | -4.854172843 |
| **A_55_P2067086** | Togaram1 | 0.063590926 | -0.260483459 | 0.387665312 | 10.3797204 | 0.476797132 | 0.649903082 | 0.847514719 | -4.854186994 |
| **A_52_P104761** | Rlim | -0.032689235 | -0.199293138 | 0.133914668 | 8.099554799 | -0.47676325 | 0.64992589 | 0.847514719 | -4.854195384 |
| **A_51_P292527** | P4htm | -0.032258748 | -0.196681733 | 0.132164237 | 7.820369119 | -0.476725267 | 0.64995146 | 0.847514719 | -4.854204788 |
| **A_51_P296866** | Msi1 | 0.074296166 | -0.304471909 | 0.453064241 | 8.919094095 | 0.476624351 | 0.650019396 | 0.847514719 | -4.854229772 |
| **A_55_P1952990** | Myl6b | -0.060617369 | -0.369664825 | 0.248430086 | 8.557827126 | -0.476601169 | 0.650035003 | 0.847514719 | -4.854235511 |
| **A_55_P1974961** | Plekhg3 | 0.120499856 | -0.4938854 | 0.734885112 | 14.13756427 | 0.476572449 | 0.650054338 | 0.847514719 | -4.85424262 |
| **A_51_P276599** | AU040320 | -0.039242057 | -0.239329154 | 0.160845039 | 10.12415938 | -0.476558121 | 0.650063984 | 0.847514719 | -4.854246166 |
| **A_51_P266546** | Zc2hc1a | -0.085552668 | -0.521779656 | 0.35067432 | 8.979631322 | -0.476545329 | 0.650072597 | 0.847514719 | -4.854249333 |
| **A_52_P409746** | Serpinc1 | -0.111716768 | -0.68157702 | 0.458143483 | 16.41126012 | -0.476357698 | 0.650198925 | 0.847622245 | -4.854295765 |
| **A_51_P309988** | Gprc5c | 0.070819544 | -0.290859348 | 0.432498436 | 9.005553366 | 0.475787636 | 0.650582815 | 0.8480157 | -4.854436737 |
| **A_55_P1955243** | Rps24 | -0.232906853 | -1.422394642 | 0.956580935 | 14.68909166 | -0.475779219 | 0.650588485 | 0.8480157 | -4.854438818 |
| **A_55_P2055137** | NA | 0.043168535 | -0.177390861 | 0.263727931 | 12.99782199 | 0.475581483 | 0.650721671 | 0.84813211 | -4.85448768 |
| **A_55_P2079560** | Lilra6 | -0.104820878 | -0.640707134 | 0.431065378 | 9.445901302 | -0.475289657 | 0.650918259 | 0.848282793 | -4.85455976 |
| **A_55_P2128969** | Mettl27 | -0.053585199 | -0.327539983 | 0.220369585 | 9.93192825 | -0.475279569 | 0.650925055 | 0.848282793 | -4.854562251 |
| **A_55_P1981291** | Spink8 | -0.031052252 | -0.189842276 | 0.127737773 | 7.658614047 | -0.475174438 | 0.650995884 | 0.848317902 | -4.854588208 |
| **A_55_P2037524** | Rps15 | -0.075598231 | -0.462427777 | 0.311231315 | 15.16901099 | -0.474870491 | 0.651200682 | 0.84847285 | -4.854663224 |
| **A_52_P305851** | 2410002F23Rik | 0.040967098 | -0.16865921 | 0.250593406 | 9.297909319 | 0.474867671 | 0.651202583 | 0.84847285 | -4.85466392 |
| **A_55_P2159289** | NA | -0.052187557 | -0.319339832 | 0.214964719 | 8.439589978 | -0.474669443 | 0.651336166 | 0.848538807 | -4.854712821 |
| **A_52_P625624** | Stau2 | -0.032606135 | -0.199522071 | 0.134309801 | 7.83184481 | -0.474662262 | 0.651341005 | 0.848538807 | -4.854714592 |
| **A_55_P2319370** | Plekha3 | -0.047516104 | -0.290902128 | 0.19586992 | 9.636544242 | -0.47438219 | 0.651529767 | 0.848689686 | -4.85478365 |
| **A_55_P2050207** | Ankrd27 | 0.070024822 | -0.288671581 | 0.428721224 | 11.72084226 | 0.474360131 | 0.651544635 | 0.848689686 | -4.854789088 |
| **A_52_P24986** | Gpat4 | 0.033639752 | -0.138806576 | 0.20608608 | 12.09467022 | 0.474004023 | 0.651784686 | 0.848937187 | -4.854876839 |
| **A_55_P1978122** | Mutyh | -0.036288906 | -0.222347589 | 0.149769777 | 8.553164052 | -0.473922274 | 0.6518398 | 0.848937187 | -4.854896975 |
| **A_55_P1971009** | Gzme | -0.03030845 | -0.185717379 | 0.125100478 | 7.681216363 | -0.473882812 | 0.651866405 | 0.848937187 | -4.854906694 |
| **A_55_P1977422** | NA | 0.047283439 | -0.195224447 | 0.289791324 | 16.50324549 | 0.473768712 | 0.651943333 | 0.848980172 | -4.854934791 |
| **A_51_P485405** | Gtsf1 | 0.03323075 | -0.1372356 | 0.2036971 | 7.67483684 | 0.47367959 | 0.652003425 | 0.849001226 | -4.854956733 |
| **A_55_P2026624** | NA | 0.042957286 | -0.17752989 | 0.263444461 | 13.68049097 | 0.473409192 | 0.652185761 | 0.849159085 | -4.855023284 |
| **A_55_P2449750** | Fam177a2 | -0.050261759 | -0.308312755 | 0.207789238 | 8.192912059 | -0.473276924 | 0.652274962 | 0.849159085 | -4.855055825 |
| **A_51_P177583** | Puf60 | 0.043374951 | -0.179342652 | 0.266092554 | 11.85420479 | 0.47322496 | 0.652310008 | 0.849159085 | -4.855068607 |
| **A_55_P2168397** | NA | 0.039425215 | -0.163020732 | 0.241871161 | 15.55905895 | 0.473203741 | 0.652324319 | 0.849159085 | -4.855073827 |

| | | | | | | | | | |
|---|---|---|---|---|---|---|---|---|---|
| **A_51_P466591** | 1600012P17Rik | -0.079014858 | -0.484776502 | 0.326746786 | 8.8746451 | -0.473174095 | 0.652344314 | 0.849159085 | -4.855081118 |
| **A_51_P208603** | Dio2 | 0.031282906 | -0.129495073 | 0.192060884 | 7.786449239 | 0.47278502 | 0.652606756 | 0.849389106 | -4.855176776 |
| **A_52_P97192** | Ergic1 | 0.088970898 | -0.36829905 | 0.546240846 | 9.692143039 | 0.472779354 | 0.652610578 | 0.849389106 | -4.855178168 |
| **A_55_P1982767** | Tmem9b | -0.049562678 | -0.304377108 | 0.205251751 | 11.20183518 | -0.472622006 | 0.652716729 | 0.849389106 | -4.855216833 |
| **A_55_P2139713** | Phf13 | -0.031134138 | -0.191219939 | 0.128951664 | 7.82008842 | -0.47257116 | 0.652751033 | 0.849389106 | -4.855229325 |
| **A_55_P2089125** | Zfp976 | 0.071688521 | -0.29695111 | 0.440328151 | 11.00959018 | 0.472531524 | 0.652777774 | 0.849389106 | -4.855239062 |
| **A_55_P1958912** | NA | 0.039171895 | -0.162263784 | 0.240607575 | 9.793758924 | 0.472521283 | 0.652784684 | 0.849389106 | -4.855241578 |
| **A_52_P10781** | Lman1 | 0.070717319 | -0.293097044 | 0.434531682 | 10.58820723 | 0.472312169 | 0.652925779 | 0.849515508 | -4.855292936 |
| **A_55_P2051566** | NA | 0.034667521 | -0.143746452 | 0.213081494 | 7.725518343 | 0.472146869 | 0.653037323 | 0.849603447 | -4.855333519 |
| **A_55_P2094034** | Nrg2 | -0.043695275 | -0.268640878 | 0.181250328 | 8.022650844 | -0.471998001 | 0.653137787 | 0.849676961 | -4.855370057 |
| **A_51_P329490** | Slc25a40 | 0.035185956 | -0.146009414 | 0.216381326 | 8.097467529 | 0.471851621 | 0.65323658 | 0.84974829 | -4.855405974 |
| **A_51_P164630** | Fitm1 | -0.033688008 | -0.207221425 | 0.139845408 | 7.658469478 | -0.471710355 | 0.653331927 | 0.84981513 | -4.855440627 |
| **A_51_P298843** | Pcbd2 | 0.071589402 | -0.29748965 | 0.440668453 | 11.33007851 | 0.471316372 | 0.653597886 | 0.849965956 | -4.855537223 |
| **A_55_P2076471** | Mon1b | 0.068451365 | -0.284450021 | 0.421352751 | 9.791848811 | 0.471315735 | 0.653598316 | 0.849965956 | -4.855537379 |
| **A_55_P2066041** | Hnrnpa1 | 0.048770145 | -0.202674355 | 0.300214646 | 9.049016215 | 0.471297457 | 0.653610657 | 0.849965956 | -4.855541858 |
| **A_55_P2117043** | Mtmr1 | 0.076700379 | -0.318760484 | 0.472161242 | 9.837213627 | 0.471278024 | 0.653623776 | 0.849965956 | -4.855546621 |
| **A_51_P447189** | 1700030C10Rik | 0.0303515 | -0.12623616 | 0.186939159 | 7.846169724 | 0.470983626 | 0.65382255 | 0.850158417 | -4.855618748 |
| **A_66_P109800** | Gcsh | 0.077455729 | -0.322196023 | 0.47710748 | 13.09276098 | 0.470928542 | 0.653859746 | 0.850158417 | -4.85563224 |
| **A_55_P2045741** | Fyb | 0.061077735 | -0.254255813 | 0.376411283 | 8.585531854 | 0.470647694 | 0.654049405 | 0.850347814 | -4.855701003 |
| **A_66_P119420** | Foxk1 | -0.036062032 | -0.222292345 | 0.150168282 | 9.360764856 | -0.470525326 | 0.65413205 | 0.850398062 | -4.855730952 |
| **A_55_P2048937** | Kif5c | 0.034049337 | -0.141900847 | 0.20999952 | 7.771249083 | 0.470221111 | 0.654337534 | 0.85060799 | -4.855805378 |
| **A_52_P125363** | Rbp2 | -0.195551915 | -1.206474741 | 0.815370911 | 10.31149006 | -0.47003182 | 0.654465409 | 0.850717007 | -4.855851666 |
| **A_52_P456059** | Pfdn1 | -0.061058084 | -0.376803715 | 0.254687548 | 11.79091847 | -0.469882215 | 0.654566483 | 0.850723285 | -4.855888237 |
| **A_55_P1963017** | Stfa1 | 0.055733785 | -0.232486932 | 0.343954503 | 16.34556639 | 0.469868718 | 0.654575602 | 0.850723285 | -4.855891536 |
| **A_55_P2129437** | Plpp2 | 0.061088828 | -0.254871536 | 0.377049193 | 12.77847754 | 0.469799312 | 0.654622496 | 0.850723285 | -4.855908499 |
| **A_55_P1966445** | Krt81 | -0.044977588 | -0.277625696 | 0.187670521 | 7.99655575 | -0.469764096 | 0.65464629 | 0.850723285 | -4.855917105 |
| **A_52_P449694** | NA | 0.036053084 | -0.150477985 | 0.222584152 | 7.989050623 | 0.469650111 | 0.65472331 | 0.85075698 | -4.855944956 |
| **A_55_P2093553** | NA | -0.030179709 | -0.186346678 | 0.12598726 | 7.66530663 | -0.469579421 | 0.654771077 | 0.85075698 | -4.855962225 |
| **A_51_P294095** | Prkar1a | -0.060414574 | -0.37306662 | 0.252237471 | 9.792631364 | -0.469530311 | 0.654804263 | 0.85075698 | -4.855974221 |
| **A_52_P191975** | Rtn3 | -0.041526213 | -0.256491893 | 0.173439468 | 9.829129103 | -0.469392756 | 0.65489722 | 0.850787397 | -4.856007815 |
| **A_55_P2018116** | Pabpc4l | -0.048743273 | -0.30108368 | 0.203597134 | 7.991654275 | -0.469365403 | 0.654915706 | 0.850787397 | -4.856014494 |
| **A_51_P520084** | Lypla2 | -0.039890235 | -0.246462957 | 0.166682486 | 12.87504026 | -0.469220325 | 0.655013756 | 0.850796701 | -4.856049915 |
| **A_55_P2134712** | NA | -0.048355958 | -0.298781437 | 0.202069521 | 11.04384905 | -0.469196397 | 0.655029928 | 0.850796701 | -4.856055756 |
| **A_55_P2170414** | Zfp180 | 0.04076734 | -0.170374897 | 0.251909577 | 9.395432411 | 0.469159424 | 0.655054917 | 0.850796701 | -4.856064781 |
| **A_51_P251129** | Stard8 | -0.044982435 | -0.278108554 | 0.188143685 | 8.858988177 | -0.468851394 | 0.655263129 | 0.851009945 | -4.856139944 |
| **A_55_P1972104** | Acsm3 | 0.111360109 | -0.465929017 | 0.688649235 | 8.721598946 | 0.468726448 | 0.655347595 | 0.85106246 | -4.856170419 |
| **A_51_P121412** | Slc35a4 | 0.043908817 | -0.183807103 | 0.271624737 | 9.082069437 | 0.468534451 | 0.6554774 | 0.851173844 | -4.838260467 |
| **A_55_P2014997** | Amz2 | -0.037850475 | -0.234298401 | 0.158597451 | 9.593179313 | -0.468173771 | 0.655721283 | 0.851373576 | -4.856305136 |
| **A_52_P625244** | Ube3a | -0.04983701 | -0.308498618 | 0.208824599 | 8.904424684 | -0.46816958 | 0.655724117 | 0.851373576 | -4.856306157 |
| **A_55_P2108334** | Med10 | -0.072374836 | -0.448057723 | 0.303308051 | 10.09103994 | -0.468111564 | 0.65576335 | 0.851373576 | -4.856320291 |
| **A_55_P2040518** | NA | 0.045122683 | -0.189208127 | 0.279453493 | 10.01429537 | 0.467895327 | 0.655909592 | 0.851506246 | -4.856372954 |
| **A_55_P2184551** | NA | 0.120688949 | -0.506474574 | 0.747852473 | 12.34859124 | 0.467595049 | 0.656112698 | 0.851712715 | -4.85644605 |
| **A_66_P119489** | Spef1l | -0.103602887 | -0.642186533 | 0.434980758 | 8.190141508 | -0.467414183 | 0.656235049 | 0.851814335 | -4.856490057 |
| **A_55_P1966420** | Scrn2 | -0.063553209 | -0.394027227 | 0.26692081 | 11.22481386 | -0.467286624 | 0.656321347 | 0.851822647 | -4.856521084 |
| **A_55_P2013268** | NA | 0.03331471 | -0.139924902 | 0.206554321 | 7.682524573 | 0.467274438 | 0.656329592 | 0.851822647 | -4.856524048 |
| **A_51_P422629** | Tbpl1 | -0.032975704 | -0.204509444 | 0.138558036 | 10.09851635 | -0.467119184 | 0.656434635 | 0.85188334 | -4.856561801 |
| **A_51_P370380** | Rdh14 | -0.044326713 | -0.274928175 | 0.186274748 | 10.52995948 | -0.467075043 | 0.656464501 | 0.85188334 | -4.856572533 |
| **A_65_P18948** | Ppm1h | -0.077780337 | -0.482563932 | 0.327003259 | 9.005569121 | -0.466906698 | 0.656578414 | 0.851973964 | -4.856613453 |
| **A_51_P338961** | Qser1 | 0.036094906 | -0.151821905 | 0.224011716 | 9.152313213 | 0.466727583 | 0.656699625 | 0.852068009 | -4.856656977 |
| **A_51_P382849** | Emb | -0.053001113 | -0.328996162 | 0.222993937 | 9.800474851 | -0.466623799 | 0.656769863 | 0.852068009 | -4.856682189 |

| | | | | | | | | | |
|---|---|---|---|---|---|---|---|---|---|
| **A_52_P404005** | NA | -0.047013206 | -0.291837456 | 0.197811043 | 11.2470247 | -0.466604185 | 0.656783138 | 0.852068009 | -4.856686953 |
| **A_52_P95759** | Sh3rf3 | -0.053153838 | -0.330077701 | 0.223770025 | 7.716373534 | -0.466398813 | 0.65692214 | 0.852191144 | -4.856736827 |
| **A_55_P1959485** | NA | 0.049382118 | -0.208021822 | 0.306786057 | 12.05896267 | 0.466162904 | 0.65708183 | 0.852341097 | -4.856794092 |
| **A_51_P322640** | Ccl24 | 0.080156434 | -0.337813489 | 0.498126356 | 9.844220411 | 0.465989944 | 0.657198921 | 0.852435777 | -4.856836061 |
| **A_55_P2064928** | Nebl | 0.040765939 | -0.171837568 | 0.253369447 | 7.691216376 | 0.465918781 | 0.6572471 | 0.852441066 | -4.856853324 |
| **A_55_P2126537** | Fam53c | 0.038539991 | -0.162609082 | 0.239689064 | 11.55258636 | 0.465561178 | 0.657489234 | 0.852697894 | -4.856940039 |
| **A_55_P1967905** | Dcbld1 | -0.038664628 | -0.240551487 | 0.163222231 | 9.004156041 | -0.465359915 | 0.65762553 | 0.852794563 | -4.856988818 |
| **A_55_P2001494** | Kitl | 0.067975718 | -0.286988883 | 0.42294032 | 8.407209566 | 0.465320246 | 0.657652396 | 0.852794563 | -4.85699843 |
| **A_55_P2179834** | Gatad2a | 0.057014925 | -0.24076422 | 0.35479407 | 10.76570014 | 0.465240453 | 0.657706437 | 0.852794563 | -4.857017762 |
| **A_55_P2013391** | Gm4402 | -0.05474425 | -0.340698303 | 0.231209804 | 9.655428926 | -0.465184718 | 0.657744185 | 0.852794563 | -4.857031263 |
| **A_55_P1952690** | Rpl12 | -0.067436458 | -0.419732606 | 0.284859691 | 15.02038997 | -0.465125385 | 0.657784373 | 0.852794563 | -4.857045635 |
| **A_51_P244969** | Rmnd5b | -0.033399 | -0.2079773 | 0.1411793 | 8.819305889 | -0.464864514 | 0.657961078 | 0.852948425 | -4.857108803 |
| **A_55_P2185651** | NA | 0.04535804 | -0.191753541 | 0.282469622 | 8.784582652 | 0.464819891 | 0.657991307 | 0.852948425 | -4.857119605 |
| **A_55_P2262593** | 4930429F24Rik | 0.039897388 | -0.168728101 | 0.248522876 | 8.162112175 | 0.464686741 | 0.658081509 | 0.852971594 | -4.857151831 |
| **A_52_P205255** | Adam11 | -0.031426699 | -0.195788187 | 0.132934789 | 7.756638829 | -0.464602871 | 0.65813833 | 0.852971594 | -4.857172126 |
| **A_55_P1953475** | NA | -0.040261021 | -0.250828135 | 0.170306094 | 15.7114273 | -0.464598094 | 0.658141567 | 0.852971594 | -4.857173282 |
| **A_55_P2103902** | Mrpl4 | -0.057718593 | -0.359671373 | 0.244234187 | 10.6828579 | -0.464472379 | 0.658226743 | 0.853024788 | -4.857203696 |
| **A_51_P311785** | Man2b2 | 0.083112152 | -0.35186647 | 0.518090774 | 9.097409007 | 0.464279815 | 0.658357221 | 0.853129041 | -4.857250268 |
| **A_55_P2077033** | Ube2e1 | -0.058740721 | -0.366231479 | 0.248750037 | 11.85394057 | -0.464184245 | 0.658421983 | 0.853129041 | -4.857273376 |
| **A_55_P1979699** | Spop | -0.059161357 | -0.368881235 | 0.250558521 | 9.862547842 | -0.464143461 | 0.658449621 | 0.853129041 | -4.857283236 |
| **A_55_P1974582** | Eif4h | 0.053425221 | -0.22629534 | 0.333145783 | 14.19163956 | 0.464093118 | 0.658483737 | 0.853129041 | -4.857295405 |
| **A_55_P1997831** | Dbndd1 | -0.064442303 | -0.401922934 | 0.273038327 | 7.813437154 | -0.463986549 | 0.658555959 | 0.853165425 | -4.857321163 |
| **A_51_P197378** | Atg12 | 0.036646812 | -0.155314549 | 0.228608173 | 11.58619259 | 0.463879915 | 0.65862823 | 0.853201867 | -4.857346931 |
| **A_66_P130813** | Samd4 | -0.051343356 | -0.320608519 | 0.217921807 | 8.179759028 | -0.463326642 | 0.65900327 | 0.85359103 | -4.857480542 |
| **A_55_P1972653** | Tada3 | -0.037494806 | -0.234163423 | 0.159173812 | 10.39628018 | -0.463254057 | 0.659052481 | 0.85359103 | -4.857498061 |
| **A_51_P521171** | Hif1an | 0.035369253 | -0.150155473 | 0.220893978 | 8.218775663 | 0.463241303 | 0.659061128 | 0.85359103 | -4.857501139 |
| **A_55_P1981090** | Thap7 | 0.041514348 | -0.176302497 | 0.259331192 | 9.785930306 | 0.463116169 | 0.659145969 | 0.853643715 | -4.857531333 |
| **A_52_P209172** | Mrpl41 | 0.036184609 | -0.153703214 | 0.226072431 | 12.76846848 | 0.463030882 | 0.659203798 | 0.85364584 | -4.857551908 |
| **A_55_P2020373** | Chd4 | -0.050250051 | -0.313991405 | 0.213491303 | 12.61177911 | -0.462957861 | 0.659253312 | 0.85364584 | -4.857569521 |
| **A_51_P395293** | Ccz1 | -0.046391327 | -0.289933613 | 0.19715096 | 11.68635367 | -0.462855683 | 0.659322599 | 0.85364584 | -4.857594164 |
| **A_51_P185593** | Synm | -0.056974882 | -0.356079644 | 0.24212988 | 7.731239757 | -0.462853224 | 0.659324266 | 0.85364584 | -4.857594756 |
| **A_51_P394788** | Actn4 | -0.061374251 | -0.383644164 | 0.260895662 | 13.12820847 | -0.462753417 | 0.65939195 | 0.853676289 | -4.857618822 |
| **A_55_P2093563** | Ufm1 | -0.039120101 | -0.244564661 | 0.166324459 | 11.56027655 | -0.46268829 | 0.659436116 | 0.853676291 | -4.857634523 |
| **A_55_P2056286** | Sf3b5 | 0.056536193 | -0.240454121 | 0.353526507 | 13.10358326 | 0.462559347 | 0.659523566 | 0.85373232 | -4.857665603 |
| **A_52_P236448** | Ngfr | 0.084762446 | -0.360935058 | 0.530459949 | 8.949512773 | 0.462111171 | 0.659827567 | 0.853983788 | -4.85062505 |
| **A_52_P640386** | Usp53 | -0.032301889 | -0.202161487 | 0.137557708 | 7.704195072 | -0.4620841 | 0.659845931 | 0.853983788 | -4.857780087 |
| **A_55_P1961436** | Prkca | -0.062179927 | -0.389157922 | 0.264798068 | 9.477051835 | -0.462077552 | 0.659850374 | 0.853983788 | -4.857781664 |
| **A_55_P1999962** | Podn | -0.066968887 | -0.419309721 | 0.285371946 | 8.085817584 | -0.461841862 | 0.660010274 | 0.854076437 | -4.8578384 |
| **A_51_P402160** | Zfp750 | 0.097988854 | -0.417585738 | 0.613563446 | 8.428741702 | 0.461815346 | 0.660028265 | 0.854076437 | -4.857844782 |
| **A_55_P2046918** | NA | -0.041661042 | -0.260881635 | 0.177559551 | 15.68297975 | -0.46177665 | 0.66005452 | 0.854076437 | -4.857854094 |
| **A_55_P1978597** | Trit1 | -0.044334268 | -0.277729705 | 0.189061168 | 10.52374092 | -0.46156235 | 0.660199931 | 0.854153422 | -4.857905652 |
| **A_55_P2102901** | Fbxl4 | -0.051240814 | -0.320997523 | 0.218515894 | 8.216077419 | -0.461558717 | 0.660202397 | 0.854153422 | -4.857906526 |
| **A_51_P302942** | Rasl10a | -0.029690764 | -0.186046066 | 0.126664539 | 7.719832984 | -0.46141525 | 0.660299754 | 0.854222203 | -4.857941031 |
| **A_52_P513177** | Sla | -0.076979549 | -0.482907906 | 0.328948808 | 9.203939913 | -0.460796489 | 0.66071973 | 0.854657564 | -4.858089735 |
| **A_55_P2060158** | Ermn | 0.030290919 | -0.129441619 | 0.190023456 | 7.936981531 | 0.460789149 | 0.660724713 | 0.854657564 | -4.858091498 |
| **A_55_P2106078** | F10 | 0.060500957 | -0.258762264 | 0.379764177 | 17.25108485 | 0.46046491 | 0.660944841 | 0.854885093 | -4.858169348 |
| **A_55_P2103603** | NA | 0.057201473 | -0.244860211 | 0.359263158 | 7.930692853 | 0.460145061 | 0.661162024 | 0.855088233 | -4.858246096 |
| **A_66_P130759** | 4631405J19Rik | 0.033492163 | -0.143389428 | 0.210373754 | 7.721628579 | 0.460091007 | 0.661198732 | 0.855088233 | -4.858259062 |
| **A_55_P2064457** | Gng7 | 0.038238589 | -0.163761575 | 0.240238753 | 8.171346028 | 0.459974042 | 0.661278164 | 0.855088233 | -4.858287113 |
| **A_55_P1968370** | Gprc5c | 0.102463316 | -0.438813249 | 0.64373988 | 11.39711362 | 0.459973032 | 0.661278851 | 0.855088233 | -4.858287355 |

| | | | | | | | | | |
|---|---|---|---|---|---|---|---|---|---|
| **A_55_P2030126** | Tfpt | 0.070836685 | -0.303444796 | 0.445118166 | 9.609727968 | 0.459878473 | 0.661343071 | 0.855114069 | -4.858310028 |
| **A_55_P2015113** | Atg5 | 0.049749187 | -0.213152314 | 0.312650689 | 10.43253725 | 0.459807566 | 0.661391229 | 0.855119135 | -4.858327027 |
| **A_51_P375509** | Spty2d1 | 0.034744812 | -0.14893036 | 0.218419984 | 8.174278167 | 0.459645168 | 0.661501534 | 0.855172667 | -4.85836595 |
| **A_55_P1983418** | Amy1 | -0.107138935 | -0.673662478 | 0.459384608 | 12.50358443 | -0.459528606 | 0.661580711 | 0.855172667 | -4.85839388 |
| **A_52_P434841** | Coa4 | -0.036124321 | -0.227143476 | 0.154894835 | 9.744472907 | -0.459521626 | 0.661585452 | 0.855172667 | -4.858395553 |
| **A_51_P377403** | Mettl14 | 0.053954735 | -0.231370646 | 0.339280116 | 8.546650177 | 0.459486071 | 0.661609605 | 0.855172667 | -4.858404071 |
| **A_51_P116407** | Pald1 | -0.050811546 | -0.319662879 | 0.218039788 | 8.683962683 | -0.459233329 | 0.661781305 | 0.855337402 | -4.858464604 |
| **A_55_P1957378** | Unc119 | 0.050152187 | -0.215363366 | 0.315667739 | 12.64552676 | 0.458968728 | 0.661961086 | 0.85547029 | -4.858527947 |
| **A_55_P1971269** | NA | -0.065301648 | -0.411034238 | 0.280430943 | 12.81845379 | -0.458951727 | 0.661972639 | 0.85547029 | -4.858532015 |
| **A_55_P2022724** | Inafm2 | -0.0701947 | -0.441929613 | 0.301540213 | 10.17242558 | -0.458832486 | 0.662053664 | 0.855517802 | -4.858560548 |
| **A_55_P2026156** | NA | 0.044566023 | -0.191554315 | 0.28068636 | 14.4970574 | 0.45862072 | 0.662197575 | 0.8556033 | -4.858611205 |
| **A_55_P2088095** | NA | 0.051785858 | -0.222673416 | 0.326245132 | 11.44035558 | 0.458475838 | 0.662296041 | 0.8556033 | -4.858645851 |
| **A_55_P1969566** | Atp9b | 0.03386167 | -0.145611814 | 0.213335154 | 8.032728984 | 0.458449254 | 0.66231411 | 0.8556033 | -4.858652207 |
| **A_55_P1978481** | Nr1h2 | -0.037927643 | -0.238993636 | 0.163138351 | 11.84067213 | -0.458353357 | 0.66237929 | 0.8556033 | -4.858675132 |
| **A_52_P620793** | Eif2a | -0.042050014 | -0.264973279 | 0.180873252 | 9.247438574 | -0.458346517 | 0.662383939 | 0.8556033 | -4.858676767 |
| **A_55_P1997429** | Gm13698 | 0.04046286 | -0.174104581 | 0.255030302 | 9.149067423 | 0.458222011 | 0.662468569 | 0.8556033 | -4.858706524 |
| **A_55_P1996086** | Gm14326 | -0.039835621 | -0.251101213 | 0.171429972 | 8.440322441 | -0.458169319 | 0.662504387 | 0.8556033 | -4.858719116 |
| **A_66_P100620** | Ensa | 0.044987921 | -0.193616332 | 0.283592174 | 11.66974447 | 0.458142872 | 0.662522365 | 0.8556033 | -4.858725435 |
| **A_55_P2087375** | Gm4987 | -0.033597595 | -0.211848756 | 0.144653565 | 16.14879569 | -0.457993189 | 0.66262412 | 0.8556033 | -4.858761195 |
| **A_55_P2084533** | Zbtb40 | 0.03078989 | -0.132579556 | 0.194159335 | 10.13547095 | 0.457952554 | 0.662651745 | 0.8556033 | -4.858770901 |
| **A_52_P325161** | Zbtb10 | -0.035385341 | -0.223149839 | 0.152379156 | 7.904089202 | -0.457923676 | 0.662671378 | 0.8556033 | -4.858777798 |
| **A_55_P2122020** | Klf4 | -0.162109395 | -1.022372452 | 0.698153662 | 8.746236372 | -0.457888884 | 0.662695031 | 0.8556033 | -4.858786107 |
| **A_52_P87843** | Aldh1a3 | 0.039722271 | -0.171111197 | 0.25055574 | 8.018232203 | 0.457802025 | 0.662754086 | 0.8556033 | -4.858806849 |
| **A_55_P2120388** | Skint7 | 0.105493826 | -0.454473273 | 0.665460924 | 15.47913037 | 0.457770414 | 0.662775578 | 0.8556033 | -4.858814397 |
| **A_52_P763938** | NA | -0.034040948 | -0.214759104 | 0.146677208 | 8.295248709 | -0.457702253 | 0.662821922 | 0.8556033 | -4.85883067 |
| **A_55_P2055257** | Maoa | -0.065385358 | -0.412585342 | 0.281814626 | 8.291341673 | -0.457597877 | 0.662892892 | 0.8556033 | -4.858855585 |
| **A_55_P1958986** | Spata7 | -0.041042183 | -0.258991497 | 0.176907131 | 7.94452586 | -0.457570622 | 0.662911425 | 0.8556033 | -4.858862091 |
| **A_55_P2024463** | Tbkbp1 | -0.058122642 | -0.366780705 | 0.250535422 | 8.525704632 | -0.457563007 | 0.662916603 | 0.8556033 | -4.858863908 |
| **A_55_P1983468** | Cflar | 0.08915608 | -0.384476153 | 0.562788313 | 9.18173553 | 0.457396572 | 0.66302978 | 0.855637582 | -4.858903624 |
| **A_51_P140321** | Mocos | 0.115254162 | -0.497025141 | 0.727533466 | 10.87028693 | 0.457393751 | 0.663031698 | 0.855637582 | -4.858904297 |
| **A_52_P585831** | Bap1 | 0.058782545 | -0.253590879 | 0.371155969 | 10.41989714 | 0.45725398 | 0.663126752 | 0.855646676 | -4.85893764 |
| **A_55_P2014189** | NA | -0.050515695 | -0.318959119 | 0.217927728 | 8.644417175 | -0.457253203 | 0.663127281 | 0.855646676 | -4.858937825 |
| **A_55_P1961247** | NA | 0.046374081 | -0.200182294 | 0.292930455 | 16.21857332 | 0.457027429 | 0.663280835 | 0.855750978 | -4.858991665 |
| **A_55_P2020035** | NA | 0.092553063 | -0.399683394 | 0.584789521 | 11.64195123 | 0.456877961 | 0.663382502 | 0.855750978 | -4.859027295 |
| **A_55_P2111870** | Rogdi | -0.065649396 | -0.414851891 | 0.2835531 | 8.970463672 | -0.456811032 | 0.66342803 | 0.855750978 | -4.859043246 |
| **A_52_P35304** | Chkb | -0.051214675 | -0.323692508 | 0.221263159 | 9.776735543 | -0.456716213 | 0.663492531 | 0.855750978 | -4.859065841 |
| **A_55_P1971804** | Hdac10 | -0.048158529 | -0.304377298 | 0.20806024 | 9.16925577 | -0.456715195 | 0.663493224 | 0.855750978 | -4.859066083 |
| **A_55_P2123962** | Micos10 | 0.058958871 | -0.25472886 | 0.372646602 | 14.63178853 | 0.456703999 | 0.66350084 | 0.855750978 | -4.859068751 |
| **A_55_P1980180** | Tardbp | 0.037056703 | -0.160112541 | 0.234225947 | 13.58115448 | 0.456678737 | 0.663518025 | 0.855750978 | -4.85907477 |
| **A_55_P2370394** | Ppp2r5e | 0.033883137 | -0.146465497 | 0.214231771 | 8.37313258 | 0.456513835 | 0.663630212 | 0.855827376 | -4.859114052 |
| **A_55_P1953361** | Luc7l2 | 0.093900718 | -0.406099436 | 0.593900871 | 15.62156066 | 0.456333091 | 0.663753186 | 0.855827376 | -4.859157092 |
| **A_52_P644354** | Zfp106 | 0.055569304 | -0.240342561 | 0.351481168 | 9.701199633 | 0.456305564 | 0.663771915 | 0.855827376 | -4.859163646 |
| **A_51_P307168** | Ddah1 | 0.136073225 | -0.58853096 | 0.860677409 | 10.40727877 | 0.45630479 | 0.663772442 | 0.855827376 | -4.85916383 |
| **A_55_P1997375** | NA | 0.029223741 | -0.126408879 | 0.184856362 | 7.829188901 | 0.45626628 | 0.663798645 | 0.855827376 | -4.859172998 |
| **A_55_P2072164** | NA | 0.054895337 | -0.237522283 | 0.347312957 | 8.42894925 | 0.4561578 | 0.663872461 | 0.855865457 | -4.85919882 |
| **A_51_P386344** | Mtmr10 | -0.081781823 | -0.517553355 | 0.35398971 | 8.972334974 | -0.45601709 | 0.663968213 | 0.855900874 | -4.859232305 |
| **A_55_P2082806** | Trib1 | -0.055432644 | -0.350872151 | 0.240006863 | 9.549741216 | -0.455911151 | 0.664040309 | 0.855900874 | -4.85925751 |
| **A_55_P1966593** | NA | 0.134485498 | -0.582465946 | 0.851436942 | 9.51399419 | 0.455794311 | 0.664119827 | 0.855900874 | -4.859285302 |
| **A_55_P2072776** | Fpgs | 0.105830832 | -0.458370117 | 0.670031781 | 12.01476855 | 0.455786636 | 0.664125051 | 0.855900874 | -4.859287127 |
| **A_55_P2149149** | NA | 0.093814986 | -0.406338399 | 0.59396837 | 9.426874801 | 0.455776777 | 0.664131761 | 0.855900874 | -4.859289472 |

| | | | | | | | | | |
|---|---|---|---|---|---|---|---|---|---|
| **A_55_P2023818** | Cysltr1 | -0.037630918 | -0.238273549 | 0.163011713 | 8.259477255 | -0.455727034 | 0.664165617 | 0.855900874 | -4.859301302 |
| **A_55_P1984635** | Parp4 | -0.043942788 | -0.278312265 | 0.190426689 | 9.184827718 | -0.455585347 | 0.664262056 | 0.855968085 | -4.859334991 |
| **A_51_P217236** | Derl1 | -0.06593199 | -0.417673917 | 0.285809938 | 11.14877089 | -0.455465233 | 0.664343816 | 0.856016373 | -4.859363543 |
| **A_52_P85864** | Eif6 | 0.071599152 | -0.310484327 | 0.453682631 | 12.11057907 | 0.455336859 | 0.664431205 | 0.856071908 | -4.859394051 |
| **A_55_P2013501** | Hmgb1 | 0.05164488 | -0.224183609 | 0.327473369 | 13.53491797 | 0.454958034 | 0.664689118 | 0.856280167 | -4.859484034 |
| **A_55_P2074085** | Gimap8 | -0.058619722 | -0.371718009 | 0.254478565 | 9.230833348 | -0.454931758 | 0.664707008 | 0.856280167 | -4.859490273 |
| **A_52_P17098** | Mpp7 | -0.039923754 | -0.253209817 | 0.173362309 | 8.633837124 | -0.454833121 | 0.664774172 | 0.856280167 | -4.85951369 |
| **A_55_P1966734** | Cyp2u1 | 0.097654155 | -0.424089538 | 0.619397848 | 15.68301226 | 0.454796075 | 0.664799398 | 0.856280167 | -4.859522484 |
| **A_51_P464064** | Zc3h13 | 0.041319917 | -0.17945355 | 0.262093384 | 8.322503277 | 0.454774126 | 0.664814344 | 0.856280167 | -4.859527694 |
| **A_55_P2017759** | Evi2a | 0.046805237 | -0.203373062 | 0.296983537 | 9.727578206 | 0.454598508 | 0.664933938 | 0.856329337 | -4.859569371 |
| **A_55_P2034345** | Terf2 | 0.037849841 | -0.164468519 | 0.2401682 | 10.01923174 | 0.454581704 | 0.664945381 | 0.856329337 | -4.859573358 |
| **A_51_P445683** | 4931406C07Rik | -0.10026915 | -0.636368476 | 0.435830176 | 11.01536487 | -0.454470043 | 0.665021427 | 0.856329337 | -4.859599848 |
| **A_52_P313217** | Adgrd1 | 0.038026372 | -0.165290992 | 0.241343736 | 7.803622776 | 0.454457849 | 0.665029732 | 0.856329337 | -4.859602741 |
| **A_51_P441898** | Krt25 | -0.029556829 | -0.187698008 | 0.128584349 | 7.687685198 | -0.454146596 | 0.665241732 | 0.85649401 | -4.85967655 |
| **A_55_P2103429** | Psmc3 | 0.039655036 | -0.172518788 | 0.251828861 | 15.11117701 | 0.454139978 | 0.66524624 | 0.85649401 | -4.859678119 |
| **A_52_P56658** | Emc3 | 0.036824616 | -0.160289371 | 0.233938602 | 9.689975701 | 0.453945767 | 0.665378538 | 0.856607283 | -4.859724149 |
| **A_52_P59318** | Rdh10 | 0.03726412 | -0.162232202 | 0.236760441 | 8.217489484 | 0.453878034 | 0.66542468 | 0.856609634 | -4.859740198 |
| **A_52_P641758** | Ggct | -0.049029885 | -0.311618914 | 0.213559144 | 9.584361209 | -0.45369865 | 0.665546894 | 0.8566631 | -4.859782693 |
| **A_55_P2154880** | NA | -0.065409375 | -0.41573095 | 0.2849122 | 10.83970981 | -0.453686968 | 0.665554854 | 0.8566631 | -4.85978546 |
| **A_66_P106238** | Gm5106 | -0.187323366 | -1.190962742 | 0.81631601 | 10.2261049 | -0.453521052 | 0.665667903 | 0.856684931 | -4.85982475 |
| **A_55_P2032192** | Tpm1 | -0.117615546 | -0.747887442 | 0.512656349 | 10.07369497 | -0.453440155 | 0.665723026 | 0.856684931 | -4.859843902 |
| **A_55_P1956472** | Cybrd1 | 0.030630886 | -0.133513452 | 0.194775224 | 8.179184397 | 0.453436885 | 0.665725254 | 0.856684931 | -4.859844677 |
| **A_55_P2084332** | Pigp | -0.031161015 | -0.198169919 | 0.135847889 | 7.780477793 | -0.453372484 | 0.665769139 | 0.856684931 | -4.859859921 |
| **A_55_P2134979** | NA | 0.050863391 | -0.221835512 | 0.323562294 | 13.68113218 | 0.453215869 | 0.665875867 | 0.856684931 | -4.859896986 |
| **A_51_P285779** | Asphd2 | -0.041940756 | -0.266834503 | 0.182952991 | 7.85370291 | -0.453150081 | 0.665920702 | 0.856684931 | -4.859912552 |
| **A_55_P2074924** | Ppp1r13l | -0.051138636 | -0.325377297 | 0.223100025 | 8.745345535 | -0.453110001 | 0.665948017 | 0.856684931 | -4.859922034 |
| **A_55_P2015277** | Cyld | -0.03343486 | -0.212736771 | 0.145867051 | 9.081815566 | -0.453103882 | 0.665952187 | 0.856684931 | -4.859923481 |
| **A_51_P132978** | Idh1 | -0.077512916 | -0.493228139 | 0.338202307 | 9.756465419 | -0.453065859 | 0.665978102 | 0.856684931 | -4.859932476 |
| **A_55_P2033250** | Fdft1 | -0.08479184 | -0.539599583 | 0.370015902 | 11.67413267 | -0.453011682 | 0.666015027 | 0.856684931 | -4.859945292 |
| **A_52_P29151** | Ankrd44 | -0.05307554 | -0.33797483 | 0.23182375 | 9.53874266 | -0.452674721 | 0.666244707 | 0.856904589 | -4.860024965 |
| **A_52_P291971** | Stau2 | -0.03152168 | -0.200740314 | 0.137696953 | 8.480159911 | -0.452631073 | 0.666274462 | 0.856904589 | -4.860035282 |
| **A_55_P2159189** | Decr1 | -0.079244273 | -0.504899846 | 0.346411299 | 13.7814819 | -0.452368914 | 0.666453187 | 0.857050599 | -4.860097227 |
| **A_52_P72434** | Khk | -0.064240118 | -0.409338837 | 0.280858601 | 12.43955758 | -0.452320396 | 0.666486266 | 0.857050599 | -4.860108687 |
| **A_55_P1965836** | Crebbp | 0.050616535 | -0.221326762 | 0.322559831 | 11.6856405 | 0.452269438 | 0.66652101 | 0.857050599 | -4.860120723 |
| **A_51_P450682** | Tepp | 0.031639735 | -0.138461009 | 0.201740479 | 7.839045534 | 0.451970213 | 0.666725044 | 0.857255929 | -4.860191372 |
| **A_55_P1988481** | NA | -0.038530785 | -0.245740146 | 0.168678575 | 9.886208296 | -0.451836882 | 0.66681597 | 0.857315809 | -4.860222839 |
| **A_52_P637353** | Zbtb43 | -0.058489332 | -0.373080056 | 0.256101391 | 9.085737585 | -0.451766415 | 0.666864027 | 0.85732057 | -4.860239467 |
| **A_52_P480064** | Tmem134 | -0.067544239 | -0.431235053 | 0.296146576 | 12.10034146 | -0.451272811 | 0.667200706 | 0.857696359 | -4.860355871 |
| **A_55_P2036280** | Psen2 | -0.08198609 | -0.523751299 | 0.359779118 | 11.88478412 | -0.450953597 | 0.667418481 | 0.857913392 | -4.860431088 |
| **A_55_P2111293** | 3110009E18Rik | 0.035820453 | -0.157253627 | 0.228894534 | 8.902190395 | 0.450806624 | 0.667518762 | 0.857913392 | -4.860465704 |
| **A_51_P426458** | Ube2d2a | 0.052645937 | -0.231129976 | 0.33642185 | 10.44153814 | 0.45078818 | 0.667531347 | 0.857913392 | -4.860470048 |
| **A_55_P1973006** | Bag1 | 0.037275939 | -0.163683785 | 0.238235663 | 13.23626275 | 0.450715773 | 0.667580753 | 0.857913392 | -4.860487097 |
| **A_55_P2109712** | Csnk1g2 | -0.060931202 | -0.389463803 | 0.267601398 | 8.555618364 | -0.450655282 | 0.667622031 | 0.857913392 | -4.860501338 |
| **A_55_P2013655** | Cd4 | 0.201812512 | -0.886500178 | 1.290125202 | 12.39170964 | 0.450585886 | 0.667669386 | 0.857913392 | -4.860517674 |
| **A_55_P2019520** | Cma2 | 0.046377073 | -0.203802127 | 0.296556273 | 8.324120441 | 0.450438315 | 0.667770093 | 0.857913392 | -4.860552405 |
| **A_55_P2015118** | NA | 0.06279694 | -0.276097054 | 0.401690934 | 11.71707992 | 0.450254226 | 0.667895732 | 0.857913392 | -4.860595716 |
| **A_66_P118772** | Tmem136 | -0.063386421 | -0.405485495 | 0.278712653 | 8.411548795 | -0.450222839 | 0.667917155 | 0.857913392 | -4.860603099 |
| **A_52_P316474** | St3gal4 | 0.042080378 | -0.185037449 | 0.269198205 | 9.401111819 | 0.45020633 | 0.667928423 | 0.857913392 | -4.860606982 |
| **A_55_P2043147** | Akr1e1 | 0.065516894 | -0.288113672 | 0.41914746 | 10.07820048 | 0.450180513 | 0.667946044 | 0.857913392 | -4.860613054 |
| **A_55_P2186282** | Agxt2 | 0.114427195 | -0.503201195 | 0.732055584 | 9.830379953 | 0.450178957 | 0.667947106 | 0.857913392 | -4.86061342 |

| | | | | | | | | | |
|---|---|---|---|---|---|---|---|---|---|
| **A_51_P341647** | Fancc | -0.062486907 | -0.399800881 | 0.274827067 | 8.914583971 | -0.450129922 | 0.667980575 | 0.857913392 | -4.860624952 |
| **A_55_P2032147** | Wnt9a | -0.088874676 | -0.568650064 | 0.390900711 | 8.002080795 | -0.450114762 | 0.667990923 | 0.857913392 | -4.860628518 |
| **A_52_P194680** | NA | 0.069167996 | -0.304299186 | 0.442635177 | 9.453903825 | 0.450024269 | 0.668052692 | 0.857935718 | -4.860649796 |
| **A_52_P54238** | Lmf2 | 0.056231363 | -0.247432805 | 0.359895531 | 9.776704143 | 0.44995414 | 0.668100563 | 0.857940193 | -4.860666284 |
| **A_51_P383270** | Fras1 | -0.143006579 | -0.915663023 | 0.629649865 | 8.994328436 | -0.449730948 | 0.668252928 | 0.857999577 | -4.860718742 |
| **A_55_P2099061** | NA | -0.046265362 | -0.296264467 | 0.203733743 | 14.20704884 | -0.449677026 | 0.668289741 | 0.857999577 | -4.860731412 |
| **A_55_P2008277** | Tcp10c | 0.059172424 | -0.260594689 | 0.378939537 | 8.023507721 | 0.449643948 | 0.668312324 | 0.857999577 | -4.860739183 |
| **A_51_P103054** | Adamts19 | -0.035242515 | -0.225704978 | 0.155219948 | 7.706225407 | -0.449614889 | 0.668332164 | 0.857999577 | -4.857217652 |
| **A_55_P2243828** | NA | -0.031070346 | -0.199005014 | 0.136864322 | 7.884181233 | -0.449561299 | 0.668368753 | 0.857999577 | -4.8607586 |
| **A_55_P2331804** | AU015791 | 0.048076586 | -0.211931423 | 0.308084594 | 7.860306368 | 0.449293444 | 0.668551646 | 0.858077058 | -4.860821502 |
| **A_52_P36218** | Upf2 | 0.033853336 | -0.149233152 | 0.216939824 | 8.997957703 | 0.44929167 | 0.668552857 | 0.858077058 | -4.860821919 |
| **A_51_P300434** | Dst | -0.043954678 | -0.28171457 | 0.193805214 | 8.623927139 | -0.449210293 | 0.668608427 | 0.858077058 | -4.860841022 |
| **A_55_P2181162** | NA | -0.039299202 | -0.251921947 | 0.173323543 | 15.61648054 | -0.449114627 | 0.668673757 | 0.858077058 | -4.860863477 |
| **A_55_P2054619** | NA | 0.050519088 | -0.222818796 | 0.323856973 | 15.91066041 | 0.449095667 | 0.668686705 | 0.858077058 | -4.860867926 |
| **A_66_P103926** | Eif3m | 0.050673171 | -0.223536157 | 0.324882498 | 12.86729731 | 0.449033813 | 0.668728947 | 0.858077058 | -4.860882441 |
| **A_55_P2028942** | Ncam1 | -0.029530336 | -0.189348395 | 0.130287723 | 7.719362504 | -0.448978682 | 0.6687666 | 0.858077058 | -4.860895377 |
| **A_52_P12806** | Gm5069 | -0.056062916 | -0.359535438 | 0.247409607 | 14.82641211 | -0.448889549 | 0.668827475 | 0.858077058 | -4.860916289 |
| **A_55_P1989986** | Fam149b | 0.04055141 | -0.178985869 | 0.260088688 | 8.829559566 | 0.448828956 | 0.66886886 | 0.858077058 | -4.860930502 |
| **A_55_P1971347** | Mta3 | 0.030879514 | -0.136326767 | 0.198085794 | 7.969979605 | 0.448746481 | 0.668925193 | 0.858077058 | -4.860949845 |
| **A_55_P2065239** | NA | -0.049808062 | -0.31952573 | 0.219909605 | 16.46350874 | -0.448717947 | 0.668944683 | 0.858077058 | -4.860956537 |
| **A_52_P424585** | Ctnnb1 | -0.087719453 | -0.562769878 | 0.387330971 | 11.5181723 | -0.448682773 | 0.668968709 | 0.858077058 | -4.860964785 |
| **A_55_P2025283** | NA | -0.034045273 | -0.218442243 | 0.150351697 | 15.52952707 | -0.448627855 | 0.669006222 | 0.858077058 | -4.860977662 |
| **A_55_P2114839** | NA | 0.056561862 | -0.249990661 | 0.363114384 | 15.10342595 | 0.448334325 | 0.669206742 | 0.858274602 | -4.861046463 |
| **A_55_P2071930** | NA | 0.034626419 | -0.153066876 | 0.222319713 | 7.874726854 | 0.448272405 | 0.669249045 | 0.858274602 | -4.861060971 |
| **A_52_P580707** | Zfp781 | 0.034929531 | -0.154517023 | 0.224376086 | 8.049554583 | 0.448011575 | 0.669427257 | 0.858446192 | -4.861122066 |
| **A_52_P498208** | H2ac15 | 0.080146201 | -0.354766966 | 0.515059368 | 14.52986005 | 0.447778847 | 0.669586289 | 0.85847096 | -4.861176552 |
| **A_55_P1985385** | Acp2 | 0.03180571 | -0.140819566 | 0.204430985 | 8.109110256 | 0.4476967 | 0.669642428 | 0.85847096 | -4.861195778 |
| **A_55_P2013414** | Dmwd | -0.065216321 | -0.419187155 | 0.288754513 | 12.01822594 | -0.447684447 | 0.669650801 | 0.85847096 | -4.861198645 |
| **A_55_P2182016** | Faf2 | 0.037921366 | -0.167936231 | 0.243778964 | 10.59775151 | 0.447610502 | 0.669701336 | 0.85847096 | -4.861215948 |
| **A_55_P2339650** | Hnrnpl | -0.049632796 | -0.319076354 | 0.219810762 | 8.71290679 | -0.447593866 | 0.669712706 | 0.85847096 | -4.86121984 |
| **A_52_P486279** | Popdc3 | -0.029763073 | -0.191339252 | 0.131813105 | 7.661487594 | -0.447593358 | 0.669713054 | 0.85847096 | -4.861219959 |
| **A_55_P2155996** | NA | 0.049058508 | -0.217315501 | 0.315432518 | 15.56729362 | 0.447513024 | 0.669767958 | 0.858484407 | -4.861238754 |
| **A_55_P1962736** | Sec23b | -0.052587031 | -0.338206946 | 0.233032884 | 11.52256383 | -0.447376695 | 0.669861138 | 0.858521529 | -4.861270642 |
| **A_51_P327418** | Rsrc2 | -0.035510324 | -0.228407763 | 0.157387115 | 10.55405297 | -0.447312842 | 0.669904783 | 0.858521529 | -4.861285574 |
| **A_52_P58283** | Kcna5 | -0.037191652 | -0.239333131 | 0.164949827 | 7.696141889 | -0.44706762 | 0.670072411 | 0.858521529 | -4.861342903 |
| **A_51_P300572** | Slc30a1 | 0.061605333 | -0.273243193 | 0.396453858 | 9.715562422 | 0.447046919 | 0.670086563 | 0.858521529 | -4.861347741 |
| **A_51_P421804** | Timm9 | 0.066311091 | -0.294132634 | 0.426754817 | 10.36980457 | 0.447025086 | 0.670101489 | 0.858521529 | -4.861352844 |
| **A_51_P135268** | Corin | -0.028832271 | -0.185563919 | 0.127899377 | 7.691550201 | -0.446997751 | 0.670120176 | 0.858521529 | -4.861359232 |
| **A_55_P2081303** | Tet2 | 0.066926279 | -0.296937217 | 0.430789776 | 10.08996298 | 0.44693193 | 0.670165175 | 0.858521529 | -4.861374612 |
| **A_52_P445253** | Dst | -0.03662047 | -0.23572088 | 0.16247994 | 8.040536612 | -0.446925308 | 0.670169702 | 0.858521529 | -4.86137616 |
| **A_55_P2073885** | Bbs5 | 0.048709172 | -0.216139205 | 0.31355755 | 8.974263253 | 0.446885869 | 0.670196666 | 0.858521529 | -4.861385375 |
| **A_55_P1987499** | Pttg1 | -0.060607034 | -0.390225611 | 0.269011543 | 12.45904907 | -0.44678084 | 0.670268474 | 0.858547244 | -4.861409911 |
| **A_52_P429364** | Ccdc69 | -0.033181626 | -0.213673397 | 0.147310144 | 8.087989307 | -0.446707713 | 0.670318473 | 0.858547244 | -4.861426991 |
| **A_51_P445912** | NA | -0.03288975 | -0.211812318 | 0.146032819 | 8.110882463 | -0.446661615 | 0.670349993 | 0.858547244 | -4.861437757 |
| **A_55_P2095909** | Lamtor3 | 0.029467242 | -0.130877614 | 0.189812098 | 8.320430942 | 0.446547484 | 0.670428033 | 0.85857987 | -4.861464408 |
| **A_55_P1996683** | Itih1 | -0.119959104 | -0.772833779 | 0.532915571 | 12.46345832 | -0.446464137 | 0.670485027 | 0.85857987 | -4.861483866 |
| **A_65_P05358** | NA | -0.107162375 | -0.690450401 | 0.47612565 | 12.92735791 | -0.446418783 | 0.670516042 | 0.85857987 | -4.861494453 |
| **A_55_P2098076** | Trim25 | 0.118596204 | -0.527117676 | 0.764310083 | 14.62690338 | 0.446286608 | 0.670606431 | 0.85857987 | -4.8615253 |
| **A_55_P2069392** | NA | 0.041056554 | -0.182488443 | 0.26460155 | 14.76001692 | 0.446273121 | 0.670615655 | 0.85857987 | -4.861528448 |
| **A_51_P463452** | Acsl1 | -0.0987362 | -0.636382093 | 0.438909693 | 10.2965104 | -0.446234625 | 0.670641982 | 0.85857987 | -4.86153743 |

| | | | | | | | | |
|---|---|---|---|---|---|---|---|---|
| **A_55_P2096121** | NA | 0.072709498 | -0.323337737 | 0.468756733 | 14.11074743 | 0.446094994 | 0.67073748 | 0.858645258 | -4.861570006 |
| **A_51_P380514** | Tjap1 | 0.047477993 | -0.211182246 | 0.306138233 | 8.793887563 | 0.446011307 | 0.67079472 | 0.858661665 | -4.861589526 |
| **A_51_P294113** | Chd2 | -0.040194713 | -0.259282869 | 0.178893443 | 8.757879807 | -0.445792971 | 0.670944066 | 0.858762564 | -4.861640436 |
| **A_55_P1952016** | R3hdm2 | -0.055846744 | -0.360267277 | 0.248573789 | 10.78823801 | -0.445766169 | 0.6709624 | 0.858762564 | -4.861646684 |
| **A_55_P2430122** | Ilf2 | 0.057471481 | -0.255861199 | 0.370804162 | 10.31292372 | 0.445686916 | 0.671016615 | 0.858775088 | -4.861665157 |
| **A_55_P2036629** | NA | -0.03761496 | -0.242754247 | 0.167524327 | 10.25777269 | -0.44554846 | 0.671111336 | 0.858839447 | -4.861697423 |
| **A_52_P292684** | Exosc4 | -0.042047013 | -0.271515901 | 0.187421875 | 10.72888627 | -0.445240364 | 0.671322133 | 0.858944538 | -4.861769189 |
| **A_55_P2187034** | NA | 0.061510526 | -0.274180895 | 0.397201948 | 11.96142509 | 0.445238168 | 0.671323636 | 0.858944538 | -4.8617697 |
| **A_52_P182592** | Ube2w | 0.040819931 | -0.181955625 | 0.263595488 | 9.251841456 | 0.445233588 | 0.67132677 | 0.858944538 | -4.861770767 |
| **A_55_P2050192** | NA | -0.072862273 | -0.470606368 | 0.324881821 | 13.46276655 | -0.445125183 | 0.671400948 | 0.858982587 | -4.861796007 |
| **A_55_P2021689** | Chrdl2 | -0.028505551 | -0.184155062 | 0.12714396 | 7.664606597 | -0.445004982 | 0.671483202 | 0.859008211 | -4.861823987 |
| **A_55_P2137937** | Fxyd1 | 0.049641914 | -0.221442688 | 0.320726515 | 14.08436234 | 0.44496603 | 0.671509859 | 0.859008211 | -4.861833053 |
| **A_55_P2114498** | Gm8801 | 0.045611661 | -0.203589555 | 0.294812876 | 9.930624413 | 0.444742791 | 0.671662639 | 0.859146791 | -4.861884996 |
| **A_55_P2158112** | Rps27 | -0.055339008 | -0.357793643 | 0.247115628 | 16.39216599 | -0.444584492 | 0.671770986 | 0.859216334 | -4.861921814 |
| **A_55_P2162452** | Mpdu1 | 0.031118069 | -0.138991317 | 0.201227455 | 12.79437222 | 0.444495689 | 0.671831771 | 0.859216334 | -4.861942464 |
| **A_55_P2097161** | Ifitm7 | 0.127860306 | -0.571140885 | 0.826861498 | 14.61069436 | 0.444468527 | 0.671850363 | 0.859216334 | -4.861948779 |
| **A_66_P109183** | Apold1 | -0.054241306 | -0.350883227 | 0.242400616 | 8.754302673 | -0.44430459 | 0.671962585 | 0.859302997 | -4.861986887 |
| **A_51_P113784** | Prpf19 | 0.053400039 | -0.238790351 | 0.345590428 | 8.933564014 | 0.444077563 | 0.672118009 | 0.859444892 | -4.86203964 |
| **A_52_P615952** | Qk | 0.04609262 | -0.206254959 | 0.298440198 | 9.592985725 | 0.443828758 | 0.672288363 | 0.859579692 | -4.862097425 |
| **A_55_P2171047** | Oaz1 | -0.039042939 | -0.252833242 | 0.174747364 | 15.65781457 | -0.443749307 | 0.672342767 | 0.859579692 | -4.862115872 |
| **A_55_P2183493** | Spcs3 | 0.055617459 | -0.248945274 | 0.360180192 | 9.663175259 | 0.443728755 | 0.67235684 | 0.859579692 | -4.862120643 |
| **A_55_P2044217** | 9930021J03Rik | -0.040722466 | -0.26375601 | 0.182311078 | 10.9505684 | -0.443656733 | 0.672406159 | 0.85958589 | -4.862137362 |
| **A_52_P320261** | Cpa2 | 0.165355021 | -0.740809416 | 1.071519458 | 8.248528227 | 0.443397828 | 0.672583466 | 0.859637771 | -4.86219744 |
| **A_55_P2094237** | NA | -0.039823204 | -0.258061227 | 0.178414819 | 16.5447055 | -0.443393136 | 0.67258668 | 0.859637771 | -4.862198529 |
| **A_51_P161037** | Cep170 | 0.038219265 | -0.171234174 | 0.247672704 | 8.197834679 | 0.44338197 | 0.672594327 | 0.859637771 | -4.862201119 |
| **A_55_P2175955** | Ano9 | -0.086312451 | -0.559424274 | 0.386799372 | 8.470025056 | -0.443295004 | 0.67265389 | 0.859637771 | -4.862221292 |
| **A_51_P199098** | Tmem107 | -0.105459063 | -0.683601855 | 0.47268373 | 10.62414738 | -0.44323289 | 0.672696433 | 0.859637771 | -4.862235698 |
| **A_55_P2148641** | Rab4a | 0.079512002 | -0.356409645 | 0.515433648 | 9.008434215 | 0.443207846 | 0.672713587 | 0.859637771 | -4.862241506 |
| **A_55_P1985850** | Timp1 | -0.098550066 | -0.639337539 | 0.442237407 | 8.977801981 | -0.442805989 | 0.672988864 | 0.859918514 | -4.862334657 |
| **A_51_P413740** | Ftcd | -0.124935272 | -0.810585433 | 0.56071489 | 12.42125787 | -0.44275724 | 0.673022261 | 0.859918514 | -4.862345952 |
| **A_55_P2028415** | Hacd2 | -0.089032224 | -0.577925924 | 0.399861476 | 12.01936698 | -0.442502744 | 0.673196627 | 0.860028867 | -4.8624049 |
| **A_51_P489413** | Luc7l | 0.044673144 | -0.200676871 | 0.290023159 | 10.04435489 | 0.442429033 | 0.673247133 | 0.860028867 | -4.862421968 |
| **A_55_P2011967** | Akirin2 | 0.046242966 | -0.2077631 | 0.300249033 | 11.83751614 | 0.442369101 | 0.6732882 | 0.860028867 | -4.862435843 |
| **A_52_P260105** | Man1a | 0.051013655 | -0.229199628 | 0.331226939 | 8.269095 | 0.442365156 | 0.673290903 | 0.860028867 | -4.862436756 |
| **A_55_P1954921** | Fbxw5 | -0.035674859 | -0.231659546 | 0.160309828 | 11.45547673 | -0.442306495 | 0.6733311 | 0.860028867 | -4.862450335 |
| **A_51_P108853** | Lyplal1 | -0.034124259 | -0.221636243 | 0.153387726 | 8.219113795 | -0.442198621 | 0.673405023 | 0.860029846 | -4.840929777 |
| **A_55_P2135725** | C8g | -0.092425925 | -0.600330709 | 0.415478859 | 9.874377513 | -0.442175517 | 0.673420856 | 0.860029846 | -4.862480649 |
| **A_55_P2137828** | Gprc5b | -0.040615875 | -0.263975264 | 0.182743514 | 8.149415235 | -0.441849929 | 0.673643998 | 0.86007531 | -4.862555968 |
| **A_55_P2125868** | NA | 0.058773988 | -0.26445843 | 0.382006406 | 11.39791898 | 0.441828206 | 0.673658888 | 0.86007531 | -4.862560992 |
| **A_51_P256827** | S100a8 | -0.07632972 | -0.496133418 | 0.343473978 | 16.45474539 | -0.441804971 | 0.673674813 | 0.86007531 | -4.862566364 |
| **A_55_P1960621** | NA | 0.037164585 | -0.167238052 | 0.241567223 | 16.12771501 | 0.441800275 | 0.673678032 | 0.86007531 | -4.86256745 |
| **A_55_P2177023** | Mafg | 0.07773209 | -0.349790285 | 0.505254465 | 10.33800525 | 0.441798953 | 0.673678938 | 0.86007531 | -4.862567756 |
| **A_55_P1998066** | Tmem121 | 0.035649106 | -0.160554165 | 0.231852378 | 8.09247518 | 0.441494802 | 0.673887427 | 0.860278211 | -4.862638063 |
| **A_55_P2149049** | Rpl32 | 0.04477927 | -0.201706141 | 0.29126468 | 16.85108319 | 0.44143725 | 0.67392688 | 0.860278211 | -4.862651362 |
| **A_51_P401907** | Cstdc4 | -0.128789614 | -0.837955502 | 0.580376274 | 15.21362348 | -0.44128198 | 0.67403333 | 0.860300417 | -4.862687233 |
| **A_55_P2008936** | Slc2a9 | 0.126027749 | -0.568008258 | 0.820063757 | 9.287194067 | 0.441232372 | 0.674067342 | 0.860300417 | -4.862698691 |
| **A_55_P1976690** | Snrnp35 | 0.061214397 | -0.275933815 | 0.39836261 | 10.60748461 | 0.441180092 | 0.674103186 | 0.860300417 | -4.862710765 |
| **A_55_P2029252** | NA | -0.07974092 | -0.518996396 | 0.359514556 | 10.34560256 | -0.441110348 | 0.674151006 | 0.860300417 | -4.86272687 |
| **A_66_P115693** | Rnf207 | -0.033685391 | -0.219252172 | 0.151881391 | 7.685897723 | -0.441087287 | 0.674166818 | 0.860300417 | -4.862732195 |
| **A_55_P1981050** | Gemin8 | 0.029514732 | -0.133161679 | 0.192191144 | 8.138549059 | 0.440856709 | 0.674324927 | 0.860445373 | -4.862785422 |

| | | | | | | | | | |
|---|---|---|---|---|---|---|---|---|---|
| **A_51_P154534** | Bloc1s5 | 0.032938901 | -0.148712612 | 0.214590415 | 10.56045718 | 0.44060891 | 0.674494865 | 0.860605402 | -4.862842595 |
| **A_51_P263591** | Pank1 | -0.113882813 | -0.742192941 | 0.514427315 | 9.36876751 | -0.440420285 | 0.674624235 | 0.860673196 | -4.862886096 |
| **A_51_P513959** | NA | 0.034198312 | -0.154492724 | 0.222889347 | 9.256298164 | 0.440389126 | 0.674645607 | 0.860673196 | -4.86289328 |
| **A_55_P2075670** | NA | -0.037563018 | -0.24491651 | 0.169790473 | 8.397259788 | -0.440182041 | 0.674787655 | 0.860673196 | -4.862941016 |
| **A_66_P110633** | Apol9b | -0.029248969 | -0.190739886 | 0.132241947 | 7.81695364 | -0.440094211 | 0.674847905 | 0.860673196 | -4.862961256 |
| **A_51_P303238** | Tmed10 | -0.047389035 | -0.309042594 | 0.214264524 | 12.1071301 | -0.440082815 | 0.674855723 | 0.860673196 | -4.862963882 |
| **A_55_P1979457** | Prdx4 | -0.056919122 | -0.371216593 | 0.257378348 | 13.54834731 | -0.440048453 | 0.674879296 | 0.860673196 | -4.862971799 |
| **A_52_P274555** | Snx14 | -0.054714581 | -0.356877485 | 0.247448323 | 9.288516626 | -0.439992345 | 0.674917788 | 0.860673196 | -4.862984726 |
| **A_55_P1989901** | Ddb1 | 0.040660424 | -0.183901446 | 0.265222293 | 14.61614602 | 0.439965956 | 0.674935892 | 0.860673196 | -4.862990805 |
| **A_66_P106478** | Cct3 | 0.036794656 | -0.166437343 | 0.240026655 | 11.38610604 | 0.439922171 | 0.674965931 | 0.860673196 | -4.863000891 |
| **A_66_P136788** | Ubald2 | 0.052801232 | -0.238868083 | 0.344470548 | 13.65807739 | 0.439882317 | 0.674993274 | 0.860673196 | -4.863010071 |
| **A_55_P2074226** | Fbxo8 | 0.056838541 | -0.257229605 | 0.370906688 | 8.876938617 | 0.439746329 | 0.675086575 | 0.860735383 | -4.863041387 |
| **A_55_P2003199** | Setd1b | -0.049527551 | -0.323440743 | 0.224385642 | 8.745608706 | -0.439356537 | 0.675354046 | 0.8608369 | -4.863131103 |
| **A_52_P238846** | Bpnt1 | -0.080643962 | -0.526677792 | 0.365389867 | 8.969768722 | -0.439326346 | 0.675374765 | 0.8608369 | -4.863138049 |
| **A_55_P1996199** | NA | -0.070114409 | -0.457955064 | 0.317726246 | 14.90905572 | -0.439275654 | 0.675409554 | 0.8608369 | -4.863149711 |
| **A_51_P409349** | Trim29 | 0.028893968 | -0.1309879 | 0.188775835 | 7.667335208 | 0.43912803 | 0.675510869 | 0.8608369 | -4.863183664 |
| **A_55_P1964812** | Tfap2a | -0.02973725 | -0.194298276 | 0.134823777 | 7.698545566 | -0.439093493 | 0.675534573 | 0.8608369 | -4.863191606 |
| **A_55_P2011436** | NA | 0.066212116 | -0.300227577 | 0.432651809 | 14.45032427 | 0.439054208 | 0.675561537 | 0.8608369 | -4.863200639 |
| **A_52_P20048** | Tcf12 | 0.05361822 | -0.243162849 | 0.35039929 | 9.90629948 | 0.438994813 | 0.675602304 | 0.8608369 | -4.863214295 |
| **A_55_P2085835** | 5330417C22Rik | -0.050875377 | -0.332480058 | 0.230729304 | 8.596828635 | -0.438986292 | 0.675608152 | 0.8608369 | -4.863216254 |
| **A_55_P2035138** | NA | 0.033840295 | -0.153496318 | 0.221176908 | 13.90313779 | 0.438929393 | 0.675647207 | 0.8608369 | -4.863229334 |
| **A_55_P2179398** | Alg1 | 0.043268325 | -0.196265492 | 0.282802143 | 9.598262661 | 0.438921078 | 0.675652915 | 0.8608369 | -4.863231245 |
| **A_51_P384113** | Acad12 | -0.03736998 | -0.244268455 | 0.169528495 | 8.305723622 | -0.438883004 | 0.675679049 | 0.8608369 | -4.863239997 |
| **A_55_P2037424** | NA | -0.046368187 | -0.303103523 | 0.210367149 | 14.40239593 | -0.438851567 | 0.675700628 | 0.8608369 | -4.863247222 |
| **A_52_P246753** | Chmp5 | -0.04584349 | -0.299794288 | 0.208107308 | 11.83717561 | -0.438643081 | 0.675843746 | 0.860962484 | -4.863295129 |
| **A_55_P2184314** | NA | 0.135874941 | -0.617559308 | 0.88930919 | 13.08792353 | 0.43820487 | 0.67614461 | 0.861249783 | -4.863395755 |
| **A_55_P2243367** | NA | 0.045352739 | -0.206142285 | 0.296847763 | 8.90358256 | 0.438184806 | 0.676158387 | 0.861249783 | -4.86340036 |
| **A_55_P2037957** | NA | 0.054037121 | -0.245698052 | 0.353772294 | 13.99284903 | 0.43806412 | 0.676241259 | 0.861298582 | -4.863428056 |
| **A_55_P2023697** | Zfp386 | -0.070152816 | -0.459339919 | 0.319034287 | 9.787967603 | -0.437995709 | 0.676288237 | 0.861301662 | -4.863443752 |
| **A_51_P384754** | Ldb1 | 0.068661926 | -0.312454907 | 0.44977876 | 10.08880774 | 0.437765005 | 0.676446675 | 0.861446684 | -4.863496669 |
| **A_55_P2069525** | Zmym6 | 0.045656445 | -0.207815535 | 0.299128425 | 8.82148448 | 0.437678613 | 0.676506009 | 0.861465489 | -4.863516478 |
| **A_55_P2064230** | Tmx3 | 0.041666364 | -0.189709177 | 0.273041906 | 8.726773177 | 0.437573837 | 0.676577974 | 0.861500373 | -4.863540498 |
| **A_55_P1971433** | Serpinb9d | -0.02769644 | -0.181580481 | 0.126187602 | 7.616228163 | -0.437334311 | 0.676742504 | 0.86165135 | -4.86359539 |
| **A_55_P1999324** | Nudt6 | -0.038732918 | -0.253967622 | 0.176501787 | 9.130117688 | -0.437271431 | 0.6767857 | 0.86165135 | -4.863609795 |
| **A_55_P1991931** | Nosip | -0.042129876 | -0.276288295 | 0.192028543 | 10.17352462 | -0.437183328 | 0.676846225 | 0.861654673 | -4.863629976 |
| **A_55_P2000938** | Ahcy | -0.129712491 | -0.850731266 | 0.591306284 | 11.97511669 | -0.437137853 | 0.676877466 | 0.861654673 | -4.863640391 |
| **A_55_P2455725** | Klhl20 | 0.041084033 | -0.187410921 | 0.269578987 | 8.955827138 | 0.436897587 | 0.677042541 | 0.861664377 | -4.863695403 |
| **A_55_P2146175** | NA | 0.031281921 | -0.14272787 | 0.205291711 | 8.859174835 | 0.436820402 | 0.677095575 | 0.861664377 | -4.863713069 |
| **A_55_P2031786** | Fcsk | -0.040025468 | -0.262695886 | 0.182644949 | 10.42713028 | -0.436774312 | 0.677127244 | 0.861664377 | -4.863723617 |
| **A_52_P316697** | Ost4 | 0.087087366 | -0.397427446 | 0.571602179 | 12.37641076 | 0.436748365 | 0.677145074 | 0.861664377 | -4.863729555 |
| **A_52_P594768** | Aprt | -0.030535948 | -0.200437344 | 0.139365447 | 13.67027721 | -0.436714543 | 0.677168314 | 0.861664377 | -4.863737294 |
| **A_55_P1973427** | Tex14 | -0.02920596 | -0.191746164 | 0.133334244 | 7.671691826 | -0.436610202 | 0.677240014 | 0.861664377 | -4.863761166 |
| **A_55_P2251082** | Qk | -0.04478481 | -0.294054154 | 0.204484535 | 8.359429672 | -0.436561114 | 0.677273747 | 0.861664377 | -4.863772395 |
| **A_55_P2163319** | Peg3 | 0.050495114 | -0.230564481 | 0.331554708 | 8.113097431 | 0.436550135 | 0.677281292 | 0.861664377 | -4.863774907 |
| **A_55_P2066792** | Sp110 | 0.088837712 | -0.405647918 | 0.583323341 | 16.03199311 | 0.436542848 | 0.6772863 | 0.861664377 | -4.863776573 |
| **A_55_P1975665** | Rhox4e | 0.028499458 | -0.130168838 | 0.187167754 | 7.690783297 | 0.436445112 | 0.677353467 | 0.861693113 | -4.863798927 |
| **A_51_P229676** | Plscr1 | -0.067369137 | -0.442504582 | 0.307766308 | 10.41595344 | -0.436371193 | 0.677404268 | 0.861701026 | -4.86381583 |
| **A_55_P2029630** | Mpdu1 | -0.052197903 | -0.342988127 | 0.23859232 | 9.505461697 | -0.436170657 | 0.677542097 | 0.861799302 | -4.863861673 |
| **A_55_P2410626** | A430061O12Rik | 0.030524802 | -0.139542583 | 0.200592187 | 8.168888625 | 0.436129047 | 0.677570697 | 0.861799302 | -4.863871183 |
| **A_65_P15689** | Mmut | -0.064784912 | -0.425830916 | 0.296261093 | 11.40248192 | -0.436008059 | 0.67765386 | 0.861848366 | -4.86389883 |

| | | | | | | | | |
|---|---|---|---|---|---|---|---|---|
| **A_51_P491051** | Ints9 | 0.038296272 | -0.175182373 | 0.251774917 | 11.44481583 | 0.435898376 | 0.677729258 | 0.861873312 | -4.863923888 |
| **A_55_P2006808** | Ntrk3 | 0.040433125 | -0.18498231 | 0.26584856 | 7.800710257 | 0.435849794 | 0.677762654 | 0.861873312 | -4.863934985 |
| **A_55_P2125564** | Lrrc58 | 0.061835498 | -0.283050865 | 0.406721862 | 10.91676702 | 0.435657336 | 0.677894965 | 0.861984855 | -4.863978934 |
| **A_55_P1987291** | Plppr5 | 0.028503358 | -0.1305306 | 0.187537316 | 7.744577075 | 0.435501194 | 0.678002318 | 0.862064649 | -4.864014577 |
| **A_51_P469951** | Srgap3 | -0.032612825 | -0.214627442 | 0.149401792 | 7.994737052 | -0.43537686 | 0.678087808 | 0.862096072 | -4.864042952 |
| **A_55_P2155834** | NA | -0.046399696 | -0.305384461 | 0.21258507 | 16.717675 | -0.43533552 | 0.678116234 | 0.862096072 | -4.864052384 |
| **A_52_P330395** | Farp1 | -0.103238263 | -0.679669241 | 0.473192714 | 10.81133856 | -0.435187655 | 0.678217911 | 0.862168629 | -4.864086116 |
| **A_55_P2183688** | NA | -0.037734664 | -0.248617715 | 0.173148386 | 15.66905869 | -0.434792461 | 0.6784897 | 0.862395012 | -4.864176218 |
| **A_55_P2054817** | Rxfp1 | 0.028308905 | -0.129904417 | 0.186522227 | 7.665123361 | 0.434773648 | 0.67850264 | 0.862395012 | -4.864180506 |
| **A_55_P1995737** | Gm14288 | 0.063233007 | -0.290244591 | 0.416710605 | 11.41397498 | 0.434675468 | 0.67857017 | 0.862395012 | -4.864202878 |
| **A_51_P392147** | Xab2 | 0.067349338 | -0.309144282 | 0.443842957 | 10.15722841 | 0.434669232 | 0.67857446 | 0.862395012 | -4.864204299 |
| **A_55_P1975185** | Sqle | 0.078255477 | -0.359318695 | 0.515829649 | 13.56294055 | 0.4345565 | 0.678652003 | 0.862436856 | -4.864229981 |
| **A_51_P249414** | Ess2 | -0.0538258 | -0.354872891 | 0.24722129 | 9.945676562 | -0.434449448 | 0.678725645 | 0.862452352 | -4.864254363 |
| **A_55_P2172956** | Cnih2 | 0.03640378 | -0.167221084 | 0.240028643 | 8.259035583 | 0.434409047 | 0.678753437 | 0.862452352 | -4.864263563 |
| **A_55_P2343648** | 1700120C14Rik | -0.027929988 | -0.184239435 | 0.128379459 | 7.680593566 | -0.434178897 | 0.678911773 | 0.862596836 | -4.86431596 |
| **A_55_P2274233** | Agap3 | -0.067227866 | -0.524075827 | 0.389620095 | 11.90966144 | -0.453777736 | 0.679193265 | 0.862840305 | -4.726143725 |
| **A_55_P2064676** | Gas2l3 | 0.045597554 | -0.209864272 | 0.30105938 | 8.610605059 | 0.433709288 | 0.679234905 | 0.862840305 | -4.864422795 |
| **A_55_P2053858** | Scamp3 | -0.036617354 | -0.241768965 | 0.168534257 | 12.60575284 | -0.433705783 | 0.679237316 | 0.862840305 | -4.864423592 |
| **A_51_P481693** | Ero1l | 0.057747289 | -0.265910166 | 0.381404743 | 8.668405026 | 0.433539999 | 0.679351409 | 0.862928526 | -4.864461282 |
| **A_55_P2036567** | Vav1 | -0.098423842 | -0.650461053 | 0.45361337 | 11.88864945 | -0.433226633 | 0.67956709 | 0.863145768 | -4.864532488 |
| **A_65_P19832** | Ttr | -0.087018585 | -0.575164123 | 0.401126953 | 16.74062352 | -0.433157543 | 0.679614648 | 0.863149453 | -4.864548181 |
| **A_55_P2155421** | Sec61g | 0.056319358 | -0.259813081 | 0.372451797 | 15.90705267 | 0.432884288 | 0.679802755 | 0.86327154 | -4.864610225 |
| **A_65_P08227** | Snx5 | 0.040650294 | -0.187588299 | 0.268888887 | 9.084391725 | 0.432770649 | 0.679880992 | 0.86327154 | -4.864636017 |
| **A_51_P298178** | Dctn3 | -0.060535382 | -0.400432533 | 0.279361769 | 13.06136082 | -0.432757849 | 0.679889804 | 0.86327154 | -4.864638922 |
| **A_52_P230688** | Rep15 | -0.028674624 | -0.189721567 | 0.132372318 | 8.202911187 | -0.432641765 | 0.679969728 | 0.86327154 | -4.864665262 |
| **A_66_P111987** | Brwd3 | 0.041796645 | -0.192966165 | 0.276559454 | 8.525356766 | 0.432608761 | 0.679992453 | 0.86327154 | -4.864672749 |
| **A_51_P323880** | COX2 | 0.044923611 | -0.207447238 | 0.297294461 | 16.82997117 | 0.432532416 | 0.68004502 | 0.86327154 | -4.864690068 |
| **A_55_P2128511** | Ociad2 | -0.029152055 | -0.192956562 | 0.134652451 | 8.394625555 | -0.432440664 | 0.680108198 | 0.86327154 | -4.864710877 |
| **A_55_P2054098** | Gm9495 | 0.040083001 | -0.185147405 | 0.265313408 | 7.986108632 | 0.432430584 | 0.680115139 | 0.86327154 | -4.864713163 |
| **A_55_P2106834** | Ndufs3 | -0.072262046 | -0.478367026 | 0.333842934 | 14.10585525 | -0.432369579 | 0.680157148 | 0.86327154 | -4.864726997 |
| **A_51_P301259** | Araf | 0.044616514 | -0.20619009 | 0.295423118 | 11.58811466 | 0.432254832 | 0.680236166 | 0.86327154 | -4.864753012 |
| **A_52_P37091** | Magi1 | 0.032104628 | -0.14842482 | 0.212634075 | 7.853874178 | 0.432118416 | 0.680330113 | 0.86327154 | -4.864783932 |
| **A_55_P2073383** | NA | 0.059522819 | -0.275202793 | 0.394248432 | 14.80226393 | 0.432093506 | 0.680347269 | 0.86327154 | -4.864789577 |
| **A_55_P2079713** | Gramd1c | -0.084529494 | -0.559909148 | 0.390850161 | 10.36065156 | -0.432066769 | 0.680365683 | 0.86327154 | -4.864795636 |
| **A_51_P141164** | Ddx50 | 0.041782758 | -0.193223475 | 0.276788992 | 9.597418886 | 0.432017078 | 0.680399907 | 0.86327154 | -4.864806895 |
| **A_51_P388002** | Snrpb2 | 0.032379629 | -0.149746869 | 0.214506126 | 11.74386361 | 0.43199818 | 0.680412922 | 0.86327154 | -4.864811177 |
| **A_51_P355301** | Cyp3a11 | -0.104294742 | -0.690948492 | 0.482359007 | 9.238466895 | -0.431980112 | 0.680425366 | 0.86327154 | -4.86481527 |
| **A_55_P2121980** | Plcb4 | 0.062587734 | -0.289602554 | 0.414778022 | 8.176573897 | 0.431812332 | 0.680540929 | 0.863313992 | -4.864853276 |
| **A_55_P2082609** | NA | -0.085372378 | -0.565855743 | 0.395110987 | 11.20715638 | -0.431739926 | 0.680590804 | 0.863313992 | -4.864869673 |
| **A_51_P438619** | Sobp | 0.040161658 | -0.185873272 | 0.266196589 | 7.896906327 | 0.431736999 | 0.680592819 | 0.863313992 | -4.864870336 |
| **A_51_P171697** | Pebp1 | -0.074361725 | -0.493184391 | 0.344460942 | 13.16255612 | -0.431422176 | 0.680809695 | 0.863532423 | -4.864941602 |
| **A_55_P2052106** | BC049762 | -0.03904099 | -0.259025719 | 0.18094374 | 8.315161798 | -0.43123249 | 0.680940382 | 0.863619852 | -4.864984518 |
| **A_55_P2003378** | Atrnl1 | -0.036024022 | -0.239027866 | 0.166979822 | 7.937925386 | -0.43119243 | 0.680967984 | 0.863619852 | -4.86499358 |
| **A_51_P463187** | Tbc1d2b | -0.043749383 | -0.290463697 | 0.202964931 | 9.486283282 | -0.430884406 | 0.681180232 | 0.863832351 | -4.865063227 |
| **A_55_P2145908** | Phb | 0.053998947 | -0.250562249 | 0.358560143 | 13.77663909 | 0.43081807 | 0.681225945 | 0.863833648 | -4.86507822 |
| **A_55_P2045916** | NA | -0.044217693 | -0.29369114 | 0.205255754 | 8.365916349 | -0.430680235 | 0.681320935 | 0.863897426 | -4.865109367 |
| **A_55_P1977149** | Susd6 | -0.038861964 | -0.258205032 | 0.180481104 | 11.45289569 | -0.430510768 | 0.681437734 | 0.863988846 | -4.86514765 |
| **A_51_P241074** | Map2k1 | -0.051297153 | -0.341018988 | 0.238424682 | 9.664586495 | -0.430224585 | 0.681634996 | 0.864087062 | -4.865212267 |
| **A_55_P2176530** | Ldah | 0.039765712 | -0.184836838 | 0.264368263 | 8.81058443 | 0.430206802 | 0.681647254 | 0.864087062 | -4.865216281 |
| **A_55_P2147427** | Prdx1 | -0.070503067 | -0.468777881 | 0.327771748 | 14.17826132 | -0.430138536 | 0.681694314 | 0.864087062 | -4.865231688 |

| | | | | | | | | | |
|---|---|---|---|---|---|---|---|---|---|
| **A_55_P2132039** | NA | 0.039367182 | -0.183039818 | 0.261774181 | 9.662168724 | 0.430099625 | 0.681721138 | 0.864087062 | -4.86524047 |
| **A_55_P2005813** | Abhd18 | -0.032575766 | -0.216625213 | 0.15147368 | 8.168590532 | -0.430074123 | 0.681738718 | 0.864087062 | -4.865246224 |
| **A_52_P450835** | Cdpf1 | 0.070481322 | -0.327818553 | 0.468781197 | 10.82005125 | 0.429978815 | 0.681804424 | 0.864113678 | -4.865267728 |
| **A_55_P2166572** | Strbp | 0.049665546 | -0.231095638 | 0.33042673 | 9.649780464 | 0.42983457 | 0.681903873 | 0.864183055 | -4.865300266 |
| **A_55_P2147616** | NA | 0.071872417 | -0.33448998 | 0.478234813 | 11.99695081 | 0.42976587 | 0.681951239 | 0.864186423 | -4.865315759 |
| **A_51_P336060** | Lcat | -0.078540623 | -0.522698159 | 0.365616913 | 9.618790717 | -0.429675406 | 0.682013615 | 0.864208808 | -4.865336157 |
| **A_55_P1972234** | NA | 0.03119968 | -0.145281727 | 0.207681087 | 16.03227813 | 0.429570421 | 0.682086005 | 0.864219914 | -4.865359824 |
| **A_51_P498772** | 1700019D03Rik | -0.085267659 | -0.567781146 | 0.397245829 | 8.987028543 | -0.429396074 | 0.682206232 | 0.864219914 | -4.865399116 |
| **A_55_P1952537** | Rrp7a | 0.057586473 | -0.268290774 | 0.383463721 | 11.348353 | 0.429387732 | 0.682211985 | 0.864219914 | -4.865400996 |
| **A_55_P2150633** | Gadd45gip1 | 0.050094464 | -0.233387138 | 0.333576066 | 13.0586504 | 0.429386162 | 0.682213068 | 0.864219914 | -4.865401349 |
| **A_51_P344878** | Thop1 | 0.060266348 | -0.280815202 | 0.401347898 | 8.924940318 | 0.429338503 | 0.682245934 | 0.864219914 | -4.865412087 |
| **A_55_P2046509** | Amot | 0.108736797 | -0.506953912 | 0.724427506 | 11.16172063 | 0.42913815 | 0.682384111 | 0.864315279 | -4.865457217 |
| **A_51_P163106** | Bdh1 | 0.1049746 | -0.489546283 | 0.699495483 | 13.61517209 | 0.429042489 | 0.682450089 | 0.864315279 | -4.865478758 |
| **A_55_P2165160** | Arhgef1 | -0.039044691 | -0.26019361 | 0.182104229 | 8.34811215 | -0.429003023 | 0.68247731 | 0.864315279 | -4.865487643 |
| **A_52_P625171** | Asph | 0.068776831 | -0.320804755 | 0.458358417 | 8.765141389 | 0.428970007 | 0.682500083 | 0.864315279 | -4.865495076 |
| **A_55_P1968240** | Cbr4 | 0.072014224 | -0.335986875 | 0.480015324 | 10.86023663 | 0.428884295 | 0.682559205 | 0.864333521 | -4.86551437 |
| **A_51_P441914** | Hsd17b2 | -0.134214577 | -0.894775536 | 0.626346383 | 11.90360984 | -0.428794144 | 0.682621391 | 0.864339572 | -4.86553466 |
| **A_51_P436447** | Zfp871 | 0.048056717 | -0.224327912 | 0.320441346 | 8.809845385 | 0.42870119 | 0.682685513 | 0.864339572 | -4.865555576 |
| **A_51_P125467** | Igf2r | 0.035629503 | -0.166326516 | 0.237585522 | 12.18776883 | 0.428682893 | 0.682698135 | 0.864339572 | -4.858265078 |
| **A_55_P2082439** | Spout1 | 0.037792665 | -0.176552982 | 0.252138313 | 11.68483304 | 0.428426219 | 0.682875213 | 0.864507138 | -4.865617424 |
| **A_52_P517224** | Trim63 | -0.034622948 | -0.231049884 | 0.161803987 | 7.69910028 | -0.428298149 | 0.682963575 | 0.864562378 | -4.865646219 |
| **A_55_P2063871** | NA | -0.056264054 | -0.375588165 | 0.263060057 | 11.09875972 | -0.428136742 | 0.683074947 | 0.86462807 | -4.865682497 |
| **A_55_P1990894** | Psmd4 | 0.033696184 | -0.157564407 | 0.224956776 | 14.1061535 | 0.428093285 | 0.683104934 | 0.86462807 | -4.865692262 |
| **A_51_P298486** | Cog6 | -0.035673105 | -0.238206488 | 0.166860278 | 9.598667417 | -0.42798395 | 0.683180382 | 0.864666946 | -4.865716827 |
| **A_55_P2036317** | NA | 0.033990684 | -0.159049787 | 0.227031156 | 16.93124739 | 0.427853145 | 0.683270651 | 0.864683337 | -4.865746208 |
| **A_55_P1968729** | Rpl10 | 0.042196575 | -0.197456791 | 0.281849941 | 16.5704335 | 0.427835538 | 0.683282802 | 0.864683337 | -4.865750163 |
| **A_55_P2057906** | Cetn2 | -0.048439266 | -0.323603659 | 0.226725128 | 11.74118887 | -0.427748509 | 0.683342865 | 0.864702732 | -4.865769706 |
| **A_55_P2120737** | NA | 0.027879746 | -0.130538994 | 0.186298485 | 7.707672295 | 0.42762732 | 0.683426507 | 0.864751961 | -4.865796914 |
| **A_55_P1959758** | Top3b | -0.030737802 | -0.205494156 | 0.144018552 | 9.762260668 | -0.427388688 | 0.683591221 | 0.86487586 | -4.865850468 |
| **A_51_P286737** | Ccl2 | 0.060196479 | -0.282071606 | 0.402464563 | 8.29457881 | 0.427354097 | 0.683615099 | 0.86487586 | -4.865858229 |
| **A_51_P362538** | Syde2 | -0.038138336 | -0.255019177 | 0.178742506 | 8.30574228 | -0.427290991 | 0.683658661 | 0.86487586 | -4.865872386 |
| **A_52_P483280** | Lamp2 | -0.062627053 | -0.41883583 | 0.293581723 | 11.9536948 | -0.427209155 | 0.683715155 | 0.864890721 | -4.865890741 |
| **A_55_P2044381** | NA | 0.035901061 | -0.168350654 | 0.240152777 | 8.042113197 | 0.427095273 | 0.683793774 | 0.864897818 | -4.86591628 |
| **A_55_P2156718** | NA | -0.027082879 | -0.181192832 | 0.127027073 | 7.674696212 | -0.427019133 | 0.68384634 | 0.864897818 | -4.865933351 |
| **A_55_P2039532** | Pax8 | -0.027829011 | -0.186189334 | 0.130531313 | 7.676489479 | -0.427006586 | 0.683855003 | 0.864897818 | -4.865936164 |
| **A_51_P361788** | Vapa | -0.053703983 | -0.359376444 | 0.251968478 | 13.01653853 | -0.426907093 | 0.683923696 | 0.864928103 | -4.865958466 |
| **A_55_P2164629** | Ptn | 0.111760385 | -0.524458913 | 0.747979682 | 9.066508021 | 0.426839143 | 0.683970612 | 0.864930845 | -4.865973695 |
| **A_55_P2002815** | Rabgap1l | 0.028715484 | -0.134896611 | 0.19232758 | 7.749153842 | 0.426465526 | 0.684228604 | 0.86520049 | -4.86605739 |
| **A_52_P246749** | Ghitm | 0.050504619 | -0.237431876 | 0.338441115 | 12.39180461 | 0.426204055 | 0.684409185 | 0.86537222 | -4.866115923 |
| **A_55_P1990299** | Ophn1 | -0.030942768 | -0.207431055 | 0.145545519 | 7.767254334 | -0.426016539 | 0.684538703 | 0.865376826 | -4.86615788 |
| **A_55_P2008061** | Itpr2 | -0.103837678 | -0.696105621 | 0.488430266 | 11.18642207 | -0.426010133 | 0.684543128 | 0.865376826 | -4.866159313 |
| **A_66_P139974** | Arl6 | -0.091249324 | -0.611723082 | 0.429224434 | 8.308150616 | -0.426004325 | 0.68454714 | 0.865376826 | -4.866160613 |
| **A_52_P663413** | P4ha2 | -0.041662976 | -0.279403458 | 0.196077506 | 10.51784422 | -0.425824199 | 0.684671565 | 0.865477515 | -4.866200899 |
| **A_51_P300297** | Srek1ip1 | 0.034570085 | -0.162753407 | 0.231893577 | 9.364281554 | 0.425701172 | 0.684756555 | 0.865528345 | -4.866228407 |
| **A_51_P324923** | Man2a1 | 0.077636676 | -0.365621993 | 0.520895345 | 12.48073898 | 0.425591439 | 0.684832365 | 0.865567566 | -4.866252935 |
| **A_51_P497821** | Zfp790 | 0.03265738 | -0.153883923 | 0.219198682 | 8.160438922 | 0.425392195 | 0.684970025 | 0.865644483 | -4.866297458 |
| **A_52_P271377** | Efr3a | -0.033052203 | -0.221871149 | 0.155766743 | 8.219645745 | -0.425341768 | 0.685004868 | 0.865644483 | -4.866308723 |
| **A_52_P748958** | Cfl1 | -0.035808936 | -0.240401341 | 0.16878347 | 13.46657995 | -0.425289923 | 0.685040691 | 0.865644483 | -4.866320304 |
| **A_55_P2074927** | Gm7932 | 0.032636336 | -0.153849691 | 0.219122364 | 7.896513931 | 0.425244092 | 0.68507236 | 0.865644483 | -4.86633054 |
| **A_55_P2112589** | Uxt | 0.045761114 | -0.21582393 | 0.307346159 | 11.30223745 | 0.425076277 | 0.685188324 | 0.865734417 | -4.866368013 |

| | | | | | | | | |
|---|---|---|---|---|---|---|---|---|
| **A_55_P2070976** | Gm6923 | 0.083115922 | -0.392326917 | 0.558558761 | 8.826103019 | 0.424784934 | 0.68538967 | 0.865932214 | -4.866433037 |
| **A_55_P2000399** | Rhox4a | -0.028661521 | -0.192641461 | 0.135318418 | 7.684244484 | -0.424709237 | 0.685441989 | 0.865941712 | -4.866449925 |
| **A_55_P2039572** | NA | -0.047180548 | -0.317334525 | 0.22297343 | 15.92424196 | -0.424360351 | 0.685683149 | 0.865986445 | -4.866527726 |
| **A_55_P1988313** | NA | 0.027761526 | -0.131210976 | 0.186734028 | 7.665394103 | 0.424330759 | 0.685703606 | 0.865986445 | -4.866534322 |
| **A_55_P2077153** | Slc27a2 | -0.13333025 | -0.896841087 | 0.630180586 | 13.79111977 | -0.424323103 | 0.685708898 | 0.865986445 | -4.866536029 |
| **A_51_P497882** | Creb3l4 | -0.049302671 | -0.331648899 | 0.233043556 | 7.978157191 | -0.424298652 | 0.685725801 | 0.865986445 | -4.866541478 |
| **A_51_P393598** | Atp6v1d | -0.034904363 | -0.234796834 | 0.164988108 | 12.18068639 | -0.424293588 | 0.685729302 | 0.865986445 | -4.866542607 |
| **A_66_P101477** | Bglap3 | 0.027956257 | -0.132154602 | 0.188067117 | 7.653231424 | 0.424269127 | 0.685746212 | 0.865986445 | -4.866548059 |
| **A_66_P126686** | NA | -0.036672557 | -0.246756087 | 0.173410973 | 15.43547459 | -0.424162603 | 0.685819856 | 0.866010224 | -4.866571797 |
| **A_51_P292943** | Ell | -0.069895527 | -0.470348693 | 0.330557638 | 10.89816972 | -0.424112277 | 0.685854649 | 0.866010224 | -4.86658301 |
| **A_55_P2105100** | Lrig1 | 0.061603037 | -0.291634362 | 0.414840437 | 9.964745732 | 0.423758709 | 0.686099116 | 0.866262318 | -4.866661754 |
| **A_55_P2064240** | Hemk1 | 0.036974648 | -0.17518133 | 0.249130626 | 8.336305961 | 0.423479082 | 0.686292487 | 0.866393159 | -4.866723987 |
| **A_52_P541161** | Rgs18 | 0.126508538 | -0.599390508 | 0.852407584 | 10.13304536 | 0.423474048 | 0.686295968 | 0.866393159 | -4.866725107 |
| **A_55_P2059337** | Fastk | 0.034244274 | -0.162283824 | 0.230772372 | 11.72638448 | 0.423395755 | 0.686350116 | 0.866393159 | -4.866742525 |
| **A_55_P2102365** | Spag7 | 0.101295894 | -0.480105251 | 0.68269704 | 15.84719338 | 0.423349595 | 0.68638204 | 0.866393159 | -4.866752793 |
| **A_52_P314129** | Pkia | -0.037652192 | -0.253845977 | 0.178541593 | 7.813971352 | -0.42318498 | 0.686495896 | 0.866404206 | -4.866789402 |
| **A_55_P2017694** | Gprc5c | 0.087208153 | -0.413617182 | 0.588033487 | 11.63149744 | 0.423110678 | 0.68654729 | 0.866404206 | -4.866805921 |
| **A_52_P674414** | Mkln1 | 0.042184882 | -0.200085692 | 0.284455456 | 9.453297673 | 0.423096427 | 0.686557147 | 0.866404206 | -4.866809089 |
| **A_55_P1962651** | NA | 0.050834882 | -0.24115435 | 0.342824115 | 8.200759365 | 0.423036828 | 0.686598373 | 0.866404206 | -4.866822338 |
| **A_51_P120830** | Mmp10 | 0.027777022 | -0.131814737 | 0.187368781 | 7.667113916 | 0.422920182 | 0.686679062 | 0.866404206 | -4.866848263 |
| **A_66_P131324** | Platr21 | -0.072866455 | -0.491549565 | 0.345816656 | 7.841679759 | -0.422888025 | 0.686701307 | 0.866404206 | -4.866855408 |
| **A_51_P329251** | Chchd1 | 0.060208431 | -0.28574683 | 0.406163692 | 13.96798527 | 0.422883323 | 0.68670456 | 0.866404206 | -4.866856453 |
| **A_55_P2121437** | NA | -0.067811912 | -0.45754195 | 0.321918125 | 13.98756285 | -0.42279058 | 0.686768719 | 0.866428599 | -4.866877059 |
| **A_52_P533792** | Sucla2 | 0.055797414 | -0.26502915 | 0.376623978 | 12.24313273 | 0.422597517 | 0.686902287 | 0.866540549 | -4.866919941 |
| **A_55_P1960972** | Lrrc59 | 0.054275203 | -0.257879129 | 0.366429536 | 8.382677579 | 0.422488877 | 0.686977453 | 0.866578816 | -4.866944064 |
| **A_51_P189899** | Olfr1134 | 0.03047516 | -0.144881289 | 0.205831609 | 7.794909237 | 0.422286741 | 0.687117319 | 0.866606459 | -4.866988932 |
| **A_52_P527842** | Reps1 | -0.028645752 | -0.193501447 | 0.136209942 | 8.401953042 | -0.42222065 | 0.687163053 | 0.866606459 | -4.867003598 |
| **A_55_P2148744** | NA | 0.063401565 | -0.301489524 | 0.428292655 | 12.08121769 | 0.422201636 | 0.687176211 | 0.866606459 | -4.867007816 |
| **A_55_P1956195** | Sap18b | -0.038396101 | -0.259430402 | 0.1826382 | 13.05518486 | -0.422095418 | 0.687249715 | 0.866606459 | -4.867031381 |
| **A_55_P2025038** | Cpe | -0.149653337 | -1.011214078 | 0.711907404 | 10.12748545 | -0.422069231 | 0.687267838 | 0.866606459 | -4.86703719 |
| **A_51_P339540** | Cdkn1c | 0.076604181 | -0.364506715 | 0.517715076 | 14.60853965 | 0.421976122 | 0.687332275 | 0.866606459 | -4.867057841 |
| **A_55_P2016981** | Nfu1 | -0.048362036 | -0.326922086 | 0.230198015 | 11.8294695 | -0.421860565 | 0.687412251 | 0.866606459 | -4.867083465 |
| **A_51_P321374** | Plpp7 | -0.034315386 | -0.231986368 | 0.163355597 | 8.166120913 | -0.421821928 | 0.687438993 | 0.866606459 | -4.867092031 |
| **A_55_P1986252** | NA | 0.029614335 | -0.140978076 | 0.200206746 | 16.42120652 | 0.421818343 | 0.687441474 | 0.866606459 | -4.867092826 |
| **A_55_P1974477** | Msh3 | 0.032492961 | -0.154685683 | 0.219671605 | 10.40945099 | 0.421809329 | 0.687447712 | 0.866606459 | -4.867094824 |
| **A_55_P2104846** | Rsl24d1 | 0.035992064 | -0.171386019 | 0.243370146 | 8.278622768 | 0.421722801 | 0.687507602 | 0.866625437 | -4.867114004 |
| **A_55_P2172058** | Trim55 | 0.02719338 | -0.12952701 | 0.183913771 | 7.706767713 | 0.421619686 | 0.687578976 | 0.866658886 | -4.867136857 |
| **A_51_P316523** | Irf2 | -0.053705525 | -0.363347159 | 0.25593611 | 9.212007705 | -0.42144684 | 0.687698624 | 0.866753176 | -4.867175153 |
| **A_55_P2061452** | NA | -0.053549523 | -0.362555374 | 0.255456327 | 15.52821183 | -0.421087255 | 0.687947567 | 0.866919376 | -4.867254775 |
| **A_55_P2138090** | U2af1l4 | -0.061107523 | -0.41374596 | 0.291530914 | 11.77847882 | -0.421064105 | 0.687963596 | 0.866919376 | -4.867259899 |
| **A_55_P2013223** | S100z | 0.026969622 | -0.128667066 | 0.18260631 | 7.700770414 | 0.421062016 | 0.687965042 | 0.866919376 | -4.867260361 |
| **A_52_P31814** | 1110038B12Rik | 0.042449934 | -0.202735502 | 0.287635371 | 12.14926542 | 0.420693249 | 0.68822039 | 0.86710117 | -4.867341948 |
| **A_55_P1965935** | Rwdd1 | 0.03018475 | -0.14417126 | 0.204540759 | 9.608767218 | 0.420662549 | 0.688241649 | 0.86710117 | -4.867348737 |
| **A_51_P297679** | Hcls1 | -0.068645641 | -0.465166142 | 0.32787486 | 11.56309959 | -0.420659314 | 0.68824389 | 0.86710117 | -4.867349453 |
| **A_51_P458778** | Hpgd | -0.124099153 | -0.841217372 | 0.593019067 | 10.79775537 | -0.420495399 | 0.688357407 | 0.867139003 | -4.867385694 |
| **A_51_P393067** | Ap4m1 | -0.040878943 | -0.277106704 | 0.195348818 | 8.527831933 | -0.420486358 | 0.688363669 | 0.867139003 | -4.867387693 |
| **A_55_P2025023** | NA | -0.029978642 | -0.20324335 | 0.143286066 | 15.99860976 | -0.420421619 | 0.688408505 | 0.867139003 | -4.867402002 |
| **A_51_P387379** | Tshz3 | -0.09654902 | -0.65476822 | 0.46167018 | 8.58879929 | -0.420267965 | 0.688514927 | 0.86719729 | -4.867435958 |
| **A_52_P74427** | Mms19 | 0.029737121 | -0.142211998 | 0.20168624 | 8.145813485 | 0.420225257 | 0.688544509 | 0.86719729 | -4.867445394 |
| **A_55_P2127258** | Dok7 | -0.027229174 | -0.184792727 | 0.130334378 | 7.65880446 | -0.419915495 | 0.688759082 | 0.867385554 | -4.867513806 |

| | | | | | | | | | |
|---|---|---|---|---|---|---|---|---|---|
| **A_52_P578436** | NA | 0.031498438 | -0.150784961 | 0.213781838 | 7.886072007 | 0.419879904 | 0.688783738 | 0.867385554 | -4.867521663 |
| **A_52_P393314** | P2rx7 | -0.033310865 | -0.226139102 | 0.159517372 | 8.221079591 | -0.419757509 | 0.688868531 | 0.86743582 | -4.86754868 |
| **A_55_P1989504** | Tmem164 | -0.03209516 | -0.21798 | 0.153789679 | 11.59527169 | -0.419545213 | 0.689015618 | 0.867564516 | -4.867595523 |
| **A_65_P13713** | Ank3 | -0.031719929 | -0.215489265 | 0.152049407 | 8.04864764 | -0.419413442 | 0.689106922 | 0.867595743 | -4.867624588 |
| **A_51_P378858** | Atg10 | -0.034070682 | -0.231474909 | 0.163333544 | 8.270268841 | -0.419379863 | 0.68913019 | 0.867595743 | -4.867631993 |
| **A_55_P2174836** | Vldlr | -0.039861883 | -0.270933041 | 0.191209276 | 7.869657182 | -0.419174886 | 0.689272231 | 0.86771805 | -4.867677185 |
| **A_51_P336833** | Fabp4 | -0.100300193 | -0.681832663 | 0.481232277 | 12.77168521 | -0.419093561 | 0.68932859 | 0.867732485 | -4.867695109 |
| **A_55_P2163669** | NA | 0.047753609 | -0.229168664 | 0.324675881 | 15.80138134 | 0.419016861 | 0.689381746 | 0.867742887 | -4.867712011 |
| **A_55_P2046398** | Eif4a1 | 0.033128665 | -0.159052875 | 0.225310205 | 14.5574509 | 0.418866336 | 0.689486071 | 0.867768588 | -4.867745174 |
| **A_55_P2063553** | Gm4234 | 0.028084615 | -0.134859179 | 0.191028409 | 7.727932625 | 0.418806911 | 0.689527258 | 0.867768588 | -4.867758263 |
| **A_55_P1996087** | Gm14326 | -0.059446974 | -0.404363061 | 0.285469113 | 10.67908999 | -0.418793075 | 0.689536848 | 0.867768588 | -4.86776131 |
| **A_55_P2102385** | Etl4 | -0.035955848 | -0.244766309 | 0.172854613 | 9.391433758 | -0.418408475 | 0.689803447 | 0.86804758 | -4.867845978 |
| **A_55_P2000454** | Tspan18 | -0.102313328 | -0.696654951 | 0.492028295 | 8.740899115 | -0.418291708 | 0.689884398 | 0.868092932 | -4.86787167 |
| **A_51_P138790** | Plekhf2 | 0.041080576 | -0.197709038 | 0.27987019 | 9.511750928 | 0.418026951 | 0.690067961 | 0.868257587 | -4.867929899 |
| **A_55_P2042993** | Arl6ip6 | 0.088697927 | -0.426943753 | 0.604339607 | 15.26242525 | 0.417973402 | 0.690105092 | 0.868257587 | -4.867941673 |
| **A_51_P240253** | Rrad | 0.062327644 | -0.300368622 | 0.42502391 | 7.775436625 | 0.417561852 | 0.690390484 | 0.86840123 | -4.868032109 |
| **A_55_P2055199** | Airn | -0.071815558 | -0.489723809 | 0.346092694 | 9.323820239 | -0.417561817 | 0.690390508 | 0.86840123 | -4.868032116 |
| **A_55_P2004927** | Dnajc25 | 0.051805651 | -0.249698975 | 0.353310277 | 9.085067909 | 0.417509486 | 0.690426802 | 0.86840123 | -4.86804361 |
| **A_51_P206225** | Uroc1 | 0.134002033 | -0.645965124 | 0.913969189 | 11.4530355 | 0.417463259 | 0.690458862 | 0.86840123 | -4.868053762 |
| **A_51_P360880** | Zfp266 | 0.060937163 | -0.293772479 | 0.415646805 | 9.13071169 | 0.417438429 | 0.690476084 | 0.86840123 | -4.868059215 |
| **A_55_P2081133** | Gapdh | 0.074441194 | -0.358893243 | 0.50777563 | 16.1601684 | 0.417420058 | 0.690488825 | 0.86840123 | -4.868063249 |
| **A_52_P647740** | Klhl41 | 0.027312321 | -0.131723081 | 0.186347724 | 7.663323556 | 0.41729962 | 0.690572359 | 0.868429578 | -4.868089691 |
| **A_55_P2148182** | NA | 0.04877981 | -0.235319787 | 0.332879407 | 10.67358168 | 0.417208048 | 0.690635877 | 0.868429578 | -4.868109791 |
| **A_66_P137152** | Zbtb8os | 0.035675051 | -0.1721082 | 0.243458302 | 12.1320832 | 0.417193234 | 0.690646152 | 0.868429578 | -4.868113042 |
| **A_55_P2097616** | Stxbp1 | -0.029964281 | -0.204537065 | 0.144608504 | 7.858941645 | -0.41707149 | 0.690730602 | 0.86843607 | -4.868139757 |
| **A_52_P472799** | Ecd | 0.034025634 | -0.16421596 | 0.232267227 | 10.6990063 | 0.41705625 | 0.690741173 | 0.86843607 | -4.868143101 |
| **A_65_P12152** | Rasa2 | 0.029328135 | -0.141632845 | 0.200289116 | 8.282067895 | 0.416841202 | 0.690890358 | 0.868564608 | -4.868190272 |
| **A_55_P2362635** | Slc9a3 | -0.028919931 | -0.197526415 | 0.139686553 | 7.653692527 | -0.41677933 | 0.690933282 | 0.868564608 | -4.86820384 |
| **A_52_P566605** | Hsd17b7 | 0.101649969 | -0.491163836 | 0.694463773 | 9.703907009 | 0.416650713 | 0.691022517 | 0.868620292 | -4.868232037 |
| **A_55_P1984103** | Gfra3 | 0.027291164 | -0.131936226 | 0.186518555 | 7.793944223 | 0.416473598 | 0.691145409 | 0.868650774 | -4.868270855 |
| **A_55_P2090441** | NA | 0.043187468 | -0.208881795 | 0.295256731 | 8.222487229 | 0.416314023 | 0.691256139 | 0.868650774 | -4.868305815 |
| **A_55_P2087622** | Il4i1 | 0.033929809 | -0.164112302 | 0.231971921 | 8.140253137 | 0.416300622 | 0.691265438 | 0.868650774 | -4.86830875 |
| **A_55_P2186562** | Hars2 | -0.035708067 | -0.244153013 | 0.172736878 | 10.58640656 | -0.416253757 | 0.69129796 | 0.868650774 | -4.868319015 |
| **A_55_P1985005** | Ghrh | 0.027722203 | -0.134108845 | 0.189553251 | 7.767833008 | 0.416245048 | 0.691304004 | 0.868650774 | -4.868320923 |
| **A_51_P398260** | Tppp | -0.037986224 | -0.259744012 | 0.183771564 | 7.85946854 | -0.416227173 | 0.691316408 | 0.868650774 | -4.868324837 |
| **A_51_P249793** | Dusp28 | -0.039901258 | -0.272984087 | 0.193181571 | 9.647616154 | -0.415967592 | 0.691496558 | 0.868820656 | -4.86838167 |
| **A_55_P2177968** | Snrpd2 | -0.051103449 | -0.349791237 | 0.24758434 | 13.93913725 | -0.41573437 | 0.691658434 | 0.868874166 | -4.868432704 |
| **A_55_P1972915** | NA | 0.100792467 | -0.488483334 | 0.690068268 | 15.12018681 | 0.415616382 | 0.691740334 | 0.868874166 | -4.868458512 |
| **A_51_P465582** | Hdhd3 | 0.04302957 | -0.208578925 | 0.294638065 | 10.00469191 | 0.41555154 | 0.691785346 | 0.868874166 | -4.868472693 |
| **A_52_P217675** | Slf1 | 0.036389247 | -0.176436148 | 0.249214642 | 8.288782053 | 0.415463434 | 0.691846509 | 0.868874166 | -4.868491958 |
| **A_55_P2007334** | Mepce | 0.05242926 | -0.254260225 | 0.359118746 | 11.70441368 | 0.415391909 | 0.691896162 | 0.868874166 | -4.868507595 |
| **A_52_P582907** | Entpd4 | -0.031863158 | -0.218287468 | 0.154561152 | 8.039076913 | -0.415307198 | 0.691954973 | 0.868874166 | -4.868526111 |
| **A_55_P1981479** | Irgm1 | -0.04538319 | -0.310923095 | 0.220156715 | 8.520151413 | -0.415287076 | 0.691968943 | 0.868874166 | -4.868530509 |
| **A_51_P375078** | Lmbrd2 | -0.037213761 | -0.254988423 | 0.1805609 | 8.552199693 | -0.415221095 | 0.692014753 | 0.868874166 | -4.868544928 |
| **A_55_P2140626** | Arl5b | 0.037442044 | -0.181676198 | 0.256560287 | 9.184796472 | 0.415206561 | 0.692024843 | 0.868874166 | -4.868548103 |
| **A_55_P1994294** | NA | -0.043599371 | -0.298793571 | 0.21159483 | 10.47467945 | -0.415138114 | 0.692072366 | 0.868874166 | -4.868563059 |
| **A_55_P2127782** | NA | -0.05495109 | -0.376692151 | 0.266789971 | 14.75993411 | -0.415004695 | 0.692165004 | 0.868874166 | -4.868592204 |
| **A_55_P2109257** | Nipal3 | 0.050339748 | -0.244423694 | 0.345103191 | 9.577034267 | 0.414973734 | 0.692186503 | 0.868874166 | -4.868598966 |
| **A_55_P1952577** | Ces1a | 0.026336986 | -0.127921584 | 0.180595556 | 7.696017451 | 0.414858469 | 0.692266541 | 0.868874166 | -4.868624136 |
| **A_55_P2344598** | E330037I15Rik | -0.057212353 | -0.392317771 | 0.277893065 | 8.250665799 | -0.414850434 | 0.692272121 | 0.868874166 | -4.868625891 |

| | | | | | | | | | |
|---|---|---|---|---|---|---|---|---|---|
| **A_55_P2010121** | Rab27b | -0.059289329 | -0.406612239 | 0.288033581 | 8.780433396 | -0.414788105 | 0.692315403 | 0.868874166 | -4.868639498 |
| **A_51_P452875** | Pigyl | -0.066509238 | -0.456137646 | 0.323119169 | 10.16199388 | -0.414776888 | 0.692323192 | 0.868874166 | -4.868641947 |
| **A_51_P191463** | H2-M2 | -0.091947961 | -0.630681434 | 0.446785512 | 11.10196984 | -0.414716516 | 0.692365117 | 0.868874166 | -4.846758459 |
| **A_55_P2058127** | Pde4dip | 0.03114074 | -0.151351978 | 0.213633457 | 7.77791613 | 0.414635586 | 0.69242132 | 0.868874166 | -4.868672789 |
| **A_66_P138999** | NA | 0.030064584 | -0.146141895 | 0.206271064 | 7.964724962 | 0.414587797 | 0.69245451 | 0.868874166 | -4.868683218 |
| **A_55_P1960896** | Fndc3c1 | 0.027008796 | -0.1313068 | 0.185324391 | 7.736417847 | 0.414538343 | 0.692488856 | 0.868874166 | -4.868694009 |
| **A_55_P2107035** | Mcmbp | 0.031896522 | -0.155099439 | 0.218892484 | 11.23972273 | 0.41447114 | 0.692535529 | 0.868874166 | -4.868708671 |
| **A_55_P2153296** | Pycrl | 0.047485423 | -0.2309412 | 0.325912046 | 8.45430709 | 0.414412398 | 0.692576328 | 0.868874166 | -4.868721485 |
| **A_55_P2145709** | NA | -0.049239378 | -0.337961353 | 0.239482597 | 7.749639603 | -0.414396335 | 0.692587485 | 0.868874166 | -4.868724988 |
| **A_55_P2155216** | Ankrd34a | -0.028924375 | -0.198575469 | 0.140726718 | 7.788703217 | -0.414276708 | 0.692670575 | 0.868874166 | -4.868751078 |
| **A_52_P637282** | Wipf1 | -0.045024897 | -0.309142123 | 0.219092329 | 9.759447622 | -0.414227753 | 0.69270458 | 0.868874166 | -4.868761753 |
| **A_52_P127206** | Gm9 | 0.026408148 | -0.128504524 | 0.18132082 | 7.679390117 | 0.414222979 | 0.692707895 | 0.868874166 | -4.868762794 |
| **A_55_P2107247** | Tssk5 | 0.210663891 | -1.025760853 | 1.447088635 | 8.523679399 | 0.414005016 | 0.692859303 | 0.869007687 | -4.868810306 |
| **A_66_P123643** | Ptbp2 | 0.030644415 | -0.149274371 | 0.210563201 | 8.310118964 | 0.413864348 | 0.692957027 | 0.869073862 | -4.868840957 |
| **A_55_P2226660** | 5033418A18Rik | 0.031139009 | -0.151718999 | 0.213997017 | 8.089925294 | 0.413784285 | 0.69301265 | 0.869087232 | -4.868858399 |
| **A_55_P1981499** | Rpl10a | -0.042327051 | -0.291022622 | 0.206368519 | 16.31741608 | -0.413554883 | 0.693172037 | 0.869230719 | -4.868908355 |
| **A_51_P413545** | Txndc11 | -0.039118085 | -0.269049588 | 0.190813418 | 11.19100575 | -0.413392221 | 0.693285064 | 0.869316056 | -4.868943763 |
| **A_65_P20249** | Prkcb | 0.085104487 | -0.415303549 | 0.585512523 | 9.127847478 | 0.413248582 | 0.69338488 | 0.869384817 | -4.868975018 |
| **A_55_P2151116** | Micu1 | 0.031660212 | -0.154585179 | 0.217905603 | 11.8951982 | 0.413058408 | 0.693517043 | 0.869494125 | -4.869016385 |
| **A_55_P2056300** | Sf3b3 | 0.043466347 | -0.21240616 | 0.299338853 | 12.30049495 | 0.412774352 | 0.693714473 | 0.869685241 | -4.86907814 |
| **A_55_P1957038** | Gstp2 | 0.067046848 | -0.327702793 | 0.461796489 | 15.59195016 | 0.412705077 | 0.693762626 | 0.869689202 | -4.869093195 |
| **A_66_P126485** | Dipk1a | -0.031970334 | -0.220268183 | 0.156327515 | 8.444714656 | -0.412557993 | 0.693864868 | 0.869760962 | -4.869125152 |
| **A_51_P476851** | Fance | -0.047751444 | -0.329065735 | 0.233562848 | 11.35529582 | -0.412456261 | 0.69393559 | 0.869793205 | -4.869147249 |
| **A_51_P325592** | 2410004B18Rik | 0.041502349 | -0.203139069 | 0.286143767 | 9.929847549 | 0.412216985 | 0.694101941 | 0.8699453 | -4.869199202 |
| **A_55_P1974412** | Kat2a | 0.038363802 | -0.187880355 | 0.264607958 | 10.0318582 | 0.412028689 | 0.694232862 | 0.86999827 | -4.869240066 |
| **A_66_P130602** | Pdzd8 | -0.038756776 | -0.267332517 | 0.189818966 | 8.157137828 | -0.412003297 | 0.694250518 | 0.86999827 | -4.869245575 |
| **A_55_P1978166** | Zfp945 | 0.035923442 | -0.175963392 | 0.247810275 | 7.978395734 | 0.411961999 | 0.694279234 | 0.86999827 | -4.869254535 |
| **A_66_P100186** | Psg28 | -0.028418798 | -0.196123314 | 0.139285717 | 7.726888053 | -0.411759985 | 0.694419711 | 0.870062713 | -4.869298351 |
| **A_55_P1965378** | Uqcrq | -0.048934183 | -0.337704716 | 0.23983635 | 15.77612946 | -0.41175858 | 0.694420688 | 0.870062713 | -4.869298656 |
| **A_55_P2170963** | Gspt1 | 0.056873088 | -0.2788521 | 0.392598276 | 12.34179323 | 0.411629102 | 0.694510732 | 0.87011913 | -4.869326729 |
| **A_52_P616392** | Sbno2 | 0.030634334 | -0.150322804 | 0.211591472 | 8.466285082 | 0.411354182 | 0.694701938 | 0.870302273 | -4.869386308 |
| **A_52_P182331** | NA | 0.043256928 | -0.212355131 | 0.298868987 | 15.27351096 | 0.411204184 | 0.694806273 | 0.870376568 | -4.8694188 |
| **A_55_P1958961** | 2010001K21Rik | -0.038837999 | -0.26854574 | 0.190869742 | 8.269499647 | -0.410832133 | 0.695065091 | 0.870623898 | -4.869499345 |
| **A_52_P562593** | Tpgs1 | 0.151153196 | -0.743002803 | 1.045309194 | 10.44562752 | 0.410759133 | 0.695115879 | 0.870623898 | -4.86951514 |
| **A_55_P1995243** | Cobl | 0.04561887 | -0.224273558 | 0.315511299 | 9.151920982 | 0.410711644 | 0.695148919 | 0.870623898 | -4.869525414 |
| **A_55_P2103944** | Mrpl30 | 0.038495456 | -0.189280973 | 0.266271885 | 12.73267205 | 0.410661394 | 0.695183881 | 0.870623898 | -4.869536285 |
| **A_52_P101982** | Tmem129 | 0.033425107 | -0.164517118 | 0.231367332 | 10.4653186 | 0.410315147 | 0.695424809 | 0.870830642 | -4.869611155 |
| **A_55_P1999314** | Dhx33 | 0.047735133 | -0.235002487 | 0.330472754 | 10.58345875 | 0.41023974 | 0.695477284 | 0.870830642 | -4.869627452 |
| **A_55_P1996419** | NA | -0.047108615 | -0.326142022 | 0.231924792 | 15.68174174 | -0.410229911 | 0.695484123 | 0.870830642 | -4.869629576 |
| **A_52_P167511** | NA | -0.041677136 | -0.288590008 | 0.205235736 | 8.836342532 | -0.410144907 | 0.69554328 | 0.8708483 | -4.869647944 |
| **A_55_P2157814** | Gmds | -0.051110255 | -0.354120844 | 0.251900335 | 10.51429271 | -0.40985801 | 0.695742955 | 0.870995621 | -4.869709913 |
| **A_55_P2060752** | Usp12 | 0.03587045 | -0.176795977 | 0.248536876 | 11.33109222 | 0.409846354 | 0.695751068 | 0.870995621 | -4.86971243 |
| **A_55_P2382741** | NA | 0.029266673 | -0.144316657 | 0.202850003 | 7.761443255 | 0.409683557 | 0.695864384 | 0.871081062 | -4.869747575 |
| **A_66_P110909** | Ppp1r12a | 0.048205655 | -0.237854935 | 0.334266246 | 11.61755614 | 0.409471001 | 0.696012348 | 0.871209861 | -4.869793442 |
| **A_51_P104077** | Usp13 | -0.028972282 | -0.200935221 | 0.142990657 | 7.709079108 | -0.40938416 | 0.696072804 | 0.871229115 | -4.842658551 |
| **A_55_P2080995** | Tgif2lx1 | 0.028016119 | -0.138300045 | 0.194332284 | 7.958517873 | 0.409314104 | 0.696121577 | 0.871233745 | -4.869827285 |
| **A_52_P402319** | Med20 | 0.037491368 | -0.185110641 | 0.260093378 | 9.561256946 | 0.409246903 | 0.696168363 | 0.871235887 | -4.869841777 |
| **A_51_P415475** | Dok4 | -0.138523721 | -0.961284036 | 0.684236594 | 9.263309019 | -0.409104261 | 0.696267677 | 0.871254828 | -4.86987253 |
| **A_55_P2034570** | Ddx19a | 0.045794062 | -0.226205032 | 0.317793157 | 9.087957826 | 0.409095686 | 0.696273647 | 0.871254828 | -4.869874378 |
| **A_55_P2117319** | Ntan1 | -0.058366205 | -0.405176432 | 0.288444022 | 11.10066876 | -0.408933557 | 0.696386538 | 0.871339681 | -4.86990932 |

| | | | | | | | | | |
|---|---|---|---|---|---|---|---|---|---|
| **A_55_P2083252** | Mtus2 | -0.065931082 | -0.458026601 | 0.326164437 | 8.545957213 | -0.408584147 | 0.696629861 | 0.871365933 | -4.869984581 |
| **A_52_P134381** | Psmd12 | -0.053350849 | -0.370650572 | 0.263948875 | 10.41443044 | -0.408559044 | 0.696647344 | 0.871365933 | -4.869989986 |
| **A_51_P465740** | Lysmd4 | -0.030535375 | -0.212210766 | 0.151140015 | 8.697562479 | -0.408404338 | 0.696755092 | 0.871365933 | -4.870023288 |
| **A_52_P37894** | Cox7a2 | -0.033537725 | -0.233080429 | 0.16600498 | 14.63354942 | -0.408395482 | 0.69676126 | 0.871365933 | -4.870025194 |
| **A_55_P2107432** | Slc18a1 | -0.037406781 | -0.25999766 | 0.185184098 | 8.196043072 | -0.408343985 | 0.696797128 | 0.871365933 | -4.870036276 |
| **A_52_P350537** | Mtmr11 | -0.053116676 | -0.369211077 | 0.262977726 | 8.564521443 | -0.408316822 | 0.696816048 | 0.871365933 | -4.870042121 |
| **A_55_P2007155** | Akr1c14 | -0.157942581 | -1.097928764 | 0.782043603 | 10.40637597 | -0.408282672 | 0.696839835 | 0.871365933 | -4.870049469 |
| **A_51_P441970** | Stox2 | -0.036401231 | -0.253042291 | 0.180239828 | 8.697717935 | -0.408280365 | 0.696841441 | 0.871365933 | -4.870049966 |
| **A_55_P2006479** | 3300002I08Rik | -0.026563968 | -0.18467659 | 0.131548653 | 8.301252432 | -0.408234403 | 0.696873456 | 0.871365933 | -4.870059854 |
| **A_55_P2157019** | Phb2 | -0.031705537 | -0.220437336 | 0.157026263 | 11.12875113 | -0.408200203 | 0.696897278 | 0.871365933 | -4.870067212 |
| **A_52_P219372** | Tmem177 | 0.035451881 | -0.175585124 | 0.246488886 | 8.347527556 | 0.408191404 | 0.696903407 | 0.871365933 | -4.870069105 |
| **A_55_P1954116** | C8b | -0.087581845 | -0.609149419 | 0.43398573 | 12.86026698 | -0.408024926 | 0.697019375 | 0.87145456 | -4.870104911 |
| **A_55_P2182324** | NA | -0.027079578 | -0.188394356 | 0.134235199 | 7.686181312 | -0.407897395 | 0.697108218 | 0.871494705 | -4.870132331 |
| **A_66_P129622** | Phldb3 | -0.056098313 | -0.390362568 | 0.278165941 | 9.399714368 | -0.407796089 | 0.697178795 | 0.871494705 | -4.870154107 |
| **A_55_P2059352** | Col18a1 | -0.080734437 | -0.561883665 | 0.400414791 | 13.43117623 | -0.407720188 | 0.697231676 | 0.871494705 | -4.870170419 |
| **A_55_P2036559** | Fbxl12os | 0.04413479 | -0.218893392 | 0.307162973 | 10.78246041 | 0.407719961 | 0.697231834 | 0.871494705 | -4.870170467 |
| **A_55_P2153721** | Ccdc127 | -0.031926003 | -0.222277855 | 0.15842585 | 8.41016431 | -0.407540363 | 0.697356968 | 0.871543174 | -4.870209054 |
| **A_55_P2024580** | Dtnbp1 | -0.043003167 | -0.299407329 | 0.213400996 | 11.93653128 | -0.407529034 | 0.697364862 | 0.871543174 | -4.870211487 |
| **A_55_P2072030** | NA | -0.030914797 | -0.215290331 | 0.153460738 | 15.47993877 | -0.407423701 | 0.697438257 | 0.871543174 | -4.87023411 |
| **A_51_P513244** | Zfp949 | -0.03826906 | -0.26651557 | 0.18997745 | 8.638071027 | -0.407405455 | 0.697450971 | 0.871543174 | -4.870238028 |
| **A_51_P119597** | Ap3b1 | -0.037113194 | -0.258568271 | 0.184341883 | 11.81187938 | -0.407216985 | 0.697582306 | 0.871650941 | -4.863143332 |
| **A_66_P134394** | Pde6c | 0.089645909 | -0.445483782 | 0.624775601 | 11.7920604 | 0.407056429 | 0.697694198 | 0.871686863 | -4.870312948 |
| **A_55_P2029046** | Fgd3 | -0.04162689 | -0.290136935 | 0.206883156 | 8.010872954 | -0.407017612 | 0.697721252 | 0.871686863 | -4.870321276 |
| **A_55_P2038117** | Tmem104 | -0.051553492 | -0.359425151 | 0.256318167 | 8.861753329 | -0.406884896 | 0.697813749 | 0.871686863 | -4.870349746 |
| **A_55_P2027819** | Rpl41 | 0.043857918 | -0.218063233 | 0.305779069 | 17.72307083 | 0.406874649 | 0.697820892 | 0.871686863 | -4.870351944 |
| **A_51_P156934** | Dnajc15 | -0.040512926 | -0.282499573 | 0.201473721 | 12.53571248 | -0.406804172 | 0.697870014 | 0.871686863 | -4.870367059 |
| **A_55_P2051176** | NA | 0.045917564 | -0.228394199 | 0.320229327 | 15.70718914 | 0.406740668 | 0.697914277 | 0.871686863 | -4.870380676 |
| **A_51_P418016** | Ccnj | 0.030049485 | -0.149504744 | 0.209603714 | 8.288673154 | 0.406653564 | 0.697974993 | 0.871686863 | -4.870399351 |
| **A_52_P556140** | Vapb | -0.05027933 | -0.35075745 | 0.250198791 | 11.7735204 | -0.406592922 | 0.698017265 | 0.871686863 | -4.87041235 |
| **A_55_P2101133** | Tomm7 | -0.064472282 | -0.449773016 | 0.320828452 | 13.91202867 | -0.40658972 | 0.698019497 | 0.871686863 | -4.870413036 |
| **A_52_P305890** | Yipf5 | -0.044130778 | -0.307905734 | 0.219644177 | 11.535833 | -0.406528706 | 0.698062029 | 0.871686863 | -4.870426113 |
| **A_55_P2419514** | Ccny | -0.032806985 | -0.228958101 | 0.163344131 | 8.19829301 | -0.406404703 | 0.698148474 | 0.871718457 | -4.870452685 |
| **A_55_P2118302** | Fam110a | -0.050155773 | -0.350137063 | 0.249825517 | 11.79926929 | -0.406265503 | 0.698245519 | 0.871718457 | -4.870482505 |
| **A_51_P190805** | Ap1m1 | 0.033968447 | -0.169224519 | 0.237161412 | 11.68301806 | 0.406209625 | 0.698284476 | 0.871718457 | -4.870494472 |
| **A_51_P464394** | Klb | 0.088752883 | -0.442195999 | 0.619701765 | 9.769234514 | 0.406174774 | 0.698308775 | 0.871718457 | -4.870501935 |
| **A_51_P507280** | Patl1 | -0.048272943 | -0.337088096 | 0.24054221 | 9.167111154 | -0.406131797 | 0.698338739 | 0.871718457 | -4.870511138 |
| **A_55_P1991416** | NA | 0.044571693 | -0.222117093 | 0.31126048 | 8.053721282 | 0.406104282 | 0.698357923 | 0.871718457 | -4.870517029 |
| **A_55_P1961414** | NA | 0.0466675 | -0.232694543 | 0.326029542 | 13.6663425 | 0.405910537 | 0.698493015 | 0.871830782 | -4.870558503 |
| **A_55_P2076448** | Tmem255a | 0.05659199 | -0.282635572 | 0.395819551 | 8.7868542 | 0.405365722 | 0.698872961 | 0.872204459 | -4.870675028 |
| **A_55_P2131274** | Zfp663 | 0.10653033 | -0.532140804 | 0.745201464 | 12.00907876 | 0.405302398 | 0.698917128 | 0.872204459 | -4.870688563 |
| **A_52_P375546** | Lrrc46 | -0.063305285 | -0.442847714 | 0.316237144 | 8.075253048 | -0.405287141 | 0.69892777 | 0.872204459 | -4.870691823 |
| **A_52_P110257** | Wdr83 | 0.03583453 | -0.179122562 | 0.250791622 | 9.317955121 | 0.40507284 | 0.699077252 | 0.87233468 | -4.870737611 |
| **A_52_P670263** | Tfpi | -0.047956143 | -0.335774894 | 0.239862609 | 8.696925218 | -0.404863242 | 0.699223467 | 0.872425046 | -4.870782371 |
| **A_52_P283628** | Rps6ka3 | -0.026832499 | -0.187907412 | 0.134242414 | 8.203171069 | -0.404777539 | 0.699283257 | 0.872425046 | -4.870800668 |
| **A_55_P2056615** | NA | -0.058956707 | -0.412874827 | 0.294961413 | 12.99428219 | -0.404774932 | 0.699285076 | 0.872425046 | -4.870801224 |
| **A_51_P496845** | Capza2 | -0.058400842 | -0.409202572 | 0.292400888 | 10.7337329 | -0.404520531 | 0.699462572 | 0.872482576 | -4.870855513 |
| **A_66_P127969** | Bcat1 | 0.025889052 | -0.129642393 | 0.181420498 | 7.654663446 | 0.404465176 | 0.699501196 | 0.872482576 | -4.870867322 |
| **A_65_P15295** | NA | 0.027284559 | -0.136643713 | 0.19121283 | 7.663418115 | 0.404432697 | 0.699523858 | 0.872482576 | -4.87087425 |
| **A_55_P2246900** | NA | 0.108644164 | -0.544170477 | 0.761458805 | 10.33366008 | 0.404389346 | 0.699554108 | 0.872482576 | -4.870883496 |
| **A_55_P2067787** | Psg29 | -0.027787892 | -0.194789708 | 0.139213925 | 7.677022744 | -0.404312892 | 0.699607457 | 0.872482576 | -4.8708998 |

| | | | | | | | | | |
|---|---|---|---|---|---|---|---|---|---|
| **A_51_P317941** | Perp | 0.049195951 | -0.246466305 | 0.344858208 | 11.84149882 | 0.40431203 | 0.699608059 | 0.872482576 | -4.870899984 |
| **A_55_P2149269** | Galt | 0.071224341 | -0.35695276 | 0.499401441 | 8.852075821 | 0.404192444 | 0.699691509 | 0.872482576 | -4.870925481 |
| **A_55_P2033997** | Ddo | -0.028388247 | -0.199049446 | 0.142272952 | 7.837186172 | -0.404191315 | 0.699692297 | 0.872482576 | -4.870925722 |
| **A_52_P88793** | Zfp933 | 0.029691425 | -0.14889601 | 0.20827886 | 8.581971631 | 0.40398322 | 0.699837523 | 0.872607372 | -4.870970073 |
| **A_51_P440568** | Stk19 | -0.063332851 | -0.444422836 | 0.317757134 | 8.651203741 | -0.40381709 | 0.699953471 | 0.872640727 | -4.871005465 |
| **A_55_P2003353** | Gemin8 | -0.028184595 | -0.197779178 | 0.141409989 | 9.047012171 | -0.403815519 | 0.699954568 | 0.872640727 | -4.871005799 |
| **A_55_P2022116** | NA | -0.062335012 | -0.437550921 | 0.312880898 | 15.93199243 | -0.403676986 | 0.700051262 | 0.872704989 | -4.871035301 |
| **A_55_P2068228** | NA | 0.098689896 | -0.495600597 | 0.692980388 | 11.80261642 | 0.403512596 | 0.700166012 | 0.872791749 | -4.871070298 |
| **A_55_P2175291** | NA | 0.030705985 | -0.154310802 | 0.215722773 | 14.36427283 | 0.403269235 | 0.700335903 | 0.872873463 | -4.871122083 |
| **A_52_P348250** | Klhl29 | -0.026582113 | -0.186794069 | 0.133629842 | 7.730370487 | -0.403160312 | 0.700411949 | 0.872873463 | -4.871145251 |
| **A_52_P355398** | Dars | -0.054613767 | -0.383798893 | 0.27457136 | 9.571534273 | -0.403129996 | 0.700433115 | 0.872873463 | -4.871151699 |
| **A_55_P2025008** | Hvcn1 | 0.082217846 | -0.413457609 | 0.577893301 | 9.574592369 | 0.403043453 | 0.70049354 | 0.872873463 | -4.871170101 |
| **A_55_P2084448** | Cybc1 | -0.030137653 | -0.211836234 | 0.151560928 | 10.82594907 | -0.403033439 | 0.700500532 | 0.872873463 | -4.871172231 |
| **A_52_P339791** | Hdgf | 0.066513288 | -0.334659712 | 0.467686287 | 9.050247595 | 0.402865343 | 0.700617903 | 0.872873463 | -4.871207963 |
| **A_55_P1960641** | Trpt1 | -0.067745974 | -0.476370506 | 0.340878559 | 9.053119024 | -0.402848954 | 0.700629347 | 0.872873463 | -4.871211447 |
| **A_51_P354433** | Ppie | 0.032291808 | -0.162492235 | 0.227075851 | 8.488757161 | 0.40283034 | 0.700642345 | 0.872873463 | -4.871215403 |
| **A_51_P315795** | Tubb4a | 0.071324964 | -0.358969227 | 0.501619155 | 10.77506108 | 0.402771996 | 0.700683086 | 0.872873463 | -4.871227801 |
| **A_51_P253633** | Mrps9 | -0.029627238 | -0.208364538 | 0.149110062 | 10.18960934 | -0.402771901 | 0.700683152 | 0.872873463 | -4.871227821 |
| **A_51_P253697** | Rpain | 0.069988859 | -0.352447125 | 0.492424844 | 10.66357787 | 0.402579075 | 0.700817808 | 0.872964994 | -4.871268785 |
| **A_55_P2160732** | Tfcp2 | 0.026413787 | -0.13303003 | 0.185857603 | 8.58339888 | 0.402537341 | 0.700846954 | 0.872964994 | -4.871277649 |
| **A_51_P107020** | Kif5a | -0.025909452 | -0.182341911 | 0.130523006 | 7.752886383 | -0.402452426 | 0.700906258 | 0.872965632 | -4.864142128 |
| **A_55_P2051479** | NA | -0.069353703 | -0.488134754 | 0.349427348 | 10.02004194 | -0.402407272 | 0.700937793 | 0.872965632 | -4.871305268 |
| **A_55_P2162765** | Dnajc24 | 0.040439949 | -0.203804157 | 0.284684055 | 10.25175125 | 0.402318215 | 0.700999994 | 0.872986849 | -4.871324174 |
| **A_55_P2054777** | Maml1 | 0.038923979 | -0.19626214 | 0.274110097 | 10.43341997 | 0.40215061 | 0.701117061 | 0.873066999 | -4.871359745 |
| **A_52_P91235** | Ccni | -0.050421328 | -0.35514118 | 0.254298523 | 12.75975893 | -0.402065421 | 0.701176566 | 0.873066999 | -4.871377819 |
| **A_55_P2179309** | P2ry10b | 0.051870201 | -0.26163187 | 0.365372272 | 8.602179725 | 0.402032075 | 0.70119986 | 0.873066999 | -4.871384893 |
| **A_55_P2153251** | Plagl1 | 0.027394376 | -0.138240309 | 0.193029061 | 8.133005382 | 0.401877148 | 0.701308085 | 0.873145506 | -4.871417752 |
| **A_55_P2100824** | LOC676689 | -0.067504148 | -0.475875042 | 0.340866745 | 10.61609773 | -0.401660267 | 0.701459603 | 0.873248413 | -4.871463731 |
| **A_52_P257426** | Eps8l2 | 0.02835301 | -0.143183573 | 0.199889593 | 8.178980458 | 0.401629502 | 0.701481096 | 0.873248413 | -4.871470251 |
| **A_55_P2055127** | Slc7a14 | 0.02596658 | -0.131293091 | 0.18322625 | 7.643468788 | 0.401218151 | 0.701768519 | 0.87342298 | -4.871557389 |
| **A_55_P2015692** | Elob | -0.069799093 | -0.492529459 | 0.352931273 | 13.41337788 | -0.401207941 | 0.701775654 | 0.87342298 | -4.87155955 |
| **A_55_P1958350** | Nipsnap3b | 0.046584576 | -0.235559086 | 0.328728239 | 11.02894792 | 0.401194563 | 0.701785002 | 0.87342298 | -4.871562383 |
| **A_51_P258281** | Gal | 0.044764994 | -0.226374714 | 0.315904701 | 8.557783513 | 0.401170132 | 0.701802075 | 0.87342298 | -4.871567555 |
| **A_51_P410744** | Dgke | -0.027968657 | -0.197439805 | 0.141502491 | 7.986160557 | -0.401013533 | 0.701911512 | 0.873445391 | -4.871600703 |
| **A_52_P614731** | Gng12 | -0.057555388 | -0.406304377 | 0.291193601 | 9.147108583 | -0.401010937 | 0.701913326 | 0.873445391 | -4.871601252 |
| **A_51_P158073** | Shfl | -0.025981528 | -0.183436814 | 0.131473758 | 9.235426559 | -0.400950381 | 0.701955647 | 0.873445391 | -4.871614067 |
| **A_55_P2160381** | NA | 0.049354902 | -0.249828128 | 0.348537932 | 11.37627379 | 0.400845049 | 0.702029263 | 0.873477214 | -4.871636353 |
| **A_51_P247481** | Adck1 | 0.037799266 | -0.191393918 | 0.266992451 | 9.184641816 | 0.400741997 | 0.702101289 | 0.873477214 | -4.871658151 |
| **A_55_P2082986** | Hmgb1 | 0.037965476 | -0.192260507 | 0.268191458 | 13.30397242 | 0.400698482 | 0.702131703 | 0.873477214 | -4.871667354 |
| **A_51_P209807** | Ccnyl1 | -0.028969642 | -0.204663095 | 0.146723811 | 7.825396291 | -0.400655163 | 0.702161982 | 0.873477214 | -4.871676515 |
| **A_51_P274947** | Atad1 | -0.071482548 | -0.505184054 | 0.362218957 | 11.22308846 | -0.400490562 | 0.702277038 | 0.873538819 | -4.871711315 |
| **A_51_P176086** | Ffar2 | 0.05500635 | -0.278759797 | 0.388772497 | 8.764842101 | 0.400455009 | 0.702301891 | 0.873538819 | -4.87171883 |
| **A_55_P1991605** | Aplp1 | 0.027766414 | -0.140803907 | 0.196336736 | 8.674347434 | 0.400241275 | 0.702451305 | 0.873663361 | -4.871763994 |
| **A_51_P202964** | Csnk2a2 | 0.027058623 | -0.137238827 | 0.191356073 | 9.580600175 | 0.400182468 | 0.702492418 | 0.873663361 | -4.871776416 |
| **A_55_P2026761** | NA | -0.069103159 | -0.488783696 | 0.350577378 | 14.85248355 | -0.400094205 | 0.702554126 | 0.873682231 | -4.871795058 |
| **A_55_P2131068** | NA | -0.038481305 | -0.272224422 | 0.195261812 | 14.41048453 | -0.400031463 | 0.702597992 | 0.873682231 | -4.871808307 |
| **A_55_P2160594** | NA | 0.036365862 | -0.184613015 | 0.257344738 | 16.21716605 | 0.399876894 | 0.702706066 | 0.873758014 | -4.87184094 |
| **A_55_P2083582** | NA | -0.072661284 | -0.514276904 | 0.368954336 | 14.96586337 | -0.399799116 | 0.702760451 | 0.873758014 | -4.871857356 |
| **A_55_P2064063** | Casp14 | 0.030054775 | -0.152632297 | 0.212741847 | 7.818292446 | 0.399750352 | 0.702794549 | 0.873758014 | -4.871867646 |
| **A_55_P2043531** | NA | 0.046922671 | -0.238394603 | 0.332239945 | 13.59698658 | 0.39961138 | 0.70289173 | 0.87382263 | -4.871896967 |

| | | | | | | | | | |
|---|---|---|---|---|---|---|---|---|---|
| A_52_P348720 | Rhobtb3 | -0.0470455 | -0.333301357 | 0.239210358 | 9.270813867 | -0.39934375 | 0.703078896 | 0.873999098 | -4.871953407 |
| A_55_P2002376 | Srm | 0.062483363 | -0.317829809 | 0.442796536 | 11.87272281 | 0.399214421 | 0.70316935 | 0.874055329 | -4.871980668 |
| A_51_P149455 | Acadl | 0.083362512 | -0.424284837 | 0.591009861 | 12.44694792 | 0.399017423 | 0.703307142 | 0.874097875 | -4.872022177 |
| A_55_P2030160 | Tcf7l2 | 0.065340709 | -0.332597792 | 0.463279209 | 8.831410295 | 0.398979948 | 0.703333355 | 0.874097875 | -4.872030071 |
| A_55_P1990629 | NA | 0.080008449 | -0.407328919 | 0.567345816 | 12.41173631 | 0.398923229 | 0.703373031 | 0.874097875 | -4.872042018 |
| A_55_P2059323 | Gm13315 | 0.088356261 | -0.449931494 | 0.626644016 | 13.05776625 | 0.398846715 | 0.703426555 | 0.874097875 | -4.872058132 |
| A_55_P2126945 | 1700066B19Rik | 0.029874374 | -0.152129598 | 0.211878347 | 8.760678524 | 0.398842236 | 0.703429688 | 0.874097875 | -4.872059075 |
| A_51_P288906 | Clip1 | 0.033791697 | -0.1721604 | 0.239743794 | 9.654267505 | 0.398682315 | 0.703541564 | 0.874169934 | -4.872092744 |
| A_52_P175066 | Cd2ap | -0.038943494 | -0.276345342 | 0.198458355 | 10.87138709 | -0.398596978 | 0.703601267 | 0.874169934 | -4.872110706 |
| A_51_P105589 | Eci1 | -0.046712822 | -0.331524038 | 0.238098393 | 12.54976234 | -0.398531093 | 0.703647362 | 0.874169934 | -4.872124571 |
| A_52_P223333 | Mib1 | -0.034084737 | -0.241917482 | 0.173748008 | 8.858450467 | -0.398500766 | 0.703668581 | 0.874169934 | -4.872130952 |
| A_55_P1997485 | NA | 0.034814235 | -0.177664957 | 0.247293426 | 11.84164306 | 0.398128837 | 0.703928824 | 0.87439574 | -4.872209176 |
| A_66_P106212 | Secisbp2 | 0.037128417 | -0.189484512 | 0.263741346 | 9.416563989 | 0.398111694 | 0.70394082 | 0.87439574 | -4.87221278 |
| A_55_P2181109 | Il10rb | -0.028969605 | -0.20582749 | 0.14788828 | 9.85387622 | -0.398016735 | 0.704007272 | 0.87442209 | -4.87223274 |
| A_55_P2100895 | Arnt | 0.042082602 | -0.214880126 | 0.299045331 | 10.70969922 | 0.397938211 | 0.704062225 | 0.874434154 | -4.872249242 |
| A_55_P2072468 | Sirt6 | -0.047969284 | -0.341092733 | 0.245154166 | 9.061437674 | -0.397645305 | 0.704267223 | 0.87463256 | -4.872310771 |
| A_55_P2150901 | Foxo4 | -0.029973758 | -0.213220624 | 0.153273109 | 7.8694451 | -0.39745487 | 0.704400519 | 0.874741897 | -4.872350752 |
| A_51_P478952 | N4bp2l1 | -0.064708276 | -0.46044319 | 0.331026637 | 8.581049494 | -0.397318378 | 0.704496064 | 0.874778196 | -4.872379397 |
| A_55_P2060101 | Tprgl | 0.035463944 | -0.181440953 | 0.252368841 | 13.55127224 | 0.397283808 | 0.704520264 | 0.874778196 | -4.872386651 |
| A_51_P191586 | Arpc1a | -0.045375313 | -0.323226389 | 0.232475763 | 11.18174429 | -0.396817428 | 0.704846782 | 0.875127406 | -4.872484452 |
| A_51_P327530 | Ciao2a | 0.060236245 | -0.308693379 | 0.429165869 | 12.03859778 | 0.396732291 | 0.704906395 | 0.875145206 | -4.872502294 |
| A_66_P127118 | Trmt112 | 0.030354732 | -0.155748327 | 0.216457791 | 8.228059949 | 0.396329205 | 0.705188666 | 0.875439417 | -4.87258672 |
| A_55_P1994339 | Dhrs7c | -0.035688795 | -0.254541729 | 0.183164139 | 7.735187569 | -0.396244 | 0.705248339 | 0.87545727 | -4.872604556 |
| A_55_P2045770 | Gmfb | -0.034584436 | -0.246760525 | 0.177591653 | 11.69790177 | -0.396065896 | 0.705373082 | 0.875482681 | -4.872641827 |
| A_52_P495372 | Twnk | -0.058434949 | -0.416998929 | 0.300129031 | 8.874364656 | -0.395994544 | 0.70542306 | 0.875482681 | -4.872656754 |
| A_55_P2015258 | Trim13 | -0.044747495 | -0.319323911 | 0.22982892 | 8.465186028 | -0.395994075 | 0.705423388 | 0.875482681 | -4.872656852 |
| A_55_P1962110 | NA | 0.053012195 | -0.272308679 | 0.378333068 | 15.18560223 | 0.395956104 | 0.705449985 | 0.875482681 | -4.872664795 |
| A_55_P2003966 | Rybp | 0.040727304 | -0.209322261 | 0.290776869 | 11.16518153 | 0.395769885 | 0.705580428 | 0.875569134 | -4.872703738 |
| A_52_P383862 | Cmc1 | -0.043175434 | -0.308284063 | 0.221933195 | 10.66742703 | -0.395727323 | 0.705610244 | 0.875569134 | -4.872712636 |
| A_55_P1993268 | Ap1s1 | -0.033667518 | -0.240573689 | 0.173238653 | 13.27048467 | -0.395385596 | 0.705849652 | 0.875755686 | -4.872784048 |
| A_66_P134690 | Spin1 | -0.029448883 | -0.210430137 | 0.151532372 | 8.031050527 | -0.395383387 | 0.705851199 | 0.875755686 | -4.872784509 |
| A_51_P341571 | Klhdc2 | 0.031676696 | -0.163175317 | 0.226528709 | 11.21040933 | 0.395019164 | 0.706106408 | 0.875965694 | -4.872860559 |
| A_55_P2077826 | Ppp1cc | 0.032162932 | -0.165727602 | 0.230053467 | 12.54348642 | 0.394924253 | 0.706172918 | 0.875965694 | -4.872880366 |
| A_55_P2104810 | Mrpl27 | -0.056454461 | -0.403810975 | 0.290902053 | 12.17196506 | -0.394917164 | 0.706177886 | 0.875965694 | -4.872881845 |
| A_55_P2036526 | Mbd2 | 0.034660881 | -0.178621119 | 0.24794288 | 9.880976709 | 0.394883126 | 0.706201739 | 0.875965694 | -4.872888947 |
| A_51_P477456 | Cnot10 | 0.051844553 | -0.2673442 | 0.371033305 | 9.744401828 | 0.394674205 | 0.706348157 | 0.876091088 | -4.872932527 |
| A_55_P1986552 | Mcam | 0.084192652 | -0.434329483 | 0.602714788 | 14.4078979 | 0.394539128 | 0.706442829 | 0.876117482 | -4.872960692 |
| A_51_P135517 | Coch | 0.026141775 | -0.134879414 | 0.187162964 | 7.665877754 | 0.394489303 | 0.706477751 | 0.876117482 | -4.872971078 |
| A_55_P2165011 | Fut10 | -0.027984701 | -0.200374707 | 0.144405305 | 7.99106023 | -0.394449835 | 0.706505416 | 0.876117482 | -4.872979305 |
| A_55_P2263098 | A130078K24Rik | 0.028250144 | -0.145833601 | 0.202333888 | 8.138478832 | 0.394317116 | 0.706598445 | 0.876126614 | -4.873006964 |
| A_55_P2056285 | Sf3b5 | -0.044108438 | -0.315955641 | 0.227738765 | 13.07276307 | -0.394257533 | 0.706640211 | 0.876126614 | -4.873019378 |
| A_52_P505234 | Stt3a | 0.053096932 | -0.27422488 | 0.380418744 | 9.460140406 | 0.394164646 | 0.706705325 | 0.876126614 | -4.873038728 |
| A_52_P282700 | Zbtb33 | -0.040805761 | -0.292379566 | 0.210768045 | 8.874304513 | -0.394129778 | 0.706729769 | 0.876126614 | -4.873045991 |
| A_55_P1958210 | Ephb1 | 0.032505961 | -0.167905297 | 0.232917219 | 7.863716804 | 0.394116017 | 0.706739416 | 0.876126614 | -4.873048857 |
| A_55_P1961265 | NA | 0.039854085 | -0.206294428 | 0.286002599 | 8.337778809 | 0.39342216 | 0.707225908 | 0.876627556 | -4.873193249 |
| A_55_P2166688 | Timm10b | 0.031249777 | -0.161762248 | 0.224261802 | 11.7818495 | 0.393410317 | 0.707234213 | 0.876627556 | -4.873195711 |
| A_66_P121288 | Tceal6 | -0.103833338 | -0.745326119 | 0.537659442 | 8.592219123 | -0.393303881 | 0.707308853 | 0.876663855 | -4.873217839 |
| A_55_P2038867 | Mcat | 0.029460166 | -0.152581446 | 0.211501779 | 11.5164395 | 0.393230964 | 0.70735999 | 0.876671021 | -4.873232995 |
| A_66_P140886 | Inafm1 | -0.032526607 | -0.233582568 | 0.168529354 | 10.60910658 | -0.393101769 | 0.707450598 | 0.876727102 | -4.873259842 |
| A_55_P1997225 | Agap2 | 0.061875634 | -0.320771202 | 0.444522471 | 8.420700366 | 0.392920535 | 0.707577712 | 0.876828414 | -4.87329749 |

| | | | | | | | | | |
|---|---|---|---|---|---|---|---|---|---|
| **A_52_P244702** | Tcf7 | -0.034773061 | -0.249995377 | 0.180449254 | 7.98940579 | -0.392589622 | 0.707809834 | 0.877052299 | -4.873366189 |
| **A_55_P1953133** | NA | -0.06071008 | -0.436519059 | 0.315098899 | 14.50291781 | -0.392533603 | 0.707849132 | 0.877052299 | -4.873377814 |
| **A_55_P2118037** | Sirpa | -0.064477292 | -0.463765786 | 0.334811203 | 10.96883945 | -0.392376672 | 0.707959227 | 0.877132485 | -4.87341037 |
| **A_55_P2076639** | Vps25 | -0.039058455 | -0.281002129 | 0.20288522 | 9.534403882 | -0.392268985 | 0.70803478 | 0.877169867 | -4.873432704 |
| **A_51_P124089** | Gtpbp2 | 0.043208749 | -0.224498663 | 0.310916161 | 9.939729104 | 0.392188167 | 0.708091484 | 0.877183893 | -4.873449461 |
| **A_55_P2046842** | Tmem253 | -0.047256856 | -0.340132004 | 0.245618291 | 7.968890931 | -0.392071699 | 0.708173203 | 0.877208491 | -4.873473605 |
| **A_55_P1959633** | Hnf4a | 0.093309665 | -0.48505006 | 0.67166939 | 13.96598518 | 0.392023206 | 0.70820723 | 0.877208491 | -4.873483655 |
| **A_51_P379069** | Dkk1 | 0.025472871 | -0.132448378 | 0.18339412 | 7.669381113 | 0.391940834 | 0.70826503 | 0.877208491 | -4.873500725 |
| **A_55_P2056348** | NA | 0.044554548 | -0.23169316 | 0.320802255 | 12.44955247 | 0.391901158 | 0.708292872 | 0.877208491 | -4.873508946 |
| **A_52_P574653** | Bid | 0.060170388 | -0.313061781 | 0.433402556 | 9.707216928 | 0.391730089 | 0.70841292 | 0.877300956 | -4.873544382 |
| **A_51_P489313** | Sri | -0.050945522 | -0.367177997 | 0.265286953 | 12.39900563 | -0.391455807 | 0.708605417 | 0.877415481 | -4.873601168 |
| **A_55_P2024376** | NA | 0.034768934 | -0.18105926 | 0.250597127 | 15.9581459 | 0.391441065 | 0.708615764 | 0.877415481 | -4.873604219 |
| **A_55_P2149821** | Srsf11 | -0.05112757 | -0.368531843 | 0.266276702 | 12.33137778 | -0.391404285 | 0.708641579 | 0.877415481 | -4.873611831 |
| **A_55_P2155827** | NA | -0.033570111 | -0.242048836 | 0.174908615 | 15.53060302 | -0.391267899 | 0.708737308 | 0.877424085 | -4.87364005 |
| **A_51_P452779** | Pygl | -0.092449021 | -0.666726722 | 0.481828681 | 15.01846773 | -0.391168207 | 0.708807286 | 0.877424085 | -4.873660672 |
| **A_55_P2292737** | 2310002F09Rik | 0.102061263 | -0.531962415 | 0.73608494 | 14.22271619 | 0.39114578 | 0.708823029 | 0.877424085 | -4.87366531 |
| **A_55_P2130399** | Rilp | 0.077008129 | -0.40139312 | 0.555409378 | 11.12397595 | 0.3911357 | 0.708830104 | 0.877424085 | -4.873667395 |
| **A_52_P330694** | Tle1 | -0.072686596 | -0.524568009 | 0.379194817 | 11.13807759 | -0.390852633 | 0.70902882 | 0.877613861 | -4.873725917 |
| **A_55_P1972923** | Polr1d | 0.072588642 | -0.379632544 | 0.524809827 | 12.18290932 | 0.390032641 | 0.709604601 | 0.878270303 | -4.873895223 |
| **A_55_P1986199** | Tut1 | 0.038115835 | -0.199413035 | 0.275644706 | 8.544407686 | 0.389917049 | 0.709685784 | 0.878274879 | -4.873919063 |
| **A_55_P1985410** | Reps2 | 0.053931458 | -0.28223347 | 0.390096386 | 9.226061078 | 0.389827928 | 0.709748379 | 0.878274879 | -4.873937439 |
| **A_55_P2009007** | NA | -0.064530595 | -0.466821923 | 0.337760734 | 14.85387411 | -0.3897698 | 0.709789207 | 0.878274879 | -4.873949423 |
| **A_66_P121976** | Slc4a11 | 0.024777792 | -0.129712281 | 0.179267865 | 7.742856465 | 0.389713285 | 0.709828903 | 0.878274879 | -4.873961072 |
| **A_55_P2146105** | NA | -0.083991826 | -0.607978091 | 0.43999444 | 16.61460561 | -0.389493583 | 0.70998323 | 0.878274879 | -4.874006345 |
| **A_55_P2048767** | Efemp1 | -0.099834605 | -0.722681963 | 0.523012753 | 8.568298935 | -0.389477762 | 0.709994344 | 0.878274879 | -4.874009604 |
| **A_51_P260639** | Elp3 | -0.027424263 | -0.198523463 | 0.143674938 | 10.2684549 | -0.389466553 | 0.710002218 | 0.878274879 | -4.874011913 |
| **A_52_P574214** | Rrp1b | 0.036651565 | -0.192046058 | 0.265349188 | 8.867505752 | 0.389416242 | 0.710037561 | 0.878274879 | -4.874022276 |
| **A_51_P198835** | Dnajb9 | -0.101235887 | -0.732925445 | 0.530453672 | 10.94228075 | -0.38941617 | 0.710037611 | 0.878274879 | -4.874022291 |
| **A_55_P2010111** | Zxdc | -0.025933472 | -0.187767438 | 0.135900494 | 8.543181555 | -0.389380486 | 0.710062679 | 0.878274879 | -4.87402964 |
| **A_51_P198645** | Parvb | 0.028852631 | -0.151412617 | 0.20911788 | 8.315523883 | 0.388916712 | 0.710388518 | 0.878509802 | -4.874125103 |
| **A_55_P2167512** | NA | 0.060417511 | -0.317088881 | 0.437923902 | 12.50750718 | 0.388885461 | 0.710410477 | 0.878509802 | -4.874131532 |
| **A_52_P526265** | Wdr73 | 0.041859349 | -0.219694307 | 0.303413005 | 11.57845802 | 0.388879333 | 0.710414783 | 0.878509802 | -4.874132792 |
| **A_51_P179258** | Kif26b | -0.032911916 | -0.238573268 | 0.172749436 | 7.715098081 | -0.388851401 | 0.71043441 | 0.878509802 | -4.874138538 |
| **A_55_P2008081** | Rhoa | -0.033299301 | -0.241573848 | 0.174975246 | 13.02250333 | -0.388492028 | 0.710686951 | 0.87876587 | -4.874212427 |
| **A_55_P2153031** | Ttn | 0.025389938 | -0.133453944 | 0.18423382 | 7.695652152 | 0.388395634 | 0.710754697 | 0.878793421 | -4.874232235 |
| **A_55_P1986406** | Ddx54 | -0.036490293 | -0.264873234 | 0.191892648 | 11.3993744 | -0.388236959 | 0.71086622 | 0.878875091 | -4.874264832 |
| **A_55_P1998374** | Ppfibp1 | 0.054893855 | -0.288777615 | 0.398565325 | 10.48867086 | 0.388117691 | 0.710950051 | 0.878922516 | -4.874289325 |
| **A_55_P1962010** | Clec1b | -0.070644648 | -0.513132995 | 0.3718437 | 9.435013032 | -0.387936473 | 0.711077435 | 0.879023774 | -4.874326527 |
| **A_51_P252859** | Ccn1 | -0.087138969 | -0.633103989 | 0.458826051 | 9.218918933 | -0.387820523 | 0.711158945 | 0.879068315 | -4.874350322 |
| **A_66_P129873** | Npm1 | 0.052444624 | -0.27622844 | 0.381117688 | 14.63995638 | 0.387721652 | 0.711228452 | 0.879098014 | -4.874370607 |
| **A_55_P2069995** | NA | 0.087255665 | -0.459844065 | 0.634355394 | 11.19682648 | 0.387534454 | 0.711360062 | 0.879204465 | -4.874409 |
| **A_55_P2085170** | 6-Sep | 0.045625327 | -0.240731322 | 0.331981977 | 8.395614038 | 0.387152361 | 0.711628726 | 0.879445622 | -4.874487311 |
| **A_55_P2083449** | Spryd7 | -0.037142654 | -0.270274743 | 0.195989434 | 8.961816905 | -0.387127543 | 0.711646178 | 0.879445622 | -4.874492395 |
| **A_55_P1998823** | NA | -0.102356362 | -0.74521623 | 0.540503507 | 11.7066409 | -0.386884841 | 0.711816857 | 0.879600308 | -4.874542097 |
| **A_55_P2060966** | Mov10 | -0.051779467 | -0.377215366 | 0.273656431 | 11.57163443 | -0.386611982 | 0.712008765 | 0.879749104 | -4.874597941 |
| **A_55_P1975690** | Bicc1 | -0.081207306 | -0.591747099 | 0.429332488 | 11.31928297 | -0.386499283 | 0.712088036 | 0.879749104 | -4.874620996 |
| **A_55_P2089223** | Ak4 | 0.103834885 | -0.548974819 | 0.756644589 | 11.87610903 | 0.386491435 | 0.712093556 | 0.879749104 | -4.874622601 |
| **A_51_P259571** | Angptl6 | -0.061834955 | -0.450657393 | 0.326987483 | 11.34359941 | -0.386425629 | 0.712139845 | 0.879749104 | -4.874636059 |
| **A_55_P2160691** | NA | 0.031315885 | -0.16563382 | 0.228265589 | 7.90112044 | 0.386360339 | 0.712185772 | 0.879749104 | -4.87464941 |
| **A_55_P2001589** | Klhl8 | -0.042532087 | -0.310071148 | 0.225006974 | 8.718461065 | -0.386289305 | 0.712235741 | 0.879749104 | -4.874663933 |

| | | | | | | | | | |
|---|---|---|---|---|---|---|---|---|---|
| A_55_P2118570 | Cd37 | -0.073943088 | -0.53910084 | 0.391214663 | 9.531800684 | -0.386260689 | 0.712255871 | 0.879749104 | -4.874669783 |
| A_51_P124798 | Tmem192 | -0.030584682 | -0.223060387 | 0.161891023 | 8.649538928 | -0.386110172 | 0.71236176 | 0.879818472 | -4.867234813 |
| A_55_P2052613 | NA | -0.098049594 | -0.715298039 | 0.519198851 | 13.22331321 | -0.385983717 | 0.712450726 | 0.879818472 | -4.874726384 |
| A_55_P2184478 | NA | 0.028957565 | -0.153343168 | 0.211258298 | 16.4856438 | 0.3859729 | 0.712458336 | 0.879818472 | -4.874728593 |
| A_55_P2032272 | Habp2 | -0.097788849 | -0.713511675 | 0.517933977 | 13.43635064 | -0.3859111 | 0.712501817 | 0.879818472 | -4.874741217 |
| A_55_P2289040 | NA | -0.039161897 | -0.285786318 | 0.207462525 | 8.050101406 | -0.38584318 | 0.712549605 | 0.879818472 | -4.874755089 |
| A_51_P140350 | Nadsyn1 | 0.071528551 | -0.378985808 | 0.52204291 | 10.10913563 | 0.385792675 | 0.71258514 | 0.879818472 | -4.874765402 |
| A_55_P2116889 | Hax1 | -0.033289425 | -0.243011424 | 0.176432574 | 11.17241591 | -0.385696319 | 0.71265294 | 0.879845981 | -4.874785075 |
| A_55_P2166743 | Sp3os | 0.068319847 | -0.362362031 | 0.499001725 | 11.00136462 | 0.385454815 | 0.712822883 | 0.879980611 | -4.874834362 |
| A_51_P159673 | Snx1 | -0.0386173 | -0.282187113 | 0.204952514 | 12.5040328 | -0.385249093 | 0.71296766 | 0.879980611 | -4.874876325 |
| A_51_P308844 | Nrn1 | -0.101106504 | -0.738877306 | 0.536664299 | 12.33249276 | -0.385210085 | 0.712995114 | 0.879980611 | -4.874884279 |
| A_52_P615375 | H2aw | -0.058688087 | -0.442860444 | 0.325484269 | 11.76692766 | -0.388656746 | 0.712997942 | 0.879980611 | -4.828483945 |
| A_55_P1957291 | Rpl39 | 0.030270985 | -0.160702818 | 0.221244787 | 16.15109382 | 0.385155355 | 0.713033633 | 0.879980611 | -4.874895438 |
| A_52_P12877 | Hspa8 | -0.072330722 | -0.52868357 | 0.384022127 | 11.30367768 | -0.385128114 | 0.713052806 | 0.879980611 | -4.874900992 |
| A_55_P2033462 | Rps23-ps1 | 0.031288596 | -0.166139379 | 0.22871657 | 16.58975163 | 0.385088521 | 0.713080672 | 0.879980611 | -4.874909063 |
| A_55_P1990653 | Comt | -0.0336636 | -0.246177028 | 0.178849828 | 12.42038784 | -0.384908385 | 0.713207464 | 0.88008089 | -4.874945775 |
| A_66_P112482 | Tmem184c | -0.04194539 | -0.306833584 | 0.222942803 | 9.190874831 | -0.384773212 | 0.713302613 | 0.880084664 | -4.874973313 |
| A_52_P153848 | Prpf4b | 0.035660861 | -0.189651034 | 0.260972755 | 8.526140357 | 0.384583714 | 0.713436012 | 0.880084664 | -4.875011903 |
| A_52_P175376 | Tfcp2l1 | 0.048782105 | -0.259501318 | 0.357065529 | 9.118052518 | 0.38449753 | 0.713496686 | 0.880084664 | -4.875029449 |
| A_52_P546552 | Pelp1 | 0.04541637 | -0.241614389 | 0.332447129 | 9.730302599 | 0.38447417 | 0.713513132 | 0.880084664 | -4.875034203 |
| A_52_P402705 | Kcnj11 | 0.026257332 | -0.139691134 | 0.192205798 | 7.660279358 | 0.384468309 | 0.713517258 | 0.880084664 | -4.875035396 |
| A_51_P415519 | Tmem39a | 0.037093702 | -0.197383982 | 0.271571385 | 10.55956582 | 0.384398647 | 0.713566302 | 0.880084664 | -4.875049574 |
| A_55_P1965772 | Atp2a3 | -0.054046995 | -0.395696393 | 0.287602402 | 9.506550409 | -0.384391774 | 0.713571142 | 0.880084664 | -4.875050973 |
| A_55_P2051476 | C8a | 0.061756591 | -0.328633017 | 0.452146199 | 8.473496426 | 0.384386612 | 0.713574776 | 0.880084664 | -4.875052024 |
| A_52_P201206 | Scrn1 | -0.028045708 | -0.205583857 | 0.149492442 | 7.778573555 | -0.38384678 | 0.71395489 | 0.880456542 | -4.875161809 |
| A_55_P2062513 | Pdzrn3 | 0.096986822 | -0.516998678 | 0.710972322 | 10.39053543 | 0.383829018 | 0.713967398 | 0.880456542 | -4.875165418 |
| A_55_P1988951 | NA | -0.0771607 | -0.565891254 | 0.411569855 | 15.25204609 | -0.383627608 | 0.714109241 | 0.88057528 | -4.87520634 |
| A_51_P240864 | Sppl3 | -0.039886052 | -0.292862356 | 0.213090251 | 12.54056844 | -0.383110806 | 0.714473257 | 0.88096795 | -4.875311252 |
| A_55_P2019776 | Gxylt1 | -0.025878202 | -0.190041508 | 0.138285105 | 8.052988247 | -0.383037407 | 0.714524963 | 0.880975506 | -4.875326141 |
| A_52_P654108 | Dync1li2 | -0.026264125 | -0.192938709 | 0.14041046 | 8.154647205 | -0.382892392 | 0.714627124 | 0.881045266 | -4.875355551 |
| A_51_P491667 | Derl3 | 0.028825953 | -0.154289163 | 0.211941069 | 7.882149955 | 0.382509892 | 0.714896621 | 0.881189811 | -4.875433073 |
| A_55_P2158483 | Nkiras2 | 0.034525526 | -0.184810947 | 0.253862 | 8.325235075 | 0.382483444 | 0.714915257 | 0.881189811 | -4.87543843 |
| A_52_P508974 | Igf2bp3 | -0.057175246 | -0.420456569 | 0.306106076 | 9.296760193 | -0.382426663 | 0.714955268 | 0.881189811 | -4.875449932 |
| A_51_P272993 | Ntm | -0.025128432 | -0.18480658 | 0.134549716 | 7.737411775 | -0.38238697 | 0.714983238 | 0.881189811 | -4.875457971 |
| A_55_P2047673 | Srr | -0.040070914 | -0.294755614 | 0.214613787 | 8.838783328 | -0.382304647 | 0.715041248 | 0.881189811 | -4.875474641 |
| A_51_P455166 | Prl4a1 | -0.025150982 | -0.185020801 | 0.134718836 | 7.642961546 | -0.382271272 | 0.715064767 | 0.881189811 | -4.875481398 |
| A_51_P210340 | 2610306M01Rik | -0.04753273 | -0.34971908 | 0.25465362 | 8.480880786 | -0.382209173 | 0.715108529 | 0.881189811 | -4.87549397 |
| A_51_P352381 | Dhrs4 | 0.087193683 | -0.467139141 | 0.641526508 | 10.1219233 | 0.382206103 | 0.715110693 | 0.881189811 | -4.875494591 |
| A_55_P2361437 | Pitpnc1 | 0.030585279 | -0.163892027 | 0.225062585 | 8.638835435 | 0.382143704 | 0.715154667 | 0.881189811 | -4.875507222 |
| A_55_P1963036 | Oxr1 | 0.043300168 | -0.23207769 | 0.318678027 | 10.7271928 | 0.382070718 | 0.715206104 | 0.881197015 | -4.875521993 |
| A_55_P2292435 | Slc8a1 | 0.025908985 | -0.138962626 | 0.190780596 | 7.667787946 | 0.381845525 | 0.715364818 | 0.881336387 | -4.875567551 |
| A_51_P245454 | Ufsp2 | 0.041551196 | -0.222945933 | 0.306048325 | 11.53685879 | 0.381720741 | 0.715452772 | 0.881388568 | -4.875592785 |
| A_55_P2031432 | Usp47 | -0.026282972 | -0.193645528 | 0.141079583 | 8.067880479 | -0.381592093 | 0.715543454 | 0.881444104 | -4.875618792 |
| A_55_P2186787 | NA | 0.028216192 | -0.151488291 | 0.207920675 | 16.79735802 | 0.381524718 | 0.715590949 | 0.881446434 | -4.875632409 |
| A_51_P389895 | Slc35f5 | -0.057641863 | -0.424972103 | 0.309688377 | 10.96938611 | -0.38129799 | 0.715750783 | 0.881569852 | -4.875678217 |
| A_51_P109449 | Ltb4r1 | -0.080408295 | -0.592880775 | 0.432064184 | 10.81797734 | -0.381253198 | 0.715782361 | 0.881569852 | -4.875687263 |
| A_55_P1964724 | NA | -0.026331264 | -0.194183099 | 0.141520571 | 8.271491927 | -0.381178859 | 0.715834771 | 0.881578229 | -4.875702276 |
| A_55_P2076797 | Ddx5 | -0.050312199 | -0.371114 | 0.270489602 | 10.56953796 | -0.381083102 | 0.715902285 | 0.881592524 | -4.875721609 |
| A_52_P577748 | Lpxn | 0.048803407 | -0.262440117 | 0.360046931 | 8.614370362 | 0.381007043 | 0.715955912 | 0.881592524 | -4.875736962 |
| A_52_P433828 | Tctn3 | -0.029497775 | -0.217649592 | 0.158654041 | 7.92236863 | -0.380946542 | 0.715998571 | 0.881592524 | -4.875749173 |

| | | | | | | | | | |
|---|---|---|---|---|---|---|---|---|---|
| **A_55_P2148046** | NA | 0.096531104 | -0.519335064 | 0.712397272 | 9.886113435 | 0.380858906 | 0.716060364 | 0.881592524 | -4.875766856 |
| **A_55_P2121225** | Cyp3a11 | 0.141509913 | -0.761486919 | 1.044506744 | 11.59909847 | 0.380788502 | 0.716110009 | 0.881592524 | -4.87578106 |
| **A_52_P113830** | 1600025M17Rik | 0.026245857 | -0.141239306 | 0.193731019 | 7.740775514 | 0.380774279 | 0.716120038 | 0.881592524 | -4.875783929 |
| **A_55_P2138960** | Tpk1 | 0.035873536 | -0.193111295 | 0.264858366 | 8.499254163 | 0.380671757 | 0.716192334 | 0.881625374 | -4.875804608 |
| **A_55_P1976470** | 2410018L13Rik | 0.028806088 | -0.155125325 | 0.212737501 | 7.866627888 | 0.380549857 | 0.716278299 | 0.881675045 | -4.875829188 |
| **A_66_P136095** | Ctnna3 | 0.026127905 | -0.140851312 | 0.193107122 | 7.647647595 | 0.3802116 | 0.716516863 | 0.881912534 | -4.875897357 |
| **A_51_P486810** | Gpx2 | -0.05141319 | -0.380182801 | 0.27735642 | 8.162304994 | -0.379984671 | 0.71667693 | 0.881976826 | -4.875943059 |
| **A_52_P440962** | Cdc73 | 0.052087587 | -0.281083319 | 0.385258493 | 8.520672655 | 0.379883434 | 0.716748344 | 0.881976826 | -4.875963439 |
| **A_55_P2462358** | Fam118b | 0.033584828 | -0.181245216 | 0.248414873 | 10.9167375 | 0.379866769 | 0.7167601 | 0.881976826 | -4.875966793 |
| **A_55_P2121491** | Bbs9 | -0.076035457 | -0.562494631 | 0.410423717 | 9.073615165 | -0.379798243 | 0.716808441 | 0.881976826 | -4.875980585 |
| **A_51_P194306** | Lrrc1 | -0.082984206 | -0.614141261 | 0.448172849 | 8.704396842 | -0.379625765 | 0.716930121 | 0.881976826 | -4.876015288 |
| **A_52_P367774** | Zfp821 | 0.037285195 | -0.201367203 | 0.275937594 | 9.044662553 | 0.379624131 | 0.716931274 | 0.881976826 | -4.876015616 |
| **A_55_P2084032** | AW146154 | -0.037306669 | -0.276104378 | 0.201491041 | 9.451110156 | -0.379611624 | 0.716940097 | 0.881976826 | -4.876018132 |
| **A_55_P2015304** | NA | 0.041343606 | -0.223338263 | 0.306025475 | 13.3625145 | 0.379548562 | 0.716984588 | 0.881976826 | -4.876030816 |
| **A_51_P301804** | St3gal1 | 0.06920282 | -0.373906726 | 0.512312365 | 12.28770229 | 0.3794861 | 0.717028658 | 0.881976826 | -4.876043377 |
| **A_55_P2093634** | Ddi2 | 0.059878188 | -0.323557717 | 0.443314092 | 9.793935864 | 0.379453934 | 0.717051353 | 0.881976826 | -4.876049846 |
| **A_51_P369516** | Nudt16l1 | 0.040606963 | -0.219492584 | 0.30070651 | 12.21299166 | 0.379353506 | 0.717122212 | 0.881976826 | -4.876070036 |
| **A_52_P579252** | NA | 0.044458411 | -0.240324247 | 0.329241068 | 13.96384528 | 0.379335585 | 0.717134856 | 0.881976826 | -4.876073639 |
| **A_55_P2130877** | Parl | 0.026770658 | -0.144784495 | 0.198325811 | 10.52791283 | 0.379173928 | 0.717248925 | 0.881976826 | -4.876106128 |
| **A_52_P637172** | Pyroxd2 | -0.043986963 | -0.325878505 | 0.237904579 | 8.602492102 | -0.379162276 | 0.717257147 | 0.881976826 | -4.876108469 |
| **A_66_P116771** | Vps8 | 0.039585129 | -0.214220439 | 0.293390697 | 9.708381357 | 0.378978094 | 0.71738712 | 0.881976826 | -4.876145469 |
| **A_55_P1980165** | NA | 0.03058936 | -0.165558522 | 0.226737242 | 15.72241248 | 0.378939564 | 0.71741431 | 0.881976826 | -4.876153206 |
| **A_51_P407904** | Trappc4 | 0.045126351 | -0.244247686 | 0.334500389 | 12.37319388 | 0.378925513 | 0.717424226 | 0.881976826 | -4.876156028 |
| **A_55_P2084812** | Sec31a | 0.050607688 | -0.27392028 | 0.375135655 | 9.473262312 | 0.378920023 | 0.7174281 | 0.881976826 | -4.876157131 |
| **A_51_P462546** | Gmppa | -0.051302561 | -0.38033567 | 0.277730549 | 9.497325718 | -0.378863379 | 0.717468076 | 0.881976826 | -4.876168504 |
| **A_66_P129444** | Pnkp | -0.050731751 | -0.37624047 | 0.274776968 | 8.803398524 | -0.378704459 | 0.717580235 | 0.881976826 | -4.876200407 |
| **A_55_P2135526** | Gzmc | -0.024308358 | -0.180281402 | 0.131664686 | 7.705784308 | -0.378694731 | 0.717587101 | 0.881976826 | -4.876202359 |
| **A_51_P348652** | Spast | -0.041009104 | -0.304164681 | 0.222146474 | 10.45854819 | -0.378661269 | 0.717610719 | 0.881976826 | -4.876209075 |
| **A_55_P2100700** | Pygo2 | -0.04716408 | -0.349825069 | 0.255496909 | 12.02608138 | -0.37865013 | 0.717618581 | 0.881976826 | -4.87621131 |
| **A_52_P455540** | Ric8b | -0.03110735 | -0.231052807 | 0.168838108 | 9.055681875 | -0.378037303 | 0.718051172 | 0.882452384 | -4.876334198 |
| **A_55_P1953708** | Cry2 | -0.025577264 | -0.190032219 | 0.138877692 | 8.407789412 | -0.377911671 | 0.718139869 | 0.882505279 | -4.876359368 |
| **A_55_P2029622** | Ppp1r3f | 0.026224791 | -0.142536367 | 0.19498595 | 7.667746826 | 0.377591955 | 0.718365612 | 0.882726568 | -4.876423386 |
| **A_55_P1992208** | Gpi1 | -0.061944165 | -0.460768488 | 0.336880158 | 14.81946979 | -0.377400053 | 0.718501124 | 0.88283696 | -4.876461787 |
| **A_51_P103865** | Ppp1r35 | 0.051359522 | -0.279442527 | 0.38216157 | 9.309248079 | 0.377255839 | 0.718602968 | 0.882905973 | -4.876490633 |
| **A_51_P164495** | Dda1 | -0.037115077 | -0.276410583 | 0.202180429 | 10.73847048 | -0.376876462 | 0.718870915 | 0.883132075 | -4.876566469 |
| **A_51_P346893** | Extl1 | -0.024210428 | -0.180332859 | 0.131912002 | 7.695880317 | -0.376808213 | 0.718919123 | 0.883132075 | -4.876580104 |
| **A_52_P446120** | Commd6 | 0.051713856 | -0.281772301 | 0.385200013 | 10.9323989 | 0.376801219 | 0.718924062 | 0.883132075 | -4.876581501 |
| **A_55_P1954604** | NA | -0.035234555 | -0.262532452 | 0.192063342 | 8.193850352 | -0.376666127 | 0.719019489 | 0.883145631 | -4.876608483 |
| **A_51_P501757** | NA | 0.061972267 | -0.337883208 | 0.461827742 | 8.038882269 | 0.376597581 | 0.719067911 | 0.883145631 | -4.876622171 |
| **A_55_P1990548** | Tbc1d22a | -0.025344111 | -0.188908492 | 0.13822027 | 11.48956007 | -0.376505665 | 0.719132844 | 0.883145631 | -4.876640521 |
| **A_51_P257684** | Stau1 | 0.038704906 | -0.211107203 | 0.288517015 | 8.981965886 | 0.376474637 | 0.719154764 | 0.883145631 | -4.876646715 |
| **A_52_P617636** | Klf12 | -0.054372054 | -0.405315837 | 0.296571729 | 8.431917692 | -0.376462202 | 0.719163549 | 0.883145631 | -4.876649197 |
| **A_55_P2081885** | NA | 0.046912404 | -0.255991959 | 0.349816766 | 16.573608 | 0.376326972 | 0.719259087 | 0.883165835 | -4.876676184 |
| **A_51_P197883** | Exo5 | 0.053025818 | -0.289367521 | 0.395419157 | 9.854288777 | 0.376309567 | 0.719271383 | 0.883165835 | -4.876679657 |
| **A_51_P334979** | Apoc2 | 0.105531603 | -0.576261427 | 0.787324633 | 16.70523771 | 0.376108516 | 0.719413434 | 0.883245896 | -4.876719761 |
| **A_55_P1965699** | NA | -0.025045247 | -0.186929144 | 0.13683865 | 7.641848681 | -0.375928159 | 0.719540874 | 0.883245896 | -4.87675572 |
| **A_55_P2096010** | NA | -0.062356419 | -0.465479503 | 0.340766665 | 12.82360655 | -0.3758605 | 0.719588684 | 0.883245896 | -4.876769206 |
| **A_55_P2091191** | Slc28a2 | -0.062905756 | -0.469584977 | 0.343773464 | 8.32321936 | -0.375856092 | 0.719591799 | 0.883245896 | -4.876770084 |
| **A_52_P75146** | Lyrm2 | -0.054511812 | -0.407063411 | 0.298039788 | 9.809897202 | -0.375708583 | 0.719696039 | 0.883245896 | -4.876799477 |
| **A_55_P1958102** | NA | -0.035940122 | -0.268383508 | 0.196503264 | 16.09182408 | -0.375703758 | 0.719699449 | 0.883245896 | -4.876800438 |

| | | | | | | | | |
|---|---|---|---|---|---|---|---|---|
| **A_55_P2092668** | Kyat3 | -0.059869935 | -0.44708307 | 0.327343201 | 8.358725536 | -0.375700602 | 0.719701679 | 0.883245896 | -4.876801067 |
| **A_55_P2024555** | Plpp1 | -0.027079247 | -0.202239753 | 0.14808126 | 11.34928147 | -0.375650155 | 0.71973733 | 0.883245896 | -4.876811117 |
| **A_55_P2125351** | Sema6d | 0.120383405 | -0.658418508 | 0.899185318 | 12.43900584 | 0.375597595 | 0.719774476 | 0.883245896 | -4.876821586 |
| **A_55_P2139341** | Zfp975 | -0.076121376 | -0.568663713 | 0.416420961 | 10.56592216 | -0.375531397 | 0.71982126 | 0.883245896 | -4.876834769 |
| **A_51_P440852** | Mtrf1 | -0.026828691 | -0.200435131 | 0.146777749 | 8.950556412 | -0.375505962 | 0.719839236 | 0.883245896 | -4.876839834 |
| **A_55_P2090219** | Mrpl48 | -0.036470875 | -0.281710157 | 0.208768408 | 12.26918834 | -0.378354154 | 0.720183666 | 0.883484062 | -4.830366109 |
| **A_55_P2027225** | Fau | -0.039891204 | -0.298375743 | 0.218593335 | 17.4164561 | -0.374995249 | 0.720200222 | 0.883484062 | -4.876941466 |
| **A_52_P533083** | Dnajc16 | 0.048693188 | -0.266857518 | 0.364243895 | 8.879274594 | 0.374957673 | 0.720226785 | 0.883484062 | -4.876948938 |
| **A_65_P19286** | Alg10b | 0.030458863 | -0.166935223 | 0.22785295 | 8.585354119 | 0.374940834 | 0.72023869 | 0.883484062 | -4.876952287 |
| **A_65_P06611** | Wtap | 0.040519016 | -0.22209466 | 0.303132693 | 9.815176347 | 0.374908032 | 0.720261878 | 0.883484062 | -4.876958809 |
| **A_52_P676359** | Fcnb | -0.07901353 | -0.591241464 | 0.433214404 | 9.477780933 | -0.374818825 | 0.720324943 | 0.883491548 | -4.876976544 |
| **A_55_P2016670** | Alpk2 | 0.024505004 | -0.134376393 | 0.183386401 | 7.642873597 | 0.374770089 | 0.720359398 | 0.883491548 | -4.876986232 |
| **A_51_P483324** | Ptpn22 | 0.068771341 | -0.377637645 | 0.515180326 | 8.515074728 | 0.374332687 | 0.720668658 | 0.883814764 | -4.877073123 |
| **A_55_P2024953** | Tbc1d4 | -0.028900956 | -0.216610743 | 0.15880883 | 8.386146776 | -0.374117899 | 0.720820543 | 0.883924997 | -4.877115757 |
| **A_55_P2138436** | NA | 0.035983161 | -0.197751103 | 0.269717426 | 15.75970821 | 0.374076239 | 0.720850004 | 0.883924997 | -4.877124023 |
| **A_55_P2002460** | Gm2399 | -0.027154977 | -0.20361996 | 0.149310007 | 7.977020569 | -0.373916033 | 0.720963303 | 0.883965142 | -4.877155805 |
| **A_55_P1970507** | Pef1 | 0.042431762 | -0.233319861 | 0.318183385 | 11.14441333 | 0.373900613 | 0.720974208 | 0.883965142 | -4.877158863 |
| **A_51_P346641** | NA | -0.03935774 | -0.295224691 | 0.216509211 | 8.138735325 | -0.373765443 | 0.721069808 | 0.884026279 | -4.877185667 |
| **A_51_P391159** | Ang | -0.031357044 | -0.235292391 | 0.172578302 | 8.536357442 | -0.373616291 | 0.721175303 | 0.884060109 | -4.877215233 |
| **A_66_P132448** | Gnpnat1 | 0.083320415 | -0.458618666 | 0.625259496 | 9.92834585 | 0.373580453 | 0.721200652 | 0.884060109 | -4.877222336 |
| **A_55_P2106801** | Bglap3 | 0.037045618 | -0.203940325 | 0.278031561 | 7.92447599 | 0.373532439 | 0.721234614 | 0.884060109 | -4.87723185 |
| **A_51_P268167** | Tasor | 0.038499416 | -0.211992187 | 0.288991019 | 9.203479772 | 0.373460057 | 0.721285814 | 0.884066804 | -4.877246192 |
| **A_55_P2177173** | Gnl1 | -0.037347873 | -0.280540925 | 0.205845179 | 9.45628395 | -0.373162421 | 0.721496366 | 0.884208852 | -4.877305136 |
| **A_51_P151968** | Eif2b5 | 0.030511934 | -0.168173754 | 0.229197622 | 9.707199807 | 0.373152484 | 0.721503396 | 0.884208852 | -4.877307103 |
| **A_66_P121249** | Ddi2 | -0.065551649 | -0.492476395 | 0.361373096 | 9.487636172 | -0.373091611 | 0.721546462 | 0.884208852 | -4.877319152 |
| **A_55_P1999576** | Josd1 | 0.039556198 | -0.218111541 | 0.297223937 | 13.56756652 | 0.373024784 | 0.721593741 | 0.884208852 | -4.877332379 |
| **A_52_P479961** | NA | -0.047586485 | -0.35760634 | 0.262433371 | 15.50104821 | -0.372972923 | 0.721630433 | 0.884208852 | -4.877342641 |
| **A_52_P639101** | Lrrc14 | 0.027003602 | -0.148969448 | 0.202976651 | 8.308060437 | 0.372871101 | 0.721702475 | 0.884241071 | -4.877362786 |
| **A_52_P63905** | Ddc | -0.039627229 | -0.298047054 | 0.218792597 | 8.132904651 | -0.372607046 | 0.721889316 | 0.884313913 | -4.877415005 |
| **A_52_P79506** | Zfp667 | 0.026408227 | -0.145818994 | 0.198635447 | 7.848554062 | 0.37258094 | 0.721907789 | 0.884313913 | -4.877420166 |
| **A_52_P478256** | Samm50 | 0.044428998 | -0.245334595 | 0.334192591 | 10.77478882 | 0.372568288 | 0.721916742 | 0.884313913 | -4.877422667 |
| **A_51_P174696** | Casq2 | -0.043359128 | -0.326175327 | 0.239457071 | 7.844090098 | -0.372528455 | 0.72194493 | 0.884313913 | -4.877430541 |
| **A_55_P2121877** | Gpr137b | -0.024123143 | -0.181522612 | 0.133276326 | 7.796379656 | -0.372403557 | 0.722033316 | 0.88435863 | -4.877455223 |
| **A_55_P2132812** | NA | -0.028790577 | -0.216691827 | 0.159110674 | 10.73673974 | -0.372309299 | 0.722100022 | 0.88435863 | -4.877473846 |
| **A_55_P2086536** | Safb2 | 0.046266543 | -0.255739891 | 0.348272978 | 8.599996542 | 0.372249436 | 0.722142388 | 0.88435863 | -4.87748567 |
| **A_55_P1987912** | Chpf2 | -0.042562197 | -0.320411648 | 0.235287254 | 9.032245926 | -0.372218267 | 0.722164448 | 0.88435863 | -4.877491826 |
| **A_55_P1973668** | Slc39a13 | 0.032434628 | -0.179354539 | 0.244223796 | 10.24239051 | 0.372124592 | 0.722230747 | 0.884383788 | -4.877510325 |
| **A_51_P150876** | Lypd3 | -0.024529739 | -0.184766383 | 0.135706905 | 7.709836786 | -0.371975449 | 0.722336309 | 0.884437733 | -4.877539768 |
| **A_55_P1996643** | Polr2j | -0.033073532 | -0.249145778 | 0.182998714 | 12.64109394 | -0.371933058 | 0.722366314 | 0.884437733 | -4.877548135 |
| **A_51_P142744** | Sulf1 | -0.090210693 | -0.680166444 | 0.499745059 | 8.912495572 | -0.371553851 | 0.722634748 | 0.884710352 | -4.877622939 |
| **A_55_P2002968** | Coro2a | 0.03981303 | -0.220619576 | 0.300245636 | 8.168544307 | 0.371460862 | 0.72270058 | 0.884734911 | -4.877641272 |
| **A_51_P119544** | Aga | -0.04190968 | -0.316158191 | 0.232338831 | 9.183189296 | -0.371324201 | 0.722797334 | 0.884752123 | -4.843419127 |
| **A_52_P236398** | NA | -0.025751302 | -0.194273413 | 0.14277081 | 7.767683423 | -0.371300456 | 0.722814146 | 0.884752123 | -4.877672886 |
| **A_55_P2092904** | Ddb2 | 0.039246443 | -0.217627754 | 0.29612064 | 8.529827198 | 0.371247048 | 0.72285196 | 0.884752123 | -4.877683409 |
| **A_55_P2172001** | Mrgpra2a | 0.039824074 | -0.220987458 | 0.300635606 | 8.506254386 | 0.371024069 | 0.723009843 | 0.884889334 | -4.877727327 |
| **A_55_P2100770** | Usp48 | -0.030551569 | -0.230830585 | 0.169727448 | 9.645399233 | -0.370664719 | 0.723264317 | 0.885144737 | -4.877798054 |
| **A_55_P1976097** | NA | -0.103846365 | -0.78486222 | 0.577169491 | 8.893523193 | -0.370524784 | 0.723363422 | 0.885160691 | -4.877825579 |
| **A_55_P2054332** | Ephb3 | 0.097499881 | -0.54190971 | 0.736909473 | 14.96863696 | 0.370516991 | 0.723368942 | 0.885160691 | -4.877827112 |
| **A_55_P1996374** | Prmt9 | 0.026816639 | -0.149105244 | 0.202738521 | 10.49194749 | 0.370397176 | 0.723453803 | 0.885208493 | -4.877850671 |
| **A_55_P2328787** | NA | -0.039847966 | -0.301338981 | 0.221643048 | 8.24680995 | -0.370281985 | 0.723535393 | 0.885252286 | -4.877873314 |

| | | | | | | | | | |
|---|---|---|---|---|---|---|---|---|---|
| **A_51_P333349** | Tmem120a | -0.036024533 | -0.27248405 | 0.200434983 | 10.29498115 | -0.370190063 | 0.723600504 | 0.885275913 | -4.877891378 |
| **A_55_P2071601** | Ppara | -0.097029358 | -0.734474219 | 0.540415504 | 11.06432399 | -0.369865413 | 0.723830485 | 0.885415706 | -4.877955144 |
| **A_55_P1991505** | Gamt | -0.092662729 | -0.701429225 | 0.516103768 | 13.54111653 | -0.369860141 | 0.72383422 | 0.885415706 | -4.877956179 |
| **A_52_P287683** | Pik3c3 | -0.023877084 | -0.180753165 | 0.132998997 | 8.213747483 | -0.369834775 | 0.72385219 | 0.885415706 | -4.877961159 |
| **A_55_P2084345** | NA | 0.026536306 | -0.147853988 | 0.200926601 | 15.22055461 | 0.369744183 | 0.723916371 | 0.885438179 | -4.877978942 |
| **A_55_P1955393** | 7-Sep | -0.048729964 | -0.369185894 | 0.271725966 | 12.35810992 | -0.369497041 | 0.724091475 | 0.885596313 | -4.878027435 |
| **A_55_P1953593** | Ece2 | -0.036256775 | -0.27478941 | 0.202275861 | 10.89256171 | -0.369338473 | 0.724203832 | 0.885677689 | -4.878058532 |
| **A_55_P1954897** | NA | -0.07035747 | -0.533474628 | 0.392759689 | 14.70133843 | -0.369149708 | 0.724337597 | 0.885785234 | -4.878095535 |
| **A_55_P2171528** | 3110082I17Rik | -0.032675853 | -0.247837571 | 0.182485865 | 8.460883733 | -0.369015933 | 0.7244324 | 0.885845123 | -4.878121748 |
| **A_55_P2140002** | Tnfaip8 | 0.037896096 | -0.211710498 | 0.28750269 | 10.13341835 | 0.368911 | 0.724506767 | 0.885857346 | -4.878142303 |
| **A_51_P499169** | Tram1 | 0.074280977 | -0.415029512 | 0.563591467 | 11.06740972 | 0.368872495 | 0.724534057 | 0.885857346 | -4.878149844 |
| **A_52_P467449** | Alox12 | -0.091997463 | -0.698175158 | 0.514180232 | 10.015287 | -0.368772977 | 0.72460459 | 0.885887547 | -4.878169331 |
| **A_55_P2067777** | Cplane1 | -0.043027454 | -0.326640875 | 0.240585966 | 8.732748974 | -0.368639664 | 0.724699081 | 0.885909843 | -4.878195429 |
| **A_55_P2096465** | NA | 0.028253448 | -0.157990457 | 0.214497353 | 12.51878226 | 0.368614635 | 0.724716822 | 0.885909843 | -4.878200327 |
| **A_52_P552550** | Agtrap | 0.034020683 | -0.190277604 | 0.258318969 | 8.681811693 | 0.368553259 | 0.724760326 | 0.885909843 | -4.878212339 |
| **A_52_P190405** | Ifnar2 | 0.041929823 | -0.234602422 | 0.318462067 | 12.08018497 | 0.368434624 | 0.724844421 | 0.885956609 | -4.87823555 |
| **A_52_P282838** | Spin2c | 0.032363695 | -0.181142237 | 0.245869627 | 8.162914715 | 0.36832512 | 0.724922047 | 0.885970644 | -4.878256969 |
| **A_55_P2034663** | Csf2rb2 | -0.040143873 | -0.305002169 | 0.224714424 | 8.359668925 | -0.368289107 | 0.724947576 | 0.885970644 | -4.878264012 |
| **A_55_P2115310** | Spata21 | -0.024891066 | -0.189148975 | 0.139366843 | 7.719375357 | -0.368214083 | 0.725000763 | 0.885979626 | -4.878278681 |
| **A_55_P2026325** | NA | -0.045086055 | -0.342683987 | 0.252511878 | 14.61446805 | -0.368125171 | 0.725063797 | 0.88600064 | -4.878296063 |
| **A_52_P270035** | Tspyl2 | -0.036916462 | -0.280760914 | 0.20692799 | 8.65827806 | -0.367866624 | 0.725247106 | 0.886038126 | -4.878346585 |
| **A_55_P2011674** | Rpl17 | -0.054677743 | -0.415876848 | 0.306521363 | 16.02386285 | -0.367829959 | 0.725273103 | 0.886038126 | -4.878353747 |
| **A_55_P2000758** | Gkap1 | 0.026875932 | -0.150667941 | 0.204419805 | 9.552611024 | 0.367824825 | 0.725276744 | 0.886038126 | -4.87835475 |
| **A_52_P563340** | Smim4 | -0.068429595 | -0.520526758 | 0.383667568 | 12.26227094 | -0.367786154 | 0.725304163 | 0.886038126 | -4.878362303 |
| **A_55_P2077510** | Zfp583 | -0.036449589 | -0.277307917 | 0.204408739 | 8.155838092 | -0.36771738 | 0.725352929 | 0.886038126 | -4.878375734 |
| **A_52_P463235** | Ankrd33b | -0.025086896 | -0.190871265 | 0.140697473 | 7.79306964 | -0.367693993 | 0.725369512 | 0.886038126 | -4.8783803 |
| **A_55_P2097478** | Gda | -0.119949299 | -0.912945044 | 0.673046445 | 11.10154193 | -0.367544601 | 0.725475448 | 0.886098383 | -4.878409465 |
| **A_51_P469789** | Agxt | -0.114424381 | -0.87099622 | 0.642147458 | 11.61226064 | -0.367495131 | 0.725510529 | 0.886098383 | -4.87841912 |
| **A_55_P2176430** | 0610010K14Rik | -0.040059634 | -0.305018679 | 0.224899411 | 10.67878429 | -0.367376538 | 0.725594631 | 0.886104155 | -4.878442261 |
| **A_51_P118742** | Drg2 | 0.039571176 | -0.22216952 | 0.301311873 | 9.454243846 | 0.36735918 | 0.725606941 | 0.886104155 | -4.878445647 |
| **A_55_P2151556** | C8g | -0.087744631 | -0.668291267 | 0.492802005 | 13.76517411 | -0.367254078 | 0.725681479 | 0.886106645 | -4.878466149 |
| **A_66_P102879** | Thra | -0.032534223 | -0.247806943 | 0.182738497 | 7.886841144 | -0.367227023 | 0.725700667 | 0.886106645 | -4.878471426 |
| **A_51_P318618** | Anks4b | 0.066471884 | -0.373743877 | 0.506687645 | 9.207666087 | 0.366906656 | 0.725927895 | 0.886328109 | -4.87853388 |
| **A_55_P2009543** | NA | -0.029186122 | -0.222688199 | 0.164315954 | 16.54974764 | -0.366499981 | 0.726216383 | 0.886508286 | -4.878613087 |
| **A_55_P1981814** | Myo3b | 0.053805525 | -0.302955402 | 0.410566452 | 8.87515301 | 0.366465233 | 0.726241035 | 0.886508286 | -4.878619851 |
| **A_51_P493270** | Mrps5 | 0.048809693 | -0.274838499 | 0.372457886 | 9.559693518 | 0.366451144 | 0.72625103 | 0.886508286 | -4.878622593 |
| **A_66_P100093** | Ccdc47 | 0.058012315 | -0.326668463 | 0.442693093 | 11.34331654 | 0.36644002 | 0.726258923 | 0.886508286 | -4.878624758 |
| **A_51_P129435** | Arid2 | -0.052478021 | -0.4005418 | 0.295585759 | 8.658321074 | -0.366354731 | 0.726319433 | 0.886526163 | -4.878641356 |
| **A_55_P2112330** | Anapc7 | -0.043605435 | -0.332931143 | 0.245720272 | 11.26385902 | -0.366215559 | 0.726418176 | 0.8865907 | -4.878668434 |
| **A_51_P121455** | Slc25a39 | 0.067263299 | -0.379203937 | 0.513730535 | 9.770766476 | 0.366076422 | 0.7265169 | 0.886655207 | -4.878695494 |
| **A_55_P1992257** | Ap1s1 | -0.029485851 | -0.225302448 | 0.166330747 | 13.4155301 | -0.365887314 | 0.72665109 | 0.886752743 | -4.878732258 |
| **A_55_P1969431** | Etfrf1 | 0.057098545 | -0.322179492 | 0.436376583 | 9.064724595 | 0.365805758 | 0.726708965 | 0.886752743 | -4.878748108 |
| **A_55_P2022519** | Tmem108 | -0.026852525 | -0.205238379 | 0.15153333 | 8.045040154 | -0.365769845 | 0.726734451 | 0.886752743 | -4.878755086 |
| **A_52_P124734** | Cybrd1 | 0.046472758 | -0.262395773 | 0.355341289 | 9.300363384 | 0.365601505 | 0.726853918 | 0.886842531 | -4.878787788 |
| **A_55_P1972118** | NA | -0.031253175 | -0.239058067 | 0.176551717 | 16.02656233 | -0.365444616 | 0.726965267 | 0.886880241 | -4.878818253 |
| **A_55_P2047375** | NA | 0.035369045 | -0.199812867 | 0.270550958 | 15.05912051 | 0.365428661 | 0.726976591 | 0.886880241 | -4.87882135 |
| **A_55_P2042202** | Pigo | -0.045482702 | -0.348209401 | 0.257243998 | 11.01846445 | -0.365072162 | 0.727229638 | 0.887132955 | -4.878890527 |
| **A_51_P293901** | Dhrs1 | -0.032387079 | -0.248009543 | 0.183235385 | 8.998446825 | -0.364973191 | 0.727299895 | 0.887162671 | -4.878909721 |
| **A_55_P2026025** | NA | -0.040590124 | -0.310912598 | 0.22973235 | 16.32319382 | -0.364855936 | 0.727383136 | 0.887208218 | -4.878932454 |
| **A_55_P1958857** | Nek6 | -0.028237094 | -0.216361488 | 0.1598873 | 9.940482004 | -0.364718733 | 0.727480543 | 0.887224556 | -4.878959046 |

| | | | | | | | | | |
|---|---|---|---|---|---|---|---|---|---|
| **A_51_P429983** | Supt5 | 0.038229356 | -0.21650737 | 0.292966082 | 11.95095554 | 0.364660403 | 0.727521955 | 0.887224556 | -4.878970348 |
| **A_55_P1969197** | NA | 0.025218717 | -0.142831145 | 0.193268578 | 16.60403351 | 0.364643109 | 0.727534234 | 0.887224556 | -4.878973699 |
| **A_55_P2080396** | Mrpl4 | -0.0273911 | -0.209967563 | 0.155185363 | 12.86017898 | -0.364542249 | 0.727605845 | 0.887255907 | -4.878993237 |
| **A_66_P129730** | Pdia3 | -0.038648089 | -0.296460635 | 0.219164458 | 12.5123723 | -0.364256375 | 0.727808833 | 0.887447447 | -4.879048588 |
| **A_55_P1996746** | NA | -0.036595411 | -0.28078764 | 0.207596818 | 7.940252947 | -0.364148042 | 0.727885762 | 0.887485264 | -4.879069553 |
| **A_51_P159284** | Bud13 | -0.030959283 | -0.237597027 | 0.17567846 | 8.376783613 | -0.364052851 | 0.727953362 | 0.887511703 | -4.87908797 |
| **A_66_P105270** | Hao1 | -0.176995741 | -1.359060944 | 1.005069461 | 11.03069563 | -0.363835082 | 0.728108021 | 0.887626593 | -4.879130086 |
| **A_55_P2180805** | Pex3 | 0.037853005 | -0.215001024 | 0.290707034 | 9.645824846 | 0.36375894 | 0.728162099 | 0.887626593 | -4.879144806 |
| **A_55_P2132472** | Prpf4b | 0.031899086 | -0.181255297 | 0.245053469 | 8.399182178 | 0.363636244 | 0.728249247 | 0.887626593 | -4.879168519 |
| **A_55_P2037897** | Brd3 | 0.044166007 | -0.250961089 | 0.339293102 | 10.45549926 | 0.363632121 | 0.728252175 | 0.887626593 | -4.879169316 |
| **A_55_P2038101** | Dock11 | -0.041227123 | -0.316772666 | 0.234318419 | 9.319855184 | -0.36355727 | 0.728305342 | 0.887626593 | -4.879183779 |
| **A_52_P270145** | Zfp329 | -0.052009572 | -0.399775151 | 0.295756007 | 9.222817055 | -0.363395777 | 0.728420055 | 0.887626593 | -4.879214973 |
| **A_51_P516615** | Ndufb10 | -0.027285506 | -0.20974176 | 0.155170749 | 14.91180558 | -0.363376168 | 0.728433985 | 0.887626593 | -4.87921876 |
| **A_51_P154997** | Brms1 | -0.055651411 | -0.427846201 | 0.316543378 | 9.098507135 | -0.363319814 | 0.728474017 | 0.887626593 | -4.879229642 |
| **A_55_P2033863** | Psmc1 | -0.028705401 | -0.220722415 | 0.163311613 | 13.93582268 | -0.363251229 | 0.728522739 | 0.887626593 | -4.879242884 |
| **A_55_P2399688** | Glipr2 | -0.037580364 | -0.289040848 | 0.21388012 | 8.905583191 | -0.36314028 | 0.728601559 | 0.887626593 | -4.8792643 |
| **A_55_P1986053** | Pcnx | -0.06012559 | -0.462481838 | 0.342230658 | 8.705959831 | -0.363104633 | 0.728626884 | 0.887626593 | -4.879271179 |
| **A_52_P229997** | Cbx8 | 0.028336767 | -0.161316608 | 0.217990142 | 8.328009588 | 0.363055405 | 0.728661858 | 0.887626593 | -4.879280678 |
| **A_52_P359819** | Ntf5 | -0.07293158 | -0.561051241 | 0.415188081 | 8.411540276 | -0.363055074 | 0.728662093 | 0.887626593 | -4.879280742 |
| **A_52_P585028** | Cnep1r1 | 0.029544115 | -0.168211811 | 0.227300041 | 10.48182881 | 0.363015082 | 0.728690506 | 0.887626593 | -4.879288458 |
| **A_55_P2076248** | NA | 0.023712582 | -0.135079299 | 0.182504464 | 8.142013919 | 0.362855531 | 0.728803865 | 0.887708735 | -4.879319235 |
| **A_55_P2126448** | Snhg16 | -0.065727606 | -0.506100565 | 0.374645352 | 10.60297202 | -0.362668941 | 0.728936446 | 0.887793786 | -4.879355211 |
| **A_52_P269461** | Ddhd2 | -0.0236215 | -0.181906516 | 0.134663516 | 8.081225201 | -0.36261925 | 0.728971754 | 0.887793786 | -4.879364788 |
| **A_55_P2143336** | Rpp21 | -0.04296682 | -0.330942922 | 0.245009281 | 9.546185928 | -0.362543361 | 0.729025681 | 0.887793786 | -4.879379414 |
| **A_55_P2000133** | Cdh2 | 0.085126148 | -0.485592159 | 0.655844454 | 11.44997404 | 0.362430189 | 0.729106103 | 0.887793786 | -4.879401218 |
| **A_55_P1967076** | Rpl11 | -0.029919431 | -0.230526839 | 0.170687978 | 16.80587224 | -0.362401132 | 0.729126752 | 0.887793786 | -4.879406816 |
| **A_51_P114028** | Bcl7c | -0.055304433 | -0.4262099 | 0.315601033 | 9.592087602 | -0.36230965 | 0.729191765 | 0.887793786 | -4.879424436 |
| **A_51_P156927** | Arid4a | 0.036762079 | -0.209790275 | 0.283314432 | 8.780001141 | 0.362304808 | 0.729195207 | 0.887793786 | -4.879425368 |
| **A_51_P507787** | Sergef | -0.035091198 | -0.270545794 | 0.200363398 | 8.683005968 | -0.362138096 | 0.729313689 | 0.887872445 | -4.879457467 |
| **A_55_P2171378** | Pdzk1ip1 | 0.041077415 | -0.234609033 | 0.316763863 | 9.162528967 | 0.362051939 | 0.729374924 | 0.887872445 | -4.87947405 |
| **A_51_P121447** | 1110038F14Rik | -0.064334992 | -0.496150133 | 0.367480148 | 11.60092818 | -0.362020009 | 0.729397618 | 0.887872445 | -4.879480195 |
| **A_55_P2061273** | Tbx6 | 0.069458162 | -0.39697451 | 0.535890835 | 8.799370692 | 0.361840782 | 0.72952501 | 0.887931108 | -4.879514677 |
| **A_66_P106418** | Prkacb | -0.035008577 | -0.270113398 | 0.200096245 | 8.902994176 | -0.36182295 | 0.729537685 | 0.887931108 | -4.879518107 |
| **A_55_P1975155** | Galnt6 | 0.06321157 | -0.361818938 | 0.488242078 | 9.653369877 | 0.361376295 | 0.729855204 | 0.888260477 | -4.879603967 |
| **A_52_P592111** | Brf1 | 0.041251851 | -0.236171527 | 0.31867523 | 9.076732013 | 0.361312992 | 0.72990021 | 0.888260477 | -4.879616127 |
| **A_51_P475995** | Thra | 0.068746872 | -0.393922415 | 0.53141616 | 11.45294481 | 0.361048431 | 0.730088315 | 0.888433457 | -4.879666928 |
| **A_66_P109720** | Jpt2 | 0.032164822 | -0.184396002 | 0.248725645 | 8.149176989 | 0.360897975 | 0.730195299 | 0.888507707 | -4.879695803 |
| **A_55_P2113026** | Foxo4 | 0.025793464 | -0.147909443 | 0.199496371 | 7.806564692 | 0.360815988 | 0.730253599 | 0.888522713 | -4.879711533 |
| **A_55_P2127971** | Gm9112 | 0.024234268 | -0.139059596 | 0.187528131 | 7.782965961 | 0.360614532 | 0.730396863 | 0.888641088 | -4.87975017 |
| **A_55_P2024841** | Gstt1 | -0.11284944 | -0.87358512 | 0.64788624 | 15.1032473 | -0.360453138 | 0.730511645 | 0.888648158 | -4.879781109 |
| **A_55_P1970825** | NA | 0.023027394 | -0.13220472 | 0.178259509 | 7.77180674 | 0.360451149 | 0.73051306 | 0.888648158 | -4.87978149 |
| **A_52_P551526** | NA | 0.036000364 | -0.206785868 | 0.278786595 | 16.37040805 | 0.360301461 | 0.730619524 | 0.888648158 | -4.879810174 |
| **A_55_P2181928** | Slc45a3 | 0.025695065 | -0.14762865 | 0.19901878 | 8.274417338 | 0.360225882 | 0.730673281 | 0.888648158 | -4.879824652 |
| **A_55_P1963712** | Cyb5b | 0.040763933 | -0.234264771 | 0.315792637 | 11.05896891 | 0.360148199 | 0.730728537 | 0.888648158 | -4.87983953 |
| **A_55_P2006869** | Fyb | 0.074385354 | -0.42764841 | 0.576419118 | 9.151895306 | 0.360029177 | 0.7308132 | 0.888648158 | -4.87986232 |
| **A_52_P217646** | Snapc3 | -0.028996052 | -0.224716616 | 0.166724512 | 9.394191421 | -0.359985989 | 0.730843922 | 0.888648158 | -4.879870588 |
| **A_55_P2160331** | Bicd2 | -0.044378628 | -0.343942316 | 0.255185059 | 10.01821884 | -0.359971315 | 0.73085436 | 0.888648158 | -4.879873397 |
| **A_55_P2150025** | Otub1 | -0.054909566 | -0.425560856 | 0.315741724 | 9.549802591 | -0.359969483 | 0.730855664 | 0.888648158 | -4.879873747 |
| **A_55_P2146555** | NA | -0.029704002 | -0.230217644 | 0.170809641 | 9.103485455 | -0.359959981 | 0.730862422 | 0.888648158 | -4.879875566 |
| **A_55_P2050602** | Ncam1 | -0.023985974 | -0.185936509 | 0.137964561 | 7.692408152 | -0.359880367 | 0.730919058 | 0.888661119 | -4.879890804 |

| | | | | | | | | | |
|---|---|---|---|---|---|---|---|---|---|
| A_55_P2005080 | NA | -0.028229102 | -0.218915538 | 0.162457335 | 16.43552418 | -0.359716572 | 0.731035583 | 0.888746889 | -4.879922143 |
| A_52_P131672 | Rbm7 | -0.028891943 | -0.224465474 | 0.166681588 | 10.39358368 | -0.358963144 | 0.731571679 | 0.889193675 | -4.880066125 |
| A_66_P140949 | NA | -0.043893006 | -0.341037702 | 0.25325169 | 13.06869181 | -0.358930638 | 0.731594811 | 0.889193675 | -4.880072331 |
| A_51_P406334 | Dnlz | -0.070452352 | -0.54740661 | 0.406501906 | 9.653119584 | -0.358923551 | 0.731599855 | 0.889193675 | -4.880073684 |
| A_55_P2091836 | Fnip1 | -0.057614889 | -0.447666193 | 0.332436415 | 10.3630295 | -0.358918782 | 0.731603249 | 0.889193675 | -4.880074594 |
| A_51_P108108 | Lrrc25 | -0.055102299 | -0.428187032 | 0.317982435 | 9.896424289 | -0.358876836 | 0.731633101 | 0.889193675 | -4.880082601 |
| A_55_P2081137 | NA | -0.060813426 | -0.472735518 | 0.351108665 | 11.04782618 | -0.358729902 | 0.731737673 | 0.889264853 | -4.880110641 |
| A_55_P2125551 | Mmadhc | 0.028383784 | -0.163972693 | 0.220740261 | 12.99470327 | 0.358547473 | 0.731867516 | 0.889357575 | -4.88014544 |
| A_55_P2170787 | Braf | 0.025128416 | -0.145201253 | 0.195458085 | 8.03777931 | 0.358474278 | 0.731919615 | 0.889357575 | -4.880159398 |
| A_51_P344376 | Gpr162 | 0.025461748 | -0.147179911 | 0.198103407 | 7.780896593 | 0.358365175 | 0.731997276 | 0.889357575 | -4.880180198 |
| A_55_P1976978 | Zfp707 | 0.051237004 | -0.296225175 | 0.398699183 | 9.017086212 | 0.358310371 | 0.732036287 | 0.889357575 | -4.880190644 |
| A_51_P163953 | Nsg2 | -0.023272674 | -0.181124731 | 0.134579384 | 7.736843388 | -0.358244246 | 0.732083359 | 0.889357575 | -4.880203245 |
| A_52_P98232 | Hira | -0.035535359 | -0.276568102 | 0.205497385 | 8.516333867 | -0.358234861 | 0.732090039 | 0.889357575 | -4.880205034 |
| A_51_P351481 | Ccnl1 | -0.0254533 | -0.198188688 | 0.147282087 | 9.347339076 | -0.358051891 | 0.732220293 | 0.889459908 | -4.880239891 |
| A_55_P2066274 | NA | -0.034843147 | -0.271366029 | 0.201679735 | 11.39358211 | -0.35795415 | 0.732289878 | 0.889465265 | -4.880258504 |
| A_55_P1972222 | NA | 0.047406191 | -0.274452364 | 0.369264745 | 15.38091227 | 0.357892989 | 0.732333422 | 0.889465265 | -4.880270149 |
| A_52_P657240 | Cdc40 | 0.02338936 | -0.135428133 | 0.182206853 | 8.230184802 | 0.35785179 | 0.732362755 | 0.889465265 | -4.880277992 |
| A_55_P2030433 | Gpi1 | -0.050857982 | -0.39632644 | 0.294610476 | 15.00617518 | -0.357712329 | 0.73246205 | 0.889529967 | -4.880304535 |
| A_55_P2025675 | Dnah8 | -0.076014056 | -0.592648992 | 0.440620879 | 8.947742213 | -0.357514226 | 0.732603108 | 0.889645377 | -4.880342223 |
| A_52_P512427 | Mkrn2os | -0.059029125 | -0.460334648 | 0.342276398 | 11.10803666 | -0.357416337 | 0.732672813 | 0.88967413 | -4.880360838 |
| A_55_P2102798 | Prl7a2 | -0.023035561 | -0.179721411 | 0.133650289 | 7.632559308 | -0.357233523 | 0.732803 | 0.889711796 | -4.880395591 |
| A_55_P2174339 | NA | 0.030317974 | -0.1759381 | 0.236574049 | 8.218670731 | 0.357171355 | 0.732847274 | 0.889711796 | -4.880407405 |
| A_55_P2287260 | Snap23 | 0.087151877 | -0.505804089 | 0.680107844 | 10.99647208 | 0.357139176 | 0.73287019 | 0.889711796 | -4.880413519 |
| A_51_P350403 | Exoc4 | -0.028829618 | -0.22502412 | 0.167364883 | 8.261508836 | -0.357055108 | 0.732930063 | 0.889711796 | -4.880429491 |
| A_51_P212630 | Chd8 | -0.034619546 | -0.270245745 | 0.201006653 | 11.93934355 | -0.357010497 | 0.732961836 | 0.889711796 | -4.880437965 |
| A_51_P319879 | Acadm | -0.109541714 | -0.85515353 | 0.636070102 | 13.35381649 | -0.356984977 | 0.732980012 | 0.889711796 | -4.880442812 |
| A_55_P2023937 | Impg1 | 0.035166709 | -0.204261255 | 0.274594672 | 9.780147358 | 0.356894652 | 0.733044345 | 0.889734012 | -4.880459965 |
| A_52_P553890 | Itgb3 | -0.085410593 | -0.667479179 | 0.496657992 | 11.93926219 | -0.356550268 | 0.73328965 | 0.889975866 | -4.880525329 |
| A_55_P2181271 | Nhsl2 | 0.054381071 | -0.316464665 | 0.425226807 | 9.097273649 | 0.356317915 | 0.733455174 | 0.890120868 | -4.880569395 |
| A_55_P1957277 | NA | 0.042089159 | -0.245090173 | 0.329268492 | 8.685027537 | 0.356123193 | 0.733593904 | 0.890200358 | -4.880606304 |
| A_66_P103999 | Rsrc1 | -0.03106468 | -0.243038922 | 0.180909562 | 8.418046552 | -0.356095905 | 0.733613345 | 0.890200358 | -4.880611475 |
| A_52_P171212 | Irs1 | 0.030223206 | -0.176061287 | 0.236507699 | 8.11775655 | 0.356005855 | 0.733677506 | 0.890200358 | -4.880628536 |
| A_55_P2133187 | NA | 0.032345034 | -0.188445555 | 0.253135623 | 8.479984721 | 0.355967415 | 0.733704895 | 0.890200358 | -4.880635818 |
| A_55_P2004179 | Col2a1 | 0.027394649 | -0.15971517 | 0.214504467 | 7.722415626 | 0.355756092 | 0.733855473 | 0.890325973 | -4.880675836 |
| A_55_P2069961 | NA | 0.026777967 | -0.156152945 | 0.20970888 | 16.69173221 | 0.355691673 | 0.733901377 | 0.890325973 | -4.88068803 |
| A_55_P1975772 | Ppp2r5a | -0.044992498 | -0.352408968 | 0.262423972 | 13.03499595 | -0.355628174 | 0.733946627 | 0.890325973 | -4.880700048 |
| A_51_P154780 | Vav1 | -0.052176422 | -0.408742673 | 0.30438983 | 8.808411974 | -0.355563559 | 0.733992673 | 0.890325973 | -4.880712276 |
| A_55_P2095019 | NA | 0.055460228 | -0.323708207 | 0.434628664 | 8.676890378 | 0.355412473 | 0.734100346 | 0.890400702 | -4.880740858 |
| A_55_P1973402 | Tada2a | -0.025689344 | -0.201465356 | 0.150086669 | 9.898788276 | -0.355121194 | 0.734307947 | 0.890596618 | -4.880795931 |
| A_52_P326214 | Cttn | 0.025237933 | -0.147538276 | 0.198014142 | 8.123741072 | 0.354938439 | 0.734438213 | 0.890698721 | -4.880830463 |
| A_55_P1972605 | Adora3 | 0.047053722 | -0.275133606 | 0.369241051 | 8.51379576 | 0.354869538 | 0.734487327 | 0.8907024 | -4.880843478 |
| A_51_P128648 | Uqcrc2 | 0.035790728 | -0.209415921 | 0.280997378 | 13.82506566 | 0.354667574 | 0.7346313 | 0.890821104 | -4.877169537 |
| A_55_P2010020 | NA | -0.067139279 | -0.527265611 | 0.392987053 | 15.53897063 | -0.354554315 | 0.734712044 | 0.890863126 | -4.88090299 |
| A_55_P2043117 | Ccdc138 | 0.081192451 | -0.475440983 | 0.637825886 | 15.31175092 | 0.354429249 | 0.734801209 | 0.890915353 | -4.880926587 |
| A_55_P2140453 | Myh6 | -0.029886469 | -0.23486153 | 0.175088592 | 7.77688786 | -0.354288266 | 0.734901727 | 0.890981339 | -4.880953179 |
| A_51_P436386 | Qrich1 | -0.06119068 | -0.481055703 | 0.358674343 | 10.4057541 | -0.35412678 | 0.735016871 | 0.891065046 | -4.880983626 |
| A_55_P2060369 | Akap9 | 0.031202407 | -0.183000471 | 0.245405285 | 10.16453577 | 0.353953315 | 0.735140565 | 0.891159108 | -4.881016317 |
| A_52_P243599 | Hsd17b12 | -0.079233705 | -0.623698119 | 0.465230709 | 10.92875493 | -0.353609281 | 0.735385912 | 0.891372516 | -4.881081108 |
| A_52_P381953 | Irf2bp1 | 0.028231698 | -0.165783621 | 0.222247017 | 8.543703468 | 0.35357713 | 0.735408842 | 0.891372516 | -4.88108716 |
| A_55_P2158152 | NA | -0.056466811 | -0.444679799 | 0.331746177 | 15.27948306 | -0.353432424 | 0.735512051 | 0.891441712 | -4.881114392 |

| | | | | | | | | |
|---|---|---|---|---|---|---|---|---|
| A_55_P2013058 | Shb | 0.06423453 | -0.3788503 | 0.507319359 | 10.47377035 | 0.352261237 | 0.736347596 | 0.892398437 | -4.881334413 |
| A_55_P1969750 | Stim1 | -0.041453195 | -0.327524861 | 0.244618471 | 11.43760508 | -0.352100267 | 0.736462465 | 0.892481692 | -4.881364599 |
| A_55_P2033084 | Zdhhc20 | -0.032527157 | -0.257099665 | 0.19204535 | 8.22163881 | -0.351943308 | 0.73657448 | 0.892561477 | -4.881394021 |
| A_55_P2012979 | Itgb2l | -0.096677042 | -0.764314399 | 0.570960315 | 10.42192574 | -0.351856773 | 0.736636238 | 0.892580357 | -4.881410237 |
| A_51_P491916 | Rassf6 | -0.05020463 | -0.397109092 | 0.296699832 | 8.850724741 | -0.351655225 | 0.73678009 | 0.892698699 | -4.88144799 |
| A_55_P2006754 | Mfsd12 | -0.030753127 | -0.243337269 | 0.181831015 | 11.79655794 | -0.35151317 | 0.736881485 | 0.89276559 | -4.881474586 |
| A_51_P314019 | Banf1 | 0.065508644 | -0.387482818 | 0.518500107 | 11.63746245 | 0.351391931 | 0.736968028 | 0.892799613 | -4.881497278 |
| A_55_P2024709 | 4933427D14Rik | -0.029778127 | -0.235721355 | 0.176165101 | 9.740162795 | -0.351344415 | 0.737001946 | 0.892799613 | -4.881506169 |
| A_51_P459128 | Serpinb9e | 0.022890652 | -0.135530139 | 0.181311442 | 7.66594752 | 0.351098657 | 0.737177389 | 0.892921076 | -4.881552138 |
| A_51_P280492 | Ndufa9 | 0.046729976 | -0.276699359 | 0.370159311 | 12.52895488 | 0.351074541 | 0.737194605 | 0.892921076 | -4.881556647 |
| A_55_P1973362 | Kcnj12 | 0.023830167 | -0.141180918 | 0.188841251 | 7.72827334 | 0.350911134 | 0.73731127 | 0.892940671 | -4.881587193 |
| A_52_P508920 | Abca6 | -0.109565317 | -0.868415188 | 0.649284553 | 9.201557368 | -0.350832991 | 0.737367062 | 0.892940671 | -4.881601796 |
| A_55_P2121456 | Fam43a | -0.054871097 | -0.434926975 | 0.325184781 | 9.760418342 | -0.35081599 | 0.737379201 | 0.892940671 | -4.881604973 |
| A_51_P208020 | Ulk3 | 0.034377746 | -0.203782855 | 0.272538347 | 9.090496706 | 0.350744338 | 0.737430361 | 0.892940671 | -4.881618359 |
| A_66_P116326 | NA | 0.086266991 | -0.511396809 | 0.683930791 | 11.01352139 | 0.350728363 | 0.737441768 | 0.892940671 | -4.881621344 |
| A_66_P100657 | Xpa | 0.03221701 | -0.191121483 | 0.255555504 | 10.71977452 | 0.350513577 | 0.737595136 | 0.893070432 | -4.881661454 |
| A_52_P496870 | 2310011J03Rik | 0.041504988 | -0.246430901 | 0.329440878 | 8.787886416 | 0.350257693 | 0.737777868 | 0.893235729 | -4.881709209 |
| A_55_P1971903 | Pla2r1 | -0.023062684 | -0.183171972 | 0.137046604 | 7.805108668 | -0.350006825 | 0.737957037 | 0.893396691 | -4.881755997 |
| A_55_P2054310 | M6pr | -0.030496753 | -0.24231166 | 0.181318154 | 10.74959359 | -0.349848706 | 0.738069973 | 0.893477455 | -4.88178547 |
| A_52_P32864 | Prps2 | -0.023123778 | -0.183867376 | 0.137619821 | 8.223108507 | -0.34954918 | 0.738283929 | 0.893564548 | -4.881841268 |
| A_55_P1957168 | Ube2s | 0.142671818 | -0.849202936 | 1.134546572 | 14.11925859 | 0.349513978 | 0.738309077 | 0.893564548 | -4.881847822 |
| A_55_P2408588 | Arntl | -0.071447619 | -0.568193661 | 0.425298423 | 9.606378533 | -0.349491433 | 0.738325182 | 0.893564548 | -4.88185202 |
| A_55_P2128526 | Pfn2 | -0.101087839 | -0.804022562 | 0.601846885 | 9.27441967 | -0.349435563 | 0.738365094 | 0.893564548 | -4.881862421 |
| A_51_P267388 | Dxo | 0.037620099 | -0.223987215 | 0.299227413 | 9.405724198 | 0.349424407 | 0.738373065 | 0.893564548 | -4.881864498 |
| A_51_P140347 | Slc4a9 | 0.125386952 | -0.747088063 | 0.997861967 | 10.13983412 | 0.349206673 | 0.738528619 | 0.893696843 | -4.881905017 |
| A_51_P323435 | Gtf2h2 | 0.025875863 | -0.154489151 | 0.206240877 | 8.41059384 | 0.348598672 | 0.738963062 | 0.894166585 | -4.882018036 |
| A_55_P2060459 | Nsd3 | 0.033052443 | -0.197496382 | 0.263601267 | 8.403560924 | 0.348356436 | 0.739136179 | 0.894263828 | -4.882063012 |
| A_55_P2028345 | Ccser2 | -0.055727449 | -0.444458864 | 0.333003965 | 10.34795401 | -0.348339494 | 0.739148288 | 0.894263828 | -4.882066157 |
| A_55_P2162651 | NA | -0.036227316 | -0.288968392 | 0.21651376 | 14.99076283 | -0.348292013 | 0.739182223 | 0.894263828 | -4.882074969 |
| A_51_P328014 | Cops3 | 0.030494748 | -0.18239878 | 0.243388277 | 10.72002529 | 0.348053323 | 0.739352826 | 0.894414243 | -4.882119251 |
| A_55_P2050488 | Cep170b | 0.081684385 | -0.48914535 | 0.652514121 | 15.28256839 | 0.347708772 | 0.739599123 | 0.894656202 | -4.882183122 |
| A_52_P269158 | Pid1 | 0.060064246 | -0.359996614 | 0.480125106 | 9.601591968 | 0.347445747 | 0.739787164 | 0.894827666 | -4.88223184 |
| A_55_P1965333 | Naa60 | -0.036999841 | -0.295826428 | 0.221826745 | 11.47701133 | -0.347355488 | 0.739851696 | 0.894849724 | -4.88224855 |
| A_51_P405423 | Smcr8 | 0.035898221 | -0.215306256 | 0.287102698 | 9.745926835 | 0.347239199 | 0.739934843 | 0.894894292 | -4.882270073 |
| A_55_P2108206 | Rrp7a | 0.048075225 | -0.288556951 | 0.384707402 | 10.76881775 | 0.347015516 | 0.740094787 | 0.89492952 | -4.882311453 |
| A_55_P2028570 | Sgsh | 0.081784311 | -0.490943625 | 0.654512248 | 15.15484686 | 0.346980304 | 0.740119966 | 0.89492952 | -4.882317965 |
| A_51_P416556 | Ssr3 | -0.053973426 | -0.432008715 | 0.324061863 | 13.48582312 | -0.346921195 | 0.740162235 | 0.89492952 | -4.882328894 |
| A_51_P519385 | Sacm1l | -0.064241222 | -0.514375697 | 0.385893252 | 9.76170973 | -0.346780532 | 0.740262826 | 0.89492952 | -4.882354897 |
| A_51_P348592 | Slc38a7 | 0.034555437 | -0.207596049 | 0.276706922 | 8.528222803 | 0.346746779 | 0.740286965 | 0.89492952 | -4.882361135 |
| A_55_P2021629 | Abcb7 | 0.039519449 | -0.23746058 | 0.316499478 | 8.339082631 | 0.346693431 | 0.740325117 | 0.89492952 | -4.882370993 |
| A_55_P2015363 | C1ra | -0.090781804 | -0.727210429 | 0.545646821 | 11.46039746 | -0.346602982 | 0.740389805 | 0.89492952 | -4.882387704 |
| A_55_P2330545 | NA | 0.026589817 | -0.159830102 | 0.213009735 | 8.590458837 | 0.34658217 | 0.740404689 | 0.89492952 | -4.882391549 |
| A_55_P2033116 | Cdcp2 | 0.069213775 | -0.416153114 | 0.554580664 | 8.474586934 | 0.346501867 | 0.740462122 | 0.89492952 | -4.882406381 |
| A_66_P125257 | Cep120 | 0.026484504 | -0.159249177 | 0.212218185 | 8.71770092 | 0.346484939 | 0.74047423 | 0.89492952 | -4.882409507 |
| A_55_P2423586 | Eya1 | 0.084497528 | -0.508115232 | 0.677110288 | 14.93821448 | 0.346462468 | 0.740490302 | 0.89492952 | -4.882413657 |
| A_52_P448466 | BC004004 | 0.053342264 | -0.320849769 | 0.427534298 | 8.908930493 | 0.346385816 | 0.740545127 | 0.89492952 | -4.88242781 |
| A_55_P2102579 | NA | -0.030700885 | -0.246083353 | 0.184681582 | 15.23617282 | -0.346356818 | 0.740565868 | 0.89492952 | -4.882433164 |
| A_55_P2062058 | Dbndd2 | -0.030748733 | -0.246540167 | 0.185042701 | 7.932146749 | -0.346239185 | 0.740650008 | 0.89493289 | -4.882454876 |
| A_55_P2081560 | Phf20 | 0.02737108 | -0.164725059 | 0.21946722 | 9.035046462 | 0.346223459 | 0.740661257 | 0.89493289 | -4.882457779 |
| A_51_P284988 | Nrbf2 | -0.03064094 | -0.245775864 | 0.184493984 | 9.646454672 | -0.346078287 | 0.740765102 | 0.895002416 | -4.882484564 |

| | | | | | | | | | |
|---|---|---|---|---|---|---|---|---|---|
| **A_55_P2122841** | NA | 0.066524043 | -0.400854796 | 0.533902882 | 16.73602482 | 0.34585398 | 0.740925566 | 0.895130698 | -4.882525929 |
| **A_55_P2065360** | Siva1 | 0.055031299 | -0.331662727 | 0.441725326 | 12.21192565 | 0.345800401 | 0.740963897 | 0.895130698 | -4.882535806 |
| **A_55_P1976926** | Tmem104 | -0.024792498 | -0.199044541 | 0.149459544 | 7.909962176 | -0.345720876 | 0.741020792 | 0.895143484 | -4.882550463 |
| **A_55_P1985591** | Rtl8a | 0.062242519 | -0.375415656 | 0.499900695 | 8.663606941 | 0.345569422 | 0.741129151 | 0.895181449 | -4.882578369 |
| **A_55_P2070347** | Rps2 | 0.028645357 | -0.172812567 | 0.230103281 | 16.21965733 | 0.345503993 | 0.741175965 | 0.895181449 | -4.882590421 |
| **A_52_P2706** | Mypn | -0.021975799 | -0.176537359 | 0.13258576 | 7.689305135 | -0.345482759 | 0.741191159 | 0.895181449 | -4.882594331 |
| **A_55_P2154097** | NA | 0.033162292 | -0.200164572 | 0.266489156 | 15.15161746 | 0.345352799 | 0.741284149 | 0.895196427 | -4.882618262 |
| **A_51_P114353** | Ctu1 | -0.029725981 | -0.238973207 | 0.179521245 | 8.522002655 | -0.34519117 | 0.741399806 | 0.895196427 | -4.882648012 |
| **A_55_P2020080** | NA | -0.052075228 | -0.41868632 | 0.314535864 | 9.622362045 | -0.345150708 | 0.74142876 | 0.895196427 | -4.882655458 |
| **A_55_P2015666** | NA | 0.02855765 | -0.172508563 | 0.229623862 | 16.81015036 | 0.34511716 | 0.741452767 | 0.895196427 | -4.882661631 |
| **A_55_P2018403** | Sycp1 | 0.02387071 | -0.144216806 | 0.191958227 | 7.658940439 | 0.345074659 | 0.741483182 | 0.895196427 | -4.88266945 |
| **A_55_P2287269** | AI415730 | 0.026414181 | -0.159642365 | 0.212470727 | 8.256787828 | 0.344965272 | 0.741561463 | 0.895196427 | -4.88268957 |
| **A_51_P384879** | Mdh2 | 0.037270654 | -0.225261924 | 0.299803232 | 15.45845204 | 0.344958617 | 0.741566226 | 0.895196427 | -4.882690794 |
| **A_55_P2051405** | Tet2 | -0.039135784 | -0.314815002 | 0.236543434 | 8.384154863 | -0.344947658 | 0.741574069 | 0.895196427 | -4.882692809 |
| **A_55_P1963084** | Rps13 | 0.024144383 | -0.146120098 | 0.194408863 | 16.08311884 | 0.344568234 | 0.741845628 | 0.895427581 | -4.88276255 |
| **A_55_P2024041** | Psg19 | -0.023181487 | -0.186664037 | 0.140301062 | 7.751292581 | -0.34455066 | 0.741858207 | 0.895427581 | -4.882765778 |
| **A_55_P2061391** | Rhobtb3 | -0.030083404 | -0.242600402 | 0.182433595 | 8.278925264 | -0.343966768 | 0.742276189 | 0.895779582 | -4.882872955 |
| **A_55_P2030805** | Actr10 | -0.026613667 | -0.214657264 | 0.161429931 | 12.93710155 | -0.343897761 | 0.742325594 | 0.895779582 | -4.88288561 |
| **A_52_P480402** | Avl9 | -0.039949595 | -0.322250499 | 0.242351309 | 9.052429821 | -0.343861299 | 0.7423517 | 0.895779582 | -4.882892296 |
| **A_51_P392090** | Tmem19 | -0.027284454 | -0.2201109 | 0.165541991 | 9.964345794 | -0.34382058 | 0.742380854 | 0.895779582 | -4.882899762 |
| **A_55_P2172396** | A530064D06Rik | -0.061330966 | -0.494774337 | 0.372112406 | 8.446268461 | -0.343819596 | 0.742381558 | 0.895779582 | -4.882899942 |
| **A_55_P2186572** | Lypla2 | -0.031376953 | -0.253192868 | 0.190438961 | 12.76121746 | -0.343717196 | 0.742454875 | 0.895812128 | -4.882918713 |
| **A_55_P1952127** | Psma3 | -0.033501204 | -0.270382248 | 0.203379841 | 13.81573563 | -0.343647581 | 0.742504721 | 0.89581635 | -4.882931471 |
| **A_51_P475298** | Zfp110 | -0.032563683 | -0.262945155 | 0.197817789 | 9.323682155 | -0.343454457 | 0.742643008 | 0.895842827 | -4.882966852 |
| **A_51_P418560** | Lnx2 | -0.039832461 | -0.321704808 | 0.242039886 | 9.660198653 | -0.343374353 | 0.74270037 | 0.895842827 | -4.882981521 |
| **A_66_P100696** | Slc26a2 | 0.024887472 | -0.15124022 | 0.201015165 | 7.895064106 | 0.343349434 | 0.742718215 | 0.895842827 | -4.882986084 |
| **A_52_P674165** | Rsbn1l | 0.025885969 | -0.157332808 | 0.209104745 | 8.24614392 | 0.343303025 | 0.742751449 | 0.895842827 | -4.882994581 |
| **A_55_P2128085** | NA | 0.052354517 | -0.31824605 | 0.422955084 | 12.02336245 | 0.343266375 | 0.742777695 | 0.895842827 | -4.88300129 |
| **A_52_P237652** | 2610035D17Rik | 0.024882278 | -0.151288708 | 0.201053264 | 8.18314161 | 0.34319341 | 0.742829948 | 0.895842827 | -4.883014646 |
| **A_55_P2178349** | Ywhaq | 0.055093291 | -0.335010644 | 0.445197225 | 12.10954554 | 0.343163882 | 0.742851095 | 0.895842827 | -4.88302005 |
| **A_51_P509223** | Arfgap1 | 0.036189346 | -0.220166161 | 0.292544853 | 8.776752946 | 0.343021445 | 0.742953106 | 0.895909949 | -4.883046111 |
| **A_51_P181523** | Tmem214 | -0.034668488 | -0.280388289 | 0.211051313 | 10.86721277 | -0.342829303 | 0.743090724 | 0.89602 | -4.883081252 |
| **A_65_P19458** | Rock2 | -0.030885172 | -0.249838582 | 0.188068238 | 9.027219211 | -0.342753184 | 0.743145245 | 0.896029845 | -4.883095167 |
| **A_55_P2179793** | NA | 0.02350167 | -0.143244308 | 0.190247649 | 7.99400498 | 0.342473168 | 0.743345825 | 0.896215784 | -4.883146334 |
| **A_52_P114889** | Upk3b | -0.078809066 | -0.638160622 | 0.48054249 | 8.457277301 | -0.3423533 | 0.743431695 | 0.896234917 | -4.883168226 |
| **A_55_P1959465** | Lrrfip1 | -0.070795968 | -0.573320714 | 0.431728778 | 9.597963798 | -0.342321566 | 0.743454429 | 0.896234917 | -4.88317402 |
| **A_55_P2069818** | Mapk10 | -0.025646641 | -0.207793009 | 0.156499727 | 7.696476473 | -0.342131581 | 0.743590539 | 0.896340324 | -4.883208699 |
| **A_51_P428742** | Pus7l | 0.03282876 | -0.200368047 | 0.266025567 | 8.418546327 | 0.342070065 | 0.743634613 | 0.896340324 | -4.883219924 |
| **A_51_P428715** | Mrpl13 | 0.032041232 | -0.195726838 | 0.259809302 | 12.47821976 | 0.341821638 | 0.743812611 | 0.896498969 | -4.883265235 |
| **A_55_P2079425** | Spred1 | -0.036700635 | -0.297662186 | 0.224260915 | 8.596321698 | -0.341727752 | 0.743879885 | 0.896524149 | -4.883282351 |
| **A_55_P1990210** | Scpep1 | -0.069441357 | -0.563544785 | 0.424662071 | 12.10647361 | -0.341494349 | 0.744047141 | 0.896669817 | -4.883324883 |
| **A_55_P2184180** | Ndufv3 | -0.023452474 | -0.190681198 | 0.143776249 | 14.60990461 | -0.340769706 | 0.744566514 | 0.897198789 | -4.883456756 |
| **A_52_P507310** | Mrps24 | -0.032923955 | -0.267790842 | 0.201942933 | 11.62952536 | -0.340622545 | 0.744672007 | 0.897198789 | -4.883483505 |
| **A_55_P2324921** | NA | 0.023475119 | -0.1439943 | 0.190944538 | 7.973671607 | 0.340608498 | 0.744682076 | 0.897198789 | -4.883486057 |
| **A_55_P2032755** | Gm10586 | 0.079746135 | -0.489194493 | 0.648686763 | 15.75000047 | 0.340585289 | 0.744698715 | 0.897198789 | -4.883490275 |
| **A_55_P2163622** | Rps27 | -0.058422195 | -0.475271961 | 0.358427571 | 16.27313877 | -0.340550479 | 0.74472367 | 0.897198789 | -4.8834966 |
| **A_51_P257541** | Plppr1 | -0.02252726 | -0.183348603 | 0.138294083 | 7.718237247 | -0.34036734 | 0.744854965 | 0.897198789 | -4.883529866 |
| **A_52_P70027** | Gm20498 | 0.045732746 | -0.280754484 | 0.372219975 | 11.71350479 | 0.340364507 | 0.744856996 | 0.897198789 | -4.88353038 |
| **A_55_P2130463** | Ptgdr2 | -0.022239392 | -0.181025486 | 0.136546702 | 7.658663588 | -0.340324833 | 0.74488544 | 0.897198789 | -4.883537585 |
| **A_55_P2043559** | NA | 0.042696247 | -0.262171857 | 0.347564351 | 9.555548146 | 0.340299181 | 0.744903832 | 0.897198789 | -4.883542242 |

| | | | | | | | | |
|---|---|---|---|---|---|---|---|---|
| **A_55_P2161390** | NA | -0.035860384 | -0.29206684 | 0.220346072 | 9.046030122 | -0.340101117 | 0.745045842 | 0.897313918 | -4.883578194 |
| **A_55_P2182159** | Ywhaq | 0.042210313 | -0.259466815 | 0.34388744 | 13.05647309 | 0.339984699 | 0.745129317 | 0.897349412 | -4.883599316 |
| **A_55_P2032538** | Pde3b | -0.093516734 | -0.7619874 | 0.574953932 | 10.42843213 | -0.339930526 | 0.745168162 | 0.897349412 | -4.883609143 |
| **A_51_P246903** | Rps27l | -0.028610065 | -0.23332307 | 0.17610294 | 14.21167511 | -0.339591334 | 0.7454114 | 0.897586404 | -4.883670636 |
| **A_51_P172155** | Hal | 0.097726786 | -0.602041459 | 0.797495031 | 10.46537503 | 0.339345876 | 0.745587441 | 0.897742457 | -4.8837151 |
| **A_51_P185693** | Slc2a2 | -0.117930047 | -0.963170283 | 0.727310188 | 11.81301451 | -0.339021702 | 0.745819962 | 0.897959612 | -4.883773776 |
| **A_52_P7183** | Tmem248 | 0.036658223 | -0.226283712 | 0.299600159 | 10.74763988 | 0.338762044 | 0.746006228 | 0.897959612 | -4.883820737 |
| **A_55_P2008437** | Cdkn2a | -0.022017155 | -0.179968274 | 0.135933964 | 7.707148506 | -0.338705063 | 0.746047106 | 0.897959612 | -4.883831037 |
| **A_55_P2143337** | Rpp21 | -0.049557045 | -0.405110165 | 0.305996075 | 12.01721793 | -0.338675747 | 0.746068138 | 0.897959612 | -4.883836336 |
| **A_51_P196590** | Hadh | 0.054264742 | -0.335090102 | 0.443619585 | 13.62324792 | 0.338653328 | 0.746084222 | 0.897959612 | -4.883840388 |
| **A_55_P2038242** | Tchhl1 | -0.021902811 | -0.179059695 | 0.135254074 | 7.668049856 | -0.338648868 | 0.746087421 | 0.897959612 | -4.883841194 |
| **A_55_P1961541** | NA | 0.076821486 | -0.474478427 | 0.628121399 | 9.065340943 | 0.338592996 | 0.746127506 | 0.897959612 | -4.883851292 |
| **A_55_P2052048** | Pih1d2 | -0.053948874 | -0.441167239 | 0.333269492 | 7.971700068 | -0.338539714 | 0.746165733 | 0.897959612 | -4.883860919 |
| **A_55_P2013765** | Raly | -0.078450313 | -0.641574672 | 0.484674045 | 13.54830865 | -0.338511603 | 0.746185901 | 0.897959612 | -4.883865998 |
| **A_65_P15875** | Chpf2 | -0.028192504 | -0.230777487 | 0.174392479 | 10.37371809 | -0.338150155 | 0.746445243 | 0.898181801 | -4.883931266 |
| **A_55_P2027052** | Prdm9 | 0.022792968 | -0.141005558 | 0.186591493 | 7.856632833 | 0.338122503 | 0.746465085 | 0.898181801 | -4.883936256 |
| **A_51_P424499** | Txn1 | 0.029660832 | -0.183532071 | 0.242853735 | 14.27878638 | 0.338059993 | 0.74650994 | 0.898181801 | -4.883947536 |
| **A_51_P446477** | Cldnd1 | -0.048028087 | -0.393575143 | 0.297518968 | 10.5050966 | -0.337731278 | 0.746745835 | 0.8983753 | -4.884006821 |
| **A_55_P2120080** | NA | -0.030979288 | -0.25395062 | 0.191992044 | 8.035807327 | -0.337602452 | 0.746838293 | 0.8983753 | -4.884030041 |
| **A_55_P2104798** | NA | -0.049873121 | -0.408838285 | 0.309092044 | 9.955360621 | -0.337596107 | 0.746842847 | 0.8983753 | -4.884031184 |
| **A_55_P2080649** | Zfp691 | -0.028245123 | -0.231552913 | 0.175062667 | 8.915411107 | -0.337576838 | 0.746856677 | 0.8983753 | -4.884034656 |
| **A_66_P124817** | Ing1 | 0.034780865 | -0.215641152 | 0.285202883 | 8.491448904 | 0.337482348 | 0.746924495 | 0.898400968 | -4.88405168 |
| **A_55_P2061159** | Ing1 | -0.063792132 | -0.523734164 | 0.3961499 | 11.71878652 | -0.337013431 | 0.747261087 | 0.898608687 | -4.884136098 |
| **A_55_P2148385** | NA | 0.039681404 | -0.246430955 | 0.325793763 | 9.217266347 | 0.337002872 | 0.747268667 | 0.898608687 | -4.884137998 |
| **A_52_P271** | Dnajc18 | -0.030110812 | -0.24727776 | 0.187056136 | 9.562599769 | -0.33690847 | 0.747336437 | 0.898608687 | -4.884154979 |
| **A_55_P2022337** | Dhcr7 | 0.059927026 | -0.372304348 | 0.4921584 | 11.05992504 | 0.336891169 | 0.747348857 | 0.898608687 | -4.88415809 |
| **A_55_P2014450** | B430105G09Rik | -0.025994027 | -0.213490836 | 0.161502783 | 7.831030956 | -0.336870409 | 0.747363761 | 0.898608687 | -4.884161824 |
| **A_55_P2170569** | Ggps1 | -0.044010802 | -0.361480165 | 0.273458561 | 10.07179984 | -0.336853178 | 0.747376131 | 0.898608687 | -4.884164922 |
| **A_55_P2346654** | AU035318 | -0.02324467 | -0.190962783 | 0.144473442 | 8.029860631 | -0.336764739 | 0.747439625 | 0.898622857 | -4.884180824 |
| **A_55_P2052470** | Uqcrh | -0.026495895 | -0.217718633 | 0.164726843 | 14.75615613 | -0.336683825 | 0.747497717 | 0.898622857 | -4.88419537 |
| **A_52_P418934** | Sec22b | -0.024220557 | -0.199043478 | 0.150602365 | 12.46808395 | -0.336642496 | 0.74752739 | 0.898622857 | -4.884202798 |
| **A_55_P2157927** | Tmem241 | -0.028145331 | -0.231385674 | 0.175095012 | 8.737404287 | -0.336495781 | 0.74763273 | 0.898649104 | -4.88422916 |
| **A_55_P1977008** | Gfi1 | 0.067635026 | -0.420783652 | 0.556053704 | 9.471097907 | 0.336482581 | 0.747642208 | 0.898649104 | -4.884231531 |
| **A_52_P528707** | Mrpl57 | 0.036136065 | -0.224978698 | 0.297250827 | 14.39554067 | 0.336273478 | 0.747792354 | 0.898773686 | -4.884269084 |
| **A_55_P2091985** | Gpr157 | 0.079267503 | -0.493757821 | 0.652292828 | 11.30044436 | 0.336127893 | 0.747896898 | 0.898843446 | -4.884295216 |
| **A_52_P110270** | Nelfe | 0.042707829 | -0.26610462 | 0.351520278 | 10.89184315 | 0.336043796 | 0.747957291 | 0.898860139 | -4.884310307 |
| **A_55_P2070913** | Rpl14 | -0.026149182 | -0.215359227 | 0.163060863 | 16.74455941 | -0.335812695 | 0.748123261 | 0.898976516 | -4.884351757 |
| **A_55_P1987231** | Fuom | -0.041995784 | -0.345898644 | 0.261907076 | 9.821194958 | -0.335779434 | 0.748147149 | 0.898976516 | -4.884357721 |
| **A_52_P136709** | Pctp | -0.043775995 | -0.361194181 | 0.273642192 | 8.498806268 | -0.335110016 | 0.748627995 | 0.899498385 | -4.884477625 |
| **A_55_P2122066** | Ppp2r3d | -0.028555689 | -0.235844858 | 0.17873348 | 8.24259016 | -0.334733535 | 0.748898477 | 0.89973421 | -4.88454496 |
| **A_55_P1960646** | Ube2w | 0.025759569 | -0.16125548 | 0.212774618 | 8.552570386 | 0.334691915 | 0.748928382 | 0.89973421 | -4.884552399 |
| **A_55_P2081690** | Tex38 | 0.024068137 | -0.15074525 | 0.198881524 | 7.772214536 | 0.334542252 | 0.749035919 | 0.89973421 | -4.884579144 |
| **A_55_P2057720** | Zfp937 | -0.030793891 | -0.254464732 | 0.192876949 | 8.305974179 | -0.334532559 | 0.749042884 | 0.89973421 | -4.884580876 |
| **A_55_P2133661** | NA | -0.083483508 | -0.689900001 | 0.522932985 | 14.07337041 | -0.334512904 | 0.749057008 | 0.89973421 | -4.884584387 |
| **A_55_P2144526** | Ripor2 | 0.037181506 | -0.233119956 | 0.307482967 | 8.193998958 | 0.334242574 | 0.749251266 | 0.899822014 | -4.884632662 |
| **A_51_P206655** | Lsm1 | -0.02549949 | -0.210888643 | 0.159889663 | 10.326675 | -0.33421841 | 0.749268631 | 0.899822014 | -4.884636976 |
| **A_51_P175871** | Got2 | 0.051912815 | -0.325511132 | 0.429336762 | 13.54249368 | 0.33421683 | 0.749269767 | 0.899822014 | -4.884637258 |
| **A_51_P233396** | Ppp2r5d | -0.043944788 | -0.363552932 | 0.275663356 | 11.10882248 | -0.334097116 | 0.7493558 | 0.899854198 | -4.884658623 |
| **A_55_P2009069** | NA | -0.049710987 | -0.411421722 | 0.311999748 | 12.99580053 | -0.333944409 | 0.749465549 | 0.899854198 | -4.884685865 |
| **A_51_P443723** | Slc35c1 | -0.030835521 | -0.255262008 | 0.193590967 | 11.50377424 | -0.333856905 | 0.749528441 | 0.899854198 | -4.884701471 |

| | | | | | | | | | |
|---|---|---|---|---|---|---|---|---|---|
| A_66_P108822 | NA | 0.032997271 | -0.207163468 | 0.273158009 | 15.90762471 | 0.333856009 | 0.749529085 | 0.899854198 | -4.884701631 |
| A_55_P1957850 | Snx9 | 0.045147075 | -0.283442718 | 0.373736868 | 10.13657146 | 0.333855657 | 0.749529338 | 0.899854198 | -4.884701693 |
| A_55_P2013396 | Gm14305 | -0.024064831 | -0.199336505 | 0.151206844 | 8.174116 | -0.333621677 | 0.749697516 | 0.899961922 | -4.884743402 |
| A_55_P2156093 | NA | 0.06386354 | -0.401303016 | 0.529030096 | 14.10702025 | 0.333601267 | 0.749712187 | 0.899961922 | -4.884747038 |
| A_52_P131548 | Ajuba | -0.051420748 | -0.426166786 | 0.32332529 | 8.843766423 | -0.333414534 | 0.749846418 | 0.900030356 | -4.884780303 |
| A_51_P196673 | Khdc1b | -0.021319364 | -0.176701726 | 0.134062999 | 7.636010632 | -0.333392408 | 0.749862324 | 0.900030356 | -4.884784243 |
| A_55_P2031671 | Gstm6 | 0.088959823 | -0.559855485 | 0.737775131 | 9.460541146 | 0.333162423 | 0.75002766 | 0.900172905 | -4.884825186 |
| A_55_P1992769 | Pafah1b1 | 0.048379136 | -0.30472581 | 0.401484083 | 12.57352164 | 0.33291817 | 0.750203268 | 0.900324211 | -4.884868639 |
| A_55_P2268221 | Arhgap10 | -0.022768205 | -0.189009987 | 0.143473576 | 7.773822945 | -0.332791178 | 0.750294578 | 0.900324211 | -4.88489122 |
| A_55_P2020926 | Trip11 | -0.033270994 | -0.27626959 | 0.209727603 | 9.160223854 | -0.332694175 | 0.750364327 | 0.900324211 | -4.884908462 |
| A_55_P2035300 | Rhot1 | -0.032140326 | -0.266900056 | 0.202619405 | 11.19904067 | -0.332667107 | 0.750383791 | 0.900324211 | -4.884913273 |
| A_52_P556337 | Txlng | 0.055837062 | -0.352080572 | 0.463754697 | 8.609191806 | 0.332608442 | 0.750425975 | 0.900324211 | -4.884923697 |
| A_51_P445662 | Hsd17b4 | 0.043830007 | -0.276419721 | 0.364079735 | 12.32512608 | 0.332556899 | 0.750463039 | 0.900324211 | -4.884932855 |
| A_55_P2043982 | Mtm1 | 0.043083598 | -0.271738096 | 0.357905292 | 8.580927494 | 0.332529747 | 0.750482564 | 0.900324211 | -4.884937679 |
| A_55_P2149540 | Srf | -0.023090872 | -0.191852062 | 0.145670317 | 9.561675459 | -0.332468844 | 0.75052636 | 0.900324211 | -4.884948497 |
| A_51_P180140 | H2bc1 | -0.041319776 | -0.343427573 | 0.26078802 | 12.88693713 | -0.332337399 | 0.750620887 | 0.900381726 | -4.884971839 |
| A_52_P239292 | AI597479 | 0.033586855 | -0.212124993 | 0.279298702 | 10.02729715 | 0.332144011 | 0.750759969 | 0.900492673 | -4.885006165 |
| A_55_P1996518 | Arhgap42 | -0.024559889 | -0.2042855 | 0.155165722 | 8.11983085 | -0.332047044 | 0.750829709 | 0.900504321 | -4.88502337 |
| A_52_P72839 | Sec23a | 0.065339885 | -0.412964576 | 0.543644347 | 9.366463671 | 0.331938115 | 0.750908056 | 0.900504321 | -4.885042691 |
| A_55_P2378827 | NA | 0.027457379 | -0.173554045 | 0.228468803 | 8.387599629 | 0.331910911 | 0.750927623 | 0.900504321 | -4.885047515 |
| A_55_P2007228 | Prkaca | 0.024118684 | -0.152472185 | 0.200709554 | 8.117275415 | 0.331870432 | 0.750956739 | 0.900504321 | -4.885054693 |
| A_51_P167232 | Cask | 0.0247698 | -0.156639334 | 0.206178934 | 8.135593708 | 0.331777201 | 0.751023799 | 0.900504321 | -4.885071222 |
| A_55_P2008879 | Cltc | 0.039123953 | -0.24749782 | 0.325745726 | 9.739737669 | 0.331678059 | 0.751095114 | 0.900504321 | -4.885088794 |
| A_51_P362881 | Fbxo38 | -0.028368248 | -0.236194401 | 0.179457905 | 11.00226584 | -0.33167711 | 0.751095797 | 0.900504321 | -4.885088962 |
| A_55_P1976017 | Cnih4 | -0.044200796 | -0.368219208 | 0.279817617 | 12.34791087 | -0.33146951 | 0.751245137 | 0.900627505 | -4.88512574 |
| A_55_P2108938 | Hps5 | -0.022910057 | -0.191044114 | 0.145224 | 9.599012896 | -0.331095796 | 0.751514003 | 0.900883332 | -4.885191893 |
| A_55_P2085142 | Spp1 | -0.06978685 | -0.582128807 | 0.442555107 | 8.412224935 | -0.330976274 | 0.7516 | 0.900883332 | -4.885213035 |
| A_55_P2097548 | NA | 0.038190983 | -0.242204569 | 0.318586535 | 16.5604538 | 0.330958018 | 0.751613136 | 0.900883332 | -4.885216264 |
| A_51_P225427 | Pkp2 | 0.056046692 | -0.355499244 | 0.467592628 | 11.32033419 | 0.330913785 | 0.751644963 | 0.900883332 | -4.885224086 |
| A_55_P1991054 | Tmf1 | 0.033870319 | -0.214918613 | 0.28265925 | 9.323110109 | 0.330804504 | 0.751723597 | 0.900921714 | -4.885243407 |
| A_55_P2100973 | Irf6 | 0.040796845 | -0.258931234 | 0.340524923 | 9.424655471 | 0.330736657 | 0.751772418 | 0.900924364 | -4.885255399 |
| A_55_P2131193 | NA | -0.060938452 | -0.509136831 | 0.387259927 | 15.30447013 | -0.330372834 | 0.752034241 | 0.901172846 | -4.885319667 |
| A_55_P2126855 | E230025N22Rik | 0.021587521 | -0.137213191 | 0.180388232 | 7.726008868 | 0.330318967 | 0.752073009 | 0.901172846 | -4.885329177 |
| A_52_P221024 | NA | 0.05264798 | -0.334755479 | 0.44005144 | 12.73356624 | 0.330218507 | 0.752145311 | 0.901185427 | -4.885346908 |
| A_55_P1995141 | Casz1 | 0.023756868 | -0.151078153 | 0.19859189 | 7.889058343 | 0.33017482 | 0.752176755 | 0.901185427 | -4.885354618 |
| A_52_P478729 | Atg2a | -0.031601728 | -0.264252642 | 0.201049185 | 8.618894499 | -0.330057239 | 0.752261384 | 0.901230959 | -4.885375362 |
| A_55_P2035023 | Zfp653 | 0.044320704 | -0.282242017 | 0.370883424 | 9.850036363 | 0.329779179 | 0.752461536 | 0.901414876 | -4.885424391 |
| A_52_P206492 | Pop4 | -0.041497522 | -0.347329632 | 0.264334588 | 11.12833905 | -0.329702524 | 0.752516717 | 0.901424735 | -4.8854379 |
| A_51_P446796 | Camta1 | -0.02529934 | -0.21178905 | 0.16119037 | 7.699295394 | -0.32963818 | 0.752563037 | 0.901424735 | -4.885449237 |
| A_55_P1995652 | Rexo4 | -0.031234105 | -0.261640832 | 0.199172623 | 12.15412493 | -0.329395071 | 0.752738057 | 0.901578504 | -4.885492054 |
| A_55_P2070296 | NA | -0.020937636 | -0.175439906 | 0.133564634 | 7.654263288 | -0.32928805 | 0.752815108 | 0.901614922 | -4.885510893 |
| A_55_P1963443 | Vps18 | -0.02552433 | -0.213958755 | 0.162910096 | 8.299246094 | -0.329137439 | 0.752923549 | 0.901684359 | -4.885537395 |
| A_52_P263251 | 3110040N11Rik | -0.024125433 | -0.202299231 | 0.154048364 | 9.041904619 | -0.329014074 | 0.753012377 | 0.901684359 | -4.885559095 |
| A_55_P1954481 | Dock9 | -0.026539716 | -0.222544286 | 0.169464853 | 7.868090275 | -0.329013162 | 0.753013034 | 0.901684359 | -4.885559255 |
| A_51_P260490 | Ralbp1 | -0.037015054 | -0.310585135 | 0.236555028 | 10.74773583 | -0.3287706 | 0.7531877 | 0.901829367 | -4.885601899 |
| A_51_P258848 | Mfsd13b | -0.021551404 | -0.18090533 | 0.137802523 | 7.678122639 | -0.328621509 | 0.753295067 | 0.901829367 | -4.885628095 |
| A_51_P468126 | Trim39 | 0.025976361 | -0.166102193 | 0.218054915 | 8.311160481 | 0.328611397 | 0.75330235 | 0.901829367 | -4.885629871 |
| A_55_P2039479 | Twistnb | 0.036915103 | -0.236131698 | 0.309961904 | 8.368538443 | 0.328511201 | 0.753374509 | 0.901829367 | -4.88564747 |
| A_52_P246889 | Chmp3 | 0.021604641 | -0.138285185 | 0.181494467 | 8.334500704 | 0.328329127 | 0.753505642 | 0.901829367 | -4.885679435 |
| A_66_P137102 | Sprydｄ7 | 0.022114505 | -0.141558477 | 0.185787488 | 8.259726722 | 0.32830948 | 0.753519793 | 0.901829367 | -4.885682884 |

| | | | | | | | | | |
|---|---|---|---|---|---|---|---|---|---|
| **A_55_P2045812** | Sigmar1 | -0.067887946 | -0.570377489 | 0.434601597 | 13.3788378 | -0.328283329 | 0.753538628 | 0.901829367 | -4.885687473 |
| **A_55_P2117081** | Trgv4 | -0.035734261 | -0.300267881 | 0.22879936 | 8.236738568 | -0.328236683 | 0.753572226 | 0.901829367 | -4.885695659 |
| **A_55_P2173438** | Arhgap4 | -0.04243947 | -0.356611769 | 0.271732829 | 8.686365965 | -0.328235275 | 0.75357324 | 0.901829367 | -4.885695906 |
| **A_55_P2077353** | Ceacam12 | -0.022581711 | -0.189769761 | 0.144606339 | 7.657578918 | -0.32819715 | 0.7536007 | 0.901829367 | -4.885702596 |
| **A_55_P1954021** | Scarb1 | -0.049150752 | -0.413260329 | 0.314958826 | 12.40514341 | -0.328005594 | 0.753738679 | 0.901938645 | -4.885736195 |
| **A_55_P2044724** | Zfp869 | 0.030659036 | -0.196585555 | 0.257903627 | 9.802368053 | 0.327829578 | 0.753865473 | 0.90196688 | -4.885767053 |
| **A_66_P110584** | Tpbpa | 0.021126022 | -0.135477564 | 0.177729608 | 7.635714874 | 0.327792649 | 0.753892076 | 0.90196688 | -4.885773525 |
| **A_52_P88983** | Dock5 | -0.043899151 | -0.369330094 | 0.281531792 | 9.327347923 | -0.327778502 | 0.753902267 | 0.90196688 | -4.885776004 |
| **A_55_P1992054** | Armc10 | -0.059963058 | -0.50470542 | 0.384779304 | 10.80962214 | -0.327610993 | 0.754022943 | 0.902055423 | -4.885805351 |
| **A_51_P279163** | Plcg2 | -0.057530535 | -0.484348986 | 0.369287916 | 11.27254773 | -0.327520445 | 0.754088178 | 0.902077633 | -4.885821209 |
| **A_55_P2119957** | Mlxipl | 0.086695032 | -0.556878313 | 0.730268378 | 10.62651341 | 0.327325133 | 0.754228898 | 0.902151631 | -4.885855401 |
| **A_55_P2166501** | Cd44 | -0.038709146 | -0.326080918 | 0.248662625 | 9.286377716 | -0.327305029 | 0.754243383 | 0.902151631 | -4.885858919 |
| **A_52_P489202** | 4732465J04Rik | -0.069840632 | -0.588675838 | 0.448994573 | 9.375440556 | -0.327085968 | 0.754401228 | 0.902284594 | -4.885897243 |
| **A_55_P2012011** | Tceal7 | -0.022213797 | -0.187508405 | 0.143080811 | 7.827213479 | -0.326548206 | 0.754788766 | 0.902692245 | -4.885991218 |
| **A_55_P2148534** | Nr1d2 | -0.032249063 | -0.272494955 | 0.207996829 | 8.745382238 | -0.326170229 | 0.755061202 | 0.902938825 | -4.886057184 |
| **A_51_P444052** | Scp2 | -0.065170912 | -0.550859546 | 0.420517723 | 13.01079908 | -0.326046126 | 0.755150661 | 0.902938825 | -4.886078826 |
| **A_55_P2014075** | Ercc2 | -0.027704698 | -0.234252906 | 0.17884351 | 10.32362947 | -0.325923126 | 0.755239329 | 0.902938825 | -4.886100269 |
| **A_55_P1989534** | Cklf | 0.024997578 | -0.161369507 | 0.211364664 | 9.797595621 | 0.325920676 | 0.755241095 | 0.902938825 | -4.886100696 |
| **A_55_P2067942** | D16Ertd472e | -0.037467699 | -0.316819678 | 0.24188428 | 9.548587276 | -0.325903075 | 0.755253783 | 0.902938825 | -4.886103764 |
| **A_55_P2037528** | Acsf3 | 0.043589221 | -0.281433323 | 0.368611765 | 11.2111975 | 0.325873325 | 0.75527523 | 0.902938825 | -4.886108949 |
| **A_51_P461452** | Ube2r2 | 0.026422172 | -0.170647871 | 0.223492216 | 12.94639177 | 0.325785011 | 0.755338897 | 0.902959091 | -4.886124338 |
| **A_55_P2430221** | Shld2 | -0.074139421 | -0.627319093 | 0.47904025 | 9.447814914 | -0.325661313 | 0.755428076 | 0.902999434 | -4.886145886 |
| **A_51_P235123** | Nfkbie | 0.029344374 | -0.189639516 | 0.248328263 | 8.557006526 | 0.325608603 | 0.755466079 | 0.902999434 | -4.886155066 |
| **A_52_P282500** | Kif21b | -0.058198573 | -0.492695556 | 0.37629841 | 10.6353854 | -0.325468349 | 0.755567201 | 0.903045965 | -4.886179484 |
| **A_52_P269016** | Nek9 | -0.029942311 | -0.253514393 | 0.193629771 | 11.08614978 | -0.325425012 | 0.755598448 | 0.903045965 | -4.886187028 |
| **A_55_P2028496** | NA | -0.03157902 | -0.267561758 | 0.204403718 | 13.38672778 | -0.325163361 | 0.755787114 | 0.903157904 | -4.88623255 |
| **A_55_P2178127** | 1-Sep | 0.053162322 | -0.344126879 | 0.450451524 | 9.14517315 | 0.325147466 | 0.755798576 | 0.903157904 | -4.886235314 |
| **A_55_P2140076** | NA | 0.03410683 | -0.22081476 | 0.28902842 | 16.55178733 | 0.325100718 | 0.755832286 | 0.903157904 | -4.886243443 |
| **A_55_P2019585** | Klhl17 | -0.044774495 | -0.379755861 | 0.290206871 | 9.297592254 | -0.324782961 | 0.756061438 | 0.903326852 | -4.886298671 |
| **A_51_P404825** | Ccdc58 | 0.032622849 | -0.211451658 | 0.276697356 | 12.25566614 | 0.32477505 | 0.756067143 | 0.903326852 | -4.886300045 |
| **A_52_P87892** | Abraxas2 | 0.038112099 | -0.247145375 | 0.323369574 | 8.756995836 | 0.324645232 | 0.75616077 | 0.903382874 | -4.886322592 |
| **A_52_P625270** | 1700024G13Rik | -0.073825436 | -0.626573919 | 0.478923047 | 7.864530034 | -0.324535083 | 0.756240215 | 0.903421948 | -4.886341717 |
| **A_55_P1960117** | Dipk2a | 0.032650573 | -0.211979035 | 0.277280181 | 8.998158422 | 0.324313464 | 0.756400067 | 0.903537496 | -4.886380177 |
| **A_55_P2031045** | Stat5b | -0.052399651 | -0.445047273 | 0.340247972 | 10.7120821 | -0.324271372 | 0.756430429 | 0.903537496 | -4.886387478 |
| **A_55_P2107957** | Abraxas1 | 0.024129063 | -0.156753443 | 0.205011568 | 8.023221493 | 0.324135854 | 0.756528185 | 0.903550283 | -4.886410981 |
| **A_51_P480119** | Prelid2 | 0.04617661 | -0.299993943 | 0.392347162 | 10.21342232 | 0.324126926 | 0.756534626 | 0.903550283 | -4.886412529 |
| **A_52_P218976** | Cyld | 0.025694788 | -0.167067499 | 0.218457075 | 8.426089111 | 0.323896431 | 0.756700906 | 0.903693037 | -4.886452481 |
| **A_55_P1984962** | NA | -0.023202641 | -0.197313172 | 0.15090789 | 7.754264974 | -0.323813959 | 0.756760404 | 0.903708257 | -4.886466769 |
| **A_55_P2092384** | Pik3r2 | 0.041624266 | -0.2707959 | 0.354044432 | 8.874936822 | 0.323735793 | 0.756816799 | 0.903719769 | -4.886480308 |
| **A_55_P2180789** | Huwe1 | 0.042298029 | -0.287415715 | 0.372011774 | 11.49515802 | 0.326381435 | 0.756915284 | 0.903765781 | -4.839148344 |
| **A_55_P2036733** | Stk36 | -0.039928587 | -0.339826688 | 0.259969514 | 8.02619994 | -0.323514251 | 0.756976643 | 0.903765781 | -4.886518665 |
| **A_51_P185259** | Gnb5 | 0.021448357 | -0.139660375 | 0.182557089 | 7.776609188 | 0.323487974 | 0.756995603 | 0.903765781 | -4.886523213 |
| **A_66_P101646** | NA | 0.170589834 | -1.111050489 | 1.452230158 | 15.5583202 | 0.323422458 | 0.757042875 | 0.903766397 | -4.88653455 |
| **A_55_P2137049** | AA467197 | -0.074890808 | -0.638532993 | 0.488751377 | 9.560352382 | -0.322855519 | 0.757451999 | 0.904198968 | -4.886632567 |
| **A_55_P2112355** | NA | 0.034501374 | -0.225236969 | 0.294239717 | 13.35758841 | 0.32276283 | 0.757518895 | 0.90422298 | -4.886648576 |
| **A_51_P290139** | Dlst | 0.030425627 | -0.198710594 | 0.259561848 | 12.18906316 | 0.322648005 | 0.75760177 | 0.904266062 | -4.886668402 |
| **A_55_P2021709** | Oxnad1 | 0.04617165 | -0.301938077 | 0.394281376 | 10.85236907 | 0.32228673 | 0.757862543 | 0.904521463 | -4.886730739 |
| **A_55_P2025443** | Oprm1 | 0.083685377 | -0.547658501 | 0.715029254 | 11.46786035 | 0.32208225 | 0.758010155 | 0.904641781 | -4.886765992 |
| **A_66_P109708** | Il1f6 | -0.020666687 | -0.176653601 | 0.135320226 | 7.640787256 | -0.321933297 | 0.75811769 | 0.904681749 | -4.886791659 |
| **A_55_P2182700** | Prkab1 | 0.036718768 | -0.240448506 | 0.313886042 | 8.579900116 | 0.321906199 | 0.758137254 | 0.904681749 | -4.886796327 |

| | | | | | | | | |
|---|---|---|---|---|---|---|---|---|
| **A_55_P2051229** | Gpr132 | -0.037624759 | -0.321749289 | 0.246499771 | 8.367596953 | -0.321771951 | 0.758234178 | 0.904730339 | -4.886819448 |
| **A_52_P77245** | Snx8 | 0.031516578 | -0.206544411 | 0.269577567 | 11.0305555 | 0.321687372 | 0.758295245 | 0.904730339 | -4.88683401 |
| **A_55_P1990217** | Zfp13 | -0.025071238 | -0.214523388 | 0.164380912 | 8.234502884 | -0.321558078 | 0.758388601 | 0.904730339 | -4.886856264 |
| **A_55_P2090924** | Tbc1d8b | 0.06113695 | -0.400869793 | 0.523143693 | 8.452658765 | 0.321542692 | 0.75839971 | 0.904730339 | -4.886858911 |
| **A_55_P2102529** | Fuca2 | -0.026834532 | -0.229631557 | 0.175962493 | 8.24057308 | -0.321525661 | 0.758412007 | 0.904730339 | -4.886861842 |
| **A_55_P2227355** | Ptpro | 0.025514386 | -0.167434635 | 0.218463407 | 8.222550624 | 0.321311102 | 0.758566939 | 0.904859317 | -4.886898749 |
| **A_51_P421182** | Sys1 | 0.028079804 | -0.18434315 | 0.240502758 | 8.525301859 | 0.321200207 | 0.75864702 | 0.904898997 | -4.886917815 |
| **A_55_P2007464** | Rps2 | 0.031604092 | -0.207529108 | 0.270737291 | 17.04969047 | 0.321134251 | 0.758694651 | 0.90489997 | -4.886929152 |
| **A_51_P345902** | Dohh | 0.024030296 | -0.157845922 | 0.205906514 | 12.43703459 | 0.321045359 | 0.758758847 | 0.904920698 | -4.886944428 |
| **A_51_P472879** | NA | 0.101373888 | -0.666111325 | 0.8688591 | 11.60046958 | 0.3209514 | 0.758826705 | 0.904945791 | -4.88696057 |
| **A_51_P475580** | Ugp2 | -0.071125144 | -0.610149311 | 0.467899022 | 10.83820405 | -0.32062554 | 0.75906206 | 0.905158174 | -4.887016518 |
| **A_55_P2139625** | Gtf2a2 | 0.037753412 | -0.248406989 | 0.323913813 | 11.78882335 | 0.320575156 | 0.759098453 | 0.905158174 | -4.887025164 |
| **A_55_P2150053** | Yipf5 | -0.033979414 | -0.291640254 | 0.223681425 | 10.12829845 | -0.320442904 | 0.759193983 | 0.905198735 | -4.887047852 |
| **A_55_P1967224** | NA | 0.058862091 | -0.387572196 | 0.505296378 | 8.703262938 | 0.320377001 | 0.759241589 | 0.905198735 | -4.887059154 |
| **A_51_P489153** | Crot | 0.086636929 | -0.570542235 | 0.743816093 | 11.36396824 | 0.32033357 | 0.759272962 | 0.905198735 | -4.887066601 |
| **A_51_P454217** | Nfkb2 | -0.030349436 | -0.260910621 | 0.200211748 | 8.485171513 | -0.319850936 | 0.759621639 | 0.90555857 | -4.887149295 |
| **A_51_P513460** | Cfhr1 | 0.091664448 | -0.605453733 | 0.788782629 | 8.87506195 | 0.319505035 | 0.75987157 | 0.90570153 | -4.887208489 |
| **A_52_P475450** | Rbm48 | 0.029477024 | -0.194710789 | 0.253664838 | 8.37396691 | 0.319488204 | 0.759883732 | 0.90570153 | -4.887211368 |
| **A_55_P2217309** | NA | 0.027544507 | -0.181949179 | 0.237038192 | 8.130723803 | 0.319482628 | 0.759887762 | 0.90570153 | -4.887212321 |
| **A_55_P2073795** | Mecr | -0.037673174 | -0.324253065 | 0.248906717 | 8.693261319 | -0.319425576 | 0.759928988 | 0.90570153 | -4.887222078 |
| **A_55_P2064004** | Rapgef2 | -0.029857064 | -0.257047151 | 0.197333023 | 9.914257614 | -0.319330875 | 0.759997423 | 0.905727245 | -4.88723827 |
| **A_51_P211573** | Wdr47 | 0.038461716 | -0.254543732 | 0.331467164 | 8.352317466 | 0.318959945 | 0.760265491 | 0.905990856 | -4.887301646 |
| **A_65_P07561** | Tpd52l2 | 0.035836634 | -0.237355091 | 0.309028359 | 9.624596087 | 0.318744618 | 0.760421123 | 0.90607214 | -4.887338404 |
| **A_55_P2013465** | L2hgdh | 0.024196412 | -0.160264022 | 0.208656846 | 8.639160805 | 0.31873586 | 0.760427454 | 0.90607214 | -4.887339898 |
| **A_55_P2076866** | Megf6 | -0.112946677 | -0.974333469 | 0.748440114 | 9.241026322 | -0.318609295 | 0.760518938 | 0.906125288 | -4.887361493 |
| **A_55_P2043702** | Tfg | 0.038751265 | -0.256922638 | 0.334425169 | 12.82001897 | 0.318460873 | 0.760626225 | 0.90619196 | -4.887386806 |
| **A_51_P488730** | Zfp612 | -0.032290614 | -0.278719891 | 0.214138663 | 8.997294436 | -0.318395696 | 0.760673341 | 0.90619196 | -4.887397918 |
| **A_55_P1956652** | Sfmbt2 | 0.033531043 | -0.222411629 | 0.289473715 | 7.78127424 | 0.318337315 | 0.760715544 | 0.90619196 | -4.887407869 |
| **A_55_P2146708** | Zcchc9 | 0.067483556 | -0.447814409 | 0.58278152 | 15.32732462 | 0.318216518 | 0.760802871 | 0.906219051 | -4.887428455 |
| **A_51_P233603** | Psma7 | -0.042958579 | -0.371027792 | 0.285110633 | 14.0432575 | -0.318176152 | 0.760832053 | 0.906219051 | -4.887435333 |
| **A_51_P196444** | Foxc2 | -0.02179875 | -0.188592886 | 0.144995386 | 7.702590071 | -0.317565998 | 0.761273212 | 0.906425312 | -4.887539186 |
| **A_51_P434776** | Jmjd8 | -0.024664341 | -0.213398109 | 0.164069428 | 8.570400834 | -0.317543399 | 0.761289554 | 0.906425312 | -4.887543029 |
| **A_55_P1967618** | Med16 | -0.02699023 | -0.233524657 | 0.179544198 | 11.10479377 | -0.317539175 | 0.761292608 | 0.906425312 | -4.887543747 |
| **A_51_P375693** | Tmem135 | -0.075287285 | -0.651404338 | 0.500829767 | 10.66168009 | -0.317536824 | 0.761294308 | 0.906425312 | -4.887544147 |
| **A_55_P2142116** | NA | 0.071493198 | -0.4757088 | 0.618695197 | 15.43235387 | 0.317468208 | 0.761343927 | 0.906425312 | -4.8875555813 |
| **A_52_P198898** | Samd5 | 0.035667448 | -0.237362364 | 0.30869726 | 8.475282947 | 0.317427942 | 0.761373045 | 0.906425312 | -4.887562658 |
| **A_55_P1967191** | Gm4632 | -0.021056642 | -0.182336618 | 0.140223333 | 7.620739416 | -0.317242865 | 0.761506887 | 0.906425312 | -4.88759411 |
| **A_55_P2082779** | Gm10240 | -0.041662178 | -0.360768561 | 0.277444205 | 15.25017801 | -0.317241263 | 0.761508045 | 0.906425312 | -4.887594382 |
| **A_55_P2100679** | Dnm2 | 0.029250706 | -0.194800699 | 0.253302111 | 8.331332566 | 0.317228264 | 0.761517446 | 0.906425312 | -4.88759659 |
| **A_55_P2144376** | NA | 0.05194443 | -0.345947403 | 0.449836264 | 8.631512131 | 0.317217509 | 0.761525224 | 0.906425312 | -4.887598417 |
| **A_51_P129464** | Scd2 | -0.050028603 | -0.433270168 | 0.333212961 | 14.93170737 | -0.317196921 | 0.761540113 | 0.906425312 | -4.887601915 |
| **A_55_P1967617** | Med16 | 0.042144758 | -0.280756069 | 0.365045585 | 16.3643195 | 0.317144798 | 0.76157781 | 0.906425312 | -4.887610768 |
| **A_51_P385598** | Slc37a4 | -0.096613955 | -0.837174727 | 0.643946816 | 13.3835377 | -0.317002253 | 0.761680904 | 0.906425312 | -4.887634972 |
| **A_55_P2144641** | Sdk1 | 0.03846295 | -0.256393987 | 0.333319888 | 8.639864238 | 0.31696728 | 0.761706198 | 0.906425312 | -4.887640909 |
| **A_55_P1959064** | Oas1b | 0.083528709 | -0.556809111 | 0.723866529 | 9.747136966 | 0.316963901 | 0.761708642 | 0.906425312 | -4.887641483 |
| **A_55_P2368355** | 2610319H10Rik | -0.027794085 | -0.240964929 | 0.185376759 | 10.32960745 | -0.316816481 | 0.761815269 | 0.906496389 | -4.887666501 |
| **A_51_P407004** | Prkrip1 | 0.05229619 | -0.348913274 | 0.453505653 | 10.55855294 | 0.3167248 | 0.761881584 | 0.906513132 | -4.887682055 |
| **A_51_P185509** | Mapre1 | -0.030950584 | -0.26846053 | 0.206559363 | 9.001249149 | -0.316643248 | 0.761940574 | 0.906513132 | -4.887695887 |
| **A_52_P337716** | NA | 0.025187499 | -0.168125808 | 0.218500805 | 11.5570589 | 0.316596768 | 0.761974195 | 0.906513132 | -4.887703768 |
| **A_51_P280158** | Sf3b2 | 0.027522759 | -0.183753033 | 0.238798551 | 12.17171871 | 0.316537682 | 0.762016937 | 0.906513132 | -4.887713786 |

| | | | | | | | | | |
|---|---|---|---|---|---|---|---|---|---|
| A_55_P1994224 | Ythdf1 | -0.033581193 | -0.291597438 | 0.224435051 | 12.38556118 | -0.316251259 | 0.762224139 | 0.906703821 | -4.887762322 |
| A_55_P2101231 | Aldh7a1 | -0.055681536 | -0.484051569 | 0.372688496 | 13.88665558 | -0.315845954 | 0.762517381 | 0.906911349 | -4.887830933 |
| A_55_P1952779 | Dcp1b | 0.022113897 | -0.148015118 | 0.192242912 | 9.888375429 | 0.315842153 | 0.762520131 | 0.906911349 | -4.887831576 |
| A_55_P2075814 | Tnni3k | 0.02670981 | -0.17883763 | 0.232257251 | 8.064792155 | 0.315748955 | 0.762587567 | 0.906911349 | -4.887847341 |
| A_55_P1983523 | Cd300ld | -0.049608571 | -0.431464134 | 0.332246991 | 8.880032917 | -0.315675435 | 0.762640765 | 0.906911349 | -4.887859774 |
| A_55_P2117202 | Mtx1 | 0.029864962 | -0.200079312 | 0.259809236 | 12.39943635 | 0.315589509 | 0.762702942 | 0.906911349 | -4.887874301 |
| A_55_P2108738 | Nck1 | 0.038498514 | -0.257933326 | 0.334930353 | 10.33454478 | 0.31557479 | 0.762713593 | 0.906911349 | -4.887876789 |
| A_55_P2415197 | 2810013P06Rik | 0.024241793 | -0.16242677 | 0.210910355 | 8.958533462 | 0.315556213 | 0.762727036 | 0.906911349 | -4.887879929 |
| A_51_P131653 | Tsacc | 0.025417326 | -0.170391596 | 0.221226248 | 9.655042033 | 0.315413715 | 0.762830156 | 0.906958682 | -4.88790401 |
| A_55_P2109439 | Lsmem1 | -0.02030042 | -0.176785382 | 0.136184543 | 7.640891892 | -0.315221332 | 0.762969383 | 0.906958682 | -4.887936505 |
| A_55_P2015700 | NA | -0.040268154 | -0.350700539 | 0.270164231 | 9.271382168 | -0.315193959 | 0.762989194 | 0.906958682 | -4.887941128 |
| A_52_P554650 | Gzmd | -0.020221399 | -0.176114806 | 0.135672007 | 7.662039955 | -0.315185804 | 0.762995095 | 0.906958682 | -4.887942504 |
| A_51_P113205 | F13b | -0.098327652 | -0.856388938 | 0.659733634 | 11.5980665 | -0.315177017 | 0.763001455 | 0.906958682 | -4.887943988 |
| A_52_P349298 | Rprd1b | 0.057980885 | -0.389203284 | 0.505165055 | 8.853747032 | 0.315051543 | 0.763092267 | 0.907003075 | -4.887965169 |
| A_55_P2302413 | BC002189 | -0.02120803 | -0.184806347 | 0.142390287 | 7.65257804 | -0.314995747 | 0.763132651 | 0.907003075 | -4.887974585 |
| A_51_P137168 | Smarcal1 | -0.02479806 | -0.216233865 | 0.166637744 | 9.59028347 | -0.31475869 | 0.763304236 | 0.907151228 | -4.888014572 |
| A_55_P2089507 | NA | -0.034098004 | -0.297383681 | 0.229187672 | 15.37156624 | -0.314691432 | 0.76335292 | 0.90715331 | -4.888025912 |
| A_51_P374900 | P2ry13 | -0.044419122 | -0.387490784 | 0.298652541 | 8.493694026 | -0.314606949 | 0.763414076 | 0.907170212 | -4.888040154 |
| A_52_P177021 | Pts | 0.032638793 | -0.21954434 | 0.284821925 | 9.7959201 | 0.314485945 | 0.76350167 | 0.907216389 | -4.888060545 |
| A_55_P2088535 | Snrpd2 | -0.043780136 | -0.38211376 | 0.294553489 | 13.99538079 | -0.314423595 | 0.763546807 | 0.907216389 | -4.888071049 |
| A_55_P2003369 | Fkbp8 | -0.042574687 | -0.371988494 | 0.28683912 | 11.97938259 | -0.314045704 | 0.763820393 | 0.907326739 | -4.888134669 |
| A_55_P2145617 | Krt80 | 0.042818104 | -0.288522686 | 0.374158894 | 8.466375039 | 0.314004387 | 0.763850307 | 0.907326739 | -4.88814162 |
| A_52_P574720 | Oit3 | 0.06990881 | -0.471168845 | 0.610986464 | 11.34284918 | 0.313946393 | 0.763892298 | 0.907326739 | -4.888151377 |
| A_55_P2046198 | Slc25a25 | -0.030341851 | -0.265254636 | 0.204570934 | 8.863151774 | -0.313847452 | 0.763963937 | 0.907326739 | -4.888168017 |
| A_55_P1954436 | NA | 0.039850501 | -0.268707347 | 0.34840835 | 8.652419998 | 0.313819827 | 0.76398394 | 0.907326739 | -4.888172662 |
| A_55_P1956147 | B4galnt2 | 0.026912706 | -0.18151307 | 0.235338481 | 7.825022812 | 0.313753893 | 0.764031683 | 0.907326739 | -4.888183747 |
| A_51_P355943 | Mvd | 0.048438285 | -0.326712392 | 0.423588963 | 10.7374657 | 0.313737344 | 0.764043666 | 0.907326739 | -4.888186529 |
| A_55_P2056021 | Snap91 | -0.019954815 | -0.174507148 | 0.134597518 | 7.649405001 | -0.313729485 | 0.764049357 | 0.907326739 | -4.88818785 |
| A_66_P102467 | Casz1 | -0.021599997 | -0.18891028 | 0.145710285 | 7.689167541 | -0.313699809 | 0.764070846 | 0.907326739 | -4.888192839 |
| A_55_P2040046 | Swi5 | 0.034951461 | -0.235847216 | 0.305750138 | 14.94932133 | 0.313618728 | 0.764129559 | 0.907326739 | -4.888206465 |
| A_66_P134149 | Snx30 | -0.044398329 | -0.388430021 | 0.299633364 | 9.113794357 | -0.313582168 | 0.764156034 | 0.907326739 | -4.888212609 |
| A_55_P2052181 | Ccdc82 | -0.031188374 | -0.272960759 | 0.21058401 | 11.41704363 | -0.313450694 | 0.764251243 | 0.907347468 | -4.888234695 |
| A_55_P2018542 | Ptges3 | 0.036688321 | -0.247739882 | 0.321116523 | 10.84902756 | 0.313428416 | 0.764267377 | 0.907347468 | -4.888238437 |
| A_52_P304858 | Gnb1 | 0.058075485 | -0.392439434 | 0.508590405 | 9.473647056 | 0.313232531 | 0.764409239 | 0.907460152 | -4.888271325 |
| A_55_P2174821 | Rab8a | -0.043493263 | -0.381307488 | 0.294320963 | 10.70077646 | -0.312843576 | 0.764690955 | 0.907738837 | -4.88833657 |
| A_51_P436068 | Gpr182 | 0.039164905 | -0.265129031 | 0.343458841 | 12.1710952 | 0.312742547 | 0.764764137 | 0.907769958 | -4.888353505 |
| A_51_P242166 | Lap3 | 0.029106309 | -0.197115028 | 0.255327646 | 12.07773864 | 0.3126344 | 0.764842476 | 0.907776452 | -4.888371627 |
| A_55_P2021908 | NA | -0.047443749 | -0.416278809 | 0.321391312 | 10.84113186 | -0.312557545 | 0.764898151 | 0.907776452 | -4.888384502 |
| A_51_P345467 | Pi4k2a | 0.031507466 | -0.213449823 | 0.276464755 | 9.444180521 | 0.312540497 | 0.764910501 | 0.907776452 | -4.888387357 |
| A_51_P497724 | Apol7a | -0.091806486 | -0.806626796 | 0.623013824 | 9.76392967 | -0.312075496 | 0.765247385 | 0.908059673 | -4.888465186 |
| A_55_P1993594 | Dcp1b | 0.024836591 | -0.168574328 | 0.218247511 | 10.21265428 | 0.312028445 | 0.765281475 | 0.908059673 | -4.888473055 |
| A_51_P390715 | Tgfb1 | 0.037813585 | -0.256664783 | 0.332291953 | 9.208108964 | 0.31201656 | 0.765290086 | 0.908059673 | -4.884693948 |
| A_52_P272054 | NA | 0.060252772 | -0.409208129 | 0.529713674 | 10.81538354 | 0.311860803 | 0.765402944 | 0.908137837 | -4.888501082 |
| A_66_P115038 | Prl3d1 | -0.019742646 | -0.173675879 | 0.134190586 | 7.64391514 | -0.311642133 | 0.765561398 | 0.908227776 | -4.88853762 |
| A_55_P2185461 | Rpl9 | -0.057584564 | -0.50660397 | 0.391434842 | 16.71737149 | -0.311619168 | 0.765578039 | 0.908227776 | -4.888541456 |
| A_51_P394833 | Tshz1 | 0.047388819 | -0.32222362 | 0.417001257 | 9.82760719 | 0.311539053 | 0.765636096 | 0.908227776 | -4.888554835 |
| A_55_P2011137 | Armc10 | 0.029071743 | -0.197733209 | 0.255876694 | 8.61939315 | 0.311459602 | 0.765693674 | 0.908227776 | -4.8885681 |
| A_55_P2408848 | Lgr4 | -0.029748164 | -0.261857476 | 0.202361148 | 8.003456941 | -0.311423082 | 0.76572014 | 0.908227776 | -4.888574197 |
| A_55_P2091741 | Lamp3 | -0.093407134 | -0.822344891 | 0.635530622 | 12.6162018 | -0.31136715 | 0.765760674 | 0.908227776 | -4.888583532 |
| A_55_P2082434 | Btbd11 | 0.024902662 | -0.169685692 | 0.219491017 | 8.180468766 | 0.310965438 | 0.766051825 | 0.908482264 | -4.888650534 |

| | | | | | | | | | |
|---|---|---|---|---|---|---|---|---|---|
| **A_55_P2018559** | Morf4l1 | -0.02468595 | -0.217595833 | 0.168223933 | 11.45155843 | -0.310941406 | 0.766069245 | 0.908482264 | -4.888654539 |
| **A_51_P155313** | Gsto1 | -0.064148527 | -0.565723018 | 0.437425964 | 13.15923496 | -0.310766672 | 0.766195901 | 0.908576723 | -4.888683655 |
| **A_52_P592101** | Washc2 | 0.031631875 | -0.215851284 | 0.279115034 | 9.346447586 | 0.310572127 | 0.766336927 | 0.908584836 | -4.888716054 |
| **A_55_P2104894** | Col5a2 | -0.063145896 | -0.557240698 | 0.430948907 | 9.156629815 | -0.310540346 | 0.766359967 | 0.908584836 | -4.888721345 |
| **A_55_P1959600** | Tmem222 | -0.028309815 | -0.249844094 | 0.193224464 | 11.84025876 | -0.310512644 | 0.766380049 | 0.908584836 | -4.888725956 |
| **A_51_P518926** | Borcs6 | -0.022994046 | -0.20293913 | 0.156951037 | 9.193880277 | -0.310497858 | 0.766390768 | 0.908584836 | -4.888728418 |
| **A_55_P1989733** | Nfkbib | 0.031585792 | -0.215812314 | 0.278983898 | 10.95075937 | 0.310226282 | 0.766587656 | 0.908758309 | -4.888773602 |
| **A_55_P1974467** | Ndufb6 | -0.026239522 | -0.231803063 | 0.179324019 | 13.22790078 | -0.310165163 | 0.766631969 | 0.908758309 | -4.888783766 |
| **A_55_P2032740** | Iars2 | 0.028254807 | -0.19314217 | 0.249651783 | 8.442343899 | 0.310101484 | 0.766678139 | 0.908758309 | -4.888794354 |
| **A_55_P2113466** | NA | 0.037706641 | -0.258069166 | 0.333482448 | 9.226491703 | 0.309769308 | 0.766918997 | 0.908941035 | -4.888849548 |
| **A_51_P376149** | Eef1akmt1 | -0.047878309 | -0.42349979 | 0.327743172 | 10.50537998 | -0.309721656 | 0.766953552 | 0.908941035 | -4.888857462 |
| **A_55_P2004811** | NA | -0.041678437 | -0.368688684 | 0.285331811 | 14.97056938 | -0.309694325 | 0.766973371 | 0.908941035 | -4.888862 |
| **A_55_P2236138** | 4930447F24Rik | 0.019733627 | -0.135203008 | 0.174670262 | 7.668742622 | 0.309482423 | 0.767127038 | 0.909067409 | -4.888897172 |
| **A_55_P2163018** | NA | -0.030380104 | -0.269070433 | 0.208310224 | 11.26249668 | -0.309269882 | 0.76728118 | 0.909194331 | -4.888932428 |
| **A_51_P337897** | Prrc1 | -0.02194865 | -0.194519136 | 0.150621835 | 8.227722467 | -0.309046969 | 0.767442857 | 0.909330165 | -4.888969379 |
| **A_55_P2106745** | Trim63 | -0.020310803 | -0.180099422 | 0.139477816 | 7.622327009 | -0.308861972 | 0.767577043 | 0.909383811 | -4.889000026 |
| **A_51_P509384** | Aldh8a1 | 0.121747065 | -0.836080426 | 1.079574556 | 11.48955816 | 0.308854825 | 0.767582227 | 0.909383811 | -4.889001209 |
| **A_55_P2126572** | Tmem25 | -0.091977813 | -0.815771745 | 0.631816119 | 9.798534227 | -0.308781541 | 0.767635387 | 0.909391051 | -4.889013344 |
| **A_66_P111090** | Sctr | 0.020110595 | -0.138218571 | 0.178439762 | 7.682923737 | 0.308636432 | 0.76774065 | 0.909460012 | -4.889037365 |
| **A_55_P2124523** | Gm13620 | 0.08194022 | -0.563672531 | 0.727552971 | 8.650855216 | 0.308395631 | 0.76791534 | 0.909611203 | -4.889077202 |
| **A_52_P437084** | NA | -0.045204767 | -0.40154689 | 0.311137357 | 14.2936188 | -0.308247993 | 0.768022452 | 0.909682332 | -4.889101612 |
| **A_55_P1956998** | Dis3l2 | -0.023966709 | -0.212963476 | 0.165030057 | 8.371235386 | -0.308132298 | 0.768106394 | 0.90972601 | -4.889120733 |
| **A_51_P418809** | Airn | 0.024154218 | -0.166447606 | 0.214756042 | 7.954786088 | 0.307927955 | 0.768254661 | 0.909794666 | -4.889154488 |
| **A_51_P301311** | Ciz1 | 0.028231566 | -0.19456707 | 0.251030201 | 11.36548234 | 0.307897122 | 0.768277034 | 0.909794666 | -4.889159579 |
| **A_55_P2338535** | Slc35a1 | 0.029830504 | -0.205616749 | 0.265277758 | 8.68289399 | 0.307857797 | 0.768305569 | 0.909794666 | -4.889166072 |
| **A_55_P1968833** | Rpgr | -0.031524012 | -0.280428648 | 0.217380624 | 8.95476231 | -0.307745483 | 0.768387068 | 0.909835435 | -4.889184612 |
| **A_51_P353392** | Cript | 0.043877898 | -0.302737779 | 0.390493574 | 9.438942043 | 0.307596081 | 0.768495485 | 0.909836195 | -4.889209264 |
| **A_55_P1997300** | Mtfr2 | 0.074214214 | -0.512047792 | 0.66047622 | 14.76081445 | 0.307594475 | 0.768496651 | 0.909836195 | -4.889209529 |
| **A_55_P2025959** | Chd9 | -0.030681174 | -0.273084975 | 0.211722627 | 9.531675696 | -0.307550004 | 0.768528923 | 0.909836195 | -4.889216864 |
| **A_51_P103929** | Met | 0.06054431 | -0.417938006 | 0.539026626 | 8.555595075 | 0.307461423 | 0.768593208 | 0.909856572 | -4.885443512 |
| **A_55_P2007496** | NA | -0.027503772 | -0.244959191 | 0.189951646 | 8.876580392 | -0.307330188 | 0.768688451 | 0.909875862 | -4.889253109 |
| **A_55_P2140423** | Arfip2 | 0.042328348 | -0.29239469 | 0.377051385 | 9.165803598 | 0.307276184 | 0.768727645 | 0.909875862 | -4.88926201 |
| **A_52_P533779** | Ankrd40 | -0.026954746 | -0.240133674 | 0.186224182 | 8.730538072 | -0.307237452 | 0.768755755 | 0.909875862 | -4.889268392 |
| **A_51_P164939** | Tmem150a | 0.070419202 | -0.486615197 | 0.6274536 | 9.934099116 | 0.30717953 | 0.768797795 | 0.909875862 | -4.889277936 |
| **A_55_P2048209** | Ganab | -0.035357553 | -0.315159421 | 0.244444315 | 12.79094532 | -0.307054014 | 0.768888896 | 0.909927966 | -4.889298611 |
| **A_55_P2011390** | Tead1 | -0.022544997 | -0.201049527 | 0.155959533 | 7.876393707 | -0.306890984 | 0.769007231 | 0.910012292 | -4.889325453 |
| **A_55_P1955778** | Adamts4 | -0.021594347 | -0.192700651 | 0.149511956 | 7.962391245 | -0.306660095 | 0.769174834 | 0.910154905 | -4.889363445 |
| **A_52_P118461** | Ipmk | 0.049224464 | -0.340914492 | 0.439363419 | 10.22404481 | 0.306580738 | 0.769232443 | 0.91016424 | -4.889376497 |
| **A_55_P1969857** | Ap2m1 | -0.029566501 | -0.263948913 | 0.20481591 | 13.64722072 | -0.306519499 | 0.769276899 | 0.91016424 | -4.889386566 |
| **A_55_P1964233** | Tnks | -0.022119524 | -0.197513413 | 0.153274364 | 8.11376048 | -0.306439339 | 0.769335094 | 0.910177379 | -4.889399744 |
| **A_55_P2144341** | Smim40 | 0.020950124 | -0.145303561 | 0.18720381 | 7.862027062 | 0.306195294 | 0.769512275 | 0.910331278 | -4.889439844 |
| **A_55_P1999648** | Plin2 | 0.047289363 | -0.328088622 | 0.422667349 | 12.88851894 | 0.306110239 | 0.769574029 | 0.910348618 | -4.889453812 |
| **A_51_P425680** | Ivd | -0.047580493 | -0.425639936 | 0.330478951 | 11.31580446 | -0.305810246 | 0.769791857 | 0.910438062 | -4.889503049 |
| **A_55_P1988168** | Proser3 | 0.071464643 | -0.496391311 | 0.639320597 | 12.22855843 | 0.305799121 | 0.769799935 | 0.910438062 | -4.889504874 |
| **A_52_P523330** | Tmem147 | 0.036007762 | -0.250119322 | 0.322134846 | 11.31557944 | 0.305787939 | 0.769808055 | 0.910438062 | -4.889506709 |
| **A_55_P2081360** | Btbd1 | -0.031971812 | -0.286084398 | 0.222140774 | 10.66477663 | -0.305720295 | 0.769857176 | 0.910438062 | -4.889517804 |
| **A_51_P147064** | 1600014C23Rik | 0.029792309 | -0.207027386 | 0.266612004 | 8.056019734 | 0.305681768 | 0.769885153 | 0.910438062 | -4.889524122 |
| **A_55_P1984283** | NA | -0.040316217 | -0.360868337 | 0.280235903 | 15.19073172 | -0.30560771 | 0.769938933 | 0.910445958 | -4.889536265 |
| **A_66_P113659** | Ripply1 | 0.022254316 | -0.154832884 | 0.199341515 | 7.709352256 | 0.305358674 | 0.770119791 | 0.910604113 | -4.889577078 |
| **A_55_P2098428** | Bloc1s4 | -0.022243754 | -0.199368092 | 0.154880584 | 9.434569366 | -0.305149756 | 0.770271525 | 0.910727814 | -4.889611292 |

| | | | | | | | | |
|---|---|---|---|---|---|---|---|---|
| A_55_P2001963 | NA | 0.022167803 | -0.154477695 | 0.198813302 | 15.56860113 | 0.304932189 | 0.770429553 | 0.910858942 | -4.889646899 |
| A_52_P385489 | Ppp1r3a | 0.020729923 | -0.144495961 | 0.185955808 | 7.750345999 | 0.304861653 | 0.770480789 | 0.910863803 | -4.889658437 |
| A_52_P357611 | Neu3 | 0.029911236 | -0.208600951 | 0.268423422 | 9.563983562 | 0.304724217 | 0.770580623 | 0.910922772 | -4.889680912 |
| A_55_P2026270 | Cfi | -0.063473127 | -0.569863015 | 0.44291676 | 15.46113907 | -0.304570658 | 0.770692175 | 0.910922772 | -4.889706013 |
| A_51_P225525 | Gm9079 | 0.032101535 | -0.224037905 | 0.288240975 | 12.34150675 | 0.304531718 | 0.770720463 | 0.910922772 | -4.889712376 |
| A_55_P2031898 | NA | -0.096093355 | -0.863217756 | 0.671031046 | 10.79318578 | -0.304376241 | 0.770833416 | 0.910922772 | -4.889737774 |
| A_51_P191611 | Cat | 0.049807619 | -0.347843889 | 0.447459127 | 9.829214113 | 0.304352125 | 0.770850936 | 0.910922772 | -4.889741712 |
| A_52_P208600 | Bms1 | 0.029186299 | -0.203833503 | 0.2622061 | 8.942411098 | 0.30434725 | 0.770854479 | 0.910922772 | -4.889742509 |
| A_66_P123155 | Ddo | -0.021953705 | -0.197234197 | 0.153326787 | 7.746500342 | -0.304338878 | 0.770860561 | 0.910922772 | -4.889743876 |
| A_55_P2068849 | Nudc | 0.038101668 | -0.266206443 | 0.34240978 | 10.8906448 | 0.304238137 | 0.770933752 | 0.910946209 | -4.889760324 |
| A_51_P520384 | Atp1b3 | -0.033770634 | -0.303546128 | 0.236004861 | 10.96175544 | -0.304172467 | 0.770981465 | 0.910946209 | -4.889771043 |
| A_52_P127925 | Tfec | 0.032173561 | -0.224890673 | 0.289237796 | 8.446484956 | 0.304116981 | 0.77102178 | 0.910946209 | -4.889780098 |
| A_52_P663585 | 10-Sep | -0.038206908 | -0.343548262 | 0.267134447 | 8.989265076 | -0.30404611 | 0.771073274 | 0.910951366 | -4.889791662 |
| A_55_P2063937 | D630033A02Rik | 0.022973713 | -0.16072753 | 0.206674956 | 8.378675388 | 0.303880115 | 0.771193888 | 0.911038177 | -4.889818737 |
| A_55_P2183955 | Olfr97 | -0.022456339 | -0.202133279 | 0.157220601 | 7.856349648 | -0.303689522 | 0.771332384 | 0.911146101 | -4.889849806 |
| A_55_P2106593 | NA | 0.023310533 | -0.163378556 | 0.209999623 | 9.202622659 | 0.303400616 | 0.771542338 | 0.911246431 | -4.889896867 |
| A_55_P2154684 | Smarca4 | 0.025774766 | -0.180775663 | 0.232325195 | 13.27707707 | 0.303215795 | 0.771676662 | 0.911246431 | -4.88992695 |
| A_51_P366061 | Fscn1 | -0.050470291 | -0.454928551 | 0.353987968 | 8.565189195 | -0.303211236 | 0.771679976 | 0.911246431 | -4.889927692 |
| A_55_P2024704 | Cpe | -0.114505886 | -1.032242824 | 0.803231052 | 9.917257568 | -0.303174584 | 0.771706615 | 0.911246431 | -4.889933656 |
| A_52_P278295 | Pear1 | -0.025606366 | -0.230880998 | 0.179668267 | 8.937969671 | -0.303106919 | 0.771755795 | 0.911246431 | -4.889944664 |
| A_55_P1965298 | Gm5640 | -0.08038529 | -0.724835977 | 0.564065398 | 9.605675363 | -0.303088932 | 0.771768869 | 0.911246431 | -4.889947589 |
| A_55_P2145942 | Atg16l1 | 0.044553397 | -0.312639012 | 0.401745805 | 9.340699554 | 0.303083147 | 0.771773074 | 0.911246431 | -4.88994853 |
| A_51_P289987 | Psmg3 | 0.028049141 | -0.196893744 | 0.252992027 | 11.53212257 | 0.302991539 | 0.771839659 | 0.911246431 | -4.889963429 |
| A_52_P565051 | Cpne3 | -0.030790974 | -0.277724453 | 0.216142504 | 8.576864254 | -0.302988848 | 0.771841615 | 0.911246431 | -4.889963866 |
| A_51_P443872 | Slu7 | 0.026533476 | -0.186327038 | 0.23939399 | 9.05376314 | 0.302888115 | 0.771914835 | 0.911267273 | -4.889980243 |
| A_51_P117618 | Ethe1 | 0.051584372 | -0.362378058 | 0.465546803 | 10.64025364 | 0.302789222 | 0.77198672 | 0.911267273 | -4.889996316 |
| A_51_P241843 | Tmem202 | 0.01936061 | -0.136017349 | 0.174738569 | 7.692390364 | 0.302769985 | 0.772000704 | 0.911267273 | -4.889999442 |
| A_55_P2425938 | 2810468N07Rik | -0.022789959 | -0.205804805 | 0.160224888 | 7.860924855 | -0.302580132 | 0.772138716 | 0.911374526 | -4.890030283 |
| A_55_P2033105 | Cdcp1 | 0.023369082 | -0.164350822 | 0.211088986 | 7.945999534 | 0.302492435 | 0.772202469 | 0.911394121 | -4.890044523 |
| A_55_P2078395 | Gm1943 | -0.020538097 | -0.185644776 | 0.144568582 | 8.14432716 | -0.302258651 | 0.772372433 | 0.911452532 | -4.890082464 |
| A_55_P2022277 | Nfe2l1 | 0.028890905 | -0.203483926 | 0.261265736 | 8.273431571 | 0.302103146 | 0.772485496 | 0.911452532 | -4.890107686 |
| A_51_P432511 | Afg1l | -0.020315508 | -0.183728163 | 0.143097146 | 8.000509803 | -0.302082242 | 0.772500695 | 0.911452532 | -4.890111075 |
| A_55_P1990225 | Fnta | -0.038693994 | -0.350017562 | 0.272629575 | 12.14312292 | -0.302005426 | 0.772556548 | 0.911452532 | -4.890123529 |
| A_51_P186476 | Slc11a1 | -0.05523403 | -0.499643665 | 0.389175606 | 9.894949204 | -0.301999658 | 0.772560741 | 0.911452532 | -4.890124464 |
| A_55_P2124976 | Grhl1 | 0.029760606 | -0.209725506 | 0.269246717 | 8.012787455 | 0.301956664 | 0.772592003 | 0.911452532 | -4.890131433 |
| A_51_P365952 | Trdmt1 | 0.025873129 | -0.182362805 | 0.234109063 | 9.512684804 | 0.301909282 | 0.772626456 | 0.911452532 | -4.890139112 |
| A_55_P2114963 | Exosc10 | 0.026710505 | -0.188267604 | 0.241688613 | 8.695643133 | 0.301905514 | 0.772629196 | 0.911452532 | -4.890139722 |
| A_52_P542794 | 2610507B11Rik | -0.023736061 | -0.214914681 | 0.167442559 | 11.96903949 | -0.301684164 | 0.772790153 | 0.911553897 | -4.89017558 |
| A_55_P1957900 | Dlg1 | -0.025705117 | -0.232761412 | 0.181351179 | 9.04027502 | -0.301657638 | 0.772809442 | 0.911553897 | -4.890179875 |
| A_55_P1963414 | NA | -0.026453203 | -0.239685794 | 0.186779387 | 14.59883753 | -0.301444857 | 0.772964182 | 0.911680783 | -4.890214318 |
| A_52_P542970 | NA | 0.022505592 | -0.159030418 | 0.204041601 | 10.63273246 | 0.301238753 | 0.773114076 | 0.911801939 | -4.890247658 |
| A_55_P2021792 | Dyrk1b | -0.026345444 | -0.238908802 | 0.186217913 | 8.235253077 | -0.3011621 | 0.773169827 | 0.911812055 | -4.890260052 |
| A_55_P2047599 | Arhgdia | -0.038358089 | -0.348005296 | 0.271289117 | 12.74984069 | -0.3010045 | 0.773284455 | 0.911887326 | -4.890285524 |
| A_55_P2180899 | Aldh3a2 | -0.0462474 | -0.419655661 | 0.327160861 | 9.191179611 | -0.300944624 | 0.773328007 | 0.911887326 | -4.890295199 |
| A_55_P2063471 | NA | 0.036055991 | -0.255246027 | 0.327358009 | 16.74845863 | 0.300757958 | 0.773463788 | 0.911991799 | -4.890325347 |
| A_55_P2341110 | Chn1os3 | -0.019027475 | -0.172923598 | 0.134868648 | 7.626925366 | -0.300425417 | 0.773705699 | 0.912221389 | -4.890379012 |
| A_52_P136914 | Nudt7 | 0.029758827 | -0.211100132 | 0.270617785 | 8.679341106 | 0.30021763 | 0.77385687 | 0.912296328 | -4.890412515 |
| A_51_P234833 | Strn4 | 0.026763309 | -0.189857562 | 0.24338418 | 11.04739889 | 0.300208304 | 0.773863655 | 0.912296328 | -4.890414018 |
| A_55_P1996314 | Amy2a5 | 0.054637901 | -0.387742515 | 0.497018317 | 12.22025931 | 0.300110578 | 0.773934758 | 0.912324507 | -4.890429767 |
| A_52_P572197 | Pank3 | -0.058077437 | -0.528670965 | 0.412516091 | 8.396420344 | -0.299878053 | 0.774103946 | 0.912452669 | -4.89046722 |

| | | | | | | | | | |
|---|---|---|---|---|---|---|---|---|---|
| A_66_P131788 | Ythdf2 | -0.042710965 | -0.38884592 | 0.30342399 | 8.31249526 | -0.299831401 | 0.774137892 | 0.912452669 | -4.890474731 |
| A_55_P1963105 | NA | -0.026738762 | -0.24351715 | 0.190039625 | 15.51232182 | -0.299715022 | 0.774222578 | 0.912496842 | -4.890493463 |
| A_55_P2288232 | Gucy1a2 | -0.035854368 | -0.326627922 | 0.254919186 | 8.438750293 | -0.299619695 | 0.774291947 | 0.912522959 | -4.890508801 |
| A_51_P363801 | Pgpep1 | -0.02342335 | -0.213436269 | 0.166589568 | 8.903634579 | -0.299536033 | 0.774352829 | 0.91253907 | -4.890522259 |
| A_66_P135882 | Rrp1 | -0.030939028 | -0.28203014 | 0.220152083 | 13.00179238 | -0.299404634 | 0.774448454 | 0.91259612 | -4.890543388 |
| A_66_P115161 | Cep85l | -0.030877838 | -0.281701519 | 0.219945843 | 8.430895787 | -0.29913108 | 0.774647545 | 0.91276104 | -4.890587347 |
| A_55_P2078940 | Nsfl1c | -0.024910001 | -0.227289179 | 0.177469178 | 12.7218134 | -0.299082569 | 0.774682853 | 0.91276104 | -4.890595139 |
| A_55_P2197914 | C030027H14Rik | -0.040396822 | -0.368768949 | 0.287975305 | 9.529969397 | -0.298926287 | 0.774796604 | 0.912839423 | -4.890620232 |
| A_65_P07062 | Serf2 | -0.048092083 | -0.439676002 | 0.343491836 | 11.20218073 | -0.298422751 | 0.775163149 | 0.91321561 | -4.890700995 |
| A_55_P2004881 | Ldlrad4 | 0.025125563 | -0.179558371 | 0.229809498 | 8.20945925 | 0.298273892 | 0.775271522 | 0.913287619 | -4.890724846 |
| A_55_P2088401 | H2-T9 | 0.021325089 | -0.152483652 | 0.19513383 | 8.729960923 | 0.298127764 | 0.775377912 | 0.913357283 | -4.890748249 |
| A_55_P2067262 | Cdv3 | 0.023145218 | -0.165672614 | 0.21196305 | 8.10541859 | 0.297852647 | 0.775578228 | 0.913464289 | -4.890792279 |
| A_55_P1982420 | Fam8a1 | -0.033749251 | -0.30913237 | 0.241633869 | 9.375021528 | -0.29778993 | 0.775623896 | 0.913464289 | -4.890802311 |
| A_55_P2011727 | Neurl1b | 0.072738434 | -0.521082711 | 0.666559579 | 14.53577246 | 0.297640123 | 0.775732982 | 0.913464289 | -4.890826265 |
| A_55_P2162910 | Rtn1 | 0.03196932 | -0.229023282 | 0.292961921 | 8.705036356 | 0.297638007 | 0.775734523 | 0.913464289 | -4.890826603 |
| A_55_P2083419 | Pkp4 | -0.02370057 | -0.217197776 | 0.169796635 | 8.988657129 | -0.297623546 | 0.775745053 | 0.913464289 | -4.890828915 |
| A_51_P145415 | Lpcat3 | 0.036251123 | -0.259721743 | 0.332223989 | 11.5531059 | 0.297613587 | 0.775752305 | 0.913464289 | -4.890830506 |
| A_55_P2327027 | Ano6 | 0.024077217 | -0.172611897 | 0.22076633 | 8.277149085 | 0.297446694 | 0.775873841 | 0.913542921 | -4.890857177 |
| A_55_P2116833 | 9430038I01Rik | 0.024382014 | -0.174833588 | 0.223597615 | 9.773458531 | 0.297392089 | 0.775913608 | 0.913542921 | -4.8908659 |
| A_51_P155323 | Hc | -0.060609006 | -0.555997878 | 0.434779865 | 14.56128678 | -0.29728576 | 0.775991044 | 0.913578444 | -4.890882881 |
| A_55_P2141415 | Ppp6r3 | 0.019660913 | -0.1411618 | 0.180483626 | 8.297139956 | 0.297056797 | 0.776157802 | 0.913693171 | -4.890919428 |
| A_52_P616591 | Eri3 | -0.028475056 | -0.261446997 | 0.204496886 | 8.185629791 | -0.29699159 | 0.776205295 | 0.913693171 | -4.890929832 |
| A_55_P2401646 | Trdn | -0.022089191 | -0.202844016 | 0.158665633 | 7.67477619 | -0.29694301 | 0.77624068 | 0.913693171 | -4.890937581 |
| A_55_P2071286 | Tmc4 | -0.054570731 | -0.501196672 | 0.392055211 | 9.98388113 | -0.296892355 | 0.776277575 | 0.913693171 | -4.89094566 |
| A_55_P1980062 | NA | -0.023512513 | -0.216049601 | 0.169024576 | 7.915883675 | -0.296734358 | 0.776392661 | 0.913772985 | -4.890970851 |
| A_55_P2089820 | Adk | 0.074070164 | -0.533319162 | 0.681459491 | 10.37439409 | 0.296318892 | 0.776695318 | 0.914073538 | -4.89103703 |
| A_51_P173735 | Dock11 | -0.040220041 | -0.370220454 | 0.289780372 | 9.659306618 | -0.296149645 | 0.776818622 | 0.914140413 | -4.891063963 |
| A_52_P302395 | Lnpep | 0.026603019 | -0.191777553 | 0.24498359 | 8.537172328 | 0.296005724 | 0.776923481 | 0.914140413 | -4.891086855 |
| A_51_P121031 | 1-Mar | -0.042992312 | -0.396014913 | 0.310030289 | 8.521881205 | -0.295918066 | 0.77698735 | 0.914140413 | -4.881694739 |
| A_55_P2163493 | Tgm3 | 0.030576596 | -0.220696685 | 0.281849877 | 9.99029728 | 0.295682764 | 0.777158804 | 0.914140413 | -4.891138186 |
| A_51_P508349 | Cdc16 | -0.038022126 | -0.350590374 | 0.274546122 | 12.23216018 | -0.295579784 | 0.777233845 | 0.914140413 | -4.891154542 |
| A_55_P2016262 | Isl2 | 0.022280364 | -0.16088389 | 0.205444618 | 7.730436643 | 0.295572992 | 0.777238794 | 0.914140413 | -4.891155621 |
| A_55_P2119458 | Slc25a44 | 0.052210846 | -0.377076655 | 0.481498346 | 8.79486347 | 0.295525984 | 0.77727305 | 0.914140413 | -4.891163085 |
| A_51_P138060 | Rnf13 | 0.02153918 | -0.155595195 | 0.198673555 | 8.075535212 | 0.295467353 | 0.777315776 | 0.914140413 | -4.891172393 |
| A_55_P2005898 | NA | -0.023138214 | -0.213456568 | 0.167180141 | 7.964523736 | -0.295414865 | 0.777354026 | 0.914140413 | -4.891180725 |
| A_55_P1986351 | Tbc1d32 | -0.027176529 | -0.250772025 | 0.196418968 | 7.95529414 | -0.29533446 | 0.777412622 | 0.914140413 | -4.891193485 |
| A_51_P370341 | Zbtb48 | -0.023323226 | -0.215256847 | 0.168610394 | 8.748580701 | -0.295270977 | 0.777458887 | 0.914140413 | -4.891203557 |
| A_51_P348433 | Rasal1 | 0.01937002 | -0.140041327 | 0.178781368 | 7.705178366 | 0.295252805 | 0.77747213 | 0.914140413 | -4.891206439 |
| A_55_P2046478 | Zfp81 | 0.02016545 | -0.145792667 | 0.186123567 | 8.136137305 | 0.295251836 | 0.777472837 | 0.914140413 | -4.891206593 |
| A_55_P2111713 | Spint2 | 0.108357325 | -0.783597653 | 1.000312302 | 10.77981648 | 0.295187885 | 0.777519444 | 0.914140413 | -4.891216737 |
| A_55_P2055411 | Rps7 | -0.03101402 | -0.286346556 | 0.224318517 | 15.63797281 | -0.295144765 | 0.77755087 | 0.914140413 | -4.891223575 |
| A_55_P2098999 | NA | 0.034086884 | -0.246606444 | 0.314780212 | 13.882451 | 0.29507907 | 0.777598751 | 0.914140413 | -4.891233992 |
| A_55_P2090880 | Stk40 | -0.030629163 | -0.282880714 | 0.221622388 | 11.43136377 | -0.295042431 | 0.777625454 | 0.914140413 | -4.891239801 |
| A_52_P344424 | Lipg | 0.02164149 | -0.156644697 | 0.199927677 | 7.701160589 | 0.29495289 | 0.777690717 | 0.914140413 | -4.891253993 |
| A_51_P427530 | Pgm1 | -0.028872776 | -0.266742396 | 0.208996844 | 8.597022076 | -0.294939374 | 0.777700568 | 0.914140413 | -4.891256135 |
| A_55_P1970806 | Mif-ps6 | -0.039256489 | -0.362693031 | 0.284180053 | 15.55692197 | -0.294920906 | 0.777714029 | 0.914140413 | -4.891259062 |
| A_52_P537571 | Trap1a | -0.019898185 | -0.183864373 | 0.144068003 | 7.682098925 | -0.294877989 | 0.77774531 | 0.914140413 | -4.891265862 |
| A_52_P212418 | Trmt13 | -0.022052704 | -0.203938402 | 0.159832994 | 8.340163956 | -0.294609325 | 0.777941143 | 0.914279198 | -4.891308411 |
| A_55_P2004930 | Hddc2 | 0.05772538 | -0.418417799 | 0.533868559 | 10.43411863 | 0.294586216 | 0.777957989 | 0.914279198 | -4.891312069 |
| A_55_P2136935 | Rgs7bp | 0.022858209 | -0.165826205 | 0.211542622 | 8.055777762 | 0.294367161 | 0.778117675 | 0.914411269 | -4.891346731 |

| | | | | | | | | | |
|---|---|---|---|---|---|---|---|---|---|
| **A_52_P264905** | Cab39l | -0.026169128 | -0.242269683 | 0.189931427 | 10.98305755 | -0.294250057 | 0.778203047 | 0.914455997 | -4.891365251 |
| **A_51_P454309** | Zfp772 | -0.024577641 | -0.227631848 | 0.178476565 | 8.019476323 | -0.294111091 | 0.778304361 | 0.914519452 | -4.891387219 |
| **A_55_P2165539** | Aebp2 | 0.047751956 | -0.347092168 | 0.442596079 | 9.093208289 | 0.293865547 | 0.778483387 | 0.914674207 | -4.891426011 |
| **A_66_P140307** | NA | 0.029086601 | -0.211700121 | 0.269873323 | 13.529569 | 0.293524011 | 0.778732426 | 0.914850044 | -4.891479916 |
| **A_51_P179498** | Fam92a | 0.040580836 | -0.295413013 | 0.376574684 | 9.829475369 | 0.293476174 | 0.77876731 | 0.914850044 | -4.891487461 |
| **A_51_P378009** | Blzf1 | 0.023235056 | -0.169149293 | 0.215619405 | 9.630098961 | 0.293465582 | 0.778775034 | 0.914850044 | -4.891489132 |
| **A_55_P2077623** | NA | 0.018833895 | -0.137160068 | 0.174827859 | 7.670544158 | 0.293369897 | 0.778844811 | 0.914873774 | -4.89150422 |
| **A_55_P2183015** | P2rx4 | 0.030695473 | -0.223597 | 0.284987946 | 12.02350026 | 0.293308072 | 0.778889897 | 0.914873774 | -4.891513967 |
| **A_52_P632423** | NA | 0.019108836 | -0.13924727 | 0.177464942 | 7.686096382 | 0.293212584 | 0.778959535 | 0.914899972 | -4.891529017 |
| **A_51_P162176** | Trappc3 | 0.069490536 | -0.506757323 | 0.645738395 | 9.873380841 | 0.293021525 | 0.779098877 | 0.915008032 | -4.891559115 |
| **A_55_P2049158** | NA | 0.035049054 | -0.255716093 | 0.325814201 | 12.30921011 | 0.292898496 | 0.779188608 | 0.915012765 | -4.891578487 |
| **A_52_P64707** | Foxa3 | 0.090261777 | -0.65866402 | 0.839187574 | 10.07861255 | 0.292852056 | 0.77922248 | 0.915012765 | -4.891585797 |
| **A_66_P112573** | Isg20 | 0.045709081 | -0.348105669 | 0.439523831 | 14.60446103 | 0.295292814 | 0.779244923 | 0.915012765 | -4.843821524 |
| **A_51_P229886** | Fpgt | 0.041200325 | -0.300771941 | 0.383172591 | 8.774000467 | 0.292747332 | 0.779298865 | 0.915020518 | -4.891602277 |
| **A_51_P134972** | Shkbp1 | -0.030778442 | -0.286352425 | 0.224795542 | 10.44186178 | -0.292626177 | 0.779387238 | 0.915023437 | -4.891621337 |
| **A_51_P170619** | Ddx54 | 0.025626514 | -0.187203258 | 0.238456286 | 11.68447433 | 0.292577107 | 0.779423032 | 0.915023437 | -4.891629054 |
| **A_55_P1963610** | NA | 0.026480317 | -0.193514214 | 0.246474848 | 15.64135066 | 0.292478863 | 0.779494697 | 0.915023437 | -4.891644501 |
| **A_55_P2010066** | Capn3 | 0.053399426 | -0.390302092 | 0.497100945 | 9.463106887 | 0.292434673 | 0.779526933 | 0.915023437 | -4.891651447 |
| **A_52_P561927** | Ncor1 | -0.029356581 | -0.273298076 | 0.214584914 | 9.28736323 | -0.292417242 | 0.779539648 | 0.915023437 | -4.891654187 |
| **A_55_P2180869** | Ocel1 | -0.026152984 | -0.243582226 | 0.191276259 | 10.22681575 | -0.292271506 | 0.779645963 | 0.915023437 | -4.891677088 |
| **A_55_P2051337** | Ppfia1 | -0.020935059 | -0.194992115 | 0.153121997 | 9.649414302 | -0.292257549 | 0.779656145 | 0.915023437 | -4.89167928 |
| **A_55_P2033947** | Fnd3c2 | -0.019380888 | -0.180559836 | 0.141798059 | 7.656817879 | -0.2921787 | 0.779713668 | 0.915023437 | -4.891691665 |
| **A_52_P381806** | Terf2ip | 0.024006529 | -0.17568152 | 0.223694578 | 8.634990688 | 0.292119463 | 0.779756885 | 0.915023437 | -4.891700968 |
| **A_55_P2021285** | Cpped1 | -0.040077034 | -0.373472896 | 0.293318829 | 11.58501723 | -0.292091253 | 0.779777465 | 0.915023437 | -4.891705397 |
| **A_55_P2040907** | Gtf2i | -0.026344918 | -0.24556885 | 0.192879013 | 8.004754484 | -0.292006208 | 0.779839512 | 0.915023437 | -4.891718748 |
| **A_51_P379687** | Gm10413 | 0.018525047 | -0.135682534 | 0.172732628 | 7.640034137 | 0.291901808 | 0.779915682 | 0.915023437 | -4.891735132 |
| **A_55_P2062787** | Mfap3 | -0.030187001 | -0.281482446 | 0.221108444 | 10.52945822 | -0.291889545 | 0.77992463 | 0.915023437 | -4.891737056 |
| **A_55_P1992045** | Msi2 | 0.035502751 | -0.260099104 | 0.331104606 | 11.28641694 | 0.291835444 | 0.779964103 | 0.915023437 | -4.891745544 |
| **A_51_P366121** | Klf14 | -0.019562275 | -0.182596034 | 0.143471483 | 7.745525046 | -0.291558034 | 0.780166518 | 0.915055403 | -4.891789043 |
| **A_55_P1970424** | Fam221a | -0.023140315 | -0.216048832 | 0.169768202 | 7.955339165 | -0.291474825 | 0.780227236 | 0.915055403 | -4.891802082 |
| **A_55_P2091631** | Coa4 | 0.032665721 | -0.239693644 | 0.305025085 | 10.04697846 | 0.29142924 | 0.780260501 | 0.915055403 | -4.891809224 |
| **A_55_P2108201** | Gnptab | -0.037711459 | -0.352211198 | 0.27678828 | 9.198529081 | -0.291364229 | 0.780307941 | 0.915055403 | -4.891819408 |
| **A_55_P1965308** | Ccdc66 | -0.02845891 | -0.265798172 | 0.208880352 | 8.263243228 | -0.291361294 | 0.780310084 | 0.915055403 | -4.891819868 |
| **A_52_P636343** | Wipi2 | 0.041350626 | -0.303599155 | 0.386300408 | 9.200392978 | 0.291279157 | 0.780370023 | 0.915055403 | -4.891832731 |
| **A_51_P224575** | Gnpat | 0.023538961 | -0.172839786 | 0.219917707 | 11.96205319 | 0.291256742 | 0.780386381 | 0.915055403 | -4.891836241 |
| **A_51_P196526** | Bscl2 | -0.02083822 | -0.194710612 | 0.153034173 | 8.452302387 | -0.291214617 | 0.780417122 | 0.915055403 | -4.891842836 |
| **A_55_P2113886** | NA | -0.018656866 | -0.174328441 | 0.137014709 | 7.670960326 | -0.291214209 | 0.78041742 | 0.915055403 | -4.8918429 |
| **A_51_P379373** | Lactb2 | 0.05018043 | -0.36890631 | 0.469267169 | 10.26174153 | 0.29094684 | 0.78061255 | 0.915132665 | -4.89188474 |
| **A_55_P2010152** | Sell | -0.091586571 | -0.856584831 | 0.673411689 | 9.598645509 | -0.290907248 | 0.780641446 | 0.915132665 | -4.891890932 |
| **A_55_P2158741** | Nos2 | 0.063611374 | -0.46772347 | 0.594946218 | 8.59550508 | 0.290903985 | 0.780643828 | 0.915132665 | -4.891891443 |
| **A_51_P446978** | Cfdp1 | 0.034090523 | -0.250700461 | 0.318881508 | 12.53966602 | 0.290864433 | 0.780672695 | 0.915132665 | -4.891897628 |
| **A_55_P2136522** | Tmem128 | -0.025909938 | -0.242462556 | 0.190642679 | 10.81750466 | -0.29072751 | 0.780772632 | 0.915179879 | -4.891919034 |
| **A_55_P1955841** | Ctsa | -0.026345865 | -0.246578278 | 0.193886547 | 13.93733963 | -0.290679511 | 0.780807667 | 0.915179879 | -4.891926536 |
| **A_55_P2168307** | Gm13213 | -0.030455381 | -0.285325645 | 0.224414883 | 16.72422001 | -0.29035416 | 0.781045156 | 0.91540273 | -4.891977354 |
| **A_55_P2136622** | Camta2 | -0.020573259 | -0.192795214 | 0.151648695 | 8.145072498 | -0.290267073 | 0.78110873 | 0.915411419 | -4.891990948 |
| **A_55_P2013680** | NA | 0.021851605 | -0.161104871 | 0.20480808 | 16.52100192 | 0.290214254 | 0.781147289 | 0.915411419 | -4.89199919 |
| **A_55_P2119802** | NA | 0.031539287 | -0.232942808 | 0.296021382 | 15.41846938 | 0.289760234 | 0.781478759 | 0.915744342 | -4.892069982 |
| **A_55_P2174085** | St8sia5 | -0.019381104 | -0.181958228 | 0.143196021 | 7.676231539 | -0.289669157 | 0.781545258 | 0.915766749 | -4.89208417 |
| **A_51_P206134** | Akna | -0.042873475 | -0.402627484 | 0.316880533 | 8.976184786 | -0.289578448 | 0.78161149 | 0.91578884 | -4.892098297 |
| **A_55_P2454896** | Amn1 | -0.029923226 | -0.281076267 | 0.221229815 | 9.517723739 | -0.289503062 | 0.781666536 | 0.915797823 | -4.892110034 |

| | | | | | | | | |
|---|---|---|---|---|---|---|---|---|
| **A_51_P444437** | Ptgr1 | -0.040566712 | -0.38114631 | 0.300012887 | 10.60296695 | -0.289423883 | 0.781724353 | 0.91581005 | -4.892122358 |
| **A_51_P338728** | Sdhaf1 | 0.040300528 | -0.298395701 | 0.378996758 | 10.74749039 | 0.289123616 | 0.781943623 | 0.915994818 | -4.892169066 |
| **A_55_P2049383** | Erp29 | -0.022516126 | -0.211777342 | 0.166745091 | 14.08621964 | -0.289078121 | 0.781976848 | 0.915994818 | -4.892176139 |
| **A_51_P465161** | Hoga1 | 0.064281506 | -0.476226724 | 0.604789736 | 10.50194917 | 0.288979423 | 0.782048928 | 0.916023738 | -4.892191479 |
| **A_52_P561671** | Msx1 | 0.019460713 | -0.144221119 | 0.183142545 | 7.902459435 | 0.288895955 | 0.782109888 | 0.91603963 | -4.892204449 |
| **A_55_P2145449** | Cep68 | -0.018741533 | -0.176429001 | 0.138945936 | 8.114778397 | -0.288795959 | 0.78218292 | 0.916069659 | -4.892219981 |
| **A_55_P1955078** | Igflr1 | 0.037297612 | -0.276693579 | 0.351288804 | 9.118159367 | 0.288633511 | 0.782301569 | 0.916099472 | -4.892245204 |
| **A_55_P1999972** | NA | -0.019336632 | -0.182150728 | 0.143477465 | 7.770691723 | -0.288583841 | 0.782337849 | 0.916099472 | -4.892252913 |
| **A_51_P180032** | Sem1 | -0.032909803 | -0.310079652 | 0.244260046 | 15.06080891 | -0.288511073 | 0.782391 | 0.916099472 | -4.892264205 |
| **A_51_P432802** | Rbm4 | -0.022695027 | -0.21390227 | 0.168512217 | 11.53623422 | -0.288409491 | 0.782465201 | 0.916099472 | -4.892279964 |
| **A_55_P2011877** | Eif2s3y | 0.355109683 | -2.636771503 | 3.346990869 | 9.779482665 | 0.288404054 | 0.782469173 | 0.916099472 | -4.892280808 |
| **A_55_P1954461** | Ptdss1 | 0.021196692 | -0.157410327 | 0.199803711 | 8.024372774 | 0.288371783 | 0.782492745 | 0.916099472 | -4.892285813 |
| **A_55_P2163729** | Tvp23a | -0.023688439 | -0.223358423 | 0.175981546 | 7.846816918 | -0.288274908 | 0.782563511 | 0.916125806 | -4.892300835 |
| **A_55_P1957624** | Insig2 | 0.023800403 | -0.176857655 | 0.224458461 | 8.870631967 | 0.288211223 | 0.782610032 | 0.916125806 | -4.892310707 |
| **A_51_P203675** | Trem3 | -0.066133604 | -0.624210987 | 0.49194378 | 9.357461234 | -0.287945937 | 0.782803834 | 0.916297179 | -4.89235181 |
| **A_55_P2068663** | Stmn1 | 0.044905733 | -0.334224142 | 0.424035608 | 14.47699402 | 0.287804095 | 0.782907463 | 0.916334927 | -4.892373772 |
| **A_55_P2088965** | Scarb1 | -0.034415803 | -0.325013742 | 0.256182136 | 8.810372584 | -0.28777202 | 0.782930897 | 0.916334927 | -4.892378737 |
| **A_55_P1959903** | Map7 | -0.022729003 | -0.214713992 | 0.169255987 | 8.109202942 | -0.287671141 | 0.783004601 | 0.916365702 | -4.892394348 |
| **A_51_P445487** | Mturn | -0.026189629 | -0.24746482 | 0.195085562 | 8.276798007 | -0.287593989 | 0.783060973 | 0.916376191 | -4.892406284 |
| **A_66_P122699** | NA | -0.062550144 | -0.591176616 | 0.466076327 | 8.724529404 | -0.287516385 | 0.783117675 | 0.916387065 | -4.892418287 |
| **A_51_P165984** | Aup1 | -0.036713797 | -0.347248799 | 0.273821205 | 10.73788794 | -0.287277711 | 0.783292075 | 0.916535657 | -4.892455182 |
| **A_51_P411909** | Sirt5 | 0.027611179 | -0.206131891 | 0.26135425 | 9.827148546 | 0.287031396 | 0.783472073 | 0.916690781 | -4.892493229 |
| **A_66_P121656** | Pcgf1 | 0.036373131 | -0.271827973 | 0.344574236 | 8.460673398 | 0.286767341 | 0.78366505 | 0.916808088 | -4.89253398 |
| **A_55_P1966843** | Gm12070 | 0.033996848 | -0.254101197 | 0.322094892 | 16.40152468 | 0.286735542 | 0.783688291 | 0.916808088 | -4.892538885 |
| **A_52_P31011** | Cep162 | 0.020483421 | -0.153120182 | 0.194087024 | 8.172261031 | 0.286699507 | 0.783714628 | 0.916808088 | -4.892544443 |
| **A_55_P1966719** | NA | -0.04163744 | -0.394789793 | 0.311514914 | 9.722923519 | -0.286487119 | 0.783869862 | 0.91693419 | -4.892577187 |
| **A_55_P1993820** | Kdm1a | -0.028812112 | -0.273367765 | 0.215743541 | 11.51869358 | -0.286273121 | 0.784026283 | 0.917020513 | -4.892610156 |
| **A_51_P244558** | Rab3gap2 | 0.036221046 | -0.2712587 | 0.343700792 | 9.917217091 | 0.286238248 | 0.784051775 | 0.917020513 | -4.892615526 |
| **A_55_P1963144** | Mtmr14 | 0.023129851 | -0.173252564 | 0.219512266 | 10.34769871 | 0.286189321 | 0.78408754 | 0.917020513 | -4.89262306 |
| **A_51_P482051** | Cyp3a16 | 0.095016584 | -0.711892209 | 0.901925377 | 11.26623599 | 0.286126546 | 0.784133428 | 0.917020513 | -4.892632724 |
| **A_55_P2223851** | NA | 0.022613443 | -0.169506022 | 0.214732907 | 8.150152335 | 0.286008212 | 0.784219933 | 0.917066192 | -4.892650935 |
| **A_55_P2137984** | NA | 0.047630288 | -0.357176622 | 0.452437198 | 9.860077715 | 0.285902847 | 0.78429696 | 0.917085617 | -4.892667145 |
| **A_51_P318640** | Rps3 | 0.04432665 | -0.332546668 | 0.421199968 | 15.27545287 | 0.285793739 | 0.784376726 | 0.917085617 | -4.892683924 |
| **A_55_P2089506** | NA | -0.042288778 | -0.40183944 | 0.317261884 | 13.71232756 | -0.28579079 | 0.784378882 | 0.917085617 | -4.892684377 |
| **A_55_P2412214** | Mhrt | 0.018266862 | -0.137266756 | 0.173800479 | 7.669924195 | 0.285379552 | 0.784679554 | 0.917381667 | -4.892747564 |
| **A_51_P104125** | Fam76b | 0.023685265 | -0.178098013 | 0.225468543 | 9.606835925 | 0.285217564 | 0.784798001 | 0.917418237 | -4.877575788 |
| **A_55_P2113031** | Foxo4 | 0.019279916 | -0.144978718 | 0.18353855 | 7.712700137 | 0.285206953 | 0.784805761 | 0.917418237 | -4.892774058 |
| **A_55_P2111153** | NA | -0.029163112 | -0.277684679 | 0.219358455 | 10.53250865 | -0.285136606 | 0.784857201 | 0.917422886 | -4.892784852 |
| **A_55_P2155438** | Rps27 | -0.037763526 | -0.359663092 | 0.28413604 | 16.52299964 | -0.285059379 | 0.784913674 | 0.917433416 | -4.892796699 |
| **A_66_P114543** | Jmjd6 | 0.025803758 | -0.194215902 | 0.245823417 | 8.401170812 | 0.284973616 | 0.78497639 | 0.917451242 | -4.892809851 |
| **A_51_P333869** | Nsmce3 | 0.02743536 | -0.206570463 | 0.261441182 | 9.344334096 | 0.284883425 | 0.785042346 | 0.917452774 | -4.892823678 |
| **A_51_P417858** | Zfp825 | 0.020740671 | -0.156204669 | 0.19768601 | 8.538591016 | 0.284817622 | 0.785090469 | 0.917452774 | -4.892833764 |
| **A_51_P254299** | Dlk2 | -0.044557416 | -0.424745153 | 0.335630321 | 8.845903929 | -0.284777111 | 0.785120096 | 0.917452774 | -4.892839972 |
| **A_51_P266640** | Nob1 | 0.020829397 | -0.157023989 | 0.198682783 | 8.19346622 | 0.284575661 | 0.785267427 | 0.917461654 | -4.89287083 |
| **A_51_P437289** | Smco4 | -0.032658521 | -0.311537428 | 0.246220385 | 9.519618863 | -0.284553571 | 0.785283584 | 0.917461654 | -4.892874213 |
| **A_55_P2021886** | Mrpl2 | -0.024760622 | -0.236232313 | 0.18671107 | 13.38851541 | -0.284506697 | 0.785317867 | 0.917461654 | -4.892881389 |
| **A_55_P2059357** | Myo7a | 0.04519491 | -0.340868252 | 0.431258071 | 10.01051035 | 0.284455508 | 0.785355306 | 0.917461654 | -4.892889225 |
| **A_55_P2020286** | Tmem179 | 0.019918106 | -0.150274699 | 0.19011091 | 7.715017532 | 0.284374089 | 0.785414858 | 0.917461654 | -4.892901686 |
| **A_52_P52156** | Isoc2a | 0.058465232 | -0.441130309 | 0.558060773 | 9.095210201 | 0.284355944 | 0.78542813 | 0.917461654 | -4.892904463 |
| **A_51_P512002** | Rer1 | 0.024279052 | -0.183221409 | 0.231779514 | 12.8187259 | 0.284312435 | 0.785459954 | 0.917461654 | -4.89291112 |

| | | | | | | | | | |
|---|---|---|---|---|---|---|---|---|---|
| **A_55_P2109744** | NA | 0.018617931 | -0.140638133 | 0.177873994 | 15.33126324 | 0.284065582 | 0.78564052 | 0.917617114 | -4.892948871 |
| **A_55_P1959576** | Slitrk4 | -0.018851095 | -0.180139497 | 0.142437307 | 7.714861814 | -0.283998886 | 0.785689309 | 0.917618649 | -4.892959065 |
| **A_55_P2023772** | Fam107b | 0.058047832 | -0.439106779 | 0.555202442 | 10.96656245 | 0.283712008 | 0.785899175 | 0.917808299 | -4.893002887 |
| **A_55_P2070241** | Preb | 0.032987546 | -0.249740651 | 0.315715742 | 12.34773357 | 0.283507186 | 0.786049025 | 0.917923577 | -4.893034149 |
| **A_52_P609109** | Pitpnc1 | 0.025976135 | -0.19675813 | 0.2487104 | 8.969419871 | 0.283380978 | 0.786141365 | 0.917923577 | -4.893053402 |
| **A_51_P484254** | Pcca | -0.067124209 | -0.642713809 | 0.50846539 | 9.286434551 | -0.283367105 | 0.786151515 | 0.917923577 | -4.893055517 |
| **A_55_P2071477** | Prl6a1 | 0.018296899 | -0.138683878 | 0.175277676 | 7.655403415 | 0.283213664 | 0.786263786 | 0.917923577 | -4.893078912 |
| **A_55_P2113633** | Bzw1 | 0.026968147 | -0.20441207 | 0.258348363 | 10.82519047 | 0.283209664 | 0.786266713 | 0.917923577 | -4.893079521 |
| **A_55_P2083499** | Shisal2a | 0.020425842 | -0.154871628 | 0.195723312 | 7.737501318 | 0.283131057 | 0.786324231 | 0.917923577 | -4.893091501 |
| **A_55_P1978502** | H2-Q1 | 0.036019886 | -0.273135061 | 0.345174834 | 8.064867158 | 0.283106165 | 0.786342445 | 0.917923577 | -4.893095294 |
| **A_52_P545810** | Lrrfip1 | 0.035234462 | -0.267303821 | 0.337772745 | 10.63906394 | 0.282989611 | 0.786427733 | 0.917923577 | -4.89311305 |
| **A_55_P1971330** | NA | 0.035012288 | -0.265625147 | 0.335649724 | 9.492875019 | 0.282983183 | 0.786432437 | 0.917923577 | -4.893114029 |
| **A_55_P2180406** | Cdkn2aipnl | 0.026604389 | -0.201887401 | 0.255096179 | 9.035905071 | 0.282921453 | 0.786477608 | 0.917923577 | -4.89312343 |
| **A_52_P386837** | Trmu | 0.025784802 | -0.195737289 | 0.247306893 | 9.542303732 | 0.282832912 | 0.786542401 | 0.917923577 | -4.89313691 |
| **A_66_P110970** | Tpd52l2 | -0.018889284 | -0.181190567 | 0.143411999 | 8.622511045 | -0.282798262 | 0.786567758 | 0.917923577 | -4.893142185 |
| **A_52_P490770** | Zfp26 | -0.025038985 | -0.240461708 | 0.190383737 | 8.28712663 | -0.282428425 | 0.786838422 | 0.918074952 | -4.893198442 |
| **A_55_P2108032** | Tmem255b | -0.018332698 | -0.176125394 | 0.139459998 | 7.706530657 | -0.282307663 | 0.786926809 | 0.918074952 | -4.893216796 |
| **A_55_P2178486** | Spin1 | 0.020581382 | -0.156574368 | 0.197737133 | 7.924604355 | 0.282294537 | 0.786936416 | 0.918074952 | -4.893218791 |
| **A_55_P2074591** | Ppm1h | -0.030440698 | -0.292465021 | 0.231583624 | 8.86477169 | -0.282290457 | 0.786939402 | 0.918074952 | -4.893219411 |
| **A_52_P22324** | Pln | -0.02012273 | -0.193335034 | 0.153089574 | 7.643675284 | -0.282287305 | 0.786941709 | 0.918074952 | -4.89321989 |
| **A_66_P127432** | Tial1 | 0.026319513 | -0.200334997 | 0.252974022 | 12.02852628 | 0.282160794 | 0.787034308 | 0.918074952 | -4.893239109 |
| **A_51_P298802** | Bfsp2 | -0.051864935 | -0.498743081 | 0.395013211 | 10.1710738 | -0.282012209 | 0.787143067 | 0.918074952 | -4.89326167 |
| **A_55_P1960231** | Txn2 | 0.03201318 | -0.243821305 | 0.307847664 | 14.06485837 | 0.282009307 | 0.787145192 | 0.918074952 | -4.893262111 |
| **A_55_P2005335** | Kcng2 | -0.019521082 | -0.187733682 | 0.148691518 | 7.744290734 | -0.281986651 | 0.787161776 | 0.918074952 | -4.89326555 |
| **A_51_P315391** | Txlnb | -0.022004757 | -0.211684088 | 0.167674573 | 7.706978084 | -0.281890063 | 0.787232479 | 0.918074952 | -4.893280208 |
| **A_52_P517896** | Man2c1 | 0.025423372 | -0.193760453 | 0.244607197 | 11.43416596 | 0.281843379 | 0.787266653 | 0.918074952 | -4.893287292 |
| **A_55_P2129219** | NA | 0.033455645 | -0.254978332 | 0.321889623 | 13.79153897 | 0.281842305 | 0.787267439 | 0.918074952 | -4.893287455 |
| **A_51_P486610** | Afg3l1 | -0.030405327 | -0.292626623 | 0.231815969 | 8.440410474 | -0.281750641 | 0.78733454 | 0.918097812 | -4.893301359 |
| **A_51_P160744** | Ndufb3 | -0.03152521 | -0.303753061 | 0.240702641 | 13.35069038 | -0.281389976 | 0.787598582 | 0.918333171 | -4.893356027 |
| **A_65_P15340** | Zfp251 | 0.035596149 | -0.271834223 | 0.343026521 | 8.769356308 | 0.281345152 | 0.787631399 | 0.918333171 | -4.893362816 |
| **A_55_P2148912** | S100a16 | -0.05496555 | -0.529838034 | 0.419906933 | 11.68459517 | -0.281252742 | 0.787699058 | 0.918348132 | -4.89337681 |
| **A_55_P2077635** | NA | 0.02230585 | -0.170442198 | 0.215053899 | 11.45193753 | 0.281197843 | 0.787739253 | 0.918348132 | -4.893385122 |
| **A_55_P2095709** | NA | 0.032357404 | -0.247315181 | 0.312029988 | 16.0658101 | 0.28112985 | 0.787789037 | 0.918350781 | -4.893395414 |
| **A_51_P455897** | Pimreg | 0.051653651 | -0.395012051 | 0.498319353 | 9.372337086 | 0.280996952 | 0.787886347 | 0.918399396 | -4.893415523 |
| **A_55_P2011425** | Atp10b | 0.018708075 | -0.143116389 | 0.180532538 | 7.683104409 | 0.280910588 | 0.787949586 | 0.918399396 | -4.893428586 |
| **A_51_P131358** | Selplg | -0.046510148 | -0.448923358 | 0.355903063 | 9.701315055 | -0.280839816 | 0.78800141 | 0.918399396 | -4.893439288 |
| **A_66_P112394** | NA | -0.021197826 | -0.20462209 | 0.162226437 | 7.855796401 | -0.28081334 | 0.788020797 | 0.918399396 | -4.89344329 |
| **A_51_P222283** | Mlycd | -0.030185925 | -0.291501223 | 0.231129372 | 11.82746386 | -0.280687356 | 0.788113054 | 0.918451539 | -4.893462333 |
| **A_51_P315595** | NA | -0.055556287 | -0.537075942 | 0.425963369 | 16.54881903 | -0.280351174 | 0.788359254 | 0.918683066 | -4.893513107 |
| **A_51_P433091** | Purb | -0.0372876 | -0.360665266 | 0.286090067 | 11.30990705 | -0.280180297 | 0.788484405 | 0.91875729 | -4.893538892 |
| **A_55_P2162573** | Art3 | 0.018849814 | -0.144652465 | 0.182352094 | 7.859551255 | 0.28013441 | 0.788518014 | 0.91875729 | -4.893545814 |
| **A_55_P1975667** | Rhox4g | 0.018166096 | -0.139545574 | 0.175877766 | 7.679307965 | 0.279885862 | 0.788700065 | 0.918914018 | -4.893583287 |
| **A_55_P2138549** | Cc2d1a | -0.020921249 | -0.202613639 | 0.160771142 | 9.657879181 | -0.279791201 | 0.788769405 | 0.918939415 | -4.89359755 |
| **A_55_P1978830** | Camk2d | -0.024958696 | -0.241943924 | 0.192026531 | 8.164068624 | -0.279495563 | 0.788985974 | 0.919136325 | -4.893642066 |
| **A_55_P1997991** | Ecpas | 0.031007214 | -0.238664634 | 0.300679062 | 8.45797599 | 0.279389656 | 0.789063561 | 0.919171312 | -4.893658003 |
| **A_52_P185907** | Crabp1 | 0.01891279 | -0.145619026 | 0.183444607 | 7.697721777 | 0.279311553 | 0.789120781 | 0.91918257 | -4.893669751 |
| **A_52_P40241** | Zfp746 | 0.02353578 | -0.18134542 | 0.22841698 | 9.101966505 | 0.279132026 | 0.789252311 | 0.919243432 | -4.893696745 |
| **A_52_P565496** | Prl8a1 | -0.01797093 | -0.174421704 | 0.138479843 | 7.628577532 | -0.279110413 | 0.789268146 | 0.919243432 | -4.893699993 |
| **A_55_P2165028** | NA | 0.026472951 | -0.204140032 | 0.257085934 | 15.31228592 | 0.27893422 | 0.789397241 | 0.919246564 | -4.893726467 |
| **A_66_P129484** | Tmed9 | 0.05248475 | -0.404975024 | 0.509944523 | 8.850206002 | 0.278781153 | 0.789509399 | 0.919246564 | -4.893749454 |

| | | | | | | | | | |
|---|---|---|---|---|---|---|---|---|---|
| **A_55_P2074197** | Zbtb24 | 0.020384492 | -0.157297513 | 0.198066496 | 8.462682764 | 0.278765889 | 0.789520584 | 0.919246564 | -4.893751745 |
| **A_51_P343889** | Slc35f6 | 0.023343318 | -0.180139146 | 0.226825782 | 8.295113246 | 0.278752512 | 0.789530386 | 0.919246564 | -4.893753753 |
| **A_55_P2042813** | Gbp11 | 0.037536149 | -0.289668318 | 0.364740616 | 9.057855764 | 0.278749233 | 0.789532789 | 0.919246564 | -4.893754245 |
| **A_55_P1993483** | NA | -0.039991563 | -0.388709917 | 0.308726791 | 10.66151381 | -0.278661365 | 0.789597175 | 0.919246564 | -4.893767434 |
| **A_55_P2112424** | Madd | 0.062936603 | -0.48596685 | 0.611840056 | 15.22775196 | 0.278606038 | 0.789637718 | 0.919246564 | -4.893775736 |
| **A_51_P469008** | Ehd1 | -0.023665312 | -0.230076769 | 0.182746144 | 13.17400661 | -0.278587505 | 0.789651299 | 0.919246564 | -4.893778516 |
| **A_55_P1953103** | Nudt7 | -0.035239144 | -0.342798787 | 0.272320498 | 10.23423929 | -0.278406389 | 0.789784024 | 0.919297727 | -4.893805681 |
| **A_52_P708507** | Myadml2 | -0.017779876 | -0.17299828 | 0.137438529 | 7.665350993 | -0.278335563 | 0.789835929 | 0.919297727 | -4.893816299 |
| **A_51_P465809** | Slc30a9 | -0.018971112 | -0.184590637 | 0.146648413 | 8.280062025 | -0.278332832 | 0.78983793 | 0.919297727 | -4.893816708 |
| **A_55_P2065586** | Gm15417 | 0.02182826 | -0.168941656 | 0.212598176 | 8.179215159 | 0.278030488 | 0.790059515 | 0.919500263 | -4.893862005 |
| **A_52_P98778** | Ang4 | 0.017756809 | -0.137486179 | 0.172999796 | 8.062765479 | 0.277930438 | 0.790132846 | 0.919530241 | -4.893876984 |
| **A_55_P1984497** | Usp48 | 0.026381537 | -0.204515525 | 0.2572786 | 9.796416101 | 0.277629033 | 0.790353771 | 0.919731971 | -4.893922078 |
| **A_55_P2115245** | Zfyve19 | -0.03202247 | -0.312475858 | 0.248430919 | 11.40293923 | -0.277445266 | 0.79048848 | 0.919762503 | -4.893949549 |
| **A_51_P206445** | Zfp773 | -0.019460228 | -0.189897035 | 0.150976579 | 7.920235727 | -0.277439137 | 0.790492973 | 0.919762503 | -4.893950465 |
| **A_55_P2017066** | NA | -0.025611499 | -0.250010972 | 0.198787974 | 11.64133664 | -0.277329678 | 0.790573215 | 0.919762503 | -4.893966819 |
| **A_55_P2463930** | Zmym5 | -0.030167327 | -0.294548577 | 0.234213924 | 9.090733474 | -0.27726137 | 0.790623292 | 0.919762503 | -4.893977021 |
| **A_52_P200359** | Aco2 | -0.029594007 | -0.288957055 | 0.22976904 | 13.45021879 | -0.277254665 | 0.790628207 | 0.919762503 | -4.893978023 |
| **A_51_P351413** | Phospho2 | -0.024251267 | -0.236845949 | 0.188343415 | 8.300744949 | -0.277182131 | 0.790681383 | 0.919762503 | -4.893988853 |
| **A_51_P248381** | Zdhhc4 | -0.018088064 | -0.176709795 | 0.140533666 | 8.35669546 | -0.277084736 | 0.790752787 | 0.919762503 | -4.894003392 |
| **A_51_P448664** | Tbxas1 | -0.046134404 | -0.4507678 | 0.358498992 | 10.08630406 | -0.277042489 | 0.79078376 | 0.919762503 | -4.894009697 |
| **A_55_P2021510** | NA | -0.026282093 | -0.256898871 | 0.204334685 | 8.656970494 | -0.276918667 | 0.790874543 | 0.919762503 | -4.894028171 |
| **A_55_P2016014** | Cpsf4l | -0.018321058 | -0.179085369 | 0.142443254 | 7.637805324 | -0.276913475 | 0.790878349 | 0.919762503 | -4.894028945 |
| **A_52_P239403** | Ipo7 | 0.024717298 | -0.192199786 | 0.241634382 | 8.403761092 | 0.276879256 | 0.790903438 | 0.919762503 | -4.894034049 |
| **A_51_P218774** | Rgs10 | 0.021319889 | -0.165971264 | 0.208611043 | 11.83922743 | 0.276599166 | 0.791108809 | 0.919894983 | -4.894075803 |
| **A_51_P153973** | Zfp273 | 0.024128166 | -0.187839553 | 0.236095885 | 8.282908959 | 0.276590832 | 0.79111492 | 0.919894983 | -4.894077045 |
| **A_66_P118759** | Safb2 | 0.019982933 | -0.155607746 | 0.195573611 | 11.66412238 | 0.276529172 | 0.791160133 | 0.919894983 | -4.894086231 |
| **A_51_P244061** | Snapc5 | -0.029329963 | -0.287114046 | 0.22845412 | 10.51738097 | -0.276464012 | 0.791207914 | 0.919895204 | -4.894095936 |
| **A_55_P1962011** | Clec1b | 0.062752643 | -0.489254446 | 0.614759733 | 11.54110583 | 0.276229818 | 0.791379653 | 0.919996559 | -4.894130801 |
| **A_55_P2112005** | Tff1 | -0.017785258 | -0.174315663 | 0.138745147 | 7.630025763 | -0.276086167 | 0.791485001 | 0.919996559 | -4.894152172 |
| **A_55_P2028491** | Hmgn2 | -0.042736759 | -0.418887636 | 0.333414117 | 10.35945096 | -0.276072214 | 0.791495234 | 0.919996559 | -4.894154247 |
| **A_52_P537545** | Smpd3 | -0.023274872 | -0.228159329 | 0.181609586 | 8.054186067 | -0.276033296 | 0.791523776 | 0.919996559 | -4.894160035 |
| **A_51_P406835** | Mei1 | -0.017988904 | -0.176370525 | 0.140392717 | 7.717985875 | -0.275983491 | 0.791560302 | 0.919996559 | -4.89416744 |
| **A_55_P2078173** | NA | -0.029028187 | -0.284651173 | 0.226594799 | 14.12067035 | -0.275932719 | 0.79159754 | 0.919996559 | -4.894174988 |
| **A_51_P370470** | Cyp20a1 | -0.031309854 | -0.30711018 | 0.244490471 | 8.875514642 | -0.275847759 | 0.791659851 | 0.919996559 | -4.894187616 |
| **A_55_P2059085** | Zfp457 | -0.038462035 | -0.377290867 | 0.300366798 | 9.081163996 | -0.275825927 | 0.791675864 | 0.919996559 | -4.894190861 |
| **A_55_P1958007** | Ddx24 | -0.03197766 | -0.313771958 | 0.249816638 | 12.54702993 | -0.275738667 | 0.791739865 | 0.920015621 | -4.894203825 |
| **A_51_P194609** | Prss34 | 0.068568758 | -0.536336501 | 0.673474017 | 8.589210525 | 0.275436894 | 0.791961213 | 0.920175209 | -4.894248631 |
| **A_55_P2169804** | Nanos3 | 0.042578779 | -0.333112293 | 0.41826985 | 9.598342513 | 0.275388319 | 0.791996845 | 0.920175209 | -4.894255839 |
| **A_51_P195483** | Ube2q2 | -0.029783112 | -0.292665392 | 0.233099169 | 9.124457494 | -0.275290957 | 0.792068265 | 0.920175209 | -4.894270282 |
| **A_55_P1955239** | Rps6 | -0.027733401 | -0.272563241 | 0.217096438 | 15.0503466 | -0.275246611 | 0.792100797 | 0.920175209 | -4.894276859 |
| **A_51_P262144** | Ddx19b | 0.033227615 | -0.260152907 | 0.326608137 | 9.674167215 | 0.275201683 | 0.792133755 | 0.920175209 | -4.894283521 |
| **A_51_P220135** | Nfatc1 | -0.023967052 | -0.235612672 | 0.187678569 | 8.237669421 | -0.275162038 | 0.792162837 | 0.920175209 | -4.894289399 |
| **A_55_P2080542** | Rbbp7 | -0.053353033 | -0.524937437 | 0.418231371 | 12.27292476 | -0.274905145 | 0.792351301 | 0.920291766 | -4.894327468 |
| **A_55_P2075213** | Kbtbd11 | -0.055551343 | -0.54660836 | 0.435505674 | 7.902782222 | -0.274881702 | 0.792368499 | 0.920291766 | -4.89433094 |
| **A_55_P2045642** | Stmn4 | -0.019554859 | -0.192446041 | 0.153336323 | 7.858146931 | -0.274830568 | 0.792406015 | 0.920291766 | -4.894338513 |
| **A_55_P2103668** | Zfp316 | 0.026071427 | -0.204671988 | 0.256814842 | 9.784990406 | 0.27454825 | 0.792613153 | 0.920477026 | -4.894380298 |
| **A_55_P2083919** | Robo2 | -0.049783038 | -0.490628351 | 0.391062275 | 8.903330005 | -0.274396367 | 0.792724598 | 0.920542944 | -4.894402761 |
| **A_51_P391445** | Ifngr1 | -0.060209015 | -0.710263787 | 0.589845756 | 12.32131502 | -0.285612442 | 0.792785373 | 0.920542944 | -4.747713813 |
| **A_51_P321734** | Bhmt2 | 0.053368457 | -0.419631153 | 0.526368068 | 12.56445986 | 0.274161871 | 0.79289667 | 0.920542944 | -4.894437419 |
| **A_51_P271336** | Wipf2 | 0.026167096 | -0.205792679 | 0.258126872 | 9.787978066 | 0.274110734 | 0.792934196 | 0.920542944 | -4.894444973 |

| | | | | | | | | | |
|---|---|---|---|---|---|---|---|---|---|
| **A_51_P286865** | Wdr33 | 0.01886476 | -0.148389398 | 0.186118918 | 10.00720561 | 0.274067507 | 0.792965918 | 0.920542944 | -4.894451357 |
| **A_55_P2065330** | NA | -0.033882019 | -0.334397507 | 0.26663347 | 15.1430574 | -0.273959022 | 0.79304553 | 0.920542944 | -4.894467376 |
| **A_55_P1954212** | Snrpd2 | -0.048014106 | -0.473918912 | 0.377890699 | 13.78164134 | -0.273929936 | 0.793066876 | 0.920542944 | -4.89447167 |
| **A_52_P660693** | Dsel | -0.038378431 | -0.378858501 | 0.30210164 | 8.265120217 | -0.273891596 | 0.793095014 | 0.920542944 | -4.894477329 |
| **A_55_P2044659** | NA | 0.018284557 | -0.14394271 | 0.180511825 | 7.768933578 | 0.273869574 | 0.793111175 | 0.920542944 | -4.894480579 |
| **A_66_P126293** | Itgb3 | 0.043468138 | -0.342264202 | 0.429200478 | 8.128744236 | 0.273821899 | 0.793146163 | 0.920542944 | -4.894487615 |
| **A_51_P356525** | Trappc11 | 0.021603438 | -0.170247602 | 0.213454478 | 9.363507193 | 0.273616256 | 0.793297091 | 0.920662832 | -4.894517948 |
| **A_66_P113760** | Acvr2a | 0.028581854 | -0.225644566 | 0.282808274 | 8.680960849 | 0.273182516 | 0.793615457 | 0.920977015 | -4.894581855 |
| **A_55_P2156274** | Plekha2 | -0.029856381 | -0.29548738 | 0.235774619 | 8.288435573 | -0.273112511 | 0.793666844 | 0.920979424 | -4.894592161 |
| **A_51_P198137** | Ubxn8 | 0.025825014 | -0.20399179 | 0.255641818 | 8.643419726 | 0.27304987 | 0.793712828 | 0.920979424 | -4.89460138 |
| **A_55_P2140118** | Qpct | 0.033765923 | -0.266831876 | 0.334363722 | 11.1308832 | 0.272945551 | 0.793789407 | 0.921006323 | -4.894616728 |
| **A_55_P2088750** | Mtif3 | -0.021300317 | -0.210963952 | 0.168363317 | 8.196046395 | -0.272888475 | 0.793831308 | 0.921006323 | -4.894625124 |
| **A_51_P514319** | Slc13a4 | -0.017631939 | -0.174861196 | 0.139597317 | 7.798960244 | -0.272489587 | 0.794124157 | 0.921243471 | -4.894683748 |
| **A_51_P328968** | Zfp322a | 0.022183317 | -0.175638732 | 0.220005367 | 11.18489627 | 0.272480223 | 0.794131032 | 0.921243471 | -4.894685124 |
| **A_52_P609334** | Klhdc8b | -0.048545297 | -0.481654825 | 0.384564231 | 8.377264199 | -0.272353284 | 0.794224235 | 0.921287311 | -4.894703762 |
| **A_55_P2019190** | NA | -0.024589818 | -0.24401809 | 0.194838454 | 15.35429941 | -0.272298923 | 0.794264149 | 0.921287311 | -4.894711741 |
| **A_51_P196726** | Itih2 | -0.070757343 | -0.702518179 | 0.561003493 | 14.12390091 | -0.272146033 | 0.794376412 | 0.921316053 | -4.894734175 |
| **A_55_P1967341** | Gm9104 | -0.028024193 | -0.278249564 | 0.222201179 | 13.51235792 | -0.272135347 | 0.794384259 | 0.921316053 | -4.894735742 |
| **A_55_P2066234** | Arfip2 | 0.053128294 | -0.421477882 | 0.52773447 | 9.022588395 | 0.27200424 | 0.794480532 | 0.921372424 | -4.894754969 |
| **A_51_P414305** | Acot12 | 0.077818806 | -0.617985904 | 0.773623517 | 9.413308221 | 0.271756781 | 0.794662253 | 0.921494805 | -4.894791234 |
| **A_55_P2073607** | Commd5 | 0.025038619 | -0.198904716 | 0.248981955 | 11.49703502 | 0.271678593 | 0.794719673 | 0.921494805 | -4.894802686 |
| **A_51_P308590** | Ppif | 0.025986683 | -0.206458378 | 0.258431743 | 8.761045354 | 0.271652505 | 0.794738832 | 0.921494805 | -4.894806506 |
| **A_51_P343913** | Parp4 | 0.022441766 | -0.178332973 | 0.223216505 | 8.455427294 | 0.271600868 | 0.794776755 | 0.921494805 | -4.894814067 |
| **A_51_P507801** | F13a1 | -0.053254367 | -0.530044561 | 0.423535827 | 10.94866497 | -0.271400786 | 0.794923703 | 0.921547674 | -4.89484335 |
| **A_51_P412595** | Eef1b2 | -0.032166101 | -0.320181403 | 0.255849201 | 15.17761309 | -0.271372633 | 0.79494438 | 0.921547674 | -4.894847468 |
| **A_55_P2038126** | Tpmt | 0.050614655 | -0.402793095 | 0.504022406 | 9.141628812 | 0.271250491 | 0.795034091 | 0.921547674 | -4.894865332 |
| **A_55_P2401403** | NA | -0.051766051 | -0.515801612 | 0.412269511 | 9.220962785 | -0.271067192 | 0.795168727 | 0.921547674 | -4.894892126 |
| **A_55_P2088018** | Fhod3 | -0.04045778 | -0.403224981 | 0.322309422 | 8.032257212 | -0.270992467 | 0.795223616 | 0.921547674 | -4.894903044 |
| **A_51_P281333** | St3gal6 | 0.025385901 | -0.202297851 | 0.253069653 | 9.92934256 | 0.270921663 | 0.795275626 | 0.921547674 | -4.894913386 |
| **A_52_P199299** | Urgcp | 0.024379999 | -0.194341286 | 0.243101284 | 9.215658805 | 0.270848114 | 0.795329653 | 0.921547674 | -4.894924127 |
| **A_55_P2065231** | Gstm3 | -0.029275125 | -0.291981706 | 0.233431456 | 9.840009699 | -0.270776514 | 0.795382251 | 0.921547674 | -4.89493458 |
| **A_52_P676461** | Lpgat1 | 0.053234467 | -0.424830631 | 0.531299566 | 8.941036272 | 0.27057587 | 0.795529649 | 0.921547674 | -4.894963859 |
| **A_55_P1972054** | NA | 0.034296198 | -0.273743985 | 0.342336381 | 8.696045892 | 0.270533964 | 0.795560435 | 0.921547674 | -4.894969972 |
| **A_55_P2322139** | C530042K13Rik | -0.017580426 | -0.175501104 | 0.140340251 | 7.684485977 | -0.270503939 | 0.795582493 | 0.921547674 | -4.894974351 |
| **A_55_P2093889** | Arhgef19 | -0.033140014 | -0.330838355 | 0.264558328 | 7.949962588 | -0.270495133 | 0.795588962 | 0.921547674 | -4.894975635 |
| **A_55_P2123822** | Rbbp7 | 0.049308362 | -0.393683478 | 0.492300201 | 14.81744925 | 0.270463109 | 0.79561249 | 0.921547674 | -4.894980305 |
| **A_66_P124179** | Atp6v0d2 | 0.017314867 | -0.138260154 | 0.172889888 | 7.74774567 | 0.270434737 | 0.795633334 | 0.921547674 | -4.894984442 |
| **A_65_P12533** | Naa10 | 0.027094257 | -0.216395434 | 0.270583947 | 11.37013259 | 0.270383294 | 0.795671129 | 0.921547674 | -4.894991941 |
| **A_55_P2097391** | Tent5a | 0.035263524 | -0.281759484 | 0.352286532 | 8.434917904 | 0.270282619 | 0.795745095 | 0.921547674 | -4.895006614 |
| **A_52_P485971** | Scap | -0.033004754 | -0.329765962 | 0.263756454 | 13.28124354 | -0.270241822 | 0.79577507 | 0.921547674 | -4.895012559 |
| **A_55_P1971430** | Serpinb9e | -0.017372346 | -0.173581695 | 0.138837002 | 7.636829701 | -0.270230674 | 0.79578326 | 0.921547674 | -4.895014183 |
| **A_55_P2083213** | Purb | 0.028634534 | -0.228977612 | 0.286246681 | 8.789229512 | 0.27008905 | 0.795887317 | 0.921547674 | -4.895034811 |
| **A_55_P1993168** | Ppargc1b | 0.025416644 | -0.203326792 | 0.254160079 | 7.928213575 | 0.269993151 | 0.795957781 | 0.921547674 | -4.895048774 |
| **A_51_P397673** | Pcsk9 | 0.061413456 | -0.491377354 | 0.614204265 | 11.71025942 | 0.269951603 | 0.79598831 | 0.921547674 | -4.895054822 |
| **A_55_P2044282** | Mpp6 | 0.046341902 | -0.370854629 | 0.463538433 | 13.11499598 | 0.26990837 | 0.796020078 | 0.921547674 | -4.895061114 |
| **A_55_P2064841** | Prss53 | 0.045078262 | -0.36084083 | 0.450997354 | 12.4719262 | 0.269842823 | 0.796068242 | 0.921547674 | -4.895070651 |
| **A_51_P292447** | Klhl36 | -0.021130695 | -0.211449248 | 0.169187859 | 8.665550472 | -0.269783781 | 0.796111628 | 0.921547674 | -4.89507924 |
| **A_55_P2011922** | 9-Mar | -0.02315362 | -0.231739565 | 0.185432325 | 9.170886308 | -0.269722423 | 0.796156715 | 0.921547674 | -4.895088165 |
| **A_55_P2129972** | NA | -0.051876036 | -0.519329332 | 0.415577259 | 10.69947454 | -0.26965703 | 0.796204769 | 0.921547674 | -4.895097673 |
| **A_55_P2088690** | NA | -0.037281813 | -0.373230473 | 0.298666848 | 8.981530506 | -0.269654265 | 0.796206801 | 0.921547674 | -4.895098076 |

| | | | | | | | | | |
|---|---|---|---|---|---|---|---|---|---|
| A_52_P485850 | Mgam | 0.044106216 | -0.353373474 | 0.441585907 | 8.357754641 | 0.269629899 | 0.796224707 | 0.921547674 | -4.895101618 |
| A_55_P2056393 | Stk11ip | 0.022547458 | -0.180687598 | 0.225782514 | 9.483943515 | 0.26957658 | 0.79626389 | 0.921547674 | -4.895109369 |
| A_55_P2082425 | Mtln | 0.041356017 | -0.331479719 | 0.414191753 | 12.01867708 | 0.269528292 | 0.796299375 | 0.921547674 | -4.895116387 |
| A_55_P1955876 | Ifitm7 | -0.019366604 | -0.193977419 | 0.155244211 | 8.93815508 | -0.269504191 | 0.796317088 | 0.921547674 | -4.89511989 |
| A_55_P2000148 | Dnaic1 | 0.062919596 | -0.504456994 | 0.630296186 | 9.60897919 | 0.269462106 | 0.796348016 | 0.921547674 | -4.895126005 |
| A_51_P520966 | Icosl | -0.037424864 | -0.375077195 | 0.300227467 | 8.529871141 | -0.26932314 | 0.796450144 | 0.921610682 | -4.89514619 |
| A_51_P482121 | Edn2 | -0.018590323 | -0.186477478 | 0.149296832 | 7.697185169 | -0.26906218 | 0.79664194 | 0.921777436 | -4.89518407 |
| A_55_P1977498 | Htr2b | -0.01705317 | -0.171210277 | 0.137103937 | 7.666636493 | -0.26879721 | 0.796836699 | 0.921947598 | -4.895222495 |
| A_52_P678895 | Dynlt3 | -0.035025499 | -0.351774283 | 0.281723286 | 9.857427585 | -0.268690652 | 0.796915027 | 0.921983034 | -4.895237937 |
| A_55_P1997421 | Csta3 | 0.067598378 | -0.543968734 | 0.679165491 | 11.60123242 | 0.268581036 | 0.796995604 | 0.92202107 | -4.895253817 |
| A_55_P2139211 | NA | -0.037549667 | -0.37751261 | 0.302413275 | 14.24564018 | -0.26838467 | 0.797139958 | 0.922132877 | -4.895282247 |
| A_55_P2045928 | NA | 0.06711327 | -0.540815549 | 0.67504209 | 11.12932922 | 0.268249462 | 0.797239358 | 0.922192672 | -4.895301812 |
| A_55_P2080168 | Dgkk | -0.020375644 | -0.204993405 | 0.164242118 | 7.784608149 | -0.268176704 | 0.797292848 | 0.922199226 | -4.895312336 |
| A_51_P199104 | Casq1 | -0.018476537 | -0.186025799 | 0.149072724 | 7.680866539 | -0.267954625 | 0.797456125 | 0.922199226 | -4.895344441 |
| A_51_P230934 | Cab39 | 0.028757143 | -0.232033908 | 0.289548195 | 10.60717867 | 0.267939193 | 0.797467472 | 0.922199226 | -4.895346671 |
| A_55_P1979397 | Palm | -0.037712137 | -0.379776059 | 0.304351786 | 10.73421689 | -0.267890341 | 0.79750339 | 0.922199226 | -4.89535373 |
| A_55_P2148002 | 2610005L07Rik | 0.044504755 | -0.359180147 | 0.448189657 | 8.954046153 | 0.267884198 | 0.797507907 | 0.922199226 | -4.895354617 |
| A_55_P2049282 | Ly6h | -0.034907699 | -0.351592505 | 0.281777106 | 8.049985096 | -0.267841081 | 0.797539609 | 0.922199226 | -4.895360846 |
| A_66_P122158 | Pisd-ps3 | 0.022292902 | -0.180108073 | 0.224693877 | 7.813133392 | 0.267631481 | 0.797693727 | 0.922199226 | -4.895391111 |
| A_55_P2070045 | Golga7b | -0.01755005 | -0.176896579 | 0.14179648 | 7.664612645 | -0.267620227 | 0.797702002 | 0.922199226 | -4.895392736 |
| A_55_P2085984 | Igf1 | -0.020649565 | -0.208149782 | 0.166850652 | 9.897062655 | -0.267603824 | 0.797714064 | 0.922199226 | -4.895395103 |
| A_55_P1996927 | 4933409K07Rik | -0.033342427 | -0.336160373 | 0.269475519 | 10.30627391 | -0.2675462 | 0.797756436 | 0.922199226 | -4.895403419 |
| A_51_P444335 | Acp6 | -0.021240603 | -0.214189787 | 0.171708582 | 10.69903581 | -0.267489709 | 0.797797977 | 0.922199226 | -4.89541157 |
| A_55_P2024220 | NA | -0.018925273 | -0.190975071 | 0.153124525 | 7.988747545 | -0.267282905 | 0.797950055 | 0.922199226 | -4.895441394 |
| A_55_P1981664 | 8-Sep | 0.030423347 | -0.246208929 | 0.307055623 | 12.14705992 | 0.267231305 | 0.797988001 | 0.922199226 | -4.895448832 |
| A_55_P2129311 | 2210016L21Rik | 0.032836203 | -0.265777061 | 0.331449468 | 9.472325511 | 0.267194202 | 0.798015287 | 0.922199226 | -4.895454179 |
| A_55_P2313163 | D10Bwg1070e | 0.059981055 | -0.485604267 | 0.605566377 | 10.4509715 | 0.267137348 | 0.798057099 | 0.922199226 | -4.895462372 |
| A_55_P2108868 | Nufip1 | -0.020964737 | -0.211678411 | 0.169748936 | 8.076493723 | -0.267110392 | 0.798076923 | 0.922199226 | -4.895466255 |
| A_66_P124715 | Ppard | -0.026791768 | -0.27051439 | 0.216930854 | 9.990860993 | -0.267109123 | 0.798077857 | 0.922199226 | -4.895466438 |
| A_55_P2022663 | NA | -0.02494291 | -0.251911149 | 0.20202533 | 15.83467457 | -0.267033172 | 0.798133715 | 0.922199226 | -4.895477379 |
| A_55_P2185489 | Zfp865 | -0.020406966 | -0.206254855 | 0.165440923 | 8.843372298 | -0.266811164 | 0.798296997 | 0.922199226 | -4.895509342 |
| A_55_P2056070 | Hfe | -0.04130913 | -0.417634821 | 0.335016561 | 10.77541557 | -0.266726012 | 0.798359627 | 0.922199226 | -4.895521595 |
| A_52_P199776 | Snhg7 | 0.024451019 | -0.198317986 | 0.247220025 | 8.667087751 | 0.266701461 | 0.798377685 | 0.922199226 | -4.895525127 |
| A_52_P162957 | Frat2 | 0.024137339 | -0.195792203 | 0.244066882 | 9.122754116 | 0.266679118 | 0.798394119 | 0.922199226 | -4.895528341 |
| A_55_P2301941 | A530030E21Rik | 0.072757103 | -0.59020745 | 0.735721657 | 12.06123908 | 0.266666389 | 0.798403481 | 0.922199226 | -4.895530172 |
| A_55_P2096599 | Tsc22d4 | -0.034421543 | -0.348126827 | 0.279283742 | 8.487199406 | -0.266619361 | 0.798438073 | 0.922199226 | -4.895536936 |
| A_51_P290921 | Sytl2 | -0.018124466 | -0.183329693 | 0.14708076 | 7.929275098 | -0.266578202 | 0.798468347 | 0.922199226 | -4.895542854 |
| A_55_P1995074 | Ubald2 | 0.029198794 | -0.237011615 | 0.295409203 | 14.46700903 | 0.266515866 | 0.798514199 | 0.922199226 | -4.895551817 |
| A_52_P76196 | NA | -0.03222183 | -0.326022271 | 0.261578611 | 13.65029532 | -0.266490019 | 0.798533211 | 0.922199226 | -4.895555532 |
| A_52_P36680 | Pggt1b | -0.019208471 | -0.194398628 | 0.155981685 | 8.31143432 | -0.266419673 | 0.798584957 | 0.922203885 | -4.895565643 |
| A_51_P138538 | Eif1b | 0.020573689 | -0.16713844 | 0.208285819 | 11.70358906 | 0.26631954 | 0.798658615 | 0.922233847 | -4.895580031 |
| A_55_P2131027 | NA | 0.085130377 | -0.691857341 | 0.862118094 | 13.23207928 | 0.266227915 | 0.798726016 | 0.922234566 | -4.895593191 |
| A_55_P1968738 | Nfkb1 | 0.020260877 | -0.164687858 | 0.205209612 | 8.61197923 | 0.266188974 | 0.798754662 | 0.922234566 | -4.895598783 |
| A_55_P2060737 | Trmt10b | 0.01902684 | -0.154776997 | 0.192830678 | 8.702120854 | 0.26600543 | 0.798889689 | 0.922335372 | -4.89562513 |
| A_51_P314153 | Nr2c2ap | 0.039103522 | -0.31826997 | 0.396477015 | 8.78359479 | 0.265874527 | 0.798985994 | 0.922391425 | -4.895643909 |
| A_66_P118989 | Gm8350 | -0.024551232 | -0.248983678 | 0.199881213 | 8.151002387 | -0.265809711 | 0.799033681 | 0.922391425 | -4.895653204 |
| A_55_P2064995 | Capzb | -0.018060578 | -0.183283783 | 0.147162627 | 7.684081271 | -0.265609618 | 0.7991809 | 0.922453704 | -4.895681886 |
| A_66_P127224 | Krt10 | -0.039223423 | -0.398054229 | 0.319607384 | 8.708679745 | -0.265606656 | 0.799183079 | 0.922453704 | -4.89568231 |
| A_55_P1959639 | NA | -0.028842186 | -0.292823572 | 0.235139201 | 13.59701826 | -0.265483816 | 0.799273463 | 0.922502942 | -4.895699908 |
| A_55_P2085127 | Ramac | -0.021910984 | -0.22255814 | 0.178736172 | 10.56809385 | -0.2653457 | 0.799375091 | 0.922565151 | -4.895719684 |

| | | | | | | | | | |
|---|---|---|---|---|---|---|---|---|---|
| **A_51_P206251** | Ahcyl1 | -0.043505549 | -0.442029914 | 0.355018816 | 9.489625213 | -0.265260732 | 0.799437615 | 0.922578647 | -4.895731846 |
| **A_55_P2190299** | Fggy | 0.08055655 | -0.65753528 | 0.81864838 | 11.47314988 | 0.265200079 | 0.799482246 | 0.922578647 | -4.895740524 |
| **A_55_P2072403** | Suv39h2 | 0.08660388 | -0.707253035 | 0.880460794 | 14.2390871 | 0.26508082 | 0.799570007 | 0.922624838 | -4.895757584 |
| **A_55_P2052709** | NA | -0.018205015 | -0.18517846 | 0.14876843 | 7.706448679 | -0.264927365 | 0.799682935 | 0.922626687 | -4.895779524 |
| **A_55_P1971938** | Atp2b2 | 0.060568527 | -0.495029729 | 0.616166782 | 9.022590024 | 0.264892294 | 0.799708745 | 0.922626687 | -4.895784536 |
| **A_66_P125071** | Ssb | 0.020850404 | -0.170424055 | 0.212124862 | 10.35720716 | 0.264874822 | 0.799721604 | 0.922626687 | -4.895787033 |
| **A_51_P211998** | Sgms2 | -0.027760673 | -0.282481067 | 0.22695972 | 8.062143239 | -0.264819197 | 0.79976254 | 0.922626687 | -4.895794982 |
| **A_55_P2048840** | Ppil2 | -0.035514361 | -0.36156786 | 0.290539138 | 9.486782398 | -0.264666106 | 0.799875211 | 0.922687856 | -4.895816849 |
| **A_51_P120717** | Lmnb1 | 0.049406269 | -0.404270774 | 0.503083313 | 10.94297337 | 0.264617429 | 0.799911036 | 0.922687856 | -4.880238082 |
| **A_51_P307082** | Hey2 | 0.019102165 | -0.156542942 | 0.194747272 | 7.679174085 | 0.26425896 | 0.800174882 | 0.922937122 | -4.895874945 |
| **A_52_P251699** | Prodh2 | 0.05614494 | -0.460291689 | 0.572581569 | 12.45062012 | 0.264165911 | 0.800243374 | 0.922961046 | -4.89588821 |
| **A_55_P1986833** | Tulp4 | 0.050653771 | -0.415687733 | 0.516995275 | 9.178052987 | 0.263931305 | 0.800416073 | 0.923105147 | -4.895921635 |
| **A_55_P1997282** | Fam135a | -0.035195219 | -0.359453909 | 0.289063471 | 8.76532865 | -0.263739535 | 0.800557248 | 0.923212878 | -4.895948937 |
| **A_55_P2132902** | Gprasp1 | -0.04286522 | -0.438058848 | 0.352328408 | 9.530998854 | -0.263559281 | 0.800689954 | 0.923310829 | -4.895974581 |
| **A_66_P108692** | NA | 0.0289473 | -0.238015103 | 0.295909704 | 15.13787508 | 0.263476057 | 0.800751226 | 0.923326401 | -4.895986415 |
| **A_51_P138760** | Abcd4 | 0.027640546 | -0.227489581 | 0.282770672 | 9.53733655 | 0.263249805 | 0.80091781 | 0.923439337 | -4.89601857 |
| **A_55_P2027127** | Triml2 | -0.017103665 | -0.174997297 | 0.140789967 | 7.656788682 | -0.263213259 | 0.800944719 | 0.923439337 | -4.896023761 |
| **A_51_P393897** | Mpi | 0.028400871 | -0.233902498 | 0.290704239 | 9.154402111 | 0.263094018 | 0.801032519 | 0.92348548 | -4.896040694 |
| **A_55_P1983046** | Plch1 | -0.02140802 | -0.219227213 | 0.176411172 | 7.856994875 | -0.262960958 | 0.801130498 | 0.923543352 | -4.896059581 |
| **A_55_P1978855** | NA | -0.03009064 | -0.308219355 | 0.248038075 | 14.85063801 | -0.262886804 | 0.801185103 | 0.923551219 | -4.896070103 |
| **A_55_P2386236** | Ceacam2 | -0.075789329 | -0.776613301 | 0.625034642 | 10.05957192 | -0.262773948 | 0.801268209 | 0.923591939 | -4.896086111 |
| **A_66_P115004** | Gabrp | -0.019256614 | -0.197548123 | 0.159034895 | 7.759550835 | -0.262441465 | 0.801513064 | 0.923655133 | -4.896133232 |
| **A_55_P1957198** | Apob | -0.080723625 | -0.828286142 | 0.666838891 | 14.11101972 | -0.262383384 | 0.80155584 | 0.923655133 | -4.896141457 |
| **A_55_P1973036** | NA | -0.018533744 | -0.19021866 | 0.153151172 | 7.748918161 | -0.262309613 | 0.801610173 | 0.923655133 | -4.896151902 |
| **A_51_P118132** | Skil | -0.029466139 | -0.302427116 | 0.243494838 | 10.03399735 | -0.262304569 | 0.801613887 | 0.923655133 | -4.888551159 |
| **A_55_P1980326** | Pcbp1 | 0.021119086 | -0.174546151 | 0.216784323 | 14.21738639 | 0.262267595 | 0.80164112 | 0.923655133 | -4.89615785 |
| **A_51_P234788** | Cxxc5 | -0.049763129 | -0.510917561 | 0.411391302 | 10.26409665 | -0.262207122 | 0.801685659 | 0.923655133 | -4.896166409 |
| **A_55_P2164258** | Prl7b1 | 0.01697231 | -0.140342329 | 0.17428695 | 7.675975798 | 0.26215311 | 0.801725442 | 0.923655133 | -4.896174052 |
| **A_55_P2020203** | Sfrp5 | 0.019055553 | -0.157584141 | 0.195695247 | 7.780318827 | 0.262129827 | 0.80174259 | 0.923655133 | -4.896177346 |
| **A_65_P13910** | Ndufb2 | -0.02519017 | -0.258708788 | 0.208328447 | 14.51772801 | -0.262115548 | 0.801753107 | 0.923655133 | -4.896179366 |
| **A_55_P1995622** | Pomp | 0.041513915 | -0.343579053 | 0.426606882 | 9.751414961 | 0.26194571 | 0.801878205 | 0.923744194 | -4.896203385 |
| **A_55_P2154690** | Smarca4 | 0.039460117 | -0.3269621 | 0.405882335 | 8.763878047 | 0.261673512 | 0.802078712 | 0.923920109 | -4.896241849 |
| **A_51_P400016** | Kalrn | -0.042209033 | -0.434257839 | 0.349839773 | 10.5448935 | -0.261606455 | 0.80212811 | 0.92392195 | -4.896251319 |
| **A_55_P1982076** | Emsy | 0.025175007 | -0.208718978 | 0.259068992 | 11.74467481 | 0.261537361 | 0.80217901 | 0.923925521 | -4.896261074 |
| **A_52_P562467** | Sync | 0.016677761 | -0.138366381 | 0.171721902 | 7.710386246 | 0.261375915 | 0.802297947 | 0.923934973 | -4.896283857 |
| **A_55_P2159870** | Tead4 | -0.030330736 | -0.312343606 | 0.251682133 | 7.790876093 | -0.261334781 | 0.802328251 | 0.923934973 | -4.89628966 |
| **A_55_P2076273** | NA | 0.035758739 | -0.296727373 | 0.368244851 | 14.268657 | 0.261331568 | 0.802330618 | 0.923934973 | -4.896290114 |
| **A_51_P370615** | Neurog2 | 0.020967531 | -0.174107805 | 0.216042867 | 7.741400369 | 0.261172905 | 0.802447513 | 0.924014534 | -4.896312488 |
| **A_55_P1969904** | Abcc6 | 0.076138293 | -0.633155257 | 0.785431843 | 10.63567001 | 0.260831665 | 0.802698938 | 0.924163336 | -4.896360564 |
| **A_51_P145433** | Lin7c | 0.017735472 | -0.147508723 | 0.182979668 | 8.280505187 | 0.260795284 | 0.802725745 | 0.924163336 | -4.896365686 |
| **A_55_P2020054** | 2610035D17Rik | 0.02060406 | -0.171377509 | 0.21258563 | 8.023588774 | 0.260781313 | 0.802736039 | 0.924163336 | -4.896367653 |
| **A_51_P461005** | Nrk | 0.017581642 | -0.146300589 | 0.181463874 | 8.106009628 | 0.260681831 | 0.802809343 | 0.924163336 | -4.896381654 |
| **A_51_P222522** | Mesd | -0.035588095 | -0.367323452 | 0.296147262 | 12.6697707 | -0.26067307 | 0.802815799 | 0.924163336 | -4.896382887 |
| **A_55_P2104617** | Kctd5 | 0.023338983 | -0.194302952 | 0.240980919 | 8.369169889 | 0.26056888 | 0.802892575 | 0.924196676 | -4.896397546 |
| **A_55_P2168312** | NA | 0.025231188 | -0.210138161 | 0.260600537 | 15.31730613 | 0.26047796 | 0.802959574 | 0.924218759 | -4.896410332 |
| **A_51_P489996** | Dst | -0.019255776 | -0.199088608 | 0.160577057 | 7.896549491 | -0.260180785 | 0.803178577 | 0.924415786 | -4.896452096 |
| **A_55_P1998697** | NA | 0.035968464 | -0.300114157 | 0.372051085 | 12.40395657 | 0.2600513 | 0.803274007 | 0.924470574 | -4.896470279 |
| **A_55_P2092671** | Nppc | -0.017504174 | -0.181103991 | 0.146095643 | 7.663111113 | -0.259981233 | 0.803325648 | 0.924474961 | -4.896480114 |
| **A_52_P211418** | G2e3 | 0.017890591 | -0.149430869 | 0.185212051 | 8.322225211 | 0.259810217 | 0.803451694 | 0.924564969 | -4.89650411 |
| **A_55_P1964960** | Il33 | -0.041177668 | -0.426428093 | 0.344072756 | 8.55868698 | -0.259717858 | 0.803519769 | 0.924588261 | -4.896517062 |

| | | | | | | | | | |
|---|---|---|---|---|---|---|---|---|---|
| **A_52_P644674** | Trappc6b | 0.020699823 | -0.173088835 | 0.214488481 | 10.76192919 | 0.259550259 | 0.803643306 | 0.924675366 | -4.896540555 |
| **A_55_P2110235** | Trip12 | 0.026454579 | -0.221653118 | 0.274562276 | 10.06433307 | 0.259085919 | 0.803985604 | 0.924968674 | -4.896605566 |
| **A_55_P2150786** | Mul1 | -0.031494454 | -0.326882147 | 0.263893239 | 10.69595749 | -0.259074624 | 0.803993931 | 0.924968674 | -4.896607146 |
| **A_55_P2144003** | NA | 0.027390512 | -0.229627208 | 0.284408232 | 13.93020531 | 0.2589526 | 0.804083891 | 0.925017113 | -4.896624211 |
| **A_55_P1958653** | Arf2 | -0.052312642 | -0.543624052 | 0.438998768 | 11.01232336 | -0.258721783 | 0.804254068 | 0.925157822 | -4.896656469 |
| **A_55_P2049061** | NA | -0.03095297 | -0.321781232 | 0.259875293 | 10.94138054 | -0.258612136 | 0.804334912 | 0.92515933 | -4.896671783 |
| **A_55_P1985025** | Pard3 | 0.027264213 | -0.228926946 | 0.283455372 | 10.17004266 | 0.258590172 | 0.804351107 | 0.92515933 | -4.89667485 |
| **A_65_P19089** | Esrrg | 0.016648615 | -0.139854061 | 0.173151291 | 7.723644841 | 0.258487492 | 0.804426817 | 0.925191357 | -4.896689184 |
| **A_55_P1959303** | Rbm47 | -0.049032086 | -0.510951407 | 0.412887235 | 8.992029914 | -0.257927371 | 0.804839858 | 0.925611329 | -4.896767281 |
| **A_55_P2030334** | D0Kist3 | 0.01883452 | -0.158647872 | 0.196316912 | 8.347839121 | 0.257859104 | 0.804890203 | 0.925614156 | -4.896776788 |
| **A_51_P439996** | Gp5 | -0.045900656 | -0.478635122 | 0.386833809 | 8.495845206 | -0.257739273 | 0.804978579 | 0.925660715 | -4.89679347 |
| **A_51_P171107** | Tmem35a | -0.021562195 | -0.22501914 | 0.18189475 | 8.099122982 | -0.257515658 | 0.805143504 | 0.925743604 | -4.896824581 |
| **A_55_P2026218** | Igsf8 | -0.047886055 | -0.499829954 | 0.404057843 | 11.14059333 | -0.257458791 | 0.805185448 | 0.925743604 | -4.896832488 |
| **A_55_P2024322** | Rps27 | 0.037710945 | -0.318262807 | 0.393684697 | 15.86900437 | 0.257414277 | 0.805218281 | 0.925743604 | -4.896838677 |
| **A_55_P1972852** | Mcrs1 | 0.032419836 | -0.273679632 | 0.338519305 | 11.44398067 | 0.257354302 | 0.805262518 | 0.925743604 | -4.896847013 |
| **A_51_P502422** | Acot13 | -0.023641193 | -0.246908105 | 0.199625719 | 13.19399538 | -0.257293145 | 0.805307628 | 0.925743604 | -4.896855512 |
| **A_51_P217047** | Gng10 | -0.029362294 | -0.30670358 | 0.247978992 | 9.756686468 | -0.257251934 | 0.805338026 | 0.925743604 | -4.896861238 |
| **A_55_P1955147** | Camk1d | 0.036985623 | -0.312538394 | 0.38650964 | 9.602791152 | 0.257121922 | 0.805433927 | 0.925798786 | -4.896879296 |
| **A_55_P2192577** | H2bfm | -0.026352806 | -0.275488465 | 0.222782853 | 8.109706058 | -0.257024287 | 0.805505949 | 0.92581863 | -4.896892851 |
| **A_51_P135357** | Sike1 | 0.018062343 | -0.152733429 | 0.188858114 | 8.063444713 | 0.256968655 | 0.805546987 | 0.92581863 | -4.896900572 |
| **A_51_P141274** | Sec24d | -0.039371017 | -0.411899267 | 0.333157232 | 10.32053388 | -0.256803309 | 0.805668964 | 0.925890453 | -4.896923512 |
| **A_51_P243134** | Adcy6 | -0.020208994 | -0.211463 | 0.171045012 | 9.29789226 | -0.256754078 | 0.805705283 | 0.925890453 | -4.896930339 |
| **A_51_P365516** | Spink1 | -0.068120942 | -0.712997553 | 0.57675567 | 15.5775125 | -0.256677148 | 0.805762037 | 0.925900625 | -4.896941006 |
| **A_55_P2420873** | NA | -0.033426475 | -0.350023492 | 0.283170543 | 9.387967123 | -0.256547004 | 0.805858053 | 0.925955909 | -4.896959043 |
| **A_55_P2002854** | Aifm2 | 0.017334037 | -0.147011412 | 0.181679487 | 7.92936333 | 0.256286205 | 0.806050471 | 0.926067493 | -4.896995161 |
| **A_55_P1981964** | Tnfrsf13b | -0.039149221 | -0.410327257 | 0.332028814 | 9.28661163 | -0.256285508 | 0.806050985 | 0.926067493 | -4.896995258 |
| **A_51_P176265** | Trip11 | 0.030627012 | -0.259928078 | 0.321182102 | 8.410017837 | 0.256129342 | 0.806166212 | 0.926144827 | -4.897016869 |
| **A_51_P335900** | Cox5a | -0.018989901 | -0.199251945 | 0.161272144 | 14.93018735 | -0.255977378 | 0.806278344 | 0.926218597 | -4.897037885 |
| **A_55_P1973723** | Etfdh | -0.058701734 | -0.616291137 | 0.49888767 | 12.1171025 | -0.25581123 | 0.806400948 | 0.926304388 | -4.89706085 |
| **A_65_P03728** | St3gal1 | -0.058049232 | -0.609953319 | 0.493854855 | 12.66385118 | -0.255573638 | 0.806576282 | 0.926442201 | -4.897093665 |
| **A_55_P2130264** | NA | -0.02334861 | -0.245383611 | 0.198686391 | 16.73580119 | -0.255518758 | 0.806616783 | 0.926442201 | -4.897101241 |
| **A_51_P417720** | Itga11 | 0.021516756 | -0.183273618 | 0.22630713 | 8.26030782 | 0.255299781 | 0.806778393 | 0.926572761 | -4.897131452 |
| **A_55_P2063646** | Krcc1 | -0.033799549 | -0.355684454 | 0.288085356 | 12.46370264 | -0.255148775 | 0.806889845 | 0.92662275 | -4.897152271 |
| **A_55_P1998801** | NA | 0.017748923 | -0.151305358 | 0.186803204 | 9.499462841 | 0.255110902 | 0.806917799 | 0.92662275 | -4.89715749 |
| **A_51_P314285** | Tmem86a | -0.037790908 | -0.398072133 | 0.322490317 | 10.00714435 | -0.254875966 | 0.807091208 | 0.926708705 | -4.897189852 |
| **A_55_P1997105** | Large2 | -0.018481843 | -0.194709586 | 0.157745901 | 10.04343125 | -0.254832125 | 0.807123569 | 0.926708705 | -4.897195888 |
| **A_66_P118863** | Ncald | -0.037065724 | -0.390518438 | 0.316386989 | 8.451142221 | -0.254814632 | 0.807136482 | 0.926708705 | -4.897198296 |
| **A_55_P2025954** | Acly | -0.05346631 | -0.563893177 | 0.456960557 | 14.22300207 | -0.254524665 | 0.807350531 | 0.926838772 | -4.89723819 |
| **A_55_P2002572** | Ephx2 | 0.091677861 | -0.783864009 | 0.96721973 | 11.37117097 | 0.254431419 | 0.807419367 | 0.926838772 | -4.897251009 |
| **A_55_P1999852** | Tchp | 0.020967618 | -0.179292756 | 0.221227993 | 8.49703177 | 0.254411811 | 0.807433842 | 0.926838772 | -4.897253704 |
| **A_55_P2153391** | NA | 0.045455175 | -0.388787345 | 0.479697696 | 7.834764344 | 0.254351428 | 0.80747842 | 0.926838772 | -4.897262002 |
| **A_51_P377508** | Srebf2 | -0.020785057 | -0.219360663 | 0.177790549 | 8.543600631 | -0.254336393 | 0.807489519 | 0.926838772 | -4.897264068 |
| **A_55_P2015681** | AU015836 | 0.019738645 | -0.16896102 | 0.20843831 | 7.79742588 | 0.254172989 | 0.807610156 | 0.926849452 | -4.897286514 |
| **A_55_P1988108** | Mrc1 | 0.060082014 | -0.514363854 | 0.634527881 | 11.11350361 | 0.254143239 | 0.80763212 | 0.926849452 | -4.897290599 |
| **A_51_P371876** | Zmynd8 | 0.034142904 | -0.292379892 | 0.360665701 | 10.13000638 | 0.254079801 | 0.807678957 | 0.926849452 | -4.897299308 |
| **A_55_P1966508** | Hoxc6 | -0.017469104 | -0.184592238 | 0.14965403 | 7.805126152 | -0.253990367 | 0.807744988 | 0.926849452 | -4.897311582 |
| **A_55_P1969042** | Cops6 | -0.019250845 | -0.203424135 | 0.164922445 | 13.05617103 | -0.253983987 | 0.807749698 | 0.926849452 | -4.897312458 |
| **A_55_P2326260** | 4930525G20Rik | 0.017218322 | -0.147542136 | 0.18197878 | 7.949265487 | 0.253934101 | 0.807786531 | 0.926849452 | -4.897319303 |
| **A_51_P117226** | NA | -0.019631848 | -0.207618814 | 0.168355118 | 8.372662198 | -0.253756182 | 0.807917899 | 0.926945158 | -4.885254008 |
| **A_55_P1957393** | NA | -0.036685382 | -0.388417255 | 0.31504649 | 13.62061329 | -0.253433787 | 0.808155961 | 0.927163258 | -4.897387877 |

| | | | | | | | | |
|---|---|---|---|---|---|---|---|---|
| **A_52_P622434** | Nop16 | -0.030986503 | -0.328198887 | 0.26622588 | 9.694155463 | -0.253331304 | 0.808231641 | 0.927195049 | -4.897401908 |
| **A_55_P1970380** | NA | 0.018473141 | -0.158798524 | 0.195744806 | 8.434399014 | 0.253212194 | 0.808319602 | 0.927231005 | -4.897418207 |
| **A_55_P2071811** | Eml2 | 0.027141189 | -0.23337901 | 0.287661388 | 8.561888144 | 0.253145757 | 0.808368666 | 0.927231005 | -4.897427296 |
| **A_51_P382253** | Rnf2 | 0.033530526 | -0.288384846 | 0.355445897 | 10.22287171 | 0.253093991 | 0.808406896 | 0.927231005 | -4.897434376 |
| **A_51_P467233** | Polr3gl | -0.029440434 | -0.312243377 | 0.253362509 | 11.00117407 | -0.252955116 | 0.808509461 | 0.927293619 | -4.897453363 |
| **A_66_P126999** | Fis1 | 0.022135384 | -0.190574137 | 0.234844906 | 14.10815873 | 0.25286186 | 0.808578336 | 0.927298302 | -4.897466107 |
| **A_51_P361201** | Atp6v0d1 | -0.03800351 | -0.403258396 | 0.327251375 | 11.00818703 | -0.252819676 | 0.808609492 | 0.927298302 | -4.89747187 |
| **A_51_P246543** | Ebf3 | -0.018024038 | -0.191353062 | 0.155304986 | 7.711263646 | -0.252675998 | 0.808715612 | 0.927307138 | -4.897491492 |
| **A_55_P2329680** | Dyrk2 | -0.036109278 | -0.383458475 | 0.31123992 | 9.465409109 | -0.252601362 | 0.80877074 | 0.927307138 | -4.897501681 |
| **A_55_P1955532** | Tfr2 | 0.067059638 | -0.578313452 | 0.712432729 | 12.71547891 | 0.252483815 | 0.808857565 | 0.927307138 | -4.897517723 |
| **A_55_P2178678** | P2rx5 | -0.017054232 | -0.181205927 | 0.147097463 | 7.835373563 | -0.252446868 | 0.808884856 | 0.927307138 | -4.897522763 |
| **A_66_P116252** | Dusp16 | -0.027012856 | -0.287130583 | 0.23310487 | 10.13984873 | -0.252338635 | 0.808964805 | 0.927307138 | -4.897537525 |
| **A_51_P208263** | NA | 0.045203665 | -0.390242825 | 0.480650154 | 15.01932615 | 0.252244694 | 0.809034199 | 0.927307138 | -4.897550332 |
| **A_55_P2177347** | NA | 0.016957771 | -0.146409413 | 0.180324955 | 7.978779124 | 0.252224421 | 0.809049174 | 0.927307138 | -4.897553095 |
| **A_66_P119034** | Pla2g7 | -0.049622388 | -0.527765828 | 0.428521051 | 11.9173869 | -0.252175361 | 0.809085416 | 0.927307138 | -4.897559781 |
| **A_51_P510722** | Rita1 | -0.018691737 | -0.198822785 | 0.161439312 | 8.447330625 | -0.252141463 | 0.809110457 | 0.927307138 | -4.897564401 |
| **A_55_P2097176** | Idua | 0.065143007 | -0.562840517 | 0.69312653 | 10.62512373 | 0.252059308 | 0.809171148 | 0.927307138 | -4.897575593 |
| **A_55_P2028883** | Rnft2 | -0.018208921 | -0.193748506 | 0.157330664 | 7.883554475 | -0.25205327 | 0.809175609 | 0.927307138 | -4.897576415 |
| **A_51_P104418** | Dusp10 | 0.019216379 | -0.166052613 | 0.204485372 | 8.152913845 | 0.252029869 | 0.809192897 | 0.927307138 | -4.897579603 |
| **A_52_P130787** | Kcna6 | -0.026759792 | -0.284841755 | 0.231322171 | 8.021718919 | -0.251946464 | 0.809254513 | 0.92730756 | -4.89759096 |
| **A_52_P127325** | AI182371 | -0.06888107 | -0.733321088 | 0.595558948 | 9.563786991 | -0.251899492 | 0.809289214 | 0.92730756 | -4.897597355 |
| **A_55_P2000102** | Mettl21a | 0.021751135 | -0.188155774 | 0.231658043 | 9.835604887 | 0.251789938 | 0.809370152 | 0.927339925 | -4.897612265 |
| **A_55_P2083954** | Scfd1 | -0.030684248 | -0.326932106 | 0.265563611 | 10.02944581 | -0.251676955 | 0.809453626 | 0.927339925 | -4.897627636 |
| **A_55_P2005753** | Manbal | 0.032610444 | -0.282365122 | 0.347586009 | 10.08087325 | 0.251572422 | 0.809530858 | 0.927339925 | -4.897641851 |
| **A_55_P2181104** | Shisa3 | 0.016922679 | -0.146561412 | 0.18040677 | 7.668163253 | 0.251522478 | 0.80956776 | 0.927339925 | -4.89764864 |
| **A_51_P463860** | Fam161b | 0.016060499 | -0.139097795 | 0.171218794 | 7.84603751 | 0.251516938 | 0.809571853 | 0.927339925 | -4.897649394 |
| **A_55_P2070039** | Zfyve9 | 0.017452685 | -0.151185695 | 0.186091065 | 8.033440253 | 0.251471643 | 0.80960532 | 0.927339925 | -4.89765555 |
| **A_55_P2142033** | Clip4 | -0.016429758 | -0.17534454 | 0.142485024 | 7.689102038 | -0.251217593 | 0.809793037 | 0.927493177 | -4.89769006 |
| **A_55_P1984333** | Emd | 0.037517197 | -0.325445428 | 0.400479822 | 8.688145419 | 0.25116069 | 0.809835084 | 0.927493177 | -4.897697785 |
| **A_55_P2078776** | Kmt2e | 0.017895032 | -0.155620836 | 0.1914109 | 10.22114324 | 0.250597349 | 0.810251392 | 0.927815362 | -4.897774171 |
| **A_55_P2172430** | Zxdb | 0.024012204 | -0.208842099 | 0.256866508 | 8.886651621 | 0.250571083 | 0.810270804 | 0.927815362 | -4.897777728 |
| **A_55_P2065501** | Rnf7 | 0.038228158 | -0.332526081 | 0.408982397 | 12.44396387 | 0.25054194 | 0.810292343 | 0.927815362 | -4.897781675 |
| **A_51_P141860** | Fbxw11 | -0.027912605 | -0.298691486 | 0.242866276 | 11.74894428 | -0.250477527 | 0.810339949 | 0.927815362 | -4.897790396 |
| **A_52_P260629** | Cpne3 | -0.031993025 | -0.342383467 | 0.278397416 | 9.154642979 | -0.250455262 | 0.810356404 | 0.927815362 | -4.897793411 |
| **A_55_P2008889** | Tmsb15b2 | 0.021659985 | -0.188582966 | 0.231902937 | 9.375520912 | 0.250334037 | 0.810446002 | 0.927839274 | -4.897809817 |
| **A_55_P2026291** | Smarce1 | 0.032606918 | -0.28393928 | 0.349153116 | 11.48832008 | 0.250297112 | 0.810473294 | 0.927839274 | -4.897814813 |
| **A_51_P276235** | Pnpla7 | 0.039850984 | -0.347235635 | 0.426937603 | 9.12207159 | 0.250157819 | 0.810576251 | 0.927902182 | -4.897833653 |
| **A_55_P2112947** | Srp9 | 0.048548574 | -0.423300032 | 0.52039718 | 9.739384478 | 0.250009776 | 0.810685679 | 0.927972491 | -4.897853665 |
| **A_55_P1973748** | Nrbp1 | -0.034828955 | -0.373442264 | 0.303784353 | 10.00177267 | -0.249930705 | 0.810744127 | 0.927975077 | -4.897864349 |
| **A_55_P2157620** | NA | 0.051924848 | -0.453006627 | 0.556856322 | 9.847381634 | 0.249876824 | 0.810783957 | 0.927975077 | -4.897871628 |
| **A_52_P287052** | Znrd2 | -0.026239569 | -0.281677794 | 0.229198655 | 9.839248189 | -0.249605414 | 0.810984594 | 0.928086121 | -4.897908268 |
| **A_51_P431737** | Cth | 0.082813975 | -0.723862221 | 0.889490172 | 11.87935445 | 0.249452339 | 0.81109776 | 0.928086121 | -4.897928915 |
| **A_55_P1965989** | Zfp329 | -0.026088396 | -0.280249876 | 0.228073083 | 8.44190299 | -0.249414011 | 0.811126096 | 0.928086121 | -4.897934083 |
| **A_55_P2019233** | Gsto2 | -0.019573604 | -0.210271897 | 0.17112469 | 7.867616054 | -0.249406156 | 0.811131903 | 0.928086121 | -4.897935142 |
| **A_55_P2131875** | Poldip3 | -0.02668075 | -0.286633345 | 0.233271846 | 12.77046681 | -0.249394606 | 0.811140442 | 0.928086121 | -4.8979367 |
| **A_55_P2055369** | NA | 0.031763447 | -0.277843501 | 0.341370394 | 8.770340191 | 0.249287277 | 0.811219793 | 0.928086121 | -4.897951166 |
| **A_51_P323299** | Trim28 | -0.023029541 | -0.24755611 | 0.201497028 | 11.26045407 | -0.249230239 | 0.811261964 | 0.928086121 | -4.897958852 |
| **A_51_P352264** | Sirt7 | 0.03083915 | -0.269832333 | 0.331510632 | 11.84041729 | 0.249225997 | 0.811265099 | 0.928086121 | -4.897959423 |
| **A_55_P2055682** | Atp5o | 0.017495859 | -0.153506985 | 0.188498703 | 13.2995653 | 0.248608008 | 0.811722049 | 0.928553914 | -4.898042584 |
| **A_51_P415945** | Psma2 | 0.022828893 | -0.200460869 | 0.246118654 | 14.76501132 | 0.248427248 | 0.811855721 | 0.92864793 | -4.89806687 |

| | | | | | | | | | |
|---|---|---|---|---|---|---|---|---|---|
| **A_66_P140452** | Map1a | -0.017880417 | -0.192811525 | 0.157050691 | 7.690533323 | -0.248366935 | 0.811900324 | 0.92864793 | -4.89807497 |
| **A_51_P107433** | Mrpl34 | 0.021733069 | -0.190949152 | 0.23441529 | 12.23984901 | 0.248297905 | 0.811951374 | 0.928651368 | -4.886517886 |
| **A_55_P2049262** | NA | -0.028046146 | -0.302626576 | 0.246534284 | 9.098305043 | -0.24819144 | 0.812030111 | 0.92868647 | -4.898098527 |
| **A_55_P2164909** | Gatd3a | -0.028234805 | -0.305013489 | 0.248543879 | 12.82511125 | -0.247876494 | 0.812263046 | 0.928897907 | -4.898140763 |
| **A_55_P2014209** | Scyl2 | 0.029125417 | -0.256831645 | 0.31508248 | 8.456645996 | 0.247488213 | 0.812550248 | 0.929082137 | -4.898192761 |
| **A_55_P1986706** | Pih1d1 | -0.024447909 | -0.26451399 | 0.215618172 | 11.88975594 | -0.247453788 | 0.812575712 | 0.929082137 | -4.898197367 |
| **A_51_P497563** | Vps18 | -0.016250383 | -0.175829622 | 0.143328856 | 9.694312527 | -0.247440275 | 0.812585708 | 0.929082137 | -4.898199175 |
| **A_55_P1953920** | Me3 | -0.01890437 | -0.204576988 | 0.166768248 | 7.692749375 | -0.24739877 | 0.812616411 | 0.929082137 | -4.898204728 |
| **A_55_P1989296** | Tnxb | -0.025070848 | -0.271571678 | 0.221429982 | 8.377345019 | -0.247134759 | 0.812811717 | 0.929250468 | -4.898240026 |
| **A_51_P399625** | Cnih1 | 0.021006046 | -0.185737602 | 0.227749695 | 13.44763614 | 0.246885354 | 0.812996231 | 0.929406443 | -4.898273338 |
| **A_52_P206526** | Gtf2ird2 | -0.020979379 | -0.227651661 | 0.185692902 | 9.327908512 | -0.246657084 | 0.81316512 | 0.929516993 | -4.898303799 |
| **A_55_P2003309** | NA | 0.031886394 | -0.28232886 | 0.346101647 | 13.84754563 | 0.246581977 | 0.813220691 | 0.929516993 | -4.898313815 |
| **A_51_P394921** | 1700020L24Rik | 0.042696273 | -0.378354691 | 0.463747237 | 10.57449138 | 0.246398743 | 0.813356271 | 0.929516993 | -4.898338239 |
| **A_55_P2071970** | Nav2 | -0.016804199 | -0.182560379 | 0.14895198 | 8.071208451 | -0.246337928 | 0.813401271 | 0.929516993 | -4.898346342 |
| **A_55_P2103055** | Gsr | 0.042323506 | -0.375193521 | 0.459840533 | 11.50668415 | 0.246314872 | 0.813418332 | 0.929516993 | -4.898349413 |
| **A_51_P190886** | Ankrd13a | 0.02198631 | -0.194919606 | 0.238892226 | 10.6694088 | 0.246299844 | 0.813429452 | 0.929516993 | -4.898351415 |
| **A_55_P2009988** | Trib3 | -0.048883637 | -0.531161512 | 0.433394238 | 10.49803824 | -0.246291465 | 0.813435652 | 0.929516993 | -4.898352531 |
| **A_55_P2120866** | NA | 0.056665527 | -0.502830157 | 0.616161212 | 8.708811527 | 0.246096469 | 0.813579946 | 0.929516993 | -4.898378492 |
| **A_55_P1953301** | Sord | 0.070229441 | -0.623220417 | 0.763679298 | 13.3432016 | 0.246086243 | 0.813587514 | 0.929516993 | -4.898379853 |
| **A_55_P2053541** | NA | -0.040168399 | -0.436868371 | 0.356531574 | 13.57124167 | -0.246039899 | 0.813621809 | 0.929516993 | -4.89838602 |
| **A_51_P438149** | Mapre2 | -0.053222422 | -0.579161138 | 0.472716293 | 10.45270302 | -0.24589103 | 0.813731978 | 0.929516993 | -4.898405822 |
| **A_52_P404461** | Tmem60 | 0.018898994 | -0.16788558 | 0.205683568 | 10.75182255 | 0.245856029 | 0.81375788 | 0.929516993 | -4.898410477 |
| **A_55_P1992174** | Tars2 | 0.027069482 | -0.240499208 | 0.294638172 | 9.544936596 | 0.245826014 | 0.813780093 | 0.929516993 | -4.898414467 |
| **A_51_P407323** | F5 | -0.062019676 | -0.675117113 | 0.551077762 | 9.961965036 | -0.245800736 | 0.8137988 | 0.929516993 | -4.898417827 |
| **A_51_P126198** | Dapk2 | 0.020315229 | -0.180528687 | 0.221159146 | 11.08911528 | 0.245779831 | 0.813814272 | 0.929516993 | -4.898420606 |
| **A_55_P1985012** | Pard3 | 0.029028735 | -0.25818286 | 0.31624033 | 8.408390189 | 0.245589241 | 0.813955326 | 0.929614927 | -4.89844593 |
| **A_55_P2177721** | Adra1b | 0.022637067 | -0.20138546 | 0.246659595 | 10.4130623 | 0.24553401 | 0.813996204 | 0.929614927 | -4.898453265 |
| **A_55_P1997061** | NA | -0.054094435 | -0.589923809 | 0.481734939 | 15.20732099 | -0.245306614 | 0.814164511 | 0.929659462 | -4.898483448 |
| **A_52_P309718** | Tasor | -0.0224028 | -0.244322558 | 0.199516959 | 8.686785981 | -0.245295461 | 0.814172767 | 0.929659462 | -4.898484928 |
| **A_55_P2093920** | Pzp | -0.030224288 | -0.329775089 | 0.269326513 | 15.81766669 | -0.245170821 | 0.814265024 | 0.929659462 | -4.898501459 |
| **A_55_P2171976** | NA | 0.027103632 | -0.24161995 | 0.295827214 | 15.34170207 | 0.245078325 | 0.81433349 | 0.929659462 | -4.898513722 |
| **A_55_P2228918** | Rslcan18 | -0.023713638 | -0.25884604 | 0.211418765 | 8.633977121 | -0.245058002 | 0.814348534 | 0.929659462 | -4.898516416 |
| **A_55_P1989125** | Rps17 | -0.020193826 | -0.220449042 | 0.180061389 | 15.71579636 | -0.245029272 | 0.814369801 | 0.929659462 | -4.898520224 |
| **A_55_P2007771** | Catsper2 | 0.016655911 | -0.148516911 | 0.181828734 | 7.860554186 | 0.245026467 | 0.814371877 | 0.929659462 | -4.898520595 |
| **A_55_P2127139** | H3c4 | 0.041596648 | -0.371092039 | 0.454285335 | 14.7898422 | 0.244917017 | 0.814452897 | 0.929697044 | -4.898535097 |
| **A_52_P1114640** | 2410003L11Rik | 0.016695603 | -0.14910255 | 0.182493756 | 7.7120301 | 0.244684026 | 0.814625375 | 0.929839014 | -4.898565947 |
| **A_55_P2084002** | Aptx | 0.016520166 | -0.147670293 | 0.180710626 | 8.080687329 | 0.244483577 | 0.814773772 | 0.929953483 | -4.898592465 |
| **A_55_P2018492** | Pgap3 | -0.016939566 | -0.185354767 | 0.151475634 | 9.35310926 | -0.244401684 | 0.814834402 | 0.929967771 | -4.898603293 |
| **A_52_P149835** | Irak1 | -0.022738768 | -0.249287855 | 0.203810319 | 8.500066602 | -0.243886522 | 0.815215836 | 0.930348167 | -4.898671326 |
| **A_55_P1967820** | AI661453 | -0.028555963 | -0.313494356 | 0.25638243 | 8.750940655 | -0.243516855 | 0.815489578 | 0.93056145 | -4.898720059 |
| **A_55_P2185504** | Masp2 | -0.048991708 | -0.537868554 | 0.439885137 | 11.51186539 | -0.243504115 | 0.815499012 | 0.93056145 | -4.898721738 |
| **A_55_P2094852** | Rabep1 | 0.03844889 | -0.345361313 | 0.422259094 | 9.653346361 | 0.243416763 | 0.815563701 | 0.930580329 | -4.898733242 |
| **A_55_P2016667** | Gstm7 | -0.043200263 | -0.474721666 | 0.388321139 | 8.91801216 | -0.243258063 | 0.815681232 | 0.930659497 | -4.898754133 |
| **A_52_P68477** | Zfp809 | 0.019999465 | -0.179830851 | 0.219829781 | 8.705739217 | 0.243186906 | 0.815733932 | 0.930664689 | -4.898763496 |
| **A_52_P413646** | Bmp6 | 0.017309836 | -0.15579747 | 0.190417141 | 8.062520959 | 0.242974523 | 0.81589123 | 0.93078921 | -4.898791425 |
| **A_55_P1967533** | Gm5069 | 0.025980261 | -0.234120447 | 0.28608097 | 12.58770035 | 0.242708604 | 0.816088192 | 0.930958963 | -4.898826361 |
| **A_55_P2059556** | NA | 0.018567429 | -0.167412288 | 0.204547146 | 15.96975017 | 0.242588038 | 0.816177499 | 0.931005894 | -4.898842189 |
| **A_55_P2067583** | Cd68 | -0.01797579 | -0.198132138 | 0.162180559 | 7.902241504 | -0.242449668 | 0.816279997 | 0.93106299 | -4.898860344 |
| **A_52_P244132** | Isoc1 | 0.028530289 | -0.257474901 | 0.314535479 | 10.84285345 | 0.242390414 | 0.816323891 | 0.93106299 | -4.898868116 |
| **A_51_P155501** | Alg3 | 0.026096932 | -0.235740788 | 0.287934652 | 8.403546326 | 0.242181204 | 0.816478874 | 0.931065741 | -4.89889554 |

| | | | | | | | | | |
|---|---|---|---|---|---|---|---|---|---|
| **A_51_P247694** | Adgrg3 | 0.040529428 | -0.366272864 | 0.447331721 | 10.09132586 | 0.242086335 | 0.816549155 | 0.931065741 | -4.898907969 |
| **A_52_P302345** | Cyp4v3 | -0.043929399 | -0.484940543 | 0.397081745 | 10.2467245 | -0.242040971 | 0.816582764 | 0.931065741 | -4.89891391 |
| **A_55_P2121701** | Mir670hg | -0.015342392 | -0.169392715 | 0.138707931 | 7.65611415 | -0.241999043 | 0.816613826 | 0.931065741 | -4.8989194 |
| **A_66_P117058** | Aasdh | 0.02311768 | -0.209010617 | 0.255245977 | 9.047411915 | 0.241991074 | 0.81661973 | 0.931065741 | -4.898920444 |
| **A_51_P267354** | Lrfn3 | -0.038462931 | -0.424900659 | 0.347974798 | 10.22330225 | -0.241849971 | 0.81672427 | 0.931065741 | -4.898938913 |
| **A_55_P2274782** | Prl7a1 | 0.015286794 | -0.138302351 | 0.168875938 | 7.619456443 | 0.241846088 | 0.816727147 | 0.931065741 | -4.898939421 |
| **A_52_P399677** | Tprkb | -0.030538354 | -0.337429739 | 0.276353031 | 8.803659193 | -0.241793218 | 0.816766318 | 0.931065741 | -4.898946339 |
| **A_55_P2005330** | Egln2 | -0.02110445 | -0.233238471 | 0.191029572 | 10.3547524 | -0.241739096 | 0.816806418 | 0.931065741 | -4.898953419 |
| **A_55_P2137887** | Mpdz | -0.019395792 | -0.214386903 | 0.175595318 | 7.894430652 | -0.241699583 | 0.816835693 | 0.931065741 | -4.898958587 |
| **A_66_P130469** | Rce1 | -0.021981535 | -0.243045193 | 0.199082124 | 8.658804691 | -0.241614973 | 0.816898384 | 0.931065741 | -4.89896965 |
| **A_51_P339822** | Arhgap4 | 0.045258711 | -0.409944817 | 0.500462239 | 9.72174456 | 0.241590444 | 0.816916558 | 0.931065741 | -4.898972857 |
| **A_55_P2144115** | 1700125H20Rik | 0.016657424 | -0.150913433 | 0.184228281 | 7.711794919 | 0.241541929 | 0.816952505 | 0.931065741 | -4.898979198 |
| **A_51_P372550** | Cgref1 | -0.041363986 | -0.457630161 | 0.374902189 | 10.35736689 | -0.241454027 | 0.817017638 | 0.931074688 | -4.898990684 |
| **A_55_P2412349** | AW111846 | -0.074640322 | -0.825946676 | 0.676666031 | 11.86976111 | -0.241401316 | 0.817056695 | 0.931074688 | -4.89899757 |
| **A_51_P488554** | Mrnip | -0.037195411 | -0.412162423 | 0.3377716 | 9.993312973 | -0.241034658 | 0.817328396 | 0.931329398 | -4.899045428 |
| **A_51_P392244** | Tbcd | 0.020883147 | -0.189869811 | 0.231636106 | 11.03526144 | 0.240771714 | 0.817523261 | 0.931404453 | -4.899079705 |
| **A_55_P2128597** | Nr1h3 | -0.025072703 | -0.278142276 | 0.227996869 | 11.11376581 | -0.240737873 | 0.817548341 | 0.931404453 | -4.899084114 |
| **A_55_P2032222** | Tm9sf2 | -0.036219853 | -0.401808344 | 0.329368638 | 10.95755515 | -0.240733957 | 0.817551243 | 0.931404453 | -4.899084624 |
| **A_55_P2156588** | Ifnk | 0.016751193 | -0.152362456 | 0.185864843 | 7.805294528 | 0.240685695 | 0.817587012 | 0.931404453 | -4.899090911 |
| **A_52_P649064** | NA | -0.019399334 | -0.215342708 | 0.17654404 | 13.47445561 | -0.24056887 | 0.817673595 | 0.931448192 | -4.899106123 |
| **A_51_P435251** | Cul4a | 0.027867198 | -0.253741945 | 0.309476341 | 10.66432544 | 0.240452747 | 0.817759662 | 0.931474573 | -4.899121237 |
| **A_55_P2073855** | Mrpl27 | -0.023300869 | -0.25880967 | 0.212207932 | 12.45593697 | -0.240407586 | 0.817793134 | 0.931474573 | -4.899127113 |
| **A_55_P2096942** | Nrn1l | 0.016959765 | -0.154572313 | 0.188491842 | 7.749993377 | 0.240246828 | 0.817912289 | 0.931555396 | -4.899148021 |
| **A_52_P531651** | Parvg | 0.037317466 | -0.34064827 | 0.415283203 | 9.069370645 | 0.239906991 | 0.818164195 | 0.931753465 | -4.899192174 |
| **A_55_P2006118** | Rbp4 | 0.048872273 | -0.446296628 | 0.544041173 | 16.7790979 | 0.239823846 | 0.81822583 | 0.931753465 | -4.899202967 |
| **A_51_P142421** | Rspo1 | 0.07997407 | -0.730337775 | 0.890285915 | 9.278724164 | 0.239817129 | 0.81823081 | 0.931753465 | -4.899203839 |
| **A_55_P2067513** | Slc10a3 | 0.017999816 | -0.164587814 | 0.200587446 | 9.461838426 | 0.239541026 | 0.818435495 | 0.931931645 | -4.899239654 |
| **A_52_P198019** | Fuz | 0.022789064 | -0.208493656 | 0.254071784 | 8.381836863 | 0.239423344 | 0.818522742 | 0.931976088 | -4.899254907 |
| **A_51_P152404** | Ikbip | -0.023407369 | -0.261056137 | 0.214241398 | 9.390161685 | -0.239331704 | 0.818590684 | 0.931998546 | -4.89926678 |
| **A_52_P505157** | Fsd2 | -0.016523836 | -0.184471162 | 0.15142349 | 7.716128239 | -0.239067736 | 0.818786399 | 0.932166468 | -4.899300954 |
| **A_55_P2005808** | Abhd18 | 0.03751131 | -0.344251838 | 0.419274459 | 9.510964423 | 0.238754421 | 0.81901872 | 0.932316202 | -4.899341469 |
| **A_55_P1971329** | Lage3 | 0.036617365 | -0.336167134 | 0.409401864 | 10.49709152 | 0.238678025 | 0.81907537 | 0.932316202 | -4.89935134 |
| **A_51_P445473** | Fut7 | 0.030726299 | -0.28211469 | 0.343567288 | 8.914179551 | 0.238654593 | 0.819092746 | 0.932316202 | -4.899354367 |
| **A_55_P1973360** | NA | -0.024885192 | -0.278280655 | 0.228510271 | 14.85332262 | -0.23863017 | 0.819110856 | 0.932316202 | -4.899357522 |
| **A_55_P2258832** | 5230400M03Rik | -0.033875264 | -0.379383095 | 0.311632567 | 8.498809985 | -0.238236327 | 0.819402927 | 0.932593722 | -4.899408352 |
| **A_52_P633379** | Plpbp | -0.019662816 | -0.220365431 | 0.181039798 | 10.94031097 | -0.238054219 | 0.819537988 | 0.932659474 | -4.899431827 |
| **A_52_P612693** | Polb | -0.027579187 | -0.309116545 | 0.253958172 | 9.520224893 | -0.238028311 | 0.819557203 | 0.932659474 | -4.899435165 |
| **A_55_P1962303** | Hba-a1 | -0.02582516 | -0.289535271 | 0.237884951 | 18.38057703 | -0.237957516 | 0.81960971 | 0.932664317 | -4.899444286 |
| **A_55_P2034110** | Tgm2 | -0.031026055 | -0.348372683 | 0.286320574 | 11.99922802 | -0.237561388 | 0.819903529 | 0.932891273 | -4.89949527 |
| **A_55_P2029294** | Cetn4 | -0.034332905 | -0.385507585 | 0.316841775 | 8.282326856 | -0.237558484 | 0.819905683 | 0.932891273 | -4.899495643 |
| **A_55_P2114110** | Cadm4 | -0.017637025 | -0.198215236 | 0.162941187 | 8.015611278 | -0.237324822 | 0.820079013 | 0.932964141 | -4.899525678 |
| **A_51_P103975** | Mpc2 | 0.035881292 | -0.331537663 | 0.403300247 | 14.14658435 | 0.237295611 | 0.820100681 | 0.932964141 | -4.899529431 |
| **A_52_P252425** | 2700062C07Rik | 0.019390577 | -0.179226117 | 0.218007272 | 8.986378527 | 0.237223758 | 0.820153984 | 0.932964141 | -4.89953866 |
| **A_55_P2010487** | Unk | -0.019997141 | -0.224884479 | 0.184890197 | 10.57586693 | -0.237157019 | 0.820203494 | 0.932964141 | -4.899547229 |
| **A_51_P169087** | Gls2 | -0.09185742 | -1.033053557 | 0.849338716 | 11.3456876 | -0.237146816 | 0.820211063 | 0.932964141 | -4.899548539 |
| **A_55_P2048110** | Tmem97 | 0.041797682 | -0.386897457 | 0.47049282 | 12.73609884 | 0.236911871 | 0.820385363 | 0.93310749 | -4.899578688 |
| **A_51_P487228** | B9d2 | -0.028889409 | -0.325333214 | 0.267554397 | 12.11972654 | -0.236798779 | 0.820469267 | 0.933148012 | -4.89959319 |
| **A_51_P443569** | Pwwp3a | -0.01912427 | -0.215593408 | 0.177344868 | 9.552172086 | -0.236523182 | 0.820673747 | 0.93318229 | -4.899628501 |
| **A_55_P2028576** | NA | 0.033094011 | -0.306979637 | 0.373167659 | 12.34931691 | 0.236461049 | 0.820719848 | 0.93318229 | -4.899636457 |
| **A_51_P363914** | Cct2 | 0.016830216 | -0.156128382 | 0.189788815 | 13.57915272 | 0.236445323 | 0.820731518 | 0.93318229 | -4.89963847 |

| | | | | | | | | | |
|---|---|---|---|---|---|---|---|---|---|
| **A_55_P2066518** | Akt2 | -0.016766279 | -0.189074215 | 0.155541657 | 12.36435434 | -0.236436538 | 0.820738036 | 0.93318229 | -4.899639594 |
| **A_51_P246653** | Clec7a | -0.021302249 | -0.240328035 | 0.197723538 | 9.84080314 | -0.236327027 | 0.820819294 | 0.93318229 | -4.89965361 |
| **A_51_P226853** | Gga1 | -0.021499852 | -0.242611419 | 0.199611714 | 8.579801686 | -0.23626925 | 0.820862165 | 0.93318229 | -4.899661001 |
| **A_55_P1975045** | Sgcg | 0.016833031 | -0.156288036 | 0.189954098 | 7.703461824 | 0.236262936 | 0.820866851 | 0.93318229 | -4.899661809 |
| **A_55_P2205650** | 2610507I01Rik | 0.016856942 | -0.15652862 | 0.190242505 | 9.488058338 | 0.236237619 | 0.820885637 | 0.93318229 | -4.899665048 |
| **A_52_P419373** | Lrrc39 | 0.015420126 | -0.143409805 | 0.174250058 | 7.786862903 | 0.235905884 | 0.821131807 | 0.933344968 | -4.899707449 |
| **A_55_P2150179** | C1s2 | 0.075112106 | -0.698597068 | 0.848821279 | 13.13754852 | 0.235893178 | 0.821141237 | 0.933344968 | -4.899709072 |
| **A_51_P342906** | Sh3bp1 | 0.024474502 | -0.227753926 | 0.27670293 | 9.199232663 | 0.235777857 | 0.821226819 | 0.933344968 | -4.899723798 |
| **A_51_P512172** | Tet1 | 0.030355477 | -0.282520052 | 0.343231005 | 8.310076024 | 0.235748343 | 0.821248722 | 0.933344968 | -4.899727565 |
| **A_52_P495031** | 9130401M01Rik | -0.019067699 | -0.215623604 | 0.177488205 | 11.26241546 | -0.235719435 | 0.821270175 | 0.933344968 | -4.899731255 |
| **A_55_P2186439** | Chtop | 0.016238438 | -0.15122559 | 0.183702466 | 10.65680046 | 0.235616617 | 0.821346482 | 0.933376809 | -4.899744375 |
| **A_52_P671062** | Uba6 | -0.019171687 | -0.216948791 | 0.178605416 | 7.922015397 | -0.235541541 | 0.821402201 | 0.933385253 | -4.899753952 |
| **A_51_P434198** | Anapc4 | -0.034367792 | -0.389070599 | 0.320335015 | 12.34113127 | -0.235434549 | 0.821481609 | 0.933420612 | -4.899767594 |
| **A_55_P2142873** | Mink1 | -0.026610052 | -0.30136475 | 0.248144646 | 10.59082822 | -0.235333517 | 0.821556596 | 0.933450943 | -4.899780471 |
| **A_55_P2061084** | Wfikkn2 | 0.016226346 | -0.151388806 | 0.183841497 | 7.816903817 | 0.235228878 | 0.821634262 | 0.933464454 | -4.899793802 |
| **A_52_P401640** | Bpgm | -0.028706227 | -0.325288583 | 0.26787613 | 12.76233291 | -0.235187366 | 0.821665074 | 0.933464454 | -4.899799089 |
| **A_51_P331385** | Paip2b | 0.021889556 | -0.204498124 | 0.248277235 | 8.643862367 | 0.234945616 | 0.821844518 | 0.933613441 | -4.899829861 |
| **A_55_P2056236** | Pdlim5 | -0.016036339 | -0.181971582 | 0.149898905 | 7.838775635 | -0.234827942 | 0.821931869 | 0.933615274 | -4.899844829 |
| **A_55_P2164778** | Ptdss1 | 0.017124021 | -0.160129979 | 0.19437802 | 10.61672167 | 0.234743129 | 0.821994828 | 0.933615274 | -4.899855612 |
| **A_55_P2153587** | Afg3l1 | 0.025068502 | -0.234508497 | 0.284645501 | 11.18185229 | 0.2346634 | 0.822054014 | 0.933615274 | -4.899865745 |
| **A_55_P2100655** | Lsm6 | 0.015339894 | -0.14352614 | 0.174205928 | 13.53033133 | 0.23462511 | 0.822082439 | 0.933615274 | -4.89987061 |
| **A_51_P379660** | Mllt11 | -0.020317005 | -0.230832641 | 0.19019863 | 9.139439874 | -0.234508466 | 0.822169032 | 0.933615274 | -4.899885427 |
| **A_51_P430973** | Paqr7 | -0.020078009 | -0.228151029 | 0.18799501 | 9.608805792 | -0.234470426 | 0.822197273 | 0.933615274 | -4.899890257 |
| **A_51_P488768** | Slc25a44 | 0.032525603 | -0.304618307 | 0.369669513 | 8.476142961 | 0.234419225 | 0.822235284 | 0.933615274 | -4.899896758 |
| **A_55_P2092156** | NA | 0.030492663 | -0.285604427 | 0.346589753 | 13.94185433 | 0.23440024 | 0.822249379 | 0.933615274 | -4.899899168 |
| **A_51_P148311** | Chic2 | -0.018446498 | -0.209703608 | 0.172810613 | 10.36032849 | -0.234357858 | 0.822280843 | 0.933615274 | -4.899904547 |
| **A_55_P2053206** | Kdm4c | 0.02480966 | -0.232989601 | 0.282608921 | 9.187174149 | 0.233841903 | 0.822663923 | 0.933945807 | -4.899969959 |
| **A_55_P2002553** | Stx16 | 0.023659172 | -0.22219189 | 0.269510234 | 8.495717537 | 0.233835608 | 0.822668597 | 0.933945807 | -4.899970757 |
| **A_55_P1953630** | E330016L19Rik | 0.015793457 | -0.148506968 | 0.180093883 | 7.652692849 | 0.233572503 | 0.822863964 | 0.934112736 | -4.900004058 |
| **A_55_P1956918** | Adamts5 | -0.015132185 | -0.172756896 | 0.142492525 | 7.753741061 | -0.233270886 | 0.823087946 | 0.934183742 | -4.900042189 |
| **A_52_P191468** | Ubfd1 | -0.021510825 | -0.245620344 | 0.202598694 | 9.411872292 | -0.233227605 | 0.823120088 | 0.934183742 | -4.900047656 |
| **A_51_P203878** | Dynll2 | -0.041853701 | -0.477955262 | 0.39424786 | 10.58572494 | -0.233200465 | 0.823140242 | 0.934183742 | -4.900051084 |
| **A_51_P335306** | Ppm1a | 0.025292956 | -0.238252973 | 0.288838885 | 9.360333121 | 0.233198884 | 0.823141417 | 0.934183742 | -4.900051284 |
| **A_52_P661722** | 2-Mar | 0.051383119 | -0.48409813 | 0.586864367 | 14.72697063 | 0.233162865 | 0.823168167 | 0.934183742 | -4.900055833 |
| **A_52_P425734** | Afm | -0.056422218 | -0.64478275 | 0.531938313 | 13.23537014 | -0.233018185 | 0.823275615 | 0.934234381 | -4.900074098 |
| **A_55_P1953931** | Fam193a | 0.023690577 | -0.223398728 | 0.270779883 | 10.03528448 | 0.232972622 | 0.823309455 | 0.934234381 | -4.900079848 |
| **A_55_P2081353** | NA | 0.025829065 | -0.243738793 | 0.295396924 | 14.37317096 | 0.23282187 | 0.823421419 | 0.934306581 | -4.900098865 |
| **A_55_P2087561** | Eno1b | -0.054395373 | -0.622790202 | 0.513999457 | 10.37819922 | -0.232538587 | 0.823631827 | 0.93442161 | -4.900134567 |
| **A_55_P1965524** | NA | -0.018292049 | -0.209459574 | 0.172875477 | 15.41546277 | -0.232504532 | 0.823657123 | 0.93442161 | -4.900138856 |
| **A_55_P2066505** | Mpo | -0.038621079 | -0.442383905 | 0.365141747 | 11.0688464 | -0.232424155 | 0.823716826 | 0.93442161 | -4.900148977 |
| **A_55_P2175381** | NA | -0.028825393 | -0.330190914 | 0.272540128 | 16.75562016 | -0.232415378 | 0.823723346 | 0.93442161 | -4.900150082 |
| **A_55_P2114482** | Rpgr | 0.018527965 | -0.175225565 | 0.212281496 | 8.704807983 | 0.232359959 | 0.823764511 | 0.93442161 | -4.900157058 |
| **A_55_P2049211** | Pisd-ps1 | 0.017262896 | -0.163422692 | 0.197948483 | 11.12144198 | 0.232152478 | 0.823918635 | 0.934541593 | -4.90018316 |
| **A_66_P112593** | Sec61b | -0.022862956 | -0.26226467 | 0.216538758 | 11.97859327 | -0.232053603 | 0.823992086 | 0.934570064 | -4.900195591 |
| **A_55_P2291776** | 4833447P13Rik | 0.059755553 | -0.566310835 | 0.685821941 | 9.72090215 | 0.231921532 | 0.8240902 | 0.934620729 | -4.900212188 |
| **A_55_P2104835** | NA | 0.019723912 | -0.186977874 | 0.226425697 | 8.057764691 | 0.231863297 | 0.824133463 | 0.934620729 | -4.900219503 |
| **A_51_P339713** | Mrps26 | 0.028966606 | -0.27488588 | 0.332819093 | 10.78133991 | 0.231642376 | 0.824297592 | 0.93466273 | -4.900247238 |
| **A_55_P2030183** | NA | 0.054494406 | -0.575040908 | 0.684029721 | 13.58995776 | 0.236340214 | 0.824304875 | 0.93466273 | -4.76955017 |
| **A_55_P2021125** | NA | 0.02600693 | -0.246827781 | 0.298841641 | 8.465363273 | 0.231618185 | 0.824315565 | 0.93466273 | -4.900250273 |
| **A_55_P1981259** | Fkbp14 | -0.023162602 | -0.266416518 | 0.220091314 | 9.652261416 | -0.231371941 | 0.824498521 | 0.934775374 | -4.900281154 |

| | | | | | | | | | |
|---|---|---|---|---|---|---|---|---|---|
| **A_55_P1970299** | Mttp | 0.065455037 | -0.622006962 | 0.752917035 | 11.14470777 | 0.231354293 | 0.824511633 | 0.934775374 | -4.900283366 |
| **A_55_P2046363** | Mapkap1 | -0.025450092 | -0.292977705 | 0.242077521 | 9.018509948 | -0.231155343 | 0.82465946 | 0.934886723 | -4.90030829 |
| **A_65_P01065** | Phf14 | -0.02486428 | -0.286305651 | 0.23657709 | 10.5260714 | -0.231091927 | 0.824706582 | 0.934886723 | -4.90031623 |
| **A_55_P1960208** | Vcan | -0.01541231 | -0.177647356 | 0.146822736 | 8.32233848 | -0.230837438 | 0.82489569 | 0.935005682 | -4.900348073 |
| **A_55_P1985360** | Man1c1 | -0.027760502 | -0.320030617 | 0.264509613 | 8.528099902 | -0.23079484 | 0.824927345 | 0.935005682 | -4.9003534 |
| **A_51_P361830** | Lgi2 | 0.019188225 | -0.182865137 | 0.221241587 | 8.199107372 | 0.230755418 | 0.824956641 | 0.935005682 | -4.900358329 |
| **A_55_P2169417** | BC021767 | -0.025642228 | -0.296008195 | 0.244723739 | 11.01353042 | -0.230455421 | 0.825179587 | 0.935203532 | -4.900395809 |
| **A_55_P2134785** | Zscan22 | 0.01958803 | -0.187067917 | 0.226243977 | 8.809642067 | 0.230317023 | 0.825282444 | 0.935265265 | -4.900413084 |
| **A_52_P313071** | Gosr1 | 0.0306736 | -0.293419712 | 0.354766912 | 10.30833047 | 0.229973539 | 0.825537739 | 0.93543323 | -4.900455914 |
| **A_55_P2423228** | AI464196 | -0.016181688 | -0.187174054 | 0.154810677 | 7.847710588 | -0.229948348 | 0.825556463 | 0.93543323 | -4.900459053 |
| **A_55_P2166349** | Dnajc19 | -0.031241483 | -0.361408577 | 0.298925611 | 9.961192014 | -0.229922276 | 0.825575843 | 0.93543323 | -4.900462301 |
| **A_55_P2068125** | Myct1 | 0.026263779 | -0.251389653 | 0.30391721 | 8.307944103 | 0.229846212 | 0.825632382 | 0.935442456 | -4.900471775 |
| **A_55_P2005420** | NA | -0.025163201 | -0.291345683 | 0.241019282 | 8.826660092 | -0.22970455 | 0.825737683 | 0.935506926 | -4.900489412 |
| **A_55_P2123101** | Cbarp | -0.043826279 | -0.507852634 | 0.420200075 | 14.27233363 | -0.229496012 | 0.825892701 | 0.935627712 | -4.900515355 |
| **A_51_P401668** | Laptm5 | -0.050593605 | -0.586436285 | 0.485249076 | 9.721575618 | -0.229425407 | 0.825945188 | 0.935632335 | -4.900524133 |
| **A_51_P205481** | Mtmr6 | -0.019842324 | -0.230135315 | 0.190450668 | 10.55441115 | -0.229271945 | 0.826059272 | 0.935706732 | -4.900543204 |
| **A_51_P130660** | Mrps30 | 0.016670549 | -0.160141195 | 0.193482292 | 11.30845291 | 0.229098356 | 0.826188325 | 0.935780341 | -4.900564762 |
| **A_66_P104365** | Ccdc167 | 0.034180421 | -0.328414868 | 0.396775711 | 10.85389854 | 0.229054296 | 0.826221082 | 0.935780341 | -4.900570231 |
| **A_55_P1968878** | Rhpn2 | 0.047650779 | -0.458270969 | 0.553572526 | 10.31965399 | 0.228859973 | 0.826365559 | 0.935814788 | -4.90059434 |
| **A_55_P1970087** | Rpl35a | -0.020860927 | -0.242405755 | 0.200683901 | 15.9881855 | -0.228799558 | 0.826410478 | 0.935814788 | -4.900601831 |
| **A_55_P1960238** | Slc2a6 | -0.019663856 | -0.228564338 | 0.189236626 | 7.965303186 | -0.228724362 | 0.826466388 | 0.935814788 | -4.900611153 |
| **A_51_P331661** | Ubiad1 | 0.029540175 | -0.284315229 | 0.343395578 | 10.63707307 | 0.22870025 | 0.826484316 | 0.935814788 | -4.900614141 |
| **A_52_P245365** | Rtf1 | 0.015671305 | -0.150840623 | 0.182183233 | 7.937413098 | 0.228687802 | 0.826493572 | 0.935814788 | -4.900615684 |
| **A_55_P2019254** | NA | 0.024794721 | -0.238844643 | 0.288434085 | 13.06648338 | 0.228524178 | 0.826615234 | 0.935894673 | -4.900635954 |
| **A_52_P671543** | Pomc | -0.036053936 | -0.419517085 | 0.347409212 | 8.075980177 | -0.228461089 | 0.826662146 | 0.935894673 | -4.900643765 |
| **A_55_P2037902** | Brd3 | -0.018064005 | -0.210242929 | 0.17411492 | 8.236868828 | -0.228397564 | 0.826709383 | 0.935894673 | -4.900651629 |
| **A_55_P1984043** | Pdcd4 | 0.053721743 | -0.518332838 | 0.625776323 | 11.31271384 | 0.228189582 | 0.826864041 | 0.936004293 | -4.90067736 |
| **A_51_P279623** | Pagr1a | -0.040774085 | -0.491799407 | 0.410251236 | 12.67590875 | -0.229998851 | 0.826903064 | 0.936004293 | -4.852182069 |
| **A_55_P2064299** | Msrb1 | 0.02173156 | -0.209800349 | 0.25326347 | 8.880386622 | 0.228067416 | 0.826954889 | 0.936008143 | -4.900692463 |
| **A_51_P502764** | Hdac10 | -0.022533838 | -0.262773287 | 0.217705612 | 9.349979336 | -0.227915586 | 0.827067801 | 0.9360464 | -4.900711222 |
| **A_55_P2029022** | Rpl34 | 0.020625675 | -0.199310702 | 0.240562052 | 16.82463105 | 0.227873751 | 0.827098914 | 0.9360464 | -4.900716389 |
| **A_51_P501550** | NA | 0.015381053 | -0.148733462 | 0.179495568 | 8.136955161 | 0.227731059 | 0.827205036 | 0.9360464 | -4.900734005 |
| **A_55_P1967483** | NA | 0.027132109 | -0.262394872 | 0.31665909 | 12.03149457 | 0.22770769 | 0.827222416 | 0.9360464 | -4.900736889 |
| **A_52_P174775** | Ripk1 | 0.030190449 | -0.292116858 | 0.352497756 | 9.229037493 | 0.227605424 | 0.827298475 | 0.9360464 | -4.900749507 |
| **A_55_P1991723** | Alkbh3 | 0.02882164 | -0.279050608 | 0.336693889 | 10.82245432 | 0.227473777 | 0.827396389 | 0.9360464 | -4.900765741 |
| **A_55_P2048434** | Hsp90ab1 | -0.024644825 | -0.287907771 | 0.23861812 | 15.69685954 | -0.227467418 | 0.827401119 | 0.9360464 | -4.900766525 |
| **A_52_P182659** | Cs | 0.025051099 | -0.242591033 | 0.292693231 | 13.83127312 | 0.227434051 | 0.827425936 | 0.9360464 | -4.900770638 |
| **A_55_P2096657** | NA | -0.028454935 | -0.332526244 | 0.275616373 | 15.3219573 | -0.227386851 | 0.827461043 | 0.9360464 | -4.900776456 |
| **A_55_P1959595** | Fmo5 | 0.054868213 | -0.531600349 | 0.641336775 | 11.37939995 | 0.227331317 | 0.827502349 | 0.9360464 | -4.900783299 |
| **A_55_P2115151** | Pxylp1 | -0.035431971 | -0.414195482 | 0.343331539 | 9.425916543 | -0.227305723 | 0.827521386 | 0.9360464 | -4.900786452 |
| **A_52_P523480** | Rfx1 | 0.018397341 | -0.178855482 | 0.215650165 | 9.123101104 | 0.226628761 | 0.82802496 | 0.936556519 | -4.900869728 |
| **A_55_P2123199** | Akirin1 | -0.020713499 | -0.242858116 | 0.201431117 | 8.443641356 | -0.226569201 | 0.82806927 | 0.936556519 | -4.900877044 |
| **A_55_P2102469** | Gm6428 | 0.017935029 | -0.174506271 | 0.21037633 | 16.1208078 | 0.226457649 | 0.82815226 | 0.936595578 | -4.900890739 |
| **A_51_P447668** | Ptpn12 | 0.025432744 | -0.247651819 | 0.298517307 | 8.874474938 | 0.226297242 | 0.828271601 | 0.936675742 | -4.900910422 |
| **A_51_P277295** | Apmap | 0.024341139 | -0.237113559 | 0.285795837 | 10.17855508 | 0.226218247 | 0.828330374 | 0.936687405 | -4.90092011 |
| **A_51_P151775** | Mief1 | 0.029072321 | -0.283554553 | 0.341699194 | 9.138260489 | 0.225962609 | 0.82852058 | 0.936793982 | -4.900951439 |
| **A_52_P358388** | Ppp4r2 | 0.021078787 | -0.205599193 | 0.247756767 | 8.945874671 | 0.225953705 | 0.828527205 | 0.936793982 | -4.900952529 |
| **A_55_P2116525** | NA | -0.014644107 | -0.172178785 | 0.142890571 | 7.675536627 | -0.225875909 | 0.828585092 | 0.936793982 | -4.900962056 |
| **A_51_P174275** | Colec11 | -0.045775793 | -0.538308931 | 0.446757344 | 11.06962041 | -0.225831029 | 0.828618486 | 0.936793982 | -4.900967551 |
| **A_55_P2154416** | Car15 | -0.015288068 | -0.179873459 | 0.149297323 | 8.245073023 | -0.22570673 | 0.828710978 | 0.93683812 | -4.900982763 |

| | | | | | | | | | |
|---|---|---|---|---|---|---|---|---|---|
| **A_51_P473576** | Apex1 | -0.019061701 | -0.224325702 | 0.1862023 | 11.38052299 | -0.225648292 | 0.828754463 | 0.93683812 | -4.900989912 |
| **A_52_P162486** | Bgn | 0.054904728 | -0.536874531 | 0.646683987 | 9.263319541 | 0.225441151 | 0.828908607 | 0.936957571 | -4.901015237 |
| **A_51_P333111** | Aox1 | -0.04974735 | -0.586984791 | 0.48749009 | 8.630783811 | -0.225002256 | 0.829235238 | 0.937271967 | -4.901068823 |
| **A_52_P622391** | Usp9x | 0.018563642 | -0.182046892 | 0.219174176 | 8.472609886 | 0.22484987 | 0.829348654 | 0.937345347 | -4.901087405 |
| **A_55_P2045903** | NA | -0.018704388 | -0.220989864 | 0.183581088 | 8.303612312 | -0.224678749 | 0.829476019 | 0.937434484 | -4.901108256 |
| **A_52_P71105** | Sertad3 | -0.024455268 | -0.289050452 | 0.240139917 | 10.58818167 | -0.224581342 | 0.829548522 | 0.937461611 | -4.901120118 |
| **A_51_P219325** | Ddx28 | 0.026697342 | -0.262304168 | 0.315698852 | 9.156350057 | 0.224466276 | 0.829634171 | 0.93750359 | -4.901134124 |
| **A_55_P2173664** | Frmpd2 | 0.060317708 | -0.593292627 | 0.713928042 | 14.66840873 | 0.224237946 | 0.829804136 | 0.937584522 | -4.901161897 |
| **A_55_P1971229** | Slc25a36 | -0.029896289 | -0.353913602 | 0.294121025 | 8.171414651 | -0.224198267 | 0.829833673 | 0.937584522 | -4.90116672 |
| **A_55_P2108609** | Srgap3 | 0.01452493 | -0.142921442 | 0.171971301 | 7.695361594 | 0.22416333 | 0.82985968 | 0.937584522 | -4.901170967 |
| **A_51_P345340** | Upk3bl | 0.018341468 | -0.180522993 | 0.217205928 | 7.856109904 | 0.224109411 | 0.829899818 | 0.937584522 | -4.901177519 |
| **A_55_P2184189** | Ncf2 | 0.059420361 | -0.585332768 | 0.70417349 | 10.11920236 | 0.223936569 | 0.83002849 | 0.937675084 | -4.901198512 |
| **A_51_P431734** | Fam185a | 0.032995317 | -0.325172566 | 0.3911632 | 8.474989094 | 0.223845748 | 0.830096103 | 0.937696662 | -4.901209537 |
| **A_55_P1970631** | Vps54 | 0.018435305 | -0.181790227 | 0.218660836 | 9.620101121 | 0.223724761 | 0.830186176 | 0.937743607 | -4.901224216 |
| **A_55_P1973259** | Gamt | -0.051786879 | -0.614459507 | 0.510885749 | 13.83091733 | -0.223638793 | 0.83025018 | 0.937761102 | -4.901234643 |
| **A_51_P418357** | Mov10l1 | 0.014384494 | -0.141956645 | 0.170725633 | 7.658869426 | 0.223565357 | 0.830304855 | 0.937768058 | -4.901243546 |
| **A_55_P2083484** | Mettl21a | 0.017735859 | -0.175152232 | 0.210623949 | 8.357279991 | 0.223424089 | 0.830410036 | 0.937832052 | -4.901260664 |
| **A_55_P2210315** | Get4 | 0.042464591 | -0.419632981 | 0.504562163 | 9.744022997 | 0.223293689 | 0.830507127 | 0.937886905 | -4.901276457 |
| **A_55_P1955059** | Rpl38 | 0.015409031 | -0.152470655 | 0.183288717 | 16.44197159 | 0.223028503 | 0.830704587 | 0.937945621 | -4.901308545 |
| **A_55_P2015632** | Gm5766 | -0.025251962 | -0.300393524 | 0.2498896 | 15.94289231 | -0.22300886 | 0.830719214 | 0.937945621 | -4.901310921 |
| **A_55_P2021991** | Zmym4 | -0.015578358 | -0.185446018 | 0.154289302 | 8.57673626 | -0.222840523 | 0.830844567 | 0.937945621 | -4.901331269 |
| **A_55_P1958447** | Prepl | 0.017279737 | -0.171169263 | 0.205728736 | 8.664886592 | 0.222805791 | 0.830870431 | 0.937945621 | -4.901335465 |
| **A_55_P2033725** | Ascl2 | -0.023629791 | -0.281346648 | 0.234087065 | 8.423641209 | -0.222792337 | 0.83088045 | 0.937945621 | -4.901337091 |
| **A_65_P08291** | Gpalpp1 | 0.023247367 | -0.230305678 | 0.276800412 | 10.89923398 | 0.222786122 | 0.830885078 | 0.937945621 | -4.901337842 |
| **A_55_P2030802** | Rpl3 | -0.019098098 | -0.227413363 | 0.189217167 | 15.75559011 | -0.222767698 | 0.830898798 | 0.937945621 | -4.901340067 |
| **A_55_P2012615** | NA | -0.031705332 | -0.377795973 | 0.314385309 | 15.27641801 | -0.222600248 | 0.831023498 | 0.937979859 | -4.901360287 |
| **A_55_P2005081** | Rpl10 | -0.022881129 | -0.272652132 | 0.226889875 | 17.43460048 | -0.222596643 | 0.831026183 | 0.937979859 | -4.901360722 |
| **A_55_P2037689** | NA | 0.049593739 | -0.492185083 | 0.591372561 | 12.14048702 | 0.222427264 | 0.831152325 | 0.938067458 | -4.901381159 |
| **A_66_P104422** | Ckap4 | -0.022774784 | -0.271722321 | 0.226172753 | 9.695048966 | -0.222294963 | 0.831250857 | 0.938087272 | -4.901397112 |
| **A_55_P2053783** | Ppp2r3a | -0.014816767 | -0.176792226 | 0.147158693 | 7.742531324 | -0.222273362 | 0.831266945 | 0.938087272 | -4.901399716 |
| **A_55_P1994519** | Tpd52l2 | 0.015967473 | -0.158723432 | 0.190658378 | 8.510143673 | 0.222100271 | 0.831395862 | 0.938132035 | -4.901420571 |
| **A_51_P292490** | Ttbk2 | -0.022132717 | -0.264285447 | 0.220020012 | 8.602530356 | -0.222089773 | 0.831403682 | 0.938132035 | -4.901421836 |
| **A_55_P1981804** | Gp1ba | 0.072288616 | -0.718974884 | 0.863552115 | 10.00980949 | 0.22198926 | 0.831478545 | 0.938157676 | -4.901433938 |
| **A_51_P341789** | Sugp1 | 0.021310479 | -0.212015167 | 0.254636125 | 11.18291146 | 0.221928935 | 0.831523478 | 0.938157676 | -4.901441199 |
| **A_55_P2037146** | Igfbpl1 | 0.014568232 | -0.145011724 | 0.174148188 | 7.657043841 | 0.22182562 | 0.831600433 | 0.938189737 | -4.901453631 |
| **A_65_P02958** | Rnf13 | -0.037783846 | -0.451968083 | 0.376400391 | 11.14464365 | -0.221664317 | 0.831720584 | 0.938270525 | -4.901473028 |
| **A_52_P456077** | Commd2 | 0.045988912 | -0.458577233 | 0.550555057 | 10.86797367 | 0.221471688 | 0.831864076 | 0.938377632 | -4.901496174 |
| **A_51_P279038** | Ppargc1a | -0.037632229 | -0.450802989 | 0.375538532 | 9.44210813 | -0.221316376 | 0.831979775 | 0.938399174 | -4.901514822 |
| **A_52_P106789** | Rbm18 | 0.019637879 | -0.195973739 | 0.235249496 | 11.03317991 | 0.221312327 | 0.831982791 | 0.938399174 | -4.901515308 |
| **A_55_P2008681** | Ktn1 | -0.023150265 | -0.277396666 | 0.231096136 | 10.72831551 | -0.221250541 | 0.832028819 | 0.938399174 | -4.901522723 |
| **A_55_P1952379** | Fkbp5 | -0.039205046 | -0.470040171 | 0.391630079 | 11.40186622 | -0.221112885 | 0.832131371 | 0.938460078 | -4.901539236 |
| **A_55_P2039842** | Frmpd1 | 0.015348293 | -0.153411491 | 0.184108078 | 7.797902576 | 0.220990863 | 0.832222279 | 0.938494102 | -4.901553865 |
| **A_52_P516091** | Pla2g15 | -0.021801111 | -0.261564815 | 0.217962593 | 9.171210469 | -0.220942048 | 0.832258648 | 0.938494102 | -4.901559715 |
| **A_55_P2135049** | NA | 0.028453125 | -0.284621987 | 0.341528238 | 13.32692675 | 0.22083333 | 0.832339647 | 0.938530687 | -4.901572739 |
| **A_55_P2046744** | Cspg5 | -0.019451438 | -0.23362852 | 0.194725643 | 7.864990705 | -0.220679459 | 0.832454291 | 0.938555043 | -4.901591162 |
| **A_52_P272534** | NA | -0.035034134 | -0.420799657 | 0.35073139 | 11.39640141 | -0.220673996 | 0.832458361 | 0.938555043 | -4.901591816 |
| **A_51_P466490** | Armc5 | 0.022614665 | -0.226650364 | 0.271879693 | 10.00418765 | 0.220450958 | 0.832624547 | 0.938605125 | -4.901618498 |
| **A_51_P185660** | Ccl9 | -0.041535267 | -0.49941212 | 0.416341587 | 13.07520438 | -0.220420256 | 0.832647425 | 0.938605125 | -4.901622168 |
| **A_52_P663526** | Nmrk1 | -0.017859578 | -0.214777743 | 0.179058588 | 8.075503192 | -0.220378182 | 0.832678775 | 0.938605125 | -4.901627198 |
| **A_55_P2130600** | Musk | 0.016857764 | -0.16906312 | 0.202778649 | 7.803784654 | 0.220320543 | 0.832721725 | 0.938605125 | -4.901634087 |

| | | | | | | | | | |
|---|---|---|---|---|---|---|---|---|---|
| **A_55_P1988882** | 9-Sep | -0.025782687 | -0.310286994 | 0.25872162 | 10.65704767 | -0.220202641 | 0.832809581 | 0.938605125 | -4.901648172 |
| **A_55_P1963109** | Prh1 | 0.052608932 | -0.528059576 | 0.63327744 | 9.610431165 | 0.220147826 | 0.832850427 | 0.938605125 | -4.901654718 |
| **A_51_P442152** | Itfg1 | 0.023527107 | -0.236202015 | 0.28325623 | 11.62127577 | 0.220105591 | 0.832881901 | 0.938605125 | -4.901659761 |
| **A_55_P2090718** | Pop5 | -0.034975943 | -0.421117305 | 0.351165418 | 12.94140955 | -0.220093035 | 0.832891257 | 0.938605125 | -4.90166126 |
| **A_51_P283016** | Slc1a6 | 0.020332572 | -0.204277686 | 0.24494283 | 8.004940806 | 0.21996112 | 0.832989561 | 0.93866118 | -4.901677004 |
| **A_52_P168338** | Fundc2 | 0.01648593 | -0.165891726 | 0.198863587 | 7.912812248 | 0.219646868 | 0.833223756 | 0.93887035 | -4.901714471 |
| **A_66_P117495** | Capza1 | 0.024166157 | -0.243486818 | 0.291819131 | 9.21173982 | 0.219390947 | 0.833414494 | 0.938945406 | -4.901744945 |
| **A_51_P464300** | Gdf1 | -0.046847882 | -0.565753532 | 0.472057767 | 8.304527488 | -0.219373798 | 0.833427275 | 0.938945406 | -4.901746985 |
| **A_55_P2124341** | Lysmd1 | -0.019070508 | -0.230428471 | 0.192287454 | 11.26744804 | -0.219243561 | 0.833524347 | 0.938945406 | -4.901762479 |
| **A_52_P64364** | Ikbkb | 0.019840902 | -0.200138023 | 0.239819827 | 8.561223287 | 0.219161127 | 0.83358579 | 0.938945406 | -4.901772281 |
| **A_55_P1984422** | Nme2 | -0.02650017 | -0.320347776 | 0.267347436 | 15.15735662 | -0.219133962 | 0.833606038 | 0.938945406 | -4.90177551 |
| **A_55_P2009572** | Ptbp3 | -0.017037741 | -0.205985193 | 0.171909711 | 8.318867252 | -0.219105944 | 0.833626922 | 0.938945406 | -4.901778841 |
| **A_52_P127362** | Vmp1 | -0.028961439 | -0.350148788 | 0.29222591 | 12.41617618 | -0.219101269 | 0.833630406 | 0.938945406 | -4.901779396 |
| **A_55_P2028788** | Slc39a11 | -0.042099176 | -0.509585604 | 0.425387253 | 11.69099025 | -0.218820382 | 0.833839781 | 0.93909365 | -4.90181276 |
| **A_55_P2274592** | Wnt7b | -0.018043221 | -0.218439862 | 0.182353421 | 7.745667658 | -0.218779592 | 0.833870187 | 0.93909365 | -4.901817601 |
| **A_55_P2019058** | Acaca | -0.04512292 | -0.546507474 | 0.456261635 | 11.66934118 | -0.218680185 | 0.83394429 | 0.93909365 | -4.901829397 |
| **A_55_P2182939** | NA | 0.028539789 | -0.288604338 | 0.345683916 | 13.7643268 | 0.218663993 | 0.833956361 | 0.93909365 | -4.901831318 |
| **A_55_P1983739** | NA | -0.033813387 | -0.409987633 | 0.34236086 | 10.50337716 | -0.21841517 | 0.834141854 | 0.939234957 | -4.901860817 |
| **A_52_P227307** | Mllt1 | 0.02260203 | -0.228903192 | 0.274107252 | 10.08245765 | 0.218365302 | 0.834179032 | 0.939234957 | -4.901866726 |
| **A_52_P253179** | Igfbp3 | -0.047006082 | -0.570320959 | 0.476308796 | 9.059058896 | -0.218260004 | 0.834257534 | 0.939268632 | -4.901879197 |
| **A_66_P114862** | Ankrd11 | -0.024920853 | -0.302746464 | 0.252904758 | 10.19424338 | -0.2179585 | 0.834482325 | 0.93938482 | -4.901914873 |
| **A_51_P374305** | Rchy1 | 0.018105793 | -0.18376938 | 0.219980965 | 10.79732656 | 0.217930402 | 0.834503275 | 0.93938482 | -4.901918195 |
| **A_55_P2067741** | Psg23 | 0.039270172 | -0.398642798 | 0.477183142 | 12.10962921 | 0.217900479 | 0.834525585 | 0.93938482 | -4.901921733 |
| **A_52_P425092** | NA | -0.058366307 | -0.709444273 | 0.592711659 | 13.91892797 | -0.21782735 | 0.834580112 | 0.93938482 | -4.901930377 |
| **A_52_P582394** | Mrps11 | 0.024435606 | -0.248317554 | 0.297188766 | 9.357015255 | 0.217689015 | 0.834683258 | 0.93938482 | -4.90194672 |
| **A_55_P2165046** | Klhl22 | 0.017778399 | -0.180671329 | 0.216228126 | 10.08931139 | 0.217683403 | 0.834687443 | 0.93938482 | -4.901947383 |
| **A_55_P2085716** | Atg12 | 0.028470605 | -0.289487255 | 0.346428465 | 9.922689779 | 0.217575667 | 0.834767777 | 0.93938482 | -4.901960104 |
| **A_52_P467930** | NA | 0.026008065 | -0.264486842 | 0.316502972 | 10.49160367 | 0.217546805 | 0.834789299 | 0.93938482 | -4.901963511 |
| **A_55_P1977066** | Klk4 | 0.014065025 | -0.14306528 | 0.17119533 | 7.677533489 | 0.217502246 | 0.834822525 | 0.93938482 | -4.901968769 |
| **A_55_P2131433** | NA | -0.015498361 | -0.188704125 | 0.157707403 | 7.820453679 | -0.21742357 | 0.834881193 | 0.93938482 | -4.901978052 |
| **A_55_P2136385** | Rps3a1 | 0.018920582 | -0.192616685 | 0.23045785 | 15.36858321 | 0.217335563 | 0.83494682 | 0.93938482 | -4.901988432 |
| **A_55_P2073015** | Lin37 | 0.033474345 | -0.340847115 | 0.407795806 | 9.561704624 | 0.217295411 | 0.834976762 | 0.93938482 | -4.901993166 |
| **A_55_P2050408** | NA | -0.017054074 | -0.207777062 | 0.173668914 | 8.178407596 | -0.217274268 | 0.834992529 | 0.93938482 | -4.901995659 |
| **A_55_P2089804** | Atf7 | -0.033546958 | -0.409126819 | 0.342032903 | 8.906305385 | -0.21703713 | 0.835169373 | 0.939408789 | -4.902023599 |
| **A_55_P1965659** | Havcr2 | 0.01504802 | -0.153430689 | 0.183526729 | 7.76960546 | 0.21702888 | 0.835175525 | 0.939408789 | -4.902024571 |
| **A_55_P2157159** | NA | 0.028236053 | -0.287909539 | 0.344381646 | 11.69120009 | 0.217020147 | 0.835182038 | 0.939408789 | -4.902025599 |
| **A_55_P2153827** | Fgfr1op | 0.017861442 | -0.182156964 | 0.217879848 | 10.13902532 | 0.216985015 | 0.835208239 | 0.939408789 | -4.902029735 |
| **A_55_P1976744** | NA | 0.016120081 | -0.164510291 | 0.196750453 | 7.739436489 | 0.216850103 | 0.835308855 | 0.93946729 | -4.902045614 |
| **A_51_P235139** | Klhl40 | -0.013788877 | -0.168447587 | 0.140869832 | 7.657931106 | -0.216639535 | 0.8354659 | 0.939576629 | -4.902070378 |
| **A_51_P411019** | Tmem127 | -0.018288339 | -0.223461355 | 0.186884677 | 11.49336429 | -0.216589404 | 0.83550329 | 0.939576629 | -4.90207627 |
| **A_55_P2115119** | Rnaseh2a | 0.0372699 | -0.381471318 | 0.456011119 | 10.37376923 | 0.216269716 | 0.835741739 | 0.93964204 | -4.902113814 |
| **A_55_P2128158** | Trappc13 | -0.018124499 | -0.221799675 | 0.185550677 | 10.54297219 | -0.216227589 | 0.835773162 | 0.93964204 | -4.902118757 |
| **A_55_P2181444** | NA | -0.033784822 | -0.413479937 | 0.345910294 | 11.94619267 | -0.216207029 | 0.835788498 | 0.93964204 | -4.90212117 |
| **A_51_P470328** | Selenop | 0.030155335 | -0.3089175 | 0.369228171 | 16.50827546 | 0.216099792 | 0.835868489 | 0.93964204 | -4.902133748 |
| **A_55_P1990092** | Snrpa | 0.018340676 | -0.18790283 | 0.224584181 | 11.69536917 | 0.216081824 | 0.835881892 | 0.93964204 | -4.902135854 |
| **A_55_P2015519** | Mtln | -0.037134557 | -0.454762796 | 0.380493682 | 10.61415391 | -0.216058609 | 0.83589921 | 0.93964204 | -4.902138576 |
| **A_52_P488612** | Btbd9 | -0.022064317 | -0.270211271 | 0.226082636 | 9.033210011 | -0.216055207 | 0.835901747 | 0.93964204 | -4.902138975 |
| **A_52_P284495** | Ankrd33b | -0.019288937 | -0.236392668 | 0.197814793 | 7.901326137 | -0.215885862 | 0.836028073 | 0.93972308 | -4.902158822 |
| **A_51_P468780** | Hmg20a | 0.020706724 | -0.212592425 | 0.254005874 | 10.00318196 | 0.215665877 | 0.836192182 | 0.93972308 | -4.90218458 |
| **A_55_P2031501** | Hnrnpa3 | 0.020857971 | -0.21417492 | 0.255890863 | 14.96623926 | 0.215638651 | 0.836212494 | 0.93972308 | -4.902187766 |

| | | | | | | | | | |
|---|---|---|---|---|---|---|---|---|---|
| **A_55_P2097759** | NA | 0.03084982 | -0.31678776 | 0.3784874 | 13.846971 | 0.215629964 | 0.836218974 | 0.93972308 | -4.902188783 |
| **A_51_P341664** | Zfp87 | 0.014017329 | -0.14401758 | 0.172052238 | 8.899778299 | 0.215523892 | 0.836298108 | 0.93972308 | -4.902201192 |
| **A_55_P2133280** | Ap4b1 | 0.031524281 | -0.323922164 | 0.386970726 | 10.50730518 | 0.215503438 | 0.836313368 | 0.93972308 | -4.902203584 |
| **A_52_P545643** | Tmem263 | 0.030966815 | -0.318232685 | 0.380166314 | 10.73173008 | 0.215479576 | 0.83633117 | 0.93972308 | -4.902206374 |
| **A_51_P488092** | Pde6d | 0.015437114 | -0.15871845 | 0.189592678 | 9.521849647 | 0.215383263 | 0.836403026 | 0.93972308 | -4.902217634 |
| **A_55_P1957204** | NA | -0.019901368 | -0.244475005 | 0.20467227 | 17.20321227 | -0.21533139 | 0.836441727 | 0.93972308 | -4.902223697 |
| **A_55_P2161595** | Hpca | -0.024510315 | -0.301243836 | 0.252223206 | 7.978384874 | -0.2152139 | 0.836529386 | 0.93972308 | -4.902237422 |
| **A_55_P2003251** | Magel2 | 0.014132814 | -0.145466101 | 0.173731728 | 7.840839936 | 0.215170082 | 0.836562079 | 0.93972308 | -4.90224254 |
| **A_52_P189957** | Dbnl | -0.040126952 | -0.49330062 | 0.413046715 | 9.037042697 | -0.215156612 | 0.836572129 | 0.93972308 | -4.902244112 |
| **A_52_P286002** | Obsl1 | -0.019607499 | -0.241166659 | 0.20195166 | 7.928356109 | -0.215038235 | 0.836660453 | 0.93972308 | -4.902257931 |
| **A_52_P515057** | Slc25a24 | -0.024177691 | -0.297453302 | 0.249097921 | 9.900270512 | -0.21497954 | 0.836704248 | 0.93972308 | -4.90226478 |
| **A_66_P117913** | NA | -0.022298015 | -0.274501443 | 0.229905414 | 15.91756095 | -0.214831718 | 0.836814547 | 0.93972308 | -4.90228202 |
| **A_51_P280085** | Zfp655 | 0.018448408 | -0.190224174 | 0.22712099 | 11.51299367 | 0.214820985 | 0.836822556 | 0.93972308 | -4.902283272 |
| **A_55_P1960873** | Mbd3 | -0.032709637 | -0.402822064 | 0.337402789 | 9.423081122 | -0.214746081 | 0.836878448 | 0.93972308 | -4.902292003 |
| **A_55_P1991377** | NA | 0.025324302 | -0.261223075 | 0.31187168 | 15.57706791 | 0.214745583 | 0.83687882 | 0.93972308 | -4.902292061 |
| **A_51_P233928** | B4galnt1 | -0.052583659 | -0.647643395 | 0.542476077 | 10.08582523 | -0.214720458 | 0.836897568 | 0.93972308 | -4.902294989 |
| **A_51_P161463** | Ccdc103 | -0.025400089 | -0.312990639 | 0.262190461 | 8.311356243 | -0.214606968 | 0.836982255 | 0.939727317 | -4.902308211 |
| **A_66_P119429** | NA | -0.019932227 | -0.245636332 | 0.205771878 | 14.32485726 | -0.214585095 | 0.836998577 | 0.939727317 | -4.902310758 |
| **A_51_P399960** | Mier2 | 0.018922957 | -0.195444889 | 0.233290803 | 8.649588518 | 0.214492717 | 0.837067512 | 0.939749428 | -4.902321514 |
| **A_51_P268145** | Naa80 | 0.018468184 | -0.190810535 | 0.227746903 | 9.200188995 | 0.2144284 | 0.837115508 | 0.939749428 | -4.902329 |
| **A_52_P193029** | Hadha | -0.029348527 | -0.362181471 | 0.303484417 | 10.69266835 | -0.214261032 | 0.837240409 | 0.939791077 | -4.902348471 |
| **A_66_P105046** | Il18 | 0.021876737 | -0.226235501 | 0.269988975 | 10.05722218 | 0.214248382 | 0.83724985 | 0.939791077 | -4.902349942 |
| **A_55_P1993836** | Dusp7 | 0.028327352 | -0.293284043 | 0.349938747 | 9.238910078 | 0.214021667 | 0.837419048 | 0.939926414 | -4.902376291 |
| **A_55_P2151751** | Matr3 | 0.017134005 | -0.177694839 | 0.211962849 | 9.426979475 | 0.213692277 | 0.837664891 | 0.940084471 | -4.902414525 |
| **A_55_P2043434** | Atmin | -0.021237581 | -0.262779964 | 0.220304802 | 9.892883985 | -0.213646059 | 0.837699387 | 0.940084471 | -4.902419885 |
| **A_66_P129795** | Ccdc59 | -0.033606485 | -0.415840021 | 0.348627052 | 11.67645763 | -0.2136375 | 0.837705776 | 0.940084471 | -4.902420877 |
| **A_51_P189733** | Sult2a8 | 0.060529873 | -0.628534613 | 0.749594359 | 9.954574432 | 0.213448488 | 0.837846857 | 0.940137123 | -4.902442785 |
| **A_51_P237585** | Btnl9 | -0.018480782 | -0.228911935 | 0.191950371 | 8.052346649 | -0.213399548 | 0.837883387 | 0.940137123 | -4.902448455 |
| **A_55_P2016682** | Alkbh7 | -0.034787651 | -0.430934587 | 0.361359285 | 12.07633132 | -0.213379155 | 0.837898609 | 0.940137123 | -4.902450817 |
| **A_52_P27332** | Ppp2r5c | -0.01870894 | -0.231926205 | 0.194508326 | 10.04504221 | -0.213211189 | 0.838023989 | 0.940220723 | -4.902470263 |
| **A_65_P09031** | Cdc42se2 | -0.0150355 | -0.186477872 | 0.156406872 | 7.934316677 | -0.213099733 | 0.838107189 | 0.940220723 | -4.902483159 |
| **A_55_P2025840** | Spred2 | 0.015295368 | -0.159195312 | 0.189786048 | 8.363620341 | 0.212995722 | 0.838184834 | 0.940220723 | -4.902495187 |
| **A_55_P1985428** | Atg16l2 | -0.022338429 | -0.277324096 | 0.232647237 | 9.220008186 | -0.212872742 | 0.838276641 | 0.940220723 | -4.902509401 |
| **A_55_P2048279** | Tlr13 | -0.026478526 | -0.328747184 | 0.275790132 | 9.45430152 | -0.212855008 | 0.838289881 | 0.940220723 | -4.902511451 |
| **A_55_P2398264** | Rnf130 | 0.016856167 | -0.175627366 | 0.2093397 | 7.917598137 | 0.212788637 | 0.83833943 | 0.940220723 | -4.902519118 |
| **A_55_P2060330** | Cyren | -0.025213679 | -0.313190462 | 0.262763104 | 10.80252806 | -0.212746259 | 0.838371067 | 0.940220723 | -4.902524013 |
| **A_66_P131091** | Rabl6 | -0.030895179 | -0.383974566 | 0.322184209 | 11.64174238 | -0.212618738 | 0.838466271 | 0.940220723 | -4.902538735 |
| **A_55_P2084303** | Nid1 | -0.014322897 | -0.178019332 | 0.149373539 | 7.887454328 | -0.212605663 | 0.838476032 | 0.940220723 | -4.902540244 |
| **A_52_P673499** | Shmt1 | 0.050306536 | -0.524975344 | 0.625588416 | 9.590191661 | 0.212484329 | 0.83856662 | 0.940220723 | -4.902554243 |
| **A_52_P937** | Spcs2 | -0.021561136 | -0.268185459 | 0.225063186 | 12.49268396 | -0.212431508 | 0.838606057 | 0.940220723 | -4.902560335 |
| **A_66_P102046** | NA | 0.033509424 | -0.349795681 | 0.41681453 | 10.81052749 | 0.212424964 | 0.838610943 | 0.940220723 | -4.90256109 |
| **A_55_P2108693** | NA | -0.05934922 | -0.73827606 | 0.619577621 | 9.378255145 | -0.212410129 | 0.838622018 | 0.940220723 | -4.9025628 |
| **A_55_P2136757** | Tomm34 | -0.027452601 | -0.341733583 | 0.28682838 | 11.12302639 | -0.212250437 | 0.838741251 | 0.940220723 | -4.902581207 |
| **A_51_P433584** | Bcas3 | 0.018669731 | -0.195084917 | 0.232424379 | 8.733644607 | 0.212229466 | 0.838756908 | 0.940220723 | -4.902583623 |
| **A_52_P532559** | Thrb | -0.031244019 | -0.389051538 | 0.3265635 | 8.628838079 | -0.212178131 | 0.838795238 | 0.940220723 | -4.902589537 |
| **A_55_P1982245** | Phactr2 | -0.016080823 | -0.200244525 | 0.168082879 | 7.87112142 | -0.212171688 | 0.838800049 | 0.940220723 | -4.902590279 |
| **A_55_P2233271** | Zfp626 | 0.018488836 | -0.193450919 | 0.230428591 | 8.905022368 | 0.211972895 | 0.838948485 | 0.940332575 | -4.902613166 |
| **A_51_P464023** | Chst12 | -0.023397841 | -0.291837013 | 0.245041332 | 9.114840855 | -0.21179372 | 0.839082277 | 0.940395311 | -4.902633775 |
| **A_51_P465281** | Lgals1 | 0.054268147 | -0.568417145 | 0.676953439 | 13.38658472 | 0.211767629 | 0.83910176 | 0.940395311 | -4.902636775 |
| **A_55_P2413458** | Tfcp2l1 | -0.026727528 | -0.33355127 | 0.280096215 | 8.615639264 | -0.21166693 | 0.839176957 | 0.940403223 | -4.902648349 |

| | | | | | | | | | |
|---|---|---|---|---|---|---|---|---|---|
| **A_55_P1988070** | Eef1g | 0.04165373 | -0.436606449 | 0.519913908 | 11.40400525 | 0.211627871 | 0.839206125 | 0.940403223 | -4.902652837 |
| **A_55_P1986531** | Mpig6b | -0.051241087 | -0.640299973 | 0.5378178 | 8.59304503 | -0.211369737 | 0.839398896 | 0.940553399 | -4.902682477 |
| **A_55_P2086128** | Fa2h | -0.013558966 | -0.169468645 | 0.142350713 | 7.68153802 | -0.211318098 | 0.839437461 | 0.940553399 | -4.902688402 |
| **A_51_P349878** | Txndc17 | 0.020250011 | -0.212682512 | 0.253182535 | 13.43613056 | 0.211241055 | 0.839494999 | 0.940563346 | -4.902697239 |
| **A_51_P480861** | Pga5 | 0.015961618 | -0.167755414 | 0.19967865 | 7.756147142 | 0.211110912 | 0.839592195 | 0.940592451 | -4.90271216 |
| **A_55_P2088692** | NA | -0.037491447 | -0.469117253 | 0.39413436 | 10.02520721 | -0.211060998 | 0.839629475 | 0.940592451 | -4.902717881 |
| **A_55_P2086265** | Gm8580 | -0.031281429 | -0.391499151 | 0.328936294 | 16.85269792 | -0.211010804 | 0.839666963 | 0.940592451 | -4.902723632 |
| **A_55_P2136426** | Prss57 | -0.030478484 | -0.381747632 | 0.320790664 | 8.434868978 | -0.210832008 | 0.839800505 | 0.940687526 | -4.902744107 |
| **A_55_P1956938** | Rpl24 | -0.016521581 | -0.207232554 | 0.174189392 | 16.60233995 | -0.210503398 | 0.840045956 | 0.940898201 | -4.902781693 |
| **A_52_P606303** | Mrpl3 | -0.016918239 | -0.212257578 | 0.1784211 | 11.87050305 | -0.210449854 | 0.840085952 | 0.940898201 | -4.902787812 |
| **A_52_P552017** | Rbsn | 0.015680027 | -0.165528961 | 0.196889016 | 8.993804285 | 0.210256904 | 0.840230085 | 0.940898201 | -4.90280985 |
| **A_55_P1964483** | Cyp2c37 | -0.09096387 | -1.142292274 | 0.960364534 | 10.44639043 | -0.210239212 | 0.840243301 | 0.940898201 | -4.90281187 |
| **A_51_P254425** | Ahrr | 0.013666499 | -0.144329909 | 0.171662907 | 7.79643939 | 0.210180902 | 0.84028686 | 0.940898201 | -4.902818525 |
| **A_51_P290074** | Fabp7 | -0.022708535 | -0.285285852 | 0.239868782 | 7.997970085 | -0.210143081 | 0.840315113 | 0.940898201 | -4.902822841 |
| **A_55_P1977314** | Ulk1 | 0.021983206 | -0.232310342 | 0.276276755 | 12.76130946 | 0.210057832 | 0.840378798 | 0.940898201 | -4.902832566 |
| **A_55_P2144597** | 9030025P20Rik | 0.017656232 | -0.18679809 | 0.222110553 | 8.547282357 | 0.209838373 | 0.840542749 | 0.940898201 | -4.902857584 |
| **A_55_P2092616** | Tbc1d25 | 0.016033671 | -0.169632441 | 0.201699782 | 9.26599127 | 0.209837693 | 0.840543257 | 0.940898201 | -4.902857662 |
| **A_55_P2005853** | Nacc2 | 0.026354554 | -0.279001498 | 0.331710606 | 10.37784241 | 0.209716367 | 0.8406339 | 0.940898201 | -4.902871482 |
| **A_51_P448203** | Cnbp | -0.024560856 | -0.309244348 | 0.260122636 | 13.2559281 | -0.209635297 | 0.840694468 | 0.940898201 | -4.902880712 |
| **A_51_P269078** | Habp4 | 0.024903487 | -0.263779539 | 0.313586513 | 8.961563007 | 0.209614879 | 0.840709723 | 0.940898201 | -4.902883036 |
| **A_52_P469956** | Zc3h6 | -0.021934456 | -0.276275749 | 0.232406836 | 8.734652545 | -0.209552666 | 0.840756205 | 0.940898201 | -4.902890116 |
| **A_51_P223575** | Tbc1d23 | -0.019934592 | -0.251090502 | 0.211221318 | 9.127606476 | -0.209548978 | 0.84075896 | 0.940898201 | -4.902890536 |
| **A_55_P2159585** | Pm20d1 | 0.041841059 | -0.443342185 | 0.527024304 | 8.798041613 | 0.209546334 | 0.840760936 | 0.940898201 | -4.902890837 |
| **A_51_P416419** | Calr | -0.018795351 | -0.236822327 | 0.199231624 | 13.08415176 | -0.209470757 | 0.840817403 | 0.940898201 | -4.902899435 |
| **A_55_P2015715** | Rab4a | 0.04675192 | -0.49560611 | 0.58910995 | 11.35289476 | 0.209457814 | 0.840827074 | 0.940898201 | -4.902900907 |
| **A_51_P208769** | Metrn | -0.022689715 | -0.285971213 | 0.240591783 | 8.916479992 | -0.209407334 | 0.84086479 | 0.940898201 | -4.902906648 |
| **A_51_P426283** | Dhx30 | -0.020120213 | -0.253809061 | 0.213568635 | 12.19652238 | -0.209207752 | 0.841013915 | 0.941001612 | -4.902929332 |
| **A_52_P246165** | Ptprk | 0.021890721 | -0.232488306 | 0.276269748 | 8.514685116 | 0.209103814 | 0.841091579 | 0.941001612 | -4.902941137 |
| **A_55_P2045040** | F7 | 0.043455385 | -0.4615515 | 0.54846227 | 9.103154226 | 0.209088185 | 0.841103257 | 0.941001612 | -4.902942912 |
| **A_55_P2060991** | Plppr3 | -0.021168714 | -0.267390124 | 0.225052696 | 9.188455132 | -0.208906451 | 0.841239056 | 0.941099069 | -4.902963537 |
| **A_55_P2018393** | NA | -0.019315856 | -0.244062653 | 0.205430941 | 15.6312919 | -0.208835167 | 0.841292324 | 0.941104192 | -4.902971623 |
| **A_55_P2051400** | Prl3d2 | -0.013182923 | -0.16662596 | 0.140260113 | 7.629166786 | -0.208760215 | 0.841348334 | 0.941112381 | -4.902980121 |
| **A_55_P2104472** | NA | 0.025398091 | -0.270482679 | 0.321278862 | 13.31531907 | 0.208577535 | 0.841484851 | 0.941157461 | -4.903000822 |
| **A_55_P2009187** | Iffo1 | -0.016765642 | -0.212082212 | 0.178550929 | 8.339001876 | -0.208575973 | 0.841486018 | 0.941157461 | -4.903000999 |
| **A_55_P1995507** | Aqp11 | -0.061148797 | -0.775095442 | 0.652797848 | 12.24676933 | -0.208115942 | 0.841829826 | 0.941487514 | -4.903053051 |
| **A_55_P2061520** | NA | -0.014839635 | -0.188462286 | 0.158783016 | 8.046226072 | -0.207682557 | 0.842153754 | 0.941772019 | -4.903101984 |
| **A_55_P2107855** | Hnrnpk | 0.020846632 | -0.223249516 | 0.26494278 | 10.26576329 | 0.207519138 | 0.842275909 | 0.941772019 | -4.90312041 |
| **A_55_P1981040** | Ddx6 | 0.017870078 | -0.191400282 | 0.227140439 | 10.56211528 | 0.20749227 | 0.842295993 | 0.941772019 | -4.903123438 |
| **A_52_P22331** | Akap10 | -0.018822423 | -0.239385194 | 0.201740348 | 8.280511027 | -0.207360724 | 0.842394327 | 0.941772019 | -4.903138258 |
| **A_52_P522379** | Map3k20 | -0.014048875 | -0.178699553 | 0.150601803 | 7.937870067 | -0.207329552 | 0.842417629 | 0.941772019 | -4.903141768 |
| **A_55_P1997895** | Rpl35a | 0.017797032 | -0.190821951 | 0.226416015 | 14.77559605 | 0.207289327 | 0.842447698 | 0.941772019 | -4.903146298 |
| **A_51_P183275** | Sympk | 0.029394966 | -0.315196415 | 0.373986346 | 11.07508685 | 0.207277297 | 0.842456692 | 0.941772019 | -4.903147652 |
| **A_66_P105074** | Enpp5 | 0.016142229 | -0.173111082 | 0.205395541 | 9.000517342 | 0.207254141 | 0.842474002 | 0.941772019 | -4.903150259 |
| **A_55_P2068167** | Nufip2 | 0.020892084 | -0.224215969 | 0.266000137 | 10.47466259 | 0.207113007 | 0.842579509 | 0.941828366 | -4.90316614 |
| **A_52_P376106** | Slc30a9 | 0.023136699 | -0.248546491 | 0.29481989 | 9.01085839 | 0.20692922 | 0.842716907 | 0.941828366 | -4.903186806 |
| **A_55_P1965902** | Pi4k2a | -0.023172109 | -0.295271665 | 0.248927446 | 9.817908339 | -0.206928794 | 0.842717226 | 0.941828366 | -4.903186854 |
| **A_51_P433733** | Nucb1 | -0.02332138 | -0.297303203 | 0.250660443 | 12.03470549 | -0.206831027 | 0.842790318 | 0.941828366 | -4.90319784 |
| **A_55_P2045224** | NA | -0.020772511 | -0.264845999 | 0.223300976 | 11.85890151 | -0.206800502 | 0.84281314 | 0.941828366 | -4.903201269 |
| **A_52_P590535** | Fbln2 | -0.062192134 | -0.792956302 | 0.668572033 | 9.531666412 | -0.206795653 | 0.842816765 | 0.941828366 | -4.903201813 |
| **A_51_P238183** | Etnk2 | -0.059786565 | -0.762965775 | 0.643392644 | 12.20740239 | -0.206595459 | 0.842966441 | 0.941903201 | -4.903224289 |

| | | | | | | | | |
|---|---|---|---|---|---|---|---|---|
| **A_55_P2285599** | 5730437C12Rik | 0.013464931 | -0.144963139 | 0.171893001 | 7.643180423 | 0.206516703 | 0.843025325 | 0.941903201 | -4.903233125 |
| **A_55_P2175060** | Plpp5 | 0.036516807 | -0.393310305 | 0.46634392 | 9.002404005 | 0.206434453 | 0.843086822 | 0.941903201 | -4.90324235 |
| **A_66_P133928** | Fundc2 | -0.039517718 | -0.504815933 | 0.425780497 | 12.44126318 | -0.20636862 | 0.843136046 | 0.941903201 | -4.903249731 |
| **A_51_P430929** | Fam20a | -0.036311798 | -0.463982315 | 0.391358719 | 10.96755602 | -0.206310641 | 0.843179398 | 0.941903201 | -4.903256229 |
| **A_55_P1972128** | Ccndbp1 | 0.027424526 | -0.29566488 | 0.350513932 | 10.20267448 | 0.206252681 | 0.843222737 | 0.941903201 | -4.903262724 |
| **A_55_P2321453** | D11Ertd726e | 0.015263481 | -0.164558638 | 0.1950856 | 8.285705189 | 0.206249864 | 0.843224843 | 0.941903201 | -4.903263039 |
| **A_55_P2178023** | NA | 0.027987794 | -0.301947776 | 0.357923364 | 12.9732285 | 0.20612123 | 0.843321028 | 0.941910581 | -4.903277446 |
| **A_55_P2300421** | NA | 0.01634305 | -0.176327701 | 0.209013801 | 8.339776724 | 0.206110689 | 0.84332891 | 0.941910581 | -4.903278626 |
| **A_52_P686091** | Phf2 | 0.017785502 | -0.191987193 | 0.227558197 | 8.647416659 | 0.206015715 | 0.843399929 | 0.941932104 | -4.903289257 |
| **A_55_P2095000** | NA | 0.017979433 | -0.194143551 | 0.230102417 | 16.21625829 | 0.20595458 | 0.843445644 | 0.941932104 | -4.903296098 |
| **A_51_P254262** | Nxph3 | 0.019555154 | -0.211323974 | 0.250434281 | 8.011325336 | 0.205806813 | 0.843556144 | 0.942001081 | -4.903312623 |
| **A_51_P181922** | Yjefn3 | -0.014811325 | -0.189864386 | 0.160241735 | 7.946261744 | -0.205592561 | 0.843716369 | 0.942125574 | -4.903336564 |
| **A_51_P149621** | Stt3b | -0.029744497 | -0.381443715 | 0.321954721 | 10.43534251 | -0.20550312 | 0.843783258 | 0.942141068 | -4.903346551 |
| **A_55_P2136776** | Sult1b1 | 0.058630327 | -0.634815872 | 0.752076526 | 8.77946752 | 0.205443656 | 0.843827729 | 0.942141068 | -4.903353188 |
| **A_55_P2024011** | NA | 0.022206746 | -0.240673574 | 0.285087066 | 13.88749164 | 0.2052627 | 0.843963064 | 0.942169945 | -4.903373375 |
| **A_55_P2083904** | Tmem185a | -0.015921081 | -0.204418132 | 0.17257597 | 8.318712901 | -0.20523488 | 0.843983872 | 0.942169945 | -4.903376477 |
| **A_55_P2042729** | NA | 0.016580333 | -0.179883673 | 0.21304434 | 12.28263024 | 0.2050659 | 0.844110256 | 0.942169945 | -4.90339531 |
| **A_55_P2080021** | Papss2 | -0.073829959 | -0.949033538 | 0.80137362 | 11.4752745 | -0.204977755 | 0.844176184 | 0.942169945 | -4.903405127 |
| **A_52_P1128182** | Tmem185a | -0.015518023 | -0.199562476 | 0.168526431 | 8.870225555 | -0.204878712 | 0.844250265 | 0.942169945 | -4.903416154 |
| **A_55_P2020706** | NA | -0.016267946 | -0.209249149 | 0.176713258 | 7.728555768 | -0.204833456 | 0.844284116 | 0.942169945 | -4.90342119 |
| **A_52_P526396** | Atl1 | 0.014447608 | -0.15704595 | 0.185941165 | 8.04293753 | 0.204706351 | 0.844379189 | 0.942169945 | -4.90343533 |
| **A_55_P2010858** | Mrps23 | 0.023327413 | -0.25357322 | 0.300228047 | 8.893732929 | 0.204703764 | 0.844381125 | 0.942169945 | -4.903435618 |
| **A_52_P373798** | Zdhhc3 | 0.01742223 | -0.189391959 | 0.224236419 | 8.215713127 | 0.204694697 | 0.844387907 | 0.942169945 | -4.903436626 |
| **A_52_P24428** | Ankrd11 | -0.015229959 | -0.196048892 | 0.165588974 | 10.87065874 | -0.20466238 | 0.84441208 | 0.942169945 | -4.90344022 |
| **A_51_P400217** | Vpreb1 | 0.027943355 | -0.303861814 | 0.359748524 | 12.62362568 | 0.204634375 | 0.844433028 | 0.942169945 | -4.903443333 |
| **A_51_P324082** | Med1 | -0.021598657 | -0.278074768 | 0.234877454 | 9.87315585 | -0.204627036 | 0.844438518 | 0.942169945 | -4.903444149 |
| **A_51_P249193** | Gsg1l | -0.014592252 | -0.187969868 | 0.158785364 | 7.713258244 | -0.204509025 | 0.844526794 | 0.942184522 | -4.903457265 |
| **A_55_P2054311** | M6pr | 0.015458295 | -0.16823595 | 0.199152539 | 10.08121281 | 0.204479242 | 0.844549073 | 0.942184522 | -4.903460574 |
| **A_55_P2040678** | Susd2 | -0.021835556 | -0.281447142 | 0.23777603 | 8.063346642 | -0.204372934 | 0.844628598 | 0.942218859 | -4.903472381 |
| **A_51_P108862** | Cops7a | -0.016142217 | -0.208170896 | 0.175886463 | 12.72999991 | -0.204258562 | 0.844714157 | 0.942259922 | -4.879619982 |
| **A_51_P464588** | Dnajc28 | -0.015800441 | -0.203850451 | 0.17224957 | 8.017447774 | -0.204163933 | 0.844784948 | 0.942284509 | -4.903495575 |
| **A_55_P2112474** | NA | -0.016146873 | -0.208383774 | 0.176090027 | 16.20950173 | -0.204096179 | 0.844835635 | 0.94228667 | -4.90350309 |
| **A_55_P2148431** | Slc4a3 | -0.014660447 | -0.189418726 | 0.160097832 | 7.670271374 | -0.203841513 | 0.84502616 | 0.942444789 | -4.903531312 |
| **A_52_P574993** | Pappa2 | 0.013181677 | -0.144036284 | 0.170399638 | 7.655966006 | 0.203728457 | 0.845110745 | 0.942461295 | -4.903543829 |
| **A_55_P2115260** | NA | -0.021352644 | -0.276072127 | 0.23336684 | 8.169449932 | -0.203691391 | 0.845138477 | 0.942461295 | -4.903547932 |
| **A_55_P2009588** | Rpl13 | -0.015331464 | -0.198333546 | 0.167670617 | 16.43274758 | -0.203568601 | 0.845230348 | 0.942509369 | -4.903561517 |
| **A_55_P2047621** | Cutal | -0.040320217 | -0.522006667 | 0.441366232 | 10.74925864 | -0.203395633 | 0.845359767 | 0.942531157 | -4.903580641 |
| **A_55_P2069935** | Fxyd2 | -0.036893054 | -0.477677048 | 0.40389094 | 8.516667968 | -0.203377068 | 0.845373658 | 0.942531157 | -4.903582692 |
| **A_51_P444994** | Tmem203 | 0.014569052 | -0.159522122 | 0.188660225 | 10.55073195 | 0.203346974 | 0.845396175 | 0.942531157 | -4.903586018 |
| **A_51_P307964** | Krt13 | 0.016931769 | -0.185542701 | 0.219406239 | 7.945892642 | 0.203196043 | 0.845509111 | 0.94259365 | -4.903602688 |
| **A_51_P349727** | Slc25a45 | -0.022180983 | -0.287559514 | 0.243197548 | 8.663761997 | -0.203094515 | 0.845585082 | 0.94259365 | -4.903613894 |
| **A_55_P1987054** | Krtap5-1 | 0.020506569 | -0.224904539 | 0.265917676 | 8.310035321 | 0.203040172 | 0.845625747 | 0.94259365 | -4.903619891 |
| **A_51_P442097** | Slc41a3 | -0.039135709 | -0.507556328 | 0.429284911 | 11.88609195 | -0.203011381 | 0.845647291 | 0.94259365 | -4.903623067 |
| **A_51_P304126** | Necap1 | 0.033262541 | -0.365468727 | 0.431993809 | 10.47714695 | 0.202702133 | 0.845878712 | 0.942782576 | -4.903657155 |
| **A_51_P311096** | Malsu1 | -0.024897055 | -0.323449662 | 0.273655552 | 12.78904885 | -0.202633057 | 0.845930406 | 0.942782576 | -4.903664762 |
| **A_55_P1962693** | Pla2g5 | -0.014723098 | -0.191355195 | 0.161908998 | 7.68550582 | -0.202540921 | 0.845999358 | 0.942782576 | -4.903674904 |
| **A_55_P2029740** | Gm10240 | -0.021289501 | -0.276792681 | 0.234213679 | 15.26078859 | -0.202466119 | 0.84605534 | 0.942782576 | -4.903683135 |
| **A_55_P2012051** | Sox1 | 0.04411763 | -0.485460636 | 0.573695896 | 8.540343822 | 0.202425495 | 0.846085743 | 0.942782576 | -4.903687604 |
| **A_52_P682990** | Timm50 | -0.025575924 | -0.332631677 | 0.28147983 | 10.51312723 | -0.202393833 | 0.846109439 | 0.942782576 | -4.903691087 |
| **A_52_P20727** | Nhlrc1 | 0.01704702 | -0.187928923 | 0.222022962 | 8.29299025 | 0.202082526 | 0.846342434 | 0.942972916 | -4.903725299 |

| | | | | | | | | | |
|---|---|---|---|---|---|---|---|---|---|
| **A_55_P2010197** | Serpina10 | -0.031703377 | -0.412998805 | 0.349592052 | 13.58561189 | -0.202035231 | 0.846377832 | 0.942972916 | -4.903730492 |
| **A_52_P301223** | Brwd1 | 0.017447388 | -0.19267857 | 0.227573347 | 9.072474005 | 0.201759458 | 0.846584249 | 0.943050859 | -4.903760749 |
| **A_55_P1965574** | A2ml1 | 0.07389516 | -0.816057833 | 0.963848152 | 10.8638484 | 0.201758627 | 0.84658487 | 0.943050859 | -4.90376084 |
| **A_66_P137462** | Nsdhl | -0.050756254 | -0.662074496 | 0.560561987 | 10.34772325 | -0.201746217 | 0.84659416 | 0.943050859 | -4.903762201 |
| **A_55_P2060255** | NA | -0.020634751 | -0.278935901 | 0.237666398 | 15.42404262 | -0.203242847 | 0.846691149 | 0.943053126 | -4.855027358 |
| **A_55_P2128792** | NA | 0.019183955 | -0.212023858 | 0.250391768 | 8.955476743 | 0.201613139 | 0.846693773 | 0.943053126 | -4.903776786 |
| **A_52_P355095** | Dstyk | -0.01838488 | -0.240458961 | 0.2036892 | 8.009254303 | -0.201162084 | 0.847031428 | 0.943272342 | -4.903826152 |
| **A_55_P2102693** | Acvr1 | -0.01954312 | -0.255684986 | 0.216598745 | 8.642899004 | -0.201096312 | 0.847080667 | 0.943272342 | -4.903833342 |
| **A_55_P2137186** | Rps25 | 0.017754919 | -0.196816917 | 0.232326754 | 15.85002881 | 0.201061611 | 0.847106646 | 0.943272342 | -4.903837134 |
| **A_51_P373619** | Mbl1 | 0.029208707 | -0.323909309 | 0.382326723 | 10.12477112 | 0.200990553 | 0.847159844 | 0.943272342 | -4.903844897 |
| **A_55_P1991109** | Gatd3a | -0.020268142 | -0.265467004 | 0.22493072 | 12.94301813 | -0.200853165 | 0.847262702 | 0.943272342 | -4.903859899 |
| **A_51_P297968** | Pdia6 | -0.034114497 | -0.446828491 | 0.378599498 | 10.1864229 | -0.200850521 | 0.847264681 | 0.943272342 | -4.903860188 |
| **A_51_P341746** | Mpl | 0.046243339 | -0.513301128 | 0.605787807 | 9.301542626 | 0.200815749 | 0.847290715 | 0.943272342 | -4.903863983 |
| **A_55_P2042229** | NA | -0.029221417 | -0.382804297 | 0.324361463 | 12.64449369 | -0.20081365 | 0.847292286 | 0.943272342 | -4.903864213 |
| **A_55_P1985569** | Tomm6 | -0.019583394 | -0.256711484 | 0.217544695 | 14.30168835 | -0.200672634 | 0.847397865 | 0.943272342 | -4.903879598 |
| **A_52_P547612** | Tmem30b | 0.056544644 | -0.628449793 | 0.741539081 | 9.919464224 | 0.200579996 | 0.847467225 | 0.943272342 | -4.903889699 |
| **A_51_P390276** | Cwc25 | 0.01984191 | -0.220621165 | 0.260304986 | 8.957856266 | 0.200501801 | 0.847525772 | 0.943272342 | -4.903898222 |
| **A_55_P2182273** | NA | -0.041052514 | -0.53870101 | 0.456595981 | 8.978290224 | -0.20044732 | 0.847566565 | 0.943272342 | -4.903904158 |
| **A_55_P1955747** | Gm12070 | -0.026904528 | -0.353097895 | 0.299288838 | 15.18397409 | -0.200416488 | 0.847589651 | 0.943272342 | -4.903907517 |
| **A_66_P112862** | Gm17821 | 0.020924248 | -0.232817052 | 0.274665549 | 11.53178545 | 0.200374244 | 0.847621281 | 0.943272342 | -4.903912118 |
| **A_55_P2062627** | Tmem238 | -0.018082771 | -0.237368367 | 0.201202825 | 8.92495905 | -0.200372478 | 0.847622604 | 0.943272342 | -4.90391231 |
| **A_55_P2060243** | Itgbl1 | 0.012665972 | -0.140985036 | 0.16631698 | 7.626133827 | 0.200302466 | 0.847675026 | 0.943276372 | -4.903919933 |
| **A_55_P2149274** | Sh3bp5l | -0.029815999 | -0.391762656 | 0.332130659 | 11.67903086 | -0.200164925 | 0.847778015 | 0.943336668 | -4.903934902 |
| **A_55_P1979743** | NA | 0.02028235 | -0.22619521 | 0.26675991 | 14.91492386 | 0.199951227 | 0.847938036 | 0.943460414 | -4.903958139 |
| **A_51_P179241** | Tssc4 | 0.019060169 | -0.212987554 | 0.251107892 | 11.35126878 | 0.199587173 | 0.848210665 | 0.94364178 | -4.903997669 |
| **A_55_P2168587** | Cdk9 | 0.01926955 | -0.215371559 | 0.25391066 | 8.833433132 | 0.199549509 | 0.848238872 | 0.94364178 | -4.904001755 |
| **A_55_P2008176** | Zscan12 | 0.014524841 | -0.162351195 | 0.191400877 | 8.799819433 | 0.199537988 | 0.8482475 | 0.94364178 | -4.904003004 |
| **A_51_P206971** | Mnat1 | -0.025151184 | -0.331691296 | 0.281388928 | 9.661187517 | -0.199367477 | 0.8483752 | 0.94370656 | -4.90402149 |
| **A_51_P399071** | Anp32a | 0.0211999 | -0.237231127 | 0.279630927 | 9.140808932 | 0.199329853 | 0.848403378 | 0.94370656 | -4.904025567 |
| **A_66_P136259** | Rbm4b | 0.018421445 | -0.206246929 | 0.243089818 | 8.345184622 | 0.19923468 | 0.848474658 | 0.943707254 | -4.904035877 |
| **A_55_P2112569** | Ern1 | -0.013129729 | -0.173289181 | 0.147029723 | 7.771384965 | -0.199198643 | 0.848501649 | 0.943707254 | -4.904039779 |
| **A_55_P1998164** | Rab28 | 0.016760261 | -0.187802923 | 0.221323445 | 8.272728008 | 0.19908406 | 0.848587469 | 0.943748399 | -4.904052182 |
| **A_52_P508089** | Psap | 0.035480352 | -0.397892586 | 0.46885329 | 11.62264635 | 0.198934139 | 0.84869976 | 0.943818978 | -4.904068401 |
| **A_55_P2113155** | Scnm1 | -0.017294106 | -0.228654605 | 0.194066394 | 9.566509277 | -0.19881881 | 0.848786144 | 0.943860739 | -4.904080868 |
| **A_55_P2172233** | NA | -0.017183701 | -0.227737427 | 0.193370026 | 15.96618297 | -0.198306504 | 0.8491699 | 0.944084038 | -4.904136167 |
| **A_55_P1992617** | Clec1b | -0.047643637 | -0.6314896 | 0.536202326 | 11.03439201 | -0.198284977 | 0.849186026 | 0.944084038 | -4.904138487 |
| **A_52_P444637** | Rcbtb2 | -0.013930539 | -0.184668949 | 0.156807871 | 8.050740578 | -0.198253049 | 0.849209945 | 0.944084038 | -4.904141929 |
| **A_51_P216496** | Tfpi2 | 0.027508719 | -0.309652142 | 0.364669579 | 8.325337574 | 0.198251463 | 0.849211133 | 0.944084038 | -4.9041421 |
| **A_51_P472249** | Slc7a7 | 0.026409914 | -0.297327094 | 0.350146921 | 11.3431373 | 0.198224724 | 0.849231164 | 0.944084038 | -4.904144981 |
| **A_51_P250551** | Dhx38 | 0.020024469 | -0.225580044 | 0.265628982 | 8.809097393 | 0.198110661 | 0.849316614 | 0.944124732 | -4.904157269 |
| **A_55_P2074636** | Adap1 | -0.018663817 | -0.248132725 | 0.210805091 | 11.36581262 | -0.197633154 | 0.849674359 | 0.944418857 | -4.904208635 |
| **A_55_P2298319** | NA | -0.051399378 | -0.683366731 | 0.580567974 | 9.115177609 | -0.197627061 | 0.849678924 | 0.944418857 | -4.90420929 |
| **A_66_P117815** | NA | -0.027742796 | -0.369181484 | 0.313695892 | 12.91593553 | -0.197433428 | 0.849824005 | 0.944493873 | -4.904230084 |
| **A_51_P360963** | Mbtd1 | -0.021622188 | -0.287947251 | 0.244702874 | 10.03937285 | -0.19727443 | 0.849943139 | 0.944493873 | -4.904247144 |
| **A_51_P267314** | Tbc1d10c | -0.022134854 | -0.294852931 | 0.250583223 | 8.982601656 | -0.197217713 | 0.849985637 | 0.944493873 | -4.904253226 |
| **A_55_P2044104** | Fgfr1op | 0.019607846 | -0.222045833 | 0.261261525 | 10.34767993 | 0.19716039 | 0.85002859 | 0.944493873 | -4.904259372 |
| **A_55_P2039110** | NA | 0.040647133 | -0.46042948 | 0.541723746 | 14.94878183 | 0.197110145 | 0.85006624 | 0.944493873 | -4.904264757 |
| **A_51_P442719** | Usp6nl | -0.020559832 | -0.274066181 | 0.232946517 | 8.448879906 | -0.197067003 | 0.850098568 | 0.944493873 | -4.90426938 |
| **A_55_P2049717** | Agrp | 0.013263984 | -0.150295151 | 0.176823119 | 7.74929045 | 0.197052692 | 0.850109291 | 0.944493873 | -4.904270913 |
| **A_55_P2051791** | Cbr4 | -0.022675764 | -0.302345329 | 0.256993802 | 8.818717312 | -0.197015277 | 0.850137328 | 0.944493873 | -4.904274921 |

| | | | | | | | | | |
|---|---|---|---|---|---|---|---|---|---|
| **A_52_P167411** | Tnk1 | -0.034117911 | -0.455104803 | 0.38686898 | 8.393423254 | -0.196923276 | 0.850206269 | 0.944516177 | -4.904284774 |
| **A_66_P135941** | Fam221a | -0.013894916 | -0.185456849 | 0.157667016 | 7.99720204 | -0.196796874 | 0.850300991 | 0.944529501 | -4.904298303 |
| **A_55_P2157033** | Bace2 | 0.025741017 | -0.292118255 | 0.343600289 | 10.36234867 | 0.196776853 | 0.850315994 | 0.944529501 | -4.904300445 |
| **A_51_P342549** | Hand1 | 0.012728453 | -0.144512505 | 0.16996941 | 7.67881267 | 0.196694909 | 0.850377402 | 0.944543432 | -4.90430921 |
| **A_55_P2187053** | Tpm1 | 0.040662581 | -0.462282151 | 0.543607313 | 10.45632146 | 0.196452639 | 0.850558963 | 0.944690812 | -4.904335104 |
| **A_52_P382886** | Gjb2 | 0.073605721 | -0.837602328 | 0.984813771 | 11.5602368 | 0.196280526 | 0.850687953 | 0.944779789 | -4.90435348 |
| **A_55_P2011300** | Fggy | -0.028661772 | -0.38401934 | 0.326695796 | 8.493921329 | -0.195984017 | 0.850910184 | 0.944914177 | -4.904385101 |
| **A_55_P2234704** | NA | -0.014997347 | -0.200962574 | 0.170967881 | 7.746206396 | -0.195959297 | 0.850928712 | 0.944914177 | -4.904387735 |
| **A_52_P476754** | Ubash3b | 0.031413735 | -0.35818402 | 0.421011489 | 10.85971433 | 0.195923404 | 0.850955615 | 0.944914177 | -4.904391559 |
| **A_55_P2156401** | NA | 0.030262576 | -0.34525033 | 0.405775481 | 14.89399955 | 0.19582323 | 0.8510307 | 0.944943267 | -4.904402228 |
| **A_52_P639567** | Ate1 | 0.016074184 | -0.183516528 | 0.215664896 | 10.06800779 | 0.195691426 | 0.851129493 | 0.94494628 | -4.904416258 |
| **A_55_P2000515** | Peli1 | -0.02633727 | -0.353366951 | 0.30069241 | 11.12608367 | -0.195689165 | 0.851131189 | 0.94494628 | -4.904416499 |
| **A_51_P304683** | Clpx | 0.049393588 | -0.56438102 | 0.663168197 | 11.80709357 | 0.19554416 | 0.851239881 | 0.945012674 | -4.904431923 |
| **A_55_P2029678** | Etfa | -0.038710998 | -0.519972473 | 0.442550477 | 13.95659628 | -0.195450353 | 0.851310198 | 0.945036459 | -4.904441895 |
| **A_55_P2083694** | Adam33 | 0.029513224 | -0.337576778 | 0.396603225 | 8.499751721 | 0.195356237 | 0.85138075 | 0.945060502 | -4.904451895 |
| **A_55_P1977681** | Vps13d | -0.020801393 | -0.279835043 | 0.238232258 | 9.131463823 | -0.195127919 | 0.851551907 | 0.945143308 | -4.904476136 |
| **A_51_P249537** | Wdr24 | -0.023324193 | -0.313932186 | 0.267283799 | 9.548970762 | -0.195021399 | 0.851631761 | 0.945143308 | -4.904487435 |
| **A_55_P1965075** | NA | 0.019130716 | -0.219230844 | 0.257492276 | 15.77814485 | 0.195019572 | 0.851633131 | 0.945143308 | -4.904487629 |
| **A_51_P232474** | Zswim1 | 0.019559694 | -0.224176458 | 0.263295847 | 7.977405976 | 0.194995819 | 0.851650939 | 0.945143308 | -4.904490148 |
| **A_55_P2044710** | Asb10 | 0.012876131 | -0.147780686 | 0.173532948 | 7.817357478 | 0.194746393 | 0.851837934 | 0.945296557 | -4.90451658 |
| **A_55_P1971951** | Cd86 | 0.039591404 | -0.454832262 | 0.534015071 | 9.697026241 | 0.194574016 | 0.851967172 | 0.945318078 | -4.904534827 |
| **A_52_P340669** | Bhlha15 | -0.034400446 | -0.463999138 | 0.395198245 | 9.249670961 | -0.194573781 | 0.851967349 | 0.945318078 | -4.904534852 |
| **A_55_P2048368** | Fggy | 0.043587878 | -0.50088201 | 0.588057766 | 9.218295335 | 0.194524833 | 0.852004047 | 0.945318078 | -4.904540031 |
| **A_55_P1998601** | Slc17a9 | -0.048991525 | -0.661474387 | 0.563491338 | 10.27230989 | -0.194361483 | 0.852126524 | 0.945348445 | -4.904557304 |
| **A_55_P2103145** | Ppp1r2 | 0.018843338 | -0.216803492 | 0.254490169 | 8.699893464 | 0.194302968 | 0.852170398 | 0.945348445 | -4.904563488 |
| **A_51_P232705** | Commd9 | -0.013150722 | -0.177705191 | 0.151403746 | 8.939434048 | -0.194188326 | 0.852256358 | 0.945348445 | -4.904575598 |
| **A_55_P2132865** | Sumo2 | 0.023166462 | -0.266791059 | 0.313123984 | 12.51837216 | 0.194137097 | 0.852294771 | 0.945348445 | -4.904581008 |
| **A_51_P365468** | Clpb | 0.026310306 | -0.303114479 | 0.35573509 | 10.5675137 | 0.194067501 | 0.852346956 | 0.945348445 | -4.904588355 |
| **A_55_P1968108** | NA | 0.026074053 | -0.300629607 | 0.352777712 | 13.81078339 | 0.193926753 | 0.852452496 | 0.945348445 | -4.904603204 |
| **A_51_P125260** | Acaa2 | -0.054708982 | -0.740300506 | 0.630882543 | 16.05516235 | -0.193899362 | 0.852473036 | 0.945348445 | -4.904606093 |
| **A_55_P1993708** | Lrp12 | 0.018137147 | -0.209163017 | 0.245437311 | 8.396414062 | 0.193888661 | 0.85248106 | 0.945348445 | -4.904607221 |
| **A_65_P05152** | Flrt2 | 0.013135816 | -0.151530739 | 0.177802371 | 7.923765421 | 0.193836177 | 0.852520417 | 0.945348445 | -4.904612755 |
| **A_55_P2272007** | Lrrfip2 | -0.016905774 | -0.228831496 | 0.195019947 | 8.380928562 | -0.193836067 | 0.8525205 | 0.945348445 | -4.904612766 |
| **A_55_P1996509** | Rpl29 | -0.025764754 | -0.349118894 | 0.297589387 | 16.71019895 | -0.193611327 | 0.852689032 | 0.945481086 | -4.904636445 |
| **A_55_P2053616** | NA | -0.033748323 | -0.45790106 | 0.390404415 | 10.21846825 | -0.19333617 | 0.852895383 | 0.945655645 | -4.904665398 |
| **A_55_P2026233** | Uba7 | 0.016334407 | -0.189220763 | 0.221889577 | 11.80076498 | 0.193089281 | 0.853080545 | 0.945713519 | -4.904691343 |
| **A_51_P302149** | Cfl2 | -0.019270539 | -0.261794938 | 0.22325386 | 8.543469736 | -0.193073031 | 0.853092733 | 0.945713519 | -4.904693049 |
| **A_55_P1956083** | Gpr68 | -0.013231296 | -0.179840811 | 0.153378219 | 7.797131287 | -0.192968206 | 0.853171354 | 0.945713519 | -4.904704054 |
| **A_55_P2064053** | NA | 0.038872009 | -0.45078871 | 0.528532727 | 11.7433179 | 0.192896748 | 0.853224949 | 0.945713519 | -4.904711553 |
| **A_55_P2080905** | NA | -0.016252441 | -0.221020812 | 0.18851593 | 16.37246561 | -0.192858557 | 0.853253594 | 0.945713519 | -4.904715559 |
| **A_51_P413539** | Nek6 | -0.025410098 | -0.345661423 | 0.294841227 | 10.84603209 | -0.192796253 | 0.853300325 | 0.945713519 | -4.904722094 |
| **A_52_P647919** | Usf2 | -0.016069032 | -0.218635385 | 0.186497321 | 9.341268022 | -0.192754975 | 0.853331285 | 0.945713519 | -4.904726422 |
| **A_55_P2163857** | Trim12c | -0.030758704 | -0.418523868 | 0.357006461 | 8.344119556 | -0.192744693 | 0.853338998 | 0.945713519 | -4.9047275 |
| **A_55_P1972991** | Pcsk6 | -0.041503894 | -0.565966507 | 0.48295872 | 10.77877412 | -0.192290353 | 0.853679799 | 0.945981367 | -4.904775076 |
| **A_55_P1995537** | Mpo | -0.039957803 | -0.544890419 | 0.464974812 | 14.80693576 | -0.192287641 | 0.853681833 | 0.945981367 | -4.904775359 |
| **A_55_P1982107** | NA | 0.020125731 | -0.234276556 | 0.274528017 | 15.88786364 | 0.192226755 | 0.853727506 | 0.945981367 | -4.904781727 |
| **A_51_P175974** | NA | 0.019365629 | -0.225565748 | 0.264297006 | 8.552514881 | 0.192119026 | 0.853808319 | 0.946016681 | -4.904792988 |
| **A_55_P1990859** | Rpl6 | 0.015964852 | -0.186047117 | 0.217976822 | 15.92438847 | 0.192030845 | 0.85387447 | 0.946031503 | -4.904802201 |
| **A_55_P2106235** | Syngr2 | -0.01938601 | -0.26476459 | 0.22599257 | 12.8344172 | -0.191970709 | 0.853919584 | 0.946031503 | -4.904808481 |
| **A_55_P2135486** | NA | -0.012843768 | -0.175495921 | 0.149808385 | 7.673996265 | -0.191873865 | 0.853992236 | 0.946057768 | -4.904818591 |

| | | | | | | | | | |
|---|---|---|---|---|---|---|---|---|---|
| **A_52_P277854** | Snrpb | -0.044308052 | -0.606280106 | 0.517664001 | 10.96362462 | -0.191580424 | 0.854212385 | 0.946211663 | -4.904849195 |
| **A_52_P674309** | Mocs2 | 0.021298413 | -0.248866967 | 0.291463793 | 12.88016937 | 0.191558197 | 0.854229061 | 0.946211663 | -4.904851511 |
| **A_55_P2186025** | NA | -0.024237887 | -0.331843767 | 0.283367993 | 14.91186971 | -0.191462319 | 0.854300995 | 0.946222143 | -4.9048615 |
| **A_55_P1998213** | NA | -0.016215428 | -0.222092616 | 0.189661761 | 16.54530507 | -0.191383005 | 0.854360503 | 0.946222143 | -4.904869759 |
| **A_52_P346033** | Trmt1l | -0.016860866 | -0.230999279 | 0.197277548 | 8.404133785 | -0.191323569 | 0.854405097 | 0.946222143 | -4.904875945 |
| **A_55_P2243431** | Gdap10 | 0.020388721 | -0.238657584 | 0.279435026 | 8.862896162 | 0.191247502 | 0.854462171 | 0.946222143 | -4.904883861 |
| **A_55_P2120278** | Ube2l3 | -0.013322963 | -0.182621097 | 0.15597517 | 12.63656407 | -0.191219355 | 0.85448329 | 0.946222143 | -4.904886789 |
| **A_55_P2182790** | NA | 0.016133372 | -0.188957803 | 0.221224546 | 14.15972223 | 0.191144301 | 0.854539605 | 0.946230294 | -4.904894595 |
| **A_55_P2020461** | Hmgn2 | -0.025106117 | -0.344476751 | 0.294264517 | 10.60694956 | -0.191015125 | 0.85463653 | 0.94628341 | -4.904908022 |
| **A_55_P2082733** | Cybb | -0.035666659 | -0.489650539 | 0.41831722 | 8.910006627 | -0.190899676 | 0.854723158 | 0.946325119 | -4.904920015 |
| **A_55_P2083559** | Hrk | -0.012027713 | -0.165378461 | 0.141323035 | 7.646374554 | -0.190581329 | 0.854962045 | 0.946535389 | -4.904953049 |
| **A_51_P107808** | Ttc19 | 0.012797857 | -0.150488725 | 0.176084438 | 7.828220135 | 0.190445158 | 0.855064231 | 0.946594303 | -4.896937873 |
| **A_51_P172532** | Nit1 | 0.029002332 | -0.341180183 | 0.399184847 | 12.0948675 | 0.190370744 | 0.855120076 | 0.946601909 | -4.90497487 |
| **A_52_P681016** | NA | 0.014352977 | -0.169201001 | 0.197906955 | 7.724352628 | 0.190003407 | 0.855395757 | 0.946852857 | -4.905012878 |
| **A_55_P2152114** | Zfyve19 | 0.022401117 | -0.264446965 | 0.3092492 | 9.435450126 | 0.189758361 | 0.855579673 | 0.946959725 | -4.905038193 |
| **A_52_P38964** | Sap25 | 0.018390901 | -0.217126384 | 0.253908187 | 8.652690985 | 0.189741973 | 0.855591973 | 0.946959725 | -4.905039885 |
| **A_55_P2090484** | Pde10a | -0.012359977 | -0.170696719 | 0.145976766 | 7.681875752 | -0.189678949 | 0.855639277 | 0.946959725 | -4.90504639 |
| **A_55_P2199717** | 9330102E08Rik | 0.012366703 | -0.146139639 | 0.170873046 | 7.734583295 | 0.18957911 | 0.855714214 | 0.946960422 | -4.905056691 |
| **A_55_P2121744** | Vars2 | -0.012287434 | -0.16980398 | 0.145229112 | 8.014149444 | -0.189547566 | 0.85573789 | 0.946960422 | -4.905059944 |
| **A_52_P435670** | Npepl1 | -0.017049381 | -0.235727986 | 0.201629224 | 10.47843338 | -0.189446032 | 0.855814102 | 0.946990542 | -4.905070412 |
| **A_52_P601688** | Abcc1 | -0.014760218 | -0.204570374 | 0.175049938 | 8.352353291 | -0.188954186 | 0.856183305 | 0.947254175 | -4.905121044 |
| **A_55_P2080870** | Cmtm7 | -0.014941552 | -0.20719362 | 0.177310517 | 11.42784667 | -0.188846033 | 0.856264496 | 0.947254175 | -4.90513216 |
| **A_55_P2066304** | Limk2 | 0.017746717 | -0.210601049 | 0.246094482 | 7.91589669 | 0.188844543 | 0.856265614 | 0.947254175 | -4.905132313 |
| **A_55_P2023652** | Vps35l | -0.015144361 | -0.21011182 | 0.179823099 | 11.31368028 | -0.188743497 | 0.856341471 | 0.947254175 | -4.905142693 |
| **A_55_P2408415** | Kcnq1ot1 | 0.022457608 | -0.266674654 | 0.31158987 | 8.85961696 | 0.188733993 | 0.856348606 | 0.947254175 | -4.905143669 |
| **A_55_P2162324** | Bmx | -0.027550823 | -0.382318447 | 0.327216802 | 8.969454537 | -0.188700817 | 0.856373512 | 0.947254175 | -4.905147076 |
| **A_55_P2007026** | Gm9112 | -0.013148202 | -0.182524917 | 0.156228513 | 7.756707506 | -0.188623519 | 0.856431543 | 0.947254175 | -4.905155011 |
| **A_55_P2107447** | Rab37 | -0.033160481 | -0.460497638 | 0.394176677 | 8.852411991 | -0.188552957 | 0.856484517 | 0.947254175 | -4.905162252 |
| **A_55_P1962344** | Trim21 | 0.016925458 | -0.201229322 | 0.235080239 | 9.160683615 | 0.188520634 | 0.856508784 | 0.947254175 | -4.905165568 |
| **A_55_P1975682** | Rhox2e | -0.012297366 | -0.170958074 | 0.146363342 | 7.767065919 | -0.188332775 | 0.856649824 | 0.947254175 | -4.905184829 |
| **A_51_P402193** | Map3k1 | 0.035495919 | -0.422493733 | 0.493485571 | 11.48207395 | 0.188324121 | 0.856656321 | 0.947254175 | -4.905185716 |
| **A_55_P2134122** | NA | -0.01462113 | -0.203337246 | 0.174094985 | 8.136810905 | -0.188258741 | 0.856705408 | 0.947254175 | -4.905192414 |
| **A_55_P2014500** | Phf24 | 0.012718356 | -0.151452384 | 0.176889096 | 7.778171921 | 0.188242818 | 0.856717363 | 0.947254175 | -4.905194045 |
| **A_55_P2045682** | Rassf5 | -0.029853212 | -0.415492361 | 0.355785937 | 10.63364413 | -0.188101881 | 0.856823181 | 0.947254175 | -4.905208476 |
| **A_52_P427818** | Zscan18 | -0.018182995 | -0.253127495 | 0.216761505 | 7.85932786 | -0.188054327 | 0.856858886 | 0.947254175 | -4.905213343 |
| **A_55_P2070546** | 1700037C18Rik | -0.032060836 | -0.446390641 | 0.382268969 | 9.020256747 | -0.188023381 | 0.856882122 | 0.947254175 | -4.90521651 |
| **A_51_P448946** | Sertad2 | 0.01716614 | -0.204681234 | 0.239013515 | 11.83319586 | 0.188018922 | 0.85688547 | 0.947254175 | -4.905216966 |
| **A_55_P1972381** | Gjb4 | 0.013207434 | -0.157764947 | 0.184179814 | 7.681876547 | 0.187704928 | 0.857121237 | 0.947454418 | -4.905249064 |
| **A_52_P683211** | NA | 0.019708521 | -0.235500086 | 0.274917128 | 8.704394749 | 0.187647122 | 0.857164644 | 0.947454418 | -4.905254968 |
| **A_51_P311038** | Adamts15 | 0.042092506 | -0.503164032 | 0.587349043 | 8.725831276 | 0.18758024 | 0.857214866 | 0.947455749 | -4.905261796 |
| **A_55_P2131506** | NA | 0.014077916 | -0.168630506 | 0.196786338 | 7.872790887 | 0.187224638 | 0.857481902 | 0.94760884 | -4.905298061 |
| **A_55_P2048650** | Dennd4a | 0.042279648 | -0.506480865 | 0.591040162 | 15.86902762 | 0.187211145 | 0.857492035 | 0.94760884 | -4.905299436 |
| **A_52_P599317** | Hs6st2 | 0.015444795 | -0.185029939 | 0.21591953 | 8.096793199 | 0.187199939 | 0.857500451 | 0.94760884 | -4.905300577 |
| **A_55_P2147816** | NA | 0.013128402 | -0.157361119 | 0.183617923 | 16.89765297 | 0.187110156 | 0.857567877 | 0.947629173 | -4.905309722 |
| **A_55_P1967978** | Nfatc2 | 0.019355256 | -0.232200178 | 0.270910689 | 8.415848676 | 0.186959869 | 0.857680742 | 0.947698262 | -4.905325019 |
| **A_51_P408703** | Rnf138rt1 | 0.012725938 | -0.152726134 | 0.17817801 | 7.691224056 | 0.186896332 | 0.85772846 | 0.947698262 | -4.905331482 |
| **A_55_P2086507** | Bglap3 | -0.013643087 | -0.191110915 | 0.163824741 | 7.739298477 | -0.186799713 | 0.857801023 | 0.947724263 | -4.905341307 |
| **A_55_P2054331** | Gm9222 | -0.012787892 | -0.179205366 | 0.153629583 | 7.875588078 | -0.186716725 | 0.857863351 | 0.947738953 | -4.905349741 |
| **A_55_P2054128** | NA | -0.025659966 | -0.359833684 | 0.308513752 | 13.39452924 | -0.186580811 | 0.857965431 | 0.947797555 | -4.905363547 |
| **A_55_P2024799** | NA | 0.026580392 | -0.320121533 | 0.373282316 | 13.24129402 | 0.186289472 | 0.858184256 | 0.947985112 | -4.905393108 |

| | | | | | | | | |
|---|---|---|---|---|---|---|---|---|
| **A_51_P114722** | Hao2 | -0.077284908 | -1.086132594 | 0.931562777 | 10.91757641 | -0.186145389 | 0.858292482 | 0.948050482 | -4.901473233 |
| **A_55_P2052116** | NA | 0.014553947 | -0.175652725 | 0.204760619 | 9.728960998 | 0.185925185 | 0.858457892 | 0.948179005 | -4.905430006 |
| **A_55_P2009783** | Sfr1 | -0.01736727 | -0.24442855 | 0.209694011 | 13.33668485 | -0.18585389 | 0.858511447 | 0.948183976 | -4.905437219 |
| **A_55_P2192633** | E330017L17Rik | 0.011739506 | -0.141928673 | 0.165407686 | 7.637042819 | 0.185630388 | 0.858679345 | 0.948263187 | -4.905459813 |
| **A_55_P1993049** | Gm4076 | -0.03460665 | -0.487698673 | 0.418485374 | 12.4665363 | -0.18559075 | 0.858709122 | 0.948263187 | -4.905463818 |
| **A_51_P137800** | Acp7 | 0.011774074 | -0.142402983 | 0.165951132 | 7.638731819 | 0.1855625 | 0.858730344 | 0.948263187 | -4.905466671 |
| **A_52_P279184** | Srp54a | -0.016404961 | -0.231501335 | 0.198691412 | 8.855311108 | -0.185321289 | 0.858911556 | 0.948386267 | -4.905491017 |
| **A_55_P2048912** | Themis2 | -0.023018968 | -0.324897523 | 0.278859586 | 8.443090398 | -0.185283516 | 0.858939934 | 0.948386267 | -4.905494827 |
| **A_55_P2001976** | Rhox12 | -0.011819179 | -0.16734708 | 0.143708722 | 7.657388615 | -0.184655474 | 0.859411804 | 0.948853074 | -4.905558058 |
| **A_55_P1988708** | Gstz1 | -0.039601536 | -0.561308307 | 0.482105236 | 13.25769125 | -0.184445788 | 0.859569363 | 0.948928743 | -4.905579122 |
| **A_55_P2074499** | Kcp | 0.013508641 | -0.164471614 | 0.191488897 | 9.014096029 | 0.184426375 | 0.859583951 | 0.948928743 | -4.905581071 |
| **A_55_P1960506** | Wdr83os | -0.024844303 | -0.35227837 | 0.302589763 | 11.7476836 | -0.184368258 | 0.859627622 | 0.948928743 | -4.905586905 |
| **A_51_P406466** | Ankrd35 | -0.0123779 | -0.175639897 | 0.150884097 | 7.771030754 | -0.184223507 | 0.859736393 | 0.948994617 | -4.905601426 |
| **A_55_P2008735** | Cdc123 | 0.027422741 | -0.334543537 | 0.389389019 | 8.981429268 | 0.184088197 | 0.859838075 | 0.949052658 | -4.90561499 |
| **A_52_P413947** | Mthfr | 0.014978027 | -0.182994717 | 0.212950772 | 10.31078408 | 0.183836782 | 0.860027012 | 0.949158554 | -4.905640167 |
| **A_55_P2079324** | Sephs2 | 0.044490164 | -0.54358276 | 0.632563089 | 9.22460756 | 0.183829843 | 0.860032227 | 0.949158554 | -4.905640861 |
| **A_51_P418526** | Sfxn1 | 0.026877207 | -0.328553411 | 0.382307824 | 11.41920503 | 0.183743711 | 0.860096957 | 0.949175796 | -4.905649478 |
| **A_51_P215540** | Rnf4 | 0.019289334 | -0.236387559 | 0.274966227 | 10.31725978 | 0.183319615 | 0.860415692 | 0.949473334 | -4.90569185 |
| **A_55_P2013598** | Ccnq | 0.014663274 | -0.179885865 | 0.209212413 | 10.40278664 | 0.183140679 | 0.860550184 | 0.949541242 | -4.905709699 |
| **A_55_P2020338** | Scml4 | 0.035798474 | -0.439333488 | 0.510930436 | 8.427656201 | 0.183076861 | 0.860598151 | 0.949541242 | -4.90571606 |
| **A_51_P146753** | Csf2rb2 | 0.039869411 | -0.48959016 | 0.569328982 | 10.07391052 | 0.182974347 | 0.860675205 | 0.949541242 | -4.905726275 |
| **A_55_P2011172** | NA | -0.015108403 | -0.215809993 | 0.185593187 | 14.00364514 | -0.182915681 | 0.860719302 | 0.949541242 | -4.905732117 |
| **A_55_P2070009** | Krt10 | -0.012461808 | -0.178010038 | 0.153086422 | 7.675960264 | -0.182910952 | 0.860722857 | 0.949541242 | -4.905732588 |
| **A_55_P2166745** | Sp3os | 0.031051849 | -0.381840593 | 0.443944291 | 10.95746612 | 0.182740041 | 0.860851327 | 0.949580448 | -4.9057496 |
| **A_55_P2021114** | Ier5 | -0.014520205 | -0.207600984 | 0.178560574 | 8.234733259 | -0.182732959 | 0.860856651 | 0.949580448 | -4.905750304 |
| **A_55_P2319468** | NA | -0.011954708 | -0.171003839 | 0.147094423 | 7.736731136 | -0.182637881 | 0.86092812 | 0.949605092 | -4.90575976 |
| **A_52_P218537** | Hnrnpc | -0.024722861 | -0.353761236 | 0.304315515 | 11.31126948 | -0.182572498 | 0.86097727 | 0.949605115 | -4.90576626 |
| **A_52_P17146** | Bmpr1a | -0.023253195 | -0.332988103 | 0.286481713 | 10.08839707 | -0.182421353 | 0.86109089 | 0.949642563 | -4.905781277 |
| **A_51_P440443** | Exosc5 | -0.021706863 | -0.3109932 | 0.267579473 | 12.45984451 | -0.182327568 | 0.861161393 | 0.949642563 | -4.905790589 |
| **A_51_P331003** | Sema4g | -0.043109606 | -0.617765564 | 0.531546352 | 10.03518019 | -0.182284326 | 0.8611939 | 0.949642563 | -4.905794881 |
| **A_52_P66199** | Fbxo11 | 0.019166705 | -0.236353476 | 0.274686886 | 8.894737342 | 0.182265911 | 0.861207744 | 0.949642563 | -4.905796709 |
| **A_55_P2151021** | NA | 0.022625119 | -0.279160833 | 0.324411072 | 12.64888833 | 0.182169238 | 0.86128042 | 0.949668525 | -4.905806299 |
| **A_55_P2174942** | NA | -0.027061772 | -0.388471849 | 0.334348306 | 14.229447 | -0.181944593 | 0.861449308 | 0.949800563 | -4.905828566 |
| **A_55_P2096807** | Rbms1 | -0.020188264 | -0.289955342 | 0.249578815 | 9.99272637 | -0.181841586 | 0.861526752 | 0.94983177 | -4.905838767 |
| **A_52_P58041** | Arpc5 | 0.025409406 | -0.31448276 | 0.365301572 | 10.33201928 | 0.18165048 | 0.861670435 | 0.949843684 | -4.905857678 |
| **A_55_P2011061** | Tspan12 | -0.025024894 | -0.359776482 | 0.309726695 | 11.09884509 | -0.181648904 | 0.861671619 | 0.949843684 | -4.905857834 |
| **A_52_P80567** | Nbas | 0.015336454 | -0.189855963 | 0.22052887 | 8.462887431 | 0.181612956 | 0.861698647 | 0.949843684 | -4.905861389 |
| **A_55_P2280237** | Zfp697 | 0.016978084 | -0.210237375 | 0.244193542 | 7.987843021 | 0.181565775 | 0.861734122 | 0.949843684 | -4.905866054 |
| **A_66_P125701** | NA | 0.016006396 | -0.198374569 | 0.230387361 | 8.224019738 | 0.181422243 | 0.861842042 | 0.949908469 | -4.905880237 |
| **A_55_P2133652** | Cep162 | 0.016935682 | -0.210115074 | 0.243986438 | 8.723915514 | 0.181243706 | 0.861976286 | 0.95000226 | -4.905897865 |
| **A_55_P2128704** | NA | 0.018907498 | -0.234711428 | 0.272526424 | 15.17422144 | 0.181148855 | 0.862047608 | 0.950026696 | -4.905907223 |
| **A_55_P2149787** | Fcnb | 0.037504046 | -0.466247534 | 0.541255626 | 9.414860818 | 0.180902634 | 0.862232757 | 0.950071019 | -4.905931492 |
| **A_55_P2336558** | Chn2 | 0.026080905 | -0.324256153 | 0.376417964 | 8.8866706 | 0.180892172 | 0.862240624 | 0.950071019 | -4.905932523 |
| **A_55_P2029711** | Cdh2 | 0.043052983 | -0.535461607 | 0.621567574 | 11.82627406 | 0.180830684 | 0.862286863 | 0.950071019 | -4.905938578 |
| **A_55_P2125466** | Ube2l3 | -0.017819015 | -0.257305024 | 0.221666995 | 11.94796001 | -0.180795131 | 0.862313598 | 0.950071019 | -4.905942079 |
| **A_55_P2070239** | Preb | 0.017866342 | -0.222291057 | 0.258023742 | 9.851878547 | 0.180768547 | 0.86233359 | 0.950071019 | -4.905944695 |
| **A_55_P2184811** | Eif2s1 | 0.015796436 | -0.196620223 | 0.228213095 | 8.600281647 | 0.180698154 | 0.862386525 | 0.950075186 | -4.905951623 |
| **A_55_P2109397** | Scn10a | -0.011990612 | -0.173441024 | 0.149459801 | 7.717339396 | -0.180461836 | 0.862564244 | 0.950170748 | -4.90597486 |
| **A_55_P1988907** | Eml4 | -0.021985366 | -0.318028507 | 0.274057775 | 10.49594283 | -0.180452079 | 0.862571582 | 0.950170748 | -4.905975819 |
| **A_55_P2164348** | Hoxa10 | 0.011630228 | -0.145080302 | 0.168340757 | 7.633294388 | 0.180332177 | 0.862661756 | 0.950215927 | -4.905987597 |

| | | | | | | | | | |
|---|---|---|---|---|---|---|---|---|---|
| **A_55_P2063626** | Matk | -0.029607723 | -0.428885649 | 0.369670203 | 9.338256851 | -0.180182617 | 0.862774237 | 0.95028567 | -4.906002277 |
| **A_52_P476904** | Chd6 | -0.011886888 | -0.172291439 | 0.148517662 | 7.951828954 | -0.180067233 | 0.862861018 | 0.950287509 | -4.906013594 |
| **A_52_P105015** | Clcn7 | -0.019245543 | -0.278974899 | 0.240483814 | 8.657605771 | -0.180049662 | 0.862874234 | 0.950287509 | -4.906015317 |
| **A_51_P355151** | Camk2n2 | -0.015573787 | -0.225831501 | 0.194683927 | 8.224220646 | -0.17998051 | 0.862926244 | 0.950290643 | -4.906022096 |
| **A_52_P297642** | Pdia5 | -0.040950083 | -0.594026386 | 0.51212622 | 10.9053324 | -0.179909013 | 0.862980019 | 0.950295721 | -4.906029102 |
| **A_55_P1993858** | Rmrp | 0.086226783 | -1.079707109 | 1.252160674 | 11.1855734 | 0.179701394 | 0.863136181 | 0.950413538 | -4.906049431 |
| **A_51_P387608** | Hif1a | -0.022284445 | -0.324172562 | 0.279603671 | 9.098418173 | -0.179365537 | 0.863388811 | 0.950586187 | -4.906082267 |
| **A_52_P442756** | Btf3l4 | -0.016740056 | -0.243522393 | 0.210042281 | 10.2830468 | -0.179362183 | 0.863391334 | 0.950586187 | -4.906082595 |
| **A_55_P2120141** | NA | 0.034548476 | -0.433812362 | 0.502909315 | 11.67779667 | 0.179238588 | 0.863484306 | 0.950587482 | -4.906094663 |
| **A_55_P1980931** | NA | -0.03322909 | -0.483725428 | 0.417267248 | 11.88799421 | -0.179229863 | 0.863490869 | 0.950587482 | -4.906095514 |
| **A_51_P129100** | Sec63 | 0.025432424 | -0.319791501 | 0.37065635 | 8.915141 | 0.179007021 | 0.863658505 | 0.950717879 | -4.902176363 |
| **A_52_P557293** | Tmprss2 | -0.019981997 | -0.291392455 | 0.251428462 | 8.511482459 | -0.178893885 | 0.863743616 | 0.950720702 | -4.906128278 |
| **A_55_P1994052** | Asxl1 | -0.013863172 | -0.202269941 | 0.174543598 | 11.82617508 | -0.178792496 | 0.863819891 | 0.950720702 | -4.906138153 |
| **A_55_P2220283** | Stk35os1 | 0.012866065 | -0.162048085 | 0.187780216 | 8.413309074 | 0.178732694 | 0.863864881 | 0.950720702 | -4.906143975 |
| **A_66_P125161** | 1700057H15Rik | -0.011853027 | -0.173053201 | 0.149347147 | 7.808823663 | -0.178668073 | 0.863913497 | 0.950720702 | -4.906150264 |
| **A_55_P1954907** | Rpl12 | -0.019935312 | -0.291113588 | 0.251242965 | 16.83341264 | -0.178628734 | 0.863943093 | 0.950720702 | -4.906154091 |
| **A_55_P2110435** | Zscan20 | 0.011398819 | -0.143673413 | 0.166471051 | 8.027610187 | 0.17861133 | 0.863956187 | 0.950720702 | -4.906155784 |
| **A_51_P418056** | Sc5d | -0.043131324 | -0.630216895 | 0.543954247 | 13.35001704 | -0.17851494 | 0.864028705 | 0.950746376 | -4.906165158 |
| **A_55_P2351193** | Snora47 | 0.050388911 | -0.636581111 | 0.737358933 | 8.840087403 | 0.178229823 | 0.864243221 | 0.95088175 | -4.906192856 |
| **A_55_P2079272** | Kars | 0.015647525 | -0.197706484 | 0.229001535 | 12.86119882 | 0.178208348 | 0.864259379 | 0.95088175 | -4.906194941 |
| **A_66_P111021** | Nlrp6 | -0.03155291 | -0.461961505 | 0.398855684 | 8.680581461 | -0.178131891 | 0.864316906 | 0.95088175 | -4.90620236 |
| **A_66_P126523** | Phb2 | 0.015899025 | -0.201130096 | 0.232928146 | 12.8505519 | 0.17800642 | 0.864411313 | 0.95088175 | -4.906214528 |
| **A_65_P19250** | Ap2m1 | -0.026941748 | -0.39477132 | 0.340887823 | 8.852421199 | -0.17797636 | 0.864433931 | 0.95088175 | -4.906217443 |
| **A_55_P2046488** | Nbeal2 | -0.023590811 | -0.345701993 | 0.29852037 | 8.517427539 | -0.177959126 | 0.864446899 | 0.95088175 | -4.906219113 |
| **A_55_P2114779** | Greb1 | 0.027158672 | -0.34393064 | 0.398247983 | 8.917497064 | 0.177833375 | 0.86454152 | 0.950931716 | -4.906231297 |
| **A_55_P2034250** | Zscan26 | -0.02074317 | -0.304302366 | 0.262816025 | 10.75641188 | -0.177752036 | 0.864602725 | 0.950932819 | -4.906239174 |
| **A_55_P2143070** | Ass1 | -0.029399456 | -0.431605358 | 0.372806445 | 13.91401222 | -0.177612715 | 0.864707562 | 0.950932819 | -4.906252657 |
| **A_55_P2160253** | 6720475J19Rik | -0.012068682 | -0.177239032 | 0.153101667 | 8.056802806 | -0.177546024 | 0.864757747 | 0.950932819 | -4.906259107 |
| **A_51_P461779** | Ppp2r2c | 0.011572716 | -0.146812802 | 0.169958234 | 7.644203383 | 0.177542779 | 0.864760189 | 0.950932819 | -4.906259421 |
| **A_55_P2138071** | NA | -0.030217629 | -0.443867247 | 0.38343199 | 13.04255637 | -0.177505144 | 0.864788509 | 0.950932819 | -4.90626306 |
| **A_55_P1964615** | Dnttip1 | 0.017947656 | -0.227868314 | 0.263763627 | 8.554483569 | 0.177411121 | 0.864859264 | 0.950956522 | -4.906272148 |
| **A_66_P138201** | Rhox10 | 0.011303515 | -0.143597497 | 0.166204526 | 7.642425983 | 0.177313758 | 0.864932532 | 0.950982987 | -4.906281553 |
| **A_51_P292057** | Tbl1xr1 | 0.019442925 | -0.247368593 | 0.286254444 | 8.913323536 | 0.177068059 | 0.865117436 | 0.951069661 | -4.906305267 |
| **A_55_P2107422** | Kif21a | -0.023419051 | -0.344824778 | 0.297986675 | 9.424525243 | -0.177051202 | 0.865130121 | 0.951069661 | -4.906306892 |
| **A_51_P370561** | Dcakd | -0.015335426 | -0.225919858 | 0.195249005 | 9.187867391 | -0.176950902 | 0.865205606 | 0.951069661 | -4.906316562 |
| **A_55_P2077501** | NA | 0.038706224 | -0.492813777 | 0.570226226 | 11.48377391 | 0.176947481 | 0.865208181 | 0.951069661 | -4.906316892 |
| **A_52_P514317** | Slc35a3 | -0.022242074 | -0.327970148 | 0.283486001 | 9.093171751 | -0.176775936 | 0.865337288 | 0.951141786 | -4.906333418 |
| **A_52_P257502** | Igfbp4 | -0.033321842 | -0.49146695 | 0.424823266 | 10.16625329 | -0.176729535 | 0.86537221 | 0.951141786 | -4.906337885 |
| **A_51_P514647** | Paip2 | 0.025071634 | -0.31985369 | 0.369996957 | 12.79833863 | 0.176620352 | 0.865454386 | 0.951141855 | -4.906348393 |
| **A_51_P400773** | Hdgfl2 | -0.022428981 | -0.33106894 | 0.286210977 | 10.91547322 | -0.176579622 | 0.865485041 | 0.951141855 | -4.906352311 |
| **A_55_P2074656** | Padi2 | -0.028138454 | -0.415446704 | 0.359169797 | 8.587668583 | -0.176533311 | 0.865519897 | 0.951141855 | -4.906356765 |
| **A_55_P2165560** | Plekhg5 | 0.048552856 | -0.621081964 | 0.718187676 | 10.01763981 | 0.176181353 | 0.865784812 | 0.951378887 | -4.906390576 |
| **A_51_P358700** | Olfr1221 | -0.011917092 | -0.176438757 | 0.152604574 | 7.741786134 | -0.176007178 | 0.865915918 | 0.951468864 | -4.906407283 |
| **A_55_P2062213** | Tmem41a | 0.024726404 | -0.316763974 | 0.366216782 | 10.75243103 | 0.17594044 | 0.865966155 | 0.951469975 | -4.906413681 |
| **A_55_P2012759** | Bicdl1 | 0.013932392 | -0.178706796 | 0.206571579 | 8.600566588 | 0.17573741 | 0.866118989 | 0.951487003 | -4.906433129 |
| **A_55_P2062538** | 1700027A15Rik | 0.011158144 | -0.143241082 | 0.165557369 | 7.953882308 | 0.175602225 | 0.866220755 | 0.951487003 | -4.906446065 |
| **A_55_P2109342** | Rnps1 | 0.019374517 | -0.248807387 | 0.28755642 | 9.98621122 | 0.175543439 | 0.86626501 | 0.951487003 | -4.906451688 |
| **A_55_P2068731** | Gm21953 | -0.034487712 | -0.512054331 | 0.443078906 | 8.977104802 | -0.175474338 | 0.86631703 | 0.951487003 | -4.906458295 |
| **A_55_P2160648** | Gm6900 | -0.012922683 | -0.191927206 | 0.166081841 | 15.66568545 | -0.175417077 | 0.866360138 | 0.951487003 | -4.906463768 |
| **A_55_P2024530** | Siah2 | -0.032513001 | -0.482997695 | 0.417971692 | 9.611286493 | -0.175371984 | 0.866394086 | 0.951487003 | -4.906468076 |

| | | | | | | | | | |
|---|---|---|---|---|---|---|---|---|---|
| **A_52_P145412** | Rictor | -0.014594397 | -0.216857535 | 0.18766874 | 9.098215546 | -0.17532854 | 0.866426792 | 0.951487003 | -4.906472226 |
| **A_51_P165098** | Gga2 | -0.021971056 | -0.326576324 | 0.282634212 | 10.93282954 | -0.175265611 | 0.866474169 | 0.951487003 | -4.906478236 |
| **A_52_P153189** | Arl2bp | 0.022798725 | -0.293303801 | 0.338901251 | 9.04233741 | 0.175253131 | 0.866483564 | 0.951487003 | -4.906479427 |
| **A_52_P163939** | Cln6 | -0.019569816 | -0.2909181 | 0.251778468 | 9.117610837 | -0.175243879 | 0.866490529 | 0.951487003 | -4.906480311 |
| **A_51_P275083** | Mrps18a | -0.015338426 | -0.228068295 | 0.197391443 | 11.58741918 | -0.175200568 | 0.866523137 | 0.951487003 | -4.906484445 |
| **A_52_P652104** | Brd3 | 0.023665796 | -0.304821004 | 0.352152595 | 9.497629037 | 0.175059779 | 0.866629133 | 0.951497539 | -4.906497877 |
| **A_55_P2094881** | Fgfr2 | 0.018093749 | -0.233139574 | 0.269327073 | 8.502417121 | 0.174998577 | 0.866675211 | 0.951497539 | -4.906503713 |
| **A_55_P2005675** | NA | 0.013387312 | -0.172504044 | 0.199278668 | 15.54402479 | 0.174991671 | 0.866680411 | 0.951497539 | -4.906504371 |
| **A_52_P89683** | NA | -0.034654415 | -0.516179077 | 0.446870247 | 12.31043111 | -0.174873189 | 0.866769617 | 0.951509825 | -4.906515663 |
| **A_51_P145785** | Ambp | 0.023268093 | -0.30011627 | 0.346652457 | 17.86396557 | 0.174833629 | 0.866799402 | 0.951509825 | -4.906519431 |
| **A_51_P208240** | Tnfsf14 | -0.024658192 | -0.367466258 | 0.318149875 | 8.467964824 | -0.174780661 | 0.866839283 | 0.951509825 | -4.906524476 |
| **A_55_P2063096** | Txnrd3 | -0.016649842 | -0.248272764 | 0.214973081 | 9.593836529 | -0.174667362 | 0.86692459 | 0.951533448 | -4.906535261 |
| **A_55_P2098688** | Mcrs1 | 0.017093877 | -0.220940971 | 0.255128725 | 8.822349683 | 0.174495094 | 0.867054301 | 0.951533448 | -4.906551645 |
| **A_51_P504522** | Derl1 | 0.016670143 | -0.215466116 | 0.248806402 | 11.62924406 | 0.174493613 | 0.867055416 | 0.951533448 | -4.906551786 |
| **A_55_P2286650** | 6720462K09Rik | 0.011728023 | -0.151718065 | 0.175174112 | 8.074740491 | 0.174354628 | 0.867160069 | 0.951533448 | -4.906564994 |
| **A_55_P2045765** | Rnf224 | 0.013781342 | -0.178346068 | 0.205908753 | 8.033982655 | 0.174295181 | 0.867204833 | 0.951533448 | -4.90657064 |
| **A_52_P179583** | Adnp2 | -0.01265127 | -0.189045511 | 0.163742971 | 8.196295452 | -0.174274127 | 0.867220687 | 0.951533448 | -4.906572639 |
| **A_55_P2018210** | NA | 0.013916277 | -0.180207782 | 0.208040337 | 16.74358392 | 0.174191476 | 0.867282924 | 0.951533448 | -4.906580485 |
| **A_51_P399614** | Zkscan14 | -0.023163921 | -0.346516293 | 0.300188451 | 8.996169882 | -0.174068112 | 0.867375821 | 0.951533448 | -4.906592188 |
| **A_55_P2104941** | Rpl30 | -0.017595747 | -0.263228433 | 0.228036939 | 15.03668385 | -0.17406231 | 0.86738019 | 0.951533448 | -4.906592739 |
| **A_66_P108188** | Car5a | -0.035341532 | -0.529128482 | 0.458445418 | 9.93029214 | -0.173911762 | 0.867493559 | 0.951533448 | -4.90660701 |
| **A_55_P2135980** | Ms4a4c | 0.038287234 | -0.496674301 | 0.573248768 | 15.58760635 | 0.173906022 | 0.867497882 | 0.951533448 | -4.906607553 |
| **A_66_P138915** | Fyn | -0.016416681 | -0.245864886 | 0.213031523 | 8.437209265 | -0.173853681 | 0.867537299 | 0.951533448 | -4.906612512 |
| **A_51_P491504** | Aldh5a1 | 0.041584229 | -0.539643762 | 0.622812221 | 11.26180785 | 0.173846249 | 0.867542895 | 0.951533448 | -4.906613216 |
| **A_55_P2028171** | Dppa5a | -0.0123509 | -0.184990343 | 0.160288544 | 7.765210785 | -0.173836814 | 0.86755 | 0.951533448 | -4.90661411 |
| **A_55_P2028046** | Cyp3a59 | -0.050118806 | -0.751170625 | 0.650933014 | 8.929841815 | -0.173713584 | 0.867642802 | 0.951581237 | -4.906625777 |
| **A_51_P514449** | Nr5a2 | -0.034481199 | -0.517368264 | 0.448405867 | 10.37172901 | -0.173508186 | 0.867797489 | 0.95165416 | -4.906645206 |
| **A_51_P403942** | A930016O22Rik | 0.011839199 | -0.153974223 | 0.177652621 | 7.654931142 | 0.173494546 | 0.867807761 | 0.95165416 | -4.906646496 |
| **A_55_P2115968** | NA | -0.011332166 | -0.170567203 | 0.14790287 | 9.337315586 | -0.172924888 | 0.868236809 | 0.952002912 | -4.906700258 |
| **A_55_P2018934** | Adgre1 | -0.019148197 | -0.288213804 | 0.249917411 | 9.849515886 | -0.172923046 | 0.868238196 | 0.952002912 | -4.906700432 |
| **A_55_P1986958** | Brms1l | 0.023959361 | -0.3128029 | 0.360721621 | 10.25096302 | 0.172876116 | 0.868273544 | 0.952002912 | -4.906704853 |
| **A_52_P80965** | Sephs1 | 0.013500565 | -0.176454964 | 0.203456094 | 7.937472389 | 0.172696364 | 0.868408939 | 0.952066535 | -4.906721776 |
| **A_55_P2282969** | Gm2a | -0.012229995 | -0.184336336 | 0.159876346 | 7.698215994 | -0.172668294 | 0.868430083 | 0.952066535 | -4.906724418 |
| **A_52_P436628** | Vegfb | 0.016555141 | -0.216623975 | 0.249734256 | 9.590627856 | 0.172514822 | 0.868545688 | 0.952139269 | -4.906738851 |
| **A_55_P2198364** | Tsen54 | -0.02097252 | -0.316612372 | 0.274667331 | 8.505096332 | -0.172373635 | 0.868652041 | 0.952201855 | -4.906752118 |
| **A_55_P2095360** | Tfpi | -0.011748741 | -0.17745262 | 0.153955138 | 9.016791434 | -0.17228277 | 0.868720489 | 0.952216247 | -4.90676065 |
| **A_66_P126153** | 1700011M02Rik | 0.0114233 | -0.149842789 | 0.172689389 | 7.650992673 | 0.172120153 | 0.868842991 | 0.952216247 | -4.906775909 |
| **A_66_P116299** | Pias1 | 0.013842591 | -0.181582004 | 0.209267187 | 10.67153032 | 0.172116048 | 0.868846084 | 0.952216247 | -4.906776294 |
| **A_52_P341449** | Pgm3 | -0.027181629 | -0.41104453 | 0.356681271 | 10.33009064 | -0.172061035 | 0.868887527 | 0.952216247 | -4.906781453 |
| **A_66_P133525** | NA | 0.010968061 | -0.143953124 | 0.165889246 | 7.687194102 | 0.172029228 | 0.868911489 | 0.952216247 | -4.906784435 |
| **A_51_P464146** | Sowahb | 0.031980761 | -0.420079943 | 0.484041464 | 8.796219061 | 0.171899741 | 0.869009039 | 0.95226916 | -4.906796569 |
| **A_51_P357561** | Fbxw9 | -0.016022221 | -0.242708555 | 0.210664112 | 8.63863157 | -0.171743605 | 0.869126667 | 0.952344068 | -4.906811188 |
| **A_55_P2049757** | NA | 0.014693239 | -0.193410128 | 0.222796607 | 14.34177186 | 0.17156221 | 0.869263331 | 0.952357262 | -4.906828156 |
| **A_52_P73848** | Clint1 | 0.027824055 | -0.366295706 | 0.421943815 | 9.09956346 | 0.171543945 | 0.869277091 | 0.952357262 | -4.906829864 |
| **A_55_P2179448** | NA | 0.014895771 | -0.196115348 | 0.22590689 | 8.052420146 | 0.171530296 | 0.869287375 | 0.952357262 | -4.90683114 |
| **A_51_P496432** | Acsl1 | -0.053756239 | -0.815825516 | 0.708313038 | 14.77544137 | -0.171402682 | 0.869383523 | 0.952357262 | -4.906843064 |
| **A_52_P554451** | 1300017J02Rik | -0.041726541 | -0.633265204 | 0.549812123 | 12.48219764 | -0.171400638 | 0.869385063 | 0.952357262 | -4.906843255 |
| **A_55_P2160204** | NA | -0.02622418 | -0.398162202 | 0.345713842 | 11.73922786 | -0.17132255 | 0.869443898 | 0.952367738 | -4.906850547 |
| **A_55_P2118362** | Tmem121 | -0.013061735 | -0.198546922 | 0.172423452 | 8.085404634 | -0.171109777 | 0.869604215 | 0.952430216 | -4.9068704 |
| **A_55_P2152901** | 2900040C04Rik | 0.017728431 | -0.234058586 | 0.269515448 | 10.17813506 | 0.171088241 | 0.869620442 | 0.952430216 | -4.906872408 |

| | | | | | | | | | |
|---|---|---|---|---|---|---|---|---|---|
| **A_55_P1964218** | Tprkb | 0.017725612 | -0.23416137 | 0.269612595 | 14.07091762 | 0.17099315 | 0.869692092 | 0.952430216 | -4.906881272 |
| **A_51_P488718** | Zfand2b | -0.035973119 | -0.54738534 | 0.475439103 | 11.58999301 | -0.170918934 | 0.869748014 | 0.952430216 | -4.906888186 |
| **A_55_P2410106** | 9130230N09Rik | -0.013744463 | -0.20915138 | 0.181662454 | 8.616515993 | -0.170911398 | 0.869753693 | 0.952430216 | -4.906888888 |
| **A_51_P243857** | Copb2 | 0.019376982 | -0.256414854 | 0.295168817 | 12.23416874 | 0.170721379 | 0.869896877 | 0.952430216 | -4.906906577 |
| **A_55_P1980611** | Trmt2b | 0.016688864 | -0.220943145 | 0.254320874 | 10.74117234 | 0.170649506 | 0.869951037 | 0.952430216 | -4.906913263 |
| **A_55_P2012101** | Bmp8b | 0.015024379 | -0.198928981 | 0.228977738 | 7.856405216 | 0.170632045 | 0.869964194 | 0.952430216 | -4.906914887 |
| **A_52_P42194** | NA | 0.013520066 | -0.179063591 | 0.206103723 | 8.129101079 | 0.170585677 | 0.869999135 | 0.952430216 | -4.906919198 |
| **A_51_P189959** | Oxsr1 | 0.012012787 | -0.159141588 | 0.183167161 | 8.926774849 | 0.170544984 | 0.8700298 | 0.952430216 | -4.906922981 |
| **A_51_P315673** | Usp11 | -0.028516653 | -0.434854475 | 0.37782117 | 9.021248534 | -0.170527524 | 0.870042958 | 0.952430216 | -4.906924604 |
| **A_55_P1979403** | Tigar | 0.020291587 | -0.269096376 | 0.30967955 | 8.363519571 | 0.170380052 | 0.870154089 | 0.952447578 | -4.906938303 |
| **A_52_P367675** | Acin1 | -0.019625061 | -0.299625528 | 0.260375405 | 8.677197343 | -0.170308162 | 0.870208265 | 0.952447578 | -4.906944977 |
| **A_51_P132459** | Rpl27a | -0.029968737 | -0.457698657 | 0.397761183 | 12.83567637 | -0.170247991 | 0.87025361 | 0.952447578 | -4.906950561 |
| **A_52_P102783** | Dync2h1 | -0.022776929 | -0.347982046 | 0.302428187 | 8.798665968 | -0.170184889 | 0.870301165 | 0.952447578 | -4.906956415 |
| **A_55_P1965507** | Trnau1ap | 0.015623935 | -0.207458956 | 0.238706826 | 12.27809646 | 0.170179541 | 0.870305195 | 0.952447578 | -4.906956911 |
| **A_55_P2082989** | Rab7b | -0.011635818 | -0.177839972 | 0.154568337 | 8.16856506 | -0.170113281 | 0.87035513 | 0.952448299 | -4.906963056 |
| **A_51_P446012** | Phf13 | -0.036074471 | -0.552051941 | 0.479902999 | 10.54011059 | -0.169883977 | 0.870527944 | 0.952569528 | -4.906984301 |
| **A_55_P2015541** | Hif3a | -0.013066358 | -0.20007899 | 0.173946274 | 13.04428068 | -0.169772285 | 0.870612124 | 0.952569528 | -4.90699464 |
| **A_66_P137516** | NA | -0.01487901 | -0.227837991 | 0.198079972 | 13.91107954 | -0.169770119 | 0.870613756 | 0.952569528 | -4.90699484 |
| **A_55_P2141876** | Ldha | -0.034497248 | -0.528595223 | 0.459600728 | 12.6701555 | -0.169650272 | 0.870704083 | 0.952589621 | -4.907005925 |
| **A_51_P370825** | Ccdc124 | 0.023213179 | -0.309367834 | 0.355794191 | 11.5577147 | 0.169597854 | 0.870743591 | 0.952589621 | -4.907010772 |
| **A_55_P2048221** | NA | -0.020343954 | -0.312010578 | 0.27132267 | 15.5007869 | -0.169485216 | 0.870828488 | 0.952589621 | -4.90702118 |
| **A_51_P446570** | Bbox1 | -0.048261481 | -0.740179445 | 0.643656484 | 10.30982722 | -0.169484203 | 0.870829251 | 0.952589621 | -4.907021274 |
| **A_55_P2138177** | Psd3 | 0.025131078 | -0.335714093 | 0.385976249 | 8.713332185 | 0.169228463 | 0.871022013 | 0.952696282 | -4.90704488 |
| **A_52_P139747** | Ndufa1 | -0.016823019 | -0.258432726 | 0.224786687 | 14.31800452 | -0.169189246 | 0.871051574 | 0.952696282 | -4.907048497 |
| **A_55_P2154943** | NA | -0.011260258 | -0.173031386 | 0.150510869 | 7.710457248 | -0.16913385 | 0.871093329 | 0.952696282 | -4.907053605 |
| **A_52_P445387** | Clk4 | -0.019344863 | -0.297330586 | 0.258640861 | 10.58806207 | -0.169093278 | 0.871123912 | 0.952696282 | -4.907057345 |
| **A_55_P2133901** | NA | -0.015178576 | -0.233454753 | 0.203097601 | 15.89467371 | -0.168969327 | 0.871217345 | 0.952710139 | -4.907068764 |
| **A_55_P2164218** | NA | 0.019534633 | -0.261423311 | 0.300492576 | 13.21058858 | 0.168945692 | 0.871235161 | 0.952710139 | -4.907070941 |
| **A_55_P2169039** | Olfr558 | -0.01102926 | -0.169839613 | 0.147781094 | 7.684590875 | -0.168752705 | 0.871380638 | 0.952815316 | -4.907088703 |
| **A_55_P1958562** | Dmrtc1b | -0.011285039 | -0.174017887 | 0.15144781 | 7.654131871 | -0.168504315 | 0.871567886 | 0.952926008 | -4.907111534 |
| **A_55_P2071683** | NA | -0.018615687 | -0.287084747 | 0.249853374 | 12.64569133 | -0.168487623 | 0.87158047 | 0.952926008 | -4.907113067 |
| **A_55_P2113703** | Spib | 0.034540529 | -0.464225626 | 0.533306684 | 9.161988406 | 0.168273295 | 0.871742048 | 0.952941556 | -4.907132738 |
| **A_51_P459108** | Insl6 | -0.015123233 | -0.233538413 | 0.203291947 | 8.346155532 | -0.168246106 | 0.871762546 | 0.952941556 | -4.907135232 |
| **A_55_P2183404** | Ptn | -0.023396729 | -0.361315277 | 0.314521818 | 8.206273991 | -0.168238857 | 0.871768011 | 0.952941556 | -4.907135897 |
| **A_55_P2187918** | Cep55 | 0.011803722 | -0.158709329 | 0.182316772 | 8.226219436 | 0.168207178 | 0.871791895 | 0.952941556 | -4.907138802 |
| **A_52_P504342** | Coa7 | -0.014297736 | -0.221454594 | 0.192859121 | 9.111696363 | -0.167706988 | 0.872169011 | 0.953299865 | -4.907184597 |
| **A_52_P92514** | Supt20 | 0.011286516 | -0.152394268 | 0.174967299 | 8.448306521 | 0.167550367 | 0.872287103 | 0.953336207 | -4.907198908 |
| **A_51_P482410** | Ceacam15 | 0.011139584 | -0.150427991 | 0.172707159 | 7.69547536 | 0.167532065 | 0.872300903 | 0.953336207 | -4.90720058 |
| **A_55_P2193512** | NA | 0.046159891 | -0.62379599 | 0.716115773 | 9.439468938 | 0.167417851 | 0.872387022 | 0.953376421 | -4.907211007 |
| **A_52_P522222** | NA | -0.017150806 | -0.266551938 | 0.232250326 | 8.337650656 | -0.167097251 | 0.872628771 | 0.953564335 | -4.907240239 |
| **A_55_P1989384** | Rnf181 | -0.01594852 | -0.247919571 | 0.21602253 | 11.31057899 | -0.167058966 | 0.87265764 | 0.953564335 | -4.907243726 |
| **A_51_P229599** | Etnk1 | -0.021446883 | -0.334000862 | 0.291107096 | 8.75327528 | -0.166733298 | 0.872903227 | 0.953737027 | -4.907273357 |
| **A_55_P2126109** | Usp16 | 0.027500223 | -0.373306989 | 0.428307434 | 11.29266257 | 0.16671853 | 0.872914364 | 0.953737027 | -4.907274699 |
| **A_55_P2320283** | Hmga2-ps1 | 0.020662743 | -0.280650498 | 0.321975983 | 8.55629834 | 0.166629901 | 0.872981202 | 0.953756142 | -4.907282753 |
| **A_52_P487686** | Cstdc5 | 0.023499695 | -0.319480904 | 0.366480295 | 16.63255428 | 0.16648529 | 0.873090261 | 0.953819526 | -4.907295884 |
| **A_52_P358076** | Ppp2ca | 0.01147474 | -0.156063918 | 0.179013399 | 11.75335443 | 0.166422107 | 0.873137912 | 0.953819526 | -4.907301617 |
| **A_55_P2088331** | Ctdsp2 | 0.02505341 | -0.341606727 | 0.391713547 | 10.18821243 | 0.166029912 | 0.873433705 | 0.954080413 | -4.90733716 |
| **A_52_P422755** | Lsm6 | 0.012657466 | -0.17264809 | 0.197963023 | 8.695094915 | 0.165974563 | 0.873475451 | 0.954080413 | -4.907342169 |
| **A_55_P2085157** | Eef1g | 0.025055019 | -0.342151901 | 0.392261939 | 12.93547582 | 0.165793336 | 0.873612141 | 0.954175797 | -4.907358559 |
| **A_52_P228031** | Tmem179b | 0.016205443 | -0.221683757 | 0.254094643 | 10.8558911 | 0.165527196 | 0.873812885 | 0.954341036 | -4.907382597 |

| | | | | | | | | | |
|---|---|---|---|---|---|---|---|---|---|
| **A_51_P464710** | Adap2 | 0.020056689 | -0.274845254 | 0.314958632 | 9.03469568 | 0.165258874 | 0.874015286 | 0.954341036 | -4.907406794 |
| **A_55_P2095177** | Usp5 | 0.012265061 | -0.168313074 | 0.192843196 | 13.02897891 | 0.165039441 | 0.874180817 | 0.954341036 | -4.907426552 |
| **A_55_P2164265** | Prl7c1 | 0.01077597 | -0.147894249 | 0.16944619 | 7.668153125 | 0.165022878 | 0.874193312 | 0.954341036 | -4.907428043 |
| **A_51_P323620** | Thyn1 | -0.01870064 | -0.294146689 | 0.256745409 | 12.96911464 | -0.164969291 | 0.874233736 | 0.954341036 | -4.907432864 |
| **A_55_P2095035** | NA | 0.014486283 | -0.198907628 | 0.227880194 | 8.236437634 | 0.164952211 | 0.874246621 | 0.954341036 | -4.9074344 |
| **A_55_P2005410** | NA | 0.017350757 | -0.2383107 | 0.273012214 | 13.55429687 | 0.16490596 | 0.874281513 | 0.954341036 | -4.907438559 |
| **A_51_P107090** | BC031181 | -0.015777836 | -0.248274665 | 0.216718993 | 10.61391888 | -0.164897318 | 0.874288032 | 0.954341036 | -4.903486503 |
| **A_55_P1994348** | Golim4 | 0.028330729 | -0.4054292 | 0.462090659 | 11.05519355 | 0.166169263 | 0.874317657 | 0.954341036 | -4.858401922 |
| **A_52_P235861** | Peg3 | -0.022650494 | -0.356823459 | 0.311522472 | 9.57029161 | -0.164698463 | 0.874438049 | 0.954341036 | -4.907457204 |
| **A_51_P517787** | Clptm1l | -0.023662998 | -0.372831783 | 0.325505788 | 11.01327579 | -0.164671158 | 0.874458649 | 0.954341036 | -4.907459656 |
| **A_55_P2244677** | 6720422M22Rik | -0.022178337 | -0.349541548 | 0.305184874 | 8.733557697 | -0.164619885 | 0.87449733 | 0.954341036 | -4.907464259 |
| **A_55_P2158030** | Glrx3 | 0.019125192 | -0.263254655 | 0.301505039 | 13.1446393 | 0.164571753 | 0.874533643 | 0.954341036 | -4.907468579 |
| **A_55_P2192492** | Slmap | 0.011433759 | -0.157385405 | 0.180252922 | 8.012257101 | 0.164569923 | 0.874535024 | 0.954341036 | -4.907468743 |
| **A_51_P151433** | Ncbp2 | 0.021340422 | -0.293787999 | 0.336468844 | 8.923038293 | 0.164550279 | 0.874549844 | 0.954341036 | -4.907470506 |
| **A_55_P2127174** | Zfp628 | 0.019537252 | -0.269081153 | 0.308155657 | 17.69227619 | 0.164483613 | 0.87460014 | 0.954341036 | -4.907476486 |
| **A_55_P1974273** | Ndufs7 | -0.012496713 | -0.19711111 | 0.172117685 | 13.65464996 | -0.164480116 | 0.874602778 | 0.954341036 | -4.9074768 |
| **A_55_P1963061** | Rps16 | -0.013634828 | -0.215211041 | 0.187941384 | 16.68271602 | -0.164359034 | 0.87469413 | 0.954386838 | -4.907487655 |
| **A_55_P1995745** | Gm14308 | -0.015903366 | -0.251300141 | 0.219493409 | 8.600583505 | -0.164161661 | 0.874843045 | 0.954452079 | -4.907505333 |
| **A_52_P194242** | Sel1l | -0.026124413 | -0.41304051 | 0.360791684 | 9.7747981 | -0.164063861 | 0.874916835 | 0.954452079 | -4.907514085 |
| **A_55_P2007210** | Abca15 | 0.01047798 | -0.144805838 | 0.165761797 | 7.657873647 | 0.163958728 | 0.87499616 | 0.954452079 | -4.907523487 |
| **A_55_P2029498** | Osbpl1a | 0.027777554 | -0.383971352 | 0.43952646 | 10.09728473 | 0.163924826 | 0.87502174 | 0.954452079 | -4.907526518 |
| **A_55_P1974352** | Fam126b | -0.024638903 | -0.389923191 | 0.340645386 | 10.07392168 | -0.163897963 | 0.875042009 | 0.954452079 | -4.907528919 |
| **A_51_P384318** | C1ra | 0.041123527 | -0.568594101 | 0.650841155 | 12.04166947 | 0.163887108 | 0.875050199 | 0.954452079 | -4.907529889 |
| **A_51_P521161** | Caml | 0.018158819 | -0.251413622 | 0.28773126 | 10.29971904 | 0.16367988 | 0.875206563 | 0.954531028 | -4.907548397 |
| **A_55_P2030859** | NA | 0.012280999 | -0.170055482 | 0.19461748 | 16.24569227 | 0.163660288 | 0.875221347 | 0.954531028 | -4.907550145 |
| **A_55_P2072666** | Cyp2d37-ps | 0.031673174 | -0.440191101 | 0.503537449 | 8.918951992 | 0.163101396 | 0.875643092 | 0.954873703 | -4.907599939 |
| **A_51_P109835** | Uqcrc1 | -0.020702729 | -0.329176523 | 0.287771065 | 12.2315297 | -0.163076924 | 0.87566156 | 0.954873703 | -4.907602116 |
| **A_55_P2150361** | NA | -0.013160413 | -0.209287928 | 0.182967102 | 13.11083593 | -0.163047515 | 0.875683753 | 0.954873703 | -4.907604731 |
| **A_51_P283649** | Brat1 | 0.011657349 | -0.162251938 | 0.185566635 | 10.02283891 | 0.162877165 | 0.875812311 | 0.95492584 | -4.90761987 |
| **A_55_P2133370** | NA | -0.010491961 | -0.167066414 | 0.146082492 | 7.662087927 | -0.162824194 | 0.875852287 | 0.95492584 | -4.907624574 |
| **A_55_P2123529** | NA | -0.016248448 | -0.258782828 | 0.226285932 | 11.04235878 | -0.162787769 | 0.875879777 | 0.95492584 | -4.907627808 |
| **A_55_P1984035** | Litaf | 0.013595618 | -0.189646336 | 0.216837573 | 8.335922691 | 0.162543193 | 0.876064361 | 0.955073211 | -4.907649504 |
| **A_55_P2043454** | NA | 0.010437119 | -0.14588531 | 0.166759547 | 8.220085615 | 0.162234238 | 0.876297545 | 0.955177843 | -4.907676865 |
| **A_51_P194377** | Pcbp4 | -0.028709623 | -0.45873931 | 0.401320065 | 9.966504709 | -0.162222928 | 0.876306082 | 0.955177843 | -4.907677866 |
| **A_51_P221625** | Stk24 | 0.018956484 | -0.265058499 | 0.302971467 | 10.00672933 | 0.162180908 | 0.876337797 | 0.955177843 | -4.907681583 |
| **A_66_P101499** | AW822252 | 0.010682297 | -0.149391331 | 0.170755924 | 7.669447627 | 0.162154136 | 0.876358004 | 0.955177843 | -4.90768395 |
| **A_55_P2085905** | Anpep | -0.05900788 | -0.943854831 | 0.825839072 | 12.44005782 | -0.162041046 | 0.876443363 | 0.95518414 | -4.907693948 |
| **A_51_P494775** | Rcor1 | 0.016304667 | -0.228228879 | 0.260838212 | 11.57011367 | 0.162015539 | 0.876462616 | 0.95518414 | -4.907696202 |
| **A_55_P1985422** | NA | 0.058837179 | -0.825203452 | 0.942877809 | 12.84281204 | 0.161719654 | 0.876685957 | 0.955373675 | -4.907722322 |
| **A_55_P2153937** | NA | 0.014089531 | -0.197834215 | 0.226013276 | 8.402557679 | 0.161547446 | 0.876815949 | 0.955434103 | -4.907737503 |
| **A_52_P68714** | NA | -0.012714651 | -0.203996678 | 0.178567375 | 8.7676756 | -0.16151523 | 0.876840268 | 0.955434103 | -4.907740341 |
| **A_52_P84353** | Homez | -0.012902083 | -0.207222355 | 0.181418189 | 9.320817701 | -0.16133363 | 0.876977356 | 0.955488583 | -4.907756329 |
| **A_51_P216550** | Trp63 | -0.010256338 | -0.164743583 | 0.144230906 | 7.633132153 | -0.161318029 | 0.876989133 | 0.955488583 | -4.907757702 |
| **A_51_P373369** | Med4 | 0.01287751 | -0.181277847 | 0.207032867 | 10.77622889 | 0.16116313 | 0.877106068 | 0.955529203 | -4.907771324 |
| **A_55_P2177278** | DXBay18 | -0.024448475 | -0.39311854 | 0.34422159 | 8.195913469 | -0.161137673 | 0.877125286 | 0.955529203 | -4.907773562 |
| **A_55_P2046699** | Tvp23b | -0.016862612 | -0.271837015 | 0.238111792 | 10.00021437 | -0.160698358 | 0.87745695 | 0.955800669 | -4.907812119 |
| **A_52_P391081** | Hinfp | -0.01746172 | -0.281623845 | 0.246700406 | 8.455673835 | -0.160620009 | 0.877516103 | 0.955800669 | -4.907818985 |
| **A_55_P1995151** | NA | -0.023898046 | -0.385449829 | 0.337653738 | 10.51380394 | -0.160610918 | 0.877522967 | 0.955800669 | -4.907819781 |
| **A_55_P1975447** | NA | 0.022256636 | -0.314702995 | 0.359216266 | 9.96927279 | 0.16049623 | 0.877609557 | 0.955800669 | -4.907829825 |
| **A_55_P2172314** | Scamp4 | 0.016995466 | -0.240337266 | 0.274328198 | 12.23552102 | 0.160480118 | 0.877621722 | 0.955800669 | -4.907831235 |

| | | | | | | | | | |
|---|---|---|---|---|---|---|---|---|---|
| **A_55_P2048800** | Cd82 | -0.025527203 | -0.412202762 | 0.361148357 | 8.827866249 | -0.160413049 | 0.877672361 | 0.955801965 | -4.907837105 |
| **A_55_P2140711** | Tufm | 0.017282338 | -0.246068244 | 0.28063292 | 11.94073306 | 0.159459868 | 0.878392104 | 0.956528478 | -4.90792026 |
| **A_52_P12001** | Erp44 | 0.021174972 | -0.301616357 | 0.343966301 | 10.37308183 | 0.15939848 | 0.878438461 | 0.956528478 | -4.907925598 |
| **A_55_P2013665** | Mybpc2 | -0.016435926 | -0.267336353 | 0.234464502 | 7.754908671 | -0.159175403 | 0.878606927 | 0.956637557 | -4.907944982 |
| **A_66_P111385** | Klhl28 | -0.012431583 | -0.202292323 | 0.177429156 | 9.549093774 | -0.159101572 | 0.878662684 | 0.956637557 | -4.907951391 |
| **A_55_P2181386** | Adm2 | -0.028371036 | -0.461888876 | 0.405146805 | 8.633976307 | -0.159019874 | 0.878724384 | 0.956637557 | -4.90795848 |
| **A_55_P2033407** | Meg3 | -0.023066843 | -0.375879475 | 0.329745789 | 8.785398045 | -0.158864631 | 0.878841628 | 0.956637557 | -4.90797194 |
| **A_52_P645862** | Agtr1a | -0.052345467 | -0.85312127 | 0.748430335 | 11.17591888 | -0.158836873 | 0.878862592 | 0.956637557 | -4.907974345 |
| **A_55_P2102419** | Lime1 | -0.018088317 | -0.294848956 | 0.258672321 | 9.373384153 | -0.158809704 | 0.878883112 | 0.956637557 | -4.907976699 |
| **A_52_P630493** | Dnajb6 | 0.013785607 | -0.197144801 | 0.224716016 | 9.053939094 | 0.158807096 | 0.878885082 | 0.956637557 | -4.907976925 |
| **A_51_P198411** | Grb2 | -0.022971281 | -0.374643866 | 0.328701304 | 11.421254 | -0.158719357 | 0.878951347 | 0.956655813 | -4.907984524 |
| **A_55_P2029366** | Abhd6 | -0.01570392 | -0.25640074 | 0.224992899 | 8.345047063 | -0.158533453 | 0.879091756 | 0.956743585 | -4.908000611 |
| **A_55_P2153555** | Zkscan3 | -0.017739644 | -0.289801263 | 0.254321975 | 9.729183472 | -0.15843853 | 0.879163452 | 0.956743585 | -4.908008818 |
| **A_55_P1984981** | NA | 0.013774063 | -0.197500034 | 0.225048159 | 7.877773937 | 0.158415981 | 0.879180483 | 0.956743585 | -4.908010766 |
| **A_55_P1980411** | Zkscan1 | -0.017282198 | -0.282569963 | 0.248005568 | 10.57190335 | -0.158294175 | 0.879272484 | 0.956789836 | -4.908021289 |
| **A_51_P285077** | Hhatl | -0.011638876 | -0.190445313 | 0.16716756 | 7.807253629 | -0.158165259 | 0.879369859 | 0.956841928 | -4.908032417 |
| **A_52_P322096** | Patl2 | -0.011172698 | -0.183066655 | 0.160721259 | 7.70537279 | -0.157935823 | 0.879543165 | 0.95693814 | -4.9080522 |
| **A_55_P2115871** | Inpp5j | -0.026730703 | -0.438132231 | 0.384670825 | 8.823527596 | -0.157880201 | 0.879585181 | 0.95693814 | -4.908056991 |
| **A_55_P2115315** | Fmr1nb | 0.0103437 | -0.148895238 | 0.169582638 | 7.663471162 | 0.157837367 | 0.879617537 | 0.95693814 | -4.90806068 |
| **A_51_P260740** | Pcdh7 | 0.011054208 | -0.159179365 | 0.18128778 | 7.729969501 | 0.157784957 | 0.879657127 | 0.95693814 | -4.908065193 |
| **A_52_P222725** | Lims1 | 0.017009994 | -0.245049087 | 0.279069074 | 11.53153875 | 0.157720496 | 0.879705821 | 0.95693814 | -4.90807074 |
| **A_55_P2015092** | E430024I08Rik | -0.011102433 | -0.182323884 | 0.160119018 | 8.731878284 | -0.157558982 | 0.879827831 | 0.956971713 | -4.908084631 |
| **A_51_P483379** | Rnaseh1 | -0.010939345 | -0.179706655 | 0.157827964 | 9.407457845 | -0.157502043 | 0.879870844 | 0.956971713 | -4.908089524 |
| **A_52_P110812** | Ggcx | -0.030355185 | -0.498717917 | 0.438007547 | 9.430753726 | -0.157483022 | 0.879885213 | 0.956971713 | -4.908091159 |
| **A_55_P1968285** | Tle5 | -0.033648235 | -0.553800929 | 0.486504458 | 14.34295071 | -0.157186278 | 0.88010939 | 0.957085477 | -4.908116631 |
| **A_51_P214663** | Tpmt | 0.03421754 | -0.495009894 | 0.563444975 | 9.316300786 | 0.157104861 | 0.880170899 | 0.957085477 | -4.908123611 |
| **A_55_P2072110** | 3-Sep | 0.040007906 | -0.578848609 | 0.65886442 | 15.08313676 | 0.157086564 | 0.880184722 | 0.957085477 | -4.908125179 |
| **A_55_P2170419** | NA | -0.014670028 | -0.24167547 | 0.212335414 | 15.92882466 | -0.15702828 | 0.880228755 | 0.957085477 | -4.908130173 |
| **A_55_P2016024** | Kif3c | 0.013025126 | -0.18861044 | 0.214660693 | 8.451617173 | 0.156963254 | 0.880277883 | 0.957085477 | -4.908135743 |
| **A_55_P2140646** | Creg1 | 0.024828706 | -0.359685118 | 0.409342531 | 14.50991702 | 0.156900859 | 0.880325023 | 0.957085477 | -4.908141085 |
| **A_55_P1960145** | Clip4 | -0.011397497 | -0.18792368 | 0.165128686 | 7.779658428 | -0.156885772 | 0.880336422 | 0.957085477 | -4.908142377 |
| **A_51_P418420** | Ddx3x | 0.017322521 | -0.251458891 | 0.286103932 | 10.14819082 | 0.156601186 | 0.880551437 | 0.957241697 | -4.908166714 |
| **A_51_P436189** | Nudt22 | -0.01179242 | -0.19481002 | 0.17122518 | 9.239999324 | -0.156564491 | 0.880579162 | 0.957241697 | -4.908169849 |
| **A_55_P1959923** | Cth | 0.057687858 | -0.839054961 | 0.954430676 | 11.38926125 | 0.156314654 | 0.880767934 | 0.957319104 | -4.908191174 |
| **A_55_P1987331** | Prl3d1 | -0.011644722 | -0.192790719 | 0.169501276 | 7.925275564 | -0.156200908 | 0.88085388 | 0.957319104 | -4.908200872 |
| **A_52_P556082** | Trmt2b | 0.011334801 | -0.165032156 | 0.187701758 | 9.316195709 | 0.156163622 | 0.880882054 | 0.957319104 | -4.908204049 |
| **A_52_P229783** | Ptgr2 | 0.026639405 | -0.387895557 | 0.441174367 | 9.55511346 | 0.156151636 | 0.880891111 | 0.957319104 | -4.908205071 |
| **A_55_P1958088** | Lyn | -0.01534228 | -0.254096983 | 0.223412424 | 8.409199221 | -0.156142509 | 0.880898008 | 0.957319104 | -4.908205848 |
| **A_52_P336768** | NA | 0.011189317 | -0.163077354 | 0.185455988 | 8.541469609 | 0.156017186 | 0.880992705 | 0.95736819 | -4.908216521 |
| **A_55_P2014169** | Ppa2 | -0.013403369 | -0.222516965 | 0.195710226 | 12.22499002 | -0.155745283 | 0.88119817 | 0.957537633 | -4.908239647 |
| **A_51_P480904** | Blmh | -0.017968127 | -0.298445747 | 0.262509493 | 10.78140681 | -0.155663854 | 0.881259704 | 0.957550667 | -4.908246566 |
| **A_55_P2137406** | Bax | 0.019508963 | -0.285388671 | 0.324406597 | 11.5919922 | 0.155475988 | 0.881401674 | 0.957651094 | -4.908262513 |
| **A_51_P138859** | Tmem68 | -0.016884688 | -0.281050783 | 0.247281407 | 9.86322736 | -0.155309903 | 0.881527188 | 0.957708743 | -4.908276595 |
| **A_55_P2038237** | Rps3 | 0.013754481 | -0.201487422 | 0.228996384 | 10.39335954 | 0.155274651 | 0.881553829 | 0.957708743 | -4.908279582 |
| **A_52_P202142** | Sv2a | -0.010209149 | -0.170151003 | 0.149732704 | 7.669673315 | -0.15509958 | 0.881686139 | 0.957791314 | -4.908294407 |
| **A_52_P399717** | Ccdc50 | 0.013120051 | -0.192500576 | 0.218740677 | 8.532131435 | 0.15504295 | 0.881728938 | 0.957791314 | -4.908299199 |
| **A_55_P2422650** | 5031425E22Rik | -0.044102469 | -0.73583275 | 0.647627812 | 15.23645188 | -0.154920647 | 0.881821372 | 0.957837892 | -4.908309543 |
| **A_55_P2149020** | Rpl35 | -0.017532539 | -0.292810836 | 0.257745759 | 17.35169778 | -0.154759039 | 0.881943514 | 0.957916734 | -4.908323197 |
| **A_55_P2074453** | Slc16a9 | -0.010443587 | -0.174619584 | 0.15373241 | 7.718349989 | -0.154569294 | 0.882086928 | 0.958012946 | -4.908339211 |
| **A_52_P515036** | Htatip2 | -0.023130024 | -0.386878128 | 0.340618081 | 9.140928734 | -0.15451069 | 0.882131223 | 0.958012946 | -4.908344154 |

| | | | | | | | | | |
|---|---|---|---|---|---|---|---|---|---|
| **A_55_P1969477** | Abcb1a | 0.010552012 | -0.155479912 | 0.176583935 | 7.924929006 | 0.154428284 | 0.882193509 | 0.958026761 | -4.9083511 |
| **A_52_P561205** | Stard4 | 0.012581951 | -0.185490257 | 0.210654159 | 8.460633479 | 0.154350353 | 0.882252414 | 0.958036905 | -4.908357665 |
| **A_51_P499530** | Ankrd10 | -0.010896558 | -0.182714232 | 0.160921117 | 9.236390506 | -0.154100723 | 0.882441104 | 0.958136124 | -4.908378675 |
| **A_52_P429876** | Tbx20 | 0.013676096 | -0.201972896 | 0.229325087 | 7.988392425 | 0.154098315 | 0.882442924 | 0.958136124 | -4.908378877 |
| **A_55_P2176300** | Slc33a1 | -0.027485977 | -0.461289938 | 0.406317983 | 8.919026047 | -0.153957503 | 0.882549365 | 0.958197869 | -4.908390713 |
| **A_55_P2335695** | Ndor1 | 0.013907551 | -0.205779933 | 0.233595035 | 8.555626339 | 0.153825567 | 0.882649098 | 0.958244746 | -4.908401793 |
| **A_55_P2394490** | NA | -0.036381878 | -0.611620523 | 0.538856768 | 9.195083502 | -0.153681023 | 0.882758366 | 0.958244746 | -4.908413922 |
| **A_55_P2128169** | Trappc13 | 0.016553021 | -0.245241127 | 0.278347168 | 8.83495242 | 0.153638662 | 0.882790388 | 0.958244746 | -4.908417474 |
| **A_66_P128313** | Mtmr2 | 0.015294865 | -0.226601883 | 0.257191614 | 10.31125053 | 0.153638061 | 0.882790843 | 0.958244746 | -4.908417524 |
| **A_66_P124958** | Ccdc47 | -0.011318317 | -0.190466925 | 0.167830291 | 8.740697866 | -0.15351528 | 0.882883661 | 0.958291683 | -4.908427815 |
| **A_51_P255329** | Riok2 | -0.017866885 | -0.30084374 | 0.26510997 | 10.16413521 | -0.153419699 | 0.882955919 | 0.958316298 | -4.90843582 |
| **A_51_P117604** | Rab5a | 0.017199923 | -0.255348825 | 0.28974867 | 11.05832343 | 0.153343542 | 0.883013492 | 0.958324974 | -4.904480305 |
| **A_55_P2002846** | Cystm1 | 0.013101712 | -0.194753059 | 0.220956482 | 9.026391271 | 0.153162071 | 0.883150686 | 0.958420056 | -4.908457372 |
| **A_55_P2022861** | Elovl1 | -0.01878959 | -0.317441949 | 0.279862769 | 9.974096876 | -0.152874322 | 0.883368237 | 0.958493032 | -4.908481402 |
| **A_55_P2003996** | H2-T24 | -0.018410024 | -0.311124308 | 0.274304259 | 8.59167558 | -0.152824729 | 0.883405732 | 0.958493032 | -4.908485539 |
| **A_55_P2025490** | Tnfrsf18 | -0.010393362 | -0.175652567 | 0.154865842 | 7.882347105 | -0.152817676 | 0.883411065 | 0.958493032 | -4.908486128 |
| **A_51_P461894** | Tnnc1 | 0.029227196 | -0.435519564 | 0.493973955 | 10.28960176 | 0.152810772 | 0.883416285 | 0.958493032 | -4.908486703 |
| **A_55_P2002426** | Rps6kb2 | -0.020428136 | -0.345403576 | 0.304547304 | 8.603045954 | -0.152743023 | 0.883467508 | 0.958494806 | -4.908492352 |
| **A_51_P244950** | Dpys | -0.057908747 | -0.980746827 | 0.864929333 | 10.70996692 | -0.152476126 | 0.883669308 | 0.958659934 | -4.908514582 |
| **A_55_P2102449** | Nxf7 | 0.009821001 | -0.146788218 | 0.16643022 | 7.700023058 | 0.152377772 | 0.883743675 | 0.958686805 | -4.908522764 |
| **A_52_P247943** | Parl | 0.013536812 | -0.20244072 | 0.229514344 | 8.593140145 | 0.152296885 | 0.883804837 | 0.958699349 | -4.90852949 |
| **A_55_P2003688** | Tmem143 | -0.023591365 | -0.400247499 | 0.35306477 | 10.64079904 | -0.152191785 | 0.883884308 | 0.95873175 | -4.908538223 |
| **A_55_P1967534** | NA | 0.020172858 | -0.302466054 | 0.34281177 | 15.05615692 | 0.15192662 | 0.884084817 | 0.958815174 | -4.908560229 |
| **A_51_P153224** | Zdhhc12 | -0.014542733 | -0.247305781 | 0.218220314 | 10.1189044 | -0.15181518 | 0.884169088 | 0.958815174 | -4.908569467 |
| **A_55_P2151143** | Dynlt1c | 0.022799505 | -0.342139701 | 0.387738712 | 13.78291516 | 0.151805704 | 0.884176254 | 0.958815174 | -4.908570252 |
| **A_51_P273684** | Gbe1 | 0.043152701 | -0.647595228 | 0.73390063 | 12.37888962 | 0.151799933 | 0.884180618 | 0.958815174 | -4.90857073 |
| **A_51_P347177** | Snx2 | 0.01595914 | -0.239563513 | 0.271481793 | 11.98519208 | 0.151762077 | 0.884209244 | 0.958815174 | -4.908573866 |
| **A_52_P460791** | Ppy | -0.010160522 | -0.17303471 | 0.152713667 | 7.678160629 | -0.151581753 | 0.884345609 | 0.95890925 | -4.908588794 |
| **A_55_P1962054** | NA | 0.014252522 | -0.214524376 | 0.24302942 | 15.88584245 | 0.151377992 | 0.884499703 | 0.959022536 | -4.908605641 |
| **A_51_P116421** | Abcc5 | -0.018428664 | -0.314443869 | 0.277586541 | 9.835466843 | -0.151273558 | 0.884578683 | 0.959054373 | -4.870345944 |
| **A_55_P2013113** | Bak1 | -0.014649722 | -0.250526012 | 0.221226569 | 11.79616028 | -0.150913565 | 0.884850945 | 0.959082019 | -4.908643956 |
| **A_52_P351669** | Gcc2 | 0.012384964 | -0.187031095 | 0.211801023 | 9.241644985 | 0.150909925 | 0.884853698 | 0.959082019 | -4.908644256 |
| **A_52_P13815** | Laptm5 | 0.034817697 | -0.525820743 | 0.595456137 | 9.257471576 | 0.150903864 | 0.884858282 | 0.959082019 | -4.908644756 |
| **A_66_P115539** | Gm11240 | 0.026430118 | -0.399161809 | 0.452022045 | 11.68196794 | 0.150899887 | 0.88486129 | 0.959082019 | -4.908645083 |
| **A_55_P2142298** | Ywhaq | -0.020411511 | -0.349093768 | 0.308270746 | 13.6336848 | -0.150897516 | 0.884863083 | 0.959082019 | -4.908645278 |
| **A_55_P1980986** | Lca5l | -0.013426697 | -0.229707511 | 0.202854118 | 7.862944165 | -0.1508462 | 0.884901895 | 0.959082019 | -4.908649504 |
| **A_55_P1990235** | Bin3 | -0.018090328 | -0.309807994 | 0.273627338 | 11.65864815 | -0.150683918 | 0.885024635 | 0.959161266 | -4.908662859 |
| **A_55_P2071329** | Rhog | 0.01575961 | -0.238566377 | 0.270085597 | 12.23244078 | 0.150569816 | 0.885110937 | 0.959169861 | -4.908672241 |
| **A_66_P124146** | Ttc16 | -0.013518907 | -0.23194784 | 0.204910026 | 8.475424388 | -0.150388492 | 0.885248087 | 0.959169861 | -4.908687135 |
| **A_55_P1971093** | NA | -0.034034858 | -0.584028879 | 0.515959162 | 12.70770572 | -0.150365833 | 0.885265226 | 0.959169861 | -4.908688995 |
| **A_55_P1990500** | Masp2 | 0.026905331 | -0.407987299 | 0.461797961 | 10.14903503 | 0.150327864 | 0.885293946 | 0.959169861 | -4.908692111 |
| **A_51_P360918** | Ehd3 | 0.033543402 | -0.5088551 | 0.575941904 | 11.30153371 | 0.150269836 | 0.885337838 | 0.959169861 | -4.908696872 |
| **A_52_P409055** | Dcaf8 | 0.012683916 | -0.192426369 | 0.217794202 | 11.74641431 | 0.150261983 | 0.885343778 | 0.959169861 | -4.908697516 |
| **A_55_P1964896** | NA | -0.010777214 | -0.185147699 | 0.163593271 | 7.861538282 | -0.1501816 | 0.885404581 | 0.959169861 | -4.908704107 |
| **A_52_P476731** | Fam110c | -0.031673162 | -0.544243368 | 0.480897044 | 8.669501241 | -0.150148587 | 0.885429553 | 0.959169861 | -4.908706813 |
| **A_51_P259415** | Tmem70 | 0.022658622 | -0.34479634 | 0.390113584 | 11.16821848 | 0.149834758 | 0.885666947 | 0.959340296 | -4.908732509 |
| **A_51_P266224** | Limd1 | 0.021517197 | -0.327486399 | 0.370520793 | 11.08864389 | 0.149809374 | 0.885686149 | 0.959340296 | -4.908734585 |
| **A_55_P2138174** | Psd3 | 0.016044852 | -0.244602656 | 0.27669236 | 8.218701751 | 0.149577175 | 0.885861803 | 0.959417583 | -4.908753559 |
| **A_55_P2130660** | Slitrk4 | 0.011591646 | -0.176800924 | 0.199984215 | 7.720456308 | 0.14950803 | 0.885914112 | 0.959417583 | -4.908759204 |
| **A_51_P233788** | Dmap1 | -0.019865752 | -0.343112044 | 0.303380541 | 10.73812757 | -0.149332604 | 0.886046825 | 0.959417583 | -4.908773513 |

| | | | | | | | | | |
|---|---|---|---|---|---|---|---|---|---|
| **A_52_P141583** | Macroh2a1 | -0.020162513 | -0.348319516 | 0.30799449 | 10.78361413 | -0.149295313 | 0.886075037 | 0.959417583 | -4.908776553 |
| **A_55_P2103011** | Sema4d | -0.026755378 | -0.462314741 | 0.408803985 | 9.09308856 | -0.149261197 | 0.886100846 | 0.959417583 | -4.908779333 |
| **A_55_P2096165** | NA | 0.011708622 | -0.178978512 | 0.202395757 | 11.60476118 | 0.149199579 | 0.886147463 | 0.959417583 | -4.908784353 |
| **A_55_P2112687** | Sipa1l1 | 0.010502682 | -0.160613504 | 0.181618868 | 9.27618268 | 0.149139375 | 0.88619301 | 0.959417583 | -4.908789255 |
| **A_51_P422208** | Dpagt1 | -0.01260535 | -0.217994823 | 0.192784123 | 11.84119757 | -0.149128252 | 0.886201426 | 0.959417583 | -4.908790161 |
| **A_55_P2081829** | Rps4x | -0.015593698 | -0.269810532 | 0.238623137 | 15.25690639 | -0.149048632 | 0.886261663 | 0.959417583 | -4.908796641 |
| **A_55_P2071596** | Ppara | -0.052601399 | -0.910191923 | 0.804989125 | 11.07453686 | -0.1490392 | 0.886268799 | 0.959417583 | -4.908797409 |
| **A_52_P441070** | Apob | 0.052764882 | -0.807809877 | 0.913339641 | 13.14092903 | 0.148983974 | 0.886310581 | 0.959417583 | -4.908801901 |
| **A_52_P642626** | Rbm20 | -0.009455209 | -0.163810289 | 0.14489987 | 7.636987514 | -0.148844713 | 0.886415944 | 0.959417583 | -4.908813222 |
| **A_51_P518586** | Emg1 | -0.013190919 | -0.228567012 | 0.202185174 | 10.57379333 | -0.148819812 | 0.886434784 | 0.959417583 | -4.908815246 |
| **A_55_P2027870** | Abhd17a | -0.014207606 | -0.246333276 | 0.217918065 | 10.33983743 | -0.14872395 | 0.886507314 | 0.959417583 | -4.908823031 |
| **A_55_P2383897** | 8030453O22Rik | 0.01074744 | -0.164989731 | 0.186484612 | 7.750656749 | 0.148601979 | 0.886599599 | 0.959417583 | -4.90883293 |
| **A_51_P441974** | Pip5k1a | -0.017478375 | -0.303374531 | 0.268417782 | 9.822953867 | -0.148551116 | 0.886638083 | 0.959417583 | -4.908837056 |
| **A_52_P467140** | Snd1 | -0.017074951 | -0.296393806 | 0.262243905 | 10.39117305 | -0.14853965 | 0.886646759 | 0.959417583 | -4.908837986 |
| **A_55_P1975415** | Cpped1 | 0.015976578 | -0.245384405 | 0.27733756 | 11.3118283 | 0.148534106 | 0.886650954 | 0.959417583 | -4.908838436 |
| **A_51_P176783** | Atxn7l3b | -0.011878574 | -0.206394346 | 0.182637197 | 11.4875032 | -0.14838592 | 0.886763077 | 0.959450613 | -4.908850446 |
| **A_52_P200925** | NA | 0.016631781 | -0.25587507 | 0.289138633 | 14.08738082 | 0.148301154 | 0.886827216 | 0.959450613 | -4.908857311 |
| **A_55_P1960775** | Rps15a | 0.016472932 | -0.253478754 | 0.286424618 | 15.90224423 | 0.148275038 | 0.886846978 | 0.959450613 | -4.908859425 |
| **A_65_P13402** | Rnf14 | 0.011759039 | -0.181000085 | 0.204518163 | 8.32140116 | 0.148231356 | 0.88688003 | 0.959450613 | -4.908862961 |
| **A_55_P2104917** | Nphp1 | 0.014571559 | -0.224486383 | 0.2536295 | 7.98393907 | 0.148110562 | 0.886971433 | 0.959495793 | -4.908872732 |
| **A_52_P381303** | Gins2 | 0.023455 | -0.361847014 | 0.408757014 | 9.15912828 | 0.147916746 | 0.887118094 | 0.95953375 | -4.908888394 |
| **A_55_P1968043** | Jmjd1c | 0.017628895 | -0.272295767 | 0.307553557 | 10.75167139 | 0.147748513 | 0.8872454 | 0.95953375 | -4.908901973 |
| **A_55_P2213418** | 4933417E11Rik | -0.009883391 | -0.172435342 | 0.152668561 | 7.72085995 | -0.147739613 | 0.887252135 | 0.95953375 | -4.908902691 |
| **A_51_P403334** | Cuedc2 | 0.019855483 | -0.306715924 | 0.346426891 | 13.1449979 | 0.147735715 | 0.887255085 | 0.95953375 | -4.908903005 |
| **A_55_P1997936** | Hsd17b7 | -0.031143358 | -0.543698331 | 0.481411615 | 9.617992802 | -0.147641409 | 0.88732645 | 0.95953375 | -4.908910609 |
| **A_55_P2047305** | Adcy5 | -0.00969917 | -0.169360955 | 0.149962614 | 7.834132245 | -0.147610343 | 0.887349959 | 0.95953375 | -4.908913113 |
| **A_55_P2054927** | Fam136a | 0.020317878 | -0.314200735 | 0.354836491 | 12.59789182 | 0.14758467 | 0.887369388 | 0.95953375 | -4.908915182 |
| **A_55_P1961750** | Dtd2 | 0.013946514 | -0.21574285 | 0.243635878 | 9.273893831 | 0.147539383 | 0.887403659 | 0.95953375 | -4.908918831 |
| **A_55_P1962354** | Lexm | -0.00964027 | -0.168558926 | 0.149278386 | 7.681712624 | -0.147399999 | 0.88750914 | 0.959594125 | -4.908930053 |
| **A_52_P571371** | Tmem144 | 0.011049932 | -0.171204023 | 0.193303887 | 8.243074923 | 0.147321397 | 0.887568626 | 0.959604764 | -4.908936377 |
| **A_55_P2009066** | Cfap74 | 0.024206734 | -0.375852724 | 0.424266192 | 14.12895689 | 0.147026238 | 0.887792005 | 0.959772979 | -4.908960095 |
| **A_55_P2138714** | Sgcd | -0.010223495 | -0.179232849 | 0.15878586 | 7.720382373 | -0.146984594 | 0.887823522 | 0.959772979 | -4.908963437 |
| **A_51_P267063** | Ugt3a2 | -0.042708885 | -0.749482047 | 0.664064277 | 8.582733722 | -0.146832224 | 0.887938843 | 0.959843962 | -4.908975659 |
| **A_52_P377416** | Disp1 | 0.011425674 | -0.177883688 | 0.200735036 | 8.43944602 | 0.146653662 | 0.88807399 | 0.959923895 | -4.908989966 |
| **A_55_P2016388** | Eif3k | 0.019390871 | -0.302002394 | 0.340784135 | 14.55162961 | 0.146603294 | 0.888112112 | 0.959923895 | -4.908993999 |
| **A_55_P1984546** | Mmp1a | 0.009683085 | -0.151151992 | 0.170518162 | 7.652086974 | 0.14629051 | 0.888348859 | 0.960107209 | -4.90901901 |
| **A_55_P2067031** | Agxt2 | 0.046117 | -0.720104574 | 0.812338574 | 11.38219441 | 0.146247974 | 0.888381056 | 0.960107209 | -4.909022407 |
| **A_52_P400425** | Pirt | -0.009662073 | -0.170622663 | 0.151298516 | 7.638554801 | -0.145859245 | 0.888675307 | 0.960334672 | -4.909053408 |
| **A_52_P262676** | NA | -0.014584684 | -0.257585403 | 0.228416036 | 9.180486846 | -0.145838654 | 0.888690893 | 0.960334672 | -4.909055048 |
| **A_55_P2150309** | Lhfpl4 | 0.012230795 | -0.191694008 | 0.216155598 | 11.01194835 | 0.145736329 | 0.888768352 | 0.960364684 | -4.909063194 |
| **A_55_P2084308** | Nid1 | -0.023021102 | -0.407173926 | 0.361131721 | 12.82230924 | -0.145614717 | 0.888860413 | 0.960410471 | -4.909072868 |
| **A_55_P2003891** | Txlna | -0.011452046 | -0.20295559 | 0.180051499 | 10.43770888 | -0.145307976 | 0.889092625 | 0.960603899 | -4.909097232 |
| **A_55_P2156741** | Stx4a | -0.01827571 | -0.324408655 | 0.287857235 | 12.50801397 | -0.1450599 | 0.889280434 | 0.960603899 | -4.9091169 |
| **A_55_P2314496** | Secisbp2l | -0.013497053 | -0.23959177 | 0.212597664 | 8.44320165 | -0.145054675 | 0.88928439 | 0.960603899 | -4.909117314 |
| **A_51_P247873** | Ndufb8 | 0.028347513 | -0.446997378 | 0.503692404 | 14.7349276 | 0.144906985 | 0.889396204 | 0.960603899 | -4.909129006 |
| **A_51_P174720** | Ccdc43 | -0.009447998 | -0.168006571 | 0.149110575 | 8.162865683 | -0.144788227 | 0.889486117 | 0.960603899 | -4.9091384 |
| **A_55_P1987938** | NA | 0.010766318 | -0.169978859 | 0.191511495 | 11.82596867 | 0.144738374 | 0.889523862 | 0.960603899 | -4.909142341 |
| **A_55_P1956497** | Serpinf2 | 0.04010433 | -0.63321658 | 0.71342524 | 15.87705787 | 0.144727931 | 0.889531769 | 0.960603899 | -4.909143166 |
| **A_55_P2419021** | 9330188P03Rik | -0.009607426 | -0.17091462 | 0.151699769 | 7.651257142 | -0.144722645 | 0.88953577 | 0.960603899 | -4.909143584 |
| **A_51_P296487** | Lss | 0.03110187 | -0.491331515 | 0.553535256 | 11.52395983 | 0.144656774 | 0.889585644 | 0.960603899 | -4.909148789 |

| | | | | | | | | | |
|---|---|---|---|---|---|---|---|---|---|
| **A_55_P2155397** | Rps8 | 0.012667908 | -0.200179123 | 0.22551494 | 17.43180862 | 0.144617384 | 0.889615467 | 0.960603899 | -4.9091519 |
| **A_52_P506344** | B3gat3 | -0.020407622 | -0.363412661 | 0.322597418 | 8.549576159 | -0.144568968 | 0.889652125 | 0.960603899 | -4.909155723 |
| **A_55_P2327958** | Mtx3 | 0.009525336 | -0.150628722 | 0.169679395 | 8.358620053 | 0.144519201 | 0.889689806 | 0.960603899 | -4.909159651 |
| **A_52_P140497** | NA | -0.015532166 | -0.276759152 | 0.24569482 | 8.975942505 | -0.144476487 | 0.889722146 | 0.960603899 | -4.909163022 |
| **A_52_P663742** | Ganc | -0.012490673 | -0.22264148 | 0.197660134 | 8.475301066 | -0.144423535 | 0.88976224 | 0.960603899 | -4.909167199 |
| **A_55_P2061294** | NA | -0.011557721 | -0.206052258 | 0.182936816 | 13.87488847 | -0.144393615 | 0.889784894 | 0.960603899 | -4.909169558 |
| **A_52_P598605** | Ptch1 | -0.012084035 | -0.215944618 | 0.191776548 | 7.928530856 | -0.144032962 | 0.890057976 | 0.960730107 | -4.909197962 |
| **A_51_P189722** | Cabin1 | 0.013463352 | -0.213705958 | 0.240632662 | 9.635024188 | 0.144008036 | 0.89007685 | 0.960730107 | -4.909199922 |
| **A_55_P2129029** | Smarcb1 | 0.015978211 | -0.253725558 | 0.28568198 | 11.41329021 | 0.143954197 | 0.890117618 | 0.960730107 | -4.909204156 |
| **A_55_P2122255** | Polm | 0.013131952 | -0.208730042 | 0.234993947 | 8.153009824 | 0.143823406 | 0.890216657 | 0.960730107 | -4.909214433 |
| **A_51_P411130** | Rpp25l | -0.019908649 | -0.356351912 | 0.316534613 | 12.27464518 | -0.143784857 | 0.890245847 | 0.960730107 | -4.909217461 |
| **A_55_P2036999** | Zic3 | -0.009151816 | -0.163864628 | 0.145560995 | 7.64290896 | -0.143735556 | 0.89028318 | 0.960730107 | -4.909221331 |
| **A_55_P2090205** | Tpm3 | -0.010751531 | -0.192600583 | 0.171097521 | 7.810697481 | -0.143662181 | 0.890338744 | 0.960730107 | -4.90922709 |
| **A_55_P1988424** | Tpi1 | -0.030884338 | -0.553313603 | 0.491544927 | 11.23910801 | -0.14364615 | 0.890350883 | 0.960730107 | -4.909228347 |
| **A_55_P2184449** | NA | 0.011037195 | -0.175847856 | 0.197922246 | 7.762115312 | 0.143505098 | 0.890457698 | 0.960730107 | -4.909239407 |
| **A_66_P125560** | Macroh2a2 | -0.020226326 | -0.362726526 | 0.322273875 | 10.64841883 | -0.143495854 | 0.890464698 | 0.960730107 | -4.909240132 |
| **A_55_P1976805** | Isca2 | 0.017607951 | -0.280670491 | 0.315886393 | 10.51593732 | 0.143439957 | 0.890507028 | 0.960730107 | -4.909244512 |
| **A_52_P267543** | Ginm1 | 0.01037695 | -0.165618007 | 0.186371907 | 7.899051841 | 0.143269154 | 0.890636376 | 0.960730107 | -4.909257884 |
| **A_55_P2043862** | Stmn1 | 0.020948285 | -0.334614267 | 0.376510838 | 13.27314082 | 0.143158009 | 0.890720548 | 0.960730107 | -4.909266577 |
| **A_55_P1998396** | Eif2ak4 | 0.015070215 | -0.240796265 | 0.270936696 | 10.43213278 | 0.14311635 | 0.890752097 | 0.960730107 | -4.909269834 |
| **A_51_P148122** | Klhdc3 | 0.019170792 | -0.306325103 | 0.344666688 | 11.87060632 | 0.143112556 | 0.89075497 | 0.960730107 | -4.909270131 |
| **A_55_P2012641** | NA | -0.012056109 | -0.216773554 | 0.192661337 | 16.91156595 | -0.143098635 | 0.890765513 | 0.960730107 | -4.909271219 |
| **A_51_P194273** | Sgcg | -0.01137729 | -0.204569851 | 0.181815271 | 7.741262655 | -0.14309736 | 0.890766479 | 0.960730107 | -4.909271318 |
| **A_55_P2180201** | Mrpl10 | 0.014879387 | -0.23787982 | 0.267638594 | 9.099776275 | 0.143041232 | 0.890808987 | 0.960730107 | -4.909275704 |
| **A_55_P2084492** | NA | 0.015398423 | -0.246459018 | 0.277255863 | 8.558737228 | 0.142887588 | 0.890925348 | 0.960730107 | -4.909287701 |
| **A_52_P137691** | Trappc5 | -0.009666175 | -0.17406169 | 0.15472934 | 8.370484871 | -0.142872251 | 0.890936963 | 0.960730107 | -4.909288897 |
| **A_51_P153557** | Hspa12a | 0.009213119 | -0.147514129 | 0.165940367 | 7.701647506 | 0.142838523 | 0.890962507 | 0.960730107 | -4.909291529 |
| **A_52_P328497** | Cdk10 | -0.012111331 | -0.21822665 | 0.194003988 | 9.916726964 | -0.142779146 | 0.891007478 | 0.960730107 | -4.90929616 |
| **A_55_P2078670** | Cbs | -0.042529143 | -0.766799921 | 0.681741634 | 13.3525831 | -0.142681892 | 0.891081135 | 0.960730107 | -4.909303741 |
| **A_55_P2090996** | Ebna1bp2 | 0.019430042 | -0.311854733 | 0.350714817 | 12.90108988 | 0.142513318 | 0.891208812 | 0.960730107 | -4.90931687 |
| **A_66_P118752** | Rabif | 0.014715439 | -0.236213675 | 0.265644554 | 8.991185605 | 0.142496885 | 0.891221259 | 0.960730107 | -4.909318149 |
| **A_51_P100034** | Mif4gd | -0.010286104 | -0.185837439 | 0.165265232 | 11.71517404 | -0.14237376 | 0.891314515 | 0.960730107 | -4.909327728 |
| **A_52_P333097** | NA | -0.013300426 | -0.240460988 | 0.213860136 | 7.687721509 | -0.142270807 | 0.891392494 | 0.960730107 | -4.909335731 |
| **A_55_P2232023** | NA | 0.009690636 | -0.155825185 | 0.175206457 | 7.677945868 | 0.14226431 | 0.891397415 | 0.960730107 | -4.909336236 |
| **A_51_P446131** | Gipc2 | 0.014184385 | -0.228208851 | 0.25657762 | 9.534597603 | 0.142191355 | 0.891452674 | 0.960730107 | -4.909341903 |
| **A_55_P2126063** | Mrps21 | 0.019166106 | -0.308447115 | 0.346779328 | 13.21118813 | 0.142152878 | 0.891481818 | 0.960730107 | -4.909344891 |
| **A_52_P528634** | Ttc27 | -0.014361946 | -0.26010764 | 0.231383749 | 9.200411256 | -0.14200726 | 0.891592117 | 0.960730107 | -4.909356191 |
| **A_55_P2171648** | Ppp1r7 | 0.022203728 | -0.357778015 | 0.402185471 | 11.33031756 | 0.141986273 | 0.891608014 | 0.960730107 | -4.909357819 |
| **A_51_P446085** | Map10 | -0.015997561 | -0.289844836 | 0.257849714 | 8.328312001 | -0.141947759 | 0.891637188 | 0.960730107 | -4.909360805 |
| **A_55_P2063598** | Tomm5 | 0.017249798 | -0.278066906 | 0.312566503 | 14.08768104 | 0.141931634 | 0.891649402 | 0.960730107 | -4.909362055 |
| **A_55_P2058348** | Pacc1 | 0.011401545 | -0.183945348 | 0.206748439 | 8.438003808 | 0.141820954 | 0.89173324 | 0.960730107 | -4.909370632 |
| **A_55_P2184264** | Timm23 | 0.030575317 | -0.493295919 | 0.554446554 | 11.92552416 | 0.141817429 | 0.89173591 | 0.960730107 | -4.909370905 |
| **A_55_P2086387** | 1700030K09Rik | -0.01495195 | -0.271147381 | 0.241243481 | 8.984673378 | -0.14181091 | 0.891740848 | 0.960730107 | -4.90937141 |
| **A_51_P396273** | Nras | 0.013625927 | -0.22033197 | 0.247583824 | 9.151087475 | 0.141517954 | 0.891962764 | 0.960915632 | -4.909394078 |
| **A_66_P105830** | 1700028B04Rik | 0.023343196 | -0.377890679 | 0.42457707 | 14.53700806 | 0.141366305 | 0.892077644 | 0.960985832 | -4.909405795 |
| **A_55_P2165414** | Serpina1a | -0.010461977 | -0.190436209 | 0.169512254 | 18.24886016 | -0.14124941 | 0.892166197 | 0.960999241 | -4.909414817 |
| **A_52_P141136** | Fggy | 0.042196205 | -0.68390851 | 0.768300919 | 10.34110152 | 0.141207357 | 0.892198055 | 0.960999241 | -4.909418061 |
| **A_55_P2156905** | NA | 0.015979957 | -0.259215674 | 0.291175589 | 13.82183511 | 0.141096836 | 0.892281783 | 0.960999241 | -4.909426582 |
| **A_55_P1998194** | Snrpn | -0.010858471 | -0.19786802 | 0.176151077 | 7.812252058 | -0.141087357 | 0.892288964 | 0.960999241 | -4.909427313 |
| **A_55_P1965363** | Serpinb6e | 0.00932161 | -0.151696198 | 0.170339419 | 7.700883909 | 0.140669591 | 0.892605466 | 0.961286553 | -4.909459461 |

| | | | | | | | | | |
|---|---|---|---|---|---|---|---|---|---|
| **A_51_P328133** | Gle1 | 0.012426168 | -0.202470828 | 0.227323165 | 8.447241763 | 0.140504449 | 0.892730584 | 0.961367734 | -4.909472143 |
| **A_65_P01783** | Med28 | 0.014471165 | -0.236419548 | 0.265361879 | 9.59516566 | 0.140152907 | 0.892996938 | 0.961533638 | -4.909499091 |
| **A_55_P1984961** | NA | 0.009776369 | -0.159790627 | 0.179343364 | 8.338857043 | 0.140093967 | 0.893041596 | 0.961533638 | -4.909503602 |
| **A_51_P358225** | Serinc1 | -0.01554454 | -0.285267107 | 0.254178028 | 13.19241386 | -0.140037314 | 0.893084522 | 0.961533638 | -4.909507937 |
| **A_51_P309733** | Cyp11a1 | -0.009658738 | -0.177325945 | 0.15800847 | 7.664904987 | -0.139976595 | 0.89313053 | 0.961533638 | -4.909512581 |
| **A_55_P2019257** | Trpc4ap | -0.009752943 | -0.179072328 | 0.159566442 | 13.30566564 | -0.139962661 | 0.893141088 | 0.961533638 | -4.909513646 |
| **A_55_P2052892** | Xiap | -0.017367388 | -0.319277117 | 0.284542341 | 11.96350834 | -0.139778568 | 0.893280581 | 0.961533638 | -4.909527712 |
| **A_52_P294253** | Zfp41 | -0.020495492 | -0.376847494 | 0.33585651 | 8.907814141 | -0.139753384 | 0.893299663 | 0.961533638 | -4.909529635 |
| **A_52_P532029** | Tbk1 | 0.019985283 | -0.327550063 | 0.367520628 | 8.709249648 | 0.139731559 | 0.893316202 | 0.961533638 | -4.909531301 |
| **A_55_P2427685** | Agl | -0.029580555 | -0.544051974 | 0.484890865 | 11.6052251 | -0.139710243 | 0.893332354 | 0.961533638 | -4.909532928 |
| **A_55_P2000521** | Lonrf1 | 0.018987302 | -0.311416337 | 0.349390941 | 9.693590088 | 0.139637358 | 0.893387582 | 0.961539539 | -4.909538489 |
| **A_51_P153170** | Cyb5r3 | 0.01868903 | -0.306705209 | 0.344083269 | 12.51541172 | 0.139559721 | 0.893446413 | 0.961549316 | -4.90954441 |
| **A_51_P131025** | Ngdn | -0.01107357 | -0.204389377 | 0.182242237 | 13.01983127 | -0.139188537 | 0.893727692 | 0.961798483 | -4.909572671 |
| **A_55_P2049122** | Tm6sf2 | -0.019286618 | -0.356390268 | 0.317817031 | 8.600308203 | -0.139019532 | 0.893855768 | 0.961831451 | -4.909585515 |
| **A_55_P2057112** | Prl8a2 | 0.009756181 | -0.16077186 | 0.180284222 | 7.697196775 | 0.139016786 | 0.893857849 | 0.961831451 | -4.909585723 |
| **A_52_P140881** | Slc26a10 | 0.010689911 | -0.176385323 | 0.197765144 | 8.157649754 | 0.138848428 | 0.893985438 | 0.961902338 | -4.909598502 |
| **A_66_P105801** | Igf1r | 0.017255491 | -0.28482707 | 0.319338052 | 10.28239239 | 0.138798528 | 0.894023255 | 0.961902338 | -4.909602286 |
| **A_55_P2177633** | NA | -0.008868999 | -0.164227299 | 0.146489301 | 7.641508423 | -0.138714974 | 0.894086578 | 0.961916925 | -4.90960862 |
| **A_51_P483617** | 0610040J01Rik | -0.022305525 | -0.413488916 | 0.368877865 | 12.17585791 | -0.138552778 | 0.894209503 | 0.961925332 | -4.909620904 |
| **A_66_P101473** | Fbxl3 | 0.016085843 | -0.266106291 | 0.298277977 | 9.445624775 | 0.138510311 | 0.894241688 | 0.961925332 | -4.909624118 |
| **A_55_P2132756** | Cacna1g | -0.019409472 | -0.359947787 | 0.321128842 | 7.934125679 | -0.138493992 | 0.894254057 | 0.961925332 | -4.909625353 |
| **A_55_P2078335** | Mapk4 | 0.009082095 | -0.150432506 | 0.168596696 | 7.678224125 | 0.138346693 | 0.894365696 | 0.961925332 | -4.909636492 |
| **A_55_P2056190** | Rps3a1 | 0.012979151 | -0.215203219 | 0.24116152 | 16.27918324 | 0.138212518 | 0.89446739 | 0.961925332 | -4.909646629 |
| **A_51_P288828** | Slc52a2 | 0.014162259 | -0.234895645 | 0.263220163 | 9.556434584 | 0.138170522 | 0.89449922 | 0.961925332 | -4.909649799 |
| **A_51_P515242** | Strip2 | -0.012161635 | -0.226046826 | 0.201723556 | 7.851242849 | -0.138163852 | 0.894504275 | 0.961925332 | -4.909650303 |
| **A_51_P115715** | Asb2 | 0.012139476 | -0.201769943 | 0.226048895 | 8.210252101 | 0.13789649 | 0.894706924 | 0.961925332 | -4.909670465 |
| **A_52_P141112** | Csnk1g3 | -0.009677472 | -0.180274154 | 0.160919211 | 7.988525545 | -0.137839761 | 0.894749922 | 0.961925332 | -4.909674738 |
| **A_55_P2124120** | Ywhae | -0.01375299 | -0.25622444 | 0.228718461 | 14.25088639 | -0.137822364 | 0.894763109 | 0.961925332 | -4.909676048 |
| **A_55_P2112379** | NA | -0.014392208 | -0.268184688 | 0.239400273 | 14.55160849 | -0.137794491 | 0.894784236 | 0.961925332 | -4.909678147 |
| **A_52_P217240** | Ppme1 | 0.016182285 | -0.269198922 | 0.301563493 | 9.158176651 | 0.137783646 | 0.894792457 | 0.961925332 | -4.909678963 |
| **A_55_P2037313** | Gm12027 | 0.029803095 | -0.495941233 | 0.555547422 | 13.15312761 | 0.137743133 | 0.894823165 | 0.961925332 | -4.909682013 |
| **A_52_P590474** | Ccnl1 | -0.017653964 | -0.32914307 | 0.293835143 | 9.021330915 | -0.137715416 | 0.894844174 | 0.961925332 | -4.909684098 |
| **A_51_P280446** | Sdf2l1 | 0.015464488 | -0.257396364 | 0.288325339 | 10.28060777 | 0.13771381 | 0.894845392 | 0.961925332 | -4.909684219 |
| **A_55_P2142008** | 4933434E20Rik | 0.009739225 | -0.162250805 | 0.181729256 | 8.304189663 | 0.137595533 | 0.894935045 | 0.961925332 | -4.909693114 |
| **A_51_P192847** | Ghdc | 0.009965469 | -0.166177473 | 0.186108411 | 8.222259227 | 0.137472453 | 0.895028341 | 0.961925332 | -4.909702363 |
| **A_51_P434240** | Mrpl11 | -0.020805929 | -0.388645796 | 0.347033938 | 12.92163987 | -0.137439479 | 0.895053336 | 0.961925332 | -4.909704839 |
| **A_55_P2064901** | NA | 0.012282238 | -0.204868065 | 0.22943254 | 15.37269836 | 0.137435916 | 0.895056037 | 0.961925332 | -4.909705107 |
| **A_66_P121474** | Chmp6 | -0.018476727 | -0.345251394 | 0.308297941 | 9.735575736 | -0.137391493 | 0.895089711 | 0.961925332 | -4.909708442 |
| **A_51_P305628** | Atp8a2 | 0.036843409 | -0.615775614 | 0.689462431 | 9.308391254 | 0.13717762 | 0.895251836 | 0.962046075 | -4.909724484 |
| **A_55_P2085156** | NA | -0.01625002 | -0.304569782 | 0.272069742 | 13.49430822 | -0.136950205 | 0.895424233 | 0.962177841 | -4.909741515 |
| **A_52_P288873** | Chmp4b | -0.020456058 | -0.383781842 | 0.342869726 | 12.16577618 | -0.136807187 | 0.895532654 | 0.962240851 | -4.909752211 |
| **A_51_P112734** | Slc7a8 | -0.025391261 | -0.476713328 | 0.425930806 | 8.844091017 | -0.136703895 | 0.895610961 | 0.962271498 | -4.897875318 |
| **A_52_P98614** | Slc6a2 | -0.009013284 | -0.169731835 | 0.151705267 | 7.637467315 | -0.136269998 | 0.895939915 | 0.962489857 | -4.909792288 |
| **A_55_P1962154** | Rarb | -0.022999579 | -0.433124568 | 0.38712541 | 8.985525741 | -0.136265791 | 0.895943104 | 0.962489857 | -4.909792601 |
| **A_51_P465473** | Btf3l4 | -0.013933528 | -0.262443333 | 0.234576277 | 8.574957964 | -0.136238787 | 0.895963578 | 0.962489857 | -4.909794611 |
| **A_51_P160083** | 1500035N22Rik | -0.012396553 | -0.233704197 | 0.20891109 | 7.762861549 | -0.136109277 | 0.896061769 | 0.962504942 | -4.909804248 |
| **A_51_P149623** | Gabrd | 0.010920912 | -0.18423906 | 0.206080885 | 7.804750904 | 0.135972589 | 0.896165405 | 0.962504942 | -4.909814409 |
| **A_55_P2068429** | Zfp958 | -0.023139153 | -0.436666475 | 0.390388168 | 10.21457353 | -0.135964786 | 0.896171321 | 0.962504942 | -4.909814988 |
| **A_55_P2140288** | Phka2 | -0.016924049 | -0.319395311 | 0.285547213 | 10.42450902 | -0.135957555 | 0.896176804 | 0.962504942 | -4.909815526 |
| **A_55_P1998186** | NA | 0.010828151 | -0.183040782 | 0.204697084 | 12.44122084 | 0.135715439 | 0.89636038 | 0.962575453 | -4.909833496 |

| | | | | | | | | | |
|---|---|---|---|---|---|---|---|---|---|
| **A_55_P2010861** | Mrps23 | -0.012428188 | -0.234964945 | 0.210108569 | 12.21920508 | -0.13570294 | 0.896369858 | 0.962575453 | -4.909834423 |
| **A_55_P2079927** | Ccdc68 | 0.009335446 | -0.157905716 | 0.176576608 | 7.809618807 | 0.135636035 | 0.896420588 | 0.962575453 | -4.909839383 |
| **A_55_P2019929** | Ythdf3 | -0.011888532 | -0.224910949 | 0.201133885 | 8.344283733 | -0.135608252 | 0.896441654 | 0.962575453 | -4.909841442 |
| **A_52_P466090** | Samhd1 | -0.010820891 | -0.204892306 | 0.183250524 | 9.378789967 | -0.135482949 | 0.896536665 | 0.962590034 | -4.909850723 |
| **A_51_P477902** | Kdm5c | -0.018099887 | -0.342887406 | 0.306687632 | 11.3743711 | -0.135412801 | 0.896589856 | 0.962590034 | -4.909855915 |
| **A_55_P2027259** | Zcchc2 | -0.020085855 | -0.380561807 | 0.340390097 | 9.846328091 | -0.135393311 | 0.896604634 | 0.962590034 | -4.909857357 |
| **A_55_P2000329** | Gskip | 0.017081406 | -0.290315457 | 0.32447827 | 8.848194357 | 0.135022884 | 0.896885526 | 0.962820642 | -4.909884725 |
| **A_52_P40293** | Hgd | -0.038344937 | -0.72888961 | 0.652199736 | 9.634603497 | -0.134927178 | 0.896958102 | 0.962820642 | -4.909891784 |
| **A_55_P2179676** | Spp2 | -0.033539169 | -0.637601518 | 0.570523179 | 14.20668458 | -0.134912977 | 0.896968871 | 0.962820642 | -4.909892831 |
| **A_51_P313337** | Smim10l1 | -0.024211761 | -0.460513264 | 0.412089742 | 10.51532315 | -0.134841278 | 0.897023243 | 0.962825536 | -4.909898116 |
| **A_55_P2083566** | NA | 0.019565025 | -0.334032135 | 0.373162184 | 16.1185958 | 0.134448147 | 0.897321377 | 0.963029207 | -4.909927042 |
| **A_51_P483220** | Impact | -0.016481119 | -0.314409951 | 0.281447712 | 9.589642263 | -0.134417979 | 0.897344256 | 0.963029207 | -4.909929258 |
| **A_52_P583973** | Lrpap1 | 0.013433895 | -0.22945371 | 0.2563215 | 11.80698979 | 0.134393971 | 0.897362463 | 0.963029207 | -4.909931022 |
| **A_52_P163795** | Tubb5 | 0.017461152 | -0.298514897 | 0.333437202 | 14.82713781 | 0.134277089 | 0.897451106 | 0.963066968 | -4.909939602 |
| **A_51_P128987** | Akr1b8 | 0.021505133 | -0.367923628 | 0.410933895 | 10.62737672 | 0.134182939 | 0.89752251 | 0.963066968 | -4.905971038 |
| **A_55_P2089304** | 2900097C17Rik | 0.009921764 | -0.16979138 | 0.189634908 | 9.007224129 | 0.134150484 | 0.897547124 | 0.963066968 | -4.909948888 |
| **A_55_P1952920** | AI854703 | -0.008889122 | -0.170065754 | 0.152287511 | 7.804774917 | -0.134010849 | 0.897653027 | 0.963104646 | -4.90995912 |
| **A_55_P1966804** | Fdps | -0.02320487 | -0.444072698 | 0.397662958 | 15.54988177 | -0.13397279 | 0.897681892 | 0.963104646 | -4.909961906 |
| **A_55_P2026562** | Cdkn1b | 0.013950519 | -0.239244365 | 0.267145402 | 12.10853687 | 0.133880899 | 0.897751586 | 0.96312596 | -4.909968632 |
| **A_52_P409731** | Scamp5 | 0.01567609 | -0.269125325 | 0.300477506 | 8.252703684 | 0.133745385 | 0.897854368 | 0.963151316 | -4.909978542 |
| **A_55_P2134148** | Mzt2 | -0.013866155 | -0.26596223 | 0.238229919 | 11.27063228 | -0.133651292 | 0.897925735 | 0.963151316 | -4.909985417 |
| **A_55_P2047800** | NA | 0.027397243 | -0.471476502 | 0.526270988 | 12.84844273 | 0.133444108 | 0.898082882 | 0.963151316 | -4.910000538 |
| **A_55_P2077884** | Kat6b | -0.008926701 | -0.171543078 | 0.153689676 | 8.014467206 | -0.133385887 | 0.898127042 | 0.963151316 | -4.910004783 |
| **A_55_P2082057** | NA | -0.015955184 | -0.306683133 | 0.274772765 | 16.83554404 | -0.133351598 | 0.898153051 | 0.963151316 | -4.910007282 |
| **A_55_P2028474** | Adam22 | -0.010803567 | -0.207719714 | 0.186112579 | 7.725166998 | -0.13331194 | 0.898183132 | 0.963151316 | -4.910010172 |
| **A_55_P2186027** | Gapdh | -0.020776674 | -0.399505745 | 0.357952397 | 16.85683571 | -0.133300129 | 0.898192091 | 0.963151316 | -4.910011033 |
| **A_51_P389531** | Prmt1 | 0.019451185 | -0.335221716 | 0.374124087 | 9.913355224 | 0.133260443 | 0.898222193 | 0.963151316 | -4.910013923 |
| **A_66_P123055** | Gm10845 | 0.035085367 | -0.604797677 | 0.674968411 | 12.21503789 | 0.133232019 | 0.898243754 | 0.963151316 | -4.910015993 |
| **A_55_P2132132** | Gm5532 | -0.008714878 | -0.167721453 | 0.150291697 | 7.657780318 | -0.133177062 | 0.898285441 | 0.963151316 | -4.910019994 |
| **A_55_P2109429** | Rfng | 0.017690116 | -0.305194551 | 0.340574783 | 11.22862175 | 0.133127092 | 0.898323345 | 0.963151316 | -4.91002363 |
| **A_51_P487175** | Acsm3 | 0.041181684 | -0.712138118 | 0.794501486 | 9.526762582 | 0.13283359 | 0.898545982 | 0.963336584 | -4.910044961 |
| **A_51_P499195** | Nkg7 | -0.020505468 | -0.396031946 | 0.35502101 | 8.917344357 | -0.132682093 | 0.898660904 | 0.963349432 | -4.910055953 |
| **A_51_P332136** | Zmat5 | -0.023237286 | -0.448820131 | 0.402345559 | 8.64834046 | -0.13267359 | 0.898667355 | 0.963349432 | -4.91005657 |
| **A_55_P1998872** | Slitrk5 | -0.008439786 | -0.163296549 | 0.146416976 | 7.659720972 | -0.132429416 | 0.898852586 | 0.963349432 | -4.910074259 |
| **A_52_P594927** | Atl3 | 0.009777585 | -0.169666353 | 0.189221523 | 8.194926518 | 0.13239939 | 0.898875364 | 0.963349432 | -4.910076432 |
| **A_55_P2122633** | Airn | -0.018441521 | -0.35701687 | 0.320133829 | 10.51087181 | -0.132350194 | 0.898912685 | 0.963349432 | -4.910079991 |
| **A_52_P394539** | Tex2 | 0.026086402 | -0.453102085 | 0.505274889 | 11.15508955 | 0.13227903 | 0.898966673 | 0.963349432 | -4.910085138 |
| **A_55_P1978927** | Srgap1 | 0.008736152 | -0.151752472 | 0.169224776 | 7.792002936 | 0.132269319 | 0.89897404 | 0.963349432 | -4.91008584 |
| **A_52_P231635** | Oxsm | -0.011139741 | -0.215868902 | 0.19358942 | 8.163448971 | -0.132214341 | 0.899015749 | 0.963349432 | -4.910089813 |
| **A_51_P427401** | Fam90a1b | 0.008793239 | -0.152915818 | 0.170502296 | 7.692756975 | 0.132128873 | 0.899080589 | 0.963349432 | -4.910095988 |
| **A_51_P317225** | Cmtm2a | -0.011552278 | -0.22405113 | 0.200946575 | 8.738552615 | -0.132097385 | 0.899104478 | 0.963349432 | -4.910098261 |
| **A_55_P2108784** | Arhgap22 | -0.03965287 | -0.769061779 | 0.689756039 | 9.379798707 | -0.132095113 | 0.899106201 | 0.963349432 | -4.910098425 |
| **A_55_P1980932** | Elob | 0.016251174 | -0.282911488 | 0.315413836 | 14.47984831 | 0.13199593 | 0.899181447 | 0.963376653 | -4.910105583 |
| **A_51_P469968** | H2-M3 | -0.01013791 | -0.196885093 | 0.176609273 | 8.49960217 | -0.131909955 | 0.899246674 | 0.963393136 | -4.910111784 |
| **A_55_P1952850** | NA | -0.018547221 | -0.360496792 | 0.32340235 | 14.36292486 | -0.131795316 | 0.89933365 | 0.963432917 | -4.910120046 |
| **A_55_P1969993** | NA | -0.01730775 | -0.336590773 | 0.301975272 | 17.11155938 | -0.131718865 | 0.899391653 | 0.963441657 | -4.910125551 |
| **A_55_P2106763** | Pxdc1 | 0.031126908 | -0.543946575 | 0.60620039 | 9.082961998 | 0.131521219 | 0.899541609 | 0.963548893 | -4.91013977 |
| **A_51_P516705** | Ern1 | -0.013061632 | -0.254764344 | 0.22864108 | 8.596323915 | -0.131310392 | 0.899701572 | 0.963666834 | -4.910154913 |
| **A_52_P117408** | Tg | 0.009805542 | -0.172036216 | 0.1916473 | 7.895207633 | 0.13102711 | 0.899916516 | 0.963843649 | -4.910175224 |
| **A_66_P125722** | Cyb5a | 0.021171617 | -0.371746739 | 0.414089973 | 15.22634723 | 0.130928715 | 0.899991177 | 0.963870204 | -4.910182268 |

| | | | | | | | | | |
|---|---|---|---|---|---|---|---|---|---|
| **A_55_P2001072** | NA | 0.021088995 | -0.370582391 | 0.412760381 | 12.85178478 | 0.130832978 | 0.900063822 | 0.963894599 | -4.910189117 |
| **A_51_P157297** | Efcab14 | -0.012399359 | -0.242818112 | 0.218019394 | 10.60508981 | -0.130756897 | 0.900121553 | 0.963903019 | -4.910194556 |
| **A_55_P1987023** | NA | 0.014401988 | -0.253437713 | 0.28224169 | 9.159050343 | 0.130656402 | 0.90019781 | 0.963931276 | -4.910201736 |
| **A_52_P244463** | D16Ertd472e | -0.013163705 | -0.258315992 | 0.231988582 | 8.692126473 | -0.130474423 | 0.900335902 | 0.964002115 | -4.910214724 |
| **A_52_P176160** | Ahcy | -0.029450268 | -0.57806701 | 0.519166474 | 9.34723478 | -0.130437776 | 0.900363711 | 0.964002115 | -4.910217337 |
| **A_55_P2103884** | Supt4b | 0.016207803 | -0.286358143 | 0.318773749 | 11.84176671 | 0.13016292 | 0.90057229 | 0.964172028 | -4.910236913 |
| **A_51_P223929** | Ash1l | -0.011447951 | -0.225847577 | 0.202951675 | 8.931192699 | -0.129743895 | 0.900890289 | 0.964459064 | -4.91026668 |
| **A_55_P2044593** | 1810037I17Rik | -0.01749278 | -0.345530818 | 0.310545257 | 13.27389576 | -0.129573989 | 0.901019237 | 0.964507134 | -4.910278722 |
| **A_51_P345927** | Ddx41 | -0.013080014 | -0.258405773 | 0.232245744 | 8.959843641 | -0.129553234 | 0.901034989 | 0.964507134 | -4.910280192 |
| **A_51_P208555** | Decr1 | 0.018036189 | -0.320682931 | 0.356755309 | 8.835084607 | 0.129386289 | 0.901161695 | 0.964589345 | -4.910292008 |
| **A_55_P1975742** | Dap3 | -0.011842278 | -0.23450113 | 0.210816574 | 11.44628166 | -0.129234499 | 0.901276899 | 0.964606317 | -4.910302737 |
| **A_55_P2057296** | Upk3bl | -0.012318011 | -0.244026759 | 0.219390737 | 7.893664019 | -0.129175859 | 0.901321407 | 0.964606317 | -4.910306879 |
| **A_55_P1966086** | NA | 0.01307171 | -0.232829228 | 0.258972648 | 9.000873695 | 0.129168141 | 0.901327264 | 0.964606317 | -4.910307424 |
| **A_55_P2102050** | Cc2d1b | -0.012052702 | -0.238962515 | 0.214857112 | 11.45481207 | -0.129066735 | 0.901404232 | 0.964635278 | -4.910314582 |
| **A_55_P1992811** | Nudc | -0.019665033 | -0.390642185 | 0.351312119 | 9.740412162 | -0.128804401 | 0.901603349 | 0.964794947 | -4.910333072 |
| **A_55_P2084652** | H2ac15 | 0.012724819 | -0.227665524 | 0.253115162 | 17.52441105 | 0.128622744 | 0.901741234 | 0.964889078 | -4.910345855 |
| **A_55_P2099520** | H2afv | -0.015904862 | -0.316662074 | 0.284852351 | 8.823299888 | -0.128498198 | 0.901835773 | 0.96493682 | -4.910354608 |
| **A_52_P189618** | Armc8 | 0.021363331 | -0.383055152 | 0.425781815 | 9.115310686 | 0.128357475 | 0.901942592 | 0.964997695 | -4.910364488 |
| **A_55_P2059164** | H3f3b | 0.014485132 | -0.260955519 | 0.289925784 | 13.76675734 | 0.127784337 | 0.902377672 | 0.965409754 | -4.910404617 |
| **A_51_P229633** | Rprd1b | 0.010974488 | -0.198076968 | 0.220025945 | 11.51741645 | 0.127559927 | 0.902548035 | 0.965506509 | -4.910420281 |
| **A_55_P2129187** | NA | 0.012226059 | -0.220765032 | 0.24521715 | 15.73859702 | 0.127505922 | 0.902589035 | 0.965506509 | -4.910424046 |
| **A_55_P2037328** | Ppp1r12b | -0.008675987 | -0.17406305 | 0.156711076 | 7.758486523 | -0.127467817 | 0.902617964 | 0.965506509 | -4.910426702 |
| **A_52_P14526** | Zyg11b | -0.012349651 | -0.247977693 | 0.223278391 | 8.386913737 | -0.127353504 | 0.902704749 | 0.965508226 | -4.910434665 |
| **A_51_P132081** | Rsl1 | -0.010285717 | -0.206564339 | 0.185992905 | 8.136949893 | -0.127334112 | 0.902719471 | 0.965508226 | -4.910436015 |
| **A_55_P2057653** | NA | -0.012399051 | -0.249382567 | 0.224584464 | 15.59471062 | -0.127131599 | 0.902873221 | 0.96552069 | -4.910450102 |
| **A_51_P240986** | Plekhg6 | -0.01897965 | -0.382598301 | 0.344639002 | 8.523381882 | -0.126830946 | 0.903101489 | 0.96552069 | -4.910470975 |
| **A_55_P2062811** | Tomm5 | -0.01493209 | -0.301188613 | 0.271324432 | 13.06796985 | -0.126750129 | 0.903162851 | 0.96552069 | -4.910476578 |
| **A_52_P561690** | BC034090 | -0.008301391 | -0.167448664 | 0.150845882 | 7.69046302 | -0.126746174 | 0.903165854 | 0.96552069 | -4.910476852 |
| **A_51_P488510** | Appl1 | 0.009528408 | -0.173234057 | 0.192290873 | 8.494694015 | 0.126682472 | 0.903214221 | 0.96552069 | -4.910481265 |
| **A_51_P121302** | Them4 | -0.011775942 | -0.237804396 | 0.214252512 | 8.632172669 | -0.12659476 | 0.903280818 | 0.96552069 | -4.873368901 |
| **A_55_P2132147** | NA | 0.025203817 | -0.458637025 | 0.50904466 | 15.67548704 | 0.126574723 | 0.903296032 | 0.96552069 | -4.910488725 |
| **A_55_P2034481** | Erdr1 | 0.044055114 | -0.801844032 | 0.88995426 | 12.79128265 | 0.126549645 | 0.903315074 | 0.96552069 | -4.91049046 |
| **A_51_P234864** | Sfxn5 | 0.021191788 | -0.386121515 | 0.428505091 | 8.838376574 | 0.126421865 | 0.903412096 | 0.96552069 | -4.910499297 |
| **A_55_P2016712** | Msl1 | -0.01654669 | -0.367082145 | 0.333988765 | 11.39854833 | -0.128879838 | 0.903419664 | 0.96552069 | -4.825360568 |
| **A_51_P261379** | Spats2 | -0.011889231 | -0.240427507 | 0.216649045 | 8.719178755 | -0.126409002 | 0.903421864 | 0.96552069 | -4.910500186 |
| **A_55_P1983958** | NA | -0.020867093 | -0.422488708 | 0.380754522 | 13.74942269 | -0.126249033 | 0.90354333 | 0.96552069 | -4.910511235 |
| **A_51_P130666** | Adk | -0.027909167 | -0.565487061 | 0.509668727 | 13.87357367 | -0.126150409 | 0.903618218 | 0.96552069 | -4.91051804 |
| **A_55_P2020636** | NA | 0.00805898 | -0.147208586 | 0.163326547 | 7.66175267 | 0.126119586 | 0.903641623 | 0.96552069 | -4.910520166 |
| **A_51_P136845** | Usp19 | 0.014737978 | -0.269221455 | 0.298697411 | 9.889340796 | 0.126114435 | 0.903645534 | 0.96552069 | -4.910520521 |
| **A_51_P144264** | Klf2 | -0.039060183 | -0.791716198 | 0.713595831 | 10.22247919 | -0.126101696 | 0.903655207 | 0.96552069 | -4.9105214 |
| **A_55_P2065929** | NA | -0.013142344 | -0.2664481 | 0.240163412 | 8.18734478 | -0.126069769 | 0.903679451 | 0.96552069 | -4.910523601 |
| **A_55_P1995126** | NA | 0.01021269 | -0.186730557 | 0.207155936 | 16.75392122 | 0.126003389 | 0.903729856 | 0.96552069 | -4.910528175 |
| **A_52_P249920** | Zscan29 | 0.011970838 | -0.218943016 | 0.242884691 | 8.443632847 | 0.125967277 | 0.903757278 | 0.96552069 | -4.910530663 |
| **A_55_P2094602** | Lrch1 | 0.009692363 | -0.177404685 | 0.196789411 | 8.594651019 | 0.125876868 | 0.903825931 | 0.96552069 | -4.910536888 |
| **A_51_P430383** | Dnajc30 | -0.01162334 | -0.236020867 | 0.212774188 | 11.18679208 | -0.125862411 | 0.903836909 | 0.96552069 | -4.910537883 |
| **A_55_P2142296** | Ywhaq | 0.014885541 | -0.27263542 | 0.302406503 | 12.91448071 | 0.125799322 | 0.903884816 | 0.96552069 | -4.910542223 |
| **A_51_P124719** | Ccdc93 | 0.009981879 | -0.182864238 | 0.202827997 | 8.336352423 | 0.125772182 | 0.903905426 | 0.96552069 | -4.91054409 |
| **A_55_P1987993** | NA | 0.027137353 | -0.497280781 | 0.551555487 | 15.51854881 | 0.12573986 | 0.903929971 | 0.96552069 | -4.910546312 |
| **A_52_P67983** | Lcmt2 | 0.011950585 | -0.219231934 | 0.243133105 | 9.877681073 | 0.125608021 | 0.904030087 | 0.965536478 | -4.910555372 |
| **A_55_P2075343** | Arfrp1 | 0.008448742 | -0.155106456 | 0.172003941 | 8.393160142 | 0.125519414 | 0.904097375 | 0.965536478 | -4.910561455 |

| | | | | | | | | | |
|---|---|---|---|---|---|---|---|---|---|
| **A_51_P290685** | Ppp6r3 | -0.009619787 | -0.196111239 | 0.176871665 | 8.685396849 | -0.125340004 | 0.904233621 | 0.965536478 | -4.91057376 |
| **A_55_P2034720** | Cfap36 | 0.021867229 | -0.402344235 | 0.446078693 | 9.764435491 | 0.125254846 | 0.904298293 | 0.965536478 | -4.910579594 |
| **A_51_P100787** | Snw1 | -0.01427709 | -0.291261495 | 0.262707315 | 10.70069711 | -0.125247068 | 0.904304199 | 0.965536478 | -4.894734526 |
| **A_51_P132013** | Cysltr2 | -0.008034634 | -0.163914299 | 0.147845031 | 7.687959962 | -0.125244827 | 0.904305901 | 0.965536478 | -4.91058028 |
| **A_55_P2041340** | Atp5a1 | -0.011911627 | -0.243060919 | 0.219237665 | 14.73082078 | -0.125216542 | 0.904327382 | 0.965536478 | -4.910582217 |
| **A_55_P2428967** | Lrrc2 | -0.008078424 | -0.164871096 | 0.148714247 | 7.697133213 | -0.125194168 | 0.904344374 | 0.965536478 | -4.910583749 |
| **A_55_P2148402** | Cplx2 | 0.016821707 | -0.310533022 | 0.344176437 | 13.16538869 | 0.124863251 | 0.904595691 | 0.965729684 | -4.910606372 |
| **A_52_P434974** | Ubqln1 | 0.009023502 | -0.166630926 | 0.184677929 | 10.70347609 | 0.124824317 | 0.90462526 | 0.965729684 | -4.91060903 |
| **A_55_P1977548** | Mapk1 | 0.013644412 | -0.25218619 | 0.279475015 | 10.66400077 | 0.124719124 | 0.904705152 | 0.965761633 | -4.910616207 |
| **A_55_P1991214** | Lair1 | -0.019824163 | -0.406527012 | 0.366878685 | 9.033482758 | -0.124566336 | 0.904821195 | 0.965832167 | -4.910626621 |
| **A_55_P2134862** | Cmip | 0.027380623 | -0.507513274 | 0.562274519 | 10.81384144 | 0.124382381 | 0.904960912 | 0.965912142 | -4.910639142 |
| **A_55_P1959718** | Cobll1 | 0.016774237 | -0.311040682 | 0.344589156 | 10.39012607 | 0.124336102 | 0.904996062 | 0.965912142 | -4.910642289 |
| **A_55_P1952882** | Cyp4f18 | -0.01364212 | -0.280495244 | 0.253211004 | 9.425224419 | -0.124220354 | 0.905083977 | 0.965952636 | -4.910650155 |
| **A_52_P44030** | Exoc3 | 0.010210957 | -0.190021885 | 0.210443799 | 8.550306721 | 0.123912278 | 0.905317981 | 0.966086394 | -4.910671056 |
| **A_52_P724940** | Ndufa12 | -0.011132155 | -0.2294667 | 0.20720239 | 13.76673577 | -0.123891064 | 0.905334095 | 0.966086394 | -4.910672493 |
| **A_55_P2052862** | Miga1 | -0.015931605 | -0.328526222 | 0.296663011 | 9.797113132 | -0.123840082 | 0.905372819 | 0.966086394 | -4.910675946 |
| **A_52_P566681** | Gpm6a | -0.01379605 | -0.284593683 | 0.257001582 | 8.099468192 | -0.123792146 | 0.905409231 | 0.966086394 | -4.910679192 |
| **A_52_P411331** | Vac14 | 0.014527826 | -0.270963065 | 0.300018718 | 8.387874854 | 0.123649275 | 0.905517756 | 0.966144559 | -4.910688858 |
| **A_66_P131205** | NA | 0.014033691 | -0.26188178 | 0.289949161 | 14.71226573 | 0.123588776 | 0.905563711 | 0.966144559 | -4.910692948 |
| **A_55_P1994913** | Ppp2cb | -0.015995443 | -0.330964614 | 0.298973728 | 12.15047955 | -0.123398936 | 0.905707919 | 0.966204176 | -4.910705768 |
| **A_55_P2016877** | Nsd1 | -0.008750478 | -0.181151547 | 0.16365059 | 11.58089926 | -0.123331786 | 0.905758928 | 0.966204176 | -4.910710299 |
| **A_51_P435321** | Ap1s2 | -0.009004281 | -0.186488529 | 0.168479968 | 8.429694852 | -0.123274259 | 0.905802629 | 0.966204176 | -4.910714178 |
| **A_52_P112888** | Atp6v1h | 0.016400735 | -0.307215135 | 0.340016605 | 8.955888878 | 0.123144972 | 0.905900842 | 0.966204176 | -4.910722889 |
| **A_51_P483922** | Tm9sf4 | -0.014047634 | -0.291410949 | 0.263315681 | 9.249686692 | -0.123065795 | 0.90596099 | 0.966204176 | -4.910728219 |
| **A_52_P65237** | Zbtb7c | 0.012398958 | -0.232473628 | 0.257271543 | 7.792742528 | 0.123034867 | 0.905984485 | 0.966204176 | -4.9107303 |
| **A_55_P2027587** | Supt4b | 0.012474253 | -0.234083267 | 0.259031773 | 10.98828396 | 0.122936116 | 0.906059504 | 0.966204176 | -4.910736942 |
| **A_55_P2047290** | Ankrd40 | -0.010818009 | -0.224694153 | 0.203058135 | 8.44790909 | -0.122904614 | 0.906083436 | 0.966204176 | -4.91073906 |
| **A_55_P1981756** | Vmn2r123 | -0.00798608 | -0.165974379 | 0.15000222 | 7.641460242 | -0.122826454 | 0.906142813 | 0.966204176 | -4.910744311 |
| **A_51_P222467** | Abcg1 | -0.021003759 | -0.436562917 | 0.394555399 | 10.64951721 | -0.122813854 | 0.906152386 | 0.966204176 | -4.910745158 |
| **A_55_P2368710** | NA | 0.013465218 | -0.253064116 | 0.279994552 | 8.614358508 | 0.122758497 | 0.906194441 | 0.966204176 | -4.910748875 |
| **A_55_P2227335** | Ptpre | 0.018473459 | -0.347572893 | 0.38451981 | 8.699192466 | 0.122629602 | 0.906292364 | 0.966204176 | -4.910757524 |
| **A_55_P2004084** | Iqce | -0.013652241 | -0.284472203 | 0.257167721 | 9.315293059 | -0.122491644 | 0.906397174 | 0.966204176 | -4.910766771 |
| **A_66_P100021** | Ncaph2 | 0.025854292 | -0.487049514 | 0.538758099 | 10.94970643 | 0.122484158 | 0.906402861 | 0.966204176 | -4.910767272 |
| **A_55_P2030736** | Scmh1 | 0.009071554 | -0.170903233 | 0.18904634 | 8.381705091 | 0.122476619 | 0.906408589 | 0.966204176 | -4.910767777 |
| **A_52_P426513** | Sh3bgrl | -0.016320223 | -0.340142325 | 0.307501879 | 11.15301624 | -0.122462407 | 0.906419386 | 0.966204176 | -4.910768729 |
| **A_51_P146633** | NA | 0.027299049 | -0.514849171 | 0.569447268 | 9.998384991 | 0.122352448 | 0.906502926 | 0.96623994 | -4.91077609 |
| **A_55_P1984178** | Pde4dip | 0.022610001 | -0.426987262 | 0.472207264 | 9.364364248 | 0.122196879 | 0.906621121 | 0.966295087 | -4.910786493 |
| **A_55_P2163093** | Nxf7 | -0.007825635 | -0.163493706 | 0.147842435 | 7.624892834 | -0.122152751 | 0.906654649 | 0.966295087 | -4.910789442 |
| **A_55_P1987514** | NA | 0.009102642 | -0.172206929 | 0.190412212 | 16.69959888 | 0.121991597 | 0.90677709 | 0.966365165 | -4.9108002 |
| **A_55_P1979973** | Nuak2 | 0.009981136 | -0.188923861 | 0.208886133 | 8.407839098 | 0.121931931 | 0.906822423 | 0.966365165 | -4.91080418 |
| **A_55_P2023523** | Aloxe3 | -0.008125064 | -0.170237406 | 0.153987277 | 7.748701466 | -0.121785033 | 0.906934036 | 0.966365165 | -4.91081397 |
| **A_55_P2171897** | Notch1 | -0.014967955 | -0.313652462 | 0.283716551 | 8.194378459 | -0.121767944 | 0.906947021 | 0.966365165 | -4.910815108 |
| **A_66_P114974** | NA | -0.028971473 | -0.607595339 | 0.549652393 | 14.09967444 | -0.121662693 | 0.907026992 | 0.966365165 | -4.910822115 |
| **A_55_P2444965** | Tmem245 | -0.008547563 | -0.179488637 | 0.162393511 | 8.022459792 | -0.121500777 | 0.907150021 | 0.966365165 | -4.910832881 |
| **A_51_P291749** | Pecr | 0.028839723 | -0.548494089 | 0.606173535 | 13.28291864 | 0.121380041 | 0.907241762 | 0.966365165 | -4.9108409 |
| **A_55_P2110758** | B430306N03Rik | 0.025843044 | -0.491604476 | 0.543290564 | 10.95070997 | 0.121355802 | 0.90726018 | 0.966365165 | -4.910842509 |
| **A_55_P1972969** | NA | -0.016668576 | -0.350425472 | 0.317088319 | 14.37106628 | -0.121353261 | 0.907262111 | 0.966365165 | -4.910842678 |
| **A_55_P2035439** | Uxt | 0.014770449 | -0.281021236 | 0.310562135 | 11.25899442 | 0.12133635 | 0.907274961 | 0.966365165 | -4.9108438 |
| **A_55_P2061025** | Hspa4 | 0.027159287 | -0.51692256 | 0.571241133 | 9.38522368 | 0.121293439 | 0.907307567 | 0.966365165 | -4.910846648 |
| **A_52_P176999** | Klhl31 | 0.007727108 | -0.14717032 | 0.162624536 | 7.620049482 | 0.121214889 | 0.907367255 | 0.966365165 | -4.910851857 |

| | | | | | | | | |
|---|---|---|---|---|---|---|---|---|
| **A_51_P266143** | Adpgk | 0.027480824 | -0.523417609 | 0.578379257 | 10.33086247 | 0.121210823 | 0.907370344 | 0.966365165 | -4.910852126 |
| **A_51_P485391** | Parn | 0.009108156 | -0.173668891 | 0.191885203 | 9.19764418 | 0.121085455 | 0.907465609 | 0.966407412 | -4.910860433 |
| **A_55_P2056639** | Bcar1 | 0.011058114 | -0.211064638 | 0.233180865 | 10.93848731 | 0.120968192 | 0.907554716 | 0.966407412 | -4.910868195 |
| **A_66_P113868** | Cdh3 | 0.011674622 | -0.222845351 | 0.246194596 | 8.345322875 | 0.120961231 | 0.907560005 | 0.966407412 | -4.910868656 |
| **A_65_P07196** | COX3 | 0.013738156 | -0.262957735 | 0.290434046 | 16.68458664 | 0.120644886 | 0.9078004 | 0.966610143 | -4.910889557 |
| **A_55_P1954001** | Snrpc | 0.01367602 | -0.262834887 | 0.290186928 | 12.16557948 | 0.120179574 | 0.908154017 | 0.966872857 | -4.910920203 |
| **A_52_P48569** | Slc38a4 | -0.038869652 | -0.824984617 | 0.747245313 | 11.7510631 | -0.120145574 | 0.908179856 | 0.966872857 | -4.910922438 |
| **A_51_P511899** | Mthfsd | 0.011403543 | -0.21926996 | 0.242077047 | 8.156352728 | 0.120122758 | 0.908197196 | 0.966872857 | -4.910923937 |
| **A_55_P2294037** | NA | -0.015650207 | -0.332637097 | 0.301336683 | 8.608447531 | -0.119967048 | 0.908315535 | 0.966945584 | -4.910934161 |
| **A_51_P257743** | Dnajc22 | 0.027330736 | -0.527241102 | 0.581902575 | 10.96573493 | 0.119750327 | 0.908480246 | 0.966995435 | -4.910948369 |
| **A_51_P469480** | Yes1 | 0.009812304 | -0.189318984 | 0.208943592 | 7.980535453 | 0.119733225 | 0.908493244 | 0.966995435 | -4.910949489 |
| **A_51_P486681** | Ap3b2 | -0.007630748 | -0.162580035 | 0.14731854 | 7.653147118 | -0.119663222 | 0.908546448 | 0.966995435 | -4.910954072 |
| **A_66_P124659** | Mark1 | 0.013647855 | -0.26362529 | 0.290921 | 8.523146621 | 0.119602372 | 0.908592697 | 0.966995435 | -4.910958054 |
| **A_51_P418820** | Tfap2c | -0.008123666 | -0.173202015 | 0.156954682 | 7.678770583 | -0.119576311 | 0.908612504 | 0.966995435 | -4.910959759 |
| **A_55_P1984173** | Pde4dip | 0.02116074 | -0.409094557 | 0.451416037 | 10.84764425 | 0.119505478 | 0.90866634 | 0.966999488 | -4.91096439 |
| **A_55_P2170978** | Ankrd13d | -0.011340099 | -0.242113277 | 0.219433079 | 8.721506709 | -0.119402858 | 0.908744338 | 0.96702925 | -4.910971096 |
| **A_55_P1985882** | Fastkd1 | -0.012047297 | -0.257401787 | 0.233307193 | 8.512948268 | -0.119310536 | 0.908814508 | 0.967050681 | -4.910977123 |
| **A_51_P196844** | Osbpl3 | -0.008142484 | -0.174104545 | 0.157819578 | 8.00118303 | -0.119215101 | 0.908887046 | 0.967074629 | -4.910983349 |
| **A_55_P2122504** | Zpr1 | 0.010501089 | -0.203970751 | 0.224972929 | 11.54722643 | 0.118972683 | 0.909071306 | 0.967173125 | -4.910999141 |
| **A_52_P378914** | AU040320 | 0.009414005 | -0.182873302 | 0.201701311 | 8.235485245 | 0.118961653 | 0.909079691 | 0.967173125 | -4.910999859 |
| **A_55_P2094613** | NA | 0.014268847 | -0.277489452 | 0.306027147 | 8.905998678 | 0.11883623 | 0.909175027 | 0.967221316 | -4.911008016 |
| **A_51_P370717** | Gsc | -0.00766362 | -0.164461454 | 0.149134213 | 7.697314004 | -0.118761891 | 0.909231534 | 0.967228196 | -4.911012847 |
| **A_66_P119771** | Nme8 | 0.007882964 | -0.153667096 | 0.169433023 | 7.651613385 | 0.118567481 | 0.909379313 | 0.967324324 | -4.911025466 |
| **A_51_P495242** | Lat | -0.023350898 | -0.502338398 | 0.455636601 | 8.940472709 | -0.11845751 | 0.909462907 | 0.967324324 | -4.911032595 |
| **A_51_P386870** | Sprr2f | -0.007793922 | -0.167683688 | 0.152095844 | 7.634659072 | -0.118445504 | 0.909472034 | 0.967324324 | -4.911033373 |
| **A_51_P140641** | Eva1a | 0.020211217 | -0.395201901 | 0.435624336 | 10.98498803 | 0.118221225 | 0.909642525 | 0.96734437 | -4.911047891 |
| **A_55_P2015767** | Yipf6 | 0.01144911 | -0.223872285 | 0.246770505 | 8.528226283 | 0.118220698 | 0.909642926 | 0.96734437 | -4.911047925 |
| **A_55_P2105638** | Ralgapa2 | -0.016468514 | -0.35499884 | 0.322061812 | 9.515027485 | -0.118206137 | 0.909653995 | 0.96734437 | -4.911048867 |
| **A_52_P518808** | Mmd | -0.017064933 | -0.368018642 | 0.333888776 | 10.62528404 | -0.118151147 | 0.909695798 | 0.96734437 | -4.911052422 |
| **A_55_P2373987** | A730009E18Rik | 0.010401026 | -0.203676405 | 0.224478457 | 8.006376758 | 0.118056115 | 0.909768041 | 0.96734437 | -4.911058562 |
| **A_55_P2337213** | Odr4 | -0.008758407 | -0.189073351 | 0.171556537 | 9.662830329 | -0.118025705 | 0.909791159 | 0.96734437 | -4.911060525 |
| **A_51_P327778** | Pfdn2 | 0.014816939 | -0.290428759 | 0.320062637 | 10.51404288 | 0.11794842 | 0.909849911 | 0.967353627 | -4.911065513 |
| **A_52_P1026830** | Tbc1d8b | 0.0171225 | -0.336315868 | 0.370560868 | 8.997504538 | 0.117716317 | 0.910026363 | 0.967488013 | -4.911080474 |
| **A_51_P260513** | Polr1d | -0.010534807 | -0.228206614 | 0.207137 | 11.46462854 | -0.117600074 | 0.910114735 | 0.96750474 | -4.911087956 |
| **A_51_P389506** | NA | 0.012420171 | -0.244286192 | 0.269126534 | 13.65404642 | 0.117563941 | 0.910142205 | 0.96750474 | -4.91109028 |
| **A_55_P2382262** | A830039N20Rik | 0.017983207 | -0.354504762 | 0.390471176 | 8.84362784 | 0.117310821 | 0.910334645 | 0.96765609 | -4.911106542 |
| **A_55_P2144210** | Snx5 | -0.013800288 | -0.299890685 | 0.272290108 | 12.59459464 | -0.117210919 | 0.910410598 | 0.96768361 | -4.91111295 |
| **A_52_P562612** | Srd5a1 | 0.00810897 | -0.160516973 | 0.176734912 | 7.776797397 | 0.116848857 | 0.910685878 | 0.96788091 | -4.911136131 |
| **A_55_P2021471** | Pwwp3a | 0.042125838 | -0.834132138 | 0.918383815 | 9.47835178 | 0.116815308 | 0.910711386 | 0.96788091 | -4.911138275 |
| **A_51_P258372** | Igsf6 | -0.024897129 | -0.54328242 | 0.493488162 | 10.9325285 | -0.116702399 | 0.910797235 | 0.96788091 | -4.911145487 |
| **A_52_P506271** | Cmtm4 | -0.007599787 | -0.165861032 | 0.150661458 | 7.794336387 | -0.116683658 | 0.910811485 | 0.96788091 | -4.911146684 |
| **A_55_P2014229** | Sema4a | 0.030351446 | -0.601950362 | 0.662653254 | 11.79931149 | 0.116637487 | 0.910846591 | 0.96788091 | -4.91114963 |
| **A_55_P2028015** | Pawr | 0.01212173 | -0.241003994 | 0.265247454 | 9.070081443 | 0.116362087 | 0.911055995 | 0.968011454 | -4.911167183 |
| **A_51_P148494** | Rnft1 | -0.017636888 | -0.386064006 | 0.35079023 | 9.025728531 | -0.116319777 | 0.911088166 | 0.968011454 | -4.911169876 |
| **A_55_P2108579** | Phkg2 | -0.009717779 | -0.212790434 | 0.193354875 | 10.59608826 | -0.116278327 | 0.911119685 | 0.968011454 | -4.911172513 |
| **A_55_P1996573** | Ttc13 | 0.016079472 | -0.320382795 | 0.352541739 | 12.0658541 | 0.116123095 | 0.911237722 | 0.968062509 | -4.911182382 |
| **A_55_P2085932** | Lamtor2 | 0.009346806 | -0.186465781 | 0.205159394 | 12.51620666 | 0.115986069 | 0.911341918 | 0.968062509 | -4.911191082 |
| **A_52_P163766** | Atp6v0a2 | 0.011490513 | -0.229297885 | 0.25227891 | 9.07090806 | 0.115954346 | 0.91136604 | 0.968062509 | -4.911193094 |
| **A_55_P2058467** | Gm4415 | -0.012264905 | -0.269332842 | 0.244803033 | 11.63349013 | -0.115930965 | 0.91138382 | 0.968062509 | -4.911194578 |
| **A_65_P17866** | Vma21 | 0.012690336 | -0.253398137 | 0.27877881 | 8.47689742 | 0.115885811 | 0.911418156 | 0.968062509 | -4.911197441 |

| | | | | | | | | | |
|---|---|---|---|---|---|---|---|---|---|
| **A_55_P2040774** | NA | -0.015069612 | -0.331268224 | 0.301129001 | 13.84574068 | -0.115804461 | 0.911480017 | 0.968075018 | -4.911202597 |
| **A_55_P2037343** | H2-Q10 | 0.025772764 | -0.515497785 | 0.567043314 | 14.82755187 | 0.115699049 | 0.911560177 | 0.968106959 | -4.911209272 |
| **A_55_P2033906** | Mrpl22 | -0.011716822 | -0.257945089 | 0.234511445 | 13.14409216 | -0.115625894 | 0.911615808 | 0.968112848 | -4.911213902 |
| **A_51_P318544** | Fdx2 | 0.014373019 | -0.288208326 | 0.316954364 | 11.64206464 | 0.11542212 | 0.911770771 | 0.968224218 | -4.911226781 |
| **A_55_P1964867** | Ndufs6 | -0.011202873 | -0.247247328 | 0.224841583 | 15.05219043 | -0.11532376 | 0.911845571 | 0.968250456 | -4.91123299 |
| **A_55_P2029051** | NA | 0.030937333 | -0.621779328 | 0.683653994 | 9.767705883 | 0.115170529 | 0.911962102 | 0.968320999 | -4.911242652 |
| **A_51_P405606** | Ndrg1 | -0.025965745 | -0.574493414 | 0.522561925 | 13.67020873 | -0.115023196 | 0.91207415 | 0.96834094 | -4.91125193 |
| **A_52_P307961** | Abca5 | 0.011586996 | -0.23320827 | 0.256382262 | 8.35420001 | 0.115014086 | 0.912081078 | 0.96834094 | -4.911252503 |
| **A_55_P2149942** | Ninj2 | -0.008674874 | -0.192166796 | 0.174817048 | 7.864302029 | -0.11487603 | 0.912186073 | 0.968353158 | -4.911261185 |
| **A_51_P358722** | Lancl3 | -0.008066531 | -0.178721068 | 0.162588005 | 7.714598202 | -0.114855601 | 0.91220161 | 0.968353158 | -4.911262469 |
| **A_65_P19933** | Zdhhc23 | -0.007844648 | -0.173910573 | 0.158221278 | 7.988795408 | -0.11478261 | 0.912257122 | 0.968353158 | -4.911267054 |
| **A_51_P244824** | Dapp1 | -0.014732173 | -0.326855964 | 0.297391617 | 8.665766927 | -0.114689361 | 0.912328042 | 0.968353158 | -4.911272908 |
| **A_52_P152219** | Rpgrip1l | 0.008711712 | -0.175966606 | 0.19339003 | 8.248920984 | 0.11462274 | 0.912378711 | 0.968353158 | -4.911277088 |
| **A_51_P406188** | Asb10 | -0.007503753 | -0.166731249 | 0.151723744 | 7.68697206 | -0.114510055 | 0.912464414 | 0.968353158 | -4.911284151 |
| **A_51_P403049** | Heatr5b | 0.009349586 | -0.189104273 | 0.207803446 | 8.390369067 | 0.114476418 | 0.912489998 | 0.968353158 | -4.911286258 |
| **A_51_P272876** | Tent5a | 0.028413672 | -0.574716926 | 0.63154427 | 11.55015616 | 0.114471979 | 0.912493374 | 0.968353158 | -4.911286536 |
| **A_55_P2014882** | Dhrs9 | -0.007569169 | -0.168570176 | 0.153431837 | 7.713095297 | -0.114235957 | 0.912672888 | 0.968440734 | -4.911301304 |
| **A_52_P116264** | Adhfe1 | -0.019622185 | -0.437013877 | 0.397769508 | 8.294253305 | -0.114231728 | 0.912676104 | 0.968440734 | -4.911301568 |
| **A_52_P664506** | Abhd2 | 0.012763728 | -0.258957894 | 0.28448535 | 8.183378919 | 0.114139645 | 0.912746143 | 0.968461887 | -4.911307321 |
| **A_51_P344046** | Nt5m | -0.010968146 | -0.245184169 | 0.223247876 | 9.82343709 | -0.11378888 | 0.913012942 | 0.96857309 | -4.911329194 |
| **A_55_P2092963** | Traf2 | 0.013796632 | -0.280895693 | 0.308488958 | 12.18652992 | 0.113759443 | 0.913035333 | 0.96857309 | -4.911331026 |
| **A_66_P121480** | Sardh | 0.030416233 | -0.619380847 | 0.680213314 | 12.23690642 | 0.113739382 | 0.913050592 | 0.96857309 | -4.911332275 |
| **A_52_P95910** | Ugcg | 0.012065411 | -0.245699392 | 0.269830213 | 9.261285622 | 0.113736976 | 0.913052422 | 0.96857309 | -4.911332425 |
| **A_55_P1995503** | Gm5643 | -0.012605357 | -0.282058385 | 0.256847672 | 12.7998059 | -0.113672459 | 0.913101497 | 0.96857309 | -4.911336439 |
| **A_55_P2029682** | Fry | 0.009458086 | -0.193131766 | 0.212047938 | 8.765084976 | 0.113440664 | 0.913277815 | 0.96868069 | -4.911350841 |
| **A_66_P110798** | NA | -0.037774737 | -0.847137986 | 0.771588513 | 13.8836974 | -0.113407337 | 0.913303166 | 0.96868069 | -4.911352909 |
| **A_55_P1964832** | Reck | 0.015996137 | -0.328617262 | 0.360609535 | 8.209986514 | 0.112788841 | 0.913773661 | 0.969126534 | -4.911391186 |
| **A_51_P275953** | Dcps | -0.010550428 | -0.238193364 | 0.217092509 | 11.18889494 | -0.112615736 | 0.91390535 | 0.96921302 | -4.911401861 |
| **A_55_P2184905** | Rpap2 | 0.007868083 | -0.162102457 | 0.177838624 | 8.413485968 | 0.112480823 | 0.914007987 | 0.969237906 | -4.91141017 |
| **A_55_P2180744** | Clstn3 | 0.025531916 | -0.526361148 | 0.577424981 | 10.03788607 | 0.112411739 | 0.914060545 | 0.969237906 | -4.911414421 |
| **A_66_P134775** | Pter | 0.025267244 | -0.521497584 | 0.572032073 | 9.991226393 | 0.112289849 | 0.914153276 | 0.969237906 | -4.911421914 |
| **A_55_P2090708** | Fam207a | -0.011643808 | -0.26393041 | 0.240642795 | 11.86215007 | -0.112146061 | 0.91426267 | 0.969237906 | -4.911430744 |
| **A_51_P499816** | Pbld1 | 0.025531824 | -0.528068048 | 0.579131695 | 10.99578506 | 0.112064756 | 0.914324527 | 0.969237906 | -4.911435732 |
| **A_51_P402458** | Anxa7 | 0.015464912 | -0.320205658 | 0.351135482 | 8.824477457 | 0.111948268 | 0.914413153 | 0.969237906 | -4.911442872 |
| **A_55_P2311780** | 4930554H23Rik | 0.007085016 | -0.146722427 | 0.160892459 | 7.637214419 | 0.111930041 | 0.91442702 | 0.969237906 | -4.911443988 |
| **A_51_P398164** | Klhl2 | -0.013749853 | -0.31226346 | 0.284763753 | 9.073539231 | -0.111922427 | 0.914432813 | 0.969237906 | -4.911444454 |
| **A_51_P331462** | Cgrrf1 | -0.010703494 | -0.243105906 | 0.221698918 | 9.437073754 | -0.1119098 | 0.91444242 | 0.969237906 | -4.911445228 |
| **A_51_P146505** | Atp11b | 0.010724845 | -0.222291547 | 0.243741236 | 8.938336255 | 0.111837571 | 0.914497374 | 0.969237906 | -4.91144965 |
| **A_55_P1963364** | NA | 0.009270434 | -0.192193067 | 0.210733934 | 9.360624415 | 0.111811599 | 0.914517134 | 0.969237906 | -4.911451239 |
| **A_51_P286748** | Frzb | 0.008302065 | -0.172207414 | 0.188811543 | 7.829468027 | 0.111755598 | 0.914559742 | 0.969237906 | -4.911454664 |
| **A_52_P508700** | Zfp951 | 0.008958409 | -0.185869778 | 0.203786596 | 7.996415152 | 0.111728066 | 0.91458069 | 0.969237906 | -4.911456348 |
| **A_55_P2048813** | NA | -0.009519516 | -0.216860251 | 0.197821218 | 11.68399459 | -0.111561266 | 0.9147076 | 0.969319254 | -4.911466539 |
| **A_51_P493649** | Sult1e1 | -0.007200655 | -0.164156674 | 0.149755363 | 7.68589752 | -0.111474935 | 0.914773286 | 0.969335719 | -4.911471807 |
| **A_51_P476601** | Ly96 | -0.012216476 | -0.278776542 | 0.254343589 | 9.305663094 | -0.11136124 | 0.914859794 | 0.969374244 | -4.911478739 |
| **A_55_P2154425** | Cd320 | 0.007245667 | -0.150954334 | 0.165445668 | 7.749720114 | 0.111289734 | 0.914914202 | 0.969378754 | -4.911483096 |
| **A_55_P1975714** | Tor1aip1 | 0.008261981 | -0.172340611 | 0.188864574 | 9.815751027 | 0.11115869 | 0.915013912 | 0.96943126 | -4.911491072 |
| **A_55_P1967330** | Gm5069 | -0.011917337 | -0.272626765 | 0.248792092 | 14.52155829 | -0.111072272 | 0.915079668 | 0.969447788 | -4.911496327 |
| **A_55_P2113210** | Cyth4 | -0.014628077 | -0.335027427 | 0.305771273 | 10.66664614 | -0.110937611 | 0.915182133 | 0.969503204 | -4.911504507 |
| **A_55_P2049448** | Phtf1 | 0.014101241 | -0.295695295 | 0.323897777 | 9.835577609 | 0.110602254 | 0.915437319 | 0.969701807 | -4.911524836 |
| **A_55_P1956267** | Syt11 | -0.009710941 | -0.223331646 | 0.203909765 | 8.019990558 | -0.110459009 | 0.915546324 | 0.969701807 | -4.911533501 |

| | | | | | | | | | |
|---|---|---|---|---|---|---|---|---|---|
| **A_55_P2131707** | Lpcat1 | 0.02803632 | -0.589153257 | 0.645225897 | 8.812931168 | 0.110378785 | 0.915607372 | 0.969701807 | -4.911538349 |
| **A_51_P407074** | Tspan31 | 0.015720271 | -0.330386067 | 0.361826608 | 12.16650584 | 0.110365581 | 0.91561742 | 0.969701807 | -4.911539146 |
| **A_51_P246166** | Wfdc18 | 0.008127904 | -0.170827067 | 0.187082875 | 7.762161597 | 0.110361599 | 0.91562045 | 0.969701807 | -4.911539387 |
| **A_55_P2096166** | NA | 0.007537902 | -0.15853052 | 0.173606324 | 10.74663901 | 0.110292661 | 0.91567291 | 0.969704234 | -4.911543549 |
| **A_52_P241484** | Rcl1 | 0.025864132 | -0.545125124 | 0.596853388 | 10.80418093 | 0.110065994 | 0.915845403 | 0.96982657 | -4.911557217 |
| **A_55_P2110837** | Rnf7 | 0.011021558 | -0.232573857 | 0.254616973 | 11.19787687 | 0.10994034 | 0.915941027 | 0.96982657 | -4.911564782 |
| **A_51_P484289** | Slc7a6 | -0.008078176 | -0.186689796 | 0.170533445 | 7.930454718 | -0.109897236 | 0.91597383 | 0.96982657 | -4.911567375 |
| **A_51_P219465** | Tmem11 | -0.011328485 | -0.261995138 | 0.239338168 | 12.70309128 | -0.109814188 | 0.916037032 | 0.96982657 | -4.911572368 |
| **A_55_P2031676** | Gstm6 | 0.019933086 | -0.421166097 | 0.461032269 | 8.569962904 | 0.109804838 | 0.916044148 | 0.96982657 | -4.91157293 |
| **A_52_P38639** | Fermt3 | 0.022415941 | -0.473895548 | 0.518727429 | 9.949543204 | 0.109745274 | 0.916089478 | 0.96982657 | -4.911576508 |
| **A_55_P2103837** | Gbp4 | -0.00730699 | -0.169282532 | 0.154668552 | 8.147080307 | -0.109615568 | 0.91618819 | 0.969853603 | -4.911584294 |
| **A_51_P283300** | Cacnb2 | -0.007675638 | -0.177961133 | 0.162609858 | 7.694164032 | -0.109526701 | 0.916255823 | 0.969853603 | -4.911589623 |
| **A_52_P243412** | Ulk2 | -0.012178016 | -0.282381135 | 0.258025103 | 8.744058566 | -0.109513932 | 0.916265541 | 0.969853603 | -4.911590388 |
| **A_55_P2112225** | NA | -0.009904249 | -0.229968255 | 0.210159757 | 7.994210304 | -0.109359299 | 0.916383227 | 0.96989465 | -4.91159965 |
| **A_51_P323812** | Slc6a12 | 0.028942125 | -0.61436906 | 0.67225331 | 9.244085126 | 0.109318207 | 0.916414501 | 0.96989465 | -4.911602109 |
| **A_51_P276063** | Phyh | -0.025510312 | -0.593089133 | 0.54206851 | 13.42077925 | -0.109212606 | 0.916494872 | 0.96989465 | -4.911608424 |
| **A_51_P509140** | U2af1l4 | -0.014124328 | -0.328556819 | 0.300308164 | 9.456634837 | -0.109149955 | 0.916542556 | 0.96989465 | -4.911612168 |
| **A_52_P561936** | Sptssa | -0.016246429 | -0.377975753 | 0.345482895 | 12.27729286 | -0.109133327 | 0.916555211 | 0.96989465 | -4.911613161 |
| **A_52_P638513** | Zbtb11os1 | -0.007713601 | -0.179573336 | 0.164146134 | 8.841688109 | -0.109060183 | 0.916610881 | 0.969900461 | -4.911617528 |
| **A_55_P1972275** | Wars | 0.010433469 | -0.222326695 | 0.243193634 | 11.77834998 | 0.108918908 | 0.916718407 | 0.969961139 | -4.911625956 |
| **A_51_P270949** | H1f5 | -0.018669832 | -0.436086011 | 0.398746346 | 14.19294489 | -0.108681177 | 0.916899351 | 0.970042111 | -4.911640112 |
| **A_55_P1995992** | NA | -0.012575856 | -0.294099796 | 0.268948084 | 8.801955995 | -0.108543897 | 0.917003841 | 0.970042111 | -4.911648273 |
| **A_55_P2035454** | Oaz1 | 0.01076878 | -0.230354861 | 0.251892421 | 15.73238134 | 0.108520025 | 0.917022011 | 0.970042111 | -4.911649691 |
| **A_55_P2081278** | Mrpl55 | 0.009591877 | -0.205208039 | 0.224391794 | 12.97078557 | 0.108505735 | 0.917032888 | 0.970042111 | -4.91165054 |
| **A_55_P2000409** | Rab44 | -0.029097981 | -0.680818777 | 0.622622815 | 10.27086566 | -0.108488689 | 0.917045863 | 0.970042111 | -4.911651552 |
| **A_51_P128075** | Tescl | 0.006982477 | -0.14963839 | 0.163603344 | 7.695849377 | 0.108328592 | 0.917167723 | 0.970117923 | -4.904914936 |
| **A_55_P2101387** | Nubpl | 0.011874834 | -0.254673819 | 0.278423487 | 8.91022233 | 0.108251583 | 0.917226341 | 0.970126837 | -4.911665615 |
| **A_52_P281209** | Kdm4a | -0.009399102 | -0.220640171 | 0.201841967 | 10.42518226 | -0.108116306 | 0.917329313 | 0.970152389 | -4.911673626 |
| **A_55_P2085880** | Aco1 | -0.015146583 | -0.355649239 | 0.325356073 | 8.419651449 | -0.108087969 | 0.917350883 | 0.970152389 | -4.911675302 |
| **A_55_P1987396** | Tbl1xr1 | 0.012685921 | -0.273065695 | 0.298437537 | 8.856436817 | 0.107873928 | 0.917513812 | 0.970183675 | -4.911687953 |
| **A_55_P2181753** | Bdh1 | 0.030019122 | -0.646512667 | 0.706550911 | 12.4727734 | 0.107818422 | 0.917556064 | 0.970183675 | -4.911691229 |
| **A_55_P2030717** | NA | -0.008633896 | -0.203246396 | 0.185978604 | 10.97846979 | -0.107800124 | 0.917569993 | 0.970183675 | -4.911692309 |
| **A_55_P2022921** | Nsa2 | 0.020780772 | -0.447692827 | 0.489254371 | 12.00667883 | 0.107785351 | 0.917581239 | 0.970183675 | -4.91169318 |
| **A_55_P1999805** | NA | 0.007203636 | -0.155449866 | 0.169857137 | 10.86694377 | 0.107614677 | 0.917711162 | 0.970267971 | -4.911703241 |
| **A_55_P2123802** | NA | -0.033327473 | -0.786308623 | 0.719653676 | 9.242056183 | -0.107547786 | 0.917762082 | 0.970268735 | -4.91170718 |
| **A_55_P2085672** | NA | -0.008212069 | -0.194582713 | 0.178158576 | 15.49220386 | -0.107067645 | 0.9181276 | 0.970552663 | -4.911735382 |
| **A_52_P301495** | Vamp2 | -0.018205426 | -0.431390174 | 0.394979321 | 9.433711582 | -0.107063087 | 0.918131069 | 0.970552663 | -4.911735649 |
| **A_55_P2005908** | Lin9 | 0.007052255 | -0.153128378 | 0.167232888 | 7.827986503 | 0.106979653 | 0.918194588 | 0.970566728 | -4.911740536 |
| **A_52_P496924** | Mgat4a | -0.007472194 | -0.177603694 | 0.162659306 | 7.821888709 | -0.10672019 | 0.918392121 | 0.970722442 | -4.911755712 |
| **A_55_P2176724** | Fbxw7 | 0.009505718 | -0.207111778 | 0.226123214 | 8.72907424 | 0.106628816 | 0.918461687 | 0.970742888 | -4.911761047 |
| **A_55_P1984337** | Emd | -0.00939725 | -0.223821808 | 0.205027308 | 11.4345676 | -0.106490154 | 0.918567256 | 0.970754221 | -4.911769135 |
| **A_55_P2007389** | Fam189b | 0.014363607 | -0.313546622 | 0.342273835 | 9.025949805 | 0.10643678 | 0.918607892 | 0.970754221 | -4.911772245 |
| **A_51_P153053** | Smpdl3a | 0.014632872 | -0.319630778 | 0.348896521 | 12.18264906 | 0.106371089 | 0.918657907 | 0.970754221 | -4.911776071 |
| **A_51_P497985** | C2 | 0.024740202 | -0.540818985 | 0.590299389 | 12.94949967 | 0.106293907 | 0.91871667 | 0.970754221 | -4.911780564 |
| **A_55_P2375121** | AI225934 | -0.00779334 | -0.185976945 | 0.170390265 | 8.382362808 | -0.106276951 | 0.91872958 | 0.970754221 | -4.91178155 |
| **A_55_P2251181** | 2900076A13Rik | 0.013964618 | -0.305530905 | 0.333460141 | 8.445416913 | 0.106205613 | 0.918783895 | 0.970754221 | -4.911785699 |
| **A_51_P321965** | Ssna1 | -0.020694396 | -0.494394793 | 0.453006002 | 11.12167575 | -0.106152981 | 0.918823968 | 0.970754221 | -4.911788758 |
| **A_55_P2085551** | Dennd4b | -0.01548384 | -0.371273613 | 0.340305934 | 9.779890874 | -0.105747082 | 0.919133019 | 0.971010263 | -4.911812299 |
| **A_55_P2095926** | NA | 0.011868944 | -0.260980803 | 0.284718691 | 8.750725941 | 0.105699204 | 0.919169474 | 0.971010263 | -4.91181507 |
| **A_55_P2047614** | Zfp809 | 0.007331621 | -0.161363013 | 0.176026255 | 7.964237497 | 0.105604379 | 0.919241676 | 0.971010263 | -4.911820554 |

| | | | | | | | | | |
|---|---|---|---|---|---|---|---|---|---|
| **A_55_P2043317** | Bag1 | 0.010639447 | -0.234243478 | 0.255522372 | 13.83711814 | 0.105570784 | 0.919267257 | 0.971010263 | -4.911822496 |
| **A_55_P2249536** | Nucks1 | 0.013119248 | -0.289187798 | 0.315426294 | 10.35524246 | 0.105449339 | 0.919359729 | 0.971024801 | -4.91182951 |
| **A_55_P2000894** | NA | -0.010436375 | -0.250987143 | 0.230114393 | 15.36904425 | -0.105420757 | 0.919381493 | 0.971024801 | -4.91183116 |
| **A_55_P2024406** | Napepld | 0.011212105 | -0.247553218 | 0.269977428 | 8.404861755 | 0.10528446 | 0.919485277 | 0.971028509 | -4.91183902 |
| **A_55_P2008704** | Gstp2 | 0.019081919 | -0.421390865 | 0.459554703 | 12.56592872 | 0.105265524 | 0.919499697 | 0.971028509 | -4.911840111 |
| **A_52_P374669** | Kdm5a | -0.00847758 | -0.204508482 | 0.187553322 | 10.27997891 | -0.105082535 | 0.919639038 | 0.971028509 | -4.911850647 |
| **A_51_P160664** | Cox7b | -0.011808044 | -0.284901443 | 0.261285355 | 15.14823554 | -0.105063039 | 0.919653883 | 0.971028509 | -4.911851768 |
| **A_55_P2023952** | Arpc3 | 0.013703901 | -0.303825398 | 0.3312332 | 9.808661828 | 0.104868148 | 0.919802291 | 0.971028509 | -4.911862967 |
| **A_52_P33800** | Fndc3a | -0.01543207 | -0.373035861 | 0.342171722 | 8.604200991 | -0.104858887 | 0.919809343 | 0.971028509 | -4.911863498 |
| **A_52_P486045** | Coq4 | 0.007827938 | -0.173687573 | 0.189343448 | 9.726147339 | 0.104789266 | 0.91986236 | 0.971028509 | -4.911867494 |
| **A_55_P2072000** | Pcmtd1 | 0.016548316 | -0.367295548 | 0.400392179 | 8.945535296 | 0.104756838 | 0.919887054 | 0.971028509 | -4.911869354 |
| **A_55_P1967414** | Asph | -0.006642562 | -0.160732472 | 0.147447349 | 7.705628058 | -0.104747722 | 0.919893996 | 0.971028509 | -4.911869876 |
| **A_55_P2137773** | NA | -0.009848358 | -0.238963583 | 0.219266866 | 16.39298623 | -0.104446307 | 0.920123531 | 0.971028509 | -4.911887135 |
| **A_55_P2137023** | NA | -0.007733084 | -0.18773635 | 0.172270182 | 8.341324026 | -0.104389193 | 0.920167026 | 0.971028509 | -4.9118904 |
| **A_55_P2171143** | Hnrnpc | 0.010627428 | -0.237009136 | 0.258263992 | 13.87413584 | 0.104278935 | 0.920250992 | 0.971028509 | -4.911896698 |
| **A_55_P2163446** | NA | 0.010183767 | -0.227371202 | 0.247738737 | 9.041866454 | 0.104166373 | 0.920336715 | 0.971028509 | -4.91190312 |
| **A_55_P2110497** | Ddc | -0.015441986 | -0.375687434 | 0.344803463 | 10.9263455 | -0.104156848 | 0.920343969 | 0.971028509 | -4.911903663 |
| **A_55_P2167803** | Abi3bp | -0.013547285 | -0.329703002 | 0.302608432 | 7.845062435 | -0.104120064 | 0.920371982 | 0.971028509 | -4.91190576 |
| **A_66_P120882** | Nfkbib | -0.009249404 | -0.225278437 | 0.206779628 | 9.13205524 | -0.104036283 | 0.920435788 | 0.971028509 | -4.911910534 |
| **A_55_P2054744** | Sh3bgrl2 | 0.009639062 | -0.215510472 | 0.234788596 | 8.0791632 | 0.104027196 | 0.920442709 | 0.971028509 | -4.911911052 |
| **A_55_P1980966** | Mettl23 | 0.011241106 | -0.251370874 | 0.273853085 | 10.70151386 | 0.104010621 | 0.920455332 | 0.971028509 | -4.911911995 |
| **A_55_P2127538** | Ppp1r21 | 0.007942535 | -0.177620549 | 0.19350562 | 8.476186212 | 0.104004171 | 0.920460244 | 0.971028509 | -4.911912363 |
| **A_55_P2107302** | Aurkaip1 | 0.014242453 | -0.318816614 | 0.34730152 | 14.1390359 | 0.10390746 | 0.920533898 | 0.971028509 | -4.911917867 |
| **A_55_P2162890** | Zfp780b | 0.008488 | -0.190047628 | 0.207023627 | 8.469495369 | 0.10388434 | 0.920551506 | 0.971028509 | -4.911919182 |
| **A_51_P401987** | Tmem37 | 0.015332336 | -0.343979553 | 0.374644225 | 14.01887574 | 0.103685954 | 0.920702596 | 0.971028509 | -4.911930454 |
| **A_51_P477121** | Pmaip1 | -0.012958395 | -0.316692527 | 0.290775738 | 8.514720768 | -0.103667062 | 0.920716984 | 0.971028509 | -4.911931526 |
| **A_52_P204618** | Rnf222 | 0.006734027 | -0.151130013 | 0.164598067 | 7.663977747 | 0.103651316 | 0.920728977 | 0.971028509 | -4.911932419 |
| **A_51_P481679** | Angptl3 | 0.028852302 | -0.647788487 | 0.705493092 | 13.81350504 | 0.103610909 | 0.920759751 | 0.971028509 | -4.911934712 |
| **A_55_P2154242** | Gfpt1 | -0.012039714 | -0.294438713 | 0.270359284 | 11.28824844 | -0.103594384 | 0.920772337 | 0.971028509 | -4.911935649 |
| **A_55_P2037918** | NA | 0.013667183 | -0.306965557 | 0.334299924 | 8.961843143 | 0.103574856 | 0.92078721 | 0.971028509 | -4.911936757 |
| **A_52_P356068** | Proz | 0.022193301 | -0.498937423 | 0.543324026 | 9.96129873 | 0.103480477 | 0.920859091 | 0.971028509 | -4.911942106 |
| **A_51_P376445** | Rhox5 | -0.014415901 | -0.353007325 | 0.324175524 | 9.342536131 | -0.103454412 | 0.920878943 | 0.971028509 | -4.911943582 |
| **A_51_P334318** | Chp2 | -0.019632542 | -0.481093756 | 0.441828672 | 8.768113555 | -0.103377142 | 0.920937795 | 0.971028509 | -4.911947957 |
| **A_52_P349844** | Obscn | 0.008332878 | -0.187603484 | 0.20426924 | 7.720046786 | 0.103338743 | 0.920967041 | 0.971028509 | -4.91195013 |
| **A_55_P2149993** | NA | 0.007213959 | -0.1624676 | 0.176895518 | 7.777501715 | 0.103305205 | 0.920992584 | 0.971028509 | -4.911952027 |
| **A_55_P2018701** | Kxd1 | 0.008326361 | -0.187910948 | 0.20456367 | 10.24090291 | 0.103099569 | 0.921149207 | 0.971140669 | -4.911963645 |
| **A_55_P2179314** | Macrod2 | -0.007095767 | -0.17455048 | 0.160358946 | 8.091142672 | -0.102963938 | 0.921252513 | 0.97116093 | -4.911971296 |
| **A_51_P139320** | Pcbd1 | -0.022957587 | -0.564852542 | 0.518937368 | 14.66342647 | -0.102942408 | 0.921268913 | 0.97116093 | -4.91197251 |
| **A_55_P2102951** | NA | 0.013180229 | -0.298606681 | 0.32496714 | 11.53591482 | 0.102718409 | 0.92143953 | 0.971287815 | -4.91198512 |
| **A_52_P256426** | NA | 0.006870556 | -0.155865131 | 0.169606243 | 8.305147432 | 0.10258698 | 0.921539639 | 0.971340369 | -4.911992507 |
| **A_55_P1966134** | Prl3c1 | -0.006542903 | -0.161668538 | 0.148582733 | 7.622640962 | -0.102487304 | 0.921615564 | 0.971367427 | -4.911998103 |
| **A_51_P302566** | Maob | -0.028691737 | -0.709413988 | 0.652030514 | 11.88111229 | -0.102416537 | 0.921669468 | 0.971371273 | -4.912002072 |
| **A_52_P105537** | Ccn3 | 0.00685593 | -0.156652131 | 0.170363991 | 7.700389071 | 0.101885035 | 0.92207434 | 0.971744993 | -4.912031798 |
| **A_55_P2089118** | Zfp976 | 0.010125593 | -0.23159942 | 0.251850605 | 9.109279701 | 0.101784593 | 0.922150854 | 0.971772645 | -4.912037398 |
| **A_52_P120740** | 0610030E20Rik | -0.008481932 | -0.21110626 | 0.194142395 | 8.595165758 | -0.101715326 | 0.92220362 | 0.97177527 | -4.912041257 |
| **A_55_P2152912** | NA | -0.006967431 | -0.173899568 | 0.159964706 | 7.721042802 | -0.101418202 | 0.92242997 | 0.971929131 | -4.91205778 |
| **A_55_P2104612** | NA | -0.009295762 | -0.232126924 | 0.2135354 | 8.3125907 | -0.101365989 | 0.922469746 | 0.971929131 | -4.912060679 |
| **A_51_P454196** | Sh2d4a | 0.008837668 | -0.203096758 | 0.220772094 | 8.645280178 | 0.101325642 | 0.922500483 | 0.971929131 | -4.912062917 |
| **A_52_P16419** | Gpd1 | -0.026864808 | -0.674153792 | 0.620424176 | 13.18061886 | -0.10084833 | 0.92286412 | 0.972259257 | -4.912089336 |
| **A_52_P405145** | NA | -0.006555751 | -0.164901285 | 0.151789783 | 7.690791597 | -0.100600425 | 0.923052994 | 0.97236556 | -4.912103008 |

| | | | | | | | | | |
|---|---|---|---|---|---|---|---|---|---|
| **A_55_P1955706** | Spg7 | -0.006566082 | -0.165187321 | 0.152055156 | 12.5054278 | -0.100583833 | 0.923065635 | 0.97236556 | -4.912103922 |
| **A_55_P1982872** | Naa10 | -0.012929726 | -0.325884019 | 0.300024567 | 11.26201209 | -0.100390266 | 0.923213113 | 0.972467917 | -4.912114572 |
| **A_55_P2170474** | Yipf2 | -0.013841089 | -0.349497668 | 0.32181549 | 10.17527483 | -0.100197823 | 0.923359738 | 0.972492211 | -4.912125141 |
| **A_51_P104747** | Gtf2h4 | 0.010647681 | -0.247583839 | 0.268879201 | 11.00468389 | 0.100191061 | 0.92336489 | 0.972492211 | -4.912125512 |
| **A_55_P2087984** | Dnaja1 | 0.009142908 | -0.212683999 | 0.230969815 | 13.52559112 | 0.10015055 | 0.923395757 | 0.972492211 | -4.912127734 |
| **A_55_P1982122** | Homer1 | 0.00760938 | -0.177111657 | 0.192330418 | 7.992345694 | 0.100095859 | 0.923437428 | 0.972492211 | -4.912130732 |
| **A_51_P408989** | L3hypdh | 0.015380272 | -0.35895103 | 0.389711574 | 9.677101403 | 0.099836892 | 0.923634745 | 0.972584228 | -4.912144907 |
| **A_55_P2068486** | Arfgap1 | -0.009813589 | -0.249273758 | 0.22964658 | 10.07809276 | -0.099581296 | 0.923829501 | 0.972584228 | -4.912158862 |
| **A_55_P2008651** | Qdpr | 0.019238751 | -0.450237269 | 0.488714771 | 13.24663895 | 0.099574161 | 0.923834937 | 0.972584228 | -4.912159251 |
| **A_51_P359485** | Zfpl1 | 0.009591993 | -0.22469415 | 0.243878136 | 10.040378 | 0.099482213 | 0.923905001 | 0.972584228 | -4.912164263 |
| **A_55_P2137496** | NA | 0.014204523 | -0.332958741 | 0.361367787 | 9.447524011 | 0.099420531 | 0.923952001 | 0.972584228 | -4.912167622 |
| **A_55_P2030030** | Adssl1 | -0.011910319 | -0.303025889 | 0.279205251 | 10.93941297 | -0.099412535 | 0.923958095 | 0.972584228 | -4.912168057 |
| **A_55_P1984825** | Bloc1s2 | -0.010740743 | -0.273412274 | 0.251930789 | 10.82263642 | -0.099358378 | 0.923999362 | 0.972584228 | -4.912171004 |
| **A_55_P2047648** | Coq2 | -0.008157561 | -0.207702773 | 0.191387651 | 11.9464503 | -0.099334979 | 0.924017192 | 0.972584228 | -4.912172277 |
| **A_55_P2169218** | NA | 0.042287024 | -0.992154802 | 1.07672885 | 10.58117117 | 0.09933087 | 0.924020323 | 0.972584228 | -4.912172501 |
| **A_55_P2121426** | NA | 0.015798568 | -0.370711307 | 0.402308444 | 14.35289274 | 0.099320825 | 0.924027977 | 0.972584228 | -4.912173047 |
| **A_55_P2046408** | NA | 0.0225124 | -0.529019786 | 0.574044586 | 11.33693825 | 0.099182289 | 0.924133542 | 0.972642376 | -4.912180576 |
| **A_52_P679105** | Prss23 | -0.013417762 | -0.342952272 | 0.316116749 | 8.955355012 | -0.098937824 | 0.92431983 | 0.972709849 | -4.912193837 |
| **A_55_P2111237** | Selenof | 0.013170283 | -0.310308474 | 0.33664904 | 13.85526131 | 0.09893103 | 0.924325007 | 0.972709849 | -4.912194205 |
| **A_55_P2156776** | NA | 0.009720636 | -0.229462349 | 0.24890362 | 7.994402215 | 0.09875238 | 0.924461147 | 0.972709849 | -4.912203874 |
| **A_55_P2019934** | Ythdf3 | 0.010246536 | -0.242175906 | 0.262668978 | 10.2766074 | 0.098635284 | 0.92455038 | 0.972709849 | -4.912210203 |
| **A_55_P2172470** | Nrgn | -0.011925275 | -0.305798323 | 0.281947772 | 8.592926972 | -0.09860339 | 0.924574685 | 0.972709849 | -4.912211925 |
| **A_55_P2161865** | Pdlim7 | -0.008922829 | -0.228823005 | 0.210977346 | 7.950867288 | -0.098596207 | 0.924580159 | 0.972709849 | -4.912212313 |
| **A_55_P2103351** | Ndufa5 | 0.012208068 | -0.288695395 | 0.313111532 | 13.07195416 | 0.098583208 | 0.924590066 | 0.972709849 | -4.912213015 |
| **A_55_P2103561** | Prdx6 | -0.008514147 | -0.218398581 | 0.201370286 | 14.35334165 | -0.098569855 | 0.924600241 | 0.972709849 | -4.912213736 |
| **A_55_P2032708** | Hmgb1 | -0.012884059 | -0.330888374 | 0.305120256 | 13.52052279 | -0.098447088 | 0.924693799 | 0.97275533 | -4.912220358 |
| **A_66_P108685** | Cideb | 0.02944111 | -0.698335212 | 0.757217432 | 13.32691281 | 0.098296815 | 0.924808319 | 0.972782 | -4.912228453 |
| **A_55_P2165854** | Gemin7 | -0.010542632 | -0.271193633 | 0.250108368 | 10.53707421 | -0.098281742 | 0.924819806 | 0.972782 | -4.912229265 |
| **A_55_P2071834** | Tln2 | 0.007478741 | -0.177655141 | 0.192612622 | 8.181361823 | 0.098158016 | 0.924914097 | 0.972828241 | -4.912235919 |
| **A_55_P2089477** | Sypl | -0.01484418 | -0.383009005 | 0.353320645 | 11.81484482 | -0.097970904 | 0.925056697 | 0.972925286 | -4.912245968 |
| **A_65_P06061** | Rpl37 | -0.010287278 | -0.266402131 | 0.245827574 | 17.43711465 | -0.097599797 | 0.92533953 | 0.973169801 | -4.91226584 |
| **A_55_P2102578** | NA | 0.009738089 | -0.233043798 | 0.252519975 | 15.15424524 | 0.097463191 | 0.925443645 | 0.973226345 | -4.912273136 |
| **A_55_P2076278** | NA | 0.007834009 | -0.187625281 | 0.203293299 | 16.69864139 | 0.097389229 | 0.925500017 | 0.973232676 | -4.912277083 |
| **A_55_P1963772** | Atp5j | 0.010161067 | -0.243568104 | 0.263890238 | 13.96620258 | 0.097308798 | 0.925561319 | 0.973243542 | -4.91228137 |
| **A_55_P2101906** | Rpl10a | -0.009492059 | -0.246674672 | 0.227690553 | 16.56330128 | -0.097243546 | 0.925611053 | 0.973243542 | -4.912284847 |
| **A_55_P2018929** | Spns2 | -0.008560162 | -0.222651683 | 0.20553136 | 9.880323899 | -0.09715512 | 0.925678449 | 0.973261463 | -4.912289553 |
| **A_51_P194249** | Stmn4 | 0.006479839 | -0.155872617 | 0.168832295 | 7.784124759 | 0.096981427 | 0.925810838 | 0.973347712 | -4.912298787 |
| **A_51_P110471** | Ddah1 | 0.021330167 | -0.513617251 | 0.556277585 | 10.3989329 | 0.096887188 | 0.925882667 | 0.973370285 | -4.892264039 |
| **A_55_P2104790** | NA | 0.010742084 | -0.25922574 | 0.280709908 | 13.81816319 | 0.096685137 | 0.926036675 | 0.973467888 | -4.912314499 |
| **A_55_P1979970** | NA | -0.015018641 | -0.392666739 | 0.362629456 | 10.84329288 | -0.096633237 | 0.926076234 | 0.973467888 | -4.912317246 |
| **A_55_P1979330** | Dapp1 | 0.016422662 | -0.397761036 | 0.430606359 | 10.11144505 | 0.096346011 | 0.926295171 | 0.973645079 | -4.912332424 |
| **A_55_P2078269** | NA | 0.010769357 | -0.261504064 | 0.283042777 | 14.02236588 | 0.096109806 | 0.926475223 | 0.97374234 | -4.912344873 |
| **A_66_P134534** | Atl2 | 0.012435642 | -0.302021997 | 0.32689328 | 8.932467014 | 0.096092444 | 0.926488457 | 0.97374234 | -4.912345786 |
| **A_55_P1969396** | NA | 0.019513814 | -0.47447687 | 0.513504498 | 13.2743186 | 0.095985711 | 0.926569818 | 0.973767231 | -4.9123514 |
| **A_52_P208521** | Khdc1a | -0.006071447 | -0.159860209 | 0.147717315 | 7.629314336 | -0.095929199 | 0.926612897 | 0.973767231 | -4.91235437 |
| **A_55_P2148400** | NA | -0.029063712 | -0.767313534 | 0.70918611 | 13.12096704 | -0.09566012 | 0.926818017 | 0.973906189 | -4.912368487 |
| **A_52_P532033** | Hltf | -0.009692244 | -0.256121119 | 0.236736632 | 9.401903972 | -0.095568752 | 0.926887668 | 0.973906189 | -4.912373272 |
| **A_55_P2030128** | Tfpt | 0.009064003 | -0.221551122 | 0.239679128 | 8.414878502 | 0.095502651 | 0.926938059 | 0.973906189 | -4.91237673 |
| **A_55_P2091359** | Padi2 | 0.013345075 | -0.326233398 | 0.352923549 | 8.39740933 | 0.095491357 | 0.926946669 | 0.973906189 | -4.912377321 |
| **A_55_P2031692** | Gstm6 | 0.025260822 | -0.619172238 | 0.669693883 | 10.90974022 | 0.095247341 | 0.927132692 | 0.97404869 | -4.912390067 |

| | | | | | | | | | |
|---|---|---|---|---|---|---|---|---|---|
| **A_51_P193863** | Gtf3c4 | -0.007314785 | -0.194075277 | 0.179445708 | 8.520880049 | -0.095169924 | 0.927191711 | 0.974051939 | -4.912394103 |
| **A_51_P396818** | Rex1bd | -0.012590322 | -0.334244292 | 0.309063648 | 9.744392298 | -0.095111081 | 0.927236571 | 0.974051939 | -4.91239717 |
| **A_51_P375533** | Polr3b | 0.010212161 | -0.252188987 | 0.272613308 | 9.035463327 | 0.094566017 | 0.927652119 | 0.97443551 | -4.912425482 |
| **A_55_P2047620** | NA | 0.009524985 | -0.235839881 | 0.25488985 | 13.44626031 | 0.094326798 | 0.927834504 | 0.974574129 | -4.912437856 |
| **A_51_P351285** | Pcsk7 | 0.006855469 | -0.169997203 | 0.183708142 | 9.468783907 | 0.094190859 | 0.927938148 | 0.97460389 | -4.912444875 |
| **A_55_P2224582** | Fam219b | 0.008037394 | -0.199379501 | 0.215454289 | 8.159327974 | 0.09415737 | 0.927963681 | 0.97460389 | -4.912446602 |
| **A_51_P493886** | Gpt2 | -0.021195979 | -0.568632227 | 0.52624027 | 10.29558141 | -0.094081257 | 0.928021713 | 0.974611883 | -4.912450526 |
| **A_55_P1975026** | Gm9104 | -0.009580709 | -0.25756855 | 0.238407132 | 15.55163741 | -0.093875111 | 0.928178891 | 0.974617266 | -4.912461137 |
| **A_52_P322421** | Mpzl2 | -0.026844143 | -0.721740257 | 0.668051971 | 11.06828364 | -0.09386698 | 0.928185091 | 0.974617266 | -4.912461555 |
| **A_51_P158584** | Poll | 0.010011073 | -0.24914652 | 0.269168666 | 9.497046462 | 0.093864178 | 0.928187227 | 0.974617266 | -4.912461699 |
| **A_51_P345593** | Sptlc1 | -0.009218198 | -0.247988361 | 0.229551965 | 9.384291728 | -0.093810009 | 0.92822853 | 0.974617266 | -4.912464483 |
| **A_55_P1989333** | NA | 0.01335828 | -0.333901737 | 0.360618296 | 12.06851014 | 0.093471443 | 0.928486681 | 0.974800699 | -4.912481848 |
| **A_55_P2115297** | Ap3s1 | 0.009462203 | -0.23665465 | 0.255579056 | 11.62654474 | 0.093418762 | 0.92852685 | 0.974800699 | -4.912484544 |
| **A_55_P2044653** | Cyp2b10 | -0.020759114 | -0.561042688 | 0.519524461 | 8.518026657 | -0.093362018 | 0.928570118 | 0.974800699 | -4.912487447 |
| **A_51_P249848** | Dsg2 | -0.013194401 | -0.35676442 | 0.330375617 | 8.778167929 | -0.093316324 | 0.92860496 | 0.974800699 | -4.912489783 |
| **A_55_P1988328** | NA | 0.006791217 | -0.170477839 | 0.184060273 | 16.17254415 | 0.093088893 | 0.928778381 | 0.974829003 | -4.912501394 |
| **A_55_P2058493** | NA | -0.008351789 | -0.226382304 | 0.209678727 | 15.96897777 | -0.093077651 | 0.928786953 | 0.974829003 | -4.912501967 |
| **A_52_P303077** | Rogdi | -0.016464937 | -0.44632537 | 0.413395495 | 9.470907855 | -0.093071305 | 0.928791792 | 0.974829003 | -4.91250229 |
| **A_55_P2000543** | Cd209f | -0.023583785 | -0.639663844 | 0.592496275 | 8.733976046 | -0.093016403 | 0.928833657 | 0.974829003 | -4.912505088 |
| **A_52_P77332** | Ireb2 | -0.006408953 | -0.174604173 | 0.161786267 | 7.939525764 | -0.092588409 | 0.929160026 | 0.975070044 | -4.912526845 |
| **A_55_P2040860** | Sun3 | -0.005986013 | -0.163169575 | 0.15119755 | 7.661124044 | -0.092536631 | 0.92919951 | 0.975070044 | -4.91252947 |
| **A_51_P156291** | F9 | -0.017570481 | -0.479705531 | 0.444564568 | 9.450321097 | -0.092384251 | 0.929315712 | 0.975070044 | -4.912537187 |
| **A_51_P315085** | Mia3 | 0.010603438 | -0.268412938 | 0.289619814 | 11.7896102 | 0.092342198 | 0.929347781 | 0.975070044 | -4.912539315 |
| **A_51_P367866** | Egr1 | 0.023090734 | -0.584549201 | 0.63073067 | 11.2559702 | 0.092336759 | 0.929351929 | 0.975070044 | -4.91253959 |
| **A_55_P2133845** | Eid2b | 0.008422343 | -0.213259796 | 0.230104482 | 7.9222585 | 0.092317793 | 0.929366393 | 0.975070044 | -4.912540549 |
| **A_55_P2030647** | Dusp13 | -0.006258704 | -0.171195527 | 0.158678118 | 7.676738247 | -0.092204048 | 0.929453134 | 0.975070044 | -4.912546297 |
| **A_55_P2007673** | Lrsam1 | -0.007298278 | -0.19966884 | 0.185072283 | 9.820405812 | -0.092186007 | 0.929466892 | 0.975070044 | -4.912547208 |
| **A_51_P442284** | Dnajc12 | -0.020497328 | -0.561589357 | 0.520594701 | 11.3507443 | -0.092046928 | 0.929572955 | 0.975074765 | -4.912554226 |
| **A_55_P2033849** | Sphkap | 0.005907461 | -0.150098672 | 0.161913595 | 7.655897916 | 0.092011561 | 0.929599927 | 0.975074765 | -4.912556009 |
| **A_51_P464576** | Psen1 | -0.007864597 | -0.215754659 | 0.200025465 | 9.108033038 | -0.091923373 | 0.929667181 | 0.975074765 | -4.912560452 |
| **A_55_P2180071** | Zfand5 | -0.007615196 | -0.209016924 | 0.193786531 | 11.66786552 | -0.091875795 | 0.929703466 | 0.975074765 | -4.912562847 |
| **A_52_P206861** | Klf8 | -0.007440649 | -0.204367347 | 0.189486049 | 7.868093611 | -0.091809881 | 0.929753734 | 0.975074765 | -4.912566163 |
| **A_55_P1981794** | Sgf29 | 0.006891203 | -0.175546654 | 0.18932906 | 8.389880236 | 0.091783216 | 0.92977407 | 0.975074765 | -4.912567504 |
| **A_51_P212801** | Ube2a | -0.008452333 | -0.23246 | 0.215555334 | 10.08451918 | -0.091684707 | 0.929849197 | 0.975100647 | -4.912572454 |
| **A_51_P396774** | Sugt1 | 0.014787119 | -0.377439359 | 0.407013597 | 12.88272437 | 0.09160725 | 0.929908269 | 0.975109691 | -4.912576342 |
| **A_51_P443693** | Rasl10b | -0.006018645 | -0.166084752 | 0.154047462 | 7.700819809 | -0.091365563 | 0.930092594 | 0.975122417 | -4.912588454 |
| **A_55_P2159506** | Gm5458 | -0.006066689 | -0.167611146 | 0.155477768 | 7.673074923 | -0.091252099 | 0.93017913 | 0.975122417 | -4.91259413 |
| **A_52_P357402** | Mlst8 | -0.00787383 | -0.217660844 | 0.201913183 | 10.5580512 | -0.091199117 | 0.930219538 | 0.975122417 | -4.912596777 |
| **A_55_P2050040** | Rpl36al | -0.008332952 | -0.230393864 | 0.21372796 | 14.91649847 | -0.091182175 | 0.93023246 | 0.975122417 | -4.912597624 |
| **A_55_P1952182** | Vmn2r111 | -0.005951848 | -0.164758874 | 0.152855178 | 7.728675383 | -0.091067898 | 0.930319618 | 0.975122417 | -4.912603328 |
| **A_51_P387868** | Tekt4 | 0.012969872 | -0.333211346 | 0.35915109 | 9.398720293 | 0.091036465 | 0.930343592 | 0.975122417 | -4.912604896 |
| **A_55_P1953082** | NA | -0.005825173 | -0.16133106 | 0.149680714 | 7.658478052 | -0.091021746 | 0.930354818 | 0.975122417 | -4.91260563 |
| **A_52_P510215** | Traf6 | -0.010453565 | -0.28974673 | 0.2688396 | 8.335383173 | -0.090946775 | 0.930411999 | 0.975122417 | -4.912609366 |
| **A_55_P2043734** | Cdkl5 | 0.006537863 | -0.168144623 | 0.181220348 | 7.697508323 | 0.090943075 | 0.930414821 | 0.975122417 | -4.912609551 |
| **A_51_P459368** | Appbp2 | -0.011150469 | -0.309190991 | 0.286890054 | 11.51412437 | -0.090907768 | 0.93044175 | 0.975122417 | -4.912611309 |
| **A_55_P2143159** | Men1 | 0.020746387 | -0.534156226 | 0.575649001 | 11.87378167 | 0.090846646 | 0.930488368 | 0.975122417 | -4.912614352 |
| **A_55_P1983235** | Tmed10 | -0.007252603 | -0.201355161 | 0.186849954 | 14.0212818 | -0.090791639 | 0.930530322 | 0.975122417 | -4.912617089 |
| **A_51_P431316** | Ube2w | 0.013434655 | -0.346357689 | 0.373226999 | 8.635009442 | 0.09073144 | 0.930576238 | 0.975122417 | -4.912620082 |
| **A_52_P139740** | Mdp1 | -0.013346224 | -0.371812775 | 0.345120327 | 9.158260222 | -0.090467578 | 0.930777494 | 0.975278506 | -4.912633178 |
| **A_55_P1974577** | Nfam1 | -0.016158385 | -0.450462824 | 0.418146054 | 10.40653147 | -0.090403839 | 0.93082611 | 0.975278506 | -4.912636335 |

| | | | | | | | | | |
|---|---|---|---|---|---|---|---|---|---|
| **A_66_P104312** | Pdcl | 0.009310274 | -0.24182745 | 0.260447999 | 9.469794136 | 0.090081105 | 0.931072279 | 0.975388955 | -4.91265229 |
| **A_51_P484842** | Asgr2 | 0.023568229 | -0.612882156 | 0.660018614 | 12.8651523 | 0.089979915 | 0.931149465 | 0.975388955 | -4.912657281 |
| **A_55_P2114785** | NA | 0.007539101 | -0.19606286 | 0.211141062 | 9.189996904 | 0.089974783 | 0.93115338 | 0.975388955 | -4.912657534 |
| **A_55_P2177426** | Abcb8 | -0.007420502 | -0.20804216 | 0.193201157 | 8.177636788 | -0.089874947 | 0.931229533 | 0.975388955 | -4.912662452 |
| **A_55_P2115732** | Kif1c | -0.006805681 | -0.190805714 | 0.177194353 | 8.152741346 | -0.089874577 | 0.931229816 | 0.975388955 | -4.91266247 |
| **A_55_P2095688** | Sh3bp5l | 0.011318509 | -0.294888215 | 0.317525233 | 11.59030225 | 0.089816823 | 0.93127387 | 0.975388955 | -4.912665313 |
| **A_51_P217395** | Cyp2d34 | 0.029649005 | -0.772591668 | 0.831889679 | 10.74956555 | 0.089802544 | 0.931284762 | 0.975388955 | -4.912666015 |
| **A_51_P509452** | Tmlhe | -0.015449553 | -0.434902182 | 0.404003075 | 8.493545582 | -0.08949859 | 0.93151662 | 0.975548509 | -4.912680943 |
| **A_55_P2092760** | Rhbdd3 | 0.008108869 | -0.212121371 | 0.228339109 | 10.98545494 | 0.089467748 | 0.931540147 | 0.975548509 | -4.912682455 |
| **A_55_P2046213** | NA | 0.007244117 | -0.189690275 | 0.20417851 | 16.43939506 | 0.089381389 | 0.931606025 | 0.975548509 | -4.912686686 |
| **A_55_P2027836** | Tnfrsf10b | -0.009189648 | -0.259245432 | 0.240866137 | 8.296080462 | -0.089298701 | 0.931669101 | 0.975548509 | -4.912690733 |
| **A_55_P2128663** | Uggt1 | -0.011010222 | -0.310694469 | 0.288674024 | 8.638307883 | -0.08927202 | 0.931689455 | 0.975548509 | -4.912692038 |
| **A_55_P1956308** | NA | -0.009112366 | -0.257699461 | 0.23947473 | 13.76213619 | -0.08907088 | 0.931842894 | 0.975614406 | -4.912701864 |
| **A_55_P2175301** | NA | -0.008056783 | -0.227881174 | 0.211767607 | 15.71193251 | -0.08905719 | 0.931853337 | 0.975614406 | -4.912702532 |
| **A_52_P444253** | Gm6900 | -0.006380959 | -0.180788792 | 0.168026873 | 14.82257113 | -0.08890029 | 0.93197303 | 0.975686871 | -4.912710181 |
| **A_51_P395842** | Timm21 | 0.009611066 | -0.253460596 | 0.272682728 | 10.2719913 | 0.088772958 | 0.932070168 | 0.975688006 | -4.912716378 |
| **A_52_P234958** | Dvl1 | -0.01266063 | -0.35928966 | 0.333968401 | 10.25162484 | -0.088751063 | 0.932086872 | 0.975688006 | -4.912717442 |
| **A_55_P2154054** | Myo18a | 0.012710983 | -0.335694516 | 0.361116483 | 10.81767594 | 0.088649713 | 0.93216419 | 0.975688006 | -4.912722367 |
| **A_51_P478098** | Epb41l5 | 0.015812817 | -0.417986586 | 0.449612221 | 8.706363562 | 0.088573436 | 0.932222381 | 0.975688006 | -4.91272607 |
| **A_51_P117865** | Fam20c | 0.020229626 | -0.534771739 | 0.57523099 | 9.757320274 | 0.088568031 | 0.932226504 | 0.975688006 | -4.912726333 |
| **A_55_P2038162** | Nt5c | 0.012030376 | -0.318274156 | 0.342334909 | 12.20884948 | 0.088500937 | 0.93227769 | 0.975688747 | -4.912729587 |
| **A_51_P421984** | Timm22 | 0.013617404 | -0.36094202 | 0.388176828 | 11.352509 | 0.088339874 | 0.932400566 | 0.975764512 | -4.912737389 |
| **A_52_P158431** | Tshz2 | -0.006566515 | -0.187618882 | 0.174485852 | 8.639430117 | -0.088127999 | 0.93256221 | 0.975880838 | -4.912747631 |
| **A_52_P437734** | Fbxo28 | -0.011266286 | -0.322274032 | 0.299741461 | 8.495729085 | -0.088022308 | 0.932642844 | 0.975886646 | -4.912752731 |
| **A_55_P1985040** | Gpbp1l1 | -0.006762626 | -0.193518069 | 0.179992818 | 8.692610939 | -0.08798837 | 0.932668737 | 0.975886646 | -4.912754368 |
| **A_55_P2043892** | Srd5a3 | -0.007979824 | -0.228592828 | 0.21263318 | 8.678120687 | -0.087891202 | 0.932742871 | 0.975911387 | -4.912759049 |
| **A_55_P2193419** | NA | 0.006059985 | -0.161727475 | 0.173847445 | 8.092558481 | 0.087759718 | 0.932843187 | 0.975963516 | -4.912765376 |
| **A_66_P136097** | Trim13 | 0.008076133 | -0.215778133 | 0.231930399 | 8.208706831 | 0.087663999 | 0.932916216 | 0.975987094 | -4.912769975 |
| **A_55_P1958133** | NA | 0.02101259 | -0.562114036 | 0.604139216 | 11.56510975 | 0.087558821 | 0.932996464 | 0.97601822 | -4.912775024 |
| **A_66_P100840** | NA | 0.012377527 | -0.331553051 | 0.356308105 | 9.038813469 | 0.087447277 | 0.93308157 | 0.976027406 | -4.912780372 |
| **A_55_P2001148** | Pdlim5 | -0.005951754 | -0.171392483 | 0.159488976 | 7.683192538 | -0.087414948 | 0.933106236 | 0.976027406 | -4.91278192 |
| **A_51_P123017** | Gmcl1 | -0.00792595 | -0.228525113 | 0.212673212 | 9.981563119 | -0.087303305 | 0.933191418 | 0.976063686 | -4.900820297 |
| **A_55_P2221647** | AI605517 | -0.006432943 | -0.185898468 | 0.173032581 | 8.016503905 | -0.087098741 | 0.933347501 | 0.976174116 | -4.912797037 |
| **A_55_P2157093** | Bcl2l14 | 0.023602897 | -0.636547361 | 0.683753155 | 7.931476338 | 0.086877177 | 0.933516557 | 0.976231084 | -4.912807597 |
| **A_55_P2106956** | Gm5523 | -0.012488951 | -0.362043181 | 0.337065279 | 17.00427009 | -0.086814966 | 0.933564026 | 0.976231084 | -4.912810557 |
| **A_52_P130912** | Pcdh7 | -0.005476412 | -0.158876721 | 0.147923897 | 7.645320533 | -0.086746722 | 0.933616099 | 0.976231084 | -4.912813802 |
| **A_52_P437850** | Fam13c | -0.039528336 | -1.146898139 | 1.067841467 | 9.589519932 | -0.086735931 | 0.933624332 | 0.976231084 | -4.912814315 |
| **A_55_P2155727** | Mafa | 0.011474546 | -0.310126642 | 0.333075733 | 9.657491875 | 0.086696396 | 0.933654499 | 0.976231084 | -4.912816193 |
| **A_52_P596541** | Kdelr1 | 0.009932537 | -0.269157068 | 0.289022141 | 9.843650065 | 0.086476818 | 0.933822047 | 0.97634827 | -4.91282661 |
| **A_51_P379750** | Myoz1 | -0.005498519 | -0.160112724 | 0.149115687 | 7.753529049 | -0.086413083 | 0.933870681 | 0.97634827 | -4.912829629 |
| **A_51_P486001** | Mob3a | -0.007357352 | -0.214389466 | 0.199674762 | 8.703358596 | -0.086350926 | 0.933918111 | 0.97634827 | -4.912832571 |
| **A_55_P1980190** | NA | -0.007297071 | -0.21294466 | 0.198350517 | 11.62186313 | -0.086220027 | 0.934017996 | 0.976356667 | -4.912838759 |
| **A_55_P2413359** | NA | 0.005913907 | -0.160776301 | 0.172604115 | 8.158752789 | 0.086208007 | 0.934027168 | 0.976356667 | -4.912839327 |
| **A_55_P2070054** | Senp3 | 0.008123192 | -0.221124157 | 0.23737054 | 11.58865255 | 0.086100479 | 0.934109221 | 0.976389635 | -4.912844403 |
| **A_55_P1964682** | Rn28s1 | 0.028009947 | -0.764768933 | 0.820788826 | 17.07776315 | 0.085850612 | 0.934299893 | 0.976536129 | -4.912856173 |
| **A_52_P367565** | Mrps2 | 0.012614068 | -0.346423718 | 0.371651854 | 10.06762772 | 0.085368608 | 0.934667721 | 0.976867761 | -4.912878783 |
| **A_52_P323658** | Zfp955a | -0.009131815 | -0.269457604 | 0.251193974 | 9.217583465 | -0.085236007 | 0.934768915 | 0.9769207 | -4.912884981 |
| **A_55_P2095108** | A430033K04Rik | 0.007311312 | -0.201500853 | 0.216123477 | 8.819844087 | 0.085079066 | 0.934888685 | 0.976944077 | -4.912892304 |
| **A_55_P2050273** | Cct6b | 0.005446205 | -0.150107049 | 0.160999459 | 7.845270506 | 0.085074239 | 0.934892369 | 0.976944077 | -4.912892529 |
| **A_55_P1981002** | NA | 0.010391854 | -0.286872487 | 0.307656196 | 9.273033425 | 0.084944139 | 0.934991657 | 0.976995012 | -4.912898589 |

| | | | | | | | | | |
|---|---|---|---|---|---|---|---|---|---|
| **A_55_P2013591** | NA | -0.009013927 | -0.267077878 | 0.249050023 | 9.339056068 | -0.084873068 | 0.935045897 | 0.976998872 | -4.912901895 |
| **A_55_P1988158** | Esd | 0.006766381 | -0.187125874 | 0.200658637 | 13.73071354 | 0.084796745 | 0.935104144 | 0.977006919 | -4.912905443 |
| **A_66_P111217** | NA | -0.009251091 | -0.27473979 | 0.256237607 | 14.2561213 | -0.084670108 | 0.935200793 | 0.977055085 | -4.912911323 |
| **A_55_P2034097** | Elavl1 | 0.011940919 | -0.331657356 | 0.355539193 | 13.14024058 | 0.084444224 | 0.935373187 | 0.977182377 | -4.912921789 |
| **A_55_P2026535** | Erc2 | -0.008970816 | -0.268154806 | 0.250213173 | 8.42008066 | -0.084102129 | 0.935634282 | 0.977282871 | -4.912937586 |
| **A_51_P452203** | Vash2 | 0.006094548 | -0.170309778 | 0.182498875 | 7.699363013 | 0.083948988 | 0.935751166 | 0.977282871 | -4.912944637 |
| **A_55_P2089310** | 2900097C17Rik | 0.005836389 | -0.163307655 | 0.174980432 | 8.404032348 | 0.083843749 | 0.93583149 | 0.977282871 | -4.912949475 |
| **A_55_P2073935** | Abcc5 | -0.006543848 | -0.196234409 | 0.183146712 | 8.162095329 | -0.083824446 | 0.935846223 | 0.977282871 | -4.912950362 |
| **A_51_P251892** | Magoh | -0.010666964 | -0.319964017 | 0.298630089 | 12.77038041 | -0.083800812 | 0.935864262 | 0.977282871 | -4.912951447 |
| **A_55_P2162582** | Rab5b | -0.009543977 | -0.28684773 | 0.267759775 | 8.780385018 | -0.083628986 | 0.93599541 | 0.977282871 | -4.912959329 |
| **A_52_P137829** | NA | -0.006087295 | -0.183165187 | 0.170990597 | 8.318029513 | -0.083530133 | 0.936070862 | 0.977282871 | -4.912963857 |
| **A_51_P105520** | Nomo1 | -0.012445809 | -0.374624909 | 0.34973329 | 12.49642428 | -0.083499323 | 0.936094379 | 0.977282871 | -4.912965267 |
| **A_55_P1975640** | Zfp217 | -0.008960263 | -0.269846156 | 0.25192563 | 10.78065693 | -0.083455193 | 0.936128063 | 0.977282871 | -4.912967285 |
| **A_51_P478003** | Papolg | 0.006544454 | -0.184222701 | 0.197311609 | 8.899624046 | 0.083359092 | 0.936201414 | 0.977282871 | -4.912971678 |
| **A_55_P2001203** | Dnm2 | 0.00830735 | -0.233858024 | 0.250472723 | 13.10141759 | 0.08335538 | 0.936204248 | 0.977282871 | -4.912971847 |
| **A_52_P495565** | Efnb3 | -0.006580985 | -0.19847073 | 0.18530876 | 7.932431457 | -0.083334022 | 0.936220551 | 0.977282871 | -4.912972823 |
| **A_51_P279854** | Ndufa11 | -0.010953704 | -0.330454724 | 0.308547315 | 10.93399041 | -0.08330517 | 0.936242573 | 0.977282871 | -4.91297414 |
| **A_51_P150430** | Mkrn2 | -0.010992137 | -0.331626755 | 0.309642481 | 8.98318627 | -0.083301903 | 0.936245067 | 0.977282871 | -4.912974289 |
| **A_51_P513072** | Rdh13 | 0.006263178 | -0.176578946 | 0.189105302 | 8.163435603 | 0.083234171 | 0.936296766 | 0.977282871 | -4.91297738 |
| **A_51_P307644** | Gtf3a | 0.010810201 | -0.305492653 | 0.327113054 | 10.57209187 | 0.083045066 | 0.936441112 | 0.977282871 | -4.912985995 |
| **A_51_P320888** | Cln5 | 0.005602206 | -0.1584115 | 0.169615913 | 10.87477786 | 0.082996952 | 0.936477837 | 0.977282871 | -4.912988184 |
| **A_55_P2108754** | Ubfd1 | 0.009374829 | -0.265286313 | 0.284035971 | 11.05711571 | 0.082937195 | 0.936523451 | 0.977282871 | -4.912990901 |
| **A_55_P1991160** | Mrrf | 0.005452603 | -0.154518335 | 0.165423541 | 7.737581557 | 0.082822054 | 0.936611342 | 0.977282871 | -4.91299613 |
| **A_55_P2020796** | Man1b1 | -0.007964629 | -0.241710372 | 0.225781115 | 11.84684621 | -0.08279517 | 0.936631863 | 0.977282871 | -4.91299735 |
| **A_55_P2151863** | NA | 0.01513527 | -0.429273671 | 0.459544211 | 11.89043139 | 0.082754302 | 0.936663059 | 0.977282871 | -4.912999204 |
| **A_52_P669005** | Lrat | -0.016380618 | -0.497371263 | 0.464610027 | 8.502857187 | -0.082751685 | 0.936665056 | 0.977282871 | -4.912999322 |
| **A_51_P121466** | Tab1 | -0.010731381 | -0.326458361 | 0.3049956 | 11.06511617 | -0.08258993 | 0.936788531 | 0.977282871 | -4.91300665 |
| **A_51_P255193** | Faf2 | 0.006259528 | -0.178053571 | 0.190572627 | 10.08339293 | 0.082521774 | 0.936840557 | 0.977282871 | -4.913009734 |
| **A_51_P197175** | Pigk | -0.00747243 | -0.22761603 | 0.212671171 | 8.417540219 | -0.082478158 | 0.936873852 | 0.977282871 | -4.913011706 |
| **A_52_P650473** | NA | 0.0053595 | -0.152616975 | 0.163335976 | 8.255299403 | 0.082435652 | 0.936906299 | 0.977282871 | -4.913013626 |
| **A_51_P480928** | Zfyve16 | -0.007687518 | -0.234404352 | 0.219029317 | 8.613873237 | -0.082392095 | 0.936939549 | 0.977282871 | -4.913015594 |
| **A_51_P483511** | Dock8 | -0.010358954 | -0.316248938 | 0.295531029 | 8.423295909 | -0.082287497 | 0.937019395 | 0.977282871 | -4.913020313 |
| **A_55_P2172227** | Ubn2 | -0.008570878 | -0.26167029 | 0.244528533 | 8.302522351 | -0.082284376 | 0.937021777 | 0.977282871 | -4.913020454 |
| **A_55_P2163353** | Sox3 | 0.005304437 | -0.151455109 | 0.162063983 | 7.672205766 | 0.082222091 | 0.937069324 | 0.977282871 | -4.913023261 |
| **A_52_P175952** | Map3k7 | 0.008334251 | -0.238248957 | 0.25491746 | 8.508946866 | 0.082127062 | 0.937141867 | 0.977282871 | -4.913027541 |
| **A_55_P2098061** | Tchp | 0.006737077 | -0.192810334 | 0.206284487 | 8.335936501 | 0.082036774 | 0.937210792 | 0.977282871 | -4.913031602 |
| **A_55_P2176565** | B3galt5 | -0.005448122 | -0.166836559 | 0.155940314 | 7.729499573 | -0.082027152 | 0.937218137 | 0.977282871 | -4.913032035 |
| **A_55_P1984751** | Ppp1r14b | 0.008295316 | -0.237504499 | 0.25409513 | 11.76497328 | 0.082003911 | 0.937235879 | 0.977282871 | -4.913033079 |
| **A_55_P1955672** | NA | 0.009887894 | -0.283116458 | 0.302892246 | 15.15060771 | 0.081999825 | 0.937238998 | 0.977282871 | -4.913033263 |
| **A_52_P318438** | Nab1 | 0.015870895 | -0.455736457 | 0.487478246 | 10.62022554 | 0.081771889 | 0.937413004 | 0.97733884 | -4.913043491 |
| **A_51_P120729** | Polr2f | 0.013459153 | -0.38676126 | 0.413679566 | 11.67323618 | 0.081714981 | 0.937456448 | 0.97733884 | -4.91304604 |
| **A_55_P2041509** | 1110004F10Rik | -0.007892826 | -0.242598474 | 0.226812823 | 11.32950931 | -0.081713189 | 0.937457816 | 0.97733884 | -4.913046121 |
| **A_55_P2034390** | Ankhd1 | -0.006632771 | -0.204150886 | 0.190885343 | 8.556695203 | -0.081596453 | 0.937546934 | 0.97733884 | -4.913051344 |
| **A_55_P2000487** | Srebf2 | 0.01033228 | -0.29756313 | 0.318227689 | 8.131091693 | 0.081541017 | 0.937589255 | 0.97733884 | -4.913053822 |
| **A_55_P1978675** | NA | 0.005209217 | -0.150039119 | 0.160457553 | 7.650985863 | 0.081532112 | 0.937596054 | 0.97733884 | -4.91305422 |
| **A_51_P382484** | Itgav | -0.010539482 | -0.325035141 | 0.303956176 | 8.892687002 | -0.081430633 | 0.937673525 | 0.977366888 | -4.913058752 |
| **A_55_P1996344** | Dcst2 | 0.005548583 | -0.160159807 | 0.171256974 | 7.674055687 | 0.081361849 | 0.937726038 | 0.977368918 | -4.91306182 |
| **A_55_P1966998** | Arhgdib | 0.008905968 | -0.257781989 | 0.275593925 | 11.06276411 | 0.081144839 | 0.937891712 | 0.977462321 | -4.913071483 |
| **A_52_P105765** | Cdc42bpg | 0.006846653 | -0.198659059 | 0.212352366 | 8.127744079 | 0.080953869 | 0.93803751 | 0.977462321 | -4.913079965 |
| **A_66_P100349** | Esyt2 | -0.013790336 | -0.428243252 | 0.400662581 | 10.20098887 | -0.080850523 | 0.938116412 | 0.977462321 | -4.913084548 |

| | | | | | | | | | |
|---|---|---|---|---|---|---|---|---|---|
| A_55_P1979356 | Morf4l1 | -0.009432339 | -0.30398859 | 0.285123911 | 13.64617724 | -0.081469217 | 0.938124314 | 0.977462321 | -4.86359752 |
| A_52_P377576 | NA | 0.006127136 | -0.178048758 | 0.19030303 | 8.330887248 | 0.080836581 | 0.938127056 | 0.977462321 | -4.913085165 |
| A_51_P438711 | Tbx18 | -0.005417283 | -0.168350205 | 0.157515639 | 7.678188391 | -0.080789675 | 0.938162867 | 0.977462321 | -4.913087243 |
| A_52_P624362 | Aamp | -0.010229493 | -0.317930664 | 0.297471677 | 10.46484301 | -0.080780804 | 0.938169641 | 0.977462321 | -4.913087635 |
| A_55_P2173892 | Isg20 | -0.011223441 | -0.34986953 | 0.327422647 | 12.93335711 | -0.080531015 | 0.938360351 | 0.977503561 | -4.913098677 |
| A_65_P10433 | Rpp14 | -0.008257375 | -0.257488889 | 0.240974139 | 9.059514037 | -0.080504886 | 0.9383803 | 0.977503561 | -4.91309983 |
| A_66_P118004 | Rab3ip | -0.007461686 | -0.232937752 | 0.218014379 | 10.89830555 | -0.080411791 | 0.938451378 | 0.977503561 | -4.913103935 |
| A_51_P467770 | Snx19 | 0.005802366 | -0.169670091 | 0.181274823 | 8.522076553 | 0.080348792 | 0.938499479 | 0.977503561 | -4.91310671 |
| A_52_P113537 | Xist | 0.052289772 | -1.529365791 | 1.633945335 | 8.183874743 | 0.080331897 | 0.938512378 | 0.977503561 | -4.913107454 |
| A_52_P373973 | Mex3c | -0.012708614 | -0.397119184 | 0.371701955 | 10.799541 | -0.080331536 | 0.938512653 | 0.977503561 | -4.91310747 |
| A_52_P285470 | Lrp2 | 0.007415088 | -0.217516313 | 0.23234649 | 8.115353918 | 0.080103117 | 0.938687056 | 0.977554297 | -4.913117512 |
| A_55_P2083997 | Zfp664 | -0.018170917 | -0.56938491 | 0.533043075 | 11.31996331 | -0.080101344 | 0.938688409 | 0.977554297 | -4.913117589 |
| A_55_P1961084 | Map3k1 | -0.01234798 | -0.38727585 | 0.362579889 | 11.65899076 | -0.080026053 | 0.938745896 | 0.977554297 | -4.913120893 |
| A_52_P277104 | Bank1 | 0.010822619 | -0.318020309 | 0.339665547 | 8.528775146 | 0.079969992 | 0.938788701 | 0.977554297 | -4.913123351 |
| A_55_P2042297 | NA | 0.006387039 | -0.187762947 | 0.200537024 | 7.863013795 | 0.079936545 | 0.938814238 | 0.977554297 | -4.913124816 |
| A_52_P590625 | Nlrp10 | 0.005125134 | -0.151053519 | 0.161303788 | 7.645272947 | 0.079738261 | 0.938965638 | 0.977567598 | -4.913133492 |
| A_55_P2048398 | Kmt5b | -0.00523486 | -0.164915898 | 0.154446178 | 8.137711269 | -0.079659014 | 0.939026147 | 0.977567598 | -4.913136954 |
| A_66_P119968 | Clec4a1 | -0.014449212 | -0.455221497 | 0.426323073 | 10.26485393 | -0.079655005 | 0.939029208 | 0.977567598 | -4.913137129 |
| A_55_P2037717 | Slc22a18 | 0.01961806 | -0.578830633 | 0.618066753 | 11.11158465 | 0.079654867 | 0.939029313 | 0.977567598 | -4.913137135 |
| A_55_P2177154 | H2bc14 | 0.005687645 | -0.168125498 | 0.179500789 | 13.31709094 | 0.079512073 | 0.939138346 | 0.977628451 | -4.913143363 |
| A_52_P632093 | Gtf3c3 | -0.006059062 | -0.192930327 | 0.180812203 | 9.108851915 | -0.078785453 | 0.939693188 | 0.978153354 | -4.913174883 |
| A_55_P2146127 | Chmp4b | 0.009136837 | -0.274608937 | 0.292882612 | 11.21828473 | 0.078243755 | 0.940106849 | 0.978531248 | -4.913198194 |
| A_66_P118600 | Lama1 | -0.01627229 | -0.522103704 | 0.489559123 | 11.16430001 | -0.078167474 | 0.940165101 | 0.978539187 | -4.913201463 |
| A_55_P1961160 | NA | -0.008446108 | -0.271452127 | 0.254559912 | 14.20163492 | -0.078032249 | 0.940268368 | 0.978593974 | -4.913207252 |
| A_55_P2167922 | Dmd | 0.00498533 | -0.15098123 | 0.16095189 | 7.657585666 | 0.077668619 | 0.940546067 | 0.978803581 | -4.913222768 |
| A_55_P2043996 | Cops7a | 0.005444622 | -0.165062251 | 0.175951494 | 8.237597739 | 0.077590581 | 0.940605665 | 0.978803581 | -4.913226088 |
| A_52_P203948 | Pla2g4e | -0.004937788 | -0.159614169 | 0.149738594 | 7.63504612 | -0.077569606 | 0.940621683 | 0.978803581 | -4.91322698 |
| A_55_P2072631 | Prr5l | -0.011304297 | -0.365733775 | 0.343125182 | 8.868926167 | -0.077499143 | 0.940675496 | 0.978806883 | -4.913229975 |
| A_52_P407280 | Adamtsl5 | -0.005077512 | -0.165216673 | 0.15506165 | 7.965071418 | -0.077043598 | 0.941023406 | 0.979099654 | -4.913249268 |
| A_55_P2070657 | Emc10 | 0.009280864 | -0.2837873 | 0.302349027 | 8.31984935 | 0.076948994 | 0.94109566 | 0.979099654 | -4.913253261 |
| A_55_P2094100 | BC085271 | 0.008646957 | -0.264465085 | 0.281759 | 10.34597088 | 0.076931757 | 0.941108825 | 0.979099654 | -4.913253988 |
| A_55_P2182423 | NA | -0.006215171 | -0.202997037 | 0.190566695 | 15.06332203 | -0.076745196 | 0.941251311 | 0.979195188 | -4.913261845 |
| A_52_P128964 | Nhej1 | 0.007175848 | -0.220229413 | 0.234581109 | 9.561304275 | 0.076675376 | 0.941304636 | 0.979197961 | -4.913264781 |
| A_55_P1958652 | Arf2 | 0.014929617 | -0.459514694 | 0.489373927 | 11.15474141 | 0.076462167 | 0.94146748 | 0.979314655 | -4.913273729 |
| A_55_P2404434 | NA | -0.007283794 | -0.238979233 | 0.224411645 | 8.52466192 | -0.076387687 | 0.941524366 | 0.979321125 | -4.913276849 |
| A_66_P128055 | Spindoc | 0.006919995 | -0.214641626 | 0.228481615 | 8.152625552 | 0.075891723 | 0.941903184 | 0.979561552 | -4.913297548 |
| A_55_P2028977 | Tcf20 | -0.005191107 | -0.171418145 | 0.161035931 | 8.102900673 | -0.075882475 | 0.941910248 | 0.979561552 | -4.913297932 |
| A_55_P2425437 | D030036P13Rik | 0.008629674 | -0.267939976 | 0.285199323 | 9.663579403 | 0.075818126 | 0.941959399 | 0.979561552 | -4.913300608 |
| A_55_P2057994 | Mif4gd | -0.005903499 | -0.195201528 | 0.18339453 | 10.27292517 | -0.075778604 | 0.941989587 | 0.979561552 | -4.91330225 |
| A_55_P1952475 | Rpl30 | 0.006054098 | -0.188259478 | 0.200367675 | 15.65937995 | 0.075705864 | 0.942045148 | 0.979561552 | -4.91330527 |
| A_55_P2157586 | NA | 0.008443079 | -0.263042084 | 0.279928241 | 8.408149812 | 0.075567999 | 0.942150453 | 0.979561552 | -4.913310986 |
| A_55_P2080163 | Carhsp1 | 0.00619203 | -0.192985634 | 0.205369694 | 13.08648135 | 0.075539757 | 0.942172026 | 0.979561552 | -4.913312155 |
| A_55_P2008056 | Cideb | 0.019595551 | -0.610984289 | 0.650175392 | 10.25723552 | 0.075509326 | 0.94219527 | 0.979561552 | -4.913313415 |
| A_52_P561020 | Slc25a32 | 0.010467023 | -0.326749217 | 0.347683262 | 8.73784673 | 0.075421969 | 0.942261998 | 0.979561552 | -4.913317029 |
| A_55_P2347756 | AU040972 | -0.021714548 | -0.721296126 | 0.677867031 | 9.775704964 | -0.075421577 | 0.942262297 | 0.979561552 | -4.913317045 |
| A_55_P1974452 | Lime1 | 0.01155402 | -0.361545284 | 0.384653324 | 9.45078088 | 0.075247466 | 0.942395293 | 0.979647123 | -4.913324235 |
| A_51_P256282 | Ppp6c | -0.007639891 | -0.254921037 | 0.239641254 | 11.55876024 | -0.075072237 | 0.942529144 | 0.979701708 | -4.913331454 |
| A_51_P346668 | Irf5 | -0.007249918 | -0.241990763 | 0.227490926 | 8.708969412 | -0.075046017 | 0.942549173 | 0.979701708 | -4.913332533 |
| A_55_P2060184 | Lamtor3 | 0.008581371 | -0.269857474 | 0.287020215 | 12.09265536 | 0.07488762 | 0.942670169 | 0.979774786 | -4.913339042 |
| A_66_P118307 | NA | -0.011829338 | -0.396728994 | 0.373070319 | 13.10751943 | -0.074678588 | 0.942829847 | 0.979888058 | -4.913347611 |

| | | | | | | | | | |
|---|---|---|---|---|---|---|---|---|---|
| **A_55_P2102816** | Rpl36a | 0.007333722 | -0.231565968 | 0.246233412 | 15.08481757 | 0.074591963 | 0.942896019 | 0.979888938 | -4.913351155 |
| **A_55_P1998545** | Ncdn | 0.004993579 | -0.157912505 | 0.167899663 | 10.29128942 | 0.074483111 | 0.942979172 | 0.979888938 | -4.913355602 |
| **A_51_P251587** | Psmd9 | 0.009657397 | -0.305416903 | 0.324731696 | 9.956378596 | 0.074478389 | 0.942982779 | 0.979888938 | -4.913355795 |
| **A_51_P327232** | Lenep | -0.005871322 | -0.197779918 | 0.186037273 | 8.335518088 | -0.074340364 | 0.943088219 | 0.979945823 | -4.913361425 |
| **A_51_P188795** | Akr7a5 | 0.011777727 | -0.373627593 | 0.397183046 | 10.10676834 | 0.074255214 | 0.943153267 | 0.979960732 | -4.913364893 |
| **A_55_P2039881** | 5-Sep | -0.004928747 | -0.1663728 | 0.156515305 | 7.866599933 | -0.074181853 | 0.94320931 | 0.97996373 | -4.913367878 |
| **A_55_P2135351** | Fam89b | 0.008478369 | -0.269590839 | 0.286547577 | 10.00329289 | 0.074087104 | 0.943281692 | 0.97996373 | -4.913371728 |
| **A_66_P130646** | H4c8 | 0.006059121 | -0.192932953 | 0.205051194 | 8.104111451 | 0.073987271 | 0.943357958 | 0.97996373 | -4.91337578 |
| **A_55_P2112030** | Tmem145 | -0.006288075 | -0.213141001 | 0.20056485 | 7.875123063 | -0.073865093 | 0.943451296 | 0.97996373 | -4.913380731 |
| **A_55_P2043762** | N6amt1 | -0.008470284 | -0.287228074 | 0.270287506 | 8.723014698 | -0.07383362 | 0.94347534 | 0.97996373 | -4.913382005 |
| **A_51_P502119** | F11 | -0.020390349 | -0.691602384 | 0.650821687 | 12.8454669 | -0.073815592 | 0.943489113 | 0.97996373 | -4.913382735 |
| **A_51_P160372** | Wbp1 | -0.016948409 | -0.575364524 | 0.541467706 | 12.25065483 | -0.073748668 | 0.94354024 | 0.97996373 | -4.913385442 |
| **A_52_P115579** | Fbxo45 | -0.005589314 | -0.189931792 | 0.178753164 | 8.121350018 | -0.073674346 | 0.943597019 | 0.97996373 | -4.913388445 |
| **A_55_P1972034** | Muc16 | 0.010603587 | -0.340343717 | 0.361550891 | 8.366850257 | 0.07341657 | 0.943793952 | 0.97996373 | -4.913398837 |
| **A_55_P2136378** | Dazap2 | 0.007451275 | -0.239281326 | 0.254183876 | 13.6717134 | 0.07338161 | 0.943820661 | 0.97996373 | -4.913400244 |
| **A_55_P2108171** | Glud1 | 0.021341931 | -0.685796259 | 0.728480122 | 15.02001157 | 0.073335222 | 0.943856101 | 0.97996373 | -4.913402109 |
| **A_51_P419246** | Mirt1 | -0.010858982 | -0.370831091 | 0.349113127 | 8.59084299 | -0.07329991 | 0.943883079 | 0.97996373 | -4.913403529 |
| **A_55_P1983544** | Tmed5 | 0.011543699 | -0.371287387 | 0.394374784 | 9.839327868 | 0.07326912 | 0.943906603 | 0.97996373 | -4.913404766 |
| **A_52_P69707** | Ergic1 | 0.010240166 | -0.329443209 | 0.349923541 | 9.720971117 | 0.073251395 | 0.943920144 | 0.97996373 | -4.913405478 |
| **A_52_P268287** | Zfp507 | 0.006232364 | -0.200605287 | 0.213070015 | 9.337011339 | 0.073216068 | 0.943947133 | 0.97996373 | -4.913406896 |
| **A_55_P2045225** | NA | 0.007821267 | -0.251842164 | 0.267484697 | 11.7764436 | 0.073189622 | 0.943967338 | 0.97996373 | -4.913407957 |
| **A_55_P2022342** | Stim1 | -0.007652085 | -0.262322499 | 0.247018328 | 10.96831747 | -0.073010363 | 0.944104293 | 0.980053269 | -4.91341514 |
| **A_55_P1999811** | NA | 0.005329091 | -0.172352705 | 0.183010888 | 8.288295428 | 0.072877489 | 0.94420581 | 0.980106015 | -4.913420454 |
| **A_66_P137157** | Arl6ip4 | 0.007995876 | -0.259056718 | 0.27504847 | 13.17441332 | 0.072753255 | 0.944300727 | 0.980151904 | -4.913425413 |
| **A_52_P462203** | Dclre1c | -0.006595171 | -0.227989637 | 0.214799295 | 8.405151282 | -0.072383983 | 0.944582864 | 0.980392106 | -4.913440103 |
| **A_55_P2109997** | Lrig1 | -0.009704038 | -0.337441265 | 0.318033189 | 9.440959069 | -0.071946536 | 0.9449171 | 0.980686353 | -4.91345741 |
| **A_55_P1984069** | NA | -0.007968673 | -0.278210851 | 0.262273504 | 13.52279395 | -0.071649981 | 0.945143694 | 0.980868857 | -4.913469082 |
| **A_55_P2128303** | Ids | 0.005866516 | -0.1935461 | 0.205279132 | 8.101623368 | 0.071484325 | 0.945270272 | 0.980902017 | -4.913475582 |
| **A_65_P01991** | Prrg4 | 0.005053534 | -0.166816957 | 0.176924024 | 7.734259597 | 0.071445857 | 0.945299665 | 0.980902017 | -4.913477089 |
| **A_55_P2041521** | Doxl2 | -0.007088572 | -0.248396533 | 0.234219388 | 7.966533908 | -0.071378978 | 0.945350769 | 0.980902017 | -4.913479707 |
| **A_55_P2149215** | Camk2a | -0.00466794 | -0.164194708 | 0.154858828 | 7.759554324 | -0.071100864 | 0.945563281 | 0.980902017 | -4.913490569 |
| **A_55_P2095663** | Pgr | 0.004538811 | -0.150586824 | 0.159664446 | 7.636373181 | 0.071095428 | 0.945567435 | 0.980902017 | -4.913490781 |
| **A_55_P2159259** | Lifr | 0.01485991 | -0.4942586 | 0.523978419 | 11.1211724 | 0.070921917 | 0.945700021 | 0.980902017 | -4.913497536 |
| **A_55_P1991016** | Basp1 | -0.00847303 | -0.299131823 | 0.282185763 | 8.523942214 | -0.070833465 | 0.945767611 | 0.980902017 | -4.913500973 |
| **A_51_P343350** | Amn | -0.013247515 | -0.46775814 | 0.44126311 | 8.255066899 | -0.070822876 | 0.945775702 | 0.980902017 | -4.913501384 |
| **A_55_P2128869** | Ccdc80 | -0.00540326 | -0.190932983 | 0.180126464 | 8.274849844 | -0.070766142 | 0.945819055 | 0.980902017 | -4.913503586 |
| **A_52_P401484** | Inha | 0.006087176 | -0.203107556 | 0.215281907 | 7.820128779 | 0.070704698 | 0.945866009 | 0.980902017 | -4.913505969 |
| **A_55_P2131520** | Vash1 | 0.009170317 | -0.30639505 | 0.324735684 | 13.88916738 | 0.070611947 | 0.945936885 | 0.980902017 | -4.913509562 |
| **A_55_P2047986** | Ankrd23 | 0.005275712 | -0.176720255 | 0.187271679 | 8.002707193 | 0.07043727 | 0.946070366 | 0.980902017 | -4.913516316 |
| **A_51_P265869** | Hspa9 | -0.016142486 | -0.573220329 | 0.540935357 | 12.7321012 | -0.070410546 | 0.946090788 | 0.980902017 | -4.913517347 |
| **A_55_P2424042** | Ttn | 0.004507761 | -0.151403086 | 0.160418609 | 7.650281217 | 0.070253465 | 0.946210826 | 0.980902017 | -4.913523404 |
| **A_65_P06572** | Smad5 | -0.004821754 | -0.171756816 | 0.162113308 | 7.884254307 | -0.070184414 | 0.946263594 | 0.980902017 | -4.913526063 |
| **A_51_P127320** | Cabs1 | -0.004513367 | -0.16103253 | 0.152005796 | 7.632271762 | -0.070067451 | 0.946352976 | 0.980902017 | -4.904389224 |
| **A_55_P2117903** | Stard5 | -0.012421753 | -0.443720884 | 0.418877378 | 9.339034534 | -0.069982185 | 0.946418136 | 0.980902017 | -4.913533833 |
| **A_55_P1953894** | Mccc2 | 0.011163351 | -0.376603909 | 0.39893061 | 9.957481571 | 0.069953044 | 0.946440406 | 0.980902017 | -4.913534951 |
| **A_52_P302463** | Arl5a | -0.007494198 | -0.268042203 | 0.253053807 | 8.935339676 | -0.069890892 | 0.946487903 | 0.980902017 | -4.913537333 |
| **A_55_P2293967** | Tbc1d5 | 0.00679615 | -0.229550234 | 0.243142535 | 9.026615402 | 0.069871041 | 0.946503073 | 0.980902017 | -4.913538094 |
| **A_55_P2085766** | NA | 0.004805578 | -0.16236733 | 0.171978486 | 7.796411653 | 0.069849437 | 0.946519583 | 0.980902017 | -4.913538921 |
| **A_55_P2074317** | Ap2b1 | 0.009137169 | -0.309012862 | 0.327287201 | 9.46064722 | 0.069785129 | 0.946568728 | 0.980902017 | -4.913541383 |
| **A_51_P400463** | Agmat | 0.018084985 | -0.611717711 | 0.647887681 | 11.7325905 | 0.069774515 | 0.946576839 | 0.980902017 | -4.913541789 |

| | | | | | | | | | |
|---|---|---|---|---|---|---|---|---|---|
| **A_51_P261443** | Mlx | -0.006333987 | -0.226983319 | 0.214315345 | 9.361103996 | -0.069752172 | 0.946593914 | 0.980902017 | -4.913542644 |
| **A_55_P2106827** | NA | -0.007093566 | -0.254311444 | 0.240124313 | 11.49789522 | -0.06972169 | 0.946617208 | 0.980902017 | -4.913543809 |
| **A_51_P476509** | Zbtb38 | 0.005042473 | -0.170745934 | 0.18083088 | 8.179640887 | 0.069700602 | 0.946633324 | 0.980902017 | -4.913544615 |
| **A_55_P2090953** | NA | -0.005357777 | -0.192308745 | 0.181593191 | 16.92570691 | -0.069637006 | 0.946681925 | 0.980902017 | -4.913547044 |
| **A_66_P120305** | Zmynd11 | -0.010063174 | -0.361241837 | 0.341115488 | 10.35038795 | -0.069628986 | 0.946688055 | 0.980902017 | -4.91354735 |
| **A_55_P2044557** | Rpl29 | -0.005388411 | -0.193541899 | 0.182765077 | 16.6362433 | -0.069587566 | 0.946719708 | 0.980902017 | -4.913548931 |
| **A_51_P108190** | Rpl23 | -0.005910817 | -0.212427432 | 0.200605798 | 11.75815114 | -0.069546567 | 0.946751041 | 0.980902017 | -4.913550495 |
| **A_51_P301964** | Arl6ip1 | -0.006160863 | -0.222040308 | 0.209718582 | 11.8060416 | -0.069344729 | 0.946905293 | 0.980902017 | -4.91355818 |
| **A_51_P235548** | Mtmr9 | 0.005950913 | -0.202667982 | 0.214569809 | 8.556182662 | 0.069312754 | 0.946929729 | 0.980902017 | -4.913559395 |
| **A_51_P263302** | Rnf24 | 0.005729879 | -0.195240006 | 0.206699764 | 8.314636494 | 0.069278364 | 0.946956011 | 0.980902017 | -4.913560702 |
| **A_55_P2161400** | Sctr | -0.004469962 | -0.161294161 | 0.152354238 | 7.656924912 | -0.06925863 | 0.946971093 | 0.980902017 | -4.913561451 |
| **A_52_P26216** | Numb | 0.004518994 | -0.154084951 | 0.163122939 | 8.038185312 | 0.069232656 | 0.946990943 | 0.980902017 | -4.913562437 |
| **A_55_P1990780** | Ptp4a1 | 0.01067893 | -0.364569045 | 0.385926905 | 13.40907201 | 0.069150063 | 0.947054065 | 0.980902017 | -4.91356557 |
| **A_52_P584415** | Cars2 | 0.005146673 | -0.175804808 | 0.186098154 | 8.248665914 | 0.069111077 | 0.94708386 | 0.980902017 | -4.913567048 |
| **A_55_P1983070** | Uhrf1bp1 | -0.007454876 | -0.269738146 | 0.254828393 | 10.3788238 | -0.06906421 | 0.947119678 | 0.980902017 | -4.913568823 |
| **A_55_P2033308** | Six2 | -0.004370362 | -0.15823425 | 0.149493526 | 7.639220724 | -0.069018246 | 0.947154806 | 0.980902017 | -4.913570563 |
| **A_55_P2075479** | Rpl37a | -0.010093054 | -0.366858556 | 0.346672448 | 17.3311096 | -0.068742124 | 0.947365838 | 0.981045304 | -4.913580991 |
| **A_55_P2139385** | Gm12070 | -0.008831317 | -0.321353586 | 0.303690951 | 15.55281013 | -0.06866378 | 0.947425714 | 0.981045304 | -4.913583942 |
| **A_55_P1980583** | Leprot | -0.005486153 | -0.199724334 | 0.188752028 | 12.45585995 | -0.068630393 | 0.947451231 | 0.981045304 | -4.913585199 |
| **A_51_P229816** | Eif2b2 | -0.014525623 | -0.529318352 | 0.500267106 | 12.35765203 | -0.068562328 | 0.947503253 | 0.981045304 | -4.913587759 |
| **A_51_P373379** | Csnk2b | 0.004699643 | -0.162069564 | 0.171468851 | 13.51089966 | 0.068475029 | 0.947569974 | 0.981045304 | -4.913591038 |
| **A_55_P2247852** | Ap2s1 | -0.010760716 | -0.39281249 | 0.371291058 | 11.90458945 | -0.06843876 | 0.947597694 | 0.981045304 | -4.9135924 |
| **A_55_P2148001** | 6820431F20Rik | -0.013725987 | -0.503270274 | 0.475818301 | 9.657546748 | -0.068129433 | 0.947834114 | 0.981207988 | -4.913603981 |
| **A_52_P568781** | Sec63 | -0.006981656 | -0.256092372 | 0.242129061 | 8.343925995 | -0.068100329 | 0.947856359 | 0.981207988 | -4.913605067 |
| **A_55_P2016976** | Zfp729a | -0.006370036 | -0.234783273 | 0.222043201 | 8.777497303 | -0.067764743 | 0.948112855 | 0.981381692 | -4.913617567 |
| **A_55_P1992099** | Ptprj | -0.014640354 | -0.53973635 | 0.510455642 | 9.795090386 | -0.067747934 | 0.948125703 | 0.981381692 | -4.913618191 |
| **A_55_P2012331** | NA | -0.009369848 | -0.346371533 | 0.327631837 | 13.38127906 | -0.06755907 | 0.94827006 | 0.981478554 | -4.913625197 |
| **A_55_P2008891** | Tmsb15b1 | -0.005407317 | -0.200145455 | 0.189330821 | 8.272400403 | -0.067470509 | 0.948337751 | 0.981496059 | -4.913628476 |
| **A_51_P165165** | Hibadh | 0.012057799 | -0.423079638 | 0.447195236 | 12.04150244 | 0.067332507 | 0.948443234 | 0.981552673 | -4.913633576 |
| **A_51_P247249** | Alox5 | 0.015087181 | -0.530475273 | 0.560649636 | 9.938038439 | 0.067196525 | 0.948547174 | 0.981607684 | -4.913638591 |
| **A_55_P2022574** | Gpank1 | 0.009482769 | -0.335361251 | 0.35432679 | 11.85243185 | 0.06681834 | 0.948836252 | 0.981763026 | -4.913652486 |
| **A_52_P326354** | Erf | -0.007041576 | -0.263250966 | 0.249167815 | 11.44681624 | -0.066781781 | 0.948864197 | 0.981763026 | -4.913653825 |
| **A_55_P2029344** | Rpl23 | -0.004920505 | -0.183994724 | 0.174153714 | 16.81445031 | -0.066766684 | 0.948875737 | 0.981763026 | -4.913654377 |
| **A_55_P1989013** | Eif6 | -0.011370253 | -0.425637442 | 0.402896937 | 11.5289684 | -0.066691849 | 0.94893294 | 0.981763026 | -4.913657116 |
| **A_51_P512783** | Rmdn1 | -0.007416776 | -0.278086576 | 0.263253024 | 10.85924958 | -0.066582259 | 0.949016712 | 0.981763026 | -4.91366112 |
| **A_55_P2027640** | NA | 0.008804397 | -0.312679217 | 0.33028801 | 17.20161739 | 0.066546305 | 0.949044195 | 0.981763026 | -4.913662432 |
| **A_55_P2128929** | Cc2d2a | -0.010735006 | -0.402779418 | 0.381309406 | 9.468282782 | -0.066535009 | 0.94905283 | 0.981763026 | -4.913662844 |
| **A_55_P2016105** | Errfi1 | -0.022459509 | -0.845002926 | 0.800083909 | 13.39511048 | -0.066347507 | 0.949196159 | 0.981858748 | -4.913669675 |
| **A_55_P2348582** | NA | -0.006027156 | -0.22789937 | 0.215845058 | 8.186963961 | -0.066007411 | 0.949456138 | 0.982060906 | -4.913682016 |
| **A_55_P2026380** | Pdcd2l | 0.009840074 | -0.352659931 | 0.372340078 | 10.47570679 | 0.065958918 | 0.949493208 | 0.982060906 | -4.913683771 |
| **A_55_P2404554** | Atcayos | -0.004299132 | -0.163063702 | 0.154465439 | 7.64963109 | -0.065797644 | 0.949616494 | 0.982090721 | -4.913689596 |
| **A_51_P158044** | NA | 0.004571291 | -0.164268011 | 0.173410594 | 8.393933331 | 0.06578828 | 0.949623652 | 0.982090721 | -4.913689934 |
| **A_52_P230904** | Wdr18 | -0.007041497 | -0.267957137 | 0.253874144 | 9.474079191 | -0.065576472 | 0.949785571 | 0.982205623 | -4.913697563 |
| **A_55_P1956687** | Rab37 | 0.010872348 | -0.394568303 | 0.416312999 | 8.999564346 | 0.065159726 | 0.950104163 | 0.982466259 | -4.9137125 |
| **A_51_P101460** | Dsp | 0.007259648 | -0.263650701 | 0.278169997 | 9.907983156 | 0.065113815 | 0.950139261 | 0.982466259 | -4.91371414 |
| **A_55_P2093720** | Dnajc5 | 0.006211641 | -0.226257952 | 0.238681233 | 8.272078943 | 0.064926705 | 0.950282307 | 0.982521295 | -4.913720811 |
| **A_51_P474053** | Erv3 | -0.004268594 | -0.164058099 | 0.155520912 | 7.720499236 | -0.064911215 | 0.950294149 | 0.982521295 | -4.913721363 |
| **A_55_P1988866** | NA | -0.005114357 | -0.196831392 | 0.186602678 | 15.10861027 | -0.064820672 | 0.950363369 | 0.982540306 | -4.913724583 |
| **A_55_P1967591** | NA | -0.022080815 | -0.852331681 | 0.808170052 | 11.49667917 | -0.064623272 | 0.950514284 | 0.982610344 | -4.913731589 |
| **A_55_P2012394** | Sfxn2 | 0.015818767 | -0.579748926 | 0.611386461 | 13.05461679 | 0.064539366 | 0.950578433 | 0.982610344 | -4.91373456 |

| | | | | | | | | | |
|---|---|---|---|---|---|---|---|---|---|
| **A_55_P2040878** | Ung | 0.01794115 | -0.658293436 | 0.694175736 | 14.86860581 | 0.064466796 | 0.950633914 | 0.982610344 | -4.913737127 |
| **A_51_P259861** | Mettl9 | -0.005764014 | -0.223022603 | 0.211494574 | 11.82974946 | -0.064466084 | 0.950634458 | 0.982610344 | -4.913737152 |
| **A_51_P432563** | Tlr9 | 0.01471779 | -0.541470536 | 0.570906116 | 11.86999287 | 0.064298957 | 0.950762233 | 0.982637582 | -4.913743053 |
| **A_51_P389957** | Rgs14 | -0.007486447 | -0.290402266 | 0.275429372 | 8.825953027 | -0.064298627 | 0.950762485 | 0.982637582 | -4.913743064 |
| **A_55_P2102350** | Stambpl1 | 0.009438565 | -0.34789729 | 0.36677442 | 9.604587178 | 0.064181899 | 0.950851728 | 0.982677274 | -4.913747176 |
| **A_52_P185044** | Adipor2 | 0.00704612 | -0.260041849 | 0.274134088 | 14.11653138 | 0.064103077 | 0.950911992 | 0.982687013 | -4.913749949 |
| **A_52_P551903** | Slc33a1 | -0.011626626 | -0.455088338 | 0.431835086 | 8.806316801 | -0.063706061 | 0.951215534 | 0.982873176 | -4.913763861 |
| **A_55_P2064442** | Pabpn1 | 0.006836963 | -0.25404022 | 0.267714145 | 13.35367249 | 0.063681064 | 0.951234646 | 0.982873176 | -4.913764734 |
| **A_55_P2035662** | Aebp1 | -0.006518456 | -0.255293689 | 0.242256778 | 11.82195258 | -0.063667934 | 0.951244685 | 0.982873176 | -4.913765193 |
| **A_55_P1980556** | NA | 0.004147073 | -0.154712022 | 0.163006168 | 7.664215847 | 0.063432643 | 0.951424584 | 0.982908783 | -4.913773393 |
| **A_55_P2033795** | Fabp6 | 0.004152073 | -0.155242621 | 0.163546766 | 7.692679394 | 0.06329571 | 0.951529282 | 0.982908783 | -4.913778151 |
| **A_51_P473940** | Slc22a15 | -0.004579511 | -0.180394992 | 0.171235969 | 8.183974454 | -0.063291476 | 0.951532519 | 0.982908783 | -4.913778298 |
| **A_55_P2042773** | Ctdnep1 | -0.004576989 | -0.180565071 | 0.171411093 | 11.07486352 | -0.063194572 | 0.951606612 | 0.982908783 | -4.913781658 |
| **A_55_P1979402** | Tigar | 0.004776909 | -0.179032517 | 0.188586334 | 8.177426037 | 0.063148399 | 0.951641916 | 0.982908783 | -4.913783258 |
| **A_55_P2010936** | Fbxo17 | -0.007963124 | -0.314403416 | 0.298477168 | 8.680320218 | -0.063142356 | 0.951646536 | 0.982908783 | -4.913783467 |
| **A_55_P2185160** | NA | 0.00826606 | -0.309978067 | 0.326510186 | 13.39950485 | 0.063113364 | 0.951668704 | 0.982908783 | -4.913784471 |
| **A_55_P2051962** | Sec61b | 0.004993799 | -0.187336736 | 0.197324334 | 14.99431211 | 0.063090799 | 0.951685957 | 0.982908783 | -4.913785251 |
| **A_52_P494380** | Dgkq | -0.006499243 | -0.257177642 | 0.244179157 | 9.319210847 | -0.06299833 | 0.95175666 | 0.982929284 | -4.913788448 |
| **A_51_P174415** | Retreg2 | -0.007555007 | -0.299410114 | 0.2843001 | 10.52332044 | -0.062900018 | 0.951831831 | 0.982949745 | -4.913791842 |
| **A_51_P322542** | Bbx | 0.00763448 | -0.287575205 | 0.302844165 | 9.215586516 | 0.062839402 | 0.951878179 | 0.982949745 | -4.913793932 |
| **A_55_P2183549** | NA | 0.006656962 | -0.251278294 | 0.264592218 | 11.10205609 | 0.062711697 | 0.951975825 | 0.982998062 | -4.913798328 |
| **A_52_P488305** | Ndel1 | 0.007721462 | -0.29219222 | 0.307635145 | 8.315419946 | 0.062558521 | 0.952092948 | 0.983066485 | -4.913803589 |
| **A_55_P2038358** | Acot1 | -0.016544008 | -0.660202252 | 0.627114237 | 11.5086722 | -0.062455196 | 0.952171954 | 0.983095545 | -4.913807131 |
| **A_55_P2159710** | Acvr1 | 0.005706567 | -0.217264124 | 0.228677257 | 11.21617966 | 0.062188531 | 0.952375859 | 0.983253551 | -4.913816245 |
| **A_51_P374464** | Gstp1 | 0.010948584 | -0.419486595 | 0.441383763 | 12.91612076 | 0.061806391 | 0.952668069 | 0.983478968 | -4.913829237 |
| **A_55_P2091551** | Arhgap9 | -0.008854848 | -0.357292113 | 0.339582417 | 11.35898366 | -0.061750389 | 0.952710893 | 0.983478968 | -4.913831135 |
| **A_52_P311297** | Als2 | -0.007589 | -0.306442803 | 0.291264803 | 8.38459765 | -0.061703379 | 0.952746841 | 0.983478968 | -4.913832726 |
| **A_55_P1961335** | Ctsk | 0.005915968 | -0.228370163 | 0.240202099 | 8.606483654 | 0.061356761 | 0.953011897 | 0.98370004 | -4.913844422 |
| **A_55_P1963491** | Ptma | -0.007728505 | -0.314550863 | 0.299093853 | 13.73451899 | -0.061205674 | 0.953127435 | 0.983754912 | -4.9138495 |
| **A_55_P1953391** | Rbfox1 | 0.003967956 | -0.153723001 | 0.161658912 | 7.706639301 | 0.061142501 | 0.953175744 | 0.983754912 | -4.913851619 |
| **A_51_P461884** | Abl2 | -0.006630309 | -0.270363038 | 0.25710242 | 9.609053689 | -0.06108758 | 0.953217742 | 0.983754912 | -4.91385346 |
| **A_55_P1962339** | Trim41 | -0.006760751 | -0.276127496 | 0.262605993 | 12.86604026 | -0.060986564 | 0.953294991 | 0.983766928 | -4.913856841 |
| **A_55_P2100495** | Synb | 0.004177434 | -0.162396001 | 0.17075087 | 7.76120818 | 0.06093784 | 0.953332252 | 0.983766928 | -4.91385847 |
| **A_51_P252363** | Stx12 | 0.006347758 | -0.247091518 | 0.259787034 | 10.62344166 | 0.060859675 | 0.953392027 | 0.983766928 | -4.913861081 |
| **A_55_P2008512** | B4galt7 | -0.004318049 | -0.176871368 | 0.16823527 | 8.359920756 | -0.060806137 | 0.953432969 | 0.983766928 | -4.913862867 |
| **A_55_P2112170** | Crtc1 | 0.004144399 | -0.162126285 | 0.170415082 | 9.39072028 | 0.060566018 | 0.953616597 | 0.983903876 | -4.913870858 |
| **A_55_P2139901** | Tsr3 | 0.005353197 | -0.210163483 | 0.220869877 | 11.6766111 | 0.060355315 | 0.953777732 | 0.983972424 | -4.913877845 |
| **A_55_P2147160** | NA | -0.011872371 | -0.489921012 | 0.46617627 | 9.047902822 | -0.06034601 | 0.953784848 | 0.983972424 | -4.913878153 |
| **A_52_P653937** | Rpl21 | 0.004601505 | -0.182171741 | 0.191374751 | 8.283974524 | 0.059864369 | 0.954153193 | 0.984299892 | -4.913894029 |
| **A_55_P2017160** | Cdc14a | 0.005381999 | -0.213678799 | 0.224442797 | 8.588362661 | 0.059698321 | 0.954280185 | 0.98432029 | -4.913899473 |
| **A_51_P357664** | Hras | 0.010099245 | -0.401193425 | 0.421391915 | 11.64632127 | 0.059665204 | 0.954305512 | 0.98432029 | -4.913900557 |
| **A_66_P127722** | NA | -0.006076085 | -0.253910148 | 0.241757978 | 12.68792836 | -0.05957253 | 0.954376389 | 0.98432029 | -4.913903587 |
| **A_55_P2161858** | Pdlim7 | -0.005563177 | -0.234420316 | 0.223293961 | 8.58665369 | -0.059066553 | 0.954763366 | 0.98432029 | -4.913920049 |
| **A_51_P403273** | Tax1bp1 | 0.006715265 | -0.269580687 | 0.283011218 | 10.9083163 | 0.059057064 | 0.954770624 | 0.98432029 | -4.913920356 |
| **A_51_P243808** | Traf3ip3 | -0.006898877 | -0.290756698 | 0.276958944 | 8.575536802 | -0.059055549 | 0.954771782 | 0.98432029 | -4.913920405 |
| **A_55_P2072980** | Sptan1 | 0.005364505 | -0.215594393 | 0.226323404 | 8.194021651 | 0.058993119 | 0.95481953 | 0.98432029 | -4.913922426 |
| **A_55_P2066897** | A230045G11Rik | -0.008554782 | -0.361056975 | 0.34394741 | 8.66012207 | -0.058969892 | 0.954837295 | 0.98432029 | -4.913923177 |
| **A_55_P2091626** | NA | 0.004435794 | -0.178344619 | 0.187216206 | 15.69539772 | 0.058969153 | 0.95483786 | 0.98432029 | -4.913923201 |
| **A_55_P2006519** | Clip1 | -0.005528832 | -0.233842414 | 0.222784751 | 9.191329124 | -0.058841644 | 0.954935383 | 0.98432029 | -4.913927321 |
| **A_51_P226417** | Rraga | 0.007402215 | -0.298562553 | 0.313366982 | 11.1261967 | 0.058785936 | 0.95497799 | 0.98432029 | -4.913929117 |

| | | | | | | | | | |
|---|---|---|---|---|---|---|---|---|---|
| **A_55_P2048655** | NA | 0.00481275 | -0.194344181 | 0.203969682 | 15.41272097 | 0.058719328 | 0.955028935 | 0.98432029 | -4.913931264 |
| **A_51_P189272** | Tmcc3 | 0.013195715 | -0.533039371 | 0.559430801 | 10.4601655 | 0.058699785 | 0.955043882 | 0.98432029 | -4.913931893 |
| **A_55_P2006499** | Esrrg | 0.004119738 | -0.166740236 | 0.174979713 | 7.693215313 | 0.058588512 | 0.95512899 | 0.98432029 | -4.913935472 |
| **A_55_P2075080** | Fat2 | 0.005594118 | -0.22641638 | 0.237604615 | 8.195468086 | 0.058587794 | 0.955129539 | 0.98432029 | -4.913935495 |
| **A_55_P2029528** | NA | 0.009532809 | -0.386386903 | 0.405452521 | 8.826116141 | 0.058505535 | 0.955192455 | 0.98432029 | -4.913938136 |
| **A_55_P1980854** | Ankrd24 | -0.006959392 | -0.2968761 | 0.282957315 | 9.356599573 | -0.058328562 | 0.955327814 | 0.98432029 | -4.913943806 |
| **A_55_P2149615** | Bex3 | -0.005125234 | -0.218907713 | 0.208657245 | 11.7078512 | -0.058253871 | 0.955384942 | 0.98432029 | -4.913946194 |
| **A_51_P103757** | Kat8 | 0.00498049 | -0.20293233 | 0.21289331 | 11.15206482 | 0.058206841 | 0.955420914 | 0.98432029 | -4.913947696 |
| **A_55_P2050483** | Abcc12 | -0.004278698 | -0.183004736 | 0.17444734 | 7.705358661 | -0.058171067 | 0.955448276 | 0.98432029 | -4.913948838 |
| **A_55_P2112934** | C030034I22Rik | 0.004766535 | -0.194414602 | 0.203947672 | 8.361433762 | 0.058148397 | 0.955465616 | 0.98432029 | -4.913949561 |
| **A_52_P50287** | Bend7 | -0.007432024 | -0.318611075 | 0.303747027 | 8.071123931 | -0.058033659 | 0.955553377 | 0.98432029 | -4.913953217 |
| **A_55_P2275057** | Nrap | -0.003726425 | -0.159754187 | 0.152301337 | 7.664648029 | -0.058032824 | 0.955554015 | 0.98432029 | -4.913953243 |
| **A_52_P223704** | Faah | -0.010110303 | -0.433681286 | 0.41346068 | 8.516523244 | -0.057999087 | 0.95557982 | 0.98432029 | -4.913954317 |
| **A_51_P222590** | Arf6 | -0.007407989 | -0.317828484 | 0.303012506 | 11.02593149 | -0.057987332 | 0.955588811 | 0.98432029 | -4.913954691 |
| **A_55_P2126014** | NA | -0.004208145 | -0.180597062 | 0.172180773 | 16.15510508 | -0.057969902 | 0.955602143 | 0.98432029 | -4.913955245 |
| **A_55_P1958097** | Eno1 | 0.007768198 | -0.317861577 | 0.333397973 | 14.5005639 | 0.057966806 | 0.955604511 | 0.98432029 | -4.913955343 |
| **A_52_P168575** | Fam169b | 0.004298433 | -0.176268622 | 0.184865489 | 7.798808151 | 0.057843542 | 0.955698794 | 0.98432029 | -4.913959258 |
| **A_51_P396297** | Hsdl1 | -0.004599445 | -0.197821237 | 0.188622348 | 8.143106168 | -0.05784056 | 0.955701074 | 0.98432029 | -4.913959353 |
| **A_55_P2022049** | Klf15 | 0.008261992 | -0.338938446 | 0.355462429 | 10.07284907 | 0.057821278 | 0.955715823 | 0.98432029 | -4.913959964 |
| **A_55_P2043267** | Cdc42se1 | -0.007770658 | -0.334587961 | 0.319046644 | 9.864823009 | -0.057774469 | 0.955751627 | 0.98432029 | -4.913961448 |
| **A_52_P417437** | Gab1 | -0.004502319 | -0.194845188 | 0.185840549 | 8.414013038 | -0.057475517 | 0.955980295 | 0.984503337 | -4.913970897 |
| **A_55_P2127557** | Abracl | -0.006640675 | -0.299362651 | 0.286081301 | 11.12317263 | -0.057716403 | 0.956135735 | 0.984544402 | -4.86441948 |
| **A_55_P1962926** | Snapc3 | -0.004730496 | -0.205788814 | 0.196327822 | 8.1412196 | -0.057169948 | 0.956214029 | 0.984544402 | -4.913980504 |
| **A_55_P2043767** | NA | -0.007241338 | -0.315837181 | 0.301354506 | 16.13801398 | -0.057017991 | 0.956330265 | 0.984544402 | -4.913985262 |
| **A_55_P1988623** | Add3 | -0.007094062 | -0.309887217 | 0.295699093 | 10.46175992 | -0.05692881 | 0.956398483 | 0.984544402 | -4.913988049 |
| **A_51_P447866** | Sash3 | 0.008837345 | -0.36861266 | 0.386287351 | 9.636602225 | 0.056891263 | 0.956427204 | 0.984544402 | -4.913989221 |
| **A_51_P390078** | NA | -0.005085779 | -0.222486684 | 0.212315126 | 8.903693486 | -0.056843306 | 0.956463889 | 0.984544402 | -4.913990717 |
| **A_51_P306017** | Dll1 | 0.003880999 | -0.162251528 | 0.170013526 | 7.88265504 | 0.056763876 | 0.956524648 | 0.984544402 | -4.913993192 |
| **A_55_P1968998** | Tmub2 | -0.00462083 | -0.202564145 | 0.193322486 | 10.20512749 | -0.056723406 | 0.956555605 | 0.984544402 | -4.913994451 |
| **A_55_P2073647** | Elp2 | -0.005573705 | -0.244545575 | 0.233398165 | 11.262803 | -0.056673546 | 0.956593745 | 0.984544402 | -4.913996002 |
| **A_52_P468343** | Bdh1 | 0.012493508 | -0.523780568 | 0.548767585 | 9.105040818 | 0.056608389 | 0.956643587 | 0.984544402 | -4.913998026 |
| **A_55_P2162503** | Tbx20 | -0.003710206 | -0.163610041 | 0.156189628 | 7.665135297 | -0.056381058 | 0.956817485 | 0.984544402 | -4.91400507 |
| **A_55_P2151907** | NA | 0.006599375 | -0.277817292 | 0.291016042 | 15.25994044 | 0.056380763 | 0.956817711 | 0.984544402 | -4.914005079 |
| **A_51_P389386** | Nsmce2 | 0.003734963 | -0.157319666 | 0.164789592 | 10.14135753 | 0.05635031 | 0.956841007 | 0.984544402 | -4.914006021 |
| **A_55_P2061134** | NA | 0.011245352 | -0.474686921 | 0.497177625 | 15.14004355 | 0.056231607 | 0.956931811 | 0.984544402 | -4.914009686 |
| **A_51_P380401** | Cars2 | 0.005295268 | -0.223784031 | 0.234374566 | 9.394248428 | 0.05616752 | 0.956980835 | 0.984544402 | -4.914011661 |
| **A_55_P2160128** | Pls1 | 0.008820834 | -0.373442667 | 0.391084335 | 8.478490238 | 0.056069928 | 0.95705549 | 0.984544402 | -4.914014665 |
| **A_55_P2114915** | Ube2i | -0.00733021 | -0.32565431 | 0.31099389 | 13.80838014 | -0.055953867 | 0.957144275 | 0.984544402 | -4.914018231 |
| **A_55_P2018944** | Snx27 | -0.003668682 | -0.163016643 | 0.155679278 | 9.459026482 | -0.055943133 | 0.957152486 | 0.984544402 | -4.91401856 |
| **A_55_P2027653** | Arhgap25 | -0.007146207 | -0.317827635 | 0.303535221 | 9.810008169 | -0.055891206 | 0.957192209 | 0.984544402 | -4.914020153 |
| **A_66_P136714** | Scpep1 | -0.008909255 | -0.396283102 | 0.378464591 | 12.43179683 | -0.055884877 | 0.957197051 | 0.984544402 | -4.914020347 |
| **A_55_P2037101** | Exoc2 | -0.006722449 | -0.299446352 | 0.286001454 | 10.51656131 | -0.055802352 | 0.957260182 | 0.984544402 | -4.914022874 |
| **A_55_P2107025** | Flad1 | -0.003831388 | -0.170886475 | 0.163223699 | 8.177315835 | -0.055728788 | 0.957316458 | 0.984544402 | -4.914025124 |
| **A_55_P2010301** | Serpina1c | -0.004500746 | -0.200784248 | 0.191782756 | 17.84295591 | -0.055716505 | 0.957325854 | 0.984544402 | -4.9140255 |
| **A_52_P407796** | Mdh1 | 0.008406935 | -0.358335915 | 0.375149786 | 13.93926917 | 0.055700519 | 0.957338083 | 0.984544402 | -4.914025988 |
| **A_55_P2107667** | Enah | -0.004023113 | -0.179795675 | 0.171749448 | 7.756527326 | -0.055615313 | 0.957403266 | 0.984544402 | -4.91402859 |
| **A_55_P2045755** | Plcxd2 | 0.013758218 | -0.588619519 | 0.616135955 | 9.787731461 | 0.05549789 | 0.957493095 | 0.984544402 | -4.914032168 |
| **A_55_P2013645** | Tubg2 | -0.004625472 | -0.207201466 | 0.197950522 | 8.147533333 | -0.055481895 | 0.957505331 | 0.984544402 | -4.914032655 |
| **A_55_P2179498** | Ppp1r8 | 0.007928665 | -0.339404913 | 0.355262243 | 9.607468238 | 0.05546723 | 0.95751655 | 0.984544402 | -4.914033101 |
| **A_65_P17492** | Med29 | 0.004776264 | -0.204607933 | 0.214160461 | 9.909227584 | 0.055427794 | 0.957546719 | 0.984544402 | -4.9140343 |

| | | | | | | | | | |
|---|---|---|---|---|---|---|---|---|---|
| **A_65_P20641** | NA | 0.011119359 | -0.476354225 | 0.498592943 | 10.00872533 | 0.055425787 | 0.957548255 | 0.984544402 | -4.914034361 |
| **A_55_P1966765** | NA | 0.007890078 | -0.340530748 | 0.356310904 | 12.19914346 | 0.05502504 | 0.957854834 | 0.984763431 | -4.9140465 |
| **A_55_P2173837** | Mpp6 | 0.011690531 | -0.505573109 | 0.528954172 | 12.52688799 | 0.054916834 | 0.957937615 | 0.984763431 | -4.914049763 |
| **A_55_P2145376** | Fastk | 0.00430013 | -0.186583681 | 0.195183941 | 8.286798085 | 0.054738846 | 0.958073783 | 0.984763431 | -4.914055116 |
| **A_55_P2125496** | Gm10731 | 0.010777006 | -0.467651024 | 0.489205036 | 11.56817774 | 0.054734946 | 0.958076766 | 0.984763431 | -4.914055233 |
| **A_52_P577384** | Il18bp | -0.012565962 | -0.571368987 | 0.546237062 | 10.62349417 | -0.054641196 | 0.958148489 | 0.984763431 | -4.914058045 |
| **A_52_P597860** | Wasf2 | -0.00555135 | -0.252659315 | 0.241556615 | 9.498450401 | -0.054587731 | 0.958189393 | 0.984763431 | -4.914059647 |
| **A_55_P2058953** | Rpl13a | -0.005609027 | -0.255407268 | 0.244189214 | 16.42165408 | -0.054560876 | 0.958209939 | 0.984763431 | -4.914060451 |
| **A_51_P462192** | Olr1 | 0.006272676 | -0.273094415 | 0.285639767 | 8.109001293 | 0.054558296 | 0.958211912 | 0.984763431 | -4.914060528 |
| **A_55_P2040362** | Bola2 | 0.00682668 | -0.297271679 | 0.310925039 | 13.77322798 | 0.05454798 | 0.958219805 | 0.984763431 | -4.914060837 |
| **A_52_P684130** | 2510002D24Rik | -0.005340127 | -0.243797352 | 0.233117099 | 8.676406956 | -0.054415704 | 0.958321003 | 0.984815071 | -4.914064789 |
| **A_52_P581390** | Kif1c | -0.003514753 | -0.160883143 | 0.153853636 | 7.895038567 | -0.05427009 | 0.958432407 | 0.984852375 | -4.91406913 |
| **A_55_P2236291** | Plpp1 | -0.00414348 | -0.189782084 | 0.181495125 | 8.115785278 | -0.054235059 | 0.958459208 | 0.984852375 | -4.914070172 |
| **A_55_P1960960** | Zdhhc9 | -0.010598407 | -0.486408568 | 0.465211755 | 12.65426699 | -0.054124021 | 0.95854416 | 0.984887309 | -4.914073472 |
| **A_52_P475886** | Rc3h1 | 0.00423843 | -0.186604719 | 0.195081578 | 8.394959947 | 0.053964928 | 0.958665878 | 0.984960015 | -4.914078189 |
| **A_55_P2016326** | NA | 0.005298016 | -0.233600863 | 0.244196895 | 10.07501673 | 0.053886797 | 0.958725655 | 0.984961496 | -4.9140805 |
| **A_52_P284958** | Ubtd2 | -0.013556268 | -0.628076096 | 0.60096356 | 9.30422807 | -0.053602796 | 0.958942942 | 0.984961496 | -4.914088872 |
| **A_51_P286357** | Adgre4 | 0.005289792 | -0.234715742 | 0.245295326 | 8.476919725 | 0.053555062 | 0.958979463 | 0.984961496 | -4.914090275 |
| **A_55_P2131899** | Sox4 | -0.004437404 | -0.205848174 | 0.196973367 | 7.956818413 | -0.053533968 | 0.958995602 | 0.984961496 | -4.914090895 |
| **A_55_P1975326** | Tmem259 | 0.005238645 | -0.232784158 | 0.243261449 | 13.12143415 | 0.05347904 | 0.959037628 | 0.984961496 | -4.914092507 |
| **A_51_P270364** | Mmaa | 0.003875359 | -0.172329994 | 0.180080712 | 10.14645446 | 0.053441181 | 0.959066594 | 0.984961496 | -4.914093617 |
| **A_51_P464918** | Mefv | 0.008017039 | -0.357126455 | 0.373160533 | 8.669331381 | 0.053349899 | 0.959136434 | 0.984961496 | -4.91409629 |
| **A_52_P172405** | Washc4 | 0.006108458 | -0.272381149 | 0.284598065 | 8.518259129 | 0.053297373 | 0.959176623 | 0.984961496 | -4.914097827 |
| **A_51_P206235** | Slc39a5 | 0.009600329 | -0.428931935 | 0.448132592 | 9.473204762 | 0.053194624 | 0.959255237 | 0.984961496 | -4.914100828 |
| **A_55_P1968523** | Btbd1 | 0.006029064 | -0.269455014 | 0.281513142 | 10.50205131 | 0.053178563 | 0.959267526 | 0.984961496 | -4.914101296 |
| **A_55_P2067960** | Cplx2 | -0.006616475 | -0.308943546 | 0.295710597 | 13.99920044 | -0.053178101 | 0.959267879 | 0.984961496 | -4.91410131 |
| **A_55_P2415372** | Mta1 | -0.006110829 | -0.286033244 | 0.273811586 | 8.368871746 | -0.053045145 | 0.959369607 | 0.984961496 | -4.914105183 |
| **A_55_P1991964** | Iqsec2 | 0.004879024 | -0.218774161 | 0.228532209 | 8.508762899 | 0.053007954 | 0.959398063 | 0.984961496 | -4.914106265 |
| **A_52_P161237** | Cpa1 | 0.024411081 | -1.094657006 | 1.143479168 | 8.73806397 | 0.053004615 | 0.959400617 | 0.984961496 | -4.914106362 |
| **A_55_P2107402** | NA | -0.005558149 | -0.261268397 | 0.250152098 | 8.422886481 | -0.052815972 | 0.959544954 | 0.984961496 | -4.914111837 |
| **A_55_P1975420** | Dhx40 | -0.004438458 | -0.208731401 | 0.199854485 | 11.32786087 | -0.052791251 | 0.959563869 | 0.984961496 | -4.914112553 |
| **A_55_P2442017** | Samd8 | -0.005455526 | -0.25668663 | 0.245775578 | 8.274890007 | -0.052765056 | 0.959583911 | 0.984961496 | -4.914113312 |
| **A_55_P1963585** | Mbd5 | 0.00472869 | -0.213262755 | 0.222720136 | 8.530985369 | 0.052708981 | 0.959626817 | 0.984961496 | -4.914114934 |
| **A_55_P1996431** | NA | -0.003352054 | -0.157952848 | 0.151248739 | 7.704486658 | -0.052684456 | 0.959645582 | 0.984961496 | -4.914115643 |
| **A_55_P1970700** | Rpsa | 0.006376611 | -0.288019049 | 0.300772272 | 16.49881734 | 0.052631013 | 0.959686474 | 0.984961496 | -4.914117186 |
| **A_55_P2153580** | Zfp655 | 0.003716768 | -0.168653157 | 0.176086692 | 7.895930547 | 0.052394668 | 0.959867313 | 0.984973911 | -4.914123994 |
| **A_55_P1979694** | NA | 0.006428645 | -0.291758496 | 0.304615787 | 14.2104867 | 0.052385819 | 0.959874083 | 0.984973911 | -4.914124249 |
| **A_52_P390114** | Eif4b | 0.006717628 | -0.305196521 | 0.318631778 | 8.558922322 | 0.05233161 | 0.959915562 | 0.984973911 | -4.914125806 |
| **A_51_P369762** | Nmrk2 | -0.003428117 | -0.162701543 | 0.15584531 | 7.677094829 | -0.052299249 | 0.959940323 | 0.984973911 | -4.914126734 |
| **A_55_P2044242** | Slc13a5 | -0.003563424 | -0.169490988 | 0.16236414 | 7.649244686 | -0.052183372 | 0.960028988 | 0.984973911 | -4.914130055 |
| **A_51_P380750** | Cbfa2t3 | -0.007590787 | -0.361198129 | 0.346016555 | 9.719209958 | -0.052161336 | 0.960045849 | 0.984973911 | -4.914130686 |
| **A_55_P2074942** | Slc6a4 | 0.008777229 | -0.401075076 | 0.418629533 | 8.358655028 | 0.052037117 | 0.960140898 | 0.984973911 | -4.914134236 |
| **A_55_P2066778** | AB124611 | -0.007008245 | -0.334379775 | 0.320363285 | 8.609603696 | -0.052017748 | 0.960155719 | 0.984973911 | -4.914134789 |
| **A_55_P2027067** | Phc2 | -0.007596313 | -0.362451037 | 0.347258411 | 12.57281935 | -0.052015818 | 0.960157195 | 0.984973911 | -4.914134844 |
| **A_55_P1979818** | Srp54b | 0.00375057 | -0.172137064 | 0.179638205 | 8.055857239 | 0.051813772 | 0.960311796 | 0.985080226 | -4.914140597 |
| **A_55_P1985804** | Thumpd2 | 0.005286461 | -0.243143168 | 0.25371609 | 8.657256508 | 0.051706461 | 0.960393909 | 0.985112177 | -4.914143644 |
| **A_51_P317542** | Slc24a2 | -0.003294905 | -0.158902203 | 0.152312394 | 7.686712903 | -0.051451269 | 0.96058918 | 0.985244878 | -4.914150865 |
| **A_55_P2030506** | Rp9 | -0.007486119 | -0.361354361 | 0.346382123 | 11.73273803 | -0.051404164 | 0.960625224 | 0.985244878 | -4.914152194 |
| **A_55_P2151273** | Elovl1 | 0.004812705 | -0.223135191 | 0.232760601 | 13.45597892 | 0.051302299 | 0.960703172 | 0.985258843 | -4.914155063 |
| **A_51_P368855** | Pfdn6 | 0.007484216 | -0.347336508 | 0.362304939 | 11.69589141 | 0.051253143 | 0.960740787 | 0.985258843 | -4.914156446 |

| | | | | | | | | | |
|---|---|---|---|---|---|---|---|---|---|
| **A_65_P19784** | Asb7 | 0.004283 | -0.199315021 | 0.207881021 | 9.665032542 | 0.051116104 | 0.96084565 | 0.985314105 | -4.914160294 |
| **A_52_P307893** | Pip4k2b | -0.006412459 | -0.313364282 | 0.300539363 | 10.29009578 | -0.050761869 | 0.961116719 | 0.985528369 | -4.914170192 |
| **A_55_P1963198** | NA | 0.003280503 | -0.154038027 | 0.160599034 | 7.67132163 | 0.050669172 | 0.961187654 | 0.985528369 | -4.914172771 |
| **A_65_P11504** | Zfp85 | 0.003615001 | -0.169833579 | 0.177063581 | 8.083420233 | 0.050643166 | 0.961207554 | 0.985528369 | -4.914173494 |
| **A_52_P253915** | Cds2 | -0.00455784 | -0.223708432 | 0.214592752 | 8.873407064 | -0.050535858 | 0.96128967 | 0.985560284 | -4.914176472 |
| **A_55_P2093014** | NA | 0.007893135 | -0.372316224 | 0.388102495 | 15.36198088 | 0.050444049 | 0.961359926 | 0.985580037 | -4.914179015 |
| **A_66_P120770** | Ywhaz | 0.003893168 | -0.184648487 | 0.192434823 | 8.613326464 | 0.05017403 | 0.961566558 | 0.985739592 | -4.914186467 |
| **A_55_P1970841** | Nr2c2 | 0.006274092 | -0.299657355 | 0.312205539 | 9.371084985 | 0.049832187 | 0.961828158 | 0.985854473 | -4.914195844 |
| **A_51_P263993** | Lipc | -0.009131262 | -0.454862514 | 0.436599991 | 8.581430504 | -0.049778396 | 0.961869324 | 0.985854473 | -4.914197313 |
| **A_55_P2162394** | Pdp1 | 0.00350029 | -0.167392264 | 0.174392844 | 8.107722086 | 0.049769594 | 0.961876059 | 0.985854473 | -4.914197554 |
| **A_55_P2185214** | NA | -0.006276563 | -0.312766359 | 0.300213232 | 16.47857441 | -0.049760998 | 0.961882638 | 0.985854473 | -4.914197788 |
| **A_55_P2025911** | Sms | 0.006039242 | -0.289624375 | 0.301702859 | 9.097286055 | 0.049632683 | 0.961980834 | 0.985896037 | -4.914201287 |
| **A_55_P2028418** | Hacd2 | 0.011016424 | -0.529409055 | 0.551441902 | 11.46410156 | 0.049532245 | 0.962057698 | 0.985896037 | -4.914204019 |
| **A_55_P1990663** | Ddx5 | 0.005035193 | -0.242201657 | 0.252272042 | 12.22635251 | 0.049486427 | 0.962092762 | 0.985896037 | -4.914205263 |
| **A_55_P1956712** | Med28 | -0.006348531 | -0.318651215 | 0.305954154 | 12.16423119 | -0.049394739 | 0.962162929 | 0.985896037 | -4.91420775 |
| **A_51_P269020** | 1700031F05Rik | -0.003127542 | -0.157042694 | 0.15078761 | 7.633954598 | -0.049374757 | 0.962178221 | 0.985896037 | -4.914208291 |
| **A_55_P1993909** | Taz | -0.003338808 | -0.167976846 | 0.161299229 | 10.1526059 | -0.049277024 | 0.962253016 | 0.98592041 | -4.914210936 |
| **A_51_P332676** | Top1mt | 0.009914098 | -0.480563332 | 0.500391529 | 10.53887216 | 0.049115366 | 0.962376733 | 0.985963758 | -4.914215299 |
| **A_55_P1953136** | NA | 0.00608623 | -0.295181297 | 0.307353756 | 14.53271739 | 0.049088437 | 0.962397342 | 0.985963758 | -4.914216024 |
| **A_65_P07361** | Trim14 | -0.008027751 | -0.405995274 | 0.389939773 | 9.187861557 | -0.049015047 | 0.962453507 | 0.985969039 | -4.914217999 |
| **A_55_P1966194** | Plek | 0.014073408 | -0.685904457 | 0.714051273 | 11.13799946 | 0.048853779 | 0.962576927 | 0.986043215 | -4.914222328 |
| **A_55_P1995623** | Pomp | -0.005202957 | -0.264587801 | 0.254181887 | 13.4916781 | -0.048740368 | 0.962663722 | 0.986051752 | -4.914225364 |
| **A_55_P2185840** | Nnmt | 0.003390814 | -0.165759495 | 0.172541123 | 7.801804862 | 0.048709575 | 0.962687289 | 0.986051752 | -4.914226187 |
| **A_51_P495131** | Prl7c1 | -0.003176234 | -0.162192983 | 0.155840515 | 7.625047002 | -0.048534748 | 0.962821089 | 0.986136542 | -4.914230851 |
| **A_55_P2022895** | Slc12a8 | -0.003236603 | -0.165687233 | 0.159214027 | 7.716573024 | -0.048411801 | 0.962915184 | 0.986180217 | -4.91423412 |
| **A_51_P201884** | Dsp | -0.003459426 | -0.177331326 | 0.170412474 | 7.747406269 | -0.048345701 | 0.962965772 | 0.986180217 | -4.914235874 |
| **A_55_P2013273** | NA | -0.008120244 | -0.417262687 | 0.401022199 | 13.92106819 | -0.048225612 | 0.963057681 | 0.986222089 | -4.914239056 |
| **A_55_P1960616** | Dnajc17 | 0.004592202 | -0.228549449 | 0.237733853 | 8.295038727 | 0.047861282 | 0.963336521 | 0.986432336 | -4.914248658 |
| **A_55_P1986660** | Anapc2 | -0.011082473 | -0.574167524 | 0.552002579 | 13.2302996 | -0.047823996 | 0.963365058 | 0.986432336 | -4.914249637 |
| **A_55_P1992199** | Psg20 | -0.003052832 | -0.158844861 | 0.152739197 | 7.663113574 | -0.047614683 | 0.963525258 | 0.986477004 | -4.914255116 |
| **A_55_P1975400** | Flnc | 0.00935583 | -0.468285163 | 0.486996822 | 8.859386252 | 0.047595282 | 0.963540107 | 0.986477004 | -4.914255623 |
| **A_55_P2113975** | Psg20 | 0.003059835 | -0.153246255 | 0.159365925 | 7.646769024 | 0.047566953 | 0.96356179 | 0.986477004 | -4.914256363 |
| **A_55_P1987483** | Pgm2 | -0.012743787 | -0.665975513 | 0.640487938 | 14.85864702 | -0.04740394 | 0.963686556 | 0.986552483 | -4.914260609 |
| **A_55_P2130109** | Nmt2 | 0.003229034 | -0.163663765 | 0.170121832 | 8.021935862 | 0.047013023 | 0.963985758 | 0.986790623 | -4.914270734 |
| **A_55_P2158151** | NA | -0.00479833 | -0.253045396 | 0.243448735 | 15.85398398 | -0.046966611 | 0.964021281 | 0.986790623 | -4.91427193 |
| **A_55_P1963817** | Rps6 | 0.003163675 | -0.160816979 | 0.167144328 | 16.05725044 | 0.046879435 | 0.964088005 | 0.986793342 | -4.914274174 |
| **A_55_P2068862** | Pigv | 0.003745322 | -0.191088306 | 0.198578949 | 9.747361562 | 0.046709841 | 0.964217812 | 0.986793342 | -4.914278528 |
| **A_52_P199614** | Wdfy4 | -0.006845601 | -0.363606318 | 0.349915116 | 11.31069098 | -0.046624884 | 0.96428284 | 0.986793342 | -4.914280703 |
| **A_66_P107038** | Prcp | -0.004515949 | -0.240120852 | 0.231088954 | 11.76038912 | -0.046574463 | 0.964321432 | 0.986793342 | -4.914281992 |
| **A_55_P2009943** | Spen | 0.006763837 | -0.346652264 | 0.360179938 | 10.51606989 | 0.046503965 | 0.964375392 | 0.986793342 | -4.914283792 |
| **A_55_P1966035** | Apoa2 | 0.006057319 | -0.31070451 | 0.322819148 | 17.0392829 | 0.046465518 | 0.96440482 | 0.986793342 | -4.914284773 |
| **A_55_P2071893** | NA | 0.006199046 | -0.31804665 | 0.330444741 | 11.96602546 | 0.046455143 | 0.964412761 | 0.986793342 | -4.914285037 |
| **A_51_P496934** | Hps3 | -0.004212386 | -0.224998633 | 0.216573862 | 8.46647616 | -0.046359558 | 0.964485924 | 0.986793342 | -4.914287471 |
| **A_55_P2006148** | Aldh3b2 | -0.003195766 | -0.170926956 | 0.164535424 | 7.712601494 | -0.046296093 | 0.964534501 | 0.986793342 | -4.914289084 |
| **A_55_P2032659** | Tox | 0.006729074 | -0.346625717 | 0.360083864 | 8.798677952 | 0.046272982 | 0.964552191 | 0.986793342 | -4.914289671 |
| **A_55_P2062981** | Abca9 | -0.00484347 | -0.260013224 | 0.250326285 | 8.308682237 | -0.046122258 | 0.964667559 | 0.986793342 | -4.914293491 |
| **A_55_P2157632** | 2510039O18Rik | -0.005008483 | -0.269377047 | 0.259360081 | 11.68217849 | -0.046034089 | 0.964735046 | 0.986793342 | -4.91429572 |
| **A_51_P381618** | Pla1a | 0.012204105 | -0.632014886 | 0.656423096 | 12.91089503 | 0.046031549 | 0.96473699 | 0.986793342 | -4.914295784 |
| **A_55_P2305294** | Ptpn21 | 0.003733741 | -0.193638381 | 0.201105863 | 8.526085449 | 0.045966514 | 0.964786771 | 0.986793342 | -4.914297425 |
| **A_55_P2007485** | Prss27 | -0.003140894 | -0.169188015 | 0.162906227 | 7.710841515 | -0.045962656 | 0.964789724 | 0.986793342 | -4.914297523 |

| | | | | | | | | | |
|---|---|---|---|---|---|---|---|---|---|
| **A_55_P2024675** | Shld2 | 0.011442052 | -0.594608269 | 0.617492372 | 10.23221064 | 0.045875246 | 0.964856631 | 0.986809557 | -4.914299725 |
| **A_55_P2027831** | Cryba4 | 0.004728266 | -0.246534686 | 0.255991218 | 8.574083333 | 0.045725305 | 0.964971402 | 0.986842194 | -4.914303492 |
| **A_55_P2115980** | Xlr5c | 0.002923281 | -0.152526852 | 0.158373414 | 7.62812207 | 0.045694363 | 0.964995086 | 0.986842194 | -4.914304268 |
| **A_55_P2001198** | NA | -0.002943492 | -0.159677297 | 0.153790313 | 8.31535377 | -0.045633456 | 0.965041707 | 0.986842194 | -4.914305794 |
| **A_51_P402994** | Ddx3y | 0.051094316 | -2.677630761 | 2.779819393 | 9.619237854 | 0.045498387 | 0.965145097 | 0.986895708 | -4.914309171 |
| **A_55_P2063312** | Mgll | 0.0074657 | -0.392889984 | 0.407821384 | 8.766909501 | 0.045311426 | 0.965288207 | 0.986989831 | -4.914313828 |
| **A_52_P339722** | Csnk1a1 | 0.005386788 | -0.285775947 | 0.296549524 | 11.06372938 | 0.044954928 | 0.965561096 | 0.987103931 | -4.914322656 |
| **A_51_P251352** | Slc25a13 | -0.015524628 | -0.855746181 | 0.824696926 | 12.42283354 | -0.044896301 | 0.965605973 | 0.987103931 | -4.914324101 |
| **A_51_P366672** | Slc36a2 | -0.007418165 | -0.408950945 | 0.394114614 | 11.76611943 | -0.044890939 | 0.965610078 | 0.987103931 | -4.914324233 |
| **A_55_P2062108** | Apold1 | 0.003936557 | -0.209195315 | 0.217068428 | 8.184523236 | 0.044879843 | 0.965618572 | 0.987103931 | -4.914324506 |
| **A_51_P404377** | Rnd2 | -0.003432918 | -0.189698873 | 0.182833036 | 7.910517876 | -0.044783001 | 0.965692702 | 0.987103931 | -4.914326888 |
| **A_55_P1971474** | NA | 0.00765859 | -0.408578352 | 0.423895532 | 9.954589638 | 0.044708634 | 0.965749629 | 0.987103931 | -4.914328714 |
| **A_55_P2114318** | NA | 0.010244889 | -0.546679379 | 0.567169158 | 11.13136945 | 0.044698642 | 0.965757278 | 0.987103931 | -4.914328959 |
| **A_55_P2075195** | Cd302 | -0.006531062 | -0.362860627 | 0.349798504 | 13.63630429 | -0.044536401 | 0.965881472 | 0.987146365 | -4.91433293 |
| **A_51_P214269** | Epha1 | -0.012349773 | -0.686558301 | 0.661858756 | 10.12077026 | -0.044509007 | 0.965902442 | 0.987146365 | -4.9143336 |
| **A_51_P104820** | Alkbh5 | 0.003741535 | -0.201099152 | 0.208582222 | 8.186473739 | 0.044383011 | 0.965998891 | 0.987146365 | -4.914336672 |
| **A_55_P1971094** | Esx1 | -0.002857149 | -0.159298956 | 0.153584658 | 7.622310189 | -0.044377542 | 0.966003078 | 0.987146365 | -4.914336805 |
| **A_55_P2026440** | Ceacam-ps1 | -0.003030986 | -0.169354175 | 0.163292204 | 7.682194821 | -0.044280676 | 0.966077229 | 0.987164292 | -4.914339161 |
| **A_66_P135804** | NA | 0.004534795 | -0.24516617 | 0.254235761 | 13.38672155 | 0.044128652 | 0.966193605 | 0.987164292 | -4.914342848 |
| **A_51_P226364** | Coa3 | 0.006392898 | -0.346669979 | 0.359455775 | 13.53410465 | 0.043997594 | 0.966293931 | 0.987164292 | -4.914346017 |
| **A_51_P455241** | Rrp36 | 0.004153987 | -0.226263568 | 0.234571542 | 10.00517079 | 0.043805916 | 0.966440664 | 0.987164292 | -4.914350633 |
| **A_66_P102062** | NA | 0.0040698 | -0.221890463 | 0.230030062 | 9.003669951 | 0.043764718 | 0.966472202 | 0.987164292 | -4.914351623 |
| **A_51_P101858** | Dlg1 | 0.005077062 | -0.277721558 | 0.287875681 | 8.399430083 | 0.043623284 | 0.966580474 | 0.987164292 | -4.914355014 |
| **A_55_P2052848** | Mif | 0.00814089 | -0.446989499 | 0.463271278 | 16.27079944 | 0.043462952 | 0.966703213 | 0.987164292 | -4.914358844 |
| **A_52_P250590** | Dnajc13 | -0.00666311 | -0.379216648 | 0.365890428 | 9.063224352 | -0.043458173 | 0.966706872 | 0.987164292 | -4.914358958 |
| **A_51_P405089** | NA | 0.014455193 | -0.793863617 | 0.822774004 | 9.800320335 | 0.043453465 | 0.966710476 | 0.987164292 | -4.91435907 |
| **A_55_P2087840** | Gpr50 | -0.002758814 | -0.157098512 | 0.151580885 | 7.654731376 | -0.043433809 | 0.966725523 | 0.987164292 | -4.914359539 |
| **A_51_P288891** | Tbc1d15 | 0.005543554 | -0.304638011 | 0.315725118 | 10.22066998 | 0.043426562 | 0.966731071 | 0.987164292 | -4.914359712 |
| **A_55_P1978681** | Tspan8 | 0.013383863 | -0.736139561 | 0.762907286 | 10.33167399 | 0.043388981 | 0.966759841 | 0.987164292 | -4.914360607 |
| **A_51_P507974** | Brf2 | -0.00378781 | -0.216740571 | 0.20916495 | 8.453611742 | -0.043220337 | 0.966888945 | 0.987164292 | -4.914364614 |
| **A_55_P1952517** | Sult2a1 | 0.01460538 | -0.810482232 | 0.839692993 | 8.661036476 | 0.043012629 | 0.967047955 | 0.987164292 | -4.914369527 |
| **A_51_P212183** | Dmac1 | -0.004755165 | -0.273937793 | 0.264427463 | 12.16518258 | -0.042924155 | 0.967115687 | 0.987164292 | -4.914371613 |
| **A_55_P2072842** | NA | 0.007841686 | -0.436197775 | 0.451881148 | 12.539124 | 0.042911243 | 0.967125572 | 0.987164292 | -4.914371917 |
| **A_55_P2054998** | NA | 0.004605344 | -0.256396515 | 0.265607203 | 10.16407834 | 0.042874758 | 0.967153503 | 0.987164292 | -4.914372776 |
| **A_55_P2065244** | Rbm26 | 0.004902271 | -0.273136446 | 0.282940988 | 9.225927176 | 0.042842545 | 0.967178164 | 0.987164292 | -4.914373533 |
| **A_55_P2065308** | Myo9b | -0.004396296 | -0.253775627 | 0.244983035 | 8.562202178 | -0.042836075 | 0.967183117 | 0.987164292 | -4.914373685 |
| **A_55_P2103682** | NA | 0.004595446 | -0.256398481 | 0.265589374 | 14.27228197 | 0.042783912 | 0.967223051 | 0.987164292 | -4.914374911 |
| **A_55_P2027698** | Ndufa5 | 0.004599906 | -0.256865606 | 0.266065418 | 14.47858134 | 0.042748189 | 0.967250399 | 0.987164292 | -4.914375749 |
| **A_55_P1983638** | Mapk6 | -0.007222142 | -0.418117879 | 0.403673594 | 10.35888799 | -0.042708822 | 0.967280537 | 0.987164292 | -4.914376672 |
| **A_55_P2162136** | Hmgn2 | 0.003624256 | -0.202573971 | 0.209822483 | 12.72175718 | 0.042708778 | 0.967280571 | 0.987164292 | -4.914376673 |
| **A_52_P502141** | Hectd2 | 0.003452253 | -0.192963521 | 0.199868028 | 8.077335062 | 0.042708024 | 0.967281149 | 0.987164292 | -4.91437669 |
| **A_52_P409769** | Mrvi1 | -0.005585493 | -0.324167425 | 0.312996439 | 9.393174157 | -0.042601369 | 0.9673628 | 0.987164292 | -4.914379186 |
| **A_66_P101423** | Ogfod3 | 0.006754111 | -0.378747011 | 0.392255233 | 10.97444377 | 0.042572165 | 0.967385158 | 0.987164292 | -4.914379869 |
| **A_55_P1954850** | Pars2 | 0.004914498 | -0.275805911 | 0.285634908 | 9.837247857 | 0.042539108 | 0.967410465 | 0.987164292 | -4.914380641 |
| **A_52_P142531** | Phc3 | -0.003855072 | -0.224332013 | 0.216621869 | 8.746660318 | -0.042486647 | 0.967450627 | 0.987164292 | -4.914381864 |
| **A_55_P2119969** | Bdh2 | -0.011170337 | -0.65117751 | 0.628836837 | 8.997814749 | -0.042409642 | 0.967509581 | 0.987172334 | -4.914383658 |
| **A_55_P2029517** | Wnk2 | 0.002683832 | -0.151349911 | 0.156717574 | 7.752759819 | 0.042337244 | 0.967565007 | 0.987176776 | -4.914385341 |
| **A_51_P273203** | Gpaa1 | 0.004740269 | -0.268487974 | 0.277968511 | 8.441330607 | 0.042156113 | 0.967703677 | 0.987208182 | -4.91438954 |
| **A_52_P59681** | Rida | -0.013458906 | -0.789823235 | 0.762905423 | 12.40366389 | -0.042123783 | 0.967728429 | 0.987208182 | -4.914390287 |
| **A_51_P165330** | Mrpl33 | 0.00561674 | -0.31913128 | 0.330364759 | 14.18672402 | 0.042026281 | 0.967803075 | 0.987208182 | -4.914392538 |

| | | | | | | | | | |
|---|---|---|---|---|---|---|---|---|---|
| **A_51_P466685** | Fbxl22 | 0.003761347 | -0.213723167 | 0.221245861 | 7.8334897 | 0.042024084 | 0.967804758 | 0.987208182 | -4.914392589 |
| **A_52_P24843** | Pten | 0.005138666 | -0.292613026 | 0.302890359 | 9.087119588 | 0.041935262 | 0.967872759 | 0.987208182 | -4.914394635 |
| **A_51_P452629** | Tlr2 | 0.003050972 | -0.173895131 | 0.179997076 | 9.091237637 | 0.041896763 | 0.967902233 | 0.987208182 | -4.885371308 |
| **A_51_P144581** | Pgrmc1 | 0.006487244 | -0.372061789 | 0.385036277 | 13.36663828 | 0.041641012 | 0.968098037 | 0.987240211 | -4.914401381 |
| **A_55_P2016351** | Eif3c | -0.003139721 | -0.186704836 | 0.180425394 | 13.77786246 | -0.041560819 | 0.968159433 | 0.987240211 | -4.914403212 |
| **A_51_P358574** | Pdzd3 | 0.002724874 | -0.15659026 | 0.162040009 | 7.707161666 | 0.041559723 | 0.968160272 | 0.987240211 | -4.914403237 |
| **A_51_P223720** | Rtcb | 0.004152878 | -0.238838352 | 0.247144107 | 11.14823233 | 0.041528068 | 0.968184507 | 0.987240211 | -4.914403958 |
| **A_52_P96847** | Smlr1 | 0.010884548 | -0.626920421 | 0.648689517 | 12.6041478 | 0.041467294 | 0.968231036 | 0.987240211 | -4.914405342 |
| **A_51_P388973** | Zhx1 | 0.004489741 | -0.258671948 | 0.26765143 | 9.094034537 | 0.04145547 | 0.968240089 | 0.987240211 | -4.914405611 |
| **A_66_P125777** | Slc43a2 | 0.005391047 | -0.311244012 | 0.322026106 | 9.59546331 | 0.041371123 | 0.968304666 | 0.987245794 | -4.914407528 |
| **A_51_P172344** | Foxn3 | 0.004290677 | -0.248058787 | 0.256640142 | 9.321914989 | 0.041314893 | 0.968347716 | 0.987245794 | -4.914408803 |
| **A_51_P500424** | Dnttip1 | 0.004876281 | -0.282938369 | 0.292690931 | 11.97112692 | 0.04116793 | 0.968460234 | 0.987308432 | -4.914412129 |
| **A_55_P2025228** | Akt2 | 0.0033763 | -0.196337827 | 0.203090427 | 12.09868599 | 0.041078583 | 0.968528641 | 0.987326095 | -4.914414145 |
| **A_51_P292368** | Tmco6 | -0.003366228 | -0.203402558 | 0.196670101 | 8.522672562 | -0.040890071 | 0.96867297 | 0.987371022 | -4.914418384 |
| **A_51_P137419** | Cst7 | 0.007944219 | -0.464165563 | 0.480054002 | 8.994603877 | 0.040887581 | 0.968674877 | 0.987371022 | -4.91441844 |
| **A_55_P1965278** | Dmrtc1a | 0.002737854 | -0.160922019 | 0.166397728 | 7.654630721 | 0.040649133 | 0.968857442 | 0.987436714 | -4.914423774 |
| **A_55_P2181071** | NA | -0.004665843 | -0.283695473 | 0.274363786 | 15.5745662 | -0.040631517 | 0.968870929 | 0.987436714 | -4.914424166 |
| **A_55_P2003720** | NA | -0.005409073 | -0.329111101 | 0.318292954 | 15.10680107 | -0.040603236 | 0.968892582 | 0.987436714 | -4.914424797 |
| **A_55_P1997898** | Tfeb | -0.003286582 | -0.203702762 | 0.197129598 | 8.197248894 | -0.039846932 | 0.969471651 | 0.987963001 | -4.914441489 |
| **A_55_P2164070** | Grip1 | 0.006454121 | -0.388051391 | 0.400959634 | 8.786379735 | 0.039752755 | 0.969543759 | 0.987963001 | -4.914443546 |
| **A_55_P2005655** | Ntsr1 | -0.002551378 | -0.158598475 | 0.15349572 | 7.688378922 | -0.039728509 | 0.969562324 | 0.987963001 | -4.914444075 |
| **A_51_P278103** | Stambp | -0.003544533 | -0.221922501 | 0.214833435 | 9.522226355 | -0.039439679 | 0.969783475 | 0.98802154 | -4.914450348 |
| **A_55_P2049473** | Sh3bp5 | -0.006857115 | -0.429419914 | 0.415705684 | 9.034333676 | -0.039430597 | 0.969790429 | 0.98802154 | -4.914450545 |
| **A_52_P48681** | Cldn1 | -0.015888162 | -0.996863419 | 0.965087095 | 10.6143152 | -0.039354897 | 0.969848391 | 0.98802154 | -4.914452181 |
| **A_66_P140533** | NA | -0.004404732 | -0.276697729 | 0.267888265 | 8.594102724 | -0.039306662 | 0.969885324 | 0.98802154 | -4.914453222 |
| **A_55_P2102295** | Tmem184c | -0.004761255 | -0.300125902 | 0.290603391 | 9.21129925 | -0.039169331 | 0.969990477 | 0.98802154 | -4.914456179 |
| **A_66_P132070** | Abra | -0.002492578 | -0.157179584 | 0.152194427 | 7.651965324 | -0.039154185 | 0.970002074 | 0.98802154 | -4.914456504 |
| **A_51_P254855** | Ptgs2 | -0.002578954 | -0.162653535 | 0.157495627 | 7.674218963 | -0.039147536 | 0.970007166 | 0.98802154 | -4.906527306 |
| **A_51_P295449** | Mfsd5 | -0.003290667 | -0.207687817 | 0.201106482 | 9.964758623 | -0.03911941 | 0.970028701 | 0.98802154 | -4.914457251 |
| **A_55_P2065897** | NA | -0.005665635 | -0.359234058 | 0.347902787 | 12.52989664 | -0.038936628 | 0.970168658 | 0.988082694 | -4.914461165 |
| **A_55_P2003648** | Bag5 | 0.003056803 | -0.187848332 | 0.193961938 | 10.2086095 | 0.038907475 | 0.97019098 | 0.988082694 | -4.914461788 |
| **A_51_P320606** | Prl5a1 | -0.002506977 | -0.160609965 | 0.155596011 | 7.641339621 | -0.038529509 | 0.970480392 | 0.98820112 | -4.914469818 |
| **A_55_P2021011** | Sfi1 | 0.002892784 | -0.179558081 | 0.185343649 | 7.785817698 | 0.038525946 | 0.97048312 | 0.98820112 | -4.914469893 |
| **A_55_P2081283** | Dlx4 | -0.002573783 | -0.16514976 | 0.160002193 | 7.71351027 | -0.038467931 | 0.970527544 | 0.98820112 | -4.914471119 |
| **A_55_P2131925** | Kprp | -0.00255893 | -0.164199599 | 0.159081738 | 7.677721796 | -0.03846724 | 0.970528072 | 0.98820112 | -4.914471133 |
| **A_52_P208222** | Pde1c | 0.002448376 | -0.152640018 | 0.157536769 | 7.643276303 | 0.038360299 | 0.97060996 | 0.98820112 | -4.914473387 |
| **A_55_P2035167** | Pcdh7 | -0.002454129 | -0.158050322 | 0.153142064 | 7.691622166 | -0.038324955 | 0.970637023 | 0.98820112 | -4.914474131 |
| **A_55_P2085616** | Nsa2 | 0.003235123 | -0.202074204 | 0.208544451 | 8.910514755 | 0.038288237 | 0.970665139 | 0.98820112 | -4.914474903 |
| **A_55_P2098558** | Slc39a1 | -0.004422385 | -0.285933163 | 0.277088393 | 13.00264981 | -0.038171984 | 0.970754157 | 0.988208026 | -4.914477342 |
| **A_55_P2144110** | Mink1 | 0.004303055 | -0.269799379 | 0.278405489 | 10.5442231 | 0.038145843 | 0.970774174 | 0.988208026 | -4.914477889 |
| **A_55_P2046178** | NA | -0.006176625 | -0.402348947 | 0.389995696 | 15.93857466 | -0.037883518 | 0.970975045 | 0.988360453 | -4.914483361 |
| **A_51_P283708** | Msh2 | 0.003985818 | -0.254760071 | 0.262731707 | 11.75932684 | 0.037430635 | 0.971321838 | 0.98866139 | -4.914492719 |
| **A_52_P354390** | Snrpn | -0.003594892 | -0.238055482 | 0.230865697 | 7.987520963 | -0.037256257 | 0.971455369 | 0.988706338 | -4.914496292 |
| **A_51_P419571** | Polr2d | -0.006021633 | -0.398933487 | 0.386890222 | 10.53766624 | -0.03723937 | 0.9714683 | 0.988706338 | -4.914496637 |
| **A_55_P2175195** | Stxbp4 | 0.002811755 | -0.181319907 | 0.186943418 | 7.95914172 | 0.037104984 | 0.971571207 | 0.988740986 | -4.914499379 |
| **A_65_P10865** | Scfd1 | -0.00592042 | -0.394377738 | 0.382536898 | 10.55018548 | -0.037033297 | 0.971626103 | 0.988740986 | -4.914500837 |
| **A_65_P10399** | Gna13 | -0.002531881 | -0.168830694 | 0.163766933 | 8.092822175 | -0.036994511 | 0.971655804 | 0.988740986 | -4.914501625 |
| **A_55_P2091245** | NA | 0.005569746 | -0.361249286 | 0.372388778 | 16.51974905 | 0.036894934 | 0.971732056 | 0.988766525 | -4.914503643 |
| **A_52_P305073** | Pias1 | 0.002943899 | -0.194463983 | 0.200351781 | 9.601529524 | 0.036236115 | 0.972236568 | 0.989175431 | -4.914516862 |
| **A_51_P363525** | Fbrsl1 | 0.003439184 | -0.227656025 | 0.234534392 | 9.753911931 | 0.036161599 | 0.972293632 | 0.989175431 | -4.914518342 |

| | | | | | | | | | |
|---|---|---|---|---|---|---|---|---|---|
| **A_51_P395111** | Nme3 | 0.006927008 | -0.459277279 | 0.473131295 | 11.96292856 | 0.036103775 | 0.972337913 | 0.989175431 | -4.914519489 |
| **A_55_P1985219** | Pnpla6 | 0.003178679 | -0.210759532 | 0.217116891 | 9.369074034 | 0.036102852 | 0.97233862 | 0.989175431 | -4.914519507 |
| **A_52_P527822** | Slc22a3 | 0.002290046 | -0.152585617 | 0.157165708 | 7.671063658 | 0.035928918 | 0.972471818 | 0.989258869 | -4.914522945 |
| **A_51_P448814** | Zfp90 | 0.00302011 | -0.202572591 | 0.208612812 | 9.288628374 | 0.035694252 | 0.972651526 | 0.989341234 | -4.914527556 |
| **A_55_P1971425** | Serpinb9f | 0.002435781 | -0.163622918 | 0.168494479 | 7.698003875 | 0.035641812 | 0.972691686 | 0.989341234 | -4.914528583 |
| **A_51_P397768** | Csnk2a1 | -0.002825418 | -0.195550999 | 0.189900162 | 8.439370371 | -0.035622679 | 0.972706338 | 0.989341234 | -4.914528957 |
| **A_55_P2115567** | Slc26a1 | 0.011284496 | -0.763959759 | 0.786528751 | 11.091001 | 0.035369328 | 0.972900358 | 0.989362272 | -4.914533892 |
| **A_55_P1999790** | Pkdcc | 0.004811488 | -0.326738966 | 0.336361941 | 12.29611876 | 0.035262486 | 0.97298218 | 0.989362272 | -4.914535962 |
| **A_66_P111285** | Pdp2 | -0.002572703 | -0.179857255 | 0.174711849 | 9.013954282 | -0.035261594 | 0.972982863 | 0.989362272 | -4.914535979 |
| **A_52_P621064** | NA | 0.003856605 | -0.262424959 | 0.270138169 | 8.719644275 | 0.035192266 | 0.973035956 | 0.989362272 | -4.91453732 |
| **A_55_P1963724** | NA | 0.010162299 | -0.693954762 | 0.71427936 | 12.40766299 | 0.035069558 | 0.973129929 | 0.989362272 | -4.914539685 |
| **A_55_P2186833** | NA | 0.007031498 | -0.480700393 | 0.494763389 | 15.1722945 | 0.035030786 | 0.973159622 | 0.989362272 | -4.914540431 |
| **A_55_P2016540** | Cry2 | 0.003800431 | -0.26011958 | 0.267720441 | 11.28641799 | 0.034989979 | 0.973190873 | 0.989362272 | -4.914541215 |
| **A_51_P190961** | Gzmf | -0.002319388 | -0.163479426 | 0.158840651 | 7.653536917 | -0.034970288 | 0.973205953 | 0.989362272 | -4.914541593 |
| **A_51_P328622** | Tlcd2 | -0.006312989 | -0.445359919 | 0.432733941 | 9.075670741 | -0.034938742 | 0.973230112 | 0.989362272 | -4.914542198 |
| **A_51_P102911** | Hoxa11 | 0.002246833 | -0.154063961 | 0.158557627 | 7.676003245 | 0.034927297 | 0.973238877 | 0.989362272 | -4.906335809 |
| **A_51_P101635** | Stub1 | 0.002808852 | -0.193556275 | 0.199173978 | 12.1524138 | 0.034757427 | 0.973368969 | 0.98938884 | -4.914545665 |
| **A_51_P336403** | Emc7 | 0.003003693 | -0.207433166 | 0.213440552 | 12.59823022 | 0.034683025 | 0.973425949 | 0.98938884 | -4.914547083 |
| **A_51_P228817** | NA | 0.003052603 | -0.210820169 | 0.216925376 | 8.418867577 | 0.034681514 | 0.973427107 | 0.98938884 | -4.914547112 |
| **A_55_P2112637** | NA | -0.007651099 | -0.544567796 | 0.529265599 | 8.595062018 | -0.034625821 | 0.973469758 | 0.98938884 | -4.914548171 |
| **A_51_P371051** | Glipr1 | 0.0058231 | -0.406119773 | 0.417765973 | 9.040008317 | 0.034347918 | 0.97368259 | 0.989553119 | -4.91455343 |
| **A_66_P111877** | Snx13 | 0.004197399 | -0.293982931 | 0.302377728 | 9.20923562 | 0.034204594 | 0.973792355 | 0.989612641 | -4.914556126 |
| **A_55_P2038504** | 6-Mar | 0.003206444 | -0.225597382 | 0.232010271 | 8.904251966 | 0.034052077 | 0.973909162 | 0.989679312 | -4.914558982 |
| **A_55_P1986630** | Gm3627 | 0.008157386 | -0.57661387 | 0.592928642 | 8.826473995 | 0.03389598 | 0.97402871 | 0.98970198 | -4.914561892 |
| **A_52_P513842** | Fubp1 | 0.002549611 | -0.18025844 | 0.185357662 | 8.236431444 | 0.033889236 | 0.974033875 | 0.98970198 | -4.914562018 |
| **A_51_P163958** | Spr | 0.003673342 | -0.261266189 | 0.268612872 | 11.54899495 | 0.033689747 | 0.974186657 | 0.989713515 | -4.914565717 |
| **A_51_P122740** | Slc37a3 | -0.002566362 | -0.188041239 | 0.182908514 | 9.14439563 | -0.033621422 | 0.974238986 | 0.989713515 | -4.906359916 |
| **A_55_P2015599** | NA | -0.005339942 | -0.391422719 | 0.380742834 | 12.42093081 | -0.033607741 | 0.974249463 | 0.989713515 | -4.914567231 |
| **A_55_P2146229** | NA | -0.002387364 | -0.175019774 | 0.170245045 | 8.27970534 | -0.033603117 | 0.974253005 | 0.989713515 | -4.914567316 |
| **A_52_P242724** | NA | -0.004976534 | -0.365509859 | 0.355556792 | 9.935361906 | -0.03354013 | 0.974301245 | 0.989713515 | -4.914568477 |
| **A_55_P1960857** | Sorbs2 | -0.002324011 | -0.171414253 | 0.166766232 | 7.791697215 | -0.03339664 | 0.974411141 | 0.989773133 | -4.914571112 |
| **A_52_P420665** | Uts2b | -0.002189181 | -0.162806029 | 0.158427666 | 7.672309219 | -0.033118737 | 0.974623982 | 0.989937307 | -4.914576184 |
| **A_55_P1955861** | Cux2 | 0.006424019 | -0.465925703 | 0.47877374 | 8.451931391 | 0.033046564 | 0.974679259 | 0.989941432 | -4.914577494 |
| **A_55_P2098771** | Taf13 | 0.003480453 | -0.253377901 | 0.260338806 | 9.793619502 | 0.032924957 | 0.974772397 | 0.989984009 | -4.914579696 |
| **A_51_P221510** | Fam81a | -0.003190925 | -0.241184966 | 0.234803115 | 7.740234663 | -0.032578699 | 0.975037595 | 0.990198113 | -4.914585919 |
| **A_51_P425798** | Mtus2 | 0.002357823 | -0.173839004 | 0.178554649 | 7.795558166 | 0.032515933 | 0.975085668 | 0.990198113 | -4.91458704 |
| **A_55_P2120161** | Myh8 | 0.00205282 | -0.152016215 | 0.156121855 | 7.654474669 | 0.03237566 | 0.975193104 | 0.990247059 | -4.914589538 |
| **A_55_P2168781** | NA | -0.006015703 | -0.459049471 | 0.447018065 | 14.16617233 | -0.032265543 | 0.975277444 | 0.990247059 | -4.914591491 |
| **A_51_P114005** | Gstm7 | -0.006022024 | -0.459717622 | 0.447673573 | 9.080630991 | -0.032252333 | 0.975287561 | 0.990247059 | -4.894446832 |
| **A_55_P2180425** | Uqcc1 | 0.002865011 | -0.213930008 | 0.21966003 | 9.113643385 | 0.032111471 | 0.97539545 | 0.990304583 | -4.914594212 |
| **A_66_P109673** | NA | 0.002556732 | -0.191593991 | 0.196707455 | 10.51672777 | 0.031998479 | 0.975481992 | 0.990332158 | -4.9145962 |
| **A_52_P615401** | Tpra1 | -0.003426795 | -0.265274482 | 0.258420893 | 8.659487754 | -0.031799666 | 0.975634268 | 0.990332158 | -4.91459968 |
| **A_55_P2064507** | Msra | 0.005386888 | -0.406760293 | 0.417534069 | 11.57216537 | 0.031759145 | 0.975665304 | 0.990332158 | -4.914600387 |
| **A_55_P2104876** | NA | -0.002750714 | -0.213287931 | 0.207786503 | 8.574823194 | -0.031746783 | 0.975674773 | 0.990332158 | -4.914600603 |
| **A_52_P141338** | Acot10 | 0.003400656 | -0.257320866 | 0.264122178 | 7.945599622 | 0.031693413 | 0.97571565 | 0.990332158 | -4.914601532 |
| **A_52_P448006** | Nit2 | -0.004627694 | -0.359633971 | 0.350378583 | 10.74765265 | -0.031674647 | 0.975730024 | 0.990332158 | -4.914601858 |
| **A_51_P519276** | Ndufb7 | 0.002477775 | -0.1885122 | 0.19346775 | 14.53283394 | 0.031523502 | 0.97584579 | 0.990381956 | -4.914604479 |
| **A_55_P2237432** | Smarca1 | -0.004179565 | -0.326823667 | 0.318464537 | 9.31136837 | -0.031476797 | 0.975881563 | 0.990381956 | -4.914605286 |
| **A_51_P143901** | Tm9sf2 | -0.004305002 | -0.343316165 | 0.334706162 | 12.78233269 | -0.030856206 | 0.976356901 | 0.990812335 | -4.914615901 |
| **A_55_P2144086** | NA | -0.002238331 | -0.180028364 | 0.175551702 | 16.40713099 | -0.030591449 | 0.976559694 | 0.990966102 | -4.914620365 |

| | | | | | | | | |
|---|---|---|---|---|---|---|---|---|
| **A_51_P432659** | Pex7 | 0.00475238 | -0.37410284 | 0.383607601 | 10.75596398 | 0.03048043 | 0.976644729 | 0.990997534 | -4.914622225 |
| **A_52_P263870** | Cdc23 | -0.002131416 | -0.172399433 | 0.168136601 | 8.068353874 | -0.030417138 | 0.976693209 | 0.990997534 | -4.914623283 |
| **A_55_P2087240** | Mipep | 0.002722399 | -0.215292726 | 0.220737525 | 8.868811688 | 0.030342292 | 0.976750538 | 0.991003683 | -4.914624531 |
| **A_55_P2001023** | Lrrc19 | 0.003753326 | -0.300903777 | 0.30841043 | 11.43097236 | 0.029935616 | 0.977062041 | 0.991188223 | -4.914631257 |
| **A_55_P1995449** | NA | 0.005611094 | -0.450184602 | 0.46140679 | 12.40858517 | 0.02991304 | 0.977079334 | 0.991188223 | -4.914631628 |
| **A_51_P234253** | Sdcbp2 | -0.001932813 | -0.159093264 | 0.155227639 | 7.806654522 | -0.029883376 | 0.977102056 | 0.991188223 | -4.914632115 |
| **A_55_P2150224** | NA | -0.003144379 | -0.259216451 | 0.252927694 | 14.18264357 | -0.029837045 | 0.977137544 | 0.991188223 | -4.914632874 |
| **A_55_P2016732** | Slc30a2 | 0.001909338 | -0.15449593 | 0.158314606 | 7.723936324 | 0.029662974 | 0.977270879 | 0.991271454 | -4.914635716 |
| **A_52_P565940** | Nsd1 | 0.002572275 | -0.209370684 | 0.214515234 | 9.587823389 | 0.02949046 | 0.977403023 | 0.991318001 | -4.914638516 |
| **A_51_P110672** | Mst1r | 0.004002694 | -0.326038304 | 0.334043692 | 8.927984481 | 0.029469154 | 0.977419343 | 0.991318001 | -4.914638861 |
| **A_55_P2087647** | Grhpr | 0.007057573 | -0.5774481 | 0.591563246 | 9.67456598 | 0.029339306 | 0.977518805 | 0.991322408 | -4.914640956 |
| **A_55_P2088699** | NA | 0.002861285 | -0.234596051 | 0.24031862 | 16.50739365 | 0.029279162 | 0.977564875 | 0.991322408 | -4.914641923 |
| **A_52_P366803** | Cyp3a44 | -0.006353874 | -0.534020401 | 0.521312652 | 8.723695849 | -0.029259188 | 0.977580176 | 0.991322408 | -4.914642244 |
| **A_66_P137219** | Elane | -0.005562132 | -0.468482348 | 0.457358083 | 12.28854999 | -0.029195664 | 0.977628834 | 0.991322408 | -4.914643263 |
| **A_51_P503625** | Gsta3 | 0.009056179 | -0.749197592 | 0.767309951 | 15.58025816 | 0.029021083 | 0.977762563 | 0.991406 | -4.914646052 |
| **A_66_P125392** | Atp8b4 | -0.003255852 | -0.27886072 | 0.272349016 | 8.41629082 | -0.028705226 | 0.978004511 | 0.991599307 | -4.914651055 |
| **A_55_P1985453** | NA | 0.002038788 | -0.17167273 | 0.175750306 | 7.891562895 | 0.028518494 | 0.978147551 | 0.991682827 | -4.914653988 |
| **A_55_P2058137** | Pde4dip | 0.005219759 | -0.44037671 | 0.450816228 | 9.50079606 | 0.028463735 | 0.978189496 | 0.991682827 | -4.914654844 |
| **A_55_P2065926** | Akap8l | 0.004007091 | -0.342068348 | 0.35008253 | 9.152478201 | 0.028134658 | 0.978441576 | 0.99188636 | -4.914659955 |
| **A_52_P401504** | Thbs4 | -0.003201902 | -0.280658263 | 0.274254459 | 7.882226479 | -0.028041195 | 0.978513171 | 0.991900233 | -4.914661395 |
| **A_51_P163233** | Khdc4 | -0.002191931 | -0.192526953 | 0.188143091 | 8.775271095 | -0.027982811 | 0.978557894 | 0.991900233 | -4.914662293 |
| **A_55_P2240478** | NA | -0.00393397 | -0.347880075 | 0.340012135 | 8.651724974 | -0.027792257 | 0.978703865 | 0.991929906 | -4.914665209 |
| **A_51_P404413** | Bckdk | 0.004145684 | -0.358568744 | 0.366860112 | 10.36893205 | 0.027772471 | 0.978719021 | 0.991929906 | -4.914665511 |
| **A_55_P2042988** | Aoah | -0.006215163 | -0.550558146 | 0.538127819 | 8.877734052 | -0.027743619 | 0.978741123 | 0.991929906 | -4.914665951 |
| **A_51_P405985** | Fig4 | -0.002385505 | -0.213457272 | 0.208686261 | 8.817468906 | -0.02746208 | 0.978956792 | 0.992037436 | -4.914670216 |
| **A_55_P2137314** | Kctd21 | 0.001964712 | -0.172410182 | 0.176339606 | 8.144100063 | 0.027377773 | 0.979021375 | 0.992037436 | -4.914671484 |
| **A_52_P987071** | Gm6803 | 0.003915374 | -0.343827621 | 0.35165837 | 12.23755667 | 0.027358864 | 0.97903586 | 0.992037436 | -4.914671768 |
| **A_55_P2093286** | Apod | -0.002077431 | -0.187028698 | 0.182873837 | 7.873393071 | -0.027293072 | 0.979086259 | 0.992037436 | -4.914672755 |
| **A_51_P188073** | Dnajb11 | 0.003537571 | -0.31192401 | 0.318999153 | 12.19639118 | 0.027248457 | 0.979120437 | 0.992037436 | -4.914673423 |
| **A_51_P417701** | Apaf1 | 0.002171563 | -0.191799444 | 0.196142569 | 8.515963565 | 0.027203121 | 0.979155166 | 0.992037436 | -4.9146741 |
| **A_52_P163515** | NA | 0.002476377 | -0.220027401 | 0.224980154 | 9.141226144 | 0.027043474 | 0.979277464 | 0.99210934 | -4.914676477 |
| **A_55_P2082235** | NA | -0.001893179 | -0.173706074 | 0.169919717 | 8.850826436 | -0.026774358 | 0.979483621 | 0.99226619 | -4.914680451 |
| **A_55_P2007495** | NA | 0.005813298 | -0.529009746 | 0.540636343 | 11.23345446 | 0.026411659 | 0.979761471 | 0.992409138 | -4.914685744 |
| **A_55_P2006375** | Dmrtc1c2 | 0.001698076 | -0.155072446 | 0.158468598 | 7.646098953 | 0.026319393 | 0.979832152 | 0.992409138 | -4.914687079 |
| **A_55_P2028655** | Slc31a1 | 0.004107126 | -0.375179302 | 0.383393554 | 12.29715012 | 0.026312002 | 0.979837814 | 0.992409138 | -4.914687186 |
| **A_51_P219505** | Slc41a2 | -0.001952447 | -0.182408201 | 0.178503307 | 7.827367251 | -0.026290065 | 0.97985462 | 0.992409138 | -4.914687503 |
| **A_52_P550843** | Desi2 | -0.004245206 | -0.397133494 | 0.388643082 | 10.25880612 | -0.026255051 | 0.979881443 | 0.992409138 | -4.914688008 |
| **A_52_P283802** | Tapt1 | 0.003494575 | -0.322078951 | 0.329068101 | 9.766302034 | 0.026081253 | 0.980014585 | 0.992491979 | -4.914690504 |
| **A_55_P2125643** | NA | 0.003770734 | -0.349790564 | 0.357332032 | 13.97199582 | 0.025914594 | 0.980142258 | 0.99251978 | -4.914692882 |
| **A_52_P599162** | Tug1 | 0.002790505 | -0.260354094 | 0.265935105 | 8.590114835 | 0.025767459 | 0.980254975 | 0.99251978 | -4.914694968 |
| **A_55_P2073397** | NA | -0.003780687 | -0.360556716 | 0.352995341 | 11.09686399 | -0.025748876 | 0.980269211 | 0.99251978 | -4.914695231 |
| **A_51_P279437** | Mfsd2a | 0.001711244 | -0.160178066 | 0.163600553 | 7.703378869 | 0.025684838 | 0.980318269 | 0.99251978 | -4.914696135 |
| **A_51_P192860** | Hint1 | 0.001800582 | -0.168653991 | 0.172255155 | 16.76122915 | 0.02566773 | 0.980331375 | 0.99251978 | -4.914696376 |
| **A_52_P289685** | Ssbp1 | 0.002608148 | -0.244531423 | 0.249747718 | 12.15040465 | 0.025643251 | 0.980350128 | 0.99251978 | -4.914696721 |
| **A_55_P2212027** | Utrn | -0.001746552 | -0.16778861 | 0.164295505 | 8.466667175 | -0.025559163 | 0.980414547 | 0.992533011 | -4.914697902 |
| **A_51_P482043** | Epm2aip1 | 0.002394876 | -0.22752688 | 0.232316632 | 9.921684857 | 0.025309648 | 0.980605697 | 0.992563139 | -4.914701385 |
| **A_52_P79385** | Magt1 | 0.004844916 | -0.460887664 | 0.470577496 | 10.36861969 | 0.025277425 | 0.980630383 | 0.992563139 | -4.914701833 |
| **A_51_P374772** | Arfgef1 | -0.003535402 | -0.34389276 | 0.336821955 | 10.40508151 | -0.025239857 | 0.980659164 | 0.992563139 | -4.914702353 |
| **A_55_P2176248** | NA | 0.001948157 | -0.185819751 | 0.189716065 | 15.85472264 | 0.02521075 | 0.980681462 | 0.992563139 | -4.914702756 |
| **A_51_P270733** | Syngr1 | 0.003344637 | -0.319346747 | 0.326036021 | 8.85607255 | 0.025185166 | 0.980701062 | 0.992563139 | -4.91470311 |

| | | | | | | | | | |
|---|---|---|---|---|---|---|---|---|---|
| **A_55_P2026547** | Gal3st2 | -0.001849412 | -0.180768628 | 0.177069805 | 7.971780418 | -0.025116538 | 0.980753637 | 0.992564378 | -4.914704057 |
| **A_51_P136337** | Galm | -0.003016051 | -0.296835839 | 0.290803736 | 8.741013389 | -0.024942552 | 0.980886928 | 0.992634566 | -4.914706447 |
| **A_55_P2162875** | NA | 0.006780629 | -0.657593717 | 0.671154974 | 11.06896483 | 0.02479935 | 0.980996635 | 0.992634566 | -4.914708402 |
| **A_51_P173622** | Ptpn11 | -0.002424719 | -0.240018074 | 0.235168635 | 13.16359557 | -0.024797642 | 0.980997944 | 0.992634566 | -4.914708425 |
| **A_55_P1962503** | Nup62cl | -0.001597634 | -0.158537733 | 0.155342465 | 7.702052462 | -0.024735833 | 0.981045295 | 0.992634566 | -4.914709266 |
| **A_55_P2063992** | Rps5 | -0.002787915 | -0.277151709 | 0.271575879 | 17.19844969 | -0.024690842 | 0.981079763 | 0.992634566 | -4.914709876 |
| **A_66_P140742** | Azin2 | -0.004434703 | -0.443746639 | 0.434877233 | 7.901692236 | -0.024528712 | 0.981203972 | 0.992708274 | -4.914712066 |
| **A_55_P2102621** | Eaf2 | 0.003346133 | -0.330293669 | 0.336985935 | 12.17346851 | 0.02436961 | 0.981325862 | 0.992710909 | -4.914714201 |
| **A_51_P508029** | Pcyox1 | -0.004443033 | -0.448232171 | 0.439346104 | 10.66234987 | -0.024326861 | 0.981358612 | 0.992710909 | -4.914714772 |
| **A_55_P2185605** | Cd48 | -0.004438881 | -0.448532718 | 0.439654955 | 9.760328277 | -0.024287453 | 0.981388803 | 0.992710909 | -4.914715298 |
| **A_55_P2117731** | Pak4 | 0.005851045 | -0.580255603 | 0.591957692 | 15.26727066 | 0.024257161 | 0.981412011 | 0.992710909 | -4.914715701 |
| **A_55_P2065040** | NA | 0.003124082 | -0.312652906 | 0.31890107 | 14.33147136 | 0.024039482 | 0.981578778 | 0.992805809 | -4.914718586 |
| **A_55_P2106800** | Bglap | 0.00156137 | -0.156837971 | 0.159960712 | 7.684423694 | 0.023951667 | 0.981646055 | 0.992805809 | -4.914719743 |
| **A_52_P240152** | Snx27 | 0.00282965 | -0.284774936 | 0.290434237 | 9.714596289 | 0.023906728 | 0.981680483 | 0.992805809 | -4.914720333 |
| **A_66_P140613** | NA | -0.002231925 | -0.229466123 | 0.225002272 | 8.577885182 | -0.023866524 | 0.981711285 | 0.992805809 | -4.91472086 |
| **A_52_P153327** | 2-Sep | -0.001670957 | -0.174100219 | 0.170758304 | 8.226102917 | -0.023547109 | 0.981955996 | 0.993001332 | -4.914725016 |
| **A_55_P1980543** | Ube2i | 0.002450826 | -0.253580011 | 0.258481663 | 10.9450557 | 0.023259663 | 0.982176219 | 0.993172071 | -4.914728708 |
| **A_55_P2418745** | D230034L24Rik | -0.002479657 | -0.262679516 | 0.257720203 | 8.452340986 | -0.02315622 | 0.98225547 | 0.99320025 | -4.914730026 |
| **A_55_P2142146** | Glul | 0.004296319 | -0.449140975 | 0.457733612 | 12.29782551 | 0.023023028 | 0.982357513 | 0.993251471 | -4.914731714 |
| **A_52_P381418** | Slc35b4 | -0.002583901 | -0.277145901 | 0.271978099 | 8.336336447 | -0.022867496 | 0.982476673 | 0.993319992 | -4.914733672 |
| **A_55_P2053848** | NA | -0.002599882 | -0.28044255 | 0.275242785 | 12.53274046 | -0.02273725 | 0.98257646 | 0.993368921 | -4.914735302 |
| **A_55_P1963339** | Washc5 | 0.001559676 | -0.166076006 | 0.169195358 | 11.38651143 | 0.022607423 | 0.982675927 | 0.99341752 | -4.914736918 |
| **A_55_P2045407** | Ppp4r1 | 0.001788136 | -0.191102427 | 0.1946787 | 11.09775159 | 0.022525419 | 0.982738754 | 0.993428452 | -4.914737934 |
| **A_55_P2008327** | Eno1b | 0.003069886 | -0.329681545 | 0.335821316 | 14.37810105 | 0.022417413 | 0.982821503 | 0.993428452 | -4.914739266 |
| **A_55_P1962603** | Cplx2 | 0.006593379 | -0.710088553 | 0.723275312 | 13.09002618 | 0.022354491 | 0.98286971 | 0.993428452 | -4.914740039 |
| **A_52_P638794** | Stk31 | 0.001432344 | -0.154465302 | 0.157329991 | 7.660730218 | 0.022324977 | 0.982892323 | 0.993428452 | -4.914740401 |
| **A_55_P2017413** | Gm4788 | -0.008590811 | -0.946450398 | 0.929268776 | 10.75906374 | -0.022257663 | 0.982943896 | 0.993428631 | -4.914741224 |
| **A_55_P2144436** | Rrp1 | 0.00235768 | -0.256619787 | 0.261335147 | 11.82527404 | 0.022121063 | 0.983048553 | 0.99344707 | -4.914742888 |
| **A_55_P2051039** | Gm11937 | -0.005600139 | -0.62133817 | 0.610137891 | 11.38238863 | -0.022099683 | 0.983064934 | 0.99344707 | -4.914743147 |
| **A_55_P2036803** | Dhx30 | 0.002003928 | -0.219283936 | 0.223291791 | 9.683954156 | 0.022004303 | 0.983138009 | 0.993468977 | -4.914744302 |
| **A_52_P453374** | Lrrn4cl | 0.001531934 | -0.170996843 | 0.174060711 | 7.887489141 | 0.021575539 | 0.983466514 | 0.993650165 | -4.914749429 |
| **A_55_P1952725** | NA | -0.001416154 | -0.161025249 | 0.158192941 | 7.691846599 | -0.021559368 | 0.983478903 | 0.993650165 | -4.914749621 |
| **A_51_P496715** | Pomk | -0.001713954 | -0.195451796 | 0.192023887 | 8.906255925 | -0.021496512 | 0.983527062 | 0.993650165 | -4.914750363 |
| **A_52_P642207** | Mkrn1 | -0.003740789 | -0.429463021 | 0.421981443 | 12.50995654 | -0.021351092 | 0.983638478 | 0.993650165 | -4.914752073 |
| **A_55_P2168098** | Plrg1 | 0.001503629 | -0.169696085 | 0.172703343 | 11.77019128 | 0.021341302 | 0.983645979 | 0.993650165 | -4.914752188 |
| **A_52_P622442** | Pdcl3 | 0.002028612 | -0.230943568 | 0.235000792 | 9.990831042 | 0.021158173 | 0.983786288 | 0.993650165 | -4.914754324 |
| **A_51_P123705** | Dus3l | -0.001868149 | -0.217886058 | 0.21414976 | 9.875699018 | -0.021013816 | 0.983896891 | 0.993650165 | -4.882232537 |
| **A_51_P491987** | Ripk3 | -0.002169945 | -0.253104347 | 0.248764458 | 9.814263731 | -0.021012209 | 0.983898123 | 0.993650165 | -4.914756013 |
| **A_51_P372992** | Vsir | -0.003193719 | -0.372600277 | 0.366212838 | 9.364441531 | -0.021007548 | 0.983901694 | 0.993650165 | -4.914756067 |
| **A_55_P2099530** | NA | -0.005238254 | -0.6133226 | 0.602846092 | 11.90128201 | -0.020931768 | 0.983959755 | 0.993650165 | -4.914756939 |
| **A_55_P2016962** | NA | -0.001346228 | -0.157714038 | 0.155021581 | 7.666350709 | -0.020919656 | 0.983969035 | 0.993650165 | -4.914757078 |
| **A_55_P2134387** | Olfr544 | 0.007204957 | -0.832863003 | 0.847272917 | 11.59105621 | 0.020840117 | 0.984029976 | 0.993650165 | -4.91475799 |
| **A_55_P2119302** | Cspp1 | 0.001739442 | -0.201629314 | 0.205108199 | 8.203216572 | 0.020783036 | 0.984073711 | 0.993650165 | -4.914758642 |
| **A_51_P287232** | Qars | -0.002876292 | -0.339309944 | 0.33355736 | 12.37868126 | -0.020773837 | 0.984080759 | 0.993650165 | -4.914758747 |
| **A_51_P213592** | Nrde2 | 0.001381349 | -0.160269861 | 0.163032559 | 8.280214493 | 0.020763835 | 0.984088422 | 0.993650165 | -4.914758861 |
| **A_51_P198694** | Gckr | -0.005264724 | -0.629366905 | 0.618837457 | 12.20573588 | -0.020497603 | 0.984292406 | 0.993804216 | -4.914761875 |
| **A_55_P1985633** | Lrp1 | 0.00129437 | -0.152835673 | 0.155424413 | 7.705939836 | 0.02040583 | 0.984362721 | 0.993823298 | -4.914762905 |
| **A_66_P105175** | Bche | 0.006291042 | -0.751509451 | 0.764091535 | 11.25958126 | 0.020172089 | 0.984541812 | 0.993952192 | -4.914765508 |
| **A_55_P2009953** | Cd300lg | 0.001743673 | -0.211162149 | 0.214649495 | 9.138602507 | 0.019900349 | 0.984750019 | 0.994110467 | -4.914768497 |
| **A_55_P2165299** | Mybpc1 | 0.001299037 | -0.160914958 | 0.163513032 | 7.814887799 | 0.019458816 | 0.985088322 | 0.994400051 | -4.914773266 |

| | | | | | | | | | |
|---|---|---|---|---|---|---|---|---|---|
| **A_51_P510939** | Chchd7 | -0.002413437 | -0.307495901 | 0.302669027 | 10.41104397 | -0.019222145 | 0.985269662 | 0.994493763 | -4.914775778 |
| **A_51_P344947** | Rab1b | 0.002470373 | -0.31011495 | 0.315055696 | 11.87027369 | 0.019203356 | 0.985284058 | 0.994493763 | -4.914775976 |
| **A_55_P2091762** | NA | 0.001277034 | -0.161540626 | 0.164094694 | 16.14169486 | 0.0190583 | 0.985395202 | 0.99449617 | -4.9147775 |
| **A_52_P411430** | Gm5914 | 0.002377108 | -0.301645208 | 0.306399424 | 8.556439084 | 0.018998821 | 0.985440776 | 0.99449617 | -4.914778121 |
| **A_52_P308507** | Chd6 | -0.001572521 | -0.202691717 | 0.199546676 | 8.120885437 | -0.018998795 | 0.985440796 | 0.99449617 | -4.914778121 |
| **A_55_P2031011** | S100pbp | 0.001547299 | -0.197764809 | 0.200859407 | 8.187812662 | 0.018863566 | 0.98554441 | 0.994546486 | -4.914779526 |
| **A_55_P2078073** | Pqlc2 | -0.003858498 | -0.502578718 | 0.494861723 | 10.79041226 | -0.01879942 | 0.985593561 | 0.994546486 | -4.914780189 |
| **A_55_P1996837** | Baz2b | 0.001375996 | -0.179526839 | 0.182278831 | 9.929725016 | 0.018482262 | 0.985836574 | 0.994703271 | -4.914783434 |
| **A_51_P121891** | Rac2 | 0.003495342 | -0.457143024 | 0.464133708 | 12.02805371 | 0.018437953 | 0.985870524 | 0.994703271 | -4.914783883 |
| **A_55_P2157616** | NA | 0.001564643 | -0.205701751 | 0.208831038 | 16.77072465 | 0.018342971 | 0.985943302 | 0.994703271 | -4.914784842 |
| **A_51_P372156** | 4930563E22Rik | -0.001219847 | -0.162943634 | 0.160503939 | 7.745582854 | -0.018327989 | 0.985954782 | 0.994703271 | -4.914784993 |
| **A_55_P1980119** | Cptp | 0.002223223 | -0.297834149 | 0.302280594 | 9.968337959 | 0.018003704 | 0.986203258 | 0.99487806 | -4.914788226 |
| **A_52_P71544** | Slc25a26 | 0.001270397 | -0.170534008 | 0.173074802 | 8.70351636 | 0.017967532 | 0.986230975 | 0.99487806 | -4.914788583 |
| **A_66_P111401** | Fbxw8 | -0.002990863 | -0.415832697 | 0.409850972 | 9.43764132 | -0.017603376 | 0.986510003 | 0.995066806 | -4.914792138 |
| **A_52_P183181** | Adar | 0.001591252 | -0.219297019 | 0.222479523 | 9.418758772 | 0.017504495 | 0.98658577 | 0.995066806 | -4.914793091 |
| **A_66_P127024** | Mbd3 | 0.001810522 | -0.250270703 | 0.253891747 | 12.97494055 | 0.017452052 | 0.986625954 | 0.995066806 | -4.914793594 |
| **A_52_P118958** | Pex19 | 0.001832108 | -0.255452486 | 0.259116701 | 9.655590504 | 0.017302959 | 0.986740195 | 0.995066806 | -4.914795016 |
| **A_55_P2001481** | Pde6c | 0.004308517 | -0.602593586 | 0.611210621 | 9.408068821 | 0.017250131 | 0.986780674 | 0.995066806 | -4.914795517 |
| **A_52_P648715** | Triml1 | 0.001100266 | -0.154621727 | 0.15682226 | 7.677622915 | 0.017168449 | 0.986843262 | 0.995066806 | -4.914796288 |
| **A_52_P202725** | Rragc | 0.001403357 | -0.197850724 | 0.200657437 | 8.514607726 | 0.017113708 | 0.986885208 | 0.995066806 | -4.914796803 |
| **A_66_P112006** | H2-T10 | 0.001298012 | -0.184312854 | 0.186908878 | 8.766220038 | 0.016992549 | 0.986978045 | 0.995066806 | -4.914797937 |
| **A_51_P294020** | Ticam1 | 0.002236106 | -0.319082109 | 0.323554321 | 9.262948474 | 0.016909869 | 0.987041399 | 0.995066806 | -4.914798707 |
| **A_52_P374292** | Tefm | -0.001192821 | -0.173290929 | 0.170905287 | 8.26985539 | -0.016841561 | 0.98709374 | 0.995066806 | -4.914799339 |
| **A_55_P2116305** | Tnfrsf25 | -0.002649312 | -0.384942317 | 0.379643693 | 8.316447435 | -0.01683914 | 0.987095595 | 0.995066806 | -4.914799362 |
| **A_52_P354744** | Slc2a3 | 0.00340279 | -0.489927712 | 0.496733291 | 9.943390979 | 0.016760244 | 0.987156048 | 0.995066806 | -4.914800089 |
| **A_55_P1960298** | Tmem80 | 0.00237776 | -0.34602593 | 0.350781449 | 8.970152965 | 0.016583204 | 0.987291706 | 0.995066806 | -4.914801709 |
| **A_55_P2154914** | Acap1 | 0.002080726 | -0.307143563 | 0.311305016 | 8.68755044 | 0.016350255 | 0.987470204 | 0.995066806 | -4.914803815 |
| **A_55_P2033435** | Stra6l | -0.0045977 | -0.68927719 | 0.68008179 | 10.01972187 | -0.016316857 | 0.987495796 | 0.995066806 | -4.914804114 |
| **A_52_P71686** | Atp6v0d2 | 0.00103569 | -0.153261116 | 0.155332496 | 7.704298072 | 0.016310078 | 0.98750099 | 0.995066806 | -4.914804175 |
| **A_52_P289256** | Arid1b | -0.001787293 | -0.268227466 | 0.264652879 | 9.817873159 | -0.01629969 | 0.987508951 | 0.995066806 | -4.914804268 |
| **A_55_P2010704** | Selenoh | -0.003195889 | -0.484652592 | 0.478260814 | 12.60915254 | -0.01612938 | 0.987639452 | 0.995066806 | -4.914805784 |
| **A_55_P2029471** | NA | 0.002383514 | -0.357288016 | 0.362055044 | 13.82217581 | 0.016102555 | 0.987660007 | 0.995066806 | -4.914806021 |
| **A_55_P2011490** | NA | -0.001607751 | -0.244235706 | 0.241020203 | 12.07291118 | -0.01610131 | 0.987660961 | 0.995066806 | -4.914806032 |
| **A_55_P2066523** | Akt2 | 0.001640561 | -0.246316238 | 0.249597359 | 8.374316779 | 0.016076794 | 0.987679747 | 0.995066806 | -4.914806248 |
| **A_55_P2094197** | Dnase1l3 | -0.002582829 | -0.393743637 | 0.388577979 | 8.351330904 | -0.016044399 | 0.987704569 | 0.995066806 | -4.914806534 |
| **A_52_P650028** | Dock10 | -0.002749195 | -0.41985855 | 0.41436016 | 9.133604215 | -0.016015438 | 0.987726761 | 0.995066806 | -4.914806789 |
| **A_55_P2138150** | Pias2 | 0.00141096 | -0.212694867 | 0.215516786 | 9.055370437 | 0.016012873 | 0.987728727 | 0.995066806 | -4.914806811 |
| **A_52_P26299** | Slc35f1 | 0.001017278 | -0.154590633 | 0.156625188 | 7.659730633 | 0.01588514 | 0.987826604 | 0.995066806 | -4.91480793 |
| **A_52_P450934** | Paqr9 | -0.003186696 | -0.49070839 | 0.484334999 | 12.06701346 | -0.015882901 | 0.987828319 | 0.995066806 | -4.914807949 |
| **A_55_P2099363** | Stac2 | -0.001038714 | -0.160149541 | 0.158072114 | 7.697651235 | -0.015862778 | 0.987843739 | 0.995066806 | -4.914808124 |
| **A_52_P606774** | Stk4 | 0.002333978 | -0.355651471 | 0.360319427 | 10.43626612 | 0.015842165 | 0.987859534 | 0.995066806 | -4.914808304 |
| **A_55_P2108983** | NA | 0.002334237 | -0.359814368 | 0.364482842 | 15.69387202 | 0.015661785 | 0.987997753 | 0.995154172 | -4.914809863 |
| **A_55_P2121541** | Gm6607 | 0.001349937 | -0.209166082 | 0.211865957 | 15.15420765 | 0.015581586 | 0.988059207 | 0.995164213 | -4.914810551 |
| **A_55_P2043622** | NA | 0.003128187 | -0.487621196 | 0.493877569 | 15.24669732 | 0.01548874 | 0.988130352 | 0.995184013 | -4.914811343 |
| **A_66_P102049** | Zfp1 | 0.001749779 | -0.274207158 | 0.277706716 | 10.48309748 | 0.015407251 | 0.988192794 | 0.995195046 | -4.914812034 |
| **A_55_P2011702** | Ppargc1a | -0.001464289 | -0.234793053 | 0.231864474 | 7.89267437 | -0.015249016 | 0.988314045 | 0.995265301 | -4.914813365 |
| **A_52_P282905** | Ces1b | -0.003540163 | -0.589517545 | 0.582437218 | 8.431142314 | -0.014679986 | 0.98875008 | 0.995536697 | -4.914818039 |
| **A_55_P1999419** | Bag5 | -0.001463585 | -0.244363398 | 0.241436229 | 9.755141971 | -0.014641107 | 0.988779872 | 0.995536697 | -4.914818352 |
| **A_52_P180335** | Mrpl28 | 0.001776341 | -0.294312659 | 0.297865341 | 10.34275457 | 0.014577641 | 0.988828504 | 0.995536697 | -4.914818861 |
| **A_55_P2071132** | Tnfrsf23 | -0.001401765 | -0.235571791 | 0.232768261 | 8.193894138 | -0.014545447 | 0.988853174 | 0.995536697 | -4.914819118 |

| | | | | | | | | | |
|---|---|---|---|---|---|---|---|---|---|
| **A_55_P2153941** | Zfp386 | 0.00199709 | -0.332583115 | 0.336577295 | 9.303334794 | 0.01450376 | 0.988885118 | 0.995536697 | -4.91481945 |
| **A_55_P1998825** | Rpl9 | -0.003252646 | -0.548546835 | 0.542041543 | 17.34015974 | -0.014494028 | 0.988892575 | 0.995536697 | -4.914819528 |
| **A_66_P127311** | Esco1 | 0.001087601 | -0.182766739 | 0.184941942 | 8.181470615 | 0.014374047 | 0.988984515 | 0.995577402 | -4.914820478 |
| **A_55_P1972406** | Tor1b | 0.001614651 | -0.274571492 | 0.277800793 | 9.684330313 | 0.014205609 | 0.989113587 | 0.995655479 | -4.914821799 |
| **A_55_P1956707** | Rpl36a | 0.000987258 | -0.173338149 | 0.175312665 | 15.29525644 | 0.013761106 | 0.989454205 | 0.995864513 | -4.914825211 |
| **A_52_P481279** | Drc1 | -0.002535758 | -0.451065215 | 0.445993699 | 9.448178947 | -0.013737256 | 0.98947248 | 0.995864513 | -4.914825391 |
| **A_55_P1961965** | Plagl1 | 0.001200866 | -0.211758199 | 0.21415993 | 7.985497125 | 0.013701921 | 0.989499558 | 0.995864513 | -4.914825657 |
| **A_52_P624149** | Hsp90b1 | 0.001643297 | -0.291015207 | 0.2943018 | 14.84371747 | 0.013643884 | 0.989544031 | 0.995864513 | -4.914826093 |
| **A_51_P443403** | Smoc1 | 0.003725984 | -0.670290814 | 0.677742783 | 10.01828151 | 0.013432396 | 0.989706093 | 0.995864513 | -4.914827664 |
| **A_51_P245368** | Abcb1b | 0.000856694 | -0.156943116 | 0.158656504 | 7.859089493 | 0.01319175 | 0.989890499 | 0.995864513 | -4.914829423 |
| **A_55_P2093949** | Zfp428 | -0.000902856 | -0.16778963 | 0.165983918 | 7.90406783 | -0.013145576 | 0.989925883 | 0.995864513 | -4.914829757 |
| **A_55_P2115401** | Bst1 | 0.001410588 | -0.259365523 | 0.262186699 | 8.104456351 | 0.01314364 | 0.989927366 | 0.995864513 | -4.914829771 |
| **A_55_P2039235** | Mat2b | -0.001144189 | -0.213676289 | 0.21138791 | 11.97235209 | -0.013081472 | 0.989975006 | 0.995864513 | -4.914830218 |
| **A_51_P371882** | Ccdc71 | -0.00123994 | -0.231795287 | 0.229315406 | 11.19304933 | -0.013067987 | 0.989985339 | 0.995864513 | -4.914830315 |
| **A_52_P652513** | Lman1 | 0.002635462 | -0.48848006 | 0.493750983 | 13.5728785 | 0.01303936 | 0.990007276 | 0.995864513 | -4.91483052 |
| **A_55_P1964375** | Gm9731 | -0.001137591 | -0.213227308 | 0.210952125 | 10.94573181 | -0.013033164 | 0.990012023 | 0.995864513 | -4.914830564 |
| **A_55_P2157788** | NA | 0.001213246 | -0.225110963 | 0.227537455 | 13.5327996 | 0.013025696 | 0.990017746 | 0.995864513 | -4.914830618 |
| **A_55_P2067905** | Scfd2 | -0.001169338 | -0.222382595 | 0.220043919 | 10.14319068 | -0.012844348 | 0.990156714 | 0.995864513 | -4.914831905 |
| **A_52_P57526** | Sin3b | -0.001056948 | -0.202102326 | 0.199988429 | 13.70976477 | -0.012774469 | 0.990210262 | 0.995864513 | -4.914832396 |
| **A_52_P109548** | Map7 | 0.001453922 | -0.276156498 | 0.279064343 | 8.486020748 | 0.012725904 | 0.990247478 | 0.995864513 | -4.914832736 |
| **A_52_P244803** | D630033O11Rik | 0.001441594 | -0.274008418 | 0.276891606 | 8.41967041 | 0.012716964 | 0.990254329 | 0.995864513 | -4.914832799 |
| **A_51_P101075** | Cnppd1 | -0.001180481 | -0.22744985 | 0.225088888 | 13.110132 | -0.012677001 | 0.990284952 | 0.995864513 | -4.914833077 |
| **A_52_P676063** | Tmem126b | -0.001100521 | -0.212377582 | 0.21017654 | 8.482637601 | -0.012656948 | 0.990300319 | 0.995864513 | -4.914833217 |
| **A_55_P2002567** | NA | -0.001316702 | -0.255613403 | 0.25298 | 15.66604974 | -0.012581426 | 0.990358191 | 0.995864513 | -4.91483374 |
| **A_51_P293989** | Senp6 | 0.000862172 | -0.167240284 | 0.168964628 | 7.851982168 | 0.012462442 | 0.99044937 | 0.995864513 | -4.914834558 |
| **A_55_P1983733** | Aldh1l1 | -0.003228465 | -0.633053186 | 0.626596257 | 13.20138925 | -0.012455453 | 0.990454726 | 0.995864513 | -4.914834605 |
| **A_55_P2026710** | 4930445E18Rik | 0.000820097 | -0.162369331 | 0.164009526 | 7.676518316 | 0.012211152 | 0.990641936 | 0.996000935 | -4.914836259 |
| **A_55_P2041700** | Hoxa11os | 0.000821415 | -0.16609202 | 0.16773485 | 7.815348797 | 0.011957891 | 0.990836012 | 0.996079333 | -4.914837939 |
| **A_55_P2120961** | NA | -0.002291074 | -0.471665498 | 0.46708335 | 10.18435994 | -0.011860497 | 0.990910647 | 0.996079333 | -4.914838576 |
| **A_55_P2417224** | NA | -0.002329103 | -0.481477239 | 0.476819032 | 9.015964822 | -0.011811422 | 0.990948254 | 0.996079333 | -4.914838895 |
| **A_55_P1986466** | Retreg2 | -0.001694917 | -0.351712048 | 0.348322213 | 8.691717942 | -0.011766367 | 0.99098278 | 0.996079333 | -4.914839186 |
| **A_51_P463120** | Hacd4 | 0.001152943 | -0.237460851 | 0.239766737 | 8.778250622 | 0.011740738 | 0.99100242 | 0.996079333 | -4.914839352 |
| **A_55_P1985565** | NA | 0.001004002 | -0.207403064 | 0.209411068 | 15.91528767 | 0.01170591 | 0.991029109 | 0.996079333 | -4.914839576 |
| **A_55_P2051899** | Atp2c1 | -0.001426261 | -0.309293109 | 0.306440587 | 9.644044761 | -0.01125691 | 0.991373186 | 0.996373352 | -4.914842405 |
| **A_55_P2010196** | Serpina10 | 0.001961985 | -0.426412837 | 0.430336808 | 9.592145047 | 0.011128971 | 0.991471227 | 0.996420077 | -4.914843191 |
| **A_55_P2162422** | NA | -0.000913521 | -0.206576593 | 0.20474955 | 16.85101569 | -0.010793081 | 0.991728628 | 0.996613329 | -4.914845212 |
| **A_52_P160936** | Gusb | 0.000771773 | -0.173781486 | 0.175325032 | 9.394388616 | 0.010743477 | 0.99176664 | 0.996613329 | -4.914845505 |
| **A_55_P1961968** | Plagl1 | 0.003602976 | -0.91846971 | 0.925675661 | 13.01156559 | 0.010668466 | 0.991978018 | 0.996773919 | -4.829169584 |
| **A_66_P103780** | Gm3646 | -0.000682047 | -0.160439424 | 0.159075331 | 7.649634112 | -0.010373766 | 0.99204996 | 0.996777469 | -4.914847648 |
| **A_55_P1975385** | Dctn1 | -0.001798662 | -0.427122242 | 0.423524918 | 12.75057596 | -0.010275744 | 0.992125076 | 0.996777469 | -4.914848204 |
| **A_51_P417839** | Hdac4 | 0.000773828 | -0.182470978 | 0.184018635 | 8.703886762 | 0.010261154 | 0.992136258 | 0.996777469 | -4.914848286 |
| **A_55_P2038318** | NA | 0.000652595 | -0.157920618 | 0.159225808 | 7.682517309 | 0.009999929 | 0.992336442 | 0.996926772 | -4.914849739 |
| **A_55_P2007991** | Tuba3b | -0.000840692 | -0.209005784 | 0.2073244 | 8.008318107 | -0.009813234 | 0.992479513 | 0.997018684 | -4.914850754 |
| **A_55_P1982524** | Cyhr1 | -0.00070505 | -0.177120329 | 0.175710229 | 8.35992068 | -0.009711067 | 0.992557806 | 0.997045517 | -4.914851301 |
| **A_55_P2015530** | NA | -0.000791574 | -0.201952892 | 0.200369743 | 7.998625921 | -0.009561597 | 0.992672351 | 0.99710876 | -4.914852092 |
| **A_55_P2179378** | Zkscan8 | -0.001098056 | -0.294390393 | 0.29219428 | 9.318986207 | -0.009097186 | 0.993028246 | 0.997414413 | -4.91485447 |
| **A_55_P2166990** | Ppcs | 0.000800164 | -0.217279666 | 0.218879994 | 10.10184806 | 0.008915523 | 0.993167462 | 0.99750241 | -4.914855368 |
| **A_52_P679205** | Slc35f6 | 0.000856282 | -0.234912073 | 0.236624636 | 9.289709495 | 0.008824995 | 0.993236837 | 0.997520255 | -4.914855809 |
| **A_55_P2083489** | Fam129a | -0.001745029 | -0.487029451 | 0.483539393 | 9.970898613 | -0.008737547 | 0.993303852 | 0.997534371 | -4.91485623 |
| **A_55_P2062190** | Gstm1 | 0.001592282 | -0.44456247 | 0.447747033 | 13.56695159 | 0.008671966 | 0.993354109 | 0.997534371 | -4.914856544 |

| | | | | | | | | | |
|---|---|---|---|---|---|---|---|---|---|
| **A_52_P650108** | Uri1 | -0.000797992 | -0.232636899 | 0.231040916 | 8.9174833 | -0.008363639 | 0.993590394 | 0.997697501 | -4.914857985 |
| **A_55_P1971360** | Ciz1 | 0.000620718 | -0.181767447 | 0.183008883 | 9.230072116 | 0.008269525 | 0.993662518 | 0.997697501 | -4.914858415 |
| **A_52_P520564** | Wasl | -0.001046426 | -0.314367596 | 0.312274744 | 8.623868976 | -0.008115251 | 0.993780745 | 0.997697501 | -4.914859109 |
| **A_55_P2177103** | Slc25a5 | 0.000738766 | -0.221567103 | 0.223044634 | 13.18286618 | 0.008074932 | 0.993811643 | 0.997697501 | -4.914859288 |
| **A_55_P2052252** | NA | -0.000843907 | -0.254891237 | 0.253203423 | 8.510421127 | -0.008071666 | 0.993814147 | 0.997697501 | -4.914859302 |
| **A_52_P117966** | Atpsckmt | 0.000942561 | -0.285649848 | 0.287534969 | 10.48333084 | 0.00799149 | 0.993875589 | 0.997697501 | -4.914859656 |
| **A_55_P2076545** | F5 | 0.002394544 | -0.732766708 | 0.737555796 | 11.61652033 | 0.007914499 | 0.993934591 | 0.997697501 | -4.914859992 |
| **A_51_P352562** | Mrpl38 | 0.001108219 | -0.340070123 | 0.34228656 | 10.44522339 | 0.007892728 | 0.993951275 | 0.997697501 | -4.914860086 |
| **A_52_P528463** | Mfsd3 | -0.000615945 | -0.193118403 | 0.191886514 | 9.522595032 | -0.007774789 | 0.994041658 | 0.997697501 | -4.914860594 |
| **A_55_P2079928** | Ccdc68 | -0.000612607 | -0.192204757 | 0.190979543 | 8.626383404 | -0.007769398 | 0.994045789 | 0.997697501 | -4.914860617 |
| **A_55_P2112504** | Kcnv2 | -0.000486794 | -0.153961631 | 0.152988043 | 7.643978253 | -0.007707101 | 0.99409353 | 0.997697501 | -4.914860881 |
| **A_55_P1977649** | Hoxd8 | 0.000497807 | -0.15758466 | 0.158580273 | 7.683343439 | 0.007651742 | 0.994135954 | 0.997697501 | -4.914861114 |
| **A_52_P581435** | Pdha1 | 0.000817809 | -0.268560726 | 0.270196345 | 10.10210812 | 0.007376872 | 0.994346601 | 0.997857092 | -4.914862248 |
| **A_55_P2300071** | 4833412C15Rik | 0.000549658 | -0.183608763 | 0.184708078 | 8.19218131 | 0.007252435 | 0.994441964 | 0.997900983 | -4.914862748 |
| **A_51_P424518** | Nin | -0.000725276 | -0.250849527 | 0.249398974 | 8.601167626 | -0.007045809 | 0.994600313 | 0.997991481 | -4.914863559 |
| **A_55_P2157384** | Xlr5b | -0.000447549 | -0.155802415 | 0.154907318 | 7.688045675 | -0.007000008 | 0.994635412 | 0.997991481 | -4.914863735 |
| **A_55_P2143946** | Slc18a2 | 0.000459387 | -0.164111573 | 0.165030347 | 8.266919889 | 0.006782795 | 0.994801875 | 0.998106693 | -4.914864557 |
| **A_51_P512061** | Doxl1 | 0.000480402 | -0.178022297 | 0.178983101 | 7.794225526 | 0.006539478 | 0.994988342 | 0.998241964 | -4.914865447 |
| **A_55_P2027521** | NA | 0.000478293 | -0.181989686 | 0.182946272 | 8.13164944 | 0.006369285 | 0.995118771 | 0.998286447 | -4.91486605 |
| **A_55_P2126333** | Cyp2a12 | 0.002350327 | -0.903752498 | 0.908453151 | 12.346495 | 0.006302806 | 0.995169718 | 0.998286447 | -4.914866281 |
| **A_55_P1980401** | Cbx7 | -0.000402042 | -0.155974818 | 0.155170734 | 7.653102306 | -0.006279442 | 0.995187622 | 0.998286447 | -4.914866361 |
| **A_66_P106789** | NA | 0.000859754 | -0.339925332 | 0.341644839 | 13.82525168 | 0.006130229 | 0.995301973 | 0.998307044 | -4.91486687 |
| **A_55_P1959500** | Ces2e | 0.002035037 | -0.811837774 | 0.815907849 | 9.943073509 | 0.006075738 | 0.995343733 | 0.998307044 | -4.914867052 |
| **A_55_P2003398** | Rmdn1 | 0.000582055 | -0.233171655 | 0.234335765 | 9.044595312 | 0.006050467 | 0.9953631 | 0.998307044 | -4.914867136 |
| **A_55_P1963508** | Slc13a5 | 0.000349575 | -0.164455789 | 0.16515494 | 7.683886615 | 0.005154097 | 0.996050043 | 0.998918854 | -4.914869893 |
| **A_55_P1982718** | Rtf2 | -0.000472427 | -0.224695545 | 0.22375069 | 10.94832832 | -0.005119621 | 0.996076465 | 0.998918854 | -4.91486999 |
| **A_52_P547662** | P2ry1 | 0.000817111 | -0.406430836 | 0.408065058 | 8.844706764 | 0.004875343 | 0.996263671 | 0.99905476 | -4.91487066 |
| **A_55_P2024933** | Zfp706 | -0.000405397 | -0.206850351 | 0.206039556 | 12.44004798 | -0.004771555 | 0.99634321 | 0.999082689 | -4.914870935 |
| **A_51_P315890** | Kcnk6 | -0.000522854 | -0.27812669 | 0.277080982 | 10.19672335 | -0.004576552 | 0.996492654 | 0.999168063 | -4.914871435 |
| **A_55_P1985673** | Zbtb37 | -0.000346924 | -0.18911277 | 0.188418922 | 7.924144472 | -0.004465746 | 0.996577572 | 0.999168063 | -4.91487171 |
| **A_55_P2111419** | Eny2 | -0.000357165 | -0.196161807 | 0.195447478 | 13.29398028 | -0.004432295 | 0.996603208 | 0.999168063 | -4.914871792 |
| **A_51_P347823** | Ipo4 | 0.000379875 | -0.209849951 | 0.2106097 | 10.11150859 | 0.004390653 | 0.996635121 | 0.999168063 | -4.914871893 |
| **A_66_P134090** | Grpel2 | 0.000348282 | -0.196722744 | 0.197419308 | 8.270952754 | 0.004294289 | 0.996708972 | 0.999190276 | -4.914872122 |
| **A_51_P468140** | Serpind1 | 0.000962113 | -0.561397578 | 0.563321805 | 14.58427891 | 0.004157145 | 0.996814074 | 0.999224339 | -4.914872441 |
| **A_51_P310030** | Naca | 0.000511148 | -0.304664377 | 0.305686672 | 15.03644101 | 0.004069864 | 0.996880963 | 0.999224339 | -4.914872638 |
| **A_52_P77764** | Slc7a2 | -0.001264696 | -0.774846717 | 0.772317325 | 9.172073864 | -0.003972492 | 0.996955587 | 0.999224339 | -4.914872853 |
| **A_52_P566390** | Zfyve26 | -0.000895654 | -0.55274962 | 0.550958312 | 9.836627052 | -0.00394366 | 0.996977683 | 0.999224339 | -4.914872915 |
| **A_51_P245533** | Tex19.1 | -0.00026212 | -0.163045209 | 0.162520969 | 7.681367441 | -0.003912676 | 0.997001428 | 0.999224339 | -4.914872982 |
| **A_51_P357735** | Inhbe | -0.000967581 | -0.640059527 | 0.638124364 | 9.372812562 | -0.00367881 | 0.997180655 | 0.999316586 | -4.914873469 |
| **A_51_P145520** | Sigirr | 0.000739263 | -0.499718747 | 0.501197273 | 10.85086294 | 0.003589338 | 0.997249224 | 0.999316586 | -4.914873648 |
| **A_55_P2052824** | Lrrc14b | -0.000233583 | -0.162794756 | 0.162327591 | 7.703680962 | -0.003491458 | 0.997324236 | 0.999316586 | -4.914873838 |
| **A_55_P2098321** | NA | 0.000414271 | -0.288739268 | 0.289567809 | 10.92376201 | 0.00348128 | 0.997332037 | 0.999316586 | -4.914873857 |
| **A_55_P2427900** | Unc13c | 0.000214708 | -0.153595607 | 0.154025023 | 7.659918561 | 0.003391926 | 0.997400515 | 0.999316586 | -4.914874026 |
| **A_55_P2000384** | Pcnx3 | -0.000394545 | -0.283378151 | 0.282589061 | 11.29138316 | -0.003387807 | 0.997403672 | 0.999316586 | -4.914874034 |
| **A_55_P2122260** | Tmem87b | -0.000318735 | -0.236560041 | 0.23592257 | 8.087613304 | -0.003278368 | 0.997487542 | 0.999348816 | -4.914874234 |
| **A_55_P2089293** | Rnf121 | -0.000420933 | -0.357753616 | 0.356911751 | 10.16385062 | -0.002862352 | 0.997806365 | 0.999499829 | -4.914874935 |
| **A_52_P514352** | Kcnk5 | -0.000212357 | -0.182574647 | 0.182149933 | 8.340434029 | -0.002829532 | 0.997831518 | 0.999499829 | -4.914874986 |
| **A_55_P2042823** | Sh3bp2 | -0.000295786 | -0.256803545 | 0.256211973 | 9.81143871 | -0.002801949 | 0.997852656 | 0.999499829 | -4.914875029 |
| **A_55_P2021585** | Tff1 | -0.000177265 | -0.159602666 | 0.159248135 | 7.659248367 | -0.002701776 | 0.997929426 | 0.999499829 | -4.91487518 |
| **A_55_P2086099** | Gm2061 | 0.000200535 | -0.181808295 | 0.182209365 | 7.797617691 | 0.002677197 | 0.997948263 | 0.999499829 | -4.914875217 |

| | | | | | | | | | |
|---|---|---|---|---|---|---|---|---|---|
| **A_55_P2095563** | Slc8a2 | 0.000191382 | -0.173532683 | 0.173915446 | 7.816488149 | 0.002676845 | 0.997948533 | 0.999499829 | -4.914875217 |
| **A_55_P1962771** | Cyfip2 | -0.00057248 | -0.547980088 | 0.546835127 | 9.865636824 | -0.002541165 | 0.998052514 | 0.999507042 | -4.914875411 |
| **A_52_P56634** | Map3k11 | 0.000190631 | -0.182715491 | 0.183096754 | 9.484441311 | 0.002532499 | 0.998059156 | 0.999507042 | -4.914875423 |
| **A_66_P139460** | Mia3 | 0.000153115 | -0.163730498 | 0.164036728 | 8.494512385 | 0.002270202 | 0.998260174 | 0.999656559 | -4.914875769 |
| **A_55_P2106922** | Arfgap3 | 0.00023145 | -0.2710983 | 0.271561201 | 10.39062534 | 0.002072734 | 0.998411507 | 0.999684956 | -4.914876005 |
| **A_55_P1976278** | Prkcb | -0.00057091 | -0.670951118 | 0.669809298 | 9.849837903 | -0.00206933 | 0.998414116 | 0.999684956 | -4.914876008 |
| **A_55_P2087399** | Tex22 | -0.00078993 | -1.018903114 | 1.017323254 | 10.10919886 | -0.001885279 | 0.998555168 | 0.999684956 | -4.914876208 |
| **A_52_P579876** | Suds3 | -0.000254205 | -0.361634195 | 0.361125786 | 9.757965802 | -0.001709238 | 0.998690082 | 0.999684956 | -4.914876382 |
| **A_55_P2046907** | Wdr83os | -0.00015643 | -0.241796514 | 0.241483654 | 11.3900854 | -0.001573022 | 0.998794474 | 0.999684956 | -4.914876505 |
| **A_55_P2335648** | Eprn | 0.000145197 | -0.238124695 | 0.23841509 | 7.816583402 | 0.00148072 | 0.998865212 | 0.999684956 | -4.914876582 |
| **A_52_P423174** | Acaa1b | 0.000279248 | -0.460862055 | 0.461420551 | 9.983597509 | 0.001471427 | 0.998872334 | 0.999684956 | -4.914876589 |
| **A_55_P1970740** | Mrps10 | -0.000200942 | -0.335582436 | 0.335180552 | 11.72206944 | -0.001455843 | 0.998884277 | 0.999684956 | -4.914876602 |
| **A_55_P1967138** | Cenpx | -0.00027335 | -0.457255608 | 0.456708907 | 11.99588832 | -0.001453462 | 0.998886102 | 0.999684956 | -4.914876604 |
| **A_51_P454103** | Manba | -0.000260416 | -0.450174033 | 0.449653201 | 9.783947489 | -0.001406441 | 0.998922138 | 0.999684956 | -4.914876641 |
| **A_55_P1994882** | Ccdc174 | -0.000147423 | -0.262321997 | 0.262027152 | 10.5093262 | -0.001366334 | 0.998952875 | 0.999684956 | -4.914876671 |
| **A_51_P170008** | Napg | 0.000147041 | -0.278825107 | 0.279119189 | 8.688451424 | 0.001280737 | 0.999018474 | 0.999684956 | -4.914876734 |
| **A_55_P2055557** | Sdsl | -0.000158126 | -0.305899813 | 0.305583562 | 11.1095166 | -0.001256697 | 0.999036898 | 0.999684956 | -4.91487675 |
| **A_55_P2042612** | C1s1 | 0.000381694 | -0.794670134 | 0.795433522 | 12.42827444 | 0.001166548 | 0.999105986 | 0.999684956 | -4.91487681 |
| **A_55_P1983959** | Ftl2-ps | 0.000133244 | -0.2880513 | 0.288317788 | 14.7130637 | 0.001123467 | 0.999139002 | 0.999684956 | -4.914876837 |
| **A_55_P1956302** | H2-Q4 | 0.000117394 | -0.288641325 | 0.288876114 | 12.89817546 | 0.000987859 | 0.999242929 | 0.999684956 | -4.914876916 |
| **A_52_P231170** | Megf9 | -8.97E-05 | -0.227045405 | 0.226865944 | 8.483702107 | -0.00096069 | 0.99926375 | 0.999684956 | -4.91487693 |
| **A_52_P496726** | Rasd1 | -8.23E-05 | -0.208686019 | 0.208521384 | 7.956186914 | -0.000958856 | 0.999265156 | 0.999684956 | -4.914876931 |
| **A_55_P2170536** | Zfp949 | -9.53E-05 | -0.266213281 | 0.266022622 | 9.038006434 | -0.000870435 | 0.999332919 | 0.999684956 | -4.914876976 |
| **A_55_P2122830** | Zfp185 | 0.000182823 | -0.525920212 | 0.526285858 | 8.411227613 | 0.000844391 | 0.999352879 | 0.999684956 | -4.914876988 |
| **A_52_P625683** | Pxmp4 | 0.000158823 | -0.472783035 | 0.473100681 | 10.98850939 | 0.000815997 | 0.99937464 | 0.999684956 | -4.914877001 |
| **A_51_P233855** | Adprhl1 | -7.68E-05 | -0.299140414 | 0.298986838 | 8.059018173 | -0.000623899 | 0.999521859 | 0.99976718 | -4.914877077 |
| **A_51_P437478** | Zfp566 | 3.2E-05 | -0.171750569 | 0.171814519 | 8.755010174 | 0.000452283 | 0.999653381 | 0.99976718 | -4.914877128 |
| **A_66_P135742** | Wdr1 | -4.06E-05 | -0.219912776 | 0.21983165 | 11.64402713 | -0.000448277 | 0.999656451 | 0.99976718 | -4.914877129 |
| **A_55_P2166446** | NA | 3.35E-05 | -0.185232051 | 0.18529896 | 8.337024693 | 0.000438776 | 0.999663733 | 0.99976718 | -4.914877131 |
| **A_52_P215539** | C030006K11Rik | -1.77E-05 | -0.324081816 | 0.324046325 | 9.401442029 | -0.000133059 | 0.999898027 | 0.999898648 | -4.914877179 |
| **A_55_P2002757** | Blnk | 1.87E-05 | -0.342851947 | 0.342889269 | 10.4038713 | 0.000132248 | 0.999898648 | 0.999898648 | -4.914877179 |

Rychlik_Arsenic_microarray_TableS3_Lung_100_vs_0_all_p_values

| | ID | logFC | CI.L | CI.R | AveExpr | t | P.Value | adj.P.Val | B |
|---|---|---|---|---|---|---|---|---|---|
| **A_55_P2008619** | Lancl2 | 1.97665165 | 1.538310644 | 2.414992656 | 11.3726537 | 13.29874195 | 0.000418605 | 0.709264768 | -4.290782617 |
| **A_55_P2279807** | NA | -0.587647689 | -0.802231054 | -0.373064323 | 9.585127373 | -6.859166186 | 0.000694267 | 0.709264768 | -3.815362811 |
| **A_55_P1997141** | Mybl2 | 0.629079759 | 0.372753269 | 0.88540625 | 10.30625876 | 6.146990846 | 0.001191761 | 0.709264768 | -3.842629892 |
| **A_66_P127175** | Ankrd16 | -0.523271455 | -0.749144132 | -0.297398778 | 10.9376435 | -5.802480508 | 0.001575772 | 0.709264768 | -3.858781857 |
| **A_55_P1962707** | NA | 0.58939521 | 0.329766024 | 0.849024397 | 11.75532544 | 5.685955227 | 0.001736882 | 0.709264768 | -3.864771618 |
| **A_51_P296036** | Nrbp2 | -0.556526364 | -0.819986261 | -0.293066468 | 9.860649177 | -5.290802757 | 0.002444122 | 0.709264768 | -3.887370335 |
| **A_51_P187901** | Nop56 | 0.617573166 | 0.319491657 | 0.915654674 | 9.89629678 | 5.189239646 | 0.002676399 | 0.709264768 | -3.893811373 |
| **A_55_P2016712** | Msl1 | 1.242137021 | 0.751441772 | 1.73283227 | 11.98963894 | 7.465349975 | 0.002939944 | 0.709264768 | -4.308057432 |
| **A_51_P438805** | Txnip | -0.646864558 | -0.966942752 | -0.326786364 | 12.3879707 | -5.061830681 | 0.003004662 | 0.709264768 | -3.902297846 |
| **A_55_P2135035** | NA | 0.538749103 | 0.271440218 | 0.806057989 | 8.581870218 | 5.048048409 | 0.003042873 | 0.709264768 | -3.903244103 |
| **A_52_P56397** | Aunip | 0.892495525 | 0.448829593 | 1.336161457 | 8.878859945 | 5.038488956 | 0.003069705 | 0.709264768 | -3.903903755 |
| **A_52_P76034** | Rcc2 | 0.544082039 | 0.269317982 | 0.818846096 | 11.9284503 | 4.959693219 | 0.003301549 | 0.709264768 | -3.909446586 |
| **A_55_P1958464** | NA | 0.630606128 | 0.311168376 | 0.950043879 | 10.84941502 | 4.944498975 | 0.003348538 | 0.709264768 | -3.910537448 |
| **A_55_P2112997** | Ddx18 | 0.475166723 | 0.233987043 | 0.716346404 | 10.82204248 | 4.934641977 | 0.003379432 | 0.709264768 | -3.911249002 |
| **A_51_P518621** | H3c1 | 0.686341696 | 0.333353803 | 1.039329589 | 8.435545501 | 4.870021957 | 0.003590272 | 0.709264768 | -3.915990532 |
| **A_66_P115053** | Anp32b | 0.582126021 | 0.281593575 | 0.882658466 | 13.26795915 | 4.851499525 | 0.003653465 | 0.709264768 | -3.917374581 |
| **A_52_P157880** | Gm1947 | 0.638773224 | 0.307754222 | 0.969792225 | 9.220662807 | 4.833304142 | 0.00371679 | 0.709264768 | -3.918745208 |
| **A_55_P2002376** | Srm | 0.629117614 | 0.296463411 | 0.961771817 | 10.43454604 | 4.736845019 | 0.004074281 | 0.709264768 | -3.926197895 |
| **A_55_P2143041** | Phgdh | 0.730927901 | 0.34296658 | 1.118889223 | 10.60518767 | 4.718853082 | 0.004145243 | 0.709264768 | -3.927623541 |
| **A_55_P2054300** | Alg8 | 0.83701536 | 0.387076 | 1.28695472 | 9.510775371 | 4.659397763 | 0.004390022 | 0.709264768 | -3.932416541 |
| **A_55_P1956847** | Nolc1 | 0.540289899 | 0.248098232 | 0.832481565 | 11.12942631 | 4.631368823 | 0.004511134 | 0.709264768 | -3.934720497 |
| **A_55_P2093770** | NA | -0.630265847 | -0.972517884 | -0.28801381 | 10.54406346 | -4.612411831 | 0.004595216 | 0.709264768 | -3.93629517 |
| **A_51_P300709** | Srm | 0.705255764 | 0.320368501 | 1.090143028 | 10.84463045 | 4.589479666 | 0.004699338 | 0.709264768 | -3.938217969 |
| **A_51_P170758** | Grwd1 | 0.510104004 | 0.22631142 | 0.793896588 | 9.607228209 | 4.502026386 | 0.005121967 | 0.709264768 | -3.945734877 |
| **A_55_P1964174** | Nme1 | 0.398268026 | 0.17221803 | 0.624318022 | 10.7138434 | 4.412871527 | 0.005598156 | 0.709264768 | -3.953708909 |
| **A_55_P1972263** | Nop56 | 0.998157447 | 0.424331425 | 1.571983468 | 11.48998499 | 4.356815393 | 0.005923407 | 0.709264768 | -3.95888978 |
| **A_51_P215077** | Mgst3 | -0.594974456 | -0.943262012 | -0.2466869 | 10.35558056 | -4.278688876 | 0.006413316 | 0.709264768 | -3.966334101 |
| **A_55_P2046411** | NA | 0.565352506 | 0.234138546 | 0.896566466 | 12.29565924 | 4.275245256 | 0.00643595 | 0.709264768 | -3.966668361 |
| **A_52_P372032** | Ddx21 | 0.471884757 | 0.187433191 | 0.756336322 | 9.421354387 | 4.155066374 | 0.007285869 | 0.709264768 | -3.9786693 |
| **A_55_P2108773** | Tedc1 | 0.781341165 | 0.308689751 | 1.253992578 | 9.349671867 | 4.140474271 | 0.007397502 | 0.709264768 | -3.980171916 |
| **A_55_P1980020** | Rsl1d1 | 0.511613035 | 0.202112402 | 0.821113669 | 12.7531786 | 4.140286592 | 0.00739895 | 0.709264768 | -3.980191308 |
| **A_51_P466162** | C1qbp | 0.694098014 | 0.272422583 | 1.115773444 | 12.65785987 | 4.122805534 | 0.007535283 | 0.709264768 | -3.982004862 |
| **A_51_P241769** | Rhd | 0.593707921 | 0.230672798 | 0.956743044 | 8.604415132 | 4.096137668 | 0.007748797 | 0.709264768 | -3.984799684 |
| **A_55_P2133187** | NA | 0.412087867 | 0.157788218 | 0.666387516 | 8.603269809 | 4.058769725 | 0.008059638 | 0.709264768 | -3.98877396 |
| **A_51_P172752** | Gpatch4 | 0.484741986 | 0.184141627 | 0.785342345 | 11.01770939 | 4.038978013 | 0.008229992 | 0.709264768 | -3.99090675 |
| **A_51_P106059** | Traf4 | 1.038859402 | 0.393462862 | 1.684255943 | 11.48813985 | 4.031628319 | 0.008294291 | 0.709264768 | -4.02424981 |
| **A_55_P2067960** | Cplx2 | -0.393873311 | -0.638930123 | -0.1488165 | 14.6206488 | -4.025688073 | 0.008346675 | 0.709264768 | -3.992349843 |
| **A_55_P2121521** | H3c11 | 0.792016557 | 0.298103855 | 1.285929259 | 8.540997027 | 4.016376444 | 0.008429548 | 0.709264768 | -3.993366224 |
| **A_55_P2002702** | Ddx18 | 0.463229217 | 0.174234941 | 0.752223494 | 10.3355364 | 4.014736514 | 0.008444239 | 0.709264768 | -3.993545678 |
| **A_55_P1974317** | Surf6 | 0.354711598 | 0.133367027 | 0.57605617 | 9.41093676 | 4.013810448 | 0.008452548 | 0.709264768 | -3.993647075 |
| **A_55_P1964658** | NA | 0.681419964 | 0.255403266 | 1.107436662 | 12.256513 | 4.006254891 | 0.008520688 | 0.709264768 | -3.994475968 |
| **A_55_P2130535** | Dnmt3b | 0.425896722 | 0.158543965 | 0.69324948 | 10.09204745 | 3.989973173 | 0.008669652 | 0.709264768 | -3.99627202 |
| **A_55_P2105512** | Nop56 | 0.969939283 | 0.35859652 | 1.581282045 | 12.07915399 | 3.973837642 | 0.008820197 | 0.709264768 | -3.998065297 |
| **A_51_P159402** | Gcat | 0.341941982 | 0.126262251 | 0.557621713 | 10.36783606 | 3.970941028 | 0.008847535 | 0.709264768 | -3.998388638 |
| **A_52_P279152** | Vamp8 | -0.48070078 | -0.784557644 | -0.176843916 | 12.47447775 | -3.962379984 | 0.008928896 | 0.709264768 | -3.999346811 |

| | | | | | | | | | |
|---|---|---|---|---|---|---|---|---|---|
| **A_55_P2158697** | H4c9 | 0.475388983 | 0.172532631 | 0.778245335 | 9.408232715 | 3.931540667 | 0.009229086 | 0.709264768 | -4.002829975 |
| **A_55_P1958160** | Sgce | -0.524698266 | -0.859054541 | -0.190341991 | 10.89293694 | -3.930524393 | 0.009239171 | 0.709264768 | -4.002945605 |
| **A_55_P2056342** | Mgst3 | -0.71729201 | -1.175448456 | -0.259135563 | 11.19423876 | -3.92132228 | 0.00933106 | 0.709264768 | -4.003995078 |
| **A_55_P2076001** | Rbm12 | 0.622087298 | 0.219524232 | 1.024650364 | 10.88387996 | 3.870506067 | 0.009857492 | 0.709264768 | -4.009871549 |
| **A_52_P131836** | Bysl | 0.381211051 | 0.133663412 | 0.628758691 | 10.41179379 | 3.857065722 | 0.010002289 | 0.709264768 | -4.01144903 |
| **A_55_P2027327** | Usp1 | 0.49340869 | 0.172875251 | 0.813942128 | 11.26517654 | 3.855527429 | 0.010019014 | 0.709264768 | -4.011630204 |
| **A_52_P569549** | Lig4 | 0.55690483 | 0.190925206 | 0.922884454 | 9.347037012 | 3.811311312 | 0.010513536 | 0.709264768 | -4.016893285 |
| **A_52_P278354** | Bmp7 | 0.732028393 | 0.250324404 | 1.213732382 | 9.135810754 | 3.806256302 | 0.010571808 | 0.709264768 | -4.017501873 |
| **A_55_P2102454** | NA | 0.419028078 | 0.142407667 | 0.695648489 | 10.50790389 | 3.794104146 | 0.01071339 | 0.709264768 | -4.018970743 |
| **A_55_P2185273** | Umps | 0.396685944 | 0.133643063 | 0.659728826 | 11.54727119 | 3.777205426 | 0.010913844 | 0.709264768 | -4.021027124 |
| **A_51_P133562** | Serpina6 | 0.524063899 | 0.172261571 | 0.875866227 | 8.092233864 | 3.731091336 | 0.011482723 | 0.709264768 | -4.02672119 |
| **A_66_P108019** | Heatr1 | 0.357303286 | 0.116674634 | 0.597931938 | 11.29529038 | 3.719118513 | 0.011635839 | 0.709264768 | -4.028219547 |
| **A_51_P314111** | Psip1 | 0.358317694 | 0.116733982 | 0.599901407 | 10.79883446 | 3.714932696 | 0.011689912 | 0.709264768 | -4.02874535 |
| **A_52_P108346** | Myc | 0.800523607 | 0.259593726 | 1.341453489 | 9.306818015 | 3.706666528 | 0.011797528 | 0.709264768 | -4.029786703 |
| **A_51_P134007** | Ncl | 0.88053816 | 0.284971719 | 1.4761046 | 13.10133744 | 3.703124466 | 0.011843983 | 0.709264768 | -4.030234144 |
| **A_55_P1958245** | Tcof1 | 0.455724827 | 0.142645008 | 0.768804646 | 10.27301761 | 3.645842982 | 0.012624576 | 0.709264768 | -4.037572844 |
| **A_51_P496751** | Nop58 | 0.751333548 | 0.235166876 | 1.26750022 | 12.07488378 | 3.64580274 | 0.012625144 | 0.709264768 | -4.037578068 |
| **A_55_P2069083** | Galnt12 | 0.73050029 | 0.227300496 | 1.233700083 | 9.667716778 | 3.636053662 | 0.012763688 | 0.709264768 | -4.038846592 |
| **A_51_P154684** | Alyref | 1.484161594 | 0.667213526 | 2.301109663 | 12.45821168 | 6.537265523 | 0.012784201 | 0.709264768 | -4.411742303 |
| **A_55_P1997791** | Fbl | 0.749959707 | 0.232152679 | 1.267766736 | 12.0813033 | 3.627607871 | 0.012885093 | 0.709264768 | -4.03995017 |
| **A_51_P185906** | Abi3 | 0.983079696 | 0.303335156 | 1.662824237 | 9.224786448 | 3.622374467 | 0.012960972 | 0.709264768 | -4.040636164 |
| **A_51_P162176** | Trappc3 | 0.620382736 | 0.190976522 | 1.04978895 | 10.21494114 | 3.618609545 | 0.013015871 | 0.709264768 | -4.041130698 |
| **A_65_P05358** | NA | 0.379325342 | 0.116080685 | 0.642569999 | 12.25425506 | 3.609130938 | 0.013155245 | 0.709264768 | -4.04237956 |
| **A_51_P393426** | Pros1 | -0.673264954 | -1.141032162 | -0.205497746 | 10.68931358 | -3.60501089 | 0.013216349 | 0.709264768 | -4.042924109 |
| **A_52_P579640** | Rrs1 | 0.572207124 | 0.17366462 | 0.970749628 | 10.03203585 | 3.596076821 | 0.013349946 | 0.709264768 | -4.044108501 |
| **A_51_P113784** | Prpf19 | 0.50316667 | 0.152567643 | 0.853765698 | 9.005695792 | 3.594607736 | 0.01337206 | 0.709264768 | -4.044303726 |
| **A_55_P1985001** | NA | -0.598307506 | -1.015972668 | -0.180642344 | 9.331035886 | -3.587951348 | 0.013472771 | 0.709264768 | -4.045189952 |
| **A_55_P2186929** | NA | 0.597414706 | 0.179441592 | 1.01538782 | 10.62449677 | 3.579957801 | 0.013594841 | 0.709264768 | -4.046257815 |
| **A_55_P2069554** | Brix1 | 0.365380642 | 0.108593115 | 0.62216817 | 11.74307731 | 3.563870832 | 0.013844289 | 0.709264768 | -4.048418879 |
| **A_55_P2131766** | Rsrp1 | -0.71531285 | -1.218142599 | -0.212483102 | 13.7222734 | -3.563078652 | 0.013856705 | 0.709264768 | -4.048525713 |
| **A_55_P2038303** | NA | 0.555390339 | 0.164638334 | 0.946142344 | 9.705740259 | 3.559979126 | 0.013905404 | 0.709264768 | -4.048944093 |
| **A_55_P2124701** | NA | 0.36109932 | 0.105686689 | 0.616511952 | 12.70052823 | 3.541071086 | 0.014206673 | 0.709264768 | -4.051509353 |
| **A_55_P2004016** | Crispld2 | -0.949140499 | -1.622754186 | -0.275526812 | 10.68270893 | -3.529148614 | 0.014400399 | 0.709264768 | -4.053138429 |
| **A_55_P2098683** | St7l | -0.333071693 | -0.569709134 | -0.096434251 | 10.55561378 | -3.525369216 | 0.014462427 | 0.709264768 | -4.053656717 |
| **A_55_P2148243** | Wdr12 | 0.446386831 | 0.129012145 | 0.763761516 | 10.22149052 | 3.522811742 | 0.01450457 | 0.709264768 | -4.05400795 |
| **A_55_P2064321** | Lama5 | -0.561736068 | -0.961332045 | -0.162140091 | 14.16701989 | -3.520963707 | 0.014535109 | 0.709264768 | -4.054262009 |
| **A_51_P237383** | Rnase4 | -0.586639214 | -1.004476787 | -0.168801641 | 9.345776014 | -3.516526913 | 0.014608721 | 0.709264768 | -4.054872845 |
| **A_55_P2056325** | Anxa3 | -0.629666661 | -1.078561664 | -0.180771658 | 10.1237707 | -3.513308396 | 0.014662382 | 0.709264768 | -4.055316737 |
| **A_51_P189269** | Mbnl2 | -0.304239562 | -0.522128838 | -0.086350287 | 9.728044653 | -3.497278178 | 0.014932957 | 0.709264768 | -4.057537439 |
| **A_52_P167278** | Mthfd1l | 0.71790709 | 0.202802626 | 1.233011554 | 10.0996106 | 3.49078639 | 0.015044121 | 0.709264768 | -4.058441442 |
| **A_52_P106739** | Ddx46 | 0.358896016 | 0.099473318 | 0.618318713 | 10.56253027 | 3.465062017 | 0.015493819 | 0.709264768 | -4.06205036 |
| **A_55_P2007500** | Tmem151 | 1.014571687 | 0.280278725 | 1.74886465 | 8.412788756 | 3.460698133 | 0.015571589 | 0.709264768 | -4.062666831 |
| **A_51_P455997** | Meg3 | -1.089508019 | -1.879046584 | -0.299969455 | 11.8647758 | -3.45626815 | 0.015650984 | 0.709264768 | -4.063293909 |
| **A_55_P1957613** | Ak2 | 0.300221683 | 0.082436239 | 0.518007126 | 13.50113613 | 3.452737426 | 0.015714587 | 0.709264768 | -4.06379461 |
| **A_55_P2118362** | Tmem121 | 0.974788565 | 0.266350882 | 1.683226247 | 8.723995779 | 3.446347912 | 0.015830424 | 0.709264768 | -4.064702796 |
| **A_51_P248044** | Lamb2 | -0.40383486 | -0.697839708 | -0.109830012 | 11.2072038 | -3.440326492 | 0.015940463 | 0.709264768 | -4.065561109 |
| **A_52_P681705** | Zfp955b | 0.739377972 | 0.198600175 | 1.28015577 | 9.849222805 | 3.424506545 | 0.016233664 | 0.709264768 | -4.0678275 |
| **A_55_P2140151** | Flcn | -0.492604137 | -0.853176297 | -0.132031977 | 10.94114293 | -3.421812504 | 0.016284193 | 0.709264768 | -4.068215098 |

| | | | | | | | | | |
|---|---|---|---|---|---|---|---|---|---|
| **A_51_P462448** | Gpx4 | -0.48549982 | -0.843026601 | -0.127973038 | 14.63566203 | -3.401189573 | 0.016676866 | 0.709264768 | -4.071198121 |
| **A_55_P2027392** | Gpr146 | -0.60502641 | -1.050917963 | -0.159134857 | 10.11968312 | -3.398563845 | 0.016727616 | 0.709264768 | -4.071579954 |
| **A_55_P2118784** | Rcor2 | 0.587317902 | 0.154073788 | 1.020562017 | 10.00774199 | 3.395399754 | 0.016789001 | 0.709264768 | -4.072040688 |
| **A_55_P2433438** | Tcf12 | 0.304173644 | 0.078782257 | 0.529565031 | 9.184656794 | 3.380139396 | 0.017088615 | 0.709264768 | -4.074272222 |
| **A_55_P2055199** | Airn | -0.385310734 | -0.671402116 | -0.099219351 | 10.22703886 | -3.373313282 | 0.017224564 | 0.709264768 | -4.075275478 |
| **A_51_P133612** | Cdt1 | 1.020776421 | 0.262396603 | 1.77915624 | 12.12233466 | 3.371275217 | 0.017265388 | 0.709264768 | -4.075575629 |
| **A_52_P18807** | Eif3c | 0.297536645 | 0.076235108 | 0.518838182 | 13.66876305 | 3.367490481 | 0.017341485 | 0.709264768 | -4.07613376 |
| **A_51_P398878** | Uck2 | 0.695080012 | 0.177628975 | 1.212531048 | 11.49758674 | 3.364463987 | 0.017402606 | 0.709264768 | -4.076580771 |
| **A_55_P2168628** | Sac3d1 | 0.340402138 | 0.086637355 | 0.594166922 | 11.70682989 | 3.359783341 | 0.017497608 | 0.709264768 | -4.077273323 |
| **A_55_P1957459** | Lilrb4a | -0.30749162 | -0.536886587 | -0.078096653 | 8.954332643 | -3.357374165 | 0.017546732 | 0.709264768 | -4.077630365 |
| **A_55_P1971159** | NA | 0.624806971 | 0.157161446 | 1.092452496 | 9.706287428 | 3.346412155 | 0.017772201 | 0.709264768 | -4.079259932 |
| **A_66_P100565** | Ccdc107 | -0.473540387 | -0.828253926 | -0.118826849 | 12.91351328 | -3.343717713 | 0.017828114 | 0.709264768 | -4.07966173 |
| **A_66_P125212** | Mapk1ip1 | 0.328646549 | 0.081759397 | 0.575533701 | 11.72475142 | 3.334117742 | 0.018028926 | 0.709264768 | -4.081097325 |
| **A_55_P1989452** | Nenf | -0.328432031 | -0.575168667 | -0.081695395 | 13.53691345 | -3.333974027 | 0.018031952 | 0.709264768 | -4.081118864 |
| **A_55_P1953169** | Saa3 | -0.726578466 | -1.273027241 | -0.180129691 | 8.447242037 | -3.330300413 | 0.018109477 | 0.709264768 | -4.081669929 |
| **A_55_P2152926** | Mybbp1a | 0.356036516 | 0.088024284 | 0.624048749 | 12.48297218 | 3.327287069 | 0.018173346 | 0.709264768 | -4.082122641 |
| **A_55_P2006525** | Adamtsl4 | -0.484101036 | -0.848560104 | -0.119641968 | 8.991064927 | -3.326883537 | 0.018181918 | 0.709264768 | -4.082183313 |
| **A_52_P399934** | Dusp2 | 0.957402176 | 0.234842632 | 1.67996172 | 10.18431271 | 3.31872383 | 0.018356221 | 0.709264768 | -4.083412555 |
| **A_51_P313761** | Shmt2 | 0.522757721 | 0.127520355 | 0.917995086 | 10.59898769 | 3.312781602 | 0.018484328 | 0.709264768 | -4.084310628 |
| **A_55_P1984113** | Nup62 | 0.304727473 | 0.074284442 | 0.535170504 | 13.18860163 | 3.312061382 | 0.018499922 | 0.709264768 | -4.084419644 |
| **A_51_P163261** | Gm6985 | 0.439760972 | 0.106862604 | 0.772659341 | 9.661704054 | 3.30868455 | 0.018573235 | 0.709264768 | -4.084931254 |
| **A_66_P131979** | Cdc6 | 0.732622867 | 0.17689787 | 1.288347864 | 8.591600569 | 3.30195297 | 0.018720347 | 0.709264768 | -4.085953486 |
| **A_55_P2054342** | 2810408I1 | 0.532064906 | 0.127927582 | 0.93620223 | 8.871771225 | 3.297509072 | 0.018818176 | 0.709264768 | -4.086630042 |
| **A_52_P295313** | Wdr46 | 0.287688933 | 0.068921058 | 0.506456808 | 11.16722146 | 3.29374485 | 0.018901487 | 0.709264768 | -4.087204197 |
| **A_51_P245796** | Ddit4 | -0.932113915 | -1.641665471 | -0.222562358 | 12.62853574 | -3.290299109 | 0.01897811 | 0.709264768 | -4.087730639 |
| **A_55_P1963309** | G3bp1 | 0.79185757 | 0.188815539 | 1.3948996 | 12.62538596 | 3.28889369 | 0.019009461 | 0.709264768 | -4.087945597 |
| **A_55_P1998066** | Tmem121 | 0.959078822 | 0.227762776 | 1.690394868 | 8.735013728 | 3.284728995 | 0.019102704 | 0.709264768 | -4.088583396 |
| **A_52_P249424** | Vegfa | -0.733360058 | -1.292927941 | -0.173792176 | 10.16651379 | -3.282576222 | 0.019151102 | 0.709264768 | -4.088913555 |
| **A_51_P430973** | Paqr7 | -0.446891966 | -0.788752994 | -0.105030938 | 9.181735939 | -3.274185631 | 0.019341042 | 0.709264768 | -4.090203473 |
| **A_51_P492765** | Wdr75 | 0.371063977 | 0.086676181 | 0.655451773 | 10.69485881 | 3.268046184 | 0.019481351 | 0.709264768 | -4.091150442 |
| **A_55_P1969396** | NA | 0.474389331 | 0.110720843 | 0.838057819 | 10.98721995 | 3.267229095 | 0.019500109 | 0.709264768 | -4.091276673 |
| **A_51_P325485** | Utp3 | 0.368976166 | 0.085879053 | 0.652073279 | 12.10036406 | 3.264474035 | 0.019563508 | 0.709264768 | -4.091702643 |
| **A_55_P2130610** | Smc3 | 0.398161971 | 0.0915359 | 0.704788042 | 12.26463978 | 3.252378226 | 0.019844578 | 0.709264768 | -4.093579153 |
| **A_51_P435333** | Dctn2 | -0.447899791 | -0.792912813 | -0.10288677 | 11.04732205 | -3.25158955 | 0.01986306 | 0.709264768 | -4.093701865 |
| **A_55_P2032659** | Tox | 0.624284705 | 0.143041193 | 1.105528216 | 8.407464266 | 3.249137969 | 0.019920633 | 0.709264768 | -4.094083594 |
| **A_52_P639774** | Gart | 0.594435439 | 0.134844797 | 1.054026082 | 8.975401041 | 3.239543838 | 0.02014773 | 0.709264768 | -4.095581565 |
| **A_55_P2111950** | Dnajc7 | 0.993486717 | 0.221198387 | 1.765775048 | 12.21690308 | 3.222054914 | 0.020569138 | 0.709264768 | -4.098329035 |
| **A_55_P2076871** | Lef1 | 1.782578976 | 0.395627614 | 3.169530339 | 9.915543258 | 3.219125395 | 0.02064068 | 0.709264768 | -4.098791391 |
| **A_51_P437144** | Dynll2 | 0.357046889 | 0.078949046 | 0.635144733 | 11.8565413 | 3.215718159 | 0.020724236 | 0.709264768 | -4.099329918 |
| **A_51_P344008** | Slc25a20 | 0.312762668 | 0.068989538 | 0.556535798 | 8.446411998 | 3.213508265 | 0.02077863 | 0.709264768 | -4.099679645 |
| **A_55_P2058628** | Hnrnpa3 | 1.450942514 | 0.311145202 | 2.590739826 | 10.45468864 | 3.188400901 | 0.021407875 | 0.709264768 | -4.103677695 |
| **A_55_P2008538** | Chmp4b | -1.965784445 | -3.511605083 | -0.419963807 | 12.91091571 | -3.185129824 | 0.0214914 | 0.709264768 | -4.104201927 |
| **A_55_P2186005** | Sphk1 | -0.558104972 | -0.99714258 | -0.119067364 | 9.026584129 | -3.183937352 | 0.021521939 | 0.709264768 | -4.10439323 |
| **A_55_P2012498** | Cdca7 | 1.535409631 | 0.326195007 | 2.744624256 | 11.61147918 | 3.18032289 | 0.0216148 | 0.709264768 | -4.104973709 |
| **A_51_P500344** | Mat2a | 0.430199848 | 0.090539797 | 0.7698599 | 13.47310372 | 3.172313706 | 0.021822154 | 0.709264768 | -4.10626336 |
| **A_55_P2059686** | Rcc1 | 0.846138239 | 0.177690632 | 1.514585846 | 11.2526059 | 3.170474721 | 0.021870075 | 0.709264768 | -4.106560135 |
| **A_51_P340402** | Eefsec | 0.401163322 | 0.084108246 | 0.718218398 | 9.49684783 | 3.169106643 | 0.021905801 | 0.709264768 | -4.106781075 |
| **A_55_P1975322** | NA | 0.485158221 | 0.101680826 | 0.868635616 | 10.59265327 | 3.168793742 | 0.021913981 | 0.709264768 | -4.106831626 |

| | | | | | | | | | |
|---|---|---|---|---|---|---|---|---|---|
| **A_51_P324228** | Satb1 | 1.799984901 | 0.375299614 | 3.224670189 | 9.50514113 | 3.164464795 | 0.0220275 | 0.709264768 | -4.107531735 |
| **A_51_P231738** | Dnajc4 | -0.296614887 | -0.531560309 | -0.061669464 | 10.65092578 | -3.16210511 | 0.022089653 | 0.709264768 | -4.107913937 |
| **A_55_P2129319** | Mre11a | 0.463733344 | 0.095665282 | 0.831801406 | 9.855133629 | 3.155662399 | 0.022260343 | 0.709264768 | -4.108959546 |
| **A_55_P1994275** | Gar1 | 0.898501217 | 0.18534427 | 1.611658163 | 12.3691936 | 3.155614174 | 0.022261626 | 0.709264768 | -4.108967384 |
| **A_55_P1978316** | Adamts14 | -0.375492505 | -0.674655613 | -0.076329397 | 10.16878272 | -3.143718005 | 0.02258067 | 0.709264768 | -4.11090609 |
| **A_55_P2084122** | Dus4l | 0.384409308 | 0.077532765 | 0.69128585 | 8.827639021 | 3.137476978 | 0.022750071 | 0.709264768 | -4.111927343 |
| **A_55_P2236382** | D8Ertd158 | -0.438974581 | -0.79184995 | -0.086099212 | 10.07582857 | -3.115791392 | 0.023349715 | 0.709264768 | -4.115498234 |
| **A_55_P2038273** | NA | 0.879446712 | 0.172085443 | 1.586807981 | 9.686222299 | 3.113999931 | 0.023400029 | 0.709264768 | -4.115794785 |
| **A_52_P515769** | Pcdh12 | -0.419539706 | -0.757185213 | -0.081894199 | 12.08107958 | -3.112163753 | 0.023451723 | 0.709264768 | -4.116098986 |
| **A_51_P214269** | Epha1 | 1.091493379 | 0.212725224 | 1.970261534 | 9.843889539 | 3.110979438 | 0.023485132 | 0.709264768 | -4.116295326 |
| **A_66_P102163** | NA | 1.267553724 | 0.246906974 | 2.288200475 | 9.673636777 | 3.110579202 | 0.023496434 | 0.709264768 | -4.116361702 |
| **A_55_P2095488** | Syncrip | 0.377744944 | 0.073411434 | 0.682078453 | 9.585589848 | 3.108846153 | 0.023545444 | 0.709264768 | -4.116649252 |
| **A_55_P1993365** | NA | 1.191416527 | 0.230739056 | 2.152093997 | 9.396192909 | 3.106249669 | 0.023619082 | 0.709264768 | -4.117080482 |
| **A_51_P514961** | Tiparp | -0.281724054 | -0.508932738 | -0.054515369 | 10.51202165 | -3.105627374 | 0.023636769 | 0.709264768 | -4.117183909 |
| **A_55_P2075359** | Eif4a3l1 | 0.40249218 | 0.077844256 | 0.727140104 | 12.301578 | 3.105239933 | 0.023647788 | 0.709264768 | -4.117248317 |
| **A_55_P1996223** | Sf3b4 | 0.290143972 | 0.055963289 | 0.524324656 | 11.77506787 | 3.103221886 | 0.023705275 | 0.709264768 | -4.117583978 |
| **A_52_P595954** | Nelfa | 0.420573431 | 0.080522858 | 0.760624004 | 10.0650663 | 3.09776642 | 0.023861457 | 0.709264768 | -4.118492904 |
| **A_55_P1966573** | Gemin4 | 0.626021106 | 0.119535528 | 1.132506683 | 10.36019061 | 3.095795307 | 0.023918166 | 0.709264768 | -4.118821855 |
| **A_55_P2005190** | Herc4 | -0.310617253 | -0.562262439 | -0.058972066 | 10.94167704 | -3.091628425 | 0.024038539 | 0.709264768 | -4.119518203 |
| **A_51_P421846** | Ech1 | -0.429602531 | -0.77883988 | -0.080365181 | 11.20744713 | -3.081034017 | 0.024347617 | 0.709264768 | -4.121294542 |
| **A_55_P1969845** | Lrpprc | 0.370058014 | 0.068858398 | 0.671257629 | 11.70344127 | 3.07727101 | 0.024458452 | 0.709264768 | -4.121927504 |
| **A_55_P1985670** | Exosc3 | 0.333286644 | 0.061251757 | 0.60532153 | 9.669671657 | 3.068623826 | 0.024715263 | 0.709264768 | -4.123386051 |
| **A_51_P169920** | Noc2l | 0.376248408 | 0.068692922 | 0.683803894 | 10.53965263 | 3.064090144 | 0.024851097 | 0.709264768 | -4.124153013 |
| **A_52_P302345** | Cyp4v3 | -0.543173505 | -0.987403531 | -0.09894348 | 8.774732884 | -3.062534673 | 0.02489789 | 0.709264768 | -4.124416509 |
| **A_52_P184149** | Mthfd2 | 0.337008661 | 0.060745152 | 0.61327217 | 8.767208857 | 3.055398659 | 0.025113813 | 0.709264768 | -4.12562769 |
| **A_55_P1989321** | Sat1 | -0.717331149 | -1.30580636 | -0.128855938 | 11.32127231 | -3.053105854 | 0.025183628 | 0.709264768 | -4.126017661 |
| **A_55_P1952379** | Fkbp5 | 1.058531768 | 0.187901125 | 1.929162411 | 11.10741518 | 3.045231359 | 0.025425036 | 0.709264768 | -4.127360025 |
| **A_51_P501248** | Sphk1 | -0.771911183 | -1.407574729 | -0.136247636 | 11.6848404 | -3.041517866 | 0.025539765 | 0.709264768 | -4.127994697 |
| **A_55_P2179020** | Mrpl15 | 0.3292786 | 0.057858361 | 0.60069884 | 10.06089472 | 3.038586632 | 0.02563073 | 0.709264768 | -4.128496414 |
| **A_66_P117659** | Nfatc3 | 0.299560369 | 0.052549861 | 0.546570877 | 9.450797446 | 3.037520889 | 0.025663892 | 0.709264768 | -4.128678991 |
| **A_51_P175232** | Fbxl4 | -0.372690112 | -0.68018796 | -0.065192263 | 9.296843372 | -3.035681014 | 0.025721252 | 0.709264768 | -4.128994392 |
| **A_51_P483329** | Sdad1 | 0.373084627 | 0.064961641 | 0.681207614 | 10.79662951 | 3.03272898 | 0.025813582 | 0.709264768 | -4.129500983 |
| **A_51_P122238** | Uchl5 | 0.625163507 | 0.108634883 | 1.141692131 | 11.16822671 | 3.031444146 | 0.025853882 | 0.709264768 | -4.172855709 |
| **A_51_P326191** | Serpina3g | 0.739060725 | 0.128315119 | 1.349806331 | 8.640914189 | 3.030889604 | 0.025871296 | 0.709264768 | -4.129816969 |
| **A_55_P2032912** | NA | -0.40530652 | -0.740248955 | -0.070364085 | 13.89035253 | -3.030845252 | 0.02587269 | 0.709264768 | -4.129824591 |
| **A_52_P432580** | NA | -0.367094613 | -0.670493163 | -0.063696064 | 12.29614135 | -3.03050397 | 0.025883415 | 0.709264768 | -4.12988325 |
| **A_52_P574190** | Psmd12 | 0.29537746 | 0.051003759 | 0.539751161 | 12.98369182 | 3.027423867 | 0.025980428 | 0.709264768 | -4.130413049 |
| **A_55_P2029592** | Dus2 | 0.275413113 | 0.047475917 | 0.50335031 | 9.311346982 | 3.026354211 | 0.026014213 | 0.709264768 | -4.130597207 |
| **A_55_P1995512** | Tmem139 | -0.487983479 | -0.89228329 | -0.083683669 | 10.56039084 | -3.023095921 | 0.026117423 | 0.709264768 | -4.131158711 |
| **A_51_P362903** | Mrps22 | 0.31083198 | 0.053037922 | 0.568626038 | 10.8034027 | 3.019973534 | 0.026216752 | 0.709264768 | -4.131697556 |
| **A_66_P118124** | Fxyd3 | -0.392857904 | -0.718768902 | -0.066946906 | 10.37809365 | -3.019164839 | 0.026242546 | 0.709264768 | -4.131837238 |
| **A_55_P2084696** | H2ac1 | 0.416056756 | 0.070667126 | 0.761446386 | 8.486959846 | 3.017127271 | 0.026307659 | 0.709264768 | -4.132189399 |
| **A_52_P625431** | Hnrnpa0 | 0.280724101 | 0.047267895 | 0.514180306 | 8.736525937 | 3.011789548 | 0.026479081 | 0.709264768 | -4.133113446 |
| **A_55_P2145136** | NA | 0.441413719 | 0.074121029 | 0.808706409 | 8.234744992 | 3.010120541 | 0.026532933 | 0.709264768 | -4.133402827 |
| **A_55_P2053923** | Rab17 | -0.350520666 | -0.642262679 | -0.058778654 | 8.3794239 | -3.009296639 | 0.026559562 | 0.709264768 | -4.133545758 |
| **A_55_P2064038** | NA | 0.894288439 | 0.149505533 | 1.639071345 | 8.949061559 | 3.007448934 | 0.026619387 | 0.709264768 | -4.133866489 |
| **A_55_P2020128** | Dhrs3 | -0.459085103 | -0.841631126 | -0.07653908 | 13.25029796 | -3.00579844 | 0.026672952 | 0.709264768 | -4.134153209 |
| **A_55_P2060567** | Ddx39 | 0.643669171 | 0.106689861 | 1.180648481 | 12.38383856 | 3.002309784 | 0.026786565 | 0.709264768 | -4.134759939 |

| | | | | | | | | | |
|---|---|---|---|---|---|---|---|---|---|
| **A_55_P2033680** | NA | 0.346481401 | 0.057320291 | 0.635642511 | 9.314317814 | 3.00116857 | 0.026823846 | 0.709264768 | -4.134958616 |
| **A_51_P126198** | Dapk2 | -0.693625644 | -1.274269053 | -0.112982234 | 10.53793985 | -2.992030286 | 0.02712444 | 0.709264768 | -4.136553145 |
| **A_51_P171772** | Bcl11b | 1.037239905 | 0.168840566 | 1.905639245 | 8.350345335 | 2.991645143 | 0.02713719 | 0.709264768 | -4.13662049 |
| **A_55_P2029417** | 2610318N | 0.971544999 | 0.153534484 | 1.789555514 | 9.031410594 | 2.974776759 | 0.027702136 | 0.709264768 | -4.13958127 |
| **A_51_P214796** | Suz12 | 0.274891964 | 0.043290771 | 0.506493158 | 10.57587873 | 2.97284042 | 0.02776781 | 0.709264768 | -4.139922551 |
| **A_51_P396708** | Med21 | -0.248846832 | -0.458536749 | -0.039156915 | 10.639661 | -2.972383995 | 0.027783316 | 0.709264768 | -4.140003038 |
| **A_51_P408946** | Ccne1 | 1.01598909 | 0.156218501 | 1.875759679 | 10.70451356 | 2.959762103 | 0.028215893 | 0.709264768 | -4.142235232 |
| **A_55_P1972018** | H4c3 | 1.192195383 | 0.181247711 | 2.203143055 | 10.50614701 | 2.95371851 | 0.028425636 | 0.709264768 | -4.143308435 |
| **A_55_P2066727** | Serpinb6a | -0.547901127 | -1.012612091 | -0.083190162 | 13.38953205 | -2.95304197 | 0.028449222 | 0.709264768 | -4.14342875 |
| **A_66_P120567** | Anxa1 | -0.340526448 | -0.629587269 | -0.051465627 | 11.84894509 | -2.950611032 | 0.028534148 | 0.709264768 | -4.14386136 |
| **A_55_P2173982** | Rrm2 | 0.462890908 | 0.069919351 | 0.855862464 | 12.39139022 | 2.950311547 | 0.028544629 | 0.709264768 | -4.143914688 |
| **A_51_P504442** | Sf3a1 | 0.287967088 | 0.043400902 | 0.532533275 | 11.07321871 | 2.949149504 | 0.028585341 | 0.709264768 | -4.144121675 |
| **A_51_P396917** | Zfyve21 | -0.480111267 | -0.887888151 | -0.072334383 | 11.11238759 | -2.948965151 | 0.028591805 | 0.709264768 | -4.144154522 |
| **A_51_P513449** | Pogz | 0.362875104 | 0.054622032 | 0.671128175 | 9.908168512 | 2.948493035 | 0.028608368 | 0.709264768 | -4.144238654 |
| **A_65_P08521** | Ssrp1 | 0.635152644 | 0.095341556 | 1.174963733 | 9.772443576 | 2.947044235 | 0.02865926 | 0.709264768 | -4.144496942 |
| **A_55_P2086475** | NA | 0.647045196 | 0.09583185 | 1.198258543 | 12.06314743 | 2.940121233 | 0.028903821 | 0.709264768 | -4.145733416 |
| **A_51_P375187** | Rpf2 | 0.722505071 | 0.106828123 | 1.338182019 | 10.91239196 | 2.939262353 | 0.028934321 | 0.709264768 | -4.145887077 |
| **A_51_P349662** | Taf5l | 0.275581691 | 0.040738495 | 0.510424888 | 10.59680092 | 2.939156583 | 0.028938079 | 0.709264768 | -4.145906004 |
| **A_55_P1957474** | Pus7 | 0.401063996 | 0.059130517 | 0.742997475 | 9.905307483 | 2.937801152 | 0.028986291 | 0.709264768 | -4.14614863 |
| **A_52_P644674** | Trappc6b | -0.272408936 | -0.504683844 | -0.040134029 | 11.08284157 | -2.937442676 | 0.028999057 | 0.709264768 | -4.146212823 |
| **A_55_P2090924** | Tbc1d8b | 0.318874774 | 0.04697508 | 0.590774469 | 8.561458011 | 2.937391207 | 0.02900089 | 0.709264768 | -4.14622204 |
| **A_66_P115323** | Utp4 | 0.295476432 | 0.043478778 | 0.547474085 | 10.24638722 | 2.936815709 | 0.029021399 | 0.709264768 | -4.146325118 |
| **A_51_P236197** | Polr1a | 0.393864667 | 0.057820927 | 0.729908408 | 10.34689739 | 2.935631628 | 0.029063644 | 0.709264768 | -4.146537281 |
| **A_51_P508191** | Vps26c | -0.420738422 | -0.77988384 | -0.061593004 | 11.53580966 | -2.934216734 | 0.029114214 | 0.709264768 | -4.146790945 |
| **A_52_P589169** | Aopep | -0.387554045 | -0.718751565 | -0.056356524 | 10.26301997 | -2.930863165 | 0.029234459 | 0.709264768 | -4.147392805 |
| **A_55_P2122633** | Airn | -0.56073028 | -1.040038965 | -0.081421596 | 11.47126715 | -2.930144626 | 0.029260293 | 0.709264768 | -4.147521874 |
| **A_55_P1957618** | Ak2 | 0.397065072 | 0.057401649 | 0.736728494 | 9.766138934 | 2.927947262 | 0.029339454 | 0.709264768 | -4.147916834 |
| **A_55_P2075048** | Atic | 0.344942589 | 0.04983 | 0.640055179 | 10.03308041 | 2.927584423 | 0.029352548 | 0.709264768 | -4.147982088 |
| **A_51_P371882** | Ccdc71 | 0.504904733 | 0.072683932 | 0.937125534 | 11.8915813 | 2.925863922 | 0.029414724 | 0.709264768 | -4.148291649 |
| **A_55_P2006975** | NA | 0.678572574 | 0.097318339 | 1.25982681 | 9.796074354 | 2.924021209 | 0.029481477 | 0.709264768 | -4.148623456 |
| **A_55_P2073789** | NA | -0.581055057 | -1.078911155 | -0.083198959 | 9.620046039 | -2.923235471 | 0.029509991 | 0.709264768 | -4.148765021 |
| **A_52_P164136** | Arrdc3 | -0.541460675 | -1.005481029 | -0.077440321 | 9.88790352 | -2.922673049 | 0.02953042 | 0.709264768 | -4.148866381 |
| **A_65_P20249** | Prkcb | 0.958951406 | 0.136785722 | 1.781117089 | 8.466248594 | 2.921376962 | 0.029577556 | 0.709264768 | -4.149100058 |
| **A_51_P420400** | Lef1 | 1.29024348 | 0.183858491 | 2.39662847 | 8.929011806 | 2.920893645 | 0.029595155 | 0.709264768 | -4.14918723 |
| **A_51_P342871** | S100a10 | -0.412229805 | -0.765794647 | -0.058664962 | 13.08075847 | -2.920254225 | 0.029618455 | 0.709264768 | -4.149302587 |
| **A_55_P1962811** | Anp32b | 0.437796361 | 0.062119394 | 0.813473327 | 13.96201107 | 2.918824107 | 0.029670641 | 0.709264768 | -4.149560708 |
| **A_55_P1989698** | Mmp11 | -0.353117564 | -0.656400925 | -0.049834204 | 11.15211405 | -2.91622536 | 0.02976573 | 0.709264768 | -4.150030166 |
| **A_55_P2014640** | Knop1 | 0.453151195 | 0.063685223 | 0.842617166 | 11.37282544 | 2.914230885 | 0.029838934 | 0.709264768 | -4.150390824 |
| **A_51_P369636** | Hat1 | 0.489795716 | 0.068641504 | 0.910949927 | 10.11838368 | 2.912890637 | 0.029888238 | 0.709264768 | -4.150633356 |
| **A_55_P2181444** | NA | -0.268144863 | -0.498837649 | -0.037452077 | 12.77958669 | -2.911292309 | 0.029947151 | 0.709264768 | -4.150922774 |
| **A_51_P405565** | Msh6 | 0.962647536 | 0.133862435 | 1.791432636 | 12.04039958 | 2.909214303 | 0.030023937 | 0.709264768 | -4.151299352 |
| **A_55_P2041397** | Ezh2 | 0.81346273 | 0.111163079 | 1.515762382 | 10.41979919 | 2.901119062 | 0.030325132 | 0.709264768 | -4.152769627 |
| **A_51_P363719** | Stx7 | -0.338722955 | -0.631260755 | -0.046185154 | 10.7383585 | -2.900100067 | 0.03036328 | 0.709264768 | -4.152955066 |
| **A_51_P483118** | Hmga1 | 0.927212235 | 0.126220156 | 1.728204313 | 14.41201713 | 2.899354225 | 0.030391235 | 0.709264768 | -4.153090847 |
| **A_66_P113884** | NA | 0.635870708 | 0.085620521 | 1.186120894 | 8.547402655 | 2.894402778 | 0.030577537 | 0.709264768 | -4.153993383 |
| **A_55_P2014337** | Bach1 | -0.324334725 | -0.605072543 | -0.043596908 | 10.78545067 | -2.89362918 | 0.030606757 | 0.709264768 | -4.154134567 |
| **A_65_P19780** | Klf9 | -0.543714967 | -1.01547288 | -0.071957054 | 10.30356963 | -2.88670524 | 0.030869653 | 0.709264768 | -4.155400322 |
| **A_51_P102257** | Tns1 | -0.460698896 | -0.860483217 | -0.060914576 | 8.96857433 | -2.886301932 | 0.030885042 | 0.709264768 | -4.155474168 |

| | | | | | | | | | |
|---|---|---|---|---|---|---|---|---|---|
| A_55_P2389528 | D130061D | -0.302014389 | -0.564184278 | -0.039844501 | 8.509117087 | -2.885327875 | 0.030922245 | 0.709264768 | -4.15565257 |
| A_55_P2202539 | BC028789 | 0.413311853 | 0.054347399 | 0.772276306 | 8.15851731 | 2.883877182 | 0.030977743 | 0.709264768 | -4.155918409 |
| A_55_P2123502 | Jam2 | -0.314553928 | -0.588016438 | -0.041091418 | 9.46221287 | -2.881029216 | 0.031087013 | 0.709264768 | -4.156440783 |
| A_55_P2075358 | NA | 0.454536873 | 0.058981437 | 0.850092309 | 12.45214023 | 2.878141277 | 0.031198247 | 0.709264768 | -4.156971147 |
| A_51_P479052 | Tars | 0.392298423 | 0.049752452 | 0.734844395 | 11.67663015 | 2.86845502 | 0.031574527 | 0.709264768 | -4.158754851 |
| A_66_P108567 | Xrcc6 | 1.108118187 | 0.139998378 | 2.076237995 | 11.75425933 | 2.866865525 | 0.031636747 | 0.709264768 | -4.159048267 |
| A_55_P2027496 | Adck5 | -0.34784727 | -0.652360905 | -0.043333635 | 11.95684272 | -2.861094488 | 0.031863782 | 0.709264768 | -4.160115273 |
| A_51_P400581 | Anp32e | 0.968842564 | 0.120501532 | 1.817183596 | 9.470802653 | 2.860441581 | 0.03188958 | 0.709264768 | -4.160236156 |
| A_55_P2064292 | Col7a1 | -0.612236025 | -1.148673041 | -0.075799009 | 8.941505246 | -2.858580903 | 0.031963224 | 0.709264768 | -4.16058084 |
| A_55_P1977875 | Arglu1 | 0.268414505 | 0.033188 | 0.503641009 | 10.7535486 | 2.858051641 | 0.031984206 | 0.709264768 | -4.160678935 |
| A_55_P2133165 | Wwc1 | -0.492375563 | -0.924264561 | -0.060486566 | 10.75513008 | -2.855450893 | 0.032087529 | 0.709264768 | -4.161161287 |
| A_55_P1955078 | Igflr1 | -0.27811551 | -0.522103961 | -0.034127058 | 8.997320224 | -2.855001124 | 0.032105434 | 0.709264768 | -4.161244758 |
| A_55_P2052016 | Crispld2 | -0.83416999 | -1.566908018 | -0.101431962 | 10.32338478 | -2.85138707 | 0.032249708 | 0.709264768 | -4.161916072 |
| A_55_P2435563 | Etl4 | -0.353986282 | -0.665392236 | -0.042580327 | 8.656894129 | -2.847146831 | 0.032419883 | 0.709264768 | -4.162705031 |
| A_55_P2151897 | NA | 0.388424417 | 0.046457235 | 0.730391599 | 15.37130142 | 2.844935591 | 0.032509018 | 0.709264768 | -4.163117036 |
| A_52_P398211 | NA | 0.729686152 | 0.08688231 | 1.372489994 | 8.638526335 | 2.843203768 | 0.032579014 | 0.709264768 | -4.163439988 |
| A_55_P2125588 | Pdgfa | -0.542088791 | -1.020444033 | -0.063733549 | 13.04832942 | -2.838378022 | 0.03277493 | 0.709264768 | -4.164341164 |
| A_55_P2173188 | Kank3 | -0.482083214 | -0.907630812 | -0.056535616 | 9.546697177 | -2.837423831 | 0.032813821 | 0.709264768 | -4.164519574 |
| A_55_P2161342 | Pkib | 0.890913048 | 0.104426955 | 1.677399141 | 11.71442189 | 2.837230483 | 0.032821708 | 0.709264768 | -4.164555734 |
| A_51_P150802 | Mpv17 | -0.387167567 | -0.729568989 | -0.044766146 | 10.6618649 | -2.832133732 | 0.033030351 | 0.709264768 | -4.165510016 |
| A_52_P620748 | Ncl | 0.330814951 | 0.037922505 | 0.623707397 | 8.961314494 | 2.828963235 | 0.03316087 | 0.709264768 | -4.16610469 |
| A_55_P2020443 | Hnrnpa3 | 0.770248171 | 0.08803948 | 1.452456862 | 9.973104437 | 2.827898212 | 0.033204841 | 0.709264768 | -4.166304632 |
| A_55_P2035205 | Nifk | 0.312701347 | 0.035728005 | 0.589674688 | 11.81567405 | 2.827757263 | 0.033210665 | 0.709264768 | -4.1663311 |
| A_55_P2031167 | Efna1 | -0.583509685 | -1.100742126 | -0.066277243 | 13.05444473 | -2.825612854 | 0.033299408 | 0.709264768 | -4.166733982 |
| A_51_P246166 | Wfdc18 | 0.534222979 | 0.060450536 | 1.007995423 | 8.26897073 | 2.824249846 | 0.033355949 | 0.709264768 | -4.16699025 |
| A_55_P2063336 | Tedc1 | 0.767190836 | 0.085516038 | 1.448865635 | 9.964938305 | 2.818879517 | 0.033579747 | 0.709264768 | -4.168001414 |
| A_55_P2141301 | Synj2bp | 0.28647447 | 0.03189353 | 0.541055409 | 10.35938584 | 2.818450328 | 0.033597703 | 0.709264768 | -4.168082325 |
| A_51_P234308 | Rpa2 | 0.60778299 | 0.067654186 | 1.147911793 | 9.341014829 | 2.818392985 | 0.033600103 | 0.709264768 | -4.168093136 |
| A_51_P196972 | Slc4a1 | 0.610939812 | 0.06793655 | 1.153943075 | 10.51256237 | 2.818034681 | 0.033615103 | 0.709264768 | -4.168160697 |
| A_55_P2091974 | Gpr137 | -0.434061923 | -0.819891992 | -0.048231855 | 10.26518566 | -2.817772757 | 0.033626072 | 0.709264768 | -4.168210091 |
| A_55_P2037962 | Trpm6 | -0.691175112 | -1.305900949 | -0.076449275 | 9.753224291 | -2.816157697 | 0.033693799 | 0.709264768 | -4.168514784 |
| A_52_P657435 | Psmc5 | 0.434245042 | 0.047968917 | 0.820521167 | 12.12055273 | 2.81570627 | 0.033712756 | 0.709264768 | -4.168599987 |
| A_55_P2048813 | NA | 0.940385206 | 0.103634228 | 1.777136183 | 12.18294013 | 2.814880129 | 0.033747479 | 0.709264768 | -4.168755956 |
| A_52_P584078 | Pop1 | 0.558462369 | 0.061508171 | 1.055416566 | 10.09726674 | 2.814672671 | 0.033756204 | 0.709264768 | -4.168795131 |
| A_55_P2065268 | NA | 0.558236004 | 0.061079695 | 1.055392312 | 9.475622038 | 2.812387987 | 0.03385246 | 0.709264768 | -4.169226786 |
| A_55_P2144210 | Snx5 | 0.327240122 | 0.035729469 | 0.618750775 | 11.90949217 | 2.811657791 | 0.033883287 | 0.709264768 | -4.169364833 |
| A_51_P341010 | Ercc8 | 0.360644683 | 0.039162314 | 0.682127052 | 9.722054391 | 2.809782601 | 0.033962593 | 0.709264768 | -4.169719545 |
| A_55_P2021821 | Prmt1 | 1.204848495 | 0.13041298 | 2.27928401 | 12.77054445 | 2.808681086 | 0.034009273 | 0.709264768 | -4.169928041 |
| A_55_P2010161 | Fastkd5 | 0.420768416 | 0.044709108 | 0.796827724 | 10.25158387 | 2.802445247 | 0.034274859 | 0.709264768 | -4.171110211 |
| A_55_P2053202 | Itsn2 | -0.264503878 | -0.500927301 | -0.028080454 | 10.52887009 | -2.802153274 | 0.034287349 | 0.709264768 | -4.171165639 |
| A_52_P552589 | Map4k1 | 0.728616069 | 0.077100668 | 1.380131469 | 9.065471738 | 2.801072852 | 0.034333613 | 0.709264768 | -4.171370806 |
| A_55_P2046262 | Phgdh | 0.58846805 | 0.061301325 | 1.115634775 | 11.30758113 | 2.795923244 | 0.034555056 | 0.709264768 | -4.17234999 |
| A_55_P2061273 | Tbx6 | -0.302867015 | -0.574571006 | -0.031163023 | 8.080845948 | -2.791941433 | 0.03472735 | 0.709264768 | -4.173108591 |
| A_55_P2112991 | Slc4a3 | -0.520029853 | -0.98681454 | -0.053245165 | 8.884695393 | -2.790371448 | 0.034795541 | 0.709264768 | -4.173408052 |
| A_55_P2033255 | NA | -0.366588734 | -0.696072065 | -0.037105404 | 14.95884076 | -2.786737184 | 0.034953953 | 0.709264768 | -4.174102021 |
| A_55_P2173457 | 1700097N | 1.446598198 | 0.146198346 | 2.746998051 | 10.71229136 | 2.786258164 | 0.034974891 | 0.709264768 | -4.17419357 |
| A_51_P500718 | Dck | 0.516456314 | 0.052177262 | 0.980735367 | 8.368391529 | 2.786152267 | 0.034979522 | 0.709264768 | -4.174213812 |
| A_55_P2145034 | Swt1 | -0.47356291 | -0.899326656 | -0.047799163 | 10.35440349 | -2.785860381 | 0.034992289 | 0.709264768 | -4.174269608 |

| | | | | | | | | | |
|---|---|---|---|---|---|---|---|---|---|
| **A_55_P2021476** | NA | -0.29318252 | -0.556862433 | -0.029502607 | 13.8916676 | -2.784911334 | 0.035033835 | 0.709264768 | -4.174451074 |
| **A_55_P1993503** | Adora2b | -0.354682605 | -0.673871903 | -0.035493308 | 8.654941765 | -2.783183972 | 0.035109592 | 0.709264768 | -4.174781547 |
| **A_52_P677147** | Prmt5 | 0.310177108 | 0.030901897 | 0.589452319 | 10.36768471 | 2.781811513 | 0.035169911 | 0.709264768 | -4.175044294 |
| **A_51_P324572** | Ctnna1 | -0.41458816 | -0.787948356 | -0.041227964 | 14.99649893 | -2.781244854 | 0.035194849 | 0.709264768 | -4.175152821 |
| **A_51_P230987** | Pi4k2b | -0.367242498 | -0.698136601 | -0.036348394 | 10.39294825 | -2.779804477 | 0.035258324 | 0.709264768 | -4.175428801 |
| **A_66_P109296** | NA | 0.420752382 | 0.041263008 | 0.800241755 | 10.67023915 | 2.777009143 | 0.035381865 | 0.709264768 | -4.175964874 |
| **A_55_P2058962** | Mcm10 | 1.029240775 | 0.100905678 | 1.957575872 | 9.552832101 | 2.776914824 | 0.035386042 | 0.709264768 | -4.175982973 |
| **A_51_P292566** | Xpot | 0.41138236 | 0.040312927 | 0.782451792 | 10.58295443 | 2.776775894 | 0.035392195 | 0.709264768 | -4.176009634 |
| **A_51_P307243** | Ppp5c | 0.374839426 | 0.036483752 | 0.713195101 | 10.59063775 | 2.77473905 | 0.03548254 | 0.709264768 | -4.176400686 |
| **A_51_P515496** | Rbm17 | 0.323978962 | 0.03148806 | 0.616469863 | 11.54101957 | 2.774308766 | 0.035501658 | 0.709264768 | -4.176483339 |
| **A_51_P151254** | Slc35b3 | -0.553011941 | -1.052290683 | -0.053733198 | 9.784060731 | -2.774225624 | 0.035505353 | 0.709264768 | -4.176499311 |
| **A_55_P2057528** | Arl4d | -0.635610937 | -1.210337272 | -0.060884602 | 9.705364635 | -2.770005358 | 0.035693478 | 0.709264768 | -4.177310804 |
| **A_51_P241861** | Tcirg1 | -0.354416373 | -0.675134858 | -0.033697888 | 8.701217622 | -2.767834579 | 0.035790667 | 0.709264768 | -4.177728776 |
| **A_52_P14666** | NA | 0.365689557 | 0.034612575 | 0.696766539 | 8.353220469 | 2.766520648 | 0.035849633 | 0.709264768 | -4.177981952 |
| **A_51_P159503** | Rnf213 | -0.369483301 | -0.704058641 | -0.034907961 | 10.6787651 | -2.765993993 | 0.035873298 | 0.709264768 | -4.178083471 |
| **A_55_P1956989** | BC00400 | -0.34640622 | -0.660280331 | -0.032532109 | 13.30140189 | -2.764270451 | 0.035950863 | 0.709264768 | -4.17841586 |
| **A_55_P1954768** | NA | 0.518923884 | 0.048585892 | 0.989261876 | 8.917446587 | 2.763401218 | 0.03599005 | 0.709264768 | -4.178583585 |
| **A_55_P2158107** | Ddx31 | 0.276099991 | 0.025628287 | 0.526571695 | 9.904893433 | 2.76094699 | 0.036100945 | 0.709264768 | -4.179057479 |
| **A_55_P2120652** | Snrpa1 | 0.667778418 | 0.061147178 | 1.274409659 | 11.90097971 | 2.757134503 | 0.036273952 | 0.709264768 | -4.179794616 |
| **A_55_P2077526** | NA | -0.328945958 | -0.628237579 | -0.029654337 | 14.45171222 | -2.752836018 | 0.036470097 | 0.709264768 | -4.180627138 |
| **A_55_P2015232** | NA | -0.25253009 | -0.482331389 | -0.02272879 | 9.886397991 | -2.752396509 | 0.036490217 | 0.709264768 | -4.180712346 |
| **A_55_P1969885** | NA | 0.299711665 | 0.026882183 | 0.572541148 | 9.107386408 | 2.751456764 | 0.036533278 | 0.709264768 | -4.180894589 |
| **A_55_P2100120** | Nme1 | 0.42251732 | 0.037782791 | 0.807251848 | 11.69270727 | 2.750639635 | 0.036570765 | 0.709264768 | -4.181053111 |
| **A_51_P389531** | Prmt1 | 0.875198312 | 0.077661705 | 1.672734919 | 9.833435047 | 2.748566132 | 0.036666079 | 0.709264768 | -4.181455612 |
| **A_55_P1971889** | F3 | -0.72483699 | -1.385386749 | -0.064287231 | 11.29839794 | -2.748433008 | 0.036672208 | 0.709264768 | -4.181481466 |
| **A_55_P2091230** | NA | 0.917617294 | 0.080859141 | 1.754375446 | 11.58825688 | 2.746704779 | 0.036751872 | 0.709264768 | -4.181817231 |
| **A_55_P2035495** | Zbtb20 | -0.290906334 | -0.556355459 | -0.025457209 | 9.08227748 | -2.744872804 | 0.036836523 | 0.709264768 | -4.182173418 |
| **A_55_P1968993** | NA | 0.33797986 | 0.029549802 | 0.646409917 | 10.65835524 | 2.744634209 | 0.036847564 | 0.709264768 | -4.182219827 |
| **A_55_P2106039** | Map3k6 | -0.284708114 | -0.544680064 | -0.024736163 | 9.847427785 | -2.742986664 | 0.036923899 | 0.709264768 | -4.182540422 |
| **A_55_P1953063** | Hspe1 | 0.533008088 | 0.046126012 | 1.019890164 | 10.97654386 | 2.741955218 | 0.036971777 | 0.709264768 | -4.182741243 |
| **A_51_P519555** | Gnb1l | 0.45540828 | 0.039305641 | 0.871510918 | 10.17804286 | 2.741263591 | 0.037003918 | 0.709264768 | -4.18287595 |
| **A_55_P2160682** | Set | 0.521146718 | 0.044819703 | 0.997473732 | 10.48881097 | 2.740344351 | 0.037046685 | 0.709264768 | -4.18305505 |
| **A_55_P2080902** | NA | 0.617684936 | 0.052937516 | 1.182432357 | 8.430461692 | 2.739448245 | 0.037088427 | 0.709264768 | -4.183229709 |
| **A_55_P1973995** | NA | 0.586954369 | 0.050084158 | 1.12382458 | 10.93791565 | 2.738327439 | 0.037140707 | 0.709264768 | -4.183448256 |
| **A_52_P113916** | NA | 0.479036088 | 0.040796179 | 0.917275997 | 9.339136618 | 2.737831103 | 0.037163885 | 0.709264768 | -4.18354507 |
| **A_51_P410451** | Tube1 | 0.609742997 | 0.05189859 | 1.167587403 | 8.73701735 | 2.737688827 | 0.037170532 | 0.709264768 | -4.183572825 |
| **A_52_P599624** | Pfas | 0.388439682 | 0.033030936 | 0.743848428 | 8.481111892 | 2.737447659 | 0.037181801 | 0.709264768 | -4.183619877 |
| **A_55_P2124085** | NA | 0.511826198 | 0.043306808 | 0.980345588 | 10.89820376 | 2.736183929 | 0.037240916 | 0.709264768 | -4.183866507 |
| **A_55_P1964154** | Asah1 | -0.239656665 | -0.459190423 | -0.020122907 | 9.133044098 | -2.734252046 | 0.037331483 | 0.709264768 | -4.184243786 |
| **A_55_P1967443** | NA | 0.479299374 | 0.039972018 | 0.91862673 | 9.908731853 | 2.732555308 | 0.037411224 | 0.709264768 | -4.184575395 |
| **A_55_P2173968** | Ctdspl | -0.516143599 | -0.989423791 | -0.042863407 | 13.70841442 | -2.731508543 | 0.037460511 | 0.709264768 | -4.184780091 |
| **A_52_P164161** | Cyp51 | 0.407348777 | 0.033671198 | 0.781026356 | 8.588001323 | 2.730358798 | 0.037514728 | 0.709264768 | -4.185005028 |
| **A_51_P518163** | Rrp9 | 0.249901407 | 0.020025845 | 0.479776969 | 8.902027401 | 2.722865837 | 0.03787016 | 0.709264768 | -4.186473607 |
| **A_51_P406020** | Abhd4 | -0.325539143 | -0.625283126 | -0.025795161 | 14.08425479 | -2.720214107 | 0.037996821 | 0.709264768 | -4.186994432 |
| **A_52_P368306** | Tmem100 | -0.545902495 | -1.048689799 | -0.043115192 | 8.685126436 | -2.719450278 | 0.038033391 | 0.709264768 | -4.187144562 |
| **A_52_P445387** | Clk4 | -0.335099647 | -0.644065327 | -0.026133966 | 10.43569894 | -2.716527299 | 0.038173689 | 0.709264768 | -4.187719513 |
| **A_51_P104747** | Gtf2h4 | 0.311349259 | 0.023894067 | 0.598804452 | 10.86738374 | 2.712864011 | 0.038350315 | 0.709264768 | -4.188441072 |
| **A_55_P2097840** | NA | 0.936885459 | 0.07148467 | 1.802286247 | 9.050160154 | 2.711562115 | 0.038413299 | 0.709264768 | -4.188697772 |

| | | | | | | | | | |
|---|---|---|---|---|---|---|---|---|---|
| **A_55_P1985554** | B4galt4 | -0.338629303 | -0.651436707 | -0.025821899 | 8.982504811 | -2.711426639 | 0.03841986 | 0.709264768 | -4.188724492 |
| **A_55_P2002703** | NA | 0.366942771 | 0.027935713 | 0.705949829 | 9.595264998 | 2.711065074 | 0.038437375 | 0.709264768 | -4.188795812 |
| **A_55_P1964068** | Slc25a38 | -0.42572938 | -0.81909483 | -0.03236393 | 10.47264662 | -2.710739322 | 0.038453163 | 0.709264768 | -4.188860076 |
| **A_55_P2095047** | Mat2a | 0.280659446 | 0.02122514 | 0.540093752 | 8.471131259 | 2.70958391 | 0.038509218 | 0.709264768 | -4.189088088 |
| **A_52_P282500** | Kif21b | 0.674230038 | 0.050937109 | 1.297522967 | 10.36853282 | 2.709357054 | 0.038520234 | 0.709264768 | -4.189132869 |
| **A_55_P2025687** | Muc4 | -0.258646622 | -0.498381347 | -0.018911897 | 9.3645725 | -2.702254272 | 0.03886689 | 0.709264768 | -4.190537083 |
| **A_55_P2148370** | NA | 1.021466117 | 0.074536301 | 1.968395933 | 9.327288001 | 2.701820667 | 0.038888162 | 0.709264768 | -4.190622941 |
| **A_66_P105075** | Mmp11 | -0.381291997 | -0.735082816 | -0.027501178 | 10.06242744 | -2.699364156 | 0.039008912 | 0.709264768 | -4.191109642 |
| **A_55_P1961210** | Trp53inp2 | -0.42979412 | -0.8286306 | -0.03095764 | 12.32208155 | -2.699081138 | 0.03902285 | 0.709264768 | -4.191165748 |
| **A_55_P2019512** | Mocs2 | -0.247268753 | -0.476905038 | -0.017632468 | 11.22370034 | -2.696988355 | 0.039126082 | 0.709264768 | -4.191580825 |
| **A_55_P2047461** | Fcho1 | 1.056514521 | 0.07521465 | 2.037814392 | 8.74118611 | 2.696646844 | 0.039142956 | 0.709264768 | -4.191648593 |
| **A_66_P135882** | Rrp1 | 0.372788935 | 0.025855288 | 0.719722582 | 12.80101775 | 2.691329904 | 0.039406686 | 0.709264768 | -4.192704908 |
| **A_66_P118696** | Nup107 | 0.626712208 | 0.043418886 | 1.210005531 | 11.94279753 | 2.691110283 | 0.039417621 | 0.709264768 | -4.19274859 |
| **A_55_P1956898** | NA | 0.32476146 | 0.02234833 | 0.62717459 | 11.8592118 | 2.689764069 | 0.039484721 | 0.709264768 | -4.193016434 |
| **A_55_P2076278** | NA | 0.267624737 | 0.018283697 | 0.516965777 | 16.75059273 | 2.688331569 | 0.039556259 | 0.709264768 | -4.19330161 |
| **A_55_P2063256** | Lgals4 | -0.456432348 | -0.881843186 | -0.031021509 | 8.29700806 | -2.687312738 | 0.039607224 | 0.709264768 | -4.193504538 |
| **A_52_P431859** | Aopep | -0.493905748 | -0.954325714 | -0.033485782 | 10.00457236 | -2.68683055 | 0.039631369 | 0.709264768 | -4.193600608 |
| **A_55_P2122709** | Arhgef3 | -0.836637348 | -1.616760995 | -0.056513702 | 10.13952571 | -2.686112214 | 0.039667368 | 0.709264768 | -4.193743763 |
| **A_66_P135534** | Zfp950 | -0.236222811 | -0.45656634 | -0.015879283 | 8.518372368 | -2.685170574 | 0.039714613 | 0.709264768 | -4.193931485 |
| **A_55_P2155931** | Tnfsfm13 | -0.304438808 | -0.588628718 | -0.020248898 | 9.178843088 | -2.683129935 | 0.039817206 | 0.709264768 | -4.194338549 |
| **A_55_P2128646** | Gmnn | 0.603040594 | 0.039815256 | 1.166265933 | 9.886292704 | 2.681727894 | 0.039887861 | 0.709264768 | -4.194618425 |
| **A_55_P2168823** | NA | 0.558136173 | 0.036778222 | 1.079494125 | 11.81997433 | 2.681356222 | 0.039906614 | 0.709264768 | -4.194692645 |
| **A_52_P635182** | Rras | -0.567956642 | -1.098530356 | -0.037382928 | 11.63624894 | -2.681141892 | 0.039917432 | 0.709264768 | -4.19473545 |
| **A_51_P514256** | Tubb2b | 1.256289507 | 0.082201593 | 2.43037742 | 10.00774488 | 2.680028798 | 0.039973667 | 0.709264768 | -4.194957816 |
| **A_55_P1986526** | Cdh26 | 0.314811638 | 0.020505778 | 0.609117498 | 9.298852079 | 2.67918199 | 0.040016507 | 0.709264768 | -4.195127052 |
| **A_51_P234174** | Pms2 | 0.313530695 | 0.020325612 | 0.606735779 | 9.219478526 | 2.678298111 | 0.040061275 | 0.709264768 | -4.195303761 |
| **A_52_P621588** | Ifnlr1 | -0.530282593 | -1.026276253 | -0.034288934 | 9.836489774 | -2.677821305 | 0.040085447 | 0.709264768 | -4.195399112 |
| **A_52_P410685** | Krt7 | -0.651094092 | -1.260258786 | -0.041929397 | 11.82625084 | -2.677067827 | 0.040123679 | 0.709264768 | -4.195549831 |
| **A_55_P2166384** | Cnot1 | 0.263964203 | 0.016965999 | 0.510962406 | 11.92217156 | 2.676711624 | 0.040141766 | 0.709264768 | -4.195621099 |
| **A_66_P114451** | Adgrg6 | -0.516168117 | -0.999260016 | -0.033076219 | 11.17953087 | -2.676158099 | 0.04016989 | 0.709264768 | -4.195731867 |
| **A_55_P2094896** | Phyhd1 | -0.312744615 | -0.605513008 | -0.019976221 | 10.66420416 | -2.675568018 | 0.040199896 | 0.709264768 | -4.195849978 |
| **A_55_P2083963** | Phgr1 | -1.100084475 | -2.130079148 | -0.070089802 | 9.065170763 | -2.675108509 | 0.040223279 | 0.709264768 | -4.195941973 |
| **A_52_P89099** | Nolc1 | 0.295009148 | 0.018757497 | 0.571260799 | 8.282102227 | 2.674736146 | 0.040242238 | 0.709264768 | -4.196016535 |
| **A_55_P2000280** | Mthfd1l | 0.343595174 | 0.021666326 | 0.665524023 | 8.481081012 | 2.673237253 | 0.040318652 | 0.709264768 | -4.196316786 |
| **A_51_P322640** | Ccl24 | 0.456708852 | 0.028726312 | 0.884691391 | 8.01721354 | 2.67278314 | 0.040341834 | 0.709264768 | -4.196407788 |
| **A_52_P514061** | Padi4 | 0.440783874 | 0.02761161 | 0.853956137 | 8.091313893 | 2.672051864 | 0.040379196 | 0.709264768 | -4.196554369 |
| **A_51_P259902** | Arl6ip5 | -0.542863318 | -1.051910117 | -0.03381652 | 11.28582051 | -2.671056855 | 0.040430092 | 0.709264768 | -4.196753884 |
| **A_51_P246066** | Slamf9 | -0.276837113 | -0.536729042 | -0.016945185 | 8.148642398 | -2.66797568 | 0.040588139 | 0.709264768 | -4.197372225 |
| **A_55_P2043798** | Mrto4 | 0.481413458 | 0.029333852 | 0.933493064 | 11.02463461 | 2.667188171 | 0.040628641 | 0.709264768 | -4.197530391 |
| **A_55_P2081615** | Timeless | 0.403737868 | 0.023982144 | 0.783493592 | 8.412949882 | 2.662842641 | 0.040852923 | 0.709264768 | -4.19840408 |
| **A_55_P2202043** | C430045I | -0.239461375 | -0.464759168 | -0.014163583 | 8.33810377 | -2.662127683 | 0.040889952 | 0.709264768 | -4.198547975 |
| **A_51_P344376** | Gpr162 | 0.566324676 | 0.032983401 | 1.09966595 | 8.169509701 | 2.659565168 | 0.041022967 | 0.709264768 | -4.199064062 |
| **A_51_P348548** | Ecsit | 0.336930237 | 0.019506678 | 0.654353797 | 11.67292515 | 2.658588836 | 0.04107377 | 0.709264768 | -4.199260836 |
| **A_55_P2122220** | Kcnk4 | 0.324506647 | 0.018273759 | 0.630739535 | 8.765737427 | 2.654129524 | 0.041306676 | 0.709264768 | -4.200160586 |
| **A_55_P1955998** | Nlrp1a | 0.662493584 | 0.037270559 | 1.287716609 | 11.97953866 | 2.653976415 | 0.041314698 | 0.709264768 | -4.200191508 |
| **A_55_P2068260** | H3c8 | 0.505620099 | 0.028336704 | 0.982903493 | 8.317567662 | 2.653373272 | 0.041346316 | 0.709264768 | -4.200313337 |
| **A_66_P133993** | Gm5093 | 1.492790987 | 0.082292294 | 2.90328968 | 11.22045999 | 2.650798168 | 0.041481602 | 0.709264768 | -4.200833821 |
| **A_55_P1970806** | Mif-ps6 | 0.813812757 | 0.044576186 | 1.583049327 | 14.63717969 | 2.649811106 | 0.041533586 | 0.709264768 | -4.201033472 |

| | | | | | | | | | |
|---|---|---|---|---|---|---|---|---|---|
| **A_55_P2039180** | Zswim8 | -0.260228773 | -0.506235521 | -0.014222024 | 12.08270258 | -2.649467753 | 0.041551685 | 0.709264768 | -4.201102941 |
| **A_51_P332602** | Surf1 | -0.391441172 | -0.761998424 | -0.020883921 | 11.072207 | -2.645827539 | 0.041744097 | 0.709264768 | -4.20184004 |
| **A_52_P47255** | Gmps | 0.354408682 | 0.018830355 | 0.689987009 | 8.92788555 | 2.645213881 | 0.041776629 | 0.709264768 | -4.201964407 |
| **A_51_P474468** | Utp11 | 0.24627543 | 0.013069832 | 0.479481027 | 12.26946458 | 2.645041323 | 0.041785781 | 0.709264768 | -4.201999383 |
| **A_52_P437084** | NA | 0.975287071 | 0.050975888 | 1.899598254 | 13.25700423 | 2.642801867 | 0.041904761 | 0.709264768 | -4.202453536 |
| **A_55_P2045066** | Eif5a | 0.614875223 | 0.032108809 | 1.197641637 | 12.49778957 | 2.642669327 | 0.041911814 | 0.709264768 | -4.202480427 |
| **A_55_P1988070** | Eef1g | 0.314809416 | 0.016421043 | 0.613197789 | 11.40911469 | 2.6425071 | 0.041920448 | 0.709264768 | -4.202513344 |
| **A_51_P127297** | Hsd11b1 | -0.526841905 | -1.026728825 | -0.026954985 | 13.03527908 | -2.639726204 | 0.042068764 | 0.709264768 | -4.233776052 |
| **A_55_P1997225** | Agap2 | 0.357179509 | 0.018133979 | 0.696225039 | 7.988849688 | 2.638632206 | 0.042127266 | 0.709264768 | -4.20330023 |
| **A_55_P2155504** | Lck | 3.251565016 | 0.160982075 | 6.342147957 | 10.30167113 | 2.63513206 | 0.042315032 | 0.709264768 | -4.204012081 |
| **A_51_P521374** | Nudt13 | -0.255117901 | -0.497679462 | -0.012556341 | 10.08546764 | -2.634324681 | 0.042358472 | 0.709264768 | -4.204176428 |
| **A_55_P2007640** | Eprs | 0.321098599 | 0.0156695 | 0.626527697 | 10.49634714 | 2.633166643 | 0.042420863 | 0.709264768 | -4.204412247 |
| **A_66_P140307** | NA | 0.578161088 | 0.027811633 | 1.128510543 | 13.26648688 | 2.631241217 | 0.042524818 | 0.709264768 | -4.20480458 |
| **A_55_P2178127** | 1-Sep | 0.936432645 | 0.04480041 | 1.828064881 | 8.715505497 | 2.630517102 | 0.042563985 | 0.709264768 | -4.204952208 |
| **A_55_P1956973** | Tln1 | -0.357709142 | -0.698330191 | -0.017088092 | 14.69427481 | -2.630321909 | 0.04257455 | 0.709264768 | -4.204992011 |
| **A_55_P2055597** | Enpp2 | -0.319754737 | -0.624454246 | -0.015055227 | 10.62903713 | -2.628424929 | 0.042677369 | 0.709264768 | -4.205378992 |
| **A_55_P2138333** | Mad2l1 | 0.355314829 | 0.016577242 | 0.694052416 | 8.107885988 | 2.627243278 | 0.042741551 | 0.709264768 | -4.205620197 |
| **A_51_P404236** | Atp6v0b | -0.660452986 | -1.290178144 | -0.030727828 | 10.49977734 | -2.6268859 | 0.042760983 | 0.709264768 | -4.20569317 |
| **A_55_P1975670** | Prps1l3 | 0.49896511 | 0.023157888 | 0.974772332 | 10.6544208 | 2.626573114 | 0.042777998 | 0.709264768 | -4.205757047 |
| **A_55_P2168761** | Sf1 | 0.328557699 | 0.01513944 | 0.641975958 | 12.80214511 | 2.625655238 | 0.042827972 | 0.709264768 | -4.20594454 |
| **A_51_P112627** | St6galnac | -0.539930733 | -1.055198506 | -0.02466296 | 11.67752668 | -2.6245534 | 0.042888044 | 0.709264768 | -4.246578233 |
| **A_55_P1978905** | NA | 0.713582033 | 0.032444912 | 1.394719154 | 8.368122897 | 2.623975058 | 0.042919612 | 0.709264768 | -4.206287928 |
| **A_51_P202430** | Pdap1 | 0.392512408 | 0.017417575 | 0.767607241 | 13.6338501 | 2.620973643 | 0.043083843 | 0.709264768 | -4.206901925 |
| **A_55_P2071691** | Kank3 | -0.514464161 | -1.006197918 | -0.022730404 | 12.18968745 | -2.62044741 | 0.043112708 | 0.709264768 | -4.207009653 |
| **A_55_P1963645** | NA | 0.846534243 | 0.036715606 | 1.656352879 | 11.7956163 | 2.618225864 | 0.043234791 | 0.709264768 | -4.207464691 |
| **A_51_P297165** | Sf3a3 | 0.51795168 | 0.021839432 | 1.014063929 | 12.96206859 | 2.614927442 | 0.043416744 | 0.709264768 | -4.208141057 |
| **A_55_P2068554** | NA | 0.543983705 | 0.022932875 | 1.065034536 | 16.71596704 | 2.614906375 | 0.043417908 | 0.709264768 | -4.20814538 |
| **A_51_P388847** | Stap2 | -0.382010943 | -0.747940198 | -0.016081689 | 9.671417512 | -2.614743061 | 0.043426939 | 0.709264768 | -4.208178893 |
| **A_52_P378084** | NA | 0.424578356 | 0.017628168 | 0.831528544 | 9.101204487 | 2.613165667 | 0.043514268 | 0.709264768 | -4.208502692 |
| **A_51_P312121** | Xdh | -0.383670141 | -0.751705823 | -0.015634459 | 9.188887838 | -2.611069434 | 0.043630614 | 0.709264768 | -4.208933315 |
| **A_55_P1993293** | Ndrg3 | 0.585710236 | 0.023795593 | 1.147624879 | 10.64960087 | 2.610735112 | 0.043649201 | 0.709264768 | -4.209002027 |
| **A_55_P2058359** | Macroh2a | 0.280982453 | 0.01138965 | 0.550575255 | 8.766952129 | 2.610485296 | 0.043663096 | 0.709264768 | -4.209053377 |
| **A_66_P116678** | Rps8 | 0.283716682 | 0.011491551 | 0.555941813 | 8.835447926 | 2.610399647 | 0.04366786 | 0.709264768 | -4.209070984 |
| **A_55_P2022699** | NA | 0.891925703 | 0.035093145 | 1.748758262 | 9.242609823 | 2.60725233 | 0.043843338 | 0.709264768 | -4.209718386 |
| **A_55_P2010962** | Surf6 | 0.24041759 | 0.009428747 | 0.471406433 | 10.3922562 | 2.606907258 | 0.043862623 | 0.709264768 | -4.209789417 |
| **A_51_P162098** | Ndufaf4 | 0.315612333 | 0.012366564 | 0.618858101 | 9.724359481 | 2.6068111 | 0.043867999 | 0.709264768 | -4.209809212 |
| **A_52_P235278** | Pms1 | 0.305013799 | 0.011782179 | 0.598245419 | 8.599480676 | 2.605307761 | 0.043952137 | 0.709264768 | -4.210118793 |
| **A_55_P1989858** | Thrap3 | 0.507845436 | 0.019165136 | 0.996525735 | 11.43288595 | 2.602897509 | 0.044087396 | 0.709264768 | -4.210615524 |
| **A_55_P2023235** | Fen1 | 1.164847184 | 0.043561606 | 2.286132761 | 12.30833925 | 2.601974665 | 0.044139303 | 0.709264768 | -4.210805841 |
| **A_65_P05252** | Cdc25a | 0.576291263 | 0.02126779 | 1.131314735 | 10.10042854 | 2.600644747 | 0.044214223 | 0.709264768 | -4.211080233 |
| **A_51_P394728** | Kti12 | 0.562099538 | 0.020485799 | 1.103713276 | 9.330343839 | 2.599404709 | 0.044284203 | 0.709264768 | -4.211336213 |
| **A_55_P1962641** | Rnf26 | 0.300264547 | 0.01091678 | 0.589612314 | 13.23919146 | 2.599167497 | 0.044297603 | 0.709264768 | -4.211385195 |
| **A_55_P2016652** | Cdk1 | 0.93418603 | 0.033809007 | 1.834563052 | 8.670281939 | 2.598718933 | 0.044322955 | 0.709264768 | -4.211477832 |
| **A_55_P2092526** | Tgif1 | -0.2303542 | -0.452373878 | -0.008334522 | 11.67829734 | -2.598693209 | 0.04432441 | 0.709264768 | -4.211483144 |
| **A_55_P2026738** | Rgs4 | -0.351026019 | -0.689838229 | -0.012213809 | 8.319632584 | -2.5949596 | 0.044536041 | 0.709264768 | -4.212254859 |
| **A_55_P2119463** | Ggh | -0.236644886 | -0.46507803 | -0.008211742 | 11.50784183 | -2.594707175 | 0.044550389 | 0.709264768 | -4.212307076 |
| **A_55_P2015687** | Phf11d | -0.661573788 | -1.300356335 | -0.02279124 | 8.92970899 | -2.594033575 | 0.0445887 | 0.709264768 | -4.212446442 |
| **A_51_P163958** | Spr | 0.701156655 | 0.023692726 | 1.378620585 | 10.78662979 | 2.592264005 | 0.044689513 | 0.709264768 | -4.212812741 |

| | | | | | | | | | |
|---|---|---|---|---|---|---|---|---|---|
| **A_52_P313080** | Hnrnpab | 0.608234966 | 0.020436157 | 1.196033775 | 12.89747084 | 2.591749522 | 0.044718869 | 0.709264768 | -4.212919287 |
| **A_51_P219721** | Ola1 | 0.476560136 | 0.015871275 | 0.937248998 | 11.1214869 | 2.590957831 | 0.044764084 | 0.709264768 | -4.213083284 |
| **A_51_P314285** | Tmem86a | -0.3276079 | -0.644717294 | -0.010498506 | 9.768812187 | -2.58759083 | 0.04495693 | 0.709264768 | -4.213781332 |
| **A_55_P1997486** | Snu13 | 0.540071227 | 0.0169867 | 1.063155755 | 12.03056444 | 2.586005905 | 0.045048017 | 0.709264768 | -4.214110244 |
| **A_52_P337427** | Izumo4 | -0.315830745 | -0.621773397 | -0.009888093 | 8.320306547 | -2.58562015 | 0.045070216 | 0.709264768 | -4.214190329 |
| **A_52_P476075** | Clspn | 0.405264045 | 0.012582163 | 0.797945928 | 8.187541284 | 2.584922675 | 0.045110385 | 0.709264768 | -4.214335162 |
| **A_55_P2046728** | C920009E | -0.494676263 | -0.974215759 | -0.015136766 | 11.48018646 | -2.583729439 | 0.045179195 | 0.709264768 | -4.214583033 |
| **A_51_P502082** | Rrm1 | 0.481035131 | 0.014299817 | 0.947770445 | 12.03393087 | 2.581406972 | 0.045313449 | 0.709264768 | -4.21506582 |
| **A_55_P2178496** | Fam53b | -0.444557847 | -0.876001108 | -0.013114587 | 9.667078041 | -2.580803489 | 0.045348405 | 0.709264768 | -4.215191343 |
| **A_52_P552547** | Lppos | -0.24236083 | -0.477877965 | -0.006843696 | 8.746934786 | -2.577450112 | 0.045543173 | 0.709264768 | -4.215889388 |
| **A_51_P453995** | Gm11961 | 0.711452543 | 0.020019504 | 1.402885581 | 10.35654778 | 2.577188317 | 0.045558416 | 0.709264768 | -4.215943923 |
| **A_66_P108474** | NA | 0.629296004 | 0.017257345 | 1.241334663 | 8.29273193 | 2.575291915 | 0.045669 | 0.709264768 | -4.216339136 |
| **A_51_P490817** | Me2 | 0.301635 | 0.008199055 | 0.595070945 | 9.079684416 | 2.574653358 | 0.045706301 | 0.709264768 | -4.216472279 |
| **A_55_P2036514** | Dis3 | 0.390389687 | 0.010134552 | 0.770644822 | 10.09609609 | 2.571423413 | 0.04589548 | 0.709264768 | -4.217146263 |
| **A_55_P2040774** | NA | -0.300050709 | -0.592592487 | -0.007508931 | 14.42133475 | -2.568958605 | 0.046040412 | 0.709264768 | -4.217661169 |
| **A_55_P2280089** | 4930570B | 0.343882548 | 0.008596683 | 0.679168412 | 10.11157723 | 2.568888395 | 0.046044548 | 0.709264768 | -4.217675844 |
| **A_55_P2010906** | Rbm45 | 0.335317886 | 0.008278265 | 0.662357507 | 9.71961481 | 2.568069038 | 0.04609284 | 0.709264768 | -4.217847127 |
| **A_52_P150114** | NA | 0.812381587 | 0.019468968 | 1.605294206 | 8.657169263 | 2.566168013 | 0.046205095 | 0.709264768 | -4.218244742 |
| **A_55_P2043702** | Tfg | 0.348861758 | 0.008327387 | 0.689396128 | 12.76608421 | 2.565917906 | 0.046219886 | 0.709264768 | -4.218297076 |
| **A_51_P403814** | Slc4a5 | -0.538100354 | -1.063490469 | -0.012710238 | 9.554759147 | -2.565261982 | 0.0462587 | 0.709264768 | -4.218434352 |
| **A_52_P175066** | Cd2ap | -0.377858761 | -0.74687812 | -0.008839402 | 11.50550502 | -2.564665275 | 0.04629404 | 0.709264768 | -4.218559265 |
| **A_52_P97192** | Ergic1 | 0.359316703 | 0.008260424 | 0.710372982 | 9.835834887 | 2.563604389 | 0.046356944 | 0.709264768 | -4.218781421 |
| **A_55_P2131520** | Vash1 | -0.255963255 | -0.506181868 | -0.005744642 | 14.56603688 | -2.562172452 | 0.046441994 | 0.709264768 | -4.219081425 |
| **A_51_P439452** | Insig2 | -0.303585565 | -0.600452268 | -0.006718863 | 10.5769122 | -2.561356178 | 0.046490552 | 0.709264768 | -4.219252518 |
| **A_55_P2097072** | Gm7219 | 0.693520602 | 0.015169132 | 1.371872071 | 8.514047084 | 2.560677834 | 0.046530946 | 0.709264768 | -4.219394743 |
| **A_51_P174681** | Bzw2 | 0.815670661 | 0.017788524 | 1.613552797 | 11.13730883 | 2.560509815 | 0.046540957 | 0.709264768 | -4.219429977 |
| **A_55_P2361547** | 2610201A | 0.331230812 | 0.007181708 | 0.655279916 | 8.654167194 | 2.560178522 | 0.046560704 | 0.709264768 | -4.219499456 |
| **A_55_P2074381** | NA | 0.371376556 | 0.007499708 | 0.735253404 | 12.38648999 | 2.55629168 | 0.046793049 | 0.709264768 | -4.220315288 |
| **A_55_P2135049** | NA | 0.568982744 | 0.01145557 | 1.126509918 | 12.85495306 | 2.556132729 | 0.046802577 | 0.709264768 | -4.220348677 |
| **A_55_P1992811** | Nudc | 0.258151379 | 0.005112927 | 0.511189831 | 9.738099765 | 2.555278692 | 0.046853807 | 0.709264768 | -4.220528115 |
| **A_55_P2126368** | Atad5 | 0.674796535 | 0.012610162 | 1.336982909 | 9.100840379 | 2.552365996 | 0.047028983 | 0.709264768 | -4.221140545 |
| **A_55_P1976278** | Prkcb | 1.110925566 | 0.02069671 | 2.201154422 | 8.763907728 | 2.552217218 | 0.04703795 | 0.709264768 | -4.221171846 |
| **A_55_P2163198** | NA | 0.476407062 | 0.008772097 | 0.944042027 | 11.29731913 | 2.551652681 | 0.047071991 | 0.709264768 | -4.221290636 |
| **A_55_P2055463** | Snapin | -0.376464742 | -0.746004252 | -0.006925232 | 11.76495446 | -2.551606942 | 0.04707475 | 0.709264768 | -4.221300261 |
| **A_55_P2161600** | Diaph2 | -0.394292913 | -0.781548262 | -0.007037565 | 9.823455037 | -2.550186207 | 0.047160541 | 0.709264768 | -4.221599333 |
| **A_55_P2046793** | Eogt | -0.379740106 | -0.752718935 | -0.006761277 | 8.55918215 | -2.550073111 | 0.047167377 | 0.709264768 | -4.221623147 |
| **A_55_P2100866** | NA | 0.491131664 | 0.00846649 | 0.973796838 | 10.2841887 | 2.548603739 | 0.047256297 | 0.709264768 | -4.221932648 |
| **A_55_P1975341** | Nasp | 0.852909245 | 0.01420659 | 1.6916119 | 12.29143467 | 2.547095036 | 0.047347783 | 0.709264768 | -4.222250619 |
| **A_55_P2168267** | NA | 0.53646054 | 0.008881277 | 1.064039804 | 12.53399962 | 2.546832662 | 0.047363713 | 0.709264768 | -4.222305936 |
| **A_51_P407165** | Abhd5 | -0.303596886 | -0.602229269 | -0.004964503 | 9.062910934 | -2.546306967 | 0.047395647 | 0.709264768 | -4.222416786 |
| **A_55_P2010758** | Npm3 | 0.537846729 | 0.008740111 | 1.066953347 | 8.595793689 | 2.546042706 | 0.047411709 | 0.709264768 | -4.222472518 |
| **A_55_P2049141** | Cops7b | 0.409096619 | 0.006643406 | 0.811549832 | 8.746690432 | 2.546014293 | 0.047413436 | 0.709264768 | -4.22247851 |
| **A_52_P380369** | Phf11d | -0.369954472 | -0.733979068 | -0.005929877 | 8.239138989 | -2.545469509 | 0.047446567 | 0.709264768 | -4.222593423 |
| **A_55_P2052848** | Mif | 0.553510404 | 0.008746227 | 1.098274582 | 15.490979 | 2.544881667 | 0.047482345 | 0.709264768 | -4.222717446 |
| **A_55_P1970536** | NA | 0.809102538 | 0.012251077 | 1.605953999 | 8.441294732 | 2.5431767 | 0.04758628 | 0.709264768 | -4.223077322 |
| **A_51_P443508** | Ppa1 | 0.344520387 | 0.005043268 | 0.683997506 | 11.22329048 | 2.541878365 | 0.04766559 | 0.709264768 | -4.223351531 |
| **A_55_P1958246** | Tcof1 | 0.524309966 | 0.007590138 | 1.041029794 | 10.99960225 | 2.541460307 | 0.047691157 | 0.709264768 | -4.223439854 |
| **A_55_P2023181** | Lsm2 | 0.584982829 | 0.008445532 | 1.161520125 | 12.51818459 | 2.541359219 | 0.047697342 | 0.709264768 | -4.223461213 |

| | | | | | | | | | |
|---|---|---|---|---|---|---|---|---|---|
| **A_51_P367285** | Rnf135 | -0.260848587 | -0.518180953 | -0.003516222 | 8.497816683 | -2.538893143 | 0.047848484 | 0.709264768 | -4.22398254 |
| **A_51_P295420** | Zc3h7a | -0.601947007 | -1.195789505 | -0.00810451 | 10.6993991 | -2.538851689 | 0.047851029 | 0.709264768 | -4.223991308 |
| **A_66_P119155** | NA | -0.38607723 | -0.767094938 | -0.005059521 | 11.73966719 | -2.537928445 | 0.047907749 | 0.709264768 | -4.224186614 |
| **A_52_P244702** | Tcf7 | 1.644704904 | 0.020850406 | 3.268559402 | 9.500642471 | 2.536829155 | 0.047975379 | 0.709264768 | -4.224419254 |
| **A_55_P2171378** | Pdzk1ip1 | -0.347357144 | -0.690348886 | -0.004365402 | 8.732055055 | -2.536547019 | 0.047992753 | 0.709264768 | -4.224478978 |
| **A_55_P2048848** | Smn1 | 0.255625546 | 0.003194824 | 0.508056269 | 10.75187055 | 2.536368719 | 0.048003736 | 0.709264768 | -4.224516724 |
| **A_55_P2060938** | NA | 0.635377629 | 0.007930928 | 1.262824329 | 11.5683832 | 2.536328054 | 0.048006242 | 0.709264768 | -4.224525334 |
| **A_55_P1992884** | NA | 0.891274612 | 0.01047623 | 1.772072993 | 8.647572053 | 2.534459603 | 0.048121506 | 0.709264768 | -4.224921057 |
| **A_55_P2003931** | NA | 0.214427036 | 0.002510867 | 0.426343206 | 7.960926649 | 2.534345343 | 0.048128565 | 0.709264768 | -4.224945266 |
| **A_55_P2060188** | NA | 0.24814642 | 0.002771329 | 0.493521511 | 8.155755429 | 2.5329574 | 0.048214393 | 0.709264768 | -4.225239422 |
| **A_55_P2027136** | H3c7 | 1.016956878 | 0.011238923 | 2.022674832 | 9.261069638 | 2.532658768 | 0.048232881 | 0.709264768 | -4.225302734 |
| **A_55_P2092310** | Phgdh | 0.582484472 | 0.005169467 | 1.159799477 | 11.115014 | 2.527096654 | 0.04857863 | 0.709264768 | -4.226483294 |
| **A_55_P2119155** | Set | 0.504683367 | 0.00443392 | 1.004932815 | 10.84319243 | 2.526868956 | 0.04859284 | 0.709264768 | -4.226531678 |
| **A_51_P125691** | Cdca4 | 0.501471103 | 0.004377902 | 0.998564303 | 9.387775714 | 2.526727659 | 0.048601661 | 0.709264768 | -4.226561704 |
| **A_55_P1978608** | NA | 1.011706576 | 0.008571645 | 2.014841506 | 9.677934929 | 2.526071069 | 0.048642672 | 0.709264768 | -4.226701255 |
| **A_51_P403760** | Braf | -0.405159417 | -0.807178103 | -0.00314073 | 10.00456166 | -2.5242365 | 0.048757456 | 0.709264768 | -4.227091362 |
| **A_55_P2146525** | Camta2 | -0.498966519 | -0.99404961 | -0.003883429 | 11.4257941 | -2.684454318 | 0.048844417 | 0.709264768 | -4.293883792 |
| **A_55_P1980951** | 1110012L | -0.242974515 | -0.484211268 | -0.001737763 | 12.0721431 | -2.522711556 | 0.04885309 | 0.709264768 | -4.227415843 |
| **A_55_P2083411** | Cyp4f18 | -0.310303067 | -0.618402419 | -0.002203714 | 9.254107656 | -2.522583948 | 0.048861102 | 0.709264768 | -4.227443004 |
| **A_55_P2073099** | Trex1 | -0.33731455 | -0.67240384 | -0.00222526 | 11.0860667 | -2.521302028 | 0.048941665 | 0.709264768 | -4.227715936 |
| **A_55_P1996751** | NA | 0.62517496 | 0.004031233 | 1.246318686 | 10.54601946 | 2.520924374 | 0.048965426 | 0.709264768 | -4.227796368 |
| **A_55_P2036673** | Ddx52 | 0.322763551 | 0.00189263 | 0.643634471 | 10.4382476 | 2.51944261 | 0.049058774 | 0.709264768 | -4.228112065 |
| **A_55_P2040988** | NA | 0.429876385 | 0.002450907 | 0.857301863 | 9.482553333 | 2.519031092 | 0.049084733 | 0.709264768 | -4.228199774 |
| **A_55_P2013043** | Serpinb6b | -1.036084744 | -2.066658083 | -0.005511404 | 12.33884875 | -2.518063751 | 0.049145812 | 0.709264768 | -4.228406002 |
| **A_52_P533198** | Dnajc2 | 0.295384721 | 0.001351079 | 0.589418362 | 9.701983071 | 2.516177938 | 0.049265118 | 0.709264768 | -4.228808264 |
| **A_51_P172251** | Ifrd2 | 0.698620126 | 0.003149009 | 1.394091242 | 10.73392988 | 2.516009868 | 0.049275766 | 0.709264768 | -4.228844129 |
| **A_55_P2108404** | NA | 0.2916426 | 0.000891868 | 0.582393332 | 10.72014139 | 2.512352018 | 0.049508121 | 0.709264768 | -4.229625275 |
| **A_51_P163953** | Nsg2 | 1.635578897 | 0.002901287 | 3.268256506 | 8.779861823 | 2.509119855 | 0.049714415 | 0.709264768 | -4.230316438 |
| **A_51_P330133** | Pspc1 | 0.503269131 | 0.000636155 | 1.005902106 | 9.170368127 | 2.507839051 | 0.049796417 | 0.709264768 | -4.230590565 |
| **A_55_P2102857** | Krtcap3 | -0.623077854 | -1.245427477 | -0.000728232 | 10.16702733 | -2.507599825 | 0.049811749 | 0.709264768 | -4.23064178 |
| **A_55_P2169149** | Washc3 | -0.253802367 | -0.507370832 | -0.000233902 | 11.18973598 | -2.506979435 | 0.049851535 | 0.709264768 | -4.230774622 |
| **A_52_P763938** | NA | -0.232295404 | -0.464385494 | -0.000205314 | 8.343639922 | -2.506884738 | 0.049857611 | 0.709264768 | -4.230794901 |
| **A_66_P118307** | NA | 0.559181577 | 0.00047001 | 1.117893144 | 12.26241396 | 2.506776056 | 0.049864585 | 0.709264768 | -4.230818177 |
| **A_52_P585233** | Ccpg1 | -0.559083027 | -1.118009598 | -0.000156456 | 10.01420111 | -2.505370142 | 0.049954897 | 0.709264768 | -4.231119361 |
| **A_55_P2117689** | Rmnd1 | -0.348973733 | -0.698106229 | 0.000158763 | 10.53747925 | -2.503530067 | 0.050073363 | 0.709264768 | -4.231513801 |
| **A_51_P369252** | Ddias | 0.416729971 | -0.000322932 | 0.833782873 | 8.342101556 | 2.502729615 | 0.050124992 | 0.709264768 | -4.231685475 |
| **A_51_P116616** | Rbm14 | 0.732033671 | -0.00064969 | 1.464717032 | 12.03025833 | 2.502448073 | 0.050143164 | 0.709264768 | -4.23174587 |
| **A_55_P2176260** | NA | 0.711402241 | -0.000701814 | 1.423506297 | 8.450650583 | 2.502200554 | 0.050159147 | 0.709264768 | -4.231798972 |
| **A_51_P405912** | Lmcd1 | -0.871564275 | -1.744466976 | 0.001338426 | 9.643327118 | -2.500828607 | 0.050247833 | 0.709264768 | -4.232093398 |
| **A_52_P105138** | Frs2 | -0.543023442 | -1.087213286 | 0.001166402 | 11.72480917 | -2.65785235 | 0.050321092 | 0.709264768 | -4.297517856 |
| **A_51_P198387** | Uckl1 | -0.327645215 | -0.656419224 | 0.001128793 | 9.96208114 | -2.496069645 | 0.050556772 | 0.709264768 | -4.233115904 |
| **A_55_P1997866** | Mtr | 0.406620768 | -0.001432241 | 0.814673777 | 10.37409112 | 2.495877796 | 0.050569269 | 0.709264768 | -4.233157164 |
| **A_52_P244463** | D16Ertd47 | 0.224547183 | -0.000829261 | 0.449923626 | 9.001881038 | 2.495453233 | 0.050596936 | 0.709264768 | -4.233248483 |
| **A_55_P1988148** | NA | 0.327176418 | -0.001211825 | 0.655564661 | 8.096544642 | 2.49542625 | 0.050598695 | 0.709264768 | -4.233254287 |
| **A_55_P2095010** | NA | 0.566608743 | -0.002333625 | 1.135551111 | 8.233694196 | 2.494395652 | 0.050665926 | 0.709264768 | -4.233476023 |
| **A_55_P2020461** | Hmgn2 | 0.265456425 | -0.001204587 | 0.532117437 | 10.96173552 | 2.493354698 | 0.05073393 | 0.709264768 | -4.233700076 |
| **A_55_P2051874** | Psmc2 | 0.311982649 | -0.001961998 | 0.625927296 | 13.04051532 | 2.489016095 | 0.051018419 | 0.709264768 | -4.234634878 |
| **A_51_P161890** | Fcgrt | -0.290202978 | -0.582247799 | 0.001841842 | 10.64539949 | -2.488872808 | 0.051027844 | 0.709264768 | -4.234665778 |

| | | | | | | | | | |
|---|---|---|---|---|---|---|---|---|---|
| **A_55_P2022337** | Dhcr7 | 0.251461267 | -0.001632822 | 0.504555356 | 8.405583475 | 2.488510296 | 0.051051696 | 0.709264768 | -4.23474396 |
| **A_51_P179907** | Appl2 | -0.308396842 | -0.619075686 | 0.002282003 | 9.154044681 | -2.486271696 | 0.051199256 | 0.709264768 | -4.235226997 |
| **A_52_P245766** | NA | 0.50875042 | -0.003919716 | 1.021420556 | 9.64144489 | 2.485519112 | 0.051248966 | 0.709264768 | -4.23538948 |
| **A_52_P417437** | Gab1 | -0.264445952 | -0.530953939 | 0.002062036 | 8.955865924 | -2.485289809 | 0.051264122 | 0.709264768 | -4.235438996 |
| **A_51_P215489** | Slc37a1 | -0.338140924 | -0.679174947 | 0.0028931 | 8.771431521 | -2.483421127 | 0.051387816 | 0.709264768 | -4.235842684 |
| **A_51_P124254** | Col4a1 | -0.31576796 | -0.634325642 | 0.002789722 | 13.68943344 | -2.482734762 | 0.051433329 | 0.709264768 | -4.244990993 |
| **A_52_P113700** | Sfxn1 | 0.465097192 | -0.004320804 | 0.934515187 | 10.99303558 | 2.481614555 | 0.051507702 | 0.709264768 | -4.236233229 |
| **A_52_P253004** | Ralb | -0.293463662 | -0.589750229 | 0.002822904 | 13.80538062 | -2.480805508 | 0.051561489 | 0.709264768 | -4.236408217 |
| **A_52_P12877** | Hspa8 | 0.36162 | -0.003503509 | 0.726743509 | 11.2393707 | 2.48063571 | 0.051572785 | 0.709264768 | -4.236444949 |
| **A_55_P2430367** | Zbtb8b | -0.618974443 | -1.244070625 | 0.006121739 | 9.330484269 | -2.480140116 | 0.05160577 | 0.709264768 | -4.236552174 |
| **A_55_P2067308** | NA | 0.589682492 | -0.006206714 | 1.185571698 | 13.25716956 | 2.478580683 | 0.051709708 | 0.709264768 | -4.236889701 |
| **A_51_P144512** | Rbm19 | 0.267152064 | -0.00282427 | 0.537128398 | 9.878598926 | 2.478467244 | 0.051717278 | 0.709264768 | -4.236914262 |
| **A_55_P2027230** | NA | 0.885871053 | -0.009759455 | 1.781501562 | 8.706065784 | 2.477376297 | 0.051790136 | 0.709264768 | -4.237150519 |
| **A_52_P460929** | BC048507 | 0.47214173 | -0.00522054 | 0.949504 | 12.71602959 | 2.477277413 | 0.051796745 | 0.709264768 | -4.237171939 |
| **A_55_P1965725** | Gapdh | 0.483281237 | -0.005354759 | 0.971917232 | 11.86619595 | 2.477221403 | 0.051800489 | 0.709264768 | -4.237184071 |
| **A_55_P1970901** | Gramd2 | -0.376270713 | -0.756727823 | 0.004186398 | 8.430433044 | -2.477108651 | 0.051808027 | 0.709264768 | -4.237208496 |
| **A_55_P2131483** | Hmgb1-ps | 1.268610021 | -0.015032818 | 2.55225286 | 12.33507188 | 2.475336699 | 0.051926643 | 0.709264768 | -4.237592482 |
| **A_55_P1971872** | NA | 0.309297878 | -0.003864757 | 0.622460514 | 8.946595284 | 2.473758771 | 0.052032516 | 0.709264768 | -4.237934642 |
| **A_51_P497100** | Lgals4 | -0.498415243 | -1.003063498 | 0.006233013 | 8.464898477 | -2.473733354 | 0.052034223 | 0.709264768 | -4.237940155 |
| **A_52_P227307** | Mllt1 | 0.469267359 | -0.006037714 | 0.944572433 | 11.14053558 | 2.472852673 | 0.052093416 | 0.709264768 | -4.238131214 |
| **A_51_P305843** | Chordc1 | 0.351610908 | -0.004588025 | 0.70780984 | 10.40492728 | 2.472407609 | 0.052123357 | 0.709264768 | -4.238227793 |
| **A_55_P1953608** | NA | 0.614791002 | -0.008064406 | 1.23764641 | 10.70688038 | 2.472239886 | 0.052134645 | 0.709264768 | -4.238264193 |
| **A_55_P2046408** | NA | 0.303156056 | -0.003980193 | 0.610292305 | 8.758513614 | 2.472210908 | 0.052136596 | 0.709264768 | -4.238270482 |
| **A_52_P370203** | Jarid2 | -0.421454062 | -0.848491381 | 0.005583258 | 10.22116472 | -2.471921979 | 0.052156048 | 0.709264768 | -4.238333193 |
| **A_51_P464981** | Map1lc3a | -0.33658435 | -0.677645325 | 0.004476626 | 10.66440043 | -2.471793775 | 0.052164682 | 0.709264768 | -4.238361022 |
| **A_51_P251357** | Ctps | 0.289762241 | -0.003917751 | 0.583442233 | 12.16368304 | 2.47125623 | 0.0522009 | 0.709264768 | -4.238477718 |
| **A_55_P2023912** | NA | 0.612513492 | -0.008419008 | 1.233445992 | 11.17468248 | 2.470709093 | 0.052237792 | 0.709264768 | -4.238596522 |
| **A_55_P2113698** | Nup205 | 0.49526134 | -0.006867165 | 0.997389845 | 11.15701877 | 2.470414898 | 0.052257641 | 0.709264768 | -4.238660412 |
| **A_66_P104118** | NA | 0.348193317 | -0.004996219 | 0.701382852 | 8.363852482 | 2.46923798 | 0.052337124 | 0.709264768 | -4.238916077 |
| **A_52_P380263** | Podxl | -0.252399168 | -0.508440846 | 0.00364251 | 8.931515894 | -2.469037007 | 0.05235071 | 0.709264768 | -4.238959746 |
| **A_51_P466964** | Pgp | 0.335459338 | -0.005003619 | 0.675922294 | 11.92596451 | 2.467859125 | 0.05243041 | 0.709264768 | -4.239215755 |
| **A_55_P1975380** | Nol11 | 0.43379634 | -0.006568589 | 0.874161269 | 10.0573511 | 2.467308784 | 0.052467692 | 0.709264768 | -4.239335409 |
| **A_51_P113274** | Hps6 | -0.22983275 | -0.46316393 | 0.00349843 | 10.48904811 | -2.467115504 | 0.052480793 | 0.709264768 | -4.239377437 |
| **A_55_P1997926** | NA | 0.610647308 | -0.009707858 | 1.231002473 | 11.23380092 | 2.465473785 | 0.052592209 | 0.709264768 | -4.239734552 |
| **A_55_P2025155** | NA | 0.977326285 | -0.015700493 | 1.970353063 | 13.71912858 | 2.465068346 | 0.052619763 | 0.709264768 | -4.239822779 |
| **A_51_P195573** | Dnmt1 | 0.320687892 | -0.005164086 | 0.64653987 | 9.685443498 | 2.46497516 | 0.052626099 | 0.709264768 | -4.239843059 |
| **A_55_P2093689** | Pold3 | 0.311574404 | -0.005196535 | 0.628345343 | 11.35425383 | 2.46358066 | 0.052721001 | 0.709264768 | -4.240146631 |
| **A_55_P2033407** | Meg3 | -0.297080601 | -0.599172306 | 0.005011104 | 8.636317583 | -2.463121521 | 0.052752288 | 0.709264768 | -4.240246617 |
| **A_55_P2110671** | NA | 1.273313446 | -0.022434029 | 2.569060921 | 12.85098152 | 2.461304239 | 0.052876317 | 0.709264768 | -4.240642535 |
| **A_55_P2064928** | Nebl | 0.593795687 | -0.010534233 | 1.198125608 | 8.049585872 | 2.461009486 | 0.052896463 | 0.709264768 | -4.240706777 |
| **A_51_P116421** | Abcc5 | -0.346374757 | -0.698907752 | 0.006158237 | 10.18888107 | -2.460916114 | 0.052902846 | 0.709264768 | -4.306670411 |
| **A_55_P1957708** | NA | 0.24092711 | -0.00440557 | 0.486259791 | 8.202690562 | 2.459691353 | 0.052986656 | 0.709264768 | -4.240994152 |
| **A_66_P111689** | NA | 1.377745335 | -0.026780248 | 2.782270917 | 11.2056409 | 2.456912222 | 0.053177359 | 0.709264768 | -4.241600521 |
| **A_55_P1954643** | 2310081J | 0.569400218 | -0.011109977 | 1.149910413 | 8.825462226 | 2.456733926 | 0.053189618 | 0.709264768 | -4.241639445 |
| **A_66_P139493** | Tmem250 | 0.841164313 | -0.016605931 | 1.698934557 | 11.04100927 | 2.4561801 | 0.053227718 | 0.709264768 | -4.241760367 |
| **A_55_P1977008** | Gfi1 | 1.310739756 | -0.025901458 | 2.647380969 | 8.421758375 | 2.456133507 | 0.053230925 | 0.709264768 | -4.241770541 |
| **A_51_P415905** | Pola1 | 1.203557187 | -0.024034379 | 2.431148753 | 9.828780232 | 2.455631412 | 0.053265494 | 0.709264768 | -4.241880192 |
| **A_55_P2017826** | Myb | 1.89421629 | -0.038171599 | 3.826604178 | 10.30463039 | 2.455192823 | 0.05329571 | 0.709264768 | -4.241975991 |

| | | | | | | | | | |
|---|---|---|---|---|---|---|---|---|---|
| **A_55_P1958480** | NA | -0.42848524 | -0.865789664 | 0.008819183 | 10.94114658 | -2.454156998 | 0.053367145 | 0.709264768 | -4.242202305 |
| **A_55_P2243828** | NA | -0.3003562 | -0.60697372 | 0.006261321 | 8.195705313 | -2.453522129 | 0.053410979 | 0.709264768 | -4.24234106 |
| **A_52_P233335** | Pphln1 | 0.29860481 | -0.006430719 | 0.603640338 | 10.36487592 | 2.451865928 | 0.053525511 | 0.709264768 | -4.242703189 |
| **A_51_P195153** | Gtse1 | 0.631182528 | -0.013794562 | 1.276159619 | 8.730825552 | 2.451099975 | 0.053578567 | 0.709264768 | -4.242870742 |
| **A_52_P5549** | Fam133b | 0.372292974 | -0.008228114 | 0.752814062 | 9.897484284 | 2.450509868 | 0.053619482 | 0.709264768 | -4.242999862 |
| **A_55_P2135023** | NA | 0.587719836 | -0.013008166 | 1.188447839 | 11.36850719 | 2.450432917 | 0.05362482 | 0.709264768 | -4.243016702 |
| **A_55_P1972688** | Wnk1 | -0.333949952 | -0.675413529 | 0.007513626 | 12.31054417 | -2.449555844 | 0.053685699 | 0.709264768 | -4.24320867 |
| **A_55_P1980190** | NA | 0.267075514 | -0.006229167 | 0.540380195 | 11.8034172 | 2.447582547 | 0.053822939 | 0.709264768 | -4.243640806 |
| **A_55_P2076592** | NA | 0.47674136 | -0.011128752 | 0.964611471 | 8.465816927 | 2.447535295 | 0.05382623 | 0.709264768 | -4.243651158 |
| **A_55_P1971769** | NA | 0.506218112 | -0.012218684 | 1.024654908 | 12.1414562 | 2.445638187 | 0.053958534 | 0.709264768 | -4.244066921 |
| **A_51_P220896** | Chmp2b | -0.217174695 | -0.439629511 | 0.005280122 | 10.23172601 | -2.445218948 | 0.053987818 | 0.709264768 | -4.244158839 |
| **A_52_P636830** | G3bp2 | 0.217331863 | -0.005355478 | 0.440019204 | 10.5473391 | 2.444433451 | 0.054042732 | 0.709264768 | -4.2443311 |
| **A_55_P2016478** | Ddx39b | 0.259634875 | -0.00682355 | 0.5260933 | 9.291199599 | 2.440528689 | 0.054316599 | 0.709264768 | -4.245188177 |
| **A_55_P1999992** | Galnt12 | 0.819927308 | -0.021901506 | 1.661756122 | 9.79233028 | 2.439506109 | 0.054388564 | 0.709264768 | -4.245412837 |
| **A_55_P2201822** | Naf1 | 0.506844704 | -0.013694 | 1.027383407 | 10.68154223 | 2.438777796 | 0.054439882 | 0.709264768 | -4.2455729 |
| **A_55_P1959828** | Slc35g1 | 0.365844891 | -0.010123627 | 0.741813409 | 8.503841713 | 2.437226323 | 0.054549373 | 0.709264768 | -4.245914015 |
| **A_55_P2340206** | E230006M | 0.410047465 | -0.011498449 | 0.831593379 | 9.030159094 | 2.436349524 | 0.054611354 | 0.709264768 | -4.246106881 |
| **A_55_P1964302** | Timm8a1 | 0.230556253 | -0.006507586 | 0.467620092 | 8.526319423 | 2.435913924 | 0.054642175 | 0.709264768 | -4.246202722 |
| **A_55_P2083023** | Sac3d1 | 0.252249186 | -0.007283196 | 0.511781569 | 10.92801653 | 2.434381092 | 0.054750778 | 0.709264768 | -4.2465401 |
| **A_51_P254045** | Traip | 0.931643459 | -0.027614397 | 1.890901314 | 9.803579947 | 2.432566491 | 0.054879643 | 0.709264768 | -4.246939747 |
| **A_51_P394515** | Tkt | 1.386685403 | -0.041400566 | 2.814771372 | 13.32558541 | 2.432058052 | 0.054915808 | 0.709264768 | -4.247051774 |
| **A_55_P2035315** | Rasgef1b | -0.738135931 | -1.498389819 | 0.022117957 | 10.81240489 | -2.431801052 | 0.054934098 | 0.709264768 | -4.247108409 |
| **A_55_P2119302** | Cspp1 | -0.210966587 | -0.428370867 | 0.006437694 | 8.470875517 | -2.430501713 | 0.055026668 | 0.709264768 | -4.247394824 |
| **A_51_P166244** | Osbpl1a | -0.350048113 | -0.710897238 | 0.010801013 | 10.67014902 | -2.429698741 | 0.055083959 | 0.709264768 | -4.247571894 |
| **A_51_P340704** | Prpf31 | 0.351286787 | -0.010880872 | 0.713454447 | 12.09850102 | 2.429419398 | 0.055103904 | 0.709264768 | -4.247633506 |
| **A_52_P658122** | Ets2 | -0.413728676 | -0.840371826 | 0.012914475 | 13.09914801 | -2.428852776 | 0.055144385 | 0.709264768 | -4.247758502 |
| **A_52_P356204** | Nostrin | -0.436665928 | -0.887042587 | 0.01371073 | 8.999630809 | -2.428419869 | 0.055175334 | 0.709264768 | -4.247854018 |
| **A_55_P1982628** | Tma16 | 0.26570223 | -0.008344569 | 0.53974903 | 9.121220542 | 2.42840328 | 0.055176521 | 0.709264768 | -4.247857679 |
| **A_55_P2062543** | Kifc5b | 0.625097948 | -0.019646344 | 1.269842239 | 9.978041383 | 2.428347937 | 0.055180479 | 0.709264768 | -4.247869891 |
| **A_55_P2415197** | 2810013P | -0.281575984 | -0.57209304 | 0.008941073 | 8.892679654 | -2.4275843 | 0.055235123 | 0.709264768 | -4.248038421 |
| **A_55_P2080703** | Cstf2 | 0.424805386 | -0.013504768 | 0.86311554 | 8.597443973 | 2.427497708 | 0.055241323 | 0.709264768 | -4.248057535 |
| **A_55_P1952882** | Cyp4f18 | -0.412307645 | -0.837921894 | 0.013306605 | 8.990365177 | -2.426361874 | 0.055322719 | 0.709264768 | -4.248308304 |
| **A_55_P1997485** | NA | 0.45318033 | -0.014705899 | 0.921066558 | 11.57389834 | 2.425946023 | 0.055352551 | 0.709264768 | -4.248400142 |
| **A_55_P1956687** | Rab37 | 0.404086141 | -0.013458556 | 0.821630837 | 8.179451303 | 2.423937007 | 0.055496916 | 0.709264768 | -4.248844022 |
| **A_55_P2036883** | H2bc18 | 1.758397036 | -0.059593925 | 3.576387998 | 11.3952553 | 2.422565728 | 0.055595684 | 0.709264768 | -4.249147189 |
| **A_55_P1964225** | Elac2 | 0.431686487 | -0.014840576 | 0.878213549 | 9.380720057 | 2.421424957 | 0.055677993 | 0.709264768 | -4.249399513 |
| **A_51_P480241** | Elf3 | -0.414656002 | -0.84383458 | 0.014522575 | 9.200385386 | -2.41991586 | 0.055787076 | 0.709264768 | -4.24973347 |
| **A_55_P2009042** | NA | -0.412367913 | -0.839321991 | 0.014586166 | 11.31325148 | -2.419101236 | 0.055846055 | 0.709264768 | -4.249913822 |
| **A_55_P2078516** | NA | 0.40187671 | -0.014402527 | 0.818155947 | 9.399867512 | 2.418011898 | 0.055925028 | 0.709264768 | -4.250155078 |
| **A_55_P1979067** | A430073D | -0.523013756 | -1.064874581 | 0.018847069 | 13.41323506 | -2.417551326 | 0.055958453 | 0.709264768 | -4.250257111 |
| **A_55_P2125351** | Sema6d | 0.524263542 | -0.018969993 | 1.067497078 | 11.0454163 | 2.417204703 | 0.055983623 | 0.709264768 | -4.250333911 |
| **A_55_P2131190** | NA | -0.259728528 | -0.528921324 | 0.009464268 | 8.185016355 | -2.416609991 | 0.056026835 | 0.709264768 | -4.250465703 |
| **A_55_P1963364** | NA | 0.316408314 | -0.011720392 | 0.644537019 | 9.271698781 | 2.415205043 | 0.056129062 | 0.709264768 | -4.250777164 |
| **A_55_P2143885** | Etl4 | -0.363702847 | -0.741012319 | 0.013606625 | 10.44160035 | -2.414345052 | 0.056191734 | 0.709264768 | -4.250967895 |
| **A_55_P1976709** | NA | 0.328975003 | -0.012443783 | 0.670393789 | 11.50785639 | 2.41338068 | 0.056262103 | 0.709264768 | -4.251181848 |
| **A_51_P387235** | Nampt | -0.268283426 | -0.546770068 | 0.010203217 | 9.903874491 | -2.412902759 | 0.05629701 | 0.709264768 | -4.251287907 |
| **A_55_P1999849** | NA | 0.448367771 | -0.017059422 | 0.913794964 | 11.92129286 | 2.412864755 | 0.056299787 | 0.709264768 | -4.251296342 |
| **A_52_P617327** | Rcan1 | -0.28201378 | -0.574818566 | 0.010791007 | 9.554327061 | -2.412362133 | 0.056336526 | 0.709264768 | -4.251407904 |

| | | | | | | | | | |
|---|---|---|---|---|---|---|---|---|---|
| **A_51_P466829** | Pheta1 | -0.692215963 | -1.411063657 | 0.026631731 | 12.32211026 | -2.411876533 | 0.056372045 | 0.709264768 | -4.251515708 |
| **A_65_P15215** | Mrpl15 | 0.297575202 | -0.011468209 | 0.606618614 | 10.9347239 | 2.411723934 | 0.056383211 | 0.709264768 | -4.251549589 |
| **A_52_P400999** | Arhgap31 | -0.331494316 | -0.67581563 | 0.012826998 | 11.14594371 | -2.41136262 | 0.05640966 | 0.709264768 | -4.251629818 |
| **A_66_P107790** | Myl12a | -0.597334716 | -1.217915414 | 0.023245982 | 14.8725695 | -2.410848046 | 0.056447351 | 0.709264768 | -4.251744097 |
| **A_51_P421804** | Timm9 | 0.288588139 | -0.011276274 | 0.588452553 | 9.70098149 | 2.410482014 | 0.056474178 | 0.709264768 | -4.2518254 |
| **A_55_P2069425** | Ttll3 | -0.60021412 | -1.2241404 | 0.02371216 | 8.619052926 | -2.409479717 | 0.056547707 | 0.709264768 | -4.252048088 |
| **A_55_P2029420** | 2610318N | 1.149739642 | -0.046001985 | 2.345481268 | 9.605230888 | 2.408310631 | 0.056633601 | 0.709264768 | -4.252307937 |
| **A_55_P2067942** | D16Ertd47 | 0.253836943 | -0.010204139 | 0.517878025 | 10.21355984 | 2.407873523 | 0.056665752 | 0.709264768 | -4.25240512 |
| **A_51_P472249** | Slc7a7 | -0.527439985 | -1.076112414 | 0.021232443 | 11.03074233 | -2.407743725 | 0.056675302 | 0.709264768 | -4.252433981 |
| **A_51_P488718** | Zfand2b | -0.261721505 | -0.534385789 | 0.010942779 | 10.91568119 | -2.40414967 | 0.056940441 | 0.709264768 | -4.253233692 |
| **A_55_P2069012** | Pidd1 | 0.638215402 | -0.026918926 | 1.303349731 | 9.144191353 | 2.403301533 | 0.057003201 | 0.709264768 | -4.253422566 |
| **A_51_P418526** | Sfxn1 | 0.303673085 | -0.012902889 | 0.62024906 | 9.870529923 | 2.402584636 | 0.057056308 | 0.709264768 | -4.253582259 |
| **A_52_P606303** | Mrpl3 | 0.280347321 | -0.012499903 | 0.573194546 | 12.02666905 | 2.397759631 | 0.05741511 | 0.709264768 | -4.25465816 |
| **A_51_P256827** | S100a8 | -0.886687664 | -1.81291954 | 0.039544211 | 11.44741708 | -2.397735588 | 0.057416904 | 0.709264768 | -4.254663527 |
| **A_52_P569038** | Cnot2 | 0.445595367 | -0.01994057 | 0.911131305 | 11.23594938 | 2.397385093 | 0.057443062 | 0.709264768 | -4.254741757 |
| **A_55_P2043382** | Llgl2 | -0.620078865 | -1.268241801 | 0.028084071 | 11.41909207 | -2.396144925 | 0.05753572 | 0.709264768 | -4.255018642 |
| **A_55_P1995633** | NA | 1.233293506 | -0.056144198 | 2.522731209 | 12.91060898 | 2.395611698 | 0.057575609 | 0.709264768 | -4.255137732 |
| **A_55_P2139027** | Plec | -0.453352701 | -0.927807889 | 0.021102487 | 12.45644728 | -2.393268107 | 0.057751273 | 0.709264768 | -4.255661421 |
| **A_55_P2187266** | Pop1 | 0.517677742 | -0.024295058 | 1.059650541 | 10.13597185 | 2.392392031 | 0.057817085 | 0.709264768 | -4.255857301 |
| **A_55_P2277014** | AA522020 | -0.408754225 | -0.836829007 | 0.019320558 | 11.66298699 | -2.391624286 | 0.057874825 | 0.709264768 | -4.256029011 |
| **A_55_P2048110** | Tmem97 | 0.285486516 | -0.013678039 | 0.584651071 | 10.33260278 | 2.390153586 | 0.057985605 | 0.709264768 | -4.256358077 |
| **A_55_P1958517** | Parp6 | -0.549713189 | -1.126075518 | 0.026649139 | 10.57642333 | -2.388861194 | 0.058083139 | 0.709264768 | -4.256647392 |
| **A_55_P1963777** | Gtf3c6 | 0.410916656 | -0.02009074 | 0.841924052 | 11.58125124 | 2.387917776 | 0.058154447 | 0.709264768 | -4.256858673 |
| **A_51_P307325** | Rbbp4 | 0.478817841 | -0.023418701 | 0.981054383 | 13.24705353 | 2.387879249 | 0.058157361 | 0.709264768 | -4.256867303 |
| **A_55_P1955587** | Hspd1 | 0.3380973 | -0.016545454 | 0.692740054 | 13.34360086 | 2.387816545 | 0.058162104 | 0.709264768 | -4.256881349 |
| **A_51_P464387** | Hspb8 | -0.80487716 | -1.649153518 | 0.039399199 | 9.989743655 | -2.387785554 | 0.058164449 | 0.709264768 | -4.256888291 |
| **A_55_P2056293** | Sf3b4 | 0.291541237 | -0.014401082 | 0.597483556 | 8.515770016 | 2.386771171 | 0.058241239 | 0.709264768 | -4.257115557 |
| **A_66_P106191** | Eef2 | 0.508832712 | -0.02517709 | 1.042842514 | 14.99802996 | 2.386580789 | 0.058255663 | 0.709264768 | -4.25715822 |
| **A_51_P308961** | Hmgcl | -0.507241586 | -1.039811931 | 0.02532876 | 11.81374877 | -2.385548312 | 0.058333954 | 0.709264768 | -4.257389643 |
| **A_55_P2047882** | Zc3h7b | 0.418086742 | -0.021126656 | 0.857300141 | 10.22452959 | 2.384191646 | 0.058436998 | 0.709264768 | -4.257693864 |
| **A_55_P2098210** | 4931406P | -0.231473118 | -0.474695961 | 0.011749725 | 12.02782756 | -2.383672283 | 0.058476497 | 0.709264768 | -4.257810367 |
| **A_52_P480544** | Tnpo2 | 0.282022815 | -0.014390323 | 0.578435954 | 8.637144358 | 2.383071863 | 0.058522196 | 0.709264768 | -4.257945079 |
| **A_66_P100561** | Bnip2 | -0.254974431 | -0.52304168 | 0.013092818 | 13.30592062 | -2.382337133 | 0.058578169 | 0.709264768 | -4.258109967 |
| **A_55_P2038452** | NA | -0.281973914 | -0.578457134 | 0.014509306 | 8.747898755 | -2.382095454 | 0.058596593 | 0.709264768 | -4.258164214 |
| **A_51_P495242** | Lat | 1.538437555 | -0.07942793 | 3.156303041 | 8.57773961 | 2.381704124 | 0.058626439 | 0.709264768 | -4.258252062 |
| **A_52_P8043** | Srsf2 | 0.697224367 | -0.036253439 | 1.430702173 | 10.28481183 | 2.380871329 | 0.058690007 | 0.709264768 | -4.258439054 |
| **A_55_P1977653** | Palld | -0.368483118 | -0.756175571 | 0.019209334 | 11.70388558 | -2.380568018 | 0.058713177 | 0.709264768 | -4.258507172 |
| **A_52_P355014** | Rbm8a | 0.415325276 | -0.021748117 | 0.852398668 | 11.84143796 | 2.380040454 | 0.058753502 | 0.709264768 | -4.258625671 |
| **A_55_P2182921** | NA | 0.463015904 | -0.024315788 | 0.950347596 | 9.097734525 | 2.379696649 | 0.058779797 | 0.709264768 | -4.258702907 |
| **A_55_P2077618** | Csgalnact | -0.479801936 | -0.984979518 | 0.025375646 | 9.240151755 | -2.378856649 | 0.058844094 | 0.709264768 | -4.258891656 |
| **A_55_P2158990** | Jun | -0.501864317 | -1.030593958 | 0.026865325 | 13.33899249 | -2.377404087 | 0.058955455 | 0.709264768 | -4.259218183 |
| **A_55_P1971614** | NA | 1.299861361 | -0.069634782 | 2.669357504 | 11.11777188 | 2.377314102 | 0.058962361 | 0.709264768 | -4.259238417 |
| **A_55_P2068330** | Cryzl1 | -0.250381236 | -0.514342826 | 0.013580353 | 9.170725174 | -2.375808271 | 0.059078057 | 0.709264768 | -4.259577112 |
| **A_51_P113722** | Fbxl14 | 0.43152815 | -0.023495221 | 0.886551521 | 10.89376991 | 2.375339953 | 0.059114088 | 0.709264768 | -4.263900714 |
| **A_51_P436719** | Eftud2 | 0.243265202 | -0.013359662 | 0.499890065 | 9.516871016 | 2.374278188 | 0.059195864 | 0.709264768 | -4.259921453 |
| **A_66_P133642** | NA | 1.31232712 | -0.072620125 | 2.697274364 | 11.46721348 | 2.373335955 | 0.059268534 | 0.709264768 | -4.260133595 |
| **A_51_P228295** | Mpzl1 | -0.430615768 | -0.885065641 | 0.023834105 | 13.85364851 | -2.373309008 | 0.059270614 | 0.709264768 | -4.260139663 |
| **A_55_P2176908** | Tdpoz8 | 0.421318728 | -0.023564978 | 0.866202434 | 10.60680426 | 2.37199959 | 0.059371765 | 0.709264768 | -4.2604346 |

| | | | | | | | | | |
|---|---|---|---|---|---|---|---|---|---|
| **A_51_P381749** | Prelid3b | 0.560238104 | -0.031340593 | 1.1518168 | 12.52544057 | 2.371976939 | 0.059373516 | 0.709264768 | -4.260439703 |
| **A_51_P105380** | Csta2 | -0.328210562 | -0.674815281 | 0.018394157 | 9.069224064 | -2.371747367 | 0.05939127 | 0.709264768 | -4.260491428 |
| **A_52_P190973** | Vcl | -0.201343517 | -0.414051801 | 0.011364766 | 10.7486698 | -2.370847362 | 0.059460925 | 0.709264768 | -4.260694247 |
| **A_55_P2081437** | NA | 0.665169287 | -0.037646812 | 1.367985386 | 8.807217184 | 2.370504765 | 0.059487462 | 0.709264768 | -4.26077147 |
| **A_55_P2020090** | Pwwp3b | -0.246336023 | -0.506694935 | 0.014022888 | 11.14383312 | -2.369767966 | 0.059544578 | 0.709264768 | -4.260937581 |
| **A_55_P2179296** | Mad2l1 | 0.626818749 | -0.0388247 | 1.292462197 | 8.196968275 | 2.508204955 | 0.059621569 | 0.709264768 | -4.318869062 |
| **A_55_P2006255** | Txnip | -0.483276035 | -0.994450101 | 0.027898031 | 8.742395983 | -2.367973254 | 0.059683944 | 0.709264768 | -4.261342382 |
| **A_55_P2102738** | NA | -0.388711055 | -0.799865258 | 0.022443149 | 14.46028704 | -2.367949861 | 0.059685763 | 0.709264768 | -4.26134766 |
| **A_55_P2072035** | Thy1 | 2.893961081 | -0.167479307 | 5.955401469 | 10.55420261 | 2.367648483 | 0.059709201 | 0.709264768 | -4.261415663 |
| **A_55_P2130256** | Polr2l | 0.346201695 | -0.020111558 | 0.712514949 | 8.987845685 | 2.367156132 | 0.059747512 | 0.709264768 | -4.261526773 |
| **A_55_P2070194** | Caprin1 | 0.443666474 | -0.025891906 | 0.913224853 | 11.96342924 | 2.366559141 | 0.059794 | 0.709264768 | -4.261661524 |
| **A_55_P1984069** | NA | 0.505214412 | -0.029483858 | 1.039912683 | 12.99098968 | 2.366558788 | 0.059794028 | 0.709264768 | -4.261661604 |
| **A_51_P149562** | Apbb2 | -0.427355249 | -0.87984132 | 0.025130821 | 11.7930492 | -2.365561122 | 0.059871803 | 0.709264768 | -4.261886859 |
| **A_55_P2094611** | NA | 0.651635492 | -0.038361689 | 1.341632672 | 8.907333718 | 2.365417251 | 0.059883027 | 0.709264768 | -4.261919349 |
| **A_55_P2064412** | NA | 0.746233087 | -0.043957399 | 1.536423573 | 8.687210411 | 2.365337137 | 0.059889278 | 0.709264768 | -4.261937442 |
| **A_51_P364786** | U2af1 | 0.45951259 | -0.027129972 | 0.946155152 | 9.052181742 | 2.365035537 | 0.059912819 | 0.709264768 | -4.262005559 |
| **A_51_P391159** | Ang | -0.280966336 | -0.578553546 | 0.016620874 | 8.556132024 | -2.364777976 | 0.059932929 | 0.709264768 | -4.262063737 |
| **A_55_P2163812** | 4-Sep | -0.557917044 | -1.148928746 | 0.033094659 | 10.41669102 | -2.364416025 | 0.059961203 | 0.709264768 | -4.262145502 |
| **A_55_P2126695** | NA | 0.323672713 | -0.019205942 | 0.666551368 | 8.384147672 | 2.364372955 | 0.059964568 | 0.709264768 | -4.262155232 |
| **A_51_P282227** | As3mt | -0.274508228 | -0.565585377 | 0.016568922 | 9.926248048 | -2.362096296 | 0.060142744 | 0.709264768 | -4.262669787 |
| **A_55_P2030428** | Sirt1 | 0.206288544 | -0.012643035 | 0.425220123 | 10.58210741 | 2.360027404 | 0.060305148 | 0.709264768 | -4.263137752 |
| **A_55_P2069856** | Tnpo3 | 0.26654274 | -0.017681853 | 0.550767333 | 11.42024321 | 2.497858947 | 0.06033235 | 0.709264768 | -4.320403166 |
| **A_51_P440892** | Dph2 | 0.378862859 | -0.023560942 | 0.781286659 | 9.301769713 | 2.358026706 | 0.060462641 | 0.709264768 | -4.263590623 |
| **A_55_P2049856** | Araf | -0.291423717 | -0.601154269 | 0.018306834 | 10.59988727 | -2.356628868 | 0.060572937 | 0.709264768 | -4.263907228 |
| **A_55_P1984690** | Ptprr | -0.295168823 | -0.608971784 | 0.018634138 | 7.991639061 | -2.355937647 | 0.060627556 | 0.709264768 | -4.264063845 |
| **A_52_P123400** | Dnaaf2 | 0.396070775 | -0.025237591 | 0.817379141 | 9.251494263 | 2.3546321 | 0.060730861 | 0.709264768 | -4.264359764 |
| **A_52_P534583** | Ahsp | 0.577683686 | -0.036930935 | 1.192298307 | 11.76950383 | 2.354168593 | 0.060767582 | 0.709264768 | -4.264464857 |
| **A_55_P2006504** | Eif4a2 | -0.349714994 | -0.721927226 | 0.022497238 | 12.96896528 | -2.353281913 | 0.060837895 | 0.709264768 | -4.264665947 |
| **A_55_P2360661** | Cbx5 | 0.260402066 | -0.016766942 | 0.537571073 | 9.80219281 | 2.353152667 | 0.060848151 | 0.709264768 | -4.264695264 |
| **A_55_P2031481** | Fam131a | -0.648187895 | -1.338207383 | 0.041831594 | 10.06945983 | -2.352826509 | 0.060874041 | 0.709264768 | -4.264769252 |
| **A_55_P2021413** | Celf4 | 0.365001183 | -0.02355752 | 0.753559886 | 9.598045404 | 2.352816068 | 0.06087487 | 0.709264768 | -4.264771621 |
| **A_55_P2136020** | Uba2 | 0.398167018 | -0.025964313 | 0.82229835 | 13.27651436 | 2.351339137 | 0.060992256 | 0.709264768 | -4.265106772 |
| **A_55_P2108143** | NA | 0.403970217 | -0.026370925 | 0.834311359 | 11.48990234 | 2.351185126 | 0.061004511 | 0.709264768 | -4.265141731 |
| **A_55_P2121364** | Tdg | 0.319151153 | -0.021053183 | 0.659355489 | 8.549061097 | 2.349670254 | 0.061125188 | 0.709264768 | -4.265485696 |
| **A_55_P2186332** | NA | 0.338908416 | -0.022423827 | 0.700240659 | 8.060451312 | 2.3492324 | 0.061160115 | 0.709264768 | -4.265585149 |
| **A_51_P158210** | Mcm2 | 1.244950582 | -0.082741091 | 2.572642254 | 10.69097341 | 2.34857929 | 0.061212252 | 0.709264768 | -4.265733525 |
| **A_55_P1968304** | NA | 0.338879629 | -0.022673553 | 0.700432812 | 12.69602454 | 2.347597401 | 0.061290724 | 0.709264768 | -4.265956658 |
| **A_55_P1999172** | Nat10 | 0.233146621 | -0.015683084 | 0.481976325 | 10.43060664 | 2.34680631 | 0.061354026 | 0.709264768 | -4.26613649 |
| **A_51_P239654** | Nr4a1 | -0.267590983 | -0.553307743 | 0.018125776 | 10.58687786 | -2.345773662 | 0.06143676 | 0.709264768 | -4.26637131 |
| **A_51_P262515** | Phf11a | -0.474167335 | -0.980633358 | 0.032298688 | 8.443408333 | -2.344939612 | 0.06150367 | 0.709264768 | -4.266561032 |
| **A_55_P2262753** | 9530053J | -0.251219437 | -0.519643851 | 0.017204977 | 8.249718449 | -2.344129339 | 0.061568746 | 0.709264768 | -4.266745401 |
| **A_55_P2121913** | Adi1 | 0.288035581 | -0.019873315 | 0.595944477 | 9.570365277 | 2.343010573 | 0.061658718 | 0.709264768 | -4.267000051 |
| **A_52_P159256** | Lpp | -0.263622021 | -0.545456202 | 0.018212159 | 9.172751405 | -2.342817011 | 0.061674299 | 0.709264768 | -4.267044119 |
| **A_52_P96782** | Wasl | -0.359701104 | -0.744338844 | 0.024936636 | 9.924208261 | -2.342287613 | 0.061716933 | 0.709264768 | -4.267164663 |
| **A_52_P431872** | Ptcd3 | 0.418270918 | -0.029155294 | 0.865697129 | 12.00437133 | 2.341459185 | 0.061783713 | 0.709264768 | -4.267353342 |
| **A_55_P2002150** | Pgs1 | -0.513738273 | -1.063328984 | 0.035852439 | 13.4596574 | -2.341277451 | 0.061798373 | 0.709264768 | -4.26739474 |
| **A_51_P294778** | Zfp954 | -0.309461701 | -0.640597155 | 0.021673754 | 9.79099078 | -2.340731343 | 0.061842447 | 0.709264768 | -4.267519158 |
| **A_55_P2005838** | Ticrr | 0.602680057 | -0.042377821 | 1.247737935 | 9.456801167 | 2.34012191 | 0.061891672 | 0.709264768 | -4.267658031 |

| | | | | | | | | | |
|---|---|---|---|---|---|---|---|---|---|
| **A_55_P1970935** | Usp7 | 0.227582956 | -0.01604793 | 0.471213842 | 12.00248728 | 2.339686853 | 0.061926838 | 0.709264768 | -4.267757187 |
| **A_52_P503208** | NA | 0.895034596 | -0.063297326 | 1.853366518 | 12.27554521 | 2.33923694 | 0.061963227 | 0.709264768 | -4.267859746 |
| **A_55_P2183237** | Ccdc191 | -0.23323674 | -0.482991888 | 0.016518409 | 9.351975118 | -2.339014199 | 0.06198125 | 0.709264768 | -4.267910526 |
| **A_55_P1971229** | Slc25a36 | -0.213401448 | -0.442119759 | 0.015316862 | 8.923737554 | -2.33693576 | 0.062149699 | 0.709264768 | -4.26838456 |
| **A_55_P2061021** | Smg6 | -0.35246953 | -0.730272227 | 0.025333167 | 11.77239538 | -2.336721048 | 0.062167129 | 0.709264768 | -4.26843355 |
| **A_55_P2176917** | NA | 1.119876726 | -0.080703942 | 2.320457394 | 10.55864942 | 2.33630328 | 0.062201056 | 0.709264768 | -4.268528881 |
| **A_65_P11853** | Arhgap28 | -0.221913912 | -0.460005216 | 0.016177391 | 9.616298101 | -2.33448637 | 0.062348836 | 0.709264768 | -4.268943649 |
| **A_52_P583458** | E2f3 | 0.557317172 | -0.040671694 | 1.155306038 | 9.91593861 | 2.334316139 | 0.062362701 | 0.709264768 | -4.268982523 |
| **A_55_P2035320** | Nfil3 | -0.356088607 | -0.738182084 | 0.02600487 | 11.10179216 | -2.334203955 | 0.06237184 | 0.709264768 | -4.269008143 |
| **A_52_P673499** | Shmt1 | 0.239099713 | -0.01755646 | 0.495755886 | 8.228888293 | 2.333338173 | 0.062442417 | 0.709264768 | -4.2692059 |
| **A_52_P646034** | Nfatc2ip | 0.359106491 | -0.026477294 | 0.744690275 | 9.020026048 | 2.33267824 | 0.062496271 | 0.709264768 | -4.269356679 |
| **A_51_P507801** | F13a1 | 1.099756455 | -0.081111425 | 2.280624336 | 10.36345591 | 2.332628382 | 0.062500342 | 0.709264768 | -4.269368072 |
| **A_55_P2021055** | Elp5 | -0.382916319 | -0.794268949 | 0.028436311 | 11.2166897 | -2.33152428 | 0.062590558 | 0.709264768 | -4.269620416 |
| **A_51_P232411** | Rag1 | 2.460392963 | -0.183151868 | 5.103937795 | 8.881423771 | 2.331138848 | 0.062622084 | 0.709264768 | -4.26970853 |
| **A_52_P480360** | Dut | 1.252946262 | -0.09360247 | 2.599494994 | 11.55213108 | 2.33056229 | 0.062669275 | 0.709264768 | -4.26984036 |
| **A_52_P684378** | Gpx1 | -0.500468764 | -1.038374963 | 0.037437435 | 14.27653017 | -2.330347961 | 0.062686827 | 0.709264768 | -4.269889374 |
| **A_55_P2079390** | 4930579G | 0.482550738 | -0.036254753 | 1.001356229 | 8.711349637 | 2.329639739 | 0.062744863 | 0.709264768 | -4.270051359 |
| **A_51_P124089** | Gtpbp2 | -0.232612237 | -0.482797422 | 0.017572948 | 9.324335196 | -2.328741669 | 0.062818538 | 0.709264768 | -4.270256824 |
| **A_55_P2118740** | NA | 0.94499175 | -0.071851084 | 1.961834583 | 13.40546261 | 2.327686728 | 0.0629052 | 0.709264768 | -4.270498263 |
| **A_51_P201661** | Xrcc5 | 0.453488719 | -0.034523959 | 0.941501396 | 9.920524176 | 2.327478776 | 0.062922298 | 0.709264768 | -4.270545867 |
| **A_51_P423091** | Flot1 | -0.476190745 | -0.98866951 | 0.03628802 | 11.15883561 | -2.327316355 | 0.062935655 | 0.709264768 | -4.27058305 |
| **A_55_P2052097** | Ip6k2 | -0.198485456 | -0.412127262 | 0.015156349 | 8.573584236 | -2.326980778 | 0.062963263 | 0.709264768 | -4.270659881 |
| **A_51_P318551** | Nle1 | 0.291651054 | -0.022295467 | 0.605597575 | 8.372110134 | 2.326795535 | 0.062978508 | 0.709264768 | -4.270702296 |
| **A_55_P2059010** | Rbp1 | -0.254146915 | -0.52777866 | 0.019484829 | 15.1413476 | -2.32631601 | 0.063017991 | 0.709264768 | -4.270812107 |
| **A_66_P125067** | Snrpd1 | 0.539772861 | -0.041499821 | 1.121045542 | 10.77342653 | 2.325848797 | 0.063056485 | 0.709264768 | -4.270919117 |
| **A_52_P361551** | Hspa14 | 0.235368744 | -0.018109523 | 0.48884701 | 12.51295302 | 2.325725239 | 0.063066669 | 0.709264768 | -4.270947419 |
| **A_51_P481821** | Spcs3 | 0.217431531 | -0.016771546 | 0.451634609 | 10.57879963 | 2.325306862 | 0.063101167 | 0.709264768 | -4.271043262 |
| **A_55_P1970120** | Vopp1 | 0.280093313 | -0.021746385 | 0.581933011 | 11.79352147 | 2.324217294 | 0.063191101 | 0.709264768 | -4.271292931 |
| **A_51_P261001** | Mtf2 | 0.65236575 | -0.050753732 | 1.355485232 | 10.45416117 | 2.323872872 | 0.063219559 | 0.709264768 | -4.271371874 |
| **A_55_P2110136** | Arhgap29 | -0.487975409 | -1.014079592 | 0.038128775 | 9.344344173 | -2.32314612 | 0.06327965 | 0.709264768 | -4.271538479 |
| **A_51_P348749** | Llgl2 | -0.565879676 | -1.176181306 | 0.044421954 | 10.53358904 | -2.322361973 | 0.063344555 | 0.709264768 | -4.27171829 |
| **A_52_P609200** | Serinc3 | -0.356109071 | -0.740198364 | 0.027980222 | 14.66502605 | -2.322208346 | 0.063357279 | 0.709264768 | -4.271753523 |
| **A_52_P496260** | Exosc8 | 0.636433078 | -0.05003014 | 1.322896296 | 10.79823637 | 2.322126193 | 0.063364084 | 0.709264768 | -4.271772365 |
| **A_51_P362959** | Fbxo36 | -0.414065428 | -0.861092775 | 0.032961919 | 9.314939282 | -2.319985252 | 0.063541709 | 0.709264768 | -4.272263594 |
| **A_51_P203501** | Vars | 0.398627255 | -0.031935077 | 0.829189586 | 9.06830288 | 2.318896163 | 0.063632269 | 0.709264768 | -4.272513622 |
| **A_55_P2045946** | Tepsin | -0.410354699 | -0.856361792 | 0.035652394 | 9.779615673 | -2.450644699 | 0.063697991 | 0.709264768 | -4.327500824 |
| **A_51_P447668** | Ptpn12 | -0.217278084 | -0.452306977 | 0.01775081 | 8.993639706 | -2.315501211 | 0.063915442 | 0.709264768 | -4.273293636 |
| **A_52_P351882** | Impad1 | 0.47133362 | -0.038703267 | 0.981370507 | 12.78834917 | 2.314606551 | 0.063990287 | 0.709264768 | -4.273499346 |
| **A_55_P2124153** | Rusc2 | -0.821982177 | -1.711490649 | 0.067526295 | 10.91313565 | -2.314529165 | 0.063996765 | 0.709264768 | -4.273517142 |
| **A_55_P1970841** | Nr2c2 | 0.23498515 | -0.019415054 | 0.489385355 | 9.616194481 | 2.313520261 | 0.064081288 | 0.709264768 | -4.273749204 |
| **A_55_P1990648** | Ndufaf4 | 0.508705572 | -0.042167453 | 1.059578597 | 9.021022518 | 2.312945149 | 0.064129522 | 0.709264768 | -4.273881525 |
| **A_51_P415247** | NA | 0.412537432 | -0.034218839 | 0.859293704 | 8.554566592 | 2.312826468 | 0.064139481 | 0.709264768 | -4.273908834 |
| **A_55_P2060404** | Pus3 | 0.349426191 | -0.029068087 | 0.72792047 | 9.678478263 | 2.312312258 | 0.064182647 | 0.709264768 | -4.274027169 |
| **A_55_P1985094** | Wrap73 | 0.422906016 | -0.035196507 | 0.881008538 | 10.46548378 | 2.312232628 | 0.064189334 | 0.709264768 | -4.274045497 |
| **A_55_P2375121** | AI225934 | -0.309629566 | -0.645045151 | 0.02578602 | 8.457877972 | -2.312115528 | 0.064199169 | 0.709264768 | -4.274072449 |
| **A_55_P2408330** | Rbm33 | -0.251932553 | -0.524934565 | 0.021069459 | 8.467822011 | -2.311366347 | 0.064262132 | 0.709264768 | -4.274244909 |
| **A_66_P132249** | Akr1c13 | 1.189199034 | -0.099521874 | 2.477919941 | 9.302751553 | 2.311245182 | 0.064272321 | 0.709264768 | -4.274272805 |
| **A_55_P2057792** | NA | 0.454952359 | -0.038357004 | 0.948261722 | 12.45234773 | 2.309919837 | 0.064383885 | 0.709264768 | -4.274578021 |

| | | | | | | | | | |
|---|---|---|---|---|---|---|---|---|---|
| **A_51_P212023** | Sart3 | 0.396332569 | -0.033632091 | 0.826297228 | 12.04788748 | 2.308752332 | 0.064482332 | 0.709264768 | -4.274847006 |
| **A_51_P502964** | Tmem63a | -0.381183478 | -0.794728861 | 0.032361904 | 9.937538206 | -2.308666699 | 0.064489559 | 0.709264768 | -4.274866739 |
| **A_55_P2113768** | NA | 1.197894873 | -0.102061041 | 2.497850787 | 12.0497769 | 2.308024569 | 0.064543779 | 0.709264768 | -4.275014732 |
| **A_51_P388298** | Mmachc | 0.306768479 | -0.026359983 | 0.639896941 | 9.137654549 | 2.30647812 | 0.064674556 | 0.709264768 | -4.275371283 |
| **A_55_P2117243** | Npm1 | 0.806264179 | -0.069711095 | 1.682239452 | 12.27181869 | 2.305344659 | 0.064770586 | 0.709264768 | -4.275632737 |
| **A_55_P2185667** | NA | 0.455609096 | -0.039407351 | 0.950625542 | 8.154679109 | 2.305276926 | 0.064776329 | 0.709264768 | -4.275648364 |
| **A_51_P450924** | Pole4 | 0.287169867 | -0.024861344 | 0.599201078 | 10.4143972 | 2.305107462 | 0.064790701 | 0.709264768 | -4.275687464 |
| **A_65_P17700** | Apobec3 | 0.290518329 | -0.025160122 | 0.60619678 | 9.976484513 | 2.305042548 | 0.064796207 | 0.709264768 | -4.275702441 |
| **A_55_P2020458** | NA | 0.626641193 | -0.054854428 | 1.308136814 | 12.59582337 | 2.303065112 | 0.064964172 | 0.709264768 | -4.27615887 |
| **A_55_P1987494** | Gabrr2 | 0.757440895 | -0.066340424 | 1.581222214 | 11.01242636 | 2.302964036 | 0.06497277 | 0.709264768 | -4.276182209 |
| **A_52_P176300** | Med1 | 0.407201824 | -0.035847082 | 0.850250731 | 9.358836816 | 2.302016285 | 0.065053447 | 0.709264768 | -4.276401087 |
| **A_51_P196590** | Hadh | -0.450000032 | -0.939816676 | 0.039816612 | 12.30005263 | -2.301067464 | 0.065134321 | 0.709264768 | -4.276620284 |
| **A_51_P267314** | Tbc1d10c | 0.696077044 | -0.061605952 | 1.453760041 | 8.275662095 | 2.301018529 | 0.065138495 | 0.709264768 | -4.276631591 |
| **A_55_P2000533** | Polk | -0.203088425 | -0.424185897 | 0.018009046 | 8.321351839 | -2.30065629 | 0.065169401 | 0.709264768 | -4.276715296 |
| **A_55_P2043337** | Colq | 0.988765832 | -0.087869855 | 2.06540152 | 8.591600313 | 2.300249922 | 0.065204091 | 0.709264768 | -4.276809212 |
| **A_52_P473953** | Ctdspl | -0.267361404 | -0.558571696 | 0.023848888 | 11.0719895 | -2.299547257 | 0.06526412 | 0.709264768 | -4.276971635 |
| **A_51_P448489** | Mea1 | -0.251223004 | -0.524904399 | 0.022458391 | 10.84266126 | -2.299135014 | 0.065299366 | 0.709264768 | -4.277066945 |
| **A_55_P1971639** | Abitram | 0.39903776 | -0.035789043 | 0.833864562 | 9.975470776 | 2.298518655 | 0.0653521 | 0.709264768 | -4.277209471 |
| **A_55_P2117081** | Trgv4 | 1.356764209 | -0.121979448 | 2.835507865 | 8.358854035 | 2.298062465 | 0.065391159 | 0.709264768 | -4.27731498 |
| **A_55_P2159600** | Herc1 | -0.203645445 | -0.425603977 | 0.018313087 | 11.30516934 | -2.298016815 | 0.065395069 | 0.709264768 | -4.277325539 |
| **A_55_P2015520** | Plekha1 | -0.539205188 | -1.126908672 | 0.048498295 | 12.33993155 | -2.297979465 | 0.065398268 | 0.709264768 | -4.277334178 |
| **A_51_P260035** | Sar1b | -0.203820024 | -0.426013182 | 0.018373134 | 10.58117125 | -2.297558158 | 0.065434367 | 0.709264768 | -4.277431637 |
| **A_55_P2032187** | NA | 0.43132719 | -0.038929676 | 0.901584056 | 9.416656186 | 2.29732287 | 0.065454536 | 0.709264768 | -4.277486071 |
| **A_55_P1999715** | Tcrb-J | 4.095668996 | -0.369808303 | 8.561146295 | 9.984345929 | 2.297244975 | 0.065461215 | 0.709264768 | -4.277504093 |
| **A_51_P478172** | Tcrb-J | 3.785461721 | -0.341980841 | 7.912904284 | 9.669035458 | 2.297143713 | 0.065469898 | 0.709264768 | -4.277527522 |
| **A_55_P2058923** | Bud23 | 0.392574267 | -0.035506863 | 0.820655398 | 11.37428634 | 2.296921165 | 0.065488986 | 0.709264768 | -4.277579016 |
| **A_55_P2057212** | Tpr | 0.44779378 | -0.040784111 | 0.93637167 | 11.07611404 | 2.295591417 | 0.065603161 | 0.709264768 | -4.277886781 |
| **A_52_P367760** | Calml4 | -0.373781062 | -0.781904208 | 0.034342083 | 9.070873986 | -2.29391021 | 0.065747815 | 0.709264768 | -4.278276094 |
| **A_55_P2153141** | NA | 0.313161668 | -0.028818466 | 0.655141803 | 12.15479902 | 2.293602031 | 0.065774367 | 0.709264768 | -4.278347483 |
| **A_55_P2041017** | Gm15365 | 0.284783884 | -0.02627273 | 0.595840499 | 13.62698243 | 2.293117526 | 0.065816135 | 0.709264768 | -4.278459732 |
| **A_55_P2110032** | Akap7 | -0.36902513 | -0.772118635 | 0.034068375 | 9.541995073 | -2.292981163 | 0.065827896 | 0.709264768 | -4.278491328 |
| **A_55_P2029235** | Btn1a1 | 0.330038673 | -0.030540233 | 0.690617579 | 13.13527528 | 2.29252912 | 0.065866897 | 0.709264768 | -4.278596079 |
| **A_51_P504314** | Prpf38b | 0.29338825 | -0.027389176 | 0.614165675 | 9.203938222 | 2.290811024 | 0.066015356 | 0.709264768 | -4.278994359 |
| **A_55_P2024595** | Atxn7l1 | 0.25098431 | -0.02360945 | 0.52557807 | 8.521104503 | 2.289318693 | 0.066144593 | 0.709264768 | -4.279340496 |
| **A_55_P1963576** | Gm11595 | 0.257110762 | -0.024304835 | 0.538526358 | 8.345552571 | 2.288349933 | 0.066228632 | 0.709264768 | -4.279565289 |
| **A_55_P2124319** | NA | 0.367598316 | -0.035235696 | 0.770432329 | 8.009165518 | 2.285586841 | 0.066468945 | 0.709264768 | -4.280206858 |
| **A_51_P189361** | Osgin1 | -0.353122357 | -0.740293941 | 0.034049226 | 8.512144208 | -2.284399653 | 0.066572481 | 0.709264768 | -4.280482701 |
| **A_55_P1967648** | Flcn | -0.406454883 | -0.852157524 | 0.039247759 | 10.39892161 | -2.284112458 | 0.066597553 | 0.709264768 | -4.280549448 |
| **A_52_P602898** | Nkrf | 0.349224805 | -0.033833503 | 0.732283112 | 8.350237211 | 2.283444937 | 0.066655866 | 0.709264768 | -4.280704612 |
| **A_66_P106478** | Cct3 | 0.28642046 | -0.028005686 | 0.600846606 | 11.25537226 | 2.281580157 | 0.066819053 | 0.709264768 | -4.281138265 |
| **A_66_P108815** | Pgls | 0.322348541 | -0.031731207 | 0.67642829 | 10.41526655 | 2.280210637 | 0.066939169 | 0.709264768 | -4.281456923 |
| **A_55_P2045622** | Setdb2 | 0.2297894 | -0.022626571 | 0.48220537 | 8.964604378 | 2.280150467 | 0.066944452 | 0.709264768 | -4.281470927 |
| **A_55_P2100008** | NA | 0.259058453 | -0.025514306 | 0.543631213 | 11.46750971 | 2.280104694 | 0.066948471 | 0.709264768 | -4.28148158 |
| **A_51_P332822** | Abce1 | 0.438490686 | -0.043366545 | 0.920347918 | 11.25883597 | 2.279251964 | 0.067023386 | 0.709264768 | -4.281680076 |
| **A_66_P124139** | Ikzf1 | 0.621194294 | -0.06157366 | 1.303962247 | 8.124748552 | 2.278791936 | 0.067063839 | 0.709264768 | -4.281787183 |
| **A_55_P2018274** | NA | 0.430224692 | -0.042765516 | 0.903214899 | 11.32619494 | 2.278208815 | 0.067115152 | 0.709264768 | -4.281922975 |
| **A_51_P386304** | Ccnl2 | -0.340027765 | -0.71410432 | 0.03404879 | 11.36161819 | -2.276691766 | 0.067248842 | 0.709264768 | -4.282276378 |
| **A_55_P2161465** | Gm10516 | -0.221726914 | -0.465706415 | 0.022252587 | 8.326424068 | -2.276226204 | 0.067289927 | 0.709264768 | -4.28238487 |

| | | | | | | | | | |
|---|---|---|---|---|---|---|---|---|---|
| **A_55_P2134032** | NA | 0.474160645 | -0.048161132 | 0.996482423 | 9.512597776 | 2.27372385 | 0.067511204 | 0.709264768 | -4.282968299 |
| **A_51_P210474** | Mrpl18 | 0.466386947 | -0.047476817 | 0.980250711 | 11.67625127 | 2.273258055 | 0.067552478 | 0.709264768 | -4.283076955 |
| **A_55_P1981814** | Myo3b | 0.343099232 | -0.034994254 | 0.721192719 | 9.503821324 | 2.272850634 | 0.067588601 | 0.709264768 | -4.283172009 |
| **A_55_P2128821** | Selenok | -0.444653296 | -0.934925575 | 0.045618983 | 13.51523878 | -2.271613929 | 0.067698376 | 0.709264768 | -4.28346062 |
| **A_55_P2164189** | Sfpq | 0.887334065 | -0.09132504 | 1.86599317 | 10.89800642 | 2.270942086 | 0.06775809 | 0.709264768 | -4.28361746 |
| **A_55_P1964433** | Zmynd12 | -0.327958843 | -0.689683439 | 0.033765753 | 9.039764 | -2.27086675 | 0.067764789 | 0.709264768 | -4.28363505 |
| **A_55_P2030183** | NA | 1.537176718 | -0.17093676 | 3.245290197 | 13.80471125 | 2.397009657 | 0.067776544 | 0.709264768 | -4.335757624 |
| **A_55_P2179069** | NA | 0.668852992 | -0.069635761 | 1.407341745 | 8.916887872 | 2.268491384 | 0.067976382 | 0.709264768 | -4.284189876 |
| **A_55_P1952743** | NA | 0.659035084 | -0.069160076 | 1.387230245 | 10.56470683 | 2.26678898 | 0.068128456 | 0.709264768 | -4.284587792 |
| **A_51_P235123** | Nfkbie | 0.501658366 | -0.05282031 | 1.056137043 | 8.331170284 | 2.266071224 | 0.06819268 | 0.709264768 | -4.284755627 |
| **A_52_P482203** | Tsen2 | 0.435849788 | -0.046325802 | 0.918025377 | 10.24623925 | 2.264028887 | 0.068375776 | 0.709264768 | -4.28523342 |
| **A_51_P468055** | Smyd1 | 0.687400332 | -0.073528931 | 1.448329595 | 9.909618769 | 2.262641754 | 0.068500428 | 0.709264768 | -4.28555812 |
| **A_51_P216702** | Eogt | -0.435310223 | -0.917268565 | 0.046648119 | 9.78266887 | -2.262245381 | 0.068536091 | 0.709264768 | -4.285650932 |
| **A_55_P2057912** | NA | 1.231933622 | -0.132408373 | 2.596275616 | 12.59113455 | 2.261592768 | 0.068594851 | 0.709264768 | -4.285803769 |
| **A_55_P1974650** | Npdc1 | -0.412313969 | -0.869353869 | 0.04472593 | 10.9621177 | -2.259562085 | 0.068778032 | 0.709264768 | -4.286279557 |
| **A_52_P200617** | Iars | 0.250536912 | -0.027200713 | 0.528274538 | 10.21006198 | 2.259369948 | 0.06879539 | 0.709264768 | -4.286324592 |
| **A_51_P352381** | Dhrs4 | 0.63297077 | -0.068815531 | 1.33475707 | 8.98395163 | 2.259067015 | 0.068822768 | 0.709264768 | -4.286395602 |
| **A_55_P1979998** | Spaca9 | -0.41180724 | -0.86844163 | 0.044827149 | 10.24150569 | -2.258789228 | 0.068847884 | 0.709264768 | -4.286460724 |
| **A_55_P2017870** | Pclaf | 0.712517837 | -0.077627176 | 1.50266285 | 13.58011239 | 2.258599787 | 0.068865017 | 0.709264768 | -4.286505138 |
| **A_55_P2092730** | Wdfy1 | -0.462698834 | -0.975838525 | 0.050440856 | 9.251256539 | -2.258463848 | 0.068877314 | 0.709264768 | -4.286537011 |
| **A_55_P2025851** | Heatr3 | 0.370107617 | -0.0403895 | 0.780604734 | 11.39562554 | 2.258230439 | 0.068898434 | 0.709264768 | -4.28659174 |
| **A_52_P495104** | NA | -0.307035842 | -0.647713791 | 0.033642107 | 11.04049554 | -2.257331792 | 0.068979812 | 0.709264768 | -4.286802493 |
| **A_55_P2060183** | Lamtor3 | -0.199435455 | -0.4207322 | 0.02186129 | 12.81872013 | -2.257239746 | 0.068988153 | 0.709264768 | -4.286824083 |
| **A_55_P1971619** | NA | 1.004826355 | -0.11044766 | 2.12010037 | 9.307725797 | 2.256626996 | 0.069043706 | 0.709264768 | -4.286967827 |
| **A_55_P1958537** | Map6 | -1.121487635 | -2.367792106 | 0.124816835 | 10.98084767 | -2.253827546 | 0.069298109 | 0.709264768 | -4.287624928 |
| **A_55_P2035981** | Mindy1 | -0.241366409 | -0.5097983 | 0.027065482 | 12.26969809 | -2.252127966 | 0.069453042 | 0.709264768 | -4.288024165 |
| **A_55_P2092826** | Anxa1 | -0.511536092 | -1.080756562 | 0.057684377 | 14.06665586 | -2.25084774 | 0.069569987 | 0.709264768 | -4.288325045 |
| **A_55_P1968178** | Smagp | -0.731795195 | -1.546136541 | 0.082546151 | 11.34107109 | -2.250781898 | 0.069576007 | 0.709264768 | -4.288340523 |
| **A_55_P2007470** | Pdgfa | -0.499495316 | -1.055357641 | 0.05636701 | 13.36072265 | -2.250684009 | 0.069584958 | 0.709264768 | -4.288363535 |
| **A_55_P2200618** | NA | -0.453942075 | -0.95928376 | 0.05139961 | 9.806044713 | -2.249912664 | 0.069655535 | 0.709264768 | -4.28854489 |
| **A_55_P2094966** | H3c14 | 1.600569753 | -0.18123603 | 3.382375536 | 11.74709863 | 2.249907104 | 0.069656043 | 0.709264768 | -4.288546197 |
| **A_55_P1984416** | NA | 0.488887498 | -0.055511253 | 1.033286248 | 10.72589182 | 2.249272932 | 0.069714126 | 0.709264768 | -4.288695337 |
| **A_55_P2135039** | Hyi | -0.50942023 | -1.076804672 | 0.057964212 | 10.66617558 | -2.248791079 | 0.069758291 | 0.709264768 | -4.288808676 |
| **A_51_P239984** | Exo1 | 1.258431882 | -0.14338658 | 2.660250343 | 9.95488113 | 2.248476137 | 0.069787174 | 0.709264768 | -4.288882766 |
| **A_55_P2127683** | 4933434E | -0.291273761 | -0.615742505 | 0.033194984 | 8.982490593 | -2.248427253 | 0.069791658 | 0.709264768 | -4.288894266 |
| **A_55_P2091473** | Bnip3 | -0.523881252 | -1.107505839 | 0.059743336 | 13.19182234 | -2.248275986 | 0.069805536 | 0.709264768 | -4.288929855 |
| **A_55_P2157200** | NA | -0.27812585 | -0.5880778 | 0.0318261 | 10.99207979 | -2.247487724 | 0.069877902 | 0.709264768 | -4.289115338 |
| **A_55_P2081656** | NA | 2.021747602 | -0.231638098 | 4.275133303 | 10.9135635 | 2.247200114 | 0.069904325 | 0.709264768 | -4.289183027 |
| **A_51_P207153** | 1110008P | -0.314638808 | -0.665408283 | 0.036130668 | 10.09098536 | -2.246678037 | 0.069952316 | 0.709264768 | -4.289305914 |
| **A_51_P369862** | Nceh1 | -0.447199964 | -0.946084797 | 0.051684869 | 9.855722735 | -2.245183309 | 0.070089909 | 0.709264768 | -4.289657865 |
| **A_52_P135873** | NA | -0.802255669 | -1.697708087 | 0.093196749 | 10.49330835 | -2.243988499 | 0.070200098 | 0.709264768 | -4.289939324 |
| **A_55_P1999847** | Nutf2 | 0.349903026 | -0.040649872 | 0.740455924 | 13.59789116 | 2.243975851 | 0.070201265 | 0.709264768 | -4.289942304 |
| **A_55_P2151082** | NA | 0.793121889 | -0.092239864 | 1.678483643 | 12.79084491 | 2.243724471 | 0.070224472 | 0.709264768 | -4.290001535 |
| **A_55_P2026400** | Capns1 | -0.256706856 | -0.543274215 | 0.029860504 | 12.5010207 | -2.243680901 | 0.070228495 | 0.709264768 | -4.290011802 |
| **A_51_P460890** | Rgl2 | -0.260047058 | -0.550430355 | 0.030336239 | 10.62802745 | -2.243007153 | 0.070290737 | 0.709264768 | -4.290170581 |
| **A_66_P125226** | NA | 0.291500235 | -0.034057954 | 0.617058424 | 12.71116265 | 2.242645512 | 0.07032417 | 0.709264768 | -4.290255823 |
| **A_51_P519251** | Nupr1 | -0.563880204 | -1.193662795 | 0.065902387 | 11.70525717 | -2.242572761 | 0.070330898 | 0.709264768 | -4.290272972 |
| **A_51_P424338** | Nqo1 | 0.416654656 | -0.048751031 | 0.882060343 | 8.360927224 | 2.242306105 | 0.070355563 | 0.709264768 | -4.290335833 |

| | | | | | | | | | |
|---|---|---|---|---|---|---|---|---|---|
| **A_52_P307893** | Pip4k2b | 0.39707107 | -0.046592378 | 0.840734518 | 10.33158078 | 2.241635219 | 0.070417658 | 0.709264768 | -4.29049401 |
| **A_52_P411601** | Ep300 | -0.239114181 | -0.506390962 | 0.0281626 | 9.029802164 | -2.24075537 | 0.070499182 | 0.709264768 | -4.29070151 |
| **A_52_P596755** | Dnph1 | 0.602736373 | -0.071070256 | 1.276543003 | 8.69340629 | 2.240487199 | 0.070524049 | 0.709264768 | -4.290764766 |
| **A_55_P2132099** | NA | 0.556148985 | -0.065605827 | 1.177903797 | 8.341739411 | 2.240383364 | 0.07053368 | 0.709264768 | -4.290789261 |
| **A_51_P407675** | Zrsr1 | -0.423222039 | -0.896384924 | 0.049940846 | 9.440978147 | -2.240309136 | 0.070540566 | 0.709264768 | -4.290806771 |
| **A_55_P1963175** | Lctl | 0.283557445 | -0.033522953 | 0.600637843 | 8.255465286 | 2.239865832 | 0.070581704 | 0.709264768 | -4.290911357 |
| **A_55_P1957249** | Pdgfrb | -0.302317103 | -0.640486399 | 0.035852194 | 11.54895732 | -2.239127831 | 0.070650245 | 0.709264768 | -4.291085503 |
| **A_55_P2041581** | NA | 0.521073279 | -0.062187112 | 1.10433367 | 9.249055028 | 2.237621695 | 0.070790344 | 0.709264768 | -4.291441038 |
| **A_55_P2063316** | Mgll | -0.343526228 | -0.728310631 | 0.041258175 | 11.5968773 | -2.23610806 | 0.070931434 | 0.709264768 | -4.291798524 |
| **A_55_P2166506** | NA | 0.279588484 | -0.03366081 | 0.592837779 | 15.32181199 | 2.235524962 | 0.070985865 | 0.709264768 | -4.291936287 |
| **A_55_P1979178** | NA | 0.340790533 | -0.041066917 | 0.722647984 | 13.70058858 | 2.23530402 | 0.071006501 | 0.709264768 | -4.291988494 |
| **A_51_P411770** | Nip7 | 0.234219201 | -0.028314081 | 0.496752484 | 11.86245176 | 2.234541749 | 0.071077746 | 0.709264768 | -4.292168641 |
| **A_55_P2156016** | NA | 0.351073537 | -0.0424469 | 0.744593973 | 8.370668563 | 2.234504063 | 0.07108127 | 0.709264768 | -4.292177549 |
| **A_55_P2034042** | Gga3 | 0.236157814 | -0.028592651 | 0.500908279 | 10.89144005 | 2.234168549 | 0.071112654 | 0.709264768 | -4.292256856 |
| **A_55_P1966790** | Impdh2 | 0.769365527 | -0.093217622 | 1.631948676 | 12.57854354 | 2.233994497 | 0.07112894 | 0.709264768 | -4.292298002 |
| **A_51_P366138** | Mertk | -0.26409483 | -0.560218297 | 0.032028636 | 8.826547078 | -2.233764686 | 0.07115045 | 0.709264768 | -4.292352332 |
| **A_55_P2063257** | Lgals4 | -0.30019189 | -0.637219806 | 0.036836026 | 8.157201027 | -2.230917063 | 0.071417551 | 0.709264768 | -4.29302589 |
| **A_55_P1957319** | Krr1 | 0.449892404 | -0.055378369 | 0.955163178 | 10.32812422 | 2.230153872 | 0.071489315 | 0.709264768 | -4.293206518 |
| **A_55_P2069787** | NA | 1.202094396 | -0.148108332 | 2.552297124 | 13.2570796 | 2.229923359 | 0.071511005 | 0.709264768 | -4.293261084 |
| **A_55_P1976682** | 4932431P | 1.251225176 | -0.155046941 | 2.657497292 | 12.37866375 | 2.22851958 | 0.071643246 | 0.709264768 | -4.293593469 |
| **A_55_P2077909** | Tpgs2 | 0.289730353 | -0.035930026 | 0.615390732 | 8.378773633 | 2.22832954 | 0.071661168 | 0.709264768 | -4.293638478 |
| **A_66_P131205** | NA | 0.384062276 | -0.047693632 | 0.815818184 | 14.50947914 | 2.227992413 | 0.071692973 | 0.709264768 | -4.293718331 |
| **A_55_P2031668** | Gstp1 | 0.271198393 | -0.033702185 | 0.576098972 | 10.20165143 | 2.227815439 | 0.071709674 | 0.709264768 | -4.293760253 |
| **A_55_P1995507** | Aqp11 | 1.307698308 | -0.162679015 | 2.778075631 | 9.141284704 | 2.227558464 | 0.071733934 | 0.709264768 | -4.29382113 |
| **A_51_P273556** | Fam83d | 0.207752812 | -0.025887152 | 0.441392776 | 9.088557259 | 2.227153376 | 0.071772193 | 0.709264768 | -4.293917106 |
| **A_55_P1980932** | Elob | -0.298250576 | -0.633669384 | 0.037168233 | 14.86658078 | -2.227123109 | 0.071775052 | 0.709264768 | -4.293924277 |
| **A_55_P2107447** | Rab37 | 0.313394739 | -0.039078486 | 0.665867964 | 8.106994973 | 2.226977939 | 0.071788769 | 0.709264768 | -4.293958675 |
| **A_55_P2197638** | 1110046J | -0.407948424 | -0.866840098 | 0.05094325 | 8.897757448 | -2.226616521 | 0.071822929 | 0.709264768 | -4.29404432 |
| **A_52_P406828** | Dkc1 | 0.825350354 | -0.103126477 | 1.753827185 | 9.073807936 | 2.226473943 | 0.071836411 | 0.709264768 | -4.294078109 |
| **A_55_P1980321** | Pcbp3 | 0.307103335 | -0.038388593 | 0.652595262 | 10.1654463 | 2.226368116 | 0.071846418 | 0.709264768 | -4.29410319 |
| **A_55_P2023449** | NA | -0.628309469 | -1.335214641 | 0.078595703 | 9.558185502 | -2.22619289 | 0.071862992 | 0.709264768 | -4.29414472 |
| **A_51_P298538** | Prmt7 | 0.344241967 | -0.043206872 | 0.731690806 | 11.91550272 | 2.225357533 | 0.071942061 | 0.709264768 | -4.294342741 |
| **A_55_P2147971** | Ranbp1 | 0.771916972 | -0.097048361 | 1.640882305 | 12.03852475 | 2.224940927 | 0.071981528 | 0.709264768 | -4.294441518 |
| **A_55_P1992839** | Dsn1 | 0.412319228 | -0.051840487 | 0.876478944 | 9.36010545 | 2.2249307 | 0.071982497 | 0.709264768 | -4.294443943 |
| **A_52_P527500** | Helb | 0.358645982 | -0.045275764 | 0.762567727 | 8.976108522 | 2.223919591 | 0.07207838 | 0.709264768 | -4.294683733 |
| **A_51_P497402** | Tonsl | 0.339945367 | -0.042956019 | 0.722846753 | 9.294097631 | 2.223681248 | 0.072101002 | 0.709264768 | -4.294740269 |
| **A_51_P481788** | Mapk1 | 0.356281912 | -0.045022704 | 0.757586529 | 10.77186674 | 2.223668092 | 0.072102251 | 0.709264768 | -4.305892545 |
| **A_55_P2156556** | NA | 0.314990456 | -0.039871436 | 0.669852348 | 11.96588129 | 2.223250391 | 0.072141914 | 0.709264768 | -4.294842481 |
| **A_55_P1977330** | Zbtb1 | 0.299738734 | -0.03796681 | 0.637444277 | 8.628086116 | 2.223079654 | 0.072158133 | 0.709264768 | -4.294882989 |
| **A_55_P2035286** | Uhrf1 | 1.731900349 | -0.219390641 | 3.683191338 | 12.02423219 | 2.223060111 | 0.07215999 | 0.709264768 | -4.294887626 |
| **A_55_P1974892** | Zbtb18 | 0.261061076 | -0.033095345 | 0.555217497 | 9.952757534 | 2.222870364 | 0.07217802 | 0.709264768 | -4.294932647 |
| **A_55_P2408415** | Kcnq1ot1 | -0.621973345 | -1.322813078 | 0.078866388 | 8.74145974 | -2.22281543 | 0.07218324 | 0.709264768 | -4.294945682 |
| **A_55_P1979217** | Abtb1 | -0.596736919 | -1.269730145 | 0.076256307 | 11.56777343 | -2.220867049 | 0.072368665 | 0.709264768 | -4.295408145 |
| **A_55_P2175813** | Ythdf2 | 0.214388094 | -0.027405995 | 0.456182183 | 10.60424909 | 2.220778938 | 0.072377063 | 0.709264768 | -4.295429066 |
| **A_55_P1961395** | Pdpn | -0.572560378 | -1.218452642 | 0.073331886 | 12.99649298 | -2.220299462 | 0.072422776 | 0.709264768 | -4.295542922 |
| **A_52_P197627** | Umps | 0.393652157 | -0.05447807 | 0.841782385 | 8.955783897 | 2.33975887 | 0.072451476 | 0.709264768 | -4.344800515 |
| **A_51_P208410** | Ppp1r12c | -0.284554075 | -0.6056004 | 0.036492251 | 11.19744345 | -2.219971767 | 0.072454036 | 0.709264768 | -4.295620747 |
| **A_55_P2159807** | Rfc4 | 1.007082753 | -0.129221129 | 2.143386634 | 11.13268452 | 2.219836631 | 0.072466931 | 0.709264768 | -4.295652843 |

| | | | | | | | | | |
|---|---|---|---|---|---|---|---|---|---|
| **A_55_P2035827** | Vma21 | 0.344975455 | -0.044341491 | 0.734292401 | 10.91117329 | 2.219398223 | 0.072508782 | 0.709264768 | -4.295756979 |
| **A_55_P2085932** | Lamtor2 | 0.803704265 | -0.103349242 | 1.710757771 | 12.15596741 | 2.219288241 | 0.072519285 | 0.709264768 | -4.295783106 |
| **A_51_P246387** | Nktr | -0.349277094 | -0.743608399 | 0.045054211 | 9.761623303 | -2.218498782 | 0.072594724 | 0.709264768 | -4.295970672 |
| **A_55_P2117741** | Nck2 | 0.209819357 | -0.027073408 | 0.446712122 | 12.384166 | 2.218421672 | 0.072602096 | 0.709264768 | -4.295988995 |
| **A_66_P132846** | Exosc1 | 0.225776234 | -0.029280819 | 0.480833288 | 9.200050771 | 2.217130382 | 0.072725679 | 0.709264768 | -4.296295904 |
| **A_52_P748958** | Cfl1 | 0.365209081 | -0.047428816 | 0.777846978 | 13.44772729 | 2.216781059 | 0.072759149 | 0.709264768 | -4.296378952 |
| **A_52_P49042** | Arfgef3 | 0.396833527 | -0.051603949 | 0.845271003 | 8.362363481 | 2.216444205 | 0.072791439 | 0.709264768 | -4.296459045 |
| **A_52_P310981** | Eif4e | 0.408225843 | -0.053377237 | 0.869828923 | 11.39490389 | 2.215042904 | 0.072925928 | 0.709264768 | -4.296792323 |
| **A_51_P242859** | Akr1c12 | 0.274900384 | -0.036082732 | 0.5858835 | 7.878169861 | 2.214057427 | 0.073020664 | 0.709264768 | -4.297026795 |
| **A_55_P2140843** | NA | 0.428149368 | -0.056431383 | 0.91273012 | 10.53918027 | 2.212990219 | 0.073123403 | 0.709264768 | -4.2972808 |
| **A_52_P397204** | Csk | 0.241765341 | -0.032189179 | 0.515719861 | 9.401656863 | 2.210374782 | 0.073375829 | 0.709264768 | -4.297903671 |
| **A_52_P555672** | Vrk1 | 0.51886014 | -0.069193378 | 1.106913658 | 8.861405752 | 2.209956887 | 0.073416247 | 0.709264768 | -4.298003243 |
| **A_55_P2099760** | NA | 0.815085073 | -0.109016067 | 1.739186213 | 9.759334632 | 2.209193615 | 0.073490128 | 0.709264768 | -4.298185143 |
| **A_51_P513541** | Rfk | -0.4353977 | -0.929218965 | 0.058423564 | 10.30638016 | -2.208343815 | 0.073572476 | 0.709264768 | -4.298387717 |
| **A_55_P2077783** | Tubb2a-ps | 0.299201134 | -0.040151166 | 0.638553435 | 8.953419725 | 2.208323955 | 0.073574402 | 0.709264768 | -4.298392453 |
| **A_51_P189746** | Pim3 | -0.35394314 | -0.755595701 | 0.047709421 | 9.694532751 | -2.207157396 | 0.073687606 | 0.709264768 | -4.298670629 |
| **A_55_P2136522** | Tmem128 | -0.427227382 | -0.912184594 | 0.057729829 | 11.46585629 | -2.206510527 | 0.073750458 | 0.709264768 | -4.298824927 |
| **A_52_P200925** | NA | 0.466143264 | -0.063097559 | 0.995384087 | 13.67550886 | 2.206055441 | 0.073794709 | 0.709264768 | -4.298933498 |
| **A_52_P136782** | Rgs5 | -0.333019279 | -0.711212627 | 0.045174069 | 8.870929641 | -2.205493771 | 0.073849362 | 0.709264768 | -4.299067519 |
| **A_51_P351970** | Hells | 0.46232503 | -0.062736355 | 0.987386414 | 8.42149815 | 2.205401535 | 0.073858341 | 0.709264768 | -4.29908953 |
| **A_55_P1968108** | NA | 0.453468962 | -0.061552087 | 0.968490012 | 13.61169851 | 2.205326687 | 0.073865628 | 0.709264768 | -4.299107392 |
| **A_51_P269084** | Chchd10 | -0.621069402 | -1.326558549 | 0.084419745 | 14.86237609 | -2.20495709 | 0.073901623 | 0.709264768 | -4.2991956 |
| **A_55_P2103796** | Arhgap11a | 0.569585632 | -0.077809451 | 1.216980715 | 8.709713853 | 2.203636586 | 0.074030377 | 0.709264768 | -4.29951084 |
| **A_55_P1972530** | Efl1 | 0.328067064 | -0.044847216 | 0.700981344 | 9.913112645 | 2.203453875 | 0.074048211 | 0.709264768 | -4.299554469 |
| **A_51_P209302** | Polr1e | 0.239810913 | -0.032963142 | 0.512584969 | 8.999479631 | 2.201994489 | 0.074190816 | 0.709264768 | -4.299903043 |
| **A_55_P1994128** | Tmem184 | -0.275635709 | -0.589253315 | 0.037981898 | 9.822881145 | -2.201331198 | 0.074255725 | 0.709264768 | -4.300061524 |
| **A_55_P2059957** | Snx7 | -0.278409899 | -0.595189743 | 0.038369945 | 9.598463327 | -2.201291101 | 0.074259651 | 0.709264768 | -4.300071106 |
| **A_51_P141071** | Cand2 | 0.381949357 | -0.052659314 | 0.816558028 | 9.006263563 | 2.20119107 | 0.074269446 | 0.709264768 | -4.300095009 |
| **A_55_P2009910** | Stxbp3 | -0.367373181 | -0.785457082 | 0.05071072 | 10.82052443 | -2.200869792 | 0.074300913 | 0.709264768 | -4.300171789 |
| **A_55_P2125774** | Ola1 | 0.425055317 | -0.058698052 | 0.908808686 | 11.52570816 | 2.200755499 | 0.074312111 | 0.709264768 | -4.300199104 |
| **A_55_P2181191** | Btg1 | 0.265286088 | -0.036655151 | 0.567227327 | 8.193383325 | 2.200606481 | 0.074326713 | 0.709264768 | -4.300234721 |
| **A_55_P2080804** | Slirp | 0.474508009 | -0.065579182 | 1.0145952 | 12.73556989 | 2.200543791 | 0.074332857 | 0.709264768 | -4.300249705 |
| **A_55_P1994224** | Ythdf1 | 0.276369831 | -0.038248533 | 0.590988196 | 12.54180351 | 2.200173404 | 0.074369169 | 0.709264768 | -4.30033824 |
| **A_51_P485240** | Adipor1 | -0.307308653 | -0.657181908 | 0.042564602 | 11.30748075 | -2.199957996 | 0.074390295 | 0.709264768 | -4.300389734 |
| **A_51_P169032** | 2810004N | 0.324585323 | -0.045236049 | 0.694406696 | 11.69370056 | 2.198301307 | 0.074552985 | 0.709264768 | -4.300785895 |
| **A_55_P1982762** | Hsd3b7 | -0.690554632 | -1.477365697 | 0.096256432 | 10.7494281 | -2.198254292 | 0.074557608 | 0.709264768 | -4.300797141 |
| **A_55_P2093874** | Pbxip1 | -0.709807463 | -1.518802286 | 0.09918736 | 12.49296475 | -2.197582381 | 0.074623701 | 0.709264768 | -4.300957877 |
| **A_55_P2135031** | NA | 0.574946966 | -0.080361158 | 1.23025509 | 8.393672044 | 2.197518703 | 0.074629968 | 0.709264768 | -4.300973112 |
| **A_55_P2149500** | Kifc2 | -0.484470261 | -1.036751816 | 0.067811294 | 8.725357225 | -2.19713595 | 0.074667649 | 0.709264768 | -4.301064693 |
| **A_55_P1986556** | Minpp1 | 0.280252402 | -0.039274895 | 0.599779699 | 10.01920466 | 2.196806091 | 0.074700139 | 0.709264768 | -4.301143627 |
| **A_55_P2114938** | Irf9 | -0.904715776 | -1.936359505 | 0.126927952 | 10.3636322 | -2.196507885 | 0.074729523 | 0.709264768 | -4.301214993 |
| **A_52_P551011** | Smim1 | -0.270926142 | -0.579899229 | 0.038046944 | 9.278090903 | -2.196244098 | 0.074755526 | 0.709264768 | -4.301278128 |
| **A_51_P270519** | Cenps | 0.41552943 | -0.058495417 | 0.889554278 | 8.539622385 | 2.195588904 | 0.074820154 | 0.709264768 | -4.301434967 |
| **A_55_P2164534** | Dtl | 1.212170765 | -0.170906685 | 2.595248215 | 9.675449341 | 2.195167431 | 0.074861759 | 0.709264768 | -4.301535876 |
| **A_55_P2044413** | Cxcl11 | 0.437483893 | -0.061715245 | 0.936683031 | 11.20166935 | 2.195020534 | 0.074876265 | 0.709264768 | -4.301571049 |
| **A_52_P659258** | Ppat | 0.235697317 | -0.03325852 | 0.504653154 | 8.583283447 | 2.19494686 | 0.074883541 | 0.709264768 | -4.30158869 |
| **A_55_P1982744** | NA | 0.79772831 | -0.112696341 | 1.708152962 | 13.36479667 | 2.194630153 | 0.07491483 | 0.709264768 | -4.30166453 |
| **A_55_P2136338** | Tomm20 | 0.29974702 | -0.042482479 | 0.641976519 | 12.88399515 | 2.193753254 | 0.075001532 | 0.709264768 | -4.301874557 |

| | | | | | | | | | |
|---|---|---|---|---|---|---|---|---|---|
| **A_52_P466090** | Samhd1 | -0.302197724 | -0.647248813 | 0.042853365 | 9.479728998 | -2.19360351 | 0.075016349 | 0.709264768 | -4.301910429 |
| **A_52_P455404** | Itpa | 0.353492547 | -0.050250808 | 0.757235902 | 12.61203381 | 2.192932278 | 0.075082801 | 0.709264768 | -4.302071244 |
| **A_51_P165704** | Mcm7 | 1.259058059 | -0.17959243 | 2.697708548 | 12.03242497 | 2.192001289 | 0.075175071 | 0.709264768 | -4.30229435 |
| **A_55_P2047778** | Krtap4-8 | 0.511971485 | -0.073034804 | 1.096977773 | 10.54270074 | 2.191974935 | 0.075177685 | 0.709264768 | -4.302300667 |
| **A_55_P2026761** | NA | 0.471837643 | -0.067314577 | 1.010989863 | 12.24670437 | 2.191954492 | 0.075179712 | 0.709264768 | -4.302305567 |
| **A_55_P1999314** | Dhx33 | 0.268488565 | -0.03833861 | 0.575315739 | 10.30758446 | 2.19170611 | 0.075204351 | 0.709264768 | -4.302365102 |
| **A_51_P383991** | 4-Sep | -0.676793466 | -1.450277075 | 0.096690142 | 12.69710694 | -2.191570209 | 0.075217836 | 0.709264768 | -4.302397679 |
| **A_55_P2072906** | Gm6788 | 0.330841116 | -0.047296898 | 0.70897913 | 13.7085774 | 2.191389032 | 0.075235817 | 0.709264768 | -4.302441111 |
| **A_55_P2115119** | Rnaseh2a | 0.217165586 | -0.031093607 | 0.465424779 | 9.629838514 | 2.190967883 | 0.075277631 | 0.709264768 | -4.302542079 |
| **A_55_P2129029** | Smarcb1 | 0.299781108 | -0.042980381 | 0.642542598 | 12.08335241 | 2.190597486 | 0.075314427 | 0.709264768 | -4.302630891 |
| **A_55_P2034097** | Elavl1 | 0.578981549 | -0.083085906 | 1.241049004 | 13.34672455 | 2.190346532 | 0.075339367 | 0.709264768 | -4.302691069 |
| **A_55_P1995165** | Mdc1 | 0.440669285 | -0.063324665 | 0.944663234 | 10.0816622 | 2.189968173 | 0.075376986 | 0.709264768 | -4.302781808 |
| **A_55_P2022840** | Csnk2a1 | 0.376699272 | -0.054156518 | 0.807555061 | 11.71970695 | 2.189844078 | 0.075389329 | 0.709264768 | -4.302811571 |
| **A_55_P2129469** | NA | 1.262139768 | -0.181748068 | 2.706027604 | 8.602787444 | 2.189396094 | 0.075433903 | 0.709264768 | -4.302919027 |
| **A_66_P140076** | NA | 0.523506451 | -0.075414603 | 1.122427505 | 11.06466535 | 2.18928753 | 0.07544471 | 0.709264768 | -4.30294507 |
| **A_55_P2125561** | Mrps36 | 0.324119198 | -0.04669348 | 0.694931877 | 12.03629212 | 2.189276053 | 0.075445852 | 0.709264768 | -4.302947823 |
| **A_52_P92514** | Supt20 | -0.291632953 | -0.625299053 | 0.042033147 | 8.580452666 | -2.189146647 | 0.075458735 | 0.709264768 | -4.302978867 |
| **A_52_P263095** | Ahnak | -0.437076175 | -0.937546392 | 0.063394042 | 13.88754858 | -2.187405206 | 0.075632331 | 0.709264768 | -4.303396757 |
| **A_66_P112164** | Rab29 | -0.286672641 | -0.614966656 | 0.041621374 | 10.38756512 | -2.187125118 | 0.075660291 | 0.709264768 | -4.303463991 |
| **A_51_P247359** | Ptprcap | 1.574926424 | -0.230457502 | 3.38031035 | 9.271420048 | 2.184947689 | 0.07587802 | 0.709264768 | -4.303986881 |
| **A_51_P268145** | Naa80 | -0.248602985 | -0.533593626 | 0.036387656 | 9.044363017 | -2.18487244 | 0.075885556 | 0.709264768 | -4.304004957 |
| **A_51_P314907** | Dbf4 | 0.203123517 | -0.02976635 | 0.436013384 | 8.788155178 | 2.184539794 | 0.07591888 | 0.709264768 | -4.304084874 |
| **A_55_P2110875** | 2700046G | -0.236975416 | -0.508763307 | 0.034812474 | 9.574923338 | -2.183853686 | 0.075987661 | 0.709264768 | -4.304249733 |
| **A_51_P395473** | Tns2 | -0.404841427 | -0.869165116 | 0.059482261 | 10.07515092 | -2.183807999 | 0.075992243 | 0.709264768 | -4.304260712 |
| **A_52_P134083** | Cyhr1 | -0.51031588 | -1.096105946 | 0.075474185 | 10.71748843 | -2.181963224 | 0.076177514 | 0.709264768 | -4.304704166 |
| **A_52_P403527** | NA | 0.897310192 | -0.132996192 | 1.927616576 | 11.03615874 | 2.181356032 | 0.076238597 | 0.709264768 | -4.304850182 |
| **A_55_P2470474** | 9530082P | -0.322621157 | -0.693068077 | 0.047825762 | 8.369453855 | -2.181309059 | 0.076243325 | 0.709264768 | -4.304861479 |
| **A_55_P2171484** | Lta4h | 0.491505903 | -0.073031645 | 1.056043451 | 11.81450466 | 2.18065143 | 0.076309544 | 0.709264768 | -4.305019659 |
| **A_51_P186053** | Rrp15 | 0.40747705 | -0.060672769 | 0.875626868 | 10.00201118 | 2.180060969 | 0.076369051 | 0.709264768 | -4.305161711 |
| **A_55_P2150294** | Synj2bp | -0.277484329 | -0.596301376 | 0.041332718 | 10.63017258 | -2.179953708 | 0.076379866 | 0.709264768 | -4.305187518 |
| **A_52_P476400** | NA | 0.220690983 | -0.032901286 | 0.474283253 | 10.22255369 | 2.179711051 | 0.076404339 | 0.709264768 | -4.305245906 |
| **A_55_P2111416** | Rcc2 | 0.524219 | -0.078211462 | 1.126649463 | 10.81382288 | 2.179496515 | 0.076425983 | 0.709264768 | -4.305297531 |
| **A_52_P627816** | Tgm1 | -1.100921191 | -2.366642098 | 0.164799716 | 8.983424404 | -2.178555474 | 0.076520996 | 0.709264768 | -4.305524021 |
| **A_55_P2026785** | Dcaf1 | 0.225078032 | -0.033758419 | 0.483914484 | 9.979672842 | 2.178000715 | 0.076577066 | 0.709264768 | -4.305657572 |
| **A_55_P1988907** | Eml4 | 0.403429561 | -0.060623338 | 0.867482459 | 11.41748099 | 2.177461942 | 0.076631561 | 0.709264768 | -4.305787298 |
| **A_55_P2068265** | Mif | 0.779044557 | -0.117152752 | 1.675241866 | 14.61017831 | 2.177253552 | 0.07665265 | 0.709264768 | -4.30583748 |
| **A_55_P2091671** | Psmg1 | 0.581549981 | -0.087540595 | 1.250640557 | 10.7774984 | 2.17697017 | 0.076681338 | 0.709264768 | -4.305905726 |
| **A_66_P108792** | Rad1 | 0.235121747 | -0.035447722 | 0.505691216 | 10.34195825 | 2.176528491 | 0.076726073 | 0.709264768 | -4.306012107 |
| **A_55_P1979694** | NA | 0.435880703 | -0.065891737 | 0.937653142 | 13.89808538 | 2.175760982 | 0.076803874 | 0.709264768 | -4.306197001 |
| **A_55_P2402134** | NA | -0.355758567 | -0.765482209 | 0.053965074 | 9.539155151 | -2.174776787 | 0.076903761 | 0.709264768 | -4.306434161 |
| **A_55_P2185160** | NA | 0.487694853 | -0.074019433 | 1.04940914 | 12.75101721 | 2.174618351 | 0.076919854 | 0.709264768 | -4.306472346 |
| **A_55_P1988009** | Gm10406 | -0.480544867 | -1.034426442 | 0.073336707 | 9.458459442 | -2.173038246 | 0.077080539 | 0.709264768 | -4.306853276 |
| **A_51_P374869** | Abca7 | -0.424089376 | -0.913330401 | 0.065151648 | 11.08839413 | -2.171125217 | 0.077275551 | 0.709264768 | -4.307314723 |
| **A_55_P2086659** | Rad54l | 0.34674843 | -0.053406922 | 0.746903782 | 8.409016618 | 2.1703822 | 0.077351432 | 0.709264768 | -4.307494024 |
| **A_52_P221024** | NA | 0.507275948 | -0.078138242 | 1.092690139 | 12.28176834 | 2.170357974 | 0.077353907 | 0.709264768 | -4.307499871 |
| **A_55_P1998072** | NA | 0.303400102 | -0.046891468 | 0.653691673 | 8.116874362 | 2.169383745 | 0.077453522 | 0.709264768 | -4.307735032 |
| **A_55_P2091284** | Ltbp3 | -0.333842003 | -0.719292175 | 0.05160817 | 11.39463853 | -2.169317291 | 0.077460322 | 0.709264768 | -4.307751076 |
| **A_55_P2106666** | Adcy9 | -0.542145691 | -1.168245069 | 0.083953687 | 9.673393368 | -2.168817874 | 0.077511444 | 0.709264768 | -4.307871657 |

| | | | | | | | | | |
|---|---|---|---|---|---|---|---|---|---|
| **A_55_P2019690** | Slc43a1 | 0.660468005 | -0.102289439 | 1.42322545 | 8.839472772 | 2.168780875 | 0.077515233 | 0.709264768 | -4.307880591 |
| **A_51_P371331** | Mpp5 | -0.335017497 | -0.72204414 | 0.052009146 | 9.788591595 | -2.168088332 | 0.077586186 | 0.709264768 | -4.308047834 |
| **A_66_P139799** | Cenpb | 0.246422018 | -0.038376912 | 0.531220947 | 8.592319293 | 2.167162622 | 0.077681133 | 0.709264768 | -4.308271442 |
| **A_51_P418725** | Plekhf1 | -0.375182193 | -0.808849008 | 0.058484621 | 8.905606026 | -2.166887549 | 0.07770937 | 0.709264768 | -4.3083379 |
| **A_55_P1974283** | Rbmx | 0.594698092 | -0.092749337 | 1.282145521 | 10.45190917 | 2.166742982 | 0.077724214 | 0.709264768 | -4.308372829 |
| **A_55_P2136832** | Gm6981 | 0.411400835 | -0.064433505 | 0.887235174 | 13.88879586 | 2.165507705 | 0.077851176 | 0.709264768 | -4.308671356 |
| **A_55_P2157922** | Ndufaf1 | 0.297702563 | -0.046725515 | 0.642130642 | 10.18865164 | 2.164882704 | 0.077915496 | 0.709264768 | -4.308822444 |
| **A_51_P465772** | Ruvbl1 | 0.534273172 | -0.084265672 | 1.152812015 | 12.39909639 | 2.163449361 | 0.078063213 | 0.709264768 | -4.309169051 |
| **A_55_P2049158** | NA | 1.24130359 | -0.196212185 | 2.678819365 | 12.3801706 | 2.162796898 | 0.078130552 | 0.709264768 | -4.30932688 |
| **A_55_P2076994** | Defa-rs10 | 0.324676991 | -0.051606085 | 0.700960066 | 8.746662041 | 2.161161254 | 0.078299627 | 0.709264768 | -4.30972268 |
| **A_55_P2068167** | Nufip2 | 0.397946387 | -0.063305063 | 0.859197838 | 10.93500269 | 2.160912428 | 0.078325382 | 0.709264768 | -4.309782909 |
| **A_55_P1963201** | Chchd7 | 0.320500319 | -0.051081835 | 0.692082472 | 11.97777269 | 2.160349239 | 0.078383706 | 0.709264768 | -4.30991925 |
| **A_55_P2243367** | NA | 0.34512387 | -0.055142695 | 0.745390434 | 9.064363094 | 2.159613479 | 0.078459971 | 0.709264768 | -4.310097404 |
| **A_55_P2130399** | Rilp | -0.35650469 | -0.769999372 | 0.056989991 | 9.176559237 | -2.159462493 | 0.078475631 | 0.709264768 | -4.310133968 |
| **A_55_P1986113** | Ppil1 | 0.583483531 | -0.093283385 | 1.260250447 | 10.33113555 | 2.159433468 | 0.078478642 | 0.709264768 | -4.310140997 |
| **A_55_P2140171** | NA | 0.786075587 | -0.126167705 | 1.698318879 | 10.85208898 | 2.158261063 | 0.078600358 | 0.709264768 | -4.31042498 |
| **A_55_P1984861** | NA | 0.513787481 | -0.082545024 | 1.110119987 | 15.01134302 | 2.15796989 | 0.078630617 | 0.709264768 | -4.310495525 |
| **A_55_P2018904** | Kbtbd6 | 0.48401256 | -0.077770021 | 1.04579514 | 8.60156707 | 2.157936736 | 0.078634063 | 0.709264768 | -4.310503558 |
| **A_51_P121275** | Cenpo | 0.419341851 | -0.067441904 | 0.906125606 | 9.130585169 | 2.157657348 | 0.07866311 | 0.709264768 | -4.310571254 |
| **A_55_P2079912** | NA | 0.436903672 | -0.070297442 | 0.944104787 | 8.954553266 | 2.157525022 | 0.078676872 | 0.709264768 | -4.310603319 |
| **A_51_P242201** | Naaa | -0.827191798 | -1.787495952 | 0.133112356 | 9.620753059 | -2.157484865 | 0.078681048 | 0.709264768 | -4.31061305 |
| **A_51_P406557** | Myorg | -0.196152916 | -0.423871735 | 0.031565904 | 8.444485432 | -2.157477077 | 0.078681858 | 0.709264768 | -4.310614937 |
| **A_51_P520936** | Bcar3 | -1.09103275 | -2.357738136 | 0.175672637 | 10.55568959 | -2.157309794 | 0.07869926 | 0.709264768 | -4.310655475 |
| **A_55_P2011551** | Anapc5 | 0.341717329 | -0.055110899 | 0.738545556 | 10.23535718 | 2.156824416 | 0.078749774 | 0.709264768 | -4.31077311 |
| **A_51_P155085** | Dennd2a | -0.434974072 | -0.940193802 | 0.070245658 | 11.63620549 | -2.156420308 | 0.078791857 | 0.709264768 | -4.310871063 |
| **A_52_P384901** | Wdr43 | 0.368492036 | -0.059533734 | 0.796517805 | 9.389450731 | 2.156296781 | 0.078804725 | 0.709264768 | -4.310901007 |
| **A_55_P1970905** | Lin28b | 0.413692089 | -0.066837805 | 0.894221983 | 7.923011285 | 2.156289913 | 0.07880544 | 0.709264768 | -4.310902672 |
| **A_51_P225771** | NA | 0.491793038 | -0.079486807 | 1.063072882 | 12.69310267 | 2.156174075 | 0.07881751 | 0.709264768 | -4.310930754 |
| **A_52_P306396** | Ppp3cb | 0.4066231 | -0.066378974 | 0.879625175 | 10.66308933 | 2.153175092 | 0.079130652 | 0.709264768 | -4.311658124 |
| **A_55_P1966114** | Epn2 | -0.627209884 | -1.356868989 | 0.102449222 | 12.48748887 | -2.152996048 | 0.079149388 | 0.709264768 | -4.311701571 |
| **A_52_P52618** | Csf2rb | -0.759590518 | -1.643444578 | 0.124263543 | 9.506037631 | -2.152530524 | 0.079198125 | 0.709264768 | -4.311814546 |
| **A_55_P2065519** | NA | 0.222153518 | -0.03642217 | 0.480729205 | 10.25502076 | 2.151869114 | 0.079267423 | 0.709264768 | -4.311975088 |
| **A_52_P207635** | Zfc3h1 | -0.216299936 | -0.468080008 | 0.035480135 | 9.415294549 | -2.151718157 | 0.079283248 | 0.709264768 | -4.312011734 |
| **A_55_P2345116** | NA | -0.26341661 | -0.570170146 | 0.043336926 | 8.262727389 | -2.150819299 | 0.079377545 | 0.709264768 | -4.312229974 |
| **A_51_P490257** | Fbxo4 | -0.244682348 | -0.529662118 | 0.040297422 | 10.18273226 | -2.150497556 | 0.079411326 | 0.709264768 | -4.312308107 |
| **A_55_P2072213** | NA | 0.612608845 | -0.101088173 | 1.326305864 | 11.69350423 | 2.14990726 | 0.079473344 | 0.709264768 | -4.312451477 |
| **A_52_P459929** | Itga1 | -0.241524868 | -0.52296283 | 0.039913095 | 9.646584281 | -2.149460757 | 0.079520288 | 0.709264768 | -4.312559941 |
| **A_55_P2064004** | Rapgef2 | -0.478876382 | -1.037187845 | 0.079435081 | 11.03490254 | -2.148311332 | 0.079641268 | 0.709264768 | -4.312839225 |
| **A_55_P2065465** | Mettl1 | 0.307135178 | -0.05100706 | 0.665277416 | 9.500236728 | 2.147950969 | 0.079679237 | 0.709264768 | -4.312926806 |
| **A_52_P392674** | Acot9 | -0.388478439 | -0.841517267 | 0.064560389 | 9.785687789 | -2.147740667 | 0.079701404 | 0.709264768 | -4.312977921 |
| **A_51_P205481** | Mtmr6 | -0.220976378 | -0.478705909 | 0.036753152 | 11.33122682 | -2.147494272 | 0.079727383 | 0.709264768 | -4.313037813 |
| **A_55_P2114074** | Acrbp | -0.333746562 | -0.723004282 | 0.055511158 | 9.027226856 | -2.147483874 | 0.079728479 | 0.709264768 | -4.31304034 |
| **A_51_P230439** | Ppfibp2 | -1.258260645 | -2.725869923 | 0.209348633 | 11.58416484 | -2.147387943 | 0.079738597 | 0.709264768 | -4.31306366 |
| **A_52_P745** | Acox3 | -0.776757249 | -1.683092132 | 0.129577633 | 9.277810511 | -2.146579441 | 0.079823917 | 0.709264768 | -4.313260224 |
| **A_51_P172853** | Cd14 | -0.207115237 | -0.448874009 | 0.034643535 | 8.538623239 | -2.14575511 | 0.079911006 | 0.709264768 | -4.313460687 |
| **A_65_P09090** | Arih2 | -0.322775513 | -0.699565169 | 0.054014143 | 10.86939109 | -2.145615779 | 0.079925736 | 0.709264768 | -4.313494575 |
| **A_55_P2008197** | Esr2 | -0.384992246 | -0.834529754 | 0.064545261 | 10.1006145 | -2.145044941 | 0.079986114 | 0.709264768 | -4.313633429 |
| **A_51_P494361** | Mbnl2 | -0.552198057 | -1.196974524 | 0.092578411 | 12.2720413 | -2.14504319 | 0.079986299 | 0.709264768 | -4.313633855 |

| | | | | | | | | | |
|---|---|---|---|---|---|---|---|---|---|
| **A_52_P656714** | Map3k5 | -0.247197913 | -0.535862337 | 0.041466511 | 8.201138579 | -2.144874482 | 0.080004153 | 0.709264768 | -4.313674897 |
| **A_52_P570240** | Kbtbd11 | 0.734337656 | -0.123274934 | 1.591950245 | 10.23666809 | 2.144642938 | 0.080028662 | 0.709264768 | -4.313731229 |
| **A_52_P608322** | Maff | -0.740319119 | -1.605001343 | 0.124363104 | 10.54510881 | -2.144434474 | 0.080050736 | 0.709264768 | -4.31378195 |
| **A_55_P2036907** | Mycbpap | -0.281727201 | -0.610815805 | 0.047361403 | 8.37644439 | -2.14420489 | 0.080075053 | 0.709264768 | -4.313837812 |
| **A_52_P592101** | Washc2 | -0.332355575 | -0.720608214 | 0.055897063 | 9.514881279 | -2.144069697 | 0.080089375 | 0.709264768 | -4.31387071 |
| **A_51_P424272** | Mt4 | 0.322150026 | -0.054277259 | 0.698577311 | 8.202870394 | 2.14351941 | 0.080147703 | 0.709264768 | -4.314004629 |
| **A_52_P656867** | Fam98b | 0.691120027 | -0.116835894 | 1.499075948 | 10.22853899 | 2.142476936 | 0.08025832 | 0.709264768 | -4.314258389 |
| **A_55_P1988168** | Proser3 | -0.483635119 | -1.049030398 | 0.081760161 | 13.15914194 | -2.142476153 | 0.080258403 | 0.709264768 | -4.31425858 |
| **A_51_P386880** | Stip1 | 0.231368781 | -0.039208648 | 0.50194621 | 11.11750354 | 2.141724174 | 0.080338295 | 0.709264768 | -4.314441679 |
| **A_55_P2049867** | Ccrl2 | -0.298094635 | -0.64672526 | 0.050535991 | 8.399499344 | -2.141603018 | 0.080351175 | 0.709264768 | -4.314471183 |
| **A_55_P2058851** | Fancf | 0.44992269 | -0.076427684 | 0.976273063 | 9.261900558 | 2.140983425 | 0.080417075 | 0.709264768 | -4.314622085 |
| **A_51_P368543** | Abhd8 | 0.303826757 | -0.051626541 | 0.659280055 | 9.133964527 | 2.140887348 | 0.080427299 | 0.709264768 | -4.314645487 |
| **A_55_P2055523** | NA | 1.35185229 | -0.229844959 | 2.933549539 | 13.31634978 | 2.140702062 | 0.080447019 | 0.709264768 | -4.31469062 |
| **A_51_P148196** | Sdf2 | -0.424204233 | -0.920666851 | 0.072258385 | 14.14587691 | -2.140123274 | 0.080508654 | 0.709264768 | -4.314831623 |
| **A_55_P2102574** | Prorsd1 | 0.361194621 | -0.06155304 | 0.783942281 | 9.96508729 | 2.139983408 | 0.080523556 | 0.709264768 | -4.3148657 |
| **A_55_P1953708** | Cry2 | -0.222500155 | -0.48292551 | 0.037925201 | 8.428004253 | -2.139919309 | 0.080530386 | 0.709264768 | -4.314881318 |
| **A_55_P2061061** | Set | 0.637390865 | -0.109288121 | 1.384069851 | 12.23048947 | 2.138071634 | 0.080727529 | 0.709264768 | -4.315331639 |
| **A_55_P2077185** | NA | 0.47389757 | -0.08126455 | 1.029059691 | 9.52059676 | 2.138035944 | 0.080731342 | 0.709264768 | -4.31534034 |
| **A_55_P1984158** | Slc44a2 | -0.291114656 | -0.632211947 | 0.049982636 | 12.7210114 | -2.137647762 | 0.080772826 | 0.709264768 | -4.315434982 |
| **A_55_P2110925** | Flrt3 | 0.340421467 | -0.058597744 | 0.739440678 | 9.201877257 | 2.136847251 | 0.080858445 | 0.709264768 | -4.31563019 |
| **A_55_P2107035** | Mcmbp | 0.446598525 | -0.076975387 | 0.970172436 | 11.54564357 | 2.13643474 | 0.080902602 | 0.709264768 | -4.315730801 |
| **A_52_P466887** | Hnrnpd | 0.545991228 | -0.09410681 | 1.186089267 | 12.46509689 | 2.136434168 | 0.080902663 | 0.709264768 | -4.315730941 |
| **A_51_P479434** | Rfc3 | 0.497669042 | -0.092603154 | 1.087941239 | 8.666699667 | 2.245696203 | 0.080921988 | 0.709264768 | -4.360175912 |
| **A_55_P1969072** | Cbx3 | 0.624744299 | -0.108611894 | 1.358100491 | 13.68130785 | 2.133721256 | 0.081193692 | 0.709264768 | -4.316392938 |
| **A_51_P516728** | Hap1 | 0.66537419 | -0.115818214 | 1.446566593 | 11.31897414 | 2.133331197 | 0.081235625 | 0.709264768 | -4.316488164 |
| **A_55_P1968863** | Synpo2l | 0.334662708 | -0.058541761 | 0.727867176 | 13.3033669 | 2.131764479 | 0.081404282 | 0.709264768 | -4.316870767 |
| **A_55_P1971579** | Acox3 | -0.708537549 | -1.541134083 | 0.124058986 | 8.769687858 | -2.131467018 | 0.081436345 | 0.709264768 | -4.316943429 |
| **A_52_P5652** | Sae1 | 0.211100571 | -0.036968964 | 0.459170105 | 9.329787327 | 2.13140667 | 0.081442852 | 0.709264768 | -4.316958172 |
| **A_55_P2088018** | Fhod3 | -0.701390836 | -1.525669871 | 0.1228882 | 9.352595939 | -2.131258745 | 0.081458802 | 0.709264768 | -4.31699431 |
| **A_51_P320614** | Txnrd1 | 0.235817967 | -0.041344154 | 0.512980088 | 11.46696813 | 2.131048631 | 0.081481465 | 0.709264768 | -4.317045643 |
| **A_51_P190765** | Rtca | 0.259295085 | -0.045471939 | 0.564062109 | 10.33470222 | 2.130966668 | 0.081490307 | 0.709264768 | -4.317065668 |
| **A_51_P155196** | Abtb2 | 0.245688725 | -0.043152862 | 0.534530312 | 9.912755167 | 2.130472091 | 0.081543682 | 0.709264768 | -4.317186515 |
| **A_55_P2054559** | NA | 0.497751564 | -0.087437917 | 1.082941046 | 9.068074382 | 2.130426069 | 0.081548651 | 0.709264768 | -4.317197761 |
| **A_51_P292016** | Uts2r | 0.269806843 | -0.047442689 | 0.587056375 | 8.883045477 | 2.130111396 | 0.081582632 | 0.709264768 | -4.31727466 |
| **A_55_P2019833** | NA | -0.475724857 | -1.035105158 | 0.083655444 | 11.75550382 | -2.130095239 | 0.081584377 | 0.709264768 | -4.317278608 |
| **A_52_P240453** | Polr3k | 0.657911397 | -0.115756379 | 1.431579173 | 10.68793435 | 2.129919779 | 0.081603331 | 0.709264768 | -4.31732149 |
| **A_55_P2157159** | NA | 0.596927805 | -0.10541571 | 1.299271321 | 11.56282415 | 2.128739789 | 0.081730922 | 0.709264768 | -4.317609935 |
| **A_55_P2004806** | Hexim2 | -0.439572232 | -0.95704796 | 0.077903496 | 12.35739146 | -2.127603084 | 0.081854028 | 0.709264768 | -4.317887898 |
| **A_52_P257502** | Igfbp4 | 0.829532163 | -0.147239266 | 1.806303592 | 9.232298157 | 2.127113321 | 0.081907129 | 0.709264768 | -4.318007692 |
| **A_55_P1959639** | NA | -0.211082865 | -0.459653774 | 0.037488044 | 14.08565405 | -2.126929162 | 0.081927106 | 0.709264768 | -4.31805274 |
| **A_51_P397564** | Snrnp40 | 0.480343965 | -0.085319888 | 1.046007817 | 10.94554342 | 2.126886218 | 0.081931765 | 0.709264768 | -4.318063246 |
| **A_66_P123014** | Twistnb | 0.364387742 | -0.064759223 | 0.793534706 | 9.967414475 | 2.126708954 | 0.081950999 | 0.709264768 | -4.318106611 |
| **A_55_P2429480** | Atad2 | 0.26919894 | -0.047976031 | 0.586373911 | 8.464121898 | 2.125811651 | 0.082048434 | 0.709264768 | -4.31832616 |
| **A_55_P2127303** | Spsb2 | -1.126652097 | -2.454500571 | 0.201196378 | 11.00100594 | -2.125160112 | 0.082119258 | 0.709264768 | -4.318485614 |
| **A_51_P293607** | Mrps14 | 0.221309488 | -0.039582002 | 0.482200979 | 8.445500862 | 2.12466501 | 0.082173119 | 0.709264768 | -4.318606803 |
| **A_55_P2037116** | Ran | 0.66286626 | -0.119046451 | 1.444778971 | 15.26820536 | 2.123332397 | 0.082318274 | 0.709264768 | -4.318933087 |
| **A_51_P407879** | Fam193b | -0.479884435 | -1.045954201 | 0.086185332 | 10.86256447 | -2.123327815 | 0.082318774 | 0.709264768 | -4.318934209 |
| **A_55_P2064906** | NA | 0.478547349 | -0.085996291 | 1.043090989 | 16.48603548 | 2.123135642 | 0.082339728 | 0.709264768 | -4.318981272 |

| | | | | | | | | | |
|---|---|---|---|---|---|---|---|---|---|
| **A_51_P252677** | Card19 | -0.304551153 | -0.664280166 | 0.055177861 | 11.21824294 | -2.120484618 | 0.082629364 | 0.709264768 | -4.319630794 |
| **A_55_P2109752** | Nans | 0.212395375 | -0.038776633 | 0.463567383 | 12.58373066 | 2.11799126 | 0.08290274 | 0.709264768 | -4.320242163 |
| **A_51_P113072** | Cdc123 | 0.287951461 | -0.052680734 | 0.628583657 | 11.86732291 | 2.117307513 | 0.082977872 | 0.709264768 | -4.320409897 |
| **A_52_P80593** | NA | 0.239423949 | -0.043976414 | 0.522824311 | 11.90987003 | 2.116009113 | 0.083120737 | 0.709264768 | -4.320728512 |
| **A_55_P2231747** | NA | -0.475510717 | -1.038374819 | 0.087353385 | 9.234313783 | -2.115958293 | 0.083126334 | 0.709264768 | -4.320740985 |
| **A_52_P486702** | G3bp1 | 0.28879627 | -0.053058191 | 0.630650732 | 9.35511768 | 2.115926966 | 0.083129784 | 0.709264768 | -4.320748674 |
| **A_55_P2139640** | Fbh1 | -0.211727678 | -0.462466438 | 0.039011083 | 10.0201309 | -2.11498117 | 0.083234025 | 0.709264768 | -4.320980848 |
| **A_52_P650855** | Myo1d | -0.535421428 | -1.169543183 | 0.098700328 | 13.03668516 | -2.114820153 | 0.083251785 | 0.709264768 | -4.321020381 |
| **A_55_P2090996** | Ebna1bp2 | 0.286241273 | -0.052777936 | 0.625260482 | 12.63531339 | 2.114746398 | 0.083259921 | 0.709264768 | -4.32103849 |
| **A_55_P1966337** | Polr2h | 0.321839959 | -0.059374032 | 0.703053949 | 11.11463127 | 2.114567138 | 0.0832797 | 0.709264768 | -4.321082505 |
| **A_55_P1956302** | H2-Q4 | -0.417279229 | -0.9116256 | 0.077067142 | 13.43185809 | -2.114198509 | 0.083320388 | 0.709264768 | -4.321173025 |
| **A_52_P522255** | Pitx1 | -1.884258317 | -4.117214363 | 0.34869773 | 8.656131756 | -2.113540683 | 0.083393049 | 0.709264768 | -4.321334585 |
| **A_55_P2064595** | NA | 0.193181484 | -0.035800357 | 0.422163325 | 8.858535051 | 2.113074455 | 0.083444586 | 0.709264768 | -4.321449109 |
| **A_51_P255657** | Misp3 | -1.189024435 | -2.598575099 | 0.220526229 | 9.692224156 | -2.112809957 | 0.083473838 | 0.709264768 | -4.321514087 |
| **A_51_P130079** | Lrmp | 0.814025258 | -0.150977128 | 1.779027643 | 8.289554254 | 2.112807059 | 0.083474159 | 0.709264768 | -4.321514799 |
| **A_51_P512820** | Dera | 0.24314119 | -0.045111405 | 0.531393785 | 9.147300755 | 2.112689419 | 0.083487173 | 0.709264768 | -4.3215437 |
| **A_55_P1973698** | Htra2 | -0.205377411 | -0.448898298 | 0.038143475 | 9.9655971 | -2.112354507 | 0.083524234 | 0.709264768 | -4.321625987 |
| **A_52_P86693** | Ifi27 | -0.618827212 | -1.352681599 | 0.115027175 | 12.49055715 | -2.112077516 | 0.083554899 | 0.709264768 | -4.321694049 |
| **A_52_P651034** | Tmx1 | 0.37029315 | -0.069076146 | 0.809662446 | 10.37697715 | 2.110893484 | 0.083686111 | 0.709264768 | -4.321985052 |
| **A_55_P2004119** | NA | 0.991691284 | -0.185018805 | 2.168401373 | 14.44409096 | 2.11084996 | 0.083690939 | 0.709264768 | -4.321995752 |
| **A_51_P418339** | Nudcd2 | 0.302280425 | -0.05652609 | 0.661086939 | 9.807011015 | 2.110085485 | 0.083775775 | 0.709264768 | -4.322183696 |
| **A_51_P130660** | Mrps30 | 0.361125795 | -0.067552529 | 0.789804119 | 10.86972424 | 2.109975113 | 0.083788031 | 0.709264768 | -4.322210835 |
| **A_52_P676271** | Tuba4a | 0.212931238 | -0.03996532 | 0.465827796 | 13.99173384 | 2.108855423 | 0.083912467 | 0.709264768 | -4.322486196 |
| **A_55_P1974533** | Eif4e | 0.277373247 | -0.052077443 | 0.606823937 | 8.955235897 | 2.108747083 | 0.083924518 | 0.709264768 | -4.322512844 |
| **A_55_P1972991** | Pcsk6 | 0.906651967 | -0.170281208 | 1.983585142 | 11.06662086 | 2.108638822 | 0.083936561 | 0.709264768 | -4.322539474 |
| **A_55_P2143204** | Pbxip1 | -0.441388903 | -0.965771373 | 0.082993566 | 10.19582835 | -2.108257198 | 0.083979029 | 0.709264768 | -4.322633353 |
| **A_51_P110371** | Wwox | 0.388685425 | -0.073412057 | 0.850782907 | 8.348975369 | 2.106759686 | 0.084145892 | 0.709264768 | -4.32300184 |
| **A_55_P1955244** | NA | 0.78475045 | -0.148258743 | 1.717759643 | 13.35585344 | 2.106667504 | 0.084156175 | 0.709264768 | -4.323024528 |
| **A_55_P2128032** | Hsf2 | 0.232555097 | -0.04394073 | 0.509050924 | 9.386374137 | 2.106626903 | 0.084160704 | 0.709264768 | -4.323034522 |
| **A_55_P2235911** | 0610009E | 0.514387637 | -0.097272148 | 1.126047422 | 8.920930722 | 2.106351955 | 0.084191384 | 0.709264768 | -4.323102198 |
| **A_55_P1962731** | Clcn4 | -0.360542975 | -0.789273156 | 0.068187206 | 12.09656134 | -2.106315026 | 0.084195505 | 0.709264768 | -4.323111288 |
| **A_55_P2021029** | Tigd3 | 0.242761532 | -0.045912106 | 0.531435171 | 8.123578738 | 2.106313879 | 0.084195633 | 0.709264768 | -4.32311157 |
| **A_52_P229959** | 6-Sep | 0.251651095 | -0.047653533 | 0.550955722 | 8.730185023 | 2.105890271 | 0.084242926 | 0.709264768 | -4.32321585 |
| **A_65_P08971** | F3 | -0.350314062 | -0.767041811 | 0.066413688 | 8.249278279 | -2.10550121 | 0.084286386 | 0.709264768 | -4.323311637 |
| **A_52_P556281** | G3bp2 | 0.344276975 | -0.065315134 | 0.753869083 | 11.01757542 | 2.105264868 | 0.084312798 | 0.709264768 | -4.323369829 |
| **A_51_P489153** | Crot | -0.195551285 | -0.428213146 | 0.037110577 | 9.596408435 | -2.105163447 | 0.084324135 | 0.709264768 | -4.323394803 |
| **A_55_P2420873** | NA | -0.265049672 | -0.580419485 | 0.05032014 | 9.754588165 | -2.105026158 | 0.084339483 | 0.709264768 | -4.32342861 |
| **A_55_P2076273** | NA | 0.579388999 | -0.110115605 | 1.268893603 | 13.94002056 | 2.104667137 | 0.084379634 | 0.709264768 | -4.323517023 |
| **A_55_P2144003** | NA | 0.427829572 | -0.081315466 | 0.93697461 | 13.46360073 | 2.104648774 | 0.084381689 | 0.709264768 | -4.323521546 |
| **A_51_P347240** | Lrr1 | 0.463810451 | -0.088158546 | 1.015779449 | 8.462814609 | 2.104632103 | 0.084383553 | 0.709264768 | -4.323525651 |
| **A_55_P2083654** | Prmt2 | -0.588987853 | -1.290000308 | 0.112024602 | 12.44841804 | -2.104412871 | 0.084408082 | 0.709264768 | -4.323579645 |
| **A_55_P2090345** | BC08527 | 0.617845273 | -0.117623366 | 1.353313911 | 11.5871824 | 2.104097765 | 0.084443351 | 0.709264768 | -4.323657257 |
| **A_51_P327232** | Lenep | -0.219328774 | -0.480472969 | 0.041815421 | 8.415918246 | -2.10361171 | 0.084497784 | 0.709264768 | -4.32377699 |
| **A_55_P2051099** | Tnfrsf1a | -0.268565069 | -0.588350661 | 0.051220524 | 11.42028271 | -2.103492546 | 0.084511134 | 0.709264768 | -4.323806347 |
| **A_51_P158876** | Dctpp1 | 0.705542844 | -0.134949287 | 1.546034975 | 12.96007806 | 2.102519757 | 0.084620202 | 0.709264768 | -4.32404604 |
| **A_51_P495269** | Lor | 0.379207893 | -0.072539656 | 0.830955442 | 8.603172818 | 2.102480171 | 0.084624644 | 0.709264768 | -4.324055795 |
| **A_66_P139333** | Atf6 | -0.286241139 | -0.627288974 | 0.054806697 | 8.570096345 | -2.102166439 | 0.084659852 | 0.709264768 | -4.324133114 |
| **A_51_P323299** | Trim28 | 0.510394273 | -0.097968264 | 1.11875681 | 11.45489293 | 2.101327169 | 0.084754113 | 0.709264768 | -4.324339986 |

| | | | | | | | | | |
|---|---|---|---|---|---|---|---|---|---|
| **A_51_P271370** | Wsb1 | -0.2029149 | -0.444840195 | 0.039010396 | 8.450182085 | -2.10079175 | 0.084814305 | 0.709264768 | -4.324471989 |
| **A_55_P2139181** | Cald1 | -0.506958079 | -1.111410363 | 0.097494204 | 9.904083478 | -2.10068228 | 0.084826617 | 0.709264768 | -4.32449898 |
| **A_51_P225224** | Htra1 | -1.253172765 | -2.74801875 | 0.241673221 | 13.15634927 | -2.099736723 | 0.084933041 | 0.709264768 | -4.324732157 |
| **A_55_P2103422** | Psmc2 | 0.216564779 | -0.041816232 | 0.474945791 | 13.37048922 | 2.099314852 | 0.084980568 | 0.709264768 | -4.324836213 |
| **A_55_P2258832** | 5230400M | -0.366319885 | -0.803442813 | 0.070803043 | 8.452150616 | -2.098974938 | 0.085018881 | 0.709264768 | -4.324920063 |
| **A_55_P2142146** | Glul | -0.468021072 | -1.026616082 | 0.090573938 | 11.64815909 | -2.098547001 | 0.085067142 | 0.709264768 | -4.325025639 |
| **A_55_P2121408** | Tpm2 | -0.490915455 | -1.076887096 | 0.095056186 | 12.46907902 | -2.098362191 | 0.085087993 | 0.709264768 | -4.325071238 |
| **A_55_P1960331** | Srbd1 | 0.399460043 | -0.077387187 | 0.876307273 | 9.792888037 | 2.098188132 | 0.085107636 | 0.709264768 | -4.325114186 |
| **A_55_P2067151** | NA | 0.518259542 | -0.100429175 | 1.136948258 | 8.658321011 | 2.098096488 | 0.08511798 | 0.709264768 | -4.325136799 |
| **A_55_P1966833** | Xaf1 | -1.076477572 | -2.361916514 | 0.208961369 | 9.164744344 | -2.097509223 | 0.085184297 | 0.709264768 | -4.325281724 |
| **A_55_P2123566** | NA | -0.493768226 | -1.083427479 | 0.095891028 | 8.507417565 | -2.097357034 | 0.085201491 | 0.709264768 | -4.325319286 |
| **A_55_P2113208** | Snrpa1 | 0.621051755 | -0.120797151 | 1.362900661 | 12.21668843 | 2.096827377 | 0.085261361 | 0.709264768 | -4.325450021 |
| **A_55_P2052555** | Dpysl5 | 0.402403377 | -0.078274327 | 0.883081081 | 12.05582653 | 2.096804711 | 0.085263924 | 0.709264768 | -4.325455617 |
| **A_52_P445307** | Cwc15 | 0.222019706 | -0.043353241 | 0.487392652 | 13.12409105 | 2.095488211 | 0.085412928 | 0.709264768 | -4.325780661 |
| **A_55_P2130178** | Fn1 | -0.763137861 | -1.675302669 | 0.149026948 | 16.06615797 | -2.095463174 | 0.085415765 | 0.709264768 | -4.325786844 |
| **A_51_P351363** | Tyms-ps | 1.229768565 | -0.240242429 | 2.699779559 | 10.39290842 | 2.09533347 | 0.08543046 | 0.709264768 | -4.325818875 |
| **A_51_P394154** | Ddx51 | 0.223064209 | -0.043662911 | 0.48979133 | 9.556984051 | 2.094657697 | 0.085507067 | 0.709264768 | -4.325985781 |
| **A_55_P1992179** | Tars2 | 0.206739327 | -0.040572963 | 0.454051617 | 11.854429 | 2.09376408 | 0.085608478 | 0.709264768 | -4.326206541 |
| **A_55_P2041075** | Ccnd3 | 0.665361684 | -0.13072909 | 1.461452458 | 13.89273924 | 2.093367814 | 0.085653487 | 0.709264768 | -4.326304454 |
| **A_51_P196973** | Chaf1a | 0.83889708 | -0.16488211 | 1.84267627 | 9.898510297 | 2.093248749 | 0.085667016 | 0.709264768 | -4.326333876 |
| **A_51_P304170** | Rtp4 | -1.226707671 | -2.694626937 | 0.241211595 | 10.43747643 | -2.093096523 | 0.085684316 | 0.709264768 | -4.326371494 |
| **A_55_P2019254** | NA | 1.173559101 | -0.230835038 | 2.57795324 | 13.00238321 | 2.092985902 | 0.08569689 | 0.709264768 | -4.326398832 |
| **A_51_P314418** | Blm | 0.563595551 | -0.111025619 | 1.238216721 | 9.250348118 | 2.092463719 | 0.085756271 | 0.709264768 | -4.326527889 |
| **A_52_P540219** | Timp2 | -0.541317778 | -1.189826598 | 0.107191042 | 14.08332508 | -2.090676073 | 0.085959878 | 0.709264768 | -4.326969857 |
| **A_55_P2004258** | Gm7102 | 0.250669443 | -0.049734849 | 0.551073736 | 9.489852639 | 2.089996735 | 0.086037383 | 0.709264768 | -4.327137874 |
| **A_52_P30877** | Gm5176 | 1.172955471 | -0.233254556 | 2.579165498 | 13.58466459 | 2.089208002 | 0.086127461 | 0.709264768 | -4.327332989 |
| **A_55_P2154982** | Cd8b1 | 1.20790617 | -0.240469123 | 2.656281462 | 8.458899658 | 2.088826831 | 0.086171028 | 0.709264768 | -4.327427299 |
| **A_55_P2113703** | Spib | 0.553156432 | -0.110128542 | 1.216441407 | 8.020384914 | 2.088806225 | 0.086173384 | 0.709264768 | -4.327432397 |
| **A_55_P2116802** | NA | 0.978742075 | -0.194920896 | 2.152405046 | 11.91952671 | 2.088695839 | 0.086186005 | 0.709264768 | -4.327459711 |
| **A_55_P1969625** | Hnrnpr | 0.540175489 | -0.107735347 | 1.188086325 | 12.40966258 | 2.088189827 | 0.086243886 | 0.709264768 | -4.32758493 |
| **A_55_P2002215** | Arfip1 | 0.391392845 | -0.07811205 | 0.86089774 | 10.07192775 | 2.087964463 | 0.086269678 | 0.709264768 | -4.327640705 |
| **A_55_P2150025** | Otub1 | 0.274518438 | -0.055196218 | 0.604233094 | 9.712941466 | 2.085372355 | 0.086566907 | 0.709264768 | -4.328282488 |
| **A_55_P1996973** | Gvin1 | -0.459162901 | -1.010850242 | 0.09252444 | 8.623663045 | -2.084606644 | 0.086654912 | 0.709264768 | -4.328472165 |
| **A_52_P255841** | Dbr1 | 0.185964849 | -0.037541142 | 0.40947084 | 9.122163393 | 2.083972763 | 0.086727835 | 0.709264768 | -4.328629217 |
| **A_52_P150683** | NA | 0.551135634 | -0.111326234 | 1.213597502 | 8.048899293 | 2.083761223 | 0.086752185 | 0.709264768 | -4.328681636 |
| **A_55_P1992676** | Ubtf | 0.377361963 | -0.076257668 | 0.830981594 | 9.031657761 | 2.08361093 | 0.08676949 | 0.709264768 | -4.328718879 |
| **A_55_P1961049** | Cep295 | 0.604531662 | -0.122372218 | 1.331435542 | 9.920169629 | 2.083015062 | 0.086838132 | 0.709264768 | -4.328866556 |
| **A_55_P2142222** | Serpina3h | 0.203074709 | -0.041108334 | 0.447257753 | 7.944484147 | 2.083006779 | 0.086839087 | 0.709264768 | -4.328868609 |
| **A_55_P2177198** | H2bc3 | 0.279284286 | -0.056669184 | 0.615237757 | 8.100119076 | 2.082177335 | 0.086934731 | 0.709264768 | -4.329074217 |
| **A_52_P656565** | Pex6 | -0.278199854 | -0.612857204 | 0.056457495 | 10.24210004 | -2.082125372 | 0.086940726 | 0.709264768 | -4.3290871 |
| **A_51_P270364** | Mmaa | -0.468094377 | -1.031428133 | 0.09523938 | 10.00831656 | -2.081220013 | 0.087045257 | 0.709264768 | -4.329311587 |
| **A_55_P1979335** | NA | -0.210116796 | -0.463012541 | 0.042778949 | 8.405578956 | -2.080988086 | 0.087072055 | 0.709264768 | -4.329369103 |
| **A_55_P2086510** | Ppid | 1.008707024 | -0.221617069 | 2.239031118 | 12.09292291 | 2.183776668 | 0.087086706 | 0.709264768 | -4.370649505 |
| **A_55_P2081360** | Btbd1 | 0.243345631 | -0.049703144 | 0.536394407 | 11.16553235 | 2.079859452 | 0.087202588 | 0.709264768 | -4.329649054 |
| **A_55_P1964664** | Fus | 0.496221706 | -0.101438628 | 1.093882039 | 13.79893874 | 2.079561028 | 0.087237136 | 0.709264768 | -4.329723091 |
| **A_51_P127756** | Ddx20 | 0.515463786 | -0.105442082 | 1.136369654 | 10.31218013 | 2.079326751 | 0.087264268 | 0.709264768 | -4.342917534 |
| **A_55_P2020572** | Psme3 | 0.408731143 | -0.083611522 | 0.901073807 | 11.21463119 | 2.079316514 | 0.087265453 | 0.709264768 | -4.329783759 |
| **A_51_P462692** | Ptbp1 | 0.559324721 | -0.114756756 | 1.233406197 | 12.78182763 | 2.078269993 | 0.087386761 | 0.709264768 | -4.330043465 |

| | | | | | | | | | |
|---|---|---|---|---|---|---|---|---|---|
| **A_55_P1965338** | Anxa8 | -0.517542995 | -1.141297372 | 0.106211382 | 9.111544823 | -2.078180063 | 0.087397193 | 0.709264768 | -4.330065786 |
| **A_51_P471779** | Phf5a | 0.472356262 | -0.097015865 | 1.041728388 | 12.23019599 | 2.07789606 | 0.087430147 | 0.709264768 | -4.33013628 |
| **A_55_P2101757** | Dscc1 | 0.685267141 | -0.140876524 | 1.511410805 | 8.788193637 | 2.077565269 | 0.087468547 | 0.709264768 | -4.330218395 |
| **A_51_P283708** | Msh2 | 0.719494761 | -0.147967642 | 1.586957163 | 12.10793182 | 2.077434408 | 0.087483743 | 0.709264768 | -4.330250881 |
| **A_55_P2049976** | NA | 0.396265346 | -0.081498722 | 0.874029414 | 8.006118534 | 2.077413532 | 0.087486167 | 0.709264768 | -4.330256064 |
| **A_52_P35304** | Chkb | -0.468392212 | -1.033217191 | 0.096432768 | 9.597941098 | -2.077046001 | 0.087528861 | 0.709264768 | -4.330347313 |
| **A_55_P2016486** | Tmem131 | 0.294103324 | -0.060607984 | 0.648814631 | 11.14295209 | 2.076707084 | 0.08756825 | 0.709264768 | -4.330431466 |
| **A_55_P2115176** | Pdhx | 0.227749377 | -0.046962101 | 0.502460854 | 9.457684261 | 2.076494276 | 0.087592992 | 0.709264768 | -4.33048431 |
| **A_55_P1978103** | Ppip5k2 | 0.303774976 | -0.062706384 | 0.670256335 | 10.12125348 | 2.076110432 | 0.087637638 | 0.709264768 | -4.330579634 |
| **A_55_P2047489** | Calm1 | -0.238559828 | -0.526370086 | 0.04925043 | 15.89294903 | -2.076067116 | 0.087642678 | 0.709264768 | -4.330590392 |
| **A_52_P267256** | LOC1008 | 0.622037449 | -0.12884772 | 1.372922618 | 13.29631147 | 2.07488175 | 0.087780707 | 0.709264768 | -4.330884838 |
| **A_55_P2055638** | Gsn | -0.542996405 | -1.198578335 | 0.112585525 | 14.09546627 | -2.074532895 | 0.087821372 | 0.709264768 | -4.330971513 |
| **A_51_P513785** | Ctcf | 0.52437132 | -0.10873764 | 1.157480279 | 11.55133726 | 2.07448747 | 0.087826669 | 0.709264768 | -4.3309828 |
| **A_55_P1967978** | Nfatc2 | 0.299522757 | -0.0622216 | 0.661267113 | 8.760086562 | 2.073855081 | 0.087900439 | 0.709264768 | -4.331139945 |
| **A_51_P268167** | Tasor | -0.209099694 | -0.461666368 | 0.04346698 | 9.16387645 | -2.07361296 | 0.0879287 | 0.709264768 | -4.331200118 |
| **A_66_P114381** | Ypel2 | -0.389414206 | -0.859810628 | 0.080982215 | 8.645185653 | -2.073471774 | 0.087945184 | 0.709264768 | -4.331235209 |
| **A_55_P1983368** | Sh3d21 | -0.726051415 | -1.603629551 | 0.151526722 | 10.36279422 | -2.072201227 | 0.08809367 | 0.709264768 | -4.331551054 |
| **A_55_P2006340** | Uba52 | 0.256872047 | -0.053747698 | 0.567491793 | 17.11153834 | 2.071276764 | 0.088201872 | 0.709264768 | -4.33178094 |
| **A_55_P2049313** | NA | 0.6816805 | -0.142736082 | 1.506097082 | 9.056650882 | 2.071020979 | 0.088231835 | 0.709264768 | -4.331844557 |
| **A_55_P2125013** | H4c11 | 1.391850632 | -0.291471325 | 3.075172588 | 11.33251697 | 2.070979444 | 0.088236701 | 0.709264768 | -4.331854888 |
| **A_55_P2078623** | Stxbp3 | -0.330460564 | -0.730167678 | 0.06924655 | 10.40863422 | -2.070752086 | 0.088263344 | 0.709264768 | -4.331911439 |
| **A_51_P424532** | Vnn1 | -1.671257564 | -3.692893734 | 0.350378606 | 9.808889402 | -2.070573885 | 0.088284232 | 0.709264768 | -4.331955766 |
| **A_55_P2059890** | Fdx1 | -0.50158511 | -1.108328858 | 0.105158637 | 13.26702498 | -2.070568829 | 0.088284824 | 0.709264768 | -4.331957023 |
| **A_55_P2009875** | NA | 0.368095115 | -0.0772115 | 0.81340173 | 10.23766393 | 2.070385672 | 0.088306299 | 0.709264768 | -4.332002586 |
| **A_51_P246773** | Sesn3 | 0.25082089 | -0.052631944 | 0.554273724 | 9.41281898 | 2.070250285 | 0.088322176 | 0.709264768 | -4.332036266 |
| **A_66_P107943** | NA | -0.261560611 | -0.578018011 | 0.054896789 | 12.48940826 | -2.070176777 | 0.088330798 | 0.709264768 | -4.332054553 |
| **A_55_P2156229** | Cd3e | 1.601806065 | -0.33671243 | 3.54032456 | 10.203399 | 2.069618655 | 0.088396288 | 0.709264768 | -4.332193415 |
| **A_66_P134728** | Psmg1 | 0.570857727 | -0.120014694 | 1.261730148 | 12.55380212 | 2.069571204 | 0.088401858 | 0.709264768 | -4.332205222 |
| **A_55_P1952312** | NA | 1.157622126 | -0.243402246 | 2.558646499 | 13.58863365 | 2.069528799 | 0.088406836 | 0.709264768 | -4.332215773 |
| **A_55_P2009857** | NA | -0.601831559 | -1.330509465 | 0.126846347 | 11.18429746 | -2.068662784 | 0.088508567 | 0.709264768 | -4.332431289 |
| **A_55_P1993544** | NA | 0.250720051 | -0.052921133 | 0.554361234 | 8.715536082 | 2.068134296 | 0.088570707 | 0.709264768 | -4.332562834 |
| **A_55_P2186672** | Rps14 | 0.265569444 | -0.056108425 | 0.587247313 | 17.60409714 | 2.067793982 | 0.088610746 | 0.709264768 | -4.332647552 |
| **A_55_P2270137** | B930036 | 0.370960225 | -0.078411369 | 0.820331819 | 10.53094885 | 2.067626436 | 0.088630465 | 0.709264768 | -4.332689264 |
| **A_55_P2079757** | NA | 0.461078204 | -0.097508072 | 1.019664481 | 9.255375456 | 2.067448389 | 0.088651425 | 0.709264768 | -4.332733592 |
| **A_55_P2049592** | NA | 0.228281136 | -0.048304377 | 0.50486665 | 9.250387671 | 2.067240201 | 0.088675939 | 0.709264768 | -4.332785428 |
| **A_55_P2107398** | NA | 0.336955706 | -0.071332127 | 0.745243539 | 12.17708312 | 2.06707732 | 0.088695124 | 0.709264768 | -4.332825985 |
| **A_51_P234692** | Neat1 | -1.743997926 | -3.858377228 | 0.370381376 | 10.03272662 | -2.065919576 | 0.088831609 | 0.709264768 | -4.333114315 |
| **A_55_P2185148** | Mettl7a3 | 0.280606201 | -0.059604085 | 0.620816487 | 8.77209982 | 2.065856589 | 0.088839041 | 0.709264768 | -4.333130005 |
| **A_52_P67995** | Eif4a3 | 0.318596806 | -0.067674167 | 0.704867779 | 10.56198262 | 2.065854309 | 0.08883931 | 0.709264768 | -4.333130573 |
| **A_55_P2026214** | NA | 0.495780303 | -0.105799629 | 1.097360235 | 12.45611039 | 2.064173861 | 0.089037821 | 0.709264768 | -4.333549258 |
| **A_52_P286098** | Epb41l4b | -0.645231838 | -1.428241937 | 0.137778261 | 11.23649648 | -2.063948095 | 0.089064525 | 0.709264768 | -4.333605523 |
| **A_55_P2033665** | NA | 0.344810859 | -0.073793242 | 0.763414959 | 9.095030109 | 2.063135733 | 0.089160683 | 0.709264768 | -4.333808009 |
| **A_51_P455906** | Pold4 | -0.42530478 | -0.941653416 | 0.091043857 | 9.315150102 | -2.063039649 | 0.089172064 | 0.709264768 | -4.333831962 |
| **A_55_P2326727** | 9430085M | 0.427447312 | -0.0915111 | 0.946405724 | 10.77030499 | 2.063005474 | 0.089176112 | 0.709264768 | -4.333840482 |
| **A_51_P152155** | Glt8d1 | -0.327677191 | -0.725781333 | 0.070426951 | 11.2629344 | -2.061578427 | 0.08934532 | 0.709264768 | -4.334196309 |
| **A_55_P1967533** | Gm5069 | 0.49330821 | -0.106197132 | 1.092813552 | 11.89911018 | 2.060988799 | 0.08941533 | 0.709264768 | -4.334343372 |
| **A_55_P2022114** | NA | 0.828386937 | -0.178780688 | 1.835554563 | 9.327324822 | 2.0600693 | 0.089524621 | 0.709264768 | -4.334572761 |
| **A_55_P1981155** | Phldb1 | -0.472943588 | -1.047980937 | 0.102093761 | 12.11983228 | -2.059982991 | 0.089534887 | 0.709264768 | -4.334594295 |

| | | | | | | | | | |
|---|---|---|---|---|---|---|---|---|---|
| **A_52_P85983** | Mrpl45 | 0.440480016 | -0.095239814 | 0.976199846 | 11.7775838 | 2.059391108 | 0.089605319 | 0.709264768 | -4.334741989 |
| **A_51_P406638** | Rfc1 | 0.567608369 | -0.122911905 | 1.258128643 | 11.35754766 | 2.058840494 | 0.089670891 | 0.709264768 | -4.334879406 |
| **A_55_P2028491** | Hmgn2 | 0.239750781 | -0.05201431 | 0.531515872 | 11.00382788 | 2.058150117 | 0.089753178 | 0.709264768 | -4.335051735 |
| **A_55_P2108784** | Arhgap22 | -0.914371235 | -2.027243919 | 0.198501449 | 10.54680563 | -2.057914932 | 0.089781228 | 0.709264768 | -4.335110448 |
| **A_55_P2424459** | Atg9a | -0.251330117 | -0.557233024 | 0.054572791 | 8.202217469 | -2.05783843 | 0.089790354 | 0.709264768 | -4.335129548 |
| **A_52_P644972** | Mzt1 | 0.199939343 | -0.0434619 | 0.443340585 | 9.316666373 | 2.057433537 | 0.089838671 | 0.709264768 | -4.33523064 |
| **A_51_P295442** | Nek7 | -0.223899751 | -0.496512452 | 0.048712949 | 10.76350347 | -2.057111691 | 0.089877097 | 0.709264768 | -4.335311006 |
| **A_55_P2128229** | NA | 0.279063494 | -0.060765121 | 0.61889211 | 8.76824197 | 2.056806459 | 0.089913555 | 0.709264768 | -4.33538723 |
| **A_51_P445683** | 4931406C | -0.174865397 | -0.387928497 | 0.038197704 | 11.22832332 | -2.055634888 | 0.090053633 | 0.709264768 | -4.335679862 |
| **A_51_P342707** | Pold2 | 0.379313391 | -0.082862345 | 0.841489126 | 8.934370951 | 2.055613111 | 0.090056239 | 0.709264768 | -4.335685302 |
| **A_51_P514712** | Parp14 | -0.846271298 | -1.87748445 | 0.184941854 | 9.449477729 | -2.055471758 | 0.090073155 | 0.709264768 | -4.335720615 |
| **A_55_P2249536** | Nucks1 | 0.47473644 | -0.103801398 | 1.053274278 | 10.65403236 | 2.055280711 | 0.090096024 | 0.709264768 | -4.335768346 |
| **A_55_P1957393** | NA | 0.437897296 | -0.095826186 | 0.971620777 | 13.4997448 | 2.054973845 | 0.090132769 | 0.709264768 | -4.335845018 |
| **A_55_P1972892** | NA | 0.740024274 | -0.161962883 | 1.64201143 | 11.59883524 | 2.054924912 | 0.09013863 | 0.709264768 | -4.335857245 |
| **A_51_P422685** | Zmat4 | -0.355616961 | -0.789150418 | 0.077916497 | 8.313949273 | -2.054519142 | 0.090187245 | 0.709264768 | -4.33595864 |
| **A_55_P2175767** | Rangrf | 0.549079115 | -0.120356079 | 1.218514309 | 9.114313455 | 2.054360849 | 0.090206218 | 0.709264768 | -4.335998198 |
| **A_52_P475040** | Etaa1 | 0.328845478 | -0.072116455 | 0.729807412 | 8.779583465 | 2.054182743 | 0.09022757 | 0.709264768 | -4.336042709 |
| **A_51_P481238** | Dop1b | -0.28335129 | -0.628877148 | 0.062174567 | 10.69957987 | -2.053974212 | 0.090252576 | 0.709264768 | -4.336094827 |
| **A_55_P1952744** | Timm8a1 | 0.643655105 | -0.141235527 | 1.428545736 | 10.76438058 | 2.053971524 | 0.090252899 | 0.709264768 | -4.336095499 |
| **A_66_P106716** | Akap8 | 0.379739787 | -0.083349757 | 0.842829331 | 9.593566018 | 2.053863005 | 0.090265915 | 0.709264768 | -4.336122622 |
| **A_55_P2094060** | Gzma | 2.812449731 | -0.617352127 | 6.242251589 | 9.869177379 | 2.053837519 | 0.090268972 | 0.709264768 | -4.336128992 |
| **A_55_P2068862** | Pigv | -0.320941432 | -0.71238516 | 0.070502296 | 10.25564806 | -2.053557143 | 0.090302611 | 0.709264768 | -4.336199074 |
| **A_52_P13815** | Laptm5 | 0.31842941 | -0.069971911 | 0.706830731 | 8.263775275 | 2.053443793 | 0.090316214 | 0.709264768 | -4.336227408 |
| **A_51_P314264** | Ssr2 | -0.180591365 | -0.400882962 | 0.039700232 | 12.10838254 | -2.053285755 | 0.090335184 | 0.709264768 | -4.336266914 |
| **A_55_P2119430** | Usp10 | 0.386571044 | -0.08499143 | 0.858133518 | 11.25592209 | 2.053243364 | 0.090340273 | 0.709264768 | -4.336277511 |
| **A_55_P2014987** | Castor2 | -0.360875171 | -0.801106562 | 0.079356221 | 10.65061464 | -2.053176761 | 0.090348269 | 0.709264768 | -4.336294161 |
| **A_55_P2184314** | NA | 0.3246289 | -0.071456437 | 0.720714237 | 12.05561959 | 2.052810025 | 0.090392313 | 0.709264768 | -4.336385848 |
| **A_51_P265406** | Ric8a | -0.250008612 | -0.555072599 | 0.055055375 | 12.18988236 | -2.052647494 | 0.090411839 | 0.709264768 | -4.336426484 |
| **A_55_P2022693** | Tcerg1 | 0.341095472 | -0.075213886 | 0.757404829 | 9.458665229 | 2.052154839 | 0.090471052 | 0.709264768 | -4.336549672 |
| **A_55_P2097578** | NA | 0.417001282 | -0.092281509 | 0.926284072 | 11.38600651 | 2.050825621 | 0.090631014 | 0.709264768 | -4.336882125 |
| **A_51_P426458** | Ube2d2a | 0.284773457 | -0.063087631 | 0.632634546 | 11.11565985 | 2.050425534 | 0.090679219 | 0.709264768 | -4.336982216 |
| **A_55_P2154228** | Ncapd3 | 0.609085282 | -0.135408181 | 1.353578746 | 10.13805591 | 2.049120801 | 0.090836604 | 0.709264768 | -4.337308704 |
| **A_55_P1983739** | NA | 0.303277767 | -0.067429733 | 0.673985267 | 10.26560633 | 2.049082986 | 0.09084117 | 0.709264768 | -4.337318168 |
| **A_51_P492830** | Cenph | 0.867128723 | -0.192904272 | 1.927161718 | 9.419749824 | 2.048870616 | 0.090866815 | 0.709264768 | -4.337371322 |
| **A_51_P146753** | Csf2rb2 | -0.469837291 | -1.044304426 | 0.104629844 | 8.91337265 | -2.048484307 | 0.090913484 | 0.709264768 | -4.337468019 |
| **A_55_P2052881** | NA | 0.617643575 | -0.137680165 | 1.372967316 | 9.410972111 | 2.048118774 | 0.090957666 | 0.709264768 | -4.337559526 |
| **A_66_P109800** | Gcsh | 0.321140897 | -0.071716492 | 0.713998287 | 11.59570977 | 2.047439303 | 0.091039853 | 0.709264768 | -4.337729648 |
| **A_55_P2059025** | U2af1 | 0.512998979 | -0.114588681 | 1.14058664 | 13.65641966 | 2.047351686 | 0.091050457 | 0.709264768 | -4.337751587 |
| **A_55_P2164139** | NA | 0.907864803 | -0.202999158 | 2.018728765 | 13.46551624 | 2.046966086 | 0.091097137 | 0.709264768 | -4.337848148 |
| **A_55_P1968381** | Cast | -0.45462228 | -1.010913596 | 0.101669035 | 9.76351384 | -2.046910159 | 0.09110391 | 0.709264768 | -4.337862154 |
| **A_51_P348119** | Ube2q1 | 0.354882008 | -0.079462709 | 0.789226726 | 10.6800432 | 2.04644362 | 0.091160426 | 0.709264768 | -4.337978999 |
| **A_55_P2086423** | Recql4 | 0.407378257 | -0.098710774 | 0.913467289 | 8.419361733 | 2.144043669 | 0.091310659 | 0.709264768 | -4.377517186 |
| **A_55_P2048928** | Cbfb | 0.597324708 | -0.134193813 | 1.328843229 | 12.17253879 | 2.045198657 | 0.091311418 | 0.709264768 | -4.338290877 |
| **A_55_P2093013** | NA | 0.403534071 | -0.090684819 | 0.897752961 | 13.86492567 | 2.045084294 | 0.091325301 | 0.709264768 | -4.338319532 |
| **A_55_P2106429** | Herc2 | -0.314039462 | -0.698680746 | 0.070601822 | 10.14187746 | -2.044931074 | 0.091343905 | 0.709264768 | -4.338357924 |
| **A_55_P2010152** | Sell | 0.615933393 | -0.138518772 | 1.370385558 | 8.189111696 | 2.044807298 | 0.091358936 | 0.709264768 | -4.33838894 |
| **A_51_P324410** | Vti1a | 0.203654775 | -0.045815394 | 0.453124944 | 9.934096142 | 2.044684575 | 0.091373842 | 0.709264768 | -4.338419692 |
| **A_55_P2076580** | Acaa1a | -0.492533021 | -1.095909157 | 0.110843116 | 14.44201242 | -2.04454921 | 0.091390287 | 0.709264768 | -4.338453614 |

| | | | | | | | | | |
|---|---|---|---|---|---|---|---|---|---|
| **A_55_P2114333** | NA | 0.275475161 | -0.062036251 | 0.612986573 | 15.24820557 | 2.044298594 | 0.09142074 | 0.709264768 | -4.338516421 |
| **A_55_P1957876** | NA | -0.282727687 | -0.629210935 | 0.063755562 | 9.985198121 | -2.043790813 | 0.091482475 | 0.709264768 | -4.33864369 |
| **A_51_P360918** | Ehd3 | -0.204498697 | -0.455285772 | 0.046288379 | 9.853149333 | -2.042376193 | 0.091654689 | 0.709264768 | -4.338998341 |
| **A_52_P222073** | Ackr2 | -0.674365438 | -1.501575211 | 0.152844335 | 9.15997182 | -2.041878954 | 0.091715301 | 0.709264768 | -4.339123034 |
| **A_55_P2110245** | Fignl1 | 1.393400604 | -0.316961612 | 3.10376282 | 11.07703799 | 2.040507738 | 0.091882664 | 0.709264768 | -4.339466985 |
| **A_52_P381303** | Gins2 | 0.385802855 | -0.087888174 | 0.859493883 | 8.874207449 | 2.039955167 | 0.091950196 | 0.709264768 | -4.339605627 |
| **A_51_P121891** | Rac2 | 1.387369805 | -0.316247324 | 3.090986933 | 9.794875899 | 2.039720148 | 0.091978935 | 0.709264768 | -4.339664601 |
| **A_51_P511250** | Tent5b | -0.640792939 | -1.427694122 | 0.146108244 | 8.859167976 | -2.039613439 | 0.091991986 | 0.709264768 | -4.339691379 |
| **A_55_P2061084** | Wfikkn2 | 0.356881349 | -0.081473053 | 0.79523575 | 8.145063306 | 2.039148366 | 0.092048892 | 0.709264768 | -4.339808095 |
| **A_55_P2007656** | Cryab | -1.167990935 | -2.60299406 | 0.267012189 | 10.47110654 | -2.038623241 | 0.092113188 | 0.709264768 | -4.3399399 |
| **A_55_P2015964** | Rif1 | 0.509920705 | -0.116584441 | 1.136425851 | 9.264561091 | 2.038582771 | 0.092118146 | 0.709264768 | -4.339950059 |
| **A_55_P2155942** | Ezh1 | -0.662663404 | -1.476863119 | 0.15153631 | 9.610363833 | -2.038507845 | 0.092127324 | 0.709264768 | -4.339968867 |
| **A_52_P59579** | Tmbim6 | -0.362799479 | -0.808656698 | 0.083057739 | 12.13552037 | -2.038079867 | 0.092179768 | 0.709264768 | -4.340076307 |
| **A_55_P1964477** | Srsf1 | 0.770782472 | -0.176820814 | 1.718385758 | 10.12721974 | 2.037302966 | 0.092275049 | 0.709264768 | -4.340271372 |
| **A_51_P117236** | Zfp354a | -0.239273883 | -0.53344237 | 0.054894605 | 8.853832804 | -2.037274242 | 0.092278574 | 0.709264768 | -4.362729725 |
| **A_55_P1952618** | Ear2 | -0.530113915 | -1.181946833 | 0.121719003 | 11.06515696 | -2.036963564 | 0.092316705 | 0.709264768 | -4.340356603 |
| **A_51_P172054** | Gas6 | 0.852692164 | -0.19616355 | 1.901547877 | 12.79695306 | 2.036230175 | 0.092406784 | 0.709264768 | -4.340540799 |
| **A_55_P2169218** | NA | 0.398743674 | -0.091753987 | 0.889241335 | 10.13317954 | 2.036138009 | 0.092418111 | 0.709264768 | -4.34056395 |
| **A_52_P125350** | Anp32e | 0.804966781 | -0.185806564 | 1.795740126 | 9.931943963 | 2.034951157 | 0.092564097 | 0.709264768 | -4.340862125 |
| **A_55_P2138529** | Mrpl40 | 0.245829336 | -0.056761849 | 0.548420522 | 11.50527176 | 2.034828359 | 0.092579215 | 0.709264768 | -4.340892981 |
| **A_52_P430274** | Dnaaf5 | 0.215654909 | -0.049931457 | 0.481241275 | 10.2574651 | 2.033779745 | 0.092708418 | 0.709264768 | -4.341156516 |
| **A_55_P2057913** | Hmgb1-ps | 1.255233612 | -0.291099512 | 2.801566736 | 13.29609976 | 2.033161357 | 0.092784698 | 0.709264768 | -4.341311964 |
| **A_51_P236175** | Rit1 | -0.247062505 | -0.55592144 | 0.061796429 | 13.21910456 | -2.130636572 | 0.09278566 | 0.709264768 | -4.379860349 |
| **A_51_P226417** | Rraga | 0.347395887 | -0.080633928 | 0.775425702 | 11.5260874 | 2.032829699 | 0.092825635 | 0.709264768 | -4.341395346 |
| **A_66_P135106** | Slco3a1 | -0.468285036 | -1.045494729 | 0.108924658 | 9.830093993 | -2.032015469 | 0.092926217 | 0.709264768 | -4.341600083 |
| **A_55_P2007273** | Pole | 0.78231313 | -0.18200515 | 1.746631411 | 10.16693018 | 2.03193853 | 0.092935727 | 0.709264768 | -4.341619431 |
| **A_51_P101573** | Klc4 | -0.337527476 | -0.753583308 | 0.078528355 | 10.57343569 | -2.03192589 | 0.09293729 | 0.709264768 | -4.34162261 |
| **A_55_P1976017** | Cnih4 | -0.360003912 | -0.803817924 | 0.083810099 | 12.60096613 | -2.031685853 | 0.092966967 | 0.709264768 | -4.341682977 |
| **A_52_P254149** | Irgq | -0.2231286 | -0.498260656 | 0.052003456 | 9.41176465 | -2.031254742 | 0.093020291 | 0.709264768 | -4.341791408 |
| **A_55_P2081137** | NA | 0.473860406 | -0.11053645 | 1.058257261 | 10.3319101 | 2.030920376 | 0.093061671 | 0.709264768 | -4.341875515 |
| **A_52_P186937** | Cmpk2 | -0.327083243 | -0.730469868 | 0.076303381 | 8.786409572 | -2.03089349 | 0.093065 | 0.709264768 | -4.341882279 |
| **A_55_P2001068** | NA | 0.51383089 | -0.119872029 | 1.147533808 | 11.04192953 | 2.030882733 | 0.093066331 | 0.709264768 | -4.341884984 |
| **A_51_P434101** | Herpud1 | -0.375924737 | -0.839587861 | 0.087738386 | 12.44964778 | -2.030713677 | 0.093087261 | 0.709264768 | -4.341927512 |
| **A_55_P2177639** | NA | -0.171358243 | -0.382755351 | 0.040038866 | 16.02327252 | -2.030281714 | 0.093140763 | 0.709264768 | -4.342036187 |
| **A_52_P488612** | Btbd9 | 0.227940586 | -0.053284653 | 0.509165825 | 9.199547274 | 2.03010131 | 0.093163116 | 0.709264768 | -4.342081577 |
| **A_65_P06061** | Rpl37 | 0.363372433 | -0.084944737 | 0.811689603 | 17.51220008 | 2.03009775 | 0.093163557 | 0.709264768 | -4.342082473 |
| **A_55_P2113912** | NA | 0.682495677 | -0.159794586 | 1.524785941 | 14.50279266 | 2.029497264 | 0.093238003 | 0.709264768 | -4.342233574 |
| **A_51_P344878** | Thop1 | 0.203751679 | -0.047732023 | 0.45523538 | 8.749901846 | 2.029278698 | 0.093265115 | 0.709264768 | -4.342288578 |
| **A_55_P2079713** | Gramd1c | -0.315054156 | -0.703995939 | 0.073887626 | 10.04415678 | -2.028854763 | 0.093317725 | 0.709264768 | -4.342395274 |
| **A_66_P111217** | NA | 0.425503499 | -0.099944208 | 0.950951207 | 13.93032094 | 2.028261654 | 0.093391381 | 0.709264768 | -4.34254457 |
| **A_55_P2097478** | Gda | -0.796997269 | -1.781325068 | 0.18733053 | 9.111771754 | -2.027997561 | 0.093424197 | 0.709264768 | -4.342611054 |
| **A_55_P1992617** | Clec1b | -0.748086908 | -1.672024044 | 0.175850228 | 8.818339544 | -2.027962765 | 0.093428521 | 0.709264768 | -4.342619814 |
| **A_55_P2072776** | Fpgs | 0.246690215 | -0.057990746 | 0.551371175 | 10.28565231 | 2.027948644 | 0.093430276 | 0.709264768 | -4.34262337 |
| **A_55_P2051229** | Gpr132 | 1.0468716 | -0.246231988 | 2.339975188 | 8.463295911 | 2.027731497 | 0.093457269 | 0.709264768 | -4.34267804 |
| **A_55_P1978471** | NA | 0.403516723 | -0.095052881 | 0.902086327 | 10.75451736 | 2.027150934 | 0.093529477 | 0.709264768 | -4.342824222 |
| **A_55_P2168321** | NA | 0.503263437 | -0.118637114 | 1.125163988 | 8.547270238 | 2.026864813 | 0.093565085 | 0.709264768 | -4.342896275 |
| **A_55_P2021721** | NA | 1.004985465 | -0.236983858 | 2.246954789 | 10.77734817 | 2.026745686 | 0.093579915 | 0.709264768 | -4.342926275 |
| **A_55_P2020612** | Arl4d | -0.556125931 | -1.243487726 | 0.131235863 | 9.421567427 | -2.026460313 | 0.093615449 | 0.709264768 | -4.342998147 |

| | | | | | | | | | |
|---|---|---|---|---|---|---|---|---|---|
| **A_55_P1974412** | Kat2a | 0.346168128 | -0.081758589 | 0.774094846 | 10.05669481 | 2.02613334 | 0.09365618 | 0.709264768 | -4.343080503 |
| **A_55_P1959550** | Carmil2 | 0.543074211 | -0.128332156 | 1.214480578 | 8.596091064 | 2.025928296 | 0.093681732 | 0.709264768 | -4.343132152 |
| **A_51_P346453** | Noc4l | 0.276620384 | -0.065404038 | 0.618644806 | 9.22974617 | 2.02571063 | 0.093708865 | 0.709264768 | -4.343186983 |
| **A_51_P367125** | Pold1 | 0.685923715 | -0.162317916 | 1.534165346 | 10.60803958 | 2.025380296 | 0.093750058 | 0.709264768 | -4.343270203 |
| **A_55_P2129925** | Mybl2 | 0.277098157 | -0.065598936 | 0.61979525 | 9.122176599 | 2.025226315 | 0.093769265 | 0.709264768 | -4.343308997 |
| **A_55_P1998651** | Entpd5 | 0.29504989 | -0.0698771 | 0.659976881 | 10.00869615 | 2.025068966 | 0.093788897 | 0.709264768 | -4.343348642 |
| **A_55_P1966583** | Bpifc | 0.332800437 | -0.078845385 | 0.74444626 | 12.74544363 | 2.024932362 | 0.093805944 | 0.709264768 | -4.343383061 |
| **A_55_P2045812** | Sigmar1 | 0.232050699 | -0.055010524 | 0.519111921 | 11.84850414 | 2.024690738 | 0.093836105 | 0.709264768 | -4.343443944 |
| **A_52_P51429** | Dennd1c | 0.261412955 | -0.062100211 | 0.58492612 | 8.196477124 | 2.02388342 | 0.093936951 | 0.709264768 | -4.343647398 |
| **A_55_P2175272** | NA | 0.352116237 | -0.083823836 | 0.788056311 | 14.05593946 | 2.023063922 | 0.094039432 | 0.709264768 | -4.343853968 |
| **A_51_P519837** | Cherp | 0.254867221 | -0.060942837 | 0.57067728 | 11.4801621 | 2.02133535 | 0.094255974 | 0.709264768 | -4.344289838 |
| **A_51_P208680** | Chtf18 | 0.756330795 | -0.180977067 | 1.693638657 | 9.677977807 | 2.021063084 | 0.094290128 | 0.709264768 | -4.34435851 |
| **A_55_P2036633** | Ssrp1 | 0.517074027 | -0.123767667 | 1.157915721 | 13.71105649 | 2.020934833 | 0.094306221 | 0.709264768 | -4.34439086 |
| **A_55_P2128734** | Pbxip1 | -0.692455048 | -1.550846893 | 0.165936796 | 12.5027766 | -2.02048834 | 0.094362268 | 0.709264768 | -4.344503492 |
| **A_55_P2179834** | Gatad2a | 0.333377319 | -0.079957252 | 0.746711889 | 11.48304499 | 2.020154866 | 0.09440415 | 0.709264768 | -4.344587622 |
| **A_52_P203691** | Arl5c | 1.172446623 | -0.281450036 | 2.626343281 | 8.700527318 | 2.019807065 | 0.094447853 | 0.709264768 | -4.344675375 |
| **A_51_P447866** | Sash3 | 1.067711583 | -0.256675035 | 2.392098202 | 8.739013498 | 2.019247324 | 0.09451823 | 0.709264768 | -4.344816621 |
| **A_55_P1964408** | Cryab | -1.44674228 | -3.241370978 | 0.347886418 | 13.84437962 | -2.019142225 | 0.09453145 | 0.709264768 | -4.344843144 |
| **A_55_P2221365** | NA | -0.231764945 | -0.519268072 | 0.055738183 | 8.530333935 | -2.019089265 | 0.094538112 | 0.709264768 | -4.344856509 |
| **A_55_P2071191** | Prss8 | -0.296186202 | -0.663631089 | 0.071258686 | 10.53457103 | -2.01893789 | 0.094557158 | 0.709264768 | -4.344894712 |
| **A_55_P2024853** | Cep57l1 | 0.177121989 | -0.042689609 | 0.396933587 | 8.717089499 | 2.018237275 | 0.094645361 | 0.709264768 | -4.345071551 |
| **A_55_P2133680** | NA | 0.513342948 | -0.123774425 | 1.15046032 | 10.89346975 | 2.018080558 | 0.094665102 | 0.709264768 | -4.345111111 |
| **A_51_P207591** | Anxa8 | -0.558196294 | -1.251336806 | 0.134944218 | 11.75382077 | -2.017046969 | 0.094795406 | 0.709264768 | -4.345372067 |
| **A_55_P1985410** | Reps2 | -0.789952864 | -1.771003825 | 0.191098096 | 9.264166265 | -2.016786643 | 0.094828254 | 0.709264768 | -4.345437804 |
| **A_55_P2103412** | Golt1a | 0.403345544 | -0.097643802 | 0.904334889 | 9.423631436 | 2.016504144 | 0.094863914 | 0.709264768 | -4.345509146 |
| **A_52_P652913** | Akip1 | -0.382583972 | -0.857892397 | 0.092724454 | 10.04697565 | -2.016051416 | 0.09492109 | 0.709264768 | -4.345623488 |
| **A_51_P148105** | Rad51 | 1.003932622 | -0.2435623 | 2.251427545 | 9.476910534 | 2.015654654 | 0.094971227 | 0.709264768 | -4.345723707 |
| **A_55_P2066041** | Hnrnpa1 | 0.551016649 | -0.133762322 | 1.235795621 | 9.076388065 | 2.015415766 | 0.095001427 | 0.709264768 | -4.345784053 |
| **A_66_P126080** | Rpl39 | 0.728265062 | -0.176863202 | 1.633393326 | 9.467506122 | 2.015253548 | 0.09502194 | 0.709264768 | -4.345825034 |
| **A_55_P2056192** | NA | 0.993637662 | -0.241311524 | 2.228586848 | 12.98504966 | 2.015251725 | 0.095022171 | 0.709264768 | -4.345825494 |
| **A_51_P452153** | Mzb1 | 0.926392944 | -0.225180446 | 2.077966335 | 8.257906454 | 2.014902164 | 0.09506639 | 0.709264768 | -4.367325246 |
| **A_55_P2077033** | Ube2e1 | 0.225981546 | -0.054981399 | 0.506944491 | 12.71809868 | 2.014532486 | 0.095113177 | 0.709264768 | -4.346007216 |
| **A_51_P202745** | Zc3h13 | 0.178378759 | -0.043403583 | 0.400161101 | 10.45695324 | 2.014496509 | 0.095117731 | 0.709264768 | -4.346016306 |
| **A_51_P516362** | Vps11 | -0.400357652 | -0.89813832 | 0.097423016 | 10.18677775 | -2.014468372 | 0.095121294 | 0.709264768 | -4.346023416 |
| **A_51_P519756** | Rusc1 | -0.431703159 | -0.968626991 | 0.105220674 | 10.69772908 | -2.013830391 | 0.0952021 | 0.709264768 | -4.34618464 |
| **A_51_P436201** | Gart | 0.199622218 | -0.048730273 | 0.447974708 | 8.969703614 | 2.013217527 | 0.095279792 | 0.709264768 | -4.346339542 |
| **A_52_P5891** | Rflnb | -0.509705295 | -1.143870702 | 0.124460112 | 8.774157138 | -2.013107389 | 0.095293761 | 0.709264768 | -4.346367382 |
| **A_55_P1970399** | NA | 0.357030991 | -0.08718238 | 0.801244362 | 14.41608645 | 2.013096686 | 0.095295119 | 0.709264768 | -4.346370088 |
| **A_55_P2119032** | Rasl2-9 | 0.862633146 | -0.210691558 | 1.93595785 | 11.10321969 | 2.013007356 | 0.09530645 | 0.709264768 | -4.346392669 |
| **A_55_P2087404** | Mapre3 | -0.304322833 | -0.683254834 | 0.074609169 | 10.43064259 | -2.0115165 | 0.095495771 | 0.709264768 | -4.346769613 |
| **A_51_P383194** | Pde9a | -0.176438124 | -0.396177333 | 0.043301085 | 8.945568349 | -2.011107197 | 0.095547815 | 0.709264768 | -4.346873127 |
| **A_51_P320852** | Cd9 | -0.563035423 | -1.264439515 | 0.138368668 | 14.46023554 | -2.010563389 | 0.095617006 | 0.709264768 | -4.347010675 |
| **A_55_P2097729** | Gtpbp4 | 0.255266631 | -0.06282783 | 0.573361091 | 9.064148785 | 2.009964031 | 0.095693326 | 0.709264768 | -4.347162296 |
| **A_52_P73369** | Hnrnpu | 0.619673374 | -0.152524522 | 1.39187127 | 12.43309954 | 2.009946823 | 0.095695518 | 0.709264768 | -4.34716665 |
| **A_55_P1989076** | Slc22a23 | -0.317807899 | -0.713878888 | 0.078263089 | 10.40978922 | -2.009749831 | 0.095720616 | 0.709264768 | -4.347216489 |
| **A_55_P2083764** | NA | 0.546336064 | -0.14514421 | 1.237816337 | 8.586635427 | 2.104470196 | 0.095739169 | 0.709264768 | -4.384470795 |
| **A_55_P2122898** | Rnf25 | 0.525573702 | -0.129733481 | 1.180880885 | 10.67654879 | 2.008810842 | 0.095840343 | 0.709264768 | -4.347454092 |
| **A_55_P2084496** | Ssbp3 | 0.590729901 | -0.14582663 | 1.327286433 | 11.16510634 | 2.008783881 | 0.095843783 | 0.709264768 | -4.347460915 |

| | | | | | | | | | |
|---|---|---|---|---|---|---|---|---|---|
| **A_55_P2025153** | Hmgb1 | 0.921302258 | -0.227657549 | 2.070262064 | 15.09445708 | 2.008388125 | 0.095894293 | 0.709264768 | -4.347561076 |
| **A_66_P117543** | NA | 0.195182484 | -0.04823578 | 0.438600747 | 8.248208675 | 2.008343643 | 0.095899972 | 0.709264768 | -4.347572334 |
| **A_55_P2016287** | Tpst1 | 0.273720663 | -0.067892011 | 0.615333338 | 8.407330853 | 2.006891778 | 0.096085516 | 0.709264768 | -4.34793988 |
| **A_52_P72587** | Prkcq | 0.506961602 | -0.125844628 | 1.139767832 | 8.176259645 | 2.006571622 | 0.09612648 | 0.709264768 | -4.348020947 |
| **A_55_P2183338** | Marveld3 | -0.464750368 | -1.0448928 | 0.115392064 | 8.813950223 | -2.006482873 | 0.096137839 | 0.709264768 | -4.348043421 |
| **A_55_P2131379** | Ybx1 | 0.983919823 | -0.244822053 | 2.212661699 | 11.74879731 | 2.00562344 | 0.096247907 | 0.709264768 | -4.348261081 |
| **A_52_P32864** | Prps2 | 0.618536728 | -0.153951067 | 1.391024523 | 8.731598297 | 2.005507138 | 0.096262812 | 0.709264768 | -4.34829054 |
| **A_51_P228193** | Ociad1 | 0.196924837 | -0.049028833 | 0.442878506 | 11.167056 | 2.005383945 | 0.096278603 | 0.709264768 | -4.348321745 |
| **A_55_P1982334** | Llph | 0.244809954 | -0.060967799 | 0.550587707 | 11.08466949 | 2.005273121 | 0.09629281 | 0.709264768 | -4.348349817 |
| **A_51_P367310** | Chaf1b | 1.213460968 | -0.30233985 | 2.729261786 | 11.09805493 | 2.005090687 | 0.096316202 | 0.709264768 | -4.348396031 |
| **A_55_P1982211** | NA | 0.985113681 | -0.246296204 | 2.216523566 | 10.470853 | 2.003706286 | 0.096493905 | 0.709264768 | -4.3487468 |
| **A_52_P97670** | Atad2b | 0.197584298 | -0.049404642 | 0.444573238 | 9.836133828 | 2.003665717 | 0.096499118 | 0.709264768 | -4.348757081 |
| **A_55_P2052081** | Eif1ad | 0.227917382 | -0.057007345 | 0.512842109 | 11.88284273 | 2.003538319 | 0.096515488 | 0.709264768 | -4.348789367 |
| **A_55_P1995195** | Fosl2 | -0.616389023 | -1.38709171 | 0.154313664 | 10.68182913 | -2.003172589 | 0.096562501 | 0.709264768 | -4.348882058 |
| **A_66_P117022** | Atp6v1e1 | -0.383896458 | -0.863904204 | 0.096111288 | 11.97628007 | -2.003162611 | 0.096563784 | 0.709264768 | -4.348884587 |
| **A_55_P2008907** | Actg1 | -0.448921564 | -1.010393213 | 0.112550085 | 15.11991722 | -2.002594324 | 0.096636881 | 0.709264768 | -4.349028634 |
| **A_55_P2404434** | NA | 0.452124968 | -0.113566904 | 1.01781684 | 8.670514901 | 2.001837857 | 0.096734272 | 0.709264768 | -4.349220413 |
| **A_66_P104340** | Atxn7l2 | 0.191444387 | -0.04809421 | 0.430982984 | 9.181800386 | 2.001785238 | 0.09674105 | 0.709264768 | -4.349233754 |
| **A_52_P486253** | Polr2b | 0.249538306 | -0.06275294 | 0.561829552 | 11.39335902 | 2.001371718 | 0.096794335 | 0.709264768 | -4.349338608 |
| **A_66_P104422** | Ckap4 | 0.500899199 | -0.126415595 | 1.128213993 | 11.22028102 | 1.99993165 | 0.096980132 | 0.709264768 | -4.349703846 |
| **A_55_P1963878** | NA | -0.287286374 | -0.647126636 | 0.072553889 | 11.81907224 | -1.999657509 | 0.097015544 | 0.709264768 | -4.34977339 |
| **A_51_P115655** | Iqcd | 0.262028978 | -0.066182957 | 0.590240912 | 9.94477331 | 1.999609998 | 0.097021682 | 0.709264768 | -4.349785444 |
| **A_51_P148675** | Dag1 | -0.298099167 | -0.671584923 | 0.075386588 | 11.96313196 | -1.999111723 | 0.097086083 | 0.709264768 | -4.349911862 |
| **A_55_P2159390** | NA | 0.626387736 | -0.158531197 | 1.411306668 | 11.32771761 | 1.99879745 | 0.097126725 | 0.709264768 | -4.349991606 |
| **A_51_P439612** | Dnajb2 | -0.622495803 | -1.403005179 | 0.158013573 | 9.794763718 | -1.997600549 | 0.097281667 | 0.709264768 | -4.350295368 |
| **A_55_P2128451** | Sfpq | 0.909186969 | -0.23100462 | 2.049378559 | 12.52061978 | 1.99721912 | 0.097331098 | 0.709264768 | -4.350392191 |
| **A_55_P2158653** | Cldn7 | 0.408253272 | -0.103809138 | 0.920315683 | 11.81483116 | 1.996903711 | 0.097371992 | 0.709264768 | -4.350472264 |
| **A_55_P1998411** | Chchd3 | 0.445457598 | -0.113290598 | 1.004205793 | 11.53583486 | 1.996827655 | 0.097381856 | 0.709264768 | -4.350491573 |
| **A_51_P332355** | Kif18a | 0.585164123 | -0.160506578 | 1.330834824 | 8.573599467 | 2.090226346 | 0.097389475 | 0.709264768 | -4.387001204 |
| **A_55_P2130930** | Nhp2 | 0.439726944 | -0.111894321 | 0.991348208 | 13.63511083 | 1.996606237 | 0.097410577 | 0.709264768 | -4.350547788 |
| **A_55_P2007535** | Edf1 | -0.325930067 | -0.734851606 | 0.082991472 | 14.9030214 | -1.996341271 | 0.097444959 | 0.709264768 | -4.350615065 |
| **A_55_P2105271** | Pdlim7 | -0.379816467 | -0.85648596 | 0.096853025 | 12.26596769 | -1.995752944 | 0.097521344 | 0.709264768 | -4.350764462 |
| **A_55_P2025840** | Spred2 | 0.237786063 | -0.06065488 | 0.536227007 | 8.920931435 | 1.995622254 | 0.09753832 | 0.709264768 | -4.350797652 |
| **A_55_P1995007** | Arhgap11a | 0.687145835 | -0.175417652 | 1.549709322 | 9.065899783 | 1.995299961 | 0.097580199 | 0.709264768 | -4.350879506 |
| **A_55_P2081105** | NA | -0.304255929 | -0.686197153 | 0.077685295 | 8.384965093 | -1.995229511 | 0.097589355 | 0.709264768 | -4.3508974 |
| **A_52_P322141** | Ccdc88b | 1.027945742 | -0.262518185 | 2.318409668 | 10.213028 | 1.995145939 | 0.097600219 | 0.709264768 | -4.350918626 |
| **A_51_P150876** | Lypd3 | -0.428837962 | -0.967253026 | 0.109577102 | 7.994604159 | -1.994924051 | 0.097629067 | 0.709264768 | -4.350974987 |
| **A_51_P268234** | Il34 | -0.568146142 | -1.281682617 | 0.145390333 | 9.084560392 | -1.994317173 | 0.097708015 | 0.709264768 | -4.351129153 |
| **A_51_P115178** | Scara3 | 0.531343366 | -0.13607644 | 1.198763172 | 9.061545385 | 1.994006263 | 0.097748485 | 0.709264768 | -4.351208143 |
| **A_55_P2053266** | St6galnac | 0.915461443 | -0.234619533 | 2.065542419 | 11.04428198 | 1.993709983 | 0.097787068 | 0.709264768 | -4.351283423 |
| **A_51_P501844** | Cyp26b1 | -1.106956359 | -2.497715735 | 0.283803017 | 10.75066801 | -1.993557877 | 0.097806882 | 0.709264768 | -4.351322072 |
| **A_51_P424256** | Gm6211 | 0.804889816 | -0.206365957 | 1.816145589 | 12.68929946 | 1.993543718 | 0.097808727 | 0.709264768 | -4.35132567 |
| **A_55_P2006869** | Fyb | 0.511754963 | -0.131296698 | 1.154806623 | 8.269494342 | 1.993271901 | 0.097844145 | 0.709264768 | -4.351394742 |
| **A_66_P131530** | Rpp30 | 0.318311337 | -0.081766209 | 0.718388884 | 8.461531866 | 1.99277504 | 0.097908922 | 0.709264768 | -4.351521013 |
| **A_55_P1980326** | Pcbp1 | 0.194800503 | -0.050055003 | 0.439656008 | 14.95162232 | 1.992647805 | 0.097925517 | 0.709264768 | -4.351553351 |
| **A_51_P141926** | Fxyd4 | -0.305863132 | -0.690369494 | 0.07864323 | 9.566119319 | -1.992388135 | 0.097959394 | 0.709264768 | -4.351619352 |
| **A_51_P399614** | Zkscan14 | -0.281916816 | -0.636390214 | 0.072556581 | 8.754010972 | -1.991992415 | 0.098011044 | 0.709264768 | -4.351719942 |
| **A_55_P2062598** | E2f7 | 0.394262861 | -0.101525536 | 0.890051259 | 10.45036554 | 1.99177307 | 0.098039685 | 0.709264768 | -4.351775702 |

| | | | | | | | | | |
|---|---|---|---|---|---|---|---|---|---|
| **A_66_P100365** | NA | 0.189922961 | -0.04892388 | 0.428769802 | 8.318023749 | 1.991628429 | 0.098058576 | 0.709264768 | -4.351812474 |
| **A_55_P2082185** | NA | 0.501371393 | -0.129329029 | 1.132071815 | 13.50072338 | 1.991071125 | 0.098131399 | 0.709264768 | -4.351954168 |
| **A_55_P1959994** | Unc5cl | 0.17471826 | -0.045083943 | 0.394520464 | 7.98373331 | 1.990932792 | 0.098149484 | 0.709264768 | -4.351989343 |
| **A_51_P483159** | Gchfr | -0.920816058 | -2.079515325 | 0.237883209 | 10.51107097 | -1.990455616 | 0.098211892 | 0.709264768 | -4.352110686 |
| **A_55_P2105002** | Pcna | 1.053935894 | -0.272488384 | 2.380360172 | 13.5709458 | 1.990132898 | 0.098254123 | 0.709264768 | -4.35219276 |
| **A_55_P1972182** | Siah1b | 0.579714055 | -0.149883796 | 1.309311905 | 10.29559724 | 1.990126256 | 0.098254992 | 0.709264768 | -4.352194449 |
| **A_65_P12359** | Pin1rt1 | 0.301671047 | -0.078063179 | 0.681405273 | 11.1049008 | 1.989776209 | 0.09830082 | 0.709264768 | -4.352283481 |
| **A_52_P391983** | Nt5c2 | 0.187421915 | -0.048570135 | 0.423413966 | 8.852115694 | 1.989176613 | 0.098379371 | 0.709264768 | -4.352436003 |
| **A_51_P368285** | Rflna | -0.537162752 | -1.213875342 | 0.139549839 | 8.995540596 | -1.988162962 | 0.098512311 | 0.709264768 | -4.352693906 |
| **A_51_P163233** | Khdc4 | -0.193509715 | -0.437302051 | 0.05028262 | 9.1396288 | -1.988076407 | 0.098523671 | 0.709264768 | -4.352715931 |
| **A_52_P89477** | Bcl9 | 0.325319059 | -0.08456415 | 0.735202268 | 8.958287522 | 1.987923763 | 0.098543708 | 0.709264768 | -4.352754775 |
| **A_55_P2024649** | Rpe | 0.408747563 | -0.106253383 | 0.923748508 | 9.804824487 | 1.987913554 | 0.098545048 | 0.709264768 | -4.352757373 |
| **A_55_P1979973** | Nuak2 | 0.341061538 | -0.088766247 | 0.770889324 | 8.711391002 | 1.987415194 | 0.098610498 | 0.709264768 | -4.352884203 |
| **A_55_P2091681** | Nudt1 | 0.499343026 | -0.130128592 | 1.128814643 | 10.5187267 | 1.986887061 | 0.098679906 | 0.709264768 | -4.353018629 |
| **A_55_P2209767** | 2610002J | 0.307383174 | -0.08015709 | 0.694923438 | 10.3242643 | 1.986614522 | 0.098715743 | 0.709264768 | -4.353088006 |
| **A_55_P2186395** | NA | 0.682329572 | -0.178105117 | 1.542764261 | 9.007664938 | 1.986216697 | 0.098768079 | 0.709264768 | -4.353189283 |
| **A_55_P1969650** | Rasgrp1 | 1.000184889 | -0.2612652 | 2.261634977 | 8.50021347 | 1.985914573 | 0.098807843 | 0.709264768 | -4.353266205 |
| **A_55_P2052172** | NA | 0.525594751 | -0.137356208 | 1.188545711 | 11.15752797 | 1.985728923 | 0.098832286 | 0.709264768 | -4.353313475 |
| **A_55_P2133632** | Sipa1l2 | -0.39571942 | -0.894854639 | 0.1034158 | 12.51450056 | -1.985726786 | 0.098832568 | 0.709264768 | -4.353314019 |
| **A_55_P2062727** | Nasp | 0.893629381 | -0.233660593 | 2.020919356 | 10.38453158 | 1.985510282 | 0.09886108 | 0.709264768 | -4.353369148 |
| **A_55_P2065754** | Krtap19-5 | 0.248358953 | -0.064958753 | 0.561676659 | 8.942616292 | 1.985387246 | 0.098877288 | 0.709264768 | -4.353400478 |
| **A_52_P144285** | Fam72a | 0.391874824 | -0.102500452 | 0.886250099 | 8.605799536 | 1.98536776 | 0.098879855 | 0.709264768 | -4.35340544 |
| **A_55_P2140118** | Qpct | -0.397726993 | -0.89951688 | 0.104062894 | 10.96034003 | -1.985242244 | 0.098896392 | 0.709264768 | -4.353437403 |
| **A_52_P660032** | Cpsf2 | 0.369977281 | -0.096818009 | 0.836772571 | 9.78760633 | 1.985175637 | 0.098905169 | 0.709264768 | -4.353454365 |
| **A_55_P2078460** | NA | 0.874197927 | -0.229144647 | 1.977540501 | 10.12963129 | 1.984493777 | 0.098995064 | 0.709264768 | -4.353628024 |
| **A_66_P132888** | Dhfr | 0.391923868 | -0.10289481 | 0.886742546 | 8.773518696 | 1.983836944 | 0.099081739 | 0.709264768 | -4.353795338 |
| **A_55_P2043723** | Rabac1 | -0.485434012 | -1.098671845 | 0.127803822 | 12.20271019 | -1.982675345 | 0.099235212 | 0.709264768 | -4.3540913 |
| **A_51_P379976** | Arl4a | -0.176750804 | -0.400038708 | 0.0465371 | 10.16774498 | -1.982652245 | 0.099238267 | 0.709264768 | -4.354097187 |
| **A_55_P2020831** | NA | -0.249779435 | -0.56535192 | 0.06579305 | 12.53963541 | -1.982475799 | 0.099261601 | 0.709264768 | -4.354142151 |
| **A_55_P2146177** | Cerk | 0.210702017 | -0.055537025 | 0.476941058 | 12.16243357 | 1.982199204 | 0.099298192 | 0.709264768 | -4.354212641 |
| **A_55_P2115003** | Prpf4 | 0.167764583 | -0.044221301 | 0.379750466 | 8.71336849 | 1.982182716 | 0.099300373 | 0.709264768 | -4.354216844 |
| **A_51_P321531** | Purg | -0.180036677 | -0.407536479 | 0.047463124 | 10.61250632 | -1.982121681 | 0.099308449 | 0.709264768 | -4.354232399 |
| **A_55_P2165719** | Pcyt1a | -0.326937463 | -0.740082928 | 0.086208001 | 12.08173213 | -1.98203831 | 0.099319482 | 0.709264768 | -4.354253648 |
| **A_55_P2122830** | Zfp185 | -0.552761449 | -1.251435159 | 0.145912261 | 9.826013793 | -1.981589492 | 0.099378898 | 0.709264768 | -4.354368044 |
| **A_55_P2014259** | Zfp82 | -0.244301677 | -0.553228157 | 0.064624803 | 9.500962927 | -1.980713484 | 0.099494972 | 0.709264768 | -4.354591363 |
| **A_55_P2120615** | Ssh3 | -0.513724333 | -1.163494213 | 0.136045547 | 9.991666702 | -1.980254035 | 0.099555905 | 0.709264768 | -4.35470851 |
| **A_55_P1987514** | NA | 0.322697489 | -0.085554191 | 0.730949169 | 16.84728504 | 1.979784642 | 0.099618197 | 0.709264768 | -4.354828207 |
| **A_55_P1994539** | Cldn12 | -0.256326529 | -0.580644566 | 0.067991508 | 10.1669405 | -1.979578824 | 0.099645523 | 0.709264768 | -4.354880695 |
| **A_51_P193794** | Lrp1 | -0.700468944 | -1.5867771 | 0.185839212 | 11.28812822 | -1.979495347 | 0.099656609 | 0.709264768 | -4.354901985 |
| **A_51_P167360** | Ptpn7 | 0.66668917 | -0.177087464 | 1.510465804 | 8.199463114 | 1.979002083 | 0.099722138 | 0.709264768 | -4.355027794 |
| **A_51_P324303** | Mylip | -0.489083156 | -1.108115638 | 0.129949326 | 9.593545838 | -1.978880705 | 0.099738269 | 0.709264768 | -4.355058754 |
| **A_55_P2030506** | Rp9 | 0.43590033 | -0.115860702 | 0.987661363 | 11.97480653 | 1.978729907 | 0.099758314 | 0.709264768 | -4.35509722 |
| **A_52_P68714** | NA | 0.237197045 | -0.06324542 | 0.53763951 | 8.836256262 | 1.977417184 | 0.099932986 | 0.709264768 | -4.355432137 |
| **A_55_P1963712** | Cyb5b | 0.171463321 | -0.04572524 | 0.388651883 | 11.40880731 | 1.977354916 | 0.099941279 | 0.709264768 | -4.355448026 |
| **A_55_P1956734** | NA | 0.865500638 | -0.230823844 | 1.96182512 | 9.872103089 | 1.977327587 | 0.099944919 | 0.709264768 | -4.355455 |
| **A_52_P468068** | Tchh | -0.283588534 | -0.642817769 | 0.075640701 | 10.31559376 | -1.977276204 | 0.099951763 | 0.709264768 | -4.355468112 |
| **A_55_P1983819** | Srsf7 | 0.725579285 | -0.193547672 | 1.644706242 | 12.46469542 | 1.977241499 | 0.099956386 | 0.709264768 | -4.355476968 |
| **A_55_P2033530** | Gm6981 | 0.518278784 | -0.138277299 | 1.174834867 | 10.62541415 | 1.977160599 | 0.099967164 | 0.709264768 | -4.355497613 |

| | | | | | | | | | |
|---|---|---|---|---|---|---|---|---|---|
| **A_55_P1991911** | NA | 1.053851925 | -0.281172447 | 2.388876297 | 14.05625314 | 1.977155123 | 0.099967893 | 0.709264768 | -4.35549901 |
| **A_55_P2056186** | Siva1 | 0.207663153 | -0.055421421 | 0.470747726 | 13.2957447 | 1.977035216 | 0.099983869 | 0.709264768 | -4.35552961 |
| **A_52_P583973** | Lrpap1 | -0.825836768 | -1.872492818 | 0.220819283 | 12.5817473 | -1.976244034 | 0.10008935 | 0.709264768 | -4.35573154 |
| **A_52_P551707** | Kdm6a | -0.27840651 | -0.631279928 | 0.074466908 | 8.662516758 | -1.976108504 | 0.100107431 | 0.709264768 | -4.355766135 |
| **A_52_P228031** | Tmem179 | -0.417651035 | -0.947104943 | 0.111802874 | 10.43840445 | -1.975767095 | 0.100152991 | 0.709264768 | -4.355853287 |
| **A_51_P467837** | Ap3m2 | -0.281389366 | -0.638171779 | 0.075393048 | 8.230824516 | -1.975397898 | 0.100202284 | 0.709264768 | -4.355947541 |
| **A_52_P329917** | Prrg2 | -0.611062708 | -1.385954629 | 0.163829213 | 10.74731847 | -1.975126851 | 0.100238488 | 0.709264768 | -4.356016744 |
| **A_51_P214306** | Haus4 | 0.464864838 | -0.125127771 | 1.054857448 | 11.92833145 | 1.973469742 | 0.100460123 | 0.709264768 | -4.356439936 |
| **A_55_P2185203** | Snrnp40 | 0.189010015 | -0.050897675 | 0.428917704 | 8.962411544 | 1.973290356 | 0.100484145 | 0.709264768 | -4.356485758 |
| **A_52_P412585** | Ear1 | -0.393211239 | -0.892562156 | 0.106139678 | 9.0721991 | -1.972288381 | 0.100618433 | 0.709264768 | -4.356741741 |
| **A_51_P105709** | Trip13 | 0.874715489 | -0.236555538 | 1.985986516 | 10.07728063 | 1.971501768 | 0.100723986 | 0.709264768 | -4.392663659 |
| **A_55_P2006792** | Ntrk2 | 0.317054574 | -0.085765285 | 0.719874432 | 9.264246264 | 1.971394296 | 0.100738416 | 0.709264768 | -4.356970216 |
| **A_51_P461761** | Zcchc8 | 0.327341339 | -0.088547938 | 0.743230617 | 9.359126005 | 1.971394211 | 0.100738427 | 0.709264768 | -4.356970238 |
| **A_55_P2064025** | A430073D | -0.630059917 | -1.430647339 | 0.170527506 | 12.38844498 | -1.971167065 | 0.100768933 | 0.709264768 | -4.357028291 |
| **A_52_P343627** | Rbp7 | 0.555703862 | -0.150528037 | 1.26193576 | 8.701321572 | 1.970817595 | 0.100815885 | 0.709264768 | -4.357117614 |
| **A_55_P1999532** | Col9a1 | 0.564605891 | -0.153267871 | 1.282479653 | 8.430422687 | 1.969915831 | 0.100937142 | 0.709264768 | -4.357348138 |
| **A_55_P2170105** | NA | -0.37233965 | -0.845754718 | 0.101075418 | 13.99436327 | -1.969915308 | 0.100937212 | 0.709264768 | -4.357348272 |
| **A_51_P494686** | Sumf1 | -0.464546948 | -1.055270754 | 0.126176858 | 10.49251002 | -1.969679133 | 0.100968994 | 0.709264768 | -4.357408656 |
| **A_52_P110270** | Nelfe | 0.211734069 | -0.057651708 | 0.481119846 | 10.61714181 | 1.968640551 | 0.101108879 | 0.709264768 | -4.357674238 |
| **A_55_P2002117** | Kbtbd4 | 0.284799339 | -0.077600721 | 0.647199399 | 10.53997227 | 1.968344275 | 0.10114882 | 0.709264768 | -4.357750013 |
| **A_55_P1974617** | Lysmd1 | -0.349134297 | -0.793405633 | 0.095137038 | 10.59946511 | -1.968314839 | 0.101152789 | 0.709264768 | -4.357757542 |
| **A_52_P557293** | Tmprss2 | -0.751615541 | -1.708063792 | 0.20483271 | 9.450512994 | -1.968269756 | 0.101158868 | 0.709264768 | -4.357769073 |
| **A_52_P497388** | NA | 0.39313825 | -0.107179402 | 0.893455901 | 15.26023303 | 1.96811205 | 0.101180138 | 0.709264768 | -4.357809411 |
| **A_55_P2076543** | NA | 0.656757403 | -0.179184555 | 1.492699362 | 10.75664825 | 1.967792034 | 0.101223311 | 0.709264768 | -4.357891269 |
| **A_51_P360492** | Mcm6 | 1.0869787 | -0.296758796 | 2.470716196 | 13.67758387 | 1.96751327 | 0.101260934 | 0.709264768 | -4.357962581 |
| **A_55_P2092085** | Tmsb10 | -0.421081661 | -0.95733424 | 0.115170918 | 15.66317113 | -1.966741489 | 0.101365172 | 0.709264768 | -4.35816004 |
| **A_55_P2050207** | Ankrd27 | 0.201069618 | -0.054998842 | 0.457138079 | 11.09852969 | 1.966711731 | 0.101369194 | 0.709264768 | -4.358167654 |
| **A_66_P127024** | Mbd3 | 0.298360952 | -0.081807529 | 0.678529433 | 13.19220668 | 1.965695405 | 0.101506634 | 0.709264768 | -4.358427741 |
| **A_51_P184849** | Pfkfb2 | -0.194367499 | -0.442055571 | 0.053320573 | 8.369681758 | -1.965481227 | 0.101535621 | 0.709264768 | -4.358482559 |
| **A_52_P491073** | Nek8 | -0.229828369 | -0.522785722 | 0.063128983 | 9.479740758 | -1.964941295 | 0.101608736 | 0.709264768 | -4.358620767 |
| **A_55_P2080243** | NA | 0.392706571 | -0.107985386 | 0.893398527 | 9.080806227 | 1.964481299 | 0.101671069 | 0.709264768 | -4.358738529 |
| **A_66_P136364** | NA | 0.387429467 | -0.106584314 | 0.881443248 | 13.02413903 | 1.964282425 | 0.10169803 | 0.709264768 | -4.358789446 |
| **A_55_P2093143** | Casd1 | 0.341011395 | -0.09402291 | 0.776045699 | 8.404652074 | 1.963340983 | 0.101825759 | 0.709264768 | -4.359030515 |
| **A_52_P616392** | Sbno2 | -0.36472399 | -0.830040854 | 0.100592873 | 8.48590421 | -1.963206053 | 0.101844079 | 0.709264768 | -4.35906507 |
| **A_55_P2123234** | NA | 1.029439111 | -0.284373817 | 2.343252038 | 10.4919337 | 1.962535307 | 0.101935198 | 0.709264768 | -4.359236865 |
| **A_51_P250445** | Zfp276 | -0.369890248 | -0.842053858 | 0.102273363 | 9.112180931 | -1.962143263 | 0.101988495 | 0.709264768 | -4.359337291 |
| **A_51_P215066** | Emc4 | -0.202445062 | -0.460908612 | 0.056018488 | 12.33179429 | -1.961815806 | 0.102033033 | 0.709264768 | -4.359421179 |
| **A_55_P2112922** | Impdh2 | 0.731608452 | -0.202468408 | 1.665685311 | 12.87624896 | 1.96176258 | 0.102040274 | 0.709264768 | -4.359434816 |
| **A_51_P269687** | Pole2 | 0.665550306 | -0.18427456 | 1.515375172 | 9.090572531 | 1.961560911 | 0.102067716 | 0.709264768 | -4.359486484 |
| **A_51_P392546** | Haus3 | 0.51777603 | -0.143424192 | 1.178976252 | 10.63587705 | 1.961368951 | 0.102093843 | 0.709264768 | -4.359535667 |
| **A_55_P2075298** | Atp8a1 | 0.347040611 | -0.096159804 | 0.790241025 | 9.980007396 | 1.961238846 | 0.102111555 | 0.709264768 | -4.359569004 |
| **A_55_P2010066** | Capn3 | 0.696713679 | -0.193157331 | 1.586584688 | 8.382616082 | 1.961000138 | 0.102144061 | 0.709264768 | -4.359630169 |
| **A_66_P139664** | Ppp2r5c | 0.235607439 | -0.065334484 | 0.536549362 | 8.561814024 | 1.960905442 | 0.102156958 | 0.709264768 | -4.359654435 |
| **A_55_P2186621** | Rbmxl1 | 1.030721317 | -0.285870903 | 2.347313537 | 13.27459346 | 1.960831698 | 0.102167004 | 0.709264768 | -4.359673332 |
| **A_51_P262171** | Irgm1 | -0.385328167 | -0.8776051 | 0.106948766 | 9.383424572 | -1.960521529 | 0.102209266 | 0.709264768 | -4.359752818 |
| **A_66_P102090** | Pkmyt1 | 0.301403438 | -0.083777317 | 0.686584193 | 10.94532026 | 1.959900244 | 0.102293972 | 0.709264768 | -4.359912052 |
| **A_51_P214747** | Parp12 | -0.1985818 | -0.452403714 | 0.055240114 | 8.605627627 | -1.959569512 | 0.102339094 | 0.709264768 | -4.359996828 |
| **A_55_P2027436** | Bri3 | -0.211650097 | -0.482212375 | 0.058912182 | 15.07982989 | -1.959302847 | 0.102375489 | 0.709264768 | -4.360065186 |

| | | | | | | | | | |
|---|---|---|---|---|---|---|---|---|---|
| **A_51_P439085** | Hilpda | -0.724386266 | -1.650416543 | 0.201644011 | 10.69014231 | -1.959274865 | 0.102379309 | 0.709264768 | -4.36007236 |
| **A_55_P2174203** | Lims2 | -1.742495043 | -3.970071094 | 0.485081008 | 10.63852309 | -1.959247751 | 0.102383011 | 0.709264768 | -4.360079311 |
| **A_55_P1979221** | Txndc9 | 0.342369957 | -0.095451152 | 0.780191067 | 10.70598016 | 1.958615995 | 0.102469295 | 0.709264768 | -4.36024128 |
| **A_51_P272553** | Bhlhe40 | -0.822826435 | -1.87512459 | 0.229471721 | 9.024180743 | -1.958482846 | 0.10248749 | 0.709264768 | -4.36027542 |
| **A_55_P2116149** | NA | -0.305784824 | -0.696903692 | 0.085334045 | 11.75459668 | -1.958202069 | 0.102525869 | 0.709264768 | -4.360347416 |
| **A_52_P117966** | Atpsckmt | -0.200654946 | -0.457347854 | 0.056037962 | 10.14087382 | -1.957881236 | 0.102569741 | 0.709264768 | -4.36042969 |
| **A_66_P133404** | Pclaf | 0.758859163 | -0.212290599 | 1.730008926 | 8.955391633 | 1.957155443 | 0.10266906 | 0.709264768 | -4.360615834 |
| **A_55_P2065113** | Gm11559 | 0.405546194 | -0.113461501 | 0.924553889 | 9.244982762 | 1.957117401 | 0.102674268 | 0.709264768 | -4.360625591 |
| **A_51_P382688** | Zmynd19 | 0.301041235 | -0.084280445 | 0.686362916 | 9.620649384 | 1.956829053 | 0.102713755 | 0.709264768 | -4.360699554 |
| **A_51_P302358** | Ltb | 1.165893559 | -0.326886686 | 2.658673805 | 8.867021766 | 1.956200518 | 0.102799883 | 0.709264768 | -4.360860796 |
| **A_55_P2341890** | NA | -0.325872105 | -0.74321398 | 0.091469771 | 8.313022506 | -1.955715006 | 0.102866462 | 0.709264768 | -4.360985364 |
| **A_55_P2120433** | Wdr12 | 0.32131095 | -0.090235627 | 0.732857528 | 8.766819007 | 1.955495756 | 0.102896543 | 0.709264768 | -4.361041622 |
| **A_55_P1994258** | Gapdh | 0.457503019 | -0.128557755 | 1.043563792 | 11.12126832 | 1.955247121 | 0.102930666 | 0.709264768 | -4.361105424 |
| **A_52_P140356** | Calm3 | 0.364820419 | -0.102618168 | 0.832259007 | 11.29021554 | 1.954811668 | 0.102990457 | 0.709264768 | -4.361217175 |
| **A_51_P333949** | Ints7 | 0.208967503 | -0.05878797 | 0.476722976 | 8.854502668 | 1.954747843 | 0.102999224 | 0.709264768 | -4.361233555 |
| **A_66_P126603** | NA | 0.79339753 | -0.223235968 | 1.810031028 | 14.19715324 | 1.954684973 | 0.10300786 | 0.709264768 | -4.361249691 |
| **A_52_P671812** | NA | 0.310707454 | -0.087700883 | 0.709115791 | 13.16805877 | 1.953320916 | 0.103195415 | 0.709264768 | -4.361599839 |
| **A_52_P546552** | Pelp1 | 0.185355301 | -0.052369888 | 0.423080491 | 9.473773023 | 1.952900677 | 0.103253268 | 0.709264768 | -4.361707737 |
| **A_55_P2022773** | Glcci1 | 0.805577747 | -0.227677343 | 1.838832837 | 9.864064381 | 1.952766218 | 0.103271785 | 0.709264768 | -4.361742263 |
| **A_55_P2062836** | Ttf2 | 0.475277206 | -0.134516111 | 1.085070523 | 10.91326434 | 1.952156682 | 0.103355771 | 0.709264768 | -4.361898788 |
| **A_55_P2032853** | Entpd1 | -0.296556415 | -0.677077277 | 0.083964446 | 8.318551537 | -1.951997235 | 0.103377753 | 0.709264768 | -4.361939737 |
| **A_55_P2152035** | Tcf19 | 0.583386487 | -0.165453072 | 1.332226045 | 9.204916448 | 1.951272537 | 0.103477719 | 0.709264768 | -4.362125875 |
| **A_55_P2031043** | Stat5b | 0.599836267 | -0.17022601 | 1.369898544 | 11.00186458 | 1.950999762 | 0.103515372 | 0.709264768 | -4.362195945 |
| **A_55_P1963873** | NA | -0.263609294 | -0.602085619 | 0.074867031 | 13.57379328 | -1.950665336 | 0.103561554 | 0.709264768 | -4.362281859 |
| **A_55_P2097612** | Zfp131 | 0.319434017 | -0.090782127 | 0.729650162 | 10.05828154 | 1.950377872 | 0.103601268 | 0.709264768 | -4.362355714 |
| **A_55_P2133385** | NA | 0.520428949 | -0.147952934 | 1.188810832 | 15.0293778 | 1.95023579 | 0.103620902 | 0.709264768 | -4.36239222 |
| **A_51_P296905** | Ppat | 0.616227011 | -0.17531759 | 1.407771612 | 9.224733978 | 1.949915025 | 0.103665243 | 0.709264768 | -4.36247464 |
| **A_55_P2092046** | NA | 0.508893594 | -0.144881579 | 1.162668766 | 9.087904981 | 1.949615214 | 0.103706705 | 0.709264768 | -4.362551681 |
| **A_65_P07561** | Tpd52l2 | -0.284807164 | -0.650721521 | 0.081107194 | 10.00094805 | -1.949493556 | 0.103723535 | 0.709264768 | -4.362582945 |
| **A_51_P484832** | Ppm1g | 0.27525179 | -0.078532467 | 0.629036047 | 9.291650144 | 1.948686581 | 0.103835237 | 0.709264768 | -4.362790346 |
| **A_55_P2129311** | 2210016L | 0.205665275 | -0.058735606 | 0.470066155 | 9.784915529 | 1.948266748 | 0.103893399 | 0.709264768 | -4.362898264 |
| **A_52_P588539** | Snapin | -0.366869983 | -0.838695939 | 0.104955972 | 8.545140366 | -1.947514489 | 0.103997698 | 0.709264768 | -4.36309166 |
| **A_55_P2017369** | NA | 0.394332953 | -0.11284258 | 0.901508485 | 11.95453265 | 1.947399804 | 0.104013608 | 0.709264768 | -4.363121147 |
| **A_55_P2080794** | Taf9 | 0.756135661 | -0.233340799 | 1.745612121 | 11.82870456 | 2.035433695 | 0.104029087 | 0.709264768 | -4.396869928 |
| **A_55_P2030775** | Fnbp1 | 0.255098247 | -0.073019172 | 0.583215666 | 10.36562729 | 1.947280583 | 0.10403015 | 0.709264768 | -4.363151802 |
| **A_51_P449911** | NA | 0.411705232 | -0.117878226 | 0.941288689 | 12.49291398 | 1.947163054 | 0.10404646 | 0.709264768 | -4.363182022 |
| **A_52_P655084** | Tbc1d17 | -0.451311385 | -1.031883583 | 0.129260813 | 10.26608438 | -1.947019942 | 0.104066324 | 0.709264768 | -4.363218821 |
| **A_52_P488427** | Sec14l2 | -0.260303838 | -0.595286053 | 0.074678376 | 8.484946148 | -1.946297248 | 0.104166691 | 0.709264768 | -4.363404673 |
| **A_51_P474454** | Rnaseh2b | 0.496581516 | -0.142464486 | 1.135627519 | 11.04417575 | 1.946295476 | 0.104166938 | 0.709264768 | -4.363405129 |
| **A_51_P508959** | Prr15 | -0.29455673 | -0.673658233 | 0.084544772 | 9.353820725 | -1.946093894 | 0.104194951 | 0.709264768 | -4.363456975 |
| **A_55_P2094100** | BC08527 | 0.425800301 | -0.122237986 | 0.973838587 | 9.961946488 | 1.946011531 | 0.104206399 | 0.709264768 | -4.363478159 |
| **A_51_P463497** | Rnh1 | -0.497632865 | -1.13835968 | 0.14309395 | 10.04505598 | -1.945299613 | 0.104305405 | 0.709264768 | -4.363661285 |
| **A_52_P349467** | Tmlhe | 0.194393445 | -0.055912623 | 0.444699514 | 8.313484623 | 1.945183533 | 0.104321557 | 0.709264768 | -4.363691147 |
| **A_51_P146303** | Mvb12a | -0.242758696 | -0.555464626 | 0.069947234 | 11.34640388 | -1.944415278 | 0.104428523 | 0.709264768 | -4.363888807 |
| **A_51_P129149** | Gatad2a | 0.258345679 | -0.074452922 | 0.591144281 | 10.30726866 | 1.944330352 | 0.104440354 | 0.709264768 | -4.363910659 |
| **A_51_P103222** | Slc39a4 | -0.983446723 | -2.250719262 | 0.283825815 | 12.80264418 | -1.943708614 | 0.104527011 | 0.709264768 | -4.364070653 |
| **A_55_P2070131** | NA | 0.668775275 | -0.193132658 | 1.530683208 | 15.29383593 | 1.943433463 | 0.104565385 | 0.709264768 | -4.364141466 |
| **A_55_P2062911** | Epha5 | 0.352195674 | -0.101761382 | 0.80615273 | 13.13094717 | 1.943209354 | 0.104596651 | 0.709264768 | -4.364199146 |

| | | | | | | | | | |
|---|---|---|---|---|---|---|---|---|---|
| **A_66_P112024** | NA | -0.345323325 | -0.790513903 | 0.099867253 | 11.27824654 | -1.942809846 | 0.104652411 | 0.709264768 | -4.364301978 |
| **A_55_P1966804** | Fdps | 0.458267399 | -0.132544946 | 1.049079744 | 13.05423392 | 1.942762656 | 0.104658999 | 0.709264768 | -4.364314126 |
| **A_55_P1967618** | Med16 | 0.395422963 | -0.114430335 | 0.905276261 | 11.4086119 | 1.942526707 | 0.104691947 | 0.709264768 | -4.364374864 |
| **A_51_P319592** | Usp42 | 0.184354991 | -0.05336966 | 0.422079641 | 10.00603529 | 1.942365819 | 0.10471442 | 0.709264768 | -4.364416281 |
| **A_55_P1979402** | Tigar | 0.169779781 | -0.049158795 | 0.388718357 | 8.751915274 | 1.94228978 | 0.104725042 | 0.709264768 | -4.364435857 |
| **A_51_P172231** | Gsdmd | -0.348431961 | -0.797753025 | 0.100889103 | 9.247373995 | -1.942278721 | 0.104726587 | 0.709264768 | -4.384048834 |
| **A_51_P345340** | Upk3bl | -1.377476744 | -3.15449935 | 0.399545863 | 9.255688591 | -1.941519104 | 0.104832768 | 0.709264768 | -4.36463428 |
| **A_55_P2038007** | Csrp1 | -0.283887533 | -0.650176266 | 0.082401199 | 16.47863083 | -1.941212629 | 0.104875639 | 0.709264768 | -4.364713198 |
| **A_51_P153013** | Map2k3os | -0.171066101 | -0.391789169 | 0.049656966 | 8.307528529 | -1.941183448 | 0.104879722 | 0.709264768 | -4.364720712 |
| **A_55_P1988010** | Gm10406 | -0.283316736 | -0.648919689 | 0.082286217 | 9.471788531 | -1.940943441 | 0.104913309 | 0.709264768 | -4.364782519 |
| **A_55_P2135025** | NA | 0.195161886 | -0.056704854 | 0.447028626 | 10.77587816 | 1.940772059 | 0.104937299 | 0.709264768 | -4.364826655 |
| **A_55_P2193419** | NA | -0.352630472 | -0.807755348 | 0.102494403 | 8.276822361 | -1.940616016 | 0.104959146 | 0.709264768 | -4.364866843 |
| **A_51_P324934** | Mcm3 | 1.575867676 | -0.459490357 | 3.611225708 | 11.77241801 | 1.939229805 | 0.105153435 | 0.709264768 | -4.36522392 |
| **A_55_P1960049** | 2810408A | 0.336134853 | -0.098195075 | 0.77046478 | 10.2588784 | 1.938403279 | 0.105269454 | 0.709264768 | -4.365436884 |
| **A_66_P109175** | Farsb | 0.259349342 | -0.075788481 | 0.594487165 | 10.97408851 | 1.938260082 | 0.105289568 | 0.709264768 | -4.365473785 |
| **A_51_P155313** | Gsto1 | -0.680684539 | -1.560345618 | 0.19897654 | 12.360737 | -1.938120857 | 0.105309127 | 0.709264768 | -4.365509663 |
| **A_55_P1965674** | Alg2 | 0.280915075 | -0.082128523 | 0.643958672 | 11.0825187 | 1.938057282 | 0.10531806 | 0.709264768 | -4.365526047 |
| **A_55_P2236818** | A430103D | 0.302912414 | -0.088579013 | 0.69440384 | 8.535117938 | 1.937961573 | 0.105331509 | 0.709264768 | -4.365550712 |
| **A_55_P2090782** | NA | 0.279231269 | -0.081770581 | 0.640233118 | 13.08896275 | 1.937336087 | 0.105419448 | 0.709264768 | -4.365711921 |
| **A_55_P2010936** | Fbxo17 | 0.28404602 | -0.083191302 | 0.651283342 | 9.593655523 | 1.937279321 | 0.105427432 | 0.709264768 | -4.365726553 |
| **A_51_P384113** | Acad12 | -0.403578975 | -0.925416482 | 0.118258532 | 8.409687559 | -1.937062296 | 0.105457964 | 0.709264768 | -4.365782495 |
| **A_55_P2128499** | Mgl2 | 0.585354404 | -0.171855877 | 1.342564685 | 13.09501715 | 1.936211225 | 0.105577783 | 0.709264768 | -4.3660019 |
| **A_55_P2430122** | Ilf2 | 0.439798914 | -0.129124693 | 1.008722521 | 10.46918635 | 1.936201462 | 0.105579158 | 0.709264768 | -4.366004417 |
| **A_55_P2161585** | Coq10a | -0.176923678 | -0.405825316 | 0.05197796 | 8.4746433 | -1.935919993 | 0.105618816 | 0.709264768 | -4.366076989 |
| **A_55_P2023607** | Ikzf1 | 0.315518227 | -0.092718205 | 0.723754659 | 7.90919829 | 1.935811379 | 0.105634123 | 0.709264768 | -4.366104995 |
| **A_51_P340699** | Rasl11a | -0.243546662 | -0.558693679 | 0.071600354 | 8.901940708 | -1.935616553 | 0.105661586 | 0.709264768 | -4.366155232 |
| **A_55_P2154914** | Acap1 | 0.430113731 | -0.126544498 | 0.986771961 | 8.067175891 | 1.935285396 | 0.105708284 | 0.709264768 | -4.366240629 |
| **A_55_P1995652** | Rexo4 | 0.318459879 | -0.093717139 | 0.730636898 | 12.08937233 | 1.935179697 | 0.105723193 | 0.709264768 | -4.366267887 |
| **A_55_P2058240** | Ipo5 | 0.758036009 | -0.223221874 | 1.739293893 | 10.27172565 | 1.934893311 | 0.1057636 | 0.709264768 | -4.366341746 |
| **A_55_P1973699** | Htra2 | -0.260752527 | -0.598354968 | 0.076849915 | 11.56963696 | -1.934520305 | 0.105816252 | 0.709264768 | -4.366437951 |
| **A_51_P185175** | Fkbp4 | 0.656560368 | -0.193647498 | 1.506768233 | 9.387747068 | 1.934193373 | 0.105862422 | 0.709264768 | -4.366522281 |
| **A_52_P472799** | Ecd | 0.224883957 | -0.066463393 | 0.516231308 | 10.18430761 | 1.933293307 | 0.105989638 | 0.709264768 | -4.366754479 |
| **A_66_P128702** | Nup43 | 0.475256491 | -0.140480741 | 1.090993723 | 9.790640447 | 1.933227605 | 0.10599893 | 0.709264768 | -4.366771431 |
| **A_55_P2157019** | Phb2 | 0.220122445 | -0.065138125 | 0.505383015 | 10.3895034 | 1.932737744 | 0.106068239 | 0.709264768 | -4.366897829 |
| **A_51_P308948** | Tamm41 | 0.566719459 | -0.167788956 | 1.301227873 | 10.98717361 | 1.932509754 | 0.106100512 | 0.709264768 | -4.366956662 |
| **A_55_P2112005** | Tff1 | -0.641832473 | -1.473818418 | 0.190153473 | 8.060451069 | -1.932217635 | 0.106141878 | 0.709264768 | -4.367032047 |
| **A_51_P281089** | S100a6 | -1.031446756 | -2.368555941 | 0.305662428 | 15.02274483 | -1.932103059 | 0.106158107 | 0.709264768 | -4.367061617 |
| **A_55_P2093439** | Mex3d | 0.448466631 | -0.132985747 | 1.029919009 | 10.68747871 | 1.931818535 | 0.106198419 | 0.709264768 | -4.36713505 |
| **A_52_P411003** | Dlgap5 | 0.304382617 | -0.09026432 | 0.699029555 | 8.178164385 | 1.93179686 | 0.106201491 | 0.709264768 | -4.367140645 |
| **A_51_P208801** | Ndufab1 | 0.250752672 | -0.074394412 | 0.575899755 | 9.60660666 | 1.931594905 | 0.106230115 | 0.709264768 | -4.367192771 |
| **A_55_P1983588** | Pmepa1 | -0.261398086 | -0.600373184 | 0.077577011 | 10.87168978 | -1.931456609 | 0.106249721 | 0.709264768 | -4.367228468 |
| **A_55_P1990944** | Casp1 | -0.266969104 | -0.613238519 | 0.079300312 | 8.43405258 | -1.931066436 | 0.106305054 | 0.709264768 | -4.367329185 |
| **A_55_P2038597** | NA | 0.704894391 | -0.209475535 | 1.619264316 | 12.6796153 | 1.930867475 | 0.106333282 | 0.709264768 | -4.367380547 |
| **A_51_P476601** | Ly96 | -0.438660898 | -1.007800363 | 0.130478566 | 8.97961908 | -1.930458938 | 0.106391267 | 0.709264768 | -4.36748602 |
| **A_55_P2081616** | Timeless | 0.7270966 | -0.216296903 | 1.670490103 | 10.68387426 | 1.930410087 | 0.106398202 | 0.709264768 | -4.367498633 |
| **A_51_P470328** | Selenop | -0.684615378 | -1.572892554 | 0.203661797 | 14.51835508 | -1.930405262 | 0.106398887 | 0.709264768 | -4.367499878 |
| **A_55_P2088331** | Ctdsp2 | 0.312251471 | -0.092942637 | 0.717445578 | 11.8195635 | 1.930152917 | 0.106434722 | 0.709264768 | -4.367565033 |
| **A_55_P2072403** | Suv39h2 | 0.327848276 | -0.097608634 | 0.753305185 | 13.14224701 | 1.930046039 | 0.106449903 | 0.709264768 | -4.36759263 |

| | | | | | | | | | |
|---|---|---|---|---|---|---|---|---|---|
| **A_55_P2025443** | Oprm1 | -0.437955483 | -1.006476561 | 0.130565596 | 12.52712326 | -1.929450948 | 0.106534472 | 0.709264768 | -4.3677463 |
| **A_55_P2004781** | Ass1 | -0.269717821 | -0.619858835 | 0.080423193 | 12.22743141 | -1.929376584 | 0.106545044 | 0.709264768 | -4.367765504 |
| **A_51_P513941** | Lrpap1 | -0.556277139 | -1.278457341 | 0.165903062 | 11.58278968 | -1.929283189 | 0.106558324 | 0.709264768 | -4.367789624 |
| **A_52_P675395** | Ackr3 | -0.324978679 | -0.746945181 | 0.096987823 | 10.12864695 | -1.928977843 | 0.106601753 | 0.709264768 | -4.367868485 |
| **A_55_P2039225** | Rap1gap | -0.551677 | -1.268123445 | 0.164769444 | 10.51841499 | -1.928641432 | 0.106649622 | 0.709264768 | -4.367955376 |
| **A_55_P1996862** | Psme2 | -0.270291557 | -0.621389409 | 0.080806294 | 14.52787976 | -1.928211435 | 0.106710838 | 0.709264768 | -4.368066449 |
| **A_51_P282508** | Rhoc | -0.725779342 | -1.668665772 | 0.217107089 | 15.06444236 | -1.927949094 | 0.106748204 | 0.709264768 | -4.36813422 |
| **A_55_P2038757** | Fblim1 | -0.704067746 | -1.618866961 | 0.210731469 | 11.66401389 | -1.927698065 | 0.106783971 | 0.709264768 | -4.368199073 |
| **A_55_P2109554** | Serinc3 | -0.316339007 | -0.727364844 | 0.094686831 | 11.33319559 | -1.927675685 | 0.10678716 | 0.709264768 | -4.368204855 |
| **A_52_P226929** | Cflar | -0.190043628 | -0.437058018 | 0.056970763 | 8.380451861 | -1.926998616 | 0.106883694 | 0.709264768 | -4.368379795 |
| **A_55_P2146620** | Atf2 | -0.241690814 | -0.555851912 | 0.072470285 | 8.474374516 | -1.926895146 | 0.106898454 | 0.709264768 | -4.368406532 |
| **A_55_P2117764** | Vangl2 | 0.172084529 | -0.051650396 | 0.395819455 | 8.560157369 | 1.926452878 | 0.106961568 | 0.709264768 | -4.368520823 |
| **A_55_P2133315** | Steap2 | -0.219480245 | -0.510021798 | 0.071061308 | 8.389489269 | -2.012101516 | 0.107002428 | 0.709264768 | -4.401136676 |
| **A_51_P493720** | Tfam | 0.341311582 | -0.102529116 | 0.785152279 | 10.60722686 | 1.926079679 | 0.107014854 | 0.709264768 | -4.368617275 |
| **A_55_P2112549** | NA | 1.079780247 | -0.324571153 | 2.484131647 | 14.14038923 | 1.925794457 | 0.107055598 | 0.709264768 | -4.368690995 |
| **A_55_P2134597** | NA | 0.197589987 | -0.059395296 | 0.45457527 | 8.365867471 | 1.925781567 | 0.107057439 | 0.709264768 | -4.368694326 |
| **A_55_P2410106** | 9130230N | -0.256120709 | -0.589303647 | 0.07706223 | 8.242507451 | -1.925361511 | 0.107117473 | 0.709264768 | -4.368802906 |
| **A_55_P1982030** | H3f3b | 0.494037784 | -0.148831089 | 1.136906658 | 12.60419004 | 1.924811091 | 0.10719619 | 0.709264768 | -4.3689452 |
| **A_52_P650379** | Strap | 0.347377595 | -0.104731296 | 0.799486485 | 10.37104035 | 1.924460945 | 0.107246296 | 0.709264768 | -4.369035729 |
| **A_55_P2020926** | Trip11 | -0.348664667 | -0.802613466 | 0.105284131 | 9.259172211 | -1.92376232 | 0.10734634 | 0.709264768 | -4.369216378 |
| **A_52_P229210** | NA | 0.461191515 | -0.139275411 | 1.061658441 | 11.54659315 | 1.923723113 | 0.107351957 | 0.709264768 | -4.369226517 |
| **A_55_P1981296** | Nup85 | 0.569773998 | -0.172211919 | 1.311759916 | 12.21798631 | 1.923345515 | 0.107406072 | 0.709264768 | -4.369324169 |
| **A_51_P285446** | Lig1 | 1.01755194 | -0.307692708 | 2.342796588 | 12.06290522 | 1.923139876 | 0.107435555 | 0.709264768 | -4.369377354 |
| **A_66_P106968** | NA | 0.458432057 | -0.138624358 | 1.055488472 | 10.89904702 | 1.923135815 | 0.107436137 | 0.709264768 | -4.369378405 |
| **A_55_P2172274** | Smc2 | 1.288440476 | -0.389653424 | 2.966534377 | 11.04420161 | 1.92308485 | 0.107443446 | 0.709264768 | -4.369391586 |
| **A_55_P1978023** | NA | 0.526853822 | -0.171942157 | 1.225649802 | 9.63370682 | 2.008179102 | 0.107511147 | 0.709264768 | -4.401857706 |
| **A_55_P2085157** | Eef1g | 0.325907696 | -0.098679171 | 0.750494563 | 13.26737908 | 1.922553372 | 0.107519688 | 0.709264768 | -4.369529057 |
| **A_55_P2040993** | NA | -0.461772162 | -1.063442823 | 0.1398985 | 8.989820752 | -1.922291556 | 0.107557267 | 0.709264768 | -4.369596785 |
| **A_55_P1991516** | E4f1 | -0.298372616 | -0.687232405 | 0.090487172 | 9.314012577 | -1.921835769 | 0.107622718 | 0.709264768 | -4.369714699 |
| **A_52_P220229** | Plekhb2 | -0.366191121 | -0.843462724 | 0.111080482 | 8.847915495 | -1.921730841 | 0.107637791 | 0.709264768 | -4.369741846 |
| **A_55_P2116054** | Caprin1 | 0.241520276 | -0.07326454 | 0.556305092 | 8.584461462 | 1.921720249 | 0.107639313 | 0.709264768 | -4.369744587 |
| **A_55_P2109658** | NA | 0.311980721 | -0.094767847 | 0.718729289 | 10.99878629 | 1.921109281 | 0.107727126 | 0.709264768 | -4.369902672 |
| **A_55_P2018825** | NA | 0.583297785 | -0.177452659 | 1.344048229 | 9.346905462 | 1.920429897 | 0.107824857 | 0.709264768 | -4.370078486 |
| **A_55_P1985159** | Mef2a | 0.370066866 | -0.112611451 | 0.852745182 | 10.87685575 | 1.920316253 | 0.107841214 | 0.709264768 | -4.370107898 |
| **A_55_P2057296** | Upk3bl | -1.408413709 | -3.245599012 | 0.428771594 | 9.332161249 | -1.920116708 | 0.10786994 | 0.709264768 | -4.370159544 |
| **A_55_P1955227** | Rabac1 | -0.797651761 | -1.838146578 | 0.242843057 | 14.39476306 | -1.920099578 | 0.107872407 | 0.709264768 | -4.370163977 |
| **A_51_P238448** | Ccnd3 | 0.635460319 | -0.193722196 | 1.464642834 | 11.92304063 | 1.919502343 | 0.107958434 | 0.709264768 | -4.370318568 |
| **A_55_P2125811** | Esam | -0.530177506 | -1.222129809 | 0.161774796 | 12.17787882 | -1.919090631 | 0.108017778 | 0.709264768 | -4.37042515 |
| **A_55_P2128969** | Mettl27 | -0.209153761 | -0.482222517 | 0.063914995 | 9.930765903 | -1.918421406 | 0.108114312 | 0.709264768 | -4.370598418 |
| **A_55_P1957875** | NA | -0.261242005 | -0.602388645 | 0.079904635 | 9.762625208 | -1.918016134 | 0.108172813 | 0.709264768 | -4.37070336 |
| **A_55_P2076941** | Lrr1 | 0.33772289 | -0.103356981 | 0.77880276 | 8.409589231 | 1.917757121 | 0.108210219 | 0.709264768 | -4.370770434 |
| **A_51_P121302** | Them4 | -0.20737698 | -0.4782506 | 0.063496641 | 8.557357637 | -1.917538876 | 0.108241748 | 0.709264768 | -4.378814596 |
| **A_55_P2177953** | Aqp11 | 0.537411995 | -0.164638702 | 1.239462692 | 8.202748529 | 1.917296265 | 0.108276807 | 0.709264768 | -4.370889788 |
| **A_55_P2102901** | Fbxl4 | -0.171570651 | -0.39572935 | 0.052588049 | 8.225794622 | -1.917069001 | 0.108309659 | 0.709264768 | -4.370948651 |
| **A_55_P1994388** | Lsm14a | 0.244848735 | -0.075054576 | 0.564752046 | 11.52164908 | 1.917032494 | 0.108314937 | 0.709264768 | -4.370958106 |
| **A_52_P255804** | NA | 0.696215413 | -0.21362238 | 1.606053205 | 11.98477667 | 1.916593481 | 0.108378431 | 0.709264768 | -4.371071823 |
| **A_51_P253697** | Rpain | -0.251731442 | -0.580768595 | 0.077305711 | 9.432832699 | -1.916208978 | 0.108434071 | 0.709264768 | -4.371171429 |
| **A_55_P2132812** | NA | -0.404520152 | -0.933276798 | 0.124236495 | 10.6260558 | -1.916172784 | 0.108439311 | 0.709264768 | -4.371180806 |

| | | | | | | | | | |
|---|---|---|---|---|---|---|---|---|---|
| **A_52_P630016** | Bet1l | -0.461907788 | -1.065760727 | 0.14194515 | 9.962942866 | -1.915907098 | 0.108477776 | 0.709264768 | -4.371249638 |
| **A_51_P238383** | B4galt4 | -0.565652906 | -1.305156477 | 0.173850664 | 10.74911938 | -1.91584378 | 0.108486945 | 0.709264768 | -4.371266043 |
| **A_51_P442964** | Knl1 | 0.326727651 | -0.100464834 | 0.753920136 | 8.154136772 | 1.91563442 | 0.108517269 | 0.709264768 | -4.371320287 |
| **A_55_P2098071** | Trim25 | -0.71574044 | -1.668760055 | 0.237279175 | 12.58861628 | -2.000398048 | 0.108527979 | 0.709264768 | -4.403291215 |
| **A_52_P257625** | Esm1 | 0.301102685 | -0.092629874 | 0.694835244 | 8.490649295 | 1.915418353 | 0.108548573 | 0.709264768 | -4.371376271 |
| **A_51_P498388** | Sbk1 | 0.411316555 | -0.126880386 | 0.949513497 | 8.72111896 | 1.914191177 | 0.10872654 | 0.709264768 | -4.371694293 |
| **A_66_P124551** | Akap13 | -0.232226223 | -0.536088574 | 0.071636129 | 11.66687493 | -1.914188529 | 0.108726924 | 0.709264768 | -4.37169498 |
| **A_55_P1963767** | Npr2 | -0.194173582 | -0.448440961 | 0.060093797 | 11.84668179 | -1.912713138 | 0.108941281 | 0.709264768 | -4.372077449 |
| **A_55_P2006519** | Clip1 | -0.45387201 | -1.048217481 | 0.140473461 | 9.387494203 | -1.912690888 | 0.108944518 | 0.709264768 | -4.372083218 |
| **A_55_P2184811** | Eif2s1 | 0.239219859 | -0.074108528 | 0.552548247 | 8.634996617 | 1.912263926 | 0.109006632 | 0.709264768 | -4.372193926 |
| **A_55_P2112240** | Cdk5rap2 | 0.458810007 | -0.142322041 | 1.059942056 | 10.43859533 | 1.911671851 | 0.109092827 | 0.709264768 | -4.372347464 |
| **A_51_P140141** | Hspbp1 | 0.179213426 | -0.05564061 | 0.414067462 | 9.493406849 | 1.911273598 | 0.109150844 | 0.709264768 | -4.372450752 |
| **A_55_P2138792** | Fdx1 | -0.484856363 | -1.120248491 | 0.150535765 | 12.90936342 | -1.911268118 | 0.109151642 | 0.709264768 | -4.372452173 |
| **A_55_P2040367** | Bola3 | 0.357606686 | -0.111082533 | 0.826295905 | 10.79797501 | 1.911045431 | 0.109184097 | 0.709264768 | -4.372509932 |
| **A_55_P1957219** | Ptma | 0.559467172 | -0.173818704 | 1.292753048 | 15.54714272 | 1.910960164 | 0.109196527 | 0.709264768 | -4.372532049 |
| **A_55_P1985079** | Mcrip1 | -0.167641266 | -0.387394508 | 0.052111977 | 14.02263904 | -1.910715318 | 0.109232226 | 0.709264768 | -4.37259556 |
| **A_52_P22617** | NA | 0.340134621 | -0.105831341 | 0.786100584 | 8.302963194 | 1.910290746 | 0.109294159 | 0.709264768 | -4.372705699 |
| **A_55_P2002472** | Zfp777 | 0.205986343 | -0.064137233 | 0.476109919 | 10.8461637 | 1.909968841 | 0.109341139 | 0.709264768 | -4.372789212 |
| **A_52_P18596** | NA | 0.877571133 | -0.273267709 | 2.028409975 | 12.79429584 | 1.909933135 | 0.109346351 | 0.709264768 | -4.372798476 |
| **A_55_P1966982** | Gsr | 0.455473152 | -0.141974137 | 1.052920441 | 12.03794204 | 1.909473051 | 0.109413537 | 0.709264768 | -4.372917849 |
| **A_55_P2086511** | Ppid | 0.740240536 | -0.230766859 | 1.711247932 | 11.98924706 | 1.909416503 | 0.109421797 | 0.709264768 | -4.372932522 |
| **A_55_P2046453** | Aarsd1 | 0.477489689 | -0.14892143 | 1.103900808 | 12.04882556 | 1.909215209 | 0.109451207 | 0.709264768 | -4.372984754 |
| **A_55_P1991075** | Lmf1 | -0.250416014 | -0.578972999 | 0.078140972 | 10.86981453 | -1.908981583 | 0.109485351 | 0.709264768 | -4.373045379 |
| **A_55_P2050439** | Dlgap5 | 0.318692611 | -0.099452214 | 0.736837435 | 8.185303243 | 1.908954662 | 0.109489286 | 0.709264768 | -4.373052365 |
| **A_55_P1981301** | Nup88 | 0.378315043 | -0.118102601 | 0.874732687 | 11.6017107 | 1.908783831 | 0.109514261 | 0.709264768 | -4.373096698 |
| **A_51_P356762** | Mcm4 | 1.35141609 | -0.422012454 | 3.124844633 | 12.30432081 | 1.908647539 | 0.10953419 | 0.709264768 | -4.373132068 |
| **A_55_P1959174** | Pmepa1 | -0.256894339 | -0.594063594 | 0.080274915 | 10.72366467 | -1.908345103 | 0.109578427 | 0.709264768 | -4.373210561 |
| **A_55_P2068172** | Nufip2 | 0.523163898 | -0.16356594 | 1.209893736 | 10.10711392 | 1.908104672 | 0.109613607 | 0.709264768 | -4.373272964 |
| **A_55_P2397599** | B930025E | -0.421259518 | -0.974759957 | 0.13224092 | 8.719929744 | -1.906259859 | 0.109883925 | 0.709264768 | -4.373751901 |
| **A_55_P2027022** | Prdm1 | -0.442559185 | -1.024594602 | 0.139476231 | 9.14574029 | -1.904461917 | 0.110148025 | 0.709264768 | -4.374218866 |
| **A_52_P109403** | Cebpz | 0.28107712 | -0.088591919 | 0.650746159 | 10.02816815 | 1.904420125 | 0.110154172 | 0.709264768 | -4.374229723 |
| **A_55_P1985890** | Tiparp | -0.196193301 | -0.45429429 | 0.061907687 | 10.11170994 | -1.903903152 | 0.110230234 | 0.709264768 | -4.37436403 |
| **A_51_P291749** | Pecr | 0.318763192 | -0.100609592 | 0.738135976 | 9.913048803 | 1.903786618 | 0.110247387 | 0.709264768 | -4.374394307 |
| **A_55_P2156905** | NA | -0.205629125 | -0.476171483 | 0.064913233 | 13.898345 | -1.903705229 | 0.110259368 | 0.709264768 | -4.374415453 |
| **A_55_P1962771** | Cyfip2 | 0.400706138 | -0.126587212 | 0.927999487 | 9.669011295 | 1.903373622 | 0.110308199 | 0.709264768 | -4.374501615 |
| **A_51_P267544** | Frg1 | -0.242500155 | -0.561610689 | 0.076610378 | 10.73277301 | -1.903361265 | 0.110310019 | 0.709264768 | -4.374504826 |
| **A_55_P2001553** | Igdcc4 | 0.621479704 | -0.196399873 | 1.43935928 | 9.614783768 | 1.903215352 | 0.110331512 | 0.709264768 | -4.374542741 |
| **A_55_P2077884** | Kat6b | -0.185162587 | -0.428876129 | 0.058550955 | 8.164722186 | -1.902934869 | 0.11037284 | 0.709264768 | -4.374615627 |
| **A_55_P1995318** | NA | 0.320346946 | -0.10145783 | 0.742151721 | 9.312892825 | 1.902214296 | 0.110479086 | 0.709264768 | -4.374802897 |
| **A_51_P111952** | Nudt19 | 0.58390082 | -0.18501893 | 1.352820569 | 10.99494063 | 1.901990813 | 0.110512059 | 0.709264768 | -4.390873727 |
| **A_51_P458866** | F11r | -0.445884201 | -1.033054583 | 0.141286181 | 12.39800878 | -1.901990261 | 0.11051214 | 0.709264768 | -4.374861128 |
| **A_52_P199928** | NA | 0.407559929 | -0.12915701 | 0.944276868 | 12.30533656 | 1.901938728 | 0.110519745 | 0.709264768 | -4.374874522 |
| **A_55_P2078735** | AI662270 | 0.283539379 | -0.08996655 | 0.657045308 | 7.919685466 | 1.901368213 | 0.11060397 | 0.709264768 | -4.375022825 |
| **A_66_P107353** | NA | 0.456567828 | -0.144904732 | 1.058040388 | 16.41523815 | 1.901252649 | 0.110621039 | 0.709264768 | -4.375052868 |
| **A_55_P1963491** | Ptma | 0.727062647 | -0.230909543 | 1.685034838 | 14.07586723 | 1.9009438 | 0.110666669 | 0.709264768 | -4.375133162 |
| **A_51_P451106** | Brca2 | 0.691728808 | -0.219742118 | 1.603199735 | 9.447451157 | 1.900830484 | 0.110683415 | 0.709264768 | -4.375162623 |
| **A_55_P2174506** | Stx17 | -0.181711439 | -0.421211931 | 0.057789053 | 8.32482085 | -1.900317657 | 0.110759234 | 0.709264768 | -4.375295962 |
| **A_52_P375493** | Tfdp1 | 0.354853787 | -0.112920354 | 0.822627927 | 9.856909419 | 1.90004366 | 0.110799765 | 0.709264768 | -4.375367211 |

| | | | | | | | | | |
|---|---|---|---|---|---|---|---|---|---|
| **A_51_P268343** | Aagab | 0.216723736 | -0.068981174 | 0.502428647 | 11.64160558 | 1.899936648 | 0.110815599 | 0.709264768 | -4.375395039 |
| **A_55_P2003492** | NA | 1.135748927 | -0.361589634 | 2.633087488 | 13.50793553 | 1.899820947 | 0.110832721 | 0.709264768 | -4.375425127 |
| **A_55_P2094602** | Lrch1 | -0.2851423 | -0.661112479 | 0.090827878 | 9.497537657 | -1.899584407 | 0.110867734 | 0.709264768 | -4.375486642 |
| **A_66_P122173** | Mrtfb | -0.297381266 | -0.689600971 | 0.094838439 | 12.01004557 | -1.899041883 | 0.110948081 | 0.709264768 | -4.375627744 |
| **A_51_P446131** | Gipc2 | -0.35579766 | -0.825175265 | 0.113579945 | 9.266317524 | -1.898589475 | 0.111015128 | 0.709264768 | -4.375745422 |
| **A_52_P39247** | Hbp1 | -0.232495064 | -0.539226654 | 0.074236525 | 12.43149962 | -1.898478039 | 0.111031649 | 0.709264768 | -4.37577441 |
| **A_55_P2108690** | Gm8709 | 0.445747143 | -0.142332317 | 1.033826604 | 11.41019225 | 1.898466345 | 0.111033382 | 0.709264768 | -4.375777452 |
| **A_55_P1970866** | Rplp0 | 0.408707542 | -0.130624508 | 0.948039592 | 16.62391236 | 1.89804615 | 0.111095702 | 0.709264768 | -4.375886765 |
| **A_55_P2093241** | Zc3h4 | 0.314224327 | -0.100505645 | 0.728954299 | 12.21262759 | 1.897687635 | 0.111148902 | 0.709264768 | -4.37598004 |
| **A_51_P293339** | Tob1 | -0.336563803 | -0.780939782 | 0.107812176 | 9.367625217 | -1.896999328 | 0.111251112 | 0.709264768 | -4.376159139 |
| **A_55_P2035087** | Arhgef15 | 0.336755369 | -0.108003688 | 0.781514425 | 11.49115576 | 1.896444221 | 0.111333611 | 0.709264768 | -4.376303599 |
| **A_55_P1957922** | Arrdc2 | -0.321115995 | -0.745343962 | 0.103111972 | 9.120965863 | -1.895889358 | 0.111416137 | 0.709264768 | -4.376448015 |
| **A_52_P212418** | Trmt13 | -0.18276632 | -0.424269845 | 0.058737205 | 8.303605483 | -1.895496729 | 0.11147457 | 0.709264768 | -4.376550216 |
| **A_55_P2094964** | NA | 0.214637933 | -0.068981734 | 0.4982576 | 10.67559053 | 1.89548556 | 0.111476233 | 0.709264768 | -4.376553124 |
| **A_55_P2179869** | NA | 0.641162624 | -0.206102141 | 1.488427389 | 13.03161604 | 1.895393544 | 0.111489932 | 0.709264768 | -4.376577077 |
| **A_51_P337089** | Gins1 | 0.96904919 | -0.31157382 | 2.249672199 | 9.727273215 | 1.895286493 | 0.111505872 | 0.709264768 | -4.376604944 |
| **A_55_P2109932** | Tmem63b | 0.46265838 | -0.18052848 | 1.105845241 | 12.5036912 | 2.121366905 | 0.111537244 | 0.709264768 | -4.466827225 |
| **A_55_P2145480** | NA | 0.680770448 | -0.219120236 | 1.580661133 | 11.61848495 | 1.894790875 | 0.111579698 | 0.709264768 | -4.376733973 |
| **A_55_P2085225** | NA | 0.38537017 | -0.124042169 | 0.894782509 | 11.6846206 | 1.894780819 | 0.111581197 | 0.709264768 | -4.376736591 |
| **A_55_P2025233** | Llph | 0.267042216 | -0.086119795 | 0.620204227 | 9.806482899 | 1.8938967 | 0.111713018 | 0.709264768 | -4.376966799 |
| **A_52_P103550** | Tfdp1 | 0.303053334 | -0.097762655 | 0.703869323 | 9.701138018 | 1.893757537 | 0.111733782 | 0.709264768 | -4.377003038 |
| **A_52_P73475** | Fam78a | 0.575092148 | -0.185731489 | 1.335915785 | 9.04746284 | 1.893231783 | 0.111812261 | 0.709264768 | -4.377139961 |
| **A_55_P2012047** | Gigyf1 | -0.235958965 | -0.548270918 | 0.076352988 | 9.821880215 | -1.892335872 | 0.111946123 | 0.709264768 | -4.377373321 |
| **A_55_P2013948** | Gsn | -0.532981634 | -1.238489352 | 0.172526085 | 14.48722722 | -1.892172907 | 0.111970489 | 0.709264768 | -4.377415774 |
| **A_51_P314397** | Crip2 | -0.321872263 | -0.74811736 | 0.104372833 | 15.95761793 | -1.891361322 | 0.112091918 | 0.709264768 | -4.377627218 |
| **A_55_P2060349** | Lzts2 | -0.431442789 | -1.003139718 | 0.14025414 | 12.44111163 | -1.890199747 | 0.112265945 | 0.709264768 | -4.377929914 |
| **A_52_P532227** | S1pr1 | -0.25058972 | -0.58265835 | 0.081478911 | 9.894610556 | -1.890104188 | 0.112280273 | 0.709264768 | -4.37795482 |
| **A_55_P1967633** | Lsm5 | 0.766819108 | -0.249357805 | 1.782996021 | 10.21389792 | 1.890052849 | 0.112287972 | 0.709264768 | -4.3779682 |
| **A_55_P2056260** | NA | 0.25956554 | -0.084454789 | 0.60358587 | 15.75312675 | 1.889788811 | 0.112327576 | 0.709264768 | -4.378037019 |
| **A_55_P2066468** | Noa1 | 0.275247981 | -0.089597197 | 0.64009316 | 9.829813842 | 1.88958258 | 0.11235852 | 0.709264768 | -4.378090775 |
| **A_52_P10732** | Map6 | -0.410260653 | -0.95422766 | 0.133706354 | 8.923286197 | -1.889024773 | 0.112442257 | 0.709264768 | -4.378236183 |
| **A_51_P125135** | Cdca5 | 1.423700927 | -0.464612175 | 3.312014028 | 10.60225854 | 1.888404849 | 0.112535393 | 0.709264768 | -4.407721029 |
| **A_55_P2011151** | Tmprss3 | 0.361570159 | -0.118011069 | 0.841151386 | 11.2745331 | 1.888342425 | 0.112544776 | 0.709264768 | -4.37841408 |
| **A_52_P84901** | Slc44a1 | -0.283647762 | -0.660011352 | 0.092715827 | 8.790994864 | -1.887652753 | 0.112648492 | 0.709264768 | -4.378593916 |
| **A_55_P2095251** | Ell3 | 0.282672137 | -0.092415505 | 0.657759779 | 8.405161892 | 1.887559244 | 0.112662562 | 0.709264768 | -4.378618301 |
| **A_55_P2115713** | Nid1 | -0.378889246 | -0.881653873 | 0.12387538 | 8.327604619 | -1.887547595 | 0.112664315 | 0.709264768 | -4.378621339 |
| **A_52_P283501** | Zc3h7b | 0.607340262 | -0.198606413 | 1.413286938 | 10.06460009 | 1.887452844 | 0.112678573 | 0.709264768 | -4.378646048 |
| **A_55_P1956160** | Gm8909 | 0.266162665 | -0.087039361 | 0.619364692 | 8.985938676 | 1.88744496 | 0.11267976 | 0.709264768 | -4.378648104 |
| **A_51_P314277** | Parp1 | 0.404523319 | -0.132369151 | 0.941415788 | 9.220620997 | 1.887150752 | 0.112724046 | 0.709264768 | -4.378724832 |
| **A_51_P455807** | Ehd4 | -0.410774793 | -0.95597647 | 0.134426883 | 13.07258366 | -1.88710884 | 0.112730356 | 0.709264768 | -4.378735763 |
| **A_51_P289881** | Txndc12 | -0.200673618 | -0.467073338 | 0.065726103 | 10.88340175 | -1.886717428 | 0.112789305 | 0.709264768 | -4.37883785 |
| **A_55_P1986552** | Mcam | 0.229766376 | -0.075261005 | 0.534793758 | 13.85816937 | 1.886678904 | 0.112795108 | 0.709264768 | -4.378847899 |
| **A_51_P468294** | Polg | -0.671246856 | -1.562383476 | 0.219889764 | 9.795706916 | -1.886636878 | 0.11280144 | 0.709264768 | -4.37885886 |
| **A_52_P365896** | Pa2g4 | 0.316241388 | -0.103668769 | 0.736151545 | 8.369919922 | 1.886308289 | 0.112850957 | 0.709264768 | -4.378944571 |
| **A_51_P498403** | Cnpy3 | 0.242213029 | -0.079402235 | 0.563828293 | 9.983589442 | 1.886301863 | 0.112851925 | 0.709264768 | -4.378946247 |
| **A_52_P152952** | Leng1 | -0.285231258 | -0.664022322 | 0.093559806 | 8.645562726 | -1.886026272 | 0.112893473 | 0.709264768 | -4.379018138 |
| **A_51_P492482** | Cstf2t | 0.483004209 | -0.158458092 | 1.12446651 | 11.1690505 | 1.885949776 | 0.112905008 | 0.709264768 | -4.379038094 |
| **A_51_P250058** | Epas1 | -1.390851201 | -3.238204546 | 0.456502143 | 13.75897262 | -1.885736661 | 0.11293715 | 0.709264768 | -4.379093691 |

| | | | | | | | | | |
|---|---|---|---|---|---|---|---|---|---|
| **A_55_P2106569** | NA | 1.159899113 | -0.381778807 | 2.701577033 | 16.61469394 | 1.884416549 | 0.11313646 | 0.709264768 | -4.379438143 |
| **A_55_P2025974** | Ruvbl2 | 0.670691072 | -0.220875917 | 1.562258061 | 11.74883808 | 1.884164821 | 0.113174507 | 0.709264768 | -4.379503836 |
| **A_52_P180408** | Bcas2 | 0.310935076 | -0.10240211 | 0.724272261 | 12.01220841 | 1.884150472 | 0.113176676 | 0.709264768 | -4.379507581 |
| **A_55_P2105873** | Ppp2r5c | 0.481921474 | -0.15874112 | 1.122584068 | 9.989148199 | 1.884070962 | 0.113188696 | 0.709264768 | -4.379528332 |
| **A_52_P240542** | Id2 | 0.672343126 | -0.221539853 | 1.566226105 | 12.72910239 | 1.883912148 | 0.113212709 | 0.709264768 | -4.37956978 |
| **A_51_P211903** | Golga5 | -0.260275152 | -0.60632849 | 0.085778186 | 10.34528485 | -1.883822641 | 0.113226244 | 0.709264768 | -4.379593141 |
| **A_51_P485391** | Parn | -0.399603958 | -0.941184836 | 0.14197692 | 9.599727749 | -1.965301405 | 0.11324361 | 0.709264768 | -4.409809143 |
| **A_51_P135340** | Panx1 | 0.742935253 | -0.244955763 | 1.730826269 | 10.6139948 | 1.883615589 | 0.113257562 | 0.709264768 | -4.379647183 |
| **A_51_P294020** | Ticam1 | -0.370308643 | -0.862715349 | 0.122098064 | 8.871925718 | -1.883606735 | 0.113258902 | 0.709264768 | -4.379649494 |
| **A_55_P2047290** | Ankrd40 | -0.204166869 | -0.475655817 | 0.067322078 | 8.624760318 | -1.883577355 | 0.113263347 | 0.709264768 | -4.379657163 |
| **A_66_P128631** | Cinp | 0.328861664 | -0.108478818 | 0.766202147 | 8.877470582 | 1.883405854 | 0.113289295 | 0.709264768 | -4.379701927 |
| **A_55_P2045163** | Ehmt1 | 0.310306147 | -0.102433787 | 0.723046082 | 12.40667574 | 1.883060328 | 0.113341593 | 0.709264768 | -4.379792121 |
| **A_55_P2052170** | NA | 0.548668904 | -0.181181935 | 1.278519743 | 12.28681908 | 1.882897078 | 0.11336631 | 0.709264768 | -4.379834737 |
| **A_66_P112065** | Sertad2 | -0.255006287 | -0.594243135 | 0.084230562 | 10.0882477 | -1.88277409 | 0.113384935 | 0.709264768 | -4.379866844 |
| **A_51_P217430** | Riox1 | 0.327174355 | -0.108115723 | 0.762464433 | 9.635143461 | 1.882568692 | 0.113416047 | 0.709264768 | -4.379920466 |
| **A_55_P1975537** | Nutf2 | 0.321777478 | -0.106457732 | 0.750012687 | 11.4040534 | 1.882017323 | 0.113499606 | 0.709264768 | -4.380064422 |
| **A_55_P1961302** | NA | 0.430464759 | -0.153555656 | 1.014485173 | 11.85913978 | 1.963234962 | 0.113527978 | 0.709264768 | -4.410195548 |
| **A_55_P1989384** | Rnf181 | -0.36578974 | -0.852818473 | 0.121238993 | 11.04158781 | -1.881166695 | 0.11362864 | 0.709264768 | -4.380286546 |
| **A_52_P80806** | Syncrip | 0.526902206 | -0.174639211 | 1.228443622 | 10.1184914 | 1.881165679 | 0.113628794 | 0.709264768 | -4.380286811 |
| **A_55_P2059412** | Ghr | -0.19388331 | -0.452100011 | 0.064333391 | 9.382622842 | -1.880643348 | 0.113708102 | 0.709264768 | -4.380423227 |
| **A_55_P1989663** | Slco3a1 | -0.364379619 | -0.849788925 | 0.121029686 | 10.25370612 | -1.880166565 | 0.113780542 | 0.709264768 | -4.380547762 |
| **A_55_P2072631** | Prr5l | -0.328575161 | -0.766479045 | 0.109328723 | 9.851343003 | -1.879343981 | 0.113905632 | 0.709264768 | -4.38076265 |
| **A_55_P1953809** | Plaat3 | 0.363159616 | -0.120887081 | 0.847206312 | 10.86556106 | 1.87914647 | 0.113935688 | 0.709264768 | -4.380814252 |
| **A_52_P465121** | NA | -0.591317118 | -1.37966073 | 0.197026494 | 10.9404461 | -1.878690522 | 0.114005102 | 0.709264768 | -4.380933384 |
| **A_55_P1988246** | NA | 1.08478556 | -0.361517602 | 2.531088722 | 10.73242334 | 1.878602541 | 0.114018501 | 0.709264768 | -4.380956374 |
| **A_55_P2169135** | NA | 1.226148562 | -0.409751962 | 2.862049087 | 13.76154887 | 1.877312393 | 0.114215171 | 0.709264768 | -4.38129354 |
| **A_55_P2117146** | Pa2g4 | 0.486238484 | -0.162742034 | 1.135219003 | 10.9984525 | 1.876584022 | 0.114326354 | 0.709264768 | -4.381483934 |
| **A_55_P2117984** | Vangl2 | 0.184938546 | -0.061960792 | 0.431837884 | 8.778558293 | 1.876108099 | 0.114399062 | 0.709264768 | -4.381608356 |
| **A_51_P416509** | H1f1 | 1.065325706 | -0.356962693 | 2.487614105 | 9.901013352 | 1.876052918 | 0.114407495 | 0.709264768 | -4.381622783 |
| **A_51_P329356** | Mcts2 | 0.486075886 | -0.162899427 | 1.135051199 | 10.71162284 | 1.87597154 | 0.114419933 | 0.709264768 | -4.381644059 |
| **A_55_P1995151** | NA | 0.465046126 | -0.155951827 | 1.08604408 | 9.974106216 | 1.87566903 | 0.11446618 | 0.709264768 | -4.381723154 |
| **A_55_P2030282** | Gm5796 | -0.336992122 | -0.787078321 | 0.113094077 | 10.16390672 | -1.875315736 | 0.114520216 | 0.709264768 | -4.381815534 |
| **A_51_P406008** | Hgh1 | 0.187748021 | -0.063052716 | 0.438548759 | 8.55176131 | 1.874981144 | 0.114571415 | 0.709264768 | -4.38190303 |
| **A_55_P1992814** | Shq1 | 0.200873266 | -0.067481088 | 0.46922762 | 8.368614036 | 1.874838403 | 0.114593264 | 0.709264768 | -4.381940359 |
| **A_55_P2157008** | Ear10 | -0.395319708 | -0.923660967 | 0.133021552 | 9.332930381 | -1.874063422 | 0.114711962 | 0.709264768 | -4.382143049 |
| **A_55_P2213348** | 5330421C | -0.178090688 | -0.416126497 | 0.059945121 | 8.12773762 | -1.873912298 | 0.114735123 | 0.709264768 | -4.382182578 |
| **A_55_P1993807** | Nudt10 | 0.353652851 | -0.119078833 | 0.826384535 | 8.41707824 | 1.873754973 | 0.11475924 | 0.709264768 | -4.38222373 |
| **A_52_P242251** | Utp25 | 0.181536765 | -0.061173162 | 0.424246693 | 8.369606897 | 1.873386554 | 0.114815735 | 0.709264768 | -4.382320106 |
| **A_51_P118742** | Drg2 | 0.205231898 | -0.069159494 | 0.47962329 | 9.147239092 | 1.873375019 | 0.114817504 | 0.709264768 | -4.382323123 |
| **A_51_P319460** | Osmr | -0.751775759 | -1.756927781 | 0.253376262 | 9.880755243 | -1.873298188 | 0.11482929 | 0.709264768 | -4.382343223 |
| **A_51_P227222** | Adamts2 | -0.570519838 | -1.333449764 | 0.192410088 | 12.4113844 | -1.872994254 | 0.114875923 | 0.709264768 | -4.382422737 |
| **A_55_P2091472** | Bnip3 | -0.441990696 | -1.033120731 | 0.149139339 | 12.89292534 | -1.872752766 | 0.114912989 | 0.709264768 | -4.382485918 |
| **A_55_P2235970** | C130068E | -0.211081147 | -0.493390729 | 0.071228436 | 8.193743485 | -1.872725701 | 0.114917144 | 0.709264768 | -4.382492999 |
| **A_55_P2029682** | Fry | -0.388320626 | -0.907706711 | 0.131065459 | 9.336384059 | -1.872623611 | 0.114932818 | 0.709264768 | -4.38251971 |
| **A_52_P654437** | NA | 0.42386297 | -0.143074081 | 0.990800022 | 11.43461994 | 1.872582593 | 0.114939116 | 0.709264768 | -4.382530442 |
| **A_55_P2102693** | Acvr1 | -0.25557799 | -0.597432051 | 0.086276071 | 9.037077023 | -1.872548404 | 0.114944366 | 0.709264768 | -4.382539388 |
| **A_55_P1965070** | Map7d1 | -0.69132642 | -1.616490566 | 0.233837727 | 13.97466317 | -1.871607194 | 0.115088987 | 0.709264768 | -4.382785681 |
| **A_55_P2035972** | Mark3 | -0.321508904 | -0.75180965 | 0.108791842 | 11.09523892 | -1.871419934 | 0.115117782 | 0.709264768 | -4.382834689 |

| | | | | | | | | | |
|---|---|---|---|---|---|---|---|---|---|
| **A_52_P546660** | 1700029I1 | -0.293098201 | -0.685475746 | 0.099279344 | 8.245379648 | -1.870937814 | 0.115191952 | 0.709264768 | -4.382960874 |
| **A_55_P2117333** | NA | 0.198017965 | -0.067078522 | 0.463114453 | 10.35415328 | 1.870901686 | 0.115197512 | 0.709264768 | -4.38297033 |
| **A_55_P1990066** | NA | 0.563733934 | -0.191031359 | 1.318499226 | 9.817204336 | 1.870736423 | 0.115222948 | 0.709264768 | -4.383013587 |
| **A_51_P148314** | Fam117a | -0.33498602 | -0.783514242 | 0.113542202 | 10.07766212 | -1.87062724 | 0.115239756 | 0.709264768 | -4.383042167 |
| **A_55_P2128944** | Bcl2l13 | 0.231517601 | -0.078590437 | 0.541625638 | 10.69357202 | 1.869912722 | 0.115349814 | 0.709264768 | -4.383229214 |
| **A_55_P2116290** | Szrd1 | 0.180180268 | -0.061184484 | 0.42154502 | 10.34283387 | 1.869750798 | 0.11537477 | 0.709264768 | -4.383271607 |
| **A_55_P2136426** | Prss57 | 0.280468803 | -0.095278294 | 0.656215901 | 7.814305004 | 1.869559419 | 0.115404272 | 0.709264768 | -4.383321713 |
| **A_55_P1992470** | NA | 0.335299756 | -0.113922292 | 0.784521804 | 13.85561325 | 1.869487303 | 0.115415391 | 0.709264768 | -4.383340594 |
| **A_55_P1977934** | Itgam | -0.373594328 | -0.874190488 | 0.127001831 | 11.74715492 | -1.869231566 | 0.115454831 | 0.709264768 | -4.383407555 |
| **A_55_P2015113** | Atg5 | -0.161101434 | -0.376990576 | 0.054787707 | 10.74926092 | -1.869041535 | 0.115484146 | 0.709264768 | -4.383457313 |
| **A_55_P2260052** | Gm17753 | 0.348980025 | -0.118689853 | 0.816649902 | 8.701767232 | 1.869009535 | 0.115489083 | 0.709264768 | -4.383465693 |
| **A_66_P117815** | NA | 0.787594594 | -0.267922253 | 1.843111441 | 13.01171366 | 1.868907912 | 0.115504764 | 0.709264768 | -4.383492303 |
| **A_55_P1959541** | NA | 0.212265088 | -0.072240739 | 0.496770914 | 10.14617254 | 1.868692101 | 0.115538071 | 0.709264768 | -4.383548816 |
| **A_51_P140797** | Dcaf6 | -0.203251635 | -0.475839989 | 0.069336719 | 8.453831142 | -1.86757089 | 0.11571127 | 0.709264768 | -4.383842464 |
| **A_55_P1956482** | NA | 0.362345979 | -0.123689713 | 0.848381672 | 10.79088995 | 1.867263591 | 0.115758786 | 0.709264768 | -4.383922959 |
| **A_55_P2028288** | Nhlrc4 | -0.251558702 | -0.59569625 | 0.092578847 | 8.070312785 | -1.947018104 | 0.11578614 | 0.709264768 | -4.413238037 |
| **A_51_P133097** | Gm4013 | 0.269961626 | -0.092207891 | 0.632131143 | 8.645008227 | 1.86698353 | 0.115802108 | 0.709264768 | -4.383996324 |
| **A_51_P209280** | Rab31 | -0.769970823 | -1.803418532 | 0.263476886 | 9.339438265 | -1.866105132 | 0.115938089 | 0.709264768 | -4.384226458 |
| **A_55_P2036978** | Mrps14 | 0.34376482 | -0.117662169 | 0.805191809 | 12.4659542 | 1.865987726 | 0.115956277 | 0.709264768 | -4.384257221 |
| **A_52_P531651** | Parvg | 0.53759537 | -0.184080237 | 1.259270977 | 8.137455629 | 1.86579463 | 0.115986195 | 0.709264768 | -4.384307818 |
| **A_51_P327828** | Ppib | -0.404283886 | -0.947247979 | 0.138680208 | 14.19581582 | -1.864943445 | 0.116118173 | 0.709264768 | -4.384530879 |
| **A_55_P2069176** | NA | 0.722101352 | -0.247800088 | 1.692002793 | 12.80655809 | 1.864751218 | 0.116147999 | 0.709264768 | -4.384581259 |
| **A_66_P128384** | Hdac1 | 0.486879725 | -0.167107109 | 1.140866559 | 12.01321948 | 1.864674495 | 0.116159906 | 0.709264768 | -4.384601368 |
| **A_55_P1977318** | Wbp11 | 0.290724458 | -0.09978773 | 0.681236647 | 11.97010823 | 1.864649986 | 0.11616371 | 0.709264768 | -4.384607792 |
| **A_55_P1953087** | Mcm3 | 1.554467879 | -0.53365795 | 3.642593709 | 12.22850902 | 1.864556005 | 0.116178297 | 0.709264768 | -4.384632425 |
| **A_55_P2078048** | BC004004 | -0.298411013 | -0.699298518 | 0.102476492 | 12.46006299 | -1.864415359 | 0.11620013 | 0.709264768 | -4.384669289 |
| **A_52_P481423** | Cttnbp2nl | -0.20868198 | -0.489067627 | 0.071703668 | 11.40397272 | -1.864144247 | 0.116242228 | 0.709264768 | -4.384740353 |
| **A_55_P1968928** | Def6 | 0.757243276 | -0.260304018 | 1.77479057 | 10.14421934 | 1.863936735 | 0.116274461 | 0.709264768 | -4.384794749 |
| **A_55_P1998801** | NA | 0.188834854 | -0.064980615 | 0.442650322 | 9.808118155 | 1.863435714 | 0.116352322 | 0.709264768 | -4.384926094 |
| **A_55_P1961968** | Plagl1 | -0.218696211 | -0.512732486 | 0.075340063 | 15.01644792 | -1.862904936 | 0.116434864 | 0.709264768 | -4.385065255 |
| **A_55_P1970636** | Unc45a | -0.47128173 | -1.104938946 | 0.162375486 | 12.01697206 | -1.862844332 | 0.116444293 | 0.709264768 | -4.385081146 |
| **A_51_P194498** | Wfdc1 | -0.615308943 | -1.44275046 | 0.212132574 | 9.346963731 | -1.862542816 | 0.116491213 | 0.709264768 | -4.385160207 |
| **A_55_P2006762** | Bzw1 | 0.309726506 | -0.106799856 | 0.726252867 | 12.16647468 | 1.862456878 | 0.116504589 | 0.709264768 | -4.385182741 |
| **A_55_P1988774** | Slc17a5 | -0.370911163 | -0.869736276 | 0.12791395 | 11.82978086 | -1.862395616 | 0.116514126 | 0.709264768 | -4.385198806 |
| **A_55_P2166572** | Strbp | 0.438157044 | -0.151117195 | 1.027431284 | 9.371664858 | 1.862355939 | 0.116520302 | 0.709264768 | -4.38520921 |
| **A_55_P2048323** | Cdk10 | -0.278874493 | -0.653951358 | 0.096202371 | 10.5202186 | -1.862253776 | 0.116536209 | 0.709264768 | -4.385236001 |
| **A_51_P259296** | Lpl | -0.27632345 | -0.647972796 | 0.095325896 | 9.34215527 | -1.862235989 | 0.116538978 | 0.709264768 | -4.385240665 |
| **A_55_P2186648** | Bard1 | 0.843150804 | -0.291073737 | 1.977375345 | 9.303138347 | 1.861900911 | 0.116591164 | 0.709264768 | -4.385328538 |
| **A_51_P192162** | NA | -0.177049822 | -0.415237911 | 0.061138266 | 9.244208589 | -1.861769029 | 0.116611711 | 0.709264768 | -4.385363125 |
| **A_55_P2082869** | Ezh1 | -0.704241043 | -1.65169779 | 0.243215704 | 12.09607519 | -1.861711085 | 0.116620739 | 0.709264768 | -4.385378322 |
| **A_51_P264053** | Pigf | 0.222782914 | -0.07696386 | 0.522529687 | 10.02548859 | 1.861562873 | 0.116643836 | 0.709264768 | -4.385417193 |
| **A_55_P2133195** | Gm4951 | 0.463431519 | -0.160100056 | 1.086963093 | 9.128111804 | 1.861561819 | 0.116644 | 0.709264768 | -4.38541747 |
| **A_55_P1978855** | NA | 0.164521397 | -0.056871358 | 0.385914151 | 15.37026116 | 1.861269797 | 0.11668952 | 0.709264768 | -4.385494062 |
| **A_55_P2023783** | Leng8 | -0.386253985 | -0.906045959 | 0.133537988 | 9.177871009 | -1.861203027 | 0.116699931 | 0.709264768 | -4.385511575 |
| **A_52_P509744** | Nup160 | 0.294227369 | -0.101734645 | 0.690189384 | 9.221772666 | 1.861143625 | 0.116709194 | 0.709264768 | -4.385527156 |
| **A_55_P2083649** | Alas1 | -0.481742277 | -1.130074593 | 0.16659004 | 12.52281763 | -1.861090262 | 0.116717515 | 0.709264768 | -4.385541153 |
| **A_51_P487690** | Ifi44 | -0.502190677 | -1.178059672 | 0.173678318 | 8.425874199 | -1.861043255 | 0.116724846 | 0.709264768 | -4.385553483 |
| **A_55_P1964403** | Agfg2 | 0.158249174 | -0.054765354 | 0.371263701 | 8.072865973 | 1.860726633 | 0.116774237 | 0.709264768 | -4.385636537 |

| | | | | | | | | | |
|---|---|---|---|---|---|---|---|---|---|
| **A_51_P260658** | Amdhd2 | -0.598341443 | -1.403822168 | 0.207139282 | 11.39297614 | -1.86056256 | 0.116799839 | 0.709264768 | -4.385679578 |
| **A_55_P2037585** | Trf | -1.255673562 | -2.946312159 | 0.434965035 | 13.71410089 | -1.860271431 | 0.116845282 | 0.709264768 | -4.385755952 |
| **A_55_P2178349** | Ywhaq | 0.368170183 | -0.127541997 | 0.863882363 | 12.35232417 | 1.86024167 | 0.116849928 | 0.709264768 | -4.38576376 |
| **A_55_P2074081** | NA | 1.002319522 | -0.347299032 | 2.351938075 | 14.57107348 | 1.860139412 | 0.116865895 | 0.709264768 | -4.385790587 |
| **A_55_P1959595** | Fmo5 | -0.248228089 | -0.582583709 | 0.086127531 | 8.970586657 | -1.859484839 | 0.116968151 | 0.709264768 | -4.385962329 |
| **A_51_P338998** | Prim2 | 0.416790237 | -0.144693855 | 0.978274329 | 9.930257985 | 1.859218478 | 0.117009787 | 0.709264768 | -4.386032222 |
| **A_52_P430247** | Nacc1 | 0.21836082 | -0.075858994 | 0.512580634 | 9.626447024 | 1.858887666 | 0.117061519 | 0.709264768 | -4.386119032 |
| **A_55_P2112797** | NA | 0.443679423 | -0.154169337 | 1.041528183 | 9.139292722 | 1.858781325 | 0.117078153 | 0.709264768 | -4.386146939 |
| **A_51_P129720** | Hectd3 | -0.334796415 | -0.786197186 | 0.116604356 | 9.569573856 | -1.857671199 | 0.117251946 | 0.709264768 | -4.386438304 |
| **A_55_P1998776** | NA | 0.462600964 | -0.161118906 | 1.086320835 | 15.96268599 | 1.857664575 | 0.117252984 | 0.709264768 | -4.386440043 |
| **A_51_P195034** | Esco2 | 0.595217033 | -0.207348368 | 1.397782434 | 8.702123984 | 1.857570319 | 0.117267752 | 0.709264768 | -4.386464784 |
| **A_55_P1976690** | Snrnp35 | 0.173008564 | -0.060397256 | 0.406414385 | 10.68062257 | 1.85654836 | 0.117427994 | 0.709264768 | -4.386733076 |
| **A_55_P1990495** | Stat5a | 0.390656827 | -0.136484581 | 0.917798235 | 9.250472571 | 1.856173773 | 0.117486784 | 0.709264768 | -4.386831429 |
| **A_55_P1960456** | Larp1 | 0.205961772 | -0.071991209 | 0.483914753 | 12.58643826 | 1.855947251 | 0.11752235 | 0.709264768 | -4.38689091 |
| **A_55_P1962906** | Als2cl | -0.365372981 | -0.85853381 | 0.127787848 | 10.3189258 | -1.855659119 | 0.117567605 | 0.709264768 | -4.386966572 |
| **A_55_P2060101** | Tprgl | -0.289930382 | -0.681264889 | 0.101404125 | 13.73216303 | -1.855649416 | 0.11756913 | 0.709264768 | -4.386969121 |
| **A_51_P414889** | Ifi35 | -0.494207387 | -1.161419375 | 0.173004601 | 9.69204114 | -1.855221368 | 0.117636394 | 0.709264768 | -4.387081534 |
| **A_52_P272534** | NA | 0.376429543 | -0.132109906 | 0.884968992 | 11.404856 | 1.853998581 | 0.11782876 | 0.709264768 | -4.387402714 |
| **A_55_P1989061** | Tsc22d3 | -0.901399258 | -2.119179406 | 0.31638089 | 12.89155587 | -1.853952708 | 0.117835982 | 0.709264768 | -4.387414765 |
| **A_55_P2068096** | Rap1gap | -0.472308302 | -1.110461032 | 0.165844428 | 9.97397756 | -1.853750554 | 0.117867817 | 0.709264768 | -4.387467872 |
| **A_55_P2157616** | NA | 0.272275168 | -0.095631834 | 0.640182171 | 16.78798223 | 1.853618377 | 0.117888637 | 0.709264768 | -4.387502597 |
| **A_52_P301495** | Vamp2 | -0.239201766 | -0.562456711 | 0.084053179 | 9.649861766 | -1.853401674 | 0.117922778 | 0.709264768 | -4.38755953 |
| **A_55_P2023136** | Rnf214 | -0.192583619 | -0.45285075 | 0.067683511 | 8.610944955 | -1.853319804 | 0.11793568 | 0.709264768 | -4.38758104 |
| **A_52_P649332** | Sdf4 | -0.669948184 | -1.575403794 | 0.235507427 | 11.37373946 | -1.853208977 | 0.117953146 | 0.709264768 | -4.387610158 |
| **A_51_P485421** | NA | 0.373759591 | -0.131516601 | 0.879035782 | 7.922453191 | 1.852737347 | 0.118027505 | 0.709264768 | -4.390441893 |
| **A_51_P213153** | Tmem208 | -0.467614377 | -1.099877719 | 0.164648965 | 13.14505267 | -1.852423144 | 0.11807707 | 0.709264768 | -4.387816646 |
| **A_52_P183038** | Impdh1 | 0.284158082 | -0.100122279 | 0.668438444 | 10.83197456 | 1.852090346 | 0.118129591 | 0.709264768 | -4.387904103 |
| **A_55_P2077079** | Rab29 | -0.214686599 | -0.505071689 | 0.075698491 | 9.640128943 | -1.851744097 | 0.11818426 | 0.709264768 | -4.387995102 |
| **A_51_P352594** | St5 | -0.245645075 | -0.577958341 | 0.086668192 | 9.277393512 | -1.851444625 | 0.118231564 | 0.709264768 | -4.388073812 |
| **A_51_P343900** | Xrcc1 | 0.375965984 | -0.132679682 | 0.884611649 | 11.31182244 | 1.851328771 | 0.118249869 | 0.709264768 | -4.388104263 |
| **A_51_P151586** | Haspin | 0.508437677 | -0.179461553 | 1.196336907 | 8.600229356 | 1.851242227 | 0.118263545 | 0.709264768 | -4.388127011 |
| **A_55_P2099700** | Klc4 | -0.320599841 | -0.754518723 | 0.11331904 | 11.48881456 | -1.850568221 | 0.118370108 | 0.709264768 | -4.388304186 |
| **A_55_P2185332** | Il17rc | -0.307329395 | -0.723317846 | 0.108659055 | 10.9376187 | -1.850432187 | 0.118391627 | 0.709264768 | -4.388339948 |
| **A_55_P1973970** | Krt73 | 0.268935131 | -0.095084503 | 0.632954765 | 8.818808125 | 1.850431762 | 0.118391695 | 0.709264768 | -4.38834006 |
| **A_55_P1987761** | Prkcz | -0.554633244 | -1.305405561 | 0.196139073 | 9.162572892 | -1.850324895 | 0.118408603 | 0.709264768 | -4.388368155 |
| **A_52_P526396** | Atl1 | -0.242258836 | -0.570250574 | 0.085732903 | 8.120947759 | -1.84998014 | 0.118463165 | 0.709264768 | -4.388458794 |
| **A_55_P1960837** | NA | 0.61817795 | -0.21878805 | 1.45514395 | 13.32620162 | 1.849933171 | 0.118470601 | 0.709264768 | -4.388471143 |
| **A_51_P162471** | Dnajc9 | 1.071991764 | -0.37957173 | 2.523555257 | 12.35192407 | 1.849718997 | 0.118504512 | 0.709264768 | -4.388527455 |
| **A_51_P233101** | Ltbp3 | -0.222825848 | -0.524557194 | 0.078905497 | 12.26385342 | -1.849675245 | 0.11851144 | 0.709264768 | -4.388538959 |
| **A_55_P2093920** | Pzp | 0.514652516 | -0.182258963 | 1.211563996 | 8.294320917 | 1.849638377 | 0.118517279 | 0.709264768 | -4.388548653 |
| **A_51_P416137** | Slc31a2 | -0.81825508 | -1.926335451 | 0.289825291 | 9.64591305 | -1.849557316 | 0.118530118 | 0.709264768 | -4.388569968 |
| **A_51_P170807** | Map3k6 | -0.200945035 | -0.473115238 | 0.071225168 | 9.874865906 | -1.849213474 | 0.118584592 | 0.709264768 | -4.388660382 |
| **A_55_P2121827** | Dgcr2 | 0.211184222 | -0.074857952 | 0.497226396 | 8.800357085 | 1.849190883 | 0.118588172 | 0.709264768 | -4.388666322 |
| **A_55_P2106150** | Cenpk | 1.080173793 | -0.383010748 | 2.543358334 | 10.05176517 | 1.849033918 | 0.118613049 | 0.709264768 | -4.388707599 |
| **A_55_P2471798** | NA | -0.189883891 | -0.447131155 | 0.067363372 | 9.735566644 | -1.848790523 | 0.118651634 | 0.709264768 | -4.388771607 |
| **A_55_P2066116** | Bcl3 | -1.069361934 | -2.518398095 | 0.379674226 | 10.45501334 | -1.848399502 | 0.118713648 | 0.709264768 | -4.388874443 |
| **A_55_P2007510** | Pef1 | -0.385293973 | -0.907416286 | 0.136828339 | 12.35570373 | -1.84829083 | 0.118730889 | 0.709264768 | -4.388903025 |
| **A_51_P502872** | 2200002D | -0.459578159 | -1.082367271 | 0.163210953 | 11.94270439 | -1.848284047 | 0.118731965 | 0.709264768 | -4.388904809 |

| | | | | | | | | | |
|---|---|---|---|---|---|---|---|---|---|
| **A_55_P2084433** | NA | 0.1858082 | -0.065989712 | 0.437606111 | 12.21900003 | 1.84826014 | 0.118735759 | 0.709264768 | -4.388911097 |
| **A_51_P417053** | Tuft1 | -0.501059573 | -1.180390558 | 0.178271412 | 9.903822745 | -1.847388712 | 0.118874108 | 0.709264768 | -4.389140316 |
| **A_55_P2150224** | NA | 1.095940016 | -0.390051511 | 2.581931544 | 14.33547378 | 1.847229251 | 0.118899441 | 0.709264768 | -4.389182265 |
| **A_55_P2383902** | 8030453C | -0.338177253 | -0.796732387 | 0.12037788 | 8.192312511 | -1.847154313 | 0.118911349 | 0.709264768 | -4.389201979 |
| **A_55_P2077879** | Kat6b | -0.1661053 | -0.39137301 | 0.05916241 | 8.271220097 | -1.846863897 | 0.118957506 | 0.709264768 | -4.389278382 |
| **A_55_P2122419** | Arl1 | -0.273068821 | -0.643439362 | 0.097301721 | 9.911345782 | -1.846656092 | 0.118990545 | 0.709264768 | -4.389333055 |
| **A_55_P1992388** | NA | -0.248302669 | -0.585116778 | 0.088511441 | 14.41405123 | -1.846466598 | 0.11902068 | 0.709264768 | -4.389382912 |
| **A_51_P362538** | Syde2 | -0.188817916 | -0.445183015 | 0.067547183 | 8.806081962 | -1.844737787 | 0.119295971 | 0.709264768 | -4.389837866 |
| **A_55_P2090798** | Mtus1 | -0.282229132 | -0.665617837 | 0.101159573 | 10.61801302 | -1.843796015 | 0.119446205 | 0.709264768 | -4.390085771 |
| **A_55_P2084626** | Rab5b | -0.247910312 | -0.584685137 | 0.088864513 | 12.01526506 | -1.843763943 | 0.119451325 | 0.709264768 | -4.390094215 |
| **A_55_P2165334** | Nde1 | 0.513267208 | -0.183997559 | 1.210531975 | 11.98823734 | 1.843724996 | 0.119457542 | 0.709264768 | -4.390104468 |
| **A_51_P430620** | Slc25a12 | 0.190791517 | -0.068418133 | 0.450001167 | 10.17565144 | 1.84356409 | 0.119483232 | 0.709264768 | -4.390146829 |
| **A_55_P1958306** | Thrap3 | 0.403389971 | -0.144672878 | 0.95145282 | 12.76432521 | 1.843508218 | 0.119492153 | 0.709264768 | -4.390161539 |
| **A_55_P2140390** | NA | 1.13924069 | -0.408891526 | 2.687372906 | 13.75512565 | 1.843137714 | 0.119551332 | 0.709264768 | -4.390259087 |
| **A_55_P2090943** | Larp4 | 0.187683407 | -0.067366881 | 0.442733695 | 11.5322592 | 1.843106397 | 0.119556336 | 0.709264768 | -4.390267332 |
| **A_55_P2017267** | Nanp | 0.511499807 | -0.183663404 | 1.206663018 | 9.265161414 | 1.842930848 | 0.119584387 | 0.709264768 | -4.390313555 |
| **A_51_P316951** | Wipf3 | -0.669660345 | -1.580231091 | 0.240910401 | 9.35551164 | -1.842006823 | 0.119732147 | 0.709264768 | -4.390556879 |
| **A_55_P2034400** | Cdk8 | 0.216343075 | -0.077852009 | 0.51053816 | 10.63484849 | 1.841865582 | 0.119754749 | 0.709264768 | -4.390594076 |
| **A_55_P2145557** | NA | -0.343142437 | -0.809836497 | 0.123551623 | 12.37574165 | -1.841588116 | 0.119799163 | 0.709264768 | -4.390667153 |
| **A_51_P251352** | Slc25a13 | 0.282717596 | -0.101844134 | 0.667279326 | 9.979794623 | 1.841353292 | 0.119836764 | 0.709264768 | -4.390729002 |
| **A_55_P2070936** | Tra2b | 0.262746171 | -0.09466219 | 0.620154533 | 8.329538689 | 1.841289316 | 0.11984701 | 0.709264768 | -4.390745853 |
| **A_52_P421713** | 9930012K | -0.412097057 | -0.972819788 | 0.148625675 | 8.756082221 | -1.840779189 | 0.119928742 | 0.709264768 | -4.390880225 |
| **A_52_P453785** | Cthrc1 | 0.309189363 | -0.111570659 | 0.729949385 | 9.252222902 | 1.840519492 | 0.119970372 | 0.709264768 | -4.390948637 |
| **A_52_P541175** | Marveld1 | 0.384105397 | -0.138609742 | 0.906820535 | 9.173489599 | 1.840499385 | 0.119973596 | 0.709264768 | -4.390953934 |
| **A_55_P2163493** | Tgm3 | 0.358555235 | -0.129459434 | 0.846569905 | 10.21051863 | 1.840236061 | 0.120015823 | 0.709264768 | -4.391023305 |
| **A_55_P1992257** | Ap1s1 | -0.203421477 | -0.480298133 | 0.073455179 | 14.09648532 | -1.840182124 | 0.120024474 | 0.709264768 | -4.391037515 |
| **A_55_P2060608** | NA | 0.645765872 | -0.233261899 | 1.524793643 | 14.43365302 | 1.840021251 | 0.120050281 | 0.709264768 | -4.391079898 |
| **A_55_P2169729** | NA | 0.686628538 | -0.248261182 | 1.621518259 | 13.96696731 | 1.839550909 | 0.120125765 | 0.709264768 | -4.391203822 |
| **A_55_P2095213** | Dusp22 | -0.306398049 | -0.723586248 | 0.11079015 | 9.116617319 | -1.839519206 | 0.120130855 | 0.709264768 | -4.391212176 |
| **A_51_P515452** | Fkbpl | -0.393710268 | -0.929816982 | 0.142396446 | 10.52918252 | -1.839398552 | 0.120150227 | 0.709264768 | -4.391243967 |
| **A_55_P2005719** | NA | 0.453323161 | -0.16402407 | 1.070670391 | 12.53834906 | 1.839199114 | 0.120182255 | 0.709264768 | -4.39129652 |
| **A_52_P81783** | Gspt1 | 0.375555842 | -0.135970561 | 0.887082244 | 9.358571239 | 1.838894495 | 0.120231191 | 0.709264768 | -4.391376792 |
| **A_55_P1973486** | Atp6v1f | -0.296877652 | -0.701256376 | 0.107501072 | 14.26497282 | -1.838821421 | 0.120242933 | 0.709264768 | -4.391396049 |
| **A_55_P2317665** | E2f1 | 0.666839412 | -0.24157988 | 1.575258703 | 9.517685467 | 1.838591537 | 0.12027988 | 0.709264768 | -4.391456632 |
| **A_55_P2141884** | Tagln2 | -0.327037401 | -0.772724724 | 0.118649923 | 11.52033055 | -1.837881415 | 0.120394084 | 0.709264768 | -4.391643792 |
| **A_55_P2045268** | Tmem59 | -0.405841728 | -0.959055669 | 0.147372213 | 13.23228075 | -1.837443221 | 0.12046461 | 0.709264768 | -4.391759296 |
| **A_55_P2097178** | Idua | -0.562364142 | -1.329167173 | 0.204438889 | 9.178959947 | -1.836894212 | 0.120553029 | 0.709264768 | -4.391904024 |
| **A_55_P2127782** | NA | 0.336738191 | -0.12246874 | 0.795945122 | 14.093385 | 1.836683338 | 0.120587009 | 0.709264768 | -4.391959619 |
| **A_66_P112862** | Gm17821 | -0.553735463 | -1.309088011 | 0.201617085 | 11.91053982 | -1.836128133 | 0.120676518 | 0.709264768 | -4.392106003 |
| **A_55_P2140031** | Fam129b | -0.52316064 | -1.237479494 | 0.191158214 | 14.73712179 | -1.834396846 | 0.120956063 | 0.709264768 | -4.392562579 |
| **A_55_P1971074** | Atp11a | -0.6267933 | -1.482753296 | 0.229166696 | 12.34476771 | -1.834092449 | 0.12100528 | 0.709264768 | -4.392642872 |
| **A_55_P2065726** | Snx29 | -0.173149853 | -0.409636563 | 0.063336858 | 8.398589334 | -1.833858114 | 0.121043182 | 0.709264768 | -4.392704687 |
| **A_55_P2058783** | Hilpda | -0.697112911 | -1.649359104 | 0.255133282 | 11.07156303 | -1.833598424 | 0.1210852 | 0.709264768 | -4.392773194 |
| **A_55_P1990324** | Ptprc | 0.361537224 | -0.132434983 | 0.85550943 | 8.005329673 | 1.833162018 | 0.121155844 | 0.709264768 | -4.392888327 |
| **A_51_P491350** | Col4a2 | -0.531722109 | -1.258265395 | 0.194821178 | 13.34496104 | -1.83304687 | 0.12117449 | 0.709264768 | -4.392918707 |
| **A_55_P2091147** | H4c12 | 1.319322669 | -0.483489796 | 3.122135134 | 12.94943084 | 1.832950844 | 0.121190042 | 0.709264768 | -4.392944043 |
| **A_55_P2131105** | NA | 0.873130692 | -0.319984114 | 2.066245498 | 12.40462211 | 1.83293627 | 0.121192403 | 0.709264768 | -4.392947888 |
| **A_51_P357561** | Fbxw9 | -0.351617213 | -0.832193646 | 0.128959219 | 8.741015809 | -1.832559162 | 0.121253499 | 0.709264768 | -4.39304739 |

| | | | | | | | | | |
|---|---|---|---|---|---|---|---|---|---|
| **A_51_P175771** | Flii | -0.258964295 | -0.612994584 | 0.095065994 | 11.47440357 | -1.8321027 | 0.121327493 | 0.709264768 | -4.39316784 |
| **A_55_P2130792** | Tuft1 | -0.29761685 | -0.704502993 | 0.109269292 | 8.535658418 | -1.832040048 | 0.121337653 | 0.709264768 | -4.393184373 |
| **A_52_P304858** | Gnb1 | 0.332920151 | -0.1222424 | 0.788082702 | 10.32888086 | 1.831993406 | 0.121345217 | 0.709264768 | -4.393196682 |
| **A_55_P2100895** | Arnt | 0.242940112 | -0.089225411 | 0.575105635 | 10.77602921 | 1.831871562 | 0.121364979 | 0.709264768 | -4.393228836 |
| **A_52_P20727** | Nhlrc1 | 0.308734869 | -0.113395765 | 0.730865503 | 8.490722282 | 1.831846831 | 0.12136899 | 0.709264768 | -4.393235363 |
| **A_55_P2131481** | NA | 0.930070359 | -0.341729237 | 2.201869955 | 14.59235329 | 1.831670988 | 0.121397517 | 0.709264768 | -4.393281769 |
| **A_55_P2000813** | Cops2 | 0.180668606 | -0.066385022 | 0.427722235 | 9.615040364 | 1.831647101 | 0.121401393 | 0.709264768 | -4.393288073 |
| **A_51_P359983** | Arhgap17 | -0.341736422 | -0.809082484 | 0.125609639 | 10.17666807 | -1.831483569 | 0.121427929 | 0.709264768 | -4.393331233 |
| **A_52_P141583** | Macroh2a | 0.574907728 | -0.211440093 | 1.361255548 | 10.93560886 | 1.831191672 | 0.12147531 | 0.709264768 | -4.393408273 |
| **A_55_P2091402** | Kpna2 | 1.209663238 | -0.444922241 | 2.864248717 | 13.36344617 | 1.831157161 | 0.121480913 | 0.709264768 | -4.393417382 |
| **A_52_P371718** | Tgs1 | 0.174930415 | -0.064343683 | 0.414204513 | 9.639107645 | 1.831133401 | 0.121484771 | 0.709264768 | -4.393423653 |
| **A_55_P2074864** | NA | 0.734700996 | -0.270281278 | 1.739683269 | 13.62153493 | 1.831059988 | 0.121496691 | 0.709264768 | -4.39344303 |
| **A_55_P2058791** | 4833420G | -0.252544552 | -0.597998563 | 0.092909458 | 8.708744568 | -1.831041181 | 0.121499745 | 0.709264768 | -4.393447994 |
| **A_52_P79187** | Ankrd12 | -0.341997345 | -0.809834202 | 0.125839511 | 8.949904356 | -1.83095912 | 0.121513071 | 0.709264768 | -4.393469654 |
| **A_55_P2048409** | Apool | 0.407901503 | -0.150137387 | 0.965940392 | 10.40620008 | 1.830801186 | 0.121538723 | 0.709264768 | -4.393511341 |
| **A_55_P2154709** | Pter | 0.327432759 | -0.120648547 | 0.775514064 | 8.654156737 | 1.830272049 | 0.121624705 | 0.709264768 | -4.393651019 |
| **A_55_P2039541** | Arhgap15 | 0.497805159 | -0.183576831 | 1.17918715 | 8.280945223 | 1.82986516 | 0.121690864 | 0.709264768 | -4.393758437 |
| **A_55_P2077671** | NA | 1.062158877 | -0.391884469 | 2.516202223 | 14.1986949 | 1.829626641 | 0.121729663 | 0.709264768 | -4.39382141 |
| **A_55_P2091247** | NA | 0.397547294 | -0.146855621 | 0.941950209 | 13.09665549 | 1.829021056 | 0.121828228 | 0.709264768 | -4.393981307 |
| **A_55_P2039324** | Ccnf | 0.67210469 | -0.248308564 | 1.592517944 | 11.38710506 | 1.828960843 | 0.121838032 | 0.709264768 | -4.393997207 |
| **A_51_P473888** | Il6st | -0.540468814 | -1.280661338 | 0.199723709 | 11.1068166 | -1.828842439 | 0.121857315 | 0.709264768 | -4.394028473 |
| **A_51_P350453** | Pdk4 | -1.026459702 | -2.43225688 | 0.379337475 | 9.154814441 | -1.82881419 | 0.121861916 | 0.709264768 | -4.394035932 |
| **A_55_P1978201** | Incenp | 1.298228556 | -0.47994044 | 3.076397551 | 12.17653119 | 1.828641071 | 0.121890115 | 0.709264768 | -4.394081647 |
| **A_51_P499195** | Nkg7 | 0.297211323 | -0.110029179 | 0.704451825 | 7.862332029 | 1.82795177 | 0.12200246 | 0.709264768 | -4.394263686 |
| **A_55_P1981992** | NA | 0.399974639 | -0.148136531 | 0.948085809 | 9.335762658 | 1.827738871 | 0.12203718 | 0.709264768 | -4.394319916 |
| **A_55_P2127139** | H3c4 | 0.483001246 | -0.178995729 | 1.14499822 | 14.4768111 | 1.827437749 | 0.122086305 | 0.709264768 | -4.394399452 |
| **A_51_P454008** | Lbp | -0.53011067 | -1.25669084 | 0.1964695 | 9.096446501 | -1.827398865 | 0.12209265 | 0.709264768 | -4.394409722 |
| **A_55_P2173623** | Samd1 | 0.268826244 | -0.099679914 | 0.637332401 | 12.87380819 | 1.827162867 | 0.122131167 | 0.709264768 | -4.39447206 |
| **A_52_P667477** | Fyco1 | -0.207040758 | -0.490897546 | 0.076816031 | 8.717301813 | -1.826866907 | 0.122179487 | 0.709264768 | -4.394550241 |
| **A_55_P2038494** | Rexo2 | -0.240421357 | -0.570084935 | 0.08924222 | 9.487541858 | -1.826637724 | 0.122216917 | 0.709264768 | -4.394610785 |
| **A_51_P476711** | Mtrex | 0.32943003 | -0.122300471 | 0.781160531 | 11.47602275 | 1.82656073 | 0.122229495 | 0.709264768 | -4.394631126 |
| **A_51_P300378** | Wwc2 | -0.441681843 | -1.047582196 | 0.164218509 | 12.73514556 | -1.825823056 | 0.122350064 | 0.709264768 | -4.394826022 |
| **A_55_P2097828** | Gmfg | 0.924798776 | -0.344283301 | 2.193880853 | 12.22801269 | 1.825189161 | 0.122453767 | 0.709264768 | -4.394993522 |
| **A_55_P2249220** | A430102J | -0.240531117 | -0.570671588 | 0.089609355 | 8.812589249 | -1.824831823 | 0.122512265 | 0.709264768 | -4.395087954 |
| **A_55_P2128668** | Ccnb1 | 0.404807778 | -0.15096822 | 0.960583776 | 8.504004527 | 1.824313228 | 0.122597212 | 0.709264768 | -4.395225013 |
| **A_55_P1952990** | Myl6b | 0.384402816 | -0.143399417 | 0.912205049 | 9.62839389 | 1.824171569 | 0.122620426 | 0.709264768 | -4.395262454 |
| **A_51_P426754** | Anxa5 | -0.517938565 | -1.229121467 | 0.193244337 | 13.62235721 | -1.824094306 | 0.122633089 | 0.709264768 | -4.395282876 |
| **A_55_P1953713** | Bop1 | 0.197222617 | -0.073594432 | 0.468039667 | 10.43590135 | 1.824026153 | 0.12264426 | 0.709264768 | -4.39530089 |
| **A_55_P1975181** | Acyp1 | 0.278405896 | -0.103941794 | 0.660753587 | 8.298557496 | 1.823770989 | 0.122686094 | 0.709264768 | -4.395368336 |
| **A_55_P1998631** | Capg | -0.736620369 | -1.748286135 | 0.275045398 | 12.85271501 | -1.823715187 | 0.122695245 | 0.709264768 | -4.395383086 |
| **A_55_P1960311** | Dnase1l1 | -1.127753369 | -2.676711489 | 0.421204752 | 10.1171788 | -1.823579926 | 0.122717428 | 0.709264768 | -4.39541884 |
| **A_65_P18167** | Hnrnpul2 | 0.296650047 | -0.110848475 | 0.704148568 | 10.17749059 | 1.823344492 | 0.12275605 | 0.709264768 | -4.395481077 |
| **A_55_P1980583** | Leprot | -0.369156107 | -0.876578691 | 0.138266477 | 13.08712675 | -1.822177208 | 0.122947718 | 0.709264768 | -4.395789688 |
| **A_51_P374464** | Gstp1 | 0.370651461 | -0.138877634 | 0.880180556 | 10.98302024 | 1.82199455 | 0.122977737 | 0.709264768 | -4.395837986 |
| **A_55_P2008599** | Pcx | -0.428619271 | -1.017944057 | 0.160705514 | 11.35095655 | -1.821660042 | 0.123032732 | 0.709264768 | -4.395926441 |
| **A_55_P2091631** | Coa4 | -0.275828877 | -0.655078985 | 0.10342123 | 9.835570704 | -1.821647595 | 0.123034779 | 0.709264768 | -4.395929733 |
| **A_51_P416689** | Ext1 | -0.253436956 | -0.601945938 | 0.095072025 | 9.990053607 | -1.82140412 | 0.123074824 | 0.709264768 | -4.395994119 |
| **A_51_P466759** | Mau2 | -0.385376408 | -0.915321367 | 0.144568551 | 8.891984608 | -1.821397369 | 0.123075935 | 0.709264768 | -4.395995905 |

| | | | | | | | | | |
|---|---|---|---|---|---|---|---|---|---|
| **A_51_P493117** | Slc16a9 | -0.886262786 | -2.105046419 | 0.332520847 | 10.02936349 | -1.821319955 | 0.12308867 | 0.709264768 | -4.396016377 |
| **A_55_P1969042** | Cops6 | 0.179107485 | -0.067326573 | 0.425541542 | 12.71411316 | 1.82038544 | 0.12324251 | 0.709264768 | -4.396263541 |
| **A_55_P2192729** | 5830469G | -0.546206918 | -1.297869178 | 0.205455341 | 9.420019718 | -1.820056195 | 0.123296757 | 0.709264768 | -4.396350632 |
| **A_52_P456750** | Aph1b | -0.218782465 | -0.519863295 | 0.082298365 | 8.82777118 | -1.820035053 | 0.123300241 | 0.709264768 | -4.396356225 |
| **A_51_P406365** | Maf1 | -0.247316944 | -0.587708242 | 0.093074353 | 12.13681924 | -1.819808835 | 0.123337528 | 0.709264768 | -4.396416067 |
| **A_51_P319879** | Acadm | -0.801467619 | -1.904684201 | 0.301748963 | 11.92879207 | -1.819598396 | 0.123372224 | 0.709264768 | -4.396471737 |
| **A_51_P282523** | Adgra2 | 0.867309444 | -0.326787954 | 2.061406841 | 11.1728151 | 1.819217684 | 0.123435018 | 0.709264768 | -4.396572458 |
| **A_55_P2065631** | Hook3 | -0.22483137 | -0.534509312 | 0.084846572 | 11.16670185 | -1.818431644 | 0.123564769 | 0.709264768 | -4.396780436 |
| **A_52_P367745** | Acads | -0.368406889 | -0.87587186 | 0.139058082 | 8.352119899 | -1.818327132 | 0.123582031 | 0.709264768 | -4.396808091 |
| **A_55_P2126805** | Wdr81 | -0.280155363 | -0.666096204 | 0.105785478 | 11.22912949 | -1.818145132 | 0.123612098 | 0.709264768 | -4.396856252 |
| **A_51_P143915** | Rnf181 | -0.337355046 | -0.802156258 | 0.127446167 | 10.66224094 | -1.817901314 | 0.123652388 | 0.709264768 | -4.396920774 |
| **A_55_P2030433** | Gpi1 | 0.271793005 | -0.102748221 | 0.64633423 | 14.53645822 | 1.817560991 | 0.123708647 | 0.709264768 | -4.397010839 |
| **A_52_P143477** | Tgoln1 | -0.863949923 | -2.054615516 | 0.32671567 | 11.60601191 | -1.817394092 | 0.123736246 | 0.709264768 | -4.39705501 |
| **A_52_P35961** | NA | -0.171236757 | -0.407274096 | 0.064800583 | 8.476189736 | -1.817048956 | 0.12379334 | 0.709264768 | -4.397146358 |
| **A_52_P450835** | Cdpf1 | -0.280686824 | -0.667722418 | 0.10634877 | 10.98624501 | -1.816441708 | 0.123893858 | 0.709264768 | -4.397307095 |
| **A_55_P2145257** | NA | 0.571316625 | -0.216514945 | 1.359148195 | 9.425385515 | 1.816326118 | 0.123913001 | 0.709264768 | -4.397337694 |
| **A_55_P1970154** | Rnf10 | 0.205137878 | -0.077742231 | 0.488017987 | 11.49059105 | 1.816325977 | 0.123913024 | 0.709264768 | -4.397337731 |
| **A_55_P1982747** | Slc8a3 | -0.401388333 | -0.954911885 | 0.152135219 | 11.49162293 | -1.816264047 | 0.123923282 | 0.709264768 | -4.397354125 |
| **A_55_P2098120** | NA | 0.62621762 | -0.237641694 | 1.490076934 | 9.315030494 | 1.815651985 | 0.124024703 | 0.709264768 | -4.397516162 |
| **A_55_P2035484** | Gucd1 | -0.308860474 | -0.734950822 | 0.117229873 | 10.90967225 | -1.815561581 | 0.124039691 | 0.709264768 | -4.397540097 |
| **A_55_P1963579** | NA | 0.349768148 | -0.133023791 | 0.832560087 | 10.05865535 | 1.814556906 | 0.124206371 | 0.709264768 | -4.397806119 |
| **A_55_P2049383** | Erp29 | -0.705083292 | -1.678522114 | 0.268355531 | 14.69083143 | -1.814187233 | 0.124267758 | 0.709264768 | -4.397904016 |
| **A_66_P102260** | Irs2 | -0.719295075 | -1.712427213 | 0.273837062 | 10.6146836 | -1.814054776 | 0.12428976 | 0.709264768 | -4.397939095 |
| **A_52_P422859** | Hspg2 | -0.638200535 | -1.519378276 | 0.242977207 | 13.1320243 | -1.81402802 | 0.124294205 | 0.709264768 | -4.397946181 |
| **A_55_P2098603** | NA | -0.42410982 | -1.009872465 | 0.161652826 | 8.299557653 | -1.813455909 | 0.124389289 | 0.709264768 | -4.398097706 |
| **A_55_P2085485** | Pidd1 | 0.239799464 | -0.091427098 | 0.571026026 | 8.154046835 | 1.813315597 | 0.124412619 | 0.709264768 | -4.398134871 |
| **A_51_P394833** | Tshz1 | 0.302045808 | -0.115214954 | 0.71930657 | 10.78157093 | 1.813074341 | 0.124452745 | 0.709264768 | -4.398198775 |
| **A_55_P2076805** | Ddx5 | -0.337678227 | -0.804247245 | 0.128890791 | 9.308946936 | -1.812748306 | 0.124506992 | 0.709264768 | -4.39828514 |
| **A_51_P423743** | Cldn6 | 0.541409783 | -0.206667143 | 1.289486708 | 8.721538741 | 1.812717741 | 0.124512078 | 0.709264768 | -4.398293237 |
| **A_52_P657123** | Gm10270 | 0.423933335 | -0.161985824 | 1.009852495 | 10.95389486 | 1.812217057 | 0.124595433 | 0.709264768 | -4.398425877 |
| **A_52_P121491** | Msrb3 | -0.307230568 | -0.731920962 | 0.117459827 | 8.610000674 | -1.811933816 | 0.124642612 | 0.709264768 | -4.398500918 |
| **A_55_P2034864** | Tubb2b | 1.264121416 | -0.483318466 | 3.011561299 | 11.66672587 | 1.811911124 | 0.124646393 | 0.709264768 | -4.398506931 |
| **A_52_P82247** | Cycs | 0.426909211 | -0.163288195 | 1.017106616 | 11.43096946 | 1.811709554 | 0.12467998 | 0.709264768 | -4.398560337 |
| **A_51_P336721** | Tipin | 1.068781538 | -0.409232008 | 2.546795084 | 11.97025355 | 1.811176916 | 0.124768776 | 0.709264768 | -4.39870147 |
| **A_52_P683441** | Capn5 | 0.258536044 | -0.099066466 | 0.616138554 | 9.123388402 | 1.810801669 | 0.124831372 | 0.709264768 | -4.398800908 |
| **A_55_P2229137** | Tmem116 | -0.389609215 | -0.928531605 | 0.149313175 | 8.03039312 | -1.810728506 | 0.12484358 | 0.709264768 | -4.398820297 |
| **A_55_P1994907** | Senp2 | -0.863773462 | -2.084545068 | 0.356998144 | 11.03324474 | -1.884638865 | 0.124925883 | 0.709264768 | -4.425105545 |
| **A_55_P1965368** | Ctc1 | 0.490319094 | -0.188099194 | 1.168737382 | 11.77433936 | 1.810221024 | 0.124928292 | 0.709264768 | -4.39895479 |
| **A_55_P2030225** | NA | 0.421671582 | -0.16196006 | 1.005303223 | 16.11499667 | 1.809613593 | 0.125029764 | 0.709264768 | -4.399115789 |
| **A_55_P2016540** | Cry2 | -0.181171807 | -0.431935113 | 0.0695915 | 11.1328975 | -1.809576605 | 0.125035946 | 0.709264768 | -4.399125593 |
| **A_52_P282838** | Spin2c | 0.23242303 | -0.089326682 | 0.554172743 | 8.782355552 | 1.809303142 | 0.125081657 | 0.709264768 | -4.399198081 |
| **A_55_P2090557** | Lrrc75a | -0.399069747 | -0.951525626 | 0.153386131 | 11.7411586 | -1.809262377 | 0.125088473 | 0.709264768 | -4.399208887 |
| **A_55_P2035400** | Ripk4 | -0.171248722 | -0.408321581 | 0.065824137 | 11.2831185 | -1.809238612 | 0.125092447 | 0.709264768 | -4.399215187 |
| **A_55_P1959628** | Pfdn4 | 0.491624089 | -0.188982272 | 1.17223045 | 9.686343593 | 1.809203821 | 0.125098264 | 0.709264768 | -4.399224409 |
| **A_55_P2162422** | NA | 0.445121671 | -0.171121225 | 1.061364567 | 17.20902295 | 1.809160755 | 0.125105465 | 0.709264768 | -4.399235825 |
| **A_55_P2086912** | NA | 1.0376046 | -0.398995675 | 2.474204874 | 12.77144442 | 1.809032166 | 0.12512697 | 0.709264768 | -4.399269913 |
| **A_55_P2149001** | NA | 0.65539456 | -0.252093635 | 1.562882755 | 14.63290216 | 1.808890149 | 0.125150724 | 0.709264768 | -4.399307562 |
| **A_55_P2182939** | NA | 1.163823993 | -0.447720936 | 2.775368922 | 13.81667504 | 1.808819511 | 0.125162541 | 0.709264768 | -4.399326288 |

| | | | | | | | | | |
|---|---|---|---|---|---|---|---|---|---|
| **A_55_P2016426** | Nova2 | 0.204481762 | -0.078666369 | 0.487629894 | 8.954801912 | 1.808802813 | 0.125165335 | 0.709264768 | -4.399330715 |
| **A_55_P1987439** | Pigt | -0.759307481 | -1.81090913 | 0.292294167 | 12.17577711 | -1.808492726 | 0.125217223 | 0.709264768 | -4.399412923 |
| **A_55_P2132507** | Bcl2l2 | -0.54683811 | -1.3042428 | 0.21056658 | 9.445744633 | -1.808344331 | 0.125242063 | 0.709264768 | -4.399452266 |
| **A_55_P2022404** | Gtf2h5 | 0.337851191 | -0.130182653 | 0.805885034 | 11.62304601 | 1.808000479 | 0.125299638 | 0.709264768 | -4.399543434 |
| **A_55_P2141415** | Ppp6r3 | -0.185954395 | -0.443572823 | 0.071664034 | 8.252170307 | -1.807922732 | 0.12531266 | 0.709264768 | -4.399564049 |
| **A_55_P2372228** | NA | 0.475196288 | -0.183151083 | 1.133543658 | 8.107430177 | 1.807874502 | 0.125320739 | 0.709264768 | -4.399576837 |
| **A_51_P183853** | Polr2m | 0.177221989 | -0.068440883 | 0.42288486 | 14.04835409 | 1.806876325 | 0.125488054 | 0.709264768 | -4.399841532 |
| **A_55_P2009329** | Gm8709 | 0.416144111 | -0.160750013 | 0.993038234 | 9.146438322 | 1.806749668 | 0.1255093 | 0.709264768 | -4.399875122 |
| **A_51_P286460** | H2ac8 | 0.185129829 | -0.071574892 | 0.441834549 | 7.986291395 | 1.806312513 | 0.125582659 | 0.709264768 | -4.399991065 |
| **A_55_P1974783** | Tsnax | 0.387653601 | -0.149876683 | 0.925183885 | 12.38820359 | 1.806305612 | 0.125583818 | 0.709264768 | -4.399992895 |
| **A_55_P2166262** | Rnf215 | -0.57466618 | -1.3717587 | 0.22242634 | 11.69825113 | -1.80574845 | 0.125677379 | 0.709264768 | -4.400140681 |
| **A_55_P1973868** | Sema3b | -0.66589803 | -1.589642623 | 0.257846564 | 10.78026568 | -1.805536057 | 0.125713063 | 0.709264768 | -4.400197021 |
| **A_55_P2057142** | Cks1brt | 0.618010837 | -0.239614904 | 1.475636579 | 9.165698397 | 1.80488123 | 0.125823144 | 0.709264768 | -4.400370739 |
| **A_55_P2089035** | Pole | 0.241063932 | -0.093468236 | 0.575596101 | 7.988906837 | 1.804864891 | 0.125825892 | 0.709264768 | -4.400375074 |
| **A_55_P1984307** | Adgrf5 | -0.67511423 | -1.61223061 | 0.262002149 | 10.09699284 | -1.804405238 | 0.125903223 | 0.709264768 | -4.400497028 |
| **A_55_P1963489** | NA | 0.801214438 | -0.311010892 | 1.913439769 | 10.19905082 | 1.804289954 | 0.125922625 | 0.709264768 | -4.400527616 |
| **A_52_P375638** | Gtpbp1 | 0.303699949 | -0.117943647 | 0.725343544 | 9.470509875 | 1.804054095 | 0.12596233 | 0.709264768 | -4.400590199 |
| **A_55_P1974542** | Gxylt2 | 0.25382484 | -0.098660197 | 0.606309876 | 11.91146873 | 1.803614762 | 0.126036321 | 0.709264768 | -4.40070678 |
| **A_55_P1979684** | Rhoc | -0.666093334 | -1.591167891 | 0.258981223 | 12.4554114 | -1.803469062 | 0.126060869 | 0.709264768 | -4.400745445 |
| **A_66_P115061** | Smc6 | 0.443315381 | -0.172392786 | 1.059023548 | 10.12306092 | 1.803384078 | 0.12607519 | 0.709264768 | -4.400767997 |
| **A_65_P01834** | Lima1 | -0.51201982 | -1.22315228 | 0.199112641 | 9.228773879 | -1.803377367 | 0.126076321 | 0.709264768 | -4.400769778 |
| **A_55_P2122555** | Csnk1e | -0.307421229 | -0.734392597 | 0.119550139 | 9.925626543 | -1.80337252 | 0.126077138 | 0.709264768 | -4.400771065 |
| **A_66_P111798** | Znrf2 | 0.16130248 | -0.062730489 | 0.385335449 | 8.887493087 | 1.803347635 | 0.126081331 | 0.709264768 | -4.400777669 |
| **A_55_P2008327** | Eno1b | 0.352174777 | -0.136970637 | 0.841320191 | 14.49823813 | 1.8033109 | 0.126087522 | 0.709264768 | -4.400787417 |
| **A_66_P133043** | Mbd2 | -0.330182982 | -0.788822363 | 0.128456399 | 11.20400241 | -1.803157608 | 0.12611336 | 0.709264768 | -4.400828099 |
| **A_55_P2140076** | NA | 0.201491446 | -0.078435123 | 0.481418015 | 16.56778741 | 1.802863463 | 0.126162954 | 0.709264768 | -4.400906165 |
| **A_51_P468464** | Gdpd5 | -0.552481032 | -1.320030014 | 0.215067951 | 9.764158615 | -1.802858399 | 0.126163808 | 0.709264768 | -4.400907509 |
| **A_55_P2115995** | Sms | 0.652390802 | -0.254454893 | 1.559236498 | 10.04121905 | 1.801875496 | 0.126329671 | 0.709264768 | -4.401168403 |
| **A_52_P274028** | Ssr1 | 0.213998678 | -0.08350313 | 0.511500487 | 9.036867029 | 1.801655811 | 0.126366772 | 0.709264768 | -4.40122672 |
| **A_52_P407692** | H2afj | 0.952056772 | -0.371569078 | 2.275682621 | 16.81247654 | 1.80155677 | 0.126383502 | 0.709264768 | -4.401253013 |
| **A_65_P02327** | Pogk | 0.348819797 | -0.136149793 | 0.833789387 | 9.021113681 | 1.801511189 | 0.126391203 | 0.709264768 | -4.401265114 |
| **A_55_P2093727** | Gemin6 | 0.354156087 | -0.138240369 | 0.846552544 | 9.847174844 | 1.801482872 | 0.126395987 | 0.709264768 | -4.401272631 |
| **A_51_P272250** | Snf8 | -0.201031456 | -0.48062486 | 0.078561948 | 10.83471827 | -1.800891063 | 0.126496011 | 0.709264768 | -4.401429751 |
| **A_55_P2089118** | Zfp976 | 0.610473792 | -0.238594972 | 1.459542555 | 9.560469414 | 1.800837418 | 0.126505082 | 0.709264768 | -4.401443994 |
| **A_55_P2131820** | A530006G | 0.346928469 | -0.135597142 | 0.829454079 | 12.13806136 | 1.800818385 | 0.1265083 | 0.709264768 | -4.401449047 |
| **A_55_P1978740** | Rpgrip1 | 0.555700528 | -0.217239956 | 1.328641012 | 11.83385711 | 1.800715488 | 0.126525701 | 0.709264768 | -4.401476368 |
| **A_51_P505696** | Inpp5k | -0.394764923 | -0.943911474 | 0.154381628 | 9.710942367 | -1.800531161 | 0.126556879 | 0.709264768 | -4.40152531 |
| **A_55_P2133679** | NA | 0.456376727 | -0.178507769 | 1.091261223 | 11.57967581 | 1.800441908 | 0.126571978 | 0.709264768 | -4.401549009 |
| **A_51_P124748** | Tgfb3 | -0.506394206 | -1.211036183 | 0.198247771 | 11.32708609 | -1.799991946 | 0.126648127 | 0.709264768 | -4.408394715 |
| **A_51_P300434** | Dst | -0.74124849 | -1.772881346 | 0.290384366 | 9.057531397 | -1.799653942 | 0.126705359 | 0.709264768 | -4.401758252 |
| **A_55_P1988678** | Znhit6 | 0.233298435 | -0.091397394 | 0.557994264 | 11.34972416 | 1.799639273 | 0.126707843 | 0.709264768 | -4.401762147 |
| **A_52_P307961** | Abca5 | -0.160877062 | -0.384825589 | 0.063071464 | 8.553325595 | -1.799269687 | 0.126770454 | 0.709264768 | -4.401860301 |
| **A_52_P135822** | Myo18a | -0.365860122 | -0.875178718 | 0.143458473 | 12.82836817 | -1.799185274 | 0.126784758 | 0.709264768 | -4.40188272 |
| **A_51_P492125** | Ciapin1 | 0.230726341 | -0.090473798 | 0.551926479 | 10.97787393 | 1.79916834 | 0.126787628 | 0.709264768 | -4.401887218 |
| **A_55_P2087528** | Tmem131 | 0.665415535 | -0.260990286 | 1.591821356 | 11.87718294 | 1.799044914 | 0.126808547 | 0.709264768 | -4.401919999 |
| **A_52_P625683** | Pxmp4 | -0.264889595 | -0.633686638 | 0.103907448 | 9.772840937 | -1.798986129 | 0.126818512 | 0.709264768 | -4.401935612 |
| **A_55_P2093994** | Maged2 | -0.350778581 | -0.839201138 | 0.137643975 | 14.58338845 | -1.798819969 | 0.126846681 | 0.709264768 | -4.401979745 |
| **A_55_P2015426** | Asnsd1 | 0.305441599 | -0.119859475 | 0.730742672 | 11.9698526 | 1.79879657 | 0.126850648 | 0.709264768 | -4.40198596 |

| | | | | | | | | | |
|---|---|---|---|---|---|---|---|---|---|
| **A_52_P869802** | Thoc3 | 0.212820589 | -0.083530045 | 0.509171223 | 9.73460766 | 1.798697478 | 0.126867451 | 0.709264768 | -4.40201228 |
| **A_51_P289987** | Psmg3 | 0.322361789 | -0.126608747 | 0.771332324 | 10.94774614 | 1.798357631 | 0.126925095 | 0.709264768 | -4.402102552 |
| **A_55_P2166528** | Mrps36 | 0.325837664 | -0.127995386 | 0.779670714 | 11.72091178 | 1.798272526 | 0.126939534 | 0.709264768 | -4.402125158 |
| **A_51_P310030** | Naca | 0.656151388 | -0.257764052 | 1.570066828 | 15.23824186 | 1.798243019 | 0.126944541 | 0.709264768 | -4.402132997 |
| **A_55_P2071802** | Slbp | 0.231539045 | -0.090960346 | 0.554038435 | 8.385430371 | 1.798231844 | 0.126946437 | 0.709264768 | -4.402135965 |
| **A_55_P2142590** | Mark3 | -0.339059807 | -0.811328989 | 0.133209375 | 11.17336938 | -1.798196093 | 0.126952504 | 0.709264768 | -4.402145462 |
| **A_52_P460703** | Rassf3 | -0.22166112 | -0.530434948 | 0.087112709 | 8.757532466 | -1.798040153 | 0.126978968 | 0.709264768 | -4.402186887 |
| **A_51_P158073** | Shfl | -0.350392756 | -0.838690225 | 0.137904714 | 9.323740135 | -1.797301721 | 0.127104362 | 0.709264768 | -4.402383063 |
| **A_51_P223404** | Plin3 | -0.39525056 | -0.946126953 | 0.155625834 | 13.80237144 | -1.797085237 | 0.127141147 | 0.709264768 | -4.402440581 |
| **A_55_P2073482** | NA | 0.954022407 | -0.375738269 | 2.283783084 | 12.29668376 | 1.79694769 | 0.127164525 | 0.709264768 | -4.402477127 |
| **A_55_P2004201** | NA | 0.769633835 | -0.303254619 | 1.842522288 | 12.63834353 | 1.79671803 | 0.127203568 | 0.709264768 | -4.40253815 |
| **A_51_P483379** | Rnaseh1 | -0.291422563 | -0.69770616 | 0.114861033 | 9.5916286 | -1.796570363 | 0.127228678 | 0.709264768 | -4.402577387 |
| **A_55_P2064538** | Zfp985 | 0.225124428 | -0.088760581 | 0.539009437 | 8.491950241 | 1.7963973 | 0.127258112 | 0.709264768 | -4.402623374 |
| **A_55_P2085181** | Chaf1b | 1.1739435 | -0.462933582 | 2.810820582 | 11.39281807 | 1.796310767 | 0.127272833 | 0.709264768 | -4.402646369 |
| **A_52_P700056** | Wfdc17 | -0.155046516 | -0.371246875 | 0.061153842 | 8.095553566 | -1.796205194 | 0.127290794 | 0.709264768 | -4.402674424 |
| **A_51_P341108** | Spint1 | -0.571478731 | -1.368475723 | 0.225518261 | 10.83464741 | -1.795947906 | 0.127334578 | 0.709264768 | -4.402742797 |
| **A_51_P225592** | Tpm4 | -0.214340029 | -0.513293253 | 0.084613195 | 15.33815889 | -1.795768666 | 0.127365088 | 0.709264768 | -4.402790431 |
| **A_51_P101660** | Ncbp3 | 0.192551261 | -0.076032591 | 0.461135113 | 8.422862618 | 1.795629841 | 0.127388725 | 0.709264768 | -4.419806034 |
| **A_51_P471520** | Stk25 | -0.231557682 | -0.554559394 | 0.091444029 | 12.23511187 | -1.795579821 | 0.127397242 | 0.709264768 | -4.40284062 |
| **A_51_P123077** | Nubp1 | 0.520350837 | -0.205500282 | 1.246201957 | 11.47625805 | 1.795556406 | 0.12740123 | 0.709264768 | -4.416361115 |
| **A_55_P2159299** | Ing1 | 0.192563594 | -0.076061966 | 0.461189153 | 11.65121866 | 1.79546604 | 0.127416619 | 0.709264768 | -4.402870859 |
| **A_55_P2043531** | NA | 1.079023916 | -0.42678444 | 2.584832273 | 14.05917699 | 1.794782034 | 0.127533167 | 0.709264768 | -4.403052663 |
| **A_51_P281333** | St3gal6 | -0.862910322 | -2.067385736 | 0.341565091 | 10.55416453 | -1.794395083 | 0.127599147 | 0.709264768 | -4.403155523 |
| **A_55_P2026535** | Erc2 | 0.205987708 | -0.081583495 | 0.493558911 | 8.800601626 | 1.794098391 | 0.12764976 | 0.709264768 | -4.403234394 |
| **A_55_P2003494** | NA | 0.547063753 | -0.216724569 | 1.310852075 | 14.65837368 | 1.793970397 | 0.127671601 | 0.709264768 | -4.403268421 |
| **A_55_P2045549** | Dpy19l1 | 0.349160694 | -0.138335885 | 0.836657274 | 10.01283747 | 1.793924334 | 0.127679462 | 0.709264768 | -4.403280667 |
| **A_55_P2004930** | Hddc2 | 0.724855062 | -0.287292671 | 1.737002796 | 11.16327313 | 1.793732244 | 0.127712249 | 0.709264768 | -4.403331735 |
| **A_51_P317941** | Perp | -0.945741978 | -2.266526626 | 0.375042671 | 12.74282536 | -1.793457127 | 0.127759222 | 0.709264768 | -4.40340488 |
| **A_55_P2077613** | Fam102a | -0.359036503 | -0.860485538 | 0.142412531 | 12.48289577 | -1.793338002 | 0.127779567 | 0.709264768 | -4.403436553 |
| **A_51_P336691** | Srsf9 | 0.530493362 | -0.210606921 | 1.271593645 | 12.34086979 | 1.792888661 | 0.127856336 | 0.709264768 | -4.403556029 |
| **A_51_P499698** | Asprv1 | -0.319061644 | -0.764819312 | 0.126696025 | 7.974045187 | -1.792776376 | 0.127875527 | 0.709264768 | -4.403585886 |
| **A_51_P473576** | Apex1 | 0.685703352 | -0.272297833 | 1.643704538 | 10.99151796 | 1.79275347 | 0.127879443 | 0.709264768 | -4.403591977 |
| **A_52_P3412** | Med17 | 0.170633122 | -0.067802343 | 0.409068587 | 9.482935727 | 1.792432585 | 0.127934304 | 0.709264768 | -4.403677306 |
| **A_55_P2163263** | NA | 0.278651593 | -0.110755473 | 0.668058658 | 9.574468321 | 1.792289037 | 0.127958853 | 0.709264768 | -4.40371548 |
| **A_55_P2060878** | NA | 0.854779789 | -0.339758996 | 2.049318574 | 11.66666683 | 1.792273714 | 0.127961474 | 0.709264768 | -4.403719555 |
| **A_52_P282762** | Myd88 | -0.313358942 | -0.75130816 | 0.124590275 | 10.08898618 | -1.792126593 | 0.12798664 | 0.709264768 | -4.417370333 |
| **A_55_P1984932** | Zc3h18 | 0.17228652 | -0.068591939 | 0.413164979 | 11.96522739 | 1.791445825 | 0.128103155 | 0.709264768 | -4.403939735 |
| **A_55_P2098801** | Gm15393 | -0.395115865 | -0.947546901 | 0.157315171 | 15.09423088 | -1.791417217 | 0.128108054 | 0.709264768 | -4.403947344 |
| **A_52_P433864** | Naa16 | 0.223475327 | -0.088980082 | 0.535930735 | 8.572410987 | 1.791397155 | 0.128111489 | 0.709264768 | -4.40395268 |
| **A_51_P291819** | 2310033P | 0.20514938 | -0.081687139 | 0.491985899 | 10.57052454 | 1.791373356 | 0.128115565 | 0.709264768 | -4.40395901 |
| **A_65_P20433** | Parpbp | 0.560775314 | -0.22341675 | 1.344967379 | 8.741886586 | 1.791087445 | 0.128164535 | 0.709264768 | -4.404035058 |
| **A_55_P2070084** | Zfp988 | 0.483558047 | -0.19268458 | 1.159800673 | 10.53220361 | 1.791003429 | 0.128178929 | 0.709264768 | -4.404057406 |
| **A_51_P367843** | Polr2g | 0.246213974 | -0.098136655 | 0.590564603 | 13.10856874 | 1.790862173 | 0.128203133 | 0.709264768 | -4.40409498 |
| **A_55_P1960748** | Arl13b | -0.29509597 | -0.707889841 | 0.117697901 | 9.190064723 | -1.790524978 | 0.128260929 | 0.709264768 | -4.404184677 |
| **A_51_P237688** | Smim14 | -0.422953918 | -1.01461825 | 0.168710415 | 10.99710967 | -1.790473957 | 0.128269677 | 0.709264768 | -4.40419825 |
| **A_55_P2059134** | Smarca5 | 0.591689103 | -0.236058504 | 1.41943671 | 11.05089781 | 1.790383153 | 0.128285246 | 0.709264768 | -4.404222406 |
| **A_52_P214630** | Sox9 | 0.797425411 | -0.318157643 | 1.913008465 | 9.491046437 | 1.79035234 | 0.12829053 | 0.709264768 | -4.408745674 |
| **A_55_P2000758** | Gkap1 | -0.373199667 | -0.89533365 | 0.148934317 | 10.0214421 | -1.790233306 | 0.128310943 | 0.709264768 | -4.40426227 |

| | | | | | | | | | |
|---|---|---|---|---|---|---|---|---|---|
| **A_55_P1960641** | Trpt1 | -0.382333536 | -0.917250514 | 0.152583442 | 9.907288985 | -1.790219802 | 0.128313259 | 0.709264768 | -4.404265863 |
| **A_52_P572702** | Gxylt1 | -0.372604844 | -0.89399388 | 0.148784192 | 11.24266102 | -1.789933711 | 0.128362336 | 0.709264768 | -4.404341975 |
| **A_55_P1953583** | Ace2 | -1.071657752 | -2.572249788 | 0.428934284 | 9.346835958 | -1.788725993 | 0.128569719 | 0.709264768 | -4.404663322 |
| **A_55_P2044369** | Hs2st1 | 0.351223398 | -0.140585163 | 0.843031959 | 9.823302168 | 1.788700801 | 0.128574049 | 0.709264768 | -4.404670025 |
| **A_51_P453856** | Csnk1a1 | 0.266605576 | -0.106793881 | 0.640005033 | 12.87987495 | 1.788322711 | 0.128639043 | 0.709264768 | -4.404770642 |
| **A_55_P2006467** | Serf2 | -0.336375436 | -0.807499708 | 0.134748836 | 14.91587027 | -1.788294906 | 0.128643824 | 0.709264768 | -4.404778042 |
| **A_55_P2172239** | NA | 0.433055317 | -0.173491031 | 1.039601664 | 16.59848119 | 1.788256157 | 0.128650488 | 0.709264768 | -4.404788354 |
| **A_52_P248604** | Cdh5 | -0.286058725 | -0.686720517 | 0.114603067 | 12.99388107 | -1.788247453 | 0.128651984 | 0.709264768 | -4.40479067 |
| **A_66_P139546** | Igfbp6 | -0.854240154 | -2.050731375 | 0.342251066 | 12.84254616 | -1.788219438 | 0.128656802 | 0.709264768 | -4.404798126 |
| **A_52_P176619** | Atat1 | -0.344814942 | -0.827784029 | 0.138154145 | 9.281399967 | -1.788204108 | 0.128659438 | 0.709264768 | -4.404802206 |
| **A_55_P1975475** | Mdm2 | -0.31633917 | -0.759460893 | 0.126782553 | 12.37207905 | -1.788052539 | 0.128685507 | 0.709264768 | -4.404842544 |
| **A_55_P1956938** | Rpl24 | 0.244217821 | -0.097980567 | 0.586416209 | 16.75819785 | 1.78751518 | 0.12877797 | 0.709264768 | -4.404985564 |
| **A_55_P1999384** | NA | -0.22528694 | -0.540977708 | 0.090403828 | 11.34555938 | -1.787411219 | 0.128795866 | 0.709264768 | -4.405013235 |
| **A_55_P2005050** | NA | 0.751559967 | -0.301605701 | 1.804725635 | 12.57236945 | 1.787381634 | 0.128800959 | 0.709264768 | -4.40502111 |
| **A_55_P2118309** | NA | 0.379222418 | -0.152285543 | 0.91073038 | 8.643888513 | 1.787041238 | 0.128859576 | 0.709264768 | -4.405111716 |
| **A_55_P1993453** | Ftsj1 | 0.667466316 | -0.268054212 | 1.602986844 | 11.33921748 | 1.787007511 | 0.128865385 | 0.709264768 | -4.405120694 |
| **A_51_P424810** | Ncapg2 | 0.948019381 | -0.380962938 | 2.277001701 | 9.740047926 | 1.786686511 | 0.128920688 | 0.709264768 | -4.405206144 |
| **A_51_P356642** | Krt19 | -0.566450487 | -1.360539951 | 0.227638976 | 13.40043135 | -1.786663917 | 0.128924582 | 0.709264768 | -4.405212159 |
| **A_51_P282092** | Urod | 0.31269808 | -0.125905295 | 0.751301454 | 12.88024508 | 1.785679822 | 0.129094279 | 0.709264768 | -4.405474156 |
| **A_55_P2059323** | Gm13315 | 0.57102407 | -0.230080326 | 1.372128466 | 11.57879053 | 1.785318256 | 0.129156684 | 0.709264768 | -4.405570428 |
| **A_55_P1988259** | Gng5 | -0.294274128 | -0.707211263 | 0.118663007 | 15.34531141 | -1.784918893 | 0.129225647 | 0.709264768 | -4.405676772 |
| **A_55_P2021585** | Tff1 | -0.594299793 | -1.428395657 | 0.23979607 | 8.073405771 | -1.784596173 | 0.129281402 | 0.709264768 | -4.405762712 |
| **A_55_P2139473** | Ilk | -0.187047819 | -0.449624732 | 0.075529095 | 12.72344803 | -1.784211991 | 0.129347807 | 0.709264768 | -4.405865026 |
| **A_55_P2061437** | Lsm5 | 0.837499558 | -0.338815264 | 2.013814379 | 11.27983377 | 1.783246428 | 0.129514853 | 0.709264768 | -4.406122204 |
| **A_55_P2156464** | Nt5c3b | 0.495212902 | -0.200463803 | 1.190889608 | 9.354059486 | 1.782932229 | 0.129569257 | 0.709264768 | -4.406205901 |
| **A_55_P2109922** | NA | 0.341635543 | -0.138305092 | 0.821576178 | 9.073660202 | 1.782895428 | 0.12957563 | 0.709264768 | -4.406215704 |
| **A_52_P492901** | Stx4a | -0.28511344 | -0.685708798 | 0.115481919 | 10.98800045 | -1.782633742 | 0.129620961 | 0.709264768 | -4.406285416 |
| **A_55_P2083252** | Mtus2 | -0.566165575 | -1.361696628 | 0.229365478 | 10.09249399 | -1.782529262 | 0.129639064 | 0.709264768 | -4.40631325 |
| **A_52_P299311** | NA | 0.44042592 | -0.178673805 | 1.059525645 | 15.2878043 | 1.781814976 | 0.129762895 | 0.709264768 | -4.406503554 |
| **A_51_P212038** | Atp6v0e2 | -0.261506529 | -0.629156167 | 0.106143109 | 9.342957277 | -1.781552968 | 0.129808347 | 0.709264768 | -4.406573365 |
| **A_55_P2086329** | Krt85 | 0.21004617 | -0.085269097 | 0.505361436 | 8.483945302 | 1.781472871 | 0.129822245 | 0.709264768 | -4.406594708 |
| **A_55_P2058122** | Kyat3 | 0.302147744 | -0.122777094 | 0.727072582 | 10.11647402 | 1.780974018 | 0.129908837 | 0.709264768 | -4.406727637 |
| **A_55_P1981994** | Krt17 | 1.283794282 | -0.521753248 | 3.089341813 | 8.836497978 | 1.780889023 | 0.129923596 | 0.709264768 | -4.406750287 |
| **A_55_P1970578** | Aamdc | -0.216844486 | -0.521858908 | 0.088169937 | 10.48473102 | -1.780649135 | 0.129965262 | 0.709264768 | -4.406814215 |
| **A_51_P414653** | Plvap | -0.353792909 | -0.851474932 | 0.143889114 | 12.00899233 | -1.780522705 | 0.129987227 | 0.709264768 | -4.406847909 |
| **A_55_P2112379** | NA | 0.336897638 | -0.137037816 | 0.810833091 | 14.3943653 | 1.780447259 | 0.130000336 | 0.709264768 | -4.406868016 |
| **A_55_P2175381** | NA | 0.241680964 | -0.098324707 | 0.581686634 | 16.995172 | 1.780355087 | 0.130016353 | 0.709264768 | -4.40689258 |
| **A_55_P2346654** | AU035318 | -0.358612455 | -0.863158991 | 0.14593408 | 8.312161696 | -1.780223323 | 0.130039253 | 0.709264768 | -4.406927698 |
| **A_55_P1996623** | Dnajb12 | -0.24992127 | -0.60155695 | 0.10171441 | 9.54791201 | -1.780166517 | 0.130049127 | 0.709264768 | -4.406942837 |
| **A_55_P2171116** | Lgals3 | -1.165851897 | -2.806385083 | 0.474681289 | 14.58818439 | -1.779953714 | 0.130086124 | 0.709264768 | -4.406999555 |
| **A_52_P316474** | St3gal4 | -0.167014698 | -0.402037054 | 0.068007658 | 8.957444852 | -1.77990106 | 0.130095279 | 0.709264768 | -4.407013589 |
| **A_51_P498631** | Gsdme | 0.161925876 | -0.065946894 | 0.389798647 | 8.311896916 | 1.779812158 | 0.130110739 | 0.709264768 | -4.407037284 |
| **A_55_P2000938** | Ahcy | 0.424019948 | -0.172728269 | 1.020768165 | 9.453742233 | 1.779694689 | 0.13013117 | 0.709264768 | -4.407068595 |
| **A_55_P2018283** | Hmgb1 | 0.363068688 | -0.148166173 | 0.874303549 | 9.284686503 | 1.77876543 | 0.130292902 | 0.709264768 | -4.407316304 |
| **A_55_P2110215** | Ckap5 | 0.58908066 | -0.24040882 | 1.418570139 | 12.38423394 | 1.778747193 | 0.130296078 | 0.709264768 | -4.407321165 |
| **A_55_P2296088** | Sp3 | 0.226448482 | -0.092471725 | 0.54536869 | 9.04573605 | 1.778433873 | 0.130350656 | 0.709264768 | -4.407404695 |
| **A_55_P2033932** | Zzz3 | 0.298245984 | -0.12180085 | 0.718292818 | 9.783848349 | 1.778390927 | 0.130358139 | 0.709264768 | -4.407416145 |
| **A_51_P405397** | Ecm1 | -1.303771253 | -3.140418295 | 0.53287579 | 10.99312352 | -1.777976607 | 0.13043035 | 0.709264768 | -4.407526609 |

| | | | | | | | | | |
|---|---|---|---|---|---|---|---|---|---|
| **A_55_P2001072** | NA | 0.432200624 | -0.176649757 | 1.041051005 | 12.35650217 | 1.777972964 | 0.130430985 | 0.709264768 | -4.40752758 |
| **A_51_P483324** | Ptpn22 | 0.405605593 | -0.165871646 | 0.977082833 | 8.039674378 | 1.77768719 | 0.130480815 | 0.709264768 | -4.407603777 |
| **A_55_P2386236** | Ceacam2 | -0.545001379 | -1.312881887 | 0.222879129 | 8.770939856 | -1.777682934 | 0.130481558 | 0.709264768 | -4.407604912 |
| **A_51_P393862** | Spryd4 | 0.209341176 | -0.08568646 | 0.504368811 | 9.170174711 | 1.777224558 | 0.130561525 | 0.709264768 | -4.407727138 |
| **A_55_P2315921** | 5330431K | -0.500037417 | -1.204877493 | 0.204802659 | 8.217807216 | -1.776897022 | 0.130618696 | 0.709264768 | -4.407814482 |
| **A_55_P2164998** | NA | 0.85784092 | -0.351502066 | 2.067183905 | 13.27364072 | 1.776673457 | 0.130657733 | 0.709264768 | -4.407874102 |
| **A_52_P641758** | Ggct | 0.294540633 | -0.120698174 | 0.70977944 | 9.050689263 | 1.776632602 | 0.130664868 | 0.709264768 | -4.407884998 |
| **A_66_P121774** | Tmem59 | -0.360470858 | -0.868685547 | 0.147743831 | 15.50626973 | -1.776533056 | 0.130682255 | 0.709264768 | -4.407911546 |
| **A_55_P2114863** | Mgll | -0.18281792 | -0.440569855 | 0.074934015 | 10.57531428 | -1.776508032 | 0.130686626 | 0.709264768 | -4.40791822 |
| **A_55_P2111005** | Pafah1b1 | 0.333962536 | -0.137021906 | 0.804946979 | 11.26668574 | 1.775994165 | 0.130776417 | 0.709264768 | -4.408055272 |
| **A_55_P2027147** | Mrps18c | 0.239224737 | -0.098213224 | 0.576662698 | 12.27525438 | 1.775670965 | 0.130832924 | 0.709264768 | -4.408141478 |
| **A_55_P2163729** | Tvp23a | 0.187137675 | -0.076852881 | 0.451128232 | 8.470108688 | 1.77551025 | 0.130861032 | 0.709264768 | -4.408184347 |
| **A_55_P2016842** | Me1 | -0.596601916 | -1.43827398 | 0.245070148 | 9.91925547 | -1.775383081 | 0.130883277 | 0.709264768 | -4.408218269 |
| **A_55_P1983858** | Scel | -1.186252131 | -2.86012418 | 0.487619919 | 8.848555284 | -1.775027532 | 0.130945491 | 0.709264768 | -4.408313114 |
| **A_55_P2014867** | Iws1 | 0.364138535 | -0.149688597 | 0.877965666 | 8.939871457 | 1.775006523 | 0.130949168 | 0.709264768 | -4.408318719 |
| **A_55_P2074080** | NA | 0.990508244 | -0.407241545 | 2.388258034 | 14.4757336 | 1.774920906 | 0.130964154 | 0.709264768 | -4.408341559 |
| **A_55_P2136501** | Midn | 0.217988004 | -0.089702577 | 0.525678586 | 12.89598231 | 1.774470316 | 0.131043052 | 0.709264768 | -4.408461768 |
| **A_55_P2147487** | Cyth3 | -0.728793412 | -1.757568843 | 0.299982019 | 9.547665325 | -1.774329199 | 0.131067771 | 0.709264768 | -4.408499417 |
| **A_52_P91402** | Zcchc17 | 0.194584139 | -0.080124489 | 0.469292766 | 12.1334858 | 1.774130178 | 0.131102642 | 0.709264768 | -4.408552516 |
| **A_55_P1993012** | NA | 0.54565808 | -0.224807971 | 1.316124132 | 11.4225769 | 1.773852191 | 0.131151363 | 0.709264768 | -4.408626687 |
| **A_51_P473919** | Mycbp2 | -0.226926659 | -0.54738723 | 0.093533912 | 9.448589249 | -1.773622798 | 0.131191581 | 0.709264768 | -4.408687895 |
| **A_66_P105689** | Trim34a | -0.594888667 | -1.435168157 | 0.245390823 | 8.732766998 | -1.773218599 | 0.131262476 | 0.709264768 | -4.408795752 |
| **A_51_P450518** | Hpf1 | 0.539206125 | -0.222635985 | 1.301048234 | 13.29412798 | 1.772720178 | 0.131349951 | 0.709264768 | -4.408928762 |
| **A_55_P2146780** | Sdhc | 0.514956888 | -0.212776433 | 1.24269021 | 12.6928606 | 1.772347826 | 0.131415337 | 0.709264768 | -4.409028137 |
| **A_55_P2125643** | NA | 0.454426545 | -0.187929411 | 1.096782501 | 13.94315378 | 1.771896226 | 0.131494684 | 0.709264768 | -4.40914867 |
| **A_51_P330213** | Asf1b | 0.988217048 | -0.409107268 | 2.385541365 | 10.07611078 | 1.771354441 | 0.131589939 | 0.709264768 | -4.409293286 |
| **A_51_P429335** | Prss16 | 1.856905396 | -0.769412401 | 4.483223193 | 9.255018524 | 1.770895145 | 0.131670745 | 0.709264768 | -4.409415894 |
| **A_55_P2008258** | Larp1b | 0.184989114 | -0.07667526 | 0.446653487 | 9.900200259 | 1.770728273 | 0.131700116 | 0.709264768 | -4.409460443 |
| **A_55_P2136289** | NA | -0.175719166 | -0.424346089 | 0.072907757 | 11.01309974 | -1.770195876 | 0.131793866 | 0.709264768 | -4.409602583 |
| **A_55_P1984070** | NA | 0.423962906 | -0.176042908 | 1.023968721 | 12.89258497 | 1.769794117 | 0.131864656 | 0.709264768 | -4.409709853 |
| **A_55_P2028961** | Idi1 | 1.119324349 | -0.465082869 | 2.703731567 | 11.30382743 | 1.769454847 | 0.131924464 | 0.709264768 | -4.409800445 |
| **A_55_P2144436** | Rrp1 | 0.288848022 | -0.12002271 | 0.697718754 | 11.59416383 | 1.769431358 | 0.131928606 | 0.709264768 | -4.409806717 |
| **A_55_P2060707** | Il15ra | 0.28022485 | -0.116463228 | 0.676912927 | 11.89285347 | 1.76932593 | 0.131947198 | 0.709264768 | -4.409834869 |
| **A_52_P608495** | Aars2 | 0.173545764 | -0.072145179 | 0.419236707 | 9.267288464 | 1.769193012 | 0.131970641 | 0.709264768 | -4.409870363 |
| **A_55_P2035613** | Tmbim1 | -0.761065108 | -1.838520753 | 0.316390537 | 12.69280833 | -1.769182995 | 0.131972408 | 0.709264768 | -4.409873038 |
| **A_55_P2325729** | Arhgap1 | 0.234276907 | -0.097407636 | 0.56596145 | 11.24607518 | 1.769109006 | 0.131985459 | 0.709264768 | -4.409892796 |
| **A_55_P1988273** | Rnf103 | -0.51531719 | -1.245036257 | 0.214401877 | 10.48511167 | -1.768761519 | 0.132046773 | 0.709264768 | -4.409985593 |
| **A_55_P2028511** | NA | 0.679745012 | -0.28288655 | 1.642376575 | 12.67302985 | 1.768627113 | 0.132070496 | 0.709264768 | -4.410021488 |
| **A_52_P217646** | Snapc3 | 0.39331774 | -0.163939984 | 0.950575465 | 9.907475338 | 1.767818945 | 0.132213231 | 0.709264768 | -4.410237337 |
| **A_55_P2067608** | Clmp | -0.427031211 | -1.032071577 | 0.178009156 | 8.6526526 | -1.767769403 | 0.132221986 | 0.709264768 | -4.41025057 |
| **A_66_P123294** | NA | 0.281322092 | -0.11742822 | 0.680072403 | 8.122529858 | 1.767067537 | 0.132346079 | 0.709264768 | -4.410438054 |
| **A_55_P2003927** | NA | 0.800038381 | -0.334253513 | 1.934330274 | 12.76344175 | 1.766592325 | 0.132430165 | 0.709264768 | -4.410565006 |
| **A_55_P2126850** | Nudt11 | 0.596398003 | -0.249200898 | 1.441996904 | 8.910769539 | 1.76653447 | 0.132440406 | 0.709264768 | -4.410580463 |
| **A_52_P95910** | Ugcg | -0.425715003 | -1.043036864 | 0.191606858 | 9.46218092 | -1.836834402 | 0.132443509 | 0.709264768 | -4.434371957 |
| **A_55_P2183055** | E4f1 | -0.218763295 | -0.528951801 | 0.091425212 | 10.1898136 | -1.766440851 | 0.132456978 | 0.709264768 | -4.410605474 |
| **A_55_P2089761** | NA | 0.298427302 | -0.124760822 | 0.721615427 | 9.290872565 | 1.766263224 | 0.132488428 | 0.709264768 | -4.410652931 |
| **A_51_P405476** | Fcer1g | -0.327316815 | -0.791586148 | 0.136952519 | 11.05986774 | -1.765829077 | 0.132565328 | 0.709264768 | -4.410768929 |
| **A_55_P2105052** | Tfb2m | 0.185351959 | -0.077563853 | 0.448267772 | 10.13067981 | 1.765756528 | 0.132578183 | 0.709264768 | -4.410788314 |

| | | | | | | | | | |
|---|---|---|---|---|---|---|---|---|---|
| **A_55_P2061025** | Hspa4 | -0.343534163 | -0.830962347 | 0.14389402 | 10.00884768 | -1.765263906 | 0.132665502 | 0.709264768 | -4.410919947 |
| **A_55_P2057519** | Rpl17 | 0.393714996 | -0.164932528 | 0.952362521 | 14.18855567 | 1.765202054 | 0.13267647 | 0.709264768 | -4.410936476 |
| **A_52_P171033** | Mcmbp | 0.276477856 | -0.115852317 | 0.66880803 | 8.969406662 | 1.765058032 | 0.132702011 | 0.709264768 | -4.410974962 |
| **A_55_P2137521** | Nmral1 | 0.402592661 | -0.168915129 | 0.974100452 | 8.422079016 | 1.764387794 | 0.132820939 | 0.709264768 | -4.411154081 |
| **A_55_P2086866** | NA | 0.583150404 | -0.244722713 | 1.41102352 | 14.13352415 | 1.764278518 | 0.132840339 | 0.709264768 | -4.411183286 |
| **A_55_P2063711** | Snu13 | 0.483592514 | -0.203108315 | 1.170293344 | 13.30389309 | 1.763852818 | 0.132915941 | 0.709264768 | -4.411297066 |
| **A_55_P2046877** | Foxq1 | -0.682337485 | -1.651269047 | 0.286594078 | 8.666088824 | -1.763828975 | 0.132920177 | 0.709264768 | -4.411303439 |
| **A_55_P2095933** | Zranb1 | -0.277748603 | -0.672196413 | 0.116699207 | 9.458814339 | -1.763651126 | 0.132951776 | 0.709264768 | -4.411350977 |
| **A_51_P287232** | Qars | 0.267066285 | -0.112277241 | 0.64640981 | 12.58008922 | 1.763342741 | 0.133006585 | 0.709264768 | -4.411433409 |
| **A_66_P100853** | NA | 1.177876123 | -0.495423242 | 2.851175489 | 16.59161204 | 1.763097451 | 0.133050196 | 0.709264768 | -4.411498979 |
| **A_55_P2037689** | NA | 0.52457286 | -0.220640628 | 1.269786348 | 10.43277015 | 1.763093956 | 0.133050818 | 0.709264768 | -4.411499913 |
| **A_55_P2087561** | Eno1b | 0.525291206 | -0.221095339 | 1.271677751 | 10.28802521 | 1.762733563 | 0.13311492 | 0.709264768 | -4.411596257 |
| **A_55_P2028243** | Dnajc19 | 0.320001621 | -0.134696334 | 0.774699576 | 10.81685071 | 1.762704541 | 0.133120083 | 0.709264768 | -4.411604015 |
| **A_55_P2011937** | Fignl1 | 1.315509769 | -0.553869445 | 3.184888982 | 11.04550398 | 1.762572597 | 0.133143561 | 0.709264768 | -4.41163929 |
| **A_52_P290725** | Arfgef2 | -0.151622413 | -0.367120159 | 0.063875333 | 8.815132189 | -1.762264194 | 0.133198452 | 0.709264768 | -4.411721742 |
| **A_52_P658974** | NA | 1.117955121 | -0.471145747 | 2.70705599 | 14.07322416 | 1.762070379 | 0.133232959 | 0.709264768 | -4.411773561 |
| **A_55_P2050799** | Nup35 | 0.477290744 | -0.201171892 | 1.155753381 | 10.73697494 | 1.76200616 | 0.133244395 | 0.709264768 | -4.411790731 |
| **A_55_P2083426** | NA | 1.038764576 | -0.437856434 | 2.515385586 | 13.9461223 | 1.761969689 | 0.13325089 | 0.709264768 | -4.411800482 |
| **A_55_P2013236** | S100g | -2.734070488 | -6.621451193 | 1.153310218 | 11.07608737 | -1.761582462 | 0.13331987 | 0.709264768 | -4.41190402 |
| **A_51_P223709** | Pask | 0.567293652 | -0.239328665 | 1.373915969 | 9.527502406 | 1.761521856 | 0.133330669 | 0.709264768 | -4.411920226 |
| **A_65_P15675** | Ipo9 | 0.281777717 | -0.118973014 | 0.682528448 | 8.915892329 | 1.76109453 | 0.133406839 | 0.709264768 | -4.412034494 |
| **A_55_P2082314** | NA | 0.277260115 | -0.11719218 | 0.67171241 | 8.490327128 | 1.760529298 | 0.133507657 | 0.709264768 | -4.412185652 |
| **A_55_P2130104** | Slc52a3 | -1.088789348 | -2.638426732 | 0.460848036 | 9.365444225 | -1.759803285 | 0.133637264 | 0.709264768 | -4.412379827 |
| **A_55_P2186619** | Rbmxl1 | 0.91553969 | -0.387694997 | 2.218774377 | 12.77832473 | 1.759563284 | 0.133680136 | 0.709264768 | -4.412444022 |
| **A_55_P2168185** | NA | 0.242560574 | -0.102817868 | 0.587939017 | 16.99115545 | 1.759038444 | 0.133773938 | 0.709264768 | -4.412584414 |
| **A_55_P1960556** | Acox1 | -0.596592835 | -1.446091839 | 0.252906169 | 12.37675969 | -1.758998645 | 0.133781054 | 0.709264768 | -4.412595061 |
| **A_55_P2128792** | NA | 0.267150689 | -0.113304023 | 0.647605402 | 8.719314024 | 1.758748243 | 0.133825832 | 0.709264768 | -4.412662047 |
| **A_55_P1953356** | Triobp | -0.359057708 | -0.870460565 | 0.15234515 | 10.21091614 | -1.758536752 | 0.133863664 | 0.709264768 | -4.412718626 |
| **A_55_P2022201** | NA | -0.165880229 | -0.402159624 | 0.070399166 | 8.73281292 | -1.758405853 | 0.133887084 | 0.709264768 | -4.412753645 |
| **A_55_P2108579** | Phkg2 | -0.291764226 | -0.707364994 | 0.123836542 | 10.59234448 | -1.758352914 | 0.133896558 | 0.709264768 | -4.412767809 |
| **A_55_P2178044** | Inppl1 | 0.25823178 | -0.109622932 | 0.626086492 | 12.53292947 | 1.758262491 | 0.13391274 | 0.709264768 | -4.412792 |
| **A_55_P2087944** | NA | 1.099802992 | -0.467058277 | 2.666664262 | 10.70833402 | 1.758064065 | 0.133948257 | 0.709264768 | -4.412845088 |
| **A_55_P2054128** | NA | 0.40158999 | -0.170610585 | 0.973790564 | 12.79802138 | 1.757862635 | 0.133984321 | 0.709264768 | -4.412898982 |
| **A_51_P450527** | Tagln | -0.394069816 | -0.955619417 | 0.167479786 | 14.00883868 | -1.75766212 | 0.134020232 | 0.709264768 | -4.412952633 |
| **A_55_P2045203** | Chp1 | -0.520215923 | -1.261671093 | 0.221239247 | 12.29927911 | -1.757312867 | 0.134082802 | 0.709264768 | -4.413046085 |
| **A_51_P212419** | Lrrc41 | -0.274464191 | -0.665654283 | 0.1167259 | 9.565932522 | -1.757309234 | 0.134083453 | 0.709264768 | -4.413047057 |
| **A_55_P1983046** | Plch1 | 0.484615318 | -0.206126604 | 1.175357239 | 8.552879993 | 1.757242379 | 0.134095434 | 0.709264768 | -4.413064946 |
| **A_66_P100840** | NA | 0.284032636 | -0.120966718 | 0.68903199 | 8.820304568 | 1.756565137 | 0.13421686 | 0.709264768 | -4.413246178 |
| **A_55_P2015304** | NA | 1.167426164 | -0.49719606 | 2.832048389 | 13.60945616 | 1.756564411 | 0.13421699 | 0.709264768 | -4.413246373 |
| **A_51_P450296** | Thap3 | -0.303420937 | -0.736117984 | 0.12927611 | 9.061052397 | -1.756353617 | 0.134254807 | 0.709264768 | -4.413302786 |
| **A_55_P2134979** | NA | 1.113299359 | -0.474356938 | 2.700955656 | 13.74398515 | 1.756328766 | 0.134259266 | 0.709264768 | -4.413309437 |
| **A_55_P2001193** | Stx18 | -0.196553173 | -0.476857023 | 0.083750677 | 10.94466285 | -1.756310681 | 0.134262511 | 0.709264768 | -4.413314277 |
| **A_55_P2040371** | Rnaset2a | -0.18297931 | -0.443929925 | 0.077971306 | 14.26766183 | -1.756281009 | 0.134267835 | 0.709264768 | -4.413322218 |
| **A_55_P2097206** | Camk4 | 0.801503162 | -0.341591727 | 1.944598052 | 9.034996391 | 1.75619729 | 0.134282859 | 0.709264768 | -4.413344624 |
| **A_65_P10357** | Srsf6 | 0.293713183 | -0.125189539 | 0.712615906 | 11.21403431 | 1.756145935 | 0.134292075 | 0.709264768 | -4.413358369 |
| **A_55_P2093291** | Coil | 0.402896832 | -0.171759752 | 0.977553415 | 9.890374011 | 1.75604569 | 0.134310068 | 0.709264768 | -4.413385198 |
| **A_55_P2092750** | Car9 | 0.280767586 | -0.119850963 | 0.681386135 | 8.432589825 | 1.755360253 | 0.13443316 | 0.709264768 | -4.41356866 |
| **A_51_P199866** | Pin1 | 0.32923405 | -0.140628911 | 0.799097011 | 11.57251016 | 1.755027308 | 0.134492991 | 0.709264768 | -4.413657783 |

| | | | | | | | | | |
|---|---|---|---|---|---|---|---|---|---|
| **A_51_P400463** | Agmat | 0.21602485 | -0.092276228 | 0.524325927 | 8.438214044 | 1.755007651 | 0.134496524 | 0.709264768 | -4.413663045 |
| **A_55_P1954718** | Cyb561 | -0.169633671 | -0.411733297 | 0.072465955 | 12.35696257 | -1.75496431 | 0.134504315 | 0.709264768 | -4.413674647 |
| **A_55_P2072980** | Sptan1 | -0.466835484 | -1.133157404 | 0.199486436 | 8.47846612 | -1.754810014 | 0.134532053 | 0.709264768 | -4.413715951 |
| **A_55_P1980311** | Pcbp2 | 0.505188714 | -0.215937894 | 1.226315322 | 9.876408898 | 1.754657938 | 0.134559398 | 0.709264768 | -4.413756662 |
| **A_55_P2152009** | Tesc | 0.603319137 | -0.258146189 | 1.464784463 | 9.97478719 | 1.754121393 | 0.13465592 | 0.709264768 | -4.413900304 |
| **A_55_P2020153** | Cycs | 0.365825635 | -0.15665675 | 0.88830802 | 13.24317051 | 1.75369001 | 0.134733573 | 0.709264768 | -4.414015802 |
| **A_51_P168632** | Prim1 | 1.283203296 | -0.549759219 | 3.116165811 | 11.29717995 | 1.753445325 | 0.134777639 | 0.709264768 | -4.414081317 |
| **A_55_P2053798** | NA | 0.570538755 | -0.244539964 | 1.385617473 | 15.12540212 | 1.753218084 | 0.134818576 | 0.709264768 | -4.414142164 |
| **A_55_P2002963** | Coro1a | 1.503046583 | -0.644297341 | 3.650390506 | 11.22748946 | 1.753158487 | 0.134829314 | 0.709264768 | -4.414158122 |
| **A_55_P2036567** | Vav1 | 1.104048951 | -0.473686019 | 2.68178392 | 9.98045753 | 1.752688041 | 0.134914109 | 0.709264768 | -4.414284099 |
| **A_55_P2077872** | Deaf1 | -0.248242866 | -0.602996966 | 0.106511235 | 10.43621983 | -1.752668164 | 0.134917693 | 0.709264768 | -4.414289422 |
| **A_52_P322181** | Adrb1 | -0.632083717 | -1.535385403 | 0.271217969 | 8.928472732 | -1.752637612 | 0.134923202 | 0.709264768 | -4.414297604 |
| **A_55_P2020338** | Scml4 | 0.181879906 | -0.078052262 | 0.441812074 | 7.960963848 | 1.752568649 | 0.134935638 | 0.709264768 | -4.414316072 |
| **A_55_P2184897** | Lbr | 0.179053095 | -0.076839917 | 0.434946106 | 8.131283954 | 1.752563457 | 0.134936574 | 0.709264768 | -4.414317462 |
| **A_55_P2121352** | Cdk5 | -0.431978152 | -1.049390345 | 0.185434042 | 9.719205309 | -1.752414852 | 0.134963375 | 0.709264768 | -4.414357259 |
| **A_55_P1962065** | NA | 0.385254467 | -0.165412157 | 0.935921091 | 14.21456792 | 1.752303282 | 0.1349835 | 0.709264768 | -4.414387138 |
| **A_66_P119999** | Atg7 | -0.322221993 | -0.782802057 | 0.138358071 | 9.511399055 | -1.752267434 | 0.134989967 | 0.709264768 | -4.414396739 |
| **A_55_P2111478** | Polq | 0.311289833 | -0.133684018 | 0.756263683 | 8.283494613 | 1.752188362 | 0.135004232 | 0.709264768 | -4.414417915 |
| **A_55_P1980621** | NA | 0.986308603 | -0.423629381 | 2.396246586 | 14.19726414 | 1.752117213 | 0.13501707 | 0.709264768 | -4.41443697 |
| **A_52_P413947** | Mthfr | -0.471709772 | -1.146045031 | 0.202625487 | 10.18029543 | -1.752061516 | 0.13502712 | 0.709264768 | -4.414451887 |
| **A_51_P343356** | Isoc2b | -0.238512959 | -0.579614249 | 0.102588332 | 9.815996317 | -1.751374262 | 0.135151193 | 0.709264768 | -4.414635958 |
| **A_52_P102559** | Dars2 | 0.160568804 | -0.069114466 | 0.390252074 | 9.021438657 | 1.750983914 | 0.135221714 | 0.709264768 | -4.414740517 |
| **A_55_P2131353** | 1110059G | 0.334784256 | -0.144124139 | 0.813692651 | 9.344233693 | 1.750906366 | 0.135235728 | 0.709264768 | -4.414761289 |
| **A_51_P401263** | Eme1 | 0.371638892 | -0.160027017 | 0.9033048 | 8.551701058 | 1.750784481 | 0.135257758 | 0.709264768 | -4.414793939 |
| **A_55_P2161390** | NA | -0.181776705 | -0.441878565 | 0.078325155 | 9.636409922 | -1.75043148 | 0.135321581 | 0.709264768 | -4.414888503 |
| **A_55_P2124528** | Capns1 | -0.346255873 | -0.841810848 | 0.149299102 | 13.96984683 | -1.750070938 | 0.135386798 | 0.709264768 | -4.414985093 |
| **A_66_P100349** | Esyt2 | -0.40049242 | -0.973695883 | 0.172711043 | 10.73174516 | -1.749991102 | 0.135401243 | 0.709264768 | -4.415006481 |
| **A_51_P370252** | Capn10 | -0.353204861 | -0.858755862 | 0.152346141 | 9.998716293 | -1.749895209 | 0.135418596 | 0.709264768 | -4.415032173 |
| **A_51_P327778** | Pfdn2 | 0.278250494 | -0.12004484 | 0.676545827 | 10.23387596 | 1.749770423 | 0.135441181 | 0.709264768 | -4.415065605 |
| **A_51_P338397** | Kctd10 | -0.487265229 | -1.184776111 | 0.210245654 | 9.523829159 | -1.749704782 | 0.135453062 | 0.709264768 | -4.415083192 |
| **A_52_P69043** | Nkap | 0.220920429 | -0.095399213 | 0.53724007 | 9.399416894 | 1.749282949 | 0.135529443 | 0.709264768 | -4.415196216 |
| **A_55_P2037111** | Ran | 0.630975015 | -0.27257316 | 1.53452319 | 10.60772011 | 1.749086126 | 0.135565095 | 0.709264768 | -4.415248954 |
| **A_55_P2145389** | Flrt3 | 0.319850672 | -0.138260607 | 0.777961951 | 8.92085148 | 1.748745576 | 0.135626805 | 0.709264768 | -4.415340208 |
| **A_51_P388374** | Ppil4 | 0.235137529 | -0.101655896 | 0.571930953 | 10.59327707 | 1.748673347 | 0.135639897 | 0.709264768 | -4.415359563 |
| **A_55_P2042612** | C1s1 | -0.368694035 | -0.896816427 | 0.159428356 | 8.900621286 | -1.748565382 | 0.135659469 | 0.709264768 | -4.415388495 |
| **A_52_P260818** | Card10 | -0.345075409 | -0.839371145 | 0.149220326 | 9.086018768 | -1.748547737 | 0.135662668 | 0.709264768 | -4.415393223 |
| **A_55_P1999408** | NA | 0.346933283 | -0.150027354 | 0.843893919 | 12.08118679 | 1.748534963 | 0.135664984 | 0.709264768 | -4.415396647 |
| **A_55_P2100700** | Pygo2 | 0.352708357 | -0.152532058 | 0.857948772 | 12.82148207 | 1.748509562 | 0.135669589 | 0.709264768 | -4.415403453 |
| **A_51_P275591** | Zfp292 | 0.306311606 | -0.132506526 | 0.745129737 | 9.639027853 | 1.748353446 | 0.135697896 | 0.709264768 | -4.41544529 |
| **A_51_P424878** | Prxl2b | 0.964075337 | -0.417110957 | 2.345261632 | 11.37997967 | 1.748272226 | 0.135712626 | 0.709264768 | -4.415467056 |
| **A_55_P1999785** | NA | 0.193614493 | -0.083776094 | 0.471005079 | 10.8954724 | 1.748221629 | 0.135721803 | 0.709264768 | -4.415480615 |
| **A_55_P2060642** | Spidr | 0.65045434 | -0.281570056 | 1.582478736 | 10.37913461 | 1.747993773 | 0.135763136 | 0.709264768 | -4.41554168 |
| **A_51_P347961** | Npc2 | -0.230181042 | -0.560005952 | 0.099643869 | 14.81978096 | -1.747979934 | 0.135765647 | 0.709264768 | -4.415545389 |
| **A_55_P2325378** | NA | -0.387390598 | -0.942503927 | 0.16772273 | 8.847957603 | -1.747904768 | 0.135779285 | 0.709264768 | -4.415565533 |
| **A_55_P2025090** | Gins3 | 0.896467999 | -0.388247329 | 2.181183327 | 13.43297102 | 1.747745655 | 0.135808159 | 0.709264768 | -4.415608177 |
| **A_51_P404193** | Sp5 | 1.006380838 | -0.436061673 | 2.448823349 | 9.467410756 | 1.747487957 | 0.135854936 | 0.709264768 | -4.415677244 |
| **A_55_P1955632** | Trp53i11 | -0.49125911 | -1.195388667 | 0.212870446 | 12.80629099 | -1.747464607 | 0.135859175 | 0.709264768 | -4.415683503 |
| **A_55_P2171303** | Bin3 | -0.169565284 | -0.412613132 | 0.073482565 | 8.231484137 | -1.747412769 | 0.135868587 | 0.709264768 | -4.415697397 |

| | | | | | | | | | |
|---|---|---|---|---|---|---|---|---|---|
| **A_52_P311245** | Scarb2 | -0.542063017 | -1.319070105 | 0.234944071 | 9.579254779 | -1.747330842 | 0.135883463 | 0.709264768 | -4.415719355 |
| **A_51_P154973** | Stard3 | -0.27345886 | -0.66547905 | 0.11856133 | 11.7503516 | -1.747164955 | 0.13591359 | 0.709264768 | -4.415763819 |
| **A_55_P1971594** | Pqbp1 | 0.30937605 | -0.134165887 | 0.752917987 | 9.924389312 | 1.747037981 | 0.135936654 | 0.709264768 | -4.415797853 |
| **A_55_P1988754** | H4f16 | 1.382150803 | -0.599597832 | 3.363899437 | 13.97574405 | 1.746856412 | 0.135969641 | 0.709264768 | -4.415846522 |
| **A_51_P257550** | Marcksl1 | 0.487507695 | -0.211581042 | 1.186596432 | 12.47302982 | 1.746624372 | 0.13601181 | 0.709264768 | -4.415908722 |
| **A_66_P139393** | Psmd10 | -0.259494432 | -0.631762952 | 0.112774088 | 9.813419777 | -1.745910901 | 0.136141551 | 0.709264768 | -4.416099987 |
| **A_51_P190111** | Mcm5 | 1.540834748 | -0.669649446 | 3.751318942 | 11.19368179 | 1.745898515 | 0.136143804 | 0.709264768 | -4.416103308 |
| **A_55_P2179604** | Vash1 | -0.241589466 | -0.588206293 | 0.105027361 | 14.5412289 | -1.745736517 | 0.13617328 | 0.709264768 | -4.416146739 |
| **A_55_P1992959** | Pfkfb4 | -0.251724743 | -0.613003436 | 0.109553951 | 9.685032972 | -1.745154578 | 0.136279218 | 0.709264768 | -4.416302764 |
| **A_55_P2065926** | Akap8l | -0.251469219 | -0.612413786 | 0.109475348 | 9.069765577 | -1.744996941 | 0.136307929 | 0.709264768 | -4.416345031 |
| **A_55_P2250424** | 9530029C | -0.391498232 | -0.953528004 | 0.170531541 | 10.09921412 | -1.74470027 | 0.136361979 | 0.709264768 | -4.41642458 |
| **A_55_P2122648** | Dtymk | 0.602460718 | -0.26276851 | 1.467689945 | 10.9671705 | 1.744005695 | 0.136488604 | 0.709264768 | -4.416610838 |
| **A_55_P2160716** | NA | -0.33493278 | -0.815962195 | 0.146096635 | 15.41691694 | -1.743959379 | 0.136497052 | 0.709264768 | -4.416623259 |
| **A_55_P2023042** | Ift43 | -0.212265483 | -0.517159106 | 0.092628141 | 12.1632861 | -1.743738599 | 0.136537329 | 0.709264768 | -4.416682468 |
| **A_55_P1979650** | S100a1 | -0.939483771 | -2.288946473 | 0.40997893 | 10.5644793 | -1.743728006 | 0.136539262 | 0.709264768 | -4.416685309 |
| **A_55_P2012789** | Impa1 | 0.522118752 | -0.227890664 | 1.272128169 | 10.50216801 | 1.743624333 | 0.136558179 | 0.709264768 | -4.416713113 |
| **A_55_P2070825** | Nudt5 | 0.394641836 | -0.172375306 | 0.961658977 | 11.00686183 | 1.743240392 | 0.136628259 | 0.709264768 | -4.416816087 |
| **A_51_P177518** | Emc6 | 0.339101614 | -0.148140088 | 0.826343316 | 11.94667512 | 1.743153978 | 0.136644038 | 0.709264768 | -4.416839264 |
| **A_52_P517001** | Rpf1 | 0.151084719 | -0.066013067 | 0.368182504 | 9.875726092 | 1.743072667 | 0.136658885 | 0.709264768 | -4.416861073 |
| **A_55_P2026635** | Asnsd1 | 0.268891718 | -0.117612741 | 0.655396177 | 11.83745144 | 1.742501908 | 0.136763155 | 0.709264768 | -4.417014167 |
| **A_55_P2103856** | Hgs | -0.226348276 | -0.551703582 | 0.099007031 | 12.70207927 | -1.74248738 | 0.13676581 | 0.709264768 | -4.417018064 |
| **A_55_P2156504** | NA | 0.377962912 | -0.165409052 | 0.921334875 | 11.75781495 | 1.742217232 | 0.136815191 | 0.709264768 | -4.41709053 |
| **A_66_P126300** | Mrpl47 | 0.217777147 | -0.095319607 | 0.530873902 | 10.45178274 | 1.742144141 | 0.136828554 | 0.709264768 | -4.417110138 |
| **A_55_P1999972** | NA | -0.412610246 | -1.005910768 | 0.180690276 | 8.214252 | -1.741869534 | 0.136878773 | 0.709264768 | -4.417183804 |
| **A_51_P332201** | Cst3 | -0.535280629 | -1.305138568 | 0.23457731 | 13.95951575 | -1.741491183 | 0.136947994 | 0.709264768 | -4.417285307 |
| **A_55_P1995537** | Mpo | 0.33099111 | -0.145089429 | 0.807071648 | 8.128039968 | 1.741350705 | 0.136973703 | 0.709264768 | -4.417322996 |
| **A_55_P1980316** | Pcbp2 | 0.250683468 | -0.109903322 | 0.611270258 | 14.29183427 | 1.741270441 | 0.136988395 | 0.709264768 | -4.41734453 |
| **A_52_P484838** | Rfxank | -0.179142871 | -0.436829866 | 0.078544125 | 9.515484031 | -1.74123494 | 0.136994894 | 0.709264768 | -4.417354055 |
| **A_55_P2072706** | Chsy1 | 0.576803094 | -0.25293435 | 1.406540537 | 12.0300708 | 1.741154212 | 0.137009673 | 0.709264768 | -4.417375714 |
| **A_52_P576442** | Dap | -0.464990679 | -1.133983999 | 0.20400264 | 11.48712328 | -1.740895939 | 0.137056966 | 0.709264768 | -4.41744501 |
| **A_55_P1997741** | Slc9a8 | -0.506560096 | -1.235389607 | 0.222269414 | 11.14765111 | -1.740826033 | 0.13706977 | 0.709264768 | -4.417463767 |
| **A_55_P2089942** | Xpo1 | 0.320009606 | -0.140450952 | 0.780470164 | 11.23186491 | 1.740687961 | 0.137095061 | 0.709264768 | -4.417500814 |
| **A_51_P246339** | Rfc5 | 0.8910085 | -0.391129493 | 2.173146493 | 11.43837317 | 1.740593764 | 0.137112319 | 0.709264768 | -4.417526089 |
| **A_51_P402458** | Anxa7 | -0.293166902 | -0.71505863 | 0.128724827 | 8.557915989 | -1.740460905 | 0.137136663 | 0.709264768 | -4.417561739 |
| **A_55_P2037712** | Smc4 | 1.32792279 | -0.583631793 | 3.239477372 | 12.03416105 | 1.739948791 | 0.13723054 | 0.709264768 | -4.41769916 |
| **A_55_P1988368** | Upp1 | -0.800705928 | -1.953466482 | 0.352054626 | 9.447123467 | -1.739739733 | 0.137268881 | 0.709264768 | -4.417755262 |
| **A_52_P39418** | Ccpg1 | -0.237785371 | -0.580120809 | 0.104550068 | 8.14566325 | -1.739737068 | 0.13726937 | 0.709264768 | -4.417755977 |
| **A_51_P417720** | Itga11 | -0.45855986 | -1.119089648 | 0.201969927 | 8.893873039 | -1.738817389 | 0.137438166 | 0.709264768 | -4.418002802 |
| **A_52_P543040** | Utp14a | 0.318681166 | -0.140367064 | 0.777729397 | 8.794663863 | 1.738795172 | 0.137442246 | 0.709264768 | -4.418008765 |
| **A_55_P2013586** | Prss8 | -0.211220102 | -0.515571309 | 0.093131105 | 10.42020803 | -1.738243306 | 0.137543636 | 0.709264768 | -4.418156893 |
| **A_55_P2094412** | Ssbp1 | 0.250232267 | -0.110344907 | 0.610809441 | 9.720214924 | 1.738182711 | 0.137554773 | 0.709264768 | -4.418173158 |
| **A_55_P1965233** | Sh3tc1 | -0.933959878 | -2.279800451 | 0.411880696 | 11.01240958 | -1.738140777 | 0.137562481 | 0.709264768 | -4.418184415 |
| **A_66_P121770** | Ilf2 | 0.599562522 | -0.264427418 | 1.463552462 | 10.56814235 | 1.738105515 | 0.137568963 | 0.709264768 | -4.41819388 |
| **A_52_P42834** | Usp25 | -0.275937439 | -0.673613864 | 0.121738986 | 8.803031177 | -1.737925396 | 0.137602076 | 0.709264768 | -4.41824223 |
| **A_55_P2157838** | NA | 0.292700721 | -0.12922711 | 0.714628551 | 12.52802108 | 1.737544616 | 0.137672107 | 0.709264768 | -4.418344449 |
| **A_51_P303620** | Whrn | 0.237457237 | -0.104867405 | 0.579781879 | 9.066240703 | 1.737391105 | 0.137700349 | 0.709264768 | -4.418385661 |
| **A_55_P2029061** | Tnfrsf9 | 0.343270313 | -0.151634663 | 0.838175288 | 11.92325402 | 1.737259804 | 0.137724511 | 0.709264768 | -4.418420911 |
| **A_55_P2089913** | Cd6 | 1.191718106 | -0.526559444 | 2.909995657 | 8.65959351 | 1.737122988 | 0.137749691 | 0.709264768 | -4.418457642 |

| | | | | | | | | | |
|---|---|---|---|---|---|---|---|---|---|
| **A_55_P2099064** | Nsd2 | 0.149291007 | -0.065965073 | 0.364547087 | 8.236709857 | 1.737114983 | 0.137751164 | 0.709264768 | -4.418459791 |
| **A_51_P392291** | Pdk3 | 0.44449327 | -0.1964078 | 1.085394339 | 9.934365176 | 1.737098874 | 0.137754129 | 0.709264768 | -4.418464116 |
| **A_55_P1968718** | Plekha6 | -0.721641607 | -1.762368487 | 0.319085273 | 8.878896799 | -1.736741328 | 0.137819958 | 0.709264768 | -4.41856011 |
| **A_55_P2150044** | Cdkl2 | -0.599692279 | -1.464556053 | 0.265171496 | 8.92043314 | -1.736725159 | 0.137822935 | 0.709264768 | -4.418564452 |
| **A_52_P679101** | Tjp2 | -0.369129491 | -0.901530589 | 0.163271607 | 12.42238118 | -1.736561413 | 0.137853094 | 0.709264768 | -4.418608417 |
| **A_51_P159610** | Ccdc120 | -0.285970516 | -0.698533809 | 0.126592777 | 8.559687151 | -1.736125117 | 0.137933483 | 0.709264768 | -4.418725565 |
| **A_51_P249127** | Pkd1 | -0.235768997 | -0.575912379 | 0.104374385 | 10.842649 | -1.736101112 | 0.137937907 | 0.709264768 | -4.418732011 |
| **A_55_P2424767** | Hecw1 | 0.235763766 | -0.104542968 | 0.5760705 | 12.78747868 | 1.735229263 | 0.138098691 | 0.709264768 | -4.418966133 |
| **A_55_P2181251** | Ccdc9b | -1.157522229 | -2.828412251 | 0.513367793 | 9.869735935 | -1.735129206 | 0.138117156 | 0.709264768 | -4.418993004 |
| **A_55_P1963631** | Yae1d1 | 0.218725281 | -0.097020085 | 0.534470647 | 10.06528795 | 1.735051392 | 0.138131517 | 0.709264768 | -4.419013902 |
| **A_51_P286563** | Gna13 | -0.258125001 | -0.63074963 | 0.114499627 | 11.4388019 | -1.735037478 | 0.138134085 | 0.709264768 | -4.419017639 |
| **A_52_P487599** | Thnsl2 | -0.203130654 | -0.496386035 | 0.090124727 | 9.107690191 | -1.734921472 | 0.138155498 | 0.709264768 | -4.419048794 |
| **A_55_P1973178** | Haus5 | 0.478370388 | -0.212277922 | 1.169018697 | 9.491133856 | 1.734833023 | 0.138171827 | 0.709264768 | -4.419072549 |
| **A_55_P2087825** | Ncoa2 | -0.25502966 | -0.623257366 | 0.113198046 | 10.74252857 | -1.734700785 | 0.138196243 | 0.709264768 | -4.419108064 |
| **A_55_P2021921** | Fkbp1a | -0.396119637 | -0.968155178 | 0.175915904 | 14.71163273 | -1.734417732 | 0.138248519 | 0.709264768 | -4.419184087 |
| **A_55_P2036813** | H2bu2 | 1.114906138 | -0.49541208 | 2.725224356 | 9.941733798 | 1.734111211 | 0.138305152 | 0.709264768 | -4.419266417 |
| **A_55_P1976212** | Selplg | 0.196650219 | -0.087384975 | 0.480685413 | 8.015019781 | 1.734093955 | 0.138308341 | 0.709264768 | -4.419271052 |
| **A_51_P402435** | Trap1 | 0.180797116 | -0.080347906 | 0.441942138 | 9.851156397 | 1.734043915 | 0.138317589 | 0.709264768 | -4.419284493 |
| **A_66_P112356** | Agt | -0.228281939 | -0.558208768 | 0.101644889 | 8.167049719 | -1.73302276 | 0.13850644 | 0.709264768 | -4.419558802 |
| **A_52_P20966** | Hnrnpab | 0.433490094 | -0.19317216 | 1.060152349 | 14.05250471 | 1.732590731 | 0.138586416 | 0.709264768 | -4.41967487 |
| **A_52_P196568** | Meg3 | -0.262614677 | -0.642306309 | 0.117076955 | 8.246721845 | -1.732360663 | 0.138629024 | 0.709264768 | -4.419736682 |
| **A_51_P357573** | Cald1 | -0.486066246 | -1.188843572 | 0.216711081 | 10.09349801 | -1.73231979 | 0.138636595 | 0.709264768 | -4.419747664 |
| **A_55_P2157788** | NA | 1.147819052 | -0.512474256 | 2.80811236 | 13.60608718 | 1.731565631 | 0.138776363 | 0.709264768 | -4.419950302 |
| **A_52_P476431** | Evc | 0.30731369 | -0.137221114 | 0.751848493 | 8.416847137 | 1.7315159 | 0.138785584 | 0.709264768 | -4.419963665 |
| **A_51_P268094** | Serpine2 | -0.289975511 | -0.709534449 | 0.129583427 | 9.935234122 | -1.731086184 | 0.138865291 | 0.709264768 | -4.420079139 |
| **A_52_P411430** | Gm5914 | 0.569377608 | -0.254443776 | 1.393198991 | 8.884112281 | 1.731082111 | 0.138866046 | 0.709264768 | -4.420080233 |
| **A_52_P268231** | NA | 0.315663691 | -0.141126107 | 0.772453489 | 9.16423643 | 1.730846603 | 0.138909749 | 0.709264768 | -4.420143522 |
| **A_55_P2036967** | Cox17 | 0.305566978 | -0.13661971 | 0.747753667 | 8.028552095 | 1.730816794 | 0.138915282 | 0.709264768 | -4.420151533 |
| **A_55_P1970424** | Fam221a | -0.187675436 | -0.459294525 | 0.083943654 | 8.583853351 | -1.730603146 | 0.138954942 | 0.709264768 | -4.42020895 |
| **A_55_P1999548** | Tma7 | 0.334848747 | -0.149817108 | 0.819514602 | 11.48865989 | 1.730440213 | 0.138985195 | 0.709264768 | -4.420252739 |
| **A_51_P100174** | Mns1 | 1.091169189 | -0.488387295 | 2.670725672 | 9.926663276 | 1.730243712 | 0.139021689 | 0.709264768 | -4.428412493 |
| **A_52_P661722** | 2-Mar | -0.354217192 | -0.867061263 | 0.158626879 | 13.77141457 | -1.729954348 | 0.139075448 | 0.709264768 | -4.420383323 |
| **A_52_P103391** | B4galt1 | -0.532288147 | -1.303125017 | 0.238548723 | 13.6750486 | -1.729556133 | 0.139149462 | 0.709264768 | -4.420490358 |
| **A_55_P1973965** | Ppp1r13b | -0.19544258 | -0.478536934 | 0.087651773 | 11.84310323 | -1.729172529 | 0.139220798 | 0.709264768 | -4.420593471 |
| **A_65_P05238** | Knop1 | 0.293977073 | -0.131920525 | 0.719874672 | 10.17207297 | 1.728855183 | 0.13927984 | 0.709264768 | -4.420678779 |
| **A_55_P2075213** | Kbtbd11 | 0.179250244 | -0.080448011 | 0.438948499 | 8.76920479 | 1.728785334 | 0.139292838 | 0.709264768 | -4.420697556 |
| **A_55_P1978481** | Nr1h2 | -0.222387364 | -0.544588655 | 0.099813928 | 12.08236595 | -1.728753908 | 0.139298687 | 0.709264768 | -4.420706004 |
| **A_52_P120612** | Grap2 | 0.189660647 | -0.085196735 | 0.464518028 | 7.791268533 | 1.72830413 | 0.139382421 | 0.709264768 | -4.420826921 |
| **A_51_P451458** | Mamdc2 | -0.149220335 | -0.365504708 | 0.067064039 | 8.272641989 | -1.728037698 | 0.139432046 | 0.709264768 | -4.420898552 |
| **A_51_P329370** | Snrnp25 | 0.403768817 | -0.18164446 | 0.989182094 | 12.80897479 | 1.727509931 | 0.139530397 | 0.709264768 | -4.421040452 |
| **A_55_P2124341** | Lysmd1 | -0.364791115 | -0.893787967 | 0.164205737 | 11.21592109 | -1.727195548 | 0.139589016 | 0.709264768 | -4.421124985 |
| **A_55_P2078403** | Rgs19 | 0.865488055 | -0.389619784 | 2.120595895 | 10.4941164 | 1.727151292 | 0.13959727 | 0.709264768 | -4.421136885 |
| **A_55_P2001891** | Prps1 | 0.847570565 | -0.381698689 | 2.076839819 | 10.40198783 | 1.726947728 | 0.139635242 | 0.709264768 | -4.421191623 |
| **A_51_P304859** | Zfand6 | -0.168523692 | -0.412952988 | 0.075905605 | 9.574571861 | -1.726863668 | 0.139650924 | 0.709264768 | -4.421214227 |
| **A_66_P107973** | Prpf38b | 0.29652967 | -0.133603279 | 0.726662619 | 8.535627353 | 1.726695623 | 0.139682281 | 0.709264768 | -4.421259416 |
| **A_51_P412348** | Scnn1b | -0.567402171 | -1.390466283 | 0.255661941 | 8.274218845 | -1.726663359 | 0.139688303 | 0.709264768 | -4.421268092 |
| **A_55_P2045903** | NA | 0.188604301 | -0.085043709 | 0.462252311 | 8.249200075 | 1.726273656 | 0.139761051 | 0.709264768 | -4.421372892 |
| **A_55_P2111163** | S100g | -2.712733808 | -6.648786025 | 1.22331841 | 11.36940862 | -1.726222107 | 0.139770677 | 0.709264768 | -4.421386755 |

| | | | | | | | | | |
|---|---|---|---|---|---|---|---|---|---|
| **A_55_P1968729** | Rpl10 | -0.180353555 | -0.442088588 | 0.081381478 | 16.81358938 | -1.725890333 | 0.139832644 | 0.709264768 | -4.421475982 |
| **A_55_P2049572** | Pld2 | -0.265777134 | -0.651537038 | 0.11998277 | 11.42570973 | -1.725642686 | 0.139878917 | 0.709264768 | -4.421542587 |
| **A_55_P2186928** | NA | 0.261726569 | -0.118304403 | 0.641757541 | 8.523758863 | 1.724960542 | 0.140006453 | 0.709264768 | -4.421726064 |
| **A_55_P1999439** | Ino80e | 0.225200434 | -0.101806991 | 0.552207859 | 10.21711465 | 1.724892188 | 0.140019239 | 0.709264768 | -4.42174445 |
| **A_55_P2149515** | Rsrc1 | 0.396770426 | -0.179371945 | 0.972912797 | 9.37242035 | 1.724883723 | 0.140020823 | 0.709264768 | -4.421746727 |
| **A_51_P181517** | Fcgr4 | -0.320151984 | -0.785140103 | 0.144836135 | 8.068971038 | -1.724505908 | 0.140091517 | 0.709264768 | -4.421848358 |
| **A_55_P1975535** | Nutf2 | 0.390219545 | -0.176628397 | 0.957067487 | 12.89192254 | 1.724220443 | 0.140144955 | 0.709264768 | -4.421925151 |
| **A_51_P140237** | Fhl2 | -0.893339308 | -2.191055833 | 0.404377216 | 11.6230407 | -1.724197277 | 0.140149293 | 0.709264768 | -4.421931383 |
| **A_55_P2185347** | NA | 0.188420373 | -0.085330638 | 0.462171384 | 7.977021661 | 1.723941293 | 0.140197231 | 0.709264768 | -4.422000248 |
| **A_52_P324754** | Nupr1l | -0.34060822 | -0.835530219 | 0.154313779 | 8.52930415 | -1.723727905 | 0.140237204 | 0.709264768 | -4.422057656 |
| **A_55_P1991693** | Pprc1 | 0.18600006 | -0.084269282 | 0.456269402 | 8.389357746 | 1.723719701 | 0.140238741 | 0.709264768 | -4.422059863 |
| **A_55_P1957080** | NA | 0.243589278 | -0.110438884 | 0.597617441 | 8.052887287 | 1.723338948 | 0.140310095 | 0.709264768 | -4.422162302 |
| **A_52_P10254** | Psmc4 | 0.240167089 | -0.108905469 | 0.589239646 | 11.94275278 | 1.723249382 | 0.140326885 | 0.709264768 | -4.4221864 |
| **A_66_P115531** | Crebzf | -0.207676131 | -0.509674651 | 0.094322388 | 10.46308336 | -1.722392465 | 0.140487624 | 0.709264768 | -4.422416973 |
| **A_52_P216539** | Fbxw8 | -0.251636037 | -0.617634493 | 0.114362418 | 8.955148718 | -1.722042756 | 0.140553274 | 0.709264768 | -4.422511079 |
| **A_55_P1969710** | NA | 0.612352026 | -0.278593111 | 1.503297163 | 14.23024002 | 1.721474298 | 0.140660053 | 0.709264768 | -4.422664061 |
| **A_66_P105307** | Adnp | 0.600748145 | -0.27344456 | 1.474940851 | 12.69225295 | 1.721216918 | 0.140708427 | 0.709264768 | -4.422733331 |
| **A_51_P243808** | Traf3ip3 | 0.270317291 | -0.123109416 | 0.663743997 | 7.93523096 | 1.720918622 | 0.14076451 | 0.709264768 | -4.422813615 |
| **A_55_P2049553** | NA | 0.86986575 | -0.396452668 | 2.136184167 | 12.14502745 | 1.720519715 | 0.140839544 | 0.709264768 | -4.422920985 |
| **A_55_P1968438** | Arxes1 | 0.524966888 | -0.239292936 | 1.289226711 | 8.819622191 | 1.720446717 | 0.140853279 | 0.709264768 | -4.422940633 |
| **A_51_P123405** | Bub1 | 0.900364545 | -0.410575615 | 2.211304706 | 9.385174216 | 1.720227406 | 0.140894553 | 0.709264768 | -4.428860162 |
| **A_55_P2067016** | Phykpl | -0.205853007 | -0.505595248 | 0.093889234 | 8.630566024 | -1.720123428 | 0.140914125 | 0.709264768 | -4.423027655 |
| **A_55_P2111907** | Noxo1 | -0.191173748 | -0.469547426 | 0.087199931 | 8.557914416 | -1.720087071 | 0.140920969 | 0.709264768 | -4.423037442 |
| **A_55_P2181743** | Elmod1 | -0.262818382 | -0.64567071 | 0.120033946 | 8.164851038 | -1.71939156 | 0.141051964 | 0.709264768 | -4.423224673 |
| **A_55_P2058467** | Gm4415 | -0.389099208 | -0.955927287 | 0.177728872 | 12.35983383 | -1.719330374 | 0.141063493 | 0.709264768 | -4.423241145 |
| **A_52_P572197** | Pank3 | 0.187785221 | -0.085814825 | 0.461385268 | 8.461777272 | 1.719078027 | 0.141111055 | 0.709264768 | -4.423309082 |
| **A_52_P309337** | Metap2 | 0.22935837 | -0.104827195 | 0.563543934 | 8.32767274 | 1.719005443 | 0.141124738 | 0.709264768 | -4.423328623 |
| **A_55_P2070546** | 1700037C | -0.208506731 | -0.512323293 | 0.09530983 | 9.544093573 | -1.718933127 | 0.141138372 | 0.709264768 | -4.423348093 |
| **A_55_P1957344** | Hnrnpa3 | 0.688608733 | -0.314985423 | 1.69220289 | 13.19839473 | 1.718560193 | 0.141208704 | 0.709264768 | -4.423448501 |
| **A_55_P2096166** | NA | 0.372494741 | -0.170641109 | 0.91563059 | 10.44718153 | 1.717758166 | 0.141360075 | 0.709264768 | -4.423664457 |
| **A_55_P1975326** | Tmem259 | -0.479260375 | -1.178078767 | 0.219558018 | 13.4558814 | -1.717740447 | 0.141363422 | 0.709264768 | -4.423669229 |
| **A_55_P2177941** | Tmem208 | -0.349770099 | -0.859799679 | 0.160259481 | 11.16458418 | -1.717661816 | 0.141378271 | 0.709264768 | -4.423690402 |
| **A_55_P1975903** | Dnmt3b | 0.159108785 | -0.07293485 | 0.39115242 | 8.688262855 | 1.717413393 | 0.141425196 | 0.709264768 | -4.4237573 |
| **A_51_P328652** | Acot7 | 0.530599001 | -0.243231368 | 1.304429371 | 10.87645444 | 1.717398201 | 0.141428066 | 0.709264768 | -4.423761391 |
| **A_52_P118958** | Pex19 | -0.31216469 | -0.767450804 | 0.143121423 | 9.375875661 | -1.717314033 | 0.141443968 | 0.709264768 | -4.423784057 |
| **A_55_P2473316** | Mapre1 | 0.176517541 | -0.080986272 | 0.434021354 | 11.76653858 | 1.716937752 | 0.141515083 | 0.709264768 | -4.423885391 |
| **A_55_P1954306** | Esrra | -0.427721748 | -1.051685502 | 0.196242007 | 11.44779598 | -1.716928918 | 0.141516753 | 0.709264768 | -4.42388777 |
| **A_55_P2147276** | Orai1 | -0.336249631 | -0.826798902 | 0.154299641 | 9.178253259 | -1.716838828 | 0.141533785 | 0.709264768 | -4.423912033 |
| **A_51_P201982** | Angpt2 | -0.552253124 | -1.357958852 | 0.253452604 | 9.957708087 | -1.716769843 | 0.141546828 | 0.709264768 | -4.423930611 |
| **A_51_P110395** | Setd2 | 0.171910442 | -0.078921768 | 0.422742651 | 12.01734367 | 1.716600748 | 0.141578805 | 0.709264768 | -4.44430499 |
| **A_51_P299805** | Slc46a3 | -0.45895796 | -1.128626723 | 0.210710802 | 9.487021996 | -1.71657669 | 0.141583355 | 0.709264768 | -4.423982632 |
| **A_52_P425706** | Rnf168 | 0.543039272 | -0.249313274 | 1.335391817 | 10.87045313 | 1.716576357 | 0.141583418 | 0.709264768 | -4.423982722 |
| **A_55_P2148288** | Tmem79 | -0.554973543 | -1.364838127 | 0.25489104 | 9.352725623 | -1.716367247 | 0.141622973 | 0.709264768 | -4.424039042 |
| **A_52_P574697** | Khdrbs1 | 0.471784728 | -0.216698704 | 1.16026816 | 12.25909972 | 1.71632975 | 0.141630067 | 0.709264768 | -4.424049141 |
| **A_55_P2030717** | NA | 0.189974138 | -0.087336691 | 0.467284967 | 11.04863823 | 1.715844782 | 0.141721849 | 0.709264768 | -4.424179765 |
| **A_52_P20048** | Tcf12 | 0.587745367 | -0.27030706 | 1.445797793 | 10.93037454 | 1.715638311 | 0.141760942 | 0.709264768 | -4.42423538 |
| **A_51_P394802** | Fam111a | 0.828928224 | -0.381245566 | 2.039102013 | 9.851490742 | 1.715613796 | 0.141765584 | 0.709264768 | -4.424241984 |
| **A_66_P122877** | NA | 0.355353563 | -0.163499397 | 0.874206523 | 16.6163089 | 1.715405195 | 0.141805093 | 0.709264768 | -4.424298174 |

| | | | | | | | | | |
|---|---|---|---|---|---|---|---|---|---|
| **A_52_P541833** | Vps37b | -0.215633528 | -0.530486407 | 0.09921935 | 10.64238933 | -1.715374567 | 0.141810894 | 0.709264768 | -4.424306425 |
| **A_55_P2075230** | BC051226 | -0.443923789 | -1.09228808 | 0.204440502 | 9.128101473 | -1.714903461 | 0.141900165 | 0.709264768 | -4.424433334 |
| **A_55_P2076989** | NA | 0.255932494 | -0.117877532 | 0.62974252 | 9.827968685 | 1.714844831 | 0.141911278 | 0.709264768 | -4.424449128 |
| **A_52_P342896** | Mrps33 | 0.354214612 | -0.163222325 | 0.871651549 | 9.503630095 | 1.714586465 | 0.141960263 | 0.709264768 | -4.424518732 |
| **A_55_P2183549** | NA | 0.836512633 | -0.385474963 | 2.058500229 | 11.34869553 | 1.714573282 | 0.141962763 | 0.709264768 | -4.424522284 |
| **A_51_P511176** | Triap1 | -0.605551564 | -1.490286183 | 0.279183055 | 10.51509027 | -1.714306432 | 0.142013375 | 0.709264768 | -4.424594177 |
| **A_51_P219849** | Alg13 | 0.289213314 | -0.133361506 | 0.711788134 | 8.333701141 | 1.714213899 | 0.14203093 | 0.709264768 | -4.424619107 |
| **A_55_P1998912** | Txnrd2 | -0.411057069 | -1.011771212 | 0.189657074 | 11.18920594 | -1.71389657 | 0.142091147 | 0.709264768 | -4.424704604 |
| **A_55_P2064811** | Nxt1 | 0.594222252 | -0.274258166 | 1.462702671 | 12.37940386 | 1.713717478 | 0.142125143 | 0.709264768 | -4.424752858 |
| **A_55_P1999637** | Gm15455 | 0.489577705 | -0.226124135 | 1.205279545 | 11.08488494 | 1.713325364 | 0.142199603 | 0.709264768 | -4.424858513 |
| **A_55_P2165869** | Cebpb | -0.455372106 | -1.12114061 | 0.210396397 | 13.75915964 | -1.713142638 | 0.142234315 | 0.709264768 | -4.42490775 |
| **A_51_P460954** | Ccl6 | -0.212075245 | -0.522141955 | 0.097991465 | 9.463386671 | -1.713109727 | 0.142240568 | 0.709264768 | -4.424916618 |
| **A_55_P2111148** | Gemin5 | 0.196938134 | -0.091008508 | 0.484884776 | 9.166099635 | 1.71304253 | 0.142253336 | 0.709264768 | -4.424934726 |
| **A_55_P1995210** | Gm11849 | 0.380997352 | -0.1761084 | 0.938103104 | 16.72257947 | 1.712910456 | 0.142278434 | 0.709264768 | -4.424970316 |
| **A_52_P482925** | Serbp1 | 0.760774388 | -0.35174432 | 1.873293095 | 13.72300363 | 1.712769443 | 0.142305236 | 0.709264768 | -4.425008316 |
| **A_55_P1987364** | Ap5z1 | -0.524032169 | -1.2904492 | 0.242384861 | 9.436497931 | -1.712549556 | 0.142347039 | 0.709264768 | -4.425067571 |
| **A_55_P2082614** | Zfp791 | 0.285768195 | -0.132178867 | 0.703715256 | 12.40976426 | 1.712548816 | 0.142347179 | 0.709264768 | -4.425067771 |
| **A_55_P2081829** | Rps4x | 0.411100303 | -0.190157645 | 1.01235825 | 15.34348848 | 1.712526546 | 0.142351414 | 0.709264768 | -4.425073772 |
| **A_55_P2009783** | Sfr1 | 0.422929225 | -0.195709728 | 1.041568178 | 14.02652694 | 1.712303639 | 0.142393805 | 0.709264768 | -4.425133844 |
| **A_51_P376857** | Det1 | 0.232546761 | -0.10761133 | 0.572704853 | 9.476110888 | 1.712299913 | 0.142394513 | 0.709264768 | -4.425134848 |
| **A_55_P2209308** | 9530083C | 0.347520456 | -0.160926745 | 0.855967656 | 11.12659946 | 1.711925489 | 0.142465747 | 0.709264768 | -4.425235757 |
| **A_55_P2063312** | Mgll | -0.317548133 | -0.782160021 | 0.147063755 | 8.977855206 | -1.71186531 | 0.142477199 | 0.709264768 | -4.425251977 |
| **A_55_P2288117** | NA | 0.219961247 | -0.101905685 | 0.541828179 | 8.827878702 | 1.711670473 | 0.142514283 | 0.709264768 | -4.425304489 |
| **A_55_P2047768** | Serpinb6c | -0.409279023 | -1.008220756 | 0.189662709 | 13.12822568 | -1.711532922 | 0.14254047 | 0.709264768 | -4.425341562 |
| **A_55_P2110713** | Anxa2 | -0.234705772 | -0.578296963 | 0.10888542 | 16.44005034 | -1.710929418 | 0.142655419 | 0.709264768 | -4.425504231 |
| **A_55_P1953796** | Fam98c | -0.313715137 | -0.773113185 | 0.145682911 | 10.25524186 | -1.710396008 | 0.142757094 | 0.709264768 | -4.425648018 |
| **A_55_P2005025** | Mbnl1 | 0.874891228 | -0.406402931 | 2.156185388 | 11.4765727 | 1.7102341 | 0.142787969 | 0.709264768 | -4.425691664 |
| **A_55_P2064451** | Pced1a | -0.26066003 | -0.64245891 | 0.12113885 | 9.758141949 | -1.709976476 | 0.142837112 | 0.709264768 | -4.425761116 |
| **A_55_P2162552** | Txndc9 | 0.287231632 | -0.133501921 | 0.707965185 | 11.6726257 | 1.709918708 | 0.142848134 | 0.709264768 | -4.42577669 |
| **A_51_P474701** | Fbp1 | 0.593386664 | -0.275808529 | 1.462581817 | 8.498591151 | 1.709900372 | 0.142851632 | 0.709264768 | -4.425781633 |
| **A_52_P428446** | Tsn | 0.50481538 | -0.234723787 | 1.244354546 | 12.17753947 | 1.709707213 | 0.142888492 | 0.709264768 | -4.425833708 |
| **A_52_P633410** | Gabarapl1 | -1.061870476 | -2.61791123 | 0.494170279 | 14.25397547 | -1.709231642 | 0.142979285 | 0.709264768 | -4.425961926 |
| **A_66_P101764** | NA | 0.481933035 | -0.224568114 | 1.188434185 | 12.03881702 | 1.708536141 | 0.143112167 | 0.709264768 | -4.426149456 |
| **A_52_P408690** | Sft2d2 | -0.154753355 | -0.381638317 | 0.072131606 | 8.770534742 | -1.708380908 | 0.143141843 | 0.709264768 | -4.426191314 |
| **A_51_P406583** | Myo1e | -0.289128081 | -0.713124487 | 0.134868326 | 10.35786788 | -1.707962938 | 0.143221775 | 0.709264768 | -4.426304024 |
| **A_51_P316801** | Wdr60 | -0.406627984 | -1.002969228 | 0.18971326 | 9.54928959 | -1.707861946 | 0.143241095 | 0.709264768 | -4.426331259 |
| **A_51_P236412** | Cdkl2 | -0.229151902 | -0.56525148 | 0.106947676 | 8.926676945 | -1.707677455 | 0.143276396 | 0.709264768 | -4.426381012 |
| **A_52_P163795** | Tubb5 | 0.45281428 | -0.211400804 | 1.117029365 | 15.16395261 | 1.707503985 | 0.143309596 | 0.709264768 | -4.426427793 |
| **A_66_P124806** | Tlr4 | -0.187960825 | -0.463717632 | 0.087795982 | 8.121495439 | -1.707227692 | 0.14336249 | 0.709264768 | -4.426502307 |
| **A_52_P244964** | Akap11 | 0.36256039 | -0.169360551 | 0.894481332 | 9.82733617 | 1.707196898 | 0.143368386 | 0.709264768 | -4.426510612 |
| **A_55_P2025328** | Tacr2 | -0.362629813 | -0.894680908 | 0.169421282 | 11.68657656 | -1.707106085 | 0.143385777 | 0.709264768 | -4.426535105 |
| **A_52_P302496** | Fah | -0.475907397 | -1.174342162 | 0.222527368 | 8.679001348 | -1.706659773 | 0.143471274 | 0.709264768 | -4.42665548 |
| **A_55_P2144556** | Flrt3 | 0.341329397 | -0.159657604 | 0.842316397 | 10.01035311 | 1.706465772 | 0.143508454 | 0.709264768 | -4.426707806 |
| **A_66_P111301** | Mplkip | 0.187625827 | -0.087772249 | 0.463023904 | 10.15256396 | 1.706404797 | 0.143520141 | 0.709264768 | -4.426724253 |
| **A_52_P659762** | Eid2 | 0.265140155 | -0.124070644 | 0.654350955 | 8.638021087 | 1.706243343 | 0.143551093 | 0.709264768 | -4.426767802 |
| **A_66_P106375** | Strap | 0.209414672 | -0.098000888 | 0.516830232 | 9.975847473 | 1.70620655 | 0.143558147 | 0.709264768 | -4.426777727 |
| **A_55_P2076757** | Znfx1 | -0.243311291 | -0.600587662 | 0.11396508 | 11.08277023 | -1.705722252 | 0.143651034 | 0.709264768 | -4.426908365 |
| **A_66_P128434** | Kit | -0.88140072 | -2.175668921 | 0.412867482 | 10.86615909 | -1.705687494 | 0.143657702 | 0.709264768 | -4.426917741 |

| | | | | | | | | | |
|---|---|---|---|---|---|---|---|---|---|
| **A_55_P2177614** | Srsf3 | 0.770996874 | -0.36140033 | 1.903394079 | 11.71076088 | 1.705313281 | 0.143729519 | 0.709264768 | -4.42701869 |
| **A_51_P466875** | Sbds | -0.405102849 | -1.000143303 | 0.189937606 | 11.43577541 | -1.705175758 | 0.14375592 | 0.709264768 | -4.42705579 |
| **A_55_P2139748** | Phf14 | 0.188235317 | -0.088273095 | 0.46474373 | 8.685482214 | 1.705073508 | 0.143775553 | 0.709264768 | -4.427083375 |
| **A_51_P182572** | Phactr1 | -1.416781875 | -3.498531259 | 0.66496751 | 10.44206986 | -1.704609454 | 0.143864689 | 0.709264768 | -4.427208572 |
| **A_55_P2061620** | Chst1 | -0.273379093 | -0.675085713 | 0.128327527 | 9.853641374 | -1.704537872 | 0.143878443 | 0.709264768 | -4.427227885 |
| **A_52_P56658** | Emc3 | -0.440126282 | -1.086947872 | 0.206695307 | 9.699334458 | -1.70428861 | 0.143926349 | 0.709264768 | -4.427295138 |
| **A_52_P539310** | Serp2 | -0.392817346 | -0.970149662 | 0.18451497 | 9.129132709 | -1.704178708 | 0.143947476 | 0.709264768 | -4.427324791 |
| **A_52_P498208** | H2ac15 | 1.445033706 | -0.679023742 | 3.569091154 | 13.56616236 | 1.703970471 | 0.143987514 | 0.709264768 | -4.427380977 |
| **A_51_P443359** | Trappc2l | -0.282969837 | -0.698934619 | 0.132994945 | 11.84787468 | -1.703860082 | 0.144008744 | 0.709264768 | -4.427410763 |
| **A_55_P2149896** | Lrrc75a | -0.349201386 | -0.862536571 | 0.164133799 | 11.56938536 | -1.703826118 | 0.144015276 | 0.709264768 | -4.427419927 |
| **A_55_P1965000** | Ddx58 | -0.523554544 | -1.293198645 | 0.246089558 | 10.33439229 | -1.703814591 | 0.144017493 | 0.709264768 | -4.427423038 |
| **A_55_P1984961** | NA | -0.283167955 | -0.699501524 | 0.133165615 | 8.147852764 | -1.703542682 | 0.144069802 | 0.709264768 | -4.427496408 |
| **A_52_P1197913** | Gadd45b | -0.578673208 | -1.429644945 | 0.272298528 | 11.36358806 | -1.703211519 | 0.144133534 | 0.709264768 | -4.427585772 |
| **A_55_P2024580** | Dtnbp1 | 0.201947677 | -0.095101073 | 0.498996427 | 12.44030371 | 1.702791522 | 0.144214403 | 0.709264768 | -4.427699112 |
| **A_52_P131548** | Ajuba | -0.208089528 | -0.514193223 | 0.098014168 | 9.836506437 | -1.702675934 | 0.144236667 | 0.709264768 | -4.427730306 |
| **A_55_P2133255** | Mad2l1 | 1.274963892 | -0.601124292 | 3.151052076 | 11.11334945 | 1.702138844 | 0.144340163 | 0.709264768 | -4.427875258 |
| **A_55_P2186558** | Tmem242 | -0.284926918 | -0.704204253 | 0.134350416 | 10.16383388 | -1.702089688 | 0.144349638 | 0.709264768 | -4.427888525 |
| **A_52_P319438** | Ankrd37 | -0.75510833 | -1.86630113 | 0.35608447 | 11.10511601 | -1.702041669 | 0.144358896 | 0.709264768 | -4.427901485 |
| **A_55_P2182705** | Kpna2 | 1.193120887 | -0.562724263 | 2.948966037 | 13.16422555 | 1.701956994 | 0.144375221 | 0.709264768 | -4.427924339 |
| **A_55_P2023441** | Elp4 | 0.400269998 | -0.188810149 | 0.989350145 | 9.278783725 | 1.70188025 | 0.144390019 | 0.709264768 | -4.427945052 |
| **A_51_P143142** | Mrpl12 | 0.320146006 | -0.151075152 | 0.791367164 | 12.6507181 | 1.701663373 | 0.144431845 | 0.709264768 | -4.428003589 |
| **A_55_P2117164** | Tmem106 | -0.467738871 | -1.156285223 | 0.220807481 | 9.283779768 | -1.701455624 | 0.144471923 | 0.709264768 | -4.428059664 |
| **A_55_P2152114** | Zfyve19 | -0.215793151 | -0.53346184 | 0.101875538 | 9.775526033 | -1.701428064 | 0.14447724 | 0.709264768 | -4.428067103 |
| **A_55_P2075769** | Ube2n | 0.516544207 | -0.24409013 | 1.277178544 | 10.30204485 | 1.70091227 | 0.144576794 | 0.709264768 | -4.428206332 |
| **A_55_P1988433** | Stard7 | 0.17649271 | -0.083458021 | 0.436443441 | 12.43724776 | 1.700536954 | 0.144649277 | 0.709264768 | -4.428307648 |
| **A_55_P2139894** | Cwc15 | 0.399662994 | -0.189193142 | 0.98851913 | 10.90504378 | 1.699945814 | 0.144763513 | 0.709264768 | -4.428467236 |
| **A_51_P394339** | Eif2b1 | 0.166189168 | -0.078685481 | 0.411063817 | 10.58343379 | 1.699844652 | 0.144783071 | 0.709264768 | -4.428494548 |
| **A_55_P2091212** | NA | 0.404956242 | -0.19186988 | 1.001782364 | 16.74310882 | 1.699458718 | 0.14485771 | 0.709264768 | -4.428598746 |
| **A_55_P1962224** | Afap1l2 | -0.833272008 | -2.06152265 | 0.394978634 | 10.64385001 | -1.699222063 | 0.144903497 | 0.709264768 | -4.428662643 |
| **A_55_P2013986** | Ddx31 | 0.169789314 | -0.080508068 | 0.420086696 | 9.389098727 | 1.699043088 | 0.144938134 | 0.709264768 | -4.428710968 |
| **A_51_P281024** | Rad9a | 0.163172231 | -0.077375077 | 0.403719539 | 8.826077608 | 1.699010628 | 0.144944417 | 0.709264768 | -4.428719732 |
| **A_66_P138976** | Lpin2 | -0.669830265 | -1.681668284 | 0.342007754 | 8.876770544 | -1.763260498 | 0.144951171 | 0.709264768 | -4.44891027 |
| **A_55_P2160204** | NA | 0.422044572 | -0.200214374 | 1.044303517 | 11.26604882 | 1.698781473 | 0.144988779 | 0.709264768 | -4.428781609 |
| **A_51_P191649** | Ndc80 | 0.541274215 | -0.256841435 | 1.339389864 | 8.821378798 | 1.698641999 | 0.145015786 | 0.709264768 | -4.42881927 |
| **A_55_P2001233** | Ifi208 | 0.234354869 | -0.111216884 | 0.579926621 | 10.99718422 | 1.698580328 | 0.145027729 | 0.709264768 | -4.428835923 |
| **A_55_P2254968** | BB212172 | 0.284806125 | -0.135163891 | 0.704776142 | 8.86166328 | 1.698561932 | 0.145031292 | 0.709264768 | -4.42884089 |
| **A_55_P2359436** | A930026E | -0.148795139 | -0.368230102 | 0.070639825 | 8.355880339 | -1.698373723 | 0.145067748 | 0.709264768 | -4.428891713 |
| **A_52_P467726** | Nsg1 | 1.200044267 | -0.57008831 | 2.970176845 | 10.44568659 | 1.698016153 | 0.145137034 | 0.709264768 | -4.428988273 |
| **A_55_P1964375** | Gm9731 | 0.157769045 | -0.074964024 | 0.390502114 | 10.84846454 | 1.697907575 | 0.145158079 | 0.709264768 | -4.429017595 |
| **A_52_P132165** | Hsd17b11 | -0.881131358 | -2.213148484 | 0.450885768 | 10.9599255 | -1.761949895 | 0.145184792 | 0.709264768 | -4.449172166 |
| **A_55_P2047621** | Cutal | -0.214380044 | -0.530686234 | 0.101926146 | 8.221858615 | -1.697567334 | 0.145224047 | 0.709264768 | -4.429109481 |
| **A_51_P120254** | Fbxl8 | -0.161632875 | -0.40016656 | 0.076900811 | 8.454633243 | -1.697189451 | 0.145297347 | 0.709264768 | -4.429211538 |
| **A_55_P2168781** | NA | 0.468316004 | -0.223017643 | 1.159649651 | 12.75970654 | 1.696686682 | 0.145394929 | 0.709264768 | -4.429347332 |
| **A_55_P1957747** | Inip | 0.182856412 | -0.087090199 | 0.452803023 | 9.418984355 | 1.696612489 | 0.145409334 | 0.709264768 | -4.429367371 |
| **A_55_P1987904** | Chpf2 | -0.357827111 | -0.886121143 | 0.170466922 | 8.382348402 | -1.696476634 | 0.145435716 | 0.709264768 | -4.429404067 |
| **A_52_P218537** | Hnrnpc | 0.417999214 | -0.199254102 | 1.03525253 | 11.85125134 | 1.696142665 | 0.145500589 | 0.709264768 | -4.429494277 |
| **A_52_P272054** | NA | 0.432083746 | -0.206120712 | 1.070288203 | 10.08536083 | 1.695736786 | 0.145579469 | 0.709264768 | -4.429603917 |
| **A_51_P168395** | Ttc30a1 | 0.39512065 | -0.188556271 | 0.97879757 | 9.264081475 | 1.695538094 | 0.145618099 | 0.709264768 | -4.429657592 |

| | | | | | | | | | |
|---|---|---|---|---|---|---|---|---|---|
| **A_51_P408071** | Kntc1 | 0.613434518 | -0.293080989 | 1.519950025 | 8.950158725 | 1.694897027 | 0.145742805 | 0.709264768 | -4.42983078 |
| **A_55_P2015430** | Gpn2 | 0.158453927 | -0.07572736 | 0.392635214 | 9.877302062 | 1.694732526 | 0.145774822 | 0.709264768 | -4.429875224 |
| **A_52_P559919** | Eif2ak2 | -0.242536371 | -0.601020813 | 0.11594807 | 8.913954237 | -1.69455984 | 0.14580844 | 0.709264768 | -4.42992188 |
| **A_55_P2076303** | 8-Mar | -0.428163868 | -1.061136295 | 0.20480856 | 11.83516275 | -1.694242484 | 0.145870241 | 0.709264768 | -4.430007625 |
| **A_55_P1976057** | Smurf2 | -0.146002225 | -0.361901844 | 0.069897393 | 8.706438534 | -1.693783689 | 0.145959631 | 0.709264768 | -4.430131593 |
| **A_66_P129852** | Fxn | 0.366902455 | -0.175701999 | 0.90950691 | 9.556370666 | 1.693626378 | 0.145990293 | 0.709264768 | -4.4301741 |
| **A_55_P2062508** | Ube2e1 | 0.195538667 | -0.093661136 | 0.48473847 | 12.45597341 | 1.693499227 | 0.146015082 | 0.709264768 | -4.430208459 |
| **A_51_P429209** | Smpd1 | -0.560971598 | -1.390696202 | 0.268753007 | 12.4077516 | -1.69339101 | 0.146036182 | 0.709264768 | -4.430237701 |
| **A_55_P1954965** | Npm1 | 0.461401786 | -0.221057827 | 1.143861399 | 14.62701344 | 1.69337312 | 0.146039671 | 0.709264768 | -4.430242536 |
| **A_52_P20357** | Aqr | 0.220322494 | -0.105566179 | 0.546211167 | 9.800346932 | 1.693323433 | 0.14604936 | 0.709264768 | -4.430255962 |
| **A_51_P228817** | NA | 0.433191406 | -0.20764717 | 1.074029982 | 9.198072181 | 1.693095795 | 0.146093759 | 0.709264768 | -4.430317477 |
| **A_55_P2110887** | NA | 0.545610729 | -0.26155796 | 1.352779419 | 11.34819676 | 1.693046712 | 0.146103334 | 0.709264768 | -4.430330741 |
| **A_55_P2176212** | NA | 0.785343834 | -0.376516712 | 1.947204381 | 13.05239717 | 1.692996965 | 0.146113039 | 0.709264768 | -4.430344185 |
| **A_51_P254471** | Birc2 | -0.19365193 | -0.480147232 | 0.092843372 | 9.288522168 | -1.692991085 | 0.146114186 | 0.709264768 | -4.430345774 |
| **A_55_P2014665** | Racgap1 | 1.04107979 | -0.499628748 | 2.581788327 | 9.949115493 | 1.692442303 | 0.146221292 | 0.709264768 | -4.430494082 |
| **A_52_P529570** | Nsl1 | 0.871534617 | -0.418631644 | 2.161700877 | 9.531174453 | 1.691956942 | 0.146316084 | 0.709264768 | -4.43062526 |
| **A_55_P2005903** | Lhx6 | -0.353065075 | -0.875800987 | 0.169670837 | 11.46626822 | -1.691697738 | 0.146366731 | 0.709264768 | -4.430695319 |
| **A_51_P259009** | Cacnb3 | -0.200427928 | -0.497182191 | 0.096326334 | 8.858927787 | -1.691654304 | 0.14637522 | 0.709264768 | -4.430707059 |
| **A_51_P477833** | Klhl21 | -0.401156046 | -0.995126404 | 0.192814311 | 9.701224466 | -1.691604828 | 0.14638489 | 0.709264768 | -4.430720431 |
| **A_52_P480402** | Avl9 | -0.149895157 | -0.371852576 | 0.072062262 | 9.768649001 | -1.69148551 | 0.146408213 | 0.709264768 | -4.430752682 |
| **A_52_P582112** | Hcfc1r1 | -0.294805462 | -0.73147836 | 0.141867437 | 12.11347604 | -1.690945584 | 0.146513798 | 0.709264768 | -4.430898627 |
| **A_55_P2130373** | Dcaf11 | -0.179747761 | -0.446029197 | 0.086533676 | 12.06973279 | -1.690724874 | 0.14655698 | 0.709264768 | -4.43095829 |
| **A_55_P2109429** | Rfng | 0.21464481 | -0.103408797 | 0.532698416 | 11.96416598 | 1.690325771 | 0.146635097 | 0.709264768 | -4.431066179 |
| **A_55_P2113920** | NA | 0.187978275 | -0.0906012 | 0.46655775 | 8.404021612 | 1.690086337 | 0.146681982 | 0.709264768 | -4.431130909 |
| **A_52_P460957** | Crem | -0.154348954 | -0.383119783 | 0.074421875 | 8.927633703 | -1.689870363 | 0.146724285 | 0.709264768 | -4.431189297 |
| **A_55_P2067351** | NA | 0.24772518 | -0.119517648 | 0.614968009 | 9.386285865 | 1.689534932 | 0.146790011 | 0.709264768 | -4.431279985 |
| **A_51_P383032** | Clec4d | -0.230294912 | -0.571819054 | 0.111229229 | 8.091011866 | -1.688936341 | 0.146907373 | 0.709264768 | -4.431441831 |
| **A_55_P2076057** | Hmga1 | 0.18593051 | -0.089806045 | 0.461667064 | 8.150749421 | 1.688910599 | 0.146912422 | 0.709264768 | -4.431448791 |
| **A_51_P114297** | Kifc3 | -0.293006413 | -0.72756448 | 0.141551654 | 8.68867324 | -1.688805577 | 0.146933023 | 0.709264768 | -4.431477189 |
| **A_55_P2028054** | Incenp | 1.257815443 | -0.607910728 | 3.123541615 | 12.68779856 | 1.688571149 | 0.146979019 | 0.709264768 | -4.431540578 |
| **A_55_P1959953** | Helz2 | -0.265108898 | -0.65836344 | 0.128145644 | 10.2534176 | -1.688499368 | 0.146993106 | 0.709264768 | -4.431559987 |
| **A_52_P446120** | Commd6 | -0.477239568 | -1.185177025 | 0.23069789 | 10.88256212 | -1.688464359 | 0.146999977 | 0.709264768 | -4.431569454 |
| **A_51_P140607** | Ints13 | 0.476870649 | -0.230552185 | 1.184293482 | 10.58328535 | 1.688386476 | 0.147015263 | 0.709264768 | -4.431590514 |
| **A_55_P1974671** | Sft2d1 | -0.193901463 | -0.481577171 | 0.093774244 | 9.206255177 | -1.688216897 | 0.147048552 | 0.709264768 | -4.431636371 |
| **A_55_P2185870** | Bcl7a | 0.608162081 | -0.294235079 | 1.51055924 | 8.944130877 | 1.687998141 | 0.147091506 | 0.709264768 | -4.431695527 |
| **A_55_P1977289** | Zbtb9 | 0.232039338 | -0.112364232 | 0.576442908 | 9.899731094 | 1.687502085 | 0.147188954 | 0.709264768 | -4.431829676 |
| **A_52_P542970** | NA | 0.435726273 | -0.211117191 | 1.082569736 | 10.92029118 | 1.68719352 | 0.147249603 | 0.709264768 | -4.431913127 |
| **A_55_P2087114** | NA | 0.184556851 | -0.08949648 | 0.458610181 | 10.83254955 | 1.686729465 | 0.147340859 | 0.709264768 | -4.432038636 |
| **A_55_P2084980** | Dcaf12l2 | 0.299246986 | -0.145162563 | 0.743656535 | 10.96667319 | 1.686540398 | 0.147378055 | 0.709264768 | -4.432089774 |
| **A_52_P142191** | Aph1b | -0.495333008 | -1.230955042 | 0.240289025 | 8.601890728 | -1.686525402 | 0.147381006 | 0.709264768 | -4.432093829 |
| **A_52_P238044** | Psmc3ip | 0.633052348 | -0.307104316 | 1.573209011 | 9.152654369 | 1.686513185 | 0.147383409 | 0.709264768 | -4.432097134 |
| **A_55_P2061170** | Tbc1d2 | -0.798519601 | -1.984603905 | 0.387564702 | 9.846327723 | -1.68624381 | 0.147436423 | 0.709264768 | -4.432169995 |
| **A_55_P2125456** | NA | 0.428396754 | -0.208147876 | 1.064941385 | 14.45517985 | 1.685650983 | 0.147553157 | 0.709264768 | -4.432330354 |
| **A_52_P584188** | Frat1 | 0.374044561 | -0.181909186 | 0.929998307 | 9.338898155 | 1.685136278 | 0.147654583 | 0.709264768 | -4.432469591 |
| **A_55_P2116664** | Hook2 | -0.200757672 | -0.499166403 | 0.09765106 | 10.86041624 | -1.68504293 | 0.147672985 | 0.709264768 | -4.432494844 |
| **A_52_P280899** | Wdr33 | 0.159637328 | -0.07767142 | 0.396946076 | 9.68975355 | 1.684888039 | 0.147703524 | 0.709264768 | -4.432536748 |
| **A_55_P2053718** | Pink1 | -0.482699545 | -1.200315423 | 0.234916332 | 11.88364922 | -1.684748957 | 0.147730952 | 0.709264768 | -4.432574375 |
| **A_55_P2009498** | Ccdc28a | -0.321549051 | -0.799589539 | 0.156491437 | 9.931344148 | -1.684740035 | 0.147732711 | 0.709264768 | -4.432576788 |

| | | | | | | | | | |
|---|---|---|---|---|---|---|---|---|---|
| **A_55_P2155826** | Rplp1 | 0.24590259 | -0.11977587 | 0.61158105 | 16.47664401 | 1.68427914 | 0.147823638 | 0.709264768 | -4.432701484 |
| **A_55_P2227355** | Ptpro | -0.212897832 | -0.52959455 | 0.103798887 | 8.209864045 | -1.683751597 | 0.147927782 | 0.709264768 | -4.43284422 |
| **A_51_P349691** | Ano10 | -0.316435629 | -0.787272465 | 0.154401207 | 9.572875213 | -1.683314599 | 0.148014105 | 0.709264768 | -4.432962465 |
| **A_52_P360650** | Abcb7 | 0.350664514 | -0.171132755 | 0.872461782 | 8.440657702 | 1.683217985 | 0.148033196 | 0.709264768 | -4.432988608 |
| **A_55_P2119145** | NA | -0.374808331 | -0.932575284 | 0.182958623 | 7.863360926 | -1.683087913 | 0.148058903 | 0.709264768 | -4.433023806 |
| **A_55_P2192662** | Lepr | -0.451997216 | -1.12469132 | 0.220696889 | 8.349059697 | -1.682939422 | 0.148088256 | 0.709264768 | -4.433063988 |
| **A_55_P2094963** | NA | 0.149662809 | -0.073102916 | 0.372428534 | 9.974739148 | 1.682735542 | 0.148128566 | 0.709264768 | -4.43311916 |
| **A_51_P420236** | Srrt | 0.228925481 | -0.111905941 | 0.569756903 | 8.525133118 | 1.682305459 | 0.148213636 | 0.709264768 | -4.43323555 |
| **A_55_P2119030** | Rasl2-9 | 0.802885664 | -0.392889079 | 1.998660407 | 13.55242834 | 1.681723809 | 0.148328763 | 0.709264768 | -4.433392967 |
| **A_55_P2069525** | Zmym6 | -0.48798007 | -1.21499821 | 0.23903807 | 9.3012249 | -1.68115278 | 0.148441873 | 0.709264768 | -4.433547522 |
| **A_52_P279599** | Ube2k | 0.223467498 | -0.109500729 | 0.556435725 | 10.50173644 | 1.68097757 | 0.148476595 | 0.709264768 | -4.433594947 |
| **A_55_P2009206** | Sumo2 | 0.574282883 | -0.281416406 | 1.429982173 | 9.621440619 | 1.680950972 | 0.148481867 | 0.709264768 | -4.433602146 |
| **A_51_P296608** | Gadd45a | -0.514987162 | -1.282480286 | 0.252505963 | 9.051508169 | -1.680630553 | 0.14854539 | 0.709264768 | -4.433688878 |
| **A_55_P1996698** | Gm16381 | 0.381676003 | -0.187230982 | 0.950582989 | 12.81494343 | 1.680366191 | 0.148597819 | 0.709264768 | -4.43376044 |
| **A_55_P1977699** | NA | 0.739435894 | -0.362844737 | 1.841716525 | 14.27021676 | 1.680191169 | 0.14863254 | 0.709264768 | -4.433807818 |
| **A_66_P120507** | Plcd3 | -0.74183846 | -1.847761297 | 0.364084376 | 8.895501025 | -1.680098968 | 0.148650835 | 0.709264768 | -4.433832778 |
| **A_55_P2009518** | NA | 0.481651819 | -0.236543467 | 1.199847105 | 15.9346731 | 1.679735885 | 0.148722898 | 0.709264768 | -4.43393107 |
| **A_52_P236448** | Ngfr | 0.374022495 | -0.18372681 | 0.9317718 | 8.317372572 | 1.679612239 | 0.148747446 | 0.709264768 | -4.441324082 |
| **A_55_P2002160** | Sumo3 | 0.222363324 | -0.109253948 | 0.553980597 | 9.858161311 | 1.679485896 | 0.148772534 | 0.709264768 | -4.433998748 |
| **A_52_P222967** | Adck2 | -0.221705019 | -0.552360569 | 0.10895053 | 8.609651675 | -1.679384169 | 0.148792738 | 0.709264768 | -4.434026288 |
| **A_55_P1991688** | Rad51ap1 | 1.083823576 | -0.532840712 | 2.700487865 | 10.14290169 | 1.679148456 | 0.148839561 | 0.709264768 | -4.434090105 |
| **A_55_P2010539** | NA | 0.214148953 | -0.105296009 | 0.533593914 | 8.652166299 | 1.679075629 | 0.148854031 | 0.709264768 | -4.434109822 |
| **A_51_P456957** | Pex11a | -0.190132248 | -0.473760782 | 0.093496287 | 10.76067912 | -1.679021302 | 0.148864825 | 0.709264768 | -4.434124531 |
| **A_55_P2001628** | Rps4l | -0.33344919 | -0.830888142 | 0.163989763 | 13.41427254 | -1.678959507 | 0.148877105 | 0.709264768 | -4.434141262 |
| **A_55_P1959137** | Usp32 | -0.43371449 | -1.080821435 | 0.213392456 | 10.90932413 | -1.678719812 | 0.148924746 | 0.709264768 | -4.43420616 |
| **A_52_P99888** | Cxcl16 | -0.189514176 | -0.472308914 | 0.093280561 | 8.283290812 | -1.678497602 | 0.148968925 | 0.709264768 | -4.434266325 |
| **A_51_P129803** | Cacybp | 0.274195419 | -0.135063829 | 0.683454666 | 11.6484993 | 1.678077593 | 0.149052465 | 0.709264768 | -4.434380052 |
| **A_55_P2095000** | NA | 0.252424527 | -0.124360272 | 0.629209327 | 16.25149529 | 1.677986735 | 0.149070543 | 0.709264768 | -4.434404654 |
| **A_66_P137297** | Otof | 0.615419829 | -0.303251234 | 1.534090892 | 10.88366715 | 1.677883464 | 0.149091093 | 0.709264768 | -4.434432619 |
| **A_51_P339822** | Arhgap4 | 0.374953417 | -0.184765701 | 0.934672534 | 8.421807212 | 1.677866953 | 0.149094379 | 0.709264768 | -4.43443709 |
| **A_55_P2073845** | Mrpl47 | 0.179378313 | -0.088455434 | 0.447212059 | 8.681797118 | 1.677470855 | 0.149173227 | 0.709264768 | -4.43454435 |
| **A_55_P2119962** | Shld1 | -0.579357701 | -1.444691332 | 0.285975931 | 10.28883621 | -1.676924642 | 0.149282024 | 0.709264768 | -4.434692271 |
| **A_51_P504588** | Tm2d2 | -0.231192925 | -0.576522261 | 0.11413641 | 10.26093931 | -1.676839181 | 0.149299054 | 0.709264768 | -4.434715415 |
| **A_55_P1978136** | Tnip2 | -0.214522948 | -0.534956655 | 0.105910759 | 9.243622905 | -1.676817923 | 0.14930329 | 0.709264768 | -4.434721172 |
| **A_51_P323521** | Snx21 | -0.603626977 | -1.505279449 | 0.298025495 | 11.21268914 | -1.676794375 | 0.149307983 | 0.709264768 | -4.43472755 |
| **A_55_P2149931** | Arap2 | -0.504536879 | -1.258191761 | 0.249118004 | 8.519425821 | -1.676759381 | 0.149314957 | 0.709264768 | -4.434737027 |
| **A_55_P2100375** | Col11a1 | 0.267225267 | -0.132061639 | 0.666512173 | 8.135338029 | 1.676265462 | 0.149413426 | 0.709264768 | -4.434870797 |
| **A_51_P248705** | Cse1l | 0.404597241 | -0.200022328 | 1.009216811 | 9.882966097 | 1.676065794 | 0.149453251 | 0.709264768 | -4.434924876 |
| **A_55_P2172956** | Cnih2 | 0.327662468 | -0.162008532 | 0.817333468 | 8.609952011 | 1.67599477 | 0.149467419 | 0.709264768 | -4.434944113 |
| **A_55_P2037584** | Trf | -1.097596211 | -2.737999312 | 0.542806889 | 14.61096816 | -1.675877859 | 0.149490745 | 0.709264768 | -4.434975779 |
| **A_55_P2168736** | Relb | -0.297235418 | -0.74150444 | 0.147033603 | 8.737316252 | -1.675733196 | 0.149519612 | 0.709264768 | -4.435014962 |
| **A_55_P2018091** | NA | -0.315835219 | -0.787974539 | 0.156304102 | 13.60989894 | -1.675485723 | 0.149569008 | 0.709264768 | -4.435081994 |
| **A_51_P320888** | Cln5 | -0.665720868 | -1.660924871 | 0.329483134 | 11.14273992 | -1.675445875 | 0.149576963 | 0.709264768 | -4.435092788 |
| **A_52_P665202** | NA | 0.624418669 | -0.309064867 | 1.557902205 | 14.75374152 | 1.675404054 | 0.149585312 | 0.709264768 | -4.435104116 |
| **A_55_P1974681** | Nsun4 | -0.295113774 | -0.736386357 | 0.14615881 | 9.479359318 | -1.675069689 | 0.149652084 | 0.709264768 | -4.435194688 |
| **A_51_P256282** | Ppp6c | 0.246991657 | -0.122387369 | 0.616370684 | 11.55306295 | 1.674790148 | 0.14970793 | 0.709264768 | -4.435270413 |
| **A_55_P1996613** | NA | -0.377269479 | -0.941895128 | 0.187356169 | 14.92677971 | -1.673560495 | 0.149953829 | 0.709264768 | -4.435603545 |
| **A_55_P1997509** | Zbtb32 | -0.184068149 | -0.459551075 | 0.091414776 | 9.004617243 | -1.673533094 | 0.149959314 | 0.709264768 | -4.435610969 |

| | | | | | | | | | |
|---|---|---|---|---|---|---|---|---|---|
| **A_55_P2052749** | Pcm1 | 0.734899792 | -0.365088777 | 1.834888361 | 10.84415053 | 1.673363523 | 0.149993256 | 0.709264768 | -4.435656912 |
| **A_55_P2307621** | 5730526G | -0.219170169 | -0.547231026 | 0.108890688 | 8.540764944 | -1.673313724 | 0.150003225 | 0.709264768 | -4.435670405 |
| **A_51_P404463** | Ecrg4 | 0.294589662 | -0.146403833 | 0.735583157 | 9.522229773 | 1.673153029 | 0.1500354 | 0.709264768 | -4.435713945 |
| **A_51_P412225** | Tbc1d16 | 0.50894233 | -0.253024383 | 1.270909044 | 11.45517506 | 1.67294984 | 0.150076092 | 0.709264768 | -4.435769 |
| **A_55_P1953231** | Map2k5 | -0.344742002 | -0.860877767 | 0.171393763 | 12.09272941 | -1.672940868 | 0.150077889 | 0.709264768 | -4.435771431 |
| **A_55_P2152836** | NA | 0.494966103 | -0.246115769 | 1.236047975 | 10.51018071 | 1.672860064 | 0.150094075 | 0.709264768 | -4.435793326 |
| **A_52_P315976** | Tpm2 | -0.480692272 | -1.200492248 | 0.239107705 | 12.40884394 | -1.672652243 | 0.150135711 | 0.709264768 | -4.435849638 |
| **A_51_P159896** | Dnajc1 | -0.664003291 | -1.658543306 | 0.330536724 | 10.70862173 | -1.672238877 | 0.150218562 | 0.709264768 | -4.43596165 |
| **A_55_P2003309** | NA | -0.195103166 | -0.487339422 | 0.09713309 | 14.29164552 | -1.672170537 | 0.150232263 | 0.709264768 | -4.435980169 |
| **A_52_P298002** | Gch1 | -0.396346972 | -0.99008714 | 0.197393196 | 8.08186802 | -1.671973766 | 0.150271721 | 0.709264768 | -4.436033491 |
| **A_55_P2176345** | Psma5 | 0.304614907 | -0.151752383 | 0.760982197 | 13.15699805 | 1.671810272 | 0.150304514 | 0.709264768 | -4.436077797 |
| **A_51_P429366** | Hes6 | 0.645466252 | -0.321585388 | 1.612517892 | 10.69887637 | 1.671761119 | 0.150314374 | 0.709264768 | -4.436091118 |
| **A_52_P372165** | Rpsa | 0.268197206 | -0.133640924 | 0.670035336 | 8.284599336 | 1.671681172 | 0.150330413 | 0.709264768 | -4.436112783 |
| **A_55_P2005315** | NA | 0.301234369 | -0.150163002 | 0.75263174 | 11.05525048 | 1.671459436 | 0.150374906 | 0.709264768 | -4.436172875 |
| **A_52_P337236** | Alyref2 | 0.457474094 | -0.228105118 | 1.143053305 | 9.526572228 | 1.671318463 | 0.1504032 | 0.709264768 | -4.43621108 |
| **A_55_P2305294** | Ptpn21 | -0.274592855 | -0.686260782 | 0.137075071 | 8.687166513 | -1.670677204 | 0.15053197 | 0.709264768 | -4.436384877 |
| **A_52_P302041** | NA | 0.733963938 | -0.366536889 | 1.834464766 | 10.99497987 | 1.670454669 | 0.150576682 | 0.709264768 | -4.436445193 |
| **A_52_P274555** | Snx14 | -0.35455697 | -0.886240556 | 0.177126615 | 9.596312306 | -1.670256307 | 0.150616549 | 0.709264768 | -4.436498958 |
| **A_52_P33800** | Fndc3a | -0.588054364 | -1.470032767 | 0.293924039 | 8.775864401 | -1.669974625 | 0.150673178 | 0.709264768 | -4.436575309 |
| **A_55_P1994947** | NA | 0.537385812 | -0.268886493 | 1.343658118 | 14.43006545 | 1.669378438 | 0.150793105 | 0.709264768 | -4.436736918 |
| **A_55_P1993493** | Efr3a | -0.153254467 | -0.383205598 | 0.076696664 | 10.80102592 | -1.669275185 | 0.150813884 | 0.709264768 | -4.436764908 |
| **A_52_P113672** | Chml | 0.308847152 | -0.154595647 | 0.772289952 | 9.365423343 | 1.669159382 | 0.150837193 | 0.709264768 | -4.4367963 |
| **A_51_P109888** | Coro1b | -0.260922146 | -0.652504213 | 0.130659921 | 9.774184993 | -1.668931428 | 0.150883085 | 0.709264768 | -4.436858096 |
| **A_55_P2064901** | NA | 0.310438035 | -0.1554647 | 0.77634077 | 15.62431686 | 1.668898836 | 0.150889648 | 0.709264768 | -4.436866932 |
| **A_51_P119544** | Aga | -0.339251321 | -0.84841442 | 0.169911778 | 9.179365611 | -1.668841042 | 0.150901285 | 0.709264768 | -4.45941135 |
| **A_51_P167971** | Bace1 | -0.194510631 | -0.486447798 | 0.097426536 | 10.62741022 | -1.668800032 | 0.150909544 | 0.709264768 | -4.436893717 |
| **A_55_P2166884** | NA | 0.427595751 | -0.214368923 | 1.069560424 | 14.34829704 | 1.668294031 | 0.15101148 | 0.709264768 | -4.437030898 |
| **A_55_P2003688** | Tmem143 | -0.246822623 | -0.617430283 | 0.123785036 | 10.15024437 | -1.668095529 | 0.151051487 | 0.709264768 | -4.437084716 |
| **A_66_P112546** | Yeats4 | 0.213179451 | -0.106948649 | 0.533307551 | 8.822905358 | 1.667907217 | 0.15108945 | 0.709264768 | -4.437135773 |
| **A_51_P185593** | Synm | 0.49177819 | -0.246730158 | 1.230286538 | 8.500164542 | 1.667877697 | 0.151095402 | 0.709264768 | -4.437143776 |
| **A_51_P276599** | AU04032C | -0.362378578 | -0.906606055 | 0.181848898 | 10.5360579 | -1.667755563 | 0.15112003 | 0.709264768 | -4.437176891 |
| **A_66_P118195** | Ciao1 | -0.220432203 | -0.551507413 | 0.110643007 | 10.28967067 | -1.667626253 | 0.151146109 | 0.709264768 | -4.437211952 |
| **A_55_P2140449** | B3galt4 | -0.376143954 | -0.941362761 | 0.189074854 | 8.450636106 | -1.666816636 | 0.151309492 | 0.709264768 | -4.437431482 |
| **A_55_P2244677** | 6720422M | -0.281511731 | -0.704559141 | 0.141535678 | 9.360546747 | -1.666701413 | 0.151332759 | 0.709264768 | -4.437462726 |
| **A_52_P528634** | Ttc27 | 0.19258071 | -0.096899873 | 0.482061293 | 8.934351013 | 1.666263539 | 0.151421209 | 0.709264768 | -4.437581468 |
| **A_55_P1980237** | Acad11 | -0.32632504 | -0.816885634 | 0.164235554 | 10.46168935 | -1.666126937 | 0.151448812 | 0.709264768 | -4.437618513 |
| **A_55_P2045481** | NA | 0.396791364 | -0.199739148 | 0.993321876 | 13.88637176 | 1.666018787 | 0.15147067 | 0.709264768 | -4.437647842 |
| **A_55_P2289800** | Fryl | 0.244902641 | -0.123291051 | 0.613096333 | 11.20735464 | 1.665971129 | 0.151480303 | 0.709264768 | -4.437660767 |
| **A_55_P1964183** | Dysf | -0.81244808 | -2.033991177 | 0.409095016 | 10.64985285 | -1.665854893 | 0.1515038 | 0.709264768 | -4.437692289 |
| **A_55_P2088425** | Tcn2 | -1.049533288 | -2.627613777 | 0.528547202 | 13.44583118 | -1.665779115 | 0.151519121 | 0.709264768 | -4.437712841 |
| **A_51_P341349** | Mtap | 0.191606487 | -0.096547642 | 0.479760615 | 8.299519037 | 1.665465753 | 0.15158249 | 0.709264768 | -4.437797826 |
| **A_66_P135941** | Fam221a | -0.172206184 | -0.431229587 | 0.086817219 | 8.324054424 | -1.665175775 | 0.151641154 | 0.709264768 | -4.437876473 |
| **A_52_P649224** | Sash1 | -0.190898927 | -0.478048534 | 0.096250679 | 11.61602528 | -1.665120272 | 0.151652386 | 0.709264768 | -4.437891527 |
| **A_51_P346871** | Ift20 | -0.231760078 | -0.580417456 | 0.116897299 | 11.95271476 | -1.664907524 | 0.151695443 | 0.709264768 | -4.43794923 |
| **A_55_P1963900** | Mib2 | -0.391692304 | -0.981056903 | 0.197672296 | 11.00097358 | -1.664605548 | 0.151756579 | 0.709264768 | -4.438031136 |
| **A_55_P2122546** | Csnk1e | -0.201735398 | -0.505325225 | 0.101854428 | 11.22854282 | -1.664352227 | 0.151807884 | 0.709264768 | -4.438099848 |
| **A_55_P2180869** | Ocel1 | -0.356096223 | -0.892026298 | 0.179833851 | 9.849549207 | -1.664215583 | 0.151835565 | 0.709264768 | -4.438136913 |
| **A_55_P1983768** | Birc5 | 1.04444793 | -0.527572328 | 2.616468188 | 9.864099443 | 1.664098391 | 0.15185931 | 0.709264768 | -4.438168702 |

| | | | | | | | | | |
|---|---|---|---|---|---|---|---|---|---|
| **A_66_P116311** | Kif5b | 0.349550632 | -0.176586297 | 0.875687561 | 9.416257021 | 1.664031916 | 0.15187278 | 0.709264768 | -4.438186734 |
| **A_55_P2042813** | Gbp11 | 0.506714934 | -0.25608246 | 1.269512328 | 8.280295519 | 1.663814287 | 0.151916888 | 0.709264768 | -4.438245769 |
| **A_51_P353592** | Commd4 | -0.236443434 | -0.592420778 | 0.119533911 | 10.55912805 | -1.663624255 | 0.151955413 | 0.709264768 | -4.438297319 |
| **A_55_P2085156** | NA | 0.22080522 | -0.111638966 | 0.553249406 | 13.5951578 | 1.663569465 | 0.151966522 | 0.709264768 | -4.438312182 |
| **A_55_P2023864** | Mfap1a | 0.157067136 | -0.079437237 | 0.393571508 | 10.00322949 | 1.663399228 | 0.152001045 | 0.709264768 | -4.438358363 |
| **A_55_P2030160** | Tcf7l2 | -0.259176157 | -0.649444201 | 0.131091886 | 8.501023113 | -1.663345245 | 0.152011994 | 0.709264768 | -4.438373008 |
| **A_51_P233396** | Ppp2r5d | 0.234714014 | -0.118838014 | 0.588266041 | 11.40323903 | 1.662784753 | 0.152125719 | 0.709264768 | -4.438525065 |
| **A_55_P2166985** | Dedd2 | -1.128647348 | -2.828840034 | 0.571545338 | 11.41625447 | -1.662686871 | 0.152145589 | 0.709264768 | -4.43855162 |
| **A_52_P229997** | Cbx8 | -0.382691863 | -0.95931462 | 0.193930893 | 8.786977273 | -1.662293842 | 0.152225395 | 0.709264768 | -4.438658253 |
| **A_55_P1958097** | Eno1 | 0.348283138 | -0.176640557 | 0.873206833 | 14.41755143 | 1.661830085 | 0.152319616 | 0.709264768 | -4.438784082 |
| **A_55_P2112111** | Ube2b | -0.214219145 | -0.537148087 | 0.108709796 | 13.26196969 | -1.661505026 | 0.152385692 | 0.709264768 | -4.438872283 |
| **A_51_P292757** | Acbd4 | -0.725100328 | -1.818195666 | 0.367995009 | 10.09146302 | -1.661461973 | 0.152394445 | 0.709264768 | -4.438883965 |
| **A_55_P2096081** | Myo5b | -0.214223572 | -0.53718146 | 0.108734317 | 11.3089562 | -1.661390427 | 0.152408993 | 0.709264768 | -4.438903378 |
| **A_52_P144297** | Tspyl3 | 0.708220152 | -0.359737626 | 1.77617793 | 9.387935499 | 1.660980534 | 0.152492366 | 0.709264768 | -4.439014604 |
| **A_51_P225793** | Prr5l | -0.449397772 | -1.12709223 | 0.228296687 | 11.25418643 | -1.660914688 | 0.152505764 | 0.709264768 | -4.439032472 |
| **A_55_P2077343** | Ahdc1 | -0.250538037 | -0.628397902 | 0.127321828 | 9.756778978 | -1.660707896 | 0.152547846 | 0.709264768 | -4.439088589 |
| **A_55_P1993909** | Taz | -0.374213157 | -0.938644877 | 0.190218563 | 10.53174593 | -1.660573054 | 0.152575293 | 0.709264768 | -4.439125181 |
| **A_55_P2001238** | Luc7l3 | 0.583595841 | -0.29668509 | 1.463876773 | 12.80238899 | 1.660509022 | 0.152588328 | 0.709264768 | -4.439142558 |
| **A_55_P2149274** | Sh3bp5l | -0.302008095 | -0.757552641 | 0.153536452 | 12.16598926 | -1.660496931 | 0.152590789 | 0.709264768 | -4.439145839 |
| **A_52_P370935** | Glcci1 | 0.764202004 | -0.388871311 | 1.917275319 | 10.01107896 | 1.659975187 | 0.152697045 | 0.709264768 | -4.439287432 |
| **A_52_P604526** | Mrpl19 | 0.214690313 | -0.109299037 | 0.538679662 | 9.687382738 | 1.659709424 | 0.152751196 | 0.709264768 | -4.439359559 |
| **A_66_P119418** | Rb1cc1 | -0.381220115 | -0.956532234 | 0.194092004 | 8.759711544 | -1.65967339 | 0.15275854 | 0.709264768 | -4.439369339 |
| **A_66_P134149** | Snx30 | -0.466108325 | -1.169579241 | 0.237362591 | 9.393848517 | -1.659552738 | 0.152783131 | 0.709264768 | -4.439402085 |
| **A_55_P2302383** | 3300002A | -0.249769177 | -0.626767973 | 0.127229619 | 8.194720553 | -1.65939289 | 0.152815717 | 0.709264768 | -4.439445469 |
| **A_55_P2104487** | Pld1 | -0.171822102 | -0.431172795 | 0.087528591 | 10.19141292 | -1.659365136 | 0.152821376 | 0.709264768 | -4.439453002 |
| **A_55_P2156462** | Nt5c3b | 0.272486746 | -0.139019033 | 0.683992526 | 8.762223189 | 1.658516474 | 0.152994502 | 0.709264768 | -4.439683352 |
| **A_51_P333159** | Rpa1 | 1.06980935 | -0.545937787 | 2.685556486 | 11.72281969 | 1.658377282 | 0.153022915 | 0.709264768 | -4.439721134 |
| **A_55_P2178862** | U2surp | 0.207006602 | -0.105686908 | 0.519700111 | 8.647670296 | 1.658118918 | 0.153075669 | 0.709264768 | -4.439791267 |
| **A_51_P145520** | Sigirr | -0.326034714 | -0.81861905 | 0.166549623 | 9.554112791 | -1.657805554 | 0.153139676 | 0.709264768 | -4.439876332 |
| **A_55_P1961463** | Bnip3l | -0.35496482 | -0.891340992 | 0.181411352 | 13.38994061 | -1.657548261 | 0.15319225 | 0.709264768 | -4.439946179 |
| **A_52_P330984** | Lin9 | 0.299480992 | -0.153072864 | 0.752034848 | 9.476501057 | 1.657483978 | 0.153205388 | 0.709264768 | -4.43996363 |
| **A_55_P2170309** | Ppfibp2 | -0.752062188 | -1.888833784 | 0.384709408 | 8.727553541 | -1.657031964 | 0.153297802 | 0.709264768 | -4.440086343 |
| **A_55_P1956195** | Sap18b | 0.295216463 | -0.15105912 | 0.741492046 | 12.95512425 | 1.65686755 | 0.153331429 | 0.709264768 | -4.44013098 |
| **A_55_P2094731** | Sox13 | 0.384663225 | -0.19688155 | 0.966208 | 9.971977638 | 1.656715197 | 0.153362596 | 0.709264768 | -4.440172344 |
| **A_55_P1957228** | Sgk3 | 0.28552125 | -0.146160739 | 0.717203238 | 11.57786156 | 1.656627447 | 0.15338055 | 0.709264768 | -4.440196168 |
| **A_55_P2084666** | H2ac10 | 0.873548877 | -0.447245719 | 2.194343473 | 16.94402165 | 1.656541316 | 0.153398175 | 0.709264768 | -4.440219553 |
| **A_55_P2047310** | Pdgfb | -0.76064392 | -1.910980238 | 0.389692398 | 12.25236608 | -1.656177622 | 0.153472619 | 0.709264768 | -4.440318299 |
| **A_51_P198775** | BC05532 | 0.558715348 | -0.286463144 | 1.40389384 | 10.86472834 | 1.655741409 | 0.153561952 | 0.709264768 | -4.440436741 |
| **A_55_P2049025** | NA | 0.197038789 | -0.101033897 | 0.495111476 | 15.03944032 | 1.655693309 | 0.153571806 | 0.709264768 | -4.440449802 |
| **A_55_P1967478** | Hbb-y | 1.283146415 | -0.658249842 | 3.224542671 | 10.29423106 | 1.655435913 | 0.153624546 | 0.709264768 | -4.440519694 |
| **A_51_P408649** | Shisa2 | 0.554821703 | -0.284694789 | 1.394338195 | 9.376475413 | 1.655291764 | 0.15365409 | 0.709264768 | -4.440558837 |
| **A_51_P345649** | Pdgfra | -0.504063417 | -1.266811631 | 0.258684797 | 11.78640551 | -1.655214665 | 0.153669894 | 0.709264768 | -4.452883165 |
| **A_55_P2107302** | Aurkaip1 | -0.230006562 | -0.578078692 | 0.118065569 | 14.46833984 | -1.655088877 | 0.153695682 | 0.709264768 | -4.44061393 |
| **A_52_P582309** | Nub1 | -0.673897771 | -1.693798989 | 0.346003447 | 10.72316231 | -1.65495525 | 0.153723082 | 0.709264768 | -4.440650217 |
| **A_55_P2005552** | Arhgef10l | 0.233226365 | -0.119752353 | 0.586205083 | 11.39904857 | 1.654929388 | 0.153728385 | 0.709264768 | -4.44065724 |
| **A_65_P19138** | 7-Mar | 0.164494474 | -0.084462431 | 0.41345138 | 11.62118132 | 1.654921823 | 0.153729937 | 0.709264768 | -4.440659294 |
| **A_55_P1992045** | Msi2 | 0.680791332 | -0.349867937 | 1.7114506 | 12.12802691 | 1.65443325 | 0.153830161 | 0.709264768 | -4.440791973 |
| **A_55_P2426272** | Scaper | -0.179277169 | -0.450717283 | 0.092162945 | 8.365460594 | -1.654250606 | 0.153867644 | 0.709264768 | -4.440841575 |

| | | | | | | | | | |
|---|---|---|---|---|---|---|---|---|---|
| **A_55_P1982524** | Cyhr1 | -0.206074444 | -0.518143321 | 0.105994434 | 8.568207733 | -1.653956268 | 0.153928069 | 0.709264768 | -4.440921512 |
| **A_55_P1965554** | Orc3 | 0.296990871 | -0.152791604 | 0.746773346 | 11.00855821 | 1.653830192 | 0.153953958 | 0.709264768 | -4.440955753 |
| **A_55_P1993268** | Ap1s1 | -0.173140347 | -0.435382755 | 0.08910206 | 13.98768674 | -1.65365804 | 0.153989316 | 0.709264768 | -4.441002508 |
| **A_55_P2219059** | Lmnb2 | 0.307895797 | -0.158478851 | 0.774270444 | 9.147210456 | 1.653557007 | 0.154010071 | 0.709264768 | -4.441029948 |
| **A_55_P2047614** | Zfp809 | -0.170684411 | -0.429224925 | 0.087856102 | 8.287885908 | -1.653543398 | 0.154012867 | 0.709264768 | -4.441033645 |
| **A_52_P26953** | Ergic2 | -0.420626662 | -1.057852572 | 0.216599248 | 9.484532295 | -1.653307809 | 0.154061274 | 0.709264768 | -4.441097632 |
| **A_55_P2020373** | Chd4 | 0.252782233 | -0.130242408 | 0.635806875 | 13.21460337 | 1.652989813 | 0.154126637 | 0.709264768 | -4.441184003 |
| **A_52_P656545** | Ogt | -0.437634933 | -1.10081466 | 0.225544795 | 8.471907572 | -1.652841026 | 0.154157229 | 0.709264768 | -4.441224417 |
| **A_55_P2068892** | Il6ra | -1.113976904 | -2.802115169 | 0.57416136 | 10.81657366 | -1.652793204 | 0.154167062 | 0.709264768 | -4.441237406 |
| **A_55_P2083153** | 1700001J | -0.396747388 | -1.013696107 | 0.220201331 | 7.879361461 | -1.712883063 | 0.154217127 | 0.709264768 | -4.459047078 |
| **A_51_P157042** | Ccn2 | -1.237539953 | -3.113314716 | 0.63823481 | 12.81100618 | -1.652452124 | 0.154237218 | 0.709264768 | -4.441330054 |
| **A_51_P469578** | Psma1 | 0.390636305 | -0.201527402 | 0.982800013 | 13.34857705 | 1.652270551 | 0.154274579 | 0.709264768 | -4.441379376 |
| **A_55_P2068233** | Phactr1 | -1.246780909 | -3.136775801 | 0.643213982 | 9.618487351 | -1.652265592 | 0.154275599 | 0.709264768 | -4.441380723 |
| **A_55_P2024391** | NA | 0.954066978 | -0.492272857 | 2.400406813 | 9.489545942 | 1.652185713 | 0.154292037 | 0.709264768 | -4.441402421 |
| **A_55_P2090239** | Casp8ap2 | 0.218547885 | -0.112806474 | 0.549902243 | 8.375263429 | 1.651978022 | 0.154334787 | 0.709264768 | -4.44145884 |
| **A_51_P170008** | Napg | -0.1526008 | -0.383970281 | 0.078768681 | 8.943917785 | -1.651965924 | 0.154337278 | 0.709264768 | -4.441462126 |
| **A_55_P2155041** | Ssbp2 | 0.60183048 | -0.310769286 | 1.514430246 | 11.86647065 | 1.651749452 | 0.154381848 | 0.709264768 | -4.441520932 |
| **A_55_P2077535** | Zfp513 | -0.3567969 | -0.897840196 | 0.184246396 | 10.97485279 | -1.651731297 | 0.154385586 | 0.709264768 | -4.441525864 |
| **A_51_P267494** | Cdc42ep3 | 0.431094616 | -0.222858184 | 1.085047416 | 10.93470906 | 1.651112027 | 0.154513162 | 0.709264768 | -4.441694099 |
| **A_51_P390775** | Ube2e1 | 0.187735513 | -0.097074151 | 0.472545177 | 11.70027109 | 1.650980931 | 0.154540182 | 0.709264768 | -4.441729715 |
| **A_55_P2138150** | Pias2 | -0.294825732 | -0.742103233 | 0.15245177 | 9.37607604 | -1.650968083 | 0.154542831 | 0.709264768 | -4.441733206 |
| **A_51_P154780** | Vav1 | 0.292756242 | -0.151398509 | 0.736910993 | 8.026933101 | 1.650905434 | 0.154555745 | 0.709264768 | -4.441750226 |
| **A_55_P1974442** | Sumf2 | 0.197881155 | -0.102364658 | 0.498126967 | 9.364896193 | 1.650736758 | 0.154590521 | 0.709264768 | -4.441796053 |
| **A_55_P2184816** | Eif2s1 | 0.193036133 | -0.099858506 | 0.485930772 | 11.70056493 | 1.650735653 | 0.154590749 | 0.709264768 | -4.441796353 |
| **A_51_P288479** | Slc25a29 | -0.521295288 | -1.312333154 | 0.269742577 | 8.872289338 | -1.650581117 | 0.154622617 | 0.709264768 | -4.441838339 |
| **A_55_P2197338** | Tnfsf13os | -0.593874376 | -1.495048894 | 0.307300143 | 8.49406 | -1.650577913 | 0.154623278 | 0.709264768 | -4.44183921 |
| **A_52_P193194** | Slc35d2 | -0.260075952 | -0.654747655 | 0.13459575 | 8.127457718 | -1.650496295 | 0.154640111 | 0.709264768 | -4.441861385 |
| **A_55_P2022845** | Csnk2a1 | 0.284476236 | -0.147287965 | 0.716240437 | 11.32010959 | 1.650249872 | 0.154690946 | 0.709264768 | -4.441928338 |
| **A_55_P1954271** | Col26a1 | 0.587029919 | -0.303978221 | 1.47803806 | 9.35896272 | 1.650170848 | 0.154707252 | 0.709264768 | -4.441949809 |
| **A_51_P244504** | LTO1 | -0.602741858 | -1.517658773 | 0.312175056 | 10.49818627 | -1.65006116 | 0.154729887 | 0.709264768 | -4.441979613 |
| **A_55_P1955656** | Ctla2a | -2.001654246 | -5.040567138 | 1.037258647 | 13.63022674 | -1.649761468 | 0.154791748 | 0.709264768 | -4.442061043 |
| **A_55_P2006940** | Sfmbt1 | 0.171602544 | -0.088948506 | 0.432153593 | 9.001136418 | 1.649609846 | 0.154823054 | 0.709264768 | -4.442102242 |
| **A_55_P2132942** | Adal | 0.268586886 | -0.139341722 | 0.676515495 | 9.357035095 | 1.64911517 | 0.154925237 | 0.709264768 | -4.442236661 |
| **A_55_P2115458** | Pot1a | 0.353363269 | -0.183335098 | 0.890061637 | 10.96165575 | 1.649079054 | 0.154932699 | 0.709264768 | -4.442246475 |
| **A_55_P1962234** | Bad | -0.265393752 | -0.668512441 | 0.137724936 | 12.56687371 | -1.648952357 | 0.154958882 | 0.709264768 | -4.442280905 |
| **A_55_P2167005** | NA | 0.298246103 | -0.154784924 | 0.75127713 | 14.55796265 | 1.648910855 | 0.15496746 | 0.709264768 | -4.442292183 |
| **A_51_P161265** | Ccl25 | 1.858829214 | -0.965214296 | 4.682872723 | 10.15313181 | 1.64861198 | 0.155029245 | 0.709264768 | -4.442373402 |
| **A_55_P1965772** | Atp2a3 | 0.357356337 | -0.185569391 | 0.900282065 | 9.014726115 | 1.648585256 | 0.155034771 | 0.709264768 | -4.442380664 |
| **A_51_P267370** | Itm2c | -0.22935354 | -0.577812888 | 0.119105807 | 11.12626951 | -1.648555888 | 0.155040843 | 0.709264768 | -4.442388645 |
| **A_55_P2016105** | Errfi1 | -0.293888932 | -0.740434761 | 0.152656896 | 12.58773222 | -1.648418728 | 0.155069208 | 0.709264768 | -4.44242592 |
| **A_52_P363216** | Gcnt2 | -0.889227741 | -2.24043997 | 0.461984488 | 8.799357939 | -1.648313369 | 0.155090999 | 0.709264768 | -4.442454552 |
| **A_55_P2157033** | Bace2 | -0.283998157 | -0.715568675 | 0.147572362 | 10.19471001 | -1.648215894 | 0.155111163 | 0.709264768 | -4.442481043 |
| **A_51_P201137** | Lbr | 0.717491633 | -0.373078045 | 1.808061312 | 10.30638206 | 1.647835172 | 0.155189944 | 0.709264768 | -4.442584513 |
| **A_55_P2038594** | Tmem234 | -0.163986738 | -0.413306604 | 0.085333128 | 12.34504695 | -1.64741186 | 0.155277583 | 0.709264768 | -4.442699563 |
| **A_51_P436020** | Tmem167 | 0.19245706 | -0.100211157 | 0.485125277 | 8.918829439 | 1.64705701 | 0.155351085 | 0.709264768 | -4.44279601 |
| **A_66_P112399** | BC029722 | -0.215563831 | -0.543482314 | 0.112354652 | 11.31795629 | -1.646494721 | 0.155467626 | 0.709264768 | -4.442948847 |
| **A_55_P2008106** | Otud6b | 0.191208309 | -0.099704377 | 0.482120995 | 8.783130909 | 1.646244917 | 0.155519428 | 0.709264768 | -4.44301675 |
| **A_51_P190805** | Ap1m1 | 0.223514304 | -0.116572283 | 0.56360089 | 11.43432462 | 1.64613771 | 0.155541665 | 0.709264768 | -4.443045892 |

| | | | | | | | | | |
|---|---|---|---|---|---|---|---|---|---|
| **A_51_P246317** | Mt2 | -1.83885713 | -4.83804048 | 1.160326221 | 14.42119117 | -1.808162638 | 0.155544152 | 0.709264768 | -4.473790698 |
| **A_52_P641216** | Galk2 | 0.196542283 | -0.102535528 | 0.495620094 | 11.32663495 | 1.645970886 | 0.155576273 | 0.709264768 | -4.44309124 |
| **A_55_P2056473** | Spc24 | 1.113291571 | -0.580821447 | 2.807404589 | 9.941799419 | 1.645950942 | 0.155580411 | 0.709264768 | -4.443096662 |
| **A_51_P428578** | Retreg1 | -0.417557185 | -1.052996123 | 0.217881752 | 8.37303491 | -1.645858459 | 0.155599601 | 0.709264768 | -4.443121802 |
| **A_51_P306017** | Dll1 | -0.179290096 | -0.452192321 | 0.093612129 | 8.169295497 | -1.645506375 | 0.155672678 | 0.709264768 | -4.443217514 |
| **A_55_P1989916** | Lin54 | 0.222817351 | -0.116342418 | 0.56197712 | 9.346816334 | 1.645489143 | 0.155676256 | 0.709264768 | -4.443222199 |
| **A_51_P367298** | NA | -0.190958915 | -0.481636185 | 0.099718355 | 8.312477801 | -1.645429242 | 0.155688692 | 0.709264768 | -4.443238483 |
| **A_55_P2081405** | BC052688 | -0.28370496 | -0.715576564 | 0.148166644 | 12.50260522 | -1.6453664 | 0.15570174 | 0.709264768 | -4.443255567 |
| **A_55_P1957754** | NA | 0.405786413 | -0.211975725 | 1.023548551 | 15.33697325 | 1.64522977 | 0.155730113 | 0.709264768 | -4.443292711 |
| **A_55_P2162136** | Hmgn2 | 0.268634441 | -0.140392933 | 0.677661816 | 13.27036699 | 1.644976368 | 0.155782748 | 0.709264768 | -4.443361602 |
| **A_55_P1961135** | Atg4d | -0.56896962 | -1.435293632 | 0.297354392 | 11.45348337 | -1.644974128 | 0.155783213 | 0.709264768 | -4.44336221 |
| **A_55_P2091145** | H4c12 | 1.228310058 | -0.642257965 | 3.098878082 | 15.71802838 | 1.644693014 | 0.155841625 | 0.709264768 | -4.443438637 |
| **A_52_P401484** | Inha | -0.179588609 | -0.453081168 | 0.09390395 | 8.115072107 | -1.644688353 | 0.155842593 | 0.709264768 | -4.443439905 |
| **A_51_P401987** | Tmem37 | -2.294704144 | -5.789760061 | 1.200351772 | 12.39932842 | -1.644458497 | 0.15589037 | 0.709264768 | -4.443502398 |
| **A_55_P2008422** | Mpc1 | -0.187268414 | -0.47258468 | 0.098047851 | 13.31041418 | -1.643949029 | 0.155996317 | 0.709264768 | -4.443640917 |
| **A_55_P2025363** | Ccnl2 | -0.164218883 | -0.414467909 | 0.086030144 | 8.578929554 | -1.643618579 | 0.156065074 | 0.709264768 | -4.443730767 |
| **A_52_P677956** | Pex11g | -0.547948419 | -1.383057611 | 0.287160774 | 10.88795238 | -1.643413157 | 0.156107831 | 0.709264768 | -4.443786624 |
| **A_55_P2032147** | Wnt9a | -0.448337625 | -1.131957426 | 0.235282176 | 9.11057125 | -1.642634343 | 0.156270039 | 0.709264768 | -4.443998404 |
| **A_51_P152990** | Grem2 | -0.225410452 | -0.569172114 | 0.118351211 | 9.204559846 | -1.642354686 | 0.156328325 | 0.709264768 | -4.444074455 |
| **A_55_P2071286** | Tmc4 | -0.771083443 | -1.947082441 | 0.404915556 | 10.96609992 | -1.642270802 | 0.156345813 | 0.709264768 | -4.444097267 |
| **A_52_P140005** | Nipal1 | -0.244509072 | -0.617465161 | 0.128447018 | 7.973771948 | -1.642054699 | 0.156390872 | 0.709264768 | -4.444156036 |
| **A_55_P2083997** | Zfp664 | 0.59892809 | -0.314699815 | 1.512555994 | 12.44027478 | 1.641933908 | 0.156416064 | 0.709264768 | -4.444188886 |
| **A_55_P2184189** | Ncf2 | -0.193071135 | -0.487606745 | 0.101464475 | 8.594594211 | -1.641836425 | 0.156436397 | 0.709264768 | -4.444215398 |
| **A_52_P456279** | Cct8 | 0.285981801 | -0.150297463 | 0.722261066 | 12.64933587 | 1.641814817 | 0.156440905 | 0.709264768 | -4.444221274 |
| **A_51_P378789** | Cxcl13 | 0.450312202 | -0.236684359 | 1.137308763 | 8.504015255 | 1.641759347 | 0.156452476 | 0.709264768 | -4.44423636 |
| **A_55_P2015666** | NA | 0.211596975 | -0.111278596 | 0.534472546 | 16.54134874 | 1.641438489 | 0.156519427 | 0.709264768 | -4.444323624 |
| **A_55_P1961436** | Prkca | 0.546244424 | -0.287286078 | 1.379774926 | 9.524952309 | 1.641405429 | 0.156526327 | 0.709264768 | -4.444332615 |
| **A_55_P2009988** | Trib3 | -0.730617083 | -1.845747276 | 0.38451311 | 10.53412115 | -1.641022718 | 0.156606225 | 0.709264768 | -4.444436705 |
| **A_52_P98778** | Ang4 | -0.161419894 | -0.40780406 | 0.084964272 | 8.101520221 | -1.640947206 | 0.156621994 | 0.709264768 | -4.444457243 |
| **A_55_P1993390** | NA | 0.506760731 | -0.266800134 | 1.280321596 | 15.16794471 | 1.640811946 | 0.156650244 | 0.709264768 | -4.444494033 |
| **A_52_P410765** | Sema7a | -0.629410302 | -1.590218967 | 0.331398363 | 10.88448012 | -1.640768393 | 0.156659342 | 0.709264768 | -4.444505879 |
| **A_55_P2101806** | Esf1 | 0.211132145 | -0.111233209 | 0.533497499 | 10.37546456 | 1.640424872 | 0.156731115 | 0.709264768 | -4.444599316 |
| **A_52_P512427** | Mkrn2os | -0.193610868 | -0.489232142 | 0.102010406 | 7.915455722 | -1.640379725 | 0.15674055 | 0.709264768 | -4.444611597 |
| **A_55_P2091626** | NA | 0.171580926 | -0.090432425 | 0.433594278 | 15.87245468 | 1.640196689 | 0.156778808 | 0.709264768 | -4.444661384 |
| **A_51_P493700** | Cep350 | -0.34558293 | -0.873395043 | 0.182229183 | 10.03789226 | -1.639922316 | 0.156836174 | 0.709264768 | -4.444736018 |
| **A_52_P266530** | Ide | 0.3268819 | -0.172370046 | 0.826133847 | 10.09015197 | 1.639915433 | 0.156837614 | 0.709264768 | -4.44473789 |
| **A_51_P123134** | Ercc6l | 0.313591123 | -0.165445174 | 0.792627419 | 8.34209128 | 1.639629354 | 0.15689745 | 0.709264768 | -4.448209143 |
| **A_51_P384640** | Mrpl10 | -0.199876783 | -0.505267629 | 0.105514063 | 9.894072755 | -1.639293365 | 0.156967753 | 0.709264768 | -4.44490711 |
| **A_51_P463440** | Elovl6 | 0.173529232 | -0.091624783 | 0.438683246 | 9.132043049 | 1.639172966 | 0.156992953 | 0.709264768 | -4.444939863 |
| **A_55_P2115691** | NA | 1.028312095 | -0.543080653 | 2.599704844 | 11.70500466 | 1.639043746 | 0.157020004 | 0.709264768 | -4.444975017 |
| **A_55_P2010014** | Map7d1 | -0.593956914 | -1.501712499 | 0.31379867 | 13.40547367 | -1.638839257 | 0.157062821 | 0.709264768 | -4.445030648 |
| **A_55_P2082203** | Baz1a | 0.353601187 | -0.186855425 | 0.894057798 | 9.562344514 | 1.638714232 | 0.157089005 | 0.709264768 | -4.445064661 |
| **A_66_P103490** | Srsf1 | 0.737425304 | -0.389860444 | 1.864711052 | 10.7512871 | 1.638454423 | 0.15714343 | 0.709264768 | -4.445135345 |
| **A_55_P2075268** | Tmem216 | -0.282083711 | -0.713321238 | 0.149153817 | 10.93817023 | -1.638369317 | 0.157161263 | 0.709264768 | -4.445158499 |
| **A_55_P2122020** | Klf4 | -0.313511388 | -0.792818184 | 0.165795408 | 10.83236309 | -1.638287355 | 0.157178438 | 0.709264768 | -4.445180798 |
| **A_52_P681310** | Plaur | -1.096103568 | -2.772071979 | 0.579864843 | 10.01231592 | -1.638083774 | 0.157221107 | 0.709264768 | -4.445236187 |
| **A_55_P2049979** | Dbi | -0.358817088 | -0.90759713 | 0.189962955 | 14.13536375 | -1.637665326 | 0.157308845 | 0.709264768 | -4.445350038 |
| **A_55_P2030938** | Trim59 | 1.004373854 | -0.531820947 | 2.540568655 | 11.45446881 | 1.637568415 | 0.157329172 | 0.709264768 | -4.445376406 |

| | | | | | | | | | |
|---|---|---|---|---|---|---|---|---|---|
| **A_55_P2184877** | NA | 0.343607208 | -0.182077104 | 0.86929152 | 10.93016895 | 1.637146691 | 0.157417656 | 0.709264768 | -4.445491155 |
| **A_51_P232901** | Cnp | 0.424924779 | -0.225171302 | 1.075020859 | 8.978987213 | 1.637136364 | 0.157419824 | 0.709264768 | -4.445493965 |
| **A_51_P324704** | Mrpl16 | 0.244737361 | -0.129708557 | 0.619183278 | 11.49898757 | 1.637048394 | 0.157438288 | 0.709264768 | -4.445517902 |
| **A_55_P2179030** | Ostf1 | -0.564629395 | -1.42855297 | 0.29929418 | 12.46403198 | -1.636961646 | 0.157456497 | 0.709264768 | -4.445541507 |
| **A_51_P325374** | Tent2 | -0.519474384 | -1.314345067 | 0.275396299 | 11.20494115 | -1.636884374 | 0.157472719 | 0.709264768 | -4.445562533 |
| **A_52_P12806** | Gm5069 | 0.30242109 | -0.160438948 | 0.765281127 | 14.19930496 | 1.636487655 | 0.15755603 | 0.709264768 | -4.445670486 |
| **A_55_P2154536** | Hagh | -0.784285573 | -1.984990206 | 0.41641906 | 10.63456479 | -1.636019159 | 0.15765447 | 0.709264768 | -4.445797977 |
| **A_65_P08022** | Hnrnpk | 0.462525846 | -0.245610354 | 1.170662046 | 11.22722976 | 1.635948228 | 0.157669379 | 0.709264768 | -4.44581728 |
| **A_55_P2126814** | NA | 0.416809519 | -0.221460416 | 1.055079454 | 9.791261641 | 1.635624423 | 0.157737457 | 0.709264768 | -4.4459054 |
| **A_55_P2060592** | Hoxa1 | -0.475834537 | -1.204493425 | 0.25282435 | 8.843178655 | -1.635618598 | 0.157738682 | 0.709264768 | -4.445906985 |
| **A_51_P100573** | Map3k3 | -0.209040811 | -0.529205844 | 0.111124221 | 8.452207361 | -1.635338004 | 0.1577977 | 0.709264768 | -4.452566902 |
| **A_51_P359549** | Fem1c | -0.372245479 | -0.942408831 | 0.197917873 | 9.315624171 | -1.635236146 | 0.157819129 | 0.709264768 | -4.44601107 |
| **A_55_P1956593** | Plekha4 | -0.449886858 | -1.139072123 | 0.239298407 | 9.234379671 | -1.63499967 | 0.15786889 | 0.709264768 | -4.44607543 |
| **A_51_P223946** | Pds5b | 0.708680019 | -0.377001461 | 1.794361498 | 9.455682393 | 1.634926014 | 0.157884392 | 0.709264768 | -4.446095477 |
| **A_55_P1965050** | Cks2 | 0.953818132 | -0.507425819 | 2.415062082 | 10.27622703 | 1.634907526 | 0.157888284 | 0.709264768 | -4.446100508 |
| **A_55_P2008651** | Qdpr | 0.440247942 | -0.234294618 | 1.114790502 | 12.09241366 | 1.634700983 | 0.157931764 | 0.709264768 | -4.446156723 |
| **A_55_P1994773** | Gm16381 | 0.394458637 | -0.20994029 | 0.998857564 | 9.780104113 | 1.634662617 | 0.157939842 | 0.709264768 | -4.446167165 |
| **A_55_P1961188** | Gm6981 | 0.340301408 | -0.181158338 | 0.861761154 | 14.53913233 | 1.634531531 | 0.157967445 | 0.709264768 | -4.446202844 |
| **A_55_P2025820** | NA | -0.165273533 | -0.418538629 | 0.087991562 | 8.223058637 | -1.634475129 | 0.157979323 | 0.709264768 | -4.446218195 |
| **A_55_P1980831** | Arpc2 | -0.476203525 | -1.205958527 | 0.253551476 | 15.50717781 | -1.634428292 | 0.157989187 | 0.709264768 | -4.446230943 |
| **A_55_P2185722** | Fermt2 | -0.396844727 | -1.005188658 | 0.211499205 | 10.31255488 | -1.633886105 | 0.158103421 | 0.709264768 | -4.44637852 |
| **A_52_P601914** | Lemd2 | -0.778434276 | -1.971777869 | 0.414909316 | 12.20616034 | -1.633829716 | 0.158115306 | 0.709264768 | -4.446393869 |
| **A_52_P632191** | Nfx1 | -0.211236167 | -0.535084541 | 0.112612206 | 8.483154625 | -1.633717287 | 0.158139006 | 0.709264768 | -4.446424472 |
| **A_55_P2050877** | Plekhg2 | -0.305764814 | -0.774546219 | 0.163016592 | 13.10151727 | -1.633681819 | 0.158146483 | 0.709264768 | -4.446434127 |
| **A_55_P2138090** | U2af1l4 | 0.174798264 | -0.093203739 | 0.442800267 | 12.11091732 | 1.633613905 | 0.158160802 | 0.709264768 | -4.446452613 |
| **A_55_P1964805** | Rpa1 | 0.895276265 | -0.477554567 | 2.268107096 | 12.74167485 | 1.63339188 | 0.15820762 | 0.709264768 | -4.44651305 |
| **A_55_P2098321** | NA | 0.180206197 | -0.09613333 | 0.456545725 | 11.31629889 | 1.633341728 | 0.158218198 | 0.709264768 | -4.446526702 |
| **A_51_P373609** | Yars2 | 0.158611111 | -0.084617509 | 0.401839731 | 9.825835584 | 1.633312463 | 0.15822437 | 0.709264768 | -4.446534668 |
| **A_55_P2044704** | NA | 0.201516806 | -0.107514718 | 0.510548329 | 11.49441549 | 1.633273189 | 0.158232654 | 0.709264768 | -4.446545359 |
| **A_55_P2021841** | Calm3 | 0.249877054 | -0.133317464 | 0.633071572 | 13.12784422 | 1.633267933 | 0.158233763 | 0.709264768 | -4.44654679 |
| **A_51_P429682** | Ebp | -0.228380555 | -0.578654768 | 0.121893659 | 12.10931989 | -1.633056846 | 0.158278294 | 0.709264768 | -4.446604251 |
| **A_52_P645862** | Agtr1a | -0.213810401 | -0.541823917 | 0.114203116 | 9.349976196 | -1.632628723 | 0.15836865 | 0.709264768 | -4.446720798 |
| **A_66_P102046** | NA | 0.367982605 | -0.19669119 | 0.9326564 | 10.18796468 | 1.632224909 | 0.158453922 | 0.709264768 | -4.446830732 |
| **A_51_P148494** | Rnft1 | -0.207353736 | -0.525593528 | 0.110886055 | 9.438172784 | -1.631953309 | 0.158511299 | 0.709264768 | -4.446904674 |
| **A_51_P377508** | Srebf2 | -0.315201143 | -0.798968887 | 0.168566601 | 8.744487031 | -1.631928853 | 0.158516467 | 0.709264768 | -4.446911332 |
| **A_65_P07062** | Serf2 | -0.275092412 | -0.697303416 | 0.147118591 | 12.25364085 | -1.631922048 | 0.158517905 | 0.709264768 | -4.446913185 |
| **A_55_P2142172** | Olfr1229 | 0.256401673 | -0.137127624 | 0.64993097 | 9.128428198 | 1.631902208 | 0.158522097 | 0.709264768 | -4.446918586 |
| **A_52_P354123** | Rhebl1 | -0.21059689 | -0.533862052 | 0.112668273 | 8.772740849 | -1.631711571 | 0.158562385 | 0.709264768 | -4.446970488 |
| **A_52_P442756** | Btf3l4 | 0.151168782 | -0.080952779 | 0.383290343 | 10.59133731 | 1.631161555 | 0.158678679 | 0.709264768 | -4.447120239 |
| **A_55_P2026049** | NA | 0.174111134 | -0.093239173 | 0.441461442 | 8.231118506 | 1.631158647 | 0.158679294 | 0.709264768 | -4.447121031 |
| **A_55_P2103626** | Slc27a4 | 0.216637435 | -0.116034272 | 0.549309142 | 8.484811692 | 1.631052667 | 0.158701712 | 0.709264768 | -4.447149887 |
| **A_51_P220993** | Pfn1 | 0.209950785 | -0.112506462 | 0.532408032 | 10.90168742 | 1.630781237 | 0.158759141 | 0.709264768 | -4.447223792 |
| **A_55_P2019719** | Oas2 | -0.406537461 | -1.031049406 | 0.217974485 | 8.009561931 | -1.630460065 | 0.158827121 | 0.709264768 | -4.447311244 |
| **A_55_P2144285** | Nnt | 0.985600535 | -0.528492828 | 2.499693899 | 11.02502452 | 1.63041672 | 0.158836297 | 0.709264768 | -4.447323047 |
| **A_51_P108459** | Gpr65 | 0.231708404 | -0.124249612 | 0.58766642 | 8.080867401 | 1.630396947 | 0.158840484 | 0.709264768 | -4.452208197 |
| **A_55_P2044593** | 1810037I1 | -0.200478095 | -0.508467967 | 0.107511778 | 13.28253042 | -1.630349956 | 0.158850433 | 0.709264768 | -4.447341226 |
| **A_55_P2114995** | NA | -0.219768949 | -0.557433829 | 0.117895932 | 15.49631791 | -1.630162064 | 0.158890221 | 0.709264768 | -4.44739239 |
| **A_66_P106509** | NA | 0.677704615 | -0.363582463 | 1.718991693 | 13.74549774 | 1.630122755 | 0.158898546 | 0.709264768 | -4.447403094 |

| | | | | | | | | | |
|---|---|---|---|---|---|---|---|---|---|
| **A_52_P401311** | NA | 0.49747275 | -0.267031139 | 1.261976639 | 10.37519815 | 1.629821127 | 0.158962443 | 0.709264768 | -4.447485229 |
| **A_52_P344290** | F2r | -0.704016941 | -1.785951056 | 0.377917174 | 13.7334324 | -1.629793721 | 0.15896825 | 0.709264768 | -4.447492692 |
| **A_51_P278550** | Tecr | 0.16727124 | -0.089845023 | 0.424387504 | 10.66533616 | 1.629453886 | 0.159040273 | 0.709264768 | -4.447585236 |
| **A_55_P2067146** | NA | 0.153244943 | -0.08244196 | 0.388931846 | 7.903955837 | 1.628549819 | 0.159232031 | 0.709264768 | -4.447831445 |
| **A_55_P1977071** | Fam110a | 0.15963131 | -0.085885603 | 0.405148223 | 9.604962955 | 1.628497171 | 0.159243205 | 0.709264768 | -4.447845784 |
| **A_52_P480351** | Loxl2 | 0.222107945 | -0.119527368 | 0.563743259 | 11.57282541 | 1.628364719 | 0.15927132 | 0.709264768 | -4.447881858 |
| **A_51_P315673** | Usp11 | 0.450372646 | -0.242383812 | 1.143129104 | 9.72010973 | 1.628327538 | 0.159279213 | 0.709264768 | -4.447891984 |
| **A_55_P1988872** | NA | 0.435326056 | -0.234518466 | 1.105170579 | 8.614012286 | 1.627762345 | 0.159399243 | 0.709264768 | -4.448045921 |
| **A_51_P336952** | Cryl1 | -0.2747499 | -0.697539545 | 0.148039745 | 10.81554409 | -1.62765946 | 0.159421102 | 0.709264768 | -4.448073944 |
| **A_55_P2058737** | Mex3d | 0.458303023 | -0.246947708 | 1.163553754 | 10.5312784 | 1.627644377 | 0.159424307 | 0.709264768 | -4.448078052 |
| **A_51_P322265** | Grap | -0.290883239 | -0.738517197 | 0.156750718 | 9.881081921 | -1.62759377 | 0.15943506 | 0.709264768 | -4.448091836 |
| **A_55_P1975685** | Nup37 | 0.552763958 | -0.29797846 | 1.403506377 | 11.73228204 | 1.627391248 | 0.1594781 | 0.709264768 | -4.448146999 |
| **A_55_P1979403** | Tigar | 0.196233333 | -0.10579975 | 0.498266416 | 8.806449542 | 1.627303697 | 0.15949671 | 0.709264768 | -4.448170846 |
| **A_65_P17827** | Cenpl | 0.453159075 | -0.244375764 | 1.150693913 | 9.003903553 | 1.627178224 | 0.159523385 | 0.709264768 | -4.448205023 |
| **A_55_P2068822** | Cdk20 | 0.427691392 | -0.230718188 | 1.086100972 | 9.425154223 | 1.626989362 | 0.159563543 | 0.709264768 | -4.448256467 |
| **A_51_P200610** | Zbtb12 | 0.965527086 | -0.521417945 | 2.452472118 | 10.30095969 | 1.626372017 | 0.159694881 | 0.709264768 | -4.448424633 |
| **A_51_P489313** | Sri | -0.767039762 | -1.948539306 | 0.414459782 | 13.21178523 | -1.626052879 | 0.159762818 | 0.709264768 | -4.44851157 |
| **A_55_P2148268** | Xxylt1 | 0.191792464 | -0.103643328 | 0.487228256 | 9.009371701 | 1.625993388 | 0.159775485 | 0.709264768 | -4.448527777 |
| **A_55_P1984284** | Gapdh | 0.271391637 | -0.146707937 | 0.689491212 | 14.53933102 | 1.625799859 | 0.159816699 | 0.709264768 | -4.448580498 |
| **A_51_P455338** | Rnase2a | -0.386449714 | -0.98180609 | 0.208906661 | 9.59004119 | -1.625797036 | 0.1598173 | 0.709264768 | -4.448581267 |
| **A_51_P173100** | Pttg1ip | -0.568366752 | -1.444053782 | 0.307320279 | 11.73309008 | -1.625661396 | 0.159846193 | 0.709264768 | -4.448618219 |
| **A_55_P2056236** | Pdlim5 | -0.213829901 | -0.54330702 | 0.115647219 | 8.164197008 | -1.625524498 | 0.159875359 | 0.709264768 | -4.448655515 |
| **A_51_P357207** | Zfp518b | 0.202455189 | -0.109495524 | 0.514405902 | 8.501217957 | 1.625523584 | 0.159875553 | 0.709264768 | -4.448655763 |
| **A_55_P2143042** | Gm8096 | 0.299007953 | -0.161751496 | 0.759767401 | 8.859814807 | 1.625394686 | 0.159903019 | 0.709264768 | -4.44869088 |
| **A_55_P2055975** | Ppip5k1 | -0.310295026 | -0.788497781 | 0.167907728 | 8.752727527 | -1.625223474 | 0.159939509 | 0.709264768 | -4.448737525 |
| **A_55_P1968235** | Rab33b | -0.165806892 | -0.421339518 | 0.089725734 | 10.39778532 | -1.625199072 | 0.15994471 | 0.709264768 | -4.448744173 |
| **A_55_P2028600** | Myh9 | -0.484720703 | -1.231747621 | 0.262306215 | 15.00259469 | -1.625195695 | 0.15994543 | 0.709264768 | -4.448745093 |
| **A_55_P2031207** | Ggnbp2 | -0.164172705 | -0.417196 | 0.08885059 | 11.82954675 | -1.625140051 | 0.159957291 | 0.709264768 | -4.448760253 |
| **A_52_P418489** | Tinagl1 | -1.1035119 | -2.804353482 | 0.597329682 | 9.776783066 | -1.625037928 | 0.159979062 | 0.709264768 | -4.448788075 |
| **A_55_P2026291** | Smarce1 | 0.718685403 | -0.389028186 | 1.826398992 | 11.99507454 | 1.625031135 | 0.159980511 | 0.709264768 | -4.448789926 |
| **A_55_P2090708** | Fam207a | 0.139489567 | -0.075561788 | 0.354540922 | 12.3042624 | 1.624612862 | 0.160069711 | 0.709264768 | -4.448903886 |
| **A_51_P192847** | Ghdc | -0.33761927 | -0.858137285 | 0.182898746 | 8.410154534 | -1.624582633 | 0.16007616 | 0.709264768 | -4.448912122 |
| **A_55_P2079089** | Fuca1 | -0.218663454 | -0.555867793 | 0.118540886 | 11.24989858 | -1.624177141 | 0.160162684 | 0.709264768 | -4.449022604 |
| **A_51_P484764** | Fam3a | -0.387598634 | -0.98542383 | 0.210226561 | 10.12560147 | -1.623896588 | 0.160222576 | 0.709264768 | -4.449099047 |
| **A_66_P108672** | Mtg1 | 0.183592673 | -0.099611981 | 0.466797327 | 10.15375858 | 1.623698183 | 0.160264944 | 0.709264768 | -4.449153108 |
| **A_55_P2051094** | Rorc | 0.753202267 | -0.408824608 | 1.915229141 | 9.02732197 | 1.623475697 | 0.160312467 | 0.709264768 | -4.449213733 |
| **A_51_P227004** | Cks1b | 1.139089866 | -0.618422054 | 2.896601785 | 12.93026724 | 1.62334211 | 0.160341008 | 0.709264768 | -4.449250134 |
| **A_55_P2137206** | Tgfbr1 | 0.180625799 | -0.098065638 | 0.459317237 | 10.34770923 | 1.623328832 | 0.160343845 | 0.709264768 | -4.449253752 |
| **A_55_P2002849** | Aifm3 | -0.65028105 | -1.653633782 | 0.353071681 | 9.395026846 | -1.62329633 | 0.16035079 | 0.709264768 | -4.449262609 |
| **A_51_P375431** | Phip | 0.322178005 | -0.174950612 | 0.819306623 | 8.621238088 | 1.623220318 | 0.160367033 | 0.709264768 | -4.449283322 |
| **A_55_P2099890** | Atxn7 | 0.260179522 | -0.141289495 | 0.661648539 | 13.0179915 | 1.623197716 | 0.160371863 | 0.709264768 | -4.449289481 |
| **A_55_P2150309** | Lhfpl4 | -0.2007335 | -0.510486272 | 0.109019273 | 11.12135239 | -1.623136335 | 0.160384981 | 0.709264768 | -4.449306207 |
| **A_55_P1971076** | Atp11a | -0.502509998 | -1.278007896 | 0.2729879 | 11.75004057 | -1.622984706 | 0.160417392 | 0.709264768 | -4.449347526 |
| **A_55_P2111444** | Trpv6 | -0.923058512 | -2.34758114 | 0.501464115 | 8.709629598 | -1.622969003 | 0.160420748 | 0.709264768 | -4.449351805 |
| **A_55_P2169721** | Rps17 | 0.352881199 | -0.191785282 | 0.89754768 | 15.94673397 | 1.62273729 | 0.16047029 | 0.709264768 | -4.449414948 |
| **A_51_P394115** | Aadac | 0.258721848 | -0.140645479 | 0.658089176 | 8.017389949 | 1.622597934 | 0.160500092 | 0.709264768 | -4.449452924 |
| **A_55_P2111293** | 3110009E | 0.279375529 | -0.151957912 | 0.71070897 | 8.956399395 | 1.62227912 | 0.160568292 | 0.709264768 | -4.449539807 |
| **A_55_P1974421** | Itga3 | -0.580760843 | -1.477506113 | 0.315984428 | 8.953276154 | -1.62210356 | 0.16060586 | 0.709264768 | -4.449587651 |

| | | | | | | | | | |
|---|---|---|---|---|---|---|---|---|---|
| **A_55_P1952256** | Ccnb1 | 0.320517377 | -0.17439497 | 0.815429723 | 8.274626842 | 1.622085107 | 0.160609809 | 0.709264768 | -4.44959268 |
| **A_55_P2060793** | NA | 1.1463465 | -0.623753378 | 2.916446378 | 13.65576062 | 1.622065857 | 0.160613929 | 0.709264768 | -4.449597926 |
| **A_55_P2053444** | Map4 | -0.273232763 | -0.695266125 | 0.148800599 | 11.75465598 | -1.621572368 | 0.16071958 | 0.709264768 | -4.449732419 |
| **A_51_P193302** | Mrps7 | 0.217631841 | -0.11855144 | 0.553815122 | 9.867959183 | 1.621424273 | 0.160751299 | 0.709264768 | -4.449772781 |
| **A_55_P2080708** | Cstf2 | 0.455271969 | -0.248043073 | 1.158587012 | 9.129513154 | 1.621329749 | 0.160771548 | 0.709264768 | -4.449798543 |
| **A_55_P2026753** | Zmat1 | -0.589662812 | -1.500595636 | 0.321270012 | 9.027695315 | -1.621316242 | 0.160774441 | 0.709264768 | -4.449802224 |
| **A_51_P252784** | Fam216a | 0.724598369 | -0.42487048 | 1.874067217 | 11.2674622 | 1.679046993 | 0.160780942 | 0.709264768 | -4.465933056 |
| **A_55_P2008891** | Tmsb15b1 | 0.247578555 | -0.134926709 | 0.630083818 | 8.589367689 | 1.621160276 | 0.160807857 | 0.709264768 | -4.449844733 |
| **A_55_P2154684** | Smarca4 | 0.191525334 | -0.104405882 | 0.48745655 | 13.41477738 | 1.621010379 | 0.160839979 | 0.709264768 | -4.449885588 |
| **A_51_P511899** | Mthfsd | -0.230207475 | -0.585934018 | 0.125519069 | 8.387998287 | -1.620889816 | 0.16086582 | 0.709264768 | -4.449918448 |
| **A_52_P116628** | Fam234b | 0.269637215 | -0.147097065 | 0.686371495 | 8.297389667 | 1.620581781 | 0.16093186 | 0.709264768 | -4.450002407 |
| **A_55_P2067533** | Foxa2 | 1.016797924 | -0.554768643 | 2.588364492 | 10.4149032 | 1.620511865 | 0.160946853 | 0.709264768 | -4.450021464 |
| **A_55_P2320263** | NA | -0.360887971 | -0.918712129 | 0.196936188 | 11.48502656 | -1.620411933 | 0.160968285 | 0.709264768 | -4.450048703 |
| **A_55_P2032760** | Ctif | -0.440826116 | -1.122370311 | 0.240718079 | 10.18371008 | -1.620032167 | 0.161049758 | 0.709264768 | -4.450152218 |
| **A_52_P462125** | Smarca5- | 0.177011568 | -0.096683563 | 0.450706699 | 10.9474882 | 1.619887759 | 0.161080749 | 0.709264768 | -4.450191581 |
| **A_51_P283344** | ccdc198 | 0.257710823 | -0.140831633 | 0.656253279 | 8.114080211 | 1.619602395 | 0.161142007 | 0.709264768 | -4.450269369 |
| **A_55_P2010469** | NA | 0.710476044 | -0.388324973 | 1.809277061 | 12.21052096 | 1.619499177 | 0.16116417 | 0.709264768 | -4.450297506 |
| **A_51_P381873** | Bub3 | 0.650075367 | -0.355362303 | 1.655513037 | 12.09600284 | 1.619417778 | 0.16118165 | 0.709264768 | -4.450319695 |
| **A_55_P2042723** | Gm2769 | 0.227107683 | -0.124197121 | 0.578412486 | 8.4875733 | 1.619191009 | 0.161230358 | 0.709264768 | -4.450381512 |
| **A_51_P196925** | Cx3cl1 | -0.545495956 | -1.389409778 | 0.298417866 | 9.459362591 | -1.618988563 | 0.161273853 | 0.709264768 | -4.450436701 |
| **A_51_P189733** | Sult2a8 | 0.199969239 | -0.109402625 | 0.509341103 | 9.278579352 | 1.618947351 | 0.161282709 | 0.709264768 | -4.450447936 |
| **A_52_P284889** | Prkcz | -0.167798842 | -0.427421483 | 0.091823798 | 9.592308004 | -1.618813225 | 0.161311534 | 0.709264768 | -4.4504845 |
| **A_66_P124755** | Fut8 | -0.595550168 | -1.517194718 | 0.326094382 | 10.01553935 | -1.618472177 | 0.16138485 | 0.709264768 | -4.450577476 |
| **A_55_P1974452** | Lime1 | -0.18542027 | -0.472384013 | 0.101543472 | 9.441723065 | -1.618380092 | 0.161404652 | 0.709264768 | -4.450602581 |
| **A_51_P389957** | Rgs14 | 0.346971473 | -0.190086743 | 0.88402969 | 8.165301314 | 1.618164803 | 0.161450955 | 0.709264768 | -4.450661275 |
| **A_52_P556140** | Vapb | -0.245641899 | -0.625863559 | 0.134579761 | 12.32218891 | -1.618139421 | 0.161456415 | 0.709264768 | -4.450668195 |
| **A_55_P2055869** | Dvl2 | 0.356680949 | -0.195417579 | 0.908779478 | 9.511213407 | 1.618130969 | 0.161458233 | 0.709264768 | -4.4506705 |
| **A_55_P2118810** | NA | 0.464826828 | -0.254706833 | 1.184360489 | 11.74718921 | 1.618044328 | 0.161476872 | 0.709264768 | -4.450694121 |
| **A_51_P483280** | Prnp | -0.676308566 | -1.723267046 | 0.370649914 | 11.71300834 | -1.617952528 | 0.161496623 | 0.709264768 | -4.450719149 |
| **A_55_P2032916** | Slc6a20a | 0.539654008 | -0.295775834 | 1.37508385 | 11.23270403 | 1.617915247 | 0.161504645 | 0.709264768 | -4.450729313 |
| **A_55_P2011943** | Gramd1a | -0.450703998 | -1.148627898 | 0.247219902 | 11.17923368 | -1.617460506 | 0.161602523 | 0.709264768 | -4.450853296 |
| **A_55_P2000409** | Rab44 | 0.353593706 | -0.193970451 | 0.901157864 | 8.12370662 | 1.617409014 | 0.16161361 | 0.709264768 | -4.450867335 |
| **A_55_P2095732** | Fbrsl1 | -0.530247122 | -1.351391379 | 0.290897135 | 8.909853196 | -1.617369339 | 0.161622153 | 0.709264768 | -4.450878153 |
| **A_51_P385697** | Mgat3 | 0.322996569 | -0.177207465 | 0.823200602 | 9.451586546 | 1.617339022 | 0.161628681 | 0.709264768 | -4.450886419 |
| **A_55_P2001438** | Coq2 | -0.272383074 | -0.694262577 | 0.149496428 | 10.54622742 | -1.617119215 | 0.16167602 | 0.709264768 | -4.450946351 |
| **A_55_P2033958** | Sltm | 0.297200737 | -0.163167786 | 0.757569261 | 11.21586375 | 1.616942611 | 0.161714065 | 0.709264768 | -4.450994504 |
| **A_55_P1962419** | Ccar1 | 0.280035521 | -0.15374545 | 0.713816491 | 9.795500036 | 1.616936524 | 0.161715376 | 0.709264768 | -4.450996164 |
| **A_55_P1961499** | Ly6c1 | -1.190365302 | -3.034357691 | 0.653627088 | 9.919765546 | -1.616856512 | 0.161732616 | 0.709264768 | -4.45101798 |
| **A_51_P336874** | Atp6v0e | -0.415672375 | -1.05970911 | 0.228364361 | 13.77212897 | -1.616556426 | 0.161797288 | 0.709264768 | -4.451099805 |
| **A_55_P2178608** | NA | 0.222681203 | -0.122345529 | 0.567707935 | 8.609255563 | 1.616520289 | 0.161805078 | 0.709264768 | -4.451109659 |
| **A_55_P2067952** | NA | 0.381786312 | -0.20981909 | 0.973391715 | 11.6823154 | 1.616361765 | 0.161839254 | 0.709264768 | -4.451152884 |
| **A_55_P1990235** | Bin3 | -0.188313343 | -0.487998552 | 0.111371866 | 11.61562725 | -1.673703365 | 0.161843412 | 0.709264768 | -4.467025978 |
| **A_55_P2095688** | Sh3bp5l | -0.343006475 | -0.874722747 | 0.188709796 | 12.40123085 | -1.615744602 | 0.161972373 | 0.709264768 | -4.451321177 |
| **A_52_P11441** | Rab6b | -0.882250058 | -2.249886179 | 0.485386062 | 11.0996665 | -1.615740009 | 0.161973364 | 0.709264768 | -4.45132243 |
| **A_55_P2093974** | Mphosph1 | 0.243012578 | -0.133813208 | 0.619838364 | 8.77421677 | 1.615245294 | 0.162080149 | 0.709264768 | -4.45145734 |
| **A_55_P2126870** | Nsf | -0.226363809 | -0.577567968 | 0.124840351 | 8.281873269 | -1.61434996 | 0.162273582 | 0.709264768 | -4.451701516 |
| **A_55_P1959703** | Zfp646 | 0.242638289 | -0.133923992 | 0.619200571 | 15.54310239 | 1.613886039 | 0.162373899 | 0.709264768 | -4.451828045 |
| **A_55_P2018549** | Gcsh | 0.350860959 | -0.193715905 | 0.895437822 | 11.84237996 | 1.613712655 | 0.162411406 | 0.709264768 | -4.451875335 |

| | | | | | | | | | |
|---|---|---|---|---|---|---|---|---|---|
| **A_55_P2440032** | Usp24 | -0.184606767 | -0.471227854 | 0.10201432 | 8.222158737 | -1.613205982 | 0.16252106 | 0.709264768 | -4.452013533 |
| **A_66_P134481** | Skp2 | 0.604731662 | -0.334525481 | 1.543988805 | 9.143786864 | 1.612607022 | 0.16265078 | 0.709264768 | -4.452176911 |
| **A_55_P2059904** | Chsy1 | 0.529284729 | -0.292932428 | 1.351501887 | 11.6515296 | 1.612327178 | 0.162711422 | 0.709264768 | -4.452253247 |
| **A_51_P425696** | Nt5c2 | 0.146973065 | -0.081377501 | 0.375323632 | 8.848313248 | 1.612077825 | 0.162765475 | 0.709264768 | -4.452321267 |
| **A_55_P2162910** | Rtn1 | -0.748361467 | -1.911145635 | 0.414422701 | 9.818461668 | -1.611991151 | 0.162784267 | 0.709264768 | -4.452344911 |
| **A_55_P2131920** | Prrx1 | 0.168266282 | -0.093182202 | 0.429714767 | 9.13485289 | 1.611986191 | 0.162785343 | 0.709264768 | -4.452346265 |
| **A_52_P400224** | Ccs | -0.162957776 | -0.41616153 | 0.090245978 | 11.81053984 | -1.611963835 | 0.162790191 | 0.709264768 | -4.452352363 |
| **A_55_P2043009** | Psmg2 | 0.163933896 | -0.090815601 | 0.418683393 | 11.04966044 | 1.611780028 | 0.162830052 | 0.709264768 | -4.452402505 |
| **A_55_P2151952** | NA | 0.494049904 | -0.273883793 | 1.2619836 | 14.37464198 | 1.611378036 | 0.162917263 | 0.709264768 | -4.45251217 |
| **A_55_P1987600** | Trmt112 | 0.438184078 | -0.242941812 | 1.119309968 | 13.15219405 | 1.611311661 | 0.162931667 | 0.709264768 | -4.452530277 |
| **A_55_P2165314** | Trio | -0.230636814 | -0.589192717 | 0.127919089 | 10.14032432 | -1.611098518 | 0.16297793 | 0.709264768 | -4.452588426 |
| **A_55_P1960566** | Ybx3 | 0.335293373 | -0.18607264 | 0.856659387 | 13.02700615 | 1.610766537 | 0.163050013 | 0.709264768 | -4.452678997 |
| **A_55_P1963109** | Prh1 | 0.177006254 | -0.098233373 | 0.452245882 | 8.565994078 | 1.610749466 | 0.16305372 | 0.709264768 | -4.452683654 |
| **A_51_P509263** | Hoxa7 | 0.475315536 | -0.263889477 | 1.21452055 | 9.493953918 | 1.61052493 | 0.163102492 | 0.709264768 | -4.452744914 |
| **A_55_P1991351** | S100a13 | -0.273483655 | -0.69882247 | 0.151855161 | 11.78737819 | -1.610447988 | 0.163119208 | 0.709264768 | -4.452765906 |
| **A_52_P141338** | Acot10 | -0.171716718 | -0.438803385 | 0.095369948 | 8.433260574 | -1.610314555 | 0.163148201 | 0.709264768 | -4.452802311 |
| **A_51_P122845** | Naa50 | 0.207641534 | -0.11535489 | 0.530637958 | 9.676603689 | 1.610151943 | 0.16318354 | 0.709264768 | -4.460701565 |
| **A_55_P2055134** | NA | 0.203185619 | -0.112889056 | 0.519260293 | 10.97152417 | 1.610102821 | 0.163194217 | 0.709264768 | -4.45286008 |
| **A_55_P1973046** | Cttnbp2nl | -0.532000648 | -1.359584607 | 0.295583312 | 10.35406895 | -1.610091073 | 0.163196771 | 0.709264768 | -4.452863286 |
| **A_55_P1957204** | NA | 0.211326593 | -0.117429558 | 0.540082744 | 17.19121195 | 1.610017551 | 0.163212753 | 0.709264768 | -4.452883345 |
| **A_51_P480233** | Nmral1 | 0.960999959 | -0.534016214 | 2.456016132 | 9.751614377 | 1.610007221 | 0.163214998 | 0.709264768 | -4.452886164 |
| **A_55_P2103706** | NA | 1.167252562 | -0.648774712 | 2.983279836 | 10.43367719 | 1.609877441 | 0.163243213 | 0.709264768 | -4.452921574 |
| **A_51_P127841** | Pdss1 | 0.211119428 | -0.117404002 | 0.539642857 | 8.52568999 | 1.60957863 | 0.163308195 | 0.709264768 | -4.460847486 |
| **A_52_P284495** | Ankrd33b | -0.94116272 | -2.445687807 | 0.563362368 | 8.460927949 | -1.666203289 | 0.163346761 | 0.709264768 | -4.468562394 |
| **A_55_P2164629** | Ptn | 0.585676174 | -0.325834534 | 1.497186882 | 10.55839542 | 1.609333781 | 0.16336146 | 0.709264768 | -4.453069914 |
| **A_52_P482897** | Areg | 0.746678303 | -0.415507038 | 1.908863644 | 8.931941199 | 1.609194295 | 0.163391812 | 0.709264768 | -4.453107974 |
| **A_51_P128775** | Zfp84 | 0.490373596 | -0.272937461 | 1.253684652 | 9.246455606 | 1.609073454 | 0.163418111 | 0.709264768 | -4.46425379 |
| **A_52_P609120** | Pdxp | 0.184541274 | -0.102722448 | 0.471804996 | 8.531684525 | 1.609026053 | 0.163428428 | 0.709264768 | -4.453153882 |
| **A_55_P2032128** | Dnase1l1 | -0.622448877 | -1.591483501 | 0.346585747 | 8.428542823 | -1.608846976 | 0.163467411 | 0.709264768 | -4.453202747 |
| **A_66_P135694** | NA | 0.349787489 | -0.194783337 | 0.894358315 | 16.50636756 | 1.608793288 | 0.1634791 | 0.709264768 | -4.453217396 |
| **A_55_P1985015** | Pard3 | -0.335925378 | -0.858952096 | 0.187101339 | 10.15382847 | -1.608678606 | 0.163504072 | 0.709264768 | -4.453248691 |
| **A_55_P2101591** | Vta1 | -0.196692154 | -0.502958821 | 0.109574514 | 9.269761503 | -1.608561417 | 0.163529593 | 0.709264768 | -4.453280669 |
| **A_55_P2129975** | Rabgef1 | -0.585193282 | -1.49646276 | 0.326076195 | 10.291858 | -1.60843255 | 0.163557662 | 0.709264768 | -4.453315834 |
| **A_52_P294253** | Zfp41 | 0.372273206 | -0.207554597 | 0.952101009 | 9.27018319 | 1.608100135 | 0.163630088 | 0.709264768 | -4.453406546 |
| **A_52_P38639** | Fermt3 | 0.258215455 | -0.143995084 | 0.660425994 | 8.075382986 | 1.607974407 | 0.163657489 | 0.709264768 | -4.453440856 |
| **A_55_P2060996** | NA | -0.144803051 | -0.370357129 | 0.080751028 | 8.394490902 | -1.607967893 | 0.163658909 | 0.709264768 | -4.453442634 |
| **A_51_P469008** | Ehd1 | -1.036416653 | -2.650921437 | 0.57808813 | 13.59413728 | -1.607849489 | 0.163684719 | 0.709264768 | -4.453474946 |
| **A_51_P344566** | Plk1 | 1.530288694 | -0.853679787 | 3.914257175 | 11.45434326 | 1.607767354 | 0.163702625 | 0.709264768 | -4.453497361 |
| **A_55_P2059864** | Igsf1 | -1.105390421 | -2.827444755 | 0.616663913 | 9.49590434 | -1.607752495 | 0.163705864 | 0.709264768 | -4.453501416 |
| **A_55_P2006703** | Bmp1 | -0.310210133 | -0.793510943 | 0.173090677 | 12.59843084 | -1.607639994 | 0.163730394 | 0.709264768 | -4.453532117 |
| **A_55_P2077497** | NA | -0.28683807 | -0.73379309 | 0.160116951 | 11.6176733 | -1.607397605 | 0.163783256 | 0.709264768 | -4.453598267 |
| **A_55_P2028773** | Slc22a17 | -0.290861699 | -0.744124764 | 0.162401366 | 10.67169263 | -1.607261533 | 0.163812939 | 0.709264768 | -4.453635402 |
| **A_55_P1963061** | Rps16 | 0.162148139 | -0.090549224 | 0.414845501 | 16.96215966 | 1.607169211 | 0.163833081 | 0.709264768 | -4.453660598 |
| **A_55_P2270412** | C230096K | 0.23644587 | -0.13204402 | 0.604935761 | 9.544180117 | 1.607150327 | 0.163837202 | 0.709264768 | -4.453665752 |
| **A_55_P2038947** | Lig3 | 0.205549814 | -0.114813256 | 0.525912884 | 9.28373861 | 1.607033708 | 0.163862649 | 0.709264768 | -4.453697579 |
| **A_55_P2044463** | NA | 0.201691067 | -0.112686456 | 0.516068589 | 8.483220308 | 1.606887682 | 0.163894518 | 0.709264768 | -4.453737433 |
| **A_55_P2084652** | H2ac15 | 1.008099783 | -0.563299931 | 2.579499497 | 16.99662345 | 1.606819883 | 0.163909317 | 0.709264768 | -4.453755937 |
| **A_55_P1995205** | Top2a | 1.313532129 | -0.734302532 | 3.36136679 | 10.46727018 | 1.606557064 | 0.163966697 | 0.709264768 | -4.453827667 |

| | | | | | | | | | |
|---|---|---|---|---|---|---|---|---|---|
| **A_55_P2436438** | Dusp22 | -0.171943698 | -0.44001432 | 0.096126923 | 8.406090974 | -1.606524628 | 0.163973779 | 0.709264768 | -4.45383652 |
| **A_51_P327632** | Igsf3 | 0.44470323 | -0.248635705 | 1.138042165 | 10.64796082 | 1.606478955 | 0.163983753 | 0.709264768 | -4.453848985 |
| **A_51_P202074** | Ncapd2 | 1.396098591 | -0.780633715 | 3.572830897 | 12.1752388 | 1.606428541 | 0.163994763 | 0.709264768 | -4.453862745 |
| **A_55_P2156002** | Sumo2 | 0.579965329 | -0.324319055 | 1.484249712 | 14.97901302 | 1.606376515 | 0.164006126 | 0.709264768 | -4.453876945 |
| **A_55_P1984366** | Cdadc1 | -0.200279075 | -0.512687454 | 0.112129304 | 9.112824422 | -1.605695721 | 0.164154884 | 0.709264768 | -4.454062763 |
| **A_55_P2049282** | Ly6h | 0.536588742 | -0.300591511 | 1.373768994 | 8.764574751 | 1.605361806 | 0.164227894 | 0.709264768 | -4.454153907 |
| **A_55_P2171413** | Me1 | -0.467623131 | -1.197394808 | 0.262148547 | 10.49846526 | -1.604941941 | 0.164319742 | 0.709264768 | -4.454268515 |
| **A_51_P369200** | Tpx2 | 1.13363005 | -0.635704084 | 2.902964185 | 10.4094842 | 1.604766461 | 0.164358144 | 0.709264768 | -4.454316416 |
| **A_55_P2102240** | Nprl3 | -0.157831713 | -0.404188899 | 0.088525472 | 8.982563928 | -1.604646536 | 0.164384393 | 0.709264768 | -4.454349152 |
| **A_51_P382253** | Rnf2 | 0.200461044 | -0.11244035 | 0.513362439 | 10.88249941 | 1.604622347 | 0.164389688 | 0.709264768 | -4.454355756 |
| **A_65_P04980** | Ilf3 | 0.639878894 | -0.359249489 | 1.639007276 | 10.44754087 | 1.604082993 | 0.164507797 | 0.709264768 | -4.454502991 |
| **A_66_P116360** | NA | 0.257573218 | -0.144625577 | 0.659772013 | 8.439946087 | 1.604021867 | 0.164521188 | 0.709264768 | -4.454519678 |
| **A_55_P2104472** | NA | 0.664381151 | -0.373066641 | 1.701828942 | 14.20264216 | 1.603989044 | 0.164528379 | 0.709264768 | -4.454528638 |
| **A_55_P2015862** | Erg | 0.286410103 | -0.160879004 | 0.733699209 | 8.628816093 | 1.603800548 | 0.16456968 | 0.709264768 | -4.454580097 |
| **A_55_P2065824** | Smtn | -0.46170792 | -1.182762058 | 0.259346219 | 12.1675633 | -1.603798474 | 0.164570135 | 0.709264768 | -4.454580663 |
| **A_55_P1973011** | Ppm1l | 0.50562948 | -0.284051856 | 1.295310815 | 9.683386822 | 1.603728543 | 0.16458546 | 0.709264768 | -4.454599754 |
| **A_55_P1967534** | NA | 0.277355494 | -0.155996173 | 0.710707162 | 14.60144596 | 1.603048446 | 0.164734574 | 0.709264768 | -4.454785426 |
| **A_52_P104824** | Diaph3 | 0.51498072 | -0.289690476 | 1.319651915 | 9.32032508 | 1.602960645 | 0.164753835 | 0.709264768 | -4.454809397 |
| **A_51_P288447** | Nup133 | 0.410880462 | -0.231192579 | 1.052953503 | 10.95849402 | 1.602807628 | 0.164787406 | 0.709264768 | -4.454851174 |
| **A_55_P2118575** | Tgs1 | 0.267762403 | -0.150733268 | 0.686258073 | 9.733343407 | 1.602540346 | 0.164846063 | 0.709264768 | -4.454924148 |
| **A_55_P2022074** | Klf10 | -0.21219615 | -0.543864711 | 0.119472412 | 10.98239751 | -1.602446498 | 0.164866663 | 0.709264768 | -4.454949772 |
| **A_52_P628067** | Cdca3 | 1.591927314 | -0.896339148 | 4.080193777 | 11.81327262 | 1.602421244 | 0.164872207 | 0.709264768 | -4.454956667 |
| **A_52_P239707** | Nfyb | 0.377941602 | -0.212833028 | 0.968716233 | 9.054460049 | 1.602334591 | 0.164891231 | 0.709264768 | -4.454980326 |
| **A_55_P1989981** | Fam129b | -0.472559333 | -1.211350167 | 0.266231502 | 14.71850773 | -1.602083661 | 0.164946332 | 0.709264768 | -4.455048839 |
| **A_51_P296416** | Orc6 | 0.404311672 | -0.227800131 | 1.036423475 | 11.47197122 | 1.602037674 | 0.164956433 | 0.709264768 | -4.455061395 |
| **A_55_P2169621** | Prorsd1 | -0.222146004 | -0.569483009 | 0.125191001 | 15.72917565 | -1.60190883 | 0.164984734 | 0.709264768 | -4.455096575 |
| **A_55_P2144126** | NA | -0.46943319 | -1.20342098 | 0.2645546 | 15.20860572 | -1.601899634 | 0.164986754 | 0.709264768 | -4.455099086 |
| **A_55_P1965079** | Gm10499 | 0.539201282 | -0.304100182 | 1.382502746 | 8.712220697 | 1.601468524 | 0.165081485 | 0.709264768 | -4.4552168 |
| **A_51_P341918** | Tsc22d1 | -0.447148863 | -1.146685956 | 0.25238823 | 13.55591363 | -1.601001462 | 0.165184175 | 0.709264768 | -4.455344335 |
| **A_51_P405606** | Ndrg1 | -1.401208709 | -3.593359265 | 0.790941847 | 12.63412287 | -1.600968531 | 0.165191418 | 0.709264768 | -4.455353328 |
| **A_55_P1979397** | Palm | 0.159777415 | -0.090199221 | 0.409754051 | 11.02293462 | 1.600907784 | 0.165204779 | 0.709264768 | -4.455369916 |
| **A_52_P346458** | Tollip | -0.238618077 | -0.611980401 | 0.134744247 | 11.16357065 | -1.60074884 | 0.165239744 | 0.709264768 | -4.455413319 |
| **A_51_P365903** | Ankzf1 | -0.218668999 | -0.560877554 | 0.123539555 | 9.645128804 | -1.600466915 | 0.165301779 | 0.709264768 | -4.455490305 |
| **A_55_P2064984** | Cenpi | 0.681175174 | -0.38484004 | 1.747190388 | 9.162412746 | 1.600463425 | 0.165302547 | 0.709264768 | -4.455491258 |
| **A_55_P2083401** | Acp1 | 0.298552338 | -0.168672802 | 0.765777477 | 11.46246895 | 1.600459244 | 0.165303467 | 0.709264768 | -4.4554924 |
| **A_51_P228171** | Cenpp | 0.90981062 | -0.514117319 | 2.333738559 | 9.746648645 | 1.600343964 | 0.16532884 | 0.709264768 | -4.455523881 |
| **A_55_P2168426** | Pced1b | -0.715504112 | -1.835491389 | 0.404483165 | 9.388175366 | -1.600108346 | 0.165380712 | 0.709264768 | -4.455588224 |
| **A_55_P2118799** | Rmi1 | 0.208523752 | -0.117902493 | 0.534949997 | 8.630491209 | 1.600003035 | 0.165403901 | 0.709264768 | -4.455616983 |
| **A_52_P231737** | Actl6a | 0.362931903 | -0.205391351 | 0.931255157 | 9.984346679 | 1.599484607 | 0.165518104 | 0.709264768 | -4.455758563 |
| **A_55_P1966109** | Epn2 | -0.412060543 | -1.057374897 | 0.233253811 | 10.37739601 | -1.599337243 | 0.165550581 | 0.709264768 | -4.455798809 |
| **A_51_P465740** | Lysmd4 | -0.160258559 | -0.411242269 | 0.09072515 | 9.310684956 | -1.599285671 | 0.165561948 | 0.709264768 | -4.455812893 |
| **A_55_P2125551** | Mmadhc | 0.161113081 | -0.091210661 | 0.413436823 | 13.00754143 | 1.599274569 | 0.165564395 | 0.709264768 | -4.455815926 |
| **A_55_P2118090** | NA | 0.714194597 | -0.40435063 | 1.832739824 | 11.20847387 | 1.599238944 | 0.165572248 | 0.709264768 | -4.455825655 |
| **A_52_P637172** | Pyroxd2 | -0.413110302 | -1.060128893 | 0.233908289 | 8.288168007 | -1.599188327 | 0.165583406 | 0.709264768 | -4.455839479 |
| **A_51_P242076** | Ubr7 | 0.159884886 | -0.090588457 | 0.41035823 | 11.08161451 | 1.59880775 | 0.165667323 | 0.709264768 | -4.455943419 |
| **A_55_P2071811** | Eml2 | 0.276568717 | -0.168631916 | 0.721769349 | 8.80373651 | 1.654664645 | 0.165687492 | 0.709264768 | -4.4709316 |
| **A_55_P2096297** | NA | 0.340856282 | -0.193155697 | 0.874868262 | 8.856065426 | 1.598713524 | 0.165688106 | 0.709264768 | -4.455969154 |
| **A_55_P1999962** | Podn | -0.344530822 | -0.884303753 | 0.19524211 | 8.750463989 | -1.598701285 | 0.165690806 | 0.709264768 | -4.455972497 |

| | | | | | | | | | |
|---|---|---|---|---|---|---|---|---|---|
| **A_55_P1959338** | Spindoc | -0.330220614 | -0.847643212 | 0.187201983 | 10.72344336 | -1.598487097 | 0.165738059 | 0.709264768 | -4.456030997 |
| **A_55_P1963610** | NA | 0.319401774 | -0.181103434 | 0.819906981 | 15.54168036 | 1.598376437 | 0.165762477 | 0.709264768 | -4.456061221 |
| **A_52_P252737** | Rangap1 | 0.239619393 | -0.135875664 | 0.61511445 | 10.3024933 | 1.598336012 | 0.165771398 | 0.709264768 | -4.456072262 |
| **A_55_P1965154** | Spc25 | 1.041577241 | -0.590695442 | 2.673849925 | 12.98090566 | 1.598266199 | 0.165786805 | 0.709264768 | -4.45609133 |
| **A_51_P270949** | H1f5 | 2.357340477 | -1.338205821 | 6.052886775 | 13.05938671 | 1.597695498 | 0.16591281 | 0.709264768 | -4.456247209 |
| **A_55_P2142439** | Echdc2 | -0.243487969 | -0.625288452 | 0.138312513 | 9.86423099 | -1.59731798 | 0.165996213 | 0.709264768 | -4.456350328 |
| **A_51_P108489** | Cfap20 | -0.166727728 | -0.428189133 | 0.094733676 | 11.37026675 | -1.597167955 | 0.166029368 | 0.709264768 | -4.456391307 |
| **A_55_P2423513** | B230369F | -0.260499211 | -0.669031272 | 0.14803285 | 10.60440875 | -1.597094497 | 0.166045605 | 0.709264768 | -4.456411373 |
| **A_55_P2170076** | Taf9-ps | 0.584301164 | -0.332057392 | 1.50065972 | 12.39760767 | 1.597061565 | 0.166052884 | 0.709264768 | -4.456420369 |
| **A_51_P165984** | Aup1 | -0.279101493 | -0.716858511 | 0.158655525 | 9.84691058 | -1.596906131 | 0.166087246 | 0.709264768 | -4.456462827 |
| **A_55_P2162160** | Anp32a | 0.846098808 | -0.481027248 | 2.173224864 | 12.71093427 | 1.596832095 | 0.166103616 | 0.709264768 | -4.456483051 |
| **A_55_P2152956** | Plekhm3 | -0.248133402 | -0.637338653 | 0.141071848 | 8.514383575 | -1.596823392 | 0.16610554 | 0.709264768 | -4.456485428 |
| **A_52_P556082** | Trmt2b | 0.463696338 | -0.263690945 | 1.191083621 | 9.229743548 | 1.596681553 | 0.166136906 | 0.709264768 | -4.456524174 |
| **A_55_P2079599** | NA | 0.350974242 | -0.199675248 | 0.901623731 | 11.76798296 | 1.596431723 | 0.166192167 | 0.709264768 | -4.45659242 |
| **A_55_P2118520** | Col1a1 | 0.478448586 | -0.272239492 | 1.229136664 | 14.54972249 | 1.596342596 | 0.166211886 | 0.709264768 | -4.456616767 |
| **A_55_P1973809** | Hbb-bt | 0.227898884 | -0.129703625 | 0.585501394 | 17.77306571 | 1.596217203 | 0.166239632 | 0.709264768 | -4.456651021 |
| **A_55_P1969530** | Zfp146 | 0.159882949 | -0.091000032 | 0.410765929 | 9.133523384 | 1.596177899 | 0.16624833 | 0.709264768 | -4.456661758 |
| **A_55_P2279977** | NA | 0.216627565 | -0.123309122 | 0.556564252 | 8.320888139 | 1.596121791 | 0.166260747 | 0.709264768 | -4.456677086 |
| **A_55_P2013780** | NA | 1.161938755 | -0.661600317 | 2.985477826 | 13.78723944 | 1.595947166 | 0.1662994 | 0.709264768 | -4.45672479 |
| **A_51_P110759** | Slc1a1 | -0.458572632 | -1.178454884 | 0.261309621 | 8.251905376 | -1.595500741 | 0.166398252 | 0.709264768 | -4.458341508 |
| **A_55_P1990785** | Zranb3 | 0.449661641 | -0.256368074 | 1.155691357 | 9.093396834 | 1.595192896 | 0.166466452 | 0.709264768 | -4.456930851 |
| **A_55_P2462377** | Nsf | -0.520298663 | -1.337272254 | 0.296674929 | 11.07834216 | -1.595126158 | 0.166481241 | 0.709264768 | -4.456949084 |
| **A_66_P124925** | Pdcd5 | 0.368474729 | -0.210174623 | 0.947124081 | 12.83576446 | 1.594933511 | 0.166523937 | 0.709264768 | -4.457001717 |
| **A_55_P1961414** | NA | 0.406355784 | -0.231832357 | 1.044543925 | 13.20162928 | 1.594806737 | 0.16655204 | 0.709264768 | -4.457036352 |
| **A_55_P1952696** | NA | 0.941424404 | -0.537294419 | 2.420143228 | 13.28121017 | 1.59459426 | 0.166599151 | 0.709264768 | -4.457094404 |
| **A_51_P376347** | Hebp1 | -0.55660348 | -1.430920595 | 0.317713636 | 10.37673902 | -1.59451013 | 0.166617808 | 0.709264768 | -4.45711739 |
| **A_55_P2028394** | Atg13 | -0.674029143 | -1.732856342 | 0.384798055 | 11.6505669 | -1.5944244 | 0.166636822 | 0.709264768 | -4.457140813 |
| **A_51_P164420** | Eef1e1 | 0.275417288 | -0.157351191 | 0.708185767 | 9.286870279 | 1.593991211 | 0.166732932 | 0.709264768 | -4.45725917 |
| **A_55_P2056285** | Sf3b5 | 0.276658167 | -0.158063664 | 0.711379997 | 13.06149379 | 1.593978246 | 0.166735809 | 0.709264768 | -4.457262713 |
| **A_51_P338443** | Angptl4 | -0.291819171 | -0.750431965 | 0.166793624 | 11.75191575 | -1.593741927 | 0.166788263 | 0.709264768 | -4.457327283 |
| **A_52_P201551** | Nav1 | -0.301347585 | -0.774961313 | 0.172266143 | 9.189555123 | -1.593653051 | 0.166807995 | 0.709264768 | -4.457351567 |
| **A_52_P575178** | Cdon | 0.410491906 | -0.234797806 | 1.055781618 | 10.13282235 | 1.593309708 | 0.166884242 | 0.709264768 | -4.457445382 |
| **A_51_P179864** | Prkx | -0.230567494 | -0.593073221 | 0.131938234 | 10.01617158 | -1.593065204 | 0.16693856 | 0.709264768 | -4.457512192 |
| **A_51_P120295** | Cyp2d26 | 0.359354989 | -0.205639283 | 0.92434926 | 8.207063456 | 1.5930521 | 0.166941471 | 0.709264768 | -4.457515773 |
| **A_55_P1967224** | NA | 0.303547075 | -0.173758404 | 0.780852553 | 8.43932134 | 1.5928687 | 0.166982227 | 0.709264768 | -4.457565887 |
| **A_55_P2068731** | Gm21953 | -0.165524691 | -0.42582736 | 0.094777977 | 9.099354315 | -1.592701951 | 0.16701929 | 0.709264768 | -4.457611452 |
| **A_55_P2151398** | Tmem116 | -0.443385417 | -1.140781673 | 0.254010839 | 8.164700758 | -1.592399891 | 0.16708645 | 0.709264768 | -4.457693994 |
| **A_55_P2106265** | Mbnl2 | -0.449449547 | -1.156424381 | 0.257525287 | 9.222628359 | -1.592308958 | 0.167106673 | 0.709264768 | -4.457718843 |
| **A_51_P331207** | Tmem218 | -0.338953115 | -0.872176809 | 0.194270579 | 10.52978902 | -1.592137369 | 0.16714484 | 0.709264768 | -4.457765733 |
| **A_55_P2185489** | Zfp865 | 0.206010698 | -0.118172778 | 0.530194174 | 9.150935443 | 1.591656123 | 0.167251929 | 0.709264768 | -4.457897246 |
| **A_51_P363630** | Snrpe | 0.655597201 | -0.376092433 | 1.687286834 | 14.50148063 | 1.591616268 | 0.167260801 | 0.709264768 | -4.457908138 |
| **A_55_P1966620** | Rab5a | -0.349330198 | -0.899136386 | 0.200475991 | 9.778421343 | -1.59139083 | 0.167310992 | 0.709264768 | -4.457969747 |
| **A_55_P1963124** | NA | 0.460268675 | -0.264233966 | 1.184771317 | 15.50219347 | 1.591189086 | 0.167355921 | 0.709264768 | -4.458024881 |
| **A_55_P2007389** | Fam189b | 0.294620865 | -0.169213666 | 0.758455395 | 9.946125086 | 1.590928891 | 0.167413883 | 0.709264768 | -4.458095991 |
| **A_55_P2153292** | Tubb4b | 0.360052455 | -0.20685622 | 0.926961131 | 14.87410807 | 1.590753983 | 0.167452857 | 0.709264768 | -4.458143793 |
| **A_52_P304323** | Isca1 | -0.369197582 | -0.950689262 | 0.212294097 | 9.668451189 | -1.590251045 | 0.167564975 | 0.709264768 | -4.45828125 |
| **A_55_P1973447** | NA | 0.490217351 | -0.281890475 | 1.262325176 | 8.627344988 | 1.59023413 | 0.167568747 | 0.709264768 | -4.458285873 |
| **A_55_P2090394** | NA | 0.23449333 | -0.134869901 | 0.603856561 | 12.78198663 | 1.590110039 | 0.167596422 | 0.709264768 | -4.458319789 |

| | | | | | | | | | |
|---|---|---|---|---|---|---|---|---|---|
| **A_66_P106585** | Sae1 | 0.43725174 | -0.251498747 | 1.126002228 | 10.53246051 | 1.590083654 | 0.167602307 | 0.709264768 | -4.458327001 |
| **A_55_P1962084** | H2ac18 | -1.57868737 | -4.065907507 | 0.908532767 | 11.73286931 | -1.589762524 | 0.167673948 | 0.709264768 | -4.458414772 |
| **A_52_P574945** | 1500026H | -0.179034318 | -0.46110626 | 0.103037625 | 8.124966915 | -1.589742341 | 0.167678452 | 0.709264768 | -4.458420289 |
| **A_66_P130601** | Foxp2 | 0.724312556 | -0.416882112 | 1.865507223 | 9.019419931 | 1.589705313 | 0.167686715 | 0.709264768 | -4.45843041 |
| **A_51_P186798** | NA | -0.558229273 | -1.437801909 | 0.321343362 | 8.261490994 | -1.589612404 | 0.167707449 | 0.709264768 | -4.458455804 |
| **A_55_P2136334** | NA | 0.803380445 | -0.462498903 | 2.069259793 | 11.86631884 | 1.589568644 | 0.167717216 | 0.709264768 | -4.458467765 |
| **A_52_P552092** | Adat1 | 0.207067636 | -0.119210248 | 0.53334552 | 8.521024848 | 1.589552708 | 0.167720773 | 0.709264768 | -4.458472121 |
| **A_51_P373461** | Brcc3 | 0.356468502 | -0.205295331 | 0.918232335 | 10.7113413 | 1.589343358 | 0.167767505 | 0.709264768 | -4.458529342 |
| **A_51_P427624** | Pgc | -0.281227808 | -0.724436165 | 0.161980548 | 8.33294084 | -1.589280913 | 0.167781447 | 0.709264768 | -4.458546411 |
| **A_55_P2124120** | Ywhae | 0.350509236 | -0.201925296 | 0.902943768 | 14.84507808 | 1.589165007 | 0.167807328 | 0.709264768 | -4.458578092 |
| **A_55_P1984168** | Casp4 | -0.698130786 | -1.79845081 | 0.402189238 | 8.620151947 | -1.589161808 | 0.167808043 | 0.709264768 | -4.458578966 |
| **A_52_P110070** | 5730416F | -0.442507612 | -1.13997815 | 0.254962927 | 8.79500251 | -1.589078031 | 0.167826752 | 0.709264768 | -4.458601866 |
| **A_55_P2082986** | Hmgb1 | 1.143157196 | -0.659492028 | 2.94580642 | 13.69957622 | 1.588345856 | 0.167990349 | 0.709264768 | -4.458802004 |
| **A_55_P1963737** | Aktip | -0.225442454 | -0.580991812 | 0.130106903 | 8.613325717 | -1.588130369 | 0.168038526 | 0.709264768 | -4.458860909 |
| **A_55_P2052897** | Arhgap30 | 0.295330438 | -0.170481059 | 0.761141935 | 8.084808763 | 1.587992152 | 0.168069435 | 0.709264768 | -4.458898692 |
| **A_55_P2048406** | NA | -0.238325638 | -0.614405686 | 0.13775441 | 11.27274017 | -1.587233483 | 0.168239192 | 0.709264768 | -4.45910609 |
| **A_51_P131358** | Selplg | 0.583596144 | -0.337443917 | 1.504636204 | 8.752568043 | 1.58702672 | 0.168285485 | 0.709264768 | -4.459162616 |
| **A_55_P2070406** | NA | -0.144779318 | -0.373275648 | 0.083717011 | 8.007549643 | -1.587002622 | 0.168290881 | 0.709264768 | -4.459169204 |
| **A_65_P09032** | Rab43 | -0.244866203 | -0.631326707 | 0.1415943 | 11.85081073 | -1.586989589 | 0.168293799 | 0.709264768 | -4.459172767 |
| **A_51_P244492** | Nbl1 | -0.910947958 | -2.348794356 | 0.526898439 | 12.64105053 | -1.586833713 | 0.168328709 | 0.709264768 | -4.459215381 |
| **A_66_P126523** | Phb2 | 0.268729059 | -0.155493161 | 0.692951279 | 12.02415396 | 1.586615033 | 0.168377696 | 0.709264768 | -4.459275166 |
| **A_55_P2062513** | Pdzrn3 | -0.272406434 | -0.702437414 | 0.157624546 | 11.07484228 | -1.586601873 | 0.168380644 | 0.709264768 | -4.459278764 |
| **A_52_P300730** | Hmga2 | 0.605548297 | -0.350597743 | 1.561694337 | 9.068878399 | 1.586261932 | 0.168456823 | 0.709264768 | -4.459371703 |
| **A_55_P2085664** | Meaf6 | 0.288779472 | -0.167256676 | 0.744815619 | 11.20267411 | 1.586051902 | 0.168503907 | 0.709264768 | -4.459429126 |
| **A_51_P431996** | Zfp367 | 0.681534863 | -0.394844175 | 1.7579139 | 9.265234457 | 1.585890474 | 0.168540103 | 0.709264768 | -4.459473261 |
| **A_52_P27020** | Sgo2a | 0.454287977 | -0.263216948 | 1.171792902 | 8.55317184 | 1.585830265 | 0.168553606 | 0.709264768 | -4.459489723 |
| **A_55_P2011322** | NA | 0.61281035 | -0.355274161 | 1.580894861 | 11.44582627 | 1.585488752 | 0.168630213 | 0.709264768 | -4.459583097 |
| **A_51_P170696** | Haus1 | 0.575629743 | -0.3337652 | 1.485024687 | 11.88285792 | 1.585407969 | 0.16864834 | 0.709264768 | -4.459605185 |
| **A_51_P341465** | Csf2ra | -0.434376071 | -1.120623276 | 0.251871134 | 11.82910816 | -1.585388303 | 0.168652752 | 0.709264768 | -4.459610562 |
| **A_66_P122362** | Aldh16a1 | 0.211524696 | -0.122651845 | 0.545701237 | 11.8194562 | 1.585387634 | 0.168652903 | 0.709264768 | -4.459610745 |
| **A_55_P2085779** | Ifi27l2b | -0.841811202 | -2.208787226 | 0.525164822 | 7.965933453 | -1.640274406 | 0.168654632 | 0.709264768 | -4.473895428 |
| **A_52_P563617** | Ssbp4 | 0.138990198 | -0.08066168 | 0.358642076 | 11.0435232 | 1.584891728 | 0.168764218 | 0.709264768 | -4.459746338 |
| **A_55_P2048563** | Cdk2ap1 | 0.744260624 | -0.431933017 | 1.920454264 | 13.96590052 | 1.584880643 | 0.168766707 | 0.709264768 | -4.459749369 |
| **A_52_P70854** | Tob2 | -0.223747295 | -0.577522973 | 0.130028384 | 9.721845873 | -1.584091141 | 0.168944073 | 0.709264768 | -4.459965249 |
| **A_66_P106789** | NA | 2.242407191 | -1.303323621 | 5.788138004 | 12.65493519 | 1.584014168 | 0.168961376 | 0.709264768 | -4.459986297 |
| **A_51_P175205** | Rps6kb1 | 0.376183949 | -0.218650963 | 0.971018861 | 8.904864675 | 1.583996279 | 0.168965397 | 0.709264768 | -4.459991189 |
| **A_51_P384609** | Ddx10 | 0.278390907 | -0.161924244 | 0.718706058 | 9.441446761 | 1.583586395 | 0.169057562 | 0.709264768 | -4.460103273 |
| **A_51_P445662** | Hsd17b4 | -0.381860032 | -0.985869849 | 0.222149784 | 12.48629902 | -1.583472603 | 0.169083158 | 0.709264768 | -4.460134391 |
| **A_55_P2012430** | NA | 0.230405048 | -0.134054036 | 0.594864132 | 8.248257784 | 1.583410628 | 0.169097099 | 0.709264768 | -4.460151338 |
| **A_55_P1970775** | Hdac1 | 0.375489912 | -0.218492942 | 0.969472766 | 13.78295933 | 1.583341922 | 0.169112557 | 0.709264768 | -4.460170127 |
| **A_55_P1979645** | S100a1 | -0.785939824 | -2.029216735 | 0.457337087 | 10.14439367 | -1.58333121 | 0.169114967 | 0.709264768 | -4.460173056 |
| **A_51_P260490** | Ralbp1 | -0.606945733 | -1.567148593 | 0.353257127 | 11.7891607 | -1.583205219 | 0.169143316 | 0.709264768 | -4.46020751 |
| **A_55_P2029846** | Fam214a | -0.815358675 | -2.105310021 | 0.47459267 | 10.27079007 | -1.583163294 | 0.16915275 | 0.709264768 | -4.460218975 |
| **A_55_P2021345** | Gm14717 | 0.218353021 | -0.136839825 | 0.573545867 | 8.804258594 | 1.63741099 | 0.169251458 | 0.709264768 | -4.474486363 |
| **A_66_P125257** | Cep120 | -0.15724629 | -0.406103679 | 0.091611099 | 9.171399706 | -1.582632982 | 0.169272132 | 0.709264768 | -4.460364001 |
| **A_55_P2057994** | Mif4gd | 0.149334303 | -0.087011999 | 0.385680604 | 9.740281298 | 1.582563384 | 0.169287805 | 0.709264768 | -4.460383034 |
| **A_55_P2056300** | Sf3b3 | 0.236108057 | -0.137580392 | 0.609796507 | 12.26876987 | 1.582528281 | 0.169295711 | 0.709264768 | -4.460392635 |
| **A_52_P258338** | Mob3b | -0.315045358 | -0.813737971 | 0.183647254 | 7.986171585 | -1.582306077 | 0.169345764 | 0.709264768 | -4.460453404 |

| | | | | | | | | | |
|---|---|---|---|---|---|---|---|---|---|
| **A_55_P2092661** | Nup210 | 0.481762274 | -0.281197746 | 1.244722294 | 11.60082426 | 1.581544269 | 0.169517472 | 0.709264768 | -4.460661752 |
| **A_51_P429472** | Mindy3 | -0.155599857 | -0.402039864 | 0.09084015 | 9.86653777 | -1.581424002 | 0.169544596 | 0.709264768 | -4.460694646 |
| **A_55_P2102210** | NA | 0.151004438 | -0.088325205 | 0.390334082 | 11.26828347 | 1.580314646 | 0.16979498 | 0.709264768 | -4.460998071 |
| **A_52_P547612** | Tmem30b | -0.179503265 | -0.464010952 | 0.105004421 | 8.784197974 | -1.580260535 | 0.169807202 | 0.709264768 | -4.461012872 |
| **A_55_P2109589** | Slc1a4 | 0.334189967 | -0.195529209 | 0.863909142 | 11.16676286 | 1.580149064 | 0.169832383 | 0.709264768 | -4.461043362 |
| **A_55_P2046398** | Eif4a1 | 0.433072174 | -0.253504697 | 1.119649045 | 15.05494879 | 1.579870379 | 0.169895352 | 0.709264768 | -4.461119591 |
| **A_52_P312102** | NA | -0.518071381 | -1.339667528 | 0.303524766 | 11.35433595 | -1.579361524 | 0.170010387 | 0.709264768 | -4.461258784 |
| **A_52_P546090** | Ssr4 | -0.77961218 | -2.016055059 | 0.4568307 | 13.78269141 | -1.579264609 | 0.170032305 | 0.709264768 | -4.461285295 |
| **A_55_P2090817** | Macrod2 | 0.555163881 | -0.325449468 | 1.43577723 | 9.14587109 | 1.579015104 | 0.170088744 | 0.709264768 | -4.461353547 |
| **A_55_P2108689** | NA | 0.308156991 | -0.180670117 | 0.796984099 | 13.83940411 | 1.578945316 | 0.170104533 | 0.709264768 | -4.461372638 |
| **A_55_P1973399** | NA | 0.230035594 | -0.134883726 | 0.594954914 | 13.39711407 | 1.578877838 | 0.170119802 | 0.709264768 | -4.461391097 |
| **A_55_P1974900** | Ndufab1 | 0.186477357 | -0.109353619 | 0.482308333 | 14.0121955 | 1.57882067 | 0.170132738 | 0.709264768 | -4.461406735 |
| **A_51_P142896** | Cd59a | -0.753576533 | -1.949097555 | 0.441944488 | 9.032914096 | -1.578775922 | 0.170142865 | 0.709264768 | -4.461418976 |
| **A_51_P448203** | Cnbp | 0.467871797 | -0.274411617 | 1.210155212 | 13.44680538 | 1.578728524 | 0.170153591 | 0.709264768 | -4.461431942 |
| **A_55_P1969776** | Pdhb | 0.155546177 | -0.09125277 | 0.402345125 | 12.7724875 | 1.578579233 | 0.170187383 | 0.709264768 | -4.461472783 |
| **A_55_P2037922** | Rpl7a | 0.524461063 | -0.307764251 | 1.356686377 | 14.55719938 | 1.578420361 | 0.17022335 | 0.709264768 | -4.461516244 |
| **A_55_P1966749** | Cyp2s1 | -0.801772947 | -2.074457853 | 0.470911958 | 12.32363072 | -1.577904995 | 0.170340074 | 0.709264768 | -4.461657233 |
| **A_55_P1989504** | Tmem164 | 0.519789992 | -0.305293318 | 1.344873302 | 11.81286284 | 1.577903563 | 0.170340398 | 0.709264768 | -4.461657625 |
| **A_55_P2093650** | NA | 0.473775625 | -0.278395255 | 1.225946505 | 12.44741039 | 1.577635039 | 0.170401247 | 0.709264768 | -4.461731087 |
| **A_55_P2038928** | NA | 0.55140155 | -0.324012771 | 1.426815872 | 12.28777113 | 1.577628275 | 0.17040278 | 0.709264768 | -4.461732938 |
| **A_55_P2159216** | NA | 0.442647122 | -0.260180187 | 1.145474431 | 11.29862475 | 1.577463657 | 0.170440093 | 0.709264768 | -4.461777974 |
| **A_55_P2114009** | Ankra2 | -0.206532498 | -0.534517164 | 0.121452168 | 10.47929393 | -1.577194319 | 0.17050116 | 0.709264768 | -4.461851662 |
| **A_51_P470724** | Eif2s2 | 0.226550029 | -0.133265967 | 0.586366026 | 10.1300387 | 1.577008381 | 0.17054333 | 0.709264768 | -4.461902533 |
| **A_55_P2124016** | Nipsnap1 | 0.707382231 | -0.416255876 | 1.831020338 | 9.339841246 | 1.576805161 | 0.170589431 | 0.709264768 | -4.461958133 |
| **A_55_P1955309** | Zfp800 | 0.1605294 | -0.094512639 | 0.415571438 | 8.469571128 | 1.576497027 | 0.170659355 | 0.709264768 | -4.462042438 |
| **A_55_P2088741** | Mtif3 | -0.220505379 | -0.570890898 | 0.129880139 | 8.705901124 | -1.57624378 | 0.170716845 | 0.709264768 | -4.462111728 |
| **A_55_P1990092** | Snrpa | 0.565883138 | -0.333580756 | 1.465347032 | 11.75319828 | 1.575771944 | 0.170824005 | 0.709264768 | -4.462240829 |
| **A_55_P2069979** | Large1 | -0.242972217 | -0.629174361 | 0.143229927 | 11.12379445 | -1.575768017 | 0.170824898 | 0.709264768 | -4.462241904 |
| **A_55_P2039903** | Wsb2 | 0.490433795 | -0.289341706 | 1.270209296 | 10.97843789 | 1.575292294 | 0.170933007 | 0.709264768 | -4.462372072 |
| **A_51_P319732** | Mrps31 | 0.230605359 | -0.136093902 | 0.597304619 | 10.95791539 | 1.575105713 | 0.170975427 | 0.709264768 | -4.462423127 |
| **A_55_P2101426** | Mrps25 | 0.396924097 | -0.234376255 | 1.02822445 | 11.53850203 | 1.574786849 | 0.171047944 | 0.709264768 | -4.462510379 |
| **A_55_P2105371** | Dlk1 | 0.855864498 | -0.505467272 | 2.217196268 | 12.83012106 | 1.574676613 | 0.171073021 | 0.709264768 | -4.462540544 |
| **A_55_P2045085** | NA | 0.352337031 | -0.208154955 | 0.912829018 | 14.90098785 | 1.574487542 | 0.171116041 | 0.709264768 | -4.462592281 |
| **A_51_P342906** | Sh3bp1 | 0.365606306 | -0.216032449 | 0.947245061 | 9.351635048 | 1.57438407 | 0.171139588 | 0.709264768 | -4.462620596 |
| **A_66_P118233** | Hnrnpa0 | 0.765892163 | -0.452789814 | 1.984574139 | 13.75303242 | 1.574082834 | 0.171208159 | 0.709264768 | -4.462703029 |
| **A_55_P1955612** | Nup93 | 0.424803271 | -0.251205538 | 1.100812079 | 12.00905049 | 1.573931554 | 0.171242605 | 0.709264768 | -4.462744427 |
| **A_51_P182257** | Catsperz | 0.474185743 | -0.280433309 | 1.228804795 | 8.619782888 | 1.573878027 | 0.171254795 | 0.709264768 | -4.462759075 |
| **A_55_P1981664** | 8-Sep | -0.791428825 | -2.051004924 | 0.468147275 | 12.66456618 | -1.573757446 | 0.171282257 | 0.709264768 | -4.462792073 |
| **A_55_P1979463** | Prdx5 | -0.229969574 | -0.595982885 | 0.136043738 | 13.44648626 | -1.573706887 | 0.171293774 | 0.709264768 | -4.462805909 |
| **A_55_P2021555** | Tbrg1 | -0.727446114 | -1.885323565 | 0.430431338 | 14.84264512 | -1.573579094 | 0.171322885 | 0.709264768 | -4.462840881 |
| **A_55_P2077313** | Ddr1 | -0.565587828 | -1.46593249 | 0.334756834 | 10.71503029 | -1.573408912 | 0.171361661 | 0.709264768 | -4.462887453 |
| **A_51_P245405** | Ppp3cc | -0.279376444 | -0.724136547 | 0.16538366 | 8.994821718 | -1.573310017 | 0.171384198 | 0.709264768 | -4.462914517 |
| **A_55_P2034420** | NA | 0.832077337 | -0.492668465 | 2.156823139 | 11.3540517 | 1.573191123 | 0.171411296 | 0.709264768 | -4.462947055 |
| **A_51_P201480** | Stat3 | -0.268941209 | -0.697168301 | 0.159285883 | 11.69849873 | -1.573017514 | 0.171450872 | 0.709264768 | -4.465890012 |
| **A_55_P1958597** | Slc27a3 | -0.321645832 | -0.833816189 | 0.190524525 | 10.9936382 | -1.572946075 | 0.17146716 | 0.709264768 | -4.463014118 |
| **A_55_P2411907** | B230213L | 0.368918498 | -0.21870285 | 0.956539846 | 11.0550771 | 1.572473053 | 0.171575045 | 0.709264768 | -4.463143573 |
| **A_55_P2070179** | Ing4 | -0.186381248 | -0.483300541 | 0.110538046 | 11.55160539 | -1.572222987 | 0.171632106 | 0.709264768 | -4.463212013 |
| **A_55_P2170681** | Cdca8 | 0.374499589 | -0.222135219 | 0.971134397 | 8.424968922 | 1.572146831 | 0.171649487 | 0.709264768 | -4.463232856 |

| | | | | | | | | | |
|---|---|---|---|---|---|---|---|---|---|
| **A_55_P2157093** | Bcl2l14 | 0.415590264 | -0.246520338 | 1.077700866 | 8.68123 | 1.572118103 | 0.171656044 | 0.709264768 | -4.463240718 |
| **A_51_P392303** | Lrrn3 | 0.812458152 | -0.48205525 | 2.106971555 | 9.516405193 | 1.571971961 | 0.171689404 | 0.709264768 | -4.463280716 |
| **A_55_P2046178** | NA | 0.206727918 | -0.122676888 | 0.536132724 | 16.02493878 | 1.571880565 | 0.17171027 | 0.709264768 | -4.463305731 |
| **A_55_P2295445** | 2600002B | 0.271891611 | -0.16136546 | 0.705148681 | 10.1319668 | 1.571811618 | 0.171726013 | 0.709264768 | -4.463324601 |
| **A_52_P234910** | Gm15032 | 0.742358028 | -0.440878516 | 1.925594573 | 13.30537603 | 1.571419654 | 0.171815536 | 0.709264768 | -4.463431881 |
| **A_52_P287052** | Znrd2 | 0.359832246 | -0.213835011 | 0.933499503 | 9.794351718 | 1.571051286 | 0.171899711 | 0.709264768 | -4.463532707 |
| **A_52_P358076** | Ppp2ca | 0.186759635 | -0.11099074 | 0.484510009 | 12.03652077 | 1.571017579 | 0.171907416 | 0.709264768 | -4.463541932 |
| **A_52_P463977** | Tmem140 | -1.997255476 | -5.182160251 | 1.187649299 | 11.10172319 | -1.570679278 | 0.171984758 | 0.709264768 | -4.46363453 |
| **A_66_P139295** | Tomm40 | 0.282167667 | -0.167789355 | 0.732124688 | 11.58625839 | 1.570675825 | 0.171985548 | 0.709264768 | -4.463635475 |
| **A_55_P1978366** | NA | 0.776040645 | -0.461471385 | 2.013552674 | 11.28919477 | 1.570671575 | 0.17198652 | 0.709264768 | -4.463636639 |
| **A_52_P480064** | Tmem134 | -0.142981078 | -0.370995913 | 0.085033758 | 11.98842555 | -1.570600773 | 0.172002711 | 0.709264768 | -4.463656018 |
| **A_51_P507622** | Itpkc | -0.265091903 | -0.687904868 | 0.157721063 | 8.770190593 | -1.570357421 | 0.172058373 | 0.709264768 | -4.463722629 |
| **A_51_P315890** | Kcnk6 | -0.412443331 | -1.0703917 | 0.245505039 | 9.905289696 | -1.570083741 | 0.172120992 | 0.709264768 | -4.463797542 |
| **A_55_P2137476** | Mpv17 | -0.325227074 | -0.844087101 | 0.193632953 | 11.2187406 | -1.569953623 | 0.172150771 | 0.709264768 | -4.46383316 |
| **A_52_P253317** | 4930481A | -0.166055515 | -0.431012441 | 0.098901411 | 9.040061093 | -1.569742344 | 0.172199136 | 0.709264768 | -4.463890994 |
| **A_51_P312748** | Oxsm | -0.182708155 | -0.482538316 | 0.117122005 | 8.811704869 | -1.623100161 | 0.172266567 | 0.709264768 | -4.477445509 |
| **A_55_P2119877** | Pnisr | 0.310138767 | -0.184851869 | 0.805129403 | 9.439817987 | 1.569312444 | 0.172297587 | 0.709264768 | -4.464008674 |
| **A_52_P536735** | Hpcal1 | 0.182209917 | -0.10861437 | 0.473034204 | 9.381539444 | 1.569248358 | 0.172312268 | 0.709264768 | -4.464026217 |
| **A_55_P1980080** | Rbms2 | -0.722542692 | -1.876064001 | 0.430978617 | 11.46279961 | -1.568874617 | 0.172397909 | 0.709264768 | -4.464128528 |
| **A_55_P2037717** | Slc22a18 | -0.997603975 | -2.590267548 | 0.595059598 | 9.336247311 | -1.568861008 | 0.172401028 | 0.709264768 | -4.464132254 |
| **A_55_P1999842** | NA | 0.381521299 | -0.227756152 | 0.990798751 | 9.899760859 | 1.568389869 | 0.172509047 | 0.709264768 | -4.464261231 |
| **A_55_P1991475** | Sesn1 | -0.443913926 | -1.152885675 | 0.265057823 | 10.6167823 | -1.568267656 | 0.172537078 | 0.709264768 | -4.464294688 |
| **A_55_P1953419** | Rab13 | 0.272109445 | -0.162660321 | 0.706879211 | 8.552622437 | 1.567597732 | 0.17269081 | 0.709264768 | -4.464478093 |
| **A_52_P598634** | 1190007I0 | 0.569539647 | -0.340540376 | 1.479619669 | 11.10437731 | 1.567453717 | 0.172723875 | 0.709264768 | -4.464517521 |
| **A_51_P273508** | Ube2g1 | 0.20084801 | -0.120166655 | 0.521862676 | 8.718632742 | 1.567086633 | 0.172808183 | 0.709264768 | -4.464618022 |
| **A_52_P541157** | Calu | -0.445263371 | -1.15698743 | 0.266460687 | 9.626591268 | -1.566951913 | 0.172839134 | 0.709264768 | -4.464654906 |
| **A_52_P512817** | Shisa5 | -0.949597298 | -2.46759361 | 0.568399013 | 14.03224882 | -1.566819975 | 0.172869452 | 0.709264768 | -4.464691029 |
| **A_51_P440460** | Hip1r | -0.361921873 | -0.940487562 | 0.216643817 | 8.505258274 | -1.566796168 | 0.172874923 | 0.709264768 | -4.464697548 |
| **A_51_P328014** | Cops3 | 0.287299625 | -0.172029857 | 0.746629108 | 10.67250042 | 1.566610682 | 0.172917554 | 0.709264768 | -4.464748332 |
| **A_55_P2071846** | NA | 1.339954015 | -0.802575538 | 3.482483567 | 12.9537611 | 1.566438753 | 0.172957079 | 0.709264768 | -4.464795406 |
| **A_55_P1996504** | Abcg2 | -0.197539182 | -0.5134109 | 0.118332535 | 11.78239878 | -1.566364585 | 0.172974132 | 0.709264768 | -4.464815712 |
| **A_55_P2046744** | Cspg5 | -0.157756071 | -0.410026458 | 0.094514315 | 8.409811924 | -1.566282717 | 0.172992958 | 0.709264768 | -4.464838128 |
| **A_55_P1958230** | Dhodh | 0.461242062 | -0.276440946 | 1.198925069 | 9.977391509 | 1.566063871 | 0.173043291 | 0.709264768 | -4.464898048 |
| **A_55_P2142451** | Echdc2 | -0.214380379 | -0.557264278 | 0.128503519 | 11.70660041 | -1.565987492 | 0.173060861 | 0.709264768 | -4.464918961 |
| **A_52_P420466** | H2ac4 | 1.465704425 | -0.878959689 | 3.810368539 | 15.15572937 | 1.565727243 | 0.173120741 | 0.709264768 | -4.464990219 |
| **A_55_P2101585** | Cd55 | -0.965634429 | -2.510593356 | 0.579324499 | 10.27486369 | -1.565475045 | 0.173178787 | 0.709264768 | -4.465059274 |
| **A_55_P1960431** | Ubn1 | 0.162915701 | -0.09774142 | 0.423572822 | 12.27970287 | 1.56546619 | 0.173180825 | 0.709264768 | -4.465061699 |
| **A_51_P459320** | Gkap1 | -0.313088236 | -0.814020511 | 0.187844038 | 11.32285256 | -1.565445966 | 0.173185481 | 0.709264768 | -4.465067236 |
| **A_52_P220879** | Tgm2 | -1.076230727 | -2.79844406 | 0.645982607 | 13.86781431 | -1.565196202 | 0.173242988 | 0.709264768 | -4.465135626 |
| **A_52_P414420** | Dcun1d1 | 0.387761769 | -0.232861644 | 1.008385183 | 10.6102596 | 1.564902119 | 0.173310722 | 0.709264768 | -4.465216153 |
| **A_51_P248644** | Tdp1 | 0.243552553 | -0.146328375 | 0.633433481 | 10.54356017 | 1.564627794 | 0.173373929 | 0.709264768 | -4.46529127 |
| **A_55_P2045127** | NA | 0.242079963 | -0.145455131 | 0.629615057 | 8.19344263 | 1.564581362 | 0.173384629 | 0.709264768 | -4.465303985 |
| **A_55_P2432388** | Prr14l | 0.144278841 | -0.086691342 | 0.375249024 | 9.983698321 | 1.564577466 | 0.173385527 | 0.709264768 | -4.465305052 |
| **A_52_P273169** | Sh3d19 | -0.169674591 | -0.441314825 | 0.101965643 | 8.466735933 | -1.564490967 | 0.173405463 | 0.709264768 | -4.465328738 |
| **A_51_P346523** | Zcchc7 | -0.196864373 | -0.512045161 | 0.118316414 | 9.20112773 | -1.564435771 | 0.173418186 | 0.709264768 | -4.465343852 |
| **A_55_P2339601** | NA | 0.175231785 | -0.105319223 | 0.455782793 | 8.107711799 | 1.564412932 | 0.17342345 | 0.709264768 | -4.465350106 |
| **A_55_P2180805** | Pex3 | -0.366486696 | -0.953351505 | 0.220378113 | 9.631415348 | -1.56412152 | 0.173490637 | 0.709264768 | -4.465429906 |
| **A_55_P2028415** | Hacd2 | -0.341965002 | -0.889579696 | 0.205649693 | 10.88366911 | -1.564072619 | 0.173501914 | 0.709264768 | -4.465443297 |

| | | | | | | | | | |
|---|---|---|---|---|---|---|---|---|---|
| **A_55_P2001714** | Blvra | -0.176099167 | -0.458119581 | 0.105921247 | 8.818888519 | -1.56396526 | 0.173526674 | 0.709264768 | -4.465472696 |
| **A_55_P1965635** | Setdb1 | 0.192170585 | -0.115623552 | 0.499964723 | 9.398141034 | 1.563784537 | 0.173568361 | 0.709264768 | -4.465522186 |
| **A_55_P2378827** | NA | -0.224299768 | -0.583711829 | 0.135112292 | 8.745206094 | -1.563099139 | 0.17372655 | 0.709264768 | -4.465709883 |
| **A_55_P2087374** | NA | 0.339338211 | -0.20442005 | 0.883096472 | 15.63515442 | 1.563065739 | 0.173734262 | 0.709264768 | -4.46571903 |
| **A_55_P1956752** | NA | 0.243540263 | -0.146741302 | 0.633821829 | 8.411039294 | 1.562942779 | 0.173762657 | 0.709264768 | -4.465752703 |
| **A_55_P2177103** | Slc25a5 | 0.719506025 | -0.433551518 | 1.872563568 | 12.89377339 | 1.562909387 | 0.173770368 | 0.709264768 | -4.465761848 |
| **A_66_P136801** | Peg13 | -0.149456582 | -0.388983281 | 0.090070118 | 11.17229798 | -1.562828993 | 0.173788937 | 0.709264768 | -4.465783865 |
| **A_55_P1967928** | Srfbp1 | 0.146910885 | -0.088593118 | 0.382414888 | 9.158898649 | 1.562449636 | 0.173876582 | 0.709264768 | -4.465887758 |
| **A_55_P2011440** | Ralgapb | -0.146966478 | -0.382560838 | 0.088627882 | 11.25762432 | -1.562441418 | 0.173878481 | 0.709264768 | -4.465890008 |
| **A_55_P2004797** | Tacc2 | -0.518069989 | -1.348612054 | 0.312472076 | 11.92868943 | -1.56234574 | 0.173900593 | 0.709264768 | -4.465916212 |
| **A_52_P658437** | Espl1 | 0.364477827 | -0.219938985 | 0.948894638 | 8.414594939 | 1.562063764 | 0.173965775 | 0.709264768 | -4.465993437 |
| **A_55_P2156052** | NA | 0.233561496 | -0.14094709 | 0.608070082 | 9.752766031 | 1.562031598 | 0.173973212 | 0.709264768 | -4.466002247 |
| **A_55_P2064771** | Ly6c1 | -1.736402509 | -4.520876845 | 1.048071827 | 12.5449132 | -1.561915486 | 0.174000061 | 0.709264768 | -4.466034047 |
| **A_55_P2060604** | Gm5797 | -0.476672028 | -1.241081717 | 0.287737662 | 9.460683286 | -1.56186621 | 0.174011456 | 0.709264768 | -4.466047543 |
| **A_55_P1956267** | Syt11 | -0.255138201 | -0.664303332 | 0.154026929 | 8.983940359 | -1.561806476 | 0.174025271 | 0.709264768 | -4.466063903 |
| **A_55_P2161829** | Hmgb1-rs | 0.652452384 | -0.393925578 | 1.698830346 | 14.1456865 | 1.561746653 | 0.174039108 | 0.709264768 | -4.466080287 |
| **A_52_P631568** | Thoc6 | 0.432616529 | -0.261221001 | 1.12645406 | 9.820967361 | 1.561693011 | 0.174051515 | 0.709264768 | -4.466094979 |
| **A_55_P2028159** | C9orf72 | -0.492795956 | -1.283234132 | 0.29764222 | 10.27389978 | -1.561527272 | 0.174089857 | 0.709264768 | -4.466140372 |
| **A_51_P489413** | Luc7l | -0.169372951 | -0.441057605 | 0.102311703 | 9.868989158 | -1.561454347 | 0.17410673 | 0.709264768 | -4.466160345 |
| **A_51_P115159** | Fam162a | 0.32127669 | -0.194141178 | 0.836694558 | 13.38659831 | 1.561241518 | 0.174155982 | 0.709264768 | -4.473177117 |
| **A_51_P462047** | Prpf38a | 0.350852936 | -0.212131319 | 0.91383719 | 10.89528605 | 1.560914848 | 0.174231605 | 0.709264768 | -4.466308109 |
| **A_55_P2041708** | Gas2 | 0.283710414 | -0.171640248 | 0.739061077 | 8.321307227 | 1.560557052 | 0.17431447 | 0.709264768 | -4.466406109 |
| **A_55_P2047698** | Pop7 | 0.144820256 | -0.087656525 | 0.377297037 | 12.49808654 | 1.560271132 | 0.174380716 | 0.709264768 | -4.466484424 |
| **A_52_P137765** | Lmna | -0.475428991 | -1.238705777 | 0.287847795 | 12.32367378 | -1.560105445 | 0.174419115 | 0.709264768 | -4.466529807 |
| **A_55_P1955279** | Specc1 | -0.397484892 | -1.035643688 | 0.240673904 | 11.30712256 | -1.560063271 | 0.174428891 | 0.709264768 | -4.466541359 |
| **A_55_P2071751** | NA | 0.364641663 | -0.220792645 | 0.950075971 | 9.410521628 | 1.560049807 | 0.174432012 | 0.709264768 | -4.466545047 |
| **A_51_P344399** | Rilpl2 | -0.399821649 | -1.041747772 | 0.242104474 | 9.549275297 | -1.560025158 | 0.174437725 | 0.709264768 | -4.466551799 |
| **A_55_P2079020** | Snhg7os | 0.572163111 | -0.346504763 | 1.490830984 | 10.29673919 | 1.559953563 | 0.174454322 | 0.709264768 | -4.466571409 |
| **A_55_P2012349** | Fermt2 | -0.175291131 | -0.456742291 | 0.106160029 | 11.98621494 | -1.559937668 | 0.174458007 | 0.709264768 | -4.466575763 |
| **A_55_P2074291** | Fbxo6 | -1.335331102 | -3.479639983 | 0.808977778 | 11.31733166 | -1.559739124 | 0.174504042 | 0.709264768 | -4.466630147 |
| **A_55_P2131027** | NA | 0.356266312 | -0.215847629 | 0.928380254 | 12.69261005 | 1.55970539 | 0.174511865 | 0.709264768 | -4.466639388 |
| **A_55_P1952877** | Snrpf | 0.686037201 | -0.415722009 | 1.78779641 | 12.85938901 | 1.559593162 | 0.174537893 | 0.709264768 | -4.466670129 |
| **A_55_P1960216** | Cd151 | -1.057603486 | -2.756549079 | 0.641342107 | 9.556155008 | -1.559170999 | 0.174635834 | 0.709264768 | -4.466785769 |
| **A_55_P2049365** | NA | 0.805786003 | -0.488788814 | 2.10036082 | 13.12443516 | 1.558988495 | 0.174678191 | 0.709264768 | -4.466835762 |
| **A_55_P2090205** | Tpm3 | -0.255669799 | -0.666480052 | 0.155140454 | 8.043499889 | -1.558793197 | 0.174723529 | 0.709264768 | -4.46688926 |
| **A_52_P236398** | NA | -0.320852906 | -0.836475762 | 0.19476995 | 7.977941717 | -1.558562285 | 0.174777148 | 0.709264768 | -4.466952515 |
| **A_51_P373369** | Med4 | 0.308539972 | -0.18733588 | 0.804415824 | 11.05876652 | 1.558435462 | 0.174806604 | 0.709264768 | -4.466987256 |
| **A_55_P2112701** | NA | 0.474163814 | -0.287975247 | 1.236302875 | 11.41113948 | 1.558276539 | 0.174843522 | 0.709264768 | -4.467030791 |
| **A_51_P190254** | NA | -0.447524096 | -1.166863013 | 0.27181482 | 8.717954574 | -1.558235928 | 0.174852957 | 0.709264768 | -4.467041917 |
| **A_55_P1962011** | Clec1b | -0.677767108 | -1.767305128 | 0.411770912 | 8.883554617 | -1.558075306 | 0.17489028 | 0.709264768 | -4.467085918 |
| **A_55_P1993758** | Nsa2 | 0.519763729 | -0.315814607 | 1.355342065 | 12.05530742 | 1.558006064 | 0.174906372 | 0.709264768 | -4.467104887 |
| **A_51_P407774** | Slc22a21 | -0.335801604 | -0.875716558 | 0.20411335 | 8.258373036 | -1.557785852 | 0.174957557 | 0.709264768 | -4.467165213 |
| **A_51_P263220** | Taf5 | 0.160318601 | -0.097456822 | 0.418094023 | 8.3564513 | 1.557732038 | 0.174970068 | 0.709264768 | -4.467179956 |
| **A_52_P407796** | Mdh1 | 0.33584601 | -0.20416033 | 0.87585235 | 13.96909895 | 1.557728191 | 0.174970962 | 0.709264768 | -4.467181009 |
| **A_51_P280532** | Supt16 | 0.159488966 | -0.097002683 | 0.415980615 | 8.334461762 | 1.557427213 | 0.17504095 | 0.709264768 | -4.467263463 |
| **A_51_P207622** | Fmod | 0.783184196 | -0.476580713 | 2.042949104 | 9.901254174 | 1.557129576 | 0.175110187 | 0.709264768 | -4.467345003 |
| **A_55_P2008692** | Eef1a1 | 0.234610544 | -0.142805029 | 0.612026117 | 16.15976612 | 1.556962154 | 0.175149145 | 0.709264768 | -4.467390871 |
| **A_66_P125437** | Rpl30 | 0.656030806 | -0.399326115 | 1.711387727 | 10.44159956 | 1.556951975 | 0.175151513 | 0.709264768 | -4.467393659 |

| | | | | | | | | | |
|---|---|---|---|---|---|---|---|---|---|
| **A_55_P1993463** | NA | 0.387992491 | -0.236239596 | 1.012224578 | 8.614552767 | 1.556781197 | 0.175191261 | 0.709264768 | -4.467440447 |
| **A_55_P2013501** | Hmgb1 | 0.635819338 | -0.387142616 | 1.658781291 | 14.2544853 | 1.556770511 | 0.175193749 | 0.709264768 | -4.467443374 |
| **A_51_P220260** | Phf1 | -0.24969013 | -0.651415924 | 0.152035663 | 9.051435969 | -1.55676122 | 0.175195911 | 0.709264768 | -4.46744592 |
| **A_52_P24783** | Pds5b | 0.583676836 | -0.35580198 | 1.523155652 | 9.49138117 | 1.556093942 | 0.175351304 | 0.709264768 | -4.467628736 |
| **A_52_P315030** | NA | 0.465972228 | -0.284112798 | 1.216057254 | 15.1609591 | 1.555965213 | 0.175381297 | 0.709264768 | -4.467664005 |
| **A_55_P2012037** | Lrrc8e | 0.278822876 | -0.17001436 | 0.727660113 | 9.454868267 | 1.555929334 | 0.175389657 | 0.709264768 | -4.467673835 |
| **A_55_P2152962** | Mtr | 0.157710941 | -0.096170937 | 0.411592818 | 8.965006765 | 1.555895646 | 0.175397508 | 0.709264768 | -4.467683065 |
| **A_51_P232858** | Rpl14 | 0.868189249 | -0.529731428 | 2.266109927 | 10.69538338 | 1.555543715 | 0.175479538 | 0.709264768 | -4.467779488 |
| **A_55_P1995127** | NA | 0.695386491 | -0.424540713 | 1.815313695 | 13.84265103 | 1.555201981 | 0.175559226 | 0.709264768 | -4.46787312 |
| **A_52_P659312** | Spsb4 | 0.274618212 | -0.167659474 | 0.716895898 | 8.563975808 | 1.555194286 | 0.175561021 | 0.709264768 | -4.467875228 |
| **A_55_P2186025** | NA | 0.27837597 | -0.170032317 | 0.726784257 | 14.67424372 | 1.554921465 | 0.175624665 | 0.709264768 | -4.46794998 |
| **A_51_P195958** | Phlda1 | 1.327810117 | -0.811039186 | 3.46665942 | 12.58250476 | 1.554913136 | 0.175626609 | 0.709264768 | -4.467952262 |
| **A_55_P2028345** | Ccser2 | -0.202300223 | -0.52821615 | 0.123615705 | 11.78059554 | -1.554680393 | 0.175680922 | 0.709264768 | -4.468016034 |
| **A_55_P2122334** | Nfic | 0.414547335 | -0.253543049 | 1.082637719 | 11.20936561 | 1.55413683 | 0.17580783 | 0.709264768 | -4.468164973 |
| **A_55_P2230638** | Ece1 | -0.247496752 | -0.646367299 | 0.151373795 | 8.554244917 | -1.554131921 | 0.175808977 | 0.709264768 | -4.468166318 |
| **A_51_P377948** | Gimap9 | 0.395369629 | -0.242144144 | 1.032883402 | 8.435115077 | 1.553331248 | 0.175996076 | 0.709264768 | -4.468385716 |
| **A_55_P2435504** | Cep295 | 0.376544872 | -0.230629273 | 0.983719017 | 9.064233805 | 1.55329453 | 0.176004661 | 0.709264768 | -4.468395778 |
| **A_51_P387379** | Tshz3 | 0.43162958 | -0.264374727 | 1.127633888 | 10.04493704 | 1.553279527 | 0.176008169 | 0.709264768 | -4.468399889 |
| **A_52_P541270** | Crebl2 | -1.031391803 | -2.694528989 | 0.631745382 | 8.764036405 | -1.553266398 | 0.176011238 | 0.709264768 | -4.468403487 |
| **A_55_P2090415** | Nat9 | -0.374530798 | -0.978531059 | 0.229469462 | 8.24329699 | -1.553104778 | 0.176049032 | 0.709264768 | -4.468447775 |
| **A_55_P2108864** | Fer | -0.326883639 | -0.8540742 | 0.200306921 | 8.353247655 | -1.553015908 | 0.176069816 | 0.709264768 | -4.468472128 |
| **A_51_P232708** | Npff | -0.138808455 | -0.362745601 | 0.085128692 | 8.75424721 | -1.552530444 | 0.176183397 | 0.709264768 | -4.468605161 |
| **A_55_P2114785** | NA | -0.197745772 | -0.51681359 | 0.121322046 | 9.124231081 | -1.552296043 | 0.176238264 | 0.709264768 | -4.468669395 |
| **A_66_P140067** | Wdyhv1 | -0.435773968 | -1.138933287 | 0.267385352 | 9.57986686 | -1.552236498 | 0.176252204 | 0.709264768 | -4.468685713 |
| **A_55_P2096485** | Bcs1l | 0.328225745 | -0.201414617 | 0.857866107 | 10.53408022 | 1.552179396 | 0.176265573 | 0.709264768 | -4.468701362 |
| **A_52_P399095** | Mettl24 | -0.257398494 | -0.672753088 | 0.1579561 | 8.199457691 | -1.552163006 | 0.176269411 | 0.709264768 | -4.468705853 |
| **A_55_P2361647** | 4831440E | -0.142417604 | -0.372257181 | 0.087421973 | 8.285727862 | -1.55199103 | 0.176309683 | 0.709264768 | -4.468752982 |
| **A_66_P137265** | Gm9933 | 0.169706192 | -0.104193576 | 0.443605959 | 8.493160361 | 1.551873688 | 0.176337166 | 0.709264768 | -4.468785139 |
| **A_55_P2117411** | NA | 0.371507025 | -0.22812009 | 0.97113414 | 11.68025833 | 1.551801303 | 0.176354121 | 0.709264768 | -4.468804976 |
| **A_55_P2106033** | Myo18b | -0.242806188 | -0.634706333 | 0.149093956 | 8.070565077 | -1.551796164 | 0.176355325 | 0.709264768 | -4.468806384 |
| **A_51_P465232** | Mcfd2 | -0.62786304 | -1.641441873 | 0.385715793 | 13.06260906 | -1.551521262 | 0.176419734 | 0.709264768 | -4.468881721 |
| **A_55_P2081050** | Kcnj13 | -0.331221583 | -0.865930065 | 0.2034869 | 9.184577949 | -1.551500428 | 0.176424616 | 0.709264768 | -4.468887431 |
| **A_55_P1981322** | Grhl2 | -0.321233569 | -0.839994586 | 0.197527447 | 10.67064265 | -1.550971924 | 0.176548509 | 0.709264768 | -4.469032271 |
| **A_51_P295967** | Proc | -1.669452433 | -4.44325094 | 1.104346073 | 8.678992405 | -1.603104543 | 0.176570724 | 0.709264768 | -4.481595782 |
| **A_51_P496804** | Adat2 | 0.238379097 | -0.146849907 | 0.623608102 | 9.26907724 | 1.549885226 | 0.176803517 | 0.709264768 | -4.469330102 |
| **A_55_P2115166** | AI661453 | -1.024853877 | -2.681054865 | 0.631347111 | 11.43387533 | -1.549884213 | 0.176803755 | 0.709264768 | -4.46933038 |
| **A_55_P2177755** | Cdk2ap1 | 0.606104277 | -0.373415548 | 1.585624103 | 10.76039174 | 1.549831429 | 0.176816151 | 0.709264768 | -4.469344847 |
| **A_55_P2333126** | Oga | -0.166002264 | -0.434279996 | 0.102275469 | 8.878414187 | -1.549814529 | 0.17682012 | 0.709264768 | -4.469349479 |
| **A_52_P126266** | Prkab2 | -0.345382609 | -0.903590232 | 0.212825014 | 8.69607566 | -1.549726459 | 0.176840804 | 0.709264768 | -4.469373617 |
| **A_55_P2075385** | Orai2-ps | 0.208090229 | -0.128247312 | 0.54442777 | 8.344858028 | 1.549625265 | 0.176864573 | 0.709264768 | -4.469401353 |
| **A_55_P2067463** | 1700110I( | 0.268992176 | -0.165856763 | 0.703841115 | 12.23658882 | 1.549357286 | 0.176927533 | 0.709264768 | -4.469474801 |
| **A_51_P298355** | Dek | 0.685346583 | -0.423113987 | 1.793807153 | 12.05749018 | 1.548603899 | 0.177104652 | 0.709264768 | -4.469681299 |
| **A_55_P2472435** | Gbp3 | -0.642014416 | -1.680392601 | 0.396363769 | 8.540114 | -1.548601124 | 0.177105305 | 0.709264768 | -4.46968206 |
| **A_52_P665675** | Abca1 | -0.456438519 | -1.194705405 | 0.281828367 | 10.86307404 | -1.548528648 | 0.177122353 | 0.709264768 | -4.469701925 |
| **A_51_P363947** | Cdkn1a | -0.579146317 | -1.515919155 | 0.35762652 | 9.133260813 | -1.54847556 | 0.177134841 | 0.709264768 | -4.473699974 |
| **A_55_P2028661** | Dusp12 | 0.325194797 | -0.200838421 | 0.851228015 | 11.18286572 | 1.548391449 | 0.177154629 | 0.709264768 | -4.469739532 |
| **A_52_P427759** | NA | 0.695315615 | -0.42947852 | 1.82010975 | 13.78117279 | 1.548314871 | 0.177172647 | 0.709264768 | -4.469760522 |
| **A_66_P124817** | Ing1 | 0.22324954 | -0.138010747 | 0.584509826 | 8.979134732 | 1.547820859 | 0.177288922 | 0.709264768 | -4.469895934 |

| | | | | | | | | | |
|---|---|---|---|---|---|---|---|---|---|
| **A_55_P1968245** | Ftl1 | -0.318392356 | -0.833689094 | 0.196904381 | 16.38312981 | -1.547588828 | 0.177343561 | 0.709264768 | -4.469959536 |
| **A_55_P1969477** | Abcb1a | -0.369163996 | -0.966702406 | 0.228374414 | 8.48425095 | -1.547404506 | 0.177386976 | 0.709264768 | -4.470010062 |
| **A_66_P132969** | NA | 0.490254967 | -0.303303441 | 1.283813375 | 15.04379819 | 1.547367426 | 0.177395711 | 0.709264768 | -4.470020226 |
| **A_55_P2045837** | NA | 0.173064906 | -0.107073883 | 0.453203695 | 8.676581719 | 1.547341269 | 0.177401874 | 0.709264768 | -4.470027396 |
| **A_55_P2091691** | Nudt3 | 0.406180503 | -0.251381065 | 1.06374207 | 13.22782669 | 1.547152047 | 0.177446457 | 0.709264768 | -4.470079265 |
| **A_51_P244690** | Ube2b | -0.161489427 | -0.422965914 | 0.099987059 | 13.00161113 | -1.546898433 | 0.17750623 | 0.709264768 | -4.470148787 |
| **A_52_P655803** | Chpt1 | -0.707787596 | -1.853964501 | 0.438389309 | 9.157870128 | -1.546684167 | 0.177556744 | 0.709264768 | -4.470207522 |
| **A_55_P2122255** | Polm | -0.369447349 | -0.967733989 | 0.228839292 | 8.451299001 | -1.546655515 | 0.1775635 | 0.709264768 | -4.470215377 |
| **A_51_P490509** | Bub1b | 0.920722517 | -0.570456873 | 2.411901907 | 9.726549455 | 1.546497483 | 0.177600766 | 0.709264768 | -4.470258698 |
| **A_55_P1990755** | Ppp3r1 | 0.279763949 | -0.173340187 | 0.732868085 | 12.02612068 | 1.546479148 | 0.177605091 | 0.709264768 | -4.470263724 |
| **A_55_P2077263** | Cenpk | 0.689861092 | -0.427480487 | 1.807202671 | 9.287055821 | 1.54641494 | 0.177620234 | 0.709264768 | -4.470281326 |
| **A_55_P2102469** | Gm6428 | -0.369068462 | -0.966877096 | 0.228740171 | 15.96628925 | -1.546304779 | 0.17764622 | 0.709264768 | -4.470311524 |
| **A_51_P180140** | H2bc1 | 0.356060514 | -0.220830034 | 0.932951062 | 11.49432208 | 1.545897649 | 0.177742286 | 0.709264768 | -4.470423133 |
| **A_52_P413855** | Slc35a3 | -0.272317627 | -0.713583804 | 0.168948549 | 8.205708455 | -1.545700902 | 0.177788728 | 0.709264768 | -4.47047707 |
| **A_55_P2171493** | BC030867 | 0.684274708 | -0.424537753 | 1.793087169 | 9.186887302 | 1.545691204 | 0.177791018 | 0.709264768 | -4.470479728 |
| **A_55_P2082156** | Rab11fip3 | -0.388389084 | -1.017823135 | 0.241044967 | 9.957002128 | -1.545493301 | 0.177837745 | 0.709264768 | -4.470533982 |
| **A_55_P2172233** | NA | 0.24461493 | -0.15183427 | 0.641064129 | 15.98686199 | 1.545417266 | 0.177855701 | 0.709264768 | -4.470554827 |
| **A_65_P19149** | Nln | -0.209958668 | -0.550248557 | 0.130331222 | 9.631012171 | -1.545379361 | 0.177864654 | 0.709264768 | -4.470565218 |
| **A_66_P100249** | Snhg12 | 0.541332273 | -0.336195955 | 1.418860501 | 11.25572412 | 1.545087822 | 0.177933521 | 0.709264768 | -4.470645144 |
| **A_55_P1955502** | Tasor2 | 0.16820722 | -0.104467969 | 0.440882409 | 9.416012775 | 1.545074257 | 0.177936726 | 0.709264768 | -4.470648862 |
| **A_55_P2134236** | Foxa2 | 1.268984076 | -0.788203293 | 3.326171445 | 12.06861454 | 1.545014889 | 0.177950754 | 0.709264768 | -4.470665138 |
| **A_55_P1977473** | Dab2 | -0.284268355 | -0.745112089 | 0.17657538 | 10.05955802 | -1.544988227 | 0.177957054 | 0.709264768 | -4.470672448 |
| **A_55_P2088315** | Vmn1r87 | 0.295157616 | -0.183348917 | 0.773664148 | 8.479617978 | 1.544957254 | 0.177964372 | 0.709264768 | -4.470680939 |
| **A_55_P2002782** | Vps26a | -0.166477817 | -0.436388188 | 0.103432555 | 11.5329849 | -1.544852941 | 0.177989024 | 0.709264768 | -4.470709537 |
| **A_65_P01783** | Med28 | 0.18002416 | -0.111855107 | 0.471903428 | 9.754795814 | 1.544820028 | 0.177996802 | 0.709264768 | -4.47071856 |
| **A_51_P385952** | Man2a2 | 0.197768643 | -0.122922968 | 0.518460253 | 8.852482359 | 1.544614757 | 0.178045323 | 0.709264768 | -4.470774837 |
| **A_55_P1994132** | Tmem184 | -0.545406159 | -1.42998096 | 0.339168642 | 8.932697363 | -1.544314752 | 0.17811626 | 0.709264768 | -4.470857086 |
| **A_51_P150394** | Vps28 | -0.170828942 | -0.447914287 | 0.106256403 | 14.13181095 | -1.544181125 | 0.178147865 | 0.709264768 | -4.470893722 |
| **A_51_P484254** | Pcca | -0.242053414 | -0.63467858 | 0.150571753 | 8.613220024 | -1.544128448 | 0.178160326 | 0.709264768 | -4.470908164 |
| **A_55_P1968841** | NA | 0.777205254 | -0.483607866 | 2.038018375 | 13.42015052 | 1.543957545 | 0.178200758 | 0.709264768 | -4.47095502 |
| **A_55_P2103055** | Gsr | 0.355425619 | -0.221183783 | 0.932035021 | 10.43151292 | 1.543893554 | 0.178215899 | 0.709264768 | -4.470972564 |
| **A_51_P240864** | Sppl3 | -0.367042656 | -0.962514813 | 0.228429501 | 13.55953552 | -1.543851147 | 0.178225934 | 0.709264768 | -4.470984191 |
| **A_55_P2008181** | Il3ra | -0.835296304 | -2.190559876 | 0.519967269 | 11.05210683 | -1.543715056 | 0.178258141 | 0.709264768 | -4.471021503 |
| **A_55_P2079064** | NA | -0.256414211 | -0.67247693 | 0.159648509 | 8.649824477 | -1.543595957 | 0.178286331 | 0.709264768 | -4.471054157 |
| **A_66_P122155** | Wdr26 | -0.201361463 | -0.528126817 | 0.125403892 | 11.18763215 | -1.543443365 | 0.178322455 | 0.709264768 | -4.471095994 |
| **A_55_P2073885** | Bbs5 | -0.16295208 | -0.427408776 | 0.101504616 | 9.092929313 | -1.543318938 | 0.178351917 | 0.709264768 | -4.471130109 |
| **A_52_P277854** | Snrpb | 0.464236476 | -0.289179754 | 1.217652707 | 10.94437713 | 1.543315205 | 0.178352801 | 0.709264768 | -4.471131132 |
| **A_51_P207403** | Pwp1 | 0.453324872 | -0.282387254 | 1.189036999 | 9.794863627 | 1.543305767 | 0.178355036 | 0.709264768 | -4.47113372 |
| **A_55_P2061159** | Ing1 | 0.152928073 | -0.095274908 | 0.401131054 | 12.78037912 | 1.543229685 | 0.178373053 | 0.709264768 | -4.47115458 |
| **A_55_P1980543** | Ube2i | 0.222953299 | -0.138912058 | 0.584818656 | 11.74939291 | 1.543182326 | 0.178384269 | 0.709264768 | -4.471167564 |
| **A_51_P311038** | Adamts15 | -0.931119855 | -2.442525095 | 0.580285385 | 9.997593599 | -1.543032274 | 0.178419811 | 0.709264768 | -4.471208706 |
| **A_51_P333518** | Sh3rf2 | -0.266673478 | -0.699588188 | 0.166241232 | 8.036230405 | -1.542864649 | 0.178459522 | 0.709264768 | -4.471254666 |
| **A_52_P381784** | Nsd2 | 0.355777668 | -0.221839268 | 0.933394603 | 9.902346092 | 1.542727109 | 0.178492113 | 0.709264768 | -4.471292377 |
| **A_55_P1968362** | Tle1 | 0.184706787 | -0.115226049 | 0.484639622 | 8.496775158 | 1.542443219 | 0.1785594 | 0.709264768 | -4.471370216 |
| **A_51_P469053** | Mfsd4a | 0.140207321 | -0.087513806 | 0.367928447 | 8.947784848 | 1.542118377 | 0.178636424 | 0.709264768 | -4.471459285 |
| **A_51_P401730** | Ints14 | 0.142671438 | -0.089070474 | 0.37441335 | 10.44206947 | 1.541994403 | 0.178665827 | 0.709264768 | -4.471493278 |
| **A_55_P2026475** | NA | 0.279626752 | -0.174723196 | 0.733976701 | 11.47942421 | 1.54148244 | 0.178787303 | 0.709264768 | -4.471633659 |
| **A_55_P1988866** | NA | 0.459222891 | -0.287274052 | 1.205719833 | 15.33371002 | 1.540798476 | 0.178949713 | 0.709264768 | -4.471821207 |

| | | | | | | | | | |
|---|---|---|---|---|---|---|---|---|---|
| **A_51_P140368** | Cmc2 | 0.561451127 | -0.351289441 | 1.474191696 | 11.36204654 | 1.540688886 | 0.178975748 | 0.709264768 | -4.471851259 |
| **A_55_P2011490** | NA | 0.380705447 | -0.238237728 | 0.999648623 | 12.82584259 | 1.540595616 | 0.17899791 | 0.709264768 | -4.471876835 |
| **A_51_P410650** | Dpp9 | 0.291953362 | -0.18275114 | 0.766657864 | 9.480640314 | 1.540424707 | 0.179038525 | 0.709264768 | -4.471923701 |
| **A_55_P1999805** | NA | 0.843463739 | -0.528014142 | 2.21494162 | 11.15913792 | 1.540380295 | 0.179049081 | 0.709264768 | -4.471935879 |
| **A_55_P2103682** | NA | 0.309203201 | -0.193584191 | 0.811990593 | 13.91443959 | 1.54031643 | 0.179064261 | 0.709264768 | -4.471953393 |
| **A_66_P130911** | Proser2 | -0.395443542 | -1.038602914 | 0.24771583 | 8.313511851 | -1.539984078 | 0.179143279 | 0.709264768 | -4.472044531 |
| **A_51_P137094** | Sall2 | 0.752189761 | -0.47122086 | 1.975600381 | 9.507866894 | 1.539946087 | 0.179152314 | 0.709264768 | -4.472054949 |
| **A_55_P2179483** | NA | -1.001083238 | -2.629311995 | 0.627145519 | 15.53198456 | -1.539944661 | 0.179152653 | 0.709264768 | -4.47205534 |
| **A_55_P2058757** | Cap1 | -0.331758184 | -0.871364234 | 0.207847865 | 11.69111139 | -1.539909439 | 0.179161029 | 0.709264768 | -4.472064999 |
| **A_55_P1982171** | H2afz | 0.896667303 | -0.562295102 | 2.355629707 | 15.67460411 | 1.539350715 | 0.179293955 | 0.709264768 | -4.472218217 |
| **A_55_P2016326** | NA | 0.318218087 | -0.199601613 | 0.836037786 | 10.46791749 | 1.53920561 | 0.179328493 | 0.709264768 | -4.47225801 |
| **A_52_P543824** | Srsf6 | 0.672070663 | -0.421578315 | 1.765719642 | 10.17778353 | 1.539172631 | 0.179336343 | 0.709264768 | -4.472267054 |
| **A_52_P425860** | Iqsec1 | -0.715056766 | -1.878982831 | 0.448869299 | 10.60473215 | -1.538740809 | 0.179439166 | 0.709264768 | -4.472385477 |
| **A_51_P514449** | Nr5a2 | 0.397740398 | -0.249695211 | 1.045176007 | 8.737448427 | 1.538698277 | 0.179449297 | 0.709264768 | -4.472397141 |
| **A_55_P2141088** | NA | 0.599258344 | -0.376289408 | 1.574806095 | 10.93534173 | 1.538565192 | 0.179480999 | 0.709264768 | -4.472433638 |
| **A_51_P267354** | Lrfn3 | 0.44252468 | -0.278014594 | 1.163063955 | 9.565326539 | 1.53826155 | 0.17955335 | 0.709264768 | -4.472516911 |
| **A_55_P1962982** | NA | 0.197023672 | -0.123863184 | 0.517910528 | 10.63131 | 1.537860093 | 0.179649052 | 0.709264768 | -4.472627011 |
| **A_55_P1985569** | Tomm6 | 0.398828077 | -0.250742516 | 1.048398671 | 13.94462244 | 1.537834906 | 0.179655058 | 0.709264768 | -4.472633919 |
| **A_55_P2089462** | Ncoa5 | 0.21585712 | -0.13574385 | 0.56745809 | 13.34664488 | 1.537682457 | 0.179691413 | 0.709264768 | -4.472675729 |
| **A_55_P2301941** | A530030E | -0.399454544 | -1.05012059 | 0.251211503 | 13.00084777 | -1.537657343 | 0.179697403 | 0.709264768 | -4.472682616 |
| **A_51_P329441** | Bsg | -0.236484255 | -0.621695621 | 0.14872711 | 12.05253783 | -1.53763581 | 0.179702539 | 0.709264768 | -4.472688522 |
| **A_55_P2000304** | Terf1 | 0.18879386 | -0.118779751 | 0.496367471 | 9.407016642 | 1.537408015 | 0.179756879 | 0.709264768 | -4.472750996 |
| **A_52_P599055** | Atg14 | -0.450938641 | -1.1857083 | 0.283831018 | 9.451797181 | -1.537151181 | 0.179818165 | 0.709264768 | -4.472821436 |
| **A_55_P1996817** | NA | 0.73342796 | -0.461647091 | 1.92850301 | 12.73997353 | 1.537137182 | 0.179821506 | 0.709264768 | -4.472825276 |
| **A_51_P491987** | Ripk3 | 0.617294612 | -0.388631688 | 1.623220912 | 9.95524966 | 1.537009914 | 0.179851883 | 0.709264768 | -4.472860181 |
| **A_51_P249360** | Suox | -0.500760375 | -1.316985128 | 0.315464379 | 10.89789061 | -1.53663436 | 0.17994155 | 0.709264768 | -4.472963183 |
| **A_55_P2005941** | Plcxd1 | 0.608183823 | -0.383176248 | 1.599543893 | 11.42258463 | 1.536575086 | 0.179955707 | 0.709264768 | -4.47297944 |
| **A_55_P2159949** | Polr1e | 0.142755625 | -0.089953292 | 0.375464543 | 8.658857719 | 1.536492875 | 0.179975343 | 0.709264768 | -4.473001988 |
| **A_52_P283910** | Hnrnpdl | 0.505600231 | -0.318688239 | 1.3298887 | 10.59726118 | 1.536308325 | 0.180019429 | 0.709264768 | -4.473052606 |
| **A_55_P1984333** | Emd | 0.16392011 | -0.103338427 | 0.431178647 | 8.682668894 | 1.536211445 | 0.180042577 | 0.709264768 | -4.473079177 |
| **A_55_P2154097** | NA | 0.517193132 | -0.326075085 | 1.36046135 | 15.44481256 | 1.53616322 | 0.180054101 | 0.709264768 | -4.473092405 |
| **A_66_P100105** | Patj | -0.499751437 | -1.314581551 | 0.315078677 | 9.67219751 | -1.536163092 | 0.180054131 | 0.709264768 | -4.47309244 |
| **A_55_P1955827** | NA | 0.778901911 | -0.491111413 | 2.048915235 | 12.8033672 | 1.536118918 | 0.180064687 | 0.709264768 | -4.473104556 |
| **A_55_P2060663** | NA | 0.183236426 | -0.115549923 | 0.482022775 | 15.77416838 | 1.536036044 | 0.180084493 | 0.709264768 | -4.473127286 |
| **A_55_P2073329** | NA | 0.295468667 | -0.186347603 | 0.777284936 | 9.895084261 | 1.535961455 | 0.180102321 | 0.709264768 | -4.473147744 |
| **A_55_P1995135** | Casz1 | -0.147455067 | -0.387912579 | 0.093002446 | 8.801771574 | -1.535930963 | 0.180109609 | 0.709264768 | -4.473156107 |
| **A_55_P2029574** | Por | -0.831277476 | -2.187077924 | 0.524522971 | 12.15513932 | -1.535679497 | 0.180169727 | 0.709264768 | -4.47322508 |
| **A_51_P356467** | Cldn4 | -0.970321262 | -2.553076701 | 0.612434177 | 8.801468834 | -1.535507984 | 0.180210741 | 0.709264768 | -4.473272123 |
| **A_55_P2003123** | Jkamp | -0.574970233 | -1.513018921 | 0.363078456 | 12.65779814 | -1.535218962 | 0.180279876 | 0.709264768 | -4.473351398 |
| **A_66_P138047** | Gfm1 | 0.278196155 | -0.175695812 | 0.732088123 | 11.87150343 | 1.53514348 | 0.180297936 | 0.709264768 | -4.473372102 |
| **A_51_P346165** | Agpat4 | -0.336689841 | -0.886037756 | 0.212658073 | 12.86621896 | -1.535086593 | 0.180311548 | 0.709264768 | -4.473387706 |
| **A_52_P639229** | Fgd4 | -0.39746148 | -1.046001095 | 0.251078134 | 8.254850006 | -1.535001775 | 0.180331844 | 0.709264768 | -4.47341097 |
| **A_51_P342549** | Hand1 | -2.080374444 | -5.574525514 | 1.413776626 | 8.522574147 | -1.585851574 | 0.180371686 | 0.709264768 | -4.485190872 |
| **A_55_P2062627** | Tmem238 | 0.40492388 | -0.255889734 | 1.065737494 | 8.498229068 | 1.534775132 | 0.180386091 | 0.709264768 | -4.473473137 |
| **A_51_P247157** | Jpt1 | 0.523430592 | -0.330810313 | 1.377671496 | 10.60788497 | 1.534719754 | 0.180399348 | 0.709264768 | -4.473488327 |
| **A_51_P302651** | Topbp1 | 0.593821492 | -0.375339809 | 1.562982792 | 9.998314159 | 1.534653002 | 0.180415329 | 0.709264768 | -4.473506637 |
| **A_51_P117604** | Rab5a | -0.165561733 | -0.435829775 | 0.10470631 | 11.33031283 | -1.534318821 | 0.180495355 | 0.709264768 | -4.473598302 |
| **A_51_P291388** | Pafah1b1 | 0.265960274 | -0.168227767 | 0.700148316 | 11.74308412 | 1.534225724 | 0.180517655 | 0.709264768 | -4.473623839 |

| | | | | | | | | | |
|---|---|---|---|---|---|---|---|---|---|
| **A_52_P19016** | Clasrp | -0.16257976 | -0.428009941 | 0.102850422 | 11.97637273 | -1.534145386 | 0.180536901 | 0.709264768 | -4.473645876 |
| **A_51_P146560** | Msln | -0.669089458 | -1.761803472 | 0.423624557 | 12.77319675 | -1.533656217 | 0.180654129 | 0.709264768 | -4.473780058 |
| **A_51_P326229** | Ddx25 | 0.240483689 | -0.15228292 | 0.633250298 | 8.544617512 | 1.53356226 | 0.180676654 | 0.709264768 | -4.473805831 |
| **A_55_P2129234** | Calm2 | -0.196044905 | -0.516243164 | 0.124153355 | 15.16667091 | -1.53351115 | 0.180688908 | 0.709264768 | -4.473819852 |
| **A_52_P24160** | Mrpl51 | 0.366059897 | -0.232003248 | 0.964123042 | 13.70375782 | 1.533046961 | 0.180800237 | 0.709264768 | -4.473947185 |
| **A_51_P379443** | Bclaf1 | 0.390521614 | -0.247597228 | 1.028640456 | 12.52415631 | 1.532829511 | 0.180852412 | 0.709264768 | -4.474006836 |
| **A_55_P1984721** | Ube2i | 0.469814058 | -0.297926544 | 1.23755466 | 13.15227696 | 1.532716542 | 0.180879524 | 0.709264768 | -4.474037825 |
| **A_55_P2021350** | NA | 0.232741853 | -0.147592878 | 0.613076584 | 16.2672154 | 1.532705965 | 0.180882062 | 0.709264768 | -4.474040727 |
| **A_51_P308814** | Toe1 | 0.153448926 | -0.097325895 | 0.404223746 | 8.97486654 | 1.532605109 | 0.18090627 | 0.709264768 | -4.474068394 |
| **A_55_P1985304** | Krtap5-3 | 0.154631952 | -0.098133935 | 0.407397838 | 8.656931523 | 1.53225526 | 0.180990267 | 0.709264768 | -4.474164366 |
| **A_55_P2148912** | S100a16 | 0.808005259 | -0.512789718 | 2.128800236 | 12.78088481 | 1.532248217 | 0.180991959 | 0.709264768 | -4.474166298 |
| **A_55_P2030524** | Vldlr | -0.332776382 | -0.876769111 | 0.211216346 | 12.9199639 | -1.532179853 | 0.181008377 | 0.709264768 | -4.474185052 |
| **A_55_P2031133** | Tmem220 | -0.313704939 | -0.826548133 | 0.199138256 | 8.386022329 | -1.532099972 | 0.181027563 | 0.709264768 | -4.474206966 |
| **A_51_P240453** | Nusap1 | 1.097030446 | -0.696701421 | 2.890762312 | 10.27933044 | 1.531833287 | 0.181091631 | 0.709264768 | -4.474280126 |
| **A_55_P1963764** | 1-Sep | 0.148186592 | -0.094116757 | 0.39048994 | 8.191011755 | 1.531792146 | 0.181101516 | 0.709264768 | -4.474291412 |
| **A_51_P154997** | Brms1 | -0.455521164 | -1.200380237 | 0.28933791 | 9.084839624 | -1.531739077 | 0.181114268 | 0.709264768 | -4.474305971 |
| **A_51_P439403** | Padi1 | 0.160625669 | -0.102034072 | 0.423285409 | 8.285053873 | 1.531693194 | 0.181125295 | 0.709264768 | -4.474318558 |
| **A_55_P2005680** | Ipo11 | 0.485890821 | -0.308672012 | 1.280453655 | 11.43462992 | 1.531654439 | 0.181134609 | 0.709264768 | -4.47432919 |
| **A_51_P425962** | Khdrbs3 | 0.6016354 | -0.382238598 | 1.585509398 | 10.42929087 | 1.531596075 | 0.181148636 | 0.709264768 | -4.474345201 |
| **A_55_P1986486** | Gtf2ird1 | 0.705205748 | -0.448174954 | 1.85858645 | 11.77975096 | 1.531417157 | 0.181191643 | 0.709264768 | -4.474394285 |
| **A_55_P2025695** | C330007F | 0.312828423 | -0.198832061 | 0.824488907 | 11.21837389 | 1.531350741 | 0.181207611 | 0.709264768 | -4.474412505 |
| **A_55_P2134712** | NA | -0.316131594 | -0.833259374 | 0.200996185 | 11.52432443 | -1.531159309 | 0.181253641 | 0.709264768 | -4.474465023 |
| **A_55_P1977149** | Susd6 | -0.362993346 | -0.956808195 | 0.230821503 | 11.74068432 | -1.531080258 | 0.181272652 | 0.709264768 | -4.47448671 |
| **A_55_P1956843** | NA | 0.62455886 | -0.397221468 | 1.646339188 | 14.53569503 | 1.530968243 | 0.181299595 | 0.709264768 | -4.47451744 |
| **A_55_P2002689** | Hmgb1 | 0.574593664 | -0.365480627 | 1.514667956 | 12.59185941 | 1.530907682 | 0.181314163 | 0.709264768 | -4.474534055 |
| **A_51_P348804** | Amotl2 | -0.409846925 | -1.080409221 | 0.260715371 | 10.15872492 | -1.530850906 | 0.181327821 | 0.709264768 | -4.474549631 |
| **A_52_P171212** | Irs1 | 0.282657835 | -0.179863419 | 0.745179088 | 8.193713736 | 1.530663333 | 0.181372952 | 0.709264768 | -4.474601091 |
| **A_51_P106859** | Lsm10 | -0.29876213 | -0.787691293 | 0.190167033 | 11.38880455 | -1.530488076 | 0.181415129 | 0.709264768 | -4.474649172 |
| **A_51_P275083** | Mrps18a | 0.164434326 | -0.104689711 | 0.433558363 | 11.63564971 | 1.530348487 | 0.18144873 | 0.709264768 | -4.474687469 |
| **A_55_P1981210** | Pex26 | 0.220834419 | -0.140603571 | 0.582272409 | 8.5159395 | 1.530323719 | 0.181454692 | 0.709264768 | -4.474694264 |
| **A_55_P2032222** | Tm9sf2 | -0.42910345 | -1.131462375 | 0.273255475 | 11.07118615 | -1.53021779 | 0.181480195 | 0.709264768 | -4.474723326 |
| **A_52_P121051** | 2610001J | 0.577392489 | -0.367922883 | 1.52270786 | 11.24830118 | 1.529835574 | 0.181572243 | 0.709264768 | -4.474828188 |
| **A_55_P2092760** | Rhbdd3 | -0.362293203 | -0.955556088 | 0.230969683 | 10.80122567 | -1.52954885 | 0.181641324 | 0.709264768 | -4.474906854 |
| **A_51_P497563** | Vps18 | -0.206988144 | -0.545963545 | 0.131987257 | 10.17938326 | -1.529423053 | 0.18167164 | 0.709264768 | -4.474941367 |
| **A_55_P1980426** | Nnat | 0.808879157 | -0.515965776 | 2.13372409 | 9.615386565 | 1.52921638 | 0.181721457 | 0.709264768 | -4.474998071 |
| **A_52_P88091** | Dsg2 | -0.738658699 | -1.984395447 | 0.507078049 | 8.055317335 | -1.57935677 | 0.181823793 | 0.709264768 | -4.486547484 |
| **A_55_P2032708** | Hmgb1 | 0.663530306 | -0.423603314 | 1.750663926 | 14.20138121 | 1.528720828 | 0.18184096 | 0.709264768 | -4.475134034 |
| **A_55_P2157627** | NA | 0.330721032 | -0.211136866 | 0.872578929 | 11.10015044 | 1.528715794 | 0.181842174 | 0.709264768 | -4.475135416 |
| **A_52_P145767** | Vcpip1 | -0.220874249 | -0.582856418 | 0.14110792 | 9.595005299 | -1.528298736 | 0.181942807 | 0.709264768 | -4.475249845 |
| **A_52_P455089** | NA | 0.244017065 | -0.156006865 | 0.644040995 | 9.141628047 | 1.52786356 | 0.182047868 | 0.709264768 | -4.475369248 |
| **A_66_P128440** | NA | 0.392774285 | -0.251190263 | 1.036738833 | 10.05401183 | 1.527676625 | 0.182093016 | 0.709264768 | -4.47542054 |
| **A_66_P126842** | Frg2f1 | -0.279515957 | -0.73781061 | 0.178778696 | 8.49484484 | -1.527608832 | 0.182109392 | 0.709264768 | -4.475439141 |
| **A_51_P295816** | Cep192 | 0.197259555 | -0.126191424 | 0.520710533 | 9.412424893 | 1.527495448 | 0.182136784 | 0.709264768 | -4.475470252 |
| **A_55_P2030334** | D0Kist3 | -0.155499795 | -0.410515427 | 0.099515836 | 8.592491229 | -1.527261362 | 0.182193348 | 0.709264768 | -4.475534482 |
| **A_52_P577748** | Lpxn | 0.293929798 | -0.188173998 | 0.776033593 | 8.21588555 | 1.52705054 | 0.182244305 | 0.709264768 | -4.475592329 |
| **A_52_P206526** | Gtf2ird2 | 0.281444448 | -0.180320088 | 0.743208985 | 9.614870413 | 1.52659015 | 0.182355631 | 0.709264768 | -4.475718657 |
| **A_55_P2124791** | Col18a1 | 1.058847482 | -0.678780318 | 2.796475281 | 13.02844073 | 1.526254641 | 0.1824368 | 0.709264768 | -4.47581072 |
| **A_55_P2035872** | Mical3 | -0.676164267 | -1.786005407 | 0.433676873 | 10.3154007 | -1.525955055 | 0.182509308 | 0.709264768 | -4.475892927 |

| | | | | | | | | | |
|---|---|---|---|---|---|---|---|---|---|
| **A_51_P191469** | Rnf31 | -0.550205563 | -1.453313454 | 0.352902329 | 9.934373559 | -1.525933772 | 0.18251446 | 0.709264768 | -4.475898767 |
| **A_55_P1986691** | Card6 | -0.209059042 | -0.55220974 | 0.134091655 | 8.410740542 | -1.525929316 | 0.182515539 | 0.709264768 | -4.475899989 |
| **A_55_P2159555** | NA | -0.869818959 | -2.297610405 | 0.557972487 | 10.45326902 | -1.525859125 | 0.182532532 | 0.709264768 | -4.47591925 |
| **A_51_P131561** | Plet1 | -1.946810107 | -5.1427215 | 1.249101285 | 10.30049026 | -1.525735348 | 0.182562501 | 0.709264768 | -4.475953215 |
| **A_55_P2068849** | Nudc | 0.219266209 | -0.140706069 | 0.579238486 | 10.87882952 | 1.525643267 | 0.182584799 | 0.709264768 | -4.475978483 |
| **A_52_P547234** | Tomm70a | 0.221854093 | -0.142371947 | 0.586080132 | 10.11637853 | 1.525621496 | 0.182590071 | 0.709264768 | -4.475984457 |
| **A_51_P393768** | Osbp | -0.240886378 | -0.636364657 | 0.154591901 | 9.167390209 | -1.525597444 | 0.182595896 | 0.709264768 | -4.475991057 |
| **A_55_P2183015** | P2rx4 | -0.904422208 | -2.389746559 | 0.580902143 | 11.34061491 | -1.525106816 | 0.182714755 | 0.709264768 | -4.47612569 |
| **A_55_P2052290** | Psat1 | 1.100884667 | -0.707187357 | 2.908956692 | 12.36620298 | 1.525023171 | 0.182735026 | 0.709264768 | -4.476148644 |
| **A_52_P526852** | Eri1 | 0.338437309 | -0.217431478 | 0.894306096 | 8.951857301 | 1.524952407 | 0.182752177 | 0.709264768 | -4.476168062 |
| **A_55_P2040048** | Swi5 | 0.364870589 | -0.234441179 | 0.964182357 | 13.82683795 | 1.524882561 | 0.182769107 | 0.709264768 | -4.476187229 |
| **A_52_P475886** | Rc3h1 | -0.211038769 | -0.557709036 | 0.135631498 | 8.421699034 | -1.524740711 | 0.182803495 | 0.709264768 | -4.476226155 |
| **A_55_P1955637** | Trp53i11 | -0.388672954 | -1.027234267 | 0.249888358 | 8.33842735 | -1.52451627 | 0.182857917 | 0.709264768 | -4.476287745 |
| **A_55_P1952235** | Spry1 | 0.826897718 | -0.53172582 | 2.185521256 | 11.89289707 | 1.524414266 | 0.182882656 | 0.709264768 | -4.476315738 |
| **A_55_P2149215** | Camk2a | 0.133250695 | -0.085720443 | 0.352221833 | 7.89727959 | 1.524168398 | 0.1829423 | 0.709264768 | -4.476383209 |
| **A_55_P2102533** | Fuca2 | -0.238992486 | -0.631834701 | 0.153849729 | 8.439355088 | -1.523759554 | 0.18304152 | 0.709264768 | -4.476495406 |
| **A_51_P123795** | Zfp637 | 0.413062868 | -0.266005399 | 1.092131135 | 12.46904899 | 1.523537208 | 0.183095501 | 0.709264768 | -4.486888608 |
| **A_55_P2079272** | Kars | 0.393374062 | -0.253391996 | 1.04014012 | 12.79200314 | 1.523382091 | 0.18313317 | 0.709264768 | -4.476598993 |
| **A_55_P2002482** | Sarnp | 0.52732353 | -0.339722947 | 1.394370007 | 12.31384973 | 1.523298864 | 0.183153383 | 0.709264768 | -4.476621834 |
| **A_52_P476699** | Gemin4 | 0.161158824 | -0.103867451 | 0.4261851 | 8.329893526 | 1.523054708 | 0.183212695 | 0.709264768 | -4.476688839 |
| **A_55_P1986296** | Tagln2 | -0.341987444 | -0.904433257 | 0.220458369 | 16.05336202 | -1.522929569 | 0.183243102 | 0.709264768 | -4.476723181 |
| **A_55_P2093173** | NA | 0.185913101 | -0.119884728 | 0.49171093 | 11.32986157 | 1.522740655 | 0.183289014 | 0.709264768 | -4.476775026 |
| **A_55_P2054350** | Fbxo44 | 0.153163054 | -0.098784272 | 0.40511038 | 8.245817307 | 1.522630794 | 0.183315719 | 0.709264768 | -4.476805177 |
| **A_55_P2174836** | Vldlr | -0.445787645 | -1.179100147 | 0.287524858 | 8.7330311 | -1.522612124 | 0.183320257 | 0.709264768 | -4.476810301 |
| **A_51_P336622** | Dexi | 0.58611531 | -0.378104583 | 1.550335203 | 12.70865464 | 1.522500081 | 0.183347497 | 0.709264768 | -4.47684105 |
| **A_55_P2080562** | Arhgap26 | 0.33978435 | -0.219228639 | 0.89879734 | 9.335224265 | 1.522410668 | 0.183369237 | 0.709264768 | -4.476865588 |
| **A_55_P2085806** | Paxx | -0.259705581 | -0.687044998 | 0.167633835 | 10.89210362 | -1.52215429 | 0.183431589 | 0.709264768 | -4.47693595 |
| **A_55_P2121980** | Plcb4 | -0.151780957 | -0.401565482 | 0.098003567 | 8.625292053 | -1.521956026 | 0.183479821 | 0.709264768 | -4.476990363 |
| **A_55_P2082050** | Ankrd12 | -0.138345165 | -0.366018691 | 0.089328361 | 8.61649993 | -1.521954949 | 0.183480083 | 0.709264768 | -4.476990658 |
| **A_55_P1958314** | Mospd1 | -0.654291127 | -1.731135253 | 0.422552999 | 11.14430858 | -1.52183838 | 0.183508446 | 0.709264768 | -4.47702265 |
| **A_55_P1980421** | Wdr81 | -0.254472593 | -0.673294046 | 0.16434886 | 11.09895758 | -1.521817035 | 0.18351364 | 0.709264768 | -4.477028509 |
| **A_51_P185108** | Dpy30 | 0.82185903 | -0.530821171 | 2.174539232 | 11.30493381 | 1.521782353 | 0.18352208 | 0.709264768 | -4.477038027 |
| **A_66_P115034** | Bend5 | -0.412411793 | -1.09137595 | 0.266552364 | 9.368711696 | -1.521369035 | 0.18362269 | 0.709264768 | -4.477151463 |
| **A_51_P255682** | Mfge8 | -0.57615291 | -1.525030891 | 0.372725072 | 15.7143645 | -1.520819723 | 0.183756484 | 0.709264768 | -4.477302226 |
| **A_52_P49378** | Kif1a | 0.383381958 | -0.24830836 | 1.015072277 | 8.944833858 | 1.520119731 | 0.183927113 | 0.709264768 | -4.477494348 |
| **A_55_P2061495** | Cks1b | 1.152648948 | -0.746719097 | 3.052016992 | 12.68319782 | 1.519981411 | 0.183960848 | 0.709264768 | -4.477532312 |
| **A_51_P284244** | Tmed3 | -0.822799872 | -2.178637239 | 0.533037495 | 13.81463418 | -1.519976811 | 0.18396197 | 0.709264768 | -4.477533575 |
| **A_52_P230688** | Rep15 | -0.160922883 | -0.426151602 | 0.104305837 | 8.358366366 | -1.519664089 | 0.184038261 | 0.709264768 | -4.477619408 |
| **A_55_P2330435** | NA | 0.558095104 | -0.361764277 | 1.477954485 | 8.73040973 | 1.519627403 | 0.184047213 | 0.709264768 | -4.477629477 |
| **A_52_P1077588** | Heatr6 | 0.217799631 | -0.141205179 | 0.57680444 | 8.384830519 | 1.51952279 | 0.184072743 | 0.709264768 | -4.47765819 |
| **A_52_P45834** | Pafah1b2 | 0.278931332 | -0.180869925 | 0.738732589 | 9.496853921 | 1.519418788 | 0.184098126 | 0.709264768 | -4.477686736 |
| **A_51_P192800** | Grn | -0.765284295 | -2.026919129 | 0.496350539 | 11.6865202 | -1.519285788 | 0.184130592 | 0.709264768 | -4.477723242 |
| **A_66_P138137** | Pnp | -0.249027459 | -0.659576932 | 0.161522013 | 12.34524894 | -1.519259934 | 0.184136904 | 0.709264768 | -4.477730338 |
| **A_65_P06453** | Nup35 | 0.418796225 | -0.271661918 | 1.109254368 | 10.81366673 | 1.519202785 | 0.184150856 | 0.709264768 | -4.477746024 |
| **A_55_P1986596** | Cacna1h | 0.501440647 | -0.325312322 | 1.328193615 | 8.664151867 | 1.519127122 | 0.18416933 | 0.709264768 | -4.477766792 |
| **A_55_P1968448** | NA | 0.586523902 | -0.380693571 | 1.553741375 | 15.23837763 | 1.518839654 | 0.184239535 | 0.709264768 | -4.477845696 |
| **A_55_P2082276** | NA | 0.224644833 | -0.145857949 | 0.595147614 | 10.24291509 | 1.51864165 | 0.184287906 | 0.709264768 | -4.477900044 |
| **A_55_P2066096** | Zfand3 | -0.277138296 | -0.734265013 | 0.179988422 | 12.19706102 | -1.518484216 | 0.184326374 | 0.709264768 | -4.477943257 |

| | | | | | | | | | |
|---|---|---|---|---|---|---|---|---|---|
| **A_55_P2039479** | Twistnb | 0.202757294 | -0.131772246 | 0.537286835 | 8.307099236 | 1.518071962 | 0.184427144 | 0.709264768 | -4.478056415 |
| **A_52_P598605** | Ptch1 | 0.4870572 | -0.316767636 | 1.290882035 | 8.777599566 | 1.517640449 | 0.184532677 | 0.709264768 | -4.478174861 |
| **A_55_P1952638** | Fgd6 | -0.616400518 | -1.633709806 | 0.400908771 | 9.129739835 | -1.517610528 | 0.184539997 | 0.709264768 | -4.478183075 |
| **A_51_P377760** | Rnasel | -0.175608229 | -0.465434086 | 0.114217629 | 8.350488882 | -1.517602657 | 0.184541923 | 0.709264768 | -4.478185235 |
| **A_52_P5901** | Saal1 | 0.293694125 | -0.191040194 | 0.778428445 | 8.973812807 | 1.517545903 | 0.184555808 | 0.709264768 | -4.478200814 |
| **A_51_P474169** | Proser2 | -1.092581175 | -2.896716496 | 0.711554146 | 10.00279227 | -1.516823156 | 0.184732717 | 0.709264768 | -4.478399206 |
| **A_55_P1967481** | Hbb-y | 1.124901475 | -0.732627638 | 2.982430588 | 9.805207914 | 1.516803083 | 0.184737633 | 0.709264768 | -4.478404716 |
| **A_52_P655265** | Bora | 0.290875106 | -0.189444747 | 0.77119496 | 8.349385814 | 1.516793165 | 0.184740061 | 0.709264768 | -4.478407438 |
| **A_51_P120093** | Snx10 | -0.30429517 | -0.806837333 | 0.198246993 | 8.797986782 | -1.516606453 | 0.184785792 | 0.709264768 | -4.479664746 |
| **A_55_P2074718** | Fh1 | 0.352938349 | -0.229944351 | 0.935821048 | 12.66476086 | 1.516589447 | 0.184789957 | 0.709264768 | -4.478463359 |
| **A_55_P2164841** | NA | -0.43561275 | -1.155111335 | 0.283885835 | 10.58977598 | -1.516425168 | 0.184830203 | 0.709264768 | -4.478508454 |
| **A_51_P225427** | Pkp2 | -0.89892692 | -2.383828339 | 0.585974498 | 10.86406899 | -1.516271981 | 0.184867738 | 0.709264768 | -4.478550505 |
| **A_52_P127465** | Ypel1 | -0.129754442 | -0.344109846 | 0.084600961 | 8.372666289 | -1.516135945 | 0.184901078 | 0.709264768 | -4.478587848 |
| **A_51_P323610** | Selenos | -0.640915687 | -1.699807415 | 0.417976041 | 11.88450226 | -1.516001712 | 0.184933981 | 0.709264768 | -4.478624696 |
| **A_51_P260169** | Gstm5 | 0.444819838 | -0.290105686 | 1.179745363 | 10.44025835 | 1.515971938 | 0.18494128 | 0.709264768 | -4.478632869 |
| **A_51_P487999** | Sgo1 | 0.966661062 | -0.630570496 | 2.56389262 | 9.950937334 | 1.515851607 | 0.184970781 | 0.709264768 | -4.478665901 |
| **A_55_P2090194** | NA | -0.409654953 | -1.086610788 | 0.267300882 | 15.02652028 | -1.515682443 | 0.185012262 | 0.709264768 | -4.478712339 |
| **A_55_P2103105** | Hacd1 | 0.403940476 | -0.263659149 | 1.071540102 | 11.94691855 | 1.515484974 | 0.185060695 | 0.709264768 | -4.478766546 |
| **A_55_P2249849** | Sema6a | -0.172965829 | -0.458836066 | 0.112904408 | 8.939159981 | -1.515450363 | 0.185069185 | 0.709264768 | -4.478776048 |
| **A_55_P2154132** | Tuba1b | 0.571880386 | -0.373399037 | 1.517159809 | 17.03612127 | 1.515288556 | 0.185108882 | 0.709264768 | -4.478820466 |
| **A_55_P2171648** | Ppp1r7 | 0.182516017 | -0.119188104 | 0.484220138 | 11.25316877 | 1.515200433 | 0.185130505 | 0.709264768 | -4.478844657 |
| **A_51_P419389** | Bmpr2 | -0.649287501 | -1.722626042 | 0.42405104 | 9.673362701 | -1.515132674 | 0.185147133 | 0.709264768 | -4.478863258 |
| **A_51_P245275** | H2afx | 0.851132297 | -0.555921265 | 2.25818586 | 13.55426201 | 1.51508426 | 0.185159015 | 0.709264768 | -4.478876549 |
| **A_52_P71544** | Slc25a26 | -0.150479216 | -0.39926445 | 0.098306018 | 9.168407376 | -1.51496383 | 0.185188573 | 0.709264768 | -4.478909609 |
| **A_55_P2045086** | NA | 0.655780474 | -0.428448141 | 1.740009089 | 12.73774364 | 1.51491394 | 0.18520082 | 0.709264768 | -4.478923305 |
| **A_55_P2061094** | Fnta | -0.175423326 | -0.465508465 | 0.114661813 | 8.437917143 | -1.514649707 | 0.185265693 | 0.709264768 | -4.478995842 |
| **A_55_P1970583** | Ttc38 | -0.154412908 | -0.409757307 | 0.100931491 | 9.330007206 | -1.514633688 | 0.185269627 | 0.709264768 | -4.47900024 |
| **A_51_P160713** | Alb | 0.710173764 | -0.464229014 | 1.884576542 | 10.19227262 | 1.514599817 | 0.185277945 | 0.709264768 | -4.479009538 |
| **A_51_P160439** | Crybg3 | -0.438297651 | -1.163115204 | 0.286519902 | 10.68160994 | -1.51457501 | 0.185284037 | 0.709264768 | -4.479016348 |
| **A_55_P2146749** | Rps13 | 0.211732081 | -0.138426823 | 0.561890984 | 16.43745391 | 1.514508924 | 0.185300267 | 0.709264768 | -4.47903449 |
| **A_52_P99848** | Pik3cd | 0.260070683 | -0.170149348 | 0.690290714 | 8.776085372 | 1.514087995 | 0.185403675 | 0.709264768 | -4.479150046 |
| **A_55_P2055742** | Adtrp | -0.597600894 | -1.616210333 | 0.421008545 | 7.903778916 | -1.562666304 | 0.185609532 | 0.709264768 | -4.490041611 |
| **A_51_P363634** | Snrpe | 0.755800456 | -0.495302636 | 2.006903547 | 12.76816857 | 1.513088735 | 0.185649381 | 0.709264768 | -4.479424375 |
| **A_55_P2144850** | Psmb5 | -0.220009068 | -0.584282071 | 0.144263935 | 15.67726918 | -1.512738783 | 0.185735503 | 0.709264768 | -4.47952045 |
| **A_55_P1985623** | Abcc3 | -0.250016977 | -0.66401273 | 0.163978776 | 9.006009966 | -1.512599522 | 0.185769785 | 0.709264768 | -4.479558683 |
| **A_55_P2081349** | Focad | 0.307207861 | -0.201594965 | 0.816010686 | 10.00382652 | 1.512283296 | 0.185847654 | 0.709264768 | -4.4796455 |
| **A_52_P585652** | Fndc3b | -0.593930735 | -1.577622009 | 0.38976054 | 10.4318564 | -1.512262999 | 0.185852653 | 0.709264768 | -4.479651073 |
| **A_51_P389751** | Relb | -0.347978668 | -0.924351628 | 0.228394293 | 9.42494798 | -1.512165649 | 0.185876632 | 0.709264768 | -4.479677799 |
| **A_55_P2127702** | Racgap1 | 1.116297475 | -0.732940892 | 2.965535843 | 10.75193635 | 1.511949872 | 0.185929791 | 0.709264768 | -4.47973704 |
| **A_55_P2034110** | Tgm2 | -1.133184036 | -3.010551903 | 0.744183831 | 12.42880028 | -1.511824618 | 0.185960656 | 0.709264768 | -4.479771428 |
| **A_55_P2443935** | Xpo7 | 0.272147797 | -0.178726885 | 0.723022478 | 8.909267947 | 1.511817331 | 0.185962452 | 0.709264768 | -4.479773429 |
| **A_55_P2131844** | NA | 0.52800777 | -0.346782145 | 1.402797684 | 9.820414447 | 1.511774067 | 0.185973114 | 0.709264768 | -4.479785307 |
| **A_66_P103926** | Eif3m | 0.518363201 | -0.340477037 | 1.377203438 | 13.12999363 | 1.511722667 | 0.185985782 | 0.709264768 | -4.479799419 |
| **A_51_P279606** | Socs1 | 0.438459703 | -0.28806472 | 1.164984126 | 9.546139232 | 1.51157539 | 0.186022084 | 0.709264768 | -4.479839854 |
| **A_55_P2078459** | Kpna2 | 1.068706796 | -0.703025827 | 2.84043942 | 13.03265063 | 1.510813075 | 0.186210095 | 0.709264768 | -4.48004915 |
| **A_51_P187716** | Zc3h11a | -0.145761423 | -0.387414701 | 0.095891854 | 13.00416198 | -1.510776627 | 0.186219088 | 0.709264768 | -4.480059157 |
| **A_51_P398260** | Tppp | -0.174265119 | -0.463174008 | 0.11464377 | 7.922799166 | -1.510775417 | 0.186219387 | 0.709264768 | -4.480059489 |
| **A_52_P609024** | Cenpw | 1.198851749 | -0.788988256 | 3.186691753 | 10.76160385 | 1.510547548 | 0.186275624 | 0.709264768 | -4.480122053 |

| | | | | | | | | | |
|---|---|---|---|---|---|---|---|---|---|
| **A_55_P2138961** | Tpk1 | 0.228785362 | -0.15072043 | 0.608291153 | 8.989417675 | 1.509941673 | 0.186425231 | 0.709264768 | -4.480288403 |
| **A_66_P110104** | Myo18b | 0.139404127 | -0.091837715 | 0.37064597 | 9.042397404 | 1.509939539 | 0.186425758 | 0.709264768 | -4.480288989 |
| **A_55_P2008407** | Ppp1r14c | 0.410446239 | -0.270445146 | 1.091337625 | 11.05790352 | 1.50983256 | 0.186452186 | 0.709264768 | -4.480318362 |
| **A_55_P2076489** | Rac1 | -0.398826906 | -1.060521434 | 0.262867621 | 13.43849015 | -1.50965341 | 0.186496451 | 0.709264768 | -4.48036755 |
| **A_55_P1978511** | H2-Q7 | -1.448133419 | -3.850812798 | 0.95454596 | 11.08462144 | -1.509604217 | 0.186508608 | 0.709264768 | -4.480381057 |
| **A_55_P2102698** | NA | 0.819723172 | -0.540572092 | 2.180018436 | 13.96881746 | 1.509330582 | 0.186576242 | 0.709264768 | -4.480456189 |
| **A_55_P1960775** | Rps15a | 0.509559921 | -0.336059102 | 1.355178945 | 16.15827537 | 1.509283635 | 0.186587848 | 0.709264768 | -4.480469079 |
| **A_52_P570543** | Sdk2 | 0.214742029 | -0.141663709 | 0.571147767 | 8.379610113 | 1.509116303 | 0.186629221 | 0.709264768 | -4.480515024 |
| **A_55_P2040046** | Swi5 | 0.169635713 | -0.111921611 | 0.451193036 | 15.37699973 | 1.509040185 | 0.186648045 | 0.709264768 | -4.480535923 |
| **A_55_P2142008** | 4933434E | -0.173732546 | -0.462089833 | 0.11462474 | 8.516508031 | -1.509039473 | 0.186648221 | 0.709264768 | -4.480536119 |
| **A_51_P490023** | Tubb2a | 0.529666225 | -0.349517365 | 1.408849814 | 13.2841946 | 1.508943757 | 0.186671893 | 0.709264768 | -4.4805624 |
| **A_52_P523330** | Tmem147 | -0.264128118 | -0.702559247 | 0.17430301 | 10.80750296 | -1.508910922 | 0.186680014 | 0.709264768 | -4.480571416 |
| **A_52_P532456** | Plagl1 | -0.219231132 | -0.583144173 | 0.144681909 | 14.90875639 | -1.508880872 | 0.186687447 | 0.709264768 | -4.480579666 |
| **A_55_P2178023** | NA | 0.78135377 | -0.515747174 | 2.078454714 | 13.32669922 | 1.508774315 | 0.186713806 | 0.709264768 | -4.480608924 |
| **A_55_P1978972** | Rp2 | -0.237298784 | -0.631259921 | 0.156662354 | 8.419321797 | -1.508663819 | 0.186741142 | 0.709264768 | -4.480639264 |
| **A_55_P1963533** | Pcdhac2 | 0.561918266 | -0.371097566 | 1.494934098 | 8.78246199 | 1.508462375 | 0.18679099 | 0.709264768 | -4.480694576 |
| **A_51_P442097** | Slc41a3 | -0.184949072 | -0.492086901 | 0.122188757 | 8.925708259 | -1.508235615 | 0.186847117 | 0.709264768 | -4.480756839 |
| **A_55_P1983095** | Adam19 | -0.874276585 | -2.326239243 | 0.577686074 | 11.69990161 | -1.508147246 | 0.186868994 | 0.709264768 | -4.480781103 |
| **A_55_P2123471** | Kpna2 | 1.053292993 | -0.69597446 | 2.802560446 | 13.95392385 | 1.508145786 | 0.186869356 | 0.709264768 | -4.480781504 |
| **A_55_P2134251** | Hmgb1 | 0.658685622 | -0.435597594 | 1.752968839 | 13.11986628 | 1.507643956 | 0.186993639 | 0.709264768 | -4.480919298 |
| **A_51_P145533** | Rnf14 | -0.514954436 | -1.370610597 | 0.340701725 | 12.64276918 | -1.507369998 | 0.18706152 | 0.709264768 | -4.480994522 |
| **A_55_P2027012** | Serpinb8 | 0.237695564 | -0.157287767 | 0.632678894 | 12.4396593 | 1.507275549 | 0.187084928 | 0.709264768 | -4.481020457 |
| **A_55_P2126945** | 1700066B | -0.171332221 | -0.456040422 | 0.11337598 | 8.853794726 | -1.507264299 | 0.187087717 | 0.709264768 | -4.481023546 |
| **A_55_P2005813** | Abhd18 | -0.403293234 | -1.073519159 | 0.266932692 | 8.221475953 | -1.507127721 | 0.187121571 | 0.709264768 | -4.481061048 |
| **A_52_P599238** | Rspry1 | 0.17492562 | -0.115830899 | 0.465682139 | 9.197654975 | 1.506864862 | 0.187186744 | 0.709264768 | -4.481133227 |
| **A_55_P2292737** | 2310002F | 0.239415307 | -0.1585981 | 0.637428713 | 13.20857697 | 1.506622875 | 0.187246762 | 0.709264768 | -4.481199674 |
| **A_66_P125560** | Macroh2a | 0.622966264 | -0.412739259 | 1.658671787 | 11.40741924 | 1.506532767 | 0.187269115 | 0.709264768 | -4.481224417 |
| **A_52_P1092823** | Irx1 | 2.408081527 | -1.715614435 | 6.53177749 | 12.04725347 | 1.555418478 | 0.187277998 | 0.709264768 | -4.491562365 |
| **A_51_P140731** | Cops5 | 0.147224353 | -0.09756724 | 0.392015946 | 13.233555 | 1.506376398 | 0.187307911 | 0.709264768 | -4.481267355 |
| **A_51_P393958** | Fbxo5 | 1.137744743 | -0.754047286 | 3.029536772 | 10.59210492 | 1.50633578 | 0.18731799 | 0.709264768 | -4.481278509 |
| **A_65_P15809** | Gm4759 | -0.260736108 | -0.694390924 | 0.172918708 | 9.197632358 | -1.505938895 | 0.1874165 | 0.709264768 | -4.481387492 |
| **A_52_P137371** | Hmgcr | 0.169505869 | -0.112454739 | 0.451466477 | 8.424187203 | 1.50572842 | 0.187468762 | 0.709264768 | -4.481445288 |
| **A_55_P2047620** | NA | -0.167079615 | -0.445010459 | 0.110851228 | 13.89881045 | -1.505695202 | 0.187477011 | 0.709264768 | -4.48145441 |
| **A_55_P2102350** | Stambpl1 | 0.13364232 | -0.088677382 | 0.355962022 | 8.864952151 | 1.505623551 | 0.187494806 | 0.709264768 | -4.481474085 |
| **A_55_P2116059** | Crybg1 | -0.856136362 | -2.280444648 | 0.568171924 | 9.099245299 | -1.505529563 | 0.187518152 | 0.709264768 | -4.481499895 |
| **A_51_P112445** | Wdr3 | 0.169375582 | -0.112456582 | 0.451207746 | 9.770396876 | 1.505256775 | 0.187585923 | 0.709264768 | -4.48881767 |
| **A_55_P2077228** | Cenpl | 0.516802845 | -0.343158007 | 1.376763697 | 9.109255227 | 1.505208147 | 0.187598007 | 0.709264768 | -4.481588156 |
| **A_55_P2048101** | Gm6762 | -0.275534326 | -0.734030871 | 0.18296222 | 8.14574005 | -1.505185367 | 0.187603668 | 0.709264768 | -4.481594412 |
| **A_55_P1964313** | Pla2g4b | 0.27763464 | -0.184420582 | 0.739689863 | 11.55317824 | 1.50497787 | 0.187655239 | 0.709264768 | -4.481651391 |
| **A_51_P236829** | Zswim7 | -0.527999964 | -1.406746921 | 0.350746993 | 9.9787825 | -1.504944224 | 0.187663602 | 0.709264768 | -4.481660631 |
| **A_55_P2056493** | NA | 0.406758035 | -0.270246604 | 1.083762674 | 8.752062223 | 1.504855643 | 0.187685623 | 0.709264768 | -4.481684955 |
| **A_55_P2161219** | Thsd7a | -0.458202905 | -1.220973573 | 0.304567763 | 8.442305888 | -1.504576242 | 0.187755097 | 0.709264768 | -4.481761681 |
| **A_51_P138790** | Plekhf2 | -0.204997805 | -0.546275009 | 0.136279398 | 9.805743518 | -1.504500296 | 0.187773985 | 0.709264768 | -4.481782537 |
| **A_55_P2055819** | Adam15 | -0.721488105 | -1.922631803 | 0.479655593 | 13.52233218 | -1.504473541 | 0.18778064 | 0.709264768 | -4.481789884 |
| **A_66_P113059** | Itgb1 | -0.43819726 | -1.167778359 | 0.291383839 | 11.46346814 | -1.504341473 | 0.187813492 | 0.709264768 | -4.481826151 |
| **A_55_P2150831** | Cdh4 | 0.250480849 | -0.166561124 | 0.667522822 | 8.974750415 | 1.50433689 | 0.187814632 | 0.709264768 | -4.481827409 |
| **A_66_P106238** | Gm5106 | 0.462688436 | -0.307708278 | 1.233085149 | 9.494324636 | 1.504265755 | 0.187832329 | 0.709264768 | -4.481846944 |
| **A_55_P2093196** | NA | 0.146976779 | -0.097750028 | 0.391703587 | 16.17608971 | 1.504241367 | 0.187838397 | 0.709264768 | -4.481853641 |

| | | | | | | | | | |
|---|---|---|---|---|---|---|---|---|---|
| **A_55_P2048588** | Cdk1 | 1.480876994 | -0.984968481 | 3.946722469 | 11.45804149 | 1.504192692 | 0.187850508 | 0.709264768 | -4.481867008 |
| **A_51_P437978** | Agtr2 | 0.708271698 | -0.471215646 | 1.887759042 | 8.994406437 | 1.504031557 | 0.187890605 | 0.709264768 | -4.481911257 |
| **A_55_P1973349** | H4c14 | 1.225677517 | -0.815589973 | 3.266945007 | 15.14013346 | 1.50392662 | 0.187916722 | 0.709264768 | -4.481940074 |
| **A_51_P506787** | Nap1l1 | 0.841517927 | -0.560325535 | 2.24336139 | 10.97493897 | 1.503537268 | 0.188013655 | 0.709264768 | -4.482046996 |
| **A_52_P106259** | Egfr | -0.674488279 | -1.79813746 | 0.449160901 | 9.620468768 | -1.503467392 | 0.188031057 | 0.709264768 | -4.482066186 |
| **A_51_P101075** | Cnppd1 | -0.89244921 | -2.379400168 | 0.594501748 | 12.99452851 | -1.50327076 | 0.188080033 | 0.709264768 | -4.483277514 |
| **A_55_P2011387** | Tifa | 0.179884681 | -0.119861069 | 0.479630431 | 8.277502235 | 1.503112519 | 0.188119456 | 0.709264768 | -4.48216364 |
| **A_51_P303749** | Depdc1b | 0.920577836 | -0.613659045 | 2.454814716 | 9.580307682 | 1.50285971 | 0.188182454 | 0.709264768 | -4.482233067 |
| **A_55_P2016604** | Glrx3 | 0.46563193 | -0.310554732 | 1.241818592 | 13.07290668 | 1.502543048 | 0.188261393 | 0.709264768 | -4.482320029 |
| **A_51_P513085** | Gins4 | 0.398332985 | -0.265707526 | 1.062373497 | 10.44637624 | 1.502456965 | 0.188282858 | 0.709264768 | -4.48234367 |
| **A_52_P116134** | Aldh2 | 1.139659914 | -0.760275372 | 3.0395952 | 11.37484331 | 1.502404272 | 0.188295998 | 0.709264768 | -4.482358141 |
| **A_65_P05152** | Flrt2 | 0.166746095 | -0.11126747 | 0.44475966 | 8.452053793 | 1.502242454 | 0.188336355 | 0.709264768 | -4.48240258 |
| **A_55_P2128794** | Clta | -0.172399716 | -0.459858236 | 0.115058804 | 13.35738464 | -1.502144483 | 0.188360794 | 0.709264768 | -4.482429485 |
| **A_55_P2145908** | Phb | 0.238468729 | -0.159166249 | 0.636103707 | 12.7781189 | 1.502094313 | 0.188373309 | 0.709264768 | -4.482443263 |
| **A_51_P151271** | Eif4b | 0.301598988 | -0.201305366 | 0.804503343 | 9.423882111 | 1.502086109 | 0.188375356 | 0.709264768 | -4.482445516 |
| **A_51_P396752** | Arl2bp | 0.43317264 | -0.289190865 | 1.155536145 | 12.76063115 | 1.501950316 | 0.188409236 | 0.709264768 | -4.482482809 |
| **A_52_P39083** | Ccbe1 | -0.411504433 | -1.097843371 | 0.274834504 | 9.069896745 | -1.50171053 | 0.188469077 | 0.709264768 | -4.482548661 |
| **A_55_P2003548** | NA | 0.451765466 | -0.301853658 | 1.20538459 | 17.32676748 | 1.501452039 | 0.188533605 | 0.709264768 | -4.482619651 |
| **A_55_P1960326** | NA | 0.234653144 | -0.156831706 | 0.626137995 | 10.12548413 | 1.501280222 | 0.188576508 | 0.709264768 | -4.482666837 |
| **A_51_P336385** | Dido1 | 0.214816019 | -0.143589253 | 0.57322129 | 10.64516576 | 1.501214048 | 0.188593035 | 0.709264768 | -4.482685011 |
| **A_55_P2088440** | Arrdc2 | -0.292124295 | -0.779617487 | 0.195368897 | 9.186506972 | -1.500892087 | 0.188673461 | 0.709264768 | -4.482773432 |
| **A_51_P351697** | Mrpl35 | 0.162484615 | -0.108697217 | 0.433666448 | 8.366534921 | 1.500728054 | 0.188714449 | 0.709264768 | -4.482818482 |
| **A_55_P2088360** | NA | 0.747723552 | -0.500262615 | 1.995709719 | 14.71363317 | 1.500657677 | 0.188732038 | 0.709264768 | -4.48283781 |
| **A_52_P190405** | Ifnar2 | -0.181918842 | -0.485555999 | 0.121718315 | 10.91108144 | -1.500628226 | 0.188739398 | 0.709264768 | -4.482845898 |
| **A_55_P2209298** | NA | -0.175850779 | -0.469403931 | 0.117702372 | 8.574759976 | -1.500402902 | 0.188795723 | 0.709264768 | -4.482907781 |
| **A_55_P2008874** | Dnmt3b | 0.131845066 | -0.088301404 | 0.351991535 | 8.235444966 | 1.500038831 | 0.188886764 | 0.709264768 | -4.48300777 |
| **A_51_P322542** | Bbx | -0.605096013 | -1.615750227 | 0.4055582 | 10.2689999 | -1.499588311 | 0.18899948 | 0.709264768 | -4.483131502 |
| **A_51_P463087** | Cenpm | 0.344619195 | -0.231019965 | 0.920258356 | 8.358035959 | 1.49947586 | 0.189027624 | 0.709264768 | -4.483162386 |
| **A_66_P125831** | Pnn | 0.511691462 | -0.343160477 | 1.366543401 | 11.41381112 | 1.499227759 | 0.189089733 | 0.709264768 | -4.483230526 |
| **A_52_P515247** | Arhgdib | 0.49448716 | -0.331643089 | 1.320617408 | 11.65250346 | 1.499190566 | 0.189099045 | 0.709264768 | -4.483240741 |
| **A_66_P138319** | Acox2 | -0.205287809 | -0.548300668 | 0.13772505 | 8.800721727 | -1.499005079 | 0.189145494 | 0.709264768 | -4.483291685 |
| **A_55_P2042590** | Polm | -0.505652463 | -1.350627696 | 0.33932277 | 9.799020159 | -1.498851108 | 0.189184059 | 0.709264768 | -4.483333973 |
| **A_55_P1956842** | NA | 0.443118381 | -0.297374091 | 1.183610853 | 15.24969379 | 1.498819944 | 0.189191866 | 0.709264768 | -4.483342532 |
| **A_55_P2057065** | Gtf3c5 | 0.161677383 | -0.108516654 | 0.431871419 | 11.19408007 | 1.498731571 | 0.189214005 | 0.709264768 | -4.483366804 |
| **A_55_P2073977** | NA | -0.454864482 | -1.215056385 | 0.305327422 | 14.58417828 | -1.498680759 | 0.189226735 | 0.709264768 | -4.483380759 |
| **A_52_P330540** | Mis18a | 0.430365987 | -0.288903219 | 1.149635193 | 10.04180965 | 1.498638269 | 0.189237381 | 0.709264768 | -4.483392429 |
| **A_55_P2048041** | Slc44a1 | -0.534975172 | -1.429077629 | 0.359127285 | 8.808582288 | -1.498637805 | 0.189237498 | 0.709264768 | -4.483392557 |
| **A_55_P2092073** | Taf11 | 0.190270071 | -0.127775885 | 0.508316028 | 9.799486825 | 1.498410984 | 0.189294338 | 0.709264768 | -4.483454853 |
| **A_55_P2110718** | Npm1 | 0.702831721 | -0.471999502 | 1.877662945 | 12.27006209 | 1.498394672 | 0.189298427 | 0.709264768 | -4.483459334 |
| **A_66_P119164** | Zfp354c | 0.797620795 | -0.535694978 | 2.130936568 | 9.629178579 | 1.498351811 | 0.189309169 | 0.709264768 | -4.483471105 |
| **A_51_P517787** | Clptm1l | -0.457619835 | -1.222685785 | 0.307446116 | 10.88518143 | -1.498153495 | 0.189358884 | 0.709264768 | -4.483525574 |
| **A_51_P493270** | Mrps5 | 0.163432233 | -0.109836675 | 0.436701142 | 9.016840702 | 1.497951798 | 0.189409458 | 0.709264768 | -4.483580971 |
| **A_55_P2181928** | Slc45a3 | -0.662210338 | -1.769586222 | 0.445165546 | 8.62883399 | -1.497791083 | 0.189449766 | 0.709264768 | -4.483625112 |
| **A_55_P2178678** | P2rx5 | -0.417937108 | -1.116834148 | 0.280959933 | 8.039797914 | -1.497780173 | 0.189452503 | 0.709264768 | -4.483628109 |
| **A_55_P2107477** | Rab5c | -0.307312784 | -0.821219042 | 0.206593474 | 13.1252244 | -1.497776688 | 0.189453377 | 0.709264768 | -4.483629066 |
| **A_55_P2145531** | Mpst | 1.070281951 | -0.71952908 | 2.860092982 | 12.14340473 | 1.49775703 | 0.189458308 | 0.709264768 | -4.483634465 |
| **A_55_P2180944** | Tmco4 | -0.320166334 | -0.855623444 | 0.215290776 | 10.9682001 | -1.497618923 | 0.189492953 | 0.709264768 | -4.483672397 |
| **A_66_P118772** | Tmem136 | -0.88123274 | -2.355275651 | 0.592810172 | 9.346530547 | -1.497375912 | 0.18955393 | 0.709264768 | -4.483739142 |

| | | | | | | | | | |
|---|---|---|---|---|---|---|---|---|---|
| **A_51_P403273** | Tax1bp1 | -0.380281673 | -1.016592245 | 0.256028898 | 10.78626001 | -1.496878681 | 0.189678753 | 0.709264768 | -4.483875712 |
| **A_51_P297968** | Pdia6 | -0.381390693 | -1.019577734 | 0.256796348 | 10.02242864 | -1.49682992 | 0.189690999 | 0.709264768 | -4.483889105 |
| **A_55_P2142808** | NA | -0.336201679 | -0.898795662 | 0.226392305 | 15.2832574 | -1.496770241 | 0.189705986 | 0.709264768 | -4.483905496 |
| **A_55_P2000090** | Arf1 | -0.132582173 | -0.354477489 | 0.089313143 | 14.99290142 | -1.496536603 | 0.189764673 | 0.709264768 | -4.483969668 |
| **A_51_P240801** | Tmem173 | 0.372652263 | -0.251107557 | 0.996412084 | 10.17921681 | 1.496362143 | 0.189808506 | 0.709264768 | -4.484017586 |
| **A_55_P2031939** | Aqp4 | 0.316165238 | -0.213069177 | 0.845399654 | 8.395798346 | 1.49629211 | 0.189826105 | 0.709264768 | -4.484036822 |
| **A_55_P2053258** | Poc1a | 0.815168568 | -0.549455913 | 2.179793049 | 11.19381609 | 1.496182645 | 0.189853615 | 0.709264768 | -4.484066888 |
| **A_55_P2133806** | Tnrc6c | 0.266958287 | -0.179955217 | 0.713871791 | 9.930522954 | 1.49613325 | 0.189866031 | 0.709264768 | -4.484080456 |
| **A_52_P488305** | Ndel1 | -0.302282277 | -0.808333495 | 0.203768942 | 8.217750191 | -1.496127325 | 0.18986752 | 0.709264768 | -4.484082083 |
| **A_55_P2171976** | NA | 0.201871136 | -0.136102497 | 0.53984477 | 15.61674501 | 1.496034994 | 0.189890729 | 0.709264768 | -4.484107444 |
| **A_51_P303180** | Fam114a1 | -0.291857774 | -0.780489719 | 0.196774172 | 10.28727318 | -1.49602811 | 0.189892459 | 0.709264768 | -4.484109334 |
| **A_52_P321627** | NA | 0.572554654 | -0.386127235 | 1.531236543 | 14.89616499 | 1.495866279 | 0.189933145 | 0.709264768 | -4.484153784 |
| **A_55_P1972799** | Eif4enif1 | 0.185059469 | -0.124803938 | 0.494922877 | 10.50816297 | 1.495861436 | 0.189934363 | 0.709264768 | -4.484155114 |
| **A_51_P272123** | Ndufa10 | 0.277043248 | -0.186870247 | 0.740956744 | 11.14976037 | 1.495756537 | 0.18996074 | 0.709264768 | -4.484183927 |
| **A_55_P2320283** | Hmga2-ps | -0.254637111 | -0.681042573 | 0.171768351 | 8.068110973 | -1.495716498 | 0.189970809 | 0.709264768 | -4.484194924 |
| **A_55_P1980854** | Ankrd24 | 0.21559412 | -0.145458089 | 0.576646329 | 9.750882924 | 1.495606183 | 0.189998553 | 0.709264768 | -4.484225225 |
| **A_55_P1965416** | Atp9a | -0.673003741 | -1.800186514 | 0.454179032 | 14.04984728 | -1.495455454 | 0.190036468 | 0.709264768 | -4.484266625 |
| **A_52_P513177** | Sla | 0.538309897 | -0.36362552 | 1.440245313 | 8.562310802 | 1.494883227 | 0.190180472 | 0.709264768 | -4.484423801 |
| **A_52_P472324** | Slpi | -1.666196752 | -4.458007152 | 1.125613648 | 9.276003791 | -1.494826225 | 0.190194822 | 0.709264768 | -4.484439458 |
| **A_51_P145415** | Lpcat3 | -0.35574051 | -0.951812521 | 0.2403315 | 10.33195244 | -1.49480637 | 0.190199821 | 0.709264768 | -4.484444911 |
| **A_66_P100093** | Ccdc47 | 0.272931274 | -0.18442352 | 0.730286068 | 11.24763511 | 1.494687533 | 0.190229743 | 0.709264768 | -4.484477553 |
| **A_55_P2045476** | Hmgxb4 | 0.341597891 | -0.230845328 | 0.91404111 | 10.01575197 | 1.494627989 | 0.190244737 | 0.709264768 | -4.484493908 |
| **A_55_P2121537** | NA | 0.194527959 | -0.131474817 | 0.520530735 | 16.55342923 | 1.494552164 | 0.190263833 | 0.709264768 | -4.484514736 |
| **A_55_P2041340** | Atp5a1 | 0.245708213 | -0.166076286 | 0.657492713 | 14.65414413 | 1.494514127 | 0.190273413 | 0.709264768 | -4.484525183 |
| **A_55_P2105152** | Trim59 | 1.03909182 | -0.703046661 | 2.781230301 | 11.44419671 | 1.493900242 | 0.190428087 | 0.709264768 | -4.484693805 |
| **A_55_P2068177** | NA | 0.354785344 | -0.24004778 | 0.949618468 | 11.90349221 | 1.493897745 | 0.190428717 | 0.709264768 | -4.484694491 |
| **A_55_P2086258** | Pan3 | -0.697240831 | -1.90171353 | 0.507231868 | 11.28332614 | -1.541872833 | 0.190436554 | 0.709264768 | -4.494409904 |
| **A_55_P2004801** | Tacc3 | 1.177225061 | -0.796573005 | 3.151023127 | 11.80218563 | 1.493850461 | 0.190440635 | 0.709264768 | -4.484707479 |
| **A_52_P114494** | Stat6 | -0.184514757 | -0.493906777 | 0.124877264 | 9.452732592 | -1.49373082 | 0.190470796 | 0.709264768 | -4.484740342 |
| **A_55_P1979605** | Eif3m | 0.490805429 | -0.332215894 | 1.313826752 | 13.46214276 | 1.49364922 | 0.190491369 | 0.709264768 | -4.484762756 |
| **A_52_P556099** | Oip5 | 0.412343781 | -0.279134812 | 1.103822374 | 8.520265921 | 1.493588823 | 0.190506598 | 0.709264768 | -4.484779346 |
| **A_55_P1984541** | NA | -0.44136509 | -1.181539312 | 0.298809131 | 8.565869767 | -1.493531441 | 0.190521067 | 0.709264768 | -4.484795107 |
| **A_55_P1988531** | Myh11 | -0.233037296 | -0.623875901 | 0.157801309 | 9.011755781 | -1.493407486 | 0.190552328 | 0.709264768 | -4.484829156 |
| **A_55_P1961385** | NA | 0.215135767 | -0.145689418 | 0.575960951 | 8.295654821 | 1.493365528 | 0.190562911 | 0.709264768 | -4.484840681 |
| **A_55_P2040743** | Zwilch | 0.416706703 | -0.282260711 | 1.115674116 | 8.895239751 | 1.493220359 | 0.190599529 | 0.709264768 | -4.484880556 |
| **A_55_P1986185** | NA | 0.616691259 | -0.418268515 | 1.651651033 | 14.97068528 | 1.492432397 | 0.190798408 | 0.709264768 | -4.485096999 |
| **A_55_P2185359** | NA | 0.328242157 | -0.222677389 | 0.879161704 | 13.65843087 | 1.492301318 | 0.190831511 | 0.709264768 | -4.485133005 |
| **A_51_P470448** | Setx | 0.384965607 | -0.261183501 | 1.031114714 | 11.49956001 | 1.492242922 | 0.19084626 | 0.709264768 | -4.485149046 |
| **A_52_P349298** | Rprd1b | -0.164428676 | -0.440418757 | 0.111561406 | 8.854257063 | -1.492225406 | 0.190850685 | 0.709264768 | -4.485153857 |
| **A_66_P131036** | NA | 0.984938225 | -0.668446151 | 2.638322601 | 10.64614637 | 1.492057324 | 0.190893145 | 0.709264768 | -4.485200028 |
| **A_55_P1982451** | Hmgn5 | 0.231579096 | -0.157230807 | 0.620389 | 11.59311031 | 1.491806113 | 0.190956621 | 0.709264768 | -4.485269033 |
| **A_55_P2045188** | NA | -0.146099795 | -0.391396921 | 0.099197331 | 8.855691882 | -1.491789313 | 0.190960866 | 0.709264768 | -4.485273648 |
| **A_55_P2161695** | Poglut2 | -0.294774287 | -0.789769227 | 0.200220654 | 9.19823775 | -1.491554691 | 0.19102017 | 0.709264768 | -4.485338097 |
| **A_55_P2032538** | Pde3b | 0.655139102 | -0.444995813 | 1.755274018 | 9.396394263 | 1.491550351 | 0.191021267 | 0.709264768 | -4.485339289 |
| **A_55_P2096165** | NA | 0.416297863 | -0.282916017 | 1.115511744 | 11.57228297 | 1.4912295 | 0.191102395 | 0.709264768 | -4.485427425 |
| **A_55_P2155727** | Mafa | 0.181439886 | -0.123330007 | 0.486209779 | 10.47349585 | 1.491114688 | 0.191131433 | 0.709264768 | -4.485458964 |
| **A_51_P322972** | Hkdc1 | -0.515627468 | -1.381763265 | 0.350508328 | 9.239924262 | -1.491078137 | 0.191140679 | 0.709264768 | -4.485469004 |
| **A_55_P1974114** | NA | 0.564020475 | -0.383464671 | 1.51150562 | 15.21376924 | 1.490983391 | 0.191164646 | 0.709264768 | -4.48549503 |

| | | | | | | | | | |
|---|---|---|---|---|---|---|---|---|---|
| **A_51_P434426** | Rwdd4a | 0.327839168 | -0.222915643 | 0.87859398 | 9.325635486 | 1.490915003 | 0.191181948 | 0.709264768 | -4.485513816 |
| **A_52_P56394** | Psmg4 | 0.622780295 | -0.423556814 | 1.669117404 | 10.51488892 | 1.490780078 | 0.191216087 | 0.709264768 | -4.48555088 |
| **A_51_P446315** | Cdc37l1 | -0.158304725 | -0.424310609 | 0.107701159 | 9.518357007 | -1.490572069 | 0.19126873 | 0.709264768 | -4.48560802 |
| **A_55_P1968664** | NA | 1.159093111 | -0.788637481 | 3.106823704 | 13.09092618 | 1.490526785 | 0.191280192 | 0.709264768 | -4.485620459 |
| **A_55_P1972034** | Muc16 | -0.390066816 | -1.045612265 | 0.265478634 | 8.995072845 | -1.490344068 | 0.191326448 | 0.709264768 | -4.485670651 |
| **A_55_P2042615** | NA | -0.250302589 | -0.671002393 | 0.170397216 | 9.314544875 | -1.490195944 | 0.191363954 | 0.709264768 | -4.485711341 |
| **A_55_P2175291** | NA | -0.187880139 | -0.503681476 | 0.127921198 | 14.74899486 | -1.490106313 | 0.191386652 | 0.709264768 | -4.485735963 |
| **A_52_P123354** | Tusc2 | -0.426015502 | -1.14213142 | 0.290100417 | 9.450132834 | -1.490021106 | 0.191408233 | 0.709264768 | -4.485759369 |
| **A_55_P2122260** | Tmem87b | -0.278493645 | -0.746634549 | 0.18964726 | 8.284956034 | -1.490009525 | 0.191411166 | 0.709264768 | -4.48576255 |
| **A_52_P660922** | Dock1 | 0.624918778 | -0.425556229 | 1.675393785 | 11.69604245 | 1.490006615 | 0.191411903 | 0.709264768 | -4.48576335 |
| **A_51_P262230** | Ggact | 0.286449242 | -0.195123996 | 0.768022479 | 8.957174107 | 1.489826443 | 0.191457544 | 0.709264768 | -4.485812843 |
| **A_52_P423174** | Acaa1b | -0.627862157 | -1.683475901 | 0.427751587 | 9.578276858 | -1.489737045 | 0.191480195 | 0.709264768 | -4.485837401 |
| **A_55_P2031898** | NA | 0.311898422 | -0.212498641 | 0.836295485 | 10.72292139 | 1.48971528 | 0.191485709 | 0.709264768 | -4.48584338 |
| **A_51_P478003** | Papolg | 0.202936389 | -0.138294958 | 0.544167737 | 9.164126276 | 1.489571497 | 0.191522145 | 0.709264768 | -4.485882878 |
| **A_55_P2058601** | Gm7097 | -0.727200481 | -1.949999444 | 0.495598481 | 9.012605626 | -1.489530641 | 0.191532499 | 0.709264768 | -4.485894101 |
| **A_55_P2100655** | Lsm6 | 0.351873607 | -0.239846776 | 0.943593991 | 13.57695762 | 1.489431411 | 0.19155765 | 0.709264768 | -4.48592136 |
| **A_66_P138999** | NA | -0.209875968 | -0.562811598 | 0.143059662 | 8.087520904 | -1.48942128 | 0.191560218 | 0.709264768 | -4.485924143 |
| **A_55_P2074453** | Slc16a9 | -0.408463493 | -1.095378389 | 0.278451402 | 8.081816083 | -1.489363339 | 0.191574905 | 0.709264768 | -4.48594006 |
| **A_66_P101600** | Lamb1 | -0.201844311 | -0.541310827 | 0.137622204 | 14.42862142 | -1.489257915 | 0.191601631 | 0.709264768 | -4.48596902 |
| **A_55_P2083904** | Tmem185 | -0.218267531 | -0.585357722 | 0.14882266 | 8.54430497 | -1.489246886 | 0.191604427 | 0.709264768 | -4.48597205 |
| **A_55_P1984283** | NA | 0.256548831 | -0.17493971 | 0.688037371 | 14.49670517 | 1.489193456 | 0.191617974 | 0.709264768 | -4.485986727 |
| **A_55_P2133405** | Dmkn | -2.042726505 | -5.478858216 | 1.393405206 | 11.35043131 | -1.488986523 | 0.191670449 | 0.709264768 | -4.486043573 |
| **A_55_P1954860** | NA | 0.723163037 | -0.493338317 | 1.93966439 | 14.10526154 | 1.488928931 | 0.191685055 | 0.709264768 | -4.486059394 |
| **A_51_P313157** | Arhgef5 | -0.674657366 | -1.809666694 | 0.460351962 | 10.6631171 | -1.48879253 | 0.191719654 | 0.709264768 | -4.486096864 |
| **A_51_P169693** | Bst2 | -1.121155473 | -3.007681167 | 0.76537022 | 11.38461158 | -1.488515847 | 0.191789854 | 0.709264768 | -4.486172871 |
| **A_51_P303160** | Arg1 | -0.196232223 | -0.526430729 | 0.133966282 | 7.901601908 | -1.488488784 | 0.191796722 | 0.709264768 | -4.486180306 |
| **A_55_P2081005** | Cox7c | 0.134387551 | -0.091751883 | 0.360526985 | 8.057359029 | 1.488446007 | 0.191807578 | 0.709264768 | -4.486192057 |
| **A_55_P2138070** | NA | 0.704594625 | -0.481187451 | 1.890376701 | 14.26636854 | 1.488280496 | 0.191849587 | 0.709264768 | -4.486237525 |
| **A_55_P2190994** | Cdsn | -0.49860475 | -1.363295822 | 0.366086323 | 7.948120745 | -1.535883028 | 0.191850154 | 0.709264768 | -4.495671233 |
| **A_52_P297332** | Aida | -0.246816503 | -0.662258362 | 0.168625356 | 8.533180987 | -1.488038909 | 0.191910921 | 0.709264768 | -4.486303891 |
| **A_51_P170946** | Hnrnpm | 0.540687815 | -0.36942977 | 1.4508054 | 13.96887221 | 1.487987978 | 0.191923853 | 0.709264768 | -4.486317882 |
| **A_55_P1993640** | Gm6994 | 0.239513236 | -0.163673652 | 0.642700124 | 13.29642428 | 1.487899042 | 0.191946438 | 0.709264768 | -4.486342314 |
| **A_55_P1992421** | Hmgb3 | 1.138746798 | -0.778218806 | 3.055712402 | 11.84201132 | 1.487863856 | 0.191955374 | 0.709264768 | -4.48635198 |
| **A_51_P230098** | Pbk | 1.233266693 | -0.842826783 | 3.309360169 | 10.28921229 | 1.487854437 | 0.191957767 | 0.709264768 | -4.486354568 |
| **A_55_P2001773** | Copg2 | 0.240703309 | -0.164593382 | 0.646 | 10.52261909 | 1.487508132 | 0.192045739 | 0.709264768 | -4.486449702 |
| **A_51_P466673** | Srsf7 | 0.369747165 | -0.253057815 | 0.992552146 | 9.387384355 | 1.486973133 | 0.192181719 | 0.709264768 | -4.486596674 |
| **A_55_P1956083** | Gpr68 | 0.231554761 | -0.158524085 | 0.621633608 | 7.980788124 | 1.486796948 | 0.192226521 | 0.709264768 | -4.486645075 |
| **A_51_P204402** | Shcbp1 | 1.167413836 | -0.79942272 | 3.134250392 | 10.34378137 | 1.486643753 | 0.192265484 | 0.709264768 | -4.48668716 |
| **A_55_P1980677** | Fam24b | 0.223893688 | -0.153325027 | 0.601112404 | 9.692825701 | 1.486616554 | 0.192272402 | 0.709264768 | -4.486694632 |
| **A_55_P2019083** | Rps6kb1 | 0.308447442 | -0.211274038 | 0.828168922 | 12.94138079 | 1.486486097 | 0.192305589 | 0.709264768 | -4.486730471 |
| **A_55_P1966883** | Rnaseh2a | 0.346187595 | -0.237305143 | 0.929680332 | 10.06718299 | 1.486025945 | 0.19242269 | 0.709264768 | -4.486856884 |
| **A_55_P2160032** | Atp5pb | 0.475343065 | -0.325846182 | 1.276532312 | 13.49546136 | 1.486012271 | 0.192426171 | 0.709264768 | -4.48686064 |
| **A_55_P2052959** | Ctbp1 | 0.266927486 | -0.182979227 | 0.716834198 | 14.40762472 | 1.486008072 | 0.19242724 | 0.709264768 | -4.486861794 |
| **A_55_P2084706** | Acaca | 0.529790638 | -0.363220812 | 1.422802088 | 10.92059332 | 1.485927423 | 0.192447771 | 0.709264768 | -4.48688395 |
| **A_55_P2016462** | Cxcl10 | 0.981840719 | -0.673195817 | 2.636877256 | 8.619613057 | 1.485880213 | 0.192459791 | 0.709264768 | -4.486896919 |
| **A_55_P2018393** | NA | 0.602421239 | -0.413084155 | 1.617926633 | 15.73119311 | 1.48582748 | 0.192473218 | 0.709264768 | -4.486911406 |
| **A_55_P2212027** | Utrn | -0.523590814 | -1.406362985 | 0.359181357 | 8.960730378 | -1.485572087 | 0.192538256 | 0.709264768 | -4.486981568 |
| **A_55_P1998023** | NA | 0.267976188 | -0.183835928 | 0.719788303 | 8.835026173 | 1.485554803 | 0.192542659 | 0.709264768 | -4.486986316 |

| | | | | | | | | | |
|---|---|---|---|---|---|---|---|---|---|
| **A_52_P346033** | Trmt1l | -0.157229428 | -0.422331838 | 0.107872983 | 8.451516883 | -1.48549263 | 0.192558495 | 0.709264768 | -4.487003397 |
| **A_55_P2129232** | Calm2 | -0.205194475 | -0.551176122 | 0.140787172 | 14.5922622 | -1.48546679 | 0.192565078 | 0.709264768 | -4.487010496 |
| **A_55_P2052754** | Pcm1 | 0.223683704 | -0.153474088 | 0.600841496 | 9.080874632 | 1.485462205 | 0.192566245 | 0.709264768 | -4.487011755 |
| **A_55_P2050483** | Abcc12 | -0.369239141 | -0.991828187 | 0.253349906 | 8.045544574 | -1.485445086 | 0.192570606 | 0.709264768 | -4.487016458 |
| **A_55_P2098057** | Sf3a2 | 0.237526368 | -0.162992796 | 0.638045533 | 12.59794959 | 1.485384447 | 0.192586054 | 0.709264768 | -4.487033117 |
| **A_52_P503334** | Erlin2 | -0.360668343 | -0.969130915 | 0.247794229 | 10.28675266 | -1.48465143 | 0.192772885 | 0.709264768 | -4.487234495 |
| **A_51_P341130** | Iqgap1 | -0.214844023 | -0.577381087 | 0.147693041 | 13.28026244 | -1.484298361 | 0.192862936 | 0.709264768 | -4.487331492 |
| **A_55_P2187918** | Cep55 | 0.201685931 | -0.149067559 | 0.552439422 | 8.387317158 | 1.531568086 | 0.192874961 | 0.709264768 | -4.496580673 |
| **A_55_P2151899** | NA | 0.7336184 | -0.504365142 | 1.971601941 | 12.78466851 | 1.484245325 | 0.192876466 | 0.709264768 | -4.487346062 |
| **A_52_P343856** | Creb3l2 | -0.681098691 | -1.83065487 | 0.468457488 | 9.46022532 | -1.483987323 | 0.1929423 | 0.709264768 | -4.487416943 |
| **A_52_P357484** | Nfrkb | 0.168942078 | -0.116229074 | 0.454113231 | 8.433984765 | 1.483824669 | 0.192983815 | 0.709264768 | -4.487461628 |
| **A_55_P1980165** | NA | 0.447537011 | -0.307925356 | 1.202999379 | 15.43846312 | 1.483769596 | 0.192997873 | 0.709264768 | -4.487476759 |
| **A_51_P159673** | Snx1 | -0.319343525 | -0.858423203 | 0.219736153 | 13.45222893 | -1.483732124 | 0.193007439 | 0.709264768 | -4.487487053 |
| **A_55_P2004126** | NA | 0.307857231 | -0.211854947 | 0.82756941 | 10.31807083 | 1.483668278 | 0.193023739 | 0.709264768 | -4.487504594 |
| **A_55_P2041060** | Rbp3 | 0.275269212 | -0.189438364 | 0.739976788 | 10.77713014 | 1.483638973 | 0.193031221 | 0.709264768 | -4.487512645 |
| **A_55_P2000618** | Gm12690 | 0.224019945 | -0.154171353 | 0.602211244 | 8.603721171 | 1.483629635 | 0.193033605 | 0.709264768 | -4.48751521 |
| **A_55_P2063146** | NA | 1.23320862 | -0.848983211 | 3.31540045 | 11.17146508 | 1.48342693 | 0.193085367 | 0.709264768 | -4.487570899 |
| **A_51_P171200** | Golm1 | -0.699312995 | -1.880533214 | 0.481907223 | 11.7422127 | -1.482829004 | 0.193238127 | 0.709264768 | -4.487735168 |
| **A_52_P213696** | Fndc5 | -0.727408712 | -1.956097473 | 0.50128005 | 10.74451652 | -1.482814955 | 0.193241718 | 0.709264768 | -4.487739028 |
| **A_55_P2137653** | NA | 0.653376976 | -0.450365804 | 1.757119755 | 13.28701614 | 1.482676133 | 0.193277201 | 0.709264768 | -4.487777167 |
| **A_55_P2181356** | Cdh11 | 0.535425083 | -0.36913792 | 1.439988087 | 9.414097585 | 1.482553032 | 0.193308672 | 0.709264768 | -4.487810987 |
| **A_51_P123047** | Smc1a | 0.776101405 | -0.53510565 | 2.08730846 | 13.31725759 | 1.482509681 | 0.193319755 | 0.709264768 | -4.487822897 |
| **A_55_P2072985** | Sptan1 | -0.189201714 | -0.508887515 | 0.130484086 | 13.13070742 | -1.482354467 | 0.193359444 | 0.709264768 | -4.48786554 |
| **A_55_P2093014** | NA | 0.333199937 | -0.229830748 | 0.896230622 | 14.84468932 | 1.482255914 | 0.193384649 | 0.709264768 | -4.487892616 |
| **A_55_P2172989** | Ptpn23 | -0.46113264 | -1.240489382 | 0.318224103 | 11.21824605 | -1.481971706 | 0.193457351 | 0.709264768 | -4.487970698 |
| **A_52_P268279** | Fbxl17 | -0.187770226 | -0.505143319 | 0.129602867 | 10.02106496 | -1.481859304 | 0.193486112 | 0.709264768 | -4.488001579 |
| **A_51_P512992** | Entpd3 | 0.468526709 | -0.323420327 | 1.260473744 | 8.845741781 | 1.481796489 | 0.193502186 | 0.709264768 | -4.488018836 |
| **A_55_P2045036** | NA | 0.160554435 | -0.110863655 | 0.431972525 | 8.647297004 | 1.481609867 | 0.19354995 | 0.709264768 | -4.488070108 |
| **A_51_P382928** | Cstf3 | 0.372565953 | -0.257281823 | 1.002413729 | 10.3297632 | 1.481555448 | 0.19356388 | 0.709264768 | -4.488085059 |
| **A_55_P2143150** | Qk | -0.284295774 | -0.764976395 | 0.196384847 | 10.12487217 | -1.481372016 | 0.193610841 | 0.709264768 | -4.488135455 |
| **A_51_P315795** | Tubb4a | 0.359672551 | -0.248507006 | 0.967852109 | 10.69391035 | 1.481241335 | 0.193644304 | 0.709264768 | -4.488171358 |
| **A_55_P2031786** | Fcsk | -0.244705991 | -0.65850508 | 0.169093098 | 10.89269416 | -1.481171739 | 0.193662127 | 0.709264768 | -4.488190479 |
| **A_55_P2008176** | Zscan12 | 0.258320395 | -0.178521829 | 0.695162618 | 8.957023456 | 1.48110017 | 0.193680458 | 0.709264768 | -4.488210141 |
| **A_52_P576778** | Tmem30a | -0.342765169 | -0.922416665 | 0.236886327 | 10.89593287 | -1.481085289 | 0.193684269 | 0.709264768 | -4.48821423 |
| **A_55_P2011235** | Rpl13 | 0.371376572 | -0.256680668 | 0.999433813 | 16.517409 | 1.481036026 | 0.193696888 | 0.709264768 | -4.488227764 |
| **A_51_P360963** | Mbtd1 | 0.511145572 | -0.353321432 | 1.375612576 | 10.74898884 | 1.480970906 | 0.193713569 | 0.709264768 | -4.488245655 |
| **A_55_P2067645** | Gm5136 | 0.247317681 | -0.170981575 | 0.665616938 | 8.730319104 | 1.48087506 | 0.193738124 | 0.709264768 | -4.488271988 |
| **A_55_P2172688** | Srbd1 | 0.175970163 | -0.121685897 | 0.473626224 | 8.82610209 | 1.480725833 | 0.19377636 | 0.709264768 | -4.488312987 |
| **A_55_P2066299** | Gpr137b | -0.822104201 | -2.212902381 | 0.568693979 | 9.566738212 | -1.48051598 | 0.193830142 | 0.709264768 | -4.488370642 |
| **A_55_P2038046** | NA | 0.577424909 | -0.399534731 | 1.554384548 | 10.02988823 | 1.480366462 | 0.19386847 | 0.709264768 | -4.488411721 |
| **A_51_P207412** | Slf1 | 0.309086297 | -0.213883744 | 0.832056337 | 9.576024356 | 1.480312092 | 0.193882409 | 0.709264768 | -4.488426658 |
| **A_55_P1960451** | Thsd7a | -0.271947178 | -0.732089087 | 0.18819473 | 8.061591316 | -1.480277415 | 0.1938913 | 0.709264768 | -4.488436186 |
| **A_51_P229633** | Rprd1b | -0.268419898 | -0.722606237 | 0.185766441 | 11.71353755 | -1.480236078 | 0.193901898 | 0.709264768 | -4.488447542 |
| **A_55_P1979699** | Spop | -0.173970558 | -0.468370384 | 0.120429267 | 9.655188324 | -1.480091465 | 0.193938982 | 0.709264768 | -4.488487274 |
| **A_55_P2058613** | NA | 0.14704785 | -0.101821222 | 0.395916921 | 8.438749227 | 1.479919513 | 0.193983085 | 0.709264768 | -4.488534517 |
| **A_55_P2094118** | Fmnl3 | 0.184924453 | -0.128062445 | 0.497911351 | 9.984364127 | 1.47985284 | 0.194000188 | 0.709264768 | -4.488552834 |
| **A_51_P235705** | Tsfm | 0.207448468 | -0.143691501 | 0.558588437 | 9.009158524 | 1.479722616 | 0.194033597 | 0.709264768 | -4.488588612 |
| **A_52_P520466** | Kif18b | 0.314302674 | -0.217793577 | 0.846398924 | 8.263738275 | 1.479477016 | 0.194096621 | 0.709264768 | -4.48865609 |

| | | | | | | | | | |
|---|---|---|---|---|---|---|---|---|---|
| **A_55_P2115235** | Bckdhb | 0.363043928 | -0.251579475 | 0.977667331 | 10.52705671 | 1.479450469 | 0.194103434 | 0.709264768 | -4.488663383 |
| **A_55_P2077558** | Sod3 | -1.295007736 | -3.487524062 | 0.89750859 | 9.287536321 | -1.479380441 | 0.194121409 | 0.709264768 | -4.488682623 |
| **A_51_P359137** | Doc2g | -0.25739046 | -0.693206507 | 0.178425588 | 8.010768467 | -1.479243173 | 0.194156646 | 0.709264768 | -4.488720337 |
| **A_51_P520857** | Selenok-p | -0.290837523 | -0.78333409 | 0.201659043 | 13.00972819 | -1.479100135 | 0.194193371 | 0.709264768 | -4.488759636 |
| **A_55_P2255449** | NA | 0.279760373 | -0.19398576 | 0.753506505 | 9.14954141 | 1.47907728 | 0.19419924 | 0.709264768 | -4.488765915 |
| **A_52_P32903** | Zzef1 | -0.24700643 | -0.66532191 | 0.17130905 | 10.00287621 | -1.478954008 | 0.194230896 | 0.709264768 | -4.488799784 |
| **A_51_P177984** | Fam234a | -0.883952863 | -2.380984829 | 0.613079102 | 11.48395783 | -1.478932589 | 0.194236397 | 0.709264768 | -4.488805668 |
| **A_55_P2170034** | Rgl1 | -0.211669547 | -0.570152597 | 0.146813502 | 10.4835579 | -1.478904402 | 0.194243636 | 0.709264768 | -4.488813413 |
| **A_52_P585448** | Ptp4a2 | 0.199964568 | -0.138714478 | 0.538643615 | 11.60479822 | 1.478819153 | 0.194265532 | 0.709264768 | -4.488836834 |
| **A_52_P364140** | Itga5 | -0.382819282 | -1.031218754 | 0.26558019 | 8.518318129 | -1.478772951 | 0.1942774 | 0.709264768 | -4.488849528 |
| **A_55_P2066274** | NA | 0.139995949 | -0.09715108 | 0.377142979 | 12.26915572 | 1.478591235 | 0.194324084 | 0.709264768 | -4.488899454 |
| **A_66_P120745** | Armc1 | 0.339127619 | -0.23535509 | 0.913610329 | 11.2565486 | 1.478551801 | 0.194334217 | 0.709264768 | -4.488910288 |
| **A_55_P1991214** | Lair1 | 0.166028834 | -0.115317334 | 0.447375001 | 8.33638933 | 1.478062706 | 0.194459928 | 0.709264768 | -4.489044666 |
| **A_51_P446825** | Trmt9b | -0.229569893 | -0.618592845 | 0.15945306 | 8.133472731 | -1.478053149 | 0.194462385 | 0.709264768 | -4.489047292 |
| **A_52_P255849** | NA | -0.332469328 | -0.895897669 | 0.230959013 | 8.628507119 | -1.477961912 | 0.194485844 | 0.709264768 | -4.489072359 |
| **A_55_P1955325** | Zfp820 | 0.332543396 | -0.23102717 | 0.896113962 | 8.669938939 | 1.477918107 | 0.194497108 | 0.709264768 | -4.489084394 |
| **A_55_P2019217** | NA | 0.193978254 | -0.13477424 | 0.522730747 | 8.194515234 | 1.477863541 | 0.194511141 | 0.709264768 | -4.489099387 |
| **A_55_P2165074** | Rnf31 | -0.51298559 | -1.382396045 | 0.356424866 | 9.583892197 | -1.477851009 | 0.194514363 | 0.709264768 | -4.48910283 |
| **A_55_P2000668** | Stxbp2 | 0.305258086 | -0.212194192 | 0.822710363 | 10.38802819 | 1.477567124 | 0.194587384 | 0.709264768 | -4.489180827 |
| **A_52_P373893** | NA | -0.546777237 | -1.473717648 | 0.380163174 | 10.48214989 | -1.477436948 | 0.194620877 | 0.709264768 | -4.489216593 |
| **A_52_P117576** | Casp3 | 0.179213548 | -0.124625326 | 0.483052422 | 8.110545486 | 1.47733112 | 0.194648109 | 0.709264768 | -4.489245669 |
| **A_55_P2161675** | Zfp133-ps | 0.252353508 | -0.175500526 | 0.680207542 | 9.746194646 | 1.477284229 | 0.194660176 | 0.709264768 | -4.489258552 |
| **A_55_P2116924** | Dbi | -0.421717615 | -1.136769645 | 0.293334416 | 14.9047103 | -1.477183483 | 0.194686105 | 0.709264768 | -4.489286233 |
| **A_55_P2148082** | Larp7 | 0.497831171 | -0.346361664 | 1.342024007 | 10.36295295 | 1.477034943 | 0.194724341 | 0.709264768 | -4.489327044 |
| **A_66_P140408** | D8Ertd738 | -0.18048609 | -0.486607516 | 0.125635336 | 10.2515424 | -1.47672747 | 0.194803511 | 0.709264768 | -4.489411523 |
| **A_52_P185044** | Adipor2 | -0.26185144 | -0.705976547 | 0.182273666 | 13.88241382 | -1.476726227 | 0.194803832 | 0.709264768 | -4.489411864 |
| **A_55_P2158030** | Glrx3 | 0.450312564 | -0.313524366 | 1.214149493 | 13.199127 | 1.47660304 | 0.194835559 | 0.709264768 | -4.48944571 |
| **A_66_P125209** | Kif2c | 0.289306535 | -0.201446581 | 0.78005965 | 8.271267241 | 1.476541043 | 0.194851528 | 0.709264768 | -4.489462744 |
| **A_55_P2100884** | Fjx1 | 0.222710525 | -0.155084273 | 0.600505322 | 8.87596629 | 1.476505652 | 0.194860645 | 0.709264768 | -4.489472468 |
| **A_55_P2050448** | Slc25a3 | 0.463390435 | -0.322719281 | 1.249500151 | 10.79224218 | 1.476434711 | 0.194878921 | 0.709264768 | -4.489491959 |
| **A_55_P2151461** | NA | 0.653743657 | -0.455364313 | 1.762851628 | 13.19924797 | 1.476331913 | 0.194905407 | 0.709264768 | -4.489520203 |
| **A_52_P188261** | Camk2d | -0.476251355 | -1.284237924 | 0.331735213 | 8.610311714 | -1.476326546 | 0.194906789 | 0.709264768 | -4.489521678 |
| **A_55_P2169010** | Ndfip1 | -0.565372113 | -1.525159466 | 0.394415241 | 11.18838277 | -1.475399748 | 0.19514573 | 0.709264768 | -4.489776319 |
| **A_51_P489522** | Ctla2b | -2.098764026 | -5.662956418 | 1.465428365 | 12.7804319 | -1.474866864 | 0.19528324 | 0.709264768 | -4.489922732 |
| **A_51_P117865** | Fam20c | -0.606512366 | -1.636591531 | 0.423566799 | 8.773535007 | -1.474753388 | 0.195312534 | 0.709264768 | -4.48995391 |
| **A_51_P100787** | Snw1 | 0.364669468 | -0.254680832 | 0.984019768 | 11.03112426 | 1.474732992 | 0.1953178 | 0.709264768 | -4.501134585 |
| **A_51_P302139** | Mastl | 0.300435813 | -0.209990621 | 0.810862247 | 8.389432471 | 1.47424237 | 0.195444507 | 0.709264768 | -4.490094317 |
| **A_55_P2134427** | NA | -0.207402922 | -0.559791144 | 0.1449853 | 9.940345016 | -1.474157315 | 0.19546648 | 0.709264768 | -4.490117686 |
| **A_52_P306351** | Asph | 0.366588483 | -0.256278016 | 0.989454983 | 10.83057644 | 1.474124586 | 0.195474937 | 0.709264768 | -4.490126679 |
| **A_55_P1962781** | NA | 0.24074619 | -0.168351319 | 0.649843698 | 8.500383883 | 1.473950618 | 0.19551989 | 0.709264768 | -4.490174478 |
| **A_55_P2143070** | Ass1 | -0.228855973 | -0.617756073 | 0.160044128 | 11.24351147 | -1.473922135 | 0.195527251 | 0.709264768 | -4.490182303 |
| **A_55_P2083449** | Spryd7 | -0.233619941 | -0.630625978 | 0.163386095 | 9.21133922 | -1.473883463 | 0.195537246 | 0.709264768 | -4.490192929 |
| **A_51_P363791** | Ap3m1 | 0.203690792 | -0.142468063 | 0.549849647 | 10.93676311 | 1.473826281 | 0.195552025 | 0.709264768 | -4.49020864 |
| **A_55_P2013645** | Tubg2 | -0.281608985 | -0.760184821 | 0.196966851 | 8.923695647 | -1.473825569 | 0.195552209 | 0.709264768 | -4.490208836 |
| **A_51_P437786** | Rbm28 | 0.208476894 | -0.145845671 | 0.562799458 | 11.87746171 | 1.473701285 | 0.195584335 | 0.709264768 | -4.490242984 |
| **A_55_P2125947** | Agpat2 | -0.329625167 | -0.889848914 | 0.230598579 | 9.079782971 | -1.473700378 | 0.19558457 | 0.709264768 | -4.490243233 |
| **A_51_P292116** | Ica1 | -0.161440646 | -0.435837391 | 0.112956099 | 9.973239293 | -1.473615824 | 0.195606429 | 0.709264768 | -4.490266465 |
| **A_55_P1967291** | Ncaph | 1.312919851 | -0.918795415 | 3.544635117 | 11.48067862 | 1.473498765 | 0.195636695 | 0.709264768 | -4.490298628 |

| | | | | | | | | | |
|---|---|---|---|---|---|---|---|---|---|
| **A_52_P533146** | Ddit3 | -0.879757842 | -2.375299699 | 0.615784015 | 9.156128398 | -1.473380507 | 0.195667276 | 0.709264768 | -4.490331121 |
| **A_55_P2066958** | Sh3glb1 | -0.312962294 | -0.84501313 | 0.219088542 | 11.25951116 | -1.473293361 | 0.195689815 | 0.709264768 | -4.490355065 |
| **A_55_P2031466** | Lrrc32 | -0.534374456 | -1.44312093 | 0.374372018 | 9.989904471 | -1.472832289 | 0.195809101 | 0.709264768 | -4.490481749 |
| **A_52_P681** | Cnot4 | 0.171965731 | -0.120513724 | 0.464445186 | 8.98497112 | 1.472641013 | 0.195858608 | 0.709264768 | -4.490534304 |
| **A_51_P215540** | Rnf4 | -0.467454131 | -1.287103553 | 0.352195292 | 10.50341283 | -1.51905525 | 0.195877695 | 0.709264768 | -4.499221634 |
| **A_55_P2005828** | Hbb-bh1 | 0.29303196 | -0.205436799 | 0.791500719 | 8.150549356 | 1.472405365 | 0.195919615 | 0.709264768 | -4.490599051 |
| **A_55_P2107120** | Kansl2 | 0.289546943 | -0.20305821 | 0.782152095 | 9.436657737 | 1.472212088 | 0.195969666 | 0.709264768 | -4.490652156 |
| **A_55_P1997097** | Bod1 | -0.198610334 | -0.536561366 | 0.139340698 | 8.958471899 | -1.471968143 | 0.196032855 | 0.709264768 | -4.490719183 |
| **A_51_P272147** | Emc9 | -0.859400172 | -2.321805268 | 0.603004924 | 10.16871764 | -1.471899271 | 0.196050698 | 0.709264768 | -4.490738107 |
| **A_55_P1970389** | Srsf10 | 0.199792909 | -0.140280875 | 0.539866694 | 9.94413436 | 1.47148982 | 0.196156811 | 0.709264768 | -4.490850608 |
| **A_55_P2055477** | Scaf11 | 0.23512069 | -0.165085958 | 0.635327337 | 10.35693172 | 1.471488574 | 0.196157134 | 0.709264768 | -4.490850951 |
| **A_52_P254795** | Ndufc1 | 0.442104761 | -0.310679865 | 1.194889387 | 14.14820951 | 1.470973322 | 0.196290743 | 0.709264768 | -4.490992523 |
| **A_51_P331462** | Cgrrf1 | -0.32910889 | -0.889786495 | 0.231568715 | 9.218836953 | -1.470201122 | 0.196491141 | 0.709264768 | -4.491204696 |
| **A_51_P158552** | Zscan12 | 0.327104427 | -0.230178542 | 0.884387395 | 10.32785594 | 1.470147793 | 0.196504988 | 0.709264768 | -4.491219349 |
| **A_52_P4046** | Baz2a | -0.510444452 | -1.380204783 | 0.359315879 | 9.520568764 | -1.469938745 | 0.196559276 | 0.709264768 | -4.491276788 |
| **A_55_P2098140** | Tmed2 | -0.411919022 | -1.113843276 | 0.290005233 | 14.15690271 | -1.469846368 | 0.19658327 | 0.709264768 | -4.49130217 |
| **A_51_P437717** | Glmp | -0.946571745 | -2.559588473 | 0.666444982 | 10.67107092 | -1.469822906 | 0.196589365 | 0.709264768 | -4.491308617 |
| **A_51_P450123** | Mrpl36 | 0.140185232 | -0.098699593 | 0.379070058 | 11.88919485 | 1.469819647 | 0.196590211 | 0.709264768 | -4.491309512 |
| **A_51_P495171** | Pbdc1 | 0.489405219 | -0.344606635 | 1.323417073 | 11.27945808 | 1.469760998 | 0.196605447 | 0.709264768 | -4.491325627 |
| **A_51_P181746** | Zfyve1 | -0.459183904 | -1.242067129 | 0.323699322 | 8.788623634 | -1.469061623 | 0.196787212 | 0.709264768 | -4.491517791 |
| **A_52_P214927** | Zmynd8 | -0.489241058 | -1.32378824 | 0.345306125 | 9.292506873 | -1.468325519 | 0.196978695 | 0.709264768 | -4.491720048 |
| **A_55_P2151951** | NA | 0.476928528 | -0.336790543 | 1.2906476 | 14.28171537 | 1.468010466 | 0.197060703 | 0.709264768 | -4.491806614 |
| **A_55_P1964078** | NA | 0.373940197 | -0.264137329 | 1.012017722 | 13.53544252 | 1.46784112 | 0.197104797 | 0.709264768 | -4.491853145 |
| **A_66_P138673** | Whamm | -0.319663402 | -0.865136564 | 0.225809761 | 8.314729982 | -1.467810108 | 0.197112873 | 0.709264768 | -4.491861666 |
| **A_51_P385598** | Slc37a4 | -0.22417135 | -0.606713556 | 0.158370855 | 10.57344882 | -1.467746643 | 0.197129401 | 0.709264768 | -4.491879104 |
| **A_55_P1975420** | Dhx40 | -0.182222384 | -0.493205017 | 0.128760248 | 11.53685377 | -1.467627818 | 0.19716035 | 0.709264768 | -4.491911753 |
| **A_55_P2152427** | Zwilch | 0.689556889 | -0.48742746 | 1.866541238 | 9.926829829 | 1.467404206 | 0.197218603 | 0.709264768 | -4.491973195 |
| **A_55_P1987650** | Ctnnd2 | 0.692429959 | -0.489761135 | 1.874621052 | 9.470784104 | 1.467028369 | 0.19731655 | 0.709264768 | -4.492076463 |
| **A_51_P220806** | Gdf9 | -0.34467909 | -0.93319759 | 0.24383941 | 8.715429102 | -1.466915725 | 0.197345914 | 0.709264768 | -4.492107414 |
| **A_55_P2081169** | NA | 0.246507924 | -0.174483188 | 0.667499036 | 16.00594716 | 1.466588593 | 0.197431217 | 0.709264768 | -4.492197299 |
| **A_66_P129873** | Npm1 | 0.549135548 | -0.388710191 | 1.486981286 | 15.11656067 | 1.466555472 | 0.197439856 | 0.709264768 | -4.4922064 |
| **A_55_P2137611** | Irgm2 | -0.434094146 | -1.175564742 | 0.30737645 | 9.197929144 | -1.46635911 | 0.197491078 | 0.709264768 | -4.492260354 |
| **A_55_P1988228** | Aspm | 1.019303558 | -0.721813146 | 2.760420262 | 9.932680378 | 1.4663107 | 0.197503708 | 0.709264768 | -4.492273655 |
| **A_55_P2184264** | Timm23 | 0.168412931 | -0.119291152 | 0.456117015 | 11.24156037 | 1.466154556 | 0.19754445 | 0.709264768 | -4.492316559 |
| **A_55_P2177026** | Mafg | -0.506865446 | -1.37279846 | 0.359067568 | 8.97051072 | -1.466083594 | 0.197562969 | 0.709264768 | -4.492336057 |
| **A_52_P426870** | Slf2 | -0.463619728 | -1.255712491 | 0.328473036 | 9.477660733 | -1.466007552 | 0.197582815 | 0.709264768 | -4.492356951 |
| **A_52_P318673** | Saa1 | -0.267845823 | -0.725480835 | 0.189789189 | 8.608023847 | -1.465939275 | 0.197600636 | 0.709264768 | -4.492375711 |
| **A_55_P2029842** | Hnrnpa1 | 0.950422885 | -0.673472384 | 2.574318154 | 13.14214143 | 1.465916435 | 0.197606598 | 0.709264768 | -4.492381987 |
| **A_51_P475501** | Ndufa8 | 0.382029546 | -0.270740718 | 1.034799809 | 12.59487597 | 1.465841238 | 0.197626227 | 0.709264768 | -4.492402649 |
| **A_55_P2140955** | NA | -0.369419232 | -1.00064423 | 0.261805767 | 16.03946613 | -1.465836921 | 0.197627354 | 0.709264768 | -4.492403835 |
| **A_55_P2066301** | Gpr137b-ן | -0.641315918 | -1.737214958 | 0.454583123 | 8.898539011 | -1.465722714 | 0.197657171 | 0.709264768 | -4.492435215 |
| **A_55_P2135880** | Fancl | 0.612751915 | -0.434455161 | 1.659958991 | 10.18672376 | 1.465556124 | 0.197700671 | 0.709264768 | -4.492480989 |
| **A_55_P2045367** | G630025F | -0.154033769 | -0.417302192 | 0.109234655 | 9.073193079 | -1.465438221 | 0.197731464 | 0.709264768 | -4.492513385 |
| **A_55_P1968129** | Eloc | 0.347258453 | -0.246302356 | 0.940819262 | 10.19051271 | 1.465338479 | 0.197757517 | 0.709264768 | -4.492540791 |
| **A_51_P249537** | Wdr24 | -0.187764441 | -0.508746711 | 0.133217829 | 9.60952516 | -1.465151893 | 0.197806262 | 0.709264768 | -4.492592059 |
| **A_55_P2063041** | Prelid3b | 0.21646615 | -0.153614021 | 0.58654632 | 9.380259801 | 1.465023267 | 0.197839872 | 0.709264768 | -4.492627402 |
| **A_55_P1989511** | Tbca | 0.432472381 | -0.306980589 | 1.17192535 | 11.79845591 | 1.464866898 | 0.197880738 | 0.709264768 | -4.492670367 |
| **A_55_P2275279** | Mid2 | -0.12689688 | -0.343935309 | 0.09014155 | 8.370917803 | -1.464416624 | 0.197998461 | 0.709264768 | -4.492794089 |

| | | | | | | | | | |
|---|---|---|---|---|---|---|---|---|---|
| **A_55_P1980636** | Aurka | 1.173811814 | -0.833924485 | 3.181548113 | 11.31139485 | 1.464340759 | 0.198018302 | 0.709264768 | -4.492814934 |
| **A_51_P155142** | Cdca8 | 1.388127593 | -0.986315563 | 3.762570748 | 11.58431372 | 1.464259181 | 0.198039639 | 0.709264768 | -4.492837349 |
| **A_55_P2145449** | Cep68 | 0.152880075 | -0.10863432 | 0.41439447 | 8.226761829 | 1.464217634 | 0.198050507 | 0.709264768 | -4.492848765 |
| **A_65_P16059** | Tgfbr3 | -0.710381187 | -1.925698078 | 0.504935704 | 11.64319421 | -1.46403771 | 0.198097577 | 0.709264768 | -4.492898202 |
| **A_55_P2090179** | Cbx5 | 0.361230412 | -0.256977743 | 0.979438566 | 8.918486671 | 1.463524248 | 0.198231963 | 0.709264768 | -4.493039286 |
| **A_66_P124952** | Hspe1 | 0.22622977 | -0.160954611 | 0.613414152 | 13.67540765 | 1.463464763 | 0.198247538 | 0.709264768 | -4.493055631 |
| **A_55_P2058933** | Kif1b | -0.16358476 | -0.443559611 | 0.116390091 | 11.12106162 | -1.46343745 | 0.198254689 | 0.709264768 | -4.493063136 |
| **A_55_P1986680** | NA | 0.170745933 | -0.121485586 | 0.462977453 | 8.519578555 | 1.463435746 | 0.198255135 | 0.709264768 | -4.493063604 |
| **A_55_P2029471** | NA | 0.663548463 | -0.47211547 | 1.799212395 | 13.80087208 | 1.463434063 | 0.198255576 | 0.709264768 | -4.493064066 |
| **A_51_P283083** | Armc1 | 0.240780244 | -0.171378125 | 0.652938612 | 11.98736295 | 1.46321139 | 0.198313888 | 0.709264768 | -4.49312525 |
| **A_55_P2000515** | Peli1 | 0.409266208 | -0.291300012 | 1.109832428 | 12.13587106 | 1.46321128 | 0.198313917 | 0.709264768 | -4.49312528 |
| **A_66_P117477** | Dlk1 | 0.493572209 | -0.351323362 | 1.33846778 | 16.13495883 | 1.463180857 | 0.198321885 | 0.709264768 | -4.493133639 |
| **A_55_P2099338** | Akt1s1 | -0.226305473 | -0.61372403 | 0.161113084 | 8.295588095 | -1.463069589 | 0.198351031 | 0.709264768 | -4.493164213 |
| **A_55_P1974554** | Eif4b | 0.1331488 | -0.094793411 | 0.361091012 | 12.20326527 | 1.463062387 | 0.198352917 | 0.709264768 | -4.493166192 |
| **A_51_P417016** | Srsf3 | 0.539505594 | -0.384094654 | 1.463105841 | 9.316486644 | 1.463060404 | 0.198353437 | 0.709264768 | -4.493166736 |
| **A_55_P2023652** | Vps35l | -0.580946121 | -1.575638545 | 0.413746304 | 12.06172538 | -1.4628419 | 0.198410684 | 0.709264768 | -4.493226775 |
| **A_55_P1993049** | Gm4076 | 0.289044851 | -0.205875717 | 0.78396542 | 11.7843141 | 1.462783593 | 0.198425963 | 0.709264768 | -4.493242796 |
| **A_51_P205820** | Klf11 | -0.175930981 | -0.477174798 | 0.125312836 | 8.243776911 | -1.462764891 | 0.198430864 | 0.709264768 | -4.493247934 |
| **A_55_P2121936** | Wiz | 0.445455466 | -0.317316715 | 1.208227647 | 11.27633157 | 1.462715261 | 0.19844387 | 0.709264768 | -4.493261571 |
| **A_55_P2077664** | Gm5458 | -0.488694891 | -1.325510624 | 0.348120841 | 8.782476471 | -1.462710262 | 0.19844518 | 0.709264768 | -4.493262945 |
| **A_55_P1994957** | NA | 0.349274514 | -0.248859799 | 0.947408826 | 15.17130578 | 1.462576278 | 0.198480297 | 0.709264768 | -4.493299759 |
| **A_55_P2099479** | H2afz | 1.138277788 | -0.81110786 | 3.087663437 | 14.10819807 | 1.462516729 | 0.198495906 | 0.709264768 | -4.493316122 |
| **A_51_P246727** | Mlxip | -0.151290449 | -0.410400015 | 0.107819117 | 8.690021153 | -1.462441186 | 0.19851571 | 0.709264768 | -4.493336879 |
| **A_55_P1969523** | Sumo2 | 0.629824816 | -0.448974282 | 1.708623914 | 11.31083836 | 1.46227663 | 0.198558855 | 0.709264768 | -4.493382094 |
| **A_55_P2146374** | Nmnat1 | -0.191430376 | -0.519331174 | 0.136470421 | 9.016534225 | -1.462240223 | 0.198568401 | 0.709264768 | -4.493392097 |
| **A_51_P286488** | Ier3 | -0.575048476 | -1.560097527 | 0.410000574 | 10.11679394 | -1.462166893 | 0.198587632 | 0.709264768 | -4.493412246 |
| **A_55_P2056348** | NA | 0.393348555 | -0.280560175 | 1.067257284 | 12.55887782 | 1.461930822 | 0.198649551 | 0.709264768 | -4.493477111 |
| **A_55_P2121275** | Gm4907 | -0.569228756 | -1.544586261 | 0.40612875 | 10.12846211 | -1.461750822 | 0.198696775 | 0.709264768 | -4.49352657 |
| **A_55_P2027822** | Sgf29 | 0.285451925 | -0.203729756 | 0.774633606 | 11.09829159 | 1.46154818 | 0.198749952 | 0.709264768 | -4.493582249 |
| **A_55_P2163143** | Cdk4 | 0.49693974 | -0.354884415 | 1.348763896 | 14.32479404 | 1.46118136 | 0.198846248 | 0.709264768 | -4.49368304 |
| **A_51_P155303** | Rbfa | -0.311901405 | -0.846593252 | 0.222790441 | 9.575390677 | -1.46104676 | 0.198881593 | 0.709264768 | -4.493720024 |
| **A_55_P2142863** | Parp9 | -0.518334443 | -1.406946916 | 0.37027803 | 9.835208709 | -1.460992576 | 0.198895823 | 0.709264768 | -4.493734912 |
| **A_55_P1984523** | NA | 0.262008445 | -0.18717863 | 0.711195521 | 11.91702616 | 1.460960199 | 0.198904327 | 0.709264768 | -4.493743808 |
| **A_51_P167843** | Timm17a | -0.143848325 | -0.390501696 | 0.102805047 | 11.92831605 | -1.460723776 | 0.198966431 | 0.709264768 | -4.49380877 |
| **A_52_P613808** | Zbtb7a | -0.201792959 | -0.547808959 | 0.144223041 | 8.254781287 | -1.460697121 | 0.198973434 | 0.709264768 | -4.493816094 |
| **A_55_P2046037** | Gm7083 | 0.851007613 | -0.608356152 | 2.310371378 | 14.04609068 | 1.46056279 | 0.19900873 | 0.709264768 | -4.493853004 |
| **A_55_P2096807** | Rbms1 | -0.416483072 | -1.130745502 | 0.297779359 | 10.89518802 | -1.460460758 | 0.199035543 | 0.709264768 | -4.493881039 |
| **A_52_P179178** | Bend3 | 0.201672483 | -0.144217437 | 0.547562402 | 10.31854013 | 1.460357163 | 0.199062771 | 0.709264768 | -4.493909504 |
| **A_55_P2022297** | Ptrh2 | 0.569224552 | -0.407080294 | 1.545529398 | 9.389259756 | 1.460321652 | 0.199072105 | 0.709264768 | -4.493919261 |
| **A_55_P2059337** | Fastk | -0.40384745 | -1.096533584 | 0.288838684 | 11.55726696 | -1.460263386 | 0.199087421 | 0.709264768 | -4.493935271 |
| **A_55_P2093649** | NA | 0.300203777 | -0.214717458 | 0.815125012 | 15.03838985 | 1.46024489 | 0.199092283 | 0.709264768 | -4.493940353 |
| **A_55_P2147220** | Ak2 | 0.251366047 | -0.17979369 | 0.682525784 | 11.44572425 | 1.460221581 | 0.19909841 | 0.709264768 | -4.493946758 |
| **A_51_P491976** | Fbxw17 | -0.16218565 | -0.440382937 | 0.116011636 | 10.81230033 | -1.460191723 | 0.19910626 | 0.709264768 | -4.493954962 |
| **A_55_P1959818** | Tmem40 | -1.289576263 | -3.501666514 | 0.922513987 | 9.258729813 | -1.460140121 | 0.199119826 | 0.709264768 | -4.49396914 |
| **A_55_P2005783** | Ifih1 | -0.762229662 | -2.069909741 | 0.545450417 | 9.253832124 | -1.459938909 | 0.199172734 | 0.709264768 | -4.494024427 |
| **A_51_P453401** | Cipc | 0.307052059 | -0.219743592 | 0.83384771 | 10.95190556 | 1.459890151 | 0.199185557 | 0.709264768 | -4.494037824 |
| **A_51_P393897** | Mpi | 0.372241549 | -0.266423517 | 1.010906616 | 9.629411578 | 1.459829148 | 0.199201601 | 0.709264768 | -4.494054586 |
| **A_51_P353095** | Zfyve27 | -0.531649843 | -1.443852673 | 0.380552987 | 10.80748549 | -1.459770626 | 0.199216994 | 0.709264768 | -4.494070666 |

| | | | | | | | | | |
|---|---|---|---|---|---|---|---|---|---|
| **A_55_P2033461** | NA | 0.578705708 | -0.414269509 | 1.571680925 | 14.82432221 | 1.459720484 | 0.199230183 | 0.709264768 | -4.494084443 |
| **A_55_P2081148** | Foxk2 | 0.451035994 | -0.323128049 | 1.225200038 | 10.23363493 | 1.459246131 | 0.199354999 | 0.709264768 | -4.49421478 |
| **A_55_P2073084** | Gtf3c2 | 0.280028994 | -0.200636433 | 0.760694421 | 10.64513066 | 1.45918535 | 0.199370997 | 0.709264768 | -4.494231481 |
| **A_51_P222657** | Tspo | -0.281049374 | -0.763504906 | 0.201406159 | 11.11231166 | -1.459068483 | 0.199401762 | 0.709264768 | -4.494263592 |
| **A_55_P2094721** | Sox11 | 0.440578401 | -0.315839636 | 1.196996438 | 9.83917002 | 1.458853466 | 0.199458375 | 0.709264768 | -4.494322672 |
| **A_66_P138289** | Rdh5 | -0.137170034 | -0.372688753 | 0.098348684 | 8.581765004 | -1.45876107 | 0.199482708 | 0.709264768 | -4.494348059 |
| **A_55_P2015022** | NA | -0.373492974 | -1.014792353 | 0.267806404 | 14.67887661 | -1.458720086 | 0.199493502 | 0.709264768 | -4.49435932 |
| **A_55_P2108803** | Tubgcp4 | 0.157735459 | -0.113122993 | 0.428593911 | 8.971750659 | 1.458603621 | 0.199524179 | 0.709264768 | -4.494391321 |
| **A_66_P111401** | Fbxw8 | -0.580666073 | -1.577818185 | 0.41648604 | 10.06664931 | -1.458530058 | 0.199543557 | 0.709264768 | -4.494411534 |
| **A_55_P2099961** | H2ac11 | 0.934721096 | -0.670478805 | 2.539920996 | 16.78182209 | 1.458489362 | 0.199554278 | 0.709264768 | -4.494422716 |
| **A_55_P2153621** | Ahnak | -0.789724353 | -2.145936699 | 0.566487993 | 9.345789652 | -1.458472292 | 0.199558776 | 0.709264768 | -4.494427406 |
| **A_55_P2034230** | Polr2c | 0.273326341 | -0.196096596 | 0.742749277 | 10.6137997 | 1.458369344 | 0.1995859 | 0.709264768 | -4.494455693 |
| **A_55_P2457153** | G2e3 | 0.362485817 | -0.26006814 | 0.985039774 | 9.564118941 | 1.458358732 | 0.199588696 | 0.709264768 | -4.494458609 |
| **A_55_P2168383** | Myo18b | -0.526575309 | -1.431027172 | 0.377876554 | 8.813533112 | -1.45822782 | 0.199623194 | 0.709264768 | -4.494494579 |
| **A_51_P294095** | Prkar1a | 0.520355558 | -0.373428461 | 1.414139578 | 10.5935378 | 1.458202902 | 0.199629761 | 0.709264768 | -4.494501426 |
| **A_66_P133204** | NA | -0.318785489 | -0.866350382 | 0.228779404 | 9.354873382 | -1.458187243 | 0.199633888 | 0.709264768 | -4.494505728 |
| **A_55_P2086143** | Cyp39a1 | 0.978364993 | -0.702159753 | 2.658889738 | 11.29353912 | 1.458163888 | 0.199640043 | 0.709264768 | -4.494512146 |
| **A_52_P22331** | Akap10 | -0.255265835 | -0.693756649 | 0.183224979 | 8.17921821 | -1.45808398 | 0.199661104 | 0.709264768 | -4.494534101 |
| **A_55_P1998389** | NA | 0.270566527 | -0.194265131 | 0.735398185 | 14.67943291 | 1.457903283 | 0.199708739 | 0.709264768 | -4.494583751 |
| **A_52_P257812** | Lpl | -0.169286608 | -0.460139593 | 0.121566377 | 8.300091397 | -1.457804962 | 0.199734662 | 0.709264768 | -4.494610767 |
| **A_51_P262101** | Ptcd2 | 0.145669977 | -0.10461267 | 0.395952624 | 9.045684101 | 1.45777218 | 0.199743306 | 0.709264768 | -4.494619774 |
| **A_51_P244558** | Rab3gap2 | -0.340641683 | -0.92595481 | 0.244671443 | 9.400475035 | -1.457672202 | 0.199769671 | 0.709264768 | -4.494647244 |
| **A_55_P1966593** | NA | 0.146352899 | -0.105145869 | 0.397851668 | 8.891116182 | 1.45752433 | 0.199808671 | 0.709264768 | -4.494687875 |
| **A_55_P2104597** | Kctd9 | -0.602921349 | -1.639389866 | 0.433547168 | 11.61492335 | -1.456984369 | 0.199951144 | 0.709264768 | -4.494836238 |
| **A_51_P361951** | Uqcrfs1 | 0.519511283 | -0.373586132 | 1.412608698 | 13.39839988 | 1.456956206 | 0.199958578 | 0.709264768 | -4.494843976 |
| **A_55_P2059342** | Lamp1 | -0.606129681 | -1.64815137 | 0.435892009 | 15.16546115 | -1.456931515 | 0.199965095 | 0.709264768 | -4.49485076 |
| **A_55_P2136622** | Camta2 | -0.194122855 | -0.527857889 | 0.13961218 | 8.287193193 | -1.456884807 | 0.199977425 | 0.709264768 | -4.494863594 |
| **A_51_P384091** | Rad21 | 0.149688516 | -0.107704394 | 0.407081426 | 13.84616982 | 1.456606517 | 0.2000509 | 0.709264768 | -4.494940058 |
| **A_55_P2363030** | Cmtm4 | -1.124190257 | -3.05729216 | 0.808911647 | 10.36465607 | -1.456583594 | 0.200056954 | 0.709264768 | -4.494946357 |
| **A_52_P355095** | Dstyk | -0.124591695 | -0.338879173 | 0.089695783 | 8.392640945 | -1.456272493 | 0.200139124 | 0.709264768 | -4.495031837 |
| **A_55_P1958857** | Nek6 | -0.273532905 | -0.744036158 | 0.196970348 | 10.07688338 | -1.456120422 | 0.200179302 | 0.709264768 | -4.49507362 |
| **A_51_P372992** | Vsir | -0.5449227 | -1.482406965 | 0.392561565 | 8.906949382 | -1.455865511 | 0.200246668 | 0.709264768 | -4.495143661 |
| **A_55_P2163009** | Mapkapk5 | -0.340319031 | -0.925831189 | 0.245193127 | 10.91720386 | -1.455796477 | 0.200264915 | 0.709264768 | -4.495162629 |
| **A_55_P2073377** | Mki67 | 1.732976278 | -1.248586196 | 4.714538752 | 12.39305883 | 1.455791066 | 0.200266346 | 0.709264768 | -4.495164116 |
| **A_55_P2019909** | Tmem35b | -0.436683163 | -1.188037918 | 0.314671591 | 9.587912548 | -1.455699573 | 0.200290532 | 0.709264768 | -4.495189254 |
| **A_52_P344376** | Eif4a2 | -0.475263988 | -1.293137128 | 0.342609153 | 9.422248177 | -1.455456759 | 0.200354734 | 0.709264768 | -4.495255971 |
| **A_55_P2139009** | NA | 0.161659673 | -0.116590727 | 0.439910074 | 15.65082409 | 1.455178414 | 0.200428354 | 0.709264768 | -4.49533245 |
| **A_51_P253803** | Mki67 | 1.863966451 | -1.344584147 | 5.07251705 | 12.82408961 | 1.455055452 | 0.200460885 | 0.709264768 | -4.495366236 |
| **A_52_P234958** | Dvl1 | -0.257326622 | -0.700298765 | 0.185645522 | 10.06005866 | -1.454985442 | 0.200479409 | 0.709264768 | -4.495385472 |
| **A_55_P2069714** | Sqstm1 | -0.73548595 | -2.001661178 | 0.530689278 | 9.712208933 | -1.454892529 | 0.200503995 | 0.709264768 | -4.495411001 |
| **A_55_P1970860** | Pemt | 0.262326215 | -0.189290303 | 0.713942733 | 9.83246042 | 1.454863407 | 0.200511702 | 0.709264768 | -4.495419002 |
| **A_55_P2034415** | Lrrc8d | -0.30514704 | -0.830537498 | 0.220243419 | 10.58171938 | -1.454713017 | 0.200551505 | 0.709264768 | -4.495460324 |
| **A_66_P114782** | Nr2f1 | 0.918401575 | -0.662894931 | 2.499698081 | 9.774005489 | 1.454687323 | 0.200558306 | 0.709264768 | -4.495467384 |
| **A_55_P2045258** | Ehd1 | -0.722126878 | -1.965935249 | 0.521681492 | 9.687511 | -1.454153928 | 0.200699542 | 0.709264768 | -4.495613941 |
| **A_55_P2139878** | NA | 0.279974594 | -0.202265157 | 0.762214344 | 8.269286215 | 1.454139136 | 0.20070346 | 0.709264768 | -4.495618005 |
| **A_52_P270164** | E2f6 | 0.243488918 | -0.175922457 | 0.662900293 | 10.61602883 | 1.454083478 | 0.200718204 | 0.709264768 | -4.495633297 |
| **A_55_P2035946** | Penk | 1.014721841 | -0.733169696 | 2.762613379 | 10.60134343 | 1.454061831 | 0.200723938 | 0.709264768 | -4.495639245 |
| **A_55_P1968606** | Synrg | -0.255028182 | -0.694389745 | 0.184333381 | 16.57839027 | -1.453839486 | 0.200782846 | 0.709264768 | -4.495700337 |

| | | | | | | | | | |
|---|---|---|---|---|---|---|---|---|---|
| **A_55_P2040996** | Klf7 | -0.347181232 | -0.945370168 | 0.251007705 | 10.60476676 | -1.453677969 | 0.200825649 | 0.709264768 | -4.495744716 |
| **A_55_P2026325** | NA | 0.265108217 | -0.191682247 | 0.721898681 | 14.40443479 | 1.453638795 | 0.200836032 | 0.709264768 | -4.495755479 |
| **A_55_P2140592** | Dmkn | -1.919415457 | -5.227179024 | 1.38834811 | 10.91813092 | -1.453399057 | 0.200899582 | 0.709264768 | -4.495821349 |
| **A_55_P2141013** | Siglech | -0.354739403 | -0.966111936 | 0.256633129 | 14.63491177 | -1.453295249 | 0.200927106 | 0.709264768 | -4.495849872 |
| **A_55_P2243431** | Gdap10 | -0.252608383 | -0.688034732 | 0.182817967 | 8.866543926 | -1.453059498 | 0.200989626 | 0.709264768 | -4.495914647 |
| **A_52_P584293** | Atrnl1 | 0.156387497 | -0.113208165 | 0.425983159 | 9.546118798 | 1.452912547 | 0.201028607 | 0.709264768 | -4.495955023 |
| **A_55_P2050226** | Ackr4 | 0.336670288 | -0.243768898 | 0.917109475 | 8.133496135 | 1.452775182 | 0.20106505 | 0.709264768 | -4.495992765 |
| **A_52_P84803** | NA | 0.191110245 | -0.138421805 | 0.520642296 | 16.04417556 | 1.45256861 | 0.201119867 | 0.709264768 | -4.496049523 |
| **A_51_P138760** | Abcd4 | -0.89897896 | -2.449492178 | 0.651534257 | 9.99603189 | -1.452193206 | 0.201219521 | 0.709264768 | -4.496152668 |
| **A_51_P122660** | Liph | -0.722495676 | -1.968851408 | 0.523860056 | 8.264364822 | -1.451922992 | 0.20129128 | 0.709264768 | -4.497197027 |
| **A_55_P2024888** | Ctss | -0.29029254 | -0.791156255 | 0.210571175 | 11.36879324 | -1.451665817 | 0.201359599 | 0.709264768 | -4.496297572 |
| **A_55_P2055137** | NA | 0.634477794 | -0.460405023 | 1.729360612 | 13.67677608 | 1.45144015 | 0.201419566 | 0.709264768 | -4.496359575 |
| **A_55_P1971674** | Golga7b | 0.309346808 | -0.224489722 | 0.843183338 | 9.651248574 | 1.451401927 | 0.201429724 | 0.709264768 | -4.496370078 |
| **A_51_P346826** | Rragc | -0.272838605 | -0.743708942 | 0.198031732 | 11.1289345 | -1.451292107 | 0.201458914 | 0.709264768 | -4.496400251 |
| **A_66_P105736** | Net1 | -0.153940166 | -0.419636391 | 0.11175606 | 10.0470453 | -1.451165384 | 0.201492602 | 0.709264768 | -4.496435069 |
| **A_55_P2083939** | Irf3 | -0.238283725 | -0.649625956 | 0.173058507 | 12.33147215 | -1.450913182 | 0.201559663 | 0.709264768 | -4.496504363 |
| **A_55_P2110355** | Zfp637 | 0.297072541 | -0.215779326 | 0.809924408 | 9.935529991 | 1.45084466 | 0.201577886 | 0.709264768 | -4.49652319 |
| **A_55_P1982249** | Phactr2 | -0.165242042 | -0.450518525 | 0.120034442 | 9.71727473 | -1.450791242 | 0.201592094 | 0.709264768 | -4.496537866 |
| **A_55_P2115151** | Pxylp1 | 0.508848172 | -0.369907341 | 1.387603684 | 9.43531952 | 1.450342259 | 0.201711548 | 0.709264768 | -4.496661226 |
| **A_55_P2162344** | Lrsam1 | -0.539492964 | -1.471244637 | 0.392258708 | 9.489780343 | -1.450226878 | 0.201742257 | 0.709264768 | -4.496692927 |
| **A_66_P128942** | Mipep | 0.174651205 | -0.126991941 | 0.47629435 | 7.990471056 | 1.4502019 | 0.201748905 | 0.709264768 | -4.49669979 |
| **A_55_P2011390** | Tead1 | -0.172187762 | -0.469608332 | 0.125232808 | 8.121224526 | -1.450045489 | 0.201790542 | 0.709264768 | -4.496742765 |
| **A_55_P1980180** | Tardbp | 0.359808993 | -0.261723701 | 0.981341686 | 14.15344915 | 1.449967878 | 0.201811205 | 0.709264768 | -4.496764089 |
| **A_51_P134972** | Shkbp1 | 0.170823423 | -0.124281889 | 0.465928735 | 10.56493683 | 1.449842204 | 0.201844669 | 0.709264768 | -4.496798618 |
| **A_51_P277718** | Cenpc1 | 0.281191682 | -0.204611512 | 0.766994876 | 9.782991164 | 1.449747768 | 0.201869819 | 0.709264768 | -4.496824564 |
| **A_66_P128586** | Cwc22 | 0.193398485 | -0.140820812 | 0.527617781 | 9.374916861 | 1.449345383 | 0.201977011 | 0.709264768 | -4.49693512 |
| **A_55_P2140941** | Tinagl1 | -0.762490114 | -2.080440632 | 0.555460403 | 8.605610018 | -1.449056963 | 0.202053877 | 0.709264768 | -4.497014364 |
| **A_51_P245090** | Aqp3 | -1.424155306 | -3.887183503 | 1.038872892 | 9.015272427 | -1.448232582 | 0.202273731 | 0.709264768 | -4.497240862 |
| **A_52_P319606** | Gpr107 | -0.593858361 | -1.621139962 | 0.433423241 | 10.18205321 | -1.447917145 | 0.202357914 | 0.709264768 | -4.497327527 |
| **A_55_P2063598** | Tomm5 | -0.282653846 | -0.771600647 | 0.206292956 | 13.98631945 | -1.447916892 | 0.202357982 | 0.709264768 | -4.497327597 |
| **A_52_P585124** | Cxcr4 | 0.442193139 | -0.322774609 | 1.207160886 | 8.920522204 | 1.447835498 | 0.202379709 | 0.709264768 | -4.497349959 |
| **A_51_P349546** | Cd109 | -0.445341331 | -1.215837697 | 0.325155034 | 9.059593195 | -1.447680598 | 0.202421065 | 0.709264768 | -4.497392518 |
| **A_55_P2020661** | Orai3 | -0.141413746 | -0.386094454 | 0.103266962 | 8.396382638 | -1.447578897 | 0.202448222 | 0.709264768 | -4.49742046 |
| **A_55_P2091461** | Casp4 | -0.689972125 | -1.883804554 | 0.503860305 | 8.717992918 | -1.447566483 | 0.202451537 | 0.709264768 | -4.49742387 |
| **A_51_P499153** | Cog3 | -0.143797629 | -0.392614087 | 0.105018828 | 9.660066943 | -1.447514653 | 0.202465378 | 0.709264768 | -4.49743811 |
| **A_55_P2049582** | Apbb3 | -0.211396607 | -0.577202066 | 0.154408851 | 9.397797485 | -1.447432024 | 0.202487447 | 0.709264768 | -4.497460812 |
| **A_55_P2158701** | H4c4 | 1.225343044 | -0.895146939 | 3.345833028 | 14.30801116 | 1.44734415 | 0.202510918 | 0.709264768 | -4.497484955 |
| **A_51_P349341** | Npc1 | -0.592982075 | -1.619190788 | 0.433226638 | 8.701677252 | -1.447292173 | 0.202524803 | 0.709264768 | -4.497499236 |
| **A_66_P107379** | Mdk | 1.124935065 | -0.822214023 | 3.072084152 | 11.7263143 | 1.447033529 | 0.202593907 | 0.709264768 | -4.497570297 |
| **A_52_P605517** | Phactr1 | -1.022338185 | -2.792083116 | 0.747406745 | 9.215372899 | -1.446885789 | 0.202633391 | 0.709264768 | -4.497610888 |
| **A_51_P392705** | Xpc | -0.339350242 | -0.926864191 | 0.248163707 | 9.362040977 | -1.446706146 | 0.202681409 | 0.709264768 | -4.497660243 |
| **A_55_P2076462** | Lnx1 | -0.638625777 | -1.744284652 | 0.467033098 | 8.969073193 | -1.446690512 | 0.202685589 | 0.709264768 | -4.497664539 |
| **A_51_P167105** | B3galt4 | -0.405472204 | -1.107533488 | 0.296589079 | 8.803945925 | -1.44655986 | 0.20272052 | 0.709264768 | -4.497700434 |
| **A_51_P281582** | Lrrc42 | 0.186320726 | -0.136310352 | 0.508951805 | 8.600255052 | 1.446456289 | 0.202748214 | 0.709264768 | -4.49772889 |
| **A_55_P2037338** | Plcd3 | -0.736273258 | -2.011354421 | 0.538807906 | 9.042412218 | -1.446277208 | 0.202796108 | 0.709264768 | -4.497778091 |
| **A_55_P1967553** | Tmem254 | -0.271990292 | -0.743110283 | 0.1991297 | 11.42427885 | -1.446013059 | 0.202866773 | 0.709264768 | -4.497850664 |
| **A_51_P301311** | Ciz1 | -0.187777003 | -0.513046916 | 0.13749291 | 11.45579601 | -1.445935287 | 0.202887582 | 0.709264768 | -4.497872031 |
| **A_55_P2002319** | Ap5s1 | -0.214470598 | -0.586035076 | 0.157093879 | 10.43043144 | -1.445719111 | 0.202945436 | 0.709264768 | -4.497931423 |

| | | | | | | | | | |
|---|---|---|---|---|---|---|---|---|---|
| **A_51_P107782** | Kdsr | -0.815385939 | -2.228134766 | 0.597362888 | 10.06311611 | -1.44560156 | 0.202976901 | 0.709264768 | -4.498910841 |
| **A_55_P1959263** | Dnpep | -0.412916519 | -1.128400043 | 0.302567005 | 13.38640759 | -1.445482923 | 0.203008662 | 0.709264768 | -4.497996313 |
| **A_55_P2082979** | Irak2 | -0.590555649 | -1.613855125 | 0.432743826 | 11.83346798 | -1.445467803 | 0.203012711 | 0.709264768 | -4.498000467 |
| **A_55_P2085596** | Nudt1 | 0.471154706 | -0.345290564 | 1.287599977 | 10.5256812 | 1.445395845 | 0.203031978 | 0.709264768 | -4.498020237 |
| **A_51_P291839** | Cox14 | -0.137914803 | -0.376909124 | 0.101079517 | 14.06721614 | -1.445352073 | 0.203043698 | 0.709264768 | -4.498032263 |
| **A_55_P1966878** | Rnaseh2a | 0.264574035 | -0.193918919 | 0.723066988 | 8.791539225 | 1.445322956 | 0.203051496 | 0.709264768 | -4.498040262 |
| **A_55_P1980416** | Zkscan1 | 0.688286255 | -0.504578599 | 1.88115111 | 11.27163721 | 1.44520082 | 0.203084205 | 0.709264768 | -4.498073818 |
| **A_55_P2020378** | Ehd4 | -0.275602865 | -0.753281613 | 0.202075883 | 9.377502196 | -1.445100839 | 0.203110984 | 0.709264768 | -4.498101286 |
| **A_51_P345046** | Metap2 | 0.346865772 | -0.254410654 | 0.948142198 | 11.46643436 | 1.444899414 | 0.203164945 | 0.709264768 | -4.498156625 |
| **A_55_P2145401** | H3c3 | 0.792605843 | -0.581383316 | 2.166595001 | 10.51933781 | 1.444855146 | 0.203176806 | 0.709264768 | -4.498168787 |
| **A_55_P2043317** | Bag1 | -0.3853596 | -1.053509994 | 0.282790794 | 14.44001781 | -1.444582334 | 0.203249917 | 0.709264768 | -4.498243739 |
| **A_55_P2054491** | Slc31a2 | -0.455959004 | -1.246536852 | 0.334618844 | 8.170025645 | -1.444546415 | 0.203259545 | 0.709264768 | -4.498253607 |
| **A_55_P1983968** | Slc8b1 | -0.624094944 | -1.706237339 | 0.458047452 | 8.401728794 | -1.444496845 | 0.203272832 | 0.709264768 | -4.498267226 |
| **A_51_P375390** | Gemin7 | -0.150904753 | -0.412567338 | 0.110757832 | 11.56198732 | -1.444480343 | 0.203277256 | 0.709264768 | -4.498271759 |
| **A_55_P2064578** | Prkci | -0.44977874 | -1.229763413 | 0.330205933 | 13.14642392 | -1.444319254 | 0.203320443 | 0.709264768 | -4.498316016 |
| **A_55_P2034175** | Usp28 | 0.527326316 | -0.387157456 | 1.441810088 | 10.19441606 | 1.44428795 | 0.203328837 | 0.709264768 | -4.498324617 |
| **A_55_P2123802** | NA | -0.327587017 | -0.895770331 | 0.240596297 | 8.171607171 | -1.444071017 | 0.203387011 | 0.709264768 | -4.498384215 |
| **A_66_P120208** | Cnnm3 | -0.293596296 | -0.802856065 | 0.215663473 | 9.085673129 | -1.443981234 | 0.203411093 | 0.709264768 | -4.498408882 |
| **A_51_P286034** | Ccdc117 | -0.387725541 | -1.060313292 | 0.284862211 | 9.337473864 | -1.443862383 | 0.203442975 | 0.709264768 | -4.498441534 |
| **A_66_P103957** | Bbs7 | 0.481391351 | -0.353845935 | 1.316628637 | 9.073271043 | 1.443573016 | 0.203520618 | 0.709264768 | -4.498521033 |
| **A_55_P2207593** | 8430437C | -0.143378628 | -0.392149945 | 0.105392689 | 8.427603704 | -1.443558741 | 0.203524449 | 0.709264768 | -4.498524955 |
| **A_55_P2011116** | L3mbtl3 | -0.167493225 | -0.45814175 | 0.123155299 | 9.767221552 | -1.443375964 | 0.203573507 | 0.709264768 | -4.498575169 |
| **A_66_P132855** | Baz1b | 0.222100927 | -0.17552695 | 0.619728805 | 9.082630311 | 1.48777156 | 0.203589358 | 0.709264768 | -4.50584699 |
| **A_65_P13402** | Rnf14 | -0.499675391 | -1.366890656 | 0.367539875 | 8.54687311 | -1.443149729 | 0.203634245 | 0.709264768 | -4.498637323 |
| **A_55_P2114915** | Ube2i | 0.180210871 | -0.13256101 | 0.492982752 | 14.78400424 | 1.443123934 | 0.203641171 | 0.709264768 | -4.49864441 |
| **A_55_P1976033** | Cnih2 | 0.306340568 | -0.22544813 | 0.838129266 | 8.775294433 | 1.442831968 | 0.203719584 | 0.709264768 | -4.498724622 |
| **A_52_P370402** | Kcmf1 | -0.283423896 | -0.775479587 | 0.208631795 | 11.98258956 | -1.442688433 | 0.203758143 | 0.709264768 | -4.498764055 |
| **A_55_P2017929** | Clu | 0.16671655 | -0.122728002 | 0.456161102 | 13.8213444 | 1.442658973 | 0.203766058 | 0.709264768 | -4.498772148 |
| **A_51_P412338** | Vps41 | -0.135036618 | -0.369528374 | 0.099455137 | 9.8938337 | -1.442362163 | 0.203845817 | 0.709264768 | -4.49885369 |
| **A_52_P215539** | C030006K | 0.24867378 | -0.183237938 | 0.680585498 | 8.726758079 | 1.442066722 | 0.203925238 | 0.709264768 | -4.498934856 |
| **A_55_P2104835** | NA | -0.254302748 | -0.695995022 | 0.187389526 | 8.001158221 | -1.442054239 | 0.203928594 | 0.709264768 | -4.498938285 |
| **A_55_P2287260** | Snap23 | -0.327508581 | -0.896381945 | 0.241364784 | 11.86448756 | -1.441973997 | 0.20395017 | 0.709264768 | -4.49896033 |
| **A_55_P2037544** | NA | 0.238004416 | -0.175437444 | 0.651446275 | 16.13655789 | 1.441852767 | 0.203982771 | 0.709264768 | -4.498993635 |
| **A_55_P2110177** | NA | 0.717616295 | -0.528970839 | 1.964203429 | 14.23114284 | 1.441849718 | 0.203983591 | 0.709264768 | -4.498994473 |
| **A_55_P2094626** | Ralgps1 | 0.30989149 | -0.228468471 | 0.848251452 | 11.83513863 | 1.441740981 | 0.204012837 | 0.709264768 | -4.499024345 |
| **A_55_P2186085** | Atp11c | 0.40257232 | -0.296823807 | 1.101968447 | 9.21158366 | 1.441687169 | 0.204027312 | 0.709264768 | -4.499039129 |
| **A_51_P341746** | Mpl | -0.132260029 | -0.362049762 | 0.097529704 | 7.887953397 | -1.441611835 | 0.204047578 | 0.709264768 | -4.499059825 |
| **A_52_P130787** | Kcna6 | 0.386544572 | -0.285051952 | 1.058141096 | 8.663752504 | 1.441589083 | 0.204053698 | 0.709264768 | -4.499066075 |
| **A_66_P118989** | Gm8350 | -0.286709671 | -0.784883448 | 0.211464106 | 8.109190047 | -1.441490634 | 0.204080186 | 0.709264768 | -4.499093122 |
| **A_51_P135268** | Corin | -0.376009856 | -1.029375935 | 0.277356223 | 8.031900668 | -1.441428123 | 0.204097005 | 0.709264768 | -4.499110295 |
| **A_52_P139650** | Ska1 | 1.052564022 | -0.776484509 | 2.881612553 | 10.01247433 | 1.441363891 | 0.20411429 | 0.709264768 | -4.499127941 |
| **A_52_P470316** | Laptm4b | -0.160945161 | -0.440642793 | 0.118752471 | 10.25250177 | -1.441250531 | 0.204144797 | 0.709264768 | -4.499159084 |
| **A_55_P1964459** | NA | -0.456386474 | -1.249543254 | 0.336770305 | 8.369414417 | -1.441199391 | 0.204158562 | 0.709264768 | -4.499173133 |
| **A_55_P2086433** | Oasl1 | -0.826072888 | -2.261794914 | 0.609649138 | 8.730546808 | -1.441114047 | 0.204181534 | 0.709264768 | -4.499196579 |
| **A_55_P1970507** | Pef1 | -0.248841793 | -0.681473815 | 0.183790229 | 11.78225823 | -1.440638466 | 0.204309592 | 0.709264768 | -4.499327231 |
| **A_55_P1996008** | Gm14399 | 0.28725556 | -0.212206814 | 0.786717933 | 9.261266288 | 1.440509118 | 0.204344433 | 0.709264768 | -4.499362766 |
| **A_55_P2053374** | Sorbs1 | -0.784364768 | -2.14825995 | 0.579530413 | 10.97135532 | -1.440414315 | 0.204369974 | 0.709264768 | -4.49938881 |
| **A_55_P2140711** | Tufm | 0.217953456 | -0.16106402 | 0.596970933 | 11.25312538 | 1.440306333 | 0.204399068 | 0.709264768 | -4.499418475 |

| | | | | | | | | | |
|---|---|---|---|---|---|---|---|---|---|
| **A_51_P290819** | Hint3 | -0.344007063 | -0.942233319 | 0.254219192 | 10.88956977 | -1.440297595 | 0.204401423 | 0.709264768 | -4.499420875 |
| **A_55_P1966528** | Pmf1 | 0.61575624 | -0.455157168 | 1.686669648 | 10.07829578 | 1.440140326 | 0.204443804 | 0.709264768 | -4.49946408 |
| **A_55_P2063163** | Agk | 0.293912939 | -0.217265307 | 0.805091185 | 9.787567973 | 1.440113387 | 0.204451065 | 0.709264768 | -4.49947148 |
| **A_55_P2108983** | NA | -0.82095744 | -2.248894441 | 0.606979562 | 15.95436188 | -1.439998173 | 0.204482119 | 0.709264768 | -4.499503132 |
| **A_55_P1976127** | Ect2 | 1.119317589 | -0.827845948 | 3.066481126 | 10.65836056 | 1.439796938 | 0.20453637 | 0.709264768 | -4.499558414 |
| **A_55_P2029549** | Dtd1 | 0.541720654 | -0.400697007 | 1.484138316 | 11.010552 | 1.439734208 | 0.204553285 | 0.709264768 | -4.499575647 |
| **A_55_P1955517** | NA | -0.217640617 | -0.596330448 | 0.161049213 | 10.45542145 | -1.43948337 | 0.204620933 | 0.709264768 | -4.499644556 |
| **A_55_P1982156** | Nelfcd | 0.149617356 | -0.110756336 | 0.409991048 | 13.72978163 | 1.439246625 | 0.204684799 | 0.709264768 | -4.499709594 |
| **A_55_P2062029** | Tsen34 | -0.277978523 | -0.761855844 | 0.205898798 | 14.77471112 | -1.43888578 | 0.204782179 | 0.709264768 | -4.499808723 |
| **A_51_P386389** | Actr6 | -0.1616321 | -0.442987391 | 0.119723192 | 9.906487783 | -1.438874358 | 0.204785263 | 0.709264768 | -4.49981186 |
| **A_52_P505501** | Nsmce2 | 0.193115347 | -0.143052506 | 0.529283199 | 8.67536522 | 1.438834869 | 0.204795922 | 0.709264768 | -4.499822708 |
| **A_65_P06611** | Wtap | 0.284492033 | -0.21078112 | 0.779765186 | 10.41788926 | 1.438717967 | 0.204827482 | 0.709264768 | -4.499854823 |
| **A_55_P2051859** | Golgb1 | -0.442058966 | -1.21164737 | 0.327529437 | 10.47606063 | -1.438705933 | 0.204830731 | 0.709264768 | -4.499858129 |
| **A_51_P228865** | Zfp72 | 0.197901594 | -0.146703662 | 0.54250685 | 8.758755353 | 1.438393593 | 0.204915075 | 0.709264768 | -4.499943932 |
| **A_55_P1984297** | Efcab14 | -0.258565251 | -0.708904196 | 0.191773695 | 10.09581747 | -1.438073216 | 0.205001623 | 0.709264768 | -4.500031942 |
| **A_55_P2168504** | Mcmbp | 0.144554277 | -0.107274555 | 0.39638311 | 9.919562064 | 1.437725052 | 0.205095717 | 0.709264768 | -4.500127586 |
| **A_55_P2012641** | NA | 0.411029801 | -0.305132045 | 1.127191648 | 17.20353408 | 1.437515302 | 0.205152423 | 0.709264768 | -4.500185205 |
| **A_52_P355084** | Metrnl | -0.73506216 | -2.0158425 | 0.545718179 | 10.71822741 | -1.437473211 | 0.205163804 | 0.709264768 | -4.500196768 |
| **A_55_P2056654** | Kif22 | 1.483162329 | -1.101209942 | 4.0675346 | 11.52958693 | 1.437420914 | 0.205177946 | 0.709264768 | -4.500211134 |
| **A_52_P66226** | Rab33a | 0.252267912 | -0.187329804 | 0.691865628 | 8.267158163 | 1.437331458 | 0.205202138 | 0.709264768 | -4.500235708 |
| **A_52_P325116** | Gramd1b | -0.648298649 | -1.778128416 | 0.481531117 | 9.840182057 | -1.437184253 | 0.205241953 | 0.709264768 | -4.500276146 |
| **A_55_P2044109** | NA | 0.288325383 | -0.214173768 | 0.790824534 | 16.32188968 | 1.437136073 | 0.205254985 | 0.709264768 | -4.500289381 |
| **A_65_P12392** | Timp2 | -1.00981869 | -2.769794216 | 0.750156836 | 10.44366064 | -1.437100436 | 0.205264626 | 0.709264768 | -4.500299171 |
| **A_52_P467266** | Mtf1 | -0.523644105 | -1.436323208 | 0.389034998 | 9.1720226 | -1.437038678 | 0.205281333 | 0.709264768 | -4.500316136 |
| **A_55_P1976109** | NA | 0.477516982 | -0.354881838 | 1.309915802 | 14.96403826 | 1.436837687 | 0.205335717 | 0.709264768 | -4.500371349 |
| **A_52_P598300** | Gtf2f2 | 0.166172324 | -0.123501617 | 0.455846264 | 9.795369681 | 1.436810891 | 0.205342969 | 0.709264768 | -4.50037871 |
| **A_55_P2089358** | Hspe1 | 0.205472664 | -0.152711631 | 0.563656959 | 13.71664607 | 1.436805088 | 0.205344539 | 0.709264768 | -4.500380304 |
| **A_55_P2010271** | Samsn1 | 0.248239571 | -0.184544695 | 0.681023838 | 8.170371547 | 1.436646419 | 0.205387483 | 0.709264768 | -4.50042389 |
| **A_55_P2062133** | Etv3 | -0.135920162 | -0.372895185 | 0.101054862 | 8.690556896 | -1.436586074 | 0.205403817 | 0.709264768 | -4.500440467 |
| **A_55_P2091676** | Fas | -0.269044338 | -0.738188325 | 0.200099649 | 9.838643532 | -1.436375694 | 0.205460774 | 0.709264768 | -4.500498259 |
| **A_55_P2022861** | Elovl1 | -0.618716037 | -1.697710223 | 0.460278148 | 9.667488445 | -1.436225437 | 0.205501462 | 0.709264768 | -4.500539534 |
| **A_51_P291460** | Gm6498 | 0.253219713 | -0.188381766 | 0.694821193 | 14.39463086 | 1.436207991 | 0.205506186 | 0.709264768 | -4.500544326 |
| **A_66_P125392** | Atp8b4 | 0.135956382 | -0.101157298 | 0.373070062 | 7.905876863 | 1.436128611 | 0.205527686 | 0.709264768 | -4.500566132 |
| **A_55_P2182412** | Adat1 | 0.133615734 | -0.09942219 | 0.366653659 | 8.192871612 | 1.436088963 | 0.205538424 | 0.709264768 | -4.500577023 |
| **A_52_P606679** | Eloa | -0.324601683 | -0.890767336 | 0.241563971 | 8.63124151 | -1.436010426 | 0.205559698 | 0.709264768 | -4.500598597 |
| **A_55_P2027077** | Shc2 | -0.214910271 | -0.589758014 | 0.159937473 | 8.434319554 | -1.435993968 | 0.205564156 | 0.709264768 | -4.500603118 |
| **A_55_P1981884** | Ercc6l2 | 0.336596607 | -0.250680002 | 0.923873216 | 9.5665767 | 1.435546868 | 0.205685306 | 0.709264768 | -4.500725935 |
| **A_51_P281593** | Plpp5 | -0.653340031 | -1.793428564 | 0.486748501 | 10.02835649 | -1.435327604 | 0.205744744 | 0.709264768 | -4.500786166 |
| **A_52_P676956** | Tirap | -0.254381717 | -0.698336917 | 0.189573484 | 8.511715942 | -1.435149326 | 0.205793084 | 0.709264768 | -4.500835137 |
| **A_55_P2138436** | NA | 0.466380563 | -0.347588594 | 1.280349721 | 15.9826539 | 1.435102231 | 0.205805855 | 0.709264768 | -4.500848074 |
| **A_55_P2129696** | Smad9 | 0.346555455 | -0.258290862 | 0.951401771 | 8.53849938 | 1.435086386 | 0.205810152 | 0.709264768 | -4.500852427 |
| **A_51_P495780** | Plin4 | -0.586850003 | -1.611267916 | 0.43756791 | 8.234759707 | -1.434829485 | 0.205879834 | 0.709264768 | -4.500922995 |
| **A_51_P143721** | Vipas39 | -0.151347523 | -0.415549808 | 0.112854763 | 10.58235214 | -1.434792481 | 0.205889872 | 0.709264768 | -4.50093316 |
| **A_51_P418560** | Lnx2 | -0.37629994 | -1.033316181 | 0.280716302 | 10.1362165 | -1.434525884 | 0.205962211 | 0.709264768 | -4.501006392 |
| **A_55_P2076134** | NA | 0.799671834 | -0.596619201 | 2.195962868 | 13.41618976 | 1.434452579 | 0.205982106 | 0.709264768 | -4.501026528 |
| **A_52_P987071** | Gm6803 | 0.355844715 | -0.265505428 | 0.977194857 | 12.0635006 | 1.434413827 | 0.205992624 | 0.709264768 | -4.501037173 |
| **A_55_P1993933** | NA | -1.407677977 | -3.865694615 | 1.05033866 | 9.073626067 | -1.434395268 | 0.205997661 | 0.709264768 | -4.501042271 |
| **A_55_P2039284** | Hspb1 | -1.062864393 | -2.919008104 | 0.793279319 | 14.77464488 | -1.434222744 | 0.206044494 | 0.709264768 | -4.501089661 |

| | | | | | | | | | |
|---|---|---|---|---|---|---|---|---|---|
| **A_66_P131110** | Zfp473 | 0.149526456 | -0.111661864 | 0.410714776 | 8.172122754 | 1.433886033 | 0.206135925 | 0.709264768 | -4.501182151 |
| **A_51_P144783** | Cct5 | 0.290619579 | -0.217072665 | 0.798311823 | 14.82818771 | 1.433754145 | 0.206171749 | 0.709264768 | -4.501218379 |
| **A_66_P137157** | Arl6ip4 | 0.33948889 | -0.253594842 | 0.932572622 | 13.11734827 | 1.433705331 | 0.206185009 | 0.709264768 | -4.501231787 |
| **A_55_P2137706** | Tmpo | 0.925893563 | -0.691720769 | 2.543507895 | 11.53178417 | 1.433627834 | 0.206206063 | 0.709264768 | -4.501253075 |
| **A_51_P301736** | Ppp2r2a | 0.203174586 | -0.151798577 | 0.558147749 | 11.52099777 | 1.433587508 | 0.206217019 | 0.709264768 | -4.501264151 |
| **A_55_P2070277** | Mfn1 | -0.370552366 | -1.017989882 | 0.276885151 | 10.31843493 | -1.433514447 | 0.20623687 | 0.709264768 | -4.50128422 |
| **A_55_P2130855** | Nap1l1 | 1.048084838 | -0.783362651 | 2.879532327 | 12.2801987 | 1.433350205 | 0.206281503 | 0.709264768 | -4.501329335 |
| **A_55_P1981739** | Donson | 0.162133553 | -0.121188535 | 0.44545564 | 10.09230064 | 1.433318846 | 0.206290026 | 0.709264768 | -4.501337948 |
| **A_51_P192589** | Pno1 | 0.218220989 | -0.163181581 | 0.59962356 | 11.15144534 | 1.433056292 | 0.206361397 | 0.709264768 | -4.501410067 |
| **A_55_P2076471** | Mon1b | 0.298873513 | -0.223500686 | 0.821247712 | 10.35122655 | 1.433032551 | 0.206367852 | 0.709264768 | -4.501416588 |
| **A_66_P125496** | NA | 0.696550305 | -0.559798578 | 1.952899189 | 13.2979187 | 1.476743045 | 0.206378888 | 0.709264768 | -4.508189779 |
| **A_66_P110343** | Snx10 | -0.252449156 | -0.693702076 | 0.188803765 | 8.043564448 | -1.432968605 | 0.206385238 | 0.709264768 | -4.501434153 |
| **A_52_P28792** | Isg20l2 | 0.408001246 | -0.305211021 | 1.121213514 | 10.24256721 | 1.432824605 | 0.206424396 | 0.709264768 | -4.501473707 |
| **A_52_P456134** | Dgat1 | -0.524481772 | -1.441314784 | 0.392351239 | 8.620348189 | -1.432816265 | 0.206426664 | 0.709264768 | -4.501475998 |
| **A_55_P2161595** | Hpca | -0.844141967 | -2.319772308 | 0.631488373 | 9.170056859 | -1.432808871 | 0.206428675 | 0.709264768 | -4.501478029 |
| **A_51_P339200** | Abhd16a | -0.346059709 | -0.951009742 | 0.258890325 | 8.556963671 | -1.432787811 | 0.206434403 | 0.709264768 | -4.501483814 |
| **A_55_P2047703** | Serac1 | -0.137599468 | -0.378142746 | 0.102943809 | 8.207927464 | -1.432761417 | 0.206441581 | 0.709264768 | -4.501491064 |
| **A_55_P2067722** | Camsap3 | -0.759108287 | -2.086268293 | 0.56805172 | 11.05226803 | -1.432619281 | 0.206480242 | 0.709264768 | -4.501530106 |
| **A_55_P2160686** | Tsc22d1 | -0.482990705 | -1.327476587 | 0.361495177 | 12.29340285 | -1.432506909 | 0.206510812 | 0.709264768 | -4.501560972 |
| **A_52_P68702** | Frmd4b | -0.246461524 | -0.677453422 | 0.184530374 | 8.198394632 | -1.432288052 | 0.206570362 | 0.709264768 | -4.501621087 |
| **A_52_P31814** | 1110038B | 0.88624229 | -0.663743833 | 2.436228413 | 12.35495431 | 1.432105476 | 0.206620053 | 0.709264768 | -4.501671236 |
| **A_55_P2015177** | NA | 0.95346767 | -0.714204919 | 2.621140259 | 12.14922154 | 1.432008272 | 0.206646513 | 0.709264768 | -4.501697935 |
| **A_52_P516091** | Pla2g15 | -0.484871863 | -1.333092548 | 0.363348823 | 8.826114681 | -1.43175421 | 0.206715687 | 0.709264768 | -4.50176772 |
| **A_55_P2165046** | Klhl22 | -0.469863128 | -1.291857476 | 0.352131219 | 10.96951409 | -1.43170282 | 0.206729681 | 0.709264768 | -4.501781835 |
| **A_55_P2062034** | Tsen34 | -0.257360031 | -0.707634149 | 0.192914087 | 14.59826795 | -1.431576174 | 0.206764173 | 0.709264768 | -4.501816621 |
| **A_55_P2101862** | Spryd3 | -0.657859574 | -1.808986534 | 0.493267386 | 9.377787531 | -1.431397715 | 0.206812786 | 0.709264768 | -4.501865639 |
| **A_52_P578562** | Slc41a1 | -0.190084569 | -0.522740253 | 0.142571116 | 8.572602938 | -1.431206362 | 0.206864923 | 0.709264768 | -4.501918198 |
| **A_55_P2143522** | Fstl3 | -1.246899553 | -3.429536739 | 0.935737632 | 10.38859881 | -1.430870286 | 0.206956522 | 0.709264768 | -4.502010508 |
| **A_55_P2176688** | NA | 0.534418738 | -0.401129046 | 1.469966523 | 14.82599663 | 1.430757556 | 0.206987255 | 0.709264768 | -4.502041471 |
| **A_51_P254805** | Kif4 | 0.599973549 | -0.450373246 | 1.650320344 | 8.967067569 | 1.430703815 | 0.207001908 | 0.709264768 | -4.502056232 |
| **A_55_P2108938** | Hps5 | -0.255883979 | -0.703955188 | 0.19218723 | 10.48220911 | -1.430363445 | 0.207094735 | 0.709264768 | -4.50214972 |
| **A_52_P32466** | Yif1a | -0.270106118 | -0.743093418 | 0.202881183 | 11.46336762 | -1.430326832 | 0.207104723 | 0.709264768 | -4.502159776 |
| **A_52_P162486** | Bgn | 0.280835296 | -0.210945778 | 0.772616369 | 9.422133163 | 1.430310164 | 0.207109269 | 0.709264768 | -4.502164354 |
| **A_55_P1963634** | Yae1d1 | 0.215688012 | -0.162034527 | 0.59341055 | 9.080295744 | 1.430221995 | 0.207133323 | 0.709264768 | -4.502188571 |
| **A_52_P526265** | Wdr73 | 0.13706874 | -0.102980841 | 0.377118322 | 11.69690095 | 1.430170494 | 0.207147374 | 0.709264768 | -4.502202717 |
| **A_55_P2068496** | NA | 0.24083418 | -0.180981182 | 0.662649541 | 11.62458598 | 1.430033058 | 0.207184876 | 0.709264768 | -4.502240465 |
| **A_51_P506748** | Arhgap35 | 0.700056922 | -0.52630736 | 1.926421204 | 12.70763314 | 1.429763502 | 0.207258448 | 0.709264768 | -4.502314502 |
| **A_55_P2052563** | Id1 | 1.045831804 | -0.78631628 | 2.877979888 | 13.14793179 | 1.429722057 | 0.207269762 | 0.709264768 | -4.502325885 |
| **A_55_P2114953** | Usp18 | -0.8627023 | -2.374115669 | 0.648711069 | 9.212979847 | -1.429644448 | 0.20729095 | 0.709264768 | -4.502347201 |
| **A_52_P150525** | Ndufaf6 | 0.239117926 | -0.179873453 | 0.658109305 | 9.530590549 | 1.429411899 | 0.20735445 | 0.709264768 | -4.502411073 |
| **A_55_P2169539** | NA | 0.133420641 | -0.100365315 | 0.367206597 | 7.992964824 | 1.429403857 | 0.207356646 | 0.709264768 | -4.502413282 |
| **A_55_P2102126** | Lamp2 | -0.789520533 | -2.17297479 | 0.593933725 | 10.60202193 | -1.429384178 | 0.207362021 | 0.709264768 | -4.502418687 |
| **A_51_P427232** | Zbed5 | -0.468254236 | -1.288964277 | 0.352455804 | 8.445772047 | -1.429033184 | 0.207457901 | 0.709264768 | -4.50251509 |
| **A_52_P127362** | Vmp1 | -0.467334757 | -1.286453948 | 0.351784435 | 11.78073097 | -1.428997028 | 0.20746778 | 0.709264768 | -4.50252502 |
| **A_55_P2124228** | Uaca | -0.777053135 | -2.139094783 | 0.584988514 | 12.95443533 | -1.42892908 | 0.207486347 | 0.709264768 | -4.502543683 |
| **A_55_P2061333** | NA | 0.96499289 | -0.726737456 | 2.656723236 | 11.67789149 | 1.428707483 | 0.207546909 | 0.709264768 | -4.502604545 |
| **A_55_P2279957** | 5730414N | 0.243185049 | -0.183151009 | 0.669521107 | 8.524433021 | 1.428680613 | 0.207554254 | 0.709264768 | -4.502611925 |
| **A_51_P271725** | Fundc1 | 0.512094946 | -0.385767734 | 1.409957626 | 11.04140153 | 1.428535097 | 0.207594033 | 0.709264768 | -4.502651892 |

| | | | | | | | | | |
|---|---|---|---|---|---|---|---|---|---|
| **A_51_P265495** | Ly6a | -1.702454995 | -4.687652141 | 1.282742152 | 10.03457189 | -1.428410282 | 0.20762816 | 0.709264768 | -4.502686172 |
| **A_51_P285770** | Fem1a | 0.242970982 | -0.183079598 | 0.669021562 | 11.45039286 | 1.428379453 | 0.20763659 | 0.709264768 | -4.50269464 |
| **A_52_P663585** | 10-Sep | 0.264149908 | -0.19904518 | 0.727344996 | 9.159698489 | 1.428357319 | 0.207642642 | 0.709264768 | -4.502700719 |
| **A_51_P164014** | Cenpe | 1.253043548 | -0.944234111 | 3.450321208 | 11.72745793 | 1.428339907 | 0.207647404 | 0.709264768 | -4.502705501 |
| **A_55_P2016089** | Uggt1 | -0.722149211 | -1.988592005 | 0.544293584 | 12.45087102 | -1.428208815 | 0.207683255 | 0.709264768 | -4.502741506 |
| **A_52_P219473** | Cdc6 | 0.32974313 | -0.248552081 | 0.908038341 | 8.198294238 | 1.428158818 | 0.20769693 | 0.709264768 | -4.502755237 |
| **A_55_P1962699** | NA | -0.300819792 | -0.828401441 | 0.226761858 | 11.53420709 | -1.42812779 | 0.207705417 | 0.709264768 | -4.502763759 |
| **A_51_P418737** | Rbm33 | -0.188829355 | -0.520023809 | 0.1423651 | 8.090347177 | -1.428028248 | 0.207732647 | 0.709264768 | -4.502791098 |
| **A_55_P2153923** | Slc17a5 | -0.430832314 | -1.186556561 | 0.324891932 | 8.179447124 | -1.427891667 | 0.207770014 | 0.709264768 | -4.50282861 |
| **A_51_P155174** | Zfp672 | 0.157845876 | -0.119042749 | 0.434734502 | 9.529529464 | 1.427836469 | 0.207785117 | 0.709264768 | -4.50284377 |
| **A_55_P2152304** | Atp5h | 0.268799244 | -0.202732704 | 0.740331192 | 15.18560295 | 1.427799633 | 0.207795197 | 0.709264768 | -4.502853887 |
| **A_55_P2129316** | NA | -0.903504756 | -2.488474053 | 0.681464542 | 8.623805928 | -1.42777553 | 0.207801792 | 0.709264768 | -4.502860507 |
| **A_51_P464308** | Gnb4 | -0.696832095 | -1.919364276 | 0.525700086 | 10.00334172 | -1.427638301 | 0.207839348 | 0.709264768 | -4.502898196 |
| **A_51_P381260** | Fxyd5 | -0.742866107 | -2.046181566 | 0.560449353 | 10.98066437 | -1.427615791 | 0.207845509 | 0.709264768 | -4.502904379 |
| **A_55_P1986863** | Ccdc126 | -0.192300869 | -0.529682693 | 0.145080955 | 9.546721333 | -1.427611082 | 0.207846798 | 0.709264768 | -4.502905672 |
| **A_55_P1962937** | Trem2 | -1.193666628 | -3.28820516 | 0.900871904 | 9.531059724 | -1.427397868 | 0.207905164 | 0.709264768 | -4.50296423 |
| **A_55_P2169069** | Gm10405 | 0.247315497 | -0.186658814 | 0.681289808 | 12.85166643 | 1.427373581 | 0.207911813 | 0.709264768 | -4.5029709 |
| **A_55_P1977583** | Anxa11 | -0.75036434 | -2.067110062 | 0.566381382 | 12.23666342 | -1.427317584 | 0.207927145 | 0.709264768 | -4.50298628 |
| **A_55_P2000783** | Axin2 | 0.419801468 | -0.316899597 | 1.156502534 | 9.018085667 | 1.427259692 | 0.207942996 | 0.709264768 | -4.503002179 |
| **A_51_P446012** | Phf13 | -0.802069035 | -2.209609297 | 0.605471227 | 12.37802578 | -1.427254002 | 0.207944554 | 0.709264768 | -4.503003742 |
| **A_51_P314186** | Syne1 | -0.170129063 | -0.468736297 | 0.128478171 | 8.097936541 | -1.427014973 | 0.208010016 | 0.709264768 | -4.50306939 |
| **A_66_P123735** | NA | -0.387554454 | -1.067867301 | 0.292758393 | 9.355463636 | -1.426837141 | 0.208058731 | 0.709264768 | -4.50311823 |
| **A_55_P2144055** | NA | 0.470301382 | -0.355511755 | 1.296114519 | 15.86415917 | 1.426411439 | 0.20817539 | 0.709264768 | -4.503235144 |
| **A_51_P467410** | Npm1 | 0.411683067 | -0.311269022 | 1.134635155 | 15.82885505 | 1.426276848 | 0.208212286 | 0.709264768 | -4.503272108 |
| **A_55_P2373987** | A730009E | -0.126965791 | -0.349959493 | 0.096027912 | 8.013313468 | -1.426081899 | 0.208265738 | 0.709264768 | -4.503325648 |
| **A_55_P1962665** | Exoc3l2 | 0.161841319 | -0.122418328 | 0.446100965 | 8.027449962 | 1.426016472 | 0.20828368 | 0.709264768 | -4.503343616 |
| **A_51_P174961** | F10 | -0.184841336 | -0.509595342 | 0.13991267 | 8.59352128 | -1.425590942 | 0.20840041 | 0.709264768 | -4.503460481 |
| **A_55_P2094852** | Rabep1 | -0.400399438 | -1.104050234 | 0.303251358 | 9.890669648 | -1.425235467 | 0.208497969 | 0.709264768 | -4.503558105 |
| **A_55_P1952676** | Gtf2a2 | 0.520163965 | -0.394035039 | 1.434362969 | 11.90119208 | 1.425114846 | 0.208531083 | 0.709264768 | -4.503591231 |
| **A_52_P275627** | Kpnb1 | 0.665341125 | -0.504042411 | 1.834724661 | 10.85774428 | 1.425075057 | 0.208542007 | 0.709264768 | -4.503602158 |
| **A_52_P606826** | Pigh | -0.207302694 | -0.571658522 | 0.157053134 | 8.265932354 | -1.425048254 | 0.208549366 | 0.709264768 | -4.503609519 |
| **A_51_P256066** | Tiam2 | -0.165115381 | -0.455331731 | 0.125100969 | 9.103660463 | -1.425003728 | 0.208561592 | 0.709264768 | -4.503621747 |
| **A_55_P2114799** | NA | -0.158414258 | -0.436882846 | 0.120054329 | 8.409591266 | -1.424847558 | 0.208604478 | 0.709264768 | -4.503664636 |
| **A_55_P1973427** | Tex14 | -0.625520512 | -1.725173798 | 0.474132774 | 8.113982094 | -1.424741664 | 0.208633562 | 0.709264768 | -4.503693717 |
| **A_66_P111660** | Mt1 | -0.88998129 | -2.454560145 | 0.674597564 | 15.21707566 | -1.424733926 | 0.208635687 | 0.709264768 | -4.503695842 |
| **A_55_P1991552** | Atraid | -0.312480055 | -0.861830087 | 0.236869977 | 12.29529916 | -1.424700228 | 0.208644944 | 0.709264768 | -4.503705096 |
| **A_55_P2102181** | NA | -1.387659004 | -3.827231152 | 1.051913144 | 12.16800049 | -1.42468692 | 0.208648599 | 0.709264768 | -4.503708751 |
| **A_51_P399889** | Ap4b1 | -0.261979177 | -0.722570371 | 0.198612017 | 10.73706624 | -1.42462804 | 0.208664774 | 0.709264768 | -4.503724921 |
| **A_52_P535012** | NA | 0.223024888 | -0.169138909 | 0.615188685 | 15.52477106 | 1.424413815 | 0.208723632 | 0.709264768 | -4.503783752 |
| **A_55_P2002893** | Pfkfb1 | -0.245439804 | -0.677057552 | 0.186177944 | 8.618362378 | -1.424282202 | 0.2087598 | 0.709264768 | -4.503819896 |
| **A_51_P313483** | Dyrk1a | -0.325028388 | -0.896640151 | 0.246583375 | 9.716108086 | -1.424198362 | 0.208782843 | 0.709264768 | -4.503842921 |
| **A_55_P1993789** | Hmgn2 | 0.424941065 | -0.322422933 | 1.172305063 | 14.98362415 | 1.424120946 | 0.208804122 | 0.709264768 | -4.503864181 |
| **A_51_P286142** | Znrd1 | 0.346379844 | -0.262831382 | 0.95559107 | 11.65205707 | 1.424082206 | 0.208814771 | 0.709264768 | -4.503874819 |
| **A_55_P1968483** | Gtpbp6 | -0.174830668 | -0.482330411 | 0.132669075 | 10.01276109 | -1.424043338 | 0.208825456 | 0.709264768 | -4.503885493 |
| **A_51_P368855** | Pfdn6 | -0.153311536 | -0.422967874 | 0.116344802 | 11.58383179 | -1.424014946 | 0.208833261 | 0.709264768 | -4.50389329 |
| **A_55_P1966209** | Fez2 | -0.447084183 | -1.233453216 | 0.33928485 | 11.06418496 | -1.424010686 | 0.208834433 | 0.709264768 | -4.50389446 |
| **A_55_P2125912** | Cct6a | 0.441517332 | -0.335121798 | 1.218156462 | 10.47769628 | 1.42389785 | 0.208865456 | 0.709264768 | -4.503925447 |
| **A_55_P2070940** | Krtap20-2 | 0.27497034 | -0.208853428 | 0.758794109 | 9.104463145 | 1.423472224 | 0.208982516 | 0.709264768 | -4.504042331 |

| | | | | | | | | | |
|---|---|---|---|---|---|---|---|---|---|
| **A_55_P2052696** | Synm | 0.745457237 | -0.566600497 | 2.05751497 | 9.403794805 | 1.42304992 | 0.209098723 | 0.709264768 | -4.504158302 |
| **A_55_P2177233** | Abhd5 | -0.765966572 | -2.114547986 | 0.582614843 | 9.59523069 | -1.422600615 | 0.209222426 | 0.709264768 | -4.504281686 |
| **A_55_P1963454** | Cenpq | 0.317279107 | -0.2413364 | 0.875894614 | 8.417696874 | 1.42258699 | 0.209226178 | 0.709264768 | -4.504285427 |
| **A_55_P2156409** | Rpl26 | 0.589504042 | -0.448463357 | 1.62747144 | 14.11605625 | 1.422503749 | 0.209249104 | 0.709264768 | -4.504308286 |
| **A_55_P2058493** | NA | 0.395809684 | -0.301132593 | 1.092751961 | 16.0880785 | 1.422459633 | 0.209261255 | 0.709264768 | -4.504320401 |
| **A_55_P2194958** | Nfia | 0.392656935 | -0.298754231 | 1.084068102 | 8.667083433 | 1.422417965 | 0.209272733 | 0.709264768 | -4.504331843 |
| **A_55_P2077009** | Acap3 | 0.290423406 | -0.220983662 | 0.801830474 | 10.58602557 | 1.42237868 | 0.209283554 | 0.709264768 | -4.504342631 |
| **A_55_P2175451** | NA | -0.297591569 | -0.821621848 | 0.226438711 | 11.64984456 | -1.422376558 | 0.209284139 | 0.709264768 | -4.504343214 |
| **A_55_P2176391** | NA | 0.260421555 | -0.19817337 | 0.719016481 | 11.95329295 | 1.422322333 | 0.209299077 | 0.709264768 | -4.504358104 |
| **A_55_P2338535** | Slc35a1 | -0.333048678 | -0.919731227 | 0.253633872 | 8.968152486 | -1.421853622 | 0.20942824 | 0.709264768 | -4.504486814 |
| **A_51_P128336** | Serinc5 | -0.468434449 | -1.293866972 | 0.356998075 | 11.70565659 | -1.4214042 | 0.209552157 | 0.709264768 | -4.51088825 |
| **A_66_P133520** | Glmn | 0.171143207 | -0.13044015 | 0.472726564 | 9.073173234 | 1.42135526 | 0.209565655 | 0.709264768 | -4.504623665 |
| **A_51_P365694** | Rala | -0.296525095 | -0.819168663 | 0.226118473 | 12.5935278 | -1.421039629 | 0.209652729 | 0.709264768 | -4.504710336 |
| **A_55_P2156401** | NA | 0.58393605 | -0.445329018 | 1.613201119 | 14.9852191 | 1.420981421 | 0.209668791 | 0.709264768 | -4.50472632 |
| **A_51_P363525** | Fbrsl1 | -0.564363968 | -1.559357301 | 0.430629365 | 10.15234008 | -1.420657712 | 0.209758134 | 0.709264768 | -4.504815209 |
| **A_55_P2004636** | Ppih | 0.84453492 | -0.644548012 | 2.333617853 | 11.19587817 | 1.420525622 | 0.209794601 | 0.709264768 | -4.504851479 |
| **A_55_P2009752** | Hlf | 0.270380457 | -0.20636707 | 0.747127983 | 8.382992987 | 1.420486773 | 0.209805328 | 0.709264768 | -4.504862147 |
| **A_52_P244193** | Cd24a | 0.744699383 | -0.568409927 | 2.057808693 | 12.55579244 | 1.420464745 | 0.20981141 | 0.709264768 | -4.504868196 |
| **A_55_P2147291** | NA | 0.357735396 | -0.273101512 | 0.988572304 | 16.20856906 | 1.42034931 | 0.209843286 | 0.709264768 | -4.504899893 |
| **A_55_P1978830** | Camk2d | -0.412975743 | -1.141498698 | 0.315547213 | 8.713100375 | -1.419814633 | 0.209990991 | 0.709264768 | -4.505046709 |
| **A_55_P2172334** | Nup155 | 0.289275198 | -0.221090924 | 0.79964132 | 9.825052699 | 1.419644836 | 0.210037918 | 0.709264768 | -4.505093333 |
| **A_55_P2037343** | H2-Q10 | 0.175268691 | -0.134055519 | 0.484592901 | 8.069820236 | 1.419190767 | 0.210163457 | 0.709264768 | -4.505218012 |
| **A_52_P594917** | Pip4k2c | -0.85058419 | -2.351844615 | 0.650676236 | 10.69933513 | -1.419095475 | 0.210189811 | 0.709264768 | -4.505244177 |
| **A_55_P2004084** | Iqce | 0.383817504 | -0.293618091 | 1.0612531 | 9.7938463 | 1.419080754 | 0.210193883 | 0.709264768 | -4.505248219 |
| **A_52_P1172945** | Ankrd34b | 0.402867631 | -0.308232302 | 1.113967564 | 8.501532238 | 1.418998979 | 0.210216502 | 0.709264768 | -4.505270673 |
| **A_51_P184969** | Rilpl1 | -0.208957934 | -0.577817256 | 0.159901387 | 9.806346195 | -1.418889087 | 0.210246902 | 0.709264768 | -4.505300847 |
| **A_55_P1971093** | NA | 0.468484075 | -0.35854795 | 1.2955161 | 11.22177906 | 1.418805459 | 0.21027004 | 0.709264768 | -4.505323809 |
| **A_52_P48777** | Copg2 | 0.281386502 | -0.215395615 | 0.778168619 | 10.77436949 | 1.418690473 | 0.210301857 | 0.709264768 | -4.505355382 |
| **A_51_P282667** | Hexa | -1.011264234 | -2.796680377 | 0.77415191 | 11.46089318 | -1.418650893 | 0.210312809 | 0.709264768 | -4.505366249 |
| **A_55_P1969831** | Ptpmt1 | -0.127849633 | -0.35360241 | 0.097903144 | 12.42012019 | -1.418458812 | 0.210365972 | 0.709264768 | -4.50541899 |
| **A_55_P1958082** | Prpsap2 | 0.228129539 | -0.174698419 | 0.630957498 | 9.663867295 | 1.418444225 | 0.210370009 | 0.709264768 | -4.505422995 |
| **A_51_P211934** | Larp7 | 0.480305444 | -0.367867781 | 1.328478669 | 10.54762744 | 1.418349618 | 0.210396198 | 0.709264768 | -4.505448972 |
| **A_51_P221428** | Cbx6 | 1.136055355 | -0.870162623 | 3.142273333 | 11.55014164 | 1.418311815 | 0.210406664 | 0.709264768 | -4.505459351 |
| **A_51_P224227** | Rlf | 0.21966977 | -0.168279392 | 0.607618933 | 10.14598615 | 1.418227241 | 0.210430079 | 0.709264768 | -4.505482573 |
| **A_55_P1982737** | Dnajc5 | -0.310939965 | -0.860110229 | 0.238230298 | 14.84309171 | -1.418142519 | 0.210453539 | 0.709264768 | -4.505505836 |
| **A_55_P2087186** | Ier3ip1 | -0.499910554 | -1.382876423 | 0.383055316 | 14.1984358 | -1.418073477 | 0.210472658 | 0.709264768 | -4.505524792 |
| **A_55_P1966124** | Erh | 0.476882177 | -0.365481584 | 1.319245938 | 14.90096963 | 1.417952759 | 0.210506091 | 0.709264768 | -4.505557938 |
| **A_51_P324871** | Cyb561a3 | -0.729327548 | -2.01762868 | 0.558973584 | 10.77567433 | -1.41793256 | 0.210511686 | 0.709264768 | -4.505563484 |
| **A_55_P1980981** | Apba3 | -0.413561947 | -1.144190529 | 0.317066636 | 9.808115515 | -1.41773238 | 0.210567138 | 0.709264768 | -4.505618447 |
| **A_55_P2182006** | Cacng6 | 0.184544753 | -0.141502334 | 0.510591839 | 8.073639936 | 1.417658816 | 0.21058752 | 0.709264768 | -4.505638645 |
| **A_55_P2183032** | Zfp787 | -0.443234827 | -1.226361575 | 0.339891921 | 9.261536573 | -1.417594977 | 0.210605209 | 0.709264768 | -4.505656173 |
| **A_55_P2272007** | Lrrfip2 | -0.416624819 | -1.152919935 | 0.319670298 | 8.500813883 | -1.417240529 | 0.210703445 | 0.709264768 | -4.505753492 |
| **A_55_P2000027** | Spink2 | -0.168268097 | -0.465672264 | 0.12913607 | 8.684835238 | -1.417114952 | 0.21073826 | 0.709264768 | -4.505787971 |
| **A_55_P2035892** | Prkcsh | -0.62896745 | -1.740762754 | 0.482827854 | 14.91569642 | -1.416947244 | 0.210784763 | 0.709264768 | -4.505834017 |
| **A_55_P2294037** | NA | -0.194171682 | -0.537424658 | 0.149081295 | 8.53395109 | -1.416843641 | 0.210813495 | 0.709264768 | -4.505862462 |
| **A_51_P184321** | Aimp1 | 0.348848683 | -0.267878126 | 0.965575492 | 12.78513167 | 1.416754517 | 0.210838215 | 0.709264768 | -4.505886932 |
| **A_52_P101982** | Tmem129 | -0.378303173 | -1.047127129 | 0.290520783 | 10.26926659 | -1.416702006 | 0.210852781 | 0.709264768 | -4.505901349 |
| **A_55_P2079860** | Asap3 | -0.228571413 | -0.632678316 | 0.17553549 | 8.618651597 | -1.416693788 | 0.210855061 | 0.709264768 | -4.505903606 |

| | | | | | | | | | |
|---|---|---|---|---|---|---|---|---|---|
| **A_55_P2091196** | Hrct1 | -0.978127381 | -2.707426359 | 0.751171597 | 9.355350696 | -1.416692768 | 0.210855344 | 0.709264768 | -4.505903886 |
| **A_51_P192149** | Slc9a6 | -0.725260559 | -2.007500336 | 0.556979219 | 8.917665585 | -1.416691083 | 0.210855811 | 0.709264768 | -4.505904348 |
| **A_55_P2034928** | BC147527 | 0.289294725 | -0.222188091 | 0.80077754 | 8.888966434 | 1.41664102 | 0.210869699 | 0.709264768 | -4.505918094 |
| **A_51_P505823** | Endod1 | -0.539271457 | -1.493064244 | 0.41452133 | 10.24259982 | -1.416132031 | 0.211010946 | 0.709264768 | -4.506057839 |
| **A_55_P2003199** | Setd1b | -0.181874806 | -0.503553732 | 0.139804119 | 8.338452514 | -1.416120729 | 0.211014084 | 0.709264768 | -4.506060942 |
| **A_55_P2154782** | NA | 0.411162826 | -0.316109965 | 1.138435617 | 9.012142278 | 1.416011721 | 0.211044346 | 0.709264768 | -4.50609087 |
| **A_55_P2038525** | C3 | -0.351795111 | -0.9740649 | 0.270474677 | 9.981326339 | -1.415994053 | 0.211049251 | 0.709264768 | -4.506095721 |
| **A_52_P162099** | Ckap2 | 0.686646411 | -0.528030947 | 1.90132377 | 9.210021452 | 1.41586734 | 0.211084434 | 0.709264768 | -4.50613051 |
| **A_55_P2007303** | Snrpg | 0.699937287 | -0.538333495 | 1.938208069 | 14.80469924 | 1.415773732 | 0.211110429 | 0.709264768 | -4.506156211 |
| **A_55_P1995012** | Plekhm2 | -0.290522971 | -0.804495137 | 0.223449195 | 12.21801423 | -1.415765165 | 0.211112808 | 0.709264768 | -4.506158563 |
| **A_51_P338728** | Sdhaf1 | 0.370106564 | -0.284667074 | 1.024880201 | 10.81660447 | 1.41574797 | 0.211117584 | 0.709264768 | -4.506163283 |
| **A_55_P2105321** | Acin1 | 0.299035939 | -0.230030184 | 0.828102063 | 12.21587311 | 1.415675702 | 0.211137655 | 0.709264768 | -4.506183125 |
| **A_55_P2147001** | Wdr55 | 0.220558976 | -0.169666283 | 0.610784236 | 10.3164642 | 1.415662426 | 0.211141342 | 0.709264768 | -4.506186769 |
| **A_55_P2014304** | Kank1 | -1.005236102 | -2.784071462 | 0.773599257 | 11.34906624 | -1.415411335 | 0.211211094 | 0.709264768 | -4.506255706 |
| **A_55_P2027278** | Gas2l1 | -0.674842698 | -1.869049718 | 0.519364322 | 11.3771985 | -1.415380731 | 0.211219597 | 0.709264768 | -4.506264108 |
| **A_55_P1961531** | Nemp1 | 0.327020737 | -0.251710798 | 0.905752273 | 9.431345775 | 1.415299949 | 0.211242043 | 0.709264768 | -4.506286286 |
| **A_55_P2098539** | Itgb3bp | 0.637353857 | -0.490613478 | 1.765321191 | 10.47919742 | 1.415254162 | 0.211254767 | 0.709264768 | -4.506298857 |
| **A_55_P2170670** | NA | 0.154301213 | -0.118904706 | 0.427507132 | 9.326806999 | 1.414586738 | 0.211440313 | 0.709264768 | -4.506482092 |
| **A_55_P1993880** | Rps24 | 0.247977371 | -0.19109922 | 0.687053963 | 15.24045354 | 1.41456241 | 0.21144708 | 0.709264768 | -4.506488771 |
| **A_51_P138923** | Peg12 | 0.604563683 | -0.465981691 | 1.675109056 | 9.446637901 | 1.414449093 | 0.211478598 | 0.709264768 | -4.506519881 |
| **A_55_P2036083** | Gm5797 | -0.608054135 | -1.684792121 | 0.46868385 | 8.636663558 | -1.414433577 | 0.211482914 | 0.709264768 | -4.506524141 |
| **A_55_P2160081** | Ccdc28b | 0.341780935 | -0.263490374 | 0.947052245 | 8.545282844 | 1.414321329 | 0.21151414 | 0.709264768 | -4.506554957 |
| **A_51_P115666** | Fam83h | -0.485072405 | -1.344138805 | 0.373993996 | 10.93949846 | -1.414263005 | 0.211530367 | 0.709264768 | -4.506570969 |
| **A_55_P2013357** | Renbp | -0.416114724 | -1.153091217 | 0.32086177 | 9.083404024 | -1.414196613 | 0.21154884 | 0.709264768 | -4.506589196 |
| **A_51_P164030** | Tcp1 | 0.574283888 | -0.442830399 | 1.591398174 | 12.50915268 | 1.414188245 | 0.211551169 | 0.709264768 | -4.506591493 |
| **A_51_P234864** | Sfxn5 | 0.142899125 | -0.11020037 | 0.395998619 | 8.713985879 | 1.414127721 | 0.21156801 | 0.709264768 | -4.506608109 |
| **A_51_P249051** | Pycr2 | 0.381801389 | -0.294466229 | 1.058069007 | 12.07500025 | 1.414064623 | 0.211585569 | 0.709264768 | -4.506625432 |
| **A_55_P2005808** | Abhd18 | -0.283362081 | -0.785344925 | 0.218620762 | 8.678367351 | -1.413849573 | 0.211645425 | 0.709264768 | -4.50668447 |
| **A_55_P1952690** | Rpl12 | 0.596834021 | -0.460482637 | 1.65415068 | 15.214583 | 1.413835369 | 0.211649379 | 0.709264768 | -4.506688369 |
| **A_66_P119421** | Prdx1 | 0.358819234 | -0.276876823 | 0.994515291 | 9.609868193 | 1.413762777 | 0.211669587 | 0.709264768 | -4.506708298 |
| **A_66_P100232** | Paxbp1 | -0.160766566 | -0.445632489 | 0.124099356 | 9.024172306 | -1.413531798 | 0.211733901 | 0.709264768 | -4.506771708 |
| **A_51_P485899** | Srsf1 | 0.71048076 | -0.548632065 | 1.969593585 | 11.40703399 | 1.413311913 | 0.211795142 | 0.709264768 | -4.506832072 |
| **A_55_P2005754** | Manbal | -0.45006621 | -1.247686658 | 0.347554238 | 10.97031487 | -1.41328736 | 0.211801982 | 0.709264768 | -4.506838813 |
| **A_51_P411019** | Tmem127 | -0.42675843 | -1.183113978 | 0.329597118 | 12.16420408 | -1.413209204 | 0.211823754 | 0.709264768 | -4.506860269 |
| **A_51_P343252** | Cd151 | -1.010736835 | -2.802122469 | 0.780648798 | 13.4612503 | -1.413186084 | 0.211830195 | 0.709264768 | -4.506866616 |
| **A_55_P2040031** | NA | 0.730040024 | -0.563900782 | 2.023980829 | 13.49558759 | 1.413131598 | 0.211845375 | 0.709264768 | -4.506881574 |
| **A_51_P438149** | Mapre2 | 0.329277224 | -0.25442518 | 0.912979628 | 11.85149148 | 1.412929703 | 0.211901633 | 0.709264768 | -4.506936998 |
| **A_51_P450021** | Higd2a | 0.300107408 | -0.231886793 | 0.832101609 | 12.50858413 | 1.412928426 | 0.211901989 | 0.709264768 | -4.506937349 |
| **A_55_P2126469** | NA | 0.223033459 | -0.172378757 | 0.618445675 | 16.08462306 | 1.412766157 | 0.211947215 | 0.709264768 | -4.506981895 |
| **A_55_P1994939** | Hmgb2 | 1.203005421 | -0.930140352 | 3.336151195 | 14.17807613 | 1.412529072 | 0.212013309 | 0.709264768 | -4.507046979 |
| **A_66_P113662** | Tmem62 | -0.585844326 | -1.624652548 | 0.452963897 | 8.922278558 | -1.412528421 | 0.212013491 | 0.709264768 | -4.507047158 |
| **A_66_P101108** | Tnrc18 | -0.132231948 | -0.366705137 | 0.10224124 | 11.01394588 | -1.412516578 | 0.212016793 | 0.709264768 | -4.507050409 |
| **A_55_P2048769** | Zfp454 | 0.490147957 | -0.379321157 | 1.359617072 | 8.782963828 | 1.411963218 | 0.212171135 | 0.709264768 | -4.507202315 |
| **A_66_P128608** | Fut11 | 0.15149914 | -0.117267411 | 0.42026569 | 8.818873438 | 1.41183939 | 0.212205687 | 0.709264768 | -4.507236307 |
| **A_55_P2143075** | Ubc | -0.435096382 | -1.207052994 | 0.33686023 | 16.10230457 | -1.411701662 | 0.212244124 | 0.709264768 | -4.507274115 |
| **A_55_P2143195** | Dyrk1a | -0.270586881 | -0.750675337 | 0.209501575 | 9.073610167 | -1.411678559 | 0.212250573 | 0.709264768 | -4.507280457 |
| **A_51_P225525** | Gm9079 | -0.257659018 | -0.714837343 | 0.199519307 | 12.28074621 | -1.411594835 | 0.212273942 | 0.709264768 | -4.50730344 |
| **A_52_P305851** | 2410002F | 0.152145977 | -0.117826296 | 0.42211825 | 9.437626165 | 1.411535017 | 0.21229064 | 0.709264768 | -4.50731986 |

| | | | | | | | | | |
|---|---|---|---|---|---|---|---|---|---|
| **A_51_P463846** | Gbp7 | -0.268651792 | -0.745374939 | 0.208071355 | 8.163165055 | -1.411477137 | 0.212306799 | 0.709264768 | -4.507335749 |
| **A_55_P2003719** | NA | 0.404593383 | -0.313432625 | 1.122619391 | 15.98812591 | 1.411331211 | 0.212347542 | 0.709264768 | -4.507375806 |
| **A_55_P2133337** | NA | 1.069288038 | -0.828382846 | 2.966958922 | 10.68270014 | 1.411315657 | 0.212351885 | 0.709264768 | -4.507380076 |
| **A_51_P359333** | Fh1 | 0.305953651 | -0.23704072 | 0.848948021 | 12.16631568 | 1.411271782 | 0.212364137 | 0.709264768 | -4.50739212 |
| **A_55_P2124751** | Col16a1 | 0.659147799 | -0.510758565 | 1.829054163 | 10.45874147 | 1.411178816 | 0.212390099 | 0.709264768 | -4.50741764 |
| **A_55_P2056190** | Rps3a1 | 0.349694152 | -0.271007247 | 0.970395551 | 16.59021854 | 1.411094149 | 0.212413746 | 0.709264768 | -4.507440881 |
| **A_55_P2184478** | NA | 0.288913161 | -0.223926952 | 0.801753275 | 16.15178982 | 1.411028172 | 0.212432175 | 0.709264768 | -4.507458992 |
| **A_55_P2136204** | Rnf38 | -0.246869861 | -0.685093093 | 0.19135337 | 11.60872072 | -1.41098703 | 0.212443667 | 0.709264768 | -4.507470286 |
| **A_55_P1989901** | Ddb1 | -0.469865857 | -1.303967197 | 0.364235483 | 15.43566951 | -1.410929828 | 0.212459647 | 0.709264768 | -4.507485988 |
| **A_55_P2227335** | Ptpre | -0.334479919 | -0.92829659 | 0.259336753 | 8.268201977 | -1.410808308 | 0.212493598 | 0.709264768 | -4.507519345 |
| **A_55_P2060854** | NA | 0.713150749 | -0.553044703 | 1.979346201 | 12.13601112 | 1.410687893 | 0.212527245 | 0.709264768 | -4.507552399 |
| **A_51_P401668** | Laptm5 | 0.304540789 | -0.236188749 | 0.845270327 | 8.486097876 | 1.41063846 | 0.21254106 | 0.709264768 | -4.507565968 |
| **A_66_P126425** | Eif1 | -0.290395065 | -0.806030452 | 0.225240322 | 14.88942832 | -1.410577211 | 0.212558177 | 0.709264768 | -4.507582781 |
| **A_55_P2126087** | NA | 0.208632258 | -0.161900486 | 0.579165001 | 14.16657355 | 1.410279558 | 0.212641383 | 0.709264768 | -4.507664486 |
| **A_55_P2012955** | NA | 0.389910343 | -0.302656379 | 1.082477065 | 16.36996188 | 1.410111589 | 0.21268835 | 0.709264768 | -4.507710592 |
| **A_55_P2130575** | NA | 0.527000352 | -0.409120165 | 1.463120869 | 14.60848662 | 1.410033682 | 0.212710137 | 0.709264768 | -4.507731977 |
| **A_55_P2008747** | Snrpd3 | 0.649332688 | -0.504145576 | 1.802810952 | 12.35975098 | 1.40996456 | 0.21272947 | 0.709264768 | -4.507750951 |
| **A_52_P284958** | Ubtd2 | 0.664812143 | -0.516248935 | 1.84587322 | 10.70911719 | 1.409863059 | 0.212757861 | 0.709264768 | -4.507778811 |
| **A_55_P2115607** | Ikbip | -0.150733276 | -0.418520826 | 0.117054274 | 10.66911414 | -1.409837645 | 0.21276497 | 0.709264768 | -4.507785788 |
| **A_55_P2083093** | Haus7 | 0.244239823 | -0.189742952 | 0.678222598 | 8.400811739 | 1.409594932 | 0.212832876 | 0.709264768 | -4.507852409 |
| **A_55_P2001143** | Ehmt2 | -0.357967867 | -0.994200588 | 0.278264854 | 14.08911733 | -1.409218672 | 0.212938187 | 0.709264768 | -4.507955687 |
| **A_51_P109258** | Cys1 | -0.399648078 | -1.10999318 | 0.310697023 | 8.729649676 | -1.409154737 | 0.212956086 | 0.709264768 | -4.507973237 |
| **A_66_P115038** | Prl3d1 | -0.946747007 | -2.629725248 | 0.736231235 | 8.110256259 | -1.408983104 | 0.213004144 | 0.709264768 | -4.508020347 |
| **A_51_P260548** | Glg1 | -0.524146326 | -1.455894135 | 0.407601483 | 13.07931809 | -1.408978971 | 0.213005301 | 0.709264768 | -4.508021481 |
| **A_51_P467110** | Dpp4 | 0.697285463 | -0.54224745 | 1.936818376 | 9.503400623 | 1.408973722 | 0.213006771 | 0.709264768 | -4.508022922 |
| **A_55_P1993733** | Gm2824 | 0.179491525 | -0.139607386 | 0.498590435 | 8.294379944 | 1.408863669 | 0.213037592 | 0.709264768 | -4.508053129 |
| **A_55_P2040280** | Dhps | -0.273830921 | -0.760648763 | 0.212986921 | 8.115838488 | -1.408855155 | 0.213039977 | 0.709264768 | -4.508055466 |
| **A_55_P2153517** | Enho | -0.822606785 | -2.285207713 | 0.639994143 | 11.16734823 | -1.4086944 | 0.213085005 | 0.709264768 | -4.50809959 |
| **A_65_P05396** | Rad18 | 0.178282835 | -0.138716943 | 0.495282612 | 8.35609887 | 1.408642928 | 0.213099424 | 0.709264768 | -4.508113718 |
| **A_55_P2129911** | NA | 0.314632706 | -0.24487608 | 0.874141492 | 15.91307672 | 1.408469025 | 0.213148148 | 0.709264768 | -4.50816145 |
| **A_55_P1969078** | Adar | -0.536510643 | -1.490589642 | 0.417568355 | 9.673846754 | -1.40845946 | 0.213150828 | 0.709264768 | -4.508164075 |
| **A_52_P876147** | Crocc2 | 0.554970138 | -0.431940369 | 1.541880646 | 8.787488818 | 1.408452444 | 0.213152794 | 0.709264768 | -4.508166001 |
| **A_51_P177583** | Puf60 | 0.19890539 | -0.154865784 | 0.552676564 | 12.1803411 | 1.408232795 | 0.213214351 | 0.709264768 | -4.508226289 |
| **A_51_P126626** | Zfp503 | 1.20834189 | -0.940836136 | 3.357519917 | 10.9363093 | 1.408211173 | 0.213220412 | 0.709264768 | -4.509843489 |
| **A_55_P1975447** | NA | 0.50357768 | -0.39217502 | 1.39933038 | 10.51001755 | 1.408084418 | 0.213255943 | 0.709264768 | -4.508267015 |
| **A_55_P1985219** | Pnpla6 | -0.585157997 | -1.626078311 | 0.455762316 | 9.761266198 | -1.408010866 | 0.213276564 | 0.709264768 | -4.508287202 |
| **A_55_P1996643** | Polr2j | 0.176071852 | -0.137143283 | 0.489286986 | 12.95845409 | 1.407983414 | 0.213284261 | 0.709264768 | -4.508294737 |
| **A_55_P2047986** | Ankrd23 | -0.569845325 | -1.583712496 | 0.444021846 | 8.210672445 | -1.407752393 | 0.213349042 | 0.709264768 | -4.508358145 |
| **A_52_P196105** | Ttyh3 | 0.684586349 | -0.533497587 | 1.902670284 | 12.01331102 | 1.407671652 | 0.213371688 | 0.709264768 | -4.508380306 |
| **A_55_P2034300** | Tmem40 | -1.239507324 | -3.444972706 | 0.965958058 | 9.129782236 | -1.407664628 | 0.213373658 | 0.709264768 | -4.508382234 |
| **A_55_P2107203** | Gpr137b | -0.737870595 | -2.050843397 | 0.575102206 | 9.092714453 | -1.407585615 | 0.213395821 | 0.709264768 | -4.50840392 |
| **A_55_P2075894** | NA | 0.898163134 | -0.700047688 | 2.496373957 | 12.34767082 | 1.407574867 | 0.213398836 | 0.709264768 | -4.50840687 |
| **A_55_P2061219** | Ces3a | 0.151411931 | -0.118040116 | 0.420863978 | 7.735084281 | 1.407436975 | 0.213437519 | 0.709264768 | -4.508444717 |
| **A_65_P19395** | H2-D1 | -1.468968266 | -4.083162123 | 1.145225591 | 12.08271069 | -1.407424056 | 0.213441144 | 0.709264768 | -4.508448263 |
| **A_55_P2019054** | Acacb | -0.630457213 | -1.752553435 | 0.491639009 | 9.117636016 | -1.407264924 | 0.213485795 | 0.709264768 | -4.508491938 |
| **A_55_P2112330** | Anapc7 | 0.180904683 | -0.14108347 | 0.502892836 | 11.40604424 | 1.407214373 | 0.213499982 | 0.709264768 | -4.508505813 |
| **A_66_P110161** | Eppk1 | -0.464337247 | -1.290829606 | 0.362155112 | 9.004403201 | -1.407165 | 0.213513838 | 0.709264768 | -4.508519364 |
| **A_51_P394676** | Mef2d | -0.238213157 | -0.66229829 | 0.185871976 | 11.77640803 | -1.406899393 | 0.213588394 | 0.709264768 | -4.508592262 |

| | | | | | | | | | |
|---|---|---|---|---|---|---|---|---|---|
| **A_55_P1955791** | Fopnl | 0.366932082 | -0.286355012 | 1.020219176 | 12.36513211 | 1.40679868 | 0.213616671 | 0.709264768 | -4.508619904 |
| **A_55_P1967568** | Chtop | 0.189897147 | -0.148205838 | 0.528000131 | 12.66050773 | 1.406759255 | 0.213627741 | 0.709264768 | -4.508630724 |
| **A_55_P2082625** | NA | 0.268432201 | -0.209501241 | 0.746365643 | 9.711603866 | 1.406751988 | 0.213629781 | 0.709264768 | -4.508632719 |
| **A_51_P168524** | NA | 0.137202156 | -0.107104893 | 0.381509205 | 8.877292809 | 1.406615126 | 0.213668215 | 0.709264768 | -4.508670281 |
| **A_55_P2174258** | Pin4 | 0.344757683 | -0.269299621 | 0.958814987 | 11.90940816 | 1.406226887 | 0.213777276 | 0.709264768 | -4.508776834 |
| **A_55_P2036317** | NA | 0.261397097 | -0.204197263 | 0.726991457 | 16.97779571 | 1.406188025 | 0.213788195 | 0.709264768 | -4.5087875 |
| **A_55_P2064598** | Cnot6 | 0.330781717 | -0.258404686 | 0.919968119 | 13.16533912 | 1.406174204 | 0.213792079 | 0.709264768 | -4.508791293 |
| **A_52_P305279** | Spata13 | 0.409673563 | -0.320048421 | 1.139395546 | 9.08431918 | 1.40614742 | 0.213799605 | 0.709264768 | -4.508798644 |
| **A_55_P2147591** | Med27 | 0.131948637 | -0.103101882 | 0.366999156 | 8.402099971 | 1.406028225 | 0.213833102 | 0.709264768 | -4.508831357 |
| **A_52_P164709** | Poc1a | 0.631387446 | -0.493368957 | 1.756143849 | 10.10547403 | 1.406008071 | 0.213838766 | 0.709264768 | -4.508836888 |
| **A_52_P106620** | Tnfrsf11b | -1.369209926 | -3.808383692 | 1.069963841 | 9.000062892 | -1.405975147 | 0.21384802 | 0.709264768 | -4.508845924 |
| **A_51_P189959** | Oxsr1 | -0.219609877 | -0.610875923 | 0.17165617 | 8.934034851 | -1.405821082 | 0.213891326 | 0.709264768 | -4.508888207 |
| **A_55_P2079927** | Ccdc68 | -0.605093608 | -1.683219284 | 0.473032068 | 8.259220762 | -1.405735207 | 0.213915468 | 0.709264768 | -4.508911775 |
| **A_52_P477431** | Mnt | -0.547002999 | -1.521631028 | 0.42762503 | 11.9609108 | -1.405727549 | 0.213917621 | 0.709264768 | -4.508913877 |
| **A_51_P391367** | Tfap4 | 0.209920593 | -0.164152142 | 0.583993329 | 8.18749396 | 1.405559826 | 0.213964781 | 0.709264768 | -4.508959907 |
| **A_55_P2038237** | Rps3 | 0.359613402 | -0.281401994 | 1.000628799 | 10.31004228 | 1.405134037 | 0.214084547 | 0.709264768 | -4.509076761 |
| **A_51_P214083** | 2310039H | 0.496991923 | -0.388939223 | 1.382923069 | 11.43276548 | 1.405075646 | 0.214100976 | 0.709264768 | -4.509092786 |
| **A_55_P2121541** | Gm6607 | 0.250315883 | -0.195904192 | 0.696535959 | 15.46760628 | 1.405043105 | 0.214110132 | 0.709264768 | -4.509101716 |
| **A_52_P629037** | Gm6987 | 0.751174556 | -0.587980175 | 2.090329287 | 12.51746939 | 1.404948661 | 0.214136709 | 0.709264768 | -4.509127635 |
| **A_55_P2026794** | Unc50 | -0.408225358 | -1.136256255 | 0.319805539 | 12.15723203 | -1.404431344 | 0.214282336 | 0.709264768 | -4.509269604 |
| **A_55_P1998596** | Nupl2 | 0.158558054 | -0.124240293 | 0.441356402 | 9.369440664 | 1.404306111 | 0.214317604 | 0.709264768 | -4.509303971 |
| **A_55_P1975021** | Rad18 | 0.226666533 | -0.177631355 | 0.63096442 | 8.734632243 | 1.404223624 | 0.214340836 | 0.709264768 | -4.509326608 |
| **A_55_P2122643** | Dtymk | 0.95000126 | -0.74464226 | 2.64464478 | 12.36955856 | 1.404093961 | 0.214377361 | 0.709264768 | -4.509362191 |
| **A_55_P2150906** | Atp5c1 | 0.453371644 | -0.355443146 | 1.262186435 | 13.1368711 | 1.403962847 | 0.2144143 | 0.709264768 | -4.509398172 |
| **A_55_P2171658** | Stx3 | -0.604744104 | -1.683691035 | 0.474202827 | 10.43832198 | -1.403853874 | 0.214445006 | 0.709264768 | -4.509428077 |
| **A_52_P26299** | Slc35f1 | 0.159263142 | -0.124910776 | 0.44343706 | 7.870114389 | 1.403722983 | 0.214481893 | 0.709264768 | -4.509463996 |
| **A_51_P187082** | G6pdx | 0.267192061 | -0.209596395 | 0.743980517 | 10.56533479 | 1.403615528 | 0.21451218 | 0.709264768 | -4.509493484 |
| **A_55_P2168990** | NA | -0.422905224 | -1.177601156 | 0.331790709 | 15.26216658 | -1.403528985 | 0.214536575 | 0.709264768 | -4.509517233 |
| **A_55_P2153620** | Ahnak | -0.325151018 | -0.905407696 | 0.25510566 | 9.536999809 | -1.403509371 | 0.214542105 | 0.709264768 | -4.509522615 |
| **A_51_P490305** | Ifi30 | -0.194220836 | -0.540829915 | 0.152388242 | 10.6044148 | -1.403480012 | 0.214550382 | 0.709264768 | -4.509530672 |
| **A_66_P114862** | Ankrd11 | -0.207762072 | -0.578541913 | 0.163017768 | 10.67826543 | -1.403461492 | 0.214555603 | 0.709264768 | -4.509535755 |
| **A_55_P1998186** | NA | 0.642316102 | -0.50400774 | 1.788639944 | 13.64561959 | 1.403433471 | 0.214563503 | 0.709264768 | -4.509543444 |
| **A_51_P511949** | Setd7 | -0.429333168 | -1.195566462 | 0.336900127 | 11.23268676 | -1.403407416 | 0.214570849 | 0.709264768 | -4.509550594 |
| **A_55_P2037424** | NA | 0.226065858 | -0.177449279 | 0.629580995 | 14.27628382 | 1.403219114 | 0.214623946 | 0.709264768 | -4.509602267 |
| **A_51_P269078** | Habp4 | -0.130627285 | -0.363791035 | 0.102536465 | 8.57573013 | -1.403211754 | 0.214626022 | 0.709264768 | -4.509604287 |
| **A_52_P413646** | Bmp6 | -0.130554615 | -0.363602466 | 0.102493236 | 8.273005166 | -1.403128586 | 0.214649477 | 0.709264768 | -4.509627109 |
| **A_55_P1999829** | Thoc7 | 0.443170753 | -0.348035959 | 1.234377465 | 12.93827418 | 1.402915372 | 0.21470962 | 0.709264768 | -4.509685618 |
| **A_55_P2130666** | Armc1 | 0.318352104 | -0.25012445 | 0.886828657 | 10.47563834 | 1.402637716 | 0.214787965 | 0.709264768 | -4.50976181 |
| **A_51_P430357** | Lrp10 | -0.422427412 | -1.176807117 | 0.331952294 | 10.76283167 | -1.40253091 | 0.214818109 | 0.709264768 | -4.509791118 |
| **A_51_P487501** | NA | 0.24313633 | -0.205293901 | 0.691566561 | 11.93236106 | 1.444167874 | 0.214839633 | 0.709264768 | -4.515128994 |
| **A_52_P88983** | Dock5 | -0.772813183 | -2.153245388 | 0.607619023 | 10.01814831 | -1.402199424 | 0.214911689 | 0.709264768 | -4.50988208 |
| **A_52_P399584** | Ckap2l | 0.461431253 | -0.362865451 | 1.285727956 | 8.661121766 | 1.40208321 | 0.214944506 | 0.709264768 | -4.509913969 |
| **A_55_P2015994** | Fgf9 | 0.547325501 | -0.430463251 | 1.525114253 | 8.872846661 | 1.402009614 | 0.21496529 | 0.709264768 | -4.509934164 |
| **A_55_P2106827** | NA | 0.765297001 | -0.601997814 | 2.132591817 | 11.48276216 | 1.401903726 | 0.214995198 | 0.709264768 | -4.50996322 |
| **A_51_P395555** | Vkorc1 | -0.208315955 | -0.580508172 | 0.163876262 | 13.39522841 | -1.401863062 | 0.215006684 | 0.709264768 | -4.509974378 |
| **A_55_P2162234** | Tm9sf3 | -0.382958353 | -1.067227214 | 0.301310507 | 8.773474656 | -1.40176469 | 0.215034474 | 0.709264768 | -4.510001371 |
| **A_51_P509669** | Uap1l1 | -0.621320885 | -1.731646189 | 0.48900442 | 9.134675553 | -1.401574089 | 0.215088328 | 0.709264768 | -4.510053671 |
| **A_55_P1981306** | Hdgfl3 | 0.519602187 | -0.409132541 | 1.448336915 | 9.523923939 | 1.401295188 | 0.215167152 | 0.709264768 | -4.5101302 |

| | | | | | | | | | |
|---|---|---|---|---|---|---|---|---|---|
| **A_55_P2172022** | B4galt1 | -0.66033612 | -1.841172186 | 0.520499947 | 8.878863649 | -1.400637629 | 0.215353102 | 0.709264768 | -4.510310626 |
| **A_55_P1964787** | Slc35e4 | -0.798661794 | -2.226955596 | 0.629632008 | 9.490999009 | -1.400540601 | 0.215380553 | 0.709264768 | -4.510337249 |
| **A_55_P2033780** | Fabp3 | -2.035054649 | -5.674507638 | 1.60439834 | 12.61179016 | -1.400523201 | 0.215385476 | 0.709264768 | -4.510342023 |
| **A_55_P2058165** | Morn4 | 0.553051903 | -0.43607894 | 1.542182745 | 9.504777773 | 1.400433504 | 0.215410856 | 0.709264768 | -4.510366635 |
| **A_51_P176783** | Atxn7l3b | 0.366163924 | -0.288735319 | 1.021063166 | 12.02991901 | 1.40039777 | 0.215420968 | 0.709264768 | -4.510376439 |
| **A_55_P2347529** | 3010003L | 0.373104349 | -0.29423041 | 1.040439108 | 9.212085871 | 1.400351014 | 0.2154342 | 0.709264768 | -4.510389268 |
| **A_66_P127160** | Eif2b3 | 0.139880075 | -0.110325584 | 0.390085733 | 8.858644625 | 1.400261262 | 0.215459601 | 0.709264768 | -4.510413894 |
| **A_51_P310649** | Arsa | -0.407866333 | -1.137495489 | 0.321762822 | 11.30560597 | -1.400122466 | 0.215498888 | 0.709264768 | -4.510451977 |
| **A_55_P1973783** | Tex9 | 0.279154931 | -0.220293147 | 0.778603009 | 8.675500667 | 1.399926723 | 0.215554305 | 0.709264768 | -4.510505684 |
| **A_55_P2001188** | Phf6 | 0.881429301 | -0.695626416 | 2.458485018 | 10.73686673 | 1.399879946 | 0.21556755 | 0.709264768 | -4.510518519 |
| **A_52_P40954** | Dubr | -0.469021659 | -1.308209587 | 0.370166268 | 8.969955571 | -1.399858108 | 0.215573734 | 0.709264768 | -4.51052451 |
| **A_55_P2152468** | NA | 0.196946854 | -0.155447368 | 0.549341075 | 9.618760791 | 1.399814906 | 0.215585968 | 0.709264768 | -4.510536364 |
| **A_51_P278018** | Vps36 | 0.511323032 | -0.403592169 | 1.426238233 | 12.52462164 | 1.399796354 | 0.215591222 | 0.709264768 | -4.510541454 |
| **A_55_P1997061** | NA | 0.358957494 | -0.283357772 | 1.00127276 | 13.35704508 | 1.399732758 | 0.215609232 | 0.709264768 | -4.510558903 |
| **A_51_P155152** | Ank | -0.565584747 | -1.577774772 | 0.446605278 | 9.789335639 | -1.399542146 | 0.215663222 | 0.709264768 | -4.510611202 |
| **A_52_P303891** | Nr1d2 | -0.127583427 | -0.35591259 | 0.100745737 | 8.241672163 | -1.399533256 | 0.21566574 | 0.709264768 | -4.510613641 |
| **A_55_P2211164** | 5330406M | -0.421685809 | -1.176357459 | 0.332985841 | 8.098911342 | -1.399527046 | 0.2156675 | 0.709264768 | -4.510615345 |
| **A_55_P1990894** | Psmd4 | -0.446862064 | -1.246643892 | 0.352919764 | 14.15055694 | -1.399433612 | 0.21569397 | 0.709264768 | -4.51064098 |
| **A_55_P1983769** | Birc5 | 0.797733884 | -0.630055726 | 2.225523494 | 9.628139337 | 1.399407404 | 0.215701395 | 0.709264768 | -4.510648171 |
| **A_55_P1968998** | Tmub2 | -0.430816641 | -1.20190247 | 0.340269189 | 10.36482435 | -1.399394276 | 0.215705114 | 0.709264768 | -4.510651773 |
| **A_51_P202596** | Sh3tc2 | -0.649471952 | -1.811965225 | 0.51302132 | 9.021229832 | -1.399330493 | 0.215723187 | 0.709264768 | -4.510669273 |
| **A_66_P134294** | Uba1y | 0.12512362 | -0.098840022 | 0.349087262 | 8.379898528 | 1.399304158 | 0.215730649 | 0.709264768 | -4.510676498 |
| **A_55_P2423228** | AI464196 | -0.162786804 | -0.454185118 | 0.128611509 | 8.061344564 | -1.39920874 | 0.215757688 | 0.709264768 | -4.510702678 |
| **A_55_P1961760** | NA | -0.466585444 | -1.301822554 | 0.368651666 | 11.76344894 | -1.399174075 | 0.215767512 | 0.709264768 | -4.510712188 |
| **A_55_P1955239** | Rps6 | 0.509392997 | -0.402488439 | 1.421274432 | 15.25656688 | 1.39915214 | 0.215773729 | 0.709264768 | -4.510718207 |
| **A_51_P372172** | Plod3 | -0.661913499 | -1.847035875 | 0.523208876 | 10.9668726 | -1.398905528 | 0.215843633 | 0.709264768 | -4.510785868 |
| **A_51_P253537** | Atf6b | 0.168548423 | -0.133237974 | 0.470334819 | 8.699358938 | 1.398863628 | 0.215855511 | 0.709264768 | -4.510797364 |
| **A_55_P1996946** | Cdc20 | 1.397876883 | -1.10534248 | 3.901096246 | 12.34908764 | 1.398686422 | 0.215905757 | 0.709264768 | -4.510845982 |
| **A_55_P2084631** | Hist1h2an | 1.11880812 | -0.884949487 | 3.122565727 | 16.57442417 | 1.398494528 | 0.21596018 | 0.709264768 | -4.51089863 |
| **A_51_P111757** | Asb1 | -0.214788015 | -0.599568615 | 0.169992584 | 9.35607278 | -1.398128935 | 0.2160639 | 0.709264768 | -4.514980538 |
| **A_52_P394385** | 1600012H | -0.316580242 | -0.883750113 | 0.25058963 | 9.750888714 | -1.398044513 | 0.216087858 | 0.709264768 | -4.511022093 |
| **A_55_P2105694** | Prr29 | 0.15139887 | -0.119848091 | 0.422645832 | 8.712994075 | 1.398002986 | 0.216099643 | 0.709264768 | -4.511033486 |
| **A_52_P26216** | Numb | -0.376744714 | -1.051781884 | 0.298292455 | 8.17862947 | -1.397879791 | 0.21613461 | 0.709264768 | -4.511067285 |
| **A_55_P1990261** | Chchd6 | 0.529081322 | -0.419019293 | 1.477181937 | 11.64361522 | 1.397714102 | 0.216181646 | 0.709264768 | -4.511112741 |
| **A_66_P102510** | Fam114a2 | -0.135741577 | -0.378996156 | 0.107513002 | 8.744163097 | -1.397662171 | 0.21619639 | 0.709264768 | -4.511126988 |
| **A_55_P2095529** | NA | 0.549613528 | -0.435461437 | 1.534688492 | 15.14256444 | 1.397457077 | 0.21625463 | 0.709264768 | -4.511183254 |
| **A_52_P641892** | Hacd3 | 0.359718456 | -0.28502288 | 1.004459793 | 13.8245861 | 1.39742192 | 0.216264614 | 0.709264768 | -4.511192899 |
| **A_51_P254262** | Nxph3 | -0.315319095 | -0.880484671 | 0.249846482 | 8.294038032 | -1.397413437 | 0.216267024 | 0.709264768 | -4.511195227 |
| **A_51_P412966** | Lrrn1 | 0.59191992 | -0.469197709 | 1.653037549 | 8.892465742 | 1.397171671 | 0.216335699 | 0.709264768 | -4.511261553 |
| **A_51_P134062** | Stxbp2 | -0.388036673 | -1.083943524 | 0.307870178 | 10.48376132 | -1.396599897 | 0.216498196 | 0.709264768 | -4.511418411 |
| **A_55_P1996927** | 4933409K | -0.358037154 | -1.000234729 | 0.284160421 | 9.457491389 | -1.396399809 | 0.216555087 | 0.709264768 | -4.511473301 |
| **A_55_P2066399** | Fcho2 | -0.234155047 | -0.65415388 | 0.185843785 | 8.072730192 | -1.396386965 | 0.216558739 | 0.709264768 | -4.511476825 |
| **A_55_P2155906** | Chmp3 | -0.249382072 | -0.696701615 | 0.197937471 | 11.16895449 | -1.396360971 | 0.216566131 | 0.709264768 | -4.511483956 |
| **A_52_P536494** | Mycn | 0.666237482 | -0.528883952 | 1.861358916 | 9.449652806 | 1.39626346 | 0.216593863 | 0.709264768 | -4.511510706 |
| **A_52_P78373** | Sorbs3 | -0.203996155 | -0.569960636 | 0.161968327 | 8.374273058 | -1.396154206 | 0.216624939 | 0.709264768 | -4.511540677 |
| **A_66_P114585** | NA | -0.452302941 | -1.263761686 | 0.359155804 | 8.273970143 | -1.396089667 | 0.216643298 | 0.709264768 | -4.511558382 |
| **A_55_P1993629** | NA | 0.278905267 | -0.221472873 | 0.779283407 | 15.27739785 | 1.396074946 | 0.216647485 | 0.709264768 | -4.51156242 |
| **A_51_P265576** | Rbm18 | -0.199623218 | -0.557780954 | 0.158534518 | 8.261344607 | -1.396005282 | 0.216667304 | 0.709264768 | -4.511581531 |

| | | | | | | | | | |
|---|---|---|---|---|---|---|---|---|---|
| **A_51_P451151** | Ube2c | 1.334701644 | -1.060546562 | 3.729949851 | 11.88504525 | 1.395674094 | 0.216761547 | 0.709264768 | -4.511672382 |
| **A_55_P1964093** | Rangrf | 0.495871245 | -0.394069009 | 1.385811499 | 9.553430374 | 1.395591833 | 0.216784961 | 0.709264768 | -4.511694948 |
| **A_51_P171616** | Wnt10a | -0.350453547 | -0.979478973 | 0.278571878 | 7.973647714 | -1.395444622 | 0.216826868 | 0.709264768 | -4.511735331 |
| **A_55_P2051337** | Ppfia1 | -0.399161564 | -1.115655597 | 0.31733247 | 10.10472964 | -1.395360684 | 0.216850766 | 0.709264768 | -4.511758356 |
| **A_55_P2098578** | Tubd1 | 0.13939438 | -0.110823247 | 0.389612006 | 8.882640187 | 1.395332494 | 0.216858792 | 0.709264768 | -4.511766089 |
| **A_66_P108692** | NA | 0.539006094 | -0.428544988 | 1.506557176 | 15.31957162 | 1.395308108 | 0.216865736 | 0.709264768 | -4.511772779 |
| **A_51_P513244** | Zfp949 | -0.134757609 | -0.376660989 | 0.107145771 | 8.32678743 | -1.395281079 | 0.216873432 | 0.709264768 | -4.511780193 |
| **A_55_P2112599** | Lgals8 | -0.215618963 | -0.602677054 | 0.171439128 | 9.705936887 | -1.395279288 | 0.216873942 | 0.709264768 | -4.511780684 |
| **A_55_P2141084** | Tenm4 | 0.571835134 | -0.454839554 | 1.598509821 | 11.95200179 | 1.39504535 | 0.216940567 | 0.709264768 | -4.511844856 |
| **A_55_P2162815** | NA | -0.300592325 | -0.840395144 | 0.239210493 | 11.26448002 | -1.394739452 | 0.217027713 | 0.709264768 | -4.511928767 |
| **A_55_P2152592** | Agpat3 | -0.267494304 | -0.747889851 | 0.212901242 | 11.65871212 | -1.394652186 | 0.21705258 | 0.709264768 | -4.511952705 |
| **A_55_P1961140** | Mettl3 | 0.173249939 | -0.137916583 | 0.484416461 | 10.84559104 | 1.394538698 | 0.217084923 | 0.709264768 | -4.511983835 |
| **A_55_P2055587** | Enpp4 | 0.451035613 | -0.359118455 | 1.26118968 | 9.631691471 | 1.39441987 | 0.217118793 | 0.709264768 | -4.51201643 |
| **A_51_P509971** | Plekho1 | -0.1770732 | -0.495150894 | 0.141004494 | 11.92245067 | -1.394344113 | 0.217140389 | 0.709264768 | -4.51203721 |
| **A_55_P2051334** | H2-K2 | -1.449670026 | -4.054225884 | 1.154885831 | 11.41406693 | -1.394074005 | 0.217217403 | 0.709264768 | -4.5121113 |
| **A_55_P2047325** | Mbtps1 | 0.192766518 | -0.15356924 | 0.539102277 | 13.47011266 | 1.394070106 | 0.217218515 | 0.709264768 | -4.51211237 |
| **A_55_P1993840** | Tmem132 | 0.807468764 | -0.643319733 | 2.25825726 | 9.576689015 | 1.394029528 | 0.217230087 | 0.709264768 | -4.5121235 |
| **A_51_P162437** | Bcr | -0.598887039 | -1.675051049 | 0.477276971 | 12.08635446 | -1.393852428 | 0.217280599 | 0.709264768 | -4.512172078 |
| **A_55_P1955528** | NA | -0.506897617 | -1.41791209 | 0.404116855 | 9.659773496 | -1.393623046 | 0.21734604 | 0.709264768 | -4.512234996 |
| **A_55_P1972653** | Tada3 | -0.187440014 | -0.524325073 | 0.149445045 | 10.93812017 | -1.393576787 | 0.217359239 | 0.709264768 | -4.512247684 |
| **A_55_P2016838** | NA | 0.553505778 | -0.441344554 | 1.54835611 | 15.10487426 | 1.39352497 | 0.217374025 | 0.709264768 | -4.512261897 |
| **A_51_P410681** | Irx5 | 0.724415315 | -0.57768173 | 2.026512361 | 9.262703615 | 1.39346035 | 0.217392466 | 0.709264768 | -4.512279622 |
| **A_55_P2141889** | NA | 0.920439449 | -0.734378304 | 2.575257202 | 10.9163598 | 1.393142041 | 0.217483324 | 0.709264768 | -4.51236693 |
| **A_55_P2019257** | Trpc4ap | -0.467805172 | -1.30891571 | 0.373305366 | 13.88755466 | -1.39303584 | 0.217513646 | 0.709264768 | -4.512396059 |
| **A_55_P2079619** | Rnf43 | 0.85554578 | -0.68282419 | 2.393915751 | 10.2076534 | 1.392941269 | 0.217540651 | 0.709264768 | -4.512421998 |
| **A_51_P251892** | Magoh | 0.450625753 | -0.359713656 | 1.260965162 | 13.18612557 | 1.392834108 | 0.217571254 | 0.709264768 | -4.51245139 |
| **A_51_P293982** | Plekho2 | -0.337017693 | -0.943142364 | 0.269106978 | 8.685654848 | -1.392647121 | 0.217624664 | 0.709264768 | -4.512502677 |
| **A_55_P1960472** | Polg | -0.539210008 | -1.509042097 | 0.430622081 | 9.067524981 | -1.392553022 | 0.217651547 | 0.709264768 | -4.512528486 |
| **A_55_P2099930** | Cers5 | -0.267042128 | -0.747351789 | 0.213267533 | 12.13238633 | -1.392543605 | 0.217654237 | 0.709264768 | -4.512531069 |
| **A_52_P366105** | Pip4p1 | -0.227059458 | -0.635460462 | 0.181341546 | 8.738776909 | -1.392525452 | 0.217659424 | 0.709264768 | -4.512536048 |
| **A_55_P1960765** | NA | 0.876079804 | -0.699837048 | 2.451996656 | 13.66670502 | 1.39238942 | 0.217698293 | 0.709264768 | -4.512573358 |
| **A_55_P2071384** | Crh | 1.643499767 | -1.312995214 | 4.599994749 | 11.18629206 | 1.392332134 | 0.217714663 | 0.709264768 | -4.51258907 |
| **A_55_P2146171** | Cox20 | 0.442523282 | -0.353612391 | 1.238658954 | 11.93788695 | 1.392192808 | 0.217754483 | 0.709264768 | -4.512627283 |
| **A_55_P2159259** | Lifr | -0.862229975 | -2.413607227 | 0.689147276 | 11.52638576 | -1.392053876 | 0.217794196 | 0.709264768 | -4.512665388 |
| **A_55_P2170474** | Yipf2 | -0.65635338 | -1.837371854 | 0.524665094 | 10.23657432 | -1.391974825 | 0.217816796 | 0.709264768 | -4.512687069 |
| **A_66_P108178** | Lin52 | 0.148093438 | -0.118385124 | 0.414572001 | 9.23806589 | 1.391950801 | 0.217823664 | 0.709264768 | -4.512693658 |
| **A_51_P386899** | Flvcr2 | -1.372706908 | -3.843119855 | 1.097706039 | 10.39492879 | -1.391741604 | 0.217883483 | 0.709264768 | -4.512751034 |
| **A_55_P1987396** | Tbl1xr1 | 0.286464436 | -0.246088689 | 0.81901756 | 9.25019796 | 1.432750209 | 0.217884781 | 0.709264768 | -4.51756735 |
| **A_55_P2013078** | Thoc5 | 0.219853492 | -0.175812116 | 0.615519099 | 9.969348527 | 1.391731354 | 0.217886415 | 0.709264768 | -4.512753845 |
| **A_55_P2024555** | Plpp1 | -0.755455668 | -2.115110906 | 0.604199571 | 11.80822574 | -1.391651619 | 0.217909219 | 0.709264768 | -4.512775714 |
| **A_51_P480119** | Prelid2 | 0.328022099 | -0.262397798 | 0.918441997 | 9.65259026 | 1.391529649 | 0.217944107 | 0.709264768 | -4.512809165 |
| **A_55_P2129309** | Foxo3 | 0.183701329 | -0.146979225 | 0.514381884 | 8.472158895 | 1.391406371 | 0.217979374 | 0.709264768 | -4.512842976 |
| **A_55_P2092156** | NA | -0.167175625 | -0.468160552 | 0.133809303 | 14.72772026 | -1.39116471 | 0.218048523 | 0.709264768 | -4.512909253 |
| **A_55_P2056140** | Flad1 | -0.391716885 | -1.096996597 | 0.313562826 | 9.422751504 | -1.39110928 | 0.218064386 | 0.709264768 | -4.512924455 |
| **A_55_P1973643** | Raf1 | -0.12426411 | -0.348005457 | 0.099477237 | 12.3154537 | -1.391072647 | 0.218074871 | 0.709264768 | -4.512934502 |
| **A_55_P2112170** | Crtc1 | 0.227733406 | -0.182384127 | 0.637850938 | 9.861814005 | 1.39081303 | 0.218149189 | 0.709264768 | -4.513005703 |
| **A_51_P468240** | Dennd4a | -0.248112027 | -0.69495548 | 0.198731427 | 8.389852693 | -1.390729805 | 0.218173018 | 0.709264768 | -4.513028527 |
| **A_51_P221337** | Ranbp10 | -0.238446818 | -0.667909259 | 0.191015623 | 10.2933346 | -1.390646313 | 0.218196926 | 0.709264768 | -4.513051425 |

| | | | | | | | | | |
|---|---|---|---|---|---|---|---|---|---|
| **A_52_P683306** | Fam13b | -0.410860402 | -1.150903934 | 0.329183129 | 8.476524345 | -1.390552421 | 0.218223815 | 0.709264768 | -4.513077174 |
| **A_55_P2065716** | Glud1 | -0.264659479 | -0.741379201 | 0.212060242 | 11.53711662 | -1.390511806 | 0.218235447 | 0.709264768 | -4.513088313 |
| **A_55_P2020743** | Abcc10 | -0.355550428 | -0.9960696 | 0.284968744 | 8.997768318 | -1.390334878 | 0.218286126 | 0.709264768 | -4.513136835 |
| **A_55_P2012394** | Sfxn2 | 0.581000838 | -0.465667435 | 1.627669112 | 11.0256324 | 1.390330483 | 0.218287385 | 0.709264768 | -4.51313804 |
| **A_55_P2001494** | Kitl | -0.472574765 | -1.323927868 | 0.378778338 | 9.347035619 | -1.390308408 | 0.218293709 | 0.709264768 | -4.513144094 |
| **A_55_P2133901** | NA | 0.455632468 | -0.365405204 | 1.276670141 | 15.39443822 | 1.389958793 | 0.218393889 | 0.709264768 | -4.513239973 |
| **A_55_P1990538** | Ndfip2 | 0.149278121 | -0.119734211 | 0.418290452 | 8.117374794 | 1.389870432 | 0.218419214 | 0.709264768 | -4.513264205 |
| **A_55_P2336943** | Dph6 | 0.425785248 | -0.341536589 | 1.193107085 | 9.658453877 | 1.389835492 | 0.21842923 | 0.709264768 | -4.513273786 |
| **A_51_P180452** | Crk | -0.206524898 | -0.57872035 | 0.165670554 | 11.32690162 | -1.389798059 | 0.21843996 | 0.709264768 | -4.513284052 |
| **A_55_P2004811** | NA | 0.729461301 | -0.585174131 | 2.044096733 | 15.05884136 | 1.389783877 | 0.218444025 | 0.709264768 | -4.513287941 |
| **A_52_P285470** | Lrp2 | -1.004401389 | -2.814749057 | 0.80594628 | 8.696397299 | -1.389618743 | 0.218491368 | 0.709264768 | -4.513333227 |
| **A_51_P193446** | Prpf40a | 0.13484789 | -0.108206388 | 0.377902168 | 9.122558391 | 1.389604562 | 0.218495434 | 0.709264768 | -4.513337115 |
| **A_52_P413492** | Ldlrap1 | -0.606847724 | -1.700658416 | 0.486962969 | 9.495447475 | -1.389593929 | 0.218498483 | 0.709264768 | -4.513340031 |
| **A_55_P2029498** | Osbpl1a | -0.568426863 | -1.593013504 | 0.456159777 | 9.408044085 | -1.38955663 | 0.218509178 | 0.709264768 | -4.51335026 |
| **A_55_P2082271** | NA | -0.158040976 | -0.44291166 | 0.126829708 | 15.73452346 | -1.389543958 | 0.218512811 | 0.709264768 | -4.513353735 |
| **A_55_P2132039** | NA | -0.156263557 | -0.437935586 | 0.125408473 | 9.653135975 | -1.389518484 | 0.218520116 | 0.709264768 | -4.513360721 |
| **A_55_P2028474** | Adam22 | 0.262774647 | -0.211018279 | 0.736567574 | 8.275171762 | 1.3891375 | 0.218629391 | 0.709264768 | -4.513465198 |
| **A_52_P381953** | Irf2bp1 | 0.310992374 | -0.24983444 | 0.871819189 | 8.748175935 | 1.38890108 | 0.218697227 | 0.709264768 | -4.51353003 |
| **A_66_P111433** | NA | 0.507567074 | -0.407865645 | 1.422999792 | 11.66298725 | 1.388728525 | 0.21874675 | 0.709264768 | -4.513577349 |
| **A_55_P1965391** | Klhl18 | -0.442384631 | -1.240425314 | 0.355656053 | 9.765586395 | -1.388434332 | 0.218831208 | 0.709264768 | -4.513658021 |
| **A_55_P2102419** | Lime1 | -0.129733172 | -0.363772709 | 0.104306364 | 9.352081192 | -1.388392167 | 0.218843315 | 0.709264768 | -4.513669584 |
| **A_55_P2074281** | Nr2f1 | 1.165307611 | -0.937026729 | 3.267641952 | 10.95973415 | 1.388318607 | 0.218864439 | 0.709264768 | -4.513689755 |
| **A_51_P155345** | Chmp2a | -0.156422025 | -0.438652203 | 0.125808153 | 11.5181238 | -1.388176859 | 0.218905149 | 0.709264768 | -4.513728624 |
| **A_55_P2156625** | NA | 0.352094945 | -0.283253577 | 0.987443468 | 13.65395785 | 1.388027631 | 0.218948014 | 0.709264768 | -4.513769544 |
| **A_52_P218379** | Kif26a | -0.138048127 | -0.387173084 | 0.11107683 | 9.95303341 | -1.387917426 | 0.218979676 | 0.709264768 | -4.513799763 |
| **A_52_P318073** | NA | 0.441495072 | -0.355268854 | 1.238258998 | 8.672298511 | 1.387862824 | 0.218995364 | 0.709264768 | -4.513814736 |
| **A_55_P1965378** | Uqcrq | -0.351271943 | -0.985221801 | 0.282677915 | 15.01996429 | -1.387838399 | 0.219002382 | 0.709264768 | -4.513821433 |
| **A_52_P251623** | Fam222a | 0.569601272 | -0.458463648 | 1.597666191 | 9.06097877 | 1.387716511 | 0.219037408 | 0.709264768 | -4.513854855 |
| **A_52_P679152** | Clk1 | -0.179596954 | -0.503749896 | 0.144555988 | 10.50020798 | -1.387712004 | 0.219038704 | 0.709264768 | -4.513856091 |
| **A_55_P2183378** | Snx11 | -0.266442478 | -0.747349448 | 0.214464491 | 11.05901205 | -1.387690896 | 0.21904477 | 0.709264768 | -4.513861879 |
| **A_66_P135804** | NA | 0.205562078 | -0.165475333 | 0.576599488 | 13.80757999 | 1.387636274 | 0.219060468 | 0.709264768 | -4.513876857 |
| **A_51_P304126** | Necap1 | -0.498032211 | -1.397059389 | 0.400994968 | 10.20284107 | -1.387506277 | 0.219097834 | 0.709264768 | -4.513912502 |
| **A_55_P2095181** | NA | 0.343180118 | -0.2763784 | 0.962738636 | 9.617977002 | 1.387363077 | 0.219139002 | 0.709264768 | -4.513951768 |
| **A_55_P2007464** | Rps2 | 0.353672815 | -0.284831174 | 0.992176805 | 16.71798103 | 1.387357575 | 0.219140584 | 0.709264768 | -4.513953277 |
| **A_55_P1980446** | Sbno1 | -0.260396506 | -0.730581792 | 0.209788781 | 10.66918786 | -1.387127761 | 0.219206666 | 0.709264768 | -4.514016291 |
| **A_51_P178872** | Ttc9c | 0.124572071 | -0.100378474 | 0.349522617 | 8.936056572 | 1.387024014 | 0.219236505 | 0.709264768 | -4.514044738 |
| **A_52_P663526** | Nmrk1 | -0.355965989 | -0.998787184 | 0.286855205 | 8.111231529 | -1.386975097 | 0.219250575 | 0.709264768 | -4.514058151 |
| **A_55_P2114839** | NA | 0.569415332 | -0.459015226 | 1.597845889 | 15.06312014 | 1.386770293 | 0.219309493 | 0.709264768 | -4.514114306 |
| **A_55_P1967332** | NA | 0.307821229 | -0.248178959 | 0.863821416 | 13.30423765 | 1.38667273 | 0.219337565 | 0.709264768 | -4.514141057 |
| **A_55_P2183692** | Rgs19 | 0.438509974 | -0.353640192 | 1.23066014 | 8.964332317 | 1.386507758 | 0.21938504 | 0.709264768 | -4.514186291 |
| **A_55_P2136880** | Ppp1r15a | -1.305654983 | -3.664342067 | 1.053032101 | 11.43376794 | -1.38646352 | 0.219397772 | 0.709264768 | -4.51419842 |
| **A_52_P740562** | Rps21 | 0.158156047 | -0.127593612 | 0.443905707 | 15.08957902 | 1.386278306 | 0.219451087 | 0.709264768 | -4.514249203 |
| **A_55_P2002712** | Mllt10 | -0.350678743 | -0.984306939 | 0.282949454 | 9.804422854 | -1.386198074 | 0.219474186 | 0.709264768 | -4.514271201 |
| **A_55_P2058861** | Mvk | 0.363370721 | -0.293214683 | 1.019956126 | 12.06466035 | 1.386146244 | 0.219489109 | 0.709264768 | -4.514285412 |
| **A_52_P404363** | Cdc42bpb | -0.217661824 | -0.610996 | 0.175672353 | 8.632444802 | -1.386024558 | 0.219524149 | 0.709264768 | -4.514318776 |
| **A_52_P91618** | Lysmd3 | -0.231196737 | -0.649002298 | 0.186608823 | 8.908605814 | -1.385982768 | 0.219536184 | 0.709264768 | -4.514330234 |
| **A_66_P133270** | Mfsd1 | -0.256344513 | -0.719622218 | 0.206933192 | 8.450532027 | -1.385903435 | 0.219559032 | 0.709264768 | -4.514351985 |
| **A_55_P2178203** | Derl2 | -0.344915103 | -0.968272265 | 0.278442058 | 10.17700744 | -1.385879926 | 0.219565803 | 0.709264768 | -4.514358431 |

| | | | | | | | | | |
|---|---|---|---|---|---|---|---|---|---|
| **A_55_P1968153** | Pcgf2 | 0.267212831 | -0.215717936 | 0.750143598 | 9.250432971 | 1.385870909 | 0.2195684 | 0.709264768 | -4.514360903 |
| **A_55_P2067976** | Gmeb1 | 0.285813373 | -0.230771079 | 0.802397825 | 9.794457642 | 1.385771294 | 0.219597094 | 0.709264768 | -4.514388215 |
| **A_55_P2009069** | NA | -1.339826719 | -3.761534987 | 1.081881549 | 13.24615223 | -1.38572533 | 0.219610335 | 0.709264768 | -4.514400817 |
| **A_55_P2138130** | NA | 0.188856513 | -0.152579135 | 0.530292161 | 12.86582512 | 1.385394471 | 0.219705668 | 0.709264768 | -4.514491529 |
| **A_55_P2038077** | Rpsa | 0.371454608 | -0.300199591 | 1.043108807 | 16.28135499 | 1.385193234 | 0.21976367 | 0.709264768 | -4.514546702 |
| **A_55_P1971194** | Timm23 | 0.124328064 | -0.100486135 | 0.349142262 | 12.40360213 | 1.385146722 | 0.219777079 | 0.709264768 | -4.514559454 |
| **A_55_P2112355** | NA | 0.459788662 | -0.371653011 | 1.291230336 | 14.37799798 | 1.385086241 | 0.219794515 | 0.709264768 | -4.514576036 |
| **A_55_P2187121** | Gpr137b-| | -0.8975559 | -2.520628158 | 0.725516358 | 10.27251464 | -1.385077252 | 0.219797106 | 0.709264768 | -4.5145785 |
| **A_55_P2014807** | Rpl4 | -0.132804631 | -0.372988678 | 0.107379416 | 15.61727163 | -1.384903161 | 0.219847303 | 0.709264768 | -4.51462623 |
| **A_55_P2136556** | Pkn2 | -0.404704139 | -1.136646551 | 0.327238274 | 9.426915055 | -1.384876603 | 0.219854962 | 0.709264768 | -4.514633511 |
| **A_51_P160673** | Kcne1l | 0.902096303 | -0.729424025 | 2.53361663 | 9.614453087 | 1.384875586 | 0.219855255 | 0.709264768 | -4.514633789 |
| **A_55_P2116496** | Tuba1b | 0.221272746 | -0.178921779 | 0.621467271 | 14.18167852 | 1.384864009 | 0.219858594 | 0.709264768 | -4.514636963 |
| **A_55_P2014506** | Amd2 | 0.318020531 | -0.257190141 | 0.893231203 | 11.96045771 | 1.384772943 | 0.219884857 | 0.709264768 | -4.51466193 |
| **A_55_P2069246** | Ddx39b | 0.468457203 | -0.37888803 | 1.315802436 | 14.17709376 | 1.384713339 | 0.219902048 | 0.709264768 | -4.514678271 |
| **A_55_P2348799** | 1700025G | -0.341575381 | -0.959417517 | 0.276266755 | 8.625199989 | -1.384711769 | 0.219902501 | 0.709264768 | -4.514678701 |
| **A_52_P1004509** | BC065397 | -0.23989857 | -0.673843859 | 0.19404672 | 8.260684434 | -1.384659616 | 0.219917544 | 0.709264768 | -4.514693 |
| **A_51_P164207** | Mta2 | 0.548559696 | -0.443754184 | 1.540873576 | 11.55651486 | 1.384602703 | 0.219933961 | 0.709264768 | -4.514708603 |
| **A_55_P2136154** | Heca | -0.570525552 | -1.602683716 | 0.461632612 | 9.805056526 | -1.384456114 | 0.219976252 | 0.709264768 | -4.514748791 |
| **A_55_P2043322** | Rnf26 | 0.170370592 | -0.137857868 | 0.478599053 | 9.088476677 | 1.384433952 | 0.219982646 | 0.709264768 | -4.514754867 |
| **A_51_P421734** | Slc12a7 | -0.656148865 | -1.843236409 | 0.530938679 | 9.322219037 | -1.384426742 | 0.219984726 | 0.709264768 | -4.514756843 |
| **A_51_P234462** | Gpn1 | 0.186988022 | -0.151326704 | 0.525302748 | 9.998989055 | 1.384341478 | 0.220009329 | 0.709264768 | -4.514780219 |
| **A_55_P1983773** | Birc5 | 1.624232176 | -1.314473868 | 4.562938219 | 13.50534835 | 1.384338537 | 0.220010178 | 0.709264768 | -4.514781025 |
| **A_55_P2036250** | Cr1l | -0.615877889 | -1.730413262 | 0.498657485 | 12.17938281 | -1.384047838 | 0.220094078 | 0.709264768 | -4.51486072 |
| **A_55_P1998506** | Aplf | 0.198200851 | -0.160482893 | 0.556884596 | 8.414273485 | 1.384025738 | 0.220100458 | 0.709264768 | -4.514866779 |
| **A_55_P2091055** | NA | 0.345781994 | -0.280132524 | 0.971696511 | 14.73555141 | 1.383686473 | 0.220198415 | 0.709264768 | -4.514959787 |
| **A_55_P2037957** | NA | 0.652178161 | -0.528689081 | 1.833045403 | 14.4228244 | 1.383297277 | 0.220310838 | 0.709264768 | -4.51506648 |
| **A_51_P187018** | Magohb | 0.588883549 | -0.477455566 | 1.655222663 | 10.89590625 | 1.38319824 | 0.220339455 | 0.709264768 | -4.51509363 |
| **A_55_P2046245** | Hes1 | 0.49347261 | -0.400108313 | 1.387053532 | 9.334969051 | 1.383182576 | 0.220343981 | 0.709264768 | -4.515097924 |
| **A_52_P530280** | Nsa2 | 0.573983499 | -0.465447211 | 1.61341421 | 11.30341041 | 1.383101999 | 0.220367267 | 0.709264768 | -4.515120013 |
| **A_55_P2124026** | Synj2 | 0.208575251 | -0.169139911 | 0.586290412 | 9.45739242 | 1.383084462 | 0.220372335 | 0.709264768 | -4.51512482 |
| **A_51_P341379** | Acad9 | 0.261349685 | -0.21194823 | 0.7346476 | 10.14094035 | 1.383049535 | 0.220382429 | 0.709264768 | -4.515134395 |
| **A_52_P594768** | Aprt | -0.749920571 | -2.108020184 | 0.608179042 | 13.61300053 | -1.3830376 | 0.220385879 | 0.709264768 | -4.515137667 |
| **A_52_P325161** | Zbtb10 | -0.361941398 | -1.017537 | 0.293654205 | 8.514456443 | -1.382778354 | 0.220460818 | 0.709264768 | -4.515208733 |
| **A_55_P1966508** | Hoxc6 | -0.348402316 | -0.979505886 | 0.282701253 | 8.002879889 | -1.382708852 | 0.220480912 | 0.709264768 | -4.515227786 |
| **A_55_P1956488** | Dmtn | 0.175545899 | -0.142459643 | 0.493551442 | 8.147675633 | 1.382631175 | 0.220503372 | 0.709264768 | -4.515249079 |
| **A_55_P2180445** | NA | -0.15608287 | -0.438847712 | 0.126681972 | 10.70942505 | -1.382547871 | 0.220527462 | 0.709264768 | -4.515271915 |
| **A_55_P1982568** | Blcap | 0.271102519 | -0.220084121 | 0.76228916 | 10.52486742 | 1.382411548 | 0.220566889 | 0.709264768 | -4.515309284 |
| **A_55_P2051254** | Pvt1 | 0.148141668 | -0.120263616 | 0.416546951 | 8.206592461 | 1.382408877 | 0.220567662 | 0.709264768 | -4.515310016 |
| **A_51_P164203** | Nme4 | 0.945108615 | -0.767331012 | 2.657548243 | 10.89815706 | 1.382346122 | 0.220585814 | 0.709264768 | -4.515327219 |
| **A_55_P2078670** | Cbs | 0.44270462 | -0.359520144 | 1.244929384 | 8.827933271 | 1.382191875 | 0.220630436 | 0.709264768 | -4.515369501 |
| **A_55_P2112302** | NA | -0.588009866 | -1.653614746 | 0.477595013 | 11.1019232 | -1.382097745 | 0.22065767 | 0.709264768 | -4.515395303 |
| **A_55_P2034067** | Myom3 | -1.033866794 | -2.908323278 | 0.840589689 | 8.828034068 | -1.381463994 | 0.220841116 | 0.709264768 | -4.515569022 |
| **A_51_P512119** | AF067063 | 0.184374379 | -0.149943344 | 0.518692102 | 8.815227803 | 1.381311145 | 0.220885381 | 0.709264768 | -4.515610918 |
| **A_51_P292008** | Gpx3 | -1.526599866 | -4.295713697 | 1.242513966 | 11.70370166 | -1.380812648 | 0.221029802 | 0.709264768 | -4.515747556 |
| **A_55_P1963443** | Vps18 | -0.256071468 | -0.720610765 | 0.20846783 | 8.421455959 | -1.380667421 | 0.221071893 | 0.709264768 | -4.515787362 |
| **A_55_P2117028** | Mtmr3 | -0.244552094 | -0.688203256 | 0.199099068 | 11.51606182 | -1.380638907 | 0.221080158 | 0.709264768 | -4.515795177 |
| **A_55_P1976207** | Arf5 | -0.155322712 | -0.437103856 | 0.126458432 | 12.43633026 | -1.380617528 | 0.221086355 | 0.709264768 | -4.515801037 |
| **A_51_P300936** | Rbbp8 | 0.131183537 | -0.106810233 | 0.369177306 | 10.88060331 | 1.380587998 | 0.221094915 | 0.709264768 | -4.515809131 |

| | | | | | | | | | |
|---|---|---|---|---|---|---|---|---|---|
| **A_55_P2058812** | Fancl | 0.552767104 | -0.450118087 | 1.555652294 | 9.515764723 | 1.380515594 | 0.221115904 | 0.709264768 | -4.515828976 |
| **A_51_P422629** | Tbpl1 | -0.392261469 | -1.103988047 | 0.319465108 | 10.5281686 | -1.380424989 | 0.221142172 | 0.709264768 | -4.51585381 |
| **A_51_P277006** | Chst8 | -0.817688214 | -2.301403281 | 0.666026854 | 9.448889736 | -1.380344777 | 0.22116543 | 0.709264768 | -4.515875795 |
| **A_51_P266224** | Limd1 | -0.25744504 | -0.724603366 | 0.209713287 | 12.46143292 | -1.380291393 | 0.22118091 | 0.709264768 | -4.515890427 |
| **A_55_P2038262** | Phf11b | -0.320290868 | -0.901583838 | 0.261002103 | 8.062182305 | -1.380065918 | 0.221246303 | 0.709264768 | -4.515952226 |
| **A_66_P104479** | NA | 0.383242876 | -0.312607093 | 1.079092844 | 17.00110559 | 1.379459085 | 0.221422387 | 0.709264768 | -4.516118545 |
| **A_55_P2042297** | NA | -0.567815665 | -1.598835629 | 0.463204298 | 8.271188851 | -1.379401332 | 0.221439152 | 0.709264768 | -4.516134374 |
| **A_55_P2001948** | Cab39l | 0.35626697 | -0.290679747 | 1.003213686 | 11.17330594 | 1.379295733 | 0.221469809 | 0.709264768 | -4.516163315 |
| **A_55_P2142200** | NA | 0.38503335 | -0.314157299 | 1.084223999 | 11.20154852 | 1.379282044 | 0.221473784 | 0.709264768 | -4.516167067 |
| **A_52_P101238** | B230354K | 0.195863664 | -0.159833959 | 0.551561287 | 8.957870771 | 1.379187325 | 0.221501286 | 0.709264768 | -4.516193027 |
| **A_55_P1968649** | Atp1b2 | 0.701445838 | -0.572500859 | 1.975392535 | 9.66479329 | 1.379091974 | 0.221528975 | 0.709264768 | -4.516219159 |
| **A_55_P2105035** | NA | 0.688824623 | -0.562455487 | 1.940104732 | 12.70668172 | 1.378810137 | 0.221610835 | 0.709264768 | -4.5162964 |
| **A_52_P592111** | Brf1 | -0.188538297 | -0.531032955 | 0.153956361 | 9.108106041 | -1.378783648 | 0.22161853 | 0.709264768 | -4.51630366 |
| **A_55_P2082989** | Rab7b | -0.315307592 | -0.888145166 | 0.257529983 | 8.268327969 | -1.378647622 | 0.221658051 | 0.709264768 | -4.516340939 |
| **A_51_P185292** | 4930581F | -0.247245269 | -0.696440586 | 0.201950048 | 7.841107256 | -1.378615368 | 0.221667423 | 0.709264768 | -4.516349779 |
| **A_55_P1956762** | Sgms1 | -0.270465751 | -0.761891019 | 0.220959517 | 8.850906458 | -1.378494828 | 0.221702451 | 0.709264768 | -4.516382813 |
| **A_51_P430033** | Ccdc13 | -0.448659245 | -1.263865067 | 0.366546577 | 8.584026284 | -1.37847754 | 0.221707476 | 0.709264768 | -4.516387551 |
| **A_55_P2151463** | NA | 0.572782265 | -0.467973299 | 1.61353783 | 10.66592835 | 1.378450473 | 0.221715342 | 0.709264768 | -4.516394969 |
| **A_55_P2031192** | Gtdc1 | 0.237260574 | -0.193850586 | 0.668371735 | 10.17692945 | 1.378436159 | 0.221719502 | 0.709264768 | -4.516398892 |
| **A_55_P2081388** | Mdm4 | -0.272350712 | -0.767365363 | 0.222663938 | 8.35218499 | -1.37803678 | 0.221835604 | 0.709264768 | -4.516508342 |
| **A_51_P310896** | Pfkl | 0.384573016 | -0.314459542 | 1.083605574 | 9.826648093 | 1.377944577 | 0.221862416 | 0.709264768 | -4.51653361 |
| **A_55_P2078633** | C4b | -0.245427322 | -0.691582719 | 0.200728076 | 8.09052243 | -1.377802922 | 0.221903614 | 0.709264768 | -4.51657243 |
| **A_55_P2158404** | Cmpk2 | -0.330215024 | -0.930528593 | 0.270098545 | 8.355796631 | -1.377745542 | 0.221920304 | 0.709264768 | -4.516588155 |
| **A_55_P2120961** | NA | 0.24355499 | -0.199217829 | 0.686327809 | 10.9332508 | 1.377737328 | 0.221922693 | 0.709264768 | -4.516590406 |
| **A_51_P203955** | Gbp2 | -0.495692386 | -1.396901593 | 0.405516821 | 9.014595002 | -1.377643901 | 0.221949871 | 0.709264768 | -4.516616009 |
| **A_55_P2045928** | NA | -0.308340665 | -0.869069125 | 0.252387795 | 11.87744101 | -1.377300009 | 0.222049934 | 0.709264768 | -4.516710248 |
| **A_55_P2153296** | Pycrl | 0.123579774 | -0.10116088 | 0.348320428 | 8.168734695 | 1.377260531 | 0.222061424 | 0.709264768 | -4.516721066 |
| **A_55_P2084308** | Nid1 | -0.581228161 | -1.638244328 | 0.475788005 | 14.07536961 | -1.377258192 | 0.222062105 | 0.709264768 | -4.516721707 |
| **A_55_P2032946** | Gsta1 | -0.145295547 | -0.409597944 | 0.119006849 | 7.909684064 | -1.376897303 | 0.222167165 | 0.709264768 | -4.516820602 |
| **A_55_P2083098** | Tatdn3 | -0.185357936 | -0.522568558 | 0.151852686 | 9.401577412 | -1.37676648 | 0.222205261 | 0.709264768 | -4.516856451 |
| **A_55_P2178343** | Rap1gds1 | 0.371235181 | -0.304142428 | 1.046612789 | 12.48174231 | 1.3767428 | 0.222212157 | 0.709264768 | -4.51686294 |
| **A_55_P2065360** | Siva1 | 0.219991135 | -0.18026402 | 0.62024629 | 12.75416667 | 1.376634321 | 0.222243751 | 0.709264768 | -4.516892666 |
| **A_51_P114634** | Amz1 | -0.894337812 | -2.521693838 | 0.733018213 | 8.721507756 | -1.376478278 | 0.222289206 | 0.709264768 | -4.516935425 |
| **A_55_P2119257** | Serpine1 | -2.591560744 | -7.308086694 | 2.124965207 | 10.63316292 | -1.376225213 | 0.222362942 | 0.709264768 | -4.51700477 |
| **A_51_P460734** | Prkcd | -0.155978493 | -0.439878215 | 0.127921229 | 10.37330227 | -1.376100329 | 0.222399337 | 0.709264768 | -4.51703899 |
| **A_51_P374101** | Pard6b | -0.297087369 | -0.837826099 | 0.24365136 | 7.989244566 | -1.376090692 | 0.222402146 | 0.709264768 | -4.517041631 |
| **A_51_P417854** | Rab20 | -0.203997824 | -0.575339876 | 0.167344228 | 8.092102407 | -1.375947134 | 0.222443991 | 0.709264768 | -4.517080967 |
| **A_51_P505795** | Tapbpl | -0.181711641 | -0.512487142 | 0.149063861 | 8.612585602 | -1.375940831 | 0.222445829 | 0.709264768 | -4.517082694 |
| **A_55_P2110351** | Eppk1 | -0.528475286 | -1.490517075 | 0.433566503 | 9.518592167 | -1.37588169 | 0.22246307 | 0.709264768 | -4.5170989 |
| **A_55_P2015742** | Cggbp1 | 0.547895639 | -0.449686508 | 1.545477786 | 10.30153754 | 1.375623293 | 0.222538413 | 0.709264768 | -4.517169703 |
| **A_52_P144925** | Taf1d | 0.417654763 | -0.3427948 | 1.178104327 | 9.755448539 | 1.375616478 | 0.222540401 | 0.709264768 | -4.51717157 |
| **A_51_P272993** | Ntm | 0.160310223 | -0.131594091 | 0.452214537 | 7.901726832 | 1.375533118 | 0.222564712 | 0.709264768 | -4.517194411 |
| **A_55_P2009351** | Prxl2c | -0.165469484 | -0.466770499 | 0.135831532 | 9.186221447 | -1.375522384 | 0.222567843 | 0.709264768 | -4.517197352 |
| **A_52_P148212** | Mis18bp1 | 0.355582177 | -0.291916383 | 1.003080738 | 8.383193953 | 1.37547127 | 0.222582751 | 0.709264768 | -4.517211358 |
| **A_55_P2081168** | Rpl7 | 0.161482216 | -0.132597872 | 0.455562304 | 16.38419613 | 1.375337947 | 0.222621642 | 0.709264768 | -4.517247888 |
| **A_55_P1986566** | Gm10318 | 0.206178073 | -0.16930806 | 0.581664205 | 8.554294098 | 1.375304673 | 0.222631349 | 0.709264768 | -4.517257005 |
| **A_55_P2150179** | C1s2 | -0.317510305 | -0.895763207 | 0.260742597 | 9.071732507 | -1.375277539 | 0.222639265 | 0.709264768 | -4.51726444 |
| **A_55_P1997977** | Rab2b | -0.196706181 | -0.554983397 | 0.161571035 | 10.32552965 | -1.375147112 | 0.22267732 | 0.709264768 | -4.517300177 |

| | | | | | | | | | |
|---|---|---|---|---|---|---|---|---|---|
| **A_55_P2153251** | Plagl1 | -0.287086394 | -0.809986691 | 0.235813903 | 8.303058431 | -1.375130982 | 0.222682027 | 0.709264768 | -4.517304596 |
| **A_55_P2151956** | NA | -0.691351742 | -1.950744698 | 0.568041214 | 8.324194986 | -1.374953931 | 0.222733696 | 0.709264768 | -4.517353108 |
| **A_51_P224593** | Arl8a | -0.171469671 | -0.483839016 | 0.140899674 | 10.35333386 | -1.374894114 | 0.222751155 | 0.709264768 | -4.517369497 |
| **A_51_P505322** | Mpnd | 0.65577933 | -0.538882565 | 1.850441225 | 12.07847349 | 1.374874501 | 0.222756879 | 0.709264768 | -4.517374871 |
| **A_55_P2049647** | H2-K1 | -1.385252927 | -3.908842935 | 1.138337082 | 11.21392138 | -1.374866793 | 0.222759129 | 0.709264768 | -4.517376983 |
| **A_55_P1972322** | Btg3 | -0.198444956 | -0.559963958 | 0.163074046 | 13.51962932 | -1.374862546 | 0.222760369 | 0.709264768 | -4.517378147 |
| **A_52_P278154** | Eif1ad | 0.138589615 | -0.113897374 | 0.391076603 | 10.0836198 | 1.374807937 | 0.22277631 | 0.709264768 | -4.517393109 |
| **A_51_P197175** | Pigk | -0.27483619 | -0.775632981 | 0.225960601 | 8.562547403 | -1.374556916 | 0.222849597 | 0.709264768 | -4.517461886 |
| **A_55_P1989333** | NA | 0.733099459 | -0.60293812 | 2.069137038 | 11.86779159 | 1.374341215 | 0.222912591 | 0.709264768 | -4.517520985 |
| **A_55_P1961160** | NA | 0.61271019 | -0.503934455 | 1.729354836 | 14.39827012 | 1.37432821 | 0.222916389 | 0.709264768 | -4.517524548 |
| **A_51_P503937** | Calm1 | -0.130828928 | -0.369260651 | 0.107602794 | 10.5156988 | -1.374327045 | 0.22291673 | 0.709264768 | -4.517524867 |
| **A_55_P1958220** | Crebl2 | -0.446242232 | -1.259517395 | 0.367032931 | 7.983540271 | -1.3743062 | 0.222922818 | 0.709264768 | -4.517530578 |
| **A_55_P1956497** | Serpinf2 | 0.303392751 | -0.249583609 | 0.856369112 | 8.310592722 | 1.37419695 | 0.222954731 | 0.709264768 | -4.517560511 |
| **A_55_P2066553** | Oacyl | -0.327988031 | -0.925800068 | 0.269824006 | 11.07357371 | -1.374180197 | 0.222959625 | 0.709264768 | -4.517565101 |
| **A_51_P235117** | Zc3h11a | -0.275145923 | -0.776656968 | 0.226365122 | 13.69532414 | -1.374146151 | 0.222969572 | 0.709264768 | -4.517574428 |
| **A_55_P2142527** | NA | 0.222883711 | -0.183381668 | 0.62914909 | 8.639280928 | 1.374101649 | 0.222982573 | 0.709264768 | -4.517586621 |
| **A_51_P504354** | Nrarp | 0.229351131 | -0.188719304 | 0.647421565 | 8.344506876 | 1.374047592 | 0.222998367 | 0.709264768 | -4.517601431 |
| **A_55_P2083859** | NA | 0.812003037 | -0.668173325 | 2.292179398 | 14.3243653 | 1.374024684 | 0.22300506 | 0.709264768 | -4.517607708 |
| **A_55_P2003005** | Capza1 | 0.174197057 | -0.143416508 | 0.491810623 | 13.34837706 | 1.37370069 | 0.223099747 | 0.709264768 | -4.517696474 |
| **A_55_P2032608** | Dipk2a | 0.597775782 | -0.492242461 | 1.687794026 | 10.88146526 | 1.373582962 | 0.223134162 | 0.709264768 | -4.517728728 |
| **A_55_P2109875** | Rpl23a | 0.587634374 | -0.483955662 | 1.659224411 | 15.27767879 | 1.373500656 | 0.223158225 | 0.709264768 | -4.517751277 |
| **A_52_P361081** | Arhgef16 | -1.004987987 | -2.837855768 | 0.827879793 | 9.836735646 | -1.373346355 | 0.223203343 | 0.709264768 | -4.51779355 |
| **A_51_P164939** | Tmem150 | -0.892264713 | -2.519587686 | 0.73505826 | 8.80100691 | -1.373315456 | 0.223212378 | 0.709264768 | -4.517802015 |
| **A_51_P514675** | Zcchc10 | 0.237661759 | -0.195811606 | 0.671135124 | 9.670257993 | 1.373242499 | 0.223233715 | 0.709264768 | -4.517822003 |
| **A_55_P1990548** | Tbc1d22a | -0.19973735 | -0.564049404 | 0.164574703 | 11.89742572 | -1.373207255 | 0.223244023 | 0.709264768 | -4.517831658 |
| **A_51_P517096** | Casp8ap2 | 0.259207161 | -0.213595822 | 0.732010144 | 9.63002198 | 1.373147317 | 0.223261554 | 0.709264768 | -4.517848079 |
| **A_55_P2159026** | Rpl22l1 | 0.326909463 | -0.269399299 | 0.923218225 | 15.09319575 | 1.373114163 | 0.223271252 | 0.709264768 | -4.517857162 |
| **A_55_P1958419** | NA | 0.21258785 | -0.175223836 | 0.600399536 | 11.60810421 | 1.372991644 | 0.223307093 | 0.709264768 | -4.517890727 |
| **A_51_P443293** | Decr2 | -0.70116922 | -1.980324214 | 0.577985774 | 9.724541341 | -1.372935131 | 0.223323627 | 0.709264768 | -4.517906209 |
| **A_55_P2170787** | Braf | -0.273190365 | -0.771604992 | 0.225224263 | 8.189583604 | -1.372855868 | 0.223346818 | 0.709264768 | -4.517927924 |
| **A_55_P2132014** | Tekt2 | 0.669060116 | -0.5516097 | 1.889729932 | 9.478655464 | 1.372831644 | 0.223353906 | 0.709264768 | -4.51793456 |
| **A_55_P1982613** | Hnrnpa3 | 0.495286761 | -0.408351093 | 1.398924616 | 14.53207575 | 1.372817004 | 0.22335819 | 0.709264768 | -4.517938571 |
| **A_55_P2070241** | Preb | -0.543143865 | -1.534156431 | 0.447868702 | 11.77447261 | -1.372732962 | 0.223382783 | 0.709264768 | -4.517961594 |
| **A_55_P2156741** | Stx4a | -0.271565104 | -0.767085232 | 0.223955024 | 13.11270991 | -1.372660094 | 0.223404109 | 0.709264768 | -4.517981557 |
| **A_55_P2081630** | Gpa33 | 0.179610577 | -0.148144627 | 0.50736578 | 9.676028797 | 1.372564169 | 0.223432185 | 0.709264768 | -4.518007836 |
| **A_55_P2008081** | Rhoa | -0.264882661 | -0.748247432 | 0.21848211 | 13.89046762 | -1.372552236 | 0.223435678 | 0.709264768 | -4.518011105 |
| **A_51_P154933** | Zdhhc7 | -0.303214424 | -0.856567377 | 0.25013853 | 10.27386952 | -1.372454545 | 0.223464275 | 0.709264768 | -4.518037867 |
| **A_55_P1966649** | Glmn | 0.157633263 | -0.130053038 | 0.445319564 | 9.619987401 | 1.372394688 | 0.223481798 | 0.709264768 | -4.518054265 |
| **A_52_P303077** | Rogdi | -0.245188809 | -0.692684147 | 0.202306529 | 8.655126001 | -1.372342376 | 0.223497114 | 0.709264768 | -4.518068595 |
| **A_51_P266774** | Mfn2 | -0.505353306 | -1.427892404 | 0.417185791 | 10.38862881 | -1.37202074 | 0.223591302 | 0.709264768 | -4.518156706 |
| **A_55_P1991585** | Blvrb | 0.266171478 | -0.219781956 | 0.752124912 | 11.07867645 | 1.371883418 | 0.223631527 | 0.709264768 | -4.518194323 |
| **A_51_P472888** | Enkd1 | 0.738888389 | -0.610215204 | 2.087991981 | 9.991084985 | 1.371778175 | 0.22366236 | 0.709264768 | -4.518223153 |
| **A_55_P2129172** | Rab40b | 0.241176112 | -0.199230562 | 0.681582786 | 8.507017274 | 1.37161032 | 0.223711544 | 0.709264768 | -4.518269134 |
| **A_52_P246749** | Ghitm | -0.124834247 | -0.352793865 | 0.103125372 | 11.72190193 | -1.371595867 | 0.223715779 | 0.709264768 | -4.518273094 |
| **A_52_P424767** | Rbbp4 | 0.23724623 | -0.195993501 | 0.67048596 | 8.317732353 | 1.37158077 | 0.223720203 | 0.709264768 | -4.518277229 |
| **A_51_P345316** | Cep76 | 0.188896008 | -0.156062459 | 0.533854474 | 8.44359279 | 1.371533173 | 0.223734152 | 0.709264768 | -4.518290267 |
| **A_55_P2139390** | NA | 0.533133335 | -0.440477644 | 1.506744315 | 16.76585181 | 1.371515503 | 0.223739331 | 0.709264768 | -4.518295108 |
| **A_51_P436068** | Gpr182 | 0.917650206 | -0.758376924 | 2.593677335 | 10.75644923 | 1.37134418 | 0.223789547 | 0.709264768 | -4.518342038 |

| | | | | | | | | | |
|---|---|---|---|---|---|---|---|---|---|
| **A_55_P2025078** | Csf1r | -1.773001837 | -5.011613802 | 1.465610129 | 12.81292647 | -1.371199401 | 0.22383199 | 0.709264768 | -4.518381697 |
| **A_66_P116154** | Cstf2 | 0.563909067 | -0.466147657 | 1.593965791 | 9.712717849 | 1.371192035 | 0.22383415 | 0.709264768 | -4.518383715 |
| **A_66_P134809** | Gas5 | 0.888689932 | -0.734777464 | 2.512157328 | 13.5631396 | 1.371061811 | 0.223872334 | 0.709264768 | -4.518419386 |
| **A_52_P593268** | Lsm6 | 0.41545831 | -0.343519323 | 1.174435943 | 12.94632637 | 1.371035873 | 0.22387994 | 0.709264768 | -4.518426491 |
| **A_55_P2052685** | Dguok | -0.158945125 | -0.44931759 | 0.13142734 | 10.85307308 | -1.371014749 | 0.223886135 | 0.709264768 | -4.518432277 |
| **A_55_P2135556** | Rpgrip1 | 0.485444564 | -0.40143053 | 1.372319657 | 11.6939972 | 1.370968666 | 0.223899649 | 0.709264768 | -4.5184449 |
| **A_51_P170911** | Ptpn9 | -0.152619401 | -0.431492115 | 0.126253314 | 8.931875888 | -1.370736777 | 0.223967665 | 0.709264768 | -4.518508419 |
| **A_55_P1980267** | Sidt2 | -0.573113736 | -1.620344616 | 0.474117144 | 11.19743251 | -1.370719916 | 0.223972611 | 0.709264768 | -4.518513038 |
| **A_52_P302544** | Col8a2 | 0.193550982 | -0.16013622 | 0.547238185 | 8.508337625 | 1.370649398 | 0.223993299 | 0.709264768 | -4.518532354 |
| **A_55_P2074193** | NA | -0.319210579 | -0.90253479 | 0.264113633 | 14.36775038 | -1.370621747 | 0.224001411 | 0.709264768 | -4.518539928 |
| **A_66_P106611** | Gna14 | -0.22433612 | -0.634297036 | 0.185624797 | 8.776891077 | -1.370588533 | 0.224011156 | 0.709264768 | -4.518549025 |
| **A_52_P206492** | Pop4 | 0.3172128 | -0.262527093 | 0.896952692 | 11.67093965 | 1.370464728 | 0.224047484 | 0.709264768 | -4.518582937 |
| **A_66_P106385** | Cd55b | -0.824973441 | -2.332819085 | 0.682872204 | 8.890399372 | -1.370356067 | 0.224079373 | 0.709264768 | -4.518612701 |
| **A_51_P421094** | NA | -0.169773987 | -0.480127865 | 0.140579891 | 10.44629314 | -1.370138018 | 0.224143376 | 0.709264768 | -4.518672426 |
| **A_55_P2079928** | Ccdc68 | -0.980043758 | -2.771614485 | 0.811526969 | 9.286730516 | -1.37013025 | 0.224145656 | 0.709264768 | -4.518674553 |
| **A_51_P108935** | Iars2 | -0.204359462 | -0.577961521 | 0.169242598 | 9.416978855 | -1.37004816 | 0.224169756 | 0.709264768 | -4.518697038 |
| **A_55_P1998442** | Sema3f | -1.200319768 | -3.394964618 | 0.994325081 | 12.26217835 | -1.369881669 | 0.224218642 | 0.709264768 | -4.51874264 |
| **A_55_P2129614** | Itprip | -0.630218606 | -1.782808302 | 0.52237109 | 12.16260185 | -1.369515127 | 0.224326303 | 0.709264768 | -4.518843035 |
| **A_55_P2006422** | Gm6900 | -0.353642774 | -1.000453546 | 0.293167998 | 15.74354975 | -1.36942386 | 0.224353118 | 0.709264768 | -4.518868032 |
| **A_55_P2173737** | Hspa4l | 0.286871595 | -0.237839695 | 0.811582885 | 9.613483515 | 1.369359517 | 0.224372024 | 0.709264768 | -4.518885655 |
| **A_51_P449048** | Cndp2 | -0.471920954 | -1.335112988 | 0.391271079 | 9.553639857 | -1.369342803 | 0.224376935 | 0.709264768 | -4.518890233 |
| **A_51_P498831** | Derl2 | -0.466775704 | -1.320644507 | 0.3870931 | 12.33703809 | -1.369201736 | 0.224418391 | 0.709264768 | -4.51892887 |
| **A_52_P535484** | Gvin1 | -0.554236949 | -1.568125283 | 0.459651385 | 8.357826492 | -1.369164703 | 0.224429274 | 0.709264768 | -4.518939012 |
| **A_55_P2046478** | Zfp81 | 0.21088396 | -0.174929855 | 0.596697775 | 8.409153259 | 1.369039943 | 0.224465945 | 0.709264768 | -4.518973182 |
| **A_55_P2128704** | NA | 0.588420995 | -0.488196206 | 1.665038197 | 15.20265691 | 1.368917235 | 0.224502019 | 0.709264768 | -4.51900679 |
| **A_55_P2014611** | Ccdc12 | 0.556320953 | -0.461649389 | 1.574291294 | 11.5370457 | 1.368802021 | 0.224535894 | 0.709264768 | -4.519038345 |
| **A_52_P286520** | Slc23a2 | -0.345718788 | -0.978439043 | 0.287001467 | 8.372876491 | -1.3685529 | 0.224609156 | 0.709264768 | -4.519106573 |
| **A_51_P454103** | Manba | -0.724366457 | -2.050378579 | 0.601645665 | 9.647627187 | -1.368236534 | 0.224702225 | 0.709264768 | -4.519193217 |
| **A_55_P1972913** | NA | 0.275048043 | -0.228465591 | 0.778561678 | 10.74204018 | 1.368193965 | 0.224714751 | 0.709264768 | -4.519204875 |
| **A_55_P2002819** | Rabgap1l | 0.282435544 | -0.234681174 | 0.799552262 | 9.114131034 | 1.367984315 | 0.224776449 | 0.709264768 | -4.519262291 |
| **A_51_P133052** | Glyr1 | 0.212262101 | -0.176438466 | 0.600962667 | 11.19315928 | 1.367752853 | 0.224844584 | 0.709264768 | -4.519325679 |
| **A_55_P2164615** | NA | 0.372910343 | -0.310038052 | 1.055858738 | 16.80071153 | 1.367624543 | 0.224882363 | 0.709264768 | -4.519360818 |
| **A_51_P223524** | Ric1 | -0.185009721 | -0.523848853 | 0.153829411 | 9.617736496 | -1.367575566 | 0.224896785 | 0.709264768 | -4.519374231 |
| **A_55_P1971249** | Eya3 | 0.186856729 | -0.155394107 | 0.529107565 | 11.49396648 | 1.367459806 | 0.224930876 | 0.709264768 | -4.519405932 |
| **A_55_P1986958** | Brms1l | -0.164420568 | -0.465614799 | 0.136773664 | 11.20944695 | -1.367287487 | 0.224981632 | 0.709264768 | -4.519453122 |
| **A_55_P2063381** | Tmem74b | 0.138325629 | -0.115102779 | 0.391754037 | 8.184775279 | 1.367091883 | 0.225039258 | 0.709264768 | -4.519506688 |
| **A_51_P213359** | Has2 | 0.234084083 | -0.194797315 | 0.662965482 | 8.367940197 | 1.367051953 | 0.225051024 | 0.709264768 | -4.519517623 |
| **A_55_P2105820** | Tbc1d31 | 0.543046473 | -0.451947299 | 1.538040244 | 9.815629192 | 1.366995172 | 0.225067755 | 0.709264768 | -4.519533172 |
| **A_55_P2176248** | NA | 0.173943274 | -0.14478046 | 0.492667008 | 15.92891345 | 1.366921521 | 0.22508946 | 0.709264768 | -4.519553341 |
| **A_51_P391926** | Ubxn6 | -0.154909612 | -0.438778732 | 0.128959508 | 12.0800501 | -1.366817597 | 0.225120089 | 0.709264768 | -4.519581799 |
| **A_55_P2134790** | Qsox1 | -0.688967767 | -1.951512202 | 0.573576667 | 12.06950308 | -1.366792472 | 0.225127494 | 0.709264768 | -4.51958868 |
| **A_55_P2011852** | Cdk2 | 0.280783545 | -0.233855521 | 0.79542261 | 8.764146601 | 1.366530243 | 0.225204799 | 0.709264768 | -4.519660488 |
| **A_51_P398887** | Ctns | -0.916652936 | -2.596901159 | 0.763595287 | 10.31381419 | -1.366412526 | 0.22523951 | 0.709264768 | -4.519692723 |
| **A_55_P1985360** | Man1c1 | -0.695995316 | -1.97183537 | 0.579844738 | 8.801863689 | -1.366345183 | 0.225259369 | 0.709264768 | -4.519711163 |
| **A_55_P2100824** | LOC67668 | -1.321251346 | -3.743267432 | 1.100764739 | 9.898912442 | -1.366339945 | 0.225260914 | 0.709264768 | -4.519712597 |
| **A_52_P161806** | Ovca2 | 0.203068058 | -0.16920227 | 0.575338385 | 9.606439715 | 1.366260584 | 0.225284319 | 0.709264768 | -4.519734329 |
| **A_55_P2006148** | Aldh3b2 | -0.683004695 | -1.935240489 | 0.569231098 | 8.231514344 | -1.366117081 | 0.225326648 | 0.709264768 | -4.519773624 |
| **A_66_P104312** | Pdcl | 0.341383132 | -0.28457088 | 0.967337144 | 9.918180218 | 1.365997726 | 0.225361859 | 0.709264768 | -4.519806307 |

| | | | | | | | | | |
|---|---|---|---|---|---|---|---|---|---|
| **A_51_P287069** | Serpinh1 | -0.232616245 | -0.659152647 | 0.193920156 | 14.68227226 | -1.365948376 | 0.22537642 | 0.709264768 | -4.51981982 |
| **A_52_P129428** | Fzd2 | 0.565796514 | -0.471737861 | 1.60333089 | 9.004715697 | 1.365866076 | 0.225400704 | 0.709264768 | -4.519842356 |
| **A_55_P2121222** | Clcc1 | 0.391594816 | -0.326506907 | 1.10969654 | 11.40222 | 1.365844665 | 0.225407021 | 0.709264768 | -4.519848218 |
| **A_55_P2169849** | Thoc6 | 0.385628459 | -0.321586619 | 1.092843536 | 9.674814503 | 1.365739627 | 0.225438019 | 0.709264768 | -4.51987698 |
| **A_55_P1992455** | Vdac3 | 0.263041846 | -0.219385676 | 0.745469369 | 14.14732974 | 1.365661648 | 0.225461034 | 0.709264768 | -4.519898332 |
| **A_55_P1972748** | Czib | 0.389205644 | -0.324624889 | 1.103036177 | 13.41432741 | 1.365634106 | 0.225469163 | 0.709264768 | -4.519905874 |
| **A_55_P2028193** | Mgat1 | -0.697537251 | -1.977000441 | 0.581925939 | 13.54539967 | -1.3654945 | 0.225510373 | 0.709264768 | -4.5199441 |
| **A_55_P2011380** | Acvrl1 | -0.85671194 | -2.428378264 | 0.714954384 | 11.7892125 | -1.365289712 | 0.225570837 | 0.709264768 | -4.520000173 |
| **A_51_P479865** | Glb1l | -0.120972062 | -0.342909998 | 0.100965873 | 8.35200955 | -1.365223921 | 0.225590264 | 0.709264768 | -4.520018187 |
| **A_51_P502054** | Gtf2b | -0.442376842 | -1.254017215 | 0.36926353 | 11.50051894 | -1.36514596 | 0.225613288 | 0.709264768 | -4.520039534 |
| **A_51_P278540** | Myg1 | 0.282965557 | -0.236219346 | 0.80215046 | 9.587439215 | 1.365091825 | 0.225629277 | 0.709264768 | -4.520054356 |
| **A_51_P296796** | Nt5dc2 | 0.459302317 | -0.383473346 | 1.30207798 | 9.054078673 | 1.365013656 | 0.225652365 | 0.709264768 | -4.52007576 |
| **A_55_P1980125** | Srsf3 | 0.90175056 | -0.753097616 | 2.556598736 | 12.10308858 | 1.364830159 | 0.225706574 | 0.709264768 | -4.520126002 |
| **A_55_P2118491** | Gm10046 | 0.301524236 | -0.251930681 | 0.854979154 | 8.628100062 | 1.364552725 | 0.225788555 | 0.709264768 | -4.520201963 |
| **A_55_P2094034** | Nrg2 | 0.298091016 | -0.249069277 | 0.845251308 | 8.358545864 | 1.364534936 | 0.225793812 | 0.709264768 | -4.520206833 |
| **A_55_P1957927** | Nono | 0.470806299 | -0.393391915 | 1.335004513 | 14.35168654 | 1.364517927 | 0.22579884 | 0.709264768 | -4.52021149 |
| **A_52_P500244** | Atrn | -0.172831091 | -0.490085921 | 0.14442374 | 10.30940182 | -1.364469941 | 0.225813023 | 0.709264768 | -4.520224628 |
| **A_55_P1998188** | Cdk4 | 0.643857864 | -0.538121575 | 1.825837303 | 13.2679256 | 1.364364556 | 0.225844174 | 0.709264768 | -4.520253482 |
| **A_55_P2071893** | NA | 0.716268881 | -0.598795679 | 2.031333441 | 12.22936824 | 1.364204115 | 0.225891608 | 0.709264768 | -4.520297409 |
| **A_52_P213889** | Tmc7 | -0.115406722 | -0.327293204 | 0.09647976 | 9.034821689 | -1.364200485 | 0.225892681 | 0.709264768 | -4.520298403 |
| **A_55_P2070838** | NA | 0.644264916 | -0.538604702 | 1.827134534 | 14.9825467 | 1.364199704 | 0.225892912 | 0.709264768 | -4.520298617 |
| **A_52_P409833** | Plat | -0.285785567 | -0.810558303 | 0.238987169 | 8.40772502 | -1.364015716 | 0.225947318 | 0.709264768 | -4.520348991 |
| **A_55_P1971448** | Snrpd1 | 0.564538034 | -0.472117457 | 1.601193524 | 14.22886534 | 1.363983446 | 0.225956862 | 0.709264768 | -4.520357825 |
| **A_52_P4698** | Rfwd3 | 0.123143291 | -0.103005636 | 0.349292218 | 11.7654674 | 1.363849884 | 0.225996366 | 0.709264768 | -4.520394392 |
| **A_55_P2042071** | Slc25a3 | 0.238479183 | -0.199510028 | 0.676468393 | 14.07614714 | 1.363758303 | 0.226023457 | 0.709264768 | -4.520419466 |
| **A_55_P2065671** | Ccnb1 | 1.353046903 | -1.132195352 | 3.838289159 | 11.35646377 | 1.363623472 | 0.226063347 | 0.709264768 | -4.520456379 |
| **A_55_P2082604** | Ndor1 | -0.212156544 | -0.601841479 | 0.177528391 | 16.6931839 | -1.363619367 | 0.226064562 | 0.709264768 | -4.520457503 |
| **A_55_P2059154** | Dtx3l | -0.379809926 | -1.077503902 | 0.317884049 | 9.455066883 | -1.363489141 | 0.226103096 | 0.709264768 | -4.520493156 |
| **A_51_P165060** | Slc22a5 | -0.734050714 | -2.082970234 | 0.614868806 | 8.588240022 | -1.362982788 | 0.226252985 | 0.709264768 | -4.520631779 |
| **A_55_P2046213** | NA | 0.239956001 | -0.201028836 | 0.680940837 | 16.48133241 | 1.362882152 | 0.226282786 | 0.709264768 | -4.52065933 |
| **A_55_P1961335** | Ctsk | -2.255780748 | -6.401823652 | 1.890262156 | 10.40048737 | -1.362741367 | 0.226324482 | 0.709264768 | -4.520697871 |
| **A_51_P371119** | Cops4 | 0.279411148 | -0.234202959 | 0.793025254 | 12.63637632 | 1.362564694 | 0.226376817 | 0.709264768 | -4.520746237 |
| **A_55_P2255628** | 9430013L | -0.326886921 | -0.927817199 | 0.274043356 | 9.02026353 | -1.362460137 | 0.226407795 | 0.709264768 | -4.52077486 |
| **A_55_P2041592** | Rdh1 | 0.134085527 | -0.112422605 | 0.380593659 | 7.964649341 | 1.36238859 | 0.226428994 | 0.709264768 | -4.520794446 |
| **A_55_P1984178** | Pde4dip | -1.138201926 | -3.231123356 | 0.954719504 | 9.713961238 | -1.362124288 | 0.226507325 | 0.709264768 | -4.520866798 |
| **A_55_P1957598** | Atad2b | 0.44827682 | -0.376015223 | 1.272568864 | 10.81621271 | 1.362120473 | 0.226508456 | 0.709264768 | -4.520867842 |
| **A_52_P431894** | Pknox1 | 0.240791234 | -0.201991837 | 0.683574305 | 10.0616007 | 1.362071825 | 0.226522876 | 0.709264768 | -4.520881159 |
| **A_55_P2003096** | Pla2g6 | -0.636812045 | -1.807973987 | 0.534349898 | 9.233581738 | -1.361898255 | 0.226574333 | 0.709264768 | -4.520928673 |
| **A_51_P174081** | Zdhhc14 | -0.74387578 | -2.112031064 | 0.624279504 | 8.280788269 | -1.361806404 | 0.226601568 | 0.709264768 | -4.520953816 |
| **A_51_P489903** | Ogg1 | 0.167930112 | -0.140941986 | 0.47680221 | 9.907875947 | 1.361758969 | 0.226615634 | 0.709264768 | -4.520966801 |
| **A_51_P513661** | Slc4a8 | -0.538544111 | -1.529085865 | 0.451997644 | 8.150295842 | -1.361754564 | 0.22661694 | 0.709264768 | -4.520968007 |
| **A_55_P1971897** | Mdk | 1.109162295 | -0.931087346 | 3.149411936 | 12.28192371 | 1.361639474 | 0.226651072 | 0.709264768 | -4.520999511 |
| **A_51_P442152** | Itfg1 | -0.277674204 | -0.788447696 | 0.233099288 | 12.13063616 | -1.361625046 | 0.226655351 | 0.709264768 | -4.521003461 |
| **A_55_P2063396** | Zfand2a | -1.391459595 | -3.951064856 | 1.168145665 | 11.19172093 | -1.361595011 | 0.226664259 | 0.709264768 | -4.521011683 |
| **A_55_P1998987** | NA | 0.299100012 | -0.251103929 | 0.849303952 | 9.216312325 | 1.361579736 | 0.22666879 | 0.709264768 | -4.521015864 |
| **A_51_P167292** | Chil3 | -0.734529188 | -2.085811823 | 0.616753448 | 8.340146541 | -1.361486087 | 0.226696569 | 0.709264768 | -4.521041499 |
| **A_55_P2023081** | Dnaja2 | 0.179598143 | -0.150804246 | 0.510000532 | 9.785960854 | 1.361472923 | 0.226700474 | 0.709264768 | -4.521045102 |
| **A_55_P2026025** | NA | 0.442866024 | -0.371864504 | 1.257596552 | 16.67715588 | 1.361472016 | 0.226700743 | 0.709264768 | -4.521045351 |

| | | | | | | | | | |
|---|---|---|---|---|---|---|---|---|---|
| **A_55_P2056496** | Tk1 | 1.069211669 | -0.897836687 | 3.036260025 | 11.40252372 | 1.361441545 | 0.226709782 | 0.709264768 | -4.521053691 |
| **A_66_P114463** | NA | 0.750152899 | -0.629956424 | 2.130262221 | 12.45406875 | 1.361402826 | 0.226721268 | 0.709264768 | -4.52106429 |
| **A_55_P2108201** | Gnptab | 0.612173357 | -0.514261979 | 1.738608693 | 10.19759423 | 1.361189229 | 0.226784644 | 0.709264768 | -4.521122757 |
| **A_52_P466147** | Rarres2 | -1.520147646 | -4.317714888 | 1.277419597 | 13.0761966 | -1.360992032 | 0.226843169 | 0.709264768 | -4.521176735 |
| **A_55_P2160451** | Stk26 | 0.132470915 | -0.111358407 | 0.376300237 | 7.975237259 | 1.360770706 | 0.226908871 | 0.709264768 | -4.521237316 |
| **A_55_P2107207** | Gpr137b-| | -1.37093271 | -3.894634038 | 1.152768619 | 9.9575085 | -1.360593926 | 0.226961362 | 0.709264768 | -4.521285704 |
| **A_51_P443872** | Slu7 | 0.135325824 | -0.113840776 | 0.384492424 | 9.550116601 | 1.360320361 | 0.227042613 | 0.709264768 | -4.521360582 |
| **A_66_P105422** | Lonrf3 | -0.715763117 | -2.033686037 | 0.602159803 | 9.169200677 | -1.36028419 | 0.227053358 | 0.709264768 | -4.521370482 |
| **A_55_P2090349** | Mettl5 | 0.126480442 | -0.106407292 | 0.359368176 | 8.246542176 | 1.360276213 | 0.227055728 | 0.709264768 | -4.521372666 |
| **A_55_P2057523** | Ppm1b | -0.170432653 | -0.484275909 | 0.143410604 | 9.435625849 | -1.360161091 | 0.22708993 | 0.709264768 | -4.521404175 |
| **A_55_P2052535** | Gm14393 | -0.251470837 | -0.714549794 | 0.21160812 | 8.776299663 | -1.360137851 | 0.227096835 | 0.709264768 | -4.521410536 |
| **A_55_P1993064** | Mbd6 | -0.263113378 | -0.747654296 | 0.22142754 | 11.43265361 | -1.360074875 | 0.227115547 | 0.709264768 | -4.521427773 |
| **A_52_P336768** | NA | -0.278998775 | -0.79281579 | 0.23481824 | 9.075187664 | -1.360016447 | 0.227132909 | 0.709264768 | -4.521443765 |
| **A_55_P2017850** | Rpsa | 0.663525599 | -0.558502643 | 1.885553841 | 13.23373883 | 1.359962036 | 0.227149079 | 0.709264768 | -4.521458657 |
| **A_55_P2103880** | Gtf2ird1 | 0.29845079 | -0.25124271 | 0.848144291 | 10.07434526 | 1.359885919 | 0.227171701 | 0.709264768 | -4.52147949 |
| **A_55_P1955715** | Gm14326 | -0.257774465 | -0.732577333 | 0.217028402 | 10.2462839 | -1.359805858 | 0.227195497 | 0.709264768 | -4.521501403 |
| **A_55_P1984635** | Parp4 | -0.651033127 | -1.850351271 | 0.548285018 | 9.918532946 | -1.359624647 | 0.227249366 | 0.709264768 | -4.521550999 |
| **A_55_P2146650** | NA | 0.170904583 | -0.143934492 | 0.485743659 | 8.560867638 | 1.359613373 | 0.227252718 | 0.709264768 | -4.521554085 |
| **A_55_P2002854** | Aifm2 | -0.281981849 | -0.801503385 | 0.237539688 | 8.077791333 | -1.359464726 | 0.227296916 | 0.709264768 | -4.521594768 |
| **A_52_P343343** | Rpl7l1 | 0.12373546 | -0.104252649 | 0.351723568 | 8.6331854 | 1.35935324 | 0.22733007 | 0.709264768 | -4.521625281 |
| **A_51_P217336** | Scamp1 | -0.196285958 | -0.557957793 | 0.165385878 | 11.89779499 | -1.359329953 | 0.227336996 | 0.709264768 | -4.521631654 |
| **A_55_P1990608** | Ube2j1 | -0.342321829 | -0.973204964 | 0.288561305 | 9.103423965 | -1.35905184 | 0.227419724 | 0.709264768 | -4.52170777 |
| **A_55_P1974204** | Fam172a | 0.446614817 | -0.376521879 | 1.269751513 | 11.24967428 | 1.358975132 | 0.227442546 | 0.709264768 | -4.521728763 |
| **A_51_P126177** | Map1lc3b | -0.188635258 | -0.53635126 | 0.159080743 | 12.7624057 | -1.358778104 | 0.227501176 | 0.709264768 | -4.521782685 |
| **A_51_P369516** | Nudt16l1 | 0.3207978 | -0.270614394 | 0.912209993 | 12.40805235 | 1.358599505 | 0.227554335 | 0.709264768 | -4.521831563 |
| **A_52_P337220** | Mgat4b | -0.621107489 | -1.766180589 | 0.523965611 | 9.085528261 | -1.358575877 | 0.227561368 | 0.709264768 | -4.521838029 |
| **A_55_P2118857** | NA | 0.178802755 | -0.150909581 | 0.508515091 | 8.330861387 | 1.358280153 | 0.227649415 | 0.709264768 | -4.521918959 |
| **A_51_P462102** | Adgra3 | -0.146467343 | -0.416572005 | 0.12363732 | 11.26854538 | -1.358185427 | 0.227677625 | 0.709264768 | -4.521944882 |
| **A_51_P502456** | H2-K2 | -1.35322498 | -3.848914189 | 1.14246423 | 10.76765348 | -1.358094062 | 0.227704837 | 0.709264768 | -4.521969886 |
| **A_55_P2042229** | NA | 0.80285198 | -0.677824432 | 2.283528391 | 13.45271127 | 1.358080991 | 0.22770873 | 0.709264768 | -4.521973463 |
| **A_55_P2024704** | Cpe | -1.010694612 | -2.874720144 | 0.853330921 | 12.54614362 | -1.358058378 | 0.227715465 | 0.709264768 | -4.521979651 |
| **A_55_P2036638** | Ints11 | 0.191444712 | -0.161685817 | 0.544575241 | 10.21919529 | 1.357870818 | 0.22777134 | 0.709264768 | -4.522030978 |
| **A_51_P214825** | Orc5 | -0.125104675 | -0.355878335 | 0.105668986 | 8.26931147 | -1.357805755 | 0.227790725 | 0.709264768 | -4.522048783 |
| **A_51_P472217** | Sapcd2 | 0.697543654 | -0.589198348 | 1.984285657 | 10.43338883 | 1.357782666 | 0.227797604 | 0.709264768 | -4.522055102 |
| **A_55_P2018457** | Hdac7 | 0.989171745 | -0.835589294 | 2.813932784 | 12.34145864 | 1.357738236 | 0.227810843 | 0.709264768 | -4.52206726 |
| **A_51_P273609** | Itpka | -0.875299696 | -2.490003178 | 0.739403787 | 10.62617652 | -1.357732898 | 0.227812434 | 0.709264768 | -4.522068721 |
| **A_55_P2281095** | 2610002M | 0.237224331 | -0.20043533 | 0.674883993 | 9.201098205 | 1.357603836 | 0.227850895 | 0.709264768 | -4.522104039 |
| **A_55_P1958350** | Nipsnap3l | 0.165179129 | -0.139578803 | 0.469937061 | 10.91350042 | 1.357533323 | 0.227871911 | 0.709264768 | -4.522123335 |
| **A_55_P2073507** | 2610528J | -0.14734131 | -0.419191278 | 0.124508657 | 8.157186083 | -1.357517971 | 0.227876487 | 0.709264768 | -4.522127536 |
| **A_55_P1998107** | NA | 0.646372553 | -0.546565983 | 1.839311089 | 12.85137983 | 1.357110416 | 0.227997992 | 0.709264768 | -4.52223906 |
| **A_51_P464822** | H1f4 | 0.987484909 | -0.835051868 | 2.810021686 | 12.70072575 | 1.357077069 | 0.228007936 | 0.709264768 | -4.522248186 |
| **A_55_P1989673** | Slco2a1 | -1.596700366 | -4.543635994 | 1.350235262 | 12.44785897 | -1.357072722 | 0.228009233 | 0.709264768 | -4.522249375 |
| **A_55_P2076048** | Nuf2 | 0.766465685 | -0.648204687 | 2.181136057 | 9.377357714 | 1.35702486 | 0.228023506 | 0.709264768 | -4.522262472 |
| **A_52_P614731** | Gng12 | -0.166220419 | -0.473019998 | 0.14057916 | 9.80817837 | -1.357000346 | 0.228030817 | 0.709264768 | -4.52226918 |
| **A_55_P2109943** | Rps15a | 0.34815571 | -0.294500457 | 0.990811877 | 16.30633356 | 1.356891707 | 0.22806322 | 0.709264768 | -4.522298907 |
| **A_55_P2052775** | Morc2a | -0.167019436 | -0.475323722 | 0.141284851 | 10.81943971 | -1.356868606 | 0.228070111 | 0.709264768 | -4.522305229 |
| **A_55_P1959305** | Rbm47 | -0.451152605 | -1.284012928 | 0.381707718 | 10.51654347 | -1.356755659 | 0.228103805 | 0.709264768 | -4.522336135 |
| **A_55_P1977633** | Irf2bpl | 0.540986043 | -0.457812787 | 1.539784873 | 12.13784503 | 1.356620518 | 0.228144125 | 0.709264768 | -4.522373113 |

| | | | | | | | | | |
|---|---|---|---|---|---|---|---|---|---|
| **A_66_P106593** | Vhl | -0.853608724 | -2.429807429 | 0.722589981 | 11.87909304 | -1.356432616 | 0.228200198 | 0.709264768 | -4.522424528 |
| **A_51_P249118** | Abcb4 | -0.71325543 | -2.030323564 | 0.603812705 | 8.540737865 | -1.356398152 | 0.228210484 | 0.709264768 | -4.522433959 |
| **A_55_P2152049** | Xrcc3 | 0.133714865 | -0.113220728 | 0.380650457 | 8.474292988 | 1.356270585 | 0.22824856 | 0.709264768 | -4.522468864 |
| **A_51_P153170** | Cyb5r3 | -1.076406051 | -3.064241565 | 0.911429463 | 11.75914855 | -1.356269611 | 0.228248851 | 0.709264768 | -4.52246913 |
| **A_55_P2123841** | Zfhx4 | 0.193210941 | -0.16360958 | 0.550031462 | 8.082364766 | 1.356226539 | 0.228261708 | 0.709264768 | -4.522480916 |
| **A_52_P937** | Spcs2 | -0.490520364 | -1.396420105 | 0.415379377 | 11.98004513 | -1.356210966 | 0.228266357 | 0.709264768 | -4.522485177 |
| **A_55_P1960083** | NA | 0.575388542 | -0.487376335 | 1.63815342 | 14.98004077 | 1.356045812 | 0.228315665 | 0.709264768 | -4.522530366 |
| **A_55_P2040628** | Gid8 | 0.15339412 | -0.1299563 | 0.436744541 | 11.94031938 | 1.355923532 | 0.228352179 | 0.709264768 | -4.522563824 |
| **A_51_P333099** | Mrps15 | 0.401768087 | -0.340424073 | 1.143960248 | 10.90186 | 1.355843053 | 0.228376214 | 0.709264768 | -4.522585843 |
| **A_51_P431903** | Pdcd6 | -0.152130479 | -0.433186853 | 0.128925894 | 13.01641657 | -1.355729797 | 0.228410041 | 0.709264768 | -4.522616832 |
| **A_55_P1994524** | Tpd52l2 | -0.18384147 | -0.523511387 | 0.155828448 | 12.36950055 | -1.355616179 | 0.228443981 | 0.709264768 | -4.522647918 |
| **A_52_P93467** | Sdc4 | -0.507702633 | -1.445933211 | 0.430527945 | 9.600668522 | -1.355345999 | 0.228524708 | 0.709264768 | -4.522721841 |
| **A_55_P1999633** | Cit | 0.197260816 | -0.167290342 | 0.561811973 | 8.097464058 | 1.355291417 | 0.228541019 | 0.709264768 | -4.522736774 |
| **A_55_P2060036** | Ulk2 | -0.377398885 | -1.074893952 | 0.320096182 | 9.438113827 | -1.35522005 | 0.228562349 | 0.709264768 | -4.5227563 |
| **A_51_P398467** | Bsdc1 | -0.418567266 | -1.192170077 | 0.355035545 | 10.38752902 | -1.355181823 | 0.228573775 | 0.709264768 | -4.522766759 |
| **A_51_P167232** | Cask | 0.396022908 | -0.33600521 | 1.128051027 | 8.654091132 | 1.355011218 | 0.228624773 | 0.709264768 | -4.522813436 |
| **A_55_P1960178** | NA | 0.534941361 | -0.453958005 | 1.523840727 | 17.32111699 | 1.35489121 | 0.228660653 | 0.709264768 | -4.522846269 |
| **A_55_P2148906** | Cltb | -0.48664381 | -1.38626673 | 0.41297911 | 9.690267103 | -1.354880642 | 0.228663813 | 0.709264768 | -4.52284916 |
| **A_55_P2016004** | Cd22 | -0.205394097 | -0.585106674 | 0.17431848 | 8.220937874 | -1.354825374 | 0.228680339 | 0.709264768 | -4.522864281 |
| **A_55_P2121081** | Cul2 | 0.194060549 | -0.164707956 | 0.552829054 | 9.084121836 | 1.354794078 | 0.228689697 | 0.709264768 | -4.522872843 |
| **A_51_P194377** | Pcbp4 | 0.436954606 | -0.370953474 | 1.244862686 | 10.92177502 | 1.354642559 | 0.228735011 | 0.709264768 | -4.522914297 |
| **A_55_P2182806** | NA | -0.50342082 | -1.434243488 | 0.427401847 | 10.49864327 | -1.354610906 | 0.228744478 | 0.709264768 | -4.522922957 |
| **A_55_P2049752** | Pamr1 | -0.403488083 | -1.149575399 | 0.342599233 | 10.02500099 | -1.354538648 | 0.228766092 | 0.709264768 | -4.522942725 |
| **A_55_P2151273** | Elovl1 | -0.758855835 | -2.162269888 | 0.644558217 | 13.96538445 | -1.35432783 | 0.228829161 | 0.709264768 | -4.523000401 |
| **A_52_P445828** | Abcf2 | 0.138466825 | -0.117629351 | 0.394563001 | 9.098524173 | 1.354231734 | 0.228857915 | 0.709264768 | -4.52302669 |
| **A_55_P2169669** | Csf1 | -0.479015223 | -1.364984239 | 0.406953793 | 8.164013045 | -1.354194754 | 0.228868981 | 0.709264768 | -4.523036807 |
| **A_55_P2134735** | Fdxacb1 | -0.300880757 | -0.857405627 | 0.255644114 | 9.233014842 | -1.354129442 | 0.228888526 | 0.709264768 | -4.523054675 |
| **A_55_P1964228** | Elac1 | -0.358772498 | -1.022460288 | 0.304915293 | 8.510160334 | -1.353959462 | 0.228939402 | 0.709264768 | -4.523101177 |
| **A_55_P2043008** | NA | 0.145235432 | -0.123434752 | 0.413905617 | 8.51488779 | 1.353952579 | 0.228941462 | 0.709264768 | -4.523103059 |
| **A_51_P286792** | Zfp275 | 0.357285263 | -0.303678735 | 1.01824926 | 9.270913178 | 1.353903292 | 0.228956216 | 0.709264768 | -4.523116543 |
| **A_55_P2008066** | Itpr1 | -0.148146411 | -0.42223449 | 0.125941668 | 10.89151471 | -1.353790096 | 0.228990105 | 0.709264768 | -4.52314751 |
| **A_52_P214752** | NA | -0.285374044 | -0.813389525 | 0.242641437 | 9.269918561 | -1.353686693 | 0.229021065 | 0.709264768 | -4.523175797 |
| **A_55_P2011341** | NA | 1.941014379 | -1.77249375 | 5.654522509 | 9.105461126 | 1.39221747 | 0.229037946 | 0.709264768 | -4.526244808 |
| **A_52_P350664** | Pygb | 0.383699431 | -0.326292979 | 1.093691842 | 9.314808523 | 1.353592049 | 0.229049406 | 0.709264768 | -4.523201688 |
| **A_66_P135700** | Cyp2d11 | 0.149440964 | -0.127102677 | 0.425984606 | 9.357722781 | 1.353493985 | 0.229078775 | 0.709264768 | -4.523228515 |
| **A_51_P241074** | Map2k1 | -0.571284146 | -1.628473796 | 0.485905504 | 9.878682547 | -1.353473057 | 0.229085043 | 0.709264768 | -4.52323424 |
| **A_55_P2059606** | Stat2 | -0.560845077 | -1.598762416 | 0.477072262 | 10.40032314 | -1.353413458 | 0.229102894 | 0.709264768 | -4.523250543 |
| **A_55_P2088401** | H2-T9 | -0.135192307 | -0.385394408 | 0.115009795 | 8.585257024 | -1.353353874 | 0.229120742 | 0.709264768 | -4.523266843 |
| **A_51_P127681** | Clic4 | -0.642528608 | -1.831836762 | 0.546779546 | 13.55978196 | -1.353157715 | 0.229179509 | 0.709264768 | -4.527858149 |
| **A_55_P2086746** | Fam3a | -0.182637265 | -0.520696748 | 0.155422219 | 8.235980213 | -1.353152102 | 0.22918119 | 0.709264768 | -4.523322039 |
| **A_55_P2101177** | Ncaph2 | 0.410844945 | -0.349624716 | 1.171314605 | 12.20499343 | 1.35315143 | 0.229181392 | 0.709264768 | -4.523322222 |
| **A_55_P1993423** | NA | -0.370213944 | -1.055478441 | 0.315050554 | 14.26087997 | -1.353146708 | 0.229182807 | 0.709264768 | -4.523323514 |
| **A_51_P224575** | Gnpat | 0.351635028 | -0.299386458 | 1.002656514 | 12.26822723 | 1.352842238 | 0.229274051 | 0.709264768 | -4.523406801 |
| **A_55_P2068734** | Ccl27a | -1.716681802 | -4.895063388 | 1.461699785 | 10.66631845 | -1.352801614 | 0.229286227 | 0.709264768 | -4.523417914 |
| **A_66_P111562** | Ccnd1 | 1.026630628 | -0.874197467 | 2.927458724 | 12.12293632 | 1.352763012 | 0.229297799 | 0.709264768 | -4.523428473 |
| **A_66_P106418** | Prkacb | 0.215897786 | -0.183897816 | 0.615693387 | 9.462797976 | 1.352572403 | 0.229354943 | 0.709264768 | -4.523480612 |
| **A_51_P497768** | Gpat2 | -0.72677172 | -2.072726408 | 0.619182968 | 8.20051335 | -1.352439747 | 0.229394721 | 0.709264768 | -4.523516899 |
| **A_55_P2005570** | Celf2 | 0.911939497 | -0.777060862 | 2.600939856 | 12.40857005 | 1.35234229 | 0.229423948 | 0.709264768 | -4.523543556 |

| | | | | | | | | | |
|---|---|---|---|---|---|---|---|---|---|
| **A_51_P317317** | Gatd1 | -0.147730384 | -0.42136049 | 0.125899722 | 8.451220269 | -1.352247831 | 0.22945228 | 0.709264768 | -4.523569394 |
| **A_52_P458000** | AI467606 | 0.605347497 | -0.515993066 | 1.72668806 | 9.106473326 | 1.352127247 | 0.229488452 | 0.709264768 | -4.523602377 |
| **A_55_P2030383** | Frmd4a | 0.214660855 | -0.182981331 | 0.612303042 | 11.54523622 | 1.352106025 | 0.229494818 | 0.709264768 | -4.523608182 |
| **A_55_P1957291** | Rpl39 | 0.470014709 | -0.400714671 | 1.340744088 | 16.15958848 | 1.352005929 | 0.229524849 | 0.709264768 | -4.523635561 |
| **A_55_P2128695** | Nckap1 | -0.149883576 | -0.427552628 | 0.127785475 | 10.41571522 | -1.352000702 | 0.229526417 | 0.709264768 | -4.523636991 |
| **A_55_P2102335** | Zfp423 | 0.321885199 | -0.27442998 | 0.918200377 | 8.969754657 | 1.351996255 | 0.229527751 | 0.709264768 | -4.523638207 |
| **A_51_P447988** | Atp6ap1 | -0.379228495 | -1.081785408 | 0.323328418 | 9.531176557 | -1.351978534 | 0.229533069 | 0.709264768 | -4.523643054 |
| **A_55_P2088028** | Mtss1 | -0.386584773 | -1.102790906 | 0.32962136 | 8.199045354 | -1.351938867 | 0.229544971 | 0.709264768 | -4.523653904 |
| **A_55_P1975947** | Fam234b | 0.296879978 | -0.253177952 | 0.846937908 | 11.22353604 | 1.351832316 | 0.229576945 | 0.709264768 | -4.523683048 |
| **A_55_P2183688** | NA | 0.348827187 | -0.297495759 | 0.995150133 | 15.83286788 | 1.351795814 | 0.229587899 | 0.709264768 | -4.523693032 |
| **A_51_P242414** | Micall1 | -0.614743542 | -1.753938888 | 0.524451803 | 10.76187346 | -1.351593577 | 0.229648601 | 0.709264768 | -4.523748348 |
| **A_55_P2106459** | Zfhx3 | 0.387628574 | -0.330706568 | 1.105963716 | 10.61046521 | 1.351571471 | 0.229655236 | 0.709264768 | -4.523754394 |
| **A_55_P1962380** | NA | 0.669746748 | -0.571566595 | 1.911060091 | 13.2734229 | 1.351386374 | 0.229710807 | 0.709264768 | -4.52380502 |
| **A_55_P1962586** | NA | 0.299394651 | -0.255513557 | 0.854302859 | 8.840019263 | 1.351366763 | 0.229716696 | 0.709264768 | -4.523810384 |
| **A_55_P1956707** | Rpl36a | 0.679533697 | -0.580046573 | 1.939113968 | 15.13739588 | 1.351249337 | 0.229751958 | 0.709264768 | -4.523842501 |
| **A_51_P162162** | Inmt | -0.219294398 | -0.625787111 | 0.187198314 | 8.277771036 | -1.35121706 | 0.229761651 | 0.709264768 | -4.523851329 |
| **A_55_P2040748** | Scrib | 0.36018675 | -0.30755326 | 1.02792676 | 12.43241505 | 1.351047689 | 0.229812522 | 0.709264768 | -4.523897653 |
| **A_55_P2056315** | Tbca | 0.361646463 | -0.308808048 | 1.032100973 | 13.82578455 | 1.351030801 | 0.229817594 | 0.709264768 | -4.523902272 |
| **A_55_P2022251** | Cdkn2d | 0.319939954 | -0.27323231 | 0.913112219 | 12.16102691 | 1.350945994 | 0.229843071 | 0.709264768 | -4.523925467 |
| **A_52_P508991** | Fmo1 | -0.239123536 | -0.682467607 | 0.204220534 | 9.094101347 | -1.350926639 | 0.229848886 | 0.709264768 | -4.523930761 |
| **A_55_P2022263** | NA | 0.387648725 | -0.331081665 | 1.106379115 | 17.27555675 | 1.35089843 | 0.229857361 | 0.709264768 | -4.523938476 |
| **A_51_P126302** | Rbmx2 | 0.169841824 | -0.145068002 | 0.484751651 | 8.183165967 | 1.35085514 | 0.229870367 | 0.709264768 | -4.525776093 |
| **A_52_P366572** | Slain2 | -0.237993262 | -0.679287477 | 0.203300953 | 8.676406172 | -1.350786693 | 0.229890932 | 0.709264768 | -4.523969036 |
| **A_55_P2004213** | Gprasp2 | 0.929612653 | -0.794170761 | 2.653396066 | 9.939617107 | 1.350733509 | 0.229906913 | 0.709264768 | -4.523983581 |
| **A_55_P2021908** | NA | 0.396102423 | -0.338408395 | 1.13061324 | 10.01803616 | 1.350702326 | 0.229916284 | 0.709264768 | -4.52399211 |
| **A_55_P2009201** | NA | 0.650337941 | -0.555710148 | 1.856386031 | 12.11289453 | 1.350593989 | 0.229948842 | 0.709264768 | -4.524021739 |
| **A_55_P2055324** | Spint1 | -0.426095577 | -1.216303166 | 0.364112011 | 9.376867306 | -1.350567131 | 0.229956914 | 0.709264768 | -4.524029085 |
| **A_55_P2097176** | Idua | -0.337934025 | -0.96467478 | 0.28880673 | 11.5164231 | -1.350499197 | 0.229977333 | 0.709264768 | -4.524047664 |
| **A_55_P1955857** | Ap5z1 | 0.235875406 | -0.201598518 | 0.67334933 | 11.75103228 | 1.35045723 | 0.229989948 | 0.709264768 | -4.524059142 |
| **A_51_P145785** | Ambp | 0.544491836 | -0.465408439 | 1.554392111 | 9.363535044 | 1.350402482 | 0.230006405 | 0.709264768 | -4.524074115 |
| **A_51_P203182** | Apobr | -1.113601252 | -3.17946157 | 0.952259066 | 8.55258008 | -1.350140928 | 0.230085044 | 0.709264768 | -4.524145646 |
| **A_55_P2083566** | NA | 0.404224679 | -0.34567946 | 1.154128818 | 16.23529176 | 1.35010461 | 0.230095966 | 0.709264768 | -4.524155578 |
| **A_55_P2012960** | Gpx7 | 0.758507287 | -0.64869427 | 2.165708843 | 9.951959819 | 1.350062257 | 0.230108703 | 0.709264768 | -4.524167161 |
| **A_51_P479786** | Smtn | -0.39969203 | -1.141236514 | 0.341852455 | 9.642199746 | -1.350015094 | 0.230122887 | 0.709264768 | -4.524180059 |
| **A_55_P1988481** | NA | 0.556747331 | -0.476207089 | 1.589701751 | 9.491742898 | 1.349979988 | 0.230133445 | 0.709264768 | -4.52418966 |
| **A_51_P193336** | Nucb2 | -0.817901689 | -2.335591737 | 0.699788358 | 10.25641876 | -1.349796708 | 0.230188576 | 0.709264768 | -4.524239782 |
| **A_55_P1953377** | NA | -0.762680246 | -2.178113459 | 0.652752966 | 12.7863899 | -1.349595004 | 0.230249262 | 0.709264768 | -4.524294943 |
| **A_52_P352563** | 1700073E | 0.531960527 | -0.455456088 | 1.519377142 | 15.31361995 | 1.349364631 | 0.230318592 | 0.709264768 | -4.524357942 |
| **A_55_P1962314** | NA | 0.299700763 | -0.256623441 | 0.856024967 | 15.90686402 | 1.34930534 | 0.230336439 | 0.709264768 | -4.524374156 |
| **A_51_P333034** | Ier3ip1 | 0.233533077 | -0.199989585 | 0.66705574 | 12.45509434 | 1.349232963 | 0.230358226 | 0.709264768 | -4.524393949 |
| **A_55_P2078845** | Qk | 0.309309064 | -0.26500257 | 0.883620699 | 11.59637526 | 1.348948528 | 0.230443865 | 0.709264768 | -4.524471729 |
| **A_55_P2185526** | 1700016P | 0.146915495 | -0.125887738 | 0.419718727 | 8.765111245 | 1.348864842 | 0.230469068 | 0.709264768 | -4.524494614 |
| **A_51_P353008** | Minpp1 | 0.238174298 | -0.204217089 | 0.680565685 | 9.981340679 | 1.348461579 | 0.230590547 | 0.709264768 | -4.524604885 |
| **A_55_P2166639** | Cdk4 | 0.600348834 | -0.514765355 | 1.715463023 | 11.70636816 | 1.348449465 | 0.230594197 | 0.709264768 | -4.524608197 |
| **A_51_P189905** | Dad1 | -0.226461489 | -0.64714528 | 0.194222302 | 13.23583305 | -1.348307421 | 0.230637001 | 0.709264768 | -4.524647038 |
| **A_52_P428521** | Fgd1 | -0.143932835 | -0.411314052 | 0.123448382 | 8.55250089 | -1.348277635 | 0.230645978 | 0.709264768 | -4.524655183 |
| **A_51_P314153** | Nr2c2ap | -0.1387811 | -0.396620301 | 0.119058101 | 8.492616263 | -1.348129845 | 0.230690523 | 0.709264768 | -4.524695594 |
| **A_51_P511386** | Zfp267 | 0.248232311 | -0.212994448 | 0.709459071 | 8.47871596 | 1.348013248 | 0.230725672 | 0.709264768 | -4.524727475 |

| | | | | | | | | | |
|---|---|---|---|---|---|---|---|---|---|
| **A_52_P123485** | Tcf25 | 0.14495992 | -0.124415642 | 0.414335483 | 13.8394987 | 1.347845438 | 0.230776268 | 0.709264768 | -4.52477336 |
| **A_51_P334289** | Zfp628 | 0.170311109 | -0.146208547 | 0.486830764 | 9.074634817 | 1.347698167 | 0.230820679 | 0.709264768 | -4.524813627 |
| **A_55_P2142728** | NA | 0.235545161 | -0.202219946 | 0.673310268 | 12.30668158 | 1.347669469 | 0.230829334 | 0.709264768 | -4.524821474 |
| **A_52_P241484** | Rcl1 | 0.167210673 | -0.14357818 | 0.477999525 | 8.989564176 | 1.347562465 | 0.230861609 | 0.709264768 | -4.524850731 |
| **A_52_P135707** | Creb3 | -0.906608803 | -2.59183273 | 0.778615125 | 11.43917085 | -1.347450004 | 0.230895533 | 0.709264768 | -4.52488148 |
| **A_55_P2313163** | D10Bwg1( | -0.255632468 | -0.730824464 | 0.219559528 | 10.89193349 | -1.347402169 | 0.230909965 | 0.709264768 | -4.524894559 |
| **A_55_P2045886** | Stk3 | 0.21995293 | -0.188948918 | 0.628854778 | 10.14500098 | 1.347289806 | 0.230943866 | 0.709264768 | -4.524925281 |
| **A_55_P2169356** | Gm1966 | -0.37410383 | -1.069609553 | 0.321401894 | 8.10724438 | -1.347230144 | 0.230961869 | 0.709264768 | -4.524941594 |
| **A_55_P2175060** | Plpp5 | -0.573018915 | -1.638395229 | 0.4923574 | 8.557255192 | -1.347150963 | 0.230985764 | 0.709264768 | -4.524963243 |
| **A_55_P2111292** | 3110009E | 0.345113948 | -0.296582101 | 0.986809997 | 9.757555817 | 1.347049304 | 0.231016445 | 0.709264768 | -4.524991037 |
| **A_52_P110812** | Ggcx | -0.196219435 | -0.561137618 | 0.168698748 | 8.905055019 | -1.346780632 | 0.23109755 | 0.709264768 | -4.525064494 |
| **A_51_P417074** | Arhgap8 | -0.693353937 | -1.982895804 | 0.59618793 | 11.86669344 | -1.346696976 | 0.231122809 | 0.709264768 | -4.525087366 |
| **A_55_P2039523** | Phf10 | 0.256356389 | -0.22043867 | 0.733151449 | 11.53473336 | 1.346674835 | 0.231129494 | 0.709264768 | -4.525093419 |
| **A_55_P2013273** | NA | 0.396439769 | -0.340919306 | 1.133798845 | 12.26089218 | 1.346630761 | 0.231142803 | 0.709264768 | -4.525105469 |
| **A_55_P2026156** | NA | 0.659667097 | -0.567471702 | 1.886805896 | 14.57519544 | 1.346422873 | 0.231205588 | 0.709264768 | -4.525162305 |
| **A_55_P2162935** | Ntn1 | -0.215548997 | -0.616534954 | 0.185436959 | 8.402938071 | -1.346378567 | 0.231218971 | 0.709264768 | -4.525174418 |
| **A_55_P1967483** | NA | 0.820791605 | -0.706323985 | 2.347907194 | 12.52772909 | 1.346205438 | 0.231271273 | 0.709264768 | -4.525221749 |
| **A_52_P585831** | Bap1 | -0.216407522 | -0.619049468 | 0.186234425 | 10.53715379 | -1.346181694 | 0.231278447 | 0.709264768 | -4.525228241 |
| **A_51_P151835** | Tmtc3 | 0.318147797 | -0.27387195 | 0.910167544 | 9.225839992 | 1.345993843 | 0.23133521 | 0.709264768 | -4.525279596 |
| **A_55_P2012296** | Gopc | 0.157557205 | -0.135675723 | 0.450790134 | 9.724468398 | 1.3457856 | 0.23139815 | 0.709264768 | -4.525336526 |
| **A_66_P134252** | NA | 0.878773932 | -0.7568041 | 2.514351963 | 13.7095474 | 1.345724758 | 0.231416542 | 0.709264768 | -4.525353159 |
| **A_52_P27122** | Ago4 | -0.145676686 | -0.416824822 | 0.125471451 | 8.439447812 | -1.345655136 | 0.231437589 | 0.709264768 | -4.525372192 |
| **A_52_P197666** | Med12 | -0.170125431 | -0.486791379 | 0.146540518 | 10.6833773 | -1.345606939 | 0.231452161 | 0.709264768 | -4.525385368 |
| **A_55_P2014169** | Ppa2 | 0.506451742 | -0.436307022 | 1.449210505 | 12.23232786 | 1.345512809 | 0.231480622 | 0.709264768 | -4.525411101 |
| **A_51_P232742** | Mfsd14b | -0.146666713 | -0.41969536 | 0.126361934 | 10.15435379 | -1.345468972 | 0.231493878 | 0.709264768 | -4.525423084 |
| **A_51_P154469** | Ccndbp1 | -0.191552754 | -0.548222875 | 0.165117368 | 11.27050109 | -1.345154024 | 0.231589135 | 0.709264768 | -4.525509181 |
| **A_55_P2069221** | Prr11 | 0.352990817 | -0.304306497 | 1.010288131 | 8.457903325 | 1.345091708 | 0.231607986 | 0.709264768 | -4.525526215 |
| **A_51_P239766** | Plcd1 | -0.38636463 | -1.105856153 | 0.333126894 | 10.67153189 | -1.344999198 | 0.231635975 | 0.709264768 | -4.525551504 |
| **A_55_P2055063** | Mcts1 | 0.342515877 | -0.295352234 | 0.980383988 | 11.56505119 | 1.34493149 | 0.231656462 | 0.709264768 | -4.525570013 |
| **A_66_P124052** | Bok | -1.095213838 | -3.134968971 | 0.944541296 | 11.46001758 | -1.344841905 | 0.231683571 | 0.709264768 | -4.525594501 |
| **A_52_P49601** | Fth1 | -0.349451489 | -1.000447882 | 0.301544903 | 15.88287963 | -1.344493353 | 0.231789072 | 0.709264768 | -4.525689778 |
| **A_52_P58524** | Neu1 | -1.152328016 | -3.299080439 | 0.994424407 | 11.421798 | -1.344449533 | 0.231802339 | 0.709264768 | -4.525701757 |
| **A_55_P2263098** | A130078K | -0.383094935 | -1.096887956 | 0.330698087 | 8.525017175 | -1.344263657 | 0.231858622 | 0.709264768 | -4.525752565 |
| **A_55_P2000548** | Cfl1 | 0.255969047 | -0.221018399 | 0.732956492 | 14.51837595 | 1.344097732 | 0.231908874 | 0.709264768 | -4.525797918 |
| **A_55_P2011436** | NA | 0.687505363 | -0.593686423 | 1.968697149 | 15.3654593 | 1.344040298 | 0.231926271 | 0.709264768 | -4.525813617 |
| **A_52_P674309** | Mocs2 | -0.167045642 | -0.478351638 | 0.144260353 | 11.83692095 | -1.343996109 | 0.231939657 | 0.709264768 | -4.525825696 |
| **A_55_P2179335** | Usp33 | -0.118886105 | -0.34046863 | 0.10269642 | 11.15228418 | -1.343834966 | 0.231988476 | 0.709264768 | -4.525869741 |
| **A_55_P2083889** | Pea15a | -0.62012549 | -1.776074104 | 0.535823125 | 10.99030887 | -1.343666223 | 0.232039608 | 0.709264768 | -4.525915864 |
| **A_55_P1955108** | Ripor1 | -0.641101885 | -1.836181986 | 0.553978216 | 13.70169279 | -1.343632142 | 0.232049936 | 0.709264768 | -4.525925179 |
| **A_51_P269320** | Adsl | 0.226878844 | -0.196056677 | 0.649814365 | 9.139116753 | 1.343600587 | 0.2320595 | 0.709264768 | -4.525933803 |
| **A_52_P121** | Osgepl1 | 0.208555248 | -0.180235728 | 0.597346223 | 8.994265871 | 1.343554512 | 0.232073464 | 0.709264768 | -4.525946397 |
| **A_55_P1968464** | Kat6a | 0.327913435 | -0.283527564 | 0.939354434 | 10.64706057 | 1.343244281 | 0.232167509 | 0.709264768 | -4.526031189 |
| **A_55_P2022921** | Nsa2 | 0.478650736 | -0.413890646 | 1.371192118 | 13.06886312 | 1.343200101 | 0.232180904 | 0.709264768 | -4.526043264 |
| **A_55_P2015178** | Nap1l1 | 0.730551719 | -0.631803456 | 2.092906894 | 13.97473077 | 1.343108096 | 0.232208803 | 0.709264768 | -4.526068411 |
| **A_51_P401683** | Tm6sf1 | 0.444114468 | -0.384252415 | 1.272481352 | 9.204009657 | 1.34283465 | 0.232291739 | 0.709264768 | -4.526143146 |
| **A_55_P1982449** | NA | 0.145473876 | -0.125877783 | 0.416825535 | 7.900711629 | 1.342773852 | 0.232310183 | 0.709264768 | -4.526159762 |
| **A_55_P2010704** | Selenoh | 0.880609099 | -0.762129486 | 2.523347685 | 13.15074644 | 1.342656925 | 0.232345658 | 0.709264768 | -4.526191719 |
| **A_55_P2089080** | Ssbp2 | 0.677042697 | -0.586044972 | 1.940130367 | 10.89587871 | 1.342557541 | 0.232375814 | 0.709264768 | -4.526218881 |

| | | | | | | | | | |
|---|---|---|---|---|---|---|---|---|---|
| **A_55_P2184449** | NA | -0.563034653 | -1.613686402 | 0.487617097 | 8.109590533 | -1.3422292 | 0.232475468 | 0.709264768 | -4.526308614 |
| **A_55_P2101920** | Eva1c | -1.177705542 | -3.375588108 | 1.020177024 | 10.76541893 | -1.342092905 | 0.232516846 | 0.709264768 | -4.526345862 |
| **A_55_P1994020** | Casc4 | 0.312751768 | -0.270929828 | 0.896433364 | 9.216912821 | 1.342066758 | 0.232524785 | 0.709264768 | -4.526353008 |
| **A_52_P99807** | Dpy19l3 | 0.176782862 | -0.153144951 | 0.506710675 | 8.592400753 | 1.342058906 | 0.232527169 | 0.709264768 | -4.526355154 |
| **A_55_P2015632** | Gm5766 | 0.40818765 | -0.35362066 | 1.16999596 | 16.17339289 | 1.342037035 | 0.23253381 | 0.709264768 | -4.526361131 |
| **A_55_P2053988** | Epas1 | -1.369079753 | -3.924281317 | 1.186121812 | 9.420987918 | -1.342004366 | 0.23254373 | 0.709264768 | -4.526370059 |
| **A_51_P133137** | Kif20a | 1.261890794 | -1.093303216 | 3.617084805 | 11.18434637 | 1.341978103 | 0.232551704 | 0.709264768 | -4.526377236 |
| **A_55_P2056286** | Sf3b5 | 0.288745753 | -0.250243446 | 0.827734952 | 13.1377614 | 1.341794132 | 0.232607575 | 0.709264768 | -4.526427512 |
| **A_52_P129697** | Stradb | -0.213748984 | -0.612792558 | 0.185294589 | 7.969039828 | -1.341634087 | 0.232656189 | 0.709264768 | -4.526471249 |
| **A_55_P2078645** | Mettl5 | 0.209978492 | -0.182054955 | 0.602011938 | 9.360808864 | 1.34153509 | 0.232686265 | 0.709264768 | -4.526498302 |
| **A_55_P2170335** | Bmi1 | 0.232009716 | -0.201177656 | 0.665197088 | 10.60907237 | 1.341469275 | 0.232706262 | 0.709264768 | -4.526516288 |
| **A_55_P2210213** | 4732423E | -0.215476094 | -0.617794079 | 0.18684189 | 9.60685504 | -1.341467002 | 0.232706952 | 0.709264768 | -4.526516909 |
| **A_51_P472726** | Pdlim2 | -0.282666418 | -0.810536263 | 0.245203428 | 13.64601065 | -1.341212854 | 0.232784185 | 0.709264768 | -4.526586359 |
| **A_55_P2373852** | 2310058N | -0.508717805 | -1.458735709 | 0.441300098 | 8.532055857 | -1.341206021 | 0.232786262 | 0.709264768 | -4.526588226 |
| **A_55_P1985768** | Ttyh2 | 0.21307659 | -0.184854236 | 0.611007416 | 9.221457495 | 1.341153541 | 0.232802213 | 0.709264768 | -4.526602567 |
| **A_55_P2124543** | Gnptg | -0.2443205 | -0.700680446 | 0.212039445 | 9.992000223 | -1.340919589 | 0.232873335 | 0.709264768 | -4.526666497 |
| **A_55_P2183597** | Tbc1d2 | -0.761341664 | -2.183483244 | 0.660799916 | 10.24910714 | -1.34087134 | 0.232888005 | 0.709264768 | -4.526679682 |
| **A_55_P2157914** | NA | -0.349620104 | -1.002773166 | 0.303532958 | 8.079999828 | -1.340700515 | 0.232939951 | 0.709264768 | -4.52672636 |
| **A_52_P637353** | Zbtb43 | -0.182565254 | -0.523634547 | 0.158504039 | 9.168040835 | -1.34068222 | 0.232945516 | 0.709264768 | -4.526731359 |
| **A_55_P1999790** | Pkdcc | 0.630470291 | -0.547552207 | 1.808492789 | 12.4403542 | 1.340483237 | 0.233006039 | 0.709264768 | -4.526785731 |
| **A_55_P2156293** | Pcyox1 | -0.314679007 | -0.902651985 | 0.273293971 | 11.74947902 | -1.340481268 | 0.233006638 | 0.709264768 | -4.526786269 |
| **A_55_P2187034** | NA | 0.733406612 | -0.637010099 | 2.103823323 | 12.5389034 | 1.340425005 | 0.233023754 | 0.709264768 | -4.526801643 |
| **A_51_P488730** | Zfp612 | 0.256258698 | -0.222578228 | 0.735095624 | 8.811093112 | 1.340421321 | 0.233024875 | 0.709264768 | -4.52680265 |
| **A_55_P2083464** | Lpin3 | -0.671597064 | -1.92659723 | 0.583403103 | 8.968385806 | -1.340341149 | 0.233049266 | 0.709264768 | -4.526824556 |
| **A_51_P122582** | Daglb | -0.408886022 | -1.173023229 | 0.355251185 | 8.847435007 | -1.340235951 | 0.233081275 | 0.709264768 | -4.530454589 |
| **A_55_P1980214** | NA | 0.226550232 | -0.196862304 | 0.649962768 | 9.852961267 | 1.340143006 | 0.233109559 | 0.709264768 | -4.526878697 |
| **A_52_P235569** | Ipo5 | 0.184734252 | -0.160599606 | 0.53006811 | 9.29978815 | 1.339857498 | 0.233196461 | 0.709264768 | -4.526956707 |
| **A_51_P484919** | Polrmt | 0.13525668 | -0.117596062 | 0.388109422 | 9.998785216 | 1.339804402 | 0.233212626 | 0.709264768 | -4.526971215 |
| **A_55_P2074154** | Dhx57 | -0.133023531 | -0.38174108 | 0.115694018 | 9.243504984 | -1.339591517 | 0.233277446 | 0.709264768 | -4.527029381 |
| **A_51_P281246** | Cep57 | 0.346807016 | -0.301728904 | 0.995342935 | 9.179633132 | 1.339381158 | 0.233341514 | 0.709264768 | -4.527086855 |
| **A_52_P155805** | 2010204K | 0.174115322 | -0.151483868 | 0.499714513 | 8.21316045 | 1.339380652 | 0.233341668 | 0.709264768 | -4.527086993 |
| **A_52_P639567** | Ate1 | -0.218048862 | -0.625813581 | 0.189715857 | 10.57890873 | -1.339351361 | 0.23335059 | 0.709264768 | -4.527094996 |
| **A_55_P2173927** | Insr | -0.130061302 | -0.373294553 | 0.113171949 | 7.980313299 | -1.339292688 | 0.233368463 | 0.709264768 | -4.527111027 |
| **A_55_P2077114** | NA | 0.617034144 | -0.53698747 | 1.771055757 | 15.16465555 | 1.339200489 | 0.233396552 | 0.709264768 | -4.527136217 |
| **A_55_P2164090** | Kcnj15 | -0.23295595 | -0.66875542 | 0.20284352 | 9.076475728 | -1.338867055 | 0.233498159 | 0.709264768 | -4.527227314 |
| **A_55_P2151609** | Sorl1 | 0.923886486 | -0.804596388 | 2.652369359 | 10.15554323 | 1.338763549 | 0.233529708 | 0.709264768 | -4.527255592 |
| **A_52_P628590** | Pvr | -0.791817446 | -2.273426488 | 0.689791597 | 9.229959571 | -1.338572171 | 0.233588052 | 0.709264768 | -4.527307877 |
| **A_51_P420547** | Clic5 | -0.428123794 | -1.229354733 | 0.373107144 | 12.28650286 | -1.338326264 | 0.233663039 | 0.709264768 | -4.527375057 |
| **A_51_P288026** | Haus6 | 0.232211224 | -0.202484518 | 0.666906967 | 8.571103023 | 1.337975521 | 0.233770033 | 0.709264768 | -4.527470875 |
| **A_55_P2065971** | Setd3 | 0.218245329 | -0.190346903 | 0.626837562 | 11.57711046 | 1.337843144 | 0.233810426 | 0.709264768 | -4.527507038 |
| **A_55_P2158741** | Nos2 | -0.668178852 | -1.919123164 | 0.58276546 | 9.713686894 | -1.337842828 | 0.233810522 | 0.709264768 | -4.527507125 |
| **A_55_P2175314** | NA | 0.423163426 | -0.369125496 | 1.215452349 | 15.66384887 | 1.337749776 | 0.23383892 | 0.709264768 | -4.527532544 |
| **A_51_P254986** | Serf1 | 0.39058531 | -0.340770188 | 1.121940807 | 13.39743075 | 1.337635298 | 0.23387386 | 0.709264768 | -4.527563817 |
| **A_55_P2106414** | Haus7 | 0.273170337 | -0.238383387 | 0.784724062 | 8.557935071 | 1.337496436 | 0.233916249 | 0.709264768 | -4.52760175 |
| **A_52_P80444** | Dpm2 | -0.386300778 | -1.109827284 | 0.337225728 | 11.59799324 | -1.337277329 | 0.233983149 | 0.709264768 | -4.527661602 |
| **A_52_P671661** | Mak16 | 0.142337509 | -0.124285473 | 0.408960491 | 8.65268479 | 1.337125358 | 0.23402956 | 0.709264768 | -4.527703115 |
| **A_51_P314019** | Banf1 | 0.645352357 | -0.563551767 | 1.854256482 | 11.31227707 | 1.337073826 | 0.234045299 | 0.709264768 | -4.527717192 |
| **A_52_P189132** | Thap1 | 0.16662671 | -0.145515249 | 0.47876867 | 10.10378974 | 1.337035117 | 0.234057123 | 0.709264768 | -4.527727765 |

| | | | | | | | | | |
|---|---|---|---|---|---|---|---|---|---|
| **A_52_P417654** | Tcea1 | 0.202102169 | -0.176567743 | 0.580772081 | 9.251547059 | 1.336781789 | 0.234134515 | 0.709264768 | -4.527796963 |
| **A_51_P417469** | Rnaseh2c | 0.775069634 | -0.677181935 | 2.227321203 | 13.56042515 | 1.336746986 | 0.234145149 | 0.709264768 | -4.527806469 |
| **A_55_P1984043** | Pdcd4 | -0.294410821 | -0.846067059 | 0.257245417 | 12.36914334 | -1.336705026 | 0.234157971 | 0.709264768 | -4.527817931 |
| **A_51_P193185** | Mb | -0.178661199 | -0.51344198 | 0.156119583 | 7.991304113 | -1.336657284 | 0.23417256 | 0.709264768 | -4.527830971 |
| **A_52_P452667** | Prom2 | -0.20714324 | -0.595408439 | 0.18112196 | 8.455897806 | -1.336265154 | 0.23429242 | 0.709264768 | -4.527938078 |
| **A_51_P158400** | Rc3h2 | -0.219333231 | -0.630447439 | 0.191780977 | 10.42687931 | -1.336264085 | 0.234292747 | 0.709264768 | -4.527938371 |
| **A_55_P1976983** | Rpl36 | 0.187077017 | -0.16362544 | 0.537779473 | 16.47880694 | 1.336078494 | 0.234349495 | 0.709264768 | -4.527989062 |
| **A_55_P2115136** | Vsir | -0.597267874 | -1.717015919 | 0.522480171 | 8.433311113 | -1.335977636 | 0.23438034 | 0.709264768 | -4.528016609 |
| **A_55_P2006281** | Ogdh | -0.366786076 | -1.054447508 | 0.320875355 | 12.20320146 | -1.335944818 | 0.234390377 | 0.709264768 | -4.528025573 |
| **A_66_P126836** | Zbtb44 | -0.142111031 | -0.408564248 | 0.124342186 | 10.70046896 | -1.335848376 | 0.234419876 | 0.709264768 | -4.528051914 |
| **A_66_P119429** | NA | 0.584400593 | -0.511340747 | 1.680141933 | 14.47119557 | 1.335835396 | 0.234423846 | 0.709264768 | -4.528055459 |
| **A_55_P2077608** | Hdac1 | 0.321502369 | -0.2813381 | 0.924342837 | 9.726422363 | 1.335771348 | 0.234443439 | 0.709264768 | -4.528072952 |
| **A_51_P171697** | Pebp1 | 0.578334647 | -0.506101247 | 1.662770541 | 13.77587473 | 1.335751506 | 0.234449509 | 0.709264768 | -4.528078371 |
| **A_55_P1985433** | Nrg1 | -0.123804608 | -0.355961897 | 0.108352681 | 8.157045805 | -1.335687409 | 0.234469118 | 0.709264768 | -4.528095878 |
| **A_52_P167189** | Isca1 | -0.250632855 | -0.720774561 | 0.219508851 | 11.72114002 | -1.33524072 | 0.234605817 | 0.709264768 | -4.528217876 |
| **A_52_P391718** | Rpl22 | 0.754843496 | -0.661107682 | 2.170794675 | 11.81621269 | 1.335238923 | 0.234606367 | 0.709264768 | -4.528218366 |
| **A_55_P1998957** | Oas1c | -0.599063185 | -1.722988989 | 0.524862619 | 8.500174412 | -1.335012508 | 0.234675685 | 0.709264768 | -4.528280202 |
| **A_55_P2148744** | NA | 0.76129482 | -0.667128914 | 2.189718554 | 12.65792402 | 1.3348921 | 0.234712555 | 0.709264768 | -4.528313086 |
| **A_55_P2004179** | Col2a1 | 0.970651756 | -0.850651045 | 2.791954558 | 9.114923186 | 1.334847445 | 0.234726231 | 0.709264768 | -4.528325281 |
| **A_52_P53906** | Ccnd2 | 0.876764275 | -0.768393743 | 2.521922293 | 10.39779067 | 1.334828814 | 0.234731937 | 0.709264768 | -4.528330369 |
| **A_52_P126825** | Cramp1l | -0.32637702 | -0.938812923 | 0.286058882 | 11.01061355 | -1.334778727 | 0.234747277 | 0.709264768 | -4.528344048 |
| **A_55_P2301063** | 9930024M | -0.255562639 | -0.735137472 | 0.224012195 | 8.192401692 | -1.334723555 | 0.234764175 | 0.709264768 | -4.528359115 |
| **A_51_P487933** | Psmd7 | 0.236395219 | -0.207266809 | 0.680057248 | 12.27877356 | 1.334555914 | 0.234815528 | 0.709264768 | -4.528404897 |
| **A_66_P104390** | Arl3 | 0.564265724 | -0.494898223 | 1.623429671 | 10.48078953 | 1.334353276 | 0.234877616 | 0.709264768 | -4.528460235 |
| **A_55_P2182736** | Limd2 | 0.233900915 | -0.20517449 | 0.672976321 | 9.607339274 | 1.334268264 | 0.234903667 | 0.709264768 | -4.52848345 |
| **A_55_P2054027** | Chst5 | 0.574832629 | -0.504339929 | 1.654005186 | 9.891983507 | 1.334138338 | 0.234943488 | 0.709264768 | -4.528518931 |
| **A_55_P2049687** | Efna2 | 0.273148117 | -0.239822161 | 0.786118395 | 8.610563312 | 1.333694482 | 0.235079571 | 0.709264768 | -4.528640137 |
| **A_55_P2131875** | Poldip3 | -0.671206691 | -1.931826366 | 0.589412983 | 13.91057666 | -1.33359065 | 0.235111416 | 0.709264768 | -4.52866849 |
| **A_55_P2171528** | 3110082I1 | 0.136638835 | -0.120038079 | 0.393315749 | 8.231758332 | 1.333330108 | 0.235191339 | 0.709264768 | -4.528739634 |
| **A_51_P484753** | Saraf | -0.597222624 | -1.719164152 | 0.524718905 | 12.63607818 | -1.333264676 | 0.235211415 | 0.709264768 | -4.5287575 |
| **A_52_P469502** | Cda | -0.667471927 | -1.921486772 | 0.586542919 | 9.364043228 | -1.33315508 | 0.235245044 | 0.709264768 | -4.528787426 |
| **A_55_P2115257** | Fer | -0.373563803 | -1.075431905 | 0.3283043 | 9.702360587 | -1.333090481 | 0.235264869 | 0.709264768 | -4.528805065 |
| **A_55_P2085574** | Nfix | 0.957863845 | -0.842062411 | 2.7577901 | 10.73528534 | 1.332905667 | 0.235321593 | 0.709264768 | -4.528855529 |
| **A_52_P577729** | Rnf169 | 0.336124589 | -0.295507494 | 0.967756672 | 10.80244551 | 1.332865872 | 0.235333809 | 0.709264768 | -4.528866394 |
| **A_52_P59228** | Zswim3 | -0.209863507 | -0.604235065 | 0.184508051 | 8.179805197 | -1.332851254 | 0.235338296 | 0.709264768 | -4.528870386 |
| **A_51_P455946** | Rac3 | 0.625784052 | -0.550183676 | 1.80175178 | 9.914391439 | 1.332844343 | 0.235340418 | 0.709264768 | -4.528872273 |
| **A_51_P502132** | Mmp23 | -0.157732855 | -0.45414397 | 0.13867826 | 9.553408347 | -1.332840023 | 0.235341744 | 0.709264768 | -4.528873453 |
| **A_55_P2130264** | NA | 0.363884315 | -0.319940066 | 1.047708695 | 17.02184511 | 1.332812632 | 0.235350153 | 0.709264768 | -4.528880931 |
| **A_55_P2064272** | NA | 0.199808712 | -0.175716079 | 0.575333503 | 8.240136371 | 1.332680837 | 0.235390616 | 0.709264768 | -4.528916917 |
| **A_55_P2009066** | Cfap74 | -1.281555228 | -3.690250512 | 1.127140056 | 14.66530688 | -1.332618413 | 0.235409784 | 0.709264768 | -4.528933961 |
| **A_55_P1994927** | Rab3il1 | 0.462212708 | -0.406672729 | 1.331098145 | 10.65794157 | 1.33238492 | 0.235481492 | 0.709264768 | -4.528997713 |
| **A_55_P1970062** | Igsf9b | 0.179486694 | -0.157947897 | 0.516921285 | 8.603519118 | 1.332272313 | 0.235516082 | 0.709264768 | -4.529028459 |
| **A_65_P19623** | Ssu72 | -0.278808483 | -0.803071035 | 0.245454069 | 12.35544224 | -1.33201002 | 0.235596669 | 0.709264768 | -4.529100071 |
| **A_51_P286665** | Rbl1 | 0.365872617 | -0.322193734 | 1.053938967 | 9.556130175 | 1.331833494 | 0.235650919 | 0.709264768 | -4.529148267 |
| **A_55_P2030486** | Srsf2 | 0.163699831 | -0.144219976 | 0.471619638 | 9.561292765 | 1.331560643 | 0.235734794 | 0.709264768 | -4.529222759 |
| **A_51_P142164** | Hspa13 | -0.253826674 | -0.731278103 | 0.223624755 | 7.93447596 | -1.331552844 | 0.235737192 | 0.709264768 | -4.529224888 |
| **A_55_P2167769** | Arid1a | 0.406152657 | -0.357883875 | 1.170189188 | 11.90933238 | 1.331452017 | 0.235768194 | 0.709264768 | -4.529252415 |
| **A_55_P2009064** | Psmc1 | 0.162615196 | -0.143315772 | 0.468546165 | 11.73873288 | 1.33133709 | 0.235803535 | 0.709264768 | -4.52928379 |

| | | | | | | | | | |
|---|---|---|---|---|---|---|---|---|---|
| **A_55_P2007733** | Epn1 | -0.565541654 | -1.629625673 | 0.498542365 | 14.85752846 | -1.331186862 | 0.23584974 | 0.709264768 | -4.529324803 |
| **A_55_P2131438** | H2ac19 | 0.715546522 | -0.6307863 | 2.061879344 | 16.6901072 | 1.331176945 | 0.23585279 | 0.709264768 | -4.52932751 |
| **A_52_P230167** | Setbp1 | 0.691842908 | -0.609948749 | 1.993634565 | 9.837808651 | 1.331117384 | 0.235871112 | 0.709264768 | -4.52934377 |
| **A_55_P1957277** | NA | 0.145930661 | -0.1286742 | 0.420535523 | 9.536109721 | 1.331032545 | 0.235897211 | 0.709264768 | -4.529366931 |
| **A_55_P2085142** | Spp1 | -0.552305082 | -1.591611121 | 0.487000956 | 7.962244926 | -1.331024148 | 0.235899794 | 0.709264768 | -4.529369224 |
| **A_55_P1953241** | Mafk | -0.70338795 | -2.02705555 | 0.620279651 | 8.866301381 | -1.330964067 | 0.235918278 | 0.709264768 | -4.529385625 |
| **A_51_P181538** | Brwd1 | -0.144676442 | -0.416972812 | 0.127619929 | 9.215430137 | -1.330780143 | 0.235974873 | 0.709264768 | -4.529435835 |
| **A_55_P2121618** | Tox4 | 0.521716212 | -0.460416325 | 1.503848749 | 8.768208528 | 1.33049908 | 0.236061381 | 0.709264768 | -4.529512561 |
| **A_55_P2104790** | NA | 0.728760601 | -0.643171984 | 2.100693186 | 13.47912536 | 1.330461954 | 0.23607281 | 0.709264768 | -4.529522696 |
| **A_52_P407280** | Adamtsl5 | -0.411608172 | -1.186550006 | 0.363333662 | 8.25420667 | -1.330347899 | 0.236107925 | 0.709264768 | -4.52955383 |
| **A_55_P1976351** | Gpcpd1 | -0.77198827 | -2.225474261 | 0.68149772 | 8.808507689 | -1.330301856 | 0.236122102 | 0.709264768 | -4.529566399 |
| **A_55_P1985143** | NA | 0.988760252 | -0.873059353 | 2.850579858 | 11.22482055 | 1.330159578 | 0.236165915 | 0.709264768 | -4.529605237 |
| **A_51_P117130** | Naga | -0.415192799 | -1.197009309 | 0.366623712 | 8.256654328 | -1.330133772 | 0.236173862 | 0.709264768 | -4.530203434 |
| **A_51_P507023** | Nt5dc3 | -0.285889935 | -0.824266986 | 0.252487115 | 10.28993573 | -1.330033785 | 0.236204658 | 0.709264768 | -4.529639575 |
| **A_66_P100789** | Kansl2 | 0.303793489 | -0.268331481 | 0.875918459 | 9.99875929 | 1.329957933 | 0.236228022 | 0.709264768 | -4.52966028 |
| **A_55_P2068607** | Rnf19b | -0.524983597 | -1.513699051 | 0.463731857 | 10.36605264 | -1.329917673 | 0.236240424 | 0.709264768 | -4.529671269 |
| **A_55_P1969857** | Ap2m1 | -0.425345023 | -1.226410865 | 0.375720819 | 13.50183407 | -1.329913785 | 0.236241622 | 0.709264768 | -4.529672331 |
| **A_55_P1970700** | Rpsa | -0.129404758 | -0.373121963 | 0.114312448 | 16.44524797 | -1.329885956 | 0.236250195 | 0.709264768 | -4.529679927 |
| **A_55_P2079485** | Ptges3 | 0.145547123 | -0.128583392 | 0.419677638 | 12.21971084 | 1.329831416 | 0.236266997 | 0.709264768 | -4.529694814 |
| **A_52_P585751** | Gns | -1.425691958 | -4.110990792 | 1.259606877 | 11.43655023 | -1.3297911 | 0.236279419 | 0.709264768 | -4.529705819 |
| **A_55_P2055232** | Sdf4 | -0.462973246 | -1.335051528 | 0.409105036 | 10.03624613 | -1.329691123 | 0.236310223 | 0.709264768 | -4.529733109 |
| **A_52_P663742** | Ganc | -0.221986595 | -0.640144362 | 0.196171172 | 8.359148537 | -1.329648745 | 0.236323282 | 0.709264768 | -4.529744676 |
| **A_55_P1994694** | Serinc2 | -0.300067539 | -0.865312473 | 0.265177396 | 8.221039522 | -1.329635747 | 0.236327287 | 0.709264768 | -4.529748224 |
| **A_55_P2091506** | Srp54c | -0.197609596 | -0.569860047 | 0.174640854 | 11.91317584 | -1.329606545 | 0.236336286 | 0.709264768 | -4.529756195 |
| **A_55_P2115245** | Zfyve19 | -0.129372259 | -0.373096126 | 0.114351609 | 11.77211712 | -1.329515624 | 0.236364307 | 0.709264768 | -4.529781012 |
| **A_55_P2171196** | Gm3411 | -0.174446874 | -0.503116872 | 0.154223124 | 8.659806935 | -1.329393266 | 0.23640202 | 0.709264768 | -4.52981441 |
| **A_55_P2060863** | NA | 0.448501186 | -0.396513215 | 1.293515587 | 12.18696575 | 1.329382113 | 0.236405458 | 0.709264768 | -4.529817454 |
| **A_55_P2162651** | NA | 0.575869069 | -0.50912634 | 1.660864478 | 15.19210537 | 1.32937099 | 0.236408887 | 0.709264768 | -4.52982049 |
| **A_55_P1994648** | Eef1d | 0.144275592 | -0.127565737 | 0.416116921 | 15.03884847 | 1.329314453 | 0.236426315 | 0.709264768 | -4.529835922 |
| **A_51_P189272** | Tmcc3 | -1.236444415 | -3.566392733 | 1.093503903 | 10.32546575 | -1.32916426 | 0.23647262 | 0.709264768 | -4.529876917 |
| **A_51_P478895** | Ppp6r1 | -0.137289609 | -0.396008653 | 0.121429434 | 9.338726261 | -1.329106 | 0.236490584 | 0.709264768 | -4.529892818 |
| **A_51_P327491** | Apoa4 | -1.089635128 | -3.143086316 | 0.96381606 | 8.214287051 | -1.329067559 | 0.236502438 | 0.709264768 | -4.529903311 |
| **A_55_P2183750** | Ncoa3 | -0.418086869 | -1.206056336 | 0.369882598 | 11.11760992 | -1.328946458 | 0.236539784 | 0.709264768 | -4.529936364 |
| **A_51_P400585** | Api5 | 0.351296395 | -0.310808745 | 1.013401535 | 10.69120566 | 1.328914619 | 0.236549604 | 0.709264768 | -4.529945054 |
| **A_52_P262219** | Fos | -0.708217727 | -2.043091083 | 0.626655629 | 8.554449108 | -1.328853406 | 0.236568484 | 0.709264768 | -4.531046005 |
| **A_51_P183894** | Fbxo15 | -0.472786333 | -1.36401278 | 0.418440114 | 8.114195005 | -1.328700791 | 0.236615561 | 0.709264768 | -4.530003415 |
| **A_66_P132777** | Cep68 | 0.437217258 | -0.386984934 | 1.261419449 | 9.774220354 | 1.328660049 | 0.236628131 | 0.709264768 | -4.530014534 |
| **A_51_P336635** | Xpo1 | 0.424491523 | -0.37573513 | 1.224718175 | 10.8546547 | 1.328637038 | 0.23663523 | 0.709264768 | -4.530020815 |
| **A_51_P297963** | Recql | 0.128482254 | -0.113731335 | 0.370695843 | 8.623935713 | 1.328602265 | 0.236645958 | 0.709264768 | -4.530030305 |
| **A_55_P2031501** | Hnrnpa3 | 0.514801539 | -0.455808695 | 1.485411774 | 15.4647259 | 1.328450314 | 0.236692845 | 0.709264768 | -4.530071777 |
| **A_55_P2146546** | NA | 0.141260965 | -0.125102126 | 0.407624057 | 8.143288351 | 1.32830702 | 0.236737068 | 0.709264768 | -4.530110885 |
| **A_55_P2156288** | NA | 0.360267644 | -0.319058338 | 1.039593626 | 8.92487942 | 1.328303693 | 0.236738095 | 0.709264768 | -4.530111793 |
| **A_55_P2046145** | Gm10693 | -0.131575959 | -0.379684479 | 0.116532561 | 8.028292232 | -1.328266476 | 0.236749582 | 0.709264768 | -4.53012195 |
| **A_52_P531175** | Fkbp3 | 0.414927763 | -0.367551843 | 1.197407369 | 12.48570444 | 1.328158215 | 0.236783 | 0.709264768 | -4.530151496 |
| **A_65_P02323** | Sec22b | -0.173814645 | -0.501675819 | 0.154046529 | 8.236049943 | -1.327842981 | 0.236880331 | 0.709264768 | -4.530237527 |
| **A_55_P2159264** | Lifr | -1.195463486 | -3.450455347 | 1.059528374 | 9.21199858 | -1.327827573 | 0.236885089 | 0.709264768 | -4.530241732 |
| **A_52_P509185** | Zfp318 | 0.130561162 | -0.115752617 | 0.376874942 | 9.111024035 | 1.327625681 | 0.236947445 | 0.709264768 | -4.530296829 |
| **A_55_P1957911** | Tom1 | 0.159441107 | -0.141358802 | 0.460241015 | 8.410462443 | 1.327617434 | 0.236949992 | 0.709264768 | -4.53029908 |

| | | | | | | | | | |
|---|---|---|---|---|---|---|---|---|---|
| **A_52_P77204** | Lrrc20 | -0.380740839 | -1.099068555 | 0.337586877 | 9.436507152 | -1.32756925 | 0.236964877 | 0.709264768 | -4.530312229 |
| **A_51_P378667** | Wrap53 | 0.336111576 | -0.298118002 | 0.970341153 | 10.85372983 | 1.327355715 | 0.237030849 | 0.709264768 | -4.530370502 |
| **A_55_P1985012** | Pard3 | -0.296376043 | -0.855646009 | 0.262893922 | 8.200554792 | -1.32730871 | 0.237045374 | 0.709264768 | -4.530383329 |
| **A_55_P2181104** | Shisa3 | -0.42196987 | -1.21824009 | 0.374300351 | 7.951814912 | -1.327306781 | 0.23704597 | 0.709264768 | -4.530383855 |
| **A_51_P384629** | Ctsd | -0.829913445 | -2.396037113 | 0.736210224 | 13.76046099 | -1.327263321 | 0.2370594 | 0.709264768 | -4.530395715 |
| **A_55_P2105242** | Arhgap39 | -0.487829872 | -1.40848545 | 0.432825706 | 10.77598869 | -1.327154694 | 0.237092971 | 0.709264768 | -4.530425358 |
| **A_55_P2132517** | Gmppa | -0.312367235 | -0.901891136 | 0.277156665 | 9.383833052 | -1.327132859 | 0.237099719 | 0.709264768 | -4.530431317 |
| **A_55_P2060111** | NA | -0.184413631 | -0.532483361 | 0.163656099 | 13.67405284 | -1.3270189 | 0.237134944 | 0.709264768 | -4.530462415 |
| **A_55_P2152225** | Ihh | -0.403170745 | -1.164234942 | 0.357893452 | 7.962593517 | -1.3268385 | 0.237190714 | 0.709264768 | -4.530511642 |
| **A_55_P2163018** | NA | 0.667309047 | -0.592416998 | 1.927035093 | 11.58375019 | 1.326787129 | 0.237206598 | 0.709264768 | -4.53052566 |
| **A_52_P650108** | Uri1 | 0.197428573 | -0.175300763 | 0.570157909 | 9.125917916 | 1.326681814 | 0.237239164 | 0.709264768 | -4.530554398 |
| **A_55_P1997696** | Plbd1 | -1.505833115 | -4.348886437 | 1.337220207 | 10.81170189 | -1.326606693 | 0.237262395 | 0.709264768 | -4.530574897 |
| **A_51_P438479** | Zfp281 | 0.412403014 | -0.366252784 | 1.191058811 | 10.84500007 | 1.326559257 | 0.237277066 | 0.709264768 | -4.530587841 |
| **A_52_P412046** | NA | 0.417621356 | -0.371018707 | 1.20626142 | 15.3293283 | 1.32633799 | 0.23734551 | 0.709264768 | -4.530648217 |
| **A_55_P2152437** | Mitf | -1.481046873 | -4.277888414 | 1.315794668 | 9.312928131 | -1.326329067 | 0.23734827 | 0.709264768 | -4.530650652 |
| **A_55_P2101431** | Mrps25 | 0.221176136 | -0.196508693 | 0.638860965 | 8.83003991 | 1.326294322 | 0.23735902 | 0.709264768 | -4.530660132 |
| **A_55_P2091494** | Tango2 | -0.254476181 | -0.735138941 | 0.226186578 | 10.01412222 | -1.326041174 | 0.237437351 | 0.709264768 | -4.530729206 |
| **A_55_P2063992** | Rps5 | 0.307936707 | -0.273720056 | 0.889593469 | 17.40094029 | 1.326004582 | 0.237448676 | 0.709264768 | -4.53073919 |
| **A_55_P1987261** | Fam214b | -1.036879559 | -2.995435269 | 0.921676151 | 9.772057217 | -1.325997573 | 0.237450845 | 0.709264768 | -4.530741103 |
| **A_51_P292943** | Ell | -0.689333352 | -1.991451779 | 0.612785075 | 10.80598789 | -1.32595612 | 0.237463675 | 0.709264768 | -4.530752413 |
| **A_55_P2124183** | NA | 0.354538671 | -0.31521467 | 1.024292011 | 9.137697238 | 1.325864276 | 0.237492103 | 0.709264768 | -4.530777473 |
| **A_51_P227777** | Top2b | 0.426576956 | -0.379292161 | 1.232446073 | 12.95120981 | 1.325815902 | 0.237507077 | 0.709264768 | -4.530790672 |
| **A_55_P2043287** | Tle5 | 0.641601061 | -0.570528094 | 1.853730215 | 10.12619961 | 1.325764915 | 0.237522861 | 0.709264768 | -4.530804583 |
| **A_55_P2119011** | Osbpl9 | -0.320602796 | -0.926300244 | 0.285094652 | 8.577899426 | -1.325750828 | 0.237527222 | 0.709264768 | -4.530808427 |
| **A_51_P314339** | Chtop | 0.308245158 | -0.274190264 | 0.89068058 | 11.03282645 | 1.325558289 | 0.237586835 | 0.709264768 | -4.53086096 |
| **A_55_P2082057** | NA | 0.370421612 | -0.329503372 | 1.070346597 | 16.90671966 | 1.325547109 | 0.237590298 | 0.709264768 | -4.530864011 |
| **A_55_P1985565** | NA | 0.446943472 | -0.397595745 | 1.29148269 | 16.25386607 | 1.325510348 | 0.237601681 | 0.709264768 | -4.53087404 |
| **A_55_P2059357** | Myo7a | -0.312304437 | -0.902440906 | 0.277832032 | 8.923640073 | -1.32548875 | 0.237608369 | 0.709264768 | -4.530879933 |
| **A_52_P335259** | Cdkal1 | 0.158533688 | -0.141051967 | 0.458119344 | 9.885029098 | 1.325411987 | 0.237632142 | 0.709264768 | -4.530900877 |
| **A_51_P320401** | Rab24 | -0.623026866 | -1.800521325 | 0.554467593 | 10.7669718 | -1.325251328 | 0.237681904 | 0.709264768 | -4.53094471 |
| **A_55_P1988373** | Cenpu | 0.377005949 | -0.335636283 | 1.08964818 | 8.629182688 | 1.325033912 | 0.237749261 | 0.709264768 | -4.531004028 |
| **A_55_P2099520** | H2afv | 0.515922373 | -0.459350046 | 1.491194791 | 9.414189779 | 1.324978297 | 0.237766494 | 0.709264768 | -4.531019201 |
| **A_51_P171107** | Tmem35a | 0.204589336 | -0.182210561 | 0.591389233 | 8.227950257 | 1.324789839 | 0.237824897 | 0.709264768 | -4.531070617 |
| **A_66_P129747** | Dnajc24 | 0.386765096 | -0.344554423 | 1.118084615 | 10.4722964 | 1.324617396 | 0.237878349 | 0.709264768 | -4.531117662 |
| **A_55_P2099560** | Apobec1 | -0.161449518 | -0.466748204 | 0.143849167 | 8.826601455 | -1.324531115 | 0.237905098 | 0.709264768 | -4.5311412 |
| **A_55_P1964108** | B230118H | 0.383027488 | -0.34132911 | 1.107384085 | 10.74532442 | 1.324426517 | 0.237937528 | 0.709264768 | -4.531169736 |
| **A_55_P1962026** | Pigg | -0.611382658 | -1.767622662 | 0.544857347 | 8.382088342 | -1.324388709 | 0.237949252 | 0.709264768 | -4.53118005 |
| **A_51_P405668** | 5-Mar | 0.139425835 | -0.124261574 | 0.403113244 | 10.87501477 | 1.324354363 | 0.237959902 | 0.709264768 | -4.53118942 |
| **A_55_P2054261** | C2cd4b | 0.399827514 | -0.356471302 | 1.156126329 | 8.659232747 | 1.324126879 | 0.238030453 | 0.709264768 | -4.531251478 |
| **A_52_P86073** | Mrpl1 | 0.165904598 | -0.147922205 | 0.4797314 | 8.859084548 | 1.32409375 | 0.23804073 | 0.709264768 | -4.531260516 |
| **A_65_P15340** | Zfp251 | 0.239349181 | -0.213461548 | 0.69215991 | 9.311094094 | 1.323931706 | 0.238090999 | 0.709264768 | -4.531304721 |
| **A_51_P123805** | Sbno1 | -0.258215451 | -0.746753497 | 0.230322595 | 8.765923026 | -1.323835979 | 0.2381207 | 0.709264768 | -4.531330835 |
| **A_52_P175242** | Irs1 | 0.971416785 | -0.866590453 | 2.809424022 | 10.76539955 | 1.323758408 | 0.238144771 | 0.709264768 | -4.531351995 |
| **A_55_P2081865** | Slc2a8 | -0.363710754 | -1.051923027 | 0.32450152 | 10.63877919 | -1.323683252 | 0.238168094 | 0.709264768 | -4.531372497 |
| **A_52_P494380** | Dgkq | -0.317101349 | -0.917309986 | 0.283107288 | 9.220505307 | -1.323263077 | 0.238298525 | 0.709264768 | -4.531487112 |
| **A_51_P479132** | Zcchc3 | 0.604099883 | -0.539399704 | 1.747599471 | 9.538856041 | 1.323192666 | 0.238320388 | 0.709264768 | -4.531506319 |
| **A_51_P518298** | Rnf187 | 0.42182767 | -0.376836179 | 1.22049152 | 13.12481719 | 1.322882839 | 0.238416615 | 0.709264768 | -4.531590829 |
| **A_52_P558699** | Ppp4r3a | 0.187650455 | -0.167684036 | 0.542984947 | 9.629048356 | 1.322703805 | 0.238472235 | 0.709264768 | -4.531639663 |

| | | | | | | | | | |
|---|---|---|---|---|---|---|---|---|---|
| **A_55_P2043554** | Slc12a4 | -0.483753141 | -1.399828741 | 0.432322459 | 11.16231019 | -1.322643579 | 0.238490948 | 0.709264768 | -4.53165609 |
| **A_55_P2029945** | NA | 0.220452404 | -0.211537926 | 0.652442734 | 15.20450343 | 1.359263075 | 0.238510565 | 0.709264768 | -4.533318872 |
| **A_52_P66371** | Nlrx1 | -0.436578952 | -1.263402448 | 0.390244544 | 8.579032871 | -1.322514158 | 0.238531166 | 0.709264768 | -4.53169139 |
| **A_55_P1985764** | NA | 0.198535316 | -0.177473813 | 0.574544444 | 13.44694123 | 1.322481874 | 0.238541199 | 0.709264768 | -4.531700196 |
| **A_52_P408757** | Fcgr3 | -0.137925474 | -0.399170683 | 0.123319736 | 8.184624242 | -1.322350225 | 0.238582117 | 0.709264768 | -4.531736103 |
| **A_55_P1974228** | NA | 0.266809764 | -0.238611341 | 0.77223087 | 15.46675786 | 1.322204683 | 0.23862736 | 0.709264768 | -4.531775799 |
| **A_55_P2100305** | Lrba | 0.144501687 | -0.129239234 | 0.418242608 | 10.88796149 | 1.322158555 | 0.238641701 | 0.709264768 | -4.53178838 |
| **A_55_P2148182** | NA | 0.833922152 | -0.746089539 | 2.413933843 | 11.43447849 | 1.321951602 | 0.238706052 | 0.709264768 | -4.531844824 |
| **A_55_P1970810** | NA | -0.423834325 | -1.226916048 | 0.379247398 | 12.24059942 | -1.321863861 | 0.238733339 | 0.709264768 | -4.531868754 |
| **A_55_P2176565** | B3galt5 | -0.29654083 | -0.858434501 | 0.26535284 | 8.174639584 | -1.321845524 | 0.238739042 | 0.709264768 | -4.531873755 |
| **A_52_P592909** | Dgat2 | -0.506219051 | -1.465474619 | 0.453036518 | 11.72310357 | -1.321765773 | 0.238763848 | 0.709264768 | -4.531895506 |
| **A_55_P2127131** | Trmt112 | 0.440133501 | -0.394048444 | 1.274315447 | 12.01548245 | 1.321520748 | 0.238840073 | 0.709264768 | -4.53196233 |
| **A_52_P484956** | Nnat | 1.01007665 | -0.904320219 | 2.92447352 | 10.5892811 | 1.321516841 | 0.238841289 | 0.709264768 | -4.531963396 |
| **A_55_P2076448** | Tmem255 | 0.862336371 | -0.772050352 | 2.496723095 | 9.708440536 | 1.321515386 | 0.238841742 | 0.709264768 | -4.531963793 |
| **A_55_P1999213** | Tdrd7 | -0.376228624 | -1.08933179 | 0.336874543 | 10.93043638 | -1.321447198 | 0.238862959 | 0.709264768 | -4.531982389 |
| **A_51_P118603** | Wdr82 | 0.197994686 | -0.177343589 | 0.573332961 | 10.56573631 | 1.321237907 | 0.238928091 | 0.709264768 | -4.532537903 |
| **A_55_P2149440** | Ehmt1 | 0.214349405 | -0.1919955 | 0.620694311 | 11.11383844 | 1.321228122 | 0.238931137 | 0.709264768 | -4.532042135 |
| **A_51_P424781** | Sf3b6 | 0.258979009 | -0.232000212 | 0.74995823 | 12.33351161 | 1.321148993 | 0.238955767 | 0.709264768 | -4.532063715 |
| **A_51_P473229** | Zbtb7b | -0.710238357 | -2.056927003 | 0.636450289 | 8.992350364 | -1.32095271 | 0.239016872 | 0.709264768 | -4.532117243 |
| **A_55_P2015118** | NA | 0.73660387 | -0.660082981 | 2.13329072 | 11.72891223 | 1.320946709 | 0.239018741 | 0.709264768 | -4.532118879 |
| **A_51_P368483** | 1700123C | 0.15347754 | -0.1375344 | 0.444489479 | 10.25169901 | 1.320943879 | 0.239019622 | 0.709264768 | -4.532119651 |
| **A_51_P424854** | Ahsg | 0.948149194 | -0.849725944 | 2.746024332 | 10.93772685 | 1.320892575 | 0.239035596 | 0.709264768 | -4.532133642 |
| **A_52_P570266** | Psmb10 | 0.556507252 | -0.498864237 | 1.611878742 | 12.68014116 | 1.320735393 | 0.239084542 | 0.709264768 | -4.532176506 |
| **A_52_P257426** | Eps8l2 | -0.616864865 | -1.786746592 | 0.553016862 | 8.371601438 | -1.320682499 | 0.239101016 | 0.709264768 | -4.53219093 |
| **A_55_P2100526** | Ptges3 | 0.206477019 | -0.185110199 | 0.598064237 | 11.90030564 | 1.320667711 | 0.239105621 | 0.709264768 | -4.532194962 |
| **A_55_P1960916** | Egln3 | -0.347563267 | -1.006785524 | 0.311658991 | 9.219691453 | -1.320542411 | 0.23914465 | 0.709264768 | -4.532229131 |
| **A_55_P1979457** | Prdx4 | -0.430946881 | -1.248342203 | 0.386448442 | 13.26277361 | -1.320510744 | 0.239154514 | 0.709264768 | -4.532237767 |
| **A_55_P2012815** | Acvr2b | -1.289233398 | -3.734599897 | 1.156133101 | 11.32286079 | -1.320498569 | 0.239158307 | 0.709264768 | -4.532241087 |
| **A_55_P1973560** | NA | 1.037296958 | -0.930272688 | 3.004866603 | 12.47918456 | 1.320454181 | 0.239172135 | 0.709264768 | -4.532253191 |
| **A_55_P2051521** | NA | 0.257794192 | -0.231201807 | 0.74679019 | 8.905036352 | 1.320438468 | 0.23917703 | 0.709264768 | -4.532257476 |
| **A_55_P2116391** | Tesk2 | -0.540987805 | -1.567206442 | 0.485230832 | 8.877910859 | -1.320376918 | 0.239196206 | 0.709264768 | -4.53227426 |
| **A_52_P267543** | Ginm1 | -0.187243867 | -0.542524058 | 0.168036323 | 8.13153603 | -1.320039585 | 0.239301327 | 0.709264768 | -4.532366245 |
| **A_51_P148612** | Cox7a1 | -0.387014078 | -1.121415155 | 0.347386999 | 11.56783767 | -1.319908433 | 0.239342209 | 0.709264768 | -4.532402007 |
| **A_51_P171288** | Gli3 | 0.595494511 | -0.53464729 | 1.725636313 | 10.1223809 | 1.319760633 | 0.239388287 | 0.709264768 | -4.532442308 |
| **A_55_P1957733** | Cysrt1 | -0.346953547 | -1.005522823 | 0.311615729 | 7.976711731 | -1.319532864 | 0.239459313 | 0.709264768 | -4.532504413 |
| **A_55_P2091616** | Krt222 | -0.338752155 | -0.981804185 | 0.304299875 | 11.58181587 | -1.319429831 | 0.239491448 | 0.709264768 | -4.532532506 |
| **A_51_P407074** | Tspan31 | -0.218694207 | -0.63384734 | 0.196458927 | 12.13119213 | -1.319408577 | 0.239498077 | 0.709264768 | -4.532538301 |
| **A_55_P2410626** | A430061C | -0.151334529 | -0.438637307 | 0.135968248 | 8.244488062 | -1.319315155 | 0.239527219 | 0.709264768 | -4.532563774 |
| **A_55_P2033876** | NA | 0.654616431 | -0.588191741 | 1.897424603 | 13.16930143 | 1.319268361 | 0.239541817 | 0.709264768 | -4.532576532 |
| **A_52_P246753** | Chmp5 | -0.121730356 | -0.35284157 | 0.109380857 | 12.86993023 | -1.319253398 | 0.239546485 | 0.709264768 | -4.532580612 |
| **A_55_P2168316** | Aldh3b2 | -0.549025092 | -1.591414687 | 0.493364504 | 8.039106456 | -1.319205554 | 0.239561412 | 0.709264768 | -4.532593657 |
| **A_55_P1961965** | Plagl1 | -0.394856449 | -1.144578446 | 0.354865548 | 8.296076367 | -1.319135256 | 0.239583345 | 0.709264768 | -4.532612824 |
| **A_55_P2168899** | NA | -0.167140602 | -0.484508458 | 0.150227254 | 8.013015549 | -1.319074636 | 0.23960226 | 0.709264768 | -4.532629352 |
| **A_52_P46085** | Mvp | -0.862346449 | -2.499790169 | 0.775097272 | 8.662689347 | -1.31906362 | 0.239605698 | 0.709264768 | -4.532632355 |
| **A_55_P1981205** | Xrcc4 | 0.211342549 | -0.189963503 | 0.612648601 | 9.005937582 | 1.319050972 | 0.239609645 | 0.709264768 | -4.532635804 |
| **A_55_P2020080** | NA | -1.390297701 | -4.03033993 | 1.249744528 | 10.12717638 | -1.319007535 | 0.2396232 | 0.709264768 | -4.532647647 |
| **A_52_P33014** | Ppp1r8 | 0.228513673 | -0.205471984 | 0.66249933 | 8.541673872 | 1.318824968 | 0.239680179 | 0.709264768 | -4.532697423 |
| **A_55_P2040962** | Gm1604b | 0.237659004 | -0.213742152 | 0.68906016 | 9.443858908 | 1.318687689 | 0.239723032 | 0.709264768 | -4.532734851 |

| | | | | | | | | | |
|---|---|---|---|---|---|---|---|---|---|
| **A_55_P1956728** | NA | 0.645986089 | -0.581037475 | 1.873009652 | 14.51590676 | 1.318622883 | 0.239743264 | 0.709264768 | -4.532752519 |
| **A_55_P2185900** | Nrg4 | -0.263568859 | -0.764303676 | 0.237165957 | 7.813388012 | -1.318368 | 0.239822853 | 0.709264768 | -4.532822008 |
| **A_55_P2028171** | Dppa5a | -0.690722232 | -2.003648499 | 0.622204035 | 8.313350106 | -1.317690585 | 0.240034497 | 0.709264768 | -4.533006683 |
| **A_55_P2009022** | NA | 0.163404963 | -0.147224067 | 0.474033994 | 8.74383896 | 1.317569544 | 0.240072332 | 0.709264768 | -4.53303968 |
| **A_66_P140533** | NA | -0.252408696 | -0.732233522 | 0.22741613 | 8.379170031 | -1.317564682 | 0.240073852 | 0.709264768 | -4.533041005 |
| **A_51_P182471** | Celf2 | 0.836610897 | -0.754090236 | 2.42731203 | 11.31233426 | 1.317301761 | 0.240156054 | 0.709264768 | -4.533112678 |
| **A_55_P2160331** | Bicd2 | -0.138098137 | -0.400682154 | 0.12448588 | 10.94154356 | -1.317255068 | 0.240170655 | 0.709264768 | -4.533125406 |
| **A_51_P491051** | Ints9 | 0.190292018 | -0.171553406 | 0.552137443 | 11.3930829 | 1.317188205 | 0.240191565 | 0.709264768 | -4.533143632 |
| **A_51_P365912** | Ciao3 | -0.130169757 | -0.377722187 | 0.117382673 | 11.01601648 | -1.317022655 | 0.240243344 | 0.709264768 | -4.533188759 |
| **A_55_P2042156** | Aatk | 0.151907728 | -0.137007117 | 0.440822574 | 8.697981132 | 1.316922921 | 0.240274543 | 0.709264768 | -4.533215945 |
| **A_65_P18858** | Btf3 | 0.661037423 | -0.596235621 | 1.918310467 | 12.79078734 | 1.316881777 | 0.240287415 | 0.709264768 | -4.533227161 |
| **A_55_P2131238** | Ttc39a | -0.95824751 | -2.780817745 | 0.864322724 | 8.823339082 | -1.316872632 | 0.240290276 | 0.709264768 | -4.533229653 |
| **A_51_P116487** | Lsm4 | 0.543722033 | -0.490446561 | 1.577890626 | 13.61176327 | 1.316848862 | 0.240297712 | 0.709264768 | -4.533236133 |
| **A_55_P2111380** | Ctnnd2 | 0.636161329 | -0.573880744 | 1.846203401 | 9.429508213 | 1.316791881 | 0.24031554 | 0.709264768 | -4.533251664 |
| **A_55_P2134148** | Mzt2 | 0.539885666 | -0.487056421 | 1.566827753 | 11.26464366 | 1.31675868 | 0.240325929 | 0.709264768 | -4.533260714 |
| **A_51_P173212** | Osbpl2 | -0.609225332 | -1.768066241 | 0.549615578 | 11.08967963 | -1.316753496 | 0.240327551 | 0.709264768 | -4.533262127 |
| **A_55_P2349198** | NA | -0.189106589 | -0.548817255 | 0.170604078 | 8.757086296 | -1.3167511 | 0.2403283 | 0.709264768 | -4.533262781 |
| **A_55_P1989885** | Smarce1 | 0.167824046 | -0.151476983 | 0.487125075 | 11.98000984 | 1.316449532 | 0.240422678 | 0.709264768 | -4.53334498 |
| **A_51_P184024** | Tsen15 | 0.370406028 | -0.33435423 | 1.075166286 | 11.31436258 | 1.316397307 | 0.240439026 | 0.709264768 | -4.533359215 |
| **A_55_P2026889** | NA | 0.137185865 | -0.123838362 | 0.398210091 | 12.60535773 | 1.316372779 | 0.240446704 | 0.709264768 | -4.5333659 |
| **A_51_P371051** | Glipr1 | -0.522234751 | -1.515912386 | 0.471442884 | 8.311518963 | -1.316347638 | 0.240454575 | 0.709264768 | -4.533372753 |
| **A_66_P122086** | 9030619P | -0.897634266 | -2.605756583 | 0.810488051 | 8.459323956 | -1.316227019 | 0.240492337 | 0.709264768 | -4.533405629 |
| **A_55_P2003673** | Mtmr12 | -0.181292486 | -0.526290822 | 0.16370585 | 10.91236391 | -1.31617352 | 0.240509088 | 0.709264768 | -4.533420211 |
| **A_51_P500541** | Flywch2 | 0.643986632 | -0.581586272 | 1.869559535 | 9.564486035 | 1.316097449 | 0.240532908 | 0.709264768 | -4.533440945 |
| **A_51_P174415** | Retreg2 | -0.334244043 | -0.970353986 | 0.3018659 | 9.804232615 | -1.316078631 | 0.240538801 | 0.709264768 | -4.533446074 |
| **A_55_P2095709** | NA | 0.319410138 | -0.288522403 | 0.927342679 | 16.77140159 | 1.315962918 | 0.240575039 | 0.709264768 | -4.533477612 |
| **A_55_P1988083** | Prc1 | 1.386905259 | -1.252927233 | 4.02673775 | 12.13975514 | 1.315893587 | 0.240596754 | 0.709264768 | -4.533496509 |
| **A_52_P474089** | Capn6 | 0.839369876 | -0.75838709 | 2.437126842 | 10.52480382 | 1.31580946 | 0.240623105 | 0.709264768 | -4.533519437 |
| **A_55_P2051479** | NA | 0.133930906 | -0.121015377 | 0.38887719 | 9.903855733 | 1.315777537 | 0.240633105 | 0.709264768 | -4.533528138 |
| **A_55_P2078238** | NA | 0.117843876 | -0.106521256 | 0.342209008 | 8.351251897 | 1.315533763 | 0.240709481 | 0.709264768 | -4.533594577 |
| **A_51_P496934** | Hps3 | -0.121051165 | -0.351525523 | 0.109423192 | 8.241348748 | -1.315517736 | 0.240714504 | 0.709264768 | -4.533598945 |
| **A_55_P2098697** | Tnfaip2 | -0.994803033 | -2.88885512 | 0.899249053 | 8.566213193 | -1.315514164 | 0.240715623 | 0.709264768 | -4.533599919 |
| **A_65_P03442** | Pgk1 | 0.428259545 | -0.38720757 | 1.24372666 | 13.39472893 | 1.315379123 | 0.240757943 | 0.709264768 | -4.533636723 |
| **A_51_P330428** | Eif4ebp1 | -0.719592727 | -2.089801286 | 0.650615832 | 12.94274701 | -1.315377579 | 0.240758427 | 0.709264768 | -4.533637143 |
| **A_51_P102507** | Vps33a | -0.122524206 | -0.355834756 | 0.110786344 | 8.656744629 | -1.315339511 | 0.240770358 | 0.709264768 | -4.53517488 |
| **A_55_P1970771** | Gm13141 | 0.739938474 | -0.66906011 | 2.148937058 | 13.73712594 | 1.315332038 | 0.2407727 | 0.709264768 | -4.533649555 |
| **A_55_P2149763** | Tapbp | -0.808521939 | -2.348261665 | 0.731217786 | 10.99456493 | -1.315209205 | 0.240811202 | 0.709264768 | -4.533683031 |
| **A_51_P519189** | Eif3i | 0.421984549 | -0.38167286 | 1.225641958 | 10.63790683 | 1.315151977 | 0.240829142 | 0.709264768 | -4.533698627 |
| **A_55_P1974367** | Fabp3 | -2.081994109 | -6.047190184 | 1.883201965 | 12.42160586 | -1.31511937 | 0.240839364 | 0.709264768 | -4.533707514 |
| **A_55_P2151907** | NA | 0.483488375 | -0.437372127 | 1.404348877 | 15.31052347 | 1.315050821 | 0.240860856 | 0.709264768 | -4.533726195 |
| **A_55_P2137023** | NA | -0.147447494 | -0.428304993 | 0.133410006 | 8.535402154 | -1.314927219 | 0.240899612 | 0.709264768 | -4.533759879 |
| **A_55_P1954820** | D1Ertd622 | 0.390603654 | -0.353439874 | 1.134647182 | 10.13719886 | 1.314886612 | 0.240912345 | 0.709264768 | -4.533770946 |
| **A_51_P270184** | Nhlrc2 | 0.276380993 | -0.250098492 | 0.802860478 | 10.25489912 | 1.314852588 | 0.240923015 | 0.709264768 | -4.533780218 |
| **A_51_P341025** | Eif3k | 0.26196286 | -0.237056924 | 0.760982643 | 11.50089782 | 1.314838174 | 0.240927536 | 0.709264768 | -4.533784146 |
| **A_55_P2135601** | Polr2i | 0.187899412 | -0.170058244 | 0.545857068 | 10.05811305 | 1.314752821 | 0.240954304 | 0.709264768 | -4.533807406 |
| **A_55_P2176838** | Mitf | -1.596282313 | -4.637314094 | 1.444749469 | 10.5728734 | -1.314737615 | 0.240959074 | 0.709264768 | -4.53381155 |
| **A_55_P2183551** | NA | 0.487588487 | -0.441324649 | 1.416501622 | 15.07103952 | 1.314706117 | 0.240968953 | 0.709264768 | -4.533820134 |
| **A_51_P449824** | NA | -0.277206643 | -0.805360345 | 0.250947059 | 8.30887043 | -1.314600071 | 0.241002218 | 0.709264768 | -4.533849033 |

| | | | | | | | | | |
|---|---|---|---|---|---|---|---|---|---|
| **A_55_P1956418** | Efr3b | -0.849458713 | -2.468038202 | 0.769120777 | 10.00171464 | -1.314493939 | 0.241035513 | 0.709264768 | -4.533877955 |
| **A_55_P2165560** | Plekhg5 | 0.516977211 | -0.468182053 | 1.502136476 | 11.11472903 | 1.314362922 | 0.241076621 | 0.709264768 | -4.533913658 |
| **A_51_P196127** | Papss1 | 0.33908698 | -0.307090449 | 0.985264409 | 10.8135372 | 1.314345903 | 0.241081962 | 0.709264768 | -4.533918295 |
| **A_52_P434841** | Coa4 | -0.218551337 | -0.635173899 | 0.198071225 | 9.543087342 | -1.313896116 | 0.241223139 | 0.709264768 | -4.534040861 |
| **A_51_P198835** | Dnajb9 | -0.559030291 | -1.624740211 | 0.506679628 | 10.384939 | -1.313852702 | 0.24123677 | 0.709264768 | -4.534052691 |
| **A_65_P12533** | Naa10 | 0.618484379 | -0.560670143 | 1.797638902 | 10.73708182 | 1.313736783 | 0.241273168 | 0.709264768 | -4.534084277 |
| **A_52_P631499** | NA | 0.262527254 | -0.238048017 | 0.763102525 | 9.658796749 | 1.313576438 | 0.241323524 | 0.709264768 | -4.534127968 |
| **A_52_P634447** | Hs1bp3 | -0.26721533 | -0.776735007 | 0.242304348 | 9.404577562 | -1.313562536 | 0.24132789 | 0.709264768 | -4.534131756 |
| **A_52_P586141** | Adcy7 | -0.917078806 | -2.665756745 | 0.831599132 | 10.8227693 | -1.313551703 | 0.241331293 | 0.709264768 | -4.534134708 |
| **A_52_P261322** | Tanc1 | -0.15711632 | -0.45671385 | 0.142481209 | 8.242153674 | -1.313510101 | 0.24134436 | 0.709264768 | -4.534146044 |
| **A_55_P2348582** | NA | -0.172269245 | -0.500762045 | 0.156223556 | 7.953473104 | -1.313506541 | 0.241345478 | 0.709264768 | -4.534147014 |
| **A_55_P2138878** | Lrfn4 | 0.539792965 | -0.489536875 | 1.569122805 | 10.98117472 | 1.313478605 | 0.241354253 | 0.709264768 | -4.534154626 |
| **A_66_P133244** | Vcl | -0.270812602 | -0.787281686 | 0.245656483 | 8.015188322 | -1.313333084 | 0.241399967 | 0.709264768 | -4.534194276 |
| **A_55_P2126557** | NA | 0.358073968 | -0.324822294 | 1.04097023 | 9.939762646 | 1.31331335 | 0.241406167 | 0.709264768 | -4.534199653 |
| **A_51_P366227** | Zfp799 | 0.334123438 | -0.303144421 | 0.971391298 | 8.466379776 | 1.313213299 | 0.241437603 | 0.709264768 | -4.534226914 |
| **A_52_P393306** | Snap23 | -0.465512683 | -1.35343713 | 0.422411764 | 9.171196211 | -1.313124336 | 0.241465557 | 0.709264768 | -4.534251153 |
| **A_51_P329490** | Slc25a40 | -0.425771925 | -1.237925144 | 0.386381294 | 8.474530051 | -1.313074588 | 0.241481191 | 0.709264768 | -4.534264708 |
| **A_55_P1981599** | Ccdc125 | 0.375470455 | -0.340737798 | 1.091678708 | 8.849427707 | 1.313066717 | 0.241483664 | 0.709264768 | -4.534266852 |
| **A_51_P387123** | Oasl2 | -0.988269182 | -2.873437917 | 0.896899552 | 9.480620035 | -1.31303218 | 0.241494519 | 0.709264768 | -4.534276263 |
| **A_55_P1962010** | Clec1b | -0.372934664 | -1.084402087 | 0.338532759 | 8.240340792 | -1.312889209 | 0.241539455 | 0.709264768 | -4.534315216 |
| **A_51_P305437** | Rcn1 | 0.264330908 | -0.24003113 | 0.768692946 | 12.92998957 | 1.312671034 | 0.241608044 | 0.709264768 | -4.534374659 |
| **A_55_P1981845** | NA | 0.363874818 | -0.330523764 | 1.058273399 | 9.489503584 | 1.312482499 | 0.241667328 | 0.709264768 | -4.534426025 |
| **A_55_P2011659** | Anapc5 | 0.32450747 | -0.294780221 | 0.943795162 | 11.12336529 | 1.312449484 | 0.241677711 | 0.709264768 | -4.53443502 |
| **A_55_P2020786** | Prkar1a | 0.192275713 | -0.174672768 | 0.559224195 | 13.08542131 | 1.312410458 | 0.241689985 | 0.709264768 | -4.534445652 |
| **A_55_P1959838** | NA | 0.474783669 | -0.431388299 | 1.380955638 | 15.36176533 | 1.312307149 | 0.241722479 | 0.709264768 | -4.534473798 |
| **A_55_P1967330** | Gm5069 | 0.268696994 | -0.244266076 | 0.781660064 | 14.26347086 | 1.311979514 | 0.241825556 | 0.709264768 | -4.534563056 |
| **A_55_P2054311** | M6pr | -0.567698552 | -1.651487154 | 0.516090051 | 9.622814517 | -1.311968936 | 0.241828885 | 0.709264768 | -4.534565938 |
| **A_55_P2016851** | Lce1a2 | 0.340429207 | -0.309524344 | 0.990382758 | 10.58051099 | 1.311882195 | 0.241856181 | 0.709264768 | -4.534589569 |
| **A_55_P2092859** | Anxa4 | -0.497437521 | -1.447209465 | 0.452334424 | 12.72462536 | -1.311805807 | 0.241880222 | 0.709264768 | -4.534610379 |
| **A_52_P350554** | Kcnb1 | -0.267167104 | -0.777278458 | 0.242944251 | 8.194576624 | -1.311802145 | 0.241881375 | 0.709264768 | -4.534611376 |
| **A_51_P497821** | Zfp790 | -0.114765501 | -0.333906125 | 0.104375123 | 8.380744174 | -1.311712954 | 0.241909448 | 0.709264768 | -4.534635674 |
| **A_55_P2000369** | Aqp4 | 0.206420055 | -0.187752892 | 0.600593003 | 8.232760555 | 1.311642322 | 0.241931682 | 0.709264768 | -4.534654916 |
| **A_51_P220682** | Retreg3 | -0.377858479 | -1.099461916 | 0.343744958 | 10.0789496 | -1.311538139 | 0.24196448 | 0.709264768 | -4.534683297 |
| **A_55_P2156490** | Gm3146 | -0.440874701 | -1.28307899 | 0.401329587 | 8.043478437 | -1.311137007 | 0.2420908 | 0.709264768 | -4.534792568 |
| **A_55_P2136209** | Ublcp1 | -0.199122625 | -0.579517013 | 0.181271763 | 9.294767614 | -1.311103127 | 0.242101472 | 0.709264768 | -4.534801797 |
| **A_55_P2002846** | Cystm1 | -0.538100147 | -1.566302448 | 0.490102154 | 8.814867263 | -1.310795329 | 0.242198445 | 0.709264768 | -4.53488564 |
| **A_55_P2006280** | Ogdh | -0.299705928 | -0.872462712 | 0.273050857 | 12.28761982 | -1.31061591 | 0.242254988 | 0.709264768 | -4.534934511 |
| **A_55_P2159710** | Acvr1 | -0.350566132 | -1.020593994 | 0.319461731 | 11.67543293 | -1.310471075 | 0.242300641 | 0.709264768 | -4.534973962 |
| **A_55_P2154208** | NA | -0.772278396 | -2.248326567 | 0.703769776 | 11.02163386 | -1.310459792 | 0.242304198 | 0.709264768 | -4.534977035 |
| **A_55_P2055881** | NA | 0.296595297 | -0.27028503 | 0.863475625 | 16.68207812 | 1.310458343 | 0.242304654 | 0.709264768 | -4.53497743 |
| **A_51_P324287** | Kif23 | 0.805183183 | -0.73386194 | 2.344228307 | 9.689696092 | 1.310369236 | 0.242332746 | 0.709264768 | -4.535001701 |
| **A_52_P563450** | Brix1 | 0.245275154 | -0.223646382 | 0.71419669 | 9.10253105 | 1.310097821 | 0.242418328 | 0.709264768 | -4.535075627 |
| **A_66_P113268** | Mme | -0.478490889 | -1.393328493 | 0.436346715 | 8.170557378 | -1.310026288 | 0.242440888 | 0.709264768 | -4.53509511 |
| **A_51_P275949** | Loxl2 | -0.217710603 | -0.63398108 | 0.198559874 | 9.400182979 | -1.309948783 | 0.242465334 | 0.709264768 | -4.535116219 |
| **A_55_P2180909** | Ccdc166 | 0.254902575 | -0.232558035 | 0.742363185 | 8.738178033 | 1.309739847 | 0.242531247 | 0.709264768 | -4.535173125 |
| **A_55_P1998374** | Ppfibp1 | -0.491354013 | -1.431037358 | 0.448329331 | 10.83578731 | -1.309674356 | 0.24255191 | 0.709264768 | -4.535190962 |
| **A_52_P317923** | Zfp948 | -0.204186275 | -0.594736261 | 0.186363711 | 9.345894044 | -1.309484207 | 0.242611914 | 0.709264768 | -4.535242749 |
| **A_55_P1975793** | Srd5a3 | -0.197271538 | -0.574600567 | 0.180057491 | 10.09241567 | -1.309467 | 0.242617345 | 0.709264768 | -4.535247436 |

| | | | | | | | | | |
|---|---|---|---|---|---|---|---|---|---|
| **A_55_P2150633** | Gadd45gi | 0.306653997 | -0.279916545 | 0.893224539 | 11.77692728 | 1.309419267 | 0.24263241 | 0.709264768 | -4.535260436 |
| **A_55_P2039771** | Itgav | -0.452327201 | -1.317644733 | 0.412990331 | 8.27199086 | -1.309264967 | 0.242681115 | 0.709264768 | -4.535302458 |
| **A_51_P378009** | Blzf1 | -0.359969968 | -1.048616624 | 0.328676688 | 10.11846873 | -1.309242734 | 0.242688133 | 0.709264768 | -4.535308513 |
| **A_55_P2049262** | NA | -0.193798286 | -0.564556222 | 0.176959649 | 9.409117023 | -1.309211534 | 0.242697983 | 0.709264768 | -4.53531701 |
| **A_51_P382152** | Procr | -1.753501933 | -5.108284824 | 1.601280958 | 10.7596815 | -1.309158335 | 0.242714779 | 0.709264768 | -4.535331498 |
| **A_55_P2112230** | Pcyt2 | 0.174255089 | -0.159206866 | 0.507717044 | 11.90292738 | 1.308848932 | 0.242812481 | 0.709264768 | -4.535415759 |
| **A_55_P2025248** | Mxd1 | -0.303282378 | -0.883688495 | 0.277123739 | 9.370118081 | -1.30877666 | 0.242835308 | 0.709264768 | -4.53543544 |
| **A_55_P2026562** | Cdkn1b | -0.167096023 | -0.486887605 | 0.152695559 | 11.74676593 | -1.308728115 | 0.242850642 | 0.709264768 | -4.53544866 |
| **A_55_P2100209** | Kif2c | 0.290473273 | -0.265487643 | 0.84643419 | 8.328201409 | 1.30861611 | 0.242886025 | 0.709264768 | -4.535479162 |
| **A_55_P2055682** | Atp5o | 0.468899056 | -0.428576668 | 1.366374781 | 13.43812066 | 1.308600235 | 0.24289104 | 0.709264768 | -4.535483485 |
| **A_55_P2184113** | Aurkaip1 | -0.291379932 | -0.849128353 | 0.266368489 | 14.70131017 | -1.308493694 | 0.242924701 | 0.709264768 | -4.535512498 |
| **A_55_P2317341** | NA | -0.168381852 | -0.49069531 | 0.153931606 | 8.28267289 | -1.308480298 | 0.242928934 | 0.709264768 | -4.535516146 |
| **A_55_P2120189** | Gas2l3 | 0.435535494 | -0.398202252 | 1.269273241 | 8.878886478 | 1.308411749 | 0.242950595 | 0.709264768 | -4.535534813 |
| **A_55_P2102579** | NA | 0.481137835 | -0.439910092 | 1.402185761 | 15.39913189 | 1.308391233 | 0.242957078 | 0.709264768 | -4.535540399 |
| **A_51_P472901** | Slc3a2 | -1.302464668 | -3.796029869 | 1.191100533 | 11.56307985 | -1.308264534 | 0.242997118 | 0.709264768 | -4.535574901 |
| **A_52_P79639** | Vat1 | -0.420972826 | -1.226947493 | 0.385001841 | 9.85190518 | -1.308226725 | 0.243009068 | 0.709264768 | -4.535585196 |
| **A_66_P124958** | Ccdc47 | -0.30622942 | -0.892547271 | 0.280088432 | 8.673795677 | -1.308169863 | 0.243027041 | 0.709264768 | -4.53560068 |
| **A_65_P16542** | NA | -0.206376686 | -0.601520041 | 0.188766668 | 11.83808069 | -1.308146242 | 0.243034507 | 0.709264768 | -4.535607112 |
| **A_55_P2012331** | NA | 0.648310881 | -0.593052386 | 1.889674147 | 13.28428583 | 1.308081388 | 0.243055009 | 0.709264768 | -4.535624773 |
| **A_55_P2230968** | NA | -0.306309431 | -0.892879022 | 0.28026016 | 8.417824307 | -1.307950082 | 0.24309652 | 0.709264768 | -4.535660527 |
| **A_55_P2006375** | Dmrtc1c2 | -1.026074861 | -2.991008889 | 0.938859167 | 8.171691298 | -1.307920719 | 0.243105804 | 0.709264768 | -4.535668523 |
| **A_51_P163778** | 2810025M | 0.969029899 | -0.88667175 | 2.824731548 | 11.8413414 | 1.307914544 | 0.243107757 | 0.709264768 | -4.535670204 |
| **A_55_P2175692** | Reep3 | -0.732360978 | -2.135055662 | 0.670333707 | 9.998069189 | -1.307712846 | 0.243171538 | 0.709264768 | -4.535725125 |
| **A_55_P1985673** | Zbtb37 | -0.165495453 | -0.482490576 | 0.15149967 | 8.340814513 | -1.307626853 | 0.243198735 | 0.709264768 | -4.53574854 |
| **A_55_P2080649** | Zfp691 | -0.138034623 | -0.40245043 | 0.126381185 | 8.80350591 | -1.307527896 | 0.243230036 | 0.709264768 | -4.535775484 |
| **A_55_P2052892** | Xiap | 0.277965741 | -0.254512894 | 0.810444377 | 12.54158972 | 1.307493179 | 0.243241018 | 0.709264768 | -4.535784937 |
| **A_55_P2098300** | Chmp6 | -0.273007992 | -0.796113882 | 0.250097898 | 8.355196426 | -1.307182113 | 0.243339439 | 0.709264768 | -4.535869634 |
| **A_52_P208600** | Bms1 | 0.146564318 | -0.134284072 | 0.427412708 | 9.091948659 | 1.307093512 | 0.243367479 | 0.709264768 | -4.535893757 |
| **A_52_P376106** | Slc30a9 | -0.16983968 | -0.495290798 | 0.155611437 | 8.854910442 | -1.307084734 | 0.243370257 | 0.709264768 | -4.535896147 |
| **A_55_P2125841** | Cog1 | -0.186101414 | -0.542729992 | 0.170527164 | 9.598978555 | -1.307024947 | 0.24338918 | 0.709264768 | -4.535912425 |
| **A_52_P623775** | Eri2 | 0.309561605 | -0.283735341 | 0.902858551 | 8.657570952 | 1.306848738 | 0.243444958 | 0.709264768 | -4.535960401 |
| **A_55_P2047598** | Arhgdia | -0.301181202 | -0.878475577 | 0.276113173 | 15.39903891 | -1.306715017 | 0.243487295 | 0.709264768 | -4.535996808 |
| **A_55_P2429225** | Psrc1 | 0.379046161 | -0.347578677 | 1.105670999 | 8.960629046 | 1.306568575 | 0.243533667 | 0.709264768 | -4.536036677 |
| **A_55_P2047225** | Uxs1 | -0.247962772 | -0.723399835 | 0.22747429 | 8.351382011 | -1.306302611 | 0.243617907 | 0.709264768 | -4.536109085 |
| **A_66_P107734** | NA | 0.464535221 | -0.426167578 | 1.355238021 | 13.94225251 | 1.306279695 | 0.243625167 | 0.709264768 | -4.536115324 |
| **A_55_P2042818** | Zfp639 | -0.150305144 | -0.438504378 | 0.137894091 | 10.6355 | -1.306265227 | 0.24362975 | 0.709264768 | -4.536119263 |
| **A_52_P182331** | NA | 0.459517286 | -0.421601648 | 1.340636219 | 15.56278417 | 1.306224017 | 0.243642806 | 0.709264768 | -4.536130482 |
| **A_51_P257675** | Tspyl4 | 0.353405084 | -0.324341678 | 1.031151846 | 8.791733744 | 1.30603762 | 0.243701866 | 0.709264768 | -4.536181226 |
| **A_52_P250578** | Tcaf1 | 0.315703714 | -0.289829172 | 0.9212366 | 8.549703317 | 1.305847019 | 0.24376227 | 0.709264768 | -4.536233114 |
| **A_52_P561690** | BC034090 | -0.13503392 | -0.394038555 | 0.123970714 | 7.960908698 | -1.305827131 | 0.243768574 | 0.709264768 | -4.536238528 |
| **A_55_P2032558** | NA | 0.550142357 | -0.505116669 | 1.605401382 | 15.00325007 | 1.305769 | 0.243787 | 0.709264768 | -4.536254353 |
| **A_66_P129508** | Hyls1 | 0.26226159 | -0.240843553 | 0.765366733 | 8.325428792 | 1.305648512 | 0.243825196 | 0.709264768 | -4.536287152 |
| **A_55_P2060991** | Plppr3 | 0.522692817 | -0.480013121 | 1.525398755 | 8.824767601 | 1.305639531 | 0.243828043 | 0.709264768 | -4.536289597 |
| **A_55_P2088692** | NA | -1.412203157 | -4.121328058 | 1.296921743 | 10.41172103 | -1.305625115 | 0.243832613 | 0.709264768 | -4.536293521 |
| **A_55_P2036526** | Mbd2 | -0.258502733 | -0.754414321 | 0.237408855 | 9.474158716 | -1.305603265 | 0.24383954 | 0.709264768 | -4.536299469 |
| **A_55_P2178803** | Zbtb5 | 0.352303621 | -0.323565594 | 1.028172837 | 10.96115065 | 1.305583905 | 0.243845678 | 0.709264768 | -4.53630474 |
| **A_55_P2286493** | 4933405E | -0.170576528 | -0.497822426 | 0.15666937 | 9.039643754 | -1.305555698 | 0.243854622 | 0.709264768 | -4.536312418 |
| **A_55_P2033445** | Tnfrsf1b | -0.344801346 | -1.006305025 | 0.316702333 | 8.20814529 | -1.305530537 | 0.2438626 | 0.709264768 | -4.536319267 |

| | | | | | | | | | |
|---|---|---|---|---|---|---|---|---|---|
| **A_55_P2064686** | Lrrc61 | -0.175814997 | -0.513142104 | 0.16151211 | 9.480691226 | -1.305434307 | 0.243893113 | 0.709264768 | -4.536345463 |
| **A_55_P2067153** | NA | 0.27173633 | -0.249645206 | 0.793117865 | 13.10199891 | 1.30539638 | 0.24390514 | 0.709264768 | -4.536355787 |
| **A_51_P392090** | Tmem19 | -0.359159687 | -1.048317169 | 0.329997795 | 10.06129976 | -1.305327399 | 0.243927017 | 0.709264768 | -4.536374564 |
| **A_55_P1968285** | Tle5 | 0.282959548 | -0.259994623 | 0.82591372 | 14.61296923 | 1.305303567 | 0.243934575 | 0.709264768 | -4.536381052 |
| **A_55_P2019233** | Gsto2 | -0.460950471 | -1.345611215 | 0.423710273 | 8.289930051 | -1.305052109 | 0.244014339 | 0.709264768 | -4.5364495 |
| **A_55_P2164724** | Gnaq | 0.213490934 | -0.196251315 | 0.623233183 | 11.02827953 | 1.305025614 | 0.244022744 | 0.709264768 | -4.536456712 |
| **A_52_P515497** | Atp1a1 | -0.434179405 | -1.267573702 | 0.399214893 | 13.9435123 | -1.304875389 | 0.244070409 | 0.709264768 | -4.536497603 |
| **A_55_P2162765** | Dnajc24 | 0.380052813 | -0.349475864 | 1.109581491 | 10.51313268 | 1.304823978 | 0.244086723 | 0.709264768 | -4.536511597 |
| **A_51_P118539** | 2700049A | 0.472723419 | -0.434827331 | 1.380274169 | 10.39971162 | 1.304627544 | 0.244149066 | 0.709264768 | -4.536565064 |
| **A_55_P1981381** | Lamc1 | -0.166472687 | -0.486083869 | 0.153138495 | 14.24257726 | -1.304581965 | 0.244163534 | 0.709264768 | -4.53657747 |
| **A_52_P605556** | Nxnl2 | -0.158681431 | -0.463348121 | 0.145985258 | 8.358770766 | -1.304522219 | 0.244182499 | 0.709264768 | -4.536593732 |
| **A_66_P126189** | Ercc5 | -0.12736103 | -0.371935113 | 0.117213052 | 8.80880036 | -1.304296944 | 0.244254022 | 0.709264768 | -4.536655048 |
| **A_51_P264527** | Dipk1b | 0.129943618 | -0.119605191 | 0.379492427 | 8.184354246 | 1.304216827 | 0.244279463 | 0.709264768 | -4.536676853 |
| **A_51_P347862** | Actn1 | 0.122312674 | -0.11258885 | 0.357214199 | 8.898587604 | 1.304175305 | 0.24429265 | 0.709264768 | -4.536688155 |
| **A_55_P2022812** | Cyp4f17 | -0.432794623 | -1.264003424 | 0.398414179 | 8.395434174 | -1.304133552 | 0.24430591 | 0.709264768 | -4.536699519 |
| **A_52_P161707** | Cand1 | 0.136574604 | -0.125748299 | 0.398897507 | 9.318969547 | 1.304019503 | 0.244342133 | 0.709264768 | -4.536730559 |
| **A_55_P2092869** | Anxa9 | -0.580821096 | -1.696456025 | 0.534813834 | 8.266372402 | -1.303979081 | 0.244354973 | 0.709264768 | -4.536741561 |
| **A_55_P2044557** | Rpl29 | 0.14703559 | -0.135405533 | 0.429476713 | 16.74518038 | 1.303901799 | 0.244379523 | 0.709264768 | -4.536762594 |
| **A_52_P517762** | Slc44a4 | -0.551720156 | -1.611561418 | 0.508121106 | 8.07452601 | -1.303852224 | 0.244395273 | 0.709264768 | -4.536776087 |
| **A_51_P242356** | Fam114a2 | -0.162204455 | -0.473819876 | 0.149410966 | 12.24313622 | -1.303749584 | 0.244427883 | 0.709264768 | -4.536804021 |
| **A_55_P2084338** | 2310009B | 0.51610031 | -0.47541313 | 1.507613751 | 10.14055948 | 1.303724599 | 0.244435822 | 0.709264768 | -4.536810821 |
| **A_55_P1960813** | NA | 0.608776835 | -0.560795884 | 1.778349554 | 15.0396805 | 1.303710713 | 0.244440234 | 0.709264768 | -4.5368146 |
| **A_51_P146760** | Pcdhb20 | 0.375742657 | -0.346140856 | 1.097626171 | 8.515492139 | 1.30368817 | 0.244447397 | 0.709264768 | -4.536820735 |
| **A_55_P2079761** | Ggps1 | -0.233424513 | -0.681906202 | 0.215057175 | 8.598818832 | -1.303623234 | 0.244468033 | 0.709264768 | -4.536838408 |
| **A_51_P103718** | Ip6k1 | -0.218890064 | -0.639464905 | 0.201684777 | 10.71011606 | -1.30356624 | 0.244486145 | 0.709264768 | -4.539134177 |
| **A_51_P114826** | Cdh13 | -0.686483615 | -2.005506872 | 0.632539641 | 8.980978966 | -1.303551125 | 0.244490949 | 0.709264768 | -4.536858032 |
| **A_55_P2128853** | Ear12 | -0.323544802 | -0.945393387 | 0.298303784 | 9.335319427 | -1.303167142 | 0.244613011 | 0.709264768 | -4.536962531 |
| **A_55_P1954418** | NA | 0.293584898 | -0.270687642 | 0.857857438 | 11.402249 | 1.303152201 | 0.244617761 | 0.709264768 | -4.536966598 |
| **A_55_P2073308** | Bin1 | 0.5238239 | -0.483032109 | 1.530679909 | 11.54767168 | 1.303071628 | 0.244643382 | 0.709264768 | -4.536988524 |
| **A_51_P176505** | Rps19bp1 | -0.160306493 | -0.468446822 | 0.147833835 | 10.88476621 | -1.30302551 | 0.244658047 | 0.709264768 | -4.537001074 |
| **A_52_P553890** | Itgb3 | -1.735570575 | -5.071768059 | 1.600626909 | 10.47796836 | -1.302989372 | 0.244669539 | 0.709264768 | -4.537010909 |
| **A_55_P2025601** | Lsm6 | 0.24917301 | -0.229829675 | 0.728175696 | 14.21762827 | 1.302906937 | 0.244695757 | 0.709264768 | -4.537033342 |
| **A_51_P193000** | Bbs4 | -0.184004326 | -0.537732904 | 0.169724253 | 9.468737185 | -1.302891436 | 0.244700687 | 0.709264768 | -4.53703756 |
| **A_51_P404084** | Psmb2 | 0.452239225 | -0.417276499 | 1.321754949 | 10.2160054 | 1.302690162 | 0.244764712 | 0.709264768 | -4.537092332 |
| **A_55_P1991906** | Acadvl | -0.146405962 | -0.427910288 | 0.135098363 | 12.99308942 | -1.302638879 | 0.244781027 | 0.709264768 | -4.537106287 |
| **A_51_P237754** | H2-T23 | -1.306264847 | -3.818129118 | 1.205599425 | 11.60017438 | -1.302523047 | 0.244817882 | 0.709264768 | -4.537137807 |
| **A_55_P2009993** | Tomm20 | 0.189488464 | -0.174897187 | 0.553874114 | 12.38833132 | 1.302482373 | 0.244830825 | 0.709264768 | -4.537148875 |
| **A_51_P159352** | Sub1 | 0.207895139 | -0.191887279 | 0.607677556 | 8.73802286 | 1.30247978 | 0.24483165 | 0.709264768 | -4.537149581 |
| **A_51_P300572** | Slc30a1 | -0.551742801 | -1.612847154 | 0.509361553 | 9.451614122 | -1.302353628 | 0.244871796 | 0.709264768 | -4.537183908 |
| **A_55_P2042202** | Pigo | -0.335396101 | -0.980469453 | 0.30967725 | 10.62538029 | -1.302264658 | 0.244900113 | 0.709264768 | -4.537208118 |
| **A_55_P2012221** | Armcx2 | 1.152707824 | -1.064435722 | 3.36985137 | 12.23462168 | 1.302194254 | 0.244922523 | 0.709264768 | -4.537227275 |
| **A_52_P380301** | Unc5c | 0.735067119 | -0.678934493 | 2.149068731 | 9.447601089 | 1.302049327 | 0.24496866 | 0.709264768 | -4.53726671 |
| **A_55_P2017526** | Dand5 | -0.179831234 | -0.525808626 | 0.166146159 | 8.329275429 | -1.301870385 | 0.245025637 | 0.709264768 | -4.5373154 |
| **A_55_P2009588** | Rpl13 | 0.38081179 | -0.351884073 | 1.113507653 | 16.44894938 | 1.301778193 | 0.245054996 | 0.709264768 | -4.537340484 |
| **A_55_P2154387** | Bmp4 | 0.399382248 | -0.369052567 | 1.167817062 | 12.05880811 | 1.301763439 | 0.245059695 | 0.709264768 | -4.537344499 |
| **A_55_P2065731** | Medag | -0.219288162 | -0.641234059 | 0.202657735 | 16.66655732 | -1.301693584 | 0.245081943 | 0.709264768 | -4.537363506 |
| **A_55_P2007738** | Mir22hg | -0.473475439 | -1.384627802 | 0.437676924 | 8.012951186 | -1.301537828 | 0.245131557 | 0.709264768 | -4.537405884 |
| **A_51_P295625** | Arf4 | 0.298654126 | -0.276079747 | 0.873387999 | 11.36109613 | 1.301523671 | 0.245136067 | 0.709264768 | -4.537409736 |

| | | | | | | | | | |
|---|---|---|---|---|---|---|---|---|---|
| **A_55_P1977917** | Cln3 | -0.735752737 | -2.15172933 | 0.680223855 | 10.1720609 | -1.301446015 | 0.245160807 | 0.709264768 | -4.537430865 |
| **A_52_P108845** | Clip3 | 0.511420467 | -0.472894841 | 1.495735776 | 9.651414074 | 1.301350283 | 0.245191309 | 0.709264768 | -4.537456911 |
| **A_55_P2033215** | Cabp5 | 0.313720589 | -0.290104662 | 0.91754584 | 10.96704413 | 1.301313984 | 0.245202876 | 0.709264768 | -4.537466788 |
| **A_51_P234253** | Sdcbp2 | -0.3395976 | -0.993236528 | 0.314041328 | 8.093165042 | -1.301298859 | 0.245207695 | 0.709264768 | -4.537470903 |
| **A_55_P2102385** | Etl4 | -0.336149238 | -0.983153994 | 0.310855518 | 10.1491095 | -1.301292731 | 0.245209648 | 0.709264768 | -4.53747257 |
| **A_52_P380329** | Tent4a | 0.175863439 | -0.162649541 | 0.514376418 | 10.37062426 | 1.301219555 | 0.245232967 | 0.709264768 | -4.537492479 |
| **A_66_P111840** | NA | 0.513883454 | -0.475278461 | 1.503045369 | 15.14069575 | 1.301210603 | 0.24523582 | 0.709264768 | -4.537494915 |
| **A_52_P180565** | 2310061I( | -0.34792581 | -1.017681888 | 0.321830267 | 9.386797875 | -1.301128919 | 0.245261853 | 0.709264768 | -4.537517138 |
| **A_55_P2164047** | NA | 0.639454327 | -0.591556192 | 1.870464846 | 13.88907293 | 1.301062358 | 0.245283068 | 0.709264768 | -4.537535247 |
| **A_51_P133792** | Prickle3 | -0.115985461 | -0.339272644 | 0.107301722 | 9.095165991 | -1.301038368 | 0.245290715 | 0.709264768 | -4.537541774 |
| **A_55_P2000553** | Slc6a8 | -0.826888285 | -2.41875686 | 0.76498029 | 11.94385328 | -1.301037982 | 0.245290838 | 0.709264768 | -4.537541879 |
| **A_51_P501897** | Pitx2 | -0.222862526 | -0.651986073 | 0.206261021 | 8.089056063 | -1.300783586 | 0.245371939 | 0.709264768 | -4.537611091 |
| **A_55_P2012780** | Rnf167 | -0.608587723 | -1.780510144 | 0.563334698 | 11.7430573 | -1.300692601 | 0.245400951 | 0.709264768 | -4.537635843 |
| **A_51_P388478** | Efnb1 | -0.158350204 | -0.463300385 | 0.146599976 | 12.87852088 | -1.300589007 | 0.245433987 | 0.709264768 | -4.537664026 |
| **A_51_P275953** | Dcps | 0.444178863 | -0.411237111 | 1.299594837 | 11.02535995 | 1.300561453 | 0.245442775 | 0.709264768 | -4.537671522 |
| **A_55_P2118609** | St6galnac | 0.282972877 | -0.26200176 | 0.827947513 | 9.518408518 | 1.300525479 | 0.245454249 | 0.709264768 | -4.537681309 |
| **A_55_P2145691** | Hlcs | -0.419041883 | -1.226083361 | 0.387999594 | 8.238997413 | -1.300504692 | 0.245460879 | 0.709264768 | -4.537686964 |
| **A_51_P454309** | Zfp772 | -0.410019886 | -1.199688746 | 0.379648974 | 8.504558058 | -1.300499692 | 0.245462473 | 0.709264768 | -4.537688324 |
| **A_51_P432520** | Wrnip1 | 0.298991324 | -0.276846714 | 0.874829362 | 10.43766256 | 1.300494686 | 0.24546407 | 0.709264768 | -4.537689686 |
| **A_55_P1996952** | NA | -1.476449128 | -4.320044721 | 1.367146464 | 9.676363327 | -1.300471985 | 0.245471311 | 0.709264768 | -4.537695862 |
| **A_52_P324767** | Tmed8 | 0.176039456 | -0.163030053 | 0.515108964 | 9.092200715 | 1.300384026 | 0.245499368 | 0.709264768 | -4.537719791 |
| **A_66_P137048** | Men1 | 0.164493679 | -0.152349937 | 0.481337294 | 11.73185242 | 1.300333043 | 0.245515632 | 0.709264768 | -4.53773366 |
| **A_55_P2005285** | Klhl22 | -0.54237289 | -1.587106535 | 0.502360756 | 12.55516216 | -1.300297531 | 0.245526961 | 0.709264768 | -4.537743321 |
| **A_51_P254912** | Rfc2 | 0.290439581 | -0.269071108 | 0.849950271 | 10.76143683 | 1.300162871 | 0.245569925 | 0.709264768 | -4.537779953 |
| **A_51_P464710** | Adap2 | -0.510019691 | -1.492614254 | 0.472574872 | 8.263847105 | -1.300058612 | 0.245603194 | 0.709264768 | -4.537808315 |
| **A_55_P2047639** | Cubn | -1.455835263 | -4.260703022 | 1.349032496 | 9.644433695 | -1.300020467 | 0.245615367 | 0.709264768 | -4.537818692 |
| **A_55_P2179106** | Ppig | 0.438321417 | -0.406303955 | 1.282946789 | 9.531278203 | 1.299807126 | 0.24568346 | 0.709264768 | -4.537876726 |
| **A_65_P03728** | St3gal1 | -0.576146506 | -1.686466285 | 0.534173273 | 11.24517434 | -1.299676307 | 0.245725223 | 0.709264768 | -4.537912312 |
| **A_52_P98614** | Slc6a2 | -1.274264865 | -3.730293543 | 1.181763813 | 8.443524255 | -1.299501008 | 0.245781195 | 0.709264768 | -4.537959996 |
| **A_52_P588881** | Iqgap3 | 0.344792499 | -0.319766516 | 1.009351515 | 8.988182488 | 1.299494965 | 0.245783125 | 0.709264768 | -4.537961639 |
| **A_51_P251129** | Stard8 | -0.632437666 | -1.851410798 | 0.586535466 | 9.694124321 | -1.299492985 | 0.245783757 | 0.709264768 | -4.537962178 |
| **A_51_P481920** | Ccna2 | 1.395508605 | -1.294558196 | 4.085575405 | 11.40517518 | 1.299330998 | 0.24583549 | 0.709264768 | -4.53800624 |
| **A_55_P2043207** | Magef1 | 0.27194652 | -0.252287244 | 0.796180284 | 9.631910508 | 1.299298276 | 0.245845941 | 0.709264768 | -4.53801514 |
| **A_55_P2090880** | Stk40 | -0.245521134 | -0.71887917 | 0.227836902 | 12.04996531 | -1.299120612 | 0.245902694 | 0.709264768 | -4.538063465 |
| **A_52_P36680** | Pggt1b | -0.132222279 | -0.387153907 | 0.122709349 | 8.57269403 | -1.299066147 | 0.245920094 | 0.709264768 | -4.53807828 |
| **A_52_P168271** | Spg20 | -0.30179291 | -0.883775593 | 0.280189774 | 8.180051473 | -1.298821039 | 0.245998416 | 0.709264768 | -4.538144947 |
| **A_52_P323830** | Zfp830 | 0.136973472 | -0.127210215 | 0.401157159 | 9.186856179 | 1.298616188 | 0.24606389 | 0.709264768 | -4.538200663 |
| **A_51_P157297** | Efcab14 | -0.201665219 | -0.590623622 | 0.187293185 | 11.53393565 | -1.298608341 | 0.246066399 | 0.709264768 | -4.538202797 |
| **A_65_P19286** | Alg10b | 0.118121776 | -0.109724378 | 0.34596793 | 8.75962352 | 1.298490011 | 0.246104227 | 0.709264768 | -4.538234981 |
| **A_51_P117581** | Cables1 | -0.691884452 | -2.026543588 | 0.642774685 | 9.498959264 | -1.298415086 | 0.246128183 | 0.709264768 | -4.538255358 |
| **A_55_P2035159** | Pcdh1 | 0.144955792 | -0.134693681 | 0.424605264 | 10.15098345 | 1.298290602 | 0.246167988 | 0.709264768 | -4.538289215 |
| **A_55_P2172121** | Mxd4 | -0.130358692 | -0.381859769 | 0.121142384 | 8.964313449 | -1.298226566 | 0.246188466 | 0.709264768 | -4.53830663 |
| **A_55_P2043837** | Cmtr1 | -0.619582917 | -1.815063238 | 0.575897404 | 12.68140804 | -1.298097606 | 0.246229711 | 0.709264768 | -4.538341703 |
| **A_52_P670026** | Rsad2 | -1.159899167 | -3.397963289 | 1.078164955 | 8.743409513 | -1.298069832 | 0.246238595 | 0.709264768 | -4.538349256 |
| **A_55_P1969993** | NA | 0.475618073 | -0.442287278 | 1.393523424 | 17.37794873 | 1.297809033 | 0.246322029 | 0.709264768 | -4.538420183 |
| **A_52_P520495** | Vcam1 | 0.225339479 | -0.209568001 | 0.660246959 | 8.272048072 | 1.297749155 | 0.246341188 | 0.709264768 | -4.538436466 |
| **A_55_P2116900** | NA | 0.205322849 | -0.191015341 | 0.60166104 | 9.010105263 | 1.29754284 | 0.246407214 | 0.709264768 | -4.538492573 |
| **A_52_P316437** | 1810013L | -0.532024411 | -1.559157383 | 0.495108561 | 8.359948145 | -1.297344259 | 0.24647078 | 0.709264768 | -4.538546575 |

| | | | | | | | | | |
|---|---|---|---|---|---|---|---|---|---|
| **A_55_P2003353** | Gemin8 | 0.462860469 | -0.430792874 | 1.356513812 | 9.173074536 | 1.29727292 | 0.24649362 | 0.709264768 | -4.538565975 |
| **A_51_P334685** | Esyt1 | -0.184219907 | -0.539919546 | 0.171479731 | 9.490735705 | -1.297189671 | 0.246520275 | 0.709264768 | -4.538588613 |
| **A_55_P2121195** | Lcorl | 0.37586754 | -0.349911083 | 1.101646162 | 8.946921544 | 1.297122506 | 0.246541781 | 0.709264768 | -4.538606877 |
| **A_55_P2048284** | Aasdhppt | 0.180692899 | -0.168223322 | 0.52960912 | 8.52432716 | 1.297090475 | 0.246552038 | 0.709264768 | -4.538615587 |
| **A_55_P2000728** | Myef2 | 0.308223786 | -0.287087033 | 0.903534606 | 8.819933753 | 1.296799161 | 0.246645343 | 0.709264768 | -4.538694802 |
| **A_51_P465161** | Hoga1 | -0.175087307 | -0.513286893 | 0.16311228 | 7.869535068 | -1.29667738 | 0.246684358 | 0.709264768 | -4.538727916 |
| **A_51_P487818** | Fabp1 | 0.287263752 | -0.26763727 | 0.842164775 | 8.054322517 | 1.296628758 | 0.246699936 | 0.709264768 | -4.538741137 |
| **A_55_P1971759** | Zfp688 | -0.232537351 | -0.681745936 | 0.216671235 | 9.563907273 | -1.29656716 | 0.246719673 | 0.709264768 | -4.538757886 |
| **A_51_P483220** | Impact | -0.534209186 | -1.566181192 | 0.497762819 | 9.801657194 | -1.296563469 | 0.246720856 | 0.709264768 | -4.538758889 |
| **A_51_P182296** | Washc3 | -0.133744373 | -0.392129701 | 0.124640956 | 11.44143869 | -1.296456692 | 0.246755073 | 0.709264768 | -4.538787923 |
| **A_52_P409746** | Serpinc1 | 0.356783229 | -0.33252714 | 1.046093598 | 8.363458111 | 1.296402818 | 0.246772339 | 0.709264768 | -4.538802571 |
| **A_51_P272817** | Map2k3 | -0.292373148 | -0.857285253 | 0.272538957 | 9.351915891 | -1.296304262 | 0.246803927 | 0.709264768 | -4.538829369 |
| **A_55_P2098558** | Slc39a1 | -0.300647335 | -0.881640967 | 0.280346297 | 13.62848045 | -1.296093498 | 0.246871491 | 0.709264768 | -4.538886675 |
| **A_51_P220062** | Mmp15 | -0.577187349 | -1.692651648 | 0.538276949 | 9.117677196 | -1.296019316 | 0.246895276 | 0.709264768 | -4.538906844 |
| **A_55_P2110915** | Flrt1 | 0.148072703 | -0.138111771 | 0.434257177 | 8.261482166 | 1.29592325 | 0.24692608 | 0.709264768 | -4.538932963 |
| **A_51_P167763** | Kcnk13 | 0.513011035 | -0.478594651 | 1.50461672 | 9.368398991 | 1.295800205 | 0.24696554 | 0.709264768 | -4.538966417 |
| **A_55_P2003720** | NA | 0.546112094 | -0.50954946 | 1.601773648 | 15.14415576 | 1.295708878 | 0.246994832 | 0.709264768 | -4.538991247 |
| **A_51_P101858** | Dlg1 | -0.262791936 | -0.770820883 | 0.245237011 | 8.501590177 | -1.295608895 | 0.247026903 | 0.709264768 | -4.53901843 |
| **A_51_P363914** | Cct2 | 0.253398885 | -0.236485695 | 0.743283465 | 14.04797371 | 1.295571171 | 0.247039005 | 0.709264768 | -4.539028686 |
| **A_51_P227165** | 2310030G | -0.769881665 | -2.258328753 | 0.718565422 | 10.31235821 | -1.295510455 | 0.247058484 | 0.709264768 | -4.539045193 |
| **A_51_P112677** | Mrpl50 | 0.577397575 | -0.539026742 | 1.693821891 | 11.16792438 | 1.295376499 | 0.247101464 | 0.709264768 | -4.540787959 |
| **A_55_P2074647** | Snap47 | -0.181387674 | -0.532135867 | 0.16936052 | 10.06371934 | -1.295277059 | 0.247133374 | 0.709264768 | -4.539108646 |
| **A_55_P1966274** | Pdzd4 | 0.517317705 | -0.483061586 | 1.517696996 | 9.110688476 | 1.295218369 | 0.247152209 | 0.709264768 | -4.539124601 |
| **A_51_P406429** | Pdk1 | 0.309319237 | -0.28888857 | 0.907527045 | 9.770443768 | 1.295105653 | 0.247188386 | 0.709264768 | -4.539155244 |
| **A_55_P1954511** | Btbd10 | -0.277866103 | -0.815296872 | 0.259564665 | 10.22417267 | -1.29498098 | 0.247228407 | 0.709264768 | -4.539189137 |
| **A_51_P103865** | Ppp1r35 | 0.146797869 | -0.137132823 | 0.430728561 | 8.904205265 | 1.294964181 | 0.2472338 | 0.709264768 | -4.539193704 |
| **A_51_P129012** | B2m | -0.688528232 | -2.020352312 | 0.643295848 | 13.65003448 | -1.294867216 | 0.247264931 | 0.709264768 | -4.539220064 |
| **A_52_P206861** | Klf8 | -0.885070171 | -2.597134361 | 0.826994018 | 8.635097503 | -1.294815849 | 0.247281424 | 0.709264768 | -4.539234028 |
| **A_52_P444253** | Gm6900 | -0.437942196 | -1.285118053 | 0.409233662 | 14.71298846 | -1.294772796 | 0.247295248 | 0.709264768 | -4.539245731 |
| **A_55_P2047673** | Srr | -0.416660367 | -1.222679961 | 0.389359227 | 9.665566675 | -1.294753037 | 0.247301593 | 0.709264768 | -4.539251103 |
| **A_52_P394539** | Tex2 | -0.714818614 | -2.097657089 | 0.66801986 | 10.53802851 | -1.294716684 | 0.247313266 | 0.709264768 | -4.539260985 |
| **A_51_P292357** | Rps3a1 | 0.401904503 | -0.375728077 | 1.179537082 | 14.97402693 | 1.29449021 | 0.247386003 | 0.709264768 | -4.539322549 |
| **A_55_P2011132** | Armc10 | 0.382116542 | -0.357239429 | 1.121472512 | 11.3903493 | 1.294471817 | 0.247391911 | 0.709264768 | -4.539327549 |
| **A_52_P127682** | Dagla | 0.481906451 | -0.450576971 | 1.414389874 | 8.985331212 | 1.294410318 | 0.247411666 | 0.709264768 | -4.539344267 |
| **A_55_P2135098** | Rplp0 | 0.317422498 | -0.296805364 | 0.931650359 | 17.41274123 | 1.294370296 | 0.247424523 | 0.709264768 | -4.539355146 |
| **A_55_P2109559** | Trim6 | -0.396326719 | -1.163273376 | 0.370619938 | 7.851425725 | -1.294310691 | 0.247443673 | 0.709264768 | -4.539371348 |
| **A_52_P173368** | NA | 0.15130887 | -0.141501255 | 0.444118996 | 8.422405693 | 1.294281202 | 0.247453147 | 0.709264768 | -4.539379364 |
| **A_55_P2068566** | NA | 0.430624165 | -0.402896407 | 1.264144738 | 14.45811495 | 1.293994468 | 0.247545287 | 0.709264768 | -4.539457304 |
| **A_55_P2011169** | NA | 0.13593211 | -0.12719128 | 0.3990555 | 8.334100827 | 1.293936453 | 0.247563934 | 0.709264768 | -4.539473074 |
| **A_55_P2038347** | Acot3 | -0.135150932 | -0.396772368 | 0.126470505 | 7.928161124 | -1.293886148 | 0.247580104 | 0.709264768 | -4.539486747 |
| **A_55_P1953758** | Serinc2 | -0.664945935 | -1.952194641 | 0.622302771 | 9.871122087 | -1.293821063 | 0.247601025 | 0.709264768 | -4.539504438 |
| **A_55_P2053933** | Foxa1 | 1.57592606 | -1.475247167 | 4.627099286 | 11.63978752 | 1.293657521 | 0.247653603 | 0.709264768 | -4.53954889 |
| **A_51_P165914** | Zfp770 | 0.39548304 | -0.370221436 | 1.161187517 | 9.409686232 | 1.293650688 | 0.2476558 | 0.709264768 | -4.539550747 |
| **A_55_P2027386** | Gpr156 | -1.071442737 | -3.145961978 | 1.003076505 | 8.529786627 | -1.29360547 | 0.247670339 | 0.709264768 | -4.539563038 |
| **A_52_P435670** | Npepl1 | -0.461480921 | -1.355037971 | 0.432076129 | 10.41376691 | -1.293545801 | 0.247689526 | 0.709264768 | -4.539579256 |
| **A_51_P482029** | Mcoln1 | -0.782542732 | -2.297856971 | 0.732771508 | 11.81315135 | -1.293468043 | 0.247714532 | 0.709264768 | -4.53960039 |
| **A_52_P281145** | Kank4 | 0.254841133 | -0.238740214 | 0.748422481 | 8.705319311 | 1.293186416 | 0.247805117 | 0.709264768 | -4.539676934 |
| **A_55_P2025611** | Psme2 | -0.152818931 | -0.448803452 | 0.14316559 | 11.65114632 | -1.293178581 | 0.247807638 | 0.709264768 | -4.539679064 |

| | | | | | | | | | |
|---|---|---|---|---|---|---|---|---|---|
| **A_55_P2425661** | Ncbp3 | 0.22881523 | -0.214366287 | 0.671996747 | 11.00742891 | 1.293164084 | 0.247812302 | 0.709264768 | -4.539683004 |
| **A_51_P472932** | Atp8b1 | -0.178949893 | -0.525656697 | 0.167756911 | 7.904111707 | -1.292764519 | 0.247940876 | 0.709264768 | -4.539791597 |
| **A_55_P2062418** | Atp6v0a1 | -0.727998503 | -2.139012647 | 0.68301564 | 10.95325025 | -1.292258701 | 0.248103728 | 0.709264768 | -4.53992906 |
| **A_55_P2072391** | Acp1 | 0.163917398 | -0.153794166 | 0.481628961 | 8.635090074 | 1.292237603 | 0.248110523 | 0.709264768 | -4.539934793 |
| **A_55_P2104422** | Patj | -0.385874974 | -1.133889404 | 0.362139456 | 8.668615424 | -1.292072797 | 0.248163606 | 0.709264768 | -4.539979579 |
| **A_51_P358906** | U2af2 | 0.128247604 | -0.120374359 | 0.376869567 | 8.687860479 | 1.291992868 | 0.248189354 | 0.709264768 | -4.5400013 |
| **A_55_P2010332** | Ewsr1 | 0.679021373 | -0.637349908 | 1.995392653 | 11.80467537 | 1.291978811 | 0.248193883 | 0.709264768 | -4.540005119 |
| **A_66_P116771** | Vps8 | -0.369697279 | -1.08641751 | 0.347022952 | 10.07991346 | -1.29195366 | 0.248201985 | 0.709264768 | -4.540011954 |
| **A_55_P2088725** | Usp14 | 0.123528183 | -0.115980312 | 0.363036679 | 11.79136063 | 1.291800585 | 0.248251305 | 0.709264768 | -4.54005355 |
| **A_51_P517495** | Cox8a | -0.136300658 | -0.400591042 | 0.127989725 | 16.10019954 | -1.291715698 | 0.24827866 | 0.709264768 | -4.540076617 |
| **A_51_P313337** | Smim10l1 | 0.633332109 | -0.594717853 | 1.861382072 | 10.47962942 | 1.291712364 | 0.248279734 | 0.709264768 | -4.540077523 |
| **A_51_P267239** | Litaf | -0.651238626 | -1.914186078 | 0.611708827 | 12.68291526 | -1.291532132 | 0.248337822 | 0.709264768 | -4.540126498 |
| **A_51_P479583** | Obsl1 | -0.378969336 | -1.113964767 | 0.356026095 | 10.50678558 | -1.291426747 | 0.248371792 | 0.709264768 | -4.540155133 |
| **A_55_P2069961** | NA | 0.220480867 | -0.207138611 | 0.648100344 | 16.63748905 | 1.291408898 | 0.248377546 | 0.709264768 | -4.540159983 |
| **A_66_P140976** | Rpl15 | 0.842709342 | -0.791807618 | 2.477226301 | 11.48565804 | 1.291334407 | 0.248401561 | 0.709264768 | -4.540180224 |
| **A_55_P2154049** | Myo18a | -0.220300525 | -0.647599186 | 0.206998137 | 8.275393754 | -1.291321389 | 0.248405758 | 0.709264768 | -4.540183761 |
| **A_51_P368496** | Tmem98 | 0.807438687 | -0.758734459 | 2.373611832 | 10.8204115 | 1.291279114 | 0.248419388 | 0.709264768 | -4.540195248 |
| **A_55_P2171143** | Hnrnpc | 0.243176881 | -0.228600714 | 0.714954476 | 14.28677122 | 1.291026979 | 0.248500694 | 0.709264768 | -4.540263756 |
| **A_51_P508349** | Cdc16 | 0.315600066 | -0.296741516 | 0.927941649 | 13.2929349 | 1.290903205 | 0.248540617 | 0.709264768 | -4.540297387 |
| **A_52_P188678** | Nectin2 | -0.274540948 | -0.807262309 | 0.258180412 | 10.50191603 | -1.290795267 | 0.248575436 | 0.709264768 | -4.540326713 |
| **A_52_P218955** | D3Ertd751 | 0.140976891 | -0.132596861 | 0.414550642 | 8.458138233 | 1.290695654 | 0.248607573 | 0.709264768 | -4.540353778 |
| **A_55_P2178658** | NA | 0.467619701 | -0.439871287 | 1.37511069 | 12.13725349 | 1.290627232 | 0.24862965 | 0.709264768 | -4.540372368 |
| **A_55_P1979728** | Atf4 | -0.389534066 | -1.145587681 | 0.366519549 | 14.86475915 | -1.290455982 | 0.248684913 | 0.709264768 | -4.540418895 |
| **A_51_P225948** | Snx33 | -0.341283161 | -1.003779667 | 0.321213345 | 9.417384646 | -1.290273014 | 0.248743969 | 0.709264768 | -4.540468604 |
| **A_66_P129469** | NA | 0.186110608 | -0.175296447 | 0.547517663 | 8.165753561 | 1.28980735 | 0.248894327 | 0.709264768 | -4.540595112 |
| **A_51_P158678** | Fam181b | 0.691568755 | -0.65142995 | 2.03456746 | 9.317061825 | 1.289763598 | 0.248908459 | 0.709264768 | -4.540606998 |
| **A_55_P2185573** | Rbms1 | -0.398010406 | -1.17098292 | 0.374962108 | 13.51171034 | -1.289676306 | 0.248936655 | 0.709264768 | -4.540630711 |
| **A_55_P2102816** | Rpl36a | 0.564525324 | -0.531994425 | 1.661045073 | 15.04562558 | 1.289488035 | 0.248997478 | 0.709264768 | -4.540681856 |
| **A_55_P2181246** | NA | -0.143280107 | -0.421589561 | 0.135029347 | 9.85595236 | -1.289461211 | 0.249006145 | 0.709264768 | -4.540689143 |
| **A_51_P312310** | Mtbp | 0.708434705 | -0.667654113 | 2.084523523 | 10.21572021 | 1.28944763 | 0.249010534 | 0.709264768 | -4.540692832 |
| **A_52_P89267** | Castor2 | -0.182130858 | -0.535927877 | 0.171666162 | 8.270175262 | -1.289376377 | 0.249033557 | 0.709264768 | -4.540712188 |
| **A_55_P2177105** | NA | 0.589141898 | -0.555410829 | 1.733694625 | 11.56219426 | 1.289242016 | 0.249076978 | 0.709264768 | -4.540748686 |
| **A_66_P121976** | Slc4a11 | -0.477769103 | -1.405969739 | 0.450431533 | 8.017116075 | -1.289218546 | 0.249084564 | 0.709264768 | -4.540755062 |
| **A_52_P367565** | Mrps2 | 0.220269367 | -0.207674172 | 0.648212906 | 9.692342031 | 1.289193109 | 0.249092785 | 0.709264768 | -4.540761971 |
| **A_55_P1991931** | Nosip | -0.306804037 | -0.902894177 | 0.289286104 | 10.4587155 | -1.289138197 | 0.249110534 | 0.709264768 | -4.540776888 |
| **A_55_P1993728** | Tmem116 | -0.541369551 | -1.593298252 | 0.510559149 | 8.680937674 | -1.28901469 | 0.249150458 | 0.709264768 | -4.540810436 |
| **A_51_P415225** | Zfp105 | 0.588052618 | -0.554588895 | 1.730694131 | 9.890314448 | 1.289010738 | 0.249151736 | 0.709264768 | -4.54081151 |
| **A_55_P2047559** | Ccp110 | 0.567311376 | -0.535046097 | 1.669668849 | 9.88354996 | 1.288989522 | 0.249158594 | 0.709264768 | -4.540817273 |
| **A_55_P1958921** | Ankrd29 | 0.314834288 | -0.296956152 | 0.926624727 | 8.797613534 | 1.288931044 | 0.2491775 | 0.709264768 | -4.540833157 |
| **A_55_P2121877** | Gpr137b | -0.441879305 | -1.300594887 | 0.416836276 | 8.157599617 | -1.288856793 | 0.249201507 | 0.709264768 | -4.540853326 |
| **A_52_P232830** | Ets1 | 0.522609505 | -0.493027035 | 1.538246045 | 10.01094166 | 1.288811292 | 0.24921622 | 0.709264768 | -4.540865685 |
| **A_51_P140321** | Mocos | -0.53106459 | -1.563313561 | 0.501184382 | 8.318074708 | -1.288585473 | 0.249289249 | 0.709264768 | -4.540927023 |
| **A_51_P345663** | Rrbp1 | -0.715849863 | -2.107376589 | 0.675676863 | 12.0747503 | -1.288489072 | 0.249320431 | 0.709264768 | -4.540953207 |
| **A_55_P1983958** | NA | -0.292034333 | -0.859758301 | 0.275689635 | 12.56196066 | -1.28838906 | 0.249352784 | 0.709264768 | -4.540980371 |
| **A_52_P281702** | Igfbp5 | 1.219759723 | -1.151555595 | 3.591075042 | 11.49088249 | 1.288354348 | 0.249364015 | 0.709264768 | -4.540989799 |
| **A_55_P2052690** | Synm | 0.35929865 | -0.339250673 | 1.057847974 | 8.466279116 | 1.288275818 | 0.249389422 | 0.709264768 | -4.541011129 |
| **A_52_P633481** | Akap11 | 0.225339739 | -0.212827493 | 0.663506972 | 8.919405761 | 1.288096016 | 0.249447605 | 0.709264768 | -4.541059963 |
| **A_66_P123804** | Fbxo30 | -0.23688704 | -0.697568719 | 0.223794638 | 8.62427493 | -1.287925396 | 0.249502827 | 0.709264768 | -4.541106303 |

| | | | | | | | | | |
|---|---|---|---|---|---|---|---|---|---|
| **A_51_P198434** | H2-K1 | -1.407211716 | -4.144129807 | 1.329706374 | 13.16277119 | -1.287798716 | 0.249543835 | 0.709264768 | -4.541926139 |
| **A_51_P518926** | Borcs6 | -0.240814849 | -0.709207618 | 0.22757792 | 9.06735295 | -1.287725888 | 0.249567413 | 0.709264768 | -4.541160487 |
| **A_55_P2038117** | Tmem104 | -1.072912331 | -3.159794575 | 1.013969913 | 9.261921218 | -1.287705761 | 0.24957393 | 0.709264768 | -4.541165953 |
| **A_55_P2007713** | Tspan4 | -0.152309107 | -0.448572613 | 0.143954399 | 10.68398581 | -1.287650672 | 0.249591767 | 0.709264768 | -4.541180915 |
| **A_65_P04472** | Atp6v1c1 | -0.396982583 | -1.169214236 | 0.375249069 | 9.295340816 | -1.287579937 | 0.249614672 | 0.709264768 | -4.541200125 |
| **A_66_P126504** | Trabd2b | 0.362376692 | -0.342636568 | 1.067389952 | 9.94021938 | 1.287399443 | 0.249673125 | 0.709264768 | -4.541249143 |
| **A_55_P2008567** | Rps6 | 0.808500184 | -0.764901625 | 2.381901992 | 12.11906997 | 1.287036381 | 0.249790742 | 0.709264768 | -4.541347738 |
| **A_55_P1999364** | Krt85 | -0.476449169 | -1.403678569 | 0.450780231 | 11.61400329 | -1.287003494 | 0.249801398 | 0.709264768 | -4.541356669 |
| **A_55_P2088690** | NA | -1.360656262 | -4.008673474 | 1.287360951 | 9.741493261 | -1.28699828 | 0.249803088 | 0.709264768 | -4.541358085 |
| **A_55_P2104689** | NA | -0.138626672 | -0.408429337 | 0.131175992 | 8.266263623 | -1.286918102 | 0.24982907 | 0.709264768 | -4.541379858 |
| **A_55_P1961265** | NA | -0.563564142 | -1.660417237 | 0.533288952 | 8.634629472 | -1.286901281 | 0.249834521 | 0.709264768 | -4.541384425 |
| **A_55_P2037156** | Pja1 | 0.518169284 | -0.490337443 | 1.526676012 | 12.23954737 | 1.28689529 | 0.249836463 | 0.709264768 | -4.541386052 |
| **A_55_P2144931** | NA | 0.516904701 | -0.489153369 | 1.522962771 | 14.76511492 | 1.286879191 | 0.249841681 | 0.709264768 | -4.541390424 |
| **A_55_P1997991** | Ecpas | -0.264021121 | -0.777909391 | 0.249867148 | 8.233775182 | -1.286827438 | 0.249858454 | 0.709264768 | -4.541404478 |
| **A_55_P2138071** | NA | 0.67490997 | -0.638787391 | 1.988607331 | 13.02573279 | 1.286769806 | 0.249877133 | 0.709264768 | -4.541420128 |
| **A_55_P2037454** | Etv5 | 1.49482132 | -1.414906533 | 4.404549172 | 13.99603732 | 1.2867295 | 0.249890198 | 0.709264768 | -4.541431073 |
| **A_55_P1970164** | Rnf11 | -0.299052478 | -0.881218522 | 0.283113566 | 10.33388849 | -1.286621725 | 0.249925134 | 0.709264768 | -4.541460338 |
| **A_55_P2073489** | NA | 0.20782459 | -0.196791595 | 0.612440776 | 9.12592069 | 1.286482928 | 0.249970134 | 0.709264768 | -4.541498028 |
| **A_51_P193563** | Cdk7 | -0.138265987 | -0.407484325 | 0.130952351 | 9.333057118 | -1.286355671 | 0.250011398 | 0.709264768 | -4.541532582 |
| **A_66_P117838** | Dhx29 | 0.156923617 | -0.148700546 | 0.46254778 | 9.131280879 | 1.28602961 | 0.250117155 | 0.709264768 | -4.541621117 |
| **A_55_P2041823** | Hyal1 | -0.185239209 | -0.546012144 | 0.175533726 | 9.278287514 | -1.286024711 | 0.250118744 | 0.709264768 | -4.541622447 |
| **A_52_P578768** | Slc38a9 | -0.22284789 | -0.656897003 | 0.211201224 | 8.900709893 | -1.285937904 | 0.250146906 | 0.709264768 | -4.541646017 |
| **A_55_P2022604** | Exoc3l4 | -1.196388246 | -3.526689146 | 1.133912654 | 10.20490085 | -1.285909724 | 0.250156049 | 0.709264768 | -4.541653668 |
| **A_51_P331021** | Ttc32 | 0.348374007 | -0.330220181 | 1.026968195 | 9.226612566 | 1.285837104 | 0.250179612 | 0.709264768 | -4.541673385 |
| **A_55_P2016216** | NA | 0.326358519 | -0.309371538 | 0.962088576 | 11.80957743 | 1.285797431 | 0.250192485 | 0.709264768 | -4.541684157 |
| **A_51_P501312** | Natd1 | -0.890297352 | -2.624581439 | 0.843986734 | 12.54269448 | -1.285775625 | 0.250199562 | 0.709264768 | -4.541690078 |
| **A_55_P2096657** | NA | 0.216998087 | -0.205754655 | 0.63975083 | 14.77619001 | 1.285641307 | 0.250243152 | 0.709264768 | -4.541726546 |
| **A_55_P2009677** | Zmiz1 | -0.46895151 | -1.382584515 | 0.444681496 | 12.49610783 | -1.285601894 | 0.250255944 | 0.709264768 | -4.541737247 |
| **A_55_P2454896** | Amn1 | -0.308609992 | -0.909872967 | 0.292652984 | 9.671114084 | -1.285570406 | 0.250266164 | 0.709264768 | -4.541745796 |
| **A_55_P1981479** | Irgm1 | -0.17924894 | -0.528489878 | 0.169991998 | 8.04772491 | -1.285528757 | 0.250279683 | 0.709264768 | -4.541757104 |
| **A_55_P1954266** | Zfp672 | 0.215127707 | -0.204203396 | 0.63445881 | 9.756456323 | 1.284960026 | 0.250464353 | 0.709264768 | -4.541911509 |
| **A_51_P282424** | Eapp | -0.218055207 | -0.643121493 | 0.20701108 | 9.84554361 | -1.284872829 | 0.250492677 | 0.709264768 | -4.541935182 |
| **A_55_P1991500** | Stn1 | 0.131877217 | -0.125202129 | 0.388956564 | 11.1226261 | 1.28485149 | 0.250499609 | 0.709264768 | -4.541940974 |
| **A_55_P2052106** | BC049762 | -0.580892159 | -1.713315837 | 0.551531518 | 8.352743361 | -1.284804115 | 0.250514999 | 0.709264768 | -4.541953836 |
| **A_55_P2035951** | Haus8 | 0.269700705 | -0.256127611 | 0.795529021 | 11.45610538 | 1.284660758 | 0.250561576 | 0.709264768 | -4.541992753 |
| **A_55_P2020341** | Camk2g | 0.184189162 | -0.174966575 | 0.5433449 | 12.10875444 | 1.284492609 | 0.250616218 | 0.709264768 | -4.5420384 |
| **A_52_P59264** | Sncaip | 0.787287283 | -0.747937681 | 2.322512247 | 9.81265771 | 1.284433305 | 0.250635491 | 0.709264768 | -4.542054498 |
| **A_55_P2169391** | Atp6v1a | -0.136937548 | -0.403986853 | 0.130111758 | 11.85841164 | -1.284344229 | 0.250664444 | 0.709264768 | -4.542078679 |
| **A_55_P2093949** | Zfp428 | 0.113694998 | -0.108063389 | 0.335453386 | 8.063200974 | 1.284137859 | 0.250731532 | 0.709264768 | -4.542134698 |
| **A_52_P527463** | Fam91a1 | -0.255999733 | -0.755320278 | 0.243320812 | 12.29384124 | -1.284134229 | 0.250732712 | 0.709264768 | -4.542135684 |
| **A_55_P1995152** | NA | 0.230280852 | -0.218883147 | 0.679444851 | 14.64390068 | 1.284112973 | 0.250739623 | 0.709264768 | -4.542141454 |
| **A_55_P2144391** | NA | 0.309529695 | -0.294244489 | 0.913303879 | 12.65020441 | 1.284038738 | 0.250763761 | 0.709264768 | -4.542161604 |
| **A_55_P1959648** | Hspg2 | -0.354144708 | -1.044955953 | 0.336666537 | 8.306083105 | -1.284019749 | 0.250769935 | 0.709264768 | -4.542166759 |
| **A_52_P448006** | Nit2 | 0.444895998 | -0.422994334 | 1.31278633 | 9.76139183 | 1.283937827 | 0.250796575 | 0.709264768 | -4.542188996 |
| **A_51_P438200** | Sil1 | -0.738666576 | -2.179641601 | 0.702308449 | 12.64437123 | -1.283932936 | 0.250798166 | 0.709264768 | -4.542190324 |
| **A_55_P2049211** | Pisd-ps1 | -0.628731896 | -1.855427935 | 0.597964142 | 11.2030656 | -1.283745328 | 0.250859183 | 0.709264768 | -4.542241248 |
| **A_51_P201884** | Dsp | -0.302562952 | -0.892884949 | 0.287759045 | 7.867256289 | -1.283740159 | 0.250860864 | 0.709264768 | -4.54224265 |
| **A_55_P2151685** | Pira11 | -0.116815866 | -0.344743742 | 0.111112009 | 7.990403536 | -1.283673975 | 0.250882393 | 0.709264768 | -4.542260615 |

| | | | | | | | | | |
|---|---|---|---|---|---|---|---|---|---|
| **A_52_P574668** | Nt5e | -1.124472457 | -3.319031296 | 1.070086382 | 8.85930557 | -1.283370164 | 0.250981241 | 0.709264768 | -4.542343077 |
| **A_51_P499061** | Ube2o | -0.79445256 | -2.344935331 | 0.75603021 | 12.2720168 | -1.283368484 | 0.250981787 | 0.709264768 | -4.542343533 |
| **A_51_P124798** | Tmem192 | -0.372303886 | -1.098920545 | 0.354312772 | 8.860403523 | -1.283342464 | 0.250990255 | 0.709264768 | -4.543521637 |
| **A_66_P108059** | Ttc39c | -0.119327608 | -0.352259229 | 0.113604012 | 8.320427782 | -1.283106882 | 0.25106693 | 0.709264768 | -4.542414535 |
| **A_55_P2073945** | Eif2b3 | 0.128191339 | -0.122046888 | 0.378429566 | 8.281796728 | 1.283084841 | 0.251074105 | 0.709264768 | -4.542420517 |
| **A_55_P1958833** | Gm9079 | -0.67382375 | -1.989188837 | 0.641541338 | 9.963806726 | -1.283069996 | 0.251078938 | 0.709264768 | -4.542424546 |
| **A_55_P2032768** | NA | -0.23449759 | -0.692269454 | 0.223274274 | 13.13085876 | -1.283038339 | 0.251089243 | 0.709264768 | -4.542433138 |
| **A_55_P2046084** | Dcun1d4 | -0.318999534 | -0.941739297 | 0.30374023 | 9.67646857 | -1.283021093 | 0.251094857 | 0.709264768 | -4.542437819 |
| **A_55_P2008896** | NA | 0.865659588 | -0.824346967 | 2.555666142 | 13.07879469 | 1.282948136 | 0.25111861 | 0.709264768 | -4.54245762 |
| **A_52_P564951** | C2cd3 | 0.258334975 | -0.24611185 | 0.762781799 | 8.797687199 | 1.282679519 | 0.251206079 | 0.709264768 | -4.542530522 |
| **A_65_P09443** | Psma4 | 0.204780653 | -0.195134614 | 0.60469592 | 12.68921056 | 1.282541085 | 0.251251168 | 0.709264768 | -4.542568091 |
| **A_55_P2087225** | Hnrnpa3 | 0.404363656 | -0.38537165 | 1.194098962 | 11.67642145 | 1.282451371 | 0.251280392 | 0.709264768 | -4.542592438 |
| **A_51_P104897** | Itpr3 | 0.434841106 | -0.414455873 | 1.284138084 | 11.85152243 | 1.282393647 | 0.251299197 | 0.709264768 | -4.545361193 |
| **A_51_P315904** | Gadd45g | -0.498611355 | -1.472716264 | 0.475493554 | 14.11954737 | -1.282055359 | 0.251409429 | 0.709264768 | -4.542699907 |
| **A_55_P2071799** | Slbp | 0.299377602 | -0.28551293 | 0.884268134 | 8.82837889 | 1.282020766 | 0.251420703 | 0.709264768 | -4.542709294 |
| **A_66_P119376** | Kctd12 | 0.304232817 | -0.290160114 | 0.898625748 | 9.827038884 | 1.281984484 | 0.251432529 | 0.709264768 | -4.54271914 |
| **A_55_P1991377** | NA | 0.429345183 | -0.40966542 | 1.268355787 | 15.66064935 | 1.281709169 | 0.25152228 | 0.709264768 | -4.542793849 |
| **A_55_P2078780** | Etv4 | 0.313674218 | -0.299322183 | 0.926670619 | 8.968315124 | 1.281655321 | 0.251539838 | 0.709264768 | -4.542808461 |
| **A_65_P10450** | Tmpo | 0.331314486 | -0.31625692 | 0.978885893 | 8.572939831 | 1.281454253 | 0.251605408 | 0.709264768 | -4.54286302 |
| **A_66_P118513** | 11-Sep | 0.425190276 | -0.405878507 | 1.256259059 | 12.99242608 | 1.281435349 | 0.251611573 | 0.709264768 | -4.54286815 |
| **A_55_P2062622** | Nono | 0.492036533 | -0.469752383 | 1.453825449 | 14.17677675 | 1.281350456 | 0.251639263 | 0.709264768 | -4.542891185 |
| **A_55_P2082439** | Spout1 | 0.317894531 | -0.303588163 | 0.939377224 | 11.74077089 | 1.281162924 | 0.251700439 | 0.709264768 | -4.542942069 |
| **A_55_P2180099** | NA | 0.56411408 | -0.538747219 | 1.666975379 | 14.2422821 | 1.281139401 | 0.251708114 | 0.709264768 | -4.542948452 |
| **A_55_P2052719** | Nadk2 | 0.503740775 | -0.481096863 | 1.488578413 | 10.12668617 | 1.281128856 | 0.251711554 | 0.709264768 | -4.542951313 |
| **A_51_P111259** | Cdk2ap2 | -0.345208476 | -1.020147102 | 0.329730149 | 11.92878946 | -1.281054227 | 0.251735904 | 0.709264768 | -4.542971562 |
| **A_55_P2049186** | Plvap | -0.239779532 | -0.708606326 | 0.229047262 | 8.863148292 | -1.281002655 | 0.251752733 | 0.709264768 | -4.542985555 |
| **A_51_P152845** | Trim24 | 0.241463252 | -0.23073466 | 0.713661164 | 10.91027438 | 1.280788232 | 0.251822711 | 0.709264768 | -4.543043733 |
| **A_52_P281209** | Kdm4a | 0.262443407 | -0.250846325 | 0.775733139 | 10.77707051 | 1.280629307 | 0.251874589 | 0.709264768 | -4.543086852 |
| **A_52_P558087** | Poc5 | 0.198695272 | -0.189919638 | 0.587310182 | 10.21035184 | 1.280614515 | 0.251879418 | 0.709264768 | -4.543090865 |
| **A_55_P2084251** | Kcna4 | 0.298417097 | -0.306165912 | 0.903000107 | 11.48822354 | 1.314711522 | 0.251914338 | 0.709264768 | -4.542898066 |
| **A_55_P2039444** | Cmss1 | 0.472671913 | -0.451898589 | 1.397242415 | 10.2750404 | 1.280472066 | 0.251925926 | 0.709264768 | -4.543129512 |
| **A_55_P2013188** | Dnajb6 | -0.764615009 | -2.260245891 | 0.731015872 | 11.8008101 | -1.280468033 | 0.251927243 | 0.709264768 | -4.543130607 |
| **A_55_P2003266** | Fam107a | -0.272858118 | -0.806590932 | 0.260874696 | 9.786128188 | -1.280452053 | 0.251932461 | 0.709264768 | -4.543134942 |
| **A_52_P134924** | Wdr20 | 0.239089252 | -0.228635694 | 0.706814198 | 8.938916327 | 1.280323937 | 0.251974297 | 0.709264768 | -4.5431697 |
| **A_55_P2067629** | Ush1g | 0.176821483 | -0.169124998 | 0.522767965 | 8.268851205 | 1.280195974 | 0.25201609 | 0.709264768 | -4.543204416 |
| **A_51_P175567** | Dact1 | 0.366211235 | -0.350385439 | 1.082807909 | 8.854362791 | 1.279991899 | 0.252082753 | 0.709264768 | -4.54325978 |
| **A_51_P128075** | Tescl | -0.274218912 | -0.810982795 | 0.26254497 | 7.876860828 | -1.279571221 | 0.252220223 | 0.709264768 | -4.543971586 |
| **A_52_P594049** | Actr5 | 0.135853294 | -0.130092926 | 0.401799514 | 9.695928513 | 1.27945995 | 0.252256596 | 0.709264768 | -4.543404085 |
| **A_55_P2072841** | NA | 0.630843844 | -0.604188448 | 1.865876137 | 10.16900117 | 1.27936334 | 0.252288179 | 0.709264768 | -4.543430292 |
| **A_52_P547491** | Lurap1 | -0.627250004 | -1.855365058 | 0.60086505 | 8.811967555 | -1.279239802 | 0.252328572 | 0.709264768 | -4.543463803 |
| **A_55_P1960941** | Wac | 0.120471503 | -0.11541437 | 0.356357376 | 11.89423625 | 1.279183183 | 0.252347086 | 0.709264768 | -4.543479161 |
| **A_55_P1954774** | NA | -0.379786679 | -1.123440604 | 0.363867247 | 11.92009136 | -1.279143294 | 0.25236013 | 0.709264768 | -4.543489981 |
| **A_51_P443403** | Smoc1 | -0.66783102 | -1.975539938 | 0.639877897 | 8.127038254 | -1.279103974 | 0.252372989 | 0.709264768 | -4.543500647 |
| **A_55_P1972128** | Ccndbp1 | -0.124986092 | -0.369738706 | 0.119766522 | 9.347825993 | -1.279041677 | 0.252393364 | 0.709264768 | -4.543517545 |
| **A_51_P196629** | Fam102b | -0.441858271 | -1.30716895 | 0.423452408 | 11.95283397 | -1.27897269 | 0.252415928 | 0.709264768 | -4.543536258 |
| **A_55_P1996941** | Ube2c | 1.034216418 | -0.991242051 | 3.059674888 | 11.11408588 | 1.278905429 | 0.252437929 | 0.709264768 | -4.543554502 |
| **A_55_P2062015** | Acyp1 | 0.204230623 | -0.195766215 | 0.60422746 | 8.31962524 | 1.2788354 | 0.252460837 | 0.709264768 | -4.543573497 |
| **A_55_P2024799** | NA | 0.685322039 | -0.657088828 | 2.027732906 | 13.60022654 | 1.278673264 | 0.252513884 | 0.709264768 | -4.543617474 |

| | | | | | | | | | |
|---|---|---|---|---|---|---|---|---|---|
| **A_55_P2084332** | Pigp | -0.412256244 | -1.219821961 | 0.395309472 | 8.110170479 | -1.278614763 | 0.252533026 | 0.709264768 | -4.543633342 |
| **A_55_P2058137** | Pde4dip | -1.087484916 | -3.218091984 | 1.043122152 | 9.558180659 | -1.278410191 | 0.252599975 | 0.709264768 | -4.543688827 |
| **A_52_P354373** | Lockd | 1.065847032 | -1.022597465 | 3.15429153 | 10.53435262 | 1.278269092 | 0.252646161 | 0.709264768 | -4.543727097 |
| **A_55_P2009582** | Nkg7 | 0.258530935 | -0.248044102 | 0.765105973 | 9.29845826 | 1.278259644 | 0.252649254 | 0.709264768 | -4.543729659 |
| **A_51_P149623** | Gabrd | -0.514765162 | -1.523430155 | 0.493899831 | 8.000177218 | -1.278240414 | 0.252655549 | 0.709264768 | -4.543734874 |
| **A_51_P440790** | Dpp7 | -1.00879872 | -2.985676686 | 0.968079246 | 9.427319115 | -1.278129937 | 0.252691718 | 0.709264768 | -4.543764838 |
| **A_51_P327796** | Itgb5 | -0.978678308 | -2.896641666 | 0.939285049 | 12.45217785 | -1.27805635 | 0.252715813 | 0.709264768 | -4.543784795 |
| **A_66_P124179** | Atp6v0d2 | -0.402496574 | -1.191388732 | 0.386395584 | 7.88317892 | -1.277894188 | 0.252768916 | 0.709264768 | -4.543828775 |
| **A_51_P365578** | Ppp2r1a | 0.217433665 | -0.208744868 | 0.643612199 | 9.726702493 | 1.277866726 | 0.25277791 | 0.709264768 | -4.543836222 |
| **A_51_P511236** | Pik3r1 | -0.270061835 | -0.799444453 | 0.259320783 | 8.768480581 | -1.277744094 | 0.252818076 | 0.709264768 | -4.54386948 |
| **A_52_P490863** | Nop10 | 0.505489127 | -0.485446583 | 1.496424838 | 12.51396722 | 1.277664078 | 0.252844287 | 0.709264768 | -4.54389118 |
| **A_55_P2225082** | Mettl4 | 0.132866334 | -0.127613978 | 0.393346646 | 8.52208226 | 1.277586735 | 0.252869625 | 0.709264768 | -4.543912155 |
| **A_51_P149960** | Letm2 | -0.268270201 | -0.794324474 | 0.257784071 | 8.044993325 | -1.27729799 | 0.252964238 | 0.709264768 | -4.543990459 |
| **A_55_P2024675** | Shld2 | -0.234281782 | -0.693742112 | 0.225178547 | 9.055601443 | -1.277146874 | 0.253013768 | 0.709264768 | -4.544031438 |
| **A_51_P442053** | Ptdss1 | -0.152832407 | -0.452587641 | 0.146922827 | 9.0867457 | -1.277023893 | 0.253054082 | 0.709264768 | -4.544064787 |
| **A_66_P122158** | Pisd-ps3 | -0.636886 | -1.886114191 | 0.612342192 | 8.128149162 | -1.276939353 | 0.253081798 | 0.709264768 | -4.544087711 |
| **A_55_P2172470** | Nrgn | -0.250001918 | -0.740384584 | 0.240380747 | 8.282400909 | -1.276904968 | 0.253093072 | 0.709264768 | -4.544097035 |
| **A_55_P2114789** | Gtf2a2 | 0.216330097 | -0.208038995 | 0.640699189 | 8.293246676 | 1.276801973 | 0.253126844 | 0.709264768 | -4.544124964 |
| **A_55_P1987186** | Ttll9 | 0.329470957 | -0.316845207 | 0.975787121 | 8.32619529 | 1.276798799 | 0.253127885 | 0.709264768 | -4.544125824 |
| **A_55_P1961888** | Twistnb | 0.129624027 | -0.124656684 | 0.383904738 | 10.07664129 | 1.276798716 | 0.253127912 | 0.709264768 | -4.544125847 |
| **A_55_P1982122** | Homer1 | -0.123933804 | -0.367067248 | 0.119199639 | 7.896468689 | -1.276719307 | 0.253153952 | 0.709264768 | -4.544147379 |
| **A_51_P494825** | Mfhas1 | -0.246079928 | -0.728842247 | 0.23668239 | 11.02459911 | -1.276712684 | 0.253156125 | 0.709264768 | -4.544149175 |
| **A_51_P465281** | Lgals1 | -0.599677221 | -1.776194779 | 0.576840338 | 14.40515247 | -1.276643048 | 0.253178963 | 0.709264768 | -4.544168057 |
| **A_55_P1959056** | Pom121 | 0.192753812 | -0.185485865 | 0.570993489 | 12.64365834 | 1.276398358 | 0.253259227 | 0.709264768 | -4.544234405 |
| **A_55_P2034517** | NA | -0.361052954 | -1.069597831 | 0.347491923 | 8.059756384 | -1.276303282 | 0.25329042 | 0.709264768 | -4.544260184 |
| **A_55_P2159852** | NA | 0.192380339 | -0.185165156 | 0.569925834 | 8.989709605 | 1.276267586 | 0.253302133 | 0.709264768 | -4.544269862 |
| **A_52_P598912** | Erc1 | -0.248619816 | -0.736567225 | 0.239327593 | 8.257158677 | -1.276183339 | 0.253329777 | 0.709264768 | -4.544292705 |
| **A_51_P289588** | Ube2e3 | 0.295162551 | -0.284132255 | 0.874457357 | 12.88196507 | 1.276180093 | 0.253330842 | 0.709264768 | -4.544293585 |
| **A_55_P2068723** | Ccl27a | -1.620026926 | -4.799660741 | 1.55960689 | 10.57218498 | -1.276131625 | 0.253346748 | 0.709264768 | -4.544306726 |
| **A_51_P462546** | Gmppa | -0.309265387 | -0.916323596 | 0.297792822 | 9.54710188 | -1.276001913 | 0.253389319 | 0.709264768 | -4.544341895 |
| **A_52_P225898** | Kcnj8 | 0.321366687 | -0.309462032 | 0.952195405 | 8.456425208 | 1.275967888 | 0.253400487 | 0.709264768 | -4.54435112 |
| **A_55_P1990210** | Scpep1 | -0.784628255 | -2.324820371 | 0.755563861 | 12.11844429 | -1.275966855 | 0.253400826 | 0.709264768 | -4.5443514 |
| **A_55_P2178152** | Ccsap | 0.158335133 | -0.152486591 | 0.469156858 | 8.522986351 | 1.275898927 | 0.253423123 | 0.709264768 | -4.544369817 |
| **A_55_P1973339** | Mapk1 | -0.206722989 | -0.612533219 | 0.19908724 | 10.18574988 | -1.275898513 | 0.253423259 | 0.709264768 | -4.544369929 |
| **A_51_P496054** | Zcchc14 | -0.124730015 | -0.369619781 | 0.120159752 | 9.055332526 | -1.275706245 | 0.25348638 | 0.709264768 | -4.544422057 |
| **A_55_P1956130** | LOC68395 | -1.307236365 | -3.873891543 | 1.259418812 | 10.95701954 | -1.275665881 | 0.253499634 | 0.709264768 | -4.544433 |
| **A_51_P248304** | Vps26b | 0.136348067 | -0.131373359 | 0.404069493 | 9.361058454 | 1.275604967 | 0.253519636 | 0.709264768 | -4.544449515 |
| **A_55_P2042277** | Pbrm1 | 0.157195147 | -0.151489915 | 0.465880209 | 9.089053644 | 1.275480629 | 0.253560468 | 0.709264768 | -4.544483224 |
| **A_55_P2162782** | Ahnak | -0.419754061 | -1.244040427 | 0.404532305 | 8.653723226 | -1.275460859 | 0.253566961 | 0.709264768 | -4.544488584 |
| **A_51_P116651** | Dpt | 0.52521203 | -0.50620717 | 1.556631229 | 8.752062288 | 1.275409945 | 0.253583683 | 0.709264768 | -4.54708073 |
| **A_51_P362209** | Commd10 | 0.285329568 | -0.275018004 | 0.84567714 | 10.32062329 | 1.275380082 | 0.253593491 | 0.709264768 | -4.544510483 |
| **A_51_P246224** | Tram2 | -1.016961229 | -3.014195541 | 0.980273083 | 9.842847094 | -1.275339242 | 0.253606906 | 0.709264768 | -4.544521554 |
| **A_55_P2157992** | NA | 0.236105891 | -0.227655844 | 0.699867627 | 14.45510263 | 1.275152882 | 0.253668127 | 0.709264768 | -4.544572076 |
| **A_51_P188126** | Gca | -0.153100192 | -0.453833949 | 0.147633565 | 8.02787172 | -1.27509899 | 0.253685834 | 0.709264768 | -4.544586686 |
| **A_52_P211223** | Cdca2 | 0.414454697 | -0.399675852 | 1.228585247 | 8.770462071 | 1.275068041 | 0.253696003 | 0.709264768 | -4.544595076 |
| **A_52_P203948** | Pla2g4e | -0.4025527 | -1.193315795 | 0.388210395 | 7.834346594 | -1.275048478 | 0.253702431 | 0.709264768 | -4.544600379 |
| **A_55_P2049757** | NA | 0.446318386 | -0.430444403 | 1.323081175 | 14.68801687 | 1.275008307 | 0.253715631 | 0.709264768 | -4.544611269 |
| **A_55_P1980611** | Trmt2b | 0.396771612 | -0.382743825 | 1.17628705 | 10.49268959 | 1.274870927 | 0.253760778 | 0.709264768 | -4.54464851 |

| | | | | | | | | | |
|---|---|---|---|---|---|---|---|---|---|
| **A_66_P117913** | NA | 0.309451891 | -0.298528641 | 0.917432424 | 16.26441142 | 1.274834516 | 0.253772745 | 0.709264768 | -4.544658381 |
| **A_55_P2150343** | Piezo1 | -0.170913829 | -0.506774816 | 0.164947158 | 12.54413145 | -1.274582611 | 0.25385555 | 0.709264768 | -4.544726666 |
| **A_55_P2183208** | Prl2c1 | -1.892839592 | -5.612572262 | 1.826893077 | 12.14345524 | -1.274536942 | 0.253870564 | 0.709264768 | -4.544739045 |
| **A_51_P140901** | Pmm2 | -0.337345314 | -1.000302319 | 0.325611691 | 8.665947611 | -1.274499484 | 0.25388288 | 0.709264768 | -4.544749199 |
| **A_55_P1990081** | Snrpg | 0.579755676 | -0.559593593 | 1.719104945 | 15.40186835 | 1.274496 | 0.253884026 | 0.709264768 | -4.544750144 |
| **A_55_P1964573** | Arhgef6 | -1.556544737 | -4.615519076 | 1.502429603 | 11.54313375 | -1.274489081 | 0.253886301 | 0.709264768 | -4.544752019 |
| **A_55_P1964525** | NA | 0.312714249 | -0.301877235 | 0.927305734 | 17.29304831 | 1.274416772 | 0.253910077 | 0.709264768 | -4.54477162 |
| **A_55_P2064240** | Hemk1 | -0.187395948 | -0.555694227 | 0.18090233 | 8.362059566 | -1.27441494 | 0.253910679 | 0.709264768 | -4.544772116 |
| **A_51_P282394** | Mrps28 | 0.327095073 | -0.315813203 | 0.970003348 | 11.99140141 | 1.274310706 | 0.253944957 | 0.709264768 | -4.54480037 |
| **A_55_P1994194** | Heph | 0.434157527 | -0.419243984 | 1.287559037 | 10.46742544 | 1.274219575 | 0.253974928 | 0.709264768 | -4.544825072 |
| **A_55_P2087872** | NA | 0.270446915 | -0.261165282 | 0.802059112 | 13.54960913 | 1.274199531 | 0.253981521 | 0.709264768 | -4.544830505 |
| **A_52_P573336** | Sbsn | -2.283874852 | -6.773406405 | 2.205656701 | 10.23458988 | -1.274153114 | 0.253996789 | 0.709264768 | -4.544843086 |
| **A_51_P102860** | Herpud2 | -0.255293287 | -0.757137561 | 0.246550987 | 9.412179031 | -1.274150611 | 0.253997612 | 0.709264768 | -4.544843764 |
| **A_52_P51548** | Pard3 | -0.140198327 | -0.415807533 | 0.135410879 | 10.13411875 | -1.274088093 | 0.254018176 | 0.709264768 | -4.54486071 |
| **A_51_P382121** | Ormdl1 | 0.18268983 | -0.176476423 | 0.541856084 | 10.44286482 | 1.273999312 | 0.254047383 | 0.709264768 | -4.544884774 |
| **A_51_P325592** | 2410004B | -0.150625148 | -0.446753221 | 0.145502926 | 10.5709664 | -1.273996542 | 0.254048294 | 0.709264768 | -4.544885525 |
| **A_55_P1960416** | Nrros | 0.622017432 | -0.600961561 | 1.844996425 | 11.76084869 | 1.273895796 | 0.25408144 | 0.709264768 | -4.544912831 |
| **A_51_P291501** | Ccno | 0.962683499 | -0.93011963 | 2.855486628 | 10.21451089 | 1.273879732 | 0.254086726 | 0.709264768 | -4.544917185 |
| **A_52_P214612** | NA | -0.168014111 | -0.498413582 | 0.162385361 | 8.676201481 | -1.273669529 | 0.254155898 | 0.709264768 | -4.544974158 |
| **A_51_P367060** | Ifrd1 | -0.61340704 | -1.819702959 | 0.592888878 | 11.68204911 | -1.273635757 | 0.254167013 | 0.709264768 | -4.544983311 |
| **A_55_P1984886** | Hcst | 0.360309193 | -0.34828012 | 1.068898506 | 8.409273427 | 1.273594254 | 0.254180673 | 0.709264768 | -4.54499456 |
| **A_51_P141818** | Mrpl32 | 0.332130065 | -0.321115971 | 0.985376101 | 11.30237648 | 1.273449576 | 0.254228297 | 0.709264768 | -4.545033772 |
| **A_55_P2121426** | NA | 0.581974281 | -0.562878709 | 1.72682727 | 14.76176585 | 1.273222824 | 0.254302953 | 0.709264768 | -4.545095226 |
| **A_66_P120901** | Atxn10 | -0.528640104 | -1.568611797 | 0.511331588 | 12.95965782 | -1.273177439 | 0.254317898 | 0.709264768 | -4.545107526 |
| **A_52_P35057** | Kctd17 | -0.513663852 | -1.524225157 | 0.496897453 | 10.53522262 | -1.273112216 | 0.254339377 | 0.709264768 | -4.545125202 |
| **A_55_P2013840** | Krt42 | -0.416777582 | -1.236892718 | 0.403337554 | 10.95368124 | -1.272857744 | 0.254423193 | 0.709264768 | -4.545194165 |
| **A_55_P2002667** | Trim26 | -0.632332099 | -1.876781707 | 0.612117509 | 11.0375355 | -1.272677186 | 0.25448268 | 0.709264768 | -4.545243096 |
| **A_51_P214423** | Tead1 | -0.217460776 | -0.645471384 | 0.210549832 | 11.16880731 | -1.272555542 | 0.254522763 | 0.709264768 | -4.54527606 |
| **A_51_P284823** | Polr1b | 0.14433762 | -0.139759544 | 0.428434785 | 8.859611011 | 1.272515223 | 0.25453605 | 0.709264768 | -4.545286986 |
| **A_55_P2110601** | Slc35a2 | -0.335491687 | -0.995860777 | 0.324877402 | 10.1007612 | -1.272463615 | 0.254553058 | 0.709264768 | -4.54530097 |
| **A_51_P110301** | C3 | -0.490158885 | -1.455022966 | 0.474705196 | 10.91412462 | -1.272392457 | 0.254576512 | 0.709264768 | -4.545994223 |
| **A_52_P425734** | Afm | -0.484181393 | -1.437362277 | 0.468999492 | 8.042712438 | -1.27228122 | 0.254613178 | 0.709264768 | -4.545350396 |
| **A_52_P203560** | Fzd10 | 0.532994561 | -0.516356449 | 1.582345572 | 9.033631751 | 1.272191056 | 0.254642901 | 0.709264768 | -4.545374828 |
| **A_52_P335064** | Mustn1 | -0.621156647 | -1.844144162 | 0.601830868 | 8.793092453 | -1.272124037 | 0.254664997 | 0.709264768 | -4.545392988 |
| **A_52_P462472** | Nexmif | 0.466465256 | -0.451989679 | 1.384920192 | 8.739228735 | 1.272072298 | 0.254682056 | 0.709264768 | -4.545407008 |
| **A_55_P1971854** | Vcpkmt | 0.24586173 | -0.238242129 | 0.72996559 | 10.44819127 | 1.27204576 | 0.254690807 | 0.709264768 | -4.545414199 |
| **A_52_P622391** | Usp9x | -0.528324414 | -1.568657378 | 0.51200855 | 8.65087594 | -1.271975265 | 0.254714052 | 0.709264768 | -4.5454333 |
| **A_51_P416585** | Hdlbp | -0.537977732 | -1.59735016 | 0.521394695 | 11.75031921 | -1.271938111 | 0.254726304 | 0.709264768 | -4.545443368 |
| **A_52_P169901** | Slc25a51 | -0.309939374 | -0.920270337 | 0.300391589 | 10.43289807 | -1.271925559 | 0.254730444 | 0.709264768 | -4.545446769 |
| **A_55_P1971815** | Clcn2 | -0.232073884 | -0.689086313 | 0.224938546 | 8.048665103 | -1.271887221 | 0.254743087 | 0.709264768 | -4.545457157 |
| **A_55_P2082653** | Zfp711 | 0.518500558 | -0.502591224 | 1.539592341 | 8.972361366 | 1.271846775 | 0.254756426 | 0.709264768 | -4.545468116 |
| **A_55_P1964233** | Tnks | -0.259004935 | -0.769135378 | 0.251125507 | 8.298255923 | -1.271677959 | 0.254812109 | 0.709264768 | -4.545513858 |
| **A_51_P335770** | Afap1 | -0.717346146 | -2.130394196 | 0.695701904 | 11.46842244 | -1.271517041 | 0.254865197 | 0.709264768 | -4.545557458 |
| **A_52_P153189** | Arl2bp | 0.175056128 | -0.169788494 | 0.519900749 | 9.54780701 | 1.271464403 | 0.254882565 | 0.709264768 | -4.54557172 |
| **A_55_P2014357** | Pdpk1 | -0.209072509 | -0.620945924 | 0.202800906 | 9.788066854 | -1.271403833 | 0.254902551 | 0.709264768 | -4.54558813 |
| **A_55_P2117649** | Slfn9 | 0.283906039 | -0.275411673 | 0.843223752 | 8.767580531 | 1.271353736 | 0.254919082 | 0.709264768 | -4.545601704 |
| **A_51_P438666** | Keap1 | -0.219050529 | -0.650606896 | 0.212505838 | 11.11083119 | -1.27132657 | 0.254928047 | 0.709264768 | -4.545609064 |
| **A_55_P1989928** | Lin52 | 0.212347712 | -0.206059113 | 0.630754536 | 8.957738621 | 1.271156935 | 0.254984034 | 0.709264768 | -4.545655024 |

| | | | | | | | | | |
|---|---|---|---|---|---|---|---|---|---|
| **A_55_P2006479** | 3300002I0 | -0.141065478 | -0.419062947 | 0.136931991 | 8.289337407 | -1.27095522 | 0.255050623 | 0.709264768 | -4.545709673 |
| **A_51_P191586** | Arpc1a | -0.121096199 | -0.359743388 | 0.11755099 | 11.52843165 | -1.270938497 | 0.255056144 | 0.709264768 | -4.545714204 |
| **A_55_P2005585** | Trps1 | 0.463955025 | -0.450442538 | 1.378352588 | 9.140559759 | 1.270840857 | 0.255088383 | 0.709264768 | -4.545740656 |
| **A_55_P2066463** | Enah | -0.330357195 | -0.981463052 | 0.320748662 | 9.914338691 | -1.270815532 | 0.255096745 | 0.709264768 | -4.545747517 |
| **A_55_P1991440** | Zscan21 | -0.115137625 | -0.342074185 | 0.111798935 | 12.21428369 | -1.270758857 | 0.25511546 | 0.709264768 | -4.545762871 |
| **A_55_P1956502** | Hnrnph1 | 0.252781092 | -0.24545832 | 0.751020503 | 14.31521432 | 1.270740443 | 0.255121541 | 0.709264768 | -4.54576786 |
| **A_55_P1985040** | Gpbp1l1 | -0.28810902 | -0.856167904 | 0.279949865 | 8.546028438 | -1.27032207 | 0.255259736 | 0.709264768 | -4.545881198 |
| **A_55_P2091039** | NA | 0.179183039 | -0.17417157 | 0.532537648 | 8.547100502 | 1.270095808 | 0.255334502 | 0.709264768 | -4.54594249 |
| **A_55_P2036942** | Gm4535 | 0.385121022 | -0.374466589 | 1.144708633 | 9.414334187 | 1.269900512 | 0.255399051 | 0.709264768 | -4.545995392 |
| **A_51_P481398** | Kif11 | 0.589499881 | -0.573191431 | 1.752191194 | 9.083850467 | 1.269900342 | 0.255399107 | 0.709264768 | -4.545995438 |
| **A_51_P408703** | Rnf138rt1 | -0.37426013 | -1.112521042 | 0.364000781 | 7.882064872 | -1.269737761 | 0.255452854 | 0.709264768 | -4.546039477 |
| **A_51_P253527** | Rffl | -0.42159029 | -1.253320361 | 0.41013978 | 8.082623425 | -1.269575528 | 0.255506497 | 0.709264768 | -4.54608342 |
| **A_52_P381430** | Tbc1d4 | -0.3945308 | -1.17295821 | 0.383896609 | 8.175869972 | -1.269442808 | 0.255550388 | 0.709264768 | -4.546119369 |
| **A_55_P1960576** | Kdm5b | -0.148331925 | -0.441006793 | 0.144342943 | 8.056876829 | -1.269403079 | 0.255563528 | 0.709264768 | -4.54613013 |
| **A_55_P2041470** | NA | 0.204876045 | -0.199369288 | 0.609121378 | 8.52879173 | 1.269394211 | 0.255566461 | 0.709264768 | -4.546132532 |
| **A_55_P2154173** | NA | -0.173642203 | -0.516273482 | 0.168989076 | 15.60855264 | -1.269341925 | 0.255583755 | 0.709264768 | -4.546146694 |
| **A_55_P2040518** | NA | 0.264592392 | -0.257542361 | 0.786727145 | 9.996871845 | 1.269243938 | 0.255616168 | 0.709264768 | -4.546173234 |
| **A_55_P1954481** | Dock9 | -0.137410318 | -0.408571215 | 0.133750579 | 8.317879153 | -1.269236719 | 0.255618557 | 0.709264768 | -4.546175189 |
| **A_51_P403693** | Uba5 | -0.37643916 | -1.119311177 | 0.366432856 | 11.25697233 | -1.269203154 | 0.25562966 | 0.709264768 | -4.54618428 |
| **A_51_P332141** | Clcn2 | -0.315692248 | -0.938721366 | 0.307336871 | 8.366887551 | -1.269129438 | 0.255654049 | 0.709264768 | -4.546204246 |
| **A_55_P1960202** | Vcan | 0.791067946 | -0.770171046 | 2.352306938 | 10.40079645 | 1.269096785 | 0.255664852 | 0.709264768 | -4.546213089 |
| **A_51_P452629** | Tlr2 | 0.359135483 | -0.349667871 | 1.067938837 | 8.904909795 | 1.269062169 | 0.255676306 | 0.709264768 | -4.547551041 |
| **A_52_P87964** | Pla2g12a | -0.215750766 | -0.641590027 | 0.210088495 | 10.02504428 | -1.268986472 | 0.255701353 | 0.709264768 | -4.546242967 |
| **A_55_P1957967** | Dhx32 | -0.493530117 | -1.467888829 | 0.480828596 | 10.65068565 | -1.268659665 | 0.255809516 | 0.709264768 | -4.546331476 |
| **A_51_P380153** | Ubr4 | -0.33991072 | -1.011107424 | 0.331285984 | 11.5972623 | -1.268426748 | 0.25588663 | 0.709264768 | -4.546394555 |
| **A_55_P2022895** | Slc12a8 | -0.21998327 | -0.654463071 | 0.214496531 | 8.032698219 | -1.268149362 | 0.255978493 | 0.709264768 | -4.546469673 |
| **A_55_P1972839** | NA | 0.285629629 | -0.27851453 | 0.849773789 | 16.90396756 | 1.268129208 | 0.255985169 | 0.709264768 | -4.546475131 |
| **A_51_P161929** | Tspo2 | 0.204478604 | -0.19944234 | 0.608399548 | 9.810972914 | 1.26794917 | 0.25604481 | 0.709264768 | -4.546523885 |
| **A_51_P214470** | Tdrp | -0.928332432 | -2.762204503 | 0.905539638 | 8.38594787 | -1.267899506 | 0.256061265 | 0.709264768 | -4.546537334 |
| **A_55_P2157620** | NA | -0.540717701 | -1.608891311 | 0.527455908 | 8.247384256 | -1.267882738 | 0.256066821 | 0.709264768 | -4.546541874 |
| **A_65_P03031** | Ncapd3 | 0.168585734 | -0.164476829 | 0.501648297 | 8.322523702 | 1.267784232 | 0.256099461 | 0.709264768 | -4.546568549 |
| **A_55_P2184246** | Rps26 | 0.539003181 | -0.525919761 | 1.603926122 | 16.88000442 | 1.267720434 | 0.256120602 | 0.709264768 | -4.546585824 |
| **A_55_P2089530** | Fkbp10 | 0.666133628 | -0.650010315 | 1.982277572 | 13.16520605 | 1.267676136 | 0.256135283 | 0.709264768 | -4.546597819 |
| **A_51_P331752** | Ccl11 | 1.098216002 | -1.071977929 | 3.268409934 | 10.23051308 | 1.267475486 | 0.256201788 | 0.709264768 | -4.546652151 |
| **A_52_P449629** | Ankrd13c | -0.352946266 | -1.050420263 | 0.344527731 | 8.722656108 | -1.267450235 | 0.256210158 | 0.709264768 | -4.546658988 |
| **A_55_P2178084** | Dnmt3a | 0.195951808 | -0.191283642 | 0.583187258 | 12.24258107 | 1.267431542 | 0.256216355 | 0.709264768 | -4.54666405 |
| **A_55_P2060303** | Mras | -0.494417269 | -1.471490694 | 0.482656156 | 9.44859537 | -1.267408969 | 0.256223838 | 0.709264768 | -4.546670162 |
| **A_55_P2000638** | Stag2 | 0.573165156 | -0.559681502 | 1.706011814 | 12.47108748 | 1.267240367 | 0.256279738 | 0.709264768 | -4.546715814 |
| **A_55_P2000107** | Atox1 | -0.341004951 | -1.015027081 | 0.333017179 | 15.25829688 | -1.26717581 | 0.256301144 | 0.709264768 | -4.546733294 |
| **A_55_P2243566** | Redrum | 0.128465652 | -0.125465165 | 0.382396469 | 7.92131453 | 1.267132295 | 0.256315574 | 0.709264768 | -4.546745076 |
| **A_55_P1989534** | Cklf | -0.35683663 | -1.06229117 | 0.34861791 | 9.782649852 | -1.266924522 | 0.256384484 | 0.709264768 | -4.546801331 |
| **A_55_P2002093** | Dph7 | 0.126938834 | -0.124062873 | 0.37794054 | 8.78036316 | 1.26668368 | 0.256464382 | 0.709264768 | -4.546866539 |
| **A_55_P2043362** | NA | -0.123829943 | -0.368690366 | 0.121030481 | 8.249147134 | -1.266652313 | 0.25647479 | 0.709264768 | -4.546875031 |
| **A_55_P2180406** | Cdkn2aipı | 0.14532068 | -0.142045284 | 0.432686644 | 8.851086872 | 1.26660862 | 0.256489287 | 0.709264768 | -4.54688686 |
| **A_51_P210603** | Cyp4f13 | -0.558318946 | -1.662413902 | 0.545776009 | 8.536480194 | -1.266561509 | 0.25650492 | 0.709264768 | -4.546899615 |
| **A_51_P135517** | Coch | -0.933686747 | -2.780230421 | 0.912856926 | 8.141909922 | -1.266461396 | 0.256538144 | 0.709264768 | -4.546926719 |
| **A_51_P519570** | 1600002K | -0.333556697 | -0.993255818 | 0.326142423 | 10.18334898 | -1.266409342 | 0.25655542 | 0.709264768 | -4.546940811 |
| **A_55_P2071532** | NA | 0.249395464 | -0.243898365 | 0.742689294 | 14.30460022 | 1.266290105 | 0.256594997 | 0.709264768 | -4.546973092 |

| | | | | | | | | | |
|---|---|---|---|---|---|---|---|---|---|
| **A_55_P2083839** | Atp13a1 | -0.33622976 | -1.001278626 | 0.328819105 | 10.61269486 | -1.266289309 | 0.256595261 | 0.709264768 | -4.546973307 |
| **A_66_P128694** | NA | 0.144758404 | -0.141572407 | 0.431089216 | 8.497640747 | 1.266269214 | 0.256601932 | 0.709264768 | -4.546978748 |
| **A_51_P413710** | Ap3d1 | -0.242082999 | -0.720946447 | 0.236780448 | 11.16264306 | -1.266201865 | 0.256624289 | 0.709264768 | -4.54699698 |
| **A_55_P1993858** | Rmrp | -0.383313454 | -1.141613655 | 0.374986747 | 10.7405212 | -1.266086091 | 0.256662726 | 0.709264768 | -4.547028323 |
| **A_51_P238523** | Shisa4 | -0.150673874 | -0.448756883 | 0.147409135 | 8.938251776 | -1.266050645 | 0.256674495 | 0.709264768 | -4.547037918 |
| **A_55_P1999648** | Plin2 | -0.375212405 | -1.117584191 | 0.367159381 | 12.45950447 | -1.265919459 | 0.256718057 | 0.709264768 | -4.547073432 |
| **A_51_P491916** | Rassf6 | -0.879665309 | -2.620173866 | 0.860843248 | 8.634001346 | -1.265877405 | 0.256732023 | 0.709264768 | -4.547084816 |
| **A_51_P219918** | Tmem125 | -1.127515313 | -3.358634325 | 1.103603699 | 10.17723417 | -1.265756183 | 0.256772284 | 0.709264768 | -4.547117631 |
| **A_55_P2000077** | Nrf1 | 0.310707865 | -0.304125638 | 0.925541368 | 10.24087236 | 1.265741639 | 0.256777114 | 0.709264768 | -4.547121568 |
| **A_52_P331212** | Cspp1 | -0.129627032 | -0.38614618 | 0.126892117 | 8.715064256 | -1.265686455 | 0.256795444 | 0.709264768 | -4.547136506 |
| **A_55_P2020395** | 2-Mar | -0.196830493 | -0.586352756 | 0.19269177 | 7.975230296 | -1.265640725 | 0.256810635 | 0.709264768 | -4.547148885 |
| **A_52_P625808** | Sf3b1 | 0.394307735 | -0.38605931 | 1.174674781 | 9.181422495 | 1.265571603 | 0.256833598 | 0.709264768 | -4.547167597 |
| **A_55_P2168189** | NA | 0.707251434 | -0.69254196 | 2.107044828 | 13.48346507 | 1.265494444 | 0.256859233 | 0.709264768 | -4.547188483 |
| **A_51_P437938** | Coa5 | -0.738875453 | -2.201594212 | 0.723843306 | 9.347247934 | -1.265204574 | 0.256955558 | 0.709264768 | -4.547266946 |
| **A_51_P141554** | Med22 | -0.194392333 | -0.579226022 | 0.190441356 | 10.50263918 | -1.265191875 | 0.256959779 | 0.709264768 | -4.547270383 |
| **A_55_P1987912** | Chpf2 | -0.747620656 | -2.227759912 | 0.732518599 | 8.965424682 | -1.265112249 | 0.256986245 | 0.709264768 | -4.547291936 |
| **A_55_P2100222** | NA | -0.304383247 | -0.907012376 | 0.298245881 | 10.85002314 | -1.265088687 | 0.256994077 | 0.709264768 | -4.547298314 |
| **A_51_P318640** | Rps3 | 0.22706709 | -0.222521 | 0.67665518 | 15.8133201 | 1.264997717 | 0.257024318 | 0.709264768 | -4.547322936 |
| **A_55_P1981670** | 8-Sep | -0.408097497 | -1.216138922 | 0.399943927 | 8.102604943 | -1.26497124 | 0.25703312 | 0.709264768 | -4.547330103 |
| **A_55_P2383897** | 8030453C | -0.214489764 | -0.639202048 | 0.210222519 | 7.839784931 | -1.264917194 | 0.257051088 | 0.709264768 | -4.547344731 |
| **A_66_P115757** | Mthfd2l | 0.555378503 | -0.54437234 | 1.655129346 | 9.409967631 | 1.264867715 | 0.257067539 | 0.709264768 | -4.547358123 |
| **A_52_P660693** | Dsel | 0.281508917 | -0.275931365 | 0.8389492 | 9.457200558 | 1.264864935 | 0.257068463 | 0.709264768 | -4.547358876 |
| **A_52_P413502** | Rtf2 | -0.215718573 | -0.642938942 | 0.211501795 | 8.348238326 | -1.264695387 | 0.257124842 | 0.709264768 | -4.547404765 |
| **A_55_P1961943** | Cd68 | -1.566144938 | -4.66796469 | 1.535674814 | 10.44067424 | -1.264636579 | 0.257144399 | 0.709264768 | -4.547420682 |
| **A_51_P457528** | Ccnb2 | 1.548023901 | -1.518190792 | 4.614238594 | 12.15285995 | 1.264519256 | 0.257183421 | 0.709264768 | -4.547452436 |
| **A_55_P2004732** | Ttc39a | -0.614817238 | -1.832611142 | 0.602976666 | 8.274295852 | -1.264510924 | 0.257186193 | 0.709264768 | -4.547454691 |
| **A_55_P2063251** | Dusp3 | -0.241705311 | -0.720496624 | 0.237086001 | 10.05475591 | -1.264416858 | 0.257217483 | 0.709264768 | -4.547480149 |
| **A_51_P165683** | Hirip3 | 0.851304109 | -0.835129773 | 2.537737992 | 10.92022308 | 1.264345468 | 0.257241233 | 0.709264768 | -4.54749947 |
| **A_55_P1983006** | Hdac5 | -0.200390601 | -0.597386149 | 0.196604946 | 12.91521952 | -1.264276479 | 0.257264186 | 0.709264768 | -4.547518141 |
| **A_52_P76196** | NA | 0.583705682 | -0.572678936 | 1.7400903 | 13.23137933 | 1.264276193 | 0.257264281 | 0.709264768 | -4.547518219 |
| **A_55_P2159870** | Tead4 | -1.142812484 | -3.407041205 | 1.121416237 | 8.801155961 | -1.26416868 | 0.257300055 | 0.709264768 | -4.547547316 |
| **A_55_P1985804** | Thumpd2 | 0.266830742 | -0.261840352 | 0.795501836 | 8.488454122 | 1.264155924 | 0.2573043 | 0.709264768 | -4.547550768 |
| **A_51_P240693** | Tecpr1 | -0.149756288 | -0.446564976 | 0.1470524 | 11.58121089 | -1.263743117 | 0.2574417 | 0.709264768 | -4.547662483 |
| **A_55_P2108868** | Nufip1 | -0.383511066 | -1.143684724 | 0.376662593 | 8.211748557 | -1.263616909 | 0.257483721 | 0.709264768 | -4.547696636 |
| **A_52_P297642** | Pdia5 | -1.234762761 | -3.682326049 | 1.212800527 | 9.816725171 | -1.263571839 | 0.257498728 | 0.709264768 | -4.547708832 |
| **A_52_P1020860** | AW11201( | -0.351774582 | -1.049194986 | 0.345645821 | 8.082844469 | -1.263339727 | 0.257576029 | 0.709264768 | -4.547771642 |
| **A_55_P2160726** | Tfcp2 | -0.190480389 | -0.568173091 | 0.187212313 | 8.064156788 | -1.263170636 | 0.257632356 | 0.709264768 | -4.547817397 |
| **A_55_P2073841** | NA | 0.550626672 | -0.541222689 | 1.642476033 | 15.03606198 | 1.263120738 | 0.257648979 | 0.709264768 | -4.547830899 |
| **A_51_P234833** | Strn4 | 0.307794465 | -0.302541645 | 0.918130576 | 11.40119767 | 1.263112655 | 0.257651672 | 0.709264768 | -4.547833086 |
| **A_66_P127224** | Krt10 | 0.449560102 | -0.441916587 | 1.341036791 | 9.294089077 | 1.263072021 | 0.25766521 | 0.709264768 | -4.547844081 |
| **A_55_P2175301** | NA | 0.435323027 | -0.427925639 | 1.298571694 | 15.87064168 | 1.263066074 | 0.257667192 | 0.709264768 | -4.54784569 |
| **A_52_P90124** | Prep | 0.174847757 | -0.171880322 | 0.521575836 | 11.82574844 | 1.263052486 | 0.257671719 | 0.709264768 | -4.547849367 |
| **A_55_P2142156** | Gm16441 | 0.14131771 | -0.138938261 | 0.421573682 | 8.69490717 | 1.262967176 | 0.257700145 | 0.709264768 | -4.54787245 |
| **A_55_P2090698** | NA | 0.454346394 | -0.446698041 | 1.355390828 | 15.00766268 | 1.262964729 | 0.25770096 | 0.709264768 | -4.547873112 |
| **A_52_P600531** | Bahd1 | -0.642438414 | -1.916697406 | 0.631820579 | 9.359144026 | -1.262769662 | 0.257765969 | 0.709264768 | -4.547925893 |
| **A_55_P1987979** | Plrg1 | 0.294554876 | -0.289747126 | 0.878856877 | 11.99614839 | 1.262638963 | 0.257809534 | 0.709264768 | -4.547961256 |
| **A_55_P1953683** | Nsmaf | 0.131322404 | -0.129193043 | 0.39183785 | 11.46167101 | 1.262570651 | 0.257832306 | 0.709264768 | -4.547979739 |
| **A_55_P2056973** | Treml2 | 0.14715273 | -0.144767208 | 0.439072669 | 7.828903242 | 1.262568387 | 0.257833061 | 0.709264768 | -4.547980351 |

| | | | | | | | | | |
|---|---|---|---|---|---|---|---|---|---|
| **A_66_P136998** | Tmem43 | -0.484558795 | -1.445852401 | 0.476734812 | 9.543660115 | -1.262527284 | 0.257846764 | 0.709264768 | -4.547991472 |
| **A_52_P118310** | Chn1 | 0.348664175 | -0.343078236 | 1.040406587 | 8.528395517 | 1.262447333 | 0.257873421 | 0.709264768 | -4.548013104 |
| **A_55_P2164218** | NA | 0.325425658 | -0.320240509 | 0.971091825 | 13.84333544 | 1.262391632 | 0.257891993 | 0.709264768 | -4.548028174 |
| **A_55_P2117574** | Ppih | 0.130657037 | -0.128576261 | 0.389890334 | 7.916355463 | 1.26238657 | 0.257893681 | 0.709264768 | -4.548029544 |
| **A_55_P2097736** | Patz1 | 0.254567196 | -0.250516146 | 0.759650539 | 8.478102248 | 1.262378936 | 0.257896227 | 0.709264768 | -4.548031609 |
| **A_55_P2103944** | Mrpl30 | 0.246788851 | -0.242955073 | 0.736532776 | 12.51314906 | 1.262137951 | 0.257976594 | 0.709264768 | -4.548096807 |
| **A_55_P1963980** | Carmil1 | -0.311803259 | -0.930613946 | 0.307007428 | 8.328074898 | -1.262040205 | 0.258009198 | 0.709264768 | -4.548123252 |
| **A_51_P450812** | Cog4 | -0.156563016 | -0.467297625 | 0.154171593 | 10.92310196 | -1.261972521 | 0.258031777 | 0.709264768 | -4.548141563 |
| **A_55_P1966445** | Krt81 | -0.372195743 | -1.110922922 | 0.366531436 | 8.97843079 | -1.261936985 | 0.258043632 | 0.709264768 | -4.548151177 |
| **A_55_P2021351** | NA | 0.430464725 | -0.423956455 | 1.284885905 | 17.24606264 | 1.261873757 | 0.258064727 | 0.709264768 | -4.548168282 |
| **A_55_P2018994** | Elk3 | 0.54928081 | -0.5409957 | 1.63955732 | 11.85592744 | 1.261851118 | 0.258072281 | 0.709264768 | -4.548174407 |
| **A_55_P2027930** | Nfx1 | -0.281054842 | -0.839064922 | 0.276955239 | 8.310798903 | -1.261535198 | 0.258177708 | 0.709264768 | -4.548259871 |
| **A_51_P183995** | Paf1 | -0.370095845 | -1.104962221 | 0.364770532 | 12.25931944 | -1.261409734 | 0.258219587 | 0.709264768 | -4.548293811 |
| **A_51_P487668** | H2afy3 | 1.17714682 | -1.160288516 | 3.514582155 | 11.15046555 | 1.261366737 | 0.258233941 | 0.709264768 | -4.548305442 |
| **A_55_P2063096** | Txnrd3 | 0.64093439 | -0.631850084 | 1.913718864 | 10.27113953 | 1.261272861 | 0.258265282 | 0.709264768 | -4.548330836 |
| **A_55_P1981964** | Tnfrsf13b | 0.378468935 | -0.373228441 | 1.130166311 | 8.589717792 | 1.261065223 | 0.258334616 | 0.709264768 | -4.548387002 |
| **A_55_P2060243** | Itgbl1 | -0.267241717 | -0.798075786 | 0.263592352 | 7.838893975 | -1.260944034 | 0.258375091 | 0.709264768 | -4.548419783 |
| **A_55_P2071858** | Mgmt | 0.553626128 | -0.546086169 | 1.653338426 | 9.896545686 | 1.260920897 | 0.258382818 | 0.709264768 | -4.548426042 |
| **A_55_P2041011** | Slc2a13 | 0.357675821 | -0.352828758 | 1.068180401 | 9.60228148 | 1.260877942 | 0.258397166 | 0.709264768 | -4.548437661 |
| **A_55_P2022364** | Nxn | 0.744454285 | -0.73445693 | 2.2233655 | 12.00663727 | 1.260800224 | 0.258423127 | 0.709264768 | -4.548458682 |
| **A_66_P125401** | NA | 0.172843948 | -0.170547668 | 0.516235564 | 9.235820467 | 1.260708946 | 0.258453621 | 0.709264768 | -4.548483371 |
| **A_66_P133450** | Dnajb9 | -0.39571975 | -1.181987653 | 0.390548153 | 8.027114415 | -1.260571616 | 0.258499506 | 0.709264768 | -4.548520516 |
| **A_66_P116252** | Dusp16 | -0.713434541 | -2.131142691 | 0.704273609 | 10.36668985 | -1.260426837 | 0.258547887 | 0.709264768 | -4.548559675 |
| **A_52_P518808** | Mmd | 0.47037054 | -0.464514074 | 1.405255154 | 9.677899391 | 1.260179605 | 0.258630525 | 0.709264768 | -4.548626542 |
| **A_55_P2028821** | Hnrnpf | 0.351745628 | -0.347370451 | 1.050861707 | 14.24628651 | 1.260171817 | 0.258633128 | 0.709264768 | -4.548628648 |
| **A_55_P2128526** | Pfn2 | 0.677963204 | -0.669538238 | 2.025464646 | 11.19762365 | 1.260164469 | 0.258635585 | 0.709264768 | -4.548630636 |
| **A_52_P146457** | NA | 0.157593101 | -0.15563526 | 0.470821461 | 15.33570779 | 1.260162261 | 0.258636323 | 0.709264768 | -4.548631233 |
| **A_55_P2107362** | Gjc1 | 0.752156436 | -0.742959728 | 2.2472726 | 10.44595198 | 1.260037832 | 0.258677923 | 0.709264768 | -4.548664885 |
| **A_51_P340653** | Slc16a4 | -0.674287213 | -2.014713611 | 0.666139185 | 8.105713565 | -1.259947059 | 0.258708275 | 0.709264768 | -4.548689435 |
| **A_55_P2262287** | 5430434G | -0.169724013 | -0.507163289 | 0.167715263 | 8.872048798 | -1.259789563 | 0.258760944 | 0.709264768 | -4.548732029 |
| **A_55_P2178808** | Zbtb5 | 0.200467448 | -0.19812651 | 0.599061405 | 9.255442101 | 1.25968946 | 0.258794426 | 0.709264768 | -4.5487591 |
| **A_55_P2150757** | Gzmm | -1.002650623 | -2.99631469 | 0.991013444 | 8.488131278 | -1.259644492 | 0.258809467 | 0.709264768 | -4.548771261 |
| **A_55_P2043734** | Cdkl5 | -0.337981849 | -1.010035647 | 0.334071949 | 7.953107111 | -1.259620394 | 0.258817528 | 0.709264768 | -4.548777778 |
| **A_55_P2102175** | Lztr1 | -0.137273429 | -0.410236317 | 0.135689458 | 10.92613771 | -1.259601663 | 0.258823794 | 0.709264768 | -4.548782844 |
| **A_51_P243514** | Macc1 | 0.389241452 | -0.384775994 | 1.163258897 | 8.6187766 | 1.259559477 | 0.258837907 | 0.709264768 | -4.548794252 |
| **A_55_P2129291** | Dclre1c | -0.288323971 | -0.861778551 | 0.285130609 | 8.422898699 | -1.259308314 | 0.258921942 | 0.709264768 | -4.548862173 |
| **A_55_P2114569** | Zfp24 | 0.358544079 | -0.354574073 | 1.071662232 | 11.63003988 | 1.259306396 | 0.258922584 | 0.709264768 | -4.548862692 |
| **A_55_P1998761** | Coq8b | -0.164772174 | -0.492498256 | 0.162953908 | 9.891094949 | -1.259282624 | 0.258930539 | 0.709264768 | -4.54886912 |
| **A_55_P1966159** | Dlg5 | -0.742153861 | -2.218499116 | 0.734191394 | 12.19825264 | -1.25908881 | 0.258995405 | 0.709264768 | -4.54892153 |
| **A_66_P111666** | Nprl3 | -0.153646809 | -0.459301182 | 0.152007564 | 9.522122737 | -1.259050868 | 0.259008106 | 0.709264768 | -4.54893179 |
| **A_55_P1981099** | NA | 0.236743062 | -0.234272954 | 0.707759077 | 14.40773715 | 1.258902022 | 0.259057933 | 0.709264768 | -4.548972038 |
| **A_66_P101519** | Abcc9 | 0.496382353 | -0.491203904 | 1.483968611 | 9.25202456 | 1.258901182 | 0.259058215 | 0.709264768 | -4.548972266 |
| **A_52_P273821** | Abhd5 | -0.485497366 | -1.451632308 | 0.480637577 | 8.518595264 | -1.258633925 | 0.259147704 | 0.709264768 | -4.549044531 |
| **A_55_P2025038** | Cpe | -0.964853226 | -2.885183985 | 0.955477534 | 12.77747083 | -1.258448827 | 0.259209699 | 0.709264768 | -4.549094578 |
| **A_55_P2062733** | NA | 0.231939387 | -0.229689992 | 0.693568766 | 14.28379758 | 1.2584368 | 0.259213728 | 0.709264768 | -4.54909783 |
| **A_55_P1963868** | Plekho2 | -0.42479197 | -1.270344542 | 0.420760601 | 11.89123351 | -1.258305312 | 0.259257776 | 0.709264768 | -4.549133382 |
| **A_51_P348227** | Stoml1 | -0.207804478 | -0.621442168 | 0.205833212 | 9.008926858 | -1.258302745 | 0.259258636 | 0.709264768 | -4.549134076 |
| **A_55_P1991600** | Aplp2 | -0.431946822 | -1.29191 | 0.428016355 | 14.1709022 | -1.258058318 | 0.259340536 | 0.709264768 | -4.549200161 |

| | | | | | | | | | |
|---|---|---|---|---|---|---|---|---|---|
| **A_55_P2259125** | D7Ertd14 | -0.953670449 | -2.852482852 | 0.945141953 | 8.472337564 | -1.257959362 | 0.2593737 | 0.709264768 | -4.549226915 |
| **A_55_P1983959** | Ftl2-ps | -0.283374095 | -0.847609005 | 0.280860814 | 13.723016 | -1.257912808 | 0.259389303 | 0.709264768 | -4.549239501 |
| **A_55_P2151855** | Acp1 | 0.268219266 | -0.265888957 | 0.80232749 | 11.60568714 | 1.257798438 | 0.25942764 | 0.709264768 | -4.549270422 |
| **A_52_P2947** | Ndufaf2 | 0.593637252 | -0.588534883 | 1.775809387 | 11.08297245 | 1.257739712 | 0.259447326 | 0.709264768 | -4.549286298 |
| **A_51_P364485** | Tnfaip2 | -1.086740451 | -3.251010818 | 1.077529915 | 8.784155068 | -1.257664104 | 0.259472674 | 0.709264768 | -4.549306739 |
| **A_55_P2155572** | NA | 0.497508442 | -0.493320776 | 1.488337659 | 13.83998137 | 1.257627413 | 0.259484976 | 0.709264768 | -4.549316658 |
| **A_55_P2003951** | Tmem192 | -0.53252745 | -1.593125922 | 0.528071021 | 11.75341265 | -1.257596581 | 0.259495314 | 0.709264768 | -4.549324993 |
| **A_55_P2059352** | Col18a1 | 1.017831631 | -1.009408445 | 3.045071706 | 13.16834916 | 1.257537966 | 0.259514968 | 0.709264768 | -4.549340839 |
| **A_55_P2008417** | Mnd1 | 0.671418022 | -0.66608221 | 2.008918255 | 9.592915118 | 1.257330568 | 0.259584521 | 0.709264768 | -4.549396907 |
| **A_52_P201206** | Scrn1 | -0.514633826 | -1.539855448 | 0.510587796 | 8.408233821 | -1.257276843 | 0.25960254 | 0.709264768 | -4.54941143 |
| **A_55_P1956284** | NA | 0.25703337 | -0.255116379 | 0.769183119 | 8.405425139 | 1.25702204 | 0.259688019 | 0.709264768 | -4.549480309 |
| **A_52_P445253** | Dst | -0.646233005 | -1.934024119 | 0.64155811 | 8.201987949 | -1.256880696 | 0.259735447 | 0.709264768 | -4.549518516 |
| **A_51_P479894** | Cpsf3 | 0.429960455 | -0.426872999 | 1.28679391 | 11.43751025 | 1.256847093 | 0.259746723 | 0.709264768 | -4.549527599 |
| **A_51_P219789** | H2-Q2 | -1.270285227 | -3.80185523 | 1.261284776 | 11.44182759 | -1.25678692 | 0.259766917 | 0.709264768 | -4.549543865 |
| **A_55_P2017845** | Ehbp1 | -0.161499065 | -0.483361505 | 0.160363375 | 8.031071309 | -1.256753373 | 0.259778176 | 0.709264768 | -4.549552933 |
| **A_55_P2121608** | NA | 0.629313869 | -0.62489111 | 1.883518847 | 12.53286639 | 1.256750678 | 0.259779081 | 0.709264768 | -4.549553661 |
| **A_52_P493091** | Fmr1 | -0.320439779 | -0.959090025 | 0.318210467 | 8.525255365 | -1.256705992 | 0.259794079 | 0.709264768 | -4.54956574 |
| **A_55_P2417569** | Prrx2 | 0.655087101 | -0.650740739 | 1.96091494 | 9.721915018 | 1.256502828 | 0.259862277 | 0.709264768 | -4.549620655 |
| **A_55_P2058497** | Tmem50a | -0.160842259 | -0.48150328 | 0.159818763 | 11.56766222 | -1.256331754 | 0.259919715 | 0.709264768 | -4.549666894 |
| **A_55_P1996988** | Tent4b | 0.127578041 | -0.126768001 | 0.381924082 | 10.46691856 | 1.256322938 | 0.259922675 | 0.709264768 | -4.549669277 |
| **A_52_P79385** | Magt1 | -0.839133941 | -2.512176228 | 0.833908347 | 9.920586698 | -1.256246067 | 0.259948489 | 0.709264768 | -4.549690054 |
| **A_52_P486279** | Popdc3 | -0.352096524 | -1.05412429 | 0.349931243 | 7.910067844 | -1.256197119 | 0.259964927 | 0.709264768 | -4.549703284 |
| **A_55_P2330545** | NA | 0.382083735 | -0.379772731 | 1.143940201 | 8.977477584 | 1.256133326 | 0.259986352 | 0.709264768 | -4.549720526 |
| **A_55_P1964752** | Slc23a3 | -1.135899951 | -3.40083032 | 1.129030419 | 8.593183514 | -1.256132844 | 0.259986514 | 0.709264768 | -4.549720656 |
| **A_55_P2127538** | Ppp1r21 | -0.154207944 | -0.461727473 | 0.153311586 | 8.542928516 | -1.255984819 | 0.260036235 | 0.709264768 | -4.549760663 |
| **A_55_P2016976** | Zfp729a | 0.203477287 | -0.202304957 | 0.609259532 | 9.216140079 | 1.255952587 | 0.260047062 | 0.709264768 | -4.549769375 |
| **A_51_P388691** | Bbx | -0.647312644 | -1.938213571 | 0.643588283 | 8.337086066 | -1.255947609 | 0.260048735 | 0.709264768 | -4.54977072 |
| **A_55_P2177023** | Mafg | -0.50852246 | -1.522735371 | 0.505690452 | 9.294900814 | -1.255831434 | 0.260087764 | 0.709264768 | -4.549802118 |
| **A_55_P2007919** | Akr1c19 | 0.531979855 | -0.529315727 | 1.593275437 | 8.978496271 | 1.255478199 | 0.260206468 | 0.709264768 | -4.549897582 |
| **A_55_P2144566** | NA | 0.21719182 | -0.216113107 | 0.650496746 | 15.5012076 | 1.255452201 | 0.260215207 | 0.709264768 | -4.549904608 |
| **A_55_P2109717** | Kif20b | 0.835115214 | -0.831190671 | 2.501421099 | 9.987289332 | 1.255284057 | 0.26027173 | 0.709264768 | -4.549950047 |
| **A_55_P1978506** | H2-Q6 | -1.230830039 | -3.686877841 | 1.225217762 | 10.54859352 | -1.255196205 | 0.260301267 | 0.709264768 | -4.549973788 |
| **A_55_P1954907** | Rpl12 | 0.518299869 | -0.515995599 | 1.552595336 | 17.22844744 | 1.255124546 | 0.260325361 | 0.709264768 | -4.549993153 |
| **A_55_P2003473** | Zc4h2 | 0.246918782 | -0.245835764 | 0.739673328 | 8.47453557 | 1.255087002 | 0.260337986 | 0.709264768 | -4.550003298 |
| **A_55_P2179398** | Alg1 | -0.354058033 | -1.060654231 | 0.352538165 | 10.08974881 | -1.255028249 | 0.260357743 | 0.709264768 | -4.550019175 |
| **A_55_P2155002** | Svbp | 0.202536804 | -0.201767738 | 0.606841346 | 12.06611283 | 1.254716699 | 0.260462534 | 0.709264768 | -4.550103362 |
| **A_55_P2114898** | Gdap2 | -0.372907583 | -1.117334208 | 0.371519041 | 9.205966291 | -1.254670432 | 0.260478099 | 0.709264768 | -4.550115864 |
| **A_66_P129622** | Phldb3 | -1.304833364 | -3.909653239 | 1.29998651 | 9.722132551 | -1.254664765 | 0.260480005 | 0.709264768 | -4.550117395 |
| **A_52_P406371** | Med14 | 0.426748169 | -0.425168963 | 1.278665302 | 9.339395241 | 1.254655978 | 0.260482962 | 0.709264768 | -4.550119769 |
| **A_65_P09657** | Zfp213 | 0.154775969 | -0.15421498 | 0.463766919 | 9.254543765 | 1.254608192 | 0.260499039 | 0.709264768 | -4.550132681 |
| **A_55_P2027052** | Prdm9 | -0.335483712 | -1.005239572 | 0.334272149 | 7.963168309 | -1.254599939 | 0.260501816 | 0.709264768 | -4.550134912 |
| **A_52_P82209** | Wnk1 | -0.489415274 | -1.46652822 | 0.487697673 | 10.45072261 | -1.254535909 | 0.26052336 | 0.709264768 | -4.550152213 |
| **A_55_P1964464** | NA | 0.287132185 | -0.286143443 | 0.860407813 | 15.93017278 | 1.254494447 | 0.260537311 | 0.709264768 | -4.550163416 |
| **A_55_P2000487** | Srebf2 | -0.432196839 | -1.295349089 | 0.43095541 | 8.061330532 | -1.254135685 | 0.260658059 | 0.709264768 | -4.550260351 |
| **A_51_P329413** | Pomgnt1 | -0.353797278 | -1.060377783 | 0.352783227 | 10.34612554 | -1.254131806 | 0.260659365 | 0.709264768 | -4.550261399 |
| **A_55_P2348815** | Casd1 | -0.152843909 | -0.458105547 | 0.152417728 | 9.626228943 | -1.25408292 | 0.260675822 | 0.709264768 | -4.550274608 |
| **A_55_P2444515** | NA | -0.649736338 | -1.947473667 | 0.64800099 | 9.870402939 | -1.254009149 | 0.260700659 | 0.709264768 | -4.550294539 |
| **A_52_P561205** | Stard4 | 0.173868483 | -0.173411201 | 0.521148167 | 8.200919157 | 1.253983532 | 0.260709284 | 0.709264768 | -4.550301461 |

| | | | | | | | | | |
|---|---|---|---|---|---|---|---|---|---|
| **A_51_P507778** | Sdr42e1 | -1.03939862 | -3.11566133 | 1.03686409 | 9.763852947 | -1.253863261 | 0.260749781 | 0.709264768 | -4.550333955 |
| **A_66_P139387** | Prlr | -0.463832806 | -1.390658893 | 0.462993282 | 8.023733524 | -1.253468885 | 0.260882615 | 0.709264768 | -4.550440501 |
| **A_51_P114028** | Bcl7c | 0.134746301 | -0.13450453 | 0.403997132 | 9.768327419 | 1.253459037 | 0.260885933 | 0.709264768 | -4.550443162 |
| **A_55_P2085672** | NA | 0.624683796 | -0.623613676 | 1.872981269 | 15.49621543 | 1.253408096 | 0.260903095 | 0.709264768 | -4.550456924 |
| **A_55_P2085335** | Mia | 1.982983662 | -1.979790481 | 5.945757805 | 12.56522987 | 1.253343638 | 0.260924813 | 0.709264768 | -4.550474337 |
| **A_55_P1978236** | Mrps7 | 0.118881236 | -0.118690653 | 0.356453124 | 12.59737944 | 1.253339149 | 0.260926326 | 0.709264768 | -4.55047555 |
| **A_55_P1996953** | NA | -1.261524488 | -3.782727203 | 1.259678227 | 9.236035052 | -1.253251591 | 0.26095583 | 0.709264768 | -4.550499204 |
| **A_51_P487298** | Vasn | -0.143978077 | -0.431754023 | 0.14379787 | 12.26853876 | -1.253118733 | 0.261000604 | 0.709264768 | -4.550535094 |
| **A_55_P1982420** | Fam8a1 | -0.417979951 | -1.253470866 | 0.417510965 | 9.858659047 | -1.253037487 | 0.261027988 | 0.709264768 | -4.550557042 |
| **A_52_P565051** | Cpne3 | -0.15537152 | -0.465981377 | 0.155238338 | 8.757760857 | -1.252871485 | 0.261083946 | 0.709264768 | -4.550601884 |
| **A_55_P2146560** | H2-Ab1 | 1.372355817 | -1.371213276 | 4.11592491 | 10.35417701 | 1.25285604 | 0.261089153 | 0.709264768 | -4.550606056 |
| **A_55_P2131173** | Sphk2 | -0.201933087 | -0.605664494 | 0.201798319 | 11.13681153 | -1.252752553 | 0.261124044 | 0.709264768 | -4.550634011 |
| **A_52_P59681** | Rida | 0.325028282 | -0.324845864 | 0.974902428 | 9.951429686 | 1.252686041 | 0.261146471 | 0.709264768 | -4.550651977 |
| **A_51_P479408** | Hyal1 | -0.231926028 | -0.695705617 | 0.231853561 | 8.163776679 | -1.252530197 | 0.261199026 | 0.709264768 | -4.550694073 |
| **A_66_P121446** | Gm7969 | -0.568719551 | -1.706154768 | 0.568715665 | 8.161883894 | -1.252338793 | 0.261263586 | 0.709264768 | -4.550745773 |
| **A_52_P207335** | Tmem246 | 0.894092378 | -0.894248674 | 2.68243343 | 10.80583107 | 1.252225064 | 0.261301954 | 0.709264768 | -4.550776491 |
| **A_52_P187132** | Ybx1 | 0.575919569 | -0.5760454 | 1.727884537 | 13.10502011 | 1.25219772 | 0.261311179 | 0.709264768 | -4.550783877 |
| **A_55_P2062777** | Mfap5 | -1.671599998 | -5.015277868 | 1.672077871 | 11.43777361 | -1.252155533 | 0.261325413 | 0.709264768 | -4.550795271 |
| **A_55_P1952076** | Npb | -0.58441582 | -1.753420751 | 0.584589111 | 8.665351391 | -1.252148871 | 0.261327661 | 0.709264768 | -4.55079707 |
| **A_52_P470401** | Tmem186 | 0.123417736 | -0.123462103 | 0.370297575 | 8.542872214 | 1.252109455 | 0.26134096 | 0.709264768 | -4.550807716 |
| **A_55_P1952793** | Son | 0.322667203 | -0.322859982 | 0.968194388 | 12.24995385 | 1.251960519 | 0.261391219 | 0.709264768 | -4.550847942 |
| **A_51_P249286** | Rgs16 | -0.595457245 | -1.786810326 | 0.595895835 | 8.345261185 | -1.251873475 | 0.261420596 | 0.709264768 | -4.550871452 |
| **A_55_P1998943** | Oas1a | -1.636010174 | -4.90941805 | 1.637397701 | 9.423977212 | -1.251803677 | 0.261444155 | 0.709264768 | -4.550890303 |
| **A_55_P2057459** | Mall | -0.348825534 | -1.046804877 | 0.349153809 | 9.18768846 | -1.251745514 | 0.261463788 | 0.709264768 | -4.550906011 |
| **A_52_P447284** | Clic6 | 1.095171855 | -1.096279026 | 3.286622736 | 11.4150842 | 1.251701807 | 0.261478543 | 0.709264768 | -4.550917815 |
| **A_51_P317181** | Pcdhb21 | 0.451387148 | -0.451847578 | 1.354621873 | 8.780945264 | 1.251696128 | 0.26148046 | 0.709264768 | -4.550919349 |
| **A_55_P2032559** | NA | 0.570361321 | -0.570971648 | 1.71169429 | 14.20890547 | 1.25166483 | 0.261491026 | 0.709264768 | -4.550927802 |
| **A_55_P2167299** | NA | 0.322619345 | -0.322978473 | 0.968217162 | 15.74276327 | 1.251637875 | 0.261500126 | 0.709264768 | -4.550935082 |
| **A_55_P2022434** | Gpi1 | 0.246545835 | -0.246826106 | 0.739917775 | 14.32905578 | 1.251623098 | 0.261505115 | 0.709264768 | -4.550939073 |
| **A_52_P36928** | Morn2 | 0.4239938 | -0.424539549 | 1.272527148 | 9.261616938 | 1.251529054 | 0.261536867 | 0.709264768 | -4.550964471 |
| **A_55_P1976754** | Ppia | 0.145924161 | -0.146132677 | 0.437980999 | 17.17669216 | 1.251440399 | 0.261566803 | 0.709264768 | -4.550988413 |
| **A_66_P116888** | Arid1a | 0.202588425 | -0.202883146 | 0.608059997 | 9.004609401 | 1.251424243 | 0.261572258 | 0.709264768 | -4.550992777 |
| **A_55_P1961466** | Dct | -0.130804787 | -0.392620845 | 0.13101127 | 7.934524875 | -1.251346852 | 0.261598393 | 0.709264768 | -4.551013677 |
| **A_66_P111572** | Fam122b | -0.529576591 | -1.589653726 | 0.530500545 | 9.372233231 | -1.251242991 | 0.261633472 | 0.709264768 | -4.551041725 |
| **A_52_P571371** | Tmem144 | -0.399425372 | -1.198999621 | 0.400148877 | 8.346299354 | -1.251201324 | 0.261647545 | 0.709264768 | -4.551052977 |
| **A_55_P1957900** | Dlg1 | -0.241079409 | -0.723728271 | 0.241569453 | 9.14408437 | -1.251062993 | 0.261694274 | 0.709264768 | -4.551090333 |
| **A_51_P358225** | Serinc1 | -0.354818587 | -1.065195683 | 0.355558508 | 13.16785327 | -1.251030097 | 0.261705387 | 0.709264768 | -4.551099216 |
| **A_51_P337918** | Aldh4a1 | 0.17582693 | -0.1761969 | 0.527850759 | 10.30912723 | 1.251018335 | 0.261709361 | 0.709264768 | -4.551102393 |
| **A_55_P2412319** | A830052D | -0.247200747 | -0.742158767 | 0.247757273 | 8.033928384 | -1.2509264 | 0.261740423 | 0.709264768 | -4.551127219 |
| **A_55_P1960145** | Clip4 | 0.186909883 | -0.187398841 | 0.561218607 | 8.068832743 | 1.250698597 | 0.261817404 | 0.709264768 | -4.551188733 |
| **A_55_P2060343** | Zfp352 | 0.270147645 | -0.270873365 | 0.811168655 | 9.919658453 | 1.250654645 | 0.261832259 | 0.709264768 | -4.551200601 |
| **A_52_P453624** | Arl6ip6 | 0.163770368 | -0.164229337 | 0.491770074 | 8.736894861 | 1.250582128 | 0.26185677 | 0.709264768 | -4.551220183 |
| **A_66_P103654** | Asxl3 | 0.313019956 | -0.313915185 | 0.939955097 | 8.545992184 | 1.250546249 | 0.261868898 | 0.709264768 | -4.551229871 |
| **A_55_P2048259** | Golph3l | 0.180162277 | -0.180719871 | 0.541044425 | 11.56154573 | 1.250399551 | 0.261918491 | 0.709264768 | -4.551269482 |
| **A_55_P2074796** | Cdkn2c | 0.640858659 | -0.642948943 | 1.924666262 | 9.411050597 | 1.250295475 | 0.261953679 | 0.709264768 | -4.551297583 |
| **A_66_P131398** | Ror2 | 0.930637155 | -0.933906941 | 2.795181252 | 10.55497818 | 1.250138339 | 0.262006816 | 0.709264768 | -4.551340011 |
| **A_51_P254855** | Ptgs2 | -0.711356799 | -2.136612952 | 0.713899353 | 8.078089467 | -1.250100441 | 0.262019633 | 0.709264768 | -4.553963843 |
| **A_52_P456977** | Deaf1 | -0.192019432 | -0.576748343 | 0.19270948 | 8.671065595 | -1.250088332 | 0.262023729 | 0.709264768 | -4.551353513 |

| | | | | | | | | | |
|---|---|---|---|---|---|---|---|---|---|
| **A_55_P1975210** | Fubp3 | 0.243115457 | -0.243993202 | 0.730224117 | 9.646620901 | 1.250077871 | 0.262027267 | 0.709264768 | -4.551356337 |
| **A_55_P2005465** | Zfp422 | 0.764606424 | -0.767768808 | 2.296981657 | 10.41563837 | 1.249750055 | 0.26213816 | 0.709264768 | -4.551444845 |
| **A_51_P327021** | Zbtb25 | 0.133885725 | -0.134511029 | 0.402282479 | 9.954867249 | 1.249416859 | 0.262250915 | 0.709264768 | -4.551534799 |
| **A_51_P415809** | Tusc1 | 0.242131202 | -0.243296788 | 0.727559193 | 9.089738582 | 1.249327469 | 0.262281172 | 0.709264768 | -4.551558931 |
| **A_55_P2161773** | Mpp3 | 0.299896318 | -0.301382852 | 0.901175489 | 9.310937885 | 1.249238385 | 0.262311329 | 0.709264768 | -4.551582981 |
| **A_55_P2073940** | Abcc5 | -0.126506347 | -0.380174197 | 0.127161504 | 8.317017364 | -1.249100065 | 0.26235816 | 0.709264768 | -4.551620321 |
| **A_55_P2012101** | Bmp8b | -0.556242518 | -1.671631929 | 0.559146894 | 8.264352228 | -1.249073545 | 0.26236714 | 0.709264768 | -4.55162748 |
| **A_55_P2054857** | Ube2v2 | 0.440728908 | -0.443049194 | 1.32450701 | 12.2582427 | 1.249046615 | 0.262376259 | 0.709264768 | -4.55163475 |
| **A_55_P1972792** | Stil | 0.278720964 | -0.28025402 | 0.837695948 | 8.414361642 | 1.248899836 | 0.262425964 | 0.709264768 | -4.551674372 |
| **A_51_P440743** | Celsr1 | 1.174121458 | -1.180700611 | 3.528943527 | 11.54399553 | 1.248835609 | 0.262447717 | 0.709264768 | -4.55169171 |
| **A_55_P2055612** | Zfp800 | 0.11662881 | -0.117290914 | 0.350548533 | 8.27938222 | 1.248789811 | 0.262463229 | 0.709264768 | -4.551704073 |
| **A_51_P144581** | Pgrmc1 | -0.175605479 | -0.527857428 | 0.17664647 | 12.91036282 | -1.248633556 | 0.262516159 | 0.709264768 | -4.551746251 |
| **A_55_P1995503** | Gm5643 | 0.79630691 | -0.801049278 | 2.393663098 | 13.45225443 | 1.248616476 | 0.262521946 | 0.709264768 | -4.551750862 |
| **A_55_P1956667** | Cyfip1 | 0.322558947 | -0.324509515 | 0.969627409 | 11.32514015 | 1.24855939 | 0.262541286 | 0.709264768 | -4.551766271 |
| **A_51_P368713** | Tmbim4 | -0.389196019 | -1.169944525 | 0.391552488 | 10.4396152 | -1.248554695 | 0.262542877 | 0.709264768 | -4.551767538 |
| **A_51_P131942** | Dph5 | 0.163344683 | -0.164349302 | 0.491038668 | 9.892603225 | 1.248495205 | 0.262563033 | 0.709264768 | -4.551783596 |
| **A_51_P131025** | Ngdn | 0.13460577 | -0.135445888 | 0.404657428 | 13.33783099 | 1.24843856 | 0.262582227 | 0.709264768 | -4.551798886 |
| **A_52_P568805** | Ubxn1 | -0.539666849 | -1.622380166 | 0.543046468 | 13.62749922 | -1.248425433 | 0.262586675 | 0.709264768 | -4.551802429 |
| **A_55_P2156435** | NA | 0.32342436 | -0.325463584 | 0.972312303 | 10.68181278 | 1.248398872 | 0.262595675 | 0.709264768 | -4.551809598 |
| **A_52_P168567** | Cebpa | 0.649695926 | -0.653929283 | 1.953321134 | 9.577267797 | 1.248267717 | 0.262640123 | 0.709264768 | -4.551844999 |
| **A_55_P2125963** | NA | 0.223858527 | -0.225368391 | 0.673085445 | 9.111490775 | 1.248125383 | 0.262688367 | 0.709264768 | -4.551883417 |
| **A_51_P212183** | Dmac1 | 0.564762758 | -0.568595584 | 1.6981211 | 11.98288253 | 1.248099332 | 0.262697197 | 0.709264768 | -4.551890448 |
| **A_55_P2013038** | Serpinb6c | -0.273848028 | -0.823439312 | 0.275743255 | 12.04768909 | -1.248015927 | 0.262725471 | 0.709264768 | -4.551912959 |
| **A_52_P506995** | Plet1 | -0.810217397 | -2.43632745 | 0.815892656 | 8.12207 | -1.247963763 | 0.262743157 | 0.709264768 | -4.551927039 |
| **A_51_P436342** | Hoxb5 | 0.770409306 | -0.775916267 | 2.316734879 | 9.592156656 | 1.24787455 | 0.262773404 | 0.709264768 | -4.551951117 |
| **A_52_P409731** | Scamp5 | -0.309171464 | -0.929727598 | 0.311384671 | 7.985213428 | -1.247868076 | 0.262775599 | 0.709264768 | -4.551952864 |
| **A_66_P116632** | Mafg | -0.572977973 | -1.723056903 | 0.577100957 | 11.94637026 | -1.247844949 | 0.262783441 | 0.709264768 | -4.551959106 |
| **A_55_P2021099** | Tmem181 | -0.187268307 | -0.563181342 | 0.188644728 | 10.38656156 | -1.24774904 | 0.262815964 | 0.709264768 | -4.551984991 |
| **A_55_P2280237** | Zfp697 | -0.35387938 | -1.064315919 | 0.35655716 | 8.214200034 | -1.24761421 | 0.262861692 | 0.709264768 | -4.55202138 |
| **A_55_P1977478** | Zfp629 | 0.724415544 | -0.729934969 | 2.178766057 | 11.45781259 | 1.247581762 | 0.262872697 | 0.709264768 | -4.552030137 |
| **A_55_P2163446** | NA | 0.117395704 | -0.118296013 | 0.353087421 | 9.093632179 | 1.247550771 | 0.262883209 | 0.709264768 | -4.552038501 |
| **A_51_P114843** | H1f10 | 0.314530136 | -0.316958715 | 0.946018987 | 8.376850854 | 1.247518289 | 0.262894227 | 0.709264768 | -4.552047267 |
| **A_55_P2113071** | Ago4 | -0.129419362 | -0.389276228 | 0.130437504 | 9.576715675 | -1.24742776 | 0.262924937 | 0.709264768 | -4.552071699 |
| **A_55_P1983754** | Pcp4l1 | -0.289403687 | -0.870497006 | 0.291689632 | 9.651962733 | -1.247407995 | 0.262931642 | 0.709264768 | -4.552077033 |
| **A_51_P489996** | Dst | -0.265578899 | -0.798850089 | 0.267692292 | 8.074063789 | -1.247371419 | 0.262944051 | 0.709264768 | -4.552086904 |
| **A_55_P2082361** | Ppt1 | -0.249733944 | -0.751198702 | 0.251730813 | 8.37535254 | -1.247347627 | 0.262952123 | 0.709264768 | -4.552093324 |
| **A_51_P268697** | Slc1a3 | 0.444572744 | -0.448181884 | 1.337327371 | 8.842208768 | 1.2472717 | 0.262977884 | 0.709264768 | -4.552113814 |
| **A_55_P2009187** | Iffo1 | -0.183969711 | -0.553491655 | 0.185552233 | 8.290351102 | -1.246971241 | 0.26307985 | 0.709264768 | -4.552194896 |
| **A_55_P1970788** | C5ar2 | -0.806486671 | -2.426412736 | 0.813439395 | 8.276006109 | -1.246959493 | 0.263083837 | 0.709264768 | -4.552198066 |
| **A_52_P505827** | Cyb5r4 | -0.358696413 | -1.079194728 | 0.361801901 | 8.785062598 | -1.246936709 | 0.263091571 | 0.709264768 | -4.552204215 |
| **A_55_P2014450** | B430105G | -0.182806202 | -0.550041959 | 0.184429555 | 8.235037443 | -1.246798614 | 0.263138449 | 0.709264768 | -4.552241479 |
| **A_55_P1968340** | Plekhh1 | -1.84731411 | -5.558649596 | 1.864021377 | 10.77516749 | -1.246696897 | 0.263172983 | 0.709264768 | -4.552268927 |
| **A_55_P2048498** | NA | 0.172054218 | -0.173667955 | 0.517776391 | 8.420570759 | 1.246488958 | 0.263243593 | 0.709264768 | -4.552325036 |
| **A_52_P171791** | Smg1 | -0.1098742 | -0.330657131 | 0.110908732 | 10.18315834 | -1.246466396 | 0.263251255 | 0.709264768 | -4.552331124 |
| **A_51_P421538** | Slc9b2 | -0.484790924 | -1.459043189 | 0.489461341 | 7.994871172 | -1.246331014 | 0.263297237 | 0.709264768 | -4.552367653 |
| **A_55_P2003581** | Scnm1 | -0.140638285 | -0.423272379 | 0.14199581 | 11.14040508 | -1.246319403 | 0.263301181 | 0.709264768 | -4.552370786 |
| **A_52_P551526** | NA | 0.375334228 | -0.379047287 | 1.129715743 | 16.53162892 | 1.246170534 | 0.263351753 | 0.709264768 | -4.552410954 |
| **A_55_P2122849** | NA | 0.136862394 | -0.138217926 | 0.411942713 | 10.72136389 | 1.2461633 | 0.263354211 | 0.709264768 | -4.552412905 |

| | | | | | | | | | |
|---|---|---|---|---|---|---|---|---|---|
| **A_52_P75465** | Gdpd1 | 0.83013569 | -0.899435294 | 2.559706674 | 10.11731849 | 1.278419542 | 0.263357474 | 0.709264768 | -4.550704638 |
| **A_55_P2185712** | Copa | -0.160235094 | -0.482394633 | 0.161924445 | 11.96097858 | -1.245767479 | 0.263488717 | 0.709264768 | -4.5525197 |
| **A_51_P497985** | C2 | -1.051204852 | -3.164810427 | 1.062400723 | 9.75866988 | -1.245700838 | 0.263511368 | 0.709264768 | -4.552537679 |
| **A_55_P2107647** | NA | 0.370653107 | -0.37465058 | 1.115956793 | 10.93565635 | 1.245617557 | 0.263539679 | 0.709264768 | -4.552560147 |
| **A_55_P2170210** | Ap1s3 | -0.651380451 | -1.961246007 | 0.658485105 | 10.26111982 | -1.245541907 | 0.263565397 | 0.709264768 | -4.552580557 |
| **A_55_P2017312** | Eapp | -0.248441958 | -0.748044053 | 0.251160137 | 11.21985145 | -1.245520954 | 0.263572521 | 0.709264768 | -4.552586209 |
| **A_66_P106536** | Atp5mpl | 0.394391849 | -0.39877669 | 1.187560388 | 14.71062146 | 1.245411285 | 0.263609809 | 0.709264768 | -4.552615796 |
| **A_55_P2025463** | Brpf3 | -0.289394132 | -0.871518168 | 0.292729905 | 9.790998531 | -1.245158203 | 0.263695878 | 0.709264768 | -4.55268407 |
| **A_51_P319070** | Retsat | -0.613916697 | -1.848884875 | 0.62105148 | 9.034137682 | -1.245099401 | 0.263715879 | 0.709264768 | -4.552699933 |
| **A_55_P1988760** | Sestd1 | -0.137219298 | -0.413253081 | 0.138814484 | 9.243027758 | -1.245097333 | 0.263716582 | 0.709264768 | -4.552700491 |
| **A_52_P339791** | Hdgf | 0.131806599 | -0.133339315 | 0.396952513 | 8.62468001 | 1.245095203 | 0.263717307 | 0.709264768 | -4.552701066 |
| **A_55_P2099695** | Msantd4 | 0.248868213 | -0.251774089 | 0.749510515 | 11.08261662 | 1.245065593 | 0.263727379 | 0.709264768 | -4.552709053 |
| **A_55_P2001290** | Ube2e3 | 0.275034737 | -0.278262927 | 0.8283324 | 12.348522 | 1.245027827 | 0.263740226 | 0.709264768 | -4.552719241 |
| **A_51_P336403** | Emc7 | -0.332107321 | -1.000222003 | 0.336007362 | 12.5291071 | -1.245024159 | 0.263741474 | 0.709264768 | -4.55272023 |
| **A_51_P423290** | Mmrn1 | 0.64695406 | -0.654641087 | 1.948549208 | 9.346191148 | 1.244938413 | 0.263770644 | 0.709264768 | -4.552743361 |
| **A_55_P2078830** | NA | -0.340987977 | -1.027062148 | 0.345086193 | 7.970771732 | -1.244853783 | 0.263799438 | 0.709264768 | -4.55276619 |
| **A_51_P301636** | Kazn | -0.870387787 | -2.621909283 | 0.881133708 | 10.11095275 | -1.244651201 | 0.263868375 | 0.709264768 | -4.552820836 |
| **A_52_P180842** | Exoc3 | 0.222073285 | -0.22481509 | 0.668961661 | 10.28196527 | 1.244651035 | 0.263868431 | 0.709264768 | -4.552820881 |
| **A_52_P611798** | Ddx5 | 0.489877529 | -0.495952566 | 1.475707623 | 13.2122204 | 1.244617182 | 0.263879953 | 0.709264768 | -4.552830012 |
| **A_51_P108978** | Myadm | -0.397548804 | -1.1976793 | 0.402581692 | 9.781253033 | -1.244457224 | 0.263934398 | 0.709264768 | -4.552873159 |
| **A_51_P170641** | Dcaf4 | -0.646446819 | -1.947642716 | 0.654749078 | 9.772123404 | -1.244344015 | 0.263972937 | 0.709264768 | -4.552903694 |
| **A_52_P219943** | Epc1 | 0.119906194 | -0.121463639 | 0.361276028 | 8.208423449 | 1.244253798 | 0.264003653 | 0.709264768 | -4.552928028 |
| **A_52_P504787** | Epha7 | 0.422341211 | -0.427837263 | 1.272519685 | 8.598596522 | 1.24423869 | 0.264008797 | 0.709264768 | -4.552932103 |
| **A_55_P2157307** | NA | 0.301389397 | -0.305321006 | 0.9080998 | 10.6964975 | 1.244219128 | 0.264015458 | 0.709264768 | -4.552937379 |
| **A_52_P471143** | Pusl1 | 0.115551429 | -0.117060071 | 0.348162928 | 8.93024844 | 1.244212284 | 0.264017788 | 0.709264768 | -4.552939225 |
| **A_51_P109369** | Fbxo32 | -1.081718046 | -3.259298342 | 1.09586225 | 8.713305117 | -1.24420013 | 0.264021927 | 0.709264768 | -4.553645869 |
| **A_52_P21550** | Gcnt1 | -0.88886424 | -2.678281652 | 0.900553173 | 9.229782825 | -1.244153945 | 0.264037654 | 0.709264768 | -4.552954961 |
| **A_55_P2002884** | Zcchc24 | 0.430732325 | -0.436504192 | 1.297968842 | 12.38781147 | 1.243999639 | 0.264090203 | 0.709264768 | -4.552996579 |
| **A_55_P2136957** | Wbp1l | -0.845321713 | -2.547428846 | 0.856785421 | 13.21138955 | -1.243900029 | 0.26412413 | 0.709264768 | -4.553023444 |
| **A_51_P447545** | Igfbp1 | -0.913791317 | -2.754017706 | 0.926435072 | 8.929488576 | -1.243730024 | 0.264182043 | 0.709264768 | -4.553069295 |
| **A_51_P429246** | Fbxl6 | -0.12856664 | -0.387495586 | 0.130362305 | 8.945399959 | -1.243649613 | 0.26420944 | 0.709264768 | -4.553090982 |
| **A_55_P2078940** | Nsfl1c | 0.151789937 | -0.153960563 | 0.457540437 | 13.29887103 | 1.243443771 | 0.264279582 | 0.709264768 | -4.553146495 |
| **A_52_P256817** | Zik1 | 0.321022681 | -0.325624246 | 0.967669607 | 8.487676014 | 1.243422852 | 0.264286712 | 0.709264768 | -4.553152136 |
| **A_55_P1975640** | Zfp217 | 0.348382236 | -0.35342849 | 1.050192961 | 10.8359969 | 1.243329811 | 0.264318422 | 0.709264768 | -4.553177227 |
| **A_55_P1967105** | Adgrb2 | 0.481680449 | -0.488679968 | 1.452040865 | 9.143858034 | 1.243301026 | 0.264328234 | 0.709264768 | -4.55318499 |
| **A_55_P2087671** | Abhd12 | -0.232935653 | -0.702234351 | 0.236363045 | 8.510173334 | -1.243188439 | 0.264366612 | 0.709264768 | -4.553215352 |
| **A_55_P2079973** | NA | 0.36401364 | -0.369421842 | 1.097449121 | 13.46482457 | 1.243100058 | 0.264396743 | 0.709264768 | -4.553239185 |
| **A_55_P2121096** | Fbn2 | 1.218011365 | -1.236136417 | 3.672159147 | 12.6295571 | 1.243085426 | 0.264401731 | 0.709264768 | -4.553243131 |
| **A_52_P430262** | Rbm15 | 0.146158387 | -0.148334517 | 0.440651291 | 9.402264801 | 1.243080499 | 0.264403411 | 0.709264768 | -4.55324446 |
| **A_55_P2008639** | NA | 0.310620847 | -0.315249151 | 0.936490846 | 12.56675667 | 1.243073511 | 0.264405794 | 0.709264768 | -4.553246344 |
| **A_51_P263591** | Pank1 | 0.300817574 | -0.305313157 | 0.906948304 | 8.373006622 | 1.243046132 | 0.264415129 | 0.709264768 | -4.553253727 |
| **A_51_P466824** | Tyk2 | -0.138277045 | -0.416911305 | 0.140357214 | 8.349981122 | -1.242985099 | 0.264435939 | 0.709264768 | -4.553270186 |
| **A_52_P95740** | Smim19 | -0.205756871 | -0.620426759 | 0.208913017 | 9.992625043 | -1.242802714 | 0.264498135 | 0.709264768 | -4.553319367 |
| **A_55_P2046838** | Amfr | -0.28252997 | -0.851940366 | 0.286880426 | 10.27486503 | -1.242766324 | 0.264510546 | 0.709264768 | -4.55332918 |
| **A_55_P1969882** | Morf4l2 | -0.379812341 | -1.145291837 | 0.385667155 | 15.05164011 | -1.242755962 | 0.26451408 | 0.709264768 | -4.553331974 |
| **A_51_P219505** | Slc41a2 | -0.931239189 | -2.808170759 | 0.945692382 | 8.348996856 | -1.242690991 | 0.26453624 | 0.709264768 | -4.553349493 |
| **A_55_P2015681** | AU01583( | -0.939068015 | -2.831867281 | 0.953731251 | 8.220638854 | -1.242632864 | 0.264556068 | 0.709264768 | -4.553365167 |
| **A_52_P579252** | NA | 0.667305515 | -0.677740066 | 2.012351096 | 14.46837596 | 1.242619195 | 0.264560731 | 0.709264768 | -4.553368853 |

| | | | | | | | | | |
|---|---|---|---|---|---|---|---|---|---|
| **A_55_P1996821** | Myo15b | -0.35248584 | -1.062970394 | 0.357998714 | 7.888543854 | -1.242617256 | 0.264561392 | 0.709264768 | -4.553369375 |
| **A_52_P610987** | Slc28a3 | -0.461257559 | -1.391015332 | 0.468500214 | 8.091749486 | -1.242579041 | 0.264574429 | 0.709264768 | -4.55337968 |
| **A_55_P2133524** | Nbr1 | -0.298296249 | -0.899588298 | 0.302995801 | 8.342480145 | -1.242546572 | 0.264585505 | 0.709264768 | -4.553388435 |
| **A_51_P288906** | Clip1 | -0.386893671 | -1.166777557 | 0.392990216 | 10.03570356 | -1.242544706 | 0.264586142 | 0.709264768 | -4.553388938 |
| **A_55_P1954021** | Scarb1 | 0.150311145 | -0.152693206 | 0.453315496 | 11.03546855 | 1.242489318 | 0.264605038 | 0.709264768 | -4.553403873 |
| **A_55_P2024302** | Spin1 | -0.644140546 | -1.942647662 | 0.65436657 | 8.846911248 | -1.24247211 | 0.264610909 | 0.709264768 | -4.553408512 |
| **A_55_P2184339** | Mdga1 | 0.5510396 | -0.560073937 | 1.662153137 | 8.952813992 | 1.242151926 | 0.264720169 | 0.709264768 | -4.553494843 |
| **A_51_P468329** | Psmb7 | 0.207042408 | -0.210455958 | 0.624540774 | 10.73126635 | 1.242095175 | 0.264739539 | 0.709264768 | -4.553510144 |
| **A_55_P2139625** | Gtf2a2 | 0.538358228 | -0.547236843 | 1.623953298 | 11.17232735 | 1.242092209 | 0.264740552 | 0.709264768 | -4.553510944 |
| **A_55_P1958652** | Arf2 | -0.77862419 | -2.348719769 | 0.79147139 | 12.80282632 | -1.242087372 | 0.264742203 | 0.709264768 | -4.553512248 |
| **A_51_P391764** | Cbx6 | 0.931638688 | -0.947058907 | 2.810336282 | 12.95335207 | 1.24205544 | 0.264753103 | 0.709264768 | -4.553520858 |
| **A_55_P2019190** | NA | 0.356747483 | -0.362666199 | 1.076161164 | 15.58771919 | 1.242031384 | 0.264761314 | 0.709264768 | -4.553527343 |
| **A_55_P2136862** | Dpy30 | 0.412223693 | -0.419121695 | 1.243569082 | 10.52303482 | 1.241943399 | 0.264791349 | 0.709264768 | -4.553551065 |
| **A_52_P201232** | Xpo5 | 0.117202536 | -0.119165791 | 0.353570863 | 8.827935557 | 1.24193273 | 0.264794992 | 0.709264768 | -4.553553942 |
| **A_52_P432153** | Yipf6 | -0.183188973 | -0.552649047 | 0.186271101 | 8.697053849 | -1.24188723 | 0.264810525 | 0.709264768 | -4.553566209 |
| **A_51_P451346** | Klf6 | -0.81873237 | -2.470019137 | 0.832554397 | 12.98499493 | -1.2418519 | 0.264822587 | 0.709264768 | -4.553575734 |
| **A_55_P2040011** | Itga7 | -0.320866913 | -0.968237714 | 0.326503887 | 10.6023369 | -1.241429827 | 0.264966728 | 0.709264768 | -4.553689523 |
| **A_51_P288828** | Slc52a2 | -0.585923294 | -1.768178547 | 0.596331958 | 9.211110448 | -1.241308868 | 0.265008049 | 0.709264768 | -4.553722132 |
| **A_52_P297457** | 0610030E | -0.144774545 | -0.436938956 | 0.147389866 | 10.02675933 | -1.241124192 | 0.265071147 | 0.709264768 | -4.553771915 |
| **A_51_P482121** | Edn2 | -1.168493285 | -3.526674033 | 1.189687463 | 8.325334144 | -1.241079143 | 0.265086541 | 0.709264768 | -4.553784059 |
| **A_55_P2029740** | Gm10240 | 0.232614417 | -0.236839808 | 0.702068642 | 15.612048 | 1.241062696 | 0.265092162 | 0.709264768 | -4.553788493 |
| **A_52_P93902** | Alg5 | -0.761899604 | -2.299568832 | 0.775769624 | 10.76856965 | -1.241038258 | 0.265100513 | 0.709264768 | -4.553795081 |
| **A_55_P1997375** | NA | -0.333512681 | -1.006635356 | 0.339609994 | 7.979441419 | -1.240990556 | 0.265116815 | 0.709264768 | -4.553807939 |
| **A_51_P157270** | Plekhd1 | -0.209819872 | -0.633395051 | 0.213755308 | 8.633403954 | -1.240699078 | 0.265216447 | 0.709264768 | -4.553886509 |
| **A_55_P2168098** | Plrg1 | 0.249435451 | -0.254115394 | 0.752986297 | 11.52772577 | 1.240695462 | 0.265217683 | 0.709264768 | -4.553887484 |
| **A_55_P2049287** | NA | 0.570391217 | -0.581152494 | 1.721934927 | 14.97834322 | 1.240631338 | 0.265239607 | 0.709264768 | -4.553904769 |
| **A_55_P2045916** | NA | -0.716481589 | -2.163030962 | 0.730067783 | 8.273651954 | -1.240572414 | 0.265259753 | 0.709264768 | -4.553920651 |
| **A_55_P1969522** | NA | 0.542496879 | -0.552819797 | 1.637813555 | 9.998264161 | 1.240531766 | 0.265273652 | 0.709264768 | -4.553931607 |
| **A_51_P418259** | Pccb | 0.310111663 | -0.316073212 | 0.936296537 | 9.403818128 | 1.240411752 | 0.265314692 | 0.709264768 | -4.553963956 |
| **A_55_P1967022** | Fgfr1 | -0.836122977 | -2.524487821 | 0.852241867 | 8.606141139 | -1.240378424 | 0.26532609 | 0.709264768 | -4.553972939 |
| **A_55_P2031217** | Vps52 | -0.186738867 | -0.563850505 | 0.19037277 | 11.41282292 | -1.240266836 | 0.265364256 | 0.709264768 | -4.554003015 |
| **A_51_P100298** | Stx3 | -0.332350714 | -1.003556178 | 0.33885475 | 8.196683738 | -1.240199292 | 0.26538736 | 0.709264768 | -4.555216723 |
| **A_52_P330694** | Tle1 | 0.383345136 | -0.390874465 | 1.157564738 | 9.992776135 | 1.240155492 | 0.265402343 | 0.709264768 | -4.554033025 |
| **A_55_P2062787** | Mfap3 | -0.272598058 | -0.823183499 | 0.277987383 | 11.60180233 | -1.24007622 | 0.265429462 | 0.709264768 | -4.554054391 |
| **A_51_P466490** | Armc5 | 0.245087045 | -0.249950519 | 0.74012461 | 10.57859046 | 1.24003101 | 0.265444929 | 0.709264768 | -4.554066576 |
| **A_55_P2083382** | Snf8 | -0.199701263 | -0.603073209 | 0.203670683 | 13.1031358 | -1.240010796 | 0.265451846 | 0.709264768 | -4.554072024 |
| **A_52_P326354** | Erf | 0.35957601 | -0.366731586 | 1.085883607 | 11.95570421 | 1.239996525 | 0.265456728 | 0.709264768 | -4.55407587 |
| **A_55_P2014124** | Gjb3 | -1.832438215 | -5.534281355 | 1.869404924 | 9.525027114 | -1.239828667 | 0.265514166 | 0.709264768 | -4.55412111 |
| **A_55_P2108599** | Tlcd1 | -0.197548562 | -0.596631053 | 0.201533928 | 8.228166932 | -1.239828299 | 0.265514292 | 0.709264768 | -4.554121209 |
| **A_55_P2083351** | Hs3st1 | -0.436705078 | -1.319018179 | 0.445608022 | 7.997931696 | -1.239697885 | 0.265558925 | 0.709264768 | -4.554156356 |
| **A_55_P1959758** | Top3b | -0.127180402 | -0.384171892 | 0.129811089 | 9.925140246 | -1.239515024 | 0.265621519 | 0.709264768 | -4.554205636 |
| **A_52_P186672** | NA | -0.452051285 | -1.365526475 | 0.461423905 | 8.695826129 | -1.239485062 | 0.265631776 | 0.709264768 | -4.554213711 |
| **A_55_P1965076** | NA | 0.621530402 | -0.634574278 | 1.877635081 | 13.00739039 | 1.239329789 | 0.265684938 | 0.709264768 | -4.554255554 |
| **A_51_P310398** | Tk2 | -0.254924321 | -0.770140021 | 0.260291379 | 8.501001372 | -1.239288809 | 0.265698971 | 0.709264768 | -4.554266598 |
| **A_52_P423364** | Fbxl7 | 0.634862965 | -0.648274293 | 1.918000222 | 10.10101703 | 1.239245136 | 0.265713926 | 0.709264768 | -4.554278367 |
| **A_51_P335969** | Des | -0.719317848 | -2.173173563 | 0.734537868 | 8.842890667 | -1.239224165 | 0.265721107 | 0.709264768 | -4.554492057 |
| **A_55_P2037732** | Foxl2 | -0.298677531 | -0.902376726 | 0.305021664 | 13.95339618 | -1.239174024 | 0.265738278 | 0.709264768 | -4.55429753 |
| **A_55_P2074062** | Sdhaf2 | 0.126997799 | -0.129699832 | 0.38369543 | 10.09513425 | 1.239152279 | 0.265745726 | 0.709264768 | -4.554303389 |

| | | | | | | | | | |
|---|---|---|---|---|---|---|---|---|---|
| **A_55_P2077213** | Cenpb | 0.131328501 | -0.134148815 | 0.396805816 | 11.00080964 | 1.239030265 | 0.265787516 | 0.709264768 | -4.554336269 |
| **A_55_P2042396** | Alkbh4 | -0.170669159 | -0.515678961 | 0.174340644 | 9.264353738 | -1.239007567 | 0.265795291 | 0.709264768 | -4.554342385 |
| **A_52_P18081** | Pacc1 | 0.56219473 | -0.574297759 | 1.698687219 | 10.52557773 | 1.23899783 | 0.265798626 | 0.709264768 | -4.554345009 |
| **A_51_P352968** | Marcks | 0.629097914 | -0.642660967 | 1.900856795 | 13.90973949 | 1.238978618 | 0.265805207 | 0.709264768 | -4.554350186 |
| **A_51_P443364** | Cbll1 | 0.119125338 | -0.121694318 | 0.359944994 | 10.26714314 | 1.23897505 | 0.265806429 | 0.709264768 | -4.554351147 |
| **A_66_P128313** | Mtmr2 | 0.189865047 | -0.193973984 | 0.573704078 | 11.081043 | 1.238928468 | 0.265822386 | 0.709264768 | -4.554363699 |
| **A_55_P2012399** | NA | -0.751040048 | -2.269548854 | 0.767468758 | 11.05722277 | -1.238785537 | 0.265871354 | 0.709264768 | -4.554402213 |
| **A_55_P1987730** | Prxl2a | 0.422718704 | -0.431974214 | 1.277411621 | 12.65316866 | 1.238772925 | 0.265875675 | 0.709264768 | -4.554405612 |
| **A_55_P2036285** | Elp4 | 0.321211921 | -0.328271267 | 0.970695109 | 8.856103364 | 1.238722672 | 0.265892894 | 0.709264768 | -4.554419153 |
| **A_55_P2094574** | Rapgef6 | 0.176437736 | -0.180332149 | 0.53320762 | 10.53231665 | 1.238664337 | 0.265912883 | 0.709264768 | -4.554434871 |
| **A_55_P1956863** | Egfr | -0.649193451 | -1.962053814 | 0.663666913 | 8.313690814 | -1.238528312 | 0.265959498 | 0.709264768 | -4.554471522 |
| **A_55_P2046363** | Mapkap1 | -0.294562856 | -0.890387481 | 0.301261769 | 9.089938163 | -1.238254399 | 0.266053389 | 0.709264768 | -4.554545324 |
| **A_51_P118132** | Skil | -0.359836206 | -1.087722179 | 0.368049768 | 10.42585768 | -1.238203004 | 0.26607101 | 0.709264768 | -4.556790286 |
| **A_55_P1990319** | Ergic3 | -0.222311618 | -0.67202161 | 0.227398375 | 12.3265225 | -1.238169116 | 0.266082629 | 0.709264768 | -4.554568302 |
| **A_55_P2058494** | NA | 0.505841932 | -0.517451299 | 1.529135163 | 14.37931018 | 1.23812665 | 0.266097189 | 0.709264768 | -4.554579743 |
| **A_66_P115693** | Rnf207 | -0.202053908 | -0.610849027 | 0.20674121 | 8.082406209 | -1.237975071 | 0.266149167 | 0.709264768 | -4.554620581 |
| **A_51_P350073** | Ndel1 | -0.488707871 | -1.47747893 | 0.500063189 | 12.35182679 | -1.237952361 | 0.266156956 | 0.709264768 | -4.5546267 |
| **A_66_P109562** | Gm12271 | 0.668500521 | -0.684082567 | 2.021083609 | 11.04867668 | 1.237907353 | 0.266172392 | 0.709264768 | -4.554638826 |
| **A_66_P137556** | Tle1 | 0.23789613 | -0.243452965 | 0.719245224 | 8.68667195 | 1.237877198 | 0.266182734 | 0.709264768 | -4.55464695 |
| **A_55_P1999473** | Ppp1r26 | 0.460772457 | -0.471642203 | 1.393187118 | 8.803199272 | 1.23773526 | 0.26623142 | 0.709264768 | -4.554685188 |
| **A_51_P114062** | Ncs1 | 0.181768698 | -0.186066956 | 0.549604351 | 8.10647867 | 1.237700651 | 0.266243292 | 0.709264768 | -4.556334232 |
| **A_55_P1963578** | NA | 0.230489937 | -0.235963783 | 0.696943656 | 15.84289225 | 1.237638337 | 0.26626467 | 0.709264768 | -4.5547113 |
| **A_51_P343913** | Parp4 | -0.709391249 | -2.145068131 | 0.726285633 | 8.707015258 | -1.237597619 | 0.266278639 | 0.709264768 | -4.554722269 |
| **A_51_P272283** | Cmbl | -0.169579181 | -0.512805538 | 0.173647176 | 9.599014653 | -1.237491569 | 0.266315026 | 0.709264768 | -4.554750838 |
| **A_55_P2061110** | Snapin | 0.183742053 | -0.18817268 | 0.555656786 | 9.308856091 | 1.237415429 | 0.266341153 | 0.709264768 | -4.554771349 |
| **A_51_P257156** | Ddrgk1 | -0.758991298 | -2.295281131 | 0.777298534 | 10.21737186 | -1.237411039 | 0.26634266 | 0.709264768 | -4.554772532 |
| **A_52_P53626** | NA | -0.198020271 | -0.598843303 | 0.20280276 | 8.580277502 | -1.23739207 | 0.266349169 | 0.709264768 | -4.554777642 |
| **A_55_P2072288** | Tmem167 | -0.295393122 | -0.893369573 | 0.30258333 | 9.823147032 | -1.237276153 | 0.266388951 | 0.709264768 | -4.554808868 |
| **A_51_P138895** | Ccdc102a | 0.729892025 | -0.747663612 | 2.207447663 | 10.97935098 | 1.237271818 | 0.266390439 | 0.709264768 | -4.554810036 |
| **A_52_P676076** | Rbx1 | 0.552658535 | -0.566258429 | 1.6715755 | 12.07138494 | 1.237112995 | 0.266444955 | 0.709264768 | -4.554852819 |
| **A_55_P2118302** | Fam110a | 0.398962822 | -0.408945264 | 1.206870909 | 12.18571618 | 1.236860778 | 0.266531548 | 0.709264768 | -4.554920758 |
| **A_55_P2018847** | Crlf2 | -0.975910435 | -2.952311213 | 1.000490342 | 12.28939289 | -1.236759603 | 0.266566291 | 0.709264768 | -4.55494801 |
| **A_66_P104527** | Tbl1x | -0.369244782 | -1.117121628 | 0.378632064 | 11.15225203 | -1.236615327 | 0.266615843 | 0.709264768 | -4.554986871 |
| **A_55_P2155499** | Tmem9 | -0.134979572 | -0.408372122 | 0.138412977 | 8.828563709 | -1.236607054 | 0.266618684 | 0.709264768 | -4.554989099 |
| **A_55_P2072041** | NA | -0.438219131 | -1.325842905 | 0.449404643 | 8.422797335 | -1.236553051 | 0.266637234 | 0.709264768 | -4.555003645 |
| **A_55_P2057777** | Fgfr1 | -1.189959859 | -3.600481194 | 1.220561476 | 11.99007034 | -1.236436102 | 0.266677408 | 0.709264768 | -4.555035144 |
| **A_55_P2007768** | Gm8709 | 0.214268183 | -0.219791772 | 0.648328137 | 14.63792568 | 1.236398051 | 0.266690481 | 0.709264768 | -4.555045392 |
| **A_51_P220135** | Nfatc1 | -0.135574796 | -0.41022165 | 0.139072057 | 8.567260696 | -1.236387708 | 0.266694034 | 0.709264768 | -4.555048178 |
| **A_52_P452689** | Atf3 | -0.970886128 | -2.937728409 | 0.995956154 | 8.515884168 | -1.236371842 | 0.266699486 | 0.709264768 | -4.555052451 |
| **A_51_P198411** | Grb2 | 0.119771221 | -0.122867971 | 0.362410412 | 11.95549239 | 1.236351248 | 0.266706561 | 0.709264768 | -4.555057998 |
| **A_55_P1994586** | Itsn1 | -0.205161576 | -0.620820486 | 0.210497334 | 8.060406655 | -1.236258464 | 0.266738441 | 0.709264768 | -4.555082988 |
| **A_55_P2025760** | Adam9 | -0.133336146 | -0.403509102 | 0.136836811 | 11.55311655 | -1.236107855 | 0.266790197 | 0.709264768 | -4.555123551 |
| **A_55_P2018666** | Thrsp | -0.269506714 | -0.815622233 | 0.276608806 | 7.969970454 | -1.236048225 | 0.266810691 | 0.709264768 | -4.55513961 |
| **A_55_P2066782** | Fam104a | -0.461950858 | -1.398056088 | 0.474154373 | 12.43407237 | -1.236008482 | 0.266824351 | 0.709264768 | -4.555150314 |
| **A_55_P2008071** | Prkcd | -0.174359992 | -0.527764855 | 0.179044872 | 9.471429945 | -1.235733054 | 0.266919035 | 0.709264768 | -4.555224489 |
| **A_51_P296448** | Casp2 | 0.689741313 | -0.708386954 | 2.087869581 | 12.88460309 | 1.235633201 | 0.266953368 | 0.709264768 | -4.55525138 |
| **A_51_P346104** | Sgta | -0.202726134 | -0.613703272 | 0.208251004 | 11.9354555 | -1.235499061 | 0.266999497 | 0.709264768 | -4.555287503 |
| **A_55_P1982902** | Tceal3 | 0.770821595 | -0.791918597 | 2.333561787 | 9.884689557 | 1.23542799 | 0.26702394 | 0.709264768 | -4.555306641 |

| | | | | | | | | | |
|---|---|---|---|---|---|---|---|---|---|
| **A_55_P1998656** | Entpd5 | 0.119769006 | -0.123056125 | 0.362594137 | 8.545041657 | 1.235381684 | 0.267039867 | 0.709264768 | -4.555319111 |
| **A_52_P657338** | Mrps17 | 0.312907323 | -0.321505292 | 0.947319937 | 11.94840406 | 1.235362069 | 0.267046614 | 0.709264768 | -4.555324393 |
| **A_51_P306151** | Sgk3 | -0.239303395 | -0.724512146 | 0.245905356 | 7.912088148 | -1.235294709 | 0.267069785 | 0.709264768 | -4.555342532 |
| **A_55_P2170515** | NA | -0.399464643 | -1.209440691 | 0.410511405 | 8.450745112 | -1.235254699 | 0.267083548 | 0.709264768 | -4.555353306 |
| **A_55_P1978860** | Trp53bp2 | 0.409208486 | -0.420535936 | 1.238952907 | 10.33220751 | 1.235237977 | 0.267089301 | 0.709264768 | -4.555357809 |
| **A_55_P1986743** | NA | -1.474884702 | -4.465848652 | 1.516079248 | 11.06337457 | -1.235086111 | 0.26714155 | 0.709264768 | -4.555398702 |
| **A_55_P1985895** | Thumpd1 | 0.19033447 | -0.195687966 | 0.576356905 | 11.53400003 | 1.234966696 | 0.26718264 | 0.709264768 | -4.555430857 |
| **A_55_P2142830** | Ninj1 | -1.01714977 | -3.080129099 | 1.045829559 | 12.38947076 | -1.234924408 | 0.267197193 | 0.709264768 | -4.555442243 |
| **A_55_P1981879** | Ercc6l2 | 0.388241445 | -0.399261692 | 1.175744583 | 10.93226559 | 1.23480946 | 0.267236754 | 0.709264768 | -4.555473194 |
| **A_51_P210956** | Vcam1 | 1.204965928 | -1.240035936 | 3.649967792 | 11.94110884 | 1.234371591 | 0.267387499 | 0.709264768 | -4.555591088 |
| **A_55_P2081925** | Zfp53 | 0.208183529 | -0.21427407 | 0.630641129 | 9.241769384 | 1.234279698 | 0.267419145 | 0.709264768 | -4.555615829 |
| **A_66_P114054** | Sco2 | -0.537614562 | -1.628666134 | 0.553437011 | 9.691533309 | -1.234173137 | 0.267455846 | 0.709264768 | -4.555644518 |
| **A_66_P139196** | Baz1a | 0.14515201 | -0.149443691 | 0.439747711 | 8.570413957 | 1.234090456 | 0.267484325 | 0.709264768 | -4.555666777 |
| **A_52_P167511** | NA | -0.115292304 | -0.349289353 | 0.118704744 | 9.116405098 | -1.234071395 | 0.267490891 | 0.709264768 | -4.555671909 |
| **A_52_P304947** | Cenpn | 0.298021985 | -0.306890736 | 0.902934706 | 8.429190695 | 1.233973779 | 0.26752452 | 0.709264768 | -4.555698189 |
| **A_55_P2042184** | NA | -0.590081548 | -1.787944205 | 0.60778111 | 8.477079416 | -1.233830079 | 0.26757403 | 0.709264768 | -4.555736875 |
| **A_52_P33453** | NA | 0.481910132 | -0.496387071 | 1.460207334 | 12.43302034 | 1.233802344 | 0.267583587 | 0.709264768 | -4.555744341 |
| **A_55_P2052181** | Ccdc82 | -0.196317927 | -0.594855513 | 0.202219658 | 11.32486001 | -1.233789358 | 0.267588062 | 0.709264768 | -4.555747837 |
| **A_51_P196895** | Ist1 | -0.27293422 | -0.82705872 | 0.281190279 | 9.861484429 | -1.233675626 | 0.267627255 | 0.709264768 | -4.555778454 |
| **A_51_P215038** | Tmem59l | -0.799728633 | -2.423432951 | 0.823975684 | 9.378364785 | -1.233633192 | 0.267641879 | 0.709264768 | -4.555789877 |
| **A_52_P550049** | Nfya | 0.221632513 | -0.228408907 | 0.671673933 | 8.969590422 | 1.233477777 | 0.267695447 | 0.709264768 | -4.555831713 |
| **A_55_P1989568** | Tasp1 | 0.166477397 | -0.171610048 | 0.504564842 | 8.278855353 | 1.233322286 | 0.267749051 | 0.709264768 | -4.555873569 |
| **A_51_P197378** | Atg12 | -0.543054593 | -1.645941439 | 0.559832254 | 12.19990214 | -1.23328338 | 0.267762465 | 0.709264768 | -4.555884042 |
| **A_55_P2122491** | Gm6222 | -0.163402547 | -0.495275093 | 0.168469999 | 15.73959794 | -1.233212279 | 0.26778698 | 0.709264768 | -4.555903181 |
| **A_51_P490171** | Nubp2 | -0.323897508 | -0.981754329 | 0.333959313 | 9.978256197 | -1.233180276 | 0.267798016 | 0.709264768 | -4.555911795 |
| **A_52_P1114640** | 2410003L | -0.289208521 | -0.876690404 | 0.298273362 | 7.797022936 | -1.233011002 | 0.267856392 | 0.709264768 | -4.555957359 |
| **A_66_P106231** | Nccrp1 | -1.395316759 | -4.229697693 | 1.439064174 | 8.689258277 | -1.233005286 | 0.267858363 | 0.709264768 | -4.555958897 |
| **A_52_P2732** | Mfn1 | -0.329725325 | -0.999524678 | 0.340074027 | 10.59637003 | -1.232985381 | 0.267865228 | 0.709264768 | -4.555964255 |
| **A_51_P324082** | Med1 | 0.349173958 | -0.360146203 | 1.058494118 | 10.43429114 | 1.232962554 | 0.267873101 | 0.709264768 | -4.555970399 |
| **A_51_P179241** | Tssc4 | 0.286178038 | -0.295210759 | 0.867566835 | 11.04120476 | 1.232877676 | 0.267902378 | 0.709264768 | -4.555993245 |
| **A_51_P132459** | Rpl27a | 0.959307215 | -0.989637865 | 2.908252294 | 13.10008868 | 1.232844935 | 0.267913672 | 0.709264768 | -4.556002057 |
| **A_55_P1953043** | E130311K | -0.295012001 | -0.894379385 | 0.304355383 | 9.31817325 | -1.232812197 | 0.267924965 | 0.709264768 | -4.556010869 |
| **A_51_P394735** | Arhgef18 | -0.920913924 | -2.791923245 | 0.950095398 | 10.16009609 | -1.232802293 | 0.267928382 | 0.709264768 | -4.556013535 |
| **A_52_P31011** | Cep162 | -0.413757922 | -1.254385309 | 0.426869465 | 8.445714377 | -1.232801439 | 0.267928677 | 0.709264768 | -4.556013765 |
| **A_55_P1976926** | Tmem104 | -0.580983643 | -1.761638886 | 0.599671601 | 8.145011176 | -1.232511983 | 0.268028548 | 0.709264768 | -4.556091671 |
| **A_51_P109171** | Os9 | -0.208674885 | -0.632760134 | 0.215410365 | 12.33080254 | -1.232444469 | 0.268051847 | 0.709264768 | -4.557619431 |
| **A_55_P2036280** | Psen2 | -0.3726626 | -1.130068128 | 0.384742928 | 10.29413108 | -1.23236026 | 0.268080911 | 0.709264768 | -4.556132505 |
| **A_55_P2127497** | Trim35 | 0.412263905 | -0.425692006 | 1.250219816 | 13.09071559 | 1.232266068 | 0.268113423 | 0.709264768 | -4.556157855 |
| **A_55_P1956718** | Rab38 | 1.057326902 | -1.091867599 | 3.206521404 | 10.27418036 | 1.232207669 | 0.268133582 | 0.709264768 | -4.556173572 |
| **A_55_P2028710** | Rnf170 | -0.209908777 | -0.636610374 | 0.216792821 | 10.56677232 | -1.232130403 | 0.268160256 | 0.709264768 | -4.556194365 |
| **A_52_P664699** | Cul1 | -0.240788714 | -0.730318044 | 0.248740616 | 8.865854005 | -1.231991624 | 0.268208171 | 0.709264768 | -4.556231713 |
| **A_55_P1997837** | Osbp2 | -0.554940906 | -1.683199563 | 0.573317752 | 8.020341537 | -1.231936746 | 0.268227121 | 0.709264768 | -4.556246482 |
| **A_55_P2183735** | Pisd | -1.047554782 | -3.177547878 | 1.082438314 | 10.53395548 | -1.231824659 | 0.268265829 | 0.709264768 | -4.556276645 |
| **A_55_P2108255** | Lgmn | -0.904369945 | -2.743288429 | 0.934548539 | 13.32106933 | -1.231782383 | 0.26828043 | 0.709264768 | -4.556288022 |
| **A_55_P2110362** | Zfp637 | 0.241049408 | -0.249125328 | 0.731224145 | 9.177912864 | 1.231701558 | 0.268308346 | 0.709264768 | -4.556309772 |
| **A_55_P2077826** | Ppp1cc | 0.50262426 | -0.519495128 | 1.524743647 | 12.46115295 | 1.231663769 | 0.268321399 | 0.709264768 | -4.556319941 |
| **A_55_P2316612** | 6720407P | 0.312433146 | -0.322997654 | 0.947863946 | 8.743690993 | 1.231513526 | 0.268373301 | 0.709264768 | -4.55636037 |
| **A_51_P429983** | Supt5 | -0.545524438 | -1.655052469 | 0.564003592 | 12.58753172 | -1.231476923 | 0.268385947 | 0.709264768 | -4.556370219 |

| | | | | | | | | | |
|---|---|---|---|---|---|---|---|---|---|
| **A_55_P1953252** | Ctnnd1 | -0.114283861 | -0.34675826 | 0.118190538 | 13.2236154 | -1.231289331 | 0.268450766 | 0.709264768 | -4.556420697 |
| **A_55_P1954925** | Endov | -0.243418004 | -0.738576355 | 0.251740348 | 16.10206912 | -1.231285978 | 0.268451925 | 0.709264768 | -4.556421599 |
| **A_55_P2183894** | Gm5617 | -0.110904547 | -0.33653195 | 0.114722856 | 10.80604612 | -1.231141165 | 0.268501972 | 0.709264768 | -4.556460564 |
| **A_55_P2122618** | Asb11 | -0.43598667 | -1.323261377 | 0.451288037 | 8.059978262 | -1.230737562 | 0.268641501 | 0.709264768 | -4.556569157 |
| **A_52_P640386** | Usp53 | -0.345466519 | -1.048529269 | 0.357596232 | 7.897015549 | -1.230728394 | 0.268644671 | 0.709264768 | -4.556571624 |
| **A_55_P2158011** | Knstrn | 0.593563971 | -0.614461124 | 1.801589065 | 9.234079516 | 1.23067087 | 0.268664563 | 0.709264768 | -4.556587101 |
| **A_52_P428745** | Camk2d | -0.222703907 | -0.675963345 | 0.23055553 | 11.22868625 | -1.23064085 | 0.268674945 | 0.709264768 | -4.556595178 |
| **A_55_P2031045** | Stat5b | 0.403746978 | -0.418007826 | 1.225501781 | 10.01018894 | 1.230601326 | 0.268688613 | 0.709264768 | -4.556605811 |
| **A_55_P2080021** | Papss2 | -0.969734989 | -2.943663959 | 1.004193981 | 9.890063629 | -1.230472438 | 0.268733191 | 0.709264768 | -4.556640487 |
| **A_55_P2419854** | NA | 0.185744064 | -0.192407715 | 0.563895844 | 8.330599415 | 1.230266339 | 0.268804488 | 0.709264768 | -4.556695934 |
| **A_55_P2166555** | NA | -0.436792823 | -1.326188906 | 0.45260326 | 11.91882324 | -1.230072266 | 0.268871639 | 0.709264768 | -4.556748143 |
| **A_55_P2124096** | NA | 0.256356095 | -0.265686786 | 0.778398975 | 13.47416239 | 1.229951016 | 0.268913601 | 0.709264768 | -4.556780761 |
| **A_55_P1971347** | Mta3 | -0.46367243 | -1.407934704 | 0.480589844 | 8.07988618 | -1.229897674 | 0.268932063 | 0.709264768 | -4.55679511 |
| **A_51_P259415** | Tmem70 | 0.175100299 | -0.18149243 | 0.531693029 | 10.6160028 | 1.229885695 | 0.268936209 | 0.709264768 | -4.556798332 |
| **A_55_P2144086** | NA | -0.165248445 | -0.501798715 | 0.171301825 | 16.16417307 | -1.229809328 | 0.268962643 | 0.709264768 | -4.556818875 |
| **A_55_P1965194** | Fbxl5 | -0.122987444 | -0.37348103 | 0.127506141 | 9.006581286 | -1.229743436 | 0.268985452 | 0.709264768 | -4.5568366 |
| **A_52_P358860** | Gss | -0.17180093 | -0.521726675 | 0.178124814 | 8.826687966 | -1.229702234 | 0.268999716 | 0.709264768 | -4.556847684 |
| **A_51_P314679** | Rrn3 | 0.23487447 | -0.243552501 | 0.713301442 | 10.97483732 | 1.229618827 | 0.269028593 | 0.709264768 | -4.556870119 |
| **A_55_P2151863** | NA | 0.449253478 | -0.465858463 | 1.364365419 | 12.62413302 | 1.229610522 | 0.269031468 | 0.709264768 | -4.556872353 |
| **A_55_P2073397** | NA | 0.855462905 | -0.88727182 | 2.59819763 | 12.02729263 | 1.229476531 | 0.269077863 | 0.709264768 | -4.556908395 |
| **A_51_P253633** | Mrps9 | 0.198289896 | -0.205696339 | 0.602276131 | 9.961233343 | 1.229374958 | 0.269113039 | 0.709264768 | -4.556935716 |
| **A_55_P2084739** | Gcnt1 | -0.675220619 | -2.051160437 | 0.700719198 | 8.182078329 | -1.229126558 | 0.269199078 | 0.709264768 | -4.557002529 |
| **A_55_P2107192** | NA | -0.642674134 | -1.952349065 | 0.667000796 | 12.8471826 | -1.229072926 | 0.269217658 | 0.709264768 | -4.557016954 |
| **A_55_P2029121** | Pex11g | -0.140681029 | -0.427377642 | 0.146015584 | 8.549266301 | -1.229032365 | 0.26923171 | 0.709264768 | -4.557027863 |
| **A_55_P2068663** | Stmn1 | 0.635861263 | -0.660074125 | 1.93179665 | 15.60664742 | 1.228936278 | 0.269265003 | 0.709264768 | -4.557053707 |
| **A_51_P465148** | Ctsb | -1.108518377 | -3.368008867 | 1.150972113 | 13.8135651 | -1.228804307 | 0.269310734 | 0.709264768 | -4.557089201 |
| **A_66_P134542** | Anln | 1.201027644 | -1.247036994 | 3.649092281 | 11.45235572 | 1.228797923 | 0.269312946 | 0.709264768 | -4.557090918 |
| **A_55_P2126644** | NA | 0.441982671 | -0.459058773 | 1.343024116 | 10.92591488 | 1.228600876 | 0.269381241 | 0.709264768 | -4.557143912 |
| **A_52_P624149** | Hsp90b1 | -0.505010158 | -1.534637134 | 0.524616818 | 15.12216511 | -1.228486948 | 0.269420734 | 0.709264768 | -4.557174551 |
| **A_52_P233441** | Gata2 | -0.960608541 | -2.919165691 | 0.997948608 | 9.449188602 | -1.228458645 | 0.269430547 | 0.709264768 | -4.557182163 |
| **A_55_P1993518** | Tbc1d14 | -0.136169887 | -0.413815278 | 0.141475504 | 11.05897924 | -1.228403242 | 0.269449755 | 0.709264768 | -4.557197062 |
| **A_55_P2035937** | Rab6a | 0.1466722 | -0.152389485 | 0.445733886 | 12.49366447 | 1.228393122 | 0.269453263 | 0.709264768 | -4.557199784 |
| **A_51_P411909** | Sirt5 | 0.417549544 | -0.433830743 | 1.268929832 | 9.681076186 | 1.228385749 | 0.26945582 | 0.709264768 | -4.557201767 |
| **A_52_P93837** | Mme | -0.776753565 | -2.360599932 | 0.807092803 | 8.389376532 | -1.228345525 | 0.269469767 | 0.709264768 | -4.557212584 |
| **A_66_P127311** | Esco1 | -0.282459339 | -0.858421109 | 0.293502431 | 8.213109977 | -1.228323121 | 0.269477535 | 0.709264768 | -4.557218609 |
| **A_52_P4846** | Zfp451 | -0.300031513 | -0.912177124 | 0.312114097 | 9.044760471 | -1.227615826 | 0.269722881 | 0.709264768 | -4.557408804 |
| **A_55_P2126572** | Tmem25 | -0.205631451 | -0.625215164 | 0.213952262 | 8.018303677 | -1.227499332 | 0.26976331 | 0.709264768 | -4.557440127 |
| **A_51_P256202** | Ctsz | -0.305257233 | -0.928135957 | 0.317621491 | 12.85247964 | -1.227475441 | 0.269771602 | 0.709264768 | -4.557446551 |
| **A_51_P435198** | Cryz | 0.42123301 | -0.438386458 | 1.280852478 | 10.95214008 | 1.227344556 | 0.269817032 | 0.709264768 | -4.557481743 |
| **A_51_P423578** | Slfn2 | -0.173093414 | -0.526332111 | 0.180145282 | 8.105260195 | -1.22733358 | 0.269820842 | 0.709264768 | -4.557484694 |
| **A_55_P2071477** | Prl6a1 | -0.422886659 | -1.285937403 | 0.440164085 | 7.82912197 | -1.227264011 | 0.269844992 | 0.709264768 | -4.557503399 |
| **A_55_P2104941** | Rpl30 | 0.3621583 | -0.376971682 | 1.101288282 | 15.41269034 | 1.227235669 | 0.269854832 | 0.709264768 | -4.55751102 |
| **A_51_P441426** | Pf4 | 0.461851456 | -0.480775186 | 1.404478099 | 12.02303199 | 1.227193234 | 0.269869564 | 0.709264768 | -4.557522429 |
| **A_55_P2048779** | Lratd2 | 0.480448682 | -0.50016143 | 1.461058793 | 8.856942082 | 1.227159417 | 0.269881305 | 0.709264768 | -4.557531521 |
| **A_55_P2052936** | NA | 0.257826121 | -0.268432536 | 0.784084779 | 16.20422299 | 1.227094493 | 0.269903848 | 0.709264768 | -4.557548976 |
| **A_55_P2173892** | Isg20 | -1.11789553 | -3.400110028 | 1.164318968 | 10.85937753 | -1.226860277 | 0.269985184 | 0.709264768 | -4.557611946 |
| **A_55_P1981681** | Syt8 | -0.496677058 | -1.510818599 | 0.517464483 | 7.924283532 | -1.226664716 | 0.270053114 | 0.709264768 | -4.557664521 |
| **A_55_P1995924** | Zfp984 | 0.164218333 | -0.171100207 | 0.499536873 | 9.630946551 | 1.226632362 | 0.270064354 | 0.709264768 | -4.557673219 |

| | | | | | | | | | |
|---|---|---|---|---|---|---|---|---|---|
| **A_52_P251366** | Neil3 | 0.441257873 | -0.459770729 | 1.342286476 | 8.68934358 | 1.226603601 | 0.270074346 | 0.709264768 | -4.557680951 |
| **A_55_P2341468** | Rbms3 | 1.106150282 | -1.152972479 | 3.365273044 | 12.07938477 | 1.226378841 | 0.270152442 | 0.709264768 | -4.557741373 |
| **A_55_P2167451** | Cnnm2 | -0.798367173 | -2.4289651 | 0.832230755 | 9.844627184 | -1.22632655 | 0.270170614 | 0.709264768 | -4.55775543 |
| **A_55_P2199717** | 9330102E | -0.281085077 | -0.855201447 | 0.293031293 | 7.873694969 | -1.226275931 | 0.270188206 | 0.709264768 | -4.557769037 |
| **A_55_P2055257** | Maoa | 0.18210141 | -0.189867884 | 0.554070703 | 9.411335298 | 1.226186593 | 0.270219257 | 0.709264768 | -4.557793052 |
| **A_55_P2007646** | Cryaa | -1.589591325 | -4.836733396 | 1.657550747 | 8.755507519 | -1.226124411 | 0.270240871 | 0.709264768 | -4.557809767 |
| **A_55_P1972699** | Rars2 | 0.122271293 | -0.127503429 | 0.372046014 | 10.95813877 | 1.226101333 | 0.270248893 | 0.709264768 | -4.557815971 |
| **A_55_P2089506** | NA | 0.599947536 | -0.625637261 | 1.825532334 | 13.86402704 | 1.226084084 | 0.27025489 | 0.709264768 | -4.557820608 |
| **A_55_P2115255** | Fer | -0.318036938 | -0.967791511 | 0.331717635 | 8.830209954 | -1.225966391 | 0.270295805 | 0.709264768 | -4.557852244 |
| **A_55_P2134616** | Med12l | -0.357119003 | -1.086756442 | 0.372518435 | 8.205827163 | -1.22590325 | 0.270317758 | 0.709264768 | -4.557869216 |
| **A_55_P2014315** | NA | -0.292780218 | -0.890969494 | 0.305409059 | 7.919393305 | -1.225895502 | 0.270320452 | 0.709264768 | -4.557871299 |
| **A_51_P494775** | Rcor1 | -0.333006142 | -1.013403986 | 0.347391701 | 12.40499273 | -1.225856576 | 0.270333987 | 0.709264768 | -4.557881762 |
| **A_55_P1994032** | Xbp1 | -0.265495773 | -0.807989066 | 0.276997519 | 13.92793218 | -1.225782972 | 0.270359582 | 0.709264768 | -4.557901546 |
| **A_51_P241843** | Tmem202 | -0.273275227 | -0.831737058 | 0.285186605 | 7.811714302 | -1.225623594 | 0.270415009 | 0.709264768 | -4.557944385 |
| **A_51_P214343** | Sephs1 | 0.611817398 | -0.63860086 | 1.862235657 | 9.830025165 | 1.225510007 | 0.270454518 | 0.709264768 | -4.557974914 |
| **A_51_P128147** | Chmp1a | -0.307076072 | -0.934678152 | 0.320526007 | 11.11616985 | -1.225496144 | 0.270459341 | 0.709264768 | -4.559087422 |
| **A_51_P158018** | Bend5 | -0.982797548 | -2.991455264 | 1.025860167 | 10.4654782 | -1.225486334 | 0.270462753 | 0.709264768 | -4.557981277 |
| **A_51_P192089** | Pinx1 | 0.25356224 | -0.26467495 | 0.771799429 | 9.572359033 | 1.225480343 | 0.270464837 | 0.709264768 | -4.557982887 |
| **A_55_P2029528** | NA | -0.786945892 | -2.395465728 | 0.821573944 | 8.485463745 | -1.225374384 | 0.270501699 | 0.709264768 | -4.558011366 |
| **A_55_P2099418** | Neil2 | -1.046250154 | -3.184823048 | 1.09232274 | 8.458982771 | -1.225354705 | 0.270508545 | 0.709264768 | -4.558016655 |
| **A_51_P411130** | Rpp25l | 0.469230042 | -0.489927931 | 1.428388014 | 12.64098213 | 1.225310102 | 0.270524064 | 0.709264768 | -4.558028643 |
| **A_55_P1964965** | Gm1673 | 0.53073109 | -0.554186439 | 1.615648619 | 11.27553889 | 1.225259698 | 0.270541601 | 0.709264768 | -4.55804219 |
| **A_55_P2153016** | Ccdc34 | 0.290456588 | -0.3033394 | 0.884252576 | 8.533679341 | 1.225164257 | 0.270574812 | 0.709264768 | -4.55806784 |
| **A_55_P1989544** | Cklf | -0.320719943 | -0.976434127 | 0.334994242 | 9.443239876 | -1.225072335 | 0.270606801 | 0.709264768 | -4.558092545 |
| **A_55_P1961089** | Npb | -0.585998538 | -1.784116882 | 0.612119807 | 8.428655147 | -1.225031229 | 0.270621108 | 0.709264768 | -4.558103592 |
| **A_51_P156800** | Mob2 | -0.197766494 | -0.602117467 | 0.206584478 | 10.47266891 | -1.225023921 | 0.270623651 | 0.709264768 | -4.558105556 |
| **A_55_P2185605** | Cd48 | 0.307429915 | -0.321168743 | 0.936028573 | 8.52207012 | 1.22496314 | 0.270644806 | 0.709264768 | -4.558121891 |
| **A_55_P2145059** | Gbp5 | -0.237486116 | -0.723090427 | 0.248118196 | 8.396470011 | -1.224915233 | 0.270661482 | 0.709264768 | -4.558134766 |
| **A_55_P2103352** | Ndufb5 | 0.37203593 | -0.388750488 | 1.132822348 | 13.29067105 | 1.224820593 | 0.270694427 | 0.709264768 | -4.5581602 |
| **A_55_P2043028** | Rexo5 | -0.142935964 | -0.435256389 | 0.149384461 | 9.078149017 | -1.22470841 | 0.270733483 | 0.709264768 | -4.558190348 |
| **A_55_P1997872** | Apip | 0.454597622 | -0.475124317 | 1.384319561 | 11.01657085 | 1.224685078 | 0.270741607 | 0.709264768 | -4.558196618 |
| **A_51_P513062** | B630019K | 0.374990374 | -0.392074296 | 1.142055044 | 8.946227896 | 1.224442753 | 0.270825991 | 0.709264768 | -4.558261736 |
| **A_51_P206445** | Zfp773 | -0.183747585 | -0.559625521 | 0.192130352 | 8.209035337 | -1.224405157 | 0.270839085 | 0.709264768 | -4.558271839 |
| **A_52_P660242** | Isy1 | 0.27744885 | -0.290137712 | 0.845035411 | 10.49712269 | 1.224337552 | 0.270862632 | 0.709264768 | -4.558290006 |
| **A_51_P184728** | Cnksr3 | 0.408587789 | -0.427277991 | 1.244453569 | 10.11377138 | 1.224331951 | 0.270864582 | 0.709264768 | -4.558291511 |
| **A_51_P234044** | 1190005I( | -0.204278864 | -0.637730478 | 0.22917275 | 9.141493974 | -1.255294179 | 0.270901604 | 0.709264768 | -4.555676039 |
| **A_55_P2136318** | Slc1a5 | -0.173276101 | -0.527793534 | 0.181241333 | 8.618837375 | -1.224197292 | 0.27091149 | 0.709264768 | -4.558327695 |
| **A_55_P2120919** | NA | -0.744989942 | -2.269221371 | 0.779241488 | 8.268871648 | -1.22419286 | 0.270913034 | 0.709264768 | -4.558328886 |
| **A_55_P2147191** | NA | 0.3731659 | -0.39036891 | 1.136700709 | 17.08962139 | 1.224118481 | 0.270938947 | 0.709264768 | -4.558348872 |
| **A_51_P245368** | Abcb1b | -0.80279893 | -2.445436807 | 0.839838947 | 8.362337768 | -1.224095489 | 0.270946958 | 0.709264768 | -4.55835505 |
| **A_55_P1998539** | C1qtnf1 | -1.739691066 | -5.299364964 | 1.819982831 | 9.228778162 | -1.224086941 | 0.270949936 | 0.709264768 | -4.558357347 |
| **A_51_P508029** | Pcyox1 | -0.300724112 | -0.916096287 | 0.314648064 | 9.735147569 | -1.223998093 | 0.270980894 | 0.709264768 | -4.55838122 |
| **A_55_P2094925** | Srgn | -1.445933586 | -4.405184488 | 1.513317317 | 11.39689404 | -1.223818185 | 0.271043589 | 0.709264768 | -4.55842956 |
| **A_55_P1985070** | Robo1 | 0.924110737 | -0.967198338 | 2.815419811 | 11.13219505 | 1.223803963 | 0.271048546 | 0.709264768 | -4.558433381 |
| **A_51_P103850** | Fbn2 | 0.850892865 | -0.890740134 | 2.592525864 | 10.80729622 | 1.223682031 | 0.271091045 | 0.709264768 | -4.558466142 |
| **A_51_P452820** | Rpl31 | 0.696757146 | -0.729534772 | 2.123049065 | 10.38814566 | 1.223554605 | 0.271135466 | 0.709264768 | -4.558500378 |
| **A_55_P2042958** | Rad50 | 0.347883469 | -0.364250286 | 1.060017225 | 9.824758479 | 1.223552379 | 0.271136242 | 0.709264768 | -4.558500976 |
| **A_55_P1961461** | Bnip3l | -0.184618294 | -0.562571977 | 0.193335389 | 13.05107483 | -1.223450766 | 0.27117167 | 0.709264768 | -4.558528277 |

| | | | | | | | | | |
|---|---|---|---|---|---|---|---|---|---|
| **A_55_P1987746** | NA | 0.397822209 | -0.416675555 | 1.212319973 | 15.62699221 | 1.223346471 | 0.271208036 | 0.709264768 | -4.558556297 |
| **A_55_P1997628** | NA | 0.238324062 | -0.249664539 | 0.726312662 | 14.74891994 | 1.223231229 | 0.271248225 | 0.709264768 | -4.558587258 |
| **A_55_P2097553** | C77080 | -0.641016608 | -1.95367454 | 0.671641323 | 11.77742617 | -1.223117164 | 0.271288008 | 0.709264768 | -4.558617901 |
| **A_55_P1953894** | Mccc2 | 0.382087447 | -0.400385191 | 1.164560084 | 9.411069383 | 1.223049277 | 0.271311687 | 0.709264768 | -4.558636139 |
| **A_55_P1968738** | Nfkb1 | 0.318900184 | -0.334244048 | 0.972044416 | 8.919661519 | 1.222914289 | 0.271358779 | 0.709264768 | -4.558672402 |
| **A_55_P1975041** | Arhgef11 | -0.284805404 | -0.868160563 | 0.298549756 | 9.673168458 | -1.222828429 | 0.271388735 | 0.709264768 | -4.558695467 |
| **A_55_P1956097** | Cyb5d2 | -0.417711656 | -1.273334891 | 0.437911579 | 9.214303833 | -1.222768861 | 0.27140952 | 0.709264768 | -4.558711469 |
| **A_55_P1975560** | Ifi204 | -0.467367218 | -1.424709016 | 0.489974579 | 8.915405295 | -1.222760982 | 0.271412269 | 0.709264768 | -4.558713586 |
| **A_55_P1998697** | NA | 0.313121137 | -0.328278871 | 0.954521145 | 12.23979326 | 1.222739014 | 0.271419935 | 0.709264768 | -4.558719487 |
| **A_66_P128431** | Fam122b | -0.542580473 | -1.654034083 | 0.568873137 | 8.166440069 | -1.222709157 | 0.271430354 | 0.709264768 | -4.558727508 |
| **A_55_P2114809** | Gtf2e2 | 0.137023845 | -0.143669173 | 0.417716863 | 11.30855037 | 1.222685845 | 0.271438489 | 0.709264768 | -4.55873377 |
| **A_55_P2024011** | NA | 0.470513388 | -0.49336073 | 1.434387506 | 14.18736466 | 1.222649605 | 0.271451135 | 0.709264768 | -4.558743504 |
| **A_55_P2036629** | NA | 0.417271376 | -0.437545942 | 1.272088693 | 10.90463179 | 1.222631632 | 0.271457408 | 0.709264768 | -4.558748333 |
| **A_66_P134674** | Far1 | 0.214771311 | -0.225209657 | 0.654752278 | 9.087213679 | 1.222623454 | 0.271460262 | 0.709264768 | -4.558750529 |
| **A_55_P2154430** | Cd320 | -1.445724422 | -4.407474119 | 1.516025275 | 12.47815279 | -1.22260878 | 0.271465383 | 0.709264768 | -4.558754471 |
| **A_51_P155843** | Igsf10 | 0.736032341 | -0.772120691 | 2.244185373 | 9.606135211 | 1.222367604 | 0.271549566 | 0.709264768 | -4.558819255 |
| **A_66_P112623** | Psmc6 | 0.463820233 | -0.486614865 | 1.414255331 | 11.65310138 | 1.222299319 | 0.271573404 | 0.709264768 | -4.558837596 |
| **A_51_P467448** | Pif1 | 0.209899501 | -0.220250658 | 0.640049661 | 9.687475987 | 1.222198268 | 0.271608686 | 0.709264768 | -4.558864739 |
| **A_66_P112573** | Isg20 | -1.230880156 | -3.753501207 | 1.291740895 | 12.29577217 | -1.2221207 | 0.271635771 | 0.709264768 | -4.558885573 |
| **A_51_P262340** | Rbm3 | 0.689883111 | -0.724106416 | 2.103872637 | 14.18628129 | 1.222023805 | 0.271669608 | 0.709264768 | -4.558911599 |
| **A_55_P2005470** | Mfap4 | 2.364275539 | -2.481665232 | 7.21021631 | 15.54837176 | 1.221997544 | 0.27167878 | 0.709264768 | -4.558918652 |
| **A_51_P506312** | Cdkn2aipı | 0.331304291 | -0.347814062 | 1.010422645 | 11.79763437 | 1.221889518 | 0.271716509 | 0.709264768 | -4.558947667 |
| **A_52_P23225** | Gpc3 | 0.400823733 | -0.420843517 | 1.222490983 | 12.48378606 | 1.221821595 | 0.271740235 | 0.709264768 | -4.558965909 |
| **A_55_P1963508** | Slc13a5 | -0.578298374 | -1.763788461 | 0.607191712 | 7.973036446 | -1.221812012 | 0.271743582 | 0.709264768 | -4.558968483 |
| **A_51_P329928** | Phlda3 | -0.330480312 | -1.007997961 | 0.347037336 | 12.55144251 | -1.221730245 | 0.271772146 | 0.709264768 | -4.558990444 |
| **A_52_P228236** | Tfrc | -0.999187699 | -3.048047769 | 1.049672371 | 10.25179838 | -1.221476528 | 0.271860795 | 0.709264768 | -4.55936716 |
| **A_55_P2105794** | Wdr61 | -0.439208543 | -1.339925461 | 0.461508374 | 12.87751392 | -1.221329381 | 0.27191222 | 0.709264768 | -4.559098101 |
| **A_51_P195483** | Ube2q2 | -0.508474549 | -1.551316244 | 0.534367146 | 9.632784799 | -1.221240444 | 0.271943305 | 0.709264768 | -4.559121985 |
| **A_55_P1980601** | Oscp1 | 0.708459996 | -0.744643512 | 2.161563504 | 9.834976083 | 1.221150318 | 0.27197481 | 0.709264768 | -4.559146188 |
| **A_66_P100601** | NA | 0.663955712 | -0.697899738 | 2.025811161 | 13.45618189 | 1.22112028 | 0.271985311 | 0.709264768 | -4.559154255 |
| **A_55_P2096942** | Nrn1l | -1.084308146 | -3.308441409 | 1.139825117 | 8.364210794 | -1.221074778 | 0.272001218 | 0.709264768 | -4.559166474 |
| **A_51_P194249** | Stmn4 | -0.480471511 | -1.466028843 | 0.505085821 | 8.800605656 | -1.22105744 | 0.27200728 | 0.709264768 | -4.55917113 |
| **A_55_P1969311** | Gramd1b | -0.756986155 | -2.309754289 | 0.795781978 | 9.104529509 | -1.221045008 | 0.272011626 | 0.709264768 | -4.559174468 |
| **A_51_P382484** | Itgav | -0.384252661 | -1.172491439 | 0.403986117 | 9.113456618 | -1.220982482 | 0.272033488 | 0.709264768 | -4.559191259 |
| **A_52_P14526** | Zyg11b | 0.161390897 | -0.169682195 | 0.492463988 | 8.786275453 | 1.220971414 | 0.272037358 | 0.709264768 | -4.559194231 |
| **A_55_P2043862** | Stmn1 | 0.748752469 | -0.787310499 | 2.284815437 | 13.84739638 | 1.220898595 | 0.27206282 | 0.709264768 | -4.559213785 |
| **A_55_P1954400** | NA | 0.705038135 | -0.741406078 | 2.151482349 | 11.89538836 | 1.220847071 | 0.272080837 | 0.709264768 | -4.559227621 |
| **A_52_P42069** | Prokr1 | 0.128292142 | -0.134964939 | 0.391549224 | 7.800240439 | 1.220591498 | 0.272170224 | 0.709264768 | -4.559296247 |
| **A_55_P2064896** | Gm8580 | 0.523397825 | -0.55064108 | 1.597436731 | 14.23371992 | 1.220568749 | 0.272178182 | 0.709264768 | -4.559302356 |
| **A_55_P1973402** | Tada2a | 0.241968337 | -0.254585982 | 0.738522655 | 10.27501911 | 1.220512189 | 0.272197968 | 0.709264768 | -4.559317542 |
| **A_55_P2050390** | Cdh22 | 0.240770715 | -0.253330166 | 0.734871596 | 8.381865459 | 1.220501675 | 0.272201646 | 0.709264768 | -4.559320366 |
| **A_51_P134627** | NA | -0.26363243 | -0.80465198 | 0.27738712 | 8.690911074 | -1.220495606 | 0.272203769 | 0.709264768 | -4.559321995 |
| **A_55_P2029151** | Snap47 | -0.13594013 | -0.414926453 | 0.143046193 | 8.50053534 | -1.220436294 | 0.272224519 | 0.709264768 | -4.559337921 |
| **A_52_P504361** | Naglu | -0.69111169 | -2.109476438 | 0.727253059 | 9.779302545 | -1.220423764 | 0.272228903 | 0.709264768 | -4.559341285 |
| **A_55_P2268221** | Arhgap10 | -0.229835165 | -0.701561135 | 0.241890805 | 7.967010652 | -1.220329292 | 0.272261958 | 0.709264768 | -4.559366651 |
| **A_55_P2060722** | Uso1 | -0.385178528 | -1.17578585 | 0.405428794 | 11.10287376 | -1.220257772 | 0.272286984 | 0.709264768 | -4.559385854 |
| **A_55_P2130388** | Mical1 | 0.256463339 | -0.269948311 | 0.782874988 | 12.00144721 | 1.220253735 | 0.272288397 | 0.709264768 | -4.559386938 |
| **A_55_P2076777** | Mdfi | -0.317865492 | -0.97031134 | 0.334580356 | 13.78726023 | -1.220251238 | 0.272289271 | 0.709264768 | -4.559387608 |

| | | | | | | | | | |
|---|---|---|---|---|---|---|---|---|---|
| **A_55_P2006677** | Gm7969 | -0.538018114 | -1.642431991 | 0.566395763 | 8.174021639 | -1.22015608 | 0.272322572 | 0.709264768 | -4.559413158 |
| **A_55_P2037787** | NA | -0.677595372 | -2.068666644 | 0.713475899 | 8.277013329 | -1.220032487 | 0.272365829 | 0.709264768 | -4.55944634 |
| **A_55_P2137586** | Nphp4 | 0.294367485 | -0.30998465 | 0.89871962 | 8.868103415 | 1.219972726 | 0.272386747 | 0.709264768 | -4.559462385 |
| **A_51_P503822** | Slitrk6 | 0.572413312 | -0.602812741 | 1.747639364 | 8.938989391 | 1.219940531 | 0.272398018 | 0.709264768 | -4.559471029 |
| **A_51_P391727** | Slc3a1 | -0.387217103 | -1.182246357 | 0.407812152 | 7.921557432 | -1.219893078 | 0.272414629 | 0.709264768 | -4.559483769 |
| **A_55_P2154870** | Sumo2 | 0.671244392 | -0.706947284 | 2.049436067 | 13.83906144 | 1.219892028 | 0.272414997 | 0.709264768 | -4.559484051 |
| **A_51_P521161** | Caml | -0.178091317 | -0.543801743 | 0.18761911 | 10.9336348 | -1.21970765 | 0.272479551 | 0.709264768 | -4.559533551 |
| **A_55_P2033610** | Fastkd3 | 0.177606039 | -0.187109207 | 0.542321285 | 9.882688259 | 1.21970318 | 0.272481116 | 0.709264768 | -4.559534751 |
| **A_51_P355943** | Mvd | 0.174721566 | -0.184133882 | 0.533577014 | 8.831744168 | 1.219487391 | 0.272556684 | 0.709264768 | -4.559592681 |
| **A_55_P2062058** | Dbndd2 | -0.144719424 | -0.441967506 | 0.152528659 | 8.182608128 | -1.219433462 | 0.272575573 | 0.709264768 | -4.559607159 |
| **A_55_P2182159** | Ywhaq | 0.373862182 | -0.394164847 | 1.141889211 | 13.80857815 | 1.219229263 | 0.272647105 | 0.709264768 | -4.559661975 |
| **A_51_P205545** | Creld1 | -0.21024378 | -0.642149458 | 0.221661898 | 9.115783349 | -1.219227048 | 0.272647881 | 0.709264768 | -4.55966257 |
| **A_55_P1962134** | NA | -0.232210181 | -0.709280587 | 0.244860226 | 8.064939825 | -1.219127492 | 0.272682761 | 0.709264768 | -4.559689294 |
| **A_55_P1985495** | Gm12005 | 0.562535947 | -0.593268074 | 1.718339969 | 15.18549425 | 1.2190357 | 0.272714925 | 0.709264768 | -4.559713934 |
| **A_52_P237159** | Spty2d1 | -0.245140941 | -0.74884313 | 0.258561247 | 9.520047579 | -1.218968146 | 0.272738598 | 0.709264768 | -4.559732067 |
| **A_55_P2055087** | Prkar2b | 0.397633249 | -0.419438605 | 1.214705103 | 11.07276521 | 1.218913219 | 0.272757848 | 0.709264768 | -4.559746811 |
| **A_55_P2057916** | Pex5 | -0.696037616 | -2.126362694 | 0.734287462 | 9.933785958 | -1.218844504 | 0.272781931 | 0.709264768 | -4.559765255 |
| **A_55_P2250249** | Taok1 | -0.212125771 | -0.64804684 | 0.223795299 | 13.63839876 | -1.218809745 | 0.272794114 | 0.709264768 | -4.559774585 |
| **A_55_P2108486** | NA | -0.24866235 | -0.759718751 | 0.26239405 | 10.97415328 | -1.21868523 | 0.272837761 | 0.709264768 | -4.559808007 |
| **A_55_P1966998** | Arhgdib | 0.18961788 | -0.200096624 | 0.579332384 | 10.39079302 | 1.21866142 | 0.272846108 | 0.709264768 | -4.559814397 |
| **A_55_P2012478** | Zfp618 | 1.129793215 | -1.192241477 | 3.451827907 | 10.7465727 | 1.218654521 | 0.272848527 | 0.709264768 | -4.559816249 |
| **A_51_P360880** | Zfp266 | 0.818470189 | -0.86371559 | 2.500655968 | 10.25773304 | 1.218650734 | 0.272849854 | 0.709264768 | -4.559817266 |
| **A_52_P315345** | Lats2 | -0.158016806 | -0.482810758 | 0.166777145 | 10.64565286 | -1.218556562 | 0.27288287 | 0.709264768 | -4.559842542 |
| **A_52_P537566** | Cenpt | -0.43096356 | -1.316812023 | 0.454884903 | 9.082999344 | -1.218516628 | 0.272896872 | 0.709264768 | -4.55985326 |
| **A_55_P2119100** | Yipf1 | -0.182051686 | -0.556312369 | 0.192208997 | 12.58639381 | -1.218346571 | 0.272956504 | 0.709264768 | -4.559898903 |
| **A_55_P2064053** | NA | 0.200774267 | -0.211998248 | 0.613546781 | 10.97090965 | 1.218281425 | 0.272979351 | 0.709264768 | -4.559916387 |
| **A_51_P196844** | Osbpl3 | -0.493322479 | -1.507586436 | 0.520941479 | 8.601502338 | -1.218232713 | 0.272996435 | 0.709264768 | -4.559929461 |
| **A_55_P2266977** | Ahcyl2 | -0.367565023 | -1.123328861 | 0.388198816 | 9.976683745 | -1.218143395 | 0.273027765 | 0.709264768 | -4.559953432 |
| **A_55_P2067563** | NA | -0.274336786 | -0.838411656 | 0.289738084 | 9.300969834 | -1.218141222 | 0.273028527 | 0.709264768 | -4.559954015 |
| **A_55_P2022574** | Gpank1 | -0.263942328 | -0.806659207 | 0.278774551 | 11.94250881 | -1.21810874 | 0.273039921 | 0.709264768 | -4.559962733 |
| **A_52_P97572** | Aplnr | 0.666524586 | -0.704017719 | 2.03706689 | 9.386128979 | 1.218075123 | 0.273051713 | 0.709264768 | -4.559971755 |
| **A_51_P475875** | Lrwd1 | 0.304175068 | -0.321403361 | 0.929753497 | 9.932190088 | 1.217845498 | 0.273132277 | 0.709264768 | -4.560033379 |
| **A_55_P2062255** | Syne3 | 0.126499904 | -0.13368061 | 0.386680419 | 7.87491688 | 1.217771413 | 0.273158274 | 0.709264768 | -4.560053261 |
| **A_55_P2175942** | NA | 0.442979012 | -0.46814907 | 1.354107094 | 12.93011086 | 1.217738575 | 0.273169797 | 0.709264768 | -4.560062073 |
| **A_55_P1974788** | Cfl1 | 0.441832973 | -0.466956742 | 1.350622687 | 11.34072801 | 1.217713344 | 0.273178652 | 0.709264768 | -4.560068844 |
| **A_51_P275350** | Pten | 0.379091421 | -0.400682781 | 1.158865623 | 11.10580481 | 1.217658315 | 0.273197965 | 0.709264768 | -4.560083611 |
| **A_51_P218774** | Rgs10 | 1.109602956 | -1.172907711 | 3.392113622 | 11.65444761 | 1.217601389 | 0.273217945 | 0.709264768 | -4.560098888 |
| **A_55_P2109744** | NA | -0.406265046 | -1.241999626 | 0.429469534 | 15.21882733 | -1.21756297 | 0.273231429 | 0.709264768 | -4.560109197 |
| **A_55_P2128663** | Uggt1 | -0.256789591 | -0.785046404 | 0.271467222 | 8.479122756 | -1.217538363 | 0.273240067 | 0.709264768 | -4.5601158 |
| **A_51_P513568** | Stx11 | -0.220977306 | -0.675596655 | 0.233642044 | 7.993961067 | -1.217447111 | 0.273272099 | 0.709264768 | -4.560140287 |
| **A_55_P2068877** | Ibtk | 0.137321238 | -0.145198588 | 0.419841065 | 10.29264924 | 1.217416336 | 0.273282903 | 0.709264768 | -4.560148546 |
| **A_55_P1987449** | Ep400 | 0.137631478 | -0.145529537 | 0.420792494 | 10.59998468 | 1.217403818 | 0.273287298 | 0.709264768 | -4.560151905 |
| **A_51_P247694** | Adgrg3 | -1.062511945 | -3.248531096 | 1.123507206 | 9.127070663 | -1.21739133 | 0.273291682 | 0.709264768 | -4.560155256 |
| **A_51_P387220** | Ddx1 | 0.231623643 | -0.244953093 | 0.708200378 | 11.12077177 | 1.217307772 | 0.273321018 | 0.709264768 | -4.560177677 |
| **A_55_P2140646** | Creg1 | -1.609908364 | -4.922634194 | 1.702817467 | 13.41580359 | -1.217211392 | 0.27335486 | 0.709264768 | -4.560203539 |
| **A_52_P30989** | Cdkn3 | 0.535595461 | -0.566518084 | 1.637709006 | 9.026809199 | 1.217197056 | 0.273359894 | 0.709264768 | -4.560207386 |
| **A_55_P2155848** | Gm14207 | 0.279777636 | -0.295976088 | 0.85553136 | 8.477550422 | 1.217100908 | 0.273393658 | 0.709264768 | -4.560233185 |
| **A_55_P2169839** | Rufy3 | -0.639426944 | -1.955445839 | 0.676591952 | 10.47204437 | -1.216967984 | 0.273440343 | 0.709264768 | -4.560268851 |

| | | | | | | | | | |
|---|---|---|---|---|---|---|---|---|---|
| **A_55_P2010641** | Slc38a5 | 1.986681916 | -2.102173271 | 6.075537103 | 12.76315484 | 1.216961824 | 0.273442507 | 0.709264768 | -4.560270504 |
| **A_55_P2286650** | 6720462K | -0.146660627 | -0.448514744 | 0.155193489 | 8.31027206 | -1.216933316 | 0.27345252 | 0.709264768 | -4.560278153 |
| **A_55_P2165011** | Fut10 | 0.170274068 | -0.180234584 | 0.520782721 | 8.227719634 | 1.216746527 | 0.273518138 | 0.709264768 | -4.56032827 |
| **A_55_P2127973** | Gm9112 | -2.359538661 | -7.216892249 | 2.497814927 | 9.247672427 | -1.216683797 | 0.273540177 | 0.709264768 | -4.5603451 |
| **A_51_P389714** | Phlpp2 | -0.774820208 | -2.370042752 | 0.820402336 | 9.915489663 | -1.21655012 | 0.273587149 | 0.709264768 | -4.560380965 |
| **A_51_P100034** | Mif4gd | 0.263493304 | -0.279012829 | 0.805999438 | 11.36819409 | 1.216508861 | 0.273601648 | 0.709264768 | -4.560392035 |
| **A_55_P2092103** | NA | -0.334568278 | -1.023432245 | 0.354295688 | 8.230011838 | -1.216470662 | 0.273615072 | 0.709264768 | -4.560402283 |
| **A_55_P1963414** | NA | 0.334513041 | -0.354295144 | 1.023321226 | 14.42278301 | 1.216368317 | 0.273651042 | 0.709264768 | -4.560429741 |
| **A_55_P2044242** | Slc13a5 | -0.654822998 | -2.00324965 | 0.693603654 | 8.020479349 | -1.216317461 | 0.273668918 | 0.709264768 | -4.560443385 |
| **A_55_P2002572** | Ephx2 | 0.152496905 | -0.16153078 | 0.466524591 | 8.813956488 | 1.216307649 | 0.273672367 | 0.709264768 | -4.560446017 |
| **A_55_P2401403** | NA | -0.109649952 | -0.33545581 | 0.116155906 | 9.08420752 | -1.216252053 | 0.27369191 | 0.709264768 | -4.560460932 |
| **A_55_P2026982** | Tdrp | -1.641860257 | -5.023336748 | 1.739616234 | 10.10850752 | -1.216130453 | 0.273734658 | 0.709264768 | -4.560493554 |
| **A_55_P2129706** | Srsf4 | 0.22429058 | -0.237664516 | 0.686245675 | 11.55300743 | 1.21607852 | 0.273752917 | 0.709264768 | -4.560507486 |
| **A_55_P1991718** | Hmmr | 0.615756746 | -0.652482007 | 1.883995498 | 9.966908004 | 1.216069803 | 0.273755982 | 0.709264768 | -4.560509824 |
| **A_55_P2106106** | C5ar2 | -0.868938572 | -2.658654879 | 0.920777734 | 8.549974364 | -1.216060625 | 0.273759209 | 0.709264768 | -4.560512287 |
| **A_55_P2014882** | Dhrs9 | -0.580065102 | -1.774811131 | 0.614680926 | 7.924109842 | -1.216050158 | 0.273762889 | 0.709264768 | -4.560515095 |
| **A_51_P329332** | Slc19a2 | -0.76025921 | -2.326152092 | 0.805633673 | 9.515245806 | -1.216045949 | 0.273764369 | 0.709264768 | -4.560516224 |
| **A_55_P2103101** | Snx9 | -0.544277599 | -1.665360842 | 0.576805644 | 9.032835559 | -1.215998235 | 0.273781146 | 0.709264768 | -4.560529024 |
| **A_51_P455266** | Ring1 | 0.123613377 | -0.131014211 | 0.378240965 | 8.403435069 | 1.215935005 | 0.27380338 | 0.709264768 | -4.560545986 |
| **A_52_P108447** | Glis2 | 0.492018777 | -0.521743871 | 1.505781426 | 11.2286478 | 1.215614123 | 0.27391624 | 0.709264768 | -4.560632062 |
| **A_51_P113182** | Lifr | -0.743857934 | -2.276697078 | 0.78898121 | 8.302187249 | -1.215468653 | 0.273967418 | 0.709264768 | -4.560671083 |
| **A_52_P612382** | Cdc25b | 0.352273724 | -0.373674041 | 1.078221489 | 10.82959141 | 1.215416767 | 0.273985674 | 0.709264768 | -4.560685 |
| **A_55_P2168664** | Atp5k | 0.509869225 | -0.540886312 | 1.560624762 | 14.42243429 | 1.215367049 | 0.274003168 | 0.709264768 | -4.560698336 |
| **A_51_P372550** | Cgref1 | -1.110917544 | -3.400359334 | 1.178524245 | 9.09568965 | -1.215353358 | 0.274007985 | 0.709264768 | -4.560702008 |
| **A_51_P345367** | Psmb8 | 0.670689025 | -0.711523796 | 2.052901845 | 10.05667205 | 1.215336744 | 0.274013832 | 0.709264768 | -4.560706464 |
| **A_52_P219372** | Tmem177 | -0.263196543 | -0.805621858 | 0.279228772 | 8.169076774 | -1.215319807 | 0.274019792 | 0.709264768 | -4.560711007 |
| **A_66_P118600** | Lama1 | -1.708598004 | -5.230140765 | 1.812944756 | 10.34120676 | -1.215226619 | 0.274052586 | 0.709264768 | -4.560736002 |
| **A_55_P1966928** | Lnx1 | -0.543610814 | -1.664032169 | 0.576810541 | 9.110324198 | -1.215226008 | 0.274052801 | 0.709264768 | -4.560736166 |
| **A_52_P354298** | Ap2a2 | -0.528680331 | -1.618457243 | 0.561096581 | 9.194029753 | -1.215082864 | 0.274103183 | 0.709264768 | -4.56077456 |
| **A_51_P212068** | Aoc1 | -2.01063431 | -6.15522245 | 2.133953831 | 8.817353015 | -1.215072116 | 0.274106966 | 0.709264768 | -4.560777442 |
| **A_55_P2141585** | NA | -0.133509739 | -0.408793278 | 0.141773801 | 11.04239138 | -1.214739204 | 0.274224172 | 0.709264768 | -4.56086673 |
| **A_52_P50417** | 1700084J | 0.121058962 | -0.128555517 | 0.370673441 | 8.327672744 | 1.214723738 | 0.274229618 | 0.709264768 | -4.560870878 |
| **A_55_P1962747** | H2-Ab1 | 1.333126083 | -1.416038877 | 4.082291043 | 10.55469751 | 1.214565026 | 0.274285511 | 0.709264768 | -4.560913442 |
| **A_55_P1999666** | Cntrl | 0.198187196 | -0.210531705 | 0.606906096 | 9.931295913 | 1.21451034 | 0.274304772 | 0.709264768 | -4.560928108 |
| **A_55_P2184364** | B4galt3 | -0.18946385 | -0.580208804 | 0.201281104 | 12.81647179 | -1.214460306 | 0.274322395 | 0.709264768 | -4.560941526 |
| **A_51_P503625** | Gsta3 | -0.197644738 | -0.605269906 | 0.209980431 | 9.068826544 | -1.214435936 | 0.27433098 | 0.709264768 | -4.560948062 |
| **A_55_P2086507** | Bglap3 | -1.258242461 | -3.853407254 | 1.336922331 | 8.333998632 | -1.214366399 | 0.274355475 | 0.709264768 | -4.56096671 |
| **A_51_P114910** | Cstb | -0.839797291 | -2.571943563 | 0.892348981 | 13.32186382 | -1.214339862 | 0.274364823 | 0.709264768 | -4.560973826 |
| **A_52_P496726** | Rasd1 | -0.478569216 | -1.46577489 | 0.508636459 | 8.046814772 | -1.214192263 | 0.274416825 | 0.709264768 | -4.561013407 |
| **A_55_P1966420** | Scrn2 | 0.501809369 | -0.533341067 | 1.536959804 | 10.70388046 | 1.214187177 | 0.274418617 | 0.709264768 | -4.561014771 |
| **A_55_P1961039** | Pnpla2 | -0.128259515 | -0.392852565 | 0.136333535 | 11.31635606 | -1.214119702 | 0.274442393 | 0.709264768 | -4.561032865 |
| **A_55_P2088716** | Ss18l2 | 0.166485387 | -0.177008339 | 0.509979114 | 11.5417214 | 1.213969169 | 0.274495442 | 0.709264768 | -4.561073231 |
| **A_55_P2139256** | Rps6ka1 | -0.322917154 | -0.989194522 | 0.343360213 | 12.63743096 | -1.213909754 | 0.274516383 | 0.709264768 | -4.561089163 |
| **A_55_P2074636** | Adap1 | -0.407236923 | -1.247620174 | 0.433146328 | 10.99222738 | -1.213724461 | 0.274581698 | 0.709264768 | -4.561138847 |
| **A_55_P2057786** | Rft1 | -0.163975684 | -0.50236126 | 0.174409892 | 9.224546833 | -1.21371845 | 0.274583817 | 0.709264768 | -4.561140459 |
| **A_55_P2077866** | Nktr | -0.468969889 | -1.436801886 | 0.498862109 | 10.76886322 | -1.21365522 | 0.274606108 | 0.709264768 | -4.561157413 |
| **A_55_P2043987** | Luzp1 | -0.486844891 | -1.491582194 | 0.517892412 | 12.04073111 | -1.213635959 | 0.274612899 | 0.709264768 | -4.561162578 |
| **A_55_P2118931** | Foxj3 | -0.172152682 | -0.527448746 | 0.183143382 | 11.52992365 | -1.213594898 | 0.274627376 | 0.709264768 | -4.561173587 |

| | | | | | | | | | |
|---|---|---|---|---|---|---|---|---|---|
| **A_55_P2097017** | Eid1 | 0.874835188 | -0.931127621 | 2.680797997 | 13.37498179 | 1.213298851 | 0.274731775 | 0.709264768 | -4.561252963 |
| **A_52_P508974** | Igf2bp3 | 0.199209568 | -0.212095867 | 0.610515004 | 9.179918251 | 1.21309857 | 0.274802423 | 0.709264768 | -4.56130666 |
| **A_66_P119801** | Optn | -0.380735558 | -1.166855911 | 0.405384794 | 8.045625594 | -1.213066877 | 0.274813604 | 0.709264768 | -4.561315157 |
| **A_52_P675617** | Agbl3 | 0.132345829 | -0.140920515 | 0.405612174 | 9.170873883 | 1.213038145 | 0.274823741 | 0.709264768 | -4.56132286 |
| **A_55_P2051716** | Tbx1 | 0.932270691 | -0.992678683 | 2.857220065 | 10.2271435 | 1.213034253 | 0.274825114 | 0.709264768 | -4.561323904 |
| **A_52_P313856** | Slc25a26 | -0.138061972 | -0.423135668 | 0.147011725 | 8.176673872 | -1.213018067 | 0.274830825 | 0.709264768 | -4.561328243 |
| **A_51_P109421** | Eif2d | -0.103600119 | -0.317550951 | 0.110350713 | 9.376130947 | -1.212820754 | 0.274900448 | 0.709264768 | -4.561381141 |
| **A_55_P2010134** | NA | 0.190439851 | -0.202866848 | 0.58374655 | 9.305397183 | 1.212765502 | 0.274919946 | 0.709264768 | -4.561395953 |
| **A_55_P2106801** | Bglap3 | -1.512182306 | -4.635832128 | 1.611467515 | 8.546389514 | -1.212529062 | 0.275003401 | 0.709264768 | -4.561459337 |
| **A_52_P198898** | Samd5 | 0.364511329 | -0.388583922 | 1.117606581 | 9.197377389 | 1.212303803 | 0.27508293 | 0.709264768 | -4.56151972 |
| **A_55_P2002460** | Gm2399 | -0.411034867 | -1.260293658 | 0.438223923 | 8.164980538 | -1.212240972 | 0.275105116 | 0.709264768 | -4.561536562 |
| **A_55_P2027653** | Arhgap25 | 0.402711871 | -0.429387205 | 1.234810947 | 9.485208545 | 1.212187322 | 0.275124062 | 0.709264768 | -4.561550943 |
| **A_51_P111164** | Rnd1 | -0.262519923 | -0.804964104 | 0.279924258 | 8.439271338 | -1.21215333 | 0.275136066 | 0.709264768 | -4.561560054 |
| **A_55_P2176145** | Rgs17 | -1.384313392 | -4.244755906 | 1.476129122 | 8.980121027 | -1.212136535 | 0.275141997 | 0.709264768 | -4.561564556 |
| **A_52_P462203** | Dclre1c | -0.263301773 | -0.807382185 | 0.280778639 | 8.192418715 | -1.21210722 | 0.275152351 | 0.709264768 | -4.561572414 |
| **A_55_P2322709** | 4633401B | -0.145754831 | -0.446942994 | 0.155433333 | 8.282253795 | -1.212091492 | 0.275157906 | 0.709264768 | -4.56157663 |
| **A_55_P2206605** | 5830444B | -0.533973234 | -1.637423145 | 0.569476678 | 8.162888401 | -1.212040715 | 0.27517584 | 0.709264768 | -4.56159024 |
| **A_55_P2093614** | NA | -0.126862486 | -0.389037467 | 0.135312494 | 7.909605717 | -1.211971254 | 0.275200375 | 0.709264768 | -4.561608859 |
| **A_52_P647919** | Usf2 | 0.214258377 | -0.228533815 | 0.657050568 | 9.29021119 | 1.211959764 | 0.275204434 | 0.709264768 | -4.561611938 |
| **A_55_P1998942** | Oas1a | -0.989851313 | -3.035796711 | 1.056094085 | 8.35450884 | -1.211786947 | 0.275265485 | 0.709264768 | -4.561658259 |
| **A_52_P550218** | Trmt12 | -0.155032936 | -0.475503064 | 0.165437193 | 8.356342796 | -1.211676714 | 0.275304433 | 0.709264768 | -4.561687804 |
| **A_55_P2213418** | 4933417E | -0.487002943 | -1.493746408 | 0.519740522 | 7.956179702 | -1.211610734 | 0.275327748 | 0.709264768 | -4.561705488 |
| **A_52_P33382** | NA | 0.415448305 | -0.443380734 | 1.274277343 | 11.67333763 | 1.211603771 | 0.275330209 | 0.709264768 | -4.561707354 |
| **A_52_P289685** | Ssbp1 | 0.420407167 | -0.448847976 | 1.289662309 | 12.26776943 | 1.211359886 | 0.275416405 | 0.709264768 | -4.561772718 |
| **A_51_P350214** | Amz2 | 0.252541355 | -0.269628343 | 0.774711054 | 10.82947637 | 1.211354303 | 0.275418378 | 0.709264768 | -4.561774214 |
| **A_55_P2047800** | NA | 0.697486351 | -0.744714232 | 2.139686934 | 12.65172393 | 1.211324197 | 0.275429021 | 0.709264768 | -4.561782283 |
| **A_55_P2103668** | Zfp316 | 0.144864395 | -0.154725983 | 0.444454772 | 9.982547052 | 1.211111538 | 0.275504203 | 0.709264768 | -4.561839274 |
| **A_55_P2116111** | Prag1 | 0.730342101 | -0.780091691 | 2.240775893 | 12.89131176 | 1.211086014 | 0.275513227 | 0.709264768 | -4.561846114 |
| **A_55_P2171185** | NA | 0.757417285 | -0.809098921 | 2.323933491 | 12.75904004 | 1.211018187 | 0.275537211 | 0.709264768 | -4.561864291 |
| **A_51_P461319** | Gatm | -1.234424395 | -3.787528734 | 1.318679944 | 8.635051537 | -1.211005952 | 0.275541537 | 0.709264768 | -4.56186757 |
| **A_51_P389895** | Slc35f5 | -0.546344431 | -1.676429591 | 0.58374073 | 10.68349582 | -1.210892791 | 0.275581556 | 0.709264768 | -4.561897894 |
| **A_55_P1987409** | Nbeal1 | -0.249334381 | -0.765085037 | 0.266416276 | 8.258514946 | -1.210856628 | 0.275594346 | 0.709264768 | -4.561907585 |
| **A_55_P2040775** | NA | -0.146247917 | -0.448808079 | 0.156312245 | 13.49900378 | -1.210676999 | 0.275657883 | 0.709264768 | -4.56195572 |
| **A_66_P132295** | Tfpi | -1.644419397 | -5.046442032 | 1.757603237 | 11.31002531 | -1.21066988 | 0.275660401 | 0.709264768 | -4.561957628 |
| **A_55_P2080895** | Rab7 | -0.442464662 | -1.357963764 | 0.473034441 | 9.774097595 | -1.210517336 | 0.275714369 | 0.709264768 | -4.561998504 |
| **A_52_P602091** | Csf1r | -1.239827307 | -3.805328329 | 1.325673714 | 8.576322212 | -1.210429086 | 0.275745594 | 0.709264768 | -4.56202215 |
| **A_55_P2448065** | Sfxn2 | 0.389775015 | -0.416917851 | 1.196467881 | 9.337358431 | 1.210197151 | 0.275827675 | 0.709264768 | -4.562084296 |
| **A_55_P2107830** | Gemin2 | 0.121377457 | -0.129837797 | 0.372592711 | 8.193481989 | 1.210158829 | 0.275841239 | 0.709264768 | -4.562094564 |
| **A_55_P2024953** | Tbc1d4 | -0.436969658 | -1.341399919 | 0.467460603 | 9.36445322 | -1.210114716 | 0.275856854 | 0.709264768 | -4.562106383 |
| **A_51_P463428** | Pik3ip1 | -0.67564249 | -2.074152396 | 0.722867416 | 9.64053335 | -1.210045644 | 0.275881304 | 0.709264768 | -4.56212489 |
| **A_52_P665386** | Ube2m | 0.131961988 | -0.141201335 | 0.405125312 | 10.18510732 | 1.209976148 | 0.275905907 | 0.709264768 | -4.56214351 |
| **A_55_P2027361** | Irak1bp1 | 0.320253452 | -0.342701987 | 0.983208891 | 12.37910965 | 1.209928837 | 0.275922657 | 0.709264768 | -4.562156185 |
| **A_55_P2139889** | Zfp783 | 0.192143579 | -0.205616455 | 0.589903613 | 9.078656847 | 1.209915604 | 0.275927342 | 0.709264768 | -4.562159731 |
| **A_55_P2004637** | NA | 0.76047659 | -0.813809079 | 2.33476226 | 11.06513161 | 1.209908849 | 0.275929734 | 0.709264768 | -4.56216154 |
| **A_51_P446417** | Ufl1 | -0.12510503 | -0.384092968 | 0.133882909 | 9.317753063 | -1.209889133 | 0.275936714 | 0.709264768 | -4.562166823 |
| **A_55_P1984173** | Pde4dip | -0.368497395 | -1.131362028 | 0.394367238 | 10.76066778 | -1.20986604 | 0.275944891 | 0.709264768 | -4.56217301 |
| **A_55_P2029046** | Fgd3 | -0.531387863 | -1.63172349 | 0.568947765 | 7.958139395 | -1.209586138 | 0.276044013 | 0.709264768 | -4.562247998 |
| **A_55_P1998461** | Rpl17 | 0.706128634 | -0.756349184 | 2.168606452 | 11.00725802 | 1.209330151 | 0.276134693 | 0.709264768 | -4.562316576 |

| | | | | | | | | | |
|---|---|---|---|---|---|---|---|---|---|
| **A_55_P2058127** | Pde4dip | -0.223756081 | -0.687211127 | 0.239698966 | 7.777057638 | -1.20925412 | 0.276161631 | 0.709264768 | -4.562336944 |
| **A_65_P15875** | Chpf2 | -0.834814668 | -2.563932669 | 0.894303333 | 10.16590932 | -1.209249133 | 0.276163398 | 0.709264768 | -4.56233828 |
| **A_55_P2100495** | Synb | -1.983953432 | -6.093316922 | 2.125410058 | 8.688437487 | -1.209225401 | 0.276171807 | 0.709264768 | -4.562344637 |
| **A_55_P2039125** | Fam219a | -0.642888257 | -1.974743228 | 0.688966714 | 8.255562811 | -1.209007243 | 0.276249116 | 0.709264768 | -4.562403076 |
| **A_55_P1975714** | Tor1aip1 | -0.789682042 | -2.425768892 | 0.846404807 | 9.966542251 | -1.208916358 | 0.276281329 | 0.709264768 | -4.562427422 |
| **A_55_P2125241** | NA | -0.195354958 | -0.600130716 | 0.209420799 | 8.215235486 | -1.208816251 | 0.276316814 | 0.709264768 | -4.562454236 |
| **A_55_P2025829** | NA | -0.166689316 | -0.512123801 | 0.178745168 | 15.78614874 | -1.208627357 | 0.276383783 | 0.709264768 | -4.562504832 |
| **A_55_P2017804** | Cox5b | 0.213966557 | -0.229460419 | 0.657393533 | 14.41103797 | 1.208576467 | 0.276401828 | 0.709264768 | -4.562518463 |
| **A_55_P2107367** | NA | 0.856880162 | -0.91901264 | 2.632772964 | 10.98140119 | 1.208519569 | 0.276422004 | 0.709264768 | -4.562533703 |
| **A_55_P2105100** | Lrig1 | 0.177480533 | -0.190361634 | 0.545322701 | 10.09709428 | 1.208480252 | 0.276435947 | 0.709264768 | -4.562544233 |
| **A_55_P1992019** | Ptpru | -0.814333249 | -2.502189175 | 0.873522677 | 10.82279459 | -1.208417874 | 0.276458069 | 0.709264768 | -4.562560941 |
| **A_55_P2032039** | Bola3 | 0.214627437 | -0.230250304 | 0.659505177 | 10.83668764 | 1.208356013 | 0.276480008 | 0.709264768 | -4.562577509 |
| **A_51_P238734** | Mfsd11 | -0.640847219 | -1.969317341 | 0.687622902 | 8.791506761 | -1.208239581 | 0.276521307 | 0.709264768 | -4.562608693 |
| **A_55_P2109114** | NA | 0.745844204 | -0.800380393 | 2.292068801 | 14.71014981 | 1.208163989 | 0.276548122 | 0.709264768 | -4.562628938 |
| **A_55_P1955483** | Grb14 | 0.889062912 | -0.954146541 | 2.732272365 | 9.926572759 | 1.208114649 | 0.276565627 | 0.709264768 | -4.562642152 |
| **A_55_P2068459** | Hspa1a | -0.330127298 | -1.014582013 | 0.354327416 | 7.939589771 | -1.208055994 | 0.276586436 | 0.709264768 | -4.562657861 |
| **A_51_P179498** | Fam92a | 0.426869114 | -0.458216742 | 1.311954969 | 10.34739146 | 1.207979815 | 0.276613466 | 0.709264768 | -4.562678263 |
| **A_55_P2025660** | Ranbp9 | -0.231200853 | -0.710582303 | 0.248180597 | 9.753259355 | -1.207976687 | 0.276614576 | 0.709264768 | -4.562679101 |
| **A_55_P2165160** | Arhgef1 | -0.387188631 | -1.190037595 | 0.415660332 | 8.448796906 | -1.207922558 | 0.276633783 | 0.709264768 | -4.562693597 |
| **A_52_P377576** | NA | -0.476168108 | -1.463580716 | 0.511244499 | 8.911817759 | -1.20784716 | 0.276660539 | 0.709264768 | -4.562713789 |
| **A_55_P2143025** | Sema3c | 0.847768203 | -0.910294572 | 2.605830977 | 10.40553834 | 1.20779462 | 0.276679185 | 0.709264768 | -4.562727859 |
| **A_52_P128691** | Gpr4 | -0.346621707 | -1.065439083 | 0.372195668 | 7.989847635 | -1.207779173 | 0.276684667 | 0.709264768 | -4.562731996 |
| **A_55_P2067682** | Atp5l | 0.39939532 | -0.428892132 | 1.227682772 | 13.52618502 | 1.20773662 | 0.27669977 | 0.709264768 | -4.562743391 |
| **A_51_P457331** | Hmgn5 | 0.20762669 | -0.222960848 | 0.638214228 | 10.93287952 | 1.207736162 | 0.276699933 | 0.709264768 | -4.562743514 |
| **A_51_P150521** | Mgat2 | -0.343648623 | -1.056377439 | 0.369080193 | 9.188536144 | -1.207648748 | 0.27673096 | 0.709264768 | -4.562766923 |
| **A_55_P2093286** | Apod | -0.543830173 | -1.67175693 | 0.584096583 | 8.390694653 | -1.207626811 | 0.276738747 | 0.709264768 | -4.562772798 |
| **A_51_P348456** | Rest | -0.399408866 | -1.227832842 | 0.42901511 | 9.20077388 | -1.207578541 | 0.276755881 | 0.709264768 | -4.562785724 |
| **A_55_P2149615** | Bex3 | -0.305267149 | -0.938468392 | 0.327934094 | 11.44891584 | -1.207504219 | 0.276782266 | 0.709264768 | -4.562805626 |
| **A_51_P168203** | Aig1 | -0.248888028 | -0.765167151 | 0.267391095 | 8.115973057 | -1.207451759 | 0.276800891 | 0.709264768 | -4.562819674 |
| **A_55_P2186446** | NA | 0.233226304 | -0.25065316 | 0.717105767 | 13.98827711 | 1.20723185 | 0.276878977 | 0.709264768 | -4.56287856 |
| **A_52_P359965** | Cpd | -0.176395926 | -0.542369368 | 0.189577517 | 10.19640485 | -1.207228069 | 0.27688032 | 0.709264768 | -4.562879572 |
| **A_52_P88793** | Zfp933 | 0.134385439 | -0.144459532 | 0.413230411 | 9.084948236 | 1.207090257 | 0.276929265 | 0.709264768 | -4.562916473 |
| **A_55_P2108476** | Mavs | -0.303349517 | -0.932809573 | 0.326110539 | 10.12407193 | -1.207050605 | 0.276943349 | 0.709264768 | -4.562927091 |
| **A_51_P239750** | Inhba | -2.204979925 | -6.780503431 | 2.370543581 | 9.405283495 | -1.207019245 | 0.276954488 | 0.709264768 | -4.562935488 |
| **A_55_P2098734** | Ttc21b | 0.575713472 | -0.619001388 | 1.770428331 | 9.975072087 | 1.206958875 | 0.276975933 | 0.709264768 | -4.562951652 |
| **A_52_P174661** | Gatc | 0.110951894 | -0.119296706 | 0.341200494 | 8.088967262 | 1.206946636 | 0.276980281 | 0.709264768 | -4.562954929 |
| **A_55_P2106803** | Bglap2 | -1.221639334 | -3.75734762 | 1.314068952 | 8.341153521 | -1.206685415 | 0.277073091 | 0.709264768 | -4.563024869 |
| **A_55_P2028532** | Ate1 | -0.533248997 | -1.640097188 | 0.573599194 | 8.586656869 | -1.206680607 | 0.2770748 | 0.709264768 | -4.563026156 |
| **A_52_P199299** | Urgcp | -0.561310383 | -1.726510379 | 0.603889614 | 9.769504472 | -1.206571177 | 0.277113688 | 0.709264768 | -4.563055454 |
| **A_55_P2065239** | NA | 0.350450747 | -0.377110729 | 1.078012223 | 16.51913248 | 1.206445312 | 0.277158423 | 0.709264768 | -4.563089152 |
| **A_52_P649296** | Nras | -0.333732779 | -1.026777665 | 0.359312108 | 9.867761213 | -1.206112578 | 0.277276713 | 0.709264768 | -4.563178229 |
| **A_55_P2094237** | NA | 0.323651236 | -0.34848092 | 0.995783392 | 16.69682967 | 1.206071187 | 0.277291431 | 0.709264768 | -4.563189309 |
| **A_51_P154842** | Oas1f | -1.274140101 | -3.920225181 | 1.371944979 | 8.871551545 | -1.206045593 | 0.277300532 | 0.709264768 | -4.563196161 |
| **A_55_P1968188** | Btf3 | 0.457786295 | -0.493101402 | 1.408673992 | 14.70537209 | 1.205823946 | 0.27737936 | 0.709264768 | -4.563255494 |
| **A_52_P516409** | Col4a6 | 0.629469231 | -0.678150335 | 1.937088796 | 10.16043412 | 1.205711606 | 0.277419321 | 0.709264768 | -4.563285566 |
| **A_51_P236267** | St8sia4 | 0.798898953 | -0.86068476 | 2.458482667 | 10.54143606 | 1.205710474 | 0.277419724 | 0.709264768 | -4.563285869 |
| **A_55_P1975510** | H6pd | -0.209471588 | -0.644638794 | 0.225695619 | 11.92933525 | -1.205644611 | 0.277443155 | 0.709264768 | -4.563303499 |
| **A_52_P152283** | Gabpa | 0.391313998 | -0.421627765 | 1.204255761 | 9.762545891 | 1.20563624 | 0.277446133 | 0.709264768 | -4.56330574 |

| | | | | | | | | | |
|---|---|---|---|---|---|---|---|---|---|
| **A_51_P413539** | Nek6 | -0.245520757 | -0.755653613 | 0.264612099 | 10.83068298 | -1.205466829 | 0.27750641 | 0.709264768 | -4.563351086 |
| **A_51_P187171** | Smim12 | -0.525226157 | -1.616666034 | 0.566213721 | 11.84802783 | -1.205304768 | 0.277564082 | 0.709264768 | -4.563394464 |
| **A_55_P2085940** | NA | -0.188283912 | -0.579550066 | 0.202982243 | 11.77822884 | -1.20528923 | 0.277569612 | 0.709264768 | -4.563398623 |
| **A_51_P468599** | Smchd1 | 0.212655227 | -0.275247139 | 0.700557593 | 11.52518844 | 1.285392192 | 0.2775812 | 0.709264768 | -4.556453158 |
| **A_51_P325318** | Serinc2 | -0.502781767 | -1.54782124 | 0.542257707 | 8.97291249 | -1.205028089 | 0.277662567 | 0.709264768 | -4.563468516 |
| **A_52_P237652** | 2610035D | -0.106266199 | -0.327144976 | 0.114612579 | 8.412395808 | -1.205012355 | 0.277668169 | 0.709264768 | -4.563472727 |
| **A_55_P2116235** | NA | 0.379620701 | -0.409470809 | 1.16871221 | 14.58963994 | 1.204960641 | 0.27768658 | 0.709264768 | -4.563486568 |
| **A_55_P2003378** | Atrnl1 | -0.277942672 | -0.855683094 | 0.29979775 | 8.304070397 | -1.204960525 | 0.277686622 | 0.709264768 | -4.563486599 |
| **A_55_P2067583** | Cd68 | -0.537112827 | -1.653579156 | 0.579353503 | 7.984129938 | -1.204953366 | 0.277689171 | 0.709264768 | -4.563488515 |
| **A_66_P107585** | Taf2 | 0.278795763 | -0.30084418 | 0.858435706 | 10.44189619 | 1.204698057 | 0.277780084 | 0.709264768 | -4.563556843 |
| **A_51_P479914** | Pik3cb | -0.951716969 | -2.93049273 | 1.027058793 | 9.325661228 | -1.204651918 | 0.277796517 | 0.709264768 | -4.563569191 |
| **A_55_P2019684** | Bspry | 0.944610215 | -1.019409631 | 2.908630061 | 10.89151411 | 1.204639533 | 0.277800928 | 0.709264768 | -4.563572505 |
| **A_55_P1972481** | NA | -1.182344557 | -3.640839534 | 1.27615042 | 11.345901 | -1.204550679 | 0.277832576 | 0.709264768 | -4.563596284 |
| **A_51_P257762** | Kat2b | -0.108561774 | -0.334304825 | 0.117181277 | 8.025909946 | -1.204516871 | 0.277844619 | 0.709264768 | -4.563605332 |
| **A_55_P2344598** | E330037I1 | -0.220949753 | -0.680399271 | 0.238499765 | 7.991759827 | -1.204497953 | 0.277851358 | 0.709264768 | -4.563610394 |
| **A_51_P399071** | Anp32a | 0.734652925 | -0.793139369 | 2.262445219 | 9.818371814 | 1.204393054 | 0.277888728 | 0.709264768 | -4.563638466 |
| **A_52_P154833** | Ankrd49 | 0.110794284 | -0.119622384 | 0.341210952 | 9.409736264 | 1.204353027 | 0.277902988 | 0.709264768 | -4.563649177 |
| **A_66_P139703** | NA | -0.723510204 | -2.228345231 | 0.781324823 | 9.374393958 | -1.204220773 | 0.277950112 | 0.709264768 | -4.563684568 |
| **A_55_P2056639** | Bcar1 | -0.452861948 | -1.394779408 | 0.489055511 | 11.77300113 | -1.204213049 | 0.277952864 | 0.709264768 | -4.563686635 |
| **A_55_P2159575** | Reep6 | -0.602534246 | -1.855815575 | 0.650747082 | 10.93317137 | -1.204158102 | 0.277972444 | 0.709264768 | -4.563701338 |
| **A_55_P1977850** | Sall1 | -0.280758647 | -0.86474381 | 0.303226516 | 7.805401856 | -1.204153004 | 0.277974261 | 0.709264768 | -4.563702702 |
| **A_51_P237514** | Eif1ax | 0.983171613 | -1.061852214 | 3.028195439 | 12.85413065 | 1.20415198 | 0.277974626 | 0.709264768 | -4.563702976 |
| **A_55_P2096422** | Inhbb | -2.025927086 | -6.240009214 | 2.188155042 | 9.663686801 | -1.204123857 | 0.277984648 | 0.709264768 | -4.563710502 |
| **A_52_P641849** | Khnyn | -0.363604391 | -1.120098332 | 0.39288955 | 8.322342379 | -1.203854528 | 0.278080644 | 0.709264768 | -4.563782569 |
| **A_51_P353882** | Zfp580 | 0.545660831 | -0.589644845 | 1.680966508 | 10.07150499 | 1.203816569 | 0.278094176 | 0.709264768 | -4.563792726 |
| **A_55_P2157330** | NA | 0.29175728 | -0.315286275 | 0.898800834 | 16.25067662 | 1.203794054 | 0.278102203 | 0.709264768 | -4.56379875 |
| **A_55_P2074924** | Ppp1r13l | -1.072411659 | -3.303802882 | 1.158979564 | 10.09912033 | -1.203749589 | 0.278118055 | 0.709264768 | -4.563810647 |
| **A_52_P563375** | Lgals2 | -0.902654671 | -2.780846347 | 0.975537005 | 8.546978729 | -1.203738269 | 0.278122091 | 0.709264768 | -4.563813676 |
| **A_55_P1989798** | NA | 0.429964973 | -0.464689765 | 1.324619712 | 15.07409583 | 1.203726874 | 0.278126154 | 0.709264768 | -4.563816725 |
| **A_55_P2115141** | Lcor | -0.429288715 | -1.322547275 | 0.463969845 | 8.123156198 | -1.203712114 | 0.278131416 | 0.709264768 | -4.563820674 |
| **A_55_P2008437** | Cdkn2a | -0.499698149 | -1.539515782 | 0.540119485 | 8.011626746 | -1.203651906 | 0.278152884 | 0.709264768 | -4.563836784 |
| **A_55_P1984035** | Litaf | -0.408452322 | -1.258439415 | 0.441534771 | 8.373073827 | -1.203592254 | 0.278174154 | 0.709264768 | -4.563852744 |
| **A_52_P122843** | Bicral | -0.109021032 | -0.335917713 | 0.117875648 | 8.649042487 | -1.20346231 | 0.278220494 | 0.709264768 | -4.563887511 |
| **A_55_P2068451** | NA | 0.227431704 | -0.245946007 | 0.700809414 | 14.87874704 | 1.203354387 | 0.278258986 | 0.709264768 | -4.563916385 |
| **A_52_P636343** | Wipi2 | -0.243502047 | -0.750343748 | 0.263339653 | 9.104620117 | -1.203318584 | 0.278271756 | 0.709264768 | -4.563925964 |
| **A_55_P1995417** | Cnppd1 | -0.447718211 | -1.379663627 | 0.484227205 | 8.85470449 | -1.203274265 | 0.278287565 | 0.709264768 | -4.563937821 |
| **A_66_P107703** | Smox | -0.229306563 | -0.706713693 | 0.248100566 | 8.557628987 | -1.20303408 | 0.278373254 | 0.709264768 | -4.564002078 |
| **A_52_P22781** | Zfp866 | 0.209450042 | -0.22665616 | 0.645556243 | 9.718701585 | 1.202924953 | 0.278412194 | 0.709264768 | -4.564031272 |
| **A_51_P347823** | Ipo4 | -0.38228338 | -1.178272684 | 0.413705925 | 10.21259279 | -1.202897245 | 0.278422082 | 0.709264768 | -4.564038684 |
| **A_52_P234354** | Hic2 | -1.114343585 | -3.434885466 | 1.206198296 | 9.843018415 | -1.202762979 | 0.278470001 | 0.709264768 | -4.564074602 |
| **A_66_P129976** | Apba1 | -0.535955448 | -1.65205265 | 0.580141753 | 8.690373192 | -1.202754572 | 0.278473002 | 0.709264768 | -4.564076851 |
| **A_55_P2132781** | Slc16a2 | 0.735701019 | -0.796401169 | 2.267803207 | 9.84401675 | 1.202718443 | 0.278485897 | 0.709264768 | -4.564086516 |
| **A_55_P2153291** | NA | 0.277319274 | -0.300225947 | 0.854864495 | 14.96409953 | 1.202664261 | 0.278505237 | 0.709264768 | -4.56410101 |
| **A_52_P161630** | St6gal1 | -0.235488386 | -0.725933647 | 0.254956874 | 10.76705355 | -1.202622423 | 0.278520172 | 0.709264768 | -4.564112202 |
| **A_52_P251183** | Tmem177 | -0.136058009 | -0.41945378 | 0.147337761 | 8.802137453 | -1.20248895 | 0.278567823 | 0.709264768 | -4.564147905 |
| **A_52_P82660** | Mcm9 | 0.14371289 | -0.155632287 | 0.443058067 | 8.368217043 | 1.202468762 | 0.27857503 | 0.709264768 | -4.564153305 |
| **A_51_P386058** | Ppp2r5b | -0.156751042 | -0.483270288 | 0.169768203 | 10.51474305 | -1.202408391 | 0.278596586 | 0.709264768 | -4.564169454 |
| **A_55_P2099526** | H2afv | 0.740796543 | -0.802380087 | 2.283973172 | 14.52277766 | 1.202357606 | 0.27861472 | 0.709264768 | -4.564183038 |

| | | | | | | | | | |
|---|---|---|---|---|---|---|---|---|---|
| **A_55_P2419296** | NA | -0.133136695 | -0.410486083 | 0.144212694 | 8.451834701 | -1.202322304 | 0.278627325 | 0.709264768 | -4.564192481 |
| **A_55_P2154240** | Rps29 | 0.267147357 | -0.289486605 | 0.82378132 | 17.10521738 | 1.202074896 | 0.278715687 | 0.709264768 | -4.564258656 |
| **A_55_P2083499** | Shisal2a | -0.369521359 | -1.139482801 | 0.400440083 | 7.888464047 | -1.202045521 | 0.278726179 | 0.709264768 | -4.564266513 |
| **A_55_P2032966** | Hmgcs1 | 0.463096828 | -0.501878864 | 1.42807252 | 13.08704315 | 1.202003628 | 0.278741144 | 0.709264768 | -4.564277718 |
| **A_55_P2110998** | Gpr153 | 0.44440713 | -0.4817126 | 1.370526861 | 10.08866482 | 1.201888632 | 0.278782226 | 0.709264768 | -4.564308475 |
| **A_55_P1968276** | Tomm22 | 0.128426321 | -0.139225286 | 0.396077929 | 12.40898297 | 1.201806456 | 0.278811587 | 0.709264768 | -4.564330453 |
| **A_51_P209930** | Rtn2 | -0.170510511 | -0.525884338 | 0.184863316 | 9.054288456 | -1.201755344 | 0.27882985 | 0.709264768 | -4.564344123 |
| **A_52_P119350** | 4732419C | -0.499225478 | -1.539891994 | 0.541441038 | 7.919730749 | -1.201532454 | 0.278909504 | 0.709264768 | -4.564403734 |
| **A_55_P1969431** | Etfrf1 | -0.117020143 | -0.360988537 | 0.126948252 | 8.904971653 | -1.201371709 | 0.278966962 | 0.709264768 | -4.564446722 |
| **A_52_P244132** | Isoc1 | 0.266397632 | -0.289016938 | 0.821812203 | 11.8220432 | 1.201333085 | 0.27898077 | 0.709264768 | -4.564457051 |
| **A_66_P115443** | Snx4 | 0.200141458 | -0.217147953 | 0.617430869 | 9.042315648 | 1.201296026 | 0.278994018 | 0.709264768 | -4.564466961 |
| **A_55_P1974961** | Plekhg3 | 0.163526636 | -0.177457487 | 0.504510759 | 13.22206668 | 1.20117059 | 0.279038866 | 0.709264768 | -4.564500505 |
| **A_55_P1986466** | Retreg2 | -0.21915128 | -0.676125187 | 0.237822626 | 8.407912127 | -1.201165793 | 0.279040581 | 0.709264768 | -4.564501788 |
| **A_55_P2105542** | Pde2a | -0.358102847 | -1.10482414 | 0.388618445 | 10.93627849 | -1.201156469 | 0.279043915 | 0.709264768 | -4.564504282 |
| **A_51_P377526** | Sts | -0.368517917 | -1.13700632 | 0.399970486 | 9.49435242 | -1.201079169 | 0.279071556 | 0.709264768 | -4.564524952 |
| **A_51_P462192** | Olr1 | -1.252369684 | -3.864020878 | 1.35928151 | 8.408159871 | -1.201068339 | 0.279075429 | 0.709264768 | -4.564527849 |
| **A_55_P2132865** | Sumo2 | 0.734090428 | -0.796820706 | 2.265001563 | 13.2757731 | 1.201019131 | 0.279093026 | 0.709264768 | -4.564541007 |
| **A_55_P2072842** | NA | 0.792561223 | -0.860434041 | 2.445556487 | 12.37054708 | 1.200913028 | 0.279130973 | 0.709264768 | -4.564569379 |
| **A_51_P302942** | Rasl10a | -0.592609721 | -1.828588164 | 0.643368722 | 8.013660498 | -1.200903805 | 0.279134272 | 0.709264768 | -4.564571846 |
| **A_51_P292877** | Snrnp200 | 0.140093676 | -0.152115267 | 0.432302618 | 10.31786987 | 1.200812979 | 0.279166759 | 0.709264768 | -4.564596132 |
| **A_55_P1970920** | Timm17b | 0.280345 | -0.304465519 | 0.865155519 | 13.06902708 | 1.200681957 | 0.279213629 | 0.709264768 | -4.564631166 |
| **A_52_P235241** | Fntb | -0.210223298 | -0.648759829 | 0.228313233 | 8.179010486 | -1.20067485 | 0.279216172 | 0.709264768 | -4.564633066 |
| **A_55_P2074942** | Slc6a4 | -0.232245185 | -0.716728746 | 0.252238375 | 7.994159311 | -1.200654408 | 0.279223485 | 0.709264768 | -4.564638532 |
| **A_55_P2085295** | Inf2 | -1.159734055 | -3.579075119 | 1.25960701 | 10.90316806 | -1.200636823 | 0.279229777 | 0.709264768 | -4.564643234 |
| **A_55_P2000943** | Nrip3 | -0.493036808 | -1.521685124 | 0.535611507 | 8.038419221 | -1.200501673 | 0.279278134 | 0.709264768 | -4.56467937 |
| **A_55_P1956652** | Sfmbt2 | -0.928411643 | -2.865416065 | 1.008592779 | 8.228334761 | -1.200494879 | 0.279280565 | 0.709264768 | -4.564681187 |
| **A_55_P2180491** | Rrbp1 | -0.672669711 | -2.076335912 | 0.730996489 | 9.86223998 | -1.200296048 | 0.279351721 | 0.709264768 | -4.564734348 |
| **A_51_P512061** | Doxl1 | -1.307625816 | -4.036400192 | 1.421148561 | 8.38892242 | -1.200234768 | 0.279373654 | 0.709264768 | -4.564750732 |
| **A_55_P2171008** | Clk4 | -0.152786731 | -0.471640054 | 0.166066593 | 9.940089694 | -1.20017627 | 0.279394594 | 0.709264768 | -4.564766372 |
| **A_55_P2117711** | Rnf185 | -0.355053764 | -1.096062281 | 0.385954753 | 11.31060398 | -1.200110586 | 0.279418107 | 0.709264768 | -4.564783933 |
| **A_52_P405145** | NA | -0.520474528 | -1.606734705 | 0.565785648 | 7.920913275 | -1.200095944 | 0.279423348 | 0.709264768 | -4.564787847 |
| **A_55_P2096121** | NA | -0.161482102 | -0.498507958 | 0.175543754 | 14.32674256 | -1.200083653 | 0.279427749 | 0.709264768 | -4.564791134 |
| **A_52_P441044** | Rfxap | 0.169803725 | -0.184596798 | 0.524204249 | 8.241620709 | 1.200060676 | 0.279435974 | 0.709264768 | -4.564797276 |
| **A_66_P137516** | NA | 0.431827088 | -0.469456637 | 1.333110813 | 14.15763242 | 1.200048223 | 0.279440432 | 0.709264768 | -4.564800605 |
| **A_66_P134785** | 2610044C | 0.18217118 | -0.198047071 | 0.56238943 | 8.663557369 | 1.200043689 | 0.279442056 | 0.709264768 | -4.564801818 |
| **A_52_P28806** | Foxm1 | 0.439394845 | -0.477742864 | 1.356532554 | 8.940886269 | 1.199971006 | 0.279468078 | 0.709264768 | -4.564821249 |
| **A_65_P15295** | NA | -0.386664508 | -1.193806451 | 0.420477434 | 7.878838733 | -1.199871507 | 0.279503705 | 0.709264768 | -4.564847849 |
| **A_51_P425048** | H2-Q2 | -0.771400029 | -2.3816848 | 0.838884741 | 8.794883635 | -1.199850981 | 0.279511055 | 0.709264768 | -4.564853337 |
| **A_55_P2130463** | Ptgdr2 | -0.819781886 | -2.531195351 | 0.891631578 | 8.152503745 | -1.199758178 | 0.279544289 | 0.709264768 | -4.564878146 |
| **A_55_P1972234** | NA | 0.20115983 | -0.218815927 | 0.621135587 | 15.9164037 | 1.199685431 | 0.279570343 | 0.709264768 | -4.564897593 |
| **A_55_P1994492** | Usp31 | -0.346637759 | -1.070395784 | 0.377120267 | 9.159942913 | -1.199589957 | 0.279604539 | 0.709264768 | -4.564923116 |
| **A_51_P114854** | Clns1a | 0.272809207 | -0.296885411 | 0.842503826 | 9.983400167 | 1.199408859 | 0.279669414 | 0.709264768 | -4.565415899 |
| **A_55_P1990285** | Rplp2 | 0.229492355 | -0.249826009 | 0.70881072 | 17.52076595 | 1.199207952 | 0.279741401 | 0.709264768 | -4.56502523 |
| **A_55_P2153631** | NA | 0.172419427 | -0.187701435 | 0.532540289 | 8.551527513 | 1.199190731 | 0.279747572 | 0.709264768 | -4.565029833 |
| **A_51_P279712** | Rell1 | -0.316878399 | -0.978722421 | 0.344965624 | 11.32881894 | -1.199188153 | 0.279748496 | 0.709264768 | -4.565030522 |
| **A_52_P891775** | Cdr2l | 0.213581871 | -0.232546892 | 0.659710634 | 10.8104316 | 1.199097529 | 0.279780974 | 0.709264768 | -4.565054745 |
| **A_51_P372702** | Il16 | 0.495038473 | -0.539014601 | 1.529091547 | 8.993002332 | 1.19907533 | 0.27978893 | 0.709264768 | -4.565060679 |
| **A_55_P1974967** | H4c14 | 0.10836277 | -0.117992412 | 0.334717952 | 7.842154532 | 1.199057475 | 0.279795329 | 0.709264768 | -4.565065451 |

| | | | | | | | | | |
|---|---|---|---|---|---|---|---|---|---|
| **A_55_P2112622** | Smad4 | -0.144921635 | -0.447658636 | 0.157815367 | 12.50795844 | -1.198996914 | 0.279817036 | 0.709264768 | -4.565081639 |
| **A_55_P2049479** | NA | 0.301625921 | -0.328564668 | 0.93181651 | 9.849130555 | 1.198800992 | 0.279887268 | 0.709264768 | -4.565134005 |
| **A_55_P2084303** | Nid1 | -0.507955306 | -1.569300837 | 0.553390226 | 8.223129898 | -1.198723587 | 0.27991502 | 0.709264768 | -4.565154693 |
| **A_51_P479818** | Lonrf3 | -0.934085362 | -2.886014786 | 1.017844063 | 8.687865063 | -1.198595936 | 0.279960792 | 0.709264768 | -4.565188809 |
| **A_55_P2058718** | Ephb4 | 0.414652387 | -0.452155737 | 1.28146051 | 9.961871127 | 1.198150966 | 0.280120396 | 0.709264768 | -4.565307727 |
| **A_51_P348325** | Poc1b | 0.455105793 | -0.496271737 | 1.406483323 | 10.14904405 | 1.198146212 | 0.280122102 | 0.709264768 | -4.565308997 |
| **A_55_P2169445** | Mfsd6 | 0.322763068 | -0.351964882 | 0.997491018 | 10.02023208 | 1.198134242 | 0.280126397 | 0.709264768 | -4.565312196 |
| **A_51_P114878** | Elp1 | 0.152263102 | -0.166053595 | 0.4705798 | 10.4207332 | 1.198079394 | 0.280146076 | 0.709264768 | -4.565549009 |
| **A_55_P2068228** | NA | 0.344027085 | -0.375285373 | 1.063339544 | 10.92668174 | 1.197913335 | 0.280205664 | 0.709264768 | -4.565371229 |
| **A_55_P2457154** | G2e3 | 0.305857277 | -0.333662076 | 0.945376631 | 9.579067574 | 1.197885952 | 0.280215491 | 0.709264768 | -4.565378546 |
| **A_55_P2065567** | Chac2 | 0.338665151 | -0.396245535 | 1.073575838 | 8.74848592 | 1.227435423 | 0.280256847 | 0.709264768 | -4.561657818 |
| **A_55_P2071389** | Erh | 0.560560847 | -0.611640091 | 1.732761785 | 14.37604899 | 1.19776341 | 0.280259472 | 0.709264768 | -4.565411291 |
| **A_51_P422124** | Fam126a | -0.114123068 | -0.352780325 | 0.12453419 | 9.786109067 | -1.197702999 | 0.280281156 | 0.709264768 | -4.565427434 |
| **A_51_P349888** | Ang2 | -0.112115809 | -0.346587882 | 0.122356265 | 8.026429088 | -1.197639401 | 0.280303986 | 0.709264768 | -4.565444428 |
| **A_51_P155873** | Ppp1r3g | -0.786493215 | -2.431396208 | 0.858409779 | 8.569434888 | -1.19758138 | 0.280324815 | 0.709264768 | -4.565459931 |
| **A_55_P2084723** | NA | -0.322818583 | -0.997980785 | 0.352343619 | 7.885537141 | -1.197569568 | 0.280329056 | 0.709264768 | -4.565463087 |
| **A_55_P1994997** | Arhgap11a | 0.172183746 | -0.187940065 | 0.532307557 | 8.098368751 | 1.197541742 | 0.280339045 | 0.709264768 | -4.565470522 |
| **A_51_P440399** | Bcap31 | -0.680198986 | -2.10304332 | 0.742645347 | 13.04810329 | -1.197371557 | 0.280400151 | 0.709264768 | -4.565515994 |
| **A_55_P1955188** | Spata21 | -1.071891149 | -3.314108593 | 1.170326294 | 8.370514984 | -1.197356024 | 0.280405729 | 0.709264768 | -4.565520145 |
| **A_55_P2054744** | Sh3bgrl2 | -0.418189876 | -1.292997338 | 0.456617587 | 8.167290279 | -1.197323154 | 0.280417533 | 0.709264768 | -4.565528927 |
| **A_51_P137611** | Card14 | -0.300442189 | -0.928969985 | 0.328085607 | 7.91575089 | -1.197255316 | 0.280441895 | 0.709264768 | -4.565547052 |
| **A_51_P161874** | Med30 | 0.658407172 | -0.719151505 | 2.035965849 | 10.9872825 | 1.19711202 | 0.280493361 | 0.709264768 | -4.565585338 |
| **A_55_P2132902** | Gprasp1 | 0.684509788 | -0.74774873 | 2.116768306 | 10.55385738 | 1.197039811 | 0.2805193 | 0.709264768 | -4.56560463 |
| **A_55_P2060255** | NA | -0.1370334 | -0.434622568 | 0.160555768 | 15.12377958 | -1.226512616 | 0.280571791 | 0.709264768 | -4.56185578 |
| **A_55_P2145706** | Thoc7 | 0.363315975 | -0.397007812 | 1.123639762 | 10.67416393 | 1.19684046 | 0.280590919 | 0.709264768 | -4.565657889 |
| **A_52_P290783** | BC016579 | -0.312540667 | -0.966607074 | 0.34152574 | 7.837499608 | -1.19683708 | 0.280592134 | 0.709264768 | -4.565658792 |
| **A_55_P2086075** | Gm2058 | -0.818093929 | -2.530243757 | 0.8940559 | 8.780169633 | -1.196772906 | 0.280615192 | 0.709264768 | -4.565675936 |
| **A_52_P199586** | Kpna6 | -0.272148402 | -0.841737518 | 0.297440715 | 11.49202457 | -1.196725241 | 0.28063232 | 0.709264768 | -4.56568867 |
| **A_52_P452019** | Usp31 | -0.200495675 | -0.620140657 | 0.219149308 | 8.698306792 | -1.19666701 | 0.280653246 | 0.709264768 | -4.565704226 |
| **A_51_P280158** | Sf3b2 | 0.228567524 | -0.249905671 | 0.707040719 | 12.52056255 | 1.196484996 | 0.280718664 | 0.709264768 | -4.56575285 |
| **A_55_P2028847** | NA | -0.189343239 | -0.585770323 | 0.207083845 | 11.75051515 | -1.196290984 | 0.280788408 | 0.709264768 | -4.565804676 |
| **A_66_P120882** | Nfkbib | -0.368306625 | -1.139435114 | 0.402821863 | 8.776256405 | -1.19628079 | 0.280792073 | 0.709264768 | -4.565807399 |
| **A_55_P2165028** | NA | 0.403979297 | -0.442035472 | 1.249994067 | 15.22427256 | 1.196000911 | 0.280892714 | 0.709264768 | -4.565882159 |
| **A_52_P375312** | Jaml | -0.808715103 | -2.50236294 | 0.884932735 | 8.331562163 | -1.195976888 | 0.280901354 | 0.709264768 | -4.565888576 |
| **A_55_P2031471** | Rufy4 | -0.796008668 | -2.463054488 | 0.871037152 | 8.4056657 | -1.195970881 | 0.280903514 | 0.709264768 | -4.56589018 |
| **A_52_P235861** | Peg3 | -0.565185434 | -1.748885814 | 0.618514947 | 9.834844145 | -1.195912812 | 0.280924399 | 0.709264768 | -4.565905691 |
| **A_51_P323880** | COX2 | -0.18973198 | -0.587175976 | 0.207712015 | 16.7384633 | -1.195679945 | 0.281008167 | 0.709264768 | -4.565967888 |
| **A_51_P518592** | Atxn7 | -0.338229075 | -1.046756133 | 0.370297983 | 9.074972505 | -1.195652134 | 0.281018173 | 0.709264768 | -4.565975316 |
| **A_55_P2087589** | Rps11 | 0.505011727 | -0.552895529 | 1.562918982 | 15.66593548 | 1.195650398 | 0.281018798 | 0.709264768 | -4.56597578 |
| **A_55_P2079471** | NA | 0.193561788 | -0.211940425 | 0.599064 | 8.882034151 | 1.195574772 | 0.281046008 | 0.709264768 | -4.565995978 |
| **A_55_P2016959** | Nme2 | -0.11498517 | -0.355876859 | 0.125906519 | 15.78516569 | -1.195557205 | 0.281052329 | 0.709264768 | -4.56600067 |
| **A_55_P2168301** | Slc46a1 | -0.721590852 | -2.233580635 | 0.790398931 | 9.708741103 | -1.195342905 | 0.281129448 | 0.709264768 | -4.566057904 |
| **A_55_P1957413** | Lsp1 | 0.723503866 | -0.792626737 | 2.23963447 | 13.27337389 | 1.195238538 | 0.281167012 | 0.709264768 | -4.566085776 |
| **A_55_P2054619** | NA | 0.416900576 | -0.456734373 | 1.290535525 | 15.40342041 | 1.195233731 | 0.281168742 | 0.709264768 | -4.56608706 |
| **A_55_P2146555** | NA | 0.107990176 | -0.118317412 | 0.334297764 | 9.738099953 | 1.195185951 | 0.281185941 | 0.709264768 | -4.56609982 |
| **A_51_P461429** | Cyp7b1 | 0.340335898 | -0.372885512 | 1.053557307 | 8.762055891 | 1.195181148 | 0.28118767 | 0.709264768 | -4.566101103 |
| **A_55_P2125982** | Lrrc4b | 0.787195459 | -0.862578544 | 2.436969463 | 9.56886382 | 1.195111623 | 0.281212698 | 0.709264768 | -4.566119669 |
| **A_55_P2122329** | Nfic | 0.212145902 | -0.232478517 | 0.656770321 | 8.648285026 | 1.195065426 | 0.28122933 | 0.709264768 | -4.566132006 |

| | | | | | | | | | |
|---|---|---|---|---|---|---|---|---|---|
| **A_52_P607683** | Ppm1e | -0.105232893 | -0.325808786 | 0.115342999 | 7.960618147 | -1.194933706 | 0.281276755 | 0.709264768 | -4.566167181 |
| **A_52_P527800** | Emilin2 | -0.953282217 | -2.951464673 | 1.044900239 | 10.7373813 | -1.194914127 | 0.281283805 | 0.709264768 | -4.56617241 |
| **A_51_P215438** | Prodh | -0.821726822 | -2.544185713 | 0.900732069 | 9.640030095 | -1.194892796 | 0.281291486 | 0.709264768 | -4.566178106 |
| **A_55_P2269497** | Pdp1 | 0.460718722 | -0.505155835 | 1.426593279 | 9.271254911 | 1.194718201 | 0.281354361 | 0.709264768 | -4.566224729 |
| **A_55_P2060459** | Nsd3 | -0.502060278 | -1.554745531 | 0.550624975 | 8.289805203 | -1.194559176 | 0.281411641 | 0.709264768 | -4.566267192 |
| **A_55_P2096465** | NA | 0.265702281 | -0.291406478 | 0.82281104 | 12.12572591 | 1.194553602 | 0.281413649 | 0.709264768 | -4.566268681 |
| **A_55_P2080168** | Dgkk | -0.997295844 | -3.088407156 | 1.093815468 | 8.433056136 | -1.194530391 | 0.28142201 | 0.709264768 | -4.566274879 |
| **A_55_P1972222** | NA | 0.52116743 | -0.571624107 | 1.613958967 | 15.36891171 | 1.194511374 | 0.281428861 | 0.709264768 | -4.566279956 |
| **A_55_P2156588** | Ifnk | -0.506317755 | -1.567974062 | 0.555338552 | 7.979846587 | -1.194509363 | 0.281429585 | 0.709264768 | -4.566280493 |
| **A_51_P394471** | Tcf3 | 0.532923225 | -0.584593926 | 1.650440376 | 10.40718591 | 1.194430256 | 0.281458084 | 0.709264768 | -4.566301616 |
| **A_55_P2064962** | Pdpn | -0.194037298 | -0.600934787 | 0.21286019 | 8.220507206 | -1.194402091 | 0.281468232 | 0.709264768 | -4.566309136 |
| **A_55_P2016316** | Xrn2 | 0.252339726 | -0.276844269 | 0.781523721 | 10.5231201 | 1.194343559 | 0.281489321 | 0.709264768 | -4.566324765 |
| **A_52_P529360** | Rab11fip5 | -0.777894249 | -2.409446241 | 0.853657743 | 11.17961869 | -1.194180537 | 0.281548065 | 0.709264768 | -4.566368292 |
| **A_51_P519791** | Ska3 | 0.247697206 | -0.271829748 | 0.767224159 | 8.258713355 | 1.194162334 | 0.281554624 | 0.709264768 | -4.566373152 |
| **A_52_P222350** | Flnb | -1.06579816 | -3.301266372 | 1.169670052 | 10.85663755 | -1.194144309 | 0.281561121 | 0.709264768 | -4.566377965 |
| **A_55_P1973683** | Morn4 | 0.632475917 | -0.694274877 | 1.95922671 | 10.37368793 | 1.194001803 | 0.281612482 | 0.709264768 | -4.566416012 |
| **A_51_P478688** | Samd10 | 0.176351344 | -0.193584677 | 0.546287365 | 10.54075406 | 1.193994972 | 0.281614945 | 0.709264768 | -4.566417836 |
| **A_52_P664506** | Abhd2 | -0.505140523 | -1.564825229 | 0.554544184 | 8.074244817 | -1.193949308 | 0.281631405 | 0.709264768 | -4.566430027 |
| **A_55_P2140201** | Dupd1 | -0.254424275 | -0.788161234 | 0.279312684 | 7.85190722 | -1.193937558 | 0.28163564 | 0.709264768 | -4.566433164 |
| **A_51_P161691** | Coq7 | 0.343381684 | -0.376990121 | 1.063753488 | 11.509953 | 1.193907732 | 0.281646392 | 0.709264768 | -4.566441127 |
| **A_55_P2024012** | Rpl5 | 0.3674195 | -0.403418055 | 1.138257054 | 15.54107574 | 1.193849776 | 0.281667285 | 0.709264768 | -4.5664566 |
| **A_52_P402897** | Cdh4 | 0.239508816 | -0.262977627 | 0.741995258 | 8.778319556 | 1.193843778 | 0.281669447 | 0.709264768 | -4.566458202 |
| **A_55_P2128263** | Phf21a | -0.18106625 | -0.560945562 | 0.198813061 | 11.08488463 | -1.19382924 | 0.281674689 | 0.709264768 | -4.566462083 |
| **A_66_P121965** | 1110032F | 0.147810938 | -0.162328165 | 0.45795004 | 7.807660258 | 1.193714288 | 0.281716134 | 0.709264768 | -4.566492772 |
| **A_52_P486045** | Coq4 | 0.193746459 | -0.21278273 | 0.600275648 | 9.160491929 | 1.193692281 | 0.281724068 | 0.709264768 | -4.566498647 |
| **A_55_P1965278** | Dmrtc1a | -1.178737976 | -3.652134522 | 1.29465857 | 8.286925302 | -1.193641394 | 0.281742417 | 0.709264768 | -4.566512232 |
| **A_55_P2070079** | Lgals9 | -0.784057891 | -2.429308324 | 0.861192542 | 9.092821731 | -1.19362103 | 0.281749761 | 0.709264768 | -4.566517668 |
| **A_51_P363103** | Blvra | -0.111807633 | -0.346423675 | 0.122808409 | 9.767587803 | -1.193614524 | 0.281752107 | 0.709264768 | -4.566519405 |
| **A_51_P514319** | Slc13a4 | -2.516183364 | -7.796576787 | 2.764210058 | 9.187654623 | -1.193510794 | 0.281789515 | 0.709264768 | -4.566547097 |
| **A_55_P2049256** | Mapkapk5 | -0.238237384 | -0.738292942 | 0.261818173 | 10.78723446 | -1.193279004 | 0.28187312 | 0.709264768 | -4.566608973 |
| **A_55_P2159895** | Zfp52 | -0.127983094 | -0.396621129 | 0.140654941 | 8.225174395 | -1.193261003 | 0.281879613 | 0.709264768 | -4.566613778 |
| **A_55_P2093535** | Rps8 | 0.38706102 | -0.425529819 | 1.199651858 | 16.70151595 | 1.193047845 | 0.281956519 | 0.709264768 | -4.566670677 |
| **A_66_P130914** | NA | 0.412269111 | -0.453244421 | 1.277782643 | 15.90623222 | 1.193046251 | 0.281957094 | 0.709264768 | -4.566671102 |
| **A_55_P2174942** | NA | 0.640654201 | -0.704399164 | 1.985707567 | 14.59931709 | 1.192983696 | 0.281979667 | 0.709264768 | -4.5666878 |
| **A_55_P2102578** | NA | 0.460481902 | -0.506336552 | 1.427300356 | 15.41739611 | 1.192938295 | 0.281996051 | 0.709264768 | -4.566699918 |
| **A_55_P1971328** | Lage3 | 0.648582909 | -0.713276192 | 2.01044201 | 11.88316641 | 1.192844049 | 0.282030064 | 0.709264768 | -4.566725074 |
| **A_51_P383140** | Slk | -0.357546481 | -1.108351951 | 0.39325899 | 8.947055683 | -1.192766478 | 0.282058062 | 0.709264768 | -4.566745779 |
| **A_51_P324442** | Wdr35 | 0.219765493 | -0.241744053 | 0.681275039 | 9.359190463 | 1.192694342 | 0.2820841 | 0.709264768 | -4.566765032 |
| **A_55_P2118674** | Tnip1 | -0.328855782 | -1.019514184 | 0.361802621 | 12.46809984 | -1.192593746 | 0.282120415 | 0.709264768 | -4.566791882 |
| **A_51_P206268** | Seh1l | 0.249736613 | -0.274759376 | 0.774232602 | 10.39290137 | 1.192587881 | 0.282122532 | 0.709264768 | -4.566793447 |
| **A_52_P543430** | NA | -0.290727987 | -0.901362769 | 0.319906794 | 11.80340302 | -1.192492481 | 0.282156975 | 0.709264768 | -4.566818909 |
| **A_55_P1961650** | Gm2066 | -0.24584673 | -0.762237368 | 0.270543909 | 7.888436567 | -1.192439685 | 0.282176038 | 0.709264768 | -4.566833 |
| **A_55_P2023523** | Aloxe3 | -0.543591291 | -1.685414571 | 0.598231988 | 8.008166191 | -1.192405424 | 0.282188409 | 0.709264768 | -4.566842143 |
| **A_55_P1984361** | NA | 0.621519999 | -0.684079099 | 1.927119097 | 14.48191282 | 1.19232764 | 0.282216498 | 0.709264768 | -4.566862903 |
| **A_55_P2108042** | Dgcr2 | 0.232794076 | -0.256227064 | 0.721815217 | 11.46616241 | 1.192324963 | 0.282217465 | 0.709264768 | -4.566863617 |
| **A_55_P2040888** | Dhx15 | 0.360426403 | -0.396733814 | 1.11758662 | 9.570825005 | 1.192282463 | 0.282232813 | 0.709264768 | -4.56687496 |
| **A_55_P2006634** | Map4k5 | -0.218693087 | -0.678193669 | 0.240807495 | 11.40206388 | -1.192063349 | 0.282311954 | 0.709264768 | -4.566933435 |
| **A_55_P2152912** | NA | -0.527550943 | -1.636274087 | 0.581172202 | 7.945888849 | -1.191767767 | 0.282418745 | 0.709264768 | -4.567012313 |

| | | | | | | | | | |
|---|---|---|---|---|---|---|---|---|---|
| **A_55_P1978311** | NA | 0.521370542 | -0.57456679 | 1.617307874 | 14.6536624 | 1.191546826 | 0.282498592 | 0.709264768 | -4.56707127 |
| **A_55_P2091368** | Cep164 | -0.133386832 | -0.413776821 | 0.147003158 | 9.103229579 | -1.191518523 | 0.282508823 | 0.709264768 | -4.567078822 |
| **A_55_P2038183** | Insc | -0.485953254 | -1.507767621 | 0.535861112 | 8.076028929 | -1.191167501 | 0.282635725 | 0.709264768 | -4.567172482 |
| **A_51_P184385** | Pla2g12b | -0.675601802 | -2.096372553 | 0.745168949 | 8.024079776 | -1.191014742 | 0.282690967 | 0.709264768 | -4.56721324 |
| **A_55_P1981050** | Gemin8 | 0.310451412 | -0.342440126 | 0.963342949 | 8.541053839 | 1.190975822 | 0.282705043 | 0.709264768 | -4.567223623 |
| **A_55_P2182077** | NA | 0.220190363 | -0.242889654 | 0.68327038 | 14.54592026 | 1.190947488 | 0.28271529 | 0.709264768 | -4.567231183 |
| **A_55_P2062642** | Magt1 | -0.772128843 | -2.396034231 | 0.851776546 | 10.24931125 | -1.190911251 | 0.282728397 | 0.709264768 | -4.567240851 |
| **A_52_P168575** | Fam169b | 0.330474718 | -0.364572422 | 1.025521859 | 8.078136375 | 1.190897343 | 0.282733428 | 0.709264768 | -4.567244561 |
| **A_51_P108190** | Rpl23 | 0.811058029 | -0.894880365 | 2.516996422 | 11.86947763 | 1.190800285 | 0.282768536 | 0.709264768 | -4.567771821 |
| **A_55_P1995045** | Zc3hav1 | -0.526187157 | -1.632945077 | 0.580570763 | 8.377157444 | -1.190797601 | 0.282769506 | 0.709264768 | -4.567271172 |
| **A_55_P2085716** | Atg12 | -0.575675501 | -1.786545407 | 0.635194404 | 10.15811559 | -1.190777467 | 0.28277679 | 0.709264768 | -4.567276543 |
| **A_55_P2375905** | 1110035E | -0.173970416 | -0.539946124 | 0.192005293 | 8.247855115 | -1.190620858 | 0.282833448 | 0.709264768 | -4.567318324 |
| **A_55_P2122619** | Asb11 | -0.50053428 | -1.553598082 | 0.552529522 | 8.107787122 | -1.19050024 | 0.282877093 | 0.709264768 | -4.567350502 |
| **A_55_P2127971** | Gm9112 | -1.481236213 | -4.59818187 | 1.635709445 | 8.521743308 | -1.190269858 | 0.28296047 | 0.709264768 | -4.567411959 |
| **A_51_P200083** | Grina | -0.714240921 | -2.217387504 | 0.788905663 | 9.349647578 | -1.190128183 | 0.283011755 | 0.709264768 | -4.567449751 |
| **A_52_P162298** | Ydjc | 0.179128649 | -0.197854508 | 0.556111805 | 8.994356313 | 1.190127387 | 0.283012043 | 0.709264768 | -4.567449963 |
| **A_52_P457411** | Ubl5 | 0.281487472 | -0.310943111 | 0.873918054 | 13.51146516 | 1.190068462 | 0.283033375 | 0.709264768 | -4.567465681 |
| **A_55_P2106394** | Knstrn | 0.226638274 | -0.250373397 | 0.703649946 | 13.3209319 | 1.190020915 | 0.28305059 | 0.709264768 | -4.567478364 |
| **A_55_P2024061** | Gtpbp10 | 0.141742804 | -0.156612055 | 0.440097664 | 9.443645974 | 1.189921334 | 0.283086646 | 0.709264768 | -4.567504926 |
| **A_55_P2133716** | Tax1bp3 | -0.141382868 | -0.438984139 | 0.156218404 | 14.13993619 | -1.189905165 | 0.283092501 | 0.709264768 | -4.567509239 |
| **A_55_P2090060** | Mab21l3 | -1.014442626 | -3.149812654 | 1.120927403 | 8.451984001 | -1.189884185 | 0.283100098 | 0.709264768 | -4.567514835 |
| **A_51_P118650** | Wdr6 | 0.499219762 | -0.55165087 | 1.550090393 | 10.21660132 | 1.189851765 | 0.283111838 | 0.709264768 | -4.567985549 |
| **A_51_P169087** | Gls2 | -0.362074897 | -1.124261889 | 0.400112096 | 8.052951527 | -1.189836338 | 0.283117424 | 0.709264768 | -4.567527597 |
| **A_55_P2135860** | Prss36 | -0.434013551 | -1.347753064 | 0.479725962 | 9.303297435 | -1.189682928 | 0.283172984 | 0.709264768 | -4.567568515 |
| **A_52_P79506** | Zfp667 | -0.372398557 | -1.156426957 | 0.411629843 | 8.02832368 | -1.189670084 | 0.283177636 | 0.709264768 | -4.567571941 |
| **A_52_P127643** | NA | -0.136000538 | -0.422339409 | 0.150338332 | 9.328579633 | -1.189626597 | 0.283193387 | 0.709264768 | -4.56758354 |
| **A_51_P148311** | Chic2 | -0.419578091 | -1.302968177 | 0.463811996 | 10.37948741 | -1.18962649 | 0.283193426 | 0.709264768 | -4.567583568 |
| **A_55_P1976539** | Nup62cl | -1.091797553 | -3.390511278 | 1.206916173 | 8.246778964 | -1.189618127 | 0.283196455 | 0.709264768 | -4.567585799 |
| **A_52_P129443** | Wdpcp | 0.386924533 | -0.427732109 | 1.201581174 | 9.054970743 | 1.189602888 | 0.283201975 | 0.709264768 | -4.567589863 |
| **A_55_P2108151** | NA | -0.422108926 | -1.310875781 | 0.466657928 | 15.2216318 | -1.189561862 | 0.283216836 | 0.709264768 | -4.567600805 |
| **A_55_P2039556** | Pak6 | -0.203579963 | -0.632227818 | 0.225067893 | 8.025780662 | -1.18955553 | 0.28321913 | 0.709264768 | -4.567602494 |
| **A_52_P387564** | Tmpo | 1.096017729 | -1.211980913 | 3.404016371 | 13.26900902 | 1.189412165 | 0.283271068 | 0.709264768 | -4.567640731 |
| **A_55_P2119985** | Baiap2 | -0.616384449 | -1.914541991 | 0.681773094 | 8.936094508 | -1.189253992 | 0.28332838 | 0.709264768 | -4.567682915 |
| **A_55_P2017636** | Thbs1 | 0.775282675 | -0.857582308 | 2.408147658 | 12.05001325 | 1.189214371 | 0.283342737 | 0.709264768 | -4.567693482 |
| **A_55_P1993404** | NA | 0.206300549 | -0.228205467 | 0.640806565 | 12.14953403 | 1.189200097 | 0.28334791 | 0.709264768 | -4.567697288 |
| **A_55_P1976908** | NA | 0.270311186 | -0.299046975 | 0.839669347 | 12.30069088 | 1.189128569 | 0.283373832 | 0.709264768 | -4.567716364 |
| **A_55_P2115401** | Bst1 | -0.19385126 | -0.602165409 | 0.214462888 | 7.885069546 | -1.189116885 | 0.283378067 | 0.709264768 | -4.56771948 |
| **A_55_P2362635** | Slc9a3 | -0.322274694 | -1.001132783 | 0.356583395 | 7.795969514 | -1.189042979 | 0.283404853 | 0.709264768 | -4.567739189 |
| **A_52_P55772** | Tbxa2r | 0.430456803 | -0.476315665 | 1.337229271 | 9.294636824 | 1.188999293 | 0.283420688 | 0.709264768 | -4.56775084 |
| **A_52_P127206** | Gm9 | -0.900874877 | -2.798893414 | 0.997143661 | 8.185707457 | -1.188815261 | 0.283487401 | 0.709264768 | -4.567799916 |
| **A_52_P16877** | Tmcc3 | -0.322979147 | -1.003569803 | 0.357611509 | 7.905066196 | -1.18860854 | 0.283562355 | 0.709264768 | -4.56785504 |
| **A_55_P1971439** | Serpinb9c | -1.179345066 | -3.664566967 | 1.305876836 | 8.233331211 | -1.188573567 | 0.283575038 | 0.709264768 | -4.567864365 |
| **A_51_P116027** | Med27 | 0.151129731 | -0.167351826 | 0.469611287 | 11.36407059 | 1.188545923 | 0.283585063 | 0.709264768 | -4.567871737 |
| **A_55_P2184508** | NA | -0.133291004 | -0.414193812 | 0.147611804 | 14.32507128 | -1.188488826 | 0.28360577 | 0.709264768 | -4.567886961 |
| **A_51_P226142** | Dennd6a | 0.127094371 | -0.140760174 | 0.394948917 | 11.60057068 | 1.188441043 | 0.283623101 | 0.709264768 | -4.567899702 |
| **A_55_P2160077** | NA | 0.268973956 | -0.297912587 | 0.835860498 | 17.39200493 | 1.188404885 | 0.283636215 | 0.709264768 | -4.567909344 |
| **A_52_P578790** | Dok3 | -0.111230219 | -0.345665594 | 0.123205155 | 7.981328148 | -1.188365392 | 0.28365054 | 0.709264768 | -4.567919874 |
| **A_55_P2137937** | Fxyd1 | 0.235087024 | -0.260425392 | 0.730599439 | 12.96803544 | 1.188295527 | 0.283675883 | 0.709264768 | -4.567938502 |

| | | | | | | | | | |
|---|---|---|---|---|---|---|---|---|---|
| **A_55_P2173438** | Arhgap4 | 0.144265144 | -0.159834718 | 0.448365007 | 8.002980344 | 1.188216382 | 0.283704595 | 0.709264768 | -4.567959605 |
| **A_55_P2115704** | Aip | 0.205110968 | -0.227261873 | 0.637483808 | 8.982803252 | 1.188176128 | 0.283719199 | 0.709264768 | -4.567970338 |
| **A_66_P116299** | Pias1 | -0.143815951 | -0.447025723 | 0.15939382 | 11.12327089 | -1.187993901 | 0.283785319 | 0.709264768 | -4.568018923 |
| **A_51_P275496** | BC026762 | -0.695951094 | -2.163377107 | 0.771474919 | 8.181702562 | -1.187880776 | 0.283826372 | 0.709264768 | -4.568049084 |
| **A_55_P2038242** | Tchhl1 | -1.248750841 | -3.881970416 | 1.384468734 | 8.273644508 | -1.187788359 | 0.283859914 | 0.709264768 | -4.568073722 |
| **A_51_P513530** | Spag5 | 0.475570666 | -0.527266311 | 1.478407642 | 9.614294433 | 1.18777742 | 0.283863885 | 0.709264768 | -4.568076639 |
| **A_55_P2315457** | Gripap1 | 0.170050889 | -0.188545888 | 0.528647667 | 10.07266747 | 1.18774407 | 0.28387599 | 0.709264768 | -4.56808553 |
| **A_55_P2113155** | Scnm1 | -0.112251776 | -0.348973582 | 0.12447003 | 9.623010803 | -1.18769602 | 0.283893431 | 0.709264768 | -4.56809834 |
| **A_51_P351285** | Pcsk7 | -0.20544468 | -0.638709379 | 0.22782002 | 9.486609804 | -1.187659478 | 0.283906696 | 0.709264768 | -4.568108082 |
| **A_51_P358354** | Jam3 | 0.533761566 | -0.59191095 | 1.659434081 | 10.49527486 | 1.187642093 | 0.283913008 | 0.709264768 | -4.568112716 |
| **A_66_P139974** | Arl6 | 0.404499203 | -0.448643776 | 1.257642183 | 9.268334812 | 1.187534389 | 0.283952109 | 0.709264768 | -4.568141429 |
| **A_55_P2042086** | Nrtn | 0.330595102 | -0.366812281 | 1.028002485 | 10.82783083 | 1.187299323 | 0.284037464 | 0.709264768 | -4.568204092 |
| **A_66_P111747** | NA | 0.434075809 | -0.481751223 | 1.349902841 | 13.44788255 | 1.187141453 | 0.284094801 | 0.709264768 | -4.568246175 |
| **A_55_P2008918** | Cbll1 | -0.109917369 | -0.341836315 | 0.122001577 | 9.17504113 | -1.187081239 | 0.284116674 | 0.709264768 | -4.568262226 |
| **A_55_P1994913** | Ppp2cb | 0.162955927 | -0.180908924 | 0.506820777 | 13.02460134 | 1.18695081 | 0.284164055 | 0.709264768 | -4.568296992 |
| **A_55_P2167035** | Zfand1 | 0.344094417 | -0.409605984 | 1.097794818 | 8.772359616 | 1.216022428 | 0.284174988 | 0.709264768 | -4.564105164 |
| **A_55_P1982872** | Naa10 | 0.348354066 | -0.38678305 | 1.083491181 | 11.11045404 | 1.186869253 | 0.284193686 | 0.709264768 | -4.568318731 |
| **A_55_P2331804** | AU015791 | -0.268103184 | -0.833899516 | 0.297693148 | 7.851773064 | -1.186840041 | 0.2842043 | 0.709264768 | -4.568326517 |
| **A_51_P456208** | Tff3 | -0.706310244 | -2.197031492 | 0.784411004 | 8.297061148 | -1.186723135 | 0.284246781 | 0.709264768 | -4.568357677 |
| **A_52_P1037027** | Apela | -0.344037879 | -1.070168622 | 0.382092864 | 8.000812144 | -1.186702298 | 0.284254353 | 0.709264768 | -4.568363231 |
| **A_55_P1978675** | NA | -0.462526914 | -1.438859742 | 0.513805914 | 7.908765472 | -1.186559341 | 0.284306308 | 0.709264768 | -4.568401333 |
| **A_55_P2027127** | Triml2 | -0.674004411 | -2.096805093 | 0.74879627 | 7.979742836 | -1.18650349 | 0.284326608 | 0.709264768 | -4.568416219 |
| **A_51_P167876** | Tfrc | -0.598636946 | -1.862410314 | 0.665136423 | 8.08171484 | -1.186436947 | 0.284350796 | 0.709264768 | -4.568433953 |
| **A_55_P2074206** | Dlgap4 | -0.506770858 | -1.576702034 | 0.563160318 | 12.40595598 | -1.186331701 | 0.284389057 | 0.709264768 | -4.568462003 |
| **A_55_P2125868** | NA | 0.485959439 | -0.54003626 | 1.511955138 | 12.23183483 | 1.186328126 | 0.284390356 | 0.709264768 | -4.568462956 |
| **A_55_P2050902** | Fetub | -1.631574872 | -5.076673592 | 1.813523848 | 8.56960276 | -1.18619389 | 0.284439162 | 0.709264768 | -4.568498731 |
| **A_52_P542794** | 2610507B | 0.207564678 | -0.230799615 | 0.645928971 | 12.32394505 | 1.185956129 | 0.284525626 | 0.709264768 | -4.568562092 |
| **A_51_P301804** | St3gal1 | -0.917589419 | -2.855538849 | 1.020360012 | 10.3943116 | -1.185922482 | 0.284537864 | 0.709264768 | -4.568571059 |
| **A_66_P125701** | NA | -0.298779054 | -0.929809784 | 0.332251676 | 8.183901977 | -1.18590523 | 0.284544139 | 0.709264768 | -4.568575656 |
| **A_51_P436189** | Nudt22 | -0.15110831 | -0.470271611 | 0.168054991 | 9.183400562 | -1.185839043 | 0.284568213 | 0.709264768 | -4.568593294 |
| **A_55_P1972648** | Pheta2 | 0.277670651 | -0.308853914 | 0.864195217 | 8.610291492 | 1.18575269 | 0.284599626 | 0.709264768 | -4.568616305 |
| **A_55_P2237432** | Smarca1 | 0.528320882 | -0.587675865 | 1.64431763 | 9.7728145 | 1.185728322 | 0.284608491 | 0.709264768 | -4.568622798 |
| **A_51_P514412** | Gsk3b | -0.318213919 | -0.990421544 | 0.353993705 | 8.618129372 | -1.185676149 | 0.284627472 | 0.709264768 | -4.568636701 |
| **A_52_P518997** | Epha2 | 0.458141524 | -0.509741759 | 1.426024808 | 10.37527291 | 1.185569486 | 0.28466628 | 0.709264768 | -4.568665123 |
| **A_55_P2083582** | NA | 0.427155808 | -0.475313028 | 1.329624645 | 15.19126421 | 1.185507887 | 0.284688695 | 0.709264768 | -4.568681537 |
| **A_55_P1997751** | Prpf8 | 0.12858492 | -0.143082407 | 0.400252246 | 15.04304659 | 1.185503865 | 0.284690158 | 0.709264768 | -4.568682608 |
| **A_55_P2013336** | Melk | 0.769302351 | -0.856133266 | 2.394737968 | 10.08600773 | 1.185434693 | 0.28471533 | 0.709264768 | -4.56870104 |
| **A_52_P58041** | Arpc5 | -0.479184486 | -1.491644207 | 0.533275235 | 10.10609458 | -1.185428433 | 0.284717608 | 0.709264768 | -4.568702708 |
| **A_55_P2026089** | Usp30 | -0.192079625 | -0.597923442 | 0.213764192 | 10.32442997 | -1.185421256 | 0.28472022 | 0.709264768 | -4.56870462 |
| **A_55_P2057469** | Caskin1 | -0.4141215 | -1.289284284 | 0.461041284 | 7.905893254 | -1.185193558 | 0.284803095 | 0.709264768 | -4.568765289 |
| **A_52_P387724** | Tcf7l1 | 0.166268303 | -0.185106522 | 0.517643128 | 10.01843203 | 1.185193244 | 0.28480321 | 0.709264768 | -4.568765373 |
| **A_55_P2137828** | Gprc5b | -0.179410423 | -0.558577869 | 0.199757023 | 9.135345575 | -1.185132677 | 0.284825258 | 0.709264768 | -4.56878151 |
| **A_55_P1955841** | Ctsa | -1.051078234 | -3.272597098 | 1.17044063 | 14.12731021 | -1.185046475 | 0.28485664 | 0.709264768 | -4.568804477 |
| **A_51_P307220** | Ankrd54 | -0.129309688 | -0.40261926 | 0.143999885 | 8.624795153 | -1.185022415 | 0.2848654 | 0.709264768 | -4.568810887 |
| **A_55_P1965659** | Havcr2 | -0.35075128 | -1.092131851 | 0.390629291 | 7.864247875 | -1.184972875 | 0.284883437 | 0.709264768 | -4.568824086 |
| **A_55_P2033287** | NA | 0.344280356 | -0.383443229 | 1.072003941 | 15.89900239 | 1.184939395 | 0.284895628 | 0.709264768 | -4.568833006 |
| **A_55_P2099269** | Ift80 | 0.604551513 | -0.673422738 | 1.882525764 | 9.793106628 | 1.184845038 | 0.284929986 | 0.709264768 | -4.568858144 |
| **A_51_P383399** | Aldh1a7 | 1.939793754 | -2.160868568 | 6.040456076 | 12.47310102 | 1.184818684 | 0.284939583 | 0.709264768 | -4.568865165 |

| | | | | | | | | | |
|---|---|---|---|---|---|---|---|---|---|
| **A_52_P537571** | Trap1a | -1.724426666 | -5.369836831 | 1.920983499 | 8.526459114 | -1.184809903 | 0.284942781 | 0.709264768 | -4.568867505 |
| **A_52_P587738** | P2ry2 | -0.207853941 | -0.647302709 | 0.231594827 | 8.52624385 | -1.184678092 | 0.284990787 | 0.709264768 | -4.568902621 |
| **A_55_P2098076** | Trim25 | 0.201058885 | -0.224026713 | 0.626144484 | 13.75532209 | 1.184669548 | 0.284993899 | 0.709264768 | -4.568904897 |
| **A_51_P150763** | Ropn1l | -0.197164553 | -0.614016774 | 0.219687669 | 9.799093126 | -1.184669105 | 0.28499406 | 0.709264768 | -4.568905015 |
| **A_52_P8903** | Tmem251 | -0.548452619 | -1.708064221 | 0.611158984 | 9.22021168 | -1.184614129 | 0.285014085 | 0.709264768 | -4.568919661 |
| **A_55_P2152305** | Atp5h | 0.317941365 | -0.35429532 | 0.99017805 | 15.16080083 | 1.18460939 | 0.285015811 | 0.709264768 | -4.568920923 |
| **A_52_P590235** | Vars2 | 0.201263864 | -0.224301159 | 0.626828886 | 8.893264808 | 1.18454135 | 0.285040595 | 0.709264768 | -4.568939049 |
| **A_52_P379277** | Enpp3 | -0.198156574 | -0.617181546 | 0.220868399 | 8.095341173 | -1.184455975 | 0.285071697 | 0.709264768 | -4.568961792 |
| **A_55_P2052939** | Dyrk2 | -0.310843449 | -0.968160672 | 0.346473774 | 8.306032038 | -1.184450873 | 0.285073556 | 0.709264768 | -4.568963151 |
| **A_51_P348022** | Nudt21 | 0.538010534 | -0.599698896 | 1.675719964 | 11.33581158 | 1.184431003 | 0.285080795 | 0.709264768 | -4.568968445 |
| **A_55_P2087182** | Car4 | -1.26982823 | -3.955176707 | 1.415520248 | 8.403595383 | -1.184389834 | 0.285095794 | 0.709264768 | -4.568979412 |
| **A_55_P1984546** | Mmp1a | -0.302453285 | -0.94211109 | 0.337204521 | 7.792214535 | -1.184297867 | 0.285129303 | 0.709264768 | -4.56900391 |
| **A_51_P513032** | Trps1 | 0.704517335 | -0.785535827 | 2.194570497 | 9.866753618 | 1.184241472 | 0.285149853 | 0.709264768 | -4.569018933 |
| **A_55_P2113026** | Foxo4 | -0.62607185 | -1.950323763 | 0.698180062 | 8.03505203 | -1.184142351 | 0.285185975 | 0.709264768 | -4.569045336 |
| **A_55_P2012439** | Tnfrsf19 | 1.155548329 | -1.288732726 | 3.599829384 | 11.11172784 | 1.184097101 | 0.285202467 | 0.709264768 | -4.569057389 |
| **A_51_P240986** | Plekhg6 | -0.466148626 | -1.452207528 | 0.519910275 | 8.144977347 | -1.184055055 | 0.285217791 | 0.709264768 | -4.569068589 |
| **A_51_P164296** | Adamdec1 | 0.517110881 | -0.576762933 | 1.610984694 | 8.935958837 | 1.184041149 | 0.285222859 | 0.709264768 | -4.569072293 |
| **A_52_P122673** | Cep41 | 0.13413968 | -0.149620026 | 0.417899386 | 8.44464961 | 1.18401413 | 0.285232708 | 0.709264768 | -4.56907949 |
| **A_55_P2113165** | Camkk1 | 0.453741611 | -0.506225244 | 1.413708466 | 9.267631213 | 1.183866459 | 0.285286537 | 0.709264768 | -4.569118824 |
| **A_55_P2096777** | Tprn | -0.284648011 | -0.886880535 | 0.317584514 | 8.581864123 | -1.183843495 | 0.285294909 | 0.709264768 | -4.56912494 |
| **A_55_P1985693** | Fhdc1 | -0.45275711 | -1.41076743 | 0.50525321 | 10.62146839 | -1.183710329 | 0.28534346 | 0.709264768 | -4.569160409 |
| **A_55_P2137314** | Kctd21 | -0.123986518 | -0.386342445 | 0.138369409 | 8.520145439 | -1.183678961 | 0.285354898 | 0.709264768 | -4.569168764 |
| **A_55_P2139385** | Gm12070 | 0.23471106 | -0.261951804 | 0.731373925 | 15.10762209 | 1.183647029 | 0.285366541 | 0.709264768 | -4.569177268 |
| **A_66_P111090** | Sctr | -1.144740162 | -3.567099187 | 1.277618863 | 8.240407572 | -1.183637603 | 0.285369978 | 0.709264768 | -4.569179779 |
| **A_55_P1965728** | Abca17 | 0.139249116 | -0.15541944 | 0.433917673 | 8.179903622 | 1.183611013 | 0.285379675 | 0.709264768 | -4.569186861 |
| **A_52_P297009** | Itpk1 | -0.905180224 | -2.820705298 | 1.01034485 | 8.62004636 | -1.183579846 | 0.28539104 | 0.709264768 | -4.569195162 |
| **A_55_P1980062** | NA | -0.277140871 | -0.863660906 | 0.309379164 | 8.058833922 | -1.183499479 | 0.285420349 | 0.709264768 | -4.569216566 |
| **A_52_P561736** | NA | 0.610865049 | -0.682179496 | 1.903909594 | 12.20079591 | 1.183265321 | 0.285505758 | 0.709264768 | -4.569278928 |
| **A_55_P2051914** | Zfp568 | -0.950171032 | -2.961501165 | 1.061159101 | 9.985121328 | -1.183228908 | 0.285519042 | 0.709264768 | -4.569288625 |
| **A_51_P217498** | Slc2a4 | 0.645793999 | -0.721282006 | 2.012870005 | 9.544914952 | 1.183182371 | 0.28553602 | 0.709264768 | -4.569414056 |
| **A_55_P2001203** | Dnm2 | -0.566769646 | -1.766604498 | 0.633065205 | 13.06788595 | -1.183138144 | 0.285552155 | 0.709264768 | -4.569312797 |
| **A_55_P1953082** | NA | -0.325569457 | -1.014833694 | 0.36369478 | 7.794125363 | -1.183064045 | 0.285579192 | 0.709264768 | -4.56933253 |
| **A_55_P2047361** | Ppih | 0.602439313 | -0.672991397 | 1.877870024 | 10.13578949 | 1.183060026 | 0.285580658 | 0.709264768 | -4.5693336 |
| **A_55_P2081353** | NA | 0.536342342 | -0.599230947 | 1.671915631 | 14.91688863 | 1.182979615 | 0.28561 | 0.709264768 | -4.569355013 |
| **A_55_P1977498** | Htr2b | -1.491464436 | -4.649716616 | 1.666787744 | 8.48017636 | -1.182813964 | 0.285670454 | 0.709264768 | -4.569399125 |
| **A_55_P2066478** | Crtc3 | -0.196170468 | -0.611582593 | 0.219241657 | 11.15878659 | -1.182782269 | 0.285682022 | 0.709264768 | -4.569407565 |
| **A_51_P327874** | Pth1r | -0.301108189 | -0.938768272 | 0.336551895 | 11.42704792 | -1.182724737 | 0.285703022 | 0.709264768 | -4.569422885 |
| **A_51_P112762** | Slc5a3 | -0.706680662 | -2.203251918 | 0.789890595 | 8.19918118 | -1.182704237 | 0.285710505 | 0.709264768 | -4.569462116 |
| **A_55_P2002113** | Rpl36 | 0.258854573 | -0.289355509 | 0.807064656 | 16.49561251 | 1.182657987 | 0.285727388 | 0.709264768 | -4.569440659 |
| **A_51_P233788** | Dmap1 | 0.149828027 | -0.167488299 | 0.467144352 | 11.1091247 | 1.18263571 | 0.28573552 | 0.709264768 | -4.569446591 |
| **A_55_P2093231** | NA | 1.647162617 | -1.841676199 | 5.136001432 | 11.98512209 | 1.182512982 | 0.285780326 | 0.709264768 | -4.56947927 |
| **A_55_P2002379** | Golga2 | -0.33128986 | -1.033011772 | 0.370432053 | 12.31673809 | -1.182479036 | 0.28579272 | 0.709264768 | -4.569488309 |
| **A_55_P2124097** | Gm6981 | 0.19787445 | -0.221320037 | 0.617068938 | 14.32180271 | 1.182291329 | 0.285861262 | 0.709264768 | -4.569538288 |
| **A_51_P497061** | Ppp2r2d | -0.281068436 | -0.876508927 | 0.314372055 | 12.32539342 | -1.182290115 | 0.285861706 | 0.709264768 | -4.569538611 |
| **A_55_P2095360** | Tfpi | -1.459198047 | -4.550840745 | 1.632444652 | 9.876803157 | -1.182157355 | 0.285910192 | 0.709264768 | -4.569573958 |
| **A_55_P2161400** | Sctr | -0.942244443 | -2.938643952 | 1.054155066 | 8.120060928 | -1.182133367 | 0.285918954 | 0.709264768 | -4.569580345 |
| **A_55_P1963150** | Creg1 | -2.316980667 | -7.226512371 | 2.592551038 | 11.49534165 | -1.182041397 | 0.285952549 | 0.709264768 | -4.569604831 |
| **A_55_P2009086** | Creb3l1 | -0.694827457 | -2.167190709 | 0.777535796 | 10.58221932 | -1.18198604 | 0.285972771 | 0.709264768 | -4.569619569 |

| | | | | | | | | | |
|---|---|---|---|---|---|---|---|---|---|
| **A_55_P2088495** | NA | -0.633740779 | -1.976698212 | 0.709216654 | 8.251283776 | -1.18195176 | 0.285985294 | 0.709264768 | -4.569628695 |
| **A_52_P348720** | Rhobtb3 | 0.768109068 | -0.859686361 | 2.395904498 | 9.809625799 | 1.181880081 | 0.286011482 | 0.709264768 | -4.569647779 |
| **A_65_P08239** | Mef2a | 0.157619424 | -0.176444336 | 0.491683184 | 8.510917347 | 1.181763896 | 0.286053934 | 0.709264768 | -4.56967871 |
| **A_52_P657317** | Tmem160 | -0.136755207 | -0.426604736 | 0.153094321 | 10.87817462 | -1.181739141 | 0.28606298 | 0.709264768 | -4.5696853 |
| **A_51_P370615** | Neurog2 | -0.541774958 | -1.690129785 | 0.606579868 | 7.923434637 | -1.181661737 | 0.286091267 | 0.709264768 | -4.569705906 |
| **A_55_P2035424** | Hpgd | -0.248461646 | -0.775115938 | 0.278192646 | 8.636984488 | -1.181636985 | 0.286100312 | 0.709264768 | -4.569712496 |
| **A_55_P1962122** | NA | 0.393199363 | -0.440258852 | 1.226657578 | 15.67445695 | 1.181624044 | 0.286105042 | 0.709264768 | -4.569715941 |
| **A_51_P216496** | Tfpi2 | -1.126205412 | -3.51364519 | 1.261234366 | 8.277267818 | -1.181504908 | 0.286148585 | 0.709264768 | -4.569747655 |
| **A_51_P441914** | Hsd17b2 | -1.523929587 | -4.75463338 | 1.706774207 | 8.446627653 | -1.181457503 | 0.286165912 | 0.709264768 | -4.569760275 |
| **A_55_P2059090** | Tagap | 3.034468735 | -3.398915603 | 9.467853074 | 11.00908439 | 1.181390612 | 0.286190364 | 0.709264768 | -4.569778081 |
| **A_51_P371867** | 3830417A | -1.827513064 | -5.702176099 | 2.047149971 | 8.56014323 | -1.181345405 | 0.286206891 | 0.709264768 | -4.569790115 |
| **A_55_P2071581** | NA | 0.123356434 | -0.138182101 | 0.384894969 | 8.326382715 | 1.181344229 | 0.286207321 | 0.709264768 | -4.569790428 |
| **A_55_P2025765** | Adam8 | -0.407716479 | -1.272170876 | 0.456737918 | 8.928021481 | -1.181317189 | 0.286217206 | 0.709264768 | -4.569797626 |
| **A_55_P1985273** | Lmcd1 | 0.157847714 | -0.176847532 | 0.49254296 | 9.932801751 | 1.181242592 | 0.286244479 | 0.709264768 | -4.569817483 |
| **A_51_P195506** | Csf1 | -0.566235118 | -1.766889331 | 0.634419096 | 9.191632944 | -1.181215662 | 0.286254326 | 0.709264768 | -4.569824651 |
| **A_55_P2016237** | Hand2 | -0.416814257 | -1.300639317 | 0.467010804 | 8.220814043 | -1.181208596 | 0.286256909 | 0.709264768 | -4.569826532 |
| **A_55_P2093232** | Myh7 | 1.540011427 | -1.725493472 | 4.805516326 | 11.71089904 | 1.18120139 | 0.286259544 | 0.709264768 | -4.56982845 |
| **A_55_P2034320** | NA | 0.349149871 | -0.391209674 | 1.089509416 | 12.61403274 | 1.181189428 | 0.286263918 | 0.709264768 | -4.569831634 |
| **A_52_P337716** | NA | 0.107304818 | -0.120245196 | 0.334854832 | 11.45038598 | 1.181116407 | 0.286290618 | 0.709264768 | -4.569851071 |
| **A_55_P2084238** | NA | 0.752463506 | -0.843316207 | 2.348243219 | 13.28723246 | 1.181035217 | 0.286320309 | 0.709264768 | -4.569872682 |
| **A_55_P2172227** | Ubn2 | -0.127219638 | -0.397041609 | 0.142602334 | 8.183957725 | -1.180938247 | 0.286355773 | 0.709264768 | -4.569898492 |
| **A_55_P2108483** | NA | -1.016527766 | -3.172539915 | 1.139484383 | 10.01499883 | -1.180914316 | 0.286364526 | 0.709264768 | -4.569904862 |
| **A_51_P295896** | 4930452B | 0.266571892 | -0.298977659 | 0.832121443 | 8.384822875 | 1.180576238 | 0.286488203 | 0.709264768 | -4.569994843 |
| **A_55_P2097161** | Ifitm7 | 0.185452064 | -0.207996737 | 0.578900865 | 13.69152275 | 1.180575566 | 0.286488449 | 0.709264768 | -4.569995022 |
| **A_66_P130469** | Rce1 | -0.302281824 | -0.943612332 | 0.339048684 | 8.842575353 | -1.180539382 | 0.286501688 | 0.709264768 | -4.570004652 |
| **A_52_P330950** | C2cd2 | -0.806154065 | -2.516610498 | 0.904302367 | 8.842723697 | -1.180473868 | 0.286525662 | 0.709264768 | -4.570022088 |
| **A_55_P1993019** | NA | 0.134187382 | -0.150527934 | 0.418902699 | 10.32667977 | 1.180459783 | 0.286530816 | 0.709264768 | -4.570025837 |
| **A_55_P2159506** | Gm5458 | -0.557450866 | -1.740256815 | 0.625355083 | 7.96612457 | -1.180438702 | 0.28653853 | 0.709264768 | -4.570031447 |
| **A_55_P2095311** | NA | -0.472717555 | -1.475820668 | 0.530385557 | 8.010577083 | -1.180338298 | 0.286575275 | 0.709264768 | -4.570058168 |
| **A_51_P486681** | Ap3b2 | -0.452312213 | -1.412165493 | 0.507541068 | 7.976314483 | -1.180276624 | 0.286597848 | 0.709264768 | -4.570074581 |
| **A_55_P2072095** | Mucl3 | -0.636799282 | -1.988159415 | 0.714560851 | 8.688004907 | -1.180271195 | 0.286599835 | 0.709264768 | -4.570076025 |
| **A_51_P315634** | Nedd1 | 0.219766589 | -0.246638651 | 0.686171829 | 9.043148811 | 1.180180928 | 0.286632876 | 0.709264768 | -4.570100048 |
| **A_55_P1999154** | Gm4987 | 0.743979269 | -0.834982405 | 2.322940943 | 14.60099735 | 1.180156469 | 0.286641829 | 0.709264768 | -4.570106557 |
| **A_51_P496432** | Acsl1 | 0.17163502 | -0.19266951 | 0.53593955 | 10.8488204 | 1.180026276 | 0.286689492 | 0.709264768 | -4.570141203 |
| **A_55_P1963547** | Prdm16 | 0.622447029 | -0.698740344 | 1.943634402 | 9.403488216 | 1.180017178 | 0.286692823 | 0.709264768 | -4.570143624 |
| **A_55_P2106175** | Gch1 | -0.706801806 | -2.207109518 | 0.793505906 | 10.46405941 | -1.179961003 | 0.28671339 | 0.709264768 | -4.570158572 |
| **A_55_P1966134** | Prl3c1 | -0.648354546 | -2.024798523 | 0.728089432 | 7.944363275 | -1.179789063 | 0.286776351 | 0.709264768 | -4.570204325 |
| **A_55_P1970159** | Rnf11 | -0.112932601 | -0.352691414 | 0.126826211 | 11.03520358 | -1.179763887 | 0.286785571 | 0.709264768 | -4.570211024 |
| **A_51_P121447** | 1110038F | 0.247080815 | -0.277484068 | 0.771645698 | 12.24977277 | 1.179750466 | 0.286790486 | 0.709264768 | -4.570214595 |
| **A_55_P1993744** | Myo10 | -0.366762731 | -1.145445191 | 0.411919728 | 8.466374793 | -1.179709704 | 0.286805415 | 0.709264768 | -4.570225442 |
| **A_52_P642207** | Mkrn1 | -0.398880274 | -1.245763705 | 0.448003157 | 10.35090087 | -1.179693725 | 0.286811267 | 0.709264768 | -4.570229694 |
| **A_52_P317040** | Edem2 | -0.442344255 | -1.381526447 | 0.496837937 | 8.14481345 | -1.179670957 | 0.286819606 | 0.709264768 | -4.570235752 |
| **A_55_P2044045** | Ybx1 | 0.597886657 | -0.671552522 | 1.867325837 | 13.87835079 | 1.179661237 | 0.286823166 | 0.709264768 | -4.570238338 |
| **A_51_P413376** | Zfp42 | -0.57962489 | -1.810410395 | 0.651160615 | 7.949664158 | -1.179546317 | 0.28686526 | 0.709264768 | -4.570268916 |
| **A_55_P2033917** | Fut10 | 0.388034066 | -0.435982157 | 1.21205029 | 9.472169444 | 1.179463318 | 0.286895665 | 0.709264768 | -4.570291 |
| **A_55_P2156583** | Cdk5r1 | -0.426102333 | -1.331012852 | 0.478808186 | 9.135830516 | -1.179393204 | 0.286921352 | 0.709264768 | -4.570309655 |
| **A_52_P597461** | Skil | -0.520309256 | -1.625311187 | 0.584692675 | 8.301143796 | -1.179366699 | 0.286931063 | 0.709264768 | -4.570316708 |
| **A_52_P302433** | Plau | -0.420892817 | -1.314784373 | 0.472998739 | 7.976820896 | -1.179334558 | 0.286942839 | 0.709264768 | -4.570325259 |

| | | | | | | | | | |
|---|---|---|---|---|---|---|---|---|---|
| **A_55_P2104876** | NA | -0.195795024 | -0.611705255 | 0.220115208 | 8.688863577 | -1.179104756 | 0.287027049 | 0.709264768 | -4.570386399 |
| **A_51_P265869** | Hspa9 | 0.18038879 | -0.202813023 | 0.563590603 | 11.40269516 | 1.179050308 | 0.287047004 | 0.709264768 | -4.570400885 |
| **A_52_P506271** | Cmtm4 | -0.292069976 | -0.912528704 | 0.328388752 | 7.991560133 | -1.179028662 | 0.287054938 | 0.709264768 | -4.570406644 |
| **A_51_P120717** | Lmnb1 | 0.560015121 | -0.629668993 | 1.749699234 | 11.41186804 | 1.179012573 | 0.287060835 | 0.709264768 | -4.570410924 |
| **A_51_P421140** | Tubb6 | 0.389553406 | -0.438033685 | 1.217140497 | 13.20676029 | 1.178972415 | 0.287075554 | 0.709264768 | -4.570421608 |
| **A_55_P1980548** | NA | -0.965289382 | -3.016240135 | 1.085661371 | 9.533455729 | -1.17883397 | 0.287126305 | 0.709264768 | -4.570458439 |
| **A_51_P327585** | Gstm4 | -0.179098101 | -0.559644063 | 0.201447861 | 8.202222471 | -1.178783937 | 0.287144647 | 0.709264768 | -4.57047175 |
| **A_52_P279687** | Lrp8 | -0.597457343 | -1.866982006 | 0.672067319 | 7.927289013 | -1.178734803 | 0.287162661 | 0.709264768 | -4.570484821 |
| **A_55_P1976108** | Rps27a | 0.396274764 | -0.445767607 | 1.238317134 | 17.44023629 | 1.178725872 | 0.287165936 | 0.709264768 | -4.570487196 |
| **A_55_P2018417** | Osbpl3 | -0.514400343 | -1.607799092 | 0.578998405 | 8.722324803 | -1.178346518 | 0.287305054 | 0.709264768 | -4.570588109 |
| **A_55_P2083894** | NA | -0.701932487 | -2.194132713 | 0.79026774 | 8.350118479 | -1.178198829 | 0.287359231 | 0.709264768 | -4.570627394 |
| **A_52_P599162** | Tug1 | -0.215040652 | -0.672193686 | 0.242112381 | 9.15954953 | -1.178173659 | 0.287368465 | 0.709264768 | -4.570634089 |
| **A_52_P112110** | Tmem82 | -0.294833578 | -0.921619777 | 0.331952622 | 7.865060867 | -1.178169735 | 0.287369905 | 0.709264768 | -4.570635132 |
| **A_51_P459128** | Serpinb9e | -0.948346069 | -2.964437654 | 1.067745516 | 8.154264417 | -1.178167225 | 0.287370826 | 0.709264768 | -4.5706358 |
| **A_55_P1974407** | Cdk14 | 0.337721886 | -0.380249186 | 1.055692957 | 9.950872847 | 1.178155472 | 0.287375138 | 0.709264768 | -4.570638926 |
| **A_52_P353417** | Gm2a | -1.170288605 | -3.658335132 | 1.317757922 | 8.94233224 | -1.178107239 | 0.287392834 | 0.709264768 | -4.570651756 |
| **A_55_P2023707** | Camta1 | -0.160738427 | -0.502536205 | 0.181059352 | 8.156036013 | -1.177879389 | 0.287476442 | 0.709264768 | -4.570712358 |
| **A_55_P2347976** | Mir17hg | -0.407205307 | -1.273157811 | 0.458747197 | 8.860380563 | -1.177794991 | 0.287507417 | 0.709264768 | -4.570734805 |
| **A_52_P263201** | Tmem33 | -0.234912967 | -0.734487089 | 0.264661156 | 8.125138433 | -1.177761629 | 0.287519662 | 0.709264768 | -4.570743678 |
| **A_51_P366061** | Fscn1 | -0.133984408 | -0.418948502 | 0.150979685 | 9.353543414 | -1.177645205 | 0.287562397 | 0.709264768 | -4.570774642 |
| **A_66_P135192** | Msl2 | 0.152378853 | -0.171741442 | 0.476499147 | 9.338913069 | 1.177521429 | 0.287607837 | 0.709264768 | -4.570807561 |
| **A_55_P2115318** | Fmr1nb | -1.459544904 | -4.56409808 | 1.645008272 | 8.449104302 | -1.177521115 | 0.287607952 | 0.709264768 | -4.570807644 |
| **A_55_P2164070** | Grip1 | 0.28661073 | -0.323104027 | 0.896325488 | 8.347087113 | 1.177378455 | 0.287660332 | 0.709264768 | -4.570845583 |
| **A_55_P2087840** | Gpr50 | -0.33666087 | -1.052870706 | 0.379548965 | 7.819352389 | -1.177342188 | 0.28767365 | 0.709264768 | -4.570855228 |
| **A_55_P2007026** | Gm9112 | -1.981181687 | -6.196037893 | 2.233674519 | 8.995150665 | -1.177312859 | 0.28768442 | 0.709264768 | -4.570863027 |
| **A_55_P1972230** | NA | 0.32877723 | -0.370692379 | 1.02824684 | 11.68784055 | 1.177289385 | 0.28769304 | 0.709264768 | -4.57086927 |
| **A_55_P1996553** | Ttc17 | -0.25034108 | -0.782961297 | 0.282279137 | 10.50932897 | -1.177239485 | 0.287711366 | 0.709264768 | -4.57088254 |
| **A_55_P2063266** | Piga | 0.542358937 | -0.61163544 | 1.696353314 | 10.21241244 | 1.177154464 | 0.287742591 | 0.709264768 | -4.570905149 |
| **A_55_P2101182** | Pcmt1 | 0.118966736 | -0.134173743 | 0.372107214 | 10.75974896 | 1.17710253 | 0.287761667 | 0.709264768 | -4.570918959 |
| **A_55_P2012974** | Sdha | 0.121285539 | -0.136820388 | 0.379391467 | 13.01241024 | 1.176959153 | 0.287814334 | 0.709264768 | -4.570957085 |
| **A_55_P1962483** | NA | 0.1747201 | -0.197100903 | 0.546541103 | 9.768392868 | 1.176953481 | 0.287816418 | 0.709264768 | -4.570958594 |
| **A_55_P2095880** | NA | 0.568339267 | -0.641164652 | 1.777843186 | 9.228134673 | 1.176930258 | 0.28782495 | 0.709264768 | -4.570964769 |
| **A_51_P361492** | Pou2f1 | 0.265452544 | -0.299484555 | 0.830389642 | 10.27960951 | 1.176893438 | 0.287838477 | 0.709264768 | -4.570974559 |
| **A_55_P1966377** | 4930502E | -0.640311381 | -2.003025418 | 0.722402656 | 7.960050532 | -1.176892614 | 0.28783878 | 0.709264768 | -4.570974779 |
| **A_51_P258848** | Mfsd13b | -0.474920724 | -1.485683279 | 0.535841831 | 7.946147899 | -1.17685328 | 0.287853231 | 0.709264768 | -4.570985238 |
| **A_55_P2121613** | Sox4 | 0.454570183 | -0.512946441 | 1.422086807 | 10.49557359 | 1.176773432 | 0.287882569 | 0.709264768 | -4.571006469 |
| **A_55_P2098305** | Coq5 | -0.2077522 | -0.649982286 | 0.234477885 | 8.086990424 | -1.176651067 | 0.287927534 | 0.709264768 | -4.571039005 |
| **A_55_P2102295** | Tmem184 | 0.107570354 | -0.121427934 | 0.336568641 | 9.959272311 | 1.176550864 | 0.28796436 | 0.709264768 | -4.571065648 |
| **A_51_P156438** | Slc25a33 | -0.497257484 | -1.55590057 | 0.561385602 | 9.199614758 | -1.17647339 | 0.287992836 | 0.709264768 | -4.571086247 |
| **A_55_P2096834** | Fbxl19 | -0.949106989 | -2.969775297 | 1.071561319 | 11.7331358 | -1.176441908 | 0.288004408 | 0.709264768 | -4.571094617 |
| **A_52_P24631** | Azin1 | 0.178804879 | -0.201903685 | 0.559513443 | 9.958511559 | 1.176351375 | 0.288037688 | 0.709264768 | -4.571118687 |
| **A_55_P2038002** | BC005624 | 0.11317204 | -0.127793533 | 0.354137614 | 10.78428465 | 1.176344405 | 0.28804025 | 0.709264768 | -4.57112054 |
| **A_66_P124420** | NA | -0.293819863 | -0.919489281 | 0.331849554 | 8.133379986 | -1.176214613 | 0.288087967 | 0.709264768 | -4.571155047 |
| **A_55_P1984103** | Gfra3 | -0.489979772 | -1.533462369 | 0.553502826 | 8.289121253 | -1.17609739 | 0.28813107 | 0.709264768 | -4.571186212 |
| **A_51_P485260** | Rgl1 | -0.455520532 | -1.425626768 | 0.514585704 | 11.30609605 | -1.1760858 | 0.288135332 | 0.709264768 | -4.571189293 |
| **A_55_P1968035** | 1810009A | 0.133190241 | -0.150529005 | 0.416909488 | 12.49004484 | 1.175801349 | 0.288239948 | 0.709264768 | -4.571264913 |
| **A_51_P427401** | Fam90a1b | -0.934637444 | -2.925628552 | 1.056353663 | 8.106618037 | -1.175774945 | 0.28824966 | 0.709264768 | -4.571271932 |
| **A_55_P2029022** | Rpl34 | 0.236751068 | -0.267583504 | 0.74108564 | 17.01746141 | 1.175773188 | 0.288250306 | 0.709264768 | -4.571272399 |

| | | | | | | | | | |
|---|---|---|---|---|---|---|---|---|---|
| **A_55_P2022678** | C1qtnf1 | -1.498011629 | -4.689132228 | 1.693108971 | 8.708102276 | -1.175769832 | 0.288251541 | 0.709264768 | -4.571273291 |
| **A_66_P118093** | NA | 0.481779924 | -0.544566977 | 1.508126826 | 9.412178358 | 1.175722607 | 0.288268913 | 0.709264768 | -4.571285845 |
| **A_55_P1991985** | Ptprs | 0.913448411 | -1.032524979 | 2.859421801 | 13.44181872 | 1.175702585 | 0.288276279 | 0.709264768 | -4.571291167 |
| **A_51_P501773** | Slc35f3 | -0.341794542 | -1.069947407 | 0.386358322 | 8.082671538 | -1.175690225 | 0.288280826 | 0.709264768 | -4.571294453 |
| **A_51_P311990** | P4hb | -0.680464504 | -2.130398744 | 0.769469736 | 13.55447211 | -1.17545908 | 0.288365872 | 0.709264768 | -4.571355895 |
| **A_55_P2063553** | Gm4234 | -0.638048802 | -1.997858771 | 0.721761167 | 8.009923814 | -1.175238533 | 0.288447038 | 0.709264768 | -4.571414517 |
| **A_55_P2239317** | Afap1l1 | -0.162602717 | -0.509212902 | 0.184007469 | 11.10667128 | -1.174997175 | 0.288535887 | 0.709264768 | -4.571478668 |
| **A_55_P1990874** | Banp | -0.287461993 | -0.900259958 | 0.325335971 | 9.858469638 | -1.17493398 | 0.288559154 | 0.709264768 | -4.571495464 |
| **A_55_P2249897** | NA | -0.189300087 | -0.592850238 | 0.214250063 | 8.770804423 | -1.174907419 | 0.288568934 | 0.709264768 | -4.571502523 |
| **A_55_P2007210** | Abca15 | -0.759277725 | -2.377950513 | 0.859395064 | 8.036450324 | -1.174875747 | 0.288580596 | 0.709264768 | -4.57151094 |
| **A_51_P275454** | Trim30a | -0.228208023 | -0.714760593 | 0.258344547 | 8.109830294 | -1.174766308 | 0.288620896 | 0.709264768 | -4.571540026 |
| **A_52_P437795** | Cdh11 | 1.269947514 | -1.437729859 | 3.977624888 | 11.42672964 | 1.17473309 | 0.288633129 | 0.709264768 | -4.571548854 |
| **A_51_P274504** | Tex264 | -1.009687255 | -3.162470752 | 1.143096241 | 11.33025303 | -1.174726767 | 0.288635458 | 0.709264768 | -4.571550534 |
| **A_51_P369371** | Ubtd1 | -0.710455107 | -2.225471667 | 0.804561453 | 9.455277213 | -1.174544852 | 0.28870246 | 0.709264768 | -4.57159888 |
| **A_52_P128068** | Kazn | -0.693299922 | -2.171741508 | 0.785141665 | 8.539690717 | -1.174538688 | 0.28870473 | 0.709264768 | -4.571600518 |
| **A_55_P2039842** | Frmpd1 | -1.498195008 | -4.693286602 | 1.696896585 | 8.676123087 | -1.174452289 | 0.288736557 | 0.709264768 | -4.571623478 |
| **A_55_P2115310** | Spata21 | -0.67980688 | -2.129641684 | 0.770027925 | 8.08824778 | -1.174403616 | 0.288754489 | 0.709264768 | -4.571636413 |
| **A_55_P2184741** | Katnal1 | -0.175455781 | -0.549665171 | 0.198753609 | 9.876095103 | -1.174365672 | 0.288768468 | 0.709264768 | -4.571646496 |
| **A_51_P309733** | Cyp11a1 | -0.937423554 | -2.93679138 | 1.061944272 | 8.12187119 | -1.174339065 | 0.288778271 | 0.709264768 | -4.571653567 |
| **A_52_P32369** | Hmgn3 | 0.642388807 | -0.727752139 | 2.012529753 | 9.835397136 | 1.174310828 | 0.288788675 | 0.709264768 | -4.57166107 |
| **A_55_P2180201** | Mrpl10 | -0.134368009 | -0.420962325 | 0.152226308 | 8.966054667 | -1.174298899 | 0.28879307 | 0.709264768 | -4.57166424 |
| **A_66_P112394** | NA | -1.394579478 | -4.369215839 | 1.580056883 | 8.818977397 | -1.174247741 | 0.28881192 | 0.709264768 | -4.571677834 |
| **A_52_P161400** | Cnpy4 | 0.144754346 | -0.164026969 | 0.453535661 | 9.69782968 | 1.174169902 | 0.288840603 | 0.709264768 | -4.571698518 |
| **A_51_P472323** | Fbh1 | -0.102456066 | -0.321012285 | 0.116100154 | 11.55124994 | -1.174153431 | 0.288846672 | 0.709264768 | -4.571702895 |
| **A_55_P1959496** | Sbno2 | -0.279634704 | -0.876166718 | 0.31689731 | 8.01633762 | -1.174106948 | 0.288863803 | 0.709264768 | -4.571715247 |
| **A_55_P1989728** | C2cd2l | -0.406143554 | -1.272556021 | 0.460268914 | 11.65411494 | -1.174100348 | 0.288866235 | 0.709264768 | -4.571717001 |
| **A_55_P2051721** | Psg22 | -1.140935051 | -3.574928516 | 1.293058414 | 8.246610931 | -1.174064239 | 0.288879542 | 0.709264768 | -4.571726595 |
| **A_51_P253207** | NA | -0.567520867 | -1.77832856 | 0.643286825 | 8.127998149 | -1.173970026 | 0.288914266 | 0.709264768 | -4.571751629 |
| **A_55_P1997320** | Zim1 | -1.02289375 | -3.205247051 | 1.159459551 | 8.644362223 | -1.173966788 | 0.28891546 | 0.709264768 | -4.57175249 |
| **A_55_P2017914** | Csf1 | -0.419889555 | -1.315728276 | 0.475949165 | 8.114253539 | -1.173966184 | 0.288915682 | 0.709264768 | -4.57175265 |
| **A_55_P2197914** | C030027H | -0.37616693 | -1.17878772 | 0.42645386 | 11.01384582 | -1.173871485 | 0.28895059 | 0.709264768 | -4.571777813 |
| **A_51_P461779** | Ppp2r2c | -0.712139534 | -2.231651165 | 0.807372097 | 8.074203133 | -1.17384678 | 0.288959697 | 0.709264768 | -4.571784377 |
| **A_55_P1962039** | Anxa7 | -0.364991208 | -1.143785251 | 0.413802835 | 10.24089635 | -1.173843306 | 0.288960977 | 0.709264768 | -4.5717853 |
| **A_52_P489295** | Adamts1 | 0.339185054 | -0.384574076 | 1.062944183 | 11.67043194 | 1.173797006 | 0.288978046 | 0.709264768 | -4.571797602 |
| **A_52_P134639** | Tgoln2 | -0.346147254 | -1.084764568 | 0.392470061 | 8.248855862 | -1.17379364 | 0.288979287 | 0.709264768 | -4.571798496 |
| **A_55_P2092671** | Nppc | -0.559052846 | -1.752038884 | 0.633933192 | 8.080502021 | -1.173729033 | 0.289003106 | 0.709264768 | -4.571815662 |
| **A_51_P466886** | Scoc | 0.20460516 | -0.232044336 | 0.641254657 | 9.121760132 | 1.173637466 | 0.289036867 | 0.709264768 | -4.571839991 |
| **A_66_P108267** | Foxn4 | 0.756510661 | -0.858037478 | 2.371058799 | 9.502807015 | 1.173584594 | 0.289056363 | 0.709264768 | -4.571854039 |
| **A_66_P140949** | NA | 0.559689325 | -0.634811659 | 1.754190309 | 13.51399328 | 1.173575022 | 0.289059892 | 0.709264768 | -4.571856582 |
| **A_55_P2040971** | Tom1 | -0.698290538 | -2.188603729 | 0.792022652 | 11.28556034 | -1.173569889 | 0.289061785 | 0.709264768 | -4.571857946 |
| **A_55_P2064776** | 4930412C | -0.391680731 | -1.227702404 | 0.444340942 | 7.879045736 | -1.173451153 | 0.289105572 | 0.709264768 | -4.571889492 |
| **A_52_P205789** | Unc119b | -0.206634935 | -0.647693258 | 0.234423388 | 9.033285 | -1.173432389 | 0.289112492 | 0.709264768 | -4.571894477 |
| **A_55_P1998848** | NA | 0.460475837 | -0.522500838 | 1.443452513 | 15.10996709 | 1.173313261 | 0.289156431 | 0.709264768 | -4.571926126 |
| **A_51_P282179** | Mtor | -0.141652017 | -0.444060837 | 0.160756804 | 8.807247364 | -1.173217826 | 0.289191634 | 0.709264768 | -4.57195148 |
| **A_55_P2096266** | Sftpb | 2.534050271 | -2.876445264 | 7.944545806 | 14.01790713 | 1.173082426 | 0.289241587 | 0.709264768 | -4.57198745 |
| **A_55_P1971956** | Ift46 | -0.162495283 | -0.509477513 | 0.184486947 | 12.49952698 | -1.172961807 | 0.289286092 | 0.709264768 | -4.572019493 |
| **A_55_P2150901** | Foxo4 | -0.742276948 | -2.327293051 | 0.842739156 | 8.1365131 | -1.172958481 | 0.28928732 | 0.709264768 | -4.572020376 |
| **A_55_P2125564** | Lrrc58 | 0.241508098 | -0.274252164 | 0.75726836 | 11.58430867 | 1.172827568 | 0.28933563 | 0.709264768 | -4.572055152 |

| | | | | | | | | | |
|---|---|---|---|---|---|---|---|---|---|
| **A_52_P574993** | Pappa2 | -0.697981732 | -2.18858072 | 0.792617256 | 7.970313284 | -1.172825985 | 0.289336214 | 0.709264768 | -4.572055573 |
| **A_51_P320606** | Prl5a1 | -0.992987989 | -3.113657142 | 1.127681165 | 8.137773508 | -1.172793152 | 0.289348332 | 0.709264768 | -4.572064294 |
| **A_55_P1971903** | Pla2r1 | -0.127574122 | -0.400028195 | 0.144879951 | 8.073909542 | -1.172788314 | 0.289350117 | 0.709264768 | -4.572065579 |
| **A_55_P2149993** | NA | -0.628109718 | -1.969751495 | 0.713532059 | 8.069517162 | -1.172598368 | 0.289420228 | 0.709264768 | -4.572116034 |
| **A_55_P2008042** | Gm15348 | -0.186064535 | -0.583523174 | 0.211394104 | 10.05966621 | -1.172524719 | 0.289447417 | 0.709264768 | -4.572135597 |
| **A_52_P581440** | Wasf2 | -0.200467257 | -0.628695414 | 0.2277609 | 13.2638337 | -1.172515448 | 0.28945084 | 0.709264768 | -4.572138059 |
| **A_52_P320193** | Clec2h | -0.760514465 | -2.385090803 | 0.864061874 | 8.018484786 | -1.172513092 | 0.28945171 | 0.709264768 | -4.572138685 |
| **A_55_P2111870** | Rogdi | -0.133311641 | -0.418104245 | 0.151480963 | 8.28963624 | -1.172437531 | 0.289479607 | 0.709264768 | -4.572158755 |
| **A_55_P2052709** | NA | -0.243965441 | -0.765181629 | 0.277250747 | 7.927424658 | -1.172359374 | 0.289508465 | 0.709264768 | -4.572179515 |
| **A_55_P1999902** | Pip5k1a | -1.021488783 | -3.204059605 | 1.161082039 | 12.10657322 | -1.172237479 | 0.289553477 | 0.709264768 | -4.57221189 |
| **A_55_P2044381** | NA | -0.782424379 | -2.454254353 | 0.889405594 | 8.291261769 | -1.172197018 | 0.28956842 | 0.709264768 | -4.572222636 |
| **A_55_P1984655** | Smtnl2 | 1.469705762 | -1.670799949 | 4.610211472 | 12.37811615 | 1.172144502 | 0.289587815 | 0.709264768 | -4.572236584 |
| **A_55_P2107502** | Fat1 | 0.357466563 | -0.40639358 | 1.121326707 | 12.16572966 | 1.172119579 | 0.28959702 | 0.709264768 | -4.572243203 |
| **A_52_P91274** | 1700018G | -0.280306113 | -0.879338526 | 0.3187263 | 7.789757423 | -1.172013441 | 0.289636224 | 0.709264768 | -4.572271392 |
| **A_52_P38157** | Lypla1 | -0.135413817 | -0.42483732 | 0.154009687 | 8.333527494 | -1.171870247 | 0.289689122 | 0.709264768 | -4.572309421 |
| **A_66_P109673** | NA | 0.497126035 | -0.565403364 | 1.559655433 | 11.13516996 | 1.171860454 | 0.289692741 | 0.709264768 | -4.572312022 |
| **A_55_P1998001** | DXBay18 | -1.568719761 | -4.921619116 | 1.784179593 | 8.493465134 | -1.171858558 | 0.289693441 | 0.709264768 | -4.572312526 |
| **A_55_P2214841** | Irs3 | -0.903903092 | -2.83586258 | 1.028056397 | 8.244443281 | -1.171855876 | 0.289694432 | 0.709264768 | -4.572313238 |
| **A_52_P338956** | Aspg | -0.519782875 | -1.630779903 | 0.591214154 | 8.529046186 | -1.171815976 | 0.289709173 | 0.709264768 | -4.572323834 |
| **A_51_P321331** | Sptlc2 | -0.375068411 | -1.176802751 | 0.42666593 | 8.437665035 | -1.171737549 | 0.289738151 | 0.709264768 | -4.572344662 |
| **A_55_P1959718** | Cobll1 | -0.48173521 | -1.5115162 | 0.548045779 | 10.66891405 | -1.171693082 | 0.289754582 | 0.709264768 | -4.572356471 |
| **A_52_P472302** | Fxyd6 | -1.494075861 | -4.687899977 | 1.699748256 | 12.47540412 | -1.171688045 | 0.289756443 | 0.709264768 | -4.572357808 |
| **A_51_P415395** | C2cd4b | 0.351300297 | -0.399660834 | 1.102261427 | 8.597287005 | 1.171686439 | 0.289757037 | 0.709264768 | -4.572358235 |
| **A_55_P2177911** | Lepr | -2.232887491 | -7.006075875 | 2.540300893 | 9.005729396 | -1.171678906 | 0.28975982 | 0.709264768 | -4.572360236 |
| **A_51_P201338** | Mtss1 | -0.529210117 | -1.660509677 | 0.602089443 | 8.811662369 | -1.171658009 | 0.289767542 | 0.709264768 | -4.572365785 |
| **A_55_P1971458** | Jak2 | -0.211453459 | -0.663484011 | 0.240577094 | 10.25057348 | -1.171648521 | 0.289771049 | 0.709264768 | -4.572368304 |
| **A_51_P520718** | Ptger3 | -0.331399026 | -1.039845791 | 0.37704774 | 8.765158712 | -1.171640434 | 0.289774037 | 0.709264768 | -4.572370452 |
| **A_55_P2025523** | Arhgap6 | 0.360363647 | -0.410018344 | 1.130745637 | 9.34572418 | 1.171615739 | 0.289783163 | 0.709264768 | -4.57237701 |
| **A_55_P2013591** | NA | 0.370549245 | -0.421673307 | 1.162771797 | 9.341337652 | 1.1715183 | 0.289819173 | 0.709264768 | -4.572402885 |
| **A_55_P2102449** | Nxf7 | -1.252867512 | -3.931645075 | 1.425910051 | 8.284479378 | -1.17143674 | 0.289849319 | 0.709264768 | -4.572424543 |
| **A_55_P2095663** | Pgr | -0.313203855 | -0.982876765 | 0.356469056 | 7.830394301 | -1.171425605 | 0.289853435 | 0.709264768 | -4.5724275 |
| **A_55_P2054310** | M6pr | -0.533804919 | -1.675201943 | 0.607592105 | 10.19930308 | -1.171375621 | 0.289871911 | 0.709264768 | -4.572440773 |
| **A_55_P2043187** | Exosc9 | 0.192766373 | -0.219418862 | 0.604951608 | 12.81175667 | 1.171356768 | 0.28987888 | 0.709264768 | -4.572445779 |
| **A_52_P155708** | Creg1 | -1.026947394 | -3.222904205 | 1.169009416 | 8.445320071 | -1.171317815 | 0.289893279 | 0.709264768 | -4.572456123 |
| **A_55_P2053850** | NA | 0.14642835 | -0.166691227 | 0.459547926 | 10.11489321 | 1.171292313 | 0.289902707 | 0.709264768 | -4.572462894 |
| **A_55_P2001718** | Pex1 | -0.274848993 | -0.862600237 | 0.312902251 | 8.746749189 | -1.171253599 | 0.289917019 | 0.709264768 | -4.572473174 |
| **A_52_P648715** | Triml1 | -0.653290653 | -2.050360124 | 0.743778818 | 7.956142232 | -1.17122083 | 0.289929134 | 0.709264768 | -4.572481875 |
| **A_51_P198645** | Parvb | -0.117726443 | -0.369487398 | 0.134034513 | 7.988175458 | -1.171213281 | 0.289931925 | 0.709264768 | -4.57248388 |
| **A_52_P367060** | Atxn2l | -0.383561101 | -1.203854218 | 0.436732015 | 12.42267751 | -1.171158934 | 0.289952019 | 0.709264768 | -4.572498311 |
| **A_55_P2086148** | NA | -0.240957098 | -0.756322707 | 0.274408511 | 11.25007889 | -1.171047833 | 0.289993099 | 0.709264768 | -4.57252781 |
| **A_55_P2133370** | NA | -0.852350057 | -2.675392371 | 0.970692257 | 8.126536196 | -1.171039626 | 0.289996134 | 0.709264768 | -4.572529989 |
| **A_55_P1985025** | Pard3 | -0.504171329 | -1.582539557 | 0.574196899 | 10.53382992 | -1.171012164 | 0.29000629 | 0.709264768 | -4.572537281 |
| **A_51_P258766** | Smo | 0.756060415 | -0.861182187 | 2.373303017 | 11.77805877 | 1.170931995 | 0.290035938 | 0.709264768 | -4.572558567 |
| **A_51_P368009** | E2f2 | 1.047168278 | -1.192790732 | 3.287127287 | 11.71121989 | 1.170918727 | 0.290040844 | 0.709264768 | -4.57256209 |
| **A_55_P2316335** | Gfm2 | -0.100720715 | -0.316186669 | 0.114745239 | 8.40349192 | -1.170820956 | 0.290077005 | 0.709264768 | -4.572588049 |
| **A_52_P189005** | Gk5 | -0.35972133 | -1.129263729 | 0.409821068 | 8.045855429 | -1.170803424 | 0.29008349 | 0.709264768 | -4.572592703 |
| **A_55_P2023953** | Arpc3 | 0.15260348 | -0.173868612 | 0.479075571 | 14.01009807 | 1.170762277 | 0.29009871 | 0.709264768 | -4.572603628 |
| **A_55_P1977144** | Cyp2c67 | 0.149855504 | -0.170742511 | 0.470453518 | 7.835354312 | 1.17074474 | 0.290105197 | 0.709264768 | -4.572608284 |

| | | | | | | | | | |
|---|---|---|---|---|---|---|---|---|---|
| **A_55_P2012592** | Ildr1 | -0.427948894 | -1.343543544 | 0.487645755 | 8.680153988 | -1.170682182 | 0.290128339 | 0.709264768 | -4.572624893 |
| **A_55_P2141395** | Ly6g6c | -1.394363208 | -4.377910806 | 1.589184391 | 9.172809509 | -1.170558949 | 0.29017393 | 0.709264768 | -4.57265761 |
| **A_51_P464576** | Psen1 | -0.594139355 | -1.865445775 | 0.677167065 | 9.393777441 | -1.17054584 | 0.29017878 | 0.709264768 | -4.572597837 |
| **A_55_P2154880** | NA | 0.597532784 | -0.681248442 | 1.87631401 | 11.2817456 | 1.170350195 | 0.290251175 | 0.709264768 | -4.57271303 |
| **A_51_P323812** | Slc6a12 | -1.717096187 | -5.391994735 | 1.95780236 | 8.525568115 | -1.170306495 | 0.290267347 | 0.709264768 | -4.572724631 |
| **A_51_P169327** | Etd | -1.227351009 | -3.854168719 | 1.399466701 | 8.269391164 | -1.170278413 | 0.29027774 | 0.709264768 | -4.572732086 |
| **A_55_P2032278** | NA | 0.149279769 | -0.170219368 | 0.468778906 | 8.514197356 | 1.170257978 | 0.290285303 | 0.709264768 | -4.572737511 |
| **A_55_P1975400** | Flnc | -0.435531301 | -1.367807252 | 0.49674465 | 10.59593703 | -1.170106083 | 0.290341526 | 0.709264768 | -4.572777833 |
| **A_55_P2023818** | Cysltr1 | 0.242721666 | -0.276839976 | 0.762283309 | 8.459119151 | 1.170096848 | 0.290344944 | 0.709264768 | -4.572780284 |
| **A_55_P2049717** | Agrp | -0.845795617 | -2.656282249 | 0.964691015 | 8.215063325 | -1.170093194 | 0.290346296 | 0.709264768 | -4.572781254 |
| **A_55_P2153580** | Zfp655 | -0.35288343 | -1.108293295 | 0.402526434 | 8.008908163 | -1.170035289 | 0.290367732 | 0.709264768 | -4.572796625 |
| **A_52_P370022** | NA | 0.402397416 | -0.45903568 | 1.263830513 | 14.83158247 | 1.169994918 | 0.290382678 | 0.709264768 | -4.572807342 |
| **A_55_P2131274** | Zfp663 | -0.35808975 | -1.124844693 | 0.408665192 | 12.84290672 | -1.169730062 | 0.290480746 | 0.709264768 | -4.572877645 |
| **A_55_P1955931** | Slc25a42 | -0.222105108 | -0.697715806 | 0.25350559 | 8.961995604 | -1.169653642 | 0.290509047 | 0.709264768 | -4.572897929 |
| **A_55_P2339379** | Sass6 | 0.125692554 | -0.143464419 | 0.394849527 | 8.178907809 | 1.169645515 | 0.290512057 | 0.709264768 | -4.572900086 |
| **A_52_P291971** | Stau2 | 0.148079658 | -0.169050418 | 0.465209735 | 8.898699997 | 1.169521787 | 0.290557884 | 0.709264768 | -4.572932926 |
| **A_52_P565496** | Prl8a1 | -1.499142234 | -4.709894548 | 1.71161008 | 8.392807281 | -1.169462716 | 0.290579765 | 0.709264768 | -4.572948604 |
| **A_52_P165654** | Star | -0.677445379 | -2.128747745 | 0.773856986 | 8.05691748 | -1.16914056 | 0.290699124 | 0.709264768 | -4.573034105 |
| **A_55_P1956862** | Egfr | -0.641748316 | -2.016617039 | 0.733120406 | 8.156701302 | -1.169105898 | 0.290711969 | 0.709264768 | -4.573043304 |
| **A_55_P2015877** | Clps | -0.554311523 | -1.741918947 | 0.6332959 | 7.949957286 | -1.169045324 | 0.290734417 | 0.709264768 | -4.573059379 |
| **A_55_P2109122** | H2bc4 | -0.912748242 | -2.868394831 | 1.042898348 | 13.57941163 | -1.168990484 | 0.290754742 | 0.709264768 | -4.573073933 |
| **A_55_P2073809** | AU01583€ | -1.873766377 | -5.888724903 | 2.14119215 | 8.604658181 | -1.168919823 | 0.290780932 | 0.709264768 | -4.573092685 |
| **A_55_P1975456** | Nfyb | 0.345796446 | -0.39520594 | 1.086798833 | 9.499453755 | 1.168829771 | 0.290814312 | 0.709264768 | -4.573116582 |
| **A_52_P566390** | Zfyve26 | -0.43772631 | -1.375724207 | 0.500271587 | 9.087301617 | -1.168829414 | 0.290814444 | 0.709264768 | -4.573116677 |
| **A_55_P2021689** | Chrdl2 | -0.533734656 | -1.677501971 | 0.610032658 | 7.944200966 | -1.168794252 | 0.290827478 | 0.709264768 | -4.573126007 |
| **A_55_P1981756** | Vmn2r123 | -0.507257095 | -1.594293255 | 0.579779065 | 7.865593949 | -1.168784612 | 0.290831052 | 0.709264768 | -4.573128566 |
| **A_55_P2116621** | Cited2 | -1.330498535 | -4.181764425 | 1.520767355 | 9.892042868 | -1.168764542 | 0.290838492 | 0.709264768 | -4.573133892 |
| **A_51_P220162** | Notch3 | 0.671875664 | -0.7679814 | 2.111732728 | 11.47315445 | 1.168745294 | 0.290845628 | 0.709264768 | -4.573138999 |
| **A_55_P2177659** | Mast1 | -0.31611264 | -0.993558064 | 0.361332785 | 7.914253361 | -1.168739958 | 0.290847606 | 0.709264768 | -4.573140415 |
| **A_55_P2097279** | Pirb | -0.197121021 | -0.619578119 | 0.225336076 | 8.824129573 | -1.168693626 | 0.290864783 | 0.709264768 | -4.57315271 |
| **A_55_P2112504** | Kcnv2 | -0.312085542 | -0.980959676 | 0.356788591 | 7.795612106 | -1.168636897 | 0.290885816 | 0.709264768 | -4.573167763 |
| **A_51_P433733** | Nucb1 | -0.496370061 | -1.560295218 | 0.567555096 | 12.30335227 | -1.168543399 | 0.290920484 | 0.709264768 | -4.573192574 |
| **A_66_P102202** | Gm648 | -1.604283002 | -5.043146048 | 1.834580044 | 8.472835683 | -1.168466989 | 0.290948819 | 0.709264768 | -4.573212849 |
| **A_55_P1962503** | Nup62cl | -1.12847856 | -3.547497566 | 1.290540445 | 8.288477849 | -1.168434515 | 0.290960862 | 0.709264768 | -4.573221465 |
| **A_52_P339996** | Slc6a6 | -0.456249609 | -1.43428906 | 0.521789842 | 11.04869099 | -1.168413261 | 0.290968745 | 0.709264768 | -4.573227105 |
| **A_55_P2274782** | Prl7a1 | -1.541208984 | -4.845038794 | 1.762620826 | 8.423023471 | -1.168407161 | 0.290971007 | 0.709264768 | -4.573228724 |
| **A_55_P1989419** | AI987944 | 0.1388419 | -0.158810133 | 0.436493933 | 8.379064292 | 1.16832062 | 0.291003104 | 0.709264768 | -4.573251686 |
| **A_51_P480861** | Pga5 | -1.02076339 | -3.209259452 | 1.167732672 | 8.221309033 | -1.168233516 | 0.291035412 | 0.709264768 | -4.573274798 |
| **A_55_P2050273** | Cct6b | -1.068761505 | -3.360321774 | 1.222798763 | 8.31473511 | -1.168153366 | 0.291065144 | 0.709264768 | -4.573296064 |
| **A_55_P1981544** | Lactb | -0.348301602 | -1.095116038 | 0.398512835 | 9.653000878 | -1.168135205 | 0.291071882 | 0.709264768 | -4.573300882 |
| **A_51_P288103** | Psmd6 | 0.124039429 | -0.141921502 | 0.390000359 | 12.00286137 | 1.16813291 | 0.291072733 | 0.709264768 | -4.573301491 |
| **A_55_P2129271** | Tead2 | 0.572086113 | -0.654633436 | 1.798805662 | 12.77666568 | 1.168063532 | 0.291098472 | 0.709264768 | -4.573319899 |
| **A_51_P258078** | Nop2 | 0.210914759 | -0.241421198 | 0.663250716 | 11.30652863 | 1.167874578 | 0.291168582 | 0.709264768 | -4.57337003 |
| **A_66_P104221** | NA | -0.212895186 | -0.669501843 | 0.24371147 | 8.564207004 | -1.167814731 | 0.291190791 | 0.709264768 | -4.573385908 |
| **A_55_P2196027** | NA | 0.180224932 | -0.206336254 | 0.566786118 | 8.109552228 | 1.167742187 | 0.291217714 | 0.709264768 | -4.573405154 |
| **A_52_P665817** | Abhd13 | 0.163652728 | -0.187365659 | 0.514671115 | 8.385184611 | 1.167733469 | 0.29122095 | 0.709264768 | -4.573407467 |
| **A_55_P2021119** | Sun2 | 0.161468578 | -0.184867083 | 0.507804239 | 13.06531288 | 1.167726552 | 0.291223517 | 0.709264768 | -4.573409302 |
| **A_55_P2067787** | Psg29 | -2.193268468 | -6.897865908 | 2.511328973 | 8.80209833 | -1.167668791 | 0.291244956 | 0.709264768 | -4.573424625 |

| | | | | | | | | | |
|---|---|---|---|---|---|---|---|---|---|
| **A_55_P2192492** | Slmap | -0.174285907 | -0.548132458 | 0.199560644 | 8.106176885 | -1.167667623 | 0.291245389 | 0.709264768 | -4.573424935 |
| **A_55_P2070913** | Rpl14 | 0.362189857 | -0.414799085 | 1.1391788 | 17.19989837 | 1.167540063 | 0.291292739 | 0.709264768 | -4.573458775 |
| **A_66_P105965** | Drp2 | -0.45385813 | -1.427513711 | 0.519797451 | 8.022432846 | -1.167522092 | 0.29129941 | 0.709264768 | -4.573463542 |
| **A_66_P123055** | Gm10845 | -0.715420759 | -2.250263014 | 0.819421497 | 12.4875045 | -1.167476468 | 0.291316347 | 0.709264768 | -4.573475645 |
| **A_55_P2174085** | St8sia5 | -1.088456333 | -3.423670388 | 1.246757721 | 8.279552178 | -1.167440245 | 0.291329796 | 0.709264768 | -4.573485255 |
| **A_51_P319449** | Acat1 | 0.486433396 | -0.557263655 | 1.530130447 | 11.02347237 | 1.167345122 | 0.291365113 | 0.709264768 | -4.573510488 |
| **A_51_P133582** | Cbx4 | -0.256900864 | -0.808149973 | 0.294348244 | 11.24419619 | -1.167261097 | 0.291396313 | 0.709264768 | -4.573532777 |
| **A_52_P382886** | Gjb2 | -2.583510006 | -8.127236857 | 2.960216846 | 9.336411311 | -1.167235989 | 0.291405636 | 0.709264768 | -4.573539437 |
| **A_51_P215995** | Zswim6 | 0.544894997 | -0.624381237 | 1.714171232 | 9.961841687 | 1.167202055 | 0.291418238 | 0.709264768 | -4.573548439 |
| **A_52_P385606** | Ckb | -0.498211466 | -1.567319238 | 0.570896305 | 9.379427869 | -1.16719274 | 0.291421697 | 0.709264768 | -4.573550909 |
| **A_55_P2108693** | NA | -1.607295076 | -5.056507437 | 1.841917285 | 8.658312277 | -1.167148257 | 0.291438217 | 0.709264768 | -4.573562709 |
| **A_55_P1992762** | Kdelr1 | -0.212960794 | -0.669994777 | 0.244073189 | 9.825706796 | -1.167082374 | 0.291462686 | 0.709264768 | -4.573580184 |
| **A_55_P2126552** | Zdhhc13 | -0.510508883 | -1.606400098 | 0.585382333 | 9.612718352 | -1.166772549 | 0.291577778 | 0.709264768 | -4.573662363 |
| **A_55_P1968703** | Gfra2 | 1.392096839 | -1.596605995 | 4.380799674 | 11.9688842 | 1.166640523 | 0.291626834 | 0.709264768 | -4.573697379 |
| **A_55_P1958275** | Bcl11a | 0.546483984 | -0.626771665 | 1.719739634 | 9.201000413 | 1.166635346 | 0.291628758 | 0.709264768 | -4.573698753 |
| **A_52_P1029978** | Lasp1 | -0.540565827 | -1.701188865 | 0.620057211 | 14.51935192 | -1.166561787 | 0.291656093 | 0.709264768 | -4.573718262 |
| **A_55_P1955276** | Specc1 | -0.314305444 | -0.989145844 | 0.360534956 | 8.09333655 | -1.166544135 | 0.291662653 | 0.709264768 | -4.573722943 |
| **A_52_P377723** | Peg10 | -1.380987155 | -4.346220372 | 1.584246062 | 8.458104548 | -1.16649029 | 0.291682664 | 0.709264768 | -4.573737224 |
| **A_51_P173622** | Ptpn11 | -0.573087188 | -1.803619109 | 0.657444733 | 13.73814861 | -1.16648232 | 0.291685626 | 0.709264768 | -4.573739337 |
| **A_55_P2082180** | Tnfrsf9 | -1.746391166 | -5.496333564 | 2.003551232 | 8.566341143 | -1.166453081 | 0.291696493 | 0.709264768 | -4.573747092 |
| **A_52_P48767** | Rassf8 | -0.960282846 | -3.022535864 | 1.101970172 | 8.408011 | -1.166292731 | 0.291756096 | 0.709264768 | -4.573789617 |
| **A_55_P1958394** | Slit2 | 1.022309652 | -1.173171236 | 3.217790539 | 10.6156483 | 1.166280853 | 0.291760511 | 0.709264768 | -4.573792767 |
| **A_55_P2019019** | Rpl36a | 0.620700699 | -0.712314623 | 1.953716022 | 15.33468782 | 1.166265527 | 0.291766208 | 0.709264768 | -4.573796832 |
| **A_55_P2004333** | Rpl24 | 0.201033762 | -0.230712129 | 0.632779652 | 15.84773433 | 1.16624859 | 0.291772504 | 0.709264768 | -4.573801323 |
| **A_55_P2057132** | Scn8a | 0.177677979 | -0.203908531 | 0.559264488 | 11.66274441 | 1.166248069 | 0.291772698 | 0.709264768 | -4.573801461 |
| **A_55_P2030284** | Trem1 | 0.311958411 | -0.358015995 | 0.981932817 | 9.456792509 | 1.166242416 | 0.2917748 | 0.709264768 | -4.57380296 |
| **A_51_P177691** | Gabrr1 | 0.104569601 | -0.120009927 | 0.32914913 | 7.958814211 | 1.166233822 | 0.291777994 | 0.709264768 | -4.573805239 |
| **A_66_P101620** | Cfap97 | -0.168447774 | -0.530219368 | 0.193323819 | 8.828025943 | -1.166221812 | 0.291782459 | 0.709264768 | -4.573808424 |
| **A_55_P2132631** | NA | 0.36156369 | -0.415005149 | 1.138132529 | 11.67647881 | 1.166152093 | 0.29180838 | 0.709264768 | -4.573826913 |
| **A_51_P134812** | Chac1 | -0.267731024 | -0.842791336 | 0.307329288 | 8.214013717 | -1.166099606 | 0.291827894 | 0.709264768 | -4.573840832 |
| **A_65_P01415** | Kremen1 | -0.543315149 | -1.710309278 | 0.62367898 | 11.04352138 | -1.166093807 | 0.291830051 | 0.709264768 | -4.57384237 |
| **A_52_P144263** | Rbms1 | -0.400733726 | -1.261524658 | 0.460057206 | 9.50101833 | -1.166026865 | 0.291854941 | 0.709264768 | -4.573860122 |
| **A_55_P2053320** | Zfr2 | -0.343302726 | -1.080755196 | 0.394149744 | 8.961796536 | -1.16598661 | 0.29186991 | 0.709264768 | -4.573870796 |
| **A_55_P1970665** | Kdm3b | -0.116314933 | -0.366183933 | 0.133554067 | 8.391326083 | -1.165932592 | 0.291889998 | 0.709264768 | -4.573885121 |
| **A_55_P2086128** | Fa2h | -1.016327897 | -3.199791301 | 1.167135507 | 8.181540005 | -1.165838183 | 0.291925109 | 0.709264768 | -4.573910155 |
| **A_55_P2027213** | Krtap20-2 | -0.358376616 | -1.128350245 | 0.411597013 | 8.391511629 | -1.165773445 | 0.291949187 | 0.709264768 | -4.573927321 |
| **A_55_P2084492** | NA | -0.219074082 | -0.689793342 | 0.251645178 | 8.191592272 | -1.165680089 | 0.291983913 | 0.709264768 | -4.573952075 |
| **A_51_P421876** | Irf7 | -0.402947362 | -1.268826525 | 0.462931802 | 7.896735534 | -1.165578085 | 0.292021859 | 0.709264768 | -4.573979122 |
| **A_55_P2152566** | Sp110 | 0.535893901 | -0.615707739 | 1.687495542 | 9.15377264 | 1.165539202 | 0.292036325 | 0.709264768 | -4.573989432 |
| **A_51_P482410** | Ceacam1! | -1.186151527 | -3.735171184 | 1.362868129 | 8.252610158 | -1.165513567 | 0.292045862 | 0.709264768 | -4.573996229 |
| **A_55_P1984481** | Mfn2 | -0.298898522 | -0.941264918 | 0.343467875 | 8.628081706 | -1.165443699 | 0.292071858 | 0.709264768 | -4.574014754 |
| **A_51_P419017** | Pde10a | -1.052293819 | -3.313875541 | 1.209287903 | 8.378486875 | -1.165400176 | 0.292088052 | 0.709264768 | -4.574026293 |
| **A_52_P202029** | Trak2 | -0.179118398 | -0.564086159 | 0.205849363 | 9.310465115 | -1.165376299 | 0.292096937 | 0.709264768 | -4.574032624 |
| **A_55_P1973036** | NA | -0.383883876 | -1.208971654 | 0.441203901 | 7.891904122 | -1.165333049 | 0.292113031 | 0.709264768 | -4.574044091 |
| **A_55_P2177278** | DXBay18 | -0.904526568 | -2.84868057 | 1.039627434 | 8.125441621 | -1.165308755 | 0.292122072 | 0.709264768 | -4.574050532 |
| **A_52_P562467** | Sync | -0.559569075 | -1.762305009 | 0.64316686 | 8.086412745 | -1.165289313 | 0.292129307 | 0.709264768 | -4.574055687 |
| **A_51_P388801** | Tbcc | -0.217393552 | -0.684664032 | 0.249876928 | 12.02233717 | -1.165275615 | 0.292134405 | 0.709264768 | -4.574059318 |
| **A_55_P2095153** | Pdf | 0.319381554 | -0.367119207 | 1.005882316 | 10.30314102 | 1.16525011 | 0.292143897 | 0.709264768 | -4.574066081 |

| | | | | | | | | | |
|---|---|---|---|---|---|---|---|---|---|
| **A_51_P299062** | Kel | 0.11760785 | -0.135204533 | 0.370420234 | 8.02872508 | 1.165167372 | 0.29217469 | 0.709264768 | -4.574088016 |
| **A_55_P1955778** | Adamts4 | -0.598975332 | -1.886579215 | 0.688628551 | 8.725009327 | -1.165137029 | 0.292185984 | 0.709264768 | -4.574096061 |
| **A_55_P2071930** | NA | -0.216554207 | -0.682091003 | 0.248982588 | 8.307191501 | -1.165099346 | 0.29220001 | 0.709264768 | -4.574106051 |
| **A_55_P2095266** | NA | 0.20935844 | -0.240729274 | 0.659446153 | 14.44491259 | 1.165047577 | 0.29221928 | 0.709264768 | -4.574119776 |
| **A_55_P2055168** | Gm2016 | -0.431177735 | -1.35818666 | 0.495831189 | 8.970864309 | -1.164991503 | 0.292240154 | 0.709264768 | -4.574134642 |
| **A_55_P2015675** | AU01583( | -1.863711451 | -5.870650976 | 2.143228074 | 8.617935419 | -1.164973996 | 0.292246672 | 0.709264768 | -4.574139283 |
| **A_55_P1967341** | Gm9104 | 0.631451957 | -0.726195557 | 1.989099471 | 13.91180566 | 1.164940195 | 0.292259255 | 0.709264768 | -4.574148244 |
| **A_66_P127722** | NA | 0.621725397 | -0.715034316 | 1.95848511 | 12.48174187 | 1.16491867 | 0.292267269 | 0.709264768 | -4.57415395 |
| **A_55_P2045642** | Stmn4 | -0.49895618 | -1.571776642 | 0.573864283 | 8.806713227 | -1.164892108 | 0.292277158 | 0.709264768 | -4.574160992 |
| **A_52_P535255** | Syf2 | -0.164882681 | -0.519439981 | 0.189674619 | 10.28994607 | -1.164766726 | 0.292323842 | 0.709264768 | -4.57419423 |
| **A_55_P1977968** | Epb41 | -0.211529782 | -0.714948957 | 0.291889393 | 13.55682966 | -1.239179693 | 0.292343766 | 0.709264768 | -4.569212781 |
| **A_51_P338961** | Qser1 | 0.400110257 | -0.460338571 | 1.260559085 | 9.74906724 | 1.164675617 | 0.292357769 | 0.709264768 | -4.574218382 |
| **A_52_P6798** | Rbm22 | 0.216194243 | -0.248760258 | 0.681148745 | 9.263821452 | 1.164619384 | 0.292378711 | 0.709264768 | -4.574233289 |
| **A_55_P2018403** | Sycp1 | -0.558713287 | -1.760349253 | 0.64292268 | 7.965516523 | -1.164572219 | 0.292396277 | 0.709264768 | -4.574245791 |
| **A_55_P1974587** | Vangl1 | -0.19616377 | -0.618066023 | 0.225738483 | 8.543517903 | -1.164547749 | 0.292405391 | 0.709264768 | -4.574252278 |
| **A_55_P1967905** | Dcbld1 | 0.294572752 | -0.339052379 | 0.928197883 | 9.222641557 | 1.164422325 | 0.292452108 | 0.709264768 | -4.574285525 |
| **A_55_P1952725** | NA | -2.220555399 | -6.997089841 | 2.555979044 | 8.821707195 | -1.164391548 | 0.292463573 | 0.709264768 | -4.574293683 |
| **A_55_P2037408** | Lrrc57 | -0.121770506 | -0.383717158 | 0.140176147 | 9.237382906 | -1.164339422 | 0.292482991 | 0.709264768 | -4.574307499 |
| **A_55_P2125557** | Irx2 | 0.94503122 | -1.088004799 | 2.978067239 | 10.56803656 | 1.164263892 | 0.29251113 | 0.709264768 | -4.57432752 |
| **A_55_P1965699** | NA | -0.212257625 | -0.668952623 | 0.244437373 | 7.750057303 | -1.164092233 | 0.292575091 | 0.709264768 | -4.574373018 |
| **A_51_P269020** | 1700031F | -0.413388095 | -1.302881579 | 0.476105389 | 7.828690205 | -1.164033664 | 0.292596917 | 0.709264768 | -4.574388542 |
| **A_55_P2017183** | NA | -0.225724021 | -0.711428019 | 0.259979977 | 10.66327298 | -1.164009289 | 0.292606001 | 0.709264768 | -4.574395002 |
| **A_55_P2341110** | Chn1os3 | -0.270034905 | -0.851097654 | 0.311027843 | 7.799821006 | -1.163984554 | 0.292615219 | 0.709264768 | -4.574401558 |
| **A_55_P1966765** | NA | 0.366383048 | -0.42207365 | 1.154839745 | 12.25825046 | 1.163879101 | 0.292654522 | 0.709264768 | -4.574429507 |
| **A_55_P2051400** | Prl3d2 | -2.127328172 | -6.70553826 | 2.450881915 | 8.71438402 | -1.163828851 | 0.292673251 | 0.709264768 | -4.574442824 |
| **A_55_P1968433** | Gpat3 | -1.14185395 | -3.599284649 | 1.315576748 | 8.980865359 | -1.163803409 | 0.292682735 | 0.709264768 | -4.574449567 |
| **A_55_P2236138** | 4930447F | -0.233534711 | -0.736188076 | 0.269118655 | 7.786289316 | -1.163678984 | 0.292729119 | 0.709264768 | -4.574482543 |
| **A_52_P355075** | Rab40c | -0.341731468 | -1.077344988 | 0.393882052 | 11.60275982 | -1.163551513 | 0.292776644 | 0.709264768 | -4.574516324 |
| **A_66_P108770** | Oxct1 | 0.608060949 | -0.701063292 | 1.91718519 | 12.7501987 | 1.163366607 | 0.292845595 | 0.709264768 | -4.574565325 |
| **A_52_P637812** | Prrg4 | -0.557512407 | -1.757861671 | 0.642836857 | 8.119303358 | -1.163314796 | 0.292864918 | 0.709264768 | -4.574579055 |
| **A_55_P1967659** | Lactb | -0.327636545 | -1.033054795 | 0.377781705 | 9.37930099 | -1.16331142 | 0.292866177 | 0.709264768 | -4.574579949 |
| **A_55_P1971840** | Muc13 | -1.519865071 | -4.792228257 | 1.752498116 | 8.626044963 | -1.163305768 | 0.292868285 | 0.709264768 | -4.574581447 |
| **A_51_P267194** | Faim | -0.303184198 | -0.956004696 | 0.3496363 | 10.28465908 | -1.163223388 | 0.292899011 | 0.709264768 | -4.574603277 |
| **A_52_P265002** | Pik3ca | -0.130627243 | -0.411919923 | 0.150665437 | 9.046390448 | -1.163123087 | 0.292936425 | 0.709264768 | -4.574629855 |
| **A_51_P474053** | Erv3 | -1.548965068 | -4.884539663 | 1.786609526 | 8.468230409 | -1.163111399 | 0.292940785 | 0.709264768 | -4.574632952 |
| **A_55_P1992199** | Psg20 | -1.781863429 | -5.619032625 | 2.055305767 | 8.55831378 | -1.163091309 | 0.29294828 | 0.709264768 | -4.574638275 |
| **A_55_P2171977** | NA | 0.634219359 | -0.731551008 | 1.999989727 | 14.69629132 | 1.163086874 | 0.292949934 | 0.709264768 | -4.574639451 |
| **A_55_P2039532** | Pax8 | -0.657618322 | -2.073794237 | 0.758557593 | 7.985769834 | -1.163073193 | 0.292955038 | 0.709264768 | -4.574643076 |
| **A_52_P555235** | Rgs19 | 0.509645471 | -0.588001755 | 1.607292698 | 9.465703881 | 1.162935774 | 0.293006307 | 0.709264768 | -4.574679488 |
| **A_66_P129188** | H2ac20 | 0.76351457 | -0.88090619 | 2.407935331 | 17.0150475 | 1.162933079 | 0.293007313 | 0.709264768 | -4.574680202 |
| **A_55_P2008704** | Gstp2 | 0.24302864 | -0.280438766 | 0.766496045 | 11.11936339 | 1.162835166 | 0.293043848 | 0.709264768 | -4.574706146 |
| **A_52_P539414** | Gtf2h3 | 0.286619154 | -0.330742657 | 0.903980964 | 9.765499373 | 1.162828839 | 0.293046209 | 0.709264768 | -4.574707822 |
| **A_55_P1956873** | Timm17a | -0.220645425 | -0.695937472 | 0.254646621 | 12.77387468 | -1.162745656 | 0.293077251 | 0.709264768 | -4.574729862 |
| **A_55_P1999838** | Cox7c | 0.321954341 | -0.371607534 | 1.015516215 | 14.25636945 | 1.162677903 | 0.293102537 | 0.709264768 | -4.574747813 |
| **A_55_P2117525** | Rbm3 | 0.685450122 | -0.791189063 | 2.162089307 | 12.80835456 | 1.162657546 | 0.293110134 | 0.709264768 | -4.574753207 |
| **A_51_P204053** | Hba-x | 0.476053873 | -0.549642413 | 1.50175016 | 8.52287639 | 1.162485824 | 0.293174232 | 0.709264768 | -4.574798703 |
| **A_52_P590474** | Ccnl1 | -0.198473034 | -0.626106459 | 0.229160391 | 8.753201979 | -1.16246587 | 0.293181681 | 0.709264768 | -4.57480399 |
| **A_55_P2088699** | NA | 0.328667949 | -0.379581078 | 1.036916976 | 16.62100493 | 1.162309303 | 0.293240134 | 0.709264768 | -4.574845469 |

| | | | | | | | | | |
|---|---|---|---|---|---|---|---|---|---|
| **A_55_P2146127** | Chmp4b | -0.349324835 | -1.102146407 | 0.403496737 | 11.93037053 | -1.162218418 | 0.293274069 | 0.709264768 | -4.574869546 |
| **A_51_P490388** | Nudt8 | -0.559839209 | -1.76639251 | 0.646714092 | 9.161747613 | -1.162163268 | 0.293294663 | 0.709264768 | -4.574884156 |
| **A_55_P1973006** | Bag1 | -0.249012052 | -0.785715335 | 0.287691231 | 13.78331172 | -1.162081163 | 0.293325325 | 0.709264768 | -4.574905907 |
| **A_51_P271984** | Tmem45b | -0.22557987 | -0.71181113 | 0.260651389 | 7.792433997 | -1.162004508 | 0.293353955 | 0.709264768 | -4.574926213 |
| **A_55_P2070045** | Golga7b | -0.621346295 | -1.960720718 | 0.718028127 | 7.986560273 | -1.161935599 | 0.293379693 | 0.709264768 | -4.574944467 |
| **A_55_P1970299** | Mttp | -0.611749539 | -1.930456385 | 0.706957307 | 8.34256286 | -1.161918684 | 0.293386011 | 0.709264768 | -4.574948948 |
| **A_51_P311694** | Ypel5 | -0.222690968 | -0.702746421 | 0.257364485 | 9.268583135 | -1.16188071 | 0.293400196 | 0.709264768 | -4.574959007 |
| **A_55_P2136385** | Rps3a1 | 0.383135543 | -0.442830072 | 1.209101159 | 15.61449108 | 1.161825276 | 0.293420904 | 0.709264768 | -4.574973691 |
| **A_51_P370341** | Zbtb48 | -0.125415676 | -0.395793339 | 0.144961986 | 8.657032978 | -1.161799971 | 0.293430357 | 0.709264768 | -4.574980394 |
| **A_55_P2059931** | Prom1 | -0.389639213 | -1.22965663 | 0.450378203 | 11.02924058 | -1.161782186 | 0.293437002 | 0.709264768 | -4.574985105 |
| **A_55_P2080905** | NA | 0.193048077 | -0.223144755 | 0.609240909 | 16.65581308 | 1.161772866 | 0.293440483 | 0.709264768 | -4.574987574 |
| **A_52_P630571** | Epg5 | -0.199428356 | -0.629384781 | 0.23052807 | 8.787500733 | -1.161750347 | 0.293448897 | 0.709264768 | -4.574993539 |
| **A_66_P131645** | Ahnak2 | -0.434520321 | -1.371345845 | 0.502305202 | 8.100055589 | -1.161720687 | 0.293459978 | 0.709264768 | -4.575001395 |
| **A_55_P2119219** | NA | 0.315532845 | -0.364772873 | 0.995838563 | 13.48144944 | 1.161691463 | 0.293470896 | 0.709264768 | -4.575009136 |
| **A_55_P2092909** | Rassf1 | -0.254871204 | -0.804421185 | 0.294678776 | 11.73817796 | -1.161619569 | 0.293497759 | 0.709264768 | -4.575028179 |
| **A_66_P134090** | Grpel2 | -0.165530873 | -0.522457778 | 0.191396033 | 8.376199643 | -1.161582509 | 0.293511607 | 0.709264768 | -4.575037995 |
| **A_52_P152219** | Rpgrip1l | 0.142664756 | -0.164976131 | 0.450305643 | 8.708916798 | 1.161510093 | 0.293538668 | 0.709264768 | -4.575057176 |
| **A_66_P136714** | Scpep1 | -0.74508742 | -2.351933195 | 0.861758355 | 12.52796381 | -1.16140417 | 0.293578254 | 0.709264768 | -4.575085231 |
| **A_55_P1984344** | Rhox2h | -0.477970284 | -1.508806164 | 0.552865596 | 7.964989203 | -1.161346234 | 0.293599908 | 0.709264768 | -4.575100577 |
| **A_55_P2051176** | NA | 0.366258985 | -0.423649792 | 1.156167762 | 15.82051332 | 1.161346175 | 0.29359993 | 0.709264768 | -4.575100592 |
| **A_66_P101477** | Bglap3 | -0.621037926 | -1.960560206 | 0.718484354 | 7.974032196 | -1.161230748 | 0.293643075 | 0.709264768 | -4.575131163 |
| **A_55_P1964812** | Tfap2a | -0.769062355 | -2.427904668 | 0.889779958 | 8.154315196 | -1.161199378 | 0.293654803 | 0.709264768 | -4.575139472 |
| **A_51_P349878** | Txndc17 | -0.141629029 | -0.447131308 | 0.16387325 | 14.12151488 | -1.161149578 | 0.29367342 | 0.709264768 | -4.575152661 |
| **A_52_P444162** | Rab11a | -0.129618635 | -0.409225613 | 0.149988343 | 11.00162522 | -1.161100424 | 0.293691796 | 0.709264768 | -4.575165679 |
| **A_55_P1981494** | H2-Bl | -0.820921013 | -2.591774178 | 0.949932151 | 8.563008957 | -1.161098773 | 0.293692413 | 0.709264768 | -4.575166116 |
| **A_55_P2368625** | Sp4 | 0.260924646 | -0.30193264 | 0.823781933 | 8.65882353 | 1.161093399 | 0.293694423 | 0.709264768 | -4.575167539 |
| **A_55_P2023617** | Kiss1r | -0.472980259 | -1.493396122 | 0.547435605 | 8.126938196 | -1.160957064 | 0.293745399 | 0.709264768 | -4.575203646 |
| **A_55_P2026440** | Ceacam-p | -0.702977767 | -2.219620855 | 0.813665321 | 7.994820578 | -1.160936713 | 0.293753008 | 0.709264768 | -4.575209035 |
| **A_55_P2069579** | NA | 0.124122391 | -0.143697475 | 0.391942258 | 8.11694978 | 1.160800777 | 0.293803844 | 0.709264768 | -4.575245034 |
| **A_51_P476851** | Fance | -0.162913429 | -0.514456389 | 0.188629531 | 11.95576629 | -1.16072363 | 0.293832698 | 0.709264768 | -4.575265464 |
| **A_51_P206134** | Akna | 0.26648625 | -0.308551916 | 0.841524417 | 8.585478641 | 1.160722708 | 0.293833042 | 0.709264768 | -4.575265708 |
| **A_51_P256246** | Tspan13 | 0.774208333 | -0.896608762 | 2.445025428 | 13.21217564 | 1.160591209 | 0.29388223 | 0.709264768 | -4.57530053 |
| **A_55_P2001976** | Rhox12 | -2.104086084 | -6.646181303 | 2.438009135 | 8.707384346 | -1.160266131 | 0.294003857 | 0.709264768 | -4.575386609 |
| **A_52_P500488** | Lars | -0.206324374 | -0.651742834 | 0.239094085 | 12.17349347 | -1.16019949 | 0.294028796 | 0.709264768 | -4.575404254 |
| **A_55_P2015878** | Clps | -0.886045996 | -2.798898536 | 1.026806544 | 8.453795611 | -1.160179323 | 0.294036343 | 0.709264768 | -4.575409594 |
| **A_55_P1977314** | Ulk1 | -0.216686122 | -0.68449301 | 0.251120765 | 12.40610886 | -1.160151838 | 0.29404663 | 0.709264768 | -4.575416871 |
| **A_55_P2218223** | Pacs2 | -0.463289851 | -1.463677066 | 0.537097363 | 8.99362925 | -1.159938597 | 0.294126447 | 0.709264768 | -4.57547333 |
| **A_52_P619248** | Prl2c5 | -2.981658044 | -9.420550207 | 3.457234119 | 9.230952504 | -1.159837185 | 0.294164413 | 0.709264768 | -4.57550018 |
| **A_55_P2124736** | Col14a1 | 1.222922206 | -1.418023667 | 3.86386808 | 12.54629937 | 1.159817551 | 0.294171763 | 0.709264768 | -4.575505378 |
| **A_55_P1963031** | Ampd2 | -0.326335807 | -1.031087333 | 0.37841572 | 8.809876412 | -1.159789169 | 0.294182389 | 0.709264768 | -4.575512892 |
| **A_52_P432780** | Mrpl50 | 0.265879025 | -0.308351522 | 0.840109572 | 8.680726988 | 1.159706608 | 0.294213303 | 0.709264768 | -4.57553475 |
| **A_55_P2009576** | Rnu12 | 0.472820325 | -0.548360885 | 1.494001536 | 13.7533267 | 1.159694688 | 0.294217766 | 0.709264768 | -4.575537905 |
| **A_55_P2025837** | NA | 0.350904071 | -0.406993132 | 1.108801273 | 14.67451803 | 1.159654046 | 0.294232985 | 0.709264768 | -4.575548665 |
| **A_55_P2002642** | Deup1 | 1.095834562 | -1.27106018 | 3.462729303 | 10.68553535 | 1.159621862 | 0.294245037 | 0.709264768 | -4.575557185 |
| **A_51_P520849** | Sfrp2 | 0.928686944 | -1.077186122 | 2.93456001 | 10.51187952 | 1.159621447 | 0.294245192 | 0.709264768 | -4.575557295 |
| **A_55_P2071028** | NA | 0.679602357 | -0.788504208 | 2.147708922 | 11.84392891 | 1.159438297 | 0.294313785 | 0.709264768 | -4.57560578 |
| **A_55_P1973362** | Kcnj12 | -0.373583558 | -1.180666299 | 0.433499183 | 8.067235856 | -1.159364611 | 0.294341386 | 0.709264768 | -4.575625287 |
| **A_55_P2158112** | Rps27 | 0.137659986 | -0.159756051 | 0.435076024 | 16.66254744 | 1.159294258 | 0.29436774 | 0.709264768 | -4.57564391 |

| | | | | | | | | | |
|---|---|---|---|---|---|---|---|---|---|
| **A_51_P233815** | Tmem126 | 0.182208322 | -0.211465508 | 0.575882152 | 9.822971125 | 1.159263089 | 0.294379417 | 0.709264768 | -4.575652161 |
| **A_55_P2145942** | Atg16l1 | -0.170002723 | -0.537319125 | 0.197313678 | 8.720238496 | -1.159220101 | 0.294395522 | 0.709264768 | -4.575663541 |
| **A_66_P100186** | Psg28 | -2.629819498 | -8.312284933 | 3.052645936 | 9.03126459 | -1.159149585 | 0.294421941 | 0.709264768 | -4.575682207 |
| **A_52_P323852** | Tnfrsf22 | -0.470037241 | -1.485694343 | 0.545619861 | 8.005244562 | -1.159138963 | 0.294425921 | 0.709264768 | -4.575685018 |
| **A_51_P499838** | Bst1 | -0.864766207 | -2.733395631 | 1.003863217 | 8.705226462 | -1.159113256 | 0.294435553 | 0.709264768 | -4.575691823 |
| **A_55_P2119441** | Vma21 | -0.392858776 | -1.241858195 | 0.456140642 | 10.42290316 | -1.158989263 | 0.294482015 | 0.709264768 | -4.575724644 |
| **A_51_P296429** | Tmem170 | -0.214734974 | -0.678801672 | 0.249331723 | 7.973568667 | -1.15897142 | 0.294488701 | 0.709264768 | -4.575729366 |
| **A_55_P2012729** | Kcnh2 | 0.186522414 | -0.216580556 | 0.589625384 | 8.156021871 | 1.158951805 | 0.294496052 | 0.709264768 | -4.575734558 |
| **A_55_P1967880** | Chtf8 | -0.239573239 | -0.757343606 | 0.278197129 | 14.06199382 | -1.158914664 | 0.294509971 | 0.709264768 | -4.575744389 |
| **A_55_P2133280** | Ap4b1 | -0.19807243 | -0.626162445 | 0.230017584 | 9.992745298 | -1.158882163 | 0.294522152 | 0.709264768 | -4.575752992 |
| **A_55_P2063126** | Patj | -0.147122395 | -0.465100895 | 0.170856105 | 10.13028057 | -1.158861074 | 0.294530056 | 0.709264768 | -4.575758573 |
| **A_55_P1977538** | Apba3 | -0.340099475 | -1.075185082 | 0.394986132 | 9.720929476 | -1.158826421 | 0.294543044 | 0.709264768 | -4.575767746 |
| **A_55_P1954001** | Snrpc | 0.366210071 | -0.425313542 | 1.157733683 | 12.05149051 | 1.158822059 | 0.294544679 | 0.709264768 | -4.5757689 |
| **A_51_P404377** | Rnd2 | -0.498626593 | -1.576371268 | 0.579118082 | 8.154987462 | -1.158803763 | 0.294551536 | 0.709264768 | -4.575773743 |
| **A_55_P2125049** | Gm11127 | -1.167616694 | -3.691428566 | 1.356195178 | 10.99270971 | -1.158760446 | 0.294567772 | 0.709264768 | -4.575785208 |
| **A_55_P1987775** | Pla2g4d | -1.00708437 | -3.184167185 | 1.169998444 | 8.20725427 | -1.158620616 | 0.29462019 | 0.709264768 | -4.575822217 |
| **A_55_P2158227** | Angpt1 | 0.735148058 | -0.854127097 | 2.324423213 | 10.09686886 | 1.158580103 | 0.294635378 | 0.709264768 | -4.575832939 |
| **A_51_P156934** | Dnajc15 | 0.400738369 | -0.465773402 | 1.26725014 | 12.48383304 | 1.158342005 | 0.294724655 | 0.709264768 | -4.575895953 |
| **A_51_P516637** | Bmp5 | 0.455535077 | -0.52949215 | 1.440562305 | 8.748012802 | 1.158307677 | 0.294737528 | 0.709264768 | -4.575905038 |
| **A_51_P469951** | Srgap3 | -0.809562947 | -2.560177458 | 0.941051564 | 8.734642854 | -1.158271697 | 0.294751022 | 0.709264768 | -4.57591456 |
| **A_51_P333622** | 2010300C | 0.676396809 | -0.786444009 | 2.139237627 | 11.24495839 | 1.15812337 | 0.294806654 | 0.709264768 | -4.575953813 |
| **A_55_P1999958** | NA | -0.642621625 | -2.032446979 | 0.747203729 | 8.071730594 | -1.158098374 | 0.294816031 | 0.709264768 | -4.575960427 |
| **A_51_P175758** | Atg101 | -0.152750256 | -0.483135202 | 0.17763469 | 11.65683968 | -1.158009285 | 0.29484945 | 0.709264768 | -4.575984003 |
| **A_55_P2163093** | Nxf7 | -0.329223501 | -1.041353047 | 0.382906045 | 7.786777272 | -1.157929636 | 0.294879331 | 0.709264768 | -4.576005079 |
| **A_55_P1968370** | Gprc5c | -0.827238438 | -2.616787172 | 0.962310296 | 10.72327447 | -1.157810601 | 0.294923994 | 0.709264768 | -4.576036578 |
| **A_52_P559975** | Cxcr2 | -0.91384715 | -2.890795403 | 1.063101104 | 8.244069994 | -1.157786831 | 0.294932913 | 0.709264768 | -4.576042868 |
| **A_66_P138201** | Rhox10 | -1.20024652 | -3.797014193 | 1.396521153 | 8.250198372 | -1.1576778 | 0.294973828 | 0.709264768 | -4.576071718 |
| **A_55_P2181665** | NA | -0.389030143 | -1.230778485 | 0.452718199 | 7.92781402 | -1.157580837 | 0.295010218 | 0.709264768 | -4.576097374 |
| **A_51_P210082** | Ercc4 | -0.104873989 | -0.331801075 | 0.122053098 | 10.11515017 | -1.157528771 | 0.29502976 | 0.709264768 | -4.57611115 |
| **A_55_P2013143** | Esrp2 | -0.109608135 | -0.346791362 | 0.127575092 | 9.726124768 | -1.157468449 | 0.295052402 | 0.709264768 | -4.57612711 |
| **A_55_P2056419** | Coq6 | 0.31226152 | -0.363475544 | 0.987998584 | 11.8034451 | 1.157420244 | 0.295070497 | 0.709264768 | -4.576139865 |
| **A_52_P87892** | Abraxas2 | -0.190146722 | -0.601634715 | 0.221341272 | 8.946449885 | -1.157396115 | 0.295079555 | 0.709264768 | -4.576146249 |
| **A_51_P450278** | 2010003K | -0.507934912 | -1.607214912 | 0.591345087 | 7.923137269 | -1.157311008 | 0.295111506 | 0.709264768 | -4.576168766 |
| **A_52_P125485** | Slc25a48 | -0.350936548 | -1.110444326 | 0.40857123 | 7.838910732 | -1.157302041 | 0.295114872 | 0.709264768 | -4.576171139 |
| **A_55_P1974005** | Tnfrsf23 | -0.515810753 | -1.632174612 | 0.600553106 | 7.959588625 | -1.157270729 | 0.295126628 | 0.709264768 | -4.576179423 |
| **A_55_P1996344** | Dcst2 | -0.531265008 | -1.681101794 | 0.618571778 | 7.921454743 | -1.157245122 | 0.295136243 | 0.709264768 | -4.576186198 |
| **A_55_P2026818** | Slc4a7 | 0.225145614 | -0.262255109 | 0.712546337 | 8.98444667 | 1.15698484 | 0.295233982 | 0.709264768 | -4.576255058 |
| **A_55_P2164265** | Prl7c1 | -1.237235671 | -3.915725133 | 1.44125379 | 8.276783024 | -1.156945328 | 0.295248822 | 0.709264768 | -4.576265511 |
| **A_55_P2092983** | NA | 0.33057095 | -0.385084692 | 1.046226593 | 14.99115642 | 1.156940254 | 0.295250727 | 0.709264768 | -4.576266854 |
| **A_52_P237077** | Esr1 | -0.254471742 | -0.805400071 | 0.296456587 | 7.804861338 | -1.15689729 | 0.295266864 | 0.709264768 | -4.576130936 |
| **A_55_P2044380** | NA | -0.638559641 | -2.021173813 | 0.744054532 | 8.282636305 | -1.156780096 | 0.295310886 | 0.709264768 | -4.576309222 |
| **A_52_P77245** | Snx8 | -0.553255596 | -1.751176014 | 0.644664822 | 10.53487793 | -1.156773135 | 0.295313501 | 0.709264768 | -4.576311064 |
| **A_51_P438701** | Atpaf2 | 0.161236832 | -0.187888987 | 0.510362651 | 9.098412643 | 1.156731695 | 0.295329068 | 0.709264768 | -4.576322026 |
| **A_55_P2219903** | AW06114 | 0.17017788 | -0.198360347 | 0.538716107 | 11.0024339 | 1.15656731 | 0.295390828 | 0.709264768 | -4.57636551 |
| **A_52_P194316** | Atp13a3 | -0.502936923 | -1.592107117 | 0.586233271 | 8.81286019 | -1.156559867 | 0.295393625 | 0.709264768 | -4.576367479 |
| **A_55_P2004071** | Gm2058 | -0.426637564 | -1.35065458 | 0.497379451 | 8.806945035 | -1.156456945 | 0.2954323 | 0.709264768 | -4.576394704 |
| **A_51_P446796** | Camta1 | -0.230601749 | -0.730069658 | 0.26886616 | 8.327959865 | -1.156392729 | 0.295456432 | 0.709264768 | -4.57641169 |
| **A_55_P1977792** | Folr1 | -2.266507317 | -7.175758629 | 2.642743995 | 9.122982465 | -1.156357725 | 0.295469587 | 0.709264768 | -4.576420948 |

| | | | | | | | | | |
|---|---|---|---|---|---|---|---|---|---|
| **A_52_P565940** | Nsd1 | -0.305684077 | -0.967933569 | 0.356565415 | 9.940923989 | -1.156116311 | 0.295560329 | 0.709264768 | -4.576484802 |
| **A_51_P162676** | Armc7 | -0.330222721 | -1.045661381 | 0.385215938 | 8.429733444 | -1.156072029 | 0.295576976 | 0.709264768 | -4.576496514 |
| **A_55_P2145709** | NA | 0.251476947 | -0.293382084 | 0.796335978 | 8.344258063 | 1.156017402 | 0.295597514 | 0.709264768 | -4.576510962 |
| **A_55_P2068199** | NA | 0.110071927 | -0.128425085 | 0.348568939 | 8.527969727 | 1.155963101 | 0.295617929 | 0.709264768 | -4.576525324 |
| **A_51_P392928** | Mthfd1 | 0.59018236 | -0.688665836 | 1.869030557 | 11.51682472 | 1.155892845 | 0.295644346 | 0.709264768 | -4.576543905 |
| **A_51_P331328** | Gpihbp1 | 0.673812651 | -0.786266059 | 2.133891361 | 11.87104524 | 1.155881301 | 0.295648687 | 0.709264768 | -4.576546958 |
| **A_51_P401958** | Taldo1 | -0.342104018 | -1.083465066 | 0.399257031 | 15.08386546 | -1.15578953 | 0.295683197 | 0.709264768 | -4.576571229 |
| **A_55_P2097820** | NA | 0.628935711 | -0.734094568 | 1.991965989 | 12.18702582 | 1.155715923 | 0.295710879 | 0.709264768 | -4.576590695 |
| **A_52_P97699** | D430019H | 0.586375799 | -0.684496188 | 1.857247786 | 10.08787187 | 1.155645351 | 0.295737421 | 0.709264768 | -4.576609359 |
| **A_55_P2282969** | Gm2a | -0.616100898 | -1.951542226 | 0.719340431 | 8.247353446 | -1.155519755 | 0.295784664 | 0.709264768 | -4.576642573 |
| **A_55_P1977353** | Ogfod2 | -0.469313891 | -1.486636048 | 0.548008267 | 10.69076441 | -1.155460891 | 0.295806807 | 0.709264768 | -4.576658139 |
| **A_55_P2216996** | Tmem87a | -0.219142102 | -0.694183055 | 0.255898851 | 8.246715864 | -1.15543393 | 0.29581695 | 0.709264768 | -4.576665269 |
| **A_55_P2360501** | NA | -0.116156461 | -0.367959889 | 0.135646966 | 8.366932833 | -1.15539925 | 0.295829997 | 0.709264768 | -4.576674439 |
| **A_51_P411345** | Mogat2 | -0.536008955 | -1.697974983 | 0.625957072 | 8.366515167 | -1.155390948 | 0.295833121 | 0.709264768 | -4.576676635 |
| **A_65_P13209** | Ctsd | -0.594221147 | -1.882481134 | 0.694038841 | 8.920254437 | -1.155300418 | 0.295867182 | 0.709264768 | -4.576700574 |
| **A_55_P2065303** | NA | 0.145630043 | -0.17009355 | 0.461353635 | 15.56952715 | 1.155298703 | 0.295867828 | 0.709264768 | -4.576701028 |
| **A_51_P502225** | Slc22a4 | -0.931521656 | -2.951056266 | 1.088012954 | 8.226237566 | -1.155292626 | 0.295870114 | 0.709264768 | -4.576702635 |
| **A_55_P2074406** | Ldoc1 | -1.123932414 | -3.560836211 | 1.312971382 | 8.241651277 | -1.15518664 | 0.295909995 | 0.709264768 | -4.57673066 |
| **A_52_P679205** | Slc35f6 | -0.483549882 | -1.531988506 | 0.564888741 | 9.248299421 | -1.155177219 | 0.295913541 | 0.709264768 | -4.576733151 |
| **A_55_P1953503** | Lrrc4b | 0.731187188 | -0.854220065 | 2.31659444 | 9.461196497 | 1.15514919 | 0.295924089 | 0.709264768 | -4.576740563 |
| **A_55_P2106763** | Pxdc1 | -0.379144928 | -1.201386674 | 0.443096819 | 8.119154858 | -1.154931091 | 0.296006175 | 0.709264768 | -4.576798231 |
| **A_55_P2100899** | Arnt | -0.183964614 | -0.582924589 | 0.21499536 | 8.442223504 | -1.154929068 | 0.296006936 | 0.709264768 | -4.576798766 |
| **A_51_P423127** | Gba | -0.769320353 | -2.437747923 | 0.899107218 | 9.286056294 | -1.154915499 | 0.296012044 | 0.709264768 | -4.576802354 |
| **A_51_P461031** | Ostm1 | -0.666929533 | -2.113371499 | 0.779512434 | 10.03473398 | -1.154859845 | 0.296032994 | 0.709264768 | -4.576817069 |
| **A_51_P135357** | Sike1 | -0.102398256 | -0.324491371 | 0.11969486 | 8.446196508 | -1.154802746 | 0.296054489 | 0.709264768 | -4.576832166 |
| **A_51_P464300** | Gdf1 | 0.532061354 | -0.621952535 | 1.686075242 | 9.245685318 | 1.154784711 | 0.296061279 | 0.709264768 | -4.576836934 |
| **A_55_P2123942** | Lmbrd1 | -0.147878022 | -0.468633469 | 0.172877425 | 9.147622859 | -1.154728642 | 0.296082388 | 0.709264768 | -4.576851759 |
| **A_55_P1985005** | Ghrh | -2.630636842 | -8.337553374 | 3.076279689 | 9.078652667 | -1.154541965 | 0.296152679 | 0.709264768 | -4.576901113 |
| **A_51_P276479** | 4930486L | -1.825705435 | -5.787075993 | 2.135665122 | 8.947350665 | -1.15434489 | 0.296226901 | 0.709264768 | -4.576953214 |
| **A_55_P2107432** | Slc18a1 | -0.31758458 | -1.00668619 | 0.371517031 | 8.191704652 | -1.15432071 | 0.296236009 | 0.709264768 | -4.576959606 |
| **A_55_P2024041** | Psg19 | -2.240772465 | -7.102931948 | 2.621387018 | 8.852482799 | -1.154300556 | 0.296243601 | 0.709264768 | -4.576964934 |
| **A_52_P506344** | B3gat3 | -0.172663396 | -0.547326694 | 0.201999902 | 8.401884102 | -1.154275484 | 0.296253045 | 0.709264768 | -4.576971562 |
| **A_52_P429876** | Tbx20 | -0.26082833 | -0.848765534 | 0.327108873 | 8.463020116 | -1.181643648 | 0.296283149 | 0.709264768 | -4.571461711 |
| **A_55_P2088625** | Fbln7 | -2.495022424 | -7.909686149 | 2.919641301 | 10.13030912 | -1.154126223 | 0.296309274 | 0.709264768 | -4.57701102 |
| **A_55_P2205156** | Aamdc | -0.211653772 | -0.670983731 | 0.247676187 | 8.881030965 | -1.154121645 | 0.296310999 | 0.709264768 | -4.57701223 |
| **A_52_P511821** | Wdfy1 | -0.363004959 | -1.150858126 | 0.424848207 | 8.015953893 | -1.154031382 | 0.296345007 | 0.709264768 | -4.577036091 |
| **A_55_P2319035** | AW01195 | -0.997019331 | -3.160920014 | 1.166881353 | 8.251649136 | -1.154028675 | 0.296346027 | 0.709264768 | -4.577036807 |
| **A_52_P523445** | Cdhr5 | -0.553017844 | -1.753274327 | 0.64723864 | 7.981743601 | -1.154025564 | 0.296347199 | 0.709264768 | -4.577037629 |
| **A_55_P1987146** | Adgrf4 | -0.315560173 | -1.000514384 | 0.369394037 | 7.852752806 | -1.153907488 | 0.296391692 | 0.709264768 | -4.577068841 |
| **A_52_P429749** | Tmem115 | -0.346382129 | -1.098256792 | 0.405492535 | 9.173843235 | -1.153879273 | 0.296402325 | 0.709264768 | -4.577076299 |
| **A_55_P2163725** | Slc7a9 | -0.876425532 | -2.778872437 | 1.026021372 | 8.119506206 | -1.153859213 | 0.296409885 | 0.709264768 | -4.577081601 |
| **A_51_P108252** | Gpsm2 | 0.40349865 | -0.472422084 | 1.279419383 | 10.18468033 | 1.15379227 | 0.296435114 | 0.709264768 | -4.576716578 |
| **A_51_P438967** | Gpnmb | -0.879907399 | -2.790198401 | 1.030383603 | 8.146405531 | -1.153686432 | 0.296475005 | 0.709264768 | -4.577127271 |
| **A_55_P2130970** | Parp10 | -0.426661291 | -1.352994581 | 0.499671999 | 11.29657191 | -1.153629404 | 0.296496501 | 0.709264768 | -4.577142345 |
| **A_51_P305547** | Snai2 | 1.222128738 | -1.431307384 | 3.87556486 | 11.54747433 | 1.153609078 | 0.296504163 | 0.709264768 | -4.577147717 |
| **A_52_P514352** | Kcnk5 | -0.850018031 | -2.695578039 | 0.995541977 | 8.698801221 | -1.153586893 | 0.296512526 | 0.709264768 | -4.577153581 |
| **A_55_P2015495** | Abat | 0.529582105 | -0.620560193 | 1.679724403 | 10.53411424 | 1.15327286 | 0.296630928 | 0.709264768 | -4.57723658 |
| **A_55_P2069729** | Ormdl2 | -0.140519788 | -0.445710373 | 0.164670797 | 11.74700475 | -1.153232041 | 0.296646321 | 0.709264768 | -4.577247367 |

| | | | | | | | | | |
|---|---|---|---|---|---|---|---|---|---|
| **A_66_P133240** | Hnrnpll | 0.463969834 | -0.543842525 | 1.471782192 | 11.67275766 | 1.153082578 | 0.296702691 | 0.709264768 | -4.577286867 |
| **A_55_P2152364** | NA | -0.209166995 | -0.663535829 | 0.245201839 | 8.029337355 | -1.153015029 | 0.296728171 | 0.709264768 | -4.577304719 |
| **A_55_P2108933** | Dao | -0.297857555 | -0.944932263 | 0.349217153 | 7.803021915 | -1.152934248 | 0.296758643 | 0.709264768 | -4.577326066 |
| **A_55_P2095035** | NA | -0.924558785 | -2.933515111 | 1.08439754 | 8.472751517 | -1.152694922 | 0.296848939 | 0.709264768 | -4.577389309 |
| **A_52_P638794** | Stk31 | -0.42140514 | -1.337069718 | 0.494259438 | 7.861073102 | -1.152693276 | 0.29684956 | 0.709264768 | -4.577389744 |
| **A_55_P2103596** | Slc30a4 | -0.616147733 | -1.954988741 | 0.722693274 | 11.29997759 | -1.152673198 | 0.296857136 | 0.709264768 | -4.57739505 |
| **A_55_P2088272** | Golga3 | -0.448723011 | -1.423844979 | 0.526398957 | 11.21080433 | -1.152576463 | 0.296893641 | 0.709264768 | -4.577420611 |
| **A_55_P1980746** | Dram2 | -0.226627071 | -0.719134736 | 0.265880595 | 10.0557009 | -1.152521767 | 0.296914284 | 0.709264768 | -4.577435064 |
| **A_55_P1973159** | Serpinf1 | 0.52396682 | -0.614728076 | 1.662661717 | 10.28219569 | 1.152515455 | 0.296916666 | 0.709264768 | -4.577436732 |
| **A_55_P1962951** | Stx5a | -0.292856092 | -0.929301589 | 0.343589404 | 12.88959509 | -1.152506521 | 0.296920038 | 0.709264768 | -4.577439092 |
| **A_55_P2042729** | NA | 0.278338391 | -0.32659134 | 0.883268122 | 12.27738831 | 1.152440543 | 0.29694494 | 0.709264768 | -4.577456526 |
| **A_55_P1980051** | Il17d | 0.152492697 | -0.178954975 | 0.48394037 | 10.37108553 | 1.152350033 | 0.296979105 | 0.709264768 | -4.577480441 |
| **A_51_P454666** | Arl6ip4 | 0.166643354 | -0.195577116 | 0.528863823 | 10.2773388 | 1.152299449 | 0.2969982 | 0.709264768 | -4.577493806 |
| **A_51_P348617** | Borcs8 | 0.321269936 | -0.377058608 | 1.019598481 | 12.59271558 | 1.152286936 | 0.297002923 | 0.709264768 | -4.577497112 |
| **A_52_P24951** | Ado | 0.277191196 | -0.325391033 | 0.879773424 | 9.731342442 | 1.152161764 | 0.29705018 | 0.709264768 | -4.577530184 |
| **A_55_P2106834** | Ndufs3 | 0.261860704 | -0.307487699 | 0.831209106 | 13.61414545 | 1.151973714 | 0.297121188 | 0.709264768 | -4.577579867 |
| **A_51_P123297** | Ppp1r7 | 0.105069946 | -0.123378978 | 0.333518871 | 8.397413508 | 1.151966204 | 0.297124024 | 0.709264768 | -4.577581851 |
| **A_55_P2183110** | Fmr1 | -0.270016097 | -0.857130923 | 0.317098729 | 8.417089741 | -1.151905771 | 0.297146847 | 0.709264768 | -4.577597817 |
| **A_51_P472621** | Zfp719 | 0.197889698 | -0.23242475 | 0.628204147 | 8.549421985 | 1.151827924 | 0.297176249 | 0.709264768 | -4.577618383 |
| **A_52_P684138** | Dpf3 | -0.390611053 | -1.240005568 | 0.458783461 | 7.899101789 | -1.15182214 | 0.297178433 | 0.709264768 | -4.577619911 |
| **A_66_P119017** | Smarca1 | 0.583579226 | -0.685524313 | 1.852682766 | 9.834625485 | 1.151736457 | 0.297210798 | 0.709264768 | -4.577642547 |
| **A_52_P584415** | Cars2 | -0.100091928 | -0.317766977 | 0.117583122 | 8.186281325 | -1.151703662 | 0.297223186 | 0.709264768 | -4.577651211 |
| **A_55_P2041521** | Doxl2 | -2.591276242 | -8.226781665 | 3.04422918 | 9.099517243 | -1.151678308 | 0.297232763 | 0.709264768 | -4.577657908 |
| **A_55_P2149983** | Foxp4 | 0.18344543 | -0.215554525 | 0.582445385 | 11.9187313 | 1.151554234 | 0.297279637 | 0.709264768 | -4.577690685 |
| **A_55_P2007088** | Rnf128 | -2.044021901 | -6.490220168 | 2.402176367 | 11.12779875 | -1.151455253 | 0.297317036 | 0.709264768 | -4.577716832 |
| **A_55_P2113886** | NA | -0.961089387 | -3.051676497 | 1.129497724 | 8.137542936 | -1.151452053 | 0.297318244 | 0.709264768 | -4.577717677 |
| **A_52_P167382** | Tmem8 | -0.299004405 | -0.949437232 | 0.351428422 | 8.075466291 | -1.151398024 | 0.29733866 | 0.709264768 | -4.577731949 |
| **A_66_P121288** | Tceal6 | 0.694117926 | -0.815838658 | 2.204074509 | 9.792727591 | 1.151381231 | 0.297345006 | 0.709264768 | -4.577736385 |
| **A_55_P2065897** | NA | 0.682909761 | -0.802674533 | 2.168494056 | 12.64310449 | 1.151373864 | 0.29734779 | 0.709264768 | -4.577738331 |
| **A_55_P2168431** | Amer1 | 0.456499708 | -0.536642877 | 1.449642292 | 10.33383774 | 1.151275454 | 0.29738498 | 0.709264768 | -4.577764326 |
| **A_55_P2010672** | Kif17 | -0.250618004 | -0.81701459 | 0.315778582 | 8.507324727 | -1.178567187 | 0.297389304 | 0.709264768 | -4.57211869 |
| **A_55_P1982727** | Dnajb5 | 0.388706816 | -0.456978431 | 1.234392063 | 9.00892272 | 1.151234371 | 0.297400507 | 0.709264768 | -4.577775178 |
| **A_52_P250555** | Dynll1 | 0.174827009 | -0.205544447 | 0.555198466 | 8.716250362 | 1.151200461 | 0.297413323 | 0.709264768 | -4.577784134 |
| **A_51_P101137** | Samd14 | 0.395231111 | -0.464721454 | 1.255183676 | 9.947491416 | 1.151136891 | 0.297437351 | 0.709264768 | -4.577800926 |
| **A_55_P2285599** | 5730437C | -0.17562728 | -0.557774657 | 0.206520097 | 7.704672862 | -1.151095717 | 0.297452914 | 0.709264768 | -4.577811801 |
| **A_55_P2021011** | Sfi1 | -0.832805919 | -2.645007026 | 0.979395189 | 8.197129846 | -1.151032952 | 0.297476641 | 0.709264768 | -4.577828379 |
| **A_55_P2102798** | Prl7a2 | -1.101952457 | -3.49998929 | 1.296084377 | 8.215359509 | -1.150952375 | 0.297507102 | 0.709264768 | -4.577849661 |
| **A_55_P2042923** | Sgk2 | -1.561409416 | -4.959725297 | 1.836906466 | 8.47440442 | -1.150809384 | 0.297561166 | 0.709264768 | -4.577887428 |
| **A_55_P1997898** | Tfeb | -0.224166862 | -0.712067948 | 0.263734224 | 8.276535715 | -1.150773822 | 0.297574613 | 0.709264768 | -4.57789682 |
| **A_52_P584335** | Folr1 | -0.546716128 | -1.736656102 | 0.643223846 | 7.989892127 | -1.150766411 | 0.297577416 | 0.709264768 | -4.577898777 |
| **A_52_P190647** | Mxd3 | 0.42529534 | -0.500435277 | 1.351025957 | 8.756132903 | 1.15068471 | 0.297608312 | 0.709264768 | -4.577920355 |
| **A_51_P229893** | Cts6 | -3.059115039 | -9.718069241 | 3.599839164 | 9.245490311 | -1.150641746 | 0.29762456 | 0.709264768 | -4.577931701 |
| **A_55_P2056521** | Rpl21 | 0.198591009 | -0.233743455 | 0.630925472 | 8.453532627 | 1.150509133 | 0.297674717 | 0.709264768 | -4.577966724 |
| **A_51_P412835** | Daxx | -0.545887985 | -1.734330079 | 0.642554108 | 11.81653496 | -1.150471477 | 0.29768896 | 0.709264768 | -4.577976668 |
| **A_51_P197321** | Clta | -0.145147556 | -0.46116736 | 0.170872249 | 14.29292116 | -1.150391786 | 0.297719105 | 0.709264768 | -4.577997713 |
| **A_55_P1966383** | NA | -0.147650309 | -0.469129999 | 0.173829381 | 10.72842365 | -1.150353091 | 0.297733744 | 0.709264768 | -4.578007932 |
| **A_55_P1962279** | Plac1 | -0.693832434 | -2.204525049 | 0.816860181 | 10.0694532 | -1.150346928 | 0.297736076 | 0.709264768 | -4.578009559 |
| **A_55_P1992640** | Hdac3 | 0.10360919 | -0.122002743 | 0.329221123 | 9.587990833 | 1.150234944 | 0.297778443 | 0.709264768 | -4.578039131 |

| | | | | | | | | | |
|---|---|---|---|---|---|---|---|---|---|
| **A_51_P515623** | Qpctl | -0.78719538 | -2.501391922 | 0.927001162 | 9.100920528 | -1.150197097 | 0.297792763 | 0.709264768 | -4.578049125 |
| **A_55_P1978466** | Pnma5 | -0.402178073 | -1.277963379 | 0.473607232 | 7.834055887 | -1.150193956 | 0.297793952 | 0.709264768 | -4.578049955 |
| **A_51_P221949** | Csnk1g3 | 0.119678755 | -0.140963554 | 0.380321063 | 11.47187957 | 1.150065284 | 0.297842642 | 0.709264768 | -4.578083932 |
| **A_55_P2275437** | Hoxb5os | 0.815785463 | -0.960927255 | 2.59249818 | 9.74964532 | 1.150029806 | 0.297856068 | 0.709264768 | -4.5780933 |
| **A_55_P2157384** | Xlr5b | -1.182256409 | -3.757286821 | 1.392774004 | 8.266099601 | -1.149951859 | 0.297885568 | 0.709264768 | -4.578113882 |
| **A_66_P140335** | Mbnl3 | -0.687414691 | -2.184651731 | 0.809822349 | 8.985392944 | -1.149949033 | 0.297886637 | 0.709264768 | -4.578114628 |
| **A_52_P327664** | Gbp5 | -0.252142056 | -0.801342349 | 0.297058237 | 7.948857493 | -1.149912713 | 0.297900384 | 0.709264768 | -4.578124218 |
| **A_55_P2009567** | Ptbp3 | -0.679783502 | -2.160466112 | 0.800899107 | 10.38170194 | -1.149897131 | 0.297906282 | 0.709264768 | -4.578128333 |
| **A_51_P479321** | Acss1 | 0.269479658 | -0.317492493 | 0.856451808 | 11.21108165 | 1.149896724 | 0.297906436 | 0.709264768 | -4.57812844 |
| **A_52_P10041** | Akr1b3 | 0.544952786 | -0.642050265 | 1.731955837 | 11.89708274 | 1.149892886 | 0.297907888 | 0.709264768 | -4.578129453 |
| **A_55_P1997106** | Large2 | -0.133020378 | -0.422813996 | 0.156773241 | 9.18366722 | -1.149687222 | 0.297985742 | 0.709264768 | -4.578183756 |
| **A_66_P126153** | 1700011M | -1.375551246 | -4.372474416 | 1.621371924 | 8.343729693 | -1.149612588 | 0.298013998 | 0.709264768 | -4.578203462 |
| **A_51_P136845** | Usp19 | -0.278290991 | -0.88461276 | 0.328030778 | 9.469494438 | -1.14959888 | 0.298019189 | 0.709264768 | -4.578207081 |
| **A_55_P2027640** | NA | 0.310088016 | -0.365511558 | 0.98568759 | 17.56105198 | 1.149597897 | 0.29801956 | 0.709264768 | -4.57820734 |
| **A_55_P2155438** | Rps27 | 0.2118287 | -0.249723855 | 0.673381255 | 17.18325481 | 1.149513266 | 0.298051606 | 0.709264768 | -4.578229685 |
| **A_55_P2083559** | Hrk | -0.771323014 | -2.451980824 | 0.909334796 | 8.096057812 | -1.149495664 | 0.298058271 | 0.709264768 | -4.578234332 |
| **A_51_P310821** | Hoxa5 | 0.984557096 | -1.160780562 | 3.129894755 | 10.16926591 | 1.149464587 | 0.298070039 | 0.709264768 | -4.578242537 |
| **A_55_P1987331** | Prl3d1 | -2.193239797 | -6.972461364 | 2.585981769 | 8.88809448 | -1.149421452 | 0.298086373 | 0.709264768 | -4.578253925 |
| **A_55_P1962756** | Ttll2 | -0.221531884 | -0.70431467 | 0.261250903 | 7.763825604 | -1.149303709 | 0.298130965 | 0.709264768 | -4.57828501 |
| **A_55_P1958887** | DXBay18 | -1.713599948 | -5.448088186 | 2.02088829 | 8.617426257 | -1.149287518 | 0.298137098 | 0.709264768 | -4.578289284 |
| **A_51_P418820** | Tfap2c | -2.036008508 | -6.47336951 | 2.401352494 | 8.786575789 | -1.149225283 | 0.29816067 | 0.709264768 | -4.578305714 |
| **A_52_P113830** | 1600025M | -2.119090517 | -6.737785654 | 2.49960462 | 8.795078472 | -1.149160148 | 0.298185343 | 0.709264768 | -4.578322909 |
| **A_51_P168792** | Xylb | -1.024332679 | -3.256943237 | 1.208277878 | 9.755053179 | -1.149154441 | 0.298187505 | 0.709264768 | -4.578324416 |
| **A_55_P2067767** | Psg25 | -2.589921913 | -8.234886439 | 3.055042613 | 9.007602091 | -1.149147559 | 0.298190112 | 0.709264768 | -4.578326233 |
| **A_51_P208263** | NA | 0.592951304 | -0.699483186 | 1.885385793 | 15.42954436 | 1.149107965 | 0.298205111 | 0.709264768 | -4.578336685 |
| **A_51_P389988** | Slc40a1 | -1.273310543 | -4.048734435 | 1.502113349 | 10.75590947 | -1.149093474 | 0.2982106 | 0.709264768 | -4.57834051 |
| **A_55_P2078123** | Rora | -0.449404213 | -1.428969139 | 0.530160712 | 8.246900885 | -1.149090566 | 0.298211702 | 0.709264768 | -4.578341278 |
| **A_55_P2115955** | Raet1e | 0.304680677 | -0.359432776 | 0.968794129 | 9.198603124 | 1.149087181 | 0.298212984 | 0.709264768 | -4.578342172 |
| **A_55_P2117630** | Snrnp25 | 0.248611767 | -0.293301089 | 0.790524624 | 11.13028834 | 1.149059643 | 0.298223417 | 0.709264768 | -4.578349441 |
| **A_55_P2153721** | Ccdc127 | -0.186640147 | -0.59347214 | 0.220191847 | 8.377457778 | -1.149053671 | 0.298225679 | 0.709264768 | -4.578351018 |
| **A_55_P2164348** | Hoxa10 | -0.61353798 | -1.951025992 | 0.723950032 | 7.951697937 | -1.148952037 | 0.298264185 | 0.709264768 | -4.578377847 |
| **A_55_P2115225** | Fap | 0.431635231 | -0.509486221 | 1.372756683 | 9.188936353 | 1.14873951 | 0.29834472 | 0.709264768 | -4.578433947 |
| **A_55_P1968878** | Rhpn2 | -0.628036192 | -1.997485804 | 0.741413419 | 9.553883504 | -1.148653289 | 0.298377397 | 0.709264768 | -4.578456706 |
| **A_55_P2109874** | Rpl23a | 0.298410163 | -0.352284236 | 0.949104561 | 16.25578794 | 1.148647805 | 0.298379476 | 0.709264768 | -4.578458153 |
| **A_55_P1962334** | Trim71 | -0.498262804 | -1.584787885 | 0.588262277 | 8.110531887 | -1.148600649 | 0.298397349 | 0.709264768 | -4.5784706 |
| **A_55_P1971094** | Esx1 | -1.188600744 | -3.78050348 | 1.403301993 | 8.228329139 | -1.148596908 | 0.298398767 | 0.709264768 | -4.578471588 |
| **A_55_P2186210** | NA | 0.135262015 | -0.159697878 | 0.430221909 | 13.44958587 | 1.148585241 | 0.29840319 | 0.709264768 | -4.578474667 |
| **A_66_P106133** | Magi3 | -0.141219425 | -0.449170879 | 0.16673203 | 9.521146546 | -1.14858337 | 0.298403899 | 0.709264768 | -4.578475161 |
| **A_51_P379710** | Rpn1 | -0.56539416 | -1.798400045 | 0.667611726 | 14.30142719 | -1.148514583 | 0.298429973 | 0.709264768 | -4.578493317 |
| **A_55_P2041700** | Hoxa11os | -0.90944728 | -2.892821156 | 1.073926596 | 8.217525965 | -1.1484796 | 0.298443234 | 0.709264768 | -4.578502551 |
| **A_52_P215176** | Opn3 | -0.29280713 | -0.931385073 | 0.345770813 | 8.323805022 | -1.148465835 | 0.298448453 | 0.709264768 | -4.578506184 |
| **A_52_P76402** | Ago1 | 0.360588228 | -0.42584152 | 1.147017976 | 9.448251105 | 1.148423199 | 0.298464616 | 0.709264768 | -4.578517437 |
| **A_52_P217437** | Htra3 | -0.523985255 | -1.666897683 | 0.618927172 | 8.153099071 | -1.148302888 | 0.298510231 | 0.709264768 | -4.578549191 |
| **A_55_P2380132** | NA | -0.498371244 | -1.585503306 | 0.588760817 | 8.016591685 | -1.148209187 | 0.29854576 | 0.709264768 | -4.578573921 |
| **A_55_P2141938** | 1810058I2 | -0.195024895 | -0.620464131 | 0.230414341 | 8.511667878 | -1.148161176 | 0.298563966 | 0.709264768 | -4.578586592 |
| **A_55_P2084758** | Agfg1 | -0.263939427 | -0.839748158 | 0.311869304 | 9.838802145 | -1.14809115 | 0.298590523 | 0.709264768 | -4.578605073 |
| **A_55_P1959409** | Deup1 | 0.358930377 | -0.454557988 | 1.172418741 | 8.369905974 | 1.175226496 | 0.298594713 | 0.709264768 | -4.572831814 |
| **A_55_P1989816** | Zbtb7b | -0.393933283 | -1.253359117 | 0.465492552 | 13.0097649 | -1.148060081 | 0.298602306 | 0.709264768 | -4.578613273 |

| | | | | | | | | | |
|---|---|---|---|---|---|---|---|---|---|
| **A_55_P2029763** | Peg10 | -1.225059403 | -3.897720165 | 1.447601359 | 8.296623954 | -1.148057542 | 0.298603269 | 0.709264768 | -4.578613943 |
| **A_51_P272806** | Ctsr | -2.879808599 | -9.162826768 | 3.403209571 | 9.17567166 | -1.148009955 | 0.298621317 | 0.709264768 | -4.578626501 |
| **A_55_P2065370** | NA | 0.842954146 | -0.996225003 | 2.682133295 | 12.85205976 | 1.147969269 | 0.298636749 | 0.709264768 | -4.578637238 |
| **A_55_P2058143** | Pclo | 0.111318863 | -0.131585746 | 0.354223472 | 8.101906946 | 1.147845275 | 0.298683783 | 0.709264768 | -4.57866996 |
| **A_52_P495553** | Zbtb10 | -0.489564044 | -1.557859443 | 0.578731355 | 8.889083274 | -1.147806028 | 0.298698672 | 0.709264768 | -4.578680317 |
| **A_51_P292196** | Prl7d1 | -3.070350284 | -9.770421537 | 3.629720969 | 9.243953806 | -1.14778052 | 0.298708349 | 0.709264768 | -4.578687049 |
| **A_52_P31641** | Tmem145 | -0.365589476 | -1.16349897 | 0.432320018 | 7.870308056 | -1.147599627 | 0.298776983 | 0.709264768 | -4.578734783 |
| **A_55_P2120577** | Cited2 | -1.253804445 | -3.990477266 | 1.482868377 | 11.34616763 | -1.147512094 | 0.298810199 | 0.709264768 | -4.578757881 |
| **A_51_P419117** | Rab15 | -1.055696996 | -3.360124744 | 1.248730751 | 13.02166719 | -1.147430885 | 0.298841018 | 0.709264768 | -4.578779309 |
| **A_55_P2017038** | Cbx3 | 0.167845664 | -0.212800194 | 0.548491522 | 11.34211892 | 1.174496491 | 0.298858708 | 0.709264768 | -4.572987606 |
| **A_55_P1972275** | Wars | -0.245459977 | -0.781316495 | 0.290396541 | 12.56187627 | -1.147314591 | 0.298885157 | 0.709264768 | -4.578809995 |
| **A_51_P359636** | Lgals3bp | -1.625149804 | -5.173006477 | 1.922706869 | 10.6350912 | -1.14730181 | 0.298890009 | 0.709264768 | -4.578813367 |
| **A_55_P2145754** | Atp5pb | 0.416966519 | -0.493355156 | 1.327288195 | 12.95388929 | 1.147246248 | 0.298911099 | 0.709264768 | -4.578828027 |
| **A_51_P502119** | F11 | -1.600837129 | -5.095813507 | 1.894139248 | 8.558883924 | -1.147237276 | 0.298914505 | 0.709264768 | -4.578830395 |
| **A_51_P369803** | Psmb9 | 0.511679681 | -0.605670747 | 1.62903011 | 9.617657161 | 1.146988642 | 0.299008901 | 0.709264768 | -4.578895995 |
| **A_55_P2060607** | NA | 0.365682611 | -0.432886987 | 1.164252209 | 15.45658803 | 1.146943124 | 0.299026184 | 0.709264768 | -4.578908003 |
| **A_51_P353502** | Sharpin | -0.109681512 | -0.349219431 | 0.129856407 | 11.97335649 | -1.146857615 | 0.299058656 | 0.709264768 | -4.578930563 |
| **A_51_P419286** | Batf3 | -0.46066709 | -1.466772374 | 0.545438193 | 8.761240535 | -1.146816951 | 0.299074099 | 0.709264768 | -4.578941291 |
| **A_55_P1955975** | Afg3l2 | 0.166357031 | -0.19697425 | 0.529688311 | 10.3951421 | 1.146802725 | 0.299079502 | 0.709264768 | -4.578945044 |
| **A_55_P1996716** | Cachd1 | 0.340876259 | -0.403613297 | 1.085365815 | 9.474477073 | 1.146802129 | 0.299079728 | 0.709264768 | -4.578945201 |
| **A_55_P2133840** | Sipa1 | 0.406172659 | -0.480981682 | 1.293327001 | 12.29954744 | 1.146731783 | 0.299106445 | 0.709264768 | -4.57896376 |
| **A_51_P245533** | Tex19.1 | -1.727504071 | -5.500930376 | 2.045922233 | 8.532202506 | -1.146657069 | 0.299134824 | 0.709264768 | -4.57898347 |
| **A_55_P1968618** | NA | 0.41161048 | -0.487527514 | 1.310748473 | 8.693800644 | 1.146595993 | 0.299158024 | 0.709264768 | -4.578999582 |
| **A_55_P1999576** | Josd1 | -0.388507413 | -1.237273187 | 0.460258361 | 12.92082916 | -1.146467629 | 0.299206789 | 0.709264768 | -4.579033445 |
| **A_51_P290626** | Gulo | -0.972338363 | -3.096891006 | 1.15221428 | 8.279336269 | -1.146305219 | 0.299268498 | 0.709264768 | -4.579076286 |
| **A_55_P1974477** | Msh3 | 0.235506231 | -0.27908633 | 0.750098793 | 10.69452376 | 1.146276118 | 0.299279557 | 0.709264768 | -4.579083963 |
| **A_51_P455166** | Prl4a1 | -1.947264617 | -6.202333738 | 2.307804504 | 8.632249813 | -1.146221892 | 0.299300163 | 0.709264768 | -4.579098266 |
| **A_55_P2121484** | Smo | -0.148403798 | -0.472693472 | 0.175885875 | 9.870209396 | -1.146204853 | 0.299306638 | 0.709264768 | -4.579102761 |
| **A_51_P455897** | Pimreg | 0.813896098 | -0.964670474 | 2.592462671 | 9.761594936 | 1.146170395 | 0.299319734 | 0.709264768 | -4.57911185 |
| **A_55_P2159289** | NA | -0.114355911 | -0.364265962 | 0.135554139 | 8.118716688 | -1.146107204 | 0.29934375 | 0.709264768 | -4.579128518 |
| **A_55_P1958562** | Dmrtc1b | -1.505463212 | -4.796629302 | 1.785702878 | 8.437166253 | -1.14569942 | 0.299498773 | 0.709264768 | -4.579236072 |
| **A_52_P72654** | Slc25a17 | 0.354158464 | -0.420100976 | 1.128417904 | 11.78539788 | 1.145675068 | 0.299508032 | 0.709264768 | -4.579242494 |
| **A_55_P2024080** | Rpl7a | 0.264202185 | -0.313408078 | 0.841812447 | 15.44014709 | 1.14564971 | 0.299517675 | 0.709264768 | -4.579249182 |
| **A_55_P2022082** | Klf13 | -0.254481694 | -0.81085828 | 0.301894893 | 10.93908373 | -1.145613298 | 0.299531522 | 0.709264768 | -4.579258785 |
| **A_66_P116412** | Flt3 | -0.769207099 | -2.451015557 | 0.912601359 | 8.09877509 | -1.145558038 | 0.299552536 | 0.709264768 | -4.579273359 |
| **A_55_P2091762** | NA | 0.308678711 | -0.36630241 | 0.983659832 | 16.17716647 | 1.145421677 | 0.299604398 | 0.709264768 | -4.57930932 |
| **A_55_P2029366** | Abhd6 | -0.950600402 | -3.029836077 | 1.128635272 | 8.50182235 | -1.145103182 | 0.299725562 | 0.709264768 | -4.579393309 |
| **A_51_P279639** | Col6a2 | 0.959480095 | -1.139347156 | 3.058307347 | 11.3853941 | 1.145010899 | 0.299760676 | 0.709264768 | -4.579417643 |
| **A_55_P1969156** | Pam | -0.182787589 | -0.582668256 | 0.217093078 | 9.125772788 | -1.144897593 | 0.299803795 | 0.709264768 | -4.57944752 |
| **A_65_P03719** | Fuca2 | -0.241095911 | -0.76860895 | 0.286417129 | 8.758346332 | -1.144740347 | 0.299863644 | 0.709264768 | -4.579488981 |
| **A_55_P2114088** | Zfp385c | -0.53662035 | -1.710776592 | 0.637535892 | 7.973359487 | -1.144699762 | 0.299879093 | 0.709264768 | -4.579499682 |
| **A_55_P2143561** | Gm6813 | 0.389452585 | -0.495889267 | 1.274794437 | 15.77167096 | 1.171672673 | 0.299881892 | 0.709264768 | -4.573590104 |
| **A_51_P480904** | Blmh | 0.513575428 | -0.610237587 | 1.637388444 | 11.29381406 | 1.144617879 | 0.299910264 | 0.709264768 | -4.579521271 |
| **A_55_P2345853** | NA | -0.933018663 | -2.974679688 | 1.108642362 | 9.343729285 | -1.144608689 | 0.299913762 | 0.709264768 | -4.579523694 |
| **A_66_P132322** | Mrpl21 | 0.351485464 | -0.417649699 | 1.120620628 | 12.53107279 | 1.144603443 | 0.299915759 | 0.709264768 | -4.579525077 |
| **A_51_P154534** | Bloc1s5 | -0.161907232 | -0.516237736 | 0.192423271 | 10.33949265 | -1.144479592 | 0.299962913 | 0.709264768 | -4.579557731 |
| **A_55_P2115846** | Gm10836 | -0.635576109 | -2.026581418 | 0.755429201 | 8.035665302 | -1.144429704 | 0.299981908 | 0.709264768 | -4.579570884 |
| **A_51_P504134** | NA | -1.741170565 | -5.551920142 | 2.069579011 | 8.559015594 | -1.144408968 | 0.299989804 | 0.709264768 | -4.579576351 |

| | | | | | | | | | |
|---|---|---|---|---|---|---|---|---|---|
| **A_51_P506755** | Tep1 | -0.136644569 | -0.435722184 | 0.162433047 | 9.93212722 | -1.144349833 | 0.300012322 | 0.709264768 | -4.579591941 |
| **A_55_P2139713** | Phf13 | -0.461098939 | -1.470338573 | 0.548140696 | 8.174705253 | -1.144327068 | 0.300020991 | 0.709264768 | -4.579597943 |
| **A_55_P2037439** | Depdc1a | 0.301535373 | -0.35849598 | 0.961566726 | 8.405209548 | 1.144258232 | 0.300047206 | 0.709264768 | -4.57961609 |
| **A_55_P2115981** | Xlr5c | -1.125674096 | -3.58969641 | 1.338348219 | 8.241166074 | -1.144243308 | 0.300052889 | 0.709264768 | -4.579620025 |
| **A_66_P110584** | Tpbpa | -0.80565062 | -2.56937682 | 0.95807558 | 8.050856585 | -1.144105109 | 0.300105526 | 0.709264768 | -4.579656457 |
| **A_52_P625277** | Accs | -0.117881316 | -0.375950069 | 0.140187437 | 9.243116192 | -1.144089234 | 0.300111573 | 0.709264768 | -4.579660642 |
| **A_51_P327904** | Yipf1 | -0.145815146 | -0.465039544 | 0.173409251 | 8.878728632 | -1.144081352 | 0.300114575 | 0.709264768 | -4.57966272 |
| **A_55_P2055404** | Slc15a4 | -0.124090929 | -0.395756329 | 0.147574472 | 12.14460718 | -1.144079095 | 0.300115435 | 0.709264768 | -4.579663315 |
| **A_52_P502577** | S1pr3 | 0.910614315 | -1.083052291 | 2.904280921 | 11.03400253 | 1.14401649 | 0.300139283 | 0.709264768 | -4.579679819 |
| **A_51_P137800** | Acp7 | -0.676397388 | -2.157475366 | 0.804680591 | 7.98945495 | -1.143863869 | 0.300197428 | 0.709264768 | -4.57972005 |
| **A_55_P2009572** | Ptbp3 | -0.424040887 | -1.352632835 | 0.504551062 | 8.245441996 | -1.143755422 | 0.300238749 | 0.709264768 | -4.579748637 |
| **A_51_P413866** | Cfb | -0.476879656 | -1.521364793 | 0.567605481 | 8.413660639 | -1.143554525 | 0.30031531 | 0.709264768 | -4.579801591 |
| **A_66_P135165** | Nfib | 0.388690324 | -0.462681352 | 1.240062001 | 8.849579103 | 1.143496599 | 0.300337388 | 0.709264768 | -4.579816859 |
| **A_51_P372312** | Mrpl54 | 0.302273115 | -0.359906986 | 0.964453217 | 12.79730475 | 1.143335639 | 0.300398745 | 0.709264768 | -4.579859283 |
| **A_51_P272106** | Cirbp | -0.104448429 | -0.333262948 | 0.124366091 | 11.70083098 | -1.143322306 | 0.300403828 | 0.709264768 | -4.579862797 |
| **A_55_P2036999** | Zic3 | -0.381771984 | -1.218120308 | 0.454576341 | 7.844371762 | -1.143318441 | 0.300405301 | 0.709264768 | -4.579863815 |
| **A_55_P2010778** | Il11ra1 | -0.315185968 | -1.005668379 | 0.375296443 | 13.51178044 | -1.143311575 | 0.300407919 | 0.709264768 | -4.579865625 |
| **A_55_P2195157** | Dbn1 | 0.457313684 | -0.544558876 | 1.459186245 | 9.913555197 | 1.143278564 | 0.300420504 | 0.709264768 | -4.579874325 |
| **A_55_P2075429** | Ttll4 | 0.099238366 | -0.118202317 | 0.316679049 | 8.41547284 | 1.143112954 | 0.300483648 | 0.709264768 | -4.579917972 |
| **A_51_P307741** | Cckar | 0.829071198 | -0.987584687 | 2.645727084 | 9.745174813 | 1.143061253 | 0.300503363 | 0.709264768 | -4.579931598 |
| **A_55_P2006906** | Armcx3 | 0.237876277 | -0.283418822 | 0.759171377 | 8.835743136 | 1.142925274 | 0.300555221 | 0.709264768 | -4.579967433 |
| **A_55_P2097656** | Rps11 | 0.270694241 | -0.322529025 | 0.863917507 | 15.64190282 | 1.142907772 | 0.300561897 | 0.709264768 | -4.579972045 |
| **A_55_P1962054** | NA | 0.376705148 | -0.448848776 | 1.202259072 | 15.88514225 | 1.142895322 | 0.300566645 | 0.709264768 | -4.579975326 |
| **A_55_P1970710** | Mllt6 | 0.637484819 | -0.759688714 | 2.034658352 | 12.7253733 | 1.142798976 | 0.300603395 | 0.709264768 | -4.580000716 |
| **A_55_P2128501** | Krt8 | -0.960589774 | -3.066017417 | 1.144837869 | 12.32804866 | -1.142741459 | 0.300625335 | 0.709264768 | -4.580015873 |
| **A_55_P2006698** | Bmp1 | -0.588619532 | -1.878806171 | 0.701567107 | 10.8340463 | -1.142700651 | 0.300640903 | 0.709264768 | -4.580026627 |
| **A_65_P01991** | Prrg4 | -0.430347739 | -1.373665501 | 0.512970023 | 7.948310991 | -1.14264641 | 0.300661596 | 0.709264768 | -4.58004092 |
| **A_51_P328001** | Sftpc | 3.414664485 | -4.070385179 | 10.89971415 | 16.3919619 | 1.142624934 | 0.300669789 | 0.709264768 | -4.580046579 |
| **A_55_P2033055** | Pdp1 | 0.380422662 | -0.453493169 | 1.214338494 | 9.152259671 | 1.142600757 | 0.300679013 | 0.709264768 | -4.58005295 |
| **A_52_P615312** | Mrps27 | 0.112038461 | -0.133566209 | 0.357643131 | 8.547040283 | 1.142564851 | 0.300692713 | 0.709264768 | -4.580062411 |
| **A_66_P107129** | 3300005D | -0.452320295 | -1.443978575 | 0.539337985 | 8.01946762 | -1.142442571 | 0.300739371 | 0.709264768 | -4.580094632 |
| **A_52_P629333** | Map3k21 | -0.247815241 | -0.791122299 | 0.295491817 | 8.103843048 | -1.142438977 | 0.300740743 | 0.709264768 | -4.580095579 |
| **A_51_P230287** | Sorcs2 | 0.524951771 | -0.626003663 | 1.675907205 | 9.390502439 | 1.142381714 | 0.300762595 | 0.709264768 | -4.580110668 |
| **A_55_P1966987** | Limk1 | -0.418841849 | -1.337175125 | 0.499491426 | 11.32470059 | -1.142352385 | 0.300773788 | 0.709264768 | -4.580118395 |
| **A_55_P1996583** | Ppdpf | 0.327070744 | -0.390064518 | 1.044206005 | 11.5923863 | 1.142328382 | 0.300782948 | 0.709264768 | -4.58012472 |
| **A_51_P156243** | Eps15 | -0.311956549 | -0.996038182 | 0.372125083 | 8.946684966 | -1.142185187 | 0.300837602 | 0.709264768 | -4.580162449 |
| **A_55_P2084448** | Cybc1 | -0.461026057 | -1.472017492 | 0.549965378 | 10.89551332 | -1.142163669 | 0.300845816 | 0.709264768 | -4.580168119 |
| **A_66_P121480** | Sardh | 0.557635674 | -0.665318842 | 1.78059019 | 9.320821107 | 1.142064388 | 0.300883715 | 0.709264768 | -4.580194276 |
| **A_55_P1982384** | NA | 0.168760335 | -0.201375414 | 0.538896085 | 12.30541067 | 1.14198314 | 0.300914733 | 0.709264768 | -4.580215682 |
| **A_51_P342652** | Cd79b | 0.496355138 | -0.592327624 | 1.585037901 | 9.76384035 | 1.14193536 | 0.300932975 | 0.709264768 | -4.580228271 |
| **A_52_P451378** | Twsg1 | -0.496221687 | -1.584710571 | 0.592267197 | 11.82284086 | -1.141831679 | 0.300972564 | 0.709264768 | -4.580255586 |
| **A_55_P2112569** | Ern1 | -0.300052009 | -0.958308819 | 0.358204801 | 7.848376795 | -1.141698746 | 0.301023328 | 0.709264768 | -4.580290606 |
| **A_55_P2028496** | NA | 0.386900319 | -0.461915382 | 1.235716021 | 14.32607282 | 1.141658013 | 0.301038885 | 0.709264768 | -4.580301337 |
| **A_66_P127463** | NA | -1.317773777 | -4.20906682 | 1.573519266 | 8.555676044 | -1.141560927 | 0.301075966 | 0.709264768 | -4.580326913 |
| **A_55_P1957154** | Mageb16 | -0.814348133 | -2.601172933 | 0.972476667 | 8.072883522 | -1.141506737 | 0.301096666 | 0.709264768 | -4.580341188 |
| **A_55_P2021285** | Cpped1 | 0.33544679 | -0.400656014 | 1.071549594 | 11.43963739 | 1.141393811 | 0.301139804 | 0.709264768 | -4.580370935 |
| **A_55_P2044659** | NA | -0.557442228 | -1.780749863 | 0.665865408 | 7.982625633 | -1.141338648 | 0.301160879 | 0.709264768 | -4.580385466 |
| **A_55_P2037797** | Tex44 | -2.234265425 | -7.137714293 | 2.669183444 | 7.264068406 | -1.141257014 | 0.30119207 | 0.709264768 | -4.580406969 |

| | | | | | | | | | |
|---|---|---|---|---|---|---|---|---|---|
| **A_55_P2033947** | Fnd3c2 | -2.002063105 | -6.396321111 | 2.392194901 | 8.67860554 | -1.141149529 | 0.301233141 | 0.709264768 | -4.580435281 |
| **A_55_P2002553** | Stx16 | -0.427882287 | -1.367134483 | 0.511369908 | 8.900979161 | -1.141017842 | 0.301283467 | 0.709264768 | -4.580469967 |
| **A_55_P2164258** | Prl7b1 | -2.180301929 | -6.966901336 | 2.606297478 | 8.807108223 | -1.140879829 | 0.301336218 | 0.709264768 | -4.580506317 |
| **A_65_P20174** | Jade1 | -0.859942646 | -2.747859864 | 1.027974572 | 8.464603199 | -1.140871905 | 0.301339247 | 0.709264768 | -4.580508404 |
| **A_52_P463235** | Ankrd33b | -0.573195652 | -1.831608138 | 0.685216834 | 8.300255512 | -1.14085438 | 0.301345946 | 0.709264768 | -4.58051302 |
| **A_55_P2070331** | NA | -0.845457463 | -2.701863133 | 1.010948206 | 8.190740365 | -1.14069417 | 0.301407193 | 0.709264768 | -4.580555215 |
| **A_55_P2073467** | Gtf2i | 1.119890356 | -1.339119385 | 3.578900097 | 13.98690752 | 1.14068466 | 0.301410829 | 0.709264768 | -4.580557719 |
| **A_52_P547662** | P2ry1 | -0.225896648 | -0.721912244 | 0.270118948 | 8.501562165 | -1.140682557 | 0.301411633 | 0.709264768 | -4.580558273 |
| **A_55_P2045278** | Gm10845 | -1.464352597 | -4.679733345 | 1.75102815 | 13.60901953 | -1.140679405 | 0.301412838 | 0.709264768 | -4.580559103 |
| **A_55_P2156712** | Gpr1 | -1.307962101 | -4.180027766 | 1.564103563 | 8.352406958 | -1.14064668 | 0.30142535 | 0.709264768 | -4.580567722 |
| **A_55_P1995233** | Lonp2 | -0.430993787 | -1.377402196 | 0.515414621 | 11.90437845 | -1.140624683 | 0.301433761 | 0.709264768 | -4.580573515 |
| **A_52_P326664** | Unc93a | -0.593397162 | -1.896509835 | 0.709715511 | 7.962057464 | -1.140548721 | 0.301462806 | 0.709264768 | -4.58059352 |
| **A_55_P2015753** | Enho | -0.766110454 | -2.448505556 | 0.916284648 | 10.584304 | -1.14054845 | 0.30146291 | 0.709264768 | -4.580593591 |
| **A_55_P2109782** | Psg17 | -2.57791784 | -8.239158276 | 3.083322596 | 9.08347273 | -1.1405329 | 0.301468856 | 0.709264768 | -4.580597687 |
| **A_55_P2166399** | Stk11 | -0.193620093 | -0.618846461 | 0.231606276 | 9.940822467 | -1.140461376 | 0.301496208 | 0.709264768 | -4.580616522 |
| **A_51_P311905** | Elp6 | 0.214983228 | -0.257178427 | 0.687144882 | 8.524179943 | 1.140418386 | 0.301512648 | 0.709264768 | -4.580627844 |
| **A_52_P539161** | Rdh11 | -0.287904003 | -0.920226355 | 0.344418349 | 9.673568801 | -1.140406055 | 0.301517364 | 0.709264768 | -4.580631091 |
| **A_55_P2022519** | Tmem108 | -0.949643028 | -3.035361735 | 1.136075679 | 8.878605629 | -1.140394183 | 0.301521905 | 0.709264768 | -4.580634218 |
| **A_51_P291682** | Tmed4 | -0.145262752 | -0.464333319 | 0.173807814 | 9.596172636 | -1.140296708 | 0.301559186 | 0.709264768 | -4.580659887 |
| **A_52_P561377** | Fam160b1 | -0.133011555 | -0.425186346 | 0.159163237 | 11.94506036 | -1.140241838 | 0.301580174 | 0.709264768 | -4.580674335 |
| **A_51_P201721** | Crtap | -0.245244738 | -0.784023715 | 0.29353424 | 12.14410708 | -1.140090695 | 0.301637993 | 0.709264768 | -4.580714135 |
| **A_52_P384574** | Stard4 | 0.506863357 | -0.606851196 | 1.620577909 | 10.86408326 | 1.139901556 | 0.301710361 | 0.709264768 | -4.580763938 |
| **A_55_P2004137** | Gnai2 | -0.205715438 | -0.657772989 | 0.246342112 | 13.95457247 | -1.139786486 | 0.301754396 | 0.709264768 | -4.580794236 |
| **A_55_P2003483** | Gldc | -0.364069108 | -1.16413002 | 0.435991803 | 8.052990738 | -1.139753995 | 0.301766831 | 0.709264768 | -4.580802791 |
| **A_52_P540434** | Ppp1cc | 0.428950139 | -0.513821425 | 1.371721703 | 10.7958084 | 1.139595389 | 0.301827537 | 0.709264768 | -4.58084455 |
| **A_55_P1993517** | Tbc1d14 | 0.149450879 | -0.179025902 | 0.477927661 | 8.151898664 | 1.139578228 | 0.301834106 | 0.709264768 | -4.580849068 |
| **A_55_P1967144** | NA | -0.183594008 | -0.587161073 | 0.219973057 | 16.71931636 | -1.139444382 | 0.301885345 | 0.709264768 | -4.580884307 |
| **A_55_P2281624** | 2610034E | -0.220981649 | -0.706734457 | 0.264771158 | 8.097205458 | -1.139439412 | 0.301887248 | 0.709264768 | -4.580885615 |
| **A_55_P1971433** | Serpinb9c | -1.325227537 | -4.238354897 | 1.587899823 | 8.346719737 | -1.139413406 | 0.301897204 | 0.709264768 | -4.580892462 |
| **A_51_P185135** | Gpr108 | -0.435283454 | -1.392208949 | 0.521642041 | 10.80573453 | -1.139316479 | 0.301934316 | 0.709264768 | -4.580917979 |
| **A_66_P128079** | Krt8 | -0.809280659 | -2.588640159 | 0.970078841 | 14.59864575 | -1.139162829 | 0.301993153 | 0.709264768 | -4.580958429 |
| **A_55_P1965298** | Gm5640 | -1.644453787 | -5.26050328 | 1.971595706 | 8.694015054 | -1.139036531 | 0.302041524 | 0.709264768 | -4.580991676 |
| **A_52_P647906** | NA | -0.166236183 | -0.531787183 | 0.199314816 | 11.89946862 | -1.139011027 | 0.302051293 | 0.709264768 | -4.58099839 |
| **A_55_P2212161** | 2310034C | -0.303612704 | -0.971270951 | 0.364045543 | 7.857563292 | -1.138979918 | 0.302063208 | 0.709264768 | -4.581006579 |
| **A_55_P2100679** | Dnm2 | -0.27145767 | -0.868410961 | 0.325495622 | 8.293449399 | -1.138969543 | 0.302067183 | 0.709264768 | -4.58100931 |
| **A_52_P133436** | Plekhm1 | -0.563755286 | -1.803520641 | 0.67601007 | 9.977253196 | -1.138941653 | 0.302077866 | 0.709264768 | -4.581016651 |
| **A_55_P2093889** | Arhgef19 | 0.182777267 | -0.219182225 | 0.584736759 | 8.540832471 | 1.138912179 | 0.302089156 | 0.709264768 | -4.58102441 |
| **A_55_P1994348** | Golim4 | 0.596828477 | -0.715765637 | 1.909422591 | 10.45296033 | 1.138857614 | 0.302110059 | 0.709264768 | -4.581038773 |
| **A_55_P2021510** | NA | 0.120632727 | -0.144697051 | 0.385962504 | 8.412674077 | 1.138752903 | 0.302150175 | 0.709264768 | -4.581066335 |
| **A_52_P340669** | Bhlha15 | -1.151087664 | -3.683014282 | 1.380838954 | 8.583382022 | -1.138695569 | 0.302172143 | 0.709264768 | -4.581081427 |
| **A_55_P1967133** | Ppp1r3b | 0.369283333 | -0.443014872 | 1.181581537 | 13.18645264 | 1.138661296 | 0.302185275 | 0.709264768 | -4.581090448 |
| **A_55_P1965592** | NA | -0.10629245 | -0.340100488 | 0.127515588 | 8.763634275 | -1.138658062 | 0.302186514 | 0.709264768 | -4.581091299 |
| **A_55_P2081318** | Gnas | -0.17409952 | -0.557064254 | 0.208865214 | 8.441356642 | -1.13864708 | 0.302190722 | 0.709264768 | -4.58109419 |
| **A_55_P2130965** | NA | -0.136325369 | -0.43620341 | 0.163552673 | 14.42720248 | -1.138629315 | 0.302197529 | 0.709264768 | -4.581098865 |
| **A_51_P346964** | Mrap | -0.173596721 | -0.555465388 | 0.208271946 | 9.021049763 | -1.138617456 | 0.302202073 | 0.709264768 | -4.581101987 |
| **A_55_P2032997** | Srp19 | 0.131889818 | -0.158252442 | 0.422032077 | 11.61051951 | 1.138546112 | 0.302229413 | 0.709264768 | -4.581120765 |
| **A_55_P1953508** | Mob3c | -0.451697218 | -1.445419049 | 0.542024612 | 8.483312077 | -1.138499727 | 0.302247189 | 0.709264768 | -4.581132974 |
| **A_55_P1954212** | Snrpd2 | 0.699034049 | -0.838835216 | 2.236903313 | 13.77242486 | 1.138490099 | 0.302250879 | 0.709264768 | -4.581135508 |

| | | | | | | | | | |
|---|---|---|---|---|---|---|---|---|---|
| **A_51_P493649** | Sult1e1 | -1.770432044 | -5.665833135 | 2.124969047 | 8.616450821 | -1.13835423 | 0.302302954 | 0.709264768 | -4.581171268 |
| **A_55_P2069515** | Dennd4a | -0.556664672 | -1.78149154 | 0.668162196 | 8.935581762 | -1.138332935 | 0.302311117 | 0.709264768 | -4.581176872 |
| **A_55_P2168692** | Cep131 | 0.235963781 | -0.283231549 | 0.755159111 | 9.820067753 | 1.138321438 | 0.302315523 | 0.709264768 | -4.581179898 |
| **A_52_P19532** | Fgd3 | -0.361843547 | -1.158165104 | 0.43447801 | 7.874088978 | -1.138105979 | 0.302398122 | 0.709264768 | -4.581236602 |
| **A_51_P375078** | Lmbrd2 | -0.288886381 | -0.92466481 | 0.346892048 | 8.436195256 | -1.138076945 | 0.302409254 | 0.709264768 | -4.581244243 |
| **A_51_P111462** | Arl15 | -0.271364387 | -0.868636018 | 0.325907245 | 10.40560357 | -1.137971301 | 0.302449762 | 0.709264768 | -4.580558176 |
| **A_55_P2329923** | Camkk2 | 0.383269833 | -0.460368841 | 1.226908506 | 11.87619228 | 1.137885341 | 0.302482726 | 0.709264768 | -4.581294666 |
| **A_55_P2019118** | NA | -0.49555972 | -1.586516281 | 0.595396841 | 8.337865262 | -1.137729151 | 0.302542629 | 0.709264768 | -4.581335767 |
| **A_55_P2096189** | Rfx8 | -0.281026951 | -0.899734608 | 0.337680705 | 7.934703011 | -1.137660887 | 0.302568814 | 0.709264768 | -4.58135373 |
| **A_55_P2040168** | Pmp22 | -0.194860916 | -0.623874347 | 0.234152514 | 10.25653613 | -1.137638281 | 0.302577486 | 0.709264768 | -4.581359678 |
| **A_52_P193029** | Hadha | 0.149463083 | -0.179602423 | 0.47852859 | 9.619676851 | 1.137632322 | 0.302579771 | 0.709264768 | -4.581361246 |
| **A_55_P2133096** | Tgds | -0.111296338 | -0.356365596 | 0.13377292 | 8.322175204 | -1.137476377 | 0.302639597 | 0.709264768 | -4.58140228 |
| **A_51_P488196** | Bmper | 0.393568475 | -0.473066777 | 1.260203727 | 10.95912946 | 1.137455195 | 0.302647724 | 0.709264768 | -4.581407853 |
| **A_55_P2071132** | Tnfrsf23 | -1.651214184 | -5.287367428 | 1.98493906 | 9.30127159 | -1.13739569 | 0.302670556 | 0.709264768 | -4.58142351 |
| **A_55_P2071626** | C530008M | 0.357582883 | -0.429945805 | 1.145111571 | 8.504010101 | 1.137262407 | 0.3027217 | 0.709264768 | -4.581458578 |
| **A_55_P2032498** | Arpin | -0.228785414 | -0.73265875 | 0.275087923 | 9.13440727 | -1.13725355 | 0.302725099 | 0.709264768 | -4.581460909 |
| **A_52_P272945** | Ccnh | 0.204911982 | -0.246399428 | 0.656223392 | 8.518099258 | 1.137211875 | 0.302741093 | 0.709264768 | -4.581471874 |
| **A_66_P121249** | Ddi2 | -0.335447503 | -1.074383321 | 0.403488315 | 8.87689443 | -1.137020228 | 0.302814651 | 0.709264768 | -4.581522295 |
| **A_55_P2048855** | Sprr2a2 | 0.913199698 | -1.098477939 | 2.924877335 | 9.377996291 | 1.136992806 | 0.302825177 | 0.709264768 | -4.581529509 |
| **A_55_P2170256** | Ptgr2 | -0.115895286 | -0.371208345 | 0.139417773 | 8.35644357 | -1.136954508 | 0.302839879 | 0.709264768 | -4.581539585 |
| **A_52_P374669** | Kdm5a | -0.246303765 | -0.788917825 | 0.296310295 | 10.75298246 | -1.136921169 | 0.302852677 | 0.709264768 | -4.581548356 |
| **A_51_P262670** | Rbm38 | 0.10265385 | -0.123513476 | 0.328821176 | 7.959497764 | 1.136830525 | 0.302887477 | 0.709264768 | -4.581572202 |
| **A_55_P2099379** | Tfe3 | -0.133049484 | -0.426189707 | 0.160090738 | 8.903225209 | -1.136810637 | 0.302895113 | 0.709264768 | -4.581577434 |
| **A_55_P2099390** | Ranbp9 | 0.124462817 | -0.149793603 | 0.398719236 | 8.935138967 | 1.13666678 | 0.30295035 | 0.709264768 | -4.581615278 |
| **A_51_P208697** | Ttl | -0.310201268 | -0.993751454 | 0.373348918 | 9.790850541 | -1.136641501 | 0.302960058 | 0.709264768 | -4.581621928 |
| **A_55_P1967500** | Nell1 | 0.322481365 | -0.388156806 | 1.033119535 | 8.574268962 | 1.136596822 | 0.302977216 | 0.709264768 | -4.581633681 |
| **A_55_P2121082** | Cul2 | 0.140608187 | -0.169264027 | 0.450480401 | 10.37988339 | 1.136523238 | 0.303005476 | 0.709264768 | -4.581653037 |
| **A_55_P2001970** | Zswim9 | 0.209751708 | -0.252523369 | 0.672026784 | 9.474073671 | 1.136463184 | 0.303028541 | 0.709264768 | -4.581668835 |
| **A_55_P2150786** | Mul1 | -0.441682518 | -1.415137983 | 0.531772947 | 10.59312489 | -1.136434652 | 0.3030395 | 0.709264768 | -4.58167634 |
| **A_66_P103092** | NA | 0.662807484 | -0.798160398 | 2.123775365 | 12.96113208 | 1.13631066 | 0.303087129 | 0.709264768 | -4.581708955 |
| **A_55_P1959393** | Hhat | 0.436605138 | -0.525783486 | 1.398993763 | 9.190146128 | 1.13628875 | 0.303095546 | 0.709264768 | -4.581714717 |
| **A_55_P1972969** | NA | 0.543071554 | -0.653997908 | 1.740141016 | 14.45489765 | 1.136287028 | 0.303096208 | 0.709264768 | -4.58171517 |
| **A_55_P2136678** | NA | 0.165448049 | -0.199268908 | 0.530165007 | 8.666633046 | 1.136203283 | 0.303128381 | 0.709264768 | -4.581737198 |
| **A_51_P257419** | Lhx2 | -1.039397802 | -3.330703595 | 1.251907991 | 8.243313219 | -1.136185092 | 0.30313537 | 0.709264768 | -4.581741982 |
| **A_55_P1959238** | Nagk | -0.317396892 | -1.017122951 | 0.382329167 | 10.89301884 | -1.136121995 | 0.303159613 | 0.709264768 | -4.581758578 |
| **A_55_P2016962** | NA | -0.725210687 | -2.324019404 | 0.87359803 | 8.025389379 | -1.136103855 | 0.303166583 | 0.709264768 | -4.58176335 |
| **A_51_P372156** | 4930563E | -0.647072418 | -2.07366912 | 0.779524284 | 8.079446993 | -1.136061959 | 0.303182682 | 0.709264768 | -4.581774369 |
| **A_51_P513224** | Cep164 | -0.141164345 | -0.452392762 | 0.170064072 | 9.045402471 | -1.136046528 | 0.303188611 | 0.709264768 | -4.581778427 |
| **A_55_P2022277** | Nfe2l1 | -0.443820778 | -1.422329445 | 0.534687889 | 8.377520759 | -1.136039153 | 0.303191445 | 0.709264768 | -4.581780367 |
| **A_52_P578043** | NA | 0.191401959 | -0.230621468 | 0.613425386 | 8.341173181 | 1.135952481 | 0.303224752 | 0.709264768 | -4.581803163 |
| **A_51_P368903** | Anapc13 | 0.313659647 | -0.377955888 | 1.005275182 | 14.40376821 | 1.135910868 | 0.303240745 | 0.709264768 | -4.581814107 |
| **A_55_P2171715** | Mrps33 | 0.351999567 | -0.424155667 | 1.128154802 | 13.01433291 | 1.135909899 | 0.303241117 | 0.709264768 | -4.581814362 |
| **A_55_P2173183** | Cdk5r1 | -0.262117154 | -0.840089166 | 0.315854857 | 8.403717421 | -1.135897077 | 0.303246045 | 0.709264768 | -4.581817734 |
| **A_55_P2030859** | NA | 0.312858502 | -0.377047527 | 1.002764531 | 16.1496847 | 1.135817005 | 0.30327682 | 0.709264768 | -4.581838793 |
| **A_51_P511015** | Fzd9 | 0.114557703 | -0.138062455 | 0.367177862 | 8.036525182 | 1.135812493 | 0.303278554 | 0.709264768 | -4.58183998 |
| **A_51_P496720** | Dnmt3l | -0.792798313 | -2.541257883 | 0.955661257 | 8.375069179 | -1.135683899 | 0.303327985 | 0.709264768 | -4.581873799 |
| **A_55_P2156718** | NA | -1.959009143 | -6.279541271 | 2.361522984 | 8.714012897 | -1.135663243 | 0.303335925 | 0.709264768 | -4.581879231 |
| **A_55_P1990046** | NA | -0.120490283 | -0.386231342 | 0.145250776 | 8.327497926 | -1.135647919 | 0.303341816 | 0.709264768 | -4.581883261 |

| | | | | | | | | | |
|---|---|---|---|---|---|---|---|---|---|
| **A_55_P1985788** | Usp2 | -0.494742163 | -1.585963463 | 0.596479137 | 9.381309476 | -1.135576599 | 0.303369235 | 0.709264768 | -4.581902017 |
| **A_55_P2180854** | Mrgprg | -1.009140518 | -3.234967671 | 1.216686635 | 8.290465165 | -1.135561222 | 0.303375147 | 0.709264768 | -4.581906061 |
| **A_55_P2162988** | Tmem179 | -0.601466492 | -1.928104468 | 0.725171484 | 8.121733663 | -1.135558096 | 0.303376349 | 0.709264768 | -4.581906883 |
| **A_52_P441070** | Apob | -1.575112868 | -5.049303352 | 1.899077616 | 8.532052101 | -1.135555588 | 0.303377313 | 0.709264768 | -4.581907542 |
| **A_55_P2090184** | Fgfr1op2 | 0.196933949 | -0.237475825 | 0.631343722 | 12.33711545 | 1.135458712 | 0.303414561 | 0.709264768 | -4.581933018 |
| **A_66_P113868** | Cdh3 | -0.904475248 | -2.89973488 | 1.090784384 | 9.464622324 | -1.135396669 | 0.303438418 | 0.709264768 | -4.581949333 |
| **A_55_P2040044** | NA | -0.204889169 | -0.65687811 | 0.247099772 | 8.368732927 | -1.135380778 | 0.303444528 | 0.709264768 | -4.581953512 |
| **A_52_P554267** | Ilf3 | 0.356261218 | -0.429685489 | 1.142207926 | 9.871960735 | 1.135339625 | 0.303460354 | 0.709264768 | -4.581964334 |
| **A_52_P470956** | NA | 0.393773498 | -0.475116218 | 1.262663214 | 16.67254167 | 1.135094899 | 0.303554478 | 0.709264768 | -4.582028684 |
| **A_51_P435339** | Epor | -0.258837774 | -0.830032227 | 0.312356679 | 8.67059975 | -1.134995189 | 0.303592835 | 0.709264768 | -4.582054902 |
| **A_55_P2442017** | Samd8 | -0.645780591 | -2.071068603 | 0.779507421 | 8.412810621 | -1.134834947 | 0.303654486 | 0.709264768 | -4.582097034 |
| **A_51_P457196** | Sfrp4 | -1.027290256 | -3.294605326 | 1.240024814 | 8.288566038 | -1.134832173 | 0.303655554 | 0.709264768 | -4.582097763 |
| **A_52_P304720** | Crlf1 | 0.296607072 | -0.358092977 | 0.951307122 | 8.42260631 | 1.134721997 | 0.303697949 | 0.709264768 | -4.58212673 |
| **A_51_P514922** | Ccar2 | 0.134190126 | -0.162009694 | 0.430389946 | 9.894703106 | 1.134713223 | 0.303701325 | 0.709264768 | -4.582129037 |
| **A_52_P496924** | Mgat4a | -1.164164241 | -3.733957011 | 1.405628528 | 8.506923151 | -1.134661968 | 0.30372105 | 0.709264768 | -4.582142513 |
| **A_51_P147034** | Ica1l | -1.023971239 | -3.284323182 | 1.236380703 | 8.649382184 | -1.134650317 | 0.303725534 | 0.709264768 | -4.582145576 |
| **A_52_P77764** | Slc7a2 | -0.159808384 | -0.512601254 | 0.192984486 | 7.844751421 | -1.134566894 | 0.30375764 | 0.709264768 | -4.582167508 |
| **A_51_P320089** | Gclm | -0.161830128 | -0.519220054 | 0.195559798 | 8.651776964 | -1.134141956 | 0.30392123 | 0.709264768 | -4.582279217 |
| **A_66_P108188** | Car5a | -0.456204181 | -1.463719153 | 0.55131079 | 7.944010879 | -1.134117622 | 0.3039306 | 0.709264768 | -4.582285614 |
| **A_55_P2045818** | NA | 0.556565484 | -0.672598757 | 1.785729725 | 15.10237518 | 1.134113964 | 0.303932009 | 0.709264768 | -4.582286575 |
| **A_55_P2002859** | Aifm2 | -0.133753579 | -0.429147161 | 0.161640002 | 7.970202203 | -1.134108756 | 0.303934014 | 0.709264768 | -4.582287944 |
| **A_52_P433828** | Tctn3 | -0.188412927 | -0.604527184 | 0.227701329 | 8.359109106 | -1.13409242 | 0.303940305 | 0.709264768 | -4.582292239 |
| **A_52_P641922** | Nhlrc3 | -0.714144714 | -2.291425355 | 0.863135927 | 8.246639649 | -1.134037977 | 0.30396127 | 0.709264768 | -4.58230655 |
| **A_55_P2016732** | Slc30a2 | -0.723695785 | -2.322114168 | 0.874722598 | 8.169139512 | -1.13400749 | 0.303973011 | 0.709264768 | -4.582314563 |
| **A_51_P130757** | Acer3 | -0.096206779 | -0.308718933 | 0.116305375 | 8.541821655 | -1.133893455 | 0.304016931 | 0.709264768 | -4.582344538 |
| **A_55_P2066007** | Crocc | 0.344061484 | -0.415962709 | 1.104085677 | 10.1484656 | 1.13385883 | 0.304030267 | 0.709264768 | -4.582353639 |
| **A_55_P2077083** | Tmem132 | -0.976938004 | -3.135384327 | 1.181508318 | 8.263122328 | -1.133642443 | 0.304113625 | 0.709264768 | -4.582410514 |
| **A_55_P2007650** | Nt5dc1 | 0.152332828 | -0.184235695 | 0.488901351 | 9.419537677 | 1.133627449 | 0.304119402 | 0.709264768 | -4.582414455 |
| **A_51_P441974** | Pip5k1a | -0.532680321 | -1.709626327 | 0.644265685 | 10.15360819 | -1.133601622 | 0.304129353 | 0.709264768 | -4.582421243 |
| **A_55_P2057112** | Prl8a2 | -2.849552717 | -9.145704032 | 3.446598599 | 9.20140291 | -1.133579242 | 0.304137975 | 0.709264768 | -4.582427125 |
| **A_51_P284608** | Cd74 | 0.735558672 | -0.889718516 | 2.360835861 | 9.54220312 | 1.133548811 | 0.3041497 | 0.709264768 | -4.582435123 |
| **A_55_P2093634** | Ddi2 | -0.373756902 | -1.199618383 | 0.452104579 | 9.127985047 | -1.133528272 | 0.304157615 | 0.709264768 | -4.582440521 |
| **A_51_P307964** | Krt13 | -0.366021599 | -1.174857785 | 0.442814587 | 8.441728068 | -1.133434654 | 0.304193689 | 0.709264768 | -4.582465125 |
| **A_55_P2000894** | NA | 0.398184117 | -0.481741082 | 1.278109317 | 15.52986144 | 1.133413872 | 0.304201698 | 0.709264768 | -4.582470587 |
| **A_55_P2083028** | NA | 0.13116258 | -0.158706537 | 0.421031697 | 11.01637139 | 1.133335122 | 0.304232047 | 0.709264768 | -4.582491283 |
| **A_52_P251450** | Cldn2 | -0.438481744 | -1.407597441 | 0.530633953 | 7.908906582 | -1.133251322 | 0.304264345 | 0.709264768 | -4.582513306 |
| **A_55_P2085060** | Vgf | -0.741202354 | -2.379415294 | 0.897010587 | 8.138929751 | -1.13322666 | 0.30427385 | 0.709264768 | -4.582519787 |
| **A_52_P364232** | Pik3c2a | -0.266737967 | -0.856288767 | 0.322812833 | 8.364478834 | -1.133219267 | 0.3042767 | 0.709264768 | -4.58252173 |
| **A_55_P1996818** | NA | 0.49669193 | -0.601147014 | 1.594530874 | 14.34773892 | 1.133179781 | 0.30429192 | 0.709264768 | -4.582532107 |
| **A_52_P458870** | Slc25a16 | 0.141077874 | -0.170767914 | 0.452923662 | 8.546038261 | 1.133102947 | 0.304321538 | 0.709264768 | -4.582552298 |
| **A_55_P1992655** | Hdac6 | -0.433670633 | -1.392282522 | 0.524941257 | 10.65105989 | -1.133098196 | 0.304323369 | 0.709264768 | -4.582553547 |
| **A_55_P1952720** | Kctd2 | -0.14205018 | -0.45605687 | 0.171956509 | 12.49394058 | -1.133060852 | 0.304337766 | 0.709264768 | -4.58256336 |
| **A_66_P107255** | Nom1 | 0.103484747 | -0.12527828 | 0.332247774 | 9.1076988 | 1.133028554 | 0.304350217 | 0.709264768 | -4.582571847 |
| **A_51_P295449** | Mfsd5 | -0.470010827 | -1.509113531 | 0.569091877 | 10.42293746 | -1.132921276 | 0.304391578 | 0.709264768 | -4.582600038 |
| **A_55_P2034481** | Erdr1 | -0.961566107 | -3.087578681 | 1.164446467 | 12.97601434 | -1.132827189 | 0.304427857 | 0.709264768 | -4.582624761 |
| **A_66_P110633** | Apol9b | -0.647298243 | -2.078730419 | 0.784133933 | 8.106983589 | -1.132619407 | 0.304507989 | 0.709264768 | -4.582679358 |
| **A_55_P1971244** | Ndufa4l2 | 0.437891964 | -0.530462931 | 1.40624686 | 12.62901569 | 1.132616199 | 0.304509227 | 0.709264768 | -4.5826802 |
| **A_55_P1973451** | Stra6 | -1.358508728 | -4.362717661 | 1.645700205 | 8.45386306 | -1.132615877 | 0.304509351 | 0.709264768 | -4.582680285 |

| | | | | | | | | | |
|---|---|---|---|---|---|---|---|---|---|
| **A_55_P2186615** | Acp1 | 0.1933222 | -0.23422388 | 0.620868281 | 10.0374504 | 1.132528517 | 0.304543047 | 0.709264768 | -4.582703239 |
| **A_55_P2056035** | Traf7 | -0.141605636 | -0.454793193 | 0.17158192 | 13.07873873 | -1.13246917 | 0.30456594 | 0.709264768 | -4.582718831 |
| **A_55_P2064333** | Lama3 | 1.205262071 | -1.460613218 | 3.87113736 | 11.07670583 | 1.13237952 | 0.304600525 | 0.709264768 | -4.582742386 |
| **A_51_P499776** | Denr | 0.105517682 | -0.127874023 | 0.338909388 | 8.444608342 | 1.132374736 | 0.304602371 | 0.709264768 | -4.582743643 |
| **A_55_P2169879** | NA | 0.455915248 | -0.552515369 | 1.464345866 | 14.38982959 | 1.132370222 | 0.304604112 | 0.709264768 | -4.582744829 |
| **A_55_P1962354** | Lexm | -0.298756163 | -0.959592552 | 0.362080226 | 7.825020718 | -1.132330666 | 0.304619374 | 0.709264768 | -4.582755221 |
| **A_66_P120305** | Zmynd11 | 0.430027137 | -0.521388904 | 1.381443177 | 11.19546543 | 1.132076405 | 0.304717486 | 0.709264768 | -4.58282202 |
| **A_55_P2013223** | S100z | -0.165481156 | -0.531604907 | 0.200642595 | 7.767409322 | -1.132064021 | 0.304722265 | 0.709264768 | -4.582825274 |
| **A_55_P2110537** | Slfn9 | 0.182174366 | -0.220918693 | 0.585267426 | 9.659898498 | 1.131963161 | 0.304761193 | 0.709264768 | -4.58285177 |
| **A_52_P77332** | Ireb2 | -0.183212425 | -0.588606938 | 0.222182088 | 8.053224136 | -1.131950411 | 0.304766114 | 0.709264768 | -4.582855119 |
| **A_55_P1971009** | Gzme | -1.384964272 | -4.449901666 | 1.679973123 | 8.422689382 | -1.131793792 | 0.30482657 | 0.709264768 | -4.582896262 |
| **A_55_P2013268** | NA | -0.431011675 | -1.384853909 | 0.522830559 | 7.884084629 | -1.131782127 | 0.304831074 | 0.709264768 | -4.582899326 |
| **A_51_P428715** | Mrpl13 | 0.593345913 | -0.719813256 | 1.906505081 | 12.68517176 | 1.131725054 | 0.304853107 | 0.709264768 | -4.582914318 |
| **A_51_P205099** | Dazap1 | 0.262208749 | -0.318192198 | 0.842609697 | 11.74675496 | 1.1315387 | 0.304925061 | 0.709264768 | -4.582963269 |
| **A_52_P117352** | Gja4 | -0.17630785 | -0.566597707 | 0.213982007 | 10.66934817 | -1.131448342 | 0.304959954 | 0.709264768 | -4.582987003 |
| **A_55_P2113031** | Foxo4 | -0.487332466 | -1.566250422 | 0.591585489 | 7.925573108 | -1.131324703 | 0.305007705 | 0.709264768 | -4.583019477 |
| **A_55_P2040026** | Itga4 | 0.488373574 | -0.592860437 | 1.569607585 | 9.952269893 | 1.131313068 | 0.305012199 | 0.709264768 | -4.583022533 |
| **A_51_P145511** | Prl2a1 | -2.928233815 | -9.411512407 | 3.555044777 | 9.317354683 | -1.131257348 | 0.305033722 | 0.709264768 | -4.583037168 |
| **A_55_P2039598** | Rpl7a | 0.60460553 | -0.734053533 | 1.943264592 | 13.60836018 | 1.131234074 | 0.305042712 | 0.709264768 | -4.583043281 |
| **A_52_P337679** | Canx | -0.4272308 | -1.373172604 | 0.518711004 | 13.51922877 | -1.131223664 | 0.305046733 | 0.709264768 | -4.583046015 |
| **A_51_P400040** | Ubxn2a | -0.373066747 | -1.199089928 | 0.452956434 | 8.679543824 | -1.131213686 | 0.305050587 | 0.709264768 | -4.583048635 |
| **A_55_P1971430** | Serpinb9e | -1.855007801 | -5.962480021 | 2.25246442 | 8.638715178 | -1.131153259 | 0.30507393 | 0.709264768 | -4.583064506 |
| **A_55_P2182106** | Rpl39 | 0.490757806 | -0.596005561 | 1.577521173 | 15.94745456 | 1.131051998 | 0.30511305 | 0.709264768 | -4.5830911 |
| **A_51_P491667** | Derl3 | -1.615156631 | -5.192285327 | 1.961972065 | 8.796258284 | -1.130916199 | 0.305165521 | 0.709264768 | -4.583126764 |
| **A_55_P2154714** | Klhl9 | 0.362232207 | -0.440131271 | 1.164595685 | 13.32293225 | 1.130749112 | 0.305230091 | 0.709264768 | -4.583170643 |
| **A_55_P2134042** | NA | -0.312865792 | -1.005886286 | 0.380154702 | 16.16362946 | -1.130738942 | 0.305234022 | 0.709264768 | -4.583173314 |
| **A_52_P126146** | Riok1 | 0.107174241 | -0.130257866 | 0.344606349 | 9.313192866 | 1.130580045 | 0.305295439 | 0.709264768 | -4.58321504 |
| **A_55_P2060376** | Gzmg | -1.105178056 | -3.553784444 | 1.343428332 | 8.239217888 | -1.130481919 | 0.305333371 | 0.709264768 | -4.583240807 |
| **A_55_P2000329** | Gskip | -0.348482375 | -1.120626148 | 0.423661397 | 8.806728442 | -1.130402191 | 0.305364195 | 0.709264768 | -4.583261742 |
| **A_51_P273679** | Dmgdh | -0.503632963 | -1.619562688 | 0.612296761 | 7.904106404 | -1.130388283 | 0.305369572 | 0.709264768 | -4.583265394 |
| **A_51_P324351** | Meltf | -0.957090541 | -3.077784817 | 1.163603735 | 8.252201397 | -1.130382188 | 0.305371929 | 0.709264768 | -4.583266995 |
| **A_52_P554650** | Gzmd | -0.429677161 | -1.381767888 | 0.522413566 | 7.854428165 | -1.130353491 | 0.305383024 | 0.709264768 | -4.58327453 |
| **A_51_P487628** | Pramef12 | -2.427748546 | -7.807296162 | 2.95179907 | 9.12524757 | -1.130337907 | 0.30538905 | 0.709264768 | -4.583278622 |
| **A_51_P109541** | E130308A | 0.207776404 | -0.252718192 | 0.668271 | 9.857381826 | 1.130113421 | 0.305475859 | 0.709264768 | -4.583337564 |
| **A_52_P359257** | NA | -0.196607487 | -0.632350181 | 0.239135206 | 7.978560088 | -1.130108873 | 0.305477617 | 0.709264768 | -4.583338758 |
| **A_51_P443475** | Bccip | 0.3995843 | -0.486044931 | 1.285213531 | 11.56359672 | 1.130073834 | 0.305491169 | 0.709264768 | -4.583347958 |
| **A_55_P2094445** | Ms4a10 | -0.68519326 | -2.203938977 | 0.833552458 | 8.059871195 | -1.129999786 | 0.305519809 | 0.709264768 | -4.583367399 |
| **A_55_P2081585** | Dennd5b | 0.194048472 | -0.236109917 | 0.624206861 | 9.532436284 | 1.129879624 | 0.305566291 | 0.709264768 | -4.583398947 |
| **A_52_P17098** | Mpp7 | -0.258350901 | -0.831101646 | 0.314399843 | 9.223332096 | -1.129782034 | 0.305604045 | 0.709264768 | -4.583424568 |
| **A_51_P470079** | Il1r2 | -1.239659645 | -3.987934366 | 1.508615075 | 8.423967925 | -1.129776837 | 0.305606056 | 0.709264768 | -4.583425932 |
| **A_55_P2094986** | Atp1a3 | -0.139525658 | -0.448866457 | 0.169815141 | 8.09800468 | -1.129710648 | 0.305631664 | 0.709264768 | -4.583443308 |
| **A_51_P174854** | Maged1 | -0.845187326 | -2.71910043 | 1.028725777 | 13.77592481 | -1.129675925 | 0.305645099 | 0.709264768 | -4.583452424 |
| **A_55_P1977431** | Cck | -0.624884632 | -2.01036875 | 0.760599485 | 8.257317549 | -1.129662308 | 0.305650368 | 0.709264768 | -4.583455999 |
| **A_52_P122696** | Arl2 | 0.229040622 | -0.278837096 | 0.736918341 | 8.962584907 | 1.129545425 | 0.305695597 | 0.709264768 | -4.583486682 |
| **A_55_P2082778** | NA | 0.580287741 | -0.706508645 | 1.867084127 | 14.789431 | 1.12949395 | 0.305715518 | 0.709264768 | -4.583500195 |
| **A_55_P2074927** | Gm7932 | -0.263744234 | -0.84863311 | 0.321144642 | 8.517837545 | -1.12943166 | 0.305739626 | 0.709264768 | -4.583516547 |
| **A_55_P2459897** | A2m | -0.327658759 | -1.05428926 | 0.398971741 | 7.878596152 | -1.12942788 | 0.305741089 | 0.709264768 | -4.583517539 |
| **A_55_P2001155** | Rpl31 | 0.25128365 | -0.305985537 | 0.808552837 | 15.24624556 | 1.129404586 | 0.305750105 | 0.709264768 | -4.583523654 |

| | | | | | | | | | |
|---|---|---|---|---|---|---|---|---|---|
| **A_55_P2102002** | Noc3l | -0.108563573 | -0.349325571 | 0.132198425 | 8.98387519 | -1.12939675 | 0.305753137 | 0.709264768 | -4.583525711 |
| **A_51_P481644** | Mbip | 0.940745531 | -1.145666485 | 3.027157548 | 11.52250154 | 1.12933408 | 0.305777394 | 0.709264768 | -4.583542161 |
| **A_55_P1965524** | NA | 0.692114082 | -0.842899662 | 2.227127826 | 15.33445952 | 1.129316733 | 0.305784109 | 0.709264768 | -4.583546715 |
| **A_55_P2022342** | Stim1 | -0.692315032 | -2.22779548 | 0.843165415 | 10.56871549 | -1.12930127 | 0.305790095 | 0.709264768 | -4.583550774 |
| **A_55_P2029775** | Gm38396 | 0.123370988 | -0.150256172 | 0.396998149 | 8.647421202 | 1.129286628 | 0.305795762 | 0.709264768 | -4.583554617 |
| **A_66_P125110** | NA | -0.291500555 | -0.938137403 | 0.355136294 | 10.37201724 | -1.129091873 | 0.305871159 | 0.709264768 | -4.583605737 |
| **A_52_P485450** | Vta1 | 0.153345203 | -0.186832576 | 0.493522981 | 10.29336108 | 1.129053702 | 0.305885938 | 0.709264768 | -4.583615756 |
| **A_55_P2085781** | Smim26 | 0.172218397 | -0.209852366 | 0.554289161 | 11.65126501 | 1.128979568 | 0.305914643 | 0.709264768 | -4.583635214 |
| **A_51_P338600** | Tmem14c | -0.262573835 | -0.845124779 | 0.319977109 | 11.99128614 | -1.128932257 | 0.305932964 | 0.709264768 | -4.583647632 |
| **A_51_P308469** | Rngtt | 0.247267288 | -0.301379929 | 0.795914505 | 9.419902704 | 1.128817753 | 0.305977307 | 0.709264768 | -4.583677684 |
| **A_52_P175086** | Usp48 | -0.428327943 | -1.37875212 | 0.522096234 | 9.431347889 | -1.128779928 | 0.305991957 | 0.709264768 | -4.583687612 |
| **A_51_P318381** | Pgf | -0.40142081 | -1.292167944 | 0.489326323 | 8.208093828 | -1.128744886 | 0.306005529 | 0.709264768 | -4.583696808 |
| **A_51_P400752** | NA | -1.041251555 | -3.351825287 | 1.269322177 | 10.5949708 | -1.128719887 | 0.306015212 | 0.709264768 | -4.583703369 |
| **A_55_P2085616** | Nsa2 | 0.186565752 | -0.227440952 | 0.600572455 | 8.951784195 | 1.128690565 | 0.30602657 | 0.709264768 | -4.583711065 |
| **A_51_P122246** | Creld2 | -0.916411862 | -2.950018462 | 1.117194738 | 13.18856021 | -1.128688512 | 0.306027365 | 0.709264768 | -4.582940628 |
| **A_55_P2150442** | NA | 0.27976251 | -0.341057988 | 0.900583009 | 13.12189038 | 1.128687755 | 0.306027658 | 0.709264768 | -4.583711802 |
| **A_55_P2142724** | NA | 0.229830073 | -0.280265796 | 0.739925942 | 8.374934768 | 1.128509954 | 0.306096535 | 0.709264768 | -4.583758464 |
| **A_55_P2061804** | NA | 0.141290972 | -0.17232335 | 0.454905294 | 9.401300434 | 1.128415055 | 0.306133303 | 0.709264768 | -4.583783368 |
| **A_55_P2107855** | Hnrnpk | 0.33608755 | -0.409939712 | 1.082114812 | 10.5881335 | 1.128361013 | 0.306154243 | 0.709264768 | -4.58379755 |
| **A_55_P2002567** | NA | 0.265252545 | -0.323577882 | 0.854082973 | 15.89423574 | 1.128287201 | 0.306182844 | 0.709264768 | -4.583816919 |
| **A_52_P271** | Dnajc18 | 0.207581781 | -0.253230961 | 0.668394523 | 10.05439193 | 1.128275352 | 0.306187436 | 0.709264768 | -4.583820029 |
| **A_55_P2095133** | Synpo2 | 0.436450904 | -0.532454236 | 1.405356044 | 9.194510044 | 1.128247769 | 0.306198125 | 0.709264768 | -4.583827267 |
| **A_55_P1967504** | Abhd11 | -0.139693311 | -0.449809751 | 0.170423129 | 10.49443962 | -1.128239153 | 0.306201464 | 0.709264768 | -4.583829528 |
| **A_55_P2157378** | Xlr5a | -0.467335549 | -1.504820809 | 0.57014971 | 7.916201563 | -1.128228923 | 0.306205429 | 0.709264768 | -4.583832212 |
| **A_55_P2139753** | Jade2 | 0.478508568 | -0.583828765 | 1.540845901 | 10.19015321 | 1.128177983 | 0.30622517 | 0.709264768 | -4.583845579 |
| **A_55_P2144341** | Smim40 | -0.526659258 | -1.695938871 | 0.642620354 | 8.024436271 | -1.128136606 | 0.306241207 | 0.709264768 | -4.583856437 |
| **A_55_P2122688** | Zmym3 | 0.475975947 | -0.580814332 | 1.532766226 | 11.42005447 | 1.128097255 | 0.306256459 | 0.709264768 | -4.583866762 |
| **A_55_P2080151** | Hspa2 | 0.105762164 | -0.129057691 | 0.34058202 | 9.989410391 | 1.128095477 | 0.306257148 | 0.709264768 | -4.583867229 |
| **A_51_P441898** | Krt25 | -0.480605183 | -1.547760618 | 0.586550252 | 7.908684779 | -1.128005235 | 0.306292127 | 0.709264768 | -4.583890908 |
| **A_51_P234714** | Ceacam14 | -2.605639369 | -8.391828744 | 3.180550006 | 9.0682688 | -1.127903669 | 0.306331499 | 0.709264768 | -4.583917558 |
| **A_55_P1952071** | Npb | -0.532140519 | -1.713836196 | 0.649555158 | 8.624379114 | -1.12790112 | 0.306332487 | 0.709264768 | -4.583918226 |
| **A_66_P118004** | Rab3ip | -0.352290559 | -1.134675581 | 0.430094463 | 11.12736012 | -1.127796708 | 0.306372968 | 0.709264768 | -4.583945622 |
| **A_52_P668715** | Nutf2 | 0.126685674 | -0.154702581 | 0.408073929 | 8.568309702 | 1.127643666 | 0.30643231 | 0.709264768 | -4.583985776 |
| **A_55_P1958133** | NA | 0.185917567 | -0.227055985 | 0.598891119 | 11.42641627 | 1.127583035 | 0.306455823 | 0.709264768 | -4.584001683 |
| **A_55_P1991109** | Gatd3a | 0.369064529 | -0.450734843 | 1.188863901 | 12.77434538 | 1.127574047 | 0.306459308 | 0.709264768 | -4.584004041 |
| **A_55_P1974827** | Pde1b | -0.199211913 | -0.641719858 | 0.243296032 | 8.677759386 | -1.127572767 | 0.306459804 | 0.709264768 | -4.584004377 |
| **A_55_P2149020** | Rpl35 | 0.190100968 | -0.232187338 | 0.612389274 | 17.37553526 | 1.127523544 | 0.306478894 | 0.709264768 | -4.584017291 |
| **A_52_P101399** | Tmem130 | -1.019894572 | -3.285640217 | 1.245851073 | 8.304201976 | -1.127442682 | 0.306510257 | 0.709264768 | -4.584038505 |
| **A_55_P2140328** | NA | 0.337569175 | -0.412360481 | 1.08749883 | 12.35592928 | 1.127437821 | 0.306512142 | 0.709264768 | -4.58403978 |
| **A_52_P390944** | Chst3 | 0.439219344 | -0.536550359 | 1.414989047 | 8.923016657 | 1.127416731 | 0.306520323 | 0.709264768 | -4.584045313 |
| **A_55_P2273811** | Msh5 | 0.172407011 | -0.210620937 | 0.555434959 | 8.240809726 | 1.127391626 | 0.306530061 | 0.709264768 | -4.584051899 |
| **A_51_P509140** | U2af1l4 | 0.181506008 | -0.221757524 | 0.584769539 | 9.892762874 | 1.127333469 | 0.30655262 | 0.709264768 | -4.584067156 |
| **A_55_P1952420** | Odf2 | 0.116183169 | -0.141970705 | 0.374337042 | 8.413630443 | 1.12723617 | 0.306590365 | 0.709264768 | -4.584092681 |
| **A_55_P2004752** | Prl2c5 | -3.064240118 | -9.873497452 | 3.745017215 | 9.343808746 | -1.1271284 | 0.306632178 | 0.709264768 | -4.584120952 |
| **A_55_P2067518** | Slc13a3 | -0.397168224 | -1.279764411 | 0.485427964 | 8.248788727 | -1.127100891 | 0.306642851 | 0.709264768 | -4.584128168 |
| **A_51_P185693** | Slc2a2 | -0.245996264 | -0.792713723 | 0.300721195 | 7.812951167 | -1.126979234 | 0.306690059 | 0.709264768 | -4.584160081 |
| **A_52_P195798** | Dmac2l | -0.179102182 | -0.577168776 | 0.218964412 | 8.4594721 | -1.126926237 | 0.306710625 | 0.709264768 | -4.584173982 |
| **A_52_P475870** | Tmem29 | 0.303817579 | -0.37146998 | 0.979105139 | 9.211825291 | 1.126871761 | 0.306731767 | 0.709264768 | -4.584188272 |

| | | | | | | | | | |
|---|---|---|---|---|---|---|---|---|---|
| **A_55_P2110595** | Slc35a2 | -0.360964668 | -1.163303147 | 0.441373811 | 8.347139497 | -1.126827453 | 0.306748964 | 0.709264768 | -4.584199894 |
| **A_51_P143031** | Eps8l3 | -0.703690787 | -2.26787011 | 0.860488536 | 8.055015217 | -1.126796969 | 0.306760796 | 0.709264768 | -4.584207889 |
| **A_51_P120830** | Mmp10 | -0.322921421 | -1.040748343 | 0.394905501 | 7.814926811 | -1.126749719 | 0.306779136 | 0.709264768 | -4.584220283 |
| **A_52_P490770** | Zfp26 | 0.100886019 | -0.123375741 | 0.32514778 | 8.61121773 | 1.126746209 | 0.306780498 | 0.709264768 | -4.584221204 |
| **A_55_P1964832** | Reck | 0.315606169 | -0.385976258 | 1.017188596 | 8.932909164 | 1.126722916 | 0.30678954 | 0.709264768 | -4.584227313 |
| **A_55_P1981724** | Nr2c1 | 0.32092607 | -0.392501227 | 1.034353367 | 9.673901292 | 1.126693065 | 0.306801127 | 0.709264768 | -4.584235143 |
| **A_51_P142421** | Rspo1 | 0.865293336 | -1.058316617 | 2.788903289 | 9.88125718 | 1.126669893 | 0.306810123 | 0.709264768 | -4.584241221 |
| **A_55_P2106800** | Bglap | -0.280937103 | -0.905552066 | 0.343677861 | 7.84156804 | -1.126541151 | 0.306860103 | 0.709264768 | -4.584274987 |
| **A_55_P1953132** | Camkv | 0.135772913 | -0.166113449 | 0.437659275 | 15.39242926 | 1.126470929 | 0.306887367 | 0.709264768 | -4.584293405 |
| **A_51_P349008** | Ptges2 | -0.097382829 | -0.313928559 | 0.119162901 | 9.822015754 | -1.12637527 | 0.306924511 | 0.709264768 | -4.584318493 |
| **A_51_P383270** | Fras1 | 1.074282453 | -1.314585412 | 3.463150318 | 10.67565046 | 1.126358652 | 0.306930964 | 0.709264768 | -4.584322851 |
| **A_55_P1977066** | Klk4 | -0.811081863 | -2.614697362 | 0.992533636 | 8.123357439 | -1.12634407 | 0.306936627 | 0.709264768 | -4.584326675 |
| **A_55_P1984144** | NA | 0.360885494 | -0.441652478 | 1.163423466 | 15.71558854 | 1.126300251 | 0.306953643 | 0.709264768 | -4.584338167 |
| **A_55_P2174953** | Akr1b3 | 0.299075351 | -0.36601262 | 0.964163322 | 9.034954557 | 1.126294266 | 0.306955968 | 0.709264768 | -4.584339737 |
| **A_55_P1953136** | NA | 0.53092921 | -0.649915413 | 1.711773833 | 15.33819626 | 1.12614473 | 0.307014045 | 0.709264768 | -4.584378953 |
| **A_55_P2143934** | NA | 0.112687985 | -0.137961595 | 0.363337565 | 8.153648679 | 1.126058567 | 0.307047513 | 0.709264768 | -4.584401548 |
| **A_55_P2151116** | Micu1 | -0.183398463 | -0.591369017 | 0.22457209 | 11.98370782 | -1.125945115 | 0.307091586 | 0.709264768 | -4.584431299 |
| **A_55_P2037425** | Gapdh | 0.209487182 | -0.256538178 | 0.675512541 | 14.72491565 | 1.125895931 | 0.307110695 | 0.709264768 | -4.584444197 |
| **A_55_P1975087** | NA | 0.564156324 | -0.690949674 | 1.819262321 | 14.05882596 | 1.125821145 | 0.307139751 | 0.709264768 | -4.584463807 |
| **A_55_P1952577** | Ces1a | -0.304163926 | -0.980878656 | 0.372550804 | 7.841568667 | -1.125777128 | 0.307156855 | 0.709264768 | -4.584475349 |
| **A_55_P2133661** | NA | 0.393137752 | -0.481587726 | 1.267863229 | 15.18151534 | 1.125701693 | 0.307186168 | 0.709264768 | -4.584495129 |
| **A_55_P2129477** | NA | 0.218632779 | -0.267850969 | 0.705116527 | 8.770665033 | 1.125634213 | 0.307212391 | 0.709264768 | -4.584512823 |
| **A_66_P101499** | AW82225 | -0.335496097 | -1.082125394 | 0.411133201 | 7.816502976 | -1.125467064 | 0.307277356 | 0.709264768 | -4.584556649 |
| **A_51_P485405** | Gtsf1 | -0.256402082 | -0.827015805 | 0.314211642 | 7.838584898 | -1.125459004 | 0.307280488 | 0.709264768 | -4.584558762 |
| **A_52_P88829** | Ctsq | -3.066699012 | -9.892158122 | 3.758760098 | 9.316542595 | -1.12535522 | 0.307320832 | 0.709264768 | -4.584585973 |
| **A_51_P337125** | Inpp5d | 0.293768199 | -0.360135893 | 0.947672291 | 9.532793244 | 1.125229402 | 0.307369746 | 0.709264768 | -4.58461896 |
| **A_51_P468073** | Ggt1 | -0.809176748 | -2.610504165 | 0.992150669 | 8.098579161 | -1.125125794 | 0.307410031 | 0.709264768 | -4.584646123 |
| **A_55_P1972381** | Gjb4 | -0.260089205 | -0.839112707 | 0.318934298 | 7.819898355 | -1.125062062 | 0.307434814 | 0.709264768 | -4.584662831 |
| **A_51_P280446** | Sdf2l1 | -0.683911013 | -2.206481277 | 0.838659251 | 10.12055005 | -1.125052008 | 0.307438723 | 0.709264768 | -4.584665467 |
| **A_55_P2036460** | Synb | -1.027342913 | -3.31485442 | 1.260168594 | 8.261983221 | -1.124870396 | 0.307509354 | 0.709264768 | -4.584713076 |
| **A_55_P2032714** | Fhit | -0.131823915 | -0.42536283 | 0.161715001 | 8.495265348 | -1.124809213 | 0.307533152 | 0.709264768 | -4.584729115 |
| **A_55_P2031949** | Bend6 | -0.387907746 | -1.25168416 | 0.475868669 | 7.935359801 | -1.124805563 | 0.307534571 | 0.709264768 | -4.584730072 |
| **A_55_P2024245** | Mthfr | -0.536765782 | -1.732075403 | 0.658543838 | 8.954548699 | -1.124746766 | 0.307557443 | 0.709264768 | -4.584745485 |
| **A_55_P2115980** | Xlr5c | -0.480250863 | -1.549711021 | 0.589209296 | 7.89666687 | -1.124744529 | 0.307558313 | 0.709264768 | -4.584746071 |
| **A_51_P417891** | Trim10 | 0.095681653 | -0.117404694 | 0.308768001 | 7.867408065 | 1.124665545 | 0.307589039 | 0.709264768 | -4.584766775 |
| **A_55_P2427150** | C430002N | -0.479423907 | -1.547141032 | 0.588293219 | 7.894013427 | -1.124640771 | 0.307598677 | 0.709264768 | -4.584773269 |
| **A_55_P1971419** | Serpinb9g | -1.910247225 | -6.164794218 | 2.344299768 | 8.686294883 | -1.12457027 | 0.307626106 | 0.709264768 | -4.584791749 |
| **A_55_P2108837** | Tuba1c | -0.351665569 | -1.134923973 | 0.431592836 | 15.28817555 | -1.124540577 | 0.307637659 | 0.709264768 | -4.584799532 |
| **A_55_P1971278** | Kdm2a | 0.206029763 | -0.252898667 | 0.664958192 | 9.434854207 | 1.124437564 | 0.307677742 | 0.709264768 | -4.584826533 |
| **A_51_P212560** | Cog7 | -0.144598572 | -0.466695089 | 0.177497945 | 9.483200403 | -1.124419375 | 0.307684821 | 0.709264768 | -4.584831301 |
| **A_55_P2176045** | NA | -0.104562447 | -0.337498157 | 0.128373263 | 7.986039858 | -1.124320191 | 0.307723419 | 0.709264768 | -4.584857297 |
| **A_51_P190961** | Gzmf | -0.874784794 | -2.823613147 | 1.074043559 | 8.10251343 | -1.124289052 | 0.307735538 | 0.709264768 | -4.584865459 |
| **A_51_P196395** | Cfap298 | 0.480624526 | -0.590122089 | 1.551371141 | 11.71669053 | 1.124267261 | 0.307744019 | 0.709264768 | -4.58487117 |
| **A_55_P2208260** | D230019N | -0.21268745 | -0.686519819 | 0.261144919 | 9.344740232 | -1.124261878 | 0.307746114 | 0.709264768 | -4.584872581 |
| **A_52_P216427** | Nkx2-1 | 1.65073616 | -2.026940168 | 5.328412488 | 11.73863406 | 1.124228281 | 0.30775919 | 0.709264768 | -4.584034765 |
| **A_51_P113003** | Cenpq | 0.56425637 | -0.693019493 | 1.821532233 | 9.223881384 | 1.124077457 | 0.307817898 | 0.709264768 | -4.584227091 |
| **A_55_P2072443** | Kansl1 | 0.131595207 | -0.161648416 | 0.424838829 | 8.873610982 | 1.123988427 | 0.307852558 | 0.709264768 | -4.584944248 |
| **A_55_P2090484** | Pde10a | -0.445369137 | -1.437880756 | 0.547142482 | 7.926287976 | -1.123918614 | 0.307879738 | 0.709264768 | -4.584962545 |

| | | | | | | | | | |
|---|---|---|---|---|---|---|---|---|---|
| **A_55_P2054942** | Iffo2 | -0.249657476 | -0.806039429 | 0.306724477 | 8.83529214 | -1.123885029 | 0.307892815 | 0.709264768 | -4.584971346 |
| **A_52_P436318** | Rps25 | 0.321733186 | -0.395308851 | 1.038775224 | 8.845623997 | 1.123832502 | 0.307913267 | 0.709264768 | -4.584985111 |
| **A_66_P110088** | NA | -0.986809171 | -3.186164497 | 1.212546155 | 9.692554217 | -1.123797659 | 0.307926835 | 0.709264768 | -4.584994242 |
| **A_51_P516705** | Ern1 | -0.641022486 | -2.069747599 | 0.787702627 | 8.495172953 | -1.123763524 | 0.307940128 | 0.709264768 | -4.585003188 |
| **A_55_P2132845** | NA | 0.118295365 | -0.14539321 | 0.381983941 | 16.29005865 | 1.123638887 | 0.307988666 | 0.709264768 | -4.585035849 |
| **A_55_P2022211** | Plxdc2 | 0.605135205 | -0.743756326 | 1.954026737 | 13.09460737 | 1.123636239 | 0.307989698 | 0.709264768 | -4.585036543 |
| **A_51_P425824** | C1qtnf12 | 0.458055568 | -0.563000829 | 1.479111965 | 11.79125919 | 1.123618244 | 0.307996706 | 0.709264768 | -4.585041259 |
| **A_51_P230734** | Slc34a3 | -0.461083654 | -1.489010768 | 0.56684346 | 7.919642415 | -1.12348622 | 0.30804813 | 0.709264768 | -4.585075854 |
| **A_51_P258698** | Phf8 | -0.485141825 | -1.566776191 | 0.596492541 | 11.28977249 | -1.123410777 | 0.308077518 | 0.709264768 | -4.585095622 |
| **A_55_P1991901** | NA | -2.678909209 | -8.654920986 | 3.297102568 | 9.115092287 | -1.12278576 | 0.308321083 | 0.709264768 | -4.585259379 |
| **A_51_P221144** | Vwa8 | -0.144745275 | -0.467651621 | 0.178161071 | 9.21740366 | -1.122737325 | 0.308339965 | 0.709264768 | -4.585272068 |
| **A_55_P2021149** | NA | -0.200755536 | -0.648644481 | 0.247133408 | 7.976649371 | -1.12265815 | 0.308370833 | 0.709264768 | -4.585292809 |
| **A_55_P1978716** | Zim1 | -0.480905486 | -1.554065987 | 0.592255015 | 8.026187824 | -1.122394157 | 0.308473772 | 0.709264768 | -4.585361964 |
| **A_52_P269461** | Ddhd2 | -0.335376812 | -1.083852704 | 0.41309908 | 8.406626736 | -1.12229121 | 0.308513923 | 0.709264768 | -4.58538893 |
| **A_52_P64763** | Gen1 | 0.118840603 | -0.146383556 | 0.384064762 | 7.91527226 | 1.122282293 | 0.308517401 | 0.709264768 | -4.585391266 |
| **A_52_P681016** | NA | -0.500114432 | -1.6163137 | 0.616084836 | 7.964356728 | -1.122219987 | 0.308541703 | 0.709264768 | -4.585407586 |
| **A_66_P119034** | Pla2g7 | -2.01734515 | -6.520328044 | 2.485637743 | 10.01878438 | -1.122096627 | 0.308589825 | 0.709264768 | -4.585439898 |
| **A_55_P2108012** | Fam78b | -0.653402601 | -2.111916435 | 0.805111232 | 8.173775675 | -1.122071811 | 0.308599506 | 0.709264768 | -4.585446398 |
| **A_51_P261999** | Ccdc166 | 0.246813103 | -0.304189735 | 0.797815941 | 8.948775835 | 1.121927315 | 0.308655882 | 0.709264768 | -4.585484243 |
| **A_52_P424840** | Mrc2 | 0.375503012 | -0.462857735 | 1.21386376 | 9.005646847 | 1.121844943 | 0.308688024 | 0.709264768 | -4.585505817 |
| **A_55_P2085979** | Igf1 | 0.174631142 | -0.215272873 | 0.564535156 | 9.076777691 | 1.121797151 | 0.308706674 | 0.709264768 | -4.585518334 |
| **A_55_P1955346** | NA | 0.307087064 | -0.378569425 | 0.992743553 | 14.88699063 | 1.12177376 | 0.308715802 | 0.709264768 | -4.58552446 |
| **A_55_P2091350** | NA | -0.314909648 | -1.018047157 | 0.388227861 | 14.12048142 | -1.121749917 | 0.308725107 | 0.709264768 | -4.585530704 |
| **A_55_P2099061** | NA | 0.60806604 | -0.749720143 | 1.965852222 | 13.86007356 | 1.121681894 | 0.308751654 | 0.709264768 | -4.585548519 |
| **A_55_P2114482** | Rpgr | 0.365942818 | -0.451197134 | 1.183082771 | 9.58547047 | 1.121675229 | 0.308754256 | 0.709264768 | -4.585550265 |
| **A_55_P2024260** | Vps33b | -0.180706311 | -0.584241036 | 0.222828415 | 8.765604198 | -1.12161227 | 0.308778829 | 0.709264768 | -4.585566753 |
| **A_51_P384879** | Mdh2 | -0.165059697 | -0.53366698 | 0.203547586 | 15.86314812 | -1.121572823 | 0.308794226 | 0.709264768 | -4.585577083 |
| **A_66_P114784** | Pla2g7 | -0.956368966 | -3.0921231 | 1.179385169 | 8.227618136 | -1.121565301 | 0.308797163 | 0.709264768 | -4.585579053 |
| **A_55_P2020203** | Sfrp5 | -1.952592535 | -6.313214253 | 2.408029184 | 8.977150535 | -1.121536873 | 0.308808259 | 0.709264768 | -4.585586498 |
| **A_52_P105765** | Cdc42bpg | -0.161718813 | -0.537148888 | 0.213711261 | 8.387175014 | -1.147345433 | 0.308828716 | 0.709264768 | -4.578770924 |
| **A_55_P2104791** | NA | 0.560245737 | -0.691170545 | 1.81166202 | 14.24611454 | 1.12131364 | 0.308895409 | 0.709264768 | -4.585644956 |
| **A_55_P2046338** | Mapk14 | -0.180820533 | -0.584725407 | 0.223084341 | 11.98632459 | -1.121292704 | 0.308903583 | 0.709264768 | -4.585650438 |
| **A_55_P2125457** | NA | 0.529679692 | -0.65349762 | 1.712857003 | 14.60398276 | 1.121279377 | 0.308908787 | 0.709264768 | -4.585653928 |
| **A_55_P2047748** | NA | -0.405716443 | -1.312049346 | 0.500616459 | 10.57724607 | -1.121205472 | 0.308937645 | 0.709264768 | -4.58567328 |
| **A_52_P402663** | Ninl | 0.156821979 | -0.193511891 | 0.507155849 | 8.142189705 | 1.121179503 | 0.308947786 | 0.709264768 | -4.58568008 |
| **A_51_P300297** | Srek1ip1 | 0.141555762 | -0.174713898 | 0.457825421 | 9.350023841 | 1.121038079 | 0.309003016 | 0.709264768 | -4.585717111 |
| **A_55_P2062714** | Zfp975 | 0.18074856 | -0.223153874 | 0.584650995 | 8.809076205 | 1.120853162 | 0.309075245 | 0.709264768 | -4.585765528 |
| **A_55_P1971425** | Serpinb9f | -2.056497689 | -6.651995938 | 2.539000559 | 8.761431009 | -1.120846052 | 0.309078022 | 0.709264768 | -4.58576739 |
| **A_51_P455866** | Elf5 | 1.400174994 | -1.728993562 | 4.529343549 | 11.26982478 | 1.120736988 | 0.30912063 | 0.709264768 | -4.585795945 |
| **A_52_P219904** | Afmid | -0.177817632 | -0.575222653 | 0.21958739 | 7.936232206 | -1.120706307 | 0.309132617 | 0.709264768 | -4.585803977 |
| **A_51_P487278** | Yipf3 | -0.27129891 | -0.877670227 | 0.335072407 | 11.21781843 | -1.120623549 | 0.309164952 | 0.709264768 | -4.585825644 |
| **A_55_P2109486** | Milr1 | -0.144751824 | -0.468287975 | 0.178784326 | 16.87661432 | -1.120602476 | 0.309173186 | 0.709264768 | -4.585831161 |
| **A_55_P2340101** | NA | 0.144566092 | -0.178598351 | 0.467730535 | 9.857794048 | 1.120451901 | 0.309232027 | 0.709264768 | -4.585870581 |
| **A_52_P381929** | Ubqln2 | 0.148206424 | -0.183099225 | 0.479512072 | 9.866254864 | 1.120439815 | 0.30923675 | 0.709264768 | -4.585873745 |
| **A_55_P2015984** | Odr4 | -0.216478621 | -0.700414511 | 0.267457269 | 10.86695968 | -1.120411418 | 0.309247848 | 0.709264768 | -4.585881179 |
| **A_51_P330369** | Ssx2ip | 0.636452152 | -0.78637985 | 2.059284154 | 10.58913904 | 1.120372602 | 0.309263019 | 0.709264768 | -4.58589134 |
| **A_55_P2066409** | Gabpb1 | 0.222769086 | -0.275308998 | 0.720847169 | 10.6977555 | 1.120231643 | 0.309318116 | 0.709264768 | -4.585928241 |
| **A_55_P2007228** | Prkaca | 0.108926999 | -0.13463712 | 0.352491119 | 8.235684618 | 1.120140693 | 0.30935367 | 0.709264768 | -4.585952049 |

| | | | | | | | | | |
|---|---|---|---|---|---|---|---|---|---|
| **A_51_P334281** | Gucy1b2 | -1.452433749 | -4.700547625 | 1.795680127 | 8.611719484 | -1.119993315 | 0.30941129 | 0.709264768 | -4.585990626 |
| **A_51_P487048** | Rhox6 | -2.863708444 | -9.26805671 | 3.540639822 | 9.226916226 | -1.119964366 | 0.309422609 | 0.709264768 | -4.585998204 |
| **A_55_P2016079** | Erich4 | -0.332611711 | -1.076521504 | 0.411298083 | 7.89068926 | -1.119869987 | 0.309459515 | 0.709264768 | -4.586022907 |
| **A_55_P1969615** | Pnck | 0.260159901 | -0.321754221 | 0.842074023 | 9.379611772 | 1.119777681 | 0.309495613 | 0.709264768 | -4.586047068 |
| **A_55_P1969745** | Pitpnm1 | -0.240628958 | -0.778860873 | 0.297602957 | 10.46188921 | -1.119769903 | 0.309498655 | 0.709264768 | -4.586049103 |
| **A_55_P2214487** | E330013F | -0.111502152 | -0.360914331 | 0.137910026 | 8.312471117 | -1.119736771 | 0.309511613 | 0.709264768 | -4.586057775 |
| **A_55_P2178510** | Mc3r | -1.48979562 | -4.822395568 | 1.842804328 | 8.697298399 | -1.119679832 | 0.309533883 | 0.709264768 | -4.586072678 |
| **A_51_P287100** | Cdh16 | 1.726043231 | -2.135131735 | 5.587218198 | 12.12889775 | 1.11965064 | 0.309545301 | 0.709264768 | -4.586080318 |
| **A_52_P466853** | Bmyc | 0.253437498 | -0.313530652 | 0.820405649 | 9.544611278 | 1.119599139 | 0.309565446 | 0.709264768 | -4.586093797 |
| **A_52_P251615** | Arhgef40 | 0.604386738 | -0.747785157 | 1.956558632 | 11.07154362 | 1.119523893 | 0.309594881 | 0.709264768 | -4.58611349 |
| **A_52_P678816** | Ctsm | -2.732029763 | -8.844996247 | 3.380936722 | 9.192786538 | -1.119396017 | 0.309644909 | 0.709264768 | -4.586146957 |
| **A_52_P480301** | Sppl3 | -0.245684171 | -0.795411834 | 0.304043491 | 8.730209445 | -1.119386155 | 0.309648768 | 0.709264768 | -4.586149538 |
| **A_55_P2337213** | Odr4 | -0.326076344 | -1.055756847 | 0.403604159 | 9.567399859 | -1.11927524 | 0.309692167 | 0.709264768 | -4.586178565 |
| **A_52_P96847** | Smlr1 | -0.465847046 | -1.508444514 | 0.576750422 | 7.93168723 | -1.11912095 | 0.309752546 | 0.709264768 | -4.586218941 |
| **A_55_P2115315** | Fmr1nb | -0.355737819 | -1.151914129 | 0.440438492 | 7.813773827 | -1.119105762 | 0.30975849 | 0.709264768 | -4.586222915 |
| **A_52_P439159** | Thap12 | 0.387599349 | -0.479953713 | 1.25515241 | 10.12380919 | 1.119018683 | 0.309792572 | 0.709264768 | -4.586245702 |
| **A_55_P2159070** | 2810025M | 0.847135703 | -1.049025747 | 2.743297153 | 11.4930335 | 1.118994671 | 0.309801971 | 0.709264768 | -4.586251985 |
| **A_52_P821** | Fli1 | 0.636728115 | -0.788556188 | 2.062012418 | 10.49647014 | 1.118929876 | 0.309827334 | 0.709264768 | -4.58626894 |
| **A_52_P112888** | Atp6v1h | -0.115085734 | -0.372700725 | 0.142529256 | 9.056069518 | -1.118924306 | 0.309829514 | 0.709264768 | -4.586270398 |
| **A_55_P2038162** | Nt5c | 0.459416193 | -0.568996444 | 1.48782883 | 12.07974885 | 1.118894759 | 0.309841081 | 0.709264768 | -4.586278129 |
| **A_55_P2095535** | NA | -0.311272463 | -1.008113703 | 0.385568777 | 7.989214473 | -1.118812225 | 0.309873391 | 0.709264768 | -4.586299725 |
| **A_51_P136699** | Ttc7b | -0.44786222 | -1.450538929 | 0.55481449 | 10.21606461 | -1.118752056 | 0.309896948 | 0.709264768 | -4.586315469 |
| **A_52_P176737** | Lcorl | 0.169617343 | -0.210124962 | 0.549359648 | 8.387427827 | 1.118746319 | 0.309899194 | 0.709264768 | -4.58631697 |
| **A_55_P2371984** | 5930436O | -0.114033562 | -0.369343104 | 0.141275981 | 8.486926301 | -1.118706049 | 0.309914961 | 0.709264768 | -4.586327506 |
| **A_55_P2112445** | Tbx4 | 1.328302905 | -1.645709752 | 4.302315562 | 12.36981626 | 1.118676862 | 0.30992639 | 0.709264768 | -4.586335143 |
| **A_51_P271503** | Il1r1 | -0.142834935 | -0.462659913 | 0.176990042 | 9.86484906 | -1.118593805 | 0.309958913 | 0.709264768 | -4.586356875 |
| **A_51_P346132** | Rhox9 | -3.003552647 | -9.729101654 | 3.72199636 | 9.338020161 | -1.118556312 | 0.309973595 | 0.709264768 | -4.586366684 |
| **A_55_P2232023** | NA | -0.248134558 | -0.803769228 | 0.307500111 | 7.951158226 | -1.118531614 | 0.309983267 | 0.709264768 | -4.586373146 |
| **A_55_P1991406** | Fbxo34 | 0.099038214 | -0.122737127 | 0.320813556 | 9.495908311 | 1.118510048 | 0.309991712 | 0.709264768 | -4.586378788 |
| **A_52_P558609** | Clec16a | -0.175930738 | -0.569896052 | 0.218034576 | 8.432573274 | -1.118495091 | 0.30999757 | 0.709264768 | -4.586382702 |
| **A_55_P2044947** | NA | 0.100160376 | -0.124164177 | 0.324484929 | 8.258731642 | 1.118328726 | 0.31006273 | 0.709264768 | -4.586426227 |
| **A_51_P114693** | Parm1 | -1.232060213 | -3.991488379 | 1.527367953 | 11.25020982 | -1.118312517 | 0.310069079 | 0.709264768 | -4.586430468 |
| **A_55_P1998688** | Gm2012 | -0.20325656 | -0.658502486 | 0.251989367 | 7.875614596 | -1.118275596 | 0.310083542 | 0.709264768 | -4.586440127 |
| **A_51_P296057** | Upf1 | 0.096291922 | -0.119403078 | 0.311986922 | 9.432232071 | 1.118150143 | 0.310132688 | 0.709264768 | -4.586472947 |
| **A_52_P276955** | Epha3 | 0.532619809 | -0.660582636 | 1.725822255 | 9.061464152 | 1.118030176 | 0.310179691 | 0.709264768 | -4.58650433 |
| **A_51_P439836** | NA | -0.272552436 | -0.883198647 | 0.338093775 | 10.70999895 | -1.117920054 | 0.310222843 | 0.709264768 | -4.586533137 |
| **A_51_P346445** | Hspb7 | -0.157140052 | -0.509242064 | 0.194961961 | 8.122755032 | -1.117811904 | 0.310265227 | 0.709264768 | -4.586351578 |
| **A_51_P379687** | Gm10413 | -0.370578305 | -1.200956458 | 0.459799847 | 7.860688571 | -1.117775079 | 0.31027966 | 0.709264768 | -4.58657106 |
| **A_51_P201971** | Kmt5a | -0.469951217 | -1.523140785 | 0.583238352 | 10.37540195 | -1.117626202 | 0.310338014 | 0.709264768 | -4.586610001 |
| **A_55_P1986541** | NA | 0.27234771 | -0.338013984 | 0.882709404 | 15.72793168 | 1.117601058 | 0.310347871 | 0.709264768 | -4.586616578 |
| **A_51_P128648** | Uqcrc2 | 0.321835197 | -0.399447444 | 1.043117838 | 13.44484564 | 1.11757944 | 0.310356346 | 0.709264768 | -4.586622232 |
| **A_52_P507305** | Iah1 | 0.195050705 | -0.242088758 | 0.632190169 | 9.75265299 | 1.117578029 | 0.310356899 | 0.709264768 | -4.586622601 |
| **A_55_P1979833** | Cited1 | -2.041603699 | -6.617675689 | 2.534468291 | 11.77120769 | -1.117452165 | 0.310406243 | 0.709264768 | -4.586655521 |
| **A_55_P2057321** | Myh7b | 0.454375583 | -0.56407685 | 1.472828016 | 8.655726152 | 1.117440946 | 0.310410642 | 0.709264768 | -4.586658456 |
| **A_55_P1961853** | Zfand3 | -0.176974838 | -0.573672794 | 0.219723119 | 8.919550458 | -1.117382601 | 0.310433518 | 0.709264768 | -4.586673716 |
| **A_55_P1979669** | Zfp946 | 0.392563464 | -0.487404249 | 1.272531176 | 9.176653302 | 1.117360939 | 0.310442012 | 0.709264768 | -4.586679381 |
| **A_52_P370311** | Plbd2 | -0.465910001 | -1.510386033 | 0.578566031 | 11.32477522 | -1.117259098 | 0.310481947 | 0.709264768 | -4.586706016 |
| **A_55_P2160691** | NA | -2.598960745 | -8.425831751 | 3.22791026 | 9.241201075 | -1.117158159 | 0.310521533 | 0.709264768 | -4.586732415 |

| | | | | | | | | | |
|---|---|---|---|---|---|---|---|---|---|
| **A_66_P140858** | NA | 0.593386314 | -0.73705241 | 1.923825038 | 12.14150464 | 1.117102424 | 0.310543393 | 0.709264768 | -4.586746991 |
| **A_55_P2031432** | Usp47 | -0.253192122 | -0.820881746 | 0.314497502 | 8.167678605 | -1.117093637 | 0.310546839 | 0.709264768 | -4.586749289 |
| **A_55_P1955921** | NA | 0.399532937 | -0.496280447 | 1.295346321 | 14.9661231 | 1.117082855 | 0.310551068 | 0.709264768 | -4.586752108 |
| **A_66_P127969** | Bcat1 | -0.482582808 | -1.564699595 | 0.599533979 | 7.938405356 | -1.116986842 | 0.310588729 | 0.709264768 | -4.586777217 |
| **A_55_P2133910** | Ttll10 | 0.145786738 | -0.181146385 | 0.472719861 | 8.638923069 | 1.116887529 | 0.310627689 | 0.709264768 | -4.586803188 |
| **A_51_P374900** | P2ry13 | 0.859466333 | -1.06821304 | 2.787145707 | 7.379693767 | 1.116720309 | 0.310693297 | 0.709264768 | -4.586846915 |
| **A_51_P199135** | Cd83 | -0.664233873 | -2.154030931 | 0.825563186 | 10.08697614 | -1.116719891 | 0.310693461 | 0.709264768 | -4.586847025 |
| **A_55_P2118268** | Chpf | -0.742800101 | -2.408815537 | 0.923215334 | 11.91132255 | -1.11671739 | 0.310694442 | 0.709264768 | -4.586847679 |
| **A_51_P197883** | Exo5 | 0.209597101 | -0.260506681 | 0.679700883 | 10.39060827 | 1.116713771 | 0.310695862 | 0.709264768 | -4.586848625 |
| **A_55_P2024525** | Prdm8 | 0.391285109 | -0.486331109 | 1.268901327 | 8.649328205 | 1.116706454 | 0.310698733 | 0.709264768 | -4.586850538 |
| **A_66_P113008** | Pla2g2e | -0.752339744 | -2.439826819 | 0.93514733 | 8.107561258 | -1.11666755 | 0.310713999 | 0.709264768 | -4.586860711 |
| **A_51_P349023** | Lsm3 | 0.470900513 | -0.585408811 | 1.527209837 | 11.42475182 | 1.116576276 | 0.310749817 | 0.709264768 | -4.586884577 |
| **A_51_P372456** | Wdr86 | 0.645771665 | -0.802907497 | 2.094450828 | 11.05758321 | 1.116495862 | 0.310781376 | 0.709264768 | -4.586905604 |
| **A_55_P2024625** | Ccl21a | 1.841671188 | -2.289863748 | 5.973206125 | 12.56519163 | 1.11648016 | 0.310787539 | 0.709264768 | -4.586909709 |
| **A_55_P1955821** | Ccl21b | 0.955053371 | -1.187772449 | 3.097879191 | 13.3001059 | 1.116326197 | 0.310847972 | 0.709264768 | -4.586949965 |
| **A_52_P531662** | Ttc9c | 0.229560321 | -0.285515736 | 0.744636378 | 9.27811806 | 1.116286843 | 0.31086342 | 0.709264768 | -4.586960254 |
| **A_52_P87843** | Aldh1a3 | -1.079435141 | -3.501632635 | 1.342762353 | 8.528888693 | -1.116187996 | 0.310902226 | 0.709264768 | -4.586986098 |
| **A_51_P494125** | Alpl | -1.313025265 | -4.259394845 | 1.633344315 | 8.901428518 | -1.116185063 | 0.310903378 | 0.709264768 | -4.586986864 |
| **A_55_P2011385** | NA | 0.120952427 | -0.150463658 | 0.392368511 | 8.209323691 | 1.116167442 | 0.310910296 | 0.709264768 | -4.586991471 |
| **A_55_P1956171** | NA | 0.505067535 | -0.628340475 | 1.638475544 | 14.87086359 | 1.116126762 | 0.310926268 | 0.709264768 | -4.587002107 |
| **A_55_P1953341** | Wfdc2 | 1.737735327 | -2.161932325 | 5.637402979 | 14.0414696 | 1.116108408 | 0.310933474 | 0.709264768 | -4.587006905 |
| **A_51_P353252** | Mal2 | 1.859426046 | -2.313401084 | 6.032253176 | 12.93574716 | 1.116089089 | 0.31094106 | 0.709264768 | -4.587011956 |
| **A_55_P2019058** | Acaca | 0.392865221 | -0.488829232 | 1.274559673 | 10.70157192 | 1.116029876 | 0.31096431 | 0.709264768 | -4.587027436 |
| **A_55_P2178486** | Spin1 | -0.140273287 | -0.455094645 | 0.17454807 | 8.535613269 | -1.115992134 | 0.310979131 | 0.709264768 | -4.587037303 |
| **A_55_P1957835** | Cpeb3 | -0.45882299 | -1.488671565 | 0.571025586 | 8.637585385 | -1.115891946 | 0.311018476 | 0.709264768 | -4.587063495 |
| **A_51_P216593** | Wfs1 | -0.94729331 | -3.073643279 | 1.179056659 | 9.817222655 | -1.115835234 | 0.311040749 | 0.709264768 | -4.58707832 |
| **A_55_P2076846** | Tead2 | 0.37832846 | -0.470911153 | 1.227568072 | 11.4559279 | 1.115807084 | 0.311051805 | 0.709264768 | -4.587085679 |
| **A_55_P2077353** | Ceacam12 | -0.838667116 | -2.721252896 | 1.043918664 | 8.08611257 | -1.115796992 | 0.311055769 | 0.709264768 | -4.587088317 |
| **A_55_P2179793** | NA | -0.871261975 | -2.827075433 | 1.084551484 | 8.821859924 | -1.11576228 | 0.311069403 | 0.709264768 | -4.587097391 |
| **A_51_P327564** | Glb1 | -0.344749925 | -1.118680723 | 0.429180872 | 10.25444696 | -1.115712753 | 0.311088857 | 0.709264768 | -4.587110338 |
| **A_55_P2419514** | Ccny | 0.124203349 | -0.154633368 | 0.403040065 | 8.467563481 | 1.115664696 | 0.311107735 | 0.709264768 | -4.587122901 |
| **A_65_P10351** | Atp2b1 | -0.222081968 | -0.720658902 | 0.276494965 | 8.805710008 | -1.115658969 | 0.311109984 | 0.709264768 | -4.587124398 |
| **A_52_P108089** | Mosmo | -0.116040722 | -0.376556232 | 0.144474788 | 10.56806176 | -1.115647982 | 0.3111143 | 0.709264768 | -4.58712727 |
| **A_51_P268559** | Idh3a | 0.162850138 | -0.202785267 | 0.528485544 | 11.15527979 | 1.115553067 | 0.311151588 | 0.709264768 | -4.58715208 |
| **A_55_P1975843** | NA | 0.737135705 | -0.917961722 | 2.392233132 | 11.58390733 | 1.115511957 | 0.31116774 | 0.709264768 | -4.587162826 |
| **A_51_P516133** | H1f2 | -0.537386609 | -1.744013462 | 0.669240244 | 11.22381685 | -1.115486194 | 0.311177862 | 0.709264768 | -4.58716956 |
| **A_51_P467505** | Tec | -0.490826972 | -1.592923403 | 0.611269459 | 8.17979048 | -1.115473275 | 0.311182938 | 0.709264768 | -4.587172937 |
| **A_55_P2056995** | Gcgr | -1.371300411 | -4.450956343 | 1.708355522 | 9.28206424 | -1.115271882 | 0.311262075 | 0.709264768 | -4.587225577 |
| **A_51_P438692** | Vti1b | -0.285007177 | -0.925113902 | 0.355099548 | 12.81465515 | -1.115202544 | 0.311289325 | 0.709264768 | -4.587243699 |
| **A_52_P198009** | Mrpl44 | 0.136517017 | -0.170091691 | 0.443125725 | 11.04541322 | 1.115199718 | 0.311290436 | 0.709264768 | -4.587244438 |
| **A_51_P379069** | Dkk1 | -0.372745345 | -1.209936297 | 0.464445608 | 7.879109276 | -1.115162219 | 0.311305175 | 0.709264768 | -4.587254239 |
| **A_52_P205255** | Adam11 | 0.125397427 | -0.156265521 | 0.407060376 | 7.94872621 | 1.115088279 | 0.311334237 | 0.709264768 | -4.587273564 |
| **A_52_P588559** | Ube2d3 | 0.318516612 | -0.396925002 | 1.033958227 | 12.24703887 | 1.115085673 | 0.311335261 | 0.709264768 | -4.587274245 |
| **A_51_P238786** | Mcl1 | -0.219017492 | -0.710988445 | 0.272953461 | 13.69380548 | -1.115038043 | 0.311353984 | 0.709264768 | -4.587286693 |
| **A_55_P2170009** | Lactb2 | -0.436464322 | -1.416945164 | 0.544016519 | 9.504899972 | -1.114961786 | 0.311383961 | 0.709264768 | -4.587306623 |
| **A_51_P408082** | Apoa1 | -1.298378464 | -4.215280735 | 1.618523807 | 8.525823283 | -1.114884225 | 0.311414453 | 0.709264768 | -4.586902434 |
| **A_55_P1966958** | Mef2d | -0.188225001 | -0.611150287 | 0.234700285 | 7.965081938 | -1.114715405 | 0.311480832 | 0.709264768 | -4.587371011 |
| **A_51_P487228** | B9d2 | 0.602883236 | -0.751833771 | 1.957600243 | 11.71417555 | 1.11464089 | 0.311510135 | 0.709264768 | -4.587390483 |

| | | | | | | | | | |
|---|---|---|---|---|---|---|---|---|---|
| **A_55_P1957909** | Mrpl49 | 0.16059643 | -0.200313144 | 0.521506004 | 10.45462236 | 1.114519903 | 0.311557717 | 0.709264768 | -4.587422098 |
| **A_51_P353392** | Cript | 0.203087711 | -0.2533314 | 0.659506822 | 9.50580256 | 1.114474587 | 0.311575541 | 0.709264768 | -4.58743394 |
| **A_55_P2053459** | Timd2 | -0.716174889 | -2.325789102 | 0.893439325 | 8.070282551 | -1.114416764 | 0.311598285 | 0.709264768 | -4.587449049 |
| **A_55_P1998578** | Ifitm2 | 0.780581819 | -0.973907054 | 2.535070692 | 12.10920506 | 1.114341126 | 0.311628038 | 0.709264768 | -4.587468813 |
| **A_51_P333349** | Tmem120 | -0.546310438 | -1.774340257 | 0.681719382 | 10.4940955 | -1.114245608 | 0.311665616 | 0.709264768 | -4.587493771 |
| **A_55_P2030647** | Dusp13 | -0.28457666 | -0.9242969 | 0.35514358 | 7.801785719 | -1.114190707 | 0.311687216 | 0.709264768 | -4.587508116 |
| **A_51_P112734** | Slc7a8 | -1.435766038 | -4.663555168 | 1.792023092 | 8.752725665 | -1.114112039 | 0.31171817 | 0.709264768 | -4.58689256 |
| **A_55_P2061268** | Rnpc3 | -0.108531918 | -0.352540512 | 0.135476676 | 8.163602525 | -1.114044917 | 0.311744582 | 0.709264768 | -4.587546207 |
| **A_52_P344424** | Lipg | -0.440841771 | -1.432125321 | 0.550441778 | 7.926757858 | -1.11387174 | 0.311812735 | 0.709264768 | -4.587591452 |
| **A_55_P1961541** | NA | -0.160949202 | -0.522866772 | 0.200968369 | 7.81638669 | -1.113857169 | 0.31181847 | 0.709264768 | -4.587595258 |
| **A_52_P58257** | Mrps6 | -0.667999091 | -2.170142362 | 0.834144181 | 11.95050172 | -1.113819611 | 0.311833253 | 0.709264768 | -4.587605071 |
| **A_55_P2062652** | Zfp941 | 0.133912374 | -0.167229538 | 0.435054286 | 8.121347391 | 1.113781118 | 0.311848404 | 0.709264768 | -4.587615127 |
| **A_55_P2125466** | Ube2l3 | 0.132993038 | -0.166097493 | 0.432083568 | 11.8298262 | 1.113721462 | 0.311871887 | 0.709264768 | -4.587630712 |
| **A_55_P2025665** | Mrpl40 | 0.191434538 | -0.239116181 | 0.621985257 | 11.43263644 | 1.113643844 | 0.311902442 | 0.709264768 | -4.587650988 |
| **A_51_P345110** | Thap4 | -0.369182678 | -1.19955223 | 0.461186874 | 11.05932621 | -1.113576982 | 0.311928765 | 0.709264768 | -4.587668455 |
| **A_51_P343350** | Amn | -0.685449149 | -2.227466593 | 0.856568295 | 8.183925078 | -1.113361759 | 0.312013511 | 0.709264768 | -4.587724676 |
| **A_55_P2067757** | Psg27 | -2.816440278 | -9.152619671 | 3.519739115 | 9.279592236 | -1.113328758 | 0.312026507 | 0.709264768 | -4.587733296 |
| **A_55_P1974645** | Entpd2 | -1.110398992 | -3.608545734 | 1.38774775 | 9.258443713 | -1.113298077 | 0.31203859 | 0.709264768 | -4.587741311 |
| **A_52_P104761** | Rlim | -0.49974734 | -1.624122546 | 0.624627866 | 8.533419777 | -1.113241984 | 0.312060681 | 0.709264768 | -4.587755962 |
| **A_55_P2043267** | Cdc42se1 | -0.459583373 | -1.493605985 | 0.574439239 | 10.23420732 | -1.113229272 | 0.312065688 | 0.709264768 | -4.587759283 |
| **A_51_P332652** | Pqlc3 | -0.522682108 | -1.698827278 | 0.653463062 | 8.660474916 | -1.113081719 | 0.312123807 | 0.709264768 | -4.587797823 |
| **A_55_P2136763** | NA | 0.307166039 | -0.38404057 | 0.998372649 | 12.7845162 | 1.113052529 | 0.312135306 | 0.709264768 | -4.587805447 |
| **A_52_P634122** | Rexo1 | -0.108680725 | -0.353256903 | 0.135895453 | 8.781725708 | -1.112983482 | 0.312162506 | 0.709264768 | -4.587823481 |
| **A_55_P1984745** | Pappa2 | -2.57774979 | -8.378783233 | 3.223283654 | 9.629067062 | -1.112975839 | 0.312165518 | 0.709264768 | -4.587825477 |
| **A_55_P1975370** | Apoc1 | -0.422087686 | -1.371972062 | 0.52779669 | 11.5481299 | -1.11296699 | 0.312169004 | 0.709264768 | -4.587827789 |
| **A_55_P2108609** | Srgap3 | -0.475919643 | -1.547433535 | 0.595594248 | 7.949979406 | -1.112464524 | 0.312367012 | 0.709264768 | -4.587959013 |
| **A_55_P2154943** | NA | -0.537849769 | -1.748833001 | 0.673133464 | 7.970403417 | -1.112431305 | 0.312380106 | 0.709264768 | -4.587967687 |
| **A_55_P2069974** | Kctd1 | 0.112867373 | -0.141260749 | 0.366995494 | 8.283307068 | 1.112412948 | 0.312387343 | 0.709264768 | -4.587972481 |
| **A_55_P1957850** | Snx9 | -0.635692557 | -2.067005211 | 0.795620097 | 10.1425278 | -1.112405074 | 0.312390447 | 0.709264768 | -4.587974537 |
| **A_55_P2122200** | Prkar2b | 0.13389743 | -0.167635755 | 0.435430616 | 8.220823832 | 1.112211732 | 0.312466671 | 0.709264768 | -4.588025024 |
| **A_55_P1972841** | Rps28 | 0.306973667 | -0.384360199 | 0.998307532 | 17.48958467 | 1.11215069 | 0.31249074 | 0.709264768 | -4.588040963 |
| **A_52_P245365** | Rtf1 | -0.259275846 | -0.843196892 | 0.3246452 | 8.105065919 | -1.112136968 | 0.312496151 | 0.709264768 | -4.588044546 |
| **A_55_P1956647** | Sfmbt2 | -1.382523139 | -4.496153955 | 1.731107676 | 8.760584385 | -1.112130209 | 0.312498816 | 0.709264768 | -4.588046311 |
| **A_51_P109050** | Paqr4 | 0.498923596 | -0.624818851 | 1.622666043 | 10.59372008 | 1.112032816 | 0.312537222 | 0.709264768 | -4.58807174 |
| **A_51_P502614** | Dusp6 | 0.551795722 | -0.691095292 | 1.794686735 | 12.47015587 | 1.111976545 | 0.312559414 | 0.709264768 | -4.588086433 |
| **A_55_P2183433** | Rab30 | -0.441239734 | -1.435267572 | 0.552788104 | 8.218025409 | -1.111799341 | 0.312629307 | 0.709264768 | -4.588132699 |
| **A_55_P2043486** | Msx2 | -0.465036801 | -1.512797752 | 0.58272415 | 8.209143098 | -1.111668908 | 0.312680761 | 0.709264768 | -4.588166752 |
| **A_55_P1968240** | Cbr4 | -0.184112968 | -0.598942204 | 0.230716268 | 10.13497319 | -1.111643079 | 0.312690951 | 0.709264768 | -4.588173496 |
| **A_55_P2182586** | Esrp1 | -0.315264811 | -1.02562362 | 0.395093999 | 9.585547245 | -1.11159881 | 0.312708417 | 0.709264768 | -4.588185053 |
| **A_55_P2042918** | Trim37 | 0.263696648 | -0.330468978 | 0.857862274 | 8.477732204 | 1.111597169 | 0.312709065 | 0.709264768 | -4.588185482 |
| **A_55_P2129217** | NA | 0.681985044 | -0.854717337 | 2.218687425 | 14.92190019 | 1.111566454 | 0.312721183 | 0.709264768 | -4.5881935 |
| **A_55_P2026380** | Pdcd2l | 0.098777106 | -0.123796639 | 0.321350851 | 10.56823055 | 1.111559486 | 0.312723933 | 0.709264768 | -4.588195319 |
| **A_55_P2156944** | Rbis | 0.217656456 | -0.292261399 | 0.727574311 | 12.40064035 | 1.136930821 | 0.312731814 | 0.709264768 | -4.580983013 |
| **A_55_P1973748** | Nrbp1 | -0.160640735 | -0.522630284 | 0.201348814 | 10.19888594 | -1.111501356 | 0.312746869 | 0.709264768 | -4.588210495 |
| **A_52_P631547** | Cytl1 | 0.640898871 | -0.860758088 | 2.14255583 | 9.039636419 | 1.136793231 | 0.312783674 | 0.709264768 | -4.581012212 |
| **A_55_P2001228** | Dctn6 | 0.207403491 | -0.260028032 | 0.674835015 | 13.17563389 | 1.111343748 | 0.312809065 | 0.709264768 | -4.588251639 |
| **A_51_P475228** | Armc6 | -0.432935822 | -1.408853551 | 0.542981907 | 9.57435719 | -1.111119218 | 0.312897687 | 0.709264768 | -4.588310249 |
| **A_52_P458697** | Higd1a | 0.318678351 | -0.399692512 | 1.037049215 | 12.46972626 | 1.111102688 | 0.312904212 | 0.709264768 | -4.588314564 |

| | | | | | | | | | |
|---|---|---|---|---|---|---|---|---|---|
| **A_51_P448391** | Nkiras1 | -0.220722658 | -0.718422491 | 0.276977174 | 8.339041107 | -1.110784394 | 0.313029882 | 0.709264768 | -4.588397643 |
| **A_55_P2075670** | NA | 0.144187608 | -0.180979908 | 0.469355125 | 8.639539686 | 1.110634421 | 0.31308911 | 0.709264768 | -4.588436785 |
| **A_51_P295312** | Gabarap | 0.236683621 | -0.297135925 | 0.770503166 | 12.67272966 | 1.110514104 | 0.313136632 | 0.709264768 | -4.588468185 |
| **A_55_P1996431** | NA | -0.314619184 | -1.0242704 | 0.395032031 | 7.855849028 | -1.11042849 | 0.313170452 | 0.709264768 | -4.588490529 |
| **A_55_P2184541** | Cdadc1 | -0.187818491 | -0.611464441 | 0.235827459 | 10.65099891 | -1.110415804 | 0.313175464 | 0.709264768 | -4.588493839 |
| **A_55_P2116650** | A130040M | -0.290950132 | -0.947405467 | 0.365505203 | 10.95110146 | -1.110104139 | 0.313298606 | 0.709264768 | -4.58857517 |
| **A_55_P2159801** | Rfc4 | 0.315983305 | -0.397011021 | 1.028977631 | 8.961233206 | 1.11001388 | 0.313334276 | 0.709264768 | -4.588598722 |
| **A_55_P2158072** | NA | -0.306083045 | -0.996867572 | 0.384701482 | 7.995636554 | -1.10980587 | 0.313416494 | 0.709264768 | -4.588652997 |
| **A_55_P2064507** | Msra | 0.372800238 | -0.468557879 | 1.214158354 | 10.53752503 | 1.109802343 | 0.313417889 | 0.709264768 | -4.588653918 |
| **A_55_P2261772** | C230098C | -0.274782719 | -0.894947638 | 0.3453822 | 9.94504862 | -1.109768943 | 0.313431092 | 0.709264768 | -4.588662632 |
| **A_66_P102954** | Gm2155 | -0.844542975 | -2.751141976 | 1.062056026 | 8.278640256 | -1.109462783 | 0.313552144 | 0.709264768 | -4.58874251 |
| **A_52_P198019** | Fuz | 0.146566304 | -0.184330722 | 0.47746333 | 8.532368002 | 1.10940883 | 0.31357348 | 0.709264768 | -4.588756586 |
| **A_52_P326214** | Cttn | -0.306276157 | -0.997837587 | 0.385285272 | 8.398390333 | -1.109258516 | 0.31363293 | 0.709264768 | -4.588795799 |
| **A_55_P1992769** | Pafah1b1 | -0.425599206 | -1.386625472 | 0.535427061 | 12.09196511 | -1.109215416 | 0.313649978 | 0.709264768 | -4.588807043 |
| **A_52_P641477** | Wars2 | -0.120507336 | -0.392622528 | 0.151607856 | 8.35251846 | -1.109203017 | 0.313654882 | 0.709264768 | -4.588810277 |
| **A_55_P2015599** | NA | 0.257765706 | -0.324335213 | 0.839866625 | 13.07921375 | 1.109116581 | 0.313689074 | 0.709264768 | -4.588832825 |
| **A_55_P2015337** | NA | 0.232162911 | -0.292127311 | 0.756453133 | 14.41065028 | 1.109101843 | 0.313694905 | 0.709264768 | -4.58883667 |
| **A_51_P465582** | Hdhd3 | 0.219620846 | -0.276385664 | 0.715627355 | 9.545639366 | 1.109012725 | 0.313730162 | 0.709264768 | -4.588859917 |
| **A_51_P517695** | Ly6f | -0.187705057 | -0.61164185 | 0.236231737 | 7.798072183 | -1.108983814 | 0.313741601 | 0.709264768 | -4.588867458 |
| **A_55_P2117119** | Efhd1 | -0.515920117 | -1.681279849 | 0.649439615 | 8.092870414 | -1.108849999 | 0.313794548 | 0.709264768 | -4.588902363 |
| **A_51_P184910** | Phpt1 | 0.692797345 | -0.872129439 | 2.25772413 | 13.23679186 | 1.108823794 | 0.313804918 | 0.709264768 | -4.588909198 |
| **A_55_P2056021** | Snap91 | -0.396137217 | -1.291082198 | 0.498807764 | 8.021420091 | -1.108663259 | 0.313868451 | 0.709264768 | -4.588951071 |
| **A_55_P2156126** | Sema6c | -0.353823536 | -1.15318107 | 0.445533999 | 9.435731646 | -1.108653904 | 0.313872153 | 0.709264768 | -4.588953511 |
| **A_55_P2176871** | NA | 0.36450694 | -0.459047546 | 1.188061425 | 16.0869182 | 1.108571756 | 0.313904668 | 0.709264768 | -4.588974936 |
| **A_55_P2163319** | Peg3 | -0.505762416 | -1.648470884 | 0.636946052 | 8.128323988 | -1.108565741 | 0.313907049 | 0.709264768 | -4.588976505 |
| **A_55_P1954436** | NA | -0.485132657 | -1.581288177 | 0.611022864 | 8.018256655 | -1.108507613 | 0.313930059 | 0.709264768 | -4.588991666 |
| **A_55_P1976224** | Ckb | -0.478885675 | -1.561000146 | 0.603228795 | 10.46673692 | -1.108431827 | 0.313960061 | 0.709264768 | -4.589011431 |
| **A_51_P306658** | Lsm8 | 0.379801575 | -0.478464377 | 1.238067527 | 11.5224583 | 1.108371175 | 0.313984073 | 0.709264768 | -4.589027249 |
| **A_51_P328300** | Pdia4 | -0.52486535 | -1.711033673 | 0.661302972 | 10.50416607 | -1.108286204 | 0.314017715 | 0.709264768 | -4.589049409 |
| **A_51_P324923** | Man2a1 | -0.495835982 | -1.616449563 | 0.624777599 | 12.23905975 | -1.108236637 | 0.314037342 | 0.709264768 | -4.589062336 |
| **A_51_P388973** | Zhx1 | 0.42669645 | -0.537660892 | 1.391053792 | 9.407666439 | 1.10823378 | 0.314038473 | 0.709264768 | -4.589063081 |
| **A_55_P2183587** | Gli1 | 1.096738197 | -1.382112354 | 3.575588748 | 10.75317898 | 1.108161278 | 0.314067183 | 0.709264768 | -4.589081988 |
| **A_52_P562612** | Srd5a1 | -0.219026957 | -0.714131329 | 0.276077415 | 7.82598597 | -1.10802907 | 0.314119541 | 0.709264768 | -4.589116465 |
| **A_51_P406466** | Ankrd35 | -0.59744164 | -1.94798372 | 0.75310044 | 8.044356107 | -1.107994778 | 0.314133123 | 0.709264768 | -4.589125407 |
| **A_55_P1977681** | Vps13d | -0.118865035 | -0.387590258 | 0.149860188 | 9.127837999 | -1.107888453 | 0.314175238 | 0.709264768 | -4.589153132 |
| **A_51_P293688** | Rab32 | -0.155996224 | -0.508669906 | 0.196677457 | 9.786465094 | -1.10787658 | 0.314179941 | 0.709264768 | -4.589156228 |
| **A_55_P2089840** | Eif2d | -0.094147915 | -0.30700542 | 0.11870959 | 9.36551034 | -1.107827351 | 0.314199442 | 0.709264768 | -4.589169065 |
| **A_51_P496381** | Ptk7 | 0.147107803 | -0.185520007 | 0.479735613 | 10.69191713 | 1.10771363 | 0.314244494 | 0.709264768 | -4.589198717 |
| **A_55_P2033906** | Mrpl22 | 0.166177583 | -0.209600635 | 0.541955802 | 12.64560516 | 1.107621 | 0.314281196 | 0.709264768 | -4.58922287 |
| **A_51_P322285** | Tecpr2 | -0.175613475 | -0.572780892 | 0.221553941 | 8.553723395 | -1.107476632 | 0.314338403 | 0.709264768 | -4.589260511 |
| **A_52_P330395** | Farp1 | 0.294257595 | -0.371241683 | 0.959756873 | 12.20918286 | 1.107466093 | 0.31434258 | 0.709264768 | -4.589263258 |
| **A_55_P1986660** | Anapc2 | -0.262616042 | -0.856572975 | 0.331340891 | 13.79295934 | -1.107430911 | 0.314356522 | 0.709264768 | -4.589272431 |
| **A_55_P1964218** | Tprkb | -0.133080138 | -0.434072482 | 0.167912205 | 14.04651379 | -1.107409233 | 0.314365114 | 0.709264768 | -4.589278083 |
| **A_52_P206613** | Adss | 0.209940432 | -0.264945436 | 0.684826299 | 10.36007496 | 1.107279313 | 0.314416607 | 0.709264768 | -4.589311954 |
| **A_55_P2152976** | NA | 0.416184498 | -0.525341776 | 1.357710771 | 15.62946995 | 1.107143212 | 0.314470558 | 0.709264768 | -4.589347436 |
| **A_55_P2146249** | NA | 0.307432429 | -0.388069404 | 1.002934263 | 13.52154516 | 1.107137964 | 0.314472639 | 0.709264768 | -4.589348804 |
| **A_51_P438821** | Pycard | 0.689842132 | -0.870797547 | 2.25048181 | 10.44438758 | 1.107126933 | 0.314477012 | 0.709264768 | -4.58935168 |
| **A_55_P2084344** | NA | 0.636273953 | -0.803292069 | 2.075839975 | 13.57056067 | 1.107038955 | 0.314511892 | 0.709264768 | -4.589374615 |

| | | | | | | | | | |
|---|---|---|---|---|---|---|---|---|---|
| **A_51_P280085** | Zfp655 | -0.79832221 | -2.604584434 | 1.007940015 | 11.9077808 | -1.107000349 | 0.314527198 | 0.709264768 | -4.589384679 |
| **A_51_P517051** | Castor1 | -0.396146989 | -1.292500302 | 0.500206324 | 8.014530463 | -1.106948654 | 0.314547696 | 0.709264768 | -4.589398155 |
| **A_55_P2056926** | Mtss2 | 0.321042772 | -0.405400036 | 1.047485581 | 11.12743306 | 1.106908733 | 0.314563525 | 0.709264768 | -4.589408561 |
| **A_55_P2029622** | Ppp1r3f | -0.517433712 | -1.688318162 | 0.653450739 | 8.059302314 | -1.106855755 | 0.314584533 | 0.709264768 | -4.589422371 |
| **A_66_P100696** | Slc26a2 | -0.503472293 | -1.642786806 | 0.63584222 | 8.211584782 | -1.10683349 | 0.314593363 | 0.709264768 | -4.589428175 |
| **A_52_P554451** | 1300017J | -0.664671123 | -2.168840806 | 0.83949856 | 8.093702634 | -1.10677751 | 0.314615563 | 0.709264768 | -4.589442767 |
| **A_51_P367866** | Egr1 | 0.349952663 | -0.442013218 | 1.141918544 | 9.498172153 | 1.106759291 | 0.314622789 | 0.709264768 | -4.589447516 |
| **A_51_P127320** | Cabs1 | -0.260249569 | -0.849255642 | 0.328756505 | 7.768974811 | -1.106676253 | 0.314655723 | 0.709264768 | -4.588938899 |
| **A_55_P2089238** | Fam76a | -0.484701715 | -1.581730527 | 0.612327097 | 10.87650449 | -1.106641284 | 0.314669593 | 0.709264768 | -4.589478276 |
| **A_52_P318955** | Ap3m1 | -0.099771309 | -0.325585635 | 0.126043017 | 9.118080013 | -1.106635311 | 0.314671962 | 0.709264768 | -4.589479832 |
| **A_55_P2036337** | Ifnar1 | 0.250838223 | -0.316890621 | 0.818567067 | 10.67531343 | 1.106631688 | 0.314673399 | 0.709264768 | -4.589480777 |
| **A_51_P211998** | Sgms2 | -0.274355056 | -0.895315585 | 0.346605473 | 7.84900538 | -1.106622048 | 0.314677223 | 0.709264768 | -4.589483289 |
| **A_52_P243391** | Sema4f | 0.366393401 | -0.462929283 | 1.195716084 | 8.596960451 | 1.106558667 | 0.314702364 | 0.709264768 | -4.58949981 |
| **A_55_P2124976** | Grhl1 | -0.192599132 | -0.628558735 | 0.243360471 | 9.057936956 | -1.106517848 | 0.314718556 | 0.709264768 | -4.589510449 |
| **A_55_P2032960** | Brap | -0.327633419 | -1.069256889 | 0.413990051 | 11.59272334 | -1.106509314 | 0.314721942 | 0.709264768 | -4.589512673 |
| **A_51_P362066** | Chil1 | -1.018371245 | -3.323569798 | 1.286827307 | 9.33092015 | -1.106491636 | 0.314728955 | 0.709264768 | -4.589517281 |
| **A_55_P1973668** | Slc39a13 | -0.544290976 | -1.776406918 | 0.687824965 | 10.81289567 | -1.106445185 | 0.314747383 | 0.709264768 | -4.589529388 |
| **A_55_P2003753** | Sh2d5 | -0.391764931 | -1.278624394 | 0.495094533 | 7.935701573 | -1.106422752 | 0.314756283 | 0.709264768 | -4.589535234 |
| **A_65_P16483** | Pdzk1 | -0.66736728 | -2.178177184 | 0.843442623 | 8.062650409 | -1.106382844 | 0.314772116 | 0.709264768 | -4.589545636 |
| **A_55_P2057906** | Cetn2 | 0.547944061 | -0.692664715 | 1.788552836 | 12.1071203 | 1.106246018 | 0.314826407 | 0.709264768 | -4.589581296 |
| **A_52_P674414** | Mkln1 | -0.184465506 | -0.602120546 | 0.233189533 | 9.257692791 | -1.106236001 | 0.314830381 | 0.709264768 | -4.589583907 |
| **A_52_P138806** | Dlgap3 | 0.137532619 | -0.173889608 | 0.448954847 | 8.082192666 | 1.106130718 | 0.314872162 | 0.709264768 | -4.589611345 |
| **A_51_P447189** | 1700030C | -0.725218162 | -2.367442367 | 0.917006042 | 8.289698352 | -1.106080074 | 0.314892261 | 0.709264768 | -4.589624543 |
| **A_55_P2097391** | Tent5a | -0.514813607 | -1.680586705 | 0.650959492 | 8.074276348 | -1.106079475 | 0.314892499 | 0.709264768 | -4.589624699 |
| **A_55_P2039572** | NA | 0.24684587 | -0.312136522 | 0.805828263 | 16.26218843 | 1.106058463 | 0.314900838 | 0.709264768 | -4.589630175 |
| **A_55_P2006876** | NA | 0.385827878 | -0.487979447 | 1.259635203 | 14.20411206 | 1.105931604 | 0.314951191 | 0.709264768 | -4.589663234 |
| **A_52_P227937** | Cystm1 | -0.319348429 | -1.042678902 | 0.403982044 | 10.45132652 | -1.105804539 | 0.315001633 | 0.709264768 | -4.589696346 |
| **A_52_P572284** | Laptm4a | -0.285804913 | -0.933226358 | 0.361616532 | 12.38063922 | -1.105688912 | 0.31504754 | 0.709264768 | -4.589726475 |
| **A_66_P114974** | NA | 0.702017675 | -0.888322189 | 2.292357538 | 14.47394828 | 1.105626519 | 0.315072314 | 0.709264768 | -4.589742733 |
| **A_51_P414637** | Col5a1 | 0.416250552 | -0.526760593 | 1.359261697 | 11.76706437 | 1.105575338 | 0.315092637 | 0.709264768 | -4.589756069 |
| **A_51_P391445** | Ifngr1 | 0.268821055 | -0.340224112 | 0.877866221 | 12.05128815 | 1.105513692 | 0.315117118 | 0.709264768 | -4.589772131 |
| **A_55_P2056241** | Pdlim5 | -0.391138872 | -1.27736583 | 0.495088085 | 8.637120996 | -1.105443037 | 0.315145178 | 0.709264768 | -4.589790541 |
| **A_52_P579876** | Suds3 | 0.192912027 | -0.244180865 | 0.630004918 | 9.734886375 | 1.105441859 | 0.315145646 | 0.709264768 | -4.589790848 |
| **A_55_P2082444** | Npepps | 0.104372044 | -0.132118029 | 0.340862117 | 11.81482267 | 1.105405496 | 0.315160088 | 0.709264768 | -4.589800322 |
| **A_66_P126293** | Itgb3 | -0.592776297 | -1.935925896 | 0.750373302 | 8.025402715 | -1.105393199 | 0.315164972 | 0.709264768 | -4.589803526 |
| **A_55_P2061737** | Tmsb4x | 0.223881199 | -0.283433394 | 0.731195792 | 16.88685662 | 1.105326583 | 0.315191431 | 0.709264768 | -4.589820883 |
| **A_51_P440047** | Lgalsl | -0.154956708 | -0.50610633 | 0.196192915 | 11.00325924 | -1.105270351 | 0.315213767 | 0.709264768 | -4.589835533 |
| **A_52_P80567** | Nbas | -0.173211064 | -0.565742229 | 0.219320101 | 8.479622746 | -1.105227881 | 0.315230638 | 0.709264768 | -4.589846598 |
| **A_55_P1987944** | Rai1 | 0.306860886 | -0.38856947 | 1.002291241 | 11.586734 | 1.105193282 | 0.315244382 | 0.709264768 | -4.589855612 |
| **A_52_P478729** | Atg2a | -0.205394635 | -0.670875773 | 0.260086502 | 8.536701038 | -1.105191039 | 0.315245273 | 0.709264768 | -4.589856197 |
| **A_55_P2052240** | Crlf1 | 1.085375794 | -1.374558021 | 3.545309609 | 13.36639801 | 1.105113934 | 0.315275905 | 0.709264768 | -4.589876285 |
| **A_51_P327295** | Akr1a1 | -0.173158201 | -0.565633917 | 0.219317515 | 14.18180386 | -1.105046668 | 0.315302631 | 0.709264768 | -4.589893809 |
| **A_55_P2132549** | Cd48 | 0.203559616 | -0.257851754 | 0.664970986 | 8.128507389 | 1.104978115 | 0.31532987 | 0.709264768 | -4.589911668 |
| **A_55_P2021709** | Oxnad1 | 0.101639567 | -0.128759008 | 0.332038142 | 10.0328002 | 1.104926431 | 0.315350407 | 0.709264768 | -4.589925132 |
| **A_55_P2166995** | NA | -0.335703801 | -1.096690898 | 0.425283297 | 15.35849446 | -1.104916122 | 0.315354503 | 0.709264768 | -4.589927817 |
| **A_52_P10781** | Lman1 | -0.395850233 | -1.293197473 | 0.501497007 | 9.990906765 | -1.104894266 | 0.315363189 | 0.709264768 | -4.589933511 |
| **A_51_P473272** | Tspan6 | 0.710636982 | -0.90031196 | 2.321585925 | 11.30778502 | 1.104883211 | 0.315367582 | 0.709264768 | -4.589936391 |
| **A_52_P204618** | Rnf222 | -0.232500553 | -0.759567389 | 0.294566283 | 7.794052418 | -1.104863549 | 0.315375395 | 0.709264768 | -4.589941513 |

| | | | | | | | | | |
|---|---|---|---|---|---|---|---|---|---|
| **A_66_P122087** | Zfp622 | -0.300878358 | -0.983056385 | 0.381299669 | 13.89462778 | -1.104698004 | 0.315441189 | 0.709264768 | -4.589984637 |
| **A_55_P2120278** | Ube2l3 | 0.187126429 | -0.237205682 | 0.61145854 | 12.93295995 | 1.104535241 | 0.315505888 | 0.709264768 | -4.590027033 |
| **A_55_P2009360** | Psg-ps1 | -2.784197057 | -9.097910505 | 3.529516391 | 9.318780429 | -1.104499309 | 0.315520173 | 0.709264768 | -4.590036393 |
| **A_55_P2018002** | Abcg4 | 0.168228915 | -0.213273089 | 0.549730919 | 8.352962923 | 1.104470613 | 0.315531581 | 0.709264768 | -4.590043867 |
| **A_55_P1985463** | NA | 0.706809635 | -0.89689319 | 2.31051246 | 14.5156139 | 1.103897914 | 0.315759338 | 0.709264768 | -4.590193023 |
| **A_52_P605834** | Tm9sf3 | -0.315393091 | -1.031074702 | 0.400288519 | 10.89884735 | -1.103780363 | 0.315806103 | 0.709264768 | -4.590223635 |
| **A_55_P2000224** | Cd63 | -0.484986279 | -1.585684872 | 0.615712314 | 15.69659738 | -1.103599223 | 0.315878179 | 0.709264768 | -4.590270805 |
| **A_55_P2097759** | NA | 0.55014817 | -0.69852659 | 1.79882293 | 14.44420449 | 1.103521211 | 0.315909224 | 0.709264768 | -4.590291118 |
| **A_55_P2057653** | NA | 0.218380284 | -0.277378068 | 0.714138637 | 15.88888765 | 1.103300292 | 0.315997154 | 0.709264768 | -4.590348641 |
| **A_55_P2102624** | Eaf2 | -0.238918432 | -0.781302104 | 0.30346524 | 8.045837149 | -1.10329943 | 0.315997497 | 0.709264768 | -4.590348865 |
| **A_55_P2033160** | Pomgnt1 | -0.298940259 | -0.977593918 | 0.3797134 | 12.09950239 | -1.103282063 | 0.31600441 | 0.709264768 | -4.590353387 |
| **A_55_P1961255** | Ppp1cc | 0.421387565 | -0.535283865 | 1.378058995 | 14.1239466 | 1.103238113 | 0.316021906 | 0.709264768 | -4.59036483 |
| **A_52_P494930** | Fnip2 | -0.413729773 | -1.35302464 | 0.525565094 | 9.365250495 | -1.103227735 | 0.316026037 | 0.709264768 | -4.590367532 |
| **A_55_P2017694** | Gprc5c | -0.745177193 | -2.437027122 | 0.946672736 | 10.66601261 | -1.103184275 | 0.316043339 | 0.709264768 | -4.590378848 |
| **A_55_P2088218** | Uqcrb | 0.252448779 | -0.3207384 | 0.825635957 | 13.28313658 | 1.103131161 | 0.316064485 | 0.709264768 | -4.590392676 |
| **A_55_P2008879** | Cltc | -0.122296698 | -0.399976671 | 0.155383275 | 9.391460324 | -1.103114311 | 0.316071194 | 0.709264768 | -4.590397063 |
| **A_55_P2277825** | E430010N | 0.240887963 | -0.306061587 | 0.787837513 | 9.950717934 | 1.103108359 | 0.316073563 | 0.709264768 | -4.590398613 |
| **A_55_P2144376** | NA | -0.096651743 | -0.316118494 | 0.122815008 | 8.027131409 | -1.103040101 | 0.316100741 | 0.709264768 | -4.590416384 |
| **A_55_P1979082** | Shc1 | 0.185917785 | -0.236250418 | 0.608085988 | 12.03931501 | 1.103026031 | 0.316106344 | 0.709264768 | -4.590420047 |
| **A_55_P1999800** | Gm14066 | 0.268647755 | -0.341401395 | 0.878696905 | 8.501410549 | 1.102982786 | 0.316123563 | 0.709264768 | -4.590431305 |
| **A_52_P550843** | Desi2 | -0.12980166 | -0.42456316 | 0.16495984 | 10.4959737 | -1.102960179 | 0.316132565 | 0.709264768 | -4.590437191 |
| **A_51_P112223** | Gsta4 | 0.654602982 | -0.831974856 | 2.141180821 | 9.496275645 | 1.102911515 | 0.316151944 | 0.709264768 | -4.589701245 |
| **A_55_P1987326** | Prl3a1 | -2.866877737 | -9.378563967 | 3.644808493 | 9.367216078 | -1.102722033 | 0.316227409 | 0.709264768 | -4.590499187 |
| **A_55_P2014555** | Lrp8 | -0.38088218 | -1.246013666 | 0.484249306 | 7.902829633 | -1.102703825 | 0.316234662 | 0.709264768 | -4.590503927 |
| **A_52_P648601** | Dusp9 | -1.878263646 | -6.144573542 | 2.388046251 | 9.069929212 | -1.102692701 | 0.316239093 | 0.709264768 | -4.590506823 |
| **A_55_P2179676** | Spp2 | -1.412545654 | -4.621151349 | 1.796060041 | 8.550324194 | -1.102646971 | 0.316257309 | 0.709264768 | -4.590518727 |
| **A_52_P52263** | Ilrun | -0.276254045 | -0.903796136 | 0.351288046 | 8.248504867 | -1.102595286 | 0.316277898 | 0.709264768 | -4.590532182 |
| **A_51_P402536** | Uvrag | -0.257577859 | -0.842711114 | 0.327555396 | 11.56425464 | -1.102564723 | 0.316290073 | 0.709264768 | -4.590540137 |
| **A_55_P2041270** | Arhgef1 | -0.457646244 | -1.497403833 | 0.582111344 | 12.48098364 | -1.102422706 | 0.316346654 | 0.709264768 | -4.590577104 |
| **A_52_P460791** | Ppy | -0.28547489 | -0.934099989 | 0.36315021 | 7.82004546 | -1.102362697 | 0.316370564 | 0.709264768 | -4.590592724 |
| **A_55_P2058646** | Nhlh1 | 0.151162259 | -0.19230053 | 0.494625048 | 8.38171288 | 1.102336091 | 0.316381166 | 0.709264768 | -4.590599649 |
| **A_55_P2133220** | Arhgef39 | 0.514530434 | -0.65461747 | 1.683678337 | 9.654350721 | 1.102280078 | 0.316403486 | 0.709264768 | -4.590614229 |
| **A_52_P322096** | Patl2 | -0.37857054 | -1.238875373 | 0.481734293 | 7.864114571 | -1.102160385 | 0.316451187 | 0.709264768 | -4.590645382 |
| **A_55_P1968450** | Gm5766 | 0.412832019 | -0.52533399 | 1.350998028 | 15.53444957 | 1.102158425 | 0.316451968 | 0.709264768 | -4.590645892 |
| **A_55_P2033250** | Fdft1 | -0.438413876 | -1.43473067 | 0.557902918 | 9.573559841 | -1.10214107 | 0.316458885 | 0.709264768 | -4.590650409 |
| **A_55_P2157102** | NA | -0.163278238 | -0.534340784 | 0.207784308 | 13.86756992 | -1.102126719 | 0.316464605 | 0.709264768 | -4.590654144 |
| **A_51_P252199** | Mmgt1 | 0.162791111 | -0.207192225 | 0.532774447 | 9.460763264 | 1.102043834 | 0.316497641 | 0.709264768 | -4.590675716 |
| **A_51_P376445** | Rhox5 | -2.32199012 | -7.599528884 | 2.955548645 | 10.18942597 | -1.10199413 | 0.316517454 | 0.709264768 | -4.590688652 |
| **A_55_P2048221** | NA | 0.231275286 | -0.294408985 | 0.756959557 | 14.8915475 | 1.101931478 | 0.316542429 | 0.709264768 | -4.590704958 |
| **A_55_P2047163** | Tmc5 | -0.85329343 | -2.793279642 | 1.086692783 | 8.191350862 | -1.101666399 | 0.316648116 | 0.709264768 | -4.590773942 |
| **A_55_P2031692** | Gstm6 | 0.206291975 | -0.262743103 | 0.675327054 | 8.281427008 | 1.101608697 | 0.316671127 | 0.709264768 | -4.590788958 |
| **A_55_P2099363** | Stac2 | -0.449855704 | -1.472675327 | 0.572963919 | 7.917810319 | -1.101601519 | 0.316673989 | 0.709264768 | -4.590790826 |
| **A_55_P2119737** | NA | -0.280946036 | -0.919752154 | 0.357860082 | 7.791221588 | -1.101549932 | 0.316694562 | 0.709264768 | -4.59080425 |
| **A_51_P273684** | Gbe1 | 0.365135951 | -0.465168613 | 1.195440514 | 10.67744104 | 1.10145692 | 0.316731658 | 0.709264768 | -4.590828453 |
| **A_55_P2113066** | Zkscan5 | -0.313141131 | -1.025367883 | 0.399085622 | 10.45819409 | -1.101215152 | 0.316828099 | 0.709264768 | -4.590891361 |
| **A_55_P1974640** | NA | -0.194023874 | -0.635347165 | 0.247299417 | 7.855266998 | -1.101155543 | 0.316851881 | 0.709264768 | -4.590906871 |
| **A_55_P1983488** | Lsr | 0.127433647 | -0.162484828 | 0.417352122 | 13.63414593 | 1.100927109 | 0.316943032 | 0.709264768 | -4.590966304 |
| **A_55_P2151868** | Mest | 0.285425877 | -0.364000102 | 0.934851856 | 9.383281965 | 1.100814222 | 0.316988085 | 0.709264768 | -4.590995673 |

| | | | | | | | | | |
|---|---|---|---|---|---|---|---|---|---|
| **A_55_P2179378** | Zkscan8 | 0.331359439 | -0.422585159 | 1.085304037 | 9.941626037 | 1.100804657 | 0.316991902 | 0.709264768 | -4.590998161 |
| **A_55_P2134852** | Cnot2 | 0.244812174 | -0.31226439 | 0.801888739 | 11.0167206 | 1.100698736 | 0.31703418 | 0.709264768 | -4.591025716 |
| **A_52_P576720** | Calm4 | -0.513121556 | -1.680790918 | 0.654547806 | 7.962363274 | -1.100653756 | 0.317052135 | 0.709264768 | -4.591037418 |
| **A_51_P183102** | Rnf139 | -0.177359165 | -0.580994143 | 0.226275813 | 10.47633297 | -1.100563704 | 0.317088085 | 0.709264768 | -4.591060844 |
| **A_51_P224517** | Jade1 | -0.94526274 | -3.096555042 | 1.206029561 | 9.513200914 | -1.100533994 | 0.317099946 | 0.709264768 | -4.591068572 |
| **A_51_P138060** | Rnf13 | -0.111258733 | -0.364500312 | 0.141982845 | 8.054577669 | -1.100397119 | 0.317154597 | 0.709264768 | -4.591104177 |
| **A_51_P212679** | Cwf19l1 | 0.104300758 | -0.133120084 | 0.341721601 | 8.439184878 | 1.100319904 | 0.31718543 | 0.709264768 | -4.591124262 |
| **A_55_P2099448** | Slc12a2 | 0.785199969 | -1.002273142 | 2.572673081 | 13.01283556 | 1.100249303 | 0.317213624 | 0.709264768 | -4.591142626 |
| **A_52_P444601** | Zfp277 | 0.525603543 | -0.671150257 | 1.722357344 | 10.3620111 | 1.10002819 | 0.317301939 | 0.709264768 | -4.591200137 |
| **A_55_P1957378** | Unc119 | -0.285709762 | -0.936274779 | 0.364855254 | 11.81855719 | -1.099979825 | 0.317321259 | 0.709264768 | -4.591212716 |
| **A_55_P2134571** | NA | 0.112728034 | -0.143957383 | 0.369413451 | 8.752599094 | 1.099970612 | 0.317324939 | 0.709264768 | -4.591215113 |
| **A_55_P1973279** | NA | -0.231974904 | -0.760216595 | 0.296266787 | 16.38337908 | -1.099913861 | 0.317347611 | 0.709264768 | -4.591229872 |
| **A_55_P1956672** | Rab34 | 0.794858914 | -1.015291839 | 2.605009666 | 12.66732051 | 1.099830222 | 0.317381027 | 0.709264768 | -4.591251625 |
| **A_52_P105015** | Clcn7 | -0.532147793 | -1.744023171 | 0.679727586 | 8.710718456 | -1.099827688 | 0.317382039 | 0.709264768 | -4.591252284 |
| **A_55_P2091858** | Hipk2 | 0.406151437 | -0.518826687 | 1.331129561 | 9.937845216 | 1.099782686 | 0.317400019 | 0.709264768 | -4.591263987 |
| **A_55_P2135271** | Zfp358 | 0.411718868 | -0.525944396 | 1.349382131 | 11.1491193 | 1.09977594 | 0.317402715 | 0.709264768 | -4.591265742 |
| **A_51_P161554** | Arid3b | -0.462374511 | -1.515432463 | 0.590683441 | 8.237039201 | -1.099744905 | 0.317415116 | 0.709264768 | -4.591273813 |
| **A_51_P282144** | Adam23 | -0.313840528 | -1.028730582 | 0.401049527 | 8.106490641 | -1.099562995 | 0.31748781 | 0.709264768 | -4.59132112 |
| **A_51_P106952** | Rbm26 | 0.323270721 | -0.413101725 | 1.059643167 | 10.22083532 | 1.099560645 | 0.317488749 | 0.709264768 | -4.589610397 |
| **A_52_P223571** | Mlec | -0.476062857 | -1.560478831 | 0.608353116 | 8.545854514 | -1.099559508 | 0.317489203 | 0.709264768 | -4.591322026 |
| **A_52_P645159** | Cpsf6 | 0.168414003 | -0.215251715 | 0.55207972 | 8.97690018 | 1.099450164 | 0.317532906 | 0.709264768 | -4.591350461 |
| **A_55_P2101191** | Arrdc4 | -0.408855542 | -1.340278063 | 0.522566979 | 9.460844402 | -1.099444978 | 0.317534979 | 0.709264768 | -4.591351809 |
| **A_55_P2108644** | NA | 0.562850855 | -0.719453282 | 1.845154991 | 13.21231013 | 1.099392151 | 0.317556094 | 0.709264768 | -4.591365546 |
| **A_52_P24843** | Pten | 0.338913691 | -0.433321454 | 1.111148837 | 9.438725145 | 1.09923335 | 0.317619577 | 0.709264768 | -4.591406839 |
| **A_51_P201904** | Ndufb5 | 0.345473705 | -0.441863821 | 1.132811231 | 10.27657172 | 1.099016952 | 0.317706102 | 0.709264768 | -4.591463104 |
| **A_52_P471502** | Dcaf13 | 0.106586688 | -0.136340944 | 0.34951432 | 9.356347813 | 1.098946108 | 0.317734432 | 0.709264768 | -4.591481524 |
| **A_55_P1996421** | NA | 0.602955052 | -0.771390206 | 1.977300309 | 14.30231625 | 1.098852589 | 0.317771834 | 0.709264768 | -4.591505838 |
| **A_51_P279163** | Plcg2 | -0.219050347 | -0.71837593 | 0.280275236 | 9.897429076 | -1.098779312 | 0.317801143 | 0.709264768 | -4.591524889 |
| **A_55_P1983426** | Abhd14a | 0.685963984 | -0.877845868 | 2.249773837 | 11.23878987 | 1.098671135 | 0.317844415 | 0.709264768 | -4.591553012 |
| **A_55_P2112693** | Sipa1l1 | 0.321608186 | -0.411573205 | 1.054789578 | 9.536403956 | 1.098666815 | 0.317846143 | 0.709264768 | -4.591554135 |
| **A_55_P2113210** | Cyth4 | 0.315738309 | -0.404167736 | 1.035644353 | 9.552668619 | 1.098504406 | 0.317911118 | 0.709264768 | -4.591596356 |
| **A_55_P2114993** | Actg1 | -0.238118619 | -0.781049305 | 0.304812067 | 15.7592996 | -1.098498105 | 0.317913639 | 0.709264768 | -4.591597994 |
| **A_55_P2055127** | Slc7a14 | -0.176350431 | -0.578453894 | 0.225753032 | 7.789457962 | -1.098472171 | 0.317924016 | 0.709264768 | -4.591604736 |
| **A_52_P156485** | Ankrd39 | -0.239769324 | -0.786513915 | 0.306975266 | 9.861384323 | -1.098397334 | 0.31795396 | 0.709264768 | -4.591624189 |
| **A_55_P1967138** | Cenpx | 0.389975599 | -0.499300663 | 1.279251862 | 11.42453259 | 1.098376115 | 0.317962452 | 0.709264768 | -4.591629705 |
| **A_51_P151897** | R3hcc1 | 0.255985665 | -0.327843086 | 0.839814417 | 9.516784393 | 1.098197658 | 0.318033869 | 0.709264768 | -4.591676093 |
| **A_52_P35064** | Tm7sf2 | 0.094499004 | -0.121054096 | 0.310052105 | 9.150407564 | 1.098053004 | 0.31809177 | 0.709264768 | -4.591713692 |
| **A_55_P1957198** | Apob | -1.948613702 | -6.393649457 | 2.496422052 | 8.924831919 | -1.097996205 | 0.318114507 | 0.709264768 | -4.591728455 |
| **A_51_P279038** | Ppargc1a | 0.408662162 | -0.523659196 | 1.34098352 | 8.990479464 | 1.097865507 | 0.318166832 | 0.709264768 | -4.591762424 |
| **A_55_P2060253** | Gm12657 | 0.800514067 | -1.025802727 | 2.626830861 | 14.8553525 | 1.097850492 | 0.318172843 | 0.709264768 | -4.591766327 |
| **A_51_P297865** | Kctd6 | -0.484280792 | -1.589145494 | 0.620583911 | 8.675167202 | -1.097838583 | 0.318177612 | 0.709264768 | -4.591769422 |
| **A_52_P530291** | Pim1 | -0.377701785 | -1.239412663 | 0.484009094 | 8.082674325 | -1.097836856 | 0.318178303 | 0.709264768 | -4.591769871 |
| **A_66_P106090** | Rpa3 | 0.639657933 | -0.819741708 | 2.099057573 | 12.41691753 | 1.097801705 | 0.318192378 | 0.709264768 | -4.591779007 |
| **A_55_P1997604** | Pla2g4a | -0.168507608 | -0.552986741 | 0.215971525 | 9.831805964 | -1.097733923 | 0.318219519 | 0.709264768 | -4.591796622 |
| **A_55_P2145109** | Rpn2 | -0.546862083 | -1.79464841 | 0.700924243 | 14.94783198 | -1.097710797 | 0.318228779 | 0.709264768 | -4.591802633 |
| **A_55_P2074776** | Gm8894 | -0.233353757 | -0.765833894 | 0.299126381 | 16.38210174 | -1.097644561 | 0.318255304 | 0.709264768 | -4.591819846 |
| **A_55_P2180292** | Dmrtc1c2 | -0.608793694 | -1.998025048 | 0.780437659 | 8.074595324 | -1.097604591 | 0.318271311 | 0.709264768 | -4.591830234 |
| **A_55_P2091836** | Fnip1 | -0.237610844 | -0.779854662 | 0.304632974 | 10.28007771 | -1.097544135 | 0.318295524 | 0.709264768 | -4.591845945 |

| | | | | | | | | | |
|---|---|---|---|---|---|---|---|---|---|
| **A_55_P2156776** | NA | 0.106631323 | -0.136708479 | 0.349971126 | 7.984027235 | 1.097544135 | 0.318295524 | 0.709264768 | -4.591845945 |
| **A_66_P118368** | Catip | 0.135948716 | -0.174318266 | 0.446215697 | 8.513569657 | 1.097463014 | 0.318328015 | 0.709264768 | -4.591867026 |
| **A_52_P536293** | NA | 0.450557737 | -0.577740602 | 1.478856076 | 14.90158165 | 1.097442217 | 0.318336346 | 0.709264768 | -4.59187243 |
| **A_55_P2183493** | Spcs3 | -0.725446036 | -2.381134568 | 0.930242495 | 9.177213122 | -1.097429972 | 0.31834125 | 0.709264768 | -4.591875612 |
| **A_55_P2097808** | P4ha1 | -0.487771198 | -1.60112403 | 0.625581633 | 9.445269211 | -1.097320974 | 0.318384913 | 0.709264768 | -4.591903936 |
| **A_51_P249313** | Herc3 | -0.25314056 | -0.830942426 | 0.324661305 | 9.532429423 | -1.097319615 | 0.318385458 | 0.709264768 | -4.59190429 |
| **A_55_P1985428** | Atg16l2 | -0.428744044 | -1.407424475 | 0.549936387 | 9.224830749 | -1.097254931 | 0.318411372 | 0.709264768 | -4.591921098 |
| **A_51_P348433** | Rasal1 | -0.561239388 | -1.842393877 | 0.719915101 | 7.996251245 | -1.097228261 | 0.318422057 | 0.709264768 | -4.591928028 |
| **A_52_P336825** | Matr3 | 0.322902983 | -0.414282759 | 1.060088725 | 10.11809679 | 1.097098133 | 0.318474196 | 0.709264768 | -4.59196184 |
| **A_51_P430563** | Mtpn | -0.207165533 | -0.680193264 | 0.265862198 | 9.160253721 | -1.096935886 | 0.318539215 | 0.709264768 | -4.592003997 |
| **A_52_P547447** | Sirt2 | -0.165341394 | -0.542877708 | 0.212194921 | 10.18467398 | -1.096915589 | 0.31854735 | 0.709264768 | -4.59200927 |
| **A_51_P269134** | Sftpb | 0.869006118 | -1.11527526 | 2.853287496 | 9.958587879 | 1.09690729 | 0.318550676 | 0.709264768 | -4.592011427 |
| **A_51_P176071** | Cmtr2 | 0.171788731 | -0.220579765 | 0.564157226 | 9.196089935 | 1.096606681 | 0.318671175 | 0.709264768 | -4.592089526 |
| **A_55_P2007001** | Ctnnbip1 | -0.531281017 | -1.744812296 | 0.682250263 | 11.09345798 | -1.096537955 | 0.318698729 | 0.709264768 | -4.592107381 |
| **A_51_P494675** | Cotl1 | 0.261810224 | -0.336221996 | 0.859842445 | 14.01819376 | 1.096509415 | 0.318710172 | 0.709264768 | -4.592114795 |
| **A_55_P2012096** | Bmp8a | -0.941741654 | -3.09302505 | 1.209541743 | 8.767850734 | -1.096439064 | 0.318738381 | 0.709264768 | -4.592133071 |
| **A_51_P403636** | Smad7 | -0.251784738 | -0.826982451 | 0.323412974 | 10.77634443 | -1.096383769 | 0.318760554 | 0.709264768 | -4.592147435 |
| **A_65_P19112** | Zfp397 | 0.174330846 | -0.223957185 | 0.572618877 | 8.314481705 | 1.096294731 | 0.318796261 | 0.709264768 | -4.592170564 |
| **A_51_P428372** | Ppbp | -0.549172132 | -1.803965061 | 0.705620798 | 8.954857271 | -1.096192366 | 0.318837316 | 0.709264768 | -4.592197155 |
| **A_52_P539632** | Plxna3 | 0.221438716 | -0.284525089 | 0.727402522 | 8.2691853 | 1.096186502 | 0.318839668 | 0.709264768 | -4.592198678 |
| **A_55_P1988623** | Add3 | 0.317903562 | -0.408476201 | 1.044283325 | 10.8183341 | 1.096180327 | 0.318842145 | 0.709264768 | -4.592200282 |
| **A_55_P1954161** | Plagl2 | -0.11701707 | -0.384391924 | 0.150357784 | 8.12750673 | -1.096172755 | 0.318845182 | 0.709264768 | -4.592202249 |
| **A_55_P2020072** | NA | 0.731716052 | -0.940277055 | 2.403709159 | 12.61304347 | 1.096120867 | 0.318865994 | 0.709264768 | -4.592215727 |
| **A_51_P516452** | Eif3f | 0.482205219 | -0.619747465 | 1.584157903 | 12.99985316 | 1.096022085 | 0.31890562 | 0.709264768 | -4.592241385 |
| **A_52_P652316** | Mapre2 | -0.253134433 | -0.831618952 | 0.325350087 | 10.2668791 | -1.095998168 | 0.318915215 | 0.709264768 | -4.592247597 |
| **A_55_P2003251** | Magel2 | -0.448361247 | -1.4731117 | 0.576389207 | 8.339514073 | -1.09587317 | 0.318965364 | 0.709264768 | -4.592280063 |
| **A_51_P194306** | Lrrc1 | -0.343067037 | -1.127251617 | 0.441117542 | 10.00960283 | -1.095748891 | 0.319015232 | 0.709264768 | -4.592312341 |
| **A_52_P42269** | Fgf10 | 0.592338114 | -0.761753755 | 1.946429983 | 9.22491207 | 1.095650128 | 0.319054865 | 0.709264768 | -4.592053787 |
| **A_55_P2115567** | Slc26a1 | -0.164075311 | -0.539161525 | 0.211010903 | 7.852054648 | -1.095626379 | 0.319064396 | 0.709264768 | -4.592344159 |
| **A_51_P422751** | Clca1 | -0.591373477 | -1.943331144 | 0.760584191 | 8.005295109 | -1.095592611 | 0.319077949 | 0.709264768 | -4.592352929 |
| **A_51_P279437** | Mfsd2a | -1.074124071 | -3.529830151 | 1.38158201 | 8.308211077 | -1.095540429 | 0.319098892 | 0.709264768 | -4.592366481 |
| **A_55_P2075731** | NA | -0.28565317 | -0.938827773 | 0.367521432 | 11.82302458 | -1.095368138 | 0.319168051 | 0.709264768 | -4.592411223 |
| **A_51_P193863** | Gtf3c4 | -0.112029392 | -0.368201928 | 0.144143144 | 8.531574487 | -1.095342041 | 0.319178527 | 0.709264768 | -4.592418 |
| **A_51_P209807** | Ccnyl1 | -0.198267515 | -0.651674592 | 0.255139563 | 7.949558334 | -1.095250885 | 0.319215124 | 0.709264768 | -4.592441671 |
| **A_55_P1953899** | Mccc1 | 0.511632449 | -0.658504743 | 1.681769641 | 10.8958025 | 1.095145047 | 0.31925762 | 0.709264768 | -4.592469154 |
| **A_66_P139310** | Cog1 | -0.145096303 | -0.47705913 | 0.186866524 | 11.04543512 | -1.094755756 | 0.319413966 | 0.709264768 | -4.592570232 |
| **A_55_P2006668** | Pi4kb | -0.253670526 | -0.834076256 | 0.326735205 | 8.952674754 | -1.094683728 | 0.319442901 | 0.709264768 | -4.592588933 |
| **A_55_P2017600** | Crmp1 | 0.287582586 | -0.370529429 | 0.945694601 | 8.468389346 | 1.094493307 | 0.319519408 | 0.709264768 | -4.592638369 |
| **A_66_P130813** | Samd4 | -0.149524513 | -0.491713346 | 0.192664319 | 8.98059815 | -1.094452484 | 0.319535811 | 0.709264768 | -4.592648967 |
| **A_51_P214107** | Tbc1d20 | -0.453962972 | -1.492865565 | 0.58493962 | 9.617955214 | -1.094450053 | 0.319536789 | 0.709264768 | -4.592649598 |
| **A_51_P401504** | Col9a2 | 0.187832266 | -0.242080085 | 0.617744617 | 8.997698391 | 1.09431064 | 0.319592813 | 0.709264768 | -4.592685789 |
| **A_55_P2106351** | NA | -0.331042514 | -1.088837246 | 0.426752218 | 9.263610479 | -1.094164288 | 0.319651636 | 0.709264768 | -4.59272378 |
| **A_55_P2020050** | 6720475J | 0.308493366 | -0.397738744 | 1.014725475 | 8.824284284 | 1.094079083 | 0.319685886 | 0.709264768 | -4.592745897 |
| **A_55_P2133047** | Tollip | -0.290154204 | -0.954446277 | 0.374137869 | 8.422416997 | -1.094007107 | 0.319714821 | 0.709264768 | -4.59276458 |
| **A_55_P2273929** | Gm20558 | -0.300796957 | -0.989456466 | 0.387862553 | 8.293122701 | -1.094004818 | 0.319715741 | 0.709264768 | -4.592765174 |
| **A_55_P2076176** | NA | 0.50765965 | -0.654664941 | 1.669984241 | 15.5133081 | 1.093945195 | 0.319739712 | 0.709264768 | -4.59278065 |
| **A_55_P2073699** | Bnip3l | -0.168468398 | -0.55419292 | 0.217256124 | 12.13900009 | -1.093935062 | 0.319743786 | 0.709264768 | -4.59278328 |
| **A_52_P592951** | Ostc | 0.120703763 | -0.155665035 | 0.397072561 | 12.27073632 | 1.093911395 | 0.319753301 | 0.709264768 | -4.592789423 |

| | | | | | | | | | |
|---|---|---|---|---|---|---|---|---|---|
| **A_55_P2148534** | Nr1d2 | -0.15728606 | -0.517427999 | 0.202855879 | 8.955203579 | -1.093872945 | 0.319768761 | 0.709264768 | -4.592799403 |
| **A_55_P1981929** | Scg5 | -0.425561258 | -1.399985707 | 0.54886319 | 8.207708342 | -1.093866338 | 0.319771417 | 0.709264768 | -4.592801117 |
| **A_55_P2096340** | Rnf170 | -0.181086546 | -0.595795565 | 0.233622473 | 8.547232194 | -1.093687002 | 0.319843532 | 0.709264768 | -4.592847663 |
| **A_55_P2092968** | Traf3 | -0.152792854 | -0.502723398 | 0.197137689 | 10.96279348 | -1.093632828 | 0.319865319 | 0.709264768 | -4.592861723 |
| **A_55_P2063495** | Dph3 | 0.215671746 | -0.278298123 | 0.709641616 | 9.001301434 | 1.093561322 | 0.319894079 | 0.709264768 | -4.592880281 |
| **A_55_P2447879** | Asxl2 | -0.241659435 | -0.795193858 | 0.311874988 | 8.64684834 | -1.093476534 | 0.319928183 | 0.709264768 | -4.592902285 |
| **A_55_P2120843** | NA | -0.45653585 | -1.502273804 | 0.589202104 | 9.012693118 | -1.093458643 | 0.31993538 | 0.709264768 | -4.592906928 |
| **A_52_P342159** | Nfatc4 | 0.46923657 | -0.605690851 | 1.544163991 | 10.34026653 | 1.093359682 | 0.31997519 | 0.709264768 | -4.59293261 |
| **A_51_P183275** | Sympk | -0.230274086 | -0.757835804 | 0.297287631 | 10.85114839 | -1.093256681 | 0.320016629 | 0.709264768 | -4.592959339 |
| **A_51_P139320** | Pcbd1 | 0.478792289 | -0.618130402 | 1.575714979 | 12.06486228 | 1.093255 | 0.320017305 | 0.709264768 | -4.592959775 |
| **A_55_P2081116** | Fam89a | -0.428268037 | -1.409467599 | 0.552931525 | 8.02821475 | -1.093222754 | 0.32003028 | 0.709264768 | -4.592968143 |
| **A_55_P1989733** | Nfkbib | -0.468677261 | -1.542484619 | 0.605130097 | 10.36237509 | -1.093195545 | 0.320041228 | 0.709264768 | -4.592975204 |
| **A_66_P126485** | Dipk1a | -0.434129689 | -1.428867422 | 0.560608045 | 8.956073716 | -1.093103387 | 0.320078311 | 0.709264768 | -4.592999117 |
| **A_55_P1979432** | Prdx2 | 0.411701225 | -0.531667979 | 1.355070429 | 14.38911512 | 1.093077135 | 0.320088875 | 0.709264768 | -4.593005929 |
| **A_55_P2115192** | Sike1 | -0.102617156 | -0.337754312 | 0.132519999 | 9.656071482 | -1.093072734 | 0.320090646 | 0.709264768 | -4.593007072 |
| **A_52_P440962** | Cdc73 | -0.347536119 | -1.143914815 | 0.448842578 | 8.635452931 | -1.093026417 | 0.320109286 | 0.709264768 | -4.59301909 |
| **A_52_P597634** | Fzd1 | 0.525409283 | -0.678664158 | 1.729482725 | 11.56509108 | 1.092936954 | 0.320145291 | 0.709264768 | -4.593042303 |
| **A_55_P2036380** | NA | -0.121692441 | -0.40059053 | 0.157205648 | 17.00655676 | -1.092869765 | 0.320172334 | 0.709264768 | -4.593059736 |
| **A_55_P2049854** | Araf | -0.104167437 | -0.342905256 | 0.134570382 | 8.252099156 | -1.092851372 | 0.320179738 | 0.709264768 | -4.593064509 |
| **A_51_P256224** | Zfp768 | 0.318047057 | -0.410925585 | 1.0470197 | 12.01017386 | 1.092774361 | 0.320210737 | 0.709264768 | -4.59308449 |
| **A_52_P253915** | Cds2 | -0.713519249 | -2.34899909 | 0.921960592 | 8.9084229 | -1.092724912 | 0.320230643 | 0.709264768 | -4.59309732 |
| **A_55_P2022139** | Pura | -0.337003018 | -1.109469336 | 0.435463301 | 11.34999834 | -1.092709159 | 0.320236985 | 0.709264768 | -4.593101407 |
| **A_52_P559817** | Fbln7 | -2.495510959 | -8.215840974 | 3.224819055 | 9.937450822 | -1.092669303 | 0.32025303 | 0.709264768 | -4.593111748 |
| **A_55_P2156731** | H2-Eb1 | 1.080936264 | -1.396957338 | 3.558829867 | 10.33235012 | 1.09261656 | 0.320274265 | 0.709264768 | -4.593125431 |
| **A_52_P177490** | Cdx2 | -0.827547573 | -2.724666591 | 1.069571445 | 8.082033109 | -1.092568656 | 0.320293552 | 0.709264768 | -4.59313786 |
| **A_51_P508115** | Trank1 | -0.275047084 | -0.905603113 | 0.355508945 | 7.927561156 | -1.092530848 | 0.320308776 | 0.709264768 | -4.593147668 |
| **A_52_P669964** | Cchcr1 | 0.317829793 | -0.410875686 | 1.046535272 | 8.593390416 | 1.092428233 | 0.320350096 | 0.709264768 | -4.59317429 |
| **A_51_P181565** | Hbegf | -0.324333778 | -1.067957827 | 0.41929027 | 9.581850042 | -1.092418636 | 0.320353961 | 0.709264768 | -4.593176779 |
| **A_55_P2066180** | Hid1 | 0.535261581 | -0.692196918 | 1.76272008 | 10.07778853 | 1.092218683 | 0.320434491 | 0.709264768 | -4.593228651 |
| **A_55_P2093221** | Rnf111 | -0.140582773 | -0.462977348 | 0.181811803 | 11.18249339 | -1.092181268 | 0.320449561 | 0.709264768 | -4.593238356 |
| **A_55_P2090995** | Ebna1bp2 | 0.107149749 | -0.138577271 | 0.352876769 | 9.424494693 | 1.092165847 | 0.320455773 | 0.709264768 | -4.593242356 |
| **A_51_P244213** | Hapln3 | -0.601652809 | -1.981510652 | 0.778205034 | 8.478117761 | -1.092098845 | 0.320482763 | 0.709264768 | -4.593259737 |
| **A_51_P370717** | Gsc | -0.339779343 | -1.119094031 | 0.439535345 | 7.963266194 | -1.092029715 | 0.320510612 | 0.709264768 | -4.593277669 |
| **A_55_P2109599** | NA | 0.453252848 | -0.586388447 | 1.492894143 | 12.50883849 | 1.091961598 | 0.320538055 | 0.709264768 | -4.593295337 |
| **A_55_P2144746** | Apip | 0.410449384 | -0.531105975 | 1.352004742 | 10.52289588 | 1.091852804 | 0.32058189 | 0.709264768 | -4.593323556 |
| **A_55_P2174987** | Creb3l3 | -0.656474593 | -2.162557773 | 0.849608587 | 8.204966478 | -1.091740219 | 0.320627258 | 0.709264768 | -4.593352757 |
| **A_55_P2011410** | Pdzk1 | -1.270947425 | -4.186757579 | 1.644862729 | 8.498406659 | -1.091738654 | 0.320627889 | 0.709264768 | -4.593353163 |
| **A_52_P441036** | Col23a1 | 0.38349235 | -0.496383088 | 1.263367787 | 8.792914843 | 1.091656126 | 0.320661149 | 0.709264768 | -4.593374568 |
| **A_51_P461877** | Thbs2 | -0.417296621 | -1.37476871 | 0.540175468 | 8.872582995 | -1.091613986 | 0.320678133 | 0.709264768 | -4.593385497 |
| **A_55_P2022181** | Arvcf | -0.406605304 | -1.339551644 | 0.526341037 | 8.526702661 | -1.091608023 | 0.320680536 | 0.709264768 | -4.593387043 |
| **A_51_P259555** | Gpatch3 | -0.225468964 | -0.742833386 | 0.291895459 | 8.500940743 | -1.091542259 | 0.320707043 | 0.709264768 | -4.593404099 |
| **A_52_P283524** | Rcan3 | -0.123135145 | -0.405689789 | 0.1594195 | 8.568986467 | -1.091515533 | 0.320717815 | 0.709264768 | -4.593411031 |
| **A_55_P1957227** | Las1l | -0.744274305 | -2.452330107 | 0.963781497 | 9.757412122 | -1.091393384 | 0.320767055 | 0.709264768 | -4.593442709 |
| **A_55_P2078955** | Aqp8 | -0.580565366 | -1.913010138 | 0.751879406 | 7.962680988 | -1.091320347 | 0.320796501 | 0.709264768 | -4.593461649 |
| **A_52_P69998** | Kdelr2 | -0.603514558 | -1.988775019 | 0.781745902 | 10.61603813 | -1.091205781 | 0.320842693 | 0.709264768 | -4.593491359 |
| **A_55_P2052769** | Mast2 | -0.395767192 | -1.304192989 | 0.512658605 | 11.46781983 | -1.091190752 | 0.320848753 | 0.709264768 | -4.593495256 |
| **A_55_P2059382** | Arl6 | 0.457831157 | -0.593072165 | 1.508734479 | 9.622597115 | 1.091171278 | 0.320856606 | 0.709264768 | -4.593500306 |
| **A_55_P1982076** | Emsy | 0.389360631 | -0.504383691 | 1.283104954 | 12.42234883 | 1.091161633 | 0.320860495 | 0.709264768 | -4.593502807 |

| | | | | | | | | | |
|---|---|---|---|---|---|---|---|---|---|
| **A_51_P118168** | Zbtb43 | -0.316892442 | -1.044310146 | 0.410525263 | 9.596095668 | -1.09113468 | 0.320871364 | 0.709264768 | -4.592928235 |
| **A_55_P2187225** | NA | 0.114988912 | -0.148982377 | 0.378960201 | 10.18843328 | 1.091062467 | 0.320900484 | 0.709264768 | -4.593528522 |
| **A_51_P391934** | Cpb1 | 0.413178119 | -0.535396412 | 1.36175265 | 8.67594082 | 1.09097852 | 0.32093434 | 0.709264768 | -4.593550289 |
| **A_55_P1964193** | Map1b | -0.115721445 | -0.381402665 | 0.149959776 | 8.196399711 | -1.090946203 | 0.320947374 | 0.709264768 | -4.593558669 |
| **A_66_P123824** | Sorl1 | 0.492222899 | -0.638010073 | 1.622455872 | 8.987839873 | 1.090797633 | 0.321007302 | 0.709264768 | -4.593597191 |
| **A_55_P1976898** | Pnp2 | -0.599264051 | -1.975373089 | 0.776844988 | 8.193585399 | -1.090726146 | 0.32103614 | 0.709264768 | -4.593615727 |
| **A_55_P2042778** | Pde5a | 0.19224434 | -0.249272039 | 0.63376072 | 8.470689422 | 1.090578894 | 0.32109555 | 0.709264768 | -4.593653904 |
| **A_51_P156857** | Smim3 | 0.587646363 | -0.761991861 | 1.937284588 | 11.04569417 | 1.090558655 | 0.321103716 | 0.709264768 | -4.593659151 |
| **A_52_P120740** | 0610030E | -0.121506945 | -0.400602523 | 0.157588634 | 8.533151189 | -1.090431755 | 0.321154923 | 0.709264768 | -4.593692051 |
| **A_55_P2047753** | Tec | -0.352055233 | -1.160786969 | 0.456676504 | 7.971663824 | -1.090326741 | 0.321197304 | 0.709264768 | -4.593719275 |
| **A_55_P2064074** | Vps37d | 0.566211652 | -0.734492658 | 1.866915961 | 9.89421219 | 1.090311439 | 0.32120348 | 0.709264768 | -4.593723242 |
| **A_55_P2184905** | Rpap2 | -0.10038826 | -0.331009805 | 0.130233285 | 8.616278413 | -1.090268321 | 0.321220883 | 0.709264768 | -4.59373442 |
| **A_51_P346923** | P3h1 | -0.192705851 | -0.635473991 | 0.25006229 | 8.369948054 | -1.090106382 | 0.32128625 | 0.709264768 | -4.593776399 |
| **A_55_P2015485** | Cfap45 | 0.700285444 | -0.908812758 | 2.309383645 | 10.2658046 | 1.090041155 | 0.321312583 | 0.709264768 | -4.593793306 |
| **A_51_P224983** | Cln8 | -0.552160289 | -1.820978872 | 0.716658294 | 10.26703621 | -1.089973613 | 0.321339852 | 0.709264768 | -4.593810814 |
| **A_55_P2150555** | Pcgf5 | -0.989432884 | -3.263198965 | 1.284333197 | 9.962697712 | -1.089910664 | 0.321365267 | 0.709264768 | -4.593827131 |
| **A_52_P660945** | Ctsf | -0.334758749 | -1.10408948 | 0.434571982 | 8.740355414 | -1.089856204 | 0.321387258 | 0.709264768 | -4.593841247 |
| **A_55_P2067112** | Gm4924 | 0.169609068 | -0.220188492 | 0.559406629 | 10.20649273 | 1.089833863 | 0.321396279 | 0.709264768 | -4.593847038 |
| **A_51_P312336** | Slc14a1 | -0.561576938 | -1.852329923 | 0.729176048 | 8.001919268 | -1.089723889 | 0.32144069 | 0.709264768 | -4.593875542 |
| **A_51_P297694** | Prpf6 | -0.201907712 | -0.665995178 | 0.262179754 | 12.02228812 | -1.089691126 | 0.321453921 | 0.709264768 | -4.593884033 |
| **A_55_P2173073** | Zfp931 | -0.097583155 | -0.321879789 | 0.126713479 | 8.62358282 | -1.08968869 | 0.321454905 | 0.709264768 | -4.593884665 |
| **A_55_P2140835** | NA | -0.13482045 | -0.444760793 | 0.175119893 | 8.671141766 | -1.089501942 | 0.321530334 | 0.709264768 | -4.593933065 |
| **A_51_P106259** | Camk1 | -0.093424101 | -0.308205528 | 0.121357326 | 8.28880486 | -1.089463163 | 0.321545999 | 0.709264768 | -4.59333063 |
| **A_51_P140742** | Islr | 0.699084187 | -0.908106633 | 2.306275007 | 10.77068955 | 1.089462739 | 0.321546171 | 0.709264768 | -4.593943225 |
| **A_55_P2065834** | Dhx8 | 0.135025842 | -0.175448476 | 0.445500161 | 10.50374176 | 1.089285089 | 0.321617942 | 0.709264768 | -4.593989264 |
| **A_51_P335981** | Narf | 0.197073146 | -0.256104595 | 0.650250887 | 8.726618906 | 1.089203993 | 0.321650709 | 0.709264768 | -4.594010279 |
| **A_55_P2148171** | Ncmap | -0.765120433 | -2.524560379 | 0.994319512 | 8.483628869 | -1.089195149 | 0.321654283 | 0.709264768 | -4.594012571 |
| **A_55_P1991896** | Ceacam5 | -2.747684396 | -9.0663264 | 3.570957609 | 9.320133854 | -1.089164413 | 0.321666703 | 0.709264768 | -4.594020536 |
| **A_55_P2013011** | NA | -0.475304094 | -1.568384268 | 0.617776081 | 8.350191237 | -1.089105329 | 0.321690579 | 0.709264768 | -4.594035847 |
| **A_55_P2138095** | NA | 0.3131248 | -0.406988417 | 1.033238017 | 16.45779029 | 1.089098172 | 0.321693471 | 0.709264768 | -4.594037701 |
| **A_52_P39157** | Bmi1 | 0.213759239 | -0.27789618 | 0.705414657 | 9.435602191 | 1.088966225 | 0.321746797 | 0.709264768 | -4.594071892 |
| **A_51_P501018** | Nek2 | 0.354410007 | -0.460764246 | 1.169584259 | 8.706467027 | 1.088944805 | 0.321755455 | 0.709264768 | -4.594077442 |
| **A_55_P2067727** | Mxra7 | -0.119279776 | -0.393637367 | 0.155077815 | 9.312210077 | -1.088930541 | 0.32176122 | 0.709264768 | -4.594081138 |
| **A_52_P46419** | Grk6 | 0.202931997 | -0.263853464 | 0.669717458 | 9.468782715 | 1.08888886 | 0.321778068 | 0.709264768 | -4.594091938 |
| **A_55_P1971010** | Gzme | -1.535991445 | -5.069251264 | 1.997268375 | 8.531132901 | -1.088838748 | 0.321798324 | 0.709264768 | -4.594104923 |
| **A_52_P382914** | Trip6 | 0.456927877 | -0.594271493 | 1.508127248 | 12.18565566 | 1.088711746 | 0.321849666 | 0.709264768 | -4.594137829 |
| **A_52_P479961** | NA | 0.35011454 | -0.455401459 | 1.155630538 | 16.05154715 | 1.088645098 | 0.321876611 | 0.709264768 | -4.594155097 |
| **A_55_P2078008** | Ddx46 | 0.257550834 | -0.335063246 | 0.850164915 | 9.844364658 | 1.088532351 | 0.321922199 | 0.709264768 | -4.594184308 |
| **A_55_P2128929** | Cc2d2a | 0.721455371 | -0.938790415 | 2.381701157 | 10.23132929 | 1.088397235 | 0.321976839 | 0.709264768 | -4.594219313 |
| **A_55_P2100155** | Mapt | 0.380933316 | -0.495710468 | 1.2575771 | 9.431799699 | 1.088368955 | 0.321988276 | 0.709264768 | -4.594226639 |
| **A_51_P412508** | Irx4 | -0.522899949 | -1.726312696 | 0.680512798 | 8.055711388 | -1.088314305 | 0.322010379 | 0.709264768 | -4.594240797 |
| **A_55_P2060072** | Clk2 | -0.192950969 | -0.637130875 | 0.251228938 | 10.63823581 | -1.088023813 | 0.322127889 | 0.709264768 | -4.594316048 |
| **A_55_P2008347** | Prelid1 | -0.181992285 | -0.600980575 | 0.236996005 | 14.73908331 | -1.087931214 | 0.322165354 | 0.709264768 | -4.594340034 |
| **A_55_P2067747** | Psg21 | -2.894921169 | -9.560113163 | 3.770270826 | 9.429276706 | -1.087863545 | 0.322192735 | 0.709264768 | -4.594357562 |
| **A_51_P140641** | Eva1a | -0.624847063 | -2.063612025 | 0.8139179 | 11.3497695 | -1.087762857 | 0.322233481 | 0.709264768 | -4.594383642 |
| **A_55_P1981291** | Spink8 | -0.528103699 | -1.744167504 | 0.687960107 | 7.939894809 | -1.087710178 | 0.3222548 | 0.709264768 | -4.594397286 |
| **A_55_P1984496** | Usp48 | -0.218442253 | -0.721594483 | 0.284709976 | 10.29808723 | -1.087395651 | 0.322382115 | 0.709264768 | -4.594478746 |
| **A_55_P2125766** | Ola1 | 0.142598225 | -0.18587851 | 0.47107496 | 8.969781874 | 1.087326195 | 0.322410235 | 0.709264768 | -4.594496733 |

| | | | | | | | | | |
|---|---|---|---|---|---|---|---|---|---|
| **A_51_P391805** | Dkkl1 | -1.30215432 | -4.302158515 | 1.697849874 | 8.633350881 | -1.087153679 | 0.32248009 | 0.709264768 | -4.594541408 |
| **A_55_P2040357** | Bola1 | 0.186938079 | -0.243752019 | 0.617628177 | 9.052709339 | 1.087134388 | 0.322487902 | 0.709264768 | -4.594546404 |
| **A_55_P2166282** | NA | -2.006455005 | -6.629379849 | 2.616469839 | 10.69395232 | -1.087083584 | 0.322508476 | 0.709264768 | -4.59455956 |
| **A_51_P111612** | Arrdc4 | -0.43371725 | -1.433015751 | 0.565581252 | 9.44718434 | -1.087080748 | 0.322509624 | 0.709264768 | -4.592284367 |
| **A_52_P289692** | NA | -0.307098896 | -1.014693765 | 0.400495972 | 10.23422449 | -1.087035999 | 0.322527748 | 0.709264768 | -4.594571882 |
| **A_51_P249848** | Dsg2 | -0.899978464 | -2.97374574 | 1.173788812 | 8.560404913 | -1.086982233 | 0.322549523 | 0.709264768 | -4.594585804 |
| **A_55_P1961870** | NA | 0.606589227 | -0.791200097 | 2.00437855 | 14.84578696 | 1.086934364 | 0.322568912 | 0.709264768 | -4.594598199 |
| **A_55_P2101526** | L1td1 | -0.759579344 | -2.509937039 | 0.990778351 | 8.53212029 | -1.086917756 | 0.322575639 | 0.709264768 | -4.5946025 |
| **A_52_P313217** | Adgrd1 | -0.416041911 | -1.374878482 | 0.542794661 | 8.158513681 | -1.086783003 | 0.322630225 | 0.709264768 | -4.594637391 |
| **A_55_P2167620** | NA | 0.412078999 | -0.537697635 | 1.361855632 | 9.210814934 | 1.086699197 | 0.322664178 | 0.709264768 | -4.59465909 |
| **A_55_P1983468** | Cflar | 0.10982106 | -0.143301098 | 0.362943219 | 8.385243653 | 1.086690356 | 0.322667759 | 0.709264768 | -4.594661379 |
| **A_55_P2128181** | NA | 0.193071748 | -0.25198654 | 0.638130036 | 9.764811581 | 1.086556166 | 0.322722131 | 0.709264768 | -4.594696122 |
| **A_55_P2179413** | Lgals3bp | -1.250048211 | -4.131731943 | 1.63163552 | 9.727666014 | -1.08650266 | 0.322743812 | 0.709264768 | -4.594709974 |
| **A_55_P2182358** | Arid3a | -0.737762603 | -2.43869368 | 0.963168474 | 10.15885993 | -1.08637626 | 0.322795037 | 0.709264768 | -4.594742698 |
| **A_55_P2324976** | 5033406C | -0.107699444 | -0.356003825 | 0.140604936 | 8.431570246 | -1.08637416 | 0.322795888 | 0.709264768 | -4.594743242 |
| **A_55_P2003981** | NA | -0.250874029 | -0.82929448 | 0.327546423 | 7.773921749 | -1.086331592 | 0.322813141 | 0.709264768 | -4.594754262 |
| **A_55_P2157632** | 2510039C | -0.303290179 | -1.002644509 | 0.39606415 | 12.31724219 | -1.08620407 | 0.32286483 | 0.709264768 | -4.594787275 |
| **A_51_P109144** | Grtp1 | -0.709604493 | -2.345882252 | 0.926673266 | 8.53780901 | -1.086199691 | 0.322866605 | 0.709264768 | -4.593013068 |
| **A_55_P2158376** | NA | 0.50215856 | -0.655777497 | 1.660094616 | 14.02263799 | 1.086192096 | 0.322869683 | 0.709264768 | -4.594790374 |
| **A_51_P275642** | NA | -0.391512253 | -1.294335217 | 0.511310712 | 8.009991221 | -1.086158253 | 0.322883402 | 0.709264768 | -4.594799135 |
| **A_52_P72237** | Actg1 | -0.171725557 | -0.567728611 | 0.224277497 | 12.92625215 | -1.08614234 | 0.322889853 | 0.709264768 | -4.594803255 |
| **A_52_P245415** | AW55491ɛ | 0.30797071 | -0.402255968 | 1.018197389 | 8.988000698 | 1.086082405 | 0.322914151 | 0.709264768 | -4.594818769 |
| **A_55_P1981734** | Tut7 | -0.208714431 | -0.69006733 | 0.272638467 | 12.24399786 | -1.086023525 | 0.322938022 | 0.709264768 | -4.594834011 |
| **A_55_P2148001** | 6820431F | -0.205149947 | -0.678291609 | 0.267991716 | 10.69117902 | -1.086001843 | 0.322946813 | 0.709264768 | -4.594839623 |
| **A_51_P208769** | Metrn | 0.413402897 | -0.54015893 | 1.366964724 | 8.987148785 | 1.085862923 | 0.323003142 | 0.709264768 | -4.594875582 |
| **A_52_P160955** | Sox11 | 0.28432264 | -0.371507111 | 0.940152392 | 9.224876207 | 1.085852098 | 0.323007531 | 0.709264768 | -4.594878384 |
| **A_55_P2179299** | Ceacam1: | -2.889596462 | -9.555264616 | 3.776071692 | 9.439952948 | -1.08578504 | 0.323034725 | 0.709264768 | -4.594895741 |
| **A_55_P2102182** | NA | -0.356627751 | -1.179303326 | 0.466047825 | 8.14201476 | -1.085767596 | 0.323041799 | 0.709264768 | -4.594900256 |
| **A_55_P2095039** | Ncmap | -1.048670267 | -3.468053138 | 1.370712604 | 9.449371232 | -1.085637156 | 0.323094702 | 0.709264768 | -4.594934017 |
| **A_51_P413111** | Ppm1l | 0.473324262 | -0.618727741 | 1.565376264 | 10.15249072 | 1.085589895 | 0.323113872 | 0.709264768 | -4.594946249 |
| **A_55_P2050083** | Gak | -0.384861734 | -1.27282537 | 0.503101903 | 11.09117929 | -1.085575158 | 0.32311985 | 0.709264768 | -4.594950063 |
| **A_51_P497113** | Igf2bp1 | -0.293381196 | -0.970280295 | 0.383517902 | 9.137287283 | -1.085572129 | 0.323121078 | 0.709264768 | -4.594950847 |
| **A_55_P2020286** | Tmem179 | -0.489462211 | -1.61884235 | 0.639917928 | 8.034424766 | -1.085498848 | 0.323150805 | 0.709264768 | -4.594969813 |
| **A_51_P290074** | Fabp7 | 0.392407857 | -0.513075078 | 1.297890793 | 8.821909943 | 1.08544487 | 0.323172702 | 0.709264768 | -4.594983783 |
| **A_51_P248865** | Foxf2 | 0.806740193 | -1.054820441 | 2.668300827 | 10.13031923 | 1.085442579 | 0.323173631 | 0.709264768 | -4.594984376 |
| **A_52_P168338** | Fundc2 | 0.16697031 | -0.218324902 | 0.552265522 | 8.119497394 | 1.085415419 | 0.32318465 | 0.709264768 | -4.594991405 |
| **A_55_P1985422** | NA | -0.935715055 | -3.09498763 | 1.22355752 | 13.2094652 | -1.085391694 | 0.323194275 | 0.709264768 | -4.594997545 |
| **A_55_P2061428** | Fads6 | -0.408596891 | -1.35158227 | 0.534388487 | 7.921333367 | -1.085276614 | 0.323240967 | 0.709264768 | -4.595027327 |
| **A_55_P2113081** | Tmem132 | 0.581810811 | -0.761073508 | 1.924695129 | 13.29979439 | 1.085159383 | 0.323288537 | 0.709264768 | -4.595057664 |
| **A_52_P582394** | Mrps11 | 0.263527836 | -0.344741084 | 0.871796756 | 9.414683433 | 1.085128612 | 0.323301024 | 0.709264768 | -4.595065627 |
| **A_66_P103301** | Dicer1 | 0.369341932 | -0.483181172 | 1.221865036 | 9.891258785 | 1.085107598 | 0.323309552 | 0.709264768 | -4.595071064 |
| **A_55_P2064741** | Nmb | -0.470349789 | -1.556035878 | 0.615336299 | 10.29556175 | -1.085093161 | 0.323315411 | 0.709264768 | -4.5950748 |
| **A_51_P324814** | Krt18 | -0.801864215 | -2.652870935 | 1.049142506 | 15.1083966 | -1.085033588 | 0.323339589 | 0.709264768 | -4.595090216 |
| **A_51_P349825** | 1700021F | -0.309457484 | -1.023816324 | 0.404901357 | 11.76942771 | -1.085012925 | 0.323347976 | 0.709264768 | -4.595095563 |
| **A_55_P1965629** | Hcfc1 | 0.1969059 | -0.257728785 | 0.651540584 | 11.5874471 | 1.084792087 | 0.323437617 | 0.709264768 | -4.595152705 |
| **A_66_P129111** | NA | 0.123290209 | -0.161376911 | 0.407957328 | 7.90211345 | 1.084779889 | 0.323442569 | 0.709264768 | -4.595155861 |
| **A_51_P232399** | Acy3 | 0.431795319 | -0.565280121 | 1.428870759 | 9.378979143 | 1.084676564 | 0.323484519 | 0.709264768 | -4.595182595 |
| **A_55_P2013730** | Islr2 | -0.833406551 | -2.758019995 | 1.091206892 | 8.440087864 | -1.084585367 | 0.323521548 | 0.709264768 | -4.595206191 |

| | | | | | | | | | |
|---|---|---|---|---|---|---|---|---|---|
| **A_51_P233027** | Rmdn3 | -0.338609559 | -1.120606406 | 0.443387288 | 8.789270976 | -1.084537461 | 0.323541001 | 0.709264768 | -4.595218585 |
| **A_51_P152404** | Ikbip | 0.12900756 | -0.1689347 | 0.42694982 | 10.07893935 | 1.084509595 | 0.323552316 | 0.709264768 | -4.595225794 |
| **A_55_P2048176** | NA | -0.233095708 | -0.771483726 | 0.30529231 | 8.117167887 | -1.084399319 | 0.3235971 | 0.709264768 | -4.595254324 |
| **A_51_P211573** | Wdr47 | -0.189284646 | -0.626498708 | 0.247929416 | 9.179622294 | -1.084355313 | 0.323614973 | 0.709264768 | -4.595265709 |
| **A_55_P2103756** | Hic2 | -0.48242141 | -1.596746136 | 0.631903316 | 8.500868948 | -1.084339185 | 0.323621524 | 0.709264768 | -4.595269881 |
| **A_55_P2112030** | Tmem145 | -0.334964671 | -1.108738793 | 0.438809451 | 7.876333056 | -1.084264275 | 0.32365195 | 0.709264768 | -4.59528926 |
| **A_52_P476560** | Tpcn1 | -0.712410516 | -2.35835772 | 0.933536688 | 11.44751107 | -1.084088572 | 0.323723326 | 0.709264768 | -4.595334712 |
| **A_52_P653937** | Rpl21 | 0.257767735 | -0.337781293 | 0.853316762 | 8.453449224 | 1.084080119 | 0.32372676 | 0.709264768 | -4.595336899 |
| **A_55_P2135331** | Evl | 0.754402476 | -0.98865139 | 2.497456342 | 13.55001823 | 1.084033348 | 0.323745762 | 0.709264768 | -4.595348998 |
| **A_55_P1979708** | Ubap1 | -0.351850033 | -1.16482926 | 0.461129194 | 10.20690039 | -1.083998031 | 0.323760111 | 0.709264768 | -4.595358133 |
| **A_52_P339742** | Cyb5r3 | -0.439082409 | -1.453637152 | 0.575472334 | 8.309799692 | -1.083979073 | 0.323767814 | 0.709264768 | -4.595363037 |
| **A_55_P2004833** | NA | 0.676956029 | -0.887255987 | 2.241168045 | 13.60902512 | 1.083964822 | 0.323773605 | 0.709264768 | -4.595366723 |
| **A_55_P2005920** | NA | 0.464226478 | -0.60848582 | 1.536938776 | 15.178223 | 1.08391941 | 0.323792057 | 0.709264768 | -4.59537847 |
| **A_51_P234627** | Nubpl | 0.171074586 | -0.224243705 | 0.566392878 | 9.560229814 | 1.083899297 | 0.323800229 | 0.709264768 | -4.595383672 |
| **A_52_P93066** | Klhl42 | -0.097658253 | -0.323340245 | 0.128023739 | 8.848594585 | -1.083833048 | 0.323827151 | 0.709264768 | -4.595400808 |
| **A_55_P2024689** | Rsbn1 | 0.222331652 | -0.291468157 | 0.73613146 | 9.017021034 | 1.083821349 | 0.323831905 | 0.709264768 | -4.595403834 |
| **A_51_P489337** | Becn1 | -0.11498326 | -0.380723291 | 0.150756772 | 8.810733358 | -1.083747179 | 0.323862047 | 0.709264768 | -4.595423018 |
| **A_52_P579933** | Slc16a6 | -0.473772606 | -1.568740369 | 0.621195157 | 8.043241153 | -1.083724666 | 0.323871197 | 0.709264768 | -4.595428841 |
| **A_51_P191463** | H2-M2 | -0.798613987 | -2.644413182 | 1.047185207 | 9.347133692 | -1.08368436 | 0.323887578 | 0.709264768 | -4.595439266 |
| **A_51_P453111** | Hexb | -0.67825149 | -2.245898446 | 0.889395466 | 10.90282119 | -1.08365949 | 0.323897687 | 0.709264768 | -4.595445699 |
| **A_55_P2008512** | B4galt7 | -0.373239076 | -1.236022462 | 0.489544309 | 8.390718361 | -1.083516871 | 0.323955658 | 0.709264768 | -4.595482585 |
| **A_55_P2184414** | Zfp330 | 0.154350551 | -0.202448718 | 0.511149819 | 11.53944201 | 1.083514114 | 0.32395678 | 0.709264768 | -4.595483298 |
| **A_55_P2111459** | Trpv2 | -0.943986385 | -3.126182872 | 1.238210101 | 9.371366789 | -1.083483306 | 0.323969304 | 0.709264768 | -4.595491266 |
| **A_55_P1981499** | Rpl10a | 0.417961153 | -0.548285269 | 1.384207575 | 16.56911439 | 1.083423785 | 0.323993501 | 0.709264768 | -4.595506659 |
| **A_51_P270733** | Syngr1 | -0.875669869 | -2.900163271 | 1.148823532 | 9.347271591 | -1.083363966 | 0.324017821 | 0.709264768 | -4.595522129 |
| **A_55_P2079561** | Lilra6 | -0.159903155 | -0.529617071 | 0.209810761 | 8.330618111 | -1.083282132 | 0.324051095 | 0.709264768 | -4.595543292 |
| **A_52_P40241** | Zfp746 | 0.246329657 | -0.32325957 | 0.815918883 | 9.936789062 | 1.083191593 | 0.32408791 | 0.709264768 | -4.595566706 |
| **A_55_P2067752** | Psg26 | -2.889520969 | -9.57136581 | 3.792323873 | 9.403718193 | -1.083128066 | 0.324113745 | 0.709264768 | -4.595583134 |
| **A_51_P509760** | Actr1a | -0.207911187 | -0.68874101 | 0.272918635 | 12.745634 | -1.083020826 | 0.324157359 | 0.709264768 | -4.595610864 |
| **A_55_P2125322** | NA | 0.354714327 | -0.46565156 | 1.175080214 | 16.06192469 | 1.08298261 | 0.324172903 | 0.709264768 | -4.595620746 |
| **A_66_P123186** | Mapk1ip1 | -0.433404999 | -1.435835276 | 0.569025279 | 9.292051159 | -1.082904318 | 0.324204749 | 0.709264768 | -4.595640991 |
| **A_55_P1957593** | Ino80b | 0.32303538 | -0.424192289 | 1.070263049 | 9.951168477 | 1.082798115 | 0.324247952 | 0.709264768 | -4.595668451 |
| **A_55_P1962339** | Trim41 | -0.653442389 | -2.164995566 | 0.858110788 | 13.78441909 | -1.082765025 | 0.324261414 | 0.709264768 | -4.595677007 |
| **A_55_P2149122** | Farp2 | -0.44744548 | -1.482570946 | 0.587679986 | 8.928585723 | -1.082673427 | 0.324298681 | 0.709264768 | -4.59570069 |
| **A_52_P477369** | Csnk1d | -0.169744839 | -0.562466229 | 0.222976552 | 8.195539174 | -1.082585899 | 0.324334295 | 0.709264768 | -4.59572332 |
| **A_55_P2142668** | NA | -1.094814927 | -3.628194034 | 1.43856418 | 8.857963578 | -1.082407695 | 0.324406815 | 0.709264768 | -4.595769392 |
| **A_55_P1980432** | Zfp488 | -0.150291771 | -0.498071989 | 0.197488448 | 9.443060247 | -1.082382274 | 0.324417162 | 0.709264768 | -4.595775964 |
| **A_51_P428742** | Pus7l | 0.093705371 | -0.123136522 | 0.310547265 | 8.303969835 | 1.082359768 | 0.324426322 | 0.709264768 | -4.595781782 |
| **A_52_P225570** | Parva | -0.096905451 | -0.32118576 | 0.127374857 | 10.60905749 | -1.082199701 | 0.324491476 | 0.709264768 | -4.595823162 |
| **A_55_P1961168** | NA | 0.491800013 | -0.646436901 | 1.630036927 | 11.86296063 | 1.082196726 | 0.324492686 | 0.709264768 | -4.595823931 |
| **A_55_P2086631** | NA | 0.524935525 | -0.690063331 | 1.739934381 | 14.97271189 | 1.082132502 | 0.324518832 | 0.709264768 | -4.595840533 |
| **A_55_P2079430** | NA | 0.643218938 | -0.845573492 | 2.132011369 | 10.27402959 | 1.082118985 | 0.324524335 | 0.709264768 | -4.595844027 |
| **A_55_P2131128** | Helq | 0.124766113 | -0.164020347 | 0.413552572 | 8.451889011 | 1.08210689 | 0.324529258 | 0.709264768 | -4.595847154 |
| **A_55_P1978441** | Kif3a | -0.110233661 | -0.365425324 | 0.144958002 | 10.43226165 | -1.081927335 | 0.324602365 | 0.709264768 | -4.595893567 |
| **A_55_P2154977** | Rpap3 | 0.245048332 | -0.32231627 | 0.812412934 | 11.7046632 | 1.081782271 | 0.324661439 | 0.709264768 | -4.595931063 |
| **A_55_P2001871** | Uros | -0.207418733 | -0.687666785 | 0.272829319 | 10.10826237 | -1.081764462 | 0.324668692 | 0.709264768 | -4.595935666 |
| **A_55_P2349265** | Ptk2 | -0.297402686 | -0.986009132 | 0.391203761 | 9.028801296 | -1.081743134 | 0.324677378 | 0.709264768 | -4.595941178 |
| **A_55_P2158351** | Rpl18a | 0.32860092 | -0.432391218 | 1.089593059 | 16.43279438 | 1.081530948 | 0.324763805 | 0.709264768 | -4.595996019 |

| | | | | | | | | | |
|---|---|---|---|---|---|---|---|---|---|
| **A_55_P2004452** | Tceal8 | 0.138468474 | -0.182227223 | 0.459164171 | 9.273107355 | 1.081454168 | 0.324795084 | 0.709264768 | -4.596015862 |
| **A_55_P2052799** | Cask | 0.527888755 | -0.694757025 | 1.750534535 | 10.44308584 | 1.081414288 | 0.324811331 | 0.709264768 | -4.596026169 |
| **A_52_P384314** | Tmem198 | 0.350331863 | -0.461076929 | 1.161740656 | 9.562092151 | 1.081409736 | 0.324813186 | 0.709264768 | -4.596027345 |
| **A_55_P2175579** | Olfr1466 | -1.635866408 | -5.42486715 | 2.153134333 | 10.01969095 | -1.08136794 | 0.324830214 | 0.709264768 | -4.596038147 |
| **A_55_P1990067** | NA | 0.460283845 | -0.605846268 | 1.526413957 | 13.57384146 | 1.081348962 | 0.324837947 | 0.709264768 | -4.596043052 |
| **A_55_P1961235** | Rpl23 | 0.124582487 | -0.163981213 | 0.413146186 | 16.42384699 | 1.081348401 | 0.324838175 | 0.709264768 | -4.596043196 |
| **A_55_P1953485** | Copz1 | 0.150021681 | -0.19747171 | 0.497515072 | 13.76238892 | 1.081328935 | 0.324846107 | 0.709264768 | -4.596048227 |
| **A_55_P2003115** | Rnf7 | 0.147768176 | -0.194528875 | 0.490065226 | 12.52355965 | 1.081254927 | 0.324876262 | 0.709264768 | -4.596067352 |
| **A_51_P209372** | Msmo1 | 0.134518823 | -0.177097502 | 0.446135148 | 11.85611124 | 1.081217841 | 0.324891375 | 0.709264768 | -4.596076936 |
| **A_55_P2024290** | Fam149a | 0.303251131 | -0.399247986 | 1.005750247 | 8.58026783 | 1.081202378 | 0.324897676 | 0.709264768 | -4.596080932 |
| **A_55_P2137203** | B4galt3 | -0.129330984 | -0.428938124 | 0.170276155 | 12.24173291 | -1.081186887 | 0.324903989 | 0.709264768 | -4.596084935 |
| **A_52_P558939** | Rab3gap1 | -0.12117441 | -0.401905597 | 0.159556777 | 8.630833535 | -1.081111773 | 0.3249346 | 0.709264768 | -4.596104346 |
| **A_55_P2454784** | Ifnar1 | -0.230331557 | -0.763991072 | 0.303327958 | 8.458443466 | -1.081034443 | 0.324966116 | 0.709264768 | -4.596124328 |
| **A_55_P2095926** | NA | 0.110346423 | -0.145327267 | 0.366020113 | 8.776877643 | 1.080992216 | 0.324983327 | 0.709264768 | -4.59613524 |
| **A_55_P1955382** | Rgma | 0.146305665 | -0.192713108 | 0.485324438 | 8.800637132 | 1.080905537 | 0.325018659 | 0.709264768 | -4.596157637 |
| **A_55_P2122066** | Ppp2r3d | -0.096508863 | -0.320145597 | 0.127127871 | 8.273127892 | -1.080872338 | 0.325032192 | 0.709264768 | -4.596166215 |
| **A_51_P409010** | Comp | -0.10179383 | -0.337692578 | 0.134104917 | 7.899290046 | -1.080802067 | 0.325060839 | 0.709264768 | -4.596184372 |
| **A_51_P388002** | Snrpb2 | 0.454158589 | -0.598370305 | 1.506687484 | 12.21476038 | 1.080746533 | 0.32508348 | 0.709264768 | -4.596198721 |
| **A_55_P2003803** | Scml2 | 0.362663749 | -0.477886854 | 1.203214352 | 8.61389113 | 1.080663861 | 0.325117186 | 0.709264768 | -4.596220081 |
| **A_51_P275123** | Psd3 | -0.159934868 | -0.530658503 | 0.210788767 | 10.07993896 | -1.080545918 | 0.325165279 | 0.709264768 | -4.596250552 |
| **A_51_P216605** | Hbp1 | -0.234870292 | -0.779297847 | 0.309557263 | 8.319587862 | -1.080533746 | 0.325170243 | 0.709264768 | -4.596253697 |
| **A_51_P444437** | Ptgr1 | 0.781125092 | -1.029582554 | 2.591832738 | 11.02708602 | 1.080494596 | 0.325186208 | 0.709264768 | -4.596263812 |
| **A_51_P199899** | Fto | 0.146313924 | -0.192858555 | 0.485486402 | 9.662362839 | 1.080476679 | 0.325193515 | 0.709264768 | -4.596268441 |
| **A_51_P380401** | Cars2 | -0.102037821 | -0.338576728 | 0.134501086 | 9.0037665 | -1.080460599 | 0.325200072 | 0.709264768 | -4.596272595 |
| **A_51_P405089** | NA | 0.874893989 | -1.153265196 | 2.903053175 | 10.86290914 | 1.080447676 | 0.325205343 | 0.709264768 | -4.596275933 |
| **A_55_P2404484** | 1700001L | 0.333012997 | -0.439117665 | 1.105143659 | 9.574414278 | 1.080241183 | 0.325289566 | 0.709264768 | -4.596329279 |
| **A_55_P1972490** | Gm4055 | -1.283115989 | -4.258187011 | 1.691955033 | 9.732724578 | -1.080236691 | 0.325291398 | 0.709264768 | -4.596330439 |
| **A_66_P111385** | Klhl28 | -0.209008031 | -0.693806939 | 0.275790876 | 10.20482024 | -1.079820796 | 0.325461087 | 0.709264768 | -4.596437868 |
| **A_66_P100050** | Ndufaf8 | 0.151362485 | -0.199734923 | 0.502459894 | 9.848053083 | 1.079794154 | 0.32547196 | 0.709264768 | -4.59644475 |
| **A_55_P2102719** | NA | 0.611077336 | -0.8063761 | 2.028530773 | 14.4606029 | 1.079786072 | 0.325475258 | 0.709264768 | -4.596446837 |
| **A_55_P2033308** | Six2 | 0.370803946 | -0.489323458 | 1.230931349 | 8.172294713 | 1.079771618 | 0.325481157 | 0.709264768 | -4.59645057 |
| **A_55_P1997747** | Cox16 | 0.302063747 | -0.398647414 | 1.002774909 | 12.22956009 | 1.079716943 | 0.325503471 | 0.709264768 | -4.596464692 |
| **A_55_P2117420** | NA | 0.693015659 | -0.915024335 | 2.301055653 | 15.71574431 | 1.079435129 | 0.325618509 | 0.709264768 | -4.596537475 |
| **A_52_P706060** | Mex3a | 1.05534324 | -1.39376945 | 3.50445593 | 10.77568469 | 1.079282933 | 0.32568065 | 0.709264768 | -4.596576779 |
| **A_55_P2040110** | Cdv3 | 0.186946157 | -0.246918248 | 0.620810562 | 8.900529034 | 1.079227161 | 0.325703424 | 0.709264768 | -4.596591182 |
| **A_55_P2025675** | Dnah8 | 0.690480902 | -0.912083139 | 2.293044942 | 9.88941538 | 1.079161946 | 0.325730056 | 0.709264768 | -4.596608022 |
| **A_55_P2114118** | Cadm1 | 0.438282099 | -0.578979941 | 1.455544139 | 9.73734575 | 1.079123723 | 0.325745665 | 0.709264768 | -4.596617893 |
| **A_66_P106861** | B3glct | 0.353995381 | -0.467639324 | 1.175630085 | 9.786679853 | 1.079118568 | 0.325747771 | 0.709264768 | -4.596619224 |
| **A_51_P276235** | Pnpla7 | -0.109319547 | -0.36305688 | 0.144417785 | 8.275991164 | -1.079105237 | 0.325753215 | 0.709264768 | -4.596622666 |
| **A_55_P2003410** | Kng1 | -0.571357701 | -1.897575551 | 0.754860149 | 8.001617219 | -1.079054951 | 0.325773753 | 0.709264768 | -4.596635651 |
| **A_55_P2081373** | Glis1 | -1.000085196 | -3.321613707 | 1.321443316 | 8.747821578 | -1.07897982 | 0.32580444 | 0.709264768 | -4.596655051 |
| **A_55_P2086692** | Nfib | 1.071209971 | -1.415439616 | 3.557859557 | 11.76442059 | 1.078972466 | 0.325807443 | 0.709264768 | -4.59665695 |
| **A_51_P343889** | Slc35f6 | -0.331087634 | -1.099834015 | 0.437658747 | 8.251506759 | -1.078723703 | 0.325909067 | 0.709264768 | -4.596721181 |
| **A_55_P1982227** | NA | -0.204482399 | -0.679307528 | 0.27034273 | 12.24064256 | -1.078630215 | 0.325947265 | 0.709264768 | -4.596745318 |
| **A_55_P2168307** | Gm13213 | 0.406049675 | -0.53683362 | 1.348932969 | 16.92289745 | 1.078627704 | 0.325948291 | 0.709264768 | -4.596745966 |
| **A_55_P2436715** | Paics | 0.521213542 | -0.689161018 | 1.731588101 | 11.95278124 | 1.078564817 | 0.325973989 | 0.709264768 | -4.596762202 |
| **A_66_P119420** | Foxk1 | -0.258111423 | -0.857519439 | 0.341296593 | 9.855236484 | -1.078536941 | 0.32598538 | 0.709264768 | -4.596769399 |
| **A_55_P2151021** | NA | 0.210917855 | -0.278899216 | 0.700734927 | 11.74700974 | 1.078523903 | 0.325990708 | 0.709264768 | -4.596772765 |

| | | | | | | | | | |
|---|---|---|---|---|---|---|---|---|---|
| **A_55_P2066799** | Sctr | -0.53246396 | -1.769060353 | 0.704132433 | 8.104364585 | -1.078481223 | 0.32600815 | 0.709264768 | -4.596783784 |
| **A_52_P54238** | Lmf2 | -0.169345082 | -0.562658371 | 0.223968208 | 9.657936237 | -1.078411008 | 0.326036846 | 0.709264768 | -4.596801911 |
| **A_52_P510877** | Bcl2l1 | -0.174868034 | -0.581020588 | 0.23128452 | 8.043821135 | -1.078379401 | 0.326049764 | 0.709264768 | -4.59681007 |
| **A_55_P2075366** | Tead3 | -0.11762099 | -0.390849325 | 0.155607345 | 8.452816411 | -1.078225107 | 0.326112832 | 0.709264768 | -4.596849901 |
| **A_55_P2067632** | Ush1c | -0.11438276 | -0.380088964 | 0.151323445 | 7.798816789 | -1.07822456 | 0.326113056 | 0.709264768 | -4.596850043 |
| **A_55_P2002898** | Ctsj | -2.993743776 | -9.948205955 | 3.960718404 | 9.412604353 | -1.078205204 | 0.326120968 | 0.709264768 | -4.596855039 |
| **A_55_P2231152** | Gm13372 | -0.202950129 | -0.674444446 | 0.268544187 | 7.851962511 | -1.078110352 | 0.326159745 | 0.709264768 | -4.596879524 |
| **A_55_P2098428** | Bloc1s4 | -0.240296 | -0.798576159 | 0.317984158 | 9.094939888 | -1.078064375 | 0.326178542 | 0.709264768 | -4.596891392 |
| **A_51_P123676** | Synpo | -0.111590405 | -0.370858804 | 0.147677994 | 8.956551433 | -1.078021978 | 0.326195876 | 0.709264768 | -4.595860729 |
| **A_55_P1997335** | Vps9d1 | -0.111577581 | -0.370822829 | 0.147667666 | 10.66308347 | -1.077994352 | 0.326207172 | 0.709264768 | -4.596909466 |
| **A_55_P1993761** | Nsa2 | 0.463780302 | -0.613802754 | 1.541363357 | 11.77278756 | 1.077982947 | 0.326211836 | 0.709264768 | -4.59691241 |
| **A_55_P2137049** | AA467197 | -1.583005348 | -5.261292383 | 2.095281687 | 8.936248647 | -1.077921443 | 0.326236985 | 0.709264768 | -4.596928286 |
| **A_55_P2111770** | Klhl13 | -1.101739835 | -3.661931875 | 1.458452206 | 12.18476433 | -1.077846348 | 0.326267693 | 0.709264768 | -4.596947668 |
| **A_55_P1976664** | Gm16430 | -0.205338968 | -0.682529909 | 0.271851973 | 7.893497626 | -1.077778536 | 0.326295426 | 0.709264768 | -4.596965171 |
| **A_51_P258409** | Hey1 | 0.82603789 | -1.093645987 | 2.745721768 | 11.61811837 | 1.077756366 | 0.326304493 | 0.709264768 | -4.596970893 |
| **A_51_P397296** | Marveld3 | -0.105004478 | -0.349034564 | 0.139025607 | 8.449245103 | -1.07774197 | 0.326310381 | 0.709264768 | -4.596974609 |
| **A_55_P2063558** | NA | 0.212649414 | -0.281575864 | 0.706874692 | 16.15808466 | 1.0776794 | 0.326335972 | 0.709264768 | -4.596990758 |
| **A_55_P1974593** | NA | 0.131563988 | -0.174232245 | 0.437360221 | 15.83494897 | 1.077594195 | 0.326370824 | 0.709264768 | -4.597012748 |
| **A_55_P2165451** | NA | 0.503437664 | -0.666719644 | 1.673594972 | 10.9587464 | 1.077585651 | 0.326374319 | 0.709264768 | -4.597014953 |
| **A_55_P2178064** | NA | -0.183916005 | -0.611422931 | 0.243590921 | 8.040494065 | -1.077523411 | 0.32639978 | 0.709264768 | -4.597031016 |
| **A_55_P2073840** | Rpl7 | 0.1137959 | -0.150720961 | 0.378312761 | 16.089896 | 1.077515681 | 0.326402942 | 0.709264768 | -4.597033011 |
| **A_55_P1953630** | E330016L | -0.264504495 | -0.879378143 | 0.350369152 | 7.804879709 | -1.077450985 | 0.326429409 | 0.709264768 | -4.597049708 |
| **A_55_P1970867** | Gm5779 | 0.542249108 | -0.71835976 | 1.802857977 | 13.94670542 | 1.077379813 | 0.326458528 | 0.709264768 | -4.597068075 |
| **A_55_P2049080** | Fam160a2 | -0.378978781 | -1.260108998 | 0.502151435 | 10.75127398 | -1.077271439 | 0.326502872 | 0.709264768 | -4.597096042 |
| **A_52_P301374** | Aaas | 0.148081439 | -0.19621255 | 0.492375428 | 9.212072428 | 1.077262471 | 0.326506541 | 0.709264768 | -4.597098356 |
| **A_55_P1992234** | Slx | -0.212436646 | -0.706402888 | 0.281529595 | 7.895576702 | -1.077165693 | 0.326546145 | 0.709264768 | -4.59712333 |
| **A_55_P1985514** | Dlg3 | -0.636703482 | -2.117199875 | 0.843792911 | 12.10397084 | -1.077159998 | 0.326548475 | 0.709264768 | -4.597124799 |
| **A_55_P2066304** | Limk2 | -0.169536431 | -0.563770934 | 0.224698071 | 7.990005883 | -1.077106762 | 0.326570262 | 0.709264768 | -4.597138536 |
| **A_55_P2148046** | NA | -0.379487961 | -1.261959044 | 0.502983122 | 8.140309421 | -1.07707976 | 0.326581314 | 0.709264768 | -4.597145504 |
| **A_51_P419571** | Polr2d | 0.387572857 | -0.513720169 | 1.288865883 | 11.09694191 | 1.077054525 | 0.326591642 | 0.709264768 | -4.597152015 |
| **A_55_P2053958** | Rabep2 | 0.310748206 | -0.411906725 | 1.033403137 | 11.21196675 | 1.077030509 | 0.326601471 | 0.709264768 | -4.597158212 |
| **A_55_P1992099** | Ptprj | -0.204829925 | -0.681191427 | 0.271531577 | 9.01340082 | -1.076978656 | 0.326622695 | 0.709264768 | -4.597171592 |
| **A_55_P2075200** | Tor3a | -0.214734229 | -0.71417873 | 0.284710273 | 10.15916778 | -1.076872746 | 0.326666049 | 0.709264768 | -4.597198919 |
| **A_55_P2022658** | Cdk11b | -0.282710978 | -0.940265435 | 0.374843478 | 12.70804266 | -1.076865079 | 0.326669187 | 0.709264768 | -4.597200897 |
| **A_51_P284577** | Hyal2 | 0.254504425 | -0.337500022 | 0.846508872 | 12.63062481 | 1.076764463 | 0.326710378 | 0.709264768 | -4.597226857 |
| **A_55_P1964347** | NA | -0.413581034 | -1.375707005 | 0.548544936 | 8.155767819 | -1.076661103 | 0.326752698 | 0.709264768 | -4.597253525 |
| **A_55_P2287910** | Rragd | -0.183500096 | -0.610385798 | 0.243385606 | 7.810253053 | -1.076651208 | 0.326756749 | 0.709264768 | -4.597256077 |
| **A_52_P515036** | Htatip2 | -0.599573941 | -1.994570702 | 0.79542282 | 8.397905348 | -1.076514528 | 0.326812718 | 0.709264768 | -4.59729134 |
| **A_55_P1966372** | Srek1 | 0.174398915 | -0.231368865 | 0.580166696 | 13.95469219 | 1.076506275 | 0.326816098 | 0.709264768 | -4.597293469 |
| **A_51_P338317** | Rhoq | 0.258153274 | -0.342625175 | 0.858931723 | 10.84399295 | 1.076251171 | 0.326920583 | 0.709264768 | -4.597359278 |
| **A_52_P615401** | Tpra1 | -0.318152964 | -1.058575744 | 0.422269816 | 8.520287215 | -1.076233601 | 0.326927781 | 0.709264768 | -4.597363811 |
| **A_55_P2109585** | Plekha7 | -0.58526592 | -1.947380485 | 0.776848646 | 8.380283238 | -1.076192458 | 0.326944635 | 0.709264768 | -4.597374424 |
| **A_55_P1953400** | Rbfox1 | 0.119602715 | -0.158758669 | 0.397964099 | 7.953904853 | 1.076173757 | 0.326952296 | 0.709264768 | -4.597379248 |
| **A_51_P502152** | Slc19a1 | 0.198223554 | -0.263156876 | 0.659603984 | 9.457691721 | 1.076084648 | 0.326988803 | 0.709264768 | -4.597402234 |
| **A_51_P394190** | Lmo4 | 0.626436138 | -0.831807552 | 2.084679827 | 13.19835902 | 1.075962271 | 0.327038944 | 0.709264768 | -4.597433799 |
| **A_52_P232637** | Dhh | 0.200942178 | -0.266845939 | 0.668730295 | 8.551461061 | 1.075900887 | 0.327064098 | 0.709264768 | -4.597449632 |
| **A_51_P441942** | Tial1 | 0.501590995 | -0.666126859 | 1.669308849 | 10.29625582 | 1.075875843 | 0.327074361 | 0.709264768 | -4.597456092 |
| **A_65_P07361** | Trim14 | -0.138401488 | -0.460611255 | 0.18380828 | 8.165962474 | -1.075851681 | 0.327084262 | 0.709264768 | -4.597462323 |

| | | | | | | | | | |
|---|---|---|---|---|---|---|---|---|---|
| **A_51_P472274** | Sox18 | 0.771216164 | -1.024352454 | 2.566784781 | 10.57764398 | 1.075782469 | 0.327112627 | 0.709264768 | -4.597480175 |
| **A_55_P2162582** | Rab5b | -0.432920535 | -1.440920978 | 0.575079907 | 8.698279918 | -1.075716449 | 0.327139685 | 0.709264768 | -4.597497202 |
| **A_55_P2004208** | Defa-rs2 | -0.434379838 | -1.445827767 | 0.577068091 | 7.975026735 | -1.075663606 | 0.327161344 | 0.709264768 | -4.597510831 |
| **A_55_P2080163** | Carhsp1 | -0.267547868 | -0.890554568 | 0.355458832 | 11.96734838 | -1.07562063 | 0.327178959 | 0.709264768 | -4.597521914 |
| **A_52_P595824** | Cpeb4 | -0.172611246 | -0.57455645 | 0.229333959 | 7.816525814 | -1.075604427 | 0.327185601 | 0.709264768 | -4.597526093 |
| **A_51_P329949** | Fam13a | 0.706443652 | -0.938900897 | 2.351788202 | 9.785322788 | 1.075402436 | 0.327268408 | 0.709264768 | -4.597578185 |
| **A_51_P222467** | Abcg1 | -1.162483317 | -3.979710053 | 1.654743419 | 11.01886473 | -1.099075768 | 0.327292408 | 0.709264768 | -4.588988781 |
| **A_55_P2093301** | NA | -0.17112725 | -0.569723172 | 0.227468672 | 8.012436223 | -1.075317381 | 0.327303282 | 0.709264768 | -4.597600119 |
| **A_55_P2173224** | Kif2a | 0.136892431 | -0.181964314 | 0.455749177 | 8.6089283 | 1.07531121 | 0.327305813 | 0.709264768 | -4.59760171 |
| **A_55_P1966659** | LOC5473 | -0.438750906 | -1.460983536 | 0.583481724 | 8.116927736 | -1.075025168 | 0.327423118 | 0.709264768 | -4.597675468 |
| **A_51_P181523** | Tmem214 | -0.388729962 | -1.294674053 | 0.517214129 | 10.62440799 | -1.074724042 | 0.327546648 | 0.709264768 | -4.597753108 |
| **A_55_P2166349** | Dnajc19 | 0.15050014 | -0.200298484 | 0.501298764 | 10.61030301 | 1.074556778 | 0.327615281 | 0.709264768 | -4.597796231 |
| **A_65_P16603** | Bad | -0.158137727 | -0.52674916 | 0.210473707 | 8.452211928 | -1.074526264 | 0.327627803 | 0.709264768 | -4.597804097 |
| **A_55_P2011087** | Gnptg | -0.286390391 | -0.954035502 | 0.381254719 | 11.09014576 | -1.074392864 | 0.32768255 | 0.709264768 | -4.597838487 |
| **A_66_P108685** | Cideb | -0.493041153 | -1.642483857 | 0.656401551 | 8.682725398 | -1.074350988 | 0.327699738 | 0.709264768 | -4.597849282 |
| **A_55_P2009792** | Hdac6 | -0.442052766 | -1.472637308 | 0.588531777 | 10.44298256 | -1.074337743 | 0.327705175 | 0.709264768 | -4.597852696 |
| **A_52_P358349** | Mtdh | 0.117840565 | -0.156915953 | 0.392597083 | 11.67237852 | 1.074229707 | 0.327749521 | 0.709264768 | -4.597880545 |
| **A_52_P246165** | Ptprk | 0.425509632 | -0.678190011 | 1.529209274 | 9.038609452 | 1.136975255 | 0.32778368 | 0.709264768 | -4.581826971 |
| **A_51_P309589** | 2700099C | 0.297305952 | -0.395975817 | 0.99058772 | 8.590601411 | 1.074098647 | 0.327803326 | 0.709264768 | -4.597914328 |
| **A_51_P246816** | Napa | -0.300082483 | -0.999988086 | 0.39982312 | 9.652054023 | -1.073869531 | 0.327897403 | 0.709264768 | -4.597973383 |
| **A_66_P125777** | Slc43a2 | -0.626928156 | -2.089220102 | 0.83536379 | 9.557274988 | -1.073826291 | 0.32791516 | 0.709264768 | -4.597984527 |
| **A_51_P257743** | Dnajc22 | -0.203330786 | -0.677630345 | 0.270968772 | 8.139796739 | -1.073744037 | 0.327948941 | 0.709264768 | -4.598005726 |
| **A_55_P1961247** | NA | 0.385692081 | -0.514049171 | 1.285433332 | 16.03574342 | 1.073676468 | 0.327976694 | 0.709264768 | -4.59802314 |
| **A_51_P517012** | Tysnd1 | 0.158330059 | -0.211112075 | 0.527772194 | 11.09360837 | 1.073414098 | 0.328084475 | 0.709264768 | -4.598090755 |
| **A_51_P101006** | Ak8 | -0.122691831 | -0.408982299 | 0.163598636 | 8.726341568 | -1.073393858 | 0.328092791 | 0.709264768 | -4.596396579 |
| **A_55_P2121603** | 2810047C | -0.368670093 | -1.228959594 | 0.491619408 | 8.608531218 | -1.073355611 | 0.328108505 | 0.709264768 | -4.598105827 |
| **A_55_P2011678** | Pdzk1ip1 | -0.396975043 | -1.323431703 | 0.529481616 | 12.1851872 | -1.073219223 | 0.328164548 | 0.709264768 | -4.598140972 |
| **A_51_P295708** | Ints1 | 0.096477987 | -0.128685709 | 0.321641684 | 9.612210929 | 1.07319888 | 0.328172908 | 0.709264768 | -4.598146214 |
| **A_52_P308507** | Chd6 | 0.225302198 | -0.300662334 | 0.75126673 | 8.742038423 | 1.072900173 | 0.32829568 | 0.709264768 | -4.598223179 |
| **A_51_P173601** | Bcam | 0.884241504 | -1.180008027 | 2.948491035 | 10.77641667 | 1.07289951 | 0.328295953 | 0.709264768 | -4.59822335 |
| **A_55_P1999294** | Gpbp1 | -0.325770909 | -1.086393062 | 0.434851245 | 12.45168753 | -1.072738023 | 0.328362342 | 0.709264768 | -4.598264955 |
| **A_55_P2133170** | Luc7l2 | -0.150759011 | -0.502806773 | 0.20128875 | 9.041922449 | -1.072585789 | 0.328424937 | 0.709264768 | -4.598304174 |
| **A_51_P217047** | Gng10 | 0.493608194 | -0.659225707 | 1.646442095 | 10.27904572 | 1.072422624 | 0.328492039 | 0.709264768 | -4.598346207 |
| **A_55_P2097869** | Frem2 | 0.564472542 | -0.753872728 | 1.882817813 | 10.60197032 | 1.07241777 | 0.328494035 | 0.709264768 | -4.598347458 |
| **A_66_P126320** | Mrpl46 | 0.114140759 | -0.152457464 | 0.380738982 | 10.13520987 | 1.07234332 | 0.328524657 | 0.709264768 | -4.598366636 |
| **A_51_P487918** | Rinl | -0.664780417 | -2.217633553 | 0.888072718 | 9.239721896 | -1.072255247 | 0.328560884 | 0.709264768 | -4.598389323 |
| **A_55_P1961152** | Pou3f1 | -0.656314897 | -2.189570662 | 0.876940868 | 8.296211476 | -1.072131364 | 0.328611847 | 0.709264768 | -4.598421233 |
| **A_55_P1968982** | Pbx3 | 0.600169754 | -0.801976014 | 2.002315521 | 9.931761881 | 1.072090099 | 0.328628824 | 0.709264768 | -4.598431862 |
| **A_55_P2019776** | Gxylt1 | -0.348521792 | -1.16276327 | 0.465719687 | 8.316648584 | -1.072079672 | 0.328633114 | 0.709264768 | -4.598434547 |
| **A_51_P410918** | Tor4a | -0.291720517 | -0.973271938 | 0.389830904 | 9.215241203 | -1.072059015 | 0.328641613 | 0.709264768 | -4.598439868 |
| **A_55_P2009940** | Xlr | -0.20939422 | -0.698612084 | 0.279823644 | 7.887817554 | -1.072044289 | 0.328647672 | 0.709264768 | -4.598443661 |
| **A_52_P552550** | Agtrap | -0.298850839 | -0.997094612 | 0.399392935 | 8.543386243 | -1.072007326 | 0.328662881 | 0.709264768 | -4.598453181 |
| **A_52_P360112** | Clip2 | -0.340969345 | -1.137684326 | 0.455745636 | 8.584521188 | -1.071920799 | 0.328698485 | 0.709264768 | -4.598475467 |
| **A_55_P2042521** | Cop1 | 0.232221097 | -0.310393539 | 0.774835732 | 11.41807358 | 1.07191541 | 0.328700702 | 0.709264768 | -4.598476855 |
| **A_51_P200477** | Hspa4 | 0.237094652 | -0.316918152 | 0.791107457 | 11.26307653 | 1.07189514 | 0.328709044 | 0.709264768 | -4.598482076 |
| **A_66_P130882** | Eif3e | 0.547298549 | -0.7316408 | 1.826237897 | 11.22043184 | 1.071827 | 0.328737085 | 0.709264768 | -4.598499625 |
| **A_51_P445487** | Mturn | -0.128252144 | -0.427981668 | 0.17147738 | 8.337016082 | -1.071730164 | 0.328776938 | 0.709264768 | -4.598524564 |
| **A_55_P2124346** | Ivns1abp | 0.203267053 | -0.271800794 | 0.6783349 | 11.59919556 | 1.071671542 | 0.328801067 | 0.709264768 | -4.598539661 |

| | | | | | | | | | |
|---|---|---|---|---|---|---|---|---|---|
| **A_51_P140942** | Rp9 | 0.323309043 | -0.432454263 | 1.079072349 | 13.27858073 | 1.071475871 | 0.328881614 | 0.709264768 | -4.598590051 |
| **A_55_P2090414** | NA | -0.380285349 | -1.26926392 | 0.508693223 | 8.890379002 | -1.071441951 | 0.328895579 | 0.709264768 | -4.598598786 |
| **A_55_P2152567** | Plekha5 | -0.263895764 | -0.880795056 | 0.353003528 | 8.23607909 | -1.071441571 | 0.328895736 | 0.709264768 | -4.598598884 |
| **A_51_P328489** | 1700025G | -0.717022563 | -2.393303879 | 0.959258753 | 10.32765952 | -1.071362061 | 0.328928472 | 0.709264768 | -4.598619358 |
| **A_51_P352264** | Sirt7 | -0.181165998 | -0.604704834 | 0.242372838 | 11.61048579 | -1.07135598 | 0.328930976 | 0.709264768 | -4.598620924 |
| **A_55_P1991920** | Dhx32 | -0.443557108 | -1.480662981 | 0.593548764 | 10.1055036 | -1.071215371 | 0.328988875 | 0.709264768 | -4.59865713 |
| **A_55_P2160322** | Anapc15 | 0.108056972 | -0.144603936 | 0.360717879 | 8.622464137 | 1.071186488 | 0.329000769 | 0.709264768 | -4.598664567 |
| **A_55_P2042526** | Apoo | 0.176494012 | -0.236324778 | 0.589312801 | 10.27428269 | 1.070830823 | 0.329147264 | 0.709264768 | -4.59875614 |
| **A_55_P1997895** | Rpl35a | 0.397819002 | -0.532720367 | 1.328358372 | 15.15022627 | 1.070782137 | 0.329167322 | 0.709264768 | -4.598768674 |
| **A_55_P2102621** | Eaf2 | 0.18454967 | -0.247139484 | 0.616238825 | 11.09329207 | 1.070760845 | 0.329176094 | 0.709264768 | -4.598774156 |
| **A_55_P2043767** | NA | 0.3979585 | -0.533040687 | 1.328957687 | 16.6581229 | 1.070628572 | 0.329230594 | 0.709264768 | -4.598808208 |
| **A_55_P2052420** | Setd5 | -0.227346575 | -0.759209794 | 0.304516643 | 12.85794065 | -1.070628511 | 0.329230619 | 0.709264768 | -4.598808224 |
| **A_55_P2160291** | Olfm1 | 0.25974694 | -0.34791634 | 0.86741022 | 8.493917982 | 1.070626015 | 0.329231648 | 0.709264768 | -4.598808866 |
| **A_52_P274496** | Tspan18 | 0.624897879 | -0.837039451 | 2.086835208 | 12.52989575 | 1.070608385 | 0.329238913 | 0.709264768 | -4.598813405 |
| **A_55_P1953998** | NA | 0.63079713 | -0.845311783 | 2.106906042 | 11.87412091 | 1.070339743 | 0.329349626 | 0.709264768 | -4.598882559 |
| **A_55_P2042062** | Mier1 | -0.230929189 | -0.771353686 | 0.309495309 | 8.145358212 | -1.070271961 | 0.329377565 | 0.709264768 | -4.598900006 |
| **A_55_P1955437** | Cmtm5 | -0.483207384 | -1.614058862 | 0.647644094 | 8.262973801 | -1.070232999 | 0.329393626 | 0.709264768 | -4.598910034 |
| **A_55_P2026863** | Trim32 | 0.33435269 | -0.448157729 | 1.11686311 | 11.4747402 | 1.070200226 | 0.329407136 | 0.709264768 | -4.59891847 |
| **A_55_P2167189** | 4921504E | 0.316926489 | -0.424845746 | 1.058698724 | 8.867371607 | 1.070134369 | 0.329434286 | 0.709264768 | -4.598935421 |
| **A_66_P105801** | Igf1r | 0.39634124 | -0.531359579 | 1.324042059 | 11.34642411 | 1.070068721 | 0.329461352 | 0.709264768 | -4.598952318 |
| **A_55_P2157586** | NA | -0.163599035 | -0.546551037 | 0.219352967 | 7.987053413 | -1.070007295 | 0.329486678 | 0.709264768 | -4.598968127 |
| **A_51_P499020** | Fbp2 | -0.166275553 | -0.555500978 | 0.222949871 | 7.777799405 | -1.069984648 | 0.329496016 | 0.709264768 | -4.598973956 |
| **A_52_P111031** | Pcdh17 | 1.216265433 | -1.631144685 | 4.06367555 | 11.32574037 | 1.069864274 | 0.329545653 | 0.709264768 | -4.599004936 |
| **A_51_P265571** | Adm | -1.105105893 | -3.692511115 | 1.482299329 | 9.190318868 | -1.069768462 | 0.329585167 | 0.709264768 | -4.599029594 |
| **A_51_P227392** | Rhou | -0.888862561 | -2.97005946 | 1.192334339 | 11.84496222 | -1.069724122 | 0.329603455 | 0.709264768 | -4.599041005 |
| **A_55_P2061525** | Cdc27 | -0.112070032 | -0.374513637 | 0.150373573 | 9.164868235 | -1.069556786 | 0.329672478 | 0.709264768 | -4.599084067 |
| **A_55_P1970027** | Itch | 0.132564042 | -0.177875575 | 0.443003659 | 10.5810624 | 1.069544708 | 0.32967746 | 0.709264768 | -4.599087175 |
| **A_51_P486618** | Akr1b10 | 0.530461035 | -0.711797872 | 1.772719943 | 10.58888451 | 1.069526906 | 0.329684804 | 0.709264768 | -4.599091756 |
| **A_55_P2029191** | Pde4a | -0.218643982 | -0.73068236 | 0.293394395 | 9.569606737 | -1.069511261 | 0.329691258 | 0.709264768 | -4.599095782 |
| **A_55_P2028798** | Nqo2 | 0.472781048 | -0.634550121 | 1.580112216 | 10.30312112 | 1.069382025 | 0.329744576 | 0.709264768 | -4.599129037 |
| **A_55_P2137431** | Rtel1 | 0.357125906 | -0.479375219 | 1.193627032 | 10.36275146 | 1.0693138 | 0.329772727 | 0.709264768 | -4.599146593 |
| **A_52_P537852** | Fam120b | 0.215602435 | -0.289477577 | 0.720682447 | 8.520123058 | 1.069162763 | 0.329835053 | 0.709264768 | -4.599185455 |
| **A_55_P1955287** | NA | 0.273143631 | -0.366737059 | 0.913024321 | 16.78209249 | 1.069159305 | 0.32983648 | 0.709264768 | -4.599186345 |
| **A_55_P2125311** | Rab3b | -0.455360469 | -1.522140288 | 0.611419351 | 8.571597834 | -1.069130894 | 0.329848205 | 0.709264768 | -4.599193654 |
| **A_55_P2016034** | Nlrc5 | -0.299697873 | -1.001828566 | 0.402432819 | 7.869943862 | -1.069094386 | 0.329863272 | 0.709264768 | -4.599203048 |
| **A_52_P528600** | Hhex | 0.666987257 | -0.895837575 | 2.22981209 | 9.905056629 | 1.068950462 | 0.329922677 | 0.709264768 | -4.599240077 |
| **A_51_P308681** | Mcur1 | -0.215094704 | -0.719100951 | 0.288911543 | 8.763189616 | -1.068917394 | 0.329936327 | 0.709264768 | -4.599248584 |
| **A_55_P1960208** | Vcan | 0.115304544 | -0.154886616 | 0.385495705 | 8.699727829 | 1.068871833 | 0.329955135 | 0.709264768 | -4.599260305 |
| **A_55_P2096933** | Cntln | 0.440499849 | -0.591784687 | 1.472784384 | 10.33988624 | 1.068800598 | 0.329984543 | 0.709264768 | -4.599278632 |
| **A_51_P348397** | Hexim1 | 0.325892015 | -0.437821134 | 1.089605165 | 9.909120789 | 1.068793483 | 0.32998748 | 0.709264768 | -4.599280462 |
| **A_52_P91235** | Ccni | 0.493474827 | -0.663063293 | 1.650012947 | 13.58945629 | 1.068698985 | 0.330026495 | 0.709264768 | -4.599304772 |
| **A_51_P102911** | Hoxa11 | -0.8567511 | -2.864931097 | 1.151428897 | 8.117742347 | -1.06856853 | 0.330080362 | 0.709264768 | -4.597881558 |
| **A_55_P2153600** | Mfsd4b3-| | -0.184505049 | -0.617045342 | 0.248035245 | 7.83568348 | -1.068395447 | 0.330151842 | 0.709264768 | -4.599382853 |
| **A_52_P423247** | Pde4b | -0.286882006 | -0.959428709 | 0.385664697 | 8.428217137 | -1.068393425 | 0.330152678 | 0.709264768 | -4.599383373 |
| **A_55_P2009973** | Ttc3 | 0.604391564 | -0.812511895 | 2.021295024 | 13.05109585 | 1.068386715 | 0.330155449 | 0.709264768 | -4.599385099 |
| **A_52_P40367** | Mrpl42 | 0.306858914 | -0.412574182 | 1.026292011 | 10.63963416 | 1.068313402 | 0.33018573 | 0.709264768 | -4.599403957 |
| **A_55_P2097048** | NA | -0.238798587 | -0.798714918 | 0.321117743 | 10.36358476 | -1.06821572 | 0.33022608 | 0.709264768 | -4.599429081 |
| **A_55_P1998309** | Sms | 0.239464049 | -0.322201652 | 0.801129751 | 11.33743311 | 1.067856175 | 0.330374635 | 0.709264768 | -4.599521552 |

| | | | | | | | | | |
|---|---|---|---|---|---|---|---|---|---|
| **A_55_P2010111** | Zxdc | -0.526857777 | -1.762987253 | 0.709271699 | 8.872701075 | -1.067529237 | 0.330509766 | 0.709264768 | -4.599605625 |
| **A_55_P2020463** | Rufy2 | -0.314732779 | -1.053176132 | 0.423710575 | 10.01583991 | -1.067517825 | 0.330514484 | 0.709264768 | -4.59960856 |
| **A_52_P193925** | Sulf2 | -0.587419178 | -1.965787955 | 0.790949599 | 10.09161842 | -1.067414357 | 0.33055726 | 0.709264768 | -4.599635165 |
| **A_55_P2019814** | Rnf20 | 0.192625683 | -0.259382004 | 0.644633369 | 11.91891204 | 1.067379153 | 0.330571815 | 0.709264768 | -4.599644217 |
| **A_51_P194628** | Tango6 | 0.148909707 | -0.200534799 | 0.498354213 | 8.539535978 | 1.067321203 | 0.330595776 | 0.709264768 | -4.599659117 |
| **A_55_P1975305** | NA | 0.117019856 | -0.157625578 | 0.39166529 | 11.66893548 | 1.067179614 | 0.330654325 | 0.709264768 | -4.599695522 |
| **A_55_P1969131** | Cidec | -0.381880687 | -1.278198887 | 0.514437514 | 7.928873897 | -1.067126305 | 0.330676371 | 0.709264768 | -4.599709227 |
| **A_52_P123693** | E130304I( | 0.329787585 | -0.44437634 | 1.10395151 | 11.69977946 | 1.066968796 | 0.330741517 | 0.709264768 | -4.599749722 |
| **A_51_P414448** | Naa15 | 0.103482741 | -0.139463483 | 0.346428964 | 9.592902238 | 1.066861675 | 0.330785829 | 0.709264768 | -4.599777261 |
| **A_66_P105791** | Prkca | 0.603470892 | -0.813355108 | 2.020296891 | 10.67884872 | 1.066817557 | 0.33080408 | 0.709264768 | -4.599788602 |
| **A_51_P175555** | Pelo | 0.184869001 | -0.298168218 | 0.667906219 | 12.80703369 | 1.128693487 | 0.330830413 | 0.709264768 | -4.5839927 |
| **A_55_P2126207** | Fthl17a | -2.807202806 | -9.398401075 | 3.783995463 | 9.511765525 | -1.066742896 | 0.330834969 | 0.709264768 | -4.599807795 |
| **A_55_P1981780** | Cul4b | -0.685391783 | -2.294796151 | 0.924012584 | 10.24353827 | -1.066655222 | 0.330871244 | 0.709264768 | -4.599830333 |
| **A_55_P2010871** | Daam2 | 0.361821481 | -0.487902162 | 1.211545125 | 10.12146654 | 1.066515055 | 0.330929245 | 0.709264768 | -4.599866363 |
| **A_55_P2052578** | Bclaf3 | -0.869506484 | -2.911648978 | 1.17263601 | 10.0694302 | -1.066441723 | 0.330959593 | 0.709264768 | -4.599885212 |
| **A_55_P2028898** | Prl2b1 | -2.894233263 | -9.692317 | 3.903850473 | 9.489135994 | -1.06634409 | 0.331000002 | 0.709264768 | -4.599910306 |
| **A_55_P2016708** | Msl1 | 0.345448821 | -0.465973395 | 1.156871037 | 10.15978934 | 1.066319045 | 0.331010368 | 0.709264768 | -4.599916743 |
| **A_51_P131408** | Tnfrsf12a | -0.706503133 | -2.366040951 | 0.953034686 | 9.983344176 | -1.066294781 | 0.331020412 | 0.709264768 | -4.59992298 |
| **A_51_P178735** | Hibch | -0.10427734 | -0.349235174 | 0.140680493 | 10.36571698 | -1.06622524 | 0.331049198 | 0.709264768 | -4.599940853 |
| **A_51_P112237** | Angel1 | 0.167058634 | -0.22546387 | 0.559581138 | 8.937188399 | 1.065993878 | 0.331144983 | 0.709264768 | -4.599228882 |
| **A_55_P2025283** | NA | -0.099290232 | -0.332592428 | 0.134011965 | 15.49322473 | -1.065952967 | 0.331161923 | 0.709264768 | -4.600010828 |
| **A_55_P2319370** | Plekha3 | -0.464737406 | -1.556757788 | 0.627282976 | 10.51327142 | -1.065926431 | 0.331172911 | 0.709264768 | -4.600017647 |
| **A_55_P2091348** | NA | -0.223797516 | -0.749685048 | 0.302090017 | 9.412242382 | -1.065890845 | 0.331187647 | 0.709264768 | -4.600026792 |
| **A_55_P2087375** | Gm4987 | 0.339123332 | -0.45776319 | 1.136009854 | 16.1776632 | 1.065887906 | 0.331188865 | 0.709264768 | -4.600027548 |
| **A_55_P2090330** | Kcnmb4 | 0.50059947 | -0.675751891 | 1.676950832 | 9.017641495 | 1.065868609 | 0.331196855 | 0.709264768 | -4.600032507 |
| **A_55_P2048937** | Kif5c | -0.630618103 | -2.112625921 | 0.851389716 | 8.478120799 | -1.065776854 | 0.331234854 | 0.709264768 | -4.600056085 |
| **A_55_P2101696** | Gnat2 | -0.299284812 | -1.002641711 | 0.404072087 | 7.886235441 | -1.065759646 | 0.331241981 | 0.709264768 | -4.600060507 |
| **A_55_P2164977** | NA | -0.300207835 | -1.005821162 | 0.405405492 | 7.84683034 | -1.065627927 | 0.331296537 | 0.709264768 | -4.600094354 |
| **A_55_P2068429** | Zfp958 | -0.185045553 | -0.619980659 | 0.249889552 | 10.94085796 | -1.065625332 | 0.331297612 | 0.709264768 | -4.600095021 |
| **A_55_P2102897** | Gm12657 | 0.858227383 | -1.159123825 | 2.875578591 | 13.94304946 | 1.065543539 | 0.331331494 | 0.709264768 | -4.600116038 |
| **A_55_P2103115** | Tor1aip2 | -0.483667564 | -1.620577669 | 0.653242541 | 11.02771091 | -1.065543496 | 0.331331511 | 0.709264768 | -4.600116049 |
| **A_52_P10017** | Tmem64 | -0.195318531 | -0.654451814 | 0.263814751 | 8.903961276 | -1.065503841 | 0.331347939 | 0.709264768 | -4.600126238 |
| **A_66_P115453** | Coro1c | -0.106438913 | -0.356666138 | 0.143788312 | 12.65549655 | -1.065408643 | 0.331387378 | 0.709264768 | -4.600150699 |
| **A_55_P2074246** | Zfp184 | 0.180358006 | -0.243701728 | 0.604417739 | 8.886042442 | 1.06526764 | 0.331445802 | 0.709264768 | -4.600186927 |
| **A_55_P2123101** | Cbarp | 0.440038286 | -0.594709534 | 1.474786106 | 14.47259716 | 1.065139009 | 0.331499106 | 0.709264768 | -4.600219974 |
| **A_51_P194853** | Sec14l4 | 1.137102512 | -1.53693227 | 3.811137295 | 10.7932362 | 1.065081675 | 0.331522868 | 0.709264768 | -4.600234704 |
| **A_55_P1996653** | D030056L | 0.211740002 | -0.286196923 | 0.709676927 | 11.60610363 | 1.065071897 | 0.33152692 | 0.709264768 | -4.600237215 |
| **A_52_P581390** | Kif1c | -0.183898304 | -0.61637042 | 0.248573812 | 8.005265088 | -1.065049906 | 0.331536035 | 0.709264768 | -4.600242865 |
| **A_55_P2181162** | NA | 0.391791475 | -0.529607982 | 1.313190932 | 15.52303218 | 1.065019048 | 0.331548824 | 0.709264768 | -4.600250793 |
| **A_51_P250807** | Spata2l | -0.187850219 | -0.629664776 | 0.253964337 | 8.261359606 | -1.06493238 | 0.331584748 | 0.709264768 | -4.600273057 |
| **A_55_P2005930** | NA | 0.597200482 | -0.807465939 | 2.001866904 | 13.28969552 | 1.064871723 | 0.331609892 | 0.709264768 | -4.600288639 |
| **A_55_P2131672** | Mical2 | -0.108381436 | -0.363316934 | 0.146554061 | 8.209101533 | -1.064816901 | 0.331632619 | 0.709264768 | -4.600302722 |
| **A_52_P399054** | Tmed2 | -0.5078948 | -1.702615871 | 0.686826272 | 8.827727415 | -1.064774369 | 0.331650252 | 0.709264768 | -4.600313648 |
| **A_51_P487308** | Pstk | 0.357967848 | -0.484095998 | 1.200031694 | 10.12452057 | 1.06475416 | 0.33165863 | 0.709264768 | -4.600318839 |
| **A_52_P202725** | Rragc | -0.240687495 | -0.806903913 | 0.325528923 | 8.592588381 | -1.064685685 | 0.33168702 | 0.709264768 | -4.600336428 |
| **A_55_P2119420** | Ndufb2 | 0.307848703 | -0.416377117 | 1.032074523 | 13.69789981 | 1.064666696 | 0.331694894 | 0.709264768 | -4.600341305 |
| **A_55_P2009953** | Cd300lg | 0.154240039 | -0.208617864 | 0.517097942 | 9.129908318 | 1.064659872 | 0.331697723 | 0.709264768 | -4.600343058 |
| **A_55_P2029643** | NA | 0.405050528 | -0.547937554 | 1.35803861 | 12.35219506 | 1.064564743 | 0.331737169 | 0.709264768 | -4.600367493 |

| | | | | | | | | | |
|---|---|---|---|---|---|---|---|---|---|
| **A_55_P1958653** | Arf2 | -0.688220545 | -2.307510681 | 0.93106959 | 12.2044724 | -1.064518734 | 0.331756248 | 0.709264768 | -4.600379311 |
| **A_55_P2037368** | Pdxdc1 | -0.263207697 | -0.882545137 | 0.356129742 | 9.533483866 | -1.064441007 | 0.331788482 | 0.709264768 | -4.600399275 |
| **A_55_P2086198** | NA | -0.19646564 | -0.658763786 | 0.265832506 | 10.1036385 | -1.06442435 | 0.331795391 | 0.709264768 | -4.600403553 |
| **A_52_P601688** | Abcc1 | -0.244768145 | -0.820739513 | 0.331203222 | 8.613598444 | -1.064398732 | 0.331806016 | 0.709264768 | -4.600410133 |
| **A_55_P2142251** | H3c15 | -0.589596858 | -1.977007504 | 0.797813788 | 8.235073087 | -1.06438926 | 0.331809944 | 0.709264768 | -4.600412565 |
| **A_55_P2008687** | Nlk | -0.127128876 | -0.426292296 | 0.172034545 | 10.48347208 | -1.064353915 | 0.331824604 | 0.709264768 | -4.600421643 |
| **A_52_P257728** | Cnnm4 | 0.09078411 | -0.122862118 | 0.304430338 | 9.379329399 | 1.064302188 | 0.331846059 | 0.709264768 | -4.600434928 |
| **A_66_P109220** | Etfbkmt | -0.198305054 | -0.665062153 | 0.268452045 | 8.314268558 | -1.064126349 | 0.331919002 | 0.709264768 | -4.600480088 |
| **A_55_P1976152** | Vrk2 | 0.138510357 | -0.18751331 | 0.464534024 | 9.413714639 | 1.064102505 | 0.331928894 | 0.709264768 | -4.600486211 |
| **A_55_P2454004** | Smim8 | 0.203195686 | -0.275097455 | 0.681488827 | 11.00204575 | 1.064071167 | 0.331941895 | 0.709264768 | -4.600494259 |
| **A_55_P2038318** | NA | -0.503113695 | -1.687447636 | 0.681220247 | 8.065767096 | -1.064001668 | 0.331970731 | 0.709264768 | -4.600512107 |
| **A_55_P2135526** | Gzmc | -0.925909395 | -3.105852525 | 1.254033735 | 8.263997541 | -1.063833525 | 0.332040502 | 0.709264768 | -4.600555285 |
| **A_52_P65494** | Iqgap2 | -0.220548486 | -0.739818215 | 0.298721243 | 7.884443637 | -1.063803514 | 0.332052957 | 0.709264768 | -4.600562991 |
| **A_55_P2157478** | NA | -0.32443008 | -1.088313251 | 0.439453091 | 8.240521881 | -1.06376211 | 0.33207014 | 0.709264768 | -4.600573623 |
| **A_51_P288876** | Tmem45a | 0.651699618 | -0.882767012 | 2.186166249 | 9.818853613 | 1.063751937 | 0.332074362 | 0.709264768 | -4.600576235 |
| **A_52_P181759** | Eif5a2 | 0.092649887 | -0.125505375 | 0.310805148 | 8.794742626 | 1.063725441 | 0.332085359 | 0.709264768 | -4.600583039 |
| **A_55_P1983235** | Tmed10 | -0.537955972 | -1.804704207 | 0.728792264 | 14.47032697 | -1.063669657 | 0.332108512 | 0.709264768 | -4.600597362 |
| **A_52_P590392** | Rab10 | 0.122201699 | -0.165590155 | 0.409993553 | 12.10185118 | 1.063528406 | 0.332167145 | 0.709264768 | -4.60063363 |
| **A_51_P297679** | Hcls1 | 0.488721137 | -0.662284598 | 1.639726873 | 10.11420243 | 1.063491396 | 0.332182509 | 0.709264768 | -4.600643133 |
| **A_55_P2072945** | NA | 0.260352714 | -0.3528623 | 0.873567727 | 16.5230097 | 1.063407393 | 0.332217383 | 0.709264768 | -4.6006647 |
| **A_66_P109708** | Il1f6 | -0.183182199 | -0.61466238 | 0.248297982 | 7.7323481 | -1.063341494 | 0.332244743 | 0.709264768 | -4.600681619 |
| **A_55_P2155996** | NA | 0.388691833 | -0.526924448 | 1.304308113 | 15.73935846 | 1.063266803 | 0.332275757 | 0.709264768 | -4.600700795 |
| **A_55_P2011672** | NA | 0.366931105 | -0.497430281 | 1.23129249 | 14.13727992 | 1.0632601 | 0.33227854 | 0.709264768 | -4.600702516 |
| **A_55_P2108248** | Art4 | 0.483688241 | -0.655724063 | 1.623100546 | 10.55115583 | 1.063248969 | 0.332283162 | 0.709264768 | -4.600705373 |
| **A_55_P1962901** | Osr1 | 0.458663372 | -0.665888264 | 1.583215007 | 8.786625316 | 1.086368996 | 0.332312827 | 0.709264768 | -4.591662277 |
| **A_55_P1980956** | Osbp | -0.37450439 | -1.256792018 | 0.507783239 | 8.077621973 | -1.063156182 | 0.332321694 | 0.709264768 | -4.600729194 |
| **A_55_P1957173** | NA | 0.195941759 | -0.265727817 | 0.657611335 | 8.455464387 | 1.063031401 | 0.332373516 | 0.709264768 | -4.600761227 |
| **A_52_P578436** | NA | -0.288214813 | -0.967362617 | 0.390932991 | 7.950076018 | -1.062924317 | 0.332417995 | 0.709264768 | -4.600788716 |
| **A_55_P2313861** | Zbed6 | -0.266268998 | -0.893746613 | 0.361208616 | 8.475022583 | -1.062851803 | 0.332448118 | 0.709264768 | -4.60080733 |
| **A_55_P2066543** | Orai2 | 0.445724436 | -0.604674081 | 1.496122953 | 9.371881123 | 1.062827272 | 0.332458308 | 0.709264768 | -4.600813627 |
| **A_55_P2077153** | Slc27a2 | -0.531291232 | -1.783405915 | 0.72082345 | 8.276971592 | -1.062769021 | 0.332482508 | 0.709264768 | -4.600828579 |
| **A_55_P1985633** | Lrp1 | -0.251254507 | -0.84349918 | 0.340990165 | 7.821696129 | -1.062583443 | 0.332559614 | 0.709264768 | -4.600876212 |
| **A_55_P2039881** | 5-Sep | -0.33554225 | -1.126542835 | 0.455458334 | 8.106184966 | -1.062479976 | 0.33260261 | 0.709264768 | -4.600902767 |
| **A_51_P406188** | Asb10 | -0.173279226 | -0.581798313 | 0.235239861 | 7.809889341 | -1.062391269 | 0.332639477 | 0.709264768 | -4.600925534 |
| **A_51_P430552** | G6pc3 | -0.3256036 | -1.093256825 | 0.442049626 | 11.09189792 | -1.062366736 | 0.332649673 | 0.709264768 | -4.60093183 |
| **A_55_P2144364** | NA | -0.423554037 | -1.422195679 | 0.575087605 | 8.269946 | -1.062305671 | 0.332675054 | 0.709264768 | -4.600947502 |
| **A_55_P1964076** | Dnajb6 | -0.354413702 | -1.190072532 | 0.481245129 | 9.380915324 | -1.06226248 | 0.332693007 | 0.709264768 | -4.600958586 |
| **A_55_P2042520** | Cop1 | 0.250144335 | -0.339670709 | 0.839959379 | 11.86404468 | 1.062246164 | 0.33269979 | 0.709264768 | -4.600962774 |
| **A_55_P2064557** | Furin | -0.461431167 | -1.549699892 | 0.626837558 | 11.52855036 | -1.0619917 | 0.332805579 | 0.709264768 | -4.601028074 |
| **A_55_P2298029** | Nsd3 | -0.662189986 | -2.224171679 | 0.899791707 | 8.957611012 | -1.061834947 | 0.33287076 | 0.709264768 | -4.601068297 |
| **A_52_P83913** | Ssx2ip | 0.130867357 | -0.177832618 | 0.439567332 | 8.022233754 | 1.061805776 | 0.332882891 | 0.709264768 | -4.601075782 |
| **A_52_P558859** | Rras2 | -0.305545844 | -1.026299923 | 0.415208236 | 9.949235462 | -1.061792409 | 0.33288845 | 0.709264768 | -4.601079211 |
| **A_65_P18433** | Ankrd28 | 0.151880816 | -0.206393441 | 0.510155074 | 9.766451898 | 1.061787635 | 0.332890435 | 0.709264768 | -4.601080436 |
| **A_51_P510418** | Aldh1b1 | 0.239243114 | -0.325124397 | 0.803610625 | 8.469010377 | 1.061763489 | 0.332900478 | 0.709264768 | -4.601086632 |
| **A_55_P1964053** | Golga4 | -0.506519962 | -1.701416268 | 0.688376344 | 11.64170138 | -1.061736366 | 0.332911758 | 0.709264768 | -4.601093591 |
| **A_51_P362738** | Dusp18 | 0.113436809 | -0.154173985 | 0.381047603 | 8.171014207 | 1.061697314 | 0.332928 | 0.709264768 | -4.601103611 |
| **A_52_P243249** | Mapk3 | 0.367732956 | -0.499800936 | 1.235266848 | 11.38168869 | 1.061686874 | 0.332932342 | 0.709264768 | -4.60110629 |
| **A_55_P1978061** | Hnrnpa3 | 0.365925238 | -0.497377441 | 1.229227917 | 9.5692992 | 1.061645738 | 0.332949452 | 0.709264768 | -4.601116844 |

| | | | | | | | | | |
|---|---|---|---|---|---|---|---|---|---|
| **A_52_P629895** | Adh1 | 1.858314514 | -2.526002943 | 6.24263197 | 13.90080607 | 1.061616285 | 0.332961703 | 0.709264768 | -4.601124401 |
| **A_55_P2151143** | Dynlt1c | 0.4455022 | -0.605745341 | 1.49674974 | 14.25822566 | 1.061439403 | 0.333035283 | 0.709264768 | -4.601169782 |
| **A_66_P112482** | Tmem184 | 0.11460849 | -0.155868804 | 0.385085783 | 10.15235843 | 1.061295499 | 0.333095156 | 0.709264768 | -4.6012067 |
| **A_55_P1988424** | Tpi1 | 0.337543502 | -0.459125618 | 1.134212622 | 10.83410248 | 1.061211905 | 0.33312994 | 0.709264768 | -4.601228145 |
| **A_52_P467690** | Sptbn1 | -0.163455187 | -0.549252172 | 0.222341798 | 10.37382217 | -1.061182852 | 0.33314203 | 0.709264768 | -4.601235598 |
| **A_55_P2038422** | Adamts17 | 1.236728216 | -1.682391474 | 4.155847906 | 11.46990232 | 1.061140066 | 0.333159836 | 0.709264768 | -4.601246574 |
| **A_55_P2166543** | Pam16 | 0.323802742 | -0.440525175 | 1.08813066 | 13.03616886 | 1.06108737 | 0.333181766 | 0.709264768 | -4.601260092 |
| **A_52_P288873** | Chmp4b | -0.387986888 | -1.303825327 | 0.52785155 | 12.97111282 | -1.061080975 | 0.333184428 | 0.709264768 | -4.601261732 |
| **A_55_P2051791** | Cbr4 | -0.089992997 | -0.302441591 | 0.122455597 | 8.419333246 | -1.060975121 | 0.333228485 | 0.709264768 | -4.601288885 |
| **A_51_P333923** | Tspan1 | -0.252423861 | -0.848406032 | 0.34355831 | 11.70792064 | -1.060834128 | 0.333287174 | 0.709264768 | -4.60132505 |
| **A_66_P113659** | Ripply1 | -0.403263904 | -1.355403643 | 0.548875836 | 7.916174372 | -1.060813417 | 0.333295796 | 0.709264768 | -4.601330362 |
| **A_55_P2031631** | Igf1 | 0.593845506 | -0.808338055 | 1.996029067 | 10.00431887 | 1.06076443 | 0.33331619 | 0.709264768 | -4.601342927 |
| **A_55_P2185548** | Cdk8 | 0.108679243 | -0.147949833 | 0.365308319 | 10.72266526 | 1.060696389 | 0.333344518 | 0.709264768 | -4.601360378 |
| **A_55_P2279685** | 2900076A | 0.169452268 | -0.230714379 | 0.569618914 | 8.405042322 | 1.060612749 | 0.333379343 | 0.709264768 | -4.60138183 |
| **A_52_P254174** | Nr3c1 | -0.155169825 | -0.521628361 | 0.211288711 | 9.129507648 | -1.060553967 | 0.33340382 | 0.709264768 | -4.601396906 |
| **A_55_P2152248** | Cpe | -0.415313926 | -1.396147784 | 0.565519933 | 8.370046146 | -1.060550589 | 0.333405226 | 0.709264768 | -4.601397772 |
| **A_52_P354682** | Elovl7 | 0.480551316 | -0.654565381 | 1.615668013 | 9.400948987 | 1.060350889 | 0.333488393 | 0.709264768 | -4.601448987 |
| **A_55_P1987264** | Gnb2 | -0.182167408 | -0.61252011 | 0.248185295 | 15.52685681 | -1.060221212 | 0.333542407 | 0.709264768 | -4.601482241 |
| **A_52_P275069** | Epp13 | 0.182503104 | -0.248689376 | 0.613695583 | 8.098772626 | 1.060106319 | 0.333590269 | 0.709264768 | -4.601511703 |
| **A_55_P2057291** | Kansl3 | -0.151883206 | -0.510763051 | 0.206996638 | 12.12589677 | -1.060012616 | 0.333629308 | 0.709264768 | -4.601535731 |
| **A_52_P123864** | Rab21 | -0.156000231 | -0.524629262 | 0.212628801 | 8.4189772 | -1.05995164 | 0.333654714 | 0.709264768 | -4.601551365 |
| **A_51_P358112** | Fads1 | 0.20679732 | -0.281865904 | 0.695460543 | 8.849088589 | 1.059950529 | 0.333655177 | 0.709264768 | -4.60155165 |
| **A_51_P174645** | Asl | -0.154094045 | -0.51822137 | 0.21003328 | 8.928748329 | -1.059944023 | 0.333657888 | 0.709264768 | -4.601553318 |
| **A_55_P1961863** | Pcdhga9 | 0.605963059 | -0.826082546 | 2.038008664 | 13.39542716 | 1.059838389 | 0.333701906 | 0.709264768 | -4.601580403 |
| **A_55_P2017710** | Igsf9 | -0.860683613 | -2.894717954 | 1.173350729 | 9.863586166 | -1.059828511 | 0.333706022 | 0.709264768 | -4.601582936 |
| **A_55_P2151822** | Plxnd1 | 0.414438582 | -0.565088156 | 1.39396532 | 13.71421512 | 1.05972756 | 0.333748093 | 0.709264768 | -4.601608819 |
| **A_52_P210338** | Ndufb6 | 0.372331105 | -0.507753683 | 1.252415893 | 12.92806474 | 1.059632209 | 0.333787834 | 0.709264768 | -4.601633265 |
| **A_51_P379478** | Nckap5 | 0.765968078 | -1.044672365 | 2.57660852 | 9.976850161 | 1.059567917 | 0.333814633 | 0.709264768 | -4.601649748 |
| **A_52_P647607** | Ubr4 | -0.326356036 | -1.097820768 | 0.445108696 | 8.345470771 | -1.059560887 | 0.333817563 | 0.709264768 | -4.60165155 |
| **A_55_P1963036** | Oxr1 | 0.179571462 | -0.244982818 | 0.604125742 | 11.04648697 | 1.059386516 | 0.333890255 | 0.709264768 | -4.601696252 |
| **A_55_P2064230** | Tmx3 | 0.27332259 | -0.372988049 | 0.919633228 | 9.114593397 | 1.059216086 | 0.333961317 | 0.709264768 | -4.601739941 |
| **A_51_P205278** | Tmem161 | -0.093198969 | -0.313593171 | 0.127195233 | 8.356481272 | -1.059159313 | 0.333984991 | 0.709264768 | -4.601754494 |
| **A_55_P2058681** | Tmc8 | 0.174571539 | -0.238259761 | 0.58740284 | 9.2947 | 1.059134633 | 0.333995283 | 0.709264768 | -4.601760821 |
| **A_51_P398723** | Flt1 | -0.650266723 | -2.188167378 | 0.887633933 | 12.62451242 | -1.059042998 | 0.334033499 | 0.709264768 | -4.601784309 |
| **A_52_P289256** | Arid1b | -0.416529031 | -1.401671491 | 0.568613429 | 10.61993109 | -1.059001521 | 0.334050798 | 0.709264768 | -4.60179494 |
| **A_52_P351816** | Itgb1 | -0.256096117 | -0.861845667 | 0.349653434 | 9.489022618 | -1.058912899 | 0.334087763 | 0.709264768 | -4.601817655 |
| **A_55_P2064841** | Prss53 | 0.294045736 | -0.401547237 | 0.98963871 | 13.11611941 | 1.058790524 | 0.334138812 | 0.709264768 | -4.60184902 |
| **A_51_P252859** | Ccn1 | -0.523674344 | -1.762509068 | 0.715160379 | 10.16766514 | -1.058761824 | 0.334150785 | 0.709264768 | -4.601856375 |
| **A_55_P1957018** | Caskin2 | -0.109347298 | -0.368035508 | 0.149340912 | 10.48191065 | -1.058721578 | 0.334167575 | 0.709264768 | -4.60186669 |
| **A_52_P227391** | Kif15 | 0.336223006 | -0.459233468 | 1.131679479 | 8.534982643 | 1.058671815 | 0.334188337 | 0.709264768 | -4.601879444 |
| **A_51_P317214** | Hpdl | -0.272517516 | -0.917368459 | 0.372333428 | 8.015557336 | -1.058486752 | 0.334265558 | 0.709264768 | -4.60192687 |
| **A_55_P1967820** | AI661453 | -0.556037377 | -1.871881952 | 0.759807199 | 9.025903165 | -1.05839977 | 0.334301858 | 0.709264768 | -4.601949161 |
| **A_55_P2074521** | Psg28 | -2.830566172 | -9.529083687 | 3.867951343 | 9.50427921 | -1.05838813 | 0.334306716 | 0.709264768 | -4.601952143 |
| **A_55_P2085994** | Zfat | -0.171143018 | -0.576155564 | 0.233869529 | 8.63502375 | -1.058378608 | 0.33431069 | 0.709264768 | -4.601954584 |
| **A_55_P2046907** | Wdr83os | 0.234608294 | -0.320614653 | 0.789831241 | 11.66231721 | 1.058342654 | 0.334325696 | 0.709264768 | -4.601963797 |
| **A_52_P484081** | Ube2v1 | 0.126370802 | -0.172699044 | 0.425440648 | 9.48595172 | 1.058338169 | 0.334327568 | 0.709264768 | -4.601964946 |
| **A_55_P2136743** | Slc36a4 | -0.31077536 | -1.046262741 | 0.424712021 | 8.411310894 | -1.05833144 | 0.334330376 | 0.709264768 | -4.60196667 |
| **A_52_P487024** | Smg7 | -0.187016492 | -0.62964472 | 0.255611735 | 9.224451184 | -1.05825699 | 0.334361451 | 0.709264768 | -4.601985748 |

| | | | | | | | | | |
|---|---|---|---|---|---|---|---|---|---|
| **A_51_P242166** | Lap3 | 0.382691903 | -0.523125032 | 1.288508838 | 12.08843315 | 1.058179109 | 0.33439396 | 0.709264768 | -4.602005704 |
| **A_51_P252410** | Cope | 0.167777977 | -0.229447037 | 0.56500299 | 11.80179049 | 1.05790997 | 0.334506326 | 0.709264768 | -4.602074664 |
| **A_55_P2020331** | Camk1g | 1.128907823 | -1.543934413 | 3.801750059 | 11.60850774 | 1.057877799 | 0.334519759 | 0.709264768 | -4.602082906 |
| **A_51_P380178** | Id3 | 0.319872222 | -0.437475076 | 1.07721952 | 12.07732433 | 1.05786876 | 0.334523534 | 0.709264768 | -4.602085222 |
| **A_55_P2107357** | Gm15421 | 0.278205312 | -0.380549983 | 0.936960607 | 14.99984775 | 1.057770972 | 0.334564369 | 0.709264768 | -4.602110275 |
| **A_55_P2130890** | Parva | -0.517630708 | -1.743438339 | 0.708176922 | 10.97432427 | -1.057664817 | 0.334608704 | 0.709264768 | -4.602137471 |
| **A_52_P106789** | Rbm18 | -0.103009224 | -0.346963161 | 0.140944712 | 11.75896358 | -1.057593157 | 0.334638635 | 0.709264768 | -4.602155829 |
| **A_55_P1956457** | 6030458C | 0.103215031 | -0.141227913 | 0.347657975 | 9.654426756 | 1.057586229 | 0.334641529 | 0.709264768 | -4.602157604 |
| **A_55_P1974984** | NA | 0.680048949 | -0.930549929 | 2.290647827 | 9.868793269 | 1.0575554 | 0.334654407 | 0.709264768 | -4.602165501 |
| **A_51_P352303** | Homer2 | -0.248029318 | -0.835542966 | 0.33948433 | 8.716693195 | -1.0573905 | 0.334723293 | 0.709264768 | -4.602207743 |
| **A_51_P200667** | Clmn | 0.39781116 | -0.544506578 | 1.340128898 | 9.479157003 | 1.057377201 | 0.334728849 | 0.709264768 | -4.60221115 |
| **A_55_P2022569** | Pak3 | 0.206491143 | -0.282685348 | 0.695667634 | 8.165402524 | 1.057270698 | 0.334773347 | 0.709264768 | -4.602238431 |
| **A_55_P2084656** | H2ac12 | 0.714679368 | -0.978758941 | 2.408117677 | 17.36130631 | 1.057041918 | 0.334868951 | 0.709264768 | -4.602297029 |
| **A_52_P284821** | Slco4c1 | 0.487106475 | -0.667168399 | 1.641381348 | 8.937934363 | 1.056975707 | 0.334896624 | 0.709264768 | -4.602313987 |
| **A_52_P264905** | Cab39l | 0.358815582 | -0.491488416 | 1.20911958 | 11.05584854 | 1.056932907 | 0.334914513 | 0.709264768 | -4.602324949 |
| **A_55_P2109857** | Rgs2 | 1.364126362 | -1.868616407 | 4.596869131 | 13.32869941 | 1.056899758 | 0.334928369 | 0.709264768 | -4.602333439 |
| **A_55_P2086885** | Slc45a4 | -1.145138991 | -3.859019675 | 1.568741693 | 9.534234392 | -1.05686082 | 0.334944645 | 0.709264768 | -4.602343411 |
| **A_51_P158120** | B3galnt2 | -0.20027975 | -0.675011854 | 0.274452354 | 9.736605313 | -1.056668556 | 0.335025022 | 0.709264768 | -4.602392649 |
| **A_55_P2117445** | NA | 0.326276663 | -0.447129115 | 1.099682442 | 13.07627965 | 1.056644618 | 0.335035031 | 0.709264768 | -4.602398779 |
| **A_66_P115467** | Sfta2 | 2.185544762 | -2.995277823 | 7.366367347 | 14.17659254 | 1.056601763 | 0.335052949 | 0.709264768 | -4.602409754 |
| **A_52_P588483** | Fbln1 | -0.640720963 | -2.15961201 | 0.878170084 | 11.46326548 | -1.056556331 | 0.335071946 | 0.709264768 | -4.602421388 |
| **A_55_P1968143** | Rbm25 | 0.128957861 | -0.176777953 | 0.434693676 | 10.45518658 | 1.056457063 | 0.335113456 | 0.709264768 | -4.602446808 |
| **A_51_P280245** | Zfp207 | 0.341829504 | -0.468606922 | 1.15226593 | 11.80225697 | 1.056430517 | 0.335124558 | 0.709264768 | -4.602453605 |
| **A_55_P2049999** | Slc9a1 | -0.166165043 | -0.560154895 | 0.227824809 | 11.59253524 | -1.056343034 | 0.335161145 | 0.709264768 | -4.602476006 |
| **A_65_P08291** | Gpalpp1 | 0.195681862 | -0.268382678 | 0.659746401 | 11.33992746 | 1.056142533 | 0.335245011 | 0.709264768 | -4.602527343 |
| **A_55_P2121961** | Psmc3 | 0.25695223 | -0.352421952 | 0.866326412 | 12.9562135 | 1.056133178 | 0.335248924 | 0.709264768 | -4.602529738 |
| **A_55_P2018210** | NA | 0.184212237 | -0.252677436 | 0.621101911 | 16.72973746 | 1.056080547 | 0.335270942 | 0.709264768 | -4.602543214 |
| **A_55_P2000117** | Ceacam1: | -2.698363442 | -9.098498101 | 3.701771217 | 9.368282714 | -1.055994553 | 0.335306919 | 0.709264768 | -4.60256523 |
| **A_55_P2039215** | Acyp2 | 0.509562028 | -0.699127916 | 1.718251973 | 9.349633107 | 1.055923594 | 0.335336608 | 0.709264768 | -4.602583397 |
| **A_55_P2061294** | NA | 0.155511538 | -0.213508829 | 0.524531906 | 13.63031238 | 1.055510666 | 0.335509423 | 0.709264768 | -4.602689104 |
| **A_55_P2055291** | Gpn2 | 0.11117408 | -0.152641148 | 0.374989308 | 9.044457733 | 1.055489775 | 0.335518168 | 0.709264768 | -4.602694451 |
| **A_52_P686785** | Lyve1 | 0.382437125 | -0.525095807 | 1.289970057 | 11.16291725 | 1.05547511 | 0.335524307 | 0.709264768 | -4.602698205 |
| **A_51_P358700** | Olfr1221 | -0.170079602 | -0.573693574 | 0.23353437 | 7.798145487 | -1.055446891 | 0.33553612 | 0.709264768 | -4.602705428 |
| **A_51_P211786** | Chst13 | -0.304502126 | -1.027145358 | 0.418141107 | 7.824262008 | -1.055399137 | 0.335556112 | 0.709264768 | -4.602717652 |
| **A_55_P2043367** | Apol9a | -0.696942973 | -2.350937786 | 0.957051839 | 8.467471505 | -1.055391146 | 0.335559457 | 0.709264768 | -4.602719697 |
| **A_52_P441974** | Evc2 | 0.331121948 | -0.454955631 | 1.117199527 | 10.24200246 | 1.055049667 | 0.335702443 | 0.709264768 | -4.602807096 |
| **A_51_P353386** | Micos13 | -0.157424865 | -0.531158739 | 0.216309009 | 12.17752317 | -1.055021265 | 0.335714338 | 0.709264768 | -4.602814365 |
| **A_52_P172014** | Ramp1 | -0.148785898 | -0.50202147 | 0.204449675 | 7.855853541 | -1.054988396 | 0.335728104 | 0.709264768 | -4.602822777 |
| **A_55_P2087985** | NA | 0.183200114 | -0.251775928 | 0.618176155 | 12.73195706 | 1.054898677 | 0.335765682 | 0.709264768 | -4.602845738 |
| **A_55_P2076772** | Hspa5 | -0.6050116 | -2.041547646 | 0.831524446 | 15.15964882 | -1.054866545 | 0.335779141 | 0.709264768 | -4.602853961 |
| **A_52_P80965** | Sephs1 | 0.242362258 | -0.3331129 | 0.817837415 | 8.411526898 | 1.054845257 | 0.335788058 | 0.709264768 | -4.602859408 |
| **A_55_P2143311** | Pdxk | 0.115865606 | -0.159252052 | 0.390983264 | 8.303916965 | 1.054839581 | 0.335790436 | 0.709264768 | -4.602860861 |
| **A_51_P165475** | C1galt1 | -0.645229628 | -2.177313478 | 0.886854222 | 8.964195649 | -1.054829124 | 0.335794817 | 0.709264768 | -4.602863537 |
| **A_55_P2031011** | S100pbp | -0.433100025 | -1.461546221 | 0.595346171 | 8.245226401 | -1.054768079 | 0.335820389 | 0.709264768 | -4.602879158 |
| **A_52_P560146** | NA | -0.319341706 | -1.07770981 | 0.439026398 | 8.386378234 | -1.054692671 | 0.33585198 | 0.709264768 | -4.602898455 |
| **A_55_P2103206** | NA | -0.246265652 | -0.83110618 | 0.338574877 | 7.95313915 | -1.054670326 | 0.335861342 | 0.709264768 | -4.602904173 |
| **A_55_P2016968** | NA | 0.122175942 | -0.168021482 | 0.412373366 | 15.96640571 | 1.054490053 | 0.335936877 | 0.709264768 | -4.602950301 |
| **A_51_P506417** | Krt14 | -0.710362458 | -2.397843701 | 0.977118786 | 8.349176175 | -1.054365999 | 0.335988864 | 0.709264768 | -4.602982043 |

| | | | | | | | | | |
|---|---|---|---|---|---|---|---|---|---|
| **A_55_P1957880** | Jpt2 | 0.203890345 | -0.280565847 | 0.688346537 | 9.434730355 | 1.054125927 | 0.33608949 | 0.709264768 | -4.603043464 |
| **A_55_P2009952** | Me1 | -0.334528626 | -1.129507605 | 0.460450353 | 11.42785388 | -1.053969363 | 0.336155127 | 0.709264768 | -4.603083518 |
| **A_55_P2028977** | Tcf20 | -0.233928801 | -0.789884714 | 0.322027113 | 8.126984101 | -1.053886122 | 0.336190029 | 0.709264768 | -4.603104812 |
| **A_51_P132013** | Cysltr2 | -0.508627323 | -1.717540366 | 0.70028572 | 7.967340345 | -1.053792174 | 0.336229423 | 0.709264768 | -4.603128845 |
| **A_52_P628117** | Abcc5 | -0.124896608 | -0.421764584 | 0.171971367 | 8.33396212 | -1.053750126 | 0.336247056 | 0.709264768 | -4.603139601 |
| **A_52_P78123** | Dcaf12 | -0.451699578 | -1.52535093 | 0.621951773 | 10.16829097 | -1.053747981 | 0.336247956 | 0.709264768 | -4.603140149 |
| **A_55_P2009013** | Dpf2 | 0.279577184 | -0.385003007 | 0.944157375 | 11.72291188 | 1.053670157 | 0.336280594 | 0.709264768 | -4.603160056 |
| **A_51_P286865** | Wdr33 | 0.12576731 | -0.173199583 | 0.424734203 | 10.12542271 | 1.053646721 | 0.336290423 | 0.709264768 | -4.603166051 |
| **A_52_P3845** | Rnf217 | 0.19414518 | -0.267366372 | 0.655656731 | 8.634258596 | 1.053645174 | 0.336291072 | 0.709264768 | -4.603166447 |
| **A_52_P263518** | Gng2 | 0.751151738 | -1.034459476 | 2.536762953 | 11.07414273 | 1.053637253 | 0.336294394 | 0.709264768 | -4.603168473 |
| **A_55_P2024868** | Psg18 | -2.836312195 | -9.579422328 | 3.906797938 | 9.518910644 | -1.053523251 | 0.336342211 | 0.709264768 | -4.603197632 |
| **A_52_P260105** | Man1a | -0.412772828 | -1.394144735 | 0.568599079 | 7.952573727 | -1.053483711 | 0.336358797 | 0.709264768 | -4.603207746 |
| **A_51_P284946** | Rnd3 | 0.453500051 | -0.624800382 | 1.531800483 | 10.98518478 | 1.053386883 | 0.336399416 | 0.709264768 | -4.603232511 |
| **A_55_P2134356** | Msl3 | 0.100573638 | -0.138565209 | 0.339712485 | 11.29929718 | 1.053378319 | 0.336403009 | 0.709264768 | -4.603234702 |
| **A_55_P2070661** | Taf7l | -1.460398618 | -4.933021302 | 2.012224067 | 10.70414536 | -1.05332929 | 0.336423578 | 0.709264768 | -4.603247241 |
| **A_52_P140881** | Slc26a10 | -0.179617222 | -0.606751391 | 0.247516947 | 7.792243982 | -1.053256158 | 0.336454262 | 0.709264768 | -4.603265945 |
| **A_51_P461005** | Nrk | -1.503722665 | -5.079901739 | 2.07245641 | 9.398024956 | -1.053170859 | 0.336490053 | 0.709264768 | -4.60328776 |
| **A_55_P1971269** | NA | 0.58753223 | -0.80983314 | 1.984897601 | 12.98169557 | 1.053105946 | 0.336517292 | 0.709264768 | -4.60330436 |
| **A_55_P2041082** | NA | -0.13270567 | -0.448361079 | 0.182949738 | 7.979828259 | -1.052995682 | 0.336563566 | 0.709264768 | -4.603332558 |
| **A_52_P183181** | Adar | -0.271752666 | -0.918213452 | 0.374708121 | 9.295764437 | -1.052887506 | 0.33660897 | 0.709264768 | -4.603360221 |
| **A_55_P2046703** | Atp5g3 | 0.149590403 | -0.206281792 | 0.505462599 | 15.47232608 | 1.052834289 | 0.336631307 | 0.709264768 | -4.603373829 |
| **A_52_P139097** | NA | 0.140512325 | -0.193780525 | 0.474805174 | 10.84566577 | 1.052780123 | 0.336654045 | 0.709264768 | -4.603387679 |
| **A_51_P455647** | Car2 | -0.152222393 | -0.514381065 | 0.209936278 | 12.50518451 | -1.052761521 | 0.336661854 | 0.709264768 | -4.603392436 |
| **A_55_P1957038** | Gstp2 | 0.252807332 | -0.348745872 | 0.854360536 | 14.29541664 | 1.052606304 | 0.336727018 | 0.709264768 | -4.603432124 |
| **A_55_P2025702** | C330007F | -0.163579508 | -0.552852551 | 0.225693535 | 8.363090283 | -1.052506806 | 0.336768796 | 0.709264768 | -4.603457563 |
| **A_51_P379660** | Mllt11 | -0.242961913 | -0.821176503 | 0.335252678 | 10.13322944 | -1.052445214 | 0.336794659 | 0.709264768 | -4.603473311 |
| **A_55_P2131522** | Arfrp1 | 0.178332887 | -0.246078733 | 0.602744506 | 11.3688282 | 1.052433152 | 0.336799724 | 0.709264768 | -4.603476394 |
| **A_55_P2041265** | Utp6 | -0.267170624 | -0.903012 | 0.368670753 | 15.21325317 | -1.052422839 | 0.336804055 | 0.709264768 | -4.603479031 |
| **A_66_P114769** | NA | 0.110566282 | -0.15257672 | 0.373709285 | 10.85976204 | 1.052400939 | 0.336813252 | 0.709264768 | -4.60348463 |
| **A_55_P2065231** | Gstm3 | 0.257929008 | -0.355947591 | 0.871805607 | 9.065924848 | 1.052372415 | 0.336825231 | 0.709264768 | -4.603491923 |
| **A_55_P1992505** | Entpd4 | -0.145184034 | -0.490848865 | 0.200480797 | 11.32759713 | -1.05199581 | 0.336983421 | 0.709264768 | -4.603588199 |
| **A_55_P1992054** | Armc10 | 0.356993446 | -0.492970023 | 1.206956914 | 12.0530835 | 1.051986891 | 0.336987169 | 0.709264768 | -4.603590479 |
| **A_52_P308465** | Plxnb1 | -0.094302076 | -0.318845327 | 0.130241175 | 8.599427322 | -1.051893066 | 0.337026589 | 0.709264768 | -4.603614462 |
| **A_55_P2181371** | NA | 0.194401657 | -0.268588386 | 0.657391699 | 8.569737259 | 1.051667993 | 0.33712117 | 0.709264768 | -4.603671991 |
| **A_55_P2181262** | NA | 0.59543123 | -0.822772563 | 2.013635024 | 13.70757233 | 1.051582408 | 0.33715714 | 0.709264768 | -4.603693866 |
| **A_51_P348665** | Ramp1 | -0.729225207 | -2.466130312 | 1.007679898 | 9.648793697 | -1.051564525 | 0.337164656 | 0.709264768 | -4.603698436 |
| **A_52_P550185** | Urm1 | -0.115495273 | -0.390601798 | 0.159611253 | 11.44457131 | -1.051510616 | 0.337187316 | 0.709264768 | -4.603712214 |
| **A_55_P2131340** | Churc1 | 0.206187299 | -0.284948635 | 0.697323232 | 11.40782213 | 1.051503068 | 0.337190488 | 0.709264768 | -4.603714143 |
| **A_55_P2051384** | Ccdc88c | 0.109317685 | -0.151116431 | 0.369751801 | 8.55930293 | 1.051339294 | 0.337259335 | 0.709264768 | -4.603755999 |
| **A_52_P276343** | Pum3 | 0.234683877 | -0.32443188 | 0.793799634 | 8.551679065 | 1.051312596 | 0.337270559 | 0.709264768 | -4.603762821 |
| **A_51_P335306** | Ppm1a | -0.207855871 | -0.703154197 | 0.287442455 | 9.065634501 | -1.051104225 | 0.337358173 | 0.709264768 | -4.603816069 |
| **A_55_P1952920** | AI854703 | -0.212647803 | -0.719444256 | 0.29414865 | 7.975043646 | -1.050939414 | 0.337427484 | 0.709264768 | -4.603858183 |
| **A_51_P472829** | Aif1l | -1.090798097 | -3.690601804 | 1.50900561 | 10.89062657 | -1.050882496 | 0.337451424 | 0.709264768 | -4.603872726 |
| **A_51_P515965** | Nfe2 | -0.161401332 | -0.546088961 | 0.223286296 | 8.672447508 | -1.050870597 | 0.337456428 | 0.709264768 | -4.603875767 |
| **A_55_P1958379** | Slit1 | -1.579797888 | -5.345223614 | 2.185627837 | 9.166758793 | -1.050842888 | 0.337468084 | 0.709264768 | -4.603882847 |
| **A_51_P509223** | Arfgap1 | -0.321365767 | -1.087358255 | 0.444626721 | 8.864186105 | -1.050813025 | 0.337480645 | 0.709264768 | -4.603890477 |
| **A_51_P451428** | Marveld2 | -0.117607707 | -0.397932841 | 0.162717428 | 8.512174994 | -1.050809733 | 0.33748203 | 0.709264768 | -4.603891318 |
| **A_55_P1965846** | Sp2 | 0.128253981 | -0.177453207 | 0.43396117 | 9.554948498 | 1.050789079 | 0.337490718 | 0.709264768 | -4.603896595 |

| | | | | | | | | | |
|---|---|---|---|---|---|---|---|---|---|
| **A_51_P218975** | Kcns3 | 0.585772406 | -0.810495517 | 1.982040328 | 9.347899098 | 1.050776845 | 0.337495864 | 0.709264768 | -4.603899721 |
| **A_52_P145349** | Stradb | -0.366284009 | -1.239385022 | 0.506817004 | 9.17079249 | -1.050760679 | 0.337502665 | 0.709264768 | -4.603903852 |
| **A_55_P2422243** | 2900072G | 0.301936905 | -0.417813565 | 1.021687376 | 10.27158113 | 1.050714166 | 0.337522231 | 0.709264768 | -4.603915736 |
| **A_65_P08768** | Fbln1 | -0.560943793 | -1.898220951 | 0.776333365 | 10.60052874 | -1.050626295 | 0.337559199 | 0.709264768 | -4.603938186 |
| **A_55_P2034245** | Meox1 | 0.622299885 | -0.861343501 | 2.105943272 | 9.381588216 | 1.050559227 | 0.337587417 | 0.709264768 | -4.603955321 |
| **A_51_P501803** | Hoxa2 | 0.400518656 | -0.554428653 | 1.355465964 | 8.727074214 | 1.050494267 | 0.33761475 | 0.709264768 | -4.603971916 |
| **A_51_P334979** | Apoc2 | -1.018710726 | -3.447759576 | 1.410338123 | 9.426854965 | -1.050424822 | 0.337643972 | 0.709264768 | -4.603989658 |
| **A_55_P1962289** | NA | -0.143346298 | -0.485217119 | 0.198524524 | 9.096799748 | -1.050206716 | 0.337735763 | 0.709264768 | -4.604045374 |
| **A_55_P2062437** | Begain | 0.344602716 | -0.477305744 | 1.166511175 | 8.658655292 | 1.050136106 | 0.337765484 | 0.709264768 | -4.604063411 |
| **A_52_P590396** | Sort1 | 0.884546993 | -1.225307558 | 2.994401545 | 11.25838363 | 1.05007118 | 0.337792815 | 0.709264768 | -4.604079995 |
| **A_51_P404077** | Fzd2 | 0.929238156 | -1.28726182 | 3.145738133 | 11.3083615 | 1.050049202 | 0.337802067 | 0.709264768 | -4.604085609 |
| **A_55_P2092989** | NA | -0.243203601 | -0.82339975 | 0.336992547 | 9.073019556 | -1.049894125 | 0.337867355 | 0.709264768 | -4.604125219 |
| **A_55_P2025959** | Chd9 | 0.326058401 | -0.451857951 | 1.103974752 | 9.854471082 | 1.049815158 | 0.337900605 | 0.709264768 | -4.604145388 |
| **A_51_P386280** | Olfr187 | 0.09125952 | -0.126478639 | 0.308997678 | 7.884698609 | 1.049769566 | 0.337919803 | 0.709264768 | -4.604157032 |
| **A_51_P147064** | 1600014C | -0.556459072 | -1.884173888 | 0.771255744 | 8.154534119 | -1.04973281 | 0.337935281 | 0.709264768 | -4.60416642 |
| **A_55_P2118630** | Lasp1 | -0.352943551 | -1.195093838 | 0.489206736 | 8.157235574 | -1.04970193 | 0.337948285 | 0.709264768 | -4.604174306 |
| **A_55_P2013716** | Napa | -0.390055057 | -1.320781276 | 0.540671163 | 12.45073426 | -1.049673684 | 0.33796018 | 0.709264768 | -4.60418152 |
| **A_55_P2010966** | Zfp688 | -0.163231087 | -0.552759031 | 0.226296858 | 9.487416052 | -1.049577708 | 0.338000601 | 0.709264768 | -4.604206031 |
| **A_55_P2054240** | Med8 | -0.225950764 | -0.765153431 | 0.313251903 | 12.90087353 | -1.049571739 | 0.338003116 | 0.709264768 | -4.604207555 |
| **A_55_P2170514** | Sfn | -0.328734031 | -1.113235797 | 0.455767736 | 8.428819265 | -1.04954505 | 0.338014357 | 0.709264768 | -4.604214371 |
| **A_55_P2009994** | NA | 0.151226267 | -0.209680659 | 0.512133192 | 11.5872013 | 1.049499802 | 0.338033415 | 0.709264768 | -4.604225927 |
| **A_55_P2005213** | Ces2c | -0.098742817 | -0.334409181 | 0.136923547 | 7.952872539 | -1.049441554 | 0.338057951 | 0.709264768 | -4.604240801 |
| **A_51_P400555** | Cts3 | -2.838910076 | -9.614984588 | 3.937164436 | 9.550527227 | -1.049358317 | 0.338093015 | 0.709264768 | -4.604262057 |
| **A_51_P461040** | Crct1 | -0.791447642 | -2.680561229 | 1.097665944 | 8.254876766 | -1.049335737 | 0.338102527 | 0.709264768 | -4.604267823 |
| **A_55_P2125496** | Gm10731 | 0.844953416 | -1.171913245 | 2.861820077 | 11.76748128 | 1.049315105 | 0.338111219 | 0.709264768 | -4.604273092 |
| **A_52_P630493** | Dnajb6 | -0.533277781 | -1.806210351 | 0.73965479 | 9.185830505 | -1.049297011 | 0.338118842 | 0.709264768 | -4.604277712 |
| **A_55_P2088601** | Fbln1 | -0.46940181 | -1.589902919 | 0.651099298 | 9.031980814 | -1.049259271 | 0.338134742 | 0.709264768 | -4.604287349 |
| **A_51_P383629** | Vps4a | 0.110954182 | -0.1539338 | 0.375842165 | 8.710532372 | 1.049135944 | 0.338186705 | 0.709264768 | -4.60431884 |
| **A_55_P1968778** | Rasa1 | 0.162342808 | -0.225275286 | 0.549960901 | 10.08079384 | 1.049009345 | 0.338240054 | 0.709264768 | -4.604351164 |
| **A_55_P2123831** | Thap6 | -0.20690343 | -0.700972029 | 0.287165169 | 8.958681463 | -1.048892025 | 0.338289498 | 0.709264768 | -4.604381118 |
| **A_51_P370163** | Ctr9 | -0.383318093 | -1.298663651 | 0.532027465 | 9.201661816 | -1.048877058 | 0.338295807 | 0.709264768 | -4.604384939 |
| **A_51_P203878** | Dynll2 | 0.101831105 | -0.141339577 | 0.345001787 | 11.19104716 | 1.048864989 | 0.338300894 | 0.709264768 | -4.604388021 |
| **A_65_P20799** | Abi2 | 0.127500437 | -0.176969523 | 0.431970397 | 8.154502962 | 1.048860112 | 0.338302949 | 0.709264768 | -4.604389266 |
| **A_52_P391081** | Hinfp | -0.168523279 | -0.571030203 | 0.233983645 | 8.341382128 | -1.048665282 | 0.338385077 | 0.709264768 | -4.604439006 |
| **A_51_P511375** | Ranbp2 | -0.157884233 | -0.535012146 | 0.21924368 | 10.53860556 | -1.048577245 | 0.338422192 | 0.709264768 | -4.60446148 |
| **A_55_P2034555** | Pdcd5 | 0.212854202 | -0.295649237 | 0.72135764 | 14.11015638 | 1.048428165 | 0.338485052 | 0.709264768 | -4.604499537 |
| **A_52_P542290** | Pdxdc1 | -0.351759955 | -1.192130181 | 0.488610271 | 9.316428348 | -1.048397763 | 0.338497872 | 0.709264768 | -4.604507297 |
| **A_51_P122649** | Degs2 | -0.896243374 | -3.037428453 | 1.244941704 | 8.376180796 | -1.048388131 | 0.338501933 | 0.709264768 | -4.602221468 |
| **A_65_P11193** | Flad1 | -0.298301792 | -1.010993632 | 0.414390048 | 9.228547287 | -1.048345468 | 0.338519925 | 0.709264768 | -4.604520646 |
| **A_51_P399625** | Cnih1 | -0.107296145 | -0.363651182 | 0.149058892 | 13.86357417 | -1.048316952 | 0.33853195 | 0.709264768 | -4.604527925 |
| **A_55_P2119478** | Ndufv2 | 0.226866839 | -0.315194041 | 0.768927718 | 13.58216443 | 1.048270345 | 0.338551606 | 0.709264768 | -4.604539822 |
| **A_52_P418665** | Ttpal | 0.172418935 | -0.23967397 | 0.584511841 | 11.19688425 | 1.047949048 | 0.338687135 | 0.709264768 | -4.604621827 |
| **A_55_P1953688** | Top1 | 0.105271932 | -0.146347844 | 0.356891708 | 11.89775554 | 1.047895962 | 0.338709532 | 0.709264768 | -4.604635376 |
| **A_51_P356525** | Trappc11 | -0.168470548 | -0.571184895 | 0.234243799 | 9.124449124 | -1.047797195 | 0.338751205 | 0.709264768 | -4.604660582 |
| **A_51_P469252** | Zfp647 | 0.325441682 | -0.452567117 | 1.103450481 | 8.832674796 | 1.047704992 | 0.338790112 | 0.709264768 | -4.604684112 |
| **A_51_P466910** | Cacna1g | 0.570561118 | -0.793446427 | 1.934568663 | 9.432314541 | 1.047697108 | 0.338793439 | 0.709264768 | -4.604686124 |
| **A_55_P1961034** | Gm13272 | 0.138611679 | -0.192773968 | 0.469997326 | 8.666702424 | 1.047650621 | 0.338813057 | 0.709264768 | -4.604697987 |
| **A_52_P389347** | Snap29 | -0.186463128 | -0.632288494 | 0.259362237 | 11.11403831 | -1.047559109 | 0.338851678 | 0.709264768 | -4.604721339 |

| | | | | | | | | | |
|---|---|---|---|---|---|---|---|---|---|
| **A_51_P170959** | NA | -0.168421618 | -0.571193686 | 0.234350451 | 7.857277501 | -1.047342759 | 0.338943001 | 0.709264768 | -4.604776545 |
| **A_55_P2060097** | Prdm5 | 0.628925427 | -0.875274792 | 2.133125647 | 9.825522837 | 1.047234284 | 0.338988796 | 0.709264768 | -4.604804222 |
| **A_55_P2155574** | NA | 0.594976833 | -0.828317448 | 2.018271115 | 14.60176606 | 1.047021735 | 0.339078544 | 0.709264768 | -4.604858451 |
| **A_55_P2044592** | NA | -0.120674814 | -0.409384696 | 0.168035068 | 13.15287151 | -1.046900324 | 0.339129818 | 0.709264768 | -4.604889425 |
| **A_51_P346893** | Extl1 | -0.372653491 | -1.264256147 | 0.518949165 | 7.95778041 | -1.046849347 | 0.339151348 | 0.709264768 | -4.604902429 |
| **A_55_P2049061** | NA | 0.105867601 | -0.147469251 | 0.359204453 | 9.84626561 | 1.046682704 | 0.339221739 | 0.709264768 | -4.60494494 |
| **A_55_P2012630** | NA | 0.398500495 | -0.555162465 | 1.352163455 | 15.54470494 | 1.046608592 | 0.339253048 | 0.709264768 | -4.604963845 |
| **A_55_P2007678** | Xntrpc | 0.372676825 | -0.519196696 | 1.264550345 | 9.038905586 | 1.046596943 | 0.339257969 | 0.709264768 | -4.604966816 |
| **A_55_P2012759** | Bicdl1 | -0.951989591 | -3.230691075 | 1.326711892 | 10.01020928 | -1.046393686 | 0.339343849 | 0.709264768 | -4.605018661 |
| **A_66_P132510** | Scamp3 | -0.138881095 | -0.471316281 | 0.19355409 | 12.30330433 | -1.046372929 | 0.339352621 | 0.709264768 | -4.605023955 |
| **A_55_P2019312** | Car12 | 0.225803141 | -0.31472183 | 0.766328111 | 8.32498306 | 1.046320085 | 0.339374952 | 0.709264768 | -4.605037433 |
| **A_51_P468140** | Serpind1 | -0.695783201 | -2.361569995 | 0.970003593 | 8.266223779 | -1.046176282 | 0.339435727 | 0.709264768 | -4.60507411 |
| **A_55_P2024376** | NA | 0.100101145 | -0.139572582 | 0.339774873 | 16.04184843 | 1.046089787 | 0.339472287 | 0.709264768 | -4.605096169 |
| **A_52_P238846** | Bpnt1 | 0.103694928 | -0.144586081 | 0.351975936 | 8.913680094 | 1.046078695 | 0.339476975 | 0.709264768 | -4.605098998 |
| **A_52_P671062** | Uba6 | -0.369943617 | -1.255748658 | 0.515861425 | 8.123739946 | -1.04603866 | 0.339493898 | 0.709264768 | -4.605109208 |
| **A_55_P2007673** | Lrsam1 | -0.40806946 | -1.385269535 | 0.569130616 | 10.00151572 | -1.04592597 | 0.339541537 | 0.709264768 | -4.605137946 |
| **A_55_P2122314** | Emc1 | -0.336705044 | -1.143052012 | 0.469641924 | 8.856078707 | -1.045870735 | 0.33956489 | 0.709264768 | -4.605152031 |
| **A_51_P500424** | Dnttip1 | 0.24735788 | -0.345064494 | 0.839780255 | 12.03767288 | 1.045790382 | 0.339598864 | 0.709264768 | -4.605172521 |
| **A_51_P161354** | Sesn2 | -0.563408256 | -1.912858032 | 0.78604152 | 9.787646323 | -1.045723401 | 0.339627186 | 0.709264768 | -4.605189601 |
| **A_51_P408561** | Oard1 | 0.291111611 | -0.406170559 | 0.98839378 | 10.58180554 | 1.045686047 | 0.339642982 | 0.709264768 | -4.605199126 |
| **A_55_P2031496** | Rufy3 | -0.16038221 | -0.544536129 | 0.22377171 | 8.419704276 | -1.045685944 | 0.339643025 | 0.709264768 | -4.605199152 |
| **A_55_P2183389** | NA | 0.462485153 | -0.645283933 | 1.57025424 | 15.33341394 | 1.045680236 | 0.339645439 | 0.709264768 | -4.605200608 |
| **A_51_P469968** | H2-M3 | -0.282086708 | -0.957771533 | 0.393598118 | 8.357849291 | -1.045655923 | 0.339655721 | 0.709264768 | -4.605206807 |
| **A_51_P104418** | Dusp10 | 0.205519052 | -0.286896787 | 0.69793489 | 8.228276047 | 1.045370931 | 0.339776258 | 0.709264768 | -4.605279472 |
| **A_65_P16081** | NA | -0.246743825 | -0.862268099 | 0.368780449 | 8.249192775 | -1.067735824 | 0.339797023 | 0.709264768 | -4.595568484 |
| **A_55_P2130600** | Musk | -0.266096057 | -0.903697239 | 0.371505126 | 7.849696219 | -1.045296919 | 0.339807567 | 0.709264768 | -4.605298342 |
| **A_51_P493886** | Gpt2 | -0.134213613 | -0.45581668 | 0.187389454 | 8.655093266 | -1.045265775 | 0.339820742 | 0.709264768 | -4.605306282 |
| **A_51_P383001** | Meox2 | 1.145131292 | -1.599284567 | 3.88954715 | 11.6921603 | 1.04509485 | 0.339893059 | 0.709264768 | -4.605349857 |
| **A_55_P1956812** | Fam83g | -0.866462363 | -2.943113934 | 1.210189208 | 9.278544176 | -1.045048421 | 0.339912705 | 0.709264768 | -4.605361693 |
| **A_55_P2046509** | Amot | -1.440622801 | -4.893399225 | 2.012153623 | 10.82648611 | -1.045038215 | 0.339917024 | 0.709264768 | -4.605364294 |
| **A_55_P2048279** | Tlr13 | -0.107937616 | -0.366654794 | 0.150779561 | 8.982338962 | -1.044955758 | 0.339951918 | 0.709264768 | -4.605385314 |
| **A_52_P615375** | H2aw | 0.157962397 | -0.220667805 | 0.536592599 | 11.03767194 | 1.044933874 | 0.339961179 | 0.709264768 | -4.605390893 |
| **A_51_P509489** | Kras | 0.223825039 | -0.312697398 | 0.760347476 | 10.20907809 | 1.044891331 | 0.339979184 | 0.709264768 | -4.605401737 |
| **A_51_P280401** | Tcta | -0.269376048 | -0.915156021 | 0.376403925 | 8.320136875 | -1.044779758 | 0.340026406 | 0.709264768 | -4.605430178 |
| **A_55_P2037478** | Kifap3 | 0.207348919 | -0.289735928 | 0.704433765 | 11.10128746 | 1.044772173 | 0.340029617 | 0.709264768 | -4.605432111 |
| **A_51_P183213** | Riiad1 | 0.446201764 | -0.623543483 | 1.515947012 | 9.034895576 | 1.044723258 | 0.340050322 | 0.709264768 | -4.605444579 |
| **A_55_P2093553** | NA | -0.309140986 | -1.050313504 | 0.432031532 | 7.902941623 | -1.044690452 | 0.340064208 | 0.709264768 | -4.605452941 |
| **A_51_P129100** | Sec63 | -0.382485017 | -1.299503739 | 0.534533704 | 8.715125508 | -1.044687916 | 0.340065282 | 0.709264768 | -4.602397527 |
| **A_55_P2054739** | Polr3c | 0.172897357 | -0.241679787 | 0.5874745 | 9.493871588 | 1.044559888 | 0.340119481 | 0.709264768 | -4.605486219 |
| **A_51_P483617** | 0610040J | 0.792578918 | -1.107966994 | 2.69312483 | 10.58785716 | 1.044514556 | 0.340138673 | 0.709264768 | -4.605497773 |
| **A_51_P382849** | Emb | -0.272580906 | -0.926218098 | 0.381056287 | 9.777239989 | -1.044501384 | 0.34014425 | 0.709264768 | -4.60550113 |
| **A_51_P503131** | Wnt11 | 0.268466151 | -0.375378907 | 0.912311209 | 8.616845925 | 1.044379926 | 0.340195677 | 0.709264768 | -4.605532085 |
| **A_55_P2083121** | Vill | -0.165351992 | -0.561912509 | 0.231208525 | 11.11451566 | -1.044360179 | 0.340204039 | 0.709264768 | -4.605537118 |
| **A_55_P1972025** | Mycl | 0.296830335 | -0.41506889 | 1.00872956 | 8.550959508 | 1.044335659 | 0.340214422 | 0.709264768 | -4.605543367 |
| **A_55_P1999841** | Cox7c | 0.416511718 | -0.582456977 | 1.415480414 | 13.39382507 | 1.044300993 | 0.340229102 | 0.709264768 | -4.605552201 |
| **A_55_P1952850** | NA | 0.533950059 | -0.746703706 | 1.814603824 | 14.46951905 | 1.044285515 | 0.340235657 | 0.709264768 | -4.605556146 |
| **A_52_P215106** | Pik3r6 | 0.382858962 | -0.53541841 | 1.301136334 | 10.39921635 | 1.044275959 | 0.340239704 | 0.709264768 | -4.605558581 |
| **A_55_P2005055** | Pepd | -0.306262108 | -1.040841907 | 0.428317692 | 12.30949012 | -1.044250355 | 0.340250547 | 0.709264768 | -4.605565106 |

| | | | | | | | | | |
|---|---|---|---|---|---|---|---|---|---|
| **A_55_P2150389** | Dnajc21 | 0.116181799 | -0.162492303 | 0.394855902 | 8.990085111 | 1.04421958 | 0.34026358 | 0.709264768 | -4.605572949 |
| **A_55_P2077994** | Ube2v2 | 0.362276587 | -0.50684266 | 1.231395833 | 9.639527771 | 1.044025834 | 0.340345642 | 0.709264768 | -4.605622321 |
| **A_51_P124535** | Mest | 0.700850913 | -0.980660423 | 2.38236225 | 14.98822522 | 1.043941565 | 0.34038134 | 0.709264768 | -4.604724532 |
| **A_55_P2006392** | NA | 0.146733149 | -0.205355145 | 0.498821443 | 16.37486668 | 1.043823326 | 0.340431433 | 0.709264768 | -4.605673922 |
| **A_55_P2033864** | Plekha8 | -0.308590146 | -1.049075334 | 0.431895042 | 9.314746903 | -1.043796952 | 0.340442608 | 0.709264768 | -4.605680642 |
| **A_55_P2135027** | NA | 0.126016726 | -0.176379993 | 0.428413444 | 9.963344878 | 1.043761955 | 0.340457436 | 0.709264768 | -4.605689559 |
| **A_55_P2115990** | 2010001K | -0.280978894 | -0.955345827 | 0.393388039 | 8.402411525 | -1.043584877 | 0.340532473 | 0.709264768 | -4.605734676 |
| **A_55_P2075274** | Rab11fip4 | -0.713157523 | -2.42479605 | 0.998481004 | 8.274429464 | -1.043575224 | 0.340536564 | 0.709264768 | -4.605737135 |
| **A_51_P315085** | Mia3 | -0.362823138 | -1.233629603 | 0.507983327 | 11.96021655 | -1.043575024 | 0.340536649 | 0.709264768 | -4.605737186 |
| **A_51_P148959** | Dhx36 | 0.250791703 | -0.351203321 | 0.852786726 | 10.42062104 | 1.043447514 | 0.34059069 | 0.709264768 | -4.605769671 |
| **A_55_P2152771** | Lhfpl2 | 0.229269018 | -0.321174667 | 0.779712703 | 9.867929855 | 1.043236619 | 0.340680087 | 0.709264768 | -4.605823396 |
| **A_55_P2082837** | Spin1 | -0.514938583 | -1.751298426 | 0.72142126 | 9.090529128 | -1.043183931 | 0.340702425 | 0.709264768 | -4.605836818 |
| **A_55_P1983733** | Aldh1l1 | 0.282174522 | -0.39540981 | 0.959758853 | 9.177014566 | 1.043049185 | 0.340759556 | 0.709264768 | -4.605871141 |
| **A_55_P2009915** | Stxbp3 | -0.263851112 | -0.897442404 | 0.36974018 | 8.224001078 | -1.043037865 | 0.340764356 | 0.709264768 | -4.605874025 |
| **A_55_P2022971** | Tspan9 | -0.276482681 | -0.940415061 | 0.387449699 | 8.167868997 | -1.043024303 | 0.340770107 | 0.709264768 | -4.605877479 |
| **A_55_P2121744** | Vars2 | 0.111747363 | -0.156601423 | 0.380096148 | 8.132899703 | 1.043008849 | 0.34077666 | 0.709264768 | -4.605881416 |
| **A_55_P2013760** | Slc39a14 | -0.834356494 | -2.838182529 | 1.169469541 | 9.212464863 | -1.042898351 | 0.340823518 | 0.709264768 | -4.605909561 |
| **A_55_P2034953** | NA | 0.17224065 | -0.241453327 | 0.585934627 | 10.46144914 | 1.042813882 | 0.340859342 | 0.709264768 | -4.605931075 |
| **A_55_P2019964** | Lancl1 | 0.299622357 | -0.420027033 | 1.019271748 | 11.84443093 | 1.042806189 | 0.340862605 | 0.709264768 | -4.605933034 |
| **A_55_P2000833** | E2f8 | 0.247226156 | -0.346596516 | 0.841048828 | 10.2316357 | 1.042768701 | 0.340878505 | 0.709264768 | -4.605942582 |
| **A_55_P2005555** | Celf2 | 0.259396362 | -0.363668929 | 0.882461653 | 8.763109161 | 1.042751126 | 0.34088596 | 0.709264768 | -4.605947058 |
| **A_51_P462918** | Ehhadh | -0.202743485 | -0.689785338 | 0.284298368 | 8.142626818 | -1.042631809 | 0.340936571 | 0.709264768 | -4.605977446 |
| **A_51_P160907** | Morc3 | 0.263188754 | -0.369070684 | 0.895448193 | 9.520057928 | 1.04261112 | 0.340945348 | 0.709264768 | -4.605982715 |
| **A_55_P2065621** | Hook2 | -0.193091448 | -0.656974925 | 0.270792029 | 10.21367337 | -1.042568217 | 0.340963548 | 0.709264768 | -4.605993642 |
| **A_55_P2093226** | Rnf121 | -0.212041546 | -0.721461177 | 0.297378085 | 8.575299842 | -1.04254697 | 0.340972562 | 0.709264768 | -4.605999052 |
| **A_66_P100268** | NA | 0.264799231 | -0.371480577 | 0.90107904 | 8.91635312 | 1.042362848 | 0.341050682 | 0.709264768 | -4.606045941 |
| **A_55_P1964960** | Il33 | 0.542621551 | -0.761276962 | 1.846520065 | 9.198143818 | 1.042326055 | 0.341066294 | 0.709264768 | -4.60605531 |
| **A_51_P508580** | Tcf15 | 0.25321077 | -0.355245197 | 0.861666737 | 8.884475505 | 1.042325505 | 0.341066528 | 0.709264768 | -4.60605545 |
| **A_55_P1979090** | Ublcp1 | -0.094841753 | -0.322750982 | 0.133067476 | 12.09312256 | -1.042288641 | 0.341082171 | 0.709264768 | -4.606064837 |
| **A_51_P317673** | Zfp764 | 0.533695507 | -0.748863785 | 1.816254798 | 10.7008828 | 1.042236889 | 0.341104133 | 0.709264768 | -4.606078015 |
| **A_55_P2131925** | Kprp | -0.348258976 | -1.185261796 | 0.488743845 | 7.850497265 | -1.042139224 | 0.341145582 | 0.709264768 | -4.606102884 |
| **A_52_P327402** | Cds1 | -0.096700338 | -0.329141203 | 0.135740528 | 8.720758692 | -1.041995521 | 0.341206576 | 0.709264768 | -4.606139473 |
| **A_51_P327751** | Ifit1 | -0.329654351 | -1.12209567 | 0.462786968 | 7.934493632 | -1.041938404 | 0.341230822 | 0.709264768 | -4.606154016 |
| **A_52_P161237** | Cpa1 | 0.449205244 | -0.630643041 | 1.529053529 | 9.181665493 | 1.041915312 | 0.341240626 | 0.709264768 | -4.606159895 |
| **A_66_P130582** | Ncoa1 | 0.477749367 | -0.670725192 | 1.626223925 | 10.16420992 | 1.041907314 | 0.341244021 | 0.709264768 | -4.606161931 |
| **A_66_P133928** | Fundc2 | 0.919771028 | -1.291363339 | 3.130905395 | 12.59097876 | 1.041873367 | 0.341258432 | 0.709264768 | -4.606170575 |
| **A_55_P2216456** | Rint1 | -0.111481381 | -0.379507001 | 0.15654424 | 9.189416438 | -1.041780862 | 0.341297706 | 0.709264768 | -4.606194126 |
| **A_55_P2178632** | Tmem80 | 0.135655815 | -0.190492551 | 0.46180418 | 9.448256209 | 1.041774093 | 0.34130058 | 0.709264768 | -4.606195849 |
| **A_55_P2119721** | NA | 0.177676166 | -0.249514517 | 0.60486685 | 17.00044196 | 1.041736181 | 0.341316677 | 0.709264768 | -4.606205501 |
| **A_55_P1975650** | Fbxo33 | -0.313659336 | -1.098954163 | 0.471635491 | 9.038508575 | -1.063869081 | 0.341368462 | 0.709264768 | -4.596376853 |
| **A_52_P328497** | Cdk10 | -0.100126868 | -0.34093043 | 0.140676693 | 9.77646293 | -1.041449156 | 0.341438565 | 0.709264768 | -4.606278569 |
| **A_55_P2094197** | Dnase1l3 | -0.285031024 | -0.970531417 | 0.400469369 | 7.818097714 | -1.041441237 | 0.341441928 | 0.709264768 | -4.606280585 |
| **A_66_P115161** | Cep85l | -0.187802987 | -0.639496296 | 0.263890321 | 8.228202668 | -1.041379886 | 0.341467986 | 0.709264768 | -4.606296202 |
| **A_55_P2072453** | 1-Mar | -0.479230182 | -1.631852176 | 0.673391811 | 8.08457173 | -1.0413761 | 0.341469594 | 0.709264768 | -4.606297166 |
| **A_55_P2010197** | Serpina10 | -0.529474655 | -1.803058898 | 0.744109588 | 8.198361944 | -1.04128076 | 0.341510092 | 0.709264768 | -4.606321433 |
| **A_55_P1953931** | Fam193a | -0.121737867 | -0.41456981 | 0.171094077 | 10.21196453 | -1.041256158 | 0.341520543 | 0.709264768 | -4.606327696 |
| **A_55_P2068486** | Arfgap1 | -0.471051837 | -1.60421963 | 0.662115955 | 10.0363984 | -1.041177622 | 0.341553907 | 0.709264768 | -4.606347685 |
| **A_51_P233947** | Tbc1d10b | -0.664325231 | -2.262479229 | 0.933828768 | 12.30860334 | -1.041147995 | 0.341566494 | 0.709264768 | -4.606355226 |

| | | | | | | | | | |
|---|---|---|---|---|---|---|---|---|---|
| **A_51_P455241** | Rrp36 | -0.193948354 | -0.660603606 | 0.272706898 | 9.945568708 | -1.040974965 | 0.341640013 | 0.709264768 | -4.606399265 |
| **A_55_P1980721** | NA | 0.384882292 | -0.541181651 | 1.310946235 | 13.36740505 | 1.040967813 | 0.341643052 | 0.709264768 | -4.606401086 |
| **A_55_P1953489** | Lrrc8b | -0.404244972 | -1.376899241 | 0.568409297 | 9.360460692 | -1.040965834 | 0.341643893 | 0.709264768 | -4.606401589 |
| **A_55_P2029051** | NA | -1.021952194 | -3.481005962 | 1.437101574 | 8.827949328 | -1.040909329 | 0.341667904 | 0.709264768 | -4.60641597 |
| **A_55_P1967592** | Ccnk | 0.106709466 | -0.150059581 | 0.363478512 | 12.2454861 | 1.040903869 | 0.341670225 | 0.709264768 | -4.606417359 |
| **A_55_P1957028** | Kat7 | -0.466176653 | -1.587989103 | 0.655635798 | 11.7870145 | -1.040831935 | 0.341700795 | 0.709264768 | -4.606435666 |
| **A_55_P1976734** | Cacna2d2 | 0.467660806 | -0.657931536 | 1.593253148 | 9.180519115 | 1.04063922 | 0.341782705 | 0.709264768 | -4.606484709 |
| **A_55_P2101910** | Rpl13a | 0.905238236 | -1.273561318 | 3.084037789 | 13.22492811 | 1.040629079 | 0.341787016 | 0.709264768 | -4.606487289 |
| **A_51_P206225** | Uroc1 | -0.20903173 | -0.712167271 | 0.294103811 | 7.783863318 | -1.040585006 | 0.341805751 | 0.709264768 | -4.606498505 |
| **A_55_P1959903** | Map7 | -0.224117652 | -0.763666993 | 0.315431688 | 8.242094705 | -1.040387783 | 0.341889599 | 0.709264768 | -4.606548689 |
| **A_66_P111430** | 2410006H | 0.42276153 | -0.595069587 | 1.440592647 | 13.20536461 | 1.040327509 | 0.341915227 | 0.709264768 | -4.606564025 |
| **A_51_P254646** | Jdp2 | -0.431345679 | -1.469956619 | 0.607265261 | 10.62876712 | -1.040214503 | 0.341963281 | 0.709264768 | -4.606592778 |
| **A_52_P622442** | Pdcl3 | 0.092690481 | -0.130496315 | 0.315877276 | 10.85608057 | 1.040200318 | 0.341969314 | 0.709264768 | -4.606596387 |
| **A_51_P394558** | Dchs1 | 0.563897924 | -0.793898092 | 1.92169394 | 9.930254294 | 1.040198711 | 0.341969997 | 0.709264768 | -4.606596796 |
| **A_55_P2170847** | Fbxo27 | -0.975965962 | -3.326044556 | 1.374112632 | 9.518359488 | -1.040165944 | 0.341983932 | 0.709264768 | -4.606605133 |
| **A_51_P375201** | Plk3 | -1.010913498 | -3.445408653 | 1.423581657 | 8.829311088 | -1.040052893 | 0.342032013 | 0.709264768 | -4.606633895 |
| **A_55_P2071137** | Asap1 | -0.380984339 | -1.298751885 | 0.536783207 | 10.14688352 | -1.03974005 | 0.342165097 | 0.709264768 | -4.606713481 |
| **A_55_P2179448** | NA | -0.183411125 | -0.625268271 | 0.25844602 | 8.946886594 | -1.039666712 | 0.342196301 | 0.709264768 | -4.606732136 |
| **A_55_P1984561** | Cpne2 | 0.210571852 | -0.296843113 | 0.717986817 | 9.064911798 | 1.039411196 | 0.342305037 | 0.709264768 | -4.606797129 |
| **A_55_P1994052** | Asxl1 | 0.476113231 | -0.671207053 | 1.623433515 | 12.42521157 | 1.039383754 | 0.342316717 | 0.709264768 | -4.606804108 |
| **A_52_P305182** | Ccdc32 | -0.190189442 | -0.648504308 | 0.268125424 | 10.66968909 | -1.039376289 | 0.342319895 | 0.709264768 | -4.606806007 |
| **A_51_P381618** | Pla1a | -1.165536994 | -3.974529513 | 1.643455526 | 8.613375528 | -1.039263861 | 0.34236775 | 0.709264768 | -4.606834602 |
| **A_51_P408989** | L3hypdh | 0.255823645 | -0.360733289 | 0.872380578 | 9.15019466 | 1.039244756 | 0.342375882 | 0.709264768 | -4.606839461 |
| **A_55_P2186439** | Chtop | 0.303460924 | -0.427991555 | 1.034913404 | 11.42033826 | 1.039123114 | 0.342427666 | 0.709264768 | -4.606870397 |
| **A_52_P655687** | Egfl6 | 0.566115244 | -0.798445935 | 1.930676422 | 9.179520322 | 1.039111577 | 0.342432578 | 0.709264768 | -4.606873331 |
| **A_51_P125679** | Rec114 | -0.231238184 | -0.788627641 | 0.326151273 | 7.820645091 | -1.03908517 | 0.342443821 | 0.709264768 | -4.606089816 |
| **A_52_P270035** | Tspyl2 | -0.107575047 | -0.366910917 | 0.151760822 | 8.512183217 | -1.038961134 | 0.342496633 | 0.709264768 | -4.606911589 |
| **A_51_P429212** | Npl | -0.328357377 | -1.119950352 | 0.463235598 | 8.191147336 | -1.038951302 | 0.34250082 | 0.709264768 | -4.60691409 |
| **A_65_P09031** | Cdc42se2 | -0.432645298 | -1.475744181 | 0.610453585 | 8.329741088 | -1.03885959 | 0.342539874 | 0.709264768 | -4.606937411 |
| **A_55_P1956712** | Med28 | 0.239904489 | -0.338542954 | 0.818351931 | 12.84032186 | 1.038782955 | 0.342572511 | 0.709264768 | -4.606956898 |
| **A_51_P458384** | Slc38a2 | -0.162442169 | -0.554162766 | 0.229278429 | 12.15736623 | -1.038658348 | 0.342625582 | 0.709264768 | -4.606988583 |
| **A_55_P2003158** | Wwp1 | -0.09643975 | -0.328999108 | 0.136119607 | 8.223802389 | -1.038658084 | 0.342625695 | 0.709264768 | -4.60698865 |
| **A_51_P206235** | Slc39a5 | -0.112534417 | -0.383921766 | 0.158852931 | 7.730879754 | -1.038594728 | 0.342652682 | 0.709264768 | -4.607004759 |
| **A_52_P145412** | Rictor | -0.12547991 | -0.42812937 | 0.177169551 | 9.099804251 | -1.038447724 | 0.342715305 | 0.709264768 | -4.607042135 |
| **A_55_P2108334** | Med10 | -0.124830872 | -0.425947264 | 0.176285521 | 10.5130411 | -1.038336092 | 0.342762867 | 0.709264768 | -4.607070516 |
| **A_55_P2183404** | Ptn | 0.349210883 | -0.493516136 | 1.191937903 | 9.419583485 | 1.037889689 | 0.342953115 | 0.709264768 | -4.607183997 |
| **A_51_P380013** | Bcl2 | 0.335020763 | -0.473544759 | 1.143586285 | 9.048540189 | 1.037783712 | 0.342998293 | 0.709264768 | -4.607210934 |
| **A_55_P2166011** | Tmem128 | -0.318075821 | -1.085761527 | 0.449609886 | 8.599992519 | -1.037761482 | 0.34300777 | 0.709264768 | -4.607216585 |
| **A_52_P207303** | Bmp3 | 0.274055679 | -0.38745298 | 0.935564339 | 8.434623671 | 1.037656518 | 0.343052522 | 0.709264768 | -4.607243263 |
| **A_55_P2086099** | Gm2061 | -0.292675932 | -0.999144437 | 0.413792574 | 7.897677346 | -1.03763485 | 0.343061761 | 0.709264768 | -4.60724877 |
| **A_55_P2089293** | Rnf121 | -0.37191998 | -1.269788573 | 0.525948614 | 9.778319703 | -1.037497537 | 0.343120314 | 0.709264768 | -4.607283669 |
| **A_51_P298178** | Dctn3 | -0.093028409 | -0.317620742 | 0.131563923 | 13.58975937 | -1.037459173 | 0.343136674 | 0.709264768 | -4.607293419 |
| **A_55_P2029187** | Pde4a | -0.14879259 | -0.508015191 | 0.210430011 | 8.710392774 | -1.037451958 | 0.343139751 | 0.709264768 | -4.607295253 |
| **A_55_P2148896** | NA | -0.960156768 | -3.278251189 | 1.357937654 | 9.147296654 | -1.037436135 | 0.343146499 | 0.709264768 | -4.607299274 |
| **A_55_P1978927** | Srgap1 | -0.266382158 | -0.909548908 | 0.376784591 | 8.02698075 | -1.037365726 | 0.343176528 | 0.709264768 | -4.607317168 |
| **A_55_P2056615** | NA | 0.622527431 | -0.880574991 | 2.125629853 | 13.39088636 | 1.037337944 | 0.343188377 | 0.709264768 | -4.607324228 |
| **A_55_P2086682** | Nfib | 1.090799675 | -1.543007525 | 3.724606875 | 13.30790619 | 1.037316689 | 0.343197443 | 0.709264768 | -4.607329629 |
| **A_55_P1996171** | Pcgf5 | -1.0464471 | -3.573344953 | 1.480450752 | 10.67916668 | -1.037241628 | 0.34322946 | 0.709264768 | -4.607348704 |

| | | | | | | | | | |
|---|---|---|---|---|---|---|---|---|---|
| **A_52_P71686** | Atp6v0d2 | -0.161756938 | -0.552358968 | 0.228845092 | 7.73826513 | -1.037238832 | 0.343230653 | 0.709264768 | -4.607349415 |
| **A_55_P2078335** | Mapk4 | -0.47310964 | -1.61559413 | 0.66937485 | 7.972637824 | -1.03719838 | 0.343247908 | 0.709264768 | -4.607359694 |
| **A_66_P103398** | Ogfrl1 | 0.301936329 | -0.427292151 | 1.031164808 | 8.996872711 | 1.037055727 | 0.343308765 | 0.709264768 | -4.607395943 |
| **A_51_P283004** | Tinag | 1.141927998 | -1.616816279 | 3.900672275 | 11.68825558 | 1.03675854 | 0.343435577 | 0.709264768 | -4.607471454 |
| **A_55_P1989469** | NA | 0.106015402 | -0.150127692 | 0.362158496 | 15.53301118 | 1.03666076 | 0.343477309 | 0.709264768 | -4.607496297 |
| **A_55_P2018116** | Pabpc4l | 0.323863865 | -0.458653305 | 1.106381035 | 8.596382389 | 1.036618522 | 0.343495337 | 0.709264768 | -4.607507028 |
| **A_51_P375509** | Spty2d1 | -0.29085683 | -0.993682459 | 0.4119688 | 8.18336015 | -1.036530346 | 0.343532975 | 0.709264768 | -4.607529429 |
| **A_55_P2133115** | NA | 0.411723814 | -0.583276398 | 1.406724027 | 14.87552247 | 1.036413734 | 0.343582756 | 0.709264768 | -4.607559053 |
| **A_66_P109395** | Rpia | -0.330768739 | -1.130165839 | 0.46862836 | 9.960769944 | -1.036363802 | 0.343604074 | 0.709264768 | -4.607571737 |
| **A_55_P1986551** | Mcam | 0.193930639 | -0.274789225 | 0.662650502 | 8.497993498 | 1.036295029 | 0.343633437 | 0.709264768 | -4.607589207 |
| **A_55_P2107155** | Rasd2 | 0.18678651 | -0.264683449 | 0.638256469 | 8.670679049 | 1.036255853 | 0.343650165 | 0.709264768 | -4.607599159 |
| **A_52_P15300** | Hoxb2 | 0.639615047 | -0.90650723 | 2.185737324 | 9.601232579 | 1.03615608 | 0.343692769 | 0.709264768 | -4.607624502 |
| **A_55_P2163404** | NA | 0.498207554 | -0.706135006 | 1.702550115 | 15.06493189 | 1.036121343 | 0.343707603 | 0.709264768 | -4.607633326 |
| **A_55_P2106235** | Syngr2 | -0.396674049 | -1.355576469 | 0.562228372 | 12.8303385 | -1.036119195 | 0.343708521 | 0.709264768 | -4.607633871 |
| **A_55_P1960074** | Pafah1b3 | 0.359594586 | -0.509692077 | 1.228881248 | 12.18154858 | 1.036097136 | 0.343717941 | 0.709264768 | -4.607639474 |
| **A_55_P2038412** | Acot8 | -0.185660402 | -0.634497193 | 0.26317639 | 9.197010305 | -1.036051114 | 0.343737596 | 0.709264768 | -4.607651164 |
| **A_51_P317695** | Trpc3 | 0.261686511 | -0.370953058 | 0.89432608 | 8.66889405 | 1.036037155 | 0.343743557 | 0.709264768 | -4.607654709 |
| **A_55_P2175400** | Zfp467 | 0.714166193 | -1.012438303 | 2.44077069 | 10.24247565 | 1.035992869 | 0.343762472 | 0.709264768 | -4.607665958 |
| **A_66_P122110** | Mitd1 | -0.106893718 | -0.365327184 | 0.151539747 | 9.144934923 | -1.035985741 | 0.343765516 | 0.709264768 | -4.607667768 |
| **A_55_P2001998** | Zfp503 | 0.671673173 | -0.952247652 | 2.295593999 | 10.07378539 | 1.035961217 | 0.343775991 | 0.709264768 | -4.607673997 |
| **A_55_P2044684** | Rsph1 | 0.762330884 | -1.080841502 | 2.605503271 | 10.2783152 | 1.035924024 | 0.343791877 | 0.709264768 | -4.607683443 |
| **A_55_P2002079** | NA | 0.381433858 | -0.540869367 | 1.303737083 | 15.76642908 | 1.035847587 | 0.343824527 | 0.709264768 | -4.607702857 |
| **A_51_P460048** | Cnrip1 | 0.491012763 | -0.696295566 | 1.678321092 | 10.01840352 | 1.035808837 | 0.343841081 | 0.709264768 | -4.607712698 |
| **A_55_P2029581** | NA | -0.203287021 | -0.694934112 | 0.28836007 | 8.73304465 | -1.035634531 | 0.343915549 | 0.709264768 | -4.607756965 |
| **A_55_P2101503** | NA | -0.243480581 | -0.832371912 | 0.34541075 | 8.342868373 | -1.035570126 | 0.343943067 | 0.709264768 | -4.607773321 |
| **A_55_P2106975** | Clcn5 | -0.490804357 | -1.677949724 | 0.69634101 | 8.514224359 | -1.035511325 | 0.343968193 | 0.709264768 | -4.607788253 |
| **A_55_P2053783** | Ppp2r3a | -0.129742696 | -0.443590496 | 0.184105103 | 7.820927798 | -1.035414344 | 0.344009637 | 0.709264768 | -4.60781288 |
| **A_51_P340355** | Tarsl2 | 0.355718005 | -0.504773417 | 1.216209426 | 8.737249516 | 1.035403546 | 0.344014251 | 0.709264768 | -4.607815622 |
| **A_55_P1961451** | Adamts7 | 0.277385677 | -0.393814168 | 0.948585523 | 8.643446487 | 1.035100529 | 0.34414377 | 0.709264768 | -4.607892562 |
| **A_51_P440443** | Exosc5 | 0.193058719 | -0.274093039 | 0.660210477 | 11.87796693 | 1.035098736 | 0.344144537 | 0.709264768 | -4.607893017 |
| **A_51_P257684** | Stau1 | -0.187986217 | -0.642959714 | 0.26698728 | 9.201486429 | -1.034880621 | 0.344237791 | 0.709264768 | -4.607948393 |
| **A_55_P2032165** | NA | 0.471811019 | -0.670110061 | 1.613732098 | 15.3125889 | 1.034861749 | 0.34424586 | 0.709264768 | -4.607953184 |
| **A_55_P2413359** | NA | -0.127910577 | -0.437497159 | 0.181676005 | 8.264355721 | -1.034843498 | 0.344253665 | 0.709264768 | -4.607957818 |
| **A_55_P1995092** | H2-Q10 | -0.411777449 | -1.40842388 | 0.584868982 | 8.230168483 | -1.034836619 | 0.344256606 | 0.709264768 | -4.607959564 |
| **A_51_P176883** | Edrf1 | -0.095755518 | -0.327522028 | 0.136010993 | 8.397753008 | -1.034816805 | 0.344265079 | 0.709264768 | -4.607964594 |
| **A_55_P2154425** | Cd320 | -0.28420674 | -0.972195781 | 0.403782301 | 7.931534172 | -1.034673196 | 0.344326494 | 0.709264768 | -4.608001051 |
| **A_66_P113487** | Pacsin2 | -0.1944555 | -0.665207968 | 0.276296969 | 13.23415484 | -1.034613093 | 0.3443522 | 0.709264768 | -4.608016308 |
| **A_55_P2069939** | Ubxn7 | -0.158617847 | -0.542614126 | 0.225378432 | 8.257725385 | -1.034606921 | 0.34435484 | 0.709264768 | -4.608017874 |
| **A_55_P2106901** | Nkain4 | 0.968528386 | -1.376479008 | 3.31353578 | 11.04421471 | 1.034471387 | 0.344412813 | 0.709264768 | -4.608052278 |
| **A_51_P265106** | Dlat | 0.103767506 | -0.147486814 | 0.355021826 | 8.94331559 | 1.034423045 | 0.344433493 | 0.709264768 | -4.608064549 |
| **A_55_P2013296** | Prkag2 | -0.116684464 | -0.399242481 | 0.165873552 | 11.31465589 | -1.034321968 | 0.344476735 | 0.709264768 | -4.608090204 |
| **A_52_P179583** | Adnp2 | -0.228719732 | -0.782578062 | 0.325138598 | 8.365393985 | -1.034320869 | 0.344477206 | 0.709264768 | -4.608090483 |
| **A_52_P330289** | Inpp4b | 0.27720235 | -0.394096207 | 0.948500908 | 8.583450464 | 1.034264314 | 0.344501403 | 0.709264768 | -4.608104837 |
| **A_55_P2089780** | Mtfr1l | -0.198586807 | -0.679531457 | 0.282357843 | 8.486717062 | -1.034202639 | 0.344527792 | 0.709264768 | -4.608120491 |
| **A_55_P1958165** | Ms4a7 | -0.156308642 | -0.534866055 | 0.222248772 | 9.319389896 | -1.034192963 | 0.344531932 | 0.709264768 | -4.608122947 |
| **A_55_P1983758** | Sdhc | 0.299410002 | -0.42575131 | 1.024571314 | 11.88446731 | 1.034146397 | 0.344551858 | 0.709264768 | -4.608134765 |
| **A_55_P1958912** | NA | 0.123004661 | -0.174940927 | 0.420950248 | 10.12562008 | 1.034034325 | 0.344599818 | 0.709264768 | -4.608163208 |
| **A_52_P447196** | Col4a6 | 0.102213264 | -0.145372586 | 0.349799115 | 8.165233926 | 1.034026774 | 0.34460305 | 0.709264768 | -4.608165124 |

| | | | | | | | | | |
|---|---|---|---|---|---|---|---|---|---|
| **A_55_P2015989** | Rpl22 | 0.320494677 | -0.455871931 | 1.096861285 | 15.98904749 | 1.033961381 | 0.344631036 | 0.709264768 | -4.60818172 |
| **A_66_P135176** | Trak2 | -0.254175502 | -0.86993738 | 0.361586375 | 8.316872481 | -1.033882628 | 0.344664744 | 0.709264768 | -4.608201705 |
| **A_52_P580707** | Zfp781 | 0.154418779 | -0.219736007 | 0.528573566 | 8.748170411 | 1.03371104 | 0.344738195 | 0.709264768 | -4.608245247 |
| **A_55_P2014214** | NA | 0.461686213 | -0.657230105 | 1.580602532 | 14.79935637 | 1.033474211 | 0.344839595 | 0.709264768 | -4.60830534 |
| **A_66_P128293** | Pheta2 | 0.287473635 | -0.409281632 | 0.984228902 | 8.832336724 | 1.033399164 | 0.344871732 | 0.709264768 | -4.608324381 |
| **A_55_P2104798** | NA | -0.386755725 | -1.324155107 | 0.550643657 | 9.524972837 | -1.033385668 | 0.344877512 | 0.709264768 | -4.608327805 |
| **A_55_P2052634** | Rpl6 | 0.228772569 | -0.325720067 | 0.783265205 | 16.03021956 | 1.033376334 | 0.344881509 | 0.709264768 | -4.608330173 |
| **A_55_P1971522** | Kif16b | -0.195758951 | -0.670272916 | 0.278755014 | 9.168320303 | -1.033291782 | 0.34491772 | 0.709264768 | -4.608351625 |
| **A_51_P243930** | Qrsl1 | 0.085973574 | -0.122427847 | 0.294374996 | 8.319217311 | 1.033271979 | 0.344926202 | 0.709264768 | -4.608356649 |
| **A_66_P106072** | Orai1 | -0.106975293 | -0.366285682 | 0.152335096 | 8.77506106 | -1.033270225 | 0.344926953 | 0.709264768 | -4.608357094 |
| **A_51_P422223** | 1700006J | 0.131111697 | -0.186706072 | 0.448929466 | 8.138792693 | 1.033269502 | 0.344927263 | 0.709264768 | -4.608357278 |
| **A_55_P1957267** | Rab9 | 0.173338993 | -0.246843772 | 0.593521758 | 11.9451671 | 1.033257057 | 0.344932593 | 0.709264768 | -4.608360435 |
| **A_52_P587948** | Zfp11 | 0.192564646 | -0.274231589 | 0.659360881 | 8.434783918 | 1.033236064 | 0.344941584 | 0.709264768 | -4.608365761 |
| **A_55_P1967538** | Hunk | 0.652732808 | -0.929639973 | 2.235105588 | 12.00130977 | 1.033182362 | 0.344964586 | 0.709264768 | -4.608379385 |
| **A_55_P2081483** | Atp5g2 | 0.234559333 | -0.334090735 | 0.803209401 | 16.15954068 | 1.033137127 | 0.344983963 | 0.709264768 | -4.608390861 |
| **A_55_P2061135** | Rpl17 | 0.531067679 | -0.756516234 | 1.818651592 | 14.84505838 | 1.033057927 | 0.34501789 | 0.709264768 | -4.608410953 |
| **A_51_P198137** | Ubxn8 | -0.270687917 | -0.927069155 | 0.385693322 | 8.478269466 | -1.032911364 | 0.345080681 | 0.709264768 | -4.608448132 |
| **A_66_P137462** | Nsdhl | 0.121996412 | -0.173839537 | 0.417832362 | 8.409259314 | 1.032871887 | 0.345097595 | 0.709264768 | -4.608458146 |
| **A_55_P1956147** | B4galnt2 | -0.653480119 | -2.238174329 | 0.931214091 | 8.127761266 | -1.03285 | 0.345106973 | 0.709264768 | -4.608463698 |
| **A_55_P2244112** | Amotl1 | -0.241257227 | -0.826437997 | 0.343923543 | 8.24063558 | -1.032620234 | 0.345205435 | 0.709264768 | -4.608521978 |
| **A_51_P193813** | Fga | -0.206504588 | -0.707400213 | 0.294391037 | 7.790424215 | -1.032601643 | 0.345213403 | 0.709264768 | -4.608526693 |
| **A_55_P1957624** | Insig2 | -0.239877364 | -0.821787002 | 0.342032275 | 9.171422616 | -1.032485739 | 0.345263082 | 0.709264768 | -4.60855609 |
| **A_55_P1984830** | Fgl1 | -0.132549943 | -0.45413069 | 0.189030803 | 8.109508167 | -1.032380642 | 0.345308133 | 0.709264768 | -4.608582744 |
| **A_55_P2060966** | Mov10 | -0.338943776 | -1.161364815 | 0.483477264 | 10.96048964 | -1.032247397 | 0.345365258 | 0.709264768 | -4.608616535 |
| **A_55_P2033795** | Fabp6 | -0.310504303 | -1.063919342 | 0.442910737 | 7.893388415 | -1.032247127 | 0.345365373 | 0.709264768 | -4.608616604 |
| **A_55_P1989846** | Tm6sf1 | 0.188270905 | -0.268569476 | 0.645111286 | 8.315688725 | 1.032212396 | 0.345380264 | 0.709264768 | -4.608625411 |
| **A_51_P412914** | Efs | -0.410157715 | -1.405427748 | 0.585112317 | 11.14948013 | -1.032191559 | 0.345389199 | 0.709264768 | -4.608630696 |
| **A_51_P329322** | Zfand1 | 0.382549607 | -0.545960525 | 1.311059739 | 10.84716818 | 1.031932899 | 0.345500119 | 0.709264768 | -4.608696286 |
| **A_55_P2065501** | Rnf7 | 0.258257666 | -0.368626619 | 0.885141951 | 13.26241278 | 1.031849087 | 0.345536066 | 0.709264768 | -4.608717537 |
| **A_55_P2103857** | Hgs | -0.105299405 | -0.360949536 | 0.150350725 | 12.14689841 | -1.031644925 | 0.345623645 | 0.709264768 | -4.608769301 |
| **A_52_P670263** | Tfpi | -0.591508304 | -2.02762818 | 0.844611572 | 9.707124373 | -1.031621771 | 0.345633578 | 0.709264768 | -4.608775172 |
| **A_55_P2076631** | Gypc | 0.865311525 | -1.235706948 | 2.966329998 | 12.03606139 | 1.031556364 | 0.34566164 | 0.709264768 | -4.608791754 |
| **A_55_P2113673** | Eml1 | 0.72801503 | -1.039752711 | 2.49578277 | 12.79963126 | 1.031491103 | 0.345689641 | 0.709264768 | -4.608808299 |
| **A_55_P2005080** | NA | 0.146461865 | -0.209184929 | 0.502108659 | 16.73598576 | 1.031468591 | 0.345699301 | 0.709264768 | -4.608814006 |
| **A_55_P2010788** | Slc6a9 | 0.598038682 | -0.854168696 | 2.05024606 | 9.979843055 | 1.031456656 | 0.345704422 | 0.709264768 | -4.608817032 |
| **A_55_P2059019** | Mapk8ip3 | -0.349412454 | -1.197936564 | 0.499111657 | 11.35505188 | -1.031393852 | 0.345731372 | 0.709264768 | -4.608832953 |
| **A_51_P286496** | Il2rb | -0.990975121 | -3.397615062 | 1.415664819 | 8.320509395 | -1.031340274 | 0.345754364 | 0.709264768 | -4.608846536 |
| **A_55_P2022805** | Rubcn | -0.403270372 | -1.382710333 | 0.576169589 | 12.42901155 | -1.03126159 | 0.345788132 | 0.709264768 | -4.608866482 |
| **A_51_P497692** | Mageh1 | 0.170065849 | -0.243003049 | 0.583134747 | 9.603110251 | 1.031204884 | 0.345812469 | 0.709264768 | -4.608880856 |
| **A_55_P2130109** | Nmt2 | -0.255748248 | -0.876953903 | 0.365457408 | 8.032486719 | -1.031163689 | 0.345830151 | 0.709264768 | -4.608891299 |
| **A_52_P67200** | Stt3b | 0.2211689 | -0.316053883 | 0.758391682 | 12.04753544 | 1.031145572 | 0.345837927 | 0.709264768 | -4.608895891 |
| **A_52_P545810** | Lrrfip1 | 0.21605957 | -0.308862466 | 0.740981606 | 11.10424599 | 1.030929692 | 0.345930599 | 0.709264768 | -4.60895061 |
| **A_52_P37757** | Eps8l1 | -0.10675417 | -0.366120821 | 0.152612481 | 8.445338562 | -1.030910731 | 0.34593874 | 0.709264768 | -4.608955416 |
| **A_55_P2088325** | Ctdsp2 | -0.296078089 | -1.015422774 | 0.423266597 | 9.889873524 | -1.030907204 | 0.345940254 | 0.709264768 | -4.60895631 |
| **A_55_P2180301** | Clcn5 | -0.485595975 | -1.665492732 | 0.694300783 | 8.481274621 | -1.030816629 | 0.345979143 | 0.709264768 | -4.608979266 |
| **A_51_P401024** | Lrrc34 | 0.436547439 | -0.624191883 | 1.49728676 | 8.952954104 | 1.030796942 | 0.345987596 | 0.709264768 | -4.608984255 |
| **A_55_P2062444** | Memo1 | 0.248357201 | -0.355161873 | 0.851876276 | 11.29422707 | 1.030709081 | 0.346025324 | 0.709264768 | -4.609006523 |
| **A_55_P1999309** | Zfp668 | 0.156533825 | -0.223893573 | 0.536961223 | 9.790004138 | 1.030591977 | 0.346075613 | 0.709264768 | -4.6090362 |

| | | | | | | | | | |
|---|---|---|---|---|---|---|---|---|---|
| **A_65_P16753** | Bex1 | -0.776338548 | -2.663119598 | 1.110442502 | 11.42618436 | -1.03057592 | 0.34608251 | 0.709264768 | -4.609040269 |
| **A_55_P2093940** | Zfp408 | -0.22650999 | -0.777046515 | 0.324026535 | 15.9568658 | -1.030508477 | 0.346111476 | 0.709264768 | -4.609057361 |
| **A_55_P2162855** | Nipa2 | -0.291900707 | -1.001449096 | 0.417647681 | 9.797323168 | -1.030394364 | 0.346160491 | 0.709264768 | -4.609086278 |
| **A_55_P1960157** | Bcat1 | -0.512720091 | -1.759153162 | 0.73371298 | 8.028122783 | -1.030295298 | 0.346203048 | 0.709264768 | -4.609111381 |
| **A_55_P1993805** | Zfp947 | 0.169928625 | -0.243202274 | 0.583059524 | 10.94339569 | 1.030218182 | 0.346236179 | 0.709264768 | -4.609130921 |
| **A_55_P2126855** | E230025N | -0.173972991 | -0.596980192 | 0.24903421 | 7.807309517 | -1.030111924 | 0.346281833 | 0.709264768 | -4.609157844 |
| **A_55_P2074414** | Gtf2h1 | -0.461740188 | -1.584620804 | 0.661140428 | 10.66935005 | -1.029945955 | 0.346353153 | 0.709264768 | -4.609199894 |
| **A_51_P104933** | Trub2 | 0.197936493 | -0.283430496 | 0.679303481 | 12.36169205 | 1.029911511 | 0.346367956 | 0.709264768 | -4.604976876 |
| **A_55_P2008870** | NA | 0.393071621 | -0.562874603 | 1.349017846 | 14.25690961 | 1.029884622 | 0.346379512 | 0.709264768 | -4.609215433 |
| **A_65_P18981** | Palb2 | 0.107088973 | -0.153353577 | 0.367531522 | 8.027441662 | 1.029871785 | 0.34638503 | 0.709264768 | -4.609218685 |
| **A_52_P308669** | Cpn1 | -0.108341769 | -0.371841589 | 0.15515805 | 7.930564412 | -1.029830966 | 0.346402573 | 0.709264768 | -4.609229026 |
| **A_66_P102047** | Tbc1d13 | -0.210149315 | -0.72133041 | 0.30103178 | 9.227664071 | -1.029682995 | 0.346466177 | 0.709264768 | -4.609266512 |
| **A_52_P672803** | Ctsa | -0.265855878 | -0.912562716 | 0.38085096 | 8.325671956 | -1.029648899 | 0.346480834 | 0.709264768 | -4.60927515 |
| **A_52_P171724** | 2310009B | 0.53612617 | -0.768143587 | 1.840395927 | 11.14872689 | 1.029555893 | 0.346520818 | 0.709264768 | -4.60929871 |
| **A_55_P2067483** | NA | 0.263691411 | -0.377870223 | 0.905253046 | 11.72238846 | 1.029456368 | 0.346563608 | 0.709264768 | -4.60932392 |
| **A_52_P661071** | Snhg3 | 0.165454842 | -0.237132878 | 0.568042562 | 10.78241326 | 1.029364779 | 0.346602991 | 0.709264768 | -4.609347119 |
| **A_55_P2112225** | NA | -0.511029064 | -1.754524655 | 0.732466528 | 8.094138408 | -1.029323045 | 0.346620937 | 0.709264768 | -4.60935769 |
| **A_55_P2358002** | 5-Mar | 0.141965267 | -0.203516845 | 0.487447379 | 8.391185051 | 1.029216841 | 0.346666611 | 0.709264768 | -4.60938459 |
| **A_51_P358574** | Pdzd3 | -0.294752352 | -1.012108025 | 0.422603322 | 7.812924664 | -1.02913675 | 0.346701057 | 0.709264768 | -4.609404874 |
| **A_51_P132718** | 2310010J | -0.098293696 | -0.337523213 | 0.140935821 | 11.38983539 | -1.029108698 | 0.346713123 | 0.709264768 | -4.609411979 |
| **A_52_P514306** | Spata2 | -0.179193307 | -0.615328415 | 0.256941801 | 9.060062402 | -1.029084607 | 0.346723485 | 0.709264768 | -4.60941808 |
| **A_51_P331003** | Sema4g | -0.242244561 | -0.831895082 | 0.34740596 | 8.198491062 | -1.028986539 | 0.346765669 | 0.709264768 | -4.609442917 |
| **A_55_P2002389** | Nelfb | -0.116466008 | -0.399981518 | 0.167049501 | 13.44085631 | -1.028898927 | 0.34680336 | 0.709264768 | -4.609465104 |
| **A_51_P414305** | Acot12 | -0.182899943 | -0.628151309 | 0.262351423 | 7.560788811 | -1.02886562 | 0.346817689 | 0.709264768 | -4.609473539 |
| **A_55_P2078073** | Pqlc2 | -0.422793272 | -1.452131838 | 0.606545293 | 9.249292845 | -1.028774449 | 0.346856915 | 0.709264768 | -4.609496626 |
| **A_51_P375987** | Fign | 0.386119226 | -0.554077652 | 1.326316103 | 8.933062136 | 1.028615271 | 0.34692541 | 0.709264768 | -4.609536933 |
| **A_51_P504522** | Derl1 | -0.677715437 | -2.328016865 | 0.972585991 | 10.7225008 | -1.028571409 | 0.346944286 | 0.709264768 | -4.60954804 |
| **A_55_P2105220** | Hscb | 0.204022716 | -0.29280845 | 0.700853881 | 11.77858871 | 1.028537283 | 0.346958972 | 0.709264768 | -4.609556681 |
| **A_55_P2220283** | Stk35os1 | 0.230507347 | -0.330861682 | 0.791876375 | 8.747792212 | 1.028458255 | 0.346992986 | 0.709264768 | -4.609576691 |
| **A_51_P116665** | 2610005L | -0.139550553 | -0.479519806 | 0.200418701 | 9.522171751 | -1.028116349 | 0.34714017 | 0.709264768 | -4.609663253 |
| **A_51_P232371** | Stab1 | 0.310474008 | -0.445900304 | 1.066848321 | 10.89907858 | 1.0281082 | 0.347143679 | 0.709264768 | -4.609665316 |
| **A_52_P283055** | Slc5a3 | -0.39766848 | -1.36649731 | 0.571160349 | 8.309873799 | -1.028074203 | 0.347158317 | 0.709264768 | -4.609673923 |
| **A_51_P365964** | Tor1a | -0.173780505 | -0.597217314 | 0.249656305 | 11.68835249 | -1.027928224 | 0.347221177 | 0.709264768 | -4.609710877 |
| **A_51_P392967** | Zmynd10 | 0.563944784 | -0.810184571 | 1.938074139 | 9.549298363 | 1.027919991 | 0.347224722 | 0.709264768 | -4.609712961 |
| **A_55_P2086835** | NA | -0.65365099 | -2.246384016 | 0.939082036 | 12.54344301 | -1.027905721 | 0.347230867 | 0.709264768 | -4.609716573 |
| **A_55_P2177273** | Kcna6 | 0.141839704 | -0.203777995 | 0.487457404 | 8.889926961 | 1.027903126 | 0.347231985 | 0.709264768 | -4.60971723 |
| **A_51_P374772** | Arfgef1 | -0.150990267 | -0.518923819 | 0.216943286 | 10.72855221 | -1.027850389 | 0.347254697 | 0.709264768 | -4.60973058 |
| **A_55_P2061701** | Osgep | 0.337330961 | -0.484706892 | 1.159368815 | 10.66292023 | 1.027814482 | 0.347270161 | 0.709264768 | -4.609739669 |
| **A_65_P15518** | NA | -0.187809388 | -0.645545812 | 0.269927035 | 8.49295466 | -1.027666434 | 0.347333929 | 0.709264768 | -4.609777142 |
| **A_52_P48569** | Slc38a4 | -0.882895845 | -3.034827824 | 1.269036135 | 10.48857287 | -1.027616996 | 0.347355225 | 0.709264768 | -4.609789655 |
| **A_55_P1984601** | Vapb | 0.192067948 | -0.276085237 | 0.660221132 | 9.348730573 | 1.027583825 | 0.347369515 | 0.709264768 | -4.609798051 |
| **A_55_P2033105** | Cdcp1 | -0.253760832 | -0.872291561 | 0.364769897 | 8.299535229 | -1.027575297 | 0.347373189 | 0.709264768 | -4.60980021 |
| **A_55_P2034081** | Hrh1 | -0.373540092 | -1.284065876 | 0.536985693 | 8.123242032 | -1.02753191 | 0.34739188 | 0.709264768 | -4.609811191 |
| **A_55_P2141699** | Rnf149 | 0.262713892 | -0.377677117 | 0.903104901 | 11.52398919 | 1.02751497 | 0.347399178 | 0.709264768 | -4.609815478 |
| **A_55_P2160594** | NA | 0.296911324 | -0.426869013 | 1.020691661 | 16.83955614 | 1.027472784 | 0.347417353 | 0.709264768 | -4.609826155 |
| **A_55_P2108122** | Arcn1 | 0.112841046 | -0.162243209 | 0.387925301 | 12.66189588 | 1.027428752 | 0.347436324 | 0.709264768 | -4.609837299 |
| **A_51_P110841** | Mrps35 | 0.201728903 | -0.290056289 | 0.693514095 | 10.42734941 | 1.027408192 | 0.347445183 | 0.709264768 | -4.606889724 |
| **A_55_P2035812** | Rnf208 | -0.626571661 | -2.154588508 | 0.901445186 | 9.017286897 | -1.027053227 | 0.347598154 | 0.709264768 | -4.609932331 |

| | | | | | | | | | |
|---|---|---|---|---|---|---|---|---|---|
| **A_52_P383862** | Cmc1 | 0.240725361 | -0.346411172 | 0.827861894 | 11.07344071 | 1.026911668 | 0.347659173 | 0.709264768 | -4.609968151 |
| **A_51_P389386** | Nsmce2 | 0.32245995 | -0.464052517 | 1.108972417 | 10.40513943 | 1.026881942 | 0.347671988 | 0.709264768 | -4.609975673 |
| **A_55_P2102951** | NA | 0.411438742 | -0.59215836 | 1.415035843 | 12.40782197 | 1.026824283 | 0.347696846 | 0.709264768 | -4.609990262 |
| **A_55_P2037513** | Pld3 | -0.457699984 | -1.574290578 | 0.65889061 | 10.63175822 | -1.026685144 | 0.347756836 | 0.709264768 | -4.610025466 |
| **A_55_P1972097** | NA | 0.321983697 | -0.463614326 | 1.10758172 | 16.80823026 | 1.026558839 | 0.3478113 | 0.709264768 | -4.610057421 |
| **A_51_P451052** | Pgm5 | 0.652077635 | -0.938921332 | 2.243076602 | 10.37857558 | 1.026549162 | 0.347815474 | 0.709264768 | -4.610059869 |
| **A_52_P86928** | Cog2 | -0.103404221 | -0.355704151 | 0.148895708 | 8.960727018 | -1.026529622 | 0.3478239 | 0.709264768 | -4.610064813 |
| **A_55_P2159475** | Sbf1 | -0.129861462 | -0.446777743 | 0.187054819 | 11.80968494 | -1.026327773 | 0.347910957 | 0.709264768 | -4.610115876 |
| **A_55_P1986833** | Tulp4 | -0.438372993 | -1.508246761 | 0.631500775 | 10.58021065 | -1.026269912 | 0.347935916 | 0.709264768 | -4.610130513 |
| **A_55_P2321453** | D11Ertd72 | -0.11024367 | -0.379314303 | 0.158826964 | 8.327930569 | -1.026213455 | 0.347960271 | 0.709264768 | -4.610144794 |
| **A_52_P57526** | Sin3b | -0.942484271 | -3.242852864 | 1.357884322 | 14.73789945 | -1.026188225 | 0.347971155 | 0.709264768 | -4.610151176 |
| **A_55_P1984675** | Abhd6 | -0.910666334 | -3.133562984 | 1.312230317 | 8.938453104 | -1.026101578 | 0.348008536 | 0.709264768 | -4.610173093 |
| **A_51_P513072** | Rdh13 | -0.173677869 | -0.597627306 | 0.250271568 | 8.319302582 | -1.02607892 | 0.348018312 | 0.709264768 | -4.610178824 |
| **A_55_P1971360** | Ciz1 | -0.106110683 | -0.365144189 | 0.152922822 | 9.360296163 | -1.026014538 | 0.34804609 | 0.709264768 | -4.610195109 |
| **A_55_P1974954** | NA | 0.349683581 | -0.503982121 | 1.203349283 | 16.15105328 | 1.025977303 | 0.348062157 | 0.709264768 | -4.610204527 |
| **A_55_P2047375** | NA | 0.268320884 | -0.386735678 | 0.923377446 | 14.94436013 | 1.025949584 | 0.348074118 | 0.709264768 | -4.610211538 |
| **A_51_P187665** | Rmi1 | -0.103847848 | -0.357379027 | 0.149683331 | 9.724483583 | -1.025927023 | 0.348083853 | 0.709264768 | -4.610217244 |
| **A_66_P112853** | Eif1a | -0.415299121 | -1.429243546 | 0.598645305 | 11.55250138 | -1.025881517 | 0.348103491 | 0.709264768 | -4.610228753 |
| **A_51_P494481** | Snx30 | -0.452275211 | -1.556603491 | 0.652053069 | 8.557910647 | -1.025781675 | 0.348146578 | 0.709264768 | -4.610254005 |
| **A_55_P2104119** | NA | 0.269616932 | -0.388857966 | 0.92809183 | 9.179768345 | 1.025553413 | 0.348245104 | 0.709264768 | -4.610311732 |
| **A_51_P366121** | Klf14 | -0.292310973 | -1.00623279 | 0.421610843 | 8.14662133 | -1.025521597 | 0.348258838 | 0.709264768 | -4.610319777 |
| **A_55_P2007485** | Prss27 | -0.143782749 | -0.494981511 | 0.207416013 | 7.758372273 | -1.025425594 | 0.348300285 | 0.709264768 | -4.610344054 |
| **A_55_P1959064** | Oas1b | -0.225254827 | -0.775541445 | 0.325031792 | 8.34145018 | -1.025263506 | 0.34837027 | 0.709264768 | -4.61038504 |
| **A_52_P432685** | Ctdsp2 | -0.231131226 | -0.795780748 | 0.333518296 | 8.865071653 | -1.025250531 | 0.348375873 | 0.709264768 | -4.610388321 |
| **A_55_P2076797** | Ddx5 | -0.274910709 | -0.946514846 | 0.396693429 | 10.49783145 | -1.025247313 | 0.348377262 | 0.709264768 | -4.610389134 |
| **A_52_P351021** | NA | -0.235485803 | -0.810801635 | 0.33983003 | 10.55915089 | -1.025200356 | 0.34839754 | 0.709264768 | -4.610401007 |
| **A_52_P58283** | Kcna5 | 0.164417303 | -0.237289524 | 0.56612413 | 8.055685644 | 1.02515292 | 0.348418025 | 0.709264768 | -4.610413001 |
| **A_52_P139747** | Ndufa1 | -0.113252733 | -0.389977367 | 0.163471902 | 14.42870779 | -1.025064549 | 0.348456191 | 0.709264768 | -4.610435345 |
| **A_51_P149373** | Fbxl20 | -0.109811801 | -0.378152119 | 0.158528517 | 10.58439431 | -1.024975374 | 0.348494707 | 0.709264768 | -4.610457891 |
| **A_55_P2135631** | Rps18 | 0.394943621 | -0.570180622 | 1.360067865 | 15.27594292 | 1.024948926 | 0.348506131 | 0.709264768 | -4.610464577 |
| **A_52_P357469** | Frk | -0.124236698 | -0.427886447 | 0.17941305 | 8.15092104 | -1.024772167 | 0.348582488 | 0.709264768 | -4.610509264 |
| **A_51_P467224** | Fbn1 | 0.703933995 | -1.016662282 | 2.424530273 | 11.44627931 | 1.024715501 | 0.34860697 | 0.709264768 | -4.610523589 |
| **A_51_P272876** | Tent5a | -1.297558085 | -4.469628727 | 1.874512557 | 9.857386812 | -1.024552702 | 0.348677313 | 0.709264768 | -4.610564742 |
| **A_55_P2062310** | NA | 0.329486232 | -0.476076197 | 1.135048661 | 15.50126031 | 1.024444452 | 0.348724093 | 0.709264768 | -4.610592104 |
| **A_51_P467751** | Hgsnat | -0.667364716 | -2.299120862 | 0.964391429 | 11.04443617 | -1.024373489 | 0.348754762 | 0.709264768 | -4.610610041 |
| **A_66_P138584** | Mnd1 | 0.378611517 | -0.547221007 | 1.30444404 | 8.914906791 | 1.024263585 | 0.348802266 | 0.709264768 | -4.610637819 |
| **A_55_P2170109** | Bivm | -0.181262538 | -0.624569577 | 0.262044502 | 10.18773187 | -1.024126899 | 0.348861352 | 0.709264768 | -4.610672365 |
| **A_55_P1955059** | Rpl38 | 0.123510222 | -0.178581222 | 0.425601665 | 16.48320103 | 1.024035053 | 0.34890106 | 0.709264768 | -4.610695577 |
| **A_51_P279232** | Dennd2d | 0.459827973 | -0.6650991 | 1.584755047 | 9.1456452 | 1.023814708 | 0.348996337 | 0.709264768 | -4.61075126 |
| **A_55_P2075479** | Rpl37a | 0.279121434 | -0.403725799 | 0.961968667 | 17.6579185 | 1.023811444 | 0.348997748 | 0.709264768 | -4.610752085 |
| **A_55_P1989653** | Slco4a1 | -1.752559737 | -6.040948997 | 2.535829522 | 9.023733316 | -1.023596933 | 0.349090523 | 0.709264768 | -4.610806289 |
| **A_52_P655948** | Mosmo | 0.248442942 | -0.359520539 | 0.856406422 | 9.902551851 | 1.0235275 | 0.349120557 | 0.709264768 | -4.610823832 |
| **A_55_P2163622** | Rps27 | 0.212877599 | -0.308063583 | 0.733818782 | 16.89230392 | 1.023508886 | 0.349128609 | 0.709264768 | -4.610828535 |
| **A_55_P2048434** | Hsp90ab1 | 0.129983135 | -0.188158787 | 0.448125057 | 16.40806909 | 1.023331756 | 0.349205239 | 0.709264768 | -4.610873288 |
| **A_52_P182659** | Cs | 0.155680669 | -0.225373323 | 0.53673466 | 14.02349354 | 1.023289502 | 0.349223521 | 0.709264768 | -4.610883963 |
| **A_52_P566681** | Gpm6a | 0.240042346 | -0.347533064 | 0.827617756 | 8.270171879 | 1.023233135 | 0.349247911 | 0.709264768 | -4.610898204 |
| **A_55_P1961873** | Akt3 | -0.133757881 | -0.461196062 | 0.193680301 | 9.866056094 | -1.023152584 | 0.349282767 | 0.709264768 | -4.610918553 |
| **A_55_P1974636** | Gm16430 | -0.180398637 | -0.622017192 | 0.261219919 | 7.867341382 | -1.023142873 | 0.349286969 | 0.709264768 | -4.610921006 |

| | | | | | | | | | |
|---|---|---|---|---|---|---|---|---|---|
| **A_55_P2085974** | Igf1 | 1.04381568 | -1.511493486 | 3.599124846 | 11.0917292 | 1.023129741 | 0.349292652 | 0.709264768 | -4.610924324 |
| **A_55_P2061909** | Tceal9 | 0.204764079 | -0.296522897 | 0.706051056 | 13.27342022 | 1.023099087 | 0.349305917 | 0.709264768 | -4.610932068 |
| **A_55_P2139211** | NA | 0.527437405 | -0.76395159 | 1.818826399 | 14.70033918 | 1.022973045 | 0.349360467 | 0.709264768 | -4.610963908 |
| **A_65_P06446** | Tfrc | -1.112926858 | -3.837947404 | 1.612093688 | 9.561607258 | -1.022932996 | 0.349377801 | 0.709264768 | -4.610974024 |
| **A_51_P456870** | Foxj1 | 0.792282512 | -1.147746487 | 2.732311512 | 10.06698449 | 1.022874128 | 0.349403282 | 0.709264768 | -4.610988894 |
| **A_51_P390078** | NA | 0.101737431 | -0.147384559 | 0.350859422 | 9.316612835 | 1.022866722 | 0.349406488 | 0.709264768 | -4.610990765 |
| **A_55_P2068977** | Armcx3 | 0.476418306 | -0.690303446 | 1.643140058 | 9.835318028 | 1.022754718 | 0.349454973 | 0.709264768 | -4.611019056 |
| **A_51_P401184** | Rarres1 | -0.43982936 | -1.517008217 | 0.637349497 | 8.439381383 | -1.022696434 | 0.349480206 | 0.709264768 | -4.611033777 |
| **A_51_P137007** | Zw10 | 0.167819919 | -0.260278595 | 0.595918434 | 9.681733986 | 1.044149038 | 0.349481291 | 0.709264768 | -4.600486632 |
| **A_55_P2318584** | Aqp8 | -2.020328914 | -6.96999601 | 2.929338182 | 9.377135097 | -1.022342546 | 0.349633445 | 0.709264768 | -4.611123154 |
| **A_55_P2128511** | Ociad2 | 0.431789631 | -0.626090931 | 1.489670193 | 9.03192167 | 1.022317788 | 0.349644168 | 0.709264768 | -4.611129406 |
| **A_55_P2142371** | Rai1 | 0.223104962 | -0.323510928 | 0.769720851 | 12.08105797 | 1.022297556 | 0.34965293 | 0.709264768 | -4.611134515 |
| **A_55_P2160563** | Mrps17 | 0.264047954 | -0.382881848 | 0.910977756 | 11.47623578 | 1.022294429 | 0.349654284 | 0.709264768 | -4.611135305 |
| **A_55_P1965892** | Gnpda1 | -0.702110863 | -2.422720799 | 1.018499074 | 11.08710827 | -1.022053456 | 0.349758666 | 0.709264768 | -4.611196156 |
| **A_52_P94401** | Rtn4rl1 | 0.113917277 | -0.165274992 | 0.393109545 | 8.614494998 | 1.021966246 | 0.349796449 | 0.709264768 | -4.611218176 |
| **A_52_P310530** | Rbfox2 | -0.231631046 | -0.799374712 | 0.33611262 | 10.0518975 | -1.02186804 | 0.349839 | 0.709264768 | -4.611242973 |
| **A_55_P2080907** | Rpl26 | 0.340792059 | -0.494660477 | 1.176244595 | 15.8691028 | 1.021687383 | 0.349917286 | 0.709264768 | -4.611288584 |
| **A_55_P2166743** | Sp3os | 0.11032421 | -0.160142608 | 0.380791028 | 11.07251774 | 1.021661859 | 0.349928348 | 0.709264768 | -4.611295028 |
| **A_55_P2067777** | Cplane1 | 0.118353928 | -0.171824743 | 0.408532598 | 9.80953519 | 1.021568593 | 0.349968771 | 0.709264768 | -4.611318574 |
| **A_55_P2031989** | Rin2 | -0.282132923 | -0.973869817 | 0.40960397 | 11.1842625 | -1.021558342 | 0.349973214 | 0.709264768 | -4.611321162 |
| **A_55_P1990314** | Elavl4 | 0.226607388 | -0.328997483 | 0.782212259 | 8.711559473 | 1.02154703 | 0.349978117 | 0.709264768 | -4.611324017 |
| **A_55_P1979705** | Spop | -0.10104151 | -0.348782145 | 0.146699126 | 7.986552255 | -1.021534232 | 0.349983665 | 0.709264768 | -4.611327248 |
| **A_52_P432570** | Chdh | -0.33291003 | -1.149240228 | 0.483420169 | 7.956350463 | -1.021436475 | 0.350026039 | 0.709264768 | -4.611351927 |
| **A_55_P2041723** | Mid1ip1 | 0.905249606 | -1.314530527 | 3.125029739 | 12.42077996 | 1.021430284 | 0.350028723 | 0.709264768 | -4.611353489 |
| **A_52_P134141** | Pyurf | 0.29142542 | -0.423187377 | 1.006038217 | 8.808899086 | 1.021426185 | 0.3500305 | 0.709264768 | -4.611354524 |
| **A_51_P292490** | Ttbk2 | 0.14737415 | -0.214038005 | 0.508786304 | 9.281685092 | 1.021336619 | 0.350069328 | 0.709264768 | -4.611377134 |
| **A_66_P120125** | Nrep | 1.29642842 | -1.882947365 | 4.475804205 | 12.76301906 | 1.021308688 | 0.350081438 | 0.709264768 | -4.611384184 |
| **A_52_P442676** | Cux1 | 0.447590062 | -0.65015827 | 1.545338395 | 12.26465498 | 1.021240419 | 0.350111037 | 0.709264768 | -4.611401417 |
| **A_55_P1962304** | Plac8 | -0.710308152 | -2.452553687 | 1.031937382 | 8.410499457 | -1.021145869 | 0.350152033 | 0.709264768 | -4.611425283 |
| **A_51_P417251** | 6330403K | 1.065978932 | -1.548838967 | 3.680796831 | 11.22441865 | 1.021074706 | 0.350182892 | 0.709264768 | -4.611443244 |
| **A_52_P424784** | Clstn2 | 0.638153117 | -0.927270554 | 2.203576787 | 9.815785842 | 1.021041381 | 0.350197344 | 0.709264768 | -4.611451655 |
| **A_55_P1965075** | NA | 0.311413327 | -0.452581499 | 1.075408154 | 16.2318391 | 1.02093272 | 0.350244469 | 0.709264768 | -4.611479081 |
| **A_66_P133202** | Nfyc | 0.139335006 | -0.202499114 | 0.481169125 | 12.41013063 | 1.020928145 | 0.350246453 | 0.709264768 | -4.611480235 |
| **A_55_P1972927** | Elk3 | 0.337382892 | -0.490339971 | 1.165105755 | 11.7166144 | 1.020912334 | 0.350253311 | 0.709264768 | -4.611484226 |
| **A_51_P469661** | Stk38 | -0.186558449 | -0.644254746 | 0.271137849 | 10.25841692 | -1.020910965 | 0.350253905 | 0.709264768 | -4.611484571 |
| **A_66_P137605** | Cgas | 0.332281875 | -0.48292922 | 1.14749297 | 9.105921215 | 1.020908726 | 0.350254876 | 0.709264768 | -4.611485136 |
| **A_55_P2001608** | Klhl4 | 0.257678986 | -0.374527107 | 0.889885079 | 8.314054821 | 1.02087054 | 0.350271438 | 0.709264768 | -4.611494774 |
| **A_51_P226429** | Rhobtb2 | 0.192115398 | -0.279252154 | 0.66348295 | 8.676994769 | 1.02082862 | 0.350289621 | 0.709264768 | -4.611505353 |
| **A_52_P466701** | Fuom | 0.326378813 | -0.474435729 | 1.127193356 | 9.233389381 | 1.020799276 | 0.350302349 | 0.709264768 | -4.611512759 |
| **A_55_P2134785** | Zscan22 | 0.159486411 | -0.231934871 | 0.550907692 | 9.266058694 | 1.020538974 | 0.350415276 | 0.709264768 | -4.611578448 |
| **A_55_P1986706** | Pih1d1 | 0.35808794 | -0.520867599 | 1.237043478 | 11.74650521 | 1.020406303 | 0.350472844 | 0.709264768 | -4.611611925 |
| **A_55_P2097657** | Rps11 | 0.427778947 | -0.622337332 | 1.477895225 | 15.49495681 | 1.020310513 | 0.350514414 | 0.709264768 | -4.611636095 |
| **A_55_P2293381** | Cldn18 | 0.461464897 | -0.671887956 | 1.59481775 | 8.918885459 | 1.019820819 | 0.350726988 | 0.709264768 | -4.61175964 |
| **A_55_P2096283** | NA | 0.603010814 | -0.878426383 | 2.08444801 | 13.93423055 | 1.019511669 | 0.350861243 | 0.709264768 | -4.611837622 |
| **A_51_P378856** | Pfkp | -0.617916954 | -2.13610202 | 0.900268112 | 8.84680861 | -1.019426085 | 0.350898417 | 0.709264768 | -4.611859209 |
| **A_55_P2016356** | Eif3b | 0.099704897 | -0.145290345 | 0.344700139 | 10.72403971 | 1.019316807 | 0.350945888 | 0.709264768 | -4.61188677 |
| **A_55_P2030030** | Adssl1 | -0.648254678 | -2.241505219 | 0.944995864 | 10.43119299 | -1.019088569 | 0.351045051 | 0.709264768 | -4.611944331 |
| **A_52_P252121** | Nfs1 | 0.109805729 | -0.160084508 | 0.379695965 | 8.035785241 | 1.019032817 | 0.351069278 | 0.709264768 | -4.611958391 |

| | | | | | | | | | |
|---|---|---|---|---|---|---|---|---|---|
| **A_52_P378914** | AU04032( | -0.217194659 | -0.751036869 | 0.316647551 | 8.179744195 | -1.019029078 | 0.351070903 | 0.709264768 | -4.611959333 |
| **A_51_P214985** | Zfp521 | 0.50171641 | -0.731576927 | 1.735009746 | 9.331556555 | 1.018925113 | 0.351116083 | 0.709264768 | -4.61198555 |
| **A_55_P2121372** | Trp53i13 | -0.356289993 | -1.232206436 | 0.519626451 | 12.44305926 | -1.018805523 | 0.35116806 | 0.709264768 | -4.612015706 |
| **A_55_P2104612** | NA | -0.326355108 | -1.12870904 | 0.475998824 | 8.050875618 | -1.018766779 | 0.3511849 | 0.709264768 | -4.612025475 |
| **A_66_P135185** | Etv1 | 1.121695863 | -1.63624488 | 3.879636606 | 11.09468978 | 1.018686459 | 0.351219814 | 0.709264768 | -4.612045728 |
| **A_55_P2138097** | NA | 0.593509476 | -0.865774612 | 2.052793564 | 15.27360033 | 1.018680883 | 0.351222238 | 0.709264768 | -4.612047134 |
| **A_51_P173961** | Pdrg1 | -0.120157838 | -0.415625464 | 0.175309788 | 12.47359179 | -1.018573908 | 0.351268743 | 0.709264768 | -4.612074105 |
| **A_51_P234359** | Sct | -2.603922862 | -9.00737054 | 3.799524816 | 10.39314132 | -1.018508353 | 0.351297244 | 0.709264768 | -4.612090634 |
| **A_52_P649285** | NA | 0.306941137 | -0.447913479 | 1.061795754 | 10.30289008 | 1.018455665 | 0.351320153 | 0.709264768 | -4.612103917 |
| **A_55_P2048867** | NA | 0.135842093 | -0.198250368 | 0.469934554 | 8.099034096 | 1.018399167 | 0.351344719 | 0.709264768 | -4.612118161 |
| **A_55_P1969746** | Pitpnm1 | -0.200118308 | -0.692311461 | 0.292074844 | 9.890221391 | -1.018360629 | 0.351361476 | 0.709264768 | -4.612127876 |
| **A_51_P143468** | Klhl26 | -0.335099813 | -1.159322117 | 0.48912249 | 9.765409883 | -1.018310376 | 0.351383329 | 0.709264768 | -4.612140545 |
| **A_55_P2044439** | Mfng | 0.655853614 | -0.957340805 | 2.269048034 | 10.83868073 | 1.018287824 | 0.351393137 | 0.709264768 | -4.612146231 |
| **A_55_P1999878** | Sec24b | -0.254031196 | -0.878908713 | 0.370846321 | 11.66756708 | -1.018222055 | 0.351421739 | 0.709264768 | -4.612162811 |
| **A_55_P2017112** | Mpnd | 0.44457023 | -0.649031378 | 1.538171838 | 10.56107105 | 1.01819646 | 0.351432871 | 0.709264768 | -4.612169263 |
| **A_55_P2015734** | Syvn1 | -0.700456438 | -2.423649609 | 1.022736733 | 13.00528667 | -1.018116585 | 0.351467611 | 0.709264768 | -4.612189398 |
| **A_55_P2033435** | Stra6l | -0.181238189 | -0.627119759 | 0.264643381 | 7.758213282 | -1.018076794 | 0.351484919 | 0.709264768 | -4.612199428 |
| **A_55_P2114776** | Greb1 | 0.183678983 | -0.268235574 | 0.635593539 | 9.183705007 | 1.018013366 | 0.351512509 | 0.709264768 | -4.612215416 |
| **A_51_P142744** | Sulf1 | 0.604384728 | -0.882779352 | 2.091548808 | 9.914521417 | 1.017899592 | 0.351562003 | 0.709264768 | -4.612244094 |
| **A_55_P1973208** | Tax1bp3 | -0.117921491 | -0.408114494 | 0.172271512 | 9.398924556 | -1.01778576 | 0.351611529 | 0.709264768 | -4.612272785 |
| **A_55_P2182423** | NA | 0.362355429 | -0.529415402 | 1.254126261 | 14.82881496 | 1.01772831 | 0.351636526 | 0.709264768 | -4.612287265 |
| **A_66_P134394** | Pde6c | -0.245461814 | -0.849617471 | 0.358693842 | 7.820859454 | -1.017619545 | 0.351683855 | 0.709264768 | -4.612314677 |
| **A_55_P1976984** | NA | 0.326179597 | -0.476809195 | 1.129168389 | 15.89115993 | 1.017413871 | 0.351773368 | 0.709264768 | -4.612366509 |
| **A_55_P2128169** | Trappc13 | 0.119305952 | -0.174432578 | 0.413044482 | 9.27862667 | 1.017305843 | 0.351820391 | 0.709264768 | -4.612393731 |
| **A_55_P2005655** | Ntsr1 | -0.277312658 | -0.960106599 | 0.405481283 | 8.029250202 | -1.017256283 | 0.351841966 | 0.709264768 | -4.61240622 |
| **A_55_P2106608** | NA | -0.175977396 | -0.609325289 | 0.257370496 | 7.77984976 | -1.017116136 | 0.35190298 | 0.709264768 | -4.612441533 |
| **A_55_P2081303** | Tet2 | 0.288902938 | -0.422533455 | 1.000339331 | 10.32394232 | 1.017106025 | 0.351907383 | 0.709264768 | -4.612444081 |
| **A_51_P302566** | Maob | 0.341222983 | -0.499064976 | 1.181510942 | 10.66170235 | 1.017092568 | 0.351913242 | 0.709264768 | -4.612447471 |
| **A_55_P2101387** | Nubpl | 0.119088821 | -0.174191739 | 0.41236938 | 8.635922781 | 1.01704007 | 0.3519361 | 0.709264768 | -4.612460699 |
| **A_51_P320304** | Plod1 | -0.423718476 | -1.467252798 | 0.619815846 | 8.378866782 | -1.017000133 | 0.35195349 | 0.709264768 | -4.612470761 |
| **A_55_P1995623** | Pomp | 0.25064894 | -0.366671315 | 0.867969196 | 14.13911213 | 1.016964261 | 0.351969111 | 0.709264768 | -4.612479799 |
| **A_52_P654043** | Msto1 | 0.113183253 | -0.165593784 | 0.391960291 | 11.01047999 | 1.016893616 | 0.351999875 | 0.709264768 | -4.612497598 |
| **A_55_P2124941** | Brinp1 | 0.74845422 | -1.095049588 | 2.591958028 | 9.755440837 | 1.016884314 | 0.352003926 | 0.709264768 | -4.612499942 |
| **A_55_P2064676** | Gas2l3 | 0.212897745 | -0.311491937 | 0.737287427 | 8.670385087 | 1.016874294 | 0.35200829 | 0.709264768 | -4.612502466 |
| **A_55_P1967683** | Bex4 | 0.688895474 | -1.007927023 | 2.38571797 | 12.91906351 | 1.016874281 | 0.352008295 | 0.709264768 | -4.612502469 |
| **A_55_P2077143** | Mcidas | -0.252620798 | -0.874858604 | 0.369617008 | 11.6432633 | -1.016864425 | 0.352012588 | 0.709264768 | -4.612504953 |
| **A_55_P1985359** | Rpl28 | 0.549685253 | -0.804289562 | 1.903660068 | 15.33062716 | 1.016842864 | 0.352021978 | 0.709264768 | -4.612510385 |
| **A_55_P1966089** | NA | 0.121136131 | -0.177267615 | 0.419539877 | 8.264458008 | 1.016763089 | 0.352056722 | 0.709264768 | -4.612530483 |
| **A_55_P2177633** | NA | -0.179475164 | -0.621683902 | 0.262733574 | 7.748229811 | -1.016546819 | 0.352150929 | 0.709264768 | -4.612584965 |
| **A_55_P2145720** | Thoc2 | 0.20147967 | -0.294996658 | 0.697955998 | 9.038372893 | 1.016443002 | 0.352196159 | 0.709264768 | -4.612611116 |
| **A_52_P229316** | Galnt4 | 0.424438504 | -0.621454664 | 1.470331671 | 9.769248532 | 1.016430749 | 0.352201497 | 0.709264768 | -4.612614203 |
| **A_55_P2043451** | NA | 0.144082342 | -0.210969685 | 0.49913437 | 11.74041577 | 1.01641042 | 0.352210354 | 0.709264768 | -4.612619323 |
| **A_66_P113245** | Mfng | 0.711230962 | -1.041409631 | 2.463871554 | 11.74372185 | 1.016408138 | 0.352211349 | 0.709264768 | -4.612619898 |
| **A_55_P2327027** | Ano6 | -0.240395176 | -0.832787224 | 0.351996872 | 8.391908511 | -1.01640519 | 0.352212633 | 0.709264768 | -4.612620641 |
| **A_55_P2123491** | B3gnt5 | 0.197146446 | -0.288688135 | 0.682981027 | 8.817843019 | 1.016367743 | 0.35222895 | 0.709264768 | -4.612630073 |
| **A_55_P2077546** | NA | 0.126771327 | -0.185636174 | 0.439178827 | 9.042055924 | 1.016365532 | 0.352229913 | 0.709264768 | -4.61263063 |
| **A_55_P2408588** | Arntl | -0.211603832 | -0.733077421 | 0.309869757 | 9.290580358 | -1.016345939 | 0.35223845 | 0.709264768 | -4.612635565 |
| **A_52_P650387** | Ccnjl | -0.344194981 | -1.192527576 | 0.504137614 | 9.887746884 | -1.016222309 | 0.352292323 | 0.709264768 | -4.612666705 |

| | | | | | | | | | |
|---|---|---|---|---|---|---|---|---|---|
| **A_55_P1986049** | Fip1l1 | -0.119391162 | -0.413653465 | 0.174871141 | 8.530921596 | -1.016220364 | 0.352293171 | 0.709264768 | -4.612667194 |
| **A_55_P1968833** | Rpgr | 0.367167573 | -0.537827936 | 1.272163083 | 9.727678356 | 1.016174379 | 0.352313211 | 0.709264768 | -4.612678776 |
| **A_55_P1988728** | Fbxl19 | -0.61424516 | -2.128354651 | 0.89986433 | 10.37147254 | -1.016096154 | 0.352347304 | 0.709264768 | -4.612698478 |
| **A_51_P144349** | Dtx4 | 0.668145009 | -0.979029034 | 2.315319051 | 11.06518176 | 1.015971638 | 0.352401576 | 0.709264768 | -4.612729837 |
| **A_66_P102581** | Ube2z | -0.239318241 | -0.829345601 | 0.350709119 | 11.6444726 | -1.015907102 | 0.352429708 | 0.709264768 | -4.61274609 |
| **A_55_P2005433** | NA | -0.346463913 | -1.20067769 | 0.507749864 | 8.214559185 | -1.015878525 | 0.352442166 | 0.709264768 | -4.612753287 |
| **A_51_P230624** | Ccdc71l | -0.324017068 | -1.123049333 | 0.475015197 | 8.959748785 | -1.015673022 | 0.352531762 | 0.709264768 | -4.612805036 |
| **A_66_P112855** | AI413582 | -0.630277034 | -2.184571156 | 0.924017088 | 11.13815929 | -1.015660642 | 0.35253716 | 0.709264768 | -4.612808154 |
| **A_55_P2032318** | 4930522L | 0.1323949 | -0.194132753 | 0.458922552 | 8.403583132 | 1.015550756 | 0.352585077 | 0.709264768 | -4.612835823 |
| **A_55_P1970825** | NA | -0.289613617 | -1.004002676 | 0.424775441 | 7.957952442 | -1.015393852 | 0.352653506 | 0.709264768 | -4.612875329 |
| **A_55_P1962831** | Ssr1 | 0.107827985 | -0.158157974 | 0.373813944 | 13.46884117 | 1.015367187 | 0.352665136 | 0.709264768 | -4.612882043 |
| **A_55_P2147160** | NA | -0.199478963 | -0.691570819 | 0.292612894 | 7.744698021 | -1.015316089 | 0.352687424 | 0.709264768 | -4.612894908 |
| **A_66_P106606** | 2900097C | 0.148525685 | -0.217870402 | 0.514921773 | 10.78755986 | 1.015315658 | 0.352687612 | 0.709264768 | -4.612895017 |
| **A_55_P2083481** | Lpin1 | -0.111799405 | -0.387598696 | 0.163999886 | 8.321579351 | -1.015305392 | 0.35269209 | 0.709264768 | -4.612897601 |
| **A_55_P2127265** | NA | 0.206936403 | -0.303565653 | 0.717438458 | 14.31421315 | 1.015289152 | 0.352699174 | 0.709264768 | -4.61290169 |
| **A_55_P2126489** | Atxn7l3b | 0.210399278 | -0.308662525 | 0.729461081 | 13.23079243 | 1.015255893 | 0.352713681 | 0.709264768 | -4.612910063 |
| **A_55_P1993879** | NA | 0.23057928 | -0.338383513 | 0.799542073 | 16.37299688 | 1.015048415 | 0.352804196 | 0.709264768 | -4.612962296 |
| **A_51_P267700** | Gkn3 | 0.311165395 | -0.456661086 | 1.078991876 | 9.00116263 | 1.015029236 | 0.352812564 | 0.709264768 | -4.612967124 |
| **A_55_P1967617** | Med16 | -0.225860709 | -0.783204387 | 0.331482968 | 16.17161697 | -1.015004469 | 0.35282337 | 0.709264768 | -4.612973359 |
| **A_55_P2026547** | Gal3st2 | -0.145743288 | -0.505408935 | 0.213922358 | 8.128227207 | -1.014939023 | 0.352851926 | 0.709264768 | -4.612989834 |
| **A_51_P354698** | Dync1h1 | -0.206654466 | -0.716695099 | 0.303386168 | 12.61706839 | -1.014823144 | 0.352902494 | 0.709264768 | -4.613019003 |
| **A_66_P126348** | Smim31 | 0.195860951 | -0.287584194 | 0.679306096 | 12.28208043 | 1.014731172 | 0.352942633 | 0.709264768 | -4.613042153 |
| **A_55_P2410007** | NA | -0.313766486 | -1.088441462 | 0.460908491 | 9.305999992 | -1.014465709 | 0.353058508 | 0.709264768 | -4.613108968 |
| **A_55_P2098100** | Arhgef9 | 0.143070244 | -0.21017815 | 0.496318638 | 8.05365302 | 1.014423887 | 0.353076767 | 0.709264768 | -4.613119493 |
| **A_55_P2088535** | Snrpd2 | 0.508488847 | -0.747031313 | 1.764009008 | 14.39472042 | 1.014397305 | 0.353088372 | 0.709264768 | -4.613126183 |
| **A_51_P340668** | Bcl9l | -0.154780155 | -0.536952218 | 0.227391908 | 8.404275848 | -1.014394034 | 0.3530898 | 0.709264768 | -4.613127006 |
| **A_52_P73848** | Clint1 | -0.376145416 | -1.304939385 | 0.552648553 | 8.37878745 | -1.014347427 | 0.353110149 | 0.709264768 | -4.613138735 |
| **A_51_P271665** | Rev3l | 0.577030309 | -0.847879803 | 2.001940421 | 11.65330748 | 1.014288503 | 0.353135876 | 0.709264768 | -4.613153564 |
| **A_66_P123448** | Pigx | -0.277528319 | -0.962903538 | 0.4078469 | 11.32586959 | -1.014213187 | 0.353168764 | 0.709264768 | -4.613172518 |
| **A_55_P2076984** | Farp1 | -0.161702094 | -0.561108436 | 0.237704248 | 8.659272937 | -1.014030537 | 0.35324853 | 0.709264768 | -4.613218479 |
| **A_55_P1996816** | NA | 0.628583521 | -0.92409861 | 2.181265652 | 14.24459512 | 1.013983252 | 0.353269182 | 0.709264768 | -4.613230377 |
| **A_55_P2046153** | NA | -0.331632508 | -1.150843488 | 0.487578473 | 14.88879168 | -1.013938644 | 0.353288666 | 0.709264768 | -4.613241601 |
| **A_52_P423810** | Mt1 | -0.270399455 | -0.938359329 | 0.397560418 | 7.952980664 | -1.013924889 | 0.353294674 | 0.709264768 | -4.613245062 |
| **A_55_P1971966** | Uba1 | 0.335911945 | -0.494051432 | 1.165875321 | 13.46351784 | 1.013717313 | 0.353385353 | 0.709264768 | -4.613297289 |
| **A_55_P2176292** | NA | 0.440352198 | -0.647687395 | 1.528391792 | 14.66600728 | 1.013691523 | 0.35339662 | 0.709264768 | -4.613303778 |
| **A_55_P2040416** | Pradc1 | 0.178081779 | -0.261939982 | 0.61810354 | 10.08038677 | 1.013667857 | 0.35340696 | 0.709264768 | -4.613309732 |
| **A_51_P390285** | Larp6 | -0.229650546 | -0.797130572 | 0.33782948 | 8.489430582 | -1.013601508 | 0.35343595 | 0.709264768 | -4.613326424 |
| **A_51_P201773** | Tubgcp3 | 0.219750466 | -0.323299547 | 0.76280048 | 10.65476134 | 1.013538667 | 0.353463409 | 0.709264768 | -4.613342233 |
| **A_55_P1987459** | Pigt | -0.19059588 | -0.66164828 | 0.280456519 | 8.009064946 | -1.01343205 | 0.35351 | 0.709264768 | -4.613369055 |
| **A_55_P2076174** | NA | 0.580272696 | -0.8539951 | 2.014540492 | 14.64998515 | 1.013333113 | 0.35355324 | 0.709264768 | -4.613393943 |
| **A_55_P2040863** | Cnn3 | 0.178505867 | -0.262737059 | 0.619748793 | 12.94300734 | 1.013269766 | 0.353580927 | 0.709264768 | -4.613409877 |
| **A_55_P2122997** | Tet1 | 0.498559786 | -0.733914131 | 1.731033703 | 9.304853951 | 1.013187572 | 0.353616855 | 0.709264768 | -4.613430552 |
| **A_55_P1967342** | NA | 0.46201537 | -0.680736248 | 1.604766988 | 15.25884877 | 1.012639641 | 0.353856435 | 0.709264768 | -4.613568359 |
| **A_55_P1978151** | NA | 0.103734543 | -0.152859237 | 0.360328324 | 16.50669958 | 1.012575976 | 0.353884281 | 0.709264768 | -4.613584368 |
| **A_55_P1979997** | Spaca9 | -0.219694937 | -0.763173851 | 0.323783976 | 9.365980387 | -1.012482898 | 0.353924995 | 0.709264768 | -4.613607774 |
| **A_52_P653902** | Eny2 | 0.311690135 | -0.459366082 | 1.082746352 | 9.424714578 | 1.012482112 | 0.353925339 | 0.709264768 | -4.613607971 |
| **A_55_P2167107** | Cdk14 | 0.481575533 | -0.759828128 | 1.722979193 | 11.69310596 | 1.033270659 | 0.354027613 | 0.709264768 | -4.602744076 |
| **A_52_P399990** | B4galt2 | 0.424091935 | -0.625365844 | 1.473549714 | 9.320822828 | 1.012151185 | 0.354070123 | 0.709264768 | -4.613691179 |

| | | | | | | | | | |
|---|---|---|---|---|---|---|---|---|---|
| **A_52_P194805** | Zfp706 | -0.35974591 | -1.250086915 | 0.530595096 | 13.11727406 | -1.012021722 | 0.354126777 | 0.709264768 | -4.613723727 |
| **A_55_P2184551** | NA | -0.119715606 | -0.416009196 | 0.176577984 | 11.80241823 | -1.011996142 | 0.354137972 | 0.709264768 | -4.613730158 |
| **A_51_P354572** | Rab35 | -0.258781102 | -0.899271727 | 0.381709524 | 12.30575775 | -1.011975797 | 0.354146877 | 0.709264768 | -4.613735273 |
| **A_55_P2054714** | NA | 0.122534482 | -0.180752591 | 0.425821554 | 9.410436643 | 1.011940003 | 0.354162542 | 0.709264768 | -4.613744271 |
| **A_55_P2015092** | E430024I( | -0.099976652 | -0.347454955 | 0.147501651 | 8.973976127 | -1.011839909 | 0.354206353 | 0.709264768 | -4.613769434 |
| **A_55_P2091985** | Gpr157 | -1.193481668 | -4.147810808 | 1.760847472 | 12.20135826 | -1.011829227 | 0.354211029 | 0.709264768 | -4.613772119 |
| **A_55_P1973838** | Slc6a9 | 0.572585666 | -0.844914923 | 1.990086255 | 9.99082637 | 1.011736852 | 0.354251466 | 0.709264768 | -4.61379534 |
| **A_55_P2015715** | Rab4a | -0.184048823 | -0.639717091 | 0.271619445 | 9.733366108 | -1.011660057 | 0.354285085 | 0.709264768 | -4.613814644 |
| **A_55_P2031242** | Sun1 | -0.201530398 | -0.700497759 | 0.297436963 | 8.270461482 | -1.011623175 | 0.354301233 | 0.709264768 | -4.613823914 |
| **A_51_P111932** | Tor1aip2 | -0.610443828 | -2.121949387 | 0.90106173 | 8.959160344 | -1.011547554 | 0.354334342 | 0.709264768 | -4.613842922 |
| **A_51_P338485** | Aldh6a1 | 0.487942655 | -0.720580392 | 1.696465703 | 11.36546506 | 1.011263177 | 0.354458874 | 0.709264768 | -4.613914396 |
| **A_51_P311785** | Man2b2 | -0.166068751 | -0.577391715 | 0.245254212 | 8.303736291 | -1.011242491 | 0.354467934 | 0.709264768 | -4.613919595 |
| **A_66_P101261** | Gm3367 | -0.192686069 | -0.669960148 | 0.284588011 | 7.78672478 | -1.01119011 | 0.354490877 | 0.709264768 | -4.613932759 |
| **A_66_P131091** | Rabl6 | -0.137935045 | -0.479598051 | 0.203727961 | 11.95390807 | -1.011176596 | 0.354496796 | 0.709264768 | -4.613936155 |
| **A_51_P116932** | Lad1 | -0.355648844 | -1.236676401 | 0.525378713 | 8.23922369 | -1.011072397 | 0.354542439 | 0.709264768 | -4.613962341 |
| **A_55_P2106255** | Mbnl3 | -0.440575768 | -1.532008231 | 0.650856694 | 11.70504089 | -1.011053381 | 0.354550769 | 0.709264768 | -4.613967119 |
| **A_51_P188073** | Dnajb11 | -0.575218504 | -2.000411552 | 0.849974543 | 12.13130777 | -1.010903032 | 0.354616637 | 0.709264768 | -4.6140049 |
| **A_55_P1985453** | NA | -0.156683827 | -0.544896829 | 0.231529175 | 7.883124873 | -1.010891256 | 0.354621797 | 0.709264768 | -4.614007859 |
| **A_55_P2052261** | Zfp280c | 0.307923159 | -0.455069994 | 1.070916311 | 8.868557111 | 1.010815885 | 0.354654821 | 0.709264768 | -4.614026798 |
| **A_51_P416419** | Calr | -0.759282169 | -2.64100053 | 1.122436192 | 13.26396786 | -1.010645681 | 0.354729406 | 0.709264768 | -4.614069563 |
| **A_51_P208020** | Ulk3 | -0.134439963 | -0.467651006 | 0.198771079 | 9.032179591 | -1.010553582 | 0.35476977 | 0.709264768 | -4.614092703 |
| **A_51_P161636** | Kng2 | -0.95186517 | -3.311333967 | 1.407603628 | 8.442092573 | -1.010442356 | 0.354818522 | 0.709264768 | -4.614120646 |
| **A_52_P582858** | Dgkd | -0.154895813 | -0.538867868 | 0.229076243 | 10.7561868 | -1.010393177 | 0.354840079 | 0.709264768 | -4.614133001 |
| **A_55_P2045225** | NA | 0.149192896 | -0.220650733 | 0.519036525 | 12.01257486 | 1.010369779 | 0.354850336 | 0.709264768 | -4.61413888 |
| **A_55_P2014224** | Sema5a | 0.830162126 | -1.227846871 | 2.888171122 | 11.2049893 | 1.010336383 | 0.354864976 | 0.709264768 | -4.614147269 |
| **A_55_P1968493** | Telo2 | 0.203944866 | -0.301650255 | 0.709539988 | 9.945646009 | 1.01032302 | 0.354870834 | 0.709264768 | -4.614150626 |
| **A_55_P1997894** | NA | 0.463884793 | -0.686144188 | 1.613913774 | 15.38121364 | 1.010303126 | 0.354879556 | 0.709264768 | -4.614155624 |
| **A_55_P2077623** | NA | -0.450564756 | -1.567624285 | 0.666494774 | 8.010156977 | -1.01025555 | 0.354900413 | 0.709264768 | -4.614167575 |
| **A_51_P134301** | Pfdn5 | 0.421789004 | -0.623972225 | 1.467550234 | 12.98406891 | 1.010213255 | 0.354918957 | 0.709264768 | -4.6141782 |
| **A_51_P345593** | Sptlc1 | -0.266674543 | -0.927932401 | 0.394583315 | 9.860347441 | -1.010092296 | 0.354971993 | 0.709264768 | -4.614208584 |
| **A_55_P2073607** | Commd5 | -0.179093718 | -0.623187133 | 0.264999697 | 11.39353489 | -1.010081379 | 0.35497678 | 0.709264768 | -4.614211326 |
| **A_51_P421958** | Sgms1 | -0.239785661 | -0.834415149 | 0.354843827 | 11.91943386 | -1.010013346 | 0.355006613 | 0.709264768 | -4.614228415 |
| **A_55_P2038484** | 4933427D | 0.148164173 | -0.219396035 | 0.515724382 | 13.2177486 | 1.009636538 | 0.355171885 | 0.709264768 | -4.614323054 |
| **A_55_P1968043** | Jmjd1c | -0.085362132 | -0.297149706 | 0.126425443 | 11.39204945 | -1.00952045 | 0.355222815 | 0.709264768 | -4.614352207 |
| **A_51_P332799** | Clptm1 | -0.371307556 | -1.29256783 | 0.549952718 | 11.69468668 | -1.009489459 | 0.355236412 | 0.709264768 | -4.614359989 |
| **A_55_P1969914** | Tcf25 | -0.722668027 | -2.51590187 | 1.070565817 | 12.78225027 | -1.009374338 | 0.355286925 | 0.709264768 | -4.614388898 |
| **A_55_P2073705** | Tfdp2 | 0.152773286 | -0.226336112 | 0.531882683 | 9.483090623 | 1.009330077 | 0.355306348 | 0.709264768 | -4.614400012 |
| **A_52_P102248** | Mex3b | 0.412416928 | -0.611017094 | 1.435850951 | 10.40099244 | 1.009315583 | 0.355312709 | 0.709264768 | -4.614403652 |
| **A_55_P1968295** | Pacsin3 | -0.205901973 | -0.716904119 | 0.305100173 | 11.2482711 | -1.009225301 | 0.355352329 | 0.709264768 | -4.614426321 |
| **A_52_P95759** | Sh3rf3 | 0.256968681 | -0.380790219 | 0.894727582 | 8.599466312 | 1.009192498 | 0.355366726 | 0.709264768 | -4.614434558 |
| **A_52_P436628** | Vegfb | 0.223765135 | -0.331591604 | 0.779121875 | 9.301795801 | 1.009184843 | 0.355370085 | 0.709264768 | -4.61443648 |
| **A_55_P2128085** | NA | 0.26129545 | -0.387225948 | 0.909816848 | 10.83094164 | 1.009155014 | 0.355383178 | 0.709264768 | -4.614443969 |
| **A_52_P296913** | Cnnm1 | -0.43516813 | -1.515317719 | 0.64498146 | 8.020466715 | -1.009075175 | 0.35541822 | 0.709264768 | -4.614464015 |
| **A_55_P2151437** | NA | -0.254317844 | -0.885596483 | 0.376960795 | 8.183418029 | -1.00903466 | 0.355436005 | 0.709264768 | -4.614474187 |
| **A_51_P478311** | Nmd3 | 0.101194941 | -0.150008005 | 0.352397887 | 8.62251215 | 1.008984324 | 0.355458101 | 0.709264768 | -4.614486825 |
| **A_66_P138929** | Lox | 0.582481504 | -0.863586496 | 2.028549504 | 10.5147495 | 1.008889888 | 0.355499558 | 0.709264768 | -4.614510534 |
| **A_66_P105244** | Tor1aip1 | -0.448382929 | -1.561565859 | 0.664800001 | 8.410581198 | -1.008864585 | 0.355510667 | 0.709264768 | -4.614516886 |
| **A_51_P389506** | NA | -0.429419226 | -1.495854063 | 0.63701561 | 11.97809488 | -1.008550171 | 0.355648727 | 0.709264768 | -4.614595815 |

| | | | | | | | | | |
|---|---|---|---|---|---|---|---|---|---|
| **A_51_P498890** | Degs1 | -0.381330464 | -1.32841045 | 0.565749522 | 12.75831551 | -1.00847512 | 0.355681689 | 0.709264768 | -4.614614653 |
| **A_55_P2176300** | Slc33a1 | -0.140931924 | -0.490962414 | 0.209098566 | 8.257781859 | -1.008448794 | 0.355693251 | 0.709264768 | -4.614621261 |
| **A_51_P463187** | Tbc1d2b | 0.397225074 | -0.589537545 | 1.383987693 | 9.4655971 | 1.008264116 | 0.355774373 | 0.709264768 | -4.614667614 |
| **A_55_P2056739** | Col16a1 | 0.594614322 | -0.882552714 | 2.071781358 | 12.26552173 | 1.008221846 | 0.355792942 | 0.709264768 | -4.614678223 |
| **A_51_P211325** | Nudt2 | 0.27449894 | -0.407459272 | 0.956457152 | 9.11353753 | 1.008168802 | 0.355816246 | 0.709264768 | -4.614691536 |
| **A_55_P1987121** | Rtraf | 0.269553014 | -0.400162995 | 0.939269023 | 14.27450533 | 1.008100562 | 0.355846228 | 0.709264768 | -4.614708662 |
| **A_52_P426768** | Cited4 | -0.350222927 | -1.220426881 | 0.519981027 | 8.360193943 | -1.008030951 | 0.355876814 | 0.709264768 | -4.614726132 |
| **A_66_P120558** | Pam16 | 0.334182337 | -0.496194869 | 1.164559543 | 12.91483848 | 1.007995095 | 0.35589257 | 0.709264768 | -4.61473513 |
| **A_55_P2028418** | Hacd2 | -0.301435125 | -1.050456838 | 0.447586589 | 10.145612 | -1.007975079 | 0.355901365 | 0.709264768 | -4.614740153 |
| **A_52_P920129** | Gnai1 | 0.474001662 | -0.703951077 | 1.651954401 | 9.689522825 | 1.007864954 | 0.35594976 | 0.709264768 | -4.614767788 |
| **A_55_P1965448** | NA | 0.271867641 | -0.403960722 | 0.947696004 | 11.6181957 | 1.007561235 | 0.356083257 | 0.709264768 | -4.614843998 |
| **A_55_P2080048** | NA | 0.357651753 | -0.531459947 | 1.246763453 | 15.51927552 | 1.00752163 | 0.356100668 | 0.709264768 | -4.614853935 |
| **A_55_P1996419** | NA | 0.139491508 | -0.207352586 | 0.486335602 | 16.02378164 | 1.007311547 | 0.356193036 | 0.709264768 | -4.614906642 |
| **A_51_P141836** | Ube3c | 0.145279678 | -0.215996662 | 0.506556018 | 9.804163139 | 1.007199948 | 0.356242111 | 0.709264768 | -4.614934639 |
| **A_51_P233928** | B4galnt1 | -0.502403306 | -1.751797906 | 0.746991295 | 8.368903755 | -1.007170992 | 0.356254845 | 0.709264768 | -4.614941903 |
| **A_52_P676461** | Lpgat1 | -0.124402476 | -0.433786112 | 0.18498116 | 8.532305258 | -1.00712188 | 0.356276444 | 0.709264768 | -4.614954223 |
| **A_51_P342481** | Gpd2 | -0.398237092 | -1.388663158 | 0.592188974 | 8.822672257 | -1.007093962 | 0.356288723 | 0.709264768 | -4.614961226 |
| **A_55_P2133225** | Mmgt2 | -0.363064666 | -1.266017449 | 0.539888116 | 10.4162056 | -1.007092334 | 0.356289439 | 0.709264768 | -4.614961634 |
| **A_55_P2000133** | Cdh2 | 0.492434702 | -0.732497664 | 1.717367068 | 9.848450176 | 1.006901264 | 0.356373483 | 0.709264768 | -4.615009562 |
| **A_51_P398406** | Kctd13 | -0.12489847 | -0.43559416 | 0.185797219 | 8.388881325 | -1.006867302 | 0.356388423 | 0.709264768 | -4.615018081 |
| **A_51_P107020** | Kif5a | -0.301325157 | -1.050900473 | 0.44825016 | 8.442904673 | -1.006863181 | 0.356390236 | 0.709264768 | -4.613854821 |
| **A_55_P1954006** | Ldhb | -0.65668708 | -2.2903715 | 0.976997339 | 13.23813113 | -1.006794074 | 0.356420639 | 0.709264768 | -4.615036448 |
| **A_52_P489202** | 4732465J | -0.130973209 | -0.456849538 | 0.19490312 | 7.764612995 | -1.006653477 | 0.356482499 | 0.709264768 | -4.615071711 |
| **A_51_P397920** | Ilrun | -0.285014423 | -0.994170091 | 0.424141245 | 10.97390598 | -1.006643294 | 0.356486979 | 0.709264768 | -4.615074265 |
| **A_55_P2086087** | NA | 0.554779236 | -0.825681462 | 1.935239935 | 14.84177875 | 1.006575829 | 0.356516666 | 0.709264768 | -4.615091185 |
| **A_55_P2050688** | Zbed4 | 0.194817525 | -0.289967451 | 0.679602501 | 11.09358528 | 1.006535775 | 0.356534292 | 0.709264768 | -4.61510123 |
| **A_51_P520966** | Icosl | 0.224513484 | -0.334189385 | 0.783216353 | 9.698603649 | 1.006495582 | 0.356551979 | 0.709264768 | -4.61511131 |
| **A_55_P2262560** | 5730471H | 0.328355696 | -0.488834735 | 1.145546127 | 8.867599867 | 1.006402315 | 0.356593026 | 0.709264768 | -4.6151347 |
| **A_55_P1976864** | Fbxl20 | -0.329933837 | -1.151078633 | 0.491210958 | 9.942162683 | -1.006369483 | 0.356607477 | 0.709264768 | -4.615142933 |
| **A_66_P120770** | Ywhaz | 0.132914283 | -0.197920181 | 0.463748746 | 9.144544317 | 1.006262418 | 0.356654602 | 0.709264768 | -4.615169781 |
| **A_55_P2048671** | Rpl27a | 0.265289112 | -0.395048123 | 0.925626348 | 16.70327017 | 1.006245579 | 0.356662015 | 0.709264768 | -4.615174003 |
| **A_55_P2044207** | Bphl | 0.183816093 | -0.273726767 | 0.641358953 | 8.230985679 | 1.006241196 | 0.356663944 | 0.709264768 | -4.615175102 |
| **A_51_P182131** | 5330417C | 0.532696296 | -0.793331196 | 1.858723787 | 9.590892266 | 1.006184202 | 0.356689033 | 0.709264768 | -4.615189394 |
| **A_51_P412270** | Rapgef6 | 0.113341305 | -0.168797994 | 0.395480604 | 8.262381593 | 1.006178359 | 0.356691606 | 0.709264768 | -4.615190859 |
| **A_55_P1958961** | 2010001K | -0.212756528 | -0.74237487 | 0.316861814 | 8.32407721 | -1.006167353 | 0.356696451 | 0.709264768 | -4.615193619 |
| **A_51_P136521** | Lypd2 | 1.034785703 | -1.541423174 | 3.610994581 | 10.88168724 | 1.006050295 | 0.356747985 | 0.709264768 | -4.61522297 |
| **A_55_P2042525** | Apoo | 0.113180554 | -0.168626771 | 0.394987879 | 10.3611322 | 1.005934916 | 0.356798787 | 0.709264768 | -4.615251898 |
| **A_51_P359570** | Ifit3 | -0.605249287 | -2.112282893 | 0.90178432 | 8.292211375 | -1.005915951 | 0.356807138 | 0.709264768 | -4.615256653 |
| **A_51_P520106** | Adprhl2 | -0.1116721 | -0.389734476 | 0.166390276 | 9.243985623 | -1.005895348 | 0.35681621 | 0.709264768 | -4.615261819 |
| **A_66_P139267** | Rbpj | 0.215509766 | -0.321171555 | 0.752191087 | 10.53386877 | 1.005774965 | 0.356869224 | 0.709264768 | -4.615292 |
| **A_55_P1996299** | Pofut1 | 0.101441407 | -0.1512246 | 0.354107414 | 11.7294911 | 1.005585015 | 0.356952886 | 0.709264768 | -4.615339619 |
| **A_51_P286814** | Ncor2 | -0.265109767 | -0.925438009 | 0.395218475 | 12.56853902 | -1.005579015 | 0.356955529 | 0.709264768 | -4.615341123 |
| **A_51_P249793** | Dusp28 | 0.09399354 | -0.140126246 | 0.328113327 | 8.931217541 | 1.005565197 | 0.356961615 | 0.709264768 | -4.615344586 |
| **A_55_P2028894** | Gata4 | -0.276996059 | -0.966956476 | 0.412964358 | 8.424478221 | -1.005540944 | 0.356972299 | 0.709264768 | -4.615350666 |
| **A_55_P2167025** | Nectin2 | -0.631877583 | -2.205821463 | 0.942066298 | 12.44896098 | -1.005527726 | 0.356978121 | 0.709264768 | -4.61535398 |
| **A_52_P483301** | Mios | 0.228439701 | -0.340635441 | 0.797514844 | 9.008341259 | 1.005431096 | 0.35702069 | 0.709264768 | -4.615378202 |
| **A_55_P2088103** | Ash2l | -0.486092443 | -1.697125228 | 0.724940343 | 12.11277349 | -1.005340815 | 0.357060465 | 0.709264768 | -4.615400832 |
| **A_55_P2172420** | Krit1 | -0.102603047 | -0.35830088 | 0.153094786 | 9.615931006 | -1.005040482 | 0.357192809 | 0.709264768 | -4.615476106 |

| | | | | | | | | | |
|---|---|---|---|---|---|---|---|---|---|
| **A_55_P2138549** | Cc2d1a | -0.282867751 | -0.987943837 | 0.422208334 | 10.09567446 | -1.004842046 | 0.357280273 | 0.709264768 | -4.615525835 |
| **A_52_P670874** | Ebag9 | 0.170788088 | -0.254991982 | 0.596568158 | 9.749627175 | 1.004668053 | 0.357356978 | 0.709264768 | -4.615569436 |
| **A_55_P2032818** | Trim2 | 0.704860533 | -1.052471798 | 2.462192865 | 10.95293695 | 1.004614959 | 0.357380387 | 0.709264768 | -4.61558274 |
| **A_51_P210928** | Tsta3 | -0.108605097 | -0.379378746 | 0.162168551 | 9.849436066 | -1.004602275 | 0.35738598 | 0.709264768 | -4.615585918 |
| **A_51_P184353** | Nr2f6 | 0.244529295 | -0.365159831 | 0.854218421 | 12.77013868 | 1.004552854 | 0.357407771 | 0.709264768 | -4.615598301 |
| **A_55_P1972772** | Gsg1 | 0.371312777 | -0.554491746 | 1.2971173 | 9.119227968 | 1.004548573 | 0.357409659 | 0.709264768 | -4.615599374 |
| **A_55_P1996082** | Zfp936 | 0.178328294 | -0.26635794 | 0.623014528 | 10.40315198 | 1.004423615 | 0.357464761 | 0.709264768 | -4.615630683 |
| **A_51_P474422** | Ift27 | 0.471494262 | -0.704265979 | 1.647254504 | 12.43085815 | 1.004402968 | 0.357473866 | 0.709264768 | -4.615635856 |
| **A_52_P147666** | Slc30a7 | -0.444113511 | -1.551696889 | 0.663469868 | 8.876873399 | -1.004310264 | 0.357514751 | 0.709264768 | -4.615659082 |
| **A_55_P1998825** | Rpl9 | 0.304665916 | -0.455360152 | 1.064691985 | 17.51560701 | 1.004027778 | 0.357639359 | 0.709264768 | -4.61572985 |
| **A_55_P2035986** | E330020D | -0.395024841 | -1.380465024 | 0.590415343 | 9.593774818 | -1.004024902 | 0.357640628 | 0.709264768 | -4.615730571 |
| **A_52_P208727** | Zmym4 | 0.136948403 | -0.204692512 | 0.478589319 | 9.324047818 | 1.004008621 | 0.357647811 | 0.709264768 | -4.615734649 |
| **A_52_P497188** | Prrg1 | 0.44069005 | -0.658776186 | 1.540156285 | 9.1282751 | 1.00392598 | 0.357684271 | 0.709264768 | -4.61575535 |
| **A_55_P2149906** | NA | 0.14449198 | -0.216004818 | 0.504988778 | 14.42696157 | 1.003905138 | 0.357693468 | 0.709264768 | -4.615760571 |
| **A_51_P397925** | Smap2 | 0.130445834 | -0.195025232 | 0.455916901 | 9.254842874 | 1.003848499 | 0.357718459 | 0.709264768 | -4.615774759 |
| **A_55_P2057441** | NA | 0.130126355 | -0.194567439 | 0.454820148 | 15.64102627 | 1.003787128 | 0.35774554 | 0.709264768 | -4.615790131 |
| **A_55_P2121491** | Bbs9 | 0.258375452 | -0.386393289 | 0.903144194 | 9.910589089 | 1.003685434 | 0.357790417 | 0.709264768 | -4.615815602 |
| **A_51_P214127** | Cpa3 | 0.383217974 | -0.573095397 | 1.339531346 | 8.578369911 | 1.003681659 | 0.357792083 | 0.709264768 | -4.615816547 |
| **A_52_P657240** | Cdc40 | -0.244384053 | -0.854295635 | 0.365527528 | 8.286173895 | -1.003590009 | 0.357832533 | 0.709264768 | -4.615839502 |
| **A_55_P2128153** | Col6a1 | 0.990517411 | -1.481881689 | 3.462916512 | 13.85006493 | 1.003445715 | 0.357896224 | 0.709264768 | -4.61587564 |
| **A_51_P432659** | Pex7 | 0.114942926 | -0.172007763 | 0.401893615 | 9.899753088 | 1.003287316 | 0.357966151 | 0.709264768 | -4.615915308 |
| **A_55_P2160253** | 6720475J | 0.185556572 | -0.277770632 | 0.648883776 | 8.261786094 | 1.003087656 | 0.35805431 | 0.709264768 | -4.615965304 |
| **A_51_P386358** | Wwp2 | 0.176745777 | -0.264591243 | 0.618082797 | 12.21236112 | 1.003064899 | 0.358064359 | 0.709264768 | -4.615971002 |
| **A_55_P2128224** | NA | 0.324546091 | -0.485922503 | 1.135014685 | 16.62766342 | 1.002975993 | 0.358103622 | 0.709264768 | -4.615993263 |
| **A_55_P1958103** | NA | 0.357141099 | -0.534737666 | 1.249019863 | 15.55557452 | 1.002961708 | 0.358109931 | 0.709264768 | -4.61599684 |
| **A_55_P2056774** | Hsd17b14 | -0.189448807 | -0.662561254 | 0.28366364 | 8.03938226 | -1.002946682 | 0.358116567 | 0.709264768 | -4.616000602 |
| **A_51_P150087** | Abhd10 | 0.149456138 | -0.223800456 | 0.522712732 | 8.584810977 | 1.00289765 | 0.358138222 | 0.709264768 | -4.616012879 |
| **A_52_P359819** | Ntf5 | -0.099541382 | -0.348164181 | 0.149081416 | 9.292486426 | -1.002797087 | 0.35818264 | 0.709264768 | -4.616038057 |
| **A_52_P258557** | Slc26a9 | 0.383717784 | -0.574797397 | 1.342232965 | 8.70509316 | 1.002682136 | 0.358233419 | 0.709264768 | -4.616066836 |
| **A_55_P1985758** | Psmd9 | -0.088160999 | -0.308399106 | 0.132077108 | 9.322581674 | -1.002615425 | 0.358262891 | 0.709264768 | -4.616083537 |
| **A_55_P1980401** | Cbx7 | -0.237015864 | -0.829283115 | 0.355251387 | 7.805413022 | -1.002328413 | 0.35838971 | 0.709264768 | -4.616155383 |
| **A_52_P467930** | NA | 0.259502844 | -0.389041718 | 0.908047405 | 9.717418334 | 1.002195955 | 0.35844825 | 0.709264768 | -4.616188538 |
| **A_52_P664404** | Zfp286 | 0.487319154 | -0.730784918 | 1.705423225 | 9.358087643 | 1.002027019 | 0.358522923 | 0.709264768 | -4.61623082 |
| **A_52_P447944** | Epcam | 0.3984031 | -0.597620064 | 1.394426265 | 10.19686117 | 1.001852107 | 0.358600251 | 0.709264768 | -4.616274595 |
| **A_51_P283499** | Drd4 | -0.412768755 | -1.444768211 | 0.619230701 | 7.931379378 | -1.001792307 | 0.358626692 | 0.709264768 | -4.61628956 |
| **A_52_P604629** | Csrnp1 | -0.777455019 | -2.721432354 | 1.166522315 | 10.5299078 | -1.001692496 | 0.358670827 | 0.709264768 | -4.616314537 |
| **A_55_P2081530** | Sh3bgrl3 | -0.184986416 | -0.64754284 | 0.277570009 | 13.89236208 | -1.001671843 | 0.358679959 | 0.709264768 | -4.616319705 |
| **A_55_P2113466** | NA | -0.151996069 | -0.532096594 | 0.228104455 | 9.105108372 | -1.001576748 | 0.358722014 | 0.709264768 | -4.6163435 |
| **A_51_P240019** | Gyg | 1.063686248 | -1.596497324 | 3.723869819 | 14.7569664 | 1.001503065 | 0.358754602 | 0.709264768 | -4.616361937 |
| **A_66_P129484** | Tmed9 | 0.221947709 | -0.333156492 | 0.77705191 | 8.914869836 | 1.001443605 | 0.358780901 | 0.709264768 | -4.616376815 |
| **A_55_P1997275** | Xpnpep1 | -0.283124743 | -0.991300328 | 0.425050841 | 13.30201386 | -1.001353042 | 0.35882096 | 0.709264768 | -4.616399474 |
| **A_51_P139678** | Sprr1a | 0.161742923 | -0.242841747 | 0.566327593 | 8.243443448 | 1.001304595 | 0.358842392 | 0.709264768 | -4.616411595 |
| **A_55_P2151546** | Adarb1 | 0.58640227 | -0.88054701 | 2.05335155 | 11.77665032 | 1.001223167 | 0.358878415 | 0.709264768 | -4.616431968 |
| **A_66_P135742** | Wdr1 | -0.450742509 | -1.578328725 | 0.676843707 | 12.18568879 | -1.001219052 | 0.358880236 | 0.709264768 | -4.616432997 |
| **A_55_P2128373** | Cops6 | 0.129718687 | -0.19484998 | 0.454287353 | 12.65780532 | 1.001028164 | 0.358964695 | 0.709264768 | -4.616480752 |
| **A_51_P196673** | Khdc1b | -0.17918621 | -0.627551924 | 0.269179504 | 7.737746628 | -1.000973395 | 0.358988931 | 0.709264768 | -4.616494453 |
| **A_55_P2018631** | NA | 0.243397694 | -0.36565463 | 0.852450019 | 13.17794587 | 1.000949577 | 0.358999472 | 0.709264768 | -4.616500411 |
| **A_52_P315111** | Mrpl52 | 0.40406784 | -0.607165333 | 1.415301012 | 14.20411755 | 1.000813888 | 0.359059523 | 0.709264768 | -4.616534353 |

| | | | | | | | | | |
|---|---|---|---|---|---|---|---|---|---|
| **A_55_P1992074** | Ptpra | -0.49270014 | -1.725845764 | 0.740445485 | 12.56757433 | -1.000734022 | 0.359094872 | 0.709264768 | -4.61655433 |
| **A_55_P1972406** | Tor1b | -0.263507118 | -0.923103599 | 0.396089364 | 9.544548861 | -1.00060891 | 0.359150254 | 0.709264768 | -4.616585623 |
| **A_55_P2185214** | NA | 0.269304759 | -0.404812477 | 0.943421996 | 16.8015034 | 1.000596415 | 0.359155786 | 0.709264768 | -4.616588748 |
| **A_55_P2177885** | Dact3 | 0.507014957 | -0.762375973 | 1.776405887 | 9.904159737 | 1.000404706 | 0.359240662 | 0.709264768 | -4.616636695 |
| **A_51_P321734** | Bhmt2 | -0.203461314 | -0.712867432 | 0.305944804 | 7.80685722 | -1.000386987 | 0.359248507 | 0.709264768 | -4.616641126 |
| **A_52_P279184** | Srp54a | -0.253791327 | -0.889323465 | 0.381740811 | 8.982372378 | -1.000206344 | 0.3593285 | 0.709264768 | -4.616686301 |
| **A_55_P2126269** | Nmb | -0.461994876 | -1.61892247 | 0.694932718 | 10.27487615 | -1.000187275 | 0.359336945 | 0.709264768 | -4.61669107 |
| **A_51_P119031** | Naa30 | 0.102243187 | -0.153798391 | 0.358284766 | 9.635148544 | 1.000170934 | 0.359344182 | 0.709264768 | -4.616695156 |
| **A_52_P450188** | Mtmr4 | -0.592686337 | -2.077154482 | 0.891781808 | 9.281083114 | -1.000010082 | 0.359415426 | 0.709264768 | -4.616735377 |
| **A_52_P463346** | Serinc3 | -0.147940679 | -0.518494021 | 0.222612663 | 10.92619237 | -0.999970569 | 0.359432929 | 0.709264768 | -4.616745257 |
| **A_55_P1974467** | Ndufb6 | 0.286976562 | -0.431865756 | 1.005818879 | 12.93363909 | 0.999915126 | 0.359457489 | 0.709264768 | -4.61675912 |
| **A_55_P2000853** | Mvb12b | 0.385362067 | -0.580269339 | 1.350993472 | 12.79003419 | 0.999557831 | 0.359615796 | 0.709264768 | -4.616848449 |
| **A_55_P1984357** | NA | 0.355054847 | -0.534699345 | 1.24480904 | 16.08573108 | 0.999483775 | 0.359648616 | 0.709264768 | -4.616866962 |
| **A_51_P190886** | Ankrd13a | -0.100300233 | -0.35165631 | 0.151055845 | 11.26740769 | -0.999454196 | 0.359661725 | 0.709264768 | -4.616874356 |
| **A_55_P1952475** | Rpl30 | 0.150061874 | -0.226002096 | 0.526125845 | 16.03632237 | 0.999445198 | 0.359665712 | 0.709264768 | -4.616876605 |
| **A_55_P2157048** | Mllt3 | 0.25668536 | -0.386586757 | 0.899957477 | 8.722201078 | 0.999439979 | 0.359668026 | 0.709264768 | -4.61687791 |
| **A_55_P2129187** | NA | 0.191482588 | -0.288412844 | 0.67137802 | 16.00774864 | 0.999385443 | 0.359692197 | 0.709264768 | -4.616891543 |
| **A_55_P2150361** | NA | 0.264658284 | -0.398654885 | 0.927971454 | 13.50274496 | 0.999349083 | 0.359708313 | 0.709264768 | -4.616900632 |
| **A_52_P70027** | Gm20498 | 0.422421348 | -0.636339473 | 1.48118217 | 12.26879074 | 0.999305647 | 0.359727566 | 0.709264768 | -4.616911489 |
| **A_55_P2005343** | NA | 0.24894131 | -0.375018444 | 0.872901065 | 16.41067395 | 0.999288152 | 0.359735321 | 0.709264768 | -4.616915862 |
| **A_55_P1964852** | Kbtbd2 | -0.08624254 | -0.302414543 | 0.129929463 | 12.61155931 | -0.999246043 | 0.359753987 | 0.709264768 | -4.616926388 |
| **A_55_P2098471** | Rps19 | 0.308554447 | -0.464862872 | 1.081971766 | 16.01571357 | 0.999236438 | 0.359758244 | 0.709264768 | -4.616928788 |
| **A_51_P298190** | Sbf2 | 0.320993326 | -0.483707118 | 1.12569377 | 10.1420823 | 0.999107243 | 0.35981552 | 0.709264768 | -4.61696108 |
| **A_55_P1959218** | Usp51 | -0.255397315 | -0.895688425 | 0.384893795 | 8.142467846 | -0.999054546 | 0.359838883 | 0.709264768 | -4.616974251 |
| **A_55_P1977910** | NA | 0.19536883 | -0.294541897 | 0.685279557 | 9.934406732 | 0.998823318 | 0.359941416 | 0.709264768 | -4.61703204 |
| **A_51_P192964** | Clec12b | -0.192298464 | -0.674522656 | 0.289925729 | 7.825001053 | -0.998796854 | 0.359953152 | 0.709264768 | -4.617038654 |
| **A_51_P222359** | Emilin1 | -0.108561001 | -0.380803424 | 0.163681421 | 8.608563862 | -0.998776662 | 0.359962107 | 0.709264768 | -4.6170437 |
| **A_51_P418317** | Mis12 | 0.088404504 | -0.133315889 | 0.310124896 | 8.265416826 | 0.998663315 | 0.360012379 | 0.709264768 | -4.617072025 |
| **A_55_P2144456** | Kmt5b | -0.086683098 | -0.304089585 | 0.13072339 | 8.426468327 | -0.998647615 | 0.360019343 | 0.709264768 | -4.617075948 |
| **A_55_P1975665** | Rhox4e | -0.580968863 | -2.03811562 | 0.876177895 | 8.060424567 | -0.998619192 | 0.36003195 | 0.709264768 | -4.617083051 |
| **A_55_P2021953** | Olfr700 | -0.235321325 | -0.825651369 | 0.35500872 | 8.067742082 | -0.99842798 | 0.360116774 | 0.709264768 | -4.617130829 |
| **A_51_P365952** | Trdmt1 | 0.35673796 | -0.538376072 | 1.251851991 | 10.05936681 | 0.998208594 | 0.360214115 | 0.709264768 | -4.617185643 |
| **A_55_P2051322** | Efhd2 | 0.166124459 | -0.250726734 | 0.582975653 | 11.60131902 | 0.998166238 | 0.360232911 | 0.709264768 | -4.617196225 |
| **A_52_P285024** | Sertad4 | 0.220659512 | -0.333048314 | 0.774367337 | 8.328343022 | 0.998142014 | 0.360243661 | 0.709264768 | -4.617202277 |
| **A_55_P2084278** | Ighmbp2 | -0.115156393 | -0.40412815 | 0.173815365 | 7.921552541 | -0.998120551 | 0.360253186 | 0.709264768 | -4.617207639 |
| **A_51_P466591** | 1600012P | -2.748893623 | -9.647356977 | 4.14956973 | 9.790929906 | -0.998058317 | 0.360280806 | 0.709264768 | -4.617223186 |
| **A_55_P2182324** | NA | -0.141486034 | -0.496558076 | 0.213586007 | 7.92668302 | -0.99803884 | 0.36028945 | 0.709264768 | -4.617228052 |
| **A_51_P506045** | Cpq | 0.695366182 | -1.049869409 | 2.440601773 | 10.98095482 | 0.997952453 | 0.360327792 | 0.709264768 | -4.617249633 |
| **A_55_P2143233** | Rasgrp2 | 0.670436044 | -1.012320034 | 2.353192121 | 10.38320748 | 0.997898873 | 0.360351575 | 0.709264768 | -4.617263017 |
| **A_51_P475249** | Rnf44 | -0.258384004 | -0.907037642 | 0.390269633 | 10.81310812 | -0.997707213 | 0.360436658 | 0.709264768 | -4.617310892 |
| **A_55_P2063366** | Commd7 | -0.135170264 | -0.474524755 | 0.204184227 | 11.74848684 | -0.997649308 | 0.360462367 | 0.709264768 | -4.617325355 |
| **A_52_P37431** | NA | 0.543758848 | -0.821388954 | 1.90890665 | 14.33660608 | 0.997647247 | 0.360463282 | 0.709264768 | -4.61732587 |
| **A_55_P2114148** | NA | 0.206874987 | -0.312500488 | 0.726250463 | 9.802067278 | 0.997646979 | 0.360463401 | 0.709264768 | -4.617325937 |
| **A_51_P443702** | Mrps34 | 0.094937569 | -0.143416684 | 0.333291822 | 12.64500692 | 0.997620922 | 0.36047497 | 0.709264768 | -4.617332445 |
| **A_55_P2118183** | Tnks2 | -0.33437872 | -1.173944442 | 0.505187002 | 8.682858896 | -0.997549092 | 0.360506865 | 0.709264768 | -4.617350386 |
| **A_51_P113906** | Gkn2 | -1.302642698 | -4.573491516 | 1.96820612 | 8.682975829 | -0.997505236 | 0.360526339 | 0.709264768 | -4.616176463 |
| **A_55_P2055854** | Ppara | 0.2822259 | -0.426462785 | 0.990914585 | 8.733638182 | 0.997451328 | 0.360550278 | 0.709264768 | -4.617374803 |
| **A_55_P2067251** | Tmpo | 0.384743435 | -0.581474928 | 1.350961799 | 9.246352056 | 0.997346979 | 0.36059662 | 0.709264768 | -4.617400863 |

| | | | | | | | | | |
|---|---|---|---|---|---|---|---|---|---|
| **A_51_P110672** | Mst1r | -0.121822335 | -0.427766159 | 0.184121488 | 8.699405013 | -0.997322405 | 0.360607535 | 0.709264768 | -4.616801624 |
| **A_55_P2147566** | Tac2 | -0.179921199 | -0.631776584 | 0.271934185 | 7.797233446 | -0.997316998 | 0.360609936 | 0.709264768 | -4.617408351 |
| **A_55_P2023762** | Epha4 | 0.369297566 | -0.558215258 | 1.29681039 | 8.8958426 | 0.997256483 | 0.360636814 | 0.709264768 | -4.617423463 |
| **A_52_P185343** | Gna13 | -0.3118692 | -1.095347756 | 0.471609355 | 9.116639644 | -0.997001285 | 0.36075018 | 0.709264768 | -4.617487189 |
| **A_55_P2050426** | Ankrd50 | -0.103921955 | -0.364999243 | 0.157155333 | 10.21144194 | -0.996984856 | 0.360757479 | 0.709264768 | -4.617491291 |
| **A_55_P2000613** | Slc6a1 | -0.153037846 | -0.537507353 | 0.23143166 | 7.905660907 | -0.99698194 | 0.360758775 | 0.709264768 | -4.61749202 |
| **A_52_P398989** | Cytip | 0.449793151 | -0.727794185 | 1.627380487 | 8.739158312 | 1.017378175 | 0.360760753 | 0.709264768 | -4.606028799 |
| **A_55_P2078710** | Morc4 | -0.758555415 | -2.664369456 | 1.147258626 | 10.40440619 | -0.996912718 | 0.360789531 | 0.709264768 | -4.617509304 |
| **A_51_P185713** | Ptdss2 | -0.203441718 | -0.714661609 | 0.307778173 | 8.281209517 | -0.996741677 | 0.360865535 | 0.709264768 | -4.617552009 |
| **A_55_P2132024** | Tekt1 | 0.809811492 | -1.225247782 | 2.844870766 | 10.5837907 | 0.996683384 | 0.360891441 | 0.709264768 | -4.617566562 |
| **A_52_P628162** | Zfp592 | 0.126495503 | -0.191411383 | 0.444402389 | 8.538145321 | 0.99661059 | 0.360923794 | 0.709264768 | -4.617584736 |
| **A_55_P1976237** | Dnajb3 | 0.105383068 | -0.159472972 | 0.370239108 | 8.886777647 | 0.996578015 | 0.360938272 | 0.709264768 | -4.617592868 |
| **A_55_P1998097** | Tmem214 | -0.359204089 | -1.26203594 | 0.543627762 | 9.042913053 | -0.996517078 | 0.360965358 | 0.709264768 | -4.617608081 |
| **A_55_P2096883** | Steap3 | -0.361653563 | -1.270644178 | 0.547337051 | 8.201281046 | -0.996514666 | 0.360966429 | 0.709264768 | -4.617608683 |
| **A_52_P88773** | Tor2a | -0.561408708 | -1.972674022 | 0.849856605 | 10.61314658 | -0.996370414 | 0.361030554 | 0.709264768 | -4.617644693 |
| **A_66_P138780** | Elob | 0.168056024 | -0.254404125 | 0.590516173 | 13.47238102 | 0.9963655 | 0.361032739 | 0.709264768 | -4.61764592 |
| **A_51_P121466** | Tab1 | 0.124884813 | -0.189065706 | 0.438835332 | 11.55842825 | 0.996319816 | 0.361053049 | 0.709264768 | -4.617657323 |
| **A_51_P284665** | Plcb1 | 0.223121197 | -0.337813605 | 0.784055998 | 8.339476079 | 0.996273987 | 0.361073424 | 0.709264768 | -4.617668763 |
| **A_51_P468126** | Trim39 | -0.14874894 | -0.522787237 | 0.225289356 | 8.383944029 | -0.996066091 | 0.361165866 | 0.709264768 | -4.617720654 |
| **A_51_P320444** | Rock2 | -0.289477727 | -1.01741563 | 0.438460176 | 8.537036574 | -0.996027126 | 0.361183194 | 0.709264768 | -4.617730379 |
| **A_52_P163939** | Cln6 | -0.255612095 | -0.898410342 | 0.387186151 | 8.932746991 | -0.995994781 | 0.361197578 | 0.709264768 | -4.617738452 |
| **A_51_P342387** | Apoh | 0.280110641 | -0.424315188 | 0.98453647 | 8.305800194 | 0.995966387 | 0.361210206 | 0.709264768 | -4.617745538 |
| **A_52_P1026830** | Tbc1d8b | 0.13428796 | -0.203460383 | 0.472036303 | 9.139444055 | 0.995850612 | 0.361261699 | 0.709264768 | -4.617774432 |
| **A_55_P2173035** | Dnajc8 | -0.15383245 | -0.54075899 | 0.233094089 | 10.39627495 | -0.995794641 | 0.361286595 | 0.709264768 | -4.617788401 |
| **A_55_P2014319** | Armcx6 | 0.537527377 | -0.814558602 | 1.889613357 | 9.517773358 | 0.995741539 | 0.361310217 | 0.709264768 | -4.617801652 |
| **A_55_P2083844** | Dis3l | 0.11120725 | -0.168523771 | 0.390938271 | 10.01444176 | 0.995732807 | 0.361314101 | 0.709264768 | -4.617803832 |
| **A_55_P2092984** | NA | 0.324019258 | -0.491105038 | 1.139143554 | 14.69128052 | 0.995628524 | 0.361360493 | 0.709264768 | -4.617829855 |
| **A_55_P2112125** | Tnk2 | -0.128039151 | -0.450153033 | 0.194074732 | 13.09605316 | -0.995597248 | 0.361374409 | 0.709264768 | -4.617837659 |
| **A_55_P2096281** | NA | 0.179269965 | -0.271769336 | 0.630309267 | 16.20835225 | 0.995505113 | 0.361415402 | 0.709264768 | -4.61786065 |
| **A_55_P2144115** | 1700125H | -0.2203878 | -0.774881273 | 0.334105673 | 7.878254151 | -0.995500442 | 0.36141748 | 0.709264768 | -4.617861815 |
| **A_52_P569348** | Dbt | 0.162409548 | -0.246271535 | 0.57109063 | 8.625040518 | 0.995353544 | 0.361482847 | 0.709264768 | -4.617898468 |
| **A_55_P1996578** | Fgfr3 | 1.101310875 | -1.670105189 | 3.872726939 | 12.01236507 | 0.995310403 | 0.361502046 | 0.709264768 | -4.617909232 |
| **A_55_P2380806** | Gm2115 | 0.283097963 | -0.429350661 | 0.995546587 | 8.556471504 | 0.995253098 | 0.361527549 | 0.709264768 | -4.617923529 |
| **A_51_P316949** | Ccdc6 | -0.375121259 | -1.319288021 | 0.569045504 | 9.776034144 | -0.995115097 | 0.361588972 | 0.709264768 | -4.617957959 |
| **A_51_P405985** | Fig4 | -0.140085674 | -0.492710176 | 0.212538829 | 8.919520927 | -0.995019477 | 0.361631536 | 0.709264768 | -4.617981813 |
| **A_51_P416613** | 1810058I2 | -0.099961223 | -0.35159581 | 0.151673364 | 8.58519916 | -0.994973634 | 0.361651945 | 0.709264768 | -4.61799325 |
| **A_55_P2040559** | NA | 0.624412933 | -0.947508796 | 2.196334663 | 15.67542893 | 0.994927232 | 0.361672602 | 0.709264768 | -4.618004825 |
| **A_66_P136899** | Nadk | -0.638014359 | -2.244324725 | 0.968296008 | 9.845632398 | -0.994835642 | 0.361713381 | 0.709264768 | -4.618027672 |
| **A_55_P1957815** | Tmub1 | -0.323577535 | -1.138293518 | 0.491138447 | 11.08615022 | -0.994769526 | 0.361742819 | 0.709264768 | -4.618044164 |
| **A_55_P2119864** | Cd2bp2 | -0.292799444 | -1.030310986 | 0.444712099 | 9.400252198 | -0.994378605 | 0.36191692 | 0.709264768 | -4.618141666 |
| **A_55_P2097665** | Arl4a | -0.127079202 | -0.447176495 | 0.193018091 | 8.559622781 | -0.994358117 | 0.361926047 | 0.709264768 | -4.618146776 |
| **A_51_P146407** | Fam104a | -0.169949087 | -0.598072637 | 0.258174464 | 9.170170148 | -0.994260216 | 0.361969659 | 0.709264768 | -4.618171191 |
| **A_55_P2154242** | Gfpt1 | -0.422508934 | -1.486893312 | 0.641875444 | 11.88979439 | -0.994232031 | 0.361982216 | 0.709264768 | -4.618178219 |
| **A_55_P1976744** | NA | -0.114110146 | -0.401600677 | 0.173380385 | 7.844012907 | -0.994148042 | 0.362019635 | 0.709264768 | -4.618199164 |
| **A_55_P2019577** | 1500011B | -0.095192289 | -0.335044256 | 0.144659678 | 8.366716978 | -0.994051376 | 0.362062707 | 0.709264768 | -4.618223268 |
| **A_55_P2145711** | Tox3 | 0.425510632 | -0.646644011 | 1.497665276 | 9.051908327 | 0.994038788 | 0.362068316 | 0.709264768 | -4.618226407 |
| **A_55_P2041626** | Ceacam12 | -2.752438211 | -9.688656201 | 4.183779778 | 9.808266058 | -0.993905721 | 0.362127614 | 0.709264768 | -4.618259587 |
| **A_55_P1969002** | Tbx3 | 0.395737071 | -0.601643229 | 1.393117372 | 12.76757718 | 0.993793827 | 0.362177484 | 0.709264768 | -4.618287486 |

| | | | | | | | | | |
|---|---|---|---|---|---|---|---|---|---|
| **A_55_P2004881** | Ldlrad4 | 0.230432997 | -0.350386738 | 0.811252731 | 8.424533515 | 0.993696246 | 0.362220978 | 0.709264768 | -4.618311814 |
| **A_55_P2048794** | Cd82 | -0.781229204 | -2.750613416 | 1.188155009 | 14.00644319 | -0.993569755 | 0.362277365 | 0.709264768 | -4.618343349 |
| **A_55_P2120737** | NA | -0.307227638 | -1.081714219 | 0.467258942 | 7.960117982 | -0.993566023 | 0.362279029 | 0.709264768 | -4.61834428 |
| **A_55_P2111990** | Tekt1 | 0.754435805 | -1.147442625 | 2.656314235 | 10.38860497 | 0.993550358 | 0.362286012 | 0.709264768 | -4.618348185 |
| **A_51_P462746** | Tex10 | 0.124736162 | -0.18978602 | 0.439258343 | 9.807584719 | 0.993325174 | 0.362386414 | 0.709264768 | -4.618404319 |
| **A_55_P2086265** | Gm8580 | 0.484777022 | -0.737701039 | 1.707255082 | 17.46030196 | 0.993233361 | 0.362427357 | 0.709264768 | -4.618427205 |
| **A_55_P2017851** | NA | 0.546988307 | -0.832459587 | 1.926436201 | 13.76115846 | 0.993168845 | 0.362456129 | 0.709264768 | -4.618443286 |
| **A_51_P288916** | Tmtc2 | 0.678489728 | -1.032809359 | 2.389788815 | 10.05161887 | 0.993042199 | 0.362512615 | 0.709264768 | -4.618474852 |
| **A_55_P1984123** | Kmt2c | -0.131168288 | -0.462013103 | 0.199676527 | 10.62940996 | -0.993012836 | 0.362525712 | 0.709264768 | -4.61848217 |
| **A_52_P156190** | Ednra | 1.090733473 | -1.660602785 | 3.84206973 | 12.57518794 | 0.992945279 | 0.362555847 | 0.709264768 | -4.618499007 |
| **A_55_P2033520** | Med6 | -0.089782006 | -0.31625621 | 0.136692198 | 10.13207066 | -0.992935201 | 0.362560342 | 0.709264768 | -4.618501519 |
| **A_55_P1952950** | Cic | 0.233212487 | -0.355123763 | 0.821548736 | 14.12246829 | 0.992833763 | 0.362605595 | 0.709264768 | -4.618526799 |
| **A_51_P119597** | Ap3b1 | -0.130069244 | -0.458216098 | 0.198077611 | 12.05345184 | -0.992788446 | 0.362625813 | 0.709264768 | -4.617282006 |
| **A_51_P484526** | Wif1 | 0.618973779 | -0.942624953 | 2.180572512 | 9.355583432 | 0.992780298 | 0.362629448 | 0.709264768 | -4.618540123 |
| **A_55_P2086687** | Nfib | 0.221272822 | -0.337026369 | 0.779572014 | 8.504063023 | 0.992684914 | 0.362672006 | 0.709264768 | -4.618563893 |
| **A_52_P216672** | Klk8 | 0.301236411 | -0.458829651 | 1.061302474 | 9.697736989 | 0.992673594 | 0.362677058 | 0.709264768 | -4.618566714 |
| **A_55_P1955691** | Alg6 | 0.149699011 | -0.228033027 | 0.52743105 | 9.262109664 | 0.992625562 | 0.36269849 | 0.709264768 | -4.618578683 |
| **A_55_P2163774** | Crip1 | -0.379502704 | -1.337269045 | 0.578263636 | 15.29671434 | -0.992443177 | 0.362779882 | 0.709264768 | -4.618624129 |
| **A_51_P151484** | Atp1b1 | -0.12464273 | -0.439242658 | 0.189957197 | 13.12646509 | -0.992335848 | 0.362827786 | 0.709264768 | -4.618650871 |
| **A_52_P515282** | 1110051M | 0.459451951 | -0.700251891 | 1.619155793 | 9.462772055 | 0.992300818 | 0.362843422 | 0.709264768 | -4.618659599 |
| **A_55_P1973387** | Srrm1 | 0.121027637 | -0.184475661 | 0.426530934 | 13.31572532 | 0.992245176 | 0.36286826 | 0.709264768 | -4.618673462 |
| **A_51_P309920** | Itga8 | 0.951899212 | -1.451079964 | 3.354878388 | 11.11814566 | 0.992181913 | 0.362896501 | 0.709264768 | -4.618689223 |
| **A_51_P493105** | Mettl15 | 0.194583537 | -0.296677945 | 0.685845019 | 8.776052405 | 0.992073217 | 0.362945028 | 0.709264768 | -4.618716302 |
| **A_51_P495131** | Prl7c1 | -0.354212785 | -1.248514023 | 0.540088453 | 7.846200036 | -0.99204357 | 0.362958265 | 0.709264768 | -4.618723688 |
| **A_51_P332676** | Top1mt | -0.233312625 | -0.822385582 | 0.355760332 | 9.053819707 | -0.99201788 | 0.362969735 | 0.709264768 | -4.618730088 |
| **A_51_P198694** | Gckr | -0.145931092 | -0.514385322 | 0.222523138 | 7.805197452 | -0.992006758 | 0.362974701 | 0.709264768 | -4.618732859 |
| **A_55_P2009543** | NA | 0.295554575 | -0.450757385 | 1.041866534 | 17.28015033 | 0.991899406 | 0.363022636 | 0.709264768 | -4.618759601 |
| **A_55_P2194064** | NA | 0.172682916 | -0.263386705 | 0.608752536 | 8.169342403 | 0.991845178 | 0.363046852 | 0.709264768 | -4.618773109 |
| **A_55_P1965114** | Spsb3 | -0.564308532 | -1.989341005 | 0.86072394 | 12.75359655 | -0.991841331 | 0.36304857 | 0.709264768 | -4.618774067 |
| **A_55_P2415372** | Mta1 | 0.168224328 | -0.256590882 | 0.593039537 | 8.581187969 | 0.991834223 | 0.363051744 | 0.709264768 | -4.618775838 |
| **A_55_P2070005** | NA | 0.725440873 | -1.106806835 | 2.55768858 | 13.75186844 | 0.991672293 | 0.363124064 | 0.709264768 | -4.618816172 |
| **A_55_P2001589** | Klhl8 | 0.208336985 | -0.317873941 | 0.734547911 | 9.341114614 | 0.991646444 | 0.36313561 | 0.709264768 | -4.618822611 |
| **A_55_P1964638** | Cxadr | 0.717312649 | -1.094460895 | 2.529086192 | 11.76199759 | 0.991642019 | 0.363137586 | 0.709264768 | -4.618823713 |
| **A_55_P1991054** | Tmf1 | -0.231599814 | -0.816595173 | 0.353395544 | 9.289727818 | -0.991599119 | 0.363156749 | 0.709264768 | -4.618834398 |
| **A_66_P123683** | Nabp1 | -0.253789688 | -0.894900611 | 0.387321235 | 8.040059425 | -0.991496397 | 0.363202635 | 0.709264768 | -4.618859982 |
| **A_55_P2001148** | Pdlim5 | -0.098382816 | -0.34692424 | 0.150158607 | 7.867738129 | -0.991449994 | 0.363223365 | 0.709264768 | -4.618871539 |
| **A_51_P399985** | Myo9b | -0.2214981 | -0.781138649 | 0.338142448 | 12.23402743 | -0.99131386 | 0.363284187 | 0.709264768 | -4.618905442 |
| **A_55_P1954633** | NA | 0.37242625 | -0.568578606 | 1.313431106 | 10.7054956 | 0.991285527 | 0.363296846 | 0.709264768 | -4.618912498 |
| **A_52_P13730** | Cbx2 | 0.117449773 | -0.17931975 | 0.414219296 | 9.194734071 | 0.991250066 | 0.363312691 | 0.709264768 | -4.618921329 |
| **A_51_P409452** | Cldn11 | -0.739128582 | -2.606771661 | 1.128514497 | 8.618305069 | -0.991234615 | 0.363319595 | 0.709264768 | -4.618925176 |
| **A_55_P2151556** | C8g | -0.093777043 | -0.330737305 | 0.143183218 | 8.544698651 | -0.991222979 | 0.363324795 | 0.709264768 | -4.618928074 |
| **A_55_P2083213** | Purb | 0.271720887 | -0.414887778 | 0.958329553 | 9.399432393 | 0.991206382 | 0.363332211 | 0.709264768 | -4.618932207 |
| **A_66_P136372** | Mrpl9 | 0.118105773 | -0.180347222 | 0.416558767 | 10.05632472 | 0.991164025 | 0.363351139 | 0.709264768 | -4.618942755 |
| **A_55_P2124582** | Xlr4a | -0.580644805 | -2.047978602 | 0.886688992 | 8.095066397 | -0.991133076 | 0.363364969 | 0.709264768 | -4.618950461 |
| **A_55_P2039566** | Arhgap23 | -0.220853588 | -0.779032859 | 0.337325684 | 12.19461215 | -0.991016989 | 0.363416851 | 0.709264768 | -4.618979368 |
| **A_55_P2030802** | Rpl3 | 0.440103466 | -0.672219606 | 1.552426538 | 16.02906856 | 0.991001219 | 0.363423899 | 0.709264768 | -4.618983294 |
| **A_52_P244064** | Ube2g2 | -0.457934207 | -1.615331823 | 0.699463408 | 10.51011415 | -0.990993597 | 0.363427305 | 0.709264768 | -4.618985192 |
| **A_55_P1981550** | Park7 | 0.468789739 | -0.716099314 | 1.653678792 | 13.1072495 | 0.99094775 | 0.363447797 | 0.709264768 | -4.618996608 |

| | | | | | | | | | |
|---|---|---|---|---|---|---|---|---|---|
| **A_55_P2016166** | Wfdc2 | 1.380294877 | -2.108569499 | 4.869159253 | 12.26768218 | 0.990918952 | 0.363460669 | 0.709264768 | -4.619003778 |
| **A_55_P1987984** | NA | -0.272821535 | -0.962458552 | 0.416815482 | 8.309469818 | -0.990851175 | 0.363490965 | 0.709264768 | -4.619020654 |
| **A_52_P625171** | Asph | -0.161216983 | -0.568772675 | 0.24633871 | 8.271074527 | -0.990773018 | 0.363525904 | 0.709264768 | -4.619040113 |
| **A_55_P1957373** | Pdcl | 0.181368009 | -0.277158147 | 0.639894165 | 10.02189897 | 0.990710842 | 0.3635537 | 0.709264768 | -4.619055592 |
| **A_51_P415688** | Gng12 | -0.709860404 | -2.504510598 | 1.084789789 | 10.76032222 | -0.990703022 | 0.363557197 | 0.709264768 | -4.619057539 |
| **A_52_P491569** | 1700017B | -0.378342527 | -1.334885525 | 0.578200471 | 9.016211103 | -0.990674556 | 0.363569923 | 0.709264768 | -4.619064626 |
| **A_51_P349961** | Gc | -0.425978473 | -1.502973835 | 0.65101689 | 8.175746793 | -0.990658943 | 0.363576904 | 0.709264768 | -4.619068513 |
| **A_51_P355753** | Hic1 | 0.197148012 | -0.301358567 | 0.69565459 | 11.78929632 | 0.990539625 | 0.363630254 | 0.709264768 | -4.619098217 |
| **A_51_P513392** | Azi2 | 0.147193155 | -0.225043211 | 0.51942952 | 9.3360346 | 0.990419448 | 0.363683994 | 0.709264768 | -4.619128134 |
| **A_55_P2074656** | Padi2 | -1.317125635 | -4.648533842 | 2.014282573 | 8.888144251 | -0.990261048 | 0.363754837 | 0.709264768 | -4.619167563 |
| **A_51_P221632** | 1700037H | 0.154090686 | -0.235697314 | 0.543878687 | 11.70256925 | 0.990143795 | 0.363807285 | 0.709264768 | -4.619196747 |
| **A_51_P453948** | Drg1 | 0.123595489 | -0.189072741 | 0.43626372 | 10.83579718 | 0.990077546 | 0.363836921 | 0.709264768 | -4.619213236 |
| **A_55_P1956627** | Gpr160 | -0.12878796 | -0.454613643 | 0.197037722 | 8.207125225 | -0.990011633 | 0.363866408 | 0.709264768 | -4.619229641 |
| **A_55_P2199118** | Bend4 | -0.481099083 | -1.698266298 | 0.736068133 | 10.29803283 | -0.989998709 | 0.36387219 | 0.709264768 | -4.619232857 |
| **A_51_P241667** | Prkdc | 0.131608414 | -0.201366057 | 0.464582886 | 8.425134454 | 0.98997235 | 0.363883983 | 0.709264768 | -4.619239418 |
| **A_51_P456465** | Cldn10 | 1.06060404 | -1.623324617 | 3.744532697 | 10.81332268 | 0.989766283 | 0.363976188 | 0.709264768 | -4.6192907 |
| **A_55_P1961270** | Cd72 | -0.216374928 | -0.763951138 | 0.331201282 | 7.871615689 | -0.98972083 | 0.363996528 | 0.709264768 | -4.619302011 |
| **A_51_P141521** | Sap130 | -0.207330359 | -0.732079599 | 0.317418881 | 11.48653731 | -0.989603966 | 0.36404883 | 0.709264768 | -4.619331091 |
| **A_66_P135987** | Ocrl | 0.298538145 | -0.457161128 | 1.054237419 | 10.66323744 | 0.989466673 | 0.364110282 | 0.709264768 | -4.619365253 |
| **A_55_P2034453** | NA | 0.102897936 | -0.157627193 | 0.363423065 | 8.167293885 | 0.98925303 | 0.364205925 | 0.709264768 | -4.619418409 |
| **A_52_P404461** | Tmem60 | -0.149222821 | -0.527040262 | 0.228594621 | 10.2887835 | -0.989244373 | 0.364209801 | 0.709264768 | -4.619420562 |
| **A_55_P2030578** | Tns1 | -0.161248548 | -0.569515525 | 0.247018429 | 13.28642721 | -0.989240536 | 0.364211518 | 0.709264768 | -4.619421517 |
| **A_55_P2061045** | Rpl29 | 0.404585272 | -0.619803341 | 1.428973885 | 15.48598173 | 0.989226341 | 0.364217874 | 0.709264768 | -4.619425048 |
| **A_55_P2038458** | Lancl2 | 0.288746578 | -0.442416752 | 1.019909908 | 8.707644362 | 0.989128669 | 0.364261608 | 0.709264768 | -4.619449348 |
| **A_51_P494863** | Vmac | -0.146935107 | -0.519004619 | 0.225134405 | 9.141301729 | -0.989126493 | 0.364262582 | 0.709264768 | -4.619449889 |
| **A_55_P2026710** | 4930445E | -0.213690417 | -0.75483322 | 0.327452385 | 7.794317268 | -0.989061978 | 0.364291471 | 0.709264768 | -4.619465938 |
| **A_51_P267346** | Topors | 0.135998336 | -0.208430901 | 0.480427572 | 11.04772248 | 0.98897185 | 0.364331833 | 0.709264768 | -4.619488359 |
| **A_55_P2137927** | Cd82 | -0.671403882 | -2.371845733 | 1.029037968 | 9.713785163 | -0.988945614 | 0.364343583 | 0.709264768 | -4.619494885 |
| **A_55_P2208579** | Onecut2 | -0.091557014 | -0.323441792 | 0.140327764 | 7.822317158 | -0.988939503 | 0.36434632 | 0.709264768 | -4.619496405 |
| **A_55_P2133943** | NA | -0.179874126 | -0.635439442 | 0.27569119 | 8.350683519 | -0.988936463 | 0.364347681 | 0.709264768 | -4.619497162 |
| **A_66_P108468** | 2610528A | -1.018388206 | -3.597737148 | 1.560960737 | 8.501191389 | -0.988902803 | 0.364362757 | 0.709264768 | -4.619505535 |
| **A_55_P2055537** | Slc16a6 | -0.97490454 | -3.444582406 | 1.494773325 | 9.390025669 | -0.988717291 | 0.364445851 | 0.709264768 | -4.619551679 |
| **A_55_P2081805** | Slc46a1 | -0.500980053 | -1.770271748 | 0.768311642 | 9.219067769 | -0.988574359 | 0.364509884 | 0.709264768 | -4.619587229 |
| **A_51_P488991** | Oaf | -0.498469751 | -1.761428965 | 0.764489463 | 10.2614399 | -0.988552705 | 0.364519585 | 0.709264768 | -4.619592614 |
| **A_51_P129464** | Scd2 | 0.442249604 | -0.678529298 | 1.563028506 | 14.75822551 | 0.988320608 | 0.364623585 | 0.709264768 | -4.619650335 |
| **A_51_P411271** | Nfe2l2 | 0.497152945 | -0.762807226 | 1.757113115 | 12.36085716 | 0.988288052 | 0.364638174 | 0.709264768 | -4.619658431 |
| **A_55_P1976012** | Cnih4 | -0.145509977 | -0.514372711 | 0.223352757 | 9.695082277 | -0.988048667 | 0.364745468 | 0.709264768 | -4.619717957 |
| **A_55_P2082826** | Fam160b1 | -0.13592573 | -0.480569217 | 0.208717757 | 11.2473663 | -0.987829391 | 0.364843771 | 0.709264768 | -4.619772477 |
| **A_55_P2044359** | Rxrg | 0.174975391 | -0.268685866 | 0.618636648 | 10.00718741 | 0.987815447 | 0.364850023 | 0.709264768 | -4.619775944 |
| **A_66_P132448** | Gnpnat1 | -0.119174658 | -0.421371602 | 0.183022287 | 9.344198263 | -0.98774352 | 0.364882273 | 0.709264768 | -4.619793826 |
| **A_51_P401964** | Aph1a | -0.101160699 | -0.357705376 | 0.155383979 | 11.3824283 | -0.987641107 | 0.364928197 | 0.709264768 | -4.619819287 |
| **A_51_P152747** | Nrip1 | -0.198091871 | -0.700458717 | 0.304274974 | 9.937004581 | -0.987633997 | 0.364931386 | 0.709264768 | -4.619821054 |
| **A_52_P597860** | Wasf2 | -0.141177552 | -0.499224285 | 0.216869181 | 9.692875036 | -0.987589076 | 0.364951531 | 0.709264768 | -4.619832222 |
| **A_55_P2022104** | Cop1 | 0.246732581 | -0.379042518 | 0.87250768 | 8.867006775 | 0.98754881 | 0.364969589 | 0.709264768 | -4.619842231 |
| **A_55_P2026218** | Igsf8 | -0.627819045 | -2.22021531 | 0.964577221 | 11.32324646 | -0.987492216 | 0.364994971 | 0.709264768 | -4.6198563 |
| **A_55_P1954393** | Susd4 | 0.484600759 | -0.744546248 | 1.713747765 | 11.90108344 | 0.987485227 | 0.364998105 | 0.709264768 | -4.619858037 |
| **A_55_P1963062** | Rps16 | 0.19362698 | -0.297493511 | 0.684747471 | 17.54693925 | 0.987479668 | 0.365000599 | 0.709264768 | -4.619859419 |
| **A_55_P2055123** | Frrs1 | -0.280825359 | -0.993120232 | 0.431469515 | 8.149262558 | -0.987476682 | 0.365001938 | 0.709264768 | -4.619860162 |

| | | | | | | | | | |
|---|---|---|---|---|---|---|---|---|---|
| **A_52_P589699** | Polr3a | -0.147540504 | -0.521824357 | 0.226743349 | 8.513302576 | -0.987325869 | 0.365069585 | 0.709264768 | -4.619897649 |
| **A_52_P603976** | Jtb | -0.099829256 | -0.353081998 | 0.153423486 | 11.65566348 | -0.987311113 | 0.365076204 | 0.709264768 | -4.619901317 |
| **A_52_P293556** | Wdr41 | 0.141886299 | -0.21810899 | 0.501881589 | 9.108544218 | 0.987174638 | 0.365137429 | 0.709264768 | -4.619935238 |
| **A_51_P293753** | Cul7 | -0.413721588 | -1.46345295 | 0.636009774 | 13.12947836 | -0.987143651 | 0.365151332 | 0.709264768 | -4.61994294 |
| **A_52_P59206** | Cst6 | -0.527772574 | -1.866929667 | 0.81138452 | 8.063978443 | -0.987110195 | 0.365166342 | 0.709264768 | -4.619951255 |
| **A_52_P303928** | Rab2a | 0.194066442 | -0.298395171 | 0.686528056 | 11.40712356 | 0.987025574 | 0.365204311 | 0.709264768 | -4.619972286 |
| **A_55_P2027999** | Hk1 | -0.543444023 | -1.922505413 | 0.835617366 | 12.11290816 | -0.987010023 | 0.365211289 | 0.709264768 | -4.619976151 |
| **A_52_P83557** | Arfgef2 | -0.261275279 | -0.924348252 | 0.401797694 | 8.259595794 | -0.986932247 | 0.36524619 | 0.709264768 | -4.61999548 |
| **A_55_P1984841** | NA | 0.321969058 | -0.495194052 | 1.139132168 | 10.87102881 | 0.986860417 | 0.365278425 | 0.709264768 | -4.620013331 |
| **A_55_P2139937** | Lrrc56 | 0.2941805 | -0.452477519 | 1.04083852 | 9.920215158 | 0.986830341 | 0.365291923 | 0.709264768 | -4.620020805 |
| **A_55_P2093794** | Usp3 | 0.303255164 | -0.466478803 | 1.072989131 | 10.64386886 | 0.986774456 | 0.365317005 | 0.709264768 | -4.620034692 |
| **A_51_P217382** | Kif1c | -0.101631772 | -0.359622683 | 0.156359139 | 11.41733337 | -0.986677983 | 0.365360307 | 0.709264768 | -4.620058665 |
| **A_55_P2096762** | Arhgdig | 0.244362352 | -0.375956802 | 0.864681505 | 8.582704803 | 0.986664382 | 0.365366412 | 0.709264768 | -4.620062045 |
| **A_55_P2091359** | Padi2 | -1.243826329 | -4.401492782 | 1.913840123 | 8.812891817 | -0.986606195 | 0.365392531 | 0.709264768 | -4.620076503 |
| **A_66_P126094** | Pycrl | 0.119524192 | -0.183982947 | 0.423031331 | 9.084889353 | 0.986364086 | 0.365501227 | 0.709264768 | -4.620136658 |
| **A_66_P121474** | Chmp6 | -0.329193683 | -1.165142537 | 0.50675517 | 10.40100939 | -0.98632975 | 0.365516644 | 0.709264768 | -4.620145189 |
| **A_52_P655285** | Zfp462 | 0.242723297 | -0.373648585 | 0.859095178 | 9.112506233 | 0.986322612 | 0.365519849 | 0.709264768 | -4.620146962 |
| **A_55_P2046852** | Gata1 | -0.672033489 | -2.378606037 | 1.03453906 | 8.423552961 | -0.98631697 | 0.365522383 | 0.709264768 | -4.620148364 |
| **A_55_P1987023** | NA | -0.414450923 | -1.46696314 | 0.638061294 | 8.890613628 | -0.986271107 | 0.365542977 | 0.709264768 | -4.620159758 |
| **A_55_P2051849** | Snap25 | -0.275990287 | -0.976955129 | 0.424974555 | 10.88440488 | -0.986161191 | 0.365592337 | 0.709264768 | -4.620187064 |
| **A_51_P143296** | Myh8 | 1.175591935 | -1.810268031 | 4.161451901 | 10.952375 | 0.986137577 | 0.365602943 | 0.709264768 | -4.620192931 |
| **A_55_P2040362** | Bola2 | 0.113359593 | -0.174574991 | 0.401294177 | 13.34359461 | 0.986085996 | 0.365626108 | 0.709264768 | -4.620205745 |
| **A_55_P2050652** | Ikbkg | -0.157070999 | -0.556035366 | 0.241893367 | 8.240758052 | -0.986080212 | 0.365628706 | 0.709264768 | -4.620207181 |
| **A_55_P2058117** | Slc35d1 | -0.361057316 | -1.278263689 | 0.556149056 | 8.778705087 | -0.985960309 | 0.365682562 | 0.709264768 | -4.620236966 |
| **A_55_P2097022** | Eid1 | 0.22925846 | -0.353169052 | 0.811685972 | 8.514088351 | 0.985902199 | 0.365708665 | 0.709264768 | -4.620251401 |
| **A_55_P2019113** | Apol7b | -0.240294495 | -0.850833097 | 0.370244107 | 8.314302542 | -0.985782353 | 0.365762504 | 0.709264768 | -4.620281169 |
| **A_55_P1993475** | Gm5168 | -0.152226092 | -0.539042855 | 0.234590671 | 7.814891379 | -0.985675945 | 0.365810313 | 0.709264768 | -4.620307598 |
| **A_55_P1978628** | Hnrnpa2b | 0.105116528 | -0.162002732 | 0.372235789 | 8.237151024 | 0.985635077 | 0.365828675 | 0.709264768 | -4.620317748 |
| **A_52_P62617** | Gm9895 | -0.098032045 | -0.347163183 | 0.151099092 | 7.741162437 | -0.985576633 | 0.365854937 | 0.709264768 | -4.620332263 |
| **A_55_P1963639** | Sycn | -0.323682368 | -1.14627291 | 0.498908175 | 8.016460808 | -0.985565915 | 0.365859753 | 0.709264768 | -4.620334925 |
| **A_55_P2011220** | Armcx1 | 0.457816937 | -0.705668066 | 1.621301939 | 9.961112568 | 0.985556238 | 0.365864102 | 0.709264768 | -4.620337328 |
| **A_55_P2349148** | 5830408B | 0.360001426 | -0.554941581 | 1.274944434 | 9.525295674 | 0.985508841 | 0.365885401 | 0.709264768 | -4.620349099 |
| **A_55_P2002133** | Ago2 | 0.147522961 | -0.227408859 | 0.522454781 | 9.109515113 | 0.985502353 | 0.365888316 | 0.709264768 | -4.62035071 |
| **A_55_P2073642** | 1600014C | -0.204935843 | -0.725789471 | 0.315917784 | 7.990818983 | -0.985490803 | 0.365893507 | 0.709264768 | -4.620353579 |
| **A_55_P2028883** | Rnft2 | -0.180289264 | -0.638640671 | 0.278062143 | 8.415796539 | -0.985193736 | 0.366027028 | 0.709264768 | -4.620427348 |
| **A_55_P2129437** | Plpp2 | -0.356357742 | -1.262344064 | 0.549628579 | 12.4719499 | -0.985178451 | 0.366033899 | 0.709264768 | -4.620431144 |
| **A_55_P2027969** | Myl6 | -0.208374507 | -0.738137645 | 0.32138863 | 16.49971974 | -0.985174578 | 0.366035639 | 0.709264768 | -4.620432105 |
| **A_52_P502771** | Rad54b | 0.218928963 | -0.337743174 | 0.7756011 | 8.445753829 | 0.985040488 | 0.366095922 | 0.709264768 | -4.6204654 |
| **A_55_P1959515** | Cep135 | 0.191242801 | -0.295101158 | 0.67758676 | 8.577007518 | 0.984899495 | 0.366159316 | 0.709264768 | -4.620500406 |
| **A_55_P2059105** | Gm9920 | 0.118715485 | -0.18324744 | 0.42067841 | 9.847602821 | 0.984700349 | 0.366248873 | 0.709264768 | -4.620549846 |
| **A_55_P1978321** | Fbxo8 | -0.111292806 | -0.394389032 | 0.17180342 | 9.80965693 | -0.984653338 | 0.366270016 | 0.709264768 | -4.620561517 |
| **A_51_P107433** | Mrpl34 | 0.157665003 | -0.243403255 | 0.558733261 | 11.87115308 | 0.984617061 | 0.366286333 | 0.709264768 | -4.619261464 |
| **A_55_P2001159** | Oser1 | -0.345953046 | -1.2260052 | 0.534099109 | 12.02284173 | -0.984598327 | 0.366294759 | 0.709264768 | -4.620575173 |
| **A_52_P312467** | Hoxb3 | 0.485470732 | -0.749650786 | 1.720592251 | 9.057608773 | 0.984472774 | 0.366351236 | 0.709264768 | -4.620606339 |
| **A_51_P246653** | Clec7a | -0.284531554 | -1.00855415 | 0.439491041 | 8.962547025 | -0.984302667 | 0.366427764 | 0.709264768 | -4.620648562 |
| **A_55_P2245920** | Lrrc8a | -0.341266091 | -1.209660812 | 0.52712863 | 12.75112192 | -0.984297334 | 0.366430164 | 0.709264768 | -4.620649885 |
| **A_51_P165934** | Phb | 0.222029724 | -0.342984231 | 0.787043679 | 11.02832895 | 0.984242899 | 0.366454657 | 0.709264768 | -4.620663396 |
| **A_51_P324651** | Adgrl1 | 0.309500821 | -0.478187147 | 1.097188789 | 9.166815405 | 0.984142392 | 0.366499882 | 0.709264768 | -4.620688341 |

| | | | | | | | | | |
|---|---|---|---|---|---|---|---|---|---|
| **A_55_P2032598** | NA | 0.129989354 | -0.200876259 | 0.460854966 | 15.86340708 | 0.984025827 | 0.366552339 | 0.709264768 | -4.620717269 |
| **A_55_P2015292** | Ltc4s | -0.319530873 | -1.132900519 | 0.493838772 | 9.218206631 | -0.983954943 | 0.366584241 | 0.709264768 | -4.620734861 |
| **A_51_P512629** | Ttll5 | -0.271440147 | -0.962461295 | 0.419581 | 8.834205655 | -0.983859513 | 0.366627194 | 0.709264768 | -4.620758542 |
| **A_51_P473240** | Zfp426 | 0.168123227 | -0.260042343 | 0.596288796 | 10.59222042 | 0.983481784 | 0.36679725 | 0.709264768 | -4.620852267 |
| **A_52_P175028** | NA | 0.489718814 | -0.757698144 | 1.737135772 | 10.00006766 | 0.983298759 | 0.366879672 | 0.709264768 | -4.620897674 |
| **A_55_P2006226** | NA | 0.154637817 | -0.239293358 | 0.548568992 | 16.59042991 | 0.983208682 | 0.366920241 | 0.709264768 | -4.62092002 |
| **A_55_P2020497** | Dcaf12l1 | -0.819738642 | -2.908234378 | 1.268757094 | 8.713622136 | -0.983087469 | 0.36697484 | 0.709264768 | -4.620950089 |
| **A_52_P566348** | Nras | 0.139899582 | -0.216557164 | 0.496356328 | 10.33036322 | 0.983014499 | 0.367007712 | 0.709264768 | -4.620968189 |
| **A_55_P2178036** | Serpina6 | 0.102139151 | -0.158120672 | 0.362398973 | 7.739584158 | 0.982959125 | 0.367032659 | 0.709264768 | -4.620981924 |
| **A_55_P1962966** | NA | -0.137876697 | -0.489301831 | 0.213548436 | 13.13586514 | -0.982671586 | 0.36716222 | 0.709264768 | -4.621053241 |
| **A_55_P2117155** | Apoc3 | -0.15363593 | -0.545279276 | 0.238007415 | 7.752387058 | -0.982544862 | 0.367219332 | 0.709264768 | -4.621084669 |
| **A_55_P2092958** | Traf2 | 0.131709814 | -0.204070098 | 0.467489725 | 8.221972544 | 0.982457496 | 0.36725871 | 0.709264768 | -4.621106334 |
| **A_55_P1986158** | Klhdc4 | -0.246295071 | -0.874282117 | 0.381691975 | 11.09443242 | -0.982325416 | 0.367318248 | 0.709264768 | -4.621139087 |
| **A_66_P132232** | Ercc1 | -0.205654353 | -0.73003987 | 0.318731164 | 9.275771127 | -0.982285115 | 0.367336416 | 0.709264768 | -4.62114908 |
| **A_51_P345836** | Prkra | 0.457757366 | -0.709503856 | 1.625018589 | 10.3385966 | 0.98224003 | 0.367356743 | 0.709264768 | -4.621160259 |
| **A_55_P2042953** | Acd | 0.278401968 | -0.4316319 | 0.988435836 | 11.53109094 | 0.982072571 | 0.367432247 | 0.709264768 | -4.62120178 |
| **A_52_P596592** | Rassf9 | 0.3162305 | -0.490373864 | 1.122834864 | 8.577855649 | 0.981959402 | 0.367483279 | 0.709264768 | -4.621229838 |
| **A_55_P2064328** | Lama2 | 0.412806691 | -0.64047293 | 1.466086312 | 9.856177067 | 0.981642588 | 0.367626174 | 0.709264768 | -4.621308377 |
| **A_52_P442710** | Ndnf | 0.679558534 | -1.054457265 | 2.413574333 | 9.559251001 | 0.981576532 | 0.367655974 | 0.709264768 | -4.621324751 |
| **A_51_P155458** | Dok7 | 0.179123291 | -0.277942259 | 0.636188841 | 8.537367441 | 0.981575966 | 0.367656229 | 0.709264768 | -4.621324891 |
| **A_52_P612137** | Runx1t1 | 0.512182149 | -0.79480878 | 1.819173078 | 9.75914858 | 0.981526908 | 0.367678361 | 0.709264768 | -4.621337051 |
| **A_51_P249414** | Ess2 | 0.099354999 | -0.154217998 | 0.352927996 | 10.33452761 | 0.98137969 | 0.367744785 | 0.709264768 | -4.621373541 |
| **A_52_P282700** | Zbtb33 | 0.229246547 | -0.355857767 | 0.814350861 | 8.985725899 | 0.981340784 | 0.367762341 | 0.709264768 | -4.621383183 |
| **A_55_P2070239** | Preb | -0.262448523 | -0.932377397 | 0.407480351 | 8.958824096 | -0.981218622 | 0.367817469 | 0.709264768 | -4.62141346 |
| **A_55_P2386256** | D130062J | -0.184742116 | -0.656320766 | 0.286836534 | 9.297849739 | -0.981210359 | 0.367821198 | 0.709264768 | -4.621415508 |
| **A_51_P118223** | Gm1943 | -0.13533109 | -0.480784243 | 0.210122063 | 8.156131729 | -0.981202766 | 0.367824625 | 0.709264768 | -4.618220929 |
| **A_55_P2052831** | 0610009B | 0.260720957 | -0.404840459 | 0.926282372 | 10.18193716 | 0.981156194 | 0.367845644 | 0.709264768 | -4.621428931 |
| **A_66_P130310** | A4galt | -0.382846613 | -1.360186361 | 0.594493134 | 8.191481269 | -0.981136864 | 0.367854368 | 0.709264768 | -4.621433722 |
| **A_51_P387608** | Hif1a | 0.128860107 | -0.200119427 | 0.457839641 | 10.20288128 | 0.981069903 | 0.367884591 | 0.709264768 | -4.621450316 |
| **A_55_P2085425** | Ophn1 | 0.345728288 | -0.537084962 | 1.228541537 | 9.570080785 | 0.980881217 | 0.367969765 | 0.709264768 | -4.621497073 |
| **A_51_P409988** | Arl13b | -0.30742045 | -1.09244132 | 0.477600421 | 9.136015996 | -0.980848418 | 0.367984572 | 0.709264768 | -4.6215052 |
| **A_55_P2003443** | Lsm7 | 0.463177046 | -0.719620627 | 1.645974719 | 14.41909427 | 0.980814579 | 0.367999849 | 0.709264768 | -4.621513585 |
| **A_52_P387009** | Egln3 | -0.307791622 | -1.093841413 | 0.478258169 | 10.31409914 | -0.980747213 | 0.368030264 | 0.709264768 | -4.621530277 |
| **A_55_P1970474** | Tmem67 | -0.103050811 | -0.366235046 | 0.160133424 | 8.715380627 | -0.980712901 | 0.368045757 | 0.709264768 | -4.621538779 |
| **A_55_P2021996** | Hbs1l | 0.092128464 | -0.143177195 | 0.327434123 | 8.135967775 | 0.98064497 | 0.36807643 | 0.709264768 | -4.62155561 |
| **A_55_P2157423** | NA | 0.101176211 | -0.157241918 | 0.359594339 | 8.450089386 | 0.980631363 | 0.368082574 | 0.709264768 | -4.621558981 |
| **A_55_P2126537** | Fam53c | -0.169115347 | -0.601088128 | 0.262857434 | 12.46983155 | -0.980566349 | 0.368111933 | 0.709264768 | -4.621575089 |
| **A_51_P157255** | Sdc2 | 0.582288417 | -0.905070234 | 2.069647068 | 10.747891 | 0.980556884 | 0.368116208 | 0.709264768 | -4.621577434 |
| **A_55_P1969367** | Col4a5 | 0.696482978 | -1.082684433 | 2.475650388 | 10.6031996 | 0.980491961 | 0.368145527 | 0.709264768 | -4.621593518 |
| **A_51_P466378** | Itgal | 0.106976867 | -0.166298663 | 0.380252397 | 7.966854785 | 0.980481663 | 0.368150178 | 0.709264768 | -4.62159607 |
| **A_51_P310548** | Osgep | 0.442451317 | -0.687998057 | 1.572900691 | 11.53795414 | 0.980312906 | 0.368226399 | 0.709264768 | -4.621637876 |
| **A_55_P1955672** | NA | 0.312380551 | -0.485779047 | 1.11054015 | 15.65692643 | 0.980267472 | 0.368246922 | 0.709264768 | -4.621649131 |
| **A_51_P357422** | Coprs | 0.211439126 | -0.328859839 | 0.751738091 | 9.253171328 | 0.980170357 | 0.368290793 | 0.709264768 | -4.621673188 |
| **A_55_P2009007** | NA | 0.417072344 | -0.648751208 | 1.482895896 | 15.3420766 | 0.980113624 | 0.368316424 | 0.709264768 | -4.621687241 |
| **A_55_P2098802** | Uqcrq | -0.133176149 | -0.473509366 | 0.207157067 | 13.74291937 | -0.980104676 | 0.368320467 | 0.709264768 | -4.621689457 |
| **A_51_P342567** | Akap12 | 0.785364226 | -1.22207854 | 2.792806991 | 13.17059575 | 0.979892172 | 0.368416485 | 0.709264768 | -4.621742091 |
| **A_55_P2170536** | Zfp949 | -0.169899516 | -0.60420357 | 0.264404538 | 8.732672877 | -0.979825198 | 0.368446751 | 0.709264768 | -4.621758679 |
| **A_55_P2063999** | Gpatch8 | 0.114758606 | -0.178600254 | 0.408117466 | 10.47342545 | 0.979797667 | 0.368459192 | 0.709264768 | -4.621765497 |

| | | | | | | | | | |
|---|---|---|---|---|---|---|---|---|---|
| **A_51_P406835** | Mei1 | -0.278584014 | -0.990761018 | 0.43359299 | 7.850211269 | -0.979757487 | 0.368477352 | 0.709264768 | -4.621775448 |
| **A_55_P2051047** | Rps10 | 0.252504721 | -0.393048507 | 0.898057949 | 17.63835369 | 0.979688006 | 0.368508754 | 0.709264768 | -4.621792655 |
| **A_55_P2108206** | Rrp7a | -0.368782282 | -1.311689856 | 0.574125292 | 10.91637952 | -0.979605621 | 0.368545992 | 0.709264768 | -4.621813057 |
| **A_51_P301930** | Lrrc17 | 0.929404065 | -1.447124795 | 3.305932925 | 11.28596529 | 0.979516645 | 0.368586212 | 0.709264768 | -4.621835091 |
| **A_55_P2152467** | NA | 0.148630326 | -0.231444135 | 0.528704788 | 8.826947013 | 0.979465374 | 0.36860939 | 0.709264768 | -4.621847786 |
| **A_51_P497724** | Apol7a | -0.120542447 | -0.428793074 | 0.187708181 | 7.739001523 | -0.979459263 | 0.368612153 | 0.709264768 | -4.6218493 |
| **A_51_P339962** | Cwf19l2 | 0.174266584 | -0.271429337 | 0.619962505 | 8.805127358 | 0.979322662 | 0.368673912 | 0.709264768 | -4.621883124 |
| **A_55_P2068137** | NA | -0.122149306 | -0.434558043 | 0.190259432 | 11.6346244 | -0.979305462 | 0.368681689 | 0.709264768 | -4.621887383 |
| **A_55_P2094292** | Erp27 | 0.121658634 | -0.189503953 | 0.432821221 | 9.507666561 | 0.979277798 | 0.368694197 | 0.709264768 | -4.621894232 |
| **A_55_P2108988** | Kcnip3 | 0.318495788 | -0.496152964 | 1.133144541 | 9.016984516 | 0.979227593 | 0.368716899 | 0.709264768 | -4.621906663 |
| **A_55_P2172747** | Mfn2 | -0.244316979 | -0.869313105 | 0.380679146 | 8.819889904 | -0.979099143 | 0.368774985 | 0.709264768 | -4.621938465 |
| **A_51_P121390** | Anapc11 | 0.296958021 | -0.462757162 | 1.056673205 | 13.47225756 | 0.979026845 | 0.368807683 | 0.709264768 | -4.620000505 |
| **A_55_P2073795** | Mecr | -0.16608608 | -0.591039068 | 0.258866909 | 8.261108113 | -0.978909836 | 0.368860606 | 0.709264768 | -4.621985331 |
| **A_51_P209327** | Apln | 0.366361418 | -0.571061981 | 1.303784818 | 9.605615085 | 0.978868352 | 0.368879371 | 0.709264768 | -4.6219956 |
| **A_55_P2000521** | Lonrf1 | -0.234744965 | -0.83541314 | 0.36592321 | 9.627799582 | -0.978840677 | 0.368891889 | 0.709264768 | -4.622002451 |
| **A_55_P1998380** | Traf6 | -0.129727801 | -0.461678637 | 0.202223035 | 8.666156787 | -0.978835325 | 0.368894311 | 0.709264768 | -4.622003776 |
| **A_55_P2143219** | Rasgrp2 | 0.830267619 | -1.294811407 | 2.955346645 | 11.26466018 | 0.978573298 | 0.369012856 | 0.709264768 | -4.622068635 |
| **A_55_P2040425** | Guf1 | 0.164225933 | -0.256112137 | 0.584564003 | 8.349684973 | 0.978573291 | 0.369012859 | 0.709264768 | -4.622068637 |
| **A_51_P402193** | Map3k1 | -0.091176804 | -0.324553345 | 0.142199738 | 11.73896719 | -0.97853758 | 0.369029018 | 0.709264768 | -4.622077476 |
| **A_52_P676063** | Tmem126 | -0.099077208 | -0.352715333 | 0.154560916 | 8.474632604 | -0.97838452 | 0.369098281 | 0.709264768 | -4.622115358 |
| **A_51_P191520** | Stard10 | -0.229867619 | -0.818357453 | 0.358622215 | 9.508097607 | -0.978338576 | 0.369119075 | 0.709264768 | -4.622126728 |
| **A_55_P2269459** | Prl8a6 | -2.684950326 | -9.559161366 | 4.189260715 | 9.843790907 | -0.978281273 | 0.36914501 | 0.709264768 | -4.62214091 |
| **A_55_P2467801** | Srp54a | 0.101318295 | -0.158094402 | 0.360730992 | 9.683740721 | 0.978243546 | 0.369162086 | 0.709264768 | -4.622150246 |
| **A_51_P175424** | Car14 | 0.283453343 | -0.442341137 | 1.009247824 | 8.660080923 | 0.978178851 | 0.369191369 | 0.709264768 | -4.622166256 |
| **A_52_P498355** | Lyrm4 | -0.130693807 | -0.465398631 | 0.204011016 | 9.748339297 | -0.978010202 | 0.369267714 | 0.709264768 | -4.622207989 |
| **A_55_P2222870** | Mylk | 0.297609226 | -0.464746398 | 1.059964849 | 9.604441893 | 0.977775446 | 0.369374007 | 0.709264768 | -4.622266075 |
| **A_51_P516870** | Itm2a | -0.176354715 | -0.628152142 | 0.275442713 | 13.36615472 | -0.977673101 | 0.369420354 | 0.709264768 | -4.622291396 |
| **A_55_P2014645** | Knop1 | -0.352850333 | -1.256871793 | 0.551171128 | 10.12916689 | -0.977602124 | 0.369452499 | 0.709264768 | -4.622308956 |
| **A_55_P2094262** | Tph1 | -0.746746323 | -2.660127739 | 1.166635093 | 8.22465317 | -0.977511526 | 0.369493534 | 0.709264768 | -4.622331369 |
| **A_55_P2027157** | NA | 0.420617274 | -0.657212205 | 1.498446754 | 9.176379509 | 0.977433889 | 0.369528701 | 0.709264768 | -4.622350574 |
| **A_66_P134690** | Spin1 | -0.290219832 | -1.033910024 | 0.453470361 | 8.29540969 | -0.977429353 | 0.369530756 | 0.709264768 | -4.622351696 |
| **A_52_P65237** | Zbtb7c | -0.143160142 | -0.510012702 | 0.223692418 | 7.842633904 | -0.977419303 | 0.369535308 | 0.709264768 | -4.622354183 |
| **A_55_P2116993** | Hnrnpdl | -0.10239424 | -0.364801164 | 0.160012683 | 12.57601208 | -0.977351052 | 0.369566226 | 0.709264768 | -4.622371066 |
| **A_51_P380986** | Epc1 | 0.275101567 | -0.42995263 | 0.980155763 | 10.48044502 | 0.977284267 | 0.369596482 | 0.709264768 | -4.622387586 |
| **A_55_P2088630** | Ap3s2 | 0.239532184 | -0.3744355 | 0.853499869 | 10.17540215 | 0.977166811 | 0.369649699 | 0.709264768 | -4.622416638 |
| **A_51_P483311** | Mpv17l2 | -0.113571835 | -0.404678823 | 0.177535153 | 8.903136225 | -0.97716602 | 0.369650057 | 0.709264768 | -4.622416834 |
| **A_55_P2102322** | Yif1b | -0.222816972 | -0.793985993 | 0.348352049 | 13.03165546 | -0.977088652 | 0.369685115 | 0.709264768 | -4.62243597 |
| **A_52_P37894** | Cox7a2 | 0.283666026 | -0.443561156 | 1.010893208 | 14.41592309 | 0.976984259 | 0.369732421 | 0.709264768 | -4.622461789 |
| **A_55_P2110435** | Zscan20 | 0.081785638 | -0.127889306 | 0.291460583 | 8.210585557 | 0.976969163 | 0.369739263 | 0.709264768 | -4.622465523 |
| **A_51_P127915** | Rnasek | -0.229534161 | -0.818018945 | 0.358950622 | 10.45748414 | -0.976927731 | 0.369758039 | 0.709264768 | -4.622475769 |
| **A_51_P456187** | Pdcd2 | 0.095653603 | -0.149585434 | 0.340892641 | 10.80632472 | 0.976926923 | 0.369758406 | 0.709264768 | -4.622475969 |
| **A_55_P2157695** | Tsc1 | -0.185569238 | -0.661440278 | 0.290301801 | 10.18392438 | -0.976713195 | 0.369855279 | 0.709264768 | -4.622528824 |
| **A_51_P204913** | Bag2 | -0.162361404 | -0.578771286 | 0.254048477 | 9.639744127 | -0.976589651 | 0.369911286 | 0.709264768 | -4.622559374 |
| **A_55_P2034027** | Nos3 | 0.650093724 | -1.017311362 | 2.31749881 | 11.14509018 | 0.976529118 | 0.36993873 | 0.709264768 | -4.622574342 |
| **A_55_P2018989** | Elk3 | 0.112076315 | -0.175385894 | 0.399538523 | 8.204275324 | 0.976525144 | 0.369940532 | 0.709264768 | -4.622575324 |
| **A_52_P151853** | Tpd52 | -0.379070791 | -1.351349887 | 0.593208305 | 9.302858755 | -0.976516799 | 0.369944316 | 0.709264768 | -4.622577388 |
| **A_52_P678895** | Dynlt3 | 0.186768118 | -0.292275001 | 0.665811237 | 10.30473505 | 0.976514016 | 0.369945577 | 0.709264768 | -4.622578076 |
| **A_55_P2032267** | Gm10767 | 0.106098603 | -0.166049675 | 0.37824688 | 11.84588253 | 0.976459916 | 0.369970107 | 0.709264768 | -4.622591453 |

| | | | | | | | | | |
|---|---|---|---|---|---|---|---|---|---|
| A_55_P2155397 | Rps8 | 0.275673141 | -0.431443692 | 0.982789975 | 17.51255944 | 0.976458128 | 0.369970917 | 0.709264768 | -4.622591895 |
| A_55_P2012406 | NA | 0.195199913 | -0.305567537 | 0.695967364 | 14.74570439 | 0.976323793 | 0.370031831 | 0.709264768 | -4.622625109 |
| A_52_P131559 | Srprb | -0.185060958 | -0.659870818 | 0.289748902 | 8.387363998 | -0.97621488 | 0.370081224 | 0.709264768 | -4.622652036 |
| A_51_P345792 | Mfsd13a | -0.239150719 | -0.852756199 | 0.374454761 | 9.160924141 | -0.976186522 | 0.370094085 | 0.709264768 | -4.622659046 |
| A_51_P267454 | Slc25a11 | 0.225163996 | -0.352707624 | 0.803035616 | 13.26286645 | 0.975928333 | 0.370211199 | 0.709264768 | -4.622722872 |
| A_55_P2067741 | Psg23 | -2.133561323 | -7.610500602 | 3.343377957 | 11.48183522 | -0.975702795 | 0.370313527 | 0.709264768 | -4.62277862 |
| A_52_P191468 | Ubfd1 | -0.552622566 | -1.971454305 | 0.866209173 | 9.767051913 | -0.975546704 | 0.37038436 | 0.709264768 | -4.622817199 |
| A_52_P582150 | Gosr2 | -0.19264266 | -0.687244525 | 0.301959204 | 11.58078361 | -0.975544452 | 0.370385382 | 0.709264768 | -4.622817755 |
| A_51_P111488 | Gfod2 | 0.255608521 | -0.400658716 | 0.911875758 | 8.84991379 | 0.975539703 | 0.370387537 | 0.709264768 | -4.622818929 |
| A_55_P2024654 | Trrap | -0.218620373 | -0.780008024 | 0.342767279 | 10.12680339 | -0.975389599 | 0.370455664 | 0.709264768 | -4.622856025 |
| A_51_P356931 | Orc1 | 0.084410852 | -0.132363554 | 0.301185258 | 7.804432276 | 0.975305392 | 0.370493886 | 0.709264768 | -4.622876834 |
| A_55_P2088905 | Hpca | 0.176088781 | -0.2761818 | 0.628359361 | 10.40155408 | 0.975177548 | 0.370551923 | 0.709264768 | -4.622908425 |
| A_55_P2108903 | Zbtb20 | -0.083582514 | -0.298266276 | 0.131101249 | 8.066322289 | -0.975139108 | 0.370569374 | 0.709264768 | -4.622917924 |
| A_55_P2070347 | Rps2 | 0.229478378 | -0.359951796 | 0.818908553 | 15.89445244 | 0.975123792 | 0.370576328 | 0.709264768 | -4.622921708 |
| A_52_P409769 | Mrvi1 | -0.167110307 | -0.596390176 | 0.262169562 | 9.015710413 | -0.975018959 | 0.370623926 | 0.709264768 | -4.622947611 |
| A_51_P221677 | Trim62 | -0.247695281 | -0.884027202 | 0.388636641 | 8.43663536 | -0.974954544 | 0.370653176 | 0.709264768 | -4.622963526 |
| A_52_P623058 | R3hdm4 | 0.093810774 | -0.147194696 | 0.334816244 | 8.684997389 | 0.974936126 | 0.370661539 | 0.709264768 | -4.622968077 |
| A_55_P2165969 | NA | 0.126278117 | -0.198150488 | 0.450706722 | 8.464488951 | 0.97489828 | 0.370678725 | 0.709264768 | -4.622977427 |
| A_55_P2033462 | Rps23-ps | 0.306328213 | -0.480701063 | 1.093357489 | 16.93936532 | 0.974869438 | 0.370691823 | 0.709264768 | -4.622984553 |
| A_55_P2171463 | Nptn | 0.113166327 | -0.17759033 | 0.403922984 | 13.10726812 | 0.974850234 | 0.370700544 | 0.709264768 | -4.622989298 |
| A_55_P1955869 | NA | 0.44844022 | -0.70383029 | 1.600710731 | 12.22359061 | 0.974766187 | 0.370738715 | 0.709264768 | -4.623010062 |
| A_55_P1991069 | Milr1 | -0.404243929 | -1.442968314 | 0.634480455 | 9.210816936 | -0.974750632 | 0.370745779 | 0.709264768 | -4.623013905 |
| A_65_P15503 | Map3k2 | -0.246361507 | -0.879437551 | 0.386714537 | 10.03701996 | -0.97469181 | 0.370772495 | 0.709264768 | -4.623028436 |
| A_51_P153423 | Fndc1 | 0.912799788 | -1.433206358 | 3.258805934 | 10.51278962 | 0.974533405 | 0.37084445 | 0.709264768 | -4.623067567 |
| A_51_P323435 | Gtf2h2 | 0.150006535 | -0.235550663 | 0.535563733 | 8.442774411 | 0.97447726 | 0.370869956 | 0.709264768 | -4.623081436 |
| A_55_P2130877 | Parl | 0.106488893 | -0.167241382 | 0.380219167 | 10.24456561 | 0.974387769 | 0.370910614 | 0.709264768 | -4.623103541 |
| A_55_P2236291 | Plpp1 | -0.265122685 | -0.946755917 | 0.416510547 | 8.442511176 | -0.974196308 | 0.370997611 | 0.709264768 | -4.62315083 |
| A_66_P123432 | NA | 0.089328497 | -0.140361162 | 0.319018156 | 7.862391712 | 0.974089655 | 0.37104608 | 0.709264768 | -4.62317717 |
| A_55_P2062981 | Abca9 | 0.139007272 | -0.218421236 | 0.49643578 | 8.24235392 | 0.974089086 | 0.371046338 | 0.709264768 | -4.623177311 |
| A_55_P2181597 | Mfap2 | 0.624689131 | -0.981774169 | 2.231152431 | 9.889830179 | 0.973965305 | 0.371102597 | 0.709264768 | -4.623207879 |
| A_55_P2067652 | Boc | 0.530988077 | -0.834831195 | 1.896807349 | 9.802157002 | 0.973737461 | 0.371206171 | 0.709264768 | -4.623264143 |
| A_55_P2079425 | Spred1 | 0.435024217 | -0.684515434 | 1.554563868 | 9.450654137 | 0.973249748 | 0.371427954 | 0.709264768 | -4.623384555 |
| A_51_P348592 | Slc38a7 | -0.351225101 | -1.255121596 | 0.552671394 | 8.463869537 | -0.973233813 | 0.371435202 | 0.709264768 | -4.623388489 |
| A_55_P2112947 | Srp9 | 0.116755711 | -0.183737651 | 0.417249072 | 9.760628902 | 0.973180942 | 0.371459252 | 0.709264768 | -4.623401541 |
| A_51_P392005 | Car8 | 0.218853454 | -0.344413441 | 0.782120349 | 8.246664597 | 0.973171817 | 0.371463403 | 0.709264768 | -4.623403793 |
| A_51_P213592 | Nrde2 | -0.215998385 | -0.771927114 | 0.339930344 | 8.543924597 | -0.973154358 | 0.371471345 | 0.709264768 | -4.623408103 |
| A_55_P1962801 | Max | -0.41784879 | -1.493297096 | 0.657599515 | 12.06206989 | -0.973150377 | 0.371473155 | 0.709264768 | -4.623409086 |
| A_55_P1966538 | Ctu2 | -0.102579096 | -0.366599487 | 0.161441295 | 11.16958084 | -0.973131976 | 0.371481526 | 0.709264768 | -4.623413628 |
| A_52_P271377 | Efr3a | -0.159215934 | -0.569015745 | 0.250583878 | 8.42729072 | -0.973117135 | 0.371488277 | 0.709264768 | -4.623417292 |
| A_51_P203192 | Mbd4 | 0.193535156 | -0.304671653 | 0.691741965 | 8.440609137 | 0.972972474 | 0.37155409 | 0.709264768 | -4.623452999 |
| A_51_P362054 | Atp5e | 0.49619663 | -0.78114288 | 1.77353614 | 15.1380609 | 0.972966327 | 0.371556886 | 0.709264768 | -4.623454517 |
| A_51_P334308 | Tspan14 | -0.446805143 | -1.597198832 | 0.703588546 | 12.75737094 | -0.972796543 | 0.371634141 | 0.709264768 | -4.623496422 |
| A_55_P2393734 | 4933421H | -0.216679151 | -0.77456653 | 0.341208229 | 9.694269974 | -0.972794111 | 0.371635247 | 0.709264768 | -4.623497023 |
| A_55_P2095578 | NA | 0.107121405 | -0.168709445 | 0.382952254 | 8.609198604 | 0.972710867 | 0.371673129 | 0.709264768 | -4.623517567 |
| A_55_P2054743 | Sh3bgrl2 | -0.314393583 | -1.123959463 | 0.495172297 | 8.15831492 | -0.972684112 | 0.371685306 | 0.709264768 | -4.62352417 |
| A_55_P1962110 | NA | 0.423497173 | -0.667033779 | 1.514028126 | 15.63061744 | 0.97266405 | 0.371694436 | 0.709264768 | -4.623529121 |
| A_55_P2144666 | Prpsap1 | 0.25145286 | -0.396108171 | 0.89901389 | 12.93061893 | 0.972581981 | 0.371731788 | 0.709264768 | -4.623549375 |
| A_52_P473966 | Kdelr3 | -0.360395541 | -1.288751921 | 0.567960839 | 9.873312651 | -0.972333006 | 0.371845123 | 0.709264768 | -4.623610813 |

| | | | | | | | | | |
|---|---|---|---|---|---|---|---|---|---|
| **A_55_P2008006** | Ap1g1 | -0.284120037 | -1.016012884 | 0.44777281 | 9.474737628 | -0.972309895 | 0.371855644 | 0.709264768 | -4.623616516 |
| **A_66_P128537** | Isg15 | -0.484449664 | -1.73243345 | 0.763534122 | 8.26922877 | -0.972277111 | 0.37187057 | 0.709264768 | -4.623624605 |
| **A_52_P507310** | Mrps24 | 0.304374779 | -0.479787931 | 1.08853749 | 11.77782126 | 0.972193746 | 0.371908526 | 0.709264768 | -4.623645175 |
| **A_52_P342620** | Csrnp2 | -0.288684391 | -1.032435517 | 0.455066736 | 8.300213067 | -0.972178498 | 0.371915469 | 0.709264768 | -4.623648937 |
| **A_55_P1953382** | Prl8a8 | -2.643419209 | -9.454962474 | 4.168124056 | 9.717169808 | -0.972010302 | 0.37199206 | 0.709264768 | -4.623690434 |
| **A_52_P663413** | P4ha2 | -0.904050735 | -3.233778984 | 1.425677513 | 11.23195092 | -0.971936482 | 0.372025679 | 0.709264768 | -4.623708646 |
| **A_52_P25812** | Prdm2 | -0.233168408 | -0.83405541 | 0.367718595 | 8.186069337 | -0.971912668 | 0.372036525 | 0.709264768 | -4.623714521 |
| **A_55_P2024431** | Gaa | -0.584631661 | -2.091281446 | 0.922018125 | 13.67409286 | -0.97189727 | 0.372043538 | 0.709264768 | -4.623718319 |
| **A_51_P251587** | Psmd9 | 0.171600818 | -0.270686355 | 0.613887991 | 9.242381156 | 0.971774178 | 0.372099604 | 0.709264768 | -4.623748685 |
| **A_55_P2008898** | NA | 0.625400781 | -0.987143562 | 2.237945124 | 13.75026095 | 0.971397763 | 0.372271096 | 0.709264768 | -4.623841531 |
| **A_51_P521176** | Rassf7 | -0.185545526 | -0.66398396 | 0.292892909 | 12.50394972 | -0.971347823 | 0.372293853 | 0.709264768 | -4.623853848 |
| **A_55_P2120277** | Gm11619 | 0.334218947 | -0.52760399 | 1.196041885 | 16.35465267 | 0.971322309 | 0.37230548 | 0.709264768 | -4.62386014 |
| **A_51_P414706** | Ddt | 0.291855816 | -0.460757693 | 1.044469325 | 12.47745149 | 0.971285015 | 0.372322476 | 0.709264768 | -4.623869338 |
| **A_55_P1976490** | Lman2l | -0.132724646 | -0.47498505 | 0.209535757 | 10.78203945 | -0.97128183 | 0.372323928 | 0.709264768 | -4.623870123 |
| **A_55_P2041828** | Tubb3 | -1.187805122 | -4.251298439 | 1.875688196 | 9.297174826 | -0.971132753 | 0.372391871 | 0.709264768 | -4.623906887 |
| **A_51_P130666** | Adk | 0.513951227 | -0.811712099 | 1.839614554 | 12.13592971 | 0.971044228 | 0.372432222 | 0.709264768 | -4.623928717 |
| **A_55_P1961014** | Selenbp1 | 1.37875047 | -2.178031693 | 4.935532634 | 14.35095051 | 0.970909502 | 0.372493639 | 0.709264768 | -4.623961938 |
| **A_55_P2029642** | NA | 0.338175819 | -0.534232673 | 1.210584311 | 13.96568119 | 0.97089667 | 0.372499489 | 0.709264768 | -4.623965102 |
| **A_51_P392147** | Xab2 | -0.200467503 | -0.717735059 | 0.316800052 | 10.14131263 | -0.970686721 | 0.372595215 | 0.709264768 | -4.624016867 |
| **A_52_P222725** | Lims1 | 0.230409416 | -0.364124659 | 0.824943491 | 11.86818321 | 0.970674941 | 0.372600587 | 0.709264768 | -4.624019772 |
| **A_55_P2024993** | Hoxb4 | 0.529230917 | -0.836708554 | 1.895170389 | 10.6688838 | 0.970429741 | 0.372712411 | 0.709264768 | -4.624080221 |
| **A_55_P2019890** | Rasef | -0.172261404 | -0.616881936 | 0.272359127 | 8.800353171 | -0.970395593 | 0.372727987 | 0.709264768 | -4.624088639 |
| **A_55_P2069995** | NA | 0.277665693 | -0.439121077 | 0.994452463 | 9.462844819 | 0.970247625 | 0.372795483 | 0.709264768 | -4.624125113 |
| **A_55_P1975690** | Bicc1 | 0.547311918 | -0.865582689 | 1.960206525 | 12.34265236 | 0.970231752 | 0.372802724 | 0.709264768 | -4.624129026 |
| **A_52_P613643** | Zfp3 | -0.128306254 | -0.459540731 | 0.202928223 | 8.072237539 | -0.970203049 | 0.372815819 | 0.709264768 | -4.624136101 |
| **A_55_P1959833** | Tmem266 | 0.192966354 | -0.305208814 | 0.691141521 | 9.12025315 | 0.970174511 | 0.372828839 | 0.709264768 | -4.624143135 |
| **A_55_P2107292** | Ptch1 | 0.176401668 | -0.279081696 | 0.631885032 | 8.441360271 | 0.970019611 | 0.372899513 | 0.709264768 | -4.624181314 |
| **A_51_P156434** | Slc25a33 | -0.159316547 | -0.570702738 | 0.252069644 | 7.92208684 | -0.969977189 | 0.37291887 | 0.709264768 | -4.624191769 |
| **A_55_P2004511** | Cd300lf | 0.21250161 | -0.336223769 | 0.761226988 | 8.547512846 | 0.96996826 | 0.372922945 | 0.709264768 | -4.62419397 |
| **A_55_P2091928** | Raet1b | 0.154047412 | -0.243740748 | 0.551835572 | 9.233292583 | 0.969957933 | 0.372927657 | 0.709264768 | -4.624196515 |
| **A_51_P275435** | Mab21l4 | 0.648606325 | -1.026277273 | 2.323489924 | 9.502815643 | 0.969944525 | 0.372933776 | 0.709264768 | -4.62419982 |
| **A_55_P2131577** | NA | 0.224069516 | -0.354544178 | 0.802683209 | 17.22563081 | 0.969938981 | 0.372936305 | 0.709264768 | -4.624201186 |
| **A_55_P2074488** | Crim1 | -0.911873033 | -3.266880519 | 1.443134454 | 9.164981753 | -0.96982288 | 0.372989288 | 0.709264768 | -4.624229798 |
| **A_55_P1963863** | Arhgef17 | 0.62717175 | -0.99265984 | 2.24700334 | 10.98910956 | 0.969766035 | 0.373015232 | 0.709264768 | -4.624243807 |
| **A_52_P8227** | Lypla1 | 0.304962647 | -0.482923577 | 1.092848872 | 9.809672022 | 0.96946802 | 0.373151268 | 0.709264768 | -4.624317242 |
| **A_55_P2153743** | Slc26a6 | -0.103130698 | -0.369583156 | 0.16332176 | 8.869290758 | -0.969434723 | 0.37316647 | 0.709264768 | -4.624325447 |
| **A_55_P2021187** | Malat1 | -0.177680735 | -0.636755407 | 0.281393937 | 8.817332489 | -0.969409687 | 0.3731779 | 0.709264768 | -4.624331615 |
| **A_55_P2000182** | Slc5a6 | -1.373999844 | -4.924153917 | 2.176154229 | 9.898418886 | -0.969370564 | 0.373195763 | 0.709264768 | -4.624341255 |
| **A_52_P456158** | Fchsd1 | 0.307732806 | -0.487425223 | 1.102890836 | 10.43553263 | 0.969327858 | 0.373215262 | 0.709264768 | -4.624351777 |
| **A_51_P192860** | Hint1 | 0.130600973 | -0.206869763 | 0.468071709 | 16.41857847 | 0.969305416 | 0.373225509 | 0.709264768 | -4.624357306 |
| **A_51_P400375** | Erg28 | -0.353038149 | -1.265288492 | 0.559212193 | 13.09918381 | -0.96929941 | 0.373228251 | 0.709264768 | -4.624358786 |
| **A_55_P2116604** | NA | 0.503967265 | -0.798488229 | 1.80642276 | 14.0101388 | 0.969147281 | 0.373297721 | 0.709264768 | -4.624396265 |
| **A_66_P124555** | Rps15 | 0.519192815 | -0.822754761 | 1.861140392 | 14.43859512 | 0.969043938 | 0.373344918 | 0.709264768 | -4.624421723 |
| **A_52_P8685** | Tmem183 | -0.090858155 | -0.325698047 | 0.143981737 | 9.431917477 | -0.969041523 | 0.373346021 | 0.709264768 | -4.624422318 |
| **A_55_P1953593** | Ece2 | 0.149686276 | -0.237222707 | 0.53659526 | 10.61150668 | 0.968999418 | 0.373365252 | 0.709264768 | -4.62443269 |
| **A_66_P101407** | Acat3 | 0.092380074 | -0.146422688 | 0.331182837 | 7.898922333 | 0.968923093 | 0.373400115 | 0.709264768 | -4.624451492 |
| **A_55_P1958447** | Prepl | 0.189750536 | -0.300772342 | 0.680273414 | 8.572897017 | 0.96888914 | 0.373415624 | 0.709264768 | -4.624459855 |
| **A_51_P341571** | Klhdc2 | -0.113065894 | -0.405412965 | 0.179281176 | 10.86994241 | -0.968686444 | 0.373508224 | 0.709264768 | -4.624509782 |

| | | | | | | | | | |
|---|---|---|---|---|---|---|---|---|---|
| **A_55_P2038573** | NA | 0.475889449 | -0.754776062 | 1.70655496 | 15.24351665 | 0.968537392 | 0.37357633 | 0.709264768 | -4.624546492 |
| **A_55_P2044364** | Mms22l | 0.151165919 | -0.239806765 | 0.542138603 | 14.02996322 | 0.968406773 | 0.37363602 | 0.709264768 | -4.62457866 |
| **A_55_P2148348** | NA | 0.380067711 | -0.602942134 | 1.363077556 | 13.95874189 | 0.96839704 | 0.373640469 | 0.709264768 | -4.624581057 |
| **A_55_P2102529** | Fuca2 | -0.211225425 | -0.757583815 | 0.335132964 | 8.383690334 | -0.968320046 | 0.373675658 | 0.709264768 | -4.624600017 |
| **A_55_P2033312** | Lgi4 | -0.152574728 | -0.547460835 | 0.242311378 | 8.409220696 | -0.967745358 | 0.373938394 | 0.709264768 | -4.624741515 |
| **A_51_P302458** | Plscr3 | 0.533985049 | -0.848065264 | 1.916035362 | 11.50662721 | 0.967733086 | 0.373944006 | 0.709264768 | -4.624744536 |
| **A_51_P511315** | Pstpip1 | -0.558799761 | -2.005150634 | 0.887551112 | 8.362313707 | -0.967682517 | 0.373967132 | 0.709264768 | -4.624756985 |
| **A_66_P104365** | Ccdc167 | 0.19633032 | -0.311882657 | 0.704543298 | 10.35416817 | 0.967591333 | 0.374008836 | 0.709264768 | -4.624779432 |
| **A_51_P361788** | Vapa | -0.192052243 | -0.689237573 | 0.305133086 | 13.79627824 | -0.967501006 | 0.374050152 | 0.709264768 | -4.624801667 |
| **A_55_P1989813** | Gcm1 | -0.196797526 | -0.706299176 | 0.312704124 | 9.671288673 | -0.96744077 | 0.374077705 | 0.709264768 | -4.624816494 |
| **A_51_P172085** | Arhgdig | 0.344948463 | -0.548138325 | 1.23803525 | 9.078087238 | 0.967410719 | 0.374091452 | 0.709264768 | -4.624823891 |
| **A_52_P58471** | Tmem199 | -0.104930619 | -0.376675258 | 0.16681402 | 8.661180847 | -0.967145011 | 0.374213018 | 0.709264768 | -4.624889287 |
| **A_51_P296528** | Cers4 | 0.449091505 | -0.713973676 | 1.612156686 | 10.4340257 | 0.967121708 | 0.374223681 | 0.709264768 | -4.624895023 |
| **A_52_P415155** | Wnt6 | -0.177294828 | -0.636519577 | 0.281929921 | 8.023271147 | -0.966988093 | 0.374284826 | 0.709264768 | -4.624927905 |
| **A_55_P1998299** | Itgb4 | -0.303548102 | -1.090018054 | 0.48292185 | 8.988676384 | -0.966708934 | 0.3744126 | 0.709264768 | -4.624996597 |
| **A_55_P1986306** | Ltv1 | 0.193183243 | -0.307348246 | 0.693714732 | 11.48381734 | 0.966692598 | 0.374420078 | 0.709264768 | -4.625000617 |
| **A_51_P436386** | Qrich1 | 0.099718741 | -0.158677535 | 0.358115016 | 10.73999511 | 0.96658685 | 0.37446849 | 0.709264768 | -4.625026636 |
| **A_55_P2125743** | NA | 0.269999664 | -0.429753559 | 0.969752888 | 16.46466669 | 0.966426127 | 0.374542079 | 0.709264768 | -4.625066178 |
| **A_52_P394395** | Abcd3 | 0.233048045 | -0.370960439 | 0.837056529 | 12.54039013 | 0.966390764 | 0.374558272 | 0.709264768 | -4.625074878 |
| **A_51_P277295** | Apmap | -0.245140885 | -0.880502116 | 0.390220346 | 9.968747102 | -0.966374329 | 0.374565798 | 0.709264768 | -4.625078921 |
| **A_55_P1973848** | Sema4b | -0.31523003 | -1.13234272 | 0.501882661 | 9.734462382 | -0.966264386 | 0.374616146 | 0.709264768 | -4.625105967 |
| **A_51_P274947** | Atad1 | 0.110536004 | -0.17599181 | 0.397063818 | 11.49294766 | 0.966245135 | 0.374624962 | 0.709264768 | -4.625110703 |
| **A_51_P477121** | Pmaip1 | -0.678705349 | -2.438267662 | 1.080856965 | 8.403920695 | -0.966110864 | 0.37468646 | 0.709264768 | -4.625143732 |
| **A_51_P190845** | Atp6v1b2 | -0.153662965 | -0.552065186 | 0.244739256 | 8.42112093 | -0.966045993 | 0.374716174 | 0.709264768 | -4.625159688 |
| **A_52_P193236** | Tedc2 | -0.425607449 | -1.529114507 | 0.677899609 | 8.923411135 | -0.966016291 | 0.37472978 | 0.709264768 | -4.625166994 |
| **A_51_P275697** | Rflnb | -0.385567891 | -1.385288002 | 0.61415222 | 8.741485401 | -0.965990324 | 0.374741675 | 0.709264768 | -4.625173381 |
| **A_51_P347177** | Snx2 | -0.10744607 | -0.386039491 | 0.17114735 | 12.23756935 | -0.965984208 | 0.374744477 | 0.709264768 | -4.625174885 |
| **A_55_P2079855** | Prl6a1 | -0.274322904 | -0.985641362 | 0.436995554 | 7.850633067 | -0.965935967 | 0.374766576 | 0.709264768 | -4.625186751 |
| **A_51_P336161** | Man2b1 | -0.186193549 | -0.669010762 | 0.296623665 | 11.13313236 | -0.965900141 | 0.374782989 | 0.709264768 | -4.625195562 |
| **A_55_P2073010** | Gm4832 | 0.414129877 | -0.659936682 | 1.488196436 | 14.56889465 | 0.96572998 | 0.374860952 | 0.709264768 | -4.625237412 |
| **A_52_P99411** | Itm2b | -0.107939494 | -0.387889093 | 0.172010105 | 13.94920836 | -0.965719215 | 0.374865885 | 0.709264768 | -4.625240059 |
| **A_51_P141535** | Oat | 0.415351467 | -0.662174224 | 1.492877157 | 11.60029854 | 0.965469282 | 0.374980422 | 0.709264768 | -4.625301521 |
| **A_51_P283473** | Fibin | 1.115655475 | -1.77888723 | 4.010198181 | 11.90496171 | 0.965384864 | 0.375019114 | 0.709264768 | -4.625322279 |
| **A_52_P445622** | 2610002M | 0.134788259 | -0.214922193 | 0.484498711 | 8.462254559 | 0.965369998 | 0.375025928 | 0.709264768 | -4.625325934 |
| **A_55_P2085905** | Anpep | -0.539099646 | -1.937861639 | 0.859662348 | 10.54960795 | -0.965329479 | 0.375044502 | 0.709264768 | -4.625335897 |
| **A_51_P227962** | Dynlrb2 | 1.601761118 | -2.554332718 | 5.757854955 | 12.4139144 | 0.965300983 | 0.375057564 | 0.709264768 | -4.625342904 |
| **A_51_P393067** | Ap4m1 | -0.098808264 | -0.355212876 | 0.157596348 | 8.988526336 | -0.965201046 | 0.375103378 | 0.709264768 | -4.625367476 |
| **A_52_P163849** | Ivns1abp | -0.081981335 | -0.294731056 | 0.130768385 | 8.063826867 | -0.965153377 | 0.375125232 | 0.709264768 | -4.625379195 |
| **A_52_P107234** | Prpf39 | -0.092030313 | -0.330863536 | 0.146802911 | 8.852775851 | -0.965131527 | 0.375135249 | 0.709264768 | -4.625384567 |
| **A_55_P2081283** | Dlx4 | -0.362195082 | -1.302155428 | 0.577765264 | 8.138628924 | -0.965124548 | 0.375138449 | 0.709264768 | -4.625386283 |
| **A_55_P2137309** | Kctd11 | -0.210211286 | -0.755751897 | 0.335329325 | 10.71999204 | -0.965115496 | 0.375142599 | 0.709264768 | -4.625388509 |
| **A_55_P2093679** | Zfp599 | 0.091626038 | -0.146173789 | 0.329425865 | 8.381318356 | 0.965067565 | 0.375164575 | 0.709264768 | -4.625400293 |
| **A_52_P742248** | Fam117b | 0.298682952 | -0.476551089 | 1.073916994 | 11.99170111 | 0.965001406 | 0.375194911 | 0.709264768 | -4.625416557 |
| **A_55_P2135486** | NA | -0.254693979 | -0.915843304 | 0.406455347 | 7.789667756 | -0.964871469 | 0.375254495 | 0.709264768 | -4.6254485 |
| **A_55_P2048011** | Larp4b | -0.324206431 | -1.165827383 | 0.517414521 | 12.62510854 | -0.964840293 | 0.375268792 | 0.709264768 | -4.625456164 |
| **A_55_P2085546** | Dnaja4 | -0.169135111 | -0.608238733 | 0.269968512 | 9.914426173 | -0.964755133 | 0.375307849 | 0.709264768 | -4.625477097 |
| **A_55_P2035138** | NA | 0.203022207 | -0.324137191 | 0.730181604 | 13.93568046 | 0.964610391 | 0.375374239 | 0.709264768 | -4.625512674 |
| **A_52_P264902** | Dnajc3 | -0.345986794 | -1.244443249 | 0.55246966 | 10.29197705 | -0.964523549 | 0.375414076 | 0.709264768 | -4.625534019 |

| | | | | | | | | | |
|---|---|---|---|---|---|---|---|---|---|
| **A_55_P1965303** | Dsc2 | -0.868987481 | -3.12567747 | 1.387702509 | 8.581667363 | -0.964477194 | 0.375435342 | 0.709264768 | -4.625545412 |
| **A_55_P2003043** | P4ha1 | -0.26430555 | -0.95075382 | 0.422142721 | 8.235169994 | -0.964381372 | 0.375479304 | 0.709264768 | -4.625568962 |
| **A_55_P2023692** | Neurl1a | 0.316939918 | -0.506244769 | 1.140124605 | 10.1445436 | 0.964339605 | 0.375498468 | 0.709264768 | -4.625579226 |
| **A_52_P675530** | Fat4 | 0.335876535 | -0.53659966 | 1.20835273 | 8.743995977 | 0.964220639 | 0.375553057 | 0.709264768 | -4.625608462 |
| **A_55_P2025403** | Gphn | 0.528994117 | -0.845257792 | 1.903246025 | 10.50533373 | 0.964128318 | 0.375595423 | 0.709264768 | -4.625631149 |
| **A_51_P226853** | Gga1 | -0.096449131 | -0.347036822 | 0.15413856 | 8.623129318 | -0.964026408 | 0.375642195 | 0.709264768 | -4.625656191 |
| **A_51_P235801** | Zfp36l1 | 0.685643483 | -1.095855093 | 2.46714206 | 12.90147109 | 0.963969335 | 0.37566839 | 0.709264768 | -4.625670214 |
| **A_51_P176474** | Kndc1 | 0.218010496 | -0.348448175 | 0.784469168 | 8.708498802 | 0.963961124 | 0.375672159 | 0.709264768 | -4.625672232 |
| **A_55_P2177968** | Snrpd2 | 0.529471869 | -0.846260964 | 1.905204702 | 13.86588983 | 0.963960269 | 0.375672551 | 0.709264768 | -4.625672442 |
| **A_55_P2186287** | Sub1 | 0.266103613 | -0.425392133 | 0.957599358 | 14.87426961 | 0.963854776 | 0.375720975 | 0.709264768 | -4.625698361 |
| **A_51_P422208** | Dpagt1 | -0.182359197 | -0.656239599 | 0.291521206 | 11.47991489 | -0.963849591 | 0.375723356 | 0.709264768 | -4.625699635 |
| **A_55_P2160128** | Pls1 | -0.130645208 | -0.470165749 | 0.208875333 | 7.953661731 | -0.963779702 | 0.375755439 | 0.709264768 | -4.625716806 |
| **A_55_P1973563** | Inava | 0.094925932 | -0.151768685 | 0.341620549 | 10.89425492 | 0.963774745 | 0.375757715 | 0.709264768 | -4.625718024 |
| **A_55_P2012799** | Rtkn | 0.686247061 | -1.09733605 | 2.469830172 | 11.44951342 | 0.963690309 | 0.37579648 | 0.709264768 | -4.625738768 |
| **A_51_P149325** | Psmd8 | -0.121982966 | -0.439031747 | 0.195065814 | 11.73067393 | -0.963659148 | 0.375810787 | 0.709264768 | -4.625746424 |
| **A_51_P278653** | Rprm | 0.620756704 | -0.992754225 | 2.234267634 | 9.590254471 | 0.963606793 | 0.375834826 | 0.709264768 | -4.625759286 |
| **A_52_P485376** | Ube2k | 0.150632934 | -0.240905101 | 0.54217097 | 12.5785934 | 0.963598962 | 0.375838421 | 0.709264768 | -4.62576121 |
| **A_55_P2143431** | Nfib | 1.076008171 | -1.721143671 | 3.873160014 | 12.89601391 | 0.963495903 | 0.375885744 | 0.709264768 | -4.625786527 |
| **A_51_P205106** | Mlf1 | 0.867900503 | -1.388345735 | 3.124146741 | 10.39340036 | 0.963460226 | 0.375902128 | 0.709264768 | -4.625795291 |
| **A_51_P204080** | Hk2 | -0.68699546 | -2.473122202 | 1.099131281 | 9.831579184 | -0.963367387 | 0.375944764 | 0.709264768 | -4.625818096 |
| **A_55_P2408355** | BB166591 | -0.321630441 | -1.157860269 | 0.514599387 | 9.745409094 | -0.963344977 | 0.375955056 | 0.709264768 | -4.6258236 |
| **A_51_P141264** | Zfp24 | 0.097551815 | -0.156113157 | 0.351216786 | 8.492592757 | 0.96321935 | 0.376012757 | 0.709264768 | -4.625854457 |
| **A_55_P2093705** | Meig1 | 0.838569203 | -1.341992377 | 3.019130783 | 10.25323147 | 0.963209813 | 0.376017138 | 0.709264768 | -4.6258568 |
| **A_55_P1981794** | Sgf29 | 0.094315906 | -0.150944085 | 0.339575897 | 8.308344164 | 0.963182486 | 0.37602969 | 0.709264768 | -4.625863512 |
| **A_52_P240152** | Snx27 | -0.270169291 | -0.972741402 | 0.432402819 | 10.11770644 | -0.963153315 | 0.37604309 | 0.709264768 | -4.625870676 |
| **A_55_P2051592** | Gfm2 | -0.152938253 | -0.550687038 | 0.244810533 | 9.250602161 | -0.963069452 | 0.376081615 | 0.709264768 | -4.625891273 |
| **A_55_P2141093** | Eya2 | 0.825904905 | -1.32217653 | 2.973986341 | 10.01295812 | 0.963007455 | 0.376110097 | 0.709264768 | -4.625906499 |
| **A_55_P2083919** | Robo2 | 0.625050576 | -1.000759594 | 2.250860747 | 9.65073642 | 0.962932111 | 0.376144714 | 0.709264768 | -4.625925002 |
| **A_55_P2007771** | Catsper2 | -0.184972524 | -0.666152865 | 0.296207817 | 7.842503405 | -0.962830175 | 0.376191552 | 0.709264768 | -4.625950035 |
| **A_55_P2082779** | Gm10240 | 0.334303633 | -0.535430769 | 1.204038035 | 15.66670559 | 0.962730639 | 0.376237292 | 0.709264768 | -4.625974477 |
| **A_55_P1970087** | Rpl35a | 0.219955128 | -0.35228853 | 0.792198787 | 16.10145744 | 0.96272766 | 0.376238661 | 0.709264768 | -4.625975209 |
| **A_55_P1975026** | Gm9104 | 0.173404928 | -0.277816305 | 0.62462616 | 15.78280689 | 0.962547682 | 0.376321377 | 0.709264768 | -4.626019401 |
| **A_55_P1966585** | NA | -0.187234909 | -0.674478622 | 0.300008805 | 16.06728719 | -0.962478249 | 0.376353292 | 0.709264768 | -4.626036449 |
| **A_55_P2159189** | Decr1 | -0.114472947 | -0.425170824 | 0.196224929 | 13.39237964 | -0.981357589 | 0.376426223 | 0.709264768 | -4.613410322 |
| **A_51_P163444** | Carhsp1 | -0.087587672 | -0.315579164 | 0.14040382 | 11.93986075 | -0.962220686 | 0.376471699 | 0.709264768 | -4.623888429 |
| **A_55_P2037883** | Ino80 | 0.09855775 | -0.157996681 | 0.355112181 | 9.019052429 | 0.962191698 | 0.376485028 | 0.709264768 | -4.626106798 |
| **A_66_P120484** | Mbtd1 | 0.358030431 | -0.57395751 | 1.290018372 | 11.3702448 | 0.962188128 | 0.376486669 | 0.709264768 | -4.626107674 |
| **A_51_P473940** | Slc22a15 | -0.158279511 | -0.57036983 | 0.253810807 | 8.153293091 | -0.962016752 | 0.376565474 | 0.709264768 | -4.626149743 |
| **A_52_P526372** | Zeb2 | 0.290447219 | -0.465862504 | 1.046756943 | 8.588771354 | 0.961873334 | 0.376631432 | 0.709264768 | -4.626184945 |
| **A_55_P2043122** | Arsg | -0.093502767 | -0.336981857 | 0.149976323 | 8.776561556 | -0.961862818 | 0.376636269 | 0.709264768 | -4.626187526 |
| **A_66_P113505** | Fgd4 | -0.24686695 | -0.889709646 | 0.395975746 | 10.29295024 | -0.961852732 | 0.376640908 | 0.709264768 | -4.626190002 |
| **A_55_P1984098** | Mst1 | -0.155205193 | -0.559371804 | 0.248961417 | 7.86905537 | -0.961825223 | 0.37665356 | 0.709264768 | -4.626196754 |
| **A_55_P2025855** | Hnrnpul1 | 0.165124291 | -0.26488856 | 0.595137142 | 9.45599794 | 0.961789156 | 0.37667015 | 0.709264768 | -4.626205606 |
| **A_55_P2129523** | NA | -0.189943633 | -0.684605857 | 0.304718592 | 7.844558745 | -0.961759178 | 0.376683939 | 0.709264768 | -4.626212964 |
| **A_51_P176365** | Gimap5 | 0.559179268 | -0.897072773 | 2.015431308 | 9.671581629 | 0.961755901 | 0.376685447 | 0.709264768 | -4.626213768 |
| **A_51_P295085** | Ogn | 1.11281601 | -1.785505582 | 4.011137602 | 12.03839211 | 0.96167237 | 0.376723871 | 0.709264768 | -4.626234269 |
| **A_51_P107808** | Ttc19 | -0.157303291 | -0.567013865 | 0.252407283 | 7.959396351 | -0.961636593 | 0.37674033 | 0.709264768 | -4.626243049 |
| **A_55_P1979818** | Srp54b | -0.137384884 | -0.495237384 | 0.220467616 | 8.273674012 | -0.961579603 | 0.376766549 | 0.709264768 | -4.626257035 |

| | | | | | | | | | |
|---|---|---|---|---|---|---|---|---|---|
| **A_51_P332136** | Zmat5 | 0.175559667 | -0.281812159 | 0.632931494 | 8.872212799 | 0.96140347 | 0.376847588 | 0.709264768 | -4.626300257 |
| **A_52_P15008** | Mapkapk2 | -0.139408408 | -0.502619387 | 0.223802571 | 10.87350842 | -0.961347379 | 0.376873399 | 0.709264768 | -4.626314021 |
| **A_51_P464738** | Slc2a1 | -1.230804523 | -4.438018566 | 1.97640952 | 13.00200298 | -0.961194959 | 0.376943544 | 0.709264768 | -4.626351421 |
| **A_55_P1968523** | Btbd1 | 0.454384451 | -0.729923333 | 1.638692235 | 11.77641442 | 0.960968657 | 0.377047708 | 0.709264768 | -4.626406943 |
| **A_51_P446978** | Cfdp1 | 0.213347474 | -0.342752448 | 0.769447397 | 13.11548518 | 0.960915099 | 0.377072364 | 0.709264768 | -4.626420082 |
| **A_51_P153995** | Gp9 | -0.229003434 | -0.825918456 | 0.367911588 | 8.808589056 | -0.960903624 | 0.377077646 | 0.709264768 | -4.626422897 |
| **A_55_P1969276** | Hhip | 1.290528521 | -2.073378354 | 4.654435395 | 12.17755814 | 0.960890696 | 0.377083598 | 0.709264768 | -4.626426069 |
| **A_51_P438527** | Cyb5r1 | -0.262942883 | -0.94845234 | 0.422566573 | 10.01316769 | -0.960723284 | 0.377160677 | 0.709264768 | -4.626467137 |
| **A_55_P2048022** | Gm6578 | 0.319261516 | -0.513101369 | 1.151624402 | 11.65225705 | 0.96069208 | 0.377175045 | 0.709264768 | -4.626474791 |
| **A_52_P515057** | Slc25a24 | -0.219054725 | -0.790182102 | 0.352072653 | 10.83177838 | -0.960660626 | 0.377189528 | 0.709264768 | -4.626482506 |
| **A_52_P181468** | Kmt5a | -0.522692916 | -1.885658226 | 0.840272395 | 9.336191844 | -0.960532706 | 0.377248436 | 0.709264768 | -4.626513883 |
| **A_66_P101646** | NA | 0.168376902 | -0.27068974 | 0.607443544 | 15.9541074 | 0.960511165 | 0.377258357 | 0.709264768 | -4.626519167 |
| **A_55_P2015403** | 9930111J | -0.293070409 | -1.057367039 | 0.471226221 | 8.755981435 | -0.960418177 | 0.377301184 | 0.709264768 | -4.626541974 |
| **A_55_P1965827** | Ubb | -0.164369314 | -0.59306405 | 0.264325422 | 16.48034955 | -0.9603354 | 0.377339312 | 0.709264768 | -4.626562275 |
| **A_51_P127464** | Ppp1r37 | -0.264643016 | -0.954865914 | 0.425579881 | 9.306844567 | -0.960332045 | 0.377340857 | 0.709264768 | -4.626563098 |
| **A_51_P432460** | Ppp1r14a | 0.907580752 | -1.459563317 | 3.274724821 | 12.24893245 | 0.960308851 | 0.377351541 | 0.709264768 | -4.626568786 |
| **A_66_P124715** | Ppard | -0.145451247 | -0.524822674 | 0.233920179 | 10.19120041 | -0.960291706 | 0.377359439 | 0.709264768 | -4.626572991 |
| **A_52_P615952** | Qk | 0.263005518 | -0.42303138 | 0.949042415 | 10.138915 | 0.960213331 | 0.377395543 | 0.709264768 | -4.626592212 |
| **A_52_P182118** | NA | 0.151781096 | -0.244156695 | 0.547718887 | 8.305838586 | 0.960154396 | 0.377422694 | 0.709264768 | -4.626606664 |
| **A_55_P2182273** | NA | -0.442209514 | -1.595814156 | 0.711395129 | 8.243575042 | -0.960110971 | 0.3774427 | 0.709264768 | -4.626617313 |
| **A_55_P2009507** | Ocrl | 0.278698016 | -0.44839696 | 1.005792993 | 11.15096659 | 0.960048291 | 0.377471579 | 0.709264768 | -4.626632684 |
| **A_55_P2180415** | Cd72 | -0.204460744 | -0.737883261 | 0.328961774 | 7.863083322 | -0.960039134 | 0.377475798 | 0.709264768 | -4.626634929 |
| **A_55_P2101392** | Tmem237 | 0.103482879 | -0.166511505 | 0.373477262 | 8.502926103 | 0.95998427 | 0.377501078 | 0.709264768 | -4.626648382 |
| **A_55_P2028571** | Sgsh | -0.081362051 | -0.293667779 | 0.130943677 | 9.687860805 | -0.959865807 | 0.377555666 | 0.709264768 | -4.626677429 |
| **A_66_P116807** | Selenot | 0.20233376 | -0.325636056 | 0.730303575 | 11.7784124 | 0.959863777 | 0.377556602 | 0.709264768 | -4.626677927 |
| **A_66_P137943** | Tctn2 | 0.24504856 | -0.394384177 | 0.884481298 | 10.80604567 | 0.959859425 | 0.377558608 | 0.709264768 | -4.626678994 |
| **A_55_P1989748** | NA | -0.210142724 | -0.758519515 | 0.338234067 | 8.127561356 | -0.95981081 | 0.377581012 | 0.709264768 | -4.626690914 |
| **A_51_P460774** | Smyd4 | 0.089945038 | -0.144820277 | 0.324710353 | 8.54035723 | 0.959607472 | 0.377674731 | 0.709264768 | -4.626740767 |
| **A_55_P2030060** | Kdm8 | 0.085965651 | -0.138415836 | 0.310347139 | 8.819921272 | 0.959595673 | 0.37768017 | 0.709264768 | -4.626743659 |
| **A_51_P146149** | Napsa | 1.181247342 | -1.901986725 | 4.264481408 | 11.32455509 | 0.959587747 | 0.377683823 | 0.709264768 | -4.626745602 |
| **A_55_P2047852** | Clcn3 | -0.224555427 | -0.810683304 | 0.36157245 | 10.7538988 | -0.959580741 | 0.377687053 | 0.709264768 | -4.62674732 |
| **A_55_P2030761** | NA | 0.151717416 | -0.24433321 | 0.547768042 | 13.30547676 | 0.959478126 | 0.377734357 | 0.709264768 | -4.626772476 |
| **A_55_P2056280** | Wdr89 | 0.1723697 | -0.277624829 | 0.622364229 | 13.45412621 | 0.95940955 | 0.377765972 | 0.709264768 | -4.626789287 |
| **A_55_P1974877** | Bcl2l1 | -0.567184302 | -2.047912727 | 0.913544123 | 14.14576039 | -0.959398719 | 0.377770965 | 0.709264768 | -4.626791942 |
| **A_52_P583155** | Ykt6 | -0.107022738 | -0.386485642 | 0.172440166 | 9.506674618 | -0.959184682 | 0.377869656 | 0.709264768 | -4.626844406 |
| **A_51_P153042** | Psg16 | -2.630954936 | -9.501083831 | 4.239173958 | 9.903323587 | -0.959177251 | 0.377873083 | 0.709264768 | -4.626846228 |
| **A_55_P2003517** | Zfp536 | 0.326957467 | -0.526853988 | 1.180768921 | 8.613829889 | 0.959134755 | 0.37789268 | 0.709264768 | -4.626856643 |
| **A_52_P110088** | Ndufaf3 | 0.181031308 | -0.291720089 | 0.653782705 | 9.987717276 | 0.95911617 | 0.37790125 | 0.709264768 | -4.626861198 |
| **A_55_P1960506** | Wdr83os | 0.220108134 | -0.354714985 | 0.794931254 | 12.05824264 | 0.959074206 | 0.377920604 | 0.709264768 | -4.626871484 |
| **A_55_P2170978** | Ankrd13d | -0.366606819 | -1.324033123 | 0.590819484 | 8.873114723 | -0.959059451 | 0.377927408 | 0.709264768 | -4.6268751 |
| **A_55_P2159994** | Atf1 | 0.103864111 | -0.167393153 | 0.375121375 | 10.79751501 | 0.959035041 | 0.377938666 | 0.709264768 | -4.626881082 |
| **A_51_P133369** | Scly | -0.172789677 | -0.624074312 | 0.278494958 | 9.810262726 | -0.958997758 | 0.377955862 | 0.709264768 | -4.62689022 |
| **A_55_P1963434** | Gpc4 | 0.180268569 | -0.29064672 | 0.651183858 | 8.68939034 | 0.958798986 | 0.378047549 | 0.709264768 | -4.626938933 |
| **A_55_P2015605** | Foxp4 | 0.156945611 | -0.253048077 | 0.566939299 | 11.88515195 | 0.95878747 | 0.378052861 | 0.709264768 | -4.626941755 |
| **A_55_P2083624** | Wbp2 | -0.145668245 | -0.526209175 | 0.234872685 | 12.8343907 | -0.958768724 | 0.378061509 | 0.709264768 | -4.626946348 |
| **A_55_P2132502** | Pkn1 | -0.143872437 | -0.51973559 | 0.231990716 | 11.27639341 | -0.958734139 | 0.378077464 | 0.709264768 | -4.626954824 |
| **A_55_P2146535** | Cnr2 | 0.33475002 | -0.539838547 | 1.209338586 | 10.57767322 | 0.958665639 | 0.378109067 | 0.709264768 | -4.626971609 |
| **A_55_P2163669** | NA | 0.392388803 | -0.632840081 | 1.417617688 | 15.53024188 | 0.95861919 | 0.378130498 | 0.709264768 | -4.626982991 |

| | | | | | | | | | |
|---|---|---|---|---|---|---|---|---|---|
| **A_55_P2043982** | Mtm1 | -0.088369094 | -0.319261389 | 0.142523202 | 8.341095811 | -0.958608564 | 0.378135401 | 0.709264768 | -4.626985594 |
| **A_55_P2053516** | Cacfd1 | -0.23287332 | -0.841366906 | 0.375620267 | 12.55719449 | -0.958548463 | 0.378163132 | 0.709264768 | -4.627000321 |
| **A_55_P2092963** | Traf2 | 0.145940024 | -0.235401077 | 0.527281125 | 12.44349671 | 0.958541992 | 0.378166118 | 0.709264768 | -4.627001906 |
| **A_55_P2003053** | Dct | -0.086066923 | -0.310961818 | 0.138827971 | 7.826139951 | -0.958532909 | 0.378170309 | 0.709264768 | -4.627004132 |
| **A_55_P1970642** | NA | 0.154142684 | -0.248686613 | 0.556971981 | 9.687077506 | 0.958411934 | 0.378226134 | 0.709264768 | -4.627033772 |
| **A_51_P159565** | Arhgef9 | 0.175475788 | -0.283122232 | 0.634073808 | 8.234904528 | 0.958374768 | 0.378243286 | 0.709264768 | -4.627042878 |
| **A_55_P1963184** | NA | 0.167080572 | -0.269585412 | 0.603746556 | 8.383145987 | 0.958356155 | 0.378251876 | 0.709264768 | -4.627047439 |
| **A_51_P261059** | Cpne2 | 0.180209953 | -0.290843192 | 0.651263099 | 8.498378817 | 0.958206718 | 0.378320847 | 0.709264768 | -4.627084049 |
| **A_66_P131324** | Platr21 | 0.355327414 | -0.573692398 | 1.284347227 | 8.754100268 | 0.9579748 | 0.378427908 | 0.709264768 | -4.627140861 |
| **A_51_P425080** | Hdgf | 0.448590809 | -0.724349366 | 1.621530983 | 13.49753523 | 0.957910326 | 0.378457675 | 0.709264768 | -4.627156654 |
| **A_55_P2076190** | NA | -0.22595208 | -0.816765945 | 0.364861786 | 15.18527133 | -0.957890817 | 0.378466683 | 0.709264768 | -4.627161433 |
| **A_51_P180001** | Ago2 | -0.304569863 | -1.100984287 | 0.491844561 | 10.7095301 | -0.957851441 | 0.378484864 | 0.709264768 | -4.627171077 |
| **A_55_P1986983** | Gne | -0.386209863 | -1.396344598 | 0.623924871 | 10.81149443 | -0.957622632 | 0.378590525 | 0.709264768 | -4.627227117 |
| **A_55_P2001048** | Rom1 | -0.378916058 | -1.370064408 | 0.612232291 | 9.677896255 | -0.957535082 | 0.37863096 | 0.709264768 | -4.627248558 |
| **A_55_P2069755** | Tymp | -0.457833313 | -1.655645255 | 0.739978628 | 9.259583036 | -0.957346375 | 0.378718128 | 0.709264768 | -4.627294769 |
| **A_55_P2179498** | Ppp1r8 | 0.141008565 | -0.227916733 | 0.509933862 | 9.584323334 | 0.957320594 | 0.378730038 | 0.709264768 | -4.627301082 |
| **A_51_P394788** | Actn4 | -0.207987408 | -0.752187154 | 0.336212337 | 13.67228986 | -0.95725811 | 0.378758905 | 0.709264768 | -4.627316382 |
| **A_51_P181365** | Atp5md | 0.251532744 | -0.406629275 | 0.909694762 | 14.55089579 | 0.957220646 | 0.378776214 | 0.709264768 | -4.627325556 |
| **A_55_P2061391** | Rhobtb3 | 0.228674859 | -0.369691246 | 0.827040964 | 8.368952839 | 0.957197998 | 0.378786677 | 0.709264768 | -4.627331101 |
| **A_55_P2158152** | NA | 0.267536738 | -0.43257238 | 0.967645856 | 15.82636765 | 0.957123632 | 0.378821038 | 0.709264768 | -4.62734931 |
| **A_55_P2086770** | Uevld | -0.086646302 | -0.313388962 | 0.140096359 | 8.536908155 | -0.957121645 | 0.378821956 | 0.709264768 | -4.627349796 |
| **A_55_P1988952** | NA | 0.481343353 | -0.77833436 | 1.741021067 | 15.20792214 | 0.957074795 | 0.378843604 | 0.709264768 | -4.627361267 |
| **A_51_P371174** | Bag3 | -0.290555581 | -1.05100339 | 0.469892227 | 11.69436558 | -0.956996072 | 0.378879982 | 0.709264768 | -4.627380541 |
| **A_55_P1991239** | NA | -0.273268471 | -0.988515747 | 0.441978804 | 8.732827786 | -0.956937691 | 0.378906962 | 0.709264768 | -4.627394834 |
| **A_55_P2115297** | Ap3s1 | 0.107079893 | -0.1731896 | 0.387349385 | 12.06187349 | 0.956935016 | 0.378908198 | 0.709264768 | -4.627395489 |
| **A_55_P2169396** | NA | 0.383447224 | -0.620185357 | 1.387079805 | 16.25728133 | 0.956932253 | 0.378909475 | 0.709264768 | -4.627396165 |
| **A_55_P1967539** | Hunk | 0.613976864 | -0.993345171 | 2.221298899 | 11.62639183 | 0.956752164 | 0.37899271 | 0.709264768 | -4.627440252 |
| **A_55_P2160381** | NA | -0.273095996 | -0.988106321 | 0.441914328 | 11.63708675 | -0.95665064 | 0.37903964 | 0.709264768 | -4.627465104 |
| **A_55_P2032740** | Iars2 | -0.23785678 | -0.86062752 | 0.38491396 | 8.320182986 | -0.95661609 | 0.379055612 | 0.709264768 | -4.627473561 |
| **A_51_P371876** | Zmynd8 | -0.205655555 | -0.744122437 | 0.332811328 | 10.24759653 | -0.956603116 | 0.379061609 | 0.709264768 | -4.627476737 |
| **A_51_P316523** | Irf2 | -0.107119007 | -0.387593563 | 0.173355548 | 9.742804017 | -0.95658467 | 0.379070137 | 0.709264768 | -4.627481252 |
| **A_51_P157083** | Gas1 | 0.209732237 | -0.33942046 | 0.758884935 | 8.890480225 | 0.956582461 | 0.379071158 | 0.709264768 | -4.627481793 |
| **A_55_P1979014** | NA | 0.121382149 | -0.19643932 | 0.439203618 | 8.322224837 | 0.956581408 | 0.379071645 | 0.709264768 | -4.62748205 |
| **A_52_P516097** | Leng9 | -0.181440387 | -0.656629916 | 0.293749141 | 9.443674976 | -0.956351291 | 0.379178041 | 0.709264768 | -4.627538373 |
| **A_55_P2164805** | Gm6749 | 0.584110943 | -0.945877944 | 2.114099831 | 14.27379321 | 0.956219095 | 0.379239174 | 0.709264768 | -4.627570726 |
| **A_55_P1961099** | Atg4c | -0.2800506 | -1.013604824 | 0.453503623 | 9.008716777 | -0.956212973 | 0.379242005 | 0.709264768 | -4.627572225 |
| **A_55_P2137376** | Dip2c | -0.097059536 | -0.351296694 | 0.157177623 | 9.649008902 | -0.956201741 | 0.379247199 | 0.709264768 | -4.627574973 |
| **A_52_P574720** | Oit3 | -0.365020306 | -1.321171266 | 0.591130653 | 8.442269957 | -0.956182752 | 0.379255981 | 0.709264768 | -4.62757962 |
| **A_55_P2153555** | Zkscan3 | 0.135313237 | -0.219154948 | 0.489781421 | 9.891615641 | 0.956122121 | 0.379284023 | 0.709264768 | -4.627594458 |
| **A_55_P1962149** | Dnah10 | -0.165994303 | -0.60084751 | 0.268858904 | 8.886299394 | -0.956094571 | 0.379296766 | 0.709264768 | -4.6276012 |
| **A_51_P116581** | Ccdc22 | 0.206435899 | -0.334477316 | 0.747349113 | 10.8346885 | 0.955890128 | 0.379391335 | 0.709264768 | -4.626207406 |
| **A_55_P2009943** | Spen | -0.169397462 | -0.613333396 | 0.274538473 | 11.0460441 | -0.955733796 | 0.379463662 | 0.709264768 | -4.627689477 |
| **A_55_P2077133** | Oxsr1 | -0.490200593 | -1.775176306 | 0.79477512 | 9.000792569 | -0.955496848 | 0.379573306 | 0.709264768 | -4.627747446 |
| **A_55_P2166414** | Uqcr10 | 0.148486109 | -0.240757846 | 0.537730065 | 14.52923117 | 0.95546393 | 0.379588541 | 0.709264768 | -4.627755499 |
| **A_55_P2098598** | Btg1 | 0.168062637 | -0.272508789 | 0.608634062 | 13.31644052 | 0.9554439 | 0.379597811 | 0.709264768 | -4.627760399 |
| **A_55_P2369474** | Nr1h4 | -0.162512127 | -0.588534666 | 0.263510413 | 7.799061779 | -0.955440271 | 0.37959949 | 0.709264768 | -4.627761287 |
| **A_55_P2018387** | Zkscan17 | 0.194307703 | -0.315103077 | 0.703718483 | 12.18020179 | 0.955371392 | 0.37963137 | 0.709264768 | -4.627778136 |
| **A_55_P2177112** | Prrg2 | -0.163545572 | -0.592325101 | 0.265233958 | 8.056789768 | -0.955333686 | 0.379648822 | 0.709264768 | -4.62778736 |

| | | | | | | | | | |
|---|---|---|---|---|---|---|---|---|---|
| **A_51_P399853** | Zfp704 | 0.373120617 | -0.605133949 | 1.351375182 | 9.684613824 | 0.955317446 | 0.379656339 | 0.709264768 | -4.627791332 |
| **A_55_P2041624** | Ceacam1 | -2.644001333 | -9.576378546 | 4.28837588 | 9.825964045 | -0.955278117 | 0.379674544 | 0.709264768 | -4.627800952 |
| **A_52_P180335** | Mrpl28 | 0.301368784 | -0.488902484 | 1.091640052 | 10.0117411 | 0.955151844 | 0.379732998 | 0.709264768 | -4.627831839 |
| **A_52_P574653** | Bid | 0.167304691 | -0.271426604 | 0.606035986 | 8.821600429 | 0.955124202 | 0.379745795 | 0.709264768 | -4.6278386 |
| **A_52_P334562** | Vdr | -0.15298693 | -0.554228035 | 0.248254176 | 8.176312138 | -0.954990949 | 0.379807489 | 0.709264768 | -4.62787119 |
| **A_51_P208580** | Nop53 | 0.205653353 | -0.333769056 | 0.745075761 | 9.720813836 | 0.954898378 | 0.379850353 | 0.709264768 | -4.62789383 |
| **A_55_P2266295** | Fzd3 | 0.111478468 | -0.180927705 | 0.403884641 | 8.49986334 | 0.954893202 | 0.37985275 | 0.709264768 | -4.627895096 |
| **A_66_P130887** | Pcdh18 | 0.700625528 | -1.137315507 | 2.538566564 | 11.52319622 | 0.954783112 | 0.379903731 | 0.709264768 | -4.627922018 |
| **A_55_P2053541** | NA | -0.125284791 | -0.454011918 | 0.203442337 | 15.10120966 | -0.954581808 | 0.379996967 | 0.709264768 | -4.627971243 |
| **A_51_P149699** | P3h3 | 0.462161895 | -0.750874169 | 1.675197959 | 10.06658732 | 0.954268895 | 0.38014193 | 0.709264768 | -4.628047749 |
| **A_52_P629451** | Gon4l | -0.286066652 | -1.037091855 | 0.464958552 | 8.683671518 | -0.954032273 | 0.38025158 | 0.709264768 | -4.628105594 |
| **A_52_P295067** | Snn | 0.577986772 | -0.939441291 | 2.095414835 | 10.68339772 | 0.95402583 | 0.380254566 | 0.709264768 | -4.628107169 |
| **A_55_P2003513** | Hsph1 | -0.331869236 | -1.203345186 | 0.539606714 | 9.387366709 | -0.953810139 | 0.380354538 | 0.709264768 | -4.628159891 |
| **A_55_P2062793** | Mfap2 | 1.120020015 | -1.821243542 | 4.061283572 | 13.32456293 | 0.95376677 | 0.380374642 | 0.709264768 | -4.628170491 |
| **A_66_P122729** | Zmat2 | 0.131436229 | -0.213767357 | 0.476639815 | 10.95875701 | 0.95365247 | 0.380427631 | 0.709264768 | -4.628198426 |
| **A_55_P1999022** | Zfp760 | 0.272597549 | -0.443419616 | 0.988614714 | 8.999609003 | 0.953561831 | 0.380469655 | 0.709264768 | -4.628220578 |
| **A_55_P2058794** | Cmip | -0.286719103 | -1.03983506 | 0.466396854 | 13.27143892 | -0.953553634 | 0.380473455 | 0.709264768 | -4.628222581 |
| **A_55_P2126470** | Rpl7 | 0.397195287 | -0.646591252 | 1.440981826 | 15.67603862 | 0.953109372 | 0.380679487 | 0.709264768 | -4.628331139 |
| **A_55_P1993594** | Dcp1b | 0.325686466 | -0.53027469 | 1.181647623 | 10.14670493 | 0.953006803 | 0.380727067 | 0.709264768 | -4.628356199 |
| **A_51_P299632** | Spef1 | 0.765522678 | -1.246460789 | 2.777506145 | 11.71208594 | 0.952980466 | 0.380739285 | 0.709264768 | -4.628362633 |
| **A_55_P2057390** | Smap1 | -0.248931001 | -0.903230312 | 0.405368311 | 12.02565768 | -0.952912156 | 0.380770977 | 0.709264768 | -4.628379322 |
| **A_51_P248122** | Bbc3 | -0.220672021 | -0.800711505 | 0.359367464 | 10.75890814 | -0.952884055 | 0.380784014 | 0.709264768 | -4.628386187 |
| **A_55_P1989405** | Elf1 | -0.134181527 | -0.486888403 | 0.218525348 | 10.66697376 | -0.952860122 | 0.380795119 | 0.709264768 | -4.628392034 |
| **A_66_P125634** | Wdr63 | 0.232467244 | -0.378627828 | 0.843562317 | 10.01517274 | 0.952803472 | 0.380821403 | 0.709264768 | -4.628405873 |
| **A_51_P193173** | Slc25a25 | -0.333664273 | -1.210871805 | 0.54354326 | 9.216651081 | -0.952703367 | 0.380867854 | 0.709264768 | -4.628430327 |
| **A_52_P555723** | Ywhag | -0.233044625 | -0.845754817 | 0.379665567 | 11.05752714 | -0.952652106 | 0.380891642 | 0.709264768 | -4.628442848 |
| **A_55_P2134122** | NA | -0.095865113 | -0.347942623 | 0.156212396 | 8.177242239 | -0.952525993 | 0.380950171 | 0.709264768 | -4.628473653 |
| **A_55_P2181114** | NA | 0.486757216 | -0.793289397 | 1.76680383 | 14.46437841 | 0.952438537 | 0.380990762 | 0.709264768 | -4.628495013 |
| **A_52_P270145** | Zfp329 | 0.27944483 | -0.455550865 | 1.014440525 | 9.980501511 | 0.952273348 | 0.381067443 | 0.709264768 | -4.628535358 |
| **A_55_P2069832** | Prpsap1 | 0.25725627 | -0.419460545 | 0.933973085 | 12.98156338 | 0.952158713 | 0.381120664 | 0.709264768 | -4.628563353 |
| **A_55_P2086203** | Kcnq1 | 0.414473451 | -0.675938388 | 1.50488529 | 9.461357416 | 0.952042869 | 0.381174452 | 0.709264768 | -4.628591642 |
| **A_52_P88808** | Vps37a | -0.333703158 | -1.211647328 | 0.544241012 | 8.887650158 | -0.952014937 | 0.381187422 | 0.709264768 | -4.628598462 |
| **A_55_P1957245** | NA | -0.225440161 | -0.818564553 | 0.367684231 | 10.61674434 | -0.951997587 | 0.381195479 | 0.709264768 | -4.628602699 |
| **A_55_P2243768** | D930030 | -0.155524228 | -0.564720911 | 0.253672455 | 8.434947868 | -0.951954728 | 0.381215381 | 0.709264768 | -4.628613165 |
| **A_55_P1955722** | Med15 | -0.200080196 | -0.726527013 | 0.32636662 | 10.86870443 | -0.951918894 | 0.381232022 | 0.709264768 | -4.628621914 |
| **A_55_P2131954** | Gm2590 | -0.122345624 | -0.444270074 | 0.199578826 | 8.683780378 | -0.951885748 | 0.381247415 | 0.709264768 | -4.628630008 |
| **A_51_P292030** | Nus1 | -0.58202927 | -2.113514568 | 0.949456027 | 9.858881147 | -0.951880302 | 0.381249945 | 0.709264768 | -4.628631337 |
| **A_51_P256304** | Mrpl37 | 0.128947027 | -0.210405893 | 0.468299947 | 8.567937458 | 0.951721957 | 0.381323487 | 0.709264768 | -4.628669999 |
| **A_51_P106799** | Pparg | -1.152703717 | -4.18665495 | 1.881247516 | 10.17529268 | -0.951610978 | 0.381375038 | 0.709264768 | -4.62503539 |
| **A_55_P2087805** | Hsd3b1 | -0.332723486 | -1.208500575 | 0.543053602 | 7.945467352 | -0.951568866 | 0.381394601 | 0.709264768 | -4.628707374 |
| **A_51_P155763** | NA | -0.188368167 | -0.684186199 | 0.307449866 | 8.539942082 | -0.951558601 | 0.381399369 | 0.709264768 | -4.62870988 |
| **A_55_P1955034** | Myo1c | -0.117982212 | -0.42853424 | 0.192569816 | 12.1905768 | -0.951551963 | 0.381402454 | 0.709264768 | -4.628711501 |
| **A_52_P413584** | NA | 0.109534493 | -0.17880468 | 0.397873666 | 8.768386883 | 0.951475478 | 0.381437986 | 0.709264768 | -4.628730172 |
| **A_55_P1995438** | NA | 0.389383676 | -0.635657712 | 1.414425064 | 15.72951956 | 0.951451566 | 0.381449096 | 0.709264768 | -4.628736009 |
| **A_55_P1967885** | Nr3c1 | 0.360943886 | -0.589318319 | 1.311206092 | 12.61717124 | 0.951363706 | 0.381489918 | 0.709264768 | -4.628757457 |
| **A_51_P377403** | Mettl14 | 0.136069232 | -0.222173035 | 0.494311498 | 8.686664427 | 0.951334956 | 0.381503277 | 0.709264768 | -4.628764475 |
| **A_51_P482633** | Rps6ka4 | 0.266396826 | -0.435047399 | 0.967841052 | 10.5230564 | 0.951231554 | 0.381551326 | 0.709264768 | -4.628789714 |
| **A_51_P140347** | Slc4a9 | -0.083587273 | -0.303693268 | 0.136518722 | 7.971490556 | -0.951171063 | 0.381579437 | 0.709264768 | -4.628804479 |

| | | | | | | | | | |
|---|---|---|---|---|---|---|---|---|---|
| **A_51_P107234** | Abcf1 | -0.415220727 | -1.508700638 | 0.678259185 | 12.25107339 | -0.95108331 | 0.38162022 | 0.709264768 | -4.628825897 |
| **A_55_P2150908** | Atp5c1 | 0.195082279 | -0.31867217 | 0.708836729 | 14.60415326 | 0.951070192 | 0.381626317 | 0.709264768 | -4.628829099 |
| **A_55_P1999823** | Zmynd19 | 0.10299963 | -0.16829458 | 0.374293841 | 8.254898642 | 0.950923291 | 0.381694598 | 0.709264768 | -4.62886495 |
| **A_55_P2072032** | NA | -0.086405308 | -0.314003009 | 0.141192392 | 16.14989518 | -0.950873841 | 0.381717585 | 0.709264768 | -4.628877018 |
| **A_51_P337771** | Pcdhb17 | 0.236701025 | -0.386802371 | 0.860204421 | 8.433171852 | 0.950849236 | 0.381729023 | 0.709264768 | -4.628883023 |
| **A_55_P2113774** | Yif1b | -0.137065806 | -0.498152454 | 0.224020842 | 10.05259259 | -0.950753735 | 0.381773421 | 0.709264768 | -4.628906328 |
| **A_55_P2058953** | Rpl13a | 0.187722692 | -0.306822803 | 0.682268187 | 16.39885402 | 0.950738038 | 0.381780719 | 0.709264768 | -4.628910158 |
| **A_55_P2085984** | Igf1 | 0.529812825 | -0.866078108 | 1.925703759 | 9.852473544 | 0.950651475 | 0.381820966 | 0.709264768 | -4.628931281 |
| **A_55_P2069306** | Ptpn18 | 0.809421158 | -1.323325441 | 2.942167757 | 12.58361898 | 0.950573363 | 0.381857287 | 0.709264768 | -4.628950341 |
| **A_55_P2353462** | 9530036C | 0.624924079 | -1.021769694 | 2.271617852 | 9.649004099 | 0.950527664 | 0.381878537 | 0.709264768 | -4.628961491 |
| **A_55_P2090406** | H13 | -0.447042653 | -1.625070293 | 0.730984986 | 9.621060226 | -0.950481848 | 0.381899843 | 0.709264768 | -4.62897267 |
| **A_51_P519592** | Srsf10 | 0.159781871 | -0.261281299 | 0.580845041 | 8.411913328 | 0.950452885 | 0.381913312 | 0.709264768 | -4.628979736 |
| **A_51_P234544** | Azin1 | -0.267446861 | -0.972375054 | 0.437481333 | 11.86639929 | -0.950261139 | 0.382002493 | 0.709264768 | -4.629026517 |
| **A_55_P2137286** | Gm10845 | -0.704561517 | -2.561768553 | 1.15264552 | 11.75483802 | -0.950186691 | 0.382037122 | 0.709264768 | -4.629044679 |
| **A_55_P1993813** | NA | 0.339691519 | -0.555756986 | 1.235140023 | 16.98366736 | 0.950154947 | 0.382051889 | 0.709264768 | -4.629052423 |
| **A_51_P245546** | Synrg | -0.085686936 | -0.311574726 | 0.140200853 | 9.056341849 | -0.950106315 | 0.382074513 | 0.709264768 | -4.629064286 |
| **A_55_P2067700** | Tas2r113 | -1.101787295 | -4.006767647 | 1.803193056 | 8.987171333 | -0.949959098 | 0.382143004 | 0.709264768 | -4.629100196 |
| **A_51_P425737** | Tbc1d7 | 0.098240209 | -0.160800861 | 0.357281279 | 8.785826258 | 0.949884934 | 0.382177512 | 0.709264768 | -4.629118286 |
| **A_51_P482503** | Tpd52l1 | 0.25251777 | -0.413325553 | 0.918361094 | 8.804327269 | 0.949883278 | 0.382178283 | 0.709264768 | -4.62911869 |
| **A_55_P1967231** | Stbd1 | 1.114236434 | -1.823883003 | 4.05235587 | 13.15042432 | 0.949857059 | 0.382190483 | 0.709264768 | -4.629125085 |
| **A_55_P1957957** | Dhx34 | -0.129923215 | -0.472539038 | 0.212692609 | 9.592830874 | -0.949794578 | 0.382219557 | 0.709264768 | -4.629140325 |
| **A_55_P2155827** | NA | 0.185072786 | -0.302992383 | 0.673137954 | 15.62400317 | 0.949762664 | 0.382234409 | 0.709264768 | -4.629148108 |
| **A_52_P116372** | Ppp1r15b | -0.338046311 | -1.229711908 | 0.553619287 | 8.403698246 | -0.949564643 | 0.382326569 | 0.709264768 | -4.629196402 |
| **A_52_P86517** | Cbwd1 | -0.331217357 | -1.204905471 | 0.542470757 | 9.472690972 | -0.949526315 | 0.382344409 | 0.709264768 | -4.629205749 |
| **A_55_P2011172** | NA | 0.349551347 | -0.572540526 | 1.271643219 | 14.51320743 | 0.949482864 | 0.382364635 | 0.709264768 | -4.629216345 |
| **A_55_P2035590** | NA | 0.449817392 | -0.73677896 | 1.636413745 | 14.38735962 | 0.949475101 | 0.382368249 | 0.709264768 | -4.629218238 |
| **A_51_P184284** | Dld | 0.099606579 | -0.16315095 | 0.362364107 | 11.10942948 | 0.94947427 | 0.382368635 | 0.709264768 | -4.62921844 |
| **A_55_P2000039** | Prf1 | 0.101271044 | -0.165903493 | 0.368445581 | 8.204294465 | 0.94938107 | 0.382412021 | 0.709264768 | -4.629241167 |
| **A_55_P2111302** | Cp | -0.47506214 | -1.728397092 | 0.778272813 | 9.623250523 | -0.949365871 | 0.382419097 | 0.709264768 | -4.629244874 |
| **A_51_P511482** | Ift57 | 0.338096732 | -0.554006379 | 1.230199844 | 10.74017935 | 0.949240512 | 0.38247746 | 0.709264768 | -4.62927544 |
| **A_55_P2167447** | Cnnm3 | -0.413779285 | -1.505660118 | 0.678101549 | 12.50731503 | -0.94916966 | 0.38251045 | 0.709264768 | -4.629292716 |
| **A_55_P1960411** | Lrrc23 | 1.167828403 | -1.913952644 | 4.249609449 | 11.1722377 | 0.949134149 | 0.382526985 | 0.709264768 | -4.629301374 |
| **A_55_P1992910** | Gyg | 1.032273627 | -1.691844513 | 3.756391766 | 14.76522587 | 0.949115879 | 0.382535493 | 0.709264768 | -4.629305828 |
| **A_55_P2102390** | Spag9 | -0.099372904 | -0.36163183 | 0.162886023 | 12.6032313 | -0.949047711 | 0.382567236 | 0.709264768 | -4.629322448 |
| **A_55_P2024591** | Atxn7l1 | -0.111491889 | -0.405785558 | 0.182801781 | 9.706049551 | -0.94888307 | 0.382643913 | 0.709264768 | -4.629362586 |
| **A_55_P1993955** | Pltp | 0.498665872 | -0.817642279 | 1.814974023 | 10.39320318 | 0.948860618 | 0.38265437 | 0.709264768 | -4.629368059 |
| **A_55_P2115507** | Egfl8 | 0.103659603 | -0.169966613 | 0.377285818 | 9.350562697 | 0.948860094 | 0.382654614 | 0.709264768 | -4.629368187 |
| **A_55_P2080542** | Rbbp7 | -0.399160467 | -1.452873883 | 0.654552949 | 11.72613702 | -0.94880149 | 0.382681911 | 0.709264768 | -4.629382473 |
| **A_66_P119643** | Atrx | 0.16760783 | -0.274893926 | 0.610109586 | 8.630462554 | 0.948701639 | 0.382728423 | 0.709264768 | -4.629406813 |
| **A_51_P185247** | Gdf10 | 0.476863567 | -0.782114738 | 1.735841871 | 9.070044854 | 0.948694193 | 0.382731892 | 0.709264768 | -4.629408628 |
| **A_52_P573255** | Cdc42ep1 | 0.660322695 | -1.083142262 | 2.403787652 | 13.43697744 | 0.948622338 | 0.382765366 | 0.709264768 | -4.629426143 |
| **A_51_P393598** | Atp6v1d | -0.207656655 | -0.75595147 | 0.34063816 | 12.57345684 | -0.948597685 | 0.382776852 | 0.709264768 | -4.629432152 |
| **A_55_P2120055** | Fgfr4 | 0.769878077 | -1.262932847 | 2.802689002 | 11.49064612 | 0.948582946 | 0.382783718 | 0.709264768 | -4.629435744 |
| **A_51_P209319** | Slc35b1 | -0.290509465 | -1.057586825 | 0.476567896 | 12.52834816 | -0.948574546 | 0.382787632 | 0.709264768 | -4.629437792 |
| **A_55_P1995449** | NA | 0.568599754 | -0.932835744 | 2.070035251 | 12.7197044 | 0.948528389 | 0.382809137 | 0.709264768 | -4.629449042 |
| **A_51_P125260** | Acaa2 | 0.844390432 | -1.385459241 | 3.074240104 | 13.73858277 | 0.948457911 | 0.382841974 | 0.709264768 | -4.629466219 |
| **A_51_P480982** | Glo1 | 0.348300602 | -0.571584017 | 1.26818522 | 11.35638433 | 0.948355601 | 0.382889648 | 0.709264768 | -4.629491154 |
| **A_52_P260555** | Vil1 | -0.232534934 | -0.846690308 | 0.381620439 | 7.944377244 | -0.948331764 | 0.382900756 | 0.709264768 | -4.629496964 |

| | | | | | | | | | |
|---|---|---|---|---|---|---|---|---|---|
| **A_51_P160744** | Ndufb3 | 0.362514972 | -0.594965975 | 1.319995918 | 13.60820828 | 0.948300877 | 0.38291515 | 0.709264768 | -4.629504491 |
| **A_55_P2058871** | Otud5 | -0.519685534 | -1.892637428 | 0.85326636 | 11.84785491 | -0.948059628 | 0.38302759 | 0.709264768 | -4.629563281 |
| **A_51_P189899** | Olfr1134 | -0.159191299 | -0.579765516 | 0.261382918 | 7.815788487 | -0.948040798 | 0.383036367 | 0.709264768 | -4.629567869 |
| **A_55_P2167530** | NA | 0.108880496 | -0.178790132 | 0.396551124 | 8.581216083 | 0.94799253 | 0.383058867 | 0.709264768 | -4.62957963 |
| **A_52_P59318** | Rdh10 | -0.181377879 | -0.660763744 | 0.298007985 | 8.440786136 | -0.947653217 | 0.383217066 | 0.709264768 | -4.629662301 |
| **A_52_P350111** | Cyth1 | -0.291693939 | -1.062756286 | 0.479368408 | 10.22356149 | -0.947519715 | 0.383279323 | 0.709264768 | -4.629694824 |
| **A_55_P2131579** | NA | 0.443856934 | -0.729444482 | 1.617158351 | 14.95090418 | 0.947509907 | 0.383283897 | 0.709264768 | -4.629697213 |
| **A_55_P2128225** | Rpl18 | 0.362480513 | -0.595739549 | 1.320700575 | 16.11340796 | 0.947479344 | 0.383298152 | 0.709264768 | -4.629704658 |
| **A_51_P285487** | Rnf113a1 | 0.137238137 | -0.22560355 | 0.500079824 | 8.860230273 | 0.947344595 | 0.383361001 | 0.709264768 | -4.629737481 |
| **A_55_P2420724** | Srek1 | 0.222595432 | -0.3659214 | 0.811112263 | 8.502163594 | 0.94734399 | 0.383361283 | 0.709264768 | -4.629737629 |
| **A_66_P121636** | Ablim3 | 1.086763055 | -1.786603815 | 3.960129925 | 11.33388212 | 0.947314384 | 0.383375093 | 0.709264768 | -4.62974484 |
| **A_55_P2069022** | Vstm4 | -0.119453821 | -0.435285979 | 0.196378336 | 8.380753093 | -0.947314197 | 0.38337518 | 0.709264768 | -4.629744885 |
| **A_51_P322989** | Ptrhd1 | 0.136480858 | -0.224389292 | 0.497351008 | 11.50766389 | 0.947264212 | 0.383398497 | 0.709264768 | -4.629757061 |
| **A_51_P205968** | Snx15 | 0.097885722 | -0.160951028 | 0.356722471 | 9.489356897 | 0.947204505 | 0.38342635 | 0.709264768 | -4.629771603 |
| **A_52_P78922** | Ythdc2 | 0.16147985 | -0.265517954 | 0.588477654 | 8.276411272 | 0.947202949 | 0.383427076 | 0.709264768 | -4.629771982 |
| **A_66_P116372** | Soat2 | -0.271410337 | -0.989144384 | 0.44632371 | 7.926544116 | -0.947137825 | 0.383457458 | 0.709264768 | -4.629787843 |
| **A_51_P433824** | Tpi1 | 0.24884625 | -0.409319266 | 0.907011765 | 13.85658916 | 0.946992025 | 0.383525484 | 0.709264768 | -4.629823352 |
| **A_55_P1967069** | Lpar6 | 0.352227598 | -0.579440651 | 1.283895847 | 11.11854673 | 0.94691813 | 0.383559966 | 0.709264768 | -4.629841347 |
| **A_52_P162509** | Ahcyl2 | -0.370104296 | -1.349099306 | 0.608890714 | 8.212162026 | -0.946877929 | 0.383578725 | 0.709264768 | -4.629851137 |
| **A_55_P1976934** | Unc13d | -0.12442539 | -0.453566698 | 0.204715919 | 8.552890883 | -0.946840798 | 0.383596053 | 0.709264768 | -4.629860179 |
| **A_55_P2047648** | Coq2 | -0.147186745 | -0.536538586 | 0.242165096 | 12.07968205 | -0.946840473 | 0.383596205 | 0.709264768 | -4.629860258 |
| **A_55_P2127020** | Nup54 | 0.266366897 | -0.438279129 | 0.971012924 | 10.61514481 | 0.946802923 | 0.383613729 | 0.709264768 | -4.629869402 |
| **A_55_P1993026** | NA | 0.381975474 | -0.628633123 | 1.392584071 | 15.6101 | 0.946679202 | 0.383671472 | 0.709264768 | -4.629899528 |
| **A_55_P2127425** | Sys1 | -0.169522026 | -0.61803548 | 0.278991428 | 11.05249005 | -0.94667521 | 0.383673335 | 0.709264768 | -4.6299005 |
| **A_55_P2027771** | Zfp260 | 0.357237196 | -0.58797623 | 1.302450621 | 12.38481983 | 0.946623181 | 0.38369762 | 0.709264768 | -4.629913169 |
| **A_55_P2021465** | Phldb2 | -0.130805348 | -0.47690316 | 0.215292464 | 11.89932707 | -0.946622871 | 0.383697765 | 0.709264768 | -4.629913244 |
| **A_55_P2070869** | Lcn2 | -0.339406004 | -1.237461389 | 0.558649381 | 9.059783849 | -0.946600532 | 0.383708192 | 0.709264768 | -4.629918683 |
| **A_52_P622434** | Nop16 | 0.099483626 | -0.163751586 | 0.362718838 | 9.22506371 | 0.946581405 | 0.38371712 | 0.709264768 | -4.62992334 |
| **A_51_P498212** | Ube2j1 | -0.348137529 | -1.269393051 | 0.573117993 | 8.491194705 | -0.946501014 | 0.383754648 | 0.709264768 | -4.629942914 |
| **A_55_P2071596** | Ppara | 0.167089623 | -0.275086464 | 0.609265711 | 8.193251652 | 0.946465032 | 0.383771445 | 0.709264768 | -4.629951674 |
| **A_55_P2038358** | Acot1 | 1.037100851 | -1.707495457 | 3.781697158 | 12.45860331 | 0.946439508 | 0.383783361 | 0.709264768 | -4.629957888 |
| **A_52_P268287** | Zfp507 | 0.208645591 | -0.343531257 | 0.760822439 | 9.769091684 | 0.946414453 | 0.383795058 | 0.709264768 | -4.629963988 |
| **A_55_P1958102** | NA | 0.240383473 | -0.395914317 | 0.876681262 | 16.04266759 | 0.946225257 | 0.383883396 | 0.709264768 | -4.630010047 |
| **A_55_P2019929** | Ythdf3 | -0.146868881 | -0.535720733 | 0.241982972 | 8.479095778 | -0.946010503 | 0.383983685 | 0.709264768 | -4.630062322 |
| **A_55_P2153570** | Zfp655 | -0.317346723 | -1.157569591 | 0.522876145 | 8.068167717 | -0.945997232 | 0.383989884 | 0.709264768 | -4.630065552 |
| **A_51_P145260** | Sall4 | 0.099351997 | -0.163715832 | 0.362419825 | 8.378747528 | 0.945930448 | 0.384021077 | 0.709264768 | -4.630081808 |
| **A_51_P288891** | Tbc1d15 | -0.104616726 | -0.381627997 | 0.172394546 | 10.21360535 | -0.94591917 | 0.384026345 | 0.709264768 | -4.630084553 |
| **A_55_P2064063** | Casp14 | -0.163219338 | -0.595423622 | 0.268984947 | 7.840463625 | -0.94587313 | 0.38404785 | 0.709264768 | -4.630095758 |
| **A_51_P365516** | Spink1 | -1.656616463 | -6.044015234 | 2.730782308 | 9.219846803 | -0.945725739 | 0.384116702 | 0.709264768 | -4.63013163 |
| **A_52_P675887** | NA | 0.30715051 | -0.506339421 | 1.120640441 | 15.53541554 | 0.945691324 | 0.38413278 | 0.709264768 | -4.630140005 |
| **A_51_P136729** | Idh3g | 0.359676976 | -0.592996448 | 1.312350399 | 13.50519262 | 0.945624974 | 0.384163779 | 0.709264768 | -4.630156152 |
| **A_51_P140819** | Wdr36 | 0.092824322 | -0.153039832 | 0.338688475 | 9.607456779 | 0.945620581 | 0.384165832 | 0.709264768 | -4.630157221 |
| **A_55_P1991016** | Basp1 | -0.605970444 | -2.211084599 | 0.999143712 | 9.458426594 | -0.945574742 | 0.38418725 | 0.709264768 | -4.630168376 |
| **A_55_P2414924** | Luc7l | -0.118869647 | -0.433766546 | 0.196027251 | 8.811871869 | -0.945481286 | 0.384230918 | 0.709264768 | -4.630191118 |
| **A_55_P2182016** | Faf2 | -0.523939274 | -1.911997305 | 0.864118757 | 10.78334822 | -0.945417585 | 0.384260685 | 0.709264768 | -4.630206618 |
| **A_51_P521052** | Ly6k | -0.583261634 | -2.128489423 | 0.961966155 | 8.157160959 | -0.945412295 | 0.384263158 | 0.709264768 | -4.630207905 |
| **A_51_P210395** | Glrx | -0.083840101 | -0.305960161 | 0.13827996 | 9.265244551 | -0.945397292 | 0.384270169 | 0.709264768 | -4.630211556 |
| **A_55_P1993820** | Kdm1a | -0.086553631 | -0.315893 | 0.142785738 | 12.21786792 | -0.945272499 | 0.384328491 | 0.709264768 | -4.63024192 |

| | | | | | | | | | |
|---|---|---|---|---|---|---|---|---|---|
| **A_55_P2053491** | Pdia6 | -0.474793863 | -1.732855425 | 0.783267698 | 15.70494697 | -0.945264939 | 0.384332025 | 0.709264768 | -4.63024376 |
| **A_52_P624362** | Aamp | -0.208927912 | -0.762532051 | 0.344676228 | 10.59688866 | -0.945251729 | 0.384338199 | 0.709264768 | -4.630246974 |
| **A_55_P2149049** | Rpl32 | 0.163144134 | -0.269145601 | 0.595433869 | 16.83388208 | 0.945250433 | 0.384338804 | 0.709264768 | -4.630247289 |
| **A_55_P2115467** | Tmem63b | -0.087772734 | -0.320352801 | 0.144807332 | 9.292311096 | -0.945229971 | 0.384348368 | 0.709264768 | -4.630252268 |
| **A_55_P2169514** | Fgf23 | -0.262807704 | -0.959203385 | 0.433587976 | 11.40254906 | -0.94521884 | 0.38435357 | 0.709264768 | -4.630254976 |
| **A_55_P2008755** | NA | 0.234387884 | -0.386798959 | 0.855574727 | 13.14106091 | 0.945068429 | 0.384423878 | 0.709264768 | -4.630291569 |
| **A_55_P2048800** | Cd82 | -0.458329251 | -1.673054346 | 0.756395844 | 8.429660363 | -0.945039403 | 0.384437447 | 0.709264768 | -4.630298631 |
| **A_52_P37498** | Dpcd | 0.288803574 | -0.476648193 | 1.05425534 | 10.35061283 | 0.945007116 | 0.384452541 | 0.709264768 | -4.630306485 |
| **A_52_P481182** | Stard5 | -0.116921156 | -0.426855327 | 0.193013015 | 7.983387994 | -0.944874189 | 0.384514688 | 0.709264768 | -4.630338822 |
| **A_55_P2004927** | Dnajc25 | -0.266082454 | -0.971418821 | 0.439253914 | 8.771444919 | -0.944866183 | 0.384518431 | 0.709264768 | -4.630340769 |
| **A_66_P102299** | Ascc1 | 0.180943819 | -0.298794304 | 0.660681943 | 8.85272807 | 0.944691193 | 0.384600257 | 0.709264768 | -4.630383334 |
| **A_52_P561936** | Sptssa | 0.251819379 | -0.415835486 | 0.919474243 | 11.43924898 | 0.944685992 | 0.384602689 | 0.709264768 | -4.630384599 |
| **A_55_P2145521** | Stk38l | -0.516256711 | -1.885229798 | 0.852716376 | 9.399792285 | -0.944541721 | 0.384670161 | 0.709264768 | -4.630419689 |
| **A_52_P209172** | Mrpl41 | 0.328815309 | -0.54314508 | 1.200775699 | 12.53225484 | 0.944507952 | 0.384685956 | 0.709264768 | -4.630427902 |
| **A_55_P2027819** | Rpl41 | 0.309502411 | -0.511350748 | 1.130355571 | 17.92005831 | 0.944384627 | 0.384743641 | 0.709264768 | -4.630457895 |
| **A_55_P2064632** | NA | 0.464552702 | -0.767635329 | 1.696740733 | 15.06337168 | 0.944296435 | 0.384784897 | 0.709264768 | -4.630479342 |
| **A_51_P381558** | Rasa4 | -0.14990283 | -0.547544803 | 0.247739143 | 8.207893414 | -0.944208615 | 0.384825982 | 0.709264768 | -4.630500697 |
| **A_51_P134452** | Lpar3 | -0.212074842 | -0.774679695 | 0.35053001 | 8.211475191 | -0.944139189 | 0.384858465 | 0.709264768 | -4.630517579 |
| **A_55_P1954302** | Esrrb | -0.28850169 | -1.053883708 | 0.476880328 | 8.245452357 | -0.944105336 | 0.384874305 | 0.709264768 | -4.630525811 |
| **A_52_P609109** | Pitpnc1 | 0.221743036 | -0.366596787 | 0.810082858 | 8.554054817 | 0.944000206 | 0.384923497 | 0.709264768 | -4.630551373 |
| **A_55_P2295951** | NA | -0.186438468 | -0.681108915 | 0.308231979 | 8.950472402 | -0.943995461 | 0.384925718 | 0.709264768 | -4.630552527 |
| **A_55_P2159432** | Fndc4 | -0.144496546 | -0.527892562 | 0.23889947 | 10.92492847 | -0.943974397 | 0.384935575 | 0.709264768 | -4.630557648 |
| **A_55_P1973941** | Slc7a5 | -0.783215781 | -2.861409887 | 1.294978325 | 9.10399923 | -0.943942774 | 0.384950374 | 0.709264768 | -4.630565337 |
| **A_55_P2400691** | Arhgap21 | -0.252278571 | -0.921748041 | 0.4171909 | 11.87286564 | -0.943843371 | 0.384996894 | 0.709264768 | -4.630589504 |
| **A_51_P411757** | Ankrd13b | 0.659870521 | -1.091507126 | 2.411248168 | 10.6868035 | 0.943689821 | 0.385068764 | 0.709264768 | -4.630626834 |
| **A_55_P2071970** | Nav2 | -0.168890583 | -0.617150635 | 0.279369469 | 8.337415052 | -0.943682157 | 0.385072352 | 0.709264768 | -4.630628697 |
| **A_51_P102789** | C1qc | 0.112918618 | -0.186785627 | 0.412622863 | 9.700638292 | 0.943676208 | 0.385075136 | 0.709264768 | -4.623891217 |
| **A_55_P2064492** | Pcmtd2 | 0.41347018 | -0.684005838 | 1.510946199 | 10.31134763 | 0.943625134 | 0.385099044 | 0.709264768 | -4.630642559 |
| **A_55_P2332731** | D130062J | -0.137124516 | -0.501103124 | 0.226854093 | 8.700867273 | -0.943603608 | 0.385109121 | 0.709264768 | -4.630647792 |
| **A_51_P206405** | Ptprz1 | 0.265472956 | -0.439248853 | 0.970194764 | 8.450645798 | 0.943523929 | 0.385146422 | 0.709264768 | -4.63066716 |
| **A_55_P2011126** | Mfap1a | 0.093650716 | -0.154998888 | 0.34230032 | 8.594212164 | 0.943351785 | 0.385227021 | 0.709264768 | -4.630709003 |
| **A_55_P1986386** | Senp3 | -0.13836205 | -0.505797687 | 0.229073588 | 9.109589416 | -0.943161484 | 0.385316136 | 0.709264768 | -4.630755254 |
| **A_51_P278266** | Gltpd2 | -0.228524116 | -0.835449107 | 0.378400874 | 7.899801892 | -0.943077457 | 0.38535549 | 0.709264768 | -4.630775675 |
| **A_55_P1960655** | NA | 0.221809157 | -0.367329152 | 0.810947466 | 11.60304612 | 0.943001868 | 0.385390895 | 0.709264768 | -4.630794044 |
| **A_55_P1969750** | Stim1 | -0.688155097 | -2.516026485 | 1.13971629 | 11.29032974 | -0.942955162 | 0.385412772 | 0.709264768 | -4.630805394 |
| **A_55_P1969660** | Pex26 | -0.120553387 | -0.44078298 | 0.199676206 | 11.43733816 | -0.942905779 | 0.385435904 | 0.709264768 | -4.630817394 |
| **A_55_P2109382** | Adora2a | 0.299669506 | -0.496366213 | 1.095705225 | 9.438492356 | 0.942888506 | 0.385443996 | 0.709264768 | -4.630821591 |
| **A_55_P2394308** | Fst | -0.596150398 | -2.180258508 | 0.987957713 | 8.593295896 | -0.942586827 | 0.385585338 | 0.709264768 | -4.630894891 |
| **A_51_P380005** | Galnt3 | 0.12500731 | -0.20722592 | 0.457240539 | 9.741066209 | 0.942416078 | 0.385665355 | 0.709264768 | -4.630936374 |
| **A_55_P2035344** | Tcf25 | -0.324774867 | -1.187967581 | 0.538417846 | 11.33332822 | -0.942377686 | 0.385683348 | 0.709264768 | -4.6309457 |
| **A_52_P343641** | Siah1a | 0.117861786 | -0.195398473 | 0.431122044 | 8.338635724 | 0.942362642 | 0.385690399 | 0.709264768 | -4.630949355 |
| **A_55_P2128283** | Nr2f2 | 0.520016346 | -0.862707054 | 1.902739747 | 11.30489178 | 0.941959062 | 0.385879588 | 0.709264768 | -4.631047384 |
| **A_52_P231635** | Oxsm | -0.178632435 | -0.653619328 | 0.296354459 | 7.996540587 | -0.941952573 | 0.385882631 | 0.709264768 | -4.63104896 |
| **A_55_P2121856** | Ier5l | 0.77272387 | -1.281994091 | 2.827441832 | 12.04295983 | 0.941938301 | 0.385889323 | 0.709264768 | -4.631052426 |
| **A_55_P2144975** | Ccdc90b | 0.102892802 | -0.170707813 | 0.376493417 | 9.130273815 | 0.941929221 | 0.38589358 | 0.709264768 | -4.631054631 |
| **A_55_P2069217** | Prr13 | -0.222368003 | -0.813700715 | 0.368964709 | 8.410417232 | -0.941869507 | 0.38592158 | 0.709264768 | -4.631069134 |
| **A_51_P284966** | Dolk | -0.246423693 | -0.901795457 | 0.408948071 | 10.54229156 | -0.941770495 | 0.385968009 | 0.709264768 | -4.631093179 |
| **A_51_P146655** | Ptgr2 | -0.157112581 | -0.574988605 | 0.260763442 | 9.865909289 | -0.941702788 | 0.385999762 | 0.709264768 | -4.631109621 |

| | | | | | | | | | |
|---|---|---|---|---|---|---|---|---|---|
| **A_55_P1967002** | NA | 0.218147244 | -0.362120681 | 0.798415168 | 9.149249426 | 0.941610974 | 0.386042823 | 0.709264768 | -4.631131916 |
| **A_51_P149621** | Stt3b | -0.290680095 | -1.063979359 | 0.482619169 | 9.49032099 | -0.941495053 | 0.386097195 | 0.709264768 | -4.631160064 |
| **A_55_P1982767** | Tmem9b | -0.21387704 | -0.782879907 | 0.355125828 | 11.94696361 | -0.941456058 | 0.386115487 | 0.709264768 | -4.631169532 |
| **A_51_P117739** | Vegfd | 0.961643934 | -1.596892034 | 3.520179903 | 11.96528518 | 0.941397662 | 0.386142882 | 0.709264768 | -4.627363313 |
| **A_55_P1960371** | NA | 0.235585696 | -0.391250829 | 0.862422222 | 15.75990721 | 0.941336654 | 0.386171503 | 0.709264768 | -4.631198523 |
| **A_55_P2083988** | Lmo2 | 0.618985169 | -1.028113251 | 2.266083588 | 10.78241956 | 0.94126311 | 0.386206006 | 0.709264768 | -4.631216378 |
| **A_55_P1987231** | Fuom | 0.197268991 | -0.327686401 | 0.722224383 | 8.838056661 | 0.94121051 | 0.386230686 | 0.709264768 | -4.631229149 |
| **A_55_P2002465** | Zfp747 | 0.344124143 | -0.571641047 | 1.259889333 | 11.36379539 | 0.941198785 | 0.386236188 | 0.709264768 | -4.631231995 |
| **A_55_P2005005** | Cdk5rap1 | -0.402236736 | -1.472698022 | 0.66822455 | 10.13673952 | -0.941154909 | 0.386256775 | 0.709264768 | -4.631242647 |
| **A_55_P2008056** | Cideb | -0.123377059 | -0.451732662 | 0.204978544 | 7.813046382 | -0.941109871 | 0.386277909 | 0.709264768 | -4.63125358 |
| **A_55_P2013601** | Ldb2 | 0.605473748 | -1.006005889 | 2.216953386 | 10.79255446 | 0.941067644 | 0.386297724 | 0.709264768 | -4.631263831 |
| **A_52_P454994** | Pi4ka | 0.083652422 | -0.139008459 | 0.306313303 | 12.39838376 | 0.940989858 | 0.386334228 | 0.709264768 | -4.631282714 |
| **A_55_P2027609** | Scgb3a2 | 1.806689199 | -3.003392431 | 6.616770828 | 12.98410823 | 0.940765423 | 0.386439567 | 0.709264768 | -4.63133719 |
| **A_55_P1963339** | Washc5 | -0.150489714 | -0.551180006 | 0.250200579 | 11.71722023 | -0.940693927 | 0.386473129 | 0.709264768 | -4.631354543 |
| **A_55_P2083841** | Atp13a1 | -0.285827067 | -1.046898111 | 0.475243977 | 12.23040355 | -0.940650952 | 0.386493304 | 0.709264768 | -4.631364973 |
| **A_52_P199614** | Wdfy4 | 0.370530214 | -0.61612086 | 1.357181287 | 9.137857535 | 0.940611707 | 0.386511728 | 0.709264768 | -4.631374498 |
| **A_55_P2135632** | Rps18 | 0.340645692 | -0.566468582 | 1.247759966 | 15.86160675 | 0.940570266 | 0.386531183 | 0.709264768 | -4.631384555 |
| **A_51_P437608** | Tulp3 | 0.309819642 | -0.515243006 | 1.134882291 | 9.404856778 | 0.940529382 | 0.386550378 | 0.709264768 | -4.631394477 |
| **A_55_P1977766** | Ttc14 | -0.112710157 | -0.412871724 | 0.18745141 | 8.569778573 | -0.94049895 | 0.386564667 | 0.709264768 | -4.631401862 |
| **A_55_P2405784** | BC023202 | 0.153851657 | -0.255892045 | 0.56359536 | 8.970322708 | 0.940459804 | 0.386583048 | 0.709264768 | -4.631411362 |
| **A_51_P229105** | Cetn3 | 0.372014774 | -0.619050127 | 1.363079676 | 11.50340616 | 0.940174435 | 0.386717059 | 0.709264768 | -4.631480608 |
| **A_51_P275101** | Chst11 | -0.578732526 | -2.120532791 | 0.963067738 | 8.58470549 | -0.940156431 | 0.386725514 | 0.709264768 | -4.631484976 |
| **A_55_P2166990** | Ppcs | -0.244560868 | -0.896235625 | 0.407113889 | 9.93133682 | -0.939953598 | 0.38682079 | 0.709264768 | -4.631534187 |
| **A_55_P2417212** | NA | 0.140858807 | -0.234507784 | 0.516225398 | 9.069810033 | 0.939893695 | 0.386848931 | 0.709264768 | -4.63154872 |
| **A_55_P2019934** | Ythdf3 | 0.307354882 | -0.511708016 | 1.126417779 | 10.45301421 | 0.93988173 | 0.386854553 | 0.709264768 | -4.631551622 |
| **A_55_P2050040** | Rpl36al | 0.198892541 | -0.331280958 | 0.729066039 | 14.41366091 | 0.939616914 | 0.386978979 | 0.709264768 | -4.631615861 |
| **A_55_P1997300** | Mtfr2 | 0.139643362 | -0.232603791 | 0.511890516 | 14.09241599 | 0.939591885 | 0.386990741 | 0.709264768 | -4.631621932 |
| **A_52_P573552** | Trib1 | 0.125733438 | -0.209438092 | 0.460904967 | 9.858662076 | 0.939580543 | 0.386996071 | 0.709264768 | -4.631624683 |
| **A_51_P229886** | Fpgt | 0.127241925 | -0.211981602 | 0.466465452 | 8.835939644 | 0.9394953 | 0.387036131 | 0.709264768 | -4.631645359 |
| **A_55_P2166049** | Vmn1r65 | -0.160323332 | -0.587750988 | 0.267104323 | 7.882671544 | -0.939473335 | 0.387046455 | 0.709264768 | -4.631650686 |
| **A_51_P183971** | Barhl1 | 0.148341959 | -0.247156063 | 0.543839982 | 8.836739545 | 0.939442148 | 0.387061112 | 0.709264768 | -4.63165825 |
| **A_55_P1968399** | Rdh7 | 0.082243931 | -0.137034398 | 0.301522261 | 7.727666654 | 0.939417167 | 0.387072853 | 0.709264768 | -4.631664309 |
| **A_55_P2000102** | Mettl21a | -0.288037911 | -1.056008457 | 0.479932635 | 9.825697393 | -0.939410551 | 0.387075963 | 0.709264768 | -4.631665914 |
| **A_55_P2140383** | Mis18a | 0.293163123 | -0.488542647 | 1.074868892 | 9.482455585 | 0.939326052 | 0.38711568 | 0.709264768 | -4.631686407 |
| **A_55_P2182222** | Rab3c | 0.24957737 | -0.415980054 | 0.915134794 | 11.9670378 | 0.939225806 | 0.387162804 | 0.709264768 | -4.631710718 |
| **A_51_P390276** | Cwc25 | -0.099456892 | -0.364683024 | 0.16576924 | 9.137910407 | -0.939223428 | 0.387163921 | 0.709264768 | -4.631711295 |
| **A_55_P2060330** | Cyren | 0.380381258 | -0.634124803 | 1.394887319 | 10.73157898 | 0.939106421 | 0.387218929 | 0.709264768 | -4.631739669 |
| **A_51_P259975** | Aspa | -0.089584934 | -0.328533034 | 0.149363166 | 7.909949635 | -0.939034917 | 0.387252548 | 0.709264768 | -4.631757008 |
| **A_51_P151433** | Ncbp2 | 0.116181843 | -0.193711146 | 0.426074832 | 9.380699751 | 0.939024353 | 0.387257515 | 0.709264768 | -4.63175957 |
| **A_51_P407004** | Prkrip1 | -0.089637171 | -0.328736526 | 0.149462184 | 10.4200853 | -0.938988088 | 0.387274566 | 0.709264768 | -4.631768363 |
| **A_51_P314161** | Qtrt1 | 0.148865182 | -0.248257691 | 0.545988056 | 8.444466008 | 0.938898351 | 0.387316763 | 0.709264768 | -4.631790122 |
| **A_55_P1975989** | Wdr63 | 0.594096365 | -0.990866987 | 2.179059718 | 9.47773241 | 0.938832285 | 0.387347831 | 0.709264768 | -4.63180614 |
| **A_55_P2040777** | Lipo3 | 0.327324227 | -0.545952847 | 1.200601302 | 9.682974445 | 0.938807256 | 0.387359602 | 0.709264768 | -4.631812209 |
| **A_55_P2030721** | Ankle1 | 0.145169201 | -0.242170843 | 0.532509245 | 8.111487319 | 0.93871214 | 0.387404335 | 0.709264768 | -4.631835269 |
| **A_55_P1952547** | Sh3rf1 | 0.236876736 | -0.395188683 | 0.868942156 | 8.815280687 | 0.938665219 | 0.387426404 | 0.709264768 | -4.631846645 |
| **A_55_P1994229** | Hrc | 0.53545497 | -0.893368314 | 1.964278253 | 9.55476188 | 0.938630756 | 0.387442614 | 0.709264768 | -4.631855 |
| **A_51_P509999** | Prl8a9 | -2.49619684 | -9.157273416 | 4.164879736 | 9.786348616 | -0.938609074 | 0.387452813 | 0.709264768 | -4.631860256 |
| **A_51_P337543** | Glce | -0.214715114 | -0.787783927 | 0.3583537 | 10.26789729 | -0.938439298 | 0.387532677 | 0.709264768 | -4.631901412 |

| | | | | | | | | | |
|---|---|---|---|---|---|---|---|---|---|
| **A_52_P524895** | Eif3a | 0.210724186 | -0.351754978 | 0.773203349 | 11.21105031 | 0.93833581 | 0.387581366 | 0.709264768 | -4.631926498 |
| **A_66_P120472** | Pglyrp2 | 0.079101109 | -0.132041164 | 0.290243381 | 7.661530249 | 0.938334587 | 0.387581941 | 0.709264768 | -4.631926794 |
| **A_55_P2157360** | Tagap1 | -0.087882609 | -0.322497097 | 0.146731879 | 8.425190667 | -0.938206553 | 0.387642185 | 0.709264768 | -4.631957827 |
| **A_55_P2048684** | Rpl27 | 0.399979108 | -0.667890549 | 1.467848764 | 14.49118132 | 0.938143787 | 0.38767172 | 0.709264768 | -4.63197304 |
| **A_55_P2184180** | Ndufv3 | -0.129226283 | -0.474254479 | 0.215801914 | 14.45760021 | -0.938094544 | 0.387694894 | 0.709264768 | -4.631984974 |
| **A_52_P354744** | Slc2a3 | -0.489776579 | -1.797970188 | 0.81841703 | 8.727283642 | -0.937726817 | 0.387867979 | 0.709264768 | -4.632074087 |
| **A_51_P443618** | Mettl26 | -0.125380061 | -0.460280871 | 0.209520749 | 11.22498228 | -0.937697214 | 0.387881915 | 0.709264768 | -4.63208126 |
| **A_55_P2036180** | Mcf2l | -0.163253811 | -0.599393054 | 0.272885432 | 10.75353397 | -0.937537197 | 0.387957254 | 0.709264768 | -4.632120031 |
| **A_51_P389864** | B3gnt9 | 0.373041662 | -0.623683234 | 1.369766558 | 9.515827004 | 0.937416033 | 0.388014308 | 0.709264768 | -4.632149386 |
| **A_52_P658034** | Tspyl1 | 0.22912795 | -0.383187723 | 0.841443623 | 11.92561453 | 0.9372448 | 0.388094951 | 0.709264768 | -4.632190868 |
| **A_55_P2147831** | Gimap4 | 0.721586756 | -1.206850554 | 2.650024067 | 10.4604046 | 0.937202361 | 0.388114939 | 0.709264768 | -4.632201149 |
| **A_66_P103792** | AI606181 | 0.169783105 | -0.283961867 | 0.623528076 | 9.976767901 | 0.937201536 | 0.388115328 | 0.709264768 | -4.632201349 |
| **A_52_P78023** | Limd2 | 0.307360398 | -0.514072573 | 1.12879337 | 14.31363802 | 0.937186717 | 0.388122307 | 0.709264768 | -4.632204938 |
| **A_51_P326393** | Gapvd1 | 0.085309638 | -0.142695344 | 0.31331462 | 10.38031294 | 0.937139204 | 0.388144687 | 0.709264768 | -4.632216448 |
| **A_55_P1988108** | Mrc1 | 0.288690513 | -0.482900199 | 1.060281225 | 9.536798794 | 0.937121424 | 0.388153062 | 0.709264768 | -4.632220754 |
| **A_66_P115004** | Gabrp | -0.254174741 | -0.933645504 | 0.425296022 | 8.052727245 | -0.93694039 | 0.388238345 | 0.709264768 | -4.632264604 |
| **A_55_P2017491** | Rbm43 | 0.454792436 | -0.761182731 | 1.670767602 | 10.43395723 | 0.936782724 | 0.388312631 | 0.709264768 | -4.632302789 |
| **A_55_P2005753** | Manbal | -0.253382255 | -0.93087076 | 0.424106251 | 10.85867596 | -0.93675196 | 0.388327126 | 0.709264768 | -4.63231024 |
| **A_66_P140613** | NA | -0.174191341 | -0.639956544 | 0.291573861 | 8.842192875 | -0.936720167 | 0.388342108 | 0.709264768 | -4.632317939 |
| **A_52_P18922** | Chid1 | -0.235086051 | -0.863688875 | 0.393516773 | 8.198421087 | -0.936700773 | 0.388351247 | 0.709264768 | -4.632322636 |
| **A_55_P2052613** | NA | 0.837478159 | -1.40213665 | 3.077092967 | 13.17683228 | 0.936592131 | 0.388402445 | 0.709264768 | -4.632348945 |
| **A_51_P381218** | Gulp1 | 0.654980357 | -1.097033003 | 2.406993717 | 10.79940473 | 0.936356452 | 0.388513529 | 0.709264768 | -4.632406013 |
| **A_55_P1954897** | NA | 0.438532962 | -0.734541982 | 1.611607905 | 15.0967695 | 0.936325452 | 0.388528142 | 0.709264768 | -4.632413519 |
| **A_51_P518586** | Emg1 | 0.26621411 | -0.44599685 | 0.97842507 | 10.02000558 | 0.936208894 | 0.388583091 | 0.709264768 | -4.632441739 |
| **A_52_P191975** | Rtn3 | -0.316160435 | -1.162076023 | 0.529755154 | 9.855085102 | -0.93611852 | 0.3886257 | 0.709264768 | -4.632463619 |
| **A_55_P2183473** | Cldn3 | 0.869728939 | -1.457396569 | 3.196854447 | 13.48513006 | 0.936083219 | 0.388642344 | 0.709264768 | -4.632472165 |
| **A_52_P209484** | Tmem88 | 0.491130711 | -0.822998159 | 1.805259581 | 11.77653329 | 0.936072488 | 0.388647404 | 0.709264768 | -4.632474762 |
| **A_52_P407854** | Dda1 | -0.242242407 | -0.890421945 | 0.405937132 | 8.394829131 | -0.936063262 | 0.388651754 | 0.709264768 | -4.632476996 |
| **A_55_P2046092** | NA | -0.382576829 | -1.406414999 | 0.64126134 | 7.937452953 | -0.935917769 | 0.388720362 | 0.709264768 | -4.632512216 |
| **A_51_P433430** | Phc1 | 0.126671213 | -0.212388928 | 0.465731354 | 8.999085612 | 0.93573212 | 0.388807919 | 0.709264768 | -4.632557154 |
| **A_55_P2143306** | Rest | -0.200920082 | -0.738810934 | 0.33697077 | 10.47615412 | -0.935576993 | 0.388881093 | 0.709264768 | -4.632594699 |
| **A_55_P2016262** | Isl2 | -0.277048862 | -1.018755239 | 0.464657515 | 7.870435175 | -0.935566588 | 0.388886001 | 0.709264768 | -4.632597218 |
| **A_52_P5213** | Chd1l | 0.103125014 | -0.172960502 | 0.37921053 | 9.223128131 | 0.935558057 | 0.388890026 | 0.709264768 | -4.632599282 |
| **A_65_P01065** | Phf14 | 0.273353588 | -0.458475705 | 1.00518288 | 11.27002123 | 0.935546407 | 0.388895521 | 0.709264768 | -4.632602102 |
| **A_52_P276794** | Tox4 | -0.141606455 | -0.520758048 | 0.237545139 | 9.166770216 | -0.93544985 | 0.388941075 | 0.709264768 | -4.632625469 |
| **A_55_P2023488** | NA | 0.171768726 | -0.288201573 | 0.631739024 | 11.02051152 | 0.935329539 | 0.38899784 | 0.709264768 | -4.632654584 |
| **A_55_P1963084** | Rps13 | 0.093213341 | -0.156411611 | 0.342838294 | 16.16939095 | 0.935277368 | 0.389022458 | 0.709264768 | -4.632667209 |
| **A_55_P2126063** | Mrps21 | 0.237243638 | -0.398170697 | 0.872657973 | 13.49743544 | 0.935164284 | 0.389075822 | 0.709264768 | -4.632694572 |
| **A_55_P2082609** | NA | 0.258811098 | -0.434388754 | 0.952010949 | 11.85382909 | 0.935136006 | 0.389089167 | 0.709264768 | -4.632701414 |
| **A_55_P1958678** | Brd2 | -0.159759951 | -0.587675441 | 0.268155539 | 12.93138222 | -0.935104737 | 0.389103924 | 0.709264768 | -4.63270898 |
| **A_51_P494541** | Abhd17b | 0.414122403 | -0.695119699 | 1.523364506 | 11.00875168 | 0.935088522 | 0.389111577 | 0.709264768 | -4.632712904 |
| **A_51_P481693** | Ero1l | -0.202505884 | -0.74496775 | 0.339955981 | 8.105201554 | -0.935015435 | 0.389146072 | 0.709264768 | -4.632730587 |
| **A_51_P482043** | Epm2aip1 | 0.282119903 | -0.473679645 | 1.037919451 | 10.15625002 | 0.934926442 | 0.389188078 | 0.709264768 | -4.632752118 |
| **A_55_P2080774** | Qdpr | 0.21005374 | -0.352733883 | 0.772841364 | 9.898301451 | 0.934837719 | 0.389229959 | 0.709264768 | -4.632773582 |
| **A_55_P2066269** | Aak1 | -0.142628277 | -0.524778587 | 0.239522033 | 8.192974196 | -0.934806591 | 0.389244654 | 0.709264768 | -4.632781113 |
| **A_51_P191782** | Olfml3 | 0.894099739 | -1.501612035 | 3.289811513 | 10.93834759 | 0.934763501 | 0.389264997 | 0.709264768 | -4.632791537 |
| **A_55_P2158873** | Ppid | 0.43195098 | -0.725746275 | 1.589648235 | 10.66791634 | 0.934522593 | 0.389378744 | 0.709264768 | -4.632849812 |
| **A_55_P2132147** | NA | -0.154469563 | -0.568502234 | 0.259563107 | 15.09797936 | -0.934455561 | 0.389410398 | 0.709264768 | -4.632866025 |

| | | | | | | | | | |
|---|---|---|---|---|---|---|---|---|---|
| **A_55_P2024933** | Zfp706 | -0.311128829 | -1.145085841 | 0.522828184 | 13.09791766 | -0.934430348 | 0.389422305 | 0.709264768 | -4.632872123 |
| **A_55_P2078325** | Smco3 | 0.257241572 | -0.432297397 | 0.946780541 | 8.473838182 | 0.93439969 | 0.389436783 | 0.709264768 | -4.632879538 |
| **A_55_P1954277** | Emid1 | 0.683194441 | -1.148122807 | 2.51451169 | 11.24367833 | 0.934396244 | 0.389438411 | 0.709264768 | -4.632880372 |
| **A_51_P142187** | Abhd11 | -0.121128628 | -0.445849983 | 0.203592728 | 10.56543324 | -0.934299876 | 0.389483925 | 0.709264768 | -4.632903679 |
| **A_55_P2084571** | NA | 0.307450738 | -0.516798895 | 1.131700372 | 16.43341768 | 0.934258641 | 0.389503401 | 0.709264768 | -4.632913651 |
| **A_55_P2024220** | NA | -0.191458034 | -0.704771052 | 0.321854985 | 8.130981847 | -0.934203868 | 0.389529273 | 0.709264768 | -4.632926898 |
| **A_55_P2019751** | Paox | 0.311354758 | -0.523574623 | 1.146284139 | 11.07261459 | 0.934019855 | 0.3896162 | 0.709264768 | -4.632971397 |
| **A_55_P2078675** | NA | -0.142089031 | -0.523141294 | 0.238963232 | 8.159556376 | -0.933955864 | 0.389646433 | 0.709264768 | -4.63298687 |
| **A_55_P2250246** | Taok3 | 0.11402495 | -0.191786186 | 0.419836085 | 8.272357611 | 0.933892611 | 0.389676318 | 0.709264768 | -4.633002165 |
| **A_55_P2056393** | Stk11ip | -0.285693369 | -1.052234441 | 0.480847703 | 9.942758257 | -0.93350162 | 0.389861094 | 0.709264768 | -4.633096694 |
| **A_52_P582261** | Ikzf5 | 0.299020195 | -0.503295815 | 1.101336205 | 9.711518629 | 0.933480838 | 0.389870918 | 0.709264768 | -4.633101718 |
| **A_55_P2162592** | Lman2 | -0.704847185 | -2.596533388 | 1.186839018 | 13.80477965 | -0.933246176 | 0.389981851 | 0.709264768 | -4.633158441 |
| **A_55_P2153904** | Gm6132 | 0.584165532 | -0.983782816 | 2.15211388 | 13.25437148 | 0.933156578 | 0.390024213 | 0.709264768 | -4.633180097 |
| **A_51_P193490** | Fbxo21 | -0.49440652 | -1.821445043 | 0.832632002 | 11.73815003 | -0.933149022 | 0.390027786 | 0.709264768 | -4.633181923 |
| **A_51_P365399** | Smg5 | -0.271038999 | -0.99870038 | 0.456622381 | 10.64229664 | -0.932938047 | 0.390127551 | 0.709264768 | -4.633232912 |
| **A_52_P206925** | Parp16 | -0.329495255 | -1.214134939 | 0.55514443 | 8.289418282 | -0.932895702 | 0.390147578 | 0.709264768 | -4.633243145 |
| **A_55_P1978615** | Rpl17 | 0.405790051 | -0.683706634 | 1.495286736 | 15.88746198 | 0.932880097 | 0.390154958 | 0.709264768 | -4.633246916 |
| **A_55_P2088530** | Htt | -0.30025729 | -1.106523191 | 0.506008612 | 10.5493695 | -0.932750762 | 0.390216131 | 0.709264768 | -4.63327817 |
| **A_51_P203710** | Taf8 | 0.283517495 | -0.510918009 | 1.077952998 | 10.92780094 | 0.950569605 | 0.390265572 | 0.709264768 | -4.619642167 |
| **A_55_P1996199** | NA | 0.419723853 | -0.707711059 | 1.547158766 | 15.35135309 | 0.932443483 | 0.390361499 | 0.709264768 | -4.633352415 |
| **A_51_P195215** | Fam149a | 0.239971547 | -0.404714745 | 0.88465784 | 9.888635539 | 0.93231283 | 0.390423321 | 0.709264768 | -4.633383979 |
| **A_55_P2118391** | Lrp6 | 0.093505358 | -0.157741612 | 0.344752327 | 12.31653653 | 0.932150441 | 0.390500171 | 0.709264768 | -4.633423207 |
| **A_52_P282035** | Rnf5 | 0.195969561 | -0.330599349 | 0.722538471 | 10.58084281 | 0.932145593 | 0.390502465 | 0.709264768 | -4.633424378 |
| **A_51_P350666** | Peli2 | 0.509386558 | -0.859335703 | 1.87810882 | 9.694134481 | 0.932142898 | 0.39050374 | 0.709264768 | -4.633425029 |
| **A_55_P2294184** | Cemip2 | 0.181397223 | -0.306072443 | 0.668866889 | 9.961532057 | 0.932037493 | 0.390553629 | 0.709264768 | -4.63345049 |
| **A_55_P2091861** | Hipk2 | -0.086567122 | -0.319205425 | 0.146071182 | 8.263445409 | -0.932013282 | 0.390565089 | 0.709264768 | -4.633456338 |
| **A_52_P154501** | Ube2q2 | -0.580911793 | -2.142123875 | 0.980300289 | 8.76574595 | -0.931962924 | 0.390588927 | 0.709264768 | -4.633468502 |
| **A_55_P1984152** | 1600029C | 0.384855404 | -0.649637416 | 1.419348224 | 15.77056499 | 0.931795168 | 0.390668343 | 0.709264768 | -4.633509018 |
| **A_55_P2169059** | Strada | -0.282518616 | -1.0419327 | 0.476895468 | 10.67594269 | -0.931791551 | 0.390670055 | 0.709264768 | -4.633509892 |
| **A_52_P533083** | Dnajc16 | -0.316602256 | -1.167682043 | 0.534477531 | 8.715942324 | -0.931738573 | 0.390695138 | 0.709264768 | -4.633522687 |
| **A_52_P440729** | Atf7ip | 0.317481101 | -0.53596398 | 1.170926182 | 10.37407716 | 0.931735501 | 0.390696592 | 0.709264768 | -4.633523429 |
| **A_51_P189985** | Ppan | 0.22334792 | -0.377078152 | 0.823773991 | 11.18930292 | 0.931692749 | 0.390716834 | 0.709264768 | -4.633533753 |
| **A_55_P1960683** | F2 | -0.890773813 | -3.285588421 | 1.504040795 | 8.767441066 | -0.931635198 | 0.390744085 | 0.709264768 | -4.633547651 |
| **A_55_P2168124** | NA | 0.534537576 | -0.902556271 | 1.971631423 | 14.46991419 | 0.93162998 | 0.390746556 | 0.709264768 | -4.633548912 |
| **A_51_P162671** | Folr2 | -0.324551253 | -1.197174652 | 0.548072146 | 8.152890168 | -0.931551311 | 0.390783808 | 0.709264768 | -4.633567909 |
| **A_55_P2029726** | NA | 0.233250886 | -0.39395192 | 0.860453692 | 15.48846366 | 0.931463099 | 0.390825582 | 0.709264768 | -4.633589209 |
| **A_55_P1996509** | Rpl29 | 0.416277889 | -0.703212717 | 1.535768496 | 17.44917823 | 0.931350679 | 0.390878826 | 0.709264768 | -4.633616354 |
| **A_55_P2144641** | Sdk1 | -0.231594212 | -0.854484335 | 0.391295912 | 8.912678991 | -0.931250677 | 0.390926194 | 0.709264768 | -4.633640499 |
| **A_52_P494372** | Zxdc | -0.357008128 | -1.317213454 | 0.603197198 | 8.358976126 | -0.931245825 | 0.390928492 | 0.709264768 | -4.633641671 |
| **A_55_P2423646** | Nf2 | -0.33800351 | -1.247118245 | 0.571111224 | 8.939531207 | -0.931221212 | 0.390940151 | 0.709264768 | -4.633647613 |
| **A_51_P197259** | Cct7 | 0.214552313 | -0.362642649 | 0.791747274 | 12.90542111 | 0.931024297 | 0.391033438 | 0.709264768 | -4.633695153 |
| **A_55_P2045741** | Fyb | 0.096801107 | -0.163622493 | 0.357224708 | 7.833769121 | 0.931001396 | 0.391044289 | 0.709264768 | -4.633700681 |
| **A_55_P1985351** | Slc35f2 | -0.773897641 | -2.855944994 | 1.308149712 | 8.8271986 | -0.930986248 | 0.391051466 | 0.709264768 | -4.633704338 |
| **A_55_P2173373** | Zfp62 | 0.091025822 | -0.15386721 | 0.335918854 | 8.729080497 | 0.930976086 | 0.391056281 | 0.709264768 | -4.633706791 |
| **A_55_P2019650** | NA | 0.140428424 | -0.253798579 | 0.534655427 | 10.66771502 | 0.948792752 | 0.391077015 | 0.709264768 | -4.619999464 |
| **A_55_P1994027** | Adra2b | -1.216683582 | -4.4903738 | 2.057006636 | 9.10279254 | -0.930872955 | 0.391105147 | 0.709264768 | -4.633731686 |
| **A_52_P205797** | Vgll4 | 0.348326846 | -0.588923408 | 1.2855771 | 8.833154449 | 0.930854336 | 0.39111397 | 0.709264768 | -4.63373618 |
| **A_52_P58181** | Suclg2 | 0.283491769 | -0.479337846 | 1.046321384 | 9.754055531 | 0.930814745 | 0.391132731 | 0.709264768 | -4.633745737 |

| | | | | | | | | | |
|---|---|---|---|---|---|---|---|---|---|
| **A_55_P2034858** | Pias4 | -0.122460773 | -0.45198805 | 0.207066504 | 8.27159416 | -0.930799139 | 0.391140127 | 0.709264768 | -4.633749504 |
| **A_55_P1968763** | Shc4 | 0.301218427 | -0.509604091 | 1.112040944 | 10.03918492 | 0.930477938 | 0.391292364 | 0.709264768 | -4.633827027 |
| **A_52_P198435** | Rasgrp3 | 0.364512541 | -0.616695516 | 1.345720599 | 10.32923901 | 0.930468585 | 0.391296797 | 0.709264768 | -4.633829284 |
| **A_55_P2138343** | Ndufaf5 | 0.158083642 | -0.267462111 | 0.583629396 | 8.7093148 | 0.930445669 | 0.391307661 | 0.709264768 | -4.633834815 |
| **A_55_P2147876** | Abhd17a | -0.114761335 | -0.423714366 | 0.194191696 | 10.18786605 | -0.930365244 | 0.391345787 | 0.709264768 | -4.633854223 |
| **A_51_P508485** | Tmtc4 | 0.290126741 | -0.490936029 | 1.071189511 | 9.594741028 | 0.930362437 | 0.391347118 | 0.709264768 | -4.6338549 |
| **A_55_P2131303** | NA | 0.4134823 | -0.699708532 | 1.526673132 | 17.42916669 | 0.930331334 | 0.391361864 | 0.709264768 | -4.633862406 |
| **A_55_P2079041** | NA | 0.179770567 | -0.304294571 | 0.663835705 | 15.7451086 | 0.930175995 | 0.391435515 | 0.709264768 | -4.63389989 |
| **A_55_P2045482** | NA | 0.222431896 | -0.376563884 | 0.821427675 | 14.591938 | 0.930087155 | 0.391477642 | 0.709264768 | -4.633921326 |
| **A_55_P2038111** | Tmem104 | -0.12580336 | -0.464587955 | 0.212981235 | 7.794365265 | -0.930077059 | 0.39148243 | 0.709264768 | -4.633923762 |
| **A_55_P2022094** | Mta3 | -0.495393046 | -1.829564621 | 0.83877853 | 10.47991789 | -0.930012033 | 0.391513267 | 0.709264768 | -4.633939451 |
| **A_55_P1957389** | Snx25 | 0.60002738 | -1.016035379 | 2.216090139 | 11.27361928 | 0.929957693 | 0.391539038 | 0.709264768 | -4.633952561 |
| **A_51_P177552** | Ndufb9 | -0.097420999 | -0.359836938 | 0.16499494 | 14.29574346 | -0.929849613 | 0.3915903 | 0.709264768 | -4.633978636 |
| **A_51_P473219** | Siae | -0.179748443 | -0.663988217 | 0.304491332 | 9.589745807 | -0.929726102 | 0.391648887 | 0.709264768 | -4.634008432 |
| **A_55_P2077964** | Vps8 | -0.152204223 | -0.562240549 | 0.257832103 | 8.208573792 | -0.929725438 | 0.391649202 | 0.709264768 | -4.634008592 |
| **A_51_P487813** | Lxn | 0.584128607 | -0.989628545 | 2.157885759 | 12.66867276 | 0.929653491 | 0.391683332 | 0.709264768 | -4.634025947 |
| **A_51_P174016** | Slc39a7 | -0.378886392 | -1.400003951 | 0.642231167 | 14.22072261 | -0.929359213 | 0.391822959 | 0.709264768 | -4.634096927 |
| **A_55_P1974288** | Rela | -0.179777037 | -0.664335781 | 0.304781707 | 13.59340837 | -0.929261895 | 0.391869142 | 0.709264768 | -4.634120398 |
| **A_51_P174720** | Ccdc43 | 0.093518591 | -0.158573776 | 0.345610957 | 8.212161767 | 0.929155931 | 0.391919433 | 0.709264768 | -4.634145952 |
| **A_51_P129866** | Atxn1l | 0.099162903 | -0.168168613 | 0.36649442 | 11.03297848 | 0.929072097 | 0.391959225 | 0.709264768 | -4.634166169 |
| **A_55_P1963300** | Vsig10 | -0.10012588 | -0.370087016 | 0.169835257 | 9.694562476 | -0.928956638 | 0.392014033 | 0.709264768 | -4.63419401 |
| **A_52_P131490** | Atg4b | -0.379291811 | -1.402006541 | 0.643422918 | 8.785724984 | -0.928900723 | 0.392040577 | 0.709264768 | -4.634207492 |
| **A_55_P2096305** | Rc3h1 | -0.116110172 | -0.42919075 | 0.196970407 | 11.15146435 | -0.928890421 | 0.392045468 | 0.709264768 | -4.634209976 |
| **A_55_P2051414** | Axl | 0.15523015 | -0.263340287 | 0.573800587 | 10.62213215 | 0.928876279 | 0.392052182 | 0.709264768 | -4.634213386 |
| **A_55_P2000127** | Tgfb1i1 | 0.668304943 | -1.133834192 | 2.470444077 | 12.34173042 | 0.928830998 | 0.39207368 | 0.709264768 | -4.634224304 |
| **A_51_P448618** | Slc16a10 | -0.654625161 | -2.420019095 | 1.110768772 | 8.564428196 | -0.928755524 | 0.392109515 | 0.709264768 | -4.634242501 |
| **A_55_P2001023** | Lrrc19 | -0.360680001 | -1.333561512 | 0.612201511 | 11.99596191 | -0.928565315 | 0.392199836 | 0.709264768 | -4.634288358 |
| **A_51_P229925** | Chst7 | 0.223348445 | -0.379223657 | 0.825920546 | 9.524147062 | 0.928376754 | 0.39228939 | 0.709264768 | -4.634333812 |
| **A_55_P1991723** | Alkbh3 | -0.08582401 | -0.317382134 | 0.145734113 | 11.32298308 | -0.928323037 | 0.392314906 | 0.709264768 | -4.63434676 |
| **A_51_P414548** | Casp7 | 0.279836256 | -0.475212982 | 1.034885493 | 9.654900784 | 0.928280128 | 0.392335288 | 0.709264768 | -4.634357103 |
| **A_55_P2083694** | Adam33 | -0.155078399 | -0.573511319 | 0.263354522 | 8.141368214 | -0.928273191 | 0.392338583 | 0.709264768 | -4.634358775 |
| **A_55_P1975682** | Rhox2e | -0.241729776 | -0.894099447 | 0.410639896 | 7.946001948 | -0.928082817 | 0.392429024 | 0.709264768 | -4.634404659 |
| **A_55_P2048730** | NA | 0.350833303 | -0.596047944 | 1.297714549 | 15.94996713 | 0.928016381 | 0.39246059 | 0.709264768 | -4.63442067 |
| **A_55_P2463930** | Zmym5 | -0.243598528 | -0.901080144 | 0.413883087 | 9.51786575 | -0.927985931 | 0.392475058 | 0.709264768 | -4.634428008 |
| **A_52_P226127** | Dido1 | -0.107457936 | -0.397529212 | 0.18261334 | 9.082733954 | -0.92786355 | 0.392533211 | 0.709264768 | -4.6344575 |
| **A_51_P453351** | Lpgat1 | -0.40816108 | -1.509961773 | 0.693639613 | 10.70815986 | -0.927852399 | 0.392538511 | 0.709264768 | -4.634460188 |
| **A_52_P348250** | Klhl29 | -0.349548676 | -1.29315664 | 0.594059288 | 8.192607824 | -0.927825724 | 0.392551187 | 0.709264768 | -4.634466616 |
| **A_55_P2086825** | Rgl3 | 0.34391392 | -0.584583537 | 1.272411376 | 10.22754011 | 0.927725258 | 0.392598934 | 0.709264768 | -4.634490824 |
| **A_55_P2141429** | Park7 | 0.424013007 | -0.720823584 | 1.568849599 | 13.68827264 | 0.927654004 | 0.3926328 | 0.709264768 | -4.634507993 |
| **A_51_P420100** | Emc2 | 0.255272564 | -0.43398908 | 0.944534207 | 10.20270538 | 0.927620578 | 0.392648688 | 0.709264768 | -4.634516047 |
| **A_51_P268903** | Mon1a | -0.249554937 | -0.923390831 | 0.424280957 | 9.437827859 | -0.927603482 | 0.392656814 | 0.709264768 | -4.634520166 |
| **A_55_P2017645** | Tap2 | 0.465555398 | -0.791621732 | 1.722732529 | 10.87288757 | 0.92752418 | 0.39269451 | 0.709264768 | -4.634539273 |
| **A_55_P2079009** | Slco2b1 | 0.124430444 | -0.211617838 | 0.460478726 | 8.472550808 | 0.927417564 | 0.392745194 | 0.709264768 | -4.634564959 |
| **A_55_P1992079** | Ptprf | 0.738939857 | -1.256779981 | 2.734659695 | 12.71911271 | 0.927384565 | 0.392760882 | 0.709264768 | -4.634572909 |
| **A_66_P138001** | Atp2b1 | -0.114770226 | -0.424740743 | 0.195200291 | 10.26484834 | -0.927383136 | 0.392761562 | 0.709264768 | -4.634573253 |
| **A_52_P272854** | NA | 0.119514884 | -0.203281825 | 0.442311593 | 8.273679914 | 0.927349072 | 0.392777757 | 0.709264768 | -4.63458146 |
| **A_55_P2043182** | Gimap6 | 0.545970442 | -0.929079755 | 2.021020639 | 10.88938465 | 0.927070319 | 0.392910306 | 0.709264768 | -4.634648608 |
| **A_55_P2208071** | Nbeal1 | -0.113922095 | -0.421712992 | 0.193868802 | 9.58628857 | -0.927048677 | 0.392920598 | 0.709264768 | -4.634653821 |

| | | | | | | | | | |
|---|---|---|---|---|---|---|---|---|---|
| **A_55_P2044119** | Klc1 | -0.099086124 | -0.36680272 | 0.168630473 | 9.063949574 | -0.92701741 | 0.392935469 | 0.709264768 | -4.634661352 |
| **A_55_P2047305** | Adcy5 | 0.146052314 | -0.248593697 | 0.540698325 | 8.110127702 | 0.926938817 | 0.392972848 | 0.709264768 | -4.634680281 |
| **A_55_P1973770** | Unc5b | -0.576868365 | -2.135706713 | 0.981969983 | 11.8389693 | -0.926885286 | 0.39299831 | 0.709264768 | -4.634693174 |
| **A_55_P1958840** | Itga6 | 0.677045107 | -1.152551612 | 2.506641826 | 11.89581929 | 0.926856664 | 0.393011924 | 0.709264768 | -4.634700068 |
| **A_55_P2144796** | Gfer | 0.135920127 | -0.231390168 | 0.503230421 | 9.112606404 | 0.92683199 | 0.39302366 | 0.709264768 | -4.63470601 |
| **A_51_P439299** | Relt | -0.177205017 | -0.656189637 | 0.301779604 | 7.97420163 | -0.926626656 | 0.393121342 | 0.709264768 | -4.634755459 |
| **A_52_P549827** | Mgst1 | 0.685089448 | -1.16671989 | 2.536898786 | 12.58886893 | 0.926619327 | 0.39312483 | 0.709264768 | -4.634757224 |
| **A_55_P2139043** | 4833439L | 0.13534038 | -0.230489636 | 0.501170397 | 11.98228221 | 0.926613028 | 0.393127827 | 0.709264768 | -4.634758741 |
| **A_52_P150236** | Apom | -1.329196873 | -4.922090508 | 2.263696761 | 9.026722885 | -0.926606401 | 0.393130979 | 0.709264768 | -4.634760336 |
| **A_51_P301994** | Trmt10c | 0.133201298 | -0.226858567 | 0.493261164 | 8.820864415 | 0.926582492 | 0.393142355 | 0.709264768 | -4.634766094 |
| **A_51_P375783** | Prap1 | -0.894374216 | -3.312031236 | 1.523282804 | 8.556556852 | -0.926562941 | 0.393151657 | 0.709264768 | -4.634770802 |
| **A_55_P1961084** | Map3k1 | -0.102822795 | -0.39289036 | 0.18724477 | 11.87781441 | -0.944175968 | 0.3931919 | 0.709264768 | -4.62092657 |
| **A_55_P2048840** | Ppil2 | -0.089373743 | -0.331004622 | 0.152257136 | 9.525720539 | -0.926419865 | 0.393219737 | 0.709264768 | -4.634805253 |
| **A_51_P386612** | Pir | 0.391361313 | -0.666733379 | 1.449456005 | 9.123806255 | 0.926410998 | 0.393223957 | 0.709264768 | -4.634807388 |
| **A_55_P2168914** | NA | -0.135168514 | -0.500645938 | 0.230308911 | 7.873456519 | -0.926329144 | 0.393262909 | 0.709264768 | -4.634827096 |
| **A_51_P486512** | Letmd1 | -0.224164704 | -0.830331714 | 0.382002305 | 10.46410583 | -0.92624373 | 0.39330356 | 0.709264768 | -4.634847661 |
| **A_55_P1969987** | Fam204a | 0.094849748 | -0.161639894 | 0.351339389 | 8.439834193 | 0.926225418 | 0.393312276 | 0.709264768 | -4.634852069 |
| **A_52_P217240** | Ppme1 | -0.168862201 | -0.625557952 | 0.28783355 | 9.093221704 | -0.926095599 | 0.393374067 | 0.709264768 | -4.634883322 |
| **A_51_P431554** | Ten1 | 0.295565728 | -0.50381038 | 1.094941836 | 9.809467525 | 0.926090132 | 0.393376669 | 0.709264768 | -4.634884638 |
| **A_52_P74427** | Mms19 | -0.129014814 | -0.477949499 | 0.219919871 | 8.181986595 | -0.926074198 | 0.393384254 | 0.709264768 | -4.634888474 |
| **A_55_P2055213** | Abcf1 | -0.175430421 | -0.649912276 | 0.299051433 | 8.297774238 | -0.926052575 | 0.393394547 | 0.709264768 | -4.63489368 |
| **A_52_P146164** | Ube2n | 0.106525957 | -0.181595064 | 0.394646979 | 8.98611986 | 0.926042343 | 0.393399418 | 0.709264768 | -4.634896143 |
| **A_55_P2064043** | Cd44 | 0.389262812 | -0.663630304 | 1.442155928 | 10.1773864 | 0.925995711 | 0.393421616 | 0.709264768 | -4.634907368 |
| **A_55_P2005410** | NA | 0.514457302 | -0.877140369 | 1.906054973 | 13.87031671 | 0.9259467 | 0.393444949 | 0.709264768 | -4.634919166 |
| **A_55_P1980883** | Ephb6 | 0.286629803 | -0.489040338 | 1.062299943 | 11.30370786 | 0.925538772 | 0.393639191 | 0.709548771 | -4.635017349 |
| **A_51_P460710** | Tdrd3 | 0.305275255 | -0.557008103 | 1.167558612 | 10.32673302 | 0.942983956 | 0.393739475 | 0.709596977 | -4.621165643 |
| **A_66_P109049** | Brd9 | 0.127034734 | -0.216844822 | 0.470914291 | 9.723688356 | 0.925265718 | 0.393769252 | 0.709596977 | -4.635083057 |
| **A_55_P1993976** | Aimp2 | 0.107947773 | -0.184272341 | 0.400167888 | 12.70757547 | 0.925238995 | 0.393781983 | 0.709596977 | -4.635089487 |
| **A_55_P1961320** | Tes | -0.141496468 | -0.524581433 | 0.241588497 | 11.69474279 | -0.925125895 | 0.393835866 | 0.709596977 | -4.6351167 |
| **A_55_P2133895** | Hnrnph2 | 0.378375568 | -0.646235689 | 1.402986825 | 9.882742916 | 0.924941591 | 0.393923684 | 0.709596977 | -4.635161041 |
| **A_52_P105840** | Fnbp1 | -0.273807252 | -1.015264747 | 0.467650243 | 8.236069376 | -0.924930354 | 0.393929039 | 0.709596977 | -4.635163745 |
| **A_52_P138205** | Plekha8 | 0.118589916 | -0.202563674 | 0.439743506 | 8.75323826 | 0.924879866 | 0.393953099 | 0.709596977 | -4.635175891 |
| **A_55_P2002155** | Sumo1 | 0.177202007 | -0.302686129 | 0.657090142 | 14.09317433 | 0.924866329 | 0.39395955 | 0.709596977 | -4.635179147 |
| **A_55_P2090914** | Tpst2 | -0.310840995 | -1.152917062 | 0.531235072 | 11.18170977 | -0.924564709 | 0.394103311 | 0.709597768 | -4.635251701 |
| **A_51_P488673** | Rack1 | -0.156109201 | -0.579020037 | 0.266801635 | 16.89950294 | -0.924549212 | 0.394110699 | 0.709597768 | -4.635255428 |
| **A_55_P1970234** | Dclk2 | 0.109564114 | -0.187342664 | 0.406470893 | 8.370500962 | 0.924269377 | 0.394244115 | 0.709597768 | -4.635322729 |
| **A_55_P2043662** | Zfp607a | 0.173991023 | -0.297514683 | 0.64549673 | 8.422407231 | 0.92425165 | 0.394252568 | 0.709597768 | -4.635326992 |
| **A_51_P193146** | Ms4a6c | -0.12584149 | -0.466923445 | 0.215240465 | 8.103619131 | -0.924092518 | 0.394328454 | 0.709597768 | -4.635365259 |
| **A_55_P2042356** | Rftn1 | 0.157542884 | -0.269463361 | 0.584549129 | 10.33535375 | 0.924091361 | 0.394329006 | 0.709597768 | -4.635365537 |
| **A_52_P337259** | Heyl | 0.675798526 | -1.156118861 | 2.507715914 | 10.19413985 | 0.92397815 | 0.394383001 | 0.709597768 | -4.635392759 |
| **A_55_P1969904** | Abcc6 | -0.085191742 | -0.316125421 | 0.145741937 | 7.689846456 | -0.923975745 | 0.394384148 | 0.709597768 | -4.635393338 |
| **A_55_P2074908** | Ube4b | -0.217167573 | -0.805868988 | 0.371533842 | 8.345937956 | -0.923953773 | 0.394394628 | 0.709597768 | -4.63539862 |
| **A_51_P181341** | Necab2 | 0.211082678 | -0.361164073 | 0.783329429 | 8.372921588 | 0.923888595 | 0.394425717 | 0.709597768 | -4.635414292 |
| **A_55_P1959303** | Rbm47 | -0.270592101 | -1.004363143 | 0.46317894 | 8.906299579 | -0.923644594 | 0.39454212 | 0.709597768 | -4.635472953 |
| **A_52_P134666** | Srpr | -0.135680966 | -0.503640289 | 0.232278358 | 9.990666789 | -0.923569238 | 0.394578075 | 0.709597768 | -4.635491068 |
| **A_55_P1997282** | Fam135a | -0.097918212 | -0.363471731 | 0.167635307 | 9.418946138 | -0.923552862 | 0.394585888 | 0.709597768 | -4.635495005 |
| **A_55_P2070721** | Rapgef1 | -0.460967203 | -1.711182363 | 0.789247957 | 10.66717374 | -0.923497261 | 0.394612419 | 0.709597768 | -4.63550837 |
| **A_55_P2071601** | Ppara | 0.105938651 | -0.1813896 | 0.393266902 | 8.834876826 | 0.923477789 | 0.394621711 | 0.709597768 | -4.635513051 |

| | | | | | | | | | |
|---|---|---|---|---|---|---|---|---|---|
| A_55_P1988313 | NA | -0.198336355 | -0.736277923 | 0.339605212 | 7.743662027 | -0.923458896 | 0.394630726 | 0.709597768 | -4.635517592 |
| A_55_P1969982 | Styx | 0.084493945 | -0.144729581 | 0.313717471 | 9.161737689 | 0.923244532 | 0.394733031 | 0.709597768 | -4.635569116 |
| A_55_P1972054 | NA | -0.096091596 | -0.356786011 | 0.16460282 | 8.244892245 | -0.923217488 | 0.394745939 | 0.709597768 | -4.635575616 |
| A_55_P1989573 | Tmem175 | -0.325683976 | -1.209289426 | 0.557921474 | 10.18916692 | -0.923184174 | 0.39476184 | 0.709597768 | -4.635583623 |
| A_55_P1978681 | Tspan8 | 0.866923455 | -1.485574334 | 3.219421244 | 11.29149392 | 0.92300037 | 0.394849581 | 0.709597768 | -4.635627795 |
| A_55_P1971649 | Ttc19 | -0.257768478 | -0.957446588 | 0.441909631 | 9.56970165 | -0.922745354 | 0.394971342 | 0.709597768 | -4.635689073 |
| A_55_P2111855 | Gale | -0.613736533 | -2.279660088 | 1.052187023 | 11.40283345 | -0.922735548 | 0.394976024 | 0.709597768 | -4.635691429 |
| A_55_P2015585 | Zcchc24 | 0.348951755 | -0.598272604 | 1.296176114 | 12.53507937 | 0.922705002 | 0.394990611 | 0.709597768 | -4.635698769 |
| A_51_P450682 | Tepp | -0.172511733 | -0.640792319 | 0.295768853 | 7.907666384 | -0.922704907 | 0.394990656 | 0.709597768 | -4.635698791 |
| A_51_P223720 | Rtcb | 0.178329631 | -0.305838888 | 0.662498151 | 11.56092718 | 0.922523226 | 0.395077423 | 0.709597768 | -4.635742441 |
| A_55_P2045158 | NA | 0.172216745 | -0.295362277 | 0.639795767 | 10.07431248 | 0.922509196 | 0.395084124 | 0.709597768 | -4.635745812 |
| A_55_P2104259 | Plin5 | 0.247758188 | -0.424933758 | 0.920450133 | 8.490011845 | 0.922490991 | 0.395092819 | 0.709597768 | -4.635750185 |
| A_55_P2133845 | Eid2b | 0.158968173 | -0.272841357 | 0.590777703 | 8.446304332 | 0.922079369 | 0.395289462 | 0.709597768 | -4.635849062 |
| A_55_P2143337 | Rpp21 | 0.144631682 | -0.248249449 | 0.537512814 | 12.82969331 | 0.92204605 | 0.395305383 | 0.709597768 | -4.635857064 |
| A_55_P1978618 | Grcc10 | 0.312544029 | -0.536484657 | 1.161572715 | 13.72919129 | 0.922017549 | 0.395319001 | 0.709597768 | -4.635863909 |
| A_51_P361830 | Lgi2 | 0.164981607 | -0.283197817 | 0.61316103 | 8.503690543 | 0.922006452 | 0.395324304 | 0.709597768 | -4.635866574 |
| A_55_P2096867 | Gap43 | 0.408354903 | -0.701024459 | 1.517734264 | 9.142058279 | 0.92195142 | 0.395350602 | 0.709597768 | -4.635879791 |
| A_55_P2026437 | Gm4802 | 0.104805921 | -0.179938395 | 0.389550237 | 16.18870897 | 0.921894239 | 0.395377929 | 0.709597768 | -4.635893524 |
| A_55_P2125823 | NA | 0.746263493 | -1.281253157 | 2.773780143 | 14.02333654 | 0.921887896 | 0.39538096 | 0.709597768 | -4.635895048 |
| A_55_P2089253 | Atg4c | -0.292105909 | -1.085768601 | 0.501556784 | 9.407474602 | -0.921838244 | 0.395404689 | 0.709597768 | -4.635906971 |
| A_55_P1999641 | Frem1 | 1.038300274 | -1.782826846 | 3.859427393 | 11.71468927 | 0.921829619 | 0.395408811 | 0.709597768 | -4.635909043 |
| A_55_P2132711 | Tmem248 | -0.605857084 | -2.252052291 | 1.040338122 | 9.551627469 | -0.921805305 | 0.395420432 | 0.709597768 | -4.635914882 |
| A_55_P2143946 | Slc18a2 | -0.190040943 | -0.706468403 | 0.326386518 | 8.237247914 | -0.921697043 | 0.395472178 | 0.709597768 | -4.635940879 |
| A_52_P565847 | AU018091 | -0.128818548 | -0.478908136 | 0.22127104 | 7.753600683 | -0.921615033 | 0.395511379 | 0.709597768 | -4.635960572 |
| A_55_P1986421 | Tcp11 | 0.315338789 | -0.541727827 | 1.172405405 | 8.630131179 | 0.921537818 | 0.395548292 | 0.709597768 | -4.635979112 |
| A_55_P2168312 | NA | 0.355150619 | -0.61025007 | 1.320551309 | 15.27454285 | 0.921415083 | 0.39560697 | 0.709597768 | -4.636008581 |
| A_55_P2007663 | Alkbh8 | 0.179178682 | -0.307948687 | 0.666306051 | 9.198421986 | 0.921285321 | 0.395669015 | 0.709597768 | -4.636039734 |
| A_66_P126686 | NA | 0.390904545 | -0.67183731 | 1.453646401 | 15.48008512 | 0.921283473 | 0.395669899 | 0.709597768 | -4.636040178 |
| A_55_P1993522 | NA | -0.388412564 | -1.444568556 | 0.667743428 | 10.6867871 | -0.921118591 | 0.395748749 | 0.709597768 | -4.63607976 |
| A_55_P2225087 | 5430439M | 0.221866452 | -0.381441264 | 0.825174168 | 9.099029974 | 0.921092199 | 0.39576137 | 0.709597768 | -4.636086095 |
| A_55_P2301312 | Wdr78 | 0.540618807 | -0.929497292 | 2.010734905 | 10.16261085 | 0.921064114 | 0.395774803 | 0.709597768 | -4.636092837 |
| A_55_P2002103 | Arhgap45 | 0.606764122 | -1.043264099 | 2.256792342 | 11.46332254 | 0.921040795 | 0.395785956 | 0.709597768 | -4.636098434 |
| A_55_P1979252 | Glod4 | 0.339565772 | -0.58386512 | 1.262996664 | 12.48708923 | 0.921021682 | 0.395795098 | 0.709597768 | -4.636103022 |
| A_55_P1960764 | Rps4x | 0.361753426 | -0.622075509 | 1.345582361 | 15.79049824 | 0.920965596 | 0.395821924 | 0.709597768 | -4.636116484 |
| A_55_P2103807 | NA | 0.375536755 | -0.645795378 | 1.396868887 | 15.08023192 | 0.920949462 | 0.395829641 | 0.709597768 | -4.636120357 |
| A_55_P1998823 | NA | 0.457591042 | -0.787347013 | 1.702529096 | 11.47979115 | 0.920619388 | 0.39598755 | 0.709597768 | -4.636199576 |
| A_51_P113403 | Slc26a11 | -0.108773768 | -0.404709574 | 0.187162038 | 9.318560676 | -0.92061279 | 0.395990707 | 0.709597768 | -4.633587592 |
| A_51_P371485 | Abraxas2 | -0.32339322 | -1.203254011 | 0.55646757 | 9.496766398 | -0.920592202 | 0.396000558 | 0.709597768 | -4.6362061 |
| A_55_P2041539 | Pigx | -0.223614233 | -0.832014617 | 0.384786151 | 11.59776443 | -0.920577401 | 0.39600764 | 0.709597768 | -4.636209652 |
| A_55_P2062926 | D430041D | 0.206451523 | -0.35539292 | 0.768295967 | 9.017225124 | 0.920348582 | 0.396117141 | 0.709597768 | -4.636264559 |
| A_55_P2266178 | Il2rb | -0.771021756 | -2.869407049 | 1.327363537 | 8.3517484 | -0.920304922 | 0.396138037 | 0.709597768 | -4.636275034 |
| A_52_P580533 | Spata33 | 0.18096133 | -0.311546967 | 0.673469628 | 8.268081306 | 0.92028549 | 0.396147338 | 0.709597768 | -4.636279697 |
| A_52_P681391 | G0s2 | 0.941381931 | -1.620729603 | 3.503493465 | 11.02718674 | 0.920276161 | 0.396151803 | 0.709597768 | -4.636281935 |
| A_55_P2096486 | Bcs1l | 0.180059013 | -0.310022212 | 0.670140239 | 9.80513799 | 0.920231608 | 0.396173128 | 0.709597768 | -4.636292625 |
| A_55_P2101216 | Aldh3a1 | 0.112180691 | -0.193179968 | 0.417541351 | 9.087526326 | 0.9201431 | 0.396215494 | 0.709597768 | -4.63631386 |
| A_55_P2162369 | Cacfd1 | -0.13443163 | -0.50048304 | 0.23161978 | 8.435220946 | -0.91983457 | 0.396363207 | 0.709597768 | -4.636387874 |
| A_51_P520927 | Ccdc97 | -0.103338713 | -0.384733594 | 0.178056168 | 10.51115272 | -0.919808037 | 0.396375911 | 0.709597768 | -4.636394238 |
| A_55_P2022663 | NA | 0.081469646 | -0.140380352 | 0.303319645 | 16.19390772 | 0.919785896 | 0.396386514 | 0.709597768 | -4.636399549 |

| | | | | | | | | | |
|---|---|---|---|---|---|---|---|---|---|
| **A_55_P1957419** | Ensa | 0.136612492 | -0.235426446 | 0.508651429 | 9.053381879 | 0.919713079 | 0.396421384 | 0.709597768 | -4.636417015 |
| **A_51_P507509** | Gimap1 | 0.469756912 | -0.809583713 | 1.749097538 | 10.00318832 | 0.919681253 | 0.396436625 | 0.709597768 | -4.636424648 |
| **A_51_P412640** | Ccdc51 | -0.234422973 | -0.872929328 | 0.404083381 | 8.908443597 | -0.919571053 | 0.396489402 | 0.709597768 | -4.636451079 |
| **A_52_P494514** | Insr | -0.091663971 | -0.341383316 | 0.158055374 | 8.61281372 | -0.919383757 | 0.396579116 | 0.709597768 | -4.636495996 |
| **A_55_P1997951** | Htatsf1 | -0.596599226 | -2.222208372 | 1.029009919 | 9.623758736 | -0.919214566 | 0.396660171 | 0.709597768 | -4.636536568 |
| **A_51_P143712** | Tdh | -0.325425685 | -1.212209364 | 0.561357995 | 7.988695257 | -0.919145957 | 0.396693043 | 0.709597768 | -4.636553019 |
| **A_51_P485594** | Tab3 | -0.389560875 | -1.451218226 | 0.672096475 | 9.776822602 | -0.919054589 | 0.396736824 | 0.709597768 | -4.636574926 |
| **A_55_P1998892** | Smox | -0.297289785 | -1.107487083 | 0.512907513 | 10.35311027 | -0.919050852 | 0.396738614 | 0.709597768 | -4.636575822 |
| **A_55_P2067086** | Togaram1 | 0.098990471 | -0.170796035 | 0.368776976 | 11.07755158 | 0.919016928 | 0.396754871 | 0.709597768 | -4.636583956 |
| **A_51_P149775** | Spcs1 | -0.088170203 | -0.328471588 | 0.152131183 | 14.02834353 | -0.919000845 | 0.396762577 | 0.709597768 | -4.636587812 |
| **A_52_P481279** | Drc1 | 0.640618107 | -1.105389502 | 2.386625715 | 10.26511586 | 0.918974421 | 0.39677524 | 0.709597768 | -4.636594147 |
| **A_66_P121459** | Cenpa | 0.731376567 | -1.262005743 | 2.724758876 | 10.72509213 | 0.918968843 | 0.396777913 | 0.709597768 | -4.636595484 |
| **A_52_P55717** | Adnp | 0.118095015 | -0.203776117 | 0.439966148 | 8.65363133 | 0.918966932 | 0.396778829 | 0.709597768 | -4.636595942 |
| **A_52_P64356** | Sparcl1 | 0.785072545 | -1.354818152 | 2.924963242 | 15.67399671 | 0.91890062 | 0.396810608 | 0.709597768 | -4.63661184 |
| **A_55_P2043147** | Akr1e1 | 0.260668678 | -0.449861658 | 0.971199015 | 9.730365895 | 0.918875284 | 0.396822751 | 0.709597768 | -4.636617914 |
| **A_55_P2028788** | Slc39a11 | -0.153646428 | -0.57255121 | 0.265258353 | 11.6099882 | -0.918665692 | 0.396923211 | 0.709711769 | -4.636668158 |
| **A_55_P2008136** | Ctr9 | -0.359478974 | -1.339788593 | 0.620830644 | 8.739880294 | -0.918460695 | 0.397021487 | 0.709821845 | -4.636717295 |
| **A_55_P1966664** | Cdc14b | -0.117118912 | -0.436541127 | 0.202303304 | 9.829490204 | -0.918358514 | 0.397070481 | 0.709843798 | -4.636741785 |
| **A_55_P1963002** | Ppfia4 | -0.148819375 | -0.554741476 | 0.257102726 | 8.504571357 | -0.918263082 | 0.397116242 | 0.709859969 | -4.636764656 |
| **A_52_P670978** | Nkain1 | -0.247149351 | -0.921588677 | 0.427289975 | 8.071556355 | -0.917839901 | 0.397319215 | 0.710157133 | -4.63686606 |
| **A_52_P357133** | Selenom | -0.172356057 | -0.64297244 | 0.298260327 | 11.59548859 | -0.917296745 | 0.397579851 | 0.710525176 | -4.636996176 |
| **A_55_P1965757** | Eme2 | -0.219919483 | -0.820432908 | 0.380593942 | 9.318506783 | -0.917257624 | 0.397598628 | 0.710525176 | -4.637005546 |
| **A_55_P2096368** | NA | 0.098291617 | -0.170131707 | 0.366714942 | 11.04415114 | 0.917163104 | 0.397644 | 0.710540581 | -4.637028184 |
| **A_52_P114889** | Upk3b | 0.284059386 | -0.49184858 | 1.059967352 | 8.892658576 | 0.916957653 | 0.397742633 | 0.710603446 | -4.637077386 |
| **A_52_P479262** | NA | 0.933501652 | -1.616415537 | 3.483418842 | 13.31669544 | 0.916936709 | 0.397752689 | 0.710603446 | -4.637082402 |
| **A_55_P2148116** | Tsr2 | 0.087614666 | -0.151771257 | 0.327000588 | 8.321955786 | 0.916702776 | 0.397865022 | 0.710726012 | -4.637138417 |
| **A_51_P105124** | P2ry6 | 0.25247383 | -0.437411713 | 0.942359374 | 9.145609165 | 0.916620719 | 0.397904431 | 0.710726012 | -4.630928337 |
| **A_51_P413740** | Ftcd | 0.09381488 | -0.162550385 | 0.350180145 | 7.785958111 | 0.916564203 | 0.397931575 | 0.710726012 | -4.637171595 |
| **A_51_P167513** | Leng1 | -0.129262962 | -0.482590073 | 0.224064149 | 10.12102262 | -0.91632068 | 0.398048555 | 0.71074028 | -4.637229894 |
| **A_55_P2003246** | Asxl2 | 0.147613196 | -0.255945557 | 0.551171949 | 9.931790647 | 0.916154582 | 0.398128358 | 0.71074028 | -4.637269653 |
| **A_55_P2064547** | Tuba1c | -0.328200424 | -1.225525213 | 0.569124366 | 14.9775091 | -0.916093532 | 0.398157692 | 0.71074028 | -4.637284265 |
| **A_55_P2065369** | H3f3a | 0.733576082 | -1.27235993 | 2.739512094 | 14.85145468 | 0.915964059 | 0.39821991 | 0.71074028 | -4.637315253 |
| **A_51_P265545** | Maco1 | 0.097736243 | -0.169530089 | 0.365002575 | 10.49576396 | 0.915928838 | 0.398236837 | 0.71074028 | -4.637323683 |
| **A_55_P2102365** | Spag7 | -0.216571456 | -0.808830348 | 0.375687437 | 15.40834548 | -0.915882943 | 0.398258895 | 0.71074028 | -4.637334667 |
| **A_55_P2127439** | Kri1 | -0.098837682 | -0.369148022 | 0.171472659 | 11.58260193 | -0.915820237 | 0.398289033 | 0.71074028 | -4.637349673 |
| **A_52_P278295** | Pear1 | 0.148964596 | -0.258471885 | 0.556401078 | 9.011570002 | 0.915742767 | 0.39832627 | 0.71074028 | -4.637368212 |
| **A_51_P266546** | Zc2hc1a | 0.223208148 | -0.387300185 | 0.83371648 | 10.01031588 | 0.915732849 | 0.398331037 | 0.71074028 | -4.637370585 |
| **A_55_P1967191** | Gm4632 | 0.232042868 | -0.402719588 | 0.866805324 | 8.024934589 | 0.915603278 | 0.398393323 | 0.71074028 | -4.63740159 |
| **A_52_P504068** | Cdk8 | -0.170520098 | -0.637020625 | 0.295980429 | 8.324507028 | -0.915532531 | 0.398427336 | 0.71074028 | -4.637418518 |
| **A_55_P2005675** | NA | 0.34957913 | -0.606862099 | 1.30602036 | 15.96786306 | 0.915456166 | 0.398464051 | 0.71074028 | -4.63743679 |
| **A_55_P1967196** | Lamb3 | 0.782499467 | -1.358458187 | 2.923457121 | 11.71696706 | 0.915432483 | 0.398475439 | 0.71074028 | -4.637442456 |
| **A_51_P349495** | Mboat1 | 0.197650014 | -0.343179584 | 0.738479613 | 10.32253658 | 0.915349068 | 0.398515547 | 0.71074028 | -4.637462413 |
| **A_51_P140803** | Slco1b2 | -0.14172182 | -0.529544562 | 0.246100922 | 7.736156693 | -0.915279624 | 0.398548941 | 0.71074028 | -4.637479027 |
| **A_51_P497882** | Creb3l4 | 0.218144145 | -0.37881019 | 0.815098481 | 8.719161976 | 0.915277522 | 0.398549952 | 0.71074028 | -4.63747953 |
| **A_55_P2179453** | Lig3 | -0.148176828 | -0.55370708 | 0.257353424 | 8.256881233 | -0.915181813 | 0.398595979 | 0.71074028 | -4.637502426 |
| **A_51_P122085** | Usp20 | -0.273449063 | -1.021833279 | 0.474935153 | 10.31114652 | -0.915170823 | 0.398601265 | 0.71074028 | -4.635814014 |
| **A_52_P83959** | Taf7 | -0.434815748 | -1.625009625 | 0.755378128 | 8.768988221 | -0.915035407 | 0.398666396 | 0.710751502 | -4.637537448 |
| **A_51_P200144** | Pak1ip1 | -0.284757028 | -1.064230592 | 0.494716535 | 11.68416227 | -0.915004874 | 0.398681082 | 0.710751502 | -4.637544751 |

| | | | | | | | | | |
|---|---|---|---|---|---|---|---|---|---|
| **A_55_P1964309** | Mlf2 | 0.184282812 | -0.3203327 | 0.688898324 | 13.84574125 | 0.914691365 | 0.398831906 | 0.710954828 | -4.637619735 |
| **A_55_P2090519** | Ppdpf | 0.317194382 | -0.551572345 | 1.185961109 | 13.50836926 | 0.914476717 | 0.398935195 | 0.710985087 | -4.637671065 |
| **A_55_P2176430** | 0610010K | 0.221122687 | -0.384536697 | 0.826782071 | 11.1731281 | 0.914439965 | 0.398952882 | 0.710985087 | -4.637679853 |
| **A_55_P2151091** | Phactr4 | -0.247060539 | -0.923838733 | 0.429717656 | 11.45532934 | -0.914339268 | 0.399001346 | 0.710985087 | -4.637703931 |
| **A_55_P2022987** | Psmd2 | -0.271879911 | -1.016676182 | 0.472916361 | 13.62584687 | -0.914302633 | 0.399018979 | 0.710985087 | -4.63771269 |
| **A_55_P1975155** | Galnt6 | -1.437183125 | -5.374612639 | 2.500246388 | 9.522345842 | -0.914217779 | 0.399059824 | 0.710985087 | -4.637732978 |
| **A_51_P469285** | Nrp1 | 0.433222954 | -0.753779037 | 1.620224944 | 9.491709167 | 0.914135042 | 0.399099652 | 0.710985087 | -4.637752759 |
| **A_51_P144868** | Mboat7 | -0.182467296 | -0.682423064 | 0.317488472 | 10.77772694 | -0.914121238 | 0.399106297 | 0.710985087 | -4.637756059 |
| **A_55_P1979814** | Abcc8 | -0.241981017 | -0.905147712 | 0.421185678 | 8.81946109 | -0.913921587 | 0.399202421 | 0.711090805 | -4.637803787 |
| **A_55_P2010773** | Tbx2 | 0.868316955 | -1.511916822 | 3.248550732 | 12.1895009 | 0.913711336 | 0.399303666 | 0.711102622 | -4.637854043 |
| **A_55_P2177426** | Abcb8 | -0.087881094 | -0.328794326 | 0.153032138 | 8.097571208 | -0.913661126 | 0.399327848 | 0.711102622 | -4.637866044 |
| **A_51_P502150** | Slc9a3r1 | -0.451845348 | -1.690629632 | 0.786938935 | 9.574353124 | -0.913575563 | 0.399369058 | 0.711102622 | -4.637886494 |
| **A_55_P2052470** | Uqcrh | 0.182334012 | -0.317661438 | 0.682329462 | 14.77196042 | 0.913381017 | 0.399462771 | 0.711102622 | -4.637932987 |
| **A_55_P2002632** | Rap1b | -0.102746973 | -0.384518624 | 0.179024678 | 13.70831136 | -0.913318143 | 0.399493061 | 0.711102622 | -4.637948011 |
| **A_55_P1978122** | Mutyh | 0.084787962 | -0.147748956 | 0.31732488 | 8.7730987 | 0.913256205 | 0.399522903 | 0.711102622 | -4.637962812 |
| **A_66_P122927** | NA | 0.226203479 | -0.394201811 | 0.846608768 | 9.118329684 | 0.913217306 | 0.399541644 | 0.711102622 | -4.637972106 |
| **A_51_P178347** | Zfp326 | 0.221124599 | -0.385429585 | 0.827678784 | 9.985881055 | 0.913098861 | 0.399598717 | 0.711102622 | -4.638000407 |
| **A_55_P2180744** | Clstn3 | -0.398924524 | -1.493244754 | 0.695395707 | 8.475677226 | -0.913054399 | 0.399620142 | 0.711102622 | -4.638011031 |
| **A_52_P616591** | Eri3 | 0.090433606 | -0.15765285 | 0.338520063 | 8.373224011 | 0.913013378 | 0.39963991 | 0.711102622 | -4.638020831 |
| **A_51_P338461** | Numa1 | -0.141002636 | -0.52783975 | 0.245834479 | 13.61113108 | -0.912955147 | 0.399667973 | 0.711102622 | -4.638034743 |
| **A_55_P2014209** | Scyl2 | -0.213923779 | -0.800875481 | 0.373027924 | 8.33658177 | -0.912866018 | 0.399710929 | 0.711102622 | -4.638056036 |
| **A_55_P2148531** | Pithd1 | -0.248380716 | -0.929894271 | 0.433132839 | 8.297289052 | -0.912838023 | 0.399724423 | 0.711102622 | -4.638062724 |
| **A_51_P111962** | Bean1 | -0.137696831 | -0.515523119 | 0.240129458 | 7.917676298 | -0.912813635 | 0.399736177 | 0.711102622 | -4.63806855 |
| **A_66_P113490** | Sh2b2 | -0.592454775 | -2.218182039 | 1.033272489 | 8.986040946 | -0.91276265 | 0.399760753 | 0.711102622 | -4.63808073 |
| **A_55_P2085127** | Ramac | -0.179895428 | -0.67360244 | 0.313811583 | 10.43809385 | -0.912643526 | 0.399818176 | 0.71113934 | -4.638109185 |
| **A_51_P300506** | Cox6b2 | 0.440928831 | -0.76928967 | 1.651147331 | 9.895924609 | 0.912546606 | 0.399864901 | 0.711157024 | -4.638132335 |
| **A_52_P218181** | Zdhhc5 | -0.214420576 | -0.803071194 | 0.374230042 | 11.48153523 | -0.912345217 | 0.399962005 | 0.711243348 | -4.638180434 |
| **A_52_P175952** | Map3k7 | -0.193572512 | -0.725055319 | 0.337910295 | 8.522522981 | -0.912230967 | 0.400017101 | 0.711243348 | -4.638207719 |
| **A_51_P376057** | Bcorl1 | -0.101618097 | -0.380637772 | 0.177401579 | 8.562963472 | -0.912192658 | 0.400035576 | 0.711243348 | -4.638216867 |
| **A_52_P198473** | NA | -0.259363393 | -0.971653578 | 0.452926792 | 8.300065595 | -0.912015175 | 0.40012118 | 0.711243348 | -4.638259247 |
| **A_55_P1962918** | Ifi211 | -0.175595065 | -0.657846005 | 0.306655874 | 8.164566361 | -0.911988936 | 0.400133836 | 0.711243348 | -4.638265513 |
| **A_52_P515826** | Med13 | 0.094417798 | -0.164889635 | 0.353725231 | 8.080595262 | 0.911988261 | 0.400134162 | 0.711243348 | -4.638265674 |
| **A_55_P2151026** | Ccdc50 | 0.098321361 | -0.171748925 | 0.368391647 | 8.276822959 | 0.911845845 | 0.400202865 | 0.711300072 | -4.638299677 |
| **A_55_P2161640** | Slc38a1 | -0.67763777 | -2.539819931 | 1.18454439 | 8.985744383 | -0.911435182 | 0.400401022 | 0.7115137 | -4.638397712 |
| **A_52_P220810** | Trib2 | 0.333255469 | -0.582603826 | 1.249114764 | 9.303880739 | 0.911378698 | 0.400428283 | 0.7115137 | -4.638411194 |
| **A_55_P1991605** | Aplp1 | -0.25964157 | -0.973210646 | 0.453927505 | 9.1825019 | -0.911357038 | 0.400438738 | 0.7115137 | -4.638416364 |
| **A_51_P230934** | Cab39 | -0.247335971 | -0.927229972 | 0.43255803 | 11.03465461 | -0.911163718 | 0.400532054 | 0.7115137 | -4.638462503 |
| **A_55_P2052744** | NA | 0.417269319 | -0.729765007 | 1.564303645 | 11.94827908 | 0.911151059 | 0.400538165 | 0.7115137 | -4.638465524 |
| **A_51_P160344** | Cenpv | 0.18288919 | -0.319862601 | 0.68564098 | 10.34466455 | 0.911139249 | 0.400543866 | 0.7115137 | -4.638468343 |
| **A_55_P1956443** | Ttc5 | -0.14987794 | -0.562038112 | 0.262282232 | 9.68124891 | -0.910797936 | 0.400708664 | 0.71162172 | -4.638549789 |
| **A_55_P2026445** | Tfam | 0.150881765 | -0.264066266 | 0.565829796 | 10.01978869 | 0.910737866 | 0.400737673 | 0.71162172 | -4.638564122 |
| **A_55_P2078285** | Uck1 | -0.18784856 | -0.704473529 | 0.328776408 | 12.82201958 | -0.910715703 | 0.400748377 | 0.71162172 | -4.63856941 |
| **A_55_P2177488** | H2bc21 | 0.132521447 | -0.231982539 | 0.497025433 | 8.178705431 | 0.910613814 | 0.400797586 | 0.71162172 | -4.638593719 |
| **A_51_P331870** | Rnf145 | 0.239733597 | -0.419696251 | 0.899163445 | 10.24868556 | 0.910564356 | 0.400821475 | 0.71162172 | -4.638605518 |
| **A_52_P14453** | Atxn1 | -0.12224552 | -0.45850663 | 0.21401559 | 8.410654802 | -0.910555989 | 0.400825516 | 0.71162172 | -4.638607514 |
| **A_55_P2009008** | NA | 0.538718523 | -0.94353202 | 2.020969066 | 14.30320961 | 0.910312771 | 0.400943009 | 0.711764955 | -4.638665534 |
| **A_52_P514317** | Slc35a3 | -0.27416925 | -1.028647988 | 0.480309487 | 9.147462996 | -0.910169095 | 0.401012428 | 0.711777025 | -4.638699804 |
| **A_55_P2130576** | NA | 0.366772443 | -0.642736216 | 1.376281102 | 15.07704493 | 0.909990786 | 0.401098593 | 0.711777025 | -4.638742331 |

| | | | | | | | | | |
|---|---|---|---|---|---|---|---|---|---|
| **A_52_P591166** | Dpysl2 | 0.50691874 | -0.888409016 | 1.902246495 | 13.47244918 | 0.909939376 | 0.401123439 | 0.711777025 | -4.638754592 |
| **A_55_P2064233** | Ipo8 | 0.107527122 | -0.188458206 | 0.40351245 | 10.30583365 | 0.90990947 | 0.401137892 | 0.711777025 | -4.638761724 |
| **A_55_P1963772** | Atp5j | 0.204288689 | -0.358071718 | 0.766649096 | 13.7444873 | 0.909871224 | 0.401156377 | 0.711777025 | -4.638770844 |
| **A_55_P2173702** | Ebpl | -0.372827169 | -1.399169188 | 0.653514849 | 11.43970866 | -0.9098416 | 0.401170696 | 0.711777025 | -4.638777909 |
| **A_55_P2028655** | Slc31a1 | 0.118660003 | -0.208064585 | 0.445384592 | 11.41183638 | 0.909646979 | 0.401264773 | 0.711824344 | -4.638824317 |
| **A_52_P119672** | Rock1 | 0.10342143 | -0.181374939 | 0.388217799 | 9.093040414 | 0.909549702 | 0.401311802 | 0.711824344 | -4.638847511 |
| **A_55_P2116833** | 9430038I0 | -0.089071771 | -0.334354294 | 0.156210751 | 9.563540533 | -0.909544243 | 0.401314441 | 0.711824344 | -4.638848812 |
| **A_55_P2136378** | Dazap2 | -0.137031061 | -0.514512253 | 0.240450131 | 14.49032446 | -0.909230611 | 0.401466098 | 0.711824344 | -4.638923583 |
| **A_55_P1960857** | Sorbs2 | -0.179237382 | -0.672994133 | 0.314519369 | 7.828246241 | -0.909213531 | 0.401474358 | 0.711824344 | -4.638927655 |
| **A_55_P2001510** | NA | 0.179433389 | -0.3148822 | 0.673748978 | 16.5801588 | 0.909178797 | 0.401491157 | 0.711824344 | -4.638935934 |
| **A_55_P2106922** | Arfgap3 | -0.248461078 | -0.932958398 | 0.436036242 | 10.54809045 | -0.909152964 | 0.401503651 | 0.711824344 | -4.638942092 |
| **A_55_P1956827** | Ogfr | -0.17378042 | -0.652563579 | 0.305002738 | 13.67449103 | -0.909101394 | 0.401528594 | 0.711824344 | -4.638954385 |
| **A_55_P2118843** | Hk1 | -0.168532392 | -0.632911722 | 0.295846937 | 8.101005635 | -0.908993653 | 0.401580708 | 0.711824344 | -4.638980065 |
| **A_55_P1957583** | Gps2 | 0.087634648 | -0.153839553 | 0.329108848 | 9.715904929 | 0.908982359 | 0.401586171 | 0.711824344 | -4.638982757 |
| **A_55_P2041758** | Scand1 | -0.14964516 | -0.56200233 | 0.262712011 | 14.48939035 | -0.908948902 | 0.401602356 | 0.711824344 | -4.638990731 |
| **A_65_P19933** | Zdhhc23 | -0.129846467 | -0.487712085 | 0.228019152 | 8.251541789 | -0.908783653 | 0.401682301 | 0.711875074 | -4.639030115 |
| **A_55_P2046158** | NA | -0.115956684 | -0.43564931 | 0.203735941 | 13.58088131 | -0.90847612 | 0.401831113 | 0.711875074 | -4.639103399 |
| **A_55_P2021530** | Clic4 | -0.182722155 | -0.68649548 | 0.321051171 | 8.191191964 | -0.908461205 | 0.401838331 | 0.711875074 | -4.639106952 |
| **A_66_P134501** | Olfm1 | 0.383062692 | -0.673119886 | 1.439245269 | 9.311000724 | 0.908408527 | 0.401863826 | 0.711875074 | -4.639119504 |
| **A_55_P2052281** | Rnf208 | -0.569386912 | -2.139591995 | 1.000818171 | 9.170944846 | -0.908241719 | 0.401944566 | 0.711875074 | -4.639159246 |
| **A_51_P218953** | Zfp536 | 0.378139493 | -0.66469424 | 1.420973225 | 8.874095783 | 0.90821216 | 0.401958875 | 0.711875074 | -4.639166288 |
| **A_55_P2451183** | Pdpk1 | -0.179931484 | -0.676160885 | 0.316297917 | 9.904835714 | -0.908186443 | 0.401971324 | 0.711875074 | -4.639172415 |
| **A_51_P458778** | Hpgd | -0.280776606 | -1.055176075 | 0.493622862 | 9.736793934 | -0.908126229 | 0.402000474 | 0.711875074 | -4.639186759 |
| **A_55_P2048767** | Efemp1 | 0.731707152 | -1.286510772 | 2.749925077 | 9.996309467 | 0.908070541 | 0.402027434 | 0.711875074 | -4.639200025 |
| **A_51_P371745** | Havcr1 | -0.143220242 | -0.538266253 | 0.25182577 | 7.733200997 | -0.908044362 | 0.402040109 | 0.711875074 | -4.639206262 |
| **A_55_P2106956** | Gm5523 | 0.172052484 | -0.302524467 | 0.646629435 | 16.62962809 | 0.908039314 | 0.402042553 | 0.711875074 | -4.639207464 |
| **A_55_P2041320** | NA | -0.937633174 | -3.524109769 | 1.648843421 | 12.94256607 | -0.907976811 | 0.402072815 | 0.711875074 | -4.639222353 |
| **A_55_P2067513** | Slc10a3 | -0.292779851 | -1.100724356 | 0.515164654 | 9.870216042 | -0.907632417 | 0.402239593 | 0.71194078 | -4.639304379 |
| **A_66_P108247** | Ucp3 | 0.227974544 | -0.401304151 | 0.85725324 | 10.18138779 | 0.907389341 | 0.402357338 | 0.71194078 | -4.639362264 |
| **A_51_P165082** | Tmem191 | -0.214718876 | -0.807422938 | 0.377985185 | 8.525420906 | -0.907366347 | 0.402368477 | 0.71194078 | -4.63936774 |
| **A_55_P2067200** | NA | 0.095118824 | -0.167446061 | 0.357683709 | 8.393517367 | 0.907361139 | 0.402371 | 0.71194078 | -4.63936898 |
| **A_52_P323658** | Zfp955a | 0.372142757 | -0.655186399 | 1.399471914 | 9.951089326 | 0.90729873 | 0.402401237 | 0.71194078 | -4.63938384 |
| **A_51_P277745** | Abcc2 | -0.130965248 | -0.492515199 | 0.230584702 | 7.714340838 | -0.907273258 | 0.402413577 | 0.71194078 | -4.639389905 |
| **A_55_P2099540** | H2afj | 0.259984223 | -0.457781467 | 0.977749912 | 9.175811964 | 0.907224238 | 0.402437328 | 0.71194078 | -4.639401576 |
| **A_51_P163378** | Zwint | -0.561560965 | -2.111955613 | 0.988833682 | 13.12151214 | -0.907204085 | 0.402447093 | 0.71194078 | -4.639406375 |
| **A_55_P2101336** | Ramp2 | 0.628577019 | -1.107070308 | 2.364224345 | 14.01697751 | 0.907083695 | 0.40250543 | 0.71194078 | -4.639435038 |
| **A_55_P2098335** | Map2k4 | -0.171417266 | -0.644743688 | 0.301909157 | 8.44001171 | -0.907077012 | 0.402508668 | 0.71194078 | -4.639436629 |
| **A_52_P633379** | Plpbp | -0.367126092 | -1.38086864 | 0.646616455 | 11.0048253 | -0.907063983 | 0.402514982 | 0.71194078 | -4.639439731 |
| **A_55_P1972719** | Pmm1 | -0.233090511 | -0.87706268 | 0.410881657 | 8.653535715 | -0.906583565 | 0.402747844 | 0.712086265 | -4.639554088 |
| **A_55_P1952166** | Arid3a | -0.575378094 | -2.165048267 | 1.014292079 | 10.40013201 | -0.906560189 | 0.402759177 | 0.712086265 | -4.639559651 |
| **A_55_P2052185** | Lamtor1 | -0.167088918 | -0.6490177 | 0.314839865 | 12.86776845 | -0.923479985 | 0.402788436 | 0.712086265 | -4.625059638 |
| **A_55_P1981709** | Rreb1 | -0.14576094 | -0.54852009 | 0.256998209 | 9.675493817 | -0.906454671 | 0.402810338 | 0.712086265 | -4.639584764 |
| **A_55_P1955717** | Hccs | -0.181816255 | -0.684242065 | 0.320609554 | 16.02686932 | -0.906381669 | 0.402845736 | 0.712086265 | -4.639602137 |
| **A_55_P2000793** | Rbm34 | 0.137879007 | -0.243174988 | 0.518933002 | 9.343929667 | 0.906279118 | 0.402895466 | 0.712086265 | -4.639626541 |
| **A_51_P141274** | Sec24d | -0.284692955 | -1.071570311 | 0.5021844 | 10.25790038 | -0.906191573 | 0.402937923 | 0.712086265 | -4.639647372 |
| **A_55_P2011425** | Atp10b | -0.122385521 | -0.460679678 | 0.215908635 | 7.889262185 | -0.906120367 | 0.402972458 | 0.712086265 | -4.639664315 |
| **A_51_P519385** | Sacm1l | -0.08197496 | -0.308578751 | 0.144628832 | 9.947716454 | -0.906075495 | 0.402994222 | 0.712086265 | -4.639674991 |
| **A_52_P15490** | Nectin3 | 0.193865167 | -0.342073525 | 0.729803858 | 8.643611674 | 0.906014226 | 0.403023942 | 0.712086265 | -4.639689569 |

| | | | | | | | | | |
|---|---|---|---|---|---|---|---|---|---|
| **A_52_P495565** | Efnb3 | 0.239765536 | -0.423068303 | 0.902599374 | 8.473350953 | 0.906008832 | 0.403026558 | 0.712086265 | -4.639690852 |
| **A_55_P2367117** | Pitpnb | 0.07947079 | -0.140249052 | 0.299190633 | 8.639545195 | 0.905917397 | 0.403070913 | 0.712086265 | -4.639712606 |
| **A_55_P2142298** | Ywhaq | 0.262187872 | -0.462822282 | 0.987198027 | 14.51278268 | 0.905771925 | 0.403141489 | 0.712086265 | -4.639747213 |
| **A_52_P227445** | Parp2 | 0.194411319 | -0.343183723 | 0.732006362 | 8.854594909 | 0.9057673 | 0.403143733 | 0.712086265 | -4.639748314 |
| **A_55_P2162890** | Zfp780b | 0.150554199 | -0.265794316 | 0.566902714 | 8.895890172 | 0.905703818 | 0.403174534 | 0.712086265 | -4.639763415 |
| **A_55_P2061452** | NA | 0.364173345 | -0.642953463 | 1.371300153 | 15.66567401 | 0.905679098 | 0.403186529 | 0.712086265 | -4.639769295 |
| **A_55_P2121466** | Ncor1 | 0.083309556 | -0.147174847 | 0.313793959 | 8.140281511 | 0.905323148 | 0.403359273 | 0.712326286 | -4.639853957 |
| **A_55_P1998601** | Slc17a9 | -0.107200418 | -0.403806396 | 0.189405559 | 7.927161704 | -0.905246652 | 0.403396404 | 0.712326795 | -4.639872149 |
| **A_55_P1967693** | Wipi2 | -0.140028731 | -0.527517675 | 0.247460214 | 8.339738882 | -0.90512421 | 0.403455843 | 0.712366692 | -4.639901266 |
| **A_51_P418168** | Manf | -0.31993707 | -1.205470187 | 0.565596046 | 11.80277248 | -0.904919823 | 0.403555078 | 0.712434734 | -4.639949865 |
| **A_51_P292368** | Tmco6 | -0.084620188 | -0.31886774 | 0.149627363 | 8.357875192 | -0.904793088 | 0.40361662 | 0.712434734 | -4.639979997 |
| **A_51_P495077** | Ift22 | 0.311971328 | -0.551676782 | 1.175619438 | 9.241424145 | 0.904749184 | 0.403637941 | 0.712434734 | -4.639990435 |
| **A_55_P1970650** | Dgkh | -0.099754588 | -0.375930996 | 0.17642182 | 8.586531743 | -0.904683453 | 0.403669864 | 0.712434734 | -4.640006062 |
| **A_51_P306066** | Eif4g2 | 0.273640583 | -0.484035187 | 1.031316352 | 12.07934683 | 0.904580983 | 0.403719633 | 0.712434734 | -4.640030421 |
| **A_55_P2008576** | Ubr2 | 0.105588667 | -0.186807758 | 0.397985092 | 8.677132638 | 0.90447297 | 0.4037721 | 0.712434734 | -4.640056097 |
| **A_51_P495825** | Apof | 0.09461841 | -0.167413263 | 0.356650084 | 7.818558921 | 0.904424257 | 0.403795764 | 0.712434734 | -4.640067675 |
| **A_52_P423128** | Arglu1 | 0.251186617 | -0.444572022 | 0.946945256 | 11.64475087 | 0.904249411 | 0.40388071 | 0.712434734 | -4.640109233 |
| **A_55_P2150737** | NA | -0.203156729 | -0.765909562 | 0.359596104 | 9.298988844 | -0.904198677 | 0.40390536 | 0.712434734 | -4.640121291 |
| **A_55_P2038737** | Slc29a1 | -0.298087141 | -1.123951747 | 0.527777465 | 11.2021356 | -0.904033934 | 0.403985414 | 0.712434734 | -4.640160442 |
| **A_52_P621617** | Dnajb4 | -0.080296654 | -0.302771102 | 0.142177794 | 8.247651967 | -0.903998383 | 0.404002691 | 0.712434734 | -4.64016889 |
| **A_55_P1999561** | Pram1 | -0.086545226 | -0.326347794 | 0.153257341 | 8.018529585 | -0.903940064 | 0.404031034 | 0.712434734 | -4.640182748 |
| **A_51_P387868** | Tekt4 | 0.529628555 | -0.938062753 | 1.997319863 | 9.715050811 | 0.903830548 | 0.404084263 | 0.712434734 | -4.640208771 |
| **A_55_P2126790** | 1500009L | 0.190708091 | -0.337810216 | 0.719226398 | 8.730040837 | 0.903773138 | 0.404112168 | 0.712434734 | -4.640222412 |
| **A_51_P238722** | Cd93 | 0.592746546 | -1.050169518 | 2.235662611 | 14.19423697 | 0.9036578 | 0.404168235 | 0.712434734 | -4.640249815 |
| **A_51_P494929** | Diablo | 0.13044091 | -0.231154078 | 0.492035899 | 10.45337963 | 0.903528308 | 0.40423119 | 0.712434734 | -4.640280579 |
| **A_55_P2080119** | NA | 0.468291143 | -0.829868752 | 1.766451038 | 14.97748287 | 0.903520688 | 0.404234895 | 0.712434734 | -4.640282389 |
| **A_55_P2143279** | Nkd2 | -0.108688974 | -0.41001194 | 0.192633992 | 10.28374153 | -0.903448913 | 0.404269793 | 0.712434734 | -4.64029944 |
| **A_52_P185485** | Arpc4 | -0.095687051 | -0.360975446 | 0.169601344 | 10.60892129 | -0.903410768 | 0.404288341 | 0.712434734 | -4.640308501 |
| **A_52_P180826** | Ndufaf7 | 0.124864064 | -0.221322184 | 0.471050312 | 10.88969087 | 0.903395659 | 0.404295687 | 0.712434734 | -4.64031209 |
| **A_51_P414396** | Mmrn2 | 0.465364554 | -0.825040121 | 1.755769228 | 12.00947983 | 0.903270274 | 0.40435666 | 0.712434734 | -4.640341874 |
| **A_52_P232648** | Slc38a1 | -0.288586496 | -1.088813741 | 0.511640749 | 8.013716267 | -0.903260496 | 0.404361416 | 0.712434734 | -4.640344197 |
| **A_55_P2130975** | Ncoa1 | 0.442746808 | -0.785485223 | 1.670978839 | 10.60765612 | 0.902870296 | 0.404551211 | 0.712434734 | -4.640436868 |
| **A_52_P563340** | Smim4 | 0.473990881 | -0.840979484 | 1.788961245 | 11.90844348 | 0.902826643 | 0.404572449 | 0.712434734 | -4.640447234 |
| **A_55_P2002115** | NA | 0.388334852 | -0.689006052 | 1.465675755 | 15.7810501 | 0.902824976 | 0.40457326 | 0.712434734 | -4.64044763 |
| **A_55_P2099750** | NA | -0.140366632 | -0.529798078 | 0.249064814 | 11.46011007 | -0.902782659 | 0.404593848 | 0.712434734 | -4.640457679 |
| **A_55_P1971709** | Dnajc8 | -0.152831644 | -0.576854061 | 0.271190773 | 9.868215856 | -0.902765207 | 0.404602339 | 0.712434734 | -4.640461823 |
| **A_51_P222283** | Mlycd | -0.096534321 | -0.36440332 | 0.171334678 | 11.35495494 | -0.90262974 | 0.404668253 | 0.712434734 | -4.640493989 |
| **A_55_P2168397** | NA | -0.301397036 | -1.137762159 | 0.534968086 | 15.38698925 | -0.90259601 | 0.404684667 | 0.712434734 | -4.640501998 |
| **A_55_P2090025** | Mest | 0.464009856 | -0.823611791 | 1.751631503 | 12.93136495 | 0.902587432 | 0.404688841 | 0.712434734 | -4.640504034 |
| **A_55_P2104846** | Rsl24d1 | 0.080058781 | -0.142120554 | 0.302238117 | 8.230132118 | 0.902517551 | 0.404722848 | 0.712434734 | -4.640520626 |
| **A_55_P2149219** | Aar2 | -0.192446334 | -0.726531086 | 0.341638417 | 10.1507513 | -0.902505401 | 0.404728761 | 0.712434734 | -4.64052351 |
| **A_66_P124549** | H13 | -0.519063256 | -1.959629939 | 0.921503427 | 12.90275924 | -0.902479334 | 0.404741446 | 0.712434734 | -4.640529699 |
| **A_55_P1967591** | NA | -0.247172806 | -0.93322062 | 0.438875008 | 11.61964458 | -0.902394933 | 0.404782524 | 0.712434734 | -4.640549736 |
| **A_51_P273433** | Prkg2 | 0.145934483 | -0.25914261 | 0.551011577 | 8.544349893 | 0.90234078 | 0.404808881 | 0.712434734 | -4.640562592 |
| **A_52_P101487** | Ddn | -0.47210023 | -1.782813819 | 0.838613359 | 8.288965082 | -0.90214585 | 0.404903769 | 0.712434734 | -4.640608864 |
| **A_55_P2186480** | NA | 0.18957485 | -0.336767178 | 0.715916878 | 9.857821624 | 0.902117311 | 0.404917662 | 0.712434734 | -4.640615639 |
| **A_55_P2043062** | Hs3st6 | 0.264365858 | -0.469905079 | 0.998636795 | 8.709508686 | 0.901777454 | 0.405083143 | 0.712434734 | -4.640696299 |
| **A_51_P516615** | Ndufb10 | 0.187679985 | -0.333617739 | 0.708977709 | 14.35154024 | 0.901742371 | 0.405100228 | 0.712434734 | -4.640704625 |

| | | | | | | | | | |
|---|---|---|---|---|---|---|---|---|---|
| **A_51_P485479** | Prrc2a | 0.087434939 | -0.155428471 | 0.33029835 | 9.589911548 | 0.901723253 | 0.405109538 | 0.712434734 | -4.640709161 |
| **A_55_P2160198** | NA | -0.273826544 | -1.034449005 | 0.486795917 | 10.24102958 | -0.901688945 | 0.405126247 | 0.712434734 | -4.640717303 |
| **A_55_P1966863** | Mad2l2 | -0.143223376 | -0.541121548 | 0.254674795 | 11.27843659 | -0.901555172 | 0.405191402 | 0.712434734 | -4.640749046 |
| **A_66_P103418** | Tmc8 | -0.246067835 | -0.929756933 | 0.437621264 | 8.346611236 | -0.901460161 | 0.405237683 | 0.712434734 | -4.640771589 |
| **A_52_P505234** | Stt3a | -0.251404522 | -0.949927038 | 0.447117995 | 8.945187337 | -0.901452858 | 0.405241241 | 0.712434734 | -4.640773322 |
| **A_51_P384441** | Tmem201 | 0.102918424 | -0.183044793 | 0.38888164 | 8.246602089 | 0.901432679 | 0.405251071 | 0.712434734 | -4.64077811 |
| **A_55_P1958140** | B9d1 | 0.494742921 | -0.879978157 | 1.869463998 | 11.6314506 | 0.901395411 | 0.405269226 | 0.712434734 | -4.640786952 |
| **A_51_P219266** | Tmprss6 | -0.13586557 | -0.513390647 | 0.241659508 | 7.805842636 | -0.90139253 | 0.40527063 | 0.712434734 | -4.640787636 |
| **A_55_P1981455** | NA | -0.260498607 | -0.984348827 | 0.463351614 | 8.100089149 | -0.901378176 | 0.405277622 | 0.712434734 | -4.640791041 |
| **A_51_P405423** | Smcr8 | -0.148514786 | -0.56121539 | 0.264185818 | 9.569470718 | -0.901332303 | 0.405299971 | 0.712434734 | -4.640801925 |
| **A_55_P2026748** | Zmat1 | -0.237268445 | -0.896782712 | 0.422245823 | 8.103775838 | -0.901085775 | 0.40542009 | 0.712581093 | -4.640860408 |
| **A_52_P262967** | Rab10 | -0.298794683 | -1.129492317 | 0.531902951 | 9.589757523 | -0.900907572 | 0.405506936 | 0.712626427 | -4.640902678 |
| **A_55_P2022585** | Pira6 | -0.079573806 | -0.300808261 | 0.141660649 | 7.991928791 | -0.900881588 | 0.4055196 | 0.712626427 | -4.640908841 |
| **A_52_P632093** | Gtf3c3 | 0.100362877 | -0.178747641 | 0.379473395 | 9.36534889 | 0.900631734 | 0.405641392 | 0.712631088 | -4.640968097 |
| **A_51_P462428** | NA | -0.152608672 | -0.577050476 | 0.271833133 | 8.015622324 | -0.900557412 | 0.405677626 | 0.712631088 | -4.640985722 |
| **A_51_P137604** | Fcna | 0.082367758 | -0.146727381 | 0.311462898 | 7.888378802 | 0.900516587 | 0.405697529 | 0.712631088 | -4.640995403 |
| **A_55_P1985385** | Acp2 | -0.18333957 | -0.693280923 | 0.326601784 | 8.167666131 | -0.900505399 | 0.405702984 | 0.712631088 | -4.640998056 |
| **A_55_P2170225** | Sh3bgrl2 | -0.408170574 | -1.543466921 | 0.727125774 | 8.344926571 | -0.900498092 | 0.405706547 | 0.712631088 | -4.640999789 |
| **A_51_P208377** | Trappc5 | 0.108281342 | -0.192975864 | 0.409538548 | 11.43699948 | 0.900257052 | 0.405824083 | 0.712648163 | -4.641056942 |
| **A_55_P2016852** | Zyx | -0.108152468 | -0.409126247 | 0.192821312 | 13.88701441 | -0.900032344 | 0.40593368 | 0.712648163 | -4.641110215 |
| **A_51_P257951** | Retnla | 1.768927829 | -3.153965818 | 6.691821477 | 13.10769531 | 0.899994813 | 0.405951987 | 0.712648163 | -4.641119112 |
| **A_55_P2117710** | Snhg1 | 0.354478295 | -0.632097933 | 1.341054522 | 9.609609389 | 0.899931279 | 0.405982979 | 0.712648163 | -4.641134172 |
| **A_51_P278489** | Itfg2 | 0.119480394 | -0.213069692 | 0.452030481 | 8.725419137 | 0.899891042 | 0.406002608 | 0.712648163 | -4.64114371 |
| **A_55_P2275054** | Nrap | 0.09243888 | -0.164867923 | 0.349745684 | 8.588576635 | 0.899816084 | 0.406039177 | 0.712648163 | -4.641161478 |
| **A_52_P243054** | Nufip2 | -0.188102525 | -0.711694314 | 0.335489265 | 8.774159725 | -0.899812749 | 0.406040804 | 0.712648163 | -4.641162268 |
| **A_55_P2020577** | Pcolce | 0.497693822 | -0.887749525 | 1.883137169 | 13.56712427 | 0.899754078 | 0.406069429 | 0.712648163 | -4.641176175 |
| **A_51_P356055** | Grp | 0.407486186 | -0.726936713 | 1.541909085 | 8.848219879 | 0.899680385 | 0.406105385 | 0.712648163 | -4.641193641 |
| **A_55_P2136786** | Zkscan1 | 0.23841749 | -0.425396361 | 0.90223134 | 14.13421714 | 0.899584878 | 0.406151989 | 0.712648163 | -4.641216276 |
| **A_55_P2059986** | Chst14 | 0.138214946 | -0.246614155 | 0.523044046 | 10.06030418 | 0.899575144 | 0.406156739 | 0.712648163 | -4.641218583 |
| **A_55_P2144526** | Ripor2 | -0.103288645 | -0.390874356 | 0.184297067 | 7.886927648 | -0.899571358 | 0.406158586 | 0.712648163 | -4.64121948 |
| **A_55_P1999760** | Lrrc51 | -0.175434506 | -0.664043794 | 0.313174782 | 10.61964478 | -0.899298036 | 0.40629198 | 0.71272244 | -4.64128425 |
| **A_66_P113749** | Il2rg | -0.416658231 | -1.577478483 | 0.744162021 | 8.848163965 | -0.899011681 | 0.406431771 | 0.71272244 | -4.641352097 |
| **A_51_P297105** | Ucp2 | 0.160137624 | -0.286010854 | 0.606286103 | 10.02125382 | 0.899009561 | 0.406432807 | 0.71272244 | -4.641352599 |
| **A_55_P2098450** | Snrnp200 | 0.194495075 | -0.347428174 | 0.736418325 | 12.16852049 | 0.898920266 | 0.406476406 | 0.71272244 | -4.641373754 |
| **A_52_P565575** | Arhgap20 | 0.384046417 | -0.686101721 | 1.454194554 | 9.230338967 | 0.898856085 | 0.406507745 | 0.71272244 | -4.641388958 |
| **A_55_P2057268** | Magi1 | -0.119713602 | -0.453329961 | 0.213902757 | 10.14664047 | -0.898765731 | 0.406551868 | 0.71272244 | -4.641410361 |
| **A_55_P2042698** | Hsd17b10 | 0.684545694 | -1.223235891 | 2.592327279 | 13.62423153 | 0.898719441 | 0.406574474 | 0.71272244 | -4.641421326 |
| **A_52_P253179** | Igfbp3 | -0.33602826 | -1.27253999 | 0.60048347 | 8.725108449 | -0.898696244 | 0.406585803 | 0.71272244 | -4.64142682 |
| **A_55_P2118675** | Tnip1 | -0.235490449 | -0.891815599 | 0.420834702 | 11.75275417 | -0.898679004 | 0.406594223 | 0.71272244 | -4.641430904 |
| **A_55_P2089223** | Ak4 | 0.257919948 | -0.460923046 | 0.976762941 | 10.57834006 | 0.898672047 | 0.40659762 | 0.71272244 | -4.641432552 |
| **A_52_P112651** | Lman2 | -0.131562493 | -0.498300134 | 0.235175147 | 9.440975213 | -0.898518357 | 0.406672687 | 0.71272244 | -4.641468953 |
| **A_51_P387845** | Fam120a | -0.110932194 | -0.420195504 | 0.198331117 | 10.23494084 | -0.898420279 | 0.406720597 | 0.71272244 | -4.641492181 |
| **A_52_P65719** | Eme2 | 0.081221586 | -0.145248944 | 0.307692116 | 8.637685879 | 0.898276661 | 0.406790759 | 0.71272244 | -4.641526192 |
| **A_55_P2037141** | Jsrp1 | -0.128737514 | -0.487705825 | 0.230230797 | 9.125776611 | -0.898254397 | 0.406801637 | 0.71272244 | -4.641531464 |
| **A_52_P598309** | Zfas1 | 0.712657444 | -1.274649825 | 2.699964714 | 12.94150795 | 0.89818573 | 0.406835188 | 0.71272244 | -4.641547725 |
| **A_55_P2056606** | NA | 0.284422282 | -0.543686877 | 1.11253144 | 8.379688001 | 0.914826494 | 0.406857361 | 0.71272244 | -4.626776247 |
| **A_55_P2159850** | NA | -0.091126577 | -0.345288678 | 0.163035525 | 13.34326476 | -0.898017104 | 0.406917587 | 0.71272244 | -4.641587652 |
| **A_51_P152612** | Nol7 | 0.161571481 | -0.289074303 | 0.612217265 | 8.467355622 | 0.898007037 | 0.406922507 | 0.71272244 | -4.641590035 |

| | | | | | | | | | |
|---|---|---|---|---|---|---|---|---|---|
| **A_55_P2119802** | NA | 0.129545394 | -0.231776148 | 0.490866936 | 15.84996437 | 0.898004403 | 0.406923794 | 0.71272244 | -4.641590659 |
| **A_51_P220723** | B230118H | 0.336433471 | -0.602008972 | 1.274875913 | 12.80837479 | 0.897928798 | 0.406960743 | 0.71272244 | -4.641608559 |
| **A_51_P108629** | Prl3b1 | -2.465854685 | -9.344296845 | 4.412587476 | 10.27287796 | -0.897899512 | 0.406975056 | 0.71272244 | -4.641615493 |
| **A_55_P1988328** | NA | 0.288628466 | -0.516770938 | 1.094027871 | 16.12873841 | 0.897590408 | 0.40712615 | 0.712862549 | -4.641688666 |
| **A_55_P2103351** | Ndufa5 | 0.341954943 | -0.612254377 | 1.296164263 | 12.71484981 | 0.897584982 | 0.407128802 | 0.712862549 | -4.64168995 |
| **A_52_P574214** | Rrp1b | -0.349111185 | -1.323647942 | 0.625425571 | 8.879165778 | -0.897254996 | 0.407290152 | 0.712926638 | -4.641768051 |
| **A_55_P2010586** | Gm6252 | -0.339988774 | -1.289142158 | 0.609164609 | 11.80489168 | -0.897177809 | 0.4073279 | 0.712926638 | -4.641786317 |
| **A_55_P2018482** | Adh5 | -0.478663314 | -1.815088463 | 0.857761836 | 7.709089515 | -0.897089655 | 0.407371016 | 0.712926638 | -4.641807178 |
| **A_55_P2109712** | Csnk1g2 | -0.117673691 | -0.44622839 | 0.210881008 | 8.546570777 | -0.89706113 | 0.407384968 | 0.712926638 | -4.641813928 |
| **A_51_P223443** | Cd81 | 0.550106554 | -0.985836926 | 2.086050033 | 15.04476228 | 0.897060905 | 0.407385078 | 0.712926638 | -4.641813981 |
| **A_51_P461578** | Fdxr | -0.458440358 | -1.738448742 | 0.821568027 | 9.644958832 | -0.897057691 | 0.40738665 | 0.712926638 | -4.641814742 |
| **A_51_P456914** | Bmt2 | 0.21636961 | -0.387868681 | 0.820607902 | 8.955776655 | 0.896888313 | 0.407469503 | 0.713007093 | -4.641854819 |
| **A_55_P1966660** | LOC5473 | -0.565613437 | -2.145652708 | 1.014425834 | 9.498819743 | -0.896607118 | 0.407607081 | 0.713174626 | -4.641921345 |
| **A_55_P2107854** | Hnrnpk | 0.177531684 | -0.318438593 | 0.673501962 | 11.07435274 | 0.896541852 | 0.407639019 | 0.713174626 | -4.641936784 |
| **A_51_P153486** | Dnajb1 | -0.343197961 | -1.302092491 | 0.61569657 | 13.83225302 | -0.896446142 | 0.407685857 | 0.713192035 | -4.641959423 |
| **A_51_P161054** | Itgb6 | -0.344636501 | -1.307696606 | 0.618423603 | 9.324201964 | -0.896309968 | 0.407752505 | 0.713244091 | -4.641991632 |
| **A_51_P386810** | Gmppb | -0.486859514 | -1.847578205 | 0.873859178 | 10.12794137 | -0.896160208 | 0.407825812 | 0.713251547 | -4.642027051 |
| **A_51_P101196** | Psme1 | 0.346147068 | -0.621306293 | 1.31360043 | 12.62855985 | 0.896150529 | 0.40783055 | 0.713251547 | -4.640225036 |
| **A_55_P1971963** | Tmem176 | 0.662934829 | -1.190176533 | 2.516046192 | 14.18691166 | 0.896024043 | 0.407892473 | 0.71328579 | -4.642059253 |
| **A_51_P477419** | Nfic | 0.380111937 | -0.682496644 | 1.442720518 | 11.40643226 | 0.895959822 | 0.407923915 | 0.71328579 | -4.642074439 |
| **A_55_P1979674** | Ikzf4 | -0.095080687 | -0.361031154 | 0.170869779 | 8.492296417 | -0.895451157 | 0.408173027 | 0.713656836 | -4.642194702 |
| **A_55_P2080603** | Hspa8 | -0.216825373 | -0.823522015 | 0.389871268 | 15.81401269 | -0.895135657 | 0.408327597 | 0.713862534 | -4.642269277 |
| **A_55_P1955209** | NA | 0.286124713 | -0.514770003 | 1.087019429 | 15.38193755 | 0.894808884 | 0.408487737 | 0.713907242 | -4.6423465 |
| **A_51_P261379** | Spats2 | -0.474522139 | -1.802764025 | 0.853719748 | 9.156527827 | -0.894807577 | 0.408488378 | 0.713907242 | -4.642346809 |
| **A_52_P652193** | Sar1a | -0.172064399 | -0.653788532 | 0.309659734 | 13.89456417 | -0.894628984 | 0.408575921 | 0.713907242 | -4.642389008 |
| **A_55_P1953475** | NA | -0.164300086 | -0.624309941 | 0.295709768 | 16.11556847 | -0.894583744 | 0.408598099 | 0.713907242 | -4.642399697 |
| **A_52_P625624** | Stau2 | -0.102569554 | -0.389770761 | 0.184631654 | 8.032735662 | -0.894504541 | 0.408636929 | 0.713907242 | -4.642418409 |
| **A_55_P1993473** | Gm5935 | -0.115157555 | -0.437627824 | 0.207312714 | 7.899013857 | -0.894443889 | 0.408666667 | 0.713907242 | -4.642432738 |
| **A_55_P2158151** | NA | 0.187003054 | -0.336659834 | 0.710665942 | 16.11767091 | 0.894431835 | 0.408672576 | 0.713907242 | -4.642435586 |
| **A_66_P136813** | Smim41 | 1.35274351 | -2.435657452 | 5.141144472 | 12.04635674 | 0.89435485 | 0.408710325 | 0.713907242 | -4.642453773 |
| **A_55_P2014249** | Sema3a | 0.627261276 | -1.129484882 | 2.384007435 | 9.92410551 | 0.894313549 | 0.408730577 | 0.713907242 | -4.642463529 |
| **A_55_P2091308** | Rpl38 | 0.106148725 | -0.191139858 | 0.403437307 | 16.6976854 | 0.894307547 | 0.40873352 | 0.713907242 | -4.642464947 |
| **A_52_P356780** | Pdzd8 | 0.15452682 | -0.278283785 | 0.587337425 | 8.948398959 | 0.894244584 | 0.408764396 | 0.713907242 | -4.64247982 |
| **A_66_P112005** | Olfr1252 | -0.387197916 | -1.471770456 | 0.697374623 | 8.341643579 | -0.894179591 | 0.408796269 | 0.713907242 | -4.642495172 |
| **A_55_P2123381** | Fga | -0.35859133 | -1.363221482 | 0.646038823 | 8.04690311 | -0.894013181 | 0.408877887 | 0.713985285 | -4.642534477 |
| **A_55_P1968723** | Noct | -1.184933246 | -4.505203199 | 2.135336707 | 10.44796919 | -0.893862741 | 0.408951683 | 0.714049656 | -4.642570006 |
| **A_55_P2024358** | Rpl34 | 0.190748716 | -0.343861377 | 0.725358809 | 16.20390059 | 0.893665137 | 0.40904863 | 0.714154436 | -4.642616669 |
| **A_55_P2021665** | Zfp638 | 0.215507563 | -0.388665696 | 0.819680821 | 10.75573838 | 0.893411137 | 0.409173272 | 0.714216563 | -4.642676642 |
| **A_52_P260994** | Fgd2 | 0.301527254 | -0.543820409 | 1.146874916 | 8.716059482 | 0.893390976 | 0.409183167 | 0.714216563 | -4.642681402 |
| **A_55_P2090718** | Pop5 | 0.185577436 | -0.334712845 | 0.705867717 | 12.75428806 | 0.893366788 | 0.409195038 | 0.714216563 | -4.642687112 |
| **A_55_P2161132** | NA | -0.114193588 | -0.434422953 | 0.206035776 | 7.932073437 | -0.893163388 | 0.409294875 | 0.714260164 | -4.642735129 |
| **A_51_P486001** | Mob3a | -0.235843839 | -0.897314644 | 0.425626966 | 8.641825784 | -0.8930262 | 0.409362222 | 0.714260164 | -4.642767512 |
| **A_55_P2025073** | Csf1r | -0.740410039 | -2.817096163 | 1.336276084 | 9.335633459 | -0.893000667 | 0.409374758 | 0.714260164 | -4.642773538 |
| **A_55_P1961690** | Tctex1d2 | 0.305007665 | -0.550583251 | 1.160598581 | 10.8759343 | 0.892883781 | 0.409432147 | 0.714260164 | -4.642801126 |
| **A_55_P2012206** | Ndufb11 | 0.358329231 | -0.647005178 | 1.363663639 | 14.69176969 | 0.892733919 | 0.409505737 | 0.714260164 | -4.642836494 |
| **A_51_P396854** | Timm10b | 0.154021318 | -0.278109875 | 0.586152512 | 13.17629092 | 0.892720617 | 0.409512269 | 0.714260164 | -4.642839633 |
| **A_51_P424641** | Sirt4 | -0.128083997 | -0.487466368 | 0.231298373 | 8.796401884 | -0.892664881 | 0.40953964 | 0.714260164 | -4.642852786 |
| **A_51_P142091** | Asb8 | -0.118507457 | -0.451030634 | 0.214015721 | 11.37962339 | -0.89263539 | 0.409554124 | 0.714260164 | -4.642859746 |

| | | | | | | | | | |
|---|---|---|---|---|---|---|---|---|---|
| **A_55_P2027737** | Il1rl1 | 0.110780522 | -0.200100503 | 0.421661548 | 8.013487375 | 0.892523249 | 0.409609202 | 0.714260164 | -4.642886208 |
| **A_51_P369897** | Zfp410 | -0.119248434 | -0.453950225 | 0.215453357 | 9.590909091 | -0.892370069 | 0.409684446 | 0.714260164 | -4.642922351 |
| **A_55_P2069597** | Tspan17 | -0.083273081 | -0.317018355 | 0.150472192 | 8.486000212 | -0.892302567 | 0.409717607 | 0.714260164 | -4.642938277 |
| **A_55_P2041131** | NA | 0.312542795 | -0.564955448 | 1.190041037 | 13.89669473 | 0.892100086 | 0.40981709 | 0.714260164 | -4.642986045 |
| **A_55_P2060184** | Lamtor3 | -0.095668262 | -0.364268731 | 0.172932206 | 12.29088717 | -0.89209574 | 0.409819225 | 0.714260164 | -4.64298707 |
| **A_55_P2156973** | Aar2 | -0.185093223 | -0.704804999 | 0.334618554 | 10.12614163 | -0.892027627 | 0.409852695 | 0.714260164 | -4.643003138 |
| **A_55_P1990041** | Nxt2 | 0.486847291 | -0.880352146 | 1.854046728 | 10.61464126 | 0.891889872 | 0.409920392 | 0.714260164 | -4.643035631 |
| **A_55_P2005873** | Zfp280d | -0.139606899 | -0.53166914 | 0.252455341 | 9.191548053 | -0.891871345 | 0.409929498 | 0.714260164 | -4.643040001 |
| **A_51_P276943** | Gpc1 | -0.709287178 | -2.701199682 | 1.282625326 | 12.78879498 | -0.891871316 | 0.409929512 | 0.714260164 | -4.643040008 |
| **A_51_P288257** | Cdk5rap3 | -0.115649867 | -0.440460827 | 0.209161094 | 11.23150616 | -0.891794535 | 0.409967249 | 0.714260164 | -4.643058118 |
| **A_55_P2100088** | Ccdc92b | 0.11104062 | -0.200833787 | 0.422915028 | 7.925157136 | 0.891769239 | 0.409979682 | 0.714260164 | -4.643064084 |
| **A_51_P432199** | Sap30 | 0.30049649 | -0.543564661 | 1.144557641 | 9.31748606 | 0.891693986 | 0.410016671 | 0.714260164 | -4.643081832 |
| **A_51_P141104** | Med18 | 0.079889282 | -0.14451626 | 0.304294825 | 8.176295383 | 0.891672323 | 0.41002732 | 0.714260164 | -4.643086941 |
| **A_55_P1986993** | Brd8 | 0.113951461 | -0.206155254 | 0.434058176 | 8.218595423 | 0.891611083 | 0.410057424 | 0.714260164 | -4.643101383 |
| **A_51_P160544** | Efemp2 | 0.481005434 | -0.87038507 | 1.832395938 | 12.90757444 | 0.891496137 | 0.410113933 | 0.714260164 | -4.64312849 |
| **A_55_P2082186** | NA | 0.271494528 | -0.491312778 | 1.034301834 | 13.22599967 | 0.891449163 | 0.410137027 | 0.714260164 | -4.643139566 |
| **A_55_P1990525** | Ube2v1 | -0.121945688 | -0.464593493 | 0.220702117 | 14.41414301 | -0.891392221 | 0.410165025 | 0.714260164 | -4.643152993 |
| **A_55_P1985464** | Gm12271 | 0.560529425 | -1.014549022 | 2.135607872 | 14.23782491 | 0.891346519 | 0.410187496 | 0.714260164 | -4.643163769 |
| **A_51_P501815** | Smdt1 | -0.109688059 | -0.41795269 | 0.198576572 | 14.58248002 | -0.891222205 | 0.410248626 | 0.714260164 | -4.64319308 |
| **A_55_P2017764** | Cep89 | 0.292392362 | -0.529350842 | 1.114135566 | 10.77311811 | 0.891210405 | 0.410254429 | 0.714260164 | -4.643195862 |
| **A_51_P432802** | Rbm4 | 0.105691364 | -0.191409892 | 0.40279262 | 11.36541737 | 0.891015712 | 0.410350181 | 0.714298798 | -4.643241762 |
| **A_52_P192426** | Tnfrsf1a | 0.18778119 | -0.340077159 | 0.715639539 | 13.06247024 | 0.891015045 | 0.410350509 | 0.714298798 | -4.643241919 |
| **A_55_P1993128** | Gas8 | 0.158824018 | -0.287771619 | 0.605419655 | 10.79935547 | 0.89074224 | 0.410484707 | 0.714468071 | -4.643306224 |
| **A_55_P2018557** | NA | 0.220387412 | -0.399459507 | 0.840234332 | 12.00415165 | 0.890538469 | 0.410584968 | 0.71457825 | -4.64335425 |
| **A_55_P2016069** | Adamtsl1 | 0.165320046 | -0.299800657 | 0.630440748 | 8.580178977 | 0.890246328 | 0.410728742 | 0.714697355 | -4.643423092 |
| **A_55_P2114218** | Zcchc7 | -0.088294109 | -0.336729889 | 0.160141672 | 10.05042505 | -0.890159698 | 0.410771384 | 0.714697355 | -4.643443504 |
| **A_51_P292550** | Chdh | -0.31902377 | -1.21678013 | 0.578732591 | 8.158769697 | -0.890051009 | 0.410824888 | 0.714697355 | -4.643469112 |
| **A_55_P2312783** | D030029J | 0.293346054 | -0.532202942 | 1.118895051 | 11.07268835 | 0.889995361 | 0.410852284 | 0.714697355 | -4.643482222 |
| **A_55_P1980501** | Znhit1 | 0.166048957 | -0.301275459 | 0.633373372 | 10.28577355 | 0.889954956 | 0.410872176 | 0.714697355 | -4.643491741 |
| **A_55_P2104894** | Col5a2 | 0.436460938 | -0.792240595 | 1.665162471 | 10.14995185 | 0.889711753 | 0.410991928 | 0.714697355 | -4.643549031 |
| **A_51_P270213** | Ceacam1 | -2.441415848 | -9.314641076 | 4.431809381 | 10.05401033 | -0.889675292 | 0.411009883 | 0.714697355 | -4.643557619 |
| **A_55_P2053810** | NA | -0.355783609 | -1.357483462 | 0.645916245 | 8.424037128 | -0.889607983 | 0.411043032 | 0.714697355 | -4.643573473 |
| **A_55_P2107347** | Ro60 | 0.305276539 | -0.554320818 | 1.164873896 | 9.499309258 | 0.889505635 | 0.41109344 | 0.714697355 | -4.643597578 |
| **A_55_P2015936** | NA | 0.208156118 | -0.37800836 | 0.794320596 | 16.28811477 | 0.889446907 | 0.411122367 | 0.714697355 | -4.64361141 |
| **A_55_P2124381** | NA | 0.087660996 | -0.159191229 | 0.334513222 | 12.15119974 | 0.889446236 | 0.411122697 | 0.714697355 | -4.643611567 |
| **A_55_P2022369** | Lbh | 0.60186002 | -1.093320785 | 2.297040826 | 10.45378965 | 0.889262165 | 0.411213372 | 0.714697355 | -4.643654915 |
| **A_55_P2140107** | Cyp21a1 | -0.380810701 | -1.453535244 | 0.691913841 | 8.214527611 | -0.889142303 | 0.411272425 | 0.714697355 | -4.643683139 |
| **A_52_P325527** | Edil3 | 0.314770763 | -0.572024056 | 1.201565581 | 8.567376487 | 0.889040581 | 0.411322547 | 0.714697355 | -4.64370709 |
| **A_55_P2173313** | Plcb1 | 0.189840903 | -0.344999283 | 0.72468109 | 8.299710302 | 0.889029364 | 0.411328074 | 0.714697355 | -4.643709731 |
| **A_55_P2383248** | 4930422N | -0.409523341 | -1.563277785 | 0.744231102 | 8.596500855 | -0.889028368 | 0.411328565 | 0.714697355 | -4.643709966 |
| **A_55_P2260094** | Tcf7l2 | -0.085612376 | -0.326813748 | 0.155588996 | 7.943973022 | -0.889010973 | 0.411337136 | 0.714697355 | -4.643714061 |
| **A_55_P2008889** | Tmsb15b2 | 0.207587144 | -0.377274271 | 0.79244856 | 9.940483592 | 0.888991951 | 0.41134651 | 0.714697355 | -4.64371854 |
| **A_55_P2137941** | Fxyd2 | -0.802272263 | -3.062662692 | 1.458118166 | 9.072159031 | -0.88897319 | 0.411355755 | 0.714697355 | -4.643722957 |
| **A_55_P2152074** | Ctps2 | 0.166846605 | -0.303444079 | 0.637137289 | 11.25848021 | 0.88858984 | 0.411544696 | 0.714908264 | -4.643813201 |
| **A_51_P124345** | Cpne5 | 0.162306596 | -0.29519385 | 0.619807042 | 8.821210614 | 0.888576848 | 0.4115511 | 0.714908264 | -4.643816259 |
| **A_55_P2013823** | Gal3st1 | 0.215272317 | -0.39160557 | 0.822150205 | 8.921782977 | 0.888458645 | 0.411609373 | 0.714931781 | -4.64384408 |
| **A_52_P561671** | Msx1 | -0.091595521 | -0.349851253 | 0.166660211 | 7.988733081 | -0.888330582 | 0.411672514 | 0.714931781 | -4.64387422 |
| **A_55_P1995996** | Zfp984 | 0.109377505 | -0.199041046 | 0.417796057 | 8.716950431 | 0.888255421 | 0.411709575 | 0.714931781 | -4.643891908 |

| | | | | | | | | | |
|---|---|---|---|---|---|---|---|---|---|
| **A_55_P2032258** | Pdlim1 | -0.903769034 | -3.452200074 | 1.644662006 | 13.20070059 | -0.888249387 | 0.411712551 | 0.714931781 | -4.643893328 |
| **A_52_P101333** | Sap18 | 0.119848528 | -0.2181449 | 0.457841957 | 8.604568718 | 0.888126431 | 0.411773185 | 0.714972856 | -4.643922262 |
| **A_55_P2047200** | NA | -0.168925735 | -0.645467701 | 0.307616232 | 8.888807945 | -0.88786106 | 0.411904072 | 0.715102824 | -4.643984702 |
| **A_55_P2174905** | Cuta | -0.085023605 | -0.324901663 | 0.154854453 | 12.1932757 | -0.88776769 | 0.411950132 | 0.715102824 | -4.644006668 |
| **A_55_P1981090** | Thap7 | -0.151367115 | -0.578502925 | 0.275768694 | 10.01377243 | -0.887597147 | 0.412034273 | 0.715102824 | -4.644046788 |
| **A_55_P1998159** | Deaf1 | -0.149546717 | -0.571558599 | 0.272465165 | 8.348020048 | -0.887569869 | 0.412047732 | 0.715102824 | -4.644053205 |
| **A_55_P1952738** | Ppp3ca | 0.331255975 | -0.603658417 | 1.266170366 | 12.71289835 | 0.887446581 | 0.412108568 | 0.715102824 | -4.644082204 |
| **A_55_P2163353** | Sox3 | -0.328288339 | -1.254868344 | 0.598291666 | 8.050154246 | -0.887407057 | 0.412128072 | 0.715102824 | -4.644091501 |
| **A_55_P2029517** | Wnk2 | -0.090471178 | -0.345829354 | 0.164886998 | 7.937710521 | -0.887382425 | 0.412140228 | 0.715102824 | -4.644097294 |
| **A_55_P2121985** | Plcb4 | -0.074883475 | -0.286273625 | 0.136506675 | 8.151380815 | -0.887261396 | 0.41219996 | 0.715102824 | -4.644125759 |
| **A_52_P195839** | Ctsc | 0.854434185 | -1.557612832 | 3.266481203 | 11.88068148 | 0.887244248 | 0.412208423 | 0.715102824 | -4.644129792 |
| **A_55_P1992208** | Gpi1 | 0.186828976 | -0.340621737 | 0.714279689 | 14.27101871 | 0.887181944 | 0.412239175 | 0.715102824 | -4.644144445 |
| **A_51_P377856** | Gstt3 | 0.380553121 | -0.694073609 | 1.455179851 | 10.53092729 | 0.886968088 | 0.412344743 | 0.715102824 | -4.644194734 |
| **A_55_P2022158** | Gc | -0.635119376 | -2.428704576 | 1.158465824 | 8.812528628 | -0.886918464 | 0.412369243 | 0.715102824 | -4.644206402 |
| **A_51_P435922** | Rsph9 | 0.70360338 | -1.283453773 | 2.690660533 | 10.3213818 | 0.886886214 | 0.412385165 | 0.715102824 | -4.644213985 |
| **A_55_P1985498** | Gm12005 | 0.408479872 | -0.745143988 | 1.562103733 | 14.8308604 | 0.886863492 | 0.412396384 | 0.715102824 | -4.644219328 |
| **A_55_P2065059** | Wnt2 | 0.306418838 | -0.55898631 | 1.171823985 | 8.722835725 | 0.886842163 | 0.412406915 | 0.715102824 | -4.644224343 |
| **A_55_P1952788** | NA | 0.095783788 | -0.174754454 | 0.36632203 | 8.834146743 | 0.886775509 | 0.412439826 | 0.715102824 | -4.644240014 |
| **A_51_P354272** | Cluh | 0.136904291 | -0.249868229 | 0.523676811 | 10.96686187 | 0.886567478 | 0.412542557 | 0.715210504 | -4.644288921 |
| **A_52_P426513** | Sh3bgrl | 0.637468283 | -1.163600169 | 2.438536735 | 11.36260998 | 0.886499935 | 0.412575915 | 0.715210504 | -4.644304799 |
| **A_55_P2000818** | Cab39l | 0.193620472 | -0.353590749 | 0.740831694 | 9.279822051 | 0.886230365 | 0.412709073 | 0.715364256 | -4.644368161 |
| **A_51_P122425** | Ctf1 | 0.123213749 | -0.225058871 | 0.47148637 | 9.225425823 | 0.886115197 | 0.412765972 | 0.715364256 | -4.644395228 |
| **A_55_P2020472** | Rufy2 | -0.295128585 | -1.129349352 | 0.539092182 | 9.017168062 | -0.88609569 | 0.41277561 | 0.715364256 | -4.644399812 |
| **A_55_P2027698** | Ndufa5 | 0.216378162 | -0.395371545 | 0.828127869 | 14.19771345 | 0.885910812 | 0.412866963 | 0.715387216 | -4.644443258 |
| **A_51_P172532** | Nit1 | -0.181338036 | -0.69402852 | 0.331352449 | 11.12897101 | -0.88589856 | 0.412873017 | 0.715387216 | -4.644446137 |
| **A_55_P2179974** | Spsb1 | -0.365853891 | -1.400308844 | 0.668601061 | 8.346109157 | -0.885821957 | 0.412910874 | 0.715387216 | -4.644464136 |
| **A_52_P595225** | Vps35 | -0.095190077 | -0.364356799 | 0.173976645 | 8.816263611 | -0.885769369 | 0.412936864 | 0.715387216 | -4.644476493 |
| **A_52_P820196** | Cxcl15 | 1.227334024 | -2.244004943 | 4.69867299 | 11.45128227 | 0.885556135 | 0.413042262 | 0.715448134 | -4.644526591 |
| **A_55_P1966615** | Actr3b | -0.173506851 | -0.664250427 | 0.317236725 | 8.073532782 | -0.885548498 | 0.413046037 | 0.715448134 | -4.644528385 |
| **A_52_P445360** | Krt20 | -0.194428324 | -0.744397866 | 0.355541218 | 7.96176677 | -0.885464674 | 0.413087475 | 0.715455437 | -4.644548077 |
| **A_52_P286928** | NA | 0.139680363 | -0.255516126 | 0.534876852 | 9.623991288 | 0.885263633 | 0.413186873 | 0.715455437 | -4.644595301 |
| **A_52_P228079** | Atf1 | -0.190887362 | -0.730976354 | 0.34920163 | 8.540832541 | -0.885242377 | 0.413197383 | 0.715455437 | -4.644600294 |
| **A_55_P2049147** | NA | 0.366033849 | -0.669609645 | 1.401677342 | 15.8212524 | 0.885240578 | 0.413198273 | 0.715455437 | -4.644600717 |
| **A_51_P113773** | Slx1b | -0.142481913 | -0.545748741 | 0.260784914 | 8.602620868 | -0.884947659 | 0.413343133 | 0.715509992 | -4.644669511 |
| **A_51_P371255** | Rwdd2b | 0.110734137 | -0.202705219 | 0.424173493 | 8.759092766 | 0.884867702 | 0.413382681 | 0.715509992 | -4.644688288 |
| **A_52_P575078** | Babam1 | 0.298192343 | -0.545889917 | 1.142274604 | 12.45683038 | 0.884834526 | 0.413399092 | 0.715509992 | -4.644696078 |
| **A_51_P341736** | Mmp2 | 0.504676688 | -0.924109801 | 1.933463177 | 11.13319426 | 0.884700464 | 0.41346541 | 0.715509992 | -4.644727557 |
| **A_55_P2021854** | NA | 0.083047524 | -0.15208526 | 0.318180308 | 8.434351489 | 0.884634449 | 0.41349807 | 0.715509992 | -4.644743058 |
| **A_55_P2104917** | Nphp1 | 0.169898403 | -0.311171668 | 0.650968474 | 8.413973854 | 0.88456816 | 0.413530867 | 0.715509992 | -4.644758621 |
| **A_51_P260639** | Elp3 | -0.139305657 | -0.533762018 | 0.255150703 | 10.43011662 | -0.884545416 | 0.413542121 | 0.715509992 | -4.644763961 |
| **A_52_P561650** | Vwa1 | 0.154133495 | -0.282316921 | 0.59058391 | 8.475239953 | 0.884529782 | 0.413549856 | 0.715509992 | -4.644767632 |
| **A_51_P423008** | Klhl7 | 0.240640879 | -0.440868731 | 0.92215049 | 8.87608519 | 0.884398031 | 0.41361505 | 0.715509992 | -4.644798562 |
| **A_55_P2159830** | Yaf2 | 0.093760646 | -0.171779266 | 0.359300557 | 8.949846204 | 0.884384513 | 0.413621739 | 0.715509992 | -4.644801735 |
| **A_55_P2168663** | Atp5k | 0.233036901 | -0.426996169 | 0.893069971 | 14.61576647 | 0.884319795 | 0.413653766 | 0.715509992 | -4.644816928 |
| **A_55_P2021886** | Mrpl2 | -0.216012804 | -0.827856443 | 0.395830834 | 13.20562533 | -0.884279164 | 0.413673875 | 0.715509992 | -4.644826466 |
| **A_51_P431329** | Car3 | 0.191360964 | -0.35075441 | 0.733476338 | 8.153225721 | 0.884121541 | 0.413751889 | 0.715551821 | -4.644863464 |
| **A_55_P2147101** | Sytl4 | 0.200462834 | -0.367504754 | 0.768430421 | 8.373998679 | 0.884017084 | 0.413803596 | 0.715551821 | -4.64488798 |
| **A_51_P426270** | Mgp | 1.033242282 | -1.894471242 | 3.960955806 | 14.686776 | 0.88394234 | 0.413840597 | 0.715551821 | -4.644905522 |

| | | | | | | | | | |
|---|---|---|---|---|---|---|---|---|---|
| **A_55_P2097269** | NA | 0.209796462 | -0.384684506 | 0.80427743 | 11.19056288 | 0.883915095 | 0.413854085 | 0.715551821 | -4.644911916 |
| **A_55_P2108678** | NA | 0.094759619 | -0.173786166 | 0.363305404 | 8.949669328 | 0.883802672 | 0.413909746 | 0.715551821 | -4.644938299 |
| **A_55_P2077055** | Itih4 | -0.625038751 | -2.396420941 | 1.146343438 | 8.794320407 | -0.883781722 | 0.413920118 | 0.715551821 | -4.644943215 |
| **A_55_P2102961** | NA | 0.147023186 | -0.269725456 | 0.563771828 | 8.859311215 | 0.883612768 | 0.41400378 | 0.715632461 | -4.64498286 |
| **A_51_P229816** | Eif2b2 | 0.270971464 | -0.497275989 | 1.039218917 | 13.04446185 | 0.883431284 | 0.41409366 | 0.715659006 | -4.645025441 |
| **A_51_P419925** | Arhgef40 | 0.324049007 | -0.594787087 | 1.242885102 | 9.12958736 | 0.883330028 | 0.414143813 | 0.715659006 | -4.645049196 |
| **A_55_P1960621** | NA | 0.251374515 | -0.461524549 | 0.96427358 | 16.47362093 | 0.883168452 | 0.414223854 | 0.715659006 | -4.645087099 |
| **A_55_P1975445** | NA | 0.352489959 | -0.647211194 | 1.352191113 | 13.60796936 | 0.883134605 | 0.414240622 | 0.715659006 | -4.645095038 |
| **A_55_P2069392** | NA | 0.697602932 | -1.280924655 | 2.676130519 | 14.29594937 | 0.883113519 | 0.414251069 | 0.715659006 | -4.645099984 |
| **A_55_P2175880** | Mgst1 | 0.651853975 | -1.197031733 | 2.500739682 | 12.10215723 | 0.883060784 | 0.414277196 | 0.715659006 | -4.645112354 |
| **A_66_P120347** | Sstr5 | 0.158868561 | -0.291739286 | 0.609476408 | 10.84437894 | 0.883058667 | 0.414278245 | 0.715659006 | -4.64511285 |
| **A_52_P24439** | Sftpa1 | 1.496832585 | -2.749337187 | 5.743002356 | 12.30743157 | 0.882929892 | 0.414342052 | 0.715705283 | -4.645143054 |
| **A_55_P1969352** | Get4 | 0.297735692 | -0.547325862 | 1.142797246 | 11.31385658 | 0.882455678 | 0.414577085 | 0.715819527 | -4.645254257 |
| **A_55_P2060278** | Fam45a | -0.178451064 | -0.684974603 | 0.328072475 | 10.06277563 | -0.882408848 | 0.4146003 | 0.715819527 | -4.645265236 |
| **A_52_P90363** | Ifi27l2a | -0.398808196 | -1.530922271 | 0.733305878 | 10.40323526 | -0.882316156 | 0.414646255 | 0.715819527 | -4.645286968 |
| **A_55_P2000022** | Ccdc151 | 0.390818108 | -0.718639076 | 1.500275291 | 9.008438721 | 0.882296337 | 0.414656081 | 0.715819527 | -4.645291614 |
| **A_55_P1954604** | NA | -0.311874072 | -1.197333599 | 0.573585456 | 9.145967622 | -0.882187501 | 0.414710045 | 0.715819527 | -4.645317129 |
| **A_51_P463452** | Acsl1 | -0.084284764 | -0.323599892 | 0.155030364 | 7.969384259 | -0.882123244 | 0.414741908 | 0.715819527 | -4.645332192 |
| **A_55_P2108655** | Mtch2 | 0.204875801 | -0.376845209 | 0.786596812 | 13.03736833 | 0.88211714 | 0.414744935 | 0.715819527 | -4.645333623 |
| **A_55_P2037912** | Cyp2d10 | 0.080703869 | -0.148456731 | 0.309864469 | 7.718162862 | 0.882073451 | 0.4147666 | 0.715819527 | -4.645343864 |
| **A_66_P111987** | Brwd3 | -0.295494506 | -1.134575803 | 0.543586792 | 8.801650074 | -0.882055098 | 0.414775702 | 0.715819527 | -4.645348166 |
| **A_55_P1955676** | Kat14 | -0.298224925 | -1.145191506 | 0.548741656 | 10.09740395 | -0.881917598 | 0.414843894 | 0.715819527 | -4.645380395 |
| **A_55_P2037613** | Elmsan1 | -0.084475044 | -0.324400974 | 0.155450887 | 9.636459488 | -0.881863938 | 0.414870509 | 0.715819527 | -4.645392972 |
| **A_55_P2041893** | NA | 0.284107383 | -0.522821602 | 1.091036369 | 15.518273 | 0.881855748 | 0.414874571 | 0.715819527 | -4.645394891 |
| **A_51_P459108** | Insl6 | -0.182875623 | -0.70230133 | 0.336550083 | 8.460681441 | -0.881825648 | 0.414889501 | 0.715819527 | -4.645401946 |
| **A_52_P131372** | NA | 0.322612932 | -0.593852935 | 1.239078799 | 17.46280062 | 0.881689814 | 0.414956882 | 0.715871898 | -4.64543378 |
| **A_55_P2111419** | Eny2 | 0.272952808 | -0.502699326 | 1.048604941 | 13.12204987 | 0.881395685 | 0.415102813 | 0.715877947 | -4.645502702 |
| **A_66_P103229** | Foxl1 | 0.259216749 | -0.477413216 | 0.995846713 | 8.424958213 | 0.881381688 | 0.415109758 | 0.715877947 | -4.645505982 |
| **A_55_P1966908** | Zfp40 | 0.166990415 | -0.30759593 | 0.64157676 | 8.452247899 | 0.881305848 | 0.415147393 | 0.715877947 | -4.645523751 |
| **A_55_P2184009** | Rnd2 | -0.459232873 | -1.764455314 | 0.845989568 | 8.824977408 | -0.881249293 | 0.41517546 | 0.715877947 | -4.645537001 |
| **A_55_P2017789** | Il4ra | -0.114913313 | -0.441527176 | 0.21170055 | 8.357495257 | -0.881223513 | 0.415188254 | 0.715877947 | -4.64554304 |
| **A_55_P1964985** | Gm10240 | 0.104476384 | -0.192490937 | 0.401443706 | 16.22109733 | 0.881170233 | 0.415214697 | 0.715877947 | -4.645555523 |
| **A_55_P2014269** | Zfp771 | -0.116693169 | -0.448415168 | 0.21502883 | 12.64553394 | -0.881092503 | 0.415253277 | 0.715877947 | -4.645573732 |
| **A_55_P2178207** | Abcc8 | -0.186216609 | -0.71557592 | 0.343142702 | 8.871090782 | -0.881085801 | 0.415256604 | 0.715877947 | -4.645575302 |
| **A_55_P1957739** | Guca2b | -0.508798399 | -1.955306036 | 0.937709238 | 8.081104856 | -0.880998869 | 0.415299754 | 0.715888503 | -4.645595666 |
| **A_55_P1973553** | Bbs12 | 0.244052146 | -0.449968138 | 0.938072431 | 8.784937824 | 0.88076655 | 0.415415088 | 0.715965355 | -4.645650081 |
| **A_51_P459944** | Tcf21 | 1.037424834 | -1.913670578 | 3.988520246 | 11.04447481 | 0.880488594 | 0.41555311 | 0.715965355 | -4.645715175 |
| **A_55_P1969543** | Pin4 | 0.262643549 | -0.484507485 | 1.009794583 | 12.09431971 | 0.880458076 | 0.415568266 | 0.715965355 | -4.645722321 |
| **A_55_P2167112** | Kank2 | 0.496099906 | -0.915177012 | 1.907376824 | 12.93029612 | 0.880455179 | 0.415569705 | 0.715965355 | -4.645723 |
| **A_51_P160372** | Wbp1 | 0.278248958 | -0.513399541 | 1.069897457 | 12.94636652 | 0.880342157 | 0.415625839 | 0.715965355 | -4.645749464 |
| **A_55_P2177154** | H2bc14 | 0.256183467 | -0.472690723 | 0.985057656 | 12.32091903 | 0.880336832 | 0.415628484 | 0.715965355 | -4.645750711 |
| **A_55_P1965655** | Mtmr14 | -0.092987489 | -0.357580672 | 0.171605695 | 8.517572979 | -0.880230093 | 0.415681503 | 0.715965355 | -4.645775703 |
| **A_52_P69020** | Slc24a5 | 0.117729938 | -0.217339925 | 0.452799802 | 8.03265609 | 0.88003895 | 0.41577646 | 0.715965355 | -4.645820452 |
| **A_55_P2005982** | NA | -0.611326191 | -2.351238991 | 1.12858661 | 10.59152915 | -0.880026732 | 0.41578253 | 0.715965355 | -4.645823312 |
| **A_51_P462516** | Senp7 | 0.130181123 | -0.24035953 | 0.500721776 | 8.540087796 | 0.879959123 | 0.415816122 | 0.715965355 | -4.645839139 |
| **A_55_P2228715** | Mm2pr | -0.395424021 | -1.521167848 | 0.730319807 | 8.204983112 | -0.879779462 | 0.415905397 | 0.715965355 | -4.645881193 |
| **A_51_P304125** | Slc29a2 | -0.192565698 | -0.740787803 | 0.355656406 | 8.035884648 | -0.87977726 | 0.41590649 | 0.715965355 | -4.645881708 |
| **A_55_P2069842** | D5Ertd579 | -0.195409235 | -0.75183426 | 0.36101579 | 7.990474924 | -0.879607201 | 0.415991007 | 0.715965355 | -4.64592151 |

| | | | | | | | | | |
|---|---|---|---|---|---|---|---|---|---|
| **A_55_P1986647** | Podxl2 | 0.116164131 | -0.214626488 | 0.44695475 | 8.066488477 | 0.879567572 | 0.416010705 | 0.715965355 | -4.645930785 |
| **A_55_P1964368** | Kctd14 | 0.38558209 | -0.712432761 | 1.48359694 | 8.920561202 | 0.879546863 | 0.416020997 | 0.715965355 | -4.645935631 |
| **A_51_P146633** | NA | 0.583760484 | -1.078631841 | 2.24615281 | 10.7810516 | 0.879531734 | 0.416028517 | 0.715965355 | -4.645939171 |
| **A_55_P2081123** | Srcin1 | 0.18232924 | -0.336961021 | 0.701619502 | 8.410335699 | 0.879420308 | 0.416083906 | 0.715965355 | -4.645965247 |
| **A_51_P154585** | Vmn1r195 | 0.11390593 | -0.210569002 | 0.438380863 | 8.188977165 | 0.879256384 | 0.4161654 | 0.715965355 | -4.646003604 |
| **A_52_P549977** | Fam32a | -0.31354036 | -1.206708658 | 0.579627939 | 9.272095311 | -0.879246195 | 0.416170465 | 0.715965355 | -4.646005988 |
| **A_55_P2055411** | Rps7 | 0.379868787 | -0.702499252 | 1.462236827 | 16.13204409 | 0.879040726 | 0.416272632 | 0.715965355 | -4.64605406 |
| **A_55_P2075631** | Slc20a2 | -0.317236928 | -1.221343776 | 0.58686992 | 10.33551721 | -0.878849121 | 0.416367921 | 0.715965355 | -4.646098883 |
| **A_51_P464238** | Atf4 | -0.147213327 | -0.566823636 | 0.272396982 | 8.63442367 | -0.878721647 | 0.416431326 | 0.715965355 | -4.646128701 |
| **A_51_P464029** | Fads3 | -0.108788769 | -0.418876277 | 0.20129874 | 13.05857038 | -0.878719238 | 0.416432525 | 0.715965355 | -4.646129264 |
| **A_55_P1970596** | NA | -0.178457886 | -0.6871428 | 0.330227028 | 16.05302456 | -0.878693129 | 0.416445512 | 0.715965355 | -4.646135371 |
| **A_55_P1974193** | Ptges | -0.09034785 | -0.347920233 | 0.167224533 | 13.64412011 | -0.878554836 | 0.416514309 | 0.715965355 | -4.646167716 |
| **A_55_P1980566** | Ndufb11 | 0.340537283 | -0.630359979 | 1.311434545 | 14.86068964 | 0.87849994 | 0.41654162 | 0.715965355 | -4.646180554 |
| **A_52_P599317** | Hs6st2 | 0.220247273 | -0.407817572 | 0.848312118 | 8.630156566 | 0.878327339 | 0.4166275 | 0.715965355 | -4.646220917 |
| **A_52_P217875** | H3f3b | 0.183570735 | -0.33991601 | 0.707057479 | 8.635695182 | 0.878310556 | 0.416635852 | 0.715965355 | -4.646224842 |
| **A_51_P176352** | Ndrg2 | 0.756860135 | -1.401729994 | 2.915450263 | 11.00481504 | 0.878204766 | 0.416688496 | 0.715965355 | -4.646249579 |
| **A_52_P418515** | Cfap299 | 0.532100869 | -0.985741291 | 2.049943028 | 9.37267373 | 0.878046876 | 0.416767077 | 0.715965355 | -4.646286495 |
| **A_51_P443569** | Pwwp3a | 0.165712158 | -0.307042509 | 0.638466826 | 10.00127528 | 0.877948203 | 0.416816192 | 0.715965355 | -4.646309564 |
| **A_55_P2049949** | NA | 0.158029505 | -0.292810108 | 0.608869118 | 8.909638747 | 0.877943277 | 0.416818644 | 0.715965355 | -4.646310715 |
| **A_52_P504302** | Pmpca | -0.245541708 | -0.946106197 | 0.45502278 | 9.33820912 | -0.877864525 | 0.416857847 | 0.715965355 | -4.646329126 |
| **A_55_P2013534** | Slc25a19 | -0.126354287 | -0.486877316 | 0.234168743 | 10.11498449 | -0.877823723 | 0.416878159 | 0.715965355 | -4.646338664 |
| **A_55_P2430221** | Shld2 | -0.135245305 | -0.521177862 | 0.250687251 | 8.347051324 | -0.877730376 | 0.416924632 | 0.715965355 | -4.646360484 |
| **A_51_P211491** | Gusb | 0.173136241 | -0.320926887 | 0.667199369 | 10.69949503 | 0.877719781 | 0.416929907 | 0.715965355 | -4.640574665 |
| **A_52_P51078** | Ctsh | 0.969285543 | -1.796692005 | 3.735263092 | 15.63724329 | 0.877714818 | 0.416932378 | 0.715965355 | -4.646364121 |
| **A_55_P2010018** | NA | 0.306306407 | -0.567862145 | 1.180474959 | 17.0373993 | 0.87762957 | 0.416974823 | 0.715965355 | -4.646384047 |
| **A_55_P1957004** | Timm10 | 0.12797117 | -0.237254568 | 0.493196908 | 11.09738279 | 0.877609085 | 0.416985023 | 0.715965355 | -4.646388835 |
| **A_51_P222522** | Mesd | 0.093747447 | -0.173820425 | 0.361315319 | 13.38054833 | 0.877558 | 0.41701046 | 0.715965355 | -4.646400775 |
| **A_52_P266320** | Akap17b | 0.112532174 | -0.208661908 | 0.433726256 | 8.350975275 | 0.877525045 | 0.41702687 | 0.715965355 | -4.646408477 |
| **A_51_P382369** | Zfp608 | 0.600251523 | -1.113070348 | 2.313573393 | 10.49659535 | 0.877495014 | 0.417041825 | 0.715965355 | -4.646415496 |
| **A_66_P135842** | Fip1l1 | 0.080223945 | -0.148791461 | 0.309239351 | 11.12236592 | 0.877383902 | 0.417097159 | 0.715965355 | -4.646441464 |
| **A_55_P2011352** | NA | -0.082562022 | -0.318271068 | 0.153147024 | 11.38686298 | -0.877312695 | 0.417132624 | 0.715965355 | -4.646458104 |
| **A_51_P185757** | Casz1 | 0.273197155 | -0.506762971 | 1.05315728 | 9.193393649 | 0.877312095 | 0.417132922 | 0.715965355 | -4.646458244 |
| **A_55_P1969566** | Atp9b | -0.209458213 | -0.834058911 | 0.415142486 | 8.211525898 | -0.893218343 | 0.417163309 | 0.715965355 | -4.63103143 |
| **A_55_P1978050** | Aven | 0.224101817 | -0.415750436 | 0.863954071 | 10.49341528 | 0.877235141 | 0.417171252 | 0.715965355 | -4.646476227 |
| **A_52_P86158** | Mtx2 | 0.252368955 | -0.468288909 | 0.973026818 | 11.34957579 | 0.877116225 | 0.417230487 | 0.715965355 | -4.646504013 |
| **A_51_P354077** | Svil | 0.718462739 | -1.333219364 | 2.770144842 | 11.90565064 | 0.877090738 | 0.417243183 | 0.715965355 | -4.646509969 |
| **A_55_P2109613** | Ahr | 0.904360854 | -1.678188985 | 3.486910693 | 11.52142562 | 0.877088445 | 0.417244326 | 0.715965355 | -4.646510504 |
| **A_55_P2073975** | NA | -0.271593701 | -1.047185821 | 0.503998419 | 15.46830347 | -0.877074836 | 0.417251106 | 0.715965355 | -4.646513684 |
| **A_51_P445912** | NA | 0.133324784 | -0.247428216 | 0.514077783 | 8.538846486 | 0.877036967 | 0.417269971 | 0.715965355 | -4.646522532 |
| **A_51_P268843** | Rasip1 | 0.554238936 | -1.028834271 | 2.137312143 | 11.24770046 | 0.876892548 | 0.417341924 | 0.716011605 | -4.646556273 |
| **A_55_P2045755** | Plcxd2 | -0.482762947 | -1.861770964 | 0.896245071 | 8.271866657 | -0.876834206 | 0.417370993 | 0.716011605 | -4.646569903 |
| **A_51_P284486** | Gstm2 | 0.643722781 | -1.19528411 | 2.482729673 | 13.03307894 | 0.876730001 | 0.417422919 | 0.716015889 | -4.646594246 |
| **A_52_P158431** | Tshz2 | 0.099809872 | -0.185382603 | 0.385002347 | 9.16436008 | 0.876568341 | 0.417503485 | 0.716015889 | -4.646632007 |
| **A_55_P2011061** | Tspan12 | 0.503080386 | -0.934408591 | 1.940569364 | 10.01243192 | 0.876563147 | 0.417506073 | 0.716015889 | -4.64663322 |
| **A_66_P109577** | Plcb3 | 0.201782658 | -0.374806455 | 0.77837177 | 11.12110151 | 0.876531941 | 0.417521627 | 0.716015889 | -4.646640509 |
| **A_55_P2042713** | Pigq | 0.080351869 | -0.149326674 | 0.310030412 | 13.15310265 | 0.876245712 | 0.417664307 | 0.716108126 | -4.646707357 |
| **A_55_P2104542** | NA | -0.15002648 | -0.578877835 | 0.278824875 | 8.17287067 | -0.876216604 | 0.417678819 | 0.716108126 | -4.646714154 |
| **A_51_P260300** | Ikbkg | -0.110071928 | -0.424718573 | 0.204574717 | 8.082097703 | -0.87620114 | 0.417686528 | 0.716108126 | -4.646717765 |

| | | | | | | | | | |
|---|---|---|---|---|---|---|---|---|---|
| **A_55_P2163128** | NA | -0.267344742 | -1.031667557 | 0.496978072 | 8.352565177 | -0.876082832 | 0.417745516 | 0.716145753 | -4.646745391 |
| **A_52_P421897** | NA | -0.125103154 | -0.48280477 | 0.232598462 | 7.93440865 | -0.875987083 | 0.417793261 | 0.716164101 | -4.646767748 |
| **A_66_P117251** | Snx6 | 0.129028203 | -0.239984406 | 0.498040812 | 12.52972945 | 0.87577751 | 0.417897777 | 0.716253535 | -4.646816676 |
| **A_55_P2051313** | Gstk1 | 0.532577893 | -0.990806009 | 2.055961796 | 12.14567059 | 0.875637029 | 0.417967847 | 0.716253535 | -4.646849471 |
| **A_55_P2135351** | Fam89b | 0.165936707 | -0.308853587 | 0.640727001 | 10.48523489 | 0.875368634 | 0.418101745 | 0.716253535 | -4.646912117 |
| **A_51_P120729** | Polr2f | 0.091928684 | -0.17111134 | 0.354968708 | 11.29694173 | 0.8753456 | 0.418113238 | 0.716253535 | -4.646917493 |
| **A_51_P160824** | Cspg4 | 0.414676266 | -0.77195253 | 1.601305062 | 9.910790048 | 0.875275237 | 0.418148347 | 0.716253535 | -4.646933914 |
| **A_55_P2031442** | Pithd1 | -0.420812137 | -1.625034086 | 0.783409812 | 11.24199199 | -0.875249888 | 0.418160996 | 0.716253535 | -4.64693983 |
| **A_55_P2137701** | Rex2 | -0.23435958 | -0.905106737 | 0.436387578 | 8.075973363 | -0.875133311 | 0.418219171 | 0.716253535 | -4.646967035 |
| **A_51_P440865** | Fam110b | -0.289969076 | -1.119884292 | 0.53994614 | 8.574968237 | -0.875121397 | 0.418225116 | 0.716253535 | -4.646969815 |
| **A_55_P2027701** | NA | 0.273000947 | -0.508462668 | 1.054464563 | 13.32904199 | 0.874995335 | 0.418288032 | 0.716253535 | -4.646999231 |
| **A_66_P129795** | Ccdc59 | 0.09323763 | -0.173659682 | 0.360134941 | 11.83692907 | 0.87497848 | 0.418296445 | 0.716253535 | -4.647003164 |
| **A_55_P1962305** | Plac8 | -1.385243444 | -5.35087514 | 2.580388252 | 11.22451823 | -0.874911393 | 0.41832993 | 0.716253535 | -4.647018817 |
| **A_55_P2148519** | Teddm2 | 0.128745712 | -0.239891105 | 0.497382528 | 8.438369141 | 0.874750927 | 0.418410033 | 0.716253535 | -4.647056255 |
| **A_55_P2258567** | D930043N | -0.103542401 | -0.400032242 | 0.192947439 | 8.563801243 | -0.874699266 | 0.418435824 | 0.716253535 | -4.647068307 |
| **A_55_P2000683** | Eci2 | 0.343511129 | -0.640167837 | 1.327190094 | 13.23383441 | 0.874656991 | 0.41845693 | 0.716253535 | -4.647078169 |
| **A_55_P1999324** | Nudt6 | -0.083788359 | -0.323728704 | 0.156151985 | 8.742219211 | -0.87464286 | 0.418463985 | 0.716253535 | -4.647081465 |
| **A_52_P281033** | Socs5 | 0.149650097 | -0.278900552 | 0.578200746 | 8.704057046 | 0.874631653 | 0.418469581 | 0.716253535 | -4.64708408 |
| **A_55_P2027107** | Rgs3 | 0.119620944 | -0.222957059 | 0.462198946 | 8.843739458 | 0.874577089 | 0.418496824 | 0.716253535 | -4.647096808 |
| **A_55_P2113975** | Psg20 | -0.216025375 | -0.834732868 | 0.402682118 | 7.785875887 | -0.874519985 | 0.418525337 | 0.716253535 | -4.647110129 |
| **A_51_P130905** | Gcm1 | -0.176192367 | -0.68092504 | 0.328540306 | 7.780249735 | -0.874331281 | 0.41861957 | 0.716253535 | -4.647154143 |
| **A_52_P229728** | Cdc42 | -0.078171258 | -0.302126064 | 0.145783547 | 11.77791591 | -0.874252862 | 0.418658735 | 0.716253535 | -4.647172432 |
| **A_55_P2157902** | Igsf10 | 0.213598402 | -0.398486554 | 0.825683358 | 8.404978733 | 0.874050727 | 0.4187597 | 0.716253535 | -4.647219571 |
| **A_55_P2136353** | NA | -0.112472248 | -0.434808579 | 0.209864082 | 9.212490584 | -0.873949755 | 0.418810142 | 0.716253535 | -4.647243116 |
| **A_66_P116195** | Psmd13 | 0.094952106 | -0.177183426 | 0.367087639 | 12.06790581 | 0.873916013 | 0.418827 | 0.716253535 | -4.647250984 |
| **A_55_P1996746** | NA | -0.104809805 | -0.405216424 | 0.195596814 | 7.72908214 | -0.87386181 | 0.418854081 | 0.716253535 | -4.647263622 |
| **A_55_P1975772** | Ppp2r5a | 0.430858799 | -0.804080378 | 1.665797976 | 13.29716782 | 0.873855741 | 0.418857113 | 0.716253535 | -4.647265037 |
| **A_55_P2039694** | NA | -0.220421068 | -0.852205212 | 0.411363077 | 15.43886086 | -0.873845642 | 0.418862158 | 0.716253535 | -4.647267391 |
| **A_51_P397548** | Mtrf1l | 0.194293381 | -0.362629747 | 0.751216509 | 10.01417737 | 0.873802128 | 0.4188839 | 0.716253535 | -4.647277537 |
| **A_55_P1957098** | Mtfr1l | -0.202624898 | -0.783442684 | 0.378192888 | 12.48502384 | -0.873782311 | 0.418893802 | 0.716253535 | -4.647282157 |
| **A_55_P2072661** | Prex1 | -0.564138517 | -2.181710238 | 1.053433204 | 10.35394096 | -0.873519396 | 0.419025186 | 0.716253535 | -4.64734345 |
| **A_52_P456059** | Pfdn1 | 0.20535801 | -0.38354019 | 0.794256211 | 11.90219514 | 0.873417253 | 0.419076238 | 0.716253535 | -4.64736726 |
| **A_55_P1953439** | Lemd3 | 0.106932647 | -0.199734128 | 0.413599422 | 9.548501868 | 0.873361287 | 0.419104212 | 0.716253535 | -4.647380305 |
| **A_55_P1969058** | Epha7 | 0.159885578 | -0.298646248 | 0.618417405 | 8.158538776 | 0.873353677 | 0.419108016 | 0.716253535 | -4.647382078 |
| **A_55_P2024461** | Tbkbp1 | 0.248182293 | -0.463575623 | 0.959940209 | 11.81358611 | 0.873351022 | 0.419109343 | 0.716253535 | -4.647382697 |
| **A_55_P1978465** | H2-Q8 | -0.693187037 | -2.681401656 | 1.295027582 | 8.748248611 | -0.873247831 | 0.419160926 | 0.716253535 | -4.647406748 |
| **A_55_P2057984** | Tbc1d19 | 0.149718714 | -0.279726051 | 0.579163478 | 9.14495283 | 0.873210844 | 0.419179416 | 0.716253535 | -4.647415369 |
| **A_51_P485161** | Cplx2 | 0.603506917 | -1.127838358 | 2.334852191 | 10.67163982 | 0.87306969 | 0.419249987 | 0.716253535 | -4.647448264 |
| **A_52_P321318** | NA | -0.165465324 | -0.640176074 | 0.309245426 | 15.15959217 | -0.873028201 | 0.419270731 | 0.716253535 | -4.647457933 |
| **A_51_P498772** | 1700019D | -0.156462487 | -0.605347257 | 0.292422283 | 7.878961557 | -0.873023037 | 0.419273313 | 0.716253535 | -4.647459136 |
| **A_55_P2077522** | NA | -0.215649109 | -0.834361979 | 0.403063762 | 9.581479437 | -0.872989184 | 0.41929024 | 0.716253535 | -4.647467025 |
| **A_55_P2002918** | Klc2 | 0.124823239 | -0.23339516 | 0.483041637 | 10.92562012 | 0.872766171 | 0.419401763 | 0.716337951 | -4.647518989 |
| **A_55_P2114680** | Car4 | -0.876884695 | -3.393442246 | 1.639672856 | 10.41810475 | -0.872742186 | 0.419413759 | 0.716337951 | -4.647524577 |
| **A_55_P2030736** | Scmh1 | 0.102693608 | -0.192242378 | 0.397629594 | 8.529449335 | 0.872099412 | 0.419735325 | 0.716636436 | -4.647674305 |
| **A_55_P1999811** | NA | -0.177549381 | -0.687549509 | 0.332450747 | 8.450261873 | -0.871965342 | 0.419802422 | 0.716636436 | -4.647705527 |
| **A_55_P2251181** | 2900076A | -0.073925116 | -0.286299993 | 0.138449762 | 7.896835757 | -0.87184487 | 0.419862719 | 0.716636436 | -4.64773358 |
| **A_55_P2068366** | Gm9292 | 0.17328169 | -0.324629867 | 0.671193248 | 8.644683124 | 0.87166742 | 0.419951547 | 0.716636436 | -4.647774897 |
| **A_55_P2359807** | NA | -0.249147167 | -0.965112165 | 0.466817831 | 9.264605597 | -0.871594554 | 0.419988027 | 0.716636436 | -4.647791862 |

| | | | | | | | | | |
|---|---|---|---|---|---|---|---|---|---|
| **A_55_P2042146** | Fech | -0.185436328 | -0.718366818 | 0.347494162 | 12.84456562 | -0.871514457 | 0.420028129 | 0.716636436 | -4.647810509 |
| **A_66_P105711** | Ap4s1 | 0.090784905 | -0.170144035 | 0.351713846 | 8.513961881 | 0.871448527 | 0.420061141 | 0.716636436 | -4.647825857 |
| **A_51_P103509** | Cyb561d2 | -0.209403864 | -0.811278046 | 0.392470317 | 10.40442934 | -0.871423613 | 0.420073616 | 0.716636436 | -4.647831657 |
| **A_55_P2069935** | Fxyd2 | -0.802231703 | -3.108048531 | 1.503585125 | 9.178607325 | -0.871415663 | 0.420077597 | 0.716636436 | -4.647833507 |
| **A_52_P31543** | Btg2 | -0.184288959 | -0.713988004 | 0.345410086 | 11.43694336 | -0.871405854 | 0.420082509 | 0.716636436 | -4.647835791 |
| **A_55_P2054511** | NA | 0.126190062 | -0.236547232 | 0.488927357 | 9.108782802 | 0.871331252 | 0.420119866 | 0.716636436 | -4.647853156 |
| **A_55_P1994887** | Zfpm1 | -0.699402307 | -2.709900983 | 1.311096368 | 11.5010438 | -0.871311839 | 0.420129587 | 0.716636436 | -4.647857675 |
| **A_55_P2123484** | Gm11627 | 0.643940545 | -1.207190119 | 2.495071209 | 9.896405714 | 0.871282601 | 0.420144229 | 0.716636436 | -4.647864481 |
| **A_55_P2056871** | Akap13 | -0.222172081 | -0.860859751 | 0.416515589 | 11.152136 | -0.871267062 | 0.420152011 | 0.716636436 | -4.647868097 |
| **A_52_P213181** | Cul4b | -0.449996756 | -1.743652962 | 0.843659451 | 9.969618035 | -0.871246109 | 0.420162505 | 0.716636436 | -4.647872974 |
| **A_52_P444457** | Dnlz | 0.139455407 | -0.261471013 | 0.540381827 | 9.498969305 | 0.871206338 | 0.420182423 | 0.716636436 | -4.647882231 |
| **A_51_P187842** | Eif4e3 | 0.11341316 | -0.212670341 | 0.439496661 | 8.667196749 | 0.871134017 | 0.420218645 | 0.716636436 | -4.647899064 |
| **A_55_P1997614** | Gm9693 | -0.099897334 | -0.387179039 | 0.18738437 | 7.89881356 | -0.870956123 | 0.420307753 | 0.716724981 | -4.647940465 |
| **A_55_P2012181** | Eps15l1 | 0.125029545 | -0.234562056 | 0.484621146 | 11.08397588 | 0.870870255 | 0.420350769 | 0.716724981 | -4.647960447 |
| **A_55_P1997948** | Ppp4r3b | 0.09169923 | -0.172083103 | 0.355481564 | 10.91336769 | 0.870703583 | 0.420434275 | 0.716724981 | -4.64799923 |
| **A_55_P2005868** | Bbs7 | 0.177868419 | -0.333811704 | 0.689548543 | 8.359204111 | 0.870664113 | 0.420454052 | 0.716724981 | -4.648008413 |
| **A_51_P145772** | Rnf113a2 | 0.088972118 | -0.166978091 | 0.344922327 | 9.602159082 | 0.870660388 | 0.420455919 | 0.716724981 | -4.64800928 |
| **A_55_P2017967** | Usf2 | 0.108213254 | -0.203135003 | 0.419561511 | 13.68810158 | 0.870531244 | 0.420520634 | 0.716732527 | -4.648039326 |
| **A_52_P453884** | Foxo1 | 0.13467262 | -0.252816118 | 0.522161358 | 10.45380546 | 0.870503599 | 0.420534488 | 0.716732527 | -4.648045758 |
| **A_52_P604613** | S1pr2 | -0.081266729 | -0.315152242 | 0.152618785 | 10.32624456 | -0.870281596 | 0.420645755 | 0.716744621 | -4.648097402 |
| **A_51_P156274** | Ltbr | 0.122695695 | -0.230429628 | 0.475821017 | 12.68664912 | 0.870263579 | 0.420654787 | 0.716744621 | -4.648101593 |
| **A_52_P427818** | Zscan18 | 0.104261402 | -0.195811416 | 0.404334221 | 8.445358932 | 0.870256449 | 0.420658361 | 0.716744621 | -4.648103251 |
| **A_52_P671543** | Pomc | -0.117818982 | -0.456963168 | 0.221325204 | 8.801943928 | -0.870124174 | 0.420724669 | 0.716744621 | -4.648134018 |
| **A_66_P139460** | Mia3 | -0.286093726 | -1.109624266 | 0.537436813 | 8.42681822 | -0.870119639 | 0.420726943 | 0.716744621 | -4.648135073 |
| **A_55_P2141955** | NA | 0.358388235 | -0.673376725 | 1.390153195 | 16.36438934 | 0.870008139 | 0.420782843 | 0.716771516 | -4.648161005 |
| **A_52_P200359** | Aco2 | -0.097193332 | -0.377082629 | 0.182695964 | 13.61190475 | -0.869762196 | 0.420906165 | 0.716771516 | -4.648218199 |
| **A_52_P674357** | Thtpa | 0.265086043 | -0.498302416 | 1.028474501 | 9.735273412 | 0.869744354 | 0.420915113 | 0.716771516 | -4.648222348 |
| **A_55_P2311208** | C130045F | -0.101402118 | -0.393460215 | 0.190655979 | 8.643036885 | -0.869617203 | 0.420978882 | 0.716771516 | -4.648251913 |
| **A_55_P2051864** | Golgb1 | 0.126071412 | -0.237067841 | 0.489210666 | 13.6761974 | 0.869548414 | 0.421013384 | 0.716771516 | -4.648267906 |
| **A_55_P2090633** | Ptp4a1 | -0.143314982 | -0.556129768 | 0.269499805 | 9.020960457 | -0.86953425 | 0.421020489 | 0.716771516 | -4.648271199 |
| **A_51_P378558** | Rpap2 | -0.086547828 | -0.335867937 | 0.162772282 | 9.418611455 | -0.869459202 | 0.421058133 | 0.716771516 | -4.648288647 |
| **A_55_P1982559** | 1700007K | 0.24269346 | -0.456452793 | 0.941839714 | 8.475732218 | 0.869441537 | 0.421066995 | 0.716771516 | -4.648292754 |
| **A_55_P2044582** | Iglon5 | 0.349261337 | -0.656970663 | 1.355493336 | 8.793529175 | 0.869366164 | 0.421104805 | 0.716771516 | -4.648310276 |
| **A_55_P1954156** | Wdr43 | 0.141251665 | -0.26571345 | 0.548216781 | 9.581670105 | 0.869334146 | 0.421120867 | 0.716771516 | -4.648317719 |
| **A_66_P105175** | Bche | 0.359634523 | -0.67659157 | 1.395860616 | 8.762509348 | 0.869275013 | 0.421150534 | 0.716771516 | -4.648331465 |
| **A_51_P278868** | H2-DMb1 | 0.299005921 | -0.562734576 | 1.160746418 | 9.253116914 | 0.869067744 | 0.421254532 | 0.716885408 | -4.648379643 |
| **A_55_P2001583** | Tspan18 | 0.158138582 | -0.297660835 | 0.613937998 | 8.30043002 | 0.868989282 | 0.421293905 | 0.716889312 | -4.648397879 |
| **A_55_P2065646** | Zfp189 | 0.212785264 | -0.400811267 | 0.826381794 | 9.12078137 | 0.868578346 | 0.421500166 | 0.717116576 | -4.648493372 |
| **A_52_P406507** | Mus81 | -0.097397163 | -0.378256668 | 0.183462342 | 8.601617409 | -0.868575403 | 0.421501643 | 0.717116576 | -4.648494055 |
| **A_55_P2087352** | NA | 0.282008548 | -0.531307101 | 1.095324197 | 13.01525448 | 0.868467337 | 0.421555897 | 0.717145774 | -4.648519163 |
| **A_51_P284466** | Tsc22d4 | 0.163518196 | -0.308162316 | 0.635198707 | 10.124513 | 0.868297397 | 0.421641226 | 0.717227825 | -4.648558643 |
| **A_51_P153224** | Zdhhc12 | -0.094873487 | -0.368640518 | 0.178893543 | 9.560107806 | -0.867988686 | 0.421796266 | 0.717279484 | -4.648630351 |
| **A_52_P645632** | Senp8 | 0.091888237 | -0.173272736 | 0.357049211 | 8.220541225 | 0.867961898 | 0.421809722 | 0.717279484 | -4.648636572 |
| **A_55_P2328787** | NA | -0.146991833 | -0.571199109 | 0.277215443 | 8.710896907 | -0.867891507 | 0.42184508 | 0.717279484 | -4.64865292 |
| **A_55_P2136962** | NA | 0.185919598 | -0.350663563 | 0.722502758 | 10.2436057 | 0.86783763 | 0.421872145 | 0.717279484 | -4.648665433 |
| **A_55_P2179221** | Man1a2 | 0.08286207 | -0.156289864 | 0.322014003 | 11.59116279 | 0.867825137 | 0.421878421 | 0.717279484 | -4.648668334 |
| **A_55_P2127959** | Zfp36l3 | -0.796453247 | -3.095215815 | 1.502309322 | 9.362928646 | -0.867793746 | 0.421894191 | 0.717279484 | -4.648675624 |
| **A_55_P2174935** | Mn1 | -0.131688993 | -0.511862319 | 0.248484332 | 9.356928555 | -0.867597281 | 0.421992899 | 0.717380971 | -4.648721245 |

| | | | | | | | | | |
|---|---|---|---|---|---|---|---|---|---|
| **A_66_P137475** | Dnajb14 | -0.341829053 | -1.328821437 | 0.645163332 | 10.85840245 | -0.867452125 | 0.42206584 | 0.717380971 | -4.648754948 |
| **A_55_P1952502** | Mapk7 | 0.279351351 | -0.527291273 | 1.085993976 | 11.72460187 | 0.867401073 | 0.422091496 | 0.717380971 | -4.648766801 |
| **A_51_P464394** | Klb | -0.133046316 | -0.517234505 | 0.251141874 | 7.738283214 | -0.867379568 | 0.422102303 | 0.717380971 | -4.648771793 |
| **A_55_P2147846** | Npcd | 0.139747309 | -0.2639568 | 0.543451418 | 9.19385257 | 0.867023022 | 0.422281519 | 0.717622475 | -4.64885456 |
| **A_52_P405206** | Tm7sf3 | -0.224666622 | -0.873910561 | 0.424577317 | 10.64923394 | -0.86672435 | 0.42243169 | 0.717649098 | -4.648923877 |
| **A_55_P2101739** | NA | 0.525274307 | -0.992965465 | 2.043514078 | 12.95628856 | 0.866555015 | 0.422516849 | 0.717649098 | -4.648963171 |
| **A_55_P2035932** | Fgf7 | 0.537687993 | -1.01653677 | 2.091912755 | 9.453007546 | 0.866496594 | 0.422546232 | 0.717649098 | -4.648976726 |
| **A_55_P2024841** | Gstt1 | 0.877421918 | -1.658981795 | 3.413825631 | 11.69296232 | 0.866443891 | 0.42257274 | 0.717649098 | -4.648988954 |
| **A_55_P2070976** | Gm6923 | -0.113358141 | -0.441083983 | 0.214367702 | 7.795603623 | -0.866347988 | 0.42262098 | 0.717649098 | -4.649011204 |
| **A_66_P124011** | Spint2 | 0.208793198 | -0.394889176 | 0.812475572 | 12.15582345 | 0.866279817 | 0.422655274 | 0.717649098 | -4.64902702 |
| **A_52_P177341** | Cox19 | 0.19964143 | -0.377635941 | 0.776918802 | 11.19226145 | 0.86619662 | 0.422697128 | 0.717649098 | -4.64904632 |
| **A_55_P2092501** | Med1 | -0.327283113 | -1.27365607 | 0.619089844 | 11.63892337 | -0.86618692 | 0.422702008 | 0.717649098 | -4.64904857 |
| **A_51_P355413** | Nisch | 0.139400614 | -0.263775636 | 0.542576865 | 15.10094685 | 0.866004388 | 0.422793849 | 0.717649098 | -4.649090911 |
| **A_55_P1987993** | NA | -0.228495306 | -0.889434209 | 0.432443596 | 14.96969328 | -0.865897158 | 0.422847808 | 0.717649098 | -4.649115782 |
| **A_55_P1963807** | Actg2 | -0.354517999 | -1.380051571 | 0.671015573 | 12.28822091 | -0.865842209 | 0.422875461 | 0.717649098 | -4.649128526 |
| **A_52_P27271** | Dhdds | -0.12170923 | -0.473795281 | 0.230376821 | 9.45078885 | -0.865814873 | 0.422889219 | 0.717649098 | -4.649134866 |
| **A_55_P2177347** | NA | -0.197197006 | -0.767701671 | 0.373307658 | 8.089928306 | -0.865747934 | 0.422922908 | 0.717649098 | -4.64915039 |
| **A_55_P2010858** | Mrps23 | -0.088541872 | -0.344728544 | 0.1676448 | 8.871684258 | -0.865650358 | 0.42297202 | 0.717649098 | -4.649173018 |
| **A_51_P233153** | Cadps2 | 0.312191687 | -0.591127997 | 1.215511372 | 8.632736429 | 0.865625829 | 0.422984367 | 0.717649098 | -4.649178706 |
| **A_55_P1970715** | NA | -0.187186778 | -0.728917076 | 0.35454352 | 8.093068766 | -0.865450811 | 0.423072471 | 0.717649098 | -4.649219289 |
| **A_55_P2181071** | NA | 0.194042548 | -0.367626938 | 0.755712034 | 15.9330448 | 0.865299562 | 0.423148621 | 0.717649098 | -4.649254356 |
| **A_55_P2173952** | Myh6 | 1.767851168 | -3.349342503 | 6.885044839 | 13.88665251 | 0.865294994 | 0.423150921 | 0.717649098 | -4.649255415 |
| **A_51_P237695** | Gnai3 | -0.268943998 | -1.047444224 | 0.509556229 | 12.28113092 | -0.86527361 | 0.423161688 | 0.717649098 | -4.649260372 |
| **A_55_P1992049** | Gucy1a1 | 0.816539571 | -1.547313802 | 3.180392944 | 12.74215242 | 0.865181147 | 0.423208247 | 0.717649098 | -4.649281808 |
| **A_55_P2018701** | Kxd1 | -0.101724323 | -0.396218291 | 0.192769645 | 9.950468369 | -0.865164617 | 0.423216571 | 0.717649098 | -4.64928564 |
| **A_55_P2047275** | Tspan7 | 0.410075675 | -0.777157687 | 1.597309037 | 12.19221864 | 0.865123803 | 0.423237124 | 0.717649098 | -4.649295101 |
| **A_52_P613977** | NA | 0.231290012 | -0.438359387 | 0.900939411 | 11.65893793 | 0.865086909 | 0.423255704 | 0.717649098 | -4.649303653 |
| **A_55_P2037313** | Gm12027 | 0.095673217 | -0.181329215 | 0.372675649 | 12.6081256 | 0.865081729 | 0.423258312 | 0.717649098 | -4.649304854 |
| **A_55_P1978795** | Pja2 | -0.098900211 | -0.385253782 | 0.187453359 | 11.73093602 | -0.865057473 | 0.423270528 | 0.717649098 | -4.649310477 |
| **A_55_P2092399** | Ccdc160 | -0.09380436 | -0.365445482 | 0.177836762 | 8.676757605 | -0.864923815 | 0.423337844 | 0.717649098 | -4.649341457 |
| **A_52_P208613** | Fancb | 0.328527759 | -0.622981169 | 1.280036687 | 8.906661399 | 0.864787789 | 0.423406362 | 0.717649098 | -4.649372984 |
| **A_52_P593037** | Acsl5 | 0.168140476 | -0.318897414 | 0.655178367 | 10.46545714 | 0.86468887 | 0.423456193 | 0.717649098 | -4.649395909 |
| **A_51_P240136** | Gng8 | 0.261983308 | -0.497022124 | 1.020988741 | 8.646471177 | 0.86452804 | 0.423537223 | 0.717649098 | -4.649433178 |
| **A_52_P478629** | Arcn1 | -0.164890684 | -0.642630477 | 0.312849109 | 8.590566009 | -0.864480196 | 0.42356133 | 0.717649098 | -4.649444264 |
| **A_55_P2012369** | Eea1 | -0.24722 | -0.963498614 | 0.469058614 | 9.719926937 | -0.864474053 | 0.423564425 | 0.717649098 | -4.649445688 |
| **A_51_P355151** | Camk2n2 | -0.23207309 | -0.904466522 | 0.440320341 | 8.710729919 | -0.864473469 | 0.42356472 | 0.717649098 | -4.649445823 |
| **A_55_P2003414** | Kng1 | -1.273041849 | -4.961577446 | 2.415493748 | 9.931619276 | -0.864448345 | 0.423577379 | 0.717649098 | -4.649451644 |
| **A_52_P670812** | Snord123 | 0.57693692 | -1.094725786 | 2.248599627 | 11.71731736 | 0.864430385 | 0.423586429 | 0.717649098 | -4.649455806 |
| **A_51_P397768** | Csnk2a1 | -0.146911565 | -0.572594443 | 0.278771313 | 8.553115155 | -0.864410731 | 0.423596333 | 0.717649098 | -4.64946036 |
| **A_52_P366932** | Gna11 | -0.148180306 | -0.57759786 | 0.281237248 | 10.58221407 | -0.864293087 | 0.423655618 | 0.717686648 | -4.649487617 |
| **A_55_P2302577** | AI426330 | 0.302457828 | -0.57425407 | 1.179169726 | 10.66271936 | 0.864088597 | 0.423758683 | 0.7177368 | -4.64953499 |
| **A_55_P1975430** | St3gal3 | 0.147319081 | -0.279723068 | 0.574361229 | 10.49877028 | 0.864049461 | 0.42377841 | 0.7177368 | -4.649544056 |
| **A_55_P2075195** | Cd302 | 0.673479328 | -1.278958774 | 2.62591743 | 10.93692848 | 0.863967371 | 0.423819791 | 0.7177368 | -4.649563071 |
| **A_55_P2189571** | NA | 0.174050524 | -0.330550514 | 0.678651562 | 9.664120135 | 0.863927982 | 0.423839648 | 0.7177368 | -4.649572195 |
| **A_51_P255193** | Faf2 | -0.156504342 | -0.610268613 | 0.29725993 | 10.40945117 | -0.863866114 | 0.423870838 | 0.7177368 | -4.649586524 |
| **A_55_P2078831** | NA | -0.132359287 | -0.516175531 | 0.251456957 | 9.92842369 | -0.86373678 | 0.423936047 | 0.717784353 | -4.649616479 |
| **A_55_P2185397** | Copz2 | -0.082074498 | -0.320185925 | 0.156036928 | 8.655803561 | -0.863333007 | 0.424139672 | 0.717796156 | -4.649709978 |
| **A_55_P2071426** | Eef2kmt | -0.108818497 | -0.424572326 | 0.206935333 | 8.422576612 | -0.863186106 | 0.424213774 | 0.717796156 | -4.649743989 |

| | | | | | | | | | |
|---|---|---|---|---|---|---|---|---|---|
| **A_55_P1969181** | Paox | 0.316508852 | -0.601894895 | 1.234912599 | 11.14034169 | 0.863182366 | 0.424215661 | 0.717796156 | -4.649744855 |
| **A_66_P101703** | Krtap5-5 | 0.109588305 | -0.208446026 | 0.427622635 | 8.315743101 | 0.863059131 | 0.424277832 | 0.717796156 | -4.649773383 |
| **A_51_P165504** | Twist2 | 0.167312599 | -0.318259744 | 0.652884942 | 9.036759584 | 0.863028322 | 0.424293376 | 0.717796156 | -4.649780515 |
| **A_51_P284988** | Nrbf2 | -0.074405394 | -0.290368599 | 0.141557812 | 9.783924228 | -0.862928871 | 0.424343555 | 0.717796156 | -4.649803536 |
| **A_55_P2366358** | 4930444F | 0.287069175 | -0.546237261 | 1.120375611 | 8.534524519 | 0.862843775 | 0.424386494 | 0.717796156 | -4.649823232 |
| **A_51_P355040** | NA | 0.097331132 | -0.185237738 | 0.379900003 | 8.348410949 | 0.862735774 | 0.424440997 | 0.717796156 | -4.649848229 |
| **A_51_P430082** | Tst | 0.540960701 | -1.029897994 | 2.111819395 | 11.71806649 | 0.862539398 | 0.42454011 | 0.717796156 | -4.649893675 |
| **A_55_P2165854** | Gemin7 | -0.098602319 | -0.384954567 | 0.187749929 | 10.64534508 | -0.862455863 | 0.424582277 | 0.717796156 | -4.649913005 |
| **A_51_P143190** | Lyl1 | 0.536016451 | -1.020954171 | 2.092987074 | 10.30888471 | 0.862279471 | 0.424671326 | 0.717796156 | -4.649953819 |
| **A_55_P2091305** | NA | 0.096215095 | -0.183291536 | 0.375721725 | 8.400659119 | 0.862186945 | 0.424718042 | 0.717796156 | -4.649975226 |
| **A_55_P2161775** | Prr3 | -0.241502973 | -0.943250868 | 0.460244923 | 9.394132687 | -0.861969121 | 0.424828036 | 0.717796156 | -4.650025616 |
| **A_55_P1967736** | Nkd1 | 0.761670328 | -1.451575179 | 2.974915834 | 11.57287853 | 0.861961348 | 0.424831962 | 0.717796156 | -4.650027414 |
| **A_51_P159453** | Serpina3n | -0.130383636 | -0.509279953 | 0.248512681 | 7.731756812 | -0.861892398 | 0.424866784 | 0.717796156 | -4.650043363 |
| **A_55_P1991618** | Ppp1r16a | -0.120575636 | -0.470976079 | 0.229824807 | 11.90318295 | -0.861876937 | 0.424874593 | 0.717796156 | -4.650046939 |
| **A_55_P1974019** | Dapk1 | 0.237273136 | -0.452268834 | 0.926815106 | 11.76072738 | 0.861862948 | 0.424881658 | 0.717796156 | -4.650050175 |
| **A_55_P1989125** | Rps17 | 0.305304965 | -0.582208616 | 1.192818547 | 15.93905634 | 0.861606974 | 0.425010956 | 0.717796156 | -4.650109377 |
| **A_55_P2169888** | Prag1 | 0.40812385 | -0.778392525 | 1.594640225 | 10.06947557 | 0.861526389 | 0.425051668 | 0.717796156 | -4.650128013 |
| **A_55_P2047571** | Rps9 | 0.409051862 | -0.780471561 | 1.598575285 | 16.67229916 | 0.861302526 | 0.425164779 | 0.717796156 | -4.650179777 |
| **A_51_P137388** | Zadh2 | 0.495679805 | -0.945917666 | 1.937277276 | 11.24222523 | 0.861207015 | 0.425213044 | 0.717796156 | -4.65020186 |
| **A_55_P2015934** | NA | 0.138996417 | -0.26526447 | 0.543257304 | 16.56736501 | 0.861176611 | 0.42522841 | 0.717796156 | -4.650208889 |
| **A_55_P2428968** | Lrrc2 | 0.233287164 | -0.445229448 | 0.911803776 | 8.849980797 | 0.861153764 | 0.425239956 | 0.717796156 | -4.650214171 |
| **A_55_P1978597** | Trit1 | -0.108655836 | -0.424684965 | 0.207373293 | 10.36207347 | -0.861145007 | 0.425244382 | 0.717796156 | -4.650216196 |
| **A_52_P444637** | Rcbtb2 | 0.138218355 | -0.263816201 | 0.540252911 | 8.366475197 | 0.861098201 | 0.425268037 | 0.717796156 | -4.650227017 |
| **A_55_P1983683** | Rtl5 | 0.24569214 | -0.469025585 | 0.960409864 | 8.762880862 | 0.861007741 | 0.425313759 | 0.717796156 | -4.650247929 |
| **A_52_P648424** | Nedd4 | 0.582683967 | -1.112420252 | 2.277788187 | 13.65586676 | 0.860967998 | 0.425333847 | 0.717796156 | -4.650257116 |
| **A_55_P1962092** | Mpc1 | -0.07706695 | -0.301291107 | 0.147157208 | 12.21373616 | -0.860867104 | 0.425384849 | 0.717796156 | -4.650280438 |
| **A_51_P403536** | Ltbp4 | 0.827396841 | -1.580043918 | 3.2348376 | 13.50593397 | 0.86081256 | 0.425412422 | 0.717796156 | -4.650293045 |
| **A_51_P439876** | Map4k4 | -0.161779171 | -0.632535392 | 0.30897705 | 12.51293594 | -0.860749707 | 0.425444198 | 0.717796156 | -4.650307573 |
| **A_55_P2048518** | Zfpm1 | -0.68361563 | -2.673038441 | 1.30580718 | 11.4234273 | -0.860667169 | 0.425485928 | 0.717796156 | -4.650326649 |
| **A_52_P596008** | Zeb2 | 0.611484758 | -1.168049832 | 2.391019349 | 11.09652447 | 0.860655895 | 0.425491628 | 0.717796156 | -4.650329255 |
| **A_55_P2126925** | Msantd3 | 0.229151582 | -0.437983614 | 0.896286778 | 10.83379653 | 0.86031868 | 0.425662154 | 0.717796156 | -4.65040718 |
| **A_55_P2177513** | Chrna2 | -0.151101954 | -0.591011578 | 0.288807669 | 8.086143482 | -0.86031395 | 0.425664546 | 0.717796156 | -4.650408273 |
| **A_55_P2098935** | Sh2d3c | 0.297594166 | -0.568881762 | 1.164070094 | 11.66534803 | 0.860237275 | 0.425703328 | 0.717796156 | -4.650425989 |
| **A_55_P1956308** | NA | 0.506149639 | -0.967662174 | 1.979961453 | 14.43105989 | 0.860175847 | 0.425734399 | 0.717796156 | -4.650440182 |
| **A_55_P2136662** | 8030462N | -0.097750013 | -0.382387948 | 0.186887921 | 8.088824435 | -0.860150395 | 0.425747273 | 0.717796156 | -4.650446062 |
| **A_55_P2173098** | Commd1 | 0.46687935 | -0.892671584 | 1.826430284 | 11.8608991 | 0.860120956 | 0.425762165 | 0.717796156 | -4.650452863 |
| **A_52_P110257** | Wdr83 | 0.177525324 | -0.339459149 | 0.694509797 | 9.205959925 | 0.860068733 | 0.425788583 | 0.717796156 | -4.650464928 |
| **A_52_P195018** | Arap3 | 0.498985129 | -0.954165454 | 1.952135713 | 12.21314637 | 0.860057185 | 0.425794425 | 0.717796156 | -4.650467596 |
| **A_52_P465886** | Gramd3 | -0.401724652 | -1.571648548 | 0.768199244 | 8.495894664 | -0.860045083 | 0.425800547 | 0.717796156 | -4.650470392 |
| **A_55_P2149719** | NA | 0.24977182 | -0.477658522 | 0.977202162 | 14.55616756 | 0.860007764 | 0.425819427 | 0.717796156 | -4.650479013 |
| **A_55_P2101906** | Rpl10a | 0.385839691 | -0.73798337 | 1.509662752 | 17.1730195 | 0.859922489 | 0.425862569 | 0.717796156 | -4.650498712 |
| **A_52_P162957** | Frat2 | 0.11091213 | -0.212149185 | 0.433973446 | 8.884576307 | 0.859893045 | 0.425877467 | 0.717796156 | -4.650505513 |
| **A_51_P202014** | Taf12 | 0.076956984 | -0.147201643 | 0.301115611 | 10.25273952 | 0.859890054 | 0.42587898 | 0.717796156 | -4.650506204 |
| **A_51_P304239** | Loxl1 | 0.626772955 | -1.198896411 | 2.452442321 | 10.94623535 | 0.859881224 | 0.425883448 | 0.717796156 | -4.650508244 |
| **A_55_P2048334** | Ubr3 | 0.136015954 | -0.260192281 | 0.532224189 | 9.51351689 | 0.859838131 | 0.425905252 | 0.717796156 | -4.650518198 |
| **A_55_P2062054** | Dbndd2 | -0.170120849 | -0.665683224 | 0.325441527 | 12.41331212 | -0.859823994 | 0.425912405 | 0.717796156 | -4.650521463 |
| **A_55_P1995622** | Pomp | 0.137350346 | -0.262764762 | 0.537465455 | 9.851302512 | 0.85979547 | 0.425926837 | 0.717796156 | -4.650528052 |
| **A_55_P1987302** | Eny2 | 0.284617411 | -0.544564459 | 1.113799282 | 12.47231823 | 0.859729863 | 0.425960036 | 0.717796156 | -4.650543205 |

| | | | | | | | | | |
|---|---|---|---|---|---|---|---|---|---|
| **A_66_P124674** | Ubxn2a | -0.357709588 | -1.399947326 | 0.684528149 | 8.480120108 | -0.859635086 | 0.426007998 | 0.717796156 | -4.650565095 |
| **A_51_P401876** | Kif1bp | 0.141828266 | -0.271457576 | 0.555114109 | 11.51243104 | 0.859533112 | 0.426059607 | 0.717796156 | -4.650588645 |
| **A_55_P2099378** | Tfe3 | -0.170026596 | -0.665514155 | 0.325460963 | 11.8260818 | -0.859477381 | 0.426087814 | 0.717796156 | -4.650601515 |
| **A_51_P167527** | Lum | -0.934461253 | -3.657797005 | 1.788874499 | 11.9108298 | -0.859429895 | 0.42611185 | 0.717796156 | -4.650612481 |
| **A_55_P1982291** | Clca3a1 | -0.173099645 | -0.677578175 | 0.331378886 | 8.990117663 | -0.85941679 | 0.426118483 | 0.717796156 | -4.650615507 |
| **A_55_P2001063** | Fcf1 | -0.093945186 | -0.367758377 | 0.179868006 | 12.58797301 | -0.859350843 | 0.426151865 | 0.717796156 | -4.650630735 |
| **A_55_P2139665** | NA | 0.285564166 | -0.546778745 | 1.117907077 | 16.57682593 | 0.859313767 | 0.426170633 | 0.717796156 | -4.650639296 |
| **A_55_P2007560** | NA | 0.083347596 | -0.159609737 | 0.326304929 | 8.434842451 | 0.859237875 | 0.426209052 | 0.717796156 | -4.650656819 |
| **A_51_P344046** | Nt5m | -0.10324692 | -0.404217884 | 0.197724044 | 9.775273666 | -0.85921698 | 0.426219631 | 0.717796156 | -4.650661644 |
| **A_55_P2020004** | NA | 0.231105979 | -0.442592713 | 0.904804671 | 12.50593164 | 0.859203077 | 0.42622667 | 0.717796156 | -4.650664854 |
| **A_55_P1976928** | Unc13d | -0.156879982 | -0.614215849 | 0.300455885 | 9.38798061 | -0.859176941 | 0.426239901 | 0.717796156 | -4.650670888 |
| **A_52_P229044** | Slc20a2 | -0.156281172 | -0.61187656 | 0.299314216 | 8.115510817 | -0.859167194 | 0.426244836 | 0.717796156 | -4.650673138 |
| **A_55_P2104761** | NA | 0.155088656 | -0.297263071 | 0.607440384 | 15.83091748 | 0.858725038 | 0.426468738 | 0.718086076 | -4.650775207 |
| **A_55_P2024575** | Sec24c | -0.187482304 | -0.734345896 | 0.359381288 | 14.06509811 | -0.858680532 | 0.42649128 | 0.718086076 | -4.650785479 |
| **A_55_P2017413** | Gm4788 | 0.157886168 | -0.302783551 | 0.618555888 | 8.158445884 | 0.85842976 | 0.426618313 | 0.718144223 | -4.650843353 |
| **A_51_P359262** | Sec11c | -0.417358858 | -1.635140384 | 0.800422668 | 10.14564251 | -0.85840176 | 0.426632498 | 0.718144223 | -4.650849814 |
| **A_51_P337523** | Adcy2 | 0.267246163 | -0.512540795 | 1.047033121 | 8.620358312 | 0.858392386 | 0.426637248 | 0.718144223 | -4.650851977 |
| **A_51_P430929** | Fam20a | -0.149579818 | -0.58607735 | 0.286917714 | 9.181265711 | -0.858304825 | 0.426681611 | 0.718156374 | -4.650872182 |
| **A_55_P2094362** | Erp29 | -0.258774932 | -1.014113188 | 0.496563323 | 9.222431842 | -0.858086498 | 0.426792243 | 0.718280052 | -4.650922556 |
| **A_55_P1986053** | Pcnx | 0.226714384 | -0.435323912 | 0.888752681 | 9.498381737 | 0.857721525 | 0.426977234 | 0.718528839 | -4.651006747 |
| **A_51_P518051** | Dusp4 | -0.270086468 | -1.058905326 | 0.51873239 | 8.155917767 | -0.857582452 | 0.42704774 | 0.718535723 | -4.651038823 |
| **A_52_P320971** | Mt3 | 0.090104535 | -0.173061001 | 0.353270071 | 9.865920357 | 0.857566845 | 0.427055653 | 0.718535723 | -4.651042423 |
| **A_55_P2070009** | Krt10 | -0.115508664 | -0.452935708 | 0.22191838 | 8.095035939 | -0.857403039 | 0.42713871 | 0.718612933 | -4.651080198 |
| **A_51_P161830** | Enpep | 0.71116128 | -1.366584157 | 2.788906717 | 10.23596909 | 0.85728675 | 0.427197682 | 0.718649612 | -4.651107014 |
| **A_55_P1967227** | Tmem158 | -0.261711353 | -1.026458813 | 0.503036107 | 10.31484125 | -0.857146124 | 0.427269004 | 0.718707058 | -4.651139438 |
| **A_55_P2075294** | NA | 0.335574801 | -0.645282734 | 1.316432336 | 8.941504573 | 0.856907126 | 0.427390237 | 0.718848443 | -4.651194537 |
| **A_55_P1999246** | Gm10318 | 0.137437263 | -0.264412256 | 0.539286783 | 11.58157625 | 0.856626273 | 0.427532735 | 0.719025565 | -4.651259273 |
| **A_55_P2397240** | B430316J | -0.111359541 | -0.437073871 | 0.214354788 | 8.309344165 | -0.856329514 | 0.427683342 | 0.719159918 | -4.651327662 |
| **A_52_P194250** | Ets1 | 0.676030827 | -1.301301208 | 2.653362863 | 10.71101913 | 0.856322279 | 0.427687014 | 0.719159918 | -4.651329329 |
| **A_55_P2093704** | Meig1 | 0.437314197 | -0.842054012 | 1.716682406 | 9.068985581 | 0.856147056 | 0.427775961 | 0.719246928 | -4.651369704 |
| **A_55_P2108451** | Galns | -0.693701261 | -2.723627827 | 1.336225304 | 11.29870264 | -0.855938384 | 0.427881904 | 0.719252945 | -4.651417778 |
| **A_55_P2133205** | Nmt2 | -0.462498151 | -1.815903353 | 0.890907051 | 9.001039559 | -0.855918681 | 0.427891908 | 0.719252945 | -4.651422317 |
| **A_55_P2162394** | Pdp1 | 0.184254562 | -0.354974914 | 0.723484039 | 8.372504065 | 0.85584471 | 0.427929469 | 0.719252945 | -4.651439357 |
| **A_51_P331661** | Ubiad1 | 0.122787489 | -0.236613394 | 0.482188372 | 10.36566262 | 0.855707472 | 0.427999162 | 0.719252945 | -4.651470969 |
| **A_55_P2132756** | Cacna1g | 0.226252729 | -0.436026325 | 0.888531782 | 8.608911056 | 0.855663786 | 0.428021349 | 0.719252945 | -4.651481031 |
| **A_51_P442284** | Dnajc12 | 0.284766208 | -0.548797541 | 1.118329957 | 9.634211828 | 0.855657534 | 0.428024524 | 0.719252945 | -4.651482471 |
| **A_51_P359272** | Alcam | 0.684355261 | -1.319169078 | 2.6878796 | 10.0647815 | 0.855534118 | 0.428087207 | 0.719252945 | -4.651510896 |
| **A_51_P477682** | Prss12 | 0.252917465 | -0.487603211 | 0.993438141 | 11.17578675 | 0.855444772 | 0.428132591 | 0.719252945 | -4.651531472 |
| **A_55_P1966204** | Cxcl12 | 0.197217249 | -0.380248284 | 0.774682782 | 9.952180397 | 0.855399861 | 0.428155405 | 0.719252945 | -4.651541814 |
| **A_52_P276607** | Ptp4a2 | -0.11971665 | -0.470275261 | 0.230841962 | 13.20998629 | -0.85535079 | 0.428180333 | 0.719252945 | -4.651553114 |
| **A_51_P357877** | Fbxo46 | 0.075868778 | -0.146297219 | 0.298034774 | 9.113234399 | 0.855334212 | 0.428188755 | 0.719252945 | -4.651556932 |
| **A_55_P2049261** | NA | -0.10548587 | -0.414435475 | 0.203463736 | 8.559060373 | -0.855178925 | 0.428267651 | 0.719264106 | -4.651592688 |
| **A_55_P2065586** | Gm15417 | -0.0930185 | -0.365454716 | 0.179417716 | 8.162013103 | -0.855174687 | 0.428269804 | 0.719264106 | -4.651593664 |
| **A_55_P1976504** | NA | 0.283888781 | -0.547659937 | 1.1154375 | 12.31664737 | 0.855088129 | 0.428313785 | 0.719264255 | -4.651613593 |
| **A_55_P2007423** | Ppp1r12a | -0.165831954 | -0.651647653 | 0.319983745 | 8.705726586 | -0.854962406 | 0.428377673 | 0.719264255 | -4.651642537 |
| **A_55_P2226660** | 5033418A | -0.084384498 | -0.331598102 | 0.162829106 | 8.040389982 | -0.854949868 | 0.428384045 | 0.719264255 | -4.651645424 |
| **A_55_P2114838** | NA | 0.229519019 | -0.442935655 | 0.901973693 | 15.199806 | 0.854881677 | 0.428418701 | 0.719264255 | -4.651661122 |
| **A_55_P1960167** | Bcat2 | 0.265091177 | -0.511665038 | 1.041847392 | 9.398821027 | 0.854792854 | 0.428463845 | 0.719277588 | -4.651681568 |

| | | | | | | | | | |
|---|---|---|---|---|---|---|---|---|---|
| **A_55_P2089304** | 2900097C | -0.249517892 | -1.013560144 | 0.514524361 | 9.100772988 | -0.869855314 | 0.428540965 | 0.719344593 | -4.63557924 |
| **A_55_P1983149** | Rps29 | 0.142720643 | -0.275623152 | 0.561064438 | 17.4364125 | 0.854483751 | 0.428620974 | 0.719416435 | -4.651752712 |
| **A_51_P130994** | NA | 0.256911 | -0.496300817 | 1.010122818 | 11.4325409 | 0.854310847 | 0.428708888 | 0.71947443 | -4.651792501 |
| **A_55_P2073647** | Elp2 | 0.198131391 | -0.382875073 | 0.779137854 | 11.70470448 | 0.85412743 | 0.42880216 | 0.71947443 | -4.651834705 |
| **A_51_P280492** | Ndufa9 | 0.169555613 | -0.327710536 | 0.666821763 | 12.11225535 | 0.854030973 | 0.428851218 | 0.71947443 | -4.651856897 |
| **A_55_P2179314** | Macrod2 | 0.166813403 | -0.322483013 | 0.656109819 | 8.250894392 | 0.853904402 | 0.428915597 | 0.71947443 | -4.651886015 |
| **A_55_P2071329** | Rhog | 0.139393896 | -0.269505494 | 0.548293286 | 11.80952011 | 0.853842246 | 0.428947215 | 0.71947443 | -4.651900314 |
| **A_55_P2143499** | Pgbd5 | 0.355914079 | -0.688219321 | 1.400047479 | 9.127897168 | 0.853767316 | 0.428985333 | 0.71947443 | -4.65191755 |
| **A_55_P2006133** | Tmc6 | 0.156924913 | -0.303442242 | 0.617292069 | 8.859928183 | 0.853764144 | 0.428986947 | 0.71947443 | -4.651918279 |
| **A_55_P1973357** | NA | 0.232464896 | -0.449604195 | 0.914533987 | 15.93715667 | 0.853649042 | 0.429045507 | 0.71947443 | -4.651944754 |
| **A_55_P2387379** | NA | -0.156495078 | -0.615699487 | 0.302709331 | 8.034826832 | -0.853581471 | 0.429079887 | 0.71947443 | -4.651960296 |
| **A_55_P2065040** | NA | 0.403763817 | -0.781011961 | 1.588539595 | 14.82802212 | 0.853574783 | 0.42908329 | 0.71947443 | -4.651961834 |
| **A_55_P1959763** | Dach1 | 0.931911748 | -1.802921544 | 3.666745039 | 10.79468594 | 0.853481819 | 0.429130594 | 0.71947443 | -4.651983214 |
| **A_55_P2003966** | Rybp | -0.247865425 | -0.975303649 | 0.4795728 | 12.50643789 | -0.853434466 | 0.429154691 | 0.71947443 | -4.651994104 |
| **A_66_P106363** | Itgb1bp1 | 0.180560682 | -0.349399884 | 0.710521248 | 10.52044218 | 0.853355469 | 0.429194893 | 0.71947443 | -4.652012271 |
| **A_55_P2014997** | Amz2 | -0.189912999 | -0.747384175 | 0.367558177 | 10.1220766 | -0.853262424 | 0.429242247 | 0.71947443 | -4.652033666 |
| **A_51_P356967** | Lgi3 | 0.929259492 | -1.798505032 | 3.657024017 | 10.68614901 | 0.853258208 | 0.429244393 | 0.71947443 | -4.652034636 |
| **A_51_P208870** | Zdhhc24 | -0.171448775 | -0.674774086 | 0.331876536 | 8.319906071 | -0.853170757 | 0.429288905 | 0.71947443 | -4.652054744 |
| **A_52_P621567** | Tcf20 | -0.266149838 | -1.047569314 | 0.515269637 | 11.46702768 | -0.85308503 | 0.429332542 | 0.71947443 | -4.652074454 |
| **A_55_P2186110** | R3hdm2 | 0.153158351 | -0.316707271 | 0.623023972 | 12.07577136 | 0.868219861 | 0.429346574 | 0.71947443 | -4.635895442 |
| **A_51_P477902** | Kdm5c | -0.235626498 | -0.927476881 | 0.456223886 | 11.2775734 | -0.853026038 | 0.429362572 | 0.71947443 | -4.652088017 |
| **A_52_P273929** | Mrpl17 | 0.091465676 | -0.177120522 | 0.360051874 | 11.06699264 | 0.852952414 | 0.429400053 | 0.719474879 | -4.652104943 |
| **A_55_P1994603** | Itsn1 | -0.207876941 | -0.818414969 | 0.402661088 | 15.12001096 | -0.85279362 | 0.429480901 | 0.719494852 | -4.652141447 |
| **A_55_P1960989** | Nlgn2 | 0.404498551 | -0.783737528 | 1.59273463 | 10.76867924 | 0.852637797 | 0.429560248 | 0.719494852 | -4.652177264 |
| **A_55_P2085835** | 5330417C | 0.17828725 | -0.345482209 | 0.702056709 | 8.708348153 | 0.852570813 | 0.429594361 | 0.719494852 | -4.652192659 |
| **A_55_P2070657** | Emc10 | -0.145539242 | -0.573132404 | 0.28205392 | 8.229620134 | -0.852510433 | 0.429625112 | 0.719494852 | -4.652206536 |
| **A_55_P2300421** | NA | -0.252485668 | -0.994404903 | 0.489433567 | 8.288697726 | -0.852374495 | 0.42969435 | 0.719494852 | -4.652237777 |
| **A_55_P2004096** | Asb8 | -0.110081659 | -0.433558105 | 0.213394787 | 8.12379984 | -0.852359192 | 0.429702145 | 0.719494852 | -4.652241293 |
| **A_55_P2015541** | Hif3a | 0.787717698 | -1.527071542 | 3.102506939 | 12.74002345 | 0.852333373 | 0.429715296 | 0.719494852 | -4.652247226 |
| **A_55_P2084812** | Sec31a | -0.128228909 | -0.505043618 | 0.2485858 | 9.339919669 | -0.852331315 | 0.429716345 | 0.719494852 | -4.652247699 |
| **A_55_P1983988** | Enox2 | 0.138994512 | -0.269484849 | 0.547473873 | 8.671002241 | 0.852271334 | 0.429746899 | 0.719494852 | -4.652261482 |
| **A_55_P1988202** | Ifi203 | -0.2092471 | -0.824246699 | 0.405752499 | 8.498439809 | -0.85218711 | 0.429789805 | 0.719504381 | -4.652280835 |
| **A_55_P2032678** | Lox | 0.227106858 | -0.440555108 | 0.894768824 | 9.173495674 | 0.851969324 | 0.429900768 | 0.719584382 | -4.652330873 |
| **A_55_P2161702** | Nrip2 | 0.080823779 | -0.156792838 | 0.318440396 | 7.850055738 | 0.85194722 | 0.429912031 | 0.719584382 | -4.65233595 |
| **A_51_P491504** | Aldh5a1 | 0.359608135 | -0.697824899 | 1.41704117 | 10.55567436 | 0.8517791 | 0.429997704 | 0.719630872 | -4.652374571 |
| **A_55_P1978176** | Abhd17c | 0.297398217 | -0.577230175 | 1.172026608 | 12.85205759 | 0.851657813 | 0.43005952 | 0.719630872 | -4.65240243 |
| **A_55_P2175260** | Tet3 | 0.077390047 | -0.150209028 | 0.304989121 | 7.917758265 | 0.851657474 | 0.430059693 | 0.719630872 | -4.652402508 |
| **A_52_P463183** | Asap1 | -0.101527817 | -0.400139732 | 0.197084098 | 9.014570368 | -0.851585508 | 0.430096374 | 0.719630872 | -4.652419037 |
| **A_55_P2148641** | Rab4a | -0.110616753 | -0.435983031 | 0.214749526 | 7.962260536 | -0.851527564 | 0.43012591 | 0.719630872 | -4.652432345 |
| **A_55_P2035439** | Uxt | 0.170934178 | -0.331954076 | 0.673822433 | 10.28838094 | 0.851349258 | 0.430216809 | 0.719677475 | -4.652473294 |
| **A_52_P445474** | Lmtk2 | -0.235269522 | -0.92753279 | 0.456993746 | 8.688909397 | -0.851225702 | 0.430279806 | 0.719677475 | -4.652501666 |
| **A_55_P2066414** | Dffa | -0.094460775 | -0.372446404 | 0.183524855 | 8.549375109 | -0.851097869 | 0.43034499 | 0.719677475 | -4.652531017 |
| **A_55_P1997501** | Gm10778 | 0.141751592 | -0.275412263 | 0.558915447 | 9.318346919 | 0.851082417 | 0.430352869 | 0.719677475 | -4.652534565 |
| **A_52_P542172** | Rb1cc1 | -0.114692187 | -0.452222402 | 0.222838028 | 9.538270076 | -0.85108223 | 0.430352965 | 0.719677475 | -4.652534608 |
| **A_51_P446085** | Map10 | 0.147044068 | -0.285719297 | 0.579807434 | 8.859225628 | 0.851034891 | 0.430377106 | 0.719677475 | -4.652545476 |
| **A_55_P2027594** | Taf3 | -0.185641967 | -0.732052626 | 0.360768693 | 8.463654988 | -0.850956466 | 0.430417102 | 0.71968211 | -4.652563481 |
| **A_52_P333097** | NA | -0.142273744 | -0.561128934 | 0.276581445 | 7.795228104 | -0.850768117 | 0.43051317 | 0.719780493 | -4.652606719 |
| **A_52_P221776** | Kif12 | 0.086903175 | -0.169016439 | 0.342822789 | 7.765794799 | 0.850515864 | 0.430641857 | 0.719933391 | -4.652664617 |

| | | | | | | | | | |
|---|---|---|---|---|---|---|---|---|---|
| **A_55_P1981714** | Rreb1 | 0.170838116 | -0.332333281 | 0.674009513 | 11.92527059 | 0.850392014 | 0.43070505 | 0.719976779 | -4.65269304 |
| **A_55_P2082425** | Mtln | 0.46753091 | -0.910096873 | 1.845158693 | 11.84397153 | 0.85001929 | 0.430895269 | 0.72004841 | -4.652778563 |
| **A_55_P1953980** | Sdhb | 0.323385899 | -0.62951089 | 1.276282687 | 12.32810357 | 0.850012986 | 0.430898487 | 0.72004841 | -4.652780009 |
| **A_55_P2148935** | NA | -0.20101672 | -0.793372946 | 0.391339506 | 11.47245606 | -0.849962119 | 0.430924452 | 0.72004841 | -4.652791679 |
| **A_55_P2159485** | NA | 0.190308997 | -0.370493897 | 0.751111891 | 9.176093578 | 0.849961824 | 0.430924602 | 0.72004841 | -4.652791747 |
| **A_55_P1962411** | Gm6813 | 0.171742995 | -0.334360666 | 0.677846656 | 16.49490022 | 0.849943191 | 0.430934114 | 0.72004841 | -4.652796021 |
| **A_52_P419162** | Cyhr1 | -0.071516159 | -0.282313916 | 0.139281598 | 8.806692433 | -0.84974485 | 0.43103537 | 0.720083151 | -4.65284152 |
| **A_55_P2098999** | NA | 0.193691187 | -0.377236305 | 0.764618679 | 13.5374154 | 0.849726672 | 0.431044651 | 0.720083151 | -4.65284569 |
| **A_52_P253748** | Zbtb45 | 0.148877722 | -0.289979067 | 0.587734511 | 9.194898045 | 0.849683608 | 0.431066638 | 0.720083151 | -4.652855568 |
| **A_55_P2062190** | Gstm1 | 0.130723748 | -0.254703825 | 0.516151321 | 11.09554776 | 0.849497404 | 0.431161719 | 0.720179756 | -4.652898275 |
| **A_51_P253732** | Il17rd | -0.116786802 | -0.461273393 | 0.22769979 | 9.356751528 | -0.849125314 | 0.431351766 | 0.720406458 | -4.652983599 |
| **A_55_P2168549** | Ap5b1 | 0.110211506 | -0.214895043 | 0.435318056 | 9.535911417 | 0.849085774 | 0.431371965 | 0.720406458 | -4.652992665 |
| **A_51_P308048** | Cmtm8 | 0.441148729 | -0.860473473 | 1.742770932 | 13.094672 | 0.848888071 | 0.431472972 | 0.720460596 | -4.65303799 |
| **A_55_P2010020** | NA | 0.166899243 | -0.325578301 | 0.659376786 | 16.16832475 | 0.848825229 | 0.431505082 | 0.720460596 | -4.653052396 |
| **A_55_P2101473** | NA | 0.206371199 | -0.402593701 | 0.8153361 | 15.64554048 | 0.848803521 | 0.431516174 | 0.720460596 | -4.653057372 |
| **A_55_P2128303** | Ids | -0.091730812 | -0.36244412 | 0.178982496 | 8.618747328 | -0.848703471 | 0.4315673 | 0.720483739 | -4.653080305 |
| **A_52_P232580** | Dyrk3 | 0.1935021 | -0.377669253 | 0.764673453 | 8.918930416 | 0.848534707 | 0.43165355 | 0.720499216 | -4.653118986 |
| **A_55_P2007495** | NA | 0.223555267 | -0.436355721 | 0.883466255 | 11.23304583 | 0.848496183 | 0.43167324 | 0.720499216 | -4.653127815 |
| **A_55_P2018181** | A730008H | 0.092052115 | -0.179711584 | 0.363815813 | 8.258265952 | 0.848384395 | 0.431730381 | 0.720499216 | -4.653153434 |
| **A_55_P2113359** | Zfp523 | 0.20061059 | -0.391659187 | 0.792880367 | 11.57685674 | 0.848368683 | 0.431738412 | 0.720499216 | -4.653157034 |
| **A_55_P1957762** | Tsr1 | 0.169539661 | -0.331037136 | 0.670116457 | 9.325636627 | 0.848302879 | 0.431772051 | 0.720499216 | -4.653172114 |
| **A_55_P2088640** | Trip4 | 0.121705657 | -0.237747699 | 0.481159013 | 9.412024874 | 0.848044353 | 0.431904228 | 0.720499216 | -4.653231349 |
| **A_51_P357664** | Hras | 0.23502232 | -0.459132613 | 0.929177253 | 11.58305447 | 0.848014035 | 0.43191973 | 0.720499216 | -4.653238295 |
| **A_55_P2176729** | Prdx6 | 0.313308358 | -0.612071132 | 1.238687849 | 12.34458062 | 0.848012896 | 0.431920313 | 0.720499216 | -4.653238556 |
| **A_52_P570652** | Flywch1 | -0.167018851 | -0.66036259 | 0.326324887 | 8.561324068 | -0.847942137 | 0.431956496 | 0.720499216 | -4.653254767 |
| **A_55_P2016685** | Csnk1g1 | 0.10862534 | -0.212253144 | 0.429503823 | 10.36371336 | 0.847892702 | 0.431981777 | 0.720499216 | -4.653266092 |
| **A_55_P2119817** | Plekhj1 | 0.078445263 | -0.153292262 | 0.310182788 | 8.351666269 | 0.847853279 | 0.432001938 | 0.720499216 | -4.653275123 |
| **A_52_P370443** | Riok3 | 0.089334993 | -0.174604025 | 0.353274012 | 10.91235079 | 0.847751092 | 0.4320542 | 0.720499216 | -4.65329853 |
| **A_55_P2087911** | Mthfsd | 0.097313197 | -0.190213485 | 0.384839879 | 8.432694082 | 0.847703419 | 0.432078584 | 0.720499216 | -4.65330945 |
| **A_65_P19250** | Ap2m1 | -0.235900559 | -0.932936421 | 0.461135303 | 8.606071289 | -0.847664884 | 0.432098294 | 0.720499216 | -4.653318277 |
| **A_55_P1995572** | Meis1 | 0.485748227 | -0.949945785 | 1.921442238 | 11.70972252 | 0.847421895 | 0.432222597 | 0.720644333 | -4.653373928 |
| **A_51_P352385** | Ism1 | 0.519874226 | -1.017082516 | 2.056830969 | 9.818254261 | 0.847202031 | 0.432335093 | 0.720645618 | -4.653424274 |
| **A_55_P2001903** | Smim4 | 0.449707975 | -0.879991132 | 1.779407083 | 12.08025363 | 0.847086105 | 0.432394416 | 0.720645618 | -4.653450817 |
| **A_52_P89425** | Pcnt | 0.331828076 | -0.64941391 | 1.313070062 | 11.42548091 | 0.847007687 | 0.432434549 | 0.720645618 | -4.653468771 |
| **A_55_P2144636** | Ccdc163 | 0.165510223 | -0.323920658 | 0.654941104 | 10.0118345 | 0.847000763 | 0.432438093 | 0.720645618 | -4.653470356 |
| **A_55_P2131068** | NA | 0.185580617 | -0.363228677 | 0.73438991 | 14.33517937 | 0.846957273 | 0.432460351 | 0.720645618 | -4.653480312 |
| **A_55_P2025468** | Tst | 0.499183656 | -0.977247106 | 1.975614418 | 10.91290671 | 0.846832697 | 0.432524116 | 0.720645618 | -4.65350883 |
| **A_66_P124497** | Imp3 | 0.22469112 | -0.439933784 | 0.889316023 | 13.8753718 | 0.846758653 | 0.432562018 | 0.720645618 | -4.653525779 |
| **A_52_P527822** | Slc22a3 | -0.161397508 | -0.638817689 | 0.316022674 | 7.74722642 | -0.846732825 | 0.43257524 | 0.720645618 | -4.653531691 |
| **A_55_P1992571** | NA | -0.100577435 | -0.398117296 | 0.196962426 | 12.99793265 | -0.846653577 | 0.432615811 | 0.720645618 | -4.653549831 |
| **A_52_P49321** | Adamts9 | 0.398512466 | -0.780673131 | 1.577698064 | 10.86604007 | 0.846467116 | 0.432711279 | 0.720645618 | -4.653592506 |
| **A_55_P2015363** | C1ra | -0.123058039 | -0.487254938 | 0.241138861 | 8.98883512 | -0.846299512 | 0.432797106 | 0.720645618 | -4.65363086 |
| **A_66_P117682** | NA | 0.088826473 | -0.174084055 | 0.351737 | 9.453369723 | 0.846222923 | 0.43283633 | 0.720645618 | -4.653648385 |
| **A_55_P1998797** | NA | -0.414440534 | -1.641214084 | 0.812333016 | 11.22475668 | -0.846151573 | 0.432872873 | 0.720645618 | -4.653664711 |
| **A_51_P279854** | Ndufa11 | 0.134810978 | -0.264308437 | 0.533930392 | 11.36613811 | 0.84600465 | 0.43294813 | 0.720645618 | -4.653698325 |
| **A_55_P2158522** | Pbx1 | -0.306452346 | -1.213783313 | 0.60087862 | 13.95941943 | -0.845955587 | 0.432973264 | 0.720645618 | -4.653709549 |
| **A_66_P100620** | Ensa | -0.079596599 | -0.31527496 | 0.156081762 | 11.5258847 | -0.845911927 | 0.43299563 | 0.720645618 | -4.653719537 |
| **A_55_P2066429** | NA | -0.409683746 | -1.623020836 | 0.803653345 | 9.367161748 | -0.845702482 | 0.433102937 | 0.720645618 | -4.653767447 |

| | | | | | | | | | |
|---|---|---|---|---|---|---|---|---|---|
| **A_55_P2006008** | Serpinb1a | -0.213244818 | -0.844801449 | 0.418311812 | 9.530377316 | -0.845700396 | 0.433104007 | 0.720645618 | -4.653767925 |
| **A_51_P483511** | Dock8 | -0.268760619 | -1.06476016 | 0.527238922 | 8.243480141 | -0.845674355 | 0.43311735 | 0.720645618 | -4.653773881 |
| **A_55_P2068988** | NA | 0.196191538 | -0.385017881 | 0.777400956 | 10.2988162 | 0.845469555 | 0.4332223 | 0.720645618 | -4.65382072 |
| **A_51_P165435** | Cox4i1 | 0.123486338 | -0.242353533 | 0.489326208 | 15.29909998 | 0.845431103 | 0.433242006 | 0.720645618 | -4.653829514 |
| **A_51_P245454** | Ufsp2 | -0.089234675 | -0.353630979 | 0.175161629 | 11.20182675 | -0.845334534 | 0.433291502 | 0.720645618 | -4.653851597 |
| **A_52_P638605** | Ap1m2 | -0.159664826 | -0.632746856 | 0.313417204 | 8.950134867 | -0.845323898 | 0.433296953 | 0.720645618 | -4.653854029 |
| **A_55_P2024366** | Raver2 | 0.805065955 | -1.580529163 | 3.190661073 | 11.4737886 | 0.845249786 | 0.433334941 | 0.720645618 | -4.653870976 |
| **A_55_P2114318** | NA | 0.200857432 | -0.394384376 | 0.796099239 | 11.27392491 | 0.845171461 | 0.433375092 | 0.720645618 | -4.653888885 |
| **A_55_P2065513** | Ebf1 | 0.182652267 | -0.358642052 | 0.723946585 | 8.587635052 | 0.845165855 | 0.433377965 | 0.720645618 | -4.653890166 |
| **A_51_P258435** | Zdhhc3 | 0.327884713 | -0.643908037 | 1.299677464 | 10.0295238 | 0.845080071 | 0.433421943 | 0.720645618 | -4.65390978 |
| **A_55_P2010048** | NA | 0.213986956 | -0.420238412 | 0.848212324 | 9.145125465 | 0.845072634 | 0.433425756 | 0.720645618 | -4.65391148 |
| **A_55_P2031999** | Dglucy | 0.19842448 | -0.389717145 | 0.786566105 | 9.043859759 | 0.845013562 | 0.433456042 | 0.720645618 | -4.653924986 |
| **A_55_P2044710** | Asb10 | -0.112886553 | -0.447492202 | 0.221719095 | 7.811620174 | -0.845005023 | 0.433460419 | 0.720645618 | -4.653926938 |
| **A_51_P299287** | Impa2 | 0.228642421 | -0.449076098 | 0.90636094 | 11.19164341 | 0.845002129 | 0.433461903 | 0.720645618 | -4.653927599 |
| **A_55_P2176963** | Hsph1 | -0.305123639 | -1.209571602 | 0.599324325 | 10.84697856 | -0.844972578 | 0.433477054 | 0.720645618 | -4.653934355 |
| **A_55_P2233271** | Zfp626 | 0.121867501 | -0.239462709 | 0.483197712 | 9.164400558 | 0.844761238 | 0.433585423 | 0.720645618 | -4.653982666 |
| **A_51_P395164** | Pex16 | -0.414182691 | -1.64231503 | 0.813949648 | 9.128557195 | -0.844689555 | 0.433622185 | 0.720645618 | -4.653999051 |
| **A_55_P2089507** | NA | 0.202491455 | -0.3980001 | 0.80298301 | 15.63644853 | 0.844598181 | 0.433669047 | 0.720645618 | -4.654019935 |
| **A_55_P2088825** | Foxp1 | 0.383906427 | -0.754588763 | 1.522401617 | 12.744988 | 0.844587263 | 0.433674647 | 0.720645618 | -4.65402243 |
| **A_52_P632742** | Shisa5 | -0.15757426 | -0.624906061 | 0.30975754 | 8.256858749 | -0.844520679 | 0.433708799 | 0.720645618 | -4.654037648 |
| **A_55_P2125786** | Lrrc8e | 0.294636504 | -0.579285098 | 1.168558106 | 9.266420714 | 0.844431496 | 0.433754545 | 0.720645618 | -4.654058029 |
| **A_52_P309737** | Cdk12 | 0.096139917 | -0.189026705 | 0.381306538 | 8.28134371 | 0.844413945 | 0.433763548 | 0.720645618 | -4.65406204 |
| **A_51_P484842** | Asgr2 | 0.073191763 | -0.143917642 | 0.290301168 | 8.012966183 | 0.844372186 | 0.43378497 | 0.720645618 | -4.654071582 |
| **A_55_P1953143** | Wt1 | -0.529705127 | -2.100991865 | 1.041581611 | 9.453566079 | -0.844362773 | 0.433789799 | 0.720645618 | -4.654073733 |
| **A_52_P356119** | Hus1 | 0.096441068 | -0.189660796 | 0.382542932 | 8.485314548 | 0.844290047 | 0.433827108 | 0.720645618 | -4.654090352 |
| **A_51_P415475** | Dok4 | 0.454822723 | -0.894559982 | 1.804205429 | 10.46466943 | 0.84422335 | 0.433861327 | 0.720645618 | -4.654105592 |
| **A_55_P1990859** | Rpl6 | 0.219711717 | -0.432139205 | 0.87156264 | 16.48439921 | 0.844219307 | 0.433863401 | 0.720645618 | -4.654106515 |
| **A_55_P2160383** | NA | -0.220897093 | -0.876436649 | 0.434642462 | 11.17686867 | -0.843998053 | 0.43397693 | 0.720726474 | -4.654157066 |
| **A_55_P2004727** | NA | -0.096665264 | -0.383537868 | 0.19020734 | 8.058904642 | -0.843979139 | 0.433986636 | 0.720726474 | -4.654161387 |
| **A_55_P2131865** | Krtap10-1 | 0.155501367 | -0.306042929 | 0.617045663 | 9.285723049 | 0.843861491 | 0.434047013 | 0.720736042 | -4.654188262 |
| **A_55_P1969874** | NA | -0.21148621 | -0.839228268 | 0.416255849 | 16.00346247 | -0.843822638 | 0.434066953 | 0.720736042 | -4.654197138 |
| **A_55_P1986103** | Suco | -0.172273015 | -0.683664882 | 0.339118852 | 8.233806051 | -0.843749993 | 0.434104239 | 0.720736054 | -4.654213731 |
| **A_52_P517896** | Man2c1 | -0.130821289 | -0.536817868 | 0.275175289 | 11.41865484 | -0.858259772 | 0.434278702 | 0.720769736 | -4.637814867 |
| **A_55_P2174601** | Plekha7 | -0.612660117 | -2.432307235 | 1.206987002 | 9.527564038 | -0.843301321 | 0.434334577 | 0.720769736 | -4.654316197 |
| **A_55_P2382070** | 4930593A | 0.106532902 | -0.209941628 | 0.423007432 | 8.2916025 | 0.843131545 | 0.43442176 | 0.720769736 | -4.654354962 |
| **A_55_P1968668** | Gpatch2l | -0.178948922 | -0.710591847 | 0.352694003 | 9.102831819 | -0.843061764 | 0.434457597 | 0.720769736 | -4.654370893 |
| **A_51_P106397** | Prkd3 | -0.086384053 | -0.343067631 | 0.170299525 | 8.432002696 | -0.842918989 | 0.43453093 | 0.720769736 | -4.654403488 |
| **A_55_P2063937** | D630033A | 0.116831912 | -0.230372338 | 0.464036162 | 8.573071639 | 0.842804404 | 0.434589789 | 0.720769736 | -4.654429644 |
| **A_55_P2023797** | Cep112 | 0.249563818 | -0.492140466 | 0.991268101 | 9.386754703 | 0.842754692 | 0.434615327 | 0.720769736 | -4.654440991 |
| **A_55_P1994294** | NA | -0.583702692 | -2.318470902 | 1.151065518 | 10.69197002 | -0.842753542 | 0.434615918 | 0.720769736 | -4.654441253 |
| **A_52_P90289** | Oaz2 | 0.16203234 | -0.319598449 | 0.643663129 | 9.570988623 | 0.842631726 | 0.434678501 | 0.720769736 | -4.654469057 |
| **A_52_P547187** | Tab2 | -0.115308681 | -0.45806559 | 0.227448228 | 11.20691884 | -0.842609074 | 0.43469014 | 0.720769736 | -4.654474227 |
| **A_55_P2181508** | Gm10653 | 0.235028545 | -0.463797553 | 0.933854644 | 15.71142119 | 0.842367966 | 0.434814032 | 0.720769736 | -4.65452925 |
| **A_55_P1988699** | Cacna2d1 | 0.411867198 | -0.812899805 | 1.636634201 | 9.313627292 | 0.842275317 | 0.434861647 | 0.720769736 | -4.65455039 |
| **A_55_P1968355** | Tle1 | 0.320021918 | -0.631637191 | 1.271681027 | 11.81972406 | 0.842264819 | 0.434867042 | 0.720769736 | -4.654552786 |
| **A_55_P2111331** | Klhl32 | 0.194076535 | -0.383114884 | 0.771267955 | 8.426076113 | 0.842177259 | 0.434912045 | 0.720769736 | -4.654572764 |
| **A_55_P2054663** | Cox4i2 | 0.38881335 | -0.767565663 | 1.545192362 | 10.1541392 | 0.842153606 | 0.434924203 | 0.720769736 | -4.65457816 |
| **A_51_P354433** | Ppie | 0.094641905 | -0.186883339 | 0.376167148 | 8.375868598 | 0.842008497 | 0.434998793 | 0.720769736 | -4.654611266 |

| | | | | | | | | | |
|---|---|---|---|---|---|---|---|---|---|
| **A_51_P509211** | Rnf34 | -0.092351157 | -0.36706459 | 0.182362275 | 11.04036601 | -0.842001357 | 0.435002464 | 0.720769736 | -4.654612895 |
| **A_55_P2058928** | Ralgds | -0.444616659 | -1.767249822 | 0.878016505 | 11.81654532 | -0.841970099 | 0.435018533 | 0.720769736 | -4.654620026 |
| **A_55_P2104327** | Pxdn | 0.529814379 | -1.046350047 | 2.105978806 | 13.5937738 | 0.841923371 | 0.435042555 | 0.720769736 | -4.654630685 |
| **A_55_P2251082** | Qk | 0.17005615 | -0.335858796 | 0.675971097 | 8.982811232 | 0.841909051 | 0.435049917 | 0.720769736 | -4.654633952 |
| **A_55_P2000658** | Gm15645 | -0.358400577 | -1.424681428 | 0.707880274 | 9.684645163 | -0.841874657 | 0.4350676 | 0.720769736 | -4.654641797 |
| **A_55_P2052479** | NA | 0.34610082 | -0.683782058 | 1.375983699 | 15.51525082 | 0.841715134 | 0.43514962 | 0.720769736 | -4.654678184 |
| **A_55_P2066017** | NA | -0.191089983 | -0.759724368 | 0.377544402 | 8.585874184 | -0.841695781 | 0.435159572 | 0.720769736 | -4.654682598 |
| **A_55_P2003211** | Igsf5 | 0.330460209 | -0.652949762 | 1.31387018 | 9.528939327 | 0.841656558 | 0.435179741 | 0.720769736 | -4.654691544 |
| **A_55_P2311780** | 4930554H | -0.118043774 | -0.469328186 | 0.233240637 | 7.701931108 | -0.841655865 | 0.435180097 | 0.720769736 | -4.654691702 |
| **A_55_P2065244** | Rbm26 | 0.204493942 | -0.404166544 | 0.813154429 | 9.320823854 | 0.841503031 | 0.435258693 | 0.720769736 | -4.654726557 |
| **A_51_P350048** | Gstt2 | 0.205734398 | -0.406732864 | 0.81820166 | 9.456105367 | 0.8413455 | 0.435339717 | 0.720769736 | -4.65476248 |
| **A_51_P475076** | Vbp1 | 0.439227863 | -0.868429215 | 1.746884941 | 10.18105823 | 0.841291224 | 0.435367635 | 0.720769736 | -4.654774856 |
| **A_51_P221014** | Cd47 | -0.183996726 | -0.731791764 | 0.363798311 | 11.56731652 | -0.841283455 | 0.435371631 | 0.720769736 | -4.654776627 |
| **A_51_P114177** | Eci3 | 0.282813605 | -0.559183363 | 1.124810572 | 11.2787036 | 0.84127913 | 0.435373856 | 0.720769736 | -4.654777613 |
| **A_52_P30451** | Ppp1r3c | 0.916790079 | -1.812820535 | 3.646400692 | 11.918284 | 0.841239298 | 0.435394346 | 0.720769736 | -4.654786695 |
| **A_51_P283016** | Slc1a6 | -0.147035626 | -0.584840069 | 0.290768816 | 8.097252009 | -0.841187398 | 0.435421044 | 0.720769736 | -4.654798529 |
| **A_55_P2153862** | Irak1 | -0.168060465 | -0.66852861 | 0.33240768 | 9.112507601 | -0.841084187 | 0.435474142 | 0.720769736 | -4.65482206 |
| **A_51_P132978** | Idh1 | 0.190436322 | -0.376682389 | 0.757555032 | 8.961853809 | 0.841058402 | 0.435487408 | 0.720769736 | -4.654827938 |
| **A_51_P159415** | Sin3a | 0.083903251 | -0.165976714 | 0.333783216 | 9.89037797 | 0.841003292 | 0.435515763 | 0.720769736 | -4.654840501 |
| **A_55_P2335695** | Ndor1 | -0.198897459 | -0.791255599 | 0.393460681 | 8.340141442 | -0.840998497 | 0.43551823 | 0.720769736 | -4.654841595 |
| **A_55_P2050508** | Spta1 | 0.071992368 | -0.142421624 | 0.286406361 | 7.828146428 | 0.840976155 | 0.435529726 | 0.720769736 | -4.654846688 |
| **A_55_P2140953** | NA | -0.283508348 | -1.128000689 | 0.560983992 | 15.19036022 | -0.840853783 | 0.435592693 | 0.720769736 | -4.654874583 |
| **A_51_P206824** | Hjv | -0.090530196 | -0.360195024 | 0.179134631 | 7.720990034 | -0.840851885 | 0.43559367 | 0.720769736 | -4.654875015 |
| **A_55_P2082688** | Ak1 | 0.432004987 | -0.854885935 | 1.71889591 | 12.09762949 | 0.840809033 | 0.435615722 | 0.720769736 | -4.654884783 |
| **A_55_P2175245** | Tusc3 | 0.280232437 | -0.554639904 | 1.115104778 | 10.67912746 | 0.840714767 | 0.435664234 | 0.720788319 | -4.654906269 |
| **A_55_P2033725** | Ascl2 | -0.219632984 | -0.874151433 | 0.434885464 | 9.687113696 | -0.840477353 | 0.435786432 | 0.720889742 | -4.654960375 |
| **A_55_P2142296** | Ywhaq | 0.318634139 | -0.631022208 | 1.268290486 | 13.89054704 | 0.840380903 | 0.435836082 | 0.720889742 | -4.654982354 |
| **A_55_P2088131** | Zfp975 | 0.486155407 | -0.962837898 | 1.935148713 | 10.5727583 | 0.840347838 | 0.435853105 | 0.720889742 | -4.654989888 |
| **A_55_P2002757** | Blnk | -0.520054197 | -2.070160647 | 1.030052254 | 8.547125406 | -0.840305929 | 0.435874681 | 0.720889742 | -4.654999437 |
| **A_55_P2068356** | 1700016K | 0.644868692 | -1.277692493 | 2.567429878 | 10.33268635 | 0.84012028 | 0.435970267 | 0.720906856 | -4.655041735 |
| **A_55_P1959633** | Hnf4a | -0.560305127 | -2.230845082 | 1.110234828 | 8.275392598 | -0.840075028 | 0.435993569 | 0.720906856 | -4.655052044 |
| **A_55_P2033500** | Ltbp4 | 0.715173017 | -1.417119214 | 2.847465248 | 11.08831306 | 0.840068579 | 0.43599689 | 0.720906856 | -4.655053513 |
| **A_51_P401501** | Tmem213 | 1.377127253 | -2.729326622 | 5.483581129 | 12.20474431 | 0.839957804 | 0.436053935 | 0.720939523 | -4.655078749 |
| **A_55_P2136572** | Cdc42se2 | -0.501066343 | -1.9955738 | 0.993441115 | 11.33174547 | -0.839745124 | 0.436163475 | 0.721015318 | -4.655127193 |
| **A_55_P1960735** | Gdf15 | -0.253430461 | -1.009353827 | 0.502492905 | 8.063859804 | -0.839713992 | 0.436179511 | 0.721015318 | -4.655134283 |
| **A_52_P204689** | Map4k2 | -0.084990704 | -0.3385335 | 0.168552093 | 8.312725514 | -0.83959626 | 0.436240158 | 0.721015318 | -4.655161096 |
| **A_55_P2098061** | Tchp | -0.102072668 | -0.406580274 | 0.202434938 | 8.803222374 | -0.839579195 | 0.436248949 | 0.721015318 | -4.655164982 |
| **A_55_P2171206** | Matn4 | 0.723909937 | -1.43617964 | 2.883999513 | 10.11825477 | 0.839388708 | 0.43634709 | 0.721042072 | -4.65520836 |
| **A_55_P2103603** | NA | -0.110718175 | -0.441152096 | 0.219715746 | 7.845025518 | -0.839237035 | 0.436425245 | 0.721042072 | -4.655242894 |
| **A_66_P132663** | Uqcrb | 0.142691366 | -0.28319444 | 0.568577172 | 14.45206596 | 0.839179518 | 0.436454885 | 0.721042072 | -4.655255989 |
| **A_51_P276939** | Gmip | -0.172941231 | -0.689115199 | 0.343232737 | 8.174875463 | -0.839175495 | 0.436456959 | 0.721042072 | -4.655256905 |
| **A_51_P160664** | Cox7b | 0.347361604 | -0.689562031 | 1.384285238 | 14.31455121 | 0.839045251 | 0.436524084 | 0.721042072 | -4.655286556 |
| **A_55_P2157770** | Neu2 | 0.263541235 | -0.523304322 | 1.050386791 | 9.150916066 | 0.838898514 | 0.436599718 | 0.721042072 | -4.655319959 |
| **A_55_P1962693** | Pla2g5 | -0.072441535 | -0.288754091 | 0.143871021 | 7.963450914 | -0.838795821 | 0.436652656 | 0.721042072 | -4.655343333 |
| **A_55_P2175615** | Csde1 | 0.165271569 | -0.328290648 | 0.658833786 | 13.36680932 | 0.838699896 | 0.43670211 | 0.721042072 | -4.655365166 |
| **A_52_P302395** | Lnpep | -0.323910079 | -1.291322084 | 0.643501926 | 8.617767659 | -0.838616369 | 0.436745175 | 0.721042072 | -4.655384175 |
| **A_51_P386344** | Mtmr10 | -0.218619477 | -0.871721157 | 0.434482203 | 8.923407454 | -0.838413758 | 0.436849651 | 0.721042072 | -4.655430281 |
| **A_55_P2120388** | Skint7 | -0.191227742 | -0.762600494 | 0.380145009 | 15.27131316 | -0.838265743 | 0.436925986 | 0.721042072 | -4.655463959 |

| | | | | | | | | | |
|---|---|---|---|---|---|---|---|---|---|
| **A_51_P160536** | Cfap161 | 0.312319958 | -0.621130566 | 1.245770482 | 8.672437683 | 0.838028482 | 0.437048368 | 0.721042072 | -4.655517936 |
| **A_51_P330630** | Usp3 | 0.194450952 | -0.386803909 | 0.775705813 | 8.838205508 | 0.837903146 | 0.437113029 | 0.721042072 | -4.655546446 |
| **A_51_P490678** | Trim68 | -0.094526991 | -0.377107479 | 0.188053498 | 8.402882946 | -0.837845625 | 0.437142706 | 0.721042072 | -4.655559529 |
| **A_55_P2006476** | NA | 0.234834277 | -0.467201144 | 0.936869698 | 11.6666414 | 0.837824022 | 0.437153852 | 0.721042072 | -4.655564443 |
| **A_55_P2219765** | Wnt5a | 0.511317166 | -1.017291097 | 2.039925429 | 9.845941574 | 0.837808024 | 0.437162107 | 0.721042072 | -4.655568081 |
| **A_55_P2175405** | Zfp467 | 0.315328847 | -0.627394875 | 1.25805257 | 9.144356758 | 0.837779276 | 0.43717694 | 0.721042072 | -4.65557462 |
| **A_55_P2021378** | Wapl | -0.383804851 | -1.531438738 | 0.763829037 | 10.65749961 | -0.837640064 | 0.437248776 | 0.721042072 | -4.655606281 |
| **A_55_P2040951** | Actc1 | 1.412276781 | -2.810789046 | 5.635342607 | 13.78448579 | 0.837610887 | 0.437263832 | 0.721042072 | -4.655612916 |
| **A_51_P111544** | Srp14 | 0.140963424 | -0.280577623 | 0.56250447 | 13.08254714 | 0.837561904 | 0.437289111 | 0.721042072 | -4.652390567 |
| **A_51_P297069** | Tmod1 | -0.13320537 | -0.531631681 | 0.26522094 | 8.347658759 | -0.837382864 | 0.437381517 | 0.721042072 | -4.655664767 |
| **A_55_P2088525** | Myh10 | 0.365252385 | -0.727280062 | 1.457784832 | 10.99040182 | 0.837353929 | 0.437396452 | 0.721042072 | -4.655671346 |
| **A_55_P2081278** | Mrpl55 | 0.07697796 | -0.153277367 | 0.307233287 | 12.94945935 | 0.837350064 | 0.437398447 | 0.721042072 | -4.655672225 |
| **A_55_P1955747** | Gm12070 | 0.156072012 | -0.310773762 | 0.622917787 | 14.72299631 | 0.837340202 | 0.437403538 | 0.721042072 | -4.655674467 |
| **A_51_P443443** | 5830417I1 | -0.258454564 | -1.031571426 | 0.514662299 | 10.78843027 | -0.837316029 | 0.437416016 | 0.721042072 | -4.655679963 |
| **A_51_P420655** | Reep4 | -0.100436589 | -0.400887161 | 0.200013982 | 9.595813748 | -0.837277204 | 0.437436057 | 0.721042072 | -4.65568879 |
| **A_52_P654965** | Eif3j1 | -0.279479862 | -1.115601874 | 0.55664215 | 11.26689035 | -0.837203835 | 0.437473932 | 0.721042072 | -4.655705471 |
| **A_66_P111089** | Ube2f | -0.217477356 | -0.897586788 | 0.462632077 | 8.641664288 | -0.851722168 | 0.437540236 | 0.721042072 | -4.639068779 |
| **A_51_P363556** | Abi2 | 0.266064383 | -0.530091084 | 1.06221985 | 9.680093661 | 0.837026495 | 0.43756549 | 0.721042072 | -4.655745785 |
| **A_51_P172502** | Cxcl12 | -0.17705087 | -0.706848259 | 0.352746519 | 12.63539743 | -0.837025324 | 0.437566094 | 0.721042072 | -4.655746052 |
| **A_55_P2088715** | Ss18l2 | 0.09186122 | -0.183025 | 0.36674744 | 9.196980496 | 0.837007956 | 0.437575062 | 0.721042072 | -4.65575 |
| **A_55_P2180899** | Aldh3a2 | -0.179507747 | -0.716775612 | 0.357760118 | 8.804761406 | -0.836840472 | 0.437661546 | 0.721042072 | -4.655788069 |
| **A_51_P256957** | Ankmy2 | 0.200671242 | -0.400006352 | 0.801348836 | 9.05245816 | 0.836746783 | 0.437709929 | 0.721042072 | -4.655809362 |
| **A_55_P2117400** | Cacng7 | 0.308010951 | -0.613993565 | 1.230015466 | 9.719994317 | 0.836726367 | 0.437720473 | 0.721042072 | -4.655814002 |
| **A_51_P141610** | Cables2 | -0.095106973 | -0.379829599 | 0.189615653 | 9.27657482 | -0.836644047 | 0.437762989 | 0.721042072 | -4.65583271 |
| **A_52_P134381** | Psmd12 | -0.160925169 | -0.642706992 | 0.320856654 | 10.13881877 | -0.836611651 | 0.437779722 | 0.721042072 | -4.655840072 |
| **A_55_P2048418** | Mdfic | 0.10695937 | -0.21328127 | 0.427200011 | 11.45860012 | 0.836551606 | 0.437810737 | 0.721042072 | -4.655853717 |
| **A_51_P247799** | Casp8 | 0.231378772 | -0.461401374 | 0.924158919 | 9.448214521 | 0.836524037 | 0.437824977 | 0.721042072 | -4.655859981 |
| **A_55_P2117830** | NA | -0.093192851 | -0.372235415 | 0.185849713 | 8.095157671 | -0.836493344 | 0.437840832 | 0.721042072 | -4.655866955 |
| **A_51_P410949** | Polr3g | -0.348505721 | -1.39203823 | 0.695026787 | 8.488996714 | -0.836477522 | 0.437849005 | 0.721042072 | -4.655870551 |
| **A_66_P110791** | Ppp1r11 | 0.116222399 | -0.231821498 | 0.464266296 | 12.29563508 | 0.836384853 | 0.437896877 | 0.721042072 | -4.655891607 |
| **A_55_P2017982** | Pde8b | 0.487529425 | -0.972633562 | 1.947692412 | 9.473194657 | 0.836276404 | 0.437952906 | 0.721042072 | -4.655916247 |
| **A_55_P2038514** | NA | 0.247382267 | -0.493619319 | 0.988383854 | 8.515717351 | 0.836179995 | 0.438002719 | 0.721042072 | -4.655938149 |
| **A_65_P17866** | Vma21 | 0.084352546 | -0.168368318 | 0.337073411 | 8.506712196 | 0.836002246 | 0.43809457 | 0.721042072 | -4.655978527 |
| **A_51_P353173** | Fam98a | 0.105995886 | -0.211577682 | 0.423569454 | 10.71238032 | 0.835978306 | 0.438106942 | 0.721042072 | -4.655983965 |
| **A_55_P2114779** | Greb1 | 0.292854523 | -0.584594688 | 1.170303734 | 9.782595664 | 0.835949983 | 0.43812158 | 0.721042072 | -4.655990398 |
| **A_51_P346884** | Snrnp27 | 0.233267373 | -0.465659144 | 0.932193889 | 12.19259554 | 0.83593561 | 0.438129008 | 0.721042072 | -4.655993663 |
| **A_51_P173384** | Tada1 | 0.171112414 | -0.341887558 | 0.684112386 | 10.35327637 | 0.835438572 | 0.438385941 | 0.721042072 | -4.656106538 |
| **A_55_P2061520** | NA | -0.130156314 | -0.520393808 | 0.260081179 | 8.173058998 | -0.835384848 | 0.438413719 | 0.721042072 | -4.656118736 |
| **A_51_P277336** | Cavin2 | 0.82312522 | -1.645291301 | 3.291541741 | 14.36396333 | 0.835214085 | 0.438502022 | 0.721042072 | -4.656157504 |
| **A_51_P376789** | Pgghg | 0.264276187 | -0.528257069 | 1.056809442 | 9.585843473 | 0.835200763 | 0.438508911 | 0.721042072 | -4.656160528 |
| **A_55_P2043627** | Fam89a | -0.807829404 | -3.230471014 | 1.614812205 | 9.865417636 | -0.835181433 | 0.438518908 | 0.721042072 | -4.656164917 |
| **A_55_P2111018** | NA | 0.078792184 | -0.157505345 | 0.315089713 | 8.84434079 | 0.835168877 | 0.438525402 | 0.721042072 | -4.656167767 |
| **A_51_P233603** | Psma7 | 0.15165358 | -0.303254382 | 0.606561542 | 14.09726975 | 0.834986539 | 0.438619708 | 0.721042072 | -4.656209157 |
| **A_51_P165098** | Gga2 | -0.334512307 | -1.337973123 | 0.668948508 | 11.40520519 | -0.834952997 | 0.438637057 | 0.721042072 | -4.65621677 |
| **A_55_P2104889** | Gm11515 | 0.3664693 | -0.733050846 | 1.465989446 | 9.063721619 | 0.834804445 | 0.438713903 | 0.721042072 | -4.656250486 |
| **A_55_P2045765** | Rnf224 | -0.192584194 | -0.770438523 | 0.385270135 | 8.074154153 | -0.834742671 | 0.438745861 | 0.721042072 | -4.656264505 |
| **A_52_P482983** | Cd164 | -0.392222314 | -1.569122746 | 0.784678118 | 14.40898698 | -0.834724038 | 0.438755501 | 0.721042072 | -4.656268733 |
| **A_55_P2173768** | Pkhd1 | -0.121681276 | -0.486821009 | 0.243458457 | 7.754082426 | -0.834670389 | 0.438783258 | 0.721042072 | -4.656280908 |

| | | | | | | | | | |
|---|---|---|---|---|---|---|---|---|---|
| **A_55_P2038831** | NA | 0.38174042 | -0.763813459 | 1.5272943 | 15.66209376 | 0.834647261 | 0.438795224 | 0.721042072 | -4.656286156 |
| **A_55_P2056014** | NA | -0.183199162 | -0.757806905 | 0.39140858 | 9.273849377 | -0.849209248 | 0.438799004 | 0.721042072 | -4.639549479 |
| **A_55_P2038747** | Ano1 | 0.4896402 | -0.979722235 | 1.959002635 | 13.25015911 | 0.834638627 | 0.438799692 | 0.721042072 | -4.656288116 |
| **A_55_P2052330** | NA | -0.880433915 | -3.522558355 | 1.761690525 | 12.92023617 | -0.834629712 | 0.438804304 | 0.721042072 | -4.656290139 |
| **A_51_P169047** | Tdp2 | 0.081963798 | -0.164042069 | 0.327969666 | 9.871827973 | 0.83450118 | 0.438870811 | 0.721042072 | -4.656319304 |
| **A_55_P1999349** | Nudt4 | 0.07652922 | -0.153167345 | 0.306225784 | 13.01494774 | 0.834493832 | 0.438874614 | 0.721042072 | -4.656320971 |
| **A_55_P2031547** | Vmn2r121 | -0.215659642 | -0.863024262 | 0.431704979 | 7.83779302 | -0.834392254 | 0.438927179 | 0.721042072 | -4.656344018 |
| **A_52_P618132** | Manea | 0.11664326 | -0.233511732 | 0.466798251 | 8.445717274 | 0.834352693 | 0.438947654 | 0.721042072 | -4.656352994 |
| **A_55_P2052759** | Scd3 | -0.126229312 | -0.505165415 | 0.25270679 | 8.444210198 | -0.834342905 | 0.438952719 | 0.721042072 | -4.656355215 |
| **A_51_P431737** | Cth | 0.193597389 | -0.387675157 | 0.774869935 | 8.86798278 | 0.834199701 | 0.439026838 | 0.721042072 | -4.656387702 |
| **A_55_P1955542** | Olfr522 | 0.080117795 | -0.160466317 | 0.320701906 | 8.104768518 | 0.834088991 | 0.439084144 | 0.721042072 | -4.656412816 |
| **A_55_P2266880** | Kirrel | 0.120939777 | -0.24228774 | 0.484167293 | 10.16114351 | 0.833951445 | 0.439155349 | 0.721042072 | -4.656444014 |
| **A_52_P242724** | NA | 0.099053282 | -0.198453751 | 0.396560315 | 10.07472748 | 0.833915369 | 0.439174027 | 0.721042072 | -4.656452196 |
| **A_66_P111285** | Pdp2 | 0.183070514 | -0.366783989 | 0.732925016 | 8.687448876 | 0.833913415 | 0.439175039 | 0.721042072 | -4.656452639 |
| **A_51_P395111** | Nme3 | 0.309590993 | -0.620431095 | 1.239613081 | 11.18730298 | 0.833768339 | 0.439250154 | 0.721042072 | -4.656485541 |
| **A_55_P2032845** | Rbm10 | 0.27037398 | -0.541870803 | 1.082618764 | 11.34691861 | 0.833735529 | 0.439267143 | 0.721042072 | -4.656492981 |
| **A_55_P2179582** | Nmur2 | 0.236667407 | -0.47453889 | 0.947873703 | 8.426759919 | 0.833476204 | 0.439401441 | 0.721042072 | -4.656551784 |
| **A_52_P515094** | Tmem176 | 0.471926474 | -0.946546908 | 1.890399856 | 10.51955773 | 0.833304057 | 0.439490608 | 0.721042072 | -4.656590812 |
| **A_51_P379373** | Lactb2 | -0.077426999 | -0.310177681 | 0.155323682 | 8.843728886 | -0.833204895 | 0.439541977 | 0.721042072 | -4.656613292 |
| **A_55_P1996314** | Amy2a5 | 0.177083946 | -0.355263963 | 0.709431854 | 7.950385011 | 0.83317069 | 0.439559697 | 0.721042072 | -4.656621045 |
| **A_55_P2062469** | Col12a1 | -0.13184268 | -0.528245413 | 0.264560053 | 9.759609768 | -0.833047431 | 0.439623557 | 0.721042072 | -4.656648984 |
| **A_52_P106463** | Jak1 | -0.175571267 | -0.703501376 | 0.352358841 | 11.79390762 | -0.83296616 | 0.439665667 | 0.721042072 | -4.656667405 |
| **A_55_P2159573** | Reep6 | -0.084222948 | -0.337478997 | 0.169033102 | 7.766261131 | -0.83295388 | 0.43967203 | 0.721042072 | -4.656670188 |
| **A_55_P2159666** | NA | 0.084288534 | -0.169197293 | 0.33777436 | 9.026202331 | 0.832846878 | 0.439727478 | 0.721042072 | -4.656694438 |
| **A_55_P2046842** | Tmem253 | -0.167151821 | -0.669854708 | 0.335551066 | 7.969989453 | -0.832817951 | 0.439742468 | 0.721042072 | -4.656700994 |
| **A_51_P351194** | Cnfn | -0.12161614 | -0.4873784 | 0.244146121 | 8.093727448 | -0.83280374 | 0.439749833 | 0.721042072 | -4.656704214 |
| **A_55_P2277171** | Gm10134 | 0.106172846 | -0.213153587 | 0.42549928 | 8.370050399 | 0.83277741 | 0.439763478 | 0.721042072 | -4.656710181 |
| **A_51_P267508** | Borcs7 | -0.119808765 | -0.480153928 | 0.240536398 | 9.170197728 | -0.832760735 | 0.43977212 | 0.721042072 | -4.65671396 |
| **A_51_P346641** | NA | 0.256693935 | -0.5154752 | 1.02886307 | 8.657273922 | 0.8326328 | 0.439838427 | 0.721042072 | -4.65674295 |
| **A_51_P117995** | Pfkm | 0.373165666 | -0.749454854 | 1.495786186 | 10.29016671 | 0.832566723 | 0.439872676 | 0.721042072 | -4.655653325 |
| **A_51_P392687** | Vim | 0.52050248 | -1.04572732 | 2.086732279 | 14.98518587 | 0.832372389 | 0.439973417 | 0.721042072 | -4.656801951 |
| **A_55_P2037919** | NA | 0.245316909 | -0.492965786 | 0.983599604 | 9.663469825 | 0.832252561 | 0.440035542 | 0.721042072 | -4.656829097 |
| **A_55_P2091741** | Lamp3 | 0.846838637 | -1.701750737 | 3.395428011 | 13.93750361 | 0.832244899 | 0.440039515 | 0.721042072 | -4.656830832 |
| **A_55_P2063490** | Csprs | -0.21042391 | -0.843703667 | 0.422855848 | 8.113954467 | -0.832242375 | 0.440040824 | 0.721042072 | -4.656831404 |
| **A_66_P120260** | Taf10 | -0.14108342 | -0.565683507 | 0.283516667 | 13.39945589 | -0.832235513 | 0.440044381 | 0.721042072 | -4.656832958 |
| **A_51_P222386** | Tbc1d10a | -0.520576948 | -2.087315811 | 1.046161914 | 9.953467114 | -0.832220985 | 0.440051914 | 0.721042072 | -4.656836249 |
| **A_55_P1957433** | Mprip | -0.111093057 | -0.445479394 | 0.22329328 | 10.6381425 | -0.832125325 | 0.440101516 | 0.721042072 | -4.656857918 |
| **A_55_P2163837** | Med14 | 0.157070536 | -0.315727397 | 0.629868469 | 8.997231825 | 0.832088468 | 0.440120629 | 0.721042072 | -4.656866266 |
| **A_52_P523712** | Tbx5 | 0.793623509 | -1.595504604 | 3.182751621 | 11.16024864 | 0.832004032 | 0.440164415 | 0.721042072 | -4.65688539 |
| **A_52_P227065** | Bud31 | 0.111447637 | -0.224072038 | 0.446967311 | 11.11292333 | 0.831961476 | 0.440186485 | 0.721042072 | -4.656895028 |
| **A_55_P2083224** | Pum2 | 0.090534253 | -0.182043984 | 0.36311249 | 10.97045236 | 0.83190185 | 0.440217409 | 0.721042072 | -4.656908532 |
| **A_51_P327530** | Ciao2a | 0.19709652 | -0.396358033 | 0.790551072 | 11.45366177 | 0.8318439 | 0.440247465 | 0.721042072 | -4.656921656 |
| **A_55_P2010768** | Dpm3 | 0.209358678 | -0.421023775 | 0.839741131 | 13.3312391 | 0.831835013 | 0.440252074 | 0.721042072 | -4.656923668 |
| **A_55_P2153381** | Ermap | -0.093831338 | -0.376403096 | 0.18874042 | 8.544977364 | -0.831705361 | 0.440319325 | 0.721042072 | -4.656953028 |
| **A_55_P1963344** | Upf3b | 0.204207792 | -0.410793248 | 0.819208831 | 9.280005348 | 0.831661896 | 0.440341872 | 0.721042072 | -4.65696287 |
| **A_55_P2069989** | Klhl12 | -0.268404749 | -1.076828631 | 0.540019132 | 11.17505848 | -0.831574967 | 0.440386969 | 0.721042072 | -4.656982552 |
| **A_65_P03022** | Cdc14b | -0.090261445 | -0.362130254 | 0.181607364 | 8.918524088 | -0.831559335 | 0.440395079 | 0.721042072 | -4.656986092 |
| **A_52_P286141** | Cisd2 | 0.106827763 | -0.214952023 | 0.428607549 | 9.837909517 | 0.831525785 | 0.440412485 | 0.721042072 | -4.656993688 |

| | | | | | | | | | |
|---|---|---|---|---|---|---|---|---|---|
| **A_55_P2134004** | Gstm2 | 0.621161482 | -1.249940339 | 2.492263303 | 11.97646368 | 0.831490787 | 0.440430643 | 0.721042072 | -4.657001611 |
| **A_51_P234025** | Rbm5 | 0.167037188 | -0.336165477 | 0.670239853 | 12.94327431 | 0.831420223 | 0.440467254 | 0.721042072 | -4.657017587 |
| **A_51_P413545** | Txndc11 | -0.249177739 | -0.999845989 | 0.50149051 | 11.35241193 | -0.831402909 | 0.440476238 | 0.721042072 | -4.657021506 |
| **A_51_P475580** | Ugp2 | 0.1197267 | -0.240979582 | 0.480432982 | 10.27776983 | 0.831357182 | 0.440499965 | 0.721042072 | -4.657031858 |
| **A_52_P520408** | Serp1 | -0.237135858 | -0.951583812 | 0.477312096 | 10.68269004 | -0.831336748 | 0.440510568 | 0.721042072 | -4.657036484 |
| **A_55_P2110978** | Atp6ap1 | -0.17893266 | -0.71815008 | 0.360284761 | 8.522560278 | -0.831143568 | 0.440610818 | 0.721042072 | -4.657080212 |
| **A_51_P101460** | Dsp | -0.440664392 | -1.768637246 | 0.887308461 | 10.73661741 | -0.831130285 | 0.440617711 | 0.721042072 | -4.648291701 |
| **A_51_P118763** | Ahctf1 | -0.208241157 | -0.83579551 | 0.419313195 | 9.781489491 | -0.831123512 | 0.440621227 | 0.721042072 | -4.642934637 |
| **A_55_P2009266** | Gm16060 | 0.409351297 | -0.824353853 | 1.643056446 | 15.05862057 | 0.831065279 | 0.44065145 | 0.721042072 | -4.657097932 |
| **A_55_P2032727** | NA | 0.253661149 | -0.510885065 | 1.018207362 | 16.05783332 | 0.83099911 | 0.440685795 | 0.721042072 | -4.657112908 |
| **A_55_P2107750** | Hectd1 | -0.076699179 | -0.307875761 | 0.154477403 | 8.794772784 | -0.830992724 | 0.440689109 | 0.721042072 | -4.657114353 |
| **A_55_P2009895** | Gm5113 | -0.071899625 | -0.288633141 | 0.144833891 | 7.990230546 | -0.830904089 | 0.440735117 | 0.721042072 | -4.657134412 |
| **A_55_P2037186** | Alkbh2 | -0.316845846 | -1.271954549 | 0.638262857 | 9.486479848 | -0.830893881 | 0.440740416 | 0.721042072 | -4.657136722 |
| **A_51_P519108** | Dzank1 | 0.195288462 | -0.393539226 | 0.78411615 | 8.724304945 | 0.830689475 | 0.440846533 | 0.721144585 | -4.657182976 |
| **A_51_P263246** | Dusp8 | -0.392479466 | -1.576208973 | 0.791250041 | 8.528279699 | -0.830452529 | 0.440969567 | 0.721144585 | -4.657236585 |
| **A_51_P477364** | Rhob | -0.180231135 | -0.723827865 | 0.363365595 | 8.298770473 | -0.830430568 | 0.440980971 | 0.721144585 | -4.657241553 |
| **A_55_P2040763** | Sec62 | -0.240448418 | -0.965706916 | 0.48481008 | 12.45381225 | -0.8303849 | 0.441004687 | 0.721144585 | -4.657251884 |
| **A_51_P342271** | Setd6 | -0.24063625 | -0.966510372 | 0.485237872 | 9.664458587 | -0.830328764 | 0.441033841 | 0.721144585 | -4.657264582 |
| **A_55_P2105180** | Bhmt | 0.085332844 | -0.172090136 | 0.342755825 | 7.801288784 | 0.830269822 | 0.441064454 | 0.721144585 | -4.657277915 |
| **A_52_P582907** | Entpd4 | -0.122917939 | -0.493731255 | 0.247895378 | 8.124548375 | -0.83025269 | 0.441073352 | 0.721144585 | -4.65728179 |
| **A_51_P283876** | Bphl | 0.620542429 | -1.251645862 | 2.492730719 | 11.65424316 | 0.830180068 | 0.441111073 | 0.721144585 | -4.657298217 |
| **A_55_P2319468** | NA | -0.07629813 | -0.306506378 | 0.153910118 | 7.830723274 | -0.830124746 | 0.441139809 | 0.721144585 | -4.657310729 |
| **A_51_P196207** | Capsl | 0.493773894 | -0.996175891 | 1.983723678 | 9.33164305 | 0.83005494 | 0.441176071 | 0.721144585 | -4.657326517 |
| **A_52_P463143** | Cdc37 | 0.105014897 | -0.211900868 | 0.421930663 | 14.90547499 | 0.829960481 | 0.441225143 | 0.721155781 | -4.65734788 |
| **A_52_P203981** | Srek1ip1 | 0.129688535 | -0.261717149 | 0.521094219 | 10.1211598 | 0.829898161 | 0.44125752 | 0.721155781 | -4.657361973 |
| **A_55_P1979743** | NA | 0.224279832 | -0.452776479 | 0.901336144 | 14.35928571 | 0.82968985 | 0.441365758 | 0.721271707 | -4.657409076 |
| **A_51_P318618** | Anks4b | -0.088729634 | -0.356634235 | 0.179174966 | 7.712621615 | -0.829542929 | 0.44144211 | 0.721294095 | -4.657442293 |
| **A_55_P2029310** | Arf4 | -0.245725577 | -0.987674303 | 0.496223149 | 11.26395066 | -0.829519913 | 0.441454072 | 0.721294095 | -4.657447497 |
| **A_55_P2062480** | NA | 0.187717635 | -0.379242436 | 0.754677705 | 14.52047095 | 0.829283349 | 0.441577032 | 0.721408077 | -4.657500973 |
| **A_66_P105074** | Enpp5 | 0.17195007 | -0.347465423 | 0.691365563 | 9.155061642 | 0.82915897 | 0.441641692 | 0.721408077 | -4.657529085 |
| **A_66_P106661** | Slc7a1 | -1.142030268 | -4.592189255 | 2.308128718 | 11.88103773 | -0.829065517 | 0.441690279 | 0.721408077 | -4.657550206 |
| **A_55_P2354531** | 2610029K | 0.291408353 | -0.588986947 | 1.171803654 | 8.891389833 | 0.829038361 | 0.441704398 | 0.721408077 | -4.657556343 |
| **A_52_P89683** | NA | 0.244094887 | -0.493389836 | 0.981579611 | 10.33639106 | 0.829002804 | 0.441722886 | 0.721408077 | -4.657564379 |
| **A_55_P2056220** | NA | 0.212085038 | -0.428725121 | 0.852895196 | 11.31352719 | 0.828955063 | 0.44174771 | 0.721408077 | -4.657575167 |
| **A_55_P2032695** | Tut4 | -0.253334531 | -1.019007517 | 0.512338454 | 11.58983295 | -0.828707773 | 0.441876309 | 0.721557142 | -4.657631045 |
| **A_55_P1988882** | 9-Sep | -0.347954398 | -1.399784788 | 0.703875992 | 11.33797694 | -0.828565719 | 0.441950195 | 0.721616845 | -4.657663138 |
| **A_52_P500002** | Tspyl5 | 0.111781129 | -0.226168881 | 0.449731139 | 7.975993428 | 0.828450135 | 0.44201032 | 0.721654072 | -4.657689249 |
| **A_55_P1993348** | NA | 0.117390573 | -0.237589391 | 0.472370537 | 15.75966176 | 0.828284868 | 0.442096299 | 0.721685308 | -4.65772658 |
| **A_55_P2160387** | Cxcl11 | -0.166040498 | -0.668175786 | 0.336094791 | 16.58515251 | -0.828216023 | 0.442132119 | 0.721685308 | -4.657742129 |
| **A_52_P267824** | Ces2g | 0.086031458 | -0.174169809 | 0.346232724 | 7.901712755 | 0.828129434 | 0.442177174 | 0.721685308 | -4.657761686 |
| **A_55_P1966444** | NA | -0.150140537 | -0.604270017 | 0.303988943 | 15.30936982 | -0.828072983 | 0.442206549 | 0.721685308 | -4.657774434 |
| **A_55_P2031112** | Dhps | -0.18732683 | -0.753955984 | 0.379302325 | 10.4952736 | -0.828040182 | 0.442223618 | 0.721685308 | -4.657781842 |
| **A_55_P2131153** | Smarcc1 | -0.68387218 | -2.752678682 | 1.384934322 | 12.07861276 | -0.827952477 | 0.442269261 | 0.721685308 | -4.657801647 |
| **A_55_P1959869** | Ppp1r2 | 0.267390911 | -0.541543346 | 1.076325168 | 11.45865589 | 0.827911202 | 0.442290742 | 0.721685308 | -4.657810967 |
| **A_55_P2131193** | NA | 0.268433783 | -0.544188882 | 1.081056448 | 15.62099172 | 0.827367745 | 0.442573653 | 0.722085992 | -4.657933658 |
| **A_52_P1042732** | Akap5 | 1.523472378 | -3.088910606 | 6.135855362 | 12.77598883 | 0.827293417 | 0.442612356 | 0.722088203 | -4.657950434 |
| **A_55_P2077510** | Zfp583 | 0.103386856 | -0.20971052 | 0.416484233 | 8.798747007 | 0.827058531 | 0.442734681 | 0.722187072 | -4.658003444 |
| **A_51_P428505** | Krtcap2 | -0.151735204 | -0.611300786 | 0.307830379 | 14.11348065 | -0.826968945 | 0.442781343 | 0.722187072 | -4.658023659 |

| | | | | | | | | | |
|---|---|---|---|---|---|---|---|---|---|
| **A_51_P143232** | Wdr59 | -0.147039712 | -0.592482783 | 0.298403359 | 8.492332673 | -0.826785365 | 0.442876973 | 0.722187072 | -4.658065081 |
| **A_65_P05316** | Zfp182 | -0.074652833 | -0.300808549 | 0.151502883 | 8.853572164 | -0.826778304 | 0.442880652 | 0.722187072 | -4.658066674 |
| **A_55_P2133427** | Vkorc1 | -0.109411863 | -0.440972843 | 0.222149118 | 11.24210951 | -0.826516145 | 0.443017243 | 0.722187072 | -4.658125815 |
| **A_51_P386870** | Sprr2f | -0.123350148 | -0.497173291 | 0.250472994 | 7.719440155 | -0.826463804 | 0.443044518 | 0.722187072 | -4.658137622 |
| **A_55_P2028362** | NA | 0.177109779 | -0.359686565 | 0.713906124 | 8.628294918 | 0.826386736 | 0.44308468 | 0.722187072 | -4.658155005 |
| **A_66_P140886** | Inafm1 | -0.105393555 | -0.424834587 | 0.214047478 | 11.26371594 | -0.82636839 | 0.443094241 | 0.722187072 | -4.658159143 |
| **A_51_P181722** | Rbks | -0.149782078 | -0.603769132 | 0.304204976 | 9.226164095 | -0.826355132 | 0.44310115 | 0.722187072 | -4.658162133 |
| **A_52_P467389** | Slc15a3 | -0.076214954 | -0.307235688 | 0.15480578 | 7.986874298 | -0.826303477 | 0.443128071 | 0.722187072 | -4.658173783 |
| **A_55_P2024337** | Tmem53 | 0.242941441 | -0.493468539 | 0.979351422 | 10.18387721 | 0.826289593 | 0.443135307 | 0.722187072 | -4.658176914 |
| **A_55_P2139673** | Rpl11 | 0.114089551 | -0.231760664 | 0.459939766 | 15.61731864 | 0.826243711 | 0.443159221 | 0.722187072 | -4.658187262 |
| **A_55_P2011727** | Neurl1b | 0.12535051 | -0.254654547 | 0.505355566 | 13.7179055 | 0.826203584 | 0.443180136 | 0.722187072 | -4.658196312 |
| **A_55_P2058157** | Ppp4r3b | -0.108755165 | -0.438501969 | 0.220991638 | 9.578953109 | -0.826075315 | 0.443246997 | 0.722187072 | -4.658225237 |
| **A_55_P1993512** | NA | -0.323752677 | -1.305378714 | 0.65787336 | 8.88702532 | -0.826071509 | 0.44324898 | 0.722187072 | -4.658226096 |
| **A_52_P504421** | Rnf138 | -0.134631157 | -0.542868313 | 0.273605998 | 9.870857178 | -0.826006368 | 0.443282939 | 0.722187072 | -4.658240784 |
| **A_55_P2124619** | Smurf2 | -0.124083306 | -0.500358463 | 0.252191851 | 9.997508001 | -0.825958365 | 0.443307964 | 0.722187072 | -4.658251608 |
| **A_52_P335508** | Tiprl | 0.075806756 | -0.154098468 | 0.305711979 | 9.416398003 | 0.825865677 | 0.443356287 | 0.722204942 | -4.658272507 |
| **A_55_P1980551** | Dennd1a | -0.334413155 | -1.34891918 | 0.68009287 | 9.148861168 | -0.82561784 | 0.443485518 | 0.722354592 | -4.658328379 |
| **A_55_P2015700** | NA | -0.132800618 | -0.535718469 | 0.270117234 | 10.22432577 | -0.825532035 | 0.443530266 | 0.722366622 | -4.658347721 |
| **A_55_P2050298** | NA | 0.33232967 | -0.676073136 | 1.340732477 | 15.42535073 | 0.825439822 | 0.44357836 | 0.722384098 | -4.658368506 |
| **A_52_P560628** | 1300002E | 0.08791458 | -0.178920762 | 0.354749923 | 8.741907041 | 0.825216495 | 0.443694852 | 0.722418744 | -4.658418837 |
| **A_55_P2178553** | Apoa2 | -1.07222532 | -4.326992816 | 2.182542176 | 11.25044791 | -0.825118708 | 0.443745867 | 0.722418744 | -4.658440873 |
| **A_55_P2136657** | Thbs3 | 0.411003636 | -0.836759481 | 1.658766753 | 10.4512485 | 0.825018839 | 0.443797973 | 0.722418744 | -4.658463376 |
| **A_55_P2176325** | Prex1 | -0.505725701 | -2.041062136 | 1.029610735 | 9.309232161 | -0.825014942 | 0.443800006 | 0.722418744 | -4.658464255 |
| **A_55_P1996087** | Gm14326 | -0.092368802 | -0.3728135 | 0.188075897 | 11.37189741 | -0.824951508 | 0.443833104 | 0.722418744 | -4.658478547 |
| **A_51_P337944** | Bmp2k | -0.12976083 | -0.523756227 | 0.264234568 | 10.27116876 | -0.824902863 | 0.443858487 | 0.722418744 | -4.658489507 |
| **A_55_P1962359** | Trim11 | 0.166335848 | -0.33891601 | 0.671587705 | 11.44098159 | 0.824571429 | 0.444031459 | 0.722418744 | -4.65856417 |
| **A_55_P2181976** | Akap11 | 0.083358264 | -0.169860097 | 0.336576624 | 8.963288606 | 0.824524971 | 0.444055709 | 0.722418744 | -4.658574634 |
| **A_51_P464911** | Msrb2 | 0.499741482 | -1.018450801 | 2.017933766 | 10.85978474 | 0.824458817 | 0.444090242 | 0.722418744 | -4.658589534 |
| **A_65_P16952** | Ywhaq | 0.395827536 | -0.806747972 | 1.598403043 | 12.60591911 | 0.82441141 | 0.444114989 | 0.722418744 | -4.658600211 |
| **A_51_P457492** | Tmem223 | 0.361391615 | -0.736594095 | 1.459377325 | 13.32530759 | 0.82438813 | 0.444127142 | 0.722418744 | -4.658605454 |
| **A_66_P133112** | Gm14635 | 0.092147408 | -0.187841079 | 0.372135896 | 8.167952551 | 0.824315177 | 0.444165228 | 0.722418744 | -4.658621884 |
| **A_55_P1989772** | Sqor | -0.157245919 | -0.635045381 | 0.320553543 | 10.26883544 | -0.824297669 | 0.444174369 | 0.722418744 | -4.658625827 |
| **A_55_P2095108** | A430033K | 0.161939245 | -0.330182646 | 0.654061136 | 8.980921867 | 0.824194612 | 0.444228176 | 0.722418744 | -4.658649034 |
| **A_55_P2032363** | Tomm40l | 0.151831794 | -0.309580085 | 0.613243674 | 8.650238336 | 0.824184227 | 0.444233598 | 0.722418744 | -4.658651373 |
| **A_51_P370286** | Med24 | 0.113587507 | -0.231610959 | 0.458785972 | 10.83916349 | 0.824160993 | 0.44424573 | 0.722418744 | -4.658656604 |
| **A_52_P438036** | Lpar4 | 0.368237709 | -0.750902551 | 1.487377969 | 9.775435952 | 0.824126894 | 0.444263535 | 0.722418744 | -4.658664283 |
| **A_55_P2058657** | NA | 0.125883727 | -0.256707144 | 0.508474598 | 16.552906 | 0.824110285 | 0.444272208 | 0.722418744 | -4.658668022 |
| **A_55_P2274592** | Wnt7b | -0.303396729 | -1.226030662 | 0.619237204 | 8.324012518 | -0.823629355 | 0.444523389 | 0.72273553 | -4.658776293 |
| **A_55_P2382741** | NA | -0.06850095 | -0.276822257 | 0.139820358 | 7.699409252 | -0.823594135 | 0.444541788 | 0.72273553 | -4.65878422 |
| **A_55_P2083184** | Gm10272 | -0.075652266 | -0.30579982 | 0.154495288 | 8.021905786 | -0.823314803 | 0.44468773 | 0.722912014 | -4.658847086 |
| **A_55_P2117903** | Stard5 | -0.184281356 | -0.744970021 | 0.37640731 | 8.594087345 | -0.82320873 | 0.44474316 | 0.72291268 | -4.658870955 |
| **A_66_P130241** | Slc25a1 | 0.074874401 | -0.152964804 | 0.302713606 | 9.824167257 | 0.82310502 | 0.444797359 | 0.72291268 | -4.65889429 |
| **A_55_P1979356** | Morf4l1 | 0.171891633 | -0.351237906 | 0.695021171 | 14.14619936 | 0.822992427 | 0.444856206 | 0.72291268 | -4.658919622 |
| **A_52_P49250** | Erbb2 | 0.351924838 | -0.719151059 | 1.423000735 | 9.419191379 | 0.822962449 | 0.444871875 | 0.72291268 | -4.65541211 |
| **A_51_P234140** | Cdkl1 | -0.09561296 | -0.399821957 | 0.208596037 | 8.068825121 | -0.837157968 | 0.444875094 | 0.72291268 | -4.641844739 |
| **A_51_P453909** | Cyp2f2 | 1.455305449 | -2.974457076 | 5.885067975 | 12.43073008 | 0.822856409 | 0.444927304 | 0.722936759 | -4.658950222 |
| **A_55_P2058903** | Sgpp1 | -0.351779726 | -1.422826748 | 0.719267296 | 12.11125532 | -0.822645287 | 0.445037676 | 0.723052282 | -4.658997711 |
| **A_51_P365656** | Tmem101 | 0.108806249 | -0.222498331 | 0.44011083 | 9.648577789 | 0.822577348 | 0.445073198 | 0.723052282 | -4.659012992 |

| | | | | | | | | | |
|---|---|---|---|---|---|---|---|---|---|
| **A_51_P292073** | Haghl | 0.331414604 | -0.678225637 | 1.341054845 | 11.04296908 | 0.822158093 | 0.445292453 | 0.723308821 | -4.659107271 |
| **A_55_P2068570** | NA | 0.082351884 | -0.168537428 | 0.333241195 | 7.982762946 | 0.822132322 | 0.445305932 | 0.723308821 | -4.659113065 |
| **A_66_P131604** | Otud4 | -0.252716185 | -1.022753158 | 0.517320789 | 12.13636648 | -0.822000011 | 0.445375144 | 0.72336047 | -4.659142812 |
| **A_55_P1979508** | Clic3 | 1.219006557 | -2.496171691 | 4.934184806 | 12.04537133 | 0.821820049 | 0.445469295 | 0.723452612 | -4.659183267 |
| **A_51_P417839** | Hdac4 | -0.368184902 | -1.490610648 | 0.754240844 | 9.066128683 | -0.821596729 | 0.445586149 | 0.723527087 | -4.659233461 |
| **A_51_P435068** | Acadsb | 0.385453307 | -0.789643643 | 1.560550257 | 10.49836122 | 0.821577283 | 0.445596325 | 0.723527087 | -4.659237831 |
| **A_51_P334104** | Dcn | -0.701918099 | -2.842030109 | 1.43819391 | 14.33475967 | -0.82148622 | 0.445643982 | 0.723527087 | -4.659258296 |
| **A_51_P149455** | Acadl | 0.340330988 | -0.697370954 | 1.378032929 | 11.23204598 | 0.821446362 | 0.445664843 | 0.723527087 | -4.659267253 |
| **A_52_P686136** | Prrc2b | 0.205422742 | -0.421260248 | 0.832105731 | 12.84408362 | 0.821014751 | 0.445890781 | 0.723786707 | -4.659364228 |
| **A_66_P120603** | Trps1 | 0.384306827 | -0.788121834 | 1.556735489 | 9.244005277 | 0.820997848 | 0.445899631 | 0.723786707 | -4.659368025 |
| **A_55_P2043892** | Srd5a3 | -0.115387546 | -0.467596601 | 0.236821508 | 8.923249711 | -0.820557024 | 0.446130483 | 0.724100634 | -4.659467036 |
| **A_55_P2013371** | Ttc29 | 0.410438714 | -0.84272905 | 1.663606479 | 9.106625749 | 0.820331615 | 0.446248559 | 0.724182777 | -4.659517651 |
| **A_55_P2022049** | Klf15 | 0.432056501 | -0.887179957 | 1.751292959 | 9.275498358 | 0.820291563 | 0.446269542 | 0.724182777 | -4.659526644 |
| **A_51_P100828** | Prdx3 | 0.165854073 | -0.340591074 | 0.672299219 | 10.90834733 | 0.820245908 | 0.446293462 | 0.724182777 | -4.659536894 |
| **A_51_P380432** | Scx | 0.887020652 | -1.821944709 | 3.595986012 | 10.99112638 | 0.820126085 | 0.446356243 | 0.724221472 | -4.659563795 |
| **A_55_P1990217** | Zfp13 | 0.177999978 | -0.390366761 | 0.746366718 | 8.426305365 | 0.83416891 | 0.446392225 | 0.724221472 | -4.642411417 |
| **A_55_P1977649** | Hoxd8 | -0.196318749 | -0.796298308 | 0.403660809 | 7.900837866 | -0.819550406 | 0.44665796 | 0.724555718 | -4.659693003 |
| **A_51_P288558** | Cfap52 | 0.568453165 | -1.168886628 | 2.305792957 | 9.574534318 | 0.81952134 | 0.446673197 | 0.724555718 | -4.659699526 |
| **A_51_P373901** | Rsph1 | 0.206411873 | -0.424533317 | 0.837357062 | 8.352672358 | 0.819395145 | 0.446739359 | 0.724571589 | -4.659727841 |
| **A_51_P510106** | B3gntl1 | -0.142535494 | -0.578247996 | 0.293177009 | 8.18688582 | -0.819357339 | 0.446759181 | 0.724571589 | -4.659736324 |
| **A_55_P1962926** | Snapc3 | 0.106018737 | -0.218094113 | 0.430131588 | 8.379270367 | 0.819288243 | 0.446795411 | 0.724571589 | -4.659751826 |
| **A_51_P319379** | Tmx4 | -0.122532692 | -0.497183059 | 0.252117675 | 9.247705236 | -0.819173995 | 0.446855321 | 0.724578244 | -4.659777457 |
| **A_55_P2043269** | Cdc42se1 | -0.191725694 | -0.777963558 | 0.39451217 | 9.047026753 | -0.819137481 | 0.446874469 | 0.724578244 | -4.659785648 |
| **A_52_P489119** | Nol6 | -0.31203551 | -1.266390081 | 0.642319062 | 9.149298519 | -0.818925901 | 0.446985437 | 0.72466692 | -4.659833107 |
| **A_51_P417747** | Kap | -0.127459338 | -0.517308215 | 0.262389539 | 7.788712781 | -0.818890279 | 0.447004121 | 0.72466692 | -4.659841097 |
| **A_52_P222096** | 4931428F | 0.231013767 | -0.475723004 | 0.937750538 | 8.854363693 | 0.818710801 | 0.447098271 | 0.724758781 | -4.659881348 |
| **A_55_P2014282** | NA | 0.249759821 | -0.514405193 | 1.013924836 | 17.42329595 | 0.818626445 | 0.447142528 | 0.724769755 | -4.659900265 |
| **A_55_P1983903** | Thap4 | -0.134509383 | -0.546123872 | 0.277105106 | 8.139592371 | -0.818487914 | 0.447215213 | 0.724826803 | -4.659931327 |
| **A_55_P2070054** | Senp3 | -0.200211351 | -0.813113629 | 0.412690927 | 11.66812201 | -0.818178017 | 0.447377843 | 0.724994575 | -4.660000803 |
| **A_55_P1986123** | Sccpdh | 0.206347251 | -0.425399828 | 0.83809433 | 11.66544542 | 0.818098867 | 0.447419388 | 0.724994575 | -4.660018545 |
| **A_51_P384718** | Efna4 | 0.178047685 | -0.367079791 | 0.723175162 | 8.578306048 | 0.818066494 | 0.44743638 | 0.724994575 | -4.660025801 |
| **A_51_P279183** | Tkfc | -0.194321435 | -0.78931896 | 0.40067609 | 8.878676909 | -0.818004882 | 0.447468722 | 0.724994575 | -4.660039611 |
| **A_55_P2143517** | Fstl1 | 0.465591629 | -0.960640989 | 1.891824248 | 15.43285342 | 0.817645676 | 0.447657311 | 0.725213731 | -4.66012011 |
| **A_55_P2102715** | NA | 0.258139667 | -0.532651648 | 1.048930982 | 16.09851462 | 0.817604364 | 0.447679005 | 0.725213731 | -4.660129367 |
| **A_55_P2162537** | Acrbp | 0.095209432 | -0.196489425 | 0.386908289 | 8.296109119 | 0.817514739 | 0.44772607 | 0.725229209 | -4.660149448 |
| **A_55_P2139396** | Unc45b | 0.202794593 | -0.418587867 | 0.824177054 | 8.373152176 | 0.817424645 | 0.447773385 | 0.725245089 | -4.660169633 |
| **A_65_P12977** | Dnaja2 | 0.09683694 | -0.19991432 | 0.393588199 | 11.84774173 | 0.817332617 | 0.44782172 | 0.725262618 | -4.66019025 |
| **A_55_P2245422** | NA | -0.102020613 | -0.414744303 | 0.210703077 | 8.927276721 | -0.817104291 | 0.447941658 | 0.725308478 | -4.660241395 |
| **A_55_P2114269** | Fam177a2 | -0.175183604 | -0.712197286 | 0.361830077 | 16.3337544 | -0.817068472 | 0.447960476 | 0.725308478 | -4.660249418 |
| **A_66_P134474** | Ang3 | -0.110907634 | -0.45091735 | 0.229102083 | 8.386958949 | -0.816996992 | 0.44799803 | 0.725308478 | -4.660265427 |
| **A_66_P125828** | NA | 0.449025231 | -0.927821426 | 1.825871887 | 10.65638602 | 0.816837215 | 0.448081981 | 0.725308478 | -4.660301209 |
| **A_55_P2419483** | 4732460I0 | -0.068965904 | -0.280457194 | 0.142525386 | 8.301160553 | -0.816755924 | 0.448124698 | 0.725308478 | -4.660319412 |
| **A_51_P416858** | Myl1 | 1.129446582 | -2.334160892 | 4.593054057 | 11.37870666 | 0.816746671 | 0.448129561 | 0.725308478 | -4.660321484 |
| **A_55_P1974134** | Mrpl58 | 0.376728132 | -0.778664772 | 1.532121036 | 12.94186607 | 0.816673949 | 0.448167778 | 0.725308478 | -4.660337767 |
| **A_51_P362881** | Fbxo38 | -0.116772337 | -0.474927424 | 0.24138275 | 11.56548269 | -0.816618461 | 0.448196939 | 0.725308478 | -4.660350191 |
| **A_55_P1991851** | Speg | -0.123504744 | -0.502319683 | 0.255310195 | 8.75680261 | -0.816595321 | 0.448209101 | 0.725308478 | -4.660355372 |
| **A_65_P05040** | Ston2 | -0.210879401 | -0.857768956 | 0.436010154 | 8.610371936 | -0.816496572 | 0.448261003 | 0.725308478 | -4.66037748 |
| **A_51_P269203** | Trpv4 | -0.085047722 | -0.345939491 | 0.175844048 | 8.28649634 | -0.816493347 | 0.448262698 | 0.725308478 | -4.660378202 |

| | | | | | | | | | |
|---|---|---|---|---|---|---|---|---|---|
| **A_66_P128055** | Spindoc | -0.172767729 | -0.70284927 | 0.357313812 | 8.650144153 | -0.816338519 | 0.448344085 | 0.725379459 | -4.660412863 |
| **A_55_P2015217** | Alg11 | -0.186578474 | -0.759219385 | 0.386062438 | 9.048930371 | -0.81607394 | 0.448483187 | 0.725474094 | -4.660472083 |
| **A_55_P2099141** | NA | 0.323256824 | -0.668999958 | 1.315513606 | 15.47004603 | 0.815969586 | 0.44853806 | 0.725474094 | -4.660495437 |
| **A_51_P245525** | NA | -0.107432765 | -0.437204777 | 0.222339247 | 15.64049679 | -0.815968336 | 0.448538717 | 0.725474094 | -4.660495717 |
| **A_52_P379337** | Rtn4 | -0.082497574 | -0.335737596 | 0.170742447 | 10.20137579 | -0.815941803 | 0.44855267 | 0.725474094 | -4.660501654 |
| **A_55_P2175356** | Uros | -0.078722039 | -0.320443144 | 0.162999066 | 7.820807651 | -0.815703088 | 0.448678216 | 0.725616447 | -4.660555069 |
| **A_52_P684130** | 2510002D | -0.079139988 | -0.322193137 | 0.163913162 | 8.580087696 | -0.815539631 | 0.448764197 | 0.725694796 | -4.660591639 |
| **A_51_P440568** | Stk19 | -0.102956463 | -0.419208046 | 0.21329512 | 9.128546233 | -0.815401023 | 0.448837116 | 0.725735254 | -4.660622646 |
| **A_51_P298843** | Pcbd2 | 0.225063988 | -0.466323781 | 0.916451756 | 11.0888839 | 0.815332328 | 0.448873258 | 0.725735254 | -4.660638012 |
| **A_51_P306527** | Ywhah | 0.144804623 | -0.300105623 | 0.589714869 | 14.40787905 | 0.815192857 | 0.448946645 | 0.725735254 | -4.660669207 |
| **A_52_P331762** | Lmo1 | 0.49270086 | -1.021294836 | 2.006696555 | 9.419312853 | 0.815096495 | 0.448997354 | 0.725735254 | -4.660690758 |
| **A_55_P1967855** | Kctd20 | -0.29441815 | -1.199132584 | 0.610296284 | 10.48111645 | -0.815085949 | 0.449002904 | 0.725735254 | -4.660693117 |
| **A_55_P2094613** | NA | 0.177409692 | -0.367765339 | 0.722584723 | 8.650933597 | 0.815064035 | 0.449014436 | 0.725735254 | -4.660698017 |
| **A_55_P2118684** | Idh2 | 0.2264542 | -0.469517873 | 0.922426272 | 15.44116236 | 0.814964914 | 0.449066602 | 0.725758897 | -4.660720183 |
| **A_65_P10195** | Myl7 | 1.276860105 | -2.647738464 | 5.201458673 | 12.12688677 | 0.814888938 | 0.449106591 | 0.725762858 | -4.660737172 |
| **A_55_P2176724** | Fbxw7 | -0.377282325 | -1.537072344 | 0.782507694 | 9.208371906 | -0.814774519 | 0.449166818 | 0.72579952 | -4.660762755 |
| **A_66_P117578** | Mtmr7 | -0.482578128 | -1.966765793 | 1.001609536 | 8.770197739 | -0.814383869 | 0.44937249 | 0.725947382 | -4.660850085 |
| **A_52_P436238** | Odc1 | -0.43174824 | -1.759679601 | 0.896183122 | 9.174747721 | -0.814339111 | 0.449396059 | 0.725947382 | -4.660860089 |
| **A_55_P1978216** | Cd300c2 | -0.185454056 | -0.755857982 | 0.38494987 | 8.900392594 | -0.814337014 | 0.449397163 | 0.725947382 | -4.660860557 |
| **A_52_P220783** | Tmem229 | -0.290303626 | -1.183295545 | 0.602688292 | 10.79661966 | -0.814245332 | 0.449445444 | 0.725947382 | -4.660881048 |
| **A_52_P194680** | NA | -0.073623032 | -0.300106237 | 0.152860173 | 9.265054406 | -0.814194273 | 0.449472334 | 0.725947382 | -4.660892459 |
| **A_55_P2431862** | Srpk2 | 0.093020808 | -0.193151252 | 0.379192869 | 10.38457284 | 0.814147746 | 0.449496839 | 0.725947382 | -4.660902857 |
| **A_51_P104392** | Rpp25 | -0.076240871 | -0.310803921 | 0.158322179 | 7.879307675 | -0.814101574 | 0.449521157 | 0.725947382 | -4.660913175 |
| **A_52_P522372** | Aard | 0.131265083 | -0.272741393 | 0.535271559 | 8.543430963 | 0.813787916 | 0.449686384 | 0.72595364 | -4.660983259 |
| **A_55_P2014229** | Sema4a | -0.097286358 | -0.396756643 | 0.202183927 | 9.849457302 | -0.813670468 | 0.449748264 | 0.72595364 | -4.661009497 |
| **A_55_P2240468** | 1110019D | -0.069988005 | -0.285429797 | 0.145453788 | 8.257810509 | -0.813661945 | 0.449752754 | 0.72595364 | -4.661011401 |
| **A_52_P528463** | Mfsd3 | -0.176414803 | -0.719559352 | 0.366729746 | 9.155242389 | -0.813523202 | 0.449825863 | 0.72595364 | -4.661042394 |
| **A_52_P555680** | Trappc12 | 0.070968545 | -0.147529881 | 0.28946697 | 11.0796773 | 0.813519437 | 0.449827847 | 0.72595364 | -4.661043235 |
| **A_51_P195240** | Gpatch11 | -0.086733563 | -0.353801899 | 0.180334773 | 10.59070863 | -0.813420532 | 0.449879969 | 0.72595364 | -4.661065326 |
| **A_52_P508700** | Zfp951 | -0.135022867 | -0.550783248 | 0.280737515 | 8.216878861 | -0.813419478 | 0.449880524 | 0.72595364 | -4.661065561 |
| **A_55_P2085641** | Lmbr1l | -0.259779205 | -1.059689266 | 0.540130856 | 10.90031236 | -0.813417609 | 0.449881509 | 0.72595364 | -4.661065979 |
| **A_55_P2016249** | Ppid | 0.429670057 | -0.893528875 | 1.752868989 | 9.870461614 | 0.81331783 | 0.449934096 | 0.72595364 | -4.661088263 |
| **A_55_P2021049** | Ugp2 | -0.137868832 | -0.581804135 | 0.306066472 | 12.12941317 | -0.827197318 | 0.449946323 | 0.72595364 | -4.643729009 |
| **A_55_P2079746** | Pls3 | 0.33418707 | -0.695165092 | 1.363539231 | 13.05038827 | 0.81316 | 0.450017287 | 0.72595364 | -4.66112351 |
| **A_55_P2027788** | Baiap2 | -0.197598008 | -0.806268345 | 0.411072329 | 8.680835651 | -0.813112748 | 0.450042195 | 0.72595364 | -4.661134061 |
| **A_55_P2104289** | NA | 0.188272531 | -0.391731778 | 0.76827684 | 8.789756405 | 0.813029093 | 0.450086296 | 0.72595364 | -4.661152741 |
| **A_51_P464630** | Rad17 | -0.112799835 | -0.460299696 | 0.234700027 | 10.36919805 | -0.813025511 | 0.450088184 | 0.72595364 | -4.661153541 |
| **A_55_P2108845** | NA | -0.143991853 | -0.587591752 | 0.299608046 | 7.925359659 | -0.813011761 | 0.450095433 | 0.72595364 | -4.661156611 |
| **A_55_P2016681** | Alkbh7 | 0.2732258 | -0.568568526 | 1.115020126 | 10.44058687 | 0.812954159 | 0.450125801 | 0.72595364 | -4.661169472 |
| **A_55_P2175752** | Mfap1b | -0.319610647 | -1.304463722 | 0.665242428 | 9.200501672 | -0.812830776 | 0.450190855 | 0.72597988 | -4.661197018 |
| **A_51_P157214** | Mfsd14a | 0.119081496 | -0.247910954 | 0.486073946 | 10.6245568 | 0.812713545 | 0.450252671 | 0.72597988 | -4.661223189 |
| **A_52_P18816** | Gosr2 | -0.286484833 | -1.169590617 | 0.59662095 | 8.8023541 | -0.812529713 | 0.450349619 | 0.72597988 | -4.661264223 |
| **A_66_P128997** | Pml | 0.089805114 | -0.187064746 | 0.366674973 | 8.688894295 | 0.812410881 | 0.450412296 | 0.72597988 | -4.661290744 |
| **A_51_P228254** | Zdhhc21 | -0.098076736 | -0.400486949 | 0.204333477 | 11.15662096 | -0.812306437 | 0.450467389 | 0.72597988 | -4.661314053 |
| **A_52_P276792** | Foxp1 | 0.257833351 | -0.537206655 | 1.052873358 | 10.50542092 | 0.81227008 | 0.450486568 | 0.72597988 | -4.661322166 |
| **A_66_P104980** | Tceal5 | 0.21789562 | -0.454010854 | 0.889802095 | 9.57552333 | 0.812250562 | 0.450496865 | 0.72597988 | -4.661326522 |
| **A_66_P109720** | Jpt2 | -0.151718392 | -0.619575075 | 0.316138291 | 8.198593065 | -0.812223851 | 0.450510956 | 0.72597988 | -4.661332482 |
| **A_52_P350616** | Mrm1 | -0.289712499 | -1.183212975 | 0.603787976 | 9.940679505 | -0.812124833 | 0.450563195 | 0.72597988 | -4.661354577 |

| | | | | | | | | | |
|---|---|---|---|---|---|---|---|---|---|
| **A_55_P2053616** | NA | 0.112483466 | -0.23442679 | 0.459393721 | 9.419786357 | 0.812123159 | 0.450564079 | 0.72597988 | -4.66135495 |
| **A_66_P132493** | Siglece | 0.144582168 | -0.301381205 | 0.590545541 | 10.00811296 | 0.812018432 | 0.450619335 | 0.72597988 | -4.661378317 |
| **A_55_P2161818** | NA | 0.135546636 | -0.282592966 | 0.553686237 | 16.75327277 | 0.811928502 | 0.450666787 | 0.72597988 | -4.66139838 |
| **A_55_P2137910** | NA | -0.122035211 | -0.498576194 | 0.254505772 | 14.40933364 | -0.811751782 | 0.450760047 | 0.72597988 | -4.661437803 |
| **A_55_P2057681** | Cdyl2 | 0.083103379 | -0.173332295 | 0.339539054 | 8.447216129 | 0.811690731 | 0.450792269 | 0.72597988 | -4.661451421 |
| **A_55_P1955393** | 7-Sep | 0.263013403 | -0.548590461 | 1.074617266 | 13.3667869 | 0.811678645 | 0.450798648 | 0.72597988 | -4.661454117 |
| **A_55_P2048454** | Rps2 | 0.368780659 | -0.769228274 | 1.506789592 | 15.5001638 | 0.811657509 | 0.450809803 | 0.72597988 | -4.661458831 |
| **A_55_P2072284** | Dnaaf3 | 0.380703417 | -0.794286223 | 1.555693057 | 9.156399359 | 0.811527205 | 0.450878583 | 0.72597988 | -4.661487894 |
| **A_55_P2001073** | Arid2 | 0.116969493 | -0.244075353 | 0.478014339 | 9.318765767 | 0.811450071 | 0.450919301 | 0.72597988 | -4.661505096 |
| **A_55_P2019014** | Tmem120 | -0.281436441 | -1.150137743 | 0.587264861 | 9.616850063 | -0.811447082 | 0.450920879 | 0.72597988 | -4.661505762 |
| **A_51_P174434** | Cenpj | 0.129224235 | -0.269678207 | 0.528126677 | 9.841651607 | 0.811386205 | 0.450953016 | 0.72597988 | -4.661519338 |
| **A_55_P2182867** | Sgcb | 0.221067293 | -0.46137003 | 0.903504617 | 9.414735343 | 0.811357152 | 0.450968355 | 0.72597988 | -4.661525817 |
| **A_55_P2054887** | Trappc1 | 0.220799394 | -0.460880358 | 0.902479146 | 12.60541161 | 0.811274505 | 0.451011989 | 0.72597988 | -4.661544246 |
| **A_55_P1982817** | NA | -0.287075106 | -1.173384356 | 0.599234145 | 10.02585603 | -0.811260997 | 0.451019121 | 0.72597988 | -4.661547258 |
| **A_51_P381360** | Ndufa13 | -0.071108581 | -0.290666656 | 0.148449494 | 15.83063679 | -0.811190662 | 0.451056258 | 0.72597988 | -4.661562941 |
| **A_51_P388737** | Slc11a2 | -0.186730293 | -0.763358504 | 0.389897918 | 8.667376838 | -0.811090357 | 0.451109223 | 0.72597988 | -4.661585305 |
| **A_55_P2148688** | NA | 0.134523701 | -0.280897995 | 0.549945396 | 10.04213427 | 0.811073064 | 0.451118355 | 0.72597988 | -4.661589161 |
| **A_55_P1956063** | Pbld1 | -0.178023547 | -0.72785379 | 0.371806696 | 16.51063714 | -0.810959512 | 0.451178322 | 0.726015953 | -4.661614476 |
| **A_51_P310254** | Fam118a | -0.071526136 | -0.292482628 | 0.149430356 | 8.673358663 | -0.810789924 | 0.451267892 | 0.726093672 | -4.661652279 |
| **A_52_P352362** | NA | -0.156366557 | -0.63946249 | 0.326729376 | 9.11618584 | -0.81070124 | 0.451314736 | 0.726093672 | -4.661672046 |
| **A_51_P495485** | Pip4p2 | 0.426276475 | -0.890784162 | 1.743337112 | 11.35497684 | 0.810654767 | 0.451339286 | 0.726093672 | -4.661682403 |
| **A_55_P1954886** | NA | 0.138211865 | -0.289063832 | 0.565487562 | 16.34572612 | 0.810191125 | 0.451584259 | 0.726408258 | -4.661785719 |
| **A_52_P317393** | Adgrg1 | -0.652573044 | -2.67023158 | 1.365085493 | 10.91918073 | -0.810087268 | 0.451639146 | 0.726408258 | -4.661808857 |
| **A_55_P1982668** | Nadk | -0.280562215 | -1.148037624 | 0.586913193 | 11.13244723 | -0.81006964 | 0.451648463 | 0.726408258 | -4.661812784 |
| **A_66_P127956** | Zfp947 | 0.079511471 | -0.166382502 | 0.325405443 | 11.79135115 | 0.809901586 | 0.45173729 | 0.726408258 | -4.661850219 |
| **A_55_P1997400** | Slc7a4 | 0.29353575 | -0.61429562 | 1.20136712 | 8.96025167 | 0.809852939 | 0.451763006 | 0.726408258 | -4.661861055 |
| **A_55_P2035023** | Zfp653 | 0.115318061 | -0.241365891 | 0.472002013 | 9.714742051 | 0.809774517 | 0.451804463 | 0.726408258 | -4.661878522 |
| **A_55_P2045859** | NA | -0.302265451 | -1.237226284 | 0.632695383 | 10.77983183 | -0.809739708 | 0.451822865 | 0.726408258 | -4.661886275 |
| **A_55_P2075237** | Ulk3 | -0.097833176 | -0.400470943 | 0.204804591 | 9.821838989 | -0.809679933 | 0.451854468 | 0.726408258 | -4.661899587 |
| **A_52_P176160** | Ahcy | 0.069822081 | -0.146183192 | 0.285827355 | 7.92299029 | 0.809615441 | 0.451888566 | 0.726408258 | -4.66191395 |
| **A_51_P368591** | Tle6 | 0.286215984 | -0.599282351 | 1.171714318 | 10.58364452 | 0.809573867 | 0.451910548 | 0.726408258 | -4.661923208 |
| **A_52_P138598** | Pum2 | 0.127182102 | -0.266511231 | 0.520875435 | 8.661356461 | 0.809129962 | 0.452145307 | 0.726725194 | -4.662022043 |
| **A_55_P2038126** | Tpmt | 0.101854072 | -0.213567774 | 0.417275917 | 8.415854695 | 0.808792235 | 0.452323974 | 0.726883583 | -4.662097215 |
| **A_51_P105520** | Nomo1 | -0.29762293 | -1.21933541 | 0.624089549 | 12.0843202 | -0.808762985 | 0.45233945 | 0.726883583 | -4.662103725 |
| **A_55_P1985519** | NA | -0.192437592 | -0.788424468 | 0.403549284 | 16.42967317 | -0.808730017 | 0.452356895 | 0.726883583 | -4.662111062 |
| **A_66_P117189** | Rad54l2 | 0.091131592 | -0.191131204 | 0.373394389 | 9.006436089 | 0.808659448 | 0.452394236 | 0.726883583 | -4.662126766 |
| **A_55_P2172624** | Gm1123 | -0.090520671 | -0.370969855 | 0.189928513 | 7.923688481 | -0.808432806 | 0.452514178 | 0.727015882 | -4.662177196 |
| **A_52_P134556** | Cd2bp2 | 0.071557946 | -0.150174293 | 0.293290186 | 9.872396857 | 0.808312637 | 0.452577783 | 0.727057652 | -4.662203931 |
| **A_55_P2167803** | Abi3bp | 0.288234506 | -0.605025581 | 1.181494592 | 8.684885511 | 0.808199146 | 0.452637859 | 0.727091248 | -4.662229178 |
| **A_55_P1967076** | Rpl11 | 0.20426405 | -0.428818716 | 0.837346816 | 17.1120495 | 0.808131048 | 0.452673909 | 0.727091248 | -4.662244326 |
| **A_51_P202498** | Sar1a | -0.383173843 | -1.571112275 | 0.804764588 | 10.44501358 | -0.807890066 | 0.452801498 | 0.727128302 | -4.662297925 |
| **A_55_P1973548** | NA | -0.291240684 | -1.194177625 | 0.611696256 | 15.07766689 | -0.807876486 | 0.452808689 | 0.727128302 | -4.662300945 |
| **A_55_P2057821** | Golga1 | -0.084410624 | -0.346110563 | 0.177289315 | 9.530448946 | -0.80787438 | 0.452809805 | 0.727128302 | -4.662301414 |
| **A_55_P2026275** | Ppp1r1b | 0.249219362 | -0.523562283 | 1.022001006 | 8.517216797 | 0.80774695 | 0.452877284 | 0.727162739 | -4.662329752 |
| **A_66_P125962** | Hipk1 | -0.09551711 | -0.391717634 | 0.200683414 | 8.076562496 | -0.80769184 | 0.45290647 | 0.727162739 | -4.662342007 |
| **A_55_P2020911** | Rps6ka6 | -0.704979018 | -2.891958759 | 1.482000724 | 9.914979835 | -0.807387045 | 0.453067911 | 0.727246255 | -4.662409773 |
| **A_55_P2050592** | NA | -0.099371225 | -0.407656747 | 0.208914296 | 7.933509105 | -0.807342587 | 0.453091462 | 0.727246255 | -4.662419657 |
| **A_55_P2037982** | Agap1 | 0.237990912 | -0.500394747 | 0.976376571 | 10.47135411 | 0.807286084 | 0.453121396 | 0.727246255 | -4.662432217 |

| | | | | | | | | | |
|---|---|---|---|---|---|---|---|---|---|
| **A_66_P129263** | Lekr1 | -0.076300096 | -0.313029633 | 0.160429441 | 8.320285399 | -0.807277748 | 0.453125812 | 0.727246255 | -4.66243407 |
| **A_55_P2053456** | Gm6548 | 0.261301809 | -0.549535334 | 1.072138952 | 16.48452513 | 0.807159062 | 0.453188694 | 0.727246255 | -4.662460452 |
| **A_52_P386837** | Trmu | -0.070161117 | -0.287882088 | 0.147559854 | 9.160100204 | -0.80713574 | 0.453201051 | 0.727246255 | -4.662465635 |
| **A_55_P1981704** | Ccdc113 | 0.597295559 | -1.256557816 | 2.451148934 | 9.766776221 | 0.806982746 | 0.453282121 | 0.727246255 | -4.662499639 |
| **A_51_P308590** | Ppif | 0.070113659 | -0.147516179 | 0.287743498 | 8.094019465 | 0.806927545 | 0.453311374 | 0.727246255 | -4.662511906 |
| **A_51_P510663** | Supt3 | 0.131237122 | -0.27611797 | 0.538592213 | 8.519120816 | 0.806926343 | 0.453312011 | 0.727246255 | -4.662512173 |
| **A_52_P571746** | Sh2d5 | -0.207358276 | -0.851024623 | 0.436308072 | 7.863174681 | -0.806883649 | 0.453334637 | 0.727246255 | -4.662521661 |
| **A_55_P2105858** | Atf5 | -0.382096984 | -1.568572778 | 0.80437881 | 13.08415531 | -0.806612732 | 0.45347823 | 0.727360182 | -4.662581859 |
| **A_55_P2090953** | NA | 0.134390261 | -0.282940749 | 0.55172127 | 17.09545842 | 0.806561497 | 0.45350539 | 0.727360182 | -4.662593241 |
| **A_52_P217474** | Ndufa6 | 0.326533379 | -0.687566418 | 1.340633176 | 13.36659821 | 0.806486741 | 0.45354502 | 0.727360182 | -4.662609849 |
| **A_55_P2078395** | Gm1943 | -0.118552783 | -0.486746317 | 0.249640752 | 8.121310536 | -0.806465771 | 0.453556137 | 0.727360182 | -4.662614508 |
| **A_66_P108345** | Sgcb | 0.249930941 | -0.526617849 | 1.026479731 | 9.738935226 | 0.806123576 | 0.45373758 | 0.727442564 | -4.662690516 |
| **A_55_P2144075** | Pofut2 | -0.097260909 | -0.399462206 | 0.204940388 | 8.327391815 | -0.806106359 | 0.45374671 | 0.727442564 | -4.66269434 |
| **A_55_P2162965** | Ptms | 0.193134917 | -0.406962474 | 0.793232308 | 14.06064398 | 0.806100898 | 0.453749606 | 0.727442564 | -4.662695552 |
| **A_55_P2027812** | Arxes2 | 0.372677957 | -0.785307705 | 1.530663619 | 9.87831226 | 0.806085056 | 0.453758008 | 0.727442564 | -4.662699071 |
| **A_51_P506937** | Mrps12 | 0.080186675 | -0.169004448 | 0.329377798 | 12.54029562 | 0.805972051 | 0.453817941 | 0.727453812 | -4.662724166 |
| **A_55_P1955289** | Slc18b1 | 0.361695547 | -0.762381856 | 1.485772949 | 9.436894225 | 0.80592994 | 0.453840275 | 0.727453812 | -4.662733517 |
| **A_52_P474636** | Stk4 | -0.105306561 | -0.432635784 | 0.222022663 | 8.307290996 | -0.805788368 | 0.453915369 | 0.727513865 | -4.662764953 |
| **A_52_P293120** | Prrg3 | 0.366224737 | -0.772339814 | 1.504789287 | 9.909485821 | 0.80563878 | 0.453994725 | 0.727580737 | -4.662798164 |
| **A_55_P2142028** | Clip4 | 0.358395274 | -0.755967938 | 1.472758486 | 8.954261611 | 0.805537669 | 0.45404837 | 0.727606397 | -4.66282061 |
| **A_55_P1999419** | Bag5 | -0.133589306 | -0.549060379 | 0.281881766 | 10.05987321 | -0.805343669 | 0.45415131 | 0.727665291 | -4.662863673 |
| **A_55_P2084611** | B4galt6 | -0.073884396 | -0.303682863 | 0.155914071 | 7.969888933 | -0.80529675 | 0.454176208 | 0.727665291 | -4.662874086 |
| **A_52_P672313** | Exoc5 | 0.128219629 | -0.27064459 | 0.527083849 | 11.31924946 | 0.805155536 | 0.454251153 | 0.727665291 | -4.662905427 |
| **A_55_P1998081** | Trmt1 | 0.083620725 | -0.176512376 | 0.343753826 | 13.06820883 | 0.805134908 | 0.454262101 | 0.727665291 | -4.662910004 |
| **A_55_P1954365** | Ttc28 | 0.38404227 | -0.810797899 | 1.578882438 | 11.08660321 | 0.805043891 | 0.454310411 | 0.727665291 | -4.662930202 |
| **A_51_P364421** | Vamp7 | 0.224283525 | -0.473557712 | 0.922124761 | 9.166029194 | 0.804991119 | 0.454338423 | 0.727665291 | -4.662941912 |
| **A_55_P2156274** | Plekha2 | -0.096523755 | -0.396878694 | 0.203831185 | 8.130756952 | -0.804914542 | 0.454379073 | 0.727665291 | -4.662958903 |
| **A_66_P121787** | Samd9l | 0.300130894 | -0.63380815 | 1.234069937 | 9.766768942 | 0.80490109 | 0.454386214 | 0.727665291 | -4.662961888 |
| **A_55_P2132358** | NA | -0.264397906 | -1.087347426 | 0.558551614 | 9.400754644 | -0.804702147 | 0.454491834 | 0.727682782 | -4.663006025 |
| **A_55_P2126950** | Zfp467 | 0.641336208 | -1.35486823 | 2.637540647 | 10.49531144 | 0.804694603 | 0.45449584 | 0.727682782 | -4.663007699 |
| **A_52_P158710** | Dcbld1 | 0.317950047 | -0.671724937 | 1.30762503 | 9.386524209 | 0.804667843 | 0.454510048 | 0.727682782 | -4.663013636 |
| **A_51_P498023** | Scyl1 | -0.087326383 | -0.359211988 | 0.184559222 | 11.98279105 | -0.804469538 | 0.45461535 | 0.727791106 | -4.663057623 |
| **A_51_P248381** | Zdhhc4 | 0.08101085 | -0.171249986 | 0.333271686 | 8.668108138 | 0.804347475 | 0.454680176 | 0.727824426 | -4.663084696 |
| **A_55_P2106241** | 4930432K | -0.142086195 | -0.584604909 | 0.30043252 | 8.300218244 | -0.804212069 | 0.454752096 | 0.727824426 | -4.663114725 |
| **A_55_P2091446** | Trim33 | 0.10857935 | -0.229744254 | 0.446902953 | 9.614024947 | 0.803831986 | 0.454954019 | 0.727824426 | -4.663198998 |
| **A_55_P2006768** | NA | 0.090683615 | -0.191914125 | 0.373281355 | 13.17982271 | 0.803730566 | 0.45500791 | 0.727824426 | -4.663221481 |
| **A_55_P2024763** | Lama3 | 0.578086086 | -1.22347129 | 2.379643463 | 10.24981539 | 0.803701473 | 0.45502337 | 0.727824426 | -4.66322793 |
| **A_55_P1956017** | Usp45 | 0.145038131 | -0.306976443 | 0.597052704 | 9.149929202 | 0.80367434 | 0.455037789 | 0.727824426 | -4.663233945 |
| **A_55_P2095196** | Akr1b10 | 0.347369671 | -0.735339871 | 1.430079213 | 9.932407504 | 0.803582145 | 0.455086785 | 0.727824426 | -4.66325438 |
| **A_55_P1964842** | Opa1 | 0.069396188 | -0.146941655 | 0.28573403 | 10.41057435 | 0.803440029 | 0.455162318 | 0.727824426 | -4.663285878 |
| **A_55_P1955427** | Cux1 | 0.264266543 | -0.559584927 | 1.088118012 | 9.666621881 | 0.803421793 | 0.455172011 | 0.727824426 | -4.66328992 |
| **A_55_P2082241** | NA | 0.182277407 | -0.385990886 | 0.750545701 | 13.95525386 | 0.803396182 | 0.455185624 | 0.727824426 | -4.663295596 |
| **A_55_P1987770** | Psenen | -0.072058333 | -0.296716529 | 0.152599862 | 13.24667236 | -0.803363863 | 0.455202803 | 0.727824426 | -4.663302758 |
| **A_55_P2161055** | Trp63 | -0.091558715 | -0.377029646 | 0.193912216 | 11.02137607 | -0.803319193 | 0.455226548 | 0.727824426 | -4.663312657 |
| **A_55_P1963105** | NA | 0.159475451 | -0.337809748 | 0.65676065 | 15.96756087 | 0.803227651 | 0.455275212 | 0.727824426 | -4.663332942 |
| **A_55_P2074337** | Ngf | 0.096886724 | -0.205321063 | 0.399094511 | 8.294045394 | 0.802987838 | 0.455402714 | 0.727824426 | -4.663386076 |
| **A_51_P444994** | Tmem203 | -0.170703476 | -0.703235257 | 0.361828306 | 10.43285189 | -0.8028736 | 0.45546346 | 0.727824426 | -4.663411383 |
| **A_55_P2088385** | Tnnt1 | 0.617262921 | -1.30841262 | 2.542938461 | 11.01607042 | 0.802855563 | 0.455473052 | 0.727824426 | -4.663415379 |

| | | | | | | | | | |
|---|---|---|---|---|---|---|---|---|---|
| **A_55_P1985945** | NA | 0.159199048 | -0.337473948 | 0.655872044 | 14.13989137 | 0.802823845 | 0.455489919 | 0.727824426 | -4.663422405 |
| **A_51_P398164** | Klhl2 | -0.17659399 | -0.727569387 | 0.374381407 | 9.307999244 | -0.802775404 | 0.455515681 | 0.727824426 | -4.663433135 |
| **A_55_P2035963** | Hipk3 | -0.172405152 | -0.710347268 | 0.365536964 | 8.376915022 | -0.802721764 | 0.455544208 | 0.727824426 | -4.663445017 |
| **A_55_P2106096** | Fam171a1 | -0.094769488 | -0.390476383 | 0.200937406 | 10.28873811 | -0.802707704 | 0.455551686 | 0.727824426 | -4.663448131 |
| **A_55_P1999197** | Clasp1 | 0.097473187 | -0.206694839 | 0.401641212 | 10.77826981 | 0.802642129 | 0.455586564 | 0.727824426 | -4.663462655 |
| **A_55_P1955891** | NA | 0.132371853 | -0.280707697 | 0.545451403 | 13.35597064 | 0.802624291 | 0.455596051 | 0.727824426 | -4.663466606 |
| **A_66_P139946** | Sfta3-ps | 1.28658229 | -2.728533768 | 5.301698348 | 12.03738697 | 0.802582732 | 0.455618157 | 0.727824426 | -4.66347581 |
| **A_51_P229599** | Etnk1 | -0.181442633 | -0.747709187 | 0.384823921 | 8.631164557 | -0.802543854 | 0.455638836 | 0.727824426 | -4.66348442 |
| **A_55_P2041050** | Zmym5 | 0.106471177 | -0.22582277 | 0.438765124 | 9.798583316 | 0.802527586 | 0.45564749 | 0.727824426 | -4.663488023 |
| **A_55_P1973573** | Spn | 0.088137526 | -0.186971368 | 0.36324642 | 7.757269292 | 0.802428915 | 0.455699978 | 0.727824426 | -4.663509874 |
| **A_51_P145631** | Svip | 0.225924719 | -0.479272409 | 0.931121848 | 8.530152681 | 0.802423358 | 0.455702935 | 0.727824426 | -4.663511105 |
| **A_51_P518893** | Ccin | -0.102260761 | -0.42147622 | 0.216954699 | 8.464375458 | -0.802371416 | 0.455730567 | 0.727824426 | -4.663522607 |
| **A_55_P2061565** | Zfp229 | -0.143396599 | -0.591038659 | 0.304245462 | 8.133253863 | -0.802339752 | 0.455747413 | 0.727824426 | -4.663529618 |
| **A_55_P2002597** | NA | -0.091765114 | -0.378249812 | 0.194719584 | 9.955756856 | -0.802281033 | 0.455778654 | 0.727824426 | -4.66354262 |
| **A_66_P121495** | Psat1 | 0.105591532 | -0.224085635 | 0.435268699 | 8.076850181 | 0.802214612 | 0.455813994 | 0.727824426 | -4.663557327 |
| **A_51_P359293** | Ubxn2b | 0.161341806 | -0.34244575 | 0.665129361 | 8.785886852 | 0.802139353 | 0.455854038 | 0.727824426 | -4.663573989 |
| **A_55_P2063465** | Thbs3 | 0.4095427 | -0.869321831 | 1.688407231 | 10.2499954 | 0.802093491 | 0.455878442 | 0.727824426 | -4.663584143 |
| **A_55_P2076533** | Pmvk | 0.177139874 | -0.376119647 | 0.730399395 | 12.23197902 | 0.801932439 | 0.455964149 | 0.72785594 | -4.663619795 |
| **A_51_P363749** | Irf6 | 0.167081965 | -0.354775171 | 0.6889391 | 10.44396925 | 0.801914918 | 0.455973474 | 0.72785594 | -4.663623674 |
| **A_55_P2148431** | Slc4a3 | -0.07494494 | -0.309077967 | 0.159188088 | 7.798049831 | -0.801733405 | 0.456070085 | 0.727876369 | -4.66366385 |
| **A_55_P2177721** | Adra1b | -0.355608766 | -1.46662274 | 0.755405208 | 10.27230064 | -0.801684121 | 0.456096318 | 0.727876369 | -4.663674757 |
| **A_55_P1976494** | Bbof1 | 0.286039007 | -0.607659161 | 1.179737175 | 8.741948044 | 0.801649894 | 0.456114539 | 0.727876369 | -4.663682332 |
| **A_55_P2141654** | NA | 0.229063842 | -0.48665898 | 0.944786665 | 16.35425053 | 0.80160796 | 0.456136862 | 0.727876369 | -4.663691612 |
| **A_55_P1987290** | Rnu1b6 | -0.415643761 | -1.715122053 | 0.88383453 | 15.03181712 | -0.80112924 | 0.45639176 | 0.728160794 | -4.663797534 |
| **A_66_P100021** | Ncaph2 | -0.101042773 | -0.416967297 | 0.214881751 | 10.70289156 | -0.80107331 | 0.456421547 | 0.728160794 | -4.663809906 |
| **A_55_P1962466** | NA | 0.232252368 | -0.493928651 | 0.958433388 | 17.18298246 | 0.801061027 | 0.456428089 | 0.728160794 | -4.663812623 |
| **A_55_P1956064** | NA | 0.133216935 | -0.283429898 | 0.549863768 | 12.24304763 | 0.800832514 | 0.456549806 | 0.72829488 | -4.663863167 |
| **A_55_P2018492** | Pgap3 | 0.099684506 | -0.212146725 | 0.411515737 | 9.870227476 | 0.800678924 | 0.456631629 | 0.728365309 | -4.663897134 |
| **A_55_P2003658** | Gnpda1 | -0.185531109 | -0.765985809 | 0.394923592 | 8.393983118 | -0.80056897 | 0.456690212 | 0.72839866 | -4.663921448 |
| **A_52_P522222** | NA | 0.087986923 | -0.187324408 | 0.363298253 | 8.598225001 | 0.800468762 | 0.456743607 | 0.728405592 | -4.663943605 |
| **A_52_P476904** | Chd6 | -0.074837846 | -0.309027333 | 0.159351642 | 8.445484713 | -0.80039474 | 0.456783052 | 0.728405592 | -4.663959971 |
| **A_51_P416546** | Ctsg | -0.126447894 | -0.522193659 | 0.269297871 | 7.878003525 | -0.80028683 | 0.456840559 | 0.728405592 | -4.663983828 |
| **A_55_P2064996** | Capzb | 0.082866391 | -0.176484583 | 0.342217364 | 11.16141765 | 0.800278014 | 0.456845257 | 0.728405592 | -4.663985777 |
| **A_52_P376337** | Sdhaf3 | 0.105438855 | -0.224659041 | 0.435536752 | 8.337213255 | 0.800033682 | 0.456975488 | 0.728542033 | -4.664039786 |
| **A_55_P2006930** | 2610005L | -0.304385607 | -1.257395607 | 0.648624393 | 11.25065723 | -0.799976078 | 0.457006195 | 0.728542033 | -4.664052517 |
| **A_55_P1958280** | Arrdc1 | 0.457847916 | -0.975867372 | 1.891563203 | 11.33477952 | 0.799850224 | 0.45707329 | 0.728588917 | -4.664080332 |
| **A_55_P2002517** | Fhl1 | 0.373850685 | -0.79693912 | 1.544640491 | 9.150466151 | 0.79977826 | 0.457111658 | 0.728590008 | -4.664096235 |
| **A_55_P2174816** | Zfp266 | 0.300594166 | -0.641327142 | 1.242515474 | 9.36344583 | 0.799311887 | 0.457360366 | 0.728827284 | -4.664199275 |
| **A_66_P108152** | Cbr1 | 0.187133917 | -0.399290574 | 0.773558408 | 11.43986719 | 0.799264924 | 0.457385416 | 0.728827284 | -4.664209649 |
| **A_55_P1991124** | Tom1l2 | -0.104245213 | -0.430928838 | 0.222438413 | 11.24160939 | -0.799243475 | 0.457396857 | 0.728827284 | -4.664214387 |
| **A_55_P2171746** | NA | 0.317454828 | -0.677418771 | 1.312328427 | 11.84620888 | 0.799216381 | 0.45741131 | 0.728827284 | -4.664220372 |
| **A_55_P2095894** | 6430548M | -0.207712283 | -0.858778466 | 0.4433539 | 9.887163112 | -0.799074711 | 0.457486885 | 0.728828865 | -4.664251662 |
| **A_55_P2073248** | Slc25a34 | -0.07054821 | -0.29167929 | 0.150582869 | 7.920753857 | -0.799073192 | 0.457487696 | 0.728828865 | -4.664251997 |
| **A_51_P493987** | Moxd1 | 0.108455093 | -0.231608909 | 0.448519095 | 8.449788352 | 0.798802905 | 0.457631909 | 0.72890909 | -4.664311686 |
| **A_55_P2160732** | Tfcp2 | -0.107884924 | -0.446174933 | 0.230405086 | 9.26222917 | -0.798770343 | 0.457649285 | 0.72890909 | -4.664318876 |
| **A_55_P1966086** | NA | -0.206147869 | -0.852589957 | 0.440294219 | 8.99656392 | -0.798729219 | 0.45767123 | 0.72890909 | -4.664327956 |
| **A_66_P113760** | Acvr2a | 0.185099113 | -0.39536192 | 0.765560146 | 9.106838171 | 0.798696189 | 0.457688857 | 0.72890909 | -4.664335249 |
| **A_66_P124788** | Tbcb | -0.072862637 | -0.301473459 | 0.155748184 | 11.37221864 | -0.79828588 | 0.457907862 | 0.728968656 | -4.664425826 |

| | | | | | | | | | |
|---|---|---|---|---|---|---|---|---|---|
| **A_66_P127484** | Abcf3 | 0.074935504 | -0.160190804 | 0.310061812 | 10.24596598 | 0.798246006 | 0.457929149 | 0.728968656 | -4.664434627 |
| **A_55_P2005778** | Uap1 | -0.162541197 | -0.672593003 | 0.347510609 | 11.74130989 | -0.798177553 | 0.457965695 | 0.728968656 | -4.664449735 |
| **A_51_P150014** | Rrp7a | -0.092514576 | -0.382835108 | 0.197805956 | 7.996751291 | -0.798146762 | 0.457982135 | 0.728968656 | -4.66445653 |
| **A_55_P2152901** | 2900040C | -0.358480868 | -1.48344115 | 0.766479415 | 10.26194203 | -0.798140113 | 0.457985685 | 0.728968656 | -4.664457998 |
| **A_52_P507498** | Plxnc1 | 0.420738233 | -0.899687417 | 1.741163883 | 9.066179893 | 0.798083573 | 0.458015873 | 0.728968656 | -4.664470475 |
| **A_66_P102733** | Ccdc28b | 0.34026305 | -0.727738341 | 1.408264441 | 12.54619828 | 0.797982409 | 0.458069891 | 0.728968656 | -4.664492799 |
| **A_51_P204740** | Cd34 | 0.539038047 | -1.153046217 | 2.231122311 | 14.55901563 | 0.79789874 | 0.458114571 | 0.728968656 | -4.664511262 |
| **A_55_P2095814** | Kalrn | 0.083844479 | -0.179351534 | 0.347040492 | 7.908416011 | 0.797894571 | 0.458116797 | 0.728968656 | -4.664512181 |
| **A_55_P2032543** | Cblc | 0.487517812 | -1.043026946 | 2.018062569 | 10.87796212 | 0.797801409 | 0.45816655 | 0.728968656 | -4.664532737 |
| **A_51_P468780** | Hmg20a | 0.197711385 | -0.423048392 | 0.818471162 | 11.06467238 | 0.797734649 | 0.458202205 | 0.728968656 | -4.664547466 |
| **A_55_P2121437** | NA | -0.080957188 | -0.33517887 | 0.173264494 | 14.08434365 | -0.797614741 | 0.458266251 | 0.728968656 | -4.664573919 |
| **A_55_P2013088** | Arpc4 | 0.111747994 | -0.239185929 | 0.462681916 | 9.207648804 | 0.797562505 | 0.458294154 | 0.728968656 | -4.664585442 |
| **A_66_P106212** | Secisbp2 | -0.076132527 | -0.315228933 | 0.162963878 | 9.141813788 | -0.797530951 | 0.458311009 | 0.728968656 | -4.664592402 |
| **A_55_P2119917** | Ikzf4 | -0.070666372 | -0.29262001 | 0.151287265 | 8.531036666 | -0.797445252 | 0.458356791 | 0.728968656 | -4.664611305 |
| **A_52_P149835** | Irak1 | -0.121457121 | -0.502969583 | 0.260055342 | 8.44853535 | -0.79737864 | 0.458392378 | 0.728968656 | -4.664625998 |
| **A_52_P556337** | Txlng | -0.125524086 | -0.5198359 | 0.268787727 | 8.414246646 | -0.79732912 | 0.458418835 | 0.728968656 | -4.66463692 |
| **A_55_P2074438** | Gtf2a1 | -0.155222291 | -0.642875636 | 0.332431054 | 10.56960397 | -0.797247611 | 0.458462385 | 0.728968656 | -4.664654896 |
| **A_52_P223333** | Mib1 | -0.137928995 | -0.571264036 | 0.295406046 | 8.996759831 | -0.7972272 | 0.458473292 | 0.728968656 | -4.664659397 |
| **A_51_P451516** | Arhgef25 | -0.070576716 | -0.292313055 | 0.151159622 | 12.96173673 | -0.797214012 | 0.458480338 | 0.728968656 | -4.664662305 |
| **A_55_P2020962** | NA | 0.24433943 | -0.523517608 | 1.012196468 | 16.99814718 | 0.797009564 | 0.458589591 | 0.729015362 | -4.664707388 |
| **A_52_P252258** | Mapk12 | 0.399840349 | -0.856726663 | 1.656407361 | 9.297672427 | 0.796987131 | 0.45860158 | 0.729015362 | -4.664712335 |
| **A_55_P2159665** | Zfp534 | 0.108640315 | -0.232797583 | 0.450078214 | 10.41593635 | 0.796947365 | 0.458622833 | 0.729015362 | -4.664721103 |
| **A_52_P236705** | Ripply3 | 0.397155123 | -0.851443214 | 1.645753461 | 9.656491733 | 0.796687059 | 0.45876197 | 0.729149848 | -4.664778489 |
| **A_51_P431316** | Ube2w | 0.140786651 | -0.301847957 | 0.583421258 | 8.933131626 | 0.796647973 | 0.458782864 | 0.729149848 | -4.664787105 |
| **A_55_P2082841** | NA | -0.288088693 | -1.194181055 | 0.61800367 | 11.58564715 | -0.796350192 | 0.458942075 | 0.729342929 | -4.664852737 |
| **A_55_P2048348** | Aig1 | -0.073813489 | -0.306052132 | 0.158425154 | 9.27292995 | -0.796070612 | 0.45909159 | 0.72946834 | -4.664914343 |
| **A_55_P2058591** | NA | 0.08882976 | -0.19066104 | 0.368320559 | 8.084259804 | 0.796051779 | 0.459101663 | 0.72946834 | -4.664918492 |
| **A_51_P427812** | Tmem108 | -0.137336756 | -0.569496908 | 0.294823397 | 8.093335762 | -0.795962137 | 0.459149611 | 0.72946834 | -4.664938241 |
| **A_51_P134142** | Cyp2c70 | -0.125088093 | -0.518759061 | 0.268582875 | 7.928761574 | -0.795853132 | 0.45920792 | 0.72946834 | -4.664962255 |
| **A_55_P1952537** | Rrp7a | -0.341901622 | -1.417998141 | 0.734194897 | 11.82836252 | -0.795793303 | 0.459239927 | 0.72946834 | -4.664975434 |
| **A_55_P1955147** | Camk1d | 0.127391742 | -0.273579025 | 0.528362509 | 8.782515093 | 0.795754147 | 0.459260874 | 0.72946834 | -4.664984059 |
| **A_55_P1963374** | Klhdc7a | 0.203763561 | -0.437627156 | 0.845154278 | 10.18872718 | 0.795708868 | 0.459285099 | 0.72946834 | -4.664994033 |
| **A_55_P1973583** | Hpn | 0.297970491 | -0.640553037 | 1.23649402 | 10.36437294 | 0.795203785 | 0.459555383 | 0.729818786 | -4.665105261 |
| **A_66_P137654** | Smpd2 | -0.083834975 | -0.347907691 | 0.180237741 | 8.455247171 | -0.795155473 | 0.459581242 | 0.729818786 | -4.665115898 |
| **A_55_P2060672** | Hoxa3 | 0.547962852 | -1.178338865 | 2.274264569 | 9.939952544 | 0.795032277 | 0.459647187 | 0.72986139 | -4.66514302 |
| **A_51_P136792** | Calcoco1 | -0.084775149 | -0.351891412 | 0.182341114 | 8.44770105 | -0.794911128 | 0.459712044 | 0.72986139 | -4.665169689 |
| **A_55_P2035657** | Psmb6 | 0.109126044 | -0.234724446 | 0.452976533 | 14.9859666 | 0.7948938 | 0.459721321 | 0.72986139 | -4.665173503 |
| **A_52_P49014** | Shh | 0.409528198 | -0.881100787 | 1.700157184 | 9.14427993 | 0.794754035 | 0.459796152 | 0.729920257 | -4.662740027 |
| **A_55_P2040907** | Gtf2i | 0.093181901 | -0.200561994 | 0.386925795 | 8.020586679 | 0.794535054 | 0.459913416 | 0.729993091 | -4.665252458 |
| **A_55_P2288232** | Gucy1a2 | 0.31301256 | -0.673884292 | 1.299909413 | 9.26665862 | 0.794402033 | 0.459984658 | 0.729993091 | -4.665281729 |
| **A_55_P2185652** | Cln3 | -0.343851517 | -1.428047696 | 0.740344662 | 9.030812556 | -0.794352776 | 0.460011041 | 0.729993091 | -4.665292567 |
| **A_55_P2053858** | Scamp3 | -0.151595534 | -0.6296105 | 0.326419432 | 12.77407162 | -0.794319561 | 0.460028832 | 0.729993091 | -4.665299875 |
| **A_51_P232207** | Hoxb6 | 0.147520227 | -0.317694545 | 0.612734999 | 8.624176173 | 0.7942339 | 0.460074717 | 0.729993091 | -4.665318721 |
| **A_55_P2106911** | Wnt1 | 0.118646732 | -0.25552195 | 0.492815414 | 8.57691323 | 0.794216108 | 0.460084247 | 0.729993091 | -4.665322635 |
| **A_55_P2020361** | Lzts3 | 0.258786062 | -0.557432319 | 1.075004444 | 10.4873775 | 0.794117666 | 0.460136984 | 0.729993091 | -4.665344291 |
| **A_55_P2090505** | NA | 0.150102287 | -0.323360437 | 0.62356501 | 11.04795998 | 0.794057336 | 0.460169305 | 0.729993091 | -4.665357562 |
| **A_51_P430259** | Cfap126 | 0.718268945 | -1.547589244 | 2.984127135 | 12.02729468 | 0.793971128 | 0.460215494 | 0.729993091 | -4.665376525 |
| **A_66_P116126** | Sdhb | 0.258904432 | -0.557845891 | 1.075654756 | 14.21212531 | 0.79396346 | 0.460219602 | 0.729993091 | -4.665378211 |

| | | | | | | | | | |
|---|---|---|---|---|---|---|---|---|---|
| **A_55_P2010622** | Sec16b | 0.089326258 | -0.192541061 | 0.371193577 | 8.266912997 | 0.793751871 | 0.460332982 | 0.730113033 | -4.665424747 |
| **A_51_P202964** | Csnk2a2 | -0.10256555 | -0.42628253 | 0.221151431 | 10.22046436 | -0.793572076 | 0.460429341 | 0.730205961 | -4.665464284 |
| **A_55_P2336173** | Mirg | -0.210538189 | -0.875212039 | 0.454135661 | 9.50793355 | -0.793364266 | 0.460540732 | 0.730322712 | -4.665509974 |
| **A_52_P24986** | Gpat4 | -0.131238865 | -0.545694724 | 0.283216994 | 12.36253124 | -0.79311201 | 0.460675973 | 0.730424777 | -4.665565426 |
| **A_51_P112308** | Tcim | 0.385328592 | -0.831627029 | 1.602284214 | 10.82131871 | 0.793061451 | 0.460703083 | 0.730424777 | -4.665576539 |
| **A_55_P2293967** | Tbc1d5 | 0.08607847 | -0.185786792 | 0.357943731 | 9.240949798 | 0.793032826 | 0.460718432 | 0.730424777 | -4.66558283 |
| **A_55_P2030209** | Cir1 | 0.108210935 | -0.233647762 | 0.450069632 | 10.07605037 | 0.79282048 | 0.460832306 | 0.730493861 | -4.665629497 |
| **A_55_P2103501** | Ankrd29 | 0.107201316 | -0.231472001 | 0.445874633 | 8.064878485 | 0.792810661 | 0.460837572 | 0.730493861 | -4.665631655 |
| **A_55_P2010038** | Tnfsf9 | -0.225603864 | -0.938466601 | 0.487258873 | 8.581920655 | -0.792667343 | 0.460914442 | 0.730555814 | -4.665663147 |
| **A_51_P290685** | Ppp6r3 | -0.093610368 | -0.389454072 | 0.202233337 | 8.752012332 | -0.792523165 | 0.460991781 | 0.730618501 | -4.665694825 |
| **A_55_P1970755** | Hdac9 | 0.172509975 | -0.37294848 | 0.71796843 | 8.560979464 | 0.792141707 | 0.461196447 | 0.730874349 | -4.665778617 |
| **A_52_P69756** | Mterf4 | 0.111970575 | -0.242103578 | 0.466044729 | 10.40000044 | 0.792063553 | 0.461238388 | 0.730874349 | -4.665795781 |
| **A_51_P317225** | Cmtm2a | -0.159335158 | -0.663242242 | 0.344571925 | 8.666589681 | -0.791975046 | 0.461285887 | 0.730874349 | -4.665815218 |
| **A_51_P147714** | Sugp2 | -0.087704541 | -0.365121911 | 0.189712829 | 8.23169241 | -0.791842445 | 0.461357058 | 0.730874349 | -4.665844336 |
| **A_55_P2017362** | Prr36 | 0.447650427 | -0.968337148 | 1.863638003 | 9.20754054 | 0.79182627 | 0.46136574 | 0.730874349 | -4.665847887 |
| **A_52_P367791** | Mri1 | 0.075080078 | -0.16241781 | 0.312577965 | 10.57585039 | 0.791799655 | 0.461380026 | 0.730874349 | -4.665853731 |
| **A_66_P102467** | Casz1 | -0.095897463 | -0.399409453 | 0.207614528 | 7.85909498 | -0.791373693 | 0.461608714 | 0.731023274 | -4.665947243 |
| **A_55_P2127934** | Palm3 | -0.277360328 | -1.155287417 | 0.600566762 | 9.314292685 | -0.791291021 | 0.461653108 | 0.731023274 | -4.665965388 |
| **A_55_P2162875** | NA | -0.556849197 | -2.319585371 | 1.205886977 | 9.900557644 | -0.791226139 | 0.46168795 | 0.731023274 | -4.665979628 |
| **A_55_P2038587** | NA | -0.090770258 | -0.378137165 | 0.196596649 | 7.895375992 | -0.791146958 | 0.461730475 | 0.731023274 | -4.665997005 |
| **A_55_P2212498** | NA | -0.132615583 | -0.552462064 | 0.287230897 | 8.313227281 | -0.791141907 | 0.461733188 | 0.731023274 | -4.665998113 |
| **A_55_P2060752** | Usp12 | -0.239156424 | -0.996350866 | 0.518038019 | 11.66358319 | -0.79108833 | 0.461761963 | 0.731023274 | -4.66600987 |
| **A_55_P2155616** | Gm9457 | 0.396813468 | -0.859746737 | 1.653373673 | 15.51064066 | 0.790958046 | 0.461831942 | 0.731023274 | -4.666038458 |
| **A_55_P2279035** | 9030419F | -0.153256791 | -0.638564169 | 0.332050587 | 8.872473598 | -0.790957555 | 0.461832206 | 0.731023274 | -4.666038566 |
| **A_51_P483473** | St3gal5 | 0.099401863 | -0.215388737 | 0.414192463 | 11.56058561 | 0.790902801 | 0.461861618 | 0.731023274 | -4.66605058 |
| **A_55_P2022149** | NA | 0.066558479 | -0.144233795 | 0.277350753 | 8.233361276 | 0.790858972 | 0.461885163 | 0.731023274 | -4.666060196 |
| **A_51_P271336** | Wipf2 | -0.089815147 | -0.374265042 | 0.194634749 | 9.989283452 | -0.790850058 | 0.461889951 | 0.731023274 | -4.666062152 |
| **A_55_P1976356** | Fgf18 | 0.53294016 | -1.155200709 | 2.221081028 | 9.906613964 | 0.79071524 | 0.46196238 | 0.731052959 | -4.666091729 |
| **A_51_P507942** | Atp13a2 | 0.149117553 | -0.32329216 | 0.621527266 | 11.6726751 | 0.790606346 | 0.462020888 | 0.731052959 | -4.666115617 |
| **A_52_P561927** | Ncor1 | -0.093220804 | -0.388587722 | 0.202146114 | 9.410230337 | -0.790499028 | 0.462078554 | 0.731052959 | -4.666139157 |
| **A_51_P391825** | Exosc10 | -0.109608512 | -0.456939748 | 0.237722725 | 9.834590382 | -0.790407009 | 0.462128003 | 0.731052959 | -4.666159339 |
| **A_55_P2247633** | Gpatch2 | 0.096414254 | -0.209106874 | 0.401935381 | 8.93741409 | 0.790406203 | 0.462128436 | 0.731052959 | -4.666159516 |
| **A_55_P2078895** | Gin1 | 0.085245514 | -0.184943487 | 0.355434515 | 8.165589414 | 0.790231277 | 0.46222245 | 0.731052959 | -4.666197878 |
| **A_51_P184197** | Smarcd2 | 0.205487246 | -0.445819828 | 0.856794319 | 10.0923746 | 0.790222555 | 0.462227137 | 0.731052959 | -4.66619979 |
| **A_55_P2089565** | Tbx22 | 0.186846612 | -0.40538717 | 0.779080394 | 8.354659398 | 0.790209771 | 0.462234009 | 0.731052959 | -4.666202593 |
| **A_51_P437478** | Zfp566 | 0.081947634 | -0.17780997 | 0.341705237 | 9.330365478 | 0.790166285 | 0.462257383 | 0.731052959 | -4.666212129 |
| **A_55_P1982533** | Elovl4 | -0.111566014 | -0.465232471 | 0.242100443 | 7.902615951 | -0.790111512 | 0.462286825 | 0.731052959 | -4.666224139 |
| **A_55_P2132512** | Ngef | 0.341213004 | -0.740733083 | 1.42315909 | 10.14542964 | 0.7898967 | 0.462402305 | 0.731175772 | -4.666271236 |
| **A_55_P1976689** | Snrnp35 | 0.07636677 | -0.165844628 | 0.318578169 | 10.55782959 | 0.789696461 | 0.462509969 | 0.731195054 | -4.666315131 |
| **A_55_P2172299** | Timm17b | 0.194818206 | -0.423130137 | 0.812766548 | 12.83819119 | 0.789637406 | 0.462541725 | 0.731195054 | -4.666328075 |
| **A_55_P2036803** | Dhx30 | -0.089341707 | -0.37274519 | 0.194061777 | 10.12799853 | -0.789585938 | 0.462569403 | 0.731195054 | -4.666339356 |
| **A_52_P472660** | Smurf2 | -0.09669268 | -0.403428759 | 0.2100434 | 9.874379756 | -0.789548984 | 0.462589276 | 0.731195054 | -4.666347455 |
| **A_66_P124017** | Map7d3 | 0.217729057 | -0.47317504 | 0.908633154 | 8.654236512 | 0.789312478 | 0.46271648 | 0.731195054 | -4.666399284 |
| **A_51_P426195** | Nppb | -0.7671309 | -3.20167088 | 1.66740908 | 9.252982159 | -0.789228775 | 0.462761505 | 0.731195054 | -4.666417625 |
| **A_55_P2033120** | Srxn1 | -0.51571491 | -2.152487855 | 1.121058036 | 9.776067123 | -0.789171867 | 0.462792118 | 0.731195054 | -4.666430093 |
| **A_55_P1986711** | NA | 0.093032701 | -0.202235547 | 0.38830095 | 8.641534415 | 0.789167569 | 0.46279443 | 0.731195054 | -4.666431035 |
| **A_55_P2037677** | Cttn | -0.092269921 | -0.385139172 | 0.20059933 | 12.93727058 | -0.789108493 | 0.462826212 | 0.731195054 | -4.666443978 |
| **A_55_P2133568** | Snhg5 | 0.154691224 | -0.336310125 | 0.645692572 | 8.29929481 | 0.789102349 | 0.462829518 | 0.731195054 | -4.666445324 |

| | | | | | | | | | |
|---|---|---|---|---|---|---|---|---|---|
| **A_55_P2122718** | Gpr19 | -0.147180134 | -0.614341823 | 0.319981554 | 9.333874382 | -0.789100506 | 0.462830509 | 0.731195054 | -4.666445728 |
| **A_55_P2185595** | Samd4b | -0.150129231 | -0.626769612 | 0.326511149 | 12.14710982 | -0.788905119 | 0.462935635 | 0.731229216 | -4.666488531 |
| **A_55_P1967216** | Adam10 | -0.139859617 | -0.583935013 | 0.304215778 | 12.42841945 | -0.788834633 | 0.462973563 | 0.731229216 | -4.666503971 |
| **A_55_P2140546** | Slc25a51 | -0.15097063 | -0.691691905 | 0.389750646 | 11.33404952 | -0.823401263 | 0.462981197 | 0.731229216 | -4.634458697 |
| **A_55_P2155062** | Naa40 | 0.1085314 | -0.236096415 | 0.453159216 | 9.867598159 | 0.788779155 | 0.463003417 | 0.731229216 | -4.666516122 |
| **A_52_P84096** | Pgd | 0.067359399 | -0.14657079 | 0.281289587 | 11.65316583 | 0.788635773 | 0.463080581 | 0.731291347 | -4.666547525 |
| **A_51_P328669** | Ncbp1 | 0.069452216 | -0.151170903 | 0.290075335 | 8.869728047 | 0.78847047 | 0.463169554 | 0.731344368 | -4.666583725 |
| **A_55_P2032458** | Gm6934 | 0.089023692 | -0.193784015 | 0.371831398 | 8.344146359 | 0.788432839 | 0.46318981 | 0.731344368 | -4.666591965 |
| **A_51_P226053** | Sra1 | -0.073818633 | -0.308422791 | 0.160785524 | 12.56186329 | -0.788098754 | 0.463369671 | 0.731497834 | -4.666665108 |
| **A_66_P112006** | H2-T10 | -0.084202145 | -0.351810914 | 0.183406624 | 8.544086372 | -0.788085181 | 0.46337698 | 0.731497834 | -4.666668079 |
| **A_51_P260871** | Sdhd | 0.283675777 | -0.617959278 | 1.185310832 | 13.56484414 | 0.788028293 | 0.463407612 | 0.731497834 | -4.666680532 |
| **A_52_P124734** | Cybrd1 | 0.528333085 | -1.151178793 | 2.207844962 | 10.04022049 | 0.787907208 | 0.463472817 | 0.731497834 | -4.666707035 |
| **A_55_P1954835** | NA | -1.146656106 | -4.791856677 | 2.498544466 | 9.281430782 | -0.787883679 | 0.463485488 | 0.731497834 | -4.666712185 |
| **A_51_P507280** | Patl1 | -0.081142573 | -0.339110798 | 0.176825652 | 9.161403733 | -0.78783071 | 0.463514015 | 0.731497834 | -4.666723778 |
| **A_55_P2172807** | NA | 0.358608762 | -0.781652966 | 1.498870491 | 8.96244401 | 0.787710608 | 0.463578701 | 0.731526863 | -4.666750062 |
| **A_55_P2004469** | NA | 0.230505478 | -0.502517872 | 0.963528828 | 16.20546986 | 0.787614654 | 0.463630386 | 0.731526863 | -4.66677106 |
| **A_55_P2036863** | Rabl2 | 0.243102691 | -0.530009714 | 1.016215097 | 11.34943746 | 0.787585061 | 0.463646326 | 0.731526863 | -4.666777535 |
| **A_55_P2162698** | Prpf40b | -0.121593449 | -0.508317666 | 0.265130767 | 8.838367627 | -0.787515582 | 0.463683754 | 0.731526863 | -4.666792737 |
| **A_52_P544555** | Hsbp1 | 0.089015874 | -0.194192333 | 0.372224081 | 12.36071278 | 0.787248735 | 0.463827523 | 0.731678333 | -4.666851118 |
| **A_66_P126492** | Brk1 | 0.185561394 | -0.40493706 | 0.776059847 | 12.64609665 | 0.787080598 | 0.463918127 | 0.731678333 | -4.666887896 |
| **A_51_P481221** | Bace2 | -0.067103574 | -0.28067415 | 0.146467003 | 7.94228507 | -0.786963473 | 0.463981249 | 0.731678333 | -4.666913513 |
| **A_65_P09523** | Scrn3 | 0.11944936 | -0.260760896 | 0.499659616 | 10.14397593 | 0.786883331 | 0.464024443 | 0.731678333 | -4.666931039 |
| **A_55_P2017418** | Cfh | 0.512517679 | -1.118894306 | 2.143929665 | 10.44476242 | 0.786856521 | 0.464038894 | 0.731678333 | -4.666936902 |
| **A_55_P1980651** | Ubr5 | -0.19566176 | -0.818483258 | 0.427159738 | 12.60968919 | -0.786851371 | 0.46404167 | 0.731678333 | -4.666938028 |
| **A_51_P137778** | NA | -0.116324298 | -0.486605065 | 0.253956469 | 7.873901372 | -0.786845798 | 0.464044673 | 0.731678333 | -4.666939247 |
| **A_55_P1955472** | Bbs1 | 0.319871593 | -0.698526887 | 1.338270074 | 9.079426908 | 0.786698418 | 0.464124118 | 0.73174392 | -4.666971475 |
| **A_55_P1998751** | Ccdc137 | -0.073344697 | -0.307022898 | 0.160333504 | 8.577279258 | -0.786141757 | 0.46442427 | 0.732063475 | -4.667093163 |
| **A_52_P8324** | Tmem178 | 0.411423229 | -0.899411123 | 1.722257581 | 9.759982813 | 0.786124515 | 0.464433569 | 0.732063475 | -4.667096932 |
| **A_55_P2008936** | Slc2a9 | 0.07089419 | -0.154995609 | 0.296783989 | 7.914506826 | 0.786075704 | 0.464459896 | 0.732063475 | -4.667107599 |
| **A_55_P2111429** | Btf3l4 | -0.075124951 | -0.314519749 | 0.164269847 | 10.61831471 | -0.785995098 | 0.464503372 | 0.732063475 | -4.667125214 |
| **A_52_P125363** | Rbp2 | -0.100143782 | -0.419294834 | 0.21900727 | 7.795445171 | -0.785919482 | 0.46454416 | 0.732063475 | -4.667141738 |
| **A_52_P480939** | Tnpo1 | 0.125525834 | -0.274561466 | 0.525613135 | 10.81691019 | 0.785830165 | 0.464592342 | 0.732063475 | -4.667161254 |
| **A_55_P2061338** | Slc16a11 | -0.074570384 | -0.312252545 | 0.163111777 | 8.022397301 | -0.785814678 | 0.464600697 | 0.732063475 | -4.667164638 |
| **A_52_P94845** | Nicn1 | 0.13918374 | -0.304474385 | 0.582841865 | 8.802093091 | 0.785760891 | 0.464629714 | 0.732063475 | -4.66717639 |
| **A_52_P533304** | Eif5 | -0.134112392 | -0.561717492 | 0.293492707 | 12.62842267 | -0.785554606 | 0.464741014 | 0.732116722 | -4.667221457 |
| **A_51_P453088** | Coa6 | 0.166217525 | -0.363791398 | 0.696226448 | 11.06118357 | 0.785495997 | 0.46477264 | 0.732116722 | -4.66723426 |
| **A_55_P2080592** | Spata31d | 0.22772024 | -0.49847335 | 0.95391383 | 8.834051178 | 0.78541568 | 0.464815982 | 0.732116722 | -4.667251803 |
| **A_55_P2151006** | C1qtnf7 | 0.583654111 | -1.277982293 | 2.445290514 | 10.41881085 | 0.785255581 | 0.464902386 | 0.732116722 | -4.667286771 |
| **A_55_P2205650** | 2610507I0 | 0.225325591 | -0.493391383 | 0.944042565 | 9.500765685 | 0.785240992 | 0.46491026 | 0.732116722 | -4.667289957 |
| **A_55_P2004501** | Immp1l | 0.320574078 | -0.702038727 | 1.343186883 | 12.29244777 | 0.785176912 | 0.464944847 | 0.732116722 | -4.667303951 |
| **A_51_P242733** | Tex261 | 0.088937144 | -0.19478907 | 0.372663359 | 11.71642536 | 0.785116423 | 0.464977498 | 0.732116722 | -4.66731716 |
| **A_66_P102879** | Thra | 0.099086545 | -0.217063127 | 0.415236217 | 8.05881698 | 0.785004769 | 0.46503777 | 0.732116722 | -4.667341541 |
| **A_55_P2128158** | Trappc13 | 0.077547484 | -0.169920302 | 0.325015269 | 11.12081196 | 0.78487299 | 0.465108913 | 0.732116722 | -4.667370313 |
| **A_55_P2105457** | Zfp7 | -0.075126897 | -0.314884757 | 0.164630964 | 8.448333265 | -0.784825201 | 0.465134715 | 0.732116722 | -4.667380747 |
| **A_51_P512561** | Zfp68 | 0.426970061 | -0.935665429 | 1.789605551 | 11.21588167 | 0.784816405 | 0.465139464 | 0.732116722 | -4.667382667 |
| **A_51_P115817** | Ralgps2 | -0.246059994 | -1.031363063 | 0.539243075 | 9.36991021 | -0.784791085 | 0.465153135 | 0.732116722 | -4.658273274 |
| **A_55_P2139611** | Acbd4 | 0.119685162 | -0.262294256 | 0.50166458 | 9.100469036 | 0.784785003 | 0.465156419 | 0.732116722 | -4.667389522 |
| **A_51_P288549** | Jmjd7 | 0.340327602 | -0.745935374 | 1.426590577 | 9.600898556 | 0.784716061 | 0.465193645 | 0.732116722 | -4.667404573 |

| | | | | | | | | | |
|---|---|---|---|---|---|---|---|---|---|
| **A_52_P72497** | Acbd3 | -0.164524428 | -0.689771598 | 0.360722742 | 10.89593974 | -0.784543474 | 0.465286843 | 0.732135079 | -4.667442245 |
| **A_55_P2174022** | Brd4 | -0.106621366 | -0.447061353 | 0.233818621 | 12.98089339 | -0.784429689 | 0.465348295 | 0.732135079 | -4.66746708 |
| **A_55_P1952414** | Odf2 | -0.160665338 | -0.673672312 | 0.352341636 | 9.643257014 | -0.784421102 | 0.465352933 | 0.732135079 | -4.667468954 |
| **A_55_P1998354** | Wbp4 | 0.068875087 | -0.15105645 | 0.288806625 | 8.619973375 | 0.784377267 | 0.465376609 | 0.732135079 | -4.66747852 |
| **A_51_P420415** | Srd5a1 | -0.641279443 | -2.689096509 | 1.406537622 | 10.3669115 | -0.784343869 | 0.465394648 | 0.732135079 | -4.667485809 |
| **A_55_P2072468** | Sirt6 | -0.12270748 | -0.514621597 | 0.269206637 | 9.712995951 | -0.784206569 | 0.465468813 | 0.732144552 | -4.66751577 |
| **A_55_P2176731** | Prdx6 | 0.224977411 | -0.493588432 | 0.943543254 | 11.27834634 | 0.784192513 | 0.465476406 | 0.732144552 | -4.667518837 |
| **A_52_P265877** | Aldh9a1 | 0.131763774 | -0.289135392 | 0.552662941 | 10.34619253 | 0.784094318 | 0.465529454 | 0.732164465 | -4.667540263 |
| **A_55_P2061809** | Ndufc2 | 0.312371321 | -0.685599088 | 1.310341731 | 14.36928578 | 0.783977928 | 0.465592337 | 0.732164465 | -4.667565656 |
| **A_55_P2253289** | NA | -0.131401485 | -0.551232032 | 0.288429062 | 9.243746292 | -0.783928738 | 0.465618915 | 0.732164465 | -4.667576388 |
| **A_51_P173564** | Ahsa1 | 0.165455547 | -0.363205447 | 0.694116542 | 13.43001982 | 0.78388871 | 0.465640543 | 0.732164465 | -4.66758512 |
| **A_55_P2071326** | Rpl36 | 0.362655485 | -0.796237561 | 1.52154853 | 15.51485409 | 0.783792744 | 0.465692399 | 0.732186456 | -4.667606054 |
| **A_55_P1973352** | NA | 0.454175732 | -0.997407827 | 1.90575929 | 11.41930943 | 0.78366821 | 0.465759699 | 0.732232723 | -4.667633217 |
| **A_55_P2074085** | Gimap8 | 0.411208801 | -0.903255978 | 1.725673581 | 9.693020963 | 0.783544729 | 0.465826437 | 0.732250811 | -4.667660148 |
| **A_55_P2164221** | Tpt1 | 0.154261132 | -0.338915473 | 0.647437737 | 17.68157698 | 0.783437568 | 0.46588436 | 0.732250811 | -4.667683517 |
| **A_55_P2457912** | Psme4 | -0.107788497 | -0.452391333 | 0.236814339 | 8.423722137 | -0.783436705 | 0.465884826 | 0.732250811 | -4.667683706 |
| **A_52_P9437** | Psmb3 | -0.094153774 | -0.395221529 | 0.206913981 | 14.48263972 | -0.78329226 | 0.46596291 | 0.732253793 | -4.667715202 |
| **A_51_P327983** | 4833427G | 0.501466231 | -1.102201342 | 2.105133803 | 9.391355805 | 0.783209038 | 0.466007902 | 0.732253793 | -4.667733347 |
| **A_55_P2185461** | Rpl9 | 0.26312256 | -0.57840634 | 1.10465146 | 17.225511 | 0.783139981 | 0.466045239 | 0.732253793 | -4.667748403 |
| **A_55_P2300071** | 4833412C | -0.084046468 | -0.352885319 | 0.184792384 | 8.37400074 | -0.78302888 | 0.466105312 | 0.732253793 | -4.667772624 |
| **A_51_P466229** | Pdgfrl | 0.369319406 | -0.812273861 | 1.550912674 | 9.635751444 | 0.782860655 | 0.466196283 | 0.732253793 | -4.667809293 |
| **A_66_P111437** | Ube2j2 | -0.233467701 | -0.980441835 | 0.513506433 | 14.09656244 | -0.78283744 | 0.466208837 | 0.732253793 | -4.667814353 |
| **A_55_P2065327** | Rpl27a | 0.274272346 | -0.603272252 | 1.151816944 | 15.83275836 | 0.782822266 | 0.466217044 | 0.732253793 | -4.66781766 |
| **A_51_P505617** | Il18r1 | 0.629131928 | -1.383917348 | 2.642181205 | 9.917012262 | 0.782776296 | 0.466241906 | 0.732253793 | -4.667827679 |
| **A_55_P1971804** | Hdac10 | -0.112849861 | -0.473984288 | 0.248284566 | 8.836898421 | -0.782676841 | 0.466295698 | 0.732253793 | -4.667849355 |
| **A_55_P2045114** | Tmem242 | -0.169652834 | -0.712589349 | 0.373283681 | 10.86054188 | -0.782640673 | 0.466315261 | 0.732253793 | -4.667857236 |
| **A_55_P2165539** | Aebp2 | -0.265440839 | -1.115053218 | 0.584171541 | 9.090747928 | -0.782523259 | 0.466378773 | 0.732253793 | -4.667882822 |
| **A_51_P285042** | Rtf1 | -0.237106257 | -0.996201974 | 0.52198946 | 10.19334523 | -0.782342312 | 0.466476665 | 0.732253793 | -4.667922248 |
| **A_55_P1963585** | Mbd5 | 0.196084366 | -0.431685073 | 0.823853805 | 8.919428863 | 0.782335692 | 0.466480247 | 0.732253793 | -4.66792369 |
| **A_66_P120736** | Bnip3 | -0.109538423 | -0.460236334 | 0.241159489 | 8.100317788 | -0.782318588 | 0.466489501 | 0.732253793 | -4.667927417 |
| **A_65_P15812** | Pum1 | -0.122195481 | -0.51345001 | 0.269059047 | 10.48990058 | -0.78225098 | 0.466526082 | 0.732253793 | -4.667942146 |
| **A_52_P179068** | Gucy1b1 | 0.647383254 | -1.425463762 | 2.720230271 | 11.29281021 | 0.782248171 | 0.466527601 | 0.732253793 | -4.667942757 |
| **A_55_P1954086** | Postn | 0.556770157 | -1.225995587 | 2.339535901 | 12.04398918 | 0.78222558 | 0.466539825 | 0.732253793 | -4.667947679 |
| **A_55_P2156093** | NA | 0.156234361 | -0.344058449 | 0.656527171 | 13.75536137 | 0.782172673 | 0.466568453 | 0.732253793 | -4.667959204 |
| **A_55_P2082961** | Fnbp1l | 0.312649043 | -0.688632573 | 1.313930659 | 11.58292354 | 0.782080049 | 0.466618575 | 0.732273014 | -4.667979381 |
| **A_55_P2115582** | Slc20a1 | -0.821598077 | -3.453490535 | 1.810294381 | 11.53560285 | -0.781882729 | 0.466725365 | 0.732300871 | -4.668022357 |
| **A_55_P2048378** | Zfp975 | 0.456545988 | -1.006281596 | 1.919373571 | 10.51201769 | 0.781702922 | 0.466822692 | 0.732300871 | -4.668061514 |
| **A_55_P2069955** | NA | 0.208933881 | -0.460530721 | 0.878398483 | 17.70201752 | 0.781684677 | 0.466832568 | 0.732300871 | -4.668065487 |
| **A_51_P328883** | Zfp810 | 0.154588318 | -0.340884134 | 0.65006077 | 8.7669476 | 0.781461353 | 0.466953473 | 0.732300871 | -4.668114111 |
| **A_55_P2094641** | NA | 0.325666466 | -0.718156382 | 1.369489314 | 9.194960516 | 0.78144171 | 0.466964108 | 0.732300871 | -4.668118387 |
| **A_51_P359237** | Tmem9 | 0.269946246 | -0.595400123 | 1.135292615 | 12.15268495 | 0.781335689 | 0.467021515 | 0.732300871 | -4.668141468 |
| **A_55_P2037902** | Brd3 | -0.235819333 | -0.991865605 | 0.520226939 | 8.508807279 | -0.781234432 | 0.467076347 | 0.732300871 | -4.668163509 |
| **A_52_P618745** | Ermp1 | -0.118867991 | -0.499969962 | 0.26223398 | 8.810068094 | -0.781221295 | 0.467083461 | 0.732300871 | -4.668166368 |
| **A_55_P2116305** | Tnfrsf25 | 0.222542576 | -0.490955028 | 0.936040179 | 8.793737935 | 0.781215654 | 0.467086516 | 0.732300871 | -4.668167596 |
| **A_55_P1965539** | Ndufb4 | 0.188978386 | -0.416915109 | 0.794871882 | 14.69410202 | 0.781207118 | 0.467091139 | 0.732300871 | -4.668169454 |
| **A_55_P2092296** | Fbxo2 | -0.211351994 | -0.889005769 | 0.466301782 | 8.384436509 | -0.781175893 | 0.467108049 | 0.732300871 | -4.668176251 |
| **A_55_P2087280** | NA | -0.253305325 | -1.06548929 | 0.55887864 | 13.52060574 | -0.781160464 | 0.467116405 | 0.732300871 | -4.668179609 |
| **A_55_P2062070** | Pdzd2 | 0.068751592 | -0.1517124 | 0.289215585 | 7.788107288 | 0.781079858 | 0.46716006 | 0.732300871 | -4.668197152 |

| | | | | | | | | | |
|---|---|---|---|---|---|---|---|---|---|
| **A_55_P1960238** | Slc2a6 | -0.145356639 | -0.611475296 | 0.320762019 | 7.964358333 | -0.781067793 | 0.467166594 | 0.732300871 | -4.668199778 |
| **A_55_P2154595** | Mrpl39 | 0.215207101 | -0.475061083 | 0.905475285 | 11.10662058 | 0.780888607 | 0.467263651 | 0.73239363 | -4.668238773 |
| **A_55_P2003133** | Ngrn | 0.087589591 | -0.1933771 | 0.368556282 | 12.23037462 | 0.780814743 | 0.467303664 | 0.732396971 | -4.668254846 |
| **A_51_P427476** | Mxra8 | -0.087980183 | -0.3702524 | 0.194292034 | 10.0276085 | -0.780669251 | 0.467382485 | 0.732461131 | -4.668286502 |
| **A_55_P2198946** | Gm10768 | -0.083392888 | -0.351004342 | 0.184218566 | 7.710193746 | -0.780503152 | 0.467472483 | 0.732542793 | -4.668322636 |
| **A_55_P1987245** | Ywhaz | -0.197899124 | -0.833034264 | 0.437236017 | 9.566490146 | -0.780419432 | 0.46751785 | 0.732554511 | -4.668340848 |
| **A_52_P817257** | Smim22 | 0.639088056 | -1.412266949 | 2.690443061 | 12.19899954 | 0.780315478 | 0.467574185 | 0.732583412 | -4.668363459 |
| **A_52_P278497** | Eif4ebp3 | -0.09040284 | -0.380671672 | 0.199865993 | 10.45262611 | -0.78006719 | 0.467708759 | 0.732661178 | -4.668417456 |
| **A_51_P424232** | Mecp2 | 0.313989897 | -0.694198001 | 1.322177796 | 10.06269355 | 0.780053771 | 0.467716033 | 0.732661178 | -4.668420374 |
| **A_51_P495232** | Gpx8 | 0.2607361 | -0.576502077 | 1.097974277 | 12.49395072 | 0.780014161 | 0.467737505 | 0.732661178 | -4.668428987 |
| **A_55_P1984497** | Usp48 | -0.136499793 | -0.574894655 | 0.301895069 | 10.10963122 | -0.779860425 | 0.467820848 | 0.732679898 | -4.668462414 |
| **A_55_P1994047** | NA | 0.58230906 | -1.287939518 | 2.452557639 | 12.81975329 | 0.779838298 | 0.467832844 | 0.732679898 | -4.668467225 |
| **A_51_P489452** | Cdo1 | 0.109550939 | -0.242366781 | 0.46146866 | 9.846231216 | 0.779696018 | 0.467909988 | 0.732679898 | -4.668498157 |
| **A_55_P2346639** | Sec14l3 | 1.318973023 | -2.918073239 | 5.556019284 | 12.26890291 | 0.779691954 | 0.467912192 | 0.732679898 | -4.66849904 |
| **A_51_P368823** | Grb7 | 0.455009987 | -1.006784573 | 1.916804547 | 10.85934563 | 0.779623522 | 0.467949299 | 0.732679898 | -4.668513916 |
| **A_66_P122758** | Tsr1 | 0.161757794 | -0.357949619 | 0.681465207 | 9.485503142 | 0.779572749 | 0.467976832 | 0.732679898 | -4.668524953 |
| **A_55_P2122605** | Cbr2 | 1.757885217 | -3.890541068 | 7.406311503 | 14.07048179 | 0.779495108 | 0.468018937 | 0.732685187 | -4.668541829 |
| **A_55_P2037525** | NA | 0.353464339 | -0.782384754 | 1.489313433 | 15.19236321 | 0.779426764 | 0.468056002 | 0.732685187 | -4.668556683 |
| **A_66_P100440** | Vezf1 | 0.310681253 | -0.687790105 | 1.309152611 | 11.18064496 | 0.779345052 | 0.46810032 | 0.732695239 | -4.668574442 |
| **A_55_P2009267** | Gm16060 | 0.367926664 | -0.814648595 | 1.550501922 | 15.31011667 | 0.779260781 | 0.468146029 | 0.732707466 | -4.668592755 |
| **A_52_P114282** | Armcx5 | 0.13604002 | -0.301389856 | 0.573469895 | 8.925782816 | 0.778948222 | 0.468315591 | 0.732872335 | -4.668660668 |
| **A_55_P2052924** | Selenow | 0.310017249 | -0.686914164 | 1.306948663 | 13.68183842 | 0.778880666 | 0.468352246 | 0.732872335 | -4.668675344 |
| **A_52_P18068** | Txlng | 0.109442888 | -0.242548216 | 0.461433992 | 8.562596248 | 0.778764602 | 0.468415225 | 0.732872335 | -4.668700557 |
| **A_52_P115579** | Fbxo45 | 0.10205383 | -0.226207471 | 0.430315132 | 8.43263992 | 0.7786817 | 0.468460213 | 0.732872335 | -4.668718564 |
| **A_55_P1982578** | Kdm3a | 0.306378803 | -0.679305994 | 1.292063601 | 11.51623639 | 0.778522203 | 0.468546776 | 0.732872335 | -4.668753205 |
| **A_51_P143901** | Tm9sf2 | -0.167180556 | -0.70504713 | 0.370686018 | 12.91436113 | -0.778505267 | 0.468555968 | 0.732872335 | -4.668756883 |
| **A_52_P127024** | Tnks | -0.167303064 | -0.705573979 | 0.37096785 | 8.572176356 | -0.778490519 | 0.468563972 | 0.732872335 | -4.668760086 |
| **A_51_P311540** | Cox6a1 | 0.093638742 | -0.207646661 | 0.394924145 | 17.35942987 | 0.778444806 | 0.468588785 | 0.732872335 | -4.668770013 |
| **A_51_P483557** | Ube2b | -0.09007073 | -0.379878512 | 0.199737052 | 10.78063002 | -0.778437922 | 0.468592521 | 0.732872335 | -4.668771508 |
| **A_55_P2048209** | Ganab | 0.080294565 | -0.178133969 | 0.3387231 | 12.44854539 | 0.778208612 | 0.468717001 | 0.733007725 | -4.6688213 |
| **A_66_P130646** | H4c8 | -0.101925198 | -0.430052904 | 0.226202509 | 8.009275682 | -0.778016855 | 0.468821115 | 0.733111245 | -4.66886293 |
| **A_55_P2065330** | NA | 0.214406972 | -0.475947837 | 0.90476178 | 15.46023834 | 0.777887681 | 0.468891258 | 0.733161632 | -4.66889097 |
| **A_66_P100159** | Cfl2 | -0.114894458 | -0.484966854 | 0.255177938 | 11.45908902 | -0.777611606 | 0.469041196 | 0.733272526 | -4.668950888 |
| **A_55_P2136765** | Uqcr10 | 0.195352893 | -0.434040086 | 0.824745873 | 12.66505043 | 0.777406736 | 0.469152484 | 0.733272526 | -4.668995342 |
| **A_55_P2123933** | D2hgdh | -0.071745874 | -0.302903006 | 0.159411257 | 8.356344823 | -0.777391845 | 0.469160574 | 0.733272526 | -4.668998573 |
| **A_65_P19832** | Ttr | -1.1566383 | -4.88325483 | 2.569978231 | 9.754203602 | -0.777379723 | 0.469167159 | 0.733272526 | -4.669001203 |
| **A_55_P2020035** | NA | -0.18328871 | -0.773852234 | 0.407274814 | 11.91415102 | -0.777355079 | 0.469180547 | 0.733272526 | -4.66900655 |
| **A_51_P243207** | Zfp574 | -0.163213299 | -0.689104494 | 0.362677897 | 9.842798144 | -0.77733816 | 0.469189739 | 0.733272526 | -4.669010221 |
| **A_55_P1973360** | NA | 0.088158339 | -0.209146677 | 0.385463354 | 15.23722734 | 0.789812254 | 0.469377169 | 0.733502456 | -4.650692509 |
| **A_55_P2122588** | Ubr2 | 0.081214134 | -0.180604963 | 0.343033232 | 9.090425281 | 0.776927769 | 0.469412739 | 0.733502456 | -4.669099243 |
| **A_55_P2166128** | NA | -0.086374468 | -0.364901207 | 0.192152271 | 8.595376892 | -0.776727778 | 0.469521438 | 0.733551015 | -4.669142614 |
| **A_55_P2027067** | Phc2 | 0.110948012 | -0.246875175 | 0.468771199 | 13.6421596 | 0.77660716 | 0.469587005 | 0.733551015 | -4.669168768 |
| **A_55_P1965402** | Uqcrb | 0.20358337 | -0.45306678 | 0.86023352 | 13.30681759 | 0.776530641 | 0.469628603 | 0.733551015 | -4.669185359 |
| **A_55_P1960053** | Hvcn1 | -0.511140151 | -2.159879375 | 1.137599073 | 8.828255056 | -0.77649448 | 0.469648263 | 0.733551015 | -4.669193199 |
| **A_51_P462299** | Aida | -0.064920823 | -0.274340722 | 0.144499075 | 9.609474542 | -0.776455233 | 0.469669601 | 0.733551015 | -4.669201708 |
| **A_55_P2134082** | Clip4 | 0.174788569 | -0.38904232 | 0.738619459 | 8.243795466 | 0.776451812 | 0.469671461 | 0.733551015 | -4.669202449 |
| **A_55_P2091736** | Rassf2 | 0.543143993 | -1.210209744 | 2.296497729 | 10.19184405 | 0.77588219 | 0.469981231 | 0.733836354 | -4.669325911 |
| **A_51_P181319** | Dcxr | 0.537083799 | -1.196749893 | 2.27091749 | 10.14406042 | 0.775862854 | 0.469991748 | 0.733836354 | -4.669330101 |

| | | | | | | | | | |
|---|---|---|---|---|---|---|---|---|---|
| **A_51_P161946** | Epop | -0.759294617 | -3.210826679 | 1.692237446 | 9.723205386 | -0.77575233 | 0.470051871 | 0.733836354 | -4.669354049 |
| **A_51_P441387** | Kansl1l | 0.312625857 | -0.69716067 | 1.322412385 | 8.988334356 | 0.775435482 | 0.47022426 | 0.733836354 | -4.669422691 |
| **A_51_P502764** | Hdac10 | -0.144364541 | -0.610766509 | 0.322037427 | 9.445693041 | -0.775265587 | 0.470316714 | 0.733836354 | -4.669459489 |
| **A_66_P126138** | Ncoa1 | 0.273299714 | -0.609711273 | 1.156310702 | 9.205802625 | 0.775217228 | 0.470343033 | 0.733836354 | -4.669469963 |
| **A_52_P367520** | Nexn | 0.437575131 | -0.976236588 | 1.85138685 | 11.89333682 | 0.77519578 | 0.470354706 | 0.733836354 | -4.669474608 |
| **A_55_P2002426** | Rps6kb2 | -0.096794094 | -0.409540334 | 0.215952147 | 8.392161446 | -0.775188116 | 0.470358877 | 0.733836354 | -4.669476268 |
| **A_52_P424585** | Ctnnb1 | 0.186404093 | -0.41589141 | 0.788699596 | 13.21256564 | 0.775168595 | 0.470369502 | 0.733836354 | -4.669480495 |
| **A_55_P2117664** | Rnf165 | -0.127664238 | -0.54017477 | 0.284846294 | 8.394037365 | -0.775147874 | 0.470380779 | 0.733836354 | -4.669484982 |
| **A_55_P2132894** | Sdcbp | -0.145632673 | -0.61624158 | 0.324976235 | 8.715413589 | -0.775084448 | 0.470415302 | 0.733836354 | -4.669498717 |
| **A_51_P411264** | Txndc16 | 0.185975174 | -0.415029675 | 0.786980023 | 9.5702822 | 0.775045757 | 0.470436362 | 0.733836354 | -4.669507096 |
| **A_51_P320481** | Mcee | 0.226453452 | -0.505469162 | 0.958376066 | 10.88884602 | 0.774932947 | 0.47049777 | 0.733836354 | -4.669531522 |
| **A_51_P265374** | Vnn3 | -0.124437892 | -0.526655538 | 0.277779754 | 7.899119038 | -0.77489324 | 0.470519385 | 0.733836354 | -4.669540119 |
| **A_55_P2091201** | Thoc2 | -0.124255635 | -0.525918854 | 0.277407584 | 8.698592112 | -0.774826335 | 0.470555808 | 0.733836354 | -4.669554604 |
| **A_66_P138829** | Ndufs8 | 0.328906138 | -0.734394358 | 1.392206633 | 11.19926729 | 0.774758424 | 0.470592782 | 0.733836354 | -4.669569307 |
| **A_52_P305995** | Ammecr1 | -0.218326921 | -0.924166777 | 0.487512936 | 9.402971183 | -0.774731933 | 0.470607205 | 0.733836354 | -4.669575041 |
| **A_51_P433091** | Purb | 0.282691348 | -0.631250425 | 1.19663312 | 12.60753381 | 0.774719226 | 0.470614123 | 0.733836354 | -4.669577792 |
| **A_55_P2040936** | Eng | -0.103507277 | -0.438153549 | 0.231138996 | 10.08403627 | -0.774702997 | 0.470622959 | 0.733836354 | -4.669581305 |
| **A_55_P1960960** | Zdhhc9 | -0.234466282 | -0.992555505 | 0.523622941 | 12.19307376 | -0.774658729 | 0.470647063 | 0.733836354 | -4.669590888 |
| **A_51_P234728** | Lpar6 | 0.342916787 | -0.765838221 | 1.451671794 | 11.9084042 | 0.774646382 | 0.470653786 | 0.733836354 | -4.669593561 |
| **A_51_P458218** | Wdr13 | 0.189890294 | -0.424194902 | 0.80397549 | 10.42198485 | 0.774505461 | 0.470730522 | 0.733836354 | -4.669624064 |
| **A_55_P1957424** | Sulf1 | 0.09647042 | -0.21553017 | 0.408471011 | 7.920597749 | 0.774442361 | 0.470764885 | 0.733836354 | -4.669637721 |
| **A_55_P1952112** | Ildr2 | 0.08519974 | -0.190369544 | 0.360769024 | 8.195836304 | 0.774386561 | 0.470795274 | 0.733836354 | -4.669649797 |
| **A_52_P400425** | Pirt | -0.068089543 | -0.288328647 | 0.152149561 | 7.716134098 | -0.774348277 | 0.470816125 | 0.733836354 | -4.669658082 |
| **A_51_P347452** | Htatsf1 | -0.413795915 | -1.752318888 | 0.924727058 | 12.12431383 | -0.774302596 | 0.470841004 | 0.733836354 | -4.669667968 |
| **A_55_P2162404** | NA | 0.076965476 | -0.172076766 | 0.326007718 | 8.345289777 | 0.774057614 | 0.470974449 | 0.733974052 | -4.669720977 |
| **A_51_P107090** | BC031181 | -0.114472094 | -0.484904043 | 0.255959856 | 10.62143717 | -0.774001022 | 0.471005279 | 0.733974052 | -4.662921211 |
| **A_55_P2071059** | Nab2 | -0.168687381 | -0.714622833 | 0.377248071 | 11.33462259 | -0.773912112 | 0.471053718 | 0.733976537 | -4.669752455 |
| **A_55_P2033393** | Slc25a21 | -0.076569552 | -0.324398841 | 0.171259736 | 7.675708991 | -0.773844719 | 0.471090436 | 0.733976537 | -4.669767034 |
| **A_55_P2140751** | NA | -0.092854725 | -0.39341513 | 0.20770568 | 13.19321645 | -0.773789061 | 0.471120763 | 0.733976537 | -4.669779074 |
| **A_51_P464182** | Mat1a | 0.114936415 | -0.257188954 | 0.487061784 | 8.020181181 | 0.773604015 | 0.4712216 | 0.734074484 | -4.669819098 |
| **A_52_P664617** | Zfp472 | 0.191826373 | -0.429304043 | 0.81295679 | 8.762976894 | 0.77352769 | 0.471263196 | 0.734080135 | -4.669835604 |
| **A_55_P1972118** | NA | 0.256312948 | -0.573753356 | 1.086379252 | 16.06066026 | 0.773407016 | 0.471328967 | 0.734123439 | -4.6698617 |
| **A_55_P2249055** | NA | -0.072952084 | -0.309250355 | 0.163346188 | 10.40485686 | -0.773263485 | 0.471407204 | 0.734186152 | -4.669892735 |
| **A_55_P1953133** | NA | 0.355935548 | -0.797190235 | 1.50906133 | 15.15064661 | 0.773116651 | 0.471487252 | 0.734251674 | -4.66992448 |
| **A_55_P1959606** | Tom1l1 | 0.231428196 | -0.518462227 | 0.981318618 | 11.17680312 | 0.77298098 | 0.471561222 | 0.734262967 | -4.669953808 |
| **A_55_P1970537** | Bcl7b | -0.159113599 | -0.674715897 | 0.3564887 | 10.32755518 | -0.77293469 | 0.471586462 | 0.734262967 | -4.669963814 |
| **A_55_P2262593** | 4930429F | 0.197294034 | -0.442088293 | 0.83667636 | 8.595427216 | 0.772865052 | 0.471624435 | 0.734262967 | -4.669978865 |
| **A_55_P2145879** | Tpt1 | 0.145461536 | -0.325968769 | 0.616891842 | 17.57172864 | 0.772824744 | 0.471646415 | 0.734262967 | -4.669987577 |
| **A_52_P269016** | Nek9 | -0.109981369 | -0.466506215 | 0.246543477 | 11.82499116 | -0.77264441 | 0.471744762 | 0.734313128 | -4.67002655 |
| **A_55_P2010191** | Serpina11 | 0.066702435 | -0.149535837 | 0.282940708 | 8.106812885 | 0.772608487 | 0.471764355 | 0.734313128 | -4.670034313 |
| **A_55_P1995774** | NA | -0.127078156 | -0.539121687 | 0.284965374 | 9.80194481 | -0.772463825 | 0.471843261 | 0.734313128 | -4.670065571 |
| **A_51_P512157** | Slc30a6 | -0.110166204 | -0.467390664 | 0.247058257 | 8.706879293 | -0.772427167 | 0.471863257 | 0.734313128 | -4.670073491 |
| **A_51_P370678** | Gfi1b | 0.074750218 | -0.167647809 | 0.317148246 | 7.810080122 | 0.772384818 | 0.471886359 | 0.734313128 | -4.670082641 |
| **A_66_P137102** | Spryd7 | -0.11936967 | -0.506504217 | 0.267764877 | 8.382233616 | -0.772293554 | 0.471936147 | 0.734313128 | -4.670102358 |
| **A_51_P153063** | Fabp12 | -0.241807329 | -1.026041939 | 0.542427281 | 8.329920521 | -0.772278246 | 0.471944498 | 0.734313128 | -4.670105665 |
| **A_55_P2147427** | Prdx1 | 0.306397214 | -0.687674122 | 1.300468549 | 13.91857244 | 0.772000544 | 0.47209602 | 0.734460993 | -4.67016565 |
| **A_52_P286002** | Obsl1 | -0.150778907 | -0.640018801 | 0.338460988 | 8.383298368 | -0.771914273 | 0.472143099 | 0.734460993 | -4.670184281 |
| **A_66_P127036** | NA | 0.194768103 | -0.437231596 | 0.826767802 | 15.6256501 | 0.771882701 | 0.472160329 | 0.734460993 | -4.6701911 |

| | | | | | | | | | |
|---|---|---|---|---|---|---|---|---|---|
| A_52_P551903 | Slc33a1 | -0.076852693 | -0.326249132 | 0.172543746 | 8.197342827 | -0.771825616 | 0.472191484 | 0.734460993 | -4.670203427 |
| A_55_P2066707 | Trim27 | 0.143765129 | -0.322833027 | 0.610363284 | 12.38589469 | 0.771722006 | 0.472248033 | 0.734489861 | -4.670225801 |
| A_55_P2007249 | Spdef | 0.157072751 | -0.35280046 | 0.666945962 | 8.963830634 | 0.771594284 | 0.472317749 | 0.734539202 | -4.670253378 |
| A_52_P185907 | Crabp1 | -0.207734101 | -0.882133224 | 0.466665021 | 8.104832488 | -0.771509262 | 0.472364162 | 0.734552297 | -4.670271734 |
| A_55_P2048096 | Mzf1 | 0.10834201 | -0.243433192 | 0.460117211 | 8.429678767 | 0.771404222 | 0.472421507 | 0.734582388 | -4.670294409 |
| A_51_P310949 | Ppox | -0.100148832 | -0.425356028 | 0.225058363 | 9.080727082 | -0.771322657 | 0.472466039 | 0.734592553 | -4.670312016 |
| A_55_P2017055 | Kctd12b | 0.248937076 | -0.559780277 | 1.05765443 | 10.13775173 | 0.770980099 | 0.4726531 | 0.734723799 | -4.670385948 |
| A_52_P451834 | Stk35 | 0.101889844 | -0.229123315 | 0.432903004 | 9.614006417 | 0.770967349 | 0.472660064 | 0.734723799 | -4.670388699 |
| A_51_P172842 | Med19 | 0.09416299 | -0.211750853 | 0.400076832 | 10.08435551 | 0.770959306 | 0.472664457 | 0.734723799 | -4.670390435 |
| A_51_P299216 | Wdr90 | 0.271261529 | -0.61032015 | 1.152843208 | 9.30557283 | 0.770683382 | 0.472815172 | 0.73489899 | -4.670449968 |
| A_55_P2136902 | Cisd3 | -0.126957761 | -0.539621422 | 0.285705901 | 11.68137865 | -0.770572262 | 0.472875878 | 0.734934262 | -4.67047394 |
| A_52_P219266 | Frmd8 | 0.189113415 | -0.425708442 | 0.803935272 | 12.24115005 | 0.770412612 | 0.472963105 | 0.73499379 | -4.670508376 |
| A_55_P2408848 | Lgr4 | -0.094686552 | -0.402539785 | 0.213166681 | 7.780186866 | -0.770362137 | 0.472990685 | 0.73499379 | -4.670519263 |
| A_55_P2013414 | Dmwd | 0.150274947 | -0.338382585 | 0.638932478 | 12.64740862 | 0.770251107 | 0.473051358 | 0.73499379 | -4.670543208 |
| A_51_P130808 | Tpbpb | -2.088677162 | -8.880786831 | 4.703432506 | 10.36250034 | -0.770223871 | 0.473066242 | 0.73499379 | -4.670549082 |
| A_55_P2034570 | Ddx19a | 0.074184066 | -0.167111577 | 0.315479709 | 8.724528881 | 0.770036834 | 0.473168463 | 0.735073235 | -4.670589413 |
| A_51_P291417 | Thbd | 0.103190356 | -0.232479215 | 0.438859926 | 12.68044576 | 0.769976521 | 0.473201429 | 0.735073235 | -4.670602417 |
| A_52_P239403 | Ipo7 | -0.147343496 | -0.626673702 | 0.331986709 | 8.273361016 | -0.769921628 | 0.473231434 | 0.735073235 | -4.670614252 |
| A_66_P124858 | Diras2 | -0.665730074 | -2.832919776 | 1.501459627 | 9.198144029 | -0.769398958 | 0.4735172 | 0.735321153 | -4.670726911 |
| A_55_P2213828 | 4933412E | -0.090091437 | -0.383396934 | 0.203214059 | 8.641266484 | -0.769331755 | 0.473553951 | 0.735321153 | -4.670741393 |
| A_52_P1026777 | Ecscr | 0.370613304 | -0.835999167 | 1.577225776 | 11.42010084 | 0.769313833 | 0.473563753 | 0.735321153 | -4.670745255 |
| A_55_P2443568 | Ube4a | -0.184731329 | -0.786237158 | 0.4167745 | 10.07723293 | -0.769220874 | 0.473614594 | 0.735321153 | -4.670765285 |
| A_51_P499441 | Prkd2 | 0.316996324 | -0.715222383 | 1.349215031 | 9.744560247 | 0.769188612 | 0.473632239 | 0.735321153 | -4.670772236 |
| A_52_P404005 | NA | 0.137753672 | -0.310852426 | 0.58635977 | 10.99647763 | 0.769109823 | 0.473675335 | 0.735321153 | -4.670789212 |
| A_55_P2016682 | Alkbh7 | 0.299947445 | -0.676984961 | 1.276879852 | 11.92812265 | 0.769008246 | 0.473730898 | 0.735321153 | -4.670811095 |
| A_51_P222092 | Igbp1 | 0.222943888 | -0.503192556 | 0.949080332 | 9.893072385 | 0.769002378 | 0.473734108 | 0.735321153 | -4.670812359 |
| A_51_P293269 | Pde7a | 0.41889766 | -0.945661735 | 1.783457055 | 10.19205227 | 0.768892874 | 0.473794013 | 0.735321153 | -4.670835948 |
| A_55_P2040873 | NA | -0.06728571 | -0.286470436 | 0.151899017 | 7.827131919 | -0.768887669 | 0.473796861 | 0.735321153 | -4.670837069 |
| A_55_P2018929 | Spns2 | -0.157137114 | -0.669118503 | 0.354844274 | 10.71486802 | -0.768731973 | 0.473882046 | 0.735321153 | -4.670870604 |
| A_55_P2040121 | Cdv3 | -0.357789424 | -1.523560592 | 0.807981744 | 9.672815158 | -0.768713546 | 0.473892129 | 0.735321153 | -4.670874573 |
| A_55_P1956472 | Cybrd1 | 0.225535073 | -0.509321983 | 0.960392129 | 8.512559826 | 0.768708293 | 0.473895003 | 0.735321153 | -4.670875704 |
| A_52_P501875 | Ccdc173 | 0.170688481 | -0.385582443 | 0.726959404 | 8.351802281 | 0.768543048 | 0.473985426 | 0.735321153 | -4.670911291 |
| A_55_P2071097 | Adrm1 | -0.281461795 | -1.198835688 | 0.635912098 | 13.62171052 | -0.768463815 | 0.474028787 | 0.735321153 | -4.670928352 |
| A_55_P2129578 | Pkp3 | -0.335581169 | -1.42939821 | 0.758235871 | 11.84208157 | -0.76842811 | 0.474048328 | 0.735321153 | -4.67093604 |
| A_55_P2173066 | NA | 0.301274803 | -0.680730497 | 1.283280103 | 9.540170135 | 0.768421176 | 0.474052123 | 0.735321153 | -4.670937533 |
| A_55_P2134487 | NA | 0.132049216 | -0.298389296 | 0.562487729 | 11.56664008 | 0.768378231 | 0.474075627 | 0.735321153 | -4.67094678 |
| A_55_P1958986 | Spata7 | 0.100384632 | -0.226873873 | 0.427643138 | 8.346324819 | 0.768292575 | 0.47412251 | 0.735334879 | -4.670965221 |
| A_55_P2035877 | Faap20 | 0.208518503 | -0.502839399 | 0.919876405 | 10.94123085 | 0.780762856 | 0.474174757 | 0.73535671 | -4.652351103 |
| A_55_P2381538 | NA | 0.106567898 | -0.240926302 | 0.454062097 | 9.801957621 | 0.768120197 | 0.474216868 | 0.73535671 | -4.67100233 |
| A_55_P2126109 | Usp16 | -0.109960633 | -0.468546944 | 0.248625679 | 11.33398661 | -0.768057739 | 0.47425106 | 0.73535671 | -4.671015775 |
| A_55_P2031021 | Bex1 | -0.775145285 | -3.30328993 | 1.75299936 | 12.92787163 | -0.767947511 | 0.474311409 | 0.73535671 | -4.6710395 |
| A_51_P315555 | Nars | 0.088912271 | -0.20113243 | 0.378956973 | 10.32194567 | 0.767798244 | 0.474393139 | 0.73535671 | -4.671071625 |
| A_55_P2114110 | Cadm4 | -0.382356556 | -1.629677912 | 0.864964801 | 8.988442103 | -0.7677866 | 0.474399515 | 0.73535671 | -4.67107413 |
| A_51_P241943 | Sap30l | -0.217150118 | -0.925541562 | 0.491241326 | 13.13005831 | -0.767780553 | 0.474402826 | 0.73535671 | -4.671075432 |
| A_51_P199168 | Cidea | -0.084458326 | -0.360140693 | 0.191224042 | 8.242891112 | -0.767332907 | 0.474647996 | 0.735677757 | -4.671171746 |
| A_55_P2159480 | Fam160a1 | -0.141625456 | -0.603985907 | 0.320734994 | 9.481242051 | -0.767204232 | 0.474718485 | 0.735728032 | -4.671199424 |
| A_55_P2101570 | Zhx3 | 0.267626546 | -0.606189436 | 1.141442528 | 10.91238423 | 0.767113368 | 0.474768267 | 0.735737394 | -4.671218967 |
| A_55_P2099080 | Irak1 | 0.090420342 | -0.204829984 | 0.385670668 | 10.20767909 | 0.767054292 | 0.474800634 | 0.735737394 | -4.671231672 |

| | | | | | | | | | |
|---|---|---|---|---|---|---|---|---|---|
| **A_55_P2054854** | Art4 | 0.087107372 | -0.19746653 | 0.371681274 | 7.992731332 | 0.76667303 | 0.475009564 | 0.735913889 | -4.671313654 |
| **A_55_P2181271** | Nhsl2 | 0.229413978 | -0.520067257 | 0.978895214 | 9.294642754 | 0.766671745 | 0.475010269 | 0.735913889 | -4.671313931 |
| **A_51_P474459** | Socs3 | -0.138453679 | -0.590793125 | 0.313885767 | 10.55432349 | -0.766638075 | 0.475028723 | 0.735913889 | -4.671321169 |
| **A_51_P173022** | Nsmce1 | 0.304633965 | -0.690913608 | 1.300181537 | 11.9489218 | 0.766419684 | 0.475148433 | 0.73593872 | -4.671368116 |
| **A_55_P2096144** | Gnao1 | 0.131949272 | -0.299297797 | 0.563196342 | 8.231189465 | 0.766357106 | 0.475182739 | 0.73593872 | -4.671381566 |
| **A_55_P2168058** | Glod4 | 0.139981696 | -0.317530394 | 0.597493786 | 10.7703453 | 0.766335635 | 0.47519451 | 0.73593872 | -4.671386181 |
| **A_51_P125183** | Coq5 | -0.116635858 | -0.497873836 | 0.264602121 | 8.396642387 | -0.766277855 | 0.475226188 | 0.73593872 | -4.666231621 |
| **A_55_P2102759** | NA | -0.146232996 | -0.624265661 | 0.331799669 | 8.225668339 | -0.766192946 | 0.475272742 | 0.73593872 | -4.671416847 |
| **A_55_P2016739** | Rbm12b2 | 0.208255028 | -0.472527031 | 0.889037088 | 9.375279378 | 0.766192222 | 0.475273139 | 0.73593872 | -4.671417003 |
| **A_51_P139651** | Nos3 | 0.232424701 | -0.527494929 | 0.99234433 | 8.731592727 | 0.76606384 | 0.475343534 | 0.73598878 | -4.671444591 |
| **A_66_P136228** | Rai14 | 0.306465729 | -0.69571813 | 1.308649588 | 12.90426305 | 0.765922552 | 0.475421015 | 0.7360498 | -4.671474949 |
| **A_55_P2051701** | Pbx1 | 0.477114783 | -1.083298966 | 2.037528532 | 10.52290169 | 0.765831896 | 0.475470734 | 0.736067834 | -4.671494426 |
| **A_51_P153004** | Phax | -0.077305466 | -0.330196213 | 0.175585281 | 9.789939927 | -0.765645277 | 0.475573094 | 0.736167351 | -4.671534515 |
| **A_51_P322994** | Mrpl22 | 0.180500884 | -0.410190573 | 0.771192341 | 9.916275352 | 0.765365687 | 0.475726479 | 0.736345829 | -4.671594564 |
| **A_55_P2078269** | NA | 0.467621901 | -1.063233333 | 1.998477134 | 13.33190917 | 0.765087428 | 0.475879169 | 0.736454034 | -4.671654312 |
| **A_52_P227267** | Atp1a2 | -0.145122179 | -0.620236917 | 0.329992558 | 8.280647601 | -0.765042629 | 0.475903755 | 0.736454034 | -4.67166393 |
| **A_55_P1957159** | NA | 0.268578549 | -0.610733807 | 1.147890905 | 15.93162413 | 0.765030047 | 0.47591066 | 0.736454034 | -4.671666631 |
| **A_55_P2038697** | Zfp235 | 0.152868655 | -0.34773587 | 0.65347318 | 8.940699036 | 0.764846034 | 0.476011657 | 0.736551371 | -4.671706133 |
| **A_55_P1975415** | Cpped1 | 0.220813266 | -0.502363456 | 0.943989988 | 11.27218315 | 0.764770396 | 0.476053177 | 0.736556668 | -4.671722369 |
| **A_51_P186476** | Slc11a1 | -0.064759348 | -0.2769023 | 0.147383604 | 8.253986805 | -0.764582244 | 0.476156468 | 0.73665753 | -4.67176275 |
| **A_55_P2087281** | NA | 0.133767811 | -0.304709504 | 0.572245126 | 8.900979647 | 0.764108155 | 0.476416803 | 0.737001317 | -4.67186447 |
| **A_55_P2015782** | Bcas1 | 0.150963943 | -0.344043344 | 0.64597123 | 8.467749339 | 0.763856862 | 0.476554835 | 0.737013037 | -4.671918369 |
| **A_55_P1967295** | Pemt | 0.065637114 | -0.1496029 | 0.280877127 | 8.336088003 | 0.763794999 | 0.47658882 | 0.737013037 | -4.671931637 |
| **A_55_P2049612** | Ube2a | -0.164116345 | -0.702363114 | 0.374130424 | 11.62126036 | -0.763696412 | 0.476642983 | 0.737013037 | -4.671952778 |
| **A_55_P2109222** | Rps3 | 0.149625932 | -0.341113091 | 0.640364955 | 15.73735398 | 0.763671564 | 0.476656635 | 0.737013037 | -4.671958107 |
| **A_51_P334942** | Aldh1a1 | 0.656124237 | -1.495870307 | 2.80811878 | 10.85897656 | 0.76365159 | 0.47666761 | 0.737013037 | -4.67196239 |
| **A_52_P48681** | Cldn1 | -0.064517349 | -0.276132479 | 0.147097781 | 8.042038222 | -0.763625013 | 0.476682213 | 0.737013037 | -4.671968088 |
| **A_55_P1982404** | Gpm6b | 0.470129428 | -1.071915261 | 2.012174118 | 11.46180233 | 0.763608621 | 0.476691219 | 0.737013037 | -4.671971603 |
| **A_55_P1957468** | She | 0.30730998 | -0.700799615 | 1.315419575 | 10.57802231 | 0.763517968 | 0.476741031 | 0.737031113 | -4.67199104 |
| **A_52_P672740** | Rala | 0.167536806 | -0.382184725 | 0.717258336 | 8.836937943 | 0.763339664 | 0.476839016 | 0.737116935 | -4.672029266 |
| **A_52_P305685** | NA | -0.171353222 | -0.733642477 | 0.390936034 | 8.965303391 | -0.763278158 | 0.476872819 | 0.737116935 | -4.672042451 |
| **A_52_P485971** | Scap | -0.276101802 | -1.182201944 | 0.62999834 | 11.74529262 | -0.763208834 | 0.476910921 | 0.737116935 | -4.672057311 |
| **A_51_P208931** | N4bp3 | 0.402505045 | -0.918824231 | 1.72383432 | 11.71473899 | 0.762975541 | 0.477039159 | 0.737256203 | -4.672107311 |
| **A_55_P2234361** | Rnf150 | 0.535127965 | -1.221781731 | 2.292037661 | 10.30359643 | 0.762884082 | 0.47708944 | 0.737274976 | -4.67212691 |
| **A_52_P527106** | Arhgap12 | -0.075229402 | -0.322287654 | 0.171828849 | 9.467364216 | -0.76267339 | 0.477205284 | 0.737345416 | -4.672172054 |
| **A_65_P11306** | NA | 0.077855359 | -0.177830369 | 0.333541087 | 8.93129457 | 0.762662458 | 0.477211296 | 0.737345416 | -4.672174397 |
| **A_66_P114102** | Zfp36l2 | 0.117788145 | -0.269106419 | 0.504682708 | 12.57832249 | 0.762534152 | 0.477281853 | 0.737395504 | -4.672201884 |
| **A_52_P364130** | Map3k14 | 0.165108719 | -0.377334864 | 0.707552303 | 8.743210676 | 0.762369965 | 0.477372151 | 0.737476082 | -4.672237053 |
| **A_52_P472319** | Plxnd1 | 0.202569091 | -0.463159682 | 0.868297865 | 9.237916046 | 0.762124983 | 0.477506908 | 0.737625324 | -4.67228952 |
| **A_55_P2006118** | Rbp4 | -1.034387407 | -4.434221691 | 2.365446877 | 11.50812262 | -0.762036584 | 0.47755554 | 0.737641512 | -4.672308449 |
| **A_51_P148037** | Sh3pxd2b | 0.306087651 | -0.700216357 | 1.312391658 | 9.354010285 | 0.761845578 | 0.477660632 | 0.737716935 | -4.672349345 |
| **A_55_P2129219** | NA | 0.41730258 | -0.954700975 | 1.789306134 | 13.59474013 | 0.76180914 | 0.477680682 | 0.737716935 | -4.672357146 |
| **A_52_P159429** | AU021092 | 0.337212721 | -0.771654575 | 1.446080016 | 9.225177664 | 0.761683803 | 0.477749654 | 0.737764522 | -4.672383978 |
| **A_51_P373890** | Guca1b | -0.122179869 | -0.524014545 | 0.279654808 | 8.654427458 | -0.761557304 | 0.477819273 | 0.7378131 | -4.672411054 |
| **A_55_P2136533** | Kmt2d | -0.233809611 | -1.003128329 | 0.535509107 | 8.516400699 | -0.761213367 | 0.478008594 | 0.737962641 | -4.672484658 |
| **A_55_P2132671** | NA | 0.085337442 | -0.195509252 | 0.366184135 | 8.376258896 | 0.761063073 | 0.47809134 | 0.737962641 | -4.672516815 |
| **A_66_P124732** | Zfp606 | 0.151394132 | -0.346846609 | 0.649634872 | 8.565230334 | 0.761062196 | 0.478091823 | 0.737962641 | -4.672517002 |
| **A_51_P221625** | Stk24 | -0.277224871 | -1.189605324 | 0.635155582 | 10.12451471 | -0.761038387 | 0.478104933 | 0.737962641 | -4.672522096 |

| | | | | | | | | | |
|---|---|---|---|---|---|---|---|---|---|
| **A_55_P2076832** | Nrd1 | 0.114165088 | -0.26159626 | 0.489926436 | 8.691058463 | 0.760977043 | 0.47813871 | 0.737962641 | -4.67253522 |
| **A_52_P428125** | Atr | 0.19150138 | -0.438828094 | 0.821830855 | 10.4949139 | 0.760947402 | 0.478155032 | 0.737962641 | -4.672541561 |
| **A_51_P261107** | Ogt | -0.170443677 | -0.731499279 | 0.390611925 | 9.680683015 | -0.760896063 | 0.478183302 | 0.737962641 | -4.672552543 |
| **A_55_P2089070** | 4933431K | 0.096287402 | -0.220754845 | 0.413329649 | 8.678536886 | 0.760681191 | 0.478301637 | 0.738086347 | -4.672598502 |
| **A_55_P2034003** | Gtf3c1 | 0.186343412 | -0.427341084 | 0.800027909 | 9.626599693 | 0.760535058 | 0.478382127 | 0.73815164 | -4.672629754 |
| **A_55_P1979620** | Eif3h | 0.451010221 | -1.034490167 | 1.936510608 | 13.87105459 | 0.760438261 | 0.478435448 | 0.738158276 | -4.672650453 |
| **A_52_P527842** | Reps1 | -0.114165457 | -0.490234299 | 0.261903385 | 8.84121317 | -0.760357288 | 0.478480056 | 0.738158276 | -4.672667767 |
| **A_55_P2121456** | Fam43a | -0.099042675 | -0.425312323 | 0.227226973 | 9.908533571 | -0.760319334 | 0.478500966 | 0.738158276 | -4.672675881 |
| **A_55_P2012146** | NA | 0.304582202 | -0.698886389 | 1.308050794 | 8.724500291 | 0.760240645 | 0.47854432 | 0.738166258 | -4.672692705 |
| **A_55_P2141860** | Aen | 0.168207313 | -0.386031837 | 0.722446463 | 11.63586377 | 0.760147759 | 0.478595498 | 0.738186308 | -4.672712562 |
| **A_55_P2159746** | Mfsd2b | -0.065483256 | -0.281294962 | 0.15032845 | 7.698887328 | -0.759986038 | 0.478684614 | 0.738244823 | -4.672747131 |
| **A_52_P82510** | Nmt1 | -0.183262122 | -0.787271336 | 0.420747092 | 11.1575054 | -0.759940329 | 0.478709804 | 0.738244823 | -4.672756901 |
| **A_51_P354126** | Reg3g | 0.906015001 | -2.080377621 | 3.892407622 | 10.69418533 | 0.75986918 | 0.478749015 | 0.738246408 | -4.672772107 |
| **A_55_P2105366** | NA | -0.134234925 | -0.576741429 | 0.30827158 | 8.524199875 | -0.759794613 | 0.478790113 | 0.7382509 | -4.672788043 |
| **A_55_P2059765** | NA | 0.698288825 | -1.604422349 | 3.000999999 | 12.95767518 | 0.759531813 | 0.478934974 | 0.738415375 | -4.672844198 |
| **A_51_P513959** | NA | 0.065246607 | -0.149940947 | 0.280434161 | 9.198622461 | 0.759435909 | 0.478987847 | 0.738421085 | -4.672864688 |
| **A_52_P139588** | Edem3 | -0.10123753 | -0.435172269 | 0.232697209 | 10.29197349 | -0.759329524 | 0.479046503 | 0.738421085 | -4.672887415 |
| **A_55_P2112510** | NA | 0.339584212 | -0.780561593 | 1.459730018 | 15.33984665 | 0.759317274 | 0.479053257 | 0.738421085 | -4.672890032 |
| **A_51_P247637** | Rnf144a | 0.497686657 | -1.144563598 | 2.139936913 | 9.861882752 | 0.759044093 | 0.4792039 | 0.738594404 | -4.67294838 |
| **A_55_P2016862** | Ncoa4 | 0.098015 | -0.22550227 | 0.42153227 | 12.31629288 | 0.758831624 | 0.479321087 | 0.738670555 | -4.672993752 |
| **A_55_P2129034** | Rheb | 0.143727868 | -0.330713651 | 0.618169387 | 10.81079076 | 0.758767361 | 0.479356536 | 0.738670555 | -4.673007473 |
| **A_55_P1973254** | Cmtm7 | -0.312497953 | -1.34429137 | 0.719295465 | 11.30880951 | -0.758585905 | 0.479456639 | 0.738670555 | -4.673046213 |
| **A_51_P383218** | Tmem161 | -0.07058666 | -0.303650192 | 0.162476872 | 8.577346161 | -0.758575222 | 0.479462533 | 0.738670555 | -4.673048494 |
| **A_52_P58059** | Usp34 | -0.074374649 | -0.319961701 | 0.171212403 | 9.137192495 | -0.758524842 | 0.479490329 | 0.738670555 | -4.673059248 |
| **A_55_P2052166** | Akap13 | -0.170899281 | -0.735214386 | 0.393415825 | 10.35923106 | -0.758523263 | 0.4794912 | 0.738670555 | -4.673059585 |
| **A_51_P108581** | Grk3 | 0.368952391 | -0.849650929 | 1.587555711 | 9.228898364 | 0.75833014 | 0.479597761 | 0.738670555 | -4.658833329 |
| **A_51_P416919** | Ube2j2 | -0.129838089 | -0.558697529 | 0.299021351 | 9.618934789 | -0.758293768 | 0.479617833 | 0.738670555 | -4.67310857 |
| **A_65_P20683** | NA | 0.203930174 | -0.469679521 | 0.87753987 | 8.911981306 | 0.758269358 | 0.479631303 | 0.738670555 | -4.673113779 |
| **A_55_P2056674** | Lrrk1 | 0.435750985 | -1.003788569 | 1.87529054 | 11.58381036 | 0.758167425 | 0.479687557 | 0.738670555 | -4.673135533 |
| **A_51_P303089** | Ttc28 | 0.280393076 | -0.645926029 | 1.20671218 | 9.446838497 | 0.758153264 | 0.479695372 | 0.738670555 | -4.673138555 |
| **A_55_P2168667** | Rbpms2 | 0.238690265 | -0.549945199 | 1.027325729 | 10.43840308 | 0.758069022 | 0.479741868 | 0.738670555 | -4.673156531 |
| **A_55_P2045693** | NA | -0.064410786 | -0.277228916 | 0.148407343 | 7.860984944 | -0.758054319 | 0.479749983 | 0.738670555 | -4.673159668 |
| **A_55_P2169332** | NA | 0.283193838 | -0.652722714 | 1.219110391 | 15.65271816 | 0.757874016 | 0.479849508 | 0.738764961 | -4.673198138 |
| **A_51_P403334** | Cuedc2 | 0.301077986 | -0.694156317 | 1.296312289 | 13.2974458 | 0.757711731 | 0.479939099 | 0.738765175 | -4.673232758 |
| **A_51_P160413** | Yod1 | 0.117051454 | -0.26992715 | 0.504030058 | 8.380690259 | 0.757600416 | 0.480000559 | 0.738765175 | -4.673256502 |
| **A_51_P124606** | Nat2 | 0.140996053 | -0.325157294 | 0.607149401 | 8.550565013 | 0.757579988 | 0.480011838 | 0.738765175 | -4.666772353 |
| **A_55_P2173837** | Mpp6 | 0.138444032 | -0.319276682 | 0.596164747 | 11.24067003 | 0.757572182 | 0.480016148 | 0.738765175 | -4.673262524 |
| **A_55_P2418745** | D230034L | 0.174769531 | -0.403083689 | 0.75262275 | 8.502259259 | 0.757527718 | 0.4800407 | 0.738765175 | -4.673272007 |
| **A_55_P2076906** | NA | 0.615826137 | -1.422598811 | 2.654251085 | 12.40238455 | 0.756682582 | 0.480507528 | 0.73942475 | -4.673452189 |
| **A_55_P1973033** | Ltbp1 | 0.251678287 | -0.58167562 | 1.085032194 | 10.31276318 | 0.756426299 | 0.480649155 | 0.739583825 | -4.673506801 |
| **A_51_P229676** | Plscr1 | 0.267123185 | -0.617501905 | 1.151748276 | 9.84606844 | 0.756314937 | 0.480710705 | 0.739619669 | -4.673530528 |
| **A_55_P2090600** | Mageb16- | -0.109406155 | -0.471831243 | 0.253018932 | 7.722623278 | -0.756090619 | 0.480834702 | 0.739670214 | -4.673578314 |
| **A_55_P1985840** | Mettl16 | 0.077562018 | -0.179409501 | 0.334533536 | 9.875941193 | 0.755987219 | 0.480891866 | 0.739670214 | -4.673600338 |
| **A_55_P2048744** | Eif2ak3 | -0.331810372 | -1.431316252 | 0.767695508 | 9.752444614 | -0.755862408 | 0.480960875 | 0.739670214 | -4.67362692 |
| **A_55_P2049072** | NA | 0.181036384 | -0.418902433 | 0.780975201 | 13.30466013 | 0.755804112 | 0.480993109 | 0.739670214 | -4.673639334 |
| **A_55_P2105181** | Bhmt | 0.069697547 | -0.161276233 | 0.300671327 | 7.806071959 | 0.75579699 | 0.480997047 | 0.739670214 | -4.673640851 |
| **A_51_P151775** | Mief1 | -0.158037597 | -0.68184445 | 0.365769257 | 8.976263357 | -0.755682884 | 0.481060145 | 0.739670214 | -4.673665149 |
| **A_55_P2168243** | NA | 0.485882116 | -1.124721127 | 2.096485359 | 11.1260066 | 0.75560129 | 0.481105269 | 0.739670214 | -4.673682522 |

| | | | | | | | | | |
|---|---|---|---|---|---|---|---|---|---|
| **A_55_P2005691** | NA | 0.198964378 | -0.46058567 | 0.858514426 | 15.85745116 | 0.755575587 | 0.481119484 | 0.739670214 | -4.673687994 |
| **A_55_P1989215** | Entpd8 | -0.141468285 | -0.632331711 | 0.349395142 | 8.179075891 | -0.767646225 | 0.481193638 | 0.739670214 | -4.654735609 |
| **A_51_P239286** | Bcl2l12 | 0.139283097 | -0.322597241 | 0.601163435 | 9.699920797 | 0.755299654 | 0.481272107 | 0.739670214 | -4.673746736 |
| **A_52_P27332** | Ppp2r5c | 0.14491413 | -0.335659036 | 0.625487296 | 10.55545649 | 0.75526883 | 0.481289159 | 0.739670214 | -4.673753297 |
| **A_65_P02321** | Usp36 | 0.091559474 | -0.212076529 | 0.395195478 | 8.813845456 | 0.755266756 | 0.481290306 | 0.739670214 | -4.673753738 |
| **A_55_P2101340** | Ramp3 | -1.170041735 | -5.050265061 | 2.710181591 | 9.534875984 | -0.755257378 | 0.481295494 | 0.739670214 | -4.673755734 |
| **A_55_P1980556** | NA | -0.068015532 | -0.293589815 | 0.157558751 | 7.670295738 | -0.75521196 | 0.48132062 | 0.739670214 | -4.673765401 |
| **A_55_P2065866** | Cygb | 0.542824929 | -1.25762464 | 2.343274498 | 11.5094012 | 0.755142944 | 0.481358801 | 0.739670214 | -4.67378009 |
| **A_66_P138046** | Gm12298 | -0.077343047 | -0.333896176 | 0.179210083 | 7.937095902 | -0.755082327 | 0.481392339 | 0.739670214 | -4.673792991 |
| **A_55_P1983558** | Tmed1 | 0.39633035 | -0.91833443 | 1.710995129 | 13.14180949 | 0.755079445 | 0.481393933 | 0.739670214 | -4.673793604 |
| **A_51_P202033** | Wls | 0.4520808 | -1.047709771 | 1.95187137 | 11.39066297 | 0.754980595 | 0.481448627 | 0.739695466 | -4.67381464 |
| **A_52_P288251** | Tmem204 | 0.39753758 | -0.921864152 | 1.716939312 | 10.08916159 | 0.754660268 | 0.481625894 | 0.739909021 | -4.673882795 |
| **A_55_P2069530** | NA | 0.305265808 | -0.708151564 | 1.31868318 | 11.13659407 | 0.754466852 | 0.481732951 | 0.739971634 | -4.673923938 |
| **A_55_P2049086** | Map3k7 | -0.148678804 | -0.64227283 | 0.344915222 | 10.57788969 | -0.754448344 | 0.481743196 | 0.739971634 | -4.673927874 |
| **A_55_P2253348** | Galnt7 | 0.17331026 | -0.402143481 | 0.748764001 | 10.25608523 | 0.754334901 | 0.481805997 | 0.740009306 | -4.673952002 |
| **A_55_P2008016** | Armc3 | 0.303622064 | -0.704713335 | 1.311957462 | 8.848952721 | 0.754186336 | 0.481888249 | 0.740076845 | -4.673983596 |
| **A_51_P414126** | Rab19 | 0.359746383 | -0.835565529 | 1.555058294 | 9.617500348 | 0.753816317 | 0.48209315 | 0.740324554 | -4.674062266 |
| **A_52_P69789** | Exosc1 | 0.069997724 | -0.162640993 | 0.302636442 | 8.515631657 | 0.753619751 | 0.482202025 | 0.740324554 | -4.674104047 |
| **A_51_P427530** | Pgm1 | -0.094496716 | -0.408566938 | 0.219573506 | 8.749038071 | -0.753598975 | 0.482213534 | 0.740324554 | -4.674108463 |
| **A_55_P2038299** | Rpl31 | 0.206839503 | -0.480642574 | 0.894321581 | 15.40855325 | 0.753568006 | 0.482230689 | 0.740324554 | -4.674115045 |
| **A_55_P2158478** | Nkiras2 | -0.074347047 | -0.321463887 | 0.172769793 | 11.95767128 | -0.7535494 | 0.482240997 | 0.740324554 | -4.674118999 |
| **A_51_P236324** | Gtf2e1 | 0.097355201 | -0.226335503 | 0.421045905 | 9.75699577 | 0.753319616 | 0.4823683 | 0.740435758 | -4.674167829 |
| **A_55_P1971962** | Tmem176 | 0.306099097 | -0.711685154 | 1.323883348 | 9.371821029 | 0.7532804 | 0.482390028 | 0.740435758 | -4.674176161 |
| **A_55_P2010957** | Gm4971 | 0.118111988 | -0.274661728 | 0.510885704 | 13.02727575 | 0.753185424 | 0.482442655 | 0.740457751 | -4.674196341 |
| **A_55_P1983036** | Tdrd3 | 0.199642048 | -0.464336621 | 0.863620718 | 10.29568895 | 0.753092349 | 0.482494232 | 0.74047813 | -4.674216114 |
| **A_55_P2025954** | Acly | 0.072712059 | -0.169155609 | 0.314579726 | 13.31799162 | 0.752972251 | 0.482560789 | 0.740488166 | -4.674241626 |
| **A_51_P195374** | Baiap2l1 | -0.066316448 | -0.286922815 | 0.154289919 | 8.566787831 | -0.752928192 | 0.482585209 | 0.740488166 | -4.674250985 |
| **A_51_P298666** | Scaf4 | -0.12056771 | -0.521677362 | 0.280541943 | 9.983530599 | -0.75286697 | 0.482619141 | 0.740488166 | -4.674263988 |
| **A_52_P375546** | Lrrc46 | 0.235416745 | -0.54784281 | 1.0186763 | 8.639828865 | 0.752804132 | 0.482653971 | 0.740488166 | -4.674277334 |
| **A_66_P137152** | Zbtb8os | 0.081272778 | -0.189243195 | 0.351788751 | 12.31235413 | 0.752493124 | 0.482826383 | 0.740548244 | -4.674343378 |
| **A_51_P323620** | Thyn1 | 0.093308247 | -0.217291743 | 0.403908238 | 13.38746737 | 0.752434915 | 0.482858657 | 0.740548244 | -4.674355737 |
| **A_55_P2035677** | Srp19 | 0.098530976 | -0.229465154 | 0.426527107 | 11.18789106 | 0.752409743 | 0.482872614 | 0.740548244 | -4.674361081 |
| **A_55_P1956807** | Gm14391 | -0.095423523 | -0.413087617 | 0.222240572 | 10.08557945 | -0.752380726 | 0.482888703 | 0.740548244 | -4.674367242 |
| **A_55_P1994355** | Ptges3 | 0.09525967 | -0.221878319 | 0.412397659 | 13.49116694 | 0.752334798 | 0.48291417 | 0.740548244 | -4.674376992 |
| **A_51_P342716** | Ppp1cb | 0.162583301 | -0.378706003 | 0.703872605 | 12.84280872 | 0.752310005 | 0.482927918 | 0.740548244 | -4.674382255 |
| **A_66_P140742** | Azin2 | -0.069455319 | -0.300711696 | 0.161801059 | 8.462391028 | -0.752249896 | 0.48296125 | 0.740548244 | -4.674395015 |
| **A_51_P433870** | Vegfc | 0.429779634 | -1.001546892 | 1.86110616 | 9.728097235 | 0.752068601 | 0.483061793 | 0.740643673 | -4.674433497 |
| **A_55_P2024749** | Hspb11 | 0.193298298 | -0.45071245 | 0.837309046 | 10.87911256 | 0.751770465 | 0.483227166 | 0.740755593 | -4.674496766 |
| **A_55_P2048607** | Hp | 1.286436264 | -3.000287215 | 5.573159742 | 13.21919978 | 0.751645653 | 0.483296411 | 0.740755593 | -4.674523248 |
| **A_66_P114627** | S100a16 | 0.574799181 | -1.340578405 | 2.490176767 | 12.69239496 | 0.751643811 | 0.483297433 | 0.740755593 | -4.674523639 |
| **A_55_P1955072** | Alad | -0.064226326 | -0.27824557 | 0.149792917 | 10.99863738 | -0.751641244 | 0.483298856 | 0.740755593 | -4.674524183 |
| **A_55_P2012615** | NA | 0.246807976 | -0.575770375 | 1.069386327 | 15.58403018 | 0.751505669 | 0.48337408 | 0.740755593 | -4.674552946 |
| **A_66_P101835** | Cd38 | 0.31435442 | -0.733364596 | 1.362073437 | 9.306750952 | 0.751493261 | 0.483380965 | 0.740755593 | -4.674555578 |
| **A_55_P1992216** | Chchd2 | -0.095741808 | -0.414858714 | 0.223375098 | 16.99723681 | -0.751453576 | 0.483402985 | 0.740755593 | -4.674563996 |
| **A_55_P2002752** | Zswim6 | 0.248930994 | -0.580880064 | 1.078742051 | 8.78945486 | 0.751363511 | 0.483452964 | 0.740773467 | -4.6745831 |
| **A_55_P1980302** | Psenen | -0.066340717 | -0.287510621 | 0.154829186 | 12.93708285 | -0.751284591 | 0.483496762 | 0.740781868 | -4.674599839 |
| **A_51_P517430** | Cd1d1 | 0.321129804 | -0.749814314 | 1.392073923 | 9.696152327 | 0.751041871 | 0.483631478 | 0.740929556 | -4.674651313 |
| **A_55_P2087622** | Il4i1 | 0.348006815 | -0.812938477 | 1.508952107 | 8.457343532 | 0.750803589 | 0.483763757 | 0.741073487 | -4.674701835 |

| | | | | | | | | | |
|---|---|---|---|---|---|---|---|---|---|
| **A_55_P1953884** | Dpm1 | -0.184427894 | -0.799871414 | 0.431015627 | 11.00661994 | -0.750565759 | 0.48389581 | 0.74121705 | -4.67475225 |
| **A_51_P214319** | Cox11 | -0.13549951 | -0.587732594 | 0.316733574 | 11.03535319 | -0.750456871 | 0.483956278 | 0.741239411 | -4.674775328 |
| **A_55_P2177910** | Lepr | -1.100568268 | -4.774681127 | 2.57354459 | 12.27585764 | -0.750265264 | 0.484062694 | 0.741239411 | -4.674815933 |
| **A_55_P2137527** | Fam183b | 0.770741795 | -1.802437876 | 3.343921465 | 11.26088046 | 0.750220874 | 0.48408735 | 0.741239411 | -4.674825339 |
| **A_51_P264495** | Pgam2 | 0.655184592 | -1.532220308 | 2.842589492 | 10.38096866 | 0.75021344 | 0.48409148 | 0.741239411 | -4.674826914 |
| **A_51_P330615** | Zfp119b | 0.170530839 | -0.398845919 | 0.739907597 | 10.38320664 | 0.750159371 | 0.484121513 | 0.741239411 | -4.674838371 |
| **A_52_P283802** | Tapt1 | 0.086893423 | -0.203243793 | 0.377030639 | 9.363868682 | 0.750125297 | 0.484140441 | 0.741239411 | -4.67484559 |
| **A_55_P1959683** | Xlr3b | -0.390116153 | -1.693339595 | 0.91310729 | 8.468701506 | -0.7497654 | 0.48434039 | 0.741390486 | -4.67492183 |
| **A_51_P234847** | 1700001L | 0.276331141 | -0.646832228 | 1.199494509 | 8.918701817 | 0.749724343 | 0.484363204 | 0.741390486 | -4.674930526 |
| **A_55_P1974567** | Plch2 | 0.371814539 | -0.870340663 | 1.613969742 | 9.545565619 | 0.74972303 | 0.484363934 | 0.741390486 | -4.674930804 |
| **A_55_P2073855** | Mrpl27 | 0.191597372 | -0.448533574 | 0.831728318 | 12.5311052 | 0.74967162 | 0.484392501 | 0.741390486 | -4.674941692 |
| **A_55_P2132844** | Rps16 | 0.084785405 | -0.198632293 | 0.368203104 | 15.83186171 | 0.749280585 | 0.484609832 | 0.741664409 | -4.675024493 |
| **A_55_P1991778** | Fam219a | 0.132228842 | -0.309858669 | 0.574316354 | 13.83599146 | 0.749149157 | 0.484682892 | 0.741717512 | -4.675052316 |
| **A_51_P329283** | Senp5 | -0.130659205 | -0.567631274 | 0.306312865 | 9.795230375 | -0.748922154 | 0.484809102 | 0.741832417 | -4.675100365 |
| **A_55_P2228327** | 4921529L | -0.16369453 | -0.711231276 | 0.383842216 | 8.708126373 | -0.748809322 | 0.484871842 | 0.741832417 | -4.675124243 |
| **A_55_P2053077** | Chuk | -0.084665139 | -0.367859778 | 0.1985295 | 11.24361864 | -0.748807085 | 0.484873087 | 0.741832417 | -4.675124717 |
| **A_51_P172155** | Hal | -0.131929911 | -0.573373096 | 0.309513275 | 7.807804754 | -0.748546521 | 0.485017996 | 0.741888736 | -4.675179851 |
| **A_51_P329251** | Chchd1 | 0.219059929 | -0.513966163 | 0.952086021 | 13.99875063 | 0.748503533 | 0.485041907 | 0.741888736 | -4.675188946 |
| **A_55_P2252184** | NA | -0.256660943 | -1.11552202 | 0.602200134 | 8.863491908 | -0.748492082 | 0.485048276 | 0.741888736 | -4.675191369 |
| **A_55_P2081378** | NA | 0.081903796 | -0.192179728 | 0.355987321 | 16.07899745 | 0.748464917 | 0.485063386 | 0.741888736 | -4.675197115 |
| **A_51_P228658** | Isyna1 | -0.133620821 | -0.580883586 | 0.313641944 | 11.54065797 | -0.748275864 | 0.485168552 | 0.741892851 | -4.675237107 |
| **A_52_P309718** | Tasor | -0.080407614 | -0.349567452 | 0.188752224 | 8.8114532 | -0.748233694 | 0.485192012 | 0.741892851 | -4.675246027 |
| **A_55_P2017066** | NA | -0.373705049 | -1.62471611 | 0.877306011 | 12.16769125 | -0.74820079 | 0.485210318 | 0.741892851 | -4.675252986 |
| **A_52_P357611** | Neu3 | 0.319175584 | -0.749316955 | 1.387668123 | 9.3311583 | 0.748184167 | 0.485219567 | 0.741892851 | -4.675256502 |
| **A_51_P374137** | Bco1 | -0.067592015 | -0.293944769 | 0.15876074 | 7.67552188 | -0.747928278 | 0.485361947 | 0.741943409 | -4.675310616 |
| **A_55_P2061979** | NA | 0.178039471 | -0.418223647 | 0.774302589 | 16.34443518 | 0.747874447 | 0.485391904 | 0.741943409 | -4.675321999 |
| **A_51_P156158** | Phlpp1 | -0.074135209 | -0.322435498 | 0.17416508 | 8.59855627 | -0.747820963 | 0.485421668 | 0.741943409 | -4.675333307 |
| **A_55_P2178378** | Gas5 | 0.171360707 | -0.402583704 | 0.745305118 | 8.718772641 | 0.747810846 | 0.485427298 | 0.741943409 | -4.675335446 |
| **A_55_P2180477** | Rpl31 | 0.097259818 | -0.228509365 | 0.423029002 | 16.24507575 | 0.747779922 | 0.485444509 | 0.741943409 | -4.675341984 |
| **A_52_P366525** | Coq10b | -0.126045226 | -0.548312586 | 0.296222135 | 8.218933704 | -0.747634326 | 0.485525544 | 0.741963088 | -4.675372765 |
| **A_55_P2134496** | Amfr | -0.140422565 | -0.610869468 | 0.330024338 | 11.69853744 | -0.747612636 | 0.485537616 | 0.741963088 | -4.67537735 |
| **A_52_P8922** | Sntb2 | 0.140014973 | -0.329105744 | 0.609135689 | 9.672376846 | 0.747549945 | 0.485572512 | 0.741963088 | -4.675390602 |
| **A_55_P1981002** | NA | -0.217971832 | -0.948403876 | 0.512460211 | 8.940713431 | -0.747430652 | 0.485638919 | 0.741972322 | -4.675415817 |
| **A_55_P1979330** | Dapp1 | 0.086728074 | -0.203937891 | 0.377394038 | 8.569494496 | 0.747335933 | 0.485691651 | 0.741972322 | -4.675435835 |
| **A_52_P99531** | Ippk | -0.391283429 | -1.702916966 | 0.920350108 | 9.783279259 | -0.747186968 | 0.48577459 | 0.741972322 | -4.675467315 |
| **A_55_P1964138** | Pign | -0.132880506 | -0.578324846 | 0.312563833 | 9.466521652 | -0.747167847 | 0.485785237 | 0.741972322 | -4.675471356 |
| **A_51_P321150** | Lyz2 | -0.301498901 | -1.312194003 | 0.709196202 | 9.109819152 | -0.747163963 | 0.4857874 | 0.741972322 | -4.675472176 |
| **A_55_P1970033** | Per1 | -0.162895852 | -0.708989105 | 0.3831974 | 12.75574184 | -0.747125504 | 0.485808815 | 0.741972322 | -4.675480303 |
| **A_55_P2155834** | NA | 0.176369514 | -0.414989759 | 0.767728788 | 17.23989548 | 0.747003184 | 0.48587693 | 0.742017738 | -4.675506147 |
| **A_51_P144319** | Efna5 | 0.203348153 | -0.478572204 | 0.88526851 | 9.193736236 | 0.746890476 | 0.485939698 | 0.742037953 | -4.675529958 |
| **A_55_P1981105** | Ap1b1 | -0.257080865 | -1.119248277 | 0.605086548 | 12.25767075 | -0.746841588 | 0.485966927 | 0.742037953 | -4.675540286 |
| **A_51_P222590** | Arf6 | -0.152585644 | -0.664452324 | 0.359281036 | 10.54632319 | -0.746632964 | 0.486083131 | 0.742156776 | -4.675584352 |
| **A_51_P213030** | Macrod1 | 0.298693626 | -0.703535173 | 1.300922426 | 9.939733508 | 0.746464954 | 0.486176728 | 0.742182363 | -4.675619834 |
| **A_52_P552017** | Rbsn | 0.103348774 | -0.243498393 | 0.45019594 | 9.737593666 | 0.746307013 | 0.486264727 | 0.742182363 | -4.675653184 |
| **A_55_P1975742** | Dap3 | 0.098972182 | -0.233200498 | 0.431144862 | 11.15483401 | 0.74627618 | 0.486281907 | 0.742182363 | -4.675659694 |
| **A_55_P2045234** | NA | 0.115809395 | -0.272939615 | 0.504558405 | 9.029846232 | 0.746147765 | 0.486353465 | 0.742182363 | -4.675686805 |
| **A_66_P139568** | Helz | -0.429304754 | -1.870608103 | 1.011998596 | 11.00106666 | -0.746037482 | 0.486414925 | 0.742182363 | -4.675710086 |
| **A_55_P2147616** | NA | 0.143058273 | -0.337231133 | 0.623347679 | 12.02562642 | 0.746036912 | 0.486415243 | 0.742182363 | -4.675710206 |

| | | | | | | | | | |
|---|---|---|---|---|---|---|---|---|---|
| **A_55_P2232754** | Smarcal1 | 0.143936399 | -0.339335843 | 0.627208641 | 8.88062376 | 0.745983338 | 0.486445101 | 0.742182363 | -4.675721515 |
| **A_55_P2183288** | Cryzl1 | 0.14042041 | -0.331050479 | 0.611891298 | 10.47586884 | 0.74597745 | 0.486448383 | 0.742182363 | -4.675722758 |
| **A_55_P2174198** | H4c9 | -0.247648521 | -1.079197093 | 0.583900051 | 10.36403673 | -0.745930666 | 0.486474458 | 0.742182363 | -4.675732633 |
| **A_55_P2004153** | NA | -0.116316855 | -0.506914992 | 0.274281282 | 9.992953523 | -0.745869465 | 0.48650857 | 0.742182363 | -4.67574555 |
| **A_51_P348652** | Spast | -0.239266932 | -1.042892115 | 0.56435825 | 10.59633722 | -0.745726351 | 0.486588346 | 0.742182363 | -4.675775753 |
| **A_51_P446106** | Lrrc38 | -0.084685235 | -0.369121669 | 0.199751199 | 8.12462126 | -0.745714895 | 0.486594732 | 0.742182363 | -4.675778171 |
| **A_65_P12152** | Rasa2 | -0.094757989 | -0.413029486 | 0.223513507 | 8.709279375 | -0.745707372 | 0.486598926 | 0.742182363 | -4.675779758 |
| **A_55_P2011862** | Mybpc3 | 1.087678363 | -2.737886552 | 4.913243279 | 10.88773673 | 0.757299411 | 0.48678456 | 0.742406932 | -4.656599869 |
| **A_52_P248483** | Zfx | -0.205804286 | -0.897799752 | 0.48619118 | 8.812567265 | -0.744906068 | 0.487045769 | 0.742652391 | -4.675948792 |
| **A_55_P2178525** | Rpl26 | 0.230338176 | -0.544185966 | 1.004862318 | 16.4316511 | 0.744871419 | 0.487065097 | 0.742652391 | -4.675956098 |
| **A_55_P2166538** | Mrps16 | 0.192092301 | -0.45387342 | 0.838058021 | 13.4452088 | 0.744819145 | 0.487094258 | 0.742652391 | -4.67596712 |
| **A_51_P117226** | NA | 0.139809129 | -0.330344896 | 0.609963155 | 8.636589282 | 0.744810376 | 0.48709915 | 0.742652391 | -4.668002275 |
| **A_55_P1967149** | Irx3 | 0.284457933 | -0.672339653 | 1.241255519 | 12.24528461 | 0.744643365 | 0.487192326 | 0.74273588 | -4.676004181 |
| **A_51_P370470** | Cyp20a1 | 0.142147358 | -0.336122196 | 0.620416912 | 8.962331062 | 0.744417204 | 0.487318522 | 0.742781132 | -4.676051856 |
| **A_51_P407690** | Sppl2b | -0.101034559 | -0.441024404 | 0.238955286 | 12.03868675 | -0.744310848 | 0.487377875 | 0.742781132 | -4.676074272 |
| **A_55_P2331974** | NA | -0.233468058 | -1.019255182 | 0.552319066 | 8.557380236 | -0.744171285 | 0.487455769 | 0.742781132 | -4.676103684 |
| **A_52_P633752** | Apc | -0.078353328 | -0.342093468 | 0.185386811 | 8.06531073 | -0.744100445 | 0.487495309 | 0.742781132 | -4.676118612 |
| **A_66_P109002** | Kcng2 | 0.070191559 | -0.166097815 | 0.306480934 | 7.765126771 | 0.744031022 | 0.487534061 | 0.742781132 | -4.67613324 |
| **A_55_P2081178** | Msl3l2 | 0.286101569 | -0.677099114 | 1.249302252 | 9.525354707 | 0.743967224 | 0.487569674 | 0.742781132 | -4.676146682 |
| **A_55_P2286503** | A930001C | 0.148093836 | -0.350549672 | 0.646737345 | 8.682754536 | 0.743870198 | 0.487623841 | 0.742781132 | -4.676167123 |
| **A_51_P352562** | Mrpl38 | 0.100906108 | -0.238881058 | 0.440693274 | 9.53877725 | 0.743807975 | 0.48765858 | 0.742781132 | -4.676180232 |
| **A_51_P453519** | Zdhhc16 | -0.118245744 | -0.516457501 | 0.279966013 | 12.25342073 | -0.743741109 | 0.487695913 | 0.742781132 | -4.676194317 |
| **A_51_P336509** | Zfpm2 | 0.261059175 | -0.618190253 | 1.140308603 | 8.774322589 | 0.743664778 | 0.487738533 | 0.742781132 | -4.676210395 |
| **A_52_P202142** | Sv2a | 0.068452171 | -0.162153431 | 0.299057773 | 7.752114555 | 0.74347731 | 0.487843219 | 0.742781132 | -4.676249879 |
| **A_55_P2171047** | Oaz1 | 0.232198594 | -0.550111285 | 1.014508474 | 15.39326786 | 0.743414653 | 0.487878211 | 0.742781132 | -4.676263074 |
| **A_51_P214197** | Stk17b | -0.135476867 | -0.591924678 | 0.320970943 | 9.038216701 | -0.743403091 | 0.487884668 | 0.742781132 | -4.676265508 |
| **A_55_P2023732** | Limch1 | 0.674121743 | -1.597125082 | 2.945368569 | 12.87025515 | 0.743403065 | 0.487884683 | 0.742781132 | -4.676265514 |
| **A_55_P2025008** | Hvcn1 | -0.42717718 | -1.866517443 | 1.012163082 | 8.763757787 | -0.743352688 | 0.487912819 | 0.742781132 | -4.676276122 |
| **A_51_P317640** | Tgfb2 | 0.242655268 | -0.575071785 | 1.060382321 | 11.60970354 | 0.743244501 | 0.487973245 | 0.742781132 | -4.676298902 |
| **A_51_P391716** | Ermap | -0.28051966 | -1.270023929 | 0.70898461 | 7.981906985 | -0.755106721 | 0.487975514 | 0.742781132 | -4.656993014 |
| **A_55_P1989703** | Galnt11 | 0.077952211 | -0.184743025 | 0.340647446 | 9.555677542 | 0.743235738 | 0.48797814 | 0.742781132 | -4.676300747 |
| **A_55_P2260648** | 6330418K | 0.065152979 | -0.154411554 | 0.284717512 | 8.846498277 | 0.743228638 | 0.487982106 | 0.742781132 | -4.676302242 |
| **A_55_P2049448** | Phtf1 | 0.112143176 | -0.265785124 | 0.490071477 | 11.40214198 | 0.743213832 | 0.487990376 | 0.742781132 | -4.676305359 |
| **A_55_P2039603** | Drosha | -0.215064702 | -0.940061918 | 0.509932514 | 11.95649893 | -0.742990299 | 0.488115248 | 0.742804548 | -4.676352418 |
| **A_66_P129048** | Faap20 | 0.16441719 | -0.389910727 | 0.718745108 | 12.71877212 | 0.742900784 | 0.48816526 | 0.742804548 | -4.67637126 |
| **A_55_P2006236** | Ugdh | 0.157611588 | -0.373778544 | 0.68900172 | 11.88691972 | 0.742890845 | 0.488170813 | 0.742804548 | -4.676373353 |
| **A_51_P144770** | 2410002F | 0.115409437 | -0.273725883 | 0.504544757 | 11.86569323 | 0.742832703 | 0.488203299 | 0.742804548 | -4.67638559 |
| **A_55_P2007334** | Mepce | 0.113162013 | -0.26840933 | 0.494733355 | 12.41680051 | 0.742805752 | 0.488218358 | 0.742804548 | -4.676391262 |
| **A_55_P1986371** | NA | 0.278175081 | -0.659924821 | 1.216274982 | 16.16851424 | 0.742710353 | 0.488271665 | 0.742804548 | -4.67641134 |
| **A_55_P2153431** | Ralgds | -0.368913244 | -1.613021829 | 0.875195341 | 10.12176048 | -0.742704927 | 0.488274697 | 0.742804548 | -4.676412482 |
| **A_51_P468249** | Phex | 0.535272039 | -1.270097225 | 2.340641303 | 9.575096488 | 0.742606692 | 0.488329594 | 0.742829612 | -4.676433154 |
| **A_52_P575771** | Mtfr1 | 0.102624242 | -0.243858062 | 0.449106546 | 10.85741703 | 0.74185538 | 0.48874959 | 0.743339636 | -4.676591196 |
| **A_55_P1958623** | Ncln | -0.183782205 | -0.804284009 | 0.436719599 | 11.70561933 | -0.741840869 | 0.488757704 | 0.743339636 | -4.676594247 |
| **A_52_P589568** | Foxo6 | 0.247987318 | -0.589405751 | 1.085380387 | 9.058942799 | 0.741737874 | 0.4888153 | 0.743339636 | -4.676615904 |
| **A_55_P2042685** | NA | -0.203133293 | -0.88928877 | 0.483022185 | 10.58309038 | -0.741496182 | 0.488950476 | 0.743339636 | -4.676666716 |
| **A_65_P01319** | Pde4b | -0.434180409 | -1.901094638 | 1.03273382 | 9.812164136 | -0.741337293 | 0.489039355 | 0.743339636 | -4.676700114 |
| **A_55_P2126448** | Snhg16 | -0.191747899 | -0.839696314 | 0.456200515 | 10.34240379 | -0.741208736 | 0.489111276 | 0.743339636 | -4.676727133 |
| **A_55_P2081313** | Gnas | -0.085044056 | -0.372430545 | 0.202342433 | 15.85833023 | -0.741187292 | 0.489123273 | 0.743339636 | -4.676731639 |

| | | | | | | | | | |
|---|---|---|---|---|---|---|---|---|---|
| **A_51_P172344** | Foxn3 | -0.142468585 | -0.623918666 | 0.338981497 | 10.01310612 | -0.741170612 | 0.489132605 | 0.743339636 | -4.676735145 |
| **A_55_P2063626** | Matk | -0.416805406 | -1.825366418 | 0.991755606 | 8.710694715 | -0.741153264 | 0.489142312 | 0.743339636 | -4.676738791 |
| **A_51_P499864** | Tubg1 | 0.130016135 | -0.309389316 | 0.569421587 | 10.75163013 | 0.74110912 | 0.48916701 | 0.743339636 | -4.676748067 |
| **A_55_P1982683** | Cib1 | -0.20732267 | -0.908032984 | 0.493387644 | 12.97216994 | -0.741068971 | 0.489189475 | 0.743339636 | -4.676756504 |
| **A_51_P451075** | Atp2a2 | 0.503892091 | -1.199198083 | 2.206982264 | 10.60303527 | 0.741054662 | 0.489197481 | 0.743339636 | -4.67675951 |
| **A_55_P2133027** | Kirrel3 | -0.112793792 | -0.494026368 | 0.268438784 | 8.073442074 | -0.741046635 | 0.489201973 | 0.743339636 | -4.676761197 |
| **A_51_P225506** | Pnrc2 | 0.230067937 | -0.547582225 | 1.007718099 | 12.92556265 | 0.741006771 | 0.489224279 | 0.743339636 | -4.676769573 |
| **A_66_P127458** | Zfp652 | 0.096873647 | -0.230581597 | 0.424328892 | 11.65928858 | 0.740975838 | 0.489241588 | 0.743339636 | -4.676776073 |
| **A_51_P256682** | Kpna1 | -0.201012908 | -0.880560065 | 0.47853425 | 9.109014399 | -0.740891634 | 0.489288708 | 0.743352812 | -4.676793764 |
| **A_55_P1979281** | Rab1a | -0.150418193 | -0.658983212 | 0.358146826 | 13.36928133 | -0.74080555 | 0.489336883 | 0.743367588 | -4.676811849 |
| **A_52_P1128182** | Tmem185 | -0.097292781 | -0.426341708 | 0.231756145 | 9.172249198 | -0.740577453 | 0.48946455 | 0.743369187 | -4.676859763 |
| **A_55_P1962035** | Coa5 | -0.122514942 | -0.536892207 | 0.291862324 | 9.377621597 | -0.740531409 | 0.489490324 | 0.743369187 | -4.676869433 |
| **A_55_P1960896** | Fndc3c1 | -0.280344719 | -1.228570727 | 0.66788129 | 8.334572613 | -0.740509887 | 0.489502371 | 0.743369187 | -4.676873953 |
| **A_55_P1973289** | Mapk9 | -0.131010023 | -0.574160501 | 0.312140454 | 10.47917134 | -0.740463487 | 0.489528346 | 0.743369187 | -4.676883698 |
| **A_51_P397673** | Pcsk9 | 0.061674587 | -0.146944898 | 0.270294073 | 7.769760814 | 0.740460211 | 0.489530179 | 0.743369187 | -4.676884386 |
| **A_55_P2035662** | Aebp1 | -0.358917486 | -1.573361167 | 0.855526196 | 12.70802131 | -0.740231535 | 0.489658205 | 0.743465991 | -4.676932406 |
| **A_55_P2033845** | NA | 0.206035737 | -0.491134042 | 0.903205517 | 16.06940659 | 0.740208978 | 0.489670835 | 0.743465991 | -4.676937142 |
| **A_52_P287683** | Pik3c3 | -0.066658139 | -0.292241657 | 0.158925378 | 8.459313356 | -0.740109822 | 0.489726357 | 0.743491903 | -4.67695796 |
| **A_55_P2127804** | Map3k7 | -0.071486946 | -0.313479057 | 0.170505165 | 8.36018914 | -0.739904865 | 0.489841134 | 0.743607765 | -4.677000986 |
| **A_52_P652104** | Brd3 | -0.37857189 | -1.660308755 | 0.903164976 | 9.721596157 | -0.739775311 | 0.489913695 | 0.743659526 | -4.677028178 |
| **A_55_P2064171** | NA | -0.197197937 | -0.864919113 | 0.470523238 | 11.51155756 | -0.739703314 | 0.489954023 | 0.743662355 | -4.677043288 |
| **A_66_P113892** | Smim6 | 0.713405772 | -1.703298502 | 3.130110046 | 10.38652579 | 0.739372773 | 0.490139197 | 0.743833692 | -4.677112646 |
| **A_55_P1955876** | Ifitm7 | -0.076696886 | -0.336515048 | 0.183121275 | 9.200863844 | -0.739364465 | 0.490143852 | 0.743833692 | -4.677114389 |
| **A_55_P1954555** | NA | -0.098438261 | -0.432005796 | 0.235129275 | 8.164425796 | -0.739146458 | 0.49026601 | 0.74391846 | -4.677160122 |
| **A_55_P2012011** | Tceal7 | -0.219234515 | -0.96231572 | 0.523846689 | 8.67494316 | -0.738963518 | 0.490368536 | 0.74391846 | -4.677198492 |
| **A_51_P245631** | Rftn2 | 0.353103033 | -0.843755576 | 1.549961641 | 9.238263116 | 0.738939607 | 0.490381937 | 0.74391846 | -4.677203506 |
| **A_55_P1982718** | Rtf2 | -0.20868095 | -0.916049262 | 0.498687362 | 10.77428019 | -0.738903205 | 0.490402341 | 0.74391846 | -4.67721114 |
| **A_55_P2182975** | P2rx2 | -0.126969621 | -0.5573828 | 0.303443558 | 8.859943665 | -0.738864171 | 0.49042422 | 0.74391846 | -4.677219326 |
| **A_55_P1990170** | NA | 0.129768826 | -0.310274111 | 0.569811762 | 14.95937366 | 0.738627828 | 0.490556706 | 0.74391846 | -4.677268882 |
| **A_55_P2138714** | Sgcd | 0.078000423 | -0.186564363 | 0.342565209 | 7.865837599 | 0.738440086 | 0.490661966 | 0.74391846 | -4.67730824 |
| **A_51_P349803** | Med11 | 0.153176122 | -0.3664189 | 0.672771144 | 10.79303676 | 0.738374065 | 0.490698986 | 0.74391846 | -4.677322079 |
| **A_55_P2034600** | Gm5523 | -0.066867048 | -0.293695025 | 0.159960929 | 8.388190492 | -0.738356115 | 0.490709051 | 0.74391846 | -4.677325841 |
| **A_55_P2025293** | Ss18 | 0.082586471 | -0.197569031 | 0.362741973 | 12.60378086 | 0.738346292 | 0.490714559 | 0.74391846 | -4.6773279 |
| **A_66_P134775** | Pter | 0.071037569 | -0.169942791 | 0.312017929 | 8.007066768 | 0.738340665 | 0.490717714 | 0.74391846 | -4.677329079 |
| **A_55_P2149540** | Srf | -0.12337919 | -0.541931266 | 0.295172885 | 10.30984436 | -0.738316818 | 0.490731087 | 0.74391846 | -4.677334078 |
| **A_51_P487547** | Ccdc91 | -0.083314868 | -0.365992211 | 0.199362475 | 9.545072807 | -0.738213283 | 0.490789148 | 0.74391846 | -4.677355777 |
| **A_51_P262701** | Cpb2 | 0.079362894 | -0.189943596 | 0.348669383 | 7.883872769 | 0.738109884 | 0.490847136 | 0.74391846 | -4.677377446 |
| **A_65_P03874** | Nav2 | 0.10160406 | -0.24318731 | 0.446395429 | 8.090156456 | 0.738082689 | 0.490862389 | 0.74391846 | -4.677383144 |
| **A_55_P2005972** | NA | -0.083917153 | -0.368710089 | 0.200875783 | 9.124616352 | -0.738026363 | 0.490893981 | 0.74391846 | -4.677394947 |
| **A_55_P2051991** | Brsk1 | 0.147930031 | -0.354164133 | 0.650024195 | 8.44158157 | 0.737940796 | 0.490941977 | 0.74391846 | -4.677412876 |
| **A_51_P131164** | Enkur | 0.288670089 | -0.691161798 | 1.268501977 | 8.94893508 | 0.737905188 | 0.490961951 | 0.74391846 | -4.677420337 |
| **A_51_P305628** | Atp8a2 | -0.102277596 | -0.449467818 | 0.244912626 | 7.924219128 | -0.737841998 | 0.490997397 | 0.74391846 | -4.677433576 |
| **A_52_P126055** | Pon2 | -0.076406921 | -0.335778365 | 0.182964524 | 8.617736106 | -0.737837768 | 0.49099977 | 0.74391846 | -4.677434462 |
| **A_52_P94830** | Galnt16 | 0.303720213 | -0.727310286 | 1.334750712 | 10.8977891 | 0.737823577 | 0.491007731 | 0.74391846 | -4.677437435 |
| **A_55_P2020602** | Pacsin1 | 0.220311778 | -0.527799319 | 0.968422876 | 11.99888506 | 0.737601794 | 0.491132158 | 0.743970847 | -4.677483893 |
| **A_52_P257547** | Dele1 | -0.148277377 | -0.651815668 | 0.355260914 | 10.41573297 | -0.737552158 | 0.491160008 | 0.743970847 | -4.67749429 |
| **A_51_P159814** | Smpd4 | 0.157757064 | -0.377976511 | 0.69349064 | 9.724142373 | 0.737548012 | 0.491162335 | 0.743970847 | -4.677495158 |
| **A_55_P2075343** | Arfrp1 | -0.069890138 | -0.307252314 | 0.167472038 | 8.640500227 | -0.737487609 | 0.491196227 | 0.743970847 | -4.677507809 |

| | | | | | | | | | |
|---|---|---|---|---|---|---|---|---|---|
| **A_55_P2021629** | Abcb7 | 0.084776183 | -0.203203371 | 0.372755736 | 8.025477988 | 0.737331094 | 0.491284058 | 0.743995005 | -4.677540586 |
| **A_52_P520397** | Timm8b | 0.228320865 | -0.547339464 | 1.003981194 | 13.27070399 | 0.737266273 | 0.491320436 | 0.743995005 | -4.677554159 |
| **A_55_P2140423** | Arfip2 | -0.073428915 | -0.322901383 | 0.176043554 | 9.845100426 | -0.737216132 | 0.491348577 | 0.743995005 | -4.677564658 |
| **A_55_P2172999** | Ptpn13 | -0.095105977 | -0.418248921 | 0.228036967 | 10.27969546 | -0.737162918 | 0.491378445 | 0.743995005 | -4.6775758 |
| **A_55_P2305010** | A730089K | 0.084517018 | -0.202708162 | 0.371742198 | 7.898574466 | 0.737007657 | 0.491465594 | 0.743995005 | -4.677608304 |
| **A_66_P124659** | Mark1 | -0.130557946 | -0.574252611 | 0.31313672 | 9.305497955 | -0.737003322 | 0.491468028 | 0.743995005 | -4.677609212 |
| **A_55_P1981949** | Klhl5 | 0.329309581 | -0.789869173 | 1.448488334 | 10.55532497 | 0.73697924 | 0.491481546 | 0.743995005 | -4.677614253 |
| **A_55_P1980591** | Hace1 | 0.238738123 | -0.572731541 | 1.050207787 | 9.155134245 | 0.736885196 | 0.49153434 | 0.74401667 | -4.677633939 |
| **A_51_P178815** | H1f8 | 0.307241548 | -0.737243855 | 1.35172695 | 8.747644881 | 0.736763182 | 0.491602842 | 0.744062106 | -4.677659477 |
| **A_66_P118863** | Ncald | 0.100459233 | -0.241127816 | 0.442046282 | 8.101172397 | 0.736612029 | 0.491687712 | 0.744132307 | -4.67769111 |
| **A_51_P469568** | Cldn13 | 0.064983589 | -0.156022846 | 0.285990025 | 7.699318083 | 0.736459929 | 0.491773124 | 0.744158042 | -4.677722936 |
| **A_55_P1990573** | Snrnp70 | 0.230975212 | -0.554577981 | 1.016528405 | 12.69008716 | 0.736444666 | 0.491781696 | 0.744158042 | -4.67772613 |
| **A_52_P527625** | Colec12 | 0.14130837 | -0.339336847 | 0.621953588 | 10.84645926 | 0.736365797 | 0.49182599 | 0.744163954 | -4.677742631 |
| **A_51_P205390** | Cabcoco1 | 0.173481407 | -0.41665076 | 0.763613575 | 8.280741923 | 0.736298633 | 0.491863712 | 0.744163954 | -4.677756682 |
| **A_55_P2085536** | Smarcal1 | 0.08324655 | -0.19998324 | 0.36647634 | 10.96991542 | 0.736169226 | 0.491936398 | 0.744163954 | -4.677783753 |
| **A_52_P491544** | Nedd9 | 0.404239714 | -0.971242042 | 1.77972147 | 10.41294601 | 0.736096054 | 0.491977502 | 0.744163954 | -4.677799058 |
| **A_55_P1967315** | Ppp1r9b | 0.080787624 | -0.19410385 | 0.355679099 | 11.70059822 | 0.736095073 | 0.491978052 | 0.744163954 | -4.677799263 |
| **A_51_P437240** | Emp2 | 0.942929067 | -2.266065913 | 4.151924047 | 13.33182981 | 0.735970372 | 0.492048106 | 0.744211694 | -4.677825345 |
| **A_66_P132890** | NA | -0.079883455 | -0.351863646 | 0.192096736 | 8.183666987 | -0.735647754 | 0.492229378 | 0.744427628 | -4.677892806 |
| **A_55_P2234744** | Spice1 | 0.163492904 | -0.393325815 | 0.720311623 | 9.252994274 | 0.735419982 | 0.492357386 | 0.744513607 | -4.677940422 |
| **A_55_P1984422** | Nme2 | -0.068407929 | -0.301441242 | 0.164625384 | 15.02276877 | -0.735256344 | 0.492449364 | 0.744513607 | -4.677974625 |
| **A_55_P2096652** | Kyat1 | -0.067971143 | -0.299524337 | 0.163582051 | 9.559625389 | -0.735231562 | 0.492463295 | 0.744513607 | -4.677979804 |
| **A_55_P2106961** | 2900097C | -0.169995862 | -0.749169051 | 0.409177327 | 8.316658491 | -0.735157253 | 0.492505068 | 0.744513607 | -4.677995334 |
| **A_55_P2157408** | NA | 0.068078472 | -0.163887975 | 0.30004492 | 7.956594227 | 0.735080627 | 0.492548146 | 0.744513607 | -4.678011346 |
| **A_55_P2140195** | Gm15517 | 0.233832593 | -0.562953706 | 1.030618893 | 10.61525303 | 0.735044334 | 0.49256855 | 0.744513607 | -4.67801893 |
| **A_51_P290576** | Plk2 | 0.442631169 | -1.065739618 | 1.951001956 | 10.87789317 | 0.73499473 | 0.492596439 | 0.744513607 | -4.678029295 |
| **A_55_P2060107** | Pkm | 0.174538362 | -0.420268505 | 0.769345229 | 13.52510208 | 0.734962647 | 0.492614478 | 0.744513607 | -4.678035998 |
| **A_51_P443514** | 1700001K | -0.359927796 | -1.586575142 | 0.866719549 | 9.025021776 | -0.734930057 | 0.492632801 | 0.744513607 | -4.678042808 |
| **A_55_P1986225** | NA | 0.135799253 | -0.327114105 | 0.598712612 | 11.98154991 | 0.734764245 | 0.492726038 | 0.744596313 | -4.678077449 |
| **A_66_P115389** | NA | 0.257851992 | -0.621532304 | 1.137236288 | 11.81176838 | 0.734416002 | 0.492921897 | 0.74479156 | -4.678150185 |
| **A_55_P2141287** | Erlin1 | -0.075468907 | -0.332878354 | 0.181940541 | 11.25569753 | -0.73433448 | 0.492967754 | 0.74479156 | -4.678167209 |
| **A_51_P407866** | 1700020D | 0.091294381 | -0.234689152 | 0.417277915 | 8.510976229 | 0.745952361 | 0.492970807 | 0.74479156 | -4.658626972 |
| **A_51_P161086** | Ppic | 0.502763967 | -1.212311522 | 2.217839456 | 14.55196817 | 0.73422852 | 0.493027363 | 0.744818802 | -4.678189334 |
| **A_55_P2023076** | Arpc1b | -0.185286748 | -0.817467906 | 0.44689441 | 13.2261767 | -0.734096507 | 0.493101635 | 0.744850224 | -4.678216897 |
| **A_52_P277139** | Ankrd17 | -0.220457442 | -0.972769849 | 0.531854964 | 11.27694059 | -0.733967595 | 0.493174169 | 0.744850224 | -4.678243808 |
| **A_51_P393761** | Ndufs2 | -0.082555162 | -0.364309738 | 0.199199415 | 10.3280828 | -0.733877543 | 0.493224842 | 0.744850224 | -4.678262605 |
| **A_51_P481832** | Rabggta | -0.30386295 | -1.340961342 | 0.733235442 | 8.957237669 | -0.733851413 | 0.493239547 | 0.744850224 | -4.678268059 |
| **A_52_P638513** | Zbtb11os1 | 0.085214694 | -0.205645243 | 0.376074632 | 8.931442027 | 0.733805444 | 0.493265416 | 0.744850224 | -4.678277654 |
| **A_66_P125173** | Aip | 0.164042542 | -0.395996944 | 0.724082028 | 9.691343356 | 0.733648759 | 0.493353599 | 0.744850224 | -4.678310354 |
| **A_55_P2044953** | Lst1 | 0.200817657 | -0.484789446 | 0.886424759 | 10.84354304 | 0.733629747 | 0.4933643 | 0.744850224 | -4.678314321 |
| **A_51_P294643** | Cdr2 | -0.196745307 | -0.868466233 | 0.47497562 | 8.857401137 | -0.733611023 | 0.493374839 | 0.744850224 | -4.678318229 |
| **A_55_P2035265** | Trpm7 | 0.075261769 | -0.181748024 | 0.332271563 | 10.71309839 | 0.733457742 | 0.493461119 | 0.744850224 | -4.678350212 |
| **A_55_P2014332** | Cntln | 0.340608109 | -0.822622508 | 1.503838726 | 9.4528539 | 0.733397634 | 0.493494956 | 0.744850224 | -4.678362753 |
| **A_55_P2113984** | Mospd2 | -0.088832919 | -0.392213575 | 0.214547737 | 8.347008983 | -0.733392378 | 0.493497915 | 0.744850224 | -4.67836385 |
| **A_55_P1981405** | Gsk3a | 0.090410802 | -0.218368 | 0.399189604 | 12.5297446 | 0.733370081 | 0.493510467 | 0.744850224 | -4.678368502 |
| **A_55_P2118103** | Zfp68 | 0.256999383 | -0.62093089 | 1.134929655 | 8.970354969 | 0.733199907 | 0.493606275 | 0.744936673 | -4.678404002 |
| **A_55_P2085186** | NA | 0.155991139 | -0.376947444 | 0.688929723 | 10.57574683 | 0.7331167 | 0.493653125 | 0.744949229 | -4.678421358 |
| **A_55_P1970929** | Hnf1a | 0.152761378 | -0.369302834 | 0.67482559 | 7.913795497 | 0.732892015 | 0.49377965 | 0.745079073 | -4.678468217 |

| | | | | | | | | | |
|---|---|---|---|---|---|---|---|---|---|
| **A_52_P373973** | Mex3c | 0.140666904 | -0.34010689 | 0.621440698 | 11.58117367 | 0.732827042 | 0.493816243 | 0.745079073 | -4.678481766 |
| **A_55_P2105888** | G6pc3 | -0.236106136 | -1.043191909 | 0.570979637 | 10.2057879 | -0.732719799 | 0.493876645 | 0.74511206 | -4.678504127 |
| **A_55_P2023727** | Limch1 | 0.858778125 | -2.077697109 | 3.795253358 | 12.60917515 | 0.732495526 | 0.494002978 | 0.74517272 | -4.678550883 |
| **A_52_P295533** | Slc25a28 | -0.139528558 | -0.616735534 | 0.337678418 | 9.637923388 | -0.732329732 | 0.494096384 | 0.74517272 | -4.678585442 |
| **A_52_P639402** | Kcnk3 | 0.28530544 | -0.690507606 | 1.261118487 | 8.914802427 | 0.732307999 | 0.494108629 | 0.74517272 | -4.678589971 |
| **A_51_P191909** | Rem2 | 0.226807133 | -0.548988296 | 1.002602562 | 8.45230207 | 0.732250772 | 0.494140873 | 0.74517272 | -4.678601898 |
| **A_55_P2038687** | Tmco1 | -0.08405636 | -0.371579372 | 0.203466651 | 12.61501553 | -0.732231349 | 0.494151818 | 0.74517272 | -4.678605946 |
| **A_55_P1992084** | Ptprg | 0.226758195 | -0.548908486 | 1.002424876 | 8.983885429 | 0.732214291 | 0.494161429 | 0.74517272 | -4.678609501 |
| **A_51_P339540** | Cdkn1c | -0.353065667 | -1.560947641 | 0.854816306 | 15.12877143 | -0.73211842 | 0.494215452 | 0.74517272 | -4.67862948 |
| **A_55_P1982996** | Ramp1 | 0.087509758 | -0.211930276 | 0.386949792 | 8.437112326 | 0.731976206 | 0.494295597 | 0.74517272 | -4.678659113 |
| **A_55_P2053206** | Kdm4c | -0.090462658 | -0.400008815 | 0.219083499 | 9.374007948 | -0.731971672 | 0.494298152 | 0.74517272 | -4.678660057 |
| **A_55_P1982049** | Icam2 | 0.475354688 | -1.151456815 | 2.10216619 | 13.83048649 | 0.731864855 | 0.494358355 | 0.74517272 | -4.678682312 |
| **A_51_P172131** | Lurap1l | 0.503699905 | -1.220154472 | 2.227554283 | 10.57864892 | 0.731849261 | 0.494367144 | 0.74517272 | -4.678685561 |
| **A_55_P2124115** | Invs | 0.078196047 | -0.189428757 | 0.34582085 | 10.1576946 | 0.731827594 | 0.494379357 | 0.74517272 | -4.678690075 |
| **A_52_P296109** | NA | 0.151682724 | -0.367603183 | 0.670968632 | 8.445921329 | 0.731610499 | 0.494501732 | 0.745214175 | -4.678735296 |
| **A_55_P2176953** | Usp51 | -0.141957711 | -0.627968415 | 0.344052993 | 7.952631266 | -0.731582821 | 0.494517336 | 0.745214175 | -4.678741061 |
| **A_52_P102783** | Dync2h1 | 0.217481085 | -0.527126216 | 0.962088387 | 9.77349663 | 0.731550897 | 0.494535333 | 0.745214175 | -4.67874771 |
| **A_52_P600318** | Scamp2 | -0.132417119 | -0.585812403 | 0.320978166 | 11.04165871 | -0.731505306 | 0.494561037 | 0.745214175 | -4.678757205 |
| **A_52_P400677** | AW20949 | -0.079065853 | -0.349821222 | 0.191689517 | 9.701194504 | -0.731412243 | 0.494613507 | 0.745235157 | -4.678776586 |
| **A_55_P1967325** | Tafa5 | -0.121072473 | -0.535798117 | 0.29365317 | 8.159793513 | -0.73119779 | 0.494734432 | 0.74535927 | -4.67882124 |
| **A_55_P2044167** | NA | -0.069906395 | -0.309391517 | 0.169578728 | 8.033568316 | -0.73112008 | 0.494778256 | 0.745367213 | -4.678837419 |
| **A_52_P459276** | Ift81 | 0.139363451 | -0.338171832 | 0.616898734 | 8.696234981 | 0.730960272 | 0.494868387 | 0.745444908 | -4.678870686 |
| **A_51_P324491** | Cmpk1 | -0.112304866 | -0.497292329 | 0.272682596 | 9.826141872 | -0.730638131 | 0.495050108 | 0.745640145 | -4.678937731 |
| **A_51_P476509** | Zbtb38 | -0.094918667 | -0.420333847 | 0.230496514 | 8.426233893 | -0.730573922 | 0.495086334 | 0.745640145 | -4.678951092 |
| **A_55_P2038540** | Hbb-b2 | -0.146495491 | -0.648771005 | 0.355780023 | 16.28171975 | -0.730520819 | 0.495116296 | 0.745640145 | -4.678962141 |
| **A_51_P281380** | Tspan5 | 0.169035035 | -0.410570296 | 0.748640366 | 10.14353377 | 0.730457077 | 0.495152262 | 0.745640145 | -4.678975403 |
| **A_55_P2171578** | Mkrn1 | -0.281008486 | -1.24483405 | 0.682817078 | 11.57096189 | -0.73024962 | 0.49526933 | 0.745751276 | -4.679018562 |
| **A_52_P73126** | Epb42 | 0.061315328 | -0.149031138 | 0.271661794 | 7.725579256 | 0.730103087 | 0.49535203 | 0.745751276 | -4.679049041 |
| **A_55_P2160648** | Gm6900 | -0.234385254 | -1.038531287 | 0.569760779 | 15.50832084 | -0.730038404 | 0.495388539 | 0.745751276 | -4.679062493 |
| **A_51_P452875** | Pigyl | 0.139032398 | -0.338075253 | 0.616140049 | 10.49411695 | 0.729877502 | 0.495479365 | 0.745751276 | -4.679095954 |
| **A_52_P260629** | Cpne3 | 0.104261741 | -0.253546497 | 0.46206998 | 9.188766343 | 0.729835498 | 0.495503077 | 0.745751276 | -4.679104688 |
| **A_55_P2136817** | Coro2b | 0.366839884 | -0.892175765 | 1.625855534 | 10.79477107 | 0.729786399 | 0.495530796 | 0.745751276 | -4.679114897 |
| **A_52_P544043** | Pcsk5 | 0.450888742 | -1.096588894 | 1.998366377 | 9.499792054 | 0.729785711 | 0.495531184 | 0.745751276 | -4.67911504 |
| **A_55_P1980411** | Zkscan1 | 0.083607441 | -0.203363081 | 0.370577963 | 11.06844265 | 0.729722924 | 0.495566632 | 0.745751276 | -4.679128095 |
| **A_66_P139876** | Psph | -0.192293527 | -0.852323948 | 0.467736894 | 8.988808649 | -0.729711277 | 0.495573207 | 0.745751276 | -4.679130516 |
| **A_52_P16419** | Gpd1 | 0.381899354 | -0.929293362 | 1.693092071 | 9.175979691 | 0.729512506 | 0.49568544 | 0.745861619 | -4.679171839 |
| **A_55_P2135220** | NA | -0.064351899 | -0.285328542 | 0.156624744 | 7.794088032 | -0.729399296 | 0.49574937 | 0.745861619 | -4.67919537 |
| **A_55_P2024322** | Rps27 | 0.082650679 | -0.201194906 | 0.366496264 | 16.218781 | 0.729314127 | 0.495797468 | 0.745861619 | -4.679213072 |
| **A_51_P239113** | Zfp35 | 0.082352439 | -0.200479445 | 0.365184323 | 10.89904859 | 0.729286956 | 0.495812814 | 0.745861619 | -4.679218719 |
| **A_51_P146044** | Ccdc92 | 0.092245561 | -0.224600598 | 0.409091719 | 9.28986396 | 0.729201199 | 0.495861249 | 0.745861619 | -4.67923654 |
| **A_52_P218976** | Cyld | -0.139663685 | -0.619401696 | 0.340074327 | 8.289460373 | -0.729171541 | 0.495878 | 0.745861619 | -4.679242703 |
| **A_52_P163766** | Atp6v0a2 | -0.257332746 | -1.141656705 | 0.626991213 | 9.211763888 | -0.728843036 | 0.496063573 | 0.74600614 | -4.679310957 |
| **A_55_P2176792** | Sh3gl3 | -0.175350112 | -0.777958371 | 0.427258148 | 8.298260385 | -0.72882173 | 0.49607561 | 0.74600614 | -4.679315383 |
| **A_55_P2125972** | Gorasp1 | 0.094319414 | -0.2298688 | 0.418507629 | 9.739465277 | 0.728709142 | 0.496139223 | 0.74600614 | -4.67933877 |
| **A_55_P1966035** | Apoa2 | -1.115843337 | -4.951397269 | 2.719710595 | 9.798755383 | -0.728660918 | 0.496166472 | 0.74600614 | -4.679348786 |
| **A_51_P155073** | Nudt14 | 0.303115615 | -0.738802876 | 1.345034106 | 9.618958629 | 0.728659966 | 0.496167009 | 0.74600614 | -4.679348984 |
| **A_55_P2078994** | Lrrc31 | -0.167333357 | -0.742628432 | 0.407961719 | 15.99605163 | -0.728521229 | 0.496245408 | 0.746049043 | -4.679377798 |
| **A_55_P2316300** | Neo1 | 0.380352128 | -0.927392047 | 1.688096302 | 10.84263151 | 0.728472902 | 0.496272719 | 0.746049043 | -4.679387834 |

| | | | | | | | | | |
|---|---|---|---|---|---|---|---|---|---|
| **A_51_P309988** | Gprc5c | -0.063422032 | -0.281551364 | 0.154707301 | 8.054823728 | -0.728243179 | 0.496402555 | 0.746186207 | -4.679435534 |
| **A_51_P446583** | Zfp445 | 0.137838563 | -0.336375612 | 0.612052738 | 9.829366403 | 0.728025433 | 0.496525644 | 0.746253971 | -4.679480737 |
| **A_55_P2078204** | Prr5 | -0.349292986 | -1.551021204 | 0.852435233 | 9.93030326 | -0.72800431 | 0.496537586 | 0.746253971 | -4.679485122 |
| **A_52_P390114** | Eif4b | 0.073439354 | -0.179248649 | 0.326127357 | 8.720969951 | 0.727938299 | 0.496574906 | 0.746253971 | -4.679498823 |
| **A_51_P358722** | Lancl3 | 0.079364262 | -0.193745989 | 0.352474512 | 7.885496414 | 0.727842357 | 0.496629151 | 0.746253971 | -4.679518736 |
| **A_51_P507290** | Klf5 | 0.185064595 | -0.451818509 | 0.821947699 | 10.69616144 | 0.727803197 | 0.496651293 | 0.746253971 | -4.679526863 |
| **A_51_P301259** | Araf | -0.088052707 | -0.391098719 | 0.214993305 | 10.88646792 | -0.727753803 | 0.496679223 | 0.746253971 | -4.679537114 |
| **A_55_P2009861** | NA | 0.165457605 | -0.404153379 | 0.735068588 | 11.49550176 | 0.727543095 | 0.496798379 | 0.746374998 | -4.679580836 |
| **A_55_P2087923** | Otud3 | -0.306694125 | -1.362725479 | 0.749337228 | 9.110138903 | -0.727409536 | 0.496873917 | 0.746424196 | -4.679608545 |
| **A_52_P149577** | Rab25 | 0.654675666 | -1.59981398 | 2.909165312 | 10.24535851 | 0.727324637 | 0.496921938 | 0.746424196 | -4.679626157 |
| **A_52_P354271** | 2310057M | -0.139920928 | -0.644448019 | 0.364606164 | 8.851048937 | -0.738687738 | 0.496961439 | 0.746424196 | -4.659915023 |
| **A_55_P2050697** | Nfkbil1 | -0.122778716 | -0.545662614 | 0.300105181 | 11.15625309 | -0.727197346 | 0.496993944 | 0.746424196 | -4.67965256 |
| **A_52_P620793** | Eif2a | 0.129438545 | -0.316515062 | 0.575392152 | 9.29060436 | 0.726983053 | 0.49711518 | 0.746424196 | -4.679697003 |
| **A_55_P2180425** | Uqcc1 | -0.093793436 | -0.416943122 | 0.22935625 | 8.755147473 | -0.726974294 | 0.497120136 | 0.746424196 | -4.679698819 |
| **A_65_P10433** | Rpp14 | -0.078130567 | -0.347318462 | 0.191057327 | 9.136032313 | -0.726968843 | 0.49712322 | 0.746424196 | -4.679699949 |
| **A_52_P28651** | Nectin1 | 0.131652703 | -0.321956674 | 0.58526208 | 10.74498594 | 0.726939222 | 0.49713998 | 0.746424196 | -4.679706091 |
| **A_55_P2096967** | Gjb1 | -0.266418739 | -1.184497275 | 0.651659797 | 8.823237811 | -0.726834076 | 0.497199476 | 0.746455557 | -4.679727894 |
| **A_55_P2059179** | Slc48a1 | 0.243940275 | -0.596842911 | 1.084723461 | 10.81616824 | 0.726691092 | 0.49728039 | 0.746519066 | -4.679757538 |
| **A_55_P1999710** | Smad2 | -0.107479624 | -0.478105354 | 0.263146106 | 9.801960301 | -0.726341599 | 0.497478204 | 0.74670643 | -4.679829979 |
| **A_52_P360330** | Map1b | -0.061902132 | -0.27537345 | 0.151569186 | 8.189814272 | -0.726300626 | 0.497501399 | 0.74670643 | -4.679838471 |
| **A_51_P297480** | Dph1 | -0.08316178 | -0.369984075 | 0.203660516 | 9.258095314 | -0.726208307 | 0.497553662 | 0.74670643 | -4.679857601 |
| **A_51_P255565** | Smarcad1 | 0.092439676 | -0.226386454 | 0.411265805 | 9.168338031 | 0.72619767 | 0.497559685 | 0.74670643 | -4.679859806 |
| **A_66_P103999** | Rsrc1 | 0.063048661 | -0.154444393 | 0.280541715 | 8.607022817 | 0.726073897 | 0.497629762 | 0.746725244 | -4.679885452 |
| **A_66_P118849** | NA | 0.203506311 | -0.498543973 | 0.905556595 | 8.701280892 | 0.726039098 | 0.497649466 | 0.746725244 | -4.679892662 |
| **A_55_P2004588** | Nabp2 | 0.088919452 | -0.217989195 | 0.395828098 | 10.77794234 | 0.725668041 | 0.497859594 | 0.74698257 | -4.679969524 |
| **A_51_P483261** | Pomt1 | -0.084686451 | -0.37705961 | 0.207686708 | 8.634926949 | -0.725482231 | 0.497964842 | 0.747082505 | -4.680008004 |
| **A_55_P1977761** | Ttc14 | -0.09160015 | -0.407974478 | 0.224774178 | 8.400682313 | -0.725179126 | 0.49813656 | 0.747282142 | -4.680070759 |
| **A_55_P2017160** | Cdc14a | 0.188973766 | -0.463777712 | 0.841725243 | 8.910310896 | 0.725110175 | 0.498175629 | 0.747282768 | -4.680085032 |
| **A_55_P2014690** | NA | 0.135545703 | -0.332729811 | 0.603821217 | 8.848650286 | 0.724994399 | 0.498241233 | 0.747323197 | -4.680108996 |
| **A_55_P2078266** | H3f3a | 0.456105697 | -1.119789434 | 2.032000828 | 13.8092567 | 0.724917408 | 0.498284864 | 0.747330662 | -4.680124931 |
| **A_51_P444052** | Scp2 | 0.270400004 | -0.663995016 | 1.204795024 | 11.75192702 | 0.724813918 | 0.498343516 | 0.747360653 | -4.680146348 |
| **A_55_P2027521** | NA | 0.196781808 | -0.483375811 | 0.876939427 | 8.547647655 | 0.72464571 | 0.498438856 | 0.747445655 | -4.680181154 |
| **A_55_P2057380** | NA | -0.066998803 | -0.298620822 | 0.164623216 | 7.965081202 | -0.724498591 | 0.498522252 | 0.747482194 | -4.680211591 |
| **A_66_P105032** | Gm13889 | 0.667786628 | -1.641245756 | 2.976819011 | 13.06482755 | 0.724365971 | 0.498597438 | 0.747482194 | -4.680239025 |
| **A_55_P2134862** | Cmip | -0.117795037 | -0.525115534 | 0.28952546 | 10.45662748 | -0.724337672 | 0.498613483 | 0.747482194 | -4.680244879 |
| **A_51_P454993** | Tmcc2 | 0.074172281 | -0.182325089 | 0.330669651 | 7.979123226 | 0.72428429 | 0.49864375 | 0.747482194 | -4.680255921 |
| **A_55_P2016618** | Neurl1a | 0.257194027 | -0.632243959 | 1.146632012 | 10.28246356 | 0.724261751 | 0.498656529 | 0.747482194 | -4.680260582 |
| **A_55_P1965571** | Rpl8 | 0.202550711 | -0.498135111 | 0.903236533 | 15.49168946 | 0.724037046 | 0.498783949 | 0.747564523 | -4.680307053 |
| **A_55_P1972720** | Pmm1 | -0.297881066 | -1.32835633 | 0.732594197 | 11.81300908 | -0.724028521 | 0.498788784 | 0.747564523 | -4.680308816 |
| **A_55_P2087726** | Cops9 | -0.107135841 | -0.477802669 | 0.263530986 | 15.04577677 | -0.723938059 | 0.498840087 | 0.747583463 | -4.680327521 |
| **A_51_P208603** | Dio2 | -0.153847125 | -0.686269272 | 0.378575022 | 7.947424535 | -0.723741736 | 0.49895144 | 0.747665081 | -4.680368109 |
| **A_55_P1952779** | Dcp1b | 0.261912807 | -0.644539791 | 1.168365406 | 10.01410772 | 0.723705682 | 0.498971891 | 0.747665081 | -4.680375563 |
| **A_52_P724940** | Ndufa12 | 0.130334702 | -0.320816239 | 0.581485642 | 13.84563334 | 0.723583308 | 0.499041311 | 0.747711153 | -4.680400858 |
| **A_51_P307644** | Gtf3a | 0.147764226 | -0.363807517 | 0.659335968 | 11.00402647 | 0.723457628 | 0.499112614 | 0.747760037 | -4.680426834 |
| **A_51_P467346** | Muc1 | -0.143606876 | -0.640922877 | 0.353709125 | 8.798933063 | -0.723257836 | 0.499225977 | 0.747871923 | -4.680468121 |
| **A_55_P1990919** | Msrb1 | -0.124230088 | -0.554543248 | 0.306083071 | 13.2973879 | -0.723090261 | 0.499321074 | 0.747876113 | -4.680502744 |
| **A_55_P2103536** | Irf2bpl | 0.211106092 | -0.520151154 | 0.942363338 | 9.905261758 | 0.723070975 | 0.499332019 | 0.747876113 | -4.680506728 |
| **A_51_P265806** | Clca3a2 | -0.16569004 | -0.739745907 | 0.408365828 | 8.709810109 | -0.722923908 | 0.499415489 | 0.747876113 | -4.680537109 |

| | | | | | | | | | |
|---|---|---|---|---|---|---|---|---|---|
| **A_66_P106760** | Adam32 | -0.067884599 | -0.30310634 | 0.167337143 | 7.849478204 | -0.722843266 | 0.499461263 | 0.747876113 | -4.680553766 |
| **A_55_P1979053** | Atp6v0c | -0.167311409 | -0.747070008 | 0.412447191 | 12.75834936 | -0.722817573 | 0.499475847 | 0.747876113 | -4.680559072 |
| **A_55_P2174288** | Snap29 | -0.27204367 | -1.214821536 | 0.670734196 | 9.897835025 | -0.722735842 | 0.499522243 | 0.747876113 | -4.680575953 |
| **A_55_P2117928** | Mxd4 | -0.139178966 | -0.621539857 | 0.343181925 | 10.55163734 | -0.722689695 | 0.49954844 | 0.747876113 | -4.680585483 |
| **A_52_P329054** | Cep70 | -0.12631545 | -0.564168451 | 0.311537552 | 9.488336336 | -0.722567604 | 0.499617755 | 0.747876113 | -4.680610695 |
| **A_52_P84275** | Derl2 | 0.109398411 | -0.269825408 | 0.48862223 | 9.839746517 | 0.722546416 | 0.499629784 | 0.747876113 | -4.68061507 |
| **A_55_P1960375** | NA | 0.359294728 | -0.886264342 | 1.604853798 | 15.24634663 | 0.722498354 | 0.499657072 | 0.747876113 | -4.680624994 |
| **A_52_P259335** | Fkbp15 | 0.071720983 | -0.176935225 | 0.320377191 | 10.79850698 | 0.722432492 | 0.499694469 | 0.747876113 | -4.680638593 |
| **A_55_P2034705** | Nmi | 0.148475107 | -0.366307124 | 0.663257338 | 10.26289097 | 0.722404504 | 0.499710361 | 0.747876113 | -4.680644371 |
| **A_55_P2070769** | Dus1l | 0.084012638 | -0.207285058 | 0.375310335 | 11.71005678 | 0.722367033 | 0.499731639 | 0.747876113 | -4.680652107 |
| **A_55_P2103698** | Isg15 | -0.801884508 | -3.584346518 | 1.980577502 | 10.60453192 | -0.721826672 | 0.500038544 | 0.74818022 | -4.680763637 |
| **A_55_P2049549** | Rab22a | -0.250395439 | -1.119317218 | 0.618526341 | 9.944111077 | -0.721765428 | 0.500073337 | 0.74818022 | -4.680776274 |
| **A_51_P352720** | Lmod2 | 0.074291112 | -0.183514588 | 0.332096812 | 7.726201949 | 0.721763125 | 0.500074645 | 0.74818022 | -4.680776749 |
| **A_52_P513842** | Fubp1 | 0.063867482 | -0.157774142 | 0.285509106 | 8.436901054 | 0.721736741 | 0.500089634 | 0.74818022 | -4.680782193 |
| **A_55_P2077628** | NA | 0.466771335 | -1.15388166 | 2.087424329 | 9.867244439 | 0.721380647 | 0.500291968 | 0.748374505 | -4.680855651 |
| **A_51_P443754** | Hivep2 | -0.106696797 | -0.477170936 | 0.263777343 | 8.479507033 | -0.721346332 | 0.500311468 | 0.748374505 | -4.680862729 |
| **A_55_P1998396** | Eif2ak4 | 0.084704091 | -0.20942399 | 0.378832172 | 10.31783948 | 0.721303837 | 0.500335618 | 0.748374505 | -4.680871493 |
| **A_55_P2081571** | Raver1 | 0.083714817 | -0.207037725 | 0.374467358 | 13.59759467 | 0.721155892 | 0.500419703 | 0.748442371 | -4.680902003 |
| **A_52_P663757** | Tnik | 0.184680581 | -0.456934898 | 0.82629606 | 8.742287676 | 0.720936055 | 0.500544664 | 0.74857136 | -4.68094733 |
| **A_52_P48689** | Bzw1 | 0.157317449 | -0.389373338 | 0.704008236 | 10.01011048 | 0.720751386 | 0.500649652 | 0.748586971 | -4.6809854 |
| **A_55_P1987914** | Map4k4 | -0.164742997 | -0.737284605 | 0.407798611 | 10.36957593 | -0.720692918 | 0.500682895 | 0.748586971 | -4.680997451 |
| **A_55_P2178247** | Fam20b | -0.202718256 | -0.907288064 | 0.501851552 | 8.607368228 | -0.72064135 | 0.500712217 | 0.748586971 | -4.68100808 |
| **A_52_P81468** | Psma4 | 0.14903956 | -0.368965876 | 0.667044996 | 11.77531475 | 0.720638711 | 0.500713717 | 0.748586971 | -4.681008624 |
| **A_55_P2083009** | Catsperd | -0.228656343 | -1.023510698 | 0.566198012 | 8.677643413 | -0.720520002 | 0.50078122 | 0.748586971 | -4.681033089 |
| **A_55_P1970695** | NA | -0.081445309 | -0.364571847 | 0.20168123 | 8.591123166 | -0.720503076 | 0.500790845 | 0.748586971 | -4.681036577 |
| **A_55_P2044217** | 9930021J | 0.09571905 | -0.237072137 | 0.428510237 | 11.34547795 | 0.720405317 | 0.500846439 | 0.748586971 | -4.681056722 |
| **A_52_P276935** | Dnajb6 | -0.492399109 | -2.204424546 | 1.219626328 | 11.3340338 | -0.72037294 | 0.500864853 | 0.748586971 | -4.681063394 |
| **A_55_P2134092** | Brsk1 | 0.130258007 | -0.32279872 | 0.583314734 | 8.332200687 | 0.720115553 | 0.501011251 | 0.748747895 | -4.681116423 |
| **A_66_P110817** | Mterf3 | 0.070413067 | -0.174558315 | 0.315384448 | 9.122463545 | 0.719926657 | 0.50111871 | 0.748850606 | -4.681155332 |
| **A_52_P614472** | Rpusd1 | 0.083352218 | -0.206665279 | 0.373369715 | 8.660994912 | 0.71985215 | 0.5011611 | 0.748856072 | -4.681170677 |
| **A_55_P2041509** | 1110004F | 0.193668575 | -0.480465544 | 0.867802694 | 11.75473652 | 0.719553674 | 0.50133094 | 0.749051962 | -4.681232139 |
| **A_55_P2102664** | NA | 0.097281418 | -0.241391506 | 0.435954342 | 8.260775819 | 0.719448584 | 0.501390748 | 0.749054188 | -4.681253775 |
| **A_55_P2064478** | Cdc27 | -0.231040104 | -1.035414584 | 0.573334377 | 9.182758317 | -0.719414907 | 0.501409915 | 0.749054188 | -4.681260708 |
| **A_55_P2093543** | Snapc4 | -0.154865607 | -0.694158656 | 0.384427442 | 10.1048211 | -0.719251046 | 0.501503183 | 0.749135636 | -4.681294437 |
| **A_52_P350950** | Dennd2c | 0.089043197 | -0.22108871 | 0.399175103 | 8.259637163 | 0.719125417 | 0.501574698 | 0.749165761 | -4.681320294 |
| **A_55_P2120807** | Dld | 0.081090826 | -0.201394521 | 0.363576174 | 10.00960334 | 0.718995457 | 0.501648685 | 0.749165761 | -4.681347038 |
| **A_55_P1998416** | Ifi47 | -0.392388828 | -1.759671132 | 0.974893476 | 9.179427766 | -0.718801187 | 0.501759298 | 0.749165761 | -4.681387011 |
| **A_52_P496870** | 2310011J | -0.066948531 | -0.300232262 | 0.1663352 | 8.432904491 | -0.718798143 | 0.501761031 | 0.749165761 | -4.681387637 |
| **A_52_P229709** | Ube2d3 | -0.165287643 | -0.74125062 | 0.410675335 | 10.93896544 | -0.718780296 | 0.501771194 | 0.749165761 | -4.681391309 |
| **A_66_P127255** | Ing3 | 0.190942148 | -0.474439138 | 0.856323434 | 10.45650212 | 0.718756139 | 0.501784949 | 0.749165761 | -4.681396279 |
| **A_55_P1964213** | Tprkb | -0.150993084 | -0.677175177 | 0.375189009 | 9.683052229 | -0.718739209 | 0.50179459 | 0.749165761 | -4.681399762 |
| **A_51_P370380** | Rdh14 | 0.176979028 | -0.439901071 | 0.793859126 | 10.46509249 | 0.718573822 | 0.501888776 | 0.749233455 | -4.681433783 |
| **A_55_P2039235** | Mat2b | -0.078135882 | -0.350532756 | 0.194260993 | 12.03942051 | -0.718453629 | 0.501957232 | 0.749233455 | -4.681458505 |
| **A_55_P2098911** | Lrrcc1 | 0.17326402 | -0.430805831 | 0.777333871 | 10.14567487 | 0.718408679 | 0.501982834 | 0.749233455 | -4.681467749 |
| **A_55_P2080308** | Gm12538 | -0.120165874 | -0.539126077 | 0.298794329 | 8.125664299 | -0.718387427 | 0.50199494 | 0.749233455 | -4.68147212 |
| **A_55_P1969273** | Zfp758 | 0.129189227 | -0.321313985 | 0.579692439 | 8.716478707 | 0.718255156 | 0.502070286 | 0.749288067 | -4.68149932 |
| **A_66_P107584** | Fam133b | 0.083457077 | -0.207618936 | 0.374533089 | 8.439082722 | 0.718136659 | 0.502137793 | 0.749309808 | -4.681523685 |
| **A_55_P2077048** | Itih5 | -0.196867775 | -0.88380784 | 0.490072289 | 9.043020682 | -0.717804427 | 0.502327095 | 0.749309808 | -4.681591983 |

| | | | | | | | | | |
|---|---|---|---|---|---|---|---|---|---|
| **A_55_P2047788** | Katnal1 | -0.093956804 | -0.421809009 | 0.233895401 | 8.311237597 | -0.717795073 | 0.502332425 | 0.749309808 | -4.681593905 |
| **A_55_P2037524** | Rps15 | 0.240512682 | -0.598736935 | 1.079762298 | 15.74489313 | 0.717789623 | 0.502335532 | 0.749309808 | -4.681595026 |
| **A_52_P30550** | Acp2 | -0.109388354 | -0.491130199 | 0.272353491 | 8.608823847 | -0.717714407 | 0.502378396 | 0.749309808 | -4.681610484 |
| **A_52_P638895** | Vegfa | -0.295521318 | -1.326887519 | 0.735844884 | 11.38253598 | -0.717672434 | 0.502402317 | 0.749309808 | -4.681619111 |
| **A_51_P337856** | Ubl7 | -0.086415874 | -0.388061729 | 0.215229981 | 9.409958208 | -0.71754065 | 0.502477427 | 0.749309808 | -4.681646192 |
| **A_51_P411917** | Gata6 | 0.460410433 | -1.146753466 | 2.067574332 | 10.90157802 | 0.717522185 | 0.502487952 | 0.749309808 | -4.681649986 |
| **A_51_P130544** | Gas5 | 0.123843648 | -0.308488857 | 0.556176152 | 8.405292708 | 0.717474042 | 0.502515393 | 0.749309808 | -4.681659879 |
| **A_52_P87503** | Hbq1b | 0.066599039 | -0.165904392 | 0.29910247 | 7.863191544 | 0.717445546 | 0.502531636 | 0.749309808 | -4.681665733 |
| **A_55_P1995447** | NA | 0.423220431 | -1.054339818 | 1.900780679 | 14.00011397 | 0.717417179 | 0.502547807 | 0.749309808 | -4.681671562 |
| **A_55_P2283207** | Maz | -0.07375611 | -0.331276876 | 0.183764656 | 8.042419478 | -0.717358244 | 0.502581402 | 0.749309808 | -4.68168367 |
| **A_55_P2011031** | Cdk5rap1 | -0.150680279 | -0.67679251 | 0.375431953 | 7.99270717 | -0.717345473 | 0.502588683 | 0.749309808 | -4.681686294 |
| **A_55_P2077053** | Itih4 | -0.080659977 | -0.362383736 | 0.201063782 | 7.712454871 | -0.717108658 | 0.502723695 | 0.749372517 | -4.681734941 |
| **A_55_P2018211** | NA | 0.260844481 | -0.650236607 | 1.17192557 | 15.72439221 | 0.717092147 | 0.502733109 | 0.749372517 | -4.681738332 |
| **A_55_P2038414** | Acot8 | -0.15347376 | -0.689558222 | 0.382610702 | 10.32927718 | -0.717053004 | 0.502755428 | 0.749372517 | -4.681746372 |
| **A_55_P1984337** | Emd | 0.123359665 | -0.307568133 | 0.554287464 | 12.18131876 | 0.716999772 | 0.502785781 | 0.749372517 | -4.681757305 |
| **A_55_P2078173** | NA | -0.146409485 | -0.65812762 | 0.36530865 | 14.06081441 | -0.716619713 | 0.50300253 | 0.749615831 | -4.681835346 |
| **A_55_P2006018** | Slc4a2 | -0.186593984 | -0.838799969 | 0.465612001 | 10.21270369 | -0.716577559 | 0.503026575 | 0.749615831 | -4.681844 |
| **A_66_P118592** | Atrx | 0.247417603 | -0.617585349 | 1.112420556 | 9.223383101 | 0.716412825 | 0.503120546 | 0.749637418 | -4.681877815 |
| **A_51_P436878** | Sertad1 | -0.222364155 | -0.999807153 | 0.555078842 | 10.79371455 | -0.716385143 | 0.503136338 | 0.749637418 | -4.681883497 |
| **A_55_P1998213** | NA | 0.232090819 | -0.579412559 | 1.043594197 | 16.8138718 | 0.716337975 | 0.503163247 | 0.749637418 | -4.681893178 |
| **A_55_P2007693** | NA | 0.170330195 | -0.425275693 | 0.765936083 | 10.5208039 | 0.7162803 | 0.503196152 | 0.749637418 | -4.681905015 |
| **A_55_P2087785** | NA | -0.099914575 | -0.449415079 | 0.249585929 | 7.912354914 | -0.716030272 | 0.503338816 | 0.749711835 | -4.681956323 |
| **A_55_P1955726** | Ptpn5 | 0.295349074 | -0.737896569 | 1.328594718 | 8.747362836 | 0.715949478 | 0.503384922 | 0.749711835 | -4.681972899 |
| **A_55_P1968133** | Ttc4 | 0.080292863 | -0.20061282 | 0.361198547 | 11.29458058 | 0.715923743 | 0.503399609 | 0.749711835 | -4.681978179 |
| **A_55_P2019068** | Flii | -0.14665058 | -0.659711656 | 0.366410497 | 8.331486077 | -0.715920934 | 0.503401213 | 0.749711835 | -4.681978756 |
| **A_55_P2002357** | Tnrc18 | -0.221031806 | -0.994440341 | 0.55237673 | 9.53388044 | -0.715807356 | 0.503466034 | 0.749735381 | -4.682002056 |
| **A_55_P2034227** | NA | -0.065650106 | -0.295381299 | 0.164081087 | 14.81485709 | -0.715757343 | 0.503494579 | 0.749735381 | -4.682012315 |
| **A_55_P2089288** | Rnf121 | -0.207281356 | -0.933102009 | 0.518539296 | 10.88569945 | -0.715288538 | 0.503762206 | 0.74990313 | -4.682108457 |
| **A_51_P386562** | Ebf2 | 0.103975736 | -0.260125412 | 0.468076883 | 8.073020944 | 0.715254007 | 0.503781923 | 0.74990313 | -4.682115537 |
| **A_51_P289889** | Npnt | 0.258210268 | -0.645990491 | 1.162411026 | 8.774748897 | 0.715251844 | 0.503783158 | 0.74990313 | -4.68211598 |
| **A_52_P271155** | Mtpap | -0.076503687 | -0.344427643 | 0.191420269 | 9.929554665 | -0.715189556 | 0.503818725 | 0.74990313 | -4.68212875 |
| **A_55_P2072164** | NA | 0.087058009 | -0.217833971 | 0.39194999 | 8.222623223 | 0.715176239 | 0.503826329 | 0.74990313 | -4.682131481 |
| **A_52_P365342** | Lrrn2 | -0.117727384 | -0.530042495 | 0.294587727 | 8.12288029 | -0.71515238 | 0.503839953 | 0.74990313 | -4.682136372 |
| **A_55_P2081133** | Gapdh | 0.156393761 | -0.391444855 | 0.704232376 | 15.89167806 | 0.715018251 | 0.50391655 | 0.7499594 | -4.682163867 |
| **A_66_P102237** | Nemf | 0.075179643 | -0.18819761 | 0.338556896 | 10.58076174 | 0.714944518 | 0.50395866 | 0.749964341 | -4.68217898 |
| **A_51_P416556** | Ssr3 | 0.079382315 | -0.198815408 | 0.357580038 | 13.47143146 | 0.714694655 | 0.504101377 | 0.750118989 | -4.682230186 |
| **A_55_P1986252** | NA | 0.252215122 | -0.631827692 | 1.136257935 | 16.95295567 | 0.714575578 | 0.504169401 | 0.750129752 | -4.682254585 |
| **A_51_P157976** | Mmab | 0.068170544 | -0.170794445 | 0.307135534 | 8.527738696 | 0.714517433 | 0.504202619 | 0.750129752 | -4.682266498 |
| **A_55_P1989328** | Rps12 | 0.260238791 | -0.652052158 | 1.17252974 | 17.5348444 | 0.714478249 | 0.504225006 | 0.750129752 | -4.682274526 |
| **A_51_P250123** | Zfp763 | 0.201795828 | -0.505798569 | 0.909390226 | 8.557841844 | 0.714295878 | 0.504329208 | 0.750163894 | -4.682311884 |
| **A_55_P2093720** | Dnajc5 | -0.127558855 | -0.574851404 | 0.319733694 | 8.20174195 | -0.714281323 | 0.504337525 | 0.750163894 | -4.682314866 |
| **A_51_P492410** | Pmvk | 0.177416254 | -0.444745066 | 0.799577575 | 10.87890945 | 0.714234369 | 0.504364356 | 0.750163894 | -4.682324483 |
| **A_51_P432511** | Afg1l | -0.062139191 | -0.280104196 | 0.155825815 | 8.068064732 | -0.714050893 | 0.504469209 | 0.75026213 | -4.682362059 |
| **A_55_P2069251** | Prr18 | 0.617346982 | -1.548508102 | 2.783202066 | 10.0135446 | 0.713921202 | 0.504543334 | 0.750275637 | -4.682388615 |
| **A_51_P336391** | Tmem18 | 0.130446849 | -0.327258335 | 0.588152034 | 11.24424085 | 0.713835443 | 0.504592353 | 0.750275637 | -4.682406174 |
| **A_51_P350403** | Exoc4 | 0.086268162 | -0.216426868 | 0.388963192 | 8.553039863 | 0.713831322 | 0.504594709 | 0.750275637 | -4.682407018 |
| **A_55_P2109544** | Trnt1 | -0.073943095 | -0.333417858 | 0.185531668 | 8.74739851 | -0.713761053 | 0.504634878 | 0.750277662 | -4.682421404 |
| **A_55_P2007404** | Vbp1 | 0.146477803 | -0.367732045 | 0.660687651 | 9.2701872 | 0.71347995 | 0.504795587 | 0.750458891 | -4.682478945 |

| | | | | | | | | | |
|---|---|---|---|---|---|---|---|---|---|
| **A_55_P1984640** | Ube2f | -0.272980374 | -1.231521697 | 0.685560949 | 10.93875983 | -0.713297875 | 0.5048997 | 0.750517251 | -4.682516206 |
| **A_55_P2012669** | NA | 0.275591771 | -0.69214951 | 1.243333051 | 17.59553594 | 0.713275528 | 0.504912479 | 0.750517251 | -4.682520779 |
| **A_55_P2007326** | Cyp2a4 | -0.105031947 | -0.473915237 | 0.263851343 | 7.814235846 | -0.713153108 | 0.504982491 | 0.750563613 | -4.682545827 |
| **A_55_P2051825** | Ephx3 | 0.086439211 | -0.217203144 | 0.390081567 | 8.542694808 | 0.713015201 | 0.505061366 | 0.750623142 | -4.682574041 |
| **A_55_P1983868** | NA | -0.092068716 | -0.415550574 | 0.231413142 | 9.002786494 | -0.712873551 | 0.505142391 | 0.750685856 | -4.682603017 |
| **A_52_P56751** | Lcp1 | 0.10832167 | -0.272341252 | 0.488984592 | 11.07683785 | 0.712730126 | 0.505224441 | 0.750721283 | -4.682632351 |
| **A_55_P2069052** | Sacs | 0.136918822 | -0.344327233 | 0.618164878 | 8.707331495 | 0.712600821 | 0.50529842 | 0.750721283 | -4.682658794 |
| **A_51_P167426** | Herc1 | -0.088113093 | -0.397826748 | 0.221600561 | 7.955190645 | -0.712574769 | 0.505313327 | 0.750721283 | -4.682664122 |
| **A_55_P1964896** | NA | -0.082662102 | -0.373236428 | 0.207912225 | 7.961195585 | -0.712524084 | 0.505342328 | 0.750721283 | -4.682674486 |
| **A_55_P2042312** | NA | 0.156190801 | -0.392876365 | 0.705257966 | 16.84066208 | 0.712492543 | 0.505360375 | 0.750721283 | -4.682680935 |
| **A_55_P2139992** | Mrps18b | -0.447476726 | -2.020761824 | 1.125808373 | 12.066972 | -0.712382706 | 0.505423228 | 0.750756968 | -4.682703392 |
| **A_51_P517094** | Sh3gl1 | -0.121857252 | -0.550418739 | 0.306704234 | 10.85100787 | -0.712178055 | 0.505540351 | 0.750819214 | -4.682745227 |
| **A_55_P2106645** | Adcy5 | 0.101460671 | -0.255369937 | 0.458291279 | 7.98226556 | 0.712173772 | 0.505542802 | 0.750819214 | -4.682746103 |
| **A_55_P2024406** | Napepld | -0.084721184 | -0.382725742 | 0.213283375 | 8.883943338 | -0.712064693 | 0.505605236 | 0.750848654 | -4.682768397 |
| **A_55_P2211341** | Map3k4 | -0.123968778 | -0.56006324 | 0.312125684 | 12.1296257 | -0.712003443 | 0.505640296 | 0.750848654 | -4.682780915 |
| **A_65_P17602** | Pgm5 | 0.351918708 | -0.886474509 | 1.590311924 | 9.323407599 | 0.711760915 | 0.505779138 | 0.750905079 | -4.682830474 |
| **A_52_P305073** | Pias1 | 0.120459257 | -0.303504891 | 0.544423406 | 10.15942737 | 0.711641708 | 0.505847391 | 0.750905079 | -4.682854829 |
| **A_52_P167411** | Tnk1 | 0.222250534 | -0.560016523 | 1.004517591 | 9.028019141 | 0.711603567 | 0.50586923 | 0.750905079 | -4.682862621 |
| **A_55_P2211319** | Fgf18 | 0.594176863 | -1.497186976 | 2.685540702 | 10.65417593 | 0.711600898 | 0.505870758 | 0.750905079 | -4.682863166 |
| **A_66_P126552** | Prr14l | -0.161926111 | -0.731870636 | 0.408018414 | 8.77164096 | -0.71159788 | 0.505872487 | 0.750905079 | -4.682863783 |
| **A_55_P2121252** | Amd2 | -0.162984538 | -0.736767784 | 0.410798708 | 9.299997544 | -0.71145738 | 0.505952942 | 0.750927355 | -4.682892483 |
| **A_55_P2102540** | Olfml3 | 0.194358484 | -0.489896545 | 0.878613513 | 8.397132844 | 0.711436021 | 0.505965173 | 0.750927355 | -4.682896845 |
| **A_52_P26626** | Fam92b | 0.461649697 | -1.16499684 | 2.088296234 | 9.38621096 | 0.710836479 | 0.506308596 | 0.751344913 | -4.683019267 |
| **A_55_P1992989** | Mrps26 | -0.128032712 | -0.579179913 | 0.323114489 | 13.03062648 | -0.710809172 | 0.506324241 | 0.751344913 | -4.683024841 |
| **A_55_P1965313** | Mctp2 | 0.17667686 | -0.445972454 | 0.799326175 | 8.847696462 | 0.710700309 | 0.506386617 | 0.751379804 | -4.683047061 |
| **A_52_P418934** | Sec22b | -0.224908999 | -1.017660166 | 0.567842168 | 12.74055256 | -0.710591959 | 0.506448704 | 0.751381761 | -4.683069175 |
| **A_51_P221495** | Nsun5 | 0.061866507 | -0.156220045 | 0.279953059 | 8.453693336 | 0.710521229 | 0.506489237 | 0.751381761 | -4.683083609 |
| **A_55_P1964757** | Stam2 | 0.092074133 | -0.232509961 | 0.416658228 | 8.773152975 | 0.710494551 | 0.506504525 | 0.751381761 | -4.683089053 |
| **A_55_P2110743** | Mettl7a2 | -0.100155931 | -0.453271977 | 0.252960114 | 13.91233853 | -0.710410819 | 0.506552513 | 0.751395296 | -4.683106139 |
| **A_51_P382970** | Itga9 | 0.45196286 | -1.141715617 | 2.045641337 | 10.47085733 | 0.71031729 | 0.506606119 | 0.751417162 | -4.683125222 |
| **A_55_P2095794** | Evi5 | -0.088405148 | -0.400259316 | 0.22344902 | 11.06376691 | -0.710029427 | 0.506771132 | 0.751547826 | -4.683183945 |
| **A_51_P151968** | Eif2b5 | 0.065119595 | -0.164608189 | 0.294847378 | 9.702735008 | 0.709983918 | 0.506797222 | 0.751547826 | -4.683193227 |
| **A_55_P2444965** | Tmem245 | -0.073763378 | -0.33399484 | 0.186468085 | 8.404889259 | -0.709955842 | 0.506813319 | 0.751547826 | -4.683198953 |
| **A_51_P285271** | Ccdc61 | 0.142111202 | -0.359444125 | 0.643666529 | 10.48002439 | 0.709675497 | 0.506974066 | 0.751547826 | -4.683256122 |
| **A_55_P2077751** | NA | 0.10694075 | -0.27050836 | 0.484389859 | 8.585171287 | 0.709635225 | 0.50699716 | 0.751547826 | -4.683264333 |
| **A_51_P496715** | Pomk | 0.137025794 | -0.346634436 | 0.620686024 | 9.026651749 | 0.709597857 | 0.50701859 | 0.751547826 | -4.683271952 |
| **A_52_P383653** | Cpne8 | -0.066965905 | -0.303382699 | 0.169450888 | 8.595652561 | -0.709456493 | 0.507099665 | 0.751547826 | -4.683300771 |
| **A_55_P1978306** | Zfp318 | 0.096338784 | -0.259883045 | 0.452560614 | 9.763286214 | 0.720349628 | 0.507139075 | 0.751547826 | -4.663131747 |
| **A_52_P316405** | Lamtor4 | 0.132371992 | -0.335005112 | 0.599749096 | 11.95449341 | 0.709380126 | 0.507143466 | 0.751547826 | -4.683316338 |
| **A_51_P105927** | Rasl12 | 0.671778265 | -1.700152987 | 3.043709517 | 11.35657651 | 0.709372252 | 0.507147982 | 0.751547826 | -4.683317943 |
| **A_55_P2086387** | 1700030K | 0.189061189 | -0.478531464 | 0.856653842 | 9.314883553 | 0.709318329 | 0.507178913 | 0.751547826 | -4.683328934 |
| **A_55_P2009091** | Mad1l1 | 0.087809559 | -0.222257709 | 0.397876827 | 9.795327599 | 0.709310221 | 0.507183563 | 0.751547826 | -4.683330586 |
| **A_55_P2174821** | Rab8a | -0.222926352 | -1.010206578 | 0.564353874 | 10.22642567 | -0.709222347 | 0.507233971 | 0.751547826 | -4.683348496 |
| **A_55_P1994917** | Wdr5 | 0.104343941 | -0.264158119 | 0.472846002 | 11.38826269 | 0.709214593 | 0.507238419 | 0.751547826 | -4.683350076 |
| **A_55_P1966322** | Eif3h | 0.172197841 | -0.43617161 | 0.780567291 | 16.04654377 | 0.708941906 | 0.507394863 | 0.751722014 | -4.683405643 |
| **A_55_P2153587** | Afg3l1 | -0.209693763 | -0.950728372 | 0.531340846 | 10.86543884 | -0.70875701 | 0.507500959 | 0.751788582 | -4.683443311 |
| **A_51_P479808** | Mapk6 | -0.275637241 | -1.249970902 | 0.69869642 | 8.459317213 | -0.708566365 | 0.507610371 | 0.751788582 | -4.683482143 |
| **A_51_P509229** | Tnrc6a | -0.075675268 | -0.343190276 | 0.191839739 | 9.951588038 | -0.708526607 | 0.507633189 | 0.751788582 | -4.68349024 |

| | | | | | | | | | |
|---|---|---|---|---|---|---|---|---|---|
| **A_55_P2061134** | NA | -0.28235096 | -1.280676175 | 0.715974254 | 14.95197277 | -0.708382093 | 0.507716139 | 0.751788582 | -4.68351967 |
| **A_51_P109835** | Uqcrc1 | 0.113711081 | -0.28835229 | 0.515774452 | 11.77518073 | 0.708367496 | 0.507724518 | 0.751788582 | -4.683522643 |
| **A_55_P1979595** | Selenon | -0.151816883 | -0.688643692 | 0.385009925 | 11.52010897 | -0.708330955 | 0.507745494 | 0.751788582 | -4.683530083 |
| **A_55_P2141724** | Gm13275 | 0.085709214 | -0.217398205 | 0.388816633 | 8.418947381 | 0.708241367 | 0.507796923 | 0.751788582 | -4.683548324 |
| **A_52_P377299** | Zbtb11 | -0.150964428 | -0.684879227 | 0.38295037 | 8.431912726 | -0.708195258 | 0.507823394 | 0.751788582 | -4.683557712 |
| **A_51_P259861** | Mettl9 | 0.135054721 | -0.342623154 | 0.612732595 | 12.44718177 | 0.708149558 | 0.507849631 | 0.751788582 | -4.683567016 |
| **A_55_P2025143** | Tlk1 | 0.089897234 | -0.228072221 | 0.407866689 | 11.19368609 | 0.708127193 | 0.507862471 | 0.751788582 | -4.683571569 |
| **A_52_P546635** | Zeb1 | 0.197093631 | -0.500088764 | 0.894276025 | 9.905419625 | 0.708070537 | 0.507895 | 0.751788582 | -4.683583103 |
| **A_51_P346938** | Lrg1 | -0.104724107 | -0.475176727 | 0.265728513 | 8.04262678 | -0.708050672 | 0.507906406 | 0.751788582 | -4.683587147 |
| **A_55_P2203051** | Gt(ROSA): | 0.148293741 | -0.376420641 | 0.673008123 | 10.29780186 | 0.707864607 | 0.508013246 | 0.75188916 | -4.683625019 |
| **A_55_P2151506** | Nudt9 | -0.112503785 | -0.510637073 | 0.285629504 | 12.44985157 | -0.707764843 | 0.508070538 | 0.751915817 | -4.683645323 |
| **A_51_P144926** | Cops8 | 0.067085751 | -0.170342712 | 0.304514213 | 9.831010763 | 0.707697805 | 0.508109038 | 0.751915817 | -4.683658965 |
| **A_52_P609972** | Eln | 0.961001667 | -2.441634957 | 4.363638291 | 11.93271229 | 0.707389997 | 0.50828584 | 0.752118605 | -4.683721592 |
| **A_55_P1982563** | Rhbdf1 | 0.131636158 | -0.334494085 | 0.597766402 | 11.42740648 | 0.707323788 | 0.508323876 | 0.752118605 | -4.68373506 |
| **A_51_P451338** | Jag2 | 0.092699394 | -0.23563159 | 0.421030377 | 9.032175286 | 0.70715623 | 0.508420142 | 0.752166843 | -4.683769141 |
| **A_55_P2117319** | Ntan1 | -0.068637671 | -0.311753163 | 0.17447782 | 11.83488718 | -0.707131614 | 0.508434285 | 0.752166843 | -4.683774148 |
| **A_51_P404825** | Ccdc58 | 0.078355567 | -0.199255099 | 0.355966232 | 11.74745836 | 0.706942439 | 0.508542988 | 0.75217304 | -4.683812618 |
| **A_55_P2186787** | NA | 0.10099263 | -0.256830995 | 0.458816255 | 16.96444417 | 0.706921216 | 0.508555184 | 0.75217304 | -4.683816933 |
| **A_55_P1990331** | Add1 | -0.075626012 | -0.343600427 | 0.192348404 | 8.493417406 | -0.706851544 | 0.508595223 | 0.75217304 | -4.683831099 |
| **A_55_P1974845** | Pde1a | 0.132376425 | -0.336712349 | 0.6014652 | 8.235033874 | 0.706815321 | 0.508616041 | 0.75217304 | -4.683838464 |
| **A_52_P633128** | Aasdhppt | 0.09110016 | -0.231753895 | 0.413954216 | 10.05327202 | 0.706745809 | 0.508655991 | 0.75217304 | -4.683852596 |
| **A_52_P141112** | Csnk1g3 | -0.112457061 | -0.51108927 | 0.286175148 | 8.243582736 | -0.706585448 | 0.508748162 | 0.75217304 | -4.683885195 |
| **A_51_P137336** | Cdh1 | 0.254385143 | -0.647541496 | 1.156311781 | 12.40498321 | 0.706432831 | 0.508835894 | 0.75217304 | -4.679618927 |
| **A_55_P1972817** | Trmt11 | 0.113545029 | -0.289052726 | 0.516142785 | 9.453507656 | 0.706394198 | 0.508858104 | 0.75217304 | -4.683924065 |
| **A_55_P2151011** | NA | 0.200913425 | -0.511521942 | 0.913348792 | 8.535017354 | 0.70634005 | 0.508889234 | 0.75217304 | -4.683935069 |
| **A_51_P336833** | Fabp4 | -0.495660775 | -2.253271513 | 1.261949963 | 11.43457341 | -0.706337396 | 0.50889076 | 0.75217304 | -4.683935609 |
| **A_55_P2122789** | Homer1 | -0.08438975 | -0.383651299 | 0.2148718 | 7.904119272 | -0.706299866 | 0.508912337 | 0.75217304 | -4.683943235 |
| **A_55_P2016024** | Kif3c | 0.091673755 | -0.233429204 | 0.416776714 | 8.860232636 | 0.706275989 | 0.508926065 | 0.75217304 | -4.683948087 |
| **A_55_P2019203** | Spire1 | -0.078555455 | -0.357149404 | 0.200038494 | 8.329634271 | -0.706244398 | 0.508944228 | 0.75217304 | -4.683954506 |
| **A_55_P2121662** | Mapk3 | 0.183986768 | -0.468680828 | 0.836654364 | 9.387230038 | 0.706065327 | 0.509047195 | 0.752267711 | -4.68399089 |
| **A_51_P499169** | Tram1 | 0.092800128 | -0.23646405 | 0.422064306 | 10.8387715 | 0.705918294 | 0.509131749 | 0.75233516 | -4.684020758 |
| **A_51_P128987** | Akr1b8 | 0.089864903 | -0.229074411 | 0.408804217 | 10.25222126 | 0.705719958 | 0.509245823 | 0.752446216 | -4.684061042 |
| **A_51_P110814** | Nudt9 | -0.126537475 | -0.575784059 | 0.322709109 | 12.74082506 | -0.70548003 | 0.50938384 | 0.75250174 | -4.681160881 |
| **A_52_P207964** | Mrps18b | -0.258032072 | -1.174228673 | 0.658164529 | 8.81371253 | -0.705399843 | 0.509429973 | 0.75250174 | -4.684126043 |
| **A_55_P1982951** | Dcaf10 | -0.08870315 | -0.403664343 | 0.226258042 | 8.405234127 | -0.705394947 | 0.50943279 | 0.75250174 | -4.684127037 |
| **A_55_P2108067** | Dgcr8 | 0.102305382 | -0.260964102 | 0.465574866 | 9.872053395 | 0.705374748 | 0.509444411 | 0.75250174 | -4.684131138 |
| **A_52_P491244** | Zfp287 | 0.135427645 | -0.345578738 | 0.616434029 | 8.734665199 | 0.705191117 | 0.50955007 | 0.75250174 | -4.684168415 |
| **A_55_P2015670** | Itga6 | -0.124645295 | -0.567382627 | 0.318092036 | 14.66568134 | -0.705147701 | 0.509575053 | 0.75250174 | -4.684177227 |
| **A_51_P119401** | Tug1 | 0.154266896 | -0.393708639 | 0.702242431 | 9.139513826 | 0.705118189 | 0.509592036 | 0.75250174 | -4.684183217 |
| **A_51_P473259** | Dpyd | 0.31353019 | -0.800254072 | 1.427314453 | 8.842798924 | 0.705064153 | 0.509623132 | 0.75250174 | -4.684194184 |
| **A_55_P2100628** | Gm12216 | 0.155939387 | -0.398033916 | 0.709912691 | 10.73201337 | 0.705045805 | 0.509633691 | 0.75250174 | -4.684197908 |
| **A_55_P2060454** | Nsd3 | -0.394184066 | -1.794936827 | 1.006568695 | 10.43476228 | -0.704835749 | 0.509754586 | 0.75257158 | -4.684240534 |
| **A_55_P2124498** | Il17rb | -0.31289315 | -1.424971779 | 0.79918548 | 8.02534806 | -0.704710765 | 0.509826528 | 0.75257158 | -4.684265893 |
| **A_52_P350786** | Phlda2 | -0.127573218 | -0.58103858 | 0.325892144 | 7.990921455 | -0.704637475 | 0.509868717 | 0.75257158 | -4.684280761 |
| **A_55_P2129786** | Slc38a10 | -0.136486033 | -0.621637681 | 0.348665615 | 8.571422657 | -0.704629865 | 0.509873098 | 0.75257158 | -4.684282305 |
| **A_55_P2339650** | Hnrnpl | 0.065131114 | -0.166384614 | 0.296646842 | 8.648755133 | 0.704625494 | 0.509875615 | 0.75257158 | -4.684283192 |
| **A_52_P117420** | Tacc1 | 0.0683141 | -0.174590533 | 0.311218732 | 10.25488654 | 0.704409003 | 0.510000254 | 0.752698084 | -4.684327105 |
| **A_55_P2030501** | Ptpa | 0.119213607 | -0.304865314 | 0.543292528 | 13.69950528 | 0.704092127 | 0.510182723 | 0.752809016 | -4.684391364 |

| | | | | | | | | | |
|---|---|---|---|---|---|---|---|---|---|
| **A_55_P2164909** | Gatd3a | 0.277376375 | -0.709370527 | 1.264123277 | 12.97656031 | 0.704067086 | 0.510197145 | 0.752809016 | -4.684396441 |
| **A_55_P2027259** | Zcchc2 | -0.247628189 | -1.128566554 | 0.633310177 | 9.45978693 | -0.704052269 | 0.510205678 | 0.752809016 | -4.684399445 |
| **A_52_P375876** | Arf3 | 0.119147975 | -0.304748112 | 0.543044062 | 12.69927064 | 0.704008015 | 0.510231165 | 0.752809016 | -4.684408417 |
| **A_55_P2190299** | Fggy | 0.08312233 | -0.212646887 | 0.378891547 | 8.789985744 | 0.703906674 | 0.510289535 | 0.752837685 | -4.684428962 |
| **A_51_P471458** | Sult5a1 | 0.404277153 | -1.034501473 | 1.843055778 | 9.323488301 | 0.703777805 | 0.510363766 | 0.752889749 | -4.684455084 |
| **A_52_P322535** | Mplkip | 0.068614661 | -0.175693705 | 0.312923027 | 8.322646158 | 0.703443025 | 0.51055664 | 0.7529475 | -4.684522929 |
| **A_55_P1968028** | Tdgf1 | 0.091590381 | -0.234561505 | 0.417742267 | 8.451387515 | 0.703364292 | 0.510602007 | 0.7529475 | -4.684538881 |
| **A_51_P209122** | Agr2 | 0.449748283 | -1.15195156 | 2.051448126 | 9.041143817 | 0.703296938 | 0.510640819 | 0.7529475 | -4.684552527 |
| **A_52_P239052** | Zfp148 | 0.121508865 | -0.311232626 | 0.554250356 | 11.53577948 | 0.703282438 | 0.510649175 | 0.7529475 | -4.684555465 |
| **A_55_P1997936** | Hsd17b7 | -0.160441062 | -0.731844637 | 0.410962512 | 8.193758599 | -0.703271344 | 0.510655568 | 0.7529475 | -4.684557712 |
| **A_55_P2165324** | Acsl3 | 0.14926994 | -0.382358433 | 0.680898313 | 9.632638547 | 0.703257793 | 0.510663377 | 0.7529475 | -4.684560458 |
| **A_52_P680870** | Ctxn1 | 0.146165894 | -0.374422949 | 0.666754738 | 8.548810665 | 0.703236716 | 0.510675523 | 0.7529475 | -4.684564727 |
| **A_55_P2095854** | Rab3c | 0.17805263 | -0.45629784 | 0.8124031 | 12.37860529 | 0.703022903 | 0.510798751 | 0.753071759 | -4.684608037 |
| **A_52_P407871** | Lsm12 | 0.068393636 | -0.175405212 | 0.312192484 | 10.98816625 | 0.702642458 | 0.511018063 | 0.753331073 | -4.684685075 |
| **A_55_P2181655** | Tns3 | -0.082327718 | -0.375821582 | 0.211166146 | 8.479639097 | -0.702582613 | 0.511052568 | 0.753331073 | -4.684697191 |
| **A_55_P2054883** | Trappc1 | 0.116505068 | -0.298982996 | 0.531993131 | 9.300037131 | 0.702322548 | 0.511202529 | 0.753435569 | -4.684749833 |
| **A_55_P2027870** | Abhd17a | 0.077001094 | -0.197631933 | 0.351634121 | 10.57270328 | 0.702254411 | 0.511241824 | 0.753435569 | -4.684763623 |
| **A_55_P2003638** | Stxbp6 | 0.10027641 | -0.257373069 | 0.457925889 | 8.949549164 | 0.702249643 | 0.511244574 | 0.753435569 | -4.684764588 |
| **A_52_P40293** | Hgd | -0.066814983 | -0.30514018 | 0.171510213 | 7.666891693 | -0.70218937 | 0.511279335 | 0.753435569 | -4.684776785 |
| **A_51_P509997** | Cox6a2 | 1.01722736 | -2.612700324 | 4.647155044 | 11.38554678 | 0.701892182 | 0.511450756 | 0.753623765 | -4.684836916 |
| **A_55_P2142033** | Clip4 | 0.087069547 | -0.223695087 | 0.397834182 | 7.912541732 | 0.701754236 | 0.511530339 | 0.753623765 | -4.684864821 |
| **A_55_P1955242** | NA | -0.093091852 | -0.425374783 | 0.239191078 | 16.2670725 | -0.701704052 | 0.511559292 | 0.753623765 | -4.684874972 |
| **A_55_P2085766** | NA | -0.084476118 | -0.386011749 | 0.217059513 | 7.824261948 | -0.701690593 | 0.511567057 | 0.753623765 | -4.684877694 |
| **A_55_P1968789** | Rasa3 | 0.393898699 | -1.012278094 | 1.800075492 | 12.38881244 | 0.7016087 | 0.511614308 | 0.753623765 | -4.684894257 |
| **A_52_P489778** | Ablim1 | 0.191082789 | -0.491117414 | 0.873282993 | 8.943202254 | 0.701552333 | 0.511646833 | 0.753623765 | -4.684905657 |
| **A_65_P16796** | Psd4 | -0.179606079 | -0.820920826 | 0.461708667 | 7.989510312 | -0.701455545 | 0.511702684 | 0.753623765 | -4.68492523 |
| **A_52_P6451** | Sec24a | -0.122163937 | -0.558401257 | 0.314073382 | 9.07001689 | -0.701407734 | 0.511730275 | 0.753623765 | -4.684934897 |
| **A_55_P2068391** | Gm9222 | 0.137772756 | -0.354260478 | 0.62980599 | 9.810209022 | 0.701324891 | 0.511778084 | 0.753623765 | -4.684951648 |
| **A_55_P2182497** | Maml3 | -0.203511594 | -0.930354188 | 0.523331 | 10.81492405 | -0.701292399 | 0.511796836 | 0.753623765 | -4.684958217 |
| **A_55_P2160043** | A330069E | 0.06614364 | -0.170186185 | 0.302473464 | 8.057991904 | 0.70100304 | 0.511963855 | 0.753677885 | -4.685016711 |
| **A_51_P478610** | Asb6 | -0.122609253 | -0.560717633 | 0.315499127 | 10.11757888 | -0.700958058 | 0.511989822 | 0.753677885 | -4.685025802 |
| **A_51_P519369** | Agtpbp1 | 0.105877504 | -0.272492927 | 0.484247935 | 9.132449706 | 0.700869001 | 0.512041236 | 0.753677885 | -4.685043801 |
| **A_66_P119247** | Gatad1 | 0.115155473 | -0.296374982 | 0.526685928 | 10.96826193 | 0.7008627 | 0.512044873 | 0.753677885 | -4.685045074 |
| **A_51_P136460** | Zmat2 | 0.083746832 | -0.215540549 | 0.383034213 | 12.28391212 | 0.700858474 | 0.512047313 | 0.753677885 | -4.685045928 |
| **A_65_P13713** | Ank3 | 0.163215857 | -0.420157184 | 0.746588897 | 8.331546615 | 0.700755216 | 0.512106929 | 0.753677885 | -4.685066794 |
| **A_55_P2034275** | Arfgap2 | 0.067664035 | -0.174189108 | 0.309517177 | 11.90565493 | 0.700739341 | 0.512116095 | 0.753677885 | -4.685070002 |
| **A_55_P2146002** | Pank4 | 0.201814765 | -0.519691817 | 0.923321348 | 10.68854633 | 0.70058847 | 0.512203211 | 0.753677885 | -4.685100486 |
| **A_55_P2072000** | Pcmtd1 | -0.066585731 | -0.304659689 | 0.171488227 | 8.627758498 | -0.700518527 | 0.5122436 | 0.753677885 | -4.685114616 |
| **A_55_P2047188** | Fgf1 | 0.6588458 | -1.697031914 | 3.014723515 | 11.1841382 | 0.700456845 | 0.512279221 | 0.753677885 | -4.685127077 |
| **A_55_P2013465** | L2hgdh | -0.114008859 | -0.521695463 | 0.293677746 | 8.023274161 | -0.700426391 | 0.512296809 | 0.753677885 | -4.685133229 |
| **A_55_P1954758** | Prickle1 | 0.259435822 | -0.668324801 | 1.187196445 | 12.2958948 | 0.700397119 | 0.512313714 | 0.753677885 | -4.685139142 |
| **A_55_P1953301** | Sord | -0.157673039 | -0.721581304 | 0.406235226 | 10.40000375 | -0.700324506 | 0.512355652 | 0.753677885 | -4.685153809 |
| **A_55_P1987839** | Erbin | 0.135795172 | -0.349896642 | 0.621486986 | 11.20186479 | 0.700283496 | 0.512379338 | 0.753677885 | -4.685162091 |
| **A_51_P123705** | Dus3l | 0.067340048 | -0.173536617 | 0.308216713 | 10.0861124 | 0.700211175 | 0.51242111 | 0.753681989 | -4.683715276 |
| **A_51_P392214** | Pcnx4 | -0.114135082 | -0.522575565 | 0.2943054 | 9.225290108 | -0.699907619 | 0.512596469 | 0.753789627 | -4.685237993 |
| **A_55_P1958210** | Ephb1 | 0.157903038 | -0.407168064 | 0.72297414 | 8.243846537 | 0.699902803 | 0.512599252 | 0.753789627 | -4.685238966 |
| **A_66_P119174** | Cmtm6 | -0.104051193 | -0.476424343 | 0.268321956 | 12.20074687 | -0.6998727 | 0.512616644 | 0.753789627 | -4.685245043 |
| **A_51_P392244** | Tbcd | -0.095857184 | -0.438938424 | 0.247224055 | 10.50916305 | -0.699806614 | 0.512654827 | 0.753789627 | -4.685258385 |

| | | | | | | | | | |
|---|---|---|---|---|---|---|---|---|---|
| **A_52_P539434** | Lbh | 0.250821045 | -0.646965781 | 1.14860787 | 8.832036911 | 0.699747072 | 0.512689231 | 0.753789627 | -4.685270404 |
| **A_55_P2044297** | Ciao2a | 0.085090039 | -0.219553308 | 0.389733387 | 12.66466483 | 0.699579978 | 0.512785787 | 0.75383463 | -4.685304131 |
| **A_55_P2027240** | Engase | -0.316774072 | -1.450973439 | 0.817425296 | 9.783559922 | -0.699536809 | 0.512810734 | 0.75383463 | -4.685312843 |
| **A_66_P132062** | Gm5608 | -0.080737138 | -0.369832509 | 0.208358233 | 7.843674336 | -0.699491689 | 0.51283681 | 0.75383463 | -4.685321949 |
| **A_51_P132081** | Rsl1 | -0.064021507 | -0.293358131 | 0.165315117 | 8.519522769 | -0.69920226 | 0.513004098 | 0.754023205 | -4.68538035 |
| **A_52_P378968** | Rgs2 | 0.275154228 | -0.710906349 | 1.261214805 | 8.674777746 | 0.698912713 | 0.513171492 | 0.754150728 | -4.685438757 |
| **A_55_P2112295** | Hmg20b | -0.107827217 | -0.494279287 | 0.278624854 | 13.29629203 | -0.698848604 | 0.513208559 | 0.754150728 | -4.685451686 |
| **A_52_P486482** | R3hcc1l | -0.061756878 | -0.283101419 | 0.159587663 | 8.439876642 | -0.698822474 | 0.513223668 | 0.754150728 | -4.685456956 |
| **A_55_P1968977** | Stk38l | -0.088013825 | -0.403484747 | 0.227457096 | 8.741505974 | -0.698782322 | 0.513246885 | 0.754150728 | -4.685465053 |
| **A_52_P413395** | Sln | 1.120790288 | -2.89751513 | 5.139095706 | 11.87520185 | 0.698605116 | 0.51334936 | 0.75424398 | -4.685500786 |
| **A_55_P1962315** | NA | 0.217720984 | -0.563107657 | 0.998549625 | 16.20752065 | 0.698384994 | 0.513476672 | 0.754364603 | -4.685545163 |
| **A_51_P321374** | Plpp7 | -0.081565147 | -0.374135763 | 0.211005469 | 8.187009618 | -0.698271417 | 0.513542369 | 0.754364603 | -4.685568056 |
| **A_55_P1956572** | Dcaf8 | -0.064616493 | -0.296451098 | 0.167218112 | 12.38141879 | -0.698096511 | 0.513643553 | 0.754364603 | -4.685603306 |
| **A_51_P394814** | Svep1 | 0.3593931 | -0.93021927 | 1.64900547 | 11.34620292 | 0.698008788 | 0.513694306 | 0.754364603 | -4.685620984 |
| **A_55_P1999601** | NA | 0.201614614 | -0.521854378 | 0.925083606 | 13.47858766 | 0.697995193 | 0.513702172 | 0.754364603 | -4.685623723 |
| **A_66_P129800** | Rab8b | 0.148939965 | -0.385579571 | 0.683459501 | 10.89024866 | 0.697907732 | 0.513752778 | 0.754364603 | -4.685641346 |
| **A_55_P2015258** | Trim13 | -0.068663778 | -0.315114389 | 0.177786834 | 8.996955949 | -0.697827593 | 0.51379915 | 0.754364603 | -4.685657491 |
| **A_55_P1956948** | Rpl27 | 0.152591983 | -0.395132245 | 0.700316211 | 16.63254059 | 0.69778256 | 0.513825209 | 0.754364603 | -4.685666564 |
| **A_55_P2210315** | Get4 | -0.127350313 | -0.584480063 | 0.329779437 | 9.364318762 | -0.697767724 | 0.513833794 | 0.754364603 | -4.685669552 |
| **A_55_P2023697** | Zfp386 | 0.142022901 | -0.367796471 | 0.651842273 | 11.08347751 | 0.697738024 | 0.513850981 | 0.754364603 | -4.685675535 |
| **A_66_P134534** | Atl2 | -0.100061434 | -0.459260402 | 0.259137533 | 8.330426759 | -0.697721315 | 0.513860651 | 0.754364603 | -4.685678901 |
| **A_51_P343309** | Cmas | -0.183119909 | -0.840630884 | 0.474391066 | 10.31624375 | -0.697562144 | 0.51395277 | 0.754419456 | -4.685710963 |
| **A_51_P481482** | Dram1 | 0.390093125 | -1.010657381 | 1.790843631 | 9.900702447 | 0.697521911 | 0.513976056 | 0.754419456 | -4.685719066 |
| **A_55_P2031871** | Plekhh3 | -0.109561516 | -0.503050762 | 0.28392773 | 11.21333613 | -0.697389672 | 0.514052599 | 0.754474528 | -4.685745697 |
| **A_52_P120925** | Metap1 | -0.149008493 | -0.684240871 | 0.386223886 | 8.295722487 | -0.697298914 | 0.514105137 | 0.754478557 | -4.685763972 |
| **A_51_P214663** | Tpmt | 0.104872611 | -0.271928355 | 0.481673578 | 8.512312731 | 0.697108566 | 0.514215335 | 0.754478557 | -4.685802296 |
| **A_51_P361220** | Fzd4 | 0.3464863 | -0.898430114 | 1.591402715 | 10.59619379 | 0.697101826 | 0.514219238 | 0.754478557 | -4.685803653 |
| **A_66_P101755** | Sp1 | 0.222285185 | -0.576390499 | 1.020960868 | 11.27438857 | 0.697092485 | 0.514224646 | 0.754478557 | -4.685805533 |
| **A_55_P1958295** | NA | 0.557424169 | -1.445541558 | 2.560389896 | 13.1301488 | 0.697047899 | 0.514250461 | 0.754478557 | -4.685814509 |
| **A_52_P160936** | Gusb | 0.081680919 | -0.211937662 | 0.375299501 | 9.615768853 | 0.696766762 | 0.514413257 | 0.754583925 | -4.685871094 |
| **A_55_P1988158** | Esd | -0.072957218 | -0.335228669 | 0.189314234 | 14.0160278 | -0.696734937 | 0.514431689 | 0.754583925 | -4.685877499 |
| **A_55_P2029711** | Cdh2 | 0.489762164 | -1.270900897 | 2.250425225 | 10.65631484 | 0.696721678 | 0.514439367 | 0.754583925 | -4.685880167 |
| **A_51_P430423** | Ada | -1.307312812 | -6.007492577 | 3.392866953 | 10.48958917 | -0.696651208 | 0.514480181 | 0.754586543 | -4.685894348 |
| **A_55_P2067842** | 5-Mar | 0.05924441 | -0.153845564 | 0.272334383 | 10.1207511 | 0.696361427 | 0.514648033 | 0.754625816 | -4.685952649 |
| **A_55_P2037398** | NA | -0.169893086 | -0.780984269 | 0.441198097 | 15.47354842 | -0.696337899 | 0.514661664 | 0.754625816 | -4.685957382 |
| **A_55_P1971330** | NA | 0.211427812 | -0.549105569 | 0.971961194 | 9.198374493 | 0.69629645 | 0.514685676 | 0.754625816 | -4.685965719 |
| **A_55_P2105286** | Lrrc56 | 0.217377597 | -0.564678371 | 0.999433566 | 10.07399963 | 0.696189222 | 0.514747799 | 0.754625816 | -4.685987286 |
| **A_51_P424054** | Btbd2 | 0.231581919 | -0.601785247 | 1.064949086 | 10.31543544 | 0.696015015 | 0.514848737 | 0.754625816 | -4.686022321 |
| **A_55_P2001342** | Cacul1 | -0.240372202 | -1.105377593 | 0.624633189 | 8.81340804 | -0.69601047 | 0.514851371 | 0.754625816 | -4.686023235 |
| **A_51_P279308** | Acbd5 | -0.080393488 | -0.369718369 | 0.208931393 | 9.6693633 | -0.695961847 | 0.514879546 | 0.754625816 | -4.686033012 |
| **A_52_P680038** | Cdk16 | -0.137848252 | -0.633957627 | 0.358261123 | 8.473810601 | -0.695943808 | 0.51489 | 0.754625816 | -4.686036639 |
| **A_51_P231687** | Ik | 0.068062866 | -0.17692166 | 0.313047392 | 13.60893625 | 0.69586008 | 0.51493852 | 0.754625816 | -4.686053474 |
| **A_55_P2184146** | NA | 0.07019727 | -0.182491391 | 0.322885931 | 8.577263937 | 0.695800623 | 0.514972978 | 0.754625816 | -4.686065428 |
| **A_55_P2398534** | Gid8 | 0.082842996 | -0.215413224 | 0.381099216 | 10.97468079 | 0.6956914 | 0.515036281 | 0.754625816 | -4.686087385 |
| **A_55_P2151138** | Dynlt1c | 0.365775947 | -0.951151637 | 1.682703531 | 13.8687065 | 0.695670512 | 0.515048388 | 0.754625816 | -4.686091584 |
| **A_55_P2030756** | Nedd4l | 0.287364732 | -0.747326625 | 1.322056088 | 9.133367842 | 0.695621491 | 0.515076801 | 0.754625816 | -4.686101438 |
| **A_55_P2280821** | Dlc1 | 0.100177549 | -0.260531337 | 0.460886435 | 9.983877275 | 0.695606939 | 0.515085236 | 0.754625816 | -4.686104363 |
| **A_55_P2018949** | Fbxo7 | -0.213819575 | -0.983732558 | 0.556093408 | 12.07067815 | -0.69559454 | 0.515092423 | 0.754625816 | -4.686106855 |

| | | | | | | | | | |
|---|---|---|---|---|---|---|---|---|---|
| **A_51_P386983** | Tnni3 | 0.840370983 | -2.186108921 | 3.866850888 | 10.76955459 | 0.695478325 | 0.515159788 | 0.754667324 | -4.686130213 |
| **A_55_P2037294** | NA | 0.131544731 | -0.342284304 | 0.605373766 | 8.449560212 | 0.695347876 | 0.515235412 | 0.754720922 | -4.686156428 |
| **A_51_P249930** | Tceal1 | 0.277175115 | -0.7214018 | 1.275752031 | 9.376000943 | 0.695221285 | 0.515308806 | 0.754771247 | -4.686181865 |
| **A_55_P2062737** | Elovl2 | 0.074860035 | -0.194868214 | 0.344588283 | 8.131187151 | 0.695142651 | 0.5153544 | 0.754780847 | -4.686197664 |
| **A_51_P448946** | Sertad2 | 0.076941935 | -0.200539537 | 0.354423407 | 11.31402957 | 0.694511529 | 0.515720435 | 0.7551466 | -4.68632442 |
| **A_55_P1955243** | Rps24 | 0.438264224 | -1.142546031 | 2.019074478 | 17.158486 | 0.694395058 | 0.515788005 | 0.7551466 | -4.686347803 |
| **A_55_P2111153** | NA | 0.21649557 | -0.564543156 | 0.997534295 | 10.59846098 | 0.694267429 | 0.515862054 | 0.7551466 | -4.686373423 |
| **A_55_P2253885** | AI503301 | 0.108107059 | -0.281912442 | 0.498126559 | 8.567968991 | 0.694253496 | 0.515870139 | 0.7551466 | -4.68637622 |
| **A_55_P1966438** | Gstm2 | 0.3120989 | -0.814045819 | 1.438243619 | 9.298911413 | 0.69414209 | 0.515934782 | 0.7551466 | -4.68639858 |
| **A_55_P2111825** | 1700001C | 0.487348117 | -1.271207385 | 2.245903618 | 9.508698821 | 0.694118402 | 0.515948528 | 0.7551466 | -4.686403334 |
| **A_55_P2102155** | Rgs20 | -0.365808991 | -1.685808229 | 0.954190246 | 11.24338599 | -0.694114379 | 0.515950862 | 0.7551466 | -4.686404142 |
| **A_51_P312497** | Atp7a | -0.130479412 | -0.601388693 | 0.340429868 | 11.33728376 | -0.693992997 | 0.516021302 | 0.7551466 | -4.686428501 |
| **A_66_P123643** | Ptbp2 | 0.168824009 | -0.440534195 | 0.778182213 | 8.562548667 | 0.693923975 | 0.516061359 | 0.7551466 | -4.686442351 |
| **A_55_P2103561** | Prdx6 | 0.331182957 | -0.864386113 | 1.526752026 | 14.41354617 | 0.69381495 | 0.516124636 | 0.7551466 | -4.686464226 |
| **A_55_P1957088** | Fam53b | 0.13402809 | -0.349815554 | 0.617871733 | 11.64231668 | 0.693810923 | 0.516126974 | 0.7551466 | -4.686465034 |
| **A_52_P301223** | Brwd1 | -0.12327876 | -0.568345189 | 0.32178767 | 9.535177249 | -0.693767202 | 0.516152351 | 0.7551466 | -4.686473805 |
| **A_55_P2083954** | Scfd1 | -0.119278999 | -0.54990571 | 0.311347711 | 9.624863331 | -0.693766569 | 0.516152718 | 0.7551466 | -4.686473932 |
| **A_55_P1968628** | NA | -0.100917287 | -0.465281512 | 0.263446938 | 8.324669011 | -0.693713558 | 0.516183489 | 0.7551466 | -4.686484567 |
| **A_51_P221362** | Tnks2 | -0.102005851 | -0.487798159 | 0.283786457 | 12.62158783 | -0.704261949 | 0.51619 | 0.7551466 | -4.665911613 |
| **A_55_P2077548** | Git1 | -0.164934102 | -0.760686336 | 0.430818132 | 11.65895889 | -0.693418024 | 0.516355054 | 0.755294474 | -4.686543845 |
| **A_55_P2096226** | Loxl1 | 0.218587823 | -0.571153555 | 1.008329201 | 8.84051793 | 0.693252456 | 0.516451188 | 0.755294474 | -4.686577047 |
| **A_66_P125734** | Tnip2 | -0.07437082 | -0.343073907 | 0.194332267 | 9.123238216 | -0.693234646 | 0.51646153 | 0.755294474 | -4.686580618 |
| **A_51_P395842** | Timm21 | 0.148023263 | -0.386810272 | 0.682856798 | 10.12645238 | 0.693205002 | 0.516478743 | 0.755294474 | -4.686586562 |
| **A_55_P2166688** | Timm10b | 0.244344482 | -0.63871403 | 1.127402994 | 11.27544793 | 0.693048137 | 0.516569838 | 0.755294474 | -4.686618011 |
| **A_55_P1990810** | 4932412D | 0.070746012 | -0.184930872 | 0.326422896 | 7.986435837 | 0.693044055 | 0.516572208 | 0.755294474 | -4.68661883 |
| **A_55_P2010116** | Rab27b | 0.296561197 | -0.775328073 | 1.368450467 | 9.739611539 | 0.692970502 | 0.516614926 | 0.755294474 | -4.686633574 |
| **A_55_P2020306** | Aasdh | -0.061360003 | -0.283151499 | 0.160431492 | 9.913707764 | -0.692932341 | 0.51663709 | 0.755294474 | -4.686641224 |
| **A_51_P226269** | Rgcc | -0.170960693 | -0.788950524 | 0.447029137 | 13.89787043 | -0.692891586 | 0.516660761 | 0.755294474 | -4.686649393 |
| **A_55_P2052252** | NA | 0.061968001 | -0.16204601 | 0.285982013 | 8.523390987 | 0.692855471 | 0.516681738 | 0.755294474 | -4.686656632 |
| **A_52_P7183** | Tmem248 | -0.273487231 | -1.262352184 | 0.715377722 | 10.50249109 | -0.692708336 | 0.516767204 | 0.755345659 | -4.68668612 |
| **A_55_P2133652** | Cep162 | -0.073211135 | -0.337966083 | 0.191543812 | 9.113152187 | -0.692601461 | 0.516829291 | 0.755345659 | -4.686707537 |
| **A_51_P237834** | Tmcc1 | -0.222874391 | -1.028885769 | 0.583136987 | 8.982817175 | -0.692579037 | 0.516842318 | 0.755345659 | -4.68671203 |
| **A_55_P2161865** | Pdlim7 | -0.071715306 | -0.331094063 | 0.187663452 | 8.107633241 | -0.692512783 | 0.51688081 | 0.755345659 | -4.686725305 |
| **A_55_P2037608** | Acp5 | -0.66292037 | -3.060746475 | 1.734905735 | 11.53668954 | -0.692458939 | 0.516912093 | 0.755345659 | -4.686736093 |
| **A_51_P122035** | Nipa1 | 0.09744663 | -0.255145178 | 0.450038438 | 9.454453287 | 0.692221289 | 0.517050183 | 0.755382924 | -4.6867837 |
| **A_66_P112436** | Prkar2b | 0.109029237 | -0.285473875 | 0.503532349 | 8.313360798 | 0.692217994 | 0.517052098 | 0.755382924 | -4.68678436 |
| **A_55_P2039250** | Mboat2 | 0.221845327 | -0.580877777 | 1.024568432 | 10.6595292 | 0.692205217 | 0.517059523 | 0.755382924 | -4.686786919 |
| **A_52_P311297** | Als2 | -0.107181854 | -0.495059661 | 0.280695953 | 8.181035781 | -0.692112479 | 0.517113417 | 0.755382924 | -4.686805493 |
| **A_55_P2086664** | Tspan33 | -0.453067041 | -2.092853335 | 1.186719254 | 10.41682997 | -0.692031021 | 0.517160759 | 0.755382924 | -4.686821806 |
| **A_55_P2079251** | Ap2m1 | -0.187789411 | -0.867474962 | 0.491896141 | 11.65771518 | -0.692011652 | 0.517172016 | 0.755382924 | -4.686825685 |
| **A_55_P1984243** | Naip2 | -0.108160053 | -0.499677472 | 0.283357365 | 8.218854985 | -0.691936356 | 0.51721578 | 0.755389779 | -4.686840763 |
| **A_55_P1957003** | Timm10 | 0.088390614 | -0.231629955 | 0.408411184 | 11.64087069 | 0.691796889 | 0.517296848 | 0.755451111 | -4.686868688 |
| **A_52_P23308** | Fam241a | 0.11784235 | -0.308874895 | 0.544559595 | 9.051316918 | 0.691690078 | 0.51735894 | 0.755484724 | -4.686890071 |
| **A_55_P2115392** | Mst1 | -0.087666026 | -0.405219066 | 0.229887015 | 7.821099986 | -0.691457337 | 0.517494256 | 0.75562525 | -4.686936658 |
| **A_55_P2148385** | NA | -0.09903701 | -0.45784713 | 0.25977311 | 8.886099634 | -0.691326464 | 0.517570355 | 0.755679297 | -4.686962849 |
| **A_52_P161087** | 2610008E | 0.228568602 | -0.599737182 | 1.056874386 | 9.649905437 | 0.691156226 | 0.517669357 | 0.7557303 | -4.686996913 |
| **A_55_P2171373** | Eloc | 0.062440064 | -0.163845096 | 0.288725224 | 12.89375034 | 0.691126605 | 0.517686584 | 0.7557303 | -4.687002839 |
| **A_52_P552665** | Fzd7 | 0.201565607 | -0.528981167 | 0.93211238 | 9.786962178 | 0.691064762 | 0.517722552 | 0.7557303 | -4.687015212 |

| | | | | | | | | | |
|---|---|---|---|---|---|---|---|---|---|
| **A_55_P2059055** | Zfp943 | 0.215422775 | -0.565648608 | 0.996494159 | 10.05812145 | 0.69079826 | 0.517877571 | 0.755817755 | -4.68706852 |
| **A_55_P2006911** | Pik3c3 | -0.065240861 | -0.301799413 | 0.171317691 | 8.488297578 | -0.690766674 | 0.517895946 | 0.755817755 | -4.687074837 |
| **A_55_P2141876** | Ldha | 0.212581381 | -0.55838473 | 0.983547493 | 10.39062569 | 0.690621798 | 0.517980233 | 0.755817755 | -4.68710381 |
| **A_51_P419971** | Tmem106 | -0.092071368 | -0.426000555 | 0.24185782 | 12.11988827 | -0.690590435 | 0.51799848 | 0.755817755 | -4.687110081 |
| **A_55_P2052380** | Gbp6 | -0.087319467 | -0.404028566 | 0.229389633 | 8.11333839 | -0.690559141 | 0.518016689 | 0.755817755 | -4.687116338 |
| **A_55_P2025730** | Hif3a | 0.172356624 | -0.452837664 | 0.797550911 | 8.454937083 | 0.690499428 | 0.518051433 | 0.755817755 | -4.687128278 |
| **A_51_P366024** | Ercc3 | -0.070934933 | -0.328242315 | 0.186372449 | 9.1113268 | -0.690491385 | 0.518056113 | 0.755817755 | -4.687129886 |
| **A_55_P1979616** | Tmem189 | -0.370056673 | -1.713029592 | 0.972916245 | 9.457474918 | -0.690162457 | 0.518247531 | 0.755951945 | -4.68719564 |
| **A_55_P2002210** | Babam2 | 0.075130229 | -0.197527696 | 0.347788155 | 11.74433209 | 0.690155468 | 0.518251598 | 0.755951945 | -4.687197037 |
| **A_51_P200561** | Shtn1 | -0.072167348 | -0.334081561 | 0.189746866 | 10.26059522 | -0.690131774 | 0.518265389 | 0.755951945 | -4.687201772 |
| **A_55_P2153887** | Tatdn3 | -0.095306544 | -0.44132985 | 0.250716762 | 9.3564579 | -0.689870725 | 0.518417342 | 0.756064477 | -4.687253939 |
| **A_51_P454691** | 4930430F | 0.139373476 | -0.366648232 | 0.645395184 | 8.705229632 | 0.68986058 | 0.518423248 | 0.756064477 | -4.687255966 |
| **A_55_P2025023** | NA | 0.166974314 | -0.439312708 | 0.773261336 | 16.12458841 | 0.689797698 | 0.518459855 | 0.756064477 | -4.68726853 |
| **A_55_P2151156** | Farsa | -0.310247305 | -1.436972282 | 0.816477672 | 11.19325745 | -0.689668581 | 0.518535028 | 0.756070147 | -4.687294325 |
| **A_55_P1982916** | Tspoap1 | 0.071721695 | -0.188754812 | 0.332198201 | 7.95010505 | 0.689655701 | 0.518542527 | 0.756070147 | -4.687296898 |
| **A_51_P206251** | Ahcyl1 | 0.103317636 | -0.271943986 | 0.478579258 | 9.630811717 | 0.689589522 | 0.51858106 | 0.756070147 | -4.687310117 |
| **A_55_P2038489** | NA | 0.102180029 | -0.269012528 | 0.473372587 | 8.352783809 | 0.689472755 | 0.518649052 | 0.756112259 | -4.68733344 |
| **A_51_P354038** | Tmc6 | -0.086501029 | -0.400792993 | 0.227790935 | 10.26456078 | -0.689347715 | 0.518721869 | 0.75612381 | -4.687358412 |
| **A_55_P2053929** | NA | -0.222113879 | -1.02916699 | 0.584939232 | 11.82347556 | -0.689324836 | 0.518735193 | 0.75612381 | -4.687362981 |
| **A_55_P2031125** | Ube2l6 | 0.119491983 | -0.314771546 | 0.553755512 | 10.70849531 | 0.689184906 | 0.51881669 | 0.756138909 | -4.687390922 |
| **A_51_P125368** | Hars | 0.074201142 | -0.195471927 | 0.343874212 | 9.632243753 | 0.689165231 | 0.51882815 | 0.756138909 | -4.68739485 |
| **A_55_P1970270** | B3gnt2 | 0.186839849 | -0.492295335 | 0.865975032 | 9.76549918 | 0.689070444 | 0.518883361 | 0.756138909 | -4.687413774 |
| **A_55_P2080870** | Cmtm7 | -0.285784908 | -1.324618942 | 0.753049126 | 11.76483948 | -0.689038465 | 0.518901988 | 0.756138909 | -4.687420159 |
| **A_55_P2017709** | Igsf9 | 0.14770307 | -0.389345175 | 0.684751315 | 12.83839857 | 0.688853018 | 0.51901002 | 0.756222876 | -4.687457176 |
| **A_55_P2109337** | Gba2 | -0.104204483 | -0.483118166 | 0.2747092 | 11.38244428 | -0.68880527 | 0.519037838 | 0.756222876 | -4.687466706 |
| **A_55_P1972575** | Tmeff1 | -0.240917532 | -1.117102767 | 0.635267703 | 11.53052386 | -0.688688484 | 0.519105882 | 0.75623874 | -4.687490014 |
| **A_66_P111773** | Gatad2a | 0.085329611 | -0.225025978 | 0.3956852 | 9.68584974 | 0.688637295 | 0.519135708 | 0.75623874 | -4.687500229 |
| **A_51_P357341** | Clic1 | 0.118915695 | -0.313629852 | 0.551461242 | 14.90732834 | 0.688585189 | 0.51916607 | 0.75623874 | -4.687510626 |
| **A_52_P489729** | Sec61a1 | -0.164584279 | -0.763400142 | 0.434231584 | 13.66481565 | -0.688407191 | 0.519269797 | 0.756332784 | -4.68754614 |
| **A_52_P576689** | Zfp944 | -0.087543752 | -0.406119803 | 0.231032299 | 8.827057307 | -0.688275604 | 0.519346487 | 0.756332784 | -4.68757239 |
| **A_55_P1997126** | Ctse | -0.087617412 | -0.406467641 | 0.231232818 | 7.819060388 | -0.688262379 | 0.519354196 | 0.756332784 | -4.687575028 |
| **A_55_P2227154** | A530041M | -0.106516128 | -0.49417285 | 0.281140594 | 9.00849802 | -0.688205909 | 0.51938711 | 0.756332784 | -4.687586292 |
| **A_51_P411297** | Nup50 | -0.235775583 | -1.094352748 | 0.622801583 | 9.468936973 | -0.687812142 | 0.519616659 | 0.756604194 | -4.687664816 |
| **A_51_P459340** | Pacrgl | 0.064596264 | -0.170651725 | 0.299844253 | 9.197545702 | 0.687751943 | 0.519651758 | 0.756604194 | -4.687676818 |
| **A_55_P1990629** | NA | 0.161782493 | -0.427484241 | 0.751049226 | 11.19626706 | 0.687653954 | 0.519708895 | 0.756619692 | -4.687696353 |
| **A_55_P2073526** | H2-Ke6 | -0.080444117 | -0.373472106 | 0.212583873 | 11.17531446 | -0.687599461 | 0.519740671 | 0.756619692 | -4.687707215 |
| **A_51_P124719** | Ccdc93 | -0.190373425 | -0.884005921 | 0.50325907 | 8.416668686 | -0.687428033 | 0.519840643 | 0.756708252 | -4.687741383 |
| **A_51_P506683** | Fbxw7 | -0.284339534 | -1.320610303 | 0.751931234 | 9.317603466 | -0.687249363 | 0.519944852 | 0.756775342 | -4.687776988 |
| **A_55_P2169415** | Cgn | 0.087188127 | -0.230612052 | 0.404988306 | 11.42493907 | 0.687153175 | 0.52000096 | 0.756775342 | -4.687796153 |
| **A_55_P1977468** | Dab2ip | -0.089262753 | -0.414627528 | 0.236102021 | 13.46447593 | -0.68714769 | 0.520004159 | 0.756775342 | -4.687797246 |
| **A_55_P2084345** | NA | 0.375991649 | -0.994733573 | 1.746716872 | 14.82318398 | 0.687033859 | 0.520070564 | 0.756788416 | -4.687819924 |
| **A_55_P1979970** | NA | 0.311668544 | -0.824860099 | 1.448197186 | 10.45020219 | 0.686851629 | 0.520176881 | 0.756788416 | -4.687856223 |
| **A_55_P2111985** | Prss35 | -0.094512517 | -0.439200343 | 0.250175309 | 8.263267542 | -0.686773817 | 0.520222282 | 0.756788416 | -4.687871721 |
| **A_55_P2067453** | Tspan17 | -0.094483937 | -0.43909016 | 0.250122286 | 10.862373 | -0.686728722 | 0.520248595 | 0.756788416 | -4.687880701 |
| **A_52_P77080** | Gypa | 0.067047185 | -0.177505871 | 0.311600241 | 7.952291261 | 0.686685381 | 0.520273886 | 0.756788416 | -4.687889333 |
| **A_55_P2143516** | NA | 0.181078787 | -0.479418308 | 0.841575883 | 10.15256659 | 0.68666832 | 0.520283842 | 0.756788416 | -4.68789273 |
| **A_51_P261351** | Mrpl49 | -0.105067958 | -0.488313583 | 0.278177666 | 10.31484731 | -0.686662663 | 0.520287143 | 0.756788416 | -4.687893857 |
| **A_55_P1996374** | Prmt9 | -0.107335512 | -0.49897659 | 0.284305566 | 10.99586231 | -0.686444673 | 0.52041436 | 0.756916515 | -4.687937261 |

| | | | | | | | | | |
|---|---|---|---|---|---|---|---|---|---|
| **A_55_P2046807** | Sparc | -0.216884472 | -1.008349333 | 0.57458039 | 14.18269426 | -0.686352415 | 0.520468207 | 0.756937891 | -4.687955628 |
| **A_55_P2034720** | Cfap36 | 0.288231211 | -0.7640787 | 1.340541123 | 10.8636836 | 0.686037241 | 0.52065219 | 0.757094721 | -4.688018359 |
| **A_55_P1991134** | Fkbp1a | 0.086014456 | -0.228019387 | 0.400048298 | 14.62618063 | 0.686033522 | 0.520654361 | 0.757094721 | -4.688019099 |
| **A_55_P2452259** | Gls | 0.117668155 | -0.312007661 | 0.54734397 | 9.535203727 | 0.685911963 | 0.520725333 | 0.757140978 | -4.688043288 |
| **A_55_P1983152** | Gramd4 | -0.155014649 | -0.721307 | 0.411277702 | 10.43481632 | -0.685618283 | 0.520896823 | 0.75728085 | -4.688101714 |
| **A_55_P2008735** | Cdc123 | -0.125375921 | -0.583396921 | 0.33264508 | 8.358404432 | -0.685613071 | 0.520899867 | 0.75728085 | -4.688102751 |
| **A_51_P467629** | Efcab10 | 0.423192199 | -1.122971223 | 1.969355622 | 9.2450092 | 0.685539691 | 0.520942722 | 0.75728621 | -4.688117347 |
| **A_66_P123527** | Ndufb4 | 0.224993138 | -0.597232718 | 1.047218993 | 13.90077449 | 0.685375362 | 0.521038702 | 0.757368789 | -4.688150029 |
| **A_55_P2048398** | Kmt5b | -0.089372687 | -0.416016602 | 0.237271228 | 8.457065549 | -0.685299776 | 0.521082853 | 0.757376025 | -4.688165059 |
| **A_51_P393078** | Tigd2 | 0.126895015 | -0.336963466 | 0.590753495 | 9.567226999 | 0.685187458 | 0.521148466 | 0.75741445 | -4.688187392 |
| **A_52_P437734** | Fbxo28 | -0.069805694 | -0.325170468 | 0.18555908 | 9.120565015 | -0.68466828 | 0.521451822 | 0.757784317 | -4.688290589 |
| **A_55_P1959425** | Slc16a3 | -0.696751011 | -3.245911683 | 1.852409661 | 11.21974649 | -0.684590304 | 0.521497394 | 0.757784317 | -4.688306084 |
| **A_55_P2145982** | NA | 0.075551733 | -0.200880802 | 0.351984268 | 8.295843434 | 0.6845507 | 0.52152054 | 0.757784317 | -4.688313953 |
| **A_51_P156631** | Glrx5 | -0.216897416 | -1.010651368 | 0.576856536 | 12.62305383 | -0.684413903 | 0.521600497 | 0.757843542 | -4.688341131 |
| **A_55_P1995442** | NA | -0.087527824 | -0.407929187 | 0.232873539 | 8.397884232 | -0.684230016 | 0.521707991 | 0.757910885 | -4.688377658 |
| **A_55_P1978089** | Dcaf17 | -0.058067371 | -0.270635979 | 0.154501236 | 9.331389545 | -0.684200495 | 0.521725249 | 0.757910885 | -4.688383522 |
| **A_52_P112958** | Ywhab | 0.192142821 | -0.511330025 | 0.895615668 | 11.64989156 | 0.68411194 | 0.521777022 | 0.757929147 | -4.68840111 |
| **A_55_P2141878** | Ldha | 0.274261954 | -0.777868515 | 1.326392423 | 11.97799934 | 0.694318398 | 0.521841111 | 0.757964961 | -4.667609547 |
| **A_55_P2036693** | Cd40 | 0.370316199 | -0.985834022 | 1.72646642 | 10.04735183 | 0.683935673 | 0.521880085 | 0.757964961 | -4.688436112 |
| **A_51_P236013** | Rnf40 | -0.099661176 | -0.464890713 | 0.26556836 | 9.899933001 | -0.683455848 | 0.522160705 | 0.758315561 | -4.688531362 |
| **A_51_P376149** | Eef1akmt1 | 0.130101958 | -0.346782851 | 0.606986766 | 11.37847659 | 0.683314582 | 0.522243342 | 0.758378607 | -4.688559396 |
| **A_55_P2060444** | Apopt1 | -0.167955865 | -0.784030761 | 0.448119031 | 16.0996104 | -0.682829079 | 0.522527414 | 0.758695354 | -4.688655709 |
| **A_51_P493857** | Katna1 | 0.065401 | -0.174502368 | 0.305304367 | 10.60157066 | 0.682807663 | 0.522539947 | 0.758695354 | -4.688659956 |
| **A_55_P2006579** | Rbbp6 | 0.065167176 | -0.174020007 | 0.304354359 | 9.9031172 | 0.682403651 | 0.52277642 | 0.758981701 | -4.688740063 |
| **A_55_P2065799** | Naa50 | 0.155571612 | -0.415642669 | 0.726785893 | 9.681010301 | 0.68215276 | 0.522923306 | 0.759137948 | -4.688789792 |
| **A_55_P2009762** | Mapkapk3 | 0.655836813 | -1.75314393 | 3.064817557 | 10.98673671 | 0.681887624 | 0.523078561 | 0.759306322 | -4.68884233 |
| **A_55_P2182790** | NA | 0.128916775 | -0.344681412 | 0.602514962 | 14.22161733 | 0.681788619 | 0.523136543 | 0.759333478 | -4.688861945 |
| **A_55_P1966721** | NA | 0.413608058 | -1.106240124 | 1.93345624 | 10.41862314 | 0.681614984 | 0.523238242 | 0.759424081 | -4.68889634 |
| **A_55_P2295671** | NA | 0.07908641 | -0.211596486 | 0.369769307 | 8.591458725 | 0.681448016 | 0.523336049 | 0.759490696 | -4.688929408 |
| **A_55_P2185651** | NA | -0.139483531 | -0.652192721 | 0.37322566 | 8.58944204 | -0.681400072 | 0.523364136 | 0.759490696 | -4.688938903 |
| **A_65_P17218** | Mndal | 0.187755517 | -0.502455774 | 0.877966808 | 8.371457037 | 0.681335461 | 0.523401988 | 0.759490696 | -4.688951697 |
| **A_51_P464023** | Chst12 | -0.136655087 | -0.639124143 | 0.36581397 | 9.279944404 | -0.681187744 | 0.523488535 | 0.759559275 | -4.688980944 |
| **A_55_P2113523** | Nsd2 | -0.114385053 | -0.535056273 | 0.306286168 | 8.147592351 | -0.681046587 | 0.523571248 | 0.759587592 | -4.689008888 |
| **A_51_P191354** | NA | 0.127640907 | -0.341824195 | 0.597106009 | 8.38958287 | 0.680984008 | 0.523607919 | 0.759587592 | -4.689021275 |
| **A_51_P282268** | Snapc1 | -0.139097417 | -0.650722799 | 0.372527964 | 8.858299382 | -0.680953302 | 0.523625914 | 0.759587592 | -4.689027353 |
| **A_55_P2054628** | Nrxn1 | 0.273404526 | -0.732459379 | 1.27926843 | 8.767358596 | 0.680795727 | 0.523718263 | 0.75962335 | -4.689058539 |
| **A_52_P42380** | Tmem106 | 0.088303743 | -0.236577596 | 0.413185081 | 8.54927515 | 0.680776711 | 0.523729409 | 0.75962335 | -4.689062302 |
| **A_55_P1993121** | NA | 0.189672892 | -0.508290422 | 0.887636206 | 16.06237474 | 0.680648693 | 0.523804445 | 0.75962335 | -4.689087634 |
| **A_55_P1965459** | Zfp521 | 0.12058989 | -0.323163617 | 0.564343397 | 8.073746037 | 0.680643102 | 0.523807722 | 0.75962335 | -4.68908874 |
| **A_55_P2138416** | NA | 0.071918388 | -0.192804798 | 0.336641574 | 13.06307705 | 0.680453277 | 0.523918999 | 0.759727738 | -4.689126296 |
| **A_55_P2077816** | Hmgxb3 | -0.195543387 | -0.915560969 | 0.524474194 | 11.65810581 | -0.680221538 | 0.524054868 | 0.759867768 | -4.689172133 |
| **A_51_P270206** | Cep83 | 0.198109721 | -0.531528131 | 0.927747574 | 10.44940931 | 0.68006242 | 0.524148172 | 0.759898599 | -4.6892036 |
| **A_51_P266958** | Nr1i2 | -0.061923177 | -0.289989916 | 0.166143561 | 7.684214996 | -0.680051223 | 0.524154738 | 0.759898599 | -4.689205814 |
| **A_55_P2054362** | NA | 0.203658962 | -0.546660226 | 0.95397815 | 10.07523201 | 0.67984173 | 0.5242776 | 0.760007684 | -4.689247234 |
| **A_52_P256426** | NA | 0.066628804 | -0.178863743 | 0.312121351 | 8.440847522 | 0.679788874 | 0.524308601 | 0.760007684 | -4.689257683 |
| **A_52_P249856** | Cc2d2a | 0.228258157 | -0.612861734 | 1.069378049 | 8.624349592 | 0.679702315 | 0.524359373 | 0.76002124 | -4.689274794 |
| **A_55_P2107402** | NA | -0.116018183 | -0.543579274 | 0.311542909 | 8.587615882 | -0.679638898 | 0.524396573 | 0.76002124 | -4.689287328 |
| **A_55_P2028248** | Dnajc19 | 0.15716329 | -0.422328183 | 0.736654764 | 11.34890162 | 0.679288728 | 0.52460201 | 0.76025438 | -4.689356527 |

| | | | | | | | | | |
|---|---|---|---|---|---|---|---|---|---|
| **A_52_P299358** | Lclat1 | 0.220378756 | -0.592489347 | 1.03324686 | 9.903218549 | 0.679047244 | 0.524743715 | 0.76025438 | -4.689404232 |
| **A_55_P1982201** | Sgsm2 | -0.188337771 | -0.883161544 | 0.506486002 | 9.488815657 | -0.678911404 | 0.524823439 | 0.76025438 | -4.689431062 |
| **A_51_P483922** | Tm9sf4 | -0.080351983 | -0.37679403 | 0.216090064 | 9.087663866 | -0.678902081 | 0.524828911 | 0.76025438 | -4.689432903 |
| **A_55_P2110837** | Rnf7 | 0.075976532 | -0.204324602 | 0.356277667 | 11.37761631 | 0.678898672 | 0.524830911 | 0.76025438 | -4.689433576 |
| **A_55_P1962094** | NA | -0.057255497 | -0.26851396 | 0.154002967 | 8.121114733 | -0.678818102 | 0.524878201 | 0.76025438 | -4.689449488 |
| **A_51_P212420** | Lama4 | 0.336783009 | -0.905895989 | 1.579462006 | 12.14784043 | 0.678799571 | 0.524889078 | 0.76025438 | -4.689453147 |
| **A_55_P1971006** | Urah | 0.192632666 | -0.518167587 | 0.903432918 | 10.24417551 | 0.678785735 | 0.5248972 | 0.76025438 | -4.689455879 |
| **A_55_P1981775** | NA | -0.081927817 | -0.38424611 | 0.220390476 | 12.95653322 | -0.678761656 | 0.524911333 | 0.76025438 | -4.689460634 |
| **A_55_P2210575** | A830073C | 0.168330384 | -0.453080167 | 0.789740936 | 13.31087525 | 0.678475605 | 0.525079259 | 0.760440628 | -4.68951711 |
| **A_65_P13270** | Csnk1g1 | 0.062322764 | -0.16787914 | 0.292524667 | 8.829586977 | 0.678091247 | 0.525304952 | 0.760680266 | -4.689592968 |
| **A_55_P1958137** | NA | 0.118473577 | -0.319193609 | 0.556140762 | 8.699388617 | 0.677997135 | 0.525360223 | 0.760680266 | -4.689611537 |
| **A_55_P1955562** | Extl3 | 0.097009888 | -0.261368042 | 0.455387819 | 11.96408172 | 0.677992816 | 0.525362759 | 0.760680266 | -4.689612389 |
| **A_51_P520639** | Mslnl | 0.306199916 | -0.82553539 | 1.437935221 | 8.832242332 | 0.677657966 | 0.525559448 | 0.760862736 | -4.689678443 |
| **A_51_P464234** | Fkbp2 | -0.096434035 | -0.452872922 | 0.260004852 | 12.25553085 | -0.677634653 | 0.525573143 | 0.760862736 | -4.689683041 |
| **A_52_P694988** | Zfp933 | 0.087262452 | -0.235303791 | 0.409828696 | 8.22170378 | 0.677577291 | 0.525606843 | 0.760862736 | -4.689694354 |
| **A_55_P2018292** | Map2k4 | 0.10631445 | -0.286764422 | 0.499393323 | 9.133695241 | 0.677427687 | 0.52569474 | 0.760933002 | -4.689723855 |
| **A_55_P2089477** | Sypl | 0.116057537 | -0.313160989 | 0.545276063 | 12.45322445 | 0.67724411 | 0.52580261 | 0.761032166 | -4.689760049 |
| **A_51_P142813** | Aspn | 0.422064075 | -1.139121275 | 1.983249424 | 9.577138498 | 0.677133446 | 0.525867644 | 0.761069319 | -4.689781865 |
| **A_51_P158388** | Cdk19 | -0.099955005 | -0.46977708 | 0.26986707 | 9.488052094 | -0.676958523 | 0.525970452 | 0.761160121 | -4.689816342 |
| **A_52_P478256** | Samm50 | -0.069019505 | -0.324408593 | 0.186369582 | 10.47308303 | -0.676892733 | 0.526009122 | 0.761160121 | -4.689829307 |
| **A_55_P2064257** | Dnaic2 | 0.307673874 | -0.831059486 | 1.446407233 | 8.805072528 | 0.676735441 | 0.526101583 | 0.761167119 | -4.689860301 |
| **A_55_P2143261** | Rpain | -0.115214679 | -0.541654323 | 0.311224965 | 11.37763009 | -0.676706874 | 0.526118377 | 0.761167119 | -4.68986593 |
| **A_52_P401504** | Thbs4 | 0.310164056 | -0.837873695 | 1.458201807 | 8.755051144 | 0.676683588 | 0.526132066 | 0.761167119 | -4.689870518 |
| **A_55_P2011747** | Lsg1 | 0.116770603 | -0.315590671 | 0.549131877 | 11.21705411 | 0.676452148 | 0.526268139 | 0.761307011 | -4.689916111 |
| **A_55_P1963718** | Lias | -0.095081087 | -0.447244951 | 0.257082778 | 10.83164541 | -0.676238187 | 0.526393956 | 0.761432047 | -4.68995825 |
| **A_55_P2005720** | NA | -0.084964765 | -0.399762784 | 0.229833255 | 10.74530743 | -0.676016369 | 0.526524413 | 0.761563777 | -4.690001927 |
| **A_55_P2012389** | Sfxn3 | -0.140868321 | -0.662912361 | 0.38117572 | 10.57159452 | -0.675859683 | 0.526616578 | 0.761626754 | -4.690032772 |
| **A_55_P2173664** | Frmpd2 | 0.116365418 | -0.314915627 | 0.547646462 | 13.9062674 | 0.67579334 | 0.526655604 | 0.761626754 | -4.690045831 |
| **A_55_P1979986** | Crcp | 0.086334059 | -0.233667393 | 0.406335511 | 10.67371235 | 0.675741446 | 0.526686133 | 0.761626754 | -4.690056045 |
| **A_55_P2162503** | Tbx20 | -0.058847646 | -0.277023006 | 0.159327714 | 7.720416056 | -0.675575262 | 0.526783905 | 0.761676332 | -4.690088751 |
| **A_52_P67983** | Lcmt2 | 0.059446629 | -0.161009137 | 0.279902395 | 9.985426555 | 0.675392321 | 0.526891548 | 0.761676332 | -4.690124747 |
| **A_55_P2221647** | AI605517 | -0.115402534 | -0.543380737 | 0.312575668 | 8.199202976 | -0.675373539 | 0.526902601 | 0.761676332 | -4.690128442 |
| **A_51_P453475** | Slc1a5 | -0.141131348 | -0.664536832 | 0.382274136 | 9.992459867 | -0.67536036 | 0.526910356 | 0.761676332 | -4.690131035 |
| **A_65_P10673** | Tmem44 | 0.297480634 | -0.805843872 | 1.400805141 | 9.433222633 | 0.675314042 | 0.526937613 | 0.761676332 | -4.690140147 |
| **A_55_P2153941** | Zfp386 | 0.12044901 | -0.326305307 | 0.567203326 | 10.0990131 | 0.675281452 | 0.526956792 | 0.761676332 | -4.690146559 |
| **A_51_P294515** | Gde1 | -0.171319993 | -0.806903398 | 0.464263412 | 10.79480202 | -0.67512757 | 0.527047356 | 0.761750186 | -4.690176828 |
| **A_55_P2021114** | Ier5 | 0.124467429 | -0.337383176 | 0.586318035 | 8.711894644 | 0.675001205 | 0.527121733 | 0.761750186 | -4.690201681 |
| **A_51_P359046** | Slurp1 | 0.302770002 | -0.820704961 | 1.426244966 | 8.887480941 | 0.674993811 | 0.527126085 | 0.761750186 | -4.690203135 |
| **A_51_P335460** | Scin | 0.111560823 | -0.30250075 | 0.525622396 | 8.030866085 | 0.674834268 | 0.527220001 | 0.761821527 | -4.690234508 |
| **A_52_P491861** | Stag1 | -0.104311704 | -0.491524251 | 0.282900843 | 10.05049786 | -0.674736126 | 0.527277779 | 0.761821527 | -4.690253805 |
| **A_52_P649064** | NA | 0.220767465 | -0.598770013 | 1.040304943 | 13.68387563 | 0.674709147 | 0.527293662 | 0.761821527 | -4.690259109 |
| **A_55_P2019349** | Txnrd2 | 0.059380547 | -0.161168848 | 0.279929942 | 7.735187749 | 0.674355136 | 0.527502112 | 0.762012539 | -4.690328694 |
| **A_55_P2146783** | Ropn1l | -0.113471478 | -0.534926484 | 0.307983528 | 8.781164147 | -0.674350745 | 0.527504697 | 0.762012539 | -4.690329557 |
| **A_51_P281806** | Cactin | 0.084184624 | -0.228597874 | 0.396967121 | 12.25101146 | 0.674125378 | 0.527637427 | 0.762088379 | -4.690373841 |
| **A_55_P2046918** | NA | 0.218541146 | -0.593568636 | 1.030650927 | 15.88654789 | 0.674013848 | 0.527703121 | 0.762088379 | -4.690395753 |
| **A_55_P2022444** | Pdss2 | -0.070040926 | -0.330325432 | 0.190243581 | 8.257139322 | -0.673990703 | 0.527716754 | 0.762088379 | -4.690400299 |
| **A_51_P428483** | Fgb | -0.705155464 | -3.325703166 | 1.915392237 | 9.578025844 | -0.673974013 | 0.527726585 | 0.762088379 | -4.690403578 |
| **A_55_P2066559** | Endou | 0.61730214 | -1.676972371 | 2.911576652 | 9.056363242 | 0.673911315 | 0.52776352 | 0.762088379 | -4.690415894 |

| | | | | | | | | | |
|---|---|---|---|---|---|---|---|---|---|
| **A_55_P1954221** | Emp1 | 0.349808733 | -0.95039456 | 1.650012026 | 14.35938986 | 0.673860083 | 0.5277937 | 0.762088379 | -4.690425957 |
| **A_52_P598835** | Ccdc71l | 0.091950133 | -0.249869936 | 0.433770202 | 9.260220762 | 0.673759885 | 0.527852731 | 0.762097001 | -4.690445637 |
| **A_51_P263965** | Hmox1 | 0.36669563 | -0.996717488 | 1.730108748 | 9.91527596 | 0.673641155 | 0.527922684 | 0.762097001 | -4.690468954 |
| **A_52_P517063** | Immp2l | 0.116169606 | -0.31583915 | 0.548178363 | 8.449297565 | 0.673519717 | 0.52799424 | 0.762097001 | -4.690492799 |
| **A_52_P502141** | Hectd2 | 0.112905733 | -0.307002963 | 0.532814428 | 8.370786169 | 0.67345948 | 0.528029736 | 0.762097001 | -4.690504626 |
| **A_51_P406077** | Fmc1 | 0.241894953 | -0.657781002 | 1.141570909 | 13.44381099 | 0.673427799 | 0.528048405 | 0.762097001 | -4.690510846 |
| **A_51_P292754** | Yaf2 | 0.078311365 | -0.212967977 | 0.369590707 | 11.33751458 | 0.673388128 | 0.528071783 | 0.762097001 | -4.690518634 |
| **A_55_P1996907** | BC067074 | 0.366296427 | -0.99615724 | 1.728750094 | 9.195259952 | 0.673381663 | 0.528075593 | 0.762097001 | -4.690519903 |
| **A_55_P1970910** | Stard13 | 0.186812527 | -0.508256004 | 0.881881059 | 10.17250318 | 0.673176141 | 0.528196721 | 0.762214913 | -4.690560246 |
| **A_55_P2012734** | Fndc10 | 0.096155505 | -0.261751064 | 0.454062075 | 9.311577365 | 0.672906664 | 0.528355569 | 0.762309627 | -4.690613128 |
| **A_66_P134739** | Clpp | -0.113165545 | -0.534394491 | 0.3080634 | 9.559737525 | -0.67289354 | 0.528363306 | 0.762309627 | -4.690615704 |
| **A_55_P2117033** | Mtmr3 | -0.19985413 | -0.943817624 | 0.544109364 | 9.524782417 | -0.672840069 | 0.528394829 | 0.762309627 | -4.690626195 |
| **A_51_P244061** | Snapc5 | 0.083159002 | -0.226422784 | 0.392740788 | 10.73932361 | 0.672797257 | 0.528420069 | 0.762309627 | -4.690634594 |
| **A_55_P2170419** | NA | 0.18083518 | -0.492474368 | 0.854144728 | 16.31233779 | 0.672695457 | 0.528480091 | 0.762339333 | -4.690654566 |
| **A_51_P108862** | Cops7a | 0.095511212 | -0.260251238 | 0.451273661 | 12.99030177 | 0.672426148 | 0.528638895 | 0.762511519 | -4.684667809 |
| **A_51_P443723** | Slc35c1 | -0.132439068 | -0.626009431 | 0.361131294 | 11.20396205 | -0.672074455 | 0.528846327 | 0.7627018 | -4.690776345 |
| **A_55_P2172314** | Scamp4 | -0.213885359 | -1.011028761 | 0.583258043 | 12.54570306 | -0.672039728 | 0.528866813 | 0.7627018 | -4.690783153 |
| **A_55_P2028942** | Ncam1 | -0.070966093 | -0.335487867 | 0.193555681 | 7.845129857 | -0.67195442 | 0.528917137 | 0.7627018 | -4.690799874 |
| **A_55_P2162678** | Dnajc10 | 0.248511473 | -0.677958724 | 1.174981671 | 11.62924538 | 0.671839194 | 0.528985117 | 0.7627018 | -4.690822459 |
| **A_55_P2022293** | Zfp579 | 0.441405288 | -1.204211583 | 2.087022159 | 11.83413953 | 0.671829618 | 0.528990767 | 0.7627018 | -4.690824335 |
| **A_55_P1959485** | NA | 0.131122776 | -0.357740795 | 0.619986347 | 11.5095197 | 0.671801244 | 0.529007507 | 0.7627018 | -4.690829896 |
| **A_51_P181851** | Eml3 | -0.063076596 | -0.298318759 | 0.172165567 | 10.4279332 | -0.671588776 | 0.529132874 | 0.762767455 | -4.69087153 |
| **A_55_P2016252** | Ctnnal1 | -0.175213102 | -0.828774937 | 0.478348732 | 8.473546247 | -0.671475609 | 0.529199657 | 0.762767455 | -4.690893701 |
| **A_51_P217236** | Derl1 | -0.174900227 | -0.827333886 | 0.477533431 | 10.66932798 | -0.671435595 | 0.529223271 | 0.762767455 | -4.69090154 |
| **A_55_P1974695** | NA | -0.087488463 | -0.413856904 | 0.238879978 | 8.284603804 | -0.671417993 | 0.529233659 | 0.762767455 | -4.690904988 |
| **A_55_P2079967** | Map3k5 | -0.260961258 | -1.234496444 | 0.712573928 | 9.35816791 | -0.671389787 | 0.529250306 | 0.762767455 | -4.690910514 |
| **A_55_P2091323** | Wdr26 | 0.111352344 | -0.30422395 | 0.526928637 | 9.590424517 | 0.671118085 | 0.529410675 | 0.762941711 | -4.690963729 |
| **A_55_P2094520** | Fam171a1 | 0.131161423 | -0.358445977 | 0.620768822 | 12.21980149 | 0.670978325 | 0.52949318 | 0.763003737 | -4.690991096 |
| **A_55_P2062108** | Apold1 | -0.068054564 | -0.322146462 | 0.186037334 | 7.852640379 | -0.670836656 | 0.52957682 | 0.763049643 | -4.691018832 |
| **A_55_P2052475** | NA | 0.098499254 | -0.269300768 | 0.466299277 | 15.56688612 | 0.670766766 | 0.529618086 | 0.763049643 | -4.691032514 |
| **A_55_P2036559** | Fbxl12os | -0.060517516 | -0.286535794 | 0.165500762 | 10.92191607 | -0.670637564 | 0.529694377 | 0.763049643 | -4.691057804 |
| **A_55_P2024679** | NA | -0.324272475 | -1.535368912 | 0.886823962 | 9.644547696 | -0.670628035 | 0.529700004 | 0.763049643 | -4.691059669 |
| **A_51_P378858** | Atg10 | 0.082953892 | -0.226880722 | 0.392788505 | 8.823106507 | 0.67059016 | 0.52972237 | 0.763049643 | -4.691067082 |
| **A_55_P2187053** | Tpm1 | -0.147635987 | -0.699344503 | 0.404072528 | 9.188375147 | -0.670243932 | 0.529926854 | 0.763287327 | -4.691134832 |
| **A_51_P311362** | 4930449E | 0.057401902 | -0.157150677 | 0.271954481 | 7.747436356 | 0.670105047 | 0.530008895 | 0.763348627 | -4.691162002 |
| **A_55_P2016014** | Cpsf4l | 0.334342115 | -0.915842362 | 1.584526593 | 8.257239975 | 0.669834218 | 0.5301689 | 0.763499389 | -4.691214971 |
| **A_51_P227108** | Bri3bp | 0.167231727 | -0.458124741 | 0.792588196 | 11.82286233 | 0.669794188 | 0.530192552 | 0.763499389 | -4.691222799 |
| **A_52_P65108** | Smad1 | 0.074652401 | -0.204623982 | 0.353928784 | 8.840756708 | 0.669514386 | 0.530357896 | 0.763563315 | -4.691277504 |
| **A_55_P2147886** | Fam118a | 0.173146897 | -0.474640775 | 0.82093457 | 11.26762309 | 0.669471942 | 0.530382981 | 0.763563315 | -4.691285801 |
| **A_51_P512172** | Tet1 | 0.062943217 | -0.172549065 | 0.298435498 | 8.018445119 | 0.669456871 | 0.530391888 | 0.763563315 | -4.691288747 |
| **A_55_P2003813** | Scn3b | 0.228671482 | -0.626873942 | 1.084216906 | 8.888169962 | 0.669451747 | 0.530394917 | 0.763563315 | -4.691289749 |
| **A_52_P51029** | Ralgapa1 | 0.113407715 | -0.311079243 | 0.537894673 | 9.718015823 | 0.669157878 | 0.530568618 | 0.763753209 | -4.691347182 |
| **A_51_P515605** | Col3a1 | 0.380159977 | -1.042918985 | 1.80323894 | 14.7099581 | 0.669094932 | 0.53060583 | 0.763753209 | -4.691359482 |
| **A_51_P109335** | Copa | -0.095727708 | -0.454127175 | 0.262671759 | 12.23210457 | -0.668991581 | 0.53066693 | 0.763768132 | -4.691379675 |
| **A_55_P2061690** | Atp5d | 0.105209886 | -0.28871851 | 0.499138282 | 13.47552057 | 0.668943759 | 0.530695204 | 0.763768132 | -4.691389018 |
| **A_51_P331288** | Akr1b7 | -0.085031312 | -0.403529016 | 0.233466391 | 7.762699515 | -0.668687064 | 0.530846987 | 0.76392971 | -4.691439159 |
| **A_55_P1953103** | Nudt7 | 0.098503632 | -0.270535855 | 0.467543119 | 9.343041774 | 0.668543622 | 0.530931816 | 0.763954685 | -4.691467172 |
| **A_55_P2070576** | Ccl27a | -0.08294024 | -0.393682298 | 0.227801819 | 8.701452251 | -0.668521828 | 0.530944706 | 0.763954685 | -4.691471428 |

| | | | | | | | | | |
|---|---|---|---|---|---|---|---|---|---|
| A_52_P651248 | Zdhhc2 | 0.117899064 | -0.323861604 | 0.559659731 | 8.534409386 | 0.668457278 | 0.530982883 | 0.763954685 | -4.691484032 |
| A_52_P311104 | Brd7 | 0.122219832 | -0.336115501 | 0.580555166 | 13.21660962 | 0.667895766 | 0.531315058 | 0.764375723 | -4.691593637 |
| A_55_P2158458 | Iqcc | 0.11034008 | -0.303526801 | 0.524206961 | 9.332423992 | 0.667763942 | 0.531393061 | 0.76443106 | -4.691619359 |
| A_51_P268529 | Csad | 0.259220227 | -0.7133383 | 1.231778754 | 10.79483169 | 0.66758026 | 0.531501763 | 0.764530547 | -4.691655193 |
| A_55_P2112589 | Uxt | 0.143524387 | -0.395062975 | 0.682111749 | 10.58321211 | 0.667451769 | 0.531577812 | 0.764583054 | -4.691680255 |
| A_55_P2035043 | Gm2436 | 0.065988959 | -0.181758831 | 0.31373675 | 10.01558068 | 0.667132096 | 0.531767043 | 0.764703848 | -4.691742593 |
| A_55_P1972907 | Cab39 | -0.106652796 | -0.507074864 | 0.293769272 | 12.64603975 | -0.667120962 | 0.531773635 | 0.764703848 | -4.691744764 |
| A_52_P351925 | Ank3 | -0.056864358 | -0.270362151 | 0.156633436 | 8.142037382 | -0.66710945 | 0.531780451 | 0.764703848 | -4.691747008 |
| A_55_P2182483 | Cd47 | -0.153235495 | -0.728659604 | 0.422188613 | 11.69346492 | -0.666993602 | 0.53184904 | 0.7647456 | -4.691769593 |
| A_66_P134453 | I830077J0 | -0.059805077 | -0.284466303 | 0.164856149 | 7.87033404 | -0.666745775 | 0.531995787 | 0.764899721 | -4.691817898 |
| A_55_P1989102 | Hmgn1 | 0.071261973 | -0.196522827 | 0.339046773 | 11.93738202 | 0.666533936 | 0.532121246 | 0.765023213 | -4.691859178 |
| A_55_P2163098 | Akr1c18 | -0.107438593 | -0.511208125 | 0.296330939 | 7.959570827 | -0.666464642 | 0.532162289 | 0.765025332 | -4.691872679 |
| A_55_P2063166 | Ogfod1 | 0.109175882 | -0.301302918 | 0.519654681 | 13.48560625 | 0.666171919 | 0.532335691 | 0.765204695 | -4.6919297 |
| A_51_P228276 | Nipsnap2 | 0.09016704 | -0.248876006 | 0.429210086 | 9.730012127 | 0.666105955 | 0.532374771 | 0.765204695 | -4.691942547 |
| A_55_P2116794 | NA | -0.246216053 | -1.172102082 | 0.679669976 | 11.4925266 | -0.6660536 | 0.532405791 | 0.765204695 | -4.691952742 |
| A_52_P182520 | Timm9 | 0.084541891 | -0.233443858 | 0.40252764 | 8.39922698 | 0.665908635 | 0.532491686 | 0.765271259 | -4.69198097 |
| A_55_P2081981 | NA | 0.064384975 | -0.177826828 | 0.306596778 | 8.055495346 | 0.66579354 | 0.532559888 | 0.765312389 | -4.692003378 |
| A_55_P2111563 | NA | -0.185551529 | -0.883867442 | 0.512764384 | 8.001796672 | -0.665522809 | 0.53272034 | 0.76546178 | -4.692056077 |
| A_55_P2032936 | Cisd1 | -0.130668996 | -0.622546408 | 0.361208416 | 11.34221147 | -0.665374298 | 0.53280837 | 0.76546178 | -4.692084978 |
| A_55_P2087796 | Eva1b | 0.300917992 | -0.831828351 | 1.433664334 | 13.00320361 | 0.665374008 | 0.532808542 | 0.76546178 | -4.692085034 |
| A_55_P2065929 | NA | 0.058007115 | -0.160396178 | 0.276410408 | 8.117119524 | 0.665230927 | 0.532893362 | 0.76546178 | -4.692112874 |
| A_66_P105270 | Hao1 | -0.06790127 | -0.323561146 | 0.187758606 | 7.685439345 | -0.66522057 | 0.532899502 | 0.76546178 | -4.692114889 |
| A_52_P551909 | Arhgap5 | -0.062844634 | -0.299466548 | 0.173777279 | 8.751530126 | -0.665217378 | 0.532901394 | 0.76546178 | -4.69211551 |
| A_52_P142208 | Akirin1 | -0.178666207 | -0.851482957 | 0.494150544 | 9.898841341 | -0.665113813 | 0.532962796 | 0.765493106 | -4.692135658 |
| A_55_P1992680 | Ubtf | -0.169558691 | -0.808179285 | 0.469061903 | 15.25385017 | -0.665008936 | 0.533024979 | 0.765525549 | -4.692156059 |
| A_51_P217535 | Actr3 | 0.069952092 | -0.193632583 | 0.333536767 | 13.72284657 | 0.664707988 | 0.533203443 | 0.765715077 | -4.692214587 |
| A_55_P2057503 | Arl4c | 0.093797509 | -0.259676465 | 0.447271482 | 10.16491776 | 0.66463653 | 0.533245824 | 0.765715077 | -4.692228481 |
| A_55_P2149269 | Galt | -0.062124139 | -0.296255453 | 0.172007175 | 8.002057152 | -0.66458606 | 0.533275758 | 0.765715077 | -4.692238294 |
| A_55_P2007699 | Nanog | -0.126232711 | -0.602073555 | 0.349608132 | 8.915221831 | -0.664447297 | 0.533358066 | 0.765776388 | -4.69226527 |
| A_55_P2128626 | Gm4349 | -0.083209808 | -0.396956694 | 0.230537079 | 8.389649886 | -0.664271223 | 0.533462517 | 0.765813585 | -4.692299493 |
| A_51_P487360 | Hpcal1 | 0.254186061 | -0.704237283 | 1.212609406 | 11.20935104 | 0.664270083 | 0.533463193 | 0.765813585 | -4.692299715 |
| A_66_P129444 | Pnkp | -0.092634552 | -0.442002356 | 0.256733253 | 8.509858896 | -0.664110687 | 0.533557761 | 0.76585779 | -4.692330691 |
| A_52_P366653 | Hnrnph3 | 0.060622733 | -0.16803114 | 0.289276606 | 8.297363546 | 0.664059962 | 0.533587859 | 0.76585779 | -4.692340547 |
| A_55_P2022509 | Mgat5b | -0.220345028 | -1.051484333 | 0.610794276 | 9.107471354 | -0.664017892 | 0.533612822 | 0.76585779 | -4.692348722 |
| A_55_P2081555 | Phf23 | -0.087962723 | -0.419824624 | 0.243899178 | 10.09849612 | -0.663883102 | 0.533692806 | 0.765915729 | -4.692374909 |
| A_55_P1954356 | Ttc23 | 0.222559434 | -0.6173698 | 1.062488669 | 10.23309837 | 0.663672247 | 0.533817942 | 0.766038454 | -4.692415866 |
| A_51_P293901 | Dhrs1 | -0.067306724 | -0.321376872 | 0.186763424 | 9.317922488 | -0.663521742 | 0.533907274 | 0.766062198 | -4.692445095 |
| A_66_P140531 | Selenok | 0.105879368 | -0.293801767 | 0.505560503 | 12.27714668 | 0.663510861 | 0.533913733 | 0.766062198 | -4.692447208 |
| A_55_P2035554 | Strip1 | 0.095870843 | -0.266110817 | 0.457852504 | 12.70890626 | 0.663361595 | 0.53400234 | 0.766077084 | -4.692476191 |
| A_55_P2111713 | Spint2 | 0.230604561 | -0.640164806 | 1.101373928 | 10.59607427 | 0.663307787 | 0.534034284 | 0.766077084 | -4.692486637 |
| A_51_P376711 | Gm15498 | -0.130745001 | -0.624458607 | 0.362968605 | 7.819828413 | -0.663285253 | 0.534047662 | 0.766077084 | -4.692491012 |
| A_52_P207614 | NA | 0.076939029 | -0.213686329 | 0.367564387 | 10.57146606 | 0.663076355 | 0.534171691 | 0.766077084 | -4.692531562 |
| A_51_P224064 | Klhl38 | -0.076730807 | -0.366580448 | 0.213118834 | 7.904459405 | -0.663051628 | 0.534186373 | 0.766077084 | -4.692536361 |
| A_55_P2117472 | Acap2 | 0.260896783 | -0.724727694 | 1.24652126 | 11.22316783 | 0.662990934 | 0.534222413 | 0.766077084 | -4.692548141 |
| A_66_P104190 | Gsk3b | 0.082267548 | -0.228527707 | 0.393062803 | 9.818305946 | 0.662986247 | 0.534225196 | 0.766077084 | -4.69254905 |
| A_51_P451588 | Plekhb1 | 0.645400561 | -1.793050534 | 3.083851656 | 10.35204047 | 0.662926888 | 0.534260445 | 0.766077084 | -4.69256057 |
| A_66_P101107 | NA | -0.181176166 | -0.865780337 | 0.503428006 | 9.430785095 | -0.662844823 | 0.53430918 | 0.766077084 | -4.692576495 |
| A_55_P2068772 | Ncbp2 | 0.149410595 | -0.415178111 | 0.713999301 | 9.285423781 | 0.66282603 | 0.534320341 | 0.766077084 | -4.692580141 |

| | | | | | | | | | |
|---|---|---|---|---|---|---|---|---|---|
| **A_55_P2086811** | Sp140 | 0.129276597 | -0.359340964 | 0.617894158 | 9.123445344 | 0.662675914 | 0.534409497 | 0.766148095 | -4.692609267 |
| **A_55_P1965427** | Uqcrh | 0.08892535 | -0.247216839 | 0.42506754 | 15.40223675 | 0.662602249 | 0.534453251 | 0.766154012 | -4.692623558 |
| **A_66_P138915** | Fyn | 0.102553869 | -0.285188144 | 0.490295882 | 8.573934315 | 0.662459807 | 0.534537863 | 0.766159094 | -4.692651188 |
| **A_52_P189618** | Armc8 | -0.074178512 | -0.354662114 | 0.20630509 | 8.91006557 | -0.662401008 | 0.534572793 | 0.766159094 | -4.692662592 |
| **A_51_P279552** | Cav2 | 0.549637011 | -1.529127343 | 2.628401365 | 10.61206843 | 0.662248607 | 0.534663334 | 0.766159094 | -4.692692146 |
| **A_55_P2069470** | 1110017D | 0.696846269 | -1.938718751 | 3.332411289 | 11.06582755 | 0.662237226 | 0.534670096 | 0.766159094 | -4.692694353 |
| **A_55_P1965507** | Trnau1ap | 0.077573278 | -0.21583309 | 0.370979646 | 12.3626603 | 0.662205761 | 0.53468879 | 0.766159094 | -4.692700455 |
| **A_51_P342535** | Rnf166 | -0.16050434 | -0.767590791 | 0.446582111 | 11.41561696 | -0.662196048 | 0.534694561 | 0.766159094 | -4.692702338 |
| **A_52_P381665** | Aff1 | 0.165360616 | -0.460260146 | 0.790981378 | 10.89538225 | 0.662020249 | 0.53479902 | 0.766202059 | -4.692736422 |
| **A_55_P2040161** | Upf3a | 0.144476658 | -0.402166885 | 0.6911202 | 9.882252579 | 0.661978385 | 0.534823897 | 0.766202059 | -4.692744538 |
| **A_55_P2187043** | Tpm1 | 0.684418673 | -1.905284316 | 3.274121661 | 12.82811447 | 0.661945505 | 0.534843436 | 0.766202059 | -4.692750911 |
| **A_55_P2063376** | Add2 | -0.092971973 | -0.4449486 | 0.259004653 | 7.916527466 | -0.661589448 | 0.535055054 | 0.766384342 | -4.692819918 |
| **A_55_P1982733** | Dnajc5 | -0.216777592 | -1.037582767 | 0.604027583 | 8.89227504 | -0.661492078 | 0.535112934 | 0.766384342 | -4.692838784 |
| **A_55_P1972257** | Sars | -0.140129792 | -0.670717122 | 0.390457539 | 12.42572097 | -0.661491011 | 0.535113569 | 0.766384342 | -4.692838991 |
| **A_55_P2289765** | Fam160a2 | 0.071095386 | -0.19811085 | 0.340301622 | 8.889189216 | 0.661464659 | 0.535129234 | 0.766384342 | -4.692844096 |
| **A_55_P2070992** | Aldoc | 0.438347905 | -1.22175098 | 2.09844679 | 11.34285267 | 0.661356038 | 0.535193808 | 0.766420049 | -4.692865139 |
| **A_55_P2050354** | Prpmp5 | 0.077710535 | -0.216749854 | 0.372170924 | 8.505593798 | 0.661002898 | 0.535403783 | 0.766663956 | -4.692933535 |
| **A_51_P129435** | Arid2 | -0.159724385 | -0.765168431 | 0.44571966 | 8.514090733 | -0.660765803 | 0.535544787 | 0.766809073 | -4.69297944 |
| **A_51_P185660** | Ccl9 | -0.131006349 | -0.6277535 | 0.365740802 | 9.178310286 | -0.660552446 | 0.535671695 | 0.766872157 | -4.693020738 |
| **A_52_P634090** | Jag1 | 0.218305446 | -0.609517705 | 1.046128597 | 9.47861093 | 0.660506884 | 0.535698799 | 0.766872157 | -4.693029556 |
| **A_55_P1981274** | NA | 0.183684966 | -0.51287194 | 0.880241872 | 17.5050576 | 0.660491687 | 0.535707839 | 0.766872157 | -4.693032497 |
| **A_55_P2071406** | NA | -0.058032321 | -0.278130983 | 0.162066341 | 7.957789268 | -0.660393645 | 0.535766165 | 0.766896077 | -4.69305147 |
| **A_55_P1960698** | Pih1d2 | 0.273653975 | -0.764330627 | 1.311638577 | 9.01575476 | 0.660330253 | 0.53580388 | 0.766896077 | -4.693063737 |
| **A_55_P2037777** | NA | -0.092645727 | -0.444152217 | 0.258860764 | 9.459515121 | -0.660149637 | 0.535911345 | 0.766993112 | -4.693098681 |
| **A_66_P103271** | Irf2bp2 | 0.266183724 | -0.743951504 | 1.276318952 | 12.71542884 | 0.66001275 | 0.535992802 | 0.767001694 | -4.69312516 |
| **A_55_P2042272** | Pbrm1 | -0.061560178 | -0.295182668 | 0.172062311 | 7.800610093 | -0.659987283 | 0.536007957 | 0.767001694 | -4.693130086 |
| **A_55_P2090055** | Phrf1 | -0.14514815 | -0.696029009 | 0.40573271 | 10.84847581 | -0.659939565 | 0.536036355 | 0.767001694 | -4.693139316 |
| **A_55_P2227967** | NA | 0.092712787 | -0.259240992 | 0.444666566 | 8.800914117 | 0.659787904 | 0.536126617 | 0.767074078 | -4.693168646 |
| **A_52_P451644** | Ndn | 0.10238916 | -0.286350965 | 0.491129285 | 8.642167257 | 0.659697679 | 0.53618032 | 0.767094148 | -4.693186092 |
| **A_55_P1953728** | Nes | 0.327395922 | -0.915962803 | 1.570754646 | 10.64333183 | 0.659518777 | 0.536286815 | 0.767139482 | -4.69322068 |
| **A_55_P2410304** | Iigp1 | 0.253658581 | -0.709677377 | 1.216994539 | 9.472197898 | 0.659511135 | 0.536291364 | 0.767139482 | -4.693222157 |
| **A_52_P528963** | Eps15l1 | -0.153754151 | -0.737780041 | 0.43027174 | 9.590425039 | -0.659394155 | 0.536361006 | 0.767154723 | -4.69324477 |
| **A_55_P2169714** | Gcn1 | -0.097672911 | -0.468701954 | 0.273356132 | 12.90015806 | -0.659350849 | 0.536386789 | 0.767154723 | -4.69325314 |
| **A_51_P433026** | Plpp6 | -0.065220242 | -0.313014099 | 0.182573615 | 9.096005567 | -0.659237971 | 0.536453997 | 0.767154723 | -4.693274956 |
| **A_55_P2066578** | Ifi204 | -0.144169737 | -0.692000799 | 0.403661325 | 8.285776528 | -0.659140199 | 0.536512216 | 0.767154723 | -4.69329385 |
| **A_51_P509384** | Aldh8a1 | -0.070026209 | -0.336127158 | 0.19607474 | 8.070729428 | -0.659120073 | 0.5365242 | 0.767154723 | -4.693297739 |
| **A_66_P124445** | Rmc1 | -0.066055969 | -0.317096245 | 0.184984308 | 8.636089589 | -0.659050974 | 0.536565348 | 0.767154723 | -4.69331109 |
| **A_55_P2044282** | Mpp6 | 0.161484278 | -0.452245998 | 0.775214555 | 12.01967784 | 0.659026752 | 0.536579773 | 0.767154723 | -4.69331577 |
| **A_52_P476029** | Nol4l | 0.17610459 | -0.493402493 | 0.845611672 | 10.52605511 | 0.658818589 | 0.536703747 | 0.767224527 | -4.693355984 |
| **A_51_P507053** | Slc38a1 | -0.756508699 | -3.632758457 | 2.119741059 | 12.112764 | -0.65877586 | 0.536729197 | 0.767224527 | -4.693364238 |
| **A_55_P1973858** | NA | 0.076740287 | -0.215062719 | 0.368543294 | 9.632616435 | 0.658694452 | 0.536777686 | 0.767224527 | -4.693379961 |
| **A_55_P2146389** | NA | 0.061252314 | -0.171663787 | 0.294168415 | 7.96732631 | 0.658678268 | 0.536787327 | 0.767224527 | -4.693383087 |
| **A_55_P1961913** | Pygm | 0.225095974 | -0.630998372 | 1.081190319 | 9.165889972 | 0.65856166 | 0.536856789 | 0.767267088 | -4.693405606 |
| **A_55_P1996837** | Baz2b | 0.181854778 | -0.509874305 | 0.873583862 | 10.47417666 | 0.658474599 | 0.536908655 | 0.767284495 | -4.693422418 |
| **A_66_P123028** | NA | -0.095624921 | -0.459513908 | 0.268264066 | 8.283212315 | -0.658191884 | 0.537077101 | 0.767458829 | -4.693476998 |
| **A_65_P20641** | NA | 0.073947995 | -0.207485689 | 0.35538168 | 8.454933706 | 0.658113312 | 0.537123921 | 0.767458829 | -4.693492164 |
| **A_52_P456077** | Commd2 | 0.266144067 | -0.74682233 | 1.279110464 | 11.63498195 | 0.658070004 | 0.537149729 | 0.767458829 | -4.693500522 |
| **A_55_P1977533** | Eno3 | 0.465388375 | -1.307245323 | 2.238022072 | 11.0387033 | 0.657577395 | 0.537443342 | 0.767714873 | -4.693595569 |

| | | | | | | | | | |
|---|---|---|---|---|---|---|---|---|---|
| **A_55_P2111277** | NA | -0.190022098 | -0.913814451 | 0.533770254 | 8.905116126 | -0.657567689 | 0.537449128 | 0.767714873 | -4.693597441 |
| **A_51_P386526** | Rcc1l | -0.140189218 | -0.674170984 | 0.393792549 | 8.657235213 | -0.657564758 | 0.537450875 | 0.767714873 | -4.693598007 |
| **A_55_P1992739** | Dync1i2 | 0.09964586 | -0.280033107 | 0.479324827 | 13.75520176 | 0.657344548 | 0.537582163 | 0.767714873 | -4.693640478 |
| **A_51_P327121** | Was | 0.261359281 | -0.734529877 | 1.257248439 | 9.0531147 | 0.657320638 | 0.537596419 | 0.767714873 | -4.693645088 |
| **A_55_P1998164** | Rab28 | -0.064417937 | -0.309906856 | 0.181070981 | 8.492913421 | -0.657241939 | 0.537643345 | 0.767714873 | -4.693660264 |
| **A_55_P2204804** | Dclk1 | 0.150119514 | -0.422042265 | 0.722281293 | 8.330390075 | 0.657156264 | 0.537694433 | 0.767714873 | -4.693676783 |
| **A_55_P2144120** | Bcl2l2 | -0.103790817 | -0.499396128 | 0.291814493 | 8.158978218 | -0.657123752 | 0.537713821 | 0.767714873 | -4.693683051 |
| **A_55_P2271742** | NA | -0.057601212 | -0.277172444 | 0.161970019 | 8.162996194 | -0.657062276 | 0.537750482 | 0.767714873 | -4.693694902 |
| **A_55_P2052048** | Pih1d2 | 0.211064017 | -0.593552303 | 1.015680337 | 8.675366039 | 0.65701564 | 0.537778294 | 0.767714873 | -4.693703893 |
| **A_51_P499530** | Ankrd10 | 0.109503045 | -0.307950686 | 0.526956775 | 9.330309399 | 0.657004272 | 0.537785074 | 0.767714873 | -4.693706084 |
| **A_52_P336290** | Rap2a | 0.38416045 | -1.080502974 | 1.848823873 | 11.51263244 | 0.656939175 | 0.537823898 | 0.767714873 | -4.693718632 |
| **A_55_P2155216** | Ankrd34a | 0.138882231 | -0.390653677 | 0.668418138 | 8.181856157 | 0.656903558 | 0.537845141 | 0.767714873 | -4.693725497 |
| **A_51_P520139** | Ccpg1os | 0.17798703 | -0.500896812 | 0.856870872 | 10.44599817 | 0.656664033 | 0.537988013 | 0.767862118 | -4.693771658 |
| **A_55_P2057686** | Cdc42ep5 | 0.290265686 | -0.817123402 | 1.397654774 | 9.88152516 | 0.65651674 | 0.538075883 | 0.767893787 | -4.693800038 |
| **A_55_P2008452** | Wtap | 0.0762516 | -0.214665166 | 0.367168365 | 11.27294124 | 0.656493687 | 0.538089636 | 0.767893787 | -4.693804479 |
| **A_55_P2057405** | Cpt1a | -0.05807316 | -0.279662964 | 0.163516643 | 7.926791051 | -0.656411276 | 0.538138804 | 0.767907273 | -4.693820355 |
| **A_51_P425833** | Lztfl1 | 0.11712785 | -0.329878971 | 0.564134671 | 10.10584191 | 0.656290877 | 0.538210643 | 0.767953105 | -4.693843547 |
| **A_52_P242973** | Sec11a | -0.141510883 | -0.681668374 | 0.398646609 | 13.2378964 | -0.656175153 | 0.538279697 | 0.767970837 | -4.693865835 |
| **A_52_P650453** | Spink4 | -0.076294908 | -0.36754726 | 0.214957445 | 7.8656202 | -0.656109696 | 0.538318759 | 0.767970837 | -4.69387844 |
| **A_55_P2089488** | Coq10b | -0.179514745 | -0.86486774 | 0.50583825 | 9.065632319 | -0.656048816 | 0.538355091 | 0.767970837 | -4.693890164 |
| **A_55_P2050192** | NA | 0.173740868 | -0.489653183 | 0.837134918 | 13.03484922 | 0.655965139 | 0.538405031 | 0.767970837 | -4.693906275 |
| **A_55_P2004777** | Micall2 | 0.130273558 | -0.367223087 | 0.627770203 | 8.966685478 | 0.655868033 | 0.538462989 | 0.767970837 | -4.693924971 |
| **A_55_P2095563** | Slc8a2 | 0.161504126 | -0.455316713 | 0.778324964 | 8.514911116 | 0.65580531 | 0.538500428 | 0.767970837 | -4.693937046 |
| **A_52_P651948** | Fam229b | 0.161556067 | -0.455464238 | 0.778576372 | 8.346714041 | 0.655804152 | 0.538501119 | 0.767970837 | -4.693937269 |
| **A_55_P1952269** | Xylt2 | 0.219585763 | -0.619363207 | 1.058534732 | 11.07221826 | 0.65556986 | 0.53864098 | 0.768070105 | -4.693982365 |
| **A_51_P371091** | Rcsd1 | 0.143632446 | -0.405164458 | 0.69242935 | 11.88849445 | 0.655528005 | 0.538665968 | 0.768070105 | -4.69399042 |
| **A_51_P148060** | Ptgfrn | 0.110811682 | -0.312608161 | 0.534231525 | 8.945283882 | 0.655487912 | 0.538689905 | 0.768070105 | -4.693998135 |
| **A_55_P2081885** | NA | 0.108023066 | -0.304787862 | 0.520833994 | 16.36838353 | 0.655413918 | 0.538734083 | 0.768076452 | -4.694012373 |
| **A_55_P2340448** | B230114F | 0.322293304 | -0.909573314 | 1.554159921 | 9.070220385 | 0.65529664 | 0.538804109 | 0.768119647 | -4.694034938 |
| **A_51_P510849** | Tmem50b | 0.30370978 | -0.857263017 | 1.464682577 | 12.51038788 | 0.6552199 | 0.538849934 | 0.768128336 | -4.694049702 |
| **A_55_P1975667** | Rhox4g | -0.740166968 | -3.57003842 | 2.089704484 | 8.593217306 | -0.655108655 | 0.538916367 | 0.768166399 | -4.694071101 |
| **A_51_P514647** | Paip2 | 0.16184022 | -0.457022422 | 0.780702862 | 13.29367664 | 0.655001867 | 0.538980143 | 0.768200669 | -4.69409164 |
| **A_52_P637730** | Ubl4a | 0.06338149 | -0.179049191 | 0.305812172 | 12.4774491 | 0.654824936 | 0.539085821 | 0.768267334 | -4.694125666 |
| **A_51_P517381** | Cers2 | -0.070114045 | -0.338384982 | 0.198156892 | 10.96788169 | -0.654608656 | 0.53921502 | 0.768267334 | -4.694167249 |
| **A_52_P601385** | Cd99l2 | -0.07016887 | -0.338681222 | 0.198343483 | 10.97735921 | -0.654531508 | 0.539261111 | 0.768267334 | -4.694182079 |
| **A_55_P1960291** | Tjp3 | 0.394106261 | -1.114118574 | 1.902331096 | 11.36926748 | 0.654481827 | 0.539290793 | 0.768267334 | -4.694191628 |
| **A_51_P413698** | Mrpl20 | 0.072728761 | -0.205609681 | 0.351067202 | 13.55432268 | 0.654460351 | 0.539303624 | 0.768267334 | -4.694195756 |
| **A_55_P2000439** | Ptprz1 | 0.200460271 | -0.566753003 | 0.967673545 | 8.553580339 | 0.654429021 | 0.539322344 | 0.768267334 | -4.694201778 |
| **A_55_P2107972** | Smad5 | 0.243325363 | -0.687971811 | 1.174622536 | 12.32846386 | 0.654409266 | 0.539334147 | 0.768267334 | -4.694205575 |
| **A_55_P2325663** | Cers3 | 0.282841638 | -0.799773476 | 1.365456752 | 9.674272641 | 0.654364308 | 0.53936101 | 0.768267334 | -4.694214216 |
| **A_55_P2134012** | Oxa1l | 0.089528834 | -0.253181441 | 0.432239108 | 12.34329781 | 0.6543139 | 0.53939113 | 0.768267334 | -4.694223903 |
| **A_55_P2000399** | Rhox4a | -0.719815651 | -3.47545457 | 2.035823267 | 8.590802313 | -0.654258421 | 0.539424282 | 0.768267334 | -4.694234564 |
| **A_52_P406036** | Cdkn2aip | -0.100001935 | -0.482972684 | 0.282968815 | 8.209136258 | -0.654023183 | 0.539564864 | 0.768324333 | -4.694279762 |
| **A_66_P117389** | Rcn2 | 0.16709219 | -0.472836842 | 0.807021221 | 10.52895991 | 0.653995384 | 0.539581478 | 0.768324333 | -4.694285103 |
| **A_55_P1973259** | Gamt | 0.186181389 | -0.526899427 | 0.899262204 | 9.890794498 | 0.653954991 | 0.539605621 | 0.768324333 | -4.694292863 |
| **A_51_P135004** | Zfp212 | 0.090732814 | -0.256792518 | 0.438258145 | 11.08183479 | 0.653925478 | 0.539623261 | 0.768324333 | -4.694298532 |
| **A_55_P2091105** | NA | 0.147842069 | -0.418562871 | 0.714247009 | 16.47566873 | 0.653764516 | 0.539719474 | 0.768404736 | -4.694329448 |
| **A_55_P2011922** | 9-Mar | -0.153033557 | -0.739430034 | 0.43336292 | 9.557617227 | -0.653650603 | 0.539787572 | 0.768445101 | -4.694351324 |

| | | | | | | | | | |
|---|---|---|---|---|---|---|---|---|---|
| **A_51_P362176** | Fgg | -0.79345049 | -3.834649206 | 2.247748226 | 9.822767562 | -0.653469587 | 0.539895795 | 0.768491837 | -4.694386081 |
| **A_51_P209518** | Galnt1 | -0.158568267 | -0.766347431 | 0.449210896 | 8.896725492 | -0.653462725 | 0.539899898 | 0.768491837 | -4.694387398 |
| **A_52_P255224** | NA | -0.184039907 | -0.889870794 | 0.521790981 | 8.334074255 | -0.653072942 | 0.540132984 | 0.768675981 | -4.694462215 |
| **A_55_P1975660** | Hemgn | -0.132529967 | -0.640870521 | 0.375810586 | 14.61309332 | -0.652994734 | 0.54017976 | 0.768675981 | -4.694477223 |
| **A_52_P796840** | Cfhr2 | 0.221921094 | -0.629340806 | 1.073182994 | 8.72870643 | 0.652958732 | 0.540201293 | 0.768675981 | -4.694484131 |
| **A_55_P2058107** | Pigyl | 0.130781242 | -0.370911501 | 0.632473985 | 12.44387722 | 0.65291701 | 0.540226248 | 0.768675981 | -4.694492137 |
| **A_51_P279465** | Farsa | -0.285491921 | -1.433345653 | 0.86236181 | 8.753521862 | -0.662475739 | 0.540228056 | 0.768675981 | -4.672938896 |
| **A_55_P2198568** | 4930506A | 0.093452662 | -0.265082587 | 0.451987912 | 8.686324823 | 0.652845123 | 0.540269248 | 0.768678022 | -4.694505929 |
| **A_52_P498985** | Zfp512 | 0.094144716 | -0.267177888 | 0.455467321 | 8.723365171 | 0.652606154 | 0.540412204 | 0.768801373 | -4.694551769 |
| **A_55_P2394490** | NA | 0.115074461 | -0.326601755 | 0.556750676 | 8.169869218 | 0.65256726 | 0.540435474 | 0.768801373 | -4.694559229 |
| **A_55_P2052485** | Ushbp1 | 0.371051313 | -1.053377405 | 1.795480031 | 10.80438351 | 0.652444534 | 0.540508903 | 0.768805667 | -4.694582765 |
| **A_52_P189957** | Dbnl | -0.136082732 | -0.658502873 | 0.386337409 | 9.171398833 | -0.652429294 | 0.540518022 | 0.768805667 | -4.694585688 |
| **A_51_P109449** | Ltb4r1 | 0.119385327 | -0.339136614 | 0.577907269 | 8.429344751 | 0.652140508 | 0.540690834 | 0.768929305 | -4.694641056 |
| **A_55_P2152245** | Rpl19 | 0.136931294 | -0.389042683 | 0.66290527 | 16.18886579 | 0.652061863 | 0.540737903 | 0.768929305 | -4.694656132 |
| **A_52_P78257** | Plekha2 | 0.083517705 | -0.237287335 | 0.404322746 | 9.256950203 | 0.652060234 | 0.540738878 | 0.768929305 | -4.694656444 |
| **A_55_P2076713** | Immp1l | 0.236988311 | -0.673396753 | 1.147373375 | 10.7039854 | 0.652006833 | 0.540770839 | 0.768929305 | -4.694666679 |
| **A_55_P2310898** | Kantr | 0.094891824 | -0.269663693 | 0.459447341 | 9.034431463 | 0.651951763 | 0.540803801 | 0.768929305 | -4.694677234 |
| **A_51_P184796** | Zfp101 | 0.211751393 | -0.601850286 | 1.025353071 | 8.710214391 | 0.651875689 | 0.540849337 | 0.768937501 | -4.694691813 |
| **A_51_P121228** | Ascc2 | -0.093281894 | -0.451747049 | 0.265183261 | 11.40321194 | -0.651779586 | 0.540906865 | 0.768962744 | -4.694710229 |
| **A_51_P409039** | Echs1 | 0.243773317 | -0.69325317 | 1.180799803 | 12.39323729 | 0.651605354 | 0.541011171 | 0.769044359 | -4.694743612 |
| **A_55_P2095342** | Rara | 0.236261257 | -0.672024925 | 1.144547438 | 10.09147636 | 0.651508594 | 0.541069104 | 0.769044359 | -4.694762148 |
| **A_51_P475995** | Thra | 0.341132553 | -0.970447079 | 1.652712186 | 11.68721951 | 0.651446638 | 0.541106201 | 0.769044359 | -4.694774016 |
| **A_55_P1997661** | Smim15 | 0.162872502 | -0.463364027 | 0.789109032 | 12.01862283 | 0.651417944 | 0.541123382 | 0.769044359 | -4.694779512 |
| **A_55_P2031671** | Gstm6 | 0.067914047 | -0.19323931 | 0.329067404 | 7.95507749 | 0.651349889 | 0.541164133 | 0.769045744 | -4.694792546 |
| **A_55_P2013680** | NA | 0.178418291 | -0.507895732 | 0.864732314 | 16.87596647 | 0.651128714 | 0.541296585 | 0.76909442 | -4.694834901 |
| **A_55_P2074972** | Zswim6 | 0.10045017 | -0.2859488 | 0.48684914 | 8.427725856 | 0.651126035 | 0.54129819 | 0.76909442 | -4.694835414 |
| **A_55_P1990879** | Banp | 0.056402846 | -0.16059311 | 0.273398802 | 8.276196145 | 0.651028088 | 0.541356853 | 0.76909442 | -4.694854167 |
| **A_55_P1962384** | Churc1 | 0.132130725 | -0.376265163 | 0.640526614 | 11.61968646 | 0.65095675 | 0.541399582 | 0.76909442 | -4.694867824 |
| **A_51_P185846** | Dnal4 | 0.198803002 | -0.566184087 | 0.963790091 | 11.53190398 | 0.650907355 | 0.541429169 | 0.76909442 | -4.69487728 |
| **A_55_P2003338** | Clint1 | 0.092672778 | -0.263936091 | 0.449281647 | 10.81034994 | 0.650894179 | 0.541437062 | 0.76909442 | -4.694879802 |
| **A_51_P259029** | Dusp26 | 0.258369963 | -0.736267873 | 1.253007798 | 8.799770058 | 0.650619975 | 0.541601327 | 0.769233902 | -4.694932283 |
| **A_55_P2166501** | Cd44 | -0.088816516 | -0.430774066 | 0.253141035 | 8.715279096 | -0.650536816 | 0.541651152 | 0.769233902 | -4.694948195 |
| **A_55_P1963724** | NA | 0.071729851 | -0.204444924 | 0.347904626 | 7.888988042 | 0.650528406 | 0.541656191 | 0.769233902 | -4.694949804 |
| **A_55_P2126391** | Srl | 0.282517374 | -0.805339646 | 1.370374393 | 8.78706345 | 0.650464633 | 0.541694402 | 0.769233902 | -4.694962006 |
| **A_55_P2007496** | NA | 0.138341987 | -0.394506167 | 0.671190142 | 8.838194284 | 0.650280739 | 0.541804597 | 0.769333877 | -4.694997187 |
| **A_52_P322927** | NA | 0.054356871 | -0.155086457 | 0.2638002 | 7.722931144 | 0.650037283 | 0.541950505 | 0.769484546 | -4.69504375 |
| **A_55_P2002212** | Babam2 | -0.063056612 | -0.306135359 | 0.180022136 | 9.330305301 | -0.649731595 | 0.542133746 | 0.769688196 | -4.695102197 |
| **A_52_P198727** | Cfap410 | -0.083233452 | -0.404164596 | 0.237697692 | 8.920889188 | -0.649585597 | 0.542221277 | 0.769689112 | -4.695130104 |
| **A_55_P2047345** | Sat2 | -0.065003413 | -0.315654976 | 0.18564815 | 9.164630129 | -0.649555239 | 0.542239479 | 0.769689112 | -4.695135906 |
| **A_52_P559498** | Clk3 | -0.078107972 | -0.379301485 | 0.22308554 | 9.899953063 | -0.649531317 | 0.542253822 | 0.769689112 | -4.695140478 |
| **A_55_P2248320** | C430003N | -0.066545621 | -0.323221559 | 0.190130316 | 8.47014412 | -0.649358717 | 0.542357318 | 0.769779503 | -4.695173463 |
| **A_51_P300207** | Rsl24d1 | 0.062122613 | -0.177526999 | 0.301772225 | 9.396096489 | 0.649267002 | 0.542412317 | 0.769801053 | -4.695190987 |
| **A_51_P216075** | Ddost | -0.183883616 | -0.893337199 | 0.525569968 | 12.7772017 | -0.649186371 | 0.542460674 | 0.769813173 | -4.695206392 |
| **A_52_P657360** | Tnni1 | 0.486923672 | -1.39233069 | 2.366178033 | 9.588086026 | 0.648971563 | 0.542589512 | 0.769939495 | -4.695247424 |
| **A_55_P2003216** | Igsf5 | 0.117798208 | -0.358896328 | 0.594492744 | 8.178533937 | 0.658205057 | 0.542727485 | 0.76996329 | -4.673640782 |
| **A_55_P2101231** | Aldh7a1 | -0.170871532 | -0.830631265 | 0.488888201 | 13.22373618 | -0.648685593 | 0.542761062 | 0.76996329 | -4.695302034 |
| **A_52_P137691** | Trappc5 | 0.094828723 | -0.271350754 | 0.4610082 | 8.543363807 | 0.648628831 | 0.542795118 | 0.76996329 | -4.695312872 |
| **A_55_P2019397** | Mtch1 | 0.113993304 | -0.326200086 | 0.554186694 | 15.65160314 | 0.648613779 | 0.542804149 | 0.76996329 | -4.695315746 |

| | | | | | | | | | |
|---|---|---|---|---|---|---|---|---|---|
| **A_55_P2034670** | Fbxl12 | 0.134983729 | -0.386267385 | 0.656234842 | 12.29814846 | 0.648611689 | 0.542805402 | 0.76996329 | -4.695316145 |
| **A_51_P479043** | Nif3l1 | 0.092181626 | -0.26390519 | 0.448268442 | 10.98968923 | 0.648393746 | 0.542936175 | 0.77009229 | -4.695357748 |
| **A_52_P235479** | Usp38 | -0.098780507 | -0.480397874 | 0.28283686 | 9.566597347 | -0.648326043 | 0.542976803 | 0.77009342 | -4.69537067 |
| **A_51_P304397** | Cpm | 0.278090676 | -0.796540184 | 1.352721536 | 11.5637558 | 0.648152895 | 0.543080716 | 0.770120159 | -4.695403713 |
| **A_51_P312437** | Dhrs7 | 0.168772458 | -0.483422407 | 0.820967322 | 13.44779396 | 0.648148538 | 0.543083331 | 0.770120159 | -4.695404544 |
| **A_55_P2082529** | Sav1 | 0.142270083 | -0.40755546 | 0.692095626 | 11.56453777 | 0.648095518 | 0.543115153 | 0.770120159 | -4.695414661 |
| **A_51_P101635** | Stub1 | -0.172292967 | -0.83827675 | 0.493690817 | 12.3297175 | -0.647969016 | 0.543191083 | 0.770168701 | -4.695438796 |
| **A_51_P185509** | Mapre1 | 0.0881371 | -0.252597397 | 0.428871596 | 9.436629908 | 0.647877646 | 0.543245931 | 0.770168701 | -4.695456226 |
| **A_55_P2073935** | Abcc5 | -0.082814405 | -0.402990482 | 0.237361671 | 8.225577618 | -0.647839397 | 0.543268891 | 0.770168701 | -4.695463522 |
| **A_52_P340073** | Efnb2 | 0.150162815 | -0.4304553 | 0.730780931 | 10.66231854 | 0.647771991 | 0.543309357 | 0.770169595 | -4.695476379 |
| **A_55_P2010341** | Rpl11 | 0.059734583 | -0.171262329 | 0.290731496 | 15.96587417 | 0.647694201 | 0.543356059 | 0.770179329 | -4.695491215 |
| **A_55_P1989797** | NA | 0.196059332 | -0.562553744 | 0.954672408 | 15.29170696 | 0.647317787 | 0.543582079 | 0.77041352 | -4.695562987 |
| **A_55_P2050268** | Nfkbid | 0.080577249 | -0.231215743 | 0.39237024 | 8.307035697 | 0.647286323 | 0.543600974 | 0.77041352 | -4.695568984 |
| **A_55_P2085551** | Dennd4b | 0.13180072 | -0.378334299 | 0.641935739 | 9.503321377 | 0.647117271 | 0.543702504 | 0.770500932 | -4.695601206 |
| **A_51_P116906** | Rapgef3 | -0.131008787 | -0.638155893 | 0.376138318 | 11.39520893 | -0.647018682 | 0.543761721 | 0.770528373 | -4.695619995 |
| **A_51_P224445** | Gpn3 | 0.096274057 | -0.276710573 | 0.469258687 | 8.72842138 | 0.646500232 | 0.544073193 | 0.770913236 | -4.695718763 |
| **A_55_P1970597** | Actg1 | -0.066116964 | -0.322357416 | 0.190123487 | 17.26621688 | -0.646272328 | 0.544210147 | 0.771050783 | -4.695762161 |
| **A_51_P112557** | Polr1c | 0.258320388 | -0.742992316 | 1.259633092 | 11.95803991 | 0.646158859 | 0.544278343 | 0.771090897 | -4.695783763 |
| **A_55_P2104332** | Pxdn | 0.130840299 | -0.37641981 | 0.638100409 | 9.703266794 | 0.646042611 | 0.544348214 | 0.771100193 | -4.695805892 |
| **A_55_P2325803** | Tnfrsf11a | 0.099670575 | -0.28676273 | 0.48610388 | 9.101867145 | 0.646015234 | 0.54436467 | 0.771100193 | -4.695811103 |
| **A_55_P2061386** | Chd3 | 0.281250165 | -0.809596401 | 1.372096731 | 13.27300265 | 0.645772375 | 0.544510664 | 0.771250489 | -4.695857323 |
| **A_55_P1962404** | Ugt1a6b | -0.22399344 | -1.093185891 | 0.645199011 | 9.729066644 | -0.645460543 | 0.544698156 | 0.771409288 | -4.695916649 |
| **A_51_P192130** | Stk10 | -0.203974005 | -0.99549131 | 0.587543301 | 9.539830203 | -0.645453193 | 0.544702576 | 0.771409288 | -4.695918047 |
| **A_52_P467046** | Wdr48 | -0.170257989 | -0.831126646 | 0.490610667 | 9.449206758 | -0.645271808 | 0.544811655 | 0.77143033 | -4.695952546 |
| **A_51_P466148** | Bckdha | 0.150661079 | -0.434202778 | 0.735524936 | 11.30533422 | 0.645203381 | 0.544852809 | 0.77143033 | -4.695965558 |
| **A_52_P680710** | Kpna3 | 0.085673225 | -0.246923094 | 0.418269544 | 9.994221632 | 0.645175731 | 0.544869439 | 0.77143033 | -4.695970816 |
| **A_55_P1966734** | Cyp2u1 | -0.17157336 | -0.837659936 | 0.494513216 | 15.13692379 | -0.645163101 | 0.544877035 | 0.77143033 | -4.695973218 |
| **A_55_P2014427** | Il17re | 0.308714735 | -0.890108504 | 1.507537974 | 9.315338349 | 0.644989361 | 0.544981538 | 0.771521788 | -4.696006251 |
| **A_66_P106814** | NA | -0.442653262 | -2.162206957 | 1.276900433 | 11.62333898 | -0.644760276 | 0.545119351 | 0.771615261 | -4.696049797 |
| **A_51_P492346** | Tmem174 | 0.133887679 | -0.386282466 | 0.654057824 | 9.146948161 | 0.644681987 | 0.545166453 | 0.771615261 | -4.696064676 |
| **A_51_P429295** | Pus10 | -0.056360806 | -0.275329963 | 0.162608351 | 9.237430413 | -0.644680588 | 0.545167295 | 0.771615261 | -4.696064942 |
| **A_51_P487389** | Drc3 | 0.187563982 | -0.541391881 | 0.916519845 | 8.401278884 | 0.644463843 | 0.545297712 | 0.771743354 | -4.696106128 |
| **A_51_P465350** | Ly86 | 0.353039456 | -1.019589589 | 1.725668501 | 10.34096621 | 0.644199534 | 0.545456775 | 0.771909392 | -4.696156338 |
| **A_52_P112178** | Srgap2 | -0.057903008 | -0.283081467 | 0.167275451 | 8.365761683 | -0.64405748 | 0.545542277 | 0.771909392 | -4.696183317 |
| **A_51_P175699** | Mtfp1 | 0.114745833 | -0.331520277 | 0.561011943 | 9.601103478 | 0.644011114 | 0.545570186 | 0.771909392 | -4.696192121 |
| **A_55_P2074736** | Prkar1b | 0.191541882 | -0.553405888 | 0.936489653 | 9.776254868 | 0.644003565 | 0.54557473 | 0.771909392 | -4.696193555 |
| **A_66_P131788** | Ythdf2 | -0.146581649 | -0.716825135 | 0.423661838 | 8.298551073 | -0.643827636 | 0.545680637 | 0.77200274 | -4.696226959 |
| **A_55_P2131707** | Lpcat1 | 0.233290837 | -0.674952533 | 1.141534207 | 9.044469587 | 0.643347756 | 0.545969586 | 0.772352108 | -4.69631804 |
| **A_55_P2169529** | NA | 0.069409702 | -0.200895683 | 0.339715088 | 8.883204425 | 0.643155265 | 0.546085518 | 0.772352108 | -4.69635456 |
| **A_55_P2160750** | Col25a1 | 0.237500183 | -0.687501314 | 1.16250168 | 8.60600081 | 0.64309015 | 0.546124738 | 0.772352108 | -4.696366912 |
| **A_51_P470311** | Cby1 | 0.230670155 | -0.667817381 | 1.129157691 | 9.020215942 | 0.643027721 | 0.546162342 | 0.772352108 | -4.696378754 |
| **A_55_P1963687** | Tsen54 | -0.082589582 | -0.404306965 | 0.239127802 | 11.63172452 | -0.642985361 | 0.546187859 | 0.772352108 | -4.696386788 |
| **A_55_P1955457** | Grb10 | -0.133699032 | -0.654527282 | 0.387129218 | 11.33195435 | -0.64296018 | 0.546203027 | 0.772352108 | -4.696391564 |
| **A_55_P2036007** | Rai2 | 0.08458465 | -0.244925769 | 0.41409507 | 8.323047412 | 0.642943412 | 0.546213128 | 0.772352108 | -4.696394744 |
| **A_51_P433584** | Bcas3 | -0.058824434 | -0.288002747 | 0.17035388 | 8.722164498 | -0.642886909 | 0.546247167 | 0.772352108 | -4.69640546 |
| **A_66_P128701** | NA | 0.223114983 | -0.646531954 | 1.092761919 | 15.74880535 | 0.642593176 | 0.546424137 | 0.772377394 | -4.696461155 |
| **A_55_P2209053** | 2900001G | 0.346946342 | -1.00543921 | 1.699331894 | 9.955671408 | 0.642557706 | 0.54644551 | 0.772377394 | -4.696467879 |
| **A_55_P2014066** | NA | -0.110439429 | -0.540954061 | 0.320075203 | 8.034304164 | -0.642519899 | 0.546468291 | 0.772377394 | -4.696475046 |

| | | | | | | | | | |
|---|---|---|---|---|---|---|---|---|---|
| **A_55_P2083794** | Tmem159 | -0.129031262 | -0.632052969 | 0.373990444 | 9.791291085 | -0.64247845 | 0.546493268 | 0.772377394 | -4.696482903 |
| **A_55_P2014927** | Zfp341 | 0.093837976 | -0.271994619 | 0.459670571 | 9.243497465 | 0.642460717 | 0.546503954 | 0.772377394 | -4.696486264 |
| **A_51_P380750** | Cbfa2t3 | 0.25515662 | -0.739587763 | 1.249901002 | 10.23730775 | 0.642459404 | 0.546504746 | 0.772377394 | -4.696486513 |
| **A_55_P2126951** | Zfp467 | 0.378611371 | -1.097971964 | 1.855194707 | 9.442035119 | 0.642223268 | 0.546647054 | 0.772429051 | -4.696531266 |
| **A_51_P448032** | Cox6b1 | 0.145967968 | -0.423318215 | 0.715254152 | 12.23785136 | 0.642210298 | 0.546654871 | 0.772429051 | -4.696533724 |
| **A_51_P144160** | Colec10 | -0.161441297 | -0.791143198 | 0.468260604 | 8.565471049 | -0.642140378 | 0.546697014 | 0.772429051 | -4.696546973 |
| **A_55_P1983638** | Mapk6 | -0.300074039 | -1.470565165 | 0.870417088 | 10.65864496 | -0.642111789 | 0.546714246 | 0.772429051 | -4.696552389 |
| **A_51_P461504** | Eef2k | -0.185795588 | -0.910641068 | 0.539049892 | 8.74068924 | -0.642007805 | 0.546776925 | 0.772429051 | -4.69657209 |
| **A_52_P597726** | Ccdc84 | -0.077260718 | -0.37868712 | 0.224165683 | 8.830544186 | -0.641989312 | 0.546788073 | 0.772429051 | -4.696575593 |
| **A_52_P485766** | Zdhhc17 | 0.114005868 | -0.330816543 | 0.558828279 | 8.904328122 | 0.641934759 | 0.546820959 | 0.772429051 | -4.696585928 |
| **A_55_P2008021** | Srpx2 | 0.247449196 | -0.718139415 | 1.213037806 | 10.60891931 | 0.641865831 | 0.546862511 | 0.772431312 | -4.696598984 |
| **A_55_P2077555** | C330021F | -0.116324335 | -0.570658575 | 0.338009906 | 8.319649062 | -0.641276691 | 0.547217754 | 0.772841552 | -4.696710537 |
| **A_55_P2043559** | NA | -0.057474084 | -0.281962512 | 0.167014343 | 8.98730072 | -0.641251582 | 0.547232898 | 0.772841552 | -4.69671529 |
| **A_55_P2151466** | Casc3 | -0.073621861 | -0.361269773 | 0.214026052 | 11.90202007 | -0.64105591 | 0.54735092 | 0.77295177 | -4.696752322 |
| **A_66_P101538** | Lrrk2 | 0.283791941 | -0.825199998 | 1.39278388 | 9.040265363 | 0.640946846 | 0.54741671 | 0.772988218 | -4.696772959 |
| **A_55_P2011967** | Akirin2 | 0.117931093 | -0.343075083 | 0.57893727 | 12.41139865 | 0.640725377 | 0.547550322 | 0.773070227 | -4.696814858 |
| **A_65_P15689** | Mmut | 0.205941291 | -0.599167638 | 1.01105022 | 10.28724701 | 0.640676999 | 0.547579511 | 0.773070227 | -4.696824009 |
| **A_55_P2117202** | Mtx1 | -0.071345002 | -0.350301027 | 0.207611023 | 12.29532565 | -0.640587051 | 0.547633785 | 0.773070227 | -4.696841022 |
| **A_55_P2027102** | Rgs3 | 0.245722945 | -0.7151728 | 1.20661869 | 11.52906253 | 0.640500963 | 0.547685732 | 0.773070227 | -4.696857302 |
| **A_52_P155124** | Zfp229 | 0.077870591 | -0.226688583 | 0.382429765 | 8.462448108 | 0.640401192 | 0.54774594 | 0.773070227 | -4.696876169 |
| **A_55_P2087647** | Grhpr | -0.109520031 | -0.537880119 | 0.318840057 | 8.114685688 | -0.640375791 | 0.547761269 | 0.773070227 | -4.696880972 |
| **A_51_P172323** | Pak2 | -0.135852596 | -0.667226087 | 0.395520896 | 11.16936637 | -0.640351455 | 0.547775956 | 0.773070227 | -4.696885573 |
| **A_51_P138879** | Zgpat | -0.098991418 | -0.486204945 | 0.288222108 | 9.152132712 | -0.640320449 | 0.547794668 | 0.773070227 | -4.696891436 |
| **A_55_P2046101** | Xlr4b | -0.672776263 | -3.306067036 | 1.96051451 | 8.679942478 | -0.63991485 | 0.548039489 | 0.773359279 | -4.696968104 |
| **A_55_P2140516** | Pex13 | -0.180891101 | -0.889013054 | 0.527230851 | 11.25637056 | -0.639822474 | 0.548095257 | 0.773376131 | -4.69698556 |
| **A_55_P2018322** | Zfp644 | 0.190152193 | -0.554296515 | 0.934600902 | 10.69372836 | 0.639759737 | 0.548133135 | 0.773376131 | -4.696997414 |
| **A_52_P72434** | Khk | 0.082309458 | -0.239965266 | 0.404584182 | 9.797244799 | 0.639696304 | 0.548171433 | 0.773376131 | -4.697009399 |
| **A_65_P19089** | Esrrg | 0.06558005 | -0.191242805 | 0.322402905 | 7.822023451 | 0.639570491 | 0.5482474 | 0.773386843 | -4.697033167 |
| **A_55_P2095384** | Glrx2 | 0.054340816 | -0.158473659 | 0.267155292 | 11.20539807 | 0.639551233 | 0.548259029 | 0.773386843 | -4.697036805 |
| **A_55_P2178317** | Gm9199 | 0.098666259 | -0.287936932 | 0.485269449 | 8.605072414 | 0.639224736 | 0.548456205 | 0.773608541 | -4.697098467 |
| **A_55_P2004447** | Fgl1 | 0.102491725 | -0.299175579 | 0.504159029 | 8.114579897 | 0.639105666 | 0.548528124 | 0.773653542 | -4.697120948 |
| **A_51_P176042** | Pklr | -0.061163499 | -0.300939425 | 0.178612427 | 7.715533087 | -0.638906181 | 0.548648628 | 0.773723804 | -4.697158606 |
| **A_51_P184100** | Capn7 | -0.079199844 | -0.389690193 | 0.231290504 | 8.946013608 | -0.638890704 | 0.548657978 | 0.773723804 | -4.697161527 |
| **A_55_P2155421** | Sec61g | -0.077876029 | -0.383313086 | 0.227561028 | 15.27177547 | -0.63860515 | 0.548830506 | 0.773910655 | -4.697215416 |
| **A_55_P2177173** | Gnl1 | -0.300716829 | -1.481000298 | 0.879566639 | 10.46379281 | -0.638148503 | 0.549106477 | 0.774243336 | -4.697301555 |
| **A_55_P2091245** | NA | 0.133723101 | -0.391259499 | 0.6587057 | 16.08136719 | 0.637987066 | 0.549204062 | 0.774324461 | -4.697331996 |
| **A_55_P2106250** | Trappc6a | 0.235969431 | -0.690618738 | 1.162557599 | 13.40420216 | 0.637851146 | 0.54928623 | 0.77438384 | -4.697357621 |
| **A_55_P2066996** | Cav2 | 0.163536743 | -0.479029493 | 0.80610298 | 8.36453396 | 0.637452441 | 0.549527308 | 0.774627521 | -4.697432765 |
| **A_51_P391380** | Mindy2 | 0.182802478 | -0.535554484 | 0.901159439 | 9.540586576 | 0.637370735 | 0.54957672 | 0.774627521 | -4.69744816 |
| **A_52_P109548** | Map7 | -0.0838188 | -0.413202834 | 0.245565234 | 8.614912474 | -0.63736651 | 0.549579275 | 0.774627521 | -4.697448956 |
| **A_55_P2027836** | Tnfrsf10b | -0.213686544 | -1.053523015 | 0.626149927 | 9.360309788 | -0.637283669 | 0.549629376 | 0.774639497 | -4.697464563 |
| **A_55_P2024327** | Kmt5c | 0.111308961 | -0.326218093 | 0.548836016 | 12.31592929 | 0.637199701 | 0.549680162 | 0.774639497 | -4.69748038 |
| **A_52_P94521** | 2510009E | 0.121129781 | -0.355042615 | 0.597302178 | 8.575761824 | 0.637143217 | 0.549714327 | 0.774639497 | -4.69749102 |
| **A_55_P1963817** | Rps6 | 0.185168007 | -0.542808262 | 0.913144276 | 16.33488497 | 0.637087488 | 0.549748036 | 0.774639497 | -4.697501517 |
| **A_55_P2107025** | Flad1 | -0.060529835 | -0.298534191 | 0.177474521 | 8.200977898 | -0.6369934 | 0.549804951 | 0.774663236 | -4.697519236 |
| **A_52_P118461** | Ipmk | -0.122754186 | -0.605519783 | 0.360011411 | 10.34507694 | -0.636869341 | 0.549880001 | 0.774712522 | -4.697542598 |
| **A_55_P2171623** | Ptbp1 | -0.102622062 | -0.506367728 | 0.301123604 | 9.203366391 | -0.636624299 | 0.55002826 | 0.774864936 | -4.697588731 |
| **A_52_P249733** | Tcap | 0.40643476 | -1.19288528 | 2.0057548 | 9.408569387 | 0.636510849 | 0.55009691 | 0.774905185 | -4.697610085 |

| | | | | | | | | | |
|---|---|---|---|---|---|---|---|---|---|
| **A_52_P408025** | Mpped2 | 0.168799133 | -0.527715472 | 0.865313738 | 9.979103364 | 0.645509575 | 0.550203832 | 0.774999335 | -4.675708772 |
| **A_66_P115406** | Pabpc1 | -0.088039559 | -0.434623878 | 0.25854476 | 15.04185077 | -0.636237546 | 0.550262311 | 0.775025244 | -4.697661516 |
| **A_55_P2033282** | Cept1 | 0.109270347 | -0.321012795 | 0.539553488 | 12.36198193 | 0.636060367 | 0.550369555 | 0.775081214 | -4.697694849 |
| **A_51_P423518** | Amph | 0.319608139 | -0.93912319 | 1.578339468 | 9.072980875 | 0.63596781 | 0.550425584 | 0.775081214 | -4.697712259 |
| **A_55_P2141479** | Rnu2-10 | -0.366936202 | -1.81207027 | 1.078197866 | 10.76434158 | -0.635964344 | 0.550427682 | 0.775081214 | -4.697712911 |
| **A_55_P2449750** | Fam177a2 | -0.081675415 | -0.403436149 | 0.240085319 | 8.76622664 | -0.635782617 | 0.550537701 | 0.775081214 | -4.697747088 |
| **A_52_P163928** | Poldip2 | -0.061440438 | -0.303531236 | 0.180650361 | 10.38334102 | -0.635662167 | 0.55061063 | 0.775081214 | -4.697769737 |
| **A_55_P2109454** | Gtf3c5 | 0.078281215 | -0.230196009 | 0.386758439 | 11.08640957 | 0.635601335 | 0.550647464 | 0.775081214 | -4.697781175 |
| **A_52_P327467** | Med7 | -0.085287464 | -0.421397001 | 0.250822073 | 10.62669043 | -0.635557301 | 0.550674128 | 0.775081214 | -4.697789453 |
| **A_51_P106144** | Zfp64 | -0.189677681 | -0.937263388 | 0.557908026 | 10.6131219 | -0.635485414 | 0.550717659 | 0.775081214 | -4.691295984 |
| **A_51_P115626** | Shank3 | 0.174371971 | -0.512894701 | 0.861638643 | 8.647687903 | 0.635479782 | 0.55072107 | 0.775081214 | -4.696185046 |
| **A_55_P2082708** | Pwp2 | 0.109522663 | -0.322163305 | 0.54120863 | 10.27834869 | 0.635457351 | 0.550734653 | 0.775081214 | -4.697808243 |
| **A_55_P2065966** | Ctdp1 | 0.101954021 | -0.299914092 | 0.503822134 | 9.477173831 | 0.635435037 | 0.550748167 | 0.775081214 | -4.697812437 |
| **A_55_P2065308** | Myo9b | -0.09856467 | -0.487108403 | 0.289979063 | 8.485018051 | -0.635377324 | 0.550783118 | 0.775081214 | -4.697823285 |
| **A_55_P2024215** | Ctnnbl1 | -0.182559169 | -0.90258118 | 0.537462842 | 12.88339753 | -0.635050443 | 0.550981105 | 0.775303397 | -4.697884713 |
| **A_55_P2008277** | Tcp10c | 0.072652557 | -0.213990766 | 0.35929588 | 8.017814486 | 0.634832889 | 0.551112899 | 0.775324309 | -4.697925583 |
| **A_51_P302377** | Fgfbp1 | 0.242222832 | -0.713450178 | 1.197895841 | 8.822045208 | 0.634828041 | 0.551115836 | 0.775324309 | -4.697926493 |
| **A_55_P2115057** | Tuba1a | 0.139803452 | -0.411781781 | 0.691388686 | 14.32566443 | 0.634827323 | 0.551116271 | 0.775324309 | -4.697926628 |
| **A_55_P2031407** | NA | 0.193273192 | -0.569437342 | 0.955983726 | 17.68103033 | 0.634690825 | 0.551198972 | 0.775384235 | -4.697952265 |
| **A_51_P367162** | Psmb1 | 0.239031734 | -0.704438083 | 1.182501552 | 13.04048026 | 0.634567605 | 0.551273635 | 0.775416669 | -4.697975404 |
| **A_55_P2138257** | Ube2d2a | -0.271041318 | -1.341339478 | 0.799256841 | 10.41775699 | -0.634280074 | 0.551447885 | 0.775416669 | -4.698029386 |
| **A_52_P12001** | Erp44 | 0.067226878 | -0.198255011 | 0.332708766 | 10.00255352 | 0.634246953 | 0.551467959 | 0.775416669 | -4.698035603 |
| **A_55_P2154307** | Peak1 | 0.283628429 | -0.836455598 | 1.403712456 | 12.67582752 | 0.634233972 | 0.551475826 | 0.775416669 | -4.69803804 |
| **A_66_P109941** | Mamdc4 | 0.126545888 | -0.37327184 | 0.626363615 | 8.364114078 | 0.634142304 | 0.551531388 | 0.775416669 | -4.698055245 |
| **A_65_P03364** | Dennd4a | -0.056417688 | -0.279290679 | 0.166455303 | 7.795363477 | -0.634027637 | 0.551600895 | 0.775416669 | -4.698076764 |
| **A_52_P154880** | Ing3 | 0.085457158 | -0.252135618 | 0.423049934 | 8.778589842 | 0.634023928 | 0.551603144 | 0.775416669 | -4.69807746 |
| **A_55_P2073253** | NA | 0.10194212 | -0.300823577 | 0.504707817 | 9.616446422 | 0.633944928 | 0.551651034 | 0.775416669 | -4.698092284 |
| **A_55_P1961391** | NA | -0.086542417 | -0.428502337 | 0.255417503 | 8.03081058 | -0.63387578 | 0.551692954 | 0.775416669 | -4.698105258 |
| **A_55_P2075636** | Epc2 | 0.085106393 | -0.25118954 | 0.421402326 | 9.648395639 | 0.633856451 | 0.551704672 | 0.775416669 | -4.698108885 |
| **A_51_P480928** | Zfyve16 | -0.075616869 | -0.374478617 | 0.223244878 | 9.218049101 | -0.633721887 | 0.551786257 | 0.775416669 | -4.698134129 |
| **A_55_P1964568** | Megf10 | 0.265993251 | -0.785380074 | 1.317366576 | 8.707619037 | 0.633671259 | 0.551816954 | 0.775416669 | -4.698143625 |
| **A_52_P502267** | Epc2 | -0.133807964 | -0.6627194 | 0.395103471 | 10.10761078 | -0.633649875 | 0.55182992 | 0.775416669 | -4.698147636 |
| **A_55_P2000148** | Dnaic1 | -0.07056204 | -0.349494058 | 0.208369978 | 8.794280078 | -0.633611575 | 0.551853143 | 0.775416669 | -4.69815482 |
| **A_51_P270899** | Zfp61 | 0.182647489 | -0.539500754 | 0.904795731 | 9.66249256 | 0.633486979 | 0.551928697 | 0.775416669 | -4.698178188 |
| **A_55_P2109983** | Mpg | -0.066056334 | -0.339899412 | 0.207786744 | 8.466815782 | -0.642503361 | 0.551984291 | 0.775416669 | -4.676194356 |
| **A_55_P2152387** | Wwox | 0.056735022 | -0.167637877 | 0.281107922 | 8.125619535 | 0.633331627 | 0.55202291 | 0.775416669 | -4.698207318 |
| **A_55_P1981259** | Fkbp14 | 0.173927871 | -0.513966738 | 0.86182248 | 10.63030906 | 0.633282695 | 0.552052587 | 0.775416669 | -4.698216492 |
| **A_51_P174275** | Colec11 | -0.062929632 | -0.311839807 | 0.185980542 | 8.393861537 | -0.633232055 | 0.552083301 | 0.775416669 | -4.698225986 |
| **A_52_P312398** | Ssr3 | -0.186161311 | -0.922531108 | 0.550208486 | 10.02550086 | -0.63320423 | 0.552100177 | 0.775416669 | -4.698231203 |
| **A_55_P2037918** | NA | 0.110803804 | -0.327493821 | 0.549101429 | 8.627688348 | 0.633192699 | 0.552107172 | 0.775416669 | -4.698233364 |
| **A_51_P171965** | Sdhaf4 | 0.17893832 | -0.528897135 | 0.886773775 | 10.72309899 | 0.633171544 | 0.552120003 | 0.775416669 | -4.69823733 |
| **A_51_P105589** | Eci1 | 0.343025218 | -1.014096268 | 1.700146703 | 11.60579457 | 0.633078651 | 0.552176349 | 0.775416669 | -4.698254743 |
| **A_55_P2124090** | Ambra1 | -0.070384017 | -0.348852493 | 0.208084459 | 10.75883384 | -0.63306508 | 0.552184581 | 0.775416669 | -4.698257287 |
| **A_55_P1962154** | Rarb | 0.212987345 | -0.629819875 | 1.055794565 | 9.203542179 | 0.632959465 | 0.552248649 | 0.775438796 | -4.698277081 |
| **A_55_P1984976** | Wnt5b | 0.193131165 | -0.571167135 | 0.957429465 | 9.897716482 | 0.632906874 | 0.552280553 | 0.775438796 | -4.698286937 |
| **A_55_P2171273** | Mier3 | -0.077859737 | -0.386171202 | 0.230451728 | 8.805689774 | -0.632519042 | 0.552515868 | 0.77571286 | -4.6983596 |
| **A_55_P2072908** | NA | 0.10494691 | -0.31070176 | 0.520595579 | 14.07237431 | 0.632402541 | 0.552586567 | 0.775717602 | -4.698381421 |
| **A_66_P122415** | Snhg6 | 0.121457597 | -0.359598859 | 0.602514054 | 13.42716627 | 0.632381247 | 0.552599489 | 0.775717602 | -4.698385409 |

| | | | | | | | | | |
|---|---|---|---|---|---|---|---|---|---|
| **A_55_P1982111** | Homer1 | -0.10944844 | -0.543046977 | 0.324150096 | 8.475058576 | -0.632225654 | 0.552693921 | 0.775753688 | -4.698414545 |
| **A_55_P2122504** | Zpr1 | -0.110553555 | -0.548567712 | 0.327460601 | 11.32087866 | -0.6321715 | 0.552726791 | 0.775753688 | -4.698424685 |
| **A_55_P2114562** | NA | 0.076076106 | -0.225352844 | 0.377505057 | 8.056071884 | 0.632140565 | 0.552745568 | 0.775753688 | -4.698430477 |
| **A_55_P1975011** | Ufc1 | 0.163696792 | -0.485569287 | 0.812962871 | 13.36654306 | 0.631491924 | 0.553139373 | 0.776201721 | -4.698551872 |
| **A_51_P396818** | Rex1bd | 0.070771866 | -0.209971337 | 0.351515069 | 10.03603368 | 0.631395879 | 0.553197699 | 0.776201721 | -4.698569839 |
| **A_55_P2065364** | Tdrkh | 0.071042112 | -0.210775458 | 0.352859682 | 8.023313529 | 0.631390646 | 0.553200877 | 0.776201721 | -4.698570818 |
| **A_55_P2064792** | Arpc3 | 0.115043222 | -0.341379374 | 0.571465818 | 12.72620654 | 0.631312292 | 0.553248463 | 0.776201721 | -4.698585474 |
| **A_55_P2060483** | Rnf141 | 0.060525174 | -0.179630818 | 0.300681165 | 8.29749896 | 0.631237751 | 0.553293735 | 0.776201721 | -4.698599416 |
| **A_51_P371750** | Marco | 0.094914752 | -0.281725767 | 0.471555271 | 8.182202066 | 0.631185512 | 0.553325464 | 0.776201721 | -4.698609185 |
| **A_51_P201785** | Morn5 | 0.559318783 | -1.660489804 | 2.77912737 | 10.30153507 | 0.631094249 | 0.553380898 | 0.776201721 | -4.698626251 |
| **A_55_P2185330** | NA | 0.109255119 | -0.324402506 | 0.542912744 | 9.214717736 | 0.631022946 | 0.55342421 | 0.776201721 | -4.698639584 |
| **A_55_P2028015** | Pawr | -0.093408633 | -0.46416992 | 0.277352653 | 9.62592384 | -0.631019793 | 0.553426126 | 0.776201721 | -4.698640173 |
| **A_55_P1958867** | Gm9790 | 0.110201243 | -0.327336743 | 0.547739229 | 14.40421577 | 0.630842691 | 0.553533715 | 0.776270619 | -4.698673283 |
| **A_55_P2082733** | Cybb | -0.076416648 | -0.379835107 | 0.227001811 | 7.915028326 | -0.630806749 | 0.553555551 | 0.776270619 | -4.698680002 |
| **A_52_P601980** | Kdm5b | -0.063083919 | -0.313839128 | 0.187671289 | 8.78347838 | -0.630113888 | 0.5539766 | 0.77679963 | -4.69880946 |
| **A_51_P310576** | Smim8 | 0.060533504 | -0.180117355 | 0.301184364 | 8.286050426 | 0.630026394 | 0.554029784 | 0.77679963 | -4.6988258 |
| **A_51_P456857** | Ift122 | 0.221153295 | -0.658095346 | 1.100401936 | 10.08618408 | 0.629987676 | 0.55405332 | 0.77679963 | -4.69883303 |
| **A_51_P491095** | Gzf1 | -0.074498901 | -0.370781869 | 0.221784067 | 9.442674414 | -0.629786761 | 0.554175464 | 0.77691454 | -4.698870544 |
| **A_55_P2150048** | Sypl | 0.140904621 | -0.419552369 | 0.701361612 | 11.19786199 | 0.629699418 | 0.554228569 | 0.776932653 | -4.69888685 |
| **A_55_P2141851** | Pls3 | 0.292792448 | -0.872100008 | 1.457684904 | 12.94931086 | 0.629541527 | 0.554324575 | 0.777010899 | -4.698916321 |
| **A_52_P654618** | Znhit3 | 0.100674871 | -0.299967108 | 0.50131685 | 8.830974595 | 0.629382953 | 0.554421006 | 0.777074502 | -4.698945914 |
| **A_55_P2025225** | Gnaz | -0.090486037 | -0.450608502 | 0.269636428 | 8.216176618 | -0.629334729 | 0.554450334 | 0.777074502 | -4.698954912 |
| **A_51_P279127** | Fam173a | 0.179664489 | -0.535841086 | 0.895170064 | 12.58594436 | 0.62892603 | 0.554698928 | 0.777191471 | -4.699031152 |
| **A_55_P2120254** | Avpi1 | 0.366715729 | -1.093831541 | 1.827262998 | 13.74273898 | 0.628874907 | 0.554730029 | 0.777191471 | -4.699040686 |
| **A_52_P451574** | Xpr1 | -0.072669975 | -0.362102411 | 0.21676246 | 11.22059743 | -0.628866065 | 0.554735408 | 0.777191471 | -4.699042335 |
| **A_52_P449871** | Id4 | 0.166883633 | -0.497853602 | 0.831620867 | 8.641014117 | 0.628802246 | 0.554774235 | 0.777191471 | -4.699054236 |
| **A_55_P2083806** | Sp8 | -0.068281263 | -0.340284116 | 0.20372159 | 8.004442214 | -0.628750625 | 0.554805642 | 0.777191471 | -4.699063861 |
| **A_66_P114501** | Zcrb1 | 0.063714051 | -0.190113646 | 0.317541748 | 12.51039087 | 0.628704476 | 0.55483372 | 0.777191471 | -4.699072466 |
| **A_52_P319123** | Vps13b | 0.076111781 | -0.227120306 | 0.379343868 | 10.20018223 | 0.628676284 | 0.554850873 | 0.777191471 | -4.699077722 |
| **A_55_P2024530** | Siah2 | -0.079434404 | -0.395907666 | 0.237038858 | 8.785016814 | -0.62866888 | 0.554855378 | 0.777191471 | -4.699079102 |
| **A_55_P1986441** | Gcdh | 0.056387319 | -0.168310526 | 0.281085163 | 8.713486761 | 0.628539945 | 0.554933832 | 0.777245053 | -4.699103139 |
| **A_51_P521171** | Hif1an | -0.087729266 | -0.43742134 | 0.261962808 | 8.282195764 | -0.628360753 | 0.555042878 | 0.777268426 | -4.699136538 |
| **A_55_P2143336** | Rpp21 | 0.102962234 | -0.307473477 | 0.513397945 | 9.710675689 | 0.628323296 | 0.555065674 | 0.777268426 | -4.699143519 |
| **A_55_P2010121** | Rab27b | 0.220766295 | -0.659281269 | 1.100813859 | 8.956036058 | 0.628314337 | 0.555071127 | 0.777268426 | -4.699145189 |
| **A_55_P2187038** | Tpm1 | 0.656071209 | -1.959525403 | 3.271667821 | 13.39772771 | 0.628247196 | 0.55511199 | 0.777269139 | -4.6991577 |
| **A_51_P135732** | Ccdc136 | 0.124107807 | -0.370805685 | 0.619021299 | 8.554425893 | 0.628087504 | 0.555209189 | 0.777269139 | -4.699187455 |
| **A_51_P414837** | Faf1 | -0.140281035 | -0.699720539 | 0.41915847 | 9.858951516 | -0.628052828 | 0.555230296 | 0.777269139 | -4.699193915 |
| **A_52_P420712** | Pja2 | -0.108010303 | -0.538756841 | 0.322736234 | 9.45792549 | -0.628049299 | 0.555232444 | 0.777269139 | -4.699194573 |
| **A_55_P1988033** | Atp6v1g1 | 0.148416674 | -0.443746641 | 0.740579989 | 13.45670262 | 0.627756967 | 0.555410409 | 0.777461978 | -4.699249024 |
| **A_55_P2032362** | NA | -0.106968975 | -0.534013403 | 0.320075452 | 8.330722456 | -0.627386432 | 0.555636033 | 0.777649904 | -4.699318013 |
| **A_66_P115996** | Kyat1 | -0.091528071 | -0.456952566 | 0.273896423 | 10.54324843 | -0.62734581 | 0.555660772 | 0.777649904 | -4.699325575 |
| **A_55_P2052116** | NA | 0.10844793 | -0.324561941 | 0.5414578 | 9.877800645 | 0.627297872 | 0.555689967 | 0.777649904 | -4.699334498 |
| **A_51_P118765** | Eef1a1 | -0.074340846 | -0.371198482 | 0.22251679 | 16.21295553 | -0.627234042 | 0.555728842 | 0.777649904 | -4.699346378 |
| **A_55_P1966760** | NA | 0.123428333 | -0.369467143 | 0.616323809 | 17.55539451 | 0.627206249 | 0.55574577 | 0.777649904 | -4.69935155 |
| **A_55_P1996202** | NA | 0.162643103 | -0.486925874 | 0.81221208 | 11.89833645 | 0.627134543 | 0.555789445 | 0.777654735 | -4.699364894 |
| **A_55_P2015692** | Elob | -0.07067174 | -0.35302316 | 0.21167968 | 13.79931789 | -0.626911377 | 0.555925386 | 0.777788654 | -4.699406418 |
| **A_55_P1983523** | Cd300ld | -0.057605387 | -0.287819467 | 0.172608693 | 8.074055834 | -0.626731559 | 0.556034937 | 0.777818002 | -4.699439867 |
| **A_55_P2004571** | Pitpnm2 | 0.07515561 | -0.225211286 | 0.375522506 | 11.36894695 | 0.626699984 | 0.556054175 | 0.777818002 | -4.69944574 |

| | | | | | | | | | |
|---|---|---|---|---|---|---|---|---|---|
| **A_55_P2058864** | Mvk | 0.119111147 | -0.356993133 | 0.595215427 | 9.727275259 | 0.626614827 | 0.556106061 | 0.777818002 | -4.699461577 |
| **A_55_P1988951** | NA | 0.177664672 | -0.532539795 | 0.887869139 | 15.40195079 | 0.626567732 | 0.556134757 | 0.777818002 | -4.699470335 |
| **A_55_P2111523** | Higd1b | 0.50592964 | -1.516563032 | 2.528422312 | 12.69202525 | 0.626546794 | 0.556147515 | 0.777818002 | -4.699474229 |
| **A_55_P2000289** | Mab21l2 | 0.063656515 | -0.190870501 | 0.318183531 | 8.166907007 | 0.626410914 | 0.556230317 | 0.777877537 | -4.699499494 |
| **A_66_P135130** | Tmem161 | -0.092246939 | -0.461355368 | 0.276861491 | 9.330174111 | -0.62596254 | 0.556503601 | 0.778187436 | -4.699582835 |
| **A_55_P2183808** | NA | -0.066781159 | -0.33402946 | 0.200467142 | 14.22332611 | -0.625877509 | 0.556555437 | 0.778187436 | -4.699598635 |
| **A_51_P491227** | Suclg1 | 0.121355654 | -0.364313614 | 0.607024922 | 12.32599362 | 0.625849252 | 0.556572663 | 0.778187436 | -4.699603885 |
| **A_55_P2159522** | Col14a1 | 0.167178867 | -0.502337707 | 0.836695441 | 8.436787356 | 0.625418022 | 0.556835598 | 0.778486918 | -4.699683985 |
| **A_55_P2113160** | Mal | 0.40377072 | -1.213484049 | 2.021025489 | 12.89748904 | 0.625326347 | 0.556891505 | 0.778486918 | -4.699701008 |
| **A_55_P2017774** | NA | 0.140264661 | -0.421572258 | 0.702101579 | 17.66398022 | 0.625299868 | 0.556907653 | 0.778486918 | -4.699705924 |
| **A_51_P479352** | Klhdc9 | 0.327590474 | -0.986348164 | 1.641529113 | 9.294383297 | 0.624462735 | 0.557418345 | 0.779144468 | -4.699861275 |
| **A_55_P2066384** | Akt1 | -0.149090953 | -0.747214721 | 0.449032815 | 13.63366086 | -0.624324784 | 0.55750253 | 0.779160006 | -4.69988686 |
| **A_55_P2135633** | NA | 0.217699938 | -0.655780437 | 1.091180313 | 16.75683666 | 0.624245612 | 0.557550848 | 0.779160006 | -4.699901542 |
| **A_55_P2008722** | Zfp984 | -0.150049267 | -0.752185295 | 0.452086762 | 8.888159192 | -0.624150912 | 0.557608648 | 0.779160006 | -4.699919101 |
| **A_55_P1964559** | Smarca5-| | 0.083914419 | -0.252859095 | 0.420687932 | 9.717911594 | 0.624092565 | 0.557644261 | 0.779160006 | -4.699929919 |
| **A_66_P134412** | Mettl25 | 0.06960988 | -0.209762528 | 0.348982288 | 10.03710944 | 0.624076342 | 0.557654163 | 0.779160006 | -4.699932926 |
| **A_55_P2046198** | Slc25a25 | -0.184401282 | -0.924569603 | 0.555767039 | 8.659612847 | -0.623998848 | 0.557701465 | 0.779160006 | -4.699947292 |
| **A_55_P2172934** | Rgs7bp | -0.105543208 | -0.529194327 | 0.318107912 | 11.41270789 | -0.623982308 | 0.557711561 | 0.779160006 | -4.699950358 |
| **A_51_P116407** | Pald1 | 0.242241186 | -0.730299313 | 1.214781684 | 9.032993382 | 0.623865018 | 0.557783161 | 0.779193274 | -4.699972099 |
| **A_55_P2046348** | Itpripl2 | -0.07789124 | -0.390662638 | 0.234880158 | 12.30988058 | -0.623751973 | 0.557852175 | 0.779193274 | -4.699993049 |
| **A_52_P304128** | Mmp14 | -0.058558265 | -0.293700855 | 0.176584324 | 9.431054436 | -0.623745252 | 0.557856279 | 0.779193274 | -4.699994295 |
| **A_55_P2454099** | Oga | 0.074194282 | -0.223811656 | 0.372200219 | 8.495283346 | 0.623585291 | 0.557953944 | 0.779251166 | -4.700023935 |
| **A_55_P2064442** | Pabpn1 | 0.063439641 | -0.191385939 | 0.31826522 | 13.29012077 | 0.623545345 | 0.557978336 | 0.779251166 | -4.700031336 |
| **A_55_P2016388** | Eif3k | 0.124402547 | -0.375397976 | 0.62420307 | 14.79561823 | 0.623423129 | 0.558052965 | 0.779299099 | -4.700053977 |
| **A_55_P2130219** | Tpr | -0.124082668 | -0.622790101 | 0.374624765 | 9.149388215 | -0.623183043 | 0.558199588 | 0.779305301 | -4.700098444 |
| **A_51_P370350** | Galnt2 | 0.13928276 | -0.420555345 | 0.699120865 | 12.32328843 | 0.623139462 | 0.558226206 | 0.779305301 | -4.700106515 |
| **A_55_P2090142** | Kcna7 | 0.084357929 | -0.254713882 | 0.423429739 | 8.654650562 | 0.623138476 | 0.558226808 | 0.779305301 | -4.700106697 |
| **A_55_P2081164** | Cd300lf | -0.152262734 | -0.764315083 | 0.459789615 | 8.649130995 | -0.623096625 | 0.558252371 | 0.779305301 | -4.700114447 |
| **A_55_P2116889** | Hax1 | -0.07893416 | -0.396256644 | 0.238388324 | 10.44622493 | -0.623037935 | 0.55828822 | 0.779305301 | -4.700125314 |
| **A_55_P2104978** | Edc4 | -0.068384448 | -0.343304102 | 0.206535206 | 11.87283693 | -0.623019874 | 0.558299252 | 0.779305301 | -4.700128658 |
| **A_55_P2172146** | Pds5a | -0.145047436 | -0.728289444 | 0.438194572 | 9.187530944 | -0.622890354 | 0.55837837 | 0.779339898 | -4.700152637 |
| **A_51_P490348** | 1110059E | 0.240417199 | -0.726457431 | 1.207291829 | 11.06714118 | 0.622795855 | 0.5584361 | 0.779339898 | -4.700170129 |
| **A_55_P2274233** | Agap3 | -0.099907923 | -0.521203455 | 0.321387608 | 13.35004939 | -0.631648959 | 0.558444965 | 0.779339898 | -4.677934354 |
| **A_55_P2011991** | St7 | 0.203650491 | -0.615509916 | 1.022810899 | 11.21647417 | 0.622682778 | 0.558505184 | 0.779367681 | -4.700191058 |
| **A_55_P2012849** | Dock10 | 0.097449887 | -0.294598581 | 0.489498356 | 8.114688774 | 0.622575356 | 0.558570819 | 0.779403017 | -4.700210938 |
| **A_51_P117109** | Ilvbl | -0.253130989 | -1.271927969 | 0.765665992 | 9.065648217 | -0.622311766 | 0.558731891 | 0.779571508 | -4.700259706 |
| **A_55_P2065953** | Btbd19 | 0.092303624 | -0.279266439 | 0.463873687 | 8.561709121 | 0.622197671 | 0.558801621 | 0.779580175 | -4.700280811 |
| **A_55_P1990440** | R3hdm1 | 0.074994702 | -0.22691155 | 0.376900954 | 10.84614948 | 0.62216965 | 0.558818746 | 0.779580175 | -4.700285994 |
| **A_65_P11092** | Tspan15 | 0.42334207 | -1.281124314 | 2.127808454 | 9.645736783 | 0.622090164 | 0.558867329 | 0.779591638 | -4.700300694 |
| **A_55_P1972019** | 4632415L | 0.074708111 | -0.226114705 | 0.375530927 | 8.235192842 | 0.622024266 | 0.558907608 | 0.779591638 | -4.70031288 |
| **A_52_P635271** | Btbd6 | 0.157279931 | -0.476303964 | 0.790863825 | 8.82675223 | 0.621755342 | 0.559072003 | 0.779748566 | -4.700362602 |
| **A_52_P678056** | Eva1b | 0.277437362 | -0.840355746 | 1.395230469 | 13.38023796 | 0.621661346 | 0.55912947 | 0.779748566 | -4.700379977 |
| **A_52_P211418** | G2e3 | 0.132500699 | -0.401359993 | 0.666361392 | 8.3080559 | 0.621642317 | 0.559141104 | 0.779748566 | -4.700383494 |
| **A_51_P259879** | Fkrp | 0.075422331 | -0.228498143 | 0.379342805 | 9.302987509 | 0.621570421 | 0.559185064 | 0.779753626 | -4.700396782 |
| **A_51_P243304** | Galk1 | 0.113929967 | -0.345281891 | 0.573141825 | 11.80519134 | 0.621405688 | 0.559285794 | 0.779787431 | -4.700427225 |
| **A_55_P2004876** | Srpx2 | 0.188487147 | -0.571246984 | 0.948221278 | 8.643878101 | 0.621398857 | 0.559289971 | 0.779787431 | -4.700428487 |
| **A_55_P1979305** | Gle1 | -0.072334467 | -0.364037302 | 0.219368367 | 9.372268694 | -0.621090639 | 0.559478469 | 0.779993995 | -4.700485428 |
| **A_51_P206971** | Mnat1 | 0.143978274 | -0.43682595 | 0.724782497 | 10.2176106 | 0.620894113 | 0.559598681 | 0.780006362 | -4.700521723 |

| | | | | | | | | | |
|---|---|---|---|---|---|---|---|---|---|
| **A_55_P2067226** | Arpc5l | 0.075089643 | -0.22782278 | 0.378002066 | 11.17565529 | 0.620888051 | 0.55960239 | 0.780006362 | -4.700522842 |
| **A_51_P511511** | Stk33 | 0.289838804 | -0.879423124 | 1.459100732 | 9.147430061 | 0.62086198 | 0.559618338 | 0.780006362 | -4.700527657 |
| **A_51_P511680** | Ttc38 | -0.053948118 | -0.271630933 | 0.163734697 | 8.181116077 | -0.620729668 | 0.559699282 | 0.780006362 | -4.700552087 |
| **A_55_P2169804** | Nanos3 | 0.084062733 | -0.255166585 | 0.423292051 | 9.6967309 | 0.620669598 | 0.559736033 | 0.780006362 | -4.700563176 |
| **A_51_P342669** | Pgam1 | 0.085143858 | -0.258462312 | 0.428750027 | 14.52323622 | 0.620644219 | 0.55975156 | 0.780006362 | -4.700567862 |
| **A_55_P2048202** | Ss18 | 0.084256074 | -0.255783645 | 0.424295793 | 12.97841327 | 0.620614496 | 0.559769746 | 0.780006362 | -4.700573348 |
| **A_55_P2105362** | Tmed5 | -0.146543913 | -0.738100874 | 0.445013049 | 8.758053173 | -0.620471102 | 0.559857485 | 0.780072399 | -4.700599816 |
| **A_51_P276039** | Txndc15 | 0.070828554 | -0.215154096 | 0.356811204 | 13.17314138 | 0.620324645 | 0.559947106 | 0.780117002 | -4.700626845 |
| **A_55_P1967776** | Slc4a4 | 0.061355071 | -0.186423601 | 0.309133744 | 8.396551336 | 0.620207324 | 0.560018906 | 0.780117002 | -4.700648492 |
| **A_66_P122559** | Myct1 | 0.187486299 | -0.569729273 | 0.94470187 | 8.636730406 | 0.620155136 | 0.560050846 | 0.780117002 | -4.700658121 |
| **A_55_P2025368** | Nkain1 | -0.224606882 | -1.131756919 | 0.682543154 | 8.861448648 | -0.62014648 | 0.560056143 | 0.780117002 | -4.700659718 |
| **A_55_P2096599** | Tsc22d4 | 0.061056521 | -0.185563473 | 0.307676515 | 8.336951384 | 0.620089132 | 0.560091243 | 0.780117002 | -4.700670298 |
| **A_55_P2044104** | Fgfr1op | 0.129206937 | -0.392837514 | 0.651251389 | 10.97813825 | 0.619910073 | 0.560200845 | 0.780133506 | -4.700703326 |
| **A_55_P2373121** | Dcaf17 | 0.073469164 | -0.223390391 | 0.37032872 | 8.890712738 | 0.619875415 | 0.56022206 | 0.780133506 | -4.700709718 |
| **A_66_P116326** | NA | -0.107315949 | -0.540939228 | 0.32630733 | 9.250030535 | -0.619872012 | 0.560224144 | 0.780133506 | -4.700710346 |
| **A_66_P123787** | Lars2 | 0.10125709 | -0.308074581 | 0.51058876 | 10.27908885 | 0.619584348 | 0.560400255 | 0.780140535 | -4.700763389 |
| **A_51_P153683** | Lrrc26 | 0.31351916 | -0.954010874 | 1.581049195 | 8.942958962 | 0.619521202 | 0.560438919 | 0.780140535 | -4.70077503 |
| **A_55_P2127692** | Timm50 | -0.093338606 | -0.470706052 | 0.284028841 | 8.77703928 | -0.619508432 | 0.560446738 | 0.780140535 | -4.700777384 |
| **A_55_P1953361** | Luc7l2 | -0.165167496 | -0.833020095 | 0.502685103 | 15.30554826 | -0.61943296 | 0.560492951 | 0.780140535 | -4.700791296 |
| **A_55_P2143802** | NA | 0.062166972 | -0.189212019 | 0.313545963 | 8.101477694 | 0.619414092 | 0.560504505 | 0.780140535 | -4.700794774 |
| **A_55_P2125229** | Arhgef2 | 0.130076329 | -0.395911308 | 0.656063965 | 12.66396709 | 0.619402681 | 0.560511492 | 0.780140535 | -4.700796877 |
| **A_55_P2061319** | Mthfs | 0.137388436 | -0.41816728 | 0.692944152 | 8.802585874 | 0.619402429 | 0.560511646 | 0.780140535 | -4.700796924 |
| **A_55_P1996963** | Mtcl1 | -0.086248219 | -0.435246426 | 0.262749988 | 10.13653315 | -0.618980953 | 0.560769774 | 0.780443623 | -4.700874589 |
| **A_52_P11174** | Hip1 | 0.241246699 | -0.73514807 | 1.217641468 | 10.01098525 | 0.618851262 | 0.560849217 | 0.780498003 | -4.700898479 |
| **A_66_P114333** | Tlr12 | 0.059801293 | -0.182325882 | 0.301928468 | 9.239371737 | 0.618610637 | 0.560996631 | 0.78064696 | -4.700942794 |
| **A_52_P338110** | Chmp1b | -0.071250007 | -0.359807736 | 0.217307721 | 9.340185736 | -0.618447087 | 0.56109684 | 0.780730213 | -4.700972906 |
| **A_51_P278887** | Hint2 | 0.169736258 | -0.517821665 | 0.857294182 | 12.25665186 | 0.618323394 | 0.561172636 | 0.780779487 | -4.700995676 |
| **A_51_P235548** | Mtmr9 | -0.069787596 | -0.352655309 | 0.213080117 | 8.532775798 | -0.617938426 | 0.561408577 | 0.781018127 | -4.70106652 |
| **A_51_P123879** | Tecrl | 0.164806508 | -0.503227037 | 0.832840052 | 8.283430808 | 0.617911719 | 0.561424947 | 0.781018127 | -4.699411782 |
| **A_55_P2004711** | Cep95 | -0.083828493 | -0.423664398 | 0.256007411 | 9.989994974 | -0.617835338 | 0.561471768 | 0.781027064 | -4.701085485 |
| **A_55_P2180764** | Gls | -0.08516756 | -0.430481576 | 0.260146457 | 8.873312437 | -0.617746568 | 0.561526186 | 0.781046567 | -4.701101814 |
| **A_55_P2156101** | Skp1a | 0.156116358 | -0.476984268 | 0.789216984 | 13.98719597 | 0.617626631 | 0.561599716 | 0.781092648 | -4.701123873 |
| **A_52_P654108** | Dync1li2 | -0.056594248 | -0.286289542 | 0.173101047 | 8.559774546 | -0.617121302 | 0.561909583 | 0.781467406 | -4.701216779 |
| **A_55_P1956918** | Adamts5 | -0.128341946 | -0.64933234 | 0.392648449 | 8.012795168 | -0.617005802 | 0.561980422 | 0.78150215 | -4.701238005 |
| **A_51_P427516** | Thsd1 | 0.382149524 | -1.169395314 | 1.933694362 | 10.30015476 | 0.616906488 | 0.562041339 | 0.78150215 | -4.701256255 |
| **A_55_P1956978** | Tln1 | -0.105076409 | -0.531707853 | 0.321555035 | 8.186834328 | -0.616882866 | 0.562055829 | 0.78150215 | -4.701260595 |
| **A_55_P1982818** | Odc1 | -0.175502972 | -0.888383907 | 0.537377963 | 8.962432825 | -0.616620304 | 0.5622169 | 0.781669894 | -4.701308829 |
| **A_55_P2076866** | Megf6 | 0.173618105 | -0.531718096 | 0.878954307 | 10.51104582 | 0.616522858 | 0.562276686 | 0.781696804 | -4.701326727 |
| **A_55_P1964628** | NA | -0.06834325 | -0.346079666 | 0.209393167 | 14.39744357 | -0.616329764 | 0.562395168 | 0.781805305 | -4.701362185 |
| **A_51_P227770** | Phykpl | -0.068515474 | -0.347130124 | 0.210099177 | 9.066359934 | -0.615935253 | 0.562637287 | 0.782044026 | -4.701434603 |
| **A_55_P2153021** | Ttn | 0.487620892 | -1.495499933 | 2.470741718 | 9.724414748 | 0.615862095 | 0.562682192 | 0.782044026 | -4.701448029 |
| **A_55_P2038247** | Tyro3 | 0.089986894 | -0.275989506 | 0.455963295 | 11.95202793 | 0.615852242 | 0.562688241 | 0.782044026 | -4.701449837 |
| **A_51_P236469** | Metap1d | -0.057477159 | -0.291276583 | 0.176322264 | 8.218871671 | -0.61574686 | 0.56275293 | 0.782062945 | -4.701469173 |
| **A_51_P216550** | Trp63 | 0.245051854 | -0.751822444 | 1.241926152 | 8.323734424 | 0.615698278 | 0.562782754 | 0.782062945 | -4.701478086 |
| **A_55_P1990633** | Iigp1 | 0.132045458 | -0.405280118 | 0.669371034 | 8.339476875 | 0.615511682 | 0.562897312 | 0.782161458 | -4.701512316 |
| **A_55_P2107288** | Tgfbr2 | -0.099751196 | -0.505703453 | 0.30620106 | 8.528123329 | -0.615451025 | 0.562934555 | 0.782161458 | -4.701523441 |
| **A_66_P102607** | Btrc | 0.08102037 | -0.248771527 | 0.410812267 | 8.689056011 | 0.615325038 | 0.563011915 | 0.782162956 | -4.701546546 |
| **A_55_P2081488** | Pglyrp1 | 0.211730391 | -0.650199708 | 1.07366049 | 8.814259267 | 0.615263991 | 0.563049402 | 0.782162956 | -4.701557741 |

| | | | | | | | | | |
|---|---|---|---|---|---|---|---|---|---|
| **A_51_P115326** | Surf4 | -0.182188357 | -0.923917905 | 0.559541192 | 13.68266168 | -0.615212829 | 0.56308082 | 0.782162956 | -4.701567122 |
| **A_55_P2024640** | Eral1 | 0.058659038 | -0.180214199 | 0.297532274 | 10.05590298 | 0.615060425 | 0.563174417 | 0.782162956 | -4.701595062 |
| **A_66_P102049** | Zfp1 | 0.065905248 | -0.202476668 | 0.334287164 | 11.01965125 | 0.615059446 | 0.563175018 | 0.782162956 | -4.701595242 |
| **A_55_P1964348** | Gdpd2 | -0.135557205 | -0.687583427 | 0.416469017 | 9.093520543 | -0.615053996 | 0.563178365 | 0.782162956 | -4.701596241 |
| **A_55_P2102065** | Gm10639 | -0.071465582 | -0.362554964 | 0.2196238 | 8.0098362 | -0.614923252 | 0.563258668 | 0.782218294 | -4.701620206 |
| **A_55_P2188464** | Uhrf1bp1l | -0.076580535 | -0.388569348 | 0.235408278 | 11.27124588 | -0.614794139 | 0.563337976 | 0.782272242 | -4.701643869 |
| **A_55_P2051962** | Sec61b | 0.13457251 | -0.413858788 | 0.683003807 | 14.92976447 | 0.614588552 | 0.563464272 | 0.782350585 | -4.701681539 |
| **A_55_P2039110** | NA | -0.271417504 | -1.377573666 | 0.834738658 | 14.30964746 | -0.614570564 | 0.563475323 | 0.782350585 | -4.701684834 |
| **A_55_P2419711** | 4932412D | 0.143898248 | -0.442673843 | 0.730470339 | 8.478118965 | 0.614447042 | 0.563551215 | 0.782363983 | -4.701707462 |
| **A_55_P2030204** | Supt6 | -0.169222576 | -0.859051005 | 0.520605853 | 12.74204723 | -0.614423135 | 0.563565904 | 0.782363983 | -4.701711841 |
| **A_55_P2088014** | Cdc42ep4 | 0.137434467 | -0.422907566 | 0.697776501 | 12.84756608 | 0.614317387 | 0.563630881 | 0.782398008 | -4.70173121 |
| **A_52_P299363** | NA | 0.059952427 | -0.184555218 | 0.304460073 | 8.880223502 | 0.614136165 | 0.563742245 | 0.782496415 | -4.701764396 |
| **A_55_P2125208** | Arid5a | 0.099662127 | -0.30700653 | 0.506330783 | 9.963514514 | 0.613818248 | 0.563937642 | 0.782561975 | -4.701822595 |
| **A_55_P2467415** | B3galt6 | 0.094493558 | -0.291132447 | 0.480119563 | 8.68913593 | 0.613742551 | 0.563984173 | 0.782561975 | -4.701836449 |
| **A_52_P353478** | Scn7a | 0.250561782 | -0.7720146 | 1.273138165 | 8.63441122 | 0.613718786 | 0.563998783 | 0.782561975 | -4.701840799 |
| **A_51_P212754** | Tgfbi | 0.500729637 | -1.542940593 | 2.544399868 | 13.84176644 | 0.613681207 | 0.564021883 | 0.782561975 | -4.701847676 |
| **A_66_P117786** | Crxos | 0.055867074 | -0.172151857 | 0.283886004 | 7.94581997 | 0.613670667 | 0.564028363 | 0.782561975 | -4.701849604 |
| **A_55_P2043434** | Atmin | -0.06102719 | -0.310109524 | 0.188055143 | 11.43851966 | -0.613664209 | 0.564032333 | 0.782561975 | -4.701850786 |
| **A_55_P2059432** | Crabp2 | -0.83541807 | -4.24637527 | 2.575539129 | 9.842636767 | -0.613448262 | 0.564165096 | 0.782690009 | -4.701890298 |
| **A_55_P2111039** | NA | -0.07980429 | -0.405682177 | 0.246073597 | 10.77717216 | -0.613368813 | 0.564213946 | 0.782701616 | -4.701904832 |
| **A_55_P2129286** | Zc3hc1 | 0.10353296 | -0.319331017 | 0.526396938 | 9.523650387 | 0.613236911 | 0.564295052 | 0.782711556 | -4.701928958 |
| **A_51_P198076** | Mien1 | -0.093535539 | -0.475573751 | 0.288502673 | 11.99678971 | -0.613225486 | 0.564302078 | 0.782711556 | -4.701931047 |
| **A_55_P2150283** | E030030I( | -0.157492331 | -0.8010186 | 0.486033939 | 11.72411508 | -0.612976006 | 0.564455504 | 0.782835598 | -4.701976668 |
| **A_51_P181002** | Vps50 | -0.082600685 | -0.420128902 | 0.254927532 | 8.894891492 | -0.612948393 | 0.564472488 | 0.782835598 | -4.701981716 |
| **A_51_P274124** | Mansc1 | 0.135424346 | -0.41817249 | 0.689021182 | 9.371943852 | 0.612707918 | 0.564620403 | 0.78298457 | -4.702025675 |
| **A_55_P2115260** | NA | -0.070625634 | -0.359380246 | 0.218128978 | 8.484607766 | -0.612609569 | 0.564680905 | 0.783012308 | -4.702043649 |
| **A_55_P1975743** | Dap3 | 0.067008223 | -0.207038854 | 0.341055301 | 13.40546642 | 0.612425513 | 0.564794141 | 0.783066249 | -4.702077281 |
| **A_55_P2027906** | Nsa2 | 0.182738484 | -0.564681092 | 0.930158059 | 13.17628374 | 0.612372802 | 0.564826573 | 0.783066249 | -4.702086911 |
| **A_55_P2054280** | Fam178b | -0.05354015 | -0.272533551 | 0.16545325 | 9.039112876 | -0.612348849 | 0.564841311 | 0.783066249 | -4.702091288 |
| **A_55_P2166446** | NA | -0.073354284 | -0.373486042 | 0.226777473 | 8.404971799 | -0.612158493 | 0.564958445 | 0.783144802 | -4.702126059 |
| **A_55_P2042273** | Pbrm1 | -0.082013356 | -0.417680814 | 0.253654102 | 8.814412451 | -0.61196374 | 0.565078299 | 0.783144802 | -4.702161625 |
| **A_52_P351669** | Gcc2 | 0.118117722 | -0.365320159 | 0.601555604 | 9.564361014 | 0.61196239 | 0.56507913 | 0.783144802 | -4.702161872 |
| **A_55_P2079560** | Lilra6 | -0.076356973 | -0.388874578 | 0.236160631 | 8.194499847 | -0.611962153 | 0.565079276 | 0.783144802 | -4.702161915 |
| **A_55_P2016967** | Gm8213 | 0.110844175 | -0.342849928 | 0.564538278 | 15.43290086 | 0.611927662 | 0.565100504 | 0.783144802 | -4.702168213 |
| **A_52_P429265** | Ppp6r2 | -0.054422587 | -0.277305578 | 0.168460404 | 8.100014421 | -0.611579046 | 0.565315093 | 0.783325675 | -4.702231852 |
| **A_55_P1989986** | Fam149b | 0.076637267 | -0.237231212 | 0.390505746 | 9.276529529 | 0.61156504 | 0.565323715 | 0.783325675 | -4.702234409 |
| **A_55_P2091751** | NA | 0.2074563 | -0.642247576 | 1.057160175 | 8.647124738 | 0.611518181 | 0.565352564 | 0.783325675 | -4.70224296 |
| **A_55_P2036894** | Med29 | -0.159085655 | -0.811033199 | 0.492861889 | 10.77380769 | -0.611179406 | 0.565561153 | 0.783487072 | -4.702304772 |
| **A_55_P2113459** | Washc1 | 0.096203916 | -0.298096941 | 0.490504773 | 11.6616957 | 0.611104349 | 0.565607373 | 0.783487072 | -4.702318463 |
| **A_55_P2053398** | NA | -0.060236235 | -0.307122283 | 0.186649812 | 17.10263994 | -0.611099067 | 0.565610626 | 0.783487072 | -4.702319426 |
| **A_55_P2040295** | Ptp4a1 | -0.113885079 | -0.580683344 | 0.352913185 | 9.761539497 | -0.611065748 | 0.565631144 | 0.783487072 | -4.702325504 |
| **A_55_P1971329** | Lage3 | 0.289540808 | -0.897564833 | 1.47664645 | 10.39051454 | 0.610900892 | 0.565732674 | 0.783530648 | -4.702355568 |
| **A_55_P1965101** | Xlr3b | -0.614757597 | -3.13531253 | 1.905797336 | 9.182701521 | -0.610883061 | 0.565743656 | 0.783530648 | -4.70235882 |
| **A_55_P2322039** | NA | 0.059762069 | -0.185296444 | 0.304820582 | 8.207424071 | 0.610810057 | 0.565788621 | 0.783536796 | -4.702372131 |
| **A_52_P137378** | Prkaa2 | -0.093762775 | -0.478309158 | 0.290783607 | 8.490454764 | -0.610705835 | 0.565852818 | 0.783569573 | -4.702391133 |
| **A_51_P385351** | Slc44a3 | 0.178724497 | -0.554428214 | 0.911877208 | 8.645107982 | 0.610576358 | 0.565932577 | 0.783572349 | -4.702414735 |
| **A_55_P1971828** | Csnk2a1 | -0.101416586 | -0.517452703 | 0.31461953 | 8.578031288 | -0.610559931 | 0.565942697 | 0.783572349 | -4.70241773 |
| **A_55_P2052472** | NA | 0.161812754 | -0.502042994 | 0.825668502 | 14.45274556 | 0.610505211 | 0.565976407 | 0.783572349 | -4.702427703 |

| | | | | | | | | | |
|---|---|---|---|---|---|---|---|---|---|
| **A_51_P434240** | Mrpl11 | -0.05853377 | -0.298737122 | 0.181669582 | 13.23558652 | -0.610348356 | 0.566073045 | 0.783650025 | -4.702456289 |
| **A_55_P1965574** | A2ml1 | 0.05863142 | -0.182146236 | 0.299409075 | 7.826856893 | 0.609908344 | 0.566344191 | 0.783969256 | -4.702536446 |
| **A_55_P2166773** | Pitpna | -0.081308325 | -0.415313027 | 0.252696377 | 12.81406165 | -0.609723282 | 0.566458255 | 0.784029472 | -4.702570146 |
| **A_51_P434670** | Qk | -0.153638828 | -0.784807964 | 0.477530309 | 11.94156166 | -0.609685092 | 0.566481795 | 0.784029472 | -4.702577099 |
| **A_55_P1993168** | Ppargc1b | -0.068540884 | -0.350150623 | 0.213068854 | 7.872413128 | -0.609610416 | 0.566527827 | 0.784029472 | -4.702590695 |
| **A_55_P2059347** | Ormdl3 | -0.2439418 | -1.246297295 | 0.758413696 | 12.40814837 | -0.60955766 | 0.566560349 | 0.784029472 | -4.702600299 |
| **A_55_P1988970** | Mkks | 0.233275665 | -0.725444126 | 1.191995455 | 10.44343318 | 0.609435977 | 0.566635364 | 0.784029472 | -4.702622448 |
| **A_55_P2072611** | 1-Mar | -0.061645096 | -0.315027345 | 0.191737153 | 7.745263146 | -0.609358242 | 0.566683291 | 0.784029472 | -4.702636596 |
| **A_51_P155501** | Alg3 | -0.081129602 | -0.414633764 | 0.252374559 | 8.135598553 | -0.609296152 | 0.566721572 | 0.784029472 | -4.702647896 |
| **A_55_P2017759** | Evi2a | 0.100827522 | -0.313699085 | 0.515354128 | 8.543400824 | 0.609224033 | 0.56676604 | 0.784029472 | -4.702661019 |
| **A_52_P358093** | Fbxo10 | -0.114977241 | -0.587704807 | 0.357750325 | 9.995658152 | -0.609187946 | 0.566788292 | 0.784029472 | -4.702667585 |
| **A_55_P2161545** | Ddx21 | 0.152842892 | -0.475577078 | 0.781262862 | 9.337676797 | 0.609179968 | 0.566793211 | 0.784029472 | -4.702669037 |
| **A_52_P634822** | Nap1l4 | 0.092083199 | -0.286615558 | 0.470781955 | 8.526352245 | 0.609027443 | 0.566887265 | 0.784103476 | -4.702696786 |
| **A_51_P136337** | Galm | 0.059321961 | -0.18470505 | 0.303348972 | 7.925543747 | 0.608874726 | 0.566981448 | 0.784177646 | -4.702724565 |
| **A_51_P286946** | Lhpp | 0.183746804 | -0.572403881 | 0.93989749 | 10.35297376 | 0.608641821 | 0.567125103 | 0.784308309 | -4.702766919 |
| **A_52_P235102** | Zfp975 | 0.268362382 | -0.836090402 | 1.372815166 | 9.770237805 | 0.608590024 | 0.567157054 | 0.784308309 | -4.702776337 |
| **A_51_P339503** | Cct4 | 0.140398103 | -0.437628753 | 0.718424959 | 11.78385644 | 0.608364088 | 0.567296436 | 0.78444495 | -4.702817409 |
| **A_51_P148355** | Zfp710 | 0.117528003 | -0.366637727 | 0.601693734 | 11.99019156 | 0.60799171 | 0.567526206 | 0.784670653 | -4.702885077 |
| **A_55_P1987291** | Plppr5 | 0.095514709 | -0.298056627 | 0.489086044 | 8.019474728 | 0.607851007 | 0.56761304 | 0.784670653 | -4.702910637 |
| **A_51_P255304** | Cav3 | 0.397585894 | -1.241105205 | 2.036276992 | 9.322428185 | 0.607692978 | 0.567710576 | 0.784670653 | -4.702939338 |
| **A_55_P2021398** | NA | -0.149419761 | -0.765287914 | 0.466448392 | 11.98305043 | -0.607673973 | 0.567722307 | 0.784670653 | -4.702942789 |
| **A_55_P2164111** | Nploc4 | -0.156276717 | -0.800459843 | 0.487906409 | 10.56307675 | -0.607624506 | 0.56775284 | 0.784670653 | -4.702951772 |
| **A_52_P387954** | Dctn5 | 0.098848134 | -0.308623382 | 0.506319649 | 11.63428903 | 0.607605317 | 0.567764685 | 0.784670653 | -4.702955257 |
| **A_52_P559808** | Arhgef12 | 0.106568252 | -0.332756605 | 0.54589311 | 11.87879854 | 0.607564532 | 0.567789861 | 0.784670653 | -4.702962662 |
| **A_51_P297586** | Birc6 | 0.054810016 | -0.171152839 | 0.280772872 | 9.898198609 | 0.607537688 | 0.567806432 | 0.784670653 | -4.702967537 |
| **A_52_P488437** | Plek2 | 0.059007016 | -0.184271309 | 0.302285342 | 7.914384514 | 0.607506018 | 0.567825982 | 0.784670653 | -4.702973287 |
| **A_55_P2022885** | Zfp866 | 0.07081493 | -0.221176678 | 0.362806538 | 8.715746876 | 0.607441986 | 0.56786551 | 0.784670653 | -4.702984912 |
| **A_66_P100932** | Tmem256 | 0.345412723 | -1.079381126 | 1.770206572 | 13.4814197 | 0.607206825 | 0.568010696 | 0.784745427 | -4.703027598 |
| **A_55_P2011525** | Rnf214 | -0.067344973 | -0.34514046 | 0.210450513 | 8.431287476 | -0.607198017 | 0.568016135 | 0.784745427 | -4.703029196 |
| **A_55_P2065449** | Kdm5b | -0.151375607 | -0.775836371 | 0.473085157 | 13.01494267 | -0.607157113 | 0.568041391 | 0.784745427 | -4.70303662 |
| **A_55_P2028300** | Ppm1d | -0.055232579 | -0.283120519 | 0.17265536 | 9.744702688 | -0.607049813 | 0.568107647 | 0.784780884 | -4.703056091 |
| **A_55_P2149942** | Ninj2 | -0.088833791 | -0.455494304 | 0.277826722 | 8.093028534 | -0.606826307 | 0.568245673 | 0.784915471 | -4.703096641 |
| **A_66_P101423** | Ogfod3 | 0.10303583 | -0.322548753 | 0.528620412 | 10.72574507 | 0.606390979 | 0.56851457 | 0.785222281 | -4.703175588 |
| **A_55_P1960972** | Lrrc59 | -0.128262008 | -0.658094775 | 0.401570759 | 8.157896775 | -0.606330713 | 0.568551801 | 0.785222281 | -4.703186514 |
| **A_55_P1997766** | Acp5 | -0.577457659 | -2.96334405 | 1.808428732 | 11.09977283 | -0.606206699 | 0.56862842 | 0.785222281 | -4.703208994 |
| **A_52_P512749** | Ears2 | -0.095272347 | -0.488911779 | 0.298367084 | 8.512347072 | -0.606203746 | 0.568630245 | 0.785222281 | -4.703209529 |
| **A_55_P2013756** | NA | 0.115102831 | -0.360826633 | 0.591032295 | 8.268307993 | 0.605750468 | 0.568910348 | 0.785507206 | -4.703291665 |
| **A_55_P2156062** | NA | -0.121007078 | -0.621359543 | 0.379345388 | 8.435603366 | -0.605738355 | 0.568917834 | 0.785507206 | -4.703293859 |
| **A_52_P264229** | Ing5 | -0.143792995 | -0.738457036 | 0.450871046 | 9.363424233 | -0.605642576 | 0.568977032 | 0.785532844 | -4.703311208 |
| **A_51_P366672** | Slc36a2 | 0.286255712 | -0.898067423 | 1.470578847 | 9.053727159 | 0.605388677 | 0.569133977 | 0.785693418 | -4.703357188 |
| **A_55_P2042753** | Cltrn | -0.055235218 | -0.283795616 | 0.17332518 | 7.709225676 | -0.605292697 | 0.569193313 | 0.785719229 | -4.703374565 |
| **A_55_P1969356** | Ccm2l | 0.181488484 | -0.569665579 | 0.932642547 | 9.10229477 | 0.605160257 | 0.569275196 | 0.785734508 | -4.70339854 |
| **A_52_P525317** | Gja5 | 0.301482544 | -0.946344131 | 1.549309219 | 9.779123115 | 0.60514333 | 0.569285662 | 0.785734508 | -4.703401604 |
| **A_55_P2029630** | Mpdu1 | -0.076074128 | -0.391042771 | 0.238894514 | 9.232609956 | -0.60495074 | 0.569404747 | 0.785801889 | -4.70343646 |
| **A_55_P1955015** | Myo1g | 0.275337744 | -0.864672829 | 1.415348317 | 10.36971332 | 0.604932917 | 0.569415769 | 0.785801889 | -4.703439686 |
| **A_55_P2026223** | St6galnac | 0.070772553 | -0.22231611 | 0.363861215 | 8.209706911 | 0.604806134 | 0.569494172 | 0.785846098 | -4.703462626 |
| **A_52_P204331** | D630039A | 0.310166334 | -0.974437952 | 1.594770621 | 8.990328582 | 0.604749665 | 0.569529095 | 0.785846098 | -4.703472843 |
| **A_66_P119489** | Spef1l | 0.223823681 | -0.703549805 | 1.151197168 | 9.043739424 | 0.604507516 | 0.569678867 | 0.785965198 | -4.703516644 |

| | | | | | | | | | |
|---|---|---|---|---|---|---|---|---|---|
| **A_55_P1967025** | NA | -0.089156582 | -0.458589752 | 0.280276588 | 8.034895643 | -0.604460419 | 0.569707999 | 0.785965198 | -4.703525162 |
| **A_51_P253547** | Ctsl | -0.56006303 | -2.881071299 | 1.76094524 | 13.94562065 | -0.604380666 | 0.569757334 | 0.785965198 | -4.703539584 |
| **A_51_P510891** | Afp | -1.105057283 | -5.684879316 | 3.47476475 | 12.84908109 | -0.604347228 | 0.569778019 | 0.785965198 | -4.701855209 |
| **A_55_P2129920** | Chd7 | -0.27083288 | -1.393688049 | 0.852022288 | 11.16804628 | -0.604126647 | 0.569914486 | 0.786023722 | -4.703585511 |
| **A_55_P2090390** | NA | -0.139731963 | -0.719147212 | 0.439683287 | 8.844804372 | -0.604026766 | 0.569976286 | 0.786023722 | -4.703603565 |
| **A_52_P160970** | Akap10 | -0.079044055 | -0.406820613 | 0.248732503 | 9.307931681 | -0.604006574 | 0.56998878 | 0.786023722 | -4.703607214 |
| **A_55_P1978424** | Bcl2a1d | 0.426266888 | -1.341489592 | 2.194023369 | 10.49645517 | 0.603961849 | 0.570016455 | 0.786023722 | -4.703615298 |
| **A_51_P239203** | Mapk13 | 0.122370446 | -0.385117609 | 0.6298585 | 10.64334713 | 0.603950108 | 0.57002372 | 0.786023722 | -4.70361742 |
| **A_55_P2109342** | Rnps1 | 0.076490714 | -0.240877021 | 0.393858448 | 9.90319033 | 0.603665404 | 0.570199909 | 0.786169428 | -4.703668864 |
| **A_55_P1996088** | NA | -0.109062154 | -0.561680847 | 0.34355654 | 12.76089926 | -0.603520364 | 0.57028968 | 0.786169428 | -4.703695065 |
| **A_55_P2053838** | Tnfaip3 | 0.204009127 | -0.642797938 | 1.050816192 | 9.43683447 | 0.603414124 | 0.570355441 | 0.786169428 | -4.703714253 |
| **A_52_P314129** | Pkia | 0.151711127 | -0.478022437 | 0.781444691 | 8.382320208 | 0.603407825 | 0.57035934 | 0.786169428 | -4.703715391 |
| **A_55_P1960281** | Med7 | -0.061071888 | -0.314584685 | 0.192440909 | 9.853532255 | -0.603381242 | 0.570375796 | 0.786169428 | -4.703720192 |
| **A_55_P2189201** | 5830426C | -0.090090007 | -0.464104152 | 0.283924138 | 8.262008706 | -0.603307798 | 0.570421261 | 0.786169428 | -4.703733455 |
| **A_55_P2131785** | Kiz | 0.19842377 | -0.625631893 | 1.022479433 | 11.18174131 | 0.603097452 | 0.570551485 | 0.786169428 | -4.703771433 |
| **A_51_P362089** | Lrrc3b | 0.076752906 | -0.24201496 | 0.395520773 | 8.269387599 | 0.603074047 | 0.570565977 | 0.786169428 | -4.703775658 |
| **A_55_P2059085** | Zfp457 | -0.068575485 | -0.367319646 | 0.230168676 | 9.571129465 | -0.611409477 | 0.570624884 | 0.786169428 | -4.681122209 |
| **A_55_P2115871** | Inpp5j | 0.144029319 | -0.454267058 | 0.742325697 | 8.268599071 | 0.602954972 | 0.570639706 | 0.786169428 | -4.703797152 |
| **A_55_P2088995** | Plscr1 | 0.195552191 | -0.616849669 | 1.007954051 | 9.643763605 | 0.602895612 | 0.570676462 | 0.786169428 | -4.703807866 |
| **A_55_P2085400** | NA | 0.081092151 | -0.255808992 | 0.417993294 | 8.23372111 | 0.602874174 | 0.570689737 | 0.786169428 | -4.703811735 |
| **A_55_P2074771** | Exosc7 | 0.32338992 | -1.020225881 | 1.667005721 | 12.00221917 | 0.602839529 | 0.570711191 | 0.786169428 | -4.703817988 |
| **A_52_P528707** | Mrpl57 | 0.139340617 | -0.439646726 | 0.718327959 | 13.92443572 | 0.602780236 | 0.570747908 | 0.786169428 | -4.703828687 |
| **A_51_P223498** | Slc39a10 | -0.062049224 | -0.319886882 | 0.195788434 | 8.572892564 | -0.60275435 | 0.57076394 | 0.786169428 | -4.703833359 |
| **A_66_P134428** | Sfrp1 | 0.078046013 | -0.246286985 | 0.40237901 | 8.018920467 | 0.602712127 | 0.570790088 | 0.786169428 | -4.703840978 |
| **A_51_P102503** | Metap1 | -0.12893555 | -0.664792235 | 0.406921135 | 10.84533689 | -0.60266278 | 0.570820649 | 0.786169428 | -4.703849881 |
| **A_55_P1986188** | Def8 | -0.208312057 | -1.074320999 | 0.657696885 | 11.61156775 | -0.602479642 | 0.570934077 | 0.786260401 | -4.70388292 |
| **A_55_P2065149** | Kdm2b | 0.1397181 | -0.441193751 | 0.72062995 | 11.43270092 | 0.602410841 | 0.570976694 | 0.786260401 | -4.70389533 |
| **A_55_P2095603** | Ccdc65 | 0.182268977 | -0.575630874 | 0.940168828 | 8.449450039 | 0.602353279 | 0.57101235 | 0.786260401 | -4.703905712 |
| **A_55_P2181888** | Tmem54 | -0.220479124 | -1.137353073 | 0.696394825 | 8.761744096 | -0.602293515 | 0.571049371 | 0.786260401 | -4.70391649 |
| **A_55_P2170094** | Lrrc27 | 0.091024539 | -0.287603857 | 0.469652935 | 8.940907975 | 0.602137471 | 0.571146042 | 0.786337505 | -4.703944628 |
| **A_52_P306537** | Emc1 | -0.051133571 | -0.263894814 | 0.161627672 | 8.498924423 | -0.601954889 | 0.571259166 | 0.786437248 | -4.703977544 |
| **A_52_P650028** | Dock10 | 0.212955656 | -0.673425731 | 1.099337042 | 8.705036908 | 0.601753877 | 0.571383725 | 0.786552719 | -4.704013774 |
| **A_55_P2023572** | NA | -0.056894261 | -0.293846953 | 0.180058431 | 7.87406566 | -0.601391326 | 0.571608424 | 0.786806014 | -4.704079094 |
| **A_51_P418809** | Airn | 0.06622802 | -0.209715744 | 0.342171784 | 8.433612637 | 0.601134335 | 0.571767733 | 0.78690832 | -4.704125377 |
| **A_51_P499816** | Pbld1 | -0.092441605 | -0.496408708 | 0.311525497 | 7.775382876 | -0.609514701 | 0.571773951 | 0.78690832 | -4.681416808 |
| **A_55_P1975095** | Timmdc1 | -0.084341564 | -0.435802734 | 0.267119607 | 9.638554672 | -0.601055592 | 0.571816552 | 0.78690832 | -4.704139556 |
| **A_55_P2020326** | Madcam1 | -0.127595312 | -0.659341296 | 0.404150673 | 8.030392194 | -0.601008818 | 0.571845552 | 0.78690832 | -4.704147977 |
| **A_55_P2053733** | NA | 0.060828638 | -0.192742189 | 0.314399464 | 15.89950499 | 0.600840434 | 0.571949956 | 0.786995172 | -4.704178289 |
| **A_51_P189814** | Cldn5 | 0.505012592 | -1.600414324 | 2.610439508 | 12.63994228 | 0.600775733 | 0.571990077 | 0.786995172 | -4.704189934 |
| **A_51_P117618** | Ethe1 | -0.106240091 | -0.549283882 | 0.336803699 | 9.302776184 | -0.600609403 | 0.572093224 | 0.787032028 | -4.704219867 |
| **A_55_P2007196** | Aldoa | -0.195602884 | -1.011363132 | 0.620157364 | 15.74917848 | -0.600569208 | 0.572118153 | 0.787032028 | -4.7042271 |
| **A_52_P603421** | Ccdc189 | 0.141029494 | -0.44721703 | 0.729276019 | 8.368552616 | 0.600483286 | 0.572171442 | 0.787032028 | -4.704242559 |
| **A_55_P2027225** | Fau | 0.101582422 | -0.322136253 | 0.525301096 | 17.59008054 | 0.600469983 | 0.572179693 | 0.787032028 | -4.704244953 |
| **A_55_P2032238** | Zbtb8os | 0.079985205 | -0.253875412 | 0.413845822 | 11.26154246 | 0.60006019 | 0.572433893 | 0.787325667 | -4.704318659 |
| **A_55_P2110081** | Cisd3 | 0.233179417 | -0.740338608 | 1.206697443 | 11.70884199 | 0.599924449 | 0.572518111 | 0.787385405 | -4.704343064 |
| **A_55_P1955237** | NA | 0.212231345 | -0.674180688 | 1.098643379 | 17.24296553 | 0.599686441 | 0.572665796 | 0.787385405 | -4.704385847 |
| **A_51_P208361** | Ak3 | 0.245404695 | -0.779647434 | 1.270456823 | 11.96270969 | 0.599635396 | 0.572697473 | 0.787385405 | -4.704395021 |
| **A_65_P16680** | B4galt5 | -0.221633514 | -1.147581336 | 0.704314309 | 11.46633476 | -0.599513908 | 0.572772868 | 0.787385405 | -4.704416852 |

| | | | | | | | | | |
|---|---|---|---|---|---|---|---|---|---|
| **A_55_P2106426** | NA | 0.117727906 | -0.374124816 | 0.609580628 | 14.39767108 | 0.59950759 | 0.572776789 | 0.787385405 | -4.704417987 |
| **A_55_P2049429** | Phtf2 | 0.072720443 | -0.231119416 | 0.376560301 | 9.727233641 | 0.599462628 | 0.572804694 | 0.787385405 | -4.704426066 |
| **A_52_P679105** | Prss23 | -0.120792049 | -0.625489894 | 0.383905797 | 9.887358475 | -0.599455903 | 0.572808868 | 0.787385405 | -4.704427274 |
| **A_52_P453374** | Lrrn4cl | 0.059355302 | -0.188647147 | 0.307357751 | 8.135670452 | 0.599451289 | 0.572811731 | 0.787385405 | -4.704428103 |
| **A_51_P243134** | Adcy6 | 0.20822572 | -0.661872578 | 1.078324018 | 9.18373799 | 0.599399531 | 0.572843856 | 0.787385405 | -4.704437402 |
| **A_51_P250551** | Dhx38 | -0.070571892 | -0.365524794 | 0.224381011 | 8.757229524 | -0.59927951 | 0.572918352 | 0.787431821 | -4.704458963 |
| **A_55_P2360427** | BC00556 | 0.08051808 | -0.256066533 | 0.417102694 | 8.568894922 | 0.599169227 | 0.57298681 | 0.787469931 | -4.704478772 |
| **A_55_P2030568** | Ifit1bl2 | -0.057407993 | -0.297437687 | 0.1826217 | 9.161508007 | -0.599042645 | 0.573065391 | 0.78747351 | -4.704501505 |
| **A_52_P513123** | Dna2 | -0.110927611 | -0.574751595 | 0.352896374 | 8.343977963 | -0.599013763 | 0.573083322 | 0.78747351 | -4.704506691 |
| **A_55_P2005833** | Adamts8 | 0.339379311 | -1.079835392 | 1.758594014 | 9.647418917 | 0.598945915 | 0.573125445 | 0.78747351 | -4.704518874 |
| **A_51_P335081** | Gnb1 | -0.080340496 | -0.416332274 | 0.255651282 | 11.91394243 | -0.598902606 | 0.573152334 | 0.78747351 | -4.704526649 |
| **A_55_P1983848** | Tc2n | 0.166464547 | -0.529863935 | 0.862793028 | 8.539169196 | 0.598767112 | 0.573236463 | 0.787533133 | -4.704550974 |
| **A_66_P129419** | Cox7a2l | 0.073818033 | -0.235065259 | 0.382701324 | 15.37828078 | 0.598574755 | 0.573355912 | 0.787641268 | -4.704585499 |
| **A_55_P2154107** | Gcm1 | -0.14581833 | -0.756289174 | 0.464652514 | 10.75305513 | -0.598270431 | 0.573544919 | 0.78777454 | -4.704640103 |
| **A_52_P661** | Plxna4 | -0.125794312 | -0.652442813 | 0.400854188 | 9.544332245 | -0.598260733 | 0.573550943 | 0.78777454 | -4.704641843 |
| **A_55_P2069765** | Kiss1 | 0.234946295 | -0.7487403 | 1.218632889 | 9.645974832 | 0.598221741 | 0.573575163 | 0.78777454 | -4.704648837 |
| **A_55_P2037101** | Exoc2 | 0.159111774 | -0.50735786 | 0.825581408 | 11.1082832 | 0.597960224 | 0.573737621 | 0.787897173 | -4.70469574 |
| **A_55_P2143030** | Timd2 | -0.061445641 | -0.318949166 | 0.196057883 | 7.694723263 | -0.597665587 | 0.573920687 | 0.787897173 | -4.704748563 |
| **A_52_P119117** | Tmem258 | 0.115590194 | -0.368888115 | 0.600068504 | 13.23377804 | 0.597581304 | 0.57397306 | 0.787897173 | -4.70476367 |
| **A_55_P2128869** | Ccdc80 | 0.143039904 | -0.456550883 | 0.74263069 | 8.555322956 | 0.597520216 | 0.574011023 | 0.787897173 | -4.704774618 |
| **A_55_P2139112** | Cdc26 | 0.151179353 | -0.482909473 | 0.78526818 | 11.1155454 | 0.597162777 | 0.574233179 | 0.787897173 | -4.704838661 |
| **A_51_P444335** | Acp6 | 0.056433812 | -0.18030112 | 0.293168743 | 10.7642184 | 0.597072935 | 0.574289026 | 0.787897173 | -4.704854754 |
| **A_51_P370050** | Nagpa | -0.119162393 | -0.619108059 | 0.380783273 | 8.618114322 | -0.596989583 | 0.574340841 | 0.787897173 | -4.704869682 |
| **A_55_P2096823** | NA | -0.074599449 | -0.387590221 | 0.238391323 | 8.51372345 | -0.596972645 | 0.574351371 | 0.787897173 | -4.704872715 |
| **A_55_P2198364** | Tsen54 | -0.050361617 | -0.2616834 | 0.160960166 | 8.470771513 | -0.596905724 | 0.574392975 | 0.787897173 | -4.704884699 |
| **A_55_P1995902** | Zfp992 | 0.076775286 | -0.261138108 | 0.414688681 | 8.873505582 | 0.605171884 | 0.574413262 | 0.787897173 | -4.682089512 |
| **A_51_P384754** | Ldb1 | 0.081938948 | -0.261924202 | 0.425802098 | 9.970791088 | 0.59683611 | 0.574436255 | 0.787897173 | -4.704897164 |
| **A_55_P2146483** | Zmynd8 | -0.104042575 | -0.540763228 | 0.332678079 | 9.901027998 | -0.596702291 | 0.574519458 | 0.787897173 | -4.704921122 |
| **A_55_P1982239** | Ctrb1 | 0.088017842 | -0.281491491 | 0.457527174 | 8.032563838 | 0.596617034 | 0.574572471 | 0.787897173 | -4.704936384 |
| **A_55_P2159740** | Mfsd2b | -0.142892337 | -0.742782814 | 0.456998141 | 15.740476 | -0.596605586 | 0.574579589 | 0.787897173 | -4.704938433 |
| **A_55_P2091846** | Tut4 | 0.067153468 | -0.214771169 | 0.349078105 | 9.833664852 | 0.596603449 | 0.574580919 | 0.787897173 | -4.704938816 |
| **A_65_P06872** | Fam210a | 0.055652528 | -0.178010477 | 0.289315532 | 8.208003002 | 0.596547849 | 0.574615493 | 0.787897173 | -4.704948768 |
| **A_55_P1973854** | Sema6b | 0.07092226 | -0.226865389 | 0.368709909 | 8.288537524 | 0.596521677 | 0.574631767 | 0.787897173 | -4.704953452 |
| **A_52_P177021** | Pts | 0.224006781 | -0.716618508 | 1.164632071 | 9.971823158 | 0.596478591 | 0.574658561 | 0.787897173 | -4.704961163 |
| **A_51_P358872** | Disc1 | -0.049936527 | -0.259626521 | 0.159753466 | 7.805621039 | -0.596473257 | 0.574661878 | 0.787897173 | -4.704962118 |
| **A_52_P799815** | Tmem171 | 0.403139467 | -1.289789011 | 2.096067945 | 9.517946876 | 0.596440399 | 0.574682312 | 0.787897173 | -4.704967998 |
| **A_55_P2041894** | NA | 0.147493077 | -0.471978965 | 0.766965119 | 17.00039733 | 0.596348693 | 0.574739346 | 0.787897173 | -4.704984409 |
| **A_55_P2096367** | NA | -0.071766408 | -0.373209328 | 0.229676512 | 13.79759455 | -0.596302276 | 0.574768214 | 0.787897173 | -4.704992714 |
| **A_55_P2145617** | Krt80 | 0.055208017 | -0.176698405 | 0.287114439 | 8.237917231 | 0.596265552 | 0.574791055 | 0.787897173 | -4.704999285 |
| **A_55_P1999078** | NA | 0.07106069 | -0.227451337 | 0.369572717 | 9.350989224 | 0.596235643 | 0.574809657 | 0.787897173 | -4.705004636 |
| **A_55_P1982593** | Rad51d | 0.059898445 | -0.191762622 | 0.311559511 | 8.316236113 | 0.596142189 | 0.574867785 | 0.787897173 | -4.705021355 |
| **A_55_P2110200** | Ubox5 | -0.14929508 | -0.776595194 | 0.478005033 | 8.65664356 | -0.59610186 | 0.574892871 | 0.787897173 | -4.705028569 |
| **A_51_P438083** | Slc6a13 | -0.149156992 | -0.775916819 | 0.477602835 | 8.044938591 | -0.596063886 | 0.574916492 | 0.787897173 | -4.705035362 |
| **A_52_P140789** | Hspa13 | -0.150895866 | -0.784966664 | 0.483174931 | 10.99259328 | -0.596059943 | 0.574918945 | 0.787897173 | -4.705036067 |
| **A_65_P01418** | Wnk2 | -0.147563814 | -0.7676675 | 0.472539873 | 9.342444905 | -0.596026958 | 0.574939464 | 0.787897173 | -4.705041967 |
| **A_51_P481679** | Angptl3 | -0.089070793 | -0.463372332 | 0.285230745 | 7.853148366 | -0.596024421 | 0.574941042 | 0.787897173 | -4.70504242 |
| **A_55_P2046488** | Nbeal2 | -0.066155692 | -0.344189833 | 0.21187845 | 8.381640914 | -0.595963183 | 0.574979136 | 0.787897173 | -4.705053373 |
| **A_55_P2181288** | Ppp1r2 | 0.177544696 | -0.56868253 | 0.923771923 | 10.15555591 | 0.595918624 | 0.575006856 | 0.787897173 | -4.705061342 |

| | | | | | | | | | |
|---|---|---|---|---|---|---|---|---|---|
| **A_55_P2138906** | Lrfn1 | 0.059612703 | -0.190967921 | 0.310193328 | 8.545447685 | 0.595856493 | 0.575045509 | 0.787897173 | -4.705072453 |
| **A_51_P357085** | Dctn6 | 0.143946793 | -0.461137589 | 0.749031176 | 12.23717322 | 0.595849249 | 0.575050016 | 0.787897173 | -4.705073748 |
| **A_51_P273538** | Syce2 | 0.212782518 | -0.681755328 | 1.107320364 | 10.685215 | 0.595782264 | 0.57509169 | 0.787898437 | -4.705085725 |
| **A_55_P2037812** | Palmd | 0.260866342 | -0.836413573 | 1.358146258 | 9.694125031 | 0.595457767 | 0.575293602 | 0.788119217 | -4.705143734 |
| **A_55_P2148708** | NA | -0.087884021 | -0.457880578 | 0.282112536 | 12.63277636 | -0.594925499 | 0.57562489 | 0.78851719 | -4.70523883 |
| **A_66_P136095** | Ctnna3 | -0.050828327 | -0.264918627 | 0.163261973 | 7.654148489 | -0.594646912 | 0.57579833 | 0.788645378 | -4.705288576 |
| **A_55_P1967672** | Tas2r109 | 0.172884235 | -0.555354202 | 0.901122672 | 8.854722086 | 0.594609907 | 0.575821371 | 0.788645378 | -4.705295183 |
| **A_52_P418644** | Plp2 | -0.058511859 | -0.305063183 | 0.188039465 | 11.52310265 | -0.594411089 | 0.575945172 | 0.788645378 | -4.705330672 |
| **A_51_P498720** | Cacna2d1 | 0.20887487 | -0.671263043 | 1.089012783 | 8.805898819 | 0.594409592 | 0.575946104 | 0.788645378 | -4.705330939 |
| **A_55_P2084741** | Zkscan6 | -0.063356969 | -0.330351147 | 0.203637208 | 9.647082625 | -0.594350935 | 0.575982633 | 0.788645378 | -4.705341408 |
| **A_51_P501656** | Poglut3 | 0.232776201 | -0.748296573 | 1.213848975 | 9.597807252 | 0.594275325 | 0.57602972 | 0.788645378 | -4.7053549 |
| **A_51_P428134** | Lrig3 | 0.22749139 | -0.731399109 | 1.186381889 | 10.51352348 | 0.594218672 | 0.576065003 | 0.788645378 | -4.705365009 |
| **A_55_P1990225** | Fnta | -0.066412823 | -0.346347083 | 0.213521437 | 12.39136805 | -0.594218588 | 0.576065056 | 0.788645378 | -4.705365024 |
| **A_51_P157982** | Tcaf1 | 0.172195758 | -0.553732439 | 0.898123956 | 8.985700131 | 0.594126781 | 0.576122235 | 0.788645378 | -4.705381405 |
| **A_55_P2005330** | Egln2 | -0.063250047 | -0.32994333 | 0.203443235 | 10.14549782 | -0.594017343 | 0.5761904 | 0.788645378 | -4.705400928 |
| **A_51_P476261** | Zfp120 | 0.097302923 | -0.312983138 | 0.507588984 | 8.866622359 | 0.594004137 | 0.576198627 | 0.788645378 | -4.705403284 |
| **A_52_P408530** | St3gal2 | 0.096081348 | -0.309085846 | 0.501248541 | 10.47148633 | 0.593957209 | 0.576227858 | 0.788645378 | -4.705411654 |
| **A_55_P2095345** | Rara | 0.318421952 | -1.024413217 | 1.661257122 | 12.19088006 | 0.593923677 | 0.576248746 | 0.788645378 | -4.705417635 |
| **A_55_P1993109** | Taf6 | 0.062668613 | -0.201726116 | 0.327063341 | 11.91305855 | 0.593673458 | 0.576404627 | 0.788750875 | -4.705462258 |
| **A_51_P293989** | Senp6 | 0.063818618 | -0.205494073 | 0.333131309 | 7.993964446 | 0.5935276 | 0.576495506 | 0.788750875 | -4.705488262 |
| **A_65_P10399** | Gna13 | -0.087568472 | -0.457143525 | 0.282006582 | 8.366758424 | -0.593465486 | 0.576534209 | 0.788750875 | -4.705499334 |
| **A_51_P425680** | Ivd | 0.173631922 | -0.559215847 | 0.906479691 | 10.82824045 | 0.593425424 | 0.576559173 | 0.788750875 | -4.705506475 |
| **A_52_P424826** | Txnl4b | -0.082367079 | -0.430020893 | 0.265286736 | 10.95975536 | -0.593412936 | 0.576566954 | 0.788750875 | -4.705508701 |
| **A_55_P2131677** | NA | -0.058863291 | -0.3073151 | 0.189588519 | 7.846966918 | -0.593407074 | 0.576570607 | 0.788750875 | -4.705509746 |
| **A_55_P1955479** | NA | -0.085016972 | -0.443940019 | 0.273906074 | 10.04011753 | -0.593273069 | 0.576654115 | 0.788809301 | -4.705533629 |
| **A_55_P2073383** | NA | 0.228650262 | -0.736879215 | 1.194179739 | 15.59756255 | 0.593139043 | 0.576737643 | 0.788867745 | -4.705557511 |
| **A_55_P2087315** | Shroom1 | -0.220630315 | -1.152667729 | 0.7114071 | 8.986568163 | -0.592901001 | 0.576886014 | 0.789014869 | -4.705599917 |
| **A_55_P1952359** | Anapc5 | 0.144568877 | -0.466327192 | 0.755464946 | 15.00187772 | 0.592731247 | 0.576991836 | 0.789103781 | -4.70563015 |
| **A_52_P508089** | Psap | -0.531593656 | -2.778317628 | 1.715130316 | 11.9409312 | -0.592625611 | 0.577057693 | 0.789138028 | -4.70564896 |
| **A_55_P2437253** | Nae1 | 0.10295483 | -0.332567624 | 0.538477285 | 9.162278057 | 0.592088358 | 0.577392709 | 0.789468103 | -4.705744585 |
| **A_55_P2153031** | Ttn | 0.089033811 | -0.287609112 | 0.465676733 | 7.911515882 | 0.592073326 | 0.577402084 | 0.789468103 | -4.70574726 |
| **A_55_P2129658** | Smad6 | 0.086284187 | -0.278746158 | 0.451314533 | 9.882546896 | 0.592042099 | 0.57742156 | 0.789468103 | -4.705752816 |
| **A_51_P141390** | Anapc1 | 0.052034464 | -0.168154506 | 0.272223433 | 11.99251781 | 0.591896633 | 0.577512292 | 0.789500648 | -4.705778694 |
| **A_55_P2009427** | Nrsn2 | 0.172586122 | -0.557758251 | 0.902930495 | 9.350457207 | 0.591873 | 0.577527034 | 0.789500648 | -4.705782898 |
| **A_55_P2061341** | Ift74 | 0.151640097 | -0.490236145 | 0.793516339 | 10.74485023 | 0.591715707 | 0.577625153 | 0.789546823 | -4.705810873 |
| **A_51_P489138** | Sptb | 0.082853394 | -0.267906323 | 0.433613112 | 7.936638002 | 0.591631024 | 0.577677983 | 0.789546823 | -4.705825932 |
| **A_52_P378287** | Lca5 | 0.181727563 | -0.587626926 | 0.951082053 | 8.875902779 | 0.591622465 | 0.577683323 | 0.789546823 | -4.705827454 |
| **A_55_P1976395** | Nsdhl | 0.054481873 | -0.176204398 | 0.285168143 | 7.808752726 | 0.591535242 | 0.57773774 | 0.789565383 | -4.705842962 |
| **A_55_P2074035** | Haao | 0.053410573 | -0.172806854 | 0.279627999 | 8.118224286 | 0.591359424 | 0.577847441 | 0.789655595 | -4.705874218 |
| **A_51_P296866** | Msi1 | 0.098348294 | -0.318243179 | 0.514939767 | 8.590212201 | 0.591298534 | 0.577885436 | 0.789655595 | -4.705885041 |
| **A_55_P2126662** | Cav1 | 0.233233551 | -0.754847617 | 1.221314718 | 9.219453286 | 0.591219497 | 0.577934757 | 0.789667179 | -4.705899088 |
| **A_55_P2099627** | Lrrc51 | -0.109498521 | -0.573583772 | 0.354586731 | 8.554978302 | -0.590963735 | 0.578094375 | 0.789762773 | -4.705944533 |
| **A_55_P1992889** | Serping1 | -0.112907973 | -0.591463451 | 0.365647506 | 12.3884838 | -0.590939014 | 0.578109805 | 0.789762773 | -4.705948925 |
| **A_55_P2151797** | Dis3l2 | 0.166313462 | -0.538702394 | 0.871329318 | 11.73905529 | 0.59085221 | 0.578163985 | 0.789762773 | -4.705964345 |
| **A_55_P2100928** | Ptgds | 0.295526343 | -0.957247109 | 1.548299795 | 9.838224951 | 0.590845598 | 0.578168112 | 0.789762773 | -4.70596552 |
| **A_55_P1989251** | Chfr | -0.076095616 | -0.398736874 | 0.246545641 | 11.03909971 | -0.590731436 | 0.578239374 | 0.789804314 | -4.705985797 |
| **A_55_P2055042** | Uqcc2 | 0.122340701 | -0.396448093 | 0.641129495 | 13.58837971 | 0.590650701 | 0.578289774 | 0.789817355 | -4.706000135 |
| **A_51_P211968** | Lcmt1 | -0.07511814 | -0.393758234 | 0.243521955 | 10.12010945 | -0.590465798 | 0.57840521 | 0.789864426 | -4.706032966 |

| | | | | | | | | | |
|---|---|---|---|---|---|---|---|---|---|
| **A_51_P208152** | 3930402G | -0.050506594 | -0.264751069 | 0.163737882 | 7.861251243 | -0.590457703 | 0.578410264 | 0.789864426 | -4.706034403 |
| **A_52_P654752** | Trub1 | 0.050334046 | -0.163226855 | 0.263894946 | 7.997280875 | 0.590324 | 0.578493745 | 0.789864426 | -4.706058138 |
| **A_51_P378051** | Aoc3 | 0.222244482 | -0.720798711 | 1.165287676 | 9.799629213 | 0.590268691 | 0.578528281 | 0.789864426 | -4.706067956 |
| **A_55_P2143790** | Gm14410 | -0.087604944 | -0.459336545 | 0.284126656 | 9.929900018 | -0.59026833 | 0.578528506 | 0.789864426 | -4.70606802 |
| **A_51_P371152** | Sos2 | -0.054223744 | -0.284406529 | 0.175959041 | 10.56382433 | -0.590020368 | 0.578683354 | 0.790020051 | -4.706112024 |
| **A_51_P273203** | Gpaa1 | 0.052299578 | -0.169788239 | 0.274387395 | 8.304275097 | 0.589825836 | 0.578804852 | 0.790104732 | -4.706146535 |
| **A_55_P2181386** | Adm2 | 0.053421618 | -0.173452621 | 0.280295857 | 7.77682002 | 0.589769345 | 0.578840137 | 0.790104732 | -4.706156556 |
| **A_66_P120277** | 9330182L | 0.228083647 | -0.740629343 | 1.196796637 | 8.587556928 | 0.589724772 | 0.57886798 | 0.790104732 | -4.706164461 |
| **A_55_P1984009** | Rsph3b | 0.113935553 | -0.370098684 | 0.59796979 | 12.465066 | 0.589567491 | 0.578966231 | 0.790158263 | -4.706192354 |
| **A_55_P2011237** | NA | 0.1350228 | -0.43865469 | 0.70870029 | 15.93296598 | 0.589507923 | 0.579003445 | 0.790158263 | -4.706202917 |
| **A_51_P145433** | Lin7c | -0.058410021 | -0.30659709 | 0.189777048 | 8.349858468 | -0.589465727 | 0.579029806 | 0.790158263 | -4.706210398 |
| **A_51_P281700** | Khsrp | -0.081490813 | -0.427902173 | 0.264920548 | 12.35732869 | -0.589205604 | 0.579192334 | 0.79032427 | -4.70625651 |
| **A_55_P2067262** | Cdv3 | -0.099118485 | -0.520605143 | 0.322368174 | 8.082401234 | -0.589007964 | 0.57931584 | 0.790437011 | -4.706291534 |
| **A_52_P154101** | Calca | 0.282848315 | -0.920282225 | 1.485978855 | 8.870692301 | 0.588831712 | 0.579425994 | 0.79050891 | -4.706322761 |
| **A_51_P103780** | Sft2d2 | 0.127255894 | -0.414078668 | 0.668590457 | 10.04682763 | 0.588792809 | 0.579450309 | 0.79050891 | -4.706329652 |
| **A_55_P2077746** | Mphosph8 | 0.06497136 | -0.211467907 | 0.341410628 | 10.80287513 | 0.588670905 | 0.579526506 | 0.790544217 | -4.706351245 |
| **A_55_P2054927** | Fam136a | 0.054844337 | -0.178526562 | 0.288215236 | 11.75508453 | 0.588620575 | 0.579557967 | 0.790544217 | -4.706360158 |
| **A_51_P352283** | Cyb561d1 | -0.13702708 | -0.720167423 | 0.446113263 | 8.946246497 | -0.588550403 | 0.579601833 | 0.790548277 | -4.706372585 |
| **A_51_P482425** | Lins1 | -0.09188734 | -0.483255212 | 0.299480532 | 8.908549316 | -0.588058938 | 0.579909113 | 0.790754275 | -4.706459586 |
| **A_55_P1985793** | Dock6 | 0.181356366 | -0.591145757 | 0.953858489 | 12.36099425 | 0.588008318 | 0.579940768 | 0.790754275 | -4.706468543 |
| **A_51_P188271** | Cd248 | 0.370055319 | -1.206249999 | 1.946360638 | 12.75027211 | 0.587999093 | 0.579946536 | 0.790754275 | -4.706470175 |
| **A_55_P2084965** | Klk13 | 0.171741962 | -0.559975793 | 0.903459718 | 8.355284621 | 0.587872537 | 0.580025682 | 0.790754275 | -4.706492568 |
| **A_55_P2131473** | Rmnd5a | 0.098515068 | -0.321222322 | 0.518252458 | 8.266876695 | 0.587861949 | 0.580032304 | 0.790754275 | -4.706494441 |
| **A_51_P208555** | Decr1 | 0.055089581 | -0.179665455 | 0.289844617 | 8.730782018 | 0.587766595 | 0.580091942 | 0.790754275 | -4.706511309 |
| **A_55_P2177539** | Sod1 | -0.06018432 | -0.316649803 | 0.196281163 | 15.48669722 | -0.587766435 | 0.580092042 | 0.790754275 | -4.706511338 |
| **A_52_P420308** | NA | -0.071319118 | -0.375284796 | 0.23264656 | 10.71589748 | -0.587667618 | 0.580153849 | 0.790754275 | -4.706528817 |
| **A_55_P1966719** | NA | -0.121359161 | -0.638735736 | 0.396017413 | 10.70584729 | -0.587511202 | 0.580251691 | 0.790754275 | -4.706556479 |
| **A_55_P1977787** | NA | 0.290616311 | -0.948403695 | 1.529636316 | 9.231939756 | 0.587478547 | 0.580272119 | 0.790754275 | -4.706562253 |
| **A_51_P435817** | Zbtb8a | 0.175092265 | -0.571404093 | 0.921588622 | 8.917368338 | 0.587475301 | 0.580274149 | 0.790754275 | -4.706562827 |
| **A_66_P122099** | Zc3h8 | 0.049733725 | -0.162319524 | 0.261786974 | 8.122512127 | 0.587430383 | 0.580302249 | 0.790754275 | -4.70657077 |
| **A_51_P247481** | Adck1 | -0.139519762 | -0.734415356 | 0.455375833 | 9.146809868 | -0.587415387 | 0.58031163 | 0.790754275 | -4.706573421 |
| **A_55_P1954081** | Ccdc115 | 0.123330281 | -0.40255509 | 0.649215653 | 11.22171975 | 0.587393285 | 0.580325458 | 0.790754275 | -4.706577329 |
| **A_55_P2109997** | Lrig1 | -0.076857443 | -0.404729321 | 0.251014435 | 9.584493076 | -0.587127073 | 0.580492015 | 0.790925484 | -4.706624388 |
| **A_55_P1989765** | NA | 0.087283778 | -0.285115719 | 0.459683275 | 8.534497652 | 0.587049599 | 0.580540493 | 0.790935797 | -4.70663808 |
| **A_55_P2040723** | Dnali1 | 0.40955782 | -1.338107977 | 2.157223617 | 9.492096747 | 0.586958209 | 0.580597682 | 0.790957975 | -4.70665423 |
| **A_55_P1954896** | Rpl17 | 0.221079674 | -0.722526971 | 1.164686319 | 15.81094663 | 0.586824409 | 0.580681415 | 0.791016309 | -4.70667787 |
| **A_55_P2052062** | Cd200 | 0.452926297 | -1.481319678 | 2.387172271 | 11.31562174 | 0.586497521 | 0.580886016 | 0.791239273 | -4.706735608 |
| **A_55_P2085412** | Ankle1 | 0.137527627 | -0.449905742 | 0.724960997 | 8.287203488 | 0.586383421 | 0.580957442 | 0.791280816 | -4.706755756 |
| **A_51_P341789** | Sugp1 | -0.112457205 | -0.592890834 | 0.367976424 | 11.68540539 | -0.586278854 | 0.581022905 | 0.791314232 | -4.706774217 |
| **A_51_P505493** | Elovl5 | 0.143160987 | -0.468939657 | 0.755261631 | 11.9424347 | 0.585803812 | 0.581320356 | 0.791663574 | -4.706858053 |
| **A_51_P126835** | Arl14ep | -0.14580194 | -0.769271727 | 0.477667848 | 8.766181858 | -0.585731033 | 0.581365935 | 0.791669882 | -4.705139153 |
| **A_51_P466371** | Pitpna | 0.11563596 | -0.379014411 | 0.610286331 | 10.6725836 | 0.585524292 | 0.581495423 | 0.791765241 | -4.706907358 |
| **A_55_P1974352** | Fam126b | -0.067100721 | -0.354151806 | 0.219950364 | 9.850501903 | -0.585488461 | 0.581517866 | 0.791765241 | -4.706913677 |
| **A_51_P424959** | Bcl6b | 0.367846779 | -1.206129235 | 1.941822794 | 9.449041058 | 0.585354813 | 0.581601584 | 0.791823465 | -4.706937243 |
| **A_51_P227077** | Mdh1b | 0.222182695 | -0.728675525 | 1.173040915 | 8.880443986 | 0.585254567 | 0.581664384 | 0.791853204 | -4.706954917 |
| **A_55_P1999240** | NA | 0.0884988 | -0.290291783 | 0.467289383 | 9.801476409 | 0.585178759 | 0.581711878 | 0.791862102 | -4.70696828 |
| **A_55_P2164502** | NA | -0.18331774 | -0.968272369 | 0.60163689 | 9.126276895 | -0.584938603 | 0.581862349 | 0.79201117 | -4.707010607 |
| **A_51_P481569** | Ddx42 | 0.060133575 | -0.1974586 | 0.31772575 | 11.59034799 | 0.584702167 | 0.582010513 | 0.792157076 | -4.707052264 |

| | | | | | | | | | |
|---|---|---|---|---|---|---|---|---|---|
| **A_52_P362033** | Polr2e | -0.059167707 | -0.312745071 | 0.194409657 | 10.25085241 | -0.58441937 | 0.58218776 | 0.792291685 | -4.707102072 |
| **A_55_P2034893** | Socs7 | -0.103954854 | -0.549482645 | 0.341572937 | 11.45765406 | -0.584413608 | 0.582191371 | 0.792291685 | -4.707103086 |
| **A_55_P2009285** | Fbxo25 | -0.060439343 | -0.319506743 | 0.198628057 | 10.59461699 | -0.584328828 | 0.582244515 | 0.792307779 | -4.707118015 |
| **A_55_P2157205** | Cdx1 | -0.387838633 | -2.05078886 | 1.275111594 | 8.282532346 | -0.584147016 | 0.582358491 | 0.792307779 | -4.707150022 |
| **A_55_P2014531** | Hes7 | 0.08208507 | -0.269874488 | 0.434044627 | 9.341798696 | 0.584146466 | 0.582358836 | 0.792307779 | -4.707150119 |
| **A_52_P682745** | Dock4 | 0.309007385 | -1.015966357 | 1.633981126 | 9.687730088 | 0.584133256 | 0.582367118 | 0.792307779 | -4.707152444 |
| **A_51_P491595** | AW54987 | -0.056090708 | -0.296659367 | 0.184477951 | 9.596981077 | -0.58398571 | 0.582459624 | 0.792377876 | -4.707178413 |
| **A_55_P2109128** | H2bc6 | 0.050767292 | -0.166999303 | 0.268533886 | 8.324525945 | 0.583906194 | 0.582509482 | 0.792389948 | -4.707192406 |
| **A_51_P359485** | Zfpl1 | -0.08068354 | -0.426843559 | 0.265476478 | 10.14795524 | -0.583792333 | 0.582580879 | 0.792431315 | -4.707212441 |
| **A_52_P376360** | Pdgfc | 0.103481211 | -0.340609195 | 0.547571617 | 8.805959726 | 0.583633831 | 0.582680276 | 0.792470554 | -4.707240325 |
| **A_51_P351860** | C1qb | 0.051203508 | -0.168543597 | 0.270950613 | 8.631297493 | 0.58361561 | 0.582691704 | 0.792470554 | -4.70724353 |
| **A_55_P1993698** | Celf1 | -0.202196853 | -1.070115534 | 0.665721828 | 9.622279736 | -0.583506505 | 0.582760131 | 0.792473108 | -4.70726272 |
| **A_55_P2034550** | 1700013F | 0.178790559 | -0.588693076 | 0.946274194 | 8.440661642 | 0.583479772 | 0.582776898 | 0.792473108 | -4.707267421 |
| **A_51_P165451** | Pbld2 | -0.055134821 | -0.291895099 | 0.181625457 | 8.020774957 | -0.583267088 | 0.582910302 | 0.792473108 | -4.70730482 |
| **A_55_P2120466** | Snx11 | -0.069248602 | -0.366661125 | 0.228163921 | 8.832380809 | -0.583179303 | 0.582965371 | 0.792473108 | -4.707320253 |
| **A_51_P377210** | Caap1 | 0.083845336 | -0.276278079 | 0.443968752 | 8.450547351 | 0.583146798 | 0.582985762 | 0.792473108 | -4.707325967 |
| **A_55_P2088615** | Fbln5 | 0.559274468 | -1.842967482 | 2.961516419 | 13.02089242 | 0.58312088 | 0.583002022 | 0.792473108 | -4.707330523 |
| **A_66_P102062** | NA | -0.140258069 | -0.742736888 | 0.46222075 | 8.973098167 | -0.583091107 | 0.5830207 | 0.792473108 | -4.707335756 |
| **A_51_P455671** | Psmd5 | 0.085166529 | -0.280667257 | 0.451000315 | 9.817139604 | 0.583089848 | 0.58302149 | 0.792473108 | -4.707335977 |
| **A_52_P399677** | Tprkb | -0.053261078 | -0.28208013 | 0.175557974 | 8.50200559 | -0.582999416 | 0.583078225 | 0.79249451 | -4.707351872 |
| **A_52_P93910** | Nrp2 | 0.332939635 | -1.097781246 | 1.763660516 | 11.68728761 | 0.58285554 | 0.583168497 | 0.792510884 | -4.707377156 |
| **A_55_P1995141** | Casz1 | -0.057385308 | -0.304042631 | 0.189272014 | 8.060653053 | -0.582716147 | 0.583255963 | 0.792510884 | -4.707401647 |
| **A_55_P2076545** | F5 | -0.048631424 | -0.257677556 | 0.160414708 | 7.825481254 | -0.582673402 | 0.583282787 | 0.792510884 | -4.707409156 |
| **A_55_P1953311** | Stx2 | -0.095977573 | -0.508548194 | 0.316593048 | 9.93855454 | -0.582668861 | 0.583285636 | 0.792510884 | -4.707409954 |
| **A_55_P2429432** | 2610301B | 0.063364959 | -0.209044603 | 0.335774521 | 8.739134848 | 0.582608959 | 0.583323228 | 0.792510884 | -4.707420477 |
| **A_55_P2118173** | Igf2 | -0.357755044 | -1.8959787 | 1.180468612 | 10.21407064 | -0.582527759 | 0.583374187 | 0.792510884 | -4.707434739 |
| **A_55_P2028444** | Ube2j2 | -0.107989253 | -0.572309738 | 0.356331231 | 12.91811299 | -0.582522948 | 0.583377206 | 0.792510884 | -4.707435584 |
| **A_52_P537381** | Dimt1 | 0.06377973 | -0.210535051 | 0.338094511 | 10.02975462 | 0.582349643 | 0.583485977 | 0.792602957 | -4.707466019 |
| **A_52_P37702** | NA | -0.063783293 | -0.338158184 | 0.210591597 | 7.739741265 | -0.582254589 | 0.583545641 | 0.792628314 | -4.707482709 |
| **A_55_P2094469** | Kdr | -0.157564536 | -0.835628471 | 0.520499399 | 12.01526942 | -0.582020357 | 0.583692679 | 0.792719698 | -4.707523827 |
| **A_55_P1952127** | Psma3 | 0.071535253 | -0.236311767 | 0.379382273 | 14.11217872 | 0.582016785 | 0.583694922 | 0.792719698 | -4.707524454 |
| **A_55_P1969820** | Slc36a1 | -0.116814424 | -0.619691071 | 0.386062223 | 8.303869635 | -0.581815585 | 0.583821243 | 0.792733818 | -4.707559762 |
| **A_52_P239292** | AI597479 | -0.061044075 | -0.323835944 | 0.201747794 | 10.48085427 | -0.581811014 | 0.583824113 | 0.792733818 | -4.707560564 |
| **A_55_P2145656** | NA | 0.322987391 | -1.06750023 | 1.713475012 | 9.403785202 | 0.581793396 | 0.583835175 | 0.792733818 | -4.707563655 |
| **A_55_P2018249** | Lmo4 | 0.186793456 | -0.617568891 | 0.991155803 | 9.455599893 | 0.581648041 | 0.583926446 | 0.792733818 | -4.707589157 |
| **A_55_P2109505** | NA | 0.122719973 | -0.405739717 | 0.651179662 | 15.7525399 | 0.581639282 | 0.583931946 | 0.792733818 | -4.707590693 |
| **A_55_P2005300** | Ranbp3 | 0.072782125 | -0.240677477 | 0.386241727 | 12.61077247 | 0.581558621 | 0.583982599 | 0.792733818 | -4.707604843 |
| **A_55_P2005898** | NA | 0.132286741 | -0.437463792 | 0.702037273 | 8.198617562 | 0.581543121 | 0.583992333 | 0.792733818 | -4.707607561 |
| **A_55_P2040245** | Piezo2 | 0.284081364 | -0.939807797 | 1.507970525 | 9.358455659 | 0.581367836 | 0.584102416 | 0.792827585 | -4.707638303 |
| **A_55_P1959228** | Slc4a4 | 0.069485153 | -0.229947137 | 0.368917443 | 8.425047852 | 0.581224259 | 0.584192596 | 0.792894324 | -4.707663477 |
| **A_55_P2039454** | Gm5088 | -0.064566209 | -0.34292795 | 0.213795533 | 7.874221606 | -0.580959803 | 0.58435872 | 0.793001403 | -4.707709834 |
| **A_55_P2176802** | Hipk1 | -0.075171037 | -0.399318301 | 0.248976227 | 8.20214509 | -0.580842688 | 0.584432298 | 0.793001403 | -4.707730358 |
| **A_51_P163173** | Rbm12b1 | 0.079993196 | -0.264950603 | 0.424936995 | 8.480434213 | 0.580838044 | 0.584435215 | 0.793001403 | -4.707731172 |
| **A_55_P2076787** | Synpo | 0.051861757 | -0.171792383 | 0.275515897 | 8.25832652 | 0.580792006 | 0.58446414 | 0.793001403 | -4.707739239 |
| **A_55_P2025103** | Snx3 | 0.053498623 | -0.177226323 | 0.28422357 | 11.29558832 | 0.580762275 | 0.584482821 | 0.793001403 | -4.707744448 |
| **A_51_P157524** | E2f4 | 0.115592946 | -0.382992599 | 0.614178492 | 8.709467505 | 0.580686855 | 0.584530209 | 0.793001403 | -4.707757662 |
| **A_52_P35048** | Serpinf1 | 0.064267459 | -0.212966842 | 0.341501759 | 7.975170883 | 0.580623369 | 0.5845701 | 0.793001403 | -4.707768784 |
| **A_55_P2077448** | Epas1 | -0.121813692 | -0.647354504 | 0.403727121 | 11.44996592 | -0.580550499 | 0.58461589 | 0.793001403 | -4.707781548 |

| | | | | | | | | | |
|---|---|---|---|---|---|---|---|---|---|
| **A_52_P121960** | NA | 0.070999834 | -0.235338043 | 0.377337712 | 14.71097835 | 0.58050636 | 0.584643627 | 0.793001403 | -4.707789279 |
| **A_51_P461724** | Atp6ap2 | 0.07490558 | -0.248317823 | 0.398128982 | 10.74439435 | 0.580445857 | 0.584681649 | 0.793001403 | -4.707799876 |
| **A_52_P684057** | Snupn | -0.116647476 | -0.620092695 | 0.386797743 | 10.13392389 | -0.580327926 | 0.584755764 | 0.793046292 | -4.707820528 |
| **A_55_P2158946** | Lama2 | 0.261082195 | -0.865894927 | 1.388059317 | 10.05589456 | 0.580246462 | 0.584806964 | 0.7930601 | -4.707834792 |
| **A_55_P2112479** | Nudt16l1 | 0.133591549 | -0.443209593 | 0.710392691 | 10.06962299 | 0.580100472 | 0.584898726 | 0.793117422 | -4.707860351 |
| **A_55_P2081510** | Atp5j2 | 0.161232034 | -0.534987424 | 0.857451492 | 14.07862093 | 0.580036765 | 0.584938771 | 0.793117422 | -4.707871502 |
| **A_55_P1954921** | Fbxw5 | 0.055806187 | -0.185193851 | 0.296806225 | 11.46745689 | 0.579983428 | 0.5849723 | 0.793117422 | -4.707880838 |
| **A_55_P2170963** | Gspt1 | 0.069093677 | -0.229359022 | 0.367546376 | 11.86859135 | 0.579846632 | 0.585058297 | 0.793158005 | -4.707904778 |
| **A_55_P2455725** | Klhl20 | -0.065622912 | -0.34918766 | 0.217941837 | 9.370926922 | -0.579633664 | 0.585192195 | 0.793158005 | -4.70794204 |
| **A_55_P2097448** | Rbpms | -0.079151994 | -0.421209178 | 0.262905189 | 9.959322477 | -0.579580135 | 0.585225853 | 0.793158005 | -4.707951404 |
| **A_55_P2409570** | Luc7l2 | -0.154051249 | -0.819891054 | 0.511788556 | 14.9280913 | -0.579489826 | 0.58528264 | 0.793158005 | -4.707967201 |
| **A_55_P1977422** | NA | -0.099993113 | -0.532188077 | 0.332201852 | 16.522455 | -0.579483041 | 0.585286906 | 0.793158005 | -4.707968387 |
| **A_55_P1992555** | Gys2 | 0.057856187 | -0.192230508 | 0.307942882 | 7.78769107 | 0.579441458 | 0.585313055 | 0.793158005 | -4.70797566 |
| **A_55_P1990299** | Ophn1 | -0.051376526 | -0.27345658 | 0.170703528 | 7.95187 | -0.579436069 | 0.585316444 | 0.793158005 | -4.707976603 |
| **A_55_P2099785** | Usp15 | -0.187498853 | -0.998034975 | 0.62303727 | 10.85613765 | -0.579397458 | 0.585340724 | 0.793158005 | -4.707983355 |
| **A_52_P85864** | Eif6 | 0.074544756 | -0.247773789 | 0.396863302 | 11.44138009 | 0.579271486 | 0.585419947 | 0.793158005 | -4.708005384 |
| **A_55_P1970002** | Sh3kbp1 | 0.0580191 | -0.192880112 | 0.308918312 | 8.107936771 | 0.579191308 | 0.585470373 | 0.793158005 | -4.708019402 |
| **A_55_P2063426** | NA | -0.066945749 | -0.356483736 | 0.222592237 | 8.35485044 | -0.579118983 | 0.585515862 | 0.793158005 | -4.708032046 |
| **A_55_P2124586** | Xlr4a | -0.623755919 | -3.321488698 | 2.07397686 | 8.795067412 | -0.57911671 | 0.585517292 | 0.793158005 | -4.708032444 |
| **A_55_P2147250** | Caprin2 | 0.079658857 | -0.264881485 | 0.424199199 | 9.293767391 | 0.579087693 | 0.585535544 | 0.793158005 | -4.708037516 |
| **A_55_P1958919** | Olfr1347 | -0.098613409 | -0.525219533 | 0.327992715 | 8.056155639 | -0.578974228 | 0.585606914 | 0.793179164 | -4.708057349 |
| **A_52_P341449** | Pgm3 | 0.055810761 | -0.185688642 | 0.297310164 | 10.09254429 | 0.578831595 | 0.585696639 | 0.793179164 | -4.708082276 |
| **A_55_P1966169** | Ppdpf | 0.201340806 | -0.670092402 | 1.072774013 | 13.5466455 | 0.578692751 | 0.585783989 | 0.793179164 | -4.708106536 |
| **A_52_P486322** | Fzr1 | 0.076643129 | -0.25511727 | 0.408403528 | 11.00333891 | 0.578627445 | 0.585825077 | 0.793179164 | -4.708117945 |
| **A_66_P106065** | Ttc3 | 0.119690212 | -0.39840632 | 0.637786744 | 9.159444453 | 0.578626471 | 0.58582569 | 0.793179164 | -4.708118115 |
| **A_55_P2077188** | Stxbp1 | 0.165382696 | -0.550519079 | 0.881284471 | 10.17975915 | 0.578611384 | 0.585835182 | 0.793179164 | -4.708120751 |
| **A_51_P279100** | Ptgs1 | 0.23029466 | -0.766648037 | 1.227237357 | 12.09426018 | 0.578580798 | 0.585854426 | 0.793179164 | -4.708126094 |
| **A_55_P2056100** | Pus1 | 0.09146488 | -0.304721121 | 0.487650881 | 8.844896598 | 0.578236617 | 0.586071007 | 0.793179164 | -4.708186203 |
| **A_51_P446558** | Timm29 | 0.053313281 | -0.177630495 | 0.284257058 | 8.110837465 | 0.578201875 | 0.586092872 | 0.793179164 | -4.708192269 |
| **A_51_P167374** | Gpatch1 | -0.134453302 | -0.71693738 | 0.448030776 | 9.519591784 | -0.578146313 | 0.58612784 | 0.793179164 | -4.708201969 |
| **A_55_P2186352** | Ttc12 | 0.243541473 | -0.811653675 | 1.298736621 | 9.598535638 | 0.578083387 | 0.586167445 | 0.793179164 | -4.708212954 |
| **A_52_P581435** | Pdha1 | 0.145757948 | -0.485783292 | 0.777299189 | 10.98945725 | 0.578070592 | 0.586175498 | 0.793179164 | -4.708215188 |
| **A_55_P2010008** | Zfp113 | 0.084854563 | -0.282860595 | 0.452569721 | 8.808890072 | 0.5779816 | 0.586231511 | 0.793179164 | -4.708230721 |
| **A_55_P2048478** | Olfml1 | 0.177872844 | -0.593036082 | 0.948781769 | 9.042663354 | 0.577905622 | 0.586279336 | 0.793179164 | -4.708243982 |
| **A_52_P220485** | Uhrf2 | -0.113679399 | -0.606381726 | 0.379022929 | 9.277026447 | -0.577893087 | 0.586287226 | 0.793179164 | -4.70824617 |
| **A_55_P2051778** | Sult2b1 | 0.056901071 | -0.189717074 | 0.303519216 | 8.401425985 | 0.577890771 | 0.586288684 | 0.793179164 | -4.708246574 |
| **A_55_P2029252** | NA | 0.094575755 | -0.315335219 | 0.504486729 | 9.795157907 | 0.577883932 | 0.586292989 | 0.793179164 | -4.708247767 |
| **A_51_P425632** | Mad2l1bp | 0.168261794 | -0.561036779 | 0.897560366 | 12.82936047 | 0.577870462 | 0.586301468 | 0.793179164 | -4.708250118 |
| **A_51_P295610** | Cyc1 | 0.205516636 | -0.685327372 | 1.096360645 | 12.10298314 | 0.577824118 | 0.586330641 | 0.793179164 | -4.708258205 |
| **A_55_P2340593** | 1810019D | -0.150226603 | -0.801843916 | 0.501390711 | 9.231807142 | -0.577436957 | 0.58657439 | 0.793400859 | -4.708325748 |
| **A_55_P1956799** | Smpx | 0.283002894 | -0.944601249 | 1.510607037 | 8.992414465 | 0.577408107 | 0.586592556 | 0.793400859 | -4.708330779 |
| **A_55_P2133356** | NA | 0.172260476 | -0.575019018 | 0.91953997 | 15.96191258 | 0.577368284 | 0.586617631 | 0.793400859 | -4.708337724 |
| **A_55_P2325698** | 2010002N | 0.076069898 | -0.254052476 | 0.406192272 | 8.686336538 | 0.577149363 | 0.586755492 | 0.793531805 | -4.708375896 |
| **A_51_P259318** | Fbrs | -0.071140722 | -0.380003486 | 0.237722041 | 8.271480965 | -0.576903351 | 0.586910437 | 0.793661521 | -4.708418778 |
| **A_55_P2123822** | Rbbp7 | -0.295521811 | -1.5787524 | 0.987708778 | 14.40057162 | -0.576813189 | 0.586967229 | 0.793661521 | -4.708434491 |
| **A_55_P2063795** | Bag6 | 0.075281203 | -0.251615449 | 0.402177855 | 12.73462508 | 0.576801555 | 0.586974557 | 0.793661521 | -4.708436518 |
| **A_55_P2087240** | Mipep | 0.119491789 | -0.399476627 | 0.638460205 | 8.812755576 | 0.576696719 | 0.587040597 | 0.793695308 | -4.708454785 |
| **A_55_P2009121** | Agrn | 0.32042733 | -1.071488457 | 1.712343117 | 14.39809103 | 0.576589775 | 0.58710797 | 0.793730893 | -4.708473416 |

| | | | | | | | | | |
|---|---|---|---|---|---|---|---|---|---|
| **A_55_P1960708** | Setd1a | -0.13482838 | -0.720608426 | 0.450951665 | 10.24741736 | -0.576497051 | 0.587166388 | 0.793754366 | -4.708489568 |
| **A_55_P2134877** | Eml1 | 0.105937605 | -0.354375506 | 0.566250716 | 8.369580266 | 0.576430764 | 0.587208152 | 0.793755325 | -4.708501113 |
| **A_52_P52128** | Leo1 | 0.086852981 | -0.290589855 | 0.464295817 | 10.97041487 | 0.576346802 | 0.587261055 | 0.79377134 | -4.708515736 |
| **A_55_P2124831** | Brdt | -0.073540269 | -0.393166137 | 0.246085598 | 8.218224544 | -0.576280125 | 0.587303069 | 0.793772634 | -4.708527346 |
| **A_55_P2351193** | Snora47 | -0.205862097 | -1.101061612 | 0.689337417 | 8.383714838 | -0.575979333 | 0.587492624 | 0.793973326 | -4.708579712 |
| **A_55_P2135759** | NA | 0.158667997 | -0.531533205 | 0.848869199 | 16.80281364 | 0.575789807 | 0.58761208 | 0.793986465 | -4.708612695 |
| **A_51_P151038** | Pmpcb | 0.153155705 | -0.513222373 | 0.819533782 | 12.04880808 | 0.575655717 | 0.587696604 | 0.793986465 | -4.708636026 |
| **A_52_P417859** | Gid4 | -0.083170241 | -0.445099854 | 0.278759372 | 9.412242907 | -0.575564749 | 0.58775395 | 0.793986465 | -4.708651851 |
| **A_55_P2170134** | Eif2b4 | -0.113597466 | -0.607941615 | 0.380746683 | 12.03498199 | -0.575558658 | 0.58775779 | 0.793986465 | -4.708652911 |
| **A_55_P2083879** | Vsig2 | 0.369450773 | -1.238346835 | 1.977248382 | 12.29533005 | 0.575540046 | 0.587769523 | 0.793986465 | -4.708656149 |
| **A_55_P1977929** | Itgam | -0.04964922 | -0.265715991 | 0.166417551 | 7.836289209 | -0.57553905 | 0.587770151 | 0.793986465 | -4.708656322 |
| **A_51_P164495** | Dda1 | -0.147505601 | -0.789465056 | 0.494453853 | 11.07223129 | -0.575507859 | 0.587789815 | 0.793986465 | -4.708661747 |
| **A_55_P2143095** | NA | -0.063460518 | -0.339678481 | 0.212757446 | 8.386970289 | -0.575442637 | 0.587830934 | 0.793986536 | -4.708673091 |
| **A_51_P488175** | Phf20l1 | 0.068390943 | -0.22933612 | 0.366118007 | 8.551150318 | 0.575348024 | 0.587890586 | 0.794007168 | -4.708689546 |
| **A_66_P111877** | Snx13 | -0.091184686 | -0.488235232 | 0.30586586 | 9.625447862 | -0.575210036 | 0.587977592 | 0.794007168 | -4.70871354 |
| **A_52_P681488** | Stx17 | 0.145563523 | -0.48841785 | 0.779544895 | 9.034802263 | 0.575077538 | 0.588061144 | 0.794007168 | -4.708736575 |
| **A_51_P210560** | Pptc7 | 0.067387409 | -0.22612798 | 0.360902797 | 11.82820408 | 0.575040226 | 0.588084674 | 0.794007168 | -4.708743061 |
| **A_52_P512575** | Hopx | 0.666858107 | -2.237910722 | 3.571626935 | 13.22551152 | 0.575005773 | 0.5881064 | 0.794007168 | -4.70874905 |
| **A_51_P153973** | Zfp273 | 0.111848198 | -0.375399271 | 0.599095667 | 8.64589677 | 0.57494956 | 0.588141851 | 0.794007168 | -4.70875882 |
| **A_55_P2035300** | Rhot1 | -0.061514612 | -0.329506074 | 0.20647685 | 11.24038029 | -0.574920344 | 0.588160277 | 0.794007168 | -4.708763898 |
| **A_55_P2134566** | NA | 0.079416465 | -0.266612556 | 0.425445485 | 8.995517955 | 0.574841842 | 0.588209788 | 0.794007168 | -4.708777541 |
| **A_52_P608132** | Snx32 | -0.048600452 | -0.260363171 | 0.163162267 | 7.964878305 | -0.574832278 | 0.58821582 | 0.794007168 | -4.708779203 |
| **A_55_P2051894** | Zfp598 | -0.097845053 | -0.524330122 | 0.328640016 | 9.354271781 | -0.574626153 | 0.588345834 | 0.794091496 | -4.708815017 |
| **A_55_P2049318** | NA | 0.265557482 | -0.892212219 | 1.423327183 | 9.54778512 | 0.574495602 | 0.588428189 | 0.794091496 | -4.708837696 |
| **A_51_P275976** | Dok1 | 0.114344738 | -0.384173851 | 0.612863327 | 9.060811525 | 0.57449357 | 0.58842947 | 0.794091496 | -4.708838048 |
| **A_55_P2058201** | Scgb2b20 | 0.13213605 | -0.444021352 | 0.708293453 | 8.552283505 | 0.57442128 | 0.588475075 | 0.794091496 | -4.708850604 |
| **A_52_P256279** | Zbtb11 | -0.110309345 | -0.591323751 | 0.370705061 | 8.816293635 | -0.574386955 | 0.58849673 | 0.794091496 | -4.708856565 |
| **A_55_P2103145** | Ppp1r2 | 0.080786537 | -0.271518071 | 0.433091145 | 9.081204714 | 0.574342581 | 0.588524726 | 0.794091496 | -4.708864272 |
| **A_55_P2073734** | Bfar | -0.061466701 | -0.329558089 | 0.206624687 | 11.38423421 | -0.574258438 | 0.588577814 | 0.794107708 | -4.708878883 |
| **A_55_P2083063** | Mob1b | -0.052152365 | -0.279735589 | 0.17543086 | 8.016539791 | -0.573963272 | 0.588764065 | 0.794117387 | -4.708930124 |
| **A_52_P646755** | Tex30 | -0.288158628 | -1.545750946 | 0.96943369 | 10.5631428 | -0.57390776 | 0.588799097 | 0.794117387 | -4.708939759 |
| **A_55_P2035038** | NA | -0.112365488 | -0.602763001 | 0.378032024 | 11.93238691 | -0.573898423 | 0.58880499 | 0.794117387 | -4.708941379 |
| **A_51_P252363** | Stx12 | -0.076413637 | -0.409927912 | 0.257100637 | 11.4030316 | -0.573861107 | 0.58882854 | 0.794117387 | -4.708947855 |
| **A_51_P224468** | Maip1 | -0.052278938 | -0.280465937 | 0.175908061 | 10.56680063 | -0.573833903 | 0.588845709 | 0.794117387 | -4.708952576 |
| **A_55_P1983450** | S100a4 | -0.195366687 | -1.048175028 | 0.657441654 | 9.782254005 | -0.573785301 | 0.588876383 | 0.794117387 | -4.70896101 |
| **A_51_P166288** | Acy1 | 0.056097686 | -0.188782689 | 0.300978062 | 8.064696435 | 0.573774592 | 0.588883142 | 0.794117387 | -4.708962868 |
| **A_52_P424272** | Gatc | 0.050655579 | -0.170499494 | 0.271810652 | 8.215797716 | 0.573694544 | 0.588933664 | 0.794117387 | -4.708976757 |
| **A_55_P2103284** | Twf2 | 0.102866595 | -0.346260736 | 0.551993926 | 9.689427475 | 0.573660868 | 0.58895492 | 0.794117387 | -4.7089826 |
| **A_51_P208121** | Klhl25 | -0.160849883 | -0.863356717 | 0.54165695 | 8.643318603 | -0.573482992 | 0.589067199 | 0.794117387 | -4.709013456 |
| **A_51_P155522** | Psmb4 | -0.053171112 | -0.285394719 | 0.179052495 | 14.65950853 | -0.573481907 | 0.589067884 | 0.794117387 | -4.709013644 |
| **A_52_P346113** | Foxn2 | 0.055678027 | -0.187501182 | 0.298857235 | 8.800876074 | 0.573466086 | 0.589077871 | 0.794117387 | -4.709016388 |
| **A_55_P2137496** | NA | -0.057902603 | -0.310999947 | 0.195194742 | 7.7870814 | -0.573008206 | 0.589366957 | 0.7944517 | -4.70909578 |
| **A_55_P2092864** | Emp2 | 0.585141661 | -1.973239203 | 3.143522526 | 11.56914787 | 0.572856926 | 0.589462487 | 0.794525079 | -4.709121999 |
| **A_51_P100289** | Srf | 0.145691086 | -0.491403251 | 0.782785422 | 10.98455784 | 0.5727691 | 0.589517952 | 0.794544446 | -4.709137218 |
| **A_55_P1958434** | Got1 | -0.375813408 | -2.019622254 | 1.267995438 | 10.87280182 | -0.572626316 | 0.58960813 | 0.79454876 | -4.709161956 |
| **A_51_P126437** | Enc1 | 0.078755569 | -0.265731758 | 0.423242897 | 10.31940807 | 0.572609265 | 0.5896189 | 0.79454876 | -4.703925783 |
| **A_55_P2422650** | 5031425E | -0.209659824 | -1.126973322 | 0.707653673 | 10.8689053 | -0.57246347 | 0.589710989 | 0.79454876 | -4.709190164 |
| **A_52_P422755** | Lsm6 | 0.15842717 | -0.534739156 | 0.851593496 | 8.739021099 | 0.572456582 | 0.589715341 | 0.79454876 | -4.709191357 |

| | | | | | | | | | |
|---|---|---|---|---|---|---|---|---|---|
| **A_55_P1964033** | Carm1 | -0.132648391 | -0.713071022 | 0.447774239 | 9.956284665 | -0.572411033 | 0.589744113 | 0.79454876 | -4.709199246 |
| **A_55_P1954476** | NA | -0.096873738 | -0.520850998 | 0.327103521 | 8.312912496 | -0.572286949 | 0.589822499 | 0.79454876 | -4.709220734 |
| **A_55_P2051899** | Atp2c1 | 0.060199409 | -0.203281439 | 0.323680257 | 9.968443543 | 0.57226017 | 0.589839417 | 0.79454876 | -4.709225371 |
| **A_51_P121179** | Erfe | 0.053530194 | -0.180802715 | 0.287863102 | 7.779651609 | 0.572157872 | 0.589904046 | 0.79454876 | -4.705774162 |
| **A_55_P2152872** | Fgfrl1 | -0.084091731 | -0.452227684 | 0.284044221 | 11.59615145 | -0.572130902 | 0.589921085 | 0.79454876 | -4.709247752 |
| **A_51_P446510** | Emp3 | 0.120866913 | -0.40827921 | 0.650013035 | 10.44362531 | 0.572113449 | 0.589932112 | 0.79454876 | -4.709250774 |
| **A_55_P2121349** | NA | 0.076691342 | -0.259196113 | 0.412578796 | 9.029087348 | 0.571877352 | 0.590081291 | 0.794676676 | -4.709291639 |
| **A_55_P2159865** | Zfp59 | 0.088137512 | -0.297990272 | 0.474265295 | 8.7156264 | 0.571715648 | 0.590183478 | 0.794676676 | -4.70931962 |
| **A_51_P503261** | Lrrc8c | 0.142113855 | -0.480521121 | 0.76474883 | 8.909821156 | 0.571680334 | 0.590205796 | 0.794676676 | -4.70932573 |
| **A_55_P2022231** | Gm826 | -0.054214584 | -0.291757288 | 0.18332812 | 7.778279218 | -0.571642856 | 0.590229482 | 0.794676676 | -4.709332213 |
| **A_51_P503542** | Zfp414 | -0.063296395 | -0.340633696 | 0.214040907 | 11.137853 | -0.571637924 | 0.590232599 | 0.794676676 | -4.709333067 |
| **A_52_P403443** | Tns3 | 0.224944232 | -0.761004401 | 1.210892865 | 12.66519858 | 0.571440368 | 0.590357462 | 0.794677293 | -4.709367239 |
| **A_51_P375969** | Ces1d | 0.675020193 | -2.283844743 | 3.633885129 | 10.74736042 | 0.57140228 | 0.590381538 | 0.794677293 | -4.709373826 |
| **A_55_P2004920** | NA | -0.078084822 | -0.420418139 | 0.264248496 | 9.009621348 | -0.571304703 | 0.590443218 | 0.794677293 | -4.7093907 |
| **A_51_P109840** | Vtn | 0.100071289 | -0.338662094 | 0.538804673 | 8.187217167 | 0.571293338 | 0.590450402 | 0.794677293 | -4.705920077 |
| **A_66_P114543** | Jmjd6 | -0.077272998 | -0.416076975 | 0.261530978 | 8.410968492 | -0.571254468 | 0.590474974 | 0.794677293 | -4.709399386 |
| **A_52_P276525** | Ankrd9 | -0.07285351 | -0.392314507 | 0.246607487 | 8.780490304 | -0.571193141 | 0.590513743 | 0.794677293 | -4.70940999 |
| **A_52_P408657** | Mycbp | -0.059044432 | -0.317957977 | 0.199869113 | 8.157341853 | -0.571182017 | 0.590520775 | 0.794677293 | -4.709411913 |
| **A_51_P338262** | Tnnt2 | 0.834933178 | -2.828010119 | 4.497876474 | 12.27431515 | 0.570915546 | 0.590689249 | 0.794848687 | -4.709457974 |
| **A_55_P2041988** | NA | -0.117355081 | -0.632427424 | 0.397717262 | 8.521032759 | -0.570668646 | 0.590845374 | 0.795003442 | -4.709500637 |
| **A_65_P02177** | Gstm4 | 0.058625025 | -0.198863558 | 0.316113608 | 8.248713378 | 0.570263285 | 0.591101754 | 0.795293063 | -4.709570648 |
| **A_55_P1958374** | NA | 0.059416984 | -0.201596854 | 0.320430823 | 11.65499751 | 0.570160882 | 0.591166532 | 0.795324871 | -4.709588328 |
| **A_51_P270286** | Dync1li1 | -0.054815599 | -0.295651869 | 0.186020671 | 8.650187149 | -0.570075818 | 0.591220344 | 0.795341925 | -4.709603013 |
| **A_52_P108502** | Slc4a7 | 0.077032093 | -0.261474834 | 0.41553902 | 8.094615605 | 0.569973265 | 0.591285225 | 0.795373864 | -4.709620714 |
| **A_52_P287992** | Tbcel | -0.053343644 | -0.287812654 | 0.181125366 | 8.359301504 | -0.569832975 | 0.591373987 | 0.79543792 | -4.709644925 |
| **A_55_P2262136** | Tek | 0.117809413 | -0.400183767 | 0.635802593 | 8.682869341 | 0.56964763 | 0.591491267 | 0.795540324 | -4.709676903 |
| **A_55_P2197998** | D4Ertd58e | 0.088252594 | -0.299936166 | 0.476441354 | 8.067208321 | 0.569422821 | 0.591633537 | 0.795676322 | -4.70971568 |
| **A_55_P2163857** | Trim12c | 0.08424044 | -0.286362606 | 0.454843487 | 8.241954448 | 0.569327274 | 0.59169401 | 0.795702302 | -4.709732157 |
| **A_55_P2114133** | Garnl3 | 0.105713551 | -0.359437093 | 0.570864195 | 9.139129357 | 0.569229478 | 0.59175591 | 0.795730197 | -4.709749019 |
| **A_66_P137495** | Btaf1 | -0.053197696 | -0.287450332 | 0.181054941 | 9.84638229 | -0.568798809 | 0.592028549 | 0.795757997 | -4.709823249 |
| **A_55_P2129000** | Cyyr1 | 0.160429796 | -0.546014594 | 0.866874186 | 9.047299048 | 0.568797129 | 0.592029613 | 0.795757997 | -4.709823538 |
| **A_55_P1981241** | Grasp | -0.072015584 | -0.389172688 | 0.245141521 | 11.21178247 | -0.568725086 | 0.592075227 | 0.795757997 | -4.709835951 |
| **A_55_P2003369** | Fkbp8 | 0.104518127 | -0.355806392 | 0.564842646 | 11.84115916 | 0.568692964 | 0.592095566 | 0.795757997 | -4.709841485 |
| **A_55_P1989578** | NA | -0.074540295 | -0.402839757 | 0.253759167 | 8.844978411 | -0.568684357 | 0.592101016 | 0.795757997 | -4.709842968 |
| **A_55_P1955568** | Extl2 | 0.066650908 | -0.226907103 | 0.360208919 | 10.18084953 | 0.56867283 | 0.592108315 | 0.795757997 | -4.709844954 |
| **A_55_P2062568** | Caprin2 | 0.09591505 | -0.326546215 | 0.518376315 | 9.627597695 | 0.568656761 | 0.59211849 | 0.795757997 | -4.709847722 |
| **A_51_P477456** | Cnot10 | 0.075347588 | -0.256543214 | 0.40723839 | 9.610696024 | 0.568623082 | 0.592139815 | 0.795757997 | -4.709853524 |
| **A_51_P374305** | Rchy1 | 0.111857011 | -0.380860218 | 0.604574239 | 11.02325952 | 0.568611719 | 0.592147011 | 0.795757997 | -4.709855481 |
| **A_55_P2055875** | Josd2 | 0.106823971 | -0.363970835 | 0.577618778 | 9.139017443 | 0.568312753 | 0.59233634 | 0.795957103 | -4.70990697 |
| **A_55_P2011286** | Hopx | 0.621584527 | -2.119256702 | 3.362425756 | 13.03713298 | 0.568023969 | 0.592519256 | 0.796106926 | -4.709956685 |
| **A_55_P2117169** | Zmym1 | -0.168400678 | -0.910976345 | 0.57417499 | 9.993518629 | -0.56800671 | 0.592530189 | 0.796106926 | -4.709959656 |
| **A_55_P2143209** | Axin1 | -0.100835114 | -0.545762325 | 0.344092097 | 9.420491372 | -0.567640236 | 0.592762364 | 0.796363528 | -4.710022714 |
| **A_55_P2030128** | Tfpt | -0.054226028 | -0.293632094 | 0.185180037 | 8.302631873 | -0.567313338 | 0.592969511 | 0.796488357 | -4.710078935 |
| **A_51_P204845** | Celf1 | 0.108197361 | -0.369501859 | 0.585896581 | 10.94439775 | 0.567299604 | 0.592978215 | 0.796488357 | -4.710081297 |
| **A_55_P2177712** | NA | 0.073275888 | -0.250299771 | 0.396851547 | 8.262260945 | 0.567199176 | 0.593041863 | 0.796488357 | -4.710098563 |
| **A_66_P105843** | Cyth2 | -0.069268915 | -0.375152762 | 0.236614931 | 12.46671078 | -0.567194734 | 0.593044678 | 0.796488357 | -4.710099327 |
| **A_51_P481159** | Cbr3 | 0.272781951 | -0.931848448 | 1.47741235 | 10.35134108 | 0.567168573 | 0.593061259 | 0.796488357 | -4.710103824 |
| **A_55_P1996469** | Zfp592 | -0.099049451 | -0.53654316 | 0.338444259 | 11.29977038 | -0.567062076 | 0.593128759 | 0.79652368 | -4.71012213 |

| | | | | | | | | | |
|---|---|---|---|---|---|---|---|---|---|
| **A_55_P1990190** | Arhgap28 | -0.102376758 | -0.554721778 | 0.349968263 | 8.333376229 | -0.566867949 | 0.593251813 | 0.796633598 | -4.710155492 |
| **A_55_P1991770** | Pdlim4 | -0.508880728 | -2.758142596 | 1.74038114 | 10.75119115 | -0.566664921 | 0.593380525 | 0.796712836 | -4.710190374 |
| **A_66_P102706** | 2700012I2 | 0.064924445 | -0.222177947 | 0.352026838 | 8.367978341 | 0.566398091 | 0.59354971 | 0.796712836 | -4.710236203 |
| **A_55_P2182700** | Prkab1 | -0.048789977 | -0.264545766 | 0.166965813 | 8.121509192 | -0.566393808 | 0.593552426 | 0.796712836 | -4.710236938 |
| **A_55_P2010552** | Hsd17b13 | 0.047673715 | -0.163174816 | 0.258522247 | 7.689374129 | 0.566315914 | 0.59360182 | 0.796712836 | -4.710250313 |
| **A_51_P284686** | Aak1 | -0.049737516 | -0.269717442 | 0.17024241 | 7.879449414 | -0.566306289 | 0.593607924 | 0.796712836 | -4.710251966 |
| **A_55_P2008884** | Cltc | -0.11325005 | -0.614156747 | 0.387656646 | 13.86373386 | -0.566280896 | 0.593624027 | 0.796712836 | -4.710256326 |
| **A_55_P2134690** | Ankrd26 | 0.049453783 | -0.169281732 | 0.268189298 | 8.341517365 | 0.56627914 | 0.593625141 | 0.796712836 | -4.710256627 |
| **A_55_P2180086** | Lrrc28 | -0.098918639 | -0.5364984 | 0.338661122 | 10.20206217 | -0.566201809 | 0.593674182 | 0.796712836 | -4.710269903 |
| **A_55_P2087984** | Dnaja1 | -0.122509166 | -0.664456093 | 0.419437762 | 14.39804666 | -0.566189967 | 0.593681692 | 0.796712836 | -4.710271936 |
| **A_52_P432969** | Pcdh19 | 0.060380994 | -0.206785279 | 0.327547267 | 8.579415559 | 0.566068476 | 0.593758743 | 0.796760935 | -4.71029279 |
| **A_51_P350332** | Rbpms | 0.157346872 | -0.538942865 | 0.853636609 | 9.545322935 | 0.566002653 | 0.593800492 | 0.796761657 | -4.710304088 |
| **A_55_P2043277** | Siglecf | 0.050247628 | -0.172143361 | 0.272638617 | 7.751753549 | 0.565911764 | 0.593858142 | 0.79677533 | -4.710319685 |
| **A_55_P2087399** | Tex22 | -0.053581366 | -0.290750249 | 0.183587517 | 7.710143997 | -0.565856645 | 0.593893105 | 0.79677533 | -4.710329143 |
| **A_51_P341177** | Ddah2 | 0.111651633 | -0.382637816 | 0.605941082 | 13.82121076 | 0.565762403 | 0.593952887 | 0.796780723 | -4.710345313 |
| **A_55_P2146136** | Gla | -0.12915743 | -0.701029856 | 0.442714995 | 15.70366301 | -0.565679689 | 0.594005359 | 0.796780723 | -4.710359502 |
| **A_51_P293069** | Flvcr1 | -0.194744455 | -1.057199546 | 0.667710636 | 9.079707593 | -0.565560351 | 0.594081069 | 0.796780723 | -4.710379972 |
| **A_66_P110490** | Polr3h | -0.114776121 | -0.62310392 | 0.393551678 | 9.002259111 | -0.565533098 | 0.59409836 | 0.796780723 | -4.710384646 |
| **A_51_P288505** | Tradd | -0.067713638 | -0.367612141 | 0.232184866 | 11.37670169 | -0.5655255 | 0.594103181 | 0.796780723 | -4.71038595 |
| **A_51_P473242** | Zfp426 | 0.084344124 | -0.28926828 | 0.457956528 | 8.926757498 | 0.565436569 | 0.594159605 | 0.796801125 | -4.710401201 |
| **A_55_P2075884** | Senp8 | 0.059897389 | -0.205537942 | 0.325332719 | 8.159796122 | 0.565196556 | 0.594311904 | 0.796950087 | -4.710442352 |
| **A_55_P2028029** | Abcb11 | -0.056310169 | -0.305897396 | 0.193277059 | 7.733218819 | -0.56508635 | 0.594381843 | 0.796988594 | -4.710461242 |
| **A_55_P1989048** | Tm9sf1 | -0.109164409 | -0.593164425 | 0.374835607 | 10.26272839 | -0.564918812 | 0.594488174 | 0.797075891 | -4.710489954 |
| **A_55_P2076792** | Anxa6 | 0.340027269 | -1.167967658 | 1.848022196 | 13.54033056 | 0.564760368 | 0.594588743 | 0.797083156 | -4.710517101 |
| **A_55_P2190152** | 4921509J | 0.078276519 | -0.268878881 | 0.425431918 | 8.082802939 | 0.564752188 | 0.594593936 | 0.797083156 | -4.710518503 |
| **A_52_P158211** | NA | 0.097410515 | -0.334721176 | 0.529542206 | 9.553019494 | 0.564598954 | 0.594691209 | 0.797083156 | -4.710544751 |
| **A_51_P424499** | Txn1 | 0.060259305 | -0.207107541 | 0.327626151 | 14.77153486 | 0.564503853 | 0.594751584 | 0.797083156 | -4.710561039 |
| **A_55_P2033521** | Med6 | -0.057837134 | -0.314458336 | 0.198784069 | 9.790043804 | -0.564500814 | 0.594753513 | 0.797083156 | -4.710561559 |
| **A_55_P2104810** | Mrpl27 | 0.136547895 | -0.469346369 | 0.742442159 | 12.21620137 | 0.564466944 | 0.594775016 | 0.797083156 | -4.710567359 |
| **A_55_P1955577** | Ccm2 | -0.146474346 | -0.796455981 | 0.503507289 | 12.67289864 | -0.564430959 | 0.594797863 | 0.797083156 | -4.710573521 |
| **A_66_P103198** | NA | 0.057356006 | -0.197194324 | 0.311906336 | 8.007564907 | 0.564359167 | 0.594843444 | 0.797083156 | -4.710585814 |
| **A_55_P1987988** | Atg2b | -0.080988355 | -0.440455408 | 0.278478699 | 8.249369866 | -0.564304911 | 0.594877893 | 0.797083156 | -4.710595103 |
| **A_55_P2059556** | NA | 0.068662927 | -0.236121472 | 0.373447327 | 16.00032673 | 0.564260862 | 0.594905863 | 0.797083156 | -4.710602644 |
| **A_55_P2088038** | Vps13d | 0.048920323 | -0.168263667 | 0.266104313 | 9.75816709 | 0.564172424 | 0.594962019 | 0.797096753 | -4.710617783 |
| **A_51_P486610** | Afg3l1 | -0.115025448 | -0.625738089 | 0.395687194 | 8.285986199 | -0.56411503 | 0.594998466 | 0.797096753 | -4.710627607 |
| **A_66_P120414** | Adcy5 | 0.090291441 | -0.310924176 | 0.491507057 | 8.158494509 | 0.563662443 | 0.59528591 | 0.797292701 | -4.710705046 |
| **A_55_P1997421** | Csta3 | -0.074963666 | -0.408072305 | 0.258144973 | 7.990935772 | -0.56365747 | 0.595289069 | 0.797292701 | -4.710705897 |
| **A_52_P64364** | Ikbkb | -0.05254846 | -0.286073142 | 0.180976223 | 8.358836864 | -0.563608517 | 0.595320165 | 0.797292701 | -4.71071427 |
| **A_55_P1989013** | Eif6 | 0.069490521 | -0.239386889 | 0.378367931 | 10.88117207 | 0.563494608 | 0.595392526 | 0.797292701 | -4.71073375 |
| **A_55_P2001861** | NA | 0.195051746 | -0.671950015 | 1.062053507 | 9.908655527 | 0.563482208 | 0.595400403 | 0.797292701 | -4.710735871 |
| **A_52_P454183** | Olfml2b | -0.102735785 | -0.559435726 | 0.353964156 | 8.497804368 | -0.563431512 | 0.59543261 | 0.797292701 | -4.71074454 |
| **A_51_P161021** | Ifit2 | -0.055621763 | -0.302882364 | 0.191638838 | 8.310912248 | -0.563430271 | 0.595433398 | 0.797292701 | -4.710744752 |
| **A_55_P2083396** | Radil | -0.119786185 | -0.653034026 | 0.413461657 | 8.242844424 | -0.56263659 | 0.595937749 | 0.797912772 | -4.710880386 |
| **A_65_P07569** | Rock1 | 0.127707869 | -0.441019467 | 0.696435205 | 9.161252458 | 0.562424073 | 0.596072837 | 0.797945574 | -4.710916677 |
| **A_66_P126313** | Rhof | -0.272674639 | -1.487008353 | 0.941659076 | 10.11073489 | -0.562415187 | 0.596078486 | 0.797945574 | -4.710918194 |
| **A_52_P238816** | Ppt2 | -0.060423573 | -0.329527068 | 0.208679922 | 9.525431888 | -0.562389769 | 0.596094644 | 0.797945574 | -4.710922534 |
| **A_55_P1993549** | Npnt | 0.266868514 | -0.921887181 | 1.455624209 | 9.014067911 | 0.562283155 | 0.596162422 | 0.797945574 | -4.710940735 |
| **A_52_P423814** | Cox8b | 0.453503418 | -1.566645333 | 2.473652169 | 9.664110386 | 0.562273429 | 0.596168606 | 0.797945574 | -4.710942395 |

| | | | | | | | | | |
|---|---|---|---|---|---|---|---|---|---|
| **A_55_P2033616** | Dhrs7b | -0.073758042 | -0.402362679 | 0.254846595 | 11.25768157 | -0.562193778 | 0.596219246 | 0.797958113 | -4.710955991 |
| **A_51_P260504** | Arhgef4 | 0.066331383 | -0.22932016 | 0.361982925 | 8.179114979 | 0.561939095 | 0.596381185 | 0.798085371 | -4.710999454 |
| **A_55_P2069460** | NA | 0.158493464 | -0.547972983 | 0.864959911 | 16.38123028 | 0.5619144 | 0.596396888 | 0.798085371 | -4.711003667 |
| **A_51_P110035** | Pik3r4 | -0.055111957 | -0.300881816 | 0.190657902 | 9.743626628 | -0.561652321 | 0.596563559 | 0.798253156 | -4.709264786 |
| **A_55_P2227321** | Ptprd | 0.155815761 | -0.53921705 | 0.850848573 | 8.3938421 | 0.561508615 | 0.596654962 | 0.79832021 | -4.711072881 |
| **A_66_P119518** | Tuba8 | 0.374094492 | -1.295336246 | 2.04352523 | 11.00008152 | 0.561258917 | 0.596813799 | 0.798451852 | -4.711115451 |
| **A_55_P2059318** | Dgkz | 0.201417979 | -0.697483851 | 1.10031981 | 11.17439373 | 0.561224105 | 0.596835946 | 0.798451852 | -4.711121384 |
| **A_55_P1975045** | Sgcg | -0.054296466 | -0.296776701 | 0.188183769 | 7.862673483 | -0.560848501 | 0.597074925 | 0.798610553 | -4.711185387 |
| **A_51_P503192** | Rbm15b | -0.140111418 | -0.765867374 | 0.485644539 | 9.210130719 | -0.560814044 | 0.597096851 | 0.798610553 | -4.711191257 |
| **A_51_P421664** | Tmem123 | -0.058989938 | -0.322451 | 0.204471123 | 11.49971027 | -0.560804967 | 0.597102628 | 0.798610553 | -4.711192803 |
| **A_66_P116804** | Smchd1 | -0.15505088 | -0.847572961 | 0.5374712 | 10.82075199 | -0.560777987 | 0.597119797 | 0.798610553 | -4.711197399 |
| **A_55_P1980986** | Lca5l | 0.104209693 | -0.361379396 | 0.569798781 | 8.6428369 | 0.560603322 | 0.597230952 | 0.798646008 | -4.711227147 |
| **A_51_P309184** | Pxmp2 | -0.157731187 | -0.862593113 | 0.547130739 | 8.945974053 | -0.560484832 | 0.597306365 | 0.798646008 | -4.711247323 |
| **A_55_P2100968** | Dnah7b | 0.291965307 | -1.012877634 | 1.596808249 | 9.095391503 | 0.560432554 | 0.597339639 | 0.798646008 | -4.711256224 |
| **A_55_P2038867** | Mcat | 0.085297198 | -0.295932862 | 0.466527258 | 11.06246918 | 0.560399801 | 0.597360487 | 0.798646008 | -4.7112618 |
| **A_55_P2002577** | Ephx1 | 0.057864524 | -0.200768356 | 0.316497405 | 8.429653198 | 0.560375313 | 0.597376074 | 0.798646008 | -4.711265969 |
| **A_55_P1970740** | Mrps10 | 0.08393273 | -0.291256755 | 0.459122215 | 11.25928285 | 0.560313435 | 0.597415461 | 0.798646008 | -4.711276503 |
| **A_55_P2021892** | Ccdc114 | 0.050419259 | -0.174973584 | 0.275812102 | 9.040965823 | 0.560282015 | 0.597435461 | 0.798646008 | -4.711281851 |
| **A_55_P1961761** | Dcc | -0.12753118 | -0.725936839 | 0.47087448 | 11.89523884 | -0.567653593 | 0.597539068 | 0.798667831 | -4.68775189 |
| **A_55_P2130660** | Slitrk4 | 0.055671288 | -0.193281634 | 0.304624211 | 7.779499187 | 0.560098473 | 0.597552303 | 0.798667831 | -4.711313088 |
| **A_52_P177847** | Tril | 0.192992439 | -0.670095618 | 1.056080496 | 8.586854845 | 0.560061261 | 0.597575994 | 0.798667831 | -4.71131942 |
| **A_51_P346575** | Rhag | 0.048333845 | -0.167846462 | 0.264514152 | 7.719745099 | 0.559996817 | 0.597617022 | 0.798667831 | -4.711330385 |
| **A_55_P2000384** | Pcnx3 | -0.073886093 | -0.404450145 | 0.256677959 | 11.77644488 | -0.559831621 | 0.597722203 | 0.798753184 | -4.711358489 |
| **A_55_P1975185** | Sqle | 0.126961895 | -0.441153146 | 0.695076935 | 11.99809077 | 0.55974143 | 0.597779633 | 0.79877472 | -4.71137383 |
| **A_55_P1954116** | C8b | 0.048588592 | -0.168928056 | 0.26610524 | 7.695835468 | 0.559489777 | 0.597939891 | 0.798911113 | -4.711416623 |
| **A_55_P2114467** | Romo1 | -0.064549566 | -0.353538578 | 0.224439445 | 13.82496943 | -0.559451375 | 0.597964348 | 0.798911113 | -4.711423152 |
| **A_55_P2004347** | NA | 0.090177133 | -0.313652922 | 0.494007189 | 12.54765008 | 0.559304267 | 0.598058044 | 0.798981082 | -4.711448159 |
| **A_52_P577662** | Ednrb | 0.20821273 | -0.724433479 | 1.140858938 | 10.10338999 | 0.55916592 | 0.598146167 | 0.799028068 | -4.711471672 |
| **A_52_P652513** | Lman1 | -0.149061587 | -0.816808071 | 0.518684896 | 13.02928378 | -0.559119293 | 0.598175869 | 0.799028068 | -4.711479595 |
| **A_52_P603654** | Stx8 | -0.059201283 | -0.324492513 | 0.206089947 | 9.567804432 | -0.55893148 | 0.598295517 | 0.799080803 | -4.711511505 |
| **A_52_P29151** | Ankrd44 | 0.126076981 | -0.43896352 | 0.691117481 | 10.2149228 | 0.558864556 | 0.598338155 | 0.799080803 | -4.711522874 |
| **A_52_P212686** | Lrrk1 | 0.114382366 | -0.398401373 | 0.627166104 | 8.668487265 | 0.558695503 | 0.598445867 | 0.799080803 | -4.711551586 |
| **A_55_P2033586** | Mmab | 0.04729632 | -0.16474621 | 0.25933885 | 8.240768851 | 0.558669188 | 0.598462635 | 0.799080803 | -4.711556055 |
| **A_55_P2151308** | G430095F | 0.091920879 | -0.320250879 | 0.504092637 | 8.155766307 | 0.558581159 | 0.598518729 | 0.799080803 | -4.711571003 |
| **A_55_P2175545** | Gys2 | 0.046579618 | -0.162318564 | 0.255477801 | 7.678870415 | 0.558485122 | 0.598579929 | 0.799080803 | -4.711587308 |
| **A_55_P2024075** | Prkacb | 0.048333235 | -0.168456434 | 0.265122903 | 12.58273203 | 0.558415709 | 0.598624165 | 0.799080803 | -4.711599092 |
| **A_55_P1990457** | Rpl15 | 0.161858183 | -0.56415376 | 0.887870126 | 16.12023475 | 0.55839464 | 0.598637592 | 0.799080803 | -4.711602668 |
| **A_55_P2019483** | Ndrg4 | -0.140522905 | -0.770852859 | 0.489807049 | 10.43291632 | -0.558379569 | 0.598647197 | 0.799080803 | -4.711605226 |
| **A_55_P2023747** | Mfsd10 | -0.082418797 | -0.452138415 | 0.28730082 | 11.13736954 | -0.558346918 | 0.598668006 | 0.799080803 | -4.711610769 |
| **A_52_P222230** | Gm40812 | 0.150138662 | -0.52356331 | 0.823840634 | 9.211773102 | 0.558181025 | 0.59877374 | 0.799080803 | -4.711638923 |
| **A_51_P372463** | Fxr1 | -0.087592729 | -0.480678821 | 0.305493363 | 11.9319356 | -0.558124036 | 0.598810065 | 0.799080803 | -4.711648594 |
| **A_55_P2039210** | Cpt1b | -0.074066186 | -0.406454646 | 0.258322274 | 8.021209392 | -0.558115895 | 0.598815254 | 0.799080803 | -4.711649975 |
| **A_55_P2033863** | Psmc1 | -0.057579778 | -0.316020666 | 0.20086111 | 13.97090288 | -0.558032001 | 0.598868731 | 0.799080803 | -4.711664209 |
| **A_51_P208240** | Tnfsf14 | 0.051344651 | -0.179124269 | 0.281813572 | 7.758544686 | 0.5579987 | 0.59888996 | 0.799080803 | -4.711669859 |
| **A_51_P345747** | E130309D | -0.155272631 | -0.852386921 | 0.541841659 | 8.903482508 | -0.557880617 | 0.598965236 | 0.799080803 | -4.71168989 |
| **A_66_P111534** | 5430431A | 0.187736977 | -0.65521056 | 1.030684514 | 8.772505923 | 0.557827114 | 0.598999346 | 0.799080803 | -4.711698965 |
| **A_51_P219444** | Plcl2 | -0.157677347 | -0.865723059 | 0.550368364 | 9.904243755 | -0.557774113 | 0.599033136 | 0.799080803 | -4.711707953 |
| **A_55_P2031979** | Sspn | 0.244046114 | -0.851836651 | 1.33992888 | 10.50236147 | 0.557773846 | 0.599033306 | 0.799080803 | -4.711707999 |

| | | | | | | | | | |
|---|---|---|---|---|---|---|---|---|---|
| **A_55_P2069350** | Psph | -0.227616239 | -1.249745931 | 0.794513453 | 10.60115557 | -0.557760281 | 0.599041955 | 0.799080803 | -4.711710299 |
| **A_55_P2425437** | D030036F | -0.052027662 | -0.285705543 | 0.181650218 | 9.751835576 | -0.557656865 | 0.599107891 | 0.799095615 | -4.711727836 |
| **A_55_P1953972** | Pdhb | -0.056139153 | -0.308303188 | 0.196024881 | 13.49458497 | -0.557613217 | 0.599135721 | 0.799095615 | -4.711735237 |
| **A_55_P1960738** | Gm4470 | 0.062790923 | -0.219331518 | 0.344913363 | 8.748694922 | 0.557454696 | 0.599236802 | 0.799169333 | -4.711762112 |
| **A_55_P1965478** | Kcnd1 | 0.086840346 | -0.303397919 | 0.477078611 | 9.27002221 | 0.557368008 | 0.599292083 | 0.799169333 | -4.711776806 |
| **A_55_P1993864** | Sh2b3 | 0.121668899 | -0.425176361 | 0.668514159 | 9.174914256 | 0.557269752 | 0.599354745 | 0.799169333 | -4.711793458 |
| **A_55_P2145029** | Gm826 | 0.05085912 | -0.177737225 | 0.279455465 | 7.800339224 | 0.557249779 | 0.599367482 | 0.799169333 | -4.711796843 |
| **A_55_P2035504** | Nt5c3 | 0.075392647 | -0.280288827 | 0.431074121 | 10.72794739 | 0.564586449 | 0.599455022 | 0.799169333 | -4.688202695 |
| **A_66_P101473** | Fbxl3 | -0.107843704 | -0.592768377 | 0.377080969 | 9.645048314 | -0.557020092 | 0.59951398 | 0.799169333 | -4.711835761 |
| **A_55_P2291776** | 4833447P | -0.079335452 | -0.436093726 | 0.277422822 | 8.306461007 | -0.556985115 | 0.59953629 | 0.799169333 | -4.711841686 |
| **A_55_P2085170** | 6-Sep | 0.111567597 | -0.390148542 | 0.613283735 | 8.339164533 | 0.556968139 | 0.599547119 | 0.799169333 | -4.711844561 |
| **A_55_P2235931** | Cacna1c | 0.340963104 | -1.192406328 | 1.874332535 | 9.840994491 | 0.556943231 | 0.599563007 | 0.799169333 | -4.711848781 |
| **A_55_P2030702** | NA | 0.19798836 | -0.692584162 | 1.088560883 | 10.30932939 | 0.556827548 | 0.599636802 | 0.799178457 | -4.711868374 |
| **A_55_P2046802** | Mlkl | -0.056389427 | -0.310046242 | 0.197267388 | 8.493303947 | -0.556802905 | 0.599652523 | 0.799178457 | -4.711872548 |
| **A_65_P19784** | Asb7 | 0.091447489 | -0.320049706 | 0.502944684 | 10.23555668 | 0.556615441 | 0.599772121 | 0.799234166 | -4.711904291 |
| **A_51_P351896** | Gask1b | 0.210959714 | -0.738333967 | 1.160253395 | 8.875501799 | 0.556607794 | 0.599777 | 0.799234166 | -4.711905586 |
| **A_55_P2163344** | NA | 0.058849052 | -0.206106044 | 0.323804147 | 8.984434625 | 0.556310859 | 0.599966468 | 0.799431543 | -4.711955847 |
| **A_55_P1973347** | A130040N | -0.124977703 | -0.687736524 | 0.437781119 | 10.65212722 | -0.556237894 | 0.60001303 | 0.79943849 | -4.711968195 |
| **A_51_P139108** | Cpxm1 | -0.305827441 | -1.683102284 | 1.071447402 | 10.53606423 | -0.556168236 | 0.600057485 | 0.799442628 | -4.711979981 |
| **A_55_P2107745** | Yy1 | 0.09295829 | -0.325846218 | 0.511762798 | 11.10400136 | 0.555938976 | 0.600203807 | 0.799513619 | -4.712018764 |
| **A_55_P2075080** | Fat2 | 0.126898259 | -0.444857343 | 0.698653861 | 8.728502129 | 0.555898599 | 0.60022958 | 0.799513619 | -4.712025593 |
| **A_55_P1993599** | Oaz2 | 0.260864752 | -0.914643398 | 1.436372902 | 15.25191456 | 0.555827592 | 0.600274905 | 0.799513619 | -4.712037602 |
| **A_55_P1952915** | Sh3bgr | 0.337022095 | -1.181678519 | 1.855722709 | 9.948147226 | 0.555823047 | 0.600277806 | 0.799513619 | -4.712038371 |
| **A_55_P2046943** | Bnip3l | 0.066650186 | -0.233742272 | 0.367042643 | 12.70134426 | 0.555728519 | 0.600338148 | 0.799513619 | -4.712054355 |
| **A_55_P2068248** | NA | 0.194105024 | -0.680777671 | 1.068987718 | 11.34463841 | 0.555696031 | 0.600358887 | 0.799513619 | -4.712059849 |
| **A_66_P109315** | Snapc2 | 0.06742669 | -0.236520122 | 0.371373502 | 11.64848657 | 0.555628602 | 0.600401934 | 0.799515875 | -4.712071249 |
| **A_52_P479500** | Npepps | -0.070650935 | -0.389244077 | 0.247942207 | 10.95589087 | -0.555433199 | 0.600526688 | 0.799599876 | -4.712104279 |
| **A_51_P199567** | Med26 | 0.067739119 | -0.237741613 | 0.373219851 | 10.34644981 | 0.555400244 | 0.600547729 | 0.799599876 | -4.712109849 |
| **A_55_P2058992** | Vamp4 | -0.047017811 | -0.259171171 | 0.165135548 | 8.653594992 | -0.555089283 | 0.600746297 | 0.799763291 | -4.712162391 |
| **A_51_P138044** | Foxo1 | 0.126303634 | -0.443665997 | 0.696273264 | 10.75030251 | 0.555027466 | 0.600785776 | 0.799763291 | -4.712172833 |
| **A_55_P2108738** | Nck1 | 0.129519781 | -0.454984768 | 0.71402433 | 11.02140906 | 0.555007117 | 0.600798772 | 0.799763291 | -4.71217627 |
| **A_55_P2031357** | Zfpm2 | 0.124295004 | -0.436749667 | 0.685339675 | 8.369518443 | 0.554889588 | 0.600873835 | 0.799763291 | -4.71219612 |
| **A_55_P2067505** | Slc16a3 | -0.557856719 | -3.075943388 | 1.96022995 | 10.79853151 | -0.554884176 | 0.600877292 | 0.799763291 | -4.712197034 |
| **A_52_P352735** | Ccdc153 | 0.622001653 | -2.186640076 | 3.430643382 | 10.59928037 | 0.554683874 | 0.601005233 | 0.799878514 | -4.712230856 |
| **A_55_P2007022** | Adgre5 | 0.264661433 | -0.930637662 | 1.459960529 | 13.0983274 | 0.55458027 | 0.601071416 | 0.799908112 | -4.712248346 |
| **A_55_P2079981** | NA | 0.099057016 | -0.348369167 | 0.546483199 | 8.555988491 | 0.554516141 | 0.601112384 | 0.799908112 | -4.71225917 |
| **A_55_P2135750** | NA | 0.077722095 | -0.273376201 | 0.428820391 | 8.379531577 | 0.55445477 | 0.601151591 | 0.799908112 | -4.712269528 |
| **A_55_P2046393** | Rcn3 | 0.14300529 | -0.503601446 | 0.789612027 | 13.93223049 | 0.553939359 | 0.601480927 | 0.800271827 | -4.712356481 |
| **A_55_P2020796** | Man1b1 | -0.078497609 | -0.433455976 | 0.276460757 | 12.33721495 | -0.553897441 | 0.601507716 | 0.800271827 | -4.71236355 |
| **A_55_P2066697** | Trim47 | 0.183495437 | -0.646354885 | 1.013345758 | 11.26554068 | 0.553829197 | 0.601551331 | 0.800274784 | -4.712375058 |
| **A_55_P2097972** | NA | -0.107957825 | -0.596307214 | 0.380391563 | 8.518205637 | -0.553699109 | 0.601634477 | 0.800294669 | -4.712396991 |
| **A_52_P580582** | Nppa | 1.228700216 | -4.329578962 | 6.786979393 | 12.79346275 | 0.553676287 | 0.601649065 | 0.800294669 | -4.704773595 |
| **A_55_P1976470** | 2410018L | -0.119623227 | -0.660978319 | 0.421731865 | 8.398769343 | -0.55345668 | 0.601789443 | 0.800425291 | -4.712437853 |
| **A_51_P196526** | Bscl2 | -0.064335202 | -0.355518625 | 0.22684822 | 8.516064654 | -0.553391354 | 0.601831205 | 0.800425291 | -4.712448861 |
| **A_55_P2124871** | Npw | 0.154575326 | -0.545186652 | 0.854337304 | 8.354009926 | 0.55327389 | 0.601906302 | 0.800425291 | -4.712468653 |
| **A_51_P407999** | 1500011B | -0.06931911 | -0.383191016 | 0.244552795 | 9.40703913 | -0.553160146 | 0.601979026 | 0.800425291 | -4.712487815 |
| **A_55_P2000158** | Pik3r3 | -0.149470237 | -0.826286348 | 0.527345873 | 10.43185066 | -0.55313913 | 0.601992463 | 0.800425291 | -4.712491355 |
| **A_55_P2073579** | Chchd5 | -0.055948361 | -0.3092904 | 0.197393678 | 9.958392669 | -0.553134125 | 0.601995664 | 0.800425291 | -4.712492198 |

| | | | | | | | | | |
|---|---|---|---|---|---|---|---|---|---|
| **A_55_P1975080** | Ugt2b5 | 0.046485556 | -0.164102647 | 0.25707376 | 7.632584631 | 0.552884403 | 0.602155347 | 0.800582553 | -4.712534254 |
| **A_51_P157462** | Rgn | 0.106126352 | -0.374787207 | 0.587039912 | 8.352439085 | 0.552721757 | 0.602259364 | 0.800665788 | -4.712561637 |
| **A_52_P477819** | Hmox2 | 0.098497135 | -0.347955239 | 0.544949509 | 12.10078135 | 0.55258464 | 0.602347063 | 0.800704656 | -4.712584717 |
| **A_55_P2244142** | Klf13 | -0.056822075 | -0.314394025 | 0.200749876 | 10.68287297 | -0.552546544 | 0.60237143 | 0.800704656 | -4.712591128 |
| **A_55_P2164957** | Slc9a3r2 | 0.116440101 | -0.411659799 | 0.644540001 | 8.807395645 | 0.55225141 | 0.602560224 | 0.800892995 | -4.712640787 |
| **A_51_P430900** | Dusp1 | -0.082998325 | -0.459498873 | 0.293502223 | 9.85119437 | -0.552146168 | 0.602627554 | 0.800892995 | -4.71265849 |
| **A_52_P247943** | Parl | 0.053740359 | -0.190065639 | 0.297546357 | 8.491041443 | 0.55208573 | 0.602666223 | 0.800892995 | -4.712668655 |
| **A_51_P292057** | Tbl1xr1 | -0.119114954 | -0.659574316 | 0.421344408 | 8.873086226 | -0.552018444 | 0.602709274 | 0.800892995 | -4.71267997 |
| **A_52_P626756** | Trnt1 | 0.056827887 | -0.201024849 | 0.314680622 | 9.884698348 | 0.552001308 | 0.602720238 | 0.800892995 | -4.712682852 |
| **A_51_P193686** | 1700012B | 0.396772895 | -1.403845379 | 2.19739117 | 9.496274892 | 0.551913082 | 0.60277669 | 0.800912964 | -4.712697687 |
| **A_55_P2078257** | NA | -0.078297283 | -0.43374876 | 0.277154195 | 8.285638681 | -0.551717441 | 0.602901885 | 0.800941438 | -4.712730577 |
| **A_66_P136259** | Rbm4b | 0.076291042 | -0.270073507 | 0.422655591 | 8.350505599 | 0.551684085 | 0.602923232 | 0.800941438 | -4.712736184 |
| **A_51_P137419** | Cst7 | 0.049729448 | -0.176049157 | 0.275508052 | 7.722698314 | 0.551672324 | 0.602930759 | 0.800941438 | -4.712738161 |
| **A_55_P1952828** | NA | -0.052828173 | -0.29269789 | 0.187041543 | 8.40200509 | -0.551620654 | 0.602963827 | 0.800941438 | -4.712746845 |
| **A_55_P2175190** | Cep250 | 0.074581742 | -0.264145356 | 0.41330884 | 8.7018816 | 0.551484011 | 0.603051282 | 0.801002575 | -4.712769808 |
| **A_51_P511060** | Csgalnact | 0.202439156 | -0.717313608 | 1.12219192 | 10.46137936 | 0.551281935 | 0.603180629 | 0.801119343 | -4.712803758 |
| **A_66_P130759** | 4631405J | -0.062939377 | -0.34900395 | 0.223125195 | 7.821690488 | -0.551072464 | 0.603314726 | 0.80120657 | -4.712838939 |
| **A_51_P518488** | Atpif1 | 0.103901441 | -0.368358388 | 0.576161271 | 15.95001908 | 0.55104988 | 0.603329185 | 0.80120657 | -4.712842732 |
| **A_51_P486068** | Zfp280b | -0.137286623 | -0.761378517 | 0.48680527 | 9.286317069 | -0.550972633 | 0.603378641 | 0.801217215 | -4.712855702 |
| **A_52_P93933** | Mcl1 | -0.0687328 | -0.381345711 | 0.243880111 | 11.24534411 | -0.55069036 | 0.603559383 | 0.801402178 | -4.712903087 |
| **A_55_P1965902** | Pi4k2a | -0.144588996 | -0.802466328 | 0.513288337 | 9.712075685 | -0.550478884 | 0.603694814 | 0.801460696 | -4.712938574 |
| **A_51_P185869** | Hbq1a | 0.052057595 | -0.184848887 | 0.288964078 | 7.896009818 | 0.550373486 | 0.603762318 | 0.801460696 | -4.712956257 |
| **A_55_P2106745** | Trim63 | 0.050717265 | -0.180096968 | 0.281531497 | 7.73281431 | 0.550355843 | 0.603773618 | 0.801460696 | -4.712959216 |
| **A_52_P485850** | Mgam | -0.048323758 | -0.268260675 | 0.171613159 | 7.684560322 | -0.550316984 | 0.603798508 | 0.801460696 | -4.712965735 |
| **A_55_P1970846** | Pemt | 0.049549576 | -0.175974244 | 0.275073396 | 7.92523648 | 0.550297915 | 0.603810722 | 0.801460696 | -4.712968933 |
| **A_51_P447976** | Tent5c | -0.191594857 | -1.063832609 | 0.680642894 | 11.47530444 | -0.550173051 | 0.603890702 | 0.80151183 | -4.712989875 |
| **A_55_P2119957** | Mlxipl | 0.064901233 | -0.230660861 | 0.360463328 | 8.179198804 | 0.549989703 | 0.604008156 | 0.801590309 | -4.71302062 |
| **A_55_P2131979** | 1-Mar | -0.163554088 | -0.908436135 | 0.581327958 | 10.64766894 | -0.549951312 | 0.604032751 | 0.801590309 | -4.713027056 |
| **A_55_P2089310** | 2900097C | -0.091144248 | -0.506341484 | 0.324052988 | 8.56408543 | -0.549825855 | 0.604113129 | 0.80159071 | -4.713048087 |
| **A_55_P2019585** | Klhl17 | 0.20227729 | -0.719256494 | 1.123811073 | 10.01588785 | 0.549776549 | 0.604144721 | 0.80159071 | -4.713056351 |
| **A_52_P117999** | Selenot | -0.067455671 | -0.374780944 | 0.239869602 | 8.455769824 | -0.549756707 | 0.604157434 | 0.80159071 | -4.713059677 |
| **A_55_P2179216** | Man1a2 | -0.120610318 | -0.670329002 | 0.429108366 | 9.149901395 | -0.549533675 | 0.604300347 | 0.801715837 | -4.71309705 |
| **A_51_P246854** | Acta1 | 0.260842565 | -0.928273459 | 1.449958588 | 9.917163279 | 0.549420141 | 0.604373105 | 0.801715837 | -4.713116071 |
| **A_52_P222624** | Hspb2 | -0.179765243 | -0.999414939 | 0.639884454 | 10.39868971 | -0.549323006 | 0.604435357 | 0.801715837 | -4.713132341 |
| **A_55_P1967613** | NA | -0.054901709 | -0.305288452 | 0.195485034 | 7.954753614 | -0.549192853 | 0.604518777 | 0.801715837 | -4.713154139 |
| **A_55_P2033041** | Sirpb1b | -0.139114301 | -0.773602172 | 0.49537357 | 8.421599987 | -0.549159875 | 0.604539914 | 0.801715837 | -4.713159661 |
| **A_55_P2182392** | Adat3 | -0.187426809 | -1.042285402 | 0.667431784 | 9.009786689 | -0.549145938 | 0.604548848 | 0.801715837 | -4.713161995 |
| **A_52_P674165** | Rsbn1l | 0.073253426 | -0.26087431 | 0.407381163 | 8.147054109 | 0.549118102 | 0.60456669 | 0.801715837 | -4.713166656 |
| **A_55_P1986276** | Tmem41b | -0.134155558 | -0.746102776 | 0.477791661 | 10.68325283 | -0.549091916 | 0.604583475 | 0.801715837 | -4.71317104 |
| **A_55_P1994074** | Mrtfa | 0.064903678 | -0.231259201 | 0.361066557 | 9.227674477 | 0.548894692 | 0.604709902 | 0.801828492 | -4.713204058 |
| **A_51_P218518** | Fiz1 | 0.057925447 | -0.206442241 | 0.322293136 | 8.379433272 | 0.548796543 | 0.604772825 | 0.801854266 | -4.713220485 |
| **A_55_P2120348** | Pdcd7 | -0.108446058 | -0.603441795 | 0.38654968 | 10.36613698 | -0.548734991 | 0.604812287 | 0.801854266 | -4.713230786 |
| **A_55_P2055284** | Chrm2 | 0.062131336 | -0.221747463 | 0.346010136 | 7.851411796 | 0.548186176 | 0.605164212 | 0.80210801 | -4.71332259 |
| **A_55_P1991802** | Kat14 | -0.195450166 | -1.088465524 | 0.697565192 | 10.56871912 | -0.548185395 | 0.605164713 | 0.80210801 | -4.71332272 |
| **A_55_P1990609** | Ube2j1 | -0.147399643 | -0.820884657 | 0.526085371 | 11.60103754 | -0.548174515 | 0.605171691 | 0.80210801 | -4.713324539 |
| **A_55_P2036764** | Prr15l | 0.17222161 | -0.614786517 | 0.959229737 | 8.576245784 | 0.548098702 | 0.605220315 | 0.80210801 | -4.713337215 |
| **A_51_P517870** | Nfatc1 | -0.06899965 | -0.384413085 | 0.246413785 | 9.495810825 | -0.547919866 | 0.605335024 | 0.80210801 | -4.71336711 |
| **A_55_P2037081** | Zfp979 | 0.09018716 | -0.322097153 | 0.502471472 | 8.718538333 | 0.547896146 | 0.60535024 | 0.80210801 | -4.713371074 |

| | | | | | | | | | |
|---|---|---|---|---|---|---|---|---|---|
| **A_52_P1055590** | Bnip1 | 0.084401703 | -0.301446244 | 0.470249651 | 9.603760861 | 0.547879893 | 0.605360666 | 0.80210801 | -4.713373791 |
| **A_55_P2009217** | Rnu3b1 | -0.185711992 | -1.03472256 | 0.663298576 | 12.08893523 | -0.547869594 | 0.605367273 | 0.80210801 | -4.713375512 |
| **A_55_P2096257** | Myl6 | -0.136832647 | -0.76240129 | 0.488735996 | 15.06971956 | -0.547854336 | 0.60537706 | 0.80210801 | -4.713378062 |
| **A_55_P2387284** | NA | 0.148982948 | -0.532463632 | 0.830429528 | 13.47295912 | 0.547589475 | 0.605546978 | 0.802230779 | -4.713422318 |
| **A_55_P2253980** | Armt1 | -0.069509323 | -0.387449483 | 0.248430837 | 8.439490052 | -0.54758055 | 0.605552704 | 0.802230779 | -4.713423809 |
| **A_55_P2157068** | Cdhr2 | -0.274404676 | -1.529701154 | 0.980891802 | 9.761526727 | -0.547514396 | 0.605595149 | 0.802232039 | -4.71343486 |
| **A_52_P354390** | Snrpn | -0.052501726 | -0.29273045 | 0.187726999 | 8.310547133 | -0.547392684 | 0.605673244 | 0.802280522 | -4.713455189 |
| **A_51_P327451** | Alas2 | -0.13937084 | -0.777711738 | 0.498970057 | 12.77892712 | -0.546851735 | 0.60602041 | 0.802685388 | -4.713545496 |
| **A_55_P2045682** | Rassf5 | 0.190880852 | -0.683718587 | 1.065480292 | 10.67302425 | 0.546642654 | 0.606154624 | 0.802776893 | -4.71358038 |
| **A_51_P487175** | Acsm3 | -0.060707838 | -0.352646493 | 0.231230818 | 7.725205811 | -0.553880234 | 0.606172539 | 0.802776893 | -4.689761585 |
| **A_55_P2118906** | Atp5j2 | 0.075929956 | -0.27205936 | 0.423919273 | 15.05010164 | 0.546509334 | 0.606240213 | 0.802811527 | -4.713602619 |
| **A_55_P1975695** | Snx18 | 0.131700449 | -0.472289859 | 0.735690758 | 11.62736168 | 0.546144586 | 0.606474413 | 0.802959017 | -4.713663437 |
| **A_51_P301215** | Knop1 | 0.094487943 | -0.338974573 | 0.527950458 | 11.28681768 | 0.54597806 | 0.606581354 | 0.802959017 | -4.713691193 |
| **A_55_P2383877** | 2810040C | 0.081830124 | -0.293571392 | 0.45723164 | 8.616414017 | 0.545968433 | 0.606587536 | 0.802959017 | -4.713692797 |
| **A_66_P122699** | NA | 0.053254387 | -0.191084375 | 0.29759315 | 7.853514778 | 0.545900342 | 0.606631267 | 0.802959017 | -4.713704144 |
| **A_55_P2084408** | Uba1 | 0.053239635 | -0.191036387 | 0.297515658 | 8.100382785 | 0.545889296 | 0.606638361 | 0.802959017 | -4.713705985 |
| **A_52_P459564** | Itga2b | -0.046707036 | -0.261066404 | 0.167652332 | 7.797403713 | -0.545745534 | 0.606730698 | 0.802959017 | -4.713729938 |
| **A_51_P446477** | Cldnd1 | -0.091610312 | -0.512065974 | 0.328845349 | 11.31940418 | -0.545725822 | 0.606743359 | 0.802959017 | -4.713733222 |
| **A_55_P2061591** | Psmd1 | -0.19603214 | -1.095885756 | 0.703821476 | 13.32136748 | -0.545639448 | 0.60679884 | 0.802959017 | -4.71374761 |
| **A_55_P1987043** | Tpp1 | -0.052285701 | -0.292297815 | 0.187726413 | 9.628274626 | -0.545632359 | 0.606803394 | 0.802959017 | -4.713748791 |
| **A_55_P2060747** | Esco1 | -0.14282288 | -0.798506549 | 0.51286079 | 9.896018668 | -0.545574124 | 0.606840802 | 0.802959017 | -4.71375849 |
| **A_55_P2112105** | NA | 0.088980137 | -0.319530455 | 0.497490729 | 10.68274256 | 0.545556953 | 0.606851833 | 0.802959017 | -4.71376135 |
| **A_55_P2086954** | NA | -0.133155142 | -0.744491572 | 0.478181288 | 11.94140353 | -0.545541775 | 0.606861583 | 0.802959017 | -4.713763878 |
| **A_55_P2061645** | NA | -0.254243174 | -1.421664855 | 0.913178507 | 9.257382438 | -0.545471284 | 0.606906867 | 0.802959017 | -4.713775617 |
| **A_55_P1992319** | Col4a3bp | -0.06221947 | -0.347939067 | 0.223500128 | 8.476669562 | -0.545426987 | 0.606935325 | 0.802959017 | -4.713782994 |
| **A_55_P2036026** | Robo4 | 0.206992738 | -0.744041875 | 1.158027351 | 12.51196468 | 0.545141357 | 0.607118841 | 0.802959017 | -4.713830546 |
| **A_51_P505967** | Spag6l | 0.243238372 | -0.874414424 | 1.360891168 | 8.786968191 | 0.545099175 | 0.607145945 | 0.802959017 | -4.713837566 |
| **A_55_P2031157** | Efna3 | -0.103287384 | -0.577995302 | 0.371420535 | 8.478849957 | -0.544968182 | 0.60723012 | 0.802959017 | -4.713859366 |
| **A_52_P421344** | Paqr5 | 0.212263534 | -0.763300215 | 1.187827284 | 9.203181853 | 0.544966847 | 0.607230978 | 0.802959017 | -4.713859588 |
| **A_55_P1982107** | NA | 0.068106593 | -0.244954204 | 0.38116739 | 16.2943355 | 0.544892481 | 0.607278768 | 0.802959017 | -4.713871962 |
| **A_51_P416822** | Hhip | 0.243411229 | -0.875507727 | 1.362330186 | 8.812634772 | 0.544869281 | 0.607293678 | 0.802959017 | -4.713875822 |
| **A_55_P2013218** | Paip1 | 0.053068169 | -0.190877222 | 0.29701356 | 8.054744352 | 0.544868666 | 0.607294073 | 0.802959017 | -4.713875924 |
| **A_55_P2098880** | Galnt10 | -0.105820116 | -0.592274217 | 0.380633985 | 11.1749051 | -0.54484969 | 0.607306268 | 0.802959017 | -4.713879081 |
| **A_52_P574759** | Rbpjl | 0.521300289 | -1.875116013 | 2.917716591 | 10.17990893 | 0.54484886 | 0.607306802 | 0.802959017 | -4.71387922 |
| **A_52_P79947** | Ndst1 | 0.141433634 | -0.508901492 | 0.79176876 | 10.66098142 | 0.544710608 | 0.607395655 | 0.802982749 | -4.713902218 |
| **A_55_P1956048** | H2-T3 | 0.045401309 | -0.163387563 | 0.25419018 | 7.692267607 | 0.544642302 | 0.607439558 | 0.802982749 | -4.713913579 |
| **A_51_P399175** | Ppm1j | -0.050781341 | -0.284385633 | 0.18282295 | 7.994851195 | -0.544469677 | 0.607550518 | 0.802982749 | -4.713942287 |
| **A_55_P2000930** | Ccnyl1 | -0.134709994 | -0.754515236 | 0.485095247 | 9.580616244 | -0.544370922 | 0.607614001 | 0.802982749 | -4.713958706 |
| **A_55_P2031100** | Ahcy | 0.046867058 | -0.168858453 | 0.262592568 | 7.848781637 | 0.544147363 | 0.607757727 | 0.802982749 | -4.713995867 |
| **A_55_P1997291** | Fam53a | 0.088878177 | -0.3203616 | 0.498117954 | 12.20538109 | 0.54396085 | 0.607877651 | 0.802982749 | -4.71402686 |
| **A_51_P247184** | Npr3 | -0.134584594 | -0.75431714 | 0.485147951 | 9.201063774 | -0.54392797 | 0.607898793 | 0.802982749 | -4.714032322 |
| **A_52_P96552** | Vkorc1l1 | -0.086345791 | -0.483970504 | 0.311278921 | 9.157015029 | -0.543898865 | 0.607917508 | 0.802982749 | -4.714037158 |
| **A_55_P2043377** | NA | -0.082306609 | -0.461429132 | 0.296815915 | 8.170539446 | -0.54375776 | 0.608008248 | 0.802982749 | -4.714060598 |
| **A_55_P2049577** | Aggf1 | -0.076688047 | -0.429956782 | 0.276580689 | 12.15857557 | -0.54371688 | 0.608034538 | 0.802982749 | -4.714067387 |
| **A_55_P1998194** | Snrpn | -0.052808169 | -0.296092372 | 0.190476034 | 8.011930342 | -0.543672726 | 0.608062935 | 0.802982749 | -4.714074721 |
| **A_55_P2064457** | Gng7 | -0.104731862 | -0.587281052 | 0.377817327 | 8.150201384 | -0.54361018 | 0.60810316 | 0.802982749 | -4.714085107 |
| **A_55_P2417224** | NA | -0.049834147 | -0.279448182 | 0.179779887 | 7.966709287 | -0.543599373 | 0.608110111 | 0.802982749 | -4.714086902 |
| **A_51_P333965** | Cisd2 | 0.060435374 | -0.218029579 | 0.338900327 | 8.199872145 | 0.543589442 | 0.608116498 | 0.802982749 | -4.714088551 |

| | | | | | | | | | |
|---|---|---|---|---|---|---|---|---|---|
| **A_55_P1986703** | Aldh3b3 | -0.186682585 | -1.046899264 | 0.673534095 | 8.687042171 | -0.543558499 | 0.608136399 | 0.802982749 | -4.714093689 |
| **A_52_P510771** | Tbl3 | 0.04679791 | -0.168847461 | 0.262443281 | 8.095413293 | 0.543546448 | 0.608144151 | 0.802982749 | -4.71409569 |
| **A_52_P467449** | Alox12 | 0.145903775 | -0.526427294 | 0.818234843 | 8.705794079 | 0.543542732 | 0.608146541 | 0.802982749 | -4.714096307 |
| **A_55_P2124233** | Vsig8 | -0.103540843 | -0.580711609 | 0.373629923 | 8.114599434 | -0.543485813 | 0.60818315 | 0.802982749 | -4.714105758 |
| **A_51_P301998** | Fmo2 | 0.610693391 | -2.203800312 | 3.425187095 | 10.9316697 | 0.543467133 | 0.608195166 | 0.802982749 | -4.714108859 |
| **A_51_P366161** | Mpg | -0.079879221 | -0.44808103 | 0.288322589 | 11.07054631 | -0.543373241 | 0.608255559 | 0.802982749 | -4.714124446 |
| **A_51_P153053** | Smpdl3a | 0.252799566 | -0.912475907 | 1.418075038 | 11.63297202 | 0.543373012 | 0.608255707 | 0.802982749 | -4.714124484 |
| **A_55_P1959624** | NA | 0.114877621 | -0.414680544 | 0.644435785 | 9.154935531 | 0.543340539 | 0.608276595 | 0.802982749 | -4.714129875 |
| **A_55_P2103011** | Sema4d | -0.282618311 | -1.585584521 | 1.0203479 | 9.389491226 | -0.543272208 | 0.608320551 | 0.802982749 | -4.714141217 |
| **A_55_P2079054** | Ndufv1 | -0.059958826 | -0.336390346 | 0.216472694 | 13.63458847 | -0.543270224 | 0.608321826 | 0.802982749 | -4.714141546 |
| **A_55_P2100550** | Mtif2 | 0.095936387 | -0.346425636 | 0.538298411 | 14.01256457 | 0.543195132 | 0.608370133 | 0.802982749 | -4.714154008 |
| **A_55_P1960936** | NA | -0.141269749 | -0.792779533 | 0.510240035 | 13.96829133 | -0.543098466 | 0.608432322 | 0.802982749 | -4.71417005 |
| **A_65_P11504** | Zfp85 | 0.100258929 | -0.362134849 | 0.562652708 | 8.476378711 | 0.54307702 | 0.608446119 | 0.802982749 | -4.714173608 |
| **A_66_P129730** | Pdia3 | -0.251352068 | -1.410958969 | 0.908254832 | 12.79290754 | -0.54290272 | 0.608558263 | 0.803049201 | -4.714202525 |
| **A_55_P2140147** | Ppp4r3b | 0.051780834 | -0.187123389 | 0.290685058 | 10.53374837 | 0.542869647 | 0.608579543 | 0.803049201 | -4.714208011 |
| **A_51_P476209** | Fscn3 | -0.054521718 | -0.306144165 | 0.197100729 | 8.214935896 | -0.54271334 | 0.608680123 | 0.803127107 | -4.714233935 |
| **A_55_P2039416** | Sgk1 | -0.402808793 | -2.26258121 | 1.456963624 | 11.28919213 | -0.542487188 | 0.608825663 | 0.803264321 | -4.714271433 |
| **A_55_P2065325** | Tmem94 | -0.064428986 | -0.362044314 | 0.233186342 | 7.894076938 | -0.54222102 | 0.60899698 | 0.803309165 | -4.714315548 |
| **A_55_P2287269** | AI415730 | -0.051398351 | -0.288823335 | 0.186026632 | 8.312867686 | -0.542216985 | 0.608999578 | 0.803309165 | -4.714316217 |
| **A_55_P2182931** | Sim2 | -0.068272641 | -0.38364806 | 0.247102777 | 8.349301984 | -0.542212108 | 0.609002717 | 0.803309165 | -4.714317025 |
| **A_51_P477676** | Mcph1 | -0.087541712 | -0.49198793 | 0.316904506 | 8.359614835 | -0.54213145 | 0.609054637 | 0.803309165 | -4.71433039 |
| **A_55_P2066484** | Crtc3 | -0.093956167 | -0.52808212 | 0.340169787 | 9.071913721 | -0.542075633 | 0.609090569 | 0.803309165 | -4.714339638 |
| **A_51_P328968** | Zfp322a | 0.114167732 | -0.413373856 | 0.641709319 | 11.38462873 | 0.542047088 | 0.609108946 | 0.803309165 | -4.714344367 |
| **A_55_P2326260** | 4930525G | -0.073758046 | -0.414720445 | 0.267204353 | 8.037936493 | -0.541817793 | 0.609256568 | 0.803425283 | -4.714382347 |
| **A_55_P2016119** | Sgcb | 0.142395434 | -0.515949883 | 0.800740752 | 9.585458502 | 0.54174219 | 0.609305246 | 0.803425283 | -4.714394867 |
| **A_55_P1984118** | Gfra4 | 0.06915985 | -0.250682602 | 0.389002302 | 8.334992975 | 0.541587064 | 0.609405134 | 0.803425283 | -4.714420551 |
| **A_55_P2419299** | NA | -0.105843214 | -0.595407482 | 0.383721054 | 9.187025145 | -0.541506474 | 0.609457031 | 0.803425283 | -4.714433892 |
| **A_55_P1973448** | Ybx2 | 0.045949505 | -0.166587609 | 0.258486619 | 8.074175366 | 0.541497433 | 0.609462853 | 0.803425283 | -4.714435388 |
| **A_51_P379597** | Ndufs1 | 0.085061641 | -0.308399861 | 0.478523143 | 12.5217565 | 0.5414793 | 0.609474531 | 0.803425283 | -4.71443839 |
| **A_52_P568781** | Sec63 | 0.056140549 | -0.203560405 | 0.315841502 | 8.60834564 | 0.541443887 | 0.609497337 | 0.803425283 | -4.714444251 |
| **A_65_P18948** | Ppm1h | -0.172061931 | -0.968086016 | 0.623962154 | 9.235130982 | -0.541388379 | 0.609533084 | 0.803425283 | -4.714453438 |
| **A_55_P1980041** | Kpna6 | -0.17419271 | -0.980231232 | 0.631845812 | 9.111698772 | -0.541283168 | 0.609600845 | 0.803425283 | -4.714470849 |
| **A_55_P1960873** | Mbd3 | 0.05867981 | -0.212857238 | 0.330216858 | 9.391001233 | 0.541265005 | 0.609612543 | 0.803425283 | -4.714473855 |
| **A_52_P111866** | Naa50 | -0.121296481 | -0.682713676 | 0.440120715 | 8.754943403 | -0.541143984 | 0.609690492 | 0.803473244 | -4.714493878 |
| **A_55_P2092668** | Kyat3 | 0.045501562 | -0.165166554 | 0.256169678 | 7.767022321 | 0.5409758 | 0.609798828 | 0.803520679 | -4.714521699 |
| **A_55_P2001481** | Pde6c | -0.051378979 | -0.289266364 | 0.186508405 | 7.659396401 | -0.540959069 | 0.609809606 | 0.803520679 | -4.714524466 |
| **A_55_P2323913** | NA | 0.084174759 | -0.305729091 | 0.474078608 | 8.970600267 | 0.540722825 | 0.609961803 | 0.80356163 | -4.714563532 |
| **A_51_P448814** | Zfp90 | -0.048147972 | -0.271174616 | 0.174878672 | 9.889837005 | -0.540718961 | 0.609964292 | 0.80356163 | -4.714564171 |
| **A_51_P459091** | Ybey | 0.066063085 | -0.239974104 | 0.372100274 | 8.336609826 | 0.540673386 | 0.609993656 | 0.80356163 | -4.714571706 |
| **A_51_P495379** | Fhod1 | 0.218723723 | -0.794634321 | 1.232081766 | 12.07317615 | 0.540609055 | 0.610035106 | 0.80356163 | -4.71458234 |
| **A_55_P2052495** | Tmx2 | -0.127690304 | -0.719356642 | 0.463976033 | 11.00848437 | -0.540544443 | 0.610076738 | 0.80356163 | -4.714593021 |
| **A_66_P124239** | Gm6557 | -0.068828764 | -0.387799884 | 0.250142356 | 8.728462657 | -0.540466717 | 0.610126822 | 0.80356163 | -4.714605867 |
| **A_51_P161342** | Abcb6 | -0.051805016 | -0.291903651 | 0.188293619 | 8.588201225 | -0.540421311 | 0.610156081 | 0.80356163 | -4.714613371 |
| **A_52_P98232** | Hira | 0.049553035 | -0.180119711 | 0.279225782 | 8.570984994 | 0.540394775 | 0.610173181 | 0.80356163 | -4.714617756 |
| **A_55_P2046374** | NA | -0.056624693 | -0.319239761 | 0.205990375 | 8.497952502 | -0.540053228 | 0.610393301 | 0.803710586 | -4.714674182 |
| **A_51_P182303** | Col1a2 | 0.117351713 | -0.426923785 | 0.661627211 | 15.92041844 | 0.540033865 | 0.610405781 | 0.803710586 | -4.71467738 |
| **A_55_P2125491** | Tnfrsf4 | -0.229611585 | -1.29456266 | 0.83533949 | 9.375800327 | -0.540025771 | 0.610410998 | 0.803710586 | -4.714678717 |
| **A_55_P2118717** | Ninl | 0.08168059 | -0.297280025 | 0.460641205 | 8.255073214 | 0.539852524 | 0.610522671 | 0.803802882 | -4.714707326 |

| | | | | | | | | | |
|---|---|---|---|---|---|---|---|---|---|
| **A_52_P479269** | Sdc1 | 0.168916201 | -0.615078755 | 0.952911157 | 11.09049808 | 0.53964528 | 0.610656274 | 0.803924037 | -4.714741539 |
| **A_55_P2068247** | NA | -0.131717681 | -0.743200301 | 0.47976494 | 11.08688287 | -0.539523421 | 0.61073484 | 0.803972724 | -4.714761651 |
| **A_51_P131800** | Cyba | 0.200924416 | -0.732085013 | 1.133933845 | 9.889940701 | 0.539382719 | 0.610825562 | 0.803994473 | -4.714784869 |
| **A_55_P2027323** | Golga7 | 0.058490616 | -0.213144661 | 0.330125893 | 12.55143381 | 0.539324776 | 0.610862925 | 0.803994473 | -4.714794429 |
| **A_55_P2054409** | Pira2 | -0.059934739 | -0.338287199 | 0.218417721 | 7.848129577 | -0.53930432 | 0.610876115 | 0.803994473 | -4.714797804 |
| **A_55_P2248330** | 2700068H | -0.070138057 | -0.395965554 | 0.255689439 | 7.96984996 | -0.539158364 | 0.610970238 | 0.804019662 | -4.71482188 |
| **A_55_P2005908** | Lin9 | 0.050932242 | -0.185679894 | 0.287544377 | 7.869516897 | 0.539145667 | 0.610978425 | 0.804019662 | -4.714823974 |
| **A_51_P416869** | Nedd8 | 0.118684739 | -0.432854624 | 0.670224101 | 13.52215249 | 0.538975111 | 0.611088423 | 0.804109682 | -4.714852101 |
| **A_55_P2007991** | Tuba3b | -0.080837475 | -0.456630557 | 0.294955607 | 7.992806082 | -0.538783521 | 0.611211999 | 0.804191929 | -4.714883688 |
| **A_55_P2223851** | NA | -0.04849657 | -0.27395927 | 0.17696613 | 8.156717912 | -0.538749234 | 0.611234116 | 0.804191929 | -4.71488934 |
| **A_55_P1987196** | Gm3448 | 0.149426932 | -0.545414279 | 0.844268143 | 8.684564392 | 0.538633867 | 0.611308536 | 0.804235114 | -4.714908355 |
| **A_52_P511381** | Tesk1 | -0.107615006 | -0.608204482 | 0.39297447 | 11.39218374 | -0.538445144 | 0.611430288 | 0.804340559 | -4.714939453 |
| **A_55_P2012166** | Spata6 | 0.132208851 | -0.482868869 | 0.74728657 | 10.48023516 | 0.538370036 | 0.611478747 | 0.804349578 | -4.714951827 |
| **A_55_P2102250** | Dzip1 | 0.050967231 | -0.186234423 | 0.288168885 | 8.43482604 | 0.538175188 | 0.611604471 | 0.804419117 | -4.714983921 |
| **A_55_P2029294** | Cetn4 | 0.17055073 | -0.623290508 | 0.964391967 | 8.780792119 | 0.53810902 | 0.611647169 | 0.804419117 | -4.714994817 |
| **A_51_P289239** | F8 | 0.065933991 | -0.24096893 | 0.372836912 | 7.819039441 | 0.538094666 | 0.611656431 | 0.804419117 | -4.714997181 |
| **A_55_P2117731** | Pak4 | -0.092508969 | -0.523249383 | 0.338231446 | 14.55786835 | -0.537921079 | 0.611768455 | 0.804425071 | -4.715025762 |
| **A_55_P1978962** | Slc9a3r2 | 0.20724915 | -0.757758954 | 1.172257254 | 10.91845047 | 0.537913128 | 0.611773586 | 0.804425071 | -4.715027071 |
| **A_55_P1995173** | Odc1 | -0.260445768 | -1.473194307 | 0.952302771 | 12.14453206 | -0.537894235 | 0.611785779 | 0.804425071 | -4.715030181 |
| **A_55_P1980966** | Mettl23 | 0.116529176 | -0.426353703 | 0.659412055 | 10.48569048 | 0.537624281 | 0.611960019 | 0.804599456 | -4.715074611 |
| **A_55_P1965150** | Pnkp | -0.065486109 | -0.370630792 | 0.239658574 | 11.53566175 | -0.537518884 | 0.612028054 | 0.80463419 | -4.715091953 |
| **A_52_P142531** | Phc3 | -0.064059858 | -0.362786954 | 0.234667238 | 9.111333897 | -0.537108101 | 0.612293261 | 0.804797859 | -4.715159515 |
| **A_55_P2013765** | Raly | -0.114643865 | -0.649323309 | 0.420035579 | 13.62234021 | -0.537041289 | 0.612336402 | 0.804797859 | -4.715170499 |
| **A_55_P2097116** | Cnep1r1 | -0.110764808 | -0.627374927 | 0.405845311 | 8.392643042 | -0.537018486 | 0.612351126 | 0.804797859 | -4.715174248 |
| **A_51_P221651** | Coq8a | -0.058585149 | -0.331831176 | 0.214660878 | 8.13533901 | -0.537012051 | 0.612355282 | 0.804797859 | -4.715175306 |
| **A_52_P429364** | Ccdc69 | 0.057780042 | -0.211715087 | 0.32727517 | 8.19698163 | 0.537003698 | 0.612360676 | 0.804797859 | -4.715176679 |
| **A_55_P2136732** | Cd99 | 0.212422241 | -0.779113311 | 1.203957794 | 13.4954252 | 0.536589341 | 0.612628274 | 0.805058631 | -4.715244775 |
| **A_52_P336958** | Jcad | -0.140699447 | -0.797516379 | 0.516117486 | 12.84254421 | -0.536535416 | 0.612663104 | 0.805058631 | -4.715253633 |
| **A_55_P2151056** | Rarg | 0.31355326 | -1.150272797 | 1.777379317 | 11.74224195 | 0.536503046 | 0.612684013 | 0.805058631 | -4.715258951 |
| **A_51_P499891** | Krit1 | -0.049169383 | -0.278830344 | 0.180491578 | 8.496845219 | -0.536238421 | 0.612854956 | 0.805228523 | -4.715302411 |
| **A_55_P1982245** | Phactr2 | -0.048571213 | -0.275480765 | 0.17833834 | 8.03992789 | -0.536137904 | 0.612919895 | 0.805259123 | -4.715318914 |
| **A_55_P2117425** | Syt5 | 0.24301776 | -0.892702493 | 1.378738014 | 9.002850049 | 0.535941008 | 0.613047111 | 0.805295736 | -4.715351235 |
| **A_51_P151820** | Urb1 | -0.080910643 | -0.459160814 | 0.297339527 | 8.745803662 | -0.53576812 | 0.613158829 | 0.805295736 | -4.715379606 |
| **A_66_P128909** | Numbl | -0.081963269 | -0.46514283 | 0.301216293 | 10.34160354 | -0.535756292 | 0.613166472 | 0.805295736 | -4.715381547 |
| **A_55_P2008895** | Tmsb15b | 0.058069548 | -0.213424646 | 0.329563741 | 8.558082934 | 0.535720473 | 0.613189619 | 0.805295736 | -4.715387423 |
| **A_55_P2069949** | Dynlrb1 | 0.097030615 | -0.356629655 | 0.550690884 | 13.81027826 | 0.535708309 | 0.61319748 | 0.805295736 | -4.715389419 |
| **A_55_P2044724** | Zfp869 | 0.061943062 | -0.227667818 | 0.351553941 | 9.8277823 | 0.53570801 | 0.613197674 | 0.805295736 | -4.715389468 |
| **A_55_P2139430** | NA | 0.15130942 | -0.556489388 | 0.859108227 | 11.0513236 | 0.535434664 | 0.613374335 | 0.805473028 | -4.715434305 |
| **A_55_P2029678** | Etfa | 0.161189884 | -0.593065851 | 0.915445619 | 12.86370078 | 0.535265814 | 0.613483476 | 0.805561635 | -4.715461992 |
| **A_52_P589119** | Rhno1 | 0.04741836 | -0.174493796 | 0.269330516 | 8.134994233 | 0.535199602 | 0.613526277 | 0.805563126 | -4.715472847 |
| **A_66_P125035** | Duxbl2 | 0.135035881 | -0.497230577 | 0.76730234 | 8.74409981 | 0.534932995 | 0.613698636 | 0.805734716 | -4.715516545 |
| **A_55_P2066778** | AB124611 | 0.047080889 | -0.173542594 | 0.267704373 | 7.779907975 | 0.534494532 | 0.613982157 | 0.806052219 | -4.71558837 |
| **A_52_P495869** | Mafb | 0.046427603 | -0.171216356 | 0.264071562 | 8.242645762 | 0.534293623 | 0.614112095 | 0.806133335 | -4.715621265 |
| **A_51_P176387** | Hook3 | -0.075676113 | -0.430447284 | 0.279095059 | 8.729527012 | -0.534270061 | 0.614127335 | 0.806133335 | -4.715625122 |
| **A_55_P2088750** | Mtif3 | 0.056900478 | -0.209987049 | 0.323788005 | 8.256857318 | 0.533995976 | 0.614304627 | 0.80630113 | -4.715669981 |
| **A_55_P2114994** | NA | -0.076990046 | -0.438141707 | 0.284161615 | 16.38692127 | -0.533943505 | 0.614338571 | 0.80630113 | -4.715678566 |
| **A_52_P130727** | Rtl6 | 0.059203499 | -0.218592615 | 0.336999612 | 8.729672692 | 0.533791376 | 0.614436991 | 0.806375563 | -4.715703454 |
| **A_55_P1971124** | Mbd1 | -0.086144474 | -0.490577676 | 0.318288727 | 10.22129238 | -0.533495756 | 0.61462827 | 0.806571844 | -4.7157518 |

| | | | | | | | | | |
|---|---|---|---|---|---|---|---|---|---|
| **A_52_P627925** | Usp36 | -0.072223208 | -0.411351006 | 0.26690459 | 8.73967448 | -0.533413165 | 0.614681716 | 0.806587234 | -4.715765303 |
| **A_55_P2154835** | NA | -0.18853839 | -1.074386474 | 0.697309694 | 11.10196496 | -0.533078158 | 0.614898531 | 0.806735059 | -4.715820056 |
| **A_55_P2162204** | Kctd15 | -0.116178204 | -0.662114978 | 0.42975857 | 11.80304331 | -0.533006685 | 0.614944794 | 0.806735059 | -4.715831734 |
| **A_52_P195922** | Exosc6 | 0.13385012 | -0.495163417 | 0.762863656 | 11.8534646 | 0.532977797 | 0.614963493 | 0.806735059 | -4.715836453 |
| **A_51_P220343** | Ccn4 | 0.18899442 | -0.699178209 | 1.07716705 | 10.34505549 | 0.532968992 | 0.614969193 | 0.806735059 | -4.715837892 |
| **A_51_P205779** | Cd5l | 0.054538574 | -0.201844145 | 0.310921294 | 7.877278015 | 0.532801425 | 0.615077665 | 0.806735059 | -4.715865264 |
| **A_55_P2018191** | Ccm2l | 0.140372026 | -0.51954436 | 0.800288411 | 8.996849253 | 0.532772748 | 0.61509623 | 0.806735059 | -4.715869947 |
| **A_55_P2062538** | 1700027A | -0.049120201 | -0.280044536 | 0.181804134 | 7.871862857 | -0.532771245 | 0.615097202 | 0.806735059 | -4.715870193 |
| **A_55_P2142191** | Rpl10 | -0.079377114 | -0.452579256 | 0.293825028 | 16.71081579 | -0.53272309 | 0.615128377 | 0.806735059 | -4.715878057 |
| **A_51_P351238** | Plgrkt | -0.149940134 | -0.854991776 | 0.555111507 | 10.88163962 | -0.532656601 | 0.615171423 | 0.806735059 | -4.715888915 |
| **A_51_P418056** | Sc5d | -0.119278444 | -0.680217447 | 0.441660559 | 10.16189764 | -0.532594495 | 0.615211633 | 0.806735059 | -4.715899055 |
| **A_55_P2175955** | Ano9 | -0.052010755 | -0.296652766 | 0.192631256 | 8.020269098 | -0.532491235 | 0.61527849 | 0.806747918 | -4.715915914 |
| **A_55_P2143494** | Doc2b | 0.079190072 | -0.29340275 | 0.451782893 | 8.235904974 | 0.532336933 | 0.615378404 | 0.806747918 | -4.7159411 |
| **A_55_P2098499** | Gnpda2 | 0.157517231 | -0.583684316 | 0.898718777 | 9.765492344 | 0.532282388 | 0.615413725 | 0.806747918 | -4.715950002 |
| **A_55_P1988766** | NA | 0.06940098 | -0.257188383 | 0.395990342 | 8.17608403 | 0.532247845 | 0.615436094 | 0.806747918 | -4.715955639 |
| **A_55_P2293351** | Slc2a4rg-| | -0.071966189 | -0.410680688 | 0.26674831 | 8.767798512 | -0.532163475 | 0.615490732 | 0.806747918 | -4.715969406 |
| **A_51_P125648** | Vwce | 0.064103158 | -0.237624164 | 0.36583048 | 7.864841405 | 0.532126799 | 0.615514485 | 0.806747918 | -4.705000209 |
| **A_55_P1961423** | Gsta3 | -0.070799089 | -0.404049487 | 0.262451309 | 8.491642142 | -0.53211725 | 0.615520668 | 0.806747918 | -4.715976948 |
| **A_55_P2084533** | Zbtb40 | 0.059056279 | -0.218948765 | 0.337061323 | 10.28393426 | 0.532063848 | 0.615555254 | 0.806747918 | -4.71598566 |
| **A_51_P490795** | Mxd1 | -0.094780086 | -0.541202576 | 0.351642404 | 9.562554352 | -0.531766993 | 0.61574753 | 0.806893041 | -4.716034077 |
| **A_51_P372895** | Zfp433 | 0.14267838 | -0.529622668 | 0.814979427 | 9.553675918 | 0.531550739 | 0.615887622 | 0.806893041 | -4.716069335 |
| **A_51_P161362** | Gnl2 | 0.074300526 | -0.275895129 | 0.424496181 | 10.72738767 | 0.531412151 | 0.615977411 | 0.806893041 | -4.716091923 |
| **A_55_P1970414** | Mccc1 | 0.125072722 | -0.464475459 | 0.714620903 | 8.71157424 | 0.531365855 | 0.616007407 | 0.806893041 | -4.716099468 |
| **A_55_P1981829** | Rhox8 | -0.139450791 | -0.796919692 | 0.518018109 | 12.06431461 | -0.53124654 | 0.616084716 | 0.806893041 | -4.71611891 |
| **A_52_P120803** | Ankrd1 | 0.558630389 | -2.075506739 | 3.192767518 | 10.13930744 | 0.531173651 | 0.616131948 | 0.806893041 | -4.716130785 |
| **A_55_P2015519** | Mtln | 0.232434812 | -0.863642293 | 1.328511917 | 10.06362665 | 0.531141716 | 0.616152641 | 0.806893041 | -4.716135987 |
| **A_55_P2020054** | 2610035D | -0.064107755 | -0.3664171 | 0.23820159 | 8.174728324 | -0.531140409 | 0.616153488 | 0.806893041 | -4.7161362 |
| **A_51_P422893** | Tmem14a | 0.154691959 | -0.57480848 | 0.884192398 | 9.443153465 | 0.531119844 | 0.616166815 | 0.806893041 | -4.71613955 |
| **A_66_P101942** | Gm9706 | -0.442541938 | -2.529607228 | 1.644523352 | 9.037520447 | -0.531090758 | 0.616185663 | 0.806893041 | -4.716144288 |
| **A_55_P2053943** | Olfr1368 | 0.051933694 | -0.193002953 | 0.296870342 | 8.230525157 | 0.531062692 | 0.616203851 | 0.806893041 | -4.71614886 |
| **A_51_P405167** | Maf | 0.260743698 | -0.96903508 | 1.490522477 | 10.6266164 | 0.531052151 | 0.616210682 | 0.806893041 | -4.716150577 |
| **A_51_P211131** | Traf3ip2 | -0.077828629 | -0.444937167 | 0.289279909 | 8.797989863 | -0.531000879 | 0.616243909 | 0.806893041 | -4.716158928 |
| **A_51_P199098** | Tmem107 | 0.080988758 | -0.301032796 | 0.463010313 | 11.50822523 | 0.53099107 | 0.616250266 | 0.806893041 | -4.716160525 |
| **A_51_P308446** | 1700094D | 0.294865599 | -1.096199612 | 1.685930809 | 9.629182169 | 0.530917406 | 0.616298007 | 0.806900905 | -4.716172522 |
| **A_55_P2080248** | Cyp2j8 | 0.114865224 | -0.427524704 | 0.657255151 | 8.400745329 | 0.530429039 | 0.616614563 | 0.807260696 | -4.71625202 |
| **A_55_P2055369** | NA | 0.053876082 | -0.200601502 | 0.308353666 | 8.262286147 | 0.53026971 | 0.616717859 | 0.807301745 | -4.716277944 |
| **A_51_P509452** | Tmlhe | 0.084652567 | -0.31520772 | 0.484512854 | 8.155515795 | 0.530251864 | 0.616729429 | 0.807301745 | -4.716280847 |
| **A_51_P249594** | Rev1 | 0.055116659 | -0.205276763 | 0.31551008 | 8.560799609 | 0.530155438 | 0.616791949 | 0.807328924 | -4.716296532 |
| **A_55_P2296118** | 9630010G | 0.269236551 | -1.002936368 | 1.54140947 | 9.7586891 | 0.530076094 | 0.616843397 | 0.807341608 | -4.716309436 |
| **A_55_P2118734** | Rnmt | -0.177821218 | -1.018409679 | 0.662767243 | 9.535380168 | -0.529847026 | 0.616991939 | 0.807481361 | -4.716346683 |
| **A_55_P2150053** | Yipf5 | -0.052233324 | -0.299219282 | 0.194752634 | 10.16824599 | -0.529694848 | 0.617090633 | 0.807527399 | -4.716371421 |
| **A_55_P2080378** | Ttc30a2 | 0.108516283 | -0.404634283 | 0.621666849 | 8.436878057 | 0.529663984 | 0.617110651 | 0.807527399 | -4.716376437 |
| **A_51_P189208** | 4933417G | 0.053118945 | -0.198112896 | 0.304350787 | 9.098068775 | 0.529572112 | 0.617170239 | 0.807545135 | -4.716391368 |
| **A_55_P1989071** | Oas1d | 0.06733206 | -0.251167395 | 0.385831515 | 7.925890119 | 0.529497059 | 0.617218921 | 0.807545135 | -4.716403563 |
| **A_55_P1997585** | NA | -0.069211371 | -0.396629651 | 0.258206909 | 7.792914355 | -0.529449903 | 0.617249509 | 0.807545135 | -4.716411225 |
| **A_55_P2023937** | Impg1 | -0.223810565 | -1.283274439 | 0.835653309 | 9.789936815 | -0.529108546 | 0.617470958 | 0.807780195 | -4.716466671 |
| **A_52_P116264** | Adhfe1 | -0.045350741 | -0.260118202 | 0.16941672 | 7.72297635 | -0.528891093 | 0.61761205 | 0.807910107 | -4.716501976 |
| **A_55_P2021570** | C87414 | 0.049679483 | -0.185705872 | 0.285064839 | 7.970823888 | 0.528625339 | 0.617784506 | 0.808081026 | -4.716545107 |

| | | | | | | | | | |
|---|---|---|---|---|---|---|---|---|---|
| **A_55_P2014189** | NA | -0.11669529 | -0.669733168 | 0.436342589 | 9.46740859 | -0.528504627 | 0.61786285 | 0.808128828 | -4.716564692 |
| **A_51_P430383** | Dnajc30 | 0.102769571 | -0.384408806 | 0.589947949 | 11.06450871 | 0.528356293 | 0.617959127 | 0.808165449 | -4.716588754 |
| **A_55_P2016877** | Nsd1 | 0.094473463 | -0.353489247 | 0.542436173 | 12.63490586 | 0.528224229 | 0.618044852 | 0.808165449 | -4.716610171 |
| **A_55_P1993851** | Gm10021 | -0.07915034 | -0.454467074 | 0.296166393 | 8.507570324 | -0.52820828 | 0.618055205 | 0.808165449 | -4.716612758 |
| **A_55_P2078138** | Otop1 | 0.157762056 | -0.590323555 | 0.905847667 | 8.925021533 | 0.528203898 | 0.618058049 | 0.808165449 | -4.716613468 |
| **A_55_P2046241** | Arl6ip6 | 0.05424707 | -0.203071218 | 0.311565359 | 8.129008903 | 0.528026817 | 0.618173007 | 0.808247298 | -4.716642179 |
| **A_51_P140350** | Nadsyn1 | -0.055965417 | -0.321458797 | 0.209527963 | 8.712569615 | -0.527978688 | 0.618204254 | 0.808247298 | -4.716649981 |
| **A_55_P2091206** | NA | 0.091008838 | -0.340839234 | 0.52285691 | 9.745102166 | 0.527840769 | 0.618293799 | 0.80830971 | -4.716672334 |
| **A_55_P1971849** | Sh3glb2 | -0.064871399 | -0.372791185 | 0.243048387 | 12.9928577 | -0.527674387 | 0.618401834 | 0.808373461 | -4.716699295 |
| **A_55_P2045802** | Nsmf | 0.202421692 | -0.758465153 | 1.163308538 | 12.08384226 | 0.527636886 | 0.618426185 | 0.808373461 | -4.716705371 |
| **A_52_P493810** | Srpk1 | -0.169120423 | -0.972214085 | 0.633973239 | 9.133179591 | -0.527448673 | 0.618548411 | 0.808478569 | -4.716735858 |
| **A_55_P2159234** | Tatdn2 | -0.119446398 | -0.686823367 | 0.447930571 | 8.250941737 | -0.52729263 | 0.618649756 | 0.808491053 | -4.716761128 |
| **A_55_P2099358** | Cars | -0.241600337 | -1.389293025 | 0.906092351 | 11.72421505 | -0.527256894 | 0.618672967 | 0.808491053 | -4.716766914 |
| **A_51_P294555** | Ifitm6 | -0.073607833 | -0.423354075 | 0.276138408 | 8.186681366 | -0.527134359 | 0.618752558 | 0.808491053 | -4.716786752 |
| **A_55_P1998406** | NA | 0.051671501 | -0.193874293 | 0.297217295 | 8.495527545 | 0.527070759 | 0.61879387 | 0.808491053 | -4.716797047 |
| **A_51_P246543** | Ebf3 | 0.080793274 | -0.303162945 | 0.464749494 | 8.358448158 | 0.527040327 | 0.618813639 | 0.808491053 | -4.716801973 |
| **A_51_P141860** | Fbxw11 | -0.061067638 | -0.351282085 | 0.229146808 | 13.23399131 | -0.527038624 | 0.618814745 | 0.808491053 | -4.716802249 |
| **A_55_P2025273** | Trabd | -0.086506481 | -0.49770377 | 0.324690808 | 9.774201678 | -0.526924933 | 0.618888601 | 0.808491053 | -4.716820649 |
| **A_55_P2003793** | Tspan3 | 0.075634486 | -0.283897772 | 0.435166744 | 13.76833988 | 0.526905029 | 0.618901531 | 0.808491053 | -4.71682387 |
| **A_55_P2006615** | Rbbp9 | 0.189979118 | -0.713182487 | 1.093140723 | 11.04721112 | 0.526854563 | 0.618934317 | 0.808491053 | -4.716832036 |
| **A_55_P2067445** | Arf5 | -0.068720903 | -0.395504043 | 0.258062237 | 12.85986766 | -0.526719698 | 0.619021938 | 0.808533058 | -4.716853856 |
| **A_52_P46310** | Dcaf7 | -0.057115911 | -0.3287371 | 0.214505279 | 8.342703307 | -0.526676335 | 0.619050111 | 0.808533058 | -4.716860871 |
| **A_55_P1989312** | Prkd3 | 0.128572139 | -0.483052137 | 0.740196414 | 12.07623503 | 0.526517123 | 0.619153562 | 0.808613549 | -4.716886622 |
| **A_51_P367780** | Adamtsl2 | 0.233167984 | -0.876295073 | 1.342631042 | 8.93692128 | 0.526388531 | 0.619237125 | 0.80863226 | -4.716907416 |
| **A_55_P2062811** | Tomm5 | -0.06864199 | -0.395302265 | 0.258018286 | 12.61027869 | -0.526312744 | 0.619286375 | 0.80863226 | -4.716919669 |
| **A_55_P2138177** | Psd3 | -0.063746394 | -0.367183141 | 0.239690353 | 8.679855039 | -0.52618419 | 0.619369922 | 0.80863226 | -4.716940451 |
| **A_51_P217706** | Ribc1 | -0.049443885 | -0.284828253 | 0.185940483 | 8.362895076 | -0.52612061 | 0.619411246 | 0.80863226 | -4.716950727 |
| **A_66_P117730** | Hapln1 | 0.296993133 | -1.116905246 | 1.710891512 | 9.105004691 | 0.526112422 | 0.619416567 | 0.80863226 | -4.71695205 |
| **A_52_P422494** | Cd300lf | -0.097044861 | -0.55905043 | 0.364960709 | 8.356853222 | -0.526108932 | 0.619418836 | 0.80863226 | -4.716952614 |
| **A_51_P255387** | Med31 | 0.113279658 | -0.426092079 | 0.652651395 | 11.47097939 | 0.526034332 | 0.619467323 | 0.808640958 | -4.71696467 |
| **A_55_P2166745** | Sp3os | -0.06163742 | -0.355168639 | 0.231893798 | 11.22419242 | -0.525945207 | 0.619525255 | 0.808661982 | -4.716979072 |
| **A_52_P474765** | Denr | 0.059962875 | -0.225678178 | 0.345603928 | 10.06074121 | 0.525789814 | 0.619626267 | 0.808680744 | -4.717004177 |
| **A_55_P1972634** | Cmc4 | 0.127712184 | -0.480665996 | 0.736090363 | 9.16299383 | 0.525786035 | 0.619628724 | 0.808680744 | -4.717004787 |
| **A_55_P1972084** | Napb | -0.077563299 | -0.44708828 | 0.291961682 | 8.413865335 | -0.525730066 | 0.619665109 | 0.808680744 | -4.717013828 |
| **A_52_P335609** | Idnk | 0.13632833 | -0.513368641 | 0.786025301 | 9.875382088 | 0.525564011 | 0.619773067 | 0.808705692 | -4.717040645 |
| **A_51_P161463** | Ccdc103 | 0.165340888 | -0.622624534 | 0.953306309 | 8.95612859 | 0.525561388 | 0.619774772 | 0.808705692 | -4.717041069 |
| **A_51_P231630** | Zfp239 | 0.080915038 | -0.304783248 | 0.466613325 | 8.27729963 | 0.525450586 | 0.619846814 | 0.808705692 | -4.71705896 |
| **A_55_P2139886** | Sft2d1 | 0.085648001 | -0.322616579 | 0.493912582 | 8.574917621 | 0.525443319 | 0.619851539 | 0.808705692 | -4.717060133 |
| **A_55_P2052834** | Lst1 | -0.108312484 | -0.62468814 | 0.408063172 | 10.41846408 | -0.525367378 | 0.619900917 | 0.808715542 | -4.717072392 |
| **A_52_P155554** | Cdc42ep2 | 0.150782579 | -0.568950361 | 0.87051552 | 11.16866181 | 0.524723039 | 0.620319972 | 0.809053642 | -4.717176354 |
| **A_55_P2119969** | Bdh2 | 0.225048667 | -0.849194254 | 1.299291588 | 10.19625924 | 0.52471598 | 0.620324564 | 0.809053642 | -4.717177492 |
| **A_51_P472481** | Dync2li1 | 0.207900114 | -0.784525228 | 1.200325456 | 9.145719013 | 0.524695364 | 0.620337974 | 0.809053642 | -4.717180816 |
| **A_55_P2096395** | Cdc42bpa | 0.078486908 | -0.296201039 | 0.453174856 | 10.48098259 | 0.524659868 | 0.620361065 | 0.809053642 | -4.71718654 |
| **A_55_P2049071** | NA | 0.107077348 | -0.404169507 | 0.618324203 | 13.98430278 | 0.524586733 | 0.620408641 | 0.809053642 | -4.717198332 |
| **A_55_P1967015** | Fgfr1 | 0.138983016 | -0.524604149 | 0.802570182 | 9.414535259 | 0.524582866 | 0.620411157 | 0.809053642 | -4.717198956 |
| **A_52_P123944** | Mospd2 | -0.056918324 | -0.328821903 | 0.214985256 | 8.342869589 | -0.524309251 | 0.620589168 | 0.809223513 | -4.71724306 |
| **A_51_P418016** | Ccnj | 0.052392727 | -0.197918085 | 0.302703539 | 8.474768584 | 0.524253983 | 0.620625129 | 0.809223513 | -4.717251967 |
| **A_55_P2038298** | Rpl31 | 0.094771581 | -0.358129842 | 0.547673004 | 16.24362244 | 0.524112824 | 0.62071698 | 0.809288698 | -4.717274711 |

| | | | | | | | | | |
|---|---|---|---|---|---|---|---|---|---|
| **A_55_P2145833** | Dmpk | 0.170005328 | -0.642552871 | 0.982563526 | 9.775273695 | 0.524032715 | 0.62076911 | 0.809302089 | -4.717287616 |
| **A_55_P2064568** | Anks1 | -0.123173837 | -0.712046483 | 0.465698809 | 11.32161981 | -0.523898838 | 0.620856235 | 0.809308782 | -4.717309179 |
| **A_51_P423880** | Smarcd3 | 0.211140045 | -0.798330969 | 1.220611058 | 9.626895486 | 0.52387431 | 0.620872197 | 0.809308782 | -4.71731313 |
| **A_55_P2126333** | Cyp2a12 | 0.044518135 | -0.168369948 | 0.257406218 | 7.693987588 | 0.523764377 | 0.620943746 | 0.809308782 | -4.717330832 |
| **A_55_P1954544** | Zfp446 | 0.074641213 | -0.282297713 | 0.431580139 | 9.265405321 | 0.523763368 | 0.620944402 | 0.809308782 | -4.717330995 |
| **A_55_P1988358** | Lrrc47 | 0.074364797 | -0.281293123 | 0.430022716 | 11.92875311 | 0.523703235 | 0.620983541 | 0.809308782 | -4.717340677 |
| **A_51_P349192** | NA | 0.162782586 | -0.615974971 | 0.941540144 | 12.75312371 | 0.523547411 | 0.621084968 | 0.80938641 | -4.717365761 |
| **A_52_P510215** | Traf6 | -0.046396857 | -0.268457622 | 0.175663908 | 8.561240461 | -0.523319688 | 0.621233212 | 0.809525034 | -4.717402409 |
| **A_55_P1977812** | Cd244a | 0.142776828 | -0.540680808 | 0.826234465 | 9.125575132 | 0.523234624 | 0.621288592 | 0.809525297 | -4.717416095 |
| **A_55_P2023314** | Casz1 | 0.16185309 | -0.613067184 | 0.936773364 | 8.982614605 | 0.523135651 | 0.621353031 | 0.809525297 | -4.717432017 |
| **A_55_P2145441** | NA | 0.103691842 | -0.392809225 | 0.600192908 | 9.739453732 | 0.523087989 | 0.621384064 | 0.809525297 | -4.717439683 |
| **A_55_P2113524** | Nsd2 | -0.082413857 | -0.477182964 | 0.31235525 | 8.224094483 | -0.522886495 | 0.621515268 | 0.809525297 | -4.717472087 |
| **A_52_P685938** | Efcab14 | 0.05773637 | -0.218875695 | 0.334348435 | 10.10784175 | 0.522791724 | 0.621576984 | 0.809525297 | -4.717487324 |
| **A_66_P102076** | Cep68 | 0.113413835 | -0.429969289 | 0.656796958 | 9.82477922 | 0.522769491 | 0.621591462 | 0.809525297 | -4.717490898 |
| **A_55_P2224582** | Fam219b | -0.059835648 | -0.346525559 | 0.226854264 | 8.338018607 | -0.522754682 | 0.621601107 | 0.809525297 | -4.717493279 |
| **A_51_P483013** | Arel1 | -0.054353359 | -0.314780059 | 0.206073341 | 10.73737347 | -0.522746613 | 0.621606361 | 0.809525297 | -4.717494576 |
| **A_55_P2154027** | Ptgis | 0.231039838 | -0.87621107 | 1.338290746 | 10.61340183 | 0.522626192 | 0.621684788 | 0.809525297 | -4.717513933 |
| **A_55_P2030638** | Dusp23 | 0.068212684 | -0.258718935 | 0.395144303 | 10.03858618 | 0.522586947 | 0.621710348 | 0.809525297 | -4.71752024 |
| **A_55_P2119458** | Slc25a44 | 0.047573476 | -0.180457266 | 0.275604219 | 8.043066283 | 0.522542757 | 0.62173913 | 0.809525297 | -4.717527342 |
| **A_55_P2147816** | NA | 0.114754996 | -0.435498433 | 0.665008425 | 17.04218365 | 0.522347103 | 0.621866571 | 0.809525297 | -4.717558779 |
| **A_52_P217675** | Slf1 | 0.056022439 | -0.212630198 | 0.324675077 | 8.287178716 | 0.522301476 | 0.621896293 | 0.809525297 | -4.717566109 |
| **A_55_P2370394** | Ppp2r5e | -0.071026181 | -0.411643578 | 0.269591217 | 8.567803792 | -0.522278298 | 0.621911392 | 0.809525297 | -4.717569832 |
| **A_66_P116635** | Igf2os | -0.420361711 | -2.436476997 | 1.595753576 | 10.11621646 | -0.522225571 | 0.62194574 | 0.809525297 | -4.717578302 |
| **A_51_P432563** | Tlr9 | -0.15440774 | -0.894979946 | 0.586164466 | 10.54433338 | -0.5222182 | 0.621950542 | 0.809525297 | -4.717579486 |
| **A_55_P1983448** | S100a4 | -0.206636815 | -1.197726135 | 0.784452505 | 10.37015041 | -0.522210078 | 0.621955833 | 0.809525297 | -4.71758079 |
| **A_55_P2175195** | Stxbp4 | 0.13689999 | -0.519771873 | 0.793571853 | 8.402550874 | 0.522162107 | 0.621987084 | 0.809525297 | -4.717588495 |
| **A_55_P2067332** | NA | 0.061468794 | -0.233534041 | 0.35647163 | 8.31214314 | 0.52188985 | 0.622164465 | 0.809701654 | -4.717632211 |
| **A_51_P120615** | Rabl3 | -0.056869293 | -0.330013296 | 0.21627471 | 8.310235228 | -0.521478618 | 0.622432446 | 0.809995888 | -4.711882549 |
| **A_51_P144543** | NA | -0.046208641 | -0.268216359 | 0.175799077 | 7.69822335 | -0.521321298 | 0.622534981 | 0.810056277 | -4.717723442 |
| **A_55_P2081482** | Atp5g2 | 0.141493013 | -0.538948618 | 0.821934643 | 15.95121516 | 0.520828166 | 0.622856447 | 0.810056277 | -4.717802503 |
| **A_51_P215887** | Aldh18a1 | -0.082037185 | -0.496209396 | 0.332135026 | 8.48342587 | -0.527585074 | 0.62286444 | 0.810056277 | -4.693491514 |
| **A_51_P338776** | Dnajc8 | -0.05255772 | -0.305326514 | 0.200211073 | 11.49323757 | -0.520790926 | 0.622880728 | 0.810056277 | -4.717808471 |
| **A_52_P374292** | Tefm | -0.04467241 | -0.259519791 | 0.17017497 | 8.139496461 | -0.520786441 | 0.622883652 | 0.810056277 | -4.71780919 |
| **A_55_P2002903** | Smoc2 | 0.106736673 | -0.406646473 | 0.620119819 | 11.56686983 | 0.520741752 | 0.622912789 | 0.810056277 | -4.717816351 |
| **A_51_P242603** | E430018J | 0.081564346 | -0.310840792 | 0.473969483 | 8.518594043 | 0.52061421 | 0.622995952 | 0.810056277 | -4.717836787 |
| **A_51_P374818** | Shd | 0.096539146 | -0.367910383 | 0.560988674 | 8.135525054 | 0.520613314 | 0.622996535 | 0.810056277 | -4.71783693 |
| **A_52_P494622** | Nr4a2 | -0.049663642 | -0.288621643 | 0.189294359 | 8.829913704 | -0.52055585 | 0.623034006 | 0.810056277 | -4.717846136 |
| **A_55_P2042973** | 1010001N | 0.07688242 | -0.293100451 | 0.446865291 | 8.230237198 | 0.520470086 | 0.623089933 | 0.810056277 | -4.717859873 |
| **A_52_P186033** | Spn | 0.348831911 | -1.330151904 | 2.027815726 | 8.923543165 | 0.520379337 | 0.623149114 | 0.810056277 | -4.717874408 |
| **A_66_P119884** | Wdr91 | 0.153409128 | -0.584995229 | 0.891813485 | 10.11279212 | 0.520364063 | 0.623159075 | 0.810056277 | -4.717876854 |
| **A_55_P2048164** | Msantd3 | 0.085451491 | -0.325855619 | 0.4967586 | 9.080521745 | 0.52035984 | 0.623161829 | 0.810056277 | -4.71787753 |
| **A_66_P105564** | Gjc2 | 0.096769118 | -0.369107365 | 0.562645601 | 9.617591722 | 0.520255091 | 0.623230144 | 0.810056277 | -4.717894303 |
| **A_51_P518246** | Fibp | -0.047511631 | -0.276314134 | 0.181290872 | 10.23726136 | -0.520103185 | 0.623329222 | 0.810056277 | -4.717918622 |
| **A_52_P299846** | Sh3rf1 | 0.125440278 | -0.478645252 | 0.729525808 | 8.891305383 | 0.520102474 | 0.623329685 | 0.810056277 | -4.717918736 |
| **A_52_P141628** | Slc23a1 | -0.046376723 | -0.269725569 | 0.176972123 | 7.725537992 | -0.520075856 | 0.623347047 | 0.810056277 | -4.717922997 |
| **A_52_P244803** | D630033C | -0.04652599 | -0.270595717 | 0.177543736 | 7.932661571 | -0.520071178 | 0.623350098 | 0.810056277 | -4.717923746 |
| **A_51_P277444** | Rnf19a | -0.064971198 | -0.377882127 | 0.24793973 | 10.62702182 | -0.520056454 | 0.623359703 | 0.810056277 | -4.717926103 |
| **A_52_P124472** | Kcnd2 | 0.047399355 | -0.180887716 | 0.275686426 | 7.731094089 | 0.520045642 | 0.623366755 | 0.810056277 | -4.717927833 |

| | | | | | | | | | |
|---|---|---|---|---|---|---|---|---|---|
| **A_52_P385615** | Rnpc3 | 0.065878057 | -0.25142848 | 0.383184593 | 8.899607162 | 0.520010493 | 0.623389682 | 0.810056277 | -4.717933459 |
| **A_51_P103054** | Adamts19 | 0.114553424 | -0.437219395 | 0.666326243 | 8.160810726 | 0.519993744 | 0.623400608 | 0.810056277 | -4.712371475 |
| **A_55_P1962479** | Gpbp1l1 | 0.050330023 | -0.192159709 | 0.292819755 | 9.391795101 | 0.519857268 | 0.623489635 | 0.810117513 | -4.71795798 |
| **A_55_P2181029** | NA | -0.045558647 | -0.265152852 | 0.174035558 | 8.098857743 | -0.519637263 | 0.623633165 | 0.810177005 | -4.717993177 |
| **A_51_P221031** | Slc16a12 | -0.063289146 | -0.368432974 | 0.241854682 | 8.767159729 | -0.519487364 | 0.623730968 | 0.810177005 | -4.718017151 |
| **A_55_P2021275** | Iqca | 0.14959896 | -0.571682405 | 0.870880325 | 8.374419408 | 0.519486431 | 0.623731577 | 0.810177005 | -4.7180173 |
| **A_55_P2059680** | Gm11938 | 0.061235604 | -0.234009487 | 0.356480696 | 8.043165679 | 0.519483392 | 0.62373356 | 0.810177005 | -4.718017786 |
| **A_55_P1974577** | Nfam1 | -0.078805446 | -0.45877561 | 0.301164717 | 8.702889683 | -0.519465948 | 0.623744942 | 0.810177005 | -4.718020576 |
| **A_55_P2057102** | Sft2d3 | -0.06995384 | -0.40736577 | 0.267458089 | 8.512376331 | -0.519279854 | 0.623866375 | 0.810280297 | -4.71805033 |
| **A_55_P2012051** | Sox1 | -0.095138645 | -0.554715757 | 0.364438467 | 8.381314219 | -0.518500185 | 0.624375282 | 0.810886794 | -4.718174891 |
| **A_51_P319453** | Zfp143 | -0.054362592 | -0.317005387 | 0.208280204 | 8.427658594 | -0.518423889 | 0.624425095 | 0.810897016 | -4.718187072 |
| **A_55_P2238079** | A830054C | 0.079263624 | -0.303788495 | 0.462315743 | 8.283696012 | 0.518282326 | 0.624517524 | 0.810962577 | -4.718209669 |
| **A_51_P249215** | Ptger2 | 0.221784775 | -0.850239324 | 1.293808873 | 8.810818684 | 0.518176277 | 0.624586772 | 0.810998029 | -4.718226593 |
| **A_52_P80944** | Zfp36 | -0.05440739 | -0.317428417 | 0.208613637 | 8.241663344 | -0.518104981 | 0.624633329 | 0.811004014 | -4.71823797 |
| **A_55_P2130725** | Rhox8 | -0.0524672 | -0.306378124 | 0.201443724 | 8.160279611 | -0.517555407 | 0.624992269 | 0.811388544 | -4.71832562 |
| **A_55_P1965989** | Zfp329 | 0.129659016 | -0.497854948 | 0.75717298 | 8.979708082 | 0.517523019 | 0.625013427 | 0.811388544 | -4.718330784 |
| **A_51_P430950** | Rdx | -0.058924386 | -0.344270859 | 0.226422087 | 11.42964838 | -0.517217133 | 0.625213262 | 0.811593475 | -4.718379532 |
| **A_55_P2143953** | Pank2 | -0.059243121 | -0.346195206 | 0.227708963 | 9.675779715 | -0.517105188 | 0.625286405 | 0.811633928 | -4.718397367 |
| **A_55_P2023484** | Osbpl5 | -0.134209265 | -0.784375977 | 0.515957446 | 13.17618208 | -0.517020919 | 0.625341468 | 0.811646082 | -4.71841079 |
| **A_52_P330214** | Fanca | 0.072659733 | -0.279379133 | 0.4246986 | 9.596341936 | 0.516955942 | 0.625383927 | 0.811646082 | -4.718421139 |
| **A_51_P488554** | Mrnip | -0.044378832 | -0.259419825 | 0.170662161 | 9.830020183 | -0.516898124 | 0.625421709 | 0.811646082 | -4.718430347 |
| **A_51_P114722** | Hao2 | -0.049652485 | -0.290370342 | 0.191065371 | 7.683858433 | -0.516634055 | 0.625594286 | 0.811815553 | -4.710923661 |
| **A_52_P84353** | Homez | 0.054600813 | -0.21019844 | 0.319400065 | 10.0342331 | 0.51645525 | 0.625711156 | 0.811912717 | -4.718500848 |
| **A_55_P2048650** | Dennd4a | -0.128926436 | -0.754793392 | 0.496940521 | 15.27539765 | -0.515953186 | 0.626039378 | 0.812284097 | -4.71858071 |
| **A_55_P2042487** | Dpysl3 | -0.113888025 | -0.667127601 | 0.439351552 | 11.89692836 | -0.51560268 | 0.626268577 | 0.812526953 | -4.718636426 |
| **A_52_P81630** | 2810001G | -0.05143054 | -0.30134187 | 0.198480791 | 8.693589009 | -0.515448737 | 0.626369256 | 0.812588856 | -4.718660886 |
| **A_55_P2123727** | Grb2 | 0.043560026 | -0.168126421 | 0.255246474 | 8.116368253 | 0.5154012 | 0.626400347 | 0.812588856 | -4.718668438 |
| **A_55_P1954092** | NA | -0.125110309 | -0.733260741 | 0.483040123 | 13.74924802 | -0.515267112 | 0.626488051 | 0.812616274 | -4.718689737 |
| **A_55_P2101776** | Adora1 | 0.239733806 | -0.925667404 | 1.405135016 | 9.856876027 | 0.515233581 | 0.626509983 | 0.812616274 | -4.718695063 |
| **A_51_P308844** | Nrn1 | -0.04987647 | -0.292373531 | 0.192620591 | 7.784793499 | -0.515156968 | 0.626560098 | 0.812616274 | -4.718707229 |
| **A_55_P2079142** | Fam124b | -0.046272943 | -0.271274962 | 0.178729075 | 8.449792895 | -0.515099413 | 0.626597748 | 0.812616274 | -4.718716368 |
| **A_55_P2109485** | Milr1 | -0.162396976 | -0.952206564 | 0.627412612 | 15.66914124 | -0.514998403 | 0.626663827 | 0.812616274 | -4.718732406 |
| **A_55_P1970105** | Ltbp2 | 0.464543906 | -1.794977867 | 2.724065679 | 12.43986443 | 0.514944688 | 0.626698969 | 0.812616274 | -4.718740933 |
| **A_51_P151126** | Cd52 | 0.240534223 | -0.929472085 | 1.410540531 | 10.83679241 | 0.51491912 | 0.626715696 | 0.812616274 | -4.718744991 |
| **A_52_P284348** | Smim20 | 0.064629534 | -0.249838831 | 0.379097899 | 11.02437616 | 0.514759544 | 0.626820101 | 0.812657679 | -4.718770319 |
| **A_52_P311031** | NA | -0.16077131 | -0.943118237 | 0.621575616 | 16.1320907 | -0.514706338 | 0.626854913 | 0.812657679 | -4.718778762 |
| **A_51_P431785** | Myom2 | 0.424866443 | -1.64283257 | 2.492565457 | 10.08184882 | 0.514654123 | 0.626889079 | 0.812657679 | -4.718787047 |
| **A_55_P2026624** | NA | -0.063324813 | -0.371538496 | 0.244888869 | 13.63172699 | -0.514603043 | 0.626922503 | 0.812657679 | -4.718795151 |
| **A_52_P981697** | Zfp874a | -0.07192676 | -0.422044411 | 0.278190891 | 8.438962647 | -0.514549115 | 0.626957792 | 0.812657679 | -4.718803707 |
| **A_51_P142515** | Antxr2 | 0.182489247 | -0.705996285 | 1.07097478 | 10.59571761 | 0.514443003 | 0.627027231 | 0.812693201 | -4.718820539 |
| **A_55_P2032723** | Dnajc10 | -0.082225549 | -0.482849275 | 0.318398177 | 8.625228851 | -0.51406787 | 0.62727275 | 0.812865917 | -4.718880021 |
| **A_55_P2041355** | Gm4750 | 0.068310597 | -0.264521014 | 0.401142209 | 12.97737546 | 0.514060059 | 0.627277863 | 0.812865917 | -4.718881259 |
| **A_51_P180032** | Sem1 | 0.082793524 | -0.320614148 | 0.486201195 | 15.48654615 | 0.514046682 | 0.627286619 | 0.812865917 | -4.718883379 |
| **A_52_P16752** | Aox3 | -0.04646557 | -0.272967146 | 0.180036006 | 7.74616925 | -0.513819269 | 0.627435485 | 0.812954265 | -4.71891942 |
| **A_55_P2176350** | 0610010F | -0.123295201 | -0.724365004 | 0.477774603 | 9.725555302 | -0.513773391 | 0.62746552 | 0.812954265 | -4.718926689 |
| **A_55_P2042016** | NA | 0.057002848 | -0.22090141 | 0.334907106 | 8.758441675 | 0.513749837 | 0.62748094 | 0.812954265 | -4.718930421 |
| **A_52_P362981** | NA | 0.167586776 | -0.649667462 | 0.984841015 | 9.373343708 | 0.513609338 | 0.627572925 | 0.813018958 | -4.718952678 |
| **A_55_P2080995** | Tgif2lx1 | 0.047281647 | -0.183369451 | 0.277932746 | 7.973421398 | 0.513437302 | 0.627685568 | 0.813109278 | -4.718979924 |

| | | | | | | | | | |
|---|---|---|---|---|---|---|---|---|---|
| **A_51_P331385** | Paip2b | 0.089237959 | -0.346139321 | 0.524615238 | 9.305330444 | 0.513374403 | 0.627726755 | 0.813109278 | -4.718989883 |
| **A_55_P2114187** | NA | -0.044001174 | -0.258805196 | 0.170802848 | 7.712562709 | -0.513064777 | 0.627929522 | 0.813289949 | -4.719038895 |
| **A_55_P2032755** | Gm10586 | -0.143429848 | -0.843666153 | 0.556806456 | 15.21027265 | -0.513032954 | 0.627950365 | 0.813289949 | -4.719043931 |
| **A_51_P167048** | Hs6st1 | -0.089335495 | -0.525636898 | 0.346965908 | 8.505905901 | -0.51284696 | 0.628072188 | 0.813361516 | -4.719073359 |
| **A_55_P2145149** | Cpn2 | -0.043340626 | -0.255081084 | 0.168399832 | 7.668986291 | -0.512674454 | 0.628185189 | 0.813361516 | -4.719100646 |
| **A_55_P1961736** | Rcan2 | 0.191719951 | -0.745139823 | 1.128579726 | 8.93088866 | 0.512558055 | 0.628261443 | 0.813361516 | -4.719119053 |
| **A_66_P108822** | NA | 0.15598113 | -0.60625896 | 0.91822122 | 16.15083917 | 0.51254337 | 0.628271063 | 0.813361516 | -4.719121375 |
| **A_55_P2044458** | Pramef8 | -0.12834096 | -0.755596492 | 0.498914572 | 10.29860775 | -0.512473164 | 0.628317059 | 0.813361516 | -4.719132475 |
| **A_55_P2123529** | NA | 0.136585554 | -0.53105056 | 0.804221669 | 11.42007689 | 0.512407284 | 0.628360222 | 0.813361516 | -4.71914289 |
| **A_55_P2024307** | Spin1 | -0.067483411 | -0.397352371 | 0.262385548 | 13.41616372 | -0.512396225 | 0.628367467 | 0.813361516 | -4.719144638 |
| **A_55_P2153827** | Fgfr1op | 0.100414327 | -0.390461709 | 0.591290363 | 10.85905843 | 0.512358795 | 0.628391991 | 0.813361516 | -4.719150555 |
| **A_55_P2129691** | AB124611 | -0.161519928 | -0.95113939 | 0.628099534 | 11.69464724 | -0.512340413 | 0.628404035 | 0.813361516 | -4.719153461 |
| **A_55_P2123002** | Tet1 | 0.072229035 | -0.280899136 | 0.425357205 | 8.092472714 | 0.512306413 | 0.628426313 | 0.813361516 | -4.719158835 |
| **A_51_P223575** | Tbc1d23 | -0.056533729 | -0.333045646 | 0.219978188 | 9.516998105 | -0.512087441 | 0.628569798 | 0.813492769 | -4.719193439 |
| **A_55_P1983418** | Amy1 | 0.072746533 | -0.283136049 | 0.428629115 | 7.839060758 | 0.511983438 | 0.628637954 | 0.81352652 | -4.71920987 |
| **A_51_P389885** | Spic | 0.050818968 | -0.197842233 | 0.299480169 | 7.793083323 | 0.511879996 | 0.628705746 | 0.813544067 | -4.719226209 |
| **A_55_P2079116** | Rpph1 | -0.253778539 | -1.495683655 | 0.988126577 | 10.47085131 | -0.511819494 | 0.6287454 | 0.813544067 | -4.719235765 |
| **A_55_P2033510** | Ppm1m | 0.085186342 | -0.331757753 | 0.502130436 | 10.93547152 | 0.511731896 | 0.628802813 | 0.813544067 | -4.719249599 |
| **A_55_P2157705** | Uqcr10 | 0.120713038 | -0.470146252 | 0.711572328 | 14.16720851 | 0.511705938 | 0.628819827 | 0.813544067 | -4.719253697 |
| **A_51_P278103** | Stambp | -0.05836033 | -0.344089156 | 0.227368496 | 10.07483616 | -0.511580555 | 0.628902013 | 0.81355556 | -4.719273493 |
| **A_55_P2247235** | Rab26os | -0.059927892 | -0.353343726 | 0.233487941 | 8.156171184 | -0.511559087 | 0.628916086 | 0.81355556 | -4.719276883 |
| **A_51_P158772** | Elof1 | 0.22172325 | -0.864077243 | 1.307523742 | 13.81856905 | 0.511459849 | 0.628981138 | 0.81355556 | -4.719292547 |
| **A_55_P2162573** | Art3 | -0.051035962 | -0.300975918 | 0.198903993 | 7.991149387 | -0.511435613 | 0.628997026 | 0.81355556 | -4.719296373 |
| **A_52_P617636** | Klf12 | 0.118360179 | -0.461379055 | 0.698099414 | 8.229808772 | 0.511355896 | 0.629049286 | 0.813568492 | -4.719308953 |
| **A_55_P2121760** | NA | 0.056715015 | -0.221126473 | 0.334556504 | 8.198000248 | 0.51127117 | 0.629104833 | 0.813568492 | -4.719322323 |
| **A_51_P121455** | Slc25a39 | -0.061879381 | -0.365070102 | 0.241311339 | 8.426465563 | -0.511187709 | 0.629159552 | 0.813568492 | -4.719335491 |
| **A_55_P2343151** | Akap1 | -0.12948054 | -0.763943648 | 0.504982567 | 9.67446788 | -0.511150128 | 0.629184193 | 0.813568492 | -4.71934142 |
| **A_52_P18765** | Hsbp1l1 | 0.053572041 | -0.20896049 | 0.316104571 | 8.034711106 | 0.511099447 | 0.629217423 | 0.813568492 | -4.719349415 |
| **A_51_P133684** | Csrp3 | 0.539778291 | -2.106538745 | 3.186095327 | 10.10927882 | 0.510885865 | 0.629357473 | 0.813609982 | -4.7193831 |
| **A_55_P1999361** | Dip2a | 0.132709481 | -0.517955947 | 0.78337491 | 10.73038581 | 0.510851374 | 0.629380092 | 0.813609982 | -4.719388539 |
| **A_51_P477941** | Fbxw4 | -0.063937096 | -0.377424473 | 0.249550281 | 10.02953782 | -0.510837996 | 0.629388865 | 0.813609982 | -4.719390649 |
| **A_55_P2093251** | NA | 0.050505264 | -0.197183554 | 0.298194082 | 9.239869847 | 0.510717327 | 0.629468 | 0.813609982 | -4.719409673 |
| **A_55_P2035454** | Oaz1 | 0.174557275 | -0.681573095 | 1.030687645 | 15.50628713 | 0.510679466 | 0.62949283 | 0.813609982 | -4.719415642 |
| **A_51_P150608** | Jagn1 | -0.112686665 | -0.665484156 | 0.440110826 | 11.35781958 | -0.510571781 | 0.629563457 | 0.813609982 | -4.719432615 |
| **A_52_P525837** | Trmt6 | -0.106515417 | -0.629041355 | 0.416010521 | 8.715459936 | -0.510569613 | 0.629564879 | 0.813609982 | -4.719432957 |
| **A_55_P2151061** | NA | -0.127664496 | -0.753979946 | 0.498650954 | 8.646766281 | -0.510537157 | 0.629586166 | 0.813609982 | -4.719438072 |
| **A_51_P159284** | Bud13 | 0.060819138 | -0.237675273 | 0.359313548 | 8.602247942 | 0.510333879 | 0.629719503 | 0.813693025 | -4.719470102 |
| **A_52_P404329** | Saa4 | -0.043168296 | -0.255043646 | 0.168707054 | 7.690817113 | -0.510310866 | 0.629734599 | 0.813693025 | -4.719473728 |
| **A_55_P1984406** | Lrrk2 | 0.403167256 | -1.577255647 | 2.38359016 | 10.50241691 | 0.509891366 | 0.630009816 | 0.813742607 | -4.719539792 |
| **A_51_P135654** | Eif4ebp2 | 0.056890819 | -0.22258723 | 0.336368868 | 11.32267564 | 0.509852823 | 0.630035106 | 0.813742607 | -4.71954586 |
| **A_66_P117289** | Cdk13 | -0.048717065 | -0.288041778 | 0.190607648 | 8.857692806 | -0.50985175 | 0.63003581 | 0.813742607 | -4.719546029 |
| **A_52_P965502** | Ssr1 | 0.045121527 | -0.176546944 | 0.266789998 | 8.712753032 | 0.509835661 | 0.630046367 | 0.813742607 | -4.719548562 |
| **A_55_P2025490** | Tnfrsf18 | -0.072468482 | -0.428487036 | 0.283550072 | 7.921255468 | -0.509831747 | 0.630048935 | 0.813742607 | -4.719549178 |
| **A_55_P2137773** | NA | -0.068587278 | -0.405599061 | 0.268424505 | 16.33653659 | -0.509740131 | 0.630109051 | 0.813742607 | -4.719563598 |
| **A_52_P316933** | Sh3bgrl2 | -0.284859992 | -1.684605356 | 1.114885373 | 9.94662318 | -0.50972128 | 0.630121421 | 0.813742607 | -4.719566565 |
| **A_55_P2172924** | NA | -0.045671765 | -0.270140805 | 0.178797274 | 8.270328063 | -0.509614407 | 0.630191554 | 0.813742607 | -4.719583384 |
| **A_55_P2002562** | Sec11a | -0.101020132 | -0.597624232 | 0.395583967 | 11.07231278 | -0.509504448 | 0.630263716 | 0.813742607 | -4.719600685 |
| **A_66_P125722** | Cyb5a | -0.080378898 | -0.475541411 | 0.314783616 | 14.55418675 | -0.509467696 | 0.630287836 | 0.813742607 | -4.719606467 |

| | | | | | | | | | |
|---|---|---|---|---|---|---|---|---|---|
| **A_55_P1971054** | Paqr9 | 0.047802099 | -0.187205046 | 0.282809243 | 7.694154099 | 0.509467219 | 0.630288149 | 0.813742607 | -4.719606542 |
| **A_55_P1959600** | Tmem222 | -0.074634679 | -0.441606951 | 0.292337594 | 11.57403589 | -0.509398618 | 0.630333173 | 0.813742607 | -4.719617333 |
| **A_55_P2170732** | Fndc10 | 0.073615124 | -0.288398659 | 0.435628908 | 9.092357736 | 0.509321829 | 0.630383572 | 0.813742607 | -4.719629412 |
| **A_55_P2079516** | Mpp1 | -0.050091497 | -0.296430417 | 0.196247423 | 8.000540763 | -0.509308966 | 0.630392015 | 0.813742607 | -4.719631435 |
| **A_55_P2111735** | Ppp1r2 | 0.104821027 | -0.410723865 | 0.620365918 | 10.12571943 | 0.509251441 | 0.630429773 | 0.813742607 | -4.719640482 |
| **A_55_P2016351** | Eif3c | -0.058116115 | -0.343995631 | 0.227763401 | 14.05834847 | -0.509171259 | 0.630482404 | 0.813742607 | -4.719653091 |
| **A_55_P1975120** | NA | 0.059687986 | -0.233962755 | 0.353338727 | 8.601352273 | 0.5091036 | 0.630526817 | 0.813742607 | -4.719663729 |
| **A_55_P2404157** | Rbm27 | -0.110051003 | -0.651481928 | 0.431379922 | 8.536568601 | -0.509097886 | 0.630530568 | 0.813742607 | -4.719664627 |
| **A_52_P653966** | Atf7ip | -0.057473897 | -0.34036697 | 0.225419175 | 10.4859056 | -0.508860431 | 0.630686453 | 0.81388946 | -4.719701953 |
| **A_55_P2114963** | Exosc10 | -0.060862445 | -0.360490855 | 0.238765964 | 8.934292266 | -0.508764446 | 0.630749472 | 0.813916458 | -4.719717037 |
| **A_66_P140452** | Map1a | 0.043933777 | -0.17242167 | 0.260289225 | 7.884263753 | 0.508605504 | 0.630853832 | 0.813946999 | -4.719742009 |
| **A_65_P07627** | Dach1 | 0.279098309 | -1.095421844 | 1.653618462 | 9.066355358 | 0.508576676 | 0.630872761 | 0.813946999 | -4.719746538 |
| **A_55_P2001347** | Rpl7a | -0.042813778 | -0.253696481 | 0.168068925 | 15.73723385 | -0.508502321 | 0.630921587 | 0.813946999 | -4.719758217 |
| **A_52_P655663** | H13 | 0.054775046 | -0.215059271 | 0.324609363 | 9.218772386 | 0.508435561 | 0.630965426 | 0.813946999 | -4.719768702 |
| **A_55_P2006754** | Mfsd12 | -0.179564301 | -1.06418707 | 0.705058468 | 11.65848637 | -0.508407831 | 0.630983636 | 0.813946999 | -4.719773057 |
| **A_51_P146813** | Mocs3 | -0.141462448 | -0.838672122 | 0.555747227 | 9.749733291 | -0.508192334 | 0.631125161 | 0.814061398 | -4.719806893 |
| **A_55_P2102050** | Cc2d1b | -0.106203497 | -0.629685641 | 0.417278647 | 12.03428887 | -0.508144573 | 0.63115653 | 0.814061398 | -4.719814391 |
| **A_55_P2121846** | NA | 0.119472707 | -0.469948084 | 0.708893499 | 16.30293926 | 0.507684144 | 0.631458979 | 0.814374134 | -4.719886638 |
| **A_55_P2010487** | Unk | 0.083484219 | -0.32843915 | 0.495407588 | 11.48213343 | 0.507619506 | 0.631501445 | 0.814374134 | -4.719896776 |
| **A_52_P54856** | Ubald1 | -0.056997793 | -0.338256825 | 0.224261239 | 9.878233072 | -0.507576969 | 0.631529392 | 0.814374134 | -4.719903448 |
| **A_51_P436483** | Kin | -0.054890583 | -0.325802757 | 0.216021591 | 10.47658169 | -0.50748086 | 0.631592538 | 0.814374134 | -4.719918519 |
| **A_51_P325836** | Hpx | -0.08243889 | -0.489336413 | 0.324458633 | 8.032853631 | -0.507454883 | 0.631609607 | 0.814374134 | -4.719922592 |
| **A_52_P417620** | Pacs1 | -0.046366973 | -0.275343264 | 0.182609318 | 8.577327371 | -0.507187541 | 0.631785279 | 0.814546318 | -4.7199645 |
| **A_51_P192751** | Mrpl14 | 0.196934841 | -0.775834629 | 1.169704311 | 14.34573497 | 0.507064224 | 0.631866321 | 0.814596483 | -4.719983825 |
| **A_55_P1996573** | Ttc13 | -0.131648066 | -0.782182019 | 0.518885888 | 12.68767219 | -0.506867983 | 0.631995299 | 0.814671932 | -4.720014569 |
| **A_55_P2047842** | Clcn3 | 0.060965816 | -0.240316464 | 0.362248096 | 11.79332443 | 0.506830974 | 0.632019624 | 0.814671932 | -4.720020366 |
| **A_51_P107362** | Socs2 | 0.336270612 | -1.325758644 | 1.998299869 | 10.73491159 | 0.50675798 | 0.632067604 | 0.814671932 | -4.718136939 |
| **A_55_P1982883** | Gys1 | -0.074033926 | -0.440026431 | 0.291958579 | 10.98032816 | -0.506651037 | 0.632137902 | 0.814671932 | -4.720048546 |
| **A_51_P257258** | Cpox | -0.116199853 | -0.69068667 | 0.458286965 | 9.944646005 | -0.506612447 | 0.632163269 | 0.814671932 | -4.720054588 |
| **A_55_P2024463** | Tbkbp1 | 0.076993316 | -0.303674611 | 0.457661243 | 8.706570967 | 0.506590546 | 0.632177666 | 0.814671932 | -4.720058017 |
| **A_55_P1961983** | Uqcr11 | 0.136742194 | -0.539602005 | 0.813086393 | 14.48979157 | 0.50639 | 0.63230951 | 0.814779253 | -4.720089411 |
| **A_51_P408471** | Cdipt | -0.06975534 | -0.414811101 | 0.275300421 | 12.72107469 | -0.506335668 | 0.632345231 | 0.814779253 | -4.720097914 |
| **A_55_P2039354** | NA | 0.07426012 | -0.293314489 | 0.441834729 | 8.207121446 | 0.506011617 | 0.632558308 | 0.814933706 | -4.720148615 |
| **A_52_P501049** | NA | 0.054239454 | -0.214253998 | 0.322732907 | 8.745778936 | 0.505978377 | 0.632580166 | 0.814933706 | -4.720153814 |
| **A_51_P293781** | Fkbp1b | -0.070682659 | -0.420584418 | 0.2792191 | 9.68778866 | -0.505961064 | 0.632591551 | 0.814933706 | -4.720156522 |
| **A_55_P2106489** | Ankrd52 | -0.0729152 | -0.434179342 | 0.288348942 | 8.564447934 | -0.505526076 | 0.632877644 | 0.815242959 | -4.720224531 |
| **A_51_P301809** | Slit3 | -0.05110122 | -0.304315431 | 0.20211299 | 8.444061313 | -0.505467855 | 0.632915941 | 0.815242959 | -4.72023363 |
| **A_55_P2087385** | Gps1 | -0.049705497 | -0.296193421 | 0.196782427 | 13.07187871 | -0.505078774 | 0.633171909 | 0.815502996 | -4.720294413 |
| **A_51_P306710** | Cldn15 | 0.076912154 | -0.304533531 | 0.45835784 | 8.19166152 | 0.505024695 | 0.633207491 | 0.815502996 | -4.720302859 |
| **A_51_P142153** | Filip1l | -0.089100105 | -0.531095032 | 0.352894822 | 10.72328618 | -0.504906868 | 0.63328502 | 0.815502996 | -4.720321256 |
| **A_55_P2018941** | Fmr1 | 0.048983865 | -0.194009327 | 0.291977058 | 9.984775345 | 0.504904557 | 0.633286541 | 0.815502996 | -4.720321617 |
| **A_55_P1959465** | Lrrfip1 | 0.140790264 | -0.557819197 | 0.839399726 | 10.84878927 | 0.504764154 | 0.633378932 | 0.815531422 | -4.720343535 |
| **A_51_P108901** | Ccdc86 | 0.099765898 | -0.395319729 | 0.594851525 | 10.37569642 | 0.504721894 | 0.633406742 | 0.815531422 | -4.720350132 |
| **A_52_P558577** | Zfp263 | 0.043508331 | -0.172419083 | 0.259435745 | 10.2578045 | 0.504678714 | 0.633435159 | 0.815531422 | -4.720356871 |
| **A_52_P69194** | Rimkla | 0.060502444 | -0.239870613 | 0.360875501 | 7.909321923 | 0.504501298 | 0.633551923 | 0.815605007 | -4.720384556 |
| **A_55_P2058433** | Cyp2c68 | 0.046439073 | -0.184142879 | 0.277021025 | 7.716267564 | 0.504438907 | 0.633592987 | 0.815605007 | -4.72039429 |
| **A_55_P2075013** | Tpt1 | 0.092398026 | -0.36641844 | 0.551214492 | 16.98879006 | 0.504398796 | 0.633619388 | 0.815605007 | -4.720400547 |
| **A_52_P92557** | Kdm2a | 0.081879233 | -0.3247556 | 0.488514067 | 9.555138638 | 0.504335496 | 0.633661054 | 0.815605007 | -4.720410421 |

| | | | | | | | | | |
|---|---|---|---|---|---|---|---|---|---|
| **A_55_P2117959** | Cyp4b1 | 0.266941029 | -1.059353778 | 1.593235837 | 9.120102247 | 0.504110342 | 0.633809266 | 0.815741469 | -4.720445533 |
| **A_52_P148678** | Plcg1 | -0.042579043 | -0.254161942 | 0.169003855 | 10.69293318 | -0.504040789 | 0.633855055 | 0.815746098 | -4.720456377 |
| **A_55_P1984962** | NA | -0.049100392 | -0.293212502 | 0.195011717 | 7.749050664 | -0.503785871 | 0.63402289 | 0.815907784 | -4.720496111 |
| **A_55_P1977418** | Ugt2b34 | -0.060467412 | -0.361280824 | 0.240346 | 7.797155698 | -0.503471083 | 0.634230176 | 0.816079261 | -4.720545152 |
| **A_51_P128065** | Prcc | -0.065127134 | -0.3891664 | 0.258912132 | 10.38006226 | -0.503401694 | 0.634275873 | 0.816079261 | -4.720555959 |
| **A_52_P485007** | Abca2 | 0.061200654 | -0.243324255 | 0.365725562 | 8.124187515 | 0.503365661 | 0.634299604 | 0.816079261 | -4.720561571 |
| **A_52_P209311** | Snx12 | 0.05673029 | -0.225572393 | 0.339032973 | 11.22089354 | 0.503327134 | 0.634324978 | 0.816079261 | -4.72056757 |
| **A_55_P1985749** | Rad23a | -0.057140872 | -0.341720208 | 0.227438464 | 7.868206386 | -0.502914141 | 0.634597011 | 0.816246548 | -4.720631856 |
| **A_52_P373798** | Zdhhc3 | 0.077709593 | -0.309354813 | 0.464773999 | 8.660574832 | 0.502853808 | 0.634636756 | 0.816246548 | -4.720641244 |
| **A_52_P150565** | Usp39 | -0.11126523 | -0.665475999 | 0.442945539 | 8.641704374 | -0.502845831 | 0.634642012 | 0.816246548 | -4.720642485 |
| **A_55_P2129407** | NA | 0.081554748 | -0.324679541 | 0.487789037 | 8.185702098 | 0.502832125 | 0.634651041 | 0.816246548 | -4.720644618 |
| **A_52_P222354** | Vps13c | -0.067725185 | -0.405176327 | 0.269725957 | 8.142914142 | -0.502677725 | 0.634752763 | 0.816246548 | -4.720668637 |
| **A_55_P2179027** | Gem | -0.072338448 | -0.432782083 | 0.288105188 | 8.350593228 | -0.502669076 | 0.634758461 | 0.816246548 | -4.720669982 |
| **A_55_P2130695** | Armc4 | 0.175169066 | -0.697724814 | 1.048062946 | 8.528273419 | 0.502627576 | 0.634785805 | 0.816246548 | -4.720676437 |
| **A_51_P227445** | NA | 0.171266607 | -0.682237037 | 1.024770251 | 8.74376328 | 0.502594417 | 0.634807652 | 0.816246548 | -4.720681594 |
| **A_55_P2003541** | Nrcam | 0.165502986 | -0.659351703 | 0.990357675 | 10.76248332 | 0.502549368 | 0.634837334 | 0.816246548 | -4.720688599 |
| **A_55_P2004427** | Fbxl15 | -0.055459603 | -0.331899451 | 0.220980245 | 9.438967997 | -0.502488882 | 0.63487719 | 0.816246548 | -4.720698005 |
| **A_51_P130095** | Fcgr2b | 0.055806004 | -0.222623567 | 0.334235575 | 7.949607088 | 0.502014097 | 0.635190078 | 0.816450256 | -4.703148002 |
| **A_55_P2093563** | Ufm1 | -0.088622592 | -0.530842219 | 0.353597036 | 11.51673772 | -0.501945744 | 0.63523513 | 0.816450256 | -4.720782421 |
| **A_55_P1980521** | Ccdc85a | 0.087700695 | -0.350006206 | 0.525407596 | 8.423025592 | 0.501845446 | 0.635301241 | 0.816450256 | -4.720798001 |
| **A_51_P490700** | Ccdc40 | 0.140638217 | -0.561398462 | 0.842674896 | 8.512817468 | 0.501757524 | 0.635359197 | 0.816450256 | -4.720811657 |
| **A_55_P2019615** | Neil1 | -0.050155834 | -0.300542276 | 0.200230607 | 9.449393629 | -0.501719519 | 0.63538425 | 0.816450256 | -4.720817559 |
| **A_55_P2076927** | Ints10 | 0.050601079 | -0.202043066 | 0.303245224 | 9.694688732 | 0.501650076 | 0.635430028 | 0.816450256 | -4.720828342 |
| **A_51_P171999** | Apoe | -0.071292927 | -0.427283615 | 0.284697762 | 13.4733504 | -0.501600719 | 0.635462567 | 0.816450256 | -4.720836006 |
| **A_52_P100252** | Fasn | 0.174060363 | -0.695396772 | 1.043517499 | 8.670803598 | 0.501420465 | 0.635581406 | 0.816450256 | -4.720863988 |
| **A_65_P12993** | Kif1b | 0.117897557 | -0.471044429 | 0.706839543 | 8.775843503 | 0.501398044 | 0.635596189 | 0.816450256 | -4.720867468 |
| **A_51_P201945** | Cfap97 | 0.07205307 | -0.287889245 | 0.431995384 | 9.061827556 | 0.501383374 | 0.635605862 | 0.816450256 | -4.720869745 |
| **A_66_P124776** | NA | 0.071227408 | -0.284607703 | 0.427062519 | 8.422116632 | 0.501358855 | 0.635622028 | 0.816450256 | -4.72087355 |
| **A_52_P484118** | Urb2 | 0.05537724 | -0.221274876 | 0.332029357 | 8.861578447 | 0.501357665 | 0.635622812 | 0.816450256 | -4.720873735 |
| **A_55_P2066613** | Lcp1 | -0.059203245 | -0.354996805 | 0.236590316 | 9.799301247 | -0.50131089 | 0.635653654 | 0.816450256 | -4.720880994 |
| **A_51_P370510** | Clba1 | -0.064266659 | -0.385373154 | 0.256839836 | 8.657658857 | -0.501287616 | 0.635669 | 0.816450256 | -4.720884606 |
| **A_55_P2241553** | 9330133C | 0.063776174 | -0.25487969 | 0.382432038 | 8.508432551 | 0.501287518 | 0.635669065 | 0.816450256 | -4.720884621 |
| **A_55_P1982454** | Eps8 | -0.068317439 | -0.410027485 | 0.273392608 | 9.951664622 | -0.500753707 | 0.636021097 | 0.81684814 | -4.720967419 |
| **A_55_P2031403** | NA | -0.049840093 | -0.299205614 | 0.199525429 | 9.301919112 | -0.500602233 | 0.63612101 | 0.816922192 | -4.720990901 |
| **A_55_P2172430** | Zxdb | -0.141473168 | -0.849624876 | 0.566678541 | 9.765043736 | -0.500377896 | 0.636268998 | 0.817057971 | -4.721025666 |
| **A_55_P2130090** | Cnn3 | -0.071032981 | -0.426798314 | 0.284732351 | 13.63255916 | -0.500088379 | 0.636460011 | 0.817248978 | -4.721070513 |
| **A_51_P396811** | Fam122a | 0.0733154 | -0.293979758 | 0.440610557 | 8.904100129 | 0.499954349 | 0.63654845 | 0.817290219 | -4.721091267 |
| **A_55_P2170414** | Zfp180 | 0.058804193 | -0.23581799 | 0.353426375 | 9.199414512 | 0.499911578 | 0.636576674 | 0.817290219 | -4.721097889 |
| **A_55_P2038767** | Fam49a | -0.095265793 | -0.572655351 | 0.382123765 | 9.069926688 | -0.499820903 | 0.63663651 | 0.817312768 | -4.721111925 |
| **A_55_P1994817** | Taf15 | 0.084444764 | -0.338841481 | 0.507731009 | 9.440823962 | 0.499676487 | 0.636731817 | 0.817380848 | -4.721134277 |
| **A_55_P2105808** | Wdr62 | -0.20809842 | -1.251466146 | 0.835269306 | 10.88529366 | -0.499553183 | 0.636813197 | 0.817394179 | -4.721153357 |
| **A_55_P2056876** | Akap13 | -0.105926409 | -0.637044049 | 0.425191231 | 9.043016355 | -0.499532638 | 0.636826757 | 0.817394179 | -4.721156536 |
| **A_55_P1974347** | Fam126b | -0.049549303 | -0.298025987 | 0.198927382 | 8.57074954 | -0.499461767 | 0.636873535 | 0.817399955 | -4.7211675 |
| **A_55_P1964682** | Rn28s1 | -0.157043492 | -0.945041841 | 0.630954857 | 16.43444309 | -0.499165983 | 0.637068784 | 0.817596275 | -4.721213246 |
| **A_55_P2000390** | Arpc5 | -0.069262892 | -0.416901772 | 0.278375989 | 14.50335205 | -0.499025366 | 0.637161618 | 0.817639873 | -4.721234986 |
| **A_51_P435321** | Ap1s2 | 0.086458288 | -0.347520257 | 0.520436834 | 8.65247912 | 0.498986412 | 0.637187337 | 0.817639873 | -4.721241007 |
| **A_66_P110909** | Ppp1r12a | -0.053726238 | -0.323564301 | 0.216111824 | 12.47797916 | -0.498693342 | 0.637380846 | 0.817801161 | -4.721286297 |
| **A_55_P2160473** | LOC63486 | -0.110857578 | -0.667666369 | 0.445951213 | 14.54307952 | -0.498665872 | 0.637398986 | 0.817801161 | -4.721290541 |

| | | | | | | | | | |
|---|---|---|---|---|---|---|---|---|---|
| **A_55_P1967983** | Use1 | 0.172495577 | -0.694012584 | 1.039003738 | 13.00436559 | 0.498603878 | 0.637439924 | 0.817801161 | -4.721300118 |
| **A_55_P2045224** | NA | 0.062814306 | -0.252767444 | 0.378396057 | 12.01663274 | 0.498536584 | 0.637484364 | 0.817803908 | -4.721310512 |
| **A_55_P2398264** | Rnf130 | 0.048966605 | -0.197086571 | 0.295019781 | 7.988980571 | 0.498449725 | 0.637541727 | 0.817823232 | -4.721323927 |
| **A_55_P2120141** | NA | -0.119388135 | -0.719505615 | 0.480729346 | 11.98908955 | -0.498282042 | 0.637652476 | 0.817861078 | -4.721349819 |
| **A_55_P2155833** | NA | -0.079715157 | -0.480416176 | 0.320985863 | 16.15052627 | -0.498276957 | 0.637655834 | 0.817861078 | -4.721350604 |
| **A_55_P2049266** | Mettl6 | -0.050966256 | -0.307497945 | 0.205565433 | 9.659262289 | -0.497613389 | 0.638094199 | 0.818319646 | -4.721452992 |
| **A_55_P2179791** | NA | 0.053223648 | -0.214678102 | 0.321125397 | 7.973222245 | 0.497598921 | 0.638103758 | 0.818319646 | -4.721455223 |
| **A_51_P124126** | Cyp2d22 | -0.042004524 | -0.253458231 | 0.169449183 | 7.717698335 | -0.49754356 | 0.638140338 | 0.818319646 | -4.72146376 |
| **A_55_P2146708** | Zcchc9 | -0.118297604 | -0.71412589 | 0.477530681 | 14.99228276 | -0.497284793 | 0.638311336 | 0.818411022 | -4.721503651 |
| **A_55_P2073814** | Gpatch8 | 0.043819345 | -0.176901215 | 0.264539904 | 8.464362576 | 0.497248449 | 0.638335355 | 0.818411022 | -4.721509253 |
| **A_52_P200207** | Rmnd5a | 0.067017761 | -0.270557876 | 0.404593399 | 9.12174613 | 0.49724356 | 0.638338586 | 0.818411022 | -4.721510006 |
| **A_55_P2018542** | Ptges3 | 0.053729417 | -0.217098309 | 0.324557142 | 10.87611486 | 0.496900404 | 0.638565392 | 0.818647523 | -4.721562876 |
| **A_51_P380309** | Ncam1 | -0.147307242 | -0.890011104 | 0.59539662 | 10.5237476 | -0.496773891 | 0.638649021 | 0.818700449 | -4.72158236 |
| **A_55_P2120545** | Wdtc1 | -0.043398873 | -0.262238698 | 0.175440952 | 8.363050264 | -0.496709469 | 0.638691608 | 0.818700759 | -4.721592279 |
| **A_55_P1981731** | Nr2c2 | 0.042147799 | -0.170443623 | 0.25473922 | 8.514789069 | 0.496568888 | 0.638784546 | 0.818717374 | -4.721613923 |
| **A_55_P2168048** | Gramd4 | -0.057061307 | -0.344879867 | 0.230757252 | 8.141129496 | -0.496561756 | 0.638789262 | 0.818717374 | -4.72161502 |
| **A_51_P254656** | Hdc | 0.45943737 | -1.860169619 | 2.779044359 | 10.39715859 | 0.496092035 | 0.63909985 | 0.81906115 | -4.721687298 |
| **A_52_P609334** | Klhdc8b | 0.174586212 | -0.707138541 | 1.056310965 | 8.760550644 | 0.495937849 | 0.639201819 | 0.819105865 | -4.72171101 |
| **A_52_P430169** | Dip2b | -0.05110792 | -0.30929656 | 0.207080721 | 9.350271183 | -0.495794173 | 0.639296845 | 0.819105865 | -4.7217331 |
| **A_55_P2274378** | AW54954: | 0.203698262 | -0.82541003 | 1.232806554 | 8.76236871 | 0.495765832 | 0.639315591 | 0.819105865 | -4.721737457 |
| **A_55_P1996212** | Zbtb2 | 0.052545331 | -0.212951083 | 0.318041745 | 10.0016643 | 0.495707875 | 0.639353926 | 0.819105865 | -4.721746366 |
| **A_55_P2176240** | Pdxk | 0.05414391 | -0.219474346 | 0.327762165 | 8.148054574 | 0.495626922 | 0.639407473 | 0.819105865 | -4.721758808 |
| **A_52_P339054** | Nap1l4 | 0.052233753 | -0.211735472 | 0.316202979 | 9.6139376 | 0.495619381 | 0.639412461 | 0.819105865 | -4.721759967 |
| **A_55_P2027312** | Cdc34 | -0.326095298 | -1.974486108 | 1.322295512 | 13.73522377 | -0.495489776 | 0.639498196 | 0.819105865 | -4.721779883 |
| **A_51_P208031** | Cnot6l | 0.049923227 | -0.2024437 | 0.302290154 | 8.912812506 | 0.49547364 | 0.63950887 | 0.819105865 | -4.721782363 |
| **A_55_P2164778** | Ptdss1 | -0.065034834 | -0.393826702 | 0.263757033 | 11.55092722 | -0.495422033 | 0.63954301 | 0.819105865 | -4.721790292 |
| **A_55_P2116165** | Pfkfb3 | -0.121419582 | -0.735362675 | 0.492523512 | 10.38988799 | -0.495348622 | 0.639591577 | 0.819105865 | -4.721801569 |
| **A_55_P2178006** | NA | 0.070688617 | -0.2867948 | 0.428172035 | 8.436982357 | 0.495272177 | 0.639642153 | 0.819105865 | -4.721813312 |
| **A_55_P2010003** | Zfp113 | 0.084501186 | -0.342835836 | 0.511838209 | 8.794447077 | 0.495270694 | 0.639643134 | 0.819105865 | -4.721813539 |
| **A_55_P2015767** | Yipf6 | 0.081258429 | -0.329737642 | 0.492254499 | 9.110287398 | 0.495200525 | 0.63968956 | 0.819111064 | -4.721824316 |
| **A_51_P395921** | Uchl4 | -0.223409363 | -1.35356346 | 0.906744735 | 10.33048258 | -0.495124084 | 0.639740137 | 0.81911355 | -4.721836055 |
| **A_51_P234788** | Cxxc5 | -0.231723457 | -1.404064984 | 0.940618069 | 9.464102667 | -0.495069529 | 0.639776235 | 0.81911355 | -4.721844432 |
| **A_51_P454217** | Nfkb2 | -0.072531399 | -0.439663218 | 0.294600421 | 8.720393006 | -0.494828119 | 0.639935983 | 0.819210047 | -4.72188149 |
| **A_52_P243412** | Ulk2 | 0.055756805 | -0.226467452 | 0.337981062 | 9.237907506 | 0.494827569 | 0.639936347 | 0.819210047 | -4.721881575 |
| **A_55_P2174339** | NA | 0.218603043 | -0.888171942 | 1.325378028 | 8.492742741 | 0.494706041 | 0.640016774 | 0.819221149 | -4.721900224 |
| **A_55_P2014511** | Amd2 | 0.071779248 | -0.291683138 | 0.435241635 | 10.07789095 | 0.494640621 | 0.640060071 | 0.819221149 | -4.721910262 |
| **A_52_P244723** | Tbcel | 0.153660372 | -0.624547475 | 0.931868218 | 10.12103839 | 0.494557304 | 0.640115216 | 0.819221149 | -4.721923044 |
| **A_51_P280437** | Slc12a6 | -0.07156141 | -0.434006078 | 0.290883258 | 9.047795886 | -0.494524167 | 0.640137148 | 0.819221149 | -4.721928127 |
| **A_55_P2115068** | Psmb5 | 0.050096706 | -0.203648692 | 0.303842103 | 12.7225519 | 0.494494356 | 0.64015688 | 0.819221149 | -4.7219327 |
| **A_55_P2207335** | Nron | 0.06929429 | -0.28177586 | 0.420364439 | 8.277161771 | 0.494372025 | 0.640237854 | 0.819270545 | -4.721951462 |
| **A_55_P2172440** | Scarf2 | 0.222226388 | -0.90390662 | 1.348359395 | 13.20770686 | 0.494260933 | 0.640311393 | 0.819310422 | -4.721968497 |
| **A_51_P373379** | Csnk2b | 0.084902117 | -0.345501387 | 0.51530562 | 13.95092989 | 0.494075211 | 0.640434345 | 0.819411209 | -4.721996969 |
| **A_51_P485756** | Nts | 0.094578205 | -0.384936831 | 0.574093241 | 8.102580188 | 0.494013917 | 0.640474925 | 0.819411209 | -4.722006363 |
| **A_51_P476018** | Sox7 | 0.213288146 | -0.868563523 | 1.295139816 | 9.8532389 | 0.493798022 | 0.640617873 | 0.819468374 | -4.722039446 |
| **A_55_P2313033** | Ptprb | 0.07443322 | -0.30312892 | 0.451995359 | 9.153004932 | 0.493774562 | 0.640633407 | 0.819468374 | -4.72204304 |
| **A_52_P148658** | Mettl22 | -0.052997838 | -0.321873589 | 0.215877912 | 8.64281425 | -0.493692882 | 0.640687494 | 0.819468374 | -4.722055552 |
| **A_52_P153848** | Prpf4b | 0.096970725 | -0.394996665 | 0.588938114 | 8.62327152 | 0.493690388 | 0.640689146 | 0.819468374 | -4.722055934 |
| **A_55_P2042481** | Gm10653 | 0.080468756 | -0.327837767 | 0.48877528 | 15.81920404 | 0.493618374 | 0.640736834 | 0.819475157 | -4.722066964 |

| | | | | | | | | | |
|---|---|---|---|---|---|---|---|---|---|
| **A_51_P488092** | Pde6d | 0.084551691 | -0.344554253 | 0.513657635 | 9.661366731 | 0.493523816 | 0.640799455 | 0.819501035 | -4.722081445 |
| **A_55_P2030899** | NA | 0.07477679 | -0.304934363 | 0.454487944 | 11.4830701 | 0.493246272 | 0.640983274 | 0.819681899 | -4.722123936 |
| **A_51_P475378** | Echdc3 | -0.109860597 | -0.667936907 | 0.448215714 | 9.63563794 | -0.493058796 | 0.641107458 | 0.819762651 | -4.722152626 |
| **A_51_P452779** | Pygl | 0.132759306 | -0.541734047 | 0.807252658 | 13.79347768 | 0.492989471 | 0.641153382 | 0.819762651 | -4.722163232 |
| **A_55_P1981674** | NA | -0.04439443 | -0.269957555 | 0.181168695 | 8.466768862 | -0.492958918 | 0.641173622 | 0.819762651 | -4.722167907 |
| **A_65_P13527** | Osbpl8 | 0.044343204 | -0.181016436 | 0.269702843 | 10.33880553 | 0.492834694 | 0.641255919 | 0.819813658 | -4.722186909 |
| **A_55_P1994175** | Ppil3 | 0.044135182 | -0.180255707 | 0.26852607 | 9.40084232 | 0.492640425 | 0.641384632 | 0.819923993 | -4.722216617 |
| **A_55_P2143139** | Myadml2 | 0.085503468 | -0.349303928 | 0.520310864 | 8.044946252 | 0.492535062 | 0.641454446 | 0.819959025 | -4.722232725 |
| **A_51_P121031** | 1-Mar | -0.05032073 | -0.306253527 | 0.205612068 | 7.878334962 | -0.49246042 | 0.641503907 | 0.819968037 | -4.719830016 |
| **A_55_P2075140** | Sirt3 | -0.059059921 | -0.359681002 | 0.241561161 | 10.72826836 | -0.49206647 | 0.641764987 | 0.820171988 | -4.722304329 |
| **A_55_P1961127** | H19 | -0.19925969 | -1.213718603 | 0.815199223 | 16.7515399 | -0.491966277 | 0.641831397 | 0.820171988 | -4.722319632 |
| **A_55_P1980336** | NA | 0.044831051 | -0.183424732 | 0.273086833 | 7.886765784 | 0.491934718 | 0.641852315 | 0.820171988 | -4.722324452 |
| **A_51_P244923** | AI837181 | 0.057550174 | -0.235472085 | 0.350572432 | 9.551585955 | 0.491922145 | 0.641860649 | 0.820171988 | -4.722326372 |
| **A_55_P2184766** | Tmc6 | -0.04869124 | -0.296623717 | 0.199241238 | 9.167660948 | -0.491889731 | 0.641882135 | 0.820171988 | -4.722331321 |
| **A_55_P2302413** | BC00218( | -0.04860637 | -0.29613392 | 0.19892118 | 7.76091304 | -0.491835632 | 0.641917996 | 0.820171988 | -4.722339582 |
| **A_55_P2049473** | Sh3bp5 | 0.042672999 | -0.174759687 | 0.260105685 | 8.365490647 | 0.491562425 | 0.642099114 | 0.820349188 | -4.722381286 |
| **A_51_P450573** | Tgfbr2 | 0.151746251 | -0.621645245 | 0.925137747 | 12.92587861 | 0.491438213 | 0.642181467 | 0.820365046 | -4.722400241 |
| **A_52_P153327** | 2-Sep | -0.087702256 | -0.534737643 | 0.35933313 | 8.319516601 | -0.491381962 | 0.642218763 | 0.820365046 | -4.722408823 |
| **A_51_P475291** | Moap1 | 0.051402022 | -0.210647874 | 0.313451918 | 8.6530935 | 0.491299767 | 0.642273264 | 0.820365046 | -4.722421362 |
| **A_55_P2141943** | NA | 0.051254171 | -0.21004838 | 0.312556722 | 8.021024367 | 0.491287722 | 0.642281251 | 0.820365046 | -4.722423199 |
| **A_51_P195100** | Vps4b | -0.049319451 | -0.300794157 | 0.202155254 | 11.71456779 | -0.491218001 | 0.642327483 | 0.820369901 | -4.722433834 |
| **A_51_P104125** | Fam76b | 0.137765664 | -0.565413282 | 0.840944611 | 9.953987485 | 0.490710643 | 0.642663965 | 0.820745433 | -4.706571735 |
| **A_51_P224771** | Tmem184 | -0.0535289 | -0.326848367 | 0.219790566 | 12.09056693 | -0.490532858 | 0.642781895 | 0.82084182 | -4.72253827 |
| **A_51_P518600** | Atp6v1c2 | 0.354376413 | -1.455333185 | 2.164086012 | 9.586567248 | 0.490463015 | 0.642828227 | 0.82084677 | -4.722548909 |
| **A_55_P2145804** | Aen | 0.094037016 | -0.386313566 | 0.574387598 | 11.73643677 | 0.490332707 | 0.642914676 | 0.820902942 | -4.722568755 |
| **A_52_P504330** | Ankib1 | -0.051868711 | -0.316885278 | 0.213147855 | 9.286384099 | -0.490210694 | 0.642995626 | 0.820952086 | -4.722587334 |
| **A_55_P2040583** | Map6d1 | -0.153886367 | -0.940455946 | 0.632683212 | 10.80172383 | -0.490019481 | 0.643122499 | 0.821059852 | -4.722616442 |
| **A_51_P302167** | Cobl | -0.044275775 | -0.270666473 | 0.182114922 | 8.076842972 | -0.489844168 | 0.643238835 | 0.821136189 | -4.722643121 |
| **A_51_P402160** | Zfp750 | -0.048439587 | -0.296142311 | 0.199263136 | 7.795304583 | -0.489801373 | 0.643267235 | 0.821136189 | -4.722649632 |
| **A_52_P144818** | Ppp1r1a | 0.048330258 | -0.198937423 | 0.295597939 | 7.799507522 | 0.48955569 | 0.643430289 | 0.821290104 | -4.722687004 |
| **A_55_P1974855** | Zfp868 | 0.067439263 | -0.277783118 | 0.412661644 | 9.268165112 | 0.489287611 | 0.643608233 | 0.821423291 | -4.722727763 |
| **A_55_P2021054** | Elp5 | -0.078923733 | -0.482949395 | 0.325101929 | 11.70921529 | -0.48927048 | 0.643619604 | 0.821423291 | -4.722730367 |
| **A_55_P2112474** | NA | -0.050166654 | -0.307034232 | 0.206700924 | 16.26406499 | -0.489165919 | 0.643689017 | 0.821436718 | -4.72274626 |
| **A_51_P326030** | Zfp959 | 0.082490621 | -0.33994236 | 0.504923603 | 8.466767806 | 0.489099369 | 0.643733198 | 0.821436718 | -4.722756373 |
| **A_51_P478952** | N4bp2l1 | -0.049424745 | -0.302566524 | 0.203717035 | 8.118584271 | -0.489024875 | 0.643782655 | 0.821436718 | -4.722767692 |
| **A_55_P1971474** | NA | -0.10113121 | -0.619128985 | 0.416866566 | 10.96176407 | -0.488998642 | 0.643800072 | 0.821436718 | -4.722771678 |
| **A_55_P2204491** | AI131754 | -0.122098161 | -0.747857763 | 0.503661441 | 8.62420764 | -0.488710811 | 0.643991186 | 0.821597513 | -4.722815397 |
| **A_52_P52156** | Isoc2a | 0.042905219 | -0.176999806 | 0.262810245 | 7.94408939 | 0.488680845 | 0.644011085 | 0.821597513 | -4.722819948 |
| **A_65_P13910** | Ndufb2 | 0.137508933 | -0.567817752 | 0.842835619 | 14.56953011 | 0.488304744 | 0.644260859 | 0.821861932 | -4.722877039 |
| **A_55_P1981366** | Lamc2 | 0.226875365 | -0.937975786 | 1.391726515 | 13.6676816 | 0.487828594 | 0.644577152 | 0.822208176 | -4.722949263 |
| **A_55_P2075814** | Tnni3k | 0.1133802 | -0.46886067 | 0.695621069 | 8.115243853 | 0.487736071 | 0.644638621 | 0.822208176 | -4.72296329 |
| **A_55_P1993609** | Slx4ip | -0.064721162 | -0.397105262 | 0.267662938 | 8.827613017 | -0.487704103 | 0.644659861 | 0.822208176 | -4.722968136 |
| **A_55_P2180096** | Gm10136 | 0.065900095 | -0.272587475 | 0.404387665 | 15.8244436 | 0.487633646 | 0.644706674 | 0.822213643 | -4.722978816 |
| **A_55_P2045136** | Ly6a2 | -0.040976226 | -0.251627728 | 0.169675276 | 7.708305733 | -0.487211733 | 0.644987038 | 0.822423137 | -4.72304274 |
| **A_52_P96748** | Ttc30b | 0.174591534 | -0.723099628 | 1.072282696 | 9.234653145 | 0.487131908 | 0.64504009 | 0.822423137 | -4.72305483 |
| **A_66_P133525** | NA | -0.089781494 | -0.551418067 | 0.371855079 | 7.941873139 | -0.487121127 | 0.645047255 | 0.822423137 | -4.723056462 |
| **A_55_P2007525** | Smu1 | -0.082320725 | -0.505624578 | 0.340983129 | 9.465664068 | -0.487087863 | 0.645069363 | 0.822423137 | -4.723061499 |
| **A_51_P465128** | Chst15 | 0.126111424 | -0.522578688 | 0.774801536 | 8.977937192 | 0.486931083 | 0.645173568 | 0.822423137 | -4.723085236 |

| | | | | | | | | | |
|---|---|---|---|---|---|---|---|---|---|
| **A_55_P2046661** | Tnfrsf14 | 0.046249695 | -0.191676242 | 0.284175632 | 7.975641173 | 0.486874953 | 0.645210878 | 0.822423137 | -4.723093733 |
| **A_51_P270666** | Uspl1 | -0.127822671 | -0.785402389 | 0.529757047 | 10.93360298 | -0.486866423 | 0.645216548 | 0.822423137 | -4.723095024 |
| **A_55_P2023424** | Trip10 | 0.130857475 | -0.542375461 | 0.804090411 | 11.357957 | 0.486836942 | 0.645236144 | 0.822423137 | -4.723099486 |
| **A_55_P1972192** | Phka1 | -0.056884845 | -0.349560812 | 0.235791123 | 9.463521272 | -0.486810412 | 0.645253779 | 0.822423137 | -4.723103501 |
| **A_55_P2143436** | NA | -0.102729438 | -0.631368487 | 0.42590961 | 7.898219796 | -0.486727651 | 0.645308795 | 0.82243904 | -4.723116026 |
| **A_55_P1960197** | P2ry14 | 0.40346344 | -1.673981934 | 2.480908814 | 9.99997202 | 0.486435116 | 0.645503277 | 0.822632678 | -4.723160283 |
| **A_55_P2074843** | Ccdc122 | 0.05872466 | -0.243720814 | 0.361170134 | 8.333228029 | 0.48632183 | 0.6455786 | 0.822660835 | -4.723177416 |
| **A_55_P2124415** | 6030468B | 0.040779335 | -0.169267773 | 0.250826444 | 7.650088645 | 0.48626586 | 0.645615815 | 0.822660835 | -4.723185879 |
| **A_55_P1982986** | Cfap410 | -0.040824539 | -0.251128686 | 0.169479607 | 8.255430128 | -0.486209906 | 0.645653021 | 0.822660835 | -4.723194339 |
| **A_55_P2145543** | Eef2k | -0.137398291 | -0.845923453 | 0.571126872 | 8.493940907 | -0.485709275 | 0.645985963 | 0.823014747 | -4.723269996 |
| **A_55_P2078088** | Pqlc1 | 0.176928596 | -0.735528079 | 1.08938527 | 11.02945064 | 0.485664236 | 0.646015921 | 0.823014747 | -4.723276799 |
| **A_66_P137383** | Ddx55 | -0.055256352 | -0.340334479 | 0.229821776 | 8.324014536 | -0.485476997 | 0.646140469 | 0.823048012 | -4.723305075 |
| **A_55_P2170813** | Pkig | 0.060132742 | -0.250172792 | 0.370438276 | 13.03800043 | 0.485368776 | 0.646212462 | 0.823048012 | -4.723321414 |
| **A_51_P196158** | Btd | -0.045339816 | -0.279329709 | 0.188650077 | 10.10063418 | -0.485325377 | 0.646241334 | 0.823048012 | -4.723327966 |
| **A_55_P2040878** | Ung | 0.092546894 | -0.385104983 | 0.570198771 | 14.01830669 | 0.485289289 | 0.646265343 | 0.823048012 | -4.723333413 |
| **A_55_P2044091** | Tdg | -0.10281012 | -0.633452607 | 0.427832367 | 8.898261935 | -0.485270835 | 0.64627762 | 0.823048012 | -4.723336199 |
| **A_51_P384946** | Ndufa2 | 0.122984029 | -0.511833369 | 0.757801428 | 14.7295633 | 0.485232902 | 0.646302857 | 0.823048012 | -4.723341924 |
| **A_55_P2060021** | Otub2 | 0.091765904 | -0.381964707 | 0.565496514 | 9.318006408 | 0.485177043 | 0.646340021 | 0.823048012 | -4.723350354 |
| **A_52_P493620** | Fgfr1op2 | -0.159673557 | -0.984294995 | 0.664947882 | 9.763475255 | -0.484985466 | 0.64646749 | 0.823075622 | -4.723379261 |
| **A_51_P146505** | Atp11b | 0.074022942 | -0.308270084 | 0.456315968 | 9.035109543 | 0.48497607 | 0.646473742 | 0.823075622 | -4.723380678 |
| **A_51_P304510** | BC017158 | -0.060171338 | -0.370942563 | 0.250599887 | 11.18444641 | -0.484952513 | 0.646489418 | 0.823075622 | -4.723384232 |
| **A_66_P134878** | D11Wsu47 | -0.085336339 | -0.526251848 | 0.355579171 | 10.96285844 | -0.484762454 | 0.646615892 | 0.823182436 | -4.723412898 |
| **A_51_P251717** | Hikeshi | 0.072819203 | -0.303532427 | 0.449170834 | 10.005592 | 0.484621264 | 0.646709855 | 0.823247849 | -4.723434188 |
| **A_55_P2045792** | Pes1 | -0.160163766 | -0.988300876 | 0.667973345 | 10.82570663 | -0.484409186 | 0.646851008 | 0.823316301 | -4.723466156 |
| **A_55_P2080758** | Slc8a1 | 0.058875221 | -0.260844186 | 0.378594629 | 7.863065542 | 0.490485597 | 0.646870182 | 0.823316301 | -4.698507044 |
| **A_52_P920158** | Rbm24 | 0.243795624 | -1.016923273 | 1.504514522 | 8.821775728 | 0.484348534 | 0.64689138 | 0.823316301 | -4.723475296 |
| **A_55_P1953030** | Tnfsf12 | -0.042270425 | -0.260919683 | 0.176378832 | 8.140346914 | -0.4842158 | 0.646979735 | 0.823374552 | -4.723495296 |
| **A_51_P321743** | NA | -0.042894781 | -0.264926311 | 0.17913675 | 7.824880392 | -0.483882754 | 0.647201458 | 0.823565765 | -4.723545456 |
| **A_51_P520384** | Atp1b3 | -0.053367737 | -0.329673449 | 0.222937975 | 11.6854739 | -0.483770373 | 0.647276284 | 0.823565765 | -4.723562376 |
| **A_55_P2133485** | Gapvd1 | 0.042400196 | -0.177133803 | 0.261934195 | 10.25327534 | 0.483744925 | 0.647293229 | 0.823565765 | -4.723566206 |
| **A_55_P2035593** | Hsf1 | -0.140355481 | -0.867103233 | 0.586392271 | 10.35034412 | -0.483722206 | 0.647308357 | 0.823565765 | -4.723569626 |
| **A_55_P2005743** | Ubxn11 | 0.090858134 | -0.37977413 | 0.561490397 | 8.389194234 | 0.48354006 | 0.647429647 | 0.823565765 | -4.723597039 |
| **A_55_P2043782** | Trpm1 | 0.059220586 | -0.247541325 | 0.365982497 | 8.215532501 | 0.483527981 | 0.647437691 | 0.823565765 | -4.723598857 |
| **A_55_P2030662** | Eda | 0.056442109 | -0.23596813 | 0.348852349 | 8.376738549 | 0.483460507 | 0.647482624 | 0.823565765 | -4.723609009 |
| **A_55_P2183438** | Runx1 | -0.177477705 | -1.09695804 | 0.742002629 | 8.79891957 | -0.483450157 | 0.647489517 | 0.823565765 | -4.723610566 |
| **A_55_P1986902** | Med23 | 0.049585304 | -0.207326334 | 0.306496943 | 9.025450545 | 0.483414364 | 0.647513354 | 0.823565765 | -4.723615952 |
| **A_55_P2167416** | NA | 0.05255333 | -0.219935559 | 0.325042219 | 8.177834845 | 0.483060791 | 0.647748848 | 0.823811092 | -4.723669127 |
| **A_55_P2175341** | NA | 0.056223278 | -0.235340263 | 0.347786819 | 9.627630806 | 0.482984609 | 0.647799594 | 0.823821439 | -4.72368058 |
| **A_55_P2102828** | NA | 0.040500571 | -0.169583227 | 0.250584369 | 7.681304953 | 0.482857447 | 0.647884304 | 0.823846604 | -4.723699694 |
| **A_55_P2142151** | Mapk1ip1 | -0.093082719 | -0.575950113 | 0.389784676 | 10.53439138 | -0.482826974 | 0.647904604 | 0.823846604 | -4.723704274 |
| **A_52_P427240** | Ltn1 | 0.065227559 | -0.273227906 | 0.403683024 | 8.328183442 | 0.482702938 | 0.647987238 | 0.823897492 | -4.723722913 |
| **A_51_P222475** | Ehbp1l1 | -0.096658832 | -0.598439039 | 0.405121376 | 9.669017712 | -0.482478938 | 0.648136484 | 0.82402384 | -4.723756562 |
| **A_66_P132446** | Usp15 | 0.083450496 | -0.349809597 | 0.516710589 | 8.105930245 | 0.482425861 | 0.648171851 | 0.82402384 | -4.723764534 |
| **A_55_P2234704** | NA | 0.104470877 | -0.438096316 | 0.64703807 | 8.248337376 | 0.482272009 | 0.648274372 | 0.824099988 | -4.723787636 |
| **A_55_P2097508** | Mcc | 0.122504034 | -0.513933972 | 0.75894204 | 11.26635105 | 0.482108323 | 0.648383456 | 0.824168001 | -4.723812208 |
| **A_51_P504065** | Spa17 | 0.154479052 | -0.648151011 | 0.957109115 | 9.635479955 | 0.482063798 | 0.64841313 | 0.824168001 | -4.72381889 |
| **A_52_P572447** | Agpat5 | -0.082000846 | -0.508144545 | 0.344142853 | 10.50512983 | -0.48196179 | 0.648481117 | 0.824179377 | -4.723834199 |
| **A_51_P507899** | Ttc8 | 0.060970133 | -0.255906584 | 0.377846851 | 9.957292427 | 0.481922451 | 0.648507337 | 0.824179377 | -4.723840101 |

| | | | | | | | | | |
|---|---|---|---|---|---|---|---|---|---|
| **A_55_P2003903** | Bfsp1 | 0.084514055 | -0.354792965 | 0.523821075 | 8.478730999 | 0.4818492 | 0.648556161 | 0.82418725 | -4.723851091 |
| **A_55_P1989276** | Romo1 | -0.055621331 | -0.344833684 | 0.233591022 | 14.14871968 | -0.481698043 | 0.648656918 | 0.824261115 | -4.723873766 |
| **A_51_P211341** | Eif3j1 | -0.108623603 | -0.673690491 | 0.456443284 | 10.50849748 | -0.481476054 | 0.648804906 | 0.824302043 | -4.723907054 |
| **A_51_P147373** | Dpp3 | -0.098524827 | -0.611082291 | 0.414032637 | 11.68312683 | -0.481452522 | 0.648820594 | 0.824302043 | -4.723910582 |
| **A_51_P375146** | Cd36 | 0.161963563 | -0.680762082 | 1.004689208 | 10.73268162 | 0.481372701 | 0.648873811 | 0.824302043 | -4.723922547 |
| **A_55_P2147126** | Zfp712 | 0.116038863 | -0.487806197 | 0.719883923 | 8.692761296 | 0.481313775 | 0.648913099 | 0.824302043 | -4.72393138 |
| **A_51_P155755** | Pld6 | 0.208482318 | -0.876521925 | 1.29348656 | 8.838674602 | 0.481269274 | 0.64894277 | 0.824302043 | -4.723938049 |
| **A_55_P1954987** | Slc10a1 | 0.065778649 | -0.276555394 | 0.408112691 | 17.49731738 | 0.481266025 | 0.648944936 | 0.824302043 | -4.723938536 |
| **A_51_P511734** | Snip1 | -0.04547038 | -0.282171585 | 0.191230824 | 8.724197317 | -0.481147755 | 0.649023796 | 0.824348054 | -4.723956259 |
| **A_55_P2033927** | NA | 0.143695612 | -0.604749975 | 0.892141198 | 16.72194682 | 0.480876572 | 0.649204635 | 0.824444275 | -4.723996882 |
| **A_52_P579531** | Pdlim3 | 0.279515004 | -1.176377702 | 1.73540771 | 10.68252567 | 0.480868247 | 0.649210187 | 0.824444275 | -4.723998129 |
| **A_55_P2008740** | Fcgr1 | -0.05736822 | -0.356194674 | 0.241458234 | 8.160558004 | -0.480842317 | 0.64922748 | 0.824444275 | -4.724002012 |
| **A_55_P1976869** | Hsdl2 | -0.048419961 | -0.300751602 | 0.203911679 | 9.604978975 | -0.480621366 | 0.649374844 | 0.82455934 | -4.724035094 |
| **A_55_P2090364** | NA | -0.106499997 | -0.661560937 | 0.448560943 | 14.7707995 | -0.480572897 | 0.649407172 | 0.82455934 | -4.72404235 |
| **A_55_P2094901** | Phyhd1 | -0.107252552 | -0.666384225 | 0.45187912 | 16.36328944 | -0.480445232 | 0.649492329 | 0.82455934 | -4.724061457 |
| **A_51_P104820** | Alkbh5 | -0.069428205 | -0.43143271 | 0.292576301 | 8.289762326 | -0.48036605 | 0.649545149 | 0.82455934 | -4.724073305 |
| **A_51_P404413** | Bckdk | 0.046751573 | -0.197033437 | 0.290536582 | 10.62344199 | 0.480329845 | 0.649569302 | 0.82455934 | -4.724078722 |
| **A_55_P2320085** | 4833415N | 0.054943043 | -0.231560384 | 0.341446469 | 8.230496313 | 0.480322833 | 0.649573979 | 0.82455934 | -4.724079771 |
| **A_52_P263870** | Cdc23 | 0.042326833 | -0.178424879 | 0.263078546 | 8.113083994 | 0.480244104 | 0.649626501 | 0.824571873 | -4.72409155 |
| **A_55_P1998872** | Slitrk5 | 0.044115086 | -0.18602503 | 0.274255202 | 7.749833787 | 0.480114861 | 0.649712726 | 0.82462718 | -4.724110881 |
| **A_55_P2282257** | NA | -0.085137152 | -0.529789639 | 0.359515334 | 9.905301276 | -0.479566393 | 0.650078705 | 0.824972709 | -4.72419287 |
| **A_55_P2060922** | Unc5a | 0.047995454 | -0.202743714 | 0.298734621 | 9.647358808 | 0.479433379 | 0.650167479 | 0.824972709 | -4.724212741 |
| **A_52_P163515** | NA | 0.060149344 | -0.254102865 | 0.374401554 | 8.646282979 | 0.47940538 | 0.650186166 | 0.824972709 | -4.724216923 |
| **A_55_P2068066** | Rapgefl1 | 0.162088353 | -0.684972372 | 1.009149078 | 8.668651444 | 0.479278127 | 0.650271103 | 0.824972709 | -4.724235929 |
| **A_55_P2115477** | Parva | -0.068096241 | -0.423992773 | 0.287800292 | 9.179978515 | -0.479236322 | 0.650299007 | 0.824972709 | -4.724242172 |
| **A_51_P268734** | Kptn | 0.194274417 | -0.821088895 | 1.209637729 | 10.6109676 | 0.479230547 | 0.650302862 | 0.824972709 | -4.724243034 |
| **A_65_P11059** | Letm1 | -0.071385224 | -0.444480996 | 0.301710549 | 11.14972637 | -0.47922376 | 0.650307392 | 0.824972709 | -4.724244047 |
| **A_52_P222593** | Cebpzos | 0.135096177 | -0.571079397 | 0.841271751 | 11.5489601 | 0.479160173 | 0.650349837 | 0.824972709 | -4.724253542 |
| **A_55_P2186668** | Cfap126 | 0.313629709 | -1.325998075 | 1.953257492 | 10.18902696 | 0.479095698 | 0.650392876 | 0.824972709 | -4.724263167 |
| **A_51_P434171** | Myo19 | 0.040801261 | -0.172516572 | 0.254119093 | 8.77386955 | 0.479067563 | 0.650411658 | 0.824972709 | -4.724267368 |
| **A_51_P156927** | Arid4a | -0.081201576 | -0.505963757 | 0.343560605 | 8.586337434 | -0.478816338 | 0.650579377 | 0.825131309 | -4.724304862 |
| **A_51_P261443** | Mlx | -0.059167739 | -0.368757512 | 0.250422034 | 9.075666821 | -0.478683782 | 0.650667881 | 0.82518221 | -4.724324638 |
| **A_52_P487362** | Ppp4r4 | 0.120844753 | -0.511537532 | 0.753227039 | 8.391484649 | 0.478628384 | 0.650704871 | 0.82518221 | -4.724332902 |
| **A_52_P501816** | Slc6a14 | 0.174607641 | -0.739323613 | 1.088538894 | 8.65503953 | 0.478519963 | 0.650777267 | 0.825216341 | -4.724349072 |
| **A_52_P43848** | Sco1 | 0.050869007 | -0.215422767 | 0.317160781 | 8.412430255 | 0.478460238 | 0.65081715 | 0.825216341 | -4.724357979 |
| **A_66_P111426** | Lce1a2 | 0.118669475 | -0.502683488 | 0.740022437 | 8.80066924 | 0.478355743 | 0.650886931 | 0.8252507 | -4.724373559 |
| **A_55_P2117726** | Stk33 | 0.128070976 | -0.542600386 | 0.798742338 | 8.360293473 | 0.478290001 | 0.650930835 | 0.825252247 | -4.72438336 |
| **A_55_P2050727** | Fcrl5 | 0.074117642 | -0.31415261 | 0.462387895 | 10.91036924 | 0.478121006 | 0.651043702 | 0.82534122 | -4.724408548 |
| **A_51_P406334** | Dnlz | 0.071064572 | -0.301510515 | 0.443639658 | 9.864846834 | 0.477737879 | 0.65129962 | 0.825472994 | -4.724465624 |
| **A_55_P2090070** | Myh14 | 0.478955954 | -2.032135046 | 2.990046954 | 13.39440412 | 0.477731052 | 0.651304181 | 0.825472994 | -4.72446664 |
| **A_51_P290018** | NA | 0.099691551 | -0.423041682 | 0.622424785 | 12.31806655 | 0.477670683 | 0.65134451 | 0.825472994 | -4.72447563 |
| **A_51_P193716** | Supv3l1 | 0.082300255 | -0.349287089 | 0.513887599 | 9.029144196 | 0.477620355 | 0.651378133 | 0.825472994 | -4.724483124 |
| **A_52_P599264** | Mdfic | -0.086817844 | -0.542132291 | 0.368496603 | 10.41067645 | -0.477581959 | 0.651403785 | 0.825472994 | -4.72448884 |
| **A_52_P345247** | Tfb1m | 0.077858285 | -0.33046798 | 0.486184551 | 9.651474977 | 0.477581906 | 0.65140382 | 0.825472994 | -4.724488848 |
| **A_51_P414243** | Pomgnt2 | 0.066584581 | -0.282688443 | 0.415857605 | 10.9336789 | 0.477484165 | 0.651469122 | 0.825501639 | -4.724503398 |
| **A_52_P506529** | Rabgap1l | -0.116849164 | -0.729925867 | 0.496227539 | 10.20710119 | -0.477376617 | 0.651540981 | 0.825538588 | -4.724519406 |
| **A_55_P2009477** | 9530068E | 0.101141761 | -0.429729197 | 0.63201272 | 13.92687859 | 0.47719061 | 0.651665271 | 0.825543383 | -4.724547084 |
| **A_51_P255456** | Cyp1b1 | 0.159960503 | -0.679646735 | 0.999567741 | 8.752507963 | 0.477185164 | 0.65166891 | 0.825543383 | -4.724547894 |

| | | | | | | | | | |
|---|---|---|---|---|---|---|---|---|---|
| **A_51_P253883** | Fam49a | 0.124610662 | -0.529458986 | 0.778680309 | 8.789287743 | 0.477179251 | 0.651672862 | 0.825543383 | -4.724548773 |
| **A_55_P2124273** | Myo1f | -0.073785704 | -0.461227368 | 0.31365596 | 8.248818912 | -0.476997662 | 0.651794213 | 0.82563523 | -4.724575784 |
| **A_52_P59457** | Nprl2 | -0.050803609 | -0.31759909 | 0.215991872 | 9.659314639 | -0.476942959 | 0.651830772 | 0.82563523 | -4.724583919 |
| **A_51_P395293** | Ccz1 | -0.111008425 | -0.69497927 | 0.47296242 | 12.45066561 | -0.47611857 | 0.652381857 | 0.826231878 | -4.724706421 |
| **A_51_P234113** | Nod1 | -0.103619574 | -0.648729935 | 0.441490787 | 10.24074383 | -0.476110446 | 0.652387289 | 0.826231878 | -4.724707627 |
| **A_55_P2079729** | NA | 0.083898265 | -0.357827484 | 0.525624014 | 16.21832075 | 0.475719122 | 0.652648967 | 0.826489778 | -4.724765712 |
| **A_52_P595717** | Smug1 | 0.062594954 | -0.267027324 | 0.392217232 | 8.500495891 | 0.47563424 | 0.652705735 | 0.826489778 | -4.724778306 |
| **A_51_P184949** | F12 | 0.043402809 | -0.185164131 | 0.271969749 | 7.720039272 | 0.475614153 | 0.65271917 | 0.826489778 | -4.724781286 |
| **A_51_P211822** | Neurl4 | 0.053241452 | -0.227460058 | 0.333942963 | 11.21169975 | 0.475067685 | 0.653084705 | 0.826898474 | -4.724862314 |
| **A_52_P707503** | Tmem212 | 0.292261158 | -1.24938277 | 1.833905085 | 9.160748879 | 0.474829146 | 0.6532443 | 0.826999266 | -4.724897659 |
| **A_66_P119350** | 2310040G | 0.09101283 | -0.389077764 | 0.571103424 | 8.557052585 | 0.474820836 | 0.65324986 | 0.826999266 | -4.72489889 |
| **A_55_P2084763** | NA | -0.054441766 | -0.341747627 | 0.232864095 | 10.49532157 | -0.474611292 | 0.653390073 | 0.827122613 | -4.724929925 |
| **A_55_P1979551** | Pitrm1 | -0.204547836 | -1.284281224 | 0.875185551 | 12.11035033 | -0.474491793 | 0.65347004 | 0.827169684 | -4.724947619 |
| **A_52_P630632** | Rcbtb2 | 0.059072202 | -0.25280591 | 0.370950315 | 10.81920158 | 0.474404294 | 0.653528597 | 0.827189649 | -4.724960572 |
| **A_55_P2095118** | Dancr | 0.239507149 | -1.025363379 | 1.504377677 | 10.62787867 | 0.474266832 | 0.653620596 | 0.827232605 | -4.724980917 |
| **A_51_P257065** | Prpf40b | -0.076473371 | -0.480374871 | 0.32742813 | 8.920083405 | -0.474225728 | 0.653648107 | 0.827232605 | -4.724987 |
| **A_52_P28057** | Gm14124 | 0.073560077 | -0.315072456 | 0.46219261 | 8.396341424 | 0.47408189 | 0.653744383 | 0.827300295 | -4.725008281 |
| **A_55_P1963529** | Abcb10 | -0.133212188 | -0.837402195 | 0.570977819 | 10.43702793 | -0.473810249 | 0.653926222 | 0.827476247 | -4.725048457 |
| **A_52_P68028** | Ace | 0.118496237 | -0.508259736 | 0.74525221 | 9.012019064 | 0.47353973 | 0.654107337 | 0.827608129 | -4.725088448 |
| **A_52_P230904** | Wdr18 | 0.044508588 | -0.190932572 | 0.279949749 | 9.033064371 | 0.473491051 | 0.654139931 | 0.827608129 | -4.725095642 |
| **A_51_P148069** | Lpgat1 | -0.143561496 | -0.941246697 | 0.654123706 | 10.45253388 | -0.479368116 | 0.654164789 | 0.827608129 | -4.699951913 |
| **A_52_P467140** | Snd1 | -0.072485502 | -0.455993388 | 0.311022384 | 10.03984198 | -0.47339885 | 0.654201668 | 0.827608129 | -4.725109266 |
| **A_51_P125467** | Igf2r | -0.060368832 | -0.379967461 | 0.259229797 | 12.94078432 | -0.47310573 | 0.654397958 | 0.827802285 | -4.717268499 |
| **A_52_P509965** | Ager | 0.582559681 | -2.502940684 | 3.668060046 | 10.95318199 | 0.472895484 | 0.65453877 | 0.827926241 | -4.725183606 |
| **A_51_P339713** | Mrps26 | -0.048201273 | -0.30353363 | 0.207131083 | 10.85566356 | -0.472827801 | 0.654584105 | 0.827929418 | -4.725193596 |
| **A_51_P343323** | Cox6c | 0.105176533 | -0.452130256 | 0.662483323 | 15.0518823 | 0.472688312 | 0.65467754 | 0.82799343 | -4.725214182 |
| **A_52_P504478** | Adprh | 0.04445159 | -0.191153502 | 0.280056681 | 11.99813836 | 0.472555662 | 0.6547664 | 0.828051647 | -4.725233754 |
| **A_51_P433789** | Agr3 | 0.153715542 | -0.661139949 | 0.968571033 | 8.452646033 | 0.472484462 | 0.654814099 | 0.828057806 | -4.725244257 |
| **A_51_P176086** | Ffar2 | 0.055050067 | -0.23693627 | 0.347036405 | 7.8628192 | 0.472221404 | 0.654990344 | 0.828226508 | -4.72528305 |
| **A_55_P2007966** | Tmem215 | -0.191517132 | -1.207587053 | 0.824552789 | 10.33785169 | -0.472100414 | 0.655071413 | 0.828274848 | -4.725300887 |
| **A_55_P2040276** | Prcd | 0.469145881 | -2.020559549 | 2.958851311 | 10.11241048 | 0.471965536 | 0.655161794 | 0.828334955 | -4.725320765 |
| **A_55_P2029750** | Ndufs5 | 0.073573603 | -0.316980299 | 0.464127505 | 14.44641883 | 0.471836344 | 0.655248372 | 0.82838673 | -4.725339802 |
| **A_51_P465473** | Btf3l4 | 0.083463487 | -0.359645452 | 0.526572426 | 9.078634483 | 0.471776561 | 0.655288438 | 0.82838673 | -4.725348609 |
| **A_55_P2050488** | Cep170b | -0.101755906 | -0.642439378 | 0.438927567 | 14.8805141 | -0.471375357 | 0.655557349 | 0.828672494 | -4.725407691 |
| **A_51_P450100** | Rab5if | 0.042547692 | -0.183655649 | 0.268751033 | 13.15675049 | 0.47111544 | 0.655731593 | 0.828786653 | -4.725445943 |
| **A_52_P175157** | NA | 0.173129118 | -0.747311128 | 1.093569364 | 10.08208858 | 0.471112755 | 0.655733393 | 0.828786653 | -4.725446338 |
| **A_55_P1979575** | Shroom2 | 0.058035605 | -0.250577883 | 0.366649093 | 10.94076907 | 0.471009817 | 0.655802408 | 0.828819699 | -4.725461483 |
| **A_52_P284778** | Rae1 | 0.061273289 | -0.26464894 | 0.387195518 | 10.6556242 | 0.470877087 | 0.655891402 | 0.828877989 | -4.725481006 |
| **A_55_P2182433** | Atg10 | -0.054216975 | -0.357316489 | 0.248882538 | 8.535850708 | -0.476444977 | 0.656090334 | 0.829075197 | -4.700327271 |
| **A_51_P344947** | Rab1b | 0.133427105 | -0.577044095 | 0.843898306 | 13.03693727 | 0.470379008 | 0.656225416 | 0.829191699 | -4.725554226 |
| **A_55_P2069277** | Mtf1 | -0.109495005 | -0.692654162 | 0.473664153 | 8.340076794 | -0.47028113 | 0.656291063 | 0.829220456 | -4.725568607 |
| **A_55_P1979027** | Tmem65 | -0.077110424 | -0.487965665 | 0.333744816 | 8.790293143 | -0.470083068 | 0.656423916 | 0.829334117 | -4.725597699 |
| **A_55_P1992450** | Vdac2 | 0.043527439 | -0.188441586 | 0.275496463 | 14.27681432 | 0.469984421 | 0.65649009 | 0.829353876 | -4.725612185 |
| **A_55_P1984981** | NA | -0.039328408 | -0.249016811 | 0.170359995 | 7.797585961 | -0.469766774 | 0.656636103 | 0.829353876 | -4.725644136 |
| **A_52_P428496** | Rhoa | 0.058986195 | -0.255541202 | 0.373513593 | 11.19555272 | 0.469723458 | 0.656665164 | 0.829353876 | -4.725650493 |
| **A_55_P2024342** | Tmem33 | 0.078255874 | -0.339026672 | 0.49553842 | 10.0270638 | 0.469717858 | 0.656668922 | 0.829353876 | -4.725651315 |
| **A_52_P402319** | Med20 | -0.052755917 | -0.334071104 | 0.228559271 | 9.427509543 | -0.469708414 | 0.656675258 | 0.829353876 | -4.725652701 |
| **A_55_P1999269** | Grk2 | -0.170535322 | -1.080069188 | 0.738998543 | 11.67627838 | -0.469619171 | 0.656735135 | 0.829353876 | -4.725665797 |

| | | | | | | | | | |
|---|---|---|---|---|---|---|---|---|---|
| **A_51_P144264** | Klf2 | -0.095252951 | -0.603292182 | 0.41278628 | 11.80713203 | -0.469603727 | 0.656745498 | 0.829353876 | -4.725668063 |
| **A_51_P126525** | D230025D | -0.092910007 | -0.588569346 | 0.402749331 | 10.19856214 | -0.46949346 | 0.656819486 | 0.829353876 | -4.723719523 |
| **A_55_P1978576** | Dtx2 | 0.075264701 | -0.326282816 | 0.476812218 | 11.379062 | 0.469466643 | 0.656837481 | 0.829353876 | -4.725688174 |
| **A_55_P1982034** | NA | 0.043699276 | -0.189465335 | 0.276863887 | 8.35371259 | 0.469420392 | 0.656868517 | 0.829353876 | -4.725694959 |
| **A_55_P2120866** | NA | -0.188198202 | -1.192619486 | 0.816223082 | 8.969535147 | -0.469299301 | 0.656949775 | 0.829402309 | -4.725712718 |
| **A_55_P2165839** | Cebpe | -0.044685562 | -0.283249987 | 0.193878863 | 7.863556455 | -0.469150184 | 0.657049849 | 0.829429538 | -4.725734582 |
| **A_55_P1959985** | Alas1 | -0.069689464 | -0.44175174 | 0.302372813 | 8.662867031 | -0.469139315 | 0.657057143 | 0.829429538 | -4.725736176 |
| **A_55_P2098995** | Cfhr2 | 0.099255154 | -0.430892103 | 0.629402411 | 9.450375216 | 0.468928788 | 0.657198444 | 0.829516449 | -4.725767033 |
| **A_55_P2131506** | NA | -0.095903952 | -0.608254411 | 0.416446506 | 8.353973435 | -0.468834672 | 0.657261617 | 0.829516449 | -4.725780824 |
| **A_55_P2015410** | 0610010K | 0.102520433 | -0.445183088 | 0.650223953 | 14.081247 | 0.468829837 | 0.657264862 | 0.829516449 | -4.725781533 |
| **A_55_P1961645** | Vcp | 0.05575841 | -0.242155431 | 0.353672252 | 13.88698101 | 0.468781051 | 0.657297611 | 0.829516449 | -4.72578868 |
| **A_55_P2089158** | NA | 0.06047368 | -0.262679754 | 0.383627114 | 8.837590485 | 0.468714044 | 0.657342591 | 0.829519068 | -4.725798497 |
| **A_51_P156263** | Zfp511 | 0.080422626 | -0.349469669 | 0.510314921 | 10.30772042 | 0.468564019 | 0.657443305 | 0.829592015 | -4.72582047 |
| **A_55_P1988689** | Rbpjl | 0.232671592 | -1.011489042 | 1.476832225 | 8.921302445 | 0.468400393 | 0.657553159 | 0.829668041 | -4.725844429 |
| **A_66_P127432** | Tial1 | 0.116596553 | -0.507022463 | 0.740215569 | 11.50219223 | 0.468291966 | 0.657625958 | 0.829668041 | -4.725860301 |
| **A_52_P161297** | Tcea3 | 0.198493028 | -0.863209433 | 1.260195489 | 9.040684926 | 0.468266165 | 0.657643283 | 0.829668041 | -4.725864078 |
| **A_55_P2207255** | Srpx | 0.196501542 | -0.854659756 | 1.247662839 | 9.732960055 | 0.468216749 | 0.657676463 | 0.829668041 | -4.72587131 |
| **A_55_P1958255** | Rnf114 | 0.06543892 | -0.284683025 | 0.415560866 | 9.023446056 | 0.468130717 | 0.657734232 | 0.829668041 | -4.7258839 |
| **A_51_P254234** | Chchd4 | 0.048595975 | -0.211432272 | 0.308624222 | 9.665342419 | 0.468090812 | 0.657761029 | 0.829668041 | -4.725889739 |
| **A_51_P216922** | Ddx59 | 0.061399161 | -0.267302182 | 0.390100504 | 8.749014652 | 0.467855032 | 0.657919367 | 0.829813624 | -4.72592423 |
| **A_51_P261517** | Tyrobp | -0.192894724 | -1.225755693 | 0.839966246 | 10.82394222 | -0.46776619 | 0.657979035 | 0.829834746 | -4.725937223 |
| **A_52_P75146** | Lyrm2 | 0.117278315 | -0.510785225 | 0.745341854 | 10.69542716 | 0.467696887 | 0.658025582 | 0.829839319 | -4.725947357 |
| **A_55_P2089362** | Wfdc13 | -0.077634757 | -0.493511006 | 0.338241492 | 8.679720133 | -0.467565463 | 0.658113856 | 0.82989651 | -4.72596657 |
| **A_55_P2110235** | Trip12 | 0.050423153 | -0.219722885 | 0.320569192 | 10.98528887 | 0.467500139 | 0.658157735 | 0.829897714 | -4.725976118 |
| **A_55_P2156425** | Upk1a | 0.134332474 | -0.585821053 | 0.854486002 | 8.933188111 | 0.467203693 | 0.65835688 | 0.830086262 | -4.726019434 |
| **A_52_P424462** | Ero1lb | -0.04293987 | -0.273166185 | 0.187286445 | 8.105711323 | -0.467149732 | 0.658393133 | 0.830086262 | -4.726027316 |
| **A_52_P682382** | Scd1 | -0.347138022 | -2.208626514 | 1.51435047 | 9.54456569 | -0.467080971 | 0.658439331 | 0.830090377 | -4.726037359 |
| **A_55_P2179796** | NA | 0.064789055 | -0.282787863 | 0.412365972 | 13.88096077 | 0.466875476 | 0.658577405 | 0.830210311 | -4.726067364 |
| **A_65_P06572** | Smad5 | -0.06074455 | -0.386812752 | 0.265323652 | 8.181598185 | -0.466604816 | 0.658759286 | 0.830301993 | -4.726106867 |
| **A_66_P119968** | Clec4a1 | -0.07244555 | -0.46137216 | 0.31648106 | 8.864072576 | -0.466545926 | 0.658798864 | 0.830301993 | -4.72611546 |
| **A_55_P2131088** | Vwa5a | -0.054872312 | -0.349486619 | 0.239741995 | 9.735777284 | -0.466497986 | 0.658831083 | 0.830301993 | -4.726122454 |
| **A_51_P263302** | Rnf24 | -0.070709904 | -0.450381855 | 0.308962048 | 8.597796206 | -0.466468236 | 0.658851077 | 0.830301993 | -4.726126794 |
| **A_51_P295237** | Lrp11 | 0.077343899 | -0.338021818 | 0.492709617 | 8.55214142 | 0.466386273 | 0.658906165 | 0.830301993 | -4.72613875 |
| **A_52_P668984** | Tmem243 | 0.276399035 | -1.2079751 | 1.76077317 | 10.00420435 | 0.466383836 | 0.658907804 | 0.830301993 | -4.726139105 |
| **A_51_P178828** | Mbl2 | -0.048733747 | -0.310504076 | 0.213036583 | 7.71715597 | -0.466293892 | 0.658968258 | 0.830324055 | -4.726152223 |
| **A_55_P2104617** | Kctd5 | 0.060011084 | -0.262395159 | 0.382417328 | 9.117632782 | 0.466206556 | 0.659026963 | 0.83034391 | -4.726164958 |
| **A_52_P727235** | Trabd2b | 0.064292982 | -0.281201277 | 0.40978724 | 8.810026394 | 0.466093534 | 0.659102936 | 0.83035436 | -4.726181436 |
| **A_51_P413916** | Pgrmc2 | 0.047355814 | -0.207137173 | 0.3018488 | 9.380749008 | 0.466066436 | 0.659121152 | 0.83035436 | -4.726185386 |
| **A_55_P2080264** | NA | 0.04048995 | -0.177173841 | 0.258153741 | 7.801243269 | 0.465920048 | 0.659219563 | 0.830424227 | -4.726206722 |
| **A_55_P2041070** | Tle3 | -0.118347046 | -0.754666941 | 0.517972848 | 11.98660973 | -0.465835163 | 0.659276632 | 0.830442009 | -4.726219091 |
| **A_55_P2078776** | Kmt2e | 0.107981534 | -0.472871461 | 0.688834528 | 10.35550939 | 0.465622121 | 0.659419871 | 0.830550726 | -4.726250126 |
| **A_51_P162955** | Serpina7 | 0.039705407 | -0.173897225 | 0.253308039 | 7.676122284 | 0.465579014 | 0.659448857 | 0.830550726 | -4.726256404 |
| **A_52_P249920** | Zscan29 | -0.041632011 | -0.265642693 | 0.18237867 | 8.359838044 | -0.465488559 | 0.65950968 | 0.830560394 | -4.726269577 |
| **A_52_P435561** | Prr15l | 0.204878842 | -0.897717399 | 1.307475084 | 8.950801005 | 0.465404898 | 0.659565939 | 0.830560394 | -4.726281758 |
| **A_51_P337269** | Aldob | -0.119610684 | -0.763372993 | 0.524151626 | 8.313053012 | -0.465366131 | 0.659592009 | 0.830560394 | -4.726287402 |
| **A_55_P1958354** | Rps29 | 0.102584932 | -0.44964837 | 0.654818233 | 17.51338935 | 0.465276723 | 0.659652135 | 0.830560394 | -4.726300416 |
| **A_55_P1993589** | NA | 0.060134193 | -0.263628712 | 0.383897099 | 15.17750975 | 0.465205399 | 0.659700103 | 0.830560394 | -4.726310797 |
| **A_51_P417701** | Apaf1 | 0.061392801 | -0.269240666 | 0.392026268 | 8.646259081 | 0.465072845 | 0.659789254 | 0.830560394 | -4.726330086 |

| | | | | | | | | | |
|---|---|---|---|---|---|---|---|---|---|
| **A_55_P1978241** | Artn | -0.172640518 | -1.102409821 | 0.757128786 | 8.528650553 | -0.465069514 | 0.659791495 | 0.830560394 | -4.726330571 |
| **A_55_P2132237** | Fastkd3 | 0.045480138 | -0.199479395 | 0.290439671 | 9.33117704 | 0.465026574 | 0.659820376 | 0.830560394 | -4.726336818 |
| **A_55_P1984289** | Cdan1 | 0.041976124 | -0.184126991 | 0.26807924 | 8.043010348 | 0.464992699 | 0.65984316 | 0.830560394 | -4.726341746 |
| **A_55_P2138816** | Mplkip | -0.048364152 | -0.308987641 | 0.212259338 | 9.320309126 | -0.464793842 | 0.659976922 | 0.830674683 | -4.72637067 |
| **A_55_P2152188** | NA | -0.176079588 | -1.125530988 | 0.773371811 | 15.46308085 | -0.464500965 | 0.660173952 | 0.830868584 | -4.726413249 |
| **A_51_P447258** | Alox12b | -0.044796792 | -0.286411217 | 0.196817634 | 7.853554733 | -0.46438095 | 0.660254699 | 0.83091612 | -4.72643069 |
| **A_55_P1954986** | Elk1 | -0.051440384 | -0.329178695 | 0.226297928 | 17.63865602 | -0.463894 | 0.660582378 | 0.831116 | -4.726501417 |
| **A_51_P416439** | Zdhhc6 | 0.060822675 | -0.267605776 | 0.389251127 | 10.69140659 | 0.463847363 | 0.660613765 | 0.831116 | -4.726508187 |
| **A_55_P2275249** | Th | 0.065242136 | -0.304831815 | 0.435316087 | 8.661516862 | 0.469572182 | 0.660629906 | 0.831116 | -4.701202285 |
| **A_52_P613498** | 4833420G | -0.060510636 | -0.387288753 | 0.266267481 | 11.27148525 | -0.463798238 | 0.660646828 | 0.831116 | -4.726515318 |
| **A_55_P2032272** | Habp2 | -0.151234855 | -0.968132015 | 0.665662305 | 8.118804386 | -0.463697606 | 0.66071456 | 0.831116 | -4.726529924 |
| **A_55_P2054456** | Adprm | -0.084741342 | -0.542572091 | 0.373089408 | 11.57127399 | -0.463597113 | 0.660782201 | 0.831116 | -4.726544507 |
| **A_51_P289414** | Spg11 | -0.054887898 | -0.35145333 | 0.241677535 | 10.53917139 | -0.463560492 | 0.660806851 | 0.831116 | -4.72654982 |
| **A_55_P2017944** | Etfb | -0.064284294 | -0.411626415 | 0.283057827 | 14.67677119 | -0.463551265 | 0.660813062 | 0.831116 | -4.726551159 |
| **A_55_P2074499** | Kcp | 0.128305589 | -0.564977556 | 0.821588734 | 9.083245565 | 0.463537931 | 0.660822037 | 0.831116 | -4.726553093 |
| **A_51_P305320** | B4gat1 | -0.046415927 | -0.297235736 | 0.204403881 | 11.203843 | -0.463506198 | 0.660843398 | 0.831116 | -4.726557697 |
| **A_55_P2034784** | Snhg17 | 0.050245531 | -0.221373279 | 0.321864341 | 8.230709209 | 0.463327359 | 0.660963788 | 0.831202072 | -4.726583637 |
| **A_55_P2147871** | NA | -0.103533136 | -0.663276757 | 0.456210486 | 8.661362715 | -0.463276808 | 0.66099782 | 0.831202072 | -4.726590968 |
| **A_51_P510764** | Agmo | -0.038967427 | -0.249687807 | 0.171752953 | 7.740561268 | -0.463175457 | 0.661066054 | 0.831233812 | -4.726605663 |
| **A_55_P2125006** | Gpatch1 | -0.109072541 | -0.699134533 | 0.48098945 | 9.958772314 | -0.462986296 | 0.661193416 | 0.831284968 | -4.726633084 |
| **A_55_P1991530** | Cep126 | 0.114837934 | -0.50652239 | 0.736198258 | 8.478648712 | 0.462905348 | 0.661247921 | 0.831284968 | -4.726644814 |
| **A_55_P2021216** | Crlf3 | 0.13072752 | -0.576633656 | 0.838088697 | 11.46785472 | 0.462888242 | 0.66125944 | 0.831284968 | -4.726647293 |
| **A_52_P100199** | Abhd10 | 0.056789811 | -0.25055992 | 0.364139541 | 8.180905365 | 0.462794225 | 0.661322749 | 0.831284968 | -4.726660915 |
| **A_51_P179921** | Igdcc3 | -0.045883401 | -0.294214269 | 0.202447468 | 8.046283364 | -0.462780696 | 0.66133186 | 0.831284968 | -4.726662876 |
| **A_66_P123155** | Ddo | 0.069913968 | -0.308525216 | 0.448353152 | 7.972747417 | 0.462719924 | 0.661372784 | 0.831284968 | -4.726671679 |
| **A_55_P2174501** | Stx17 | -0.063850341 | -0.409531421 | 0.281830739 | 8.605807916 | -0.462634438 | 0.661430354 | 0.831284968 | -4.726684062 |
| **A_55_P2005420** | NA | -0.073170903 | -0.469338703 | 0.322996897 | 8.438509495 | -0.462604218 | 0.661450706 | 0.831284968 | -4.726688439 |
| **A_55_P2105843** | NA | 0.110852683 | -0.489729267 | 0.711434633 | 12.19518663 | 0.462300411 | 0.661655325 | 0.831488077 | -4.726732425 |
| **A_55_P2154416** | Car15 | 0.049136928 | -0.21715669 | 0.315430545 | 8.38845636 | 0.462165567 | 0.661746156 | 0.831548172 | -4.726751941 |
| **A_51_P114353** | Ctu1 | 0.067218454 | -0.29716259 | 0.431599497 | 8.698932329 | 0.462043738 | 0.661828225 | 0.831555697 | -4.726769568 |
| **A_55_P2153391** | NA | 0.152018231 | -0.67216991 | 0.976206371 | 8.580577564 | 0.46197626 | 0.661873683 | 0.831555697 | -4.72677933 |
| **A_51_P176517** | Ntmt1 | -0.08046944 | -0.516756289 | 0.355817408 | 10.86482573 | -0.46196514 | 0.661881174 | 0.831555697 | -4.726780938 |
| **A_55_P1954755** | NA | -0.069350594 | -0.44550391 | 0.306802722 | 15.83814027 | -0.46178055 | 0.662005537 | 0.831657898 | -4.726807635 |
| **A_52_P618187** | Mrs2 | -0.136435749 | -0.876839701 | 0.603968203 | 9.045137377 | -0.461540479 | 0.662167297 | 0.831807064 | -4.726842342 |
| **A_55_P2159585** | Pm20d1 | -0.040012264 | -0.257218941 | 0.177194413 | 7.772937661 | -0.461392254 | 0.662267181 | 0.831878487 | -4.726863763 |
| **A_55_P2022116** | NA | 0.138029149 | -0.612170161 | 0.88822846 | 16.390392 | 0.460833982 | 0.662643452 | 0.832220018 | -4.726944389 |
| **A_55_P2046184** | Ccdc90b | 0.117956582 | -0.523210262 | 0.759123427 | 10.4708448 | 0.460788329 | 0.662674227 | 0.832220018 | -4.726950978 |
| **A_51_P453246** | Uqcc3 | -0.0546334 | -0.35161587 | 0.242349071 | 12.8074346 | -0.460763168 | 0.662691188 | 0.832220018 | -4.72695461 |
| **A_55_P2018919** | Spns1 | -0.071796428 | -0.462213345 | 0.31862049 | 13.40485054 | -0.460600657 | 0.662800744 | 0.832220018 | -4.72697806 |
| **A_51_P280893** | Pla2g1b | 0.194666936 | -0.864013862 | 1.253347734 | 8.713058517 | 0.46055076 | 0.662834384 | 0.832220018 | -4.726985259 |
| **A_51_P438619** | Sobp | -0.046918849 | -0.302099226 | 0.208261528 | 8.489405295 | -0.460522041 | 0.662853747 | 0.832220018 | -4.726989402 |
| **A_52_P174915** | Gja1 | -0.044500245 | -0.286603368 | 0.197602879 | 9.648704847 | -0.460375658 | 0.662952442 | 0.832220018 | -4.725029425 |
| **A_55_P2008609** | Rxrb | 0.057396422 | -0.25490771 | 0.369700554 | 12.50915336 | 0.460317449 | 0.662991691 | 0.832220018 | -4.727018911 |
| **A_66_P129893** | 1700047M | -0.041877504 | -0.269743656 | 0.185988647 | 7.889066949 | -0.460310964 | 0.662996063 | 0.832220018 | -4.727019846 |
| **A_52_P199776** | Snhg7 | 0.060474932 | -0.268624229 | 0.389574094 | 8.59901615 | 0.460255472 | 0.66303348 | 0.832220018 | -4.727027847 |
| **A_55_P2132467** | Nfx1 | -0.071979688 | -0.463686724 | 0.319727348 | 9.027365716 | -0.460255444 | 0.663033499 | 0.832220018 | -4.727027852 |
| **A_51_P380514** | Tjap1 | 0.066218095 | -0.294160538 | 0.426596728 | 9.249969732 | 0.460222654 | 0.663055609 | 0.832220018 | -4.727032579 |
| **A_51_P176377** | Usp25 | -0.041465957 | -0.267269312 | 0.184337398 | 9.924964906 | -0.459951088 | 0.663238742 | 0.832395835 | -4.727071723 |

| | | | | | | | | | |
|---|---|---|---|---|---|---|---|---|---|
| **A_52_P552832** | Ndufa4 | 0.203728006 | -0.905939717 | 1.313395729 | 13.58738165 | 0.45984146 | 0.663312677 | 0.832422152 | -4.727087519 |
| **A_52_P385206** | Harbi1 | -0.042764699 | -0.275731885 | 0.190202487 | 8.622394641 | -0.459770404 | 0.663360601 | 0.832422152 | -4.727097755 |
| **A_55_P2086153** | Bcdin3d | 0.040536442 | -0.18031211 | 0.261384994 | 10.141445 | 0.459728484 | 0.663388875 | 0.832422152 | -4.727103794 |
| **A_55_P2066827** | Hexim2 | -0.077377659 | -0.499361677 | 0.344606359 | 8.651341135 | -0.459271956 | 0.663696832 | 0.832702167 | -4.727169525 |
| **A_55_P2161535** | Ddx23 | -0.068096065 | -0.439566766 | 0.303374636 | 8.160619897 | -0.45914282 | 0.663783956 | 0.832702167 | -4.727188107 |
| **A_51_P483373** | Txndc5 | 0.096203574 | -0.428702265 | 0.621109413 | 10.24628286 | 0.459050166 | 0.66384647 | 0.832702167 | -4.727201437 |
| **A_55_P2157814** | Gmds | -0.179501065 | -1.15906506 | 0.80006293 | 10.74612447 | -0.458970277 | 0.663900374 | 0.832702167 | -4.727212929 |
| **A_55_P2045414** | Hipk4 | -0.080151788 | -0.517567349 | 0.357263773 | 8.899105102 | -0.458954182 | 0.663911234 | 0.832702167 | -4.727215244 |
| **A_55_P2047932** | Cyb5r4 | -0.06551865 | -0.423106562 | 0.292069261 | 9.937809995 | -0.458915218 | 0.663937526 | 0.832702167 | -4.727220848 |
| **A_51_P194479** | 1110059G | 0.056502281 | -0.251876256 | 0.364880819 | 9.709552487 | 0.458914927 | 0.663937722 | 0.832702167 | -4.72722089 |
| **A_55_P2379480** | NA | 0.068296956 | -0.304477343 | 0.441071255 | 8.478593987 | 0.458886974 | 0.663956584 | 0.832702167 | -4.72722491 |
| **A_55_P1996983** | Btg1c | 0.081192991 | -0.362164476 | 0.524550458 | 9.11179188 | 0.45868534 | 0.66409265 | 0.83281879 | -4.727253902 |
| **A_55_P2073218** | Hps4 | -0.04382477 | -0.2831988 | 0.195549261 | 11.02875048 | -0.458556609 | 0.664179527 | 0.832848792 | -4.727272406 |
| **A_55_P2087865** | Tinf2 | -0.067273075 | -0.434838074 | 0.300291925 | 9.142886604 | -0.458413579 | 0.664276061 | 0.832848792 | -4.727292959 |
| **A_55_P2031436** | Ly6e | -0.128192488 | -0.828608073 | 0.572223098 | 12.76357016 | -0.458413205 | 0.664276313 | 0.832848792 | -4.727293013 |
| **A_55_P2127825** | Bcas3 | 0.056120329 | -0.25054525 | 0.362785908 | 8.98684364 | 0.458358745 | 0.664313071 | 0.832848792 | -4.727300838 |
| **A_55_P2152547** | Myl3 | -0.260820388 | -1.686142011 | 1.164501235 | 9.929746681 | -0.458330764 | 0.664331958 | 0.832848792 | -4.727304858 |
| **A_51_P407683** | Ints3 | -0.093321018 | -0.603561887 | 0.416919851 | 10.14293333 | -0.458093967 | 0.664491799 | 0.83299212 | -4.727338868 |
| **A_55_P2123673** | Trnp1 | -0.11769837 | -0.761309164 | 0.525912424 | 8.937072749 | -0.458033745 | 0.664532454 | 0.83299212 | -4.727347515 |
| **A_55_P2028570** | Sgsh | -0.109202945 | -0.706465112 | 0.488059221 | 14.47845442 | -0.457951717 | 0.66458783 | 0.833007527 | -4.727359292 |
| **A_55_P2073294** | Hadhb | -0.057618886 | -0.373143601 | 0.25790583 | 12.22124507 | -0.457384893 | 0.664970555 | 0.833393844 | -4.727440619 |
| **A_55_P1972426** | Pdzd11 | 0.04353799 | -0.194896308 | 0.281972288 | 9.439824972 | 0.457351384 | 0.664993184 | 0.833393844 | -4.727445424 |
| **A_55_P1974582** | Eif4h | 0.050475048 | -0.225978896 | 0.326928992 | 14.91195436 | 0.457303255 | 0.665025687 | 0.833393844 | -4.727452325 |
| **A_55_P2046563** | Cym | 0.366712817 | -1.642105863 | 2.375531498 | 9.580259906 | 0.457231031 | 0.665074463 | 0.833393844 | -4.72746268 |
| **A_52_P504743** | Cdyl | 0.048384599 | -0.216693582 | 0.31346278 | 8.72386502 | 0.457176095 | 0.665111565 | 0.833393844 | -4.727470555 |
| **A_51_P461452** | Ube2r2 | -0.098734053 | -0.639975393 | 0.442507287 | 13.12444506 | -0.456905462 | 0.665294359 | 0.833568864 | -4.727509339 |
| **A_55_P2092329** | Adrm1 | -0.110865405 | -0.718768229 | 0.497037419 | 11.93425636 | -0.456785418 | 0.665375448 | 0.833599383 | -4.727526535 |
| **A_55_P2305525** | C81189 | -0.05983811 | -0.388097988 | 0.268421769 | 8.265542927 | -0.456573191 | 0.66551882 | 0.833599383 | -4.727556928 |
| **A_55_P1973930** | Krt79 | -0.074077704 | -0.480464263 | 0.332308855 | 8.511205926 | -0.456560698 | 0.66552726 | 0.833599383 | -4.727558717 |
| **A_55_P2115281** | Chchd2-p | -0.069835637 | -0.45300395 | 0.313332675 | 15.59367555 | -0.45649693 | 0.665570342 | 0.833599383 | -4.727567846 |
| **A_55_P2012523** | Twf1 | -0.040003482 | -0.25949462 | 0.179487656 | 10.992262 | -0.456489879 | 0.665575106 | 0.833599383 | -4.727568856 |
| **A_55_P1994042** | Zbp1 | -0.187604513 | -1.21708953 | 0.841880504 | 8.381401385 | -0.456429385 | 0.665615978 | 0.833599383 | -4.727577515 |
| **A_51_P273755** | Exoc7 | -0.056224052 | -0.364759666 | 0.252311561 | 9.564140285 | -0.456422652 | 0.665620527 | 0.833599383 | -4.727578479 |
| **A_55_P1988260** | NA | -0.174887589 | -1.134798298 | 0.78502312 | 13.87566897 | -0.456329451 | 0.6656835 | 0.833624249 | -4.727591818 |
| **A_52_P266132** | Fgl2 | 0.068168161 | -0.306234899 | 0.442571221 | 8.211942042 | 0.456029075 | 0.665886473 | 0.833756161 | -4.727634794 |
| **A_55_P2043612** | Prdm6 | 0.161703957 | -0.726997012 | 1.050404926 | 9.674997076 | 0.455738103 | 0.666083123 | 0.833756161 | -4.7276764 |
| **A_51_P219465** | Tmem11 | -0.042506966 | -0.276132587 | 0.191118656 | 12.97251314 | -0.455711492 | 0.666101108 | 0.833756161 | -4.727680204 |
| **A_52_P602996** | Cdip1 | -0.041401556 | -0.26895208 | 0.186148968 | 8.170998152 | -0.455710643 | 0.666101683 | 0.833756161 | -4.727680325 |
| **A_51_P342082** | Cnpy2 | 0.101339845 | -0.455704906 | 0.658384596 | 12.89663088 | 0.45565957 | 0.666136203 | 0.833756161 | -4.727687625 |
| **A_55_P2151601** | Samd9l | 0.18658212 | -0.83910102 | 1.212265259 | 9.9364694 | 0.455624587 | 0.666159849 | 0.833756161 | -4.727692625 |
| **A_65_P17492** | Med29 | -0.112435241 | -0.730581492 | 0.50571101 | 9.752083508 | -0.455576759 | 0.666192177 | 0.833756161 | -4.72769946 |
| **A_55_P2125149** | Tulp1 | 0.228557696 | -1.028184179 | 1.485299571 | 11.50240054 | 0.455512301 | 0.666235748 | 0.833756161 | -4.727708671 |
| **A_55_P1961620** | Vax2 | -0.188561679 | -1.225404542 | 0.848281185 | 9.898143539 | -0.455502577 | 0.666242321 | 0.833756161 | -4.72771006 |
| **A_55_P1978696** | NA | 0.043257779 | -0.194605893 | 0.281121452 | 7.735624701 | 0.455497973 | 0.666245434 | 0.833756161 | -4.727710718 |
| **A_55_P2102569** | Cox7a2l | 0.106586706 | -0.479540654 | 0.692714065 | 12.19751131 | 0.455471692 | 0.666263199 | 0.833756161 | -4.727714473 |
| **A_51_P202050** | Dtx1 | -0.632005488 | -4.109624573 | 2.845613597 | 9.653320253 | -0.455186302 | 0.666456131 | 0.833906581 | -4.727755238 |
| **A_55_P2250296** | 6330566A | -0.095603136 | -0.621686087 | 0.430479815 | 8.54445502 | -0.455164368 | 0.66647096 | 0.833906581 | -4.72775837 |
| **A_55_P1985077** | Mcrip1 | -0.042186779 | -0.274362874 | 0.189989316 | 8.59663407 | -0.45510249 | 0.666512796 | 0.833906581 | -4.727767205 |

| | | | | | | | | | |
|---|---|---|---|---|---|---|---|---|---|
| **A_55_P2044843** | NA | 0.070745621 | -0.318722321 | 0.460213563 | 15.35002578 | 0.454965212 | 0.666605614 | 0.833968742 | -4.727786803 |
| **A_55_P2155546** | Nlrc3 | 0.049577206 | -0.223576323 | 0.322730735 | 8.950946576 | 0.454595964 | 0.666855308 | 0.834227145 | -4.72783949 |
| **A_52_P282987** | Yeats2 | 0.121671137 | -0.548836345 | 0.79217862 | 10.40985244 | 0.454500415 | 0.666919929 | 0.834254005 | -4.727853117 |
| **A_55_P1976574** | Tmod4 | 0.047224816 | -0.213165675 | 0.307615307 | 8.476950876 | 0.454250589 | 0.667088902 | 0.834348694 | -4.727888737 |
| **A_55_P2157676** | Wdr34 | -0.101999837 | -0.664420801 | 0.460421128 | 15.26169147 | -0.454243081 | 0.66709398 | 0.834348694 | -4.727889807 |
| **A_55_P2067772** | Cplane1 | -0.048142293 | -0.313644991 | 0.217360405 | 8.861525898 | -0.454159264 | 0.667150676 | 0.834348694 | -4.727901753 |
| **A_51_P399960** | Mier2 | -0.08278015 | -0.539345913 | 0.373785614 | 8.702659573 | -0.4541227 | 0.66717541 | 0.834348694 | -4.727906964 |
| **A_51_P240723** | 1700022A | 0.084292867 | -0.380697574 | 0.549283308 | 8.172634938 | 0.454043168 | 0.667229211 | 0.834348694 | -4.727918297 |
| **A_51_P381321** | Ddx17 | -0.081057232 | -0.528235614 | 0.36612115 | 14.78057823 | -0.454005708 | 0.667254552 | 0.834348694 | -4.727923634 |
| **A_66_P103477** | Nkapd1 | 0.042160443 | -0.190488131 | 0.274809017 | 9.405283768 | 0.453894707 | 0.667329646 | 0.834382828 | -4.727939447 |
| **A_55_P2132132** | Gm5532 | -0.041808401 | -0.272543285 | 0.188926483 | 7.845434866 | -0.453837775 | 0.667368162 | 0.834382828 | -4.727947557 |
| **A_51_P155747** | Scyl3 | 0.040571831 | -0.183371058 | 0.264514719 | 8.735763388 | 0.453771977 | 0.667412679 | 0.834384529 | -4.727956927 |
| **A_55_P2107957** | Abraxas1 | 0.046502015 | -0.210225199 | 0.303229228 | 8.143874331 | 0.453680598 | 0.667474505 | 0.834407868 | -4.72796994 |
| **A_55_P2122604** | Cbr2 | 0.600897526 | -2.72090075 | 3.922695802 | 10.95107617 | 0.453082727 | 0.667879092 | 0.834837065 | -4.728055019 |
| **A_55_P2186027** | Gapdh | 0.085853292 | -0.388787811 | 0.560494394 | 16.40394546 | 0.453045637 | 0.667904196 | 0.834837065 | -4.728060294 |
| **A_55_P2181109** | Il10rb | -0.081830041 | -0.534326461 | 0.370666379 | 9.761883367 | -0.452947602 | 0.66797055 | 0.83486603 | -4.728074234 |
| **A_55_P2010843** | Fahd1 | -0.052772565 | -0.344697336 | 0.239152206 | 8.798727477 | -0.45278038 | 0.668083742 | 0.834953527 | -4.728098007 |
| **A_55_P2041090** | Tyw5 | -0.039712479 | -0.259459356 | 0.180034397 | 10.4974182 | -0.452641778 | 0.668177568 | 0.835016812 | -4.728117704 |
| **A_52_P407697** | NA | -0.043094899 | -0.281621938 | 0.195432139 | 7.882866346 | -0.452520858 | 0.66825943 | 0.835065138 | -4.728134885 |
| **A_55_P2372902** | 5330438D | 0.060249658 | -0.273309603 | 0.393808918 | 9.0800297 | 0.452409721 | 0.668334673 | 0.835068448 | -4.728150672 |
| **A_51_P305532** | Eif2s3x | 0.228169218 | -1.035097408 | 1.491435845 | 9.594186478 | 0.452389355 | 0.668348462 | 0.835068448 | -4.728153565 |
| **A_51_P298802** | Bfsp2 | 0.040646292 | -0.184450805 | 0.26574339 | 7.892887799 | 0.452273756 | 0.668426732 | 0.835112274 | -4.728169982 |
| **A_55_P2061381** | Ghitm | -0.114105143 | -0.746212031 | 0.518001744 | 12.33183983 | -0.45213179 | 0.668522861 | 0.835131333 | -4.728190138 |
| **A_55_P1998545** | Ncdn | 0.052201201 | -0.237049661 | 0.341452063 | 10.67162438 | 0.452018466 | 0.6685996 | 0.835131333 | -4.728206224 |
| **A_51_P425373** | Smg9 | 0.051860076 | -0.235507833 | 0.339227984 | 9.51601003 | 0.452007067 | 0.668607319 | 0.835131333 | -4.728207842 |
| **A_51_P437021** | Trak1 | 0.054189333 | -0.2460928 | 0.354471465 | 10.24126521 | 0.451996069 | 0.668614767 | 0.835131333 | -4.728209403 |
| **A_55_P2105638** | Ralgapa2 | 0.041540738 | -0.188767646 | 0.271849121 | 8.983726955 | 0.45176731 | 0.668769691 | 0.8352449 | -4.728241862 |
| **A_51_P277795** | Resf1 | -0.050405514 | -0.329891575 | 0.229080547 | 10.5408293 | -0.451718879 | 0.668802493 | 0.8352449 | -4.728248732 |
| **A_55_P2017684** | Macf1 | 0.172248682 | -0.783041275 | 1.127538639 | 12.95660037 | 0.451617791 | 0.668870961 | 0.8352449 | -4.728263069 |
| **A_55_P2158281** | NA | -0.107163312 | -0.701504909 | 0.487178284 | 16.29500402 | -0.45160667 | 0.668878493 | 0.8352449 | -4.728264646 |
| **A_51_P364210** | Ms4a3 | 0.038026237 | -0.172919492 | 0.248971966 | 7.667890195 | 0.451505407 | 0.668947084 | 0.835276603 | -4.728279005 |
| **A_55_P2144476** | NA | 0.046006367 | -0.209325284 | 0.301338018 | 8.773404828 | 0.451298235 | 0.669087423 | 0.835397884 | -4.728308374 |
| **A_51_P375533** | Polr3b | -0.039818545 | -0.260928554 | 0.181291465 | 9.211928153 | -0.45105274 | 0.669253742 | 0.835483641 | -4.72834316 |
| **A_55_P2018345** | Amz2 | 0.051480245 | -0.234395179 | 0.33735567 | 9.793514516 | 0.451039038 | 0.669263025 | 0.835483641 | -4.728345101 |
| **A_55_P2052145** | Ly6g6d | 0.268926045 | -1.224560691 | 1.762412781 | 9.380637408 | 0.451005502 | 0.669285747 | 0.835483641 | -4.728349851 |
| **A_51_P512002** | Rer1 | -0.079749998 | -0.522720866 | 0.363220871 | 12.87434268 | -0.450926603 | 0.669339205 | 0.835496429 | -4.728361027 |
| **A_51_P116088** | Rexo2 | -0.081462059 | -0.534330646 | 0.371406527 | 9.928100681 | -0.450540187 | 0.669601055 | 0.83563564 | -4.714463224 |
| **A_55_P2065537** | NA | 0.094554116 | -0.43110979 | 0.620218023 | 8.763744255 | 0.450528871 | 0.669608724 | 0.83563564 | -4.728417335 |
| **A_55_P2236607** | Ccnc | 0.041047616 | -0.187180529 | 0.26927576 | 8.321262504 | 0.450473327 | 0.669646367 | 0.83563564 | -4.728425195 |
| **A_55_P2162920** | NA | -0.067858502 | -0.445200274 | 0.30948327 | 16.04760904 | -0.450422139 | 0.669681059 | 0.83563564 | -4.728432438 |
| **A_55_P2161126** | NA | -0.113095523 | -0.742080254 | 0.515889207 | 8.851948402 | -0.450355694 | 0.669726093 | 0.83563564 | -4.728441839 |
| **A_52_P214891** | Stx6 | 0.113428594 | -0.517434844 | 0.744292032 | 10.88130055 | 0.450336902 | 0.669738829 | 0.83563564 | -4.728444497 |
| **A_51_P424518** | Nin | 0.08875122 | -0.404885775 | 0.582388215 | 9.01095458 | 0.450315585 | 0.669753278 | 0.83563564 | -4.728447513 |
| **A_55_P2169963** | Zfp982 | -0.061140698 | -0.401262364 | 0.278980968 | 8.085744166 | -0.450242451 | 0.669802848 | 0.83564356 | -4.728457858 |
| **A_51_P313581** | Fabp2 | -0.041432735 | -0.271967956 | 0.189102485 | 7.82261535 | -0.450149389 | 0.669865927 | 0.835667305 | -4.728471019 |
| **A_55_P2025902** | NA | -0.054937055 | -0.360694934 | 0.250820823 | 8.979320859 | -0.450026477 | 0.669949244 | 0.835667305 | -4.728488398 |
| **A_55_P2043454** | NA | 0.049791635 | -0.227339354 | 0.326922625 | 8.531970343 | 0.450009459 | 0.66996078 | 0.835667305 | -4.728490804 |
| **A_55_P2019557** | Mrgpra2b | -0.045162639 | -0.296565198 | 0.206239921 | 7.745031184 | -0.449945547 | 0.670004107 | 0.835667305 | -4.72849984 |

| | | | | | | | | | |
|---|---|---|---|---|---|---|---|---|---|
| **A_55_P1969441** | Eef1akmt2 | -0.05387783 | -0.353827735 | 0.246072076 | 10.25246548 | -0.449895561 | 0.670037993 | 0.835667305 | -4.728506905 |
| **A_52_P594927** | Atl3 | 0.081173795 | -0.370949572 | 0.533297162 | 8.849536685 | 0.449685872 | 0.670180155 | 0.835786697 | -4.728536537 |
| **A_55_P2045040** | F7 | -0.041576574 | -0.273180983 | 0.190027835 | 7.757036582 | -0.449626839 | 0.67022018 | 0.835786697 | -4.728544877 |
| **A_55_P2013138** | NA | -0.072921457 | -0.479557002 | 0.333714088 | 12.05795876 | -0.449159247 | 0.670537255 | 0.836128171 | -4.728610904 |
| **A_55_P2072193** | Firre | -0.069732211 | -0.458732984 | 0.319268563 | 8.13716459 | -0.448986533 | 0.670654393 | 0.836194496 | -4.728635277 |
| **A_51_P315666** | Nid2 | -0.077669286 | -0.51097908 | 0.355640509 | 11.1182775 | -0.448953283 | 0.670676944 | 0.836194496 | -4.728639968 |
| **A_51_P396297** | Hsdl1 | 0.049194789 | -0.225475187 | 0.323864765 | 8.197907313 | 0.448598955 | 0.670917291 | 0.836440218 | -4.728689941 |
| **A_51_P484054** | F7 | -0.041047686 | -0.270295733 | 0.18820036 | 7.798511422 | -0.448469987 | 0.671004783 | 0.836495356 | -4.728708122 |
| **A_55_P2117130** | Zfp937 | 0.056144524 | -0.257583169 | 0.369872217 | 9.062810067 | 0.448234099 | 0.671164825 | 0.836640924 | -4.728741363 |
| **A_55_P2035018** | Clec4n | 0.114364968 | -0.524865015 | 0.753594952 | 9.041482302 | 0.448111638 | 0.671247918 | 0.836690558 | -4.728758614 |
| **A_52_P390918** | Trip4 | -0.049737868 | -0.327817456 | 0.22834172 | 8.496754028 | -0.447990083 | 0.671330401 | 0.836730127 | -4.728775733 |
| **A_51_P247614** | Smim10l2 | -0.083056329 | -0.547470909 | 0.381358251 | 8.581164143 | -0.447937303 | 0.671366218 | 0.836730127 | -4.728783165 |
| **A_52_P509710** | Zwint | -0.177737258 | -1.17189771 | 0.816423193 | 11.05166433 | -0.447787885 | 0.671467618 | 0.836771919 | -4.728804201 |
| **A_55_P2403453** | BC037032 | 0.08351881 | -0.38369123 | 0.550728849 | 9.534133928 | 0.447736474 | 0.671502509 | 0.836771919 | -4.728811437 |
| **A_55_P2302285** | Tarbp2 | -0.047233561 | -0.311484944 | 0.217017821 | 11.56893649 | -0.447696572 | 0.67152959 | 0.836771919 | -4.728817053 |
| **A_51_P221256** | Slc34a2 | 0.302378626 | -1.390409354 | 1.995166606 | 9.520762394 | 0.447402976 | 0.671728866 | 0.836966289 | -4.728858361 |
| **A_52_P175679** | Gm13023 | -0.050470066 | -0.333070743 | 0.232130611 | 8.157878266 | -0.447312485 | 0.671790292 | 0.836988232 | -4.728871088 |
| **A_55_P2022595** | Gm10653 | 0.07857585 | -0.361591087 | 0.518742787 | 15.47898104 | 0.447117858 | 0.671922417 | 0.836988232 | -4.728898454 |
| **A_55_P2095394** | Dhx37 | -0.073961999 | -0.48830529 | 0.340381291 | 8.403809767 | -0.447093832 | 0.671938728 | 0.836988232 | -4.728901831 |
| **A_55_P2012201** | App | -0.061434722 | -0.40569446 | 0.282825016 | 14.59067277 | -0.44696962 | 0.672023058 | 0.836988232 | -4.72891929 |
| **A_55_P1966102** | Nme5 | 0.167734468 | -0.772287227 | 1.107756163 | 10.47847484 | 0.446925139 | 0.672053258 | 0.836988232 | -4.728925541 |
| **A_55_P1978676** | Mospd3 | 0.062629242 | -0.288408529 | 0.413667014 | 10.23787812 | 0.446862235 | 0.672095968 | 0.836988232 | -4.72893438 |
| **A_55_P2144110** | Mink1 | -0.05831421 | -0.385176089 | 0.268547669 | 10.22343683 | -0.446848671 | 0.672105178 | 0.836988232 | -4.728936286 |
| **A_52_P587648** | Phf3 | -0.086231014 | -0.569623516 | 0.397161487 | 11.04388688 | -0.446800788 | 0.67213769 | 0.836988232 | -4.728943013 |
| **A_52_P476754** | Ubash3b | 0.053427425 | -0.246108291 | 0.352963141 | 9.937734606 | 0.446751452 | 0.67217119 | 0.836988232 | -4.728949944 |
| **A_55_P1964282** | Sbk3 | 0.121407802 | -0.559417286 | 0.802232891 | 9.207531378 | 0.446643885 | 0.672244232 | 0.836988232 | -4.728965053 |
| **A_51_P296100** | Rhot2 | 0.043063223 | -0.198453553 | 0.284579999 | 10.63930961 | 0.446590594 | 0.67228042 | 0.836988232 | -4.728972537 |
| **A_51_P441782** | Micu2 | -0.063943653 | -0.422569776 | 0.29468247 | 11.15319553 | -0.446586784 | 0.672283007 | 0.836988232 | -4.728973072 |
| **A_55_P1968053** | NA | 0.047041272 | -0.216811179 | 0.310893723 | 15.84817321 | 0.446548125 | 0.672309261 | 0.836988232 | -4.728978501 |
| **A_55_P2061874** | Prelid1 | -0.0604426 | -0.399814233 | 0.278929033 | 14.30635012 | -0.446085334 | 0.672623574 | 0.837325618 | -4.729043457 |
| **A_55_P2151071** | Cox7a2l | 0.05175484 | -0.239042067 | 0.342551746 | 15.60712575 | 0.445770711 | 0.672837299 | 0.83744527 | -4.729087582 |
| **A_55_P2083609** | Dkk3 | 0.202707716 | -0.936525452 | 1.341940884 | 12.1073431 | 0.445664464 | 0.672909481 | 0.83744527 | -4.729102477 |
| **A_52_P683211** | NA | -0.064730073 | -0.428556062 | 0.299095916 | 8.418247763 | -0.445618001 | 0.672941048 | 0.83744527 | -4.72910899 |
| **A_51_P513163** | Tra2a | 0.075594442 | -0.349325686 | 0.50051457 | 12.86003821 | 0.445587405 | 0.672961835 | 0.83744527 | -4.729113278 |
| **A_55_P2018559** | Morf4l1 | 0.117328417 | -0.542215119 | 0.776871953 | 11.78424944 | 0.445563995 | 0.672977741 | 0.83744527 | -4.729116559 |
| **A_55_P2025911** | Sms | 0.106980285 | -0.49439648 | 0.70835705 | 9.572388591 | 0.44556129 | 0.672979578 | 0.83744527 | -4.729116938 |
| **A_51_P179131** | Vasp | 0.047858283 | -0.22125325 | 0.316969815 | 10.04843438 | 0.445425573 | 0.673071792 | 0.837506116 | -4.729135956 |
| **A_51_P139400** | Taf1 | -0.039193361 | -0.259627487 | 0.181240765 | 8.897847902 | -0.445332126 | 0.673135288 | 0.837531222 | -4.729149048 |
| **A_66_P102395** | Elac2 | 0.040709884 | -0.188340097 | 0.269759864 | 8.340752851 | 0.44516391 | 0.673249598 | 0.837619545 | -4.729172609 |
| **A_55_P2022539** | NA | -0.106462451 | -0.705779671 | 0.492854769 | 8.941899247 | -0.444928319 | 0.673409708 | 0.837747001 | -4.729205592 |
| **A_55_P2063871** | NA | 0.057829611 | -0.267746263 | 0.383405486 | 11.57663764 | 0.444885656 | 0.673438704 | 0.837747001 | -4.729211564 |
| **A_55_P2089820** | Adk | 0.139441516 | -0.645999596 | 0.924882627 | 9.098840735 | 0.444660765 | 0.673591563 | 0.837830755 | -4.729243033 |
| **A_55_P2026014** | Mc5r | -0.040157409 | -0.266355459 | 0.18604064 | 7.891061994 | -0.444659093 | 0.673592699 | 0.837830755 | -4.729243267 |
| **A_55_P2026233** | Uba7 | -0.086282495 | -0.572538219 | 0.399973229 | 10.0787772 | -0.444435064 | 0.67374499 | 0.837914881 | -4.729274601 |
| **A_55_P1957439** | NA | -0.075977504 | -0.504161032 | 0.352206025 | 8.313692071 | -0.444432089 | 0.673747012 | 0.837914881 | -4.729275017 |
| **A_55_P1952905** | Dnm1l | -0.070003351 | -0.464629838 | 0.324623136 | 11.3770737 | -0.444306784 | 0.6738322 | 0.837966923 | -4.729292537 |
| **A_55_P2031032** | NA | -0.125130569 | -0.830650519 | 0.580389381 | 10.44073024 | -0.444226503 | 0.673886781 | 0.837975797 | -4.729303759 |
| **A_51_P103975** | Mpc2 | 0.101858429 | -0.472521531 | 0.676238388 | 12.54104929 | 0.44416879 | 0.67392602 | 0.837975797 | -4.727302339 |

| | | | | | | | | | |
|---|---|---|---|---|---|---|---|---|---|
| **A_55_P2050408** | NA | -0.116515342 | -0.773657481 | 0.540626798 | 9.122254338 | -0.444093218 | 0.673977403 | 0.837985795 | -4.729322387 |
| **A_55_P2132472** | Prpf4b | -0.055428109 | -0.368095566 | 0.257239347 | 8.320975868 | -0.444015089 | 0.674030528 | 0.837986626 | -4.729333304 |
| **A_51_P472518** | Capn1 | -0.058888048 | -0.391110529 | 0.273334433 | 8.231832454 | -0.44396475 | 0.674064757 | 0.837986626 | -4.729340337 |
| **A_55_P2144095** | NA | 0.202133421 | -0.938405064 | 1.342671906 | 10.13236282 | 0.443893236 | 0.674113386 | 0.837993197 | -4.729350327 |
| **A_51_P199104** | Casq1 | 0.050533075 | -0.234673355 | 0.335739506 | 7.879508257 | 0.443779016 | 0.674191058 | 0.83803587 | -4.729366281 |
| **A_52_P289213** | Notch2 | 0.049796385 | -0.231297266 | 0.330890037 | 11.30539523 | 0.443707867 | 0.674239444 | 0.838042135 | -4.729376216 |
| **A_55_P2121141** | Tbc1d22b | -0.090420467 | -0.601003955 | 0.42016302 | 10.02220954 | -0.44355791 | 0.674341429 | 0.838115016 | -4.729397152 |
| **A_52_P447424** | NA | 0.103055986 | -0.479111874 | 0.685223846 | 9.409574925 | 0.44337923 | 0.674462959 | 0.838212177 | -4.72942209 |
| **A_51_P340816** | Ipp | -0.044442913 | -0.295615955 | 0.20673013 | 9.002885945 | -0.443179673 | 0.674598701 | 0.838326988 | -4.729449932 |
| **A_55_P2139814** | Fgf13 | 0.041938275 | -0.195132512 | 0.279009062 | 7.878638975 | 0.443080732 | 0.674666008 | 0.838356745 | -4.729463732 |
| **A_51_P237575** | Akirin1 | 0.066082181 | -0.307599669 | 0.439764031 | 10.95342133 | 0.442927565 | 0.674770209 | 0.83843234 | -4.729485089 |
| **A_55_P2154817** | Ccdc28a | -0.133055734 | -0.885871039 | 0.619759571 | 9.227061478 | -0.442685707 | 0.674934764 | 0.838525799 | -4.7295188 |
| **A_66_P133383** | NA | 0.186669377 | -0.869653951 | 1.242992705 | 8.674770769 | 0.442615433 | 0.674982581 | 0.838525799 | -4.729528592 |
| **A_51_P304683** | Clpx | -0.062564369 | -0.41661319 | 0.291484452 | 10.8055029 | -0.442602905 | 0.674991106 | 0.838525799 | -4.729530338 |
| **A_52_P393589** | Clmp | 0.045376291 | -0.211447807 | 0.302200389 | 8.794682454 | 0.442530864 | 0.675040127 | 0.838525799 | -4.729540375 |
| **A_51_P390538** | Mpeg1 | -0.184274568 | -1.227364088 | 0.858814951 | 9.930232151 | -0.442480531 | 0.675074378 | 0.838525799 | -4.729547386 |
| **A_55_P2066548** | Baiap2l2 | 0.097744881 | -0.45563728 | 0.651127043 | 9.485402528 | 0.442404172 | 0.67512634 | 0.838525799 | -4.729558022 |
| **A_55_P2064351** | Vipr1 | 0.070943751 | -0.330770342 | 0.472657845 | 8.044178384 | 0.442331049 | 0.675176103 | 0.838525799 | -4.729568205 |
| **A_55_P1971991** | Kdf1 | -0.090509486 | -0.60305965 | 0.422040679 | 10.93607081 | -0.442290962 | 0.675203384 | 0.838525799 | -4.729573787 |
| **A_52_P614960** | Skiv2l | -0.054760835 | -0.364901744 | 0.255380074 | 8.986024951 | -0.442243392 | 0.675235759 | 0.838525799 | -4.729580411 |
| **A_52_P519538** | Uba6 | -0.119639972 | -0.79739955 | 0.558119607 | 8.578208869 | -0.442131018 | 0.67531224 | 0.838566914 | -4.729596054 |
| **A_52_P540470** | Zfp407 | -0.058492664 | -0.389966157 | 0.272980829 | 10.25187517 | -0.441980329 | 0.675414804 | 0.838609519 | -4.729617027 |
| **A_51_P303238** | Tmed10 | -0.080410929 | -0.536148686 | 0.375326828 | 12.56018163 | -0.441926875 | 0.675451189 | 0.838609519 | -4.729624465 |
| **A_51_P434697** | Ube2e2 | 0.202757446 | -0.946668824 | 1.352183717 | 11.90807223 | 0.44182068 | 0.675523477 | 0.838609519 | -4.729639239 |
| **A_55_P2138366** | Xpo4 | 0.064803585 | -0.302570585 | 0.432177756 | 8.552209642 | 0.441815309 | 0.675527132 | 0.838609519 | -4.729639987 |
| **A_51_P129317** | Mob4 | 0.048734905 | -0.227578604 | 0.325048413 | 11.26848783 | 0.441761995 | 0.675563425 | 0.838609519 | -4.717651941 |
| **A_51_P196444** | Foxc2 | 0.07114583 | -0.332415802 | 0.474707462 | 8.199117464 | 0.441560204 | 0.675700799 | 0.838625992 | -4.729675465 |
| **A_55_P2059860** | Igsf1 | -0.038945349 | -0.25985781 | 0.181967112 | 7.733189068 | -0.441555941 | 0.675703701 | 0.838625992 | -4.729676058 |
| **A_55_P2145227** | Slmap | -0.046115916 | -0.30770521 | 0.215473377 | 11.3995305 | -0.441551357 | 0.675706822 | 0.838625992 | -4.729676695 |
| **A_55_P2077501** | NA | -0.117549543 | -0.784817581 | 0.549718495 | 11.73018462 | -0.441236028 | 0.675921519 | 0.838794279 | -4.729720524 |
| **A_55_P1952220** | Pnpo | -0.267814806 | -1.788182561 | 1.252552948 | 12.15780599 | -0.441200788 | 0.675945515 | 0.838794279 | -4.72972542 |
| **A_55_P2011674** | Rpl17 | 0.195128421 | -0.912703076 | 1.302959918 | 16.75251193 | 0.441161056 | 0.67597257 | 0.838794279 | -4.72973094 |
| **A_55_P2002314** | Zmiz2 | -0.061247454 | -0.409043901 | 0.286548994 | 10.464645 | -0.441075811 | 0.676030618 | 0.838812474 | -4.729742782 |
| **A_55_P1962284** | Klhl24 | -0.045677029 | -0.305230524 | 0.213876467 | 7.966011831 | -0.440779419 | 0.676232469 | 0.839009084 | -4.72978394 |
| **A_51_P279623** | Pagr1a | -0.05605093 | -0.374718578 | 0.262616718 | 13.15374549 | -0.44054999 | 0.676388737 | 0.839149117 | -4.729815783 |
| **A_55_P2007377** | Elf2 | -0.049383049 | -0.330217098 | 0.231451001 | 10.73891456 | -0.440431611 | 0.676469374 | 0.839163554 | -4.729832207 |
| **A_55_P2028576** | NA | -0.054182557 | -0.362328986 | 0.253963872 | 12.72616996 | -0.44040547 | 0.676487181 | 0.839163554 | -4.729835833 |
| **A_55_P2107667** | Enah | -0.037973345 | -0.253980395 | 0.178033706 | 8.023925665 | -0.440312761 | 0.676550336 | 0.839182081 | -4.729848693 |
| **A_55_P1984825** | Bloc1s2 | -0.06642012 | -0.444328183 | 0.311487942 | 10.78418731 | -0.440213995 | 0.67661762 | 0.839182081 | -4.729862389 |
| **A_55_P2347384** | NA | -0.03774163 | -0.25249311 | 0.17700985 | 7.811581956 | -0.440184593 | 0.676637652 | 0.839182081 | -4.729866466 |
| **A_66_P117495** | Capza1 | -0.138993521 | -0.929973001 | 0.65198596 | 9.300611512 | -0.440128695 | 0.676675734 | 0.839182081 | -4.729874216 |
| **A_52_P71105** | Sertad3 | 0.142097145 | -0.666818729 | 0.951013018 | 9.777377245 | 0.439979396 | 0.676777456 | 0.839223017 | -4.729894912 |
| **A_52_P138110** | 1110032A | 0.042141312 | -0.197770858 | 0.282053481 | 8.178300244 | 0.439952831 | 0.676795556 | 0.839223017 | -4.729898594 |
| **A_55_P1991151** | Rpl13 | 0.115000551 | -0.539873076 | 0.769874178 | 15.16660259 | 0.43983802 | 0.676873786 | 0.839266195 | -4.729914504 |
| **A_51_P487105** | Bud31 | 0.054101739 | -0.254133834 | 0.362337312 | 10.36072309 | 0.439621389 | 0.677021406 | 0.8393954 | -4.729944515 |
| **A_55_P1953226** | H2al1e | -0.03698791 | -0.247785793 | 0.173809972 | 7.769245191 | -0.43948484 | 0.677114464 | 0.839451033 | -4.729963424 |
| **A_55_P2055419** | Arid4a | -0.040270104 | -0.269803161 | 0.189262954 | 8.670484268 | -0.439428129 | 0.677153114 | 0.839451033 | -4.729971276 |
| **A_51_P251205** | Psd | -0.119880977 | -0.803533902 | 0.563771948 | 9.271565493 | -0.439202641 | 0.677306801 | 0.839551912 | -4.730002487 |

| | | | | | | | | | |
|---|---|---|---|---|---|---|---|---|---|
| **A_51_P388696** | Ndufs4 | 0.135410539 | -0.636841308 | 0.907662385 | 12.93984904 | 0.439181316 | 0.677321337 | 0.839551912 | -4.730005438 |
| **A_55_P2083149** | NA | 0.084156747 | -0.39590366 | 0.564217154 | 9.505019432 | 0.439079739 | 0.677390576 | 0.839583909 | -4.730019492 |
| **A_55_P2069415** | Fxr1 | -0.08474631 | -0.568473487 | 0.398980867 | 10.87948085 | -0.438804077 | 0.677578496 | 0.839717446 | -4.73005762 |
| **A_55_P1959748** | Asns | -0.224767352 | -1.508160924 | 1.058626219 | 11.17786513 | -0.438655638 | 0.677679699 | 0.839717446 | -4.730078142 |
| **A_55_P2079579** | Pira7 | -0.05856942 | -0.393052325 | 0.275913485 | 8.181944697 | -0.438578503 | 0.677732291 | 0.839717446 | -4.730088804 |
| **A_51_P436727** | Arrb1 | -0.115882042 | -0.777670646 | 0.545906563 | 11.76641166 | -0.43857836 | 0.677732388 | 0.839717446 | -4.730088823 |
| **A_52_P161495** | Bcl6 | 0.099293922 | -0.467809057 | 0.6663969 | 10.46514714 | 0.438541887 | 0.677757257 | 0.839717446 | -4.730093864 |
| **A_51_P262144** | Ddx19b | 0.068976892 | -0.324983296 | 0.46293708 | 9.308986663 | 0.438532345 | 0.677763763 | 0.839717446 | -4.730095183 |
| **A_55_P2050236** | Hikeshi | 0.040942362 | -0.192929322 | 0.274814046 | 11.59033742 | 0.438475767 | 0.677802341 | 0.839717446 | -4.730103001 |
| **A_51_P223929** | Ash1l | -0.072170857 | -0.484564215 | 0.340222501 | 9.385213477 | -0.438329346 | 0.677902184 | 0.839787328 | -4.730123231 |
| **A_52_P282741** | Sdc3 | -0.046798146 | -0.314298263 | 0.220701971 | 9.60499506 | -0.438182486 | 0.678002334 | 0.839857581 | -4.730143515 |
| **A_51_P501757** | NA | 0.053329074 | -0.251742724 | 0.358400873 | 8.605646896 | 0.43783687 | 0.678238053 | 0.840057011 | -4.730191229 |
| **A_51_P496845** | Capza2 | 0.086950834 | -0.410476303 | 0.58437797 | 11.28205337 | 0.437819017 | 0.67825023 | 0.840057011 | -4.730193692 |
| **A_55_P1964509** | Lcn12 | 0.051937516 | -0.245309536 | 0.349184569 | 9.427417533 | 0.437636933 | 0.678374433 | 0.840157023 | -4.730218815 |
| **A_51_P196726** | Itih2 | 0.044049314 | -0.208341763 | 0.296440391 | 8.03661892 | 0.437134917 | 0.678716927 | 0.840527355 | -4.730288032 |
| **A_51_P262208** | Itgb2 | 0.079634338 | -0.376880627 | 0.536149302 | 8.467395289 | 0.436913737 | 0.678867851 | 0.840641373 | -4.730318506 |
| **A_55_P1978451** | Ttc1 | -0.063272814 | -0.426027252 | 0.299481625 | 10.57029446 | -0.436872552 | 0.678895956 | 0.840641373 | -4.730324179 |
| **A_66_P106098** | Kif3a | -0.047315258 | -0.318949078 | 0.224318561 | 9.48288802 | -0.436282428 | 0.679298722 | 0.841044074 | -4.730405411 |
| **A_55_P2092384** | Pik3r2 | 0.057706245 | -0.273613513 | 0.389026003 | 9.372844512 | 0.436240343 | 0.679327451 | 0.841044074 | -4.730411201 |
| **A_55_P2097474** | Creb5 | -0.11857385 | -0.799419661 | 0.562271962 | 14.88617785 | -0.436204855 | 0.679351676 | 0.841044074 | -4.730416082 |
| **A_51_P417858** | Zfp825 | 0.055907771 | -0.265307816 | 0.377123357 | 8.493009586 | 0.435939186 | 0.679533044 | 0.841135947 | -4.730452615 |
| **A_55_P1996683** | Itih1 | -0.043035488 | -0.290326237 | 0.204255261 | 7.751460944 | -0.435882274 | 0.679571901 | 0.841135947 | -4.730460439 |
| **A_55_P2045050** | Zfp953 | 0.048429223 | -0.229879877 | 0.326738322 | 8.348692541 | 0.435843363 | 0.679598467 | 0.841135947 | -4.730465787 |
| **A_51_P380807** | Ckm | 0.2022724 | -0.960135731 | 1.364680531 | 8.698140646 | 0.435841252 | 0.679599908 | 0.841135947 | -4.730466077 |
| **A_55_P2071536** | NA | -0.044421027 | -0.299767413 | 0.210925358 | 7.996202607 | -0.43572174 | 0.679681509 | 0.841183094 | -4.730482502 |
| **A_51_P345699** | Tnnt3 | -0.127428058 | -0.860055427 | 0.605199311 | 8.499973038 | -0.435644535 | 0.679734226 | 0.84119449 | -4.73049311 |
| **A_51_P106428** | Nsmce4a | -0.086860326 | -0.58637077 | 0.412650117 | 12.98859172 | -0.435539181 | 0.679806167 | 0.841229674 | -4.730507584 |
| **A_51_P101545** | Hgfac | -0.044100451 | -0.29780051 | 0.209599608 | 7.829687353 | -0.435384349 | 0.679911901 | 0.841306668 | -4.730528849 |
| **A_51_P135389** | Zfp474 | 0.133240513 | -0.633883902 | 0.900364928 | 8.364731347 | 0.435031631 | 0.680152799 | 0.841499522 | -4.730577266 |
| **A_55_P2197847** | Lhx1os | -0.061505857 | -0.415680925 | 0.292669212 | 8.435479753 | -0.434959511 | 0.680202061 | 0.841499522 | -4.730587162 |
| **A_55_P2348676** | B130052F | -0.064050463 | -0.43289678 | 0.304795853 | 8.256804992 | -0.434937817 | 0.680216879 | 0.841499522 | -4.730590139 |
| **A_55_P2041910** | NA | 0.152737883 | -0.726905124 | 1.03238089 | 9.579351948 | 0.434901253 | 0.680241855 | 0.841499522 | -4.730595155 |
| **A_55_P2018479** | Carnmt1 | 0.051698774 | -0.246299334 | 0.349696883 | 10.88776246 | 0.434527316 | 0.680497308 | 0.841761674 | -4.730646434 |
| **A_52_P376214** | Cstf3 | -0.073795974 | -0.499410708 | 0.35181876 | 9.267149164 | -0.434276531 | 0.680668658 | 0.841919764 | -4.730680803 |
| **A_55_P2048759** | Mgarp | 0.095518133 | -0.455880002 | 0.646916268 | 8.128438994 | 0.433881244 | 0.680938782 | 0.8422 | -4.73073494 |
| **A_55_P2078680** | Tpm1 | -0.094389097 | -0.639435634 | 0.45065744 | 10.07486706 | -0.433749105 | 0.681029092 | 0.842257818 | -4.730753028 |
| **A_51_P405227** | Nradd | 0.075297139 | -0.360101424 | 0.510695701 | 8.748057273 | 0.433153499 | 0.681436233 | 0.842687398 | -4.730834495 |
| **A_55_P2079158** | Saxo2 | 0.132334635 | -0.63316261 | 0.89783188 | 8.405261614 | 0.432992365 | 0.681546401 | 0.842687398 | -4.730856518 |
| **A_55_P1990196** | NA | -0.042275971 | -0.286837829 | 0.202285887 | 10.90416387 | -0.432967416 | 0.681563459 | 0.842687398 | -4.730859927 |
| **A_55_P2016193** | Prxl2c | 0.140607505 | -0.672810396 | 0.954025406 | 10.34093866 | 0.432957354 | 0.681570339 | 0.842687398 | -4.730861302 |
| **A_55_P2047626** | Cutal | -0.093093231 | -0.63171536 | 0.445528899 | 8.591176382 | -0.43289668 | 0.681611825 | 0.842687398 | -4.730869593 |
| **A_52_P35377** | Plagl2 | 0.059002243 | -0.282405174 | 0.400409661 | 9.214243113 | 0.432858466 | 0.681637955 | 0.842687398 | -4.730874813 |
| **A_52_P102002** | Srrd | 0.042292639 | -0.202547051 | 0.287132329 | 8.477466034 | 0.432646617 | 0.681782819 | 0.842735903 | -4.730903749 |
| **A_51_P428754** | Grik5 | 0.080959316 | -0.387741007 | 0.549659639 | 8.641496609 | 0.432635272 | 0.681790578 | 0.842735903 | -4.730905298 |
| **A_55_P2144248** | Zfp790 | 0.18344095 | -0.878621911 | 1.245503811 | 8.808866221 | 0.432609861 | 0.681807955 | 0.842735903 | -4.730908768 |
| **A_55_P2040893** | Tnni2 | -0.282718566 | -1.920665171 | 1.355228039 | 10.31420875 | -0.43231961 | 0.68200646 | 0.842882622 | -4.730948389 |
| **A_55_P2156394** | Btbd16 | -0.085192324 | -0.578771383 | 0.408386735 | 9.193465598 | -0.432308808 | 0.682013848 | 0.842882622 | -4.730949863 |
| **A_55_P2054777** | Maml1 | 0.040878199 | -0.196198669 | 0.277955067 | 10.93642059 | 0.431869878 | 0.682314092 | 0.843199786 | -4.731009733 |

| | | | | | | | | | |
|---|---|---|---|---|---|---|---|---|---|
| **A_55_P2069885** | Slc50a1 | -0.050120807 | -0.340890479 | 0.240648865 | 9.031515485 | -0.431737024 | 0.682404981 | 0.843258207 | -4.731027843 |
| **A_66_P138567** | NA | 0.042220341 | -0.202762279 | 0.287202962 | 9.910913745 | 0.431655033 | 0.682461077 | 0.843273629 | -4.731039018 |
| **A_51_P120066** | 9330151L | -0.040881045 | -0.278226617 | 0.196464526 | 8.233072421 | -0.431410991 | 0.682628056 | 0.84342605 | -4.723036237 |
| **A_51_P165087** | Snai1 | 0.11625206 | -0.559197278 | 0.791701399 | 11.20978696 | 0.431080347 | 0.682854322 | 0.84360559 | -4.731117289 |
| **A_51_P439857** | Dnajc21 | -0.052991225 | -0.360888165 | 0.254905715 | 9.159075173 | -0.431071126 | 0.682860633 | 0.84360559 | -4.731118544 |
| **A_51_P424221** | Irf2bp2 | 0.244629177 | -1.177008349 | 1.666266702 | 10.78652291 | 0.430992508 | 0.682914439 | 0.843618156 | -4.731129244 |
| **A_51_P286106** | NA | 0.048990644 | -0.235775485 | 0.333756772 | 8.874786997 | 0.430898677 | 0.682978659 | 0.843643585 | -4.731142013 |
| **A_52_P1016836** | NA | 0.176474775 | -0.84979785 | 1.202747399 | 8.686474387 | 0.430695404 | 0.683117793 | 0.843755798 | -4.731169665 |
| **A_55_P2083814** | Zzz3 | 0.046370574 | -0.223328853 | 0.316070002 | 11.81497673 | 0.430638442 | 0.683156784 | 0.843755798 | -4.731177412 |
| **A_55_P2171158** | Hmgn1 | -0.074623203 | -0.508751534 | 0.359505129 | 15.41599853 | -0.43053266 | 0.683229197 | 0.843791331 | -4.731191797 |
| **A_55_P2140453** | Myh6 | 0.087297053 | -0.420775286 | 0.595369391 | 8.008149515 | 0.430352546 | 0.683352501 | 0.843841726 | -4.731216281 |
| **A_52_P60353** | Greb1l | 0.044757574 | -0.215737533 | 0.305252681 | 8.026360719 | 0.430345547 | 0.683357293 | 0.843841726 | -4.731217232 |
| **A_51_P465809** | Slc30a9 | -0.056414642 | -0.384834705 | 0.272005421 | 8.158048256 | -0.430241702 | 0.68342839 | 0.843875622 | -4.731231344 |
| **A_55_P2101666** | Vamp5 | -0.082053127 | -0.560058339 | 0.395952086 | 10.06697616 | -0.429944946 | 0.683631581 | 0.844072609 | -4.731271656 |
| **A_55_P2060415** | NA | 0.121646959 | -0.587289965 | 0.830583884 | 16.64463709 | 0.429777827 | 0.683746022 | 0.84412138 | -4.731294347 |
| **A_55_P2070823** | Ino80d | 0.067746901 | -0.327201368 | 0.462695169 | 10.44640206 | 0.429634911 | 0.683843896 | 0.84412138 | -4.731313745 |
| **A_52_P339722** | Csnk1a1 | 0.043357976 | -0.209461594 | 0.296177546 | 11.72509182 | 0.429544989 | 0.683905482 | 0.84412138 | -4.731325948 |
| **A_51_P467770** | Snx19 | 0.037564767 | -0.181496599 | 0.256626133 | 9.174508845 | 0.429502059 | 0.683934885 | 0.84412138 | -4.731331772 |
| **A_55_P2044653** | Cyp2b10 | -0.089954255 | -0.614561171 | 0.434652662 | 8.188707514 | -0.429475154 | 0.683953312 | 0.84412138 | -4.731335422 |
| **A_51_P232474** | Zswim1 | -0.09304678 | -0.635821896 | 0.449728337 | 8.153417234 | -0.429370065 | 0.684025291 | 0.84412138 | -4.731349678 |
| **A_55_P1985298** | Krtap5-2 | 0.050692057 | -0.245029393 | 0.346413507 | 8.191300208 | 0.429346012 | 0.684041766 | 0.84412138 | -4.731352941 |
| **A_55_P2028310** | Trappc1 | -0.049810189 | -0.340394642 | 0.240774263 | 8.385776632 | -0.429334872 | 0.684049397 | 0.84412138 | -4.731354451 |
| **A_51_P296487** | Lss | -0.044316403 | -0.302863863 | 0.214231058 | 8.286786865 | -0.429313522 | 0.684064021 | 0.84412138 | -4.731357347 |
| **A_55_P2006662** | Zfp967 | -0.077461529 | -0.529514614 | 0.374591556 | 8.650034702 | -0.429187411 | 0.684150406 | 0.844174098 | -4.731374448 |
| **A_55_P2042870** | Aoc2 | -0.135910148 | -0.929215588 | 0.657395291 | 14.98163229 | -0.429103246 | 0.684208061 | 0.844179538 | -4.731385859 |
| **A_55_P1953459** | Yap1 | 0.056053952 | -0.271170037 | 0.383277941 | 13.10942377 | 0.4290535 | 0.68424214 | 0.844179538 | -4.731392602 |
| **A_55_P2279997** | 4930549C | -0.036734517 | -0.25138844 | 0.177919406 | 7.856169002 | -0.428633246 | 0.684530068 | 0.844480879 | -4.731449543 |
| **A_55_P2277019** | Loxl3 | 0.076589817 | -0.371123479 | 0.524303112 | 8.474649117 | 0.428470951 | 0.684641277 | 0.844564184 | -4.731471519 |
| **A_55_P2111439** | Gpsm1 | -0.035862452 | -0.245574785 | 0.173849881 | 7.901408925 | -0.428318028 | 0.684746072 | 0.844639566 | -4.731492219 |
| **A_55_P2070766** | NA | -0.09966244 | -0.682949344 | 0.483624464 | 12.28200847 | -0.427956508 | 0.684993846 | 0.844812165 | -4.73154113 |
| **A_55_P2061991** | Pnldc1 | 0.058616536 | -0.284463977 | 0.401697049 | 8.443487493 | 0.427931683 | 0.685010862 | 0.844812165 | -4.731544487 |
| **A_51_P235878** | Guk1 | -0.068934774 | -0.472416132 | 0.334546584 | 11.0389946 | -0.427922604 | 0.685017085 | 0.844812165 | -4.731545715 |
| **A_55_P1995768** | Zfp967 | -0.079397773 | -0.544374488 | 0.385578942 | 9.645809495 | -0.427688391 | 0.685177633 | 0.844956267 | -4.731577381 |
| **A_51_P439996** | Gp5 | -0.0359304 | -0.246496153 | 0.174635353 | 7.716353759 | -0.427390304 | 0.685381991 | 0.845154373 | -4.731617659 |
| **A_55_P2452384** | Mga | -0.054893219 | -0.376743893 | 0.266957454 | 8.981164839 | -0.427183651 | 0.685523683 | 0.845235565 | -4.731645569 |
| **A_52_P377416** | Disp1 | 0.058044151 | -0.282294573 | 0.398382876 | 8.637359541 | 0.427166754 | 0.685535269 | 0.845235565 | -4.73164785 |
| **A_55_P2090617** | Pilra | -0.054300121 | -0.372799024 | 0.264198782 | 7.932863933 | -0.427015069 | 0.685639283 | 0.845309902 | -4.731668327 |
| **A_52_P110052** | Ackr1 | 0.037377079 | -0.181908198 | 0.256662356 | 7.782171146 | 0.426919735 | 0.685704659 | 0.845336598 | -4.731681194 |
| **A_55_P2033135** | Sh3bgr | 0.256207569 | -1.247433118 | 1.759848255 | 9.783730622 | 0.426774274 | 0.685804416 | 0.845405673 | -4.731700821 |
| **A_55_P2093483** | NA | 0.095906245 | -0.467057707 | 0.658870198 | 16.06169479 | 0.42669411 | 0.685859395 | 0.845419544 | -4.731711635 |
| **A_52_P525070** | Vamp4 | -0.035916905 | -0.246828421 | 0.174994611 | 8.524425263 | -0.426529386 | 0.685972376 | 0.845451992 | -4.73173385 |
| **A_55_P2017714** | Kat5 | -0.041768164 | -0.28704016 | 0.203503831 | 11.85348342 | -0.426528218 | 0.685973177 | 0.845451992 | -4.731734008 |
| **A_55_P2074371** | Lin28a | -0.087890219 | -0.60449481 | 0.428714372 | 8.23718905 | -0.426120697 | 0.686252726 | 0.845630955 | -4.731788935 |
| **A_55_P2075303** | Zfp609 | 0.042850843 | -0.209044357 | 0.294746044 | 8.563703932 | 0.426078704 | 0.686281535 | 0.845630955 | -4.731794592 |
| **A_52_P357402** | Mlst8 | -0.044497035 | -0.306104339 | 0.217110269 | 10.34067487 | -0.426021537 | 0.686320755 | 0.845630955 | -4.731802293 |
| **A_51_P312485** | Fpr1 | -0.121181395 | -0.833897542 | 0.591534753 | 8.290771734 | -0.425862788 | 0.686429673 | 0.845630955 | -4.731823672 |
| **A_66_P115378** | Orc2 | -0.041281413 | -0.284082419 | 0.201519592 | 8.06034095 | -0.425847814 | 0.686439947 | 0.845630955 | -4.731825688 |
| **A_51_P173709** | Gprc5b | 0.068841841 | -0.336080733 | 0.473764415 | 9.200857652 | 0.425824686 | 0.686455816 | 0.845630955 | -4.731828803 |

| | | | | | | | | | |
|---|---|---|---|---|---|---|---|---|---|
| **A_51_P409429** | Aars | -0.172769822 | -1.189066197 | 0.843526553 | 11.08088949 | -0.425792349 | 0.686478004 | 0.845630955 | -4.731833156 |
| **A_52_P273594** | Zc3h14 | 0.065249788 | -0.318587654 | 0.44908723 | 10.38665555 | 0.425776917 | 0.686488593 | 0.845630955 | -4.731835234 |
| **A_55_P2006277** | Ogdh | -0.167822221 | -1.155129933 | 0.819485491 | 9.228194191 | -0.42574277 | 0.686512024 | 0.845630955 | -4.731839831 |
| **A_52_P325443** | Arhgap29 | -0.072495679 | -0.499108214 | 0.354116856 | 10.79856865 | -0.42562669 | 0.686591676 | 0.845646584 | -4.731855456 |
| **A_51_P502348** | Tfip11 | -0.081282226 | -0.559634219 | 0.397069766 | 9.38986443 | -0.425596796 | 0.68661219 | 0.845646584 | -4.731859479 |
| **A_55_P2014284** | Serpina11 | 0.038968403 | -0.190563729 | 0.268500535 | 7.922578072 | 0.425225655 | 0.686866896 | 0.8459064 | -4.731909407 |
| **A_55_P2039196** | Gfra4 | -0.168659456 | -1.162486255 | 0.825167343 | 13.93166224 | -0.425060098 | 0.68698053 | 0.845990716 | -4.731931666 |
| **A_55_P2054998** | NA | -0.057546155 | -0.396686418 | 0.281594108 | 10.57369404 | -0.424998409 | 0.687022874 | 0.845990716 | -4.731939958 |
| **A_55_P2127688** | 4933434E | -0.035603393 | -0.245462517 | 0.17425573 | 12.24002954 | -0.424926565 | 0.687072189 | 0.845997561 | -4.731949614 |
| **A_52_P312371** | Zfp345 | 0.069438889 | -0.339940933 | 0.478818712 | 8.345617278 | 0.42484125 | 0.687130754 | 0.846015793 | -4.731961079 |
| **A_55_P2248651** | Aak1 | 0.047367513 | -0.232006495 | 0.326741521 | 10.45296528 | 0.424663494 | 0.687252784 | 0.846090641 | -4.731984959 |
| **A_55_P2096204** | Kdm1b | -0.131852645 | -0.909676351 | 0.645971061 | 10.18689583 | -0.424578519 | 0.687311123 | 0.846090641 | -4.731996371 |
| **A_52_P350537** | Mtmr11 | -0.03658189 | -0.252394051 | 0.179230271 | 8.216318235 | -0.424561466 | 0.68732283 | 0.846090641 | -4.731998661 |
| **A_55_P2059640** | Abhd14b | 0.237587256 | -1.164346334 | 1.639520847 | 10.57477893 | 0.424469067 | 0.68738627 | 0.846114863 | -4.732011068 |
| **A_55_P2123331** | Cog2 | -0.052564322 | -0.36289491 | 0.257766267 | 8.133775977 | -0.424245091 | 0.687540059 | 0.846247836 | -4.732041132 |
| **A_55_P1967708** | NA | -0.09455928 | -0.652900848 | 0.463782289 | 8.599808529 | -0.424184249 | 0.687581838 | 0.846247836 | -4.732049296 |
| **A_55_P1960298** | Tmem80 | 0.095960586 | -0.470935004 | 0.662856177 | 9.251297696 | 0.423974912 | 0.687725595 | 0.846305966 | -4.732077379 |
| **A_55_P1974362** | Hp1bp3 | 0.057752322 | -0.283472416 | 0.39897706 | 13.65953289 | 0.423915491 | 0.687766403 | 0.846305966 | -4.732085348 |
| **A_55_P2011877** | Eif2s3y | 0.492225881 | -2.416071252 | 3.400523013 | 9.6222651 | 0.423912297 | 0.687768597 | 0.846305966 | -4.732085777 |
| **A_55_P2187261** | NA | 0.041517767 | -0.203850232 | 0.286885766 | 8.504643969 | 0.423805324 | 0.687842065 | 0.846305966 | -4.732100121 |
| **A_55_P2024909** | Dnah17 | -0.040037582 | -0.276662533 | 0.196587368 | 7.824729408 | -0.42379678 | 0.687847934 | 0.846305966 | -4.732101266 |
| **A_55_P2133067** | Ube2v1 | 0.036402178 | -0.178909358 | 0.251713713 | 8.35858515 | 0.423458067 | 0.688080587 | 0.846538344 | -4.732146662 |
| **A_66_P127411** | Acat2 | 0.040561886 | -0.19942987 | 0.280553642 | 8.213057939 | 0.423323286 | 0.688173175 | 0.846598382 | -4.732164717 |
| **A_55_P2060680** | Acsf2 | 0.106700411 | -0.524809736 | 0.738210559 | 11.36070193 | 0.423190691 | 0.688264268 | 0.846656573 | -4.732182474 |
| **A_55_P2057946** | Hsp90aa1 | -0.115004823 | -0.795985602 | 0.565975956 | 14.2052064 | -0.422991408 | 0.688401186 | 0.846681179 | -4.732209152 |
| **A_51_P108108** | Lrrc25 | -0.039494784 | -0.273376838 | 0.19438727 | 8.160882827 | -0.422954051 | 0.688426854 | 0.846681179 | -4.732214152 |
| **A_55_P2167512** | NA | 0.048039525 | -0.236459941 | 0.33253899 | 12.94773523 | 0.422929123 | 0.688443982 | 0.846681179 | -4.732217488 |
| **A_55_P2117043** | Mtmr1 | 0.048607142 | -0.239269177 | 0.336483461 | 9.84236294 | 0.422906626 | 0.68845944 | 0.846681179 | -4.732220499 |
| **A_55_P2127884** | Btbd1 | 0.037025861 | -0.182358539 | 0.256410261 | 8.115759606 | 0.422717053 | 0.688589704 | 0.846729497 | -4.732245862 |
| **A_55_P2124461** | NA | 0.11940173 | -0.588237619 | 0.827041079 | 12.95497067 | 0.422618973 | 0.688657104 | 0.846729497 | -4.732258981 |
| **A_51_P481210** | Ofd1 | -0.039206547 | -0.271565828 | 0.193152735 | 10.03759389 | -0.422618897 | 0.688657157 | 0.846729497 | -4.732258991 |
| **A_55_P2110290** | Tmem254 | -0.063743206 | -0.441541842 | 0.31405543 | 10.37264489 | -0.422594521 | 0.688673908 | 0.846729497 | -4.732262251 |
| **A_55_P2062213** | Tmem41a | -0.125258621 | -0.867857219 | 0.617339976 | 10.73261722 | -0.422477757 | 0.688754153 | 0.846774309 | -4.732277864 |
| **A_55_P2092831** | NA | -0.060511431 | -0.419427955 | 0.298405093 | 12.13074385 | -0.422273974 | 0.688894211 | 0.846864599 | -4.732305104 |
| **A_55_P2066878** | Kcnj13 | -0.077222437 | -0.535378856 | 0.380933982 | 8.756162816 | -0.422162907 | 0.688970553 | 0.846864599 | -4.732319946 |
| **A_55_P2033666** | NA | 0.103436747 | -0.510288539 | 0.717162033 | 16.88490583 | 0.422134826 | 0.688989854 | 0.846864599 | -4.732323698 |
| **A_51_P415519** | Tmem39a | -0.102071185 | -0.707721147 | 0.503578778 | 10.81386751 | -0.422115992 | 0.6890028 | 0.846864599 | -4.732326214 |
| **A_55_P2308835** | E130110C | -0.03665349 | -0.25419365 | 0.18088667 | 8.253485959 | -0.422013396 | 0.689073325 | 0.846897442 | -4.732339919 |
| **A_55_P2089020** | Slc35g2 | 0.110719531 | -0.546511727 | 0.76795079 | 8.921130592 | 0.421945513 | 0.689119989 | 0.846900957 | -4.732348985 |
| **A_55_P1984023** | Arrb2 | -0.050831324 | -0.35302804 | 0.251365392 | 8.693531718 | -0.42130055 | 0.689563428 | 0.847392062 | -4.732435061 |
| **A_52_P669922** | Hamp | -0.039228324 | -0.272509264 | 0.194052615 | 7.70521518 | -0.421183012 | 0.689644255 | 0.847437525 | -4.732450735 |
| **A_55_P2051727** | Rnf8 | 0.062878743 | -0.311141812 | 0.436899297 | 11.37106863 | 0.421074289 | 0.689719024 | 0.847438759 | -4.73246523 |
| **A_55_P2253746** | Nip7 | -0.051812512 | -0.36002279 | 0.256397765 | 8.856530223 | -0.421054081 | 0.689732922 | 0.847438759 | -4.732467924 |
| **A_66_P106289** | Vps54 | 0.043387647 | -0.214747593 | 0.301522887 | 9.643191011 | 0.420987449 | 0.689778747 | 0.847441209 | -4.732476805 |
| **A_55_P2090441** | NA | 0.039199935 | -0.194299992 | 0.272699861 | 8.052652858 | 0.420483481 | 0.690125395 | 0.847633925 | -4.732543938 |
| **A_55_P2060498** | Zfp553 | -0.050595992 | -0.351981852 | 0.250789868 | 9.89553474 | -0.4204783 | 0.690128959 | 0.847633925 | -4.732544627 |
| **A_52_P677891** | Fam174b | 0.235071016 | -1.165228011 | 1.635370044 | 9.922354027 | 0.420463831 | 0.690138912 | 0.847633925 | -4.732546554 |
| **A_55_P2109354** | Dot1l | -0.055429837 | -0.385644199 | 0.274784524 | 10.47681649 | -0.420434158 | 0.690159325 | 0.847633925 | -4.732550504 |

| | | | | | | | | | |
|---|---|---|---|---|---|---|---|---|---|
| **A_55_P2147402** | NA | -0.036630832 | -0.254904543 | 0.181642878 | 8.07404227 | -0.420335143 | 0.690227443 | 0.847633925 | -4.732563684 |
| **A_52_P522977** | Ssbp2 | 0.037448494 | -0.185698091 | 0.260595079 | 7.921004774 | 0.420333936 | 0.690228273 | 0.847633925 | -4.732563844 |
| **A_51_P262533** | Cxxc1 | 0.124784994 | -0.619022104 | 0.868592092 | 9.606131777 | 0.42019646 | 0.690322856 | 0.847633925 | -4.732582139 |
| **A_55_P2078271** | Rbis | 0.084962385 | -0.421499224 | 0.591423995 | 12.5095411 | 0.420175293 | 0.690337419 | 0.847633925 | -4.732584955 |
| **A_55_P2142873** | Mink1 | -0.052162446 | -0.363134214 | 0.258809321 | 10.21620218 | -0.420133522 | 0.690366159 | 0.847633925 | -4.732590513 |
| **A_55_P2065991** | S100a11 | -0.074561631 | -0.519080534 | 0.369957271 | 12.89351368 | -0.420122087 | 0.690374026 | 0.847633925 | -4.732592034 |
| **A_55_P2000628** | NA | 0.054036079 | -0.268226015 | 0.376298174 | 9.338416765 | 0.41997646 | 0.690474227 | 0.847703118 | -4.732611404 |
| **A_51_P295215** | Alkbh5 | -0.08180663 | -0.570557529 | 0.406944269 | 11.10118749 | -0.419228963 | 0.690988663 | 0.848280831 | -4.732710738 |
| **A_55_P2068148** | Pias1 | -0.045136643 | -0.314857821 | 0.224584535 | 8.574387171 | -0.419145256 | 0.691046282 | 0.848297703 | -4.732721851 |
| **A_55_P1958349** | Nipsnap3l | 0.076389406 | -0.380163635 | 0.532942446 | 8.770070829 | 0.419075468 | 0.691094323 | 0.848302815 | -4.732731116 |
| **A_52_P320261** | Cpa2 | 0.083680083 | -0.417010927 | 0.584371093 | 8.538066142 | 0.418603306 | 0.691419389 | 0.848644386 | -4.732793759 |
| **A_55_P1977548** | Mapk1 | 0.069436754 | -0.346089884 | 0.484963391 | 11.05921561 | 0.418543772 | 0.691460381 | 0.848644386 | -4.732801653 |
| **A_51_P409349** | Trim29 | 0.108464238 | -0.540822602 | 0.757751078 | 8.081733056 | 0.418408321 | 0.69155365 | 0.848689039 | -4.732819609 |
| **A_55_P2219657** | Tmf1 | -0.038324417 | -0.267766023 | 0.191117189 | 8.435736449 | -0.418363441 | 0.691584556 | 0.848689039 | -4.732825558 |
| **A_55_P2026109** | Rpap1 | 0.075263925 | -0.375400298 | 0.525928148 | 11.57714086 | 0.418296402 | 0.691630721 | 0.848691823 | -4.732834442 |
| **A_55_P2154252** | Gfpt2 | -0.065924523 | -0.460763234 | 0.328914188 | 8.343414589 | -0.418193823 | 0.691701363 | 0.848697592 | -4.732848034 |
| **A_55_P2005081** | Rpl10 | 0.11108421 | -0.554277825 | 0.776446244 | 17.78019247 | 0.418162092 | 0.691723216 | 0.848697592 | -4.732852238 |
| **A_52_P642488** | Kcnk1 | -0.143333874 | -1.002016924 | 0.715349176 | 9.790602925 | -0.418086645 | 0.691775176 | 0.848707484 | -4.732862233 |
| **A_55_P2107422** | Kif21a | -0.057802811 | -0.404209231 | 0.28860361 | 9.188388427 | -0.417939452 | 0.691876553 | 0.848778 | -4.732881727 |
| **A_55_P1960496** | Ppp1r2 | 0.04427998 | -0.221127129 | 0.309687088 | 8.844313101 | 0.417873862 | 0.691921729 | 0.848779565 | -4.732890411 |
| **A_55_P2005345** | Epb41 | -0.037243923 | -0.26054785 | 0.186060005 | 7.768667799 | -0.417743209 | 0.692011724 | 0.848836104 | -4.732907707 |
| **A_52_P561020** | Slc25a32 | -0.066335136 | -0.464215352 | 0.331545079 | 8.835965136 | -0.417581863 | 0.692122867 | 0.848904206 | -4.73292906 |
| **A_51_P335900** | Cox5a | -0.056907401 | -0.398277968 | 0.284463166 | 14.92542065 | -0.417535133 | 0.692155059 | 0.848904206 | -4.732935242 |
| **A_55_P1986351** | Tbc1d32 | 0.06775131 | -0.338850615 | 0.474353234 | 8.43185916 | 0.417348263 | 0.692283799 | 0.848924728 | -4.732959961 |
| **A_55_P2079684** | NA | -0.050066951 | -0.350548463 | 0.250414561 | 14.35200178 | -0.417333968 | 0.692293647 | 0.848924728 | -4.732961851 |
| **A_55_P2112967** | Diaph1 | -0.097683945 | -0.683974143 | 0.488606253 | 8.763945489 | -0.417312028 | 0.692308763 | 0.848924728 | -4.732964753 |
| **A_52_P566605** | Hsd17b7 | -0.119459946 | -0.836544158 | 0.597624265 | 8.446442218 | -0.417255913 | 0.692347425 | 0.848924728 | -4.732972173 |
| **A_52_P393314** | P2rx7 | 0.088467528 | -0.442734423 | 0.619669478 | 8.428587399 | 0.417133025 | 0.692432095 | 0.848932192 | -4.73298842 |
| **A_52_P420394** | Apeh | 0.043870537 | -0.219570374 | 0.307311449 | 11.48191741 | 0.417099892 | 0.692454925 | 0.848932192 | -4.732992799 |
| **A_51_P210340** | 2610306M | 0.075475112 | -0.377797882 | 0.528748106 | 8.839541391 | 0.417055899 | 0.692485238 | 0.848932192 | -4.732998614 |
| **A_66_P119123** | A1cf | 0.040064087 | -0.200598375 | 0.280726549 | 7.669380223 | 0.416962737 | 0.692549433 | 0.848957059 | -4.733010926 |
| **A_55_P1959152** | Gm10324 | -0.130568384 | -0.915237662 | 0.654100895 | 8.809631409 | -0.416775061 | 0.692678763 | 0.849061764 | -4.73303572 |
| **A_51_P168630** | Klf1 | 0.035572334 | -0.178249602 | 0.24939427 | 7.725085544 | 0.416687474 | 0.692739125 | 0.849081922 | -4.733047288 |
| **A_52_P367294** | Fsd1l | -0.048892664 | -0.342920237 | 0.245134908 | 8.420391385 | -0.416491356 | 0.692874291 | 0.849193758 | -4.733073183 |
| **A_55_P2107901** | Dlk2 | -0.044390914 | -0.311436231 | 0.222654402 | 8.555508376 | -0.416350863 | 0.692971126 | 0.849253139 | -4.733091726 |
| **A_66_P130647** | NA | -0.035789579 | -0.251120896 | 0.179541738 | 7.706711913 | -0.416293609 | 0.693010591 | 0.849253139 | -4.733099281 |
| **A_55_P2125623** | NA | 0.052734257 | -0.264657117 | 0.37012563 | 8.124420988 | 0.416148232 | 0.693110804 | 0.849322112 | -4.733118461 |
| **A_51_P446469** | Dok2 | -0.198410524 | -1.392904233 | 0.996083184 | 9.630839348 | -0.41603626 | 0.693187994 | 0.849362866 | -4.73313323 |
| **A_51_P413785** | Commd3 | 0.075443684 | -0.378974606 | 0.529861974 | 11.90211532 | 0.415831541 | 0.693329131 | 0.849376181 | -4.733160222 |
| **A_51_P385773** | Eif2ak1 | -0.042829942 | -0.300890315 | 0.215230431 | 8.42185829 | -0.415696639 | 0.693422143 | 0.849376181 | -4.733178002 |
| **A_55_P2182810** | Arl16 | -0.03639935 | -0.255714943 | 0.182916244 | 8.727138573 | -0.415694674 | 0.693423498 | 0.849376181 | -4.733178261 |
| **A_55_P2122841** | NA | 0.085345215 | -0.428923591 | 0.59961402 | 14.06389652 | 0.415661059 | 0.693446675 | 0.849376181 | -4.73318269 |
| **A_55_P2073015** | Lin37 | -0.066710617 | -0.468706462 | 0.335285227 | 9.479247026 | -0.415646129 | 0.69345697 | 0.849376181 | -4.733184658 |
| **A_55_P2146050** | Dennd5b | -0.175060017 | -1.229985726 | 0.879865692 | 11.48124774 | -0.415638181 | 0.693462451 | 0.849376181 | -4.733185705 |
| **A_55_P1960479** | NA | -0.119403568 | -0.8393619 | 0.600554764 | 11.48121071 | -0.415394066 | 0.693630782 | 0.84952193 | -4.733217862 |
| **A_55_P2062262** | Aspscr1 | -0.041010268 | -0.288319937 | 0.206299402 | 9.004815799 | -0.415338177 | 0.693669324 | 0.84952193 | -4.733225223 |
| **A_55_P2368710** | NA | -0.105135964 | -0.739359985 | 0.529088057 | 9.081535038 | -0.415201545 | 0.693763551 | 0.849583512 | -4.733243212 |
| **A_55_P2259500** | D130012F | 0.042204986 | -0.212685591 | 0.297095563 | 8.234866663 | 0.414725107 | 0.694092169 | 0.849785915 | -4.733305898 |

| | | | | | | | | | |
|---|---|---|---|---|---|---|---|---|---|
| **A_55_P2100108** | Zfp148 | -0.064247414 | -0.452358528 | 0.323863699 | 8.344942828 | -0.414619687 | 0.694164891 | 0.849785915 | -4.73331976 |
| **A_55_P1981804** | Gp1ba | -0.03579925 | -0.252068445 | 0.180469944 | 7.877555478 | -0.414600303 | 0.694178263 | 0.849785915 | -4.733322309 |
| **A_51_P246705** | Nop14 | -0.05132123 | -0.361435859 | 0.258793399 | 8.391941214 | -0.414500582 | 0.694247058 | 0.849785915 | -4.733335418 |
| **A_55_P2151442** | C330007F | 0.061169571 | -0.308462638 | 0.43080178 | 10.74282035 | 0.414491829 | 0.694253097 | 0.849785915 | -4.733336568 |
| **A_55_P2152292** | Map2k2 | -0.064470427 | -0.454132319 | 0.325191464 | 9.83532936 | -0.414403066 | 0.694314335 | 0.849785915 | -4.733348234 |
| **A_52_P667913** | Pcdhga9 | -0.07647864 | -0.538901268 | 0.385943987 | 8.938422883 | -0.414239421 | 0.694427243 | 0.849785915 | -4.733369736 |
| **A_51_P293938** | Rasl11b | 0.24625662 | -1.242810327 | 1.735323567 | 12.31606798 | 0.414213296 | 0.694445268 | 0.849785915 | -4.733373168 |
| **A_55_P2008681** | Ktn1 | 0.064109686 | -0.323605192 | 0.451824564 | 11.48528532 | 0.414153685 | 0.694486399 | 0.849785915 | -4.733380998 |
| **A_52_P173442** | Wscd2 | -0.078122184 | -0.550598381 | 0.394354013 | 10.55061281 | -0.414137719 | 0.694497416 | 0.849785915 | -4.733383095 |
| **A_51_P251209** | Cacna1d | 0.114718691 | -0.57917025 | 0.808607631 | 8.478315126 | 0.414089827 | 0.694530463 | 0.849785915 | -4.733389384 |
| **A_55_P1980817** | Gm6548 | 0.066276198 | -0.33460862 | 0.467161016 | 15.65526572 | 0.414083879 | 0.694534567 | 0.849785915 | -4.733390166 |
| **A_52_P469381** | Comtd1 | -0.109372631 | -0.770941421 | 0.55219616 | 10.12700034 | -0.414079751 | 0.694537415 | 0.849785915 | -4.733390708 |
| **A_55_P2056808** | Tanc1 | -0.038562055 | -0.271820186 | 0.194696076 | 8.396982187 | -0.414069958 | 0.694544173 | 0.849785915 | -4.733391994 |
| **A_55_P2004152** | NA | -0.08953143 | -0.631243442 | 0.452180581 | 12.14046706 | -0.413959071 | 0.69462069 | 0.849820935 | -4.733406554 |
| **A_55_P2157023** | Sox17 | 0.135720293 | -0.685614082 | 0.957054667 | 11.03880664 | 0.413880661 | 0.6946748 | 0.849820935 | -4.733416847 |
| **A_55_P2003996** | H2-T24 | -0.03657422 | -0.258039299 | 0.184890859 | 7.780277984 | -0.413637746 | 0.694842444 | 0.849820935 | -4.733448725 |
| **A_55_P2141709** | Gabarapl2 | 0.073507863 | -0.371636586 | 0.518652311 | 13.02405708 | 0.413602523 | 0.694866754 | 0.849820935 | -4.733453346 |
| **A_55_P1991728** | Zfp956 | 0.054703457 | -0.276586998 | 0.385993912 | 9.53456943 | 0.413576826 | 0.69488449 | 0.849820935 | -4.733456717 |
| **A_55_P2167000** | Dmac2 | 0.051966466 | -0.262836903 | 0.366769835 | 11.20413867 | 0.413460627 | 0.694964692 | 0.849820935 | -4.733471959 |
| **A_52_P238858** | Shisa9 | 0.037310397 | -0.188714002 | 0.263334796 | 7.862308063 | 0.413451802 | 0.694970783 | 0.849820935 | -4.733473116 |
| **A_55_P2061421** | NA | 0.061511392 | -0.311153643 | 0.434176426 | 14.19177709 | 0.413415972 | 0.694995515 | 0.849820935 | -4.733477815 |
| **A_51_P304527** | Cluap1 | 0.103192904 | -0.522093102 | 0.72847891 | 10.78500659 | 0.413353359 | 0.695038734 | 0.849820935 | -4.733486025 |
| **A_51_P279571** | Syap1 | -0.047088896 | -0.33242669 | 0.238248899 | 11.57048401 | -0.413342012 | 0.695046566 | 0.849820935 | -4.733487513 |
| **A_66_P121117** | Serpina1d | -0.091884255 | -0.648681023 | 0.464912513 | 7.997708188 | -0.413327915 | 0.695056297 | 0.849820935 | -4.733489361 |
| **A_66_P111011** | Gata3 | -0.465882861 | -3.290091256 | 2.358325535 | 9.178440732 | -0.413171483 | 0.695164282 | 0.849843454 | -4.733509867 |
| **A_52_P141488** | Grk5 | 0.17043165 | -0.862752751 | 1.203616051 | 9.352643921 | 0.413164266 | 0.695169264 | 0.849843454 | -4.733510813 |
| **A_52_P575668** | NA | -0.08558044 | -0.604450882 | 0.433290003 | 9.131917091 | -0.413110208 | 0.695206583 | 0.849843454 | -4.733517898 |
| **A_52_P294510** | Fgl1 | -0.034675203 | -0.245019131 | 0.175668725 | 7.700097252 | -0.412894763 | 0.695355322 | 0.849970364 | -4.733546125 |
| **A_51_P108266** | Actn2 | 0.245178325 | -1.242466708 | 1.732823358 | 9.0174812 | 0.41279374 | 0.695425072 | 0.849970364 | -4.733559356 |
| **A_52_P517224** | Trim63 | 0.077345297 | -0.391983701 | 0.546674295 | 8.169789198 | 0.412768806 | 0.695442288 | 0.849970364 | -4.733562621 |
| **A_52_P253567** | Hsd3b6 | -0.039876011 | -0.281983555 | 0.202231532 | 7.690817625 | -0.41252829 | 0.695608363 | 0.850119601 | -4.733594108 |
| **A_55_P2426549** | NA | 0.062741049 | -0.318282006 | 0.443764105 | 8.647897072 | 0.41243059 | 0.695675831 | 0.85014228 | -4.733606894 |
| **A_51_P259571** | Angptl6 | -0.048420439 | -0.342515491 | 0.245674613 | 9.099946743 | -0.412374069 | 0.695714863 | 0.85014228 | -4.73361429 |
| **A_55_P2170737** | Igf2bp2 | -0.096883243 | -0.685508901 | 0.491742415 | 9.83889573 | -0.412249202 | 0.695801097 | 0.850193921 | -4.733630625 |
| **A_51_P482711** | Dhcr24 | -0.174988964 | -1.238404977 | 0.88842705 | 12.23597571 | -0.412152377 | 0.695867969 | 0.850221897 | -4.733643289 |
| **A_55_P2144781** | Erich5 | -0.160279987 | -1.134553473 | 0.813993499 | 9.341827838 | -0.4120489 | 0.695939437 | 0.850224516 | -4.733656819 |
| **A_66_P121196** | Katnb1 | -0.076303026 | -0.540204709 | 0.387598656 | 10.43368238 | -0.41197054 | 0.695993561 | 0.850224516 | -4.733667064 |
| **A_55_P2103452** | Npy1r | 0.046524056 | -0.236380274 | 0.329428387 | 7.878174778 | 0.411896709 | 0.696044559 | 0.850224516 | -4.733676715 |
| **A_52_P596541** | Kdelr1 | -0.050857699 | -0.360115708 | 0.258400309 | 9.897547359 | -0.411894602 | 0.696046014 | 0.850224516 | -4.73367699 |
| **A_51_P352738** | Mpv17l | 0.042091867 | -0.213910532 | 0.298094265 | 8.487152675 | 0.411817214 | 0.696099471 | 0.850236096 | -4.733687104 |
| **A_55_P2324921** | NA | -0.040860999 | -0.289471801 | 0.207749803 | 8.053541651 | -0.411660628 | 0.696207639 | 0.850305285 | -4.733707563 |
| **A_55_P2014041** | Rab14 | 0.06605143 | -0.335877147 | 0.467980007 | 13.72331003 | 0.411607885 | 0.696244076 | 0.850305285 | -4.733714453 |
| **A_55_P2102095** | 1810010H | 0.169453673 | -0.862000864 | 1.200908211 | 8.878949009 | 0.41148238 | 0.696330782 | 0.850324412 | -4.733730844 |
| **A_52_P231170** | Megf9 | 0.037422214 | -0.190378155 | 0.265222582 | 8.203986232 | 0.411457893 | 0.696347699 | 0.850324412 | -4.733734042 |
| **A_55_P1996872** | Polr2k | 0.058679208 | -0.298605667 | 0.415964083 | 12.29565659 | 0.411358011 | 0.696416709 | 0.850354973 | -4.733747083 |
| **A_51_P382805** | Zpbp2 | -0.077924885 | -0.552552748 | 0.396702978 | 7.90735745 | -0.411219108 | 0.696512682 | 0.850383274 | -4.733765213 |
| **A_66_P125862** | A930005H | 0.070147358 | -0.357184245 | 0.49747896 | 8.497205284 | 0.41114655 | 0.696562818 | 0.850383274 | -4.733774682 |
| **A_52_P572045** | Pfkp | -0.060725942 | -0.43071662 | 0.309264736 | 7.966699376 | -0.411087074 | 0.696603916 | 0.850383274 | -4.733782443 |

| | | | | | | | | | |
|---|---|---|---|---|---|---|---|---|---|
| **A_55_P2054842** | Clpb | 0.056724156 | -0.289002129 | 0.402450441 | 10.80132905 | 0.410947165 | 0.696700598 | 0.850383274 | -4.733800694 |
| **A_51_P151902** | Spon1 | -0.10358264 | -0.734987613 | 0.527822334 | 8.570775765 | -0.410893548 | 0.69673765 | 0.850383274 | -4.733807687 |
| **A_55_P2026315** | Gys1 | -0.053220589 | -0.377660561 | 0.271219383 | 9.041140591 | -0.410861709 | 0.696759653 | 0.850383274 | -4.733811839 |
| **A_66_P130777** | Ephb4 | 0.072351293 | -0.36873673 | 0.513439315 | 8.682763539 | 0.410838727 | 0.696775536 | 0.850383274 | -4.733814836 |
| **A_55_P2026530** | Erc2 | 0.083754315 | -0.426881151 | 0.594389781 | 8.4996319 | 0.410815255 | 0.696791757 | 0.850383274 | -4.733817897 |
| **A_51_P312846** | Fbxo9 | -0.059976988 | -0.425853052 | 0.305899075 | 10.69261673 | -0.410583035 | 0.696952252 | 0.850525459 | -4.73384817 |
| **A_55_P2370096** | NA | -0.036097451 | -0.256629473 | 0.18443457 | 8.15380307 | -0.409972974 | 0.697373969 | 0.850966412 | -4.733927625 |
| **A_55_P2037221** | Klhdc10 | 0.143658406 | -0.734086908 | 1.021403721 | 12.35104851 | 0.40993299 | 0.697401613 | 0.850966412 | -4.733932829 |
| **A_52_P450934** | Paqr9 | -0.041737553 | -0.296866109 | 0.213391002 | 7.754328495 | -0.409749341 | 0.697528591 | 0.851059309 | -4.733956725 |
| **A_55_P1952182** | Vmn2r111 | -0.057624362 | -0.409960549 | 0.294711824 | 8.029933611 | -0.409637048 | 0.697606237 | 0.851059309 | -4.733971332 |
| **A_55_P2175284** | Ei24 | 0.044880166 | -0.229536854 | 0.319297186 | 9.298531504 | 0.409631891 | 0.697609803 | 0.851059309 | -4.733972002 |
| **A_51_P417321** | Zfp236 | -0.062049901 | -0.441556157 | 0.317456355 | 9.004260676 | -0.409517533 | 0.697688882 | 0.851102078 | -4.733986874 |
| **A_55_P2096530** | Icmt | 0.061978931 | -0.317184245 | 0.441142106 | 9.856953175 | 0.409419263 | 0.697756839 | 0.851131276 | -4.73399965 |
| **A_66_P138765** | Trappc10 | -0.063247395 | -0.450252348 | 0.323757558 | 9.923408967 | -0.409332725 | 0.697816685 | 0.85113529 | -4.734010898 |
| **A_55_P2149293** | Agfg2 | 0.042536868 | -0.217835992 | 0.302909729 | 7.935854921 | 0.409185414 | 0.697918566 | 0.85113529 | -4.734030042 |
| **A_55_P2035167** | Pcdh7 | 0.038160901 | -0.195429289 | 0.271751091 | 7.798249407 | 0.409179968 | 0.697922333 | 0.85113529 | -4.734030749 |
| **A_55_P2060284** | Itpripl2 | -0.043349683 | -0.308742561 | 0.222043196 | 8.304499941 | -0.409116506 | 0.697966225 | 0.85113529 | -4.734038994 |
| **A_55_P2107499** | Galnt18 | 0.221054626 | -1.132340064 | 1.574449316 | 10.35237472 | 0.409096238 | 0.697980244 | 0.85113529 | -4.734041627 |
| **A_55_P2006808** | Ntrk3 | 0.074620403 | -0.382321855 | 0.53156266 | 8.137677969 | 0.409021945 | 0.69803163 | 0.851140841 | -4.734051278 |
| **A_51_P266640** | Nob1 | 0.039168821 | -0.200718613 | 0.279056254 | 8.228489063 | 0.408962363 | 0.698072842 | 0.851140841 | -4.734059016 |
| **A_55_P2355345** | A130019F | -0.064295538 | -0.458240941 | 0.329649864 | 8.363142392 | -0.408785182 | 0.698195403 | 0.851236594 | -4.734082021 |
| **A_55_P2158483** | Nkiras2 | -0.046508249 | -0.331606395 | 0.238589898 | 8.234251721 | -0.408588312 | 0.698331595 | 0.851348954 | -4.734107573 |
| **A_51_P400366** | Rhbg | -0.068142365 | -0.485963277 | 0.349678546 | 7.923239029 | -0.40848619 | 0.698402247 | 0.851381402 | -4.734120823 |
| **A_55_P2068134** | NA | -0.037393911 | -0.266911646 | 0.192123825 | 7.856928715 | -0.408070295 | 0.698690013 | 0.851618984 | -4.734174753 |
| **A_55_P1987645** | Unc13b | 0.128939751 | -0.662616315 | 0.920495817 | 10.30408079 | 0.407995611 | 0.698741694 | 0.851618984 | -4.734184432 |
| **A_55_P2065732** | Medag | -0.091967448 | -0.65663107 | 0.472696174 | 15.12373894 | -0.407938477 | 0.698781233 | 0.851618984 | -4.734191835 |
| **A_51_P149422** | Gng13 | 0.101440997 | -0.521466931 | 0.724348925 | 8.501535665 | 0.407887123 | 0.698816772 | 0.851618984 | -4.734198489 |
| **A_55_P1962602** | NA | 0.299527229 | -1.539751683 | 2.13880614 | 13.01921646 | 0.407886247 | 0.698817378 | 0.851618984 | -4.734198603 |
| **A_55_P1972104** | Acsm3 | -0.036553583 | -0.261161447 | 0.18805428 | 7.704592395 | -0.407619871 | 0.699001734 | 0.851789961 | -4.734233104 |
| **A_55_P2129685** | Pkig | 0.037134254 | -0.191092498 | 0.265361006 | 8.186160191 | 0.407528984 | 0.699064642 | 0.851792757 | -4.734244872 |
| **A_55_P1984770** | Fam220a | -0.088783887 | -0.634501964 | 0.45693419 | 11.96050812 | -0.407489254 | 0.699092141 | 0.851792757 | -4.734250015 |
| **A_55_P2054728** | Ldlrad3 | 0.067547389 | -0.348629383 | 0.483724161 | 14.2538508 | 0.406519214 | 0.699763732 | 0.852526427 | -4.734375449 |
| **A_55_P2051622** | Gm8479 | -0.050372145 | -0.360748501 | 0.260004211 | 8.275667042 | -0.406492146 | 0.699782476 | 0.852526427 | -4.734378945 |
| **A_55_P1995892** | NA | -0.05263335 | -0.377021272 | 0.271754572 | 9.302528113 | -0.406393436 | 0.699850834 | 0.852555985 | -4.734391693 |
| **A_55_P1959923** | Cth | 0.067928481 | -0.351492579 | 0.487349541 | 8.525276117 | 0.405650499 | 0.700365431 | 0.85312911 | -4.734487553 |
| **A_66_P102719** | Spop | 0.098529124 | -0.510073079 | 0.707131326 | 11.93197085 | 0.405491212 | 0.700475784 | 0.853209778 | -4.734508085 |
| **A_55_P2335903** | NA | -0.172792993 | -1.240531094 | 0.894945108 | 10.34350389 | -0.405332785 | 0.70058555 | 0.853252711 | -4.734528498 |
| **A_51_P342773** | Vipr2 | 0.126404737 | -0.654725084 | 0.907534557 | 8.534291013 | 0.405312946 | 0.700599296 | 0.853252711 | -4.734531054 |
| **A_55_P2054098** | Gm9495 | -0.054562976 | -0.391830996 | 0.282705045 | 8.324731668 | -0.405203537 | 0.700675104 | 0.853282939 | -4.734545147 |
| **A_55_P2130905** | NA | 0.223196069 | -1.156620401 | 1.603012539 | 12.68680415 | 0.405149738 | 0.700712383 | 0.853282939 | -4.734552076 |
| **A_55_P2175494** | NA | -0.075251216 | -0.540813579 | 0.390311148 | 8.772559063 | -0.40484241 | 0.700925356 | 0.853488527 | -4.734591639 |
| **A_55_P2001920** | Flna | 0.135917503 | -0.70539313 | 0.977228136 | 14.81311491 | 0.404640505 | 0.701065289 | 0.853605157 | -4.734617617 |
| **A_55_P2161077** | NA | -0.073754587 | -0.530546408 | 0.383037235 | 9.059466893 | -0.404409231 | 0.701225593 | 0.853746575 | -4.734647359 |
| **A_55_P2181222** | Ifi208 | -0.036247971 | -0.260783459 | 0.188287518 | 8.799080634 | -0.404342182 | 0.701272069 | 0.853749398 | -4.734655978 |
| **A_52_P594756** | Asb4 | 0.04897811 | -0.254854388 | 0.352810607 | 8.927190787 | 0.40375521 | 0.701679009 | 0.854191031 | -4.734731382 |
| **A_55_P2087539** | Clpsl2 | -0.049084007 | -0.35396661 | 0.255798596 | 8.031210283 | -0.40323453 | 0.702040083 | 0.854563295 | -4.734798188 |
| **A_51_P321341** | Sult1a1 | -0.243237665 | -1.754273963 | 1.267798633 | 10.67307525 | -0.403186771 | 0.702073206 | 0.854563295 | -4.734804312 |
| **A_51_P390967** | Slc43a3 | 0.144286456 | -0.752330413 | 1.040903325 | 11.19587918 | 0.403059355 | 0.70216158 | 0.8545676 | -4.734820647 |

| | | | | | | | | | |
|---|---|---|---|---|---|---|---|---|---|
| **A_55_P2013118** | NA | -0.049606313 | -0.357871033 | 0.258658407 | 8.166008718 | -0.403054217 | 0.702165144 | 0.8545676 | -4.734821305 |
| **A_55_P2146384** | Filip1 | 0.098785309 | -0.515442975 | 0.713013593 | 8.406107906 | 0.402821736 | 0.702326403 | 0.854659485 | -4.734851096 |
| **A_55_P2005440** | Dennd5a | 0.142975258 | -0.872764576 | 1.158715091 | 12.51942221 | 0.415117387 | 0.702367469 | 0.854659485 | -4.68969759 |
| **A_51_P348183** | Tmem141 | 0.143897366 | -0.751107332 | 1.038902064 | 11.47669258 | 0.402696519 | 0.702413266 | 0.854659485 | -4.734867136 |
| **A_55_P1978226** | Prkn | -0.044040153 | -0.31796272 | 0.229882414 | 8.70061374 | -0.402690469 | 0.702417462 | 0.854659485 | -4.734867911 |
| **A_51_P108659** | Pon1 | -0.06726699 | -0.485906672 | 0.351372692 | 7.880348717 | -0.402450015 | 0.702584281 | 0.854808664 | -4.724257107 |
| **A_55_P1986531** | Mpig6b | -0.034991065 | -0.252976752 | 0.182994622 | 7.78895993 | -0.402049501 | 0.702862183 | 0.855092968 | -4.734949944 |
| **A_55_P2030373** | Atp1a2 | 0.056251862 | -0.294312931 | 0.406816655 | 8.080160259 | 0.401900874 | 0.702965323 | 0.855164636 | -4.734968948 |
| **A_66_P139785** | Plpp3 | 0.06627641 | -0.346941199 | 0.479494019 | 12.47409317 | 0.401726517 | 0.703086327 | 0.855258025 | -4.734991235 |
| **A_52_P482124** | Fam32a | -0.064088451 | -0.463829551 | 0.335652649 | 10.4557351 | -0.401560806 | 0.70320134 | 0.855344115 | -4.735012409 |
| **A_51_P415607** | Fkbp7 | 0.138654078 | -0.72662658 | 1.003934736 | 10.90873866 | 0.401352523 | 0.703345913 | 0.855466148 | -4.735039011 |
| **A_55_P2016623** | S100a11- | -0.075601812 | -0.547522901 | 0.396319277 | 11.79703018 | -0.401248263 | 0.703418287 | 0.855500356 | -4.735052322 |
| **A_52_P621418** | NA | 0.058558315 | -0.307211035 | 0.424327665 | 12.68962754 | 0.400988212 | 0.703598821 | 0.855666097 | -4.735085511 |
| **A_55_P2014361** | Pdpk1 | -0.087661609 | -0.635455079 | 0.46013186 | 11.41504027 | -0.400814047 | 0.703719743 | 0.855724781 | -4.735107728 |
| **A_55_P2008790** | Zfp30 | 0.124156351 | -0.651735322 | 0.900048023 | 8.631828088 | 0.400791215 | 0.703735596 | 0.855724781 | -4.735110639 |
| **A_51_P352687** | Rnf216 | -0.125778226 | -0.911970348 | 0.660413897 | 10.78011513 | -0.400707177 | 0.703793948 | 0.855741915 | -4.735121356 |
| **A_55_P1978166** | Zfp945 | 0.082325518 | -0.432401853 | 0.597052889 | 8.280241695 | 0.400596871 | 0.703870541 | 0.85576662 | -4.735135419 |
| **A_52_P660385** | Trmt6 | 0.051073349 | -0.268291772 | 0.37043847 | 8.592151598 | 0.40055043 | 0.703902791 | 0.85576662 | -4.735141338 |
| **A_66_P138053** | Gm9917 | 0.089287511 | -0.469213792 | 0.647788813 | 8.463817038 | 0.400421022 | 0.703992655 | 0.855822057 | -4.73515783 |
| **A_51_P125205** | Aqp1 | -0.159135656 | -1.154760422 | 0.83648911 | 13.694343 | -0.400333702 | 0.704053295 | 0.855841962 | -4.735168956 |
| **A_55_P2193512** | NA | -0.034205506 | -0.248261508 | 0.179850496 | 7.795615253 | -0.400238588 | 0.704119352 | 0.855868449 | -4.735181072 |
| **A_51_P290921** | Sytl2 | 0.052367512 | -0.275450104 | 0.380185128 | 8.273936631 | 0.400110548 | 0.704208279 | 0.855874802 | -4.735197378 |
| **A_55_P2162152** | NA | -0.08344006 | -0.605821742 | 0.438941623 | 9.614878129 | -0.400070945 | 0.704235786 | 0.855874802 | -4.735202421 |
| **A_55_P2174582** | Urah | 0.099612316 | -0.524111293 | 0.723335925 | 9.345511568 | 0.400010324 | 0.704277891 | 0.855874802 | -4.735210139 |
| **A_55_P2047967** | Bfar | -0.082030952 | -0.595712584 | 0.43165068 | 14.12398101 | -0.399976118 | 0.70430165 | 0.855874802 | -4.735214493 |
| **A_55_P2171993** | Mrgbp | -0.048851683 | -0.354856961 | 0.257153595 | 8.337741287 | -0.399853556 | 0.704386783 | 0.855921803 | -4.735230093 |
| **A_55_P2074762** | Lect2 | -0.03913043 | -0.284279254 | 0.206018394 | 7.725402316 | -0.399792971 | 0.704428868 | 0.855921803 | -4.735237802 |
| **A_55_P2004037** | NA | 0.034062303 | -0.179376207 | 0.247500813 | 7.707281031 | 0.399716035 | 0.704482312 | 0.85593295 | -4.735247591 |
| **A_52_P475450** | Rbm48 | -0.049146599 | -0.357280532 | 0.258987335 | 8.212853872 | -0.399488504 | 0.704640381 | 0.856032793 | -4.73527653 |
| **A_52_P58145** | Aldh1a2 | 0.187419395 | -0.98775529 | 1.36259408 | 9.447141832 | 0.399450022 | 0.704667116 | 0.856032793 | -4.735281423 |
| **A_55_P2034998** | Tjp1 | 0.127221038 | -0.670579076 | 0.925021151 | 12.43165461 | 0.399406552 | 0.704697317 | 0.856032793 | -4.73528695 |
| **A_55_P1990663** | Ddx5 | 0.035750325 | -0.188506635 | 0.260007286 | 12.63002799 | 0.399286301 | 0.704780867 | 0.85604582 | -4.735302236 |
| **A_51_P442206** | Ankrd42 | 0.073389979 | -0.387001548 | 0.533781507 | 8.555928441 | 0.399263665 | 0.704796595 | 0.85604582 | -4.735305113 |
| **A_55_P2074591** | Ppm1h | -0.187292583 | -1.362429362 | 0.987844197 | 9.050253025 | -0.399192621 | 0.704845958 | 0.856051998 | -4.735314141 |
| **A_52_P174775** | Ripk1 | 0.036916884 | -0.194946976 | 0.268780743 | 8.75001269 | 0.39878822 | 0.705126979 | 0.856321898 | -4.735365505 |
| **A_55_P1959061** | NA | -0.038727707 | -0.281990945 | 0.204535532 | 11.64244857 | -0.398745358 | 0.705156767 | 0.856321898 | -4.735370946 |
| **A_55_P1984314** | Trim43c | -0.075840092 | -0.552519053 | 0.40083887 | 9.230821893 | -0.398495307 | 0.705330558 | 0.856449902 | -4.735402679 |
| **A_66_P131708** | Znhit1 | 0.082373338 | -0.435406928 | 0.600153605 | 13.22199383 | 0.398466228 | 0.70535077 | 0.856449902 | -4.735406368 |
| **A_55_P2083489** | Fam129a | -0.035690685 | -0.260116513 | 0.188735143 | 9.003757985 | -0.398320255 | 0.705452235 | 0.856519312 | -4.735424884 |
| **A_52_P342131** | Zfp532 | 0.083724241 | -0.443134581 | 0.610583064 | 8.918976129 | 0.398022213 | 0.705659424 | 0.856717068 | -4.735462669 |
| **A_51_P312997** | Zfp346 | -0.044905674 | -0.342058075 | 0.252246727 | 8.278348835 | -0.402517423 | 0.705792345 | 0.856823203 | -4.709169614 |
| **A_55_P2036952** | NA | -0.046813208 | -0.341586295 | 0.247959879 | 8.565479522 | -0.397768989 | 0.705835479 | 0.856823203 | -4.735494752 |
| **A_55_P2026270** | Cfi | -0.251747113 | -1.837268636 | 1.33377441 | 9.688791525 | -0.397688198 | 0.705891654 | 0.856837597 | -4.735504984 |
| **A_55_P2000643** | NA | 0.125761752 | -0.666929434 | 0.918452938 | 9.192048198 | 0.397369834 | 0.706113035 | 0.857052509 | -4.735545286 |
| **A_51_P256384** | Atp2b2 | 0.035687712 | -0.189434868 | 0.260810291 | 7.735307617 | 0.397054381 | 0.706332424 | 0.857183988 | -4.735585192 |
| **A_51_P363338** | Prkd1 | 0.076229419 | -0.404648026 | 0.557106864 | 8.262454856 | 0.397043919 | 0.706339701 | 0.857183988 | -4.735586515 |
| **A_52_P134023** | Rtp3 | 0.03895611 | -0.206803462 | 0.284715681 | 7.831880784 | 0.397022834 | 0.706354366 | 0.857183988 | -4.735589181 |
| **A_51_P473953** | Arhgef26 | 0.187092605 | -0.993620774 | 1.367805984 | 9.373476533 | 0.396882988 | 0.706451636 | 0.857248223 | -4.735606861 |

| | | | | | | | | | |
|---|---|---|---|---|---|---|---|---|---|
| **A_55_P1966327** | NA | 0.170094883 | -0.903662468 | 1.243852235 | 10.88640856 | 0.39676691 | 0.706532379 | 0.857292395 | -4.735621532 |
| **A_55_P2121945** | Cnot6l | 0.064457291 | -0.342679886 | 0.471594468 | 12.29856247 | 0.396535096 | 0.70669364 | 0.857387223 | -4.735650819 |
| **A_55_P2092611** | Slc45a3 | -0.147239662 | -1.077347804 | 0.782868481 | 9.112115543 | -0.396498647 | 0.706718998 | 0.857387223 | -4.735655423 |
| **A_55_P2028837** | Tspan2 | -0.132261069 | -0.967888777 | 0.703366638 | 9.451629138 | -0.396432767 | 0.706764831 | 0.857387223 | -4.735663743 |
| **A_55_P2013029** | NA | 0.039853107 | -0.212013041 | 0.291719255 | 8.48973356 | 0.396317028 | 0.706845355 | 0.857387223 | -4.735678356 |
| **A_55_P2005549** | Tnn | 0.115862467 | -0.61645645 | 0.848181383 | 8.28905082 | 0.396271523 | 0.706877016 | 0.857387223 | -4.7356841 |
| **A_52_P456271** | Tmem167 | -0.098829996 | -0.723510983 | 0.52585099 | 11.99412852 | -0.39626055 | 0.70688465 | 0.857387223 | -4.735685485 |
| **A_51_P503303** | C1galt1c1 | 0.113581791 | -0.604479936 | 0.831643518 | 10.79387259 | 0.396184316 | 0.706937693 | 0.857387223 | -4.735695107 |
| **A_55_P1956179** | Git2 | -0.063614747 | -0.465884014 | 0.33865452 | 10.38584535 | -0.396087646 | 0.707004957 | 0.857387223 | -4.735707306 |
| **A_55_P2038640** | Gcn1 | -0.049693965 | -0.3639402 | 0.264552271 | 9.623544414 | -0.396080907 | 0.707009646 | 0.857387223 | -4.735708157 |
| **A_51_P257885** | Mmd2 | 0.033184443 | -0.176702027 | 0.243070914 | 7.7012582 | 0.396004788 | 0.707062613 | 0.857397677 | -4.73571776 |
| **A_55_P1981836** | Ccm2l | 0.16508612 | -0.879752168 | 1.209924409 | 10.69572736 | 0.395741711 | 0.707245687 | 0.85756589 | -4.735750938 |
| **A_55_P2149951** | Prx | -0.207451863 | -1.520911959 | 1.106008234 | 11.71094209 | -0.395595007 | 0.707347788 | 0.857571087 | -4.735769431 |
| **A_55_P1982186** | Sgsm1 | -0.066423937 | -0.487047848 | 0.354199974 | 8.510004503 | -0.395531434 | 0.707392034 | 0.857571087 | -4.735777443 |
| **A_52_P519882** | Etf1 | -0.134206274 | -0.984138746 | 0.715726198 | 9.94045931 | -0.395492947 | 0.707418821 | 0.857571087 | -4.735782293 |
| **A_55_P1985984** | Actg1 | -0.043908731 | -0.322050859 | 0.234233397 | 13.453524 | -0.39539799 | 0.707484914 | 0.857571087 | -4.735794257 |
| **A_65_P02958** | Rnf13 | -0.080783006 | -0.592577463 | 0.431011451 | 11.127746 | -0.39534366 | 0.707522731 | 0.857571087 | -4.735801101 |
| **A_52_P272811** | Cpsf1 | 0.058521618 | -0.331175985 | 0.44821922 | 9.54806456 | 0.39999199 | 0.707522943 | 0.857571087 | -4.709448922 |
| **A_55_P2075474** | Gid8 | -0.046151315 | -0.338579482 | 0.246276851 | 8.775330793 | -0.395289455 | 0.707560462 | 0.857571087 | -4.735807928 |
| **A_55_P2098688** | Mcrs1 | 0.063692816 | -0.340262118 | 0.46764775 | 8.993179949 | 0.394918865 | 0.707818444 | 0.857829988 | -4.735854584 |
| **A_55_P2063594** | Gm5801 | 0.07449422 | -0.398171896 | 0.547160336 | 11.35844287 | 0.39474665 | 0.707938345 | 0.857921522 | -4.735876251 |
| **A_55_P1968085** | NA | -0.044652469 | -0.328446372 | 0.239141434 | 8.167008418 | -0.394087592 | 0.708397284 | 0.858351311 | -4.735959092 |
| **A_55_P2037702** | Gm10638 | -0.04236017 | -0.311593421 | 0.226873082 | 9.056184583 | -0.394075414 | 0.708405766 | 0.858351311 | -4.735960622 |
| **A_55_P1997415** | Exoc6b | -0.038811472 | -0.285508221 | 0.207885278 | 10.7310976 | -0.394046095 | 0.708426185 | 0.858351311 | -4.735964304 |
| **A_51_P430327** | Gmpr2 | 0.05448682 | -0.291953148 | 0.400926787 | 9.109691808 | 0.393925246 | 0.708510356 | 0.8583995 | -4.735979479 |
| **A_55_P2036377** | NA | 0.037358666 | -0.200240249 | 0.274957582 | 16.65104388 | 0.393819537 | 0.708583985 | 0.858434912 | -4.73599275 |
| **A_55_P2055442** | Pafah2 | -0.044514451 | -0.327821185 | 0.238792282 | 9.152507359 | -0.393545068 | 0.708775177 | 0.858612737 | -4.736027191 |
| **A_55_P2167160** | Hdhd2 | -0.048039167 | -0.353948516 | 0.257870181 | 8.354388514 | -0.393326374 | 0.708927534 | 0.858695961 | -4.736054618 |
| **A_51_P240594** | Zfp7 | 0.0598938 | -0.32151204 | 0.44129964 | 9.543926201 | 0.393318951 | 0.708932705 | 0.858695961 | -4.736055549 |
| **A_55_P2350022** | Akap6 | 0.053184244 | -0.303042739 | 0.409411226 | 7.938491962 | 0.397666384 | 0.709118434 | 0.858843372 | -4.709704764 |
| **A_51_P321886** | Cmtm3 | -0.040003346 | -0.294955957 | 0.214949264 | 11.9994333 | -0.392995163 | 0.709158306 | 0.858843372 | -4.73609613 |
| **A_55_P2017998** | NA | 0.053613665 | -0.288117988 | 0.395345317 | 8.757773108 | 0.392953021 | 0.709187671 | 0.858843372 | -4.73610141 |
| **A_55_P2060962** | Tm2d3 | 0.045969432 | -0.247209738 | 0.339148603 | 11.46618855 | 0.392723034 | 0.709347939 | 0.858943232 | -4.736130214 |
| **A_52_P258218** | Proser1 | 0.044349202 | -0.238545196 | 0.3272436 | 8.58826467 | 0.392655608 | 0.709394928 | 0.858943232 | -4.736138655 |
| **A_55_P2110567** | Proca1 | 0.10489255 | -0.5642161 | 0.7740012 | 9.131515221 | 0.392643438 | 0.70940341 | 0.858943232 | -4.736140179 |
| **A_65_P19319** | Soga1 | 0.068734524 | -0.370065755 | 0.507534803 | 10.91442834 | 0.392336197 | 0.709617545 | 0.859040729 | -4.736178627 |
| **A_52_P183524** | Tmem86b | -0.032604664 | -0.240806742 | 0.175597413 | 7.689089115 | -0.392233805 | 0.709688916 | 0.859040729 | -4.736191434 |
| **A_55_P2032600** | Rps24 | 0.114296018 | -0.615653054 | 0.844245091 | 17.40687129 | 0.39218311 | 0.709724252 | 0.859040729 | -4.736197774 |
| **A_55_P2175469** | Chd2 | -0.039014329 | -0.288231876 | 0.210203217 | 9.8427394 | -0.392099126 | 0.709782796 | 0.859040729 | -4.736208275 |
| **A_65_P20699** | Tlk2 | -0.051136497 | -0.377793807 | 0.275520813 | 9.694247226 | -0.392092865 | 0.70978716 | 0.859040729 | -4.736209058 |
| **A_51_P256093** | Map2k6 | -0.089728699 | -0.66292075 | 0.483463352 | 9.03544441 | -0.392086201 | 0.709791805 | 0.859040729 | -4.736209891 |
| **A_51_P461475** | Atn1 | 0.040225292 | -0.216739128 | 0.297189711 | 9.761908796 | 0.392081685 | 0.709794954 | 0.859040729 | -4.736210456 |
| **A_55_P2090219** | Mrpl48 | 0.103918766 | -0.560317192 | 0.768154724 | 12.53636703 | 0.391851888 | 0.709955151 | 0.859135023 | -4.736239178 |
| **A_55_P2008244** | NA | 0.071706071 | -0.386641359 | 0.530053501 | 9.626069877 | 0.391842439 | 0.709961739 | 0.859135023 | -4.736240358 |
| **A_52_P464003** | Atg3 | 0.054268317 | -0.292701701 | 0.401238335 | 10.19957095 | 0.391746164 | 0.710028861 | 0.859162473 | -4.736252387 |
| **A_55_P2027337** | NA | -0.117805098 | -0.871358747 | 0.635748551 | 11.49689671 | -0.391561743 | 0.710157444 | 0.859264285 | -4.736275421 |
| **A_52_P302463** | Arl5a | -0.035604991 | -0.26339983 | 0.192189849 | 8.548055882 | -0.391486997 | 0.710209561 | 0.859273571 | -4.736284754 |
| **A_55_P2063785** | Ank1 | 0.072736546 | -0.39286002 | 0.538333112 | 8.037368246 | 0.391285048 | 0.710350383 | 0.859314112 | -4.736309961 |

| | | | | | | | | | |
|---|---|---|---|---|---|---|---|---|---|
| **A_55_P2084686** | Pdik1l | -0.036043882 | -0.266804033 | 0.194716269 | 8.618058988 | -0.391220036 | 0.710395719 | 0.859314112 | -4.736318074 |
| **A_55_P2062936** | Abca3 | 0.151710381 | -0.819577107 | 1.122997869 | 13.86146315 | 0.391217119 | 0.710397754 | 0.859314112 | -4.736318438 |
| **A_51_P175871** | Got2 | 0.056361395 | -0.30450879 | 0.417231581 | 11.84528997 | 0.391183995 | 0.710420853 | 0.859314112 | -4.73632257 |
| **A_51_P112355** | Igtp | -0.156754277 | -1.160844594 | 0.84733604 | 9.229782583 | -0.391018193 | 0.710536483 | 0.859377165 | -4.72804826 |
| **A_55_P1967720** | Polr2i | 0.130263937 | -0.704220083 | 0.964747957 | 13.40568014 | 0.390981781 | 0.710561879 | 0.859377165 | -4.736347793 |
| **A_52_P225260** | Ywhaz | -0.041021495 | -0.303885748 | 0.221842758 | 8.79467767 | -0.390868165 | 0.710641121 | 0.859419243 | -4.73636196 |
| **A_55_P2183672** | Pebp1 | -0.072641352 | -0.538331476 | 0.393048773 | 16.79060154 | -0.390694444 | 0.710762292 | 0.859460547 | -4.736383614 |
| **A_55_P2108032** | Tmem255 | 0.03559076 | -0.192619895 | 0.263801415 | 7.791486306 | 0.390617495 | 0.710815966 | 0.859460547 | -4.736393202 |
| **A_55_P2082045** | Vps13a | -0.034005699 | -0.252066333 | 0.184054936 | 8.10016792 | -0.390593288 | 0.710832852 | 0.859460547 | -4.736396218 |
| **A_52_P532033** | Hltf | -0.04339702 | -0.321732316 | 0.234938276 | 9.53509672 | -0.390518822 | 0.710884798 | 0.859460547 | -4.736405495 |
| **A_51_P323842** | Ciao2b | 0.068655664 | -0.371873404 | 0.509184732 | 13.10361051 | 0.39034817 | 0.711003846 | 0.859460547 | -4.736426749 |
| **A_52_P494982** | Dhx40 | 0.033086271 | -0.179240281 | 0.245412823 | 8.507043622 | 0.390295784 | 0.711040393 | 0.859460547 | -4.736433272 |
| **A_55_P2066440** | Myh3 | 0.36721601 | -1.989396666 | 2.723828686 | 9.440025473 | 0.390286693 | 0.711046735 | 0.859460547 | -4.736434404 |
| **A_51_P422369** | Odf3b | 0.25584678 | -1.38613594 | 1.897829499 | 9.415066088 | 0.3902669 | 0.711060544 | 0.859460547 | -4.736436868 |
| **A_55_P2028660** | Dusp12 | 0.038651956 | -0.20942346 | 0.286727372 | 8.153891575 | 0.39024567 | 0.711075355 | 0.859460547 | -4.736439511 |
| **A_51_P221510** | Fam81a | 0.069762462 | -0.378066566 | 0.51759149 | 8.076022427 | 0.390175417 | 0.71112437 | 0.85946606 | -4.736448256 |
| **A_55_P2053177** | NA | 0.052940873 | -0.287128146 | 0.393009893 | 11.93935622 | 0.389918982 | 0.711303292 | 0.859521613 | -4.736480166 |
| **A_52_P16563** | Lsm14b | 0.074291512 | -0.402963193 | 0.551546217 | 10.89149487 | 0.38988751 | 0.711325252 | 0.859521613 | -4.73648408 |
| **A_52_P217710** | Fzd6 | 0.086050414 | -0.466838748 | 0.638939576 | 8.638358965 | 0.389821003 | 0.711371661 | 0.859521613 | -4.736492352 |
| **A_52_P305890** | Yipf5 | 0.040589508 | -0.22023004 | 0.301409055 | 11.93026016 | 0.389783988 | 0.71139749 | 0.859521613 | -4.736496956 |
| **A_55_P1987499** | Pttg1 | -0.033248567 | -0.24691261 | 0.180415476 | 12.63941807 | -0.389755126 | 0.711417631 | 0.859521613 | -4.736500545 |
| **A_55_P1982084** | NA | 0.086932951 | -0.471761014 | 0.645626917 | 10.31645658 | 0.389727264 | 0.711437074 | 0.859521613 | -4.736504009 |
| **A_52_P602669** | Serpinb6c | 0.089776753 | -0.487687709 | 0.667241215 | 8.651532387 | 0.38939375 | 0.711669827 | 0.859742009 | -4.736545463 |
| **A_66_P104134** | Homer1 | -0.039240811 | -0.291682475 | 0.213200854 | 7.886207855 | -0.389338435 | 0.711708434 | 0.859742009 | -4.736552335 |
| **A_66_P121583** | Tmem267 | 0.048865335 | -0.265656532 | 0.363387202 | 8.457004234 | 0.389135077 | 0.711850374 | 0.859859749 | -4.736577592 |
| **A_52_P429534** | Elavl1 | -0.036616105 | -0.272511854 | 0.199279644 | 8.615824733 | -0.388778617 | 0.712099209 | 0.860106585 | -4.736621835 |
| **A_55_P2072656** | Ckmt1 | 0.067932061 | -0.369819657 | 0.505683778 | 10.8616688 | 0.388684548 | 0.712164883 | 0.860132174 | -4.736633505 |
| **A_52_P300416** | Dcp1a | -0.046127523 | -0.343463828 | 0.251208783 | 8.948599145 | -0.388563979 | 0.712249061 | 0.860152365 | -4.736648458 |
| **A_55_P2017769** | B020031M | 0.062901302 | -0.342590322 | 0.468392926 | 8.395860374 | 0.388533161 | 0.712270578 | 0.860152365 | -4.736652279 |
| **A_55_P1979341** | Cyba | 0.163222935 | -0.889233188 | 1.215679057 | 13.10515249 | 0.388443205 | 0.712333387 | 0.860174487 | -4.736663432 |
| **A_55_P1987938** | NA | -0.064318733 | -0.47916797 | 0.350530505 | 11.94097983 | -0.388326945 | 0.712414566 | 0.860189642 | -4.736677843 |
| **A_52_P66313** | Psmf1 | 0.041908097 | -0.228415128 | 0.312231323 | 8.498050696 | 0.388297797 | 0.712434919 | 0.860189642 | -4.736681456 |
| **A_55_P2122380** | Zfp14 | 0.033642193 | -0.183454082 | 0.250738468 | 8.614216387 | 0.38813452 | 0.712548936 | 0.860273581 | -4.736701686 |
| **A_52_P17112** | Znrf1 | 0.073731717 | -0.402218589 | 0.549682023 | 10.520819 | 0.388010146 | 0.712635792 | 0.860324721 | -4.736717091 |
| **A_55_P2069592** | Pitpnm3 | 0.188068568 | -1.027101592 | 1.403238727 | 8.864443995 | 0.387640787 | 0.712893762 | 0.860551696 | -4.736762815 |
| **A_55_P2023777** | Tmx2 | -0.053118964 | -0.396473148 | 0.29023522 | 8.437161191 | -0.387487412 | 0.713000896 | 0.860551696 | -4.736781789 |
| **A_51_P330044** | Cyp2j9 | 0.096134459 | -0.525321354 | 0.717590272 | 8.38451017 | 0.387453133 | 0.71302484 | 0.860551696 | -4.736786029 |
| **A_55_P1955548** | Ezr | -0.149284038 | -1.114354829 | 0.815786754 | 12.00453164 | -0.387440081 | 0.713033958 | 0.860551696 | -4.736787644 |
| **A_51_P134551** | Coq9 | -0.057709647 | -0.430800075 | 0.31538078 | 9.299865132 | -0.387422339 | 0.713046352 | 0.860551696 | -4.736789838 |
| **A_55_P2142758** | Slc9a7 | 0.033201605 | -0.181620772 | 0.248023982 | 7.746408396 | 0.387106006 | 0.713267342 | 0.860764671 | -4.736828945 |
| **A_52_P229972** | Slc22a1 | 0.137479738 | -0.752356201 | 1.027315676 | 8.588874987 | 0.386971604 | 0.713361244 | 0.86082426 | -4.736845552 |
| **A_52_P633489** | C1qtnf4 | 0.049089269 | -0.268991229 | 0.367169767 | 8.251551192 | 0.386544829 | 0.713659457 | 0.861130371 | -4.736898251 |
| **A_51_P349015** | Pacrg | 0.100864534 | -0.553428025 | 0.755157093 | 8.465690497 | 0.386115158 | 0.71395975 | 0.861438954 | -4.736951254 |
| **A_55_P2278604** | A330009N | 0.126419965 | -0.694044253 | 0.946884182 | 9.281249056 | 0.385928068 | 0.714090523 | 0.861507201 | -4.736974316 |
| **A_55_P2084002** | Aptx | 0.059884383 | -0.328786103 | 0.44855487 | 8.300669711 | 0.385906741 | 0.714105431 | 0.861507201 | -4.736976944 |
| **A_55_P2180551** | Sinhcaf | 0.045590642 | -0.250426828 | 0.341608113 | 8.936053076 | 0.385752469 | 0.714213274 | 0.861583543 | -4.736995953 |
| **A_55_P2087265** | NA | -0.180631105 | -1.353850492 | 0.992588283 | 10.98580418 | -0.385623642 | 0.714303336 | 0.861598037 | -4.73701182 |
| **A_51_P290139** | Dlst | 0.062536418 | -0.343661362 | 0.468734198 | 11.95808266 | 0.385607792 | 0.714314417 | 0.861598037 | -4.737013772 |

| | | | | | | | | | |
|---|---|---|---|---|---|---|---|---|---|
| **A_55_P2174967** | NA | 0.049553541 | -0.272377977 | 0.371485058 | 9.119929545 | 0.385532983 | 0.714366718 | 0.861607369 | -4.737022984 |
| **A_55_P2175502** | M1ap | 0.044490194 | -0.244793145 | 0.333773534 | 8.054840757 | 0.385204387 | 0.714596469 | 0.861810467 | -4.737063427 |
| **A_55_P2023942** | Ccdc77 | 0.061532703 | -0.338605413 | 0.461670819 | 9.188144435 | 0.385164645 | 0.714624259 | 0.861810467 | -4.737068316 |
| **A_55_P1989594** | Mynn | -0.035220519 | -0.264307857 | 0.193866819 | 9.405415665 | -0.385074719 | 0.714687141 | 0.861829715 | -4.737079378 |
| **A_51_P499854** | Ghr | 0.095855194 | -0.527720391 | 0.719430779 | 11.08849237 | 0.385014329 | 0.714729371 | 0.861829715 | -4.737086805 |
| **A_51_P333071** | Anapc16 | 0.050875561 | -0.280268512 | 0.382019634 | 11.10663596 | 0.38480665 | 0.714874607 | 0.861951085 | -4.737112338 |
| **A_66_P134704** | Gm4952 | 0.032192121 | -0.177417933 | 0.241802174 | 7.682394886 | 0.384669564 | 0.714970483 | 0.862012928 | -4.737129185 |
| **A_55_P2280868** | 0610005C | -0.032455801 | -0.243895998 | 0.178984396 | 7.66605226 | -0.384463511 | 0.715114604 | 0.862132927 | -4.737154497 |
| **A_52_P464129** | Gon7 | 0.042076128 | -0.23209077 | 0.316243026 | 9.633694844 | 0.384389125 | 0.715166635 | 0.862141896 | -4.737163632 |
| **A_65_P07196** | COX3 | -0.03876832 | -0.291500106 | 0.213963465 | 16.31816404 | -0.38420894 | 0.715292679 | 0.862228337 | -4.737185753 |
| **A_66_P125660** | Nde1 | 0.033882408 | -0.187026624 | 0.254791441 | 8.016038164 | 0.384159116 | 0.715327533 | 0.862228337 | -4.737191868 |
| **A_66_P105288** | Rplp0 | -0.041817665 | -0.314550414 | 0.230915085 | 16.04890859 | -0.384036789 | 0.715413111 | 0.862277732 | -4.737206879 |
| **A_55_P2164469** | Prcd | 0.386928581 | -2.138099357 | 2.911956519 | 10.13043478 | 0.383808836 | 0.715572596 | 0.862390657 | -4.737234839 |
| **A_55_P2057446** | Ccdc134 | -0.08581337 | -0.645865182 | 0.474238442 | 9.943922414 | -0.383775368 | 0.715596012 | 0.862390657 | -4.737238943 |
| **A_55_P2011300** | Fggy | -0.033135944 | -0.249557034 | 0.183285145 | 7.825733704 | -0.383486534 | 0.715798118 | 0.862570034 | -4.737274346 |
| **A_55_P2006554** | Eif4e2 | -0.041535926 | -0.312901362 | 0.229829511 | 11.91682833 | -0.383371397 | 0.715878689 | 0.862570034 | -4.737288452 |
| **A_55_P2123962** | Micos10 | 0.044454898 | -0.246044332 | 0.334954128 | 15.23231712 | 0.383287785 | 0.715937203 | 0.862570034 | -4.737298694 |
| **A_55_P2012939** | Arid4b | 0.042160547 | -0.23336575 | 0.317686845 | 9.561352693 | 0.383260031 | 0.715956626 | 0.862570034 | -4.737302093 |
| **A_55_P2158448** | Gdf3 | 0.050909577 | -0.2818139 | 0.383633054 | 8.137986277 | 0.383236081 | 0.715973387 | 0.862570034 | -4.737305026 |
| **A_55_P1953693** | Pex2 | 0.072819816 | -0.403205443 | 0.548845074 | 10.67799077 | 0.383150965 | 0.716032957 | 0.862570034 | -4.737315448 |
| **A_55_P2157610** | Rpusd3 | -0.072375518 | -0.545717686 | 0.40096665 | 10.03002547 | -0.382971835 | 0.716158331 | 0.862570034 | -4.737337375 |
| **A_55_P1991964** | Iqsec2 | -0.057740316 | -0.435388444 | 0.319907813 | 8.879772446 | -0.382950079 | 0.716173559 | 0.862570034 | -4.737340037 |
| **A_55_P2034350** | Terf2 | -0.062811993 | -0.473631633 | 0.348007647 | 9.281656797 | -0.382949688 | 0.716173832 | 0.862570034 | -4.737340085 |
| **A_65_P15245** | Nrp2 | 0.108366433 | -0.600539313 | 0.817272179 | 8.869455254 | 0.382874664 | 0.716226345 | 0.862570034 | -4.737349265 |
| **A_55_P2013391** | Gm4402 | -0.061609553 | -0.46465757 | 0.341438465 | 8.824232398 | -0.382861425 | 0.716235612 | 0.862570034 | -4.737350885 |
| **A_51_P168695** | Mast4 | 0.085267327 | -0.472656303 | 0.643190958 | 9.008714871 | 0.382787934 | 0.716287054 | 0.862578256 | -4.737359876 |
| **A_55_P2099530** | NA | 0.05792079 | -0.321310831 | 0.437152411 | 10.72180792 | 0.382543018 | 0.716458501 | 0.862705329 | -4.737389827 |
| **A_55_P2091153** | BC051142 | 0.043415875 | -0.240870757 | 0.327702507 | 8.389462847 | 0.382509712 | 0.716481818 | 0.862705329 | -4.737393899 |
| **A_51_P138772** | Ddx27 | -0.11997958 | -0.905754749 | 0.665795589 | 10.26556068 | -0.382436542 | 0.716533043 | 0.86271328 | -4.737402843 |
| **A_51_P363642** | 1700088E | 0.114105826 | -0.633968297 | 0.86217995 | 9.364785685 | 0.382044132 | 0.716807791 | 0.862949724 | -4.737450783 |
| **A_55_P2161170** | Srp68 | 0.074729024 | -0.415212013 | 0.564670061 | 11.96278827 | 0.382028566 | 0.716818691 | 0.862949724 | -4.737452684 |
| **A_55_P2135980** | Ms4a4c | -0.105046719 | -0.794305594 | 0.584212156 | 14.9400439 | -0.381724884 | 0.71703135 | 0.863136697 | -4.737489752 |
| **A_55_P1974098** | Gm12063 | -0.079037793 | -0.597702345 | 0.439626759 | 10.26722667 | -0.381679278 | 0.717063289 | 0.863136697 | -4.737495317 |
| **A_55_P2144305** | Gm3211 | 0.057584957 | -0.320569335 | 0.435739249 | 8.399019329 | 0.381408492 | 0.71725294 | 0.863311233 | -4.737528343 |
| **A_52_P111963** | 1810030C | 0.036409215 | -0.202728344 | 0.275546773 | 9.806821035 | 0.381341322 | 0.717299987 | 0.863314115 | -4.737536532 |
| **A_51_P467233** | Polr3gl | 0.105636107 | -0.588631274 | 0.799903488 | 11.34558719 | 0.381097389 | 0.717470855 | 0.863466014 | -4.737566261 |
| **A_55_P1959738** | Zfp511 | -0.039062659 | -0.295966617 | 0.217841299 | 9.582861015 | -0.380838944 | 0.717651908 | 0.86363015 | -4.737597739 |
| **A_51_P184806** | Elmod2 | 0.03833372 | -0.214095995 | 0.290763434 | 9.455055636 | 0.380356493 | 0.717989944 | 0.863983168 | -4.737656449 |
| **A_55_P1956223** | Chka | -0.05789021 | -0.439227688 | 0.323447268 | 9.238310899 | -0.380229648 | 0.71807883 | 0.864036352 | -4.737671873 |
| **A_55_P2021540** | Myocd | 0.076992426 | -0.430366837 | 0.584351689 | 8.641993297 | 0.380086773 | 0.718178957 | 0.864103053 | -4.737689242 |
| **A_66_P126332** | Zfp703 | 0.05568072 | -0.311350699 | 0.422712138 | 12.8093461 | 0.379972305 | 0.718259181 | 0.8641458 | -4.737703152 |
| **A_55_P2066657** | Vps13a | 0.040792259 | -0.228281112 | 0.309865631 | 8.224439055 | 0.379714678 | 0.71843975 | 0.86430926 | -4.737734446 |
| **A_55_P2047509** | Ufd1 | -0.084444954 | -0.641712896 | 0.472822989 | 12.07741569 | -0.379542127 | 0.718560701 | 0.864400983 | -4.737755395 |
| **A_55_P2077666** | Porcn | 0.057145424 | -0.320129882 | 0.434420731 | 9.476329338 | 0.379379121 | 0.71867497 | 0.86440221 | -4.737775178 |
| **A_55_P2045911** | Spata32 | -0.043094872 | -0.327637632 | 0.241447888 | 7.895116071 | -0.379339795 | 0.718702539 | 0.86440221 | -4.737779949 |
| **A_52_P74524** | NA | -0.047555365 | -0.361609634 | 0.266498905 | 8.233237762 | -0.379267091 | 0.718753509 | 0.86440221 | -4.737788769 |
| **A_52_P89717** | NA | 0.039481859 | -0.22125907 | 0.300222788 | 8.25208993 | 0.379261472 | 0.718757448 | 0.86440221 | -4.737789451 |
| **A_55_P2126259** | Slc35e2 | -0.057700234 | -0.438796448 | 0.32339598 | 8.554790963 | -0.379221794 | 0.718785266 | 0.86440221 | -4.737794263 |

| | | | | | | | | | |
|---|---|---|---|---|---|---|---|---|---|
| **A_51_P246924** | Tppp3 | 0.240007508 | -1.345862437 | 1.825877453 | 9.671336857 | 0.379059691 | 0.718898917 | 0.864485114 | -4.737813921 |
| **A_55_P1994000** | Snx16 | -0.044913464 | -0.341923269 | 0.252096341 | 9.80846008 | -0.378753028 | 0.719113944 | 0.864628238 | -4.737851088 |
| **A_52_P546513** | NA | 0.061165494 | -0.343416084 | 0.465747072 | 8.136785756 | 0.378661132 | 0.719178385 | 0.864628238 | -4.73786222 |
| **A_51_P116813** | Cyp17a1 | -0.155500443 | -1.184078519 | 0.873077634 | 11.92705885 | -0.378655886 | 0.719182064 | 0.864628238 | -4.737862855 |
| **A_55_P2119892** | NA | 0.056260952 | -0.315905177 | 0.42842708 | 7.910478996 | 0.378634843 | 0.71919682 | 0.864628238 | -4.737865404 |
| **A_52_P830312** | Iba57 | -0.068987393 | -0.525425407 | 0.387450622 | 8.789528162 | -0.378563092 | 0.719247137 | 0.864634966 | -4.737874094 |
| **A_51_P512783** | Rmdn1 | -0.040716779 | -0.31023606 | 0.228802502 | 9.849022849 | -0.378385011 | 0.719372028 | 0.864731335 | -4.737895654 |
| **A_55_P2086978** | Col23a1 | 0.046895834 | -0.263660351 | 0.35745202 | 8.803586731 | 0.378219945 | 0.7194878 | 0.864816731 | -4.737915631 |
| **A_66_P136097** | Trim13 | -0.039746353 | -0.303007985 | 0.223515279 | 8.98021576 | -0.378146481 | 0.719539328 | 0.864824901 | -4.737924519 |
| **A_51_P245789** | Pcolce2 | 0.270764878 | -1.523427735 | 2.064957491 | 9.46520441 | 0.377984169 | 0.71965318 | 0.864907974 | -4.737944151 |
| **A_55_P2122666** | Pnma3 | -0.054903842 | -0.419110722 | 0.309303038 | 8.41576441 | -0.377576483 | 0.719939181 | 0.865163524 | -4.737993427 |
| **A_51_P377376** | Gnl3 | 0.073145959 | -0.41211716 | 0.558409077 | 9.215701505 | 0.377540369 | 0.719964519 | 0.865163524 | -4.73799779 |
| **A_55_P2092616** | Tbc1d25 | -0.083481444 | -0.637483615 | 0.470520726 | 9.621299706 | -0.37742341 | 0.720046579 | 0.865163524 | -4.738011916 |
| **A_55_P2143015** | Sema6c | -0.057302834 | -0.437668154 | 0.323062485 | 8.516920787 | -0.377333649 | 0.72010956 | 0.865163524 | -4.738022755 |
| **A_55_P2044788** | NA | -0.034631153 | -0.264507997 | 0.195245691 | 8.844188131 | -0.377330635 | 0.720111674 | 0.865163524 | -4.738023119 |
| **A_55_P2042655** | NA | 0.120033911 | -0.676802835 | 0.916870656 | 12.05948344 | 0.377298387 | 0.720134302 | 0.865163524 | -4.738027012 |
| **A_55_P2106434** | Zfp114 | 0.067219755 | -0.379161795 | 0.513601304 | 8.427481349 | 0.377173379 | 0.720222019 | 0.865215143 | -4.738042102 |
| **A_66_P107680** | Med8 | 0.03605392 | -0.203437869 | 0.275545709 | 8.472442927 | 0.377061515 | 0.720300517 | 0.865246172 | -4.738055601 |
| **A_52_P451073** | Tnfrsf21 | -0.034996246 | -0.267541981 | 0.197549488 | 8.790625536 | -0.37693237 | 0.720391147 | 0.865246172 | -4.738071181 |
| **A_52_P497424** | Aanat | -0.033051468 | -0.252724131 | 0.186621194 | 7.907362007 | -0.376847025 | 0.720451041 | 0.865246172 | -4.738081475 |
| **A_51_P173906** | Mto1 | 0.033929775 | -0.191649304 | 0.259508854 | 8.202062603 | 0.376731999 | 0.72053177 | 0.865246172 | -4.738095344 |
| **A_51_P423509** | Faap100 | 0.033295556 | -0.188074924 | 0.254666037 | 8.091733656 | 0.37671847 | 0.720541265 | 0.865246172 | -4.738096976 |
| **A_55_P1986406** | Ddx54 | 0.066129447 | -0.373747613 | 0.506006507 | 11.08946199 | 0.376542431 | 0.720664823 | 0.865246172 | -4.738118194 |
| **A_51_P335480** | 1810055G | -0.073343807 | -0.561250997 | 0.414563382 | 9.276576587 | -0.376510056 | 0.720687547 | 0.865246172 | -4.738122095 |
| **A_55_P2156895** | Tmem39b | -0.098160161 | -0.751154445 | 0.554834124 | 9.083384244 | -0.376509749 | 0.720687762 | 0.865246172 | -4.738122132 |
| **A_51_P177171** | Tie1 | 0.210882549 | -1.192393779 | 1.614158878 | 10.34271522 | 0.376398418 | 0.720765909 | 0.865246172 | -4.738135546 |
| **A_51_P394394** | Tspan2 | 0.229012505 | -1.295036826 | 1.753061837 | 11.01760622 | 0.376366117 | 0.720788583 | 0.865246172 | -4.738139437 |
| **A_66_P126726** | Cwc27 | 0.036132531 | -0.204333453 | 0.276598515 | 8.913232518 | 0.376352738 | 0.720797974 | 0.865246172 | -4.738141048 |
| **A_51_P461108** | Osbpl10 | 0.051805892 | -0.292969576 | 0.39658136 | 9.458134648 | 0.376351059 | 0.720799153 | 0.865246172 | -4.73814125 |
| **A_55_P2097329** | Sgsm3 | -0.12699801 | -0.972327565 | 0.718331546 | 12.34789887 | -0.376288726 | 0.720842908 | 0.865246172 | -4.738148758 |
| **A_52_P237669** | Nup54 | 0.03352356 | -0.18969596 | 0.256743081 | 8.377312128 | 0.376156276 | 0.720935889 | 0.865246172 | -4.738164707 |
| **A_55_P2122425** | NA | 0.040403399 | -0.228715005 | 0.309521804 | 12.89282163 | 0.376032042 | 0.721023106 | 0.865246172 | -4.738179661 |
| **A_55_P2069485** | Ptpn13 | 0.079479005 | -0.449933434 | 0.608891443 | 10.92654313 | 0.376017991 | 0.72103297 | 0.865246172 | -4.738181353 |
| **A_51_P302520** | Myom1 | 0.120601708 | -0.682762309 | 0.923965724 | 8.617391958 | 0.376003102 | 0.721043423 | 0.865246172 | -4.738183144 |
| **A_52_P554703** | Gprin3 | 0.104339737 | -0.590773798 | 0.799453273 | 8.608883484 | 0.375962336 | 0.721072044 | 0.865246172 | -4.73818805 |
| **A_55_P2039712** | Gigyf2 | -0.108047376 | -0.827932467 | 0.611837715 | 11.3701428 | -0.375925157 | 0.721098147 | 0.865246172 | -4.738192524 |
| **A_52_P209514** | Cacnb2 | 0.035960269 | -0.20375745 | 0.275677988 | 7.854281082 | 0.375727639 | 0.721236828 | 0.865358871 | -4.738216285 |
| **A_55_P2024155** | Zbtb16 | 0.17868105 | -1.013425863 | 1.370787963 | 10.48519895 | 0.375416741 | 0.721455141 | 0.865567092 | -4.738253663 |
| **A_65_P11455** | Pphln1 | -0.070155719 | -0.538499746 | 0.398188308 | 8.413337473 | -0.375187569 | 0.721616084 | 0.865624422 | -4.738281196 |
| **A_55_P2077635** | NA | 0.045765824 | -0.259761465 | 0.351293113 | 11.18023863 | 0.375181679 | 0.72162022 | 0.865624422 | -4.738281904 |
| **A_66_P134181** | Aym1 | -0.059326344 | -0.455407667 | 0.336754979 | 8.414394053 | -0.375157443 | 0.721637242 | 0.865624422 | -4.738284815 |
| **A_55_P1955706** | Spg7 | -0.062114882 | -0.476940482 | 0.352710718 | 12.71584107 | -0.375042479 | 0.721717986 | 0.865667569 | -4.73829862 |
| **A_55_P2163138** | Tm4sf5 | 0.043635274 | -0.248066601 | 0.33533715 | 8.211493424 | 0.374669926 | 0.721979672 | 0.865906071 | -4.738343332 |
| **A_66_P100780** | Katnbl1 | -0.049266362 | -0.37864552 | 0.280112795 | 10.03016614 | -0.374631876 | 0.722006402 | 0.865906071 | -4.738347896 |
| **A_55_P1953414** | Rab12 | 0.054831836 | -0.311936903 | 0.421600575 | 12.49025803 | 0.374447403 | 0.722135997 | 0.865977771 | -4.738370018 |
| **A_55_P2056344** | Unc93a2 | 0.228058704 | -1.297534746 | 1.753652155 | 12.56659259 | 0.374419262 | 0.722155767 | 0.865977771 | -4.738373392 |
| **A_55_P1985850** | Timp1 | 0.04124777 | -0.234775158 | 0.317270698 | 10.17429756 | 0.374287794 | 0.722248132 | 0.866034817 | -4.738389151 |
| **A_55_P1961715** | Nme5 | 0.143833892 | -0.819746442 | 1.107414226 | 8.957239116 | 0.373872611 | 0.722539861 | 0.866330894 | -4.738438884 |

| | | | | | | | | | |
|---|---|---|---|---|---|---|---|---|---|
| **A_55_P1977563** | Galk2 | -0.032114258 | -0.247372842 | 0.183144326 | 8.839285912 | -0.373669596 | 0.722682529 | 0.866448221 | -4.738463184 |
| **A_55_P2065334** | Gm14326 | 0.059331833 | -0.33847167 | 0.457135337 | 9.109804691 | 0.373567863 | 0.722754025 | 0.866470906 | -4.738475357 |
| **A_55_P2169276** | 4732491K | -0.039875463 | -0.307267138 | 0.227516211 | 8.080428614 | -0.373515138 | 0.722791082 | 0.866470906 | -4.738481665 |
| **A_55_P2143324** | Dyrk1b | -0.106658658 | -0.822720289 | 0.609402973 | 8.832898324 | -0.373074922 | 0.723100504 | 0.866708739 | -4.738534295 |
| **A_52_P110291** | Phf12 | 0.037683506 | -0.215351448 | 0.290718461 | 9.092514545 | 0.373010562 | 0.723145747 | 0.866708739 | -4.738541985 |
| **A_55_P2050807** | Mtg2 | -0.03403307 | -0.262566177 | 0.194500036 | 10.15559317 | -0.372994435 | 0.723157084 | 0.866708739 | -4.738543912 |
| **A_55_P2002557** | Srebf1 | -0.040772956 | -0.314576828 | 0.233030915 | 8.475365987 | -0.372977783 | 0.72316879 | 0.866708739 | -4.738545901 |
| **A_55_P1954598** | NA | -0.072723691 | -0.561360698 | 0.415913317 | 11.06762846 | -0.372769097 | 0.7233155 | 0.866830836 | -4.738570826 |
| **A_55_P2067895** | Krba1 | 0.060577351 | -0.346587055 | 0.467741757 | 10.24842588 | 0.372641155 | 0.723405451 | 0.866884901 | -4.7385861 |
| **A_55_P2034390** | Ankhd1 | -0.033042323 | -0.255203707 | 0.189119061 | 8.434278705 | -0.37252236 | 0.723488977 | 0.86693126 | -4.738600278 |
| **A_55_P2005859** | Fn3k | 0.033196368 | -0.190078102 | 0.256470838 | 7.794423508 | 0.372393292 | 0.72357973 | 0.866970774 | -4.738615677 |
| **A_55_P2163659** | Rspo3 | -0.047469598 | -0.36678287 | 0.271843674 | 8.195426311 | -0.372347916 | 0.723611637 | 0.866970774 | -4.73862109 |
| **A_55_P2023200** | Miox | -0.032152261 | -0.248483174 | 0.184178652 | 7.776688544 | -0.372257354 | 0.723675319 | 0.866982586 | -4.738631891 |
| **A_52_P411331** | Vac14 | -0.042235884 | -0.32648626 | 0.242014493 | 8.14421112 | -0.372161018 | 0.723743065 | 0.866982586 | -4.738643378 |
| **A_51_P192344** | Ash2l | -0.088123831 | -0.681398673 | 0.50515101 | 11.25153744 | -0.372038415 | 0.723829285 | 0.866982586 | -4.738657993 |
| **A_51_P130727** | Fkbp11 | -0.172178934 | -1.331383352 | 0.987025484 | 8.724225912 | -0.372023464 | 0.723839799 | 0.866982586 | -4.738659775 |
| **A_55_P2146254** | Ifitm1 | -0.172026716 | -1.330232475 | 0.986179042 | 11.18481866 | -0.372015063 | 0.723845707 | 0.866982586 | -4.738660776 |
| **A_55_P1998947** | Trim16 | 0.141624003 | -0.812291954 | 1.095539959 | 9.144313328 | 0.371857972 | 0.72395619 | 0.867061201 | -4.738679495 |
| **A_66_P117695** | Zpr1 | 0.045604139 | -0.261708199 | 0.352916477 | 11.94890367 | 0.371684634 | 0.724078106 | 0.8671535 | -4.738700141 |
| **A_55_P2079636** | Scgb3a2 | 0.164343294 | -0.943880726 | 1.272567314 | 8.624389906 | 0.371428115 | 0.724258546 | 0.86729475 | -4.738730679 |
| **A_55_P2020552** | Iscu | 0.053740649 | -0.30868901 | 0.416170309 | 12.03253617 | 0.371389418 | 0.724285768 | 0.86729475 | -4.738735284 |
| **A_55_P2059164** | H3f3b | 0.047120316 | -0.270827209 | 0.365067841 | 14.63948336 | 0.371195835 | 0.724421952 | 0.86734359 | -4.738758315 |
| **A_51_P270891** | 1700056E | -0.042886549 | -0.332286891 | 0.246513793 | 8.744366146 | -0.371169606 | 0.724440405 | 0.86734359 | -4.738761435 |
| **A_66_P111714** | Utp20 | 0.034825039 | -0.200309481 | 0.269959559 | 8.292815732 | 0.370958704 | 0.724588788 | 0.86734359 | -4.738786512 |
| **A_52_P389230** | Smarcc2 | -0.032453139 | -0.25158599 | 0.186679713 | 8.369637834 | -0.370936494 | 0.724604414 | 0.86734359 | -4.738789152 |
| **A_52_P356068** | Proz | -0.031266572 | -0.242391651 | 0.179858506 | 7.65958722 | -0.370929006 | 0.724609683 | 0.86734359 | -4.738790042 |
| **A_51_P183051** | Upb1 | 0.088971062 | -0.511861228 | 0.689803351 | 8.302931303 | 0.370890624 | 0.724636689 | 0.86734359 | -4.738794604 |
| **A_51_P434758** | Fam83e | 0.057390613 | -0.330180886 | 0.444962112 | 8.563678824 | 0.370885091 | 0.724640582 | 0.86734359 | -4.738795261 |
| **A_55_P2148400** | NA | 0.251878003 | -1.449940536 | 1.953696542 | 14.00569276 | 0.370704055 | 0.724767967 | 0.867442359 | -4.738816773 |
| **A_51_P265219** | Ppm1k | 0.134633958 | -0.776015832 | 1.045283747 | 9.122904655 | 0.370299876 | 0.725052401 | 0.867729069 | -4.738864764 |
| **A_51_P339444** | Rab9 | 0.070819048 | -0.408564952 | 0.550203048 | 12.6119221 | 0.370012926 | 0.725254366 | 0.867794616 | -4.738898807 |
| **A_52_P239536** | Ppp1r9a | 0.116977307 | -0.674870585 | 0.9088252 | 10.1486472 | 0.370007221 | 0.725258381 | 0.867794616 | -4.738899484 |
| **A_51_P365019** | Gclc | -0.070561781 | -0.548281381 | 0.407157819 | 10.40260981 | -0.369953228 | 0.725296387 | 0.867794616 | -4.738905886 |
| **A_55_P2040815** | Lrrc45 | 0.11586393 | -0.668590027 | 0.900317887 | 11.0963202 | 0.369939872 | 0.725305788 | 0.867794616 | -4.73890747 |
| **A_55_P1971498** | Exoc1 | -0.060269013 | -0.468359355 | 0.34782133 | 11.41510708 | -0.369903213 | 0.725331592 | 0.867794616 | -4.738911817 |
| **A_55_P2113733** | NA | 0.039758974 | -0.229748528 | 0.309266477 | 10.57529629 | 0.369500185 | 0.725615313 | 0.86805436 | -4.738959577 |
| **A_52_P642626** | Rbm20 | -0.034005691 | -0.264534912 | 0.19652353 | 7.691159277 | -0.369467264 | 0.72563849 | 0.86805436 | -4.738963476 |
| **A_55_P2010347** | Zdhhc6 | -0.033834058 | -0.263278188 | 0.195610072 | 8.593673446 | -0.369340974 | 0.725727406 | 0.868089317 | -4.738978431 |
| **A_52_P476731** | Fam110c | 0.051518521 | -0.297921087 | 0.40095813 | 8.031416132 | 0.369267942 | 0.725778827 | 0.868089317 | -4.738987077 |
| **A_55_P2058746** | Mterf1a | 0.033332135 | -0.192844805 | 0.259509076 | 8.88348414 | 0.369117941 | 0.725884447 | 0.868089317 | -4.73900483 |
| **A_51_P117881** | Leap2 | 0.033939808 | -0.196389593 | 0.264269208 | 7.724933062 | 0.369071362 | 0.725917245 | 0.868089317 | -4.739010341 |
| **A_52_P708507** | Myadml2 | 0.03204225 | -0.185423976 | 0.249508476 | 7.681872147 | 0.369046879 | 0.725934485 | 0.868089317 | -4.739013238 |
| **A_55_P2095727** | Pml | 0.048106912 | -0.278472339 | 0.374686163 | 10.36415082 | 0.368951462 | 0.726001677 | 0.868089317 | -4.739024525 |
| **A_55_P2050997** | Serinc4 | -0.042018783 | -0.327307916 | 0.243270351 | 8.279911643 | -0.368899937 | 0.726037961 | 0.868089317 | -4.73903062 |
| **A_52_P9703** | Cyp2a4 | -0.036243933 | -0.282347337 | 0.209859471 | 7.719722777 | -0.368865503 | 0.72606221 | 0.868089317 | -4.739034692 |
| **A_51_P186899** | Egln1 | 0.033269083 | -0.192667938 | 0.259206104 | 7.917882177 | 0.36881093 | 0.726100642 | 0.868089317 | -4.739041145 |
| **A_55_P1967266** | Plxnb2 | -0.064202765 | -0.5002436 | 0.371838069 | 13.02972518 | -0.368788116 | 0.726116708 | 0.868089317 | -4.739043843 |
| **A_55_P2052210** | Gdi2 | 0.038285835 | -0.221808177 | 0.298379848 | 13.98275781 | 0.36868725 | 0.726187744 | 0.868120562 | -4.739055767 |

| | | | | | | | | | |
|---|---|---|---|---|---|---|---|---|---|
| **A_55_P2184128** | Ndufa3 | 0.053352798 | -0.309403422 | 0.416109018 | 14.85462651 | 0.368377147 | 0.726406157 | 0.868307328 | -4.739092409 |
| **A_55_P1983268** | 4930444P | 0.182160283 | -1.056719117 | 1.421039683 | 9.04935374 | 0.368277347 | 0.726476454 | 0.868307328 | -4.739104195 |
| **A_55_P2043762** | N6amt1 | 0.078711166 | -0.456611284 | 0.614033616 | 9.038619881 | 0.368274148 | 0.726478708 | 0.868307328 | -4.739104573 |
| **A_55_P2119927** | NA | 0.121564501 | -0.706324464 | 0.949453467 | 9.058414511 | 0.367777388 | 0.726828662 | 0.868671902 | -4.739163196 |
| **A_51_P159711** | Car7 | 0.075581484 | -0.439360678 | 0.590523646 | 9.130764369 | 0.367626923 | 0.726934676 | 0.868744902 | -4.739180938 |
| **A_55_P2029687** | Hmox1 | -0.051648509 | -0.403812749 | 0.300515731 | 7.918853216 | -0.367335482 | 0.727140036 | 0.868936613 | -4.739215284 |
| **A_51_P517120** | Zfp930 | -0.076048744 | -0.59492944 | 0.442831951 | 11.74331039 | -0.367091966 | 0.727311646 | 0.869087971 | -4.739243964 |
| **A_55_P1999182** | Cdk12 | 0.041406771 | -0.24116898 | 0.323982521 | 8.598185672 | 0.367017537 | 0.727364101 | 0.869096936 | -4.739252726 |
| **A_51_P488768** | Slc25a44 | -0.042257464 | -0.330914454 | 0.246399526 | 8.29347585 | -0.366666892 | 0.727611245 | 0.869316723 | -4.739293984 |
| **A_55_P2020532** | Selenov | -0.085572252 | -0.670257398 | 0.499112893 | 8.868809993 | -0.366573654 | 0.727676968 | 0.869316723 | -4.739304948 |
| **A_55_P2042988** | Aoah | -0.032821071 | -0.257081102 | 0.191438959 | 7.851583807 | -0.366565191 | 0.727682934 | 0.869316723 | -4.739305943 |
| **A_51_P173285** | Nkx2-5 | 0.154138469 | -0.899401429 | 1.207678367 | 8.403595529 | 0.36644635 | 0.727766709 | 0.86934628 | -4.739319915 |
| **A_55_P2002723** | Inpp5a | 0.094325819 | -0.550470388 | 0.739122026 | 11.69787012 | 0.366402522 | 0.727797605 | 0.86934628 | -4.739325066 |
| **A_55_P2076360** | NA | -0.049619701 | -0.389042106 | 0.289802703 | 8.206396574 | -0.36615417 | 0.727972692 | 0.869467449 | -4.739354247 |
| **A_55_P1992582** | Hmgcs2 | 0.030873445 | -0.180328964 | 0.242075854 | 7.672543561 | 0.366131059 | 0.727988987 | 0.869467449 | -4.739356961 |
| **A_55_P1990490** | Recql | 0.054239673 | -0.316953368 | 0.425432713 | 8.618687253 | 0.365988619 | 0.728089416 | 0.869533681 | -4.739373688 |
| **A_55_P1952901** | Gm833 | 0.04481299 | -0.261981773 | 0.351607753 | 7.97703043 | 0.365852752 | 0.728185216 | 0.869594377 | -4.739389638 |
| **A_55_P1956464** | Stam | 0.107534457 | -0.62895758 | 0.844026494 | 9.528518317 | 0.36570419 | 0.728289974 | 0.869665762 | -4.739407072 |
| **A_66_P128839** | Gm13262 | -0.045236186 | -0.355166428 | 0.264694056 | 8.031993891 | -0.365571534 | 0.728383521 | 0.869723752 | -4.739422633 |
| **A_51_P398235** | Dnajc11 | 0.046649585 | -0.273046989 | 0.366346158 | 8.720970004 | 0.365477079 | 0.728450132 | 0.869723843 | -4.735082336 |
| **A_55_P2113683** | Raph1 | 0.123303276 | -0.721789328 | 0.96839588 | 11.25207107 | 0.365443852 | 0.728473565 | 0.869723843 | -4.739437607 |
| **A_55_P2117614** | Tnfrsf13c | -0.031877601 | -0.250544551 | 0.186789349 | 7.682538521 | -0.365134463 | 0.728691774 | 0.869791886 | -4.739473869 |
| **A_55_P1955861** | Cux2 | 0.034931658 | -0.204725142 | 0.274588459 | 7.802028168 | 0.365073066 | 0.728735081 | 0.869791886 | -4.739481061 |
| **A_55_P2031337** | Trmt2a | 0.037091413 | -0.217388613 | 0.291571439 | 10.78055611 | 0.365064854 | 0.728740873 | 0.869791886 | -4.739482023 |
| **A_55_P2212908** | Pon2 | -0.038960445 | -0.306269103 | 0.228348214 | 8.028002889 | -0.365057459 | 0.728746089 | 0.869791886 | -4.73948289 |
| **A_55_P2136906** | Vpreb3 | -0.096319378 | -0.757193363 | 0.564554606 | 8.379688185 | -0.365044123 | 0.728755496 | 0.869791886 | -4.739484452 |
| **A_51_P103594** | Fam162b | 0.183360575 | -1.075074429 | 1.441795579 | 10.62601459 | 0.364943404 | 0.728826541 | 0.869822984 | -4.739496248 |
| **A_55_P2054420** | Tcf4 | 0.09971287 | -0.584754787 | 0.784180527 | 11.93363526 | 0.3648788 | 0.728872113 | 0.869823679 | -4.739503813 |
| **A_55_P2170139** | Rheb | 0.08436939 | -0.495082979 | 0.663821758 | 11.30272141 | 0.364684673 | 0.729009058 | 0.86986872 | -4.739526536 |
| **A_55_P2174067** | Cerk | 0.048739964 | -0.28604603 | 0.383525958 | 9.146360687 | 0.364643325 | 0.729038228 | 0.86986872 | -4.739531375 |
| **A_52_P67212** | 2900052N | 0.035242959 | -0.206865945 | 0.277351863 | 7.881151563 | 0.364596044 | 0.729071584 | 0.86986872 | -4.739536907 |
| **A_55_P2021565** | Ntf3 | 0.064437043 | -0.378258132 | 0.507132218 | 8.170161719 | 0.364570194 | 0.729089822 | 0.86986872 | -4.739539932 |
| **A_51_P361580** | Rab11b | 0.034074213 | -0.200246576 | 0.268395002 | 12.27294782 | 0.364221312 | 0.729335977 | 0.870108712 | -4.73958073 |
| **A_52_P625244** | Ube3a | -0.03479592 | -0.274122138 | 0.204530298 | 8.715193724 | -0.364156771 | 0.729381518 | 0.870109353 | -4.739588274 |
| **A_52_P577384** | Il18bp | -0.046610225 | -0.367298209 | 0.27407776 | 7.858236578 | -0.36403979 | 0.729464065 | 0.8701229 | -4.739601944 |
| **A_51_P160083** | 1500035N | -0.056638943 | -0.446355129 | 0.333077244 | 8.005080997 | -0.364013122 | 0.729482885 | 0.8701229 | -4.73960506 |
| **A_51_P239924** | Polr2c | 0.094428427 | -0.555615454 | 0.744472309 | 12.90354567 | 0.363839987 | 0.729605065 | 0.870214949 | -4.739625282 |
| **A_55_P2030080** | Kdm6b | 0.038166138 | -0.224635363 | 0.300967639 | 10.86653194 | 0.363748088 | 0.729669922 | 0.87023862 | -4.739636013 |
| **A_52_P124173** | Zdhhc8 | 0.032098885 | -0.189032796 | 0.253230567 | 7.9214453 | 0.363571079 | 0.72979485 | 0.870243855 | -4.739656674 |
| **A_51_P434198** | Anapc4 | 0.030594622 | -0.180182711 | 0.241371955 | 13.31699162 | 0.36355618 | 0.729805366 | 0.870243855 | -4.739658413 |
| **A_55_P2116385** | Gpr183 | 0.03072183 | -0.180943784 | 0.242387444 | 7.892407929 | 0.363535748 | 0.729819787 | 0.870243855 | -4.739660797 |
| **A_55_P1974273** | Ndufs7 | -0.051981697 | -0.410170654 | 0.30620726 | 13.70938689 | -0.363486771 | 0.729854355 | 0.870243855 | -4.739666512 |
| **A_55_P2156345** | Zfp51 | 0.04188728 | -0.246864125 | 0.330638686 | 9.203309486 | 0.363335977 | 0.729960793 | 0.870275022 | -4.739684102 |
| **A_55_P2062148** | Arl1 | -0.04586019 | -0.362175684 | 0.270455305 | 10.56744136 | -0.363133006 | 0.73010407 | 0.870275022 | -4.739707769 |
| **A_55_P1988789** | Acss2 | 0.042554132 | -0.250979515 | 0.336087779 | 9.816753886 | 0.363106639 | 0.730122684 | 0.870275022 | -4.739710842 |
| **A_52_P178904** | Seh1l | 0.042264952 | -0.249314542 | 0.333844445 | 8.35336598 | 0.363056106 | 0.730158357 | 0.870275022 | -4.739716732 |
| **A_51_P108334** | Slc25a4 | 0.097266226 | -0.573762901 | 0.768295353 | 14.99333894 | 0.363053845 | 0.730159954 | 0.870275022 | -4.722485389 |
| **A_55_P2093665** | NA | -0.080888117 | -0.638961288 | 0.477185055 | 9.010663282 | -0.363031179 | 0.730175955 | 0.870275022 | -4.739719637 |

| | | | | | | | | | |
|---|---|---|---|---|---|---|---|---|---|
| **A_55_P2139763** | Trim56 | -0.076941642 | -0.607869721 | 0.453986438 | 11.04088331 | -0.362974485 | 0.730215979 | 0.870275022 | -4.739726244 |
| **A_51_P321965** | Ssna1 | 0.04423457 | -0.261030967 | 0.349500106 | 11.15839031 | 0.362939616 | 0.730240596 | 0.870275022 | -4.739730307 |
| **A_55_P1987201** | Capn2 | -0.081798333 | -0.646534987 | 0.48293832 | 13.16494872 | -0.362784585 | 0.73035005 | 0.870351816 | -4.739748366 |
| **A_55_P2142116** | NA | -0.09131019 | -0.721918803 | 0.539298423 | 14.79045817 | -0.36266838 | 0.730432096 | 0.870395941 | -4.739761898 |
| **A_55_P1981217** | Aatf | 0.04258278 | -0.251660406 | 0.336825966 | 8.740037582 | 0.362474902 | 0.73056871 | 0.870505081 | -4.73978442 |
| **A_55_P2075829** | NA | 0.031410848 | -0.185774428 | 0.248596124 | 8.715386007 | 0.362242689 | 0.73073269 | 0.870646813 | -4.739811436 |
| **A_55_P2170569** | Ggps1 | 0.03598825 | -0.213135835 | 0.285112335 | 10.84599663 | 0.36182232 | 0.731029578 | 0.870890392 | -4.739860302 |
| **A_55_P2025174** | Rps27a | 0.050878745 | -0.301354359 | 0.403111849 | 15.8435759 | 0.361790006 | 0.731052402 | 0.870890392 | -4.739864056 |
| **A_55_P2027117** | Rgs3 | 0.135754962 | -0.804148482 | 1.075658405 | 10.31445906 | 0.361761892 | 0.73107226 | 0.870890392 | -4.739867322 |
| **A_55_P2102936** | Pnrc1 | 0.109035317 | -0.64641051 | 0.864481143 | 10.59278795 | 0.361504917 | 0.73125378 | 0.871006971 | -4.739897164 |
| **A_55_P2184106** | Gpn2 | 0.03650732 | -0.216470404 | 0.289485045 | 8.551515757 | 0.361449825 | 0.731292698 | 0.871006971 | -4.739903559 |
| **A_51_P519364** | Fam83f | 0.08240055 | -0.488784948 | 0.653586049 | 9.144516079 | 0.361329387 | 0.731377781 | 0.871006971 | -4.739917536 |
| **A_51_P306067** | Lpp | 0.125999232 | -0.747488088 | 0.999486553 | 11.31751735 | 0.361294741 | 0.731402258 | 0.871006971 | -4.739921556 |
| **A_55_P2083629** | Tle2 | 0.10304232 | -0.611329881 | 0.817414521 | 8.653229029 | 0.361277926 | 0.731414137 | 0.871006971 | -4.739923507 |
| **A_52_P24308** | Uchl1 | -0.042828601 | -0.339819142 | 0.254161939 | 8.443641981 | -0.36119491 | 0.731472787 | 0.871006971 | -4.739933138 |
| **A_52_P399717** | Ccdc50 | 0.03727021 | -0.22118909 | 0.295729511 | 8.61147804 | 0.36117695 | 0.731485476 | 0.871006971 | -4.739935221 |
| **A_55_P2096827** | Art3 | 0.056142956 | -0.333466043 | 0.445751956 | 8.37702278 | 0.360924734 | 0.731663678 | 0.87116551 | -4.739964467 |
| **A_52_P577388** | Epdr1 | 0.068632658 | -0.407976839 | 0.545242155 | 10.51514532 | 0.36067702 | 0.731838717 | 0.87119398 | -4.739993172 |
| **A_52_P359621** | Luzp1 | -0.178393303 | -1.417341283 | 1.060554678 | 10.7585851 | -0.360641598 | 0.731863748 | 0.87119398 | -4.739997276 |
| **A_51_P452798** | Zfp524 | 0.088735649 | -0.527556099 | 0.705027397 | 10.50679923 | 0.360630224 | 0.731871786 | 0.87119398 | -4.739998593 |
| **A_51_P421182** | Sys1 | 0.033476606 | -0.19908751 | 0.266040722 | 8.395055653 | 0.360536349 | 0.731938126 | 0.87119398 | -4.740009465 |
| **A_51_P502422** | Acot13 | 0.084300643 | -0.501377939 | 0.669979225 | 12.88760571 | 0.360513799 | 0.731954062 | 0.87119398 | -4.740012077 |
| **A_55_P2073709** | Tfdp2 | 0.043563224 | -0.259096823 | 0.346223271 | 8.948861407 | 0.360508297 | 0.731957951 | 0.87119398 | -4.740012714 |
| **A_51_P453043** | Aacs | -0.095903401 | -0.762411323 | 0.570604522 | 10.42860612 | -0.360395233 | 0.732037855 | 0.871235451 | -4.740025804 |
| **A_55_P1984386** | Depdc5 | -0.041218776 | -0.327780874 | 0.245343322 | 8.584135161 | -0.360268832 | 0.73212719 | 0.871288138 | -4.740040434 |
| **A_55_P2035667** | Gk | 0.048846909 | -0.290934564 | 0.388628382 | 8.048989146 | 0.360070663 | 0.732267257 | 0.87132609 | -4.740063361 |
| **A_55_P2171406** | Tm7sf2 | -0.031154377 | -0.247887496 | 0.185578741 | 7.934630242 | -0.360034519 | 0.732292805 | 0.87132609 | -4.740067542 |
| **A_51_P400773** | Hdgfl2 | 0.041957265 | -0.249930501 | 0.33384503 | 10.75339392 | 0.360032429 | 0.732294282 | 0.87132609 | -4.740067783 |
| **A_55_P2032453** | Mrpl23 | 0.049909893 | -0.297564283 | 0.397384069 | 14.05039362 | 0.359761306 | 0.732485935 | 0.871352663 | -4.740099129 |
| **A_66_P116998** | Tro | 0.063146306 | -0.376544136 | 0.502836749 | 10.69111465 | 0.359708974 | 0.73252293 | 0.871352663 | -4.740105177 |
| **A_55_P1996852** | NA | 0.109896016 | -0.655398288 | 0.87519032 | 10.06151784 | 0.359669667 | 0.732550718 | 0.871352663 | -4.740109719 |
| **A_55_P2123199** | Akirin1 | -0.063872953 | -0.508672883 | 0.380926977 | 8.439163548 | -0.359668688 | 0.73255141 | 0.871352663 | -4.740109832 |
| **A_55_P2018944** | Snx27 | -0.051640561 | -0.411277047 | 0.307995925 | 9.556651255 | -0.359647919 | 0.732566093 | 0.871352663 | -4.740112232 |
| **A_55_P2174743** | Akap7 | -0.041216066 | -0.328309695 | 0.245877564 | 9.190610974 | -0.359578175 | 0.7326154 | 0.871352663 | -4.740120289 |
| **A_51_P377452** | Ncf4 | 0.090062142 | -0.537318897 | 0.717443181 | 9.288570129 | 0.359551602 | 0.732634187 | 0.871352663 | -4.740123359 |
| **A_55_P1975090** | NA | 0.082357711 | -0.491450573 | 0.656165995 | 10.6798581 | 0.359490816 | 0.732677162 | 0.871352663 | -4.74013038 |
| **A_55_P2136813** | Rab10os | -0.127124226 | -1.013511174 | 0.759262723 | 9.638356155 | -0.359215703 | 0.732871679 | 0.871530387 | -4.740162142 |
| **A_55_P2074055** | NA | 0.061490624 | -0.367722796 | 0.490704045 | 9.19497752 | 0.358827695 | 0.733146055 | 0.871768016 | -4.740206901 |
| **A_51_P434503** | Dtwd1 | 0.0361412 | -0.216184363 | 0.288466763 | 8.258511114 | 0.358749797 | 0.733201146 | 0.871768016 | -4.740215882 |
| **A_55_P2123982** | Rhoh | 0.03537948 | -0.211633488 | 0.282392448 | 8.223333404 | 0.358741844 | 0.733206771 | 0.871768016 | -4.740216798 |
| **A_55_P2102679** | NA | 0.031172824 | -0.186574728 | 0.248920377 | 8.232891102 | 0.358569395 | 0.733328735 | 0.871794314 | -4.740236673 |
| **A_52_P434974** | Ubqln1 | -0.06619867 | -0.52868951 | 0.39629217 | 10.92407297 | -0.358506038 | 0.733373547 | 0.871794314 | -4.740243972 |
| **A_55_P1983209** | Rcbtb1 | 0.057221985 | -0.342575781 | 0.457019752 | 10.54046204 | 0.358486582 | 0.733387308 | 0.871794314 | -4.740246213 |
| **A_51_P203675** | Trem3 | -0.032073521 | -0.256186943 | 0.192039901 | 7.733899102 | -0.35845044 | 0.733412871 | 0.871794314 | -4.740250377 |
| **A_51_P431734** | Fam185a | 0.033165939 | -0.198618662 | 0.264950541 | 8.099836376 | 0.358391804 | 0.733454345 | 0.871794314 | -4.74025713 |
| **A_55_P2075592** | Clstn1 | 0.04457847 | -0.267184404 | 0.356341344 | 8.445132623 | 0.358138579 | 0.733633467 | 0.871930446 | -4.740286284 |
| **A_51_P360004** | Ocstamp | 0.065109089 | -0.390346846 | 0.520565024 | 8.313820577 | 0.358051582 | 0.73369501 | 0.871930446 | -4.740296296 |
| **A_55_P2013928** | Dhrs13 | -0.099505734 | -0.795600813 | 0.596589344 | 9.447492186 | -0.358038634 | 0.73370417 | 0.871930446 | -4.740297786 |

| | | | | | | | | | |
|---|---|---|---|---|---|---|---|---|---|
| **A_51_P217395** | Cyp2d34 | 0.03031815 | -0.181904086 | 0.242540386 | 7.639862404 | 0.357817979 | 0.733860275 | 0.872010103 | -4.740323168 |
| **A_55_P2009111** | NA | 0.030147789 | -0.180882887 | 0.241178464 | 7.759923805 | 0.357816384 | 0.733861404 | 0.872010103 | -4.740323352 |
| **A_51_P267278** | Slc15a2 | -0.087639386 | -0.701635635 | 0.526356863 | 9.767539735 | -0.35750651 | 0.734080652 | 0.872157692 | -4.740358972 |
| **A_55_P2128042** | NA | 0.124853933 | -0.749871305 | 0.999579172 | 10.16225566 | 0.35750401 | 0.734082421 | 0.872157692 | -4.74035926 |
| **A_55_P2127952** | NA | -0.075835745 | -0.607221025 | 0.455549536 | 8.084487991 | -0.357449572 | 0.734120941 | 0.872157692 | -4.740365515 |
| **A_51_P454196** | Sh2d4a | 0.101609063 | -0.610604199 | 0.813822325 | 8.798407746 | 0.35733268 | 0.734203656 | 0.872202365 | -4.740378942 |
| **A_55_P2104653** | Vma21 | 0.031761589 | -0.190918958 | 0.254442136 | 8.321310267 | 0.357248396 | 0.734263299 | 0.872219626 | -4.740388622 |
| **A_55_P2088202** | Hcn2 | 0.07892466 | -0.474590237 | 0.632439558 | 13.9128542 | 0.357136102 | 0.734342768 | 0.872260433 | -4.740401515 |
| **A_51_P433127** | Rdm1 | -0.057964119 | -0.464613844 | 0.348685606 | 12.2465289 | -0.357017168 | 0.734426939 | 0.872306821 | -4.740415166 |
| **A_55_P2003824** | Asgr1 | -0.045297172 | -0.363406624 | 0.27281228 | 7.909357495 | -0.356652164 | 0.734685282 | 0.872487162 | -4.740457034 |
| **A_55_P2019214** | NA | 0.033469112 | -0.201640903 | 0.268579126 | 8.038562379 | 0.356552433 | 0.734755876 | 0.872487162 | -4.740468467 |
| **A_55_P1972448** | NA | 0.059438702 | -0.358137468 | 0.477014873 | 8.531571115 | 0.35652005 | 0.7347788 | 0.872487162 | -4.740472179 |
| **A_55_P2172375** | NA | 0.033420851 | -0.201382531 | 0.268224234 | 8.196553738 | 0.356503259 | 0.734790685 | 0.872487162 | -4.740474104 |
| **A_55_P2005475** | Sult1a1 | -0.206453263 | -1.657216204 | 1.244309678 | 10.40968759 | -0.356431144 | 0.734841735 | 0.872487162 | -4.740482368 |
| **A_51_P459368** | Appbp2 | 0.043775905 | -0.263886307 | 0.351438117 | 12.39994584 | 0.356378357 | 0.734879103 | 0.872487162 | -4.740488416 |
| **A_55_P1965467** | 2300009A | 0.038491805 | -0.232049814 | 0.309033423 | 9.999045872 | 0.356356376 | 0.734894664 | 0.872487162 | -4.740490935 |
| **A_52_P468033** | Setd1b | -0.05008081 | -0.402215662 | 0.302054041 | 10.36749924 | -0.356215392 | 0.734994472 | 0.872530576 | -4.740507084 |
| **A_55_P1991505** | Gamt | 0.1161902 | -0.700869317 | 0.933249717 | 9.949230246 | 0.356177229 | 0.73502149 | 0.872530576 | -4.740511454 |
| **A_51_P103209** | Ttc7 | 0.071011882 | -0.428503284 | 0.570527049 | 9.89928522 | 0.356067791 | 0.735098971 | 0.872568978 | -4.740523984 |
| **A_55_P1965931** | Nat8f1 | -0.029935866 | -0.240701562 | 0.18082983 | 8.934208263 | -0.355747815 | 0.735325531 | 0.872784322 | -4.7405606 |
| **A_55_P2042121** | Slc25a22 | -0.10225552 | -0.822560495 | 0.618049456 | 8.878154318 | -0.355566381 | 0.735454009 | 0.87288323 | -4.740581349 |
| **A_55_P2172396** | A530064D | -0.030880393 | -0.248503564 | 0.186742778 | 7.735225861 | -0.355408677 | 0.735565691 | 0.872962192 | -4.740599375 |
| **A_55_P2073617** | NA | -0.077766515 | -0.625960686 | 0.470427656 | 8.761789797 | -0.355310931 | 0.735634916 | 0.872972634 | -4.740610545 |
| **A_55_P2107528** | NA | -0.034381855 | -0.276776289 | 0.208012579 | 7.719990756 | -0.355268744 | 0.735664794 | 0.872972634 | -4.740615365 |
| **A_51_P229911** | Adcy4 | -0.076278406 | -0.615009105 | 0.462452293 | 10.08092366 | -0.354633886 | 0.736114484 | 0.873452647 | -4.740687833 |
| **A_55_P1990336** | Add1 | 0.060383824 | -0.366292668 | 0.487060315 | 14.14156103 | 0.354464087 | 0.736234777 | 0.873541773 | -4.740707195 |
| **A_55_P2124712** | Ces2c | -0.039587984 | -0.334158904 | 0.254982935 | 8.154707803 | -0.357961422 | 0.736620883 | 0.873946253 | -4.713867852 |
| **A_55_P2240478** | NA | -0.044806079 | -0.36218301 | 0.272570851 | 8.29315209 | -0.353599737 | 0.736847253 | 0.87409741 | -4.740805624 |
| **A_52_P533779** | Ankrd40 | 0.038329369 | -0.233180889 | 0.309839627 | 9.137127274 | 0.353586578 | 0.736856579 | 0.87409741 | -4.740807121 |
| **A_51_P231320** | Mmp8 | -0.048904383 | -0.395423891 | 0.297615124 | 8.025204565 | -0.353484556 | 0.736928886 | 0.87409741 | -4.740818724 |
| **A_55_P2037528** | Acsf3 | -0.059749196 | -0.483111696 | 0.363613304 | 10.50747682 | -0.353484214 | 0.736929129 | 0.87409741 | -4.740818763 |
| **A_55_P2007964** | Cx3cr1 | 0.126730806 | -0.772625159 | 1.02608677 | 10.00519701 | 0.352940033 | 0.737314863 | 0.874478493 | -4.7408806 |
| **A_55_P1969166** | Fxr2 | -0.065925289 | -0.533818357 | 0.401967779 | 11.95873831 | -0.352903347 | 0.737340871 | 0.874478493 | -4.740884765 |
| **A_51_P459350** | Dstn | 0.115024738 | -0.701559881 | 0.931609357 | 12.82999326 | 0.352809607 | 0.737407327 | 0.874493422 | -4.740895407 |
| **A_55_P1981830** | NA | -0.086416617 | -0.700093934 | 0.527260701 | 10.01074117 | -0.352701684 | 0.737483841 | 0.874493422 | -4.740907656 |
| **A_55_P2120931** | Dleu2 | 0.03531739 | -0.215503376 | 0.286138155 | 8.287263172 | 0.352675632 | 0.737502311 | 0.874493422 | -4.740910613 |
| **A_55_P2055107** | Agpat1 | -0.106161089 | -0.860203826 | 0.647881649 | 11.4786083 | -0.352630398 | 0.737534382 | 0.874493422 | -4.740915745 |
| **A_51_P198675** | Ttc36 | -0.029675535 | -0.240604885 | 0.181253815 | 7.712545388 | -0.352380517 | 0.737711557 | 0.874513315 | -4.740944087 |
| **A_55_P1955417** | Atad3a | 0.070470463 | -0.430535749 | 0.571476676 | 12.21175222 | 0.352301393 | 0.737767664 | 0.874513315 | -4.740953058 |
| **A_55_P2176492** | Tollip | -0.042512866 | -0.344803662 | 0.25977793 | 8.226938534 | -0.352245785 | 0.737807095 | 0.874513315 | -4.740959361 |
| **A_55_P2139587** | NA | -0.02963995 | -0.240428829 | 0.181148929 | 7.72109004 | -0.352192511 | 0.737844873 | 0.874513315 | -4.740965399 |
| **A_51_P364984** | Gabpa | -0.032992052 | -0.267650257 | 0.201666153 | 8.391765876 | -0.35214695 | 0.737877182 | 0.874513315 | -4.740970563 |
| **A_55_P1979156** | Arap3 | 0.066277073 | -0.405236285 | 0.53779043 | 8.297584343 | 0.352062415 | 0.73793713 | 0.874513315 | -4.740980141 |
| **A_55_P2097259** | Zup1 | -0.037602054 | -0.305126521 | 0.229922413 | 9.718201979 | -0.352045181 | 0.737949352 | 0.874513315 | -4.740982093 |
| **A_55_P2040838** | Gm14548 | -0.060441579 | -0.490487157 | 0.369603998 | 8.173515348 | -0.352023506 | 0.737964724 | 0.874513315 | -4.740984548 |
| **A_51_P309534** | Pex14 | -0.120845538 | -0.980849207 | 0.73915813 | 10.05417194 | -0.351949752 | 0.738017029 | 0.874513315 | -4.740992903 |
| **A_51_P258372** | Igsf6 | -0.075772334 | -0.615105731 | 0.463561063 | 8.621716493 | -0.351887384 | 0.738061261 | 0.874513315 | -4.740999966 |
| **A_66_P123526** | 4930504D | 0.034153072 | -0.208961211 | 0.277267356 | 7.805560638 | 0.351859798 | 0.738080826 | 0.874513315 | -4.741003089 |

| | | | | | | | | | |
|---|---|---|---|---|---|---|---|---|---|
| **A_52_P674759** | Btbd3 | 0.074952126 | -0.458613053 | 0.608517305 | 11.14315724 | 0.351841304 | 0.738093942 | 0.874513315 | -4.741005183 |
| **A_51_P375862** | Srpk3 | -0.053020935 | -0.430598843 | 0.324556972 | 7.926623948 | -0.351715215 | 0.738183371 | 0.874565678 | -4.741019457 |
| **A_55_P2180056** | NA | -0.099467779 | -0.808021034 | 0.609085475 | 9.51409273 | -0.351609233 | 0.738258542 | 0.874601143 | -4.741031452 |
| **A_55_P1959119** | NA | -0.089759377 | -0.729322231 | 0.549803477 | 13.16714805 | -0.351517494 | 0.738323613 | 0.874624639 | -4.741041831 |
| **A_55_P1953920** | Me3 | -0.040791061 | -0.331719524 | 0.250137402 | 7.958269237 | -0.351179485 | 0.738563388 | 0.874841844 | -4.741080053 |
| **A_51_P385635** | Mrpl53 | 0.166304265 | -1.020102225 | 1.352710755 | 13.18806983 | 0.351091422 | 0.738625863 | 0.874841844 | -4.741090005 |
| **A_55_P2108883** | NA | -0.048470763 | -0.394401074 | 0.297459548 | 9.317050629 | -0.35094704 | 0.738728297 | 0.874841844 | -4.741106318 |
| **A_55_P1992403** | Simc1 | -0.065179868 | -0.530367575 | 0.400007838 | 9.165589538 | -0.350942202 | 0.73873173 | 0.874841844 | -4.741106864 |
| **A_55_P2025615** | Ammecr1l | 0.048618519 | -0.29846979 | 0.395706827 | 13.11624521 | 0.350842408 | 0.738802534 | 0.874841844 | -4.741118135 |
| **A_55_P1987483** | Pgm2 | 0.069435798 | -0.426346823 | 0.565218419 | 13.53104464 | 0.350786182 | 0.738842429 | 0.874841844 | -4.741124485 |
| **A_55_P2156186** | BC028528 | 0.167269224 | -1.027290246 | 1.361828694 | 9.701096716 | 0.35071845 | 0.738890487 | 0.874841844 | -4.741132132 |
| **A_66_P105764** | Tor1b | -0.067858931 | -0.552593064 | 0.416875202 | 9.536855727 | -0.350633782 | 0.738950565 | 0.874841844 | -4.741141689 |
| **A_51_P202801** | Abcb9 | 0.231788359 | -1.424073912 | 1.88765063 | 10.06618954 | 0.350604718 | 0.738971188 | 0.874841844 | -4.741144969 |
| **A_55_P2052490** | Ushbp1 | 0.058238513 | -0.357843559 | 0.474320584 | 8.290673062 | 0.350575545 | 0.738991889 | 0.874841844 | -4.741148261 |
| **A_55_P2029344** | Rpl23 | 0.045805703 | -0.281467661 | 0.373079067 | 16.93085975 | 0.350557482 | 0.739004707 | 0.874841844 | -4.7411503 |
| **A_55_P2094686** | Smim10l1 | -0.041967802 | -0.342090999 | 0.258155395 | 8.401007389 | -0.350241018 | 0.739229284 | 0.875054122 | -4.741185996 |
| **A_51_P190437** | Ccdc181 | -0.087048285 | -0.709794234 | 0.535697665 | 10.80187596 | -0.350106079 | 0.739325052 | 0.875113907 | -4.741201207 |
| **A_55_P2040773** | NA | 0.071273763 | -0.43902554 | 0.581573065 | 9.107199633 | 0.349828396 | 0.739522144 | 0.875223969 | -4.741232493 |
| **A_51_P272586** | Dym | 0.030656002 | -0.18885816 | 0.250170164 | 9.459674957 | 0.349786721 | 0.739551725 | 0.875223969 | -4.741237187 |
| **A_55_P2097372** | Tctex1d4 | 0.180244977 | -1.110596053 | 1.471086007 | 8.889365821 | 0.349736335 | 0.73958749 | 0.875223969 | -4.74124286 |
| **A_55_P2184567** | NA | -0.052515392 | -0.428681053 | 0.323650268 | 8.09679055 | -0.349669548 | 0.739634899 | 0.875223969 | -4.74125038 |
| **A_52_P680495** | Pdcd10 | -0.086184548 | -0.703544612 | 0.531175515 | 11.96951414 | -0.349656192 | 0.73964438 | 0.875223969 | -4.741251884 |
| **A_55_P2113723** | Pecr | -0.03325447 | -0.271557604 | 0.205048665 | 7.966609166 | -0.349518861 | 0.739741868 | 0.875269071 | -4.741267341 |
| **A_51_P128463** | Grrp1 | 0.148254352 | -0.914531417 | 1.211040121 | 10.66531354 | 0.349391283 | 0.739832438 | 0.875269071 | -4.712944315 |
| **A_55_P1962603** | Cplx2 | 0.227084783 | -1.400840661 | 1.855010227 | 13.8366309 | 0.349384688 | 0.739837121 | 0.875269071 | -4.741282437 |
| **A_55_P1976376** | Gimap4 | 0.087996038 | -0.5429229 | 0.718914977 | 8.555778695 | 0.349333231 | 0.739873652 | 0.875269071 | -4.741288225 |
| **A_52_P551743** | Aff4 | 0.062805969 | -0.387567714 | 0.513179652 | 10.30557315 | 0.349283654 | 0.73990885 | 0.875269071 | -4.741293802 |
| **A_52_P113537** | Xist | -0.168211244 | -1.374839862 | 1.038417373 | 8.085020821 | -0.34916584 | 0.739992496 | 0.875306447 | -4.741307049 |
| **A_51_P163578** | Ugt2b35 | 0.029320466 | -0.181048166 | 0.239689098 | 7.625341499 | 0.349092265 | 0.740044734 | 0.875306447 | -4.74131532 |
| **A_55_P2023703** | Zfp386 | 0.047344235 | -0.292397242 | 0.387085711 | 8.485618747 | 0.349034917 | 0.740085453 | 0.875306447 | -4.741321766 |
| **A_55_P2010076** | Miga2 | 0.055846323 | -0.344964363 | 0.456657009 | 10.5796544 | 0.348984098 | 0.740121537 | 0.875306447 | -4.741327478 |
| **A_51_P307168** | Ddah1 | 0.038046717 | -0.235180662 | 0.311274096 | 8.55556438 | 0.34877337 | 0.740271172 | 0.875429863 | -4.741351151 |
| **A_55_P1968774** | Ptk2b | 0.046273936 | -0.286343314 | 0.378891185 | 8.233685981 | 0.348451241 | 0.740499934 | 0.875646834 | -4.741387315 |
| **A_51_P223036** | Fbxo30 | -0.058365616 | -0.477998395 | 0.361267162 | 8.9815045 | -0.348367808 | 0.74055919 | 0.875663347 | -4.741396677 |
| **A_55_P2064567** | Anks1 | -0.0566582 | -0.464180325 | 0.350863924 | 9.256838841 | -0.34822659 | 0.74065949 | 0.875665788 | -4.741412517 |
| **A_52_P552879** | Cracr2b | 0.108327771 | -0.670918424 | 0.887573967 | 9.096077397 | 0.348189334 | 0.740685952 | 0.875665788 | -4.741416695 |
| **A_52_P159885** | Ddx49 | -0.040358605 | -0.330726369 | 0.25000916 | 9.122951775 | -0.348127305 | 0.740730011 | 0.875665788 | -4.741423651 |
| **A_65_P02543** | Srp72 | 0.036820889 | -0.22810739 | 0.301749168 | 9.177871363 | 0.348109834 | 0.740742421 | 0.875665788 | -4.741425609 |
| **A_55_P2025626** | NA | -0.063270527 | -0.518731992 | 0.392190939 | 8.125704062 | -0.347936632 | 0.740865451 | 0.87572758 | -4.741445023 |
| **A_55_P1970670** | Scp2 | 0.085939857 | -0.532759498 | 0.704639212 | 13.47426519 | 0.347908716 | 0.740885281 | 0.87572758 | -4.741448151 |
| **A_55_P2090794** | NA | -0.123166754 | -1.010080442 | 0.763746935 | 9.271983585 | -0.347826352 | 0.74094379 | 0.875743198 | -4.74145738 |
| **A_55_P2088571** | Repin1 | -0.040485053 | -0.332109971 | 0.251139865 | 10.25105972 | -0.3477126 | 0.7410246 | 0.875785171 | -4.741470122 |
| **A_55_P2106180** | Dnah2 | 0.141062674 | -0.87575863 | 1.157883978 | 8.962349117 | 0.347470406 | 0.741196667 | 0.875934985 | -4.741497238 |
| **A_55_P1972852** | Mcrs1 | 0.060125852 | -0.373446915 | 0.493698619 | 12.17227873 | 0.347335837 | 0.741292279 | 0.875948971 | -4.741512296 |
| **A_55_P2240823** | Epb41 | -0.031608604 | -0.259557475 | 0.196340267 | 7.81945785 | -0.347310738 | 0.741310113 | 0.875948971 | -4.741515104 |
| **A_55_P2145307** | Ints11 | 0.042537611 | -0.264269634 | 0.349344857 | 9.774838019 | 0.347262455 | 0.74134442 | 0.875948971 | -4.741520506 |
| **A_55_P1990870** | Mylk3 | -0.081067894 | -0.665924021 | 0.503788232 | 8.331846375 | -0.3471764 | 0.741405567 | 0.875953604 | -4.741530131 |
| **A_55_P2126890** | NA | 0.043306157 | -0.269171028 | 0.355783342 | 8.19358985 | 0.347121633 | 0.741444483 | 0.875953604 | -4.741536256 |

| | | | | | | | | | |
|---|---|---|---|---|---|---|---|---|---|
| **A_52_P516021** | Ptpn1 | 0.044863226 | -0.278901259 | 0.368627712 | 9.087732043 | 0.347065655 | 0.74148426 | 0.875953604 | -4.741542515 |
| **A_52_P119187** | NA | 0.10342403 | -0.643171968 | 0.850020027 | 10.31451291 | 0.34696538 | 0.741555517 | 0.875984259 | -4.741553724 |
| **A_51_P427828** | Mlip | 0.058978738 | -0.366890289 | 0.484847765 | 7.969445501 | 0.346872417 | 0.741621581 | 0.876008776 | -4.741564114 |
| **A_55_P2066976** | Acsm1 | -0.033797769 | -0.277906235 | 0.210310696 | 8.162717375 | -0.346781197 | 0.741686408 | 0.87603183 | -4.741574306 |
| **A_55_P2155764** | NA | 0.042698943 | -0.265761803 | 0.351159688 | 10.45538509 | 0.346710954 | 0.74173633 | 0.876037276 | -4.741582153 |
| **A_51_P476579** | NA | 0.039444867 | -0.245893797 | 0.324783531 | 9.406515669 | 0.346242376 | 0.742069381 | 0.876330686 | -4.741634459 |
| **A_55_P2017729** | Gm2138 | -0.029938988 | -0.246518757 | 0.18664078 | 7.97033255 | -0.346233893 | 0.74207541 | 0.876330686 | -4.741635405 |
| **A_55_P2159595** | NA | -0.031612245 | -0.260552566 | 0.197328076 | 7.97323613 | -0.345846512 | 0.742350798 | 0.876590026 | -4.741678597 |
| **A_55_P2366273** | Amn1 | -0.038741708 | -0.319354433 | 0.241871016 | 8.491893037 | -0.345797422 | 0.742385698 | 0.876590026 | -4.741684067 |
| **A_66_P109368** | Prex2 | -0.059642553 | -0.491937296 | 0.37265219 | 8.582016657 | -0.345562508 | 0.742552721 | 0.876732955 | -4.741710234 |
| **A_51_P507851** | Gcc1 | -0.043622803 | -0.359890215 | 0.272644609 | 9.113530729 | -0.345469304 | 0.742618993 | 0.876732955 | -4.741720612 |
| **A_55_P2021266** | Hpse | 0.035811615 | -0.223894593 | 0.295517823 | 9.340169975 | 0.345375818 | 0.742685469 | 0.876732955 | -4.741731018 |
| **A_51_P311904** | Ptk2b | -0.035644908 | -0.294144953 | 0.222855138 | 8.073447057 | -0.345372073 | 0.742688131 | 0.876732955 | -4.741731435 |
| **A_55_P1999541** | Ccdc42 | -0.060563242 | -0.499999703 | 0.378873219 | 8.295295961 | -0.345194108 | 0.742814684 | 0.87681228 | -4.741751237 |
| **A_51_P290904** | Raver2 | -0.039186933 | -0.323556886 | 0.24518302 | 8.00652276 | -0.345150029 | 0.74284603 | 0.87681228 | -4.74175614 |
| **A_51_P188772** | Arhgef28 | 0.145370128 | -0.910307668 | 1.201047925 | 9.742740852 | 0.344900745 | 0.743023317 | 0.876968 | -4.741783859 |
| **A_55_P2108808** | Tubgcp4 | 0.032624896 | -0.204363884 | 0.269613676 | 8.371643255 | 0.344803522 | 0.743092465 | 0.876992734 | -4.741794665 |
| **A_51_P121412** | Slc35a4 | 0.035552265 | -0.222745854 | 0.293850383 | 9.213147704 | 0.344743729 | 0.743134993 | 0.876992734 | -4.726348471 |
| **A_66_P136099** | Coasy | 0.04455498 | -0.279548406 | 0.368658366 | 12.18967962 | 0.34432062 | 0.74343596 | 0.877204362 | -4.741848294 |
| **A_55_P2165655** | Zbtb4 | 0.059802828 | -0.375268238 | 0.494873895 | 10.29157076 | 0.344280056 | 0.743464816 | 0.877204362 | -4.741852796 |
| **A_55_P2088296** | Fam160b2 | 0.051499967 | -0.323188013 | 0.426187947 | 14.04046027 | 0.344260771 | 0.743478536 | 0.877204362 | -4.741854936 |
| **A_51_P470989** | Paip1 | 0.036674928 | -0.230209372 | 0.303559227 | 9.096239825 | 0.344188684 | 0.74352982 | 0.877204362 | -4.741862934 |
| **A_55_P2043466** | Eif5 | -0.054617747 | -0.452155406 | 0.342919912 | 12.74437658 | -0.344116779 | 0.743580974 | 0.877204362 | -4.741870911 |
| **A_55_P2029106** | Bmf | 0.134845212 | -0.846653903 | 1.116344328 | 10.70215503 | 0.344108947 | 0.743586547 | 0.877204362 | -4.74187178 |
| **A_55_P2053083** | Gm6098 | -0.081463892 | -0.674623685 | 0.5116959 | 10.16078382 | -0.3439884 | 0.743672311 | 0.877252011 | -4.741885149 |
| **A_55_P2062632** | Smc5 | -0.126280034 | -1.046129196 | 0.793569128 | 10.53089681 | -0.343849517 | 0.743771126 | 0.877315048 | -4.741900546 |
| **A_55_P1999852** | Tchp | -0.04378323 | -0.362787888 | 0.275221427 | 8.834672519 | -0.343764576 | 0.743831564 | 0.877332812 | -4.74190996 |
| **A_55_P2116105** | Ap2a1 | -0.051641339 | -0.428047207 | 0.324764529 | 10.96633067 | -0.343630302 | 0.743927108 | 0.877391978 | -4.741924837 |
| **A_51_P292527** | P4htm | -0.028879192 | -0.239451658 | 0.181693275 | 8.09699415 | -0.343505581 | 0.744015859 | 0.877443126 | -4.741938651 |
| **A_55_P2109377** | Camk2b | 0.044426393 | -0.279707893 | 0.368560679 | 8.430679837 | 0.343294172 | 0.744166306 | 0.877523521 | -4.741962056 |
| **A_55_P2180071** | Zfand5 | -0.061914581 | -0.513658124 | 0.389828963 | 12.30723363 | -0.343282233 | 0.744174803 | 0.877523521 | -4.741963377 |
| **A_55_P2134387** | Olfr544 | -0.09325771 | -0.773907374 | 0.587391954 | 11.39935973 | -0.343171694 | 0.744253473 | 0.877562764 | -4.741975609 |
| **A_55_P2047599** | Arhgdia | 0.044340248 | -0.280159518 | 0.368840013 | 13.19132173 | 0.342242605 | 0.744914835 | 0.878289024 | -4.742078275 |
| **A_55_P2143159** | Men1 | -0.109773737 | -0.913445409 | 0.693897934 | 11.53277806 | -0.34211344 | 0.745006799 | 0.87834389 | -4.742092527 |
| **A_52_P117408** | Tg | -0.032392936 | -0.269677052 | 0.20489118 | 7.771202945 | -0.341925893 | 0.745140338 | 0.878447762 | -4.742113213 |
| **A_55_P2045512** | NA | 0.044410494 | -0.281180563 | 0.370001551 | 15.18471263 | 0.341635883 | 0.745346855 | 0.87863765 | -4.742145179 |
| **A_55_P2169586** | Ccdc80 | 0.168463111 | -1.06717935 | 1.404105573 | 10.52774974 | 0.341477694 | 0.745459512 | 0.878683447 | -4.742162605 |
| **A_55_P2160712** | NA | 0.040238967 | -0.254926364 | 0.335404298 | 8.60431987 | 0.3414537 | 0.7454766 | 0.878683447 | -4.742165247 |
| **A_51_P501107** | Cul5 | -0.040680123 | -0.339254486 | 0.25789424 | 9.089971548 | -0.341255837 | 0.745617522 | 0.878795974 | -4.742187031 |
| **A_51_P503866** | Edem1 | -0.087841807 | -0.733180176 | 0.557496562 | 10.04684384 | -0.340929139 | 0.745850227 | 0.879016659 | -4.742222974 |
| **A_51_P225083** | Snx17 | -0.039869629 | -0.332850034 | 0.253110777 | 13.004173 | -0.340842672 | 0.745911821 | 0.879024894 | -4.742232482 |
| **A_52_P121342** | Nrxn1 | 0.10025435 | -0.636571042 | 0.837079742 | 8.41060431 | 0.340791682 | 0.745948145 | 0.879024894 | -4.742238087 |
| **A_55_P2050513** | Pgbd1 | 0.042170611 | -0.268010272 | 0.352351493 | 8.190045793 | 0.340522025 | 0.746140254 | 0.879097492 | -4.742267719 |
| **A_55_P2401084** | 4930558N | 0.054372227 | -0.345653514 | 0.454397968 | 8.947913248 | 0.340439175 | 0.746199281 | 0.879097492 | -4.742276819 |
| **A_55_P1971938** | Atp2b2 | -0.029691852 | -0.248154616 | 0.188770911 | 7.895825316 | -0.340416195 | 0.746215655 | 0.879097492 | -4.742279343 |
| **A_55_P2114288** | Mrpl24 | 0.034442237 | -0.218995876 | 0.287880349 | 13.18171711 | 0.340384492 | 0.746238242 | 0.879097492 | -4.742282824 |
| **A_55_P1968231** | Rbpms | 0.089029856 | -0.566191593 | 0.744251305 | 8.986930144 | 0.34032818 | 0.746278365 | 0.879097492 | -4.742289007 |
| **A_51_P462385** | G6pc | -0.028715504 | -0.240052922 | 0.182621914 | 7.715900623 | -0.340322283 | 0.746282566 | 0.879097492 | -4.742289654 |

| | | | | | | | | | |
|---|---|---|---|---|---|---|---|---|---|
| **A_55_P2058270** | Gm6507 | -0.040396311 | -0.338089288 | 0.257296667 | 7.90843703 | -0.339878317 | 0.746598925 | 0.879373843 | -4.742338367 |
| **A_52_P589550** | Nr2c1 | -0.029468233 | -0.246715101 | 0.187778635 | 8.345557528 | -0.339743314 | 0.746695135 | 0.879373843 | -4.742353168 |
| **A_55_P2024600** | Zfp658 | -0.093241167 | -0.780677197 | 0.594194863 | 14.39592186 | -0.33972363 | 0.746709164 | 0.879373843 | -4.742355325 |
| **A_51_P511707** | Ntpcr | 0.06329199 | -0.403394284 | 0.529978264 | 10.1661269 | 0.3396832 | 0.746737977 | 0.879373843 | -4.742359756 |
| **A_51_P262721** | 0610009L | 0.090637175 | -0.57769826 | 0.758972611 | 10.76257885 | 0.339673932 | 0.746744582 | 0.879373843 | -4.742360772 |
| **A_55_P2052425** | Setd4 | -0.082250687 | -0.688988616 | 0.524487242 | 10.0236324 | -0.339538274 | 0.746841268 | 0.879429234 | -4.742375637 |
| **A_55_P2158498** | Btk | 0.041474584 | -0.264522873 | 0.347472042 | 8.079879804 | 0.339480295 | 0.746882591 | 0.879429234 | -4.742381988 |
| **A_55_P2066081** | Prmt6 | 0.062910855 | -0.401814965 | 0.527636675 | 8.381831733 | 0.339062011 | 0.747180747 | 0.879726725 | -4.742427778 |
| **A_55_P2147141** | Spata2 | -0.097302031 | -0.817274278 | 0.622670217 | 11.04462723 | -0.338498301 | 0.747582639 | 0.880146313 | -4.742489406 |
| **A_55_P1989865** | Thrap3 | -0.064755446 | -0.544167134 | 0.414656243 | 9.173337695 | -0.338312484 | 0.747715136 | 0.880210517 | -4.7425097 |
| **A_55_P2214625** | 4930519A | 0.053632663 | -0.343453952 | 0.450719278 | 8.222080041 | 0.338294126 | 0.747728227 | 0.880210517 | -4.742511704 |
| **A_55_P1953003** | Atp6v0a4 | 0.220958148 | -1.415297741 | 1.857214036 | 10.55526126 | 0.338227678 | 0.74777561 | 0.880212703 | -4.742518958 |
| **A_51_P260513** | Polr1d | -0.034752351 | -0.292163679 | 0.222658978 | 11.57940153 | -0.33814804 | 0.747832401 | 0.880225961 | -4.742527651 |
| **A_66_P106654** | Camsap1 | 0.043970569 | -0.281920119 | 0.369861257 | 10.71717078 | 0.337940684 | 0.747980276 | 0.88026398 | -4.742550274 |
| **A_55_P2230506** | 4931402H | 0.158141726 | -1.014251636 | 1.330535087 | 14.77267076 | 0.337849646 | 0.748045204 | 0.88026398 | -4.742560203 |
| **A_55_P2046253** | Tmem239 | 0.055054864 | -0.353166669 | 0.463276397 | 14.14790364 | 0.337792598 | 0.748085891 | 0.88026398 | -4.742566423 |
| **A_55_P2062603** | 4932438A | -0.069808405 | -0.587524439 | 0.44790763 | 9.284073655 | -0.337727515 | 0.74813231 | 0.88026398 | -4.742573518 |
| **A_55_P2041634** | Vps51 | 0.035699318 | -0.229056804 | 0.300455439 | 10.70029343 | 0.337725808 | 0.748133527 | 0.88026398 | -4.742573704 |
| **A_55_P2039289** | Hspb6 | 0.088385304 | -0.567381142 | 0.744151751 | 9.53173557 | 0.337583504 | 0.748235027 | 0.88026398 | -4.742589214 |
| **A_51_P254299** | Dlk2 | -0.077605097 | -0.653427844 | 0.49821765 | 9.275397082 | -0.337560618 | 0.748251351 | 0.88026398 | -4.742591708 |
| **A_51_P501922** | Plxna1 | 0.066476461 | -0.426899531 | 0.559852454 | 8.956802414 | 0.337473927 | 0.748313187 | 0.88026398 | -4.742601152 |
| **A_55_P2044385** | Fgfbp3 | -0.040891684 | -0.344520903 | 0.262737534 | 8.837731851 | -0.337319762 | 0.748423158 | 0.88026398 | -4.742617942 |
| **A_51_P266143** | Adpgk | -0.040675696 | -0.342719118 | 0.261367726 | 9.627542324 | -0.337299703 | 0.748437467 | 0.88026398 | -4.742620126 |
| **A_55_P2061278** | Sik1 | 0.102153942 | -0.656498575 | 0.860806459 | 10.62087758 | 0.337258242 | 0.748467043 | 0.88026398 | -4.74262464 |
| **A_55_P2050857** | D11Wsu4 | -0.057270897 | -0.482644636 | 0.368102843 | 9.829845598 | -0.337220255 | 0.748494143 | 0.88026398 | -4.742628776 |
| **A_51_P166023** | Hip1 | 0.057448876 | -0.369323447 | 0.4842212 | 8.424767491 | 0.337159682 | 0.748537354 | 0.88026398 | -4.742635369 |
| **A_55_P2117943** | NA | 0.071737499 | -0.461273901 | 0.604748899 | 9.77062242 | 0.33710103 | 0.748579197 | 0.88026398 | -4.742641752 |
| **A_52_P297075** | Heatr5a | 0.052182631 | -0.335599005 | 0.439964267 | 10.65693377 | 0.337045922 | 0.748618512 | 0.88026398 | -4.742647749 |
| **A_55_P1973462** | Pex11b | -0.037159682 | -0.313336573 | 0.239017209 | 10.31330222 | -0.337003953 | 0.748648454 | 0.88026398 | -4.742652315 |
| **A_55_P2113598** | Ikzf3 | 0.028108519 | -0.180824523 | 0.237041561 | 7.777818153 | 0.336962195 | 0.748678246 | 0.88026398 | -4.742656858 |
| **A_55_P2010196** | Serpina10 | -0.029722326 | -0.250656435 | 0.191211783 | 7.663575073 | -0.336953809 | 0.74868423 | 0.88026398 | -4.74265777 |
| **A_51_P160113** | Copb1 | -0.042160519 | -0.355865849 | 0.271544811 | 12.00455913 | -0.336615721 | 0.748925455 | 0.880319282 | -4.742694529 |
| **A_55_P2013058** | Shb | -0.055845236 | -0.471403771 | 0.359713298 | 12.02929073 | -0.336592373 | 0.748942115 | 0.880319282 | -4.742697066 |
| **A_52_P522864** | Zfp442 | -0.0511656 | -0.431925475 | 0.329594274 | 12.23279471 | -0.336571428 | 0.748957061 | 0.880319282 | -4.742699342 |
| **A_52_P88818** | Clybl | -0.03344465 | -0.282336863 | 0.215447562 | 8.024597188 | -0.336562478 | 0.748963447 | 0.880319282 | -4.742700315 |
| **A_55_P2016902** | Entr1 | -0.053787204 | -0.454128528 | 0.346554121 | 9.654527458 | -0.33651071 | 0.749000387 | 0.880319282 | -4.742705939 |
| **A_51_P172573** | Sod2 | 0.096190912 | -0.619775849 | 0.812157673 | 13.05517751 | 0.336505005 | 0.749004458 | 0.880319282 | -4.742706559 |
| **A_55_P2240116** | 2310005E | -0.060855334 | -0.513927643 | 0.392216974 | 8.00756333 | -0.336419747 | 0.749065297 | 0.880337272 | -4.742715821 |
| **A_51_P121915** | Rdh16f2 | -0.030882055 | -0.26085975 | 0.199095641 | 7.674730521 | -0.336334034 | 0.749126463 | 0.880355643 | -4.728295831 |
| **A_55_P2108850** | 2-Mar | 0.065911873 | -0.425089345 | 0.556913091 | 9.347843321 | 0.336226104 | 0.749203486 | 0.880367926 | -4.742736848 |
| **A_52_P420665** | Uts2b | 0.077284668 | -0.498630863 | 0.653200198 | 7.912006338 | 0.336112682 | 0.749284432 | 0.880367926 | -4.742749159 |
| **A_55_P1978875** | Kmt2b | -0.046485843 | -0.39293537 | 0.299963684 | 10.78740134 | -0.336071037 | 0.749314154 | 0.880367926 | -4.742753678 |
| **A_51_P224311** | Popdc2 | 0.174497227 | -1.126021654 | 1.475016108 | 9.077959776 | 0.336064172 | 0.749319054 | 0.880367926 | -4.742754423 |
| **A_55_P2039699** | Camp | 0.164439326 | -1.062895638 | 1.39177429 | 9.112528217 | 0.335577572 | 0.749666373 | 0.880722468 | -4.742807189 |
| **A_55_P2148002** | 2610005L | -0.064131654 | -0.54306118 | 0.414797872 | 9.64031964 | -0.335390822 | 0.749799687 | 0.880772595 | -4.74282742 |
| **A_52_P89567** | Rhob | -0.037119063 | -0.314321483 | 0.240083357 | 8.847245596 | -0.33539017 | 0.749800152 | 0.880772595 | -4.742827491 |
| **A_55_P2066505** | Mpo | 0.028543151 | -0.184684692 | 0.241770993 | 7.666344246 | 0.335280536 | 0.74987842 | 0.880811019 | -4.742839363 |
| **A_55_P2072801** | Lypd5 | -0.043789704 | -0.37108744 | 0.283508032 | 8.19649353 | -0.335103802 | 0.750004598 | 0.880878329 | -4.742858494 |

| | | | | | | | | | |
|---|---|---|---|---|---|---|---|---|---|
| **A_55_P2039038** | Zfp664 | 0.054257615 | -0.351318225 | 0.459833455 | 10.70071928 | 0.33507264 | 0.750026847 | 0.880878329 | -4.742861867 |
| **A_55_P2062707** | Mthfsl | 0.098929199 | -0.640737073 | 0.838595471 | 9.66053297 | 0.334995539 | 0.750081896 | 0.880889472 | -4.742870209 |
| **A_55_P2108708** | Kcne3 | -0.03322531 | -0.281867407 | 0.215416786 | 8.001585388 | -0.334691539 | 0.750298964 | 0.881090875 | -4.742903084 |
| **A_55_P2071300** | Rnf126 | -0.082781191 | -0.702954188 | 0.537391805 | 10.50293186 | -0.33432524 | 0.75056055 | 0.881303256 | -4.74294266 |
| **A_55_P2060445** | Apopt1 | 0.052768092 | -0.342572631 | 0.448108815 | 14.30733843 | 0.334310629 | 0.750570984 | 0.881303256 | -4.742944238 |
| **A_52_P117313** | Azin1 | 0.030047529 | -0.195197207 | 0.255292265 | 7.894896344 | 0.334121532 | 0.75070604 | 0.881358618 | -4.742964652 |
| **A_55_P2452914** | N4bp2 | -0.157061715 | -1.334456641 | 1.020333211 | 9.966527305 | -0.33411696 | 0.750709306 | 0.881358618 | -4.742965145 |
| **A_52_P274238** | Maea | 0.043108089 | -0.296995031 | 0.383211208 | 11.85418111 | 0.337606545 | 0.750904451 | 0.881518752 | -4.715849111 |
| **A_51_P512591** | Cdc42se1 | -0.070694698 | -0.601241003 | 0.459851607 | 10.83401368 | -0.33374433 | 0.750975472 | 0.881518752 | -4.74300534 |
| **A_55_P2176361** | Zcchc9 | -0.034080103 | -0.289850411 | 0.221690204 | 8.4133365 | -0.333734512 | 0.750982485 | 0.881518752 | -4.743006399 |
| **A_55_P2072110** | 3-Sep | -0.057535352 | -0.489707373 | 0.374636668 | 14.35539583 | -0.333448276 | 0.751186969 | 0.881663774 | -4.743037246 |
| **A_55_P1958730** | NA | -0.120001444 | -1.021421175 | 0.781418286 | 11.43563366 | -0.333433905 | 0.751197235 | 0.881663774 | -4.743038794 |
| **A_55_P1967643** | Plb1 | 0.136178685 | -0.887852694 | 1.160210064 | 11.13587976 | 0.333078205 | 0.751451375 | 0.881866643 | -4.74307709 |
| **A_55_P2139809** | Fgf13 | 0.029707679 | -0.193696342 | 0.253111701 | 7.779299322 | 0.333064302 | 0.751461309 | 0.881866643 | -4.743078586 |
| **A_55_P2126281** | NA | -0.048430119 | -0.413047683 | 0.316187445 | 8.291066193 | -0.332681229 | 0.751735048 | 0.882134341 | -4.743119786 |
| **A_51_P510939** | Chchd7 | 0.055525004 | -0.362687886 | 0.473737895 | 9.455879737 | 0.332538192 | 0.751837269 | 0.88220075 | -4.743135158 |
| **A_51_P109939** | Naxe | 0.07121234 | -0.465540745 | 0.607965424 | 13.32901263 | 0.332300543 | 0.752007119 | 0.882346501 | -4.743160685 |
| **A_52_P437850** | Fam13c | -0.158232123 | -1.351458445 | 1.034994199 | 9.034721923 | -0.33214076 | 0.752121326 | 0.882381587 | -4.743177838 |
| **A_55_P2075794** | Tti2 | -0.041000419 | -0.350233669 | 0.268232831 | 11.36176452 | -0.332087442 | 0.752159438 | 0.882381587 | -4.743183561 |
| **A_52_P220573** | Serpini1 | -0.029005423 | -0.247800455 | 0.189789609 | 7.935513368 | -0.33204129 | 0.752192428 | 0.882381587 | -4.743188513 |
| **A_55_P1993708** | Lrp12 | -0.076492963 | -0.653598612 | 0.500612686 | 9.296552439 | -0.331983505 | 0.752233733 | 0.882381587 | -4.743194713 |
| **A_52_P509906** | Ift172 | 0.090888205 | -0.595141547 | 0.776917956 | 9.382310804 | 0.331829444 | 0.752343864 | 0.882381587 | -4.743211237 |
| **A_55_P1954724** | A130040M | 0.115758929 | -0.758170502 | 0.98968836 | 12.49635922 | 0.331763406 | 0.752391073 | 0.882381587 | -4.743218318 |
| **A_55_P1954486** | Mapt | 0.12679486 | -0.830627985 | 1.084217705 | 9.836408685 | 0.331702091 | 0.752434907 | 0.882381587 | -4.743224892 |
| **A_55_P2024709** | 4933427D | 0.061798942 | -0.40509281 | 0.528690693 | 10.63718707 | 0.331524159 | 0.752562116 | 0.882381587 | -4.743243961 |
| **A_51_P492446** | Ldah | -0.047513389 | -0.406484627 | 0.31145785 | 10.96140613 | -0.331517681 | 0.752566747 | 0.882381587 | -4.743244655 |
| **A_51_P289697** | Nek4 | -0.040873246 | -0.349678528 | 0.267932036 | 9.364382767 | -0.3315162 | 0.752567807 | 0.882381587 | -4.743244814 |
| **A_51_P103929** | Met | -0.028555447 | -0.244311128 | 0.187200233 | 7.855112628 | -0.33149507 | 0.752582914 | 0.882381587 | -4.736747763 |
| **A_55_P2030752** | Nedd4l | 0.115721018 | -0.758796059 | 0.990238095 | 12.58544343 | 0.331431892 | 0.752628084 | 0.882381587 | -4.743253845 |
| **A_55_P2003648** | Bag5 | -0.091015832 | -0.778839481 | 0.596807817 | 11.13252933 | -0.331428755 | 0.752630327 | 0.882381587 | -4.743254182 |
| **A_51_P324640** | C330018D | -0.030127326 | -0.257881433 | 0.197626781 | 9.429122501 | -0.331317754 | 0.752709692 | 0.882397392 | -4.74326607 |
| **A_52_P66130** | Myo5b | 0.055062253 | -0.361436714 | 0.47156122 | 9.010335794 | 0.331123797 | 0.752848379 | 0.882397392 | -4.743286833 |
| **A_52_P676198** | Fam210b | 0.120357108 | -0.790533755 | 1.031247972 | 9.416573453 | 0.330944939 | 0.752976278 | 0.882397392 | -4.74330597 |
| **A_55_P2462358** | Fam118b | 0.037572179 | -0.246810068 | 0.321954425 | 11.16952635 | 0.330913315 | 0.752998893 | 0.882397392 | -4.743309353 |
| **A_55_P2066309** | Slc39a8 | 0.095357675 | -0.626471149 | 0.817186499 | 10.85466655 | 0.330880962 | 0.75302203 | 0.882397392 | -4.743312813 |
| **A_51_P502798** | Nxpe4 | 0.095877357 | -0.62991025 | 0.821664964 | 8.507409564 | 0.330869588 | 0.753030164 | 0.882397392 | -4.74331403 |
| **A_55_P2388197** | 1010001B | 0.06306003 | -0.414304447 | 0.540424508 | 9.489858432 | 0.330867738 | 0.753031487 | 0.882397392 | -4.743314228 |
| **A_55_P2030269** | Dnah6 | 0.184579221 | -1.21330048 | 1.582458923 | 9.209334785 | 0.330722206 | 0.753135564 | 0.882397392 | -4.743329788 |
| **A_55_P2074536** | Ppm1b | 0.031087931 | -0.204357983 | 0.266533844 | 12.63825402 | 0.330712799 | 0.753142292 | 0.882397392 | -4.743330794 |
| **A_65_P08227** | Snx5 | -0.045578071 | -0.390791007 | 0.299634865 | 8.550294326 | -0.330688601 | 0.753159599 | 0.882397392 | -4.743333381 |
| **A_51_P285779** | Asphd2 | 0.068300187 | -0.449080024 | 0.585680398 | 8.354694537 | 0.330645356 | 0.753190527 | 0.882397392 | -4.743338003 |
| **A_51_P237752** | Cavin1 | 0.127207979 | -0.836408079 | 1.090824036 | 15.42116601 | 0.330644017 | 0.753191484 | 0.882397392 | -4.743338146 |
| **A_55_P2293013** | Ces2a | 0.030602625 | -0.201334235 | 0.262539486 | 7.688393052 | 0.330475492 | 0.753312016 | 0.882485126 | -4.743356154 |
| **A_55_P2014978** | Dvl3 | -0.116218797 | -0.997287395 | 0.7648498 | 12.96158852 | -0.330382473 | 0.753378549 | 0.882509595 | -4.743366089 |
| **A_52_P429774** | Cnst | -0.027888563 | -0.239726325 | 0.183949198 | 9.399612122 | -0.329741121 | 0.753837344 | 0.88299353 | -4.743434524 |
| **A_51_P235726** | Icam4 | -0.033132097 | -0.285017941 | 0.218753746 | 8.1143194 | -0.329454554 | 0.754042378 | 0.883161493 | -4.743465061 |
| **A_51_P486091** | Ift52 | 0.038358249 | -0.253334218 | 0.330050717 | 10.32744097 | 0.329369899 | 0.754102951 | 0.883161493 | -4.743474078 |
| **A_55_P2037897** | Brd3 | 0.043255467 | -0.285698225 | 0.37220916 | 11.52609557 | 0.329349181 | 0.754117776 | 0.883161493 | -4.743476284 |

| | | | | | | | | | |
|---|---|---|---|---|---|---|---|---|---|
| **A_51_P258620** | Rab4b | 0.028196565 | -0.186428616 | 0.242821746 | 8.481565409 | 0.329053018 | 0.754329708 | 0.883306067 | -4.743507809 |
| **A_66_P139647** | Vwa3a | 0.108828545 | -0.719557212 | 0.937214303 | 8.497290999 | 0.329048978 | 0.754332599 | 0.883306067 | -4.743508239 |
| **A_55_P2039463** | Timmdc1 | -0.115577868 | -0.995771563 | 0.764615826 | 11.61425401 | -0.328887049 | 0.754448484 | 0.88330803 | -4.743525463 |
| **A_55_P1980035** | Tbc1d9b | -0.034297954 | -0.295507806 | 0.226911898 | 11.12118104 | -0.328873594 | 0.754458114 | 0.88330803 | -4.743526894 |
| **A_52_P432919** | Rab3d | 0.113392092 | -0.750239474 | 0.977023659 | 9.650066602 | 0.328855119 | 0.754471336 | 0.88330803 | -4.743528859 |
| **A_55_P2029612** | Dhx35 | -0.03917569 | -0.337729882 | 0.259378502 | 9.711236565 | -0.32865771 | 0.754612624 | 0.883371393 | -4.743549845 |
| **A_55_P1980863** | NA | -0.032456912 | -0.279812373 | 0.214898548 | 8.09087416 | -0.328651825 | 0.754616837 | 0.883371393 | -4.743550471 |
| **A_51_P131653** | Tsacc | 0.044629165 | -0.295616635 | 0.384874964 | 9.235104727 | 0.328530983 | 0.75470333 | 0.883387687 | -4.743563312 |
| **A_51_P237668** | Bex2 | 0.169045453 | -1.120030836 | 1.458121742 | 12.08633192 | 0.328454503 | 0.754758074 | 0.883387687 | -4.743571436 |
| **A_55_P2000909** | Atp5l | 0.055892927 | -0.370347571 | 0.482133425 | 15.09288688 | 0.328437311 | 0.754770379 | 0.883387687 | -4.743573262 |
| **A_55_P2078474** | Fam50a | -0.038421491 | -0.331525467 | 0.254682484 | 12.76002894 | -0.32832417 | 0.754851367 | 0.883387687 | -4.743585277 |
| **A_55_P2427685** | Agl | -0.051359759 | -0.443312138 | 0.340592621 | 11.0095498 | -0.328201086 | 0.754939477 | 0.883387687 | -4.743598344 |
| **A_51_P333111** | Aox1 | -0.038271226 | -0.330373182 | 0.25383073 | 7.984479102 | -0.328161973 | 0.754967477 | 0.883387687 | -4.743602495 |
| **A_66_P114768** | Bach2 | 0.042550466 | -0.282231372 | 0.367332304 | 9.321060932 | 0.328142842 | 0.754981172 | 0.883387687 | -4.743604525 |
| **A_55_P1953053** | NA | 0.049274967 | -0.327026121 | 0.425576055 | 11.32990579 | 0.327975358 | 0.755101074 | 0.883387687 | -4.743622295 |
| **A_55_P2026669** | Atp5l | 0.0531563 | -0.352787149 | 0.459099749 | 14.76523909 | 0.327974103 | 0.755101973 | 0.883387687 | -4.743622428 |
| **A_55_P1967335** | NA | -0.059385965 | -0.512915828 | 0.394143899 | 10.90810246 | -0.327965585 | 0.755108071 | 0.883387687 | -4.743623332 |
| **A_55_P1963017** | Stfa1 | 0.217754374 | -1.445412128 | 1.880920876 | 10.67135596 | 0.327930268 | 0.755133356 | 0.883387687 | -4.743627078 |
| **A_55_P1965836** | Crebbp | -0.033354585 | -0.288170618 | 0.221461447 | 11.63104358 | -0.327852985 | 0.755188687 | 0.883398853 | -4.743635274 |
| **A_51_P470851** | Cplx3 | -0.036560848 | -0.315925741 | 0.242804046 | 8.025289998 | -0.327789299 | 0.755234284 | 0.883398853 | -4.743642026 |
| **A_52_P75777** | NA | 0.087418186 | -0.581141991 | 0.755978362 | 8.281778699 | 0.327500246 | 0.75544125 | 0.883587485 | -4.743672658 |
| **A_55_P2149288** | Cmya5 | 0.035659947 | -0.237282896 | 0.30860279 | 8.055061001 | 0.327234682 | 0.755631418 | 0.883756448 | -4.743700779 |
| **A_55_P2087013** | Hipk1 | -0.032506318 | -0.281376274 | 0.216363638 | 7.869255018 | -0.327149044 | 0.755692746 | 0.883774713 | -4.743709842 |
| **A_51_P411355** | Taf1c | -0.050542849 | -0.437615232 | 0.336529533 | 8.738805546 | -0.327052808 | 0.755761667 | 0.883801856 | -4.743720025 |
| **A_55_P1992650** | Bloc1s1 | 0.122181846 | -0.81441645 | 1.058780143 | 13.82970178 | 0.326741024 | 0.755984973 | 0.884009524 | -4.743752995 |
| **A_55_P2137840** | Etfbkmt | 0.031930689 | -0.212936589 | 0.276797967 | 9.196678551 | 0.326608804 | 0.75607968 | 0.8840668 | -4.743766969 |
| **A_55_P1962214** | Kpna3 | -0.080605795 | -0.699132381 | 0.537920792 | 9.233165025 | -0.326406078 | 0.756224897 | 0.884183126 | -4.743788383 |
| **A_55_P2023118** | Prss28 | -0.038396352 | -0.333144174 | 0.256351471 | 7.985734261 | -0.326279433 | 0.756315622 | 0.884225162 | -4.743801754 |
| **A_66_P130654** | Abcg4 | 0.038483619 | -0.256980497 | 0.333947736 | 8.044709241 | 0.326228209 | 0.756352318 | 0.884225162 | -4.743807161 |
| **A_51_P357685** | Dhx16 | -0.03649656 | -0.316928729 | 0.243935609 | 10.89838632 | -0.325967605 | 0.756539024 | 0.884389957 | -4.743834656 |
| **A_51_P314763** | Rabggtb | -0.041858454 | -0.363597948 | 0.27988104 | 9.617071049 | -0.325858576 | 0.756617142 | 0.884427801 | -4.743846153 |
| **A_55_P2010386** | Tspan32 | 0.049613867 | -0.331907707 | 0.431135441 | 8.178390148 | 0.325712425 | 0.756721862 | 0.884478253 | -4.743861559 |
| **A_55_P2074226** | Fbxo8 | -0.037516737 | -0.326050597 | 0.251017123 | 8.5229478 | -0.325670647 | 0.756751798 | 0.884478253 | -4.743865962 |
| **A_55_P2247852** | Ap2s1 | 0.029652476 | -0.198913762 | 0.258218715 | 11.68886112 | 0.32493705 | 0.757277531 | 0.885039218 | -4.743943185 |
| **A_55_P1977593** | NA | 0.069861337 | -0.468754172 | 0.608476845 | 11.55726356 | 0.324869082 | 0.757326248 | 0.885042654 | -4.743950331 |
| **A_51_P395743** | Spopl | -0.033572771 | -0.292539189 | 0.225393646 | 8.741050959 | -0.324708824 | 0.75744112 | 0.885107092 | -4.743967176 |
| **A_65_P11603** | Ldb3 | 0.174021199 | -1.168489639 | 1.516532036 | 9.022373223 | 0.324664424 | 0.757472946 | 0.885107092 | -4.743971842 |
| **A_55_P2009146** | Gtpbp3 | -0.028667227 | -0.249928767 | 0.192594313 | 7.897802841 | -0.324511503 | 0.757582567 | 0.885181685 | -4.743987906 |
| **A_51_P464064** | Zc3h13 | 0.035764375 | -0.240371934 | 0.311900685 | 8.430740316 | 0.324397482 | 0.757664306 | 0.885223694 | -4.743999879 |
| **A_55_P2139913** | Ssb | 0.031170162 | -0.209569598 | 0.271909923 | 11.90117366 | 0.324295994 | 0.757737064 | 0.885255205 | -4.744010533 |
| **A_55_P2184385** | NA | 0.028678498 | -0.192929088 | 0.250286084 | 7.840466646 | 0.32413216 | 0.757854523 | 0.885338934 | -4.744027725 |
| **A_52_P612803** | Ccng1 | -0.032630125 | -0.284836869 | 0.219576619 | 10.03851781 | -0.324050271 | 0.757913236 | 0.885354027 | -4.744036315 |
| **A_55_P2167984** | NA | 0.084781756 | -0.570702004 | 0.740265515 | 10.41742791 | 0.323959571 | 0.757978268 | 0.8853765 | -4.744045827 |
| **A_55_P2176585** | Donson | 0.027578074 | -0.185709821 | 0.240865969 | 7.994256635 | 0.323853108 | 0.758054606 | 0.885412176 | -4.744056989 |
| **A_55_P2161858** | Pdlim7 | -0.039458479 | -0.344830084 | 0.265913126 | 9.056391846 | -0.323639882 | 0.758207504 | 0.885537264 | -4.744079334 |
| **A_55_P1964359** | Cnot8 | 0.04527898 | -0.305237818 | 0.395795777 | 12.07915425 | 0.32354757 | 0.758273702 | 0.885561084 | -4.744089004 |
| **A_55_P2070799** | Tyw1 | -0.069915404 | -0.611558475 | 0.471727666 | 9.40518128 | -0.323303218 | 0.758448941 | 0.88567899 | -4.744114587 |
| **A_51_P239673** | Hprt | -0.095388368 | -0.834428821 | 0.643652086 | 12.51146603 | -0.323279042 | 0.758466279 | 0.88567899 | -4.744117117 |

| | | | | | | | | | |
|---|---|---|---|---|---|---|---|---|---|
| **A_52_P55190** | BC005537 | -0.030722955 | -0.268857249 | 0.20741134 | 9.303755146 | -0.323140491 | 0.758565651 | 0.885741532 | -4.744131614 |
| **A_55_P2060487** | Zfp593 | 0.054886958 | -0.370663409 | 0.480437325 | 9.677234506 | 0.323049102 | 0.758631199 | 0.885764575 | -4.744141173 |
| **A_55_P1990866** | Mylk3 | -0.026915124 | -0.235744127 | 0.181913878 | 7.778440229 | -0.322816644 | 0.758797939 | 0.885905757 | -4.744165476 |
| **A_51_P243122** | Sec14l3 | 0.12180253 | -0.823542923 | 1.067147982 | 8.537526323 | 0.322712743 | 0.75887247 | 0.885939276 | -4.744176334 |
| **A_55_P1970785** | NA | -0.106357162 | -0.932774057 | 0.720059734 | 10.97386012 | -0.322342743 | 0.759137908 | 0.886195647 | -4.744214971 |
| **A_55_P2230913** | Etv6 | -0.041696191 | -0.366537719 | 0.283145337 | 9.191362494 | -0.321495709 | 0.759745708 | 0.886827473 | -4.744303267 |
| **A_55_P1972039** | Nox4 | 0.072366599 | -0.491479679 | 0.636212877 | 8.860602937 | 0.32146063 | 0.759770884 | 0.886827473 | -4.744306919 |
| **A_51_P390715** | Tgfb1 | -0.036038141 | -0.316991552 | 0.244915271 | 8.534907509 | -0.321276094 | 0.759903326 | 0.886928519 | -4.744326125 |
| **A_55_P1977741** | NA | -0.035593285 | -0.313216565 | 0.242029994 | 9.306328622 | -0.32111644 | 0.760017919 | 0.886994282 | -4.744342732 |
| **A_55_P2107686** | Ttll1 | -0.031846682 | -0.280323848 | 0.216630485 | 8.146572078 | -0.321017009 | 0.76008929 | 0.886994282 | -4.744353071 |
| **A_55_P2007413** | Agps | 0.03297462 | -0.224312003 | 0.290261243 | 11.75396069 | 0.321005845 | 0.760097303 | 0.886994282 | -4.744354232 |
| **A_55_P2091476** | NA | 0.069153017 | -0.470865122 | 0.609171156 | 9.386831491 | 0.320740003 | 0.760288136 | 0.887163427 | -4.744381859 |
| **A_55_P1991792** | Slc25a46 | -0.06645772 | -0.585911619 | 0.452996179 | 11.61735437 | -0.320441512 | 0.760502428 | 0.88732747 | -4.744412854 |
| **A_52_P277104** | Bank1 | -0.026881316 | -0.237010431 | 0.183247799 | 7.756629538 | -0.32041633 | 0.760520508 | 0.88732747 | -4.744415468 |
| **A_55_P2120956** | NA | -0.069036681 | -0.608827324 | 0.470753963 | 8.936006129 | -0.320335371 | 0.760578634 | 0.88734174 | -4.744423869 |
| **A_55_P2018480** | Carnmt1 | 0.030374734 | -0.207170864 | 0.267920332 | 10.55961447 | 0.320269694 | 0.76062579 | 0.88734321 | -4.744430683 |
| **A_51_P257541** | Plppr1 | 0.03270605 | -0.223320946 | 0.288733046 | 7.714758449 | 0.319957785 | 0.760849755 | 0.887550933 | -4.744463027 |
| **A_55_P2001250** | Cpz | 0.046791457 | -0.319806799 | 0.413389713 | 9.231568063 | 0.319688139 | 0.761043394 | 0.887670344 | -4.744490963 |
| **A_51_P142923** | Chka | -0.059629406 | -0.526810591 | 0.40755178 | 8.826596535 | -0.319687372 | 0.761043944 | 0.887670344 | -4.744491043 |
| **A_51_P413461** | Parg | 0.028818031 | -0.197097637 | 0.254733698 | 10.67912107 | 0.319498111 | 0.761179869 | 0.887775326 | -4.744510638 |
| **A_52_P461343** | Upk3a | 0.212771159 | -1.455810259 | 1.881352576 | 9.281650512 | 0.319385872 | 0.761260482 | 0.887815789 | -4.744522254 |
| **A_52_P208222** | Pde1c | -0.027073267 | -0.239901952 | 0.185755419 | 7.652156467 | -0.318611057 | 0.761817064 | 0.88841131 | -4.744602335 |
| **A_55_P2158866** | Nme6 | -0.03339813 | -0.296022668 | 0.229226408 | 10.828512 | -0.318520358 | 0.761882227 | 0.888433714 | -4.744611697 |
| **A_55_P2008933** | Slc17a3 | -0.032420486 | -0.28743129 | 0.222590319 | 7.684780595 | -0.318428022 | 0.761948569 | 0.88843534 | -4.744621226 |
| **A_55_P2149149** | NA | -0.046929185 | -0.41610501 | 0.322246639 | 8.276148279 | -0.318390505 | 0.761975525 | 0.88843534 | -4.744625097 |
| **A_52_P600625** | Rad23b | 0.030106889 | -0.206798352 | 0.26701213 | 8.693121973 | 0.318303608 | 0.762037962 | 0.888451461 | -4.744634061 |
| **A_52_P130912** | Pcdh7 | -0.026458098 | -0.234719766 | 0.181803571 | 7.709050625 | -0.318199591 | 0.762112702 | 0.888451461 | -4.744644788 |
| **A_55_P2059650** | Atpaf2 | -0.061817402 | -0.548455503 | 0.4248207 | 9.954604231 | -0.31816689 | 0.7621362 | 0.888451461 | -4.74464816 |
| **A_55_P2036240** | NA | 0.105230614 | -0.723340537 | 0.933801766 | 10.54627708 | 0.318099248 | 0.762184805 | 0.888451461 | -4.744655133 |
| **A_55_P1972277** | Samd11 | -0.026757222 | -0.237471476 | 0.183957032 | 7.671302828 | -0.318051501 | 0.762219116 | 0.888451461 | -4.744660054 |
| **A_51_P117794** | Bik | -0.063332336 | -0.562352123 | 0.435687451 | 8.293775412 | -0.317876255 | 0.762345051 | 0.888544684 | -4.744678112 |
| **A_55_P2064766** | NA | 0.042537407 | -0.292929625 | 0.37800444 | 11.16022437 | 0.317593434 | 0.762548308 | 0.88866091 | -4.744707234 |
| **A_55_P2078300** | Fndc7 | -0.091192067 | -0.81044122 | 0.628057087 | 8.463496123 | -0.317561646 | 0.762571155 | 0.88866091 | -4.744710506 |
| **A_52_P179729** | Txnl4a | -0.038401056 | -0.341363842 | 0.264561731 | 11.24687956 | -0.317471117 | 0.762636221 | 0.88866091 | -4.744719822 |
| **A_55_P2083128** | Sec16a | -0.068226089 | -0.60651169 | 0.470059512 | 10.72174234 | -0.317459303 | 0.762644712 | 0.88866091 | -4.744721038 |
| **A_55_P2017082** | Hnrnpa2b | 0.034071177 | -0.234910167 | 0.30305252 | 12.48918671 | 0.31726 | 0.762787967 | 0.88866091 | -4.744741537 |
| **A_51_P173735** | Dock11 | -0.06131392 | -0.545378927 | 0.422751087 | 9.904218106 | -0.317253 | 0.762792998 | 0.88866091 | -4.744742257 |
| **A_55_P2108424** | NA | -0.028149788 | -0.25039393 | 0.194094354 | 10.2961411 | -0.317245266 | 0.762798558 | 0.88866091 | -4.744743052 |
| **A_55_P2171091** | Sap30bp | 0.028577298 | -0.197055833 | 0.254210428 | 9.965816269 | 0.3172259 | 0.762812478 | 0.88866091 | -4.744745044 |
| **A_55_P2047155** | Meis2 | -0.042122112 | -0.374781361 | 0.290537137 | 10.47608941 | -0.3171472 | 0.76286905 | 0.888668246 | -4.744753134 |
| **A_51_P315042** | Avpr1a | 0.028456051 | -0.196345242 | 0.253257343 | 7.704211471 | 0.317048841 | 0.762939755 | 0.888668246 | -4.744763243 |
| **A_55_P2116988** | 1700024P | 0.051872302 | -0.357946545 | 0.46169115 | 9.513302873 | 0.317025314 | 0.762956667 | 0.888668246 | -4.744765661 |
| **A_65_P11773** | Adi1 | 0.046605631 | -0.321919616 | 0.415130878 | 9.989986717 | 0.316753552 | 0.763152038 | 0.888815027 | -4.744793575 |
| **A_55_P2162269** | Bri3 | -0.073442702 | -0.654272066 | 0.507386662 | 12.3948003 | -0.316701724 | 0.763189299 | 0.888815027 | -4.744798896 |
| **A_51_P328622** | Tlcd2 | -0.037030068 | -0.329926531 | 0.255866396 | 8.084870534 | -0.316658188 | 0.7632206 | 0.888815027 | -4.744803365 |
| **A_51_P486239** | Clec3b | 0.187540091 | -1.296677297 | 1.671757479 | 9.036160729 | 0.316480498 | 0.763348357 | 0.888893751 | -4.744821599 |
| **A_52_P550147** | NA | 0.13807223 | -0.954802588 | 1.230947049 | 10.11485592 | 0.316436278 | 0.763380152 | 0.888893751 | -4.744826135 |
| **A_51_P349727** | Slc25a45 | -0.032160305 | -0.286889931 | 0.22256932 | 8.167681933 | -0.316221249 | 0.763534768 | 0.88895759 | -4.744848185 |

| | | | | | | | | | |
|---|---|---|---|---|---|---|---|---|---|
| **A_51_P170562** | Cpn2 | -0.027030371 | -0.24115343 | 0.187092688 | 7.669671839 | -0.316183289 | 0.763562065 | 0.88895759 | -4.744852076 |
| **A_66_P106113** | Rhoj | 0.066877429 | -0.462922291 | 0.596677149 | 11.28973197 | 0.316168203 | 0.763572913 | 0.88895759 | -4.744853622 |
| **A_51_P230977** | Cdc42 | 0.032466417 | -0.224969857 | 0.289902691 | 14.27217798 | 0.315874789 | 0.763783916 | 0.889149701 | -4.744883683 |
| **A_55_P1959989** | D430042( | -0.035119389 | -0.313717989 | 0.243479211 | 10.46692989 | -0.31573183 | 0.76388673 | 0.889215849 | -4.74489832 |
| **A_55_P2105200** | Abraxas1 | 0.030144156 | -0.20908162 | 0.269369932 | 8.409534483 | 0.315606182 | 0.763977099 | 0.889267504 | -4.744911179 |
| **A_51_P419439** | Gnmt | 0.082393015 | -0.571709659 | 0.736495689 | 8.502664603 | 0.315496696 | 0.764055848 | 0.889272999 | -4.74492238 |
| **A_55_P2018076** | Arap1 | 0.09582449 | -0.665040032 | 0.856689011 | 12.87280122 | 0.315442006 | 0.764095185 | 0.889272999 | -4.744927974 |
| **A_55_P2412214** | Mhrt | -0.026411153 | -0.236143449 | 0.183321143 | 7.679627003 | -0.315407776 | 0.764119806 | 0.889272999 | -4.744931474 |
| **A_51_P396273** | Nras | 0.03849788 | -0.267323182 | 0.344318941 | 9.723631771 | 0.315296947 | 0.764199526 | 0.889312245 | -4.744942806 |
| **A_55_P2112360** | Arhgap17 | -0.034556407 | -0.309142041 | 0.240029227 | 11.02655781 | -0.315210815 | 0.764261484 | 0.889330817 | -4.744951611 |
| **A_55_P1967677** | Cped1 | 0.08852712 | -0.615243864 | 0.792298105 | 8.511219774 | 0.315061492 | 0.764368901 | 0.889402283 | -4.744966869 |
| **A_55_P1988048** | Emcn | 0.158399721 | -1.101376669 | 1.418176112 | 9.887217709 | 0.314928013 | 0.764464926 | 0.889412526 | -4.744980502 |
| **A_52_P239023** | Zfp955b | -0.034025897 | -0.3046446 | 0.236592805 | 8.337797179 | -0.314921365 | 0.764469709 | 0.889412526 | -4.744981181 |
| **A_52_P578634** | Cela1 | 0.062031705 | -0.431513123 | 0.555576534 | 8.161143237 | 0.314801984 | 0.764555597 | 0.889430785 | -4.744993369 |
| **A_55_P2115732** | Kif1c | 0.029566225 | -0.205720392 | 0.264852843 | 8.171272178 | 0.314737871 | 0.764601723 | 0.889430785 | -4.744999913 |
| **A_65_P20104** | Mical2 | -0.026573785 | -0.238066966 | 0.184919395 | 8.108520298 | -0.314707724 | 0.764623414 | 0.889430785 | -4.74500299 |
| **A_55_P2054332** | Ephb3 | 0.056959752 | -0.396510488 | 0.510429992 | 14.26743518 | 0.314607914 | 0.764695226 | 0.889460805 | -4.745013174 |
| **A_52_P139740** | Mdp1 | 0.03640104 | -0.253610955 | 0.326413034 | 9.533944745 | 0.314375123 | 0.764862727 | 0.889602115 | -4.745036915 |
| **A_55_P2056896** | Kif2a | -0.063495189 | -0.569524601 | 0.442534222 | 9.5863179 | -0.314279034 | 0.76493187 | 0.889608711 | -4.74504671 |
| **A_55_P1970655** | Kdm3b | -0.030478311 | -0.273454259 | 0.212497636 | 8.368621211 | -0.314179586 | 0.765003434 | 0.889608711 | -4.745056844 |
| **A_55_P1990970** | Casp9 | -0.041881509 | -0.375801631 | 0.292038613 | 11.4188863 | -0.314144945 | 0.765028362 | 0.889608711 | -4.745060374 |
| **A_51_P210877** | Pcnp | 0.067456499 | -0.470448828 | 0.605361827 | 11.87856073 | 0.314100263 | 0.765060516 | 0.889608711 | -4.745064925 |
| **A_55_P1952324** | NA | -0.076616219 | -0.687787411 | 0.534554973 | 8.74220614 | -0.313984483 | 0.765143838 | 0.889608711 | -4.745076717 |
| **A_55_P2011545** | Pyroxd1 | 0.042822905 | -0.298778387 | 0.384424196 | 8.845127673 | 0.313983599 | 0.765144474 | 0.889608711 | -4.745076807 |
| **A_55_P2006708** | Bmp1 | -0.08253675 | -0.741251353 | 0.576177852 | 9.221507906 | -0.313834309 | 0.765251916 | 0.889649834 | -4.745092006 |
| **A_51_P189916** | Nol12 | -0.063283632 | -0.56838634 | 0.441819076 | 10.87586248 | -0.31380658 | 0.765271873 | 0.889649834 | -4.745094829 |
| **A_55_P2041121** | Rn4.5s | -0.049228832 | -0.442889625 | 0.344431961 | 16.83454964 | -0.31321872 | 0.765695009 | 0.890088222 | -4.745154606 |
| **A_51_P114005** | Gstm7 | 0.030841859 | -0.21586796 | 0.277551677 | 7.784570334 | 0.313115418 | 0.765769374 | 0.890093878 | -4.73387001 |
| **A_55_P2100770** | Usp48 | -0.096674999 | -0.870073935 | 0.676723936 | 9.443404208 | -0.313084058 | 0.765791951 | 0.890093878 | -4.745168285 |
| **A_65_P12530** | NA | 0.034060555 | -0.238549934 | 0.306671044 | 8.140806832 | 0.312938864 | 0.765896479 | 0.890112718 | -4.745183027 |
| **A_51_P294807** | Stom | 0.064244878 | -0.449996111 | 0.578485866 | 9.965285818 | 0.312911961 | 0.765915848 | 0.890112718 | -4.745185758 |
| **A_51_P276063** | Phyh | 0.135667257 | -0.950440387 | 1.221774902 | 12.59423719 | 0.312861786 | 0.765951971 | 0.890112718 | -4.74519085 |
| **A_55_P2064876** | Fam89b | 0.090673792 | -0.635360934 | 0.816708518 | 12.65479746 | 0.312805752 | 0.765992314 | 0.890112718 | -4.745196536 |
| **A_51_P465273** | Mettl17 | -0.030819126 | -0.277670046 | 0.216031794 | 8.291388445 | -0.312705787 | 0.766064288 | 0.890142853 | -4.745206678 |
| **A_55_P2112424** | Madd | -0.067337307 | -0.607052605 | 0.472377991 | 14.81542653 | -0.312493767 | 0.766216948 | 0.890183341 | -4.745228179 |
| **A_55_P2081223** | Sptbn1 | -0.045117974 | -0.406757653 | 0.316521706 | 14.93211806 | -0.312481173 | 0.766226017 | 0.890183341 | -4.745229456 |
| **A_55_P1993636** | Rab9 | -0.037409528 | -0.337277732 | 0.262458676 | 10.75585967 | -0.31246556 | 0.766237259 | 0.890183341 | -4.745231038 |
| **A_55_P2051405** | Tet2 | 0.07057117 | -0.495293389 | 0.636435729 | 8.591000685 | 0.312367017 | 0.766308218 | 0.890212286 | -4.745241026 |
| **A_55_P2095325** | AI182371 | 0.026728253 | -0.187714873 | 0.241171379 | 7.651298433 | 0.312182668 | 0.76644097 | 0.890311232 | -4.745259702 |
| **A_66_P123582** | Mog | 0.036230611 | -0.254508347 | 0.32696957 | 8.179817783 | 0.312120849 | 0.766485489 | 0.890311232 | -4.745265963 |
| **A_66_P133413** | NA | 0.082890171 | -0.582739312 | 0.748519653 | 16.48035178 | 0.311903918 | 0.766641719 | 0.89037184 | -4.745287923 |
| **A_51_P438952** | Bag4 | -0.03433652 | -0.310071141 | 0.241398102 | 8.505095194 | -0.311899959 | 0.76664457 | 0.89037184 | -4.745288323 |
| **A_55_P1968969** | NA | -0.035632861 | -0.321817389 | 0.250551667 | 15.20396898 | -0.311856564 | 0.766675824 | 0.89037184 | -4.745292714 |
| **A_52_P15212** | Ctbp2 | 0.068491324 | -0.482889529 | 0.619872177 | 11.37711085 | 0.311124511 | 0.767203137 | 0.890930714 | -4.745366703 |
| **A_52_P604195** | Mbp | -0.060366909 | -0.546871538 | 0.42613772 | 11.33891069 | -0.310786617 | 0.767446576 | 0.891159886 | -4.745400799 |
| **A_66_P126563** | Ptges3l | 0.098206334 | -0.694033919 | 0.890446587 | 9.376363103 | 0.310479507 | 0.767667862 | 0.891318071 | -4.745431758 |
| **A_55_P1995874** | Gm14410 | 0.083928965 | -0.593155949 | 0.761013879 | 8.905361244 | 0.310469596 | 0.767675004 | 0.891318071 | -4.745432757 |
| **A_51_P483351** | Lix1l | 0.067164863 | -0.474925952 | 0.609255677 | 8.77489537 | 0.310327618 | 0.767777313 | 0.891341953 | -4.745447059 |

| | | | | | | | | | |
|---|---|---|---|---|---|---|---|---|---|
| **A_55_P1994204** | Ythdc1 | 0.04233544 | -0.299413786 | 0.384084665 | 12.10987177 | 0.31027507 | 0.767815181 | 0.891341953 | -4.745452351 |
| **A_55_P1987222** | NA | 0.033477786 | -0.236791386 | 0.303746958 | 8.224668253 | 0.310249123 | 0.76783388 | 0.891341953 | -4.745454964 |
| **A_55_P2056542** | Nin | 0.036970405 | -0.261628768 | 0.335569578 | 9.523643428 | 0.310110131 | 0.767934045 | 0.891367013 | -4.745468956 |
| **A_55_P2057370** | Slc22a19 | 0.103783424 | -0.734627649 | 0.942194497 | 8.41464906 | 0.310042575 | 0.767982732 | 0.891367013 | -4.745475755 |
| **A_55_P2082434** | Btbd11 | 0.08289659 | -0.586845284 | 0.752638463 | 8.361121404 | 0.310012752 | 0.768004226 | 0.891367013 | -4.745478756 |
| **A_55_P1978351** | Apobec2 | 0.029556705 | -0.209291057 | 0.268404467 | 7.809177181 | 0.309945396 | 0.76805277 | 0.891367013 | -4.745485533 |
| **A_55_P2160286** | Olfm1 | 0.072771942 | -0.515385641 | 0.660929526 | 8.537086032 | 0.309899311 | 0.768085985 | 0.891367013 | -4.745490168 |
| **A_55_P2184925** | Tmem185 | -0.122626075 | -1.114514613 | 0.869262463 | 10.45530786 | -0.309649441 | 0.768266083 | 0.891522504 | -4.745515292 |
| **A_55_P2039274** | Hspb3 | 0.084630277 | -0.6000664 | 0.769326953 | 8.177846126 | 0.309583557 | 0.768313572 | 0.891524103 | -4.745521913 |
| **A_55_P2182458** | Kcnh4 | 0.034947476 | -0.248284192 | 0.318179143 | 8.175230677 | 0.309046871 | 0.768700462 | 0.891919506 | -4.7455758 |
| **A_55_P1998811** | NA | 0.071002851 | -0.504856172 | 0.646861874 | 10.53781758 | 0.308823228 | 0.768861704 | 0.892011762 | -4.745598229 |
| **A_55_P2023952** | Arpc3 | -0.059348316 | -0.540707594 | 0.422010963 | 9.694380042 | -0.308808607 | 0.768872247 | 0.892011762 | -4.745599695 |
| **A_51_P256665** | Pon3 | 0.169442004 | -1.205382886 | 1.544266893 | 11.3499591 | 0.30869105 | 0.768957009 | 0.892027214 | -4.745611478 |
| **A_51_P418908** | Larp1 | -0.070287588 | -0.64093629 | 0.500361115 | 9.004300224 | -0.308503539 | 0.769092218 | 0.892027214 | -4.745630264 |
| **A_55_P2148402** | Cplx2 | -0.044594108 | -0.406669609 | 0.317481394 | 13.49292744 | -0.308481188 | 0.769108335 | 0.892027214 | -4.745632503 |
| **A_51_P415945** | Psma2 | 0.036296336 | -0.258442227 | 0.331034899 | 15.03428426 | 0.308443891 | 0.76913523 | 0.892027214 | -4.745636238 |
| **A_55_P1954910** | Fbxw2 | -0.029487087 | -0.268954136 | 0.209979963 | 8.480998613 | -0.308415679 | 0.769155574 | 0.892027214 | -4.745639063 |
| **A_55_P2111399** | Pcp2 | -0.038389122 | -0.350159888 | 0.273381644 | 8.044191571 | -0.308406225 | 0.769162392 | 0.892027214 | -4.74564001 |
| **A_52_P181709** | Crkl | -0.030226471 | -0.275758885 | 0.215305944 | 8.531179387 | -0.308339349 | 0.769210618 | 0.892029636 | -4.745646705 |
| **A_55_P2167840** | Gm14151 | -0.026344509 | -0.240457465 | 0.187768447 | 7.815992444 | -0.308175073 | 0.769329087 | 0.892040044 | -4.745663146 |
| **A_55_P1967761** | Foxc1 | 0.040846738 | -0.291204547 | 0.372898024 | 8.574923145 | 0.308107706 | 0.769377671 | 0.892040044 | -4.745669886 |
| **A_55_P2132403** | Szt2 | 0.033484771 | -0.238730018 | 0.30569956 | 9.929474456 | 0.308095935 | 0.76938616 | 0.892040044 | -4.745671064 |
| **A_52_P239787** | Ccdc25 | -0.026412186 | -0.241225573 | 0.188401201 | 8.537608405 | -0.307959318 | 0.769484691 | 0.892040044 | -4.745684727 |
| **A_52_P365011** | Fes | -0.059915976 | -0.54723066 | 0.427398708 | 9.038809051 | -0.307952324 | 0.769489735 | 0.892040044 | -4.745685427 |
| **A_55_P2130716** | Ube2g2 | -0.058801781 | -0.537074917 | 0.419471355 | 8.592122153 | -0.307939101 | 0.769499272 | 0.892040044 | -4.745686749 |
| **A_52_P327615** | Rab18 | 0.041319067 | -0.294821303 | 0.377459437 | 11.78073689 | 0.30787908 | 0.769542562 | 0.892040044 | -4.745692749 |
| **A_55_P2040170** | Pmp22 | 0.086874405 | -0.620035867 | 0.793784677 | 12.92468123 | 0.307806578 | 0.769594856 | 0.892047179 | -4.745699996 |
| **A_52_P478745** | Paxbp1 | 0.051171477 | -0.36534486 | 0.467687814 | 8.562297437 | 0.307713292 | 0.769662142 | 0.89207169 | -4.745709319 |
| **A_65_P01643** | Chd2 | -0.037781741 | -0.345390947 | 0.269827465 | 9.833105011 | -0.307633042 | 0.769720027 | 0.892085301 | -4.745717336 |
| **A_66_P132037** | Jmjd4 | -0.031982461 | -0.292666585 | 0.228701662 | 8.349348012 | -0.307289449 | 0.769967883 | 0.89231907 | -4.74575164 |
| **A_55_P1992174** | Tars2 | -0.031621807 | -0.289480781 | 0.226237168 | 8.844228139 | -0.307153008 | 0.770066314 | 0.892379653 | -4.745765252 |
| **A_55_P2006093** | Nebl | 0.052525249 | -0.376132113 | 0.48118261 | 7.976098092 | 0.306907977 | 0.770243099 | 0.89253102 | -4.745789684 |
| **A_52_P306357** | NA | -0.027403372 | -0.251280018 | 0.196473274 | 7.677576602 | -0.30658123 | 0.770478863 | 0.892736464 | -4.745822235 |
| **A_52_P600535** | Zfp39 | -0.033804016 | -0.31001414 | 0.242406109 | 8.169943486 | -0.306534278 | 0.770512743 | 0.892736464 | -4.745826909 |
| **A_52_P487615** | Otulinl | -0.038836949 | -0.356289938 | 0.278616041 | 8.500957753 | -0.306419235 | 0.77059576 | 0.892744831 | -4.74583836 |
| **A_51_P258281** | Gal | -0.032247152 | -0.295854923 | 0.231360618 | 8.93165562 | -0.306396295 | 0.770612315 | 0.892744831 | -4.745840643 |
| **A_55_P2403325** | NA | 0.028646983 | -0.205589035 | 0.262883002 | 7.883534694 | 0.306320149 | 0.770667265 | 0.892754997 | -4.74584822 |
| **A_55_P1968553** | Btbd9 | 0.110219288 | -0.792053698 | 1.012492274 | 10.72980665 | 0.305963762 | 0.770924471 | 0.892999445 | -4.745883658 |
| **A_55_P2167763** | NA | -0.033781271 | -0.310495984 | 0.242933443 | 7.817782212 | -0.305769437 | 0.77106473 | 0.8930835 | -4.745902964 |
| **A_55_P1971184** | Ube2l3 | -0.069315131 | -0.637164191 | 0.498533929 | 12.91592157 | -0.305735229 | 0.771089421 | 0.8930835 | -4.745906362 |
| **A_51_P382925** | Fadd | 0.028256468 | -0.203426484 | 0.259939419 | 8.898724359 | 0.30547392 | 0.771278045 | 0.893248456 | -4.745932302 |
| **A_55_P2186693** | NA | 0.037039262 | -0.266957714 | 0.341036237 | 8.481612628 | 0.3051711 | 0.771496655 | 0.893412992 | -4.745962338 |
| **A_51_P211305** | Alpk3 | 0.038667927 | -0.278719069 | 0.356054923 | 8.034038517 | 0.305149107 | 0.771512533 | 0.893412992 | -4.745964518 |
| **A_55_P2179726** | Mettl18 | -0.037722813 | -0.347428336 | 0.27198271 | 9.149441465 | -0.305074188 | 0.771566623 | 0.893422117 | -4.745971944 |
| **A_55_P2180789** | Huwe1 | -0.034842752 | -0.321042465 | 0.251356962 | 11.94654317 | -0.304925397 | 0.77167405 | 0.89347015 | -4.745986687 |
| **A_51_P113195** | Upk1b | -0.051761112 | -0.477006426 | 0.373484203 | 8.349935481 | -0.304869798 | 0.771714193 | 0.89347015 | -4.745992195 |
| **A_55_P2004230** | Emc6 | 0.035733819 | -0.257949533 | 0.32941717 | 11.16464652 | 0.304754724 | 0.771797281 | 0.89347015 | -4.74600359 |
| **A_55_P1994564** | Sap30bp | 0.028332267 | -0.204559101 | 0.261223635 | 8.657570487 | 0.30470409 | 0.771833843 | 0.89347015 | -4.746008603 |

| | | | | | | | | | |
|---|---|---|---|---|---|---|---|---|---|
| **A_55_P2134581** | Papola | -0.064518025 | -0.594886027 | 0.465849977 | 9.108131072 | -0.304687121 | 0.771846095 | 0.89347015 | -4.746010283 |
| **A_55_P2084797** | Aff3 | -0.081499198 | -0.751706705 | 0.58870831 | 9.072306233 | -0.304575097 | 0.771926987 | 0.89347015 | -4.74602137 |
| **A_51_P263993** | Lipc | -0.025277293 | -0.233149108 | 0.182594522 | 7.64114847 | -0.304568721 | 0.771931592 | 0.89347015 | -4.746022001 |
| **A_55_P1979242** | Kcne2 | 0.158564124 | -1.145912783 | 1.463041031 | 8.911858669 | 0.304452036 | 0.772015852 | 0.893499116 | -4.746033546 |
| **A_55_P1992834** | Socs2 | 0.189478883 | -1.369563245 | 1.748521012 | 10.24829471 | 0.304406072 | 0.772049045 | 0.893499116 | -4.746038092 |
| **A_55_P2122987** | Dlx3 | 0.102787752 | -0.743284258 | 0.948859762 | 10.54705068 | 0.304287691 | 0.772134535 | 0.893509508 | -4.746049798 |
| **A_52_P557129** | Slc12a5 | -0.092567141 | -0.854565912 | 0.66943163 | 8.442807524 | -0.304265649 | 0.772150453 | 0.893509508 | -4.746051978 |
| **A_52_P507382** | Unc93b1 | -0.06383396 | -0.58964225 | 0.461974331 | 9.202318343 | -0.304070789 | 0.772291184 | 0.893618873 | -4.746071236 |
| **A_55_P2021659** | Abcb8 | 0.04099073 | -0.296901842 | 0.378883303 | 10.83874918 | 0.303848681 | 0.772451604 | 0.893751006 | -4.746093173 |
| **A_55_P1952016** | R3hdm2 | 0.052306107 | -0.378954307 | 0.483566521 | 10.99368209 | 0.303782778 | 0.772499205 | 0.893752596 | -4.746099679 |
| **A_55_P2141038** | Gnpnat1 | -0.031744064 | -0.293559358 | 0.230071231 | 8.630013881 | -0.303681165 | 0.772572603 | 0.893768136 | -4.746109708 |
| **A_51_P180974** | Cavin3 | 0.125254235 | -0.907957272 | 1.158465741 | 11.70838279 | 0.303636187 | 0.772605092 | 0.893768136 | -4.746114147 |
| **A_55_P2053848** | NA | 0.038009701 | -0.275624701 | 0.351644102 | 11.99597488 | 0.303543615 | 0.772671962 | 0.893792013 | -4.746123279 |
| **A_55_P2033849** | Sphkap | 0.037370083 | -0.27107239 | 0.345812556 | 7.806983163 | 0.303459143 | 0.772732983 | 0.893809123 | -4.746131611 |
| **A_55_P1997145** | Nlrx1 | 0.038387201 | -0.278553572 | 0.355327974 | 8.39147617 | 0.303360257 | 0.772804419 | 0.893838277 | -4.746141361 |
| **A_52_P529486** | Wdr77 | 0.041684617 | -0.302740081 | 0.386109315 | 9.20356403 | 0.303132063 | 0.772969277 | 0.893960493 | -4.746163849 |
| **A_55_P1968943** | NA | 0.029784679 | -0.216369782 | 0.27593914 | 15.87740643 | 0.303064846 | 0.773017839 | 0.893960493 | -4.74617047 |
| **A_52_P485891** | Dnah5 | 0.094090264 | -0.683625459 | 0.871805987 | 8.388258642 | 0.303021995 | 0.773048799 | 0.893960493 | -4.74617469 |
| **A_55_P1959973** | Dcaf12l2 | 0.105550737 | -0.767467117 | 0.978568591 | 11.97387413 | 0.302822744 | 0.773192764 | 0.894073498 | -4.746194307 |
| **A_55_P2042217** | Fam161a | 0.105912273 | -0.770508454 | 0.982332999 | 8.635499422 | 0.302680187 | 0.773295771 | 0.894139133 | -4.746208334 |
| **A_55_P2123421** | Dnal4 | 0.047280585 | -0.344228816 | 0.438789985 | 8.852960532 | 0.302476048 | 0.773443285 | 0.894185387 | -4.74622841 |
| **A_52_P595642** | Smim7 | -0.036455909 | -0.338346446 | 0.265434629 | 10.28576657 | -0.302460574 | 0.773454467 | 0.894185387 | -4.746229931 |
| **A_55_P2016064** | NA | -0.040267093 | -0.373748564 | 0.293214377 | 8.43425707 | -0.302432821 | 0.773474523 | 0.894185387 | -4.746232659 |
| **A_51_P286737** | Ccl2 | -0.094558995 | -0.878080123 | 0.688962133 | 8.856481026 | -0.302275174 | 0.77358845 | 0.894263622 | -4.741725638 |
| **A_51_P453167** | Naxd | 0.064550625 | -0.47076088 | 0.59986213 | 12.937782 | 0.302025922 | 0.77376859 | 0.894375504 | -4.746272632 |
| **A_52_P206881** | Prpf18 | 0.043696743 | -0.318690936 | 0.406084422 | 11.22026152 | 0.302013246 | 0.773777752 | 0.894375504 | -4.746273877 |
| **A_55_P2043117** | Ccdc138 | -0.068465016 | -0.636449515 | 0.499519482 | 14.76497557 | -0.301913531 | 0.773849823 | 0.894401542 | -4.746283664 |
| **A_55_P2005229** | NA | 0.092622226 | -0.676030695 | 0.861275146 | 11.40932564 | 0.301811148 | 0.773923825 | 0.894401542 | -4.746293711 |
| **A_55_P2137973** | Bnip2 | 0.035229161 | -0.257153934 | 0.327612255 | 8.85954043 | 0.301786899 | 0.773941353 | 0.894401542 | -4.74629609 |
| **A_55_P2109397** | Scn10a | 0.030889949 | -0.225531705 | 0.287311602 | 7.745316023 | 0.30172607 | 0.773985321 | 0.894401542 | -4.746302056 |
| **A_51_P177432** | NA | 0.029030382 | -0.212064573 | 0.270125338 | 8.012407003 | 0.301588639 | 0.774084663 | 0.894462878 | -4.746315533 |
| **A_55_P2045876** | NA | 0.055529314 | -0.406355742 | 0.51741437 | 8.589254106 | 0.301119404 | 0.774423885 | 0.894801374 | -4.746361504 |
| **A_51_P148122** | Klhdc3 | -0.039969772 | -0.372545375 | 0.292605832 | 11.95351669 | -0.301017416 | 0.774497622 | 0.894833096 | -4.746371486 |
| **A_52_P425839** | Retnlg | -0.064428154 | -0.600738112 | 0.471881804 | 8.290527856 | -0.300891677 | 0.774588535 | 0.894884657 | -4.746383789 |
| **A_55_P2133637** | Etfa | 0.062387676 | -0.457484472 | 0.582259824 | 9.323993759 | 0.300574823 | 0.774817646 | 0.895007913 | -4.746414771 |
| **A_55_P2023391** | Grhl3 | 0.050844958 | -0.372847617 | 0.474537533 | 8.038715505 | 0.300571214 | 0.774820256 | 0.895007913 | -4.746415124 |
| **A_66_P128918** | Hacl1 | 0.035869659 | -0.263077035 | 0.334816354 | 9.485979671 | 0.300527241 | 0.774852055 | 0.895007913 | -4.746419421 |
| **A_55_P2120714** | NA | -0.032611487 | -0.30444274 | 0.239219766 | 8.121297997 | -0.300484143 | 0.77488322 | 0.895007913 | -4.746423632 |
| **A_55_P1993836** | Dusp7 | 0.049347127 | -0.362065416 | 0.460759669 | 9.96184342 | 0.300424043 | 0.774926682 | 0.895007913 | -4.746429503 |
| **A_55_P2170345** | Gm6729 | -0.035637749 | -0.332992307 | 0.26171681 | 8.07214384 | -0.30018294 | 0.775101046 | 0.895155822 | -4.746453046 |
| **A_51_P309740** | Usp40 | -0.030520691 | -0.285793948 | 0.224752566 | 8.906292094 | -0.299460387 | 0.775623678 | 0.89566299 | -4.746523494 |
| **A_51_P256945** | Opa3 | 0.027242017 | -0.200618252 | 0.255102287 | 8.337533446 | 0.299447714 | 0.775632846 | 0.89566299 | -4.746524728 |
| **A_51_P384618** | Dhdds | 0.044561142 | -0.328336525 | 0.41745881 | 9.933194554 | 0.299307083 | 0.775734581 | 0.895722564 | -4.74653842 |
| **A_55_P2043932** | Tmem8b | 0.054279098 | -0.400040946 | 0.508599142 | 8.612668154 | 0.299240982 | 0.775782402 | 0.895722564 | -4.746544854 |
| **A_55_P2089233** | Pou2af1 | 0.031005612 | -0.228584994 | 0.290596218 | 7.798226459 | 0.299158728 | 0.775841911 | 0.895722564 | -4.746552858 |
| **A_55_P1963563** | Zfp689 | 0.039296565 | -0.28986487 | 0.368458001 | 8.865247989 | 0.299017077 | 0.775944395 | 0.895722564 | -4.746566637 |
| **A_51_P386670** | Dse | 0.038592369 | -0.284776561 | 0.361961298 | 8.431635937 | 0.29891898 | 0.776015371 | 0.895722564 | -4.746576175 |
| **A_55_P2147641** | Cstf1 | 0.048108796 | -0.355045583 | 0.451263175 | 10.92847004 | 0.29888454 | 0.77604029 | 0.895722564 | -4.746579523 |

| | | | | | | | | | |
|---|---|---|---|---|---|---|---|---|---|
| **A_55_P1991960** | Nwd1 | 0.031223389 | -0.230438729 | 0.292885506 | 7.921749734 | 0.298874959 | 0.776047222 | 0.895722564 | -4.746580455 |
| **A_51_P370561** | Dcakd | -0.066455749 | -0.623414144 | 0.490502647 | 9.819552123 | -0.298854739 | 0.776061853 | 0.895722564 | -4.74658242 |
| **A_55_P2107045** | Myl4 | -0.180771974 | -1.696175721 | 1.334631772 | 14.87169049 | -0.298781078 | 0.776115151 | 0.895722564 | -4.746589579 |
| **A_55_P2143657** | Gm5784 | -0.081492338 | -0.76474208 | 0.601757403 | 8.513518326 | -0.298736061 | 0.776147725 | 0.895722564 | -4.746593954 |
| **A_51_P278427** | 3830406C | 0.055340014 | -0.408800061 | 0.519480089 | 9.617986179 | 0.298634889 | 0.776220933 | 0.895753582 | -4.746603782 |
| **A_55_P2044547** | NA | 0.091340963 | -0.676728571 | 0.859410497 | 9.395528371 | 0.2978622 | 0.776780135 | 0.896345397 | -4.746678745 |
| **A_55_P2024808** | Abl1 | 0.061681338 | -0.457375008 | 0.580737684 | 11.80539577 | 0.297638858 | 0.776941798 | 0.896430185 | -4.746700378 |
| **A_51_P355301** | Cyp3a11 | 0.024710123 | -0.183255121 | 0.232675367 | 7.690147778 | 0.297601074 | 0.776969148 | 0.896430185 | -4.746704037 |
| **A_55_P2115127** | Mphosph1 | 0.05321588 | -0.394860265 | 0.501292024 | 8.453176354 | 0.297467667 | 0.77706572 | 0.896430185 | -4.74671695 |
| **A_52_P83623** | Rapgef6 | -0.028035657 | -0.264195922 | 0.208124607 | 8.6458917 | -0.297340631 | 0.777157684 | 0.896430185 | -4.746729241 |
| **A_52_P109010** | Hipk3 | 0.027449861 | -0.203790409 | 0.25869013 | 12.17321266 | 0.297321985 | 0.777171182 | 0.896430185 | -4.746731045 |
| **A_55_P2106690** | Tal1 | 0.109830729 | -0.815573036 | 1.035234495 | 9.00716361 | 0.297264433 | 0.777212847 | 0.896430185 | -4.746736612 |
| **A_52_P141136** | Fggy | 0.028404362 | -0.210928823 | 0.267737547 | 8.244417846 | 0.297257256 | 0.777218043 | 0.896430185 | -4.746737306 |
| **A_55_P2063471** | NA | 0.066363685 | -0.492846448 | 0.625573818 | 15.89588327 | 0.29723901 | 0.777231252 | 0.896430185 | -4.74673907 |
| **A_51_P209818** | Prtn3 | -0.026355636 | -0.248495663 | 0.195784392 | 7.741657039 | -0.297164563 | 0.77728515 | 0.896430185 | -4.746746269 |
| **A_55_P2044932** | Gpr84 | -0.033148622 | -0.312704298 | 0.246407053 | 7.86047132 | -0.296993891 | 0.777408717 | 0.896430185 | -4.746762765 |
| **A_55_P2023772** | Fam107b | 0.058795502 | -0.437058803 | 0.554649808 | 10.80453555 | 0.296988999 | 0.777412259 | 0.896430185 | -4.746763238 |
| **A_51_P365008** | Tns1 | -0.093943234 | -0.88637918 | 0.698492712 | 13.5207039 | -0.296928365 | 0.77745616 | 0.896430185 | -4.746769096 |
| **A_55_P2036733** | Stk36 | 0.049854695 | -0.37078648 | 0.470495869 | 8.475365364 | 0.296855175 | 0.777509153 | 0.896430185 | -4.746776166 |
| **A_55_P2327177** | Exoc8 | 0.033485491 | -0.249057164 | 0.316028147 | 9.153635088 | 0.296840395 | 0.777519855 | 0.896430185 | -4.746777594 |
| **A_55_P2068882** | Ccng2 | 0.063070108 | -0.469172951 | 0.595313167 | 8.596046459 | 0.296800011 | 0.777549095 | 0.896430185 | -4.746781494 |
| **A_51_P318830** | Syt10 | 0.03613214 | -0.268853505 | 0.341117784 | 7.791260791 | 0.296732167 | 0.77759822 | 0.896433366 | -4.746788045 |
| **A_55_P2041961** | Bmp2 | 0.025111625 | -0.186963741 | 0.237186991 | 8.561017327 | 0.296575267 | 0.777711832 | 0.896510885 | -4.74680319 |
| **A_52_P109319** | E2f5 | 0.051575502 | -0.384673909 | 0.487824913 | 8.388505895 | 0.296114012 | 0.778045867 | 0.896793756 | -4.746847668 |
| **A_51_P428364** | Psma6 | 0.052327484 | -0.390292016 | 0.494946985 | 12.1601297 | 0.296107672 | 0.778050459 | 0.896793756 | -4.746848279 |
| **A_55_P1999334** | Inpp5b | -0.051707245 | -0.48928682 | 0.38587233 | 11.51287336 | -0.295967963 | 0.778151645 | 0.896793756 | -4.746861738 |
| **A_55_P1960023** | NA | 0.02826613 | -0.210957677 | 0.267489937 | 8.511718252 | 0.295945883 | 0.778167637 | 0.896793756 | -4.746863865 |
| **A_55_P1962274** | Ccnt1 | 0.033626841 | -0.25099465 | 0.318248333 | 9.030555543 | 0.295916193 | 0.778189141 | 0.896793756 | -4.746866724 |
| **A_55_P2152946** | NA | -0.07513463 | -0.712027115 | 0.561757855 | 8.746345292 | -0.295477471 | 0.778506928 | 0.897065279 | -4.746908942 |
| **A_51_P396570** | Plod2 | 0.06016509 | -0.449860652 | 0.570190833 | 9.453228323 | 0.295462808 | 0.77851755 | 0.897065279 | -4.746910352 |
| **A_55_P2020200** | Snx6 | -0.038453242 | -0.364551546 | 0.287645063 | 8.982513377 | -0.295348495 | 0.778600361 | 0.897097398 | -4.746921342 |
| **A_55_P2186808** | NA | -0.054130894 | -0.51326298 | 0.405001192 | 10.03338946 | -0.295296229 | 0.778638224 | 0.897097398 | -4.746926366 |
| **A_55_P2048912** | Themis2 | -0.025451719 | -0.241405079 | 0.190501641 | 7.814996487 | -0.295193978 | 0.778712302 | 0.897129284 | -4.746936191 |
| **A_55_P2002815** | Rabgap1l | -0.026725141 | -0.253635547 | 0.200185264 | 7.739018561 | -0.294995877 | 0.778855825 | 0.897192674 | -4.746955218 |
| **A_55_P2009732** | Prr9 | -0.041033694 | -0.389526444 | 0.307459056 | 7.956531461 | -0.294915236 | 0.778914252 | 0.897192674 | -4.74696296 |
| **A_55_P1974976** | Sprr2a2 | 0.033569855 | -0.251578318 | 0.318718027 | 16.15362233 | 0.294869066 | 0.778947705 | 0.897192674 | -4.746967391 |
| **A_55_P2029533** | Zfp57 | -0.026554951 | -0.252122851 | 0.199012948 | 7.905092897 | -0.294861834 | 0.778952945 | 0.897192674 | -4.746968085 |
| **A_51_P169880** | Zbtb3 | 0.026737299 | -0.200432581 | 0.253907178 | 8.375875194 | 0.29479297 | 0.779002841 | 0.897196695 | -4.746974693 |
| **A_55_P2046550** | Adamts8 | -0.024502397 | -0.232882107 | 0.183877313 | 7.883551801 | -0.294512335 | 0.779206191 | 0.897331658 | -4.747001609 |
| **A_55_P2079787** | Prss50 | 0.025351799 | -0.190258342 | 0.240961939 | 7.845986273 | 0.294503149 | 0.779212848 | 0.897331658 | -4.747002489 |
| **A_55_P2172599** | Fam174a | -0.050391071 | -0.479104355 | 0.378322212 | 10.91994614 | -0.294399452 | 0.779287993 | 0.897364744 | -4.747012428 |
| **A_52_P662013** | Plg | 0.040439737 | -0.303804123 | 0.384683597 | 7.951250247 | 0.294233734 | 0.779408087 | 0.897449584 | -4.747028305 |
| **A_55_P2092884** | NA | -0.062804515 | -0.597651788 | 0.472042757 | 8.546625727 | -0.294111109 | 0.779496957 | 0.897498461 | -4.747040048 |
| **A_52_P157170** | Rnf157 | -0.056955451 | -0.542201738 | 0.428290836 | 8.264699885 | -0.293983815 | 0.779589214 | 0.897551233 | -4.747052233 |
| **A_66_P112593** | Sec61b | 0.056874313 | -0.428394072 | 0.542142698 | 11.43974617 | 0.293551641 | 0.779902466 | 0.897858418 | -4.747093564 |
| **A_52_P576049** | Inhbc | -0.024964078 | -0.23835309 | 0.188424933 | 7.660113335 | -0.293017682 | 0.780289558 | 0.898217874 | -4.74714455 |
| **A_52_P437230** | Rbx1 | 0.036794913 | -0.277801785 | 0.351391611 | 14.45207263 | 0.292943567 | 0.780343293 | 0.898217874 | -4.74715162 |
| **A_55_P2137591** | Rnf220 | -0.059306107 | -0.566399369 | 0.447787155 | 9.151021899 | -0.2929287 | 0.780354072 | 0.898217874 | -4.747153038 |

| | | | | | | | | | |
|---|---|---|---|---|---|---|---|---|---|
| **A_55_P1995243** | Cobl | 0.061948123 | -0.468091631 | 0.591987876 | 10.30077086 | 0.2927319 | 0.780496763 | 0.898328636 | -4.747171803 |
| **A_55_P2037862** | NA | 0.029130556 | -0.220173553 | 0.278434665 | 7.889040543 | 0.292664254 | 0.780545813 | 0.898331612 | -4.74717825 |
| **A_51_P328932** | Tmem165 | 0.034445224 | -0.260500771 | 0.329391218 | 12.30621671 | 0.292507395 | 0.780659555 | 0.898367223 | -4.747193194 |
| **A_55_P1980539** | Cetn4 | 0.106242045 | -0.803525963 | 1.016010052 | 9.484275633 | 0.292493423 | 0.780669686 | 0.898367223 | -4.747194525 |
| **A_55_P1957213** | NA | 0.103520996 | -0.783169719 | 0.990211711 | 12.47646643 | 0.292419699 | 0.780723147 | 0.898375272 | -4.747201546 |
| **A_55_P2183668** | NA | -0.126706891 | -1.213325376 | 0.959911594 | 12.86885588 | -0.292060949 | 0.780983315 | 0.898538365 | -4.747235686 |
| **A_52_P483129** | Utp23 | -0.043209364 | -0.413778608 | 0.32735988 | 9.385794121 | -0.292051103 | 0.780990455 | 0.898538365 | -4.747236623 |
| **A_55_P2370384** | 1600029I1 | 0.140543468 | -1.064854788 | 1.345941725 | 9.447294087 | 0.292032007 | 0.781004305 | 0.898538365 | -4.747238439 |
| **A_52_P537545** | Smpd3 | 0.03815122 | -0.289635314 | 0.365937754 | 8.197555469 | 0.291519537 | 0.781376009 | 0.898795517 | -4.747287135 |
| **A_66_P105046** | Il18 | 0.080010084 | -0.607450878 | 0.767471046 | 8.601250283 | 0.291505687 | 0.781386056 | 0.898795517 | -4.74728845 |
| **A_51_P110471** | Ddah1 | -0.025642831 | -0.246140394 | 0.194854731 | 8.477931887 | -0.291281251 | 0.781548866 | 0.898795517 | -4.740661068 |
| **A_55_P2155654** | Pvt1 | 0.025575281 | -0.194341867 | 0.245492428 | 8.04943498 | 0.291280667 | 0.781549289 | 0.898795517 | -4.747309805 |
| **A_55_P2021923** | Fkbp1a | 0.133907668 | -1.017820598 | 1.285635933 | 12.11439517 | 0.291209652 | 0.781600807 | 0.898795517 | -4.747316541 |
| **A_51_P453010** | Mphosph6 | 0.044400589 | -0.337529106 | 0.426330285 | 10.13243023 | 0.291176054 | 0.781625181 | 0.898795517 | -4.747319728 |
| **A_51_P326685** | Lrtm1 | 0.024896204 | -0.189304518 | 0.239096926 | 7.67333751 | 0.291113634 | 0.781670465 | 0.898795517 | -4.747325647 |
| **A_51_P136888** | Rb1 | -0.067871137 | -0.651869648 | 0.516127374 | 9.725598037 | -0.291087618 | 0.781689339 | 0.898795517 | -4.738335567 |
| **A_55_P2329680** | Dyrk2 | -0.03463672 | -0.332708596 | 0.263435155 | 9.501751082 | -0.291048997 | 0.781717358 | 0.898795517 | -4.747331775 |
| **A_55_P2167050** | Paip2 | 0.078461779 | -0.596798964 | 0.753722522 | 12.74369197 | 0.291029487 | 0.781731513 | 0.898795517 | -4.747333624 |
| **A_55_P2140002** | Tnfaip8 | -0.092837923 | -0.891852219 | 0.706176373 | 9.792060024 | -0.291018912 | 0.781739186 | 0.898795517 | -4.747334627 |
| **A_55_P1991199** | Nos1ap | -0.031381973 | -0.301570875 | 0.23880693 | 8.462543761 | -0.290912965 | 0.781816052 | 0.898830442 | -4.747344668 |
| **A_55_P2120161** | Myh8 | 0.026365154 | -0.200829869 | 0.253560178 | 7.721776073 | 0.290657712 | 0.782001254 | 0.898989907 | -4.747368844 |
| **A_51_P248666** | Cd274 | -0.136725136 | -1.315395776 | 1.041945504 | 8.269344939 | -0.290540208 | 0.782086517 | 0.899034468 | -4.747379967 |
| **A_51_P292447** | Klhl36 | 0.041641716 | -0.317608498 | 0.40089193 | 8.732660373 | 0.290323324 | 0.782243899 | 0.899161922 | -4.747400486 |
| **A_55_P2043996** | Cops7a | -0.024759143 | -0.238413157 | 0.18889487 | 8.272156955 | -0.290251789 | 0.78229581 | 0.899168135 | -4.74740725 |
| **A_55_P1962454** | NA | 0.038674114 | -0.295293165 | 0.372641392 | 8.25084896 | 0.290045944 | 0.782445196 | 0.899286376 | -4.747426707 |
| **A_65_P14951** | Cblb | 0.058868639 | -0.449665997 | 0.567403276 | 10.01303405 | 0.289943786 | 0.782519338 | 0.899318129 | -4.747436358 |
| **A_55_P2021991** | Zmym4 | -0.033286263 | -0.320986705 | 0.254414179 | 8.944549194 | -0.289784304 | 0.782635088 | 0.899350216 | -4.747451418 |
| **A_55_P2130448** | NA | -0.156110928 | -1.50544515 | 1.193223294 | 11.34479079 | -0.289777136 | 0.782640291 | 0.899350216 | -4.747452095 |
| **A_55_P2107247** | Tssk5 | -0.083296557 | -0.803691043 | 0.63709793 | 8.313604898 | -0.28960564 | 0.782764768 | 0.899439797 | -4.74746828 |
| **A_55_P2103952** | NA | -0.035734834 | -0.345033651 | 0.273563984 | 8.046632186 | -0.289376895 | 0.78293081 | 0.899577125 | -4.747489855 |
| **A_51_P248857** | Inka1 | -0.035010569 | -0.338267299 | 0.26824616 | 8.281441773 | -0.289160569 | 0.783087849 | 0.899662277 | -4.747510243 |
| **A_51_P138141** | Angel2 | 0.050479129 | -0.38678525 | 0.487743507 | 10.60599656 | 0.289146604 | 0.783097986 | 0.899662277 | -4.747511558 |
| **A_55_P2110027** | NA | 0.1358753 | -1.041501565 | 1.313252165 | 13.47320798 | 0.289051591 | 0.783166964 | 0.899687357 | -4.747520508 |
| **A_52_P90684** | Klhl31 | 0.035926761 | -0.275492207 | 0.34734573 | 7.87884501 | 0.288950436 | 0.783240402 | 0.899687357 | -4.747530033 |
| **A_55_P2150976** | NA | -0.051319642 | -0.49626245 | 0.393623166 | 15.11190826 | -0.288888181 | 0.783285601 | 0.899687357 | -4.747535894 |
| **A_66_P130960** | Psmd11 | 0.041455538 | -0.318000217 | 0.400911292 | 10.05936141 | 0.288860032 | 0.783306038 | 0.899687357 | -4.747538544 |
| **A_66_P117058** | Aasdh | -0.025465765 | -0.246325126 | 0.195393596 | 9.559401377 | -0.288796057 | 0.783352486 | 0.899687357 | -4.747544565 |
| **A_55_P2013655** | Cd4 | 0.118768224 | -0.912707587 | 1.150244035 | 9.737633706 | 0.288397545 | 0.783641845 | 0.899966228 | -4.747582041 |
| **A_55_P2019520** | Cma2 | 0.027654782 | -0.212575872 | 0.267885436 | 8.216354871 | 0.288331547 | 0.78368977 | 0.899967808 | -4.747588242 |
| **A_52_P44030** | Exoc3 | -0.041308511 | -0.40040148 | 0.317784458 | 8.584808484 | -0.288126356 | 0.783838778 | 0.899979941 | -4.747607515 |
| **A_55_P1976805** | Isca2 | -0.029575456 | -0.286673644 | 0.227522731 | 9.647435886 | -0.288126222 | 0.783838876 | 0.899979941 | -4.747607528 |
| **A_51_P202623** | Mterf2 | 0.034404422 | -0.264673345 | 0.33348219 | 9.424050885 | 0.288124697 | 0.783839983 | 0.899979941 | -4.747607671 |
| **A_55_P2055299** | NA | 0.04567313 | -0.351457662 | 0.442803923 | 8.416255741 | 0.288056422 | 0.783889566 | 0.899983424 | -4.747614081 |
| **A_55_P1958804** | Hsd3b3 | 0.024385229 | -0.187723722 | 0.236494181 | 7.684071165 | 0.287950737 | 0.783966319 | 0.900000705 | -4.747624 |
| **A_51_P369971** | Fra10ac1 | 0.028405723 | -0.218711607 | 0.275523052 | 10.77346769 | 0.287907504 | 0.783997717 | 0.900000705 | -4.747628057 |
| **A_51_P170619** | Ddx54 | 0.037676473 | -0.290287675 | 0.365640622 | 11.96881682 | 0.287736013 | 0.78412227 | 0.900090243 | -4.747644143 |
| **A_55_P1997429** | Gm13698 | -0.028255565 | -0.27435876 | 0.217847631 | 9.49904312 | -0.287565699 | 0.784245973 | 0.900123982 | -4.747660109 |
| **A_51_P463860** | Fam161b | 0.034320143 | -0.264693953 | 0.333334239 | 8.205115564 | 0.287480087 | 0.784308159 | 0.900123982 | -4.747668132 |

| | | | | | | | | | |
|---|---|---|---|---|---|---|---|---|---|
| **A_51_P407323** | F5 | 0.024537392 | -0.189268349 | 0.238343134 | 7.69896829 | 0.287448066 | 0.784331418 | 0.900123982 | -4.747671132 |
| **A_51_P149349** | Nfxl1 | -0.040921044 | -0.397533888 | 0.3156918 | 8.668039208 | -0.287408804 | 0.784359937 | 0.900123982 | -4.74767481 |
| **A_55_P2003789** | Taf4 | 0.025020012 | -0.193083333 | 0.243123357 | 10.02945664 | 0.2873264 | 0.784419796 | 0.900123982 | -4.747682528 |
| **A_52_P598714** | Afg3l1 | -0.050928034 | -0.494899482 | 0.393043414 | 9.969249063 | -0.287310974 | 0.784431001 | 0.900123982 | -4.747683973 |
| **A_55_P2008991** | Syp | -0.09118391 | -0.886552966 | 0.704185146 | 9.773840254 | -0.28714408 | 0.784552239 | 0.900209673 | -4.747699597 |
| **A_51_P345927** | Ddx41 | -0.029556536 | -0.28748023 | 0.228367159 | 8.759347873 | -0.287020313 | 0.784642151 | 0.900259412 | -4.747711178 |
| **A_51_P406306** | Hecw2 | 0.034975804 | -0.270338023 | 0.340289631 | 8.199319096 | 0.286927107 | 0.784709865 | 0.900283677 | -4.747719896 |
| **A_55_P1956517** | Polr2l | 0.050447149 | -0.390070284 | 0.490964582 | 14.27923622 | 0.286829537 | 0.784780751 | 0.900311579 | -4.74772902 |
| **A_55_P2100050** | NA | 0.062294266 | -0.482561686 | 0.607150218 | 10.13758854 | 0.286362878 | 0.78511982 | 0.900586241 | -4.747772616 |
| **A_55_P2000047** | 9930104L | -0.030173004 | -0.294091887 | 0.233745879 | 8.417899384 | -0.286350824 | 0.785128579 | 0.900586241 | -4.747773741 |
| **A_55_P1960703** | NA | -0.063447376 | -0.618495925 | 0.491601173 | 9.189754613 | -0.286307708 | 0.785159909 | 0.900586241 | -4.747777766 |
| **A_55_P2025790** | Fanci | 0.045415519 | -0.352251492 | 0.443082529 | 9.246672359 | 0.28604546 | 0.785350481 | 0.900751392 | -4.747802231 |
| **A_51_P439824** | Gipc1 | 0.055144222 | -0.428194333 | 0.538482777 | 10.49600405 | 0.285758344 | 0.785559144 | 0.90093727 | -4.747828992 |
| **A_51_P239036** | Ubn2 | -0.046340509 | -0.473805328 | 0.38112431 | 11.04372147 | -0.288750756 | 0.785656395 | 0.900995358 | -4.72016841 |
| **A_55_P2158350** | Gm15427 | 0.039613324 | -0.30793987 | 0.387166519 | 16.47726508 | 0.285476493 | 0.785764 | 0.901065313 | -4.747855238 |
| **A_52_P68477** | Zfp809 | 0.055212156 | -0.429363852 | 0.539788164 | 9.110908772 | 0.285379744 | 0.785834324 | 0.901092511 | -4.747864241 |
| **A_55_P2145696** | P3h4 | 0.035123408 | -0.273436028 | 0.343682844 | 11.93522296 | 0.285107185 | 0.78603245 | 0.901266243 | -4.74788959 |
| **A_51_P425798** | Mtus2 | -0.030019148 | -0.293885075 | 0.233846779 | 8.086233754 | -0.284947857 | 0.786148277 | 0.901309743 | -4.747904397 |
| **A_55_P1992004** | NA | 0.026561945 | -0.206957422 | 0.260081312 | 10.93386402 | 0.284896631 | 0.786185518 | 0.901309743 | -4.747909157 |
| **A_55_P2000823** | Rbm39 | 0.056586049 | -0.440949651 | 0.554121749 | 13.39151887 | 0.284862623 | 0.786210242 | 0.901309743 | -4.747912316 |
| **A_55_P2183498** | Spcs2 | -0.092151022 | -0.903713547 | 0.719411503 | 9.865914598 | -0.284399295 | 0.786547109 | 0.901588885 | -4.747955318 |
| **A_66_P123989** | Ikzf5 | 0.02631083 | -0.205449441 | 0.2580711 | 8.370848914 | 0.284345199 | 0.786586444 | 0.901588885 | -4.747960335 |
| **A_55_P2153541** | Sik3 | 0.033342737 | -0.26036866 | 0.327054135 | 12.08756355 | 0.284335312 | 0.786593633 | 0.901588885 | -4.747961252 |
| **A_51_P436447** | Zfp871 | 0.068506061 | -0.535295816 | 0.672307938 | 9.065605436 | 0.284174356 | 0.786710673 | 0.90166958 | -4.747976172 |
| **A_52_P334796** | Taar4 | -0.027817928 | -0.273289116 | 0.21765326 | 7.902693713 | -0.283840657 | 0.786953343 | 0.901894246 | -4.748007079 |
| **A_51_P158638** | Ggnbp1 | 0.044511784 | -0.348435365 | 0.437458934 | 8.126971435 | 0.28372082 | 0.787040496 | 0.901894624 | -4.74801817 |
| **A_51_P219061** | Oxa1l | 0.074163505 | -0.580903054 | 0.729230064 | 9.203442515 | 0.283566656 | 0.787152619 | 0.901894624 | -4.748032431 |
| **A_55_P2138306** | NA | -0.037164808 | -0.365454987 | 0.29112537 | 11.9984336 | -0.28354654 | 0.787167251 | 0.901894624 | -4.748034292 |
| **A_66_P128446** | Ttpa | 0.024814309 | -0.194431473 | 0.244060092 | 7.740447651 | 0.283479261 | 0.787216185 | 0.901894624 | -4.748040513 |
| **A_55_P2014326** | NA | -0.029357245 | -0.288757184 | 0.230042695 | 9.062805982 | -0.283462599 | 0.787228303 | 0.901894624 | -4.748042053 |
| **A_55_P2051039** | Gm11937 | -0.063754966 | -0.627106753 | 0.499596821 | 8.581114063 | -0.28345537 | 0.787233561 | 0.901894624 | -4.748042721 |
| **A_55_P2337138** | Ago2 | 0.051732935 | -0.405818359 | 0.509284229 | 11.16008571 | 0.283189847 | 0.787426696 | 0.902045734 | -4.748067258 |
| **A_55_P2055217** | NA | 0.106922035 | -0.838895828 | 1.052739898 | 10.65381454 | 0.283145752 | 0.787458772 | 0.902045734 | -4.748071331 |
| **A_55_P1953236** | Epb42 | 0.0241348 | -0.18943879 | 0.237708391 | 7.653879013 | 0.283039146 | 0.78753632 | 0.90208112 | -4.748081174 |
| **A_51_P103397** | Vwf | 0.055361159 | -0.43467665 | 0.545398967 | 11.09731336 | 0.282960575 | 0.787593477 | 0.902093145 | -4.748088427 |
| **A_55_P2115320** | Metap1d | -0.027445071 | -0.270485099 | 0.215594956 | 8.057325151 | -0.282837444 | 0.787683052 | 0.902116459 | -4.748099789 |
| **A_55_P1974028** | Stab2 | -0.076837918 | -0.757356358 | 0.603680522 | 9.703391493 | -0.282804319 | 0.787707151 | 0.902116459 | -4.748102845 |
| **A_55_P2106058** | Gm11110 | 0.032998976 | -0.259362935 | 0.325360888 | 8.164847423 | 0.282702742 | 0.787781049 | 0.902147652 | -4.748112213 |
| **A_51_P361448** | Scara5 | 0.057057662 | -0.448818766 | 0.562934091 | 8.597745996 | 0.282500927 | 0.78792788 | 0.902262356 | -4.748130818 |
| **A_55_P2167123** | Cyp4b1-p | 0.025567296 | -0.201228015 | 0.252362607 | 7.873167384 | 0.28235864 | 0.788031407 | 0.902327463 | -4.748143927 |
| **A_51_P141164** | Ddx50 | 0.027607005 | -0.217372519 | 0.272586529 | 9.802576824 | 0.282253835 | 0.788107665 | 0.90236134 | -4.748153578 |
| **A_51_P507172** | 2310022A | -0.042618678 | -0.420942474 | 0.335705118 | 10.44566436 | -0.282154292 | 0.788180097 | 0.902390832 | -4.748162742 |
| **A_55_P2020453** | NA | 0.031629847 | -0.249279386 | 0.31253908 | 8.499111809 | 0.282020987 | 0.788277099 | 0.902414291 | -4.74817501 |
| **A_55_P2159890** | Zbtb7a | 0.054570504 | -0.430117773 | 0.539258781 | 9.570817189 | 0.281997849 | 0.788293936 | 0.902414291 | -4.748177138 |
| **A_55_P2120080** | NA | -0.024408084 | -0.241466395 | 0.192650228 | 8.318469683 | -0.281648606 | 0.788548093 | 0.902651796 | -4.748209249 |
| **A_51_P244950** | Dpys | -0.02357533 | -0.233427073 | 0.186276413 | 7.66326474 | -0.281381504 | 0.788742492 | 0.902820872 | -4.748233782 |
| **A_55_P2067031** | Agxt2 | -0.024893968 | -0.246567552 | 0.196779615 | 7.733585376 | -0.281274614 | 0.788820293 | 0.902830369 | -4.748243593 |
| **A_55_P2075941** | Nf1 | 0.070510546 | -0.557438731 | 0.698459822 | 9.405864872 | 0.281241792 | 0.788844183 | 0.902830369 | -4.748246605 |

| | | | | | | | | | |
|---|---|---|---|---|---|---|---|---|---|
| **A_51_P409893** | Prkar2a | 0.028686266 | -0.226884808 | 0.28425734 | 8.062806252 | 0.281133546 | 0.788922974 | 0.902867099 | -4.748256536 |
| **A_52_P208521** | Khdc1a | -0.13142215 | -1.302970643 | 1.040126344 | 8.054647866 | -0.280969153 | 0.789042639 | 0.902894781 | -4.748271611 |
| **A_66_P110742** | Gm9949 | 0.030870666 | -0.244373019 | 0.306114351 | 7.890052363 | 0.280917624 | 0.789080149 | 0.902894781 | -4.748276335 |
| **A_55_P2326517** | Ube2i | -0.040969299 | -0.406268987 | 0.324330388 | 9.77205137 | -0.280905072 | 0.789089286 | 0.902894781 | -4.748277486 |
| **A_51_P224175** | 2210016F | -0.029931838 | -0.296897594 | 0.237033918 | 8.339223608 | -0.280820087 | 0.789151152 | 0.902894781 | -4.748285274 |
| **A_52_P105537** | Ccn3 | -0.027487093 | -0.272683259 | 0.217709074 | 8.179957637 | -0.280779552 | 0.789180661 | 0.902894781 | -4.748288988 |
| **A_55_P2036220** | Chadl | -0.079474725 | -0.788755179 | 0.629805729 | 9.825720232 | -0.280647636 | 0.789276696 | 0.902951221 | -4.748301071 |
| **A_55_P2095271** | Pkn3 | 0.095296943 | -0.755485247 | 0.946079133 | 9.263403953 | 0.28055042 | 0.789347472 | 0.90297876 | -4.748309973 |
| **A_55_P2144512** | NA | 0.041544899 | -0.329512129 | 0.412601927 | 8.309373468 | 0.280431887 | 0.78943377 | 0.902988811 | -4.748320822 |
| **A_51_P507787** | Sergef | 0.042989001 | -0.341054175 | 0.427032177 | 8.597076156 | 0.280367485 | 0.78948066 | 0.902988811 | -4.748326715 |
| **A_55_P2038952** | Ubb | -0.03828419 | -0.380451059 | 0.30388268 | 16.18837627 | -0.280241112 | 0.789572672 | 0.902988811 | -4.748338274 |
| **A_55_P2413458** | Tfcp2l1 | -0.056261773 | -0.559140238 | 0.446616692 | 8.615402437 | -0.280221026 | 0.789587297 | 0.902988811 | -4.748340111 |
| **A_52_P200265** | Mocs1 | 0.043380103 | -0.344397734 | 0.43115794 | 9.445296726 | 0.280193427 | 0.789607392 | 0.902988811 | -4.748342635 |
| **A_55_P2114908** | Nek3 | -0.025145057 | -0.249950575 | 0.19966046 | 8.898020825 | -0.28015347 | 0.789636486 | 0.902988811 | -4.748346288 |
| **A_55_P1995074** | Ubald2 | 0.032325476 | -0.256792697 | 0.321443649 | 14.45630496 | 0.28003988 | 0.789719197 | 0.903029983 | -4.748356671 |
| **A_55_P1969197** | NA | 0.050778125 | -0.403614103 | 0.505170354 | 16.85470644 | 0.279895628 | 0.789824237 | 0.903054745 | -4.748369851 |
| **A_52_P446457** | Ly6g6f | -0.023474606 | -0.233563015 | 0.186613803 | 7.675223225 | -0.279863699 | 0.789847489 | 0.903054745 | -4.748372767 |
| **A_55_P2015941** | NA | -0.025890731 | -0.257705886 | 0.205924424 | 8.271115607 | -0.27973888 | 0.789938384 | 0.903054745 | -4.748384165 |
| **A_55_P2171033** | NA | 0.046969641 | -0.373681351 | 0.467620632 | 8.775476834 | 0.279669861 | 0.789988647 | 0.903054745 | -4.748390466 |
| **A_51_P249608** | Trdn | 0.104350409 | -0.830298874 | 1.038999691 | 8.424408848 | 0.279637765 | 0.79001202 | 0.903054745 | -4.748393395 |
| **A_55_P2087384** | Gps1 | -0.030002819 | -0.298792129 | 0.238786491 | 13.35193661 | -0.279576338 | 0.790056755 | 0.903054745 | -4.748399001 |
| **A_52_P257523** | 1110065P | -0.050133406 | -0.499293046 | 0.399026233 | 11.74376092 | -0.27956116 | 0.790067809 | 0.903054745 | -4.748400386 |
| **A_55_P2068812** | Pkm | 0.056411696 | -0.449139955 | 0.561963347 | 14.93269974 | 0.279482083 | 0.7901254 | 0.903062561 | -4.7484076 |
| **A_55_P2036615** | Lrif1 | 0.029418255 | -0.234278215 | 0.293114724 | 9.221791936 | 0.279423503 | 0.790168064 | 0.903062561 | -4.748412943 |
| **A_52_P87210** | Dcun1d5 | 0.025539317 | -0.203631827 | 0.254710462 | 11.31984676 | 0.279125617 | 0.790385026 | 0.903230658 | -4.748440095 |
| **A_55_P2103884** | Supt4b | 0.063192174 | -0.504000848 | 0.630385196 | 11.85839303 | 0.279050471 | 0.790439762 | 0.903230658 | -4.748446941 |
| **A_51_P241995** | Col5a3 | -0.121611649 | -1.213242903 | 0.970019604 | 9.086656666 | -0.279029142 | 0.790455298 | 0.903230658 | -4.748448883 |
| **A_55_P2414524** | NA | 0.055034552 | -0.439430623 | 0.549499726 | 8.265350958 | 0.278772588 | 0.79064218 | 0.90339081 | -4.748472239 |
| **A_51_P416825** | Tspan15 | 0.060138478 | -0.481058352 | 0.601335308 | 8.408767678 | 0.278321998 | 0.790970442 | 0.903661274 | -4.748513209 |
| **A_51_P118885** | Amacr | -0.026286722 | -0.262859828 | 0.210286383 | 9.862495963 | -0.278305259 | 0.790982637 | 0.903661274 | -4.730395878 |
| **A_55_P2034260** | Ctdsp1 | -0.032573173 | -0.325775296 | 0.260628951 | 11.8155013 | -0.278255204 | 0.791019106 | 0.903661274 | -4.748519277 |
| **A_51_P353856** | Ubac2 | -0.055779444 | -0.558039286 | 0.446480397 | 8.959920729 | -0.278160895 | 0.791087819 | 0.903686375 | -4.748527842 |
| **A_55_P2015238** | Dctn4 | -0.029707134 | -0.297330308 | 0.237916041 | 11.76554063 | -0.278027258 | 0.79118519 | 0.903715033 | -4.748539975 |
| **A_51_P130332** | Hspa12b | 0.068122017 | -0.545634219 | 0.681878253 | 10.27023857 | 0.277998162 | 0.791206391 | 0.903715033 | -4.748542615 |
| **A_55_P2064243** | Rhot2 | -0.024458585 | -0.244873702 | 0.195956532 | 8.073622189 | -0.277933121 | 0.791253783 | 0.903715775 | -4.748548517 |
| **A_55_P2162324** | Bmx | 0.044564025 | -0.357199436 | 0.446327486 | 8.522977359 | 0.277820521 | 0.791335831 | 0.903756097 | -4.748558732 |
| **A_55_P2055557** | Sdsl | 0.029259065 | -0.234894911 | 0.293413041 | 9.949244171 | 0.277430137 | 0.791620317 | 0.904027597 | -4.748594116 |
| **A_51_P460332** | Apoc4 | 0.024358497 | -0.195625569 | 0.244342563 | 7.743322798 | 0.277338144 | 0.791687361 | 0.904050761 | -4.748602448 |
| **A_55_P1987156** | Crhr2 | -0.023817352 | -0.239044095 | 0.191409392 | 7.984352458 | -0.277170866 | 0.791809276 | 0.904136579 | -4.74861759 |
| **A_55_P1996086** | Gm14326 | 0.032691806 | -0.262804259 | 0.328187871 | 8.744501917 | 0.277100657 | 0.791860448 | 0.904141611 | -4.748623943 |
| **A_51_P515242** | Strip2 | -0.037091297 | -0.372682101 | 0.298499507 | 8.443023116 | -0.276829466 | 0.792058116 | 0.904200898 | -4.748648468 |
| **A_55_P2186961** | NA | 0.026677484 | -0.214694595 | 0.268049562 | 8.972327511 | 0.27682683 | 0.792060037 | 0.904200898 | -4.748648706 |
| **A_55_P2032294** | Zfp932 | 0.026529671 | -0.213538316 | 0.266597659 | 9.241904168 | 0.276788448 | 0.792088015 | 0.904200898 | -4.748652176 |
| **A_55_P1976958** | NA | 0.03166087 | -0.254947041 | 0.318268781 | 9.908418221 | 0.276684617 | 0.792163702 | 0.904200898 | -4.748661558 |
| **A_52_P15197** | Dcun1d2 | -0.035045718 | -0.352304965 | 0.28221353 | 10.65410627 | -0.276675697 | 0.792170205 | 0.904200898 | -4.748662364 |
| **A_55_P2106358** | NA | 0.031426501 | -0.253146983 | 0.315999984 | 8.00558492 | 0.27659985 | 0.792225494 | 0.904200898 | -4.748669215 |
| **A_55_P1977776** | Rs5-8s1 | -0.224211489 | -2.255999552 | 1.807576575 | 10.83878023 | -0.276394759 | 0.792375005 | 0.904200898 | -4.748687733 |
| **A_55_P2006915** | Myo5c | 0.178914797 | -1.442997523 | 1.800827118 | 9.342903795 | 0.276292587 | 0.792449492 | 0.904200898 | -4.748696953 |

| | | | | | | | | | |
|---|---|---|---|---|---|---|---|---|---|
| **A_51_P259214** | Slc39a6 | -0.087715627 | -0.882887282 | 0.707456028 | 10.51196437 | -0.276290801 | 0.792450794 | 0.904200898 | -4.748697114 |
| **A_55_P2033650** | Cpt2 | -0.043757679 | -0.440629866 | 0.353114508 | 11.43490596 | -0.276155665 | 0.792549317 | 0.904200898 | -4.748709304 |
| **A_51_P354165** | Apcs | 0.023178252 | -0.187162643 | 0.233519147 | 7.695103102 | 0.275998873 | 0.792663634 | 0.904200898 | -4.74872344 |
| **A_55_P2105833** | Chchd5 | -0.031492859 | -0.317359297 | 0.254373578 | 10.42763067 | -0.275930222 | 0.792713688 | 0.904200898 | -4.748729627 |
| **A_51_P181451** | C1qa | 0.030891254 | -0.24951604 | 0.311298548 | 8.719891573 | 0.275928511 | 0.792714936 | 0.904200898 | -4.748729781 |
| **A_51_P357735** | Inhbe | -0.030165964 | -0.304014803 | 0.243682874 | 7.860727352 | -0.275903149 | 0.792733428 | 0.904200898 | -4.748732066 |
| **A_55_P2032034** | Tbp | 0.032296262 | -0.260910715 | 0.32550324 | 9.26400473 | 0.275885139 | 0.79274656 | 0.904200898 | -4.748733689 |
| **A_55_P2110497** | Ddc | -0.108233823 | -1.090869862 | 0.874402216 | 10.13521461 | -0.275880278 | 0.792750104 | 0.904200898 | -4.748734127 |
| **A_55_P2038207** | Zfp687 | -0.034256944 | -0.345274172 | 0.276760285 | 9.88646322 | -0.275876377 | 0.792752949 | 0.904200898 | -4.748734478 |
| **A_51_P203474** | Tsg101 | -0.038151098 | -0.384591338 | 0.308289142 | 10.48859197 | -0.275822098 | 0.792792527 | 0.904200898 | -4.748739368 |
| **A_51_P219028** | Wdr53 | 0.023896579 | -0.193209167 | 0.241002324 | 10.74050784 | 0.275686026 | 0.792891747 | 0.904200898 | -4.748751622 |
| **A_55_P1965308** | Ccdc66 | -0.037120769 | -0.374375255 | 0.300133717 | 8.58933248 | -0.275682738 | 0.792894144 | 0.904200898 | -4.748751918 |
| **A_51_P480499** | Emc8 | 0.039828174 | -0.322024605 | 0.401680953 | 10.52183259 | 0.275682267 | 0.792894487 | 0.904200898 | -4.748751961 |
| **A_51_P105408** | Alg14 | -0.028922877 | -0.291771309 | 0.233925555 | 10.39203825 | -0.275604591 | 0.792951129 | 0.904201217 | -4.737563808 |
| **A_51_P164630** | Fitm1 | 0.032304618 | -0.261450933 | 0.326060169 | 7.804358863 | 0.275441181 | 0.793070292 | 0.904201217 | -4.748773658 |
| **A_51_P351481** | Ccnl1 | -0.037724528 | -0.38078832 | 0.305339263 | 9.160325157 | -0.275422415 | 0.793083977 | 0.904201217 | -4.748775346 |
| **A_66_P100220** | NA | -0.027714589 | -0.279762614 | 0.224333436 | 8.084889984 | -0.275407326 | 0.793094981 | 0.904201217 | -4.748776703 |
| **A_55_P2086983** | Atxn2 | -0.062400881 | -0.629995529 | 0.505193767 | 11.64572545 | -0.275361219 | 0.793128605 | 0.904201217 | -4.74878085 |
| **A_55_P2038972** | Erich1 | 0.024888747 | -0.201593188 | 0.251370682 | 8.408773119 | 0.275245235 | 0.793213189 | 0.904244328 | -4.748791279 |
| **A_52_P641367** | Desi1 | 0.083358572 | -0.675446437 | 0.842163581 | 8.94638849 | 0.275150573 | 0.793282227 | 0.904269711 | -4.748799788 |
| **A_52_P495031** | 9130401M | 0.035903954 | -0.291106299 | 0.362914207 | 11.88082027 | 0.274999088 | 0.79339271 | 0.904342333 | -4.748813398 |
| **A_55_P2035717** | Pgap2 | 0.057700705 | -0.467986218 | 0.583387629 | 11.51587371 | 0.274918708 | 0.793451335 | 0.90435584 | -4.748820617 |
| **A_55_P2011702** | Ppargc1a | 0.029386523 | -0.238552694 | 0.29732574 | 7.822785967 | 0.274702283 | 0.793609194 | 0.904482443 | -4.748840044 |
| **A_55_P1965333** | Naa60 | -0.035787161 | -0.362320802 | 0.290746479 | 11.45469359 | -0.274504623 | 0.793753376 | 0.904491871 | -4.748857774 |
| **A_51_P286357** | Adgre4 | -0.023775911 | -0.240746147 | 0.193194326 | 7.739309422 | -0.274465234 | 0.793782109 | 0.904491871 | -4.748861306 |
| **A_52_P344978** | NA | -0.041062812 | -0.415818432 | 0.333692808 | 8.692957889 | -0.274442188 | 0.793798921 | 0.904491871 | -4.748863372 |
| **A_55_P2099840** | NA | 0.025604825 | -0.208081659 | 0.259291309 | 7.843439579 | 0.274434408 | 0.793804597 | 0.904491871 | -4.748864069 |
| **A_55_P2040860** | Sun3 | -0.023166532 | -0.234812073 | 0.188479009 | 7.696058938 | -0.274158835 | 0.794005631 | 0.90459904 | -4.748888762 |
| **A_55_P2169198** | NA | -0.025195493 | -0.255413044 | 0.205022058 | 7.842123592 | -0.274116248 | 0.7940367 | 0.90459904 | -4.748892576 |
| **A_52_P182592** | Ube2w | 0.040449232 | -0.329212727 | 0.410111191 | 9.201820703 | 0.274066445 | 0.794073034 | 0.90459904 | -4.748897035 |
| **A_51_P491635** | Ly6g6e | -0.074238467 | -0.752850916 | 0.604373981 | 8.734053755 | -0.274004389 | 0.794118309 | 0.90459904 | -4.74890259 |
| **A_55_P2067221** | Iqsec3 | 0.114838083 | -0.935062301 | 1.164738467 | 11.29124508 | 0.273960648 | 0.794150222 | 0.90459904 | -4.748906505 |
| **A_55_P2040490** | NA | -0.035914258 | -0.364306144 | 0.292477628 | 8.65182432 | -0.273920685 | 0.794179379 | 0.90459904 | -4.748910082 |
| **A_55_P2440441** | Wdhd1 | 0.029069238 | -0.236853619 | 0.294992095 | 8.553382326 | 0.273796773 | 0.794269787 | 0.904648723 | -4.748921168 |
| **A_51_P121818** | Prkag1 | -0.027358356 | -0.277819888 | 0.223103176 | 11.0081865 | -0.273589427 | 0.794421078 | 0.904767738 | -4.748939708 |
| **A_55_P1990864** | NA | -0.023566735 | -0.239407873 | 0.192274403 | 7.723645263 | -0.273473685 | 0.794505534 | 0.904803222 | -4.748950051 |
| **A_55_P2232988** | Scube2 | -0.105391481 | -1.070837548 | 0.860054587 | 10.46670871 | -0.273418461 | 0.794545832 | 0.904803222 | -4.748954985 |
| **A_51_P336827** | Cyb5b | 0.034914928 | -0.285203074 | 0.355032931 | 9.201564412 | 0.273181574 | 0.794718699 | 0.904946777 | -4.748976137 |
| **A_51_P456721** | Azgp1 | 0.024635141 | -0.201359172 | 0.250629454 | 7.744898592 | 0.273028437 | 0.794830457 | 0.905020734 | -4.748989802 |
| **A_55_P1958808** | Stoml2 | 0.029543877 | -0.241631554 | 0.300719309 | 12.1594736 | 0.272877355 | 0.794940721 | 0.90509298 | -4.749003276 |
| **A_55_P2079568** | NA | 0.033677813 | -0.275889002 | 0.343244627 | 8.574563912 | 0.272483259 | 0.795228368 | 0.905367169 | -4.74903839 |
| **A_55_P2041310** | NA | -0.048955513 | -0.499346656 | 0.40143563 | 8.483701682 | -0.272246379 | 0.795401282 | 0.905414973 | -4.749059473 |
| **A_51_P421984** | Timm22 | 0.023491852 | -0.192708417 | 0.23969212 | 11.6705643 | 0.272151897 | 0.795470254 | 0.905414973 | -4.749067877 |
| **A_51_P117477** | Slc27a1 | -0.042462781 | -0.433281328 | 0.348355766 | 8.794302899 | -0.272134506 | 0.79548295 | 0.905414973 | -4.749069423 |
| **A_65_P09285** | 2610524H | -0.044308471 | -0.452207406 | 0.363590465 | 9.260326952 | -0.272072429 | 0.795528267 | 0.905414973 | -4.749074943 |
| **A_55_P1961894** | Fbxw11 | -0.029504304 | -0.301119733 | 0.242111126 | 12.72518956 | -0.272070391 | 0.795529755 | 0.905414973 | -4.749075124 |
| **A_55_P1967867** | Xpa | 0.038362162 | -0.314836727 | 0.39156105 | 9.061974165 | 0.272040828 | 0.795551337 | 0.905414973 | -4.749077753 |
| **A_55_P2066523** | Akt2 | -0.040774684 | -0.416382038 | 0.334832671 | 8.363230617 | -0.271898529 | 0.795655223 | 0.905479903 | -4.7490904 |

| | | | | | | | | | |
|---|---|---|---|---|---|---|---|---|---|
| **A_51_P283754** | Cpsf7 | 0.089534533 | -0.735855383 | 0.914924449 | 11.86775605 | 0.271695071 | 0.795803767 | 0.905595646 | -4.749108472 |
| **A_55_P2084512** | NA | -0.023752628 | -0.242981075 | 0.195475818 | 7.897461216 | -0.271372049 | 0.796039624 | 0.905810728 | -4.749137138 |
| **A_55_P1979863** | Syne2 | 0.145263329 | -1.196471025 | 1.486997684 | 12.0743787 | 0.27116885 | 0.796188003 | 0.90592625 | -4.749155153 |
| **A_51_P243857** | Copb2 | -0.043818508 | -0.449412766 | 0.361775751 | 12.33818189 | -0.270592735 | 0.796608744 | 0.906351642 | -4.749206162 |
| **A_51_P309158** | Snx20 | -0.034975747 | -0.358855165 | 0.288903671 | 9.069602165 | -0.270479276 | 0.796691612 | 0.906392588 | -4.749216195 |
| **A_66_P120249** | Nr1h3 | 0.102763694 | -0.849069172 | 1.05459656 | 10.63605345 | 0.270414115 | 0.796739207 | 0.906393399 | -4.749221956 |
| **A_55_P2098771** | Taf13 | 0.02572498 | -0.212782228 | 0.264232188 | 10.35235003 | 0.270149324 | 0.796932622 | 0.906469434 | -4.74924535 |
| **A_55_P2144386** | Adgre1 | 0.042538871 | -0.351857833 | 0.436935575 | 8.647009669 | 0.270148791 | 0.796933011 | 0.906469434 | -4.749245398 |
| **A_55_P2175206** | Rbbp6 | -0.026105722 | -0.268160218 | 0.215948773 | 8.567111427 | -0.270130056 | 0.796946697 | 0.906469434 | -4.749247052 |
| **A_55_P2026420** | Pou6f1 | 0.060475768 | -0.500417186 | 0.621368723 | 8.51389335 | 0.270054709 | 0.797001737 | 0.906478709 | -4.749253705 |
| **A_55_P1982019** | Rbfox2 | 0.043747658 | -0.362253977 | 0.449749292 | 10.54176428 | 0.269884142 | 0.797126339 | 0.906567096 | -4.749268758 |
| **A_55_P2034853** | 2610028H | 0.079219665 | -0.656313269 | 0.814752598 | 9.55667984 | 0.269762279 | 0.797215366 | 0.906615016 | -4.749279507 |
| **A_51_P278034** | Ufsp1 | -0.095829875 | -0.986109749 | 0.794449998 | 8.676069082 | -0.269602996 | 0.797331736 | 0.906694023 | -4.74929355 |
| **A_55_P1987534** | Tnrc6b | 0.035373283 | -0.29333623 | 0.364082795 | 8.597118533 | 0.269533925 | 0.7973822 | 0.90669808 | -4.749299638 |
| **A_51_P271865** | Clec4f | 0.022727132 | -0.188566235 | 0.2340205 | 7.685127839 | 0.269407153 | 0.797474823 | 0.906750072 | -4.749310806 |
| **A_55_P2067142** | NA | 0.026481863 | -0.219816743 | 0.27278047 | 17.15272751 | 0.269300359 | 0.797552853 | 0.906785467 | -4.74932021 |
| **A_55_P1976978** | Zfp707 | 0.039189712 | -0.325397727 | 0.403777152 | 8.945700242 | 0.269228305 | 0.797605502 | 0.906792001 | -4.749326554 |
| **A_52_P420504** | Acta2 | 0.107670887 | -0.894933846 | 1.110275619 | 13.86157815 | 0.268979316 | 0.797787442 | 0.906816585 | -4.749348461 |
| **A_55_P2045407** | Ppp4r1 | -0.032376392 | -0.333870972 | 0.269118188 | 12.09454494 | -0.268967177 | 0.797796312 | 0.906816585 | -4.749349528 |
| **A_55_P2045952** | Mysm1 | 0.032446888 | -0.269728285 | 0.334622061 | 9.079097357 | 0.268945709 | 0.797812 | 0.906816585 | -4.749351416 |
| **A_52_P623751** | Fem1b | -0.046252817 | -0.477007456 | 0.384501822 | 12.54455061 | -0.268941965 | 0.797814736 | 0.906816585 | -4.749351745 |
| **A_55_P1963483** | Vps16 | -0.034365924 | -0.354511642 | 0.285779794 | 11.87039282 | -0.268862774 | 0.797872606 | 0.90682905 | -4.749358708 |
| **A_55_P2091297** | NA | 0.024613595 | -0.204859928 | 0.254087118 | 7.849190835 | 0.268653689 | 0.798025404 | 0.906858243 | -4.749377082 |
| **A_55_P2058384** | NA | 0.046954001 | -0.390933187 | 0.484841189 | 8.908787032 | 0.268571987 | 0.798085115 | 0.906858243 | -4.749384257 |
| **A_55_P2016049** | Gp6 | -0.025631172 | -0.264688514 | 0.213426171 | 7.834101784 | -0.268544781 | 0.798104998 | 0.906858243 | -4.749386647 |
| **A_51_P285077** | Hhatl | -0.023758844 | -0.245374473 | 0.197856786 | 7.797504252 | -0.268519146 | 0.798123733 | 0.906858243 | -4.749388897 |
| **A_55_P2326582** | 5730408K | -0.027257499 | -0.281529981 | 0.227014983 | 7.814700747 | -0.268495489 | 0.798141023 | 0.906858243 | -4.749390974 |
| **A_51_P209736** | Atoh8 | 0.059684658 | -0.497233289 | 0.616602606 | 8.43238272 | 0.268424309 | 0.798193045 | 0.906858243 | -4.749397223 |
| **A_55_P1959379** | BC080695 | -0.040311682 | -0.416754093 | 0.336130729 | 8.219102408 | -0.268214788 | 0.798346184 | 0.906858243 | -4.749415606 |
| **A_55_P2011887** | Kif13a | -0.031465915 | -0.325384741 | 0.26245291 | 11.03375814 | -0.268141054 | 0.798400078 | 0.906858243 | -4.749422072 |
| **A_55_P2092776** | Apoo | 0.027073826 | -0.225848147 | 0.279995799 | 9.68522524 | 0.268110254 | 0.798422591 | 0.906858243 | -4.749424773 |
| **A_51_P300337** | Csrp2 | -0.067459177 | -0.697714453 | 0.562796098 | 13.72537776 | -0.26808647 | 0.798439976 | 0.906858243 | -4.749426858 |
| **A_55_P1962661** | Cyp2c67 | 0.024192733 | -0.201867705 | 0.250253172 | 7.748981732 | 0.268046857 | 0.798468931 | 0.906858243 | -4.74943033 |
| **A_55_P2101441** | Gpbar1 | -0.033530848 | -0.346930001 | 0.279868304 | 7.957110371 | -0.267976722 | 0.798520197 | 0.906858243 | -4.749436477 |
| **A_55_P1974189** | Ptges | 0.096395065 | -0.804668792 | 0.997458921 | 8.574694037 | 0.267947417 | 0.798541618 | 0.906858243 | -4.749439045 |
| **A_55_P2106525** | Nmnat3 | 0.046504914 | -0.388294992 | 0.48130482 | 9.705722495 | 0.267892003 | 0.798582125 | 0.906858243 | -4.7494439 |
| **A_55_P2049597** | Tpp2 | -0.023234894 | -0.240492541 | 0.194022753 | 10.06528076 | -0.267865001 | 0.798601864 | 0.906858243 | -4.749446265 |
| **A_51_P283649** | Brat1 | -0.033773298 | -0.349803253 | 0.282256657 | 10.34209162 | -0.267667456 | 0.798746274 | 0.906968959 | -4.749463564 |
| **A_52_P1130990** | Trim67 | 0.091710178 | -0.767055267 | 0.950475623 | 8.362525751 | 0.267481236 | 0.798882413 | 0.907070271 | -4.749479859 |
| **A_55_P1965363** | Serpinb6e | -0.029492622 | -0.305789401 | 0.246804156 | 7.837321772 | -0.267354754 | 0.798974884 | 0.907120603 | -4.749490921 |
| **A_52_P548202** | NA | -0.022557169 | -0.23398396 | 0.188869623 | 7.703123951 | -0.267223662 | 0.79907073 | 0.907120603 | -4.749502381 |
| **A_55_P2154690** | Smarca4 | 0.028071337 | -0.235065302 | 0.291207976 | 8.978810708 | 0.267197333 | 0.79908998 | 0.907120603 | -4.749504682 |
| **A_55_P2136340** | NA | -0.030686368 | -0.318371912 | 0.256999176 | 14.32123155 | -0.267163912 | 0.799114416 | 0.907120603 | -4.749507602 |
| **A_55_P2065874** | D430042C | -0.028006863 | -0.290779974 | 0.234766248 | 9.032874491 | -0.266952435 | 0.799269044 | 0.907220307 | -4.749526073 |
| **A_51_P232439** | Ice1 | 0.030223108 | -0.253383128 | 0.313829345 | 10.07053969 | 0.266915441 | 0.799296095 | 0.907220307 | -4.749529303 |
| **A_55_P2123220** | Slc35e1 | -0.092046304 | -0.956107783 | 0.772015175 | 9.851409126 | -0.266816115 | 0.799368725 | 0.907249482 | -4.749537973 |
| **A_55_P1966843** | Gm12070 | -0.047608154 | -0.495186754 | 0.399970447 | 16.03823098 | -0.266417269 | 0.799660395 | 0.907482682 | -4.749572755 |
| **A_55_P2140780** | Clip4 | 0.073528391 | -0.617788774 | 0.764845557 | 8.339560367 | 0.266396227 | 0.799675783 | 0.907482682 | -4.749574589 |

| | | | | | | | | | |
|---|---|---|---|---|---|---|---|---|---|
| **A_55_P2076683** | Sort1 | 0.035700799 | -0.30010034 | 0.371501937 | 8.20937423 | 0.266284639 | 0.799757393 | 0.907482682 | -4.74958431 |
| **A_51_P397375** | Gatb | 0.048798477 | -0.410209934 | 0.507806888 | 9.26291782 | 0.266278417 | 0.799761943 | 0.907482682 | -4.749584852 |
| **A_55_P2033075** | NA | 0.034124144 | -0.286983445 | 0.355231734 | 16.2635853 | 0.266171496 | 0.799840143 | 0.907518152 | -4.749594163 |
| **A_51_P445841** | Deptor | 0.036896812 | -0.310499066 | 0.384292689 | 8.247789051 | 0.266020143 | 0.799950843 | 0.907590493 | -4.749607338 |
| **A_52_P494686** | Kansl1l | 0.056307397 | -0.474191661 | 0.586806454 | 8.431263437 | 0.265846641 | 0.800077749 | 0.907681212 | -4.749622431 |
| **A_52_P192307** | Rbak | 0.061178716 | -0.515547148 | 0.637904581 | 9.282845028 | 0.265693713 | 0.800189613 | 0.907754854 | -4.749635727 |
| **A_51_P419300** | Pdcd2l | 0.040161815 | -0.338802088 | 0.419125719 | 11.5143978 | 0.265439673 | 0.800375448 | 0.907912398 | -4.749657798 |
| **A_51_P328133** | Gle1 | -0.029936031 | -0.312544189 | 0.252672128 | 8.397261849 | -0.265313814 | 0.800467523 | 0.907951877 | -4.749668725 |
| **A_55_P2209046** | NA | 0.034491266 | -0.291181643 | 0.360164175 | 8.133715817 | 0.265263716 | 0.800504174 | 0.907951877 | -4.749673073 |
| **A_51_P371675** | Slc7a6os | -0.048249943 | -0.504110523 | 0.407610638 | 9.207799647 | -0.265103284 | 0.800621548 | 0.907967209 | -4.749686992 |
| **A_52_P627269** | Ces2b | -0.028554443 | -0.298342908 | 0.241234021 | 8.143904858 | -0.265094471 | 0.800627995 | 0.907967209 | -4.749687756 |
| **A_55_P2110507** | NA | -0.026590651 | -0.277864409 | 0.224683106 | 7.984516652 | -0.265052671 | 0.800658578 | 0.907967209 | -4.749691381 |
| **A_55_P2006261** | Krt15 | 0.177222404 | -1.499902435 | 1.854347243 | 11.42720742 | 0.264669306 | 0.80093908 | 0.908172167 | -4.749724603 |
| **A_55_P2047867** | NA | -0.04609235 | -0.482314167 | 0.390129467 | 10.48589145 | -0.264649949 | 0.800953244 | 0.908172167 | -4.749726279 |
| **A_52_P402705** | Kcnj11 | 0.02389023 | -0.202274619 | 0.250055079 | 7.708535474 | 0.26457303 | 0.801009528 | 0.908172167 | -4.749732938 |
| **A_55_P2184526** | NA | 0.028688013 | -0.242961012 | 0.300337039 | 8.023506671 | 0.264510348 | 0.801055396 | 0.908172167 | -4.749738364 |
| **A_66_P115949** | Myo9a | 0.026865319 | -0.227549157 | 0.281279794 | 8.660649074 | 0.264484683 | 0.801074177 | 0.908172167 | -4.749740585 |
| **A_55_P2335648** | Eprn | -0.024323939 | -0.255038075 | 0.206390197 | 8.01419111 | -0.26406452 | 0.801381659 | 0.908433321 | -4.749776918 |
| **A_51_P196509** | Plekhj1 | 0.025186265 | -0.213727916 | 0.264100447 | 8.589712161 | 0.2640415 | 0.801398506 | 0.908433321 | -4.749778907 |
| **A_55_P1953650** | Zdhhc18 | -0.028697861 | -0.30099589 | 0.243600168 | 12.70190547 | -0.263970487 | 0.801450479 | 0.908438973 | -4.749785042 |
| **A_55_P2094352** | Zfp523 | 0.027791418 | -0.23603063 | 0.291613465 | 8.111613413 | 0.263845663 | 0.801541837 | 0.908489265 | -4.749795822 |
| **A_55_P2043668** | Zfp617 | 0.072721874 | -0.618297515 | 0.763741264 | 9.585219596 | 0.263587722 | 0.801730635 | 0.908636714 | -4.749818082 |
| **A_55_P2067181** | Usp21 | 0.052961152 | -0.450448385 | 0.556370689 | 9.841976198 | 0.263503463 | 0.801792311 | 0.908636714 | -4.749825349 |
| **A_55_P2007457** | Muc5b | 0.249975004 | -2.126397528 | 2.626347537 | 9.651997799 | 0.263470749 | 0.801816257 | 0.908636714 | -4.74982817 |
| **A_55_P1988113** | Cyth2 | 0.034606711 | -0.294454426 | 0.363667849 | 10.56223719 | 0.263411106 | 0.801859916 | 0.908636714 | -4.749833312 |
| **A_55_P1957937** | NA | 0.022946521 | -0.19533282 | 0.241225862 | 7.82461978 | 0.263302246 | 0.801939604 | 0.908673757 | -4.749842694 |
| **A_55_P1971174** | Cd1d2 | -0.024279887 | -0.255330976 | 0.206771202 | 8.074714364 | -0.263201883 | 0.802013074 | 0.908686342 | -4.749851341 |
| **A_52_P273120** | Myl1 | 0.055503293 | -0.472761193 | 0.583767779 | 8.045629387 | 0.263158668 | 0.802044711 | 0.908686342 | -4.749855063 |
| **A_66_P129672** | 4930453N | 0.028729507 | -0.245027525 | 0.30248654 | 10.50230879 | 0.262853182 | 0.802268358 | 0.908817023 | -4.749881358 |
| **A_51_P206585** | Runx1 | 0.042941213 | -0.366240312 | 0.452122739 | 8.597923678 | 0.262850399 | 0.802270395 | 0.908817023 | -4.749881598 |
| **A_55_P1993054** | Psip1 | 0.039383476 | -0.336054976 | 0.414821928 | 9.266442223 | 0.262739663 | 0.80235147 | 0.908817023 | -4.749891122 |
| **A_52_P401640** | Bpgm | -0.034564878 | -0.364078465 | 0.294948708 | 10.99087843 | -0.262731444 | 0.802357488 | 0.908817023 | -4.749891829 |
| **A_55_P2335529** | 4921531C | -0.026358416 | -0.277687352 | 0.22497052 | 8.359629547 | -0.262680093 | 0.802395085 | 0.908817023 | -4.749896244 |
| **A_52_P42194** | NA | -0.037104633 | -0.391107961 | 0.316898695 | 8.111010118 | -0.262525285 | 0.802508435 | 0.908849498 | -4.74990955 |
| **A_55_P2071354** | Togaram2 | 0.097487236 | -0.832659114 | 1.027633586 | 8.578395824 | 0.262510583 | 0.8025192 | 0.908849498 | -4.749910814 |
| **A_55_P2156068** | NA | 0.049155433 | -0.419958198 | 0.518269063 | 9.042625086 | 0.262448332 | 0.802564781 | 0.908849498 | -4.749916162 |
| **A_52_P250590** | Dnajc13 | -0.022842881 | -0.240901017 | 0.195215255 | 9.355660253 | -0.262378916 | 0.80261561 | 0.908853825 | -4.749922124 |
| **A_66_P106836** | Kmo | 0.022024542 | -0.188390914 | 0.232439999 | 7.802484574 | 0.26216795 | 0.802770093 | 0.908960782 | -4.749940235 |
| **A_55_P2102471** | Gm16378 | -0.080147821 | -0.845990177 | 0.685694535 | 13.43444446 | -0.262121523 | 0.802804092 | 0.908960782 | -4.749944219 |
| **A_55_P2091191** | Slc28a2 | -0.049902183 | -0.527556358 | 0.427751992 | 7.849579208 | -0.261671433 | 0.803133715 | 0.909263204 | -4.749982804 |
| **A_51_P365409** | Rpusd4 | -0.03727968 | -0.394244923 | 0.319685563 | 10.20224787 | -0.261575213 | 0.803204188 | 0.909263204 | -4.749991045 |
| **A_55_P1968568** | Rapgef5 | 0.046152678 | -0.395880421 | 0.488185777 | 9.074920255 | 0.261512507 | 0.803250115 | 0.909263204 | -4.749996414 |
| **A_55_P2056070** | Hfe | 0.065521283 | -0.562082629 | 0.693125196 | 9.556476631 | 0.261485192 | 0.803270122 | 0.909263204 | -4.749998752 |
| **A_66_P127567** | Nxf1 | -0.030583899 | -0.323667165 | 0.262499366 | 13.10092786 | -0.261367859 | 0.803356063 | 0.909263204 | -4.750008794 |
| **A_55_P2092953** | Traf5 | 0.074241483 | -0.637287574 | 0.78577054 | 9.314777433 | 0.261339071 | 0.803377149 | 0.909263204 | -4.750011257 |
| **A_55_P2425938** | 2810468N | -0.024606308 | -0.260470359 | 0.211257743 | 8.011350883 | -0.261297373 | 0.803407692 | 0.909263204 | -4.750014824 |
| **A_51_P480881** | Thap11 | 0.026078868 | -0.223952337 | 0.276110073 | 11.88958222 | 0.261243125 | 0.803447429 | 0.909263204 | -4.750019464 |
| **A_52_P541161** | Rgs18 | 0.022037556 | -0.189367458 | 0.23344257 | 7.813269249 | 0.26109496 | 0.803555961 | 0.909290684 | -4.750032132 |

| | | | | | | | | | |
|---|---|---|---|---|---|---|---|---|---|
| **A_55_P2401646** | Trdn | 0.0468827 | -0.402883432 | 0.496648833 | 7.98553036 | 0.261081568 | 0.803565772 | 0.909290684 | -4.750033277 |
| **A_55_P2085915** | Tnfrsf14 | 0.022301431 | -0.191878845 | 0.236481706 | 7.712907826 | 0.260797605 | 0.803773793 | 0.909472845 | -4.750057536 |
| **A_52_P545643** | Tmem263 | -0.072851736 | -0.772834943 | 0.62713147 | 11.34869576 | -0.260676951 | 0.803862185 | 0.909498554 | -4.750067835 |
| **A_51_P336060** | Lcat | -0.021845417 | -0.231794226 | 0.188103392 | 7.657150837 | -0.260613711 | 0.803908517 | 0.909498554 | -4.750073232 |
| **A_66_P118256** | Strn3 | -0.030112746 | -0.319655122 | 0.259429629 | 11.22742298 | -0.260488515 | 0.804000242 | 0.909498554 | -4.750083912 |
| **A_55_P2028804** | Uchl3 | 0.061299673 | -0.528161394 | 0.65076074 | 8.299126166 | 0.260467401 | 0.804015711 | 0.909498554 | -4.750085713 |
| **A_55_P2039637** | Ift88 | 0.056429733 | -0.486247252 | 0.599106719 | 10.70317357 | 0.26044555 | 0.804031721 | 0.909498554 | -4.750087576 |
| **A_55_P2224830** | D230044E | 0.042418289 | -0.366092893 | 0.450929471 | 8.518503342 | 0.26007556 | 0.804302818 | 0.909751985 | -4.750119106 |
| **A_55_P2063216** | Dusp14 | -0.059143694 | -0.629017071 | 0.510729683 | 9.555791973 | -0.259944376 | 0.804398946 | 0.90975835 | -4.750130274 |
| **A_55_P2106424** | NA | 0.045061385 | -0.389131483 | 0.479254253 | 16.27814171 | 0.259939452 | 0.804402554 | 0.90975835 | -4.750130693 |
| **A_51_P333274** | Gzmb | -0.050955107 | -0.54283688 | 0.440926666 | 8.401666289 | -0.259464133 | 0.804750887 | 0.91000288 | -4.750171115 |
| **A_55_P1990038** | Mark2 | 0.027857904 | -0.241087164 | 0.296802971 | 8.506974881 | 0.259438961 | 0.804769336 | 0.91000288 | -4.750173254 |
| **A_51_P440327** | Efhc1 | -0.060869913 | -0.648582159 | 0.526842333 | 9.624018173 | -0.25941094 | 0.804789873 | 0.91000288 | -4.750175634 |
| **A_55_P2028922** | Stoml2 | 0.032075152 | -0.277645378 | 0.341795682 | 11.95577026 | 0.259387521 | 0.804807037 | 0.91000288 | -4.750177624 |
| **A_52_P259402** | Dsg3 | -0.022052007 | -0.235140385 | 0.191036371 | 7.620202692 | -0.259202212 | 0.804942855 | 0.910002974 | -4.750193359 |
| **A_55_P1975396** | Ighmbp2 | 0.024297346 | -0.210503345 | 0.259098037 | 9.25994294 | 0.259184969 | 0.804955494 | 0.910002974 | -4.750194822 |
| **A_55_P1983162** | Pstpip2 | -0.030433966 | -0.324587252 | 0.26371932 | 8.14260045 | -0.259140441 | 0.804988131 | 0.910002974 | -4.750198602 |
| **A_55_P1987504** | Ccnc | -0.022071316 | -0.23541229 | 0.191269659 | 8.273255025 | -0.259122003 | 0.805001645 | 0.910002974 | -4.750200166 |
| **A_55_P2280581** | A130088E | 0.043403603 | -0.376255603 | 0.46306281 | 8.20397324 | 0.259047483 | 0.805056267 | 0.910002974 | -4.750206489 |
| **A_55_P2121886** | Map3k9 | -0.02455776 | -0.26213755 | 0.213022031 | 8.531352448 | -0.258898536 | 0.805165445 | 0.910002974 | -4.750219122 |
| **A_55_P2102060** | Gstm3 | 0.0221849 | -0.19245708 | 0.23682688 | 7.694567681 | 0.258876819 | 0.805181364 | 0.910002974 | -4.750220963 |
| **A_55_P1998962** | Oas1b | -0.025400421 | -0.271155978 | 0.220355136 | 13.43600816 | -0.258873687 | 0.80518366 | 0.910002974 | -4.750221228 |
| **A_55_P2086949** | Cntf | -0.026604387 | -0.284312217 | 0.231103443 | 8.342687723 | -0.258568723 | 0.805407216 | 0.910202427 | -4.750247069 |
| **A_55_P2172852** | Hacd4 | -0.026394707 | -0.28243858 | 0.229649166 | 8.273970996 | -0.258197958 | 0.805679036 | 0.910456396 | -4.750278446 |
| **A_51_P501550** | NA | -0.047250682 | -0.506246925 | 0.411745561 | 8.343554419 | -0.257839408 | 0.80594193 | 0.91061176 | -4.750308748 |
| **A_52_P566840** | Adgrf1 | -0.021418168 | -0.229496226 | 0.18665989 | 7.652082031 | -0.257813935 | 0.805960608 | 0.91061176 | -4.750310899 |
| **A_55_P1967835** | NA | -0.073802626 | -0.790997981 | 0.643392729 | 8.872178449 | -0.257741703 | 0.806013573 | 0.91061176 | -4.750316998 |
| **A_55_P2149511** | Mff | 0.041379058 | -0.360808137 | 0.443566254 | 12.27123345 | 0.257693052 | 0.806049248 | 0.91061176 | -4.750321105 |
| **A_51_P163252** | Schip1 | -0.028992355 | -0.310790083 | 0.252805374 | 8.15380061 | -0.257689279 | 0.806052015 | 0.91061176 | -4.750321424 |
| **A_55_P2046593** | Gm15226 | 0.032808527 | -0.286383358 | 0.352000413 | 9.867034212 | 0.257445463 | 0.806230809 | 0.910760529 | -4.750341995 |
| **A_55_P2091364** | Cep164 | 0.027350031 | -0.238876962 | 0.293577024 | 11.00545821 | 0.257309654 | 0.806330404 | 0.91081982 | -4.750353445 |
| **A_55_P2001218** | Cd80 | 0.025753806 | -0.225023412 | 0.276531023 | 8.417619859 | 0.257219374 | 0.806396614 | 0.910841393 | -4.750361053 |
| **A_55_P2080333** | NA | 0.026055343 | -0.227873713 | 0.2799844 | 8.334415505 | 0.257000962 | 0.806556801 | 0.910969108 | -4.75037945 |
| **A_51_P438711** | Tbx18 | 0.024010175 | -0.210157336 | 0.258177687 | 7.854873644 | 0.256814203 | 0.806693781 | 0.911026876 | -4.750395168 |
| **A_52_P434820** | Gnpda2 | 0.045825339 | -0.401122022 | 0.492772699 | 8.883169274 | 0.256802739 | 0.806702189 | 0.911026876 | -4.750396133 |
| **A_52_P66199** | Fbxo11 | -0.034977359 | -0.37626321 | 0.306308491 | 9.616530511 | -0.256695986 | 0.806780491 | 0.911062089 | -4.750405112 |
| **A_55_P2035539** | D3Ertd751 | -0.026215248 | -0.282122646 | 0.22969215 | 8.605724413 | -0.256579214 | 0.806866146 | 0.911079796 | -4.75041493 |
| **A_51_P255441** | Mrpl43 | -0.022045905 | -0.237280866 | 0.193189056 | 13.0709352 | -0.256546125 | 0.806890418 | 0.911079796 | -4.750417711 |
| **A_55_P2172619** | NA | 0.050002759 | -0.438333638 | 0.538339156 | 16.56768046 | 0.256463298 | 0.806951176 | 0.91109519 | -4.750424672 |
| **A_52_P11833** | NA | -0.038473444 | -0.414499612 | 0.337552723 | 9.987934937 | -0.25626739 | 0.807094889 | 0.911171897 | -4.750441127 |
| **A_51_P209527** | Bcl10 | 0.058667482 | -0.544665002 | 0.661999966 | 10.4115609 | 0.259002196 | 0.807113371 | 0.911171897 | -4.72248932 |
| **A_52_P172563** | Dffa | -0.024440342 | -0.26347888 | 0.214598197 | 8.285310156 | -0.256088276 | 0.80722629 | 0.911246167 | -4.750456161 |
| **A_55_P2055632** | Rtf2 | -0.030627755 | -0.330678456 | 0.269422946 | 8.428805616 | -0.255664754 | 0.807537022 | 0.911543717 | -4.750491669 |
| **A_51_P421303** | Caly | 0.056678827 | -0.499593171 | 0.612950825 | 8.395608827 | 0.255201958 | 0.807876614 | 0.911806728 | -4.750530405 |
| **A_66_P136857** | Phrf1 | 0.04655229 | -0.410411333 | 0.503515912 | 10.84686517 | 0.255158337 | 0.807908625 | 0.911806728 | -4.750534052 |
| **A_55_P2105256** | Trim47 | 0.07302451 | -0.643805069 | 0.789854089 | 11.24301465 | 0.255154411 | 0.807911506 | 0.911806728 | -4.750534381 |
| **A_51_P133422** | Nck2 | 0.026666599 | -0.235367434 | 0.288700633 | 7.981350459 | 0.254894393 | 0.808102326 | 0.911968852 | -4.750556111 |
| **A_55_P2082478** | Kidins220 | -0.070829672 | -0.767063166 | 0.625403823 | 9.085822125 | -0.254806593 | 0.808166762 | 0.911988337 | -4.750563444 |

| | | | | | | | | | |
|---|---|---|---|---|---|---|---|---|---|
| **A_51_P317272** | Bod1 | -0.030842313 | -0.334233684 | 0.272549057 | 12.37229847 | -0.254620912 | 0.808303042 | 0.91204553 | -4.750578943 |
| **A_55_P2042879** | Mbtd1 | -0.021814973 | -0.236415744 | 0.192785799 | 7.94506517 | -0.254608991 | 0.808311791 | 0.91204553 | -4.750579938 |
| **A_55_P2068125** | Myct1 | 0.046893231 | -0.414535307 | 0.508321769 | 8.279688504 | 0.254540007 | 0.808362423 | 0.912049433 | -4.750585693 |
| **A_52_P613241** | Icam1 | 0.067242469 | -0.594602083 | 0.729087021 | 8.486435004 | 0.254470825 | 0.808413202 | 0.912053501 | -4.750591464 |
| **A_52_P553316** | Snrk | 0.121695799 | -1.076991857 | 1.320383455 | 11.59698761 | 0.254284506 | 0.808549962 | 0.912154567 | -4.750606997 |
| **A_55_P2071466** | Ncf1 | 0.1035924 | -0.917317985 | 1.124502784 | 8.70743794 | 0.254150294 | 0.80864848 | 0.912212481 | -4.750618179 |
| **A_55_P2069917** | Uba52 | 0.035067615 | -0.310763927 | 0.380899156 | 17.27383783 | 0.253975585 | 0.808776731 | 0.912303927 | -4.750632727 |
| **A_55_P2100973** | Irf6 | 0.054400184 | -0.483099771 | 0.59190014 | 9.052070347 | 0.253496685 | 0.809128317 | 0.912573624 | -4.750672555 |
| **A_52_P15388** | Ltf | -0.067374783 | -0.733209002 | 0.598459437 | 8.361601612 | -0.253443764 | 0.809167172 | 0.912573624 | -4.750676952 |
| **A_55_P2044533** | NA | -0.076446835 | -0.832035727 | 0.679142058 | 9.119853704 | -0.253410315 | 0.809191731 | 0.912573624 | -4.750679731 |
| **A_52_P263251** | 3110040N | -0.027279553 | -0.296925197 | 0.24236609 | 8.890298857 | -0.253392755 | 0.809204624 | 0.912573624 | -4.750681189 |
| **A_55_P2158473** | 1700028P | 0.069606654 | -0.618810151 | 0.758023459 | 8.816800027 | 0.253250109 | 0.809309361 | 0.912606377 | -4.750693034 |
| **A_55_P2089125** | Zfp976 | -0.052318782 | -0.570012136 | 0.465374573 | 11.96845779 | -0.253125198 | 0.80940108 | 0.912606377 | -4.750703401 |
| **A_55_P1953578** | Top3a | -0.040375053 | -0.439889336 | 0.35913923 | 9.624293305 | -0.253122727 | 0.809402894 | 0.912606377 | -4.750703606 |
| **A_51_P298486** | Cog6 | 0.02718199 | -0.241885485 | 0.296249464 | 9.98366823 | 0.253029052 | 0.809471679 | 0.912606377 | -4.750711377 |
| **A_51_P221132** | L2hgdh | 0.023834187 | -0.21216735 | 0.259835724 | 9.042820819 | 0.25295068 | 0.809529229 | 0.912606377 | -4.750717877 |
| **A_55_P2081560** | Phf20 | -0.052426055 | -0.571869227 | 0.467017118 | 9.439563126 | -0.252789762 | 0.809647398 | 0.912606377 | -4.750731216 |
| **A_55_P2145376** | Fastk | 0.036933745 | -0.329045469 | 0.402912959 | 8.533771287 | 0.252765195 | 0.809665439 | 0.912606377 | -4.750733252 |
| **A_55_P2153797** | Slc38a6 | -0.043385684 | -0.473311025 | 0.386539658 | 9.087254831 | -0.252757322 | 0.80967122 | 0.912606377 | -4.750733904 |
| **A_55_P2138174** | Psd3 | -0.026699914 | -0.291343808 | 0.237943979 | 8.160058073 | -0.252695983 | 0.809716266 | 0.912606377 | -4.750738986 |
| **A_55_P1991416** | NA | -0.027930953 | -0.304827654 | 0.248965748 | 7.883773037 | -0.25264943 | 0.809750454 | 0.912606377 | -4.750742842 |
| **A_51_P240614** | Tm4sf1 | 0.16293657 | -1.452531148 | 1.778404289 | 12.75510497 | 0.252621688 | 0.809770828 | 0.912606377 | -4.750745139 |
| **A_51_P296249** | Foxs1 | -0.041809348 | -0.456430566 | 0.372811869 | 8.957272261 | -0.252564451 | 0.809812862 | 0.912606377 | -4.750749879 |
| **A_51_P386911** | Taco1 | 0.024493028 | -0.218448208 | 0.267434265 | 8.266814291 | 0.252517566 | 0.809847295 | 0.912606377 | -4.750753761 |
| **A_52_P449694** | NA | 0.052877088 | -0.472061711 | 0.577815886 | 8.512303704 | 0.252295323 | 0.81001052 | 0.912701019 | -4.750772151 |
| **A_55_P2135261** | Ccdc174 | -0.028865433 | -0.315451319 | 0.257720452 | 10.61860613 | -0.252274661 | 0.810025695 | 0.912701019 | -4.75077386 |
| **A_52_P556290** | Arih1 | -0.039558884 | -0.432552964 | 0.353435195 | 9.286078277 | -0.252120624 | 0.810138833 | 0.91272337 | -4.750786596 |
| **A_51_P244824** | Dapp1 | 0.034350659 | -0.307087877 | 0.375789195 | 8.065367017 | 0.251983951 | 0.810239223 | 0.91272337 | -4.75079789 |
| **A_51_P506297** | Cst9 | -0.029436297 | -0.322033009 | 0.263160415 | 7.77672881 | -0.251978845 | 0.810242973 | 0.91272337 | -4.750798312 |
| **A_51_P403049** | Heatr5b | 0.023074878 | -0.206290094 | 0.25243985 | 8.361863593 | 0.251978024 | 0.810243576 | 0.91272337 | -4.75079838 |
| **A_51_P142989** | Zfp429 | -0.030164477 | -0.330102327 | 0.269773374 | 9.337113227 | -0.251892285 | 0.810306555 | 0.91272337 | -4.750805462 |
| **A_52_P676359** | Fcnb | 0.021091781 | -0.1886577 | 0.230841261 | 7.667886756 | 0.251862028 | 0.81032878 | 0.91272337 | -4.75080796 |
| **A_55_P2336558** | Chn2 | 0.030233203 | -0.270650215 | 0.331116622 | 8.484400048 | 0.251672785 | 0.810467795 | 0.912826771 | -4.750823582 |
| **A_55_P2070686** | Wdfy1 | -0.082502552 | -0.903925989 | 0.738920886 | 9.047347465 | -0.251565242 | 0.810546797 | 0.912862572 | -4.750832454 |
| **A_52_P585028** | Cnep1r1 | 0.039355241 | -0.352613177 | 0.431323659 | 11.08336637 | 0.251479069 | 0.810610103 | 0.912880692 | -4.750839561 |
| **A_55_P1982494** | H2al1m | -0.02139168 | -0.23454902 | 0.191765659 | 7.742283136 | -0.251359297 | 0.810698094 | 0.912894666 | -4.750849434 |
| **A_55_P2002122** | Rgs12 | 0.033527391 | -0.300590316 | 0.367645099 | 11.78207003 | 0.251333637 | 0.810716945 | 0.912894666 | -4.750851549 |
| **A_55_P2121543** | Rps20 | 0.024362124 | -0.21874228 | 0.267466528 | 17.07527389 | 0.250999391 | 0.810962519 | 0.913038987 | -4.750879077 |
| **A_55_P2039071** | Dhdh | 0.071934182 | -0.645918996 | 0.789787359 | 9.279175045 | 0.250986306 | 0.810972133 | 0.913038987 | -4.750880154 |
| **A_55_P2063505** | NA | -0.064905408 | -0.712772585 | 0.582961769 | 11.82470805 | -0.250925762 | 0.811016619 | 0.913038987 | -4.750885136 |
| **A_55_P2141786** | Pard6g | -0.086343253 | -0.948278163 | 0.775591657 | 12.05224161 | -0.250902091 | 0.811034011 | 0.913038987 | -4.750887084 |
| **A_66_P110798** | NA | -0.034570798 | -0.379785389 | 0.310643793 | 13.37703646 | -0.250824874 | 0.811090748 | 0.913049695 | -4.750893436 |
| **A_51_P439626** | Map2k2 | 0.031943863 | -0.287617433 | 0.351505159 | 11.00425244 | 0.250370758 | 0.811424447 | 0.913372162 | -4.750930754 |
| **A_55_P1997105** | Large2 | 0.030403664 | -0.273914323 | 0.33472165 | 10.19973176 | 0.25023534 | 0.811523965 | 0.913428662 | -4.75094187 |
| **A_55_P1973334** | Mapk1 | 0.026495023 | -0.238968122 | 0.291958168 | 12.69304434 | 0.249982963 | 0.811709447 | 0.913428662 | -4.750962571 |
| **A_55_P2102424** | Msto1 | 0.028707743 | -0.258932877 | 0.316348363 | 10.04079982 | 0.249976494 | 0.811714201 | 0.913428662 | -4.750963101 |
| **A_55_P2011712** | NA | -0.031447002 | -0.346541191 | 0.283647187 | 9.020298772 | -0.249970754 | 0.81171842 | 0.913428662 | -4.750963572 |
| **A_55_P2016387** | Eif3k | -0.022415097 | -0.24704786 | 0.202217667 | 15.22505956 | -0.249929698 | 0.811748595 | 0.913428662 | -4.750966937 |

| | | | | | | | | | |
|---|---|---|---|---|---|---|---|---|---|
| **A_55_P2082235** | NA | 0.031334527 | -0.28270052 | 0.345369574 | 9.447120884 | 0.249916753 | 0.811758109 | 0.913428662 | -4.750967998 |
| **A_55_P2079103** | Pan2 | -0.034965192 | -0.385876248 | 0.315945863 | 11.21407649 | -0.249568182 | 0.812014314 | 0.913663781 | -4.750996549 |
| **A_55_P1997831** | Dbndd1 | 0.046105477 | -0.416871752 | 0.509082706 | 8.375894989 | 0.249426869 | 0.812118189 | 0.913727483 | -4.751008113 |
| **A_51_P479769** | Ampd2 | -0.092171131 | -1.018441509 | 0.834099247 | 10.75409597 | -0.249234113 | 0.812259885 | 0.91378586 | -4.751023877 |
| **A_55_P2038187** | BC051019 | 0.05715132 | -0.517203544 | 0.631506184 | 10.71723326 | 0.249227698 | 0.812264601 | 0.91378586 | -4.751024401 |
| **A_55_P2010699** | Map11 | 0.044763679 | -0.405356967 | 0.494884324 | 9.169985271 | 0.249084774 | 0.81236967 | 0.913819173 | -4.751036081 |
| **A_51_P116601** | Cfap69 | 0.080106261 | -0.725575087 | 0.885787609 | 8.892214044 | 0.249031049 | 0.812409167 | 0.913819173 | -4.744254784 |
| **A_66_P119680** | Polr2e | -0.022290267 | -0.246518511 | 0.201937976 | 13.03470679 | -0.248986217 | 0.812442127 | 0.913819173 | -4.751044132 |
| **A_55_P2218334** | 9430011C | 0.024058044 | -0.218007506 | 0.266123595 | 7.75661308 | 0.248930251 | 0.812483272 | 0.913819173 | -4.751048702 |
| **A_55_P2023384** | NA | -0.022709218 | -0.251285958 | 0.205867522 | 8.44589659 | -0.248840169 | 0.8125495 | 0.9138405 | -4.751056056 |
| **A_55_P2073915** | Nos1 | -0.04184205 | -0.463208221 | 0.379524121 | 7.99444155 | -0.24871595 | 0.812640829 | 0.913888318 | -4.751066193 |
| **A_52_P352525** | Qser1 | 0.032121398 | -0.308308222 | 0.372551017 | 8.21404484 | 0.25132159 | 0.812686555 | 0.913888318 | -4.723049787 |
| **A_55_P2069768** | Kiss1 | 0.064744703 | -0.587604613 | 0.717094019 | 8.551592769 | 0.24858469 | 0.812737338 | 0.91389227 | -4.751076899 |
| **A_66_P116373** | Pwwp2b | -0.0663741 | -0.735324046 | 0.602575846 | 10.58113393 | -0.248516579 | 0.812787418 | 0.913895431 | -4.751082453 |
| **A_55_P2425235** | NA | -0.027590472 | -0.305997663 | 0.25081672 | 9.274800158 | -0.248215571 | 0.813008752 | 0.914065538 | -4.751106977 |
| **A_55_P2007630** | Sez6l2 | 0.023517719 | -0.21382441 | 0.260859847 | 8.048692361 | 0.248182242 | 0.813033261 | 0.914065538 | -4.751109691 |
| **A_52_P152631** | Tmem17 | 0.071320823 | -0.649625915 | 0.792267561 | 9.252420925 | 0.247778437 | 0.813330216 | 0.914179892 | -4.751142541 |
| **A_66_P119074** | Tbc1d22b | 0.02715365 | -0.247430009 | 0.301737309 | 8.504598187 | 0.247687377 | 0.813397186 | 0.914179892 | -4.751149942 |
| **A_51_P284793** | Ints12 | -0.021324969 | -0.237066276 | 0.194416339 | 8.744782162 | -0.247574234 | 0.813480399 | 0.914179892 | -4.751159134 |
| **A_55_P2157927** | Tmem241 | 0.021388699 | -0.195004391 | 0.237781788 | 8.848645023 | 0.247566181 | 0.813486321 | 0.914179892 | -4.751159788 |
| **A_55_P2004741** | Cyb5rl | 0.033256995 | -0.303270117 | 0.369784106 | 8.652079059 | 0.247521707 | 0.813519032 | 0.914179892 | -4.7511634 |
| **A_52_P682990** | Timm50 | -0.030540491 | -0.33959108 | 0.278510098 | 9.942750486 | -0.247512299 | 0.813525951 | 0.914179892 | -4.751164164 |
| **A_51_P302856** | Ugt2b37 | 0.02096696 | -0.191209273 | 0.233143193 | 7.67914078 | 0.247507909 | 0.81352918 | 0.914179892 | -4.751164521 |
| **A_66_P134040** | Vps25 | -0.028562269 | -0.317672381 | 0.260547844 | 9.699913692 | -0.247445616 | 0.813574997 | 0.914179892 | -4.751169579 |
| **A_51_P440210** | Prkg1 | -0.058724465 | -0.653234315 | 0.535785386 | 8.495521278 | -0.247406073 | 0.81360408 | 0.914179892 | -4.751172789 |
| **A_51_P418935** | Neurl2 | 0.053544383 | -0.488658006 | 0.595746772 | 9.651297534 | 0.24734483 | 0.813649126 | 0.914179892 | -4.751177759 |
| **A_65_P10865** | Scfd1 | -0.033831737 | -0.376630771 | 0.308967298 | 10.71923151 | -0.247192361 | 0.813761274 | 0.914179892 | -4.751190129 |
| **A_55_P2010298** | Serpina1d | -0.135080658 | -1.503959319 | 1.233798002 | 8.770238853 | -0.247160213 | 0.813784921 | 0.914179892 | -4.751192736 |
| **A_55_P1976789** | Dgcr6 | 0.045466292 | -0.415284259 | 0.506216844 | 11.09975708 | 0.24715763 | 0.813786821 | 0.914179892 | -4.751192946 |
| **A_55_P2073219** | Hps4 | -0.032305356 | -0.359702612 | 0.295091901 | 11.553103 | -0.247143867 | 0.813796944 | 0.914179892 | -4.751194062 |
| **A_55_P2127243** | Tpsb2 | -0.023655848 | -0.263507152 | 0.216195457 | 8.404386531 | -0.247028337 | 0.813881927 | 0.914222239 | -4.751203428 |
| **A_66_P127262** | NA | -0.066782876 | -0.744406687 | 0.610840935 | 9.087420643 | -0.246846404 | 0.814015759 | 0.91431945 | -4.751218169 |
| **A_51_P464146** | Sowahb | -0.021818483 | -0.243278283 | 0.199641317 | 8.022936625 | -0.246762971 | 0.814077137 | 0.914326368 | -4.751224925 |
| **A_55_P1983999** | Desi1 | -0.123459856 | -1.377305653 | 1.130385941 | 8.723941686 | -0.246622095 | 0.814180774 | 0.914326368 | -4.751236329 |
| **A_55_P2038504** | 6-Mar | -0.023084308 | -0.257565274 | 0.211396658 | 8.812027368 | -0.246580999 | 0.814211008 | 0.914326368 | -4.751239654 |
| **A_55_P2261882** | D230040J | 0.044545273 | -0.407927598 | 0.497018145 | 8.293164438 | 0.246580898 | 0.814211083 | 0.914326368 | -4.751239662 |
| **A_55_P2089710** | Ednrb | -0.0338073 | -0.377449899 | 0.309835299 | 9.024438757 | -0.246407453 | 0.814338687 | 0.914360919 | -4.751253691 |
| **A_51_P185939** | Gper1 | 0.045995139 | -0.421738369 | 0.513728646 | 8.365961887 | 0.24629965 | 0.814418002 | 0.914360919 | -4.751262406 |
| **A_55_P1983177** | Acsl4 | 0.031056197 | -0.284804971 | 0.346917364 | 11.9129291 | 0.246264821 | 0.814443628 | 0.914360919 | -4.75126522 |
| **A_55_P2078433** | Mcoln2 | -0.104944278 | -1.172343289 | 0.962454733 | 8.331934264 | -0.246253444 | 0.814451998 | 0.914360919 | -4.75126614 |
| **A_51_P205215** | Klhl30 | 0.064466371 | -0.591322948 | 0.72025569 | 8.324170474 | 0.246217677 | 0.814478315 | 0.914360919 | -4.75126903 |
| **A_55_P2058290** | Cluh | 0.022560221 | -0.207016468 | 0.25213691 | 8.029460125 | 0.246130769 | 0.81454226 | 0.914379612 | -4.751276051 |
| **A_51_P322473** | Perm1 | 0.021348675 | -0.196019822 | 0.238717173 | 7.749768325 | 0.245994091 | 0.814642828 | 0.914439412 | -4.751287087 |
| **A_55_P1986813** | Ibsp | 0.100950626 | -0.927766283 | 1.129667535 | 8.538448257 | 0.245789589 | 0.814793307 | 0.914484763 | -4.751303589 |
| **A_55_P2160862** | Wrn | -0.026449928 | -0.295991653 | 0.243091796 | 8.220088387 | -0.245781302 | 0.814799406 | 0.914484763 | -4.751304257 |
| **A_51_P440852** | Mtrf1 | -0.025246177 | -0.282557503 | 0.232065148 | 8.806063804 | -0.245746348 | 0.814825127 | 0.914484763 | -4.751307077 |
| **A_51_P202633** | Ebi3 | -0.046150784 | -0.51683358 | 0.424532011 | 8.770497345 | -0.245584587 | 0.814944165 | 0.914545134 | -4.751320118 |
| **A_55_P2187206** | NA | 0.045410169 | -0.417814791 | 0.508635128 | 8.819360478 | 0.245533926 | 0.814981446 | 0.914545134 | -4.751324201 |

| | | | | | | | | | |
|---|---|---|---|---|---|---|---|---|---|
| **A_55_P2061492** | Pnldc1 | 0.026731197 | -0.246095928 | 0.299558322 | 8.135859982 | 0.245403756 | 0.815077241 | 0.914545134 | -4.751334687 |
| **A_55_P2327518** | 1500032P | -0.038045735 | -0.426499792 | 0.350408322 | 8.419658881 | -0.245310796 | 0.815145656 | 0.914545134 | -4.751342173 |
| **A_51_P359603** | Itgb7 | 0.069786189 | -0.642778836 | 0.782351214 | 8.29720788 | 0.245298746 | 0.815154524 | 0.914545134 | -4.751343143 |
| **A_55_P2149787** | Fcnb | -0.020886822 | -0.234165352 | 0.192391707 | 7.720408043 | -0.245287591 | 0.815162734 | 0.914545134 | -4.751344041 |
| **A_55_P2131393** | Eef1akmt2 | -0.033862637 | -0.379803285 | 0.31207801 | 10.17103899 | -0.245171244 | 0.815248362 | 0.91458813 | -4.751353405 |
| **A_55_P2049289** | NA | 0.033096178 | -0.305133316 | 0.371325673 | 16.37056578 | 0.245084992 | 0.815311844 | 0.914606277 | -4.751360344 |
| **A_55_P2121225** | Cyp3a11 | 0.020814608 | -0.192142696 | 0.233771912 | 7.67232781 | 0.244808247 | 0.815515538 | 0.914781702 | -4.751382593 |
| **A_51_P429276** | Tmod3 | 0.080205937 | -0.740641353 | 0.901053228 | 11.06214479 | 0.244734106 | 0.815570112 | 0.914789844 | -4.75138855 |
| **A_55_P1994898** | NA | -0.063156512 | -0.709868113 | 0.583555089 | 12.42934962 | -0.244600777 | 0.815668255 | 0.914824514 | -4.751399257 |
| **A_51_P296878** | Triqk | 0.075907911 | -0.701494082 | 0.853309904 | 9.125587833 | 0.244563553 | 0.815695655 | 0.914824514 | -4.751402245 |
| **A_52_P88695** | Dgka | -0.056641665 | -0.636929356 | 0.523646027 | 10.18296674 | -0.244479807 | 0.815757303 | 0.914840586 | -4.751408967 |
| **A_55_P1971025** | Paqr6 | 0.111249832 | -1.029349887 | 1.251849551 | 8.990438011 | 0.244296053 | 0.815892574 | 0.91491381 | -4.751423707 |
| **A_55_P2039190** | Gfra4 | -0.042926684 | -0.48309705 | 0.397243682 | 8.643023614 | -0.244262549 | 0.815917239 | 0.91491381 | -4.751426394 |
| **A_55_P2103902** | Mrpl4 | -0.021146433 | -0.23816144 | 0.195868574 | 10.6417168 | -0.244060616 | 0.816065903 | 0.914985716 | -4.751442578 |
| **A_51_P426739** | Gpt | 0.053314947 | -0.493859831 | 0.600489726 | 8.708212696 | 0.244046879 | 0.816076016 | 0.914985716 | -4.751443678 |
| **A_55_P2130565** | Prdm11 | -0.024139365 | -0.272151322 | 0.223872592 | 8.346036794 | -0.243783085 | 0.816270236 | 0.915061528 | -4.7514648 |
| **A_55_P2155593** | NA | 0.039874301 | -0.369848973 | 0.449597574 | 9.830533369 | 0.243754583 | 0.816291221 | 0.915061528 | -4.751467081 |
| **A_55_P2121142** | Tbc1d22b | 0.03028756 | -0.280977599 | 0.341552719 | 9.905157432 | 0.243716045 | 0.816319596 | 0.915061528 | -4.751470164 |
| **A_55_P1998384** | Wsb2 | -0.053344795 | -0.601624217 | 0.494934628 | 11.34694741 | -0.243691537 | 0.816337642 | 0.915061528 | -4.751472125 |
| **A_51_P436817** | Cbarp | 0.03326765 | -0.308739531 | 0.37527483 | 9.797322127 | 0.243633632 | 0.816380278 | 0.915061528 | -4.751476757 |
| **A_52_P538470** | Spats2l | 0.059576404 | -0.55340623 | 0.672559038 | 8.817944423 | 0.243431324 | 0.816529244 | 0.91508284 | -4.751492931 |
| **A_66_P119170** | Phf21a | 0.057287059 | -0.532180037 | 0.646754154 | 9.256680573 | 0.24341498 | 0.816541279 | 0.91508284 | -4.751494237 |
| **A_55_P1989539** | Cklf | 0.046156652 | -0.428781823 | 0.521095128 | 8.315280192 | 0.243414976 | 0.816541282 | 0.91508284 | -4.751494237 |
| **A_52_P455540** | Ric8b | 0.024265003 | -0.225598402 | 0.274128408 | 9.566886208 | 0.243236109 | 0.816672995 | 0.915177401 | -4.751508526 |
| **A_51_P142046** | Oxld1 | 0.047701585 | -0.443826163 | 0.539229332 | 10.32143392 | 0.243072099 | 0.816793774 | 0.915259698 | -4.751521618 |
| **A_55_P2009375** | Cdk18 | -0.029338307 | -0.331786934 | 0.273110319 | 10.70528892 | -0.242959444 | 0.816876739 | 0.915287257 | -4.751530606 |
| **A_51_P506674** | Rpp38 | -0.026379252 | -0.298378168 | 0.245619665 | 9.526036567 | -0.242910139 | 0.81691305 | 0.915287257 | -4.751534539 |
| **A_55_P1971513** | NA | 0.037585354 | -0.350367551 | 0.425538259 | 16.12888479 | 0.242655414 | 0.817100652 | 0.915361584 | -4.751554843 |
| **A_51_P243914** | Skap2 | 0.063433567 | -0.59136262 | 0.718229754 | 11.46427871 | 0.242640525 | 0.817111618 | 0.915361584 | -4.751556029 |
| **A_55_P2028064** | Kdm4b | 0.031501458 | -0.293691768 | 0.356694684 | 10.44208099 | 0.242627214 | 0.817121422 | 0.915361584 | -4.75155709 |
| **A_51_P290207** | Insig1 | -0.022318671 | -0.25329908 | 0.208661739 | 8.177793482 | -0.242015692 | 0.817571862 | 0.915774275 | -4.751605746 |
| **A_55_P2151209** | Krtap19-4 | -0.092387266 | -1.048589723 | 0.863815191 | 11.68123047 | -0.241998462 | 0.817584554 | 0.915774275 | -4.751607115 |
| **A_52_P367675** | Acin1 | 0.020637147 | -0.19309188 | 0.234366174 | 8.477269741 | 0.241844653 | 0.817697861 | 0.915808069 | -4.751619333 |
| **A_55_P2010223** | Gabarap | 0.026429686 | -0.24738833 | 0.300247702 | 14.33759284 | 0.241757709 | 0.817761912 | 0.915808069 | -4.751626236 |
| **A_51_P247873** | Ndufb8 | 0.061777304 | -0.578275517 | 0.701830124 | 14.34082382 | 0.241748328 | 0.817768823 | 0.915808069 | -4.751626981 |
| **A_51_P169745** | Tuba1a | 0.047623896 | -0.445893009 | 0.541140801 | 14.89734447 | 0.241698098 | 0.817805829 | 0.915808069 | -4.751630968 |
| **A_55_P1992475** | NA | -0.120205946 | -1.366507515 | 1.126095622 | 9.690539286 | -0.24157565 | 0.81789604 | 0.915808069 | -4.751640684 |
| **A_51_P505521** | NA | -0.089771853 | -1.02054626 | 0.841002554 | 10.42538933 | -0.241571725 | 0.817898931 | 0.915808069 | -4.751640996 |
| **A_55_P2091116** | Kbtbd12 | 0.026856762 | -0.251822688 | 0.305536211 | 7.796429976 | 0.241378757 | 0.818041103 | 0.915914217 | -4.751656297 |
| **A_55_P2094049** | Hectd4 | -0.05405388 | -0.615650826 | 0.507543065 | 11.06990909 | -0.241075171 | 0.81826479 | 0.916069806 | -4.751680347 |
| **A_52_P318438** | Nab1 | 0.05071089 | -0.476183583 | 0.577605364 | 9.920144446 | 0.241061546 | 0.818274829 | 0.916069806 | -4.751681426 |
| **A_52_P241508** | Zbtb46 | 0.021493784 | -0.201976906 | 0.244964474 | 7.905543176 | 0.240903245 | 0.818391477 | 0.916147346 | -4.751693954 |
| **A_55_P2088965** | Scarb1 | 0.020156721 | -0.189557606 | 0.229871048 | 8.198734434 | 0.240736604 | 0.818514274 | 0.916231761 | -4.751707133 |
| **A_55_P2424042** | Ttn | -0.020678061 | -0.235874901 | 0.194518778 | 7.697656222 | -0.240671286 | 0.818562409 | 0.916232595 | -4.751712297 |
| **A_55_P1988795** | Acss2 | -0.020070245 | -0.229217217 | 0.189076727 | 7.88257499 | -0.240354048 | 0.818796202 | 0.916360256 | -4.751737356 |
| **A_66_P139116** | Rpl8 | 0.046268681 | -0.435939637 | 0.528477 | 16.83689233 | 0.240327113 | 0.818816053 | 0.916360256 | -4.751739483 |
| **A_55_P2048036** | Zfx | -0.024646176 | -0.281534972 | 0.23224262 | 11.08592053 | -0.240300527 | 0.818835647 | 0.916360256 | -4.751741581 |
| **A_51_P394014** | Qprt | -0.026496055 | -0.302713683 | 0.249721573 | 7.850274628 | -0.240259279 | 0.818866047 | 0.916360256 | -4.751744836 |

| | | | | | | | | | |
|---|---|---|---|---|---|---|---|---|---|
| **A_55_P2015007** | Zfp46 | 0.08426822 | -0.794864771 | 0.963401212 | 12.30840983 | 0.240081993 | 0.818996713 | 0.916453435 | -4.751758821 |
| **A_52_P560728** | Serhl | -0.052650406 | -0.602219255 | 0.496918444 | 10.06030039 | -0.239955087 | 0.819090251 | 0.916505059 | -4.751768826 |
| **A_52_P648573** | Gorab | 0.026829516 | -0.25331505 | 0.306974081 | 8.731390176 | 0.239872783 | 0.819150916 | 0.916515063 | -4.751775312 |
| **A_55_P1988498** | Naa20 | -0.035715993 | -0.428212547 | 0.356780561 | 11.28266735 | -0.242375982 | 0.819194001 | 0.916515063 | -4.723682318 |
| **A_66_P127118** | Trmt112 | -0.024927255 | -0.285544814 | 0.235690303 | 8.235358057 | -0.239563769 | 0.8193787 | 0.91666866 | -4.751799644 |
| **A_55_P2041457** | NA | 0.084893653 | -0.803028946 | 0.972816252 | 10.71443306 | 0.239469637 | 0.819448091 | 0.916693247 | -4.75180705 |
| **A_55_P1956819** | Gm4832 | 0.050464179 | -0.477772606 | 0.578700963 | 15.67949879 | 0.239279181 | 0.819588495 | 0.916759054 | -4.751822026 |
| **A_52_P637282** | Wipf1 | 0.037589992 | -0.355915045 | 0.431095029 | 10.30000026 | 0.2392612 | 0.819601751 | 0.916759054 | -4.751823439 |
| **A_51_P362661** | Spin4 | 0.02734499 | -0.259012854 | 0.313702834 | 8.415143101 | 0.239176788 | 0.819663982 | 0.916775624 | -4.751830072 |
| **A_55_P2274553** | NA | 0.020049612 | -0.190021404 | 0.230120628 | 7.656478363 | 0.239050793 | 0.819756872 | 0.916826481 | -4.751839969 |
| **A_55_P2053157** | Zc3h3 | -0.027818392 | -0.319519274 | 0.263882491 | 10.80093289 | -0.238860657 | 0.819897057 | 0.916930224 | -4.751854895 |
| **A_52_P605812** | Ptrh1 | -0.137419978 | -1.579316512 | 1.304476555 | 9.716156662 | -0.238707532 | 0.820009959 | 0.916997684 | -4.751866907 |
| **A_51_P401343** | Cldn14 | -0.020657309 | -0.237458818 | 0.1961442 | 7.761969309 | -0.238650194 | 0.820052237 | 0.916997684 | -4.751871403 |
| **A_55_P1965935** | Rwdd1 | -0.026448369 | -0.304245335 | 0.251348597 | 9.702669472 | -0.238463405 | 0.820189969 | 0.917023058 | -4.751886042 |
| **A_52_P535907** | Sde2 | -0.023328725 | -0.268364788 | 0.221707339 | 9.141745001 | -0.238457694 | 0.820194181 | 0.917023058 | -4.751886489 |
| **A_55_P2159934** | Rnf186 | 0.115784035 | -1.10052685 | 1.332094919 | 9.107871288 | 0.238426449 | 0.82021722 | 0.917023058 | -4.751888937 |
| **A_55_P2257341** | 2410004I( | -0.082000952 | -0.943858059 | 0.779856154 | 8.267423403 | -0.23830545 | 0.820306446 | 0.917069783 | -4.751898413 |
| **A_55_P2039379** | Slc27a6 | 0.020227902 | -0.192810779 | 0.233266583 | 10.03122537 | 0.237816909 | 0.820666731 | 0.917413844 | -4.751936624 |
| **A_55_P2071683** | NA | 0.050542977 | -0.481900331 | 0.582986286 | 13.19127752 | 0.237759454 | 0.820709105 | 0.917413844 | -4.751941113 |
| **A_55_P1984622** | Smarcc1 | -0.040985714 | -0.472901703 | 0.390930275 | 8.194305611 | -0.237675035 | 0.820771368 | 0.9174304 | -4.751947707 |
| **A_52_P298548** | Nbn | 0.02903846 | -0.27708747 | 0.33516439 | 8.659512358 | 0.237587622 | 0.82083584 | 0.917449423 | -4.751954532 |
| **A_52_P183152** | Txnl1 | 0.02186662 | -0.208734081 | 0.252467322 | 8.300539217 | 0.237504252 | 0.820897332 | 0.917465114 | -4.751961039 |
| **A_55_P2137887** | Mpdz | -0.038237634 | -0.441658807 | 0.36518354 | 8.314076578 | -0.237401066 | 0.820973441 | 0.917497139 | -4.75196909 |
| **A_52_P644354** | Zfp106 | 0.026796682 | -0.256002314 | 0.309595678 | 10.04818953 | 0.237330469 | 0.821025514 | 0.9175023 | -4.751974596 |
| **A_55_P2111790** | Gem | -0.059588614 | -0.688995085 | 0.569817858 | 9.245351463 | -0.237127774 | 0.821175031 | 0.917537298 | -4.751990397 |
| **A_51_P351394** | Vps45 | -0.021161418 | -0.244699761 | 0.202376924 | 9.230501143 | -0.237106296 | 0.821190874 | 0.917537298 | -4.75199207 |
| **A_52_P536481** | Mogs | 0.062165635 | -0.594551455 | 0.718882726 | 13.23347501 | 0.237095003 | 0.821199204 | 0.917537298 | -4.75199295 |
| **A_55_P2087061** | Tmem240 | 0.088890874 | -0.850870949 | 1.028652697 | 8.825524203 | 0.236913455 | 0.82133313 | 0.917633906 | -4.752007089 |
| **A_51_P183630** | Bcor | -0.030529019 | -0.35371792 | 0.292659882 | 10.26357956 | -0.236595651 | 0.821567586 | 0.917841548 | -4.752031815 |
| **A_66_P110251** | Tmem91 | -0.031168137 | -0.361210547 | 0.298874273 | 8.704126754 | -0.236532839 | 0.821613928 | 0.917841548 | -4.752036698 |
| **A_52_P522379** | Map3k20 | 0.027190557 | -0.260912687 | 0.315293801 | 8.023559312 | 0.236385211 | 0.821722847 | 0.917910188 | -4.75204817 |
| **A_55_P2092364** | Ermard | -0.028949436 | -0.336128131 | 0.278229259 | 9.523675317 | -0.236047477 | 0.821972044 | 0.918135507 | -4.752074388 |
| **A_51_P169516** | Ppp1r3d | 0.053864455 | -0.51792149 | 0.625650401 | 9.446718621 | 0.235949544 | 0.822044307 | 0.918163179 | -4.752081984 |
| **A_51_P136303** | Cyp4f15 | -0.01959381 | -0.227646632 | 0.188459011 | 7.64701274 | -0.235882453 | 0.822093814 | 0.918165432 | -4.752087186 |
| **A_55_P1983262** | Gm4026 | 0.038479222 | -0.370288326 | 0.447246771 | 8.541297007 | 0.23577634 | 0.822172118 | 0.918199846 | -4.752095411 |
| **A_55_P2112185** | Nhsl1 | -0.025107328 | -0.292061428 | 0.241846772 | 9.305681112 | -0.23556689 | 0.822326683 | 0.918306954 | -4.752111634 |
| **A_51_P427934** | Megf8 | -0.027269509 | -0.31734643 | 0.262807411 | 9.948503493 | -0.235458565 | 0.822406626 | 0.918306954 | -4.752120019 |
| **A_52_P282905** | Ces1b | -0.020591379 | -0.239684657 | 0.198501899 | 7.707933641 | -0.235400144 | 0.822449742 | 0.918306954 | -4.75212454 |
| **A_51_P335419** | Csl | 0.025709328 | -0.247852204 | 0.29927086 | 8.560419509 | 0.235388936 | 0.822458013 | 0.918306954 | -4.752125407 |
| **A_55_P2060369** | Akap9 | 0.036757711 | -0.354701003 | 0.428216425 | 10.56548287 | 0.235186745 | 0.822607238 | 0.918420531 | -4.752141044 |
| **A_51_P262878** | Tmed9 | -0.061451243 | -0.716913256 | 0.59401077 | 14.80539423 | -0.234819139 | 0.822878566 | 0.918626985 | -4.75216944 |
| **A_51_P284426** | Cstad | 0.070596425 | -0.682448955 | 0.823641804 | 8.643089472 | 0.234807468 | 0.822887181 | 0.918626985 | -4.752170341 |
| **A_51_P383857** | Prp2 | 0.030651972 | -0.296428236 | 0.35773218 | 8.320301926 | 0.23472238 | 0.822949988 | 0.918644057 | -4.752176907 |
| **A_55_P2171788** | NA | -0.056304844 | -0.657489936 | 0.544880247 | 11.81523327 | -0.234578339 | 0.823056314 | 0.918709704 | -4.752188017 |
| **A_55_P2115442** | Clec2d | 0.060864057 | -0.589205378 | 0.710933492 | 9.680162831 | 0.234504671 | 0.823110695 | 0.918717364 | -4.752193697 |
| **A_55_P1976694** | 11-Sep | -0.026875298 | -0.314140924 | 0.260390327 | 8.622651381 | -0.234325729 | 0.823242792 | 0.918811762 | -4.752207486 |
| **A_55_P2048119** | Slc29a4 | 0.025439345 | -0.246711579 | 0.297590269 | 8.772174546 | 0.234124281 | 0.823391512 | 0.9189247 | -4.752222998 |
| **A_55_P2128597** | Nr1h3 | 0.087878232 | -0.852983122 | 1.028739587 | 10.2993913 | 0.233940831 | 0.823526952 | 0.918973819 | -4.752237112 |

| | | | | | | | | | |
|---|---|---|---|---|---|---|---|---|---|
| **A_55_P2120906** | NA | 0.035309554 | -0.342737355 | 0.413356462 | 8.978938957 | 0.233935905 | 0.823530588 | 0.918973819 | -4.752237491 |
| **A_51_P486155** | Ocln | 0.045655778 | -0.443307348 | 0.534618903 | 8.650208342 | 0.233867559 | 0.82358105 | 0.918977089 | -4.752242746 |
| **A_51_P275514** | Mprip | -0.044479685 | -0.521632198 | 0.432672827 | 12.4509796 | -0.233482771 | 0.823865163 | 0.919241059 | -4.752272307 |
| **A_55_P2140288** | Phka2 | 0.049258655 | -0.479382422 | 0.577899733 | 9.321639953 | 0.23338449 | 0.823937735 | 0.919268981 | -4.75227985 |
| **A_52_P630867** | Abcc4 | -0.024098827 | -0.282844224 | 0.23464657 | 8.683645055 | -0.233277915 | 0.824016434 | 0.919298995 | -4.752288026 |
| **A_55_P2024858** | Fars2 | -0.045052771 | -0.528899047 | 0.438793505 | 11.93067702 | -0.23321928 | 0.824059733 | 0.919298995 | -4.752292523 |
| **A_51_P106527** | Mcrip2 | -0.04090188 | -0.480924685 | 0.399120924 | 9.82994855 | -0.232819008 | 0.824355332 | 0.919575698 | -4.75232319 |
| **A_52_P636151** | Golph3 | 0.020879695 | -0.203894892 | 0.245654281 | 11.28932247 | 0.23266298 | 0.824470567 | 0.919618717 | -4.75233513 |
| **A_52_P316644** | Bet1 | 0.041648423 | -0.406754341 | 0.490051188 | 10.77487388 | 0.232637986 | 0.824489026 | 0.919618717 | -4.752337042 |
| **A_55_P2008691** | NA | 0.029043939 | -0.283860852 | 0.341948729 | 15.26780203 | 0.232484307 | 0.824602532 | 0.919618833 | -4.752348794 |
| **A_55_P2050044** | NA | 0.034803807 | -0.340163002 | 0.409770615 | 10.62917836 | 0.232479289 | 0.824606239 | 0.919618833 | -4.752349177 |
| **A_55_P2026125** | Srrm2 | -0.039451496 | -0.464554591 | 0.3856516 | 8.543071925 | -0.232444647 | 0.824631826 | 0.919618833 | -4.752351825 |
| **A_55_P1967414** | Asph | -0.020416679 | -0.24048875 | 0.199655391 | 7.72340029 | -0.2323649 | 0.824690728 | 0.919631476 | -4.75235792 |
| **A_55_P1992445** | Vdac1 | -0.045832288 | -0.540205661 | 0.448541086 | 14.50929249 | -0.232202456 | 0.824810717 | 0.919712233 | -4.752370328 |
| **A_55_P1990735** | Ift122 | 0.035244559 | -0.345239413 | 0.415728531 | 9.130792193 | 0.232009657 | 0.824953133 | 0.919778461 | -4.752385043 |
| **A_55_P1980262** | Rtl8b | 0.049841577 | -0.488263118 | 0.587946272 | 13.90180757 | 0.231993244 | 0.824965258 | 0.919778461 | -4.752386295 |
| **A_55_P2159094** | P3h1 | -0.058461494 | -0.690073783 | 0.573150796 | 9.63688774 | -0.231830024 | 0.825085831 | 0.919859846 | -4.752398743 |
| **A_55_P2021515** | Rpl3 | 0.077853726 | -0.763930514 | 0.919637965 | 16.83340856 | 0.231648213 | 0.825220145 | 0.919921344 | -4.752412599 |
| **A_55_P2007802** | Smarca2 | 0.077994456 | -0.765390315 | 0.921379228 | 10.23002714 | 0.231626544 | 0.825236154 | 0.919921344 | -4.752414249 |
| **A_55_P2183777** | NA | -0.040450947 | -0.478029583 | 0.397127689 | 9.65440214 | -0.231538346 | 0.825301313 | 0.919940939 | -4.752420966 |
| **A_55_P2357303** | 5430401H | -0.020059732 | -0.237171636 | 0.197052172 | 7.732103945 | -0.231415179 | 0.825392309 | 0.919989329 | -4.752430343 |
| **A_52_P487436** | Nags | 0.02012724 | -0.197954413 | 0.238208893 | 7.741837735 | 0.231161469 | 0.825579761 | 0.920112567 | -4.752449641 |
| **A_55_P1960616** | Dnajc17 | -0.025297416 | -0.299428064 | 0.248833231 | 8.44438182 | -0.231136708 | 0.825598056 | 0.920112567 | -4.752451524 |
| **A_55_P2072666** | Cyp2d37- | -0.01939326 | -0.229610786 | 0.190824267 | 7.626165077 | -0.231063975 | 0.825651797 | 0.920119422 | -4.752457052 |
| **A_55_P2102135** | Rdh12 | -0.069727723 | -0.826083448 | 0.686628002 | 9.885696173 | -0.230903083 | 0.825770682 | 0.92018846 | -4.752469275 |
| **A_55_P1984730** | Sgpl1 | 0.038439562 | -0.378690076 | 0.455569201 | 11.32966837 | 0.230811654 | 0.825838241 | 0.92018846 | -4.752476217 |
| **A_51_P143470** | Timm44 | -0.044985884 | -0.533341995 | 0.443370227 | 11.50174087 | -0.230722514 | 0.825904111 | 0.92018846 | -4.752482982 |
| **A_52_P349844** | Obscn | 0.038897604 | -0.383380128 | 0.461175336 | 7.896017981 | 0.230714567 | 0.825909984 | 0.92018846 | -4.752483585 |
| **A_55_P2168296** | Rnu7 | 0.021506219 | -0.212275216 | 0.255287654 | 8.08897679 | 0.23041163 | 0.826133851 | 0.92018846 | -4.752506559 |
| **A_55_P1990964** | Osbpl11 | 0.032796038 | -0.323712524 | 0.389304601 | 10.70990517 | 0.230410234 | 0.826134882 | 0.92018846 | -4.752506664 |
| **A_65_P20371** | Bclaf1 | -0.032040426 | -0.380426178 | 0.316345325 | 10.70263891 | -0.230350017 | 0.826179384 | 0.92018846 | -4.752511227 |
| **A_55_P1981011** | Slc35c2 | -0.038162171 | -0.453206932 | 0.37688259 | 9.353161666 | -0.230297108 | 0.826218486 | 0.92018846 | -4.752515236 |
| **A_55_P2396375** | 1810073C | -0.052615468 | -0.624980093 | 0.519749158 | 8.312158611 | -0.230245418 | 0.826256687 | 0.92018846 | -4.752519151 |
| **A_51_P479902** | Mars | 0.03001237 | -0.296587365 | 0.356612106 | 9.827402877 | 0.230162632 | 0.826317871 | 0.92018846 | -4.752525419 |
| **A_55_P2162452** | Mpdu1 | -0.054805306 | -0.651239425 | 0.541628812 | 12.86331353 | -0.230149733 | 0.826327404 | 0.92018846 | -4.752526396 |
| **A_66_P128963** | Cyp2d9 | -0.019219283 | -0.22840649 | 0.189967924 | 7.653495657 | -0.230118964 | 0.826350145 | 0.92018846 | -4.752528725 |
| **A_55_P2149382** | Gm3448 | 0.03158146 | -0.312218849 | 0.37538177 | 8.19237549 | 0.230078635 | 0.826379951 | 0.92018846 | -4.752531778 |
| **A_55_P2133993** | NA | -0.034897207 | -0.414793474 | 0.344999059 | 10.15269692 | -0.230078477 | 0.826380068 | 0.92018846 | -4.75253179 |
| **A_55_P2106079** | NA | -0.03575344 | -0.425094821 | 0.353587941 | 8.08682778 | -0.230005178 | 0.826434242 | 0.920195787 | -4.752537336 |
| **A_51_P431894** | Radil | -0.056694288 | -0.674342901 | 0.560954326 | 8.452940634 | -0.229904874 | 0.826508378 | 0.920225338 | -4.752544923 |
| **A_55_P1973779** | Pde4c | -0.075276708 | -0.895816448 | 0.745263033 | 15.02021169 | -0.229779533 | 0.826601022 | 0.92027549 | -4.7525544 |
| **A_55_P1958088** | Lyn | 0.021664886 | -0.214811189 | 0.258140961 | 8.170621967 | 0.229466634 | 0.826832309 | 0.920420722 | -4.752578035 |
| **A_55_P2032445** | Slc25a37 | 0.037989807 | -0.376754426 | 0.452734039 | 9.117219857 | 0.229423062 | 0.826864518 | 0.920420722 | -4.752581324 |
| **A_51_P442719** | Usp6nl | 0.029738104 | -0.294973826 | 0.354450034 | 9.288487194 | 0.229385187 | 0.826892516 | 0.920420722 | -4.752584182 |
| **A_51_P104527** | Atrip | -0.028477183 | -0.339474946 | 0.282520579 | 9.840872773 | -0.229345443 | 0.826921896 | 0.920420722 | -4.752587181 |
| **A_55_P1975922** | Olfr629 | 0.072208122 | -0.717726088 | 0.862142331 | 9.017906418 | 0.228952543 | 0.827212355 | 0.920691018 | -4.7526168 |
| **A_52_P639101** | Lrrc14 | -0.027891022 | -0.33322337 | 0.277441325 | 8.329946124 | -0.228792595 | 0.827330609 | 0.920769629 | -4.752628843 |
| **A_51_P122740** | Slc37a3 | -0.042576111 | -0.508925103 | 0.423772881 | 9.743582962 | -0.228667947 | 0.827422768 | 0.920800935 | -4.752638223 |

| | | | | | | | | | |
|---|---|---|---|---|---|---|---|---|---|
| **A_55_P1971951** | Cd86 | 0.024873646 | -0.247625229 | 0.297372522 | 7.975641332 | 0.228625719 | 0.82745399 | 0.920800935 | -4.7526414 |
| **A_55_P2167645** | NA | 0.02945617 | -0.293950205 | 0.352862544 | 8.394668813 | 0.22812771 | 0.827822229 | 0.921157696 | -4.75267882 |
| **A_55_P2110101** | Tm2d3 | 0.025407596 | -0.254000903 | 0.304816096 | 11.63789317 | 0.227758351 | 0.828095374 | 0.921373686 | -4.752706522 |
| **A_55_P2140626** | Arl5b | 0.027779116 | -0.277813443 | 0.333371676 | 9.097111342 | 0.227680585 | 0.828152886 | 0.921373686 | -4.752712349 |
| **A_51_P125842** | Tmem51 | 0.033369788 | -0.333738631 | 0.400478208 | 10.41088962 | 0.227671911 | 0.8281593 | 0.921373686 | -4.752712999 |
| **A_55_P2176385** | St6galnac | 0.038681296 | -0.386986268 | 0.46434886 | 9.333293552 | 0.22760448 | 0.82820917 | 0.921376149 | -4.75271805 |
| **A_55_P1960366** | Fam47e | -0.034771737 | -0.4190855 | 0.349542025 | 7.984146046 | -0.22661612 | 0.828940234 | 0.922136391 | -4.752791916 |
| **A_55_P1953402** | Mlana | 0.026648094 | -0.268062441 | 0.321358629 | 7.854913645 | 0.226475295 | 0.829044414 | 0.922156851 | -4.752802415 |
| **A_51_P114049** | Tmem109 | -0.031728673 | -0.38272989 | 0.319272543 | 13.78756199 | -0.226408974 | 0.829093478 | 0.922156851 | -4.732045775 |
| **A_55_P2013396** | Gm14305 | -0.028004211 | -0.337869033 | 0.281860612 | 8.523000537 | -0.226360897 | 0.829129046 | 0.922156851 | -4.75281094 |
| **A_55_P2093595** | Mmd | 0.066106681 | -0.665725971 | 0.797939333 | 10.95993747 | 0.226247566 | 0.829212892 | 0.922156851 | -4.752819381 |
| **A_51_P482051** | Cyp3a16 | 0.019198276 | -0.193421497 | 0.231818049 | 7.65623402 | 0.226156425 | 0.829280323 | 0.922156851 | -4.752826166 |
| **A_55_P1985882** | Fastkd1 | -0.021104119 | -0.254843493 | 0.212635255 | 8.402716384 | -0.226144327 | 0.829289274 | 0.922156851 | -4.752827066 |
| **A_55_P2062549** | Gm6524 | 0.029583858 | -0.298168374 | 0.357336091 | 9.950057615 | 0.226078624 | 0.829337886 | 0.922156851 | -4.752831956 |
| **A_52_P326657** | Fam167b | 0.05282788 | -0.532447187 | 0.638102947 | 8.790741892 | 0.226075503 | 0.829340195 | 0.922156851 | -4.752832188 |
| **A_51_P285413** | Rbbp6 | 0.027973467 | -0.28235337 | 0.338300304 | 8.339703051 | 0.225775757 | 0.829561979 | 0.922291373 | -4.752854477 |
| **A_55_P1972157** | Espnl | 0.032910477 | -0.332384948 | 0.398205901 | 8.429181203 | 0.22565257 | 0.82965313 | 0.922291373 | -4.752863628 |
| **A_55_P1966194** | Plek | 0.046036983 | -0.465022245 | 0.557096211 | 9.075530056 | 0.22562435 | 0.829674012 | 0.922291373 | -4.752865724 |
| **A_55_P2064995** | Capzb | 0.024787598 | -0.250442257 | 0.300017452 | 7.836759361 | 0.225574105 | 0.829711192 | 0.922291373 | -4.752869455 |
| **A_51_P233160** | Lysmd2 | -0.028206529 | -0.341461114 | 0.285048056 | 9.778992183 | -0.225529084 | 0.829744507 | 0.922291373 | -4.752872798 |
| **A_55_P2433218** | Bloc1s6 | -0.056561444 | -0.684729425 | 0.571606537 | 10.76320648 | -0.22552518 | 0.829747395 | 0.922291373 | -4.752873087 |
| **A_55_P2083406** | Myh3 | 0.165622585 | -1.674643321 | 2.00588849 | 8.968180587 | 0.225418379 | 0.829826427 | 0.922326194 | -4.752881013 |
| **A_55_P2162472** | NA | -0.077648571 | -0.941221229 | 0.785924088 | 9.826691397 | -0.225208578 | 0.829981686 | 0.922348273 | -4.752896573 |
| **A_55_P2054157** | Hspa4 | 0.068734192 | -0.695743561 | 0.833211944 | 12.90272473 | 0.225194782 | 0.829991895 | 0.922348273 | -4.752897596 |
| **A_55_P2064150** | NA | -0.028980806 | -0.351317965 | 0.293356353 | 15.80162161 | -0.225190692 | 0.829994922 | 0.922348273 | -4.752897899 |
| **A_55_P2182452** | Tnfsf15 | -0.070618535 | -0.85626758 | 0.71503051 | 8.370379497 | -0.225133677 | 0.830037116 | 0.922348273 | -4.752902125 |
| **A_55_P1977589** | Nfu1 | 0.019017331 | -0.192626645 | 0.230661308 | 11.66264526 | 0.225057766 | 0.830093296 | 0.922357688 | -4.752907749 |
| **A_51_P309854** | Kcnn2 | 0.02976992 | -0.30173936 | 0.3612792 | 7.95904428 | 0.2249222 | 0.830193626 | 0.922416158 | -4.75061818 |
| **A_51_P194609** | Prss34 | 0.020478209 | -0.207747475 | 0.248703893 | 7.827531134 | 0.224738669 | 0.830329461 | 0.922492092 | -4.752931372 |
| **A_55_P1996329** | Kat5 | -0.022063921 | -0.268131913 | 0.224004071 | 9.444351986 | -0.224583538 | 0.830444282 | 0.922492092 | -4.752942845 |
| **A_55_P1952434** | Tm2d1 | 0.022563482 | -0.229127967 | 0.274254932 | 11.242277 | 0.22453705 | 0.830478691 | 0.922492092 | -4.752946282 |
| **A_66_P132832** | Myzap | 0.063469197 | -0.644926249 | 0.771864644 | 8.53117574 | 0.224407616 | 0.830574497 | 0.922492092 | -4.752955846 |
| **A_55_P1992849** | Adrb3 | 0.056318652 | -0.572273489 | 0.684910794 | 8.513175328 | 0.224405581 | 0.830576003 | 0.922492092 | -4.752955997 |
| **A_52_P103977** | Ints8 | 0.033715805 | -0.342681933 | 0.410113543 | 8.541897426 | 0.224355579 | 0.830613015 | 0.922492092 | -4.75295969 |
| **A_55_P2096187** | Hax1 | -0.021642617 | -0.263281872 | 0.219996639 | 12.78862946 | -0.224332721 | 0.830629935 | 0.922492092 | -4.752961378 |
| **A_52_P651078** | Fyttd1 | 0.019388396 | -0.197164496 | 0.235941289 | 8.086061265 | 0.224247826 | 0.830692776 | 0.922492092 | -4.752967647 |
| **A_55_P2079167** | Dnm1 | -0.039894527 | -0.485707984 | 0.40591893 | 10.03933403 | -0.224135421 | 0.830775983 | 0.922492092 | -4.752975943 |
| **A_55_P2162811** | NA | -0.072000882 | -0.876604011 | 0.732602247 | 15.09257804 | -0.22413333 | 0.830777531 | 0.922492092 | -4.752976097 |
| **A_52_P532029** | Tbk1 | -0.029200701 | -0.355534073 | 0.297132671 | 8.614288183 | -0.224120787 | 0.830786817 | 0.922492092 | -4.752977022 |
| **A_55_P2199040** | NA | -0.051675173 | -0.629680643 | 0.526330298 | 8.393915376 | -0.22392384 | 0.830932612 | 0.922502854 | -4.752991548 |
| **A_51_P114817** | Ccna1 | 0.073189456 | -0.745735029 | 0.892113942 | 8.444684665 | 0.223848923 | 0.830988073 | 0.922502854 | -4.746139592 |
| **A_55_P2056729** | Igfbp2 | -0.122687277 | -1.49554273 | 1.250168175 | 10.58863725 | -0.223833488 | 0.8309995 | 0.922502854 | -4.752998207 |
| **A_51_P345902** | Dohh | -0.021875218 | -0.266667322 | 0.222916886 | 11.84109284 | -0.223823317 | 0.831007029 | 0.922502854 | -4.752998957 |
| **A_55_P2062797** | 4921524J | 0.019197364 | -0.195696499 | 0.234091227 | 8.758244656 | 0.22375252 | 0.831059442 | 0.922502854 | -4.753004173 |
| **A_52_P382876** | Bloc1s3 | 0.027607544 | -0.281524286 | 0.336739373 | 8.371489562 | 0.223683725 | 0.831110373 | 0.922502854 | -4.75300924 |
| **A_55_P2032668** | NA | 0.039464175 | -0.402484452 | 0.481412802 | 11.81357616 | 0.22365653 | 0.831130507 | 0.922502854 | -4.753011242 |
| **A_51_P158922** | Prkaa1 | 0.041818837 | -0.427094672 | 0.510732346 | 10.186944 | 0.223372422 | 0.831340854 | 0.92264868 | -4.75303215 |
| **A_55_P2039601** | Drosha | 0.040967192 | -0.41857075 | 0.500505134 | 11.48653239 | 0.223287891 | 0.831403441 | 0.92264868 | -4.753038366 |

| | | | | | | | | | |
|---|---|---|---|---|---|---|---|---|---|
| **A_52_P328078** | Atp5b | -0.024003349 | -0.293256782 | 0.245250084 | 15.93868668 | -0.223285713 | 0.831405054 | 0.92264868 | -4.753038526 |
| **A_51_P485862** | Eef1a2 | 0.020025052 | -0.204797446 | 0.24484755 | 7.767085136 | 0.22309212 | 0.831548398 | 0.922717752 | -4.753052752 |
| **A_55_P2056212** | Gpr3 | -0.041547429 | -0.508043499 | 0.424948641 | 8.756372412 | -0.223072744 | 0.831562745 | 0.922717752 | -4.753054175 |
| **A_55_P2053032** | Xkr8 | 0.022583529 | -0.231157543 | 0.276324601 | 8.610508566 | 0.222921208 | 0.831674954 | 0.922742628 | -4.753065302 |
| **A_55_P2099019** | NA | -0.047953444 | -0.586760979 | 0.490854091 | 11.1404645 | -0.222913561 | 0.831680616 | 0.922742628 | -4.753065863 |
| **A_55_P2089804** | Atf7 | 0.040478932 | -0.438565356 | 0.519523219 | 8.221898007 | 0.225069207 | 0.831831666 | 0.922857258 | -4.724843803 |
| **A_55_P2147310** | Mrtfa | 0.049466178 | -0.507366146 | 0.606298502 | 12.15409293 | 0.222502177 | 0.831985259 | 0.922974696 | -4.753096031 |
| **A_55_P2106844** | NA | -0.02707157 | -0.331922605 | 0.277779466 | 8.165831192 | -0.222421166 | 0.832045254 | 0.922976832 | -4.753101965 |
| **A_51_P378550** | Oplah | -0.065399339 | -0.802023955 | 0.671225277 | 9.853162637 | -0.222370656 | 0.832082662 | 0.922976832 | -4.753105664 |
| **A_55_P2154235** | 1700008J | 0.035122228 | -0.360987306 | 0.431231762 | 8.38748521 | 0.222083918 | 0.832295027 | 0.923072863 | -4.753126647 |
| **A_51_P315682** | Igf2bp2 | 0.05113927 | -0.525699043 | 0.627977583 | 10.09217953 | 0.222049998 | 0.83232015 | 0.923072863 | -4.753129128 |
| **A_55_P2037657** | NA | 0.026653841 | -0.274099723 | 0.327407405 | 8.242196798 | 0.2219726 | 0.832377476 | 0.923072863 | -4.753134787 |
| **A_55_P2040227** | Ptchd4 | -0.047519687 | -0.583780802 | 0.488741429 | 9.052717876 | -0.221946145 | 0.83239707 | 0.923072863 | -4.75313672 |
| **A_51_P337195** | Pipox | 0.024117441 | -0.248096338 | 0.296331219 | 7.827721508 | 0.221907234 | 0.832425891 | 0.923072863 | -4.753139564 |
| **A_51_P158584** | Poll | -0.033341624 | -0.409777262 | 0.343094013 | 9.685769105 | -0.221843327 | 0.832473227 | 0.923072863 | -4.753144233 |
| **A_55_P2075996** | Cpne1 | 0.028931299 | -0.297770686 | 0.355633284 | 12.08716422 | 0.221802537 | 0.83250344 | 0.923072863 | -4.753147213 |
| **A_55_P1996046** | Zfp931 | -0.030847496 | -0.379400139 | 0.317705148 | 9.373174181 | -0.221667424 | 0.832603521 | 0.923097412 | -4.753157079 |
| **A_51_P481676** | Zfp787 | 0.029938525 | -0.308505921 | 0.368382971 | 9.768553718 | 0.221561018 | 0.83268234 | 0.923097412 | -4.753164845 |
| **A_55_P2014500** | Phf24 | 0.018606441 | -0.191757739 | 0.228970622 | 7.796860151 | 0.221534753 | 0.832701796 | 0.923097412 | -4.753166762 |
| **A_55_P1974780** | NA | 0.04347032 | -0.448285009 | 0.53522565 | 9.294076737 | 0.221408408 | 0.832795388 | 0.923097412 | -4.753175977 |
| **A_55_P2053511** | Mrfap1 | 0.019399646 | -0.200064433 | 0.238863725 | 14.80328825 | 0.221401576 | 0.83280045 | 0.923097412 | -4.753176475 |
| **A_55_P2074197** | Zbtb24 | -0.021637321 | -0.266433049 | 0.223158408 | 9.031198615 | -0.221385917 | 0.832812049 | 0.923097412 | -4.753177617 |
| **A_51_P308254** | Surf2 | 0.018439981 | -0.190517796 | 0.227397758 | 8.480559911 | 0.221030536 | 0.833075324 | 0.923201289 | -4.753203508 |
| **A_55_P1959500** | Ces2e | 0.03085923 | -0.318831749 | 0.380550209 | 7.916617104 | 0.221029887 | 0.833075804 | 0.923201289 | -4.753203555 |
| **A_55_P1990134** | Cox18 | -0.033009426 | -0.407198036 | 0.341179183 | 9.476004741 | -0.22095191 | 0.833133574 | 0.923201289 | -4.75320923 |
| **A_51_P382764** | Akr1c20 | -0.018887783 | -0.233041328 | 0.195265763 | 7.658167935 | -0.220905258 | 0.833168137 | 0.923201289 | -4.753212625 |
| **A_66_P111021** | Nlrp6 | 0.018404239 | -0.19031483 | 0.227123307 | 7.715090935 | 0.220854408 | 0.833205812 | 0.923201289 | -4.753216324 |
| **A_55_P1985559** | Gatad1 | 0.022452213 | -0.232210557 | 0.277114984 | 8.599140048 | 0.220822869 | 0.833229179 | 0.923201289 | -4.753218618 |
| **A_55_P2039436** | Ilkap | 0.019871434 | -0.205533952 | 0.245276821 | 12.09728505 | 0.220808239 | 0.833240018 | 0.923201289 | -4.753219682 |
| **A_51_P249193** | Gsg1l | -0.028675266 | -0.354055318 | 0.296704786 | 8.073399301 | -0.220732801 | 0.83329591 | 0.92321031 | -4.753225168 |
| **A_65_P15412** | Usp13 | 0.032528405 | -0.337213058 | 0.402269868 | 8.049511682 | 0.220350967 | 0.833578829 | 0.923470839 | -4.753252905 |
| **A_52_P6828** | Xk | 0.020698498 | -0.214787295 | 0.256184291 | 8.1150601 | 0.220152927 | 0.833725578 | 0.923523813 | -4.753267272 |
| **A_51_P254583** | Foxred1 | -0.037371007 | -0.462645625 | 0.387903611 | 9.701816499 | -0.220097791 | 0.833766435 | 0.923523813 | -4.75327127 |
| **A_55_P2033010** | Mrrf | -0.02123724 | -0.262918007 | 0.220443527 | 7.963565694 | -0.220093052 | 0.833769947 | 0.923523813 | -4.753271613 |
| **A_66_P133328** | Tpd52 | -0.032798378 | -0.406429482 | 0.340832726 | 11.75792357 | -0.219866817 | 0.8339376 | 0.923625909 | -4.753288007 |
| **A_55_P1968908** | Hypk | 0.024020693 | -0.249650873 | 0.29769226 | 11.36674351 | 0.219839742 | 0.833957665 | 0.923625909 | -4.753289968 |
| **A_55_P2096055** | NA | 0.033194965 | -0.345796592 | 0.412186522 | 13.78161565 | 0.219377977 | 0.834299891 | 0.923952005 | -4.753323373 |
| **A_51_P186856** | Krt5 | -0.15236045 | -1.894302277 | 1.589581377 | 8.880421693 | -0.219073045 | 0.834525906 | 0.924080317 | -4.753345395 |
| **A_55_P2064299** | Msrb1 | 0.019068406 | -0.198982618 | 0.237119431 | 8.219264772 | 0.219031516 | 0.83455669 | 0.924080317 | -4.753348392 |
| **A_55_P2150299** | NA | 0.062571048 | -0.652952161 | 0.778094258 | 9.494893659 | 0.219028209 | 0.834559141 | 0.924080317 | -4.753348631 |
| **A_55_P2131607** | Trnau1ap | -0.029855815 | -0.371467984 | 0.311756355 | 10.06822905 | -0.21890009 | 0.834654109 | 0.924125612 | -4.753357873 |
| **A_55_P2279575** | 5430400D | 0.03081324 | -0.321844123 | 0.383470603 | 8.48737591 | 0.218844058 | 0.834695643 | 0.924125612 | -4.753361913 |
| **A_55_P2415472** | 4931408D | 0.018506093 | -0.193431028 | 0.230443214 | 7.693369949 | 0.218704673 | 0.834798968 | 0.924187085 | -4.75337196 |
| **A_52_P366803** | Cyp3a44 | -0.018242564 | -0.227335507 | 0.190850378 | 7.664521888 | -0.218522849 | 0.834933756 | 0.924209384 | -4.753385055 |
| **A_55_P2181419** | Tcf4 | 0.021977581 | -0.230221712 | 0.274176874 | 8.118241892 | 0.218266144 | 0.835124067 | 0.924209384 | -4.753403527 |
| **A_51_P232142** | Thumpd3 | -0.04634172 | -0.578144926 | 0.485461485 | 10.57160408 | -0.218258692 | 0.835129592 | 0.924209384 | -4.753404063 |
| **A_51_P165165** | Hibadh | -0.027589652 | -0.344240826 | 0.289061522 | 11.7993474 | -0.218230508 | 0.835150487 | 0.924209384 | -4.753406089 |
| **A_55_P2066533** | NA | 0.045944815 | -0.481584948 | 0.573474578 | 15.15628138 | 0.218142297 | 0.835215886 | 0.924209384 | -4.75341243 |

| | | | | | | | | | |
|---|---|---|---|---|---|---|---|---|---|
| **A_52_P253230** | Ankrd46 | 0.038816672 | -0.406909802 | 0.484543146 | 10.82399684 | 0.218122375 | 0.835230657 | 0.924209384 | -4.753413862 |
| **A_55_P1960646** | Ube2w | -0.0232824 | -0.290655068 | 0.244090267 | 8.550824128 | -0.218102724 | 0.835245226 | 0.924209384 | -4.753415275 |
| **A_55_P2045931** | Btnl10 | 0.018524953 | -0.194262047 | 0.231311953 | 7.625753524 | 0.218053149 | 0.835281982 | 0.924209384 | -4.753418837 |
| **A_66_P122219** | Kcnd3 | 0.02436533 | -0.255534984 | 0.304265644 | 8.055091886 | 0.218031505 | 0.835298029 | 0.924209384 | -4.753420392 |
| **A_55_P2151986** | Gngt2 | 0.078844432 | -0.827180304 | 0.984869167 | 10.93329552 | 0.217962267 | 0.835349364 | 0.924209384 | -4.753425365 |
| **A_55_P2007756** | Krt16 | -0.066561628 | -0.831460197 | 0.69833694 | 8.54991548 | -0.217956807 | 0.835353413 | 0.924209384 | -4.753425757 |
| **A_55_P2066897** | A230045G | -0.031255105 | -0.390513208 | 0.328002997 | 8.876912307 | -0.217903769 | 0.835392738 | 0.924209384 | -4.753429565 |
| **A_55_P2393483** | 6430584L | 0.030063098 | -0.31566548 | 0.375791677 | 8.344672399 | 0.21779545 | 0.835473052 | 0.924245351 | -4.753437341 |
| **A_55_P2135967** | NA | -0.035361749 | -0.442382949 | 0.371659451 | 13.59738614 | -0.217604091 | 0.835614943 | 0.924315155 | -4.753451067 |
| **A_55_P2074836** | Mapk8ip1 | -0.036154823 | -0.452546873 | 0.380237227 | 11.13479457 | -0.217477411 | 0.835708878 | 0.924315155 | -4.753460148 |
| **A_52_P367774** | Zfp821 | -0.021933087 | -0.274627399 | 0.230761225 | 9.22021511 | -0.217397547 | 0.835768101 | 0.924315155 | -4.75346587 |
| **A_52_P359502** | Pdxk | -0.075345079 | -0.943564404 | 0.792874246 | 11.12250038 | -0.217358082 | 0.835797366 | 0.924315155 | -4.753468697 |
| **A_51_P343613** | Ehd2 | 0.060123898 | -0.632909055 | 0.753156851 | 8.808094395 | 0.217291925 | 0.835846425 | 0.924315155 | -4.753473435 |
| **A_52_P194242** | Sel1l | -0.030282749 | -0.37935362 | 0.318788122 | 10.10522096 | -0.217286144 | 0.835850712 | 0.924315155 | -4.753473849 |
| **A_55_P2144511** | NA | -0.031894803 | -0.399593764 | 0.335804158 | 8.346106999 | -0.217259047 | 0.835870807 | 0.924315155 | -4.753475789 |
| **A_52_P361993** | Zc2hc1a | 0.029189003 | -0.307662176 | 0.366040183 | 9.468687281 | 0.21703588 | 0.836036306 | 0.924445292 | -4.753491757 |
| **A_55_P2073452** | Glp2r | 0.018427079 | -0.194379675 | 0.231233833 | 7.760830384 | 0.216880968 | 0.836151194 | 0.92449824 | -4.753502833 |
| **A_55_P2014570** | Zfp961 | 0.024801288 | -0.261720515 | 0.31132309 | 10.29914979 | 0.216803805 | 0.836208421 | 0.92449824 | -4.753508347 |
| **A_55_P2137932** | NA | -0.02429549 | -0.305007516 | 0.256416536 | 8.066847332 | -0.21677789 | 0.836227642 | 0.92449824 | -4.753510198 |
| **A_55_P2048224** | NA | -0.018608302 | -0.233727062 | 0.196510458 | 7.723912122 | -0.216660035 | 0.836315052 | 0.92454201 | -4.753518616 |
| **A_55_P2083929** | Robo2 | 0.086221313 | -0.911901345 | 1.084343972 | 8.66944724 | 0.216362039 | 0.83653608 | 0.92473348 | -4.753539878 |
| **A_55_P1988183** | Cdkn2b | 0.14208927 | -1.503992368 | 1.788170909 | 10.41350524 | 0.216202275 | 0.836654586 | 0.92479618 | -4.753551266 |
| **A_51_P392867** | Pex12 | 0.025675861 | -0.271858992 | 0.323210714 | 9.008869194 | 0.216141179 | 0.836699906 | 0.92479618 | -4.753555619 |
| **A_55_P2106063** | NA | -0.022142665 | -0.278846143 | 0.234560813 | 12.22151627 | -0.216047118 | 0.836769679 | 0.92479618 | -4.753562318 |
| **A_55_P2208682** | B230334C | -0.02434498 | -0.306605701 | 0.257915742 | 8.140253669 | -0.216027637 | 0.83678413 | 0.92479618 | -4.753563705 |
| **A_55_P2027212** | NA | -0.044715206 | -0.563805862 | 0.474375451 | 8.513220386 | -0.215755745 | 0.836985828 | 0.924966219 | -4.753583051 |
| **A_55_P2042426** | Rpl6 | 0.026942691 | -0.28604379 | 0.339929172 | 16.47773965 | 0.215608429 | 0.837095117 | 0.925034122 | -4.753593523 |
| **A_55_P2132681** | Fzr1 | 0.025074945 | -0.266471054 | 0.316620944 | 9.791792954 | 0.215418626 | 0.837235933 | 0.925061718 | -4.753607006 |
| **A_51_P327418** | Rsrc2 | 0.063051391 | -0.670604538 | 0.796707319 | 11.10003747 | 0.215254671 | 0.837357576 | 0.925061718 | -4.753618643 |
| **A_55_P2080451** | NA | -0.054035528 | -0.682859852 | 0.574788797 | 12.23911082 | -0.215228812 | 0.837376762 | 0.925061718 | -4.753620477 |
| **A_51_P244969** | Rmnd5b | 0.019195797 | -0.204195762 | 0.242587356 | 8.919746075 | 0.215223519 | 0.83738069 | 0.925061718 | -4.753620853 |
| **A_51_P181922** | Yjefn3 | 0.022127496 | -0.235455694 | 0.279710685 | 8.131980894 | 0.215161764 | 0.83742651 | 0.925061718 | -4.753625233 |
| **A_55_P1983708** | Insrr | -0.019829558 | -0.250665008 | 0.211005891 | 7.932145049 | -0.215159676 | 0.837428058 | 0.925061718 | -4.753625381 |
| **A_51_P216219** | Ptk2 | 0.029270331 | -0.311522491 | 0.370063153 | 10.77432419 | 0.215123346 | 0.837455014 | 0.925061718 | -4.753627957 |
| **A_52_P468343** | Bdh1 | -0.018757502 | -0.237226005 | 0.199711002 | 7.729141674 | -0.215048542 | 0.837510518 | 0.925070175 | -4.753633261 |
| **A_51_P271439** | Garem1 | -0.01974796 | -0.249970086 | 0.210474167 | 8.133099684 | -0.214845133 | 0.837661449 | 0.925147292 | -4.753647673 |
| **A_55_P1983414** | NA | 0.035108928 | -0.374294588 | 0.444512443 | 8.725933286 | 0.21479113 | 0.837701521 | 0.925147292 | -4.753651497 |
| **A_55_P2121344** | Nudcd3 | 0.029766456 | -0.317443494 | 0.376976405 | 12.79374512 | 0.214726336 | 0.837749601 | 0.925147292 | -4.753656084 |
| **A_51_P353232** | Tnnc2 | 0.258318018 | -2.756669742 | 3.273305778 | 10.33865384 | 0.214594947 | 0.837847099 | 0.925147292 | -4.753665381 |
| **A_55_P2108171** | Glud1 | -0.030771257 | -0.389948058 | 0.328405545 | 12.30562697 | -0.21457904 | 0.837858903 | 0.925147292 | -4.753666506 |
| **A_55_P2037146** | Igfbpl1 | 0.018865118 | -0.201404983 | 0.239135219 | 7.688479959 | 0.214513351 | 0.83790765 | 0.925147292 | -4.753671152 |
| **A_55_P2146520** | Carns1 | -0.018774292 | -0.237994424 | 0.20044584 | 8.033674751 | -0.214503055 | 0.837915291 | 0.925147292 | -4.75367188 |
| **A_55_P2001064** | NA | -0.021845167 | -0.277124901 | 0.233434567 | 12.00984791 | -0.214333168 | 0.838041365 | 0.925233654 | -4.753683888 |
| **A_55_P1955532** | Tfr2 | 0.023192206 | -0.24802883 | 0.294413241 | 7.827885444 | 0.214175124 | 0.838158655 | 0.925290317 | -4.75369505 |
| **A_51_P186547** | Pah | 0.017972909 | -0.192404993 | 0.228350811 | 7.684148897 | 0.213977745 | 0.838305144 | 0.925290317 | -4.75370898 |
| **A_55_P2192633** | E330017L | -0.020351648 | -0.258586365 | 0.217883069 | 7.752166762 | -0.213966053 | 0.838313822 | 0.925290317 | -4.753709805 |
| **A_55_P1990127** | Dolpp1 | 0.029647677 | -0.317488306 | 0.37678366 | 11.74452449 | 0.213915068 | 0.838351662 | 0.925290317 | -4.753713401 |
| **A_55_P2133624** | NA | 0.082817445 | -0.887092597 | 1.052727486 | 12.08093503 | 0.213865492 | 0.838388458 | 0.925290317 | -4.753716896 |

| | | | | | | | | | |
|---|---|---|---|---|---|---|---|---|---|
| **A_55_P2131348** | Loxl2 | -0.033844486 | -0.430324472 | 0.3626355 | 9.680889681 | -0.213804576 | 0.838433671 | 0.925290317 | -4.753721191 |
| **A_51_P361426** | Arl8b | 0.02282566 | -0.244733657 | 0.290384978 | 11.15779314 | 0.213674954 | 0.838529881 | 0.925290317 | -4.753730325 |
| **A_55_P2135725** | C8g | -0.019269571 | -0.245188116 | 0.206648973 | 7.847164043 | -0.213634073 | 0.838560225 | 0.925290317 | -4.753733204 |
| **A_55_P2129373** | Capn11 | 0.047588736 | -0.510450308 | 0.60562778 | 8.898555795 | 0.213594432 | 0.838589648 | 0.925290317 | -4.753735996 |
| **A_55_P2108754** | Ubfd1 | -0.03263871 | -0.415453504 | 0.350176085 | 11.57218192 | -0.213547559 | 0.83862444 | 0.925290317 | -4.753739296 |
| **A_55_P2055763** | Gm5878 | 0.073955302 | -0.793688087 | 0.941598691 | 10.97431313 | 0.213490424 | 0.83866685 | 0.925290317 | -4.753743318 |
| **A_51_P410744** | Dgke | 0.035864891 | -0.38490227 | 0.456632052 | 8.54442191 | 0.21349024 | 0.838666986 | 0.925290317 | -4.753743331 |
| **A_52_P451888** | Tlk2 | -0.018074732 | -0.230294166 | 0.194144702 | 8.384319195 | -0.213322694 | 0.838791355 | 0.925295353 | -4.753755119 |
| **A_51_P497317** | Tcp11l2 | -0.045544242 | -0.580457591 | 0.489369107 | 10.12711237 | -0.213255572 | 0.838841181 | 0.925295353 | -4.75375984 |
| **A_55_P2135311** | Nol3 | 0.021725267 | -0.233455203 | 0.276905737 | 7.852264634 | 0.213239687 | 0.838852973 | 0.925295353 | -4.753760956 |
| **A_55_P1979893** | Tef | 0.086378564 | -0.928270261 | 1.101027388 | 13.61351493 | 0.2132262 | 0.838862985 | 0.925295353 | -4.753761905 |
| **A_55_P1978726** | NA | -0.023877819 | -0.304828306 | 0.257072668 | 10.62255635 | -0.212870371 | 0.839127138 | 0.925493826 | -4.753786899 |
| **A_55_P2148478** | Klhl25 | -0.020607103 | -0.263128191 | 0.221913984 | 11.6238668 | -0.212822622 | 0.839162587 | 0.925493826 | -4.75379025 |
| **A_55_P2012899** | Vsig4 | 0.018740771 | -0.201864303 | 0.239345844 | 7.677677752 | 0.21277583 | 0.839197325 | 0.925493826 | -4.753793533 |
| **A_52_P179238** | Macroh2a | 0.029687812 | -0.319909047 | 0.379284671 | 8.8494898 | 0.212696826 | 0.83925598 | 0.925493826 | -4.753799075 |
| **A_52_P398925** | Stfa2l1 | 0.116663052 | -1.257940533 | 1.491266636 | 9.604288846 | 0.212572072 | 0.839348602 | 0.925493826 | -4.753807822 |
| **A_55_P2121231** | Nipbl | -0.018686133 | -0.238883965 | 0.2015117 | 8.53157794 | -0.212547858 | 0.83936658 | 0.925493826 | -4.753809519 |
| **A_66_P110607** | Sart1 | -0.05743614 | -0.734594458 | 0.619722179 | 9.884970107 | -0.212444441 | 0.839443363 | 0.925493826 | -4.753816765 |
| **A_55_P2121076** | Cul2 | 0.037882187 | -0.408798825 | 0.484563199 | 8.696932314 | 0.212416327 | 0.839464237 | 0.925493826 | -4.753818734 |
| **A_51_P498257** | NA | 0.046550047 | -0.502420086 | 0.595520179 | 9.918627375 | 0.212383977 | 0.839488256 | 0.925493826 | -4.753821 |
| **A_66_P117417** | NA | 0.089598591 | -0.967271561 | 1.146468743 | 9.551460475 | 0.212339061 | 0.839521606 | 0.925493826 | -4.753824145 |
| **A_51_P125050** | Wdr74 | 0.03494909 | -0.377623312 | 0.447521492 | 9.359387697 | 0.21217101 | 0.839646385 | 0.925541445 | -4.744519423 |
| **A_55_P2141311** | AU022252 | 0.034768921 | -0.375834697 | 0.445372538 | 9.13874681 | 0.212089312 | 0.839707049 | 0.925541445 | -4.753841622 |
| **A_51_P108226** | Wfdc21 | 0.046498307 | -0.502628278 | 0.595624893 | 8.23166192 | 0.212087473 | 0.839708414 | 0.925541445 | -4.739877858 |
| **A_55_P2031676** | Gstm6 | 0.018875909 | -0.204191797 | 0.241943614 | 7.741334556 | 0.211944188 | 0.83981481 | 0.925605948 | -4.753851768 |
| **A_55_P2039611** | Eed | 0.023457119 | -0.253899131 | 0.30081337 | 12.13960728 | 0.211829804 | 0.839899749 | 0.925626769 | -4.75385976 |
| **A_55_P2003129** | Nemp1 | -0.0198601 | -0.254743933 | 0.215023734 | 8.256964053 | -0.211776928 | 0.839939015 | 0.925626769 | -4.753863453 |
| **A_55_P1983284** | Flot2 | 0.020607223 | -0.223172172 | 0.264386619 | 12.23908382 | 0.211725334 | 0.839977328 | 0.925626769 | -4.753867056 |
| **A_51_P349073** | Endog | -0.029886579 | -0.383739147 | 0.323965988 | 9.359375813 | -0.211545701 | 0.840110728 | 0.925695904 | -4.753879592 |
| **A_51_P437289** | Smco4 | 0.047479105 | -0.515091542 | 0.610049751 | 9.703778844 | 0.211385794 | 0.840229484 | 0.925695904 | -4.753890744 |
| **A_51_P271425** | Lhfpl4 | -0.053762577 | -0.690954349 | 0.583429194 | 9.827164241 | -0.211329569 | 0.84027124 | 0.925695904 | -4.753894663 |
| **A_55_P2157551** | NA | 0.035592142 | -0.386249045 | 0.457433329 | 9.594143317 | 0.211327248 | 0.840272964 | 0.925695904 | -4.753894825 |
| **A_55_P2002602** | NA | -0.026813916 | -0.344713883 | 0.291086051 | 15.28311356 | -0.211261377 | 0.840321886 | 0.925695904 | -4.753899415 |
| **A_52_P140497** | NA | 0.020979594 | -0.227797518 | 0.269756707 | 9.125342097 | 0.211220959 | 0.840351904 | 0.925695904 | -4.75390223 |
| **A_51_P365468** | Clpb | 0.023161549 | -0.251691036 | 0.298014134 | 10.30468116 | 0.211065924 | 0.840467051 | 0.925695904 | -4.753913026 |
| **A_52_P521507** | Trio | -0.025382233 | -0.326606202 | 0.275841737 | 11.1417749 | -0.211052567 | 0.840476971 | 0.925695904 | -4.753913956 |
| **A_51_P328089** | Crb3 | -0.021425011 | -0.275709402 | 0.232859381 | 9.624170495 | -0.211033639 | 0.840491029 | 0.925695904 | -4.753915273 |
| **A_55_P2050353** | Prpmp5 | 0.02966144 | -0.322440159 | 0.381763038 | 8.36117605 | 0.210996172 | 0.840518857 | 0.925695904 | -4.75391788 |
| **A_51_P227345** | Dpep1 | 0.16120606 | -1.753811383 | 2.076223502 | 9.949072926 | 0.210842897 | 0.840632703 | 0.925768551 | -4.753928542 |
| **A_51_P477440** | Sec23ip | -0.022896566 | -0.295117942 | 0.249324811 | 10.11273392 | -0.210667947 | 0.840762653 | 0.925844395 | -4.753940703 |
| **A_55_P2028721** | Rnf130 | 0.021896282 | -0.238523074 | 0.282315638 | 11.31174747 | 0.210594709 | 0.840817055 | 0.925844395 | -4.753945791 |
| **A_55_P2113061** | NA | -0.031334663 | -0.404074799 | 0.341405472 | 8.581176058 | -0.210556775 | 0.840845233 | 0.925844395 | -4.753948425 |
| **A_55_P2020248** | NA | 0.025084573 | -0.273512188 | 0.323681333 | 13.97353074 | 0.210412705 | 0.840952253 | 0.925849706 | -4.753958427 |
| **A_55_P1953846** | Abca8b | 0.021223931 | -0.231479235 | 0.273927096 | 7.943637349 | 0.210361123 | 0.840990571 | 0.925849706 | -4.753962006 |
| **A_55_P2007981** | Fmn2 | -0.029466177 | -0.380312878 | 0.321380524 | 8.947730652 | -0.210356887 | 0.840993718 | 0.925849706 | -4.7539623 |
| **A_52_P197095** | Cdk17 | 0.031206926 | -0.34070688 | 0.403120732 | 8.392125573 | 0.210164344 | 0.841136753 | 0.925954449 | -4.753975653 |
| **A_55_P2098354** | Shf | 0.020850851 | -0.228060324 | 0.269762026 | 8.928147912 | 0.209811717 | 0.841398728 | 0.926184098 | -4.754000077 |
| **A_55_P2143832** | Col4a4 | 0.103693627 | -1.134506803 | 1.341894057 | 8.878381644 | 0.209754585 | 0.841441176 | 0.926184098 | -4.75400403 |

| | | | | | | | | | |
|---|---|---|---|---|---|---|---|---|---|
| **A_52_P454295** | Ttn | 0.066888531 | -0.732118516 | 0.865895578 | 8.342337124 | 0.209677289 | 0.841498605 | 0.926194582 | -4.754009377 |
| **A_55_P2018747** | Cox8a | -0.034399294 | -0.445472613 | 0.376674026 | 15.55975161 | -0.209594839 | 0.841559864 | 0.926196015 | -4.754015079 |
| **A_55_P2041693** | Gm5834 | 0.018492964 | -0.202557163 | 0.239543091 | 7.766057125 | 0.209539592 | 0.841600913 | 0.926196015 | -4.754018898 |
| **A_55_P1962174** | Rab8a | -0.022397061 | -0.290291166 | 0.245497044 | 9.880111223 | -0.209400743 | 0.84170408 | 0.926196015 | -4.754028492 |
| **A_52_P586004** | Hk3 | -0.017444782 | -0.226126692 | 0.191237129 | 7.783422526 | -0.209378017 | 0.841720966 | 0.926196015 | -4.754030061 |
| **A_55_P2008427** | Dnd1 | 0.01965195 | -0.215460955 | 0.254764855 | 7.969546589 | 0.209353166 | 0.841739432 | 0.926196015 | -4.754031778 |
| **A_51_P300655** | Tbccd1 | 0.027423878 | -0.300888863 | 0.355736618 | 9.02451039 | 0.209214352 | 0.841842578 | 0.926256795 | -4.754041361 |
| **A_55_P2065819** | Fbxo22 | -0.035723289 | -0.463744198 | 0.39229762 | 10.79191894 | -0.209043562 | 0.841969489 | 0.926335404 | -4.754053143 |
| **A_55_P2399688** | Glipr2 | 0.034195197 | -0.375623258 | 0.444013653 | 8.288150997 | 0.208989251 | 0.842009848 | 0.926335404 | -4.754056888 |
| **A_51_P501069** | Spta1 | -0.017423085 | -0.226304723 | 0.191458553 | 7.722997061 | -0.20891765 | 0.842063055 | 0.92634123 | -4.754061824 |
| **A_51_P459661** | Lipa | -0.030519631 | -0.396753573 | 0.335714312 | 10.16559556 | -0.208723344 | 0.842207451 | 0.926348656 | -4.754075209 |
| **A_55_P2295786** | Mindy2 | -0.017479292 | -0.227286013 | 0.192327429 | 7.879034008 | -0.208667484 | 0.842248964 | 0.926348656 | -4.754079055 |
| **A_52_P288614** | Rars | 0.022087987 | -0.243061183 | 0.287237158 | 8.160097935 | 0.208648958 | 0.842262732 | 0.926348656 | -4.75408033 |
| **A_66_P104442** | Cog6 | -0.023653635 | -0.307653713 | 0.260346443 | 8.152977776 | -0.20860743 | 0.842293595 | 0.926348656 | -4.754083188 |
| **A_66_P110970** | Tpd52l2 | -0.029067178 | -0.378101095 | 0.31996674 | 8.882117113 | -0.208586202 | 0.842309371 | 0.926348656 | -4.754084649 |
| **A_55_P2096630** | Ssxb9 | -0.018761481 | -0.244165676 | 0.206642714 | 7.839107591 | -0.208475715 | 0.842391484 | 0.926374215 | -4.75409225 |
| **A_55_P2032081** | Dbp | 0.044351115 | -0.488619193 | 0.577321424 | 9.667015957 | 0.20842599 | 0.842428439 | 0.926374215 | -4.75409567 |
| **A_55_P1957168** | Ube2s | 0.078622171 | -0.867769375 | 1.025013717 | 13.79433003 | 0.208077215 | 0.842687664 | 0.926606568 | -4.754119632 |
| **A_65_P09177** | Strn3 | -0.025769425 | -0.336164579 | 0.284625729 | 10.00109937 | -0.207941009 | 0.842788904 | 0.926665188 | -4.75412898 |
| **A_51_P431870** | Map1s | 0.033686589 | -0.372388596 | 0.439761774 | 10.24832176 | 0.207778656 | 0.842909583 | 0.926745173 | -4.754140114 |
| **A_52_P591153** | Pld4 | 0.026914055 | -0.29779891 | 0.351627021 | 8.027021185 | 0.207601197 | 0.843041496 | 0.926837501 | -4.754152274 |
| **A_55_P1964615** | Dnttip1 | -0.018665598 | -0.244044029 | 0.206712833 | 8.298932833 | -0.207433984 | 0.843165798 | 0.926900245 | -4.754163723 |
| **A_55_P2008591** | Kras | -0.033033515 | -0.431972002 | 0.365904971 | 8.152389739 | -0.20739544 | 0.843194451 | 0.926900245 | -4.754166361 |
| **A_55_P2253284** | NA | -0.032043014 | -0.419563883 | 0.355477855 | 8.406443312 | -0.207104058 | 0.843411072 | 0.927073033 | -4.754186287 |
| **A_55_P2119907** | Dnah11 | 0.084374653 | -0.936274776 | 1.105024081 | 8.673786251 | 0.207055012 | 0.843447535 | 0.927073033 | -4.754189638 |
| **A_51_P427768** | Diaph1 | -0.025054684 | -0.328378881 | 0.278269513 | 8.457662401 | -0.206886532 | 0.843572795 | 0.927158003 | -4.754201145 |
| **A_55_P2087766** | Dnaja3 | 0.025123505 | -0.279266175 | 0.329513185 | 11.43667459 | 0.206728639 | 0.843690189 | 0.927234317 | -4.754211919 |
| **A_51_P387188** | Vrk3 | 0.05071315 | -0.564513812 | 0.665940111 | 11.35421645 | 0.206459832 | 0.843890059 | 0.927333949 | -4.754230245 |
| **A_55_P2002449** | NA | 0.031574757 | -0.351507738 | 0.414657252 | 9.865972315 | 0.206441998 | 0.843903319 | 0.927333949 | -4.75423146 |
| **A_66_P108276** | Gsg1l | -0.02168102 | -0.284763902 | 0.241401863 | 7.915290681 | -0.206413197 | 0.843924735 | 0.927333949 | -4.754233422 |
| **A_51_P357744** | Jund | 0.05827283 | -0.903787435 | 1.020333095 | 12.57792901 | 0.217958203 | 0.844142239 | 0.927393178 | -4.670970032 |
| **A_55_P2082070** | Ak6 | -0.027820226 | -0.365932527 | 0.310292075 | 10.19080852 | -0.206086729 | 0.844167497 | 0.927393178 | -4.754255642 |
| **A_55_P1968365** | NA | 0.018536834 | -0.20680619 | 0.243879859 | 8.057637043 | 0.206035378 | 0.844205684 | 0.927393178 | -4.754259134 |
| **A_51_P314323** | Naa38 | 0.052930607 | -0.590561184 | 0.696422398 | 13.60073721 | 0.20602229 | 0.844215417 | 0.927393178 | -4.754260024 |
| **A_55_P1956557** | Rab3a | 0.056832844 | -0.634113283 | 0.747778971 | 9.665835672 | 0.206018182 | 0.844218471 | 0.927393178 | -4.754260303 |
| **A_51_P441687** | Lrrc10 | -0.035720781 | -0.470650696 | 0.399209134 | 7.957019219 | -0.205708395 | 0.844448852 | 0.927593552 | -4.75428135 |
| **A_55_P2000623** | Stag1 | -0.023615951 | -0.311537793 | 0.264305891 | 10.8249977 | -0.205438185 | 0.844649814 | 0.927719215 | -4.754299682 |
| **A_55_P2046109** | NA | 0.018953954 | -0.212194688 | 0.250102596 | 7.803197005 | 0.205380317 | 0.844692854 | 0.927719215 | -4.754303605 |
| **A_55_P2068414** | 6030426L | -0.038721388 | -0.510983646 | 0.43354087 | 8.310710008 | -0.205361024 | 0.844707203 | 0.927719215 | -4.754304913 |
| **A_51_P469902** | Ggps1 | -0.020061614 | -0.26483452 | 0.224711292 | 9.081224971 | -0.205282943 | 0.844765277 | 0.927730296 | -4.754310204 |
| **A_52_P672453** | Prpf3 | -0.025343421 | -0.334947917 | 0.284261075 | 10.65758346 | -0.205025713 | 0.844956605 | 0.927887709 | -4.754327621 |
| **A_51_P247249** | Alox5 | -0.021084689 | -0.279074652 | 0.236905273 | 8.142023749 | -0.204698538 | 0.845199975 | 0.92810225 | -4.754349744 |
| **A_55_P2044202** | Vps29 | -0.023523894 | -0.311885688 | 0.2648379 | 11.3106441 | -0.204325156 | 0.84547774 | 0.928312565 | -4.754374949 |
| **A_52_P323820** | Ufm1 | 0.018818096 | -0.21201901 | 0.249655201 | 8.144703265 | 0.204183387 | 0.84558321 | 0.928312565 | -4.754384507 |
| **A_66_P121361** | C330021F | 0.034392529 | -0.38750785 | 0.456292909 | 8.688133092 | 0.20417593 | 0.845588758 | 0.928312565 | -4.754385009 |
| **A_52_P516034** | Ptp4a1 | 0.020518316 | -0.231230138 | 0.27226677 | 9.054301478 | 0.204138656 | 0.845616489 | 0.928312565 | -4.754387521 |
| **A_55_P2018706** | Cep19 | 0.021466743 | -0.241944557 | 0.284878043 | 11.13190772 | 0.204118377 | 0.845631576 | 0.928312565 | -4.754388888 |
| **A_51_P185259** | Gnb5 | 0.022126589 | -0.250075373 | 0.294328551 | 7.895372724 | 0.203598028 | 0.846018733 | 0.928661664 | -4.754423904 |

| | | | | | | | | | |
|---|---|---|---|---|---|---|---|---|---|
| **A_51_P317191** | Eepd1 | -0.029219099 | -0.388737206 | 0.330299007 | 8.09509138 | -0.203561856 | 0.846045647 | 0.928661664 | -4.754426335 |
| **A_51_P307316** | Syde1 | -0.031135361 | -0.414499942 | 0.35222922 | 10.46003454 | -0.20341935 | 0.846151686 | 0.928704069 | -4.754435908 |
| **A_55_P2085880** | Aco1 | -0.018691835 | -0.248957631 | 0.21157396 | 8.116623036 | -0.203316611 | 0.846228136 | 0.928704069 | -4.754442805 |
| **A_51_P345467** | Pi4k2a | -0.040017618 | -0.533080781 | 0.453045544 | 9.2364183 | -0.203282046 | 0.846253856 | 0.928704069 | -4.754445125 |
| **A_51_P434776** | Jmjd8 | 0.027592446 | -0.312428999 | 0.367613891 | 8.517218549 | 0.203251725 | 0.846276419 | 0.928704069 | -4.75444716 |
| **A_55_P1960117** | Dipk2a | -0.024048143 | -0.320547459 | 0.272451173 | 8.994100844 | -0.203145963 | 0.846355122 | 0.928737722 | -4.754454255 |
| **A_51_P309501** | Praf2 | -0.025733631 | -0.343127024 | 0.291659762 | 8.792752437 | -0.203073627 | 0.846408952 | 0.928744079 | -4.754459105 |
| **A_55_P2427900** | Unc13c | 0.017466657 | -0.198130291 | 0.233063604 | 7.707414917 | 0.202916573 | 0.846525829 | 0.928811183 | -4.75446963 |
| **A_55_P2004046** | NA | -0.019143823 | -0.255505786 | 0.217218139 | 7.94302199 | -0.202862343 | 0.846566188 | 0.928811183 | -4.754473263 |
| **A_51_P240253** | Rrad | -0.035345074 | -0.472362818 | 0.401672669 | 8.07013467 | -0.202572354 | 0.846782007 | 0.928959184 | -4.754492671 |
| **A_55_P2191002** | 2610027K | 0.025786117 | -0.293115118 | 0.344687351 | 8.365937743 | 0.202525676 | 0.846816748 | 0.928959184 | -4.754495792 |
| **A_51_P108853** | Lyplal1 | 0.041986819 | -0.477369339 | 0.561342976 | 8.375968885 | 0.202487412 | 0.846845227 | 0.928959184 | -4.745314466 |
| **A_55_P2012984** | Ldha | 0.043911836 | -0.499607694 | 0.587431366 | 16.07809224 | 0.202356327 | 0.846942791 | 0.929013499 | -4.754507111 |
| **A_51_P472329** | Meak7 | 0.019161274 | -0.218599975 | 0.256922524 | 8.669107035 | 0.20185228 | 0.847317975 | 0.929372312 | -4.754540746 |
| **A_51_P152826** | Golt1b | -0.051009941 | -0.686070971 | 0.58405109 | 10.32418864 | -0.201182268 | 0.847816763 | 0.929866649 | -4.754585329 |
| **A_55_P1958902** | NA | -0.020576135 | -0.276971338 | 0.235819068 | 10.68860942 | -0.201003795 | 0.84794964 | 0.929894311 | -4.754597181 |
| **A_52_P63709** | Fam49b | -0.017427253 | -0.234596781 | 0.199742275 | 7.945030804 | -0.200992751 | 0.847957863 | 0.929894311 | -4.754597914 |
| **A_55_P2042823** | Sh3bp2 | -0.026186842 | -0.352575858 | 0.300202174 | 9.376232252 | -0.200954593 | 0.847986273 | 0.929894311 | -4.754600446 |
| **A_52_P559545** | Cercam | 0.018392868 | -0.210951983 | 0.24773772 | 8.089946196 | 0.200868028 | 0.848050725 | 0.929912246 | -4.754606189 |
| **A_55_P2121701** | Mir670hg | 0.01925172 | -0.220980191 | 0.259483631 | 7.707462105 | 0.200719326 | 0.848161445 | 0.929980909 | -4.754616049 |
| **A_51_P452203** | Vash2 | -0.019323865 | -0.260994597 | 0.222346868 | 8.044484378 | -0.20027202 | 0.84849452 | 0.930293357 | -4.754645666 |
| **A_65_P10913** | Tgfb2 | 0.040015002 | -0.461734254 | 0.541764258 | 12.31879345 | 0.199749844 | 0.84888339 | 0.930666938 | -4.754680159 |
| **A_51_P172688** | Stard3nl | -0.047637371 | -0.645386254 | 0.550111511 | 10.23986207 | -0.199608652 | 0.848988545 | 0.930698966 | -4.754689471 |
| **A_55_P1986998** | Enoph1 | -0.029179686 | -0.395549396 | 0.337190024 | 11.09953826 | -0.199485527 | 0.849080248 | 0.930698966 | -4.754697585 |
| **A_51_P379775** | Nrm | -0.033092725 | -0.44863896 | 0.38245351 | 9.005438117 | -0.19946354 | 0.849096624 | 0.930698966 | -4.754699034 |
| **A_55_P2035102** | Odc1 | -0.109072141 | -1.479240733 | 1.261096451 | 11.44126619 | -0.199383941 | 0.84915591 | 0.930698966 | -4.754704277 |
| **A_55_P2104769** | Utf1 | 0.022532324 | -0.260527995 | 0.305592643 | 8.048006355 | 0.199378051 | 0.849160297 | 0.930698966 | -4.754704665 |
| **A_55_P2046699** | Tvp23b | 0.029796729 | -0.344728259 | 0.404321717 | 10.52131006 | 0.199268267 | 0.849242068 | 0.930698966 | -4.754711892 |
| **A_51_P400659** | Bcap29 | 0.021436302 | -0.248088532 | 0.290961136 | 9.914513626 | 0.199205546 | 0.849288785 | 0.930698966 | -4.75471602 |
| **A_55_P1954331** | Cavin4 | -0.054373275 | -0.738065316 | 0.629318766 | 8.64099273 | -0.199193569 | 0.849297706 | 0.930698966 | -4.754716808 |
| **A_66_P104309** | Myl2 | 0.110798169 | -1.283407609 | 1.505003947 | 8.740067825 | 0.19904719 | 0.849406739 | 0.930759434 | -4.754726435 |
| **A_66_P118759** | Safb2 | -0.021074184 | -0.286365335 | 0.244216967 | 11.7923864 | -0.19896576 | 0.849467395 | 0.930759434 | -4.754731788 |
| **A_55_P2045026** | Mgrn1 | 0.0193552 | -0.224345814 | 0.263056215 | 13.15743104 | 0.198925601 | 0.849497309 | 0.930759434 | -4.754734427 |
| **A_55_P2062039** | Tsen34 | 0.017461394 | -0.202810675 | 0.237733462 | 12.99272943 | 0.19854997 | 0.849777129 | 0.930970139 | -4.754759085 |
| **A_51_P369762** | Nmrk2 | 0.023780071 | -0.276218709 | 0.323778851 | 7.785640847 | 0.198538167 | 0.849785922 | 0.930970139 | -4.754759859 |
| **A_55_P2109033** | Hmgcs2 | -0.018345054 | -0.250108242 | 0.213418134 | 7.98593465 | -0.198255333 | 0.849996631 | 0.931148216 | -4.754778395 |
| **A_55_P2006499** | Esrrg | 0.0213197 | -0.248309065 | 0.290948465 | 7.805362483 | 0.198045607 | 0.850152884 | 0.93118696 | -4.754792123 |
| **A_55_P1972541** | Zfp960 | 0.024023748 | -0.27991339 | 0.327960885 | 10.06370769 | 0.197973625 | 0.850206515 | 0.93118696 | -4.754796831 |
| **A_55_P2068121** | Sun5 | 0.02316537 | -0.269933556 | 0.316264297 | 8.056989519 | 0.197959052 | 0.850217373 | 0.93118696 | -4.754797784 |
| **A_55_P2038987** | Gas7 | -0.024535892 | -0.335036635 | 0.285964852 | 8.906595835 | -0.197919938 | 0.850246516 | 0.93118696 | -4.754800342 |
| **A_55_P2058152** | Ppp4r3b | -0.031726849 | -0.433300494 | 0.369846795 | 9.518704032 | -0.197884641 | 0.850272814 | 0.93118696 | -4.75480265 |
| **A_55_P2103097** | Ccr8 | -0.020200144 | -0.276135976 | 0.235735688 | 7.754711755 | -0.197685 | 0.850421566 | 0.931297115 | -4.754815694 |
| **A_52_P322421** | Mpzl2 | -0.042190485 | -0.577175029 | 0.49279406 | 10.08043317 | -0.197525707 | 0.85054026 | 0.931374342 | -4.754826093 |
| **A_51_P394921** | 1700020L | -0.039176868 | -0.536342224 | 0.457988487 | 8.453044409 | -0.19736912 | 0.850656942 | 0.931429944 | -4.754836307 |
| **A_55_P2010861** | Mrps23 | 0.024035992 | -0.281050582 | 0.329122565 | 12.37779278 | 0.197328263 | 0.850687387 | 0.931429944 | -4.754838971 |
| **A_55_P2073557** | Slc27a5 | 0.017202231 | -0.201359536 | 0.235763998 | 7.770591388 | 0.197133721 | 0.850832359 | 0.931535921 | -4.754851647 |
| **A_52_P32081** | Crbn | 0.024484266 | -0.28695925 | 0.335927783 | 8.912275383 | 0.196905635 | 0.851002335 | 0.931630539 | -4.754866494 |
| **A_66_P118579** | 3300002A | 0.140493136 | -1.646809323 | 1.927795595 | 9.690651335 | 0.196882628 | 0.851019481 | 0.931630539 | -4.75486799 |

| | | | | | | | | | |
|---|---|---|---|---|---|---|---|---|---|
| **A_51_P369784** | Ces1e | -0.01892136 | -0.259703968 | 0.221861249 | 7.743385142 | -0.196823781 | 0.851063338 | 0.931630539 | -4.754871818 |
| **A_66_P123938** | Smim13 | 0.016781694 | -0.196958572 | 0.230521961 | 11.51770952 | 0.196652652 | 0.851190876 | 0.931717398 | -4.754882941 |
| **A_55_P2063646** | Krcc1 | 0.030501162 | -0.358333061 | 0.419335384 | 12.81215301 | 0.196472714 | 0.851324985 | 0.931806107 | -4.754894627 |
| **A_51_P426283** | Dhx30 | -0.021366431 | -0.293830089 | 0.251097227 | 12.64027824 | -0.196414588 | 0.851368309 | 0.931806107 | -4.7548984 |
| **A_55_P2140745** | NA | 0.032667884 | -0.384466945 | 0.449802714 | 8.091164476 | 0.196152976 | 0.851563303 | 0.931915989 | -4.754915367 |
| **A_52_P381806** | Terf2ip | -0.024475326 | -0.337233077 | 0.288282425 | 8.920500198 | -0.196006624 | 0.851672393 | 0.931915989 | -4.754924849 |
| **A_55_P2045770** | Gmfb | -0.041372389 | -0.570109668 | 0.48736489 | 12.50995323 | -0.195984179 | 0.851689123 | 0.931915989 | -4.754926302 |
| **A_55_P1983070** | Uhrf1bp1 | -0.02760476 | -0.380461669 | 0.325252148 | 10.23013442 | -0.195945684 | 0.851717818 | 0.931915989 | -4.754928795 |
| **A_51_P226932** | Tubgcp2 | 0.022994407 | -0.270981857 | 0.316970672 | 8.996065967 | 0.195911666 | 0.851743176 | 0.931915989 | -4.754930997 |
| **A_55_P1958007** | Ddx24 | -0.043544486 | -0.600702087 | 0.513613114 | 12.80513757 | -0.195751663 | 0.851862447 | 0.931915989 | -4.754941351 |
| **A_52_P47645** | Cul3 | 0.021628643 | -0.255114834 | 0.29837212 | 9.512607821 | 0.195750208 | 0.851863532 | 0.931915989 | -4.754941445 |
| **A_55_P2364315** | D030002E | -0.021516033 | -0.296884376 | 0.253852309 | 8.407024254 | -0.195703479 | 0.851898366 | 0.931915989 | -4.754944467 |
| **A_55_P2109257** | Nipal3 | 0.044877869 | -0.529498251 | 0.619253989 | 10.43165508 | 0.195697915 | 0.851902513 | 0.931915989 | -4.754944827 |
| **A_51_P505868** | Lhfp | -0.043880839 | -0.606473005 | 0.518711326 | 9.275274236 | -0.195358176 | 0.852155784 | 0.932140307 | -4.750384571 |
| **A_52_P127325** | AI182371 | 0.016565155 | -0.196146539 | 0.22927685 | 7.694311602 | 0.195053834 | 0.852382684 | 0.932335755 | -4.75498641 |
| **A_55_P2138718** | H2al1a | -0.016200492 | -0.224583806 | 0.192182822 | 7.703296433 | -0.194722263 | 0.852629901 | 0.932549782 | -4.755007765 |
| **A_51_P153557** | Hspa12a | -0.016848527 | -0.233634443 | 0.199937389 | 7.850092542 | -0.194662016 | 0.852674824 | 0.932549782 | -4.755011641 |
| **A_51_P290601** | Znhit2 | -0.024937601 | -0.3460537 | 0.296178499 | 10.69351137 | -0.194510446 | 0.852787841 | 0.93262063 | -4.755021388 |
| **A_55_P2020706** | NA | 0.024522969 | -0.29136396 | 0.340409899 | 7.940658347 | 0.194442744 | 0.852838324 | 0.932623086 | -4.75502574 |
| **A_55_P2414479** | NA | -0.02234309 | -0.310555938 | 0.265869759 | 7.991949613 | -0.194169152 | 0.85304234 | 0.932793428 | -4.755043309 |
| **A_55_P2149873** | Mapk1ip1 | -0.052079002 | -0.724576919 | 0.620418914 | 8.577920836 | -0.193964414 | 0.853195021 | 0.932838765 | -4.755056441 |
| **A_55_P1995126** | NA | -0.020701834 | -0.288130979 | 0.246727311 | 16.65076926 | -0.193887778 | 0.853252173 | 0.932838765 | -4.755061353 |
| **A_55_P2000603** | Slc50a1 | 0.031426461 | -0.395868908 | 0.458721831 | 10.75225046 | 0.195897999 | 0.853265809 | 0.932838765 | -4.726613879 |
| **A_51_P123920** | Mydgf | -0.041060421 | -0.571574932 | 0.48945409 | 11.50996125 | -0.193854761 | 0.853276796 | 0.932838765 | -4.741181884 |
| **A_55_P2112934** | C030034I | -0.020652074 | -0.28769511 | 0.246390963 | 8.421738605 | -0.193701396 | 0.853391173 | 0.932897764 | -4.755073291 |
| **A_52_P369123** | Fam189a2 | 0.021446155 | -0.255934101 | 0.298826411 | 7.939246665 | 0.193653002 | 0.853427265 | 0.932897764 | -4.755076389 |
| **A_52_P57308** | Cand1 | -0.033462607 | -0.466841167 | 0.399915954 | 12.83505255 | -0.193393864 | 0.853620538 | 0.93305628 | -4.755092964 |
| **A_55_P1992329** | Gmpr | 0.01622554 | -0.194419192 | 0.226870271 | 8.011254021 | 0.192929612 | 0.853966818 | 0.933376704 | -4.755122606 |
| **A_55_P2169829** | Armh4 | 0.017603571 | -0.211000135 | 0.246207276 | 8.218926591 | 0.192871406 | 0.854010236 | 0.933376704 | -4.755126317 |
| **A_55_P2101357** | Aftph | 0.019763667 | -0.236986522 | 0.276513855 | 11.53342111 | 0.192800029 | 0.854063479 | 0.933382133 | -4.755130867 |
| **A_66_P104473** | Polg2 | 0.022174743 | -0.266039031 | 0.310388517 | 9.478823325 | 0.192705543 | 0.854133962 | 0.933406399 | -4.755136887 |
| **A_55_P2050602** | Ncam1 | 0.020107196 | -0.24142501 | 0.281639402 | 7.805774575 | 0.192564701 | 0.854239027 | 0.933438257 | -4.755145855 |
| **A_52_P622230** | Gm4890 | -0.027813992 | -0.389639704 | 0.33401172 | 8.144428325 | -0.192537018 | 0.854259678 | 0.933438257 | -4.755147617 |
| **A_55_P2038101** | Dock11 | -0.034529269 | -0.483863472 | 0.414804934 | 9.704745888 | -0.192472307 | 0.854307953 | 0.933438257 | -4.755151735 |
| **A_52_P537466** | Zfp330 | -0.022803344 | -0.319924533 | 0.274317844 | 9.7606324 | -0.19222739 | 0.854490667 | 0.933585136 | -4.755167308 |
| **A_55_P2159869** | Zfp59 | 0.027806811 | -0.334801139 | 0.390414761 | 9.036949176 | 0.192072066 | 0.854606548 | 0.933658982 | -4.755177175 |
| **A_52_P381484** | Spon2 | -0.059886459 | -0.841867374 | 0.722094457 | 10.37022449 | -0.191815113 | 0.854798259 | 0.933815659 | -4.755193479 |
| **A_52_P443167** | Naa35 | 0.020900564 | -0.252172782 | 0.293973909 | 11.33938477 | 0.191703054 | 0.854881869 | 0.933854233 | -4.755200583 |
| **A_51_P311096** | Malsu1 | 0.027435334 | -0.331150512 | 0.38602118 | 13.04991152 | 0.191631744 | 0.854935077 | 0.933859592 | -4.755205102 |
| **A_52_P316697** | Ost4 | -0.034324028 | -0.483350206 | 0.41470215 | 12.11602776 | -0.191459506 | 0.855063594 | 0.933914201 | -4.755216009 |
| **A_55_P1976097** | NA | -0.028748358 | -0.404968705 | 0.347471989 | 10.22505066 | -0.191390822 | 0.855114845 | 0.933914201 | -4.755220356 |
| **A_55_P2002466** | Zfp747 | -0.037500402 | -0.528307912 | 0.453307108 | 10.26756518 | -0.191370534 | 0.855129984 | 0.933914201 | -4.755221639 |
| **A_66_P121059** | Mirg | -0.044784917 | -0.632271208 | 0.542701374 | 10.60824777 | -0.190934489 | 0.855455372 | 0.934150835 | -4.755249197 |
| **A_55_P2096737** | Elmod3 | 0.031643242 | -0.384023335 | 0.447309818 | 10.85654007 | 0.190671688 | 0.855651497 | 0.934150835 | -4.755265776 |
| **A_52_P412452** | Cntn6 | 0.016285866 | -0.197682201 | 0.230253933 | 7.775208266 | 0.190639215 | 0.855675731 | 0.934150835 | -4.755267823 |
| **A_55_P2144597** | 9030025P | 0.017025955 | -0.206752365 | 0.240804274 | 8.51883677 | 0.190565296 | 0.855730899 | 0.934150835 | -4.755272481 |
| **A_51_P323878** | Coro7 | -0.026574037 | -0.375881303 | 0.322733229 | 10.5742786 | -0.190546186 | 0.855745161 | 0.934150835 | -4.755273685 |
| **A_51_P418357** | Mov10l1 | -0.01655267 | -0.234215513 | 0.201110172 | 7.688536214 | -0.190473305 | 0.855799554 | 0.934150835 | -4.755278276 |

| | | | | | | | | | |
|---|---|---|---|---|---|---|---|---|---|
| **A_66_P121053** | Sftpd | -0.204089762 | -2.88838364 | 2.480204117 | 10.51937454 | -0.19043269 | 0.855829867 | 0.934150835 | -4.755280834 |
| **A_55_P2013598** | Ccnq | 0.019916524 | -0.242105701 | 0.281938748 | 10.44298264 | 0.190381941 | 0.855867744 | 0.934150835 | -4.755284029 |
| **A_51_P407606** | Ndc1 | -0.061424331 | -0.869650388 | 0.746801727 | 9.49188386 | -0.190352213 | 0.855889932 | 0.934150835 | -4.7552859 |
| **A_66_P107174** | NA | -0.018742306 | -0.265440876 | 0.227956263 | 8.254070644 | -0.190285958 | 0.855939382 | 0.934150835 | -4.755290069 |
| **A_51_P215922** | Casp6 | 0.067729561 | -0.823845432 | 0.959304555 | 10.8953361 | 0.19027018 | 0.855951159 | 0.934150835 | -4.755291062 |
| **A_51_P233855** | Adprhl1 | 0.063955501 | -0.778239442 | 0.906150443 | 8.508649824 | 0.190202237 | 0.85600187 | 0.934150835 | -4.755295336 |
| **A_55_P2096010** | NA | -0.032699699 | -0.463362971 | 0.397963573 | 13.14445654 | -0.190176243 | 0.856021271 | 0.934150835 | -4.75529697 |
| **A_55_P2050988** | NA | 0.064274428 | -0.7822468 | 0.910795656 | 9.571195632 | 0.190173812 | 0.856023086 | 0.934150835 | -4.755297123 |
| **A_55_P2112687** | Sipa1l1 | 0.0282846 | -0.344493046 | 0.401062246 | 10.29821775 | 0.190042409 | 0.856121165 | 0.934160769 | -4.755305383 |
| **A_51_P123017** | Gmcl1 | 0.027266273 | -0.332191602 | 0.386724147 | 10.77925671 | 0.189988851 | 0.856161142 | 0.934160769 | -4.746085297 |
| **A_55_P2134804** | Cinp | 0.018061824 | -0.220078441 | 0.25620209 | 8.67259925 | 0.189967418 | 0.85617714 | 0.934160769 | -4.755310095 |
| **A_55_P2157388** | Skint9 | 0.02952325 | -0.359987778 | 0.419034278 | 8.229418777 | 0.18984307 | 0.856269958 | 0.934184663 | -4.755317903 |
| **A_51_P126337** | Fgf12 | -0.040583543 | -0.576114259 | 0.494947172 | 8.001411254 | -0.189808616 | 0.856295676 | 0.934184663 | -4.748377407 |
| **A_52_P321140** | Defb1 | -0.016601048 | -0.235763027 | 0.202560932 | 7.66314338 | -0.18972328 | 0.856359376 | 0.934201443 | -4.755325421 |
| **A_55_P2146175** | NA | -0.027694588 | -0.393722875 | 0.3383337 | 9.10510033 | -0.189509331 | 0.856519085 | 0.934322951 | -4.755338836 |
| **A_55_P2210301** | Gm12992 | -0.039638696 | -0.563896689 | 0.484619298 | 8.834428504 | -0.189375869 | 0.856618716 | 0.934378915 | -4.755347197 |
| **A_55_P2050932** | Tlr7 | 0.031433443 | -0.385119797 | 0.447986684 | 8.32877869 | 0.189004344 | 0.85689608 | 0.93456095 | -4.755370441 |
| **A_55_P2023446** | NA | 0.018328477 | -0.224614007 | 0.261270962 | 11.41170408 | 0.188961472 | 0.856928088 | 0.93456095 | -4.75537312 |
| **A_51_P419226** | S100a14 | 0.137957067 | -1.691529131 | 1.967443265 | 11.47250643 | 0.188870948 | 0.856995673 | 0.93456095 | -4.755378776 |
| **A_51_P208919** | Acbd6 | -0.024476301 | -0.349137011 | 0.30018441 | 10.65567681 | -0.188827999 | 0.857027739 | 0.93456095 | -4.755381458 |
| **A_51_P302149** | Cfl2 | -0.026506037 | -0.378132345 | 0.325120271 | 9.034410926 | -0.18880513 | 0.857044813 | 0.93456095 | -4.755382886 |
| **A_55_P1976112** | C130074C | 0.028537949 | -0.350355866 | 0.407431764 | 10.53306216 | 0.188649467 | 0.857161037 | 0.93456095 | -4.755392602 |
| **A_55_P2115968** | NA | 0.044578335 | -0.547314323 | 0.636470994 | 9.923633552 | 0.188638892 | 0.857168932 | 0.93456095 | -4.755393262 |
| **A_51_P115471** | Apol8 | 0.016043597 | -0.197100731 | 0.229187924 | 7.677547686 | 0.18852906 | 0.857250939 | 0.93456095 | -4.750807427 |
| **A_51_P358243** | Tstd3 | -0.019123449 | -0.273193904 | 0.234947005 | 8.445891293 | -0.188522161 | 0.85725609 | 0.93456095 | -4.755400542 |
| **A_55_P2071236** | Aqp5 | -0.073735648 | -1.053462942 | 0.905991645 | 8.388217252 | -0.188504899 | 0.857268979 | 0.93456095 | -4.755401618 |
| **A_51_P299934** | BC048403 | -0.026484659 | -0.378686207 | 0.325716888 | 9.240695887 | -0.18834473 | 0.857388574 | 0.934633777 | -4.755411599 |
| **A_51_P418420** | Ddx3x | -0.03442807 | -0.492406588 | 0.423550448 | 10.62840718 | -0.188285949 | 0.857432466 | 0.934633777 | -4.75541526 |
| **A_51_P213691** | Scnn1a | -0.122156374 | -1.748641781 | 1.504329032 | 11.25116956 | -0.188111917 | 0.857562419 | 0.934722731 | -4.755426093 |
| **A_55_P2178578** | Tmprss4 | 0.044174031 | -0.545234262 | 0.633582323 | 9.110557499 | 0.187715931 | 0.857858127 | 0.934961579 | -4.755450704 |
| **A_55_P2058942** | Aldh3b1 | -0.064393448 | -0.923710225 | 0.794923329 | 9.234981675 | -0.187688963 | 0.857878267 | 0.934961579 | -4.755452378 |
| **A_52_P464629** | Nme7 | -0.018768307 | -0.269527182 | 0.231990568 | 8.818416261 | -0.187464541 | 0.85804587 | 0.935091531 | -4.755466302 |
| **A_66_P118934** | Plaa | -0.040575629 | -0.582893538 | 0.50174228 | 10.61315941 | -0.187396579 | 0.858096627 | 0.935094138 | -4.755470515 |
| **A_55_P1962303** | Hba-a1 | 0.038842915 | -0.480786204 | 0.558472034 | 18.16819909 | 0.187227088 | 0.858223214 | 0.935179374 | -4.755481016 |
| **A_55_P2015032** | Kazald1 | 0.033540176 | -0.415492124 | 0.482572475 | 8.656844926 | 0.187084624 | 0.858329619 | 0.935201419 | -4.755489836 |
| **A_51_P103650** | Trim8 | -0.022475717 | -0.323498307 | 0.278546873 | 12.04464943 | -0.187009992 | 0.858385362 | 0.935201419 | -4.748545402 |
| **A_51_P128876** | Ifitm3 | 0.06756618 | -0.837685938 | 0.972818298 | 12.12194126 | 0.18694341 | 0.858435094 | 0.935201419 | -4.755498571 |
| **A_51_P356283** | Fbxo31 | -0.023263081 | -0.334946081 | 0.28841992 | 10.96647148 | -0.186940954 | 0.858436929 | 0.935201419 | -4.755498723 |
| **A_55_P2150058** | Zfp593 | -0.031693225 | -0.456706728 | 0.393320278 | 9.811781826 | -0.186772981 | 0.858562394 | 0.935285403 | -4.755509105 |
| **A_55_P2003891** | Txlna | -0.032384597 | -0.467160888 | 0.402391693 | 10.91355849 | -0.186561916 | 0.858720054 | 0.935324144 | -4.755522138 |
| **A_52_P50329** | Stimate | 0.043632406 | -0.542199263 | 0.629464076 | 9.429994005 | 0.186546312 | 0.85873171 | 0.935324144 | -4.755523101 |
| **A_55_P2016667** | Gstm7 | 0.017697657 | -0.219939814 | 0.255335128 | 7.768151378 | 0.186531079 | 0.858743089 | 0.935324144 | -4.755524041 |
| **A_55_P1976764** | Polr3e | 0.038095145 | -0.473965275 | 0.550155566 | 9.743145426 | 0.186336859 | 0.858888172 | 0.935408316 | -4.755536019 |
| **A_51_P238183** | Etnk2 | 0.028066012 | -0.34926513 | 0.405397155 | 8.594152787 | 0.186298092 | 0.858917132 | 0.935408316 | -4.755538409 |
| **A_51_P492595** | Fbxo42 | -0.051476168 | -0.743906452 | 0.640954116 | 9.943220958 | -0.186200354 | 0.858990146 | 0.935435141 | -4.755544431 |
| **A_55_P2120993** | Zfp282 | -0.070256238 | -1.016122047 | 0.875609572 | 9.673108492 | -0.186039733 | 0.85911014 | 0.93551312 | -4.75555432 |
| **A_55_P1989921** | Eml2 | 0.034254732 | -0.42719928 | 0.495708744 | 11.89142638 | 0.185927013 | 0.859194351 | 0.935551752 | -4.755561256 |
| **A_51_P235088** | Dusp27 | 0.019060715 | -0.237803547 | 0.275924977 | 7.787240017 | 0.185859963 | 0.859244444 | 0.935551752 | -4.755565379 |

| | | | | | | | | | |
|---|---|---|---|---|---|---|---|---|---|
| **A_52_P141161** | Hnrnpf | 0.032464064 | -0.405230505 | 0.470158634 | 9.507033242 | 0.185772778 | 0.859309581 | 0.935551752 | -4.755570739 |
| **A_55_P2123362** | NA | 0.029083297 | -0.363113924 | 0.421280517 | 8.553261395 | 0.185733171 | 0.859339172 | 0.935551752 | -4.755573173 |
| **A_51_P464918** | Mefv | -0.016502287 | -0.239187606 | 0.206183032 | 7.731915732 | -0.185610653 | 0.859430709 | 0.935598724 | -4.755580698 |
| **A_55_P1999289** | Ovol2 | 0.029894951 | -0.374167565 | 0.433957466 | 11.77609838 | 0.185310327 | 0.859655103 | 0.93573426 | -4.755599126 |
| **A_55_P2154797** | 2810006K | 0.016422102 | -0.205630287 | 0.238474491 | 8.29118512 | 0.185235248 | 0.859711202 | 0.93573426 | -4.755603728 |
| **A_55_P2122429** | Cym | 0.061676983 | -0.772397478 | 0.895751445 | 8.992992891 | 0.185211797 | 0.859728725 | 0.93573426 | -4.755605165 |
| **A_55_P2370250** | Syn3 | 0.018648234 | -0.233573449 | 0.270869916 | 7.932068623 | 0.185184923 | 0.859748805 | 0.93573426 | -4.755606811 |
| **A_51_P103706** | Cyp2c29 | 0.016661524 | -0.208834481 | 0.242157528 | 7.702043025 | 0.185065817 | 0.859837804 | 0.935754537 | -4.743996677 |
| **A_55_P1994882** | Ccdc174 | -0.020065986 | -0.291698336 | 0.251566365 | 10.84509802 | -0.185024548 | 0.859868641 | 0.935754537 | -4.755616633 |
| **A_65_P16688** | Taok2 | 0.028570156 | -0.358323565 | 0.415463877 | 8.647681859 | 0.184957217 | 0.859918955 | 0.935754537 | -4.755620754 |
| **A_55_P2102515** | Daam1 | -0.024485677 | -0.356168831 | 0.307197478 | 9.895496738 | -0.184900906 | 0.859961033 | 0.935754537 | -4.7556242 |
| **A_51_P179258** | Kif26b | -0.018244268 | -0.265584113 | 0.229095577 | 8.091666339 | -0.184749256 | 0.860074356 | 0.935825179 | -4.755633473 |
| **A_51_P386539** | Rnf125 | -0.021523795 | -0.313779411 | 0.270731821 | 8.691742647 | -0.184461749 | 0.860289213 | 0.936006282 | -4.755651035 |
| **A_55_P2255399** | F630048H | 0.020747434 | -0.261813781 | 0.303308648 | 8.191210553 | 0.18390866 | 0.860702577 | 0.936321157 | -4.755684742 |
| **A_55_P2098226** | C1rl | 0.020415712 | -0.257776126 | 0.29860755 | 8.18227971 | 0.183810577 | 0.860775887 | 0.936321157 | -4.75569071 |
| **A_55_P1982414** | NA | 0.020331551 | -0.256734311 | 0.297397413 | 8.160801375 | 0.183796755 | 0.860786218 | 0.936321157 | -4.75569155 |
| **A_55_P2094881** | Fgfr2 | -0.02976247 | -0.456698144 | 0.397173204 | 8.932503446 | -0.185681756 | 0.860808712 | 0.936321157 | -4.727177563 |
| **A_51_P160482** | Vsig2 | 0.056185619 | -0.709729245 | 0.822100482 | 8.740042921 | 0.18373632 | 0.86083139 | 0.936321157 | -4.755695225 |
| **A_51_P520084** | Lypla2 | -0.028510153 | -0.417314476 | 0.360294171 | 13.35152493 | -0.183661786 | 0.860887101 | 0.936321157 | -4.755699756 |
| **A_55_P1999923** | Nyap1 | 0.055084994 | -0.696298063 | 0.806468051 | 9.137909776 | 0.183620959 | 0.860917617 | 0.936321157 | -4.755702237 |
| **A_51_P329158** | Zc3h10 | -0.018732438 | -0.274527434 | 0.237062557 | 8.916024276 | -0.183422504 | 0.861065959 | 0.936376071 | -4.755714289 |
| **A_55_P1974233** | Dtd2 | 0.01642451 | -0.20792995 | 0.240778971 | 8.403945845 | 0.183361464 | 0.861111586 | 0.936376071 | -4.755717993 |
| **A_51_P200776** | Grsf1 | 0.034747161 | -0.439967971 | 0.509462294 | 10.51942958 | 0.183331293 | 0.861134139 | 0.936376071 | -4.755719824 |
| **A_51_P214251** | Rnls | 0.023237109 | -0.294312472 | 0.34078669 | 8.04552168 | 0.183282454 | 0.861170647 | 0.936376071 | -4.755722787 |
| **A_55_P1998209** | NA | 0.029886237 | -0.378739477 | 0.43851195 | 15.90793592 | 0.18318752 | 0.861241613 | 0.936376071 | -4.755728543 |
| **A_55_P2160416** | Acoxl | -0.064688412 | -0.949342824 | 0.819966001 | 8.333351132 | -0.183148423 | 0.861270839 | 0.936376071 | -4.755730913 |
| **A_55_P2034740** | Exosc2 | 0.016561486 | -0.210024697 | 0.243147668 | 17.5024273 | 0.183069595 | 0.861329767 | 0.936376071 | -4.75573569 |
| **A_51_P423825** | Morn1 | 0.023076946 | -0.292710102 | 0.338863994 | 8.402242839 | 0.183035091 | 0.861355561 | 0.936376071 | -4.75573778 |
| **A_55_P2001963** | NA | 0.035909656 | -0.455740664 | 0.527559976 | 15.56301086 | 0.182938564 | 0.861427721 | 0.936401865 | -4.755743625 |
| **A_55_P2126627** | Phf20l1 | -0.016629292 | -0.244577809 | 0.211319224 | 8.125612446 | -0.182720526 | 0.861590725 | 0.936491331 | -4.755756818 |
| **A_51_P315595** | NA | -0.024150901 | -0.355246515 | 0.306944713 | 16.44686112 | -0.182696512 | 0.861608678 | 0.936491331 | -4.75575827 |
| **A_55_P2061465** | Ppp1r21 | 0.021476169 | -0.273215214 | 0.316167552 | 9.484656557 | 0.182532297 | 0.86173145 | 0.936491331 | -4.755768194 |
| **A_52_P89064** | Dusp18 | -0.02263861 | -0.333321785 | 0.288044564 | 8.316061109 | -0.182508195 | 0.86174947 | 0.936491331 | -4.75576965 |
| **A_55_P2165234** | Slurp2 | 0.031590736 | -0.402071078 | 0.465252551 | 8.385933375 | 0.182456321 | 0.861788253 | 0.936491331 | -4.755772783 |
| **A_51_P175974** | NA | -0.021414538 | -0.315472507 | 0.272643431 | 8.356467582 | -0.182400533 | 0.861829964 | 0.936491331 | -4.755776151 |
| **A_55_P2010611** | Slc30a5 | -0.018889403 | -0.278455062 | 0.240676256 | 10.41087859 | -0.18227258 | 0.861925631 | 0.936491331 | -4.755783873 |
| **A_51_P469480** | Yes1 | 0.026659468 | -0.358925397 | 0.412244333 | 8.353430161 | 0.184159533 | 0.861934118 | 0.936491331 | -4.727259039 |
| **A_52_P73559** | NA | 0.019925711 | -0.253921058 | 0.293772479 | 8.560484177 | 0.182245387 | 0.861945963 | 0.936491331 | -4.755785513 |
| **A_55_P1996518** | Arhgap42 | -0.02214654 | -0.326661969 | 0.282368888 | 8.49068052 | -0.18215745 | 0.862011712 | 0.936495655 | -4.755790816 |
| **A_66_P109183** | Apold1 | 0.019978257 | -0.254833746 | 0.29479026 | 8.363715716 | 0.182084192 | 0.862066488 | 0.936495655 | -4.755795231 |
| **A_55_P1994339** | Dhrs7c | 0.030449151 | -0.388484329 | 0.449382632 | 8.011038006 | 0.182045718 | 0.862095256 | 0.936495655 | -4.75579755 |
| **A_55_P2088375** | Tnnt3 | -0.066714062 | -0.986206573 | 0.852778448 | 8.852623342 | -0.181727033 | 0.86233355 | 0.936673131 | -4.755816734 |
| **A_55_P2080396** | Mrpl4 | 0.021321158 | -0.272587126 | 0.315229441 | 12.98227642 | 0.181697645 | 0.862355526 | 0.936673131 | -4.755818502 |
| **A_55_P2044468** | Marf1 | -0.054976691 | -0.813733803 | 0.703780421 | 8.964386378 | -0.181478912 | 0.862519093 | 0.936789507 | -4.755831648 |
| **A_51_P446570** | Bbox1 | -0.017438295 | -0.258183571 | 0.22330698 | 7.772176365 | -0.18142478 | 0.862559574 | 0.936789507 | -4.7558349 |
| **A_51_P110931** | Znrd1as | -0.023356027 | -0.346591267 | 0.299879213 | 9.411406457 | -0.180980012 | 0.862892195 | 0.937098113 | -4.751257734 |
| **A_55_P2036459** | Med24 | 0.015061131 | -0.193673067 | 0.223795328 | 8.607739036 | 0.180723372 | 0.863084138 | 0.937253916 | -4.755876939 |
| **A_55_P2026823** | Uspl1 | -0.054646919 | -0.812416355 | 0.703122517 | 10.4528456 | -0.180625451 | 0.863157377 | 0.937280805 | -4.755882796 |

| | | | | | | | | | |
|---|---|---|---|---|---|---|---|---|---|
| **A_51_P288138** | Fpr2 | 0.061482949 | -0.791627626 | 0.914593523 | 8.470671921 | 0.180509352 | 0.863244213 | 0.937322454 | -4.755889735 |
| **A_51_P406165** | Nxpe2 | 0.015549845 | -0.200490537 | 0.231590227 | 7.788270128 | 0.18027748 | 0.86341765 | 0.937349384 | -4.755903582 |
| **A_55_P1964038** | Gm10032 | -0.028044885 | -0.417719446 | 0.361629677 | 8.528203085 | -0.180261071 | 0.863429924 | 0.937349384 | -4.755904561 |
| **A_55_P2032079** | Dbp | 0.043147521 | -0.556482051 | 0.642777094 | 9.526906732 | 0.180228371 | 0.863454384 | 0.937349384 | -4.755906512 |
| **A_66_P105830** | 1700028B | 0.04546206 | -0.586518929 | 0.677443048 | 14.57597206 | 0.180175377 | 0.863494024 | 0.937349384 | -4.755909673 |
| **A_55_P2042993** | Arl6ip6 | -0.041650855 | -0.620725974 | 0.537424263 | 14.68924753 | -0.180152115 | 0.863511424 | 0.937349384 | -4.755911061 |
| **A_55_P2376423** | Igkv4-72 | -0.024077767 | -0.359469276 | 0.311313741 | 8.301268708 | -0.179810271 | 0.86376714 | 0.937574325 | -4.755931428 |
| **A_51_P343833** | Traf1 | -0.125904802 | -1.881225877 | 1.629416273 | 9.391362211 | -0.179653662 | 0.863884297 | 0.937623749 | -4.755940747 |
| **A_55_P2065529** | Mok | 0.05949694 | -0.770145357 | 0.889139238 | 9.854718187 | 0.179619752 | 0.863909665 | 0.937623749 | -4.755942764 |
| **A_55_P2018611** | Igfbp7 | -0.062999164 | -0.943696489 | 0.817698161 | 8.717526705 | -0.179167178 | 0.864248254 | 0.937871032 | -4.755969643 |
| **A_55_P2122180** | Snta1 | -0.045772976 | -0.685773249 | 0.594227298 | 10.30049295 | -0.17913454 | 0.864272673 | 0.937871032 | -4.755971578 |
| **A_51_P401527** | Mrm3 | -0.024279094 | -0.363776969 | 0.315218782 | 8.263238837 | -0.179120691 | 0.864283034 | 0.937871032 | -4.7559724 |
| **A_52_P400857** | Zfp595 | 0.017805237 | -0.231448619 | 0.267059093 | 8.616053576 | 0.178918902 | 0.864434013 | 0.937872316 | -4.75598436 |
| **A_55_P2071952** | Wdr92 | 0.057947149 | -0.753806908 | 0.869701205 | 12.33516776 | 0.178796062 | 0.864525925 | 0.937872316 | -4.755991634 |
| **A_52_P494126** | Usp46 | 0.024176462 | -0.314664105 | 0.363017028 | 9.575346698 | 0.178709518 | 0.86459068 | 0.937872316 | -4.755996756 |
| **A_55_P2032388** | Trim24 | 0.02809432 | -0.365730703 | 0.421919342 | 9.853654015 | 0.178675728 | 0.864615964 | 0.937872316 | -4.755998755 |
| **A_51_P184062** | Hbs1l | 0.040240737 | -0.524041417 | 0.604522891 | 10.63505411 | 0.178615833 | 0.864660781 | 0.937872316 | -4.756002298 |
| **A_51_P176265** | Trip11 | -0.023141426 | -0.347668965 | 0.301386114 | 8.682277878 | -0.178603062 | 0.864670337 | 0.937872316 | -4.756003053 |
| **A_51_P423976** | Crem | -0.017083823 | -0.25670789 | 0.222540245 | 8.401877297 | -0.178568549 | 0.864696162 | 0.937872316 | -4.756005094 |
| **A_52_P78770** | Chpt1 | 0.021860909 | -0.28480146 | 0.328523278 | 8.432580071 | 0.178549267 | 0.86471059 | 0.937872316 | -4.756006234 |
| **A_52_P49406** | Pde12 | 0.038259988 | -0.49863089 | 0.575150866 | 8.897545102 | 0.178488051 | 0.864756397 | 0.937872316 | -4.756009852 |
| **A_55_P2289040** | NA | -0.026193612 | -0.393796207 | 0.341408983 | 8.410982233 | -0.178470797 | 0.864769307 | 0.937872316 | -4.756010872 |
| **A_55_P2041843** | Nde1 | 0.015508988 | -0.202534039 | 0.233552016 | 8.368711451 | 0.178152371 | 0.865007589 | 0.93807812 | -4.756029672 |
| **A_55_P2142393** | Cacna1a | 0.050613788 | -0.661255972 | 0.762483548 | 9.018373826 | 0.178081433 | 0.865060675 | 0.938083071 | -4.756033855 |
| **A_51_P372141** | Pnkd | -0.016272154 | -0.245249124 | 0.212704817 | 7.92456412 | -0.177993267 | 0.865126654 | 0.938102004 | -4.756039053 |
| **A_55_P1997544** | NA | -0.015162898 | -0.228733448 | 0.198407652 | 7.674002875 | -0.177824338 | 0.865253076 | 0.93816036 | -4.756049005 |
| **A_66_P118165** | NA | -0.064945613 | -0.979937767 | 0.85004654 | 9.855683906 | -0.177779958 | 0.86528629 | 0.93816036 | -4.756051617 |
| **A_52_P14456** | Srrm4 | -0.014991914 | -0.226382459 | 0.19639863 | 7.803812393 | -0.177632274 | 0.865396817 | 0.93816036 | -4.756060308 |
| **A_55_P2106078** | F10 | -0.029197195 | -0.440991615 | 0.382597224 | 17.14531499 | -0.177586939 | 0.865430747 | 0.93816036 | -4.756062974 |
| **A_55_P2096887** | Steap3 | -0.02962307 | -0.447490245 | 0.388244105 | 8.39477057 | -0.177558781 | 0.865451821 | 0.93816036 | -4.75606463 |
| **A_55_P2158319** | Ntan1 | -0.022614394 | -0.341663726 | 0.296434938 | 10.88614636 | -0.177532335 | 0.865471614 | 0.93816036 | -4.756066184 |
| **A_52_P106537** | Chrac1 | 0.037475346 | -0.4920678 | 0.567018492 | 10.01942561 | 0.177253429 | 0.865680362 | 0.938318014 | -4.756082568 |
| **A_55_P2217309** | NA | 0.016638381 | -0.218529109 | 0.251805872 | 7.982256608 | 0.177208331 | 0.865714117 | 0.938318014 | -4.756085215 |
| **A_66_P130602** | Pdzd8 | -0.016604109 | -0.251448751 | 0.218240533 | 8.146655696 | -0.177086425 | 0.865805363 | 0.938350964 | -4.756092366 |
| **A_55_P2004606** | Serpina4- | -0.014800247 | -0.224188701 | 0.194588207 | 7.656104202 | -0.177038033 | 0.865841585 | 0.938350964 | -4.756095204 |
| **A_55_P1961059** | Usp16 | 0.023798218 | -0.31310392 | 0.360700355 | 11.02256099 | 0.176925735 | 0.865925642 | 0.93838946 | -4.756101785 |
| **A_55_P2115028** | Fastkd1 | -0.027720035 | -0.420483897 | 0.365043827 | 8.676523727 | -0.176771645 | 0.866040984 | 0.938461853 | -4.75611081 |
| **A_51_P407248** | Antxr1 | 0.065616236 | -0.864539889 | 0.995772361 | 11.73200944 | 0.176687494 | 0.866103976 | 0.938477513 | -4.756115735 |
| **A_55_P2412349** | AW11184 | -0.014981439 | -0.227494441 | 0.197531563 | 7.69861851 | -0.17657059 | 0.866191487 | 0.938519739 | -4.756122573 |
| **A_55_P1964902** | NA | -0.016982402 | -0.258148281 | 0.224183478 | 7.977946899 | -0.176373604 | 0.866338951 | 0.938626914 | -4.756134085 |
| **A_51_P224564** | Ppm1f | 0.034170271 | -0.451768957 | 0.520109499 | 12.19276358 | 0.176123298 | 0.866526338 | 0.938777329 | -4.756148696 |
| **A_51_P414115** | Naprt | -0.047189611 | -0.718571629 | 0.624192408 | 9.129126094 | -0.176046354 | 0.866583942 | 0.938787132 | -4.756153183 |
| **A_55_P2070352** | Csf3r | -0.019126916 | -0.291659035 | 0.253405203 | 14.43126712 | -0.175783298 | 0.866780888 | 0.938902199 | -4.756168509 |
| **A_51_P308912** | Mybphl | 0.055141423 | -0.730585827 | 0.840868674 | 8.253522462 | 0.175774756 | 0.866787284 | 0.938902199 | -4.756169007 |
| **A_52_P504268** | B3galnt1 | 0.024664067 | -0.327051289 | 0.376379423 | 10.38002138 | 0.175640114 | 0.866888092 | 0.938923888 | -4.756176842 |
| **A_55_P2174632** | Rpl7 | 0.028279841 | -0.375147565 | 0.431707247 | 16.24321839 | 0.175574689 | 0.866937078 | 0.938923888 | -4.756180647 |
| **A_66_P126459** | Gm13429 | -0.021846023 | -0.333591473 | 0.289899427 | 7.869836573 | -0.175518384 | 0.866979237 | 0.938923888 | -4.756183921 |
| **A_51_P478098** | Epb41l5 | -0.022589604 | -0.345000817 | 0.299821608 | 8.480987454 | -0.175488568 | 0.867001561 | 0.938923888 | -4.756185654 |

| | | | | | | | | | |
|---|---|---|---|---|---|---|---|---|---|
| **A_55_P2027831** | Cryba4 | 0.016830633 | -0.223554176 | 0.257215441 | 8.216157344 | 0.175365342 | 0.867093827 | 0.938971214 | -4.756192814 |
| **A_52_P200458** | Fut4 | 0.032880614 | -0.437321645 | 0.503082872 | 8.569145521 | 0.175148148 | 0.86725646 | 0.939094729 | -4.756205422 |
| **A_55_P2028046** | Cyp3a59 | -0.014489579 | -0.221831958 | 0.192852799 | 7.658250362 | -0.175032238 | 0.867343254 | 0.939136115 | -4.756212144 |
| **A_51_P166740** | Kcnj4 | -0.015033151 | -0.230380712 | 0.20031441 | 7.759343162 | -0.174847895 | 0.867481297 | 0.939190444 | -4.756222826 |
| **A_55_P2185840** | Nnmt | -0.016733745 | -0.256522978 | 0.223055488 | 7.971065384 | -0.174788886 | 0.867525486 | 0.939190444 | -4.756226243 |
| **A_55_P2203996** | 4932433N | -0.016537568 | -0.253589109 | 0.220513973 | 8.929612157 | -0.17473472 | 0.867566048 | 0.939190444 | -4.756229378 |
| **A_55_P2064712** | NA | 0.031597943 | -0.421405963 | 0.484601849 | 15.85930323 | 0.174705756 | 0.867587739 | 0.939190444 | -4.756231055 |
| **A_55_P2071834** | Tln2 | 0.020715905 | -0.276410055 | 0.317841864 | 8.335851949 | 0.174627909 | 0.867646036 | 0.939200967 | -4.756235558 |
| **A_55_P2016670** | Alpk2 | 0.014590911 | -0.19483384 | 0.224015662 | 7.66287187 | 0.174503743 | 0.867739024 | 0.939249036 | -4.756242738 |
| **A_51_P136781** | Pnliprp1 | 0.023285565 | -0.311333434 | 0.357904563 | 7.910515945 | 0.17429564 | 0.867894876 | 0.939336685 | -4.75625476 |
| **A_51_P250358** | Prpf39 | -0.024113791 | -0.370809393 | 0.322581812 | 9.066383079 | -0.17420776 | 0.867960692 | 0.939336685 | -4.756259833 |
| **A_55_P1973521** | Rnf7 | -0.017671029 | -0.271778156 | 0.236436098 | 10.3272775 | -0.174178818 | 0.867982369 | 0.939336685 | -4.756261503 |
| **A_55_P2019463** | Tnks1bp1 | 0.0444355 | -0.594811334 | 0.683682333 | 11.14569146 | 0.174105234 | 0.86803748 | 0.939336685 | -4.756265748 |
| **A_55_P2057720** | Zfp937 | -0.016491738 | -0.253787336 | 0.22080386 | 8.943814541 | -0.174071264 | 0.868062922 | 0.939336685 | -4.756267707 |
| **A_52_P262676** | NA | -0.04610199 | -0.710704827 | 0.618500848 | 9.842679131 | -0.173743202 | 0.868308638 | 0.939529332 | -4.756286606 |
| **A_51_P283292** | Gm14326 | 0.026657596 | -0.357750357 | 0.411065549 | 9.159068209 | 0.17369166 | 0.868347244 | 0.939529332 | -4.756289573 |
| **A_51_P249867** | Snrnp48 | -0.020815367 | -0.321110258 | 0.279479524 | 9.600070789 | -0.17361469 | 0.868404897 | 0.939529332 | -4.756294 |
| **A_55_P2163928** | Rtl8c | 0.015211927 | -0.204295767 | 0.234719621 | 8.369471712 | 0.173574061 | 0.868435331 | 0.939529332 | -4.756296337 |
| **A_55_P2228918** | Rslcan18 | -0.016357192 | -0.252779443 | 0.220065059 | 8.550696712 | -0.1732889 | 0.868648935 | 0.93966431 | -4.756312721 |
| **A_66_P135194** | Yrdc | -0.021483273 | -0.332131161 | 0.289164616 | 8.417809028 | -0.173213755 | 0.868705225 | 0.93966431 | -4.756317034 |
| **A_66_P125071** | Ssb | 0.019221299 | -0.258719922 | 0.29716252 | 10.37420428 | 0.173212854 | 0.868705901 | 0.93966431 | -4.756317086 |
| **A_52_P471395** | Ifngr2 | 0.063334561 | -0.853262321 | 0.979931443 | 11.21879297 | 0.173066389 | 0.868815619 | 0.939730415 | -4.756325487 |
| **A_51_P105017** | Rad52 | -0.017095683 | -0.265003253 | 0.230811888 | 8.382205201 | -0.172721739 | 0.869073813 | 0.939957099 | -4.756345229 |
| **A_55_P1962299** | Hba-a2 | 0.018346894 | -0.247888553 | 0.284582342 | 17.74073574 | 0.172602479 | 0.869163161 | 0.940001149 | -4.756352051 |
| **A_55_P2016346** | Eif3d | 0.01610061 | -0.217640001 | 0.249841221 | 14.14765905 | 0.172527564 | 0.869219287 | 0.940009268 | -4.756356334 |
| **A_51_P125446** | Lzic | 0.021600784 | -0.292277335 | 0.335478903 | 9.69924407 | 0.172368864 | 0.869338187 | 0.940085268 | -4.746923175 |
| **A_55_P2207055** | NA | -0.015006583 | -0.233204292 | 0.203191127 | 7.732720009 | -0.172259018 | 0.869420488 | 0.940121684 | -4.756371673 |
| **A_55_P2201676** | Cyp2r1 | 0.044398428 | -0.601792366 | 0.690589223 | 9.06735511 | 0.17209061 | 0.869546669 | 0.940165937 | -4.75638128 |
| **A_51_P268193** | Slc7a10 | 0.056250305 | -0.7625511 | 0.87505171 | 8.577387051 | 0.172066628 | 0.869564638 | 0.940165937 | -4.756382647 |
| **A_55_P2140212** | NA | -0.054633493 | -0.850639604 | 0.741372618 | 13.62384072 | -0.171906743 | 0.869684437 | 0.940165937 | -4.756391759 |
| **A_55_P1970954** | Pknox2 | 0.019810485 | -0.268853971 | 0.30847494 | 9.087768187 | 0.171890605 | 0.86969653 | 0.940165937 | -4.756392678 |
| **A_51_P181400** | Efcab2 | 0.019090427 | -0.259125116 | 0.297305971 | 8.260879447 | 0.171863876 | 0.869716558 | 0.940165937 | -4.7563942 |
| **A_55_P2124523** | Gm13620 | -0.01470754 | -0.229109772 | 0.199694692 | 7.772891947 | -0.171815004 | 0.869753178 | 0.940165937 | -4.756396983 |
| **A_51_P341664** | Zfp87 | 0.021557404 | -0.292974619 | 0.336089428 | 9.013592884 | 0.171665073 | 0.869865525 | 0.94023481 | -4.756405514 |
| **A_55_P1965219** | Syt6 | -0.016221758 | -0.253167128 | 0.220723612 | 8.466569485 | -0.171474694 | 0.870008185 | 0.940336441 | -4.756416337 |
| **A_51_P113205** | F13b | -0.01524451 | -0.238502207 | 0.208013186 | 7.672092036 | -0.171024128 | 0.870345838 | 0.940648803 | -4.74477354 |
| **A_55_P2000968** | Ppp2r3c | 0.014420088 | -0.197028375 | 0.225868551 | 9.952989295 | 0.170810171 | 0.870506188 | 0.940769516 | -4.756454023 |
| **A_55_P2076524** | 1700028I1 | -0.050344791 | -0.789267282 | 0.688577701 | 10.02449016 | -0.170649885 | 0.870626318 | 0.940830996 | -4.756463092 |
| **A_55_P2110935** | Mlip | -0.018387133 | -0.288331397 | 0.251557131 | 7.832663308 | -0.170604412 | 0.8706604 | 0.940830996 | -4.756465663 |
| **A_55_P1992838** | Socs2 | 0.095644049 | -1.309461384 | 1.500749482 | 9.608589108 | 0.170490187 | 0.870746012 | 0.940870921 | -4.756472118 |
| **A_51_P466288** | Frmd5 | 0.041730413 | -0.573209768 | 0.656670594 | 8.309910912 | 0.169969171 | 0.87113654 | 0.941240297 | -4.756501511 |
| **A_51_P173086** | Eif3l | 0.017788913 | -0.244534588 | 0.280112415 | 15.04069212 | 0.169848832 | 0.871226746 | 0.941285158 | -4.756508287 |
| **A_55_P2038519** | NA | 0.021472367 | -0.295587736 | 0.338532471 | 9.042213287 | 0.169624538 | 0.871394882 | 0.941414207 | -4.756520905 |
| **A_55_P2063151** | NA | -0.02289911 | -0.361397882 | 0.315599662 | 10.83889477 | -0.169438403 | 0.871534419 | 0.94148929 | -4.756531363 |
| **A_51_P509098** | Asah1 | -0.052138801 | -0.82315152 | 0.718873918 | 10.43388177 | -0.169375208 | 0.871581795 | 0.94148929 | -4.756534911 |
| **A_51_P106488** | Speg | -0.039103497 | -0.617484711 | 0.539277718 | 10.53727892 | -0.16933696 | 0.871610469 | 0.94148929 | -4.756537058 |
| **A_55_P1961750** | Dtd2 | 0.017370032 | -0.239762645 | 0.274502709 | 9.14968028 | 0.169197403 | 0.871715095 | 0.9415497 | -4.756544887 |
| **A_52_P496655** | Leprotl1 | 0.015716864 | -0.217207888 | 0.248641616 | 8.468156966 | 0.169005406 | 0.871859038 | 0.941570588 | -4.756555648 |

| | | | | | | | | | |
|---|---|---|---|---|---|---|---|---|---|
| **A_55_P2203478** | Scn5a | -0.027979312 | -0.442654367 | 0.386695742 | 8.367223573 | -0.168997185 | 0.871865202 | 0.941570588 | -4.756556108 |
| **A_55_P2386592** | Cep63 | -0.027207118 | -0.430486459 | 0.376072222 | 11.47337301 | -0.168976735 | 0.871880533 | 0.941570588 | -4.756557254 |
| **A_55_P2172196** | Thnsl1 | -0.014655578 | -0.232245333 | 0.202934177 | 7.860972334 | -0.168699913 | 0.872088084 | 0.941716578 | -4.756572745 |
| **A_55_P2039439** | Ppp1ca | 0.025858421 | -0.358134759 | 0.409851601 | 15.35651077 | 0.168666503 | 0.872113134 | 0.941716578 | -4.756574613 |
| **A_55_P2093764** | Eif2s3x | 0.025434142 | -0.352859401 | 0.403727686 | 9.075813236 | 0.168398615 | 0.872313998 | 0.941880869 | -4.756589578 |
| **A_55_P1972923** | Polr1d | 0.02106187 | -0.292827212 | 0.334950952 | 12.88250318 | 0.168062595 | 0.872565962 | 0.942100314 | -4.756608315 |
| **A_52_P98625** | Sowahc | 0.033920011 | -0.472274761 | 0.540114783 | 8.770367633 | 0.167837376 | 0.872734852 | 0.942168218 | -4.756620854 |
| **A_55_P2055708** | Vmn1r48 | 0.01549557 | -0.215831503 | 0.246822644 | 7.857061042 | 0.167776624 | 0.872780411 | 0.942168218 | -4.756624233 |
| **A_55_P2021455** | NA | 0.062196723 | -0.86662544 | 0.991018886 | 11.6241825 | 0.167720164 | 0.87282275 | 0.942168218 | -4.756627372 |
| **A_52_P2706** | Mypn | 0.014054945 | -0.195908365 | 0.224018255 | 7.687801469 | 0.167662555 | 0.872865953 | 0.942168218 | -4.756630575 |
| **A_55_P2111355** | Egfl7 | -0.077661782 | -1.238159937 | 1.082836374 | 14.96350252 | -0.167615138 | 0.872901513 | 0.942168218 | -4.75663321 |
| **A_55_P2254779** | NA | -0.021113197 | -0.336741993 | 0.294515599 | 8.606634848 | -0.167543555 | 0.872955197 | 0.942168218 | -4.756637186 |
| **A_65_P03606** | Rpap2 | 0.017725185 | -0.247375822 | 0.282826191 | 8.135397355 | 0.167467191 | 0.873012467 | 0.942168218 | -4.756641427 |
| **A_55_P2091486** | Tmem219 | -0.020245676 | -0.323058634 | 0.282567283 | 10.88787367 | -0.167458873 | 0.873018704 | 0.942168218 | -4.756641888 |
| **A_51_P475938** | Mfsd6l | 0.037736283 | -0.52708172 | 0.602554286 | 8.402115229 | 0.167340453 | 0.873107517 | 0.942211472 | -4.75664846 |
| **A_51_P101228** | Gmeb2 | 0.016506826 | -0.231267017 | 0.26428067 | 8.233492183 | 0.166862394 | 0.873466068 | 0.942545793 | -4.756674942 |
| **A_51_P115085** | Nup153 | -0.018802822 | -0.301277021 | 0.263671377 | 10.38168593 | -0.166722643 | 0.87357089 | 0.942606295 | -4.741979709 |
| **A_55_P2074005** | Tctn1 | -0.022571164 | -0.362265153 | 0.317122824 | 8.505559666 | -0.166424188 | 0.87379476 | 0.942795239 | -4.756699151 |
| **A_51_P463765** | Timp3 | -0.053729347 | -0.863168331 | 0.755709638 | 13.57316184 | -0.166256176 | 0.87392079 | 0.942878603 | -4.756708417 |
| **A_52_P493519** | Igll1 | -0.014008765 | -0.225485707 | 0.197468176 | 7.684392614 | -0.165915587 | 0.874176289 | 0.943101633 | -4.756727171 |
| **A_55_P2011802** | Pdk2 | 0.035079256 | -0.495001243 | 0.565159755 | 9.576833723 | 0.165752044 | 0.874298979 | 0.943142541 | -4.756736163 |
| **A_51_P265348** | Ldb3 | -0.016844478 | -0.271447891 | 0.237758934 | 8.082418225 | -0.16570808 | 0.874331962 | 0.943142541 | -4.756738579 |
| **A_55_P2086500** | Dennd5a | 0.02433256 | -0.343632897 | 0.392298017 | 12.65036279 | 0.165626986 | 0.8743928 | 0.943142541 | -4.756743033 |
| **A_55_P1955970** | Cbfa2t2 | 0.019147239 | -0.27044251 | 0.308736987 | 10.84857647 | 0.165604949 | 0.874409334 | 0.943142541 | -4.756744243 |
| **A_66_P119191** | Inpp5e | -0.022315839 | -0.360520026 | 0.315888347 | 8.937747355 | -0.165266409 | 0.874663326 | 0.943363871 | -4.756762813 |
| **A_55_P2082658** | NA | -0.030935856 | -0.500016459 | 0.438144746 | 8.38013453 | -0.165182872 | 0.874726004 | 0.943378845 | -4.75676739 |
| **A_51_P491470** | Ddx47 | -0.029959982 | -0.484830767 | 0.424910804 | 9.914529887 | -0.164969572 | 0.874886046 | 0.943463582 | -4.756779065 |
| **A_55_P1970380** | NA | -0.01606019 | -0.260016285 | 0.227895906 | 8.619067257 | -0.164888111 | 0.874947169 | 0.943463582 | -4.75678352 |
| **A_55_P2010312** | Serpina1a | -0.093942267 | -1.520979754 | 1.333095219 | 9.05080365 | -0.164883046 | 0.874950969 | 0.943463582 | -4.756783797 |
| **A_55_P1955039** | Myo1b | 0.038898002 | -0.552442412 | 0.630238416 | 9.578828333 | 0.16475556 | 0.875046628 | 0.94351411 | -4.756790764 |
| **A_52_P136709** | Pctp | 0.013959921 | -0.198387808 | 0.22630765 | 7.844378878 | 0.164659084 | 0.87511902 | 0.943539546 | -4.756796033 |
| **A_66_P126984** | Gm10523 | -0.019800576 | -0.321308489 | 0.281707336 | 8.122237312 | -0.1644862 | 0.875248751 | 0.943614299 | -4.756805468 |
| **A_51_P484111** | Matn2 | 0.033038991 | -0.470205019 | 0.536283002 | 10.9833143 | 0.164436608 | 0.875285965 | 0.943614299 | -4.756808172 |
| **A_51_P137168** | Smarcal1 | 0.022853176 | -0.325660186 | 0.371366539 | 10.04841879 | 0.164239449 | 0.875433917 | 0.943721179 | -4.756818916 |
| **A_55_P2076782** | 1700028P | 0.038535126 | -0.550147093 | 0.627217345 | 8.64243818 | 0.163955584 | 0.875646945 | 0.943866356 | -4.756834363 |
| **A_65_P10180** | Rad23b | -0.053265965 | -0.867153285 | 0.760621356 | 10.76024999 | -0.163921478 | 0.87567254 | 0.943866356 | -4.756836217 |
| **A_55_P1990653** | Comt | 0.023895524 | -0.341443255 | 0.389234303 | 11.94724177 | 0.163821587 | 0.875747508 | 0.943866356 | -4.756841646 |
| **A_55_P2026945** | Zranb2 | -0.017572915 | -0.286327555 | 0.251181726 | 10.88915861 | -0.163771442 | 0.875785142 | 0.943866356 | -4.75684437 |
| **A_55_P2000958** | Sdr39u1 | 0.022434668 | -0.320818339 | 0.365687675 | 10.31718725 | 0.163702624 | 0.87583679 | 0.943866356 | -4.756848106 |
| **A_55_P2014706** | Naip5 | -0.023217327 | -0.378657487 | 0.332222833 | 14.29562402 | -0.16360481 | 0.875910201 | 0.943866356 | -4.756853415 |
| **A_55_P2013113** | Bak1 | 0.02107775 | -0.301820726 | 0.343976227 | 12.12076574 | 0.163496556 | 0.87599145 | 0.943866356 | -4.756859286 |
| **A_55_P2146229** | NA | -0.034870489 | -0.569474635 | 0.499733657 | 8.426009594 | -0.163371411 | 0.876085377 | 0.943866356 | -4.756866069 |
| **A_55_P2144716** | Hecw2 | 0.043634825 | -0.625428103 | 0.712697753 | 9.71114405 | 0.163349051 | 0.87610216 | 0.943866356 | -4.756867281 |
| **A_55_P1958200** | P2rx1 | 0.055168322 | -0.791386985 | 0.901723628 | 8.663674044 | 0.163224287 | 0.876195804 | 0.943866356 | -4.756874037 |
| **A_51_P267796** | Ddr2 | 0.045736195 | -0.65608887 | 0.74756126 | 10.1414254 | 0.163223054 | 0.87619673 | 0.943866356 | -4.756874104 |
| **A_55_P2170509** | Yipf4 | -0.020106488 | -0.32873243 | 0.288519455 | 11.14979915 | -0.163175192 | 0.876232654 | 0.943866356 | -4.756876694 |
| **A_51_P435968** | Tnfaip8 | 0.029569404 | -0.424354275 | 0.483493082 | 10.09718442 | 0.163158639 | 0.876245079 | 0.943866356 | -4.75687759 |
| **A_55_P2326091** | 9330175M | -0.040989727 | -0.670365778 | 0.588386324 | 9.84686335 | -0.163122985 | 0.876271841 | 0.943866356 | -4.756879519 |

| | | | | | | | | | |
|---|---|---|---|---|---|---|---|---|---|
| **A_55_P1959576** | Slitrk4 | 0.018215543 | -0.276242907 | 0.312673992 | 7.706449184 | 0.164771007 | 0.876300874 | 0.943866356 | -4.728239528 |
| **A_55_P2412369** | Ubxn7 | -0.01767868 | -0.289444166 | 0.254086805 | 8.847179824 | -0.162931813 | 0.876415337 | 0.943937058 | -4.756889855 |
| **A_51_P337675** | Cd53 | 0.017413592 | -0.250585052 | 0.285412236 | 7.843581692 | 0.16274442 | 0.876556001 | 0.944018289 | -4.756899975 |
| **A_52_P425092** | NA | 0.049793808 | -0.716746183 | 0.816333799 | 8.716838923 | 0.162701243 | 0.876588412 | 0.944018289 | -4.756902306 |
| **A_52_P127925** | Tfec | -0.017886385 | -0.293568817 | 0.257796046 | 7.910889888 | -0.162503919 | 0.876736537 | 0.94412522 | -4.756912947 |
| **A_52_P67643** | Mtor | 0.014079901 | -0.20302912 | 0.231188922 | 8.016517944 | 0.162432183 | 0.876790389 | 0.944130625 | -4.756916813 |
| **A_52_P45738** | Trim35 | -0.01514557 | -0.248853065 | 0.218561926 | 7.942159026 | -0.162316744 | 0.876877049 | 0.944161441 | -4.756923029 |
| **A_55_P2181753** | Bdh1 | -0.064203451 | -1.055233102 | 0.9268262 | 8.483311502 | -0.16226396 | 0.876916675 | 0.944161441 | -4.756925871 |
| **A_51_P163797** | Bdp1 | -0.0158224 | -0.260803516 | 0.229158716 | 8.948407646 | -0.161767064 | 0.877289722 | 0.944468668 | -4.756952573 |
| **A_55_P1994751** | NA | 0.023735453 | -0.34379639 | 0.391267295 | 8.766420197 | 0.161753203 | 0.877300129 | 0.944468668 | -4.756953316 |
| **A_55_P2273705** | NA | 0.014385234 | -0.208451945 | 0.237222413 | 8.820702827 | 0.161688682 | 0.877348571 | 0.944468668 | -4.756956777 |
| **A_51_P427663** | Cnn2 | -0.058603943 | -0.967161255 | 0.849953369 | 13.03122091 | -0.161556656 | 0.877447698 | 0.944522788 | -4.756963855 |
| **A_55_P2114034** | NA | -0.030564858 | -0.504956815 | 0.443827099 | 8.426043417 | -0.161374687 | 0.877584327 | 0.944537271 | -4.756973601 |
| **A_55_P1975837** | NA | -0.061212365 | -1.011416384 | 0.888991655 | 12.6470005 | -0.161351364 | 0.877601839 | 0.944537271 | -4.756974849 |
| **A_52_P409055** | Dcaf8 | 0.038554532 | -0.559979049 | 0.637088112 | 11.7562557 | 0.161338218 | 0.877611709 | 0.944537271 | -4.756975553 |
| **A_66_P113578** | Washc1 | 0.023874437 | -0.346897688 | 0.394646563 | 11.49572382 | 0.161278476 | 0.877656567 | 0.944537271 | -4.75697875 |
| **A_55_P2058348** | Pacc1 | -0.014814441 | -0.245050614 | 0.215421732 | 8.57549628 | -0.16116178 | 0.87774419 | 0.944578992 | -4.75698499 |
| **A_55_P1994668** | Tti1 | -0.019670618 | -0.325701311 | 0.286360075 | 10.01997884 | -0.160991657 | 0.877871934 | 0.944629703 | -4.75699408 |
| **A_52_P128964** | Nhej1 | 0.023200969 | -0.337805112 | 0.384207049 | 9.009803764 | 0.160968889 | 0.87788903 | 0.944629703 | -4.756995296 |
| **A_51_P337662** | Ints6l | -0.056118827 | -0.930699862 | 0.818462209 | 9.240591296 | -0.16071591 | 0.878078998 | 0.944729809 | -4.757008794 |
| **A_51_P461894** | Tnnc1 | 0.084774528 | -1.236398519 | 1.405947575 | 9.671604983 | 0.160714855 | 0.87807979 | 0.944729809 | -4.75700885 |
| **A_55_P2169417** | BC021767 | -0.018873363 | -0.313317 | 0.275570274 | 10.58202422 | -0.160545251 | 0.878207155 | 0.944814263 | -4.757017887 |
| **A_55_P2003823** | Asgr1 | -0.013364246 | -0.22225044 | 0.195521949 | 7.654003614 | -0.160245211 | 0.878432481 | 0.945001463 | -4.757033852 |
| **A_52_P573366** | Hnrnpul1 | 0.013902656 | -0.20348288 | 0.231288191 | 8.027894765 | 0.160183386 | 0.878478912 | 0.945001463 | -4.757037138 |
| **A_55_P2015299** | NA | 0.058431423 | -0.856500203 | 0.97336305 | 16.42411032 | 0.159958812 | 0.878647575 | 0.945117083 | -4.757049064 |
| **A_55_P2066792** | Sp110 | -0.042679557 | -0.711168692 | 0.625809579 | 15.51073262 | -0.159910098 | 0.878684161 | 0.945117083 | -4.757051648 |
| **A_55_P2192577** | H2bfm | 0.017934829 | -0.263209868 | 0.299079526 | 8.012669219 | 0.159778261 | 0.878783179 | 0.945138633 | -4.757058639 |
| **A_51_P389636** | Kcnn4 | 0.036579927 | -0.536933364 | 0.610093219 | 8.011064977 | 0.159753248 | 0.878801966 | 0.945138633 | -4.757059965 |
| **A_55_P2016675** | Ogfr | -0.015232874 | -0.254567958 | 0.224102209 | 10.6270576 | -0.15941377 | 0.879056949 | 0.945360276 | -4.757077938 |
| **A_51_P492298** | Aqp2 | 0.031407609 | -0.462713679 | 0.525528898 | 8.717579257 | 0.159203151 | 0.879215153 | 0.945424297 | -4.75708907 |
| **A_52_P646783** | Fbxl22 | -0.029763415 | -0.498167031 | 0.438640201 | 8.233607984 | -0.159152279 | 0.879253366 | 0.945424297 | -4.757091757 |
| **A_51_P226364** | Coa3 | 0.02891767 | -0.426339423 | 0.484174764 | 13.72977557 | 0.159095146 | 0.879296282 | 0.945424297 | -4.757094773 |
| **A_52_P175376** | Tfcp2l1 | 0.068867935 | -1.015732524 | 1.153468393 | 9.28448953 | 0.159036797 | 0.879340112 | 0.945424297 | -4.757097853 |
| **A_55_P1973723** | Etfdh | -0.014659514 | -0.245575948 | 0.21625692 | 11.28179842 | -0.159006575 | 0.879362814 | 0.945424297 | -4.757099447 |
| **A_51_P206655** | Lsm1 | 0.015511779 | -0.22892582 | 0.259949378 | 10.34648605 | 0.158943925 | 0.879409877 | 0.945424297 | -4.757102752 |
| **A_51_P210143** | Syne2 | 0.021142637 | -0.312622527 | 0.3549078 | 9.561441216 | 0.158660379 | 0.879622879 | 0.945592839 | -4.757117691 |
| **A_51_P255329** | Riok2 | -0.017788354 | -0.298699629 | 0.263122921 | 10.24187012 | -0.15860502 | 0.879664466 | 0.945592839 | -4.757120605 |
| **A_55_P2009813** | NA | -0.014257806 | -0.239565723 | 0.211050112 | 7.87424047 | -0.158499021 | 0.879744098 | 0.945625863 | -4.757126181 |
| **A_51_P484289** | Slc7a6 | -0.035429653 | -0.595728149 | 0.524868843 | 8.571048163 | -0.15837907 | 0.879834213 | 0.94562631 | -4.757132487 |
| **A_51_P400016** | Kalrn | 0.018184881 | -0.269417626 | 0.305787389 | 10.06350062 | 0.15836826 | 0.879842334 | 0.94562631 | -4.757133055 |
| **A_55_P2119716** | Mapk1ip1 | 0.025979853 | -0.385229213 | 0.437188919 | 9.62939685 | 0.158242944 | 0.879936482 | 0.945674927 | -4.757139638 |
| **A_51_P326994** | Calhm2 | 0.015440958 | -0.229254487 | 0.260136402 | 8.624620381 | 0.158051527 | 0.880080294 | 0.945776911 | -4.757149682 |
| **A_52_P381418** | Slc35b4 | -0.016167811 | -0.27268621 | 0.240350589 | 8.401388539 | -0.157863976 | 0.880221206 | 0.945875767 | -4.757159512 |
| **A_55_P2012689** | Stk16 | -0.018434271 | -0.311282525 | 0.274413984 | 9.949735632 | -0.157664409 | 0.880371152 | 0.945984319 | -4.75716996 |
| **A_51_P200517** | Rasgef1a | -0.085922108 | -1.452289338 | 1.280445122 | 10.27114129 | -0.157502638 | 0.880492704 | 0.946025011 | -4.757178419 |
| **A_55_P2101001** | Smad3 | 0.019402533 | -0.289181157 | 0.327986223 | 12.02950179 | 0.157483768 | 0.880506883 | 0.946025011 | -4.757179405 |
| **A_55_P2125714** | Atp2b4 | -0.015819264 | -0.267723915 | 0.236085386 | 8.090457364 | -0.157289759 | 0.880652663 | 0.946129061 | -4.757189537 |
| **A_55_P2100241** | Rn18s | 0.027236145 | -0.407502663 | 0.461974953 | 17.24811116 | 0.156916125 | 0.88093343 | 0.946316008 | -4.757209016 |

| | | | | | | | | | |
|---|---|---|---|---|---|---|---|---|---|
| **A_52_P247788** | Ino80c | -0.015385632 | -0.261001956 | 0.230230693 | 8.208312702 | -0.156894764 | 0.880949482 | 0.946316008 | -4.757210128 |
| **A_52_P513347** | Phkb | 0.021090528 | -0.315907116 | 0.358088171 | 8.375472931 | 0.15675122 | 0.881057354 | 0.946316008 | -4.757217599 |
| **A_51_P209183** | Cxcl14 | -0.037671608 | -0.639706589 | 0.564363373 | 9.945539223 | -0.156726625 | 0.881075837 | 0.946316008 | -4.757218878 |
| **A_55_P2430472** | Dido1 | -0.040540995 | -0.688460482 | 0.607378492 | 8.921550172 | -0.156719743 | 0.881081008 | 0.946316008 | -4.757219236 |
| **A_66_P115539** | Gm11240 | -0.037741122 | -0.641358606 | 0.565876361 | 10.48709297 | -0.156604179 | 0.881167855 | 0.946316008 | -4.757225244 |
| **A_51_P267388** | Dxo | 0.01559307 | -0.23379974 | 0.26498588 | 9.240593124 | 0.156602267 | 0.881169292 | 0.946316008 | -4.757225343 |
| **A_51_P448664** | Tbxas1 | 0.020699158 | -0.310602009 | 0.352000324 | 8.214124864 | 0.156487644 | 0.881255435 | 0.946354615 | -4.757231298 |
| **A_52_P303176** | 1810037I1 | 0.022429217 | -0.337015465 | 0.381873899 | 9.387922566 | 0.156290431 | 0.88140365 | 0.946354615 | -4.757241534 |
| **A_51_P219325** | Ddx28 | 0.014075331 | -0.211504383 | 0.239655046 | 8.917559706 | 0.15628199 | 0.881409994 | 0.946354615 | -4.757241971 |
| **A_51_P384148** | Ryk | -0.033140724 | -0.56436297 | 0.498081522 | 11.2284623 | -0.156255776 | 0.881429695 | 0.946354615 | -4.757243331 |
| **A_51_P354744** | Cox15 | -0.022891603 | -0.38989112 | 0.344107914 | 9.357275883 | -0.156228787 | 0.881449979 | 0.946354615 | -4.75724473 |
| **A_51_P141772** | Uhrf2 | 0.02276381 | -0.342574408 | 0.388102027 | 11.08886547 | 0.156063085 | 0.881574518 | 0.946435767 | -4.757253317 |
| **A_55_P2200628** | 4733401D | -0.039588601 | -0.676002245 | 0.596825043 | 9.353390367 | -0.155804866 | 0.881768598 | 0.946483034 | -4.75726668 |
| **A_55_P2017640** | Phf7 | 0.034131074 | -0.514724098 | 0.582986246 | 9.889375901 | 0.155755194 | 0.881805933 | 0.946483034 | -4.757269249 |
| **A_51_P194273** | Sgcg | -0.019232187 | -0.328552943 | 0.290088569 | 8.00209365 | -0.155729167 | 0.881825496 | 0.946483034 | -4.757270594 |
| **A_55_P2163961** | Kcnj5 | 0.07206684 | -1.087269164 | 1.231402845 | 8.93347327 | 0.155695659 | 0.881850682 | 0.946483034 | -4.757272326 |
| **A_55_P2089219** | NA | 0.241355252 | -3.642633246 | 4.12534375 | 10.87793302 | 0.155642846 | 0.881890378 | 0.946483034 | -4.757275054 |
| **A_55_P2024175** | Map4k3 | -0.040988356 | -0.700713388 | 0.618736676 | 11.91464618 | -0.155613719 | 0.881912272 | 0.946483034 | -4.757276559 |
| **A_55_P2113291** | Sik3 | -0.020594463 | -0.352249045 | 0.311060119 | 12.01541654 | -0.155530228 | 0.881975029 | 0.946497847 | -4.75728087 |
| **A_55_P1961129** | Egfem1 | 0.057701711 | -0.871959098 | 0.987362519 | 8.532010458 | 0.155458513 | 0.882028935 | 0.946503159 | -4.757284571 |
| **A_51_P114616** | Batf | 0.051746113 | -0.784890599 | 0.888382825 | 8.642093271 | 0.154914176 | 0.88243812 | 0.9468897 | -4.750296932 |
| **A_52_P558368** | Zdhhc1 | -0.054289345 | -0.933649699 | 0.825071009 | 9.351830076 | -0.154631534 | 0.882650602 | 0.947065139 | -4.757327128 |
| **A_55_P2382262** | A830039N | 0.027962815 | -0.425748885 | 0.481674515 | 8.743213258 | 0.154365859 | 0.882850339 | 0.947226884 | -4.757340752 |
| **A_55_P2133266** | Rtl8c | 0.021592761 | -0.329494287 | 0.372679809 | 14.04678796 | 0.154043621 | 0.883092612 | 0.947434247 | -4.757357246 |
| **A_52_P246889** | Chmp3 | -0.017501065 | -0.302843677 | 0.267841548 | 8.594523241 | -0.153620149 | 0.88341102 | 0.947705995 | -4.757378871 |
| **A_55_P2072228** | Cdk9 | -0.026386654 | -0.456725235 | 0.403951926 | 12.25258597 | -0.153576368 | 0.883443941 | 0.947705995 | -4.757381103 |
| **A_55_P2171897** | Notch1 | 0.026603766 | -0.408238618 | 0.46144615 | 8.159285332 | 0.153236281 | 0.883699671 | 0.947878356 | -4.757398422 |
| **A_66_P129029** | Taf10 | -0.01565477 | -0.27155561 | 0.24024607 | 11.96468974 | -0.153223483 | 0.883709295 | 0.947878356 | -4.757399073 |
| **A_55_P1962876** | Adam11 | 0.023349906 | -0.35876107 | 0.405460883 | 8.689429726 | 0.153054454 | 0.883836404 | 0.947878356 | -4.757407666 |
| **A_52_P497534** | Wipi1 | 0.021125538 | -0.324787796 | 0.367038872 | 10.09085224 | 0.152964561 | 0.883904004 | 0.947878356 | -4.757412232 |
| **A_55_P2069730** | NA | -0.014557965 | -0.252947704 | 0.223831775 | 8.056258904 | -0.152954921 | 0.883911254 | 0.947878356 | -4.757412722 |
| **A_66_P126415** | NA | 0.016794579 | -0.25823706 | 0.291826217 | 7.874166195 | 0.152945534 | 0.883918312 | 0.947878356 | -4.757413198 |
| **A_55_P2033997** | Ddo | 0.016991 | -0.26133394 | 0.29531594 | 7.890982256 | 0.15290341 | 0.883949991 | 0.947878356 | -4.757415337 |
| **A_55_P2087567** | Olfr373 | 0.022912244 | -0.352559899 | 0.398384387 | 8.518385423 | 0.152841132 | 0.883996826 | 0.947878356 | -4.757418498 |
| **A_52_P343690** | Kpna4 | 0.025987917 | -0.400686184 | 0.452662019 | 10.92967691 | 0.152554682 | 0.884212252 | 0.94805677 | -4.757433019 |
| **A_55_P2020256** | Sik2 | -0.023542762 | -0.410509762 | 0.363424238 | 8.793534682 | -0.152382052 | 0.884342083 | 0.948143395 | -4.757441757 |
| **A_55_P2081055** | 1700056E | 0.016151264 | -0.249588207 | 0.281890735 | 8.331076583 | 0.152230192 | 0.884456298 | 0.94819294 | -4.757449436 |
| **A_55_P1963198** | NA | 0.012719839 | -0.19661682 | 0.222056499 | 7.697637026 | 0.152190196 | 0.88448638 | 0.94819294 | -4.757451457 |
| **A_55_P2148873** | Cat | -0.019867293 | -0.347158014 | 0.307423429 | 10.52883858 | -0.152039118 | 0.88460001 | 0.948199652 | -4.757459087 |
| **A_55_P2112085** | Vwa8 | 0.021346748 | -0.330495953 | 0.373189449 | 8.688539877 | 0.151961484 | 0.884658403 | 0.948199652 | -4.757463005 |
| **A_55_P1955733** | NA | 0.056495717 | -0.874825384 | 0.987816817 | 11.47358751 | 0.151938007 | 0.884676061 | 0.948199652 | -4.757464189 |
| **A_52_P372151** | Ctnnal1 | -0.018889302 | -0.330402362 | 0.292623758 | 10.44324095 | -0.151876295 | 0.884722478 | 0.948199652 | -4.757467301 |
| **A_55_P2131288** | Ank2 | 0.020746543 | -0.321441292 | 0.362934378 | 8.905535553 | 0.151855849 | 0.884737857 | 0.948199652 | -4.757468332 |
| **A_51_P461884** | Abl2 | 0.021338129 | -0.330857242 | 0.373533499 | 10.20739892 | 0.151748019 | 0.884818964 | 0.948233603 | -4.757473767 |
| **A_55_P2043582** | 4930480E | 0.017268913 | -0.267981581 | 0.302519407 | 8.124819845 | 0.15163133 | 0.884906736 | 0.948233603 | -4.757479644 |
| **A_52_P638459** | Ccl5 | 0.030544659 | -0.474040536 | 0.535129854 | 8.210672245 | 0.151618123 | 0.88491667 | 0.948233603 | -4.757480309 |
| **A_55_P2018017** | Tnfsf10 | 0.044499303 | -0.692184401 | 0.781183007 | 9.501993705 | 0.151294274 | 0.885160276 | 0.948371713 | -4.757496594 |
| **A_55_P2000219** | Ubap2 | -0.034183216 | -0.600119334 | 0.531752902 | 12.20737838 | -0.151284994 | 0.885167257 | 0.948371713 | -4.75749706 |

| | | | | | | | | | |
|---|---|---|---|---|---|---|---|---|---|
| **A_55_P2006861** | Trio | -0.021428906 | -0.37640995 | 0.333552139 | 8.217179365 | -0.151197697 | 0.885232925 | 0.948371713 | -4.757501444 |
| **A_55_P2424988** | C78228 | 0.014197619 | -0.221099719 | 0.249494957 | 7.958830047 | 0.151129365 | 0.885284329 | 0.948371713 | -4.757504874 |
| **A_51_P334789** | Nudt16 | -0.014302016 | -0.251343049 | 0.222739017 | 8.506735287 | -0.151120739 | 0.885290818 | 0.948371713 | -4.757505307 |
| **A_55_P1977985** | Pigp | 0.037819154 | -0.589402921 | 0.665041229 | 11.99641076 | 0.151022209 | 0.885364939 | 0.948377152 | -4.757510249 |
| **A_52_P671769** | Arhgap18 | 0.04523011 | -0.705092916 | 0.795553136 | 10.21049039 | 0.150983579 | 0.885394 | 0.948377152 | -4.757512186 |
| **A_55_P2054352** | Rbm6 | 0.016493211 | -0.257481945 | 0.290468368 | 10.61318791 | 0.150780224 | 0.885546983 | 0.948395995 | -4.757522374 |
| **A_55_P2153937** | NA | 0.02254738 | -0.352017677 | 0.397112437 | 8.675728541 | 0.150771471 | 0.885553568 | 0.948395995 | -4.757522812 |
| **A_55_P2035122** | NA | 0.042412555 | -0.662299224 | 0.747124334 | 10.25209314 | 0.150741645 | 0.885576007 | 0.948395995 | -4.757524305 |
| **A_52_P376135** | Mknk2 | -0.01985429 | -0.349838563 | 0.310129983 | 13.25149287 | -0.150699379 | 0.885607804 | 0.948395995 | -4.75752642 |
| **A_55_P2155312** | Dock3 | -0.031132119 | -0.548893038 | 0.486628801 | 9.733629213 | -0.150601659 | 0.885681322 | 0.948422192 | -4.757531309 |
| **A_55_P1971179** | Cd1d1 | 0.022176922 | -0.347037801 | 0.391391644 | 8.110079743 | 0.150443213 | 0.885800528 | 0.948455265 | -4.757539228 |
| **A_51_P432851** | Dhrs11 | 0.012763821 | -0.199754334 | 0.225281976 | 7.91171296 | 0.150430197 | 0.88581032 | 0.948455265 | -4.757539878 |
| **A_51_P445473** | Fut7 | 0.013795775 | -0.216088344 | 0.243679894 | 7.82064589 | 0.150309866 | 0.885900853 | 0.948460873 | -4.757545886 |
| **A_55_P2053551** | Mast4 | -0.018267742 | -0.322749015 | 0.286213532 | 9.120411558 | -0.150270807 | 0.88593024 | 0.948460873 | -4.757547836 |
| **A_52_P487686** | Cstdc5 | 0.09180376 | -1.438793749 | 1.622401269 | 11.12556889 | 0.150227628 | 0.885962727 | 0.948460873 | -4.75754999 |
| **A_55_P1993483** | NA | 0.036586231 | -0.574757644 | 0.647930106 | 11.32383074 | 0.14989338 | 0.886214217 | 0.948677575 | -4.757566646 |
| **A_55_P2157872** | Srsf5 | 0.024716975 | -0.388597142 | 0.438031092 | 11.72899162 | 0.149784 | 0.886296519 | 0.948679706 | -4.757572088 |
| **A_55_P1990086** | Mks1 | -0.020945612 | -0.371250881 | 0.329359657 | 9.509192501 | -0.14976031 | 0.886314344 | 0.948679706 | -4.757573266 |
| **A_55_P1968488** | Cblif | -0.067601795 | -1.199115006 | 1.063911416 | 8.557006031 | -0.149640429 | 0.886404549 | 0.948698733 | -4.757579226 |
| **A_55_P2113439** | Caln1 | 0.013836302 | -0.217956639 | 0.245629244 | 7.974132111 | 0.149509981 | 0.886502708 | 0.948698733 | -4.757585705 |
| **A_55_P1959006** | Imp4 | 0.033288734 | -0.524405504 | 0.590982971 | 10.91847473 | 0.149503536 | 0.886507557 | 0.948698733 | -4.757586025 |
| **A_52_P289893** | Prdm4 | -0.015168402 | -0.269641571 | 0.239304768 | 9.096967589 | -0.149295997 | 0.886663729 | 0.948698733 | -4.757596322 |
| **A_55_P2128606** | Nr1h5 | 0.012532824 | -0.19793059 | 0.222996238 | 7.642352354 | 0.149149801 | 0.886773744 | 0.948698733 | -4.757603567 |
| **A_55_P2177463** | B130024G | 0.065001402 | -1.026879929 | 1.156882733 | 9.041387611 | 0.149106861 | 0.886806058 | 0.948698733 | -4.757605694 |
| **A_51_P314521** | Smim11 | 0.032862218 | -0.519152544 | 0.58487698 | 12.28703567 | 0.149106484 | 0.886806341 | 0.948698733 | -4.757605712 |
| **A_55_P2120354** | Cib2 | -0.029707507 | -0.528769274 | 0.46935426 | 10.70184698 | -0.149094715 | 0.886815198 | 0.948698733 | -4.757606295 |
| **A_55_P2258261** | 1810008I1 | 0.01348354 | -0.213042895 | 0.240009974 | 7.762304394 | 0.149085487 | 0.886822142 | 0.948698733 | -4.757606752 |
| **A_55_P1991164** | Mlc1 | 0.129783867 | -2.050626656 | 2.31019439 | 10.5163022 | 0.149084601 | 0.886822809 | 0.948698733 | -4.757606796 |
| **A_55_P2120526** | NA | -0.026940559 | -0.480002125 | 0.426121007 | 8.329170226 | -0.148936013 | 0.886934629 | 0.948760659 | -4.757614149 |
| **A_55_P2048655** | NA | -0.025460755 | -0.453805308 | 0.402883798 | 15.21930369 | -0.148877263 | 0.886978841 | 0.948760659 | -4.757617054 |
| **A_55_P1973526** | Fam216b | 0.043438407 | -0.687925991 | 0.774802806 | 8.391855559 | 0.148761457 | 0.887065993 | 0.948801389 | -4.757622778 |
| **A_66_P118752** | Rabif | -0.013870633 | -0.247533088 | 0.219791823 | 8.883762036 | -0.148681757 | 0.887125974 | 0.948813054 | -4.757626715 |
| **A_55_P2128113** | Tbc1d12 | 0.048289392 | -0.76582324 | 0.862402023 | 8.841109865 | 0.148565369 | 0.887213567 | 0.948854248 | -4.757632459 |
| **A_55_P2136776** | Sult1b1 | -0.01237963 | -0.221220949 | 0.196461689 | 7.633528523 | -0.148470986 | 0.887284601 | 0.94887773 | -4.757637115 |
| **A_52_P72839** | Sec23a | 0.017403899 | -0.276425183 | 0.311232981 | 9.290784981 | 0.148354977 | 0.887371912 | 0.948918615 | -4.757642833 |
| **A_55_P2015277** | Cyld | -0.015195326 | -0.273474203 | 0.243083551 | 9.600390984 | -0.147357243 | 0.888122904 | 0.94965389 | -4.75769183 |
| **A_51_P308298** | Myl9 | 0.081605744 | -1.305903802 | 1.46911529 | 14.69553819 | 0.147310972 | 0.888157735 | 0.94965389 | -4.757694094 |
| **A_55_P2014635** | Ccdc82 | 0.029624578 | -0.47473678 | 0.533985936 | 10.13545067 | 0.147116272 | 0.888304301 | 0.94975808 | -4.757703615 |
| **A_51_P124568** | Mpp1 | -0.023907917 | -0.431487575 | 0.383671741 | 11.18787487 | -0.146919548 | 0.888452397 | 0.949809738 | -4.757713222 |
| **A_55_P2048751** | Prox1 | 0.012482714 | -0.200372624 | 0.225338053 | 7.700351576 | 0.146884113 | 0.888479073 | 0.949809738 | -4.757714951 |
| **A_55_P2038121** | Rbx1 | 0.01985166 | -0.31872306 | 0.35842638 | 14.05255418 | 0.14685632 | 0.888499996 | 0.949809738 | -4.757716307 |
| **A_51_P415220** | Zmat3 | -0.025147606 | -0.454604853 | 0.404309641 | 9.472378283 | -0.146665194 | 0.888643883 | 0.949883325 | -4.757725624 |
| **A_55_P2020143** | Slc39a3 | -0.015392833 | -0.2783186 | 0.247532933 | 9.491667125 | -0.146634364 | 0.888667094 | 0.949883325 | -4.757727126 |
| **A_51_P188795** | Akr7a5 | 0.013467763 | -0.217101825 | 0.244037351 | 9.532944384 | 0.14629982 | 0.888918965 | 0.950057665 | -4.757743402 |
| **A_55_P1993940** | Sh3bgrl3 | -0.019177642 | -0.347529334 | 0.30917405 | 11.53482985 | -0.146287187 | 0.888928476 | 0.950057665 | -4.757744016 |
| **A_55_P2028214** | NA | -0.012967801 | -0.235181551 | 0.209245949 | 7.870178497 | -0.146165795 | 0.889019873 | 0.950102826 | -4.757749913 |
| **A_52_P417825** | Osgin2 | -0.012831656 | -0.233156119 | 0.207492807 | 8.786636379 | -0.145871462 | 0.889241488 | 0.950287139 | -4.75776419 |
| **A_51_P384673** | Lrrc59 | 0.046634652 | -0.755639806 | 0.84890911 | 10.51258738 | 0.145591535 | 0.889452266 | 0.950367691 | -4.757777742 |

| | | | | | | | | | |
|---|---|---|---|---|---|---|---|---|---|
| **A_51_P341647** | Fancc | -0.037069775 | -0.674999761 | 0.600860211 | 9.106455146 | -0.145544995 | 0.88948731 | 0.950367691 | -4.757779992 |
| **A_55_P2043682** | Ubac1 | -0.02330075 | -0.424366773 | 0.377765273 | 11.25150321 | -0.145513862 | 0.889510753 | 0.950367691 | -4.757781497 |
| **A_55_P1972605** | Adora3 | 0.019271874 | -0.31279022 | 0.351333967 | 7.960390389 | 0.14536337 | 0.889624075 | 0.950367691 | -4.757788769 |
| **A_55_P2148248** | NA | 0.025226898 | -0.409452588 | 0.459906385 | 8.634822394 | 0.145360049 | 0.889626576 | 0.950367691 | -4.757788929 |
| **A_55_P2081690** | Tex38 | -0.012663767 | -0.231173077 | 0.205845544 | 7.803829955 | -0.145158775 | 0.889778142 | 0.950367691 | -4.757798642 |
| **A_51_P150678** | Tnfaip8l2 | 0.025739123 | -0.41847535 | 0.469953596 | 8.191219232 | 0.145128057 | 0.889801274 | 0.950367691 | -4.757800123 |
| **A_52_P275408** | Ppp4c | -0.015472569 | -0.282537043 | 0.251591906 | 10.8765912 | -0.145109767 | 0.889815048 | 0.950367691 | -4.757801005 |
| **A_55_P2155901** | Snhg10 | 0.022823976 | -0.371274121 | 0.416922073 | 8.936675291 | 0.145056538 | 0.889855132 | 0.950367691 | -4.757803571 |
| **A_55_P1990500** | Masp2 | 0.012145328 | -0.197569224 | 0.221859881 | 7.662781554 | 0.145054444 | 0.889856709 | 0.950367691 | -4.757803671 |
| **A_52_P325237** | Rgs6 | 0.016144577 | -0.262627594 | 0.294916749 | 7.851633011 | 0.145053297 | 0.889857573 | 0.950367691 | -4.757803727 |
| **A_55_P2027587** | Supt4b | 0.03244452 | -0.528693005 | 0.593582044 | 10.78614664 | 0.144817947 | 0.89003481 | 0.950452471 | -4.757815059 |
| **A_52_P429450** | Ngp | 0.017421307 | -0.283976768 | 0.318819382 | 7.866064772 | 0.144774011 | 0.890067898 | 0.950452471 | -4.757817173 |
| **A_55_P1964867** | Ndufs6 | 0.016007555 | -0.260973856 | 0.292988967 | 15.06095998 | 0.144752054 | 0.890084433 | 0.950452471 | -4.757818229 |
| **A_55_P2230688** | Pigm | -0.022169454 | -0.40655655 | 0.362217643 | 9.002770334 | -0.144456315 | 0.89030716 | 0.950586909 | -4.757832437 |
| **A_55_P2068673** | Stmn2 | 0.04187143 | -0.684542434 | 0.768285294 | 9.376801543 | 0.144372346 | 0.8903704 | 0.950586909 | -4.757836466 |
| **A_55_P2029746** | NA | -0.044978275 | -0.82552385 | 0.735567299 | 13.79169279 | -0.144329424 | 0.890402727 | 0.950586909 | -4.757838524 |
| **A_55_P2062891** | Atf2 | 0.021020168 | -0.343774694 | 0.385815029 | 11.3003957 | 0.144323751 | 0.890407 | 0.950586909 | -4.757838796 |
| **A_55_P2139901** | Tsr3 | -0.024463825 | -0.449488237 | 0.400560587 | 11.77601406 | -0.14416533 | 0.890526317 | 0.950661798 | -4.757846388 |
| **A_55_P2074393** | NA | 0.026126635 | -0.428576388 | 0.480829658 | 8.720618503 | 0.143914973 | 0.890714884 | 0.950810598 | -4.75785837 |
| **A_55_P2017724** | NA | 0.035585121 | -0.584198733 | 0.655368975 | 9.314807749 | 0.143806507 | 0.890796582 | 0.950845311 | -4.757863554 |
| **A_55_P2137461** | NA | 0.038572179 | -0.633906705 | 0.711051063 | 16.38492163 | 0.143663309 | 0.890904443 | 0.950907944 | -4.757870393 |
| **A_55_P2097518** | Zdhhc22 | 0.04472347 | -0.735639222 | 0.825086162 | 13.86981574 | 0.143545419 | 0.890993243 | 0.950950227 | -4.757876018 |
| **A_55_P1978866** | NA | -0.038305889 | -0.707106062 | 0.630494283 | 12.32320548 | -0.143456265 | 0.891060399 | 0.950969406 | -4.757880269 |
| **A_52_P37681** | Fkbp9 | 0.020869516 | -0.3441382 | 0.385877233 | 12.53638401 | 0.143205824 | 0.891249052 | 0.951118241 | -4.757892196 |
| **A_51_P104077** | Usp13 | 0.015122942 | -0.24950643 | 0.279752314 | 7.933279832 | 0.143135905 | 0.891301722 | 0.95112195 | -4.748377278 |
| **A_51_P286321** | Trir | -0.018757307 | -0.347221616 | 0.309707001 | 13.09404358 | -0.143031817 | 0.891380133 | 0.951153126 | -4.757900471 |
| **A_55_P1973926** | Ipo13 | 0.012350726 | -0.204136098 | 0.22883755 | 10.77477088 | 0.142893137 | 0.891484605 | 0.951163151 | -4.757907059 |
| **A_55_P2118570** | Cd37 | 0.02102942 | -0.347591222 | 0.389650063 | 8.207148082 | 0.142888734 | 0.891487921 | 0.951163151 | -4.757907268 |
| **A_55_P1991783** | NA | 0.030080908 | -0.497642856 | 0.557804671 | 12.21650896 | 0.142769235 | 0.891577945 | 0.95120671 | -4.75791294 |
| **A_55_P2116525** | NA | -0.011988449 | -0.22260371 | 0.198626813 | 7.691930193 | -0.142568473 | 0.891729193 | 0.951257987 | -4.757922458 |
| **A_55_P2133365** | 1110017D | 0.067224652 | -1.114106254 | 1.248555559 | 8.880694102 | 0.142530348 | 0.891757916 | 0.951257987 | -4.757924264 |
| **A_55_P2185821** | Cyp3a13 | -0.013278834 | -0.246660337 | 0.220102669 | 7.807934472 | -0.142509513 | 0.891773612 | 0.951257987 | -4.75792525 |
| **A_55_P2028365** | Lrrc36 | 0.03969202 | -0.658513982 | 0.737898023 | 8.414569849 | 0.14238688 | 0.891866003 | 0.951304055 | -4.757931055 |
| **A_66_P140599** | NA | 0.012236203 | -0.203198222 | 0.227670628 | 8.377480049 | 0.142259711 | 0.891961813 | 0.951353765 | -4.757937069 |
| **A_55_P2292435** | Slc8a1 | 0.012047327 | -0.200197676 | 0.224292331 | 7.676000301 | 0.142168567 | 0.892030484 | 0.951374524 | -4.757941377 |
| **A_55_P2050773** | LOC1000 | -0.022672132 | -0.422498072 | 0.377153809 | 7.947013022 | -0.142027269 | 0.892136942 | 0.951409093 | -4.757948049 |
| **A_66_P140711** | Rps10 | -0.014675395 | -0.273536813 | 0.244186024 | 16.32110705 | -0.141994922 | 0.892161314 | 0.951409093 | -4.757949576 |
| **A_55_P1995055** | Prr7 | -0.019308024 | -0.360046061 | 0.321430012 | 8.465508296 | -0.141927831 | 0.892211864 | 0.951410523 | -4.757952741 |
| **A_55_P2100963** | Hmces | 0.031743664 | -0.52940175 | 0.592889078 | 11.61135178 | 0.141687646 | 0.892392838 | 0.951448294 | -4.757964059 |
| **A_55_P2161690** | Tomm40 | 0.016758505 | -0.279507643 | 0.313024652 | 8.431873381 | 0.141678379 | 0.89239982 | 0.951448294 | -4.757964496 |
| **A_55_P2133751** | Txnl4a | 0.031317738 | -0.522421568 | 0.585057044 | 13.0103283 | 0.141656133 | 0.892416582 | 0.951448294 | -4.757965543 |
| **A_55_P2009604** | NA | 0.015922393 | -0.265774207 | 0.297618993 | 8.165385782 | 0.141571906 | 0.892480047 | 0.951448294 | -4.757969507 |
| **A_55_P1980931** | NA | 0.016644004 | -0.277855925 | 0.311143932 | 11.96922789 | 0.141554263 | 0.892493341 | 0.951448294 | -4.757970337 |
| **A_55_P2021125** | NA | 0.024373337 | -0.407450507 | 0.456197181 | 8.410433945 | 0.141370476 | 0.892631828 | 0.951543462 | -4.757978977 |
| **A_55_P2018507** | Gorasp2 | 0.018781932 | -0.314329631 | 0.351893496 | 8.763861658 | 0.14122153 | 0.892744065 | 0.951610638 | -4.757985971 |
| **A_55_P2097616** | Stxbp1 | -0.012412511 | -0.232759194 | 0.207934172 | 7.995941153 | -0.14109235 | 0.892841409 | 0.951661933 | -4.757992031 |
| **A_55_P2413924** | Gm10404 | -0.029694513 | -0.557181517 | 0.497792492 | 8.750937843 | -0.140998594 | 0.892912061 | 0.951684773 | -4.757996426 |
| **A_55_P2165414** | Serpina1a | -0.122179263 | -2.293801548 | 2.049443022 | 10.99455212 | -0.140917055 | 0.892973507 | 0.951694952 | -4.758000246 |

| | | | | | | | | | |
|---|---|---|---|---|---|---|---|---|---|
| **A_55_P2072315** | NA | -0.052927514 | -0.994077215 | 0.888222186 | 11.90699125 | -0.140855282 | 0.893020059 | 0.951694952 | -4.758003138 |
| **A_55_P2110758** | B430306N | -0.0120828 | -0.227286743 | 0.203121143 | 7.958263318 | -0.140626678 | 0.893192336 | 0.951797578 | -4.758013831 |
| **A_55_P2167323** | Cort | 0.01193363 | -0.200703479 | 0.22457074 | 7.922967901 | 0.14056716 | 0.893237191 | 0.951797578 | -4.758016612 |
| **A_51_P155323** | Hc | -0.178544252 | -3.36070748 | 3.003618977 | 10.71648353 | -0.140531527 | 0.893264045 | 0.951797578 | -4.758018277 |
| **A_55_P1974243** | Dbh | 0.027062642 | -0.455813284 | 0.509938568 | 8.082159313 | 0.140373452 | 0.893383177 | 0.951862714 | -4.758025656 |
| **A_52_P296632** | NA | -0.071799776 | -1.353405952 | 1.2098064 | 10.31738462 | -0.140319763 | 0.89342364 | 0.951862714 | -4.75802816 |
| **A_55_P2021358** | Klhl15 | -0.015691858 | -0.296698209 | 0.265314494 | 8.193748706 | -0.139864844 | 0.893766508 | 0.952152916 | -4.758049342 |
| **A_55_P2124425** | Zfp661 | 0.026395388 | -0.446413087 | 0.499203863 | 8.750175681 | 0.139827678 | 0.893794521 | 0.952152916 | -4.75805107 |
| **A_51_P320105** | Smyd2 | -0.037934566 | -0.718204076 | 0.642334944 | 12.20691222 | -0.139670428 | 0.893913045 | 0.952203375 | -4.758058374 |
| **A_55_P2139407** | Myl12b | -0.029574869 | -0.560070138 | 0.500920401 | 13.76694944 | -0.139634152 | 0.893940387 | 0.952203375 | -4.758060058 |
| **A_55_P2419021** | 9330188P | -0.015689007 | -0.297377425 | 0.265999412 | 7.825427394 | -0.139500832 | 0.894040878 | 0.952257952 | -4.758066243 |
| **A_55_P2180884** | Psmd10 | 0.019132655 | -0.324797397 | 0.363062707 | 9.529871367 | 0.139333472 | 0.89416703 | 0.952327878 | -4.758073998 |
| **A_55_P2017075** | Zfp54 | -0.013434188 | -0.255015606 | 0.22814723 | 8.153423421 | -0.139283042 | 0.894205043 | 0.952327878 | -4.758076333 |
| **A_55_P2161923** | Rabgap1 | 0.016446149 | -0.279682157 | 0.312574456 | 8.479921187 | 0.139102408 | 0.894341205 | 0.952420427 | -4.758084691 |
| **A_55_P2051476** | C8a | 0.011713201 | -0.199594868 | 0.22302127 | 7.640673583 | 0.138838479 | 0.894540162 | 0.952579836 | -4.758096883 |
| **A_52_P481686** | Wtip | -0.021994024 | -0.419479903 | 0.375491856 | 12.7516404 | -0.138590459 | 0.894727135 | 0.95267606 | -4.758108319 |
| **A_66_P109595** | Ube4b | 0.020875631 | -0.356405141 | 0.398156402 | 10.70887004 | 0.138587888 | 0.894729073 | 0.95267606 | -4.758108437 |
| **A_51_P196862** | Amdhd1 | -0.011815933 | -0.225693695 | 0.20206183 | 7.694445659 | -0.138373435 | 0.894890747 | 0.952756844 | -4.758118309 |
| **A_55_P2179309** | P2ry10b | 0.014403854 | -0.246349218 | 0.275156925 | 7.978491279 | 0.138356517 | 0.894903501 | 0.952756844 | -4.758119087 |
| **A_51_P459070** | Aplf | 0.019429551 | -0.332548362 | 0.371407465 | 9.382797349 | 0.138260368 | 0.894975989 | 0.952763829 | -4.758123508 |
| **A_55_P2029558** | Adrb2 | 0.017193081 | -0.294454348 | 0.32884051 | 7.923183982 | 0.138178511 | 0.895037702 | 0.952763829 | -4.758127269 |
| **A_51_P378381** | Crnde | 0.056002999 | -0.959962605 | 1.071968602 | 8.762771425 | 0.138064691 | 0.895123515 | 0.952763829 | -4.758132495 |
| **A_51_P303906** | Actr2 | -0.016129945 | -0.308755763 | 0.276495872 | 12.33177281 | -0.138060869 | 0.895126396 | 0.952763829 | -4.75813267 |
| **A_55_P2041045** | Csnk1g2 | -0.015216399 | -0.291348614 | 0.260915816 | 12.95273295 | -0.138020997 | 0.895156457 | 0.952763829 | -4.7581345 |
| **A_55_P2028259** | Rhbdf2 | 0.044861964 | -0.769725435 | 0.859449364 | 10.06979045 | 0.137940229 | 0.895217352 | 0.952764577 | -4.758138205 |
| **A_51_P154753** | Klc3 | 0.044731647 | -0.767789236 | 0.85725253 | 10.29778249 | 0.137889342 | 0.895255719 | 0.952764577 | -4.758140538 |
| **A_55_P2003398** | Rmdn1 | -0.013519428 | -0.259621029 | 0.232582173 | 8.498111042 | -0.137592326 | 0.895479663 | 0.952773701 | -4.758154138 |
| **A_66_P119735** | BC049265 | -0.016637141 | -0.319535308 | 0.286261027 | 8.049934023 | -0.137572739 | 0.895494432 | 0.952773701 | -4.758155034 |
| **A_51_P391616** | Etnppl | -0.011695752 | -0.22470287 | 0.201311366 | 7.662066054 | -0.137525861 | 0.895529778 | 0.952773701 | -4.758157178 |
| **A_52_P191537** | NA | 0.020635582 | -0.355344609 | 0.396615774 | 9.721190805 | 0.137468158 | 0.895573287 | 0.952773701 | -4.758159815 |
| **A_55_P1972040** | Nox4 | -0.029638172 | -0.569777764 | 0.510501419 | 9.363621354 | -0.137434496 | 0.895598668 | 0.952773701 | -4.758161353 |
| **A_52_P494115** | Gatad2b | -0.020684844 | -0.397673257 | 0.356303569 | 8.620039431 | -0.1374278 | 0.895603717 | 0.952773701 | -4.758161659 |
| **A_55_P2379599** | Asph | 0.01899302 | -0.3271798 | 0.36516584 | 8.800924876 | 0.137420462 | 0.89560925 | 0.952773701 | -4.758161995 |
| **A_55_P1958250** | Rnf114 | -0.033476091 | -0.644206866 | 0.577254683 | 13.85453417 | -0.137288855 | 0.895708485 | 0.952785512 | -4.758168004 |
| **A_51_P443693** | Rasl10b | 0.013863021 | -0.239176159 | 0.266902201 | 7.807902269 | 0.137220958 | 0.895759682 | 0.952785512 | -4.758171102 |
| **A_55_P1952512** | Sult2a2 | 0.012173253 | -0.210064615 | 0.234411122 | 7.669296289 | 0.13719521 | 0.895779098 | 0.952785512 | -4.758172277 |
| **A_51_P412595** | Eef1b2 | 0.011889811 | -0.205254011 | 0.229033633 | 15.80379752 | 0.137144317 | 0.895817473 | 0.952785512 | -4.758174598 |
| **A_66_P134383** | NA | -0.05031436 | -0.971041431 | 0.870412712 | 10.60850747 | -0.136870982 | 0.896023587 | 0.952952309 | -4.758187048 |
| **A_55_P2156785** | Ppfia3 | -0.040583457 | -0.784150243 | 0.702983329 | 9.544552345 | -0.136703427 | 0.896149939 | 0.953034264 | -4.758194668 |
| **A_55_P2040341** | NA | 0.029434712 | -0.510490048 | 0.569359473 | 9.17778728 | 0.136545344 | 0.896269151 | 0.953108618 | -4.758201848 |
| **A_51_P267740** | Zc3h15 | -0.016152614 | -0.312856084 | 0.280550857 | 10.12072232 | -0.13635483 | 0.896412825 | 0.953208974 | -4.758210491 |
| **A_55_P2014075** | Ercc2 | -0.016817455 | -0.327080544 | 0.293445634 | 10.24572141 | -0.135762715 | 0.896859389 | 0.953543449 | -4.758237277 |
| **A_51_P138538** | Eif1b | 0.013789911 | -0.240640266 | 0.268220087 | 11.93452474 | 0.135751044 | 0.896868192 | 0.953543449 | -4.758237804 |
| **A_66_P132070** | Abra | 0.011801683 | -0.205959958 | 0.229563323 | 7.694545473 | 0.135741579 | 0.89687533 | 0.953543449 | -4.758238231 |
| **A_51_P244856** | Atp2a1 | -0.011398397 | -0.22199008 | 0.199193286 | 7.798485427 | -0.135566661 | 0.897007259 | 0.953619434 | -4.758246121 |
| **A_55_P2026487** | NA | -0.013008306 | -0.25343374 | 0.227417129 | 7.791945927 | -0.135516032 | 0.897045445 | 0.953619434 | -4.758248402 |
| **A_51_P317433** | Mesp2 | 0.011552306 | -0.202169886 | 0.225274497 | 7.845685893 | 0.135384643 | 0.897144547 | 0.953672349 | -4.75825432 |
| **A_52_P75384** | B230219D | -0.03096601 | -0.604448403 | 0.542516382 | 12.03185173 | -0.135243222 | 0.897251217 | 0.953733302 | -4.758260683 |

| | | | | | | | | | |
|---|---|---|---|---|---|---|---|---|---|
| **A_55_P2026492** | NA | -0.011747411 | -0.229505485 | 0.206010662 | 7.814637303 | -0.135119571 | 0.897344486 | 0.953769863 | -4.75826624 |
| **A_55_P2000062** | Irf1 | -0.043164565 | -0.843605043 | 0.757275912 | 11.82186991 | -0.135066821 | 0.897384275 | 0.953769863 | -4.75826861 |
| **A_55_P2070296** | NA | -0.011753819 | -0.229860001 | 0.206352363 | 7.688087331 | -0.134977498 | 0.897451652 | 0.953789041 | -4.75827262 |
| **A_55_P2256686** | NA | -0.059245351 | -1.159318752 | 1.040828051 | 8.426517566 | -0.134890994 | 0.897516904 | 0.953805959 | -4.758276502 |
| **A_55_P2136413** | NA | 0.028619183 | -0.503242277 | 0.560480643 | 11.06574879 | 0.13477491 | 0.897604469 | 0.95382344 | -4.758281706 |
| **A_51_P333869** | Nsmce3 | -0.014420759 | -0.282618834 | 0.253777315 | 9.172616589 | -0.13467371 | 0.89768081 | 0.95382344 | -4.75828624 |
| **A_55_P2001168** | NA | 0.021812364 | -0.383857359 | 0.427482087 | 11.33414113 | 0.134672987 | 0.897681355 | 0.95382344 | -4.758286272 |
| **A_51_P174696** | Casq2 | 0.063507498 | -1.118267241 | 1.245282238 | 8.58722901 | 0.134598633 | 0.897737444 | 0.953830618 | -4.758289601 |
| **A_51_P488510** | Appl1 | -0.012811081 | -0.25181052 | 0.226188359 | 8.92766708 | -0.134257708 | 0.897994631 | 0.954041012 | -4.758304841 |
| **A_51_P191611** | Cat | 0.022534791 | -0.398031221 | 0.443100803 | 8.422444233 | 0.134205315 | 0.898034157 | 0.954041012 | -4.758307179 |
| **A_51_P519276** | Ndufb7 | 0.021420335 | -0.379059128 | 0.421899799 | 14.21504181 | 0.133966546 | 0.898214289 | 0.954179948 | -4.758317826 |
| **A_51_P306047** | Sec13 | 0.011599779 | -0.205650536 | 0.228850095 | 10.11558703 | 0.133733328 | 0.89839024 | 0.954314427 | -4.758328207 |
| **A_55_P1999202** | Vmn1r217 | -0.026836272 | -0.529745815 | 0.476073271 | 8.61734449 | -0.133654212 | 0.89844993 | 0.954325401 | -4.758331724 |
| **A_52_P69558** | Gm8221 | -0.011699778 | -0.231286697 | 0.207887142 | 7.676309404 | -0.133450895 | 0.89860333 | 0.954417802 | -4.758340754 |
| **A_55_P2062246** | Tgtp2 | 0.094642683 | -1.682225972 | 1.871511338 | 9.121573422 | 0.133408058 | 0.898635651 | 0.954417802 | -4.758342655 |
| **A_55_P1999082** | Clec14a | -0.029079732 | -0.575539602 | 0.517380137 | 8.935734693 | -0.133285368 | 0.898728221 | 0.954458959 | -4.758348096 |
| **A_55_P2087414** | Sema3e | -0.016527564 | -0.327319219 | 0.294264092 | 8.823933386 | -0.133195586 | 0.898795963 | 0.954458959 | -4.758352074 |
| **A_55_P2046378** | Eif4g1 | -0.032840997 | -0.650560828 | 0.584878834 | 13.16840419 | -0.133160413 | 0.898822502 | 0.954458959 | -4.758353632 |
| **A_55_P2161958** | Gm10324 | -0.054416796 | -1.078798029 | 0.969964437 | 9.791001506 | -0.13305209 | 0.898904236 | 0.954493328 | -4.758358427 |
| **A_55_P2117843** | Coasy | 0.02001207 | -0.357523917 | 0.397548058 | 12.1462776 | 0.132765126 | 0.899120768 | 0.954670818 | -4.758371111 |
| **A_52_P275354** | Osbpl7 | 0.01487558 | -0.265992186 | 0.295743346 | 10.45623383 | 0.132654613 | 0.899204159 | 0.954706931 | -4.758375989 |
| **A_55_P1995737** | Gm14288 | 0.023251039 | -0.4173903 | 0.463892378 | 11.79306769 | 0.132162267 | 0.899575693 | 0.955048952 | -4.75839767 |
| **A_55_P2005248** | Meis3 | 0.011909923 | -0.2141171 | 0.237936946 | 8.922256301 | 0.131977206 | 0.899715351 | 0.955144773 | -4.758405799 |
| **A_52_P635338** | Fes | 0.034077575 | -0.613019827 | 0.681174977 | 10.1003825 | 0.131901389 | 0.899772568 | 0.955153069 | -4.758409126 |
| **A_51_P424561** | Zfp93 | 0.030756954 | -0.554157362 | 0.61567127 | 9.371484278 | 0.131704743 | 0.899920975 | 0.95525816 | -4.758417747 |
| **A_55_P2113356** | Miip | 0.013071082 | -0.235631791 | 0.261773955 | 8.203963814 | 0.131637944 | 0.899971389 | 0.955259228 | -4.758420672 |
| **A_66_P116098** | Ptpn2 | -0.023096373 | -0.463319761 | 0.417127015 | 8.812790046 | -0.131407761 | 0.900145114 | 0.955363877 | -4.758430742 |
| **A_55_P2040115** | Cdv3 | 0.038950882 | -0.703641336 | 0.7815431 | 13.28790621 | 0.131376366 | 0.900168808 | 0.955363877 | -4.758432114 |
| **A_52_P64707** | Foxa3 | -0.011371717 | -0.228567757 | 0.205824322 | 7.727257015 | -0.131136773 | 0.900349642 | 0.955503348 | -4.758442574 |
| **A_55_P2105436** | Foxp1 | 0.022385995 | -0.405590985 | 0.450362975 | 8.514841821 | 0.13101057 | 0.900444898 | 0.955551987 | -4.758448076 |
| **A_55_P2431118** | Itsn1 | 0.012158869 | -0.220429371 | 0.244747108 | 10.4615951 | 0.130935003 | 0.900501934 | 0.955560066 | -4.758451369 |
| **A_51_P375693** | Tmem135 | 0.016967677 | -0.307974745 | 0.341910098 | 9.547364574 | 0.130787523 | 0.900613252 | 0.955625741 | -4.758457788 |
| **A_55_P2061456** | Nfyc | 0.015002217 | -0.272640571 | 0.302645004 | 9.366186776 | 0.130632817 | 0.900730028 | 0.955697199 | -4.758464514 |
| **A_51_P374782** | Epn3 | 0.048205776 | -0.879318614 | 0.975730166 | 11.52439179 | 0.130173951 | 0.901076404 | 0.955944323 | -4.758484419 |
| **A_51_P215097** | Klf16 | -0.020257132 | -0.410138117 | 0.369623853 | 8.506660803 | -0.130135639 | 0.901105326 | 0.955944323 | -4.758486078 |
| **A_55_P2102628** | Usp4 | 0.017420789 | -0.317890524 | 0.352732103 | 10.96398013 | 0.130127765 | 0.901111269 | 0.955944323 | -4.758486418 |
| **A_55_P2072556** | Ncoa6 | -0.014489467 | -0.29418185 | 0.265202915 | 11.82704419 | -0.12975441 | 0.901393118 | 0.956109815 | -4.758502557 |
| **A_55_P2327958** | Mtx3 | 0.021970894 | -0.402315968 | 0.446257756 | 8.867558976 | 0.129699558 | 0.901434527 | 0.956109815 | -4.758504924 |
| **A_55_P2022499** | Pnisr | 0.034422235 | -0.630473026 | 0.699317496 | 9.556923424 | 0.129669003 | 0.901457595 | 0.956109815 | -4.758506242 |
| **A_55_P2089677** | Synrg | 0.040553793 | -0.742837701 | 0.823945286 | 10.09748602 | 0.12965909 | 0.901465078 | 0.956109815 | -4.75850667 |
| **A_55_P2044252** | Zfp512b | -0.017282627 | -0.351860845 | 0.317295591 | 10.78422012 | -0.129378598 | 0.901676837 | 0.956281951 | -4.758518755 |
| **A_52_P212214** | Pomt2 | 0.01552939 | -0.285310683 | 0.316369463 | 8.333526844 | 0.129291226 | 0.901742801 | 0.956299453 | -4.758522514 |
| **A_52_P533792** | Sucla2 | 0.018309243 | -0.355358165 | 0.391976651 | 12.23246603 | 0.130511227 | 0.901819858 | 0.956328715 | -4.729710534 |
| **A_55_P2037930** | NA | 0.016985779 | -0.313612752 | 0.34758431 | 8.0572123 | 0.128687066 | 0.902198953 | 0.956624455 | -4.75854844 |
| **A_51_P386270** | Cyp2d12 | -0.011089305 | -0.226983231 | 0.204804622 | 7.733177958 | -0.128651318 | 0.902225944 | 0.956624455 | -4.75854997 |
| **A_51_P384318** | C1ra | -0.018112757 | -0.370821005 | 0.33459549 | 9.21395771 | -0.128623196 | 0.902247178 | 0.956624455 | -4.758551174 |
| **A_55_P1983135** | NA | -0.016604064 | -0.340280288 | 0.30707216 | 8.302999951 | -0.128485445 | 0.902351189 | 0.956682271 | -4.758557065 |
| **A_55_P2138960** | Tpk1 | -0.011849919 | -0.243337724 | 0.219637886 | 8.40921817 | -0.128214643 | 0.90255567 | 0.956846593 | -4.758568629 |

| | | | | | | | | | |
|---|---|---|---|---|---|---|---|---|---|
| **A_51_P423549** | Shbg | 0.010666608 | -0.197852405 | 0.219185621 | 7.655500946 | 0.128124153 | 0.902624 | 0.956866565 | -4.758572488 |
| **A_51_P260740** | Pcdh7 | 0.010919106 | -0.20265276 | 0.224490972 | 7.822054668 | 0.12805407 | 0.90267692 | 0.9568702 | -4.758575474 |
| **A_55_P2371801** | A430046D | 0.013725844 | -0.255143522 | 0.28259521 | 8.036601088 | 0.127863943 | 0.902820492 | 0.956962623 | -4.758583568 |
| **A_66_P136834** | Usp7 | -0.011531707 | -0.237623247 | 0.214559832 | 8.303983934 | -0.127749629 | 0.902906816 | 0.956962623 | -4.758588429 |
| **A_51_P427674** | Cpt1a | -0.021470831 | -0.442467346 | 0.399525684 | 10.14362551 | -0.12773817 | 0.90291547 | 0.956962623 | -4.751526493 |
| **A_55_P2099952** | Car1 | 0.010596628 | -0.197280783 | 0.218474039 | 7.660672353 | 0.12767643 | 0.902962094 | 0.956962623 | -4.75859154 |
| **A_51_P316173** | Vmn1r192 | -0.010925224 | -0.225492862 | 0.203642414 | 7.807742518 | -0.127531204 | 0.903071765 | 0.957026392 | -4.758597705 |
| **A_55_P2134246** | Serpinb9 | 0.055730304 | -1.039956715 | 1.151417323 | 9.841183076 | 0.127395838 | 0.903173992 | 0.957053919 | -4.758603446 |
| **A_55_P2052862** | Miga1 | -0.01212109 | -0.250484451 | 0.22624227 | 10.37104207 | -0.127365713 | 0.903196742 | 0.957053919 | -4.758604723 |
| **A_55_P1994608** | Itsn1 | -0.01296648 | -0.268348392 | 0.242415432 | 8.796771065 | -0.127169308 | 0.90334507 | 0.957158634 | -4.75861304 |
| **A_55_P2128324** | Ubap2l | -0.019748979 | -0.408957367 | 0.36945941 | 8.644831874 | -0.127090415 | 0.903404653 | 0.957169309 | -4.758616377 |
| **A_55_P2043622** | NA | -0.039339138 | -0.815278273 | 0.736599997 | 12.84039741 | -0.12698357 | 0.903485346 | 0.957182557 | -4.758620893 |
| **A_55_P2029021** | NA | -0.031499911 | -0.653015116 | 0.590015293 | 15.41543352 | -0.126942755 | 0.903516172 | 0.957182557 | -4.758622617 |
| **A_55_P2172058** | Trim55 | -0.011129432 | -0.231132945 | 0.20887408 | 7.71455417 | -0.126704997 | 0.903695742 | 0.957320337 | -4.758632651 |
| **A_52_P253044** | Syt13 | 0.081694003 | -1.54009803 | 1.703486036 | 9.533800567 | 0.126166879 | 0.904102186 | 0.957669372 | -4.75865529 |
| **A_55_P2006250** | Myo5a | -0.0153956 | -0.321100464 | 0.290309265 | 9.557696339 | -0.126137614 | 0.904124291 | 0.957669372 | -4.758656518 |
| **A_55_P1961009** | Apbb1ip | -0.059191894 | -1.236509582 | 1.118125794 | 8.680643225 | -0.125927016 | 0.904283367 | 0.957717465 | -4.75866535 |
| **A_55_P2060333** | S100a5 | 0.033980066 | -0.642050311 | 0.710010443 | 8.725616989 | 0.125894963 | 0.904307579 | 0.957717465 | -4.758666693 |
| **A_55_P2075000** | NA | -0.017811587 | -0.372211468 | 0.336588295 | 12.54824377 | -0.125880767 | 0.904318302 | 0.957717465 | -4.758667288 |
| **A_55_P1981415** | Gsk3b | -0.027216954 | -0.569538023 | 0.515104115 | 12.29868349 | -0.125699453 | 0.904455263 | 0.957810049 | -4.758674878 |
| **A_55_P1999301** | Cyp2e1 | 0.030023604 | -0.569447021 | 0.62949423 | 8.199518406 | 0.125442664 | 0.904649243 | 0.957905904 | -4.758685608 |
| **A_55_P2149575** | Sdr39u1 | 0.012258588 | -0.232519102 | 0.257036278 | 8.12857943 | 0.125435065 | 0.904654983 | 0.957905904 | -4.758685925 |
| **A_55_P2011912** | Ecm2 | 0.02074567 | -0.393798018 | 0.435289358 | 8.055531801 | 0.125345141 | 0.904722914 | 0.957905904 | -4.758689677 |
| **A_66_P112886** | Ubl3 | -0.034078533 | -0.715193259 | 0.647036193 | 10.8749683 | -0.125317283 | 0.904743959 | 0.957905904 | -4.758690839 |
| **A_52_P176999** | Klhl31 | 0.010499246 | -0.199667137 | 0.220665628 | 7.642967424 | 0.125125318 | 0.904888977 | 0.958006981 | -4.758698838 |
| **A_55_P2088070** | Spata5 | -0.02327712 | -0.490057755 | 0.443503515 | 9.448740936 | -0.124901241 | 0.90505826 | 0.958060004 | -4.758708161 |
| **A_55_P2126827** | NA | -0.027468484 | -0.578304424 | 0.523367457 | 13.35861301 | -0.124900094 | 0.905059127 | 0.958060004 | -4.758708208 |
| **A_51_P223458** | Polr3d | 0.024189166 | -0.461032398 | 0.509410729 | 8.836843069 | 0.124862246 | 0.90508772 | 0.958060004 | -4.758709781 |
| **A_51_P267063** | Ugt3a2 | -0.010461228 | -0.220653147 | 0.199730692 | 7.703354427 | -0.124657089 | 0.905242715 | 0.95807965 | -4.758718299 |
| **A_55_P2128326** | Ubap2l | -0.015902744 | -0.335789578 | 0.30398409 | 9.042879726 | -0.124516254 | 0.905349118 | 0.95807965 | -4.758724138 |
| **A_52_P362111** | Fam3c | -0.019186942 | -0.405305482 | 0.366931599 | 11.21238188 | -0.124461619 | 0.905390397 | 0.95807965 | -4.758726402 |
| **A_66_P100657** | Xpa | 0.029393643 | -0.562448285 | 0.621235571 | 10.92379759 | 0.1243936 | 0.905441787 | 0.95807965 | -4.758729218 |
| **A_55_P2175550** | Atf2 | -0.015890687 | -0.335889617 | 0.304108243 | 8.669749702 | -0.124378263 | 0.905453375 | 0.95807965 | -4.758729853 |
| **A_55_P1988102** | Hmbs | -0.014361568 | -0.303572565 | 0.27484943 | 8.353088247 | -0.124376229 | 0.905454911 | 0.95807965 | -4.758729938 |
| **A_55_P2363442** | Zfp867 | -0.033469479 | -0.707887054 | 0.640948096 | 9.011141995 | -0.124299798 | 0.905512658 | 0.95807965 | -4.7587331 |
| **A_52_P238468** | Gipr | -0.035649748 | -0.754549262 | 0.683249765 | 8.32655841 | -0.12420487 | 0.905584381 | 0.95807965 | -4.758737026 |
| **A_51_P376238** | Serping1 | -0.025308874 | -0.535747802 | 0.485130054 | 11.53451834 | -0.124187928 | 0.905597182 | 0.95807965 | -4.758737726 |
| **A_55_P2068104** | NA | -0.035679975 | -0.75552813 | 0.684168179 | 9.864190943 | -0.124146362 | 0.905628588 | 0.95807965 | -4.758739444 |
| **A_52_P412321** | Pcgf6 | 0.013690348 | -0.262644004 | 0.290024701 | 8.874553695 | 0.124088053 | 0.905672644 | 0.95807965 | -4.758741852 |
| **A_55_P2150962** | Cpne7 | -0.012662829 | -0.268334239 | 0.243008581 | 8.618851924 | -0.124050617 | 0.905700929 | 0.95807965 | -4.758743398 |
| **A_55_P2404554** | Atcayos | -0.012597108 | -0.267469869 | 0.242275653 | 7.767265416 | -0.123793477 | 0.905895221 | 0.9582048 | -4.758754003 |
| **A_51_P373619** | Mbl1 | 0.010873655 | -0.209183543 | 0.230930854 | 7.703641058 | 0.123762857 | 0.905918358 | 0.9582048 | -4.758755265 |
| **A_55_P2008061** | Itpr2 | -0.037293006 | -0.793374434 | 0.718788421 | 11.06368857 | -0.123540448 | 0.906086414 | 0.958330128 | -4.758764417 |
| **A_51_P428086** | Sptbn1 | 0.036140403 | -0.697193612 | 0.769474417 | 9.395534825 | 0.123435905 | 0.906165411 | 0.958361252 | -4.758768714 |
| **A_55_P2069721** | Hspa8 | -0.029042198 | -0.618755068 | 0.560670672 | 15.30389077 | -0.12335002 | 0.906230309 | 0.958377464 | -4.758772241 |
| **A_55_P1966332** | Plac9a | 0.043406286 | -0.839289883 | 0.926102454 | 15.05434204 | 0.123166253 | 0.906369175 | 0.958471893 | -4.75877978 |
| **A_51_P509643** | Snca | -0.025816033 | -0.551604789 | 0.499972722 | 11.48624289 | -0.122978323 | 0.90651119 | 0.958491129 | -4.758787478 |
| **A_52_P370473** | Msl1 | -0.021816567 | -0.466178998 | 0.422545865 | 11.9118036 | -0.122970069 | 0.906517428 | 0.958491129 | -4.758787816 |

| | | | | | | | | | |
|---|---|---|---|---|---|---|---|---|---|
| **A_51_P316935** | Depp1 | 0.05799138 | -1.123421233 | 1.239403992 | 10.78049784 | 0.122945371 | 0.906536091 | 0.958491129 | -4.758788827 |
| **A_51_P462814** | Sars2 | 0.017815509 | -0.345729328 | 0.381360347 | 8.83710567 | 0.122741269 | 0.906690333 | 0.958601788 | -4.758797172 |
| **A_52_P648304** | Pigs | 0.01316217 | -0.255627652 | 0.281951992 | 12.31535417 | 0.122649287 | 0.906759846 | 0.95862286 | -4.758800928 |
| **A_55_P1999057** | Cps1 | 0.010216108 | -0.198741924 | 0.21917414 | 7.686253048 | 0.122455065 | 0.906906627 | 0.958723655 | -4.758808851 |
| **A_55_P2206461** | A930006K | -0.020469665 | -0.439367774 | 0.398428444 | 8.178691694 | -0.122391901 | 0.906954363 | 0.958723655 | -4.758811425 |
| **A_52_P590625** | Nlrp10 | 0.012481246 | -0.243503395 | 0.268465886 | 7.755968128 | 0.122122129 | 0.907158249 | 0.958886752 | -4.758822403 |
| **A_55_P2002864** | Aifm1 | 0.014260075 | -0.278418997 | 0.306939147 | 11.31229874 | 0.122033899 | 0.907224932 | 0.958904813 | -4.758825988 |
| **A_51_P462972** | Ap5m1 | -0.010430314 | -0.224772037 | 0.203911408 | 9.44580314 | -0.12188241 | 0.907339428 | 0.958973405 | -4.758832138 |
| **A_55_P2135701** | BC024978 | -0.017229855 | -0.372049111 | 0.337589402 | 8.269236835 | -0.121625538 | 0.907533578 | 0.959126173 | -4.758842548 |
| **A_51_P102438** | Ugt2b36 | 0.010085655 | -0.19810383 | 0.218275139 | 7.678824759 | 0.121337668 | 0.907751167 | 0.959173451 | -4.758854189 |
| **A_55_P2021471** | Pwwp3a | 0.010765136 | -0.211461074 | 0.232991346 | 7.840146759 | 0.121331787 | 0.907755612 | 0.959173451 | -4.758854427 |
| **A_55_P2074473** | Gtf2f1 | -0.023292578 | -0.504285272 | 0.457700117 | 13.24606353 | -0.121291234 | 0.907786265 | 0.959173451 | -4.758856064 |
| **A_55_P2036854** | Pnkd | -0.021901394 | -0.474314574 | 0.430511787 | 11.80734461 | -0.121251424 | 0.907816356 | 0.959173451 | -4.758857671 |
| **A_55_P1971199** | Slc4a1ap | 0.011938188 | -0.234693778 | 0.258570155 | 10.77343392 | 0.12123818 | 0.907826367 | 0.959173451 | -4.758858206 |
| **A_55_P2112120** | Tnk2 | -0.01317764 | -0.285661192 | 0.259305912 | 8.410355063 | -0.121128876 | 0.907908988 | 0.959208327 | -4.758862615 |
| **A_66_P139167** | Safb | -0.01596514 | -0.346298549 | 0.31436827 | 11.12173805 | -0.12105161 | 0.907967393 | 0.959217616 | -4.75886573 |
| **A_55_P2171912** | Gja3 | -0.012414572 | -0.269540128 | 0.244710985 | 8.097147853 | -0.120930775 | 0.908058734 | 0.959261696 | -4.758870597 |
| **A_52_P31490** | Vamp3 | -0.019680428 | -0.427694567 | 0.388333711 | 9.161016883 | -0.120811887 | 0.908148604 | 0.959304218 | -4.75887538 |
| **A_55_P2176894** | NA | 0.010539951 | -0.208618734 | 0.229698636 | 7.852649962 | 0.120456502 | 0.908417255 | 0.959496397 | -4.758889652 |
| **A_55_P2229351** | NA | -0.014889718 | -0.324536355 | 0.294756918 | 8.440169545 | -0.120439922 | 0.90842979 | 0.959496397 | -4.758890317 |
| **A_55_P2172001** | Mrgpra2a | 0.010317521 | -0.204491582 | 0.225126624 | 7.695264348 | 0.120302052 | 0.908534016 | 0.959554062 | -4.758895842 |
| **A_55_P2174401** | NA | 0.01517587 | -0.301975295 | 0.332327035 | 8.273748661 | 0.119849887 | 0.908875855 | 0.959862663 | -4.758913917 |
| **A_55_P1966644** | Trpc6 | 0.035997655 | -0.719270465 | 0.791265775 | 8.606001594 | 0.119377754 | 0.909232814 | 0.960074519 | -4.758932718 |
| **A_55_P2082570** | NA | 0.015434309 | -0.308511203 | 0.339379822 | 8.272726078 | 0.11933438 | 0.909265609 | 0.960074519 | -4.758934441 |
| **A_51_P128575** | Scgb1a1 | -0.172105302 | -3.784843613 | 3.440633008 | 11.57059425 | -0.119318584 | 0.909277551 | 0.960074519 | -4.75185997 |
| **A_55_P2137186** | Rps25 | 0.026825847 | -0.536288692 | 0.589940385 | 16.22289837 | 0.119318297 | 0.909277769 | 0.960074519 | -4.75893508 |
| **A_55_P2127557** | Abracl | 0.023848123 | -0.477152544 | 0.52484879 | 11.68667437 | 0.1192247 | 0.909348536 | 0.960074519 | -4.758938797 |
| **A_55_P2012599** | NA | -0.014730906 | -0.324286159 | 0.294824348 | 8.149091873 | -0.119190492 | 0.909374401 | 0.960074519 | -4.758940155 |
| **A_55_P1985259** | C2cd3 | -0.013580401 | -0.299253356 | 0.272092555 | 8.287726507 | -0.119067656 | 0.909467279 | 0.960120147 | -4.758945027 |
| **A_55_P2127174** | Zfp628 | 0.014394946 | -0.288948788 | 0.317738681 | 17.73503124 | 0.118857164 | 0.909626437 | 0.960235737 | -4.758953364 |
| **A_51_P400543** | Aif1 | 0.020510657 | -0.412184894 | 0.453206208 | 9.186512195 | 0.11872645 | 0.909725275 | 0.960241362 | -4.758958533 |
| **A_52_P54261** | Tmem56 | 0.011617325 | -0.233481071 | 0.25671572 | 7.88909204 | 0.118717845 | 0.909731782 | 0.960241362 | -4.758958874 |
| **A_52_P234729** | Pkd2 | -0.018215053 | -0.402719892 | 0.366289786 | 10.92287193 | -0.118653069 | 0.909780762 | 0.960241362 | -4.758961433 |
| **A_55_P2120773** | Dctn4 | -0.013181904 | -0.291655657 | 0.265291849 | 11.78878747 | -0.118561648 | 0.909849892 | 0.960261903 | -4.758965043 |
| **A_55_P2180846** | Wdsub1 | -0.015790055 | -0.349678678 | 0.318098568 | 8.476487222 | -0.118449264 | 0.909934873 | 0.960299173 | -4.758969478 |
| **A_55_P2037328** | Ppp1r12b | -0.009864136 | -0.218728101 | 0.198999829 | 7.744685266 | -0.118289408 | 0.910055754 | 0.960357437 | -4.758975778 |
| **A_66_P133498** | Pdcd6ip | -0.023369711 | -0.51838812 | 0.471648697 | 11.43682716 | -0.11824488 | 0.910089426 | 0.960357437 | -4.758977531 |
| **A_52_P223704** | Faah | 0.011648112 | -0.235233049 | 0.258529273 | 7.810495127 | 0.11817291 | 0.910143851 | 0.960362452 | -4.758980364 |
| **A_51_P486810** | Gpx2 | -0.012690384 | -0.282015481 | 0.256634713 | 8.746532538 | -0.118018009 | 0.910260989 | 0.960433637 | -4.758986455 |
| **A_55_P2092904** | Ddb2 | -0.011352353 | -0.252480242 | 0.229775536 | 8.693529021 | -0.117920355 | 0.910334838 | 0.960459141 | -4.75899029 |
| **A_51_P115005** | Edn1 | 0.020488404 | -0.415393117 | 0.456369925 | 8.610746206 | 0.117730779 | 0.910478204 | 0.960557983 | -4.758997728 |
| **A_52_P160518** | Sfmbt1 | 0.017525837 | -0.356022975 | 0.391074649 | 8.599454846 | 0.117511876 | 0.910643753 | 0.960675429 | -4.759006301 |
| **A_55_P1958532** | Hr | -0.021487449 | -0.479707554 | 0.436732656 | 8.929382956 | -0.117452175 | 0.910688904 | 0.960675429 | -4.759008636 |
| **A_55_P2072030** | NA | 0.02433222 | -0.496419825 | 0.545084265 | 16.05093424 | 0.117031046 | 0.911007406 | 0.960958982 | -4.759025077 |
| **A_55_P2183955** | Olfr97 | 0.010107765 | -0.206382016 | 0.226597546 | 7.801391479 | 0.116941346 | 0.911075249 | 0.960978116 | -4.759028571 |
| **A_52_P178998** | Fam168b | -0.011514375 | -0.258617539 | 0.235588788 | 13.36574363 | -0.116711169 | 0.911249343 | 0.961109312 | -4.759037526 |
| **A_51_P377390** | Ndst2 | 0.01189141 | -0.243790255 | 0.267573076 | 9.693524448 | 0.116488789 | 0.911417546 | 0.96123428 | -4.75904616 |
| **A_55_P1995992** | NA | 0.020217715 | -0.415307978 | 0.455743409 | 9.348477425 | 0.116270258 | 0.911582841 | 0.961356169 | -4.759054629 |

| | | | | | | | | | |
|---|---|---|---|---|---|---|---|---|---|
| **A_55_P1992487** | NA | -0.017338006 | -0.392090197 | 0.357414186 | 10.08221346 | -0.115879151 | 0.911878686 | 0.961615715 | -4.759069747 |
| **A_55_P2051596** | NA | 0.02120081 | -0.437363668 | 0.479765288 | 9.707139598 | 0.115798357 | 0.911939802 | 0.961615723 | -4.759072863 |
| **A_55_P1954850** | Pars2 | -0.016842089 | -0.381288903 | 0.347604725 | 9.773578683 | -0.11574764 | 0.911978168 | 0.961615723 | -4.759074819 |
| **A_51_P163106** | Bdh1 | 0.070664779 | -1.459650845 | 1.600980402 | 9.328922228 | 0.11565711 | 0.912046651 | 0.961635488 | -4.759078307 |
| **A_55_P2072373** | Msn | 0.030188899 | -0.62460642 | 0.684984218 | 14.41491282 | 0.115476086 | 0.912183592 | 0.961718442 | -4.759085274 |
| **A_51_P221248** | Tspan32 | 0.009929265 | -0.205537533 | 0.225396063 | 7.735217736 | 0.115421598 | 0.912224812 | 0.961718442 | -4.759087369 |
| **A_55_P2017677** | Cap2 | 0.023402779 | -0.485463597 | 0.532269155 | 8.060212897 | 0.1151898 | 0.912400169 | 0.961850865 | -4.75909627 |
| **A_52_P114620** | Elmo3 | -0.028788901 | -0.656078295 | 0.598500493 | 10.49772144 | -0.114949605 | 0.912581883 | 0.961953468 | -4.759105475 |
| **A_51_P232705** | Commd9 | -0.010740288 | -0.245074568 | 0.223593991 | 8.877088335 | -0.114796981 | 0.912697351 | 0.961953468 | -4.759111314 |
| **A_52_P219966** | Spg21 | -0.011160778 | -0.254778943 | 0.232457387 | 8.566766165 | -0.114745361 | 0.912736405 | 0.961953468 | -4.759113287 |
| **A_55_P1981959** | Trafd1 | -0.010663467 | -0.243467171 | 0.222140238 | 12.03841961 | -0.114725214 | 0.912751647 | 0.961953468 | -4.759114057 |
| **A_55_P2026119** | Dlec1 | -0.014161871 | -0.323483736 | 0.295159995 | 9.046653448 | -0.114672784 | 0.912791315 | 0.961953468 | -4.759116059 |
| **A_55_P2127765** | Crp | 0.009493675 | -0.198011264 | 0.216998614 | 7.652279162 | 0.114592516 | 0.912852043 | 0.961953468 | -4.759119124 |
| **A_51_P483658** | Trim3 | -0.027512548 | -0.629071641 | 0.574046544 | 9.781540618 | -0.114552051 | 0.912882659 | 0.961953468 | -4.759120668 |
| **A_55_P2143000** | NA | -0.011156209 | -0.255121825 | 0.232809407 | 8.036940764 | -0.114535035 | 0.912895532 | 0.961953468 | -4.759121317 |
| **A_55_P2022724** | Inafm2 | -0.022140648 | -0.507771057 | 0.463489761 | 9.898622575 | -0.114191767 | 0.91315525 | 0.962174702 | -4.759134392 |
| **A_55_P2139341** | Zfp975 | -0.012329726 | -0.283166458 | 0.258507006 | 12.08564617 | -0.114023986 | 0.913282198 | 0.96220639 | -4.759140768 |
| **A_51_P336599** | Kcne3 | -0.011998618 | -0.275570304 | 0.251573068 | 8.032046287 | -0.11402047 | 0.913284859 | 0.96220639 | -4.759140901 |
| **A_51_P426633** | Dnttip2 | -0.021699596 | -0.500358269 | 0.456959077 | 12.34210973 | -0.113547104 | 0.913643037 | 0.962531304 | -4.759158841 |
| **A_55_P2153382** | Ermap | 0.009543008 | -0.201210965 | 0.220296981 | 7.685082249 | 0.113412217 | 0.913745105 | 0.962550597 | -4.759163939 |
| **A_55_P1960097** | Epb41l3 | 0.020000724 | -0.421789666 | 0.461791113 | 10.83599361 | 0.113391315 | 0.913760922 | 0.962550597 | -4.759164728 |
| **A_51_P350817** | Cnn1 | 0.059457772 | -1.255348493 | 1.374264037 | 11.7318424 | 0.113265387 | 0.913856213 | 0.96259853 | -4.759169482 |
| **A_52_P77227** | Rnf212 | -0.009521533 | -0.220525633 | 0.201482567 | 7.756351935 | -0.113022868 | 0.914039734 | 0.962681179 | -4.759178621 |
| **A_51_P413059** | Mcpt8 | 0.012487426 | -0.264365466 | 0.289340318 | 7.839774975 | 0.112972883 | 0.914077559 | 0.962681179 | -4.759180503 |
| **A_55_P2091120** | Kbtbd12 | -0.010118985 | -0.234604564 | 0.214366594 | 7.703933968 | -0.112901273 | 0.91413175 | 0.962681179 | -4.759183196 |
| **A_55_P2027412** | NA | 0.027031168 | -0.572659066 | 0.626721403 | 8.893322157 | 0.112898504 | 0.914133845 | 0.962681179 | -4.759183301 |
| **A_55_P2034250** | Zscan26 | 0.019467441 | -0.413242691 | 0.452177573 | 10.61939706 | 0.11268397 | 0.914296197 | 0.962799709 | -4.75919136 |
| **A_55_P2077068** | Malt1 | 0.01136389 | -0.24147006 | 0.26419784 | 8.983303209 | 0.112575007 | 0.914378658 | 0.962834103 | -4.759195448 |
| **A_55_P2425190** | Nhsl2 | 0.012388128 | -0.263829383 | 0.288605639 | 8.091507222 | 0.112332343 | 0.914562305 | 0.962975035 | -4.759204538 |
| **A_51_P485458** | Txlna | 0.016011929 | -0.341643068 | 0.373666926 | 8.41577362 | 0.112132035 | 0.914713902 | 0.963082208 | -4.759212026 |
| **A_55_P2091756** | Nxnl1 | 0.021423372 | -0.457886863 | 0.500733607 | 8.42496271 | 0.111949323 | 0.914852185 | 0.963175351 | -4.759218845 |
| **A_55_P2091551** | Arhgap9 | 0.078052915 | -1.670137808 | 1.826243638 | 10.0889829 | 0.111828027 | 0.914943989 | 0.963219552 | -4.759223366 |
| **A_51_P365980** | 2310022B | -0.022699117 | -0.531512481 | 0.486114246 | 11.10548695 | -0.111737978 | 0.915012144 | 0.963238854 | -4.759226719 |
| **A_65_P19458** | Rock2 | -0.01153175 | -0.270208097 | 0.247144597 | 9.528551827 | -0.111657741 | 0.915072873 | 0.963250336 | -4.759229705 |
| **A_52_P37091** | Magi1 | 0.016090383 | -0.345767369 | 0.377948136 | 8.062606602 | 0.111372726 | 0.915288598 | 0.963424965 | -4.759240292 |
| **A_51_P329966** | D6Wsu16 | -0.014701025 | -0.345756362 | 0.316354313 | 9.071425925 | -0.111223704 | 0.915401394 | 0.963491238 | -4.759245817 |
| **A_66_P107038** | Prcp | 0.013993769 | -0.301514623 | 0.329502162 | 12.37691394 | 0.11108979 | 0.915502758 | 0.963545472 | -4.759250776 |
| **A_55_P2051429** | NA | 0.014349531 | -0.309691118 | 0.33839018 | 8.28908852 | 0.110914557 | 0.915635399 | 0.963608059 | -4.759257256 |
| **A_55_P2157966** | Map1a | 0.010161863 | -0.219385479 | 0.239709205 | 7.904872348 | 0.11087954 | 0.915661905 | 0.963608059 | -4.75925855 |
| **A_55_P2144766** | Pigc | -0.010083349 | -0.23803636 | 0.217869663 | 8.594552778 | -0.110792355 | 0.915727901 | 0.96362506 | -4.759261769 |
| **A_51_P140211** | Ndufv3 | -0.011121561 | -0.263223598 | 0.240980476 | 9.344771878 | -0.110494265 | 0.915953548 | 0.963810052 | -4.759272756 |
| **A_51_P156291** | F9 | -0.015037062 | -0.37420429 | 0.344130166 | 8.018477841 | -0.111513897 | 0.91603271 | 0.963840893 | -4.730381031 |
| **A_55_P2070039** | Zfyve9 | 0.009753957 | -0.211958753 | 0.231466667 | 8.185263145 | 0.110189595 | 0.916184185 | 0.963947815 | -4.759283956 |
| **A_52_P612693** | Polb | 0.015720673 | -0.342662068 | 0.374103414 | 9.361384531 | 0.109868803 | 0.916427037 | 0.964150859 | -4.759295715 |
| **A_51_P191893** | 2410022M | -0.012217197 | -0.291141205 | 0.266706812 | 11.36302 | -0.109707421 | 0.916549213 | 0.964180769 | -4.759301618 |
| **A_52_P576100** | Sh2b1 | 0.016884365 | -0.368620986 | 0.402389716 | 11.23184987 | 0.109699505 | 0.916555206 | 0.964180769 | -4.759301907 |
| **A_52_P639551** | Tinf2 | -0.015647844 | -0.373277506 | 0.341981817 | 9.194078622 | -0.109590102 | 0.916638032 | 0.964215436 | -4.759305903 |
| **A_55_P2003643** | Tstd2 | 0.015040595 | -0.330875554 | 0.360956743 | 10.43954907 | 0.108904171 | 0.91715736 | 0.964709233 | -4.75933087 |

| | | | | | | | | | |
|---|---|---|---|---|---|---|---|---|---|
| **A_55_P1985288** | Bhlha9 | -0.011651073 | -0.279849364 | 0.256547219 | 7.882427808 | -0.108807855 | 0.917230286 | 0.964733454 | -4.759334363 |
| **A_52_P286360** | Otc | 0.009072093 | -0.200395587 | 0.218539773 | 7.66157964 | 0.108477786 | 0.917480206 | 0.964943823 | -4.759346311 |
| **A_55_P2041584** | NA | -0.010834197 | -0.261445583 | 0.239777189 | 7.973466588 | -0.108279508 | 0.917630342 | 0.965002351 | -4.759353471 |
| **A_55_P2154054** | Myo18a | -0.017844657 | -0.430645485 | 0.39495617 | 11.12857003 | -0.108272459 | 0.917635679 | 0.965002351 | -4.759353725 |
| **A_55_P1961567** | Ypel3 | -0.025749552 | -0.621898013 | 0.570398909 | 13.05207665 | -0.108184638 | 0.917702178 | 0.965019793 | -4.759356892 |
| **A_66_P103780** | Gm3646 | -0.010226044 | -0.247123957 | 0.22667187 | 7.748561126 | -0.108117688 | 0.917752875 | 0.965020617 | -4.759359304 |
| **A_51_P119429** | Nckap1l | -0.038384888 | -0.928639204 | 0.851869428 | 8.860601585 | -0.10799323 | 0.917847118 | 0.965028147 | -4.745164429 |
| **A_55_P2088912** | Gapdh | -0.027224128 | -0.658727215 | 0.60427896 | 14.29922414 | -0.107976399 | 0.917859863 | 0.965028147 | -4.759364391 |
| **A_55_P2048249** | NA | 0.013205414 | -0.293723311 | 0.320134139 | 12.98665195 | 0.107761801 | 0.918022367 | 0.965146516 | -4.759372104 |
| **A_55_P2065829** | Cfap300 | 0.032774294 | -0.729695263 | 0.79524385 | 13.51797678 | 0.107661687 | 0.918098179 | 0.965173737 | -4.759375697 |
| **A_55_P2358679** | NA | -0.031234216 | -0.759489766 | 0.697021334 | 11.2112713 | -0.10742297 | 0.918278955 | 0.965213554 | -4.759384251 |
| **A_52_P469956** | Zc3h6 | -0.009207995 | -0.223931991 | 0.205516 | 9.144518576 | -0.107407561 | 0.918290624 | 0.965213554 | -4.759384803 |
| **A_55_P2095177** | Usp5 | 0.016250049 | -0.362837724 | 0.395337822 | 13.0597592 | 0.107365623 | 0.918322383 | 0.965213554 | -4.759386303 |
| **A_51_P286373** | Ttll12 | -0.030934735 | -0.752727204 | 0.690857733 | 10.25380965 | -0.107345639 | 0.918337518 | 0.965213554 | -4.759387018 |
| **A_51_P161037** | Cep170 | 0.009605629 | -0.214865288 | 0.234076546 | 8.358525684 | 0.107180571 | 0.918462524 | 0.965213554 | -4.759392918 |
| **A_51_P288341** | Hpd | -0.008983402 | -0.218959712 | 0.200992908 | 7.667321062 | -0.107157081 | 0.918480314 | 0.965213554 | -4.759393757 |
| **A_52_P118560** | Kynu | -0.008885404 | -0.216584446 | 0.198813638 | 7.681994902 | -0.107150212 | 0.918485516 | 0.965213554 | -4.759394002 |
| **A_66_P111449** | 1700018A | -0.009429818 | -0.230028556 | 0.21116892 | 8.111911072 | -0.107065765 | 0.918549469 | 0.965228297 | -4.759397016 |
| **A_55_P2087087** | Pecam1 | -0.012479943 | -0.304698284 | 0.279738399 | 9.120340322 | -0.106968391 | 0.918623214 | 0.96525073 | -4.759400488 |
| **A_55_P2095909** | Lamtor3 | 0.011273706 | -0.252855277 | 0.275402689 | 8.431799617 | 0.106905732 | 0.918670668 | 0.96525073 | -4.759402721 |
| **A_55_P2185543** | Tle2 | -0.012203328 | -0.298430716 | 0.274024059 | 11.47429309 | -0.106786773 | 0.918760761 | 0.965260553 | -4.759406956 |
| **A_55_P2015530** | NA | -0.022289722 | -0.545215626 | 0.500636182 | 8.304569326 | -0.106761545 | 0.918779867 | 0.965260553 | -4.759407854 |
| **A_55_P2117375** | NA | 0.010671812 | -0.239971638 | 0.261315261 | 8.534888332 | 0.106642948 | 0.918869688 | 0.965302464 | -4.759412071 |
| **A_52_P313102** | Yipf7 | 0.012160758 | -0.273693841 | 0.298015356 | 7.785011221 | 0.106553028 | 0.918937791 | 0.965321556 | -4.759415265 |
| **A_51_P489229** | Eif3g | 0.011234272 | -0.253296546 | 0.275765089 | 14.39337362 | 0.10636996 | 0.919076443 | 0.965414753 | -4.75942176 |
| **A_55_P1956998** | Dis3l2 | 0.012216018 | -0.275892848 | 0.300324883 | 8.460970784 | 0.10619972 | 0.919205383 | 0.965497739 | -4.759427789 |
| **A_55_P1992720** | NA | -0.00888109 | -0.218620419 | 0.200858239 | 7.7016069 | -0.106056366 | 0.919313961 | 0.965559329 | -4.759432859 |
| **A_55_P2359610** | AI646519 | 0.03589266 | -0.812990186 | 0.884775507 | 9.411679304 | 0.105902993 | 0.91943013 | 0.965628884 | -4.759438276 |
| **A_52_P624434** | Clec1a | 0.049914709 | -1.133245975 | 1.233075392 | 9.695437634 | 0.105665974 | 0.91960966 | 0.965662032 | -4.759446631 |
| **A_55_P1962651** | NA | -0.021640066 | -0.561830878 | 0.518550746 | 8.054482869 | -0.106702376 | 0.919638528 | 0.965662032 | -4.730534314 |
| **A_55_P2114402** | Notum | -0.026553731 | -0.656356806 | 0.603249344 | 10.39435868 | -0.10560175 | 0.919658307 | 0.965662032 | -4.759448892 |
| **A_55_P1959164** | Gm10324 | -0.028885335 | -0.7140167 | 0.65624603 | 8.752987785 | -0.105597565 | 0.919661477 | 0.965662032 | -4.759449039 |
| **A_52_P27103** | Necap2 | -0.009218942 | -0.22828027 | 0.209842386 | 9.400009452 | -0.105406089 | 0.919806515 | 0.965719523 | -4.759455771 |
| **A_52_P269158** | Pid1 | 0.034847399 | -0.793299364 | 0.862994163 | 8.659641074 | 0.105393398 | 0.919816128 | 0.965719523 | -4.759456217 |
| **A_52_P279557** | NA | -0.008799368 | -0.218073273 | 0.200474538 | 7.660450075 | -0.10531415 | 0.919876157 | 0.965730106 | -4.759458999 |
| **A_51_P246471** | Pbx2 | 0.014212853 | -0.324439633 | 0.352865339 | 10.96180992 | 0.105118061 | 0.920024695 | 0.965763317 | -4.759465875 |
| **A_51_P291078** | Sel1l3 | 0.023927768 | -0.546614412 | 0.594469947 | 8.640239135 | 0.105042433 | 0.920081984 | 0.965763317 | -4.759468523 |
| **A_51_P140871** | Ankhd1 | 0.009072309 | -0.207378064 | 0.225522682 | 11.14438373 | 0.104980787 | 0.920128683 | 0.965763317 | -4.759470681 |
| **A_55_P2136752** | Ermn | -0.026771725 | -0.66566266 | 0.612119209 | 10.64673454 | -0.104954238 | 0.920148794 | 0.965763317 | -4.75947161 |
| **A_55_P2131899** | Sox4 | -0.009789893 | -0.243819379 | 0.224239592 | 8.084366615 | -0.104775014 | 0.920284562 | 0.965763317 | -4.759477873 |
| **A_55_P2021792** | Dyrk1b | -0.010295268 | -0.256498985 | 0.235908449 | 8.598486039 | -0.104735378 | 0.920314588 | 0.965763317 | -4.759479257 |
| **A_55_P1986088** | Prr14 | 0.011579507 | -0.265378095 | 0.288537109 | 12.11834967 | 0.104719396 | 0.920326695 | 0.965763317 | -4.759479815 |
| **A_55_P2279690** | 2900076A | 0.009081262 | -0.208202893 | 0.226365418 | 7.794717084 | 0.104681155 | 0.920355665 | 0.965763317 | -4.759481149 |
| **A_51_P456366** | Pard6a | 0.008819168 | -0.202198458 | 0.219836795 | 8.447959296 | 0.104678923 | 0.920357356 | 0.965763317 | -4.759481227 |
| **A_55_P2246900** | NA | -0.013886267 | -0.347687711 | 0.319915177 | 8.756400786 | -0.104195184 | 0.920723823 | 0.96609543 | -4.759498064 |
| **A_55_P1995745** | Gm14308 | 0.012378285 | -0.28571504 | 0.31047161 | 9.170914647 | 0.104006045 | 0.920867116 | 0.966193348 | -4.759504626 |
| **A_55_P2067905** | Scfd2 | 0.017742139 | -0.409833089 | 0.445317367 | 10.31500183 | 0.103930683 | 0.920924211 | 0.96620082 | -4.759507238 |
| **A_55_P2185504** | Masp2 | -0.008621866 | -0.217198586 | 0.199954855 | 7.669745128 | -0.103534659 | 0.921224255 | 0.96646317 | -4.759520929 |

| | | | | | | | | | |
|---|---|---|---|---|---|---|---|---|---|
| **A_52_P507214** | Mmp9 | 0.009901326 | -0.230134877 | 0.249937528 | 7.731629742 | 0.103315848 | 0.921390041 | 0.966584648 | -4.750099146 |
| **A_55_P2347756** | AU040972 | 0.008905146 | -0.207380177 | 0.22519047 | 8.474496361 | 0.103125095 | 0.921534573 | 0.966683817 | -4.759535035 |
| **A_52_P60194** | C4bp | 0.008633472 | -0.201560638 | 0.218827583 | 7.650516156 | 0.102876292 | 0.921723092 | 0.966788861 | -4.759543576 |
| **A_55_P2066234** | Arfip2 | -0.010440575 | -0.26468941 | 0.243808259 | 9.434139639 | -0.102852726 | 0.921740948 | 0.966788861 | -4.759544384 |
| **A_52_P238242** | Sertad2 | -0.01271185 | -0.322444723 | 0.297021024 | 9.663543776 | -0.102794953 | 0.921784724 | 0.966788861 | -4.759546364 |
| **A_55_P2170355** | NA | -0.014657728 | -0.372866207 | 0.343550752 | 8.232573685 | -0.102489914 | 0.922015864 | 0.966941948 | -4.7595568 |
| **A_51_P330090** | Trappc9 | 0.01015037 | -0.237953831 | 0.258254571 | 11.110292 | 0.102470322 | 0.92203071 | 0.966941948 | -4.75955747 |
| **A_55_P2158384** | Aanat | -0.016475365 | -0.419846773 | 0.386896042 | 8.516340769 | -0.102301097 | 0.922158943 | 0.967023973 | -4.759563245 |
| **A_55_P2069337** | NA | 0.014084141 | -0.331693499 | 0.359861781 | 8.942491823 | 0.102019645 | 0.922372223 | 0.967133718 | -4.759572829 |
| **A_55_P2081183** | Slc28a1 | 0.015577456 | -0.367170703 | 0.398325615 | 8.324901102 | 0.101937449 | 0.922434511 | 0.967133718 | -4.759575623 |
| **A_55_P2136757** | Tomm34 | 0.019586633 | -0.461895047 | 0.501068314 | 11.2920257 | 0.10188972 | 0.92247068 | 0.967133718 | -4.759577244 |
| **A_55_P2011137** | Armc10 | 0.010488232 | -0.247417279 | 0.268393743 | 9.175153936 | 0.101857262 | 0.922495277 | 0.967133718 | -4.759578347 |
| **A_55_P1957845** | Tnfaip1 | 0.009064904 | -0.214163131 | 0.23229294 | 13.47506444 | 0.101710277 | 0.922606666 | 0.967133718 | -4.759583334 |
| **A_55_P2423586** | Eya1 | 0.018847674 | -0.445572131 | 0.483267478 | 14.44125325 | 0.101647656 | 0.922654121 | 0.967133718 | -4.759585456 |
| **A_51_P441970** | Stox2 | 0.010278709 | -0.24316579 | 0.263723208 | 8.635748715 | 0.101579495 | 0.922705776 | 0.967133718 | -4.759587765 |
| **A_55_P2032643** | Ajap1 | 0.011272454 | -0.266945799 | 0.289490707 | 7.915976845 | 0.101480641 | 0.922780692 | 0.967133718 | -4.75959111 |
| **A_55_P2071486** | Suv39h1 | -0.016778803 | -0.430984559 | 0.397426952 | 9.181246473 | -0.101460078 | 0.922796275 | 0.967133718 | -4.759591806 |
| **A_55_P2126780** | Pgam5 | -0.033914682 | -0.871423079 | 0.803593714 | 11.68464932 | -0.101425914 | 0.922822166 | 0.967133718 | -4.759592961 |
| **A_55_P1975285** | Lrig2 | -0.024395247 | -0.627145571 | 0.578355076 | 8.816773396 | -0.101372025 | 0.922863006 | 0.967133718 | -4.759594783 |
| **A_51_P322273** | Ptp4a3 | -0.01745945 | -0.44884702 | 0.413928121 | 10.4281625 | -0.101370892 | 0.922863865 | 0.967133718 | -4.759594821 |
| **A_66_P128593** | Lims1 | 0.041893364 | -1.003545736 | 1.087332463 | 10.82847883 | 0.100368362 | 0.923623683 | 0.967877523 | -4.759628531 |
| **A_51_P321651** | Fam192a | -0.009687131 | -0.251852574 | 0.232478312 | 10.61168966 | -0.100192073 | 0.923757302 | 0.96796508 | -4.759634424 |
| **A_55_P1960231** | Txn2 | -0.012063572 | -0.314064869 | 0.289937725 | 13.08772113 | -0.100050087 | 0.923864924 | 0.96796683 | -4.759639163 |
| **A_66_P106583** | NA | 0.011520852 | -0.276957301 | 0.299999004 | 13.96464979 | 0.100028095 | 0.923881593 | 0.96796683 | -4.759639897 |
| **A_55_P1986199** | Tut1 | -0.012664329 | -0.329985325 | 0.304656667 | 8.39954236 | -0.099961719 | 0.923931905 | 0.96796683 | -4.75964211 |
| **A_55_P2052824** | Lrrc14b | -0.009865208 | -0.257139824 | 0.237409409 | 7.835627496 | -0.099925663 | 0.923959235 | 0.96796683 | -4.759643311 |
| **A_55_P2068733** | Gm21953 | -0.011192614 | -0.292171282 | 0.269786054 | 8.42833819 | -0.099771963 | 0.924075738 | 0.967996249 | -4.759648427 |
| **A_55_P2087162** | Specc1l | 0.012642794 | -0.304790275 | 0.330075864 | 8.604538992 | 0.099756511 | 0.924087451 | 0.967996249 | -4.759648941 |
| **A_55_P2012819** | Sec61a2 | -0.015027037 | -0.392655396 | 0.362601322 | 9.64155339 | -0.099668768 | 0.92415396 | 0.967997405 | -4.759651858 |
| **A_52_P650473** | NA | -0.010175761 | -0.266009508 | 0.245657986 | 8.446967742 | -0.099622954 | 0.924188688 | 0.967997405 | -4.75965338 |
| **A_55_P2086536** | Safb2 | 0.008791633 | -0.212889784 | 0.230473049 | 8.391483273 | 0.099332323 | 0.924408994 | 0.968125534 | -4.759663018 |
| **A_52_P239726** | Larp4 | 0.021456023 | -0.519574174 | 0.56248622 | 9.018755296 | 0.099329458 | 0.924411166 | 0.968125534 | -4.759663113 |
| **A_55_P2108389** | D830030K | 0.017869048 | -0.433158832 | 0.468896927 | 9.58429744 | 0.099231229 | 0.924485628 | 0.968137547 | -4.759666364 |
| **A_55_P2299524** | Scube1 | -0.011130912 | -0.292222143 | 0.26996032 | 8.203387053 | -0.099182212 | 0.924522785 | 0.968137547 | -4.759667985 |
| **A_51_P463120** | Hacd4 | -0.022222647 | -0.58492203 | 0.540476737 | 8.709283201 | -0.098916715 | 0.924724049 | 0.96829586 | -4.759676753 |
| **A_55_P2070316** | Ankrd36 | 0.008176469 | -0.199348314 | 0.215701253 | 7.679500855 | 0.098683871 | 0.924900564 | 0.968427222 | -4.759684422 |
| **A_66_P107818** | Gdf3 | -0.009341698 | -0.246646607 | 0.227963211 | 7.672467862 | -0.098598304 | 0.924965432 | 0.968427222 | -4.759687236 |
| **A_51_P466685** | Fbxl22 | 0.009692481 | -0.236715319 | 0.256100282 | 7.787004585 | 0.098521467 | 0.925023684 | 0.968427222 | -4.759689761 |
| **A_51_P278994** | Swap70 | 0.020560978 | -0.502335177 | 0.543457132 | 10.05751927 | 0.098486943 | 0.925049857 | 0.968427222 | -4.759690895 |
| **A_55_P1997931** | Map9 | 0.012542754 | -0.306783204 | 0.331868712 | 10.14178768 | 0.098380499 | 0.925130553 | 0.968459263 | -4.759694388 |
| **A_55_P2176530** | Ldah | -0.011557412 | -0.306046139 | 0.282931315 | 8.746790637 | -0.098297455 | 0.925193511 | 0.968472731 | -4.759697111 |
| **A_51_P507974** | Brf2 | -0.008777814 | -0.232876847 | 0.215321218 | 8.393611164 | -0.098106268 | 0.925338457 | 0.968543738 | -4.759703371 |
| **A_51_P115715** | Asb2 | 0.026815161 | -0.65805585 | 0.711686173 | 8.211438921 | 0.098066794 | 0.925368385 | 0.968543738 | -4.759704662 |
| **A_55_P1964364** | Kctd14 | -0.01261197 | -0.334914688 | 0.309690749 | 17.34085589 | -0.098009753 | 0.925411631 | 0.968543738 | -4.759706526 |
| **A_55_P1964483** | Cyp2c37 | 0.008161403 | -0.201030125 | 0.217352931 | 7.673185944 | 0.097717212 | 0.925633426 | 0.968696234 | -4.759716072 |
| **A_55_P2127258** | Dok7 | 0.008539327 | -0.210483439 | 0.227562093 | 7.724450006 | 0.097652806 | 0.925682258 | 0.968696234 | -4.759718169 |
| **A_52_P447971** | NA | 0.026653309 | -0.657658681 | 0.710965299 | 10.81511565 | 0.097554506 | 0.925756788 | 0.968696234 | -4.759721368 |
| **A_51_P320539** | Otulin | 0.012117575 | -0.299145984 | 0.323381134 | 10.52209593 | 0.097507445 | 0.92579247 | 0.968696234 | -4.759722899 |

| | | | | | | | | | |
|---|---|---|---|---|---|---|---|---|---|
| **A_55_P2129348** | Pilra | 0.029847361 | -0.736999907 | 0.796694628 | 8.596859217 | 0.097487156 | 0.925807852 | 0.968696234 | -4.759723558 |
| **A_55_P1984751** | Ppp1r14b | 0.023734844 | -0.587302267 | 0.634771956 | 12.35096484 | 0.097290211 | 0.925957178 | 0.968800048 | -4.759729953 |
| **A_51_P150120** | Mgst2 | -0.035552312 | -0.952064259 | 0.880959635 | 10.22480919 | -0.097158335 | 0.92605717 | 0.968852236 | -4.759734228 |
| **A_52_P48546** | Rnf41 | 0.008984119 | -0.22285256 | 0.240820797 | 8.955544991 | 0.097060759 | 0.926131156 | 0.968857822 | -4.759737388 |
| **A_55_P2086094** | Slc35b2 | -0.010254042 | -0.275135757 | 0.254627674 | 11.72560491 | -0.096960189 | 0.926207412 | 0.968857822 | -4.759740641 |
| **A_55_P1977245** | Slc22a13 | -0.012845306 | -0.344731348 | 0.319040737 | 8.19164924 | -0.096940623 | 0.926222248 | 0.968857822 | -4.759741273 |
| **A_52_P168097** | Atp2a2 | 0.008529708 | -0.211975721 | 0.229035137 | 7.959484385 | 0.096886936 | 0.926262956 | 0.968857822 | -4.759743008 |
| **A_55_P1981040** | Ddx6 | -0.012207769 | -0.328185555 | 0.303770017 | 11.09916187 | -0.09676763 | 0.926353421 | 0.968899174 | -4.75974686 |
| **A_52_P607507** | Olfm4 | 0.008020814 | -0.199769466 | 0.215811095 | 7.682850537 | 0.096681545 | 0.926418697 | 0.968899174 | -4.759749637 |
| **A_52_P440709** | Ryr2 | 0.024136683 | -0.601448737 | 0.649722104 | 8.201857205 | 0.096636528 | 0.926452831 | 0.968899174 | -4.759751087 |
| **A_55_P2173005** | Dnase2a | 0.019191974 | -0.479935953 | 0.5183199 | 12.45302791 | 0.096307058 | 0.926702666 | 0.969058013 | -4.759761686 |
| **A_52_P484405** | Twf1 | -0.010641827 | -0.287595022 | 0.266311367 | 11.35516003 | -0.096241011 | 0.926752749 | 0.969058013 | -4.759763806 |
| **A_55_P2047009** | NA | -0.020285501 | -0.548231811 | 0.507660809 | 10.71034931 | -0.096237941 | 0.926755077 | 0.969058013 | -4.759763905 |
| **A_51_P288618** | Dusp11 | 0.00821458 | -0.205863172 | 0.222292332 | 8.276624155 | 0.096109027 | 0.926852835 | 0.969058501 | -4.759768039 |
| **A_55_P2065050** | NA | 0.009456471 | -0.236995808 | 0.255908749 | 8.466787772 | 0.096105132 | 0.926855788 | 0.969058501 | -4.759768164 |
| **A_55_P2170834** | NA | -0.033321974 | -0.903451561 | 0.836807614 | 10.44197831 | -0.09591734 | 0.926998197 | 0.969103915 | -4.759774176 |
| **A_51_P337897** | Prrc1 | 0.01032452 | -0.259282202 | 0.279931242 | 8.555012341 | 0.095915657 | 0.926999473 | 0.969103915 | -4.75977423 |
| **A_55_P2007566** | Trmt61b | 0.007984811 | -0.200816592 | 0.216786214 | 8.205108701 | 0.095781483 | 0.927101223 | 0.969157883 | -4.759778518 |
| **A_52_P338180** | Timm13 | 0.020250408 | -0.510432774 | 0.550933589 | 13.63763331 | 0.095575987 | 0.927257062 | 0.969172764 | -4.759785074 |
| **A_52_P162184** | Snx12 | 0.008916632 | -0.224764336 | 0.242597599 | 8.706293712 | 0.095571373 | 0.927260561 | 0.969172764 | -4.759785221 |
| **A_55_P2090535** | Thrap3 | -0.012953077 | -0.352458238 | 0.326552085 | 8.168664181 | -0.095560166 | 0.92726906 | 0.969172764 | -4.759785578 |
| **A_55_P1958457** | Nbeal2 | -0.015728271 | -0.428601353 | 0.397144812 | 8.672854097 | -0.095414582 | 0.927379467 | 0.969172764 | -4.759790214 |
| **A_51_P390377** | Rcor3 | -0.010310519 | -0.281028234 | 0.260407195 | 9.416284739 | -0.095392496 | 0.927396217 | 0.969172764 | -4.759790917 |
| **A_52_P490272** | Nup98 | -0.016096589 | -0.439012915 | 0.406819736 | 8.959428096 | -0.095330036 | 0.927443585 | 0.969172764 | -4.759792903 |
| **A_51_P244531** | NA | -0.009731893 | -0.265504894 | 0.246041108 | 9.335007333 | -0.095300014 | 0.927466354 | 0.969172764 | -4.759793857 |
| **A_51_P137029** | Plce1 | -0.00984905 | -0.268956839 | 0.249258738 | 7.879205631 | -0.095205982 | 0.927537667 | 0.969194901 | -4.759796844 |
| **A_52_P64687** | Camk2n1 | -0.023859481 | -0.652255078 | 0.604536116 | 9.553135261 | -0.095099492 | 0.92761843 | 0.969226909 | -4.759800223 |
| **A_55_P1953028** | Tnfsf12 | -0.018519294 | -0.506701826 | 0.469663239 | 9.064974441 | -0.095015079 | 0.92768245 | 0.96924142 | -4.759802899 |
| **A_51_P328400** | Cops2 | 0.011994443 | -0.304741519 | 0.328730405 | 10.77871503 | 0.094849066 | 0.927808357 | 0.969320587 | -4.759808154 |
| **A_52_P652212** | Psmd14 | 0.010658326 | -0.271210588 | 0.29252724 | 9.622261492 | 0.094709201 | 0.927914436 | 0.96937903 | -4.759812575 |
| **A_51_P149714** | Ms4a6d | 0.012359812 | -0.315423529 | 0.340143153 | 8.347391492 | 0.094444212 | 0.928115417 | 0.969536605 | -4.759820933 |
| **A_51_P310164** | 2810459M | 0.008473338 | -0.216590542 | 0.233537218 | 7.778080406 | 0.094297262 | 0.928226874 | 0.969546573 | -4.759825557 |
| **A_51_P254425** | Ahrr | -0.008206962 | -0.226261983 | 0.20984806 | 7.822253155 | -0.094268512 | 0.928248681 | 0.969546573 | -4.759826461 |
| **A_51_P496400** | Med9 | -0.012154426 | -0.335212424 | 0.310903571 | 11.09603577 | -0.094233282 | 0.928275401 | 0.969546573 | -4.759827569 |
| **A_55_P2059372** | Ppip5k2 | 0.007817023 | -0.200113093 | 0.215747139 | 7.88396311 | 0.094161714 | 0.928329685 | 0.969550893 | -4.759829817 |
| **A_51_P361201** | Atp6v0d1 | 0.014579414 | -0.373571725 | 0.402730553 | 10.76912189 | 0.09407832 | 0.928392939 | 0.96956458 | -4.759832434 |
| **A_51_P318544** | Fdx2 | 0.008953266 | -0.229621875 | 0.247528406 | 11.4733771 | 0.093995407 | 0.928455827 | 0.969577885 | -4.759835034 |
| **A_55_P1956882** | Cpt1b | 0.023342643 | -0.600100122 | 0.646785409 | 8.282631868 | 0.09377861 | 0.92862027 | 0.969697235 | -4.759841823 |
| **A_66_P139618** | Stfa2 | 0.061953299 | -1.594216653 | 1.71812325 | 9.926411927 | 0.09369359 | 0.92868476 | 0.969712203 | -4.75984448 |
| **A_51_P139069** | Wdr45 | -0.035453079 | -0.987181101 | 0.916274942 | 12.45462227 | -0.093302107 | 0.928981716 | 0.969843361 | -4.759856687 |
| **A_52_P13897** | Hook1 | -0.010158706 | -0.282907841 | 0.26259043 | 8.084314968 | -0.093287905 | 0.92899249 | 0.969843361 | -4.759857129 |
| **A_55_P2031966** | Fbxo39 | 0.018679834 | -0.482853003 | 0.520212671 | 8.237127637 | 0.093287614 | 0.92899271 | 0.969843361 | -4.759857138 |
| **A_55_P2018934** | Adgre1 | 0.009082127 | -0.234826011 | 0.252990264 | 8.128550743 | 0.093263477 | 0.929011019 | 0.969843361 | -4.759857889 |
| **A_55_P1976007** | Psme4 | -0.008019429 | -0.223608519 | 0.207569661 | 9.106574643 | -0.093168051 | 0.929083406 | 0.969866561 | -4.759860856 |
| **A_55_P1962946** | Syk | 0.010616042 | -0.275573089 | 0.296805172 | 9.271328884 | 0.092909437 | 0.929279586 | 0.969986223 | -4.759868881 |
| **A_55_P2075627** | Apc2 | 0.010334091 | -0.268328452 | 0.288996634 | 8.476220994 | 0.092884667 | 0.929298376 | 0.969986223 | -4.759869648 |
| **A_55_P2054331** | Gm9222 | -0.013449941 | -0.376609856 | 0.349709974 | 8.266108573 | -0.092762582 | 0.92939099 | 0.970030523 | -4.759873427 |
| **A_52_P375873** | Srp9 | -0.008784872 | -0.247052529 | 0.229482785 | 12.05550502 | -0.092346553 | 0.929706598 | 0.970259126 | -4.75988627 |

| | | | | | | | | | |
|---|---|---|---|---|---|---|---|---|---|
| **A_51_P418116** | Tmem119 | -0.023861931 | -0.671329329 | 0.623605467 | 10.53707895 | -0.092307722 | 0.929736057 | 0.970259126 | -4.759887466 |
| **A_55_P2391065** | Cep128 | 0.009685039 | -0.253199912 | 0.27256999 | 8.173184074 | 0.092275413 | 0.929760568 | 0.970259126 | -4.75988846 |
| **A_55_P2343648** | 1700120C | -0.007757286 | -0.218695987 | 0.203181415 | 7.700476175 | -0.092109389 | 0.929886522 | 0.970338192 | -4.759893565 |
| **A_55_P1953919** | Me3 | -0.013290363 | -0.375311969 | 0.348731243 | 8.145467475 | -0.091950203 | 0.930007291 | 0.970411839 | -4.759898451 |
| **A_52_P355057** | Wdr4 | -0.01244348 | -0.351925679 | 0.327038718 | 10.35079225 | -0.091806874 | 0.930116032 | 0.970472928 | -4.759902843 |
| **A_55_P2004099** | NA | 0.010964284 | -0.2894093 | 0.311337867 | 8.142408888 | 0.091425821 | 0.930405136 | 0.97072219 | -4.759914488 |
| **A_55_P2151751** | Matr3 | -0.017721845 | -0.505240655 | 0.469796966 | 9.566794209 | -0.091047473 | 0.9306922 | 0.970926919 | -4.759926001 |
| **A_66_P101862** | NA | 0.008852615 | -0.2347631 | 0.252468329 | 7.968318118 | 0.091015762 | 0.93071626 | 0.970926919 | -4.759926964 |
| **A_55_P2062495** | Smyd1 | -0.007720407 | -0.220288874 | 0.20484806 | 7.717073913 | -0.090968635 | 0.930752018 | 0.970926919 | -4.759928394 |
| **A_55_P2139089** | Slc22a28 | 0.01286889 | -0.34347538 | 0.369213159 | 8.387695075 | 0.090452723 | 0.931143477 | 0.971255341 | -4.759944004 |
| **A_55_P2149821** | Srsf11 | -0.012228261 | -0.350982427 | 0.326525905 | 12.04599844 | -0.0904129 | 0.931173695 | 0.971255341 | -4.759945206 |
| **A_52_P368262** | Bcl7c | 0.010855719 | -0.290146413 | 0.311857851 | 8.867977067 | 0.090331531 | 0.931235438 | 0.971255341 | -4.759947658 |
| **A_51_P283300** | Cacnb2 | 0.011272891 | -0.301534097 | 0.324079879 | 7.896542664 | 0.090262884 | 0.931287528 | 0.971255341 | -4.759949726 |
| **A_51_P354706** | Lefty1 | -0.00764469 | -0.219868684 | 0.204579303 | 7.865664993 | -0.090222689 | 0.931318028 | 0.971255341 | -4.759950936 |
| **A_55_P1959403** | Ccdc77 | -0.016676224 | -0.48049788 | 0.447145432 | 9.356157306 | -0.090052762 | 0.931446972 | 0.97127087 | -4.759956045 |
| **A_51_P410927** | NA | -0.009733162 | -0.280467231 | 0.261000906 | 9.19236734 | -0.090045375 | 0.931452578 | 0.97127087 | -4.759956267 |
| **A_52_P669005** | Lrat | 0.007737869 | -0.207593708 | 0.223069446 | 7.735594613 | 0.090004456 | 0.931483628 | 0.97127087 | -4.759957496 |
| **A_55_P2183172** | Rdx | 0.01607654 | -0.432104043 | 0.464257123 | 12.30720702 | 0.089844169 | 0.93160526 | 0.97134531 | -4.759962304 |
| **A_52_P151320** | Tnfaip8l1 | 0.014525807 | -0.391201227 | 0.42025284 | 11.18423978 | 0.089671961 | 0.93173594 | 0.97138814 | -4.75996746 |
| **A_52_P597775** | Gprc5a | 0.057209091 | -1.542099676 | 1.656517859 | 13.92408816 | 0.089594856 | 0.931794451 | 0.97138814 | -4.759969765 |
| **A_51_P396983** | Ssu2 | 0.018725888 | -0.504807107 | 0.542258882 | 8.377287124 | 0.089587765 | 0.931799833 | 0.97138814 | -4.759969977 |
| **A_52_P644690** | Flt3l | -0.013872217 | -0.401978703 | 0.374234269 | 8.296016274 | -0.089525205 | 0.931847307 | 0.97138814 | -4.759971846 |
| **A_55_P2025712** | Immt | -0.009799002 | -0.284301999 | 0.264703995 | 9.953307545 | -0.089409793 | 0.93193489 | 0.971427063 | -4.759975291 |
| **A_66_P137219** | Elane | -0.025754265 | -0.748392128 | 0.696883598 | 8.766640987 | -0.089264503 | 0.932045148 | 0.971489616 | -4.75997962 |
| **A_51_P393518** | Hmcn1 | 0.027879771 | -0.757081619 | 0.81284116 | 8.387567717 | 0.088959278 | 0.932276784 | 0.97167867 | -4.759988694 |
| **A_55_P2061724** | NA | 0.008172532 | -0.222320586 | 0.238665651 | 8.19767182 | 0.088807373 | 0.932392068 | 0.971682295 | -4.759993198 |
| **A_55_P2279140** | F830014C | 0.008412567 | -0.228999311 | 0.245824445 | 8.002063032 | 0.088751651 | 0.932434356 | 0.971682295 | -4.759994848 |
| **A_55_P2076248** | NA | -0.015808623 | -0.462204369 | 0.430587124 | 8.503028741 | -0.088700146 | 0.932473445 | 0.971682295 | -4.759996372 |
| **A_55_P1966144** | Slfn14 | -0.007405453 | -0.216914403 | 0.202103497 | 7.665436838 | -0.088531817 | 0.932601197 | 0.971682295 | -4.760001349 |
| **A_55_P1953157** | Mettl18 | -0.008366173 | -0.245131172 | 0.228398825 | 9.415668971 | -0.08850335 | 0.932622802 | 0.971682295 | -4.760002189 |
| **A_55_P2088203** | Hcn2 | 0.016562764 | -0.452178054 | 0.485303581 | 8.969267793 | 0.088501448 | 0.932624245 | 0.971682295 | -4.760002245 |
| **A_51_P407193** | Clp1 | 0.007597242 | -0.207436455 | 0.22263094 | 8.15810285 | 0.088491144 | 0.932632066 | 0.971682295 | -4.76000255 |
| **A_55_P2015405** | 9930111J | -0.008592111 | -0.25212761 | 0.234943387 | 7.891293003 | -0.088366564 | 0.932726617 | 0.971728441 | -4.760006225 |
| **A_55_P1973829** | NA | 0.010930245 | -0.299333114 | 0.321193604 | 8.469638427 | 0.088236799 | 0.932825103 | 0.971765972 | -4.760010047 |
| **A_52_P662244** | Slc25a47 | -0.010088333 | -0.296872825 | 0.276696159 | 8.612414904 | -0.088107744 | 0.932923053 | 0.971765972 | -4.760013843 |
| **A_51_P287198** | Krt23 | 0.013831435 | -0.379538223 | 0.407201093 | 9.013348366 | 0.088067714 | 0.932953435 | 0.971765972 | -4.76001502 |
| **A_55_P2032849** | NA | -0.020078108 | -0.591637331 | 0.551481115 | 9.000428673 | -0.08798566 | 0.933015714 | 0.971765972 | -4.760017429 |
| **A_55_P1980826** | Kcnrg | 0.03217944 | -0.884078542 | 0.948437421 | 8.503297536 | 0.087965232 | 0.933031218 | 0.971765972 | -4.760018029 |
| **A_55_P2145005** | NA | 0.020421727 | -0.56183045 | 0.602673904 | 8.65337032 | 0.087847962 | 0.933120225 | 0.971765972 | -4.760021468 |
| **A_51_P371001** | Tm4sf4 | -0.007609967 | -0.224584934 | 0.209365001 | 7.813085168 | -0.087846298 | 0.933121489 | 0.971765972 | -4.760021517 |
| **A_51_P295192** | Nfkbia | -0.025600897 | -0.756006091 | 0.704804297 | 13.32348532 | -0.087789317 | 0.933164737 | 0.971765972 | -4.760023186 |
| **A_55_P2109439** | Lsmem1 | 0.008358045 | -0.231001986 | 0.247718076 | 7.754512095 | 0.087458779 | 0.933415622 | 0.971974882 | -4.760032848 |
| **A_51_P396774** | Sugt1 | -0.008833019 | -0.262639758 | 0.244973721 | 13.79050402 | -0.087167852 | 0.933636448 | 0.972107849 | -4.760041323 |
| **A_51_P128667** | Lynx1 | 0.011402448 | -0.316336197 | 0.339141093 | 8.785573015 | 0.087140649 | 0.933657097 | 0.972107849 | -4.760042114 |
| **A_55_P2147186** | A430005L | -0.017602132 | -0.524287697 | 0.489083433 | 12.4698487 | -0.087011586 | 0.933755064 | 0.972107849 | -4.760045863 |
| **A_55_P2034663** | Csf2rb2 | 0.014513115 | -0.403462565 | 0.432488796 | 8.203330198 | 0.086968098 | 0.933788074 | 0.972107849 | -4.760047125 |
| **A_52_P228318** | Smu1 | -0.01559101 | -0.46465506 | 0.433473041 | 13.31275142 | -0.086959353 | 0.933794712 | 0.972107849 | -4.760047379 |
| **A_55_P2074811** | Zfp609 | 0.016591362 | -0.461683581 | 0.494866306 | 8.804464179 | 0.086886992 | 0.933849639 | 0.9721105 | -4.760049478 |

| | | | | | | | | | |
|---|---|---|---|---|---|---|---|---|---|
| **A_51_P163694** | Rufy1 | 0.013325017 | -0.371072616 | 0.397722649 | 9.480406325 | 0.086823522 | 0.933897819 | 0.9721105 | -4.760051317 |
| **A_52_P355398** | Dars | -0.013338172 | -0.398869152 | 0.372192809 | 9.187256437 | -0.086653751 | 0.93402669 | 0.972192302 | -4.76005623 |
| **A_51_P225179** | Smndc1 | 0.012684811 | -0.354893163 | 0.380262785 | 8.839052338 | 0.086434052 | 0.934193464 | 0.972313546 | -4.760062573 |
| **A_55_P2113650** | Clybl | 0.009907317 | -0.277726272 | 0.297540907 | 8.249977681 | 0.086271394 | 0.934316941 | 0.972389715 | -4.760067259 |
| **A_51_P279062** | Treml1 | -0.028182311 | -0.847042252 | 0.790677631 | 9.380340114 | -0.086201996 | 0.934369623 | 0.9723922 | -4.760069256 |
| **A_52_P157274** | Abi1 | 0.00847606 | -0.238178472 | 0.255130593 | 8.635321462 | 0.08607069 | 0.934469301 | 0.972402745 | -4.76007303 |
| **A_52_P647740** | Klhl41 | -0.007981136 | -0.240326845 | 0.224364572 | 7.726349135 | -0.086036043 | 0.934495604 | 0.972402745 | -4.760074024 |
| **A_52_P661587** | Ccdc141 | 0.008036076 | -0.226034599 | 0.242106752 | 7.78109103 | 0.08598989 | 0.93453064 | 0.972402745 | -4.760075349 |
| **A_55_P2047837** | NA | -0.010135749 | -0.305595231 | 0.285323732 | 7.985143828 | -0.085922773 | 0.934581592 | 0.972403428 | -4.760077274 |
| **A_55_P2036495** | NA | 0.009150303 | -0.258052483 | 0.276353088 | 13.56753782 | 0.08577186 | 0.934696159 | 0.972425768 | -4.760081596 |
| **A_51_P480328** | Adgrl4 | -0.009472748 | -0.286123261 | 0.267177766 | 8.362053387 | -0.085761986 | 0.934703655 | 0.972425768 | -4.760081879 |
| **A_55_P2181738** | Lyz1 | -0.087419292 | -2.64657012 | 2.471731535 | 13.18871466 | -0.085558222 | 0.934858347 | 0.972509505 | -4.760087702 |
| **A_55_P2169039** | Olfr558 | -0.007448781 | -0.225596329 | 0.210698767 | 7.807497783 | -0.085523449 | 0.934884745 | 0.972509505 | -4.760088695 |
| **A_65_P11718** | Nol8 | -0.008979564 | -0.272580816 | 0.254621688 | 8.354878778 | -0.085321431 | 0.935038116 | 0.972616717 | -4.760094453 |
| **A_51_P209782** | Cyp2c23 | 0.008272558 | -0.234957818 | 0.251502934 | 7.882752523 | 0.085186811 | 0.93514032 | 0.972670698 | -4.760098282 |
| **A_55_P2138422** | Obox1 | -0.007384785 | -0.224792341 | 0.210022771 | 7.826186979 | -0.085077276 | 0.93522348 | 0.972704866 | -4.760101393 |
| **A_52_P632423** | NA | 0.007736536 | -0.220589582 | 0.236062655 | 7.766943334 | 0.084867483 | 0.93538276 | 0.972818197 | -4.760107341 |
| **A_51_P205385** | Uox | 0.009285035 | -0.265863104 | 0.284433174 | 7.929941096 | 0.08452152 | 0.935645431 | 0.973039038 | -4.760117118 |
| **A_55_P2182690** | Orai3 | 0.011873507 | -0.340795924 | 0.364542938 | 11.35130507 | 0.084326009 | 0.935793876 | 0.973141071 | -4.760122626 |
| **A_55_P2181084** | Rpl17 | 0.018017526 | -0.519219815 | 0.555254868 | 15.19713631 | 0.084000007 | 0.936041404 | 0.973346125 | -4.760131781 |
| **A_55_P2008349** | Prelid1 | -0.010165696 | -0.313661974 | 0.293330582 | 13.90236538 | -0.083894614 | 0.936121429 | 0.973364105 | -4.760134733 |
| **A_51_P490296** | Slx4 | 0.011902672 | -0.343662798 | 0.367468142 | 10.18343062 | 0.083844627 | 0.936159384 | 0.973364105 | -4.760136132 |
| **A_66_P135391** | Igfbp2 | 0.050738315 | -1.471065399 | 1.572542029 | 11.51021488 | 0.083507936 | 0.936415042 | 0.973577566 | -4.760145533 |
| **A_55_P2088490** | Acad10 | -0.027306226 | -0.847621799 | 0.793009348 | 11.17652929 | -0.083374082 | 0.936516682 | 0.973630883 | -4.76014926 |
| **A_55_P2166123** | Atp9a | 0.014040553 | -0.409685621 | 0.437766727 | 8.359892986 | 0.08299449 | 0.936804929 | 0.973878186 | -4.760159796 |
| **A_55_P2083967** | Trmt5 | 0.012434062 | -0.364018443 | 0.388886567 | 8.676689265 | 0.082728124 | 0.937007203 | 0.974036091 | -4.760167162 |
| **A_55_P2096535** | Icmt | -0.01918349 | -0.601090213 | 0.562723233 | 8.714327157 | -0.082570438 | 0.937126949 | 0.974083059 | -4.760171511 |
| **A_52_P137829** | NA | 0.009872665 | -0.289727216 | 0.309472545 | 8.780259139 | 0.082535939 | 0.937153149 | 0.974083059 | -4.760172461 |
| **A_55_P1978052** | Gatb | -0.007109596 | -0.223403266 | 0.209184073 | 8.053043162 | -0.082328744 | 0.937310495 | 0.974194233 | -4.760178161 |
| **A_55_P2131279** | Atp6v1a | -0.01123799 | -0.354466721 | 0.33199074 | 10.73699693 | -0.082007837 | 0.937554203 | 0.974369455 | -4.76018696 |
| **A_55_P2009449** | Pnma2 | -0.018792432 | -0.593144304 | 0.55555944 | 8.943322519 | -0.081951196 | 0.937597219 | 0.974369455 | -4.76018851 |
| **A_51_P254895** | Cyp4a10 | -0.00711476 | -0.224678245 | 0.210448725 | 7.832313581 | -0.081907672 | 0.937630273 | 0.974369455 | -4.7601897 |
| **A_55_P2100715** | Auh | -0.008107374 | -0.256727734 | 0.240512985 | 8.440377427 | -0.081675892 | 0.937806301 | 0.974500001 | -4.760196026 |
| **A_55_P2362601** | D4Ertd681 | 0.018693598 | -0.556417184 | 0.59380438 | 8.413468438 | 0.08141262 | 0.938006251 | 0.97461856 | -4.76020319 |
| **A_65_P01641** | Rnf6 | 0.009313221 | -0.277937478 | 0.29656392 | 12.68784278 | 0.081206193 | 0.938163032 | 0.97461856 | -4.760208792 |
| **A_55_P2112637** | NA | -0.006783306 | -0.216027823 | 0.202461211 | 7.632927862 | -0.081196567 | 0.938170343 | 0.97461856 | -4.760209053 |
| **A_51_P475298** | Zfp110 | 0.017224331 | -0.514158334 | 0.548606996 | 9.664410599 | 0.081186782 | 0.938177775 | 0.97461856 | -4.760209318 |
| **A_52_P532559** | Thrb | -0.007044838 | -0.224403538 | 0.210313861 | 7.874025033 | -0.081179123 | 0.938183592 | 0.97461856 | -4.760209525 |
| **A_55_P2093418** | Zmpste24 | -0.016627936 | -0.529986699 | 0.496730826 | 11.40044239 | -0.08112743 | 0.938222853 | 0.97461856 | -4.760210925 |
| **A_55_P1990780** | Ptp4a1 | -0.011164523 | -0.356446652 | 0.334117605 | 13.15836919 | -0.080987209 | 0.938329353 | 0.974676824 | -4.760214718 |
| **A_51_P378825** | Actr8 | -0.007233567 | -0.23131265 | 0.216845516 | 10.69488736 | -0.080854004 | 0.938430526 | 0.974729547 | -4.760218316 |
| **A_55_P2085288** | Acoxl | 0.025216153 | -0.7567233 | 0.807155606 | 8.27519104 | 0.080771109 | 0.938493487 | 0.974742578 | -4.760220551 |
| **A_55_P1967168** | Lrp4 | 0.034177944 | -1.02728054 | 1.095636427 | 9.238806398 | 0.080647937 | 0.938587042 | 0.97478738 | -4.760223869 |
| **A_51_P369628** | Ptov1 | 0.026302995 | -0.793573878 | 0.846179868 | 11.93354331 | 0.08035389 | 0.938810387 | 0.974880535 | -4.760231769 |
| **A_55_P2174895** | Mrpl43 | -0.007800111 | -0.251127867 | 0.235527645 | 12.27734712 | -0.080289635 | 0.938859193 | 0.974880535 | -4.760233492 |
| **A_55_P2132207** | 1810037I1 | 0.010296346 | -0.311094589 | 0.331687281 | 8.674753884 | 0.080241651 | 0.938895641 | 0.974880535 | -4.760234777 |
| **A_51_P426276** | Pdk2 | -0.007467217 | -0.240926053 | 0.225991619 | 8.232179969 | -0.080112226 | 0.938993949 | 0.974880535 | -4.76023824 |
| **A_55_P2235059** | 6230415J | -0.023420715 | -0.755908491 | 0.709067062 | 9.052183853 | -0.080084802 | 0.93901478 | 0.974880535 | -4.760238973 |

| | | | | | | | | | |
|---|---|---|---|---|---|---|---|---|---|
| **A_52_P549166** | Arsj | 0.009180434 | -0.277957288 | 0.296318155 | 7.997505392 | 0.080079857 | 0.939018536 | 0.974880535 | -4.760239105 |
| **A_66_P137660** | Fam166b | 0.02975788 | -0.901154385 | 0.960670145 | 8.532029535 | 0.080065161 | 0.939029699 | 0.974880535 | -4.760239498 |
| **A_55_P2166421** | Rpusd3 | 0.016205919 | -0.491722204 | 0.524134041 | 9.065137334 | 0.079913793 | 0.939144677 | 0.974883834 | -4.760243539 |
| **A_55_P1989296** | Tnxb | -0.027858194 | -0.901344745 | 0.845628357 | 8.745863113 | -0.07988166 | 0.939169085 | 0.974883834 | -4.760244396 |
| **A_55_P2069818** | Mapk10 | 0.018307972 | -0.556867366 | 0.59348331 | 8.456244474 | 0.079724228 | 0.939288671 | 0.974883834 | -4.760248589 |
| **A_55_P2079388** | Snora74a | 0.008041946 | -0.24464822 | 0.260732111 | 7.964201857 | 0.079711894 | 0.93929804 | 0.974883834 | -4.760248917 |
| **A_55_P1973213** | Trmt61a | -0.023378381 | -0.758305987 | 0.711549226 | 9.293046712 | -0.079674659 | 0.939326325 | 0.974883834 | -4.760249908 |
| **A_51_P237585** | Btnl9 | -0.006891465 | -0.223728479 | 0.209945549 | 7.74113507 | -0.079602827 | 0.939380889 | 0.974883834 | -4.760251817 |
| **A_52_P313071** | Gosr1 | -0.009186899 | -0.298306044 | 0.279932245 | 10.52772246 | -0.07958706 | 0.939392867 | 0.974883834 | -4.760252236 |
| **A_55_P2090209** | Tpm3 | -0.008940652 | -0.290512292 | 0.272630989 | 15.73204519 | -0.07952993 | 0.939436263 | 0.974883834 | -4.760253753 |
| **A_55_P2083419** | Pkp4 | 0.00940174 | -0.287187175 | 0.305990655 | 9.391172301 | 0.079396924 | 0.939537299 | 0.974936354 | -4.76025728 |
| **A_55_P2095365** | Meaf6 | 0.007951964 | -0.243456111 | 0.259360038 | 8.43061037 | 0.079221945 | 0.93967022 | 0.975021952 | -4.760261912 |
| **A_55_P2127204** | Amigo1 | 0.006961545 | -0.213832307 | 0.227755397 | 8.006512339 | 0.078971247 | 0.939860665 | 0.975156925 | -4.76026853 |
| **A_55_P2095301** | NA | -0.006877803 | -0.225206121 | 0.211450515 | 7.832140749 | -0.078902362 | 0.939912995 | 0.975156925 | -4.760270345 |
| **A_55_P2100065** | Fsd1l | 0.013725404 | -0.422711073 | 0.450161882 | 8.18901993 | 0.078768841 | 0.940014427 | 0.975156925 | -4.760273858 |
| **A_51_P139030** | Slc38a3 | 0.006940604 | -0.214204922 | 0.228086129 | 7.980557917 | 0.078608488 | 0.940136245 | 0.975156925 | -4.76027807 |
| **A_55_P2060198** | NA | -0.063535682 | -2.089532193 | 1.962460829 | 10.25423004 | -0.078546954 | 0.940182992 | 0.975156925 | -4.760279684 |
| **A_52_P22324** | Pln | -0.010712983 | -0.35246063 | 0.331034664 | 7.815562255 | -0.078515467 | 0.940206913 | 0.975156925 | -4.760280509 |
| **A_55_P2069875** | Trim44 | 0.009164191 | -0.283257074 | 0.301585455 | 13.45906465 | 0.078493828 | 0.940223352 | 0.975156925 | -4.760281076 |
| **A_51_P150430** | Mkrn2 | -0.007314788 | -0.240822016 | 0.226192441 | 9.064569967 | -0.078460622 | 0.940248578 | 0.975156925 | -4.760281946 |
| **A_55_P2111237** | Selenof | 0.014438631 | -0.446524113 | 0.475401374 | 14.30631314 | 0.078453175 | 0.940254236 | 0.975156925 | -4.760282141 |
| **A_55_P1953093** | Pabpc4 | -0.014503158 | -0.479643895 | 0.450637578 | 10.77740128 | -0.078095957 | 0.94052562 | 0.975370631 | -4.760291473 |
| **A_51_P466221** | Amhr2 | -0.031287429 | -1.035579444 | 0.973004586 | 9.241771189 | -0.078029749 | 0.94057592 | 0.975370631 | -4.760293199 |
| **A_55_P2088095** | NA | -0.010134197 | -0.335626872 | 0.315358479 | 11.01916669 | -0.077982734 | 0.940611638 | 0.975370631 | -4.760294423 |
| **A_52_P501721** | Ccdc127 | -0.009582856 | -0.318484415 | 0.299318703 | 8.462992539 | -0.077700748 | 0.940825876 | 0.975540464 | -4.760301749 |
| **A_66_P101631** | Zfp27 | 0.007958851 | -0.248905016 | 0.264822719 | 8.071133703 | 0.077606434 | 0.940897532 | 0.975562444 | -4.760304194 |
| **A_55_P2329313** | C530014F | -0.008303756 | -0.277881759 | 0.261274248 | 8.242099646 | -0.077150803 | 0.94124371 | 0.975798868 | -4.760315962 |
| **A_55_P2014928** | Zfp341 | 0.010657022 | -0.335403034 | 0.356717078 | 8.89369354 | 0.077132024 | 0.941257978 | 0.975798868 | -4.760316445 |
| **A_55_P2114498** | Gm8801 | -0.01138114 | -0.381074904 | 0.358312624 | 9.843414559 | -0.077107031 | 0.941276968 | 0.975798868 | -4.760317089 |
| **A_55_P2039586** | Grk3 | 0.016415197 | -0.518160147 | 0.550990542 | 8.19245153 | 0.076910836 | 0.941426037 | 0.975851259 | -4.760322132 |
| **A_51_P487913** | NA | -0.00651587 | -0.218719679 | 0.205687939 | 7.761448502 | -0.076907659 | 0.941428451 | 0.975851259 | -4.760322214 |
| **A_52_P215170** | Acot4 | -0.006824791 | -0.22938346 | 0.215733879 | 7.711759573 | -0.076806001 | 0.941505692 | 0.975857519 | -4.760324822 |
| **A_55_P1965945** | NA | -0.015995811 | -0.538082007 | 0.506090385 | 16.78832606 | -0.076738693 | 0.941556834 | 0.975857519 | -4.760326547 |
| **A_55_P2068152** | Anapc15 | -0.015620963 | -0.527460387 | 0.496218461 | 10.99734835 | -0.076440657 | 0.941783291 | 0.975857519 | -4.760334166 |
| **A_55_P2099105** | Rhobtb1 | -0.020266291 | -0.68451236 | 0.643979778 | 9.213198885 | -0.076417993 | 0.941800513 | 0.975857519 | -4.760334744 |
| **A_55_P2085460** | 5530601H | -0.016152481 | -0.545735709 | 0.513430747 | 8.670057242 | -0.076393316 | 0.941819264 | 0.975857519 | -4.760335374 |
| **A_55_P1959076** | Zfp930 | 0.00744305 | -0.236677035 | 0.251563135 | 8.181309368 | 0.076365603 | 0.941840321 | 0.975857519 | -4.76033608 |
| **A_51_P219385** | Six5 | 0.019483227 | -0.619927105 | 0.658893559 | 9.350189548 | 0.076318809 | 0.941875877 | 0.975857519 | -4.760337273 |
| **A_55_P2175275** | Hmbox1 | 0.01274882 | -0.405737049 | 0.431234689 | 9.724483452 | 0.076302636 | 0.941888166 | 0.975857519 | -4.760337685 |
| **A_55_P1971574** | Acox3 | 0.026087825 | -0.83026568 | 0.882441331 | 10.30427713 | 0.076301863 | 0.941888754 | 0.975857519 | -4.760337704 |
| **A_51_P136355** | Gng11 | -0.03183302 | -1.077716384 | 1.014050345 | 14.3206107 | -0.076233337 | 0.941940823 | 0.975859173 | -4.760339449 |
| **A_55_P2145606** | Kmo | -0.00641907 | -0.217798169 | 0.204960029 | 7.680831766 | -0.076060718 | 0.94207199 | 0.975876982 | -4.760343837 |
| **A_51_P110341** | Scgb3a1 | 0.120567712 | -3.850679505 | 4.091814929 | 11.08822117 | 0.07604216 | 0.942086092 | 0.975876982 | -4.75546889 |
| **A_55_P2100149** | Mapt | -0.029311373 | -0.996700996 | 0.93807825 | 10.39900245 | -0.075890092 | 0.942201645 | 0.975876982 | -4.760348164 |
| **A_55_P2042486** | Dpysl3 | 0.006350297 | -0.203257613 | 0.215958206 | 7.820686287 | 0.075881639 | 0.942208067 | 0.975876982 | -4.760348378 |
| **A_52_P147803** | Syne4 | 0.019357987 | -0.619627952 | 0.658343926 | 9.259475042 | 0.075878587 | 0.942210387 | 0.975876982 | -4.760348455 |
| **A_55_P2051949** | Elmod3 | -0.0137825 | -0.469364495 | 0.441799494 | 10.63267381 | -0.075772533 | 0.942290976 | 0.97590817 | -4.76035114 |
| **A_51_P116755** | Cap2 | 0.023203684 | -0.745308188 | 0.791715557 | 8.381659947 | 0.075623489 | 0.942404233 | 0.975947101 | -4.753227828 |

| | | | | | | | | | |
|---|---|---|---|---|---|---|---|---|---|
| **A_52_P523480** | Rfx1 | 0.007838259 | -0.251881166 | 0.267557684 | 9.405776397 | 0.075590209 | 0.942429522 | 0.975947101 | -4.760355746 |
| **A_52_P229320** | Rptor | 0.012983978 | -0.417941433 | 0.443909389 | 10.64965484 | 0.075466812 | 0.942523292 | 0.975957559 | -4.760358857 |
| **A_55_P2079659** | NA | 0.006370675 | -0.205207954 | 0.217949304 | 7.65866878 | 0.075416084 | 0.942561841 | 0.975957559 | -4.760360135 |
| **A_66_P111781** | 4930432M | -0.01393404 | -0.476938182 | 0.449070102 | 9.237474275 | -0.075377637 | 0.942591057 | 0.975957559 | -4.760361102 |
| **A_55_P1987054** | Krtap5-1 | 0.008688549 | -0.28096945 | 0.298346548 | 8.199417142 | 0.075129772 | 0.942779415 | 0.976080185 | -4.760367329 |
| **A_52_P410636** | Trp73 | -0.010080616 | -0.34646354 | 0.326302309 | 8.206244128 | -0.075059119 | 0.942833107 | 0.976080185 | -4.7603691 |
| **A_51_P130110** | Idh3b | 0.014487904 | -0.469525922 | 0.49850173 | 10.59780326 | 0.074971833 | 0.942899439 | 0.976080185 | -4.760371286 |
| **A_55_P1994477** | Prrc2c | 0.007508007 | -0.243524291 | 0.258540306 | 9.598127703 | 0.074910973 | 0.94294569 | 0.976080185 | -4.760372809 |
| **A_55_P1965620** | Bud31 | 0.008249346 | -0.267735416 | 0.284234109 | 13.13176198 | 0.074866027 | 0.942979846 | 0.976080185 | -4.760373932 |
| **A_51_P372473** | Auh | -0.011470827 | -0.396089566 | 0.373147912 | 10.96734869 | -0.074698974 | 0.943106799 | 0.976080185 | -4.760378103 |
| **A_52_P295104** | Smim5 | 0.016818479 | -0.547309991 | 0.580946949 | 8.352873868 | 0.074672217 | 0.943127133 | 0.976080185 | -4.76037877 |
| **A_51_P416059** | Slc49a4 | -0.007052017 | -0.243733935 | 0.229629901 | 7.902685609 | -0.074627451 | 0.943161154 | 0.976080185 | -4.760379886 |
| **A_55_P2002968** | Coro2a | -0.009874389 | -0.34129741 | 0.321548632 | 8.192222998 | -0.074623894 | 0.943163857 | 0.976080185 | -4.760379974 |
| **A_51_P444447** | Cebpd | -0.029453117 | -1.019939943 | 0.961033709 | 13.59902431 | -0.074478841 | 0.943274093 | 0.976142018 | -4.760383584 |
| **A_55_P1974830** | Pde1c | 0.006221244 | -0.203381899 | 0.215824387 | 7.692771285 | 0.074341239 | 0.943378667 | 0.976197985 | -4.760387002 |
| **A_55_P2063715** | Cfap53 | 0.01846563 | -0.605492604 | 0.642423864 | 8.410953865 | 0.074124018 | 0.943543753 | 0.97631656 | -4.760392385 |
| **A_55_P2097548** | NA | 0.013454605 | -0.442033322 | 0.468942532 | 16.27195262 | 0.073985127 | 0.943649311 | 0.97637353 | -4.760395819 |
| **A_52_P621064** | NA | -0.006931591 | -0.242277892 | 0.22841471 | 9.16022112 | -0.073769341 | 0.943813311 | 0.97649096 | -4.760401141 |
| **A_55_P2113837** | Tbc1d1 | 0.011786524 | -0.389322347 | 0.412895394 | 11.75838861 | 0.073599321 | 0.943942531 | 0.976544691 | -4.760405324 |
| **A_55_P2322139** | C530042K | -0.006207562 | -0.217547641 | 0.205132518 | 7.734862129 | -0.073568097 | 0.943966263 | 0.976544691 | -4.760406091 |
| **A_52_P570700** | Dpp8 | -0.006586723 | -0.23227409 | 0.219100644 | 8.934396223 | -0.073099176 | 0.944322667 | 0.976861126 | -4.760417571 |
| **A_52_P69109** | Slc10a1 | 0.006078062 | -0.202761826 | 0.21491795 | 7.63656238 | 0.072895713 | 0.944477314 | 0.976884 | -4.76042253 |
| **A_55_P2060737** | Trmt10b | 0.006453544 | -0.215291407 | 0.228198495 | 9.074205257 | 0.072894523 | 0.944478218 | 0.976884 | -4.760422559 |
| **A_66_P104713** | Lrp5 | 0.009874064 | -0.329511673 | 0.3492598 | 9.827047423 | 0.072870656 | 0.944496359 | 0.976884 | -4.760423139 |
| **A_55_P2000798** | Ccdc144b | -0.01256433 | -0.445918479 | 0.420789819 | 8.342900368 | -0.072618408 | 0.94468809 | 0.976925445 | -4.760429266 |
| **A_55_P1978642** | Wasf1 | -0.020705452 | -0.735567805 | 0.694156901 | 9.246820039 | -0.07254586 | 0.944743233 | 0.976925445 | -4.760431024 |
| **A_55_P2079344** | Zfp362 | -0.012911425 | -0.458720498 | 0.432897648 | 10.58593228 | -0.072539677 | 0.944747933 | 0.976925445 | -4.760431173 |
| **A_55_P2013833** | Ddx3x | 0.011814254 | -0.396124403 | 0.41975291 | 12.43750097 | 0.072537366 | 0.94474969 | 0.976925445 | -4.760431229 |
| **A_51_P122723** | Grpel1 | -0.007332479 | -0.260759928 | 0.246094971 | 12.71032804 | -0.072468207 | 0.944802258 | 0.976925445 | -4.753302971 |
| **A_51_P165330** | Mrpl33 | -0.012834801 | -0.456736199 | 0.431066597 | 14.33638814 | -0.072419075 | 0.944839604 | 0.976925445 | -4.760434092 |
| **A_51_P246677** | Rec8 | 0.016384087 | -0.551745961 | 0.584514136 | 11.14386508 | 0.072231201 | 0.94498241 | 0.976978218 | -4.760438628 |
| **A_51_P387302** | Zmym2 | -0.009178845 | -0.3276069 | 0.30924921 | 10.10000953 | -0.072198314 | 0.945007408 | 0.976978218 | -4.760439421 |
| **A_55_P2062678** | Hmox2 | 0.009166311 | -0.309125291 | 0.327457913 | 10.1907697 | 0.072130637 | 0.945058851 | 0.976978218 | -4.760441052 |
| **A_55_P1968600** | NA | 0.009774319 | -0.32988664 | 0.349435278 | 8.781085606 | 0.072076091 | 0.945100314 | 0.976978218 | -4.760442365 |
| **A_52_P131672** | Rbm7 | -0.014013202 | -0.501359187 | 0.473332784 | 10.19960466 | -0.072019537 | 0.945143302 | 0.976978218 | -4.760443725 |
| **A_51_P435251** | Cul4a | 0.008636911 | -0.292225536 | 0.309499359 | 10.50669447 | 0.071901975 | 0.945232666 | 0.976982225 | -4.76044655 |
| **A_55_P2001198** | NA | 0.006376215 | -0.216271687 | 0.229024116 | 8.44263114 | 0.071728981 | 0.945364167 | 0.976982225 | -4.760450698 |
| **A_51_P427964** | Lmod3 | 0.011764431 | -0.399564732 | 0.423093594 | 8.320692184 | 0.071636072 | 0.945434792 | 0.976982225 | -4.760452921 |
| **A_55_P2051566** | NA | -0.012444864 | -0.44760195 | 0.422712221 | 7.838764657 | -0.071629919 | 0.94543947 | 0.976982225 | -4.760453069 |
| **A_51_P180724** | Mlh1 | -0.008739218 | -0.314482721 | 0.297004286 | 9.913542451 | -0.071592195 | 0.945468147 | 0.976982225 | -4.760453971 |
| **A_51_P513254** | Rac1 | -0.007056676 | -0.254153501 | 0.240040148 | 13.12006554 | -0.071529203 | 0.945516031 | 0.976982225 | -4.760455476 |
| **A_51_P251736** | Reep5 | 0.015162397 | -0.515832284 | 0.546157077 | 9.521620863 | 0.071520086 | 0.945522962 | 0.976982225 | -4.760455693 |
| **A_55_P1985825** | NA | -0.00737442 | -0.26603379 | 0.251284949 | 7.991961278 | -0.071408518 | 0.945607772 | 0.976982225 | -4.760458355 |
| **A_51_P504815** | Stfa3 | 0.032171795 | -1.096730723 | 1.161074313 | 9.215257815 | 0.071378792 | 0.945630369 | 0.976982225 | -4.760459064 |
| **A_55_P2096917** | Mreg | -0.045882487 | -1.658060983 | 1.56629601 | 8.426717886 | -0.071282704 | 0.945703413 | 0.976982225 | -4.760461352 |
| **A_51_P132170** | Ccdc141 | -0.010968015 | -0.396672622 | 0.374736593 | 7.958681877 | -0.071223537 | 0.945748391 | 0.976982225 | -4.76046276 |
| **A_55_P2078428** | Tstd1 | -0.010565906 | -0.382165324 | 0.361033511 | 8.389573774 | -0.071216738 | 0.94575356 | 0.976982225 | -4.760462922 |
| **A_51_P165293** | Ckmt2 | -0.013416421 | -0.486648583 | 0.459815741 | 8.050360713 | -0.071008899 | 0.945911558 | 0.977093234 | -4.760467857 |

| | | | | | | | | | |
|---|---|---|---|---|---|---|---|---|---|
| **A_55_P2045658** | Nme6 | 0.006671727 | -0.228895679 | 0.242239134 | 9.006550413 | 0.070937098 | 0.945966141 | 0.977097413 | -4.760469558 |
| **A_55_P2076560** | Nr5a1 | -0.006686061 | -0.243193483 | 0.229821361 | 7.986234093 | -0.070806953 | 0.946065079 | 0.977147403 | -4.760472638 |
| **A_55_P2171348** | Grm2 | -0.01258507 | -0.458835321 | 0.433665181 | 8.682275228 | -0.07063623 | 0.946194866 | 0.977177767 | -4.760476669 |
| **A_51_P513460** | Cfhr1 | 0.005851419 | -0.201719413 | 0.213422251 | 7.620732328 | 0.070606586 | 0.946217402 | 0.977177767 | -4.760477368 |
| **A_51_P493016** | Cmya5 | 0.016888638 | -0.583617313 | 0.617394589 | 8.268808252 | 0.070441348 | 0.946343022 | 0.977177767 | -4.760481259 |
| **A_55_P2353079** | NA | 0.006481512 | -0.22400394 | 0.236966963 | 7.735023989 | 0.070434128 | 0.946348511 | 0.977177767 | -4.760481429 |
| **A_55_P2247608** | 4931428A | 0.009133742 | -0.315887518 | 0.334155001 | 8.578531757 | 0.070386165 | 0.946384974 | 0.977177767 | -4.760482557 |
| **A_55_P1975110** | Pnpt1 | 0.008887583 | -0.307573722 | 0.325348889 | 10.26703492 | 0.070341791 | 0.946418709 | 0.977177767 | -4.760483599 |
| **A_51_P120461** | Riox2 | 0.011736671 | -0.407284386 | 0.430757728 | 9.955742781 | 0.07015513 | 0.946560618 | 0.977177767 | -4.753356029 |
| **A_65_P00881** | NA | -0.009991093 | -0.366735557 | 0.346753372 | 8.874223712 | -0.070146513 | 0.946567169 | 0.977177767 | -4.760488179 |
| **A_55_P2153067** | 2610028H | 0.008497111 | -0.295053288 | 0.31204751 | 17.29984755 | 0.070111756 | 0.946593593 | 0.977177767 | -4.760488993 |
| **A_52_P520607** | Ankrd22 | -0.007675769 | -0.281942632 | 0.266591093 | 7.777350101 | -0.070096917 | 0.946604874 | 0.977177767 | -4.76048934 |
| **A_52_P84027** | Cyp7a1 | -0.005820454 | -0.213972063 | 0.202331154 | 7.641125904 | -0.070036986 | 0.946650438 | 0.977177767 | -4.760490743 |
| **A_55_P2005853** | Nacc2 | 0.012316226 | -0.428698293 | 0.453330744 | 10.33889623 | 0.069947966 | 0.946718116 | 0.977195455 | -4.760492823 |
| **A_55_P2114044** | Commd8 | -0.006705631 | -0.247475101 | 0.234063839 | 10.21494989 | -0.069757126 | 0.946863207 | 0.977293041 | -4.760497274 |
| **A_55_P2167294** | NA | -0.005785045 | -0.213892942 | 0.202322853 | 7.661529308 | -0.069625528 | 0.946963259 | 0.97732991 | -4.760500337 |
| **A_51_P248444** | Poglut1 | 0.007300413 | -0.255663458 | 0.270264283 | 8.796400974 | 0.069534713 | 0.947032305 | 0.97732991 | -4.760502446 |
| **A_51_P212801** | Ube2a | 0.011489533 | -0.402511273 | 0.42549034 | 10.59228429 | 0.06951068 | 0.947050578 | 0.97732991 | -4.760503004 |
| **A_55_P2353004** | Pitpnm2os | 0.013719295 | -0.481913888 | 0.509352479 | 8.293486759 | 0.069330092 | 0.947187879 | 0.977371716 | -4.76050719 |
| **A_52_P221756** | Spock2 | 0.028999719 | -1.019711255 | 1.077710692 | 8.616813445 | 0.06926093 | 0.947240463 | 0.977371716 | -4.760508791 |
| **A_55_P1970229** | Csnk1d | -0.006151689 | -0.228623606 | 0.216320228 | 14.15391964 | -0.069257931 | 0.947242744 | 0.977371716 | -4.76050886 |
| **A_55_P2023114** | Hsd3b2 | -0.005868553 | -0.218465775 | 0.206728668 | 7.673516023 | -0.069139116 | 0.947333081 | 0.977410968 | -4.760511606 |
| **A_55_P2041673** | Ttc38 | -0.008086695 | -0.301311741 | 0.285138351 | 17.31454867 | -0.069074915 | 0.947381894 | 0.977410968 | -4.760513087 |
| **A_51_P212630** | Chd8 | -0.008633106 | -0.322122078 | 0.304855866 | 12.55095528 | -0.068975544 | 0.947457448 | 0.977436759 | -4.760515377 |
| **A_55_P2066230** | Hck | 0.065691468 | -2.324309793 | 2.455692729 | 10.77319238 | 0.068843221 | 0.947558058 | 0.977488395 | -4.760518422 |
| **A_55_P2028595** | AW55198 | 0.006734052 | -0.239268815 | 0.252736918 | 8.431361209 | 0.068562496 | 0.947771506 | 0.977565869 | -4.760524862 |
| **A_52_P412529** | Fbxo3 | 0.013140492 | -0.467235853 | 0.493516837 | 8.821797139 | 0.068514165 | 0.947808255 | 0.977565869 | -4.760525969 |
| **A_55_P2357860** | Atxn7l1 | 0.007889777 | -0.281001478 | 0.296781032 | 8.336913681 | 0.068403871 | 0.947892118 | 0.977565869 | -4.76052849 |
| **A_51_P206346** | Reep3 | -0.008885449 | -0.33426166 | 0.316490762 | 12.09967378 | -0.06839808 | 0.947896522 | 0.977565869 | -4.760528622 |
| **A_55_P1971298** | Lonp1 | 0.012083198 | -0.430686996 | 0.454853392 | 12.74158716 | 0.068352413 | 0.947931245 | 0.977565869 | -4.760529665 |
| **A_51_P508444** | St13 | -0.008748849 | -0.329369572 | 0.311871874 | 11.90786279 | -0.06834546 | 0.947936532 | 0.977565869 | -4.760529824 |
| **A_55_P2103429** | Psmc3 | 0.01290967 | -0.461181029 | 0.48700037 | 15.13617248 | 0.068203092 | 0.948044785 | 0.977577694 | -4.760533069 |
| **A_55_P1995698** | Thumpd1 | 0.01281745 | -0.458322658 | 0.483957558 | 10.65164124 | 0.068139963 | 0.948092787 | 0.977577694 | -4.760534506 |
| **A_51_P365859** | Sowaha | -0.005823173 | -0.220115013 | 0.208468667 | 7.690551277 | -0.068061949 | 0.948152108 | 0.977577694 | -4.760536281 |
| **A_52_P243516** | Rmdn2 | 0.015600749 | -0.55979985 | 0.591001349 | 9.09058626 | 0.067908712 | 0.948268627 | 0.977577694 | -4.760539759 |
| **A_51_P510722** | Rita1 | -0.006652877 | -0.252093213 | 0.238787459 | 8.23891279 | -0.067891267 | 0.948281893 | 0.977577694 | -4.760540155 |
| **A_51_P158044** | NA | -0.007966433 | -0.301955113 | 0.286022247 | 8.697771112 | -0.067870904 | 0.948297376 | 0.977577694 | -4.760540617 |
| **A_51_P181891** | Amer2 | 0.005995877 | -0.215598147 | 0.227589901 | 7.894284858 | 0.067771175 | 0.94837321 | 0.977577694 | -4.760542875 |
| **A_55_P2109263** | Ppp1r14b | 0.021787551 | -0.783885707 | 0.82746081 | 13.16726105 | 0.067732923 | 0.948402298 | 0.977577694 | -4.760543741 |
| **A_55_P2080850** | Ptpn2 | -0.012970279 | -0.492975946 | 0.467035387 | 10.93243443 | -0.067678903 | 0.948443375 | 0.977577694 | -4.760544962 |
| **A_55_P2018307** | NA | 0.027143033 | -0.977569726 | 1.031855792 | 11.91419451 | 0.067665423 | 0.948453625 | 0.977577694 | -4.760545267 |
| **A_66_P125161** | 1700057H | -0.008496298 | -0.323827464 | 0.306834869 | 8.164130702 | -0.067485919 | 0.948590123 | 0.977604428 | -4.760549317 |
| **A_55_P1977747** | Picalm | -0.015137497 | -0.577118786 | 0.546843792 | 13.81955433 | -0.067465627 | 0.948605553 | 0.977604428 | -4.760549775 |
| **A_55_P2057803** | Stum | 0.006228815 | -0.225434971 | 0.237892601 | 7.729034619 | 0.067343798 | 0.948698194 | 0.977604428 | -4.760552517 |
| **A_52_P430934** | Dcun1d1 | 0.006418555 | -0.232610096 | 0.245447207 | 8.961344848 | 0.067257028 | 0.948764177 | 0.977604428 | -4.760554467 |
| **A_55_P1973818** | Mmel1 | -0.005660666 | -0.216550175 | 0.205228843 | 7.690885002 | -0.067229971 | 0.948784752 | 0.977604428 | -4.760555075 |
| **A_55_P2033420** | Sppl2a | 0.00935297 | -0.339754882 | 0.358460823 | 9.970420543 | 0.067102746 | 0.9488815 | 0.977604428 | -4.760557928 |
| **A_55_P2031382** | Crebzf | -0.007042469 | -0.269969322 | 0.255884384 | 8.537833587 | -0.067087303 | 0.948893243 | 0.977604428 | -4.760558274 |

| | | | | | | | | | |
|---|---|---|---|---|---|---|---|---|---|
| **A_55_P1953851** | Stk39 | -0.011153246 | -0.427998568 | 0.405692075 | 10.09572859 | -0.067015722 | 0.948947676 | 0.977604428 | -4.760559877 |
| **A_55_P2113857** | Serpina1e | -0.056226178 | -2.159428787 | 2.046976431 | 11.02342932 | -0.066958821 | 0.948990947 | 0.977604428 | -4.76056115 |
| **A_51_P104891** | Selenoi | -0.007065193 | -0.271536729 | 0.257406343 | 9.11499781 | -0.06691068 | 0.949027556 | 0.977604428 | -4.748640509 |
| **A_55_P2258366** | C030009H | 0.027912755 | -1.017984799 | 1.073810309 | 11.80858165 | 0.066844226 | 0.949078091 | 0.977604428 | -4.76056371 |
| **A_55_P2007384** | Sgca | 0.006193956 | -0.225932655 | 0.238320566 | 7.751229797 | 0.066833391 | 0.949086331 | 0.977604428 | -4.760563952 |
| **A_51_P269792** | Rad51b | -0.005861641 | -0.226297803 | 0.214574522 | 8.055952268 | -0.066601912 | 0.949262364 | 0.977693327 | -4.760569109 |
| **A_55_P2095019** | NA | 0.00627188 | -0.229645131 | 0.242188891 | 8.286164089 | 0.066586908 | 0.949273773 | 0.977693327 | -4.760569443 |
| **A_55_P2173358** | NA | -0.006212939 | -0.240842937 | 0.228417059 | 8.008603951 | -0.066322961 | 0.949474501 | 0.977721392 | -4.760575301 |
| **A_55_P2007155** | Akr1c14 | 0.044576691 | -1.639721951 | 1.728875334 | 9.359645742 | 0.066288635 | 0.949500605 | 0.977721392 | -4.760576061 |
| **A_52_P625270** | 1700024G | -0.019605901 | -0.760645086 | 0.721433283 | 8.281269131 | -0.066266798 | 0.949517212 | 0.977721392 | -4.760576544 |
| **A_55_P1985808** | Ranbp9 | -0.006158525 | -0.239005993 | 0.226688943 | 11.21088911 | -0.066245372 | 0.949533507 | 0.977721392 | -4.760577018 |
| **A_51_P472879** | NA | 0.005643957 | -0.207834533 | 0.219122447 | 7.704201598 | 0.066218592 | 0.949553873 | 0.977721392 | -4.760577611 |
| **A_55_P2093822** | Ube2d2a | 0.00875786 | -0.323070997 | 0.340586716 | 12.9937078 | 0.06610498 | 0.949640275 | 0.977751817 | -4.760580121 |
| **A_66_P126999** | Fis1 | -0.015630074 | -0.609010161 | 0.577750013 | 14.27559065 | -0.065974849 | 0.94973924 | 0.977751817 | -4.760582991 |
| **A_55_P2001683** | Atp5g1 | -0.008586267 | -0.335074274 | 0.31790174 | 14.38321758 | -0.065869973 | 0.949819 | 0.977751817 | -4.7605853 |
| **A_55_P1972997** | A530016L | -0.005492188 | -0.214394861 | 0.203410486 | 7.736304821 | -0.065849384 | 0.949834658 | 0.977751817 | -4.760585753 |
| **A_55_P2034372** | Eif4g3 | -0.009913934 | -0.387015812 | 0.367187944 | 10.81393159 | -0.065847252 | 0.94983628 | 0.977751817 | -4.7605858 |
| **A_55_P2009345** | Ovch2 | -0.006422595 | -0.25114359 | 0.2382984 | 7.995931334 | -0.065733939 | 0.949922457 | 0.977788467 | -4.760588289 |
| **A_55_P2003586** | Unkl | -0.006007528 | -0.236253291 | 0.224238235 | 8.906931769 | -0.065351338 | 0.950213439 | 0.977987838 | -4.760596664 |
| **A_55_P2033922** | Tsga8 | -0.006094133 | -0.239804323 | 0.227616057 | 7.998676913 | -0.06531074 | 0.950244316 | 0.977987838 | -4.76059755 |
| **A_55_P2096127** | NA | -0.027926649 | -1.100414621 | 1.044561323 | 11.23083724 | -0.065219392 | 0.95031379 | 0.977987838 | -4.760599541 |
| **A_55_P1988708** | Gstz1 | -0.022634689 | -0.891973766 | 0.846704388 | 10.71062737 | -0.065213225 | 0.950318481 | 0.977987838 | -4.760599675 |
| **A_51_P465292** | Hnmt | 0.009463218 | -0.354413204 | 0.373339639 | 7.914626845 | 0.065138126 | 0.950375598 | 0.977994561 | -4.76060131 |
| **A_52_P498193** | Aldh1l2 | -0.006246412 | -0.246700736 | 0.234207911 | 7.973026909 | -0.065065146 | 0.950431104 | 0.977999625 | -4.760602897 |
| **A_52_P136914** | Nudt7 | -0.005954142 | -0.236229898 | 0.224321613 | 8.318104814 | -0.064762162 | 0.950661546 | 0.97818469 | -4.760609465 |
| **A_55_P2016981** | Nfu1 | 0.006610031 | -0.249935503 | 0.263155565 | 11.81043091 | 0.064534122 | 0.95083499 | 0.978311091 | -4.760614389 |
| **A_51_P317443** | Cd3eap | 0.013611792 | -0.515544442 | 0.542768026 | 9.179958797 | 0.06442905 | 0.950914908 | 0.978321427 | -4.760616652 |
| **A_52_P331901** | Trp53rkb | -0.00672873 | -0.268474363 | 0.255016904 | 8.231141897 | -0.064387858 | 0.950946239 | 0.978321427 | -4.760617538 |
| **A_55_P2021065** | E130308A | -0.007220147 | -0.28858647 | 0.274146176 | 8.76778621 | -0.064272362 | 0.951034086 | 0.978332656 | -4.760620019 |
| **A_51_P416191** | Hectd1 | 0.006701097 | -0.254806181 | 0.268208376 | 10.58952259 | 0.06418189 | 0.951102901 | 0.978332656 | -4.76062196 |
| **A_55_P2046709** | NA | -0.056325367 | -2.254669947 | 2.142019214 | 11.02175647 | -0.064173926 | 0.951108958 | 0.978332656 | -4.760622131 |
| **A_55_P2103963** | Bhmt | 0.005365934 | -0.20432422 | 0.215056088 | 7.678163965 | 0.064094039 | 0.951169721 | 0.978343107 | -4.760623842 |
| **A_55_P2065434** | Rprl2 | 0.019317141 | -0.738453844 | 0.777088125 | 9.954611191 | 0.063849164 | 0.95135598 | 0.978482632 | -4.760629074 |
| **A_55_P1963080** | Rps12 | -0.009919029 | -0.400054036 | 0.380215978 | 16.44735964 | -0.063680225 | 0.951484482 | 0.978532842 | -4.760632671 |
| **A_55_P2050628** | Cyp4a31 | -0.005585009 | -0.225525843 | 0.214355825 | 7.835173367 | -0.063601651 | 0.951544249 | 0.978532842 | -4.760634342 |
| **A_55_P2151666** | NA | -0.006826516 | -0.275727427 | 0.262074394 | 8.345073276 | -0.063585369 | 0.951556634 | 0.978532842 | -4.760634687 |
| **A_55_P2025228** | Akt2 | -0.009710666 | -0.393066057 | 0.373644725 | 12.30342418 | -0.063445054 | 0.951663365 | 0.978577304 | -4.760637664 |
| **A_55_P1975874** | Bcl2l15 | 0.00561334 | -0.216189896 | 0.227416577 | 7.753823359 | 0.063387532 | 0.95170712 | 0.978577304 | -4.760638882 |
| **A_51_P357433** | Ap1g2 | 0.007236494 | -0.279042688 | 0.293515676 | 8.446797678 | 0.0633124 | 0.95176427 | 0.978577304 | -4.760640472 |
| **A_55_P1963144** | Mtmr14 | 0.005538729 | -0.213749356 | 0.224826814 | 10.38993065 | 0.063262368 | 0.951802328 | 0.978577304 | -4.76064153 |
| **A_52_P292684** | Exosc4 | 0.005993503 | -0.233143764 | 0.24513077 | 10.27693 | 0.06277458 | 0.95217338 | 0.978816209 | -4.760651797 |
| **A_51_P204247** | C8a | -0.005325478 | -0.217991889 | 0.207340933 | 7.639526606 | -0.062720574 | 0.952214462 | 0.978816209 | -4.760652929 |
| **A_52_P365005** | Zbtb22 | 0.006463301 | -0.251663833 | 0.264590435 | 9.061112051 | 0.06271495 | 0.95221874 | 0.978816209 | -4.760653046 |
| **A_51_P248638** | Myoz2 | -0.021053747 | -0.862828136 | 0.820720643 | 8.505730743 | -0.062644657 | 0.952272213 | 0.978816209 | -4.760654518 |
| **A_51_P493558** | Rnpep | -0.012528764 | -0.513951683 | 0.488894154 | 11.37254878 | -0.062582715 | 0.952319332 | 0.978816209 | -4.760655813 |
| **A_55_P2138222** | NA | -0.010536853 | -0.432409477 | 0.411335772 | 8.978793116 | -0.062557575 | 0.952338456 | 0.978816209 | -4.760656339 |
| **A_51_P400217** | Vpreb1 | 0.007072809 | -0.276617599 | 0.290763216 | 8.450345292 | 0.06244499 | 0.952424101 | 0.978852199 | -4.760658689 |
| **A_55_P2314496** | Secisbp2l | 0.010433087 | -0.408608732 | 0.429474906 | 9.023693567 | 0.062359958 | 0.952488786 | 0.978866645 | -4.760660461 |

| | | | | | | | | | |
|---|---|---|---|---|---|---|---|---|---|
| **A_51_P500550** | Mfsd1 | -0.008287387 | -0.342164381 | 0.325589607 | 9.130164071 | -0.062170089 | 0.952633224 | 0.978948451 | -4.76066441 |
| **A_52_P562593** | Tpgs1 | -0.017414965 | -0.719929613 | 0.685099683 | 10.04926416 | -0.062089415 | 0.952694596 | 0.978948451 | -4.760666084 |
| **A_55_P2044005** | NA | -0.008042829 | -0.332695074 | 0.316609417 | 9.560190039 | -0.062049853 | 0.952724692 | 0.978948451 | -4.760666905 |
| **A_66_P122372** | Ppp4r2 | -0.005479316 | -0.2269847 | 0.216026068 | 8.971339185 | -0.061957286 | 0.952795111 | 0.978948451 | -4.760668822 |
| **A_51_P145662** | Clec4g | -0.006719767 | -0.278755411 | 0.265315877 | 7.959880512 | -0.061869803 | 0.952861664 | 0.978948451 | -4.760670631 |
| **A_55_P2101133** | Tomm7 | 0.009790801 | -0.386657853 | 0.406239454 | 14.74105118 | 0.061855967 | 0.952872189 | 0.978948451 | -4.760670917 |
| **A_55_P1980341** | Romo1 | -0.007244719 | -0.301180397 | 0.28669096 | 13.83472633 | -0.061733311 | 0.9529655 | 0.978992295 | -4.760673448 |
| **A_55_P1996998** | Polr2a | 0.007566356 | -0.300202733 | 0.315335444 | 9.003128872 | 0.061576088 | 0.953085109 | 0.979063147 | -4.760676686 |
| **A_55_P2144771** | NA | 0.007183215 | -0.285500223 | 0.299866652 | 8.276806369 | 0.061471108 | 0.953164974 | 0.979093168 | -4.760678843 |
| **A_52_P38964** | Sap25 | 0.012591383 | -0.503624068 | 0.528806834 | 8.538165489 | 0.061093186 | 0.95345249 | 0.979266999 | -4.760686579 |
| **A_51_P164835** | Ppl | -0.022516526 | -0.945736528 | 0.900703476 | 10.2908345 | -0.061086681 | 0.953457439 | 0.979266999 | -4.760686712 |
| **A_65_P11425** | Bmp2k | -0.008782822 | -0.369117697 | 0.351552053 | 8.564678723 | -0.061048939 | 0.953486152 | 0.979266999 | -4.760687482 |
| **A_55_P2125200** | Mterf1b | 0.005945884 | -0.23848245 | 0.250374218 | 8.357687712 | 0.060927764 | 0.953578341 | 0.979273871 | -4.760689951 |
| **A_55_P2104583** | C87436 | -0.007304608 | -0.30797487 | 0.293365654 | 9.3921712 | -0.060849468 | 0.953637909 | 0.979273871 | -4.760691543 |
| **A_51_P225832** | 2700097C | 0.007458416 | -0.299793968 | 0.3147108 | 10.07021601 | 0.060799733 | 0.953675748 | 0.979273871 | -4.760692553 |
| **A_55_P2071271** | Abracl | -0.010263238 | -0.433241505 | 0.412715028 | 11.75832915 | -0.060773845 | 0.953695444 | 0.979273871 | -4.760693079 |
| **A_55_P2083484** | Mettl21a | 0.005645161 | -0.227853116 | 0.239143438 | 8.389830986 | 0.06055402 | 0.95386269 | 0.979393587 | -4.760697534 |
| **A_55_P2045602** | Gpkow | 0.008481066 | -0.342977271 | 0.359939403 | 9.617929132 | 0.060440345 | 0.953949176 | 0.979426795 | -4.760699831 |
| **A_55_P2084032** | AW14615 | -0.009632953 | -0.409279515 | 0.390013609 | 9.428167991 | -0.060371739 | 0.954001373 | 0.979426795 | -4.760701215 |
| **A_55_P2076288** | Zfp661 | 0.012842485 | -0.520489244 | 0.546174215 | 8.663243313 | 0.06031176 | 0.954047007 | 0.979426795 | -4.760702424 |
| **A_51_P392861** | Med25 | 0.015690264 | -0.638785845 | 0.670166374 | 10.61155321 | 0.060046377 | 0.954248921 | 0.979550834 | -4.760707759 |
| **A_55_P2019293** | NA | 0.0049894 | -0.203221898 | 0.213200698 | 7.635732854 | 0.060019775 | 0.954269161 | 0.979550834 | -4.760708292 |
| **A_55_P2076639** | Vps25 | -0.006599014 | -0.282618764 | 0.269420737 | 9.436678927 | -0.059881022 | 0.954374732 | 0.979607192 | -4.760711071 |
| **A_55_P2129972** | NA | 0.022568221 | -0.923020019 | 0.968156462 | 11.31058964 | 0.059778583 | 0.954452674 | 0.979635186 | -4.760713118 |
| **A_55_P2001593** | Rnf180 | -0.008885287 | -0.381675969 | 0.363905396 | 8.671194111 | -0.059697581 | 0.954514305 | 0.979646437 | -4.760714735 |
| **A_52_P464437** | Trp53bp1 | 0.007563375 | -0.310409371 | 0.32553612 | 9.161686004 | 0.05957665 | 0.954606318 | 0.979688866 | -4.760717144 |
| **A_52_P267106** | Ksr2 | -0.005793158 | -0.250137698 | 0.238551382 | 8.002579339 | -0.059383125 | 0.954753567 | 0.979758715 | -4.760720989 |
| **A_52_P606774** | Stk4 | -0.012263602 | -0.5297732 | 0.505245996 | 10.46225979 | -0.059353998 | 0.954775729 | 0.979758715 | -4.760721566 |
| **A_55_P2099232** | Bdp1 | 0.010603708 | -0.438255617 | 0.459463033 | 10.2870067 | 0.059169492 | 0.954916117 | 0.97985077 | -4.760725219 |
| **A_52_P334670** | Clock | -0.007087046 | -0.308902958 | 0.294728865 | 9.007490499 | -0.058813021 | 0.955187357 | 0.980018126 | -4.760732244 |
| **A_55_P2035804** | Fzd8 | -0.005419536 | -0.236318861 | 0.225479789 | 8.041566938 | -0.05878815 | 0.955206282 | 0.980018126 | -4.760732732 |
| **A_55_P2185826** | Cyp3a13 | -0.005201124 | -0.227067179 | 0.21666493 | 7.874201352 | -0.058716034 | 0.955261156 | 0.980018126 | -4.760734148 |
| **A_51_P283266** | Zfp874b | -0.009344621 | -0.408146943 | 0.389457701 | 8.504736881 | -0.058688681 | 0.95528197 | 0.980018126 | -4.760734684 |
| **A_55_P2089655** | Eif2s2 | -0.010998025 | -0.482394187 | 0.460398138 | 13.3884858 | -0.0584358 | 0.955474394 | 0.980063669 | -4.760739631 |
| **A_55_P2368355** | 2610319H | -0.011440072 | -0.502044174 | 0.47916403 | 10.75058505 | -0.058404717 | 0.955498046 | 0.980063669 | -4.760740238 |
| **A_51_P194004** | Glyctk | -0.004899815 | -0.215094047 | 0.205294417 | 7.76496207 | -0.058386066 | 0.955512238 | 0.980063669 | -4.760740602 |
| **A_55_P2065751** | Krtap19-5 | -0.015633646 | -0.686547157 | 0.655279865 | 9.294323568 | -0.05836387 | 0.955529128 | 0.980063669 | -4.760741034 |
| **A_51_P237040** | Nog | 0.009891435 | -0.416247305 | 0.436030175 | 8.197181896 | 0.058137805 | 0.955701151 | 0.980079376 | -4.760745433 |
| **A_55_P2167999** | Ldlr | -0.005793999 | -0.255611819 | 0.244023821 | 11.55689192 | -0.058090531 | 0.955737124 | 0.980079376 | -4.760746351 |
| **A_51_P269173** | Alg9 | -0.006410229 | -0.283472487 | 0.270652029 | 9.114071929 | -0.057949076 | 0.955844765 | 0.980079376 | -4.760749093 |
| **A_55_P2005135** | Tardbp | -0.007421842 | -0.328430256 | 0.313586571 | 9.579292066 | -0.057908946 | 0.955875302 | 0.980079376 | -4.76074987 |
| **A_55_P2145800** | NA | 0.008291291 | -0.350544294 | 0.367126877 | 8.450210384 | 0.057873137 | 0.955902551 | 0.980079376 | -4.760750562 |
| **A_55_P2048210** | Ganab | 0.007561597 | -0.319807209 | 0.334930403 | 12.38304665 | 0.057853095 | 0.955917802 | 0.980079376 | -4.760750949 |
| **A_55_P2169659** | Vwf | -0.006222368 | -0.275943668 | 0.263498932 | 8.293507762 | -0.057781762 | 0.955972084 | 0.980079376 | -4.760752327 |
| **A_55_P1975385** | Dctn1 | 0.006044921 | -0.256174642 | 0.268264484 | 13.61963957 | 0.057739883 | 0.956003952 | 0.980079376 | -4.760753135 |
| **A_52_P505157** | Fsd2 | 0.004970528 | -0.21069199 | 0.220633046 | 7.700508898 | 0.057726895 | 0.956013836 | 0.980079376 | -4.760753386 |
| **A_65_P12104** | Tmtc1 | 0.019082906 | -0.810010427 | 0.84817624 | 9.272030999 | 0.057648955 | 0.956073146 | 0.980079376 | -4.760754888 |
| **A_55_P2104662** | Ttc33 | -0.006446241 | -0.28670017 | 0.273807688 | 10.60039126 | -0.057610967 | 0.956102054 | 0.980079376 | -4.760755619 |

| | | | | | | | | | |
|---|---|---|---|---|---|---|---|---|---|
| **A_52_P17146** | Bmpr1a | -0.00991564 | -0.441544823 | 0.421713544 | 11.32265493 | -0.05753873 | 0.956157025 | 0.980083762 | -4.760757009 |
| **A_55_P2139615** | Rtl6 | 0.007336321 | -0.314552969 | 0.32922561 | 8.570134588 | 0.057085015 | 0.956502298 | 0.980378998 | -4.760765696 |
| **A_55_P2118037** | Sirpa | 0.011022488 | -0.474596317 | 0.496641292 | 9.7992092 | 0.056850524 | 0.956680748 | 0.980378998 | -4.760770158 |
| **A_55_P2048368** | Fggy | 0.004961471 | -0.213841014 | 0.223763955 | 7.946849132 | 0.056794795 | 0.956723159 | 0.980378998 | -4.760771216 |
| **A_52_P449417** | Vangl1 | -0.008875879 | -0.400559847 | 0.38280809 | 10.67280542 | -0.056757845 | 0.956751278 | 0.980378998 | -4.760771917 |
| **A_55_P1994302** | NA | 0.005066136 | -0.218688334 | 0.228820606 | 7.918318549 | 0.056709456 | 0.956788104 | 0.980378998 | -4.760772834 |
| **A_51_P265008** | Ppig | -0.006833279 | -0.308699271 | 0.295032714 | 11.09127141 | -0.056697681 | 0.956797064 | 0.980378998 | -4.760773057 |
| **A_55_P2160323** | NA | -0.006183711 | -0.279373125 | 0.267005702 | 8.197734056 | -0.056693816 | 0.956800005 | 0.980378998 | -4.760773131 |
| **A_55_P1959800** | NA | 0.008214092 | -0.35572256 | 0.372150745 | 8.844393412 | 0.056530668 | 0.956924166 | 0.980403057 | -4.760776216 |
| **A_55_P2063970** | Ube2l3 | 0.012691838 | -0.549647166 | 0.575030843 | 12.88347122 | 0.056529699 | 0.956924904 | 0.980403057 | -4.760776234 |
| **A_51_P282837** | St14 | 0.010330056 | -0.449736713 | 0.470396824 | 9.700729149 | 0.056238294 | 0.957146674 | 0.980578306 | -4.760781724 |
| **A_51_P506309** | Hrg | -0.00486582 | -0.2218319 | 0.21210026 | 7.77148601 | -0.056171308 | 0.957197653 | 0.980578574 | -4.760782981 |
| **A_55_P1992031** | Dydc2 | -0.018573568 | -0.848395776 | 0.811248639 | 8.394449808 | -0.056060975 | 0.957281623 | 0.980612636 | -4.76078505 |
| **A_55_P2125820** | Gm14288 | -0.007266265 | -0.336156509 | 0.321623979 | 10.41312017 | -0.055336359 | 0.957833109 | 0.981121874 | -4.760798534 |
| **A_51_P134262** | 1700052K | -0.009690045 | -0.449125476 | 0.429745386 | 10.06489699 | -0.055230766 | 0.957913475 | 0.981121874 | -4.760800484 |
| **A_55_P2059958** | Snx7 | -0.007626132 | -0.353829267 | 0.338577003 | 9.762946759 | -0.055172628 | 0.957957724 | 0.981121874 | -4.760801556 |
| **A_52_P56634** | Map3k11 | -0.006041628 | -0.280469974 | 0.268386719 | 9.641227225 | -0.055141089 | 0.957981728 | 0.981121874 | -4.760802138 |
| **A_55_P1982096** | Drap1 | 0.012043813 | -0.536385718 | 0.560473344 | 14.6121343 | 0.055003903 | 0.95808614 | 0.981152434 | -4.760804662 |
| **A_55_P1993719** | Tmem216 | -0.013525746 | -0.630431243 | 0.603379752 | 9.284885332 | -0.054915245 | 0.95815362 | 0.981152434 | -4.76080629 |
| **A_52_P653825** | Keg1 | 0.004589259 | -0.204776614 | 0.213955132 | 7.630134037 | 0.054901856 | 0.95816381 | 0.981152434 | -4.760806535 |
| **A_51_P449935** | Ftsj3 | 0.010991306 | -0.492238406 | 0.514221018 | 9.070269761 | 0.054705799 | 0.958313032 | 0.981253266 | -4.760810125 |
| **A_55_P2048656** | Atp5l | 0.010315762 | -0.466721065 | 0.487352589 | 14.66582614 | 0.05416263 | 0.958726458 | 0.981583993 | -4.760820004 |
| **A_55_P2107182** | Angptl8 | 0.007095895 | -0.32171077 | 0.335902559 | 8.384124361 | 0.054052637 | 0.958810179 | 0.981583993 | -4.760821992 |
| **A_55_P1980292** | Purb | 0.00761432 | -0.346179948 | 0.361408588 | 10.73310986 | 0.053905202 | 0.958922401 | 0.981583993 | -4.760824651 |
| **A_55_P1962344** | Trim21 | 0.00663369 | -0.301919187 | 0.315186568 | 9.017033429 | 0.053848788 | 0.958965341 | 0.981583993 | -4.760825667 |
| **A_55_P2186562** | Hars2 | 0.007079465 | -0.322360839 | 0.336519768 | 10.68541741 | 0.053823761 | 0.958984391 | 0.981583993 | -4.760826117 |
| **A_55_P2001558** | Zfp128 | -0.011965439 | -0.569074363 | 0.545143485 | 8.713838613 | -0.053794624 | 0.959006569 | 0.981583993 | -4.760826641 |
| **A_52_P23379** | Depdc7 | 0.004695892 | -0.2141254 | 0.223517185 | 7.96593031 | 0.053750052 | 0.959040496 | 0.981583993 | -4.760827441 |
| **A_52_P252322** | Pdcd11 | 0.005414229 | -0.247459488 | 0.258287946 | 8.190155194 | 0.05362697 | 0.959134182 | 0.981583993 | -4.760829649 |
| **A_51_P273326** | Sec31a | -0.011015524 | -0.553113246 | 0.531082197 | 11.82777491 | -0.054124051 | 0.959167001 | 0.981583993 | -4.731771113 |
| **A_51_P159612** | Hebp2 | 0.009876616 | -0.452193355 | 0.471946587 | 8.74316205 | 0.053536599 | 0.95920297 | 0.981583993 | -4.760831267 |
| **A_55_P1957918** | Asap2 | 0.005868412 | -0.268881012 | 0.280617837 | 9.666268785 | 0.053497586 | 0.959232666 | 0.981583993 | -4.760831965 |
| **A_55_P2112617** | Smad4 | 0.008994601 | -0.412248149 | 0.430237351 | 10.30557389 | 0.053481034 | 0.959245265 | 0.981583993 | -4.760832261 |
| **A_55_P2134800** | Cinp | -0.010962484 | -0.528240623 | 0.506315655 | 8.730458736 | -0.053080524 | 0.959550131 | 0.981843992 | -4.76083939 |
| **A_55_P2156340** | NA | -0.008933698 | -0.431071424 | 0.413204028 | 16.41055783 | -0.053006296 | 0.959606633 | 0.981849843 | -4.760840706 |
| **A_52_P251699** | Prodh2 | 0.004521555 | -0.210829685 | 0.219872796 | 7.738281881 | 0.052588506 | 0.959924661 | 0.982123268 | -4.760848076 |
| **A_51_P175580** | Trp53inp1 | -0.010794156 | -0.530733866 | 0.509145555 | 10.88935321 | -0.051997927 | 0.960374233 | 0.982514044 | -4.760858395 |
| **A_52_P216226** | Masp1 | -0.004348313 | -0.213980721 | 0.205284094 | 7.676167917 | -0.051953254 | 0.96040824 | 0.982514044 | -4.760859171 |
| **A_55_P2042773** | Ctdnep1 | -0.009499641 | -0.474220613 | 0.455221332 | 11.28680726 | -0.051199444 | 0.960982095 | 0.983049091 | -4.760872162 |
| **A_55_P2068027** | NA | -0.004336516 | -0.217020206 | 0.208347175 | 7.740674305 | -0.051068968 | 0.961081425 | 0.983081364 | -4.760874391 |
| **A_55_P2006713** | NA | 0.004481745 | -0.215709956 | 0.224673445 | 16.77516693 | 0.050979609 | 0.961149454 | 0.983081364 | -4.760875915 |
| **A_55_P1990678** | NA | 0.007905424 | -0.380661888 | 0.396472736 | 8.32469078 | 0.050957632 | 0.961166185 | 0.983081364 | -4.760876289 |
| **A_52_P337126** | Bves | -0.01975037 | -0.99194536 | 0.952444619 | 9.054397736 | -0.050882942 | 0.961223047 | 0.983087516 | -4.76087756 |
| **A_51_P484158** | Steap1 | -0.004322717 | -0.218670882 | 0.210025448 | 8.158126845 | -0.050511167 | 0.961506084 | 0.983324974 | -4.760883858 |
| **A_52_P24428** | Ankrd11 | 0.008155916 | -0.398700248 | 0.41501208 | 11.08428788 | 0.050209074 | 0.961736076 | 0.983508161 | -4.760888941 |
| **A_55_P1953866** | Sema4c | -0.010491755 | -0.535813594 | 0.514830083 | 14.57516371 | -0.05002338 | 0.961877452 | 0.983590785 | -4.760892051 |
| **A_66_P115116** | Wwtr1 | 0.016431758 | -0.807987669 | 0.840851184 | 11.72750243 | 0.04992133 | 0.961955147 | 0.983590785 | -4.760893755 |
| **A_55_P2005335** | Kcng2 | 0.005339926 | -0.2626777 | 0.273357552 | 7.776384708 | 0.049902489 | 0.961969492 | 0.983590785 | -4.760894069 |

| | | | | | | | | | |
|---|---|---|---|---|---|---|---|---|---|
| **A_55_P2117681** | Rnf115 | -0.011467777 | -0.588417821 | 0.565482268 | 9.936186675 | -0.04978418 | 0.962059567 | 0.983630865 | -4.76089604 |
| **A_55_P1972842** | Klf3 | 0.013773657 | -0.684330252 | 0.711877565 | 12.2499537 | 0.049417359 | 0.96233885 | 0.983802124 | -4.76090212 |
| **A_55_P1993762** | NA | 0.004337372 | -0.215896498 | 0.224571243 | 7.913752465 | 0.049327937 | 0.962406934 | 0.983802124 | -4.760903595 |
| **A_55_P1970017** | Zer1 | -0.004976623 | -0.258436912 | 0.248483665 | 12.07257053 | -0.049178491 | 0.962520719 | 0.983802124 | -4.760906055 |
| **A_55_P1980119** | Cptp | -0.00484794 | -0.251759774 | 0.242063893 | 10.09372294 | -0.049177416 | 0.962521538 | 0.983802124 | -4.760906072 |
| **A_52_P50478** | Rnf123 | -0.005325986 | -0.276648504 | 0.265996531 | 8.563547697 | -0.049165965 | 0.962530256 | 0.983802124 | -4.760906261 |
| **A_55_P2120682** | Golga7 | 0.004351018 | -0.217316097 | 0.226018133 | 10.19179723 | 0.049163179 | 0.962532377 | 0.983802124 | -4.760906306 |
| **A_55_P2129354** | Pilra | 0.0188251 | -0.951958208 | 0.989608407 | 8.718137927 | 0.04856969 | 0.962984256 | 0.984166076 | -4.760915998 |
| **A_55_P1970631** | Vps54 | 0.007301639 | -0.3692946 | 0.383897878 | 10.23320981 | 0.048561796 | 0.962990267 | 0.984166076 | -4.760916126 |
| **A_55_P2066518** | Akt2 | 0.006004216 | -0.30430432 | 0.316312752 | 12.71381438 | 0.048463297 | 0.963065265 | 0.984190699 | -4.760917723 |
| **A_51_P325165** | Il27 | -0.007075013 | -0.374526295 | 0.360376269 | 8.299875444 | -0.048225621 | 0.963246236 | 0.984323611 | -4.760921562 |
| **A_55_P2127659** | Mtmr2 | -0.004845178 | -0.257481926 | 0.24779157 | 8.511007791 | -0.04803564 | 0.963390893 | 0.984352526 | -4.760924618 |
| **A_52_P460537** | Lpar2 | -0.004797441 | -0.255192175 | 0.245597293 | 8.33747768 | -0.047988237 | 0.963426987 | 0.984352526 | -4.760925379 |
| **A_51_P340947** | Selenoo | 0.007396195 | -0.3789482 | 0.39374059 | 9.795458971 | 0.047949499 | 0.963456484 | 0.984352526 | -4.760926 |
| **A_51_P316553** | Kdr | 0.010556012 | -0.541973416 | 0.56308544 | 12.6725857 | 0.047851417 | 0.963531166 | 0.984352526 | -4.73683667 |
| **A_55_P2147842** | Zglp1 | -0.010415202 | -0.556670628 | 0.535840224 | 9.5067168 | -0.047755377 | 0.963604295 | 0.984352526 | -4.760929104 |
| **A_55_P2010301** | Serpina1c | -0.041882435 | -2.241373028 | 2.157608158 | 11.10732032 | -0.047693606 | 0.96365133 | 0.984352526 | -4.760930089 |
| **A_55_P2029902** | Gab3 | -0.01093873 | -0.585495045 | 0.563617585 | 11.67940452 | -0.047685312 | 0.963657646 | 0.984352526 | -4.760930221 |
| **A_55_P2019699** | Samhd1 | 0.020191951 | -1.041096482 | 1.081480384 | 11.75168333 | 0.047653543 | 0.963681836 | 0.984352526 | -4.760930727 |
| **A_51_P230904** | Copz2 | 0.005130544 | -0.265917617 | 0.276178704 | 9.800834579 | 0.047409708 | 0.963867505 | 0.984490166 | -4.7609346 |
| **A_55_P2186833** | NA | -0.015676493 | -0.845049473 | 0.813696486 | 16.10407194 | -0.047342304 | 0.963918831 | 0.98449058 | -4.760935668 |
| **A_52_P283628** | Rps6ka3 | -0.004291744 | -0.231691024 | 0.223107535 | 8.317547741 | -0.047271036 | 0.963973099 | 0.984493999 | -4.760936794 |
| **A_52_P686091** | Phf2 | 0.004511927 | -0.235098608 | 0.244122462 | 9.32409361 | 0.047163554 | 0.964054943 | 0.984525579 | -4.76093849 |
| **A_55_P1961094** | Tmem230 | -0.004351099 | -0.236124895 | 0.227422697 | 10.7369316 | -0.047020258 | 0.964164058 | 0.984585005 | -4.760940745 |
| **A_55_P2143693** | Sytl1 | 0.013521816 | -0.708501436 | 0.735545068 | 8.769108055 | 0.046906625 | 0.964250587 | 0.984586791 | -4.760942529 |
| **A_55_P1978013** | Obp1a | -0.006528283 | -0.355404441 | 0.342347876 | 8.205208351 | -0.046868171 | 0.964279869 | 0.984586791 | -4.760943131 |
| **A_51_P487869** | Cyp4f16 | -0.005942959 | -0.323883893 | 0.311997974 | 8.22797231 | -0.046817331 | 0.964318583 | 0.984586791 | -4.760943927 |
| **A_52_P508920** | Abca6 | 0.003913548 | -0.206362377 | 0.214189472 | 7.670576522 | 0.046615615 | 0.964472187 | 0.984691623 | -4.760947076 |
| **A_55_P2409306** | Vps35l | -0.008859832 | -0.488340826 | 0.470621161 | 8.371920702 | -0.046281183 | 0.964726857 | 0.984899622 | -4.760952268 |
| **A_55_P2074317** | Ap2b1 | 0.009067841 | -0.484050607 | 0.502186288 | 9.658910844 | 0.046057777 | 0.964896983 | 0.98498376 | -4.760955715 |
| **A_51_P229602** | Plxna2 | 0.010761041 | -0.574672092 | 0.596194175 | 11.25751437 | 0.046039156 | 0.964911163 | 0.98498376 | -4.760956001 |
| **A_66_P136788** | Ubald2 | 0.005919885 | -0.317563232 | 0.329403003 | 13.91045512 | 0.045836559 | 0.965065444 | 0.985043351 | -4.760959111 |
| **A_51_P379750** | Myoz1 | 0.00599328 | -0.321556626 | 0.333543186 | 7.92321913 | 0.04582869 | 0.965071437 | 0.985043351 | -4.760959232 |
| **A_66_P121745** | Arhgef28 | -0.012358421 | -0.689700742 | 0.664983901 | 8.560371125 | -0.045698833 | 0.965170326 | 0.985092282 | -4.760961218 |
| **A_55_P2113633** | Bzw1 | 0.00661069 | -0.357240548 | 0.370461928 | 11.11273643 | 0.045506483 | 0.965316807 | 0.985189777 | -4.76096415 |
| **A_51_P294113** | Chd2 | -0.004276262 | -0.241666266 | 0.233113742 | 8.666378097 | -0.045118249 | 0.965612465 | 0.985427629 | -4.760970029 |
| **A_52_P676798** | Gigyf1 | 0.006759302 | -0.368902807 | 0.382421411 | 8.971366695 | 0.0450666 | 0.965651798 | 0.985427629 | -4.760970807 |
| **A_51_P301964** | Arl6ip1 | 0.006188931 | -0.339473675 | 0.351851538 | 12.07633737 | 0.044844956 | 0.965820594 | 0.985508146 | -4.760974137 |
| **A_66_P112816** | B4galt6 | 0.004894358 | -0.268560538 | 0.278349253 | 8.489061536 | 0.044829133 | 0.965832644 | 0.985508146 | -4.760974374 |
| **A_55_P2072150** | Vpreb2 | -0.003917676 | -0.224225476 | 0.216390124 | 7.763230316 | -0.044539873 | 0.966052936 | 0.985608377 | -4.760978694 |
| **A_55_P2096043** | Acot11 | -0.004484091 | -0.256763791 | 0.247795609 | 8.336562403 | -0.044518699 | 0.966069061 | 0.985608377 | -4.760979009 |
| **A_55_P2246650** | 9330159N | -0.00473569 | -0.271460209 | 0.261988829 | 8.620537583 | -0.044470364 | 0.966105873 | 0.985608377 | -4.760979728 |
| **A_55_P2052623** | Pgap2 | -0.01111919 | -0.637911608 | 0.615673229 | 11.31427827 | -0.044432398 | 0.966134787 | 0.985608377 | -4.760980292 |
| **A_55_P1957584** | Gps2 | -0.00521941 | -0.300705182 | 0.290266361 | 9.686814974 | -0.044242045 | 0.966279757 | 0.985682371 | -4.760983113 |
| **A_55_P2106593** | NA | -0.004499463 | -0.259450401 | 0.250451474 | 9.391724489 | -0.044203277 | 0.966309282 | 0.985682371 | -4.760983686 |
| **A_55_P1978047** | Aven | 0.004034998 | -0.226219242 | 0.234289238 | 8.027822402 | 0.043892065 | 0.9665463 | 0.985821601 | -4.760988267 |
| **A_52_P358388** | Ppp4r2 | 0.006516648 | -0.36536736 | 0.378400656 | 8.814107695 | 0.043890156 | 0.966547754 | 0.985821601 | -4.760988295 |
| **A_51_P363801** | Pgpep1 | 0.005512018 | -0.310054101 | 0.321078137 | 8.742805955 | 0.043749248 | 0.96665507 | 0.985856993 | -4.760990359 |

| | | | | | | | | | |
|---|---|---|---|---|---|---|---|---|---|
| **A_52_P149391** | Usp40 | 0.007673442 | -0.432361587 | 0.44770847 | 9.134942589 | 0.043677049 | 0.966710058 | 0.985856993 | -4.760991413 |
| **A_51_P108020** | Zmynd15 | -0.004049794 | -0.236710261 | 0.228610672 | 7.961590333 | -0.043597415 | 0.966770708 | 0.985856993 | -4.760992575 |
| **A_55_P2428967** | Lrrc2 | -0.003919584 | -0.229456434 | 0.221617266 | 7.787730186 | -0.043528412 | 0.966823262 | 0.985856993 | -4.760993579 |
| **A_52_P883557** | Slc30a10 | 0.00380324 | -0.215132457 | 0.222738936 | 7.671513155 | 0.043509838 | 0.966837408 | 0.985856993 | -4.76099385 |
| **A_55_P1985591** | Rtl8a | 0.003974843 | -0.226208182 | 0.234157867 | 9.092540655 | 0.043251083 | 0.967034482 | 0.985958554 | -4.760997601 |
| **A_55_P2044197** | Vps29 | -0.004873676 | -0.287318966 | 0.277571615 | 11.3697483 | -0.043218792 | 0.967059076 | 0.985958554 | -4.760998068 |
| **A_52_P460836** | Lrrn4 | -0.010219509 | -0.60303007 | 0.582591052 | 8.307941756 | -0.043178191 | 0.967089998 | 0.985958554 | -4.760998654 |
| **A_55_P2030126** | Tfpt | 0.004585845 | -0.261962192 | 0.271133882 | 9.456640251 | 0.04309176 | 0.967155827 | 0.985973676 | -4.7609999 |
| **A_52_P496202** | Pcsk1n | -0.007051003 | -0.417796077 | 0.403694071 | 8.344092358 | -0.042996084 | 0.967228697 | 0.985995974 | -4.761001276 |
| **A_55_P2040529** | Mtfr1 | 0.00405588 | -0.233275361 | 0.241387121 | 8.854652782 | 0.042803621 | 0.967375285 | 0.986063053 | -4.761004036 |
| **A_55_P2080756** | Olfr704 | -0.003803786 | -0.226528628 | 0.218921056 | 7.944728858 | -0.042775763 | 0.967396503 | 0.986063053 | -4.761004434 |
| **A_55_P2076635** | Vps25 | -0.004308496 | -0.257576753 | 0.248959761 | 9.221482481 | -0.042608406 | 0.96752397 | 0.98614099 | -4.761006822 |
| **A_55_P2136935** | Rgs7bp | 0.005120117 | -0.297089625 | 0.307329858 | 8.035463479 | 0.04243476 | 0.967656228 | 0.986223802 | -4.76100929 |
| **A_52_P434052** | Wdr37 | -0.004976128 | -0.299401728 | 0.289449471 | 8.990083286 | -0.042331762 | 0.967734678 | 0.986227536 | -4.761010749 |
| **A_51_P409724** | Secisbp2 | 0.003606912 | -0.209985857 | 0.217199681 | 8.621402046 | 0.042296006 | 0.967761912 | 0.986227536 | -4.761011255 |
| **A_55_P2297485** | Ago2 | -0.00771562 | -0.465809544 | 0.450378305 | 11.35917659 | -0.042185832 | 0.967845828 | 0.986261068 | -4.76101281 |
| **A_51_P297336** | Chmp7 | -0.005865708 | -0.355728191 | 0.343996774 | 11.09621696 | -0.041992665 | 0.967992958 | 0.98635901 | -4.761015528 |
| **A_55_P1954461** | Ptdss1 | -0.003651088 | -0.222925702 | 0.215623526 | 8.374114292 | -0.041704632 | 0.968212348 | 0.986498415 | -4.761019557 |
| **A_55_P2177386** | Cradd | -0.008566794 | -0.524535189 | 0.507401601 | 9.651226017 | -0.041585849 | 0.968302823 | 0.986498415 | -4.76102121 |
| **A_51_P103757** | Kat8 | -0.005326312 | -0.326394545 | 0.315741921 | 11.27497897 | -0.041550821 | 0.968329504 | 0.986498415 | -4.75862297 |
| **A_55_P2229645** | NA | 0.011272507 | -0.669554369 | 0.692099383 | 8.777155021 | 0.041470013 | 0.968391055 | 0.986498415 | -4.761022818 |
| **A_51_P105068** | Lypd6b | 0.01174348 | -0.69754715 | 0.72103411 | 8.380449655 | 0.041468941 | 0.968391871 | 0.986498415 | -4.756130466 |
| **A_52_P177293** | Snx24 | 0.005842028 | -0.347831202 | 0.359515259 | 8.672328509 | 0.041372506 | 0.968465326 | 0.986498415 | -4.761024168 |
| **A_51_P264695** | Crym | 0.005074232 | -0.302327805 | 0.312476268 | 8.651176045 | 0.041344134 | 0.968486936 | 0.986498415 | -4.76102456 |
| **A_55_P2094378** | NA | 0.008046641 | -0.482526474 | 0.498619755 | 8.609829899 | 0.041082911 | 0.968685911 | 0.986517468 | -4.761028158 |
| **A_55_P2142580** | NA | -0.00524028 | -0.325371149 | 0.314890588 | 10.774812 | -0.040999383 | 0.968749535 | 0.986517468 | -4.761029304 |
| **A_55_P2061076** | Gm3219 | 0.003528649 | -0.212070622 | 0.21912792 | 7.849691288 | 0.040993171 | 0.968754267 | 0.986517468 | -4.761029389 |
| **A_52_P508317** | Erlec1 | -0.007850005 | -0.488065152 | 0.472365143 | 8.813272604 | -0.040943447 | 0.968792143 | 0.986517468 | -4.76103007 |
| **A_51_P370825** | Ccdc124 | 0.004476754 | -0.269907514 | 0.278861023 | 11.50465094 | 0.040865274 | 0.968851688 | 0.986517468 | -4.761031139 |
| **A_55_P1962736** | Sec23b | -0.003856765 | -0.240405845 | 0.232692314 | 11.48639631 | -0.040836856 | 0.968873335 | 0.986517468 | -4.761031527 |
| **A_51_P186487** | NA | 0.00379559 | -0.229253232 | 0.236844411 | 7.758651539 | 0.040792724 | 0.968906951 | 0.986517468 | -4.761032129 |
| **A_55_P2078765** | Kmt2e | -0.011377493 | -0.710109323 | 0.687354337 | 10.85178045 | -0.040783679 | 0.968913841 | 0.986517468 | -4.761032252 |
| **A_51_P494006** | Scaf8 | -0.005131573 | -0.32384799 | 0.313584843 | 11.29696612 | -0.04032705 | 0.969261669 | 0.986659976 | -4.761038442 |
| **A_55_P2172832** | Kcne1l | 0.003425458 | -0.209744846 | 0.216595761 | 7.941731069 | 0.040247808 | 0.969322031 | 0.986659976 | -4.761039509 |
| **A_55_P1998471** | S100a9 | -0.026297735 | -1.663724827 | 1.611129358 | 12.0813968 | -0.040225987 | 0.969338652 | 0.986659976 | -4.761039803 |
| **A_51_P419959** | Eaf1 | 0.007824746 | -0.479443103 | 0.495092595 | 9.529473167 | 0.040220997 | 0.969342453 | 0.986659976 | -4.76103987 |
| **A_52_P495372** | Twnk | -0.003514653 | -0.222387655 | 0.215358348 | 8.296286197 | -0.04021987 | 0.969343312 | 0.986659976 | -4.761039885 |
| **A_55_P1994519** | Tpd52l2 | 0.004915996 | -0.301391656 | 0.311223648 | 8.720866085 | 0.040197962 | 0.96936 | 0.986659976 | -4.761040179 |
| **A_55_P1972187** | Slc25a14 | 0.003648662 | -0.224574307 | 0.23187163 | 10.30788552 | 0.040042817 | 0.969478181 | 0.986728319 | -4.76104226 |
| **A_51_P325856** | Mcemp1 | -0.012513477 | -0.799831765 | 0.774804811 | 9.059061565 | -0.039808701 | 0.969656519 | 0.98685788 | -4.761045384 |
| **A_55_P2092622** | NA | -0.00440833 | -0.283398109 | 0.274581448 | 8.09811521 | -0.03957639 | 0.969833485 | 0.986964762 | -4.761048466 |
| **A_51_P470715** | Cish | -0.021394234 | -1.376725488 | 1.33393702 | 10.96163726 | -0.039536811 | 0.969863635 | 0.986964762 | -4.76104899 |
| **A_66_P121656** | Pcgf1 | 0.003748766 | -0.234944756 | 0.242442288 | 8.437981335 | 0.039336709 | 0.970016067 | 0.987058944 | -4.761051627 |
| **A_55_P2142218** | NA | -0.011103686 | -0.719101459 | 0.696894087 | 15.10810508 | -0.03928128 | 0.97005829 | 0.987058944 | -4.761052356 |
| **A_51_P189082** | Akr1c6 | 0.003257155 | -0.205315979 | 0.211830289 | 7.718947592 | 0.039113838 | 0.970185844 | 0.987116736 | -4.761054549 |
| **A_55_P2149363** | 6430548M | -0.003417133 | -0.222465027 | 0.21563076 | 8.581729612 | -0.039072678 | 0.970217199 | 0.987116736 | -4.761055087 |
| **A_55_P2128672** | Ikbip | -0.005350149 | -0.349992137 | 0.33929184 | 10.34557795 | -0.038881949 | 0.970362494 | 0.987204428 | -4.761057572 |
| **A_51_P230507** | Ell2 | 0.004470475 | -0.283924155 | 0.292865105 | 8.283595413 | 0.03882548 | 0.970405511 | 0.987204428 | -4.761058305 |

| | | | | | | | | | |
|---|---|---|---|---|---|---|---|---|---|
| **A_52_P267391** | Trim12a | 0.00988649 | -0.63127964 | 0.651052621 | 9.227854776 | 0.038620859 | 0.970561391 | 0.987293789 | -4.761060954 |
| **A_55_P2034345** | Terf2 | 0.003537577 | -0.226150142 | 0.233225295 | 10.50735186 | 0.038576109 | 0.970595482 | 0.987293789 | -4.761061531 |
| **A_55_P1966432** | Gstm1 | -0.005940201 | -0.393520649 | 0.381640247 | 9.428440665 | -0.038387479 | 0.97073918 | 0.987388011 | -4.761063958 |
| **A_52_P317653** | Car1 | 0.0035549 | -0.229229838 | 0.236339638 | 8.021133025 | 0.03824928 | 0.970844461 | 0.987408532 | -4.761065728 |
| **A_52_P438082** | Cebpg | -0.005878484 | -0.391483935 | 0.379726968 | 9.003983959 | -0.038183214 | 0.970894791 | 0.987408532 | -4.761066572 |
| **A_55_P2097244** | Bptf | 0.003797439 | -0.245451959 | 0.253046836 | 13.04720776 | 0.038159879 | 0.970912568 | 0.987408532 | -4.76106687 |
| **A_66_P132613** | C230057M | 0.004350858 | -0.283944508 | 0.292646224 | 9.018116518 | 0.037799633 | 0.971187011 | 0.987594953 | -4.761071443 |
| **A_55_P2127421** | NA | 0.003189653 | -0.208567896 | 0.214947202 | 7.895766412 | 0.037727224 | 0.971242174 | 0.987594953 | -4.761072357 |
| **A_55_P2002578** | Ephx1 | -0.008621299 | -0.581118194 | 0.563875596 | 11.08518338 | -0.03771811 | 0.971249118 | 0.987594953 | -4.761072472 |
| **A_55_P2084646** | Csrnp3 | -0.004436226 | -0.300248077 | 0.291375626 | 8.270759022 | -0.037561974 | 0.971368067 | 0.987630924 | -4.761074436 |
| **A_52_P392509** | Gin1 | -0.003808832 | -0.258303294 | 0.25068563 | 8.533451212 | -0.037485546 | 0.971426292 | 0.987630924 | -4.761075395 |
| **A_55_P2167922** | Dmd | 0.003519939 | -0.231765882 | 0.23880576 | 7.714273658 | 0.037470519 | 0.971437741 | 0.987630924 | -4.761075583 |
| **A_55_P1969645** | Sdcbp | -0.003447581 | -0.234387956 | 0.227492794 | 11.19117685 | -0.037390815 | 0.971498463 | 0.987640723 | -4.761076581 |
| **A_52_P473975** | Cpsf4 | 0.003979381 | -0.264049399 | 0.272008161 | 10.14978964 | 0.037186427 | 0.971654174 | 0.987747084 | -4.761079128 |
| **A_52_P298088** | Itgb1bp2 | -0.00610387 | -0.418640267 | 0.406432526 | 8.000239739 | -0.037058972 | 0.971751275 | 0.98776551 | -4.76108071 |
| **A_66_P130908** | Cyb5r4 | 0.005294874 | -0.353487297 | 0.364077045 | 9.335044403 | 0.036963673 | 0.971823879 | 0.98776551 | -4.761081889 |
| **A_55_P2007871** | Ptpdc1 | 0.007298288 | -0.487767661 | 0.502364237 | 10.03172409 | 0.036923962 | 0.971854133 | 0.98776551 | -4.761082379 |
| **A_51_P317542** | Slc24a2 | -0.003422494 | -0.235767086 | 0.228922099 | 7.769279938 | -0.036894397 | 0.971876658 | 0.98776551 | -4.761082744 |
| **A_55_P1973813** | NA | 0.003054644 | -0.205055624 | 0.211164912 | 7.684499167 | 0.036763545 | 0.971976348 | 0.987814903 | -4.761084355 |
| **A_55_P2074831** | Fam168a | -0.008247643 | -0.571505722 | 0.555010435 | 11.20828459 | -0.036675226 | 0.972043636 | 0.987831362 | -4.761085439 |
| **A_55_P2068289** | Slc17a2 | -0.003080231 | -0.214661031 | 0.208500569 | 7.657820937 | -0.036463416 | 0.972205007 | 0.987943425 | -4.761088028 |
| **A_55_P1999102** | Pi16 | 0.014259481 | -0.976099697 | 1.004618658 | 9.828665722 | 0.036062957 | 0.972510108 | 0.988201526 | -4.761092882 |
| **A_55_P2068701** | NA | 0.004661684 | -0.32267729 | 0.332000658 | 8.044534923 | 0.03566937 | 0.972809978 | 0.988454284 | -4.761097601 |
| **A_52_P504342** | Coa7 | 0.004457152 | -0.311957289 | 0.320871593 | 8.861578607 | 0.035281863 | 0.973105221 | 0.988668527 | -4.761102197 |
| **A_55_P2165299** | Mybpc1 | -0.00980422 | -0.708252823 | 0.688644383 | 8.241225584 | -0.035158386 | 0.9731993 | 0.988668527 | -4.761103651 |
| **A_55_P2010292** | Serpina1e | 0.018676373 | -1.312425216 | 1.349777961 | 9.257925746 | 0.035142421 | 0.973211463 | 0.988668527 | -4.761103838 |
| **A_55_P1992299** | Cep295 | -0.00495113 | -0.358011261 | 0.348109002 | 8.124308261 | -0.035124162 | 0.973225375 | 0.988668527 | -4.761104053 |
| **A_55_P2030046** | Barx2 | 0.010862375 | -0.767180096 | 0.788904847 | 9.015801516 | 0.034968084 | 0.973344294 | 0.98871085 | -4.761105881 |
| **A_51_P462271** | Acan | 0.003399349 | -0.240352475 | 0.247151172 | 7.837597716 | 0.034929967 | 0.973373336 | 0.98871085 | -4.761106326 |
| **A_55_P2148609** | NA | -0.004811574 | -0.350941407 | 0.34131826 | 8.649526272 | -0.03481757 | 0.973458974 | 0.98871085 | -4.761107637 |
| **A_51_P389664** | Scrt2 | -0.003640554 | -0.265655459 | 0.25837435 | 7.927210878 | -0.034801011 | 0.973471591 | 0.98871085 | -4.761107829 |
| **A_51_P503654** | Tmem205 | -0.006320338 | -0.464620671 | 0.451979995 | 9.31321885 | -0.034541443 | 0.973669362 | 0.988781613 | -4.761110838 |
| **A_51_P394585** | Snx22 | -0.003525378 | -0.259224624 | 0.252173868 | 8.064211154 | -0.034532385 | 0.973676264 | 0.988781613 | -4.761110942 |
| **A_52_P252425** | 2700062C | -0.003584907 | -0.264277659 | 0.257107844 | 9.328607186 | -0.034442867 | 0.973744471 | 0.988781613 | -4.761111975 |
| **A_55_P2132651** | Ccn4 | -0.006424936 | -0.474159571 | 0.4613097 | 8.417984633 | -0.034404844 | 0.973773442 | 0.988781613 | -4.761112412 |
| **A_55_P2037538** | NA | 0.004277367 | -0.307394248 | 0.315948981 | 9.49819254 | 0.03437396 | 0.973796974 | 0.988781613 | -4.761112767 |
| **A_55_P2133057** | Pdia2 | 0.00378295 | -0.275235698 | 0.282801599 | 7.918835295 | 0.033958442 | 0.974113575 | 0.989051143 | -4.761117513 |
| **A_55_P1995497** | Atad3a | -0.004611597 | -0.3458767 | 0.336653507 | 8.43375333 | -0.03384619 | 0.974199106 | 0.989086043 | -4.761118785 |
| **A_55_P2021490** | Gm16386 | -0.004942073 | -0.372152787 | 0.362268642 | 8.823985726 | -0.033708864 | 0.974303742 | 0.989140336 | -4.761120336 |
| **A_66_P111216** | 4632427E | 0.004687203 | -0.345064602 | 0.354439008 | 8.271475999 | 0.033566355 | 0.974412329 | 0.989198633 | -4.761121938 |
| **A_55_P2165544** | Aebp2 | -0.00518482 | -0.39398902 | 0.383619381 | 9.500863803 | -0.033400506 | 0.9745387 | 0.989274978 | -4.761123795 |
| **A_55_P1983358** | Gm4593 | -0.003603272 | -0.274931514 | 0.26772497 | 7.893805727 | -0.033262306 | 0.974644004 | 0.98932993 | -4.761125335 |
| **A_55_P2032192** | Tpm1 | -0.00505525 | -0.388414724 | 0.378304223 | 11.20353524 | -0.033028344 | 0.974822277 | 0.989458941 | -4.761127927 |
| **A_52_P53470** | NA | -0.002955502 | -0.228576985 | 0.222665982 | 8.015005065 | -0.032809614 | 0.974988945 | 0.98957616 | -4.761130334 |
| **A_51_P469789** | Agxt | 0.002728972 | -0.206465726 | 0.21192367 | 7.688637381 | 0.032673733 | 0.975092485 | 0.989617908 | -4.761131822 |
| **A_51_P120398** | Asb17 | 0.002882622 | -0.218445656 | 0.224210899 | 7.678392582 | 0.032621287 | 0.975132448 | 0.989617908 | -4.745184594 |
| **A_66_P121369** | Gars | 0.006711432 | -0.518868573 | 0.532291438 | 14.24143884 | 0.031983555 | 0.9756184 | 0.99005911 | -4.761139281 |
| **A_51_P419246** | Mirt1 | -0.003356044 | -0.266963328 | 0.26025124 | 7.820850497 | -0.031887512 | 0.975691586 | 0.990074258 | -4.761140306 |

| | | | | | | | | | |
|---|---|---|---|---|---|---|---|---|---|
| **A_52_P114722** | Ptpn6 | 0.008251285 | -0.641042565 | 0.657545135 | 9.79112599 | 0.031829561 | 0.975735746 | 0.990074258 | -4.761140923 |
| **A_55_P2131964** | Pdia2 | 0.002999886 | -0.233967425 | 0.239967196 | 7.837863567 | 0.031707835 | 0.975828503 | 0.990116414 | -4.761142216 |
| **A_55_P1952202** | NA | 0.003188516 | -0.252462583 | 0.258839615 | 8.583167188 | 0.031238579 | 0.976186087 | 0.990427257 | -4.761147154 |
| **A_55_P2151672** | NA | -0.004209908 | -0.34254553 | 0.334125714 | 8.868948435 | -0.031165582 | 0.976241713 | 0.99043172 | -4.761147915 |
| **A_51_P475613** | Fahd2a | 0.00337682 | -0.269761947 | 0.276515588 | 9.742269613 | 0.030965277 | 0.976394353 | 0.990497115 | -4.761149995 |
| **A_55_P2011867** | Kif19a | -0.002785917 | -0.228481396 | 0.222909562 | 8.182866418 | -0.030916877 | 0.976431236 | 0.990497115 | -4.761150496 |
| **A_55_P2085445** | Acsm1 | -0.0025435 | -0.210194034 | 0.205107033 | 7.690566071 | -0.030679559 | 0.976612082 | 0.990497115 | -4.76115294 |
| **A_55_P2203141** | 2900011F | 0.005676086 | -0.458586132 | 0.469938303 | 8.911859672 | 0.030622168 | 0.976655816 | 0.990497115 | -4.761153528 |
| **A_55_P2121185** | Omp | 0.002600347 | -0.210157305 | 0.215358 | 7.801044678 | 0.030612338 | 0.976663308 | 0.990497115 | -4.761153628 |
| **A_55_P2046499** | Vmn1r70 | -0.00430739 | -0.356750684 | 0.348135903 | 8.280307121 | -0.030610846 | 0.976664444 | 0.990497115 | -4.761153644 |
| **A_51_P246903** | Rps27l | -0.005952726 | -0.493029347 | 0.481123895 | 14.45461519 | -0.030610396 | 0.976664788 | 0.990497115 | -4.761153648 |
| **A_55_P2163873** | Uox | 0.002572038 | -0.208933404 | 0.214077481 | 7.671224173 | 0.03045834 | 0.976780662 | 0.990562671 | -4.7611552 |
| **A_55_P1994117** | Ifnl3 | -0.002570558 | -0.216041278 | 0.210900161 | 7.983765294 | -0.030160567 | 0.977007582 | 0.990740826 | -4.761158218 |
| **A_55_P1961458** | Speer1 | 0.00396194 | -0.328251331 | 0.336175211 | 8.093763912 | 0.029870415 | 0.977228696 | 0.990913075 | -4.761161129 |
| **A_55_P2093696** | NA | -0.00254091 | -0.216988058 | 0.211906237 | 7.949373851 | -0.029676961 | 0.977376121 | 0.990960058 | -4.761163055 |
| **A_55_P2113539** | NA | 0.002538602 | -0.211727125 | 0.216804328 | 7.943037724 | 0.029675101 | 0.977377539 | 0.990960058 | -4.761163073 |
| **A_51_P143805** | Tmem42 | 0.008174366 | -0.684161504 | 0.700510235 | 11.23074413 | 0.029572468 | 0.977455753 | 0.99098739 | -4.76116409 |
| **A_51_P402994** | Ddx3y | 0.033827426 | -2.855107822 | 2.922762674 | 9.512194668 | 0.029327935 | 0.977642107 | 0.99112435 | -4.761166498 |
| **A_55_P2166069** | 2310079G | 0.002680515 | -0.227994384 | 0.233355414 | 7.868702276 | 0.02910504 | 0.977811972 | 0.991244582 | -4.761168675 |
| **A_55_P2013665** | Mybpc2 | -0.004177253 | -0.365173393 | 0.356818887 | 7.946466648 | -0.028982681 | 0.97790522 | 0.991287135 | -4.761169863 |
| **A_55_P2000543** | Cd209f | 0.002407066 | -0.206925174 | 0.211739307 | 7.686101841 | 0.028800651 | 0.978043944 | 0.99137578 | -4.761171621 |
| **A_51_P307082** | Hey2 | 0.002800734 | -0.241856038 | 0.247457506 | 7.779735892 | 0.02867246 | 0.978141639 | 0.991422828 | -4.761172853 |
| **A_55_P2012979** | Itgb2l | -0.002533151 | -0.224736918 | 0.219670616 | 7.687922426 | -0.028553545 | 0.978232264 | 0.991451503 | -4.76117399 |
| **A_55_P1998115** | Klkb1 | 0.002410473 | -0.209423805 | 0.214244751 | 7.744055858 | 0.028500761 | 0.97827249 | 0.991451503 | -4.761174494 |
| **A_55_P2168587** | Cdk9 | -0.003154984 | -0.281167691 | 0.274857722 | 8.696948336 | -0.02842385 | 0.978331105 | 0.991458936 | -4.761175226 |
| **A_51_P138859** | Tmem68 | 0.002759722 | -0.243260283 | 0.248779727 | 10.29271265 | 0.028096049 | 0.978580926 | 0.991655179 | -4.761178323 |
| **A_52_P642453** | Obscn | -0.002748801 | -0.248355373 | 0.242857771 | 7.728562978 | -0.028031972 | 0.97862976 | 0.991655179 | -4.761178924 |
| **A_55_P2054913** | Olfr406 | -0.002396125 | -0.217278503 | 0.212486253 | 7.76133371 | -0.027929234 | 0.978708058 | 0.991655179 | -4.761179885 |
| **A_55_P2004168** | NA | 0.002726963 | -0.242076027 | 0.247529954 | 9.829274034 | 0.027900558 | 0.978729912 | 0.991655179 | -4.761180153 |
| **A_55_P2215880** | Ephb2 | -0.002615345 | -0.239776032 | 0.234545342 | 8.064623243 | -0.027620821 | 0.978943106 | 0.991819212 | -4.761182749 |
| **A_55_P1952517** | Sult2a1 | -0.002292625 | -0.211396961 | 0.206811711 | 7.654117447 | -0.02746125 | 0.979064721 | 0.991890448 | -4.761184218 |
| **A_52_P520564** | Wasl | -0.004498923 | -0.418613812 | 0.409615966 | 8.799889149 | -0.027210598 | 0.979255751 | 0.992031998 | -4.761186509 |
| **A_55_P2116744** | Xirp1 | -0.002778197 | -0.259986408 | 0.254430013 | 7.831059303 | -0.027053818 | 0.979375238 | 0.992101062 | -4.761187931 |
| **A_55_P1973886** | Socs4 | -0.003467152 | -0.325887521 | 0.318953216 | 8.398176171 | -0.026933997 | 0.97946656 | 0.992105258 | -4.761189013 |
| **A_51_P340829** | AA986860 | -0.011696038 | -1.100163091 | 1.076771015 | 10.96881141 | -0.026913726 | 0.979482009 | 0.992105258 | -4.761189195 |
| **A_55_P1983016** | NA | -0.003142041 | -0.298639655 | 0.292355572 | 8.486992593 | -0.026632276 | 0.979696515 | 0.992270545 | -4.761191715 |
| **A_51_P493037** | Dclre1a | 0.002663313 | -0.249495859 | 0.254822484 | 8.157071729 | 0.02645439 | 0.979832091 | 0.992324288 | -4.761193293 |
| **A_55_P2020636** | NA | 0.002224595 | -0.208607892 | 0.213057082 | 7.67975714 | 0.026427969 | 0.979852228 | 0.992324288 | -4.761193527 |
| **A_55_P1983314** | Crat | 0.002211447 | -0.207919059 | 0.212341952 | 7.730357862 | 0.026359532 | 0.979904388 | 0.992325133 | -4.761194131 |
| **A_55_P1990121** | Aqp5 | 0.03618755 | -3.430599613 | 3.502974713 | 10.38370439 | 0.026144621 | 0.980068185 | 0.992439024 | -4.761196017 |
| **A_55_P2048660** | NA | 0.006958432 | -0.661466299 | 0.675383163 | 11.20093759 | 0.026074094 | 0.980121938 | 0.992441476 | -4.761196633 |
| **A_51_P343517** | Ly6d | -0.015315574 | -1.498974536 | 1.468343387 | 8.563240328 | -0.025855298 | 0.980288697 | 0.992558348 | -4.761198532 |
| **A_52_P249588** | Cnot11 | -0.003646688 | -0.360299848 | 0.353006472 | 8.841729522 | -0.025609606 | 0.980475956 | 0.992585082 | -4.761200646 |
| **A_55_P2101246** | Aldh6a1 | -0.002922526 | -0.290200077 | 0.284355025 | 7.99936363 | -0.025480447 | 0.980574399 | 0.992585082 | -4.761201749 |
| **A_55_P1960924** | Egln2 | 0.0028639 | -0.278848178 | 0.284575978 | 13.55221409 | 0.025462598 | 0.980588002 | 0.992585082 | -4.761201901 |
| **A_55_P2107785** | Gm13105 | -0.008623511 | -0.85840937 | 0.841162348 | 9.651156195 | -0.02541704 | 0.980622726 | 0.992585082 | -4.761202289 |
| **A_55_P2020592** | Trim3 | 0.003894358 | -0.380677582 | 0.388466299 | 8.407301726 | 0.025363471 | 0.980663555 | 0.992585082 | -4.761202743 |
| **A_55_P2183854** | Sctr | 0.002810742 | -0.275024863 | 0.280646347 | 7.954087871 | 0.025338651 | 0.980682473 | 0.992585082 | -4.761202954 |

| | | | | | | | | | |
|---|---|---|---|---|---|---|---|---|---|
| **A_55_P2030805** | Actr10 | 0.002209964 | -0.216608528 | 0.221028457 | 13.24098519 | 0.025295983 | 0.980714993 | 0.992585082 | -4.761203315 |
| **A_52_P468041** | Alkbh5 | -0.003531337 | -0.353381181 | 0.346318507 | 10.43034134 | -0.025281789 | 0.980725811 | 0.992585082 | -4.761203435 |
| **A_55_P2097309** | Nudt4 | -0.003422454 | -0.347528452 | 0.340683543 | 10.83481125 | -0.024911263 | 0.981008221 | 0.992818936 | -4.761206544 |
| **A_55_P2100864** | Tma7 | 0.002454807 | -0.247829643 | 0.252739258 | 14.44694143 | 0.024565969 | 0.981271403 | 0.992996748 | -4.7612094 |
| **A_51_P286748** | Frzb | 0.00216577 | -0.219046205 | 0.223377744 | 7.737016277 | 0.024521892 | 0.981304998 | 0.992996748 | -4.761209762 |
| **A_55_P2083725** | NA | 0.005523748 | -0.559670432 | 0.570717928 | 8.289332202 | 0.024478596 | 0.981337998 | 0.992996748 | -4.761210117 |
| **A_52_P63905** | Ddc | -0.00209792 | -0.217524101 | 0.213328262 | 7.773213747 | -0.024391623 | 0.981404289 | 0.993011855 | -4.761210827 |
| **A_52_P347591** | Ndufa7 | 0.005318016 | -0.544173806 | 0.554809839 | 15.04502356 | 0.024240344 | 0.981519594 | 0.993070155 | -4.761212057 |
| **A_55_P2088416** | Lamtor5 | 0.003268475 | -0.335276736 | 0.341813685 | 12.81014357 | 0.024181252 | 0.981564635 | 0.993070155 | -4.761212535 |
| **A_51_P215475** | Ptprb | -0.006427663 | -0.675276342 | 0.662421016 | 12.15512866 | -0.024069972 | 0.981649454 | 0.993104 | -4.761213433 |
| **A_55_P2181542** | Sult1d1 | -0.001976772 | -0.209874071 | 0.205920527 | 7.660830038 | -0.02381541 | 0.981843483 | 0.993197369 | -4.761215471 |
| **A_55_P2079520** | Mpp1 | -0.003962935 | -0.420768244 | 0.412842374 | 9.72837365 | -0.023814093 | 0.981844487 | 0.993197369 | -4.761215481 |
| **A_55_P2094706** | Sox17 | 0.005533053 | -0.580758517 | 0.591824623 | 10.17093684 | 0.023637499 | 0.98197909 | 0.993272289 | -4.761216882 |
| **A_55_P1957633** | Gpsm3 | 0.009172764 | -0.965071034 | 0.983416561 | 11.26383233 | 0.023582123 | 0.982021299 | 0.993272289 | -4.761217319 |
| **A_51_P320980** | Sfxn4 | 0.002532129 | -0.267365869 | 0.272430127 | 8.263442607 | 0.023498302 | 0.982085189 | 0.993284842 | -4.761217979 |
| **A_55_P2416494** | 8430426J | 0.01282397 | -1.357998963 | 1.383646904 | 9.771531529 | 0.023431036 | 0.982136461 | 0.993284842 | -4.761218507 |
| **A_51_P464588** | Dnajc28 | -0.001934483 | -0.213635663 | 0.209766697 | 7.851398308 | -0.022887162 | 0.982551017 | 0.993626407 | -4.761222718 |
| **A_52_P587831** | Ddx56 | 0.003217821 | -0.349450941 | 0.355886582 | 9.881351406 | 0.022853104 | 0.982576978 | 0.993626407 | -4.761222979 |
| **A_55_P2062851** | 7-Mar | 0.00423592 | -0.462013555 | 0.470485396 | 11.70235111 | 0.022755155 | 0.982651638 | 0.993649936 | -4.761223726 |
| **A_55_P1958697** | Rbm33 | -0.002881327 | -0.322187548 | 0.316424893 | 10.36974546 | -0.022601412 | 0.982768826 | 0.993716463 | -4.761224891 |
| **A_55_P2161347** | Acmsd | -0.001871573 | -0.210093958 | 0.206350811 | 7.685471 | -0.022512813 | 0.98283636 | 0.993732778 | -4.76122556 |
| **A_55_P2174576** | Nova1 | 0.001992333 | -0.223138142 | 0.227122807 | 7.900154001 | 0.022165522 | 0.983101081 | 0.993831108 | -4.761228154 |
| **A_51_P205740** | Ube3b | -0.002803358 | -0.320085514 | 0.314478797 | 12.35005667 | -0.022130097 | 0.983128084 | 0.993831108 | -4.761228416 |
| **A_52_P418734** | Rnf220 | -0.003638684 | -0.418629519 | 0.41135215 | 12.71120715 | -0.021961209 | 0.983256819 | 0.993831108 | -4.761229661 |
| **A_55_P2304507** | Noxa1 | -0.008197628 | -0.943249258 | 0.926854002 | 8.598725703 | -0.021958515 | 0.983258873 | 0.993831108 | -4.761229681 |
| **A_55_P1966189** | Il4ra | 0.001934691 | -0.218790296 | 0.222659678 | 7.919305181 | 0.02195384 | 0.983262436 | 0.993831108 | -4.761229715 |
| **A_55_P2070174** | NA | -0.005680369 | -0.656800555 | 0.645439818 | 8.81608481 | -0.021850717 | 0.983341042 | 0.993831108 | -4.76123047 |
| **A_55_P2145496** | Wdr45 | -0.009691033 | -1.120782055 | 1.101399988 | 12.80331187 | -0.021845943 | 0.983344681 | 0.993831108 | -4.761230505 |
| **A_66_P102628** | 1700016J | -0.004915953 | -0.569968457 | 0.560136551 | 8.38899619 | -0.021790604 | 0.983386864 | 0.993831108 | -4.761230909 |
| **A_51_P130427** | Psmd3 | 0.003827636 | -0.437589034 | 0.445244305 | 14.10797077 | 0.021718621 | 0.983441733 | 0.993831108 | -4.761231433 |
| **A_55_P2051840** | NA | -0.003122037 | -0.364024387 | 0.357780313 | 9.164577698 | -0.021666997 | 0.983481084 | 0.993831108 | -4.761231807 |
| **A_51_P139184** | Fank1 | -0.005242209 | -0.611892765 | 0.601408346 | 8.343732822 | -0.02164343 | 0.983499048 | 0.993831108 | -4.761231978 |
| **A_52_P232314** | NA | 0.005333918 | -0.613894266 | 0.624562101 | 8.782850896 | 0.021574758 | 0.983551394 | 0.99383206 | -4.761232474 |
| **A_55_P2023537** | NA | 0.001839174 | -0.213568584 | 0.217246931 | 7.808221927 | 0.021385125 | 0.983695944 | 0.993926176 | -4.761233836 |
| **A_55_P2033660** | Crls1 | 0.002399787 | -0.280376162 | 0.285175736 | 9.867073465 | 0.021255952 | 0.983794408 | 0.993930129 | -4.761234757 |
| **A_52_P69707** | Ergic1 | -0.003355337 | -0.398929102 | 0.392218429 | 8.790546486 | -0.021245109 | 0.983802673 | 0.993930129 | -4.761234834 |
| **A_55_P2033116** | Cdcp2 | 0.001733269 | -0.205899915 | 0.209366454 | 7.666206329 | 0.020908342 | 0.984059381 | 0.994134354 | -4.761237207 |
| **A_55_P2046812** | Sparc | 0.003378403 | -0.402559561 | 0.409316367 | 17.23591021 | 0.020845013 | 0.984107655 | 0.994134354 | -4.761237649 |
| **A_51_P516850** | Rbbp5 | 0.001898396 | -0.227061244 | 0.230858037 | 8.387863888 | 0.020767215 | 0.984166959 | 0.994142319 | -4.76123819 |
| **A_55_P1957855** | NA | 0.008036364 | -0.975986938 | 0.992059666 | 10.40940801 | 0.020455239 | 0.984404771 | 0.994308342 | -4.76124034 |
| **A_51_P195875** | Notch4 | 0.005084365 | -0.618653629 | 0.62882236 | 10.19666657 | 0.020416669 | 0.984434172 | 0.994308342 | -4.761240604 |
| **A_55_P2131433** | NA | -0.003712892 | -0.461397243 | 0.45397146 | 8.388138751 | -0.02031873 | 0.98450883 | 0.994331803 | -4.761241271 |
| **A_51_P407904** | Trappc4 | -0.003512017 | -0.439265503 | 0.432241469 | 12.9727096 | -0.020186735 | 0.984609447 | 0.99438148 | -4.761242165 |
| **A_51_P408363** | Cfp | -0.007965007 | -1.003709229 | 0.987779215 | 9.862355887 | -0.020034971 | 0.984725135 | 0.994446371 | -4.761243185 |
| **A_55_P2150108** | Pgm2l1 | -0.003374001 | -0.428859411 | 0.422111409 | 8.8554564 | -0.019861446 | 0.984857412 | 0.994528006 | -4.761244343 |
| **A_52_P82765** | NA | -0.004002613 | -0.51122914 | 0.503223915 | 8.946004001 | -0.019764778 | 0.984931101 | 0.994550473 | -4.761244983 |
| **A_55_P2001078** | Mmel1 | -0.001637331 | -0.211546648 | 0.208271985 | 7.691834128 | -0.019536879 | 0.985104828 | 0.994656725 | -4.761246481 |
| **A_51_P505229** | Gtpbp8 | 0.002436953 | -0.311746992 | 0.316620898 | 8.854901314 | 0.019427348 | 0.985188323 | 0.994656725 | -4.761247194 |

| | | | | | | | | | |
|---|---|---|---|---|---|---|---|---|---|
| **A_51_P334318** | Chp2 | -0.002793054 | -0.362944235 | 0.357358126 | 7.915679983 | -0.019424279 | 0.985190663 | 0.994656725 | -4.761247214 |
| **A_55_P2158102** | Camp | 0.006121077 | -0.794964344 | 0.807206498 | 8.608343656 | 0.019138124 | 0.9854088 | 0.994781746 | -4.761249059 |
| **A_55_P2152246** | Rpl19 | -0.002135268 | -0.282008434 | 0.277737898 | 17.05738717 | -0.019109154 | 0.985430883 | 0.994781746 | -4.761249244 |
| **A_51_P199624** | Sfswap | -0.001762591 | -0.233392064 | 0.229866883 | 8.70059675 | -0.019059347 | 0.985468852 | 0.994781746 | -4.761249562 |
| **A_52_P462350** | Dr1 | -0.003138808 | -0.423910105 | 0.417632489 | 9.919220393 | -0.018683963 | 0.98575501 | 0.994782075 | -4.761251931 |
| **A_55_P2046867** | Gata5 | 0.004614981 | -0.616317147 | 0.62554711 | 9.964889339 | 0.018615563 | 0.985807153 | 0.994782075 | -4.761252358 |
| **A_55_P1963389** | NA | 0.002740202 | -0.366478591 | 0.371958995 | 10.48897069 | 0.018588702 | 0.985827629 | 0.994782075 | -4.761252525 |
| **A_55_P2110512** | Lmo7 | 0.008778571 | -1.174659393 | 1.192216535 | 10.43058435 | 0.018579272 | 0.985834818 | 0.994782075 | -4.761252583 |
| **A_55_P2024021** | Atp5l | -0.003813684 | -0.518894439 | 0.511267071 | 15.08134316 | -0.018544695 | 0.985861177 | 0.994782075 | -4.761252798 |
| **A_55_P1978502** | H2-Q1 | -0.001706154 | -0.232175936 | 0.228763627 | 8.096823192 | -0.018541918 | 0.985863293 | 0.994782075 | -4.761252815 |
| **A_55_P2186282** | Agxt2 | 0.00154544 | -0.207884803 | 0.210975683 | 7.702330049 | 0.018482606 | 0.985908508 | 0.994782075 | -4.761253182 |
| **A_55_P2116853** | Brpf1 | 0.003093938 | -0.416380628 | 0.422568504 | 11.30729649 | 0.018473803 | 0.985915218 | 0.994782075 | -4.761253237 |
| **A_55_P2127179** | NA | -0.001795939 | -0.245673435 | 0.242081558 | 7.789654824 | -0.018444636 | 0.985937453 | 0.994782075 | -4.761253417 |
| **A_55_P2054708** | NA | -0.003503249 | -0.480792965 | 0.473786467 | 8.413754783 | -0.018383967 | 0.985983702 | 0.994782075 | -4.76125379 |
| **A_55_P1963206** | NA | -0.002110889 | -0.290795297 | 0.28657352 | 8.202145176 | -0.018314384 | 0.986036747 | 0.994783681 | -4.761254217 |
| **A_55_P1964724** | NA | 0.001871444 | -0.255125228 | 0.258868117 | 8.471354497 | 0.018238946 | 0.986094254 | 0.994789789 | -4.761254678 |
| **A_55_P2120303** | NA | 0.002354942 | -0.33229343 | 0.337003314 | 8.11061307 | 0.017625514 | 0.986561889 | 0.995038828 | -4.761258356 |
| **A_51_P324838** | Evpl | 0.003229141 | -0.45576943 | 0.462227712 | 11.22028546 | 0.017620814 | 0.986565472 | 0.995038828 | -4.761258384 |
| **A_51_P235687** | Alox5ap | -0.006874286 | -0.989909433 | 0.976160862 | 9.763198122 | -0.017514949 | 0.986646177 | 0.995038828 | -4.761259006 |
| **A_51_P398191** | Auts2 | 0.002245723 | -0.319198521 | 0.323689966 | 9.142754069 | 0.017498499 | 0.986658717 | 0.995038828 | -4.761259102 |
| **A_55_P2071496** | Fcf1 | 0.001792723 | -0.256281263 | 0.259866709 | 12.20087333 | 0.017398803 | 0.986734719 | 0.995038828 | -4.761259684 |
| **A_55_P1978191** | C2cd4c | 0.001865168 | -0.266700171 | 0.270430508 | 8.01723942 | 0.01739476 | 0.986737801 | 0.995038828 | -4.761259708 |
| **A_51_P215559** | Gstp3 | 0.001457977 | -0.208712713 | 0.211628666 | 7.723744602 | 0.017375159 | 0.986752743 | 0.995038828 | -4.761259821 |
| **A_55_P2051124** | Scaf1 | 0.002619923 | -0.376329665 | 0.381569511 | 13.31820305 | 0.017316395 | 0.986797541 | 0.995038828 | -4.761260162 |
| **A_55_P2097126** | Cnep1r1 | 0.00295459 | -0.424994531 | 0.430903711 | 10.2011315 | 0.017292407 | 0.986815828 | 0.995038828 | -4.761260301 |
| **A_51_P295582** | Cutc | 0.001588044 | -0.229126765 | 0.232302853 | 8.225636671 | 0.017240008 | 0.986855774 | 0.995038828 | -4.761260604 |
| **A_51_P211506** | Muc20 | 0.002848399 | -0.413905196 | 0.419601995 | 8.268010237 | 0.017118742 | 0.986948219 | 0.995080146 | -4.7612613 |
| **A_52_P365768** | Nsun2 | -0.003146918 | -0.469698082 | 0.463404246 | 11.52688334 | -0.016894155 | 0.987119431 | 0.995134067 | -4.761262577 |
| **A_55_P2103837** | Gbp4 | -0.006760358 | -1.015514977 | 1.001994261 | 8.399997624 | -0.016785509 | 0.987202256 | 0.995134067 | -4.761263189 |
| **A_55_P2104572** | Tmem268 | 0.002456584 | -0.366647479 | 0.371560648 | 9.26929811 | 0.016669907 | 0.987290385 | 0.995134067 | -4.761263836 |
| **A_51_P303725** | Tmem38a | 0.004418229 | -0.661777296 | 0.670613755 | 9.848400357 | 0.016611042 | 0.98733526 | 0.995134067 | -4.761264163 |
| **A_55_P1960491** | Snta1 | -0.004651888 | -0.707781998 | 0.698478223 | 11.06240825 | -0.016570815 | 0.987365927 | 0.995134067 | -4.761264387 |
| **A_66_P124146** | Ttc16 | 0.001886404 | -0.283889571 | 0.287662379 | 8.731478984 | 0.016533294 | 0.987394532 | 0.995134067 | -4.761264594 |
| **A_55_P2024515** | Crtc2 | -0.003436941 | -0.524563241 | 0.517689359 | 10.00655803 | -0.016518836 | 0.987405554 | 0.995134067 | -4.761264674 |
| **A_51_P214487** | 2-Sep | -0.00150284 | -0.229514266 | 0.226508586 | 14.06069902 | -0.016508458 | 0.987413465 | 0.995134067 | -4.761264731 |
| **A_52_P109304** | Tbl2 | 0.002978078 | -0.45477442 | 0.460730576 | 8.762848748 | 0.016295049 | 0.987576157 | 0.995246152 | -4.761265902 |
| **A_52_P448466** | BC00400 | -0.001431437 | -0.225850724 | 0.222987851 | 8.191383242 | -0.015975787 | 0.987819547 | 0.995439546 | -4.761267624 |
| **A_55_P2155921** | Hace1 | 0.001363995 | -0.218509114 | 0.221237104 | 8.186545081 | 0.015537851 | 0.988153411 | 0.995635252 | -4.761269932 |
| **A_55_P2110037** | Akap7 | -0.00272615 | -0.44247035 | 0.437018049 | 10.0228663 | -0.015527446 | 0.988161344 | 0.995635252 | -4.761269986 |
| **A_55_P2010931** | Cnn2 | -0.003671584 | -0.597708476 | 0.590365308 | 10.85901578 | -0.015480693 | 0.988196987 | 0.995635252 | -4.761270228 |
| **A_55_P1976839** | Spata5 | -0.002651912 | -0.432541838 | 0.427238014 | 9.093774972 | -0.015450845 | 0.988219742 | 0.995635252 | -4.761270383 |
| **A_51_P346668** | Irf5 | -0.002431427 | -0.407309314 | 0.402446461 | 8.040397886 | -0.015041371 | 0.98853191 | 0.995884798 | -4.76127247 |
| **A_51_P468762** | Alkbh6 | -0.002540958 | -0.431260663 | 0.426178746 | 10.91567193 | -0.014844802 | 0.988681769 | 0.995884798 | -4.761273453 |
| **A_51_P468418** | Ice2 | -0.00206067 | -0.349932848 | 0.345811509 | 8.606841237 | -0.014836758 | 0.988687901 | 0.995884798 | -4.761273492 |
| **A_55_P1983544** | Tmed5 | 0.002960032 | -0.498807 | 0.504727065 | 9.759942488 | 0.014775585 | 0.988734538 | 0.995884798 | -4.761273795 |
| **A_52_P632691** | Zfp395 | -0.001619593 | -0.290859397 | 0.287620211 | 9.112539065 | -0.01491456 | 0.988741747 | 0.995884798 | -4.732167912 |
| **A_55_P2000454** | Tspan18 | -0.00289639 | -0.49571041 | 0.489917629 | 9.546411887 | -0.014720561 | 0.988776487 | 0.995884798 | -4.761274067 |
| **A_51_P218653** | Jph2 | -0.001255376 | -0.216406144 | 0.213895391 | 7.718307993 | -0.014614414 | 0.98885741 | 0.995907414 | -4.761274587 |

| | | | | | | | | | |
|---|---|---|---|---|---|---|---|---|---|
| **A_55_P2361437** | Pitpnc1 | -0.001675616 | -0.292150768 | 0.288799536 | 8.61716359 | -0.01444827 | 0.988984075 | 0.995907414 | -4.761275395 |
| **A_55_P2039105** | Rbck1 | 0.002906494 | -0.501139275 | 0.506952262 | 11.42114359 | 0.014442745 | 0.988988287 | 0.995907414 | -4.761275421 |
| **A_55_P2112737** | Actb | 0.001925223 | -0.333956924 | 0.337807369 | 17.65792651 | 0.014356362 | 0.989054143 | 0.995907414 | -4.761275837 |
| **A_55_P2007601** | Sftpd | -0.019080151 | -3.374486502 | 3.3363262 | 12.47056415 | -0.014242527 | 0.989140929 | 0.995907414 | -4.761276382 |
| **A_55_P1957954** | Rsu1 | 0.003133581 | -0.550263771 | 0.556530934 | 11.72892797 | 0.014182547 | 0.989186656 | 0.995907414 | -4.761276667 |
| **A_55_P2040678** | Susd2 | -0.001366975 | -0.244111338 | 0.241377388 | 8.648790506 | -0.014104635 | 0.989246054 | 0.995907414 | -4.761277035 |
| **A_55_P2173333** | Plcb4 | -0.001817939 | -0.324685967 | 0.321050089 | 9.708527342 | -0.014102774 | 0.989247473 | 0.995907414 | -4.761277044 |
| **A_55_P1959753** | NA | -0.002597137 | -0.464800591 | 0.459606317 | 8.857127706 | -0.014073822 | 0.989269546 | 0.995907414 | -4.76127718 |
| **A_52_P193233** | Trnau1ap | 0.001687295 | -0.299845271 | 0.303219861 | 9.956251803 | 0.01401545 | 0.989314047 | 0.995907414 | -4.761277454 |
| **A_52_P172405** | Washc4 | 0.001491899 | -0.269833551 | 0.27281735 | 8.624819521 | 0.013772071 | 0.989499595 | 0.996022411 | -4.761278585 |
| **A_55_P2066453** | NA | 0.005191662 | -0.94558617 | 0.955969494 | 8.671781533 | 0.013676586 | 0.989572392 | 0.996022411 | -4.761279023 |
| **A_55_P2092286** | NA | -0.003133958 | -0.577648332 | 0.571380416 | 8.329795091 | -0.013662891 | 0.989582833 | 0.996022411 | -4.761279086 |
| **A_55_P2128493** | Rap1a | 0.001470034 | -0.271834446 | 0.274774514 | 9.419005598 | 0.013471964 | 0.989728394 | 0.996117062 | -4.761279952 |
| **A_55_P2079324** | Sephs2 | 0.001289986 | -0.240203359 | 0.242783332 | 7.923383127 | 0.013379206 | 0.989799111 | 0.996136381 | -4.761280369 |
| **A_55_P2126014** | NA | 0.001178461 | -0.223057938 | 0.22541486 | 15.92266507 | 0.01316314 | 0.989963838 | 0.996250304 | -4.761281328 |
| **A_51_P199608** | Ttll1 | 0.002031762 | -0.392732203 | 0.396795727 | 9.072195027 | 0.01289097 | 0.990171338 | 0.996407258 | -4.761282513 |
| **A_55_P2186572** | Lypla2 | 0.001946837 | -0.3826776 | 0.386571275 | 13.34292483 | 0.012677778 | 0.990333875 | 0.996518952 | -4.761283425 |
| **A_66_P139250** | Kctd3 | -0.001492049 | -0.304868331 | 0.301884233 | 11.19116923 | -0.01231833 | 0.990607919 | 0.996656495 | -4.761284927 |
| **A_51_P126563** | Otc | 0.00102973 | -0.208410523 | 0.210469984 | 7.651006133 | 0.012314413 | 0.990610905 | 0.996656495 | -4.758883 |
| **A_55_P2160094** | Plscr2 | -0.003051297 | -0.626424101 | 0.620321507 | 8.371813844 | -0.012259902 | 0.990652464 | 0.996656495 | -4.761285167 |
| **A_55_P1996456** | Srrm1 | 0.001999492 | -0.407556595 | 0.411555578 | 9.165023498 | 0.012228031 | 0.990676763 | 0.996656495 | -4.761285297 |
| **A_55_P2427377** | NA | -0.000978853 | -0.208778343 | 0.206820637 | 7.669709758 | -0.011798402 | 0.991004314 | 0.996926325 | -4.761287023 |
| **A_55_P1991160** | Mrrf | 0.000987725 | -0.209720966 | 0.211696416 | 7.755453259 | 0.011740971 | 0.9910481 | 0.996926325 | -4.761287249 |
| **A_55_P1976310** | Tbx20 | -0.00100717 | -0.222362912 | 0.220348571 | 7.728032452 | -0.011396262 | 0.991310909 | 0.99697794 | -4.761288583 |
| **A_55_P2060158** | Ermn | 0.000967886 | -0.212719195 | 0.214654968 | 7.937413975 | 0.011344788 | 0.991350154 | 0.99697794 | -4.761288779 |
| **A_55_P2137984** | NA | -0.001755049 | -0.389832812 | 0.386322713 | 9.819136371 | -0.011327157 | 0.991363596 | 0.99697794 | -4.761288845 |
| **A_55_P1986630** | Gm3627 | -0.000965911 | -0.214875712 | 0.212943889 | 7.773396583 | -0.011309853 | 0.991376789 | 0.99697794 | -4.761288911 |
| **A_51_P315391** | Txlnb | -0.002736652 | -0.609135731 | 0.603662428 | 8.180752435 | -0.011303458 | 0.991381665 | 0.99697794 | -4.761288935 |
| **A_52_P385489** | Ppp1r3a | 0.00116164 | -0.257069948 | 0.259393228 | 7.763412331 | 0.01126711 | 0.991409376 | 0.99697794 | -4.761289072 |
| **A_55_P2139402** | Amy2a5 | -0.000950829 | -0.213581719 | 0.211680061 | 7.657140818 | -0.011200221 | 0.991460373 | 0.99697794 | -4.761289324 |
| **A_55_P1953387** | Fabp5 | -0.002168411 | -0.502894791 | 0.498557968 | 14.88139926 | -0.010846549 | 0.991730019 | 0.997133202 | -4.761290627 |
| **A_55_P1959313** | Rbm42 | 0.001094266 | -0.251921477 | 0.254110009 | 13.10848343 | 0.010832426 | 0.991740787 | 0.997133202 | -4.761290679 |
| **A_55_P2132800** | Zfp367 | -0.002430868 | -0.566993235 | 0.562131498 | 8.661438122 | -0.010784496 | 0.991777329 | 0.997133202 | -4.761290852 |
| **A_55_P2114596** | Zbtb14 | -0.001079271 | -0.255897758 | 0.253739217 | 9.904841686 | -0.010608401 | 0.991911588 | 0.997133202 | -4.761291482 |
| **A_55_P2033084** | Zdhhc20 | 0.001944429 | -0.457531544 | 0.461420401 | 8.602364437 | 0.010599357 | 0.991918482 | 0.997133202 | -4.761291514 |
| **A_55_P1975832** | 1810009N | -0.001278745 | -0.303665954 | 0.301108463 | 10.21556296 | -0.010591831 | 0.991924221 | 0.997133202 | -4.76129154 |
| **A_66_P131389** | Ugt2b38 | -0.000882487 | -0.211055703 | 0.209290728 | 7.689853522 | -0.010516748 | 0.991981465 | 0.997138902 | -4.761291805 |
| **A_55_P2129856** | Mtfmt | 0.000987419 | -0.24147106 | 0.243445897 | 10.14420182 | 0.010200332 | 0.992222708 | 0.997329547 | -4.761292901 |
| **A_51_P220407** | Rnpepl1 | 0.001146011 | -0.288982661 | 0.291274682 | 12.93820804 | 0.009893463 | 0.992456672 | 0.997512856 | -4.761293932 |
| **A_52_P419373** | Lrrc39 | 0.000830387 | -0.211553177 | 0.213213952 | 7.678826337 | 0.009792875 | 0.992533364 | 0.997522092 | -4.761294264 |
| **A_52_P610808** | Hexdc | 0.001102204 | -0.282156128 | 0.284360535 | 8.168338511 | 0.009746069 | 0.99256905 | 0.997522092 | -4.761294416 |
| **A_55_P1975732** | Sipa1l3 | 0.002117181 | -0.546979551 | 0.551213912 | 10.79218772 | 0.009657383 | 0.992636666 | 0.997538194 | -4.761294704 |
| **A_55_P2171086** | NA | -0.001150543 | -0.309704993 | 0.307403907 | 8.693912592 | -0.009339452 | 0.992879067 | 0.997691088 | -4.761295714 |
| **A_51_P282616** | Maats1 | -0.001580152 | -0.426119754 | 0.422959451 | 8.557678893 | -0.009322469 | 0.992892015 | 0.997691088 | -4.761295767 |
| **A_52_P525183** | Acot2 | -0.00115787 | -0.315476654 | 0.313160915 | 8.18108535 | -0.009226556 | 0.992965142 | 0.997712714 | -4.761296064 |
| **A_55_P1954258** | Zfp652 | -0.001391143 | -0.391931771 | 0.389149486 | 9.183021582 | -0.008921869 | 0.993197446 | 0.997842933 | -4.761296989 |
| **A_51_P483544** | Aass | 0.001591477 | -0.447218407 | 0.450401361 | 8.97218303 | 0.00888154 | 0.993228193 | 0.997842933 | -4.761297109 |
| **A_51_P395736** | Srp72 | 0.000781561 | -0.221427377 | 0.222990499 | 11.09989601 | 0.008809509 | 0.993283113 | 0.997842933 | -4.761297322 |

| | | | | | | | | | |
|---|---|---|---|---|---|---|---|---|---|
| **A_52_P36218** | Upf2 | -0.001043879 | -0.298634256 | 0.296546498 | 8.987232897 | -0.008785807 | 0.993301184 | 0.997842933 | -4.761297392 |
| **A_66_P133269** | Hesx1 | 0.000727948 | -0.210204983 | 0.211660878 | 7.708205869 | 0.008643827 | 0.993409435 | 0.997899829 | -4.761297806 |
| **A_51_P492087** | Pole3 | 0.001731258 | -0.506237198 | 0.509699715 | 9.765311536 | 0.008536414 | 0.99349133 | 0.997929817 | -4.761298114 |
| **A_55_P2275057** | Nrap | 0.000695801 | -0.208911579 | 0.21030318 | 7.662319333 | 0.008314359 | 0.993660633 | 0.997929817 | -4.76129874 |
| **A_51_P173678** | Slc10a6 | 0.001366792 | -0.418939435 | 0.421673019 | 8.429934715 | 0.008144925 | 0.993789817 | 0.997929817 | -4.761299206 |
| **A_51_P304109** | Cyp2c39 | -0.000677115 | -0.209392822 | 0.208038592 | 7.640558667 | -0.008125637 | 0.993804523 | 0.997929817 | -4.761299259 |
| **A_55_P2117487** | NA | 0.001071345 | -0.331980526 | 0.334123216 | 14.38917387 | 0.008056898 | 0.993856932 | 0.997929817 | -4.761299445 |
| **A_52_P346581** | Colgalt1 | 0.001330198 | -0.415964497 | 0.418624893 | 10.56362652 | 0.007984059 | 0.993912467 | 0.997929817 | -4.76129964 |
| **A_51_P272563** | Naa25 | -0.001239629 | -0.39165713 | 0.389177873 | 9.385627914 | -0.007952664 | 0.993936404 | 0.997929817 | -4.761299724 |
| **A_52_P590535** | Fbln2 | -0.002586732 | -0.821477542 | 0.816304078 | 11.44343406 | -0.007911809 | 0.993967554 | 0.997929817 | -4.761299832 |
| **A_55_P2123716** | Tug1 | 0.001542711 | -0.488922099 | 0.492007521 | 8.929878447 | 0.007878199 | 0.993993179 | 0.997929817 | -4.761299921 |
| **A_55_P1996674** | Itih3 | -0.001560358 | -0.498318158 | 0.495197441 | 8.193158802 | -0.007867378 | 0.99400143 | 0.997929817 | -4.761299949 |
| **A_51_P245989** | Ccr2 | 0.00253917 | -0.80659332 | 0.811671661 | 8.583257199 | 0.00786 | 0.994007055 | 0.997929817 | -4.761299969 |
| **A_51_P351413** | Phospho2 | 0.000658748 | -0.211127199 | 0.212444694 | 8.357837045 | 0.007790623 | 0.994059951 | 0.997931102 | -4.76130015 |
| **A_55_P2068154** | Anapc15 | 0.00144261 | -0.467945327 | 0.470830546 | 9.871637674 | 0.007697811 | 0.994130715 | 0.997932387 | -4.761300391 |
| **A_52_P644534** | Dhrsx | -0.000751839 | -0.246880446 | 0.245376767 | 8.611723769 | -0.007650912 | 0.994166473 | 0.997932387 | -4.761300511 |
| **A_52_P222709** | Pigu | 0.000924362 | -0.304278038 | 0.306126762 | 10.0295508 | 0.007585853 | 0.994216077 | 0.997932387 | -4.761300677 |
| **A_55_P2101944** | NA | 0.001787465 | -0.594907758 | 0.598482688 | 12.39061784 | 0.007503006 | 0.994279244 | 0.99794398 | -4.761300886 |
| **A_52_P483280** | Lamp2 | 0.001006375 | -0.341486089 | 0.343498839 | 12.73264763 | 0.007359686 | 0.994388517 | 0.998001847 | -4.761301242 |
| **A_52_P604515** | Zfp319 | 0.000835277 | -0.288722959 | 0.290393514 | 8.715250888 | 0.007225118 | 0.994491118 | 0.998009614 | -4.76130157 |
| **A_66_P137134** | NA | -0.001184338 | -0.415978092 | 0.413609415 | 8.685016864 | -0.007151447 | 0.994547288 | 0.998009614 | -4.761301747 |
| **A_55_P2146185** | Rs1 | 0.002210792 | -0.774039139 | 0.778460723 | 12.73117348 | 0.007133402 | 0.994561047 | 0.998009614 | -4.76130179 |
| **A_52_P610403** | Sf1 | 0.000623122 | -0.219856516 | 0.22110276 | 8.44136572 | 0.007078727 | 0.994602733 | 0.998009614 | -4.76130192 |
| **A_55_P2099650** | Slc24a3 | 0.001107934 | -0.402560788 | 0.404776655 | 9.880654212 | 0.006874468 | 0.994758471 | 0.998083803 | -4.761302397 |
| **A_55_P1953391** | Rbfox1 | -0.000572745 | -0.210105978 | 0.208960488 | 7.764579299 | -0.006846341 | 0.994779916 | 0.998083803 | -4.761302461 |
| **A_51_P366770** | Rims1 | 0.002246275 | -0.832675425 | 0.837167976 | 8.44779596 | 0.006738568 | 0.994862087 | 0.998114452 | -4.761302706 |
| **A_51_P343429** | Slc25a37 | -0.001899819 | -0.718799017 | 0.714999379 | 12.30598711 | -0.0066375 | 0.994939147 | 0.998139968 | -4.761302932 |
| **A_55_P2087322** | Rapsn | 0.001222106 | -0.484903733 | 0.487347945 | 12.89764513 | 0.006296664 | 0.995199017 | 0.998298497 | -4.761303669 |
| **A_55_P1999818** | NA | -0.003198201 | -1.280832262 | 1.27443586 | 13.81526923 | -0.006269741 | 0.995219545 | 0.998298497 | -4.761303726 |
| **A_55_P2127357** | NA | -0.001061887 | -0.432820692 | 0.430696919 | 10.89323329 | -0.006160094 | 0.995303146 | 0.998298497 | -4.761303954 |
| **A_55_P1962139** | Mff | 0.001064894 | -0.431968204 | 0.434097993 | 11.31042763 | 0.006159361 | 0.995303704 | 0.998298497 | -4.761303955 |
| **A_55_P2136101** | Meis3 | -0.000884225 | -0.376384741 | 0.37461629 | 8.750827643 | -0.005897973 | 0.995503001 | 0.998315113 | -4.761304482 |
| **A_66_P112305** | Myo1f | 0.001788728 | -0.761307484 | 0.76488494 | 9.034800616 | 0.005871044 | 0.995523533 | 0.998315113 | -4.761304535 |
| **A_55_P2146105** | NA | 0.002219789 | -0.945619516 | 0.950059093 | 16.80309836 | 0.0058658 | 0.995527531 | 0.998315113 | -4.761304545 |
| **A_51_P367563** | Rsph3a | -0.000499459 | -0.213938236 | 0.212939318 | 8.383491888 | -0.005861074 | 0.995531135 | 0.998315113 | -4.761304555 |
| **A_52_P503730** | Fchsd2 | 0.000859854 | -0.372627282 | 0.37434699 | 9.45573374 | 0.005766329 | 0.995603373 | 0.998315113 | -4.761304739 |
| **A_55_P1976242** | Arhgef7 | -0.001460888 | -0.642582666 | 0.639660891 | 13.19810582 | -0.005707247 | 0.995648421 | 0.998315113 | -4.761304852 |
| **A_55_P2082806** | Trib1 | -0.001214317 | -0.53999233 | 0.537563696 | 9.570930629 | -0.005645112 | 0.995695796 | 0.998315113 | -4.76130497 |
| **A_55_P2298319** | NA | 0.000465031 | -0.207679819 | 0.208609881 | 7.669854587 | 0.005595855 | 0.995733352 | 0.998315113 | -4.761305063 |
| **A_55_P2053497** | Poli | -0.000648665 | -0.299366042 | 0.298068713 | 9.596199396 | -0.005438887 | 0.995853033 | 0.998316687 | -4.761305352 |
| **A_55_P2360266** | 4833411C | 0.000465585 | -0.217468205 | 0.218399374 | 7.722318519 | 0.005350871 | 0.995920142 | 0.998316687 | -4.761305511 |
| **A_55_P1977096** | Elk4 | 0.003331164 | -1.563511574 | 1.570173903 | 10.79137114 | 0.005325017 | 0.995939855 | 0.998316687 | -4.761305557 |
| **A_52_P609868** | Timd4 | -0.000474579 | -0.224547623 | 0.223598464 | 7.718036287 | -0.005304804 | 0.995955266 | 0.998316687 | -4.761305593 |
| **A_55_P2090324** | Zfp956 | 0.000868159 | -0.412904322 | 0.41464064 | 9.594092999 | 0.005255184 | 0.995993099 | 0.998316687 | -4.761305681 |
| **A_51_P315646** | Nfs1 | -0.000504176 | -0.24869835 | 0.247689998 | 10.14208255 | -0.005087931 | 0.996120623 | 0.998392748 | -4.76130597 |
| **A_55_P2049122** | Tm6sf2 | 0.000501624 | -0.253199146 | 0.254202394 | 8.402106914 | 0.004952301 | 0.996224035 | 0.998424802 | -4.761306198 |
| **A_51_P245156** | Gdf2 | -0.000425015 | -0.218166103 | 0.217316074 | 7.68731808 | -0.004888932 | 0.996272351 | 0.998424802 | -4.761306302 |
| **A_51_P236864** | Parp8 | -0.000936089 | -0.486276704 | 0.484404527 | 8.601638759 | -0.004830819 | 0.99631666 | 0.998424802 | -4.761306397 |

| | | | | | | | | | |
|---|---|---|---|---|---|---|---|---|---|
| **A_52_P229783** | Ptgr2 | -0.000709489 | -0.372858092 | 0.371439114 | 9.16684594 | -0.00477507 | 0.996359167 | 0.998424802 | -4.761306486 |
| **A_52_P243599** | Hsd17b12 | 0.001103734 | -0.591123156 | 0.593330624 | 9.827150668 | 0.004667956 | 0.996440837 | 0.998454892 | -4.761306655 |
| **A_55_P1970309** | Arhgap44 | 0.000827245 | -0.465479728 | 0.467134217 | 11.45940069 | 0.004443369 | 0.996612076 | 0.998574724 | -4.761306997 |
| **A_66_P140658** | Olfr787 | -0.000416576 | -0.24090193 | 0.240068779 | 7.98916446 | -0.004338661 | 0.996691912 | 0.998602965 | -4.761307151 |
| **A_52_P50287** | Bend7 | 0.00058407 | -0.349881076 | 0.351049216 | 8.525720567 | 0.004174173 | 0.996817327 | 0.998676868 | -4.761307385 |
| **A_51_P363400** | Prg2 | 0.000335446 | -0.208969555 | 0.209640446 | 7.674441235 | 0.004014145 | 0.996939343 | 0.998747358 | -4.761307604 |
| **A_55_P2043833** | Srgap2 | 0.000975286 | -0.625949195 | 0.627899767 | 8.842081795 | 0.003896432 | 0.997029095 | 0.998785519 | -4.76130776 |
| **A_55_P2043684** | Ubac1 | 0.000639016 | -0.421863211 | 0.423141243 | 11.77073057 | 0.003788203 | 0.997111616 | 0.998788609 | -4.761307899 |
| **A_55_P2094242** | NA | -0.000343092 | -0.229078912 | 0.228392727 | 8.090115802 | -0.003756879 | 0.997135499 | 0.998788609 | -4.761307939 |
| **A_55_P1970676** | Sh3yl1 | 0.000778283 | -0.552433033 | 0.553989599 | 9.308665931 | 0.003523683 | 0.997313303 | 0.998870846 | -4.761308222 |
| **A_51_P289742** | Slc27a5 | -0.000291606 | -0.209176432 | 0.20859322 | 7.671918033 | -0.003496549 | 0.997333991 | 0.998870846 | -4.761308254 |
| **A_51_P401907** | Cstdc4 | 0.001739078 | -1.262309538 | 1.265787694 | 9.624019761 | 0.003445924 | 0.997372591 | 0.998870846 | -4.761308313 |
| **A_51_P487678** | Dtnbp1 | -0.00032471 | -0.245956217 | 0.245306796 | 9.595126839 | -0.003311026 | 0.997475446 | 0.998922111 | -4.761308466 |
| **A_55_P2158547** | Cyp2c54 | 0.000269655 | -0.208832305 | 0.209371615 | 7.640165055 | 0.003229985 | 0.997537237 | 0.998932249 | -4.761308554 |
| **A_66_P120572** | Gm14488 | 0.000325513 | -0.261255841 | 0.261906867 | 8.330120666 | 0.003116822 | 0.99762352 | 0.998957952 | -4.761308674 |
| **A_55_P1964332** | NA | 0.000315492 | -0.262420649 | 0.263051633 | 10.58174835 | 0.003007591 | 0.997706805 | 0.998957952 | -4.761308786 |
| **A_55_P2067712** | Hmcn2 | -0.000673257 | -0.564078173 | 0.562731659 | 9.951015713 | -0.002993028 | 0.997717909 | 0.998957952 | -4.761308801 |
| **A_55_P1972915** | NA | 0.000573078 | -0.508618307 | 0.509764462 | 14.52970911 | 0.00281892 | 0.997850661 | 0.999018304 | -4.76130897 |
| **A_55_P1960999** | Pigr | 0.000740663 | -0.66694314 | 0.668424467 | 12.71695832 | 0.002778435 | 0.997881529 | 0.999018304 | -4.761309008 |
| **A_55_P2088720** | NA | 0.000797747 | -0.775503985 | 0.777099479 | 12.72400677 | 0.00257386 | 0.99803751 | 0.999031228 | -4.761309191 |
| **A_55_P2011146** | Reep1 | -0.000279964 | -0.274316878 | 0.273756951 | 9.593224873 | -0.00255884 | 0.998048962 | 0.999031228 | -4.761309204 |
| **A_55_P2138739** | NA | 0.000474512 | -0.46410928 | 0.465058304 | 8.928829683 | 0.002558194 | 0.998049455 | 0.999031228 | -4.761309204 |
| **A_52_P354823** | Irf8 | -0.000448039 | -0.479957426 | 0.479061348 | 10.34662319 | -0.002340288 | 0.998215602 | 0.999104769 | -4.761309383 |
| **A_66_P119771** | Nme8 | -0.000198285 | -0.214994302 | 0.214597732 | 7.656313633 | -0.002312143 | 0.998237061 | 0.999104769 | -4.761309405 |
| **A_51_P162116** | Mblac1 | 0.000347778 | -0.385334167 | 0.386029722 | 8.746285964 | 0.002258513 | 0.998277952 | 0.999104769 | -4.761309446 |
| **A_51_P189722** | Cabin1 | -0.000430044 | -0.547087203 | 0.546227116 | 9.984174237 | -0.001970371 | 0.998497651 | 0.999272923 | -4.761309649 |
| **A_51_P332932** | NA | 0.000219091 | -0.304716045 | 0.305154228 | 8.085420713 | 0.001799569 | 0.998627883 | 0.999351526 | -4.761309757 |
| **A_55_P2054817** | Rxfp1 | 0.000161895 | -0.238260275 | 0.238584065 | 7.76648777 | 0.001700735 | 0.99870324 | 0.999375211 | -4.761309815 |
| **A_51_P394172** | Ces1c | -0.000123574 | -0.211593048 | 0.2113459 | 7.666075529 | -0.001463621 | 0.998884033 | 0.999504394 | -4.76130994 |
| **A_51_P431046** | St8sia3 | 0.000115202 | -0.215282385 | 0.21551279 | 7.69724811 | 0.001339586 | 0.998978605 | 0.999521207 | -4.761309999 |
| **A_55_P2054062** | Chst2 | -0.00028732 | -0.566515565 | 0.565940925 | 8.967329278 | -0.001270939 | 0.999030947 | 0.999521207 | -4.761310029 |
| **A_55_P2143968** | Gps1 | 0.000114176 | -0.249774476 | 0.250002828 | 13.0516049 | 0.001144405 | 0.999127425 | 0.999521207 | -4.76131008 |
| **A_55_P2110513** | Lmo7 | 0.000550285 | -1.211202057 | 1.212302627 | 11.49209203 | 0.001137429 | 0.999132744 | 0.999521207 | -4.761310082 |
| **A_52_P456640** | Fgr | -0.000115236 | -0.263739407 | 0.263508935 | 7.907889981 | -0.001094844 | 0.999165214 | 0.999521207 | -4.761310098 |
| **A_55_P2137406** | Bax | 0.000144252 | -0.349019831 | 0.349308334 | 11.94632264 | 0.001034767 | 0.999211021 | 0.999521207 | -4.76131012 |
| **A_66_P136186** | Wee1 | -0.000186738 | -0.567461821 | 0.567088345 | 9.536377522 | -0.000824498 | 0.999371344 | 0.99962986 | -4.761310185 |
| **A_51_P403942** | A930016C | 4.89E-05 | -0.207611944 | 0.20770974 | 7.658519508 | 0.000589776 | 0.999550313 | 0.999757153 | -4.76131024 |
| **A_51_P144249** | Ubr1 | 9.56E-05 | -0.505410807 | 0.505601942 | 8.54508233 | 0.000473513 | 0.99963896 | 0.999794095 | -4.761310261 |
| **A_55_P2112892** | Dab1 | 4.29E-05 | -0.32417334 | 0.324259097 | 8.106024087 | 0.000331246 | 0.999747434 | 0.999850864 | -4.76131028 |
| **A_51_P235139** | Klhl40 | 1.86E-05 | -0.209975486 | 0.210012647 | 7.69383055 | 0.000221613 | 0.999831027 | 0.999882743 | -4.76131029 |
| **A_55_P2032099** | Zfp715 | -3.81E-06 | -0.264285596 | 0.264277967 | 8.147247622 | -3.62E-05 | 0.999972436 | 0.999972436 | -4.761310298 |

Rychlik_Arsenic_microarray_TableS4_Placenta_100_vs_0_all_p_values

| | ID | logFC | CI.L | CI.R | AveExpr | t | P.Value | adj.P.Val | B |
|---|---|---|---|---|---|---|---|---|---|
| **A_55_P1974780** | NA | 0.383081841 | 0.268389721 | 0.497773961 | 9.614858909 | 8.217262278 | 0.000196413 | 0.594255995 | -2.015519346 |
| **A_55_P2137927** | Cd82 | -0.401739763 | -0.531793337 | -0.271686188 | 10.93896799 | -7.599616713 | 0.000300166 | 0.594255995 | -2.082031972 |
| **A_55_P1997951** | Htatsf1 | -0.328771083 | -0.456646113 | -0.200896052 | 11.29191221 | -6.325240242 | 0.000795192 | 0.594255995 | -2.268306324 |
| **A_55_P2472435** | Gbp3 | -0.429681825 | -0.598085173 | -0.261278477 | 9.535067169 | -6.277192203 | 0.0008274 | 0.594255995 | -2.276990038 |
| **A_51_P105520** | Nomo1 | 0.343634467 | 0.206307068 | 0.480961865 | 12.86052651 | 6.156142843 | 0.00091541 | 0.594255995 | -2.299511966 |
| **A_51_P487228** | B9d2 | 0.276009626 | 0.160669888 | 0.391349365 | 10.01786429 | 5.887276434 | 0.001152227 | 0.594255995 | -2.353041638 |
| **A_55_P2155421** | Sec61g | 0.37917056 | 0.215460796 | 0.542880324 | 15.63278957 | 5.698088521 | 0.00136121 | 0.594255995 | -2.393843139 |
| **A_55_P1958697** | Rbm33 | -0.286397279 | -0.410369974 | -0.162424584 | 10.41621354 | -5.68344954 | 0.001379114 | 0.594255995 | -2.397115325 |
| **A_55_P2052848** | Mif | 0.44263203 | 0.249973694 | 0.635290367 | 15.11947549 | 5.652286864 | 0.001418132 | 0.594255995 | -2.404137783 |
| **A_55_P2170813** | Pkig | -0.395083836 | -0.567554127 | -0.222613546 | 12.74880601 | -5.635651509 | 0.001439476 | 0.594255995 | -2.407918447 |
| **A_51_P345367** | Psmb8 | -0.332944351 | -0.478886018 | -0.187002685 | 9.21397659 | -5.61256667 | 0.001469709 | 0.594255995 | -2.413202023 |
| **A_51_P317941** | Perp | -0.474692677 | -0.684517634 | -0.264867721 | 14.41330966 | -5.565761003 | 0.001533263 | 0.594255995 | -2.424048865 |
| **A_55_P1975560** | Ifi204 | -0.361892504 | -0.522100864 | -0.201684144 | 9.897812164 | -5.557296595 | 0.001545089 | 0.594255995 | -2.42602981 |
| **A_55_P2151506** | Nudt9 | -0.30625077 | -0.443724727 | -0.168776813 | 13.00412813 | -5.480572413 | 0.001657176 | 0.594255995 | -2.44426147 |
| **A_52_P493810** | Srpk1 | -0.259860144 | -0.376713042 | -0.143007246 | 9.624030585 | -5.471033197 | 0.001671751 | 0.594255995 | -2.446563395 |
| **A_66_P125035** | Duxbl2 | 0.242848106 | 0.133575401 | 0.352120811 | 8.232057947 | 5.467542775 | 0.00167712 | 0.594255995 | -2.447407642 |
| **A_52_P370402** | Kcmf1 | -0.31302885 | -0.454983611 | -0.171074089 | 12.56895662 | -5.425047765 | 0.0017441 | 0.594255995 | -2.457771278 |
| **A_55_P2097126** | Cnep1r1 | -0.242958653 | -0.353575251 | -0.132342055 | 10.1798922 | -5.403575657 | 0.001779102 | 0.594255995 | -2.463068263 |
| **A_55_P2025790** | Fanci | -0.346391221 | -0.504583232 | -0.188199211 | 9.313582654 | -5.387056046 | 0.001806578 | 0.594255995 | -2.467171405 |
| **A_51_P262171** | Irgm1 | -0.402286706 | -0.589260595 | -0.215312817 | 9.804891823 | -5.293268 | 0.001972064 | 0.594255995 | -2.490934526 |
| **A_55_P2114110** | Cadm4 | -0.265032513 | -0.390351235 | -0.139713791 | 9.848673493 | -5.202982435 | 0.002147899 | 0.594255995 | -2.514582594 |
| **A_51_P338776** | Dnajc8 | -0.243121999 | -0.358665019 | -0.127578979 | 11.55356192 | -5.176660758 | 0.00220248 | 0.594255995 | -2.5216236 |
| **A_52_P452689** | Atf3 | -0.307714788 | -0.456047188 | -0.159382387 | 10.42908396 | -5.103657312 | 0.002362325 | 0.594255995 | -2.541507284 |
| **A_51_P223524** | Ric1 | -0.286299735 | -0.42486983 | -0.14772964 | 9.820986572 | -5.083005686 | 0.002409912 | 0.594255995 | -2.547228402 |
| **A_52_P346113** | Foxn2 | -0.242865831 | -0.360474744 | -0.125256918 | 8.873509711 | -5.080370011 | 0.002416064 | 0.594255995 | -2.547961658 |
| **A_55_P1995233** | Lonp2 | -0.361659639 | -0.537015151 | -0.186304127 | 12.93232334 | -5.073992084 | 0.002431024 | 0.594255995 | -2.549738935 |
| **A_51_P134007** | Ncl | -0.422439032 | -0.628097299 | -0.216780765 | 12.39650444 | -5.053439029 | 0.002479955 | 0.594255995 | -2.555494393 |
| **A_55_P1994032** | Xbp1 | -0.21798003 | -0.326023772 | -0.109936288 | 14.6546033 | -4.963479734 | 0.002707742 | 0.594255995 | -2.581198726 |
| **A_55_P2103596** | Slc30a4 | -0.24430638 | -0.365719955 | -0.122892805 | 13.13169107 | -4.950359352 | 0.002742914 | 0.594255995 | -2.585018588 |
| **A_51_P304170** | Rtp4 | -0.293825057 | -0.44240887 | -0.145241243 | 12.42171823 | -4.865040855 | 0.002984711 | 0.594255995 | -2.610309609 |
| **A_52_P463977** | Tmem140 | -0.378055467 | -0.57066226 | -0.185448674 | 15.02385417 | -4.828954185 | 0.003094204 | 0.594255995 | -2.621246422 |
| **A_51_P472901** | Slc3a2 | -0.382742891 | -0.578028893 | -0.187456889 | 14.48427463 | -4.821755472 | 0.003116588 | 0.594255995 | -2.623445467 |
| **A_52_P617327** | Rcan1 | -0.262727633 | -0.397855879 | -0.127599387 | 9.483709246 | -4.783312727 | 0.003239279 | 0.594255995 | -2.635287443 |
| **A_55_P1960281** | Med7 | -0.262390827 | -0.397529033 | -0.12725262 | 10.120033 | -4.776828616 | 0.003260509 | 0.594255995 | -2.637301298 |
| **A_51_P112355** | Igtp | -0.332162747 | -0.504145855 | -0.160179639 | 10.08286215 | -4.751538946 | 0.003344834 | 0.594255995 | -2.712398645 |
| **A_55_P1979397** | Palm | 0.264484785 | 0.126966529 | 0.40200304 | 11.0786458 | 4.731616119 | 0.003413009 | 0.594255995 | -2.651477255 |
| **A_55_P2062310** | NA | -0.298065805 | -0.453202537 | -0.142929074 | 14.60598367 | -4.726794072 | 0.003429745 | 0.594255995 | -2.653003066 |
| **A_55_P1978830** | Camk2d | -0.204783678 | -0.311734286 | -0.09783307 | 9.357678878 | -4.710654415 | 0.003486445 | 0.594255995 | -2.658129696 |
| **A_55_P2137611** | Irgm2 | -0.291540387 | -0.444359128 | -0.138721645 | 9.953342993 | -4.693440002 | 0.003548095 | 0.594255995 | -2.663631237 |
| **A_55_P2108644** | NA | 0.484512284 | 0.228078323 | 0.740946245 | 12.12703551 | 4.64834565 | 0.003715534 | 0.594255995 | -2.678208578 |
| **A_55_P2116165** | Pfkfb3 | -0.252149456 | -0.385939403 | -0.118359509 | 10.67906666 | -4.636643625 | 0.003760435 | 0.594255995 | -2.682030963 |
| **A_55_P2088741** | Mtif3 | -0.254407384 | -0.389479479 | -0.119335288 | 8.854545119 | -4.633756812 | 0.003771607 | 0.594255995 | -2.682976446 |
| **A_55_P2078048** | BC004004 | 0.225254624 | 0.105591133 | 0.344918115 | 12.82030551 | 4.631068548 | 0.003782043 | 0.594255995 | -2.6838578 |
| **A_51_P288891** | Tbc1d15 | -0.247187726 | -0.380834585 | -0.113540867 | 10.77644002 | -4.5502715 | 0.004111524 | 0.594255995 | -2.71075696 |
| **A_52_P574668** | Nt5e | -0.33993092 | -0.524017114 | -0.155844726 | 11.4039504 | -4.542956879 | 0.004142923 | 0.594255995 | -2.713231741 |
| **A_55_P2047489** | Calm1 | -0.396550452 | -0.611430929 | -0.181669976 | 16.13704738 | -4.540154729 | 0.004155023 | 0.594255995 | -2.714181562 |
| **A_55_P1980951** | 1110012L19Rik | -0.32729225 | -0.49480274 | -0.159781761 | 12.08975262 | -5.063418176 | 0.004189188 | 0.594255995 | -3.018370963 |
| **A_51_P413785** | Commd3 | -0.233496594 | -0.360536769 | -0.106456418 | 11.5973461 | -4.521771665 | 0.004235402 | 0.594255995 | -2.720436983 |

| | | | | | | | | | |
|---|---|---|---|---|---|---|---|---|---|
| **A_51_P359570** | Ifit3 | -0.404506278 | -0.624602349 | -0.184410208 | 9.695730723 | -4.52149589 | 0.004236621 | 0.594255995 | -2.720531145 |
| **A_51_P268559** | Idh3a | -0.256253981 | -0.39576382 | -0.116744143 | 11.00319378 | -4.518923076 | 0.004248014 | 0.594255995 | -2.721410085 |
| **A_51_P180452** | Crk | -0.302847543 | -0.468221443 | -0.137473642 | 11.44578635 | -4.505326263 | 0.004308801 | 0.594255995 | -2.726068905 |
| **A_51_P374772** | Arfgef1 | -0.274707942 | -0.424973724 | -0.12444216 | 11.1960766 | -4.497594349 | 0.004343807 | 0.594255995 | -2.728728552 |
| **A_51_P238786** | Mcl1 | -0.324345379 | -0.501881298 | -0.146809459 | 13.35982339 | -4.494595823 | 0.004357469 | 0.594255995 | -2.729762025 |
| **A_55_P1994947** | NA | 0.543574361 | 0.245904955 | 0.841243767 | 13.89762937 | 4.492560414 | 0.004366771 | 0.594255995 | -2.730464197 |
| **A_55_P2161773** | Mpp3 | 0.235389477 | 0.106300086 | 0.364478868 | 9.871533862 | 4.486065964 | 0.0043966 | 0.594255995 | -2.732708151 |
| **A_55_P2095894** | 6430548M08Rik | 0.282118801 | 0.126560959 | 0.437676642 | 10.71590097 | 4.461791609 | 0.00451015 | 0.594255995 | -2.741142867 |
| **A_55_P2062246** | Tgtp2 | -0.611858679 | -0.950434496 | -0.273282863 | 9.924752511 | -4.445947745 | 0.00458605 | 0.594255995 | -2.74668883 |
| **A_55_P2013780** | NA | -0.381396197 | -0.59342304 | -0.169369355 | 12.39048179 | -4.425421846 | 0.004686535 | 0.594255995 | -2.753921748 |
| **A_52_P186937** | Cmpk2 | -0.285271485 | -0.444029291 | -0.12651368 | 9.184394258 | -4.420714207 | 0.004709931 | 0.594255995 | -2.755588311 |
| **A_51_P496845** | Capza2 | -0.312666741 | -0.486812651 | -0.138520831 | 10.96373588 | -4.417103455 | 0.004727964 | 0.594255995 | -2.756868513 |
| **A_55_P2103837** | Gbp4 | -0.261000323 | -0.406501057 | -0.11549959 | 9.074219416 | -4.413112771 | 0.004747986 | 0.594255995 | -2.758285392 |
| **A_51_P189746** | Pim3 | -0.294404807 | -0.458836027 | -0.129973587 | 9.588062581 | -4.404836387 | 0.004789816 | 0.594255995 | -2.761230508 |
| **A_55_P2066578** | Ifi204 | -0.328413419 | -0.511860267 | -0.144966571 | 8.96732398 | -4.404329498 | 0.004792392 | 0.594255995 | -2.761411173 |
| **A_51_P369636** | Hat1 | -0.283818434 | -0.44238365 | -0.125253217 | 9.758546475 | -4.403538904 | 0.004796412 | 0.594255995 | -2.761693023 |
| **A_55_P1987331** | Prl3d1 | -0.854340893 | -1.333115126 | -0.37556666 | 13.86607711 | -4.390051228 | 0.004865583 | 0.594255995 | -2.766514024 |
| **A_55_P2048800** | Cd82 | -0.208675221 | -0.325643357 | -0.091707084 | 9.153078285 | -4.389069662 | 0.00487066 | 0.594255995 | -2.766865803 |
| **A_55_P1992505** | Entpd4 | 0.307778099 | 0.13482467 | 0.480731527 | 11.44329106 | 4.37801959 | 0.004928238 | 0.594255995 | -2.770834731 |
| **A_55_P2154536** | Hagh | -0.260318837 | -0.406862317 | -0.113775358 | 11.89525435 | -4.370269904 | 0.004969076 | 0.594255995 | -2.773627836 |
| **A_52_P8685** | Tmem183a | -0.193396052 | -0.302546349 | -0.084245754 | 9.501292786 | -4.359049928 | 0.005028881 | 0.594255995 | -2.777685758 |
| **A_55_P2051400** | Prl3d2 | -0.824649093 | -1.291098639 | -0.358199546 | 13.43308044 | -4.349443297 | 0.005080732 | 0.594255995 | -2.781173455 |
| **A_55_P2170737** | Igf2bp2 | -0.222901018 | -0.349003095 | -0.096798941 | 10.4295796 | -4.348695249 | 0.005084795 | 0.594255995 | -2.78144555 |
| **A_52_P48689** | Bzw1 | -0.238057084 | -0.373454575 | -0.102659593 | 9.751394122 | -4.325533268 | 0.005212423 | 0.594255995 | -2.789907455 |
| **A_55_P1959833** | Tmem266 | 0.24902565 | 0.107047677 | 0.391003622 | 9.706814172 | 4.315114003 | 0.005271011 | 0.594255995 | -2.793737439 |
| **A_55_P2024012** | Rpl5 | -0.331418148 | -0.521247645 | -0.14158865 | 14.77323435 | -4.295183777 | 0.00538516 | 0.594255995 | -2.80110432 |
| **A_55_P2145496** | Wdr45 | -0.262698119 | -0.41359604 | -0.111800198 | 11.59100857 | -4.28294868 | 0.005456615 | 0.594255995 | -2.805653493 |
| **A_51_P110395** | Setd2 | -0.215204798 | -0.338879586 | -0.09153001 | 11.85033723 | -4.280947858 | 0.005468401 | 0.594255995 | -2.896499028 |
| **A_52_P108502** | Slc4a7 | 0.374670484 | 0.159338828 | 0.59000214 | 8.725073642 | 4.280660291 | 0.005470098 | 0.594255995 | -2.806506609 |
| **A_55_P2112355** | NA | -0.340734391 | -0.53660257 | -0.144866212 | 13.36387959 | -4.279778053 | 0.005475306 | 0.594255995 | -2.806835699 |
| **A_51_P121390** | Anapc11 | 0.347126399 | 0.147104424 | 0.547148374 | 12.85878428 | 4.269520424 | 0.00553627 | 0.594255995 | -2.900460348 |
| **A_66_P115038** | Prl3d1 | -0.67084572 | -1.059049371 | -0.282642068 | 9.576473611 | -4.251402272 | 0.00564583 | 0.594255995 | -2.817477154 |
| **A_65_P01319** | Pde4b | -0.442533054 | -0.699497682 | -0.185568426 | 8.607216516 | -4.236834824 | 0.005735692 | 0.594255995 | -2.822983224 |
| **A_55_P2059957** | Snx7 | -0.21897883 | -0.346767952 | -0.091189708 | 9.833985091 | -4.215774846 | 0.005868465 | 0.594255995 | -2.830995229 |
| **A_55_P1961302** | NA | 0.386591177 | 0.160926651 | 0.612255704 | 12.35340779 | 4.214614008 | 0.005875884 | 0.594255995 | -2.831438648 |
| **A_51_P159673** | Snx1 | -0.4253526 | -0.673791866 | -0.176913334 | 14.05231918 | -4.212093642 | 0.005892027 | 0.594255995 | -2.83240203 |
| **A_55_P2110743** | Mettl7a2 | 0.726543118 | 0.301245442 | 1.151840794 | 15.05574817 | 4.202788613 | 0.00595206 | 0.594255995 | -2.835966455 |
| **A_55_P2313033** | Ptprb | -0.274643746 | -0.435452949 | -0.113834543 | 8.890783363 | -4.201728404 | 0.005958944 | 0.594255995 | -2.836373352 |
| **A_55_P2069989** | Klhl12 | -0.284101585 | -0.450523368 | -0.117679802 | 11.82893746 | -4.199839066 | 0.005971234 | 0.594255995 | -2.837098849 |
| **A_55_P2005941** | Plcxd1 | 0.378247425 | 0.156211286 | 0.600283564 | 10.95095646 | 4.191036818 | 0.006028866 | 0.594255995 | -2.840485469 |
| **A_51_P272806** | Ctsr | -0.214337198 | -0.340306319 | -0.088368077 | 16.60714039 | -4.18603267 | 0.006061909 | 0.594255995 | -2.842415641 |
| **A_66_P126348** | Smim31 | 1.166957579 | 0.480359025 | 1.853556133 | 13.31128452 | 4.181396466 | 0.006092705 | 0.594255995 | -2.844207037 |
| **A_52_P480544** | Tnpo2 | -0.178584215 | -0.283757503 | -0.073410928 | 8.876207002 | -4.177406939 | 0.006119346 | 0.594255995 | -2.845750986 |
| **A_52_P478256** | Samm50 | -0.257725658 | -0.409692628 | -0.105758687 | 9.894790426 | -4.172320043 | 0.006153505 | 0.594255995 | -2.847722875 |
| **A_55_P1965219** | Syt6 | 0.283864599 | 0.116176037 | 0.451553162 | 8.67279741 | 4.164635108 | 0.006205518 | 0.594255995 | -2.85070881 |
| **A_55_P2082986** | Hmgb1 | -0.413760126 | -0.658425785 | -0.169094467 | 12.33702978 | -4.160492862 | 0.006233758 | 0.594255995 | -2.852321725 |
| **A_55_P1981845** | NA | 0.551195124 | 0.225141263 | 0.877248986 | 9.512307515 | 4.158964205 | 0.006244216 | 0.594255995 | -2.852917571 |
| **A_52_P47255** | Gmps | -0.179720254 | -0.286117983 | -0.073322525 | 8.671228992 | -4.155600867 | 0.006267296 | 0.594255995 | -2.854229714 |
| **A_55_P2166501** | Cd44 | -0.286940675 | -0.457473932 | -0.116407417 | 8.328932579 | -4.139541013 | 0.006378833 | 0.594255995 | -2.860517365 |
| **A_55_P2115057** | Tuba1a | 0.25573596 | 0.103637469 | 0.407834451 | 14.15885181 | 4.136528856 | 0.0064 | 0.594255995 | -2.861700763 |

| | | | | | | | | | |
|---|---|---|---|---|---|---|---|---|---|
| **A_55_P2094262** | Tph1 | -0.66135844 | -1.054720577 | -0.267996303 | 10.1968429 | -4.136314083 | 0.006401512 | 0.594255995 | -2.861785192 |
| **A_55_P2176492** | Tollip | 0.258056145 | 0.104171442 | 0.411940848 | 8.563082888 | 4.125607594 | 0.006477411 | 0.594255995 | -2.866002335 |
| **A_55_P1961391** | NA | 0.350675422 | 0.141407027 | 0.559943816 | 8.661235533 | 4.122596678 | 0.006498937 | 0.594255995 | -2.867191256 |
| **A_55_P2163143** | Cdk4 | 0.326750698 | 0.131388292 | 0.522113104 | 13.39215624 | 4.114761722 | 0.00655533 | 0.594255995 | -2.870291153 |
| **A_52_P6798** | Rbm22 | -0.193177778 | -0.30899334 | -0.077362216 | 9.083864414 | -4.103546976 | 0.006637009 | 0.594255995 | -2.874743653 |
| **A_55_P2056876** | Akap13 | 0.427386678 | 0.170967732 | 0.683805624 | 9.364772743 | 4.100530378 | 0.006659174 | 0.594255995 | -2.875944408 |
| **A_55_P2185148** | Mettl7a3 | 0.285561186 | 0.114141906 | 0.456980466 | 8.996614477 | 4.098346502 | 0.006675273 | 0.594255995 | -2.87681452 |
| **A_55_P2147871** | NA | 0.661157924 | 0.264219482 | 1.058096366 | 9.80333032 | 4.097804258 | 0.006679277 | 0.594255995 | -2.877030671 |
| **A_55_P1972322** | Btg3 | -0.286161298 | -0.458084302 | -0.114238294 | 13.52181627 | -4.094926105 | 0.006700574 | 0.594255995 | -2.878178682 |
| **A_55_P2134004** | Gstm2 | 0.309730707 | 0.123443208 | 0.496018207 | 9.96242609 | 4.090437048 | 0.006733944 | 0.594255995 | -2.879971632 |
| **A_52_P565051** | Cpne3 | -0.290125804 | -0.464837091 | -0.115414517 | 8.742581261 | -4.085399739 | 0.006771612 | 0.594255995 | -2.881987038 |
| **A_55_P2043723** | Rabac1 | -0.265997022 | -0.426551885 | -0.10544216 | 12.83871264 | -4.075890187 | 0.006843369 | 0.594255995 | -2.885801831 |
| **A_55_P2154714** | Klhl9 | 0.298333171 | 0.11809341 | 0.478572931 | 12.55197878 | 4.072115669 | 0.006872087 | 0.594255995 | -2.887319646 |
| **A_55_P2035344** | Tcf25 | -0.336998661 | -0.540853522 | -0.133143801 | 11.73900223 | -4.067019623 | 0.006911075 | 0.594255995 | -2.889372176 |
| **A_55_P2061421** | NA | -0.35820316 | -0.575436249 | -0.140970071 | 14.47808734 | -4.056697256 | 0.00699081 | 0.594255995 | -2.893541348 |
| **A_55_P2016426** | Nova2 | 0.609319413 | 0.23943351 | 0.979205315 | 9.526203304 | 4.052718125 | 0.007021822 | 0.594255995 | -2.895152678 |
| **A_55_P1998987** | NA | 0.269041007 | 0.105550785 | 0.432531229 | 10.10168098 | 4.048515998 | 0.00705474 | 0.594255995 | -2.896856833 |
| **A_51_P484832** | Ppm1g | -0.217492069 | -0.349849186 | -0.085134952 | 8.753352199 | -4.042642048 | 0.007101045 | 0.594255995 | -2.899243342 |
| **A_55_P2108255** | Lgmn | -0.207580916 | -0.334272513 | -0.080889319 | 15.42479986 | -4.030962685 | 0.007194131 | 0.594255995 | -2.904003625 |
| **A_51_P280245** | Zfp207 | -0.255192144 | -0.41131512 | -0.099069168 | 10.97996833 | -4.021329341 | 0.007271941 | 0.594255995 | -2.907945165 |
| **A_55_P2042813** | Gbp11 | -0.199726858 | -0.321947952 | -0.077505764 | 8.343122699 | -4.020309296 | 0.007280235 | 0.594255995 | -2.908363327 |
| **A_55_P2010858** | Mrps23 | -0.199833717 | -0.322136481 | -0.077530953 | 8.851937573 | -4.019774181 | 0.007284591 | 0.594255995 | -2.908582757 |
| **A_55_P2146002** | Pank4 | 0.211536326 | 0.082051892 | 0.34102076 | 9.847186331 | 4.019171689 | 0.007289498 | 0.594255995 | -2.908829865 |
| **A_55_P2070546** | 1700037C18Rik | 0.226713965 | 0.087923077 | 0.365504854 | 9.426485593 | 4.01870826 | 0.007293275 | 0.594255995 | -2.909019975 |
| **A_52_P686785** | Lyve1 | -0.33544823 | -0.541188 | -0.129708459 | 10.14537101 | -4.011219392 | 0.007354617 | 0.594255995 | -2.912096511 |
| **A_52_P350554** | Kcnb1 | -0.197955188 | -0.319695476 | -0.076214899 | 8.436990252 | -4.000384378 | 0.007444395 | 0.594255995 | -2.916562463 |
| **A_52_P410685** | Krt7 | -0.249165938 | -0.402546723 | -0.095785154 | 12.12610861 | -3.996565003 | 0.007476334 | 0.594255995 | -2.918140899 |
| **A_55_P2025023** | NA | 0.311149457 | 0.119609799 | 0.502689116 | 15.41519223 | 3.996497106 | 0.007476903 | 0.594255995 | -2.918168978 |
| **A_55_P2075000** | NA | -0.292997273 | -0.473673 | -0.112321547 | 12.54347556 | -3.989632855 | 0.007534697 | 0.594255995 | -2.921011321 |
| **A_55_P2166773** | Pitpna | -0.197039162 | -0.318556056 | -0.075522268 | 12.73048285 | -3.989193061 | 0.007538417 | 0.594255995 | -2.921193671 |
| **A_51_P326191** | Serpina3g | -0.302166961 | -0.488664705 | -0.115669218 | 8.10852133 | -3.986048312 | 0.007565076 | 0.594255995 | -2.922498408 |
| **A_51_P152845** | Trim24 | -0.238887919 | -0.386448546 | -0.091327292 | 10.44670821 | -3.982839974 | 0.007592383 | 0.594255995 | -2.923831056 |
| **A_51_P446583** | Zfp445 | -0.271306181 | -0.438974848 | -0.103637515 | 9.032877323 | -3.980860334 | 0.007609288 | 0.594255995 | -2.92465411 |
| **A_52_P456977** | Deaf1 | -0.203555697 | -0.329382617 | -0.077728776 | 8.676655081 | -3.979961376 | 0.007616978 | 0.594255995 | -2.925028054 |
| **A_55_P2054559** | NA | 0.565167787 | 0.215724247 | 0.914611327 | 9.697112597 | 3.978959763 | 0.007625557 | 0.594255995 | -2.925444843 |
| **A_51_P319070** | Retsat | -0.360374223 | -0.583387431 | -0.137361014 | 10.14287994 | -3.975504925 | 0.00765523 | 0.594255995 | -2.92688362 |
| **A_52_P319606** | Gpr107 | -0.259712116 | -0.420549856 | -0.098874377 | 11.04965724 | -3.972586966 | 0.007680393 | 0.594255995 | -2.92810021 |
| **A_52_P496726** | Rasd1 | -0.243870609 | -0.395342595 | -0.092398624 | 8.765105543 | -3.960922162 | 0.007781915 | 0.594255995 | -2.93297645 |
| **A_51_P125050** | Wdr74 | -0.194569285 | -0.315450465 | -0.073688105 | 8.936365868 | -3.959904898 | 0.007790839 | 0.594255995 | -2.93340267 |
| **A_55_P2165560** | Plekhg5 | 0.210649994 | 0.079162581 | 0.342137407 | 9.743603783 | 3.941362982 | 0.007955534 | 0.594255995 | -2.941198926 |
| **A_52_P380369** | Phf11d | -0.255828665 | -0.415537082 | -0.096120248 | 8.411762122 | -3.940855978 | 0.007960091 | 0.594255995 | -2.941412836 |
| **A_55_P2005838** | Ticrr | -0.231050034 | -0.375580465 | -0.086519604 | 8.693396259 | -3.932927166 | 0.008031751 | 0.594255995 | -2.944763171 |
| **A_51_P211903** | Golga5 | -0.220799789 | -0.359089597 | -0.082509981 | 10.81233235 | -3.928055758 | 0.008076135 | 0.594255995 | -2.946826339 |
| **A_65_P11059** | Letm1 | -0.213329483 | -0.34698131 | -0.079677657 | 11.39390023 | -3.926857537 | 0.008087095 | 0.594255995 | -2.947334371 |
| **A_65_P07561** | Tpd52l2 | -0.263652237 | -0.428946094 | -0.098358381 | 10.00239648 | -3.924134745 | 0.008112061 | 0.594255995 | -2.948489619 |
| **A_55_P2141287** | Erlin1 | 0.235097229 | 0.087681174 | 0.382513284 | 11.76371669 | 3.923484021 | 0.00811804 | 0.594255995 | -2.94876588 |
| **A_52_P194805** | Zfp706 | -0.283452436 | -0.461644953 | -0.105259919 | 14.14128364 | -3.913450942 | 0.008210859 | 0.594255995 | -2.953033566 |
| **A_55_P2118173** | Igf2 | -0.354330474 | -0.577442475 | -0.131218472 | 10.93676827 | -3.907101883 | 0.008270209 | 0.594255995 | -2.955742172 |
| **A_55_P2096823** | NA | 0.328013256 | 0.121332268 | 0.534694244 | 9.067106686 | 3.90445147 | 0.008295127 | 0.594255995 | -2.956874708 |
| **A_52_P140356** | Calm3 | 0.325732307 | 0.120131574 | 0.53133304 | 10.70431311 | 3.897672458 | 0.008359242 | 0.594255995 | -2.959776325 |

| | | | | | | | | | |
|---|---|---|---|---|---|---|---|---|---|
| **A_55_P2038111** | Tmem104 | 0.844875501 | 0.311544544 | 1.378206458 | 8.852627522 | 3.897311214 | 0.008362674 | 0.594255995 | -2.959931146 |
| **A_51_P257419** | Lhx2 | -0.446794896 | -0.72885656 | -0.164733232 | 10.75198135 | -3.89702599 | 0.008365385 | 0.594255995 | -2.960053401 |
| **A_55_P2009217** | Rnu3b1 | 0.828387624 | 0.305244262 | 1.351530986 | 11.84546246 | 3.895668967 | 0.008378297 | 0.594255995 | -2.96063523 |
| **A_51_P298355** | Dek | -0.19340458 | -0.315606653 | -0.071202507 | 10.79637609 | -3.8936539 | 0.008397511 | 0.594255995 | -2.961499722 |
| **A_55_P2051566** | NA | 0.315641901 | 0.115630079 | 0.515653724 | 8.401089649 | 3.882470337 | 0.008505048 | 0.594255995 | -2.966309 |
| **A_55_P2089125** | Zfp976 | -0.249106932 | -0.407192142 | -0.091021723 | 12.44469204 | -3.876713769 | 0.008561001 | 0.594255995 | -2.968792031 |
| **A_51_P366770** | Rims1 | 1.096176914 | 0.400495699 | 1.791858129 | 9.621969662 | 3.876497767 | 0.008563109 | 0.594255995 | -2.968885301 |
| **A_55_P2074636** | Adap1 | 0.33718703 | 0.123117258 | 0.551256802 | 11.89454912 | 3.875115735 | 0.008576607 | 0.594255995 | -2.969482234 |
| **A_55_P2017998** | NA | 0.456517706 | 0.166445424 | 0.746589987 | 9.290877054 | 3.871868138 | 0.008608419 | 0.594255995 | -2.970886114 |
| **A_55_P2000090** | Arf1 | -0.271056745 | -0.443306607 | -0.098806882 | 15.2472406 | -3.871421858 | 0.008612801 | 0.594255995 | -2.971079161 |
| **A_55_P2177273** | Kcna6 | 0.508119502 | 0.185097565 | 0.831141439 | 9.503315692 | 3.869929116 | 0.008627476 | 0.594255995 | -2.971725098 |
| **A_52_P366105** | Pip4p1 | -0.242939435 | -0.397523511 | -0.088355359 | 8.994981015 | -3.866361222 | 0.008662665 | 0.594255995 | -2.973270394 |
| **A_51_P317443** | Cd3eap | -0.255261301 | -0.417844297 | -0.092678304 | 9.442234012 | -3.862593605 | 0.008699998 | 0.594255995 | -2.974904335 |
| **A_55_P2071486** | Suv39h1 | 0.542178914 | 0.19628622 | 0.888071609 | 9.372658017 | 3.856296314 | 0.0087628 | 0.594255995 | -2.977640272 |
| **A_55_P2088825** | Foxp1 | -0.329178872 | -0.539214219 | -0.119143525 | 11.1908809 | -3.855748748 | 0.008768285 | 0.594255995 | -2.97787846 |
| **A_51_P440399** | Bcap31 | -0.26725938 | -0.437989266 | -0.096529493 | 14.52251453 | -3.851168921 | 0.008814309 | 0.594255995 | -2.979872489 |
| **A_55_P1977071** | Fam110a | 0.181858155 | 0.065619336 | 0.298096973 | 9.797025179 | 3.849025212 | 0.008835944 | 0.594255995 | -2.98080697 |
| **A_51_P329283** | Senp5 | -0.278788214 | -0.457979829 | -0.099596599 | 10.28338137 | -3.827594208 | 0.009055532 | 0.594255995 | -2.990188584 |
| **A_51_P393761** | Ndufs2 | -0.191332518 | -0.314530063 | -0.068134973 | 9.891947539 | -3.820813994 | 0.009126268 | 0.594255995 | -2.993171664 |
| **A_55_P2039903** | Wsb2 | -0.49202292 | -0.808958007 | -0.175087833 | 11.26423618 | -3.819303542 | 0.00914211 | 0.594255995 | -2.993837198 |
| **A_51_P386810** | Gmppb | 0.239055144 | 0.085048516 | 0.393061772 | 11.35608267 | 3.818808178 | 0.009147312 | 0.594255995 | -2.994055544 |
| **A_55_P1977812** | Cd244a | 0.670655239 | 0.238114288 | 1.103196191 | 10.20822409 | 3.814531921 | 0.009192358 | 0.594255995 | -2.995942024 |
| **A_55_P2025615** | Ammecr1l | -0.283040387 | -0.465698467 | -0.100382306 | 13.05091316 | -3.812226379 | 0.009216747 | 0.594255995 | -2.996960314 |
| **A_55_P1995512** | Tmem139 | -0.18645076 | -0.307072099 | -0.065829422 | 10.96902658 | -3.802849847 | 0.009316685 | 0.594255995 | -3.001110281 |
| **A_55_P2072706** | Chsy1 | 0.238728446 | 0.0841498 | 0.393307091 | 11.14883841 | 3.799477146 | 0.009352927 | 0.594255995 | -3.0026064 |
| **A_55_P1965827** | Ubb | -0.188034237 | -0.309905321 | -0.066163153 | 16.8279036 | -3.795818354 | 0.009392422 | 0.594255995 | -3.004231461 |
| **A_52_P277854** | Snrpb | -0.251120714 | -0.414096967 | -0.088144461 | 9.688747117 | -3.790769447 | 0.009447228 | 0.594255995 | -3.006477423 |
| **A_55_P1986833** | Tulp4 | -0.264498646 | -0.436267135 | -0.092730157 | 11.86052384 | -3.788341488 | 0.009473711 | 0.594255995 | -3.007558917 |
| **A_55_P1957739** | Guca2b | 1.572811512 | 0.550565118 | 2.595057905 | 10.76030231 | 3.785212411 | 0.009507963 | 0.594255995 | -3.00895409 |
| **A_55_P2110362** | Zfp637 | -0.168585624 | -0.278297155 | -0.058874092 | 8.459931154 | -3.7803971 | 0.009560945 | 0.594255995 | -3.011104145 |
| **A_51_P517094** | Sh3gl1 | -0.307362733 | -0.507431786 | -0.10729368 | 11.1049361 | -3.779553142 | 0.009570264 | 0.594255995 | -3.011481355 |
| **A_55_P2163128** | NA | 0.215351601 | 0.075128581 | 0.35557462 | 9.063391173 | 3.778313166 | 0.009583975 | 0.594255995 | -3.012035771 |
| **A_52_P869802** | Thoc3 | -0.26517159 | -0.437907157 | -0.092436023 | 9.190150709 | -3.776716507 | 0.009601663 | 0.594255995 | -3.012750027 |
| **A_66_P116098** | Ptpn2 | -0.173210116 | -0.286041136 | -0.060379096 | 9.151635828 | -3.776712292 | 0.00960171 | 0.594255995 | -3.012751913 |
| **A_52_P95740** | Smim19 | -0.175290886 | -0.2896105 | -0.060971272 | 10.45650991 | -3.772313269 | 0.00965063 | 0.594255995 | -3.01472189 |
| **A_55_P2157384** | Xlr5b | -0.242626306 | -0.4010203 | -0.084232311 | 11.27721204 | -3.768498416 | 0.00969328 | 0.594255995 | -3.016432757 |
| **A_66_P100657** | Xpa | -0.216831573 | -0.358764577 | -0.074898568 | 10.59428998 | -3.758445763 | 0.009806677 | 0.594255995 | -3.020952234 |
| **A_51_P109888** | Coro1b | -0.359017724 | -0.594071252 | -0.123964195 | 9.499797925 | -3.757666881 | 0.009815525 | 0.594255995 | -3.021303077 |
| **A_55_P2182921** | NA | 0.274223083 | 0.094432247 | 0.454013919 | 9.505524998 | 3.752369695 | 0.009875934 | 0.594255995 | -3.02369174 |
| **A_55_P2026437** | Gm4802 | -0.358002013 | -0.592747798 | -0.123256229 | 15.83007372 | -3.75194817 | 0.009880759 | 0.594255995 | -3.023882011 |
| **A_55_P1952725** | NA | -0.421347559 | -0.698058978 | -0.144636141 | 14.09056057 | -3.746127595 | 0.009947652 | 0.594255995 | -3.026512256 |
| **A_55_P1995417** | Cnppd1 | -0.243387672 | -0.403239153 | -0.083536191 | 9.536737653 | -3.74585597 | 0.009950786 | 0.594255995 | -3.026635132 |
| **A_55_P2115980** | Xlr5c | -0.238314314 | -0.395058616 | -0.081570012 | 9.09632742 | -3.74048165 | 0.010013021 | 0.594255995 | -3.029068774 |
| **A_51_P320105** | Smyd2 | 0.27951881 | 0.095557956 | 0.463479665 | 12.61005621 | 3.738133304 | 0.010040352 | 0.594255995 | -3.030133625 |
| **A_55_P2009576** | Rnu12 | 0.37892478 | 0.129309746 | 0.628539813 | 12.67836654 | 3.734663108 | 0.01008089 | 0.594255995 | -3.031708797 |
| **A_51_P295192** | Nfkbia | -0.387306858 | -0.642524266 | -0.13208945 | 12.53687314 | -3.733481959 | 0.01009473 | 0.594255995 | -3.03224538 |
| **A_55_P2083009** | Catsperd | 0.425301338 | 0.144773185 | 0.705829491 | 9.731073525 | 3.729833621 | 0.010137612 | 0.594255995 | -3.033904193 |
| **A_55_P2112360** | Arhgap17 | 0.379291851 | 0.128954005 | 0.629629696 | 10.9671243 | 3.727487235 | 0.010165298 | 0.594255995 | -3.034972171 |
| **A_55_P2326582** | 5730408K05Rik | 0.436014795 | 0.147882158 | 0.724147433 | 8.45810475 | 3.722870546 | 0.01022002 | 0.594255995 | -3.037076085 |
| **A_55_P2079116** | Rpph1 | 0.359010934 | 0.121653183 | 0.596368685 | 9.179998093 | 3.721117816 | 0.010240881 | 0.594255995 | -3.037875738 |

| | | | | | | | | | |
|---|---|---|---|---|---|---|---|---|---|
| **A_55_P2149615** | Bex3 | -0.20414069 | -0.339142698 | -0.069138682 | 11.72286089 | -3.720133387 | 0.010252618 | 0.594255995 | -3.038325083 |
| **A_51_P483118** | Hmga1 | 0.209543061 | 0.070782799 | 0.348303322 | 13.19885351 | 3.715158404 | 0.010312165 | 0.594255995 | -3.040598322 |
| **A_55_P1994229** | Hrc | 0.243784991 | 0.082256204 | 0.405313778 | 8.16837651 | 3.713010655 | 0.010337991 | 0.594255995 | -3.041580936 |
| **A_55_P2013760** | Slc39a14 | 0.312565022 | 0.105419593 | 0.519710451 | 11.766745 | 3.71222409 | 0.010347467 | 0.594255995 | -3.041940983 |
| **A_55_P2085806** | Paxx | 0.289295541 | 0.097426504 | 0.481164579 | 10.92512955 | 3.709419979 | 0.010381329 | 0.594255995 | -3.043225368 |
| **A_55_P2160691** | NA | -0.372199841 | -0.619193395 | -0.125206288 | 15.79450696 | -3.707317048 | 0.010406804 | 0.594255995 | -3.04418942 |
| **A_51_P143712** | Tdh | -0.358145463 | -0.595950971 | -0.120339954 | 8.506712177 | -3.705157774 | 0.010433035 | 0.594255995 | -3.045180047 |
| **A_51_P130110** | Idh3b | -0.282625789 | -0.470347519 | -0.094904059 | 9.938313955 | -3.703960981 | 0.010447605 | 0.594255995 | -3.045729434 |
| **A_66_P123028** | NA | 0.372475613 | 0.125068274 | 0.619882952 | 8.940516075 | 3.70385885 | 0.010448849 | 0.594255995 | -3.045776328 |
| **A_55_P2152567** | Plekha5 | 0.744494773 | 0.249043883 | 1.239945663 | 9.553159008 | 3.69683616 | 0.010534815 | 0.594255995 | -3.049004882 |
| **A_52_P126825** | Cramp1l | -0.35953896 | -0.599213972 | -0.119863947 | 11.79468325 | -3.690560778 | 0.010612299 | 0.594255995 | -3.051896643 |
| **A_55_P2089219** | NA | 0.230375686 | 0.076571631 | 0.38417974 | 7.801959623 | 3.68500448 | 0.010681435 | 0.594255995 | -3.054462382 |
| **A_51_P478098** | Epb41l5 | -0.206276078 | -0.344165017 | -0.068387139 | 8.515255534 | -3.680345568 | 0.010739791 | 0.594255995 | -3.056617604 |
| **A_55_P2010539** | NA | 0.256632721 | 0.08504583 | 0.428219612 | 8.748157644 | 3.679570141 | 0.010749538 | 0.594255995 | -3.056976662 |
| **A_51_P416869** | Nedd8 | -0.211158502 | -0.352360919 | -0.069956084 | 12.82233678 | -3.679049255 | 0.010756091 | 0.594255995 | -3.05721791 |
| **A_66_P115116** | Wwtr1 | -0.202632259 | -0.338172372 | -0.067092146 | 11.4871594 | -3.677984599 | 0.010769499 | 0.594255995 | -3.057711143 |
| **A_55_P2064841** | Prss53 | 0.327668453 | 0.108274229 | 0.547062676 | 12.86558154 | 3.674333772 | 0.010815618 | 0.594255995 | -3.059403899 |
| **A_52_P87892** | Abraxas2 | -0.201370076 | -0.336237924 | -0.066502228 | 9.327176555 | -3.673293824 | 0.010828795 | 0.594255995 | -3.059886483 |
| **A_55_P2112170** | Crtc1 | 0.307870616 | 0.101595839 | 0.514145393 | 9.412209638 | 3.671903655 | 0.010846438 | 0.594255995 | -3.06053186 |
| **A_66_P138976** | Lpin2 | -0.299217548 | -0.499714632 | -0.098720463 | 9.215149814 | -3.671539124 | 0.01085107 | 0.594255995 | -3.060701144 |
| **A_51_P331288** | Akr1b7 | 0.2662702 | 0.087688251 | 0.444852149 | 8.178222263 | 3.668209827 | 0.010893473 | 0.594255995 | -3.062248232 |
| **A_52_P101487** | Ddn | 0.552428731 | 0.181747534 | 0.923109929 | 10.72842336 | 3.666442375 | 0.010916059 | 0.594255995 | -3.063070283 |
| **A_55_P2065434** | Rprl2 | 0.690402817 | 0.226973487 | 1.153832148 | 9.273669771 | 3.665120026 | 0.010932991 | 0.594255995 | -3.063685648 |
| **A_51_P286034** | Ccdc117 | -0.293183044 | -0.49005072 | -0.096315368 | 9.922389005 | -3.663815528 | 0.010949723 | 0.594255995 | -3.064292986 |
| **A_52_P222725** | Lims1 | -0.297392657 | -0.497170803 | -0.097614511 | 11.37316816 | -3.662278992 | 0.010969468 | 0.594255995 | -3.065008711 |
| **A_51_P249867** | Snrnp48 | -0.246307699 | -0.411949525 | -0.080665873 | 9.292397705 | -3.65828166 | 0.01102102 | 0.594255995 | -3.066872496 |
| **A_55_P2032912** | NA | -0.501154786 | -0.838389828 | -0.163919744 | 13.89439064 | -3.656018207 | 0.011050331 | 0.594255995 | -3.067929006 |
| **A_55_P2022139** | Pura | 0.395116728 | 0.128983756 | 0.661249699 | 12.14354166 | 3.6525477 | 0.011095441 | 0.594255995 | -3.069550563 |
| **A_55_P2060198** | NA | 0.986804348 | 0.321873852 | 1.651734844 | 11.44645081 | 3.651102001 | 0.011114292 | 0.594255995 | -3.070226633 |
| **A_52_P658974** | NA | -0.437490876 | -0.732282246 | -0.142699505 | 12.74141889 | -3.651097214 | 0.011114355 | 0.594255995 | -3.070228872 |
| **A_55_P2143261** | Rpain | 0.364010342 | 0.11861529 | 0.609405393 | 11.49259584 | 3.649363172 | 0.011137014 | 0.594255995 | -3.071040236 |
| **A_55_P2046877** | Foxq1 | -0.243342254 | -0.407488416 | -0.079196091 | 9.56523618 | -3.647169563 | 0.011165751 | 0.594255995 | -3.072067341 |
| **A_55_P1966615** | Actr3b | 0.22767545 | 0.073995595 | 0.381355304 | 8.713198565 | 3.644755702 | 0.01119747 | 0.594255995 | -3.073198485 |
| **A_51_P414889** | Ifi35 | -0.174264611 | -0.291909576 | -0.056619645 | 9.819003322 | -3.6442236 | 0.011204475 | 0.594255995 | -3.073447959 |
| **A_55_P1975732** | Sipa1l3 | 0.307403393 | 0.099754663 | 0.515052122 | 10.58658403 | 3.642072117 | 0.01123285 | 0.594255995 | -3.074457147 |
| **A_55_P2042086** | Nrtn | 0.282835934 | 0.091781785 | 0.473890082 | 10.13716267 | 3.642061502 | 0.011232991 | 0.594255995 | -3.074462128 |
| **A_66_P112886** | Ubl3 | -0.21489088 | -0.360048839 | -0.06973292 | 10.67883347 | -3.642053259 | 0.011233099 | 0.594255995 | -3.074465996 |
| **A_66_P106585** | Sae1 | -0.198906678 | -0.333476044 | -0.064337311 | 9.507588038 | -3.636405602 | 0.011307967 | 0.594255995 | -3.077118759 |
| **A_51_P230934** | Cab39 | -0.21091097 | -0.353878262 | -0.067943679 | 11.57151171 | -3.629373319 | 0.011401966 | 0.594255995 | -3.08042923 |
| **A_51_P371485** | Abraxas2 | -0.280138933 | -0.470085574 | -0.090192293 | 9.99732626 | -3.628365404 | 0.011415509 | 0.594255995 | -3.080904377 |
| **A_52_P584293** | Atrnl1 | -0.43265867 | -0.72636164 | -0.138955701 | 9.875917255 | -3.6241506 | 0.011472339 | 0.594255995 | -3.082893117 |
| **A_55_P1965114** | Spsb3 | -0.3879212 | -0.651314709 | -0.124527691 | 13.78996801 | -3.623327665 | 0.011483472 | 0.594255995 | -3.083281758 |
| **A_51_P489413** | Luc7l | -0.207772276 | -0.348847426 | -0.066697125 | 9.82176095 | -3.623316222 | 0.011483626 | 0.594255995 | -3.083287163 |
| **A_55_P2173005** | Dnase2a | 0.628599076 | 0.200704183 | 1.056493968 | 13.51381307 | 3.614146883 | 0.011608485 | 0.594255995 | -3.087625073 |
| **A_55_P2038077** | Rpsa | 0.225669089 | 0.071758257 | 0.379579921 | 15.16536922 | 3.607215214 | 0.011703875 | 0.594255995 | -3.090913594 |
| **A_55_P2015878** | Clps | 0.425098386 | 0.135059002 | 0.71513777 | 10.58936891 | 3.605800093 | 0.011723456 | 0.594255995 | -3.091585933 |
| **A_66_P108059** | Ttc39c | -0.268872385 | -0.452499535 | -0.085245235 | 8.461056074 | -3.602288306 | 0.011772206 | 0.594255995 | -3.093255854 |
| **A_55_P2209053** | 2900001G08Rik | 0.367013457 | 0.116225263 | 0.617801651 | 8.93468754 | 3.600344539 | 0.011799286 | 0.594255995 | -3.094181031 |
| **A_52_P579640** | Rrs1 | -0.167726811 | -0.282450098 | -0.053003523 | 9.568982684 | -3.596831288 | 0.011848407 | 0.594255995 | -3.095854827 |
| **A_52_P304858** | Gnb1 | -0.328892728 | -0.554158359 | -0.103627098 | 9.799190299 | -3.591935251 | 0.011917241 | 0.594255995 | -3.098190832 |

| | | | | | | | | |
|---|---|---|---|---|---|---|---|---|
| **A_55_P2068731** | Gm21953 | 0.374244289 | 0.117653387 | 0.630835191 | 9.295265027 | 3.588253283 | 0.011969299 | 0.594255995 | -3.099950203 |
| **A_51_P384609** | Ddx10 | -0.279706801 | -0.471527821 | -0.08788578 | 9.198304291 | -3.587368513 | 0.011981846 | 0.594255995 | -3.100373312 |
| **A_66_P120347** | Sstr5 | 0.662633889 | 0.207906994 | 1.117360784 | 11.81231392 | 3.585024806 | 0.012015153 | 0.594255995 | -3.101494735 |
| **A_52_P497534** | Wipi1 | -0.213533962 | -0.360079623 | -0.0669883 | 10.37916585 | -3.584785344 | 0.012018562 | 0.594255995 | -3.101609365 |
| **A_55_P2139430** | NA | 0.52269935 | 0.163962737 | 0.881435964 | 11.16879284 | 3.584638951 | 0.012020646 | 0.594255995 | -3.101679448 |
| **A_55_P2187038** | Tpm1 | -0.273780805 | -0.46178749 | -0.08577412 | 10.6310972 | -3.582604504 | 0.012049656 | 0.594255995 | -3.102653766 |
| **A_51_P146303** | Mvb12a | 0.233100816 | 0.073006044 | 0.393195588 | 11.53803522 | 3.58208434 | 0.012057086 | 0.594255995 | -3.102902989 |
| **A_66_P111401** | Fbxw8 | -0.27985328 | -0.472085406 | -0.087621153 | 10.74864348 | -3.581571238 | 0.01206442 | 0.594255995 | -3.103148873 |
| **A_51_P448203** | Cnbp | -0.332019082 | -0.560160386 | -0.103877778 | 12.54574554 | -3.580373125 | 0.012081564 | 0.594255995 | -3.103723191 |
| **A_55_P2280089** | 4930570B17Rik | 0.426191968 | 0.133338616 | 0.719045321 | 10.26030349 | 3.580339616 | 0.012082044 | 0.594255995 | -3.103739257 |
| **A_55_P2148082** | Larp7 | -0.186710579 | -0.315069295 | -0.058351862 | 9.320185162 | -3.578596399 | 0.012107037 | 0.594255995 | -3.104575309 |
| **A_55_P2156588** | Ifnk | -0.329929909 | -0.556777732 | -0.103082087 | 9.165021063 | -3.57813099 | 0.01211372 | 0.594255995 | -3.104798606 |
| **A_55_P1972257** | Sars | 0.214149057 | 0.06671234 | 0.361585774 | 12.75299137 | 3.573383921 | 0.012182115 | 0.594255995 | -3.107078249 |
| **A_51_P520139** | Ccpg1os | -0.324514395 | -0.548160604 | -0.100868185 | 10.00918847 | -3.569781111 | 0.01223431 | 0.594255995 | -3.108810904 |
| **A_55_P1952690** | Rpl12 | 0.319782409 | 0.099359996 | 0.540204821 | 14.01402109 | 3.569176142 | 0.012243098 | 0.594255995 | -3.109102057 |
| **A_55_P1982996** | Ramp1 | 0.312050039 | 0.096577709 | 0.52752237 | 8.855642059 | 3.56288575 | 0.012334898 | 0.594255995 | -3.11213305 |
| **A_55_P2145706** | Thoc7 | -0.311227193 | -0.526145777 | -0.096308609 | 9.893990254 | -3.562646471 | 0.012338405 | 0.594255995 | -3.112248476 |
| **A_52_P192426** | Tnfrsf1a | -0.332953503 | -0.562888874 | -0.103018131 | 12.48984488 | -3.562435139 | 0.012341503 | 0.594255995 | -3.123291632 |
| **A_55_P2021841** | Calm3 | 0.380300309 | 0.117508222 | 0.643092396 | 12.91500478 | 3.560275104 | 0.012373221 | 0.594255995 | -3.113392918 |
| **A_51_P203474** | Tsg101 | -0.283205587 | -0.479024212 | -0.087386963 | 10.7113454 | -3.558090457 | 0.012405392 | 0.594255995 | -3.11444808 |
| **A_55_P2000329** | Gskip | 0.172189573 | 0.053119451 | 0.291259694 | 9.783723032 | 3.557731875 | 0.012410681 | 0.594255995 | -3.114621348 |
| **A_51_P327751** | Ifit1 | -0.251504257 | -0.425612573 | -0.077395941 | 8.695771126 | -3.55381641 | 0.0124686 | 0.594255995 | -3.116514711 |
| **A_55_P2048119** | Slc29a4 | 0.300918699 | 0.09247788 | 0.509359517 | 8.70525952 | 3.551694217 | 0.012500117 | 0.594255995 | -3.117541993 |
| **A_55_P2139341** | Zfp975 | 0.455947558 | 0.139713808 | 0.772181308 | 12.95375013 | 3.547119658 | 0.012568357 | 0.594255995 | -3.119758949 |
| **A_51_P442053** | Ptdss1 | -0.163539948 | -0.277041153 | -0.050038743 | 9.296427067 | -3.544806595 | 0.012603018 | 0.594255995 | -3.120881257 |
| **A_55_P2008327** | Eno1b | 0.54191302 | 0.165767511 | 0.918058529 | 14.56558977 | 3.544401144 | 0.012609105 | 0.594255995 | -3.121078077 |
| **A_55_P2173664** | Frmpd2 | -0.237560202 | -0.40248054 | -0.072639864 | 13.99584976 | -3.543795342 | 0.012618205 | 0.594255995 | -3.121372204 |
| **A_55_P1958597** | Slc27a3 | 0.536084571 | 0.223082321 | 0.849086822 | 11.4551523 | 5.593953166 | 0.012697185 | 0.594255995 | -3.870592447 |
| **A_55_P2056280** | Wdr89 | -0.308440218 | -0.522975976 | -0.093904461 | 13.70155033 | -3.53704412 | 0.012720117 | 0.594255995 | -3.124654212 |
| **A_55_P2136101** | Meis3 | 0.209049407 | 0.063290473 | 0.354808341 | 8.871850112 | 3.528441483 | 0.012851302 | 0.594255995 | -3.128847359 |
| **A_55_P2061874** | Prelid1 | 0.314882739 | 0.095281521 | 0.534483956 | 14.50288253 | 3.527631999 | 0.012863723 | 0.594255995 | -3.129242563 |
| **A_55_P1959313** | Rbm42 | -0.184384376 | -0.313300961 | -0.055467791 | 12.71845611 | -3.518718217 | 0.013001384 | 0.594255995 | -3.133601731 |
| **A_55_P2014361** | Pdpk1 | 0.319926927 | 0.102901614 | 0.53695224 | 12.00008677 | 3.820238579 | 0.013004181 | 0.594255995 | -3.317290587 |
| **A_55_P2103011** | Sema4d | 0.419075221 | 0.12605379 | 0.712096653 | 10.58554913 | 3.518534066 | 0.013004245 | 0.594255995 | -3.133691929 |
| **A_66_P133204** | NA | -0.220369832 | -0.374539049 | -0.066200615 | 9.400347811 | -3.516605385 | 0.013034253 | 0.594255995 | -3.134636948 |
| **A_55_P2176260** | NA | 0.572321105 | 0.171461157 | 0.973181054 | 8.733382512 | 3.512499395 | 0.013098392 | 0.594255995 | -3.1366509 |
| **A_55_P2040583** | Map6d1 | 0.625345851 | 0.18613865 | 1.064553051 | 11.73338402 | 3.50283745 | 0.013250705 | 0.594255995 | -3.141401236 |
| **A_55_P2052380** | Gbp6 | -0.218557571 | -0.372126578 | -0.064988565 | 8.152013093 | -3.501317099 | 0.01327485 | 0.594255995 | -3.142150161 |
| **A_55_P2009266** | Gm16060 | -0.206693854 | -0.352065601 | -0.061322108 | 13.80664433 | -3.497975133 | 0.013328097 | 0.594255995 | -3.143797787 |
| **A_51_P499153** | Cog3 | -0.317228397 | -0.540417482 | -0.094039312 | 9.677946201 | -3.496779663 | 0.013347201 | 0.594255995 | -3.144387627 |
| **A_55_P2063366** | Commd7 | -0.220508181 | -0.375682684 | -0.065333679 | 11.85925056 | -3.496016779 | 0.013359408 | 0.594255995 | -3.144764157 |
| **A_55_P2055557** | Sdsl | 0.260448616 | 0.07708831 | 0.443808923 | 9.955439782 | 3.494506817 | 0.013383607 | 0.594255995 | -3.145509706 |
| **A_51_P392861** | Med25 | -0.181606883 | -0.309509532 | -0.053704234 | 9.879163518 | -3.493187744 | 0.013404785 | 0.594255995 | -3.14616132 |
| **A_52_P321318** | NA | -0.202804471 | -0.345726106 | -0.059882836 | 15.31703608 | -3.490990862 | 0.013440141 | 0.594255995 | -3.14724722 |
| **A_55_P2150224** | NA | -0.441852346 | -0.753552663 | -0.130152028 | 12.98451287 | -3.487458778 | 0.0134972 | 0.594255995 | -3.148994816 |
| **A_51_P153170** | Cyb5r3 | -0.409428374 | -0.698583651 | -0.120273096 | 13.05809189 | -3.483501292 | 0.013561451 | 0.594255995 | -3.150955405 |
| **A_51_P386539** | Rnf125 | -0.177150536 | -0.302298122 | -0.05200295 | 8.529319678 | -3.482484223 | 0.013578018 | 0.594255995 | -3.151459703 |
| **A_51_P219061** | Oxa1l | -0.185363301 | -0.316489194 | -0.054237408 | 8.613333666 | -3.477798851 | 0.013654628 | 0.594255995 | -3.153785145 |
| **A_52_P26216** | Numb | -0.188456865 | -0.321776135 | -0.055137595 | 8.403538846 | -3.477668639 | 0.013656764 | 0.594255995 | -3.153849825 |
| **A_51_P300709** | Srm | 0.298256977 | 0.086918892 | 0.509595062 | 10.19633233 | 3.472018311 | 0.013749805 | 0.594255995 | -3.156659282 |

| | | | | | | | | | |
|---|---|---|---|---|---|---|---|---|---|
| **A_55_P2034570** | Ddx19a | -0.166783675 | -0.284963569 | -0.048603781 | 8.594427945 | -3.47199511 | 0.013750189 | 0.594255995 | -3.156670829 |
| **A_55_P1977593** | NA | 0.254223749 | 0.0740853 | 0.434362197 | 12.28415782 | 3.471993177 | 0.013750221 | 0.594255995 | -3.156671791 |
| **A_55_P2013840** | Krt42 | -0.195208961 | -0.333531368 | -0.056886554 | 11.42721133 | -3.471973288 | 0.01375055 | 0.594255995 | -3.15668169 |
| **A_52_P552665** | Fzd7 | -0.206403758 | -0.352689402 | -0.060118114 | 9.035893091 | -3.471243298 | 0.013762622 | 0.594255995 | -3.157045057 |
| **A_55_P2037962** | Trpm6 | -0.313173506 | -0.535434689 | -0.090912322 | 10.24703932 | -3.466494816 | 0.013841437 | 0.594255995 | -3.159410926 |
| **A_55_P2077448** | Epas1 | 0.207087246 | 0.059941241 | 0.354233251 | 11.6760127 | 3.462374487 | 0.013910231 | 0.594255995 | -3.161466937 |
| **A_65_P20174** | Jade1 | -0.314179347 | -0.537601213 | -0.090757482 | 10.25642861 | -3.459562039 | 0.013957405 | 0.594255995 | -3.162871988 |
| **A_51_P509223** | Arfgap1 | -0.222338263 | -0.380465805 | -0.064210721 | 9.024101454 | -3.459201421 | 0.013963467 | 0.594255995 | -3.163052244 |
| **A_55_P2020004** | NA | 1.04250163 | 0.3007272 | 1.78427606 | 13.86445377 | 3.457594409 | 0.013990514 | 0.594255995 | -3.163855784 |
| **A_51_P248381** | Zdhhc4 | -0.190587599 | -0.326207342 | -0.054967857 | 8.476480795 | -3.457330411 | 0.013994963 | 0.594255995 | -3.163987831 |
| **A_55_P1985359** | Rpl28 | -0.176639598 | -0.302589295 | -0.050689902 | 13.71462614 | -3.4503258 | 0.014113578 | 0.594255995 | -3.167495748 |
| **A_55_P2113837** | Tbc1d1 | 0.161795797 | 0.046397344 | 0.277194249 | 11.33727983 | 3.449343252 | 0.014130305 | 0.594255995 | -3.167988479 |
| **A_52_P515247** | Arhgdib | 0.314949432 | 0.090259316 | 0.539639549 | 11.34909839 | 3.448466598 | 0.014145248 | 0.594255995 | -3.168428246 |
| **A_52_P110812** | Ggcx | -0.198387432 | -0.340019959 | -0.056754904 | 9.276525161 | -3.446039991 | 0.014186702 | 0.594255995 | -3.169646218 |
| **A_51_P217430** | Riox1 | -0.167238601 | -0.286675134 | -0.047802068 | 9.045579676 | -3.444835591 | 0.014207326 | 0.594255995 | -3.170251109 |
| **A_55_P2177154** | H2bc14 | -0.367510651 | -0.630098188 | -0.104923114 | 12.54755884 | -3.443221679 | 0.014235015 | 0.594255995 | -3.17106206 |
| **A_55_P2003951** | Tmem192 | 0.267335885 | 0.076187636 | 0.458484135 | 12.90203209 | 3.440773605 | 0.01427713 | 0.594255995 | -3.172293008 |
| **A_52_P682990** | Timm50 | -0.20126028 | -0.345217177 | -0.057303383 | 9.604406767 | -3.439495612 | 0.01429917 | 0.594255995 | -3.172936019 |
| **A_55_P2057446** | Ccdc134 | 0.300983545 | 0.085550671 | 0.516416419 | 10.43167508 | 3.437161514 | 0.014339521 | 0.594255995 | -3.174111122 |
| **A_52_P47645** | Cul3 | -0.182385198 | -0.312939259 | -0.051831137 | 9.500997138 | -3.436911635 | 0.014343848 | 0.594255995 | -3.174236978 |
| **A_51_P143721** | Vipas39 | -0.147770009 | -0.253572739 | -0.041967279 | 10.50504055 | -3.436042984 | 0.014358902 | 0.594255995 | -3.174674577 |
| **A_55_P2147427** | Prdx1 | -0.358439456 | -0.615158029 | -0.101720883 | 13.10999278 | -3.43500744 | 0.014376871 | 0.594255995 | -3.175196419 |
| **A_55_P2055107** | Agpat1 | 0.210160534 | 0.059629767 | 0.3606913 | 12.15885781 | 3.434749376 | 0.014381353 | 0.594255995 | -3.175326494 |
| **A_55_P1954258** | Zfp652 | -0.174795541 | -0.300058969 | -0.049532114 | 9.462406248 | -3.433011211 | 0.014411581 | 0.594255995 | -3.176202898 |
| **A_55_P2138529** | Mrpl40 | 0.30168692 | 0.085012706 | 0.518361135 | 11.41192621 | 3.425456159 | 0.014543786 | 0.594255995 | -3.180018263 |
| **A_51_P130727** | Fkbp11 | 0.629632362 | 0.176822228 | 1.082442496 | 10.93525694 | 3.420897545 | 0.014624205 | 0.594255995 | -3.182325138 |
| **A_55_P2035159** | Pcdh1 | 0.284551098 | 0.079795069 | 0.489307126 | 10.25178743 | 3.418949858 | 0.014658715 | 0.594255995 | -3.183311847 |
| **A_51_P490171** | Nubp2 | -0.184071603 | -0.316597263 | -0.051545943 | 10.31006212 | -3.417086572 | 0.014691813 | 0.594255995 | -3.184256408 |
| **A_55_P1974019** | Dapk1 | -0.254520415 | -0.437771843 | -0.071268986 | 11.26805003 | -3.416995567 | 0.014693431 | 0.594255995 | -3.184302557 |
| **A_55_P1964805** | Rpa1 | 0.29048294 | 0.08120261 | 0.49976327 | 11.05980846 | 3.414769418 | 0.014733088 | 0.594255995 | -3.185431881 |
| **A_55_P2010586** | Gm6252 | 0.595080994 | 0.166163752 | 1.023998235 | 13.1873937 | 3.413278605 | 0.014759711 | 0.594255995 | -3.186188646 |
| **A_55_P2133027** | Kirrel3 | 0.201039343 | 0.055943539 | 0.346135147 | 8.803226415 | 3.408751691 | 0.014840881 | 0.594255995 | -3.188488935 |
| **A_55_P2142200** | NA | -0.291510717 | -0.501958653 | -0.08106278 | 10.29818971 | -3.407838584 | 0.014857313 | 0.594255995 | -3.188953345 |
| **A_51_P418725** | Plekhf1 | -0.185322106 | -0.31917635 | -0.051467863 | 9.021787777 | -3.406153743 | 0.014887686 | 0.594255995 | -3.189810639 |
| **A_55_P2156556** | NA | -0.29441818 | -0.507157775 | -0.081678584 | 11.97142203 | -3.404751796 | 0.014913012 | 0.594255995 | -3.19052436 |
| **A_52_P581440** | Wasf2 | -0.349769506 | -0.602563919 | -0.096975092 | 12.70174806 | -3.403953622 | 0.014927452 | 0.594255995 | -3.190930856 |
| **A_55_P1965478** | Kcnd1 | 0.361924801 | 0.100316907 | 0.623532694 | 9.879227312 | 3.403585488 | 0.014934117 | 0.594255995 | -3.191118377 |
| **A_51_P440568** | Stk19 | -0.17523068 | -0.301902752 | -0.048558609 | 8.956261348 | -3.403285928 | 0.014939543 | 0.594255995 | -3.191270985 |
| **A_55_P2109544** | Trnt1 | -0.143941848 | -0.248010977 | -0.03987272 | 8.819999742 | -3.402783531 | 0.014948648 | 0.594255995 | -3.191526961 |
| **A_55_P2176894** | NA | 0.62903716 | 0.173917147 | 1.084157173 | 8.461834655 | 3.400317982 | 0.01499342 | 0.594255995 | -3.192783811 |
| **A_52_P444457** | Dnlz | -0.187732027 | -0.323584558 | -0.051879495 | 9.186478923 | -3.399693751 | 0.015004779 | 0.594255995 | -3.193102187 |
| **A_52_P456271** | Tmem167 | 0.239624245 | 0.066019193 | 0.413229297 | 12.93022499 | 3.395764474 | 0.015076498 | 0.594255995 | -3.195107781 |
| **A_66_P110607** | Sart1 | 0.396675476 | 0.109158506 | 0.684192447 | 10.71868471 | 3.394228152 | 0.015104643 | 0.594255995 | -3.195892678 |
| **A_51_P463846** | Gbp7 | -0.312235239 | -0.53870795 | -0.085762528 | 8.489491887 | -3.391838768 | 0.01514853 | 0.594255995 | -3.197114208 |
| **A_55_P1995165** | Mdc1 | 0.163485952 | 0.044848247 | 0.282123658 | 9.865959059 | 3.390212084 | 0.015178489 | 0.594255995 | -3.197946385 |
| **A_51_P513163** | Tra2a | -0.268935992 | -0.464147292 | -0.073724692 | 12.71194974 | -3.389324395 | 0.015194865 | 0.594255995 | -3.1984007 |
| **A_55_P1976682** | 4932431P20Rik | -0.30180825 | -0.520957883 | -0.082658618 | 11.08713608 | -3.388125771 | 0.015217009 | 0.594255995 | -3.199014366 |
| **A_51_P340653** | Slc16a4 | -0.24995611 | -0.431484513 | -0.068427707 | 9.943946236 | -3.387570429 | 0.01522728 | 0.594255995 | -3.199298771 |
| **A_52_P20048** | Tcf12 | -0.376647976 | -0.650262676 | -0.103033275 | 10.0888965 | -3.386611458 | 0.015245035 | 0.594255995 | -3.199790011 |
| **A_55_P2275054** | Nrap | 0.57373733 | 0.156937816 | 0.990536843 | 8.996347903 | 3.386531093 | 0.015246524 | 0.594255995 | -3.199831186 |

| | | | | | | | | | |
|---|---|---|---|---|---|---|---|---|---|
| **A_66_P117838** | Dhx29 | -0.239411239 | -0.413450333 | -0.065372146 | 8.819890824 | -3.384284657 | 0.01528821 | 0.594255995 | -3.200982591 |
| **A_51_P483922** | Tm9sf4 | -0.242829559 | -0.419371818 | -0.0662873 | 9.298222633 | -3.383935175 | 0.015294707 | 0.594255995 | -3.201161796 |
| **A_55_P2012201** | App | -0.287430358 | -0.496424647 | -0.078436068 | 14.58683873 | -3.383509333 | 0.015302627 | 0.594255995 | -3.201380184 |
| **A_55_P2169879** | NA | 0.261025029 | 0.071143084 | 0.450906974 | 13.22386652 | 3.381953595 | 0.0153316 | 0.594255995 | -3.202178293 |
| **A_52_P91274** | 1700018G05Rik | 0.35069908 | 0.09549551 | 0.60590265 | 8.670396023 | 3.380781037 | 0.015353477 | 0.594255995 | -3.202780103 |
| **A_52_P676798** | Gigyf1 | 0.300239778 | 0.081678817 | 0.518800738 | 9.493235441 | 3.379596116 | 0.015375619 | 0.594255995 | -3.203388499 |
| **A_55_P2086099** | Gm2061 | 0.20556239 | 0.055861015 | 0.355263764 | 9.025908304 | 3.378213022 | 0.01540151 | 0.594255995 | -3.204098955 |
| **A_52_P327664** | Gbp5 | -0.330913605 | -0.572073145 | -0.089754065 | 8.322101052 | -3.375820443 | 0.015446411 | 0.594255995 | -3.205328733 |
| **A_55_P2043322** | Rnf26 | -0.211946751 | -0.366425987 | -0.057467515 | 8.516373863 | -3.375404409 | 0.015454233 | 0.594255995 | -3.205542675 |
| **A_66_P138289** | Rdh5 | 0.158536061 | 0.042935681 | 0.27413644 | 8.659058712 | 3.373944743 | 0.015481712 | 0.594255995 | -3.20629353 |
| **A_55_P1956171** | NA | -0.230100239 | -0.39791469 | -0.062285787 | 13.30051629 | -3.373314817 | 0.015493588 | 0.594255995 | -3.206617678 |
| **A_52_P59318** | Rdh10 | -0.172886112 | -0.299082847 | -0.046689378 | 9.212846095 | -3.370397728 | 0.015548713 | 0.594255995 | -3.208119656 |
| **A_51_P443514** | 1700001K19Rik | 0.911850336 | 0.246047333 | 1.577653339 | 11.24484586 | 3.369356586 | 0.01556844 | 0.594255995 | -3.208656085 |
| **A_55_P2161545** | Ddx21 | 0.213515795 | 0.057558988 | 0.369472601 | 9.889768138 | 3.368176467 | 0.015590834 | 0.594255995 | -3.209264347 |
| **A_51_P368009** | E2f2 | 0.379898884 | 0.102007247 | 0.657790522 | 9.233238765 | 3.363269767 | 0.015684325 | 0.594255995 | -3.211795961 |
| **A_66_P115389** | NA | 0.24963381 | 0.067028261 | 0.432239359 | 11.23637998 | 3.363245711 | 0.015684784 | 0.594255995 | -3.211808383 |
| **A_55_P2071601** | Ppara | 0.278589946 | 0.07480311 | 0.482376781 | 9.175426239 | 3.363244779 | 0.015684802 | 0.594255995 | -3.211808864 |
| **A_51_P417016** | Srsf3 | 0.172960086 | 0.046396228 | 0.299523945 | 8.443489479 | 3.362059161 | 0.015707486 | 0.594255995 | -3.212421215 |
| **A_55_P2121141** | Tbc1d22b | 0.362110719 | 0.097082894 | 0.627138543 | 10.19795912 | 3.361391301 | 0.01572028 | 0.594255995 | -3.21276626 |
| **A_55_P2325803** | Tnfrsf11a | 0.168504959 | 0.04511927 | 0.291890648 | 8.964606817 | 3.359828115 | 0.01575027 | 0.594255995 | -3.213574171 |
| **A_55_P2064150** | NA | -0.214127154 | -0.371020842 | -0.057233465 | 15.59331295 | -3.357650086 | 0.015792161 | 0.594255995 | -3.214700561 |
| **A_66_P111562** | Ccnd1 | -0.339046538 | -0.587512373 | -0.090580703 | 9.54351274 | -3.357080818 | 0.01580313 | 0.594255995 | -3.214995099 |
| **A_55_P2002846** | Cystm1 | 0.363159584 | 0.096986857 | 0.629332312 | 10.29510606 | 3.356627268 | 0.015811876 | 0.594255995 | -3.215229805 |
| **A_55_P2000062** | Irf1 | -0.295827515 | -0.512667251 | -0.078987779 | 11.09225438 | -3.356362495 | 0.015816984 | 0.594255995 | -3.215366838 |
| **A_55_P1984297** | Efcab14 | -0.318190723 | -0.551720991 | -0.084660454 | 10.30850346 | -3.352073229 | 0.015899985 | 0.594255995 | -3.217588437 |
| **A_52_P463183** | Asap1 | -0.254936959 | -0.442246208 | -0.067627709 | 9.175728724 | -3.348441687 | 0.015970634 | 0.594255995 | -3.219471866 |
| **A_55_P2013218** | Paip1 | 0.394482236 | 0.104547985 | 0.684416487 | 8.510817588 | 3.347318915 | 0.015992546 | 0.594255995 | -3.220054633 |
| **A_55_P2090345** | BC085271 | -0.183417991 | -0.31828526 | -0.048550723 | 10.44002569 | -3.345835107 | 0.016021556 | 0.594255995 | -3.220825129 |
| **A_55_P2112424** | Madd | -0.32517026 | -0.564508221 | -0.0858323 | 15.20573171 | -3.342476763 | 0.01608743 | 0.594255995 | -3.222570426 |
| **A_51_P197321** | Clta | -0.282423037 | -0.49034076 | -0.074505315 | 14.37673348 | -3.341779365 | 0.016101146 | 0.594255995 | -3.222933102 |
| **A_51_P262515** | Phf11a | -0.309677758 | -0.537678238 | -0.081677278 | 9.073531402 | -3.341515002 | 0.016106349 | 0.594255995 | -3.223070603 |
| **A_55_P2143436** | NA | 0.181216763 | 0.047763152 | 0.314670375 | 8.610259591 | 3.340697889 | 0.016122443 | 0.594255995 | -3.223495681 |
| **A_51_P371331** | Mpp5 | -0.364183895 | -0.632468845 | -0.095898945 | 10.32896693 | -3.339593362 | 0.016144225 | 0.594255995 | -3.224070462 |
| **A_52_P561690** | BC034090 | -0.150857006 | -0.262006902 | -0.03970711 | 7.98316787 | -3.339070477 | 0.016154548 | 0.594255995 | -3.224342638 |
| **A_55_P2139640** | Fbh1 | -0.274093853 | -0.476237535 | -0.071950171 | 9.787919841 | -3.335863619 | 0.01621802 | 0.594255995 | -3.226012935 |
| **A_55_P2081690** | Tex38 | 0.477143663 | 0.125103074 | 0.829184252 | 8.27091261 | 3.334460557 | 0.016245876 | 0.594255995 | -3.226744283 |
| **A_55_P2081183** | Slc28a1 | 0.486217367 | 0.127476788 | 0.844957946 | 9.01782349 | 3.334410963 | 0.016246862 | 0.594255995 | -3.226770141 |
| **A_66_P111430** | 2410006H16Rik | 0.457888891 | 0.120043262 | 0.79573452 | 12.62174245 | 3.334348318 | 0.016248107 | 0.594255995 | -3.226802803 |
| **A_55_P2125766** | Ola1 | 0.273797287 | 0.071691467 | 0.475903107 | 9.332987336 | 3.332878507 | 0.016277349 | 0.594255995 | -3.227569341 |
| **A_55_P1999805** | NA | 0.240565408 | 0.062967195 | 0.418163621 | 9.798168048 | 3.332450823 | 0.016285869 | 0.594255995 | -3.227792458 |
| **A_51_P412595** | Eef1b2 | -0.192706804 | -0.335002308 | -0.0504113 | 15.78489847 | -3.331770146 | 0.016299439 | 0.594255995 | -3.228147623 |
| **A_55_P2116993** | Hnrnpdl | -0.359449912 | -0.624966684 | -0.093933139 | 12.42659632 | -3.330547132 | 0.016323852 | 0.594255995 | -3.228785973 |
| **A_55_P2229645** | NA | 0.247378588 | 0.06461916 | 0.430138016 | 9.474152256 | 3.33005555 | 0.016333676 | 0.594255995 | -3.229042627 |
| **A_55_P2102322** | Yif1b | 0.234768582 | 0.061232949 | 0.408304214 | 13.0621389 | 3.328284751 | 0.016369119 | 0.594255995 | -3.229967505 |
| **A_52_P230904** | Wdr18 | -0.182404129 | -0.317286313 | -0.047521945 | 8.649405988 | -3.326972716 | 0.016395434 | 0.594255995 | -3.230653125 |
| **A_55_P1966719** | NA | 0.647888419 | 0.168713025 | 1.127063813 | 12.14335131 | 3.32640306 | 0.016406874 | 0.594255995 | -3.230950899 |
| **A_55_P1990859** | Rpl6 | -0.271488262 | -0.472341306 | -0.070635218 | 15.65758958 | -3.325383993 | 0.016427361 | 0.594255995 | -3.231483733 |
| **A_51_P484753** | Saraf | -0.341890168 | -0.595112036 | -0.088668299 | 13.53334917 | -3.321655326 | 0.016502562 | 0.594255995 | -3.23343486 |
| **A_55_P1981775** | NA | 0.321407941 | 0.083102317 | 0.559713566 | 12.57905288 | 3.318115311 | 0.016574309 | 0.594255995 | -3.235289511 |
| **A_55_P2065330** | NA | 0.288449775 | 0.074491281 | 0.50240827 | 15.00983586 | 3.316727825 | 0.016602524 | 0.594255995 | -3.236017024 |

| | | | | | | | | | |
|---|---|---|---|---|---|---|---|---|---|
| **A_51_P293901** | Dhrs1 | -0.216720004 | -0.377492281 | -0.055947727 | 9.077659522 | -3.316324349 | 0.016610738 | 0.594255995 | -3.236228645 |
| **A_55_P2064632** | NA | -0.364066511 | -0.634168007 | -0.093965015 | 13.53145612 | -3.316064007 | 0.016616041 | 0.594255995 | -3.236365209 |
| **A_52_P535255** | Syf2 | -0.199104251 | -0.346820602 | -0.0513879 | 10.49924052 | -3.31605042 | 0.016616318 | 0.594255995 | -3.236372336 |
| **A_55_P2109263** | Ppp1r14b | 0.26473754 | 0.068265786 | 0.461209295 | 13.24898831 | 3.315007908 | 0.016637572 | 0.594255995 | -3.23691931 |
| **A_55_P2086265** | Gm8580 | -0.247472756 | -0.431176303 | -0.063769209 | 16.16937598 | -3.314202319 | 0.016654016 | 0.594255995 | -3.237342107 |
| **A_55_P1973563** | Inava | -0.277972305 | -0.484421617 | -0.071522992 | 10.4976016 | -3.312510401 | 0.01668861 | 0.594255995 | -3.238230445 |
| **A_65_P19623** | Ssu72 | 0.258687348 | 0.071106727 | 0.44626797 | 13.13397834 | 3.57385807 | 0.016713279 | 0.594255995 | -3.397260072 |
| **A_51_P267278** | Slc15a2 | -0.520876341 | -0.908130707 | -0.133621975 | 9.046064249 | -3.309081565 | 0.016758962 | 0.594255995 | -3.240032279 |
| **A_55_P2063312** | Mgll | -0.288023203 | -0.502253725 | -0.073792681 | 8.723068922 | -3.30761759 | 0.016789099 | 0.594255995 | -3.240802214 |
| **A_55_P2210315** | Get4 | -0.150844957 | -0.26309891 | -0.038591004 | 9.333186682 | -3.305965479 | 0.016823181 | 0.594255995 | -3.241671543 |
| **A_55_P2048498** | NA | 0.34668938 | 0.088603735 | 0.604775024 | 8.621331575 | 3.304805106 | 0.016847164 | 0.594255995 | -3.242282408 |
| **A_55_P1966109** | Epn2 | -0.349615825 | -0.610308541 | -0.088923109 | 11.27413367 | -3.299372486 | 0.01695995 | 0.594255995 | -3.245145471 |
| **A_52_P79506** | Zfp667 | -0.25415317 | -0.443674631 | -0.064631708 | 8.949510892 | -3.299182171 | 0.016963916 | 0.594255995 | -3.245245863 |
| **A_65_P07569** | Rock1 | -0.159960024 | -0.279329082 | -0.040590966 | 8.941081457 | -3.296771565 | 0.017014241 | 0.594255995 | -3.246518015 |
| **A_52_P480402** | Avl9 | -0.222691294 | -0.388899663 | -0.056482926 | 10.14020754 | -3.296245246 | 0.01702525 | 0.594255995 | -3.246795904 |
| **A_51_P355753** | Hic1 | 0.364378218 | 0.092229363 | 0.636527073 | 11.27155041 | 3.293935259 | 0.017073664 | 0.594255995 | -3.248016122 |
| **A_55_P1965448** | NA | 0.447708026 | 0.11309667 | 0.782319382 | 11.84199117 | 3.291723648 | 0.017120157 | 0.594255995 | -3.249185246 |
| **A_52_P220879** | Tgm2 | -0.255003809 | -0.445617543 | -0.064390075 | 15.68657403 | -3.29125583 | 0.01713001 | 0.594255995 | -3.249432659 |
| **A_51_P292943** | Ell | -0.349531008 | -0.611011097 | -0.088050919 | 11.80031458 | -3.288639348 | 0.017185229 | 0.594255995 | -3.250817128 |
| **A_51_P207403** | Pwp1 | -0.184714492 | -0.322900562 | -0.046528422 | 9.040857904 | -3.288560735 | 0.017186891 | 0.594255995 | -3.250858743 |
| **A_55_P2087087** | Pecam1 | -0.198127168 | -0.3465129 | -0.049741437 | 8.547683565 | -3.284891678 | 0.017264664 | 0.594255995 | -3.252802232 |
| **A_55_P2128042** | NA | 0.348852326 | 0.087575167 | 0.610129484 | 11.34086733 | 3.284803097 | 0.017266546 | 0.594255995 | -3.252849182 |
| **A_55_P2090209** | Tpm3 | -0.2144883 | -0.375177615 | -0.053798986 | 15.59853827 | -3.28386861 | 0.017286419 | 0.594255995 | -3.253344568 |
| **A_51_P442719** | Usp6nl | -0.317739677 | -0.555841781 | -0.079637574 | 9.321708539 | -3.283049121 | 0.017303867 | 0.594255995 | -3.253779117 |
| **A_55_P2035439** | Uxt | 0.24828886 | 0.062205297 | 0.434372422 | 9.959411535 | 3.282603174 | 0.017313369 | 0.594255995 | -3.254015639 |
| **A_55_P1978013** | Obp1a | 0.325136767 | 0.081399186 | 0.568874347 | 8.833585665 | 3.281804833 | 0.017330396 | 0.594255995 | -3.254439149 |
| **A_55_P2107901** | Dlk2 | 0.265740834 | 0.066425449 | 0.46505622 | 8.982413186 | 3.2800966 | 0.017366889 | 0.594255995 | -3.255345721 |
| **A_66_P100932** | Tmem256 | 0.593030632 | 0.147814552 | 1.038246713 | 12.40713574 | 3.276992507 | 0.01743342 | 0.594255995 | -3.256994393 |
| **A_66_P115996** | Kyat1 | 0.217890453 | 0.054246559 | 0.381534348 | 10.22885825 | 3.275725934 | 0.017460647 | 0.594255995 | -3.25766759 |
| **A_55_P1968085** | NA | 0.348372169 | 0.086677153 | 0.610067185 | 8.497715237 | 3.275044193 | 0.017475321 | 0.594255995 | -3.258030059 |
| **A_55_P2133485** | Gapvd1 | -0.144344271 | -0.252795504 | -0.035893038 | 10.3246896 | -3.274418776 | 0.017488795 | 0.594255995 | -3.258362653 |
| **A_51_P266224** | Limd1 | -0.240918814 | -0.42200083 | -0.059836797 | 13.22596532 | -3.273139811 | 0.017516385 | 0.594255995 | -3.259043014 |
| **A_55_P2104472** | NA | -0.324520458 | -0.568477754 | -0.080563162 | 12.76579211 | -3.272633962 | 0.01752731 | 0.594255995 | -3.259312186 |
| **A_51_P372172** | Plod3 | -0.294525034 | -0.516040365 | -0.073009704 | 11.83585337 | -3.271053075 | 0.017561503 | 0.594255995 | -3.260153695 |
| **A_55_P2182939** | NA | -0.25938129 | -0.45447311 | -0.06428947 | 12.5722477 | -3.27091116 | 0.017564576 | 0.594255995 | -3.260229258 |
| **A_55_P2012599** | NA | 0.396755178 | 0.098120208 | 0.695390148 | 8.709830805 | 3.268519685 | 0.01761645 | 0.594255995 | -3.261503135 |
| **A_55_P2083694** | Adam33 | 0.473019731 | 0.116956506 | 0.829082956 | 8.970088037 | 3.268295368 | 0.017621324 | 0.594255995 | -3.261622674 |
| **A_52_P74524** | NA | 0.281924792 | 0.06965492 | 0.494194665 | 8.604555977 | 3.267488542 | 0.017638869 | 0.594255995 | -3.262052707 |
| **A_55_P2055523** | NA | -0.258669033 | -0.453448623 | -0.063889444 | 12.10260304 | -3.26715815 | 0.017646059 | 0.594255995 | -3.262228837 |
| **A_55_P2108678** | NA | -0.221343772 | -0.388168849 | -0.054518694 | 8.816786088 | -3.26418778 | 0.017710844 | 0.594255995 | -3.26381318 |
| **A_55_P1961853** | Zfand3 | 0.331782235 | 0.081525381 | 0.582039089 | 9.500327883 | 3.261642152 | 0.017766573 | 0.594255995 | -3.265172204 |
| **A_55_P2022693** | Tcerg1 | -0.289858612 | -0.508610678 | -0.071106545 | 9.020332564 | -3.259891951 | 0.017805001 | 0.594255995 | -3.266107235 |
| **A_55_P2168761** | Sf1 | 0.255166489 | 0.062468654 | 0.447864325 | 12.43018167 | 3.257736589 | 0.01785245 | 0.594255995 | -3.267259459 |
| **A_55_P2124319** | NA | 0.444818885 | 0.108880796 | 0.780756974 | 8.405415338 | 3.257565318 | 0.017856227 | 0.594255995 | -3.267351053 |
| **A_55_P2065537** | NA | 0.144901268 | 0.035457312 | 0.254345225 | 8.283830135 | 3.257238535 | 0.017863434 | 0.594255995 | -3.267525827 |
| **A_55_P1955188** | Spata21 | 0.605367603 | 0.148035153 | 1.062700053 | 11.06007849 | 3.256539277 | 0.017878868 | 0.594255995 | -3.267899876 |
| **A_51_P328652** | Acot7 | 0.209723677 | 0.051167431 | 0.368279923 | 9.843291821 | 3.254117804 | 0.017932429 | 0.594255995 | -3.269195839 |
| **A_55_P1973521** | Rnf7 | -0.193938165 | -0.340583258 | -0.047293071 | 10.01549433 | -3.253605608 | 0.017943781 | 0.594255995 | -3.269470096 |
| **A_51_P317214** | Hpdl | -0.168386175 | -0.295855329 | -0.04091702 | 8.55806059 | -3.249903672 | 0.018026062 | 0.594255995 | -3.27145368 |
| **A_55_P2058681** | Tmc8 | 0.247684473 | 0.060101423 | 0.435267523 | 9.83785965 | 3.248436277 | 0.018058792 | 0.594255995 | -3.272240609 |

| | | | | | | | | | |
|---|---|---|---|---|---|---|---|---|---|
| **A_51_P493270** | Mrps5 | -0.15239762 | -0.267820798 | -0.036974442 | 8.825680238 | -3.248286454 | 0.018062137 | 0.594255995 | -3.272320978 |
| **A_55_P2143209** | Axin1 | -0.225388826 | -0.396153187 | -0.054624465 | 9.500785434 | -3.247164626 | 0.018087208 | 0.594255995 | -3.272922872 |
| **A_55_P1968153** | Pcgf2 | 0.206621788 | 0.049957628 | 0.363285949 | 8.886211869 | 3.244708027 | 0.018142243 | 0.594255995 | -3.274241684 |
| **A_55_P1980237** | Acad11 | -0.186827924 | -0.325488214 | -0.048167634 | 11.06414898 | -3.491722855 | 0.018209837 | 0.594255995 | -3.42578994 |
| **A_55_P2126269** | Nmb | -0.307335743 | -0.540587967 | -0.074083518 | 11.44500403 | -3.241577729 | 0.018212638 | 0.594255995 | -3.275923703 |
| **A_55_P2076866** | Megf6 | 0.326581223 | 0.07868218 | 0.574480266 | 9.721505308 | 3.24104846 | 0.01822457 | 0.594255995 | -3.276208268 |
| **A_55_P2081510** | Atp5j2 | 0.239726856 | 0.057735442 | 0.421718269 | 13.466196 | 3.240671075 | 0.018233083 | 0.594255995 | -3.276411201 |
| **A_55_P1984561** | Cpne2 | 0.425152325 | 0.102242018 | 0.748062632 | 9.115064666 | 3.239155625 | 0.018267313 | 0.594255995 | -3.277226365 |
| **A_55_P2140955** | NA | -0.181943206 | -0.320163233 | -0.043723179 | 16.71721492 | -3.238426412 | 0.018283809 | 0.594255995 | -3.277618755 |
| **A_55_P1956728** | NA | 0.393757878 | 0.094448658 | 0.693067098 | 13.58251183 | 3.236520215 | 0.018327007 | 0.594255995 | -3.278644919 |
| **A_55_P2015782** | Bcas1 | 0.342470064 | 0.082048615 | 0.602891513 | 8.418863737 | 3.235303496 | 0.018354639 | 0.594255995 | -3.279300251 |
| **A_55_P2178578** | Tmprss4 | 0.244755596 | 0.058541891 | 0.430969301 | 8.798715537 | 3.233628706 | 0.01839275 | 0.594255995 | -3.280202727 |
| **A_55_P1968028** | Tdgf1 | 0.239963844 | 0.057351195 | 0.422576494 | 8.509857162 | 3.232839295 | 0.018410743 | 0.594255995 | -3.28062828 |
| **A_51_P108629** | Prl3b1 | -0.254895674 | -0.44891213 | -0.060879217 | 17.93639105 | -3.23216155 | 0.018426206 | 0.594255995 | -3.280993723 |
| **A_55_P2337213** | Odr4 | -0.273840756 | -0.482332487 | -0.065349024 | 10.0542197 | -3.23130841 | 0.018445692 | 0.594255995 | -3.281453855 |
| **A_55_P1990894** | Psmd4 | -0.231954468 | -0.408821297 | -0.05508764 | 14.86111592 | -3.226454294 | 0.018556991 | 0.594255995 | -3.284074307 |
| **A_55_P2022114** | NA | 0.607382188 | 0.143928348 | 1.070836027 | 9.48766307 | 3.224220466 | 0.018608457 | 0.594255995 | -3.285281612 |
| **A_55_P2184551** | NA | 0.63370054 | 0.150117085 | 1.117283995 | 12.24207035 | 3.223901812 | 0.018615811 | 0.594255995 | -3.285453905 |
| **A_55_P2095035** | NA | -0.206813324 | -0.364646846 | -0.048979802 | 10.60199214 | -3.223654058 | 0.018621532 | 0.594255995 | -3.285587875 |
| **A_52_P137829** | NA | -0.185002708 | -0.326274973 | -0.043730443 | 8.854270287 | -3.221738896 | 0.018665815 | 0.594255995 | -3.286623842 |
| **A_55_P1987914** | Map4k4 | -0.176488264 | -0.311278614 | -0.041697915 | 10.72127787 | -3.221263094 | 0.018676835 | 0.594255995 | -3.286881318 |
| **A_52_P246889** | Chmp3 | -0.189462997 | -0.334238811 | -0.044687183 | 8.54798352 | -3.219567619 | 0.018716161 | 0.594255995 | -3.28779913 |
| **A_55_P2180415** | Cd72 | -0.29107883 | -0.513669509 | -0.068488151 | 8.185988688 | -3.217161103 | 0.018772135 | 0.594255995 | -3.289102719 |
| **A_55_P2025840** | Spred2 | -0.155269762 | -0.274020619 | -0.036518905 | 8.95992134 | -3.216764723 | 0.018781372 | 0.594255995 | -3.289317533 |
| **A_55_P2087567** | Olfr373 | 0.548387913 | 0.128660414 | 0.968115412 | 9.437167838 | 3.214323721 | 0.018838367 | 0.594255995 | -3.290641014 |
| **A_51_P163778** | 2810025M15Rik | 0.39638055 | 0.092960237 | 0.699800862 | 10.16871229 | 3.213933212 | 0.018847503 | 0.594255995 | -3.29085284 |
| **A_55_P2086433** | Oasl1 | -0.398975389 | -0.70448596 | -0.093464819 | 10.32063828 | -3.21283949 | 0.018873115 | 0.594255995 | -3.291446259 |
| **A_52_P88829** | Ctsq | -0.19385591 | -0.342341655 | -0.045370164 | 17.337602 | -3.211910572 | 0.018894898 | 0.594255995 | -3.291950425 |
| **A_55_P1995318** | NA | 0.477135381 | 0.111522339 | 0.842748423 | 9.681440774 | 3.210621469 | 0.018925174 | 0.594255995 | -3.292650332 |
| **A_55_P2096485** | Bcs1l | 0.454877968 | 0.106145499 | 0.803610437 | 10.30817193 | 3.209014514 | 0.018962988 | 0.594255995 | -3.293523223 |
| **A_55_P2074414** | Gtf2h1 | -0.314220281 | -0.555243995 | -0.073196566 | 11.60084118 | -3.207329657 | 0.019002725 | 0.594255995 | -3.29443892 |
| **A_55_P2055257** | Maoa | 0.36943197 | 0.085798608 | 0.653065332 | 9.715317671 | 3.204397068 | 0.019072108 | 0.594255995 | -3.296033933 |
| **A_55_P2026119** | Dlec1 | 0.362347635 | 0.084078628 | 0.640616643 | 9.628657686 | 3.203537081 | 0.019092507 | 0.594255995 | -3.296501961 |
| **A_52_P420712** | Pja2 | -0.210701651 | -0.372514393 | -0.048888908 | 9.321436034 | -3.203497451 | 0.019093448 | 0.594255995 | -3.296523532 |
| **A_55_P1979863** | Syne2 | 0.159753378 | 0.037008885 | 0.282497871 | 11.05383535 | 3.20196955 | 0.019129752 | 0.594255995 | -3.297355388 |
| **A_55_P1977875** | Arglu1 | -0.255788856 | -0.452334548 | -0.059243163 | 10.29669187 | -3.201748774 | 0.019135004 | 0.594255995 | -3.297475623 |
| **A_55_P2162965** | Ptms | 0.615388445 | 0.142425714 | 1.088351176 | 13.84523316 | 3.201043534 | 0.019151791 | 0.594255995 | -3.297859753 |
| **A_55_P2016064** | NA | 0.307760625 | 0.071182378 | 0.544338872 | 8.984326539 | 3.200423194 | 0.019166571 | 0.594255995 | -3.298197711 |
| **A_55_P1962303** | Hba-a1 | -0.146430178 | -0.259032882 | -0.033827474 | 18.1138498 | -3.199270297 | 0.019194072 | 0.594255995 | -3.298825986 |
| **A_55_P2097279** | Pirb | -0.272799309 | -0.482666846 | -0.062931773 | 8.651696485 | -3.197916829 | 0.019226413 | 0.594255995 | -3.299563861 |
| **A_52_P119672** | Rock1 | -0.400390664 | -0.708616543 | -0.092164786 | 8.694314333 | -3.195832481 | 0.019276336 | 0.594255995 | -3.300700824 |
| **A_55_P2075627** | Apc2 | 0.173001348 | 0.039765936 | 0.30623676 | 8.804185109 | 3.194471224 | 0.019309016 | 0.594255995 | -3.301443772 |
| **A_55_P1975456** | Nfyb | -0.206746466 | -0.366066034 | -0.047426897 | 8.734532263 | -3.192553144 | 0.019355168 | 0.594255995 | -3.302491176 |
| **A_52_P400677** | AW209491 | -0.173537956 | -0.307319068 | -0.039756844 | 9.629294212 | -3.191308876 | 0.019385171 | 0.594255995 | -3.30317098 |
| **A_55_P1995497** | Atad3a | -0.155261092 | -0.274998092 | -0.035524092 | 8.276504851 | -3.190093598 | 0.019414525 | 0.594255995 | -3.303835209 |
| **A_55_P2018542** | Ptges3 | -0.15943778 | -0.282399532 | -0.036476028 | 10.68624126 | -3.189997622 | 0.019416845 | 0.594255995 | -3.303887677 |
| **A_55_P2084739** | Gcnt1 | -0.29958162 | -0.530739076 | -0.068424165 | 9.626736396 | -3.188426601 | 0.019454868 | 0.594255995 | -3.304746754 |
| **A_51_P164030** | Tcp1 | -0.227383653 | -0.402857783 | -0.051909523 | 11.32659547 | -3.18797792 | 0.019465743 | 0.594255995 | -3.304992185 |
| **A_55_P2074393** | NA | 0.433745565 | 0.0987277 | 0.768763429 | 9.383779747 | 3.185196607 | 0.019533301 | 0.594255995 | -3.306514373 |
| **A_55_P2012974** | Sdha | -0.370542856 | -0.656821301 | -0.084264411 | 12.61648102 | -3.184336528 | 0.019554244 | 0.594255995 | -3.306985362 |

| | | | | | | | | | |
|---|---|---|---|---|---|---|---|---|---|
| **A_51_P384879** | Mdh2 | -0.269549879 | -0.477873387 | -0.06122637 | 16.18709816 | -3.183244683 | 0.019580866 | 0.594255995 | -3.307583458 |
| **A_55_P1960074** | Pafah1b3 | 0.182450694 | 0.041369729 | 0.32353166 | 11.12336034 | 3.181605004 | 0.019620921 | 0.594255995 | -3.308482043 |
| **A_51_P389988** | Slc40a1 | -0.412066405 | -0.730787318 | -0.093345492 | 14.22661632 | -3.180722722 | 0.019642511 | 0.594255995 | -3.308965752 |
| **A_52_P189005** | Gk5 | -0.232452901 | -0.412338052 | -0.052567749 | 8.349205574 | -3.179133878 | 0.019681457 | 0.594255995 | -3.309837178 |
| **A_55_P2077143** | Mcidas | 0.593471072 | 0.133932812 | 1.053009331 | 12.54547108 | 3.177218208 | 0.019728526 | 0.594255995 | -3.310888449 |
| **A_55_P2022699** | NA | 0.300729073 | 0.067855224 | 0.533602922 | 9.490442165 | 3.177048594 | 0.0197327 | 0.594255995 | -3.31098156 |
| **A_55_P2104769** | Utf1 | 0.279788689 | 0.063061434 | 0.496515944 | 8.484975506 | 3.176038701 | 0.019757569 | 0.594255995 | -3.311536055 |
| **A_55_P2008244** | NA | -0.234510784 | -0.41620013 | -0.052821439 | 9.80647288 | -3.175429836 | 0.019772579 | 0.594255995 | -3.311870447 |
| **A_55_P2178525** | Rpl26 | -0.199918041 | -0.354816692 | -0.04501939 | 15.75487675 | -3.175217766 | 0.019777811 | 0.594255995 | -3.311986933 |
| **A_55_P2173224** | Kif2a | -0.174129729 | -0.30921461 | -0.039044848 | 8.405330675 | -3.171285517 | 0.019875082 | 0.594255995 | -3.31414827 |
| **A_55_P2050488** | Cep170b | -0.306488663 | -0.544360733 | -0.068616594 | 15.26213736 | -3.169860331 | 0.019910466 | 0.594255995 | -3.314932289 |
| **A_55_P2127952** | NA | 0.249194624 | 0.055743542 | 0.442645707 | 8.501927158 | 3.169105288 | 0.019929239 | 0.594255995 | -3.315347796 |
| **A_55_P1974542** | Gxylt2 | -0.267551127 | -0.475298622 | -0.059803632 | 11.95680287 | -3.168401069 | 0.019946767 | 0.594255995 | -3.315735426 |
| **A_51_P106488** | Speg | 0.203384782 | 0.045452879 | 0.361316685 | 9.956416974 | 3.168237632 | 0.019950837 | 0.594255995 | -3.3158254 |
| **A_52_P265002** | Pik3ca | -0.22564731 | -0.400899168 | -0.050395452 | 9.046294795 | -3.167646377 | 0.01996557 | 0.594255995 | -3.316150934 |
| **A_55_P2154173** | NA | 0.241638773 | 0.053918924 | 0.429358622 | 15.75375708 | 3.166836115 | 0.019985778 | 0.594255995 | -3.31659715 |
| **A_55_P2042016** | NA | 0.446391254 | 0.098760216 | 0.794022292 | 8.772912119 | 3.159121248 | 0.020179321 | 0.594255995 | -3.320851578 |
| **A_52_P207964** | Mrps18b | -0.201156446 | -0.357817365 | -0.044495526 | 9.774876305 | -3.158947761 | 0.020183697 | 0.594255995 | -3.320947369 |
| **A_55_P2171033** | NA | 0.400778436 | 0.088466664 | 0.713090209 | 9.523666507 | 3.157076856 | 0.020230952 | 0.594255995 | -3.321980737 |
| **A_52_P648601** | Dusp9 | -0.319543433 | -0.568721062 | -0.070365805 | 13.28432811 | -3.154932065 | 0.020285275 | 0.594255995 | -3.323166142 |
| **A_55_P1952112** | Ildr2 | 0.177138287 | 0.038992424 | 0.315284151 | 8.538064279 | 3.154595789 | 0.020293806 | 0.594255995 | -3.323352072 |
| **A_51_P399071** | Anp32a | -0.137816705 | -0.245309743 | -0.030323668 | 8.115633802 | -3.154210625 | 0.020303583 | 0.594255995 | -3.323565058 |
| **A_55_P2098057** | Sf3a2 | 0.261017476 | 0.057312095 | 0.464722856 | 12.994996 | 3.152363195 | 0.020350548 | 0.594255995 | -3.324587002 |
| **A_55_P1993733** | Gm2824 | 0.36996753 | 0.081073272 | 0.658861789 | 8.709409546 | 3.150604311 | 0.020395373 | 0.594255995 | -3.325560525 |
| **A_55_P2066017** | NA | 0.348544273 | 0.076300028 | 0.620788517 | 9.21468268 | 3.149694315 | 0.020418607 | 0.594255995 | -3.326064413 |
| **A_55_P1990299** | Ophn1 | 0.559543541 | 0.122232194 | 0.996854888 | 8.606279851 | 3.147837494 | 0.020466105 | 0.594255995 | -3.327093036 |
| **A_55_P2179834** | Gatad2a | 0.278336053 | 0.064874219 | 0.491797887 | 10.99218835 | 3.379086511 | 0.020518016 | 0.594255995 | -3.466515738 |
| **A_51_P140347** | Slc4a9 | 0.226070274 | 0.049238244 | 0.402902303 | 8.263164677 | 3.1452248 | 0.020533142 | 0.594255995 | -3.328541421 |
| **A_51_P341465** | Csf2ra | -0.257497894 | -0.458950405 | -0.056045382 | 12.69292994 | -3.144634369 | 0.020548325 | 0.594255995 | -3.328868903 |
| **A_55_P2011867** | Kif19a | 0.155162756 | 0.03370875 | 0.276616761 | 8.283330027 | 3.143003074 | 0.020590337 | 0.594255995 | -3.329774018 |
| **A_55_P2004071** | Gm2058 | -0.248366105 | -0.442879876 | -0.053852334 | 9.496983906 | -3.141312455 | 0.020633977 | 0.594255995 | -3.330712544 |
| **A_55_P2214841** | Irs3 | 0.191957745 | 0.041614504 | 0.342300987 | 10.44718261 | 3.14116604 | 0.020637761 | 0.594255995 | -3.330793849 |
| **A_55_P2076543** | NA | 0.481082324 | 0.104280651 | 0.857883996 | 11.30580433 | 3.141056448 | 0.020640594 | 0.594255995 | -3.330854708 |
| **A_55_P2095709** | NA | -0.274064233 | -0.488764222 | -0.059364243 | 15.99368371 | -3.140432766 | 0.020656724 | 0.594255995 | -3.331201093 |
| **A_52_P5891** | Rflnb | -0.183462429 | -0.327335091 | -0.039589768 | 9.380922776 | -3.137169795 | 0.020741339 | 0.594255995 | -3.333014429 |
| **A_52_P505234** | Stt3a | -0.226932012 | -0.40490538 | -0.048958644 | 9.49553531 | -3.136966691 | 0.020746618 | 0.594255995 | -3.333127363 |
| **A_51_P388973** | Zhx1 | -0.217418346 | -0.387967891 | -0.046868801 | 8.554256913 | -3.136279689 | 0.020764487 | 0.594255995 | -3.333509416 |
| **A_55_P2034415** | Lrrc8d | -0.193647127 | -0.345550162 | -0.041744092 | 11.33682697 | -3.136272355 | 0.020764678 | 0.594255995 | -3.333513495 |
| **A_55_P2008991** | Syp | 0.581737091 | 0.125244917 | 1.038229265 | 10.91399825 | 3.135180732 | 0.020793105 | 0.594255995 | -3.334120738 |
| **A_55_P2081116** | Fam89a | -0.180412516 | -0.321987653 | -0.038837379 | 8.696669406 | -3.135081506 | 0.020795691 | 0.594255995 | -3.334175945 |
| **A_55_P2085994** | Zfat | 0.187062567 | 0.040264064 | 0.33386107 | 9.064161738 | 3.134977489 | 0.020798403 | 0.594255995 | -3.33423382 |
| **A_55_P2021351** | NA | -0.225617614 | -0.402886406 | -0.048348822 | 16.36971807 | -3.131193302 | 0.020897306 | 0.594255995 | -3.336340638 |
| **A_55_P2040962** | Gm1604b | 0.25953022 | 0.055531641 | 0.463528799 | 9.548674943 | 3.129896346 | 0.020931322 | 0.594255995 | -3.337063291 |
| **A_55_P2129217** | NA | -0.158834075 | -0.283714385 | -0.033953765 | 13.49459411 | -3.129095905 | 0.020952345 | 0.594255995 | -3.337509439 |
| **A_66_P104527** | Tbl1x | -0.179798685 | -0.321183568 | -0.038413802 | 11.66149729 | -3.128619144 | 0.020964878 | 0.594255995 | -3.337775228 |
| **A_55_P1998823** | NA | 0.348111013 | 0.074209586 | 0.62201244 | 10.13559251 | 3.126746206 | 0.021014194 | 0.594255995 | -3.338819763 |
| **A_52_P519882** | Etf1 | -0.210208992 | -0.375625781 | -0.044792202 | 10.50043566 | -3.126373528 | 0.021024022 | 0.594255995 | -3.339027679 |
| **A_55_P2040416** | Pradc1 | -0.218290273 | -0.390100039 | -0.046480507 | 9.505427064 | -3.125760508 | 0.021040198 | 0.594255995 | -3.339369734 |
| **A_55_P2038928** | NA | -0.331224614 | -0.591928995 | -0.070520232 | 11.43773137 | -3.125672232 | 0.021042529 | 0.594255995 | -3.339418996 |
| **A_55_P2007525** | Smu1 | -0.177239357 | -0.316758109 | -0.037720606 | 9.570381668 | -3.125336238 | 0.021051402 | 0.594255995 | -3.339606509 |

| | | | | | | | | | |
|---|---|---|---|---|---|---|---|---|---|
| **A_55_P2079927** | Ccdc68 | -0.220032321 | -0.393287809 | -0.046776833 | 9.279091384 | -3.124414491 | 0.021075766 | 0.594255995 | -3.340121023 |
| **A_55_P2067332** | NA | 0.211565726 | 0.044958895 | 0.378172558 | 8.633302974 | 3.124076578 | 0.021084706 | 0.594255995 | -3.340309682 |
| **A_51_P193716** | Supv3l1 | -0.15092318 | -0.269851307 | -0.031995053 | 8.796159819 | -3.122054864 | 0.021138277 | 0.594255995 | -3.34143884 |
| **A_55_P2230688** | Pigm | 0.177312258 | 0.037504233 | 0.317120283 | 9.702557107 | 3.120152515 | 0.021188821 | 0.594255995 | -3.342501991 |
| **A_55_P1963634** | Yae1d1 | 0.22758964 | 0.048054796 | 0.407124485 | 8.987526955 | 3.118694993 | 0.021227636 | 0.594255995 | -3.343316979 |
| **A_52_P380263** | Podxl | -0.245514562 | -0.439194233 | -0.051834892 | 8.748530661 | -3.118619441 | 0.02122965 | 0.594255995 | -3.343359234 |
| **A_55_P1981992** | NA | 0.824790437 | 0.173976975 | 1.4756039 | 10.11518126 | 3.117857429 | 0.021249976 | 0.594255995 | -3.34378548 |
| **A_51_P121891** | Rac2 | -0.153917019 | -0.275396083 | -0.032437955 | 8.293145785 | -3.117126079 | 0.021269504 | 0.594255995 | -3.381424003 |
| **A_55_P2061492** | Pnldc1 | 0.633052056 | 0.133154703 | 1.132949409 | 8.854755919 | 3.115499959 | 0.021312995 | 0.594255995 | -3.345104828 |
| **A_66_P138047** | Gfm1 | -0.241906825 | -0.43307421 | -0.050739441 | 11.48367658 | -3.113174657 | 0.021375353 | 0.594255995 | -3.346407136 |
| **A_55_P1989885** | Smarce1 | -0.204962231 | -0.367032045 | -0.042892417 | 11.65094425 | -3.111292695 | 0.021425969 | 0.594255995 | -3.347461847 |
| **A_51_P301964** | Arl6ip1 | -0.485128729 | -0.868831067 | -0.101426391 | 11.54333812 | -3.110510788 | 0.021447037 | 0.594255995 | -3.347900237 |
| **A_55_P2025368** | Nkain1 | 0.163615913 | 0.03420207 | 0.293029757 | 9.965963998 | 3.110383704 | 0.021450463 | 0.594255995 | -3.347971499 |
| **A_51_P394190** | Lmo4 | -0.247888119 | -0.444085698 | -0.051690539 | 11.39230318 | -3.108359408 | 0.021505122 | 0.594255995 | -3.349107005 |
| **A_51_P315595** | NA | -0.186020067 | -0.333290162 | -0.038749971 | 16.53690871 | -3.107523279 | 0.021527743 | 0.594255995 | -3.349576234 |
| **A_52_P257426** | Eps8l2 | -0.196586798 | -0.352315288 | -0.040858308 | 9.185043771 | -3.105670942 | 0.021577949 | 0.594255995 | -3.350616191 |
| **A_55_P2061701** | Osgep | -0.171088862 | -0.306652062 | -0.035525662 | 9.350072532 | -3.104910513 | 0.021598597 | 0.594255995 | -3.351043295 |
| **A_55_P2044005** | NA | 0.210374405 | 0.043553199 | 0.377195612 | 9.817222233 | 3.102492974 | 0.021664384 | 0.594255995 | -3.352401812 |
| **A_55_P2046807** | Sparc | 0.668570124 | 0.138362606 | 1.198777643 | 15.41884816 | 3.102203238 | 0.021672283 | 0.594255995 | -3.352564696 |
| **A_55_P1976689** | Snrnp35 | -0.212067448 | -0.380326181 | -0.043808714 | 10.33632321 | -3.100741468 | 0.021712184 | 0.594255995 | -3.353386704 |
| **A_55_P2065268** | NA | 0.244179748 | 0.050416304 | 0.437943193 | 9.50828961 | 3.100323115 | 0.021723618 | 0.594255995 | -3.353622029 |
| **A_55_P2084797** | Aff3 | 0.40411191 | 0.083431464 | 0.724792356 | 9.609549988 | 3.100261553 | 0.021725301 | 0.594255995 | -3.35365666 |
| **A_52_P494372** | Zxdc | -0.191018151 | -0.342616522 | -0.03941978 | 9.141005584 | -3.099911297 | 0.02173488 | 0.594255995 | -3.353853708 |
| **A_51_P255682** | Mfge8 | -0.2630242 | -0.471779534 | -0.054268866 | 16.68398473 | -3.099754439 | 0.021739171 | 0.594255995 | -3.35394196 |
| **A_55_P2155827** | NA | -0.423900759 | -0.760530539 | -0.087270979 | 14.68157819 | -3.097995794 | 0.021787347 | 0.594255995 | -3.35493172 |
| **A_66_P107943** | NA | 0.182032124 | 0.037450792 | 0.326613456 | 12.75325393 | 3.097454799 | 0.021802191 | 0.594255995 | -3.355236301 |
| **A_51_P487628** | Pramef12 | -0.242486656 | -0.435095613 | -0.049877698 | 15.2392824 | -3.097280415 | 0.021806978 | 0.594255995 | -3.35533449 |
| **A_55_P2168321** | NA | 0.245897128 | 0.050504542 | 0.441289713 | 8.705181387 | 3.096096863 | 0.021839497 | 0.594255995 | -3.356001048 |
| **A_55_P2181928** | Slc45a3 | 0.199493438 | 0.040901583 | 0.358085294 | 10.27703095 | 3.09468819 | 0.021878271 | 0.594255995 | -3.356794713 |
| **A_55_P2142728** | NA | 0.370658226 | 0.075951155 | 0.665365296 | 12.28504721 | 3.094227562 | 0.021890967 | 0.594255995 | -3.357054313 |
| **A_51_P435251** | Cul4a | -0.281855054 | -0.50607299 | -0.057637119 | 10.27796268 | -3.092606329 | 0.021935713 | 0.594255995 | -3.357968305 |
| **A_51_P378825** | Actr8 | -0.15354819 | -0.275760293 | -0.031336087 | 10.40436144 | -3.09100457 | 0.02198002 | 0.594255995 | -3.358871774 |
| **A_55_P2117146** | Pa2g4 | -0.220247718 | -0.395630454 | -0.044864982 | 9.753707021 | -3.08953941 | 0.022020635 | 0.594255995 | -3.359698594 |
| **A_55_P1997061** | NA | 0.152106138 | 0.030960885 | 0.273251391 | 12.87700344 | 3.088940206 | 0.022037268 | 0.594255995 | -3.360036846 |
| **A_55_P2122419** | Arl1 | 0.2356528 | 0.047962466 | 0.423343134 | 10.39309125 | 3.088871634 | 0.022039173 | 0.594255995 | -3.360075559 |
| **A_66_P116412** | Flt3 | -0.296434695 | -0.532594874 | -0.060274517 | 10.03474502 | -3.088101223 | 0.022060581 | 0.594255995 | -3.360510562 |
| **A_55_P1959379** | BC080695 | 0.553979884 | 0.112588705 | 0.995371062 | 8.905972561 | 3.087731361 | 0.022070867 | 0.594255995 | -3.360719436 |
| **A_55_P2131979** | 1-Mar | 0.569633485 | 0.115263091 | 1.024003879 | 11.49818835 | 3.08428604 | 0.022166934 | 0.594255995 | -3.362666301 |
| **A_66_P124715** | Ppard | 0.297157634 | 0.06003391 | 0.534281358 | 10.43116809 | 3.083053402 | 0.022201415 | 0.594255995 | -3.363363345 |
| **A_51_P517787** | Clptm1l | -0.222160034 | -0.399530224 | -0.044789844 | 11.26244945 | -3.081445324 | 0.022246486 | 0.594255995 | -3.3642731 |
| **A_55_P1986088** | Prr14 | -0.231076864 | -0.415578741 | -0.046574986 | 12.09454666 | -3.0812351 | 0.022252385 | 0.594255995 | -3.364392066 |
| **A_51_P301259** | Araf | 0.188105281 | 0.037913007 | 0.338297556 | 10.6510128 | 3.081218931 | 0.022252839 | 0.594255995 | -3.364401216 |
| **A_55_P1960873** | Mbd3 | -0.278740085 | -0.501317968 | -0.056162202 | 8.611637127 | -3.08096372 | 0.022260004 | 0.594255995 | -3.364545652 |
| **A_55_P2002465** | Zfp747 | 0.248507489 | 0.049950375 | 0.447064603 | 10.60527698 | 3.079095731 | 0.02231252 | 0.594255995 | -3.365603184 |
| **A_65_P08022** | Hnrnpk | -0.208685785 | -0.375463076 | -0.041908494 | 10.39478282 | -3.078400458 | 0.022332101 | 0.594255995 | -3.365996959 |
| **A_66_P129484** | Tmed9 | -0.259448261 | -0.466830715 | -0.052065808 | 9.119904622 | -3.07785332 | 0.022347524 | 0.594255995 | -3.366306897 |
| **A_55_P1968483** | Gtpbp6 | 0.164167235 | 0.032929135 | 0.295405334 | 9.999874603 | 3.077483362 | 0.022357959 | 0.594255995 | -3.366516498 |
| **A_55_P1984751** | Ppp1r14b | 0.271690626 | 0.054287177 | 0.489094076 | 12.43634998 | 3.074520456 | 0.022441721 | 0.594255995 | -3.368196012 |
| **A_55_P2027102** | Rgs3 | 0.306955106 | 0.061193819 | 0.552716393 | 10.4150806 | 3.072773521 | 0.022491268 | 0.594255995 | -3.369186986 |
| **A_55_P2175813** | Ythdf2 | -0.245938519 | -0.442865054 | -0.049011984 | 10.35732429 | -3.072497071 | 0.02249912 | 0.594255995 | -3.369343855 |

| | | | | | | | | | |
|---|---|---|---|---|---|---|---|---|---|
| **A_52_P533198** | Dnajc2 | -0.242776643 | -0.437264966 | -0.04828832 | 9.530066475 | -3.071019214 | 0.022541144 | 0.594255995 | -3.370182685 |
| **A_55_P1982737** | Dnajc5 | 0.323917239 | 0.064033145 | 0.583801334 | 15.76027435 | 3.066362708 | 0.022674119 | 0.594255995 | -3.372828241 |
| **A_51_P403334** | Cuedc2 | 0.221559806 | 0.043692008 | 0.399427605 | 11.94502107 | 3.064522468 | 0.022726906 | 0.594255995 | -3.373874817 |
| **A_55_P2056186** | Siva1 | 0.161945048 | 0.031859218 | 0.292030877 | 13.15890708 | 3.062716829 | 0.022778831 | 0.594255995 | -3.374902298 |
| **A_55_P2246650** | 9330159M07Rik | 0.191678622 | 0.037686009 | 0.345671236 | 8.614248069 | 3.062266279 | 0.022791808 | 0.594255995 | -3.375158769 |
| **A_51_P311540** | Cox6a1 | -0.259870628 | -0.468650804 | -0.051090452 | 17.36601622 | -3.062225008 | 0.022792997 | 0.594255995 | -3.375182264 |
| **A_55_P1981455** | NA | -0.217163634 | -0.391778858 | -0.04254841 | 8.44567141 | -3.059666977 | 0.02286683 | 0.594255995 | -3.376639097 |
| **A_55_P1958097** | Eno1 | 0.529114037 | 0.103490394 | 0.95473768 | 14.48057385 | 3.058388851 | 0.022903819 | 0.594255995 | -3.377367441 |
| **A_55_P2031125** | Ube2l6 | -0.153459316 | -0.276942661 | -0.02997597 | 10.64799545 | -3.05741248 | 0.022932119 | 0.594255995 | -3.377924024 |
| **A_55_P1991728** | Zfp956 | 0.43625586 | 0.085186938 | 0.787324782 | 10.00400174 | 3.057159173 | 0.022939467 | 0.594255995 | -3.37806845 |
| **A_51_P517120** | Zfp930 | -0.237117091 | -0.42803442 | -0.046199762 | 12.20510711 | -3.055530847 | 0.022986766 | 0.594255995 | -3.378997131 |
| **A_55_P2072656** | Ckmt1 | 0.536311315 | 0.104392304 | 0.968230327 | 11.48185578 | 3.054807178 | 0.02300782 | 0.594255995 | -3.37941001 |
| **A_55_P2003443** | Lsm7 | 0.249891746 | 0.048623967 | 0.451159525 | 13.38609278 | 3.054547064 | 0.023015393 | 0.594255995 | -3.379558438 |
| **A_55_P2065586** | Gm15417 | 0.263355263 | 0.051233428 | 0.475477097 | 8.612404814 | 3.054399151 | 0.023019701 | 0.594255995 | -3.379642845 |
| **A_55_P1962586** | NA | 0.48712625 | 0.094742489 | 0.879510011 | 9.281221979 | 3.054215456 | 0.023025052 | 0.594255995 | -3.379747678 |
| **A_55_P1975120** | NA | 0.359359769 | 0.069849556 | 0.648869982 | 9.023594739 | 3.053758791 | 0.02303836 | 0.594255995 | -3.380008318 |
| **A_51_P507851** | Gcc1 | -0.20471401 | -0.369677092 | -0.039750928 | 9.283245929 | -3.053022236 | 0.023059842 | 0.594255995 | -3.380428781 |
| **A_52_P315111** | Mrpl52 | 0.261890811 | 0.050807343 | 0.472974279 | 13.47344406 | 3.052356126 | 0.023079289 | 0.594255995 | -3.380809114 |
| **A_55_P2059179** | Slc48a1 | 0.184955728 | 0.035840184 | 0.334071271 | 9.917524279 | 3.051504629 | 0.023104174 | 0.594255995 | -3.381295413 |
| **A_52_P318673** | Saa1 | -0.166297641 | -0.300381002 | -0.03221428 | 9.075655126 | -3.051268059 | 0.023111093 | 0.594255995 | -3.381430543 |
| **A_55_P2061979** | NA | -0.246506749 | -0.445338841 | -0.047674656 | 15.63401674 | -3.050081852 | 0.023145819 | 0.594255995 | -3.382108261 |
| **A_51_P195100** | Vps4b | -0.244074267 | -0.440997076 | -0.047151459 | 11.69874313 | -3.049264775 | 0.023169773 | 0.594255995 | -3.382575229 |
| **A_66_P125437** | Rpl30 | -0.19550722 | -0.353251008 | -0.037763433 | 9.178348868 | -3.049156395 | 0.023172952 | 0.594255995 | -3.382637178 |
| **A_55_P2164778** | Ptdss1 | -0.225583344 | -0.404664993 | -0.046501695 | 11.62802068 | -3.264419983 | 0.023216712 | 0.594255995 | -3.50994208 |
| **A_66_P138137** | Pnp | -0.298661825 | -0.539859111 | -0.05746454 | 12.58370989 | -3.046326594 | 0.023256132 | 0.594255995 | -3.384255405 |
| **A_55_P2120303** | NA | 0.152911713 | 0.02939487 | 0.276428556 | 8.40927251 | 3.045676222 | 0.023275296 | 0.594255995 | -3.384627522 |
| **A_51_P214107** | Tbc1d20 | -0.280515675 | -0.507289495 | -0.053741855 | 10.1108691 | -3.043220228 | 0.023347817 | 0.594255995 | -3.386033419 |
| **A_55_P2038867** | Mcat | 0.210270183 | 0.040237868 | 0.380302498 | 10.6909299 | 3.042393517 | 0.023372284 | 0.594255995 | -3.386506897 |
| **A_52_P67983** | Lcmt2 | 0.208350719 | 0.039727465 | 0.376973974 | 9.909767603 | 3.039811823 | 0.023448871 | 0.594255995 | -3.387986278 |
| **A_55_P1972852** | Mcrs1 | 0.158145046 | 0.030005945 | 0.286284148 | 11.69147291 | 3.036288833 | 0.023553823 | 0.594255995 | -3.390006952 |
| **A_55_P2150309** | Lhfpl4 | 0.720230243 | 0.136309684 | 1.304150802 | 11.66004893 | 3.034497266 | 0.023607391 | 0.594255995 | -3.39103538 |
| **A_55_P2003581** | Scnm1 | 0.169544792 | 0.032043369 | 0.307046215 | 11.02072777 | 3.033516944 | 0.023636759 | 0.594255995 | -3.391598363 |
| **A_55_P2016851** | Lce1a2 | 0.492539004 | 0.092961498 | 0.89211651 | 11.33841419 | 3.032555506 | 0.0236656 | 0.594255995 | -3.392150666 |
| **A_55_P1993880** | Rps24 | 0.26689652 | 0.050335532 | 0.483457509 | 15.37629988 | 3.032018518 | 0.023681725 | 0.594255995 | -3.392459213 |
| **A_52_P208600** | Bms1 | -0.183471946 | -0.332366179 | -0.034577712 | 8.758094216 | -3.031523586 | 0.023696598 | 0.594255995 | -3.39274364 |
| **A_52_P482925** | Serbp1 | -0.307314685 | -0.556719229 | -0.057910142 | 12.20189314 | -3.031433989 | 0.023699292 | 0.594255995 | -3.392795135 |
| **A_55_P1955393** | 7-Sep | -0.350525971 | -0.635018972 | -0.066032969 | 12.6661186 | -3.031222226 | 0.023705659 | 0.594255995 | -3.392916847 |
| **A_55_P2053456** | Gm6548 | -0.185079182 | -0.335346547 | -0.034811817 | 15.46608464 | -3.030135601 | 0.023738362 | 0.594255995 | -3.393541519 |
| **A_51_P324572** | Ctnna1 | -0.318658378 | -0.577380053 | -0.059936702 | 15.32623146 | -3.030132982 | 0.023738441 | 0.594255995 | -3.393543024 |
| **A_55_P2182975** | P2rx2 | -0.162720801 | -0.294870747 | -0.030570854 | 8.906221642 | -3.029320631 | 0.023762921 | 0.594255995 | -3.394010159 |
| **A_51_P346132** | Rhox9 | -0.207009768 | -0.375183935 | -0.0388356 | 16.95324782 | -3.02831268 | 0.023793335 | 0.594255995 | -3.394589935 |
| **A_55_P2011087** | Gnptg | 0.255710242 | 0.04797146 | 0.463449024 | 11.96879313 | 3.028305637 | 0.023793547 | 0.594255995 | -3.394593987 |
| **A_55_P1983284** | Flot2 | 0.468856378 | 0.087937262 | 0.849775494 | 12.39284649 | 3.028141908 | 0.023798491 | 0.594255995 | -3.394688182 |
| **A_55_P2173188** | Kank3 | -0.207117047 | -0.375414771 | -0.038819323 | 9.538797024 | -3.027657644 | 0.023813122 | 0.594255995 | -3.39496681 |
| **A_55_P2095794** | Evi5 | -0.285092079 | -0.516855019 | -0.053329139 | 11.48059222 | -3.026288471 | 0.02385454 | 0.594255995 | -3.395754809 |
| **A_52_P164136** | Arrdc3 | -0.374744921 | -0.679551473 | -0.06993837 | 10.5597183 | -3.024688269 | 0.023903046 | 0.594255995 | -3.396676192 |
| **A_51_P128876** | Ifitm3 | -0.363512319 | -0.659203021 | -0.067821618 | 11.28489768 | -3.024479389 | 0.023909386 | 0.594255995 | -3.396796497 |
| **A_55_P2022595** | Gm10653 | 0.26947915 | 0.050269293 | 0.488689006 | 15.17715344 | 3.024365351 | 0.023912848 | 0.594255995 | -3.39686218 |
| **A_66_P117815** | NA | -0.333675465 | -0.605539611 | -0.061811319 | 11.56118237 | -3.019544969 | 0.024059682 | 0.594255995 | -3.39964073 |
| **A_55_P2030805** | Actr10 | 0.235845928 | 0.043683539 | 0.428008318 | 13.49636767 | 3.01945913 | 0.024062306 | 0.594255995 | -3.399690246 |

| | | | | | | | | | |
|---|---|---|---|---|---|---|---|---|---|
| **A_55_P2059904** | Chsy1 | 0.237110124 | 0.043916474 | 0.430303774 | 10.81221191 | 3.019440069 | 0.024062889 | 0.594255995 | -3.399701242 |
| **A_55_P2124543** | Gnptg | 0.174438285 | 0.032242198 | 0.316634372 | 10.65070425 | 3.018028426 | 0.024106081 | 0.594255995 | -3.400515741 |
| **A_52_P305995** | Ammecr1 | -0.380162952 | -0.69012383 | -0.070202074 | 9.678145854 | -3.01739434 | 0.02412551 | 0.594255995 | -3.400881715 |
| **A_51_P281333** | St3gal6 | -0.261407085 | -0.47462544 | -0.04818873 | 12.46316369 | -3.016212458 | 0.024161769 | 0.594255995 | -3.40156405 |
| **A_51_P191893** | 2410022M11Rik | -0.220039813 | -0.399524157 | -0.040555468 | 11.09967369 | -3.016086945 | 0.024165623 | 0.594255995 | -3.401636527 |
| **A_55_P2041624** | Ceacam12 | -0.15868408 | -0.288129406 | -0.029238755 | 17.40834078 | -3.015894535 | 0.024171532 | 0.594255995 | -3.401747638 |
| **A_66_P138673** | Whamm | -0.217644232 | -0.395217022 | -0.040071442 | 8.555139923 | -3.015365121 | 0.024187801 | 0.594255995 | -3.402053394 |
| **A_55_P2146708** | Zcchc9 | -0.345976294 | -0.628345774 | -0.063606813 | 15.29561608 | -3.014378269 | 0.024218158 | 0.594255995 | -3.40262347 |
| **A_55_P1953475** | NA | -0.202336182 | -0.367503131 | -0.037169233 | 16.45732366 | -3.013835668 | 0.024234866 | 0.594255995 | -3.402936987 |
| **A_55_P2115151** | Pxylp1 | 0.273914789 | 0.050314086 | 0.497515492 | 8.626222804 | 3.013779537 | 0.024236596 | 0.594255995 | -3.402969423 |
| **A_55_P2219059** | Lmnb2 | 0.243005167 | 0.044617042 | 0.441393292 | 8.984046088 | 3.013484747 | 0.024245679 | 0.594255995 | -3.403139779 |
| **A_55_P2043531** | NA | -0.613219165 | -1.114276606 | -0.112161725 | 12.6192315 | -3.010907179 | 0.024325263 | 0.594255995 | -3.404629984 |
| **A_55_P2048650** | Dennd4a | -0.176321008 | -0.320405047 | -0.032236968 | 15.90544224 | -3.010629736 | 0.024333846 | 0.594255995 | -3.404790457 |
| **A_55_P1982524** | Cyhr1 | -0.172570427 | -0.313617955 | -0.031522899 | 8.502941764 | -3.010024688 | 0.024352576 | 0.594255995 | -3.405140462 |
| **A_52_P289893** | Prdm4 | -0.144399405 | -0.26250787 | -0.026290939 | 8.931622046 | -3.007831751 | 0.024420591 | 0.594255995 | -3.406409563 |
| **A_51_P178815** | H1f8 | 0.785624988 | 0.142876665 | 1.428373311 | 9.078658041 | 3.007069732 | 0.024444273 | 0.594255995 | -3.406850759 |
| **A_52_P67200** | Stt3b | -0.220601972 | -0.401107694 | -0.04009625 | 11.55498903 | -3.006682576 | 0.024456315 | 0.594255995 | -3.407074955 |
| **A_55_P2037528** | Acsf3 | 0.178105224 | 0.032336504 | 0.323873943 | 10.14382793 | 3.005948119 | 0.024479177 | 0.594255995 | -3.407500341 |
| **A_55_P2125149** | Tulp1 | 0.578781098 | 0.104647557 | 1.05291464 | 10.96168579 | 3.003190097 | 0.024565237 | 0.594255995 | -3.409098596 |
| **A_52_P286002** | Obsl1 | -0.142263087 | -0.258807747 | -0.025718426 | 8.616259367 | -3.003094572 | 0.024568224 | 0.594255995 | -3.409153976 |
| **A_55_P2103884** | Supt4b | -0.139535653 | -0.253853474 | -0.025217833 | 11.36488516 | -3.002896875 | 0.024574406 | 0.594255995 | -3.409268595 |
| **A_52_P464129** | Gon7 | -0.22046089 | -0.401181115 | -0.039740666 | 9.542619381 | -3.001193258 | 0.024627751 | 0.594255995 | -3.41025659 |
| **A_55_P2125623** | NA | 0.492809557 | 0.088656506 | 0.896962609 | 8.562776961 | 2.999869913 | 0.024669275 | 0.594255995 | -3.411024404 |
| **A_52_P552017** | Rbsn | -0.178187135 | -0.324368455 | -0.032005816 | 9.710493023 | -2.998842319 | 0.024701572 | 0.594255995 | -3.411620835 |
| **A_52_P326354** | Erf | 0.17791785 | 0.031845691 | 0.323990009 | 10.64567333 | 2.996547978 | 0.024773848 | 0.594255995 | -3.41295318 |
| **A_55_P2082837** | Spin1 | -0.239421504 | -0.436095468 | -0.042747539 | 10.20763493 | -2.994921513 | 0.024825224 | 0.594255995 | -3.413898246 |
| **A_52_P16877** | Tmcc3 | -0.162005597 | -0.295103645 | -0.02890755 | 8.607908893 | -2.994521929 | 0.024837864 | 0.594255995 | -3.414130498 |
| **A_66_P123804** | Fbxo30 | -0.196176976 | -0.357396576 | -0.034957376 | 8.935861743 | -2.99363853 | 0.024865833 | 0.594255995 | -3.41464406 |
| **A_55_P2046363** | Mapkap1 | -0.181843968 | -0.331381983 | -0.032305953 | 9.639063992 | -2.99168904 | 0.024927676 | 0.594255995 | -3.41577788 |
| **A_55_P2145982** | NA | -0.178607628 | -0.325540499 | -0.031674756 | 8.253316322 | -2.99054401 | 0.024964077 | 0.594255995 | -3.416444142 |
| **A_55_P1971619** | NA | 0.956802036 | 0.169365859 | 1.744238214 | 9.859517346 | 2.989343969 | 0.025002289 | 0.594255995 | -3.41714266 |
| **A_52_P419162** | Cyhr1 | -0.150133257 | -0.2737005 | -0.026566013 | 8.488209939 | -2.989115593 | 0.025009568 | 0.594255995 | -3.417275622 |
| **A_66_P107255** | Nom1 | -0.166206667 | -0.303025727 | -0.029387607 | 9.253857498 | -2.988622091 | 0.025025306 | 0.594255995 | -3.417562973 |
| **A_55_P1962134** | NA | 0.195952273 | 0.034639852 | 0.357264694 | 9.166502564 | 2.988488992 | 0.025029552 | 0.594255995 | -3.41764048 |
| **A_51_P514712** | Parp14 | -0.545452071 | -0.994514787 | -0.096389356 | 10.48008162 | -2.988262605 | 0.025036777 | 0.594255995 | -3.417772317 |
| **A_66_P121636** | Ablim3 | 0.216436942 | 0.038246115 | 0.394627769 | 8.240776526 | 2.988238167 | 0.025037557 | 0.594255995 | -3.41778655 |
| **A_55_P2116235** | NA | 0.317984442 | 0.056167912 | 0.579800973 | 14.14275452 | 2.987981965 | 0.025045735 | 0.594255995 | -3.417935763 |
| **A_55_P2097176** | Idua | 0.504227058 | 0.08867859 | 0.919775526 | 12.35471501 | 2.985201317 | 0.025134691 | 0.594255995 | -3.419555968 |
| **A_55_P1959600** | Tmem222 | 0.182380213 | 0.032048746 | 0.33271168 | 11.57894602 | 2.984674579 | 0.02515158 | 0.594255995 | -3.419863037 |
| **A_51_P183971** | Barhl1 | 0.343870171 | 0.060378979 | 0.627361364 | 9.209076815 | 2.984173691 | 0.025167652 | 0.594255995 | -3.420155082 |
| **A_51_P365019** | Gclc | -0.398343586 | -0.726823043 | -0.069864128 | 10.62080697 | -2.983450091 | 0.02519089 | 0.594255995 | -3.420577059 |
| **A_55_P1999213** | Tdrd7 | -0.275758734 | -0.503153451 | -0.048364017 | 11.48249072 | -2.983445226 | 0.025191046 | 0.594255995 | -3.420579896 |
| **A_55_P2019233** | Gsto2 | 0.182867235 | 0.032040971 | 0.3336935 | 9.515740454 | 2.982827151 | 0.025210914 | 0.594255995 | -3.420940409 |
| **A_51_P480328** | Adgrl4 | -0.18972042 | -0.346224488 | -0.033216352 | 8.734155196 | -2.982343141 | 0.025226484 | 0.594255995 | -3.42122277 |
| **A_51_P314679** | Rrn3 | -0.147385706 | -0.26909032 | -0.025681092 | 11.11709071 | -2.979322279 | 0.025323899 | 0.594255995 | -3.422986015 |
| **A_55_P2043582** | 4930480E11Rik | 0.29688195 | 0.051666522 | 0.542097378 | 8.588462154 | 2.978551858 | 0.025348809 | 0.594255995 | -3.42343596 |
| **A_51_P332602** | Surf1 | -0.161033165 | -0.294187726 | -0.027878603 | 11.32882861 | -2.975284104 | 0.02545476 | 0.594255995 | -3.425345573 |
| **A_51_P476018** | Sox7 | -0.205031315 | -0.374610436 | -0.035452194 | 8.952524492 | -2.974520593 | 0.025479585 | 0.594255995 | -3.425792026 |
| **A_55_P1999240** | NA | 0.250674301 | 0.043294647 | 0.458053955 | 10.39988188 | 2.973807351 | 0.025502799 | 0.594255995 | -3.426209178 |
| **A_55_P2141013** | Siglech | 0.745460506 | 0.128672337 | 1.362248674 | 15.62751668 | 2.97343034 | 0.025515079 | 0.594255995 | -3.426429715 |

| | | | | | | | | | |
|---|---|---|---|---|---|---|---|---|---|
| **A_55_P1983638** | Mapk6 | 0.207368588 | 0.035750654 | 0.378986522 | 11.88471966 | 2.972688912 | 0.025539247 | 0.594255995 | -3.426863497 |
| **A_55_P2041011** | Slc2a13 | -0.254666242 | -0.465457887 | -0.043874596 | 8.905531467 | -2.972262406 | 0.025553161 | 0.594255995 | -3.427113074 |
| **A_55_P2032849** | NA | 0.375413198 | 0.064224327 | 0.686602068 | 9.59951545 | 2.967936757 | 0.025694747 | 0.594255995 | -3.429646109 |
| **A_51_P421303** | Caly | 0.307026083 | 0.052422755 | 0.561629411 | 8.740172358 | 2.966745953 | 0.025733873 | 0.594255995 | -3.430344006 |
| **A_55_P2169135** | NA | -0.200738923 | -0.367233856 | -0.034243989 | 12.51658444 | -2.966195216 | 0.025751991 | 0.594255995 | -3.430666862 |
| **A_55_P2065874** | D430042O09Rik | 0.40214675 | 0.068388765 | 0.735904735 | 9.408151814 | 2.964299372 | 0.025814464 | 0.594255995 | -3.431778664 |
| **A_55_P2423513** | B230369F24Rik | 0.358019289 | 0.060875855 | 0.655162723 | 11.24205962 | 2.964213293 | 0.025817305 | 0.594255995 | -3.43182916 |
| **A_52_P559808** | Arhgef12 | -0.197668088 | -0.36175285 | -0.033583325 | 11.54722923 | -2.963722052 | 0.025833521 | 0.594255995 | -3.432117355 |
| **A_55_P2016967** | Gm8213 | 0.259923107 | 0.043990479 | 0.475855736 | 15.54642016 | 2.961391205 | 0.025910617 | 0.594255995 | -3.433485368 |
| **A_55_P2170105** | NA | -0.313521338 | -0.574240098 | -0.052802578 | 14.66377475 | -2.958448309 | 0.026008315 | 0.594255995 | -3.435213971 |
| **A_55_P2116900** | NA | 0.359731117 | 0.060551313 | 0.658910922 | 9.580934766 | 2.958113927 | 0.026019441 | 0.594255995 | -3.435410478 |
| **A_55_P2021570** | C87414 | 0.354579367 | 0.059581977 | 0.649576757 | 8.368620161 | 2.957089364 | 0.026053564 | 0.594255995 | -3.436012706 |
| **A_52_P300730** | Hmga2 | -0.197466278 | -0.361765706 | -0.033166851 | 8.135042227 | -2.956827939 | 0.026062278 | 0.594255995 | -3.436166399 |
| **A_55_P2015687** | Phf11d | -0.274932859 | -0.503747387 | -0.046118331 | 9.653523607 | -2.956053 | 0.026088129 | 0.594255995 | -3.43662206 |
| **A_66_P108692** | NA | -0.268512148 | -0.492030855 | -0.04499344 | 13.96079334 | -2.95542011 | 0.026109262 | 0.594255995 | -3.436994275 |
| **A_55_P1999601** | NA | 0.743857713 | 0.124476503 | 1.363238924 | 14.86939191 | 2.954615742 | 0.026136148 | 0.594255995 | -3.437467442 |
| **A_55_P2325729** | Arhgap1 | 0.520479239 | 0.086886828 | 0.95407165 | 11.73370389 | 2.953186685 | 0.026183988 | 0.594255995 | -3.438308361 |
| **A_52_P116134** | Aldh2 | -0.21380301 | -0.391924349 | -0.03568167 | 8.861933888 | -2.953024328 | 0.026189429 | 0.594255995 | -3.438403922 |
| **A_55_P2157788** | NA | -0.3172926 | -0.581801729 | -0.052783471 | 12.29037648 | -2.951130678 | 0.026252983 | 0.594255995 | -3.439518837 |
| **A_55_P1957433** | Mprip | -0.360682875 | -0.661492539 | -0.059873211 | 9.911636415 | -2.949870165 | 0.026295381 | 0.594255995 | -3.440261334 |
| **A_52_P532029** | Tbk1 | -0.163080838 | -0.299110475 | -0.027051201 | 8.514377196 | -2.949433159 | 0.026310097 | 0.594255995 | -3.440518815 |
| **A_55_P2000618** | Gm12690 | 0.531535 | 0.088103688 | 0.974966311 | 9.2788425 | 2.948999325 | 0.026324715 | 0.594255995 | -3.44077446 |
| **A_51_P415519** | Tmem39a | 0.203159109 | 0.033669034 | 0.372649185 | 11.88604569 | 2.948907768 | 0.026327801 | 0.594255995 | -3.440828416 |
| **A_66_P135192** | Msl2 | -0.153260927 | -0.281198548 | -0.025323306 | 8.991963763 | -2.947150683 | 0.026387105 | 0.594255995 | -3.441864181 |
| **A_55_P2071751** | NA | 0.615964056 | 0.101669709 | 1.130258403 | 9.594487269 | 2.946542993 | 0.026407649 | 0.594255995 | -3.442222528 |
| **A_55_P2326260** | 4930525G20Rik | 0.188104265 | 0.031016674 | 0.345191856 | 8.559196235 | 2.945953763 | 0.026427585 | 0.594255995 | -3.442570052 |
| **A_55_P2074193** | NA | -0.261773982 | -0.480441717 | -0.043106247 | 13.91258896 | -2.945173853 | 0.026453998 | 0.594255995 | -3.443030131 |
| **A_52_P424585** | Ctnnb1 | -0.485578005 | -0.891244441 | -0.079911569 | 12.68106995 | -2.944822301 | 0.026465914 | 0.594255995 | -3.443237551 |
| **A_52_P29151** | Ankrd44 | 0.185101655 | 0.030344277 | 0.339859034 | 10.13273907 | 2.942578805 | 0.026542092 | 0.594255995 | -3.444561757 |
| **A_55_P2074055** | NA | -0.144680245 | -0.265784663 | -0.023575827 | 8.562314882 | -2.939127412 | 0.026659749 | 0.594255995 | -3.446600645 |
| **A_55_P2169356** | Gm1966 | -0.261427783 | -0.480319748 | -0.042535817 | 8.416274876 | -2.938265809 | 0.026689209 | 0.594255995 | -3.447109959 |
| **A_55_P2088038** | Vps13d | -0.259988687 | -0.477719487 | -0.042257886 | 9.870338964 | -2.937674975 | 0.026709431 | 0.594255995 | -3.44745929 |
| **A_55_P2082235** | NA | -0.222916464 | -0.409799813 | -0.036033115 | 9.372153245 | -2.9345445 | 0.026816855 | 0.594255995 | -3.449311214 |
| **A_55_P2137527** | Fam183b | 0.144335982 | 0.023275758 | 0.265396205 | 8.620655758 | 2.933204232 | 0.02686299 | 0.594255995 | -3.450104616 |
| **A_51_P172323** | Pak2 | -0.199031877 | -0.36606045 | -0.032003305 | 11.79524351 | -2.931575154 | 0.026919183 | 0.594255995 | -3.451069412 |
| **A_55_P2179483** | NA | -0.155856995 | -0.286694501 | -0.025019489 | 17.42719445 | -2.930644932 | 0.026951327 | 0.594255995 | -3.451620531 |
| **A_52_P656565** | Pex6 | -0.247026738 | -0.454441334 | -0.039612141 | 9.814323224 | -2.930041762 | 0.026972192 | 0.594255995 | -3.451977966 |
| **A_55_P2161829** | Hmgb1-rs17 | -0.346713608 | -0.637857279 | -0.055569938 | 12.57212554 | -2.929764406 | 0.026981792 | 0.594255995 | -3.452142347 |
| **A_55_P1954476** | NA | 0.412666658 | 0.066059726 | 0.759273591 | 9.38926706 | 2.929080446 | 0.027005482 | 0.594255995 | -3.452547769 |
| **A_52_P311245** | Scarb2 | -0.220376927 | -0.405494368 | -0.035259486 | 10.28503884 | -2.928788074 | 0.027015616 | 0.594255995 | -3.452721099 |
| **A_55_P2083844** | Dis3l | 0.161239375 | 0.025792828 | 0.296685923 | 9.842331823 | 2.928682758 | 0.027019267 | 0.594255995 | -3.452783539 |
| **A_55_P2077009** | Acap3 | 0.168995748 | 0.027017274 | 0.310974221 | 10.07929319 | 2.928346317 | 0.027030935 | 0.594255995 | -3.45298302 |
| **A_52_P67995** | Eif4a3 | -0.212950458 | -0.391903316 | -0.0339976 | 9.752579689 | -2.927582232 | 0.027057454 | 0.594255995 | -3.453436133 |
| **A_55_P2036952** | NA | 0.242711442 | 0.038734867 | 0.446688017 | 9.009974901 | 2.927380111 | 0.027064474 | 0.594255995 | -3.45355601 |
| **A_55_P2097161** | Ifitm7 | -0.223290192 | -0.410948378 | -0.035632006 | 13.81023767 | -2.927327395 | 0.027066305 | 0.594255995 | -3.453587277 |
| **A_55_P2294184** | Cemip2 | -0.209222584 | -0.385086675 | -0.033358492 | 8.99972782 | -2.926850576 | 0.027082874 | 0.594255995 | -3.453870111 |
| **A_55_P1955577** | Ccm2 | 0.167618276 | 0.026721012 | 0.30851554 | 13.28386758 | 2.926765816 | 0.02708582 | 0.594255995 | -3.453920392 |
| **A_52_P572284** | Laptm4a | -0.171558709 | -0.315784075 | -0.027333342 | 13.04785831 | -2.926444426 | 0.027096996 | 0.594255995 | -3.454111057 |
| **A_55_P2002150** | Pgs1 | -0.342106373 | -0.629841903 | -0.054370844 | 13.59888142 | -2.925073855 | 0.02714471 | 0.594255995 | -3.454924355 |
| **A_55_P2012799** | Rtkn | 0.268571955 | 0.042654208 | 0.494489701 | 9.716227355 | 2.924687549 | 0.027158176 | 0.594255995 | -3.45515365 |

| | | | | | | | | | |
|---|---|---|---|---|---|---|---|---|---|
| **A_55_P2043267** | Cdc42se1 | 0.200416912 | 0.031814386 | 0.369019439 | 11.1700799 | 2.924418026 | 0.027167574 | 0.594255995 | -3.455313642 |
| **A_55_P2095010** | NA | 0.439693102 | 0.069598666 | 0.809787539 | 8.54882107 | 2.922848192 | 0.027222388 | 0.594255995 | -3.45624577 |
| **A_55_P1962918** | Ifi211 | -0.29909395 | -0.550852721 | -0.04733518 | 8.813276941 | -2.922753457 | 0.0272257 | 0.594255995 | -3.456302035 |
| **A_55_P1987326** | Prl3a1 | -0.352623361 | -0.649447307 | -0.055799415 | 16.7656321 | -2.922680285 | 0.027228258 | 0.594255995 | -3.456345495 |
| **A_52_P452019** | Usp31 | -0.152634132 | -0.281164201 | -0.024104064 | 9.243122956 | -2.92156858 | 0.027267157 | 0.594255995 | -3.457005893 |
| **A_55_P1997931** | Map9 | 0.47169669 | 0.074157785 | 0.869235596 | 10.83740529 | 2.919122727 | 0.027352951 | 0.594255995 | -3.458459594 |
| **A_55_P2083859** | NA | 0.205717458 | 0.032269276 | 0.379165641 | 13.05942505 | 2.917901045 | 0.027395913 | 0.594255995 | -3.459186099 |
| **A_55_P2166985** | Dedd2 | -0.250071708 | -0.461057421 | -0.039085995 | 13.55715311 | -2.915954044 | 0.027464535 | 0.594255995 | -3.460344476 |
| **A_55_P2050354** | Prpmp5 | 0.545027055 | 0.0851516 | 1.00490251 | 9.240939896 | 2.915727906 | 0.027472517 | 0.594255995 | -3.460479061 |
| **A_55_P2156583** | Cdk5r1 | -0.184719664 | -0.340662146 | -0.028777182 | 10.24432258 | -2.914189918 | 0.027526872 | 0.594255995 | -3.461394625 |
| **A_55_P2118570** | Cd37 | 0.383857422 | 0.059797559 | 0.707917284 | 8.70687749 | 2.914163149 | 0.027527819 | 0.594255995 | -3.461410565 |
| **A_55_P2007088** | Rnf128 | -0.37864275 | -0.698449423 | -0.058836076 | 16.21768352 | -2.912804163 | 0.027575946 | 0.594255995 | -3.462219922 |
| **A_55_P2091039** | NA | 0.234216632 | 0.036244532 | 0.432188731 | 8.986861218 | 2.910602546 | 0.027654109 | 0.594255995 | -3.463531805 |
| **A_55_P2028595** | AW551984 | 0.659924611 | 0.102056986 | 1.217792236 | 8.985729318 | 2.910263614 | 0.027666163 | 0.594255995 | -3.463733841 |
| **A_55_P2156229** | Cd3e | 0.695974711 | 0.107502377 | 1.284447045 | 10.1670784 | 2.909622002 | 0.027688997 | 0.594255995 | -3.464116358 |
| **A_55_P2012780** | Rnf167 | -0.236711947 | -0.436907972 | -0.036515923 | 12.84502451 | -2.908934166 | 0.0277135 | 0.594255995 | -3.464526513 |
| **A_66_P111689** | NA | 0.756620539 | 0.11646956 | 1.396771518 | 11.63956255 | 2.907802299 | 0.027753871 | 0.594255995 | -3.465201625 |
| **A_55_P2050727** | Fcrl5 | 0.620806777 | 0.095455196 | 1.146158357 | 11.88974641 | 2.907204412 | 0.027775222 | 0.594255995 | -3.465558331 |
| **A_51_P461578** | Fdxr | -0.174453165 | -0.32209896 | -0.026807369 | 10.61008173 | -2.906878848 | 0.027786855 | 0.594255995 | -3.465752591 |
| **A_51_P387123** | Oasl2 | -0.384854037 | -0.710609592 | -0.059098482 | 11.00890048 | -2.906520426 | 0.027799669 | 0.594255995 | -3.46596648 |
| **A_52_P82209** | Wnk1 | -0.451642338 | -0.833988599 | -0.069296076 | 10.56897588 | -2.90607596 | 0.027815568 | 0.594255995 | -3.466231747 |
| **A_55_P2038452** | NA | 0.25159954 | 0.038522181 | 0.464676898 | 9.079791675 | 2.904970329 | 0.02785516 | 0.594255995 | -3.466891763 |
| **A_51_P167971** | Bace1 | -0.313667132 | -0.579371844 | -0.047962421 | 10.28279992 | -2.904282927 | 0.027879806 | 0.594255995 | -3.467302221 |
| **A_55_P2016838** | NA | 0.469832215 | 0.071626317 | 0.868038113 | 14.37307204 | 2.902714126 | 0.027936143 | 0.594255995 | -3.468239287 |
| **A_51_P487501** | NA | -0.249142188 | -0.460346042 | -0.037938335 | 11.76785195 | -2.90211488 | 0.027957695 | 0.594255995 | -3.468597338 |
| **A_55_P2173035** | Dnajc8 | -0.240791579 | -0.444924181 | -0.036658977 | 10.62719561 | -2.902004527 | 0.027961666 | 0.594255995 | -3.468663281 |
| **A_55_P1961894** | Fbxw11 | -0.183683231 | -0.339440686 | -0.027925777 | 13.01991148 | -2.90128126 | 0.027987706 | 0.594255995 | -3.469095534 |
| **A_55_P2148531** | Pithd1 | -0.143038216 | -0.264340454 | -0.021735979 | 8.703944819 | -2.901031451 | 0.027996707 | 0.594255995 | -3.469244851 |
| **A_52_P581435** | Pdha1 | -0.196644788 | -0.363415031 | -0.029874545 | 10.66986703 | -2.90090189 | 0.028001376 | 0.594255995 | -3.469322297 |
| **A_55_P2065926** | Akap8l | -0.170248246 | -0.31466892 | -0.025827572 | 8.642441458 | -2.900163235 | 0.028028011 | 0.594255995 | -3.46976389 |
| **A_55_P1977910** | NA | -0.371282038 | -0.686314699 | -0.056249377 | 10.41589802 | -2.899462662 | 0.028053299 | 0.594255995 | -3.470182805 |
| **A_55_P2136154** | Heca | -0.216147752 | -0.399565091 | -0.032730413 | 11.15291852 | -2.899208819 | 0.028062468 | 0.594255995 | -3.470334614 |
| **A_51_P493558** | Rnpep | -0.22602065 | -0.417909263 | -0.034132036 | 10.7801385 | -2.897797795 | 0.028113493 | 0.594255995 | -3.471178675 |
| **A_52_P467389** | Slc15a3 | -0.13390968 | -0.247608185 | -0.020211174 | 7.929600297 | -2.89751957 | 0.028123566 | 0.594255995 | -3.471345147 |
| **A_51_P348227** | Stoml1 | -0.163036455 | -0.301528468 | -0.024544442 | 8.945716438 | -2.896204066 | 0.028171246 | 0.594255995 | -3.472132447 |
| **A_55_P2141876** | Ldha | 0.160381019 | 0.024095006 | 0.296667032 | 10.14434417 | 2.8951485 | 0.028209568 | 0.594255995 | -3.4727644 |
| **A_52_P281145** | Kank4 | -0.141556702 | -0.261863694 | -0.021249709 | 8.139214522 | -2.894734443 | 0.028224616 | 0.594255995 | -3.473012343 |
| **A_51_P342387** | Apoh | -0.319040557 | -0.590260941 | -0.047820174 | 8.335455566 | -2.893961389 | 0.028252733 | 0.594255995 | -3.473475339 |
| **A_55_P1986566** | Gm10318 | 0.238753316 | 0.035619377 | 0.441887255 | 8.864719709 | 2.891585808 | 0.028339328 | 0.594255995 | -3.474898775 |
| **A_55_P2007673** | Lrsam1 | 0.45879525 | 0.068316161 | 0.84927434 | 11.37329637 | 2.89061528 | 0.028374789 | 0.594255995 | -3.475480595 |
| **A_55_P1952071** | Npb | 0.669581906 | 0.099507514 | 1.239656297 | 10.42365405 | 2.889624029 | 0.028411056 | 0.594255995 | -3.476075009 |
| **A_55_P1992834** | Socs2 | -0.13985956 | -0.258979917 | -0.020739202 | 8.7269948 | -2.888519688 | 0.02845152 | 0.594255995 | -3.476737441 |
| **A_55_P1954302** | Esrrb | 0.173426496 | 0.025708744 | 0.321144248 | 9.148801 | 2.888363997 | 0.02845723 | 0.594255995 | -3.476830848 |
| **A_55_P1968362** | Tle1 | 0.209031369 | 0.030933936 | 0.387128801 | 8.649220344 | 2.887506489 | 0.0284887 | 0.594255995 | -3.47734539 |
| **A_55_P1962479** | Gpbp1l1 | 0.149489359 | 0.022098822 | 0.276879895 | 9.419085915 | 2.886970023 | 0.028508407 | 0.594255995 | -3.477667358 |
| **A_52_P20966** | Hnrnpab | -0.442727076 | -0.82003093 | -0.065423221 | 12.83880114 | -2.886782103 | 0.028515314 | 0.594255995 | -3.477780152 |
| **A_55_P2091476** | NA | 0.146834179 | 0.02165408 | 0.272014278 | 9.574591355 | 2.885765465 | 0.02855271 | 0.594255995 | -3.478390474 |
| **A_55_P2185722** | Fermt2 | 0.327898631 | 0.048354574 | 0.607442688 | 11.36723601 | 2.885748658 | 0.028553329 | 0.594255995 | -3.478400566 |
| **A_55_P2059154** | Dtx3l | -0.19860453 | -0.3679421 | -0.02926696 | 9.963230659 | -2.88539311 | 0.02856642 | 0.594255995 | -3.478614057 |
| **A_55_P2047275** | Tspan7 | 0.198100889 | 0.029049876 | 0.367151903 | 11.30983323 | 2.882954646 | 0.028656381 | 0.594255995 | -3.480078847 |

| | | | | | | | | | |
|---|---|---|---|---|---|---|---|---|---|
| **A_55_P1971614** | NA | -0.2073447 | -0.384288787 | -0.030400613 | 10.16020467 | -2.882876561 | 0.028659267 | 0.594255995 | -3.480125769 |
| **A_51_P267544** | Frg1 | -0.151728103 | -0.281294219 | -0.022161987 | 10.53785085 | -2.881003259 | 0.028728593 | 0.594255995 | -3.481251797 |
| **A_52_P522864** | Zfp442 | -0.246542422 | -0.457128408 | -0.035956436 | 12.46142035 | -2.880257751 | 0.028756233 | 0.594255995 | -3.481700088 |
| **A_55_P2060592** | Hoxa1 | 0.166651036 | 0.024173105 | 0.309128967 | 9.823839331 | 2.877594413 | 0.028855213 | 0.594255995 | -3.483302406 |
| **A_51_P227345** | Dpep1 | 0.18681979 | 0.027078391 | 0.346561189 | 8.181393625 | 2.877229861 | 0.02886879 | 0.594255995 | -3.483521825 |
| **A_55_P2178152** | Ccsap | 0.222639819 | 0.032216987 | 0.413062651 | 8.481630651 | 2.876424527 | 0.028898807 | 0.594255995 | -3.484006626 |
| **A_52_P1020860** | AW112010 | -0.313910317 | -0.582468279 | -0.045352355 | 8.828908595 | -2.875654597 | 0.028927536 | 0.594255995 | -3.48447022 |
| **A_55_P2052081** | Eif1ad | 0.184469699 | 0.02663519 | 0.342304209 | 11.97458821 | 2.875360075 | 0.028938534 | 0.594255995 | -3.484647586 |
| **A_52_P585028** | Cnep1r1 | -0.171971035 | -0.319123969 | -0.024818101 | 11.00645287 | -2.875117129 | 0.02894761 | 0.594255995 | -3.484793905 |
| **A_55_P1973970** | Krt73 | 0.544636871 | 0.07816791 | 1.011105832 | 9.222257029 | 2.872456435 | 0.029047204 | 0.594255995 | -3.486397024 |
| **A_55_P2075343** | Arfrp1 | -0.133946465 | -0.248686621 | -0.019206309 | 8.532847622 | -2.872003631 | 0.02906419 | 0.594255995 | -3.486669971 |
| **A_55_P2121945** | Cnot6l | -0.653881881 | -1.17829122 | -0.129472543 | 12.00362771 | -4.072507232 | 0.029068137 | 0.594255995 | -3.961684297 |
| **A_55_P1983262** | Gm4026 | 0.479944675 | 0.068631534 | 0.891257816 | 8.982888864 | 2.870699642 | 0.029113167 | 0.594255995 | -3.487456205 |
| **A_51_P405606** | Ndrg1 | -0.232263776 | -0.431475056 | -0.033052495 | 15.13156035 | -2.868380211 | 0.029200503 | 0.594255995 | -3.488855429 |
| **A_55_P2075200** | Tor3a | -0.205061751 | -0.38102779 | -0.029095712 | 9.850845093 | -2.866982005 | 0.029253289 | 0.594255995 | -3.489699368 |
| **A_55_P2088716** | Ss18l2 | 0.201848418 | 0.02858188 | 0.375114955 | 11.35984009 | 2.866023956 | 0.029289517 | 0.594255995 | -3.49027783 |
| **A_51_P301215** | Knop1 | -0.399519795 | -0.7425903 | -0.056449291 | 11.297798 | -2.864996427 | 0.029328427 | 0.594255995 | -3.490898423 |
| **A_55_P2117525** | Rbm3 | -0.161404128 | -0.300219643 | -0.022588613 | 10.93012738 | -2.86052529 | 0.029498388 | 0.594255995 | -3.493600985 |
| **A_55_P1964902** | NA | 0.13217858 | 0.018337729 | 0.246019432 | 7.992821912 | 2.856486072 | 0.029652847 | 0.594255995 | -3.496045475 |
| **A_55_P2093251** | NA | 0.264485044 | 0.036658497 | 0.49231159 | 9.363373134 | 2.856050977 | 0.029669537 | 0.594255995 | -3.496308959 |
| **A_52_P370473** | Msl1 | -0.235763515 | -0.43887822 | -0.03264881 | 11.93001357 | -2.855646168 | 0.029685074 | 0.594255995 | -3.496554133 |
| **A_55_P2076639** | Vps25 | -0.144339113 | -0.268779625 | -0.0198986 | 9.149086968 | -2.85358887 | 0.029764174 | 0.594255995 | -3.497800582 |
| **A_55_P1968245** | Ftl1 | 0.16260242 | 0.022374756 | 0.302830085 | 17.10428791 | 2.852741818 | 0.029796808 | 0.594255995 | -3.498313997 |
| **A_55_P2158351** | Rpl18a | -0.161641479 | -0.301068866 | -0.022214091 | 15.50633788 | -2.852160022 | 0.029819245 | 0.594255995 | -3.498666708 |
| **A_55_P1952990** | Myl6b | 0.156237297 | 0.021462467 | 0.291012127 | 9.072016748 | 2.851970891 | 0.029826543 | 0.594255995 | -3.498781381 |
| **A_51_P191520** | Stard10 | -0.154490607 | -0.287762995 | -0.021218218 | 9.549564397 | -2.851878777 | 0.029830098 | 0.594255995 | -3.498837233 |
| **A_52_P103391** | B4galt1 | -0.246510104 | -0.459243182 | -0.033777026 | 14.50752815 | -2.85081388 | 0.029871229 | 0.594255995 | -3.499483027 |
| **A_55_P2064412** | NA | 0.691578349 | 0.094659039 | 1.28849766 | 9.186154994 | 2.85032855 | 0.029889995 | 0.594255995 | -3.499777414 |
| **A_55_P2080850** | Ptpn2 | -0.215634334 | -0.401769905 | -0.029498762 | 11.20631078 | -2.850084245 | 0.029899446 | 0.594255995 | -3.499925619 |
| **A_55_P1957439** | NA | 0.191255255 | 0.026153619 | 0.35635689 | 8.76344231 | 2.849910122 | 0.029906184 | 0.594255995 | -3.500031254 |
| **A_55_P2003658** | Gnpda1 | 0.229256854 | 0.031310452 | 0.427203255 | 9.34163649 | 2.849337406 | 0.029928359 | 0.594255995 | -3.500378743 |
| **A_51_P300506** | Cox6b2 | 0.16851606 | 0.022996543 | 0.314035577 | 8.641579206 | 2.848978785 | 0.029942253 | 0.594255995 | -3.500596361 |
| **A_55_P2095384** | Glrx2 | 0.202513986 | 0.027608723 | 0.377419249 | 11.55470331 | 2.848533336 | 0.029959521 | 0.594255995 | -3.500866699 |
| **A_51_P422893** | Tmem14a | 0.209430522 | 0.028466773 | 0.390394271 | 8.869754257 | 2.847197229 | 0.030011379 | 0.594255995 | -3.501677775 |
| **A_52_P677956** | Pex11g | 0.276236443 | 0.037531669 | 0.514941216 | 11.86531306 | 2.847010258 | 0.030018644 | 0.594255995 | -3.501791299 |
| **A_51_P396273** | Nras | -0.273303129 | -0.509486294 | -0.037119964 | 9.583938338 | -2.846851599 | 0.03002481 | 0.594255995 | -3.501887638 |
| **A_55_P2165451** | NA | -0.172890762 | -0.322313642 | -0.023467881 | 9.751764397 | -2.846582965 | 0.030035253 | 0.594255995 | -3.502050764 |
| **A_55_P1996613** | NA | 0.415971182 | 0.056347443 | 0.77559492 | 15.9221273 | 2.845666857 | 0.030070897 | 0.594255995 | -3.50260716 |
| **A_55_P2074371** | Lin28a | 0.433308475 | 0.058634037 | 0.807982914 | 8.946234495 | 2.845196603 | 0.030089212 | 0.594255995 | -3.502892825 |
| **A_55_P2235059** | 6230415J03Rik | 0.475157496 | 0.063984779 | 0.886330214 | 9.923224034 | 2.843036648 | 0.030173488 | 0.594255995 | -3.504205422 |
| **A_51_P444437** | Ptgr1 | 0.22002285 | 0.029594612 | 0.410451087 | 9.300188475 | 2.842533556 | 0.030193153 | 0.594255995 | -3.504511266 |
| **A_55_P2098305** | Coq5 | -0.161546621 | -0.301377186 | -0.021716056 | 8.326967808 | -2.84226737 | 0.030203564 | 0.594255995 | -3.504673106 |
| **A_55_P2133679** | NA | -0.223424919 | -0.41687215 | -0.029977689 | 10.98735396 | -2.841438472 | 0.030236008 | 0.594255995 | -3.505177152 |
| **A_55_P1967342** | NA | -0.214498457 | -0.400245338 | -0.028751577 | 13.83073349 | -2.841003756 | 0.030253039 | 0.594255995 | -3.505441547 |
| **A_51_P461504** | Eef2k | -0.190567999 | -0.355637053 | -0.025498945 | 9.366924849 | -2.840229759 | 0.030283386 | 0.594255995 | -3.505912374 |
| **A_51_P269020** | 1700031F05Rik | -0.23509784 | -0.438756596 | -0.031439083 | 8.924097018 | -2.839976286 | 0.030293332 | 0.594255995 | -3.506066586 |
| **A_55_P2056212** | Gpr3 | 0.139530843 | 0.018650089 | 0.260411598 | 9.306345015 | 2.839763802 | 0.030301672 | 0.594255995 | -3.506195869 |
| **A_52_P680038** | Cdk16 | -0.137245468 | -0.256164043 | -0.018326894 | 8.658252166 | -2.839340531 | 0.030318293 | 0.594255995 | -3.506453426 |
| **A_55_P2074521** | Psg28 | -0.288407508 | -0.538393085 | -0.038421931 | 17.22982512 | -2.838316172 | 0.030358558 | 0.594255995 | -3.507076868 |
| **A_55_P2175752** | Mfap1b | -0.157151135 | -0.293411163 | -0.020891108 | 9.252968853 | -2.837384639 | 0.030395224 | 0.594255995 | -3.507643974 |

| | | | | | | | | |
|---|---|---|---|---|---|---|---|---|
| **A_55_P2117943** | NA | 0.187642852 | 0.024922488 | 0.350363216 | 10.13143057 | 2.83699965 | 0.030410392 | 0.594255995 | -3.507878395 |
| **A_52_P93467** | Sdc4 | -0.242178372 | -0.452302661 | -0.032054083 | 10.02530529 | -2.835490921 | 0.030469911 | 0.594255995 | -3.508797311 |
| **A_51_P320444** | Rock2 | -0.242823558 | -0.453661555 | -0.031985561 | 8.767575516 | -2.833420947 | 0.030551775 | 0.594255995 | -3.510058706 |
| **A_55_P1982211** | NA | 0.324748975 | 0.042719978 | 0.606777971 | 9.387872899 | 2.832847461 | 0.030574498 | 0.594255995 | -3.510408308 |
| **A_55_P2082070** | Ak6 | -0.290601466 | -0.542987718 | -0.038215214 | 10.40626811 | -2.832704483 | 0.030580166 | 0.594255995 | -3.510495477 |
| **A_55_P1992884** | NA | 0.703848233 | 0.092227705 | 1.315468761 | 9.008253386 | 2.831171215 | 0.030641019 | 0.594255995 | -3.511430487 |
| **A_55_P1979014** | NA | 0.183886533 | 0.024023855 | 0.343749211 | 8.688211808 | 2.829905967 | 0.030691333 | 0.594255995 | -3.512202363 |
| **A_55_P1958652** | Arf2 | -0.335195213 | -0.626604799 | -0.043785626 | 14.87910768 | -2.829848174 | 0.030693634 | 0.594255995 | -3.512237626 |
| **A_51_P178872** | Ttc9c | -0.182251537 | -0.340717475 | -0.0237856 | 8.803555623 | -2.829465742 | 0.03070886 | 0.594255995 | -3.51247099 |
| **A_55_P2183668** | NA | -0.318093977 | -0.594790676 | -0.041397278 | 12.56937174 | -2.82826846 | 0.030756583 | 0.594255995 | -3.51320175 |
| **A_55_P2062926** | D430041D05Rik | 0.15410506 | 0.020038462 | 0.288171659 | 8.494208186 | 2.827909277 | 0.030770916 | 0.594255995 | -3.513421026 |
| **A_52_P634822** | Nap1l4 | 0.18701871 | 0.024287711 | 0.349749708 | 8.712334387 | 2.827378362 | 0.030792114 | 0.594255995 | -3.513745182 |
| **A_55_P1961270** | Cd72 | -0.259781642 | -0.485833207 | -0.033730077 | 8.334630259 | -2.827288343 | 0.03079571 | 0.594255995 | -3.513800149 |
| **A_55_P1967672** | Tas2r109 | 1.002884881 | 0.129834856 | 1.875934905 | 10.36860617 | 2.826058309 | 0.030844889 | 0.594255995 | -3.514551367 |
| **A_55_P2074337** | Ngf | 0.796472281 | 0.102839946 | 1.490104616 | 9.347478283 | 2.824948265 | 0.030889344 | 0.594255995 | -3.515229529 |
| **A_52_P313080** | Hnrnpab | -0.218154315 | -0.408157535 | -0.028151095 | 11.95417587 | -2.824697921 | 0.03089938 | 0.594255995 | -3.515382502 |
| **A_55_P2322039** | NA | 0.193940108 | 0.024992882 | 0.362887333 | 8.503795186 | 2.824136811 | 0.030921885 | 0.594255995 | -3.515725408 |
| **A_55_P2029945** | NA | 0.561500364 | 0.072357742 | 1.050642986 | 15.09548403 | 2.824123588 | 0.030922416 | 0.594255995 | -3.515733489 |
| **A_55_P1993026** | NA | 0.281160547 | 0.036215641 | 0.526105452 | 14.28987601 | 2.823937826 | 0.03092987 | 0.594255995 | -3.515847025 |
| **A_55_P2038262** | Phf11b | -0.290574299 | -0.543803755 | -0.037344844 | 8.652195506 | -2.823008208 | 0.030967205 | 0.594255995 | -3.516415286 |
| **A_55_P2020443** | Hnrnpa3 | -0.186475416 | -0.34898678 | -0.023964053 | 9.116271923 | -2.822974886 | 0.030968545 | 0.594255995 | -3.516435658 |
| **A_55_P1994608** | Itsn1 | 0.605424434 | 0.077677342 | 1.133171526 | 9.521679789 | 2.822300539 | 0.030995659 | 0.594255995 | -3.516847974 |
| **A_52_P600531** | Bahd1 | -0.262520649 | -0.491378493 | -0.033662804 | 10.7076767 | -2.822063808 | 0.031005184 | 0.594255995 | -3.516992737 |
| **A_55_P2003673** | Mtmr12 | -0.177102292 | -0.331556723 | -0.02264786 | 11.30446336 | -2.820934291 | 0.031050673 | 0.594255995 | -3.517683581 |
| **A_55_P1959238** | Nagk | 0.268454613 | 0.03431155 | 0.502597677 | 11.80544588 | 2.820711947 | 0.031059636 | 0.594255995 | -3.517819599 |
| **A_55_P2156052** | NA | 0.179530633 | 0.022945443 | 0.336115823 | 9.629242843 | 2.820700932 | 0.03106008 | 0.594255995 | -3.517826337 |
| **A_52_P337679** | Canx | -0.298443767 | -0.558808122 | -0.038079413 | 14.67630647 | -2.820006767 | 0.031088082 | 0.594255995 | -3.518251045 |
| **A_52_P301374** | Aaas | -0.145887263 | -0.273161827 | -0.0186127 | 8.869177268 | -2.819972836 | 0.031089451 | 0.594255995 | -3.518271807 |
| **A_66_P121053** | Sftpd | -0.207744542 | -0.389078065 | -0.026411018 | 8.300486734 | -2.818517167 | 0.031148261 | 0.594255995 | -3.519162703 |
| **A_55_P2153517** | Enho | 0.175962027 | 0.022318909 | 0.329605144 | 12.84285797 | 2.817571903 | 0.031186514 | 0.594255995 | -3.519741419 |
| **A_55_P1990879** | Banp | 0.510458684 | 0.064715658 | 0.95620171 | 8.579865683 | 2.81737854 | 0.031194346 | 0.594255995 | -3.519859819 |
| **A_55_P2100526** | Ptges3 | 0.171303485 | 0.021715185 | 0.320891785 | 11.88473614 | 2.8173301 | 0.031196308 | 0.594255995 | -3.519889481 |
| **A_51_P259555** | Gpatch3 | -0.1536296 | -0.287806216 | -0.019452983 | 8.666287781 | -2.81687273 | 0.031214842 | 0.594255995 | -3.52016957 |
| **A_55_P2078257** | NA | 0.134206621 | 0.016966323 | 0.251446919 | 8.758333673 | 2.816217488 | 0.031241414 | 0.594255995 | -3.520570897 |
| **A_51_P159711** | Car7 | 0.470693094 | 0.059406062 | 0.881980125 | 10.26051745 | 2.815541764 | 0.031268843 | 0.594255995 | -3.520984846 |
| **A_51_P386670** | Dse | -0.142237774 | -0.266538307 | -0.017937242 | 8.231071869 | -2.815212117 | 0.031282233 | 0.594255995 | -3.521186817 |
| **A_51_P184353** | Nr2f6 | 0.256251482 | 0.032137727 | 0.480365236 | 12.13886347 | 2.812982544 | 0.031372961 | 0.594255995 | -3.522553345 |
| **A_55_P2396375** | 1810073O08Rik | 0.480972928 | 0.060161752 | 0.901784104 | 9.256134152 | 2.81191721 | 0.031416413 | 0.594255995 | -3.523206603 |
| **A_55_P1970860** | Pemt | 0.300877911 | 0.037590651 | 0.564165172 | 9.800356435 | 2.811445199 | 0.031435686 | 0.594255995 | -3.5234961 |
| **A_51_P215475** | Ptprb | -0.392091682 | -0.735233099 | -0.048950265 | 11.30839173 | -2.811147622 | 0.031447843 | 0.594255995 | -3.523678633 |
| **A_55_P2024600** | Zfp658 | 0.227391186 | 0.028213093 | 0.426569278 | 15.41096526 | 2.808673269 | 0.031549125 | 0.594255995 | -3.525196982 |
| **A_55_P2121537** | NA | -0.180519026 | -0.33870279 | -0.022335262 | 16.14298418 | -2.807567718 | 0.031594492 | 0.594255995 | -3.525875729 |
| **A_55_P1975305** | NA | -0.173002335 | -0.323155777 | -0.022848894 | 11.91251865 | -2.985841559 | 0.03162152 | 0.594255995 | -3.624109781 |
| **A_51_P485240** | Adipor1 | -0.333615399 | -0.626146239 | -0.04108456 | 12.05929986 | -2.805715261 | 0.031670667 | 0.594255995 | -3.527013509 |
| **A_52_P191975** | Rtn3 | -0.155177764 | -0.291263399 | -0.01909213 | 10.22414842 | -2.805345594 | 0.031685892 | 0.594255995 | -3.52724063 |
| **A_55_P2128501** | Krt8 | -0.442749737 | -0.831196854 | -0.05430262 | 13.84230097 | -2.804113401 | 0.031736696 | 0.594255995 | -3.527997851 |
| **A_51_P270184** | Nhlrc2 | -0.239488487 | -0.449605578 | -0.029371395 | 9.894174102 | -2.804093062 | 0.031737536 | 0.594255995 | -3.528010352 |
| **A_52_P423128** | Arglu1 | -0.264831675 | -0.497458462 | -0.032204888 | 10.71350988 | -2.800782401 | 0.031874478 | 0.594255995 | -3.530046167 |
| **A_52_P366653** | Hnrnph3 | -0.210188019 | -0.394843685 | -0.025532353 | 8.093466124 | -2.80036398 | 0.031891831 | 0.594255995 | -3.5303036 |
| **A_55_P2085766** | NA | 0.399698536 | 0.048524259 | 0.750872812 | 8.435418923 | 2.800135345 | 0.031901317 | 0.594255995 | -3.530444281 |

| | | | | | | | | | |
|---|---|---|---|---|---|---|---|---|---|
| **A_51_P279183** | Tkfc | 0.214182851 | 0.025901525 | 0.402464177 | 9.689617036 | 2.798637248 | 0.03196355 | 0.594255995 | -3.531366291 |
| **A_66_P103198** | NA | 0.283588202 | 0.034189659 | 0.532986745 | 8.451345806 | 2.797456942 | 0.032012674 | 0.594255995 | -3.532092988 |
| **A_52_P659762** | Eid2 | -0.15214827 | -0.28597327 | -0.01832327 | 8.119431344 | -2.797041621 | 0.032029979 | 0.594255995 | -3.532348752 |
| **A_51_P133052** | Glyr1 | -0.169326951 | -0.318264125 | -0.020389776 | 11.18261317 | -2.796997894 | 0.032031802 | 0.594255995 | -3.532375682 |
| **A_51_P306067** | Lpp | -0.193349795 | -0.363432837 | -0.023266753 | 11.02011628 | -2.796738516 | 0.032042615 | 0.594255995 | -3.53253543 |
| **A_66_P125634** | Wdr63 | 0.392298086 | 0.047114036 | 0.737482136 | 9.886642461 | 2.79598368 | 0.032074105 | 0.594255995 | -3.533000391 |
| **A_55_P1965699** | NA | 0.487451813 | 0.058417067 | 0.916486559 | 9.055490817 | 2.795171039 | 0.032108044 | 0.594255995 | -3.533501069 |
| **A_55_P2054456** | Adprm | -0.255846899 | -0.48107733 | -0.030616467 | 11.21611596 | -2.794616711 | 0.032131217 | 0.594255995 | -3.533842662 |
| **A_55_P2086631** | NA | -0.21429837 | -0.403024898 | -0.025571843 | 13.38822624 | -2.79354121 | 0.032176229 | 0.594255995 | -3.534505569 |
| **A_51_P347961** | Npc2 | -0.247370784 | -0.465263952 | -0.029477616 | 14.80939042 | -2.793019418 | 0.032198092 | 0.594255995 | -3.534827258 |
| **A_55_P2079761** | Ggps1 | 0.155168132 | 0.018485024 | 0.29185124 | 9.320510386 | 2.792909402 | 0.032202703 | 0.594255995 | -3.53489509 |
| **A_55_P2168189** | NA | 0.26378971 | 0.031374739 | 0.496204682 | 12.55931118 | 2.792305388 | 0.032228035 | 0.594255995 | -3.535267539 |
| **A_55_P2131274** | Zfp663 | 0.595257859 | 0.070656694 | 1.119859023 | 13.72954668 | 2.791547685 | 0.032259842 | 0.594255995 | -3.535734846 |
| **A_55_P2046499** | Vmn1r70 | 0.514473721 | 0.060971038 | 0.967976404 | 9.054413435 | 2.79095284 | 0.032284837 | 0.594255995 | -3.536101781 |
| **A_55_P2101216** | Aldh3a1 | 1.019558042 | 0.120636287 | 1.918479798 | 10.21942341 | 2.790353504 | 0.032310041 | 0.594255995 | -3.536471548 |
| **A_55_P2090142** | Kcna7 | 0.135358336 | 0.015985093 | 0.254731579 | 8.962141733 | 2.789633676 | 0.03234034 | 0.594255995 | -3.536915735 |
| **A_65_P11455** | Pphln1 | -0.181066249 | -0.340812688 | -0.021319809 | 8.844948914 | -2.788531021 | 0.032386813 | 0.594255995 | -3.537596329 |
| **A_55_P2126644** | NA | -0.167377437 | -0.315056539 | -0.019698334 | 10.0645592 | -2.788348367 | 0.032394519 | 0.594255995 | -3.537709089 |
| **A_51_P215540** | Rnf4 | -0.167860129 | -0.315967931 | -0.019752326 | 11.2659575 | -2.788295364 | 0.032396755 | 0.594255995 | -3.537741811 |
| **A_51_P145260** | Sall4 | 0.575140575 | 0.067656211 | 1.08262494 | 8.861881177 | 2.788178053 | 0.032401705 | 0.594255995 | -3.537814236 |
| **A_52_P363216** | Gcnt2 | -0.302641855 | -0.569716881 | -0.03556683 | 10.07577385 | -2.787820936 | 0.032416779 | 0.594255995 | -3.538034726 |
| **A_52_P81468** | Psma4 | -0.252618221 | -0.47562619 | -0.029610253 | 10.97391898 | -2.786849107 | 0.032457839 | 0.594255995 | -3.538634863 |
| **A_55_P2076989** | NA | 0.164495556 | 0.019260534 | 0.309730578 | 9.827674124 | 2.786454633 | 0.032474521 | 0.594255995 | -3.53887851 |
| **A_66_P106593** | Vhl | -0.260501981 | -0.490600048 | -0.030403913 | 13.74912993 | -2.785269482 | 0.032524698 | 0.594255995 | -3.53961068 |
| **A_66_P137134** | NA | 0.225084257 | 0.026181556 | 0.423986957 | 9.405117505 | 2.78402803 | 0.032577347 | 0.594255995 | -3.540377892 |
| **A_55_P2045911** | Spata32 | 0.306195218 | 0.035372104 | 0.577018333 | 8.43072758 | 2.781517774 | 0.032684087 | 0.594255995 | -3.541930028 |
| **A_55_P2015700** | NA | 0.204766912 | 0.023623268 | 0.385910557 | 10.49142722 | 2.781031083 | 0.032704826 | 0.594255995 | -3.542231084 |
| **A_55_P2144716** | Hecw2 | -0.157219345 | -0.296301168 | -0.018137522 | 9.268512529 | -2.781024225 | 0.032705118 | 0.594255995 | -3.542235326 |
| **A_66_P132613** | C230057M02Rik | 0.414503096 | 0.047564434 | 0.781441758 | 9.336283615 | 2.779095486 | 0.032787444 | 0.594255995 | -3.5434288 |
| **A_55_P2122425** | NA | 0.160255986 | 0.018382284 | 0.302129689 | 13.18601097 | 2.778955044 | 0.032793447 | 0.594255995 | -3.543515729 |
| **A_55_P2085835** | 5330417C22Rik | 0.236462874 | 0.027117722 | 0.445808026 | 8.446964267 | 2.778876332 | 0.032796813 | 0.594255995 | -3.54356445 |
| **A_52_P120740** | 0610030E20Rik | -0.1446851 | -0.272821334 | -0.016548866 | 8.568787479 | -2.777928195 | 0.032837377 | 0.594255995 | -3.544151413 |
| **A_51_P155303** | Rbfa | -0.264964838 | -0.499657999 | -0.030271678 | 9.72690154 | -2.777518594 | 0.032854918 | 0.594255995 | -3.544405033 |
| **A_51_P296866** | Msi1 | 0.357453293 | 0.04079582 | 0.674110766 | 9.078018612 | 2.777146071 | 0.032870879 | 0.594255995 | -3.544635719 |
| **A_52_P622434** | Nop16 | -0.140333456 | -0.264660214 | -0.016006698 | 8.880166211 | -2.776935392 | 0.03287991 | 0.594255995 | -3.544766193 |
| **A_55_P1960738** | Gm4470 | 0.723522959 | 0.082286617 | 1.364759302 | 9.484907167 | 2.775897027 | 0.032924459 | 0.594255995 | -3.545409368 |
| **A_55_P2172956** | Cnih2 | 0.399328729 | 0.045365224 | 0.753292233 | 8.436515207 | 2.775500003 | 0.03294151 | 0.594255995 | -3.545655338 |
| **A_55_P1992838** | Socs2 | -0.13254349 | -0.250029737 | -0.015057244 | 8.553741843 | -2.775495563 | 0.0329417 | 0.594255995 | -3.545658089 |
| **A_52_P665675** | Abca1 | -0.19360825 | -0.365290226 | -0.021926273 | 11.9529943 | -2.77439515 | 0.032989009 | 0.594255995 | -3.546339975 |
| **A_55_P1974098** | Gm12063 | 0.684917179 | 0.077451808 | 1.292382549 | 11.53366029 | 2.773867403 | 0.033011724 | 0.594255995 | -3.546667075 |
| **A_51_P368713** | Tmbim4 | -0.154732143 | -0.291973127 | -0.01749116 | 11.50480487 | -2.773740851 | 0.033017173 | 0.594255995 | -3.546745519 |
| **A_55_P2166349** | Dnajc19 | 0.222880162 | 0.025147624 | 0.420612701 | 10.56409217 | 2.773080242 | 0.033045635 | 0.594255995 | -3.547155049 |
| **A_52_P672453** | Prpf3 | -0.185957001 | -0.35096718 | -0.020946821 | 10.67482369 | -2.772496198 | 0.03307082 | 0.594255995 | -3.547517176 |
| **A_51_P228193** | Ociad1 | -0.208262521 | -0.393238261 | -0.023286781 | 11.07160993 | -2.769909054 | 0.033182632 | 0.594255995 | -3.549121999 |
| **A_52_P232648** | Slc38a1 | -0.125292303 | -0.236599388 | -0.013985218 | 8.992911002 | -2.769304731 | 0.033208808 | 0.594255995 | -3.54949703 |
| **A_55_P2019257** | Trpc4ap | -0.250635644 | -0.473347161 | -0.027924127 | 14.66665618 | -2.768658055 | 0.033236844 | 0.594255995 | -3.549898413 |
| **A_55_P2103952** | NA | 0.187601679 | 0.020898298 | 0.35430506 | 8.471345762 | 2.768608043 | 0.033239013 | 0.594255995 | -3.549929458 |
| **A_55_P2133637** | Etfa | -0.168316767 | -0.318084768 | -0.018548767 | 8.783865588 | -2.764887805 | 0.033400805 | 0.594255995 | -3.552239988 |
| **A_55_P2095177** | Usp5 | 0.185901317 | 0.0204632 | 0.351339434 | 13.00936637 | 2.764496543 | 0.03341787 | 0.594255995 | -3.552483128 |
| **A_55_P1990919** | Msrb1 | 0.353289572 | 0.038882694 | 0.667696451 | 13.24743458 | 2.764444932 | 0.033420122 | 0.594255995 | -3.552515202 |

| | | | | | | | | | |
|---|---|---|---|---|---|---|---|---|---|
| **A_51_P282179** | Mtor | -0.311323946 | -0.588412506 | -0.034235386 | 8.980708641 | -2.76415946 | 0.033432579 | 0.594255995 | -3.55269262 |
| **A_55_P2020831** | NA | 0.352442836 | 0.038664469 | 0.666221204 | 12.98461548 | 2.763343357 | 0.033468221 | 0.594255995 | -3.553199897 |
| **A_55_P1952324** | NA | 0.214208987 | 0.023476561 | 0.404941414 | 9.409864223 | 2.763009109 | 0.033482831 | 0.594255995 | -3.553407693 |
| **A_52_P274238** | Maea | -0.402167897 | -0.760382504 | -0.04395329 | 12.50595987 | -2.762061039 | 0.033524307 | 0.594255995 | -3.553997196 |
| **A_66_P136097** | Trim13 | -0.167379713 | -0.316468786 | -0.01829064 | 9.043789882 | -2.762015851 | 0.033526285 | 0.594255995 | -3.554025297 |
| **A_55_P1977934** | Itgam | 0.407784276 | 0.044539152 | 0.771029401 | 12.61805558 | 2.761848461 | 0.033533614 | 0.594255995 | -3.554129397 |
| **A_55_P2112549** | NA | -0.400713276 | -0.757660587 | -0.043765966 | 12.87906758 | -2.761841691 | 0.033533911 | 0.594255995 | -3.554133607 |
| **A_52_P84803** | NA | -0.280462815 | -0.530372479 | -0.030553152 | 15.26851832 | -2.760968185 | 0.033572185 | 0.594255995 | -3.554676916 |
| **A_55_P2033308** | Six2 | 0.155188957 | 0.016854788 | 0.293523126 | 8.353415355 | 2.759945539 | 0.033617055 | 0.594255995 | -3.555313153 |
| **A_51_P327418** | Rsrc2 | -0.359764193 | -0.680499268 | -0.039029117 | 10.39950622 | -2.759565088 | 0.033633764 | 0.594255995 | -3.555549896 |
| **A_52_P278497** | Eif4ebp3 | -0.204928824 | -0.387704176 | -0.022153472 | 10.26414229 | -2.758383087 | 0.033685733 | 0.594255995 | -3.556285576 |
| **A_55_P2131785** | Kiz | -0.162373693 | -0.307224674 | -0.017522711 | 10.35851763 | -2.757803948 | 0.033711228 | 0.594255995 | -3.556646121 |
| **A_51_P403636** | Smad7 | -0.233903415 | -0.442587053 | -0.025219776 | 11.45547138 | -2.757511445 | 0.033724113 | 0.594255995 | -3.556828241 |
| **A_55_P2064741** | Nmb | -0.244420276 | -0.462490615 | -0.026349938 | 11.5280987 | -2.757463652 | 0.033726218 | 0.594255995 | -3.556858 |
| **A_55_P2158458** | Iqcc | 0.208242706 | 0.022405486 | 0.394079925 | 9.145662204 | 2.756806359 | 0.033755192 | 0.594255995 | -3.557267307 |
| **A_51_P182131** | 5330417C22Rik | 0.299496986 | 0.032174244 | 0.566819728 | 8.33669103 | 2.756295094 | 0.033777748 | 0.594255995 | -3.557585732 |
| **A_55_P1959174** | Pmepa1 | -0.345647696 | -0.654188337 | -0.037107056 | 10.60904438 | -2.756071329 | 0.033787625 | 0.594255995 | -3.55772511 |
| **A_55_P2162988** | Tmem179 | -0.589311491 | -1.115492061 | -0.063130922 | 8.801996986 | -2.755365815 | 0.033818787 | 0.594255995 | -3.558164617 |
| **A_55_P1974593** | NA | 0.312827661 | 0.033497316 | 0.592158005 | 15.83657274 | 2.755219434 | 0.033825257 | 0.594255995 | -3.558255817 |
| **A_52_P372151** | Ctnnal1 | -0.178671245 | -0.338277831 | -0.01906466 | 10.79813376 | -2.754057607 | 0.033876651 | 0.594255995 | -3.558979802 |
| **A_55_P2050353** | Prpmp5 | 0.35438664 | 0.037723106 | 0.671050174 | 8.946357085 | 2.753267768 | 0.033911639 | 0.594255995 | -3.559472117 |
| **A_52_P489119** | Nol6 | 0.274727155 | 0.028990981 | 0.52046333 | 9.914991702 | 2.750436728 | 0.034037365 | 0.594255995 | -3.56123761 |
| **A_55_P2165234** | Slurp2 | 0.876224889 | 0.092454106 | 1.659995671 | 9.778307622 | 2.750398852 | 0.03403905 | 0.594255995 | -3.561261239 |
| **A_52_P426768** | Cited4 | -0.273888106 | -0.518905643 | -0.028870569 | 9.119523545 | -2.750078973 | 0.034053289 | 0.594255995 | -3.561460811 |
| **A_55_P2118684** | Idh2 | 0.27776932 | 0.029274524 | 0.526264117 | 14.78592343 | 2.750021823 | 0.034055833 | 0.594255995 | -3.561496468 |
| **A_51_P464308** | Gnb4 | -0.380164774 | -0.720475911 | -0.039853637 | 10.63330159 | -2.748304657 | 0.034132381 | 0.594255995 | -3.562568115 |
| **A_66_P128839** | Gm13262 | 0.64247613 | 0.066923369 | 1.21802889 | 8.909298696 | 2.746256 | 0.034223949 | 0.594255995 | -3.563847298 |
| **A_55_P2144086** | NA | -0.211076674 | -0.400209931 | -0.021943418 | 16.69048598 | -2.7456268 | 0.034252125 | 0.594255995 | -3.564240315 |
| **A_51_P140942** | Rp9 | -0.285048397 | -0.540488074 | -0.02960872 | 12.31431744 | -2.745360687 | 0.034264049 | 0.594255995 | -3.564406557 |
| **A_55_P2020532** | Selenov | 0.20932323 | 0.021732881 | 0.396913578 | 9.67011408 | 2.745213336 | 0.034270653 | 0.594255995 | -3.564498613 |
| **A_55_P1964138** | Pign | 0.30036082 | 0.031128488 | 0.569593152 | 10.09351085 | 2.744639008 | 0.034296409 | 0.594255995 | -3.564857454 |
| **A_55_P2119864** | Cd2bp2 | -0.203064352 | -0.385136393 | -0.020992312 | 9.655176465 | -2.743845128 | 0.034332044 | 0.594255995 | -3.565353565 |
| **A_55_P2157408** | NA | 0.35984245 | 0.03693037 | 0.682754529 | 8.282622368 | 2.741556916 | 0.03443498 | 0.594255995 | -3.566784112 |
| **A_55_P1996088** | NA | -0.260660507 | -0.494593423 | -0.026727591 | 12.97397477 | -2.741277892 | 0.034447555 | 0.594255995 | -3.566958614 |
| **A_55_P2159094** | P3h1 | 0.405842446 | 0.041554903 | 0.77012999 | 10.36106347 | 2.740831325 | 0.03446769 | 0.594255995 | -3.567237925 |
| **A_55_P2015022** | NA | 0.748068664 | 0.076574245 | 1.419563082 | 15.81874518 | 2.740742402 | 0.034471701 | 0.594255995 | -3.567293547 |
| **A_66_P110251** | Tmem91 | 0.384254232 | 0.039324694 | 0.72918377 | 9.245995443 | 2.740674351 | 0.034474771 | 0.594255995 | -3.567336114 |
| **A_66_P133413** | NA | -0.218117315 | -0.413951148 | -0.022283483 | 15.84366255 | -2.740132623 | 0.03449922 | 0.594255995 | -3.567675004 |
| **A_55_P2002449** | NA | 0.236673095 | 0.024144452 | 0.449201737 | 9.9993907 | 2.739684698 | 0.034519449 | 0.594255995 | -3.567955252 |
| **A_55_P1994603** | Itsn1 | -0.230061362 | -0.436717688 | -0.023405036 | 15.7288968 | -2.738824129 | 0.03455835 | 0.594255995 | -3.568493768 |
| **A_55_P2033922** | Tsga8 | 0.564410341 | 0.057377778 | 1.071442904 | 8.600485052 | 2.738597918 | 0.034568583 | 0.594255995 | -3.568635345 |
| **A_55_P1957754** | NA | 0.299310918 | 0.030427338 | 0.568194497 | 15.31525019 | 2.738592665 | 0.034568821 | 0.594255995 | -3.568638633 |
| **A_55_P2166884** | NA | 0.283917995 | 0.028862319 | 0.538973672 | 14.21618699 | 2.738590449 | 0.034568921 | 0.594255995 | -3.56864002 |
| **A_51_P466371** | Pitpna | -0.267023095 | -0.507022782 | -0.027023408 | 10.18498334 | -2.737204926 | 0.034631672 | 0.594255995 | -3.569507356 |
| **A_55_P2372902** | 5330438D12Rik | -0.194615769 | -0.369590378 | -0.01964116 | 8.77323302 | -2.736353229 | 0.034670306 | 0.594255995 | -3.570040681 |
| **A_55_P2149983** | Foxp4 | 0.242854285 | 0.02448052 | 0.46122805 | 11.80026547 | 2.735989787 | 0.034686807 | 0.594255995 | -3.570268302 |
| **A_55_P2038494** | Rexo2 | -0.198842327 | -0.377691999 | -0.019992656 | 9.508070847 | -2.735204832 | 0.034722473 | 0.594255995 | -3.570759991 |
| **A_55_P1982668** | Nadk | -0.145490114 | -0.276362827 | -0.014617401 | 12.19511139 | -2.734976141 | 0.034732871 | 0.594255995 | -3.57090326 |
| **A_55_P2013188** | Dnajb6 | -0.247566391 | -0.470280776 | -0.024852006 | 13.5680614 | -2.734718198 | 0.034744604 | 0.594255995 | -3.571064867 |
| **A_51_P177583** | Puf60 | -0.147574167 | -0.280337153 | -0.014811182 | 11.86880284 | -2.734654693 | 0.034747493 | 0.594255995 | -3.571104655 |

| | | | | | | | | | |
|---|---|---|---|---|---|---|---|---|---|
| **A_55_P2083997** | Zfp664 | -0.186203474 | -0.353744941 | -0.018662008 | 11.32073923 | -2.734227398 | 0.03476694 | 0.594255995 | -3.571372393 |
| **A_55_P2095000** | NA | 0.235519743 | 0.023595262 | 0.447444223 | 15.68089637 | 2.734106034 | 0.034772465 | 0.594255995 | -3.571448444 |
| **A_51_P187901** | Nop56 | -0.16216549 | -0.308129172 | -0.016201809 | 9.112644583 | -2.73327163 | 0.03481048 | 0.594255995 | -3.571971377 |
| **A_55_P2077671** | NA | -0.480337075 | -0.912718129 | -0.047956021 | 12.8235922 | -2.733056539 | 0.034820287 | 0.594255995 | -3.572106198 |
| **A_55_P1970389** | Srsf10 | 0.213182016 | 0.021200316 | 0.405163715 | 9.958608231 | 2.731869164 | 0.034874476 | 0.594255995 | -3.572850591 |
| **A_55_P1970806** | Mif-ps6 | 0.205672157 | 0.020429559 | 0.390914756 | 13.75573739 | 2.731516341 | 0.034890596 | 0.594255995 | -3.573071831 |
| **A_55_P1979432** | Prdx2 | 0.29090582 | 0.028870248 | 0.552941392 | 13.42896077 | 2.73124916 | 0.034902808 | 0.594255995 | -3.573239382 |
| **A_51_P389714** | Phlpp2 | -0.286554536 | -0.544690444 | -0.028418628 | 11.82647049 | -2.731039802 | 0.03491238 | 0.594255995 | -3.57337068 |
| **A_52_P168271** | Spg20 | -0.140614307 | -0.267293448 | -0.013935166 | 8.362652904 | -2.730823086 | 0.034922292 | 0.594255995 | -3.573506601 |
| **A_55_P2021721** | NA | 1.017771038 | 0.10034629 | 1.935195786 | 10.57013598 | 2.729284445 | 0.034992751 | 0.594255995 | -3.574471842 |
| **A_55_P2102250** | Dzip1 | 0.27485244 | 0.027070489 | 0.522634391 | 8.943474337 | 2.72897207 | 0.035007075 | 0.594255995 | -3.574667854 |
| **A_55_P2098911** | Lrrcc1 | -0.267187294 | -0.508105987 | -0.026268601 | 9.536877923 | -2.728440362 | 0.035031469 | 0.594255995 | -3.575001534 |
| **A_66_P102733** | Ccdc28b | 0.210361806 | 0.020565853 | 0.40015776 | 11.4933578 | 2.726773752 | 0.035108052 | 0.594255995 | -3.576047746 |
| **A_51_P129100** | Sec63 | -0.153343147 | -0.291703889 | -0.014982405 | 9.57318192 | -2.726595118 | 0.035116271 | 0.594255995 | -3.60858741 |
| **A_66_P101473** | Fbxl3 | -0.213091156 | -0.40548371 | -0.020698601 | 9.459988568 | -2.72487338 | 0.035195595 | 0.594255995 | -3.577241277 |
| **A_51_P469480** | Yes1 | -0.132586792 | -0.252375648 | -0.012797936 | 8.309411503 | -2.723033644 | 0.035280569 | 0.594255995 | -3.578397309 |
| **A_55_P2107855** | Hnrnpk | -0.148177301 | -0.282055661 | -0.014298942 | 9.829786675 | -2.722954983 | 0.035284207 | 0.594255995 | -3.57844675 |
| **A_55_P2184508** | NA | -0.165352906 | -0.314763518 | -0.015942294 | 14.6397251 | -2.722698482 | 0.035296073 | 0.594255995 | -3.578607976 |
| **A_55_P1978972** | Rp2 | -0.176824945 | -0.336665675 | -0.016984215 | 9.063580949 | -2.721605912 | 0.035346665 | 0.594255995 | -3.579294846 |
| **A_55_P2101910** | Rpl13a | 0.337181807 | 0.032317302 | 0.642046312 | 10.72789604 | 2.720986791 | 0.035375368 | 0.594255995 | -3.579684161 |
| **A_66_P100220** | NA | 0.525311277 | 0.050304033 | 1.000318521 | 8.684977475 | 2.720731308 | 0.03538722 | 0.594255995 | -3.579844832 |
| **A_55_P2253348** | Galnt7 | 0.179741212 | 0.017179028 | 0.342303397 | 10.71147005 | 2.720177874 | 0.035412908 | 0.594255995 | -3.580192921 |
| **A_51_P384618** | Dhdds | -0.180793046 | -0.344401724 | -0.017184367 | 9.966493762 | -2.718595214 | 0.03548648 | 0.594255995 | -3.581188639 |
| **A_51_P379710** | Rpn1 | -0.272310748 | -0.518744869 | -0.025876627 | 15.27162627 | -2.718523595 | 0.035489813 | 0.594255995 | -3.581233708 |
| **A_55_P2169804** | Nanos3 | 0.757601211 | 0.071961218 | 1.443241203 | 10.50286864 | 2.718402005 | 0.035495472 | 0.594255995 | -3.581310224 |
| **A_55_P2120388** | Skint7 | -0.229465602 | -0.437151593 | -0.021779611 | 15.8219597 | -2.718188378 | 0.035505418 | 0.594255995 | -3.581444665 |
| **A_55_P2034003** | Gtf3c1 | 0.142117999 | 0.01346233 | 0.270773668 | 8.9119543 | 2.71762369 | 0.035531723 | 0.594255995 | -3.581800076 |
| **A_55_P2070686** | Wdfy1 | 0.750276814 | 0.070995963 | 1.429557665 | 10.5709474 | 2.717323326 | 0.035545723 | 0.594255995 | -3.581989145 |
| **A_55_P2175260** | Tet3 | 0.35098244 | 0.033153543 | 0.668811337 | 8.308123723 | 2.716821908 | 0.035569109 | 0.594255995 | -3.582304805 |
| **A_52_P78770** | Chpt1 | -0.16010962 | -0.305128275 | -0.015090964 | 8.520738935 | -2.716206065 | 0.035597853 | 0.594255995 | -3.582692557 |
| **A_52_P181759** | Eif5a2 | 0.400483985 | 0.037689969 | 0.763278002 | 9.232518853 | 2.715777599 | 0.035617866 | 0.594255995 | -3.582962369 |
| **A_55_P2133751** | Txnl4a | -0.341784486 | -0.651446383 | -0.032122589 | 13.38277353 | -2.715399529 | 0.035635535 | 0.594255995 | -3.583200472 |
| **A_55_P2068096** | Rap1gap | 0.149584894 | 0.014019236 | 0.285150551 | 10.64438974 | 2.714608572 | 0.035672531 | 0.594255995 | -3.583698683 |
| **A_55_P1968993** | NA | 0.206377387 | 0.019241488 | 0.393513285 | 10.49110873 | 2.713152252 | 0.035740757 | 0.594255995 | -3.584616273 |
| **A_66_P124925** | Pdcd5 | 0.165416261 | 0.015374622 | 0.3154579 | 12.6480803 | 2.7122865 | 0.035781382 | 0.594255995 | -3.58516193 |
| **A_66_P110104** | Myo18b | 0.710549743 | 0.0660245 | 1.355074987 | 9.737653215 | 2.712212534 | 0.035784855 | 0.594255995 | -3.585208555 |
| **A_51_P248304** | Vps26b | -0.142365864 | -0.271521007 | -0.013210721 | 9.300889058 | -2.711835401 | 0.035802569 | 0.594255995 | -3.585446295 |
| **A_52_P142531** | Phc3 | -0.151453643 | -0.288881353 | -0.014025933 | 8.896342036 | -2.711281297 | 0.035828612 | 0.594255995 | -3.585795638 |
| **A_55_P2104289** | NA | -0.131231237 | -0.250340731 | -0.012121744 | 8.245599712 | -2.710566142 | 0.035862256 | 0.594255995 | -3.586246596 |
| **A_55_P2083411** | Cyp4f18 | 0.372260486 | 0.034305721 | 0.71021525 | 9.892737747 | 2.709926608 | 0.03589237 | 0.594255995 | -3.586649943 |
| **A_55_P2029366** | Abhd6 | -0.202570413 | -0.386499614 | -0.018641212 | 10.76857887 | -2.709533301 | 0.035910904 | 0.594255995 | -3.586898031 |
| **A_51_P429276** | Tmod3 | -0.163881628 | -0.312695724 | -0.015067533 | 10.63367953 | -2.709289201 | 0.035922412 | 0.594255995 | -3.587052016 |
| **A_52_P288873** | Chmp4b | -0.136201597 | -0.259888897 | -0.012514297 | 13.58367801 | -2.709107597 | 0.035930976 | 0.594255995 | -3.587166584 |
| **A_52_P637730** | Ubl4a | 0.123447906 | 0.011279726 | 0.235616085 | 12.52482411 | 2.707591975 | 0.036002535 | 0.594255995 | -3.588122954 |
| **A_55_P2247608** | 4931428A05Rik | 0.564319969 | 0.051496745 | 1.077143194 | 9.378109408 | 2.707240862 | 0.036019134 | 0.594255995 | -3.588344565 |
| **A_55_P2132147** | NA | 0.323325956 | 0.029502344 | 0.617149568 | 15.79396262 | 2.707216822 | 0.036020271 | 0.594255995 | -3.588359738 |
| **A_55_P1965363** | Serpinb6e | 0.332647113 | 0.030007971 | 0.635286254 | 8.548804462 | 2.704131607 | 0.036166488 | 0.594255995 | -3.590307927 |
| **A_55_P2243367** | NA | -0.130702148 | -0.249627742 | -0.011776553 | 8.931432722 | -2.703812405 | 0.036181652 | 0.594255995 | -3.590509582 |
| **A_51_P253732** | Il17rd | -0.305395541 | -0.583274688 | -0.027516394 | 9.208421799 | -2.703808308 | 0.036181847 | 0.594255995 | -3.59051217 |
| **A_55_P1981959** | Trafd1 | 0.174272568 | 0.015688098 | 0.332857039 | 12.21183204 | 2.703569498 | 0.036193196 | 0.594255995 | -3.590663049 |

| | | | | | | | | | |
|---|---|---|---|---|---|---|---|---|---|
| **A_55_P2079020** | Snhg7os | 1.017874651 | 0.091376833 | 1.94437247 | 10.58928814 | 2.702832056 | 0.036228268 | 0.594255995 | -3.591129022 |
| **A_55_P2143324** | Dyrk1b | 0.570495144 | 0.051136235 | 1.089854052 | 10.32843772 | 2.702424248 | 0.036247678 | 0.594255995 | -3.591386745 |
| **A_55_P2007560** | NA | 0.552449697 | 0.049432268 | 1.055467125 | 9.082996621 | 2.701959717 | 0.036269801 | 0.594255995 | -3.59168035 |
| **A_55_P2128794** | Clta | -0.120124924 | -0.229516217 | -0.01073363 | 13.27523814 | -2.701590562 | 0.036287393 | 0.594255995 | -3.5919137 |
| **A_55_P2326091** | 9330175M20Rik | 0.314835776 | 0.028078414 | 0.601593138 | 10.38174505 | 2.701087565 | 0.036311377 | 0.594255995 | -3.592231689 |
| **A_55_P2134979** | NA | -0.191417556 | -0.365852897 | -0.016982215 | 12.54994319 | -2.699705868 | 0.036377349 | 0.594255995 | -3.593105404 |
| **A_51_P487690** | Ifi44 | -0.217146539 | -0.415061245 | -0.019231833 | 9.159458813 | -2.699255594 | 0.036398875 | 0.594255995 | -3.593390204 |
| **A_55_P2041310** | NA | 0.231410287 | 0.020452603 | 0.44236797 | 8.943434961 | 2.698711592 | 0.036424901 | 0.594255995 | -3.593734332 |
| **A_55_P1976007** | Psme4 | -0.15607714 | -0.298365165 | -0.013789116 | 9.086529268 | -2.698609917 | 0.036429768 | 0.594255995 | -3.593798656 |
| **A_55_P1977741** | NA | 0.849653517 | 0.074744475 | 1.624562559 | 10.06253888 | 2.697492799 | 0.036483283 | 0.594255995 | -3.594505506 |
| **A_51_P506674** | Rpp38 | 0.194276834 | 0.017062749 | 0.371490919 | 9.881571797 | 2.697068295 | 0.036503641 | 0.594255995 | -3.594774163 |
| **A_55_P2087322** | Rapsn | 1.006935744 | 0.08842199 | 1.925449498 | 14.12775886 | 2.697026767 | 0.036505633 | 0.594255995 | -3.594800447 |
| **A_52_P642207** | Mkrn1 | -0.171364605 | -0.327867902 | -0.014861307 | 11.01489787 | -2.69380896 | 0.036660354 | 0.594255995 | -3.596837919 |
| **A_55_P2000968** | Ppp2r3c | 0.270193642 | 0.023387011 | 0.517000273 | 10.23250628 | 2.69331695 | 0.036684073 | 0.594255995 | -3.597149607 |
| **A_55_P1988882** | 9-Sep | -0.211139611 | -0.404140866 | -0.018138357 | 12.32912208 | -2.6914031 | 0.036776493 | 0.594255995 | -3.598362414 |
| **A_51_P149960** | Letm2 | -0.161410888 | -0.308958984 | -0.013862793 | 8.382746631 | -2.691338861 | 0.0367796 | 0.594255995 | -3.598403133 |
| **A_55_P2087061** | Tmem240 | 0.439999269 | 0.037749584 | 0.842248955 | 10.39928102 | 2.691072545 | 0.036792481 | 0.594255995 | -3.59857195 |
| **A_51_P322285** | Tecpr2 | -0.241087274 | -0.461513796 | -0.020660753 | 8.741569573 | -2.690788747 | 0.036806214 | 0.594255995 | -3.598751861 |
| **A_55_P2067505** | Slc16a3 | 0.314915194 | 0.026952299 | 0.602878089 | 13.43320226 | 2.690458186 | 0.036822216 | 0.594255995 | -3.598961434 |
| **A_51_P429983** | Supt5 | -0.323732636 | -0.619766928 | -0.027698345 | 13.08911027 | -2.690380001 | 0.036826002 | 0.594255995 | -3.599011005 |
| **A_51_P138044** | Foxo1 | -0.241155981 | -0.461703255 | -0.020608707 | 10.27488263 | -2.690081928 | 0.036840439 | 0.594255995 | -3.599200001 |
| **A_66_P111426** | Lce1a2 | 0.354439355 | 0.030191095 | 0.678687615 | 9.616939442 | 2.689264047 | 0.036880085 | 0.594255995 | -3.599718661 |
| **A_55_P1976934** | Unc13d | 0.431976663 | 0.03678602 | 0.827167305 | 9.51986582 | 2.689198099 | 0.036883284 | 0.594255995 | -3.599760487 |
| **A_55_P2186332** | NA | 0.136138056 | 0.011571036 | 0.260705075 | 7.962433063 | 2.688720352 | 0.036906465 | 0.594255995 | -3.600063509 |
| **A_51_P148314** | Fam117a | -0.270490034 | -0.518046599 | -0.02293347 | 10.44684507 | -2.68810349 | 0.03693642 | 0.594255995 | -3.600454823 |
| **A_55_P2043454** | NA | -0.152160195 | -0.291420743 | -0.012899647 | 8.537781003 | -2.688079558 | 0.036937583 | 0.594255995 | -3.600470006 |
| **A_51_P295449** | Mfsd5 | -0.145603292 | -0.278872788 | -0.012333797 | 11.13122137 | -2.68787832 | 0.036947361 | 0.594255995 | -3.600597679 |
| **A_52_P412046** | NA | -0.442435462 | -0.847418083 | -0.03745284 | 14.50267658 | -2.687711841 | 0.036955452 | 0.594255995 | -3.600703304 |
| **A_55_P1996862** | Psme2 | -0.223108847 | -0.427344148 | -0.018873547 | 14.95259054 | -2.687541299 | 0.036963743 | 0.594255995 | -3.600811512 |
| **A_66_P119034** | Pla2g7 | -0.736868682 | -1.411664793 | -0.062072571 | 14.84292866 | -2.686499047 | 0.037014454 | 0.594255995 | -3.601472921 |
| **A_55_P2055232** | Sdf4 | -0.153837464 | -0.294734107 | -0.012940822 | 10.51040503 | -2.686152302 | 0.037031341 | 0.594255995 | -3.601693005 |
| **A_55_P1993820** | Kdm1a | -0.21329797 | -0.408693986 | -0.017901953 | 12.3909164 | -2.68559289 | 0.037058604 | 0.594255995 | -3.602048113 |
| **A_52_P344978** | NA | -0.135959896 | -0.260559032 | -0.011360759 | 8.698387208 | -2.684509553 | 0.03711146 | 0.594255995 | -3.602735953 |
| **A_55_P2049158** | NA | -0.309222788 | -0.592624841 | -0.025820735 | 11.01230789 | -2.684340864 | 0.037119698 | 0.594255995 | -3.602843076 |
| **A_55_P1961188** | Gm6981 | -0.342401921 | -0.656368861 | -0.028434981 | 14.23644204 | -2.683004713 | 0.037185017 | 0.594255995 | -3.603691742 |
| **A_55_P2417212** | NA | 0.251496916 | 0.020870342 | 0.48212349 | 9.463597213 | 2.682825834 | 0.037193771 | 0.594255995 | -3.603805381 |
| **A_55_P1968928** | Def6 | 0.647244308 | 0.053569861 | 1.240918755 | 10.54089831 | 2.682186959 | 0.037225054 | 0.594255995 | -3.604211292 |
| **A_55_P1953169** | Saa3 | -0.553352633 | -1.061111674 | -0.045593592 | 8.644692059 | -2.681102876 | 0.037278202 | 0.594255995 | -3.604900224 |
| **A_55_P1960924** | Egln2 | 0.169507974 | 0.013899586 | 0.325116361 | 13.14889243 | 2.679947537 | 0.037334932 | 0.594255995 | -3.605634655 |
| **A_52_P350664** | Pygb | -0.136418802 | -0.261670407 | -0.011167198 | 8.173319312 | -2.679539097 | 0.03735501 | 0.594255995 | -3.605894347 |
| **A_55_P1967776** | Slc4a4 | -0.180500404 | -0.346239781 | -0.014761026 | 8.733616964 | -2.67930181 | 0.03736668 | 0.594255995 | -3.606045231 |
| **A_51_P257258** | Cpox | 0.2025763 | 0.01656172 | 0.388590881 | 10.76535383 | 2.679235017 | 0.037369966 | 0.594255995 | -3.606087705 |
| **A_55_P2052613** | NA | 0.398166877 | 0.032331309 | 0.764002445 | 10.75163293 | 2.677616433 | 0.037449679 | 0.594255995 | -3.607117182 |
| **A_55_P2047626** | Cutal | 0.328773112 | 0.026647669 | 0.630898554 | 9.735802033 | 2.677183594 | 0.037471026 | 0.594255995 | -3.607392557 |
| **A_55_P2155312** | Dock3 | 0.64527403 | 0.052058849 | 1.238489212 | 10.87514512 | 2.67609233 | 0.037524905 | 0.594255995 | -3.608086966 |
| **A_55_P2036894** | Med29 | 0.384181767 | 0.030819015 | 0.737544518 | 11.33990857 | 2.674762001 | 0.0375907 | 0.594255995 | -3.608933769 |
| **A_55_P2034893** | Socs7 | -0.26720766 | -0.513128304 | -0.021287016 | 11.81330074 | -2.673148462 | 0.037670668 | 0.594255995 | -3.609961242 |
| **A_55_P2091305** | NA | 0.278491767 | 0.022086732 | 0.534896802 | 8.985642778 | 2.672113954 | 0.037722034 | 0.594255995 | -3.610620227 |
| **A_52_P133436** | Plekhm1 | -0.246318053 | -0.473139076 | -0.019497029 | 11.19314714 | -2.671665606 | 0.037744319 | 0.594255995 | -3.610905882 |
| **A_55_P2053943** | Olfr1368 | 0.523419534 | 0.0413705 | 1.005468569 | 8.808331995 | 2.671332035 | 0.037760909 | 0.594255995 | -3.611118431 |

| | | | | | | | | | |
|---|---|---|---|---|---|---|---|---|---|
| **A_55_P2047648** | Coq2 | 0.225661614 | 0.017834826 | 0.433488402 | 12.82128151 | 2.671316382 | 0.037761687 | 0.594255995 | -3.611128405 |
| **A_51_P518893** | Ccin | 0.336068404 | 0.026555332 | 0.645581477 | 9.029987729 | 2.671270395 | 0.037763975 | 0.594255995 | -3.611157709 |
| **A_55_P2143306** | Rest | -0.275059509 | -0.528409277 | -0.021709741 | 10.31084908 | -2.671008778 | 0.037776992 | 0.594255995 | -3.611324425 |
| **A_66_P110970** | Tpd52l2 | -0.13136954 | -0.252373021 | -0.010366059 | 8.508881961 | -2.670951348 | 0.037779851 | 0.594255995 | -3.611361024 |
| **A_55_P2003927** | NA | 0.451659109 | 0.035602865 | 0.867715352 | 11.48915365 | 2.670717133 | 0.037791509 | 0.594255995 | -3.61151029 |
| **A_52_P317040** | Edem2 | -0.209900304 | -0.403276092 | -0.016524516 | 8.711906048 | -2.670423461 | 0.037806134 | 0.594255995 | -3.611697461 |
| **A_55_P2159740** | Mfsd2b | 0.34148915 | 0.026873697 | 0.656104604 | 16.58736118 | 2.670336685 | 0.037810456 | 0.594255995 | -3.611752771 |
| **A_55_P1970536** | NA | 0.485956092 | 0.038219016 | 0.933693168 | 8.636327405 | 2.670195966 | 0.037817467 | 0.594255995 | -3.611842464 |
| **A_51_P464710** | Adap2 | -0.181252169 | -0.348258408 | -0.01424593 | 8.932920303 | -2.67005175 | 0.037824653 | 0.594255995 | -3.611934391 |
| **A_65_P20699** | Tlk2 | -0.208756262 | -0.401202279 | -0.016310246 | 9.587269108 | -2.66869996 | 0.037892082 | 0.594255995 | -3.61279622 |
| **A_55_P2109875** | Rpl23a | -0.191243902 | -0.367566246 | -0.014921558 | 14.07036022 | -2.668390645 | 0.037907529 | 0.594255995 | -3.612993465 |
| **A_55_P1993789** | Hmgn2 | -0.34755155 | -0.668027079 | -0.02707602 | 14.20650197 | -2.668047264 | 0.037924686 | 0.594255995 | -3.613212453 |
| **A_55_P2114785** | NA | -0.185489096 | -0.356532574 | -0.014445617 | 9.115630424 | -2.667970523 | 0.037928521 | 0.594255995 | -3.613261396 |
| **A_51_P512119** | AF067063 | 0.406132168 | 0.031626584 | 0.780637752 | 9.192120957 | 2.667953493 | 0.037929372 | 0.594255995 | -3.613272257 |
| **A_51_P185108** | Dpy30 | -0.23300551 | -0.447874119 | -0.018136902 | 9.64854574 | -2.667855996 | 0.037934246 | 0.594255995 | -3.61333444 |
| **A_55_P2174905** | Cuta | 0.229085026 | 0.017756339 | 0.440413712 | 12.27534197 | 2.666904175 | 0.037981858 | 0.594255995 | -3.613941586 |
| **A_55_P2173358** | NA | 0.506719571 | 0.039242474 | 0.974196667 | 8.701259891 | 2.666714383 | 0.037991359 | 0.594255995 | -3.614062669 |
| **A_55_P2007485** | Prss27 | 0.483476295 | 0.037433197 | 0.929519392 | 8.760817668 | 2.666659255 | 0.03799412 | 0.594255995 | -3.61409784 |
| **A_51_P520384** | Atp1b3 | -0.154847646 | -0.297736748 | -0.011958544 | 12.18172493 | -2.666089129 | 0.038022679 | 0.594255995 | -3.614461607 |
| **A_55_P1964154** | Asah1 | -0.201014715 | -0.38650846 | -0.015520969 | 9.083171649 | -2.666046637 | 0.038024809 | 0.594255995 | -3.614488721 |
| **A_52_P552092** | Adat1 | 0.292875771 | 0.022600108 | 0.563151434 | 8.672494327 | 2.665911126 | 0.038031601 | 0.594255995 | -3.614575193 |
| **A_55_P2011712** | NA | 0.389118664 | 0.029997362 | 0.748239966 | 9.427528733 | 2.665692549 | 0.038042559 | 0.594255995 | -3.614714676 |
| **A_55_P1957319** | Krr1 | -0.144388471 | -0.27765917 | -0.011117773 | 9.767868898 | -2.665428325 | 0.03805581 | 0.594255995 | -3.614883298 |
| **A_55_P1970270** | B3gnt2 | 0.191811715 | 0.014704152 | 0.368919278 | 9.548491323 | 2.664447599 | 0.038105037 | 0.594255995 | -3.615509281 |
| **A_55_P2080995** | Tgif2lx1 | 0.489660942 | 0.037338123 | 0.94198376 | 8.464459474 | 2.663275718 | 0.038163949 | 0.594255995 | -3.616257486 |
| **A_55_P1987222** | NA | 0.132167056 | 0.010040643 | 0.254293469 | 8.57731972 | 2.662458072 | 0.038205111 | 0.594255995 | -3.61677966 |
| **A_52_P350111** | Cyth1 | -0.144560339 | -0.278152073 | -0.010968605 | 10.88637058 | -2.662188079 | 0.038218713 | 0.594255995 | -3.616952109 |
| **A_66_P138001** | Atp2b1 | -0.156559588 | -0.301294948 | -0.011824227 | 10.46459026 | -2.661179592 | 0.038269567 | 0.594255995 | -3.617596358 |
| **A_55_P2029533** | Zfp57 | 0.826959027 | 0.062412927 | 1.591505127 | 8.716109184 | 2.661028207 | 0.038277207 | 0.594255995 | -3.617693081 |
| **A_55_P2096367** | NA | -0.177162829 | -0.340975953 | -0.013349706 | 13.48143564 | -2.660682656 | 0.038294652 | 0.594255995 | -3.617913876 |
| **A_55_P1967002** | NA | 0.326003932 | 0.024533143 | 0.627474721 | 8.933146387 | 2.660398926 | 0.038308983 | 0.594255995 | -3.618095184 |
| **A_55_P2040362** | Bola2 | 0.257788624 | 0.019378098 | 0.496199149 | 13.18906899 | 2.660158328 | 0.038321139 | 0.594255995 | -3.618248941 |
| **A_51_P229633** | Rprd1b | -0.236732298 | -0.455752943 | -0.017711653 | 11.8539706 | -2.65914257 | 0.038372507 | 0.594255995 | -3.618898177 |
| **A_51_P269320** | Adsl | -0.119964138 | -0.230983379 | -0.008944898 | 8.400243762 | -2.658412371 | 0.03840948 | 0.594255995 | -3.619365 |
| **A_55_P1966209** | Fez2 | -0.156123926 | -0.300615318 | -0.011632535 | 11.8499705 | -2.658255097 | 0.038417448 | 0.594255995 | -3.619465558 |
| **A_51_P156800** | Mob2 | -0.149156442 | -0.287337493 | -0.010975392 | 10.43298325 | -2.655600183 | 0.038552226 | 0.594255995 | -3.621163686 |
| **A_55_P2027012** | Serpinb8 | -0.213463055 | -0.411266505 | -0.015659605 | 12.50519911 | -2.654960238 | 0.038584789 | 0.594255995 | -3.62157318 |
| **A_55_P1985015** | Pard3 | -0.228280874 | -0.439828634 | -0.016733113 | 10.33528973 | -2.654790513 | 0.038593431 | 0.594255995 | -3.621681797 |
| **A_55_P2131193** | NA | 0.23996512 | 0.017545944 | 0.462384296 | 15.20627001 | 2.654269733 | 0.038619958 | 0.594255995 | -3.622015103 |
| **A_55_P2174198** | H4c9 | 0.406650688 | 0.029711916 | 0.78358946 | 11.20624788 | 2.654115773 | 0.038627804 | 0.594255995 | -3.622113649 |
| **A_55_P1990319** | Ergic3 | 0.232702519 | 0.017001058 | 0.44840398 | 12.78032574 | 2.654099263 | 0.038628646 | 0.594255995 | -3.622124216 |
| **A_55_P2163018** | NA | 0.749628443 | 0.05448428 | 1.444772607 | 11.03205237 | 2.653018869 | 0.038683754 | 0.594255995 | -3.622815858 |
| **A_51_P209527** | Bcl10 | -0.123457998 | -0.237958358 | -0.008957639 | 10.17646541 | -2.652659731 | 0.038702091 | 0.594255995 | -3.623045812 |
| **A_51_P115655** | Iqcd | 0.654645881 | 0.047153308 | 1.262138453 | 10.52658245 | 2.651152012 | 0.038779176 | 0.594255995 | -3.624011429 |
| **A_55_P2311780** | 4930554H23Rik | 0.437194789 | 0.031410783 | 0.842978794 | 8.622549973 | 2.650630647 | 0.03880587 | 0.594255995 | -3.624345425 |
| **A_55_P1989858** | Thrap3 | 0.225129574 | 0.016166704 | 0.434092444 | 11.32973989 | 2.650529165 | 0.038811068 | 0.594255995 | -3.624410441 |
| **A_55_P1954835** | NA | -0.605997482 | -1.168483247 | -0.043511717 | 12.70097179 | -2.650503893 | 0.038812363 | 0.594255995 | -3.624426632 |
| **A_55_P2393483** | 6430584L05Rik | 0.837608363 | 0.060099363 | 1.615117363 | 9.489601764 | 2.650359235 | 0.038819774 | 0.594255995 | -3.624519314 |
| **A_51_P203955** | Gbp2 | -0.443987142 | -0.856165083 | -0.031809201 | 9.771291929 | -2.650054528 | 0.038835391 | 0.594255995 | -3.624714549 |
| **A_51_P460710** | Tdrd3 | -0.277188599 | -0.534643376 | -0.019733823 | 9.498477405 | -2.648765879 | 0.038901508 | 0.594255995 | -3.625540395 |

| | | | | | | | | | |
|---|---|---|---|---|---|---|---|---|---|
| **A_55_P2003638** | Stxbp6 | 0.285174272 | 0.020046546 | 0.550301999 | 9.053130818 | 2.646210315 | 0.039032983 | 0.594255995 | -3.627178973 |
| **A_52_P476904** | Chd6 | -0.145347229 | -0.2805031 | -0.010191359 | 8.508179041 | -2.645702488 | 0.039059166 | 0.594255995 | -3.62750471 |
| **A_55_P1962094** | NA | 0.192940742 | 0.013462732 | 0.372418751 | 8.379191326 | 2.644733171 | 0.039109194 | 0.594255995 | -3.628126581 |
| **A_51_P310398** | Tk2 | -0.181825098 | -0.350987666 | -0.01266253 | 8.84690853 | -2.644348701 | 0.039129057 | 0.594255995 | -3.628373283 |
| **A_55_P2034110** | Tgm2 | -0.35177542 | -0.679084475 | -0.024466365 | 13.89122254 | -2.644092429 | 0.039142302 | 0.594255995 | -3.628537738 |
| **A_66_P116326** | NA | -0.182330854 | -0.352048079 | -0.01261363 | 9.633908801 | -2.643038013 | 0.03919685 | 0.594255995 | -3.629214492 |
| **A_51_P222467** | Abcg1 | -0.507238193 | -0.97941793 | -0.035058457 | 12.79478245 | -2.642857605 | 0.039206191 | 0.594255995 | -3.629330301 |
| **A_55_P2156718** | NA | -0.245840446 | -0.474815352 | -0.01686554 | 13.33565413 | -2.641402642 | 0.039281613 | 0.594255995 | -3.630264482 |
| **A_55_P2132844** | Rps16 | -0.178534406 | -0.344830672 | -0.012238141 | 15.55322531 | -2.641244404 | 0.039289825 | 0.594255995 | -3.630366103 |
| **A_55_P2102135** | Rdh12 | 1.168192397 | 0.079850976 | 2.256533818 | 11.56381375 | 2.640695833 | 0.039318308 | 0.594255995 | -3.630718426 |
| **A_52_P431872** | Ptcd3 | -0.131836983 | -0.254672341 | -0.009001624 | 11.55484271 | -2.640480837 | 0.039329477 | 0.594255995 | -3.630856522 |
| **A_66_P126686** | NA | -0.413609604 | -0.799139812 | -0.028079396 | 13.80191483 | -2.639376616 | 0.039386895 | 0.594255995 | -3.631565906 |
| **A_55_P2117731** | Pak4 | -0.273692668 | -0.528932867 | -0.01845247 | 15.09299617 | -2.638051398 | 0.039455924 | 0.594255995 | -3.632417529 |
| **A_55_P1953693** | Pex2 | -0.147695362 | -0.28546625 | -0.009924473 | 10.57707724 | -2.637415535 | 0.039489091 | 0.594255995 | -3.632826256 |
| **A_51_P421876** | Irf7 | -0.131602262 | -0.254366931 | -0.008837592 | 8.518922761 | -2.637297467 | 0.039495253 | 0.594255995 | -3.632902156 |
| **A_55_P2083213** | Purb | -0.166755968 | -0.322321669 | -0.011190267 | 8.654452197 | -2.637161324 | 0.039502359 | 0.594255995 | -3.632989679 |
| **A_55_P2152114** | Zfyve19 | -0.155435074 | -0.300575808 | -0.010294341 | 9.380538676 | -2.634686076 | 0.0396318 | 0.594255995 | -3.634581482 |
| **A_55_P2032187** | NA | -0.147118997 | -0.284499249 | -0.009738745 | 8.608949634 | -2.634593454 | 0.039636652 | 0.594255995 | -3.634641066 |
| **A_55_P1969874** | NA | 0.441534593 | 0.029207657 | 0.853861529 | 16.92096017 | 2.634463535 | 0.039643459 | 0.594255995 | -3.634724645 |
| **A_55_P2149293** | Agfg2 | 0.164584148 | 0.010886296 | 0.318282 | 8.09533695 | 2.634446405 | 0.039644357 | 0.594255995 | -3.634735665 |
| **A_55_P1962641** | Rnf26 | 0.204692396 | 0.013499431 | 0.395885361 | 12.74058288 | 2.633898072 | 0.039673103 | 0.594255995 | -3.63508845 |
| **A_55_P1978740** | Rpgrip1 | 0.152607148 | 0.010053998 | 0.295160297 | 11.06386606 | 2.633705565 | 0.0396832 | 0.594255995 | -3.635212316 |
| **A_52_P656545** | Ogt | -0.292603122 | -0.566029349 | -0.019176894 | 8.943885869 | -2.632739856 | 0.039733894 | 0.594255995 | -3.635833782 |
| **A_55_P2112589** | Uxt | 0.213705232 | 0.013934644 | 0.413475821 | 10.17732838 | 2.631799312 | 0.039783333 | 0.594255995 | -3.6364392 |
| **A_55_P2104889** | Gm11515 | 0.138172968 | 0.008957144 | 0.267388792 | 8.090916788 | 2.630731637 | 0.039839535 | 0.594255995 | -3.637126628 |
| **A_66_P104713** | Lrp5 | 0.441720807 | 0.028606729 | 0.854834885 | 10.40719251 | 2.630552808 | 0.039848956 | 0.594255995 | -3.637241786 |
| **A_55_P2164221** | Tpt1 | -0.163340345 | -0.316127381 | -0.010553308 | 17.30487238 | -2.630123392 | 0.03987159 | 0.594255995 | -3.637518333 |
| **A_55_P2053491** | Pdia6 | -0.257928226 | -0.499265552 | -0.0165909 | 17.05144854 | -2.629320561 | 0.039913943 | 0.594255995 | -3.638035442 |
| **A_51_P384946** | Ndufa2 | 0.127775769 | 0.008190511 | 0.247361027 | 14.2128 | 2.628693934 | 0.039947034 | 0.594255995 | -3.638439132 |
| **A_55_P2008567** | Rps6 | 0.231589346 | 0.014815529 | 0.448363163 | 10.56104683 | 2.628336186 | 0.039965939 | 0.594255995 | -3.638669631 |
| **A_51_P458866** | F11r | -0.167886231 | -0.325077758 | -0.010694704 | 12.94749423 | -2.627574923 | 0.040006199 | 0.594255995 | -3.639160188 |
| **A_55_P1986371** | NA | -0.327129747 | -0.633429797 | -0.020829697 | 14.78827363 | -2.627496421 | 0.040010353 | 0.594255995 | -3.639210779 |
| **A_55_P1968969** | NA | 0.222736281 | 0.014177028 | 0.431295534 | 15.4666107 | 2.627427033 | 0.040014025 | 0.594255995 | -3.639255498 |
| **A_55_P1995774** | NA | -0.170980335 | -0.331124051 | -0.010836618 | 10.03906432 | -2.626669453 | 0.040054142 | 0.594255995 | -3.639743794 |
| **A_55_P2059606** | Stat2 | -0.430123282 | -0.833531896 | -0.026714668 | 11.45949599 | -2.623112678 | 0.040243062 | 0.594255995 | -3.642037561 |
| **A_51_P501922** | Plxna1 | 0.435620565 | 0.027043588 | 0.844197541 | 9.773124014 | 2.623032348 | 0.04024734 | 0.594255995 | -3.64208939 |
| **A_55_P2051849** | Snap25 | 0.630415869 | 0.039136177 | 1.221695561 | 11.93339174 | 2.62303048 | 0.04024744 | 0.594255995 | -3.642090596 |
| **A_55_P2019254** | NA | -0.280186882 | -0.543012957 | -0.017360808 | 11.80320659 | -2.622699376 | 0.040265077 | 0.594255995 | -3.642304235 |
| **A_66_P108474** | NA | 0.276586824 | 0.017073983 | 0.536099665 | 8.270996042 | 2.622055008 | 0.040299425 | 0.594255995 | -3.642720054 |
| **A_55_P2172934** | Rgs7bp | 0.619318564 | 0.03818715 | 1.200449977 | 12.10039597 | 2.621856443 | 0.040310015 | 0.594255995 | -3.642848204 |
| **A_51_P247614** | Smim10l2a | 0.126500986 | 0.007766445 | 0.245235527 | 9.013506926 | 2.621114518 | 0.040349613 | 0.594255995 | -3.643327086 |
| **A_55_P2049553** | NA | 0.474533 | 0.029091982 | 0.919974018 | 11.7804874 | 2.620869378 | 0.040362706 | 0.594255995 | -3.643485333 |
| **A_55_P2030046** | Barx2 | 0.606941632 | 0.037129363 | 1.1767539 | 9.907010713 | 2.620500792 | 0.0403824 | 0.594255995 | -3.643723288 |
| **A_55_P2159234** | Tatdn2 | -0.239110135 | -0.463596488 | -0.014623782 | 8.917128078 | -2.620458001 | 0.040384687 | 0.594255995 | -3.643750915 |
| **A_55_P2091756** | Nxnl1 | 0.61381248 | 0.037491623 | 1.190133337 | 9.303919671 | 2.620236793 | 0.040396513 | 0.594255995 | -3.643893737 |
| **A_51_P179921** | Igdcc3 | 0.334046673 | 0.020379325 | 0.64771402 | 8.411703587 | 2.620034452 | 0.040407333 | 0.594255995 | -3.644024385 |
| **A_55_P2163138** | Tm4sf5 | 0.707402472 | 0.043110189 | 1.371694754 | 9.100532272 | 2.61985062 | 0.040417165 | 0.594255995 | -3.644143087 |
| **A_51_P137007** | Zw10 | -0.127451576 | -0.247178279 | -0.007724873 | 9.344071711 | -2.618926738 | 0.040466621 | 0.594255995 | -3.644739732 |
| **A_51_P200776** | Grsf1 | -0.143999599 | -0.279274538 | -0.00872466 | 10.37049765 | -2.618864918 | 0.040469933 | 0.594255995 | -3.64477966 |
| **A_55_P1991377** | NA | 0.282069792 | 0.017061375 | 0.54707821 | 15.04447573 | 2.618581363 | 0.040485126 | 0.594255995 | -3.644962811 |

| | | | | | | | | | |
|---|---|---|---|---|---|---|---|---|---|
| **A_55_P1981814** | Myo3b | 0.49796828 | 0.029941223 | 0.965995337 | 9.403533055 | 2.617579507 | 0.040538854 | 0.594255995 | -3.645610026 |
| **A_55_P2131379** | Ybx1 | 0.278779638 | 0.016695935 | 0.540863341 | 11.18093019 | 2.616918498 | 0.040574346 | 0.594255995 | -3.646037137 |
| **A_55_P2129685** | Pkig | -0.124631373 | -0.24186839 | -0.007394356 | 8.20404994 | -2.615361835 | 0.040658058 | 0.594255995 | -3.64704326 |
| **A_55_P2094292** | Erp27 | 0.397600194 | 0.023416824 | 0.771783565 | 10.19335354 | 2.614154601 | 0.040723106 | 0.594255995 | -3.647823807 |
| **A_55_P2014506** | Amd2 | -0.145049455 | -0.281571955 | -0.008526954 | 11.69469219 | -2.613852213 | 0.040739416 | 0.594255995 | -3.648019355 |
| **A_66_P123294** | NA | 0.310364751 | 0.01822785 | 0.602501652 | 8.549744405 | 2.613696382 | 0.040747824 | 0.594255995 | -3.648120134 |
| **A_55_P2097548** | NA | 0.179078422 | 0.010508879 | 0.347647966 | 16.83146831 | 2.613565034 | 0.040754913 | 0.594255995 | -3.648205082 |
| **A_55_P1960708** | Setd1a | 0.321522681 | 0.018757726 | 0.624287637 | 10.82854996 | 2.61261354 | 0.040806302 | 0.594255995 | -3.648820538 |
| **A_55_P2080895** | Rab7 | -0.157088556 | -0.305075062 | -0.00910205 | 10.07885511 | -2.611509399 | 0.040866022 | 0.594255995 | -3.649534915 |
| **A_55_P2040295** | Ptp4a1 | -0.162753878 | -0.316110431 | -0.009397325 | 10.16120644 | -2.610947681 | 0.04089644 | 0.594255995 | -3.649898422 |
| **A_55_P2038952** | Ubb | -0.168712827 | -0.327721288 | -0.009704367 | 16.51502419 | -2.610339729 | 0.040929389 | 0.594255995 | -3.650291907 |
| **A_52_P199928** | NA | 0.476004119 | 0.02736707 | 0.924641168 | 12.46838603 | 2.61026583 | 0.040933396 | 0.594255995 | -3.65033974 |
| **A_55_P2083481** | Lpin1 | -0.140155354 | -0.272267533 | -0.008043174 | 8.363004016 | -2.609972893 | 0.040949284 | 0.594255995 | -3.650529362 |
| **A_55_P2065754** | Krtap19-5 | 1.16226566 | 0.066567553 | 2.257963768 | 9.965208634 | 2.609658373 | 0.040966349 | 0.594255995 | -3.650732971 |
| **A_55_P1963374** | Klhdc7a | 0.743617372 | 0.042583867 | 1.444650877 | 11.55871824 | 2.609635881 | 0.04096757 | 0.594255995 | -3.650747533 |
| **A_55_P1976709** | NA | 0.330132936 | 0.018824813 | 0.641441059 | 11.47969332 | 2.608960866 | 0.041004223 | 0.594255995 | -3.65118457 |
| **A_52_P455540** | Ric8b | -0.138393395 | -0.268919405 | -0.007867385 | 9.328485793 | -2.608479711 | 0.04103037 | 0.594255995 | -3.651496139 |
| **A_55_P2062054** | Dbndd2 | 0.240526693 | 0.013644 | 0.467409386 | 12.35980323 | 2.608141052 | 0.041048785 | 0.594255995 | -3.651715457 |
| **A_55_P2034453** | NA | 0.194026093 | 0.010965882 | 0.377086304 | 8.47609885 | 2.607566198 | 0.041080063 | 0.594255995 | -3.652087781 |
| **A_55_P2102698** | NA | 0.174103828 | 0.009781022 | 0.338426634 | 12.85659554 | 2.606631506 | 0.041130974 | 0.594255995 | -3.65269328 |
| **A_52_P606826** | Pigh | -0.120667284 | -0.234570338 | -0.006764231 | 8.525752666 | -2.606293582 | 0.041149397 | 0.594255995 | -3.652912225 |
| **A_65_P10865** | Scfd1 | -0.232718858 | -0.45244468 | -0.012993035 | 10.72809543 | -2.605671366 | 0.041183342 | 0.594255995 | -3.653315413 |
| **A_55_P2112330** | Anapc7 | 0.164556156 | 0.00916686 | 0.319945453 | 10.94748658 | 2.605326742 | 0.041202155 | 0.594255995 | -3.653538752 |
| **A_52_P356780** | Pdzd8 | -0.123029022 | -0.239208959 | -0.006849086 | 8.704205883 | -2.605227191 | 0.041207592 | 0.594255995 | -3.653603271 |
| **A_55_P2177278** | DXBay18 | -0.181344222 | -0.352648748 | -0.010039697 | 9.820750642 | -2.604378188 | 0.041253986 | 0.594255995 | -3.654153578 |
| **A_55_P2143159** | Men1 | 0.69154751 | 0.038210469 | 1.344884551 | 12.2258527 | 2.604077493 | 0.041270431 | 0.594255995 | -3.654348511 |
| **A_55_P2093650** | NA | -0.323378006 | -0.628911349 | -0.017844663 | 11.77904025 | -2.603880386 | 0.041281214 | 0.594255995 | -3.654476298 |
| **A_51_P241861** | Tcirg1 | -0.119497366 | -0.232409859 | -0.006584872 | 8.758262702 | -2.603667349 | 0.041292873 | 0.594255995 | -3.65461442 |
| **A_55_P2125311** | Rab3b | 0.291024839 | 0.015988009 | 0.56606167 | 9.892544017 | 2.603204979 | 0.041318188 | 0.594255995 | -3.654914221 |
| **A_51_P167292** | Chil3 | 0.548102359 | 0.029979003 | 1.066225715 | 10.45689559 | 2.602541507 | 0.041354543 | 0.594255995 | -3.655344478 |
| **A_55_P2093940** | Zfp408 | 0.196129348 | 0.010722465 | 0.38153623 | 16.67689079 | 2.602470978 | 0.04135841 | 0.594255995 | -3.65539022 |
| **A_66_P140599** | NA | 0.264737964 | 0.014407587 | 0.515068341 | 8.560230795 | 2.601787566 | 0.041395896 | 0.594255995 | -3.655833491 |
| **A_51_P275642** | NA | 0.564625461 | 0.030648879 | 1.098602043 | 9.943559197 | 2.601401643 | 0.041417081 | 0.594255995 | -3.656083839 |
| **A_55_P2182458** | Kcnh4 | 0.60719747 | 0.032863875 | 1.181531066 | 9.030852001 | 2.600967322 | 0.041440937 | 0.594255995 | -3.656365613 |
| **A_55_P2184146** | NA | 0.269694901 | 0.014545662 | 0.524844141 | 8.950918178 | 2.600444692 | 0.041469662 | 0.594255995 | -3.656704719 |
| **A_55_P2064292** | Col7a1 | -0.451080847 | -0.877951114 | -0.024210579 | 9.023208739 | -2.599726398 | 0.041509176 | 0.594255995 | -3.657170853 |
| **A_55_P2359610** | AI646519 | 0.391890305 | 0.021014543 | 0.762766068 | 10.84819128 | 2.599592232 | 0.041516561 | 0.594255995 | -3.657257928 |
| **A_51_P253547** | Ctsl | -0.160782558 | -0.312949563 | -0.008615553 | 16.23382561 | -2.599486732 | 0.041522369 | 0.594255995 | -3.657326401 |
| **A_55_P2047837** | NA | 0.569475545 | 0.03043765 | 1.10851344 | 8.564736513 | 2.599111701 | 0.041543023 | 0.594255995 | -3.657569824 |
| **A_55_P2089223** | Ak4 | 0.291911945 | 0.015595382 | 0.568228508 | 10.42843517 | 2.599046869 | 0.041546595 | 0.594255995 | -3.657611907 |
| **A_51_P383857** | Prp2 | 0.288343238 | 0.015368681 | 0.561317794 | 8.876574344 | 2.598703699 | 0.041565505 | 0.594255995 | -3.657834673 |
| **A_55_P2317665** | E2f1 | 0.194670845 | 0.010363229 | 0.37897846 | 8.359178245 | 2.598524377 | 0.041575391 | 0.594255995 | -3.657951086 |
| **A_55_P1969820** | Slc36a1 | 0.277640058 | 0.014748657 | 0.540531459 | 9.01361243 | 2.598213937 | 0.04159251 | 0.594255995 | -3.658152631 |
| **A_55_P2008692** | Eef1a1 | -0.172417929 | -0.335724974 | -0.009110885 | 16.00655508 | -2.597446847 | 0.041634843 | 0.594255995 | -3.658650712 |
| **A_55_P2253289** | NA | 0.720735346 | 0.037743458 | 1.403727233 | 10.3027908 | 2.59614795 | 0.04170663 | 0.594255995 | -3.659494317 |
| **A_51_P309740** | Usp40 | -0.158901325 | -0.309541472 | -0.008261178 | 8.484587088 | -2.595111079 | 0.04176403 | 0.594255995 | -3.660167939 |
| **A_55_P2041910** | NA | 0.602857964 | 0.031277182 | 1.174438745 | 10.58279523 | 2.594815882 | 0.041780388 | 0.594255995 | -3.660359751 |
| **A_66_P119489** | Spef1l | 0.160090357 | 0.008246189 | 0.311934525 | 8.274117125 | 2.593798412 | 0.041836819 | 0.594255995 | -3.661020986 |
| **A_51_P131942** | Dph5 | -0.186075894 | -0.362624959 | -0.009526829 | 10.02757905 | -2.592948283 | 0.041884032 | 0.594255995 | -3.661573597 |
| **A_55_P2025233** | Llph | 0.204884036 | 0.010423654 | 0.399344417 | 9.708466006 | 2.59206656 | 0.041933059 | 0.594255995 | -3.66214687 |

| | | | | | | | | | |
|---|---|---|---|---|---|---|---|---|---|
| **A_55_P2098558** | Slc39a1 | 0.272555239 | 0.013756492 | 0.531353986 | 14.37530572 | 2.590964896 | 0.041994402 | 0.594255995 | -3.662863319 |
| **A_55_P2082961** | Fnbp1l | -0.316028922 | -0.616237967 | -0.015819877 | 10.52297783 | -2.589835744 | 0.042057375 | 0.594255995 | -3.663597847 |
| **A_55_P2113208** | Snrpa1 | -0.227117172 | -0.442872162 | -0.011362182 | 11.34816301 | -2.589752655 | 0.042062013 | 0.594255995 | -3.663651906 |
| **A_55_P2174743** | Akap7 | 0.145198375 | 0.007247786 | 0.283148964 | 9.385636209 | 2.589448981 | 0.042078968 | 0.594255995 | -3.663849489 |
| **A_51_P177691** | Gabrr1 | 0.159618102 | 0.00782433 | 0.311411873 | 8.191814524 | 2.587005507 | 0.04221566 | 0.594255995 | -3.665439858 |
| **A_55_P2171976** | NA | 0.44139442 | 0.021579476 | 0.861209364 | 15.6402643 | 2.586652618 | 0.04223544 | 0.594255995 | -3.665669621 |
| **A_55_P2067351** | NA | 0.273631713 | 0.013361491 | 0.533901935 | 9.603470086 | 2.586491826 | 0.042244456 | 0.594255995 | -3.665774317 |
| **A_51_P156158** | Phlpp1 | -0.148653398 | -0.290048686 | -0.007258111 | 8.838649095 | -2.586479671 | 0.042245138 | 0.594255995 | -3.665782232 |
| **A_52_P26626** | Fam92b | 0.152902752 | 0.007425377 | 0.298380128 | 7.807701523 | 2.585764734 | 0.042285252 | 0.594255995 | -3.666247801 |
| **A_51_P246066** | Slamf9 | -0.217853731 | -0.425135996 | -0.010571467 | 8.277395176 | -2.585663594 | 0.04229093 | 0.594255995 | -3.66631367 |
| **A_55_P1985519** | NA | 0.23355932 | 0.01131888 | 0.455799759 | 17.05987137 | 2.585492472 | 0.042300539 | 0.594255995 | -3.666425121 |
| **A_51_P301998** | Fmo2 | -0.202781941 | -0.395784701 | -0.009779182 | 7.936628462 | -2.584847475 | 0.042336777 | 0.594255995 | -3.666845244 |
| **A_51_P322265** | Grap | -0.263478208 | -0.514250801 | -0.012705614 | 9.5520626 | -2.58484075 | 0.042337155 | 0.594255995 | -3.666849625 |
| **A_55_P1957598** | Atad2b | -0.194945914 | -0.380561129 | -0.0093307 | 9.981803436 | -2.58386447 | 0.04239207 | 0.594255995 | -3.66748566 |
| **A_51_P205820** | Klf11 | -0.140442239 | -0.274182452 | -0.006702025 | 8.738602336 | -2.583478756 | 0.042413787 | 0.594255995 | -3.667736991 |
| **A_55_P1984655** | Smtnl2 | 0.249545133 | 0.011813149 | 0.487277117 | 9.040971261 | 2.58244245 | 0.042472194 | 0.594255995 | -3.668412364 |
| **A_55_P1994042** | Zbp1 | -0.32121455 | -0.627224257 | -0.015204842 | 9.387891346 | -2.582433623 | 0.042472691 | 0.594255995 | -3.668418117 |
| **A_66_P127722** | NA | 0.325667621 | 0.015390926 | 0.635944315 | 11.1062279 | 2.582228014 | 0.04248429 | 0.594255995 | -3.668552136 |
| **A_55_P2130399** | Rilp | 0.493316526 | 0.023313892 | 0.96331916 | 9.824031923 | 2.582227701 | 0.042484308 | 0.594255995 | -3.66855234 |
| **A_52_P276792** | Foxp1 | -0.182540622 | -0.356516948 | -0.008564295 | 9.341212461 | -2.581300297 | 0.042536665 | 0.594255995 | -3.66915692 |
| **A_51_P320401** | Rab24 | -0.192091914 | -0.375184929 | -0.008998898 | 11.46227417 | -2.581109739 | 0.042547431 | 0.594255995 | -3.669281163 |
| **A_55_P2113920** | NA | 0.118747628 | 0.005545005 | 0.231950252 | 8.324337761 | 2.580700551 | 0.04257056 | 0.594255995 | -3.669547971 |
| **A_55_P2159390** | NA | 0.48101151 | 0.022446312 | 0.939576709 | 11.2121381 | 2.580616916 | 0.042575289 | 0.594255995 | -3.669602508 |
| **A_55_P2114269** | Fam177a2 | 0.27378702 | 0.012755714 | 0.534818326 | 17.06232582 | 2.580414195 | 0.042586754 | 0.594255995 | -3.669734703 |
| **A_55_P2104487** | Pld1 | 0.206207473 | 0.009583032 | 0.402831913 | 10.34409204 | 2.580097165 | 0.042604691 | 0.594255995 | -3.669941454 |
| **A_55_P2089462** | Ncoa5 | 0.701380506 | 0.032241467 | 1.370519545 | 13.8544256 | 2.57873364 | 0.042681925 | 0.594255995 | -3.670830858 |
| **A_55_P1968943** | NA | 0.336781817 | 0.01537368 | 0.658189954 | 15.97582752 | 2.577869522 | 0.042730949 | 0.594255995 | -3.671394662 |
| **A_51_P358700** | Olfr1221 | 0.367820979 | 0.016762757 | 0.7188792 | 8.636592033 | 2.577665209 | 0.042742549 | 0.594255995 | -3.671527986 |
| **A_55_P2055419** | Arid4a | -0.191336737 | -0.374007615 | -0.008665859 | 8.43746698 | -2.57690382 | 0.042785807 | 0.594255995 | -3.672024887 |
| **A_51_P298190** | Sbf2 | -0.312845604 | -0.611553641 | -0.014137567 | 9.370411565 | -2.576631502 | 0.04280129 | 0.594255995 | -3.672202631 |
| **A_55_P2009345** | Ovch2 | 0.245507252 | 0.011036577 | 0.479977926 | 8.372301876 | 2.575994634 | 0.042837523 | 0.594255995 | -3.672618367 |
| **A_55_P2182690** | Orai3 | 0.225929671 | 0.010155778 | 0.441703565 | 11.28153353 | 2.575986219 | 0.042838002 | 0.594255995 | -3.67262386 |
| **A_52_P306396** | Ppp3cb | 0.224035883 | 0.010052113 | 0.438019652 | 10.39662808 | 2.575763072 | 0.042850705 | 0.594255995 | -3.672769542 |
| **A_52_P330214** | Fanca | -0.174593678 | -0.341380322 | -0.007807034 | 9.680605132 | -2.575350874 | 0.042874182 | 0.594255995 | -3.673038667 |
| **A_52_P305279** | Spata13 | -0.12220728 | -0.238961058 | -0.005453503 | 8.311377514 | -2.575107139 | 0.04288807 | 0.594255995 | -3.673197815 |
| **A_55_P1961983** | Uqcr11 | 0.250533339 | 0.011110968 | 0.489955709 | 13.99853029 | 2.574364055 | 0.04293044 | 0.594255995 | -3.673683074 |
| **A_51_P365694** | Rala | -0.202810382 | -0.396659039 | -0.008961725 | 12.89309325 | -2.573928915 | 0.042955272 | 0.594255995 | -3.673967276 |
| **A_66_P133993** | Gm5093 | 1.454094907 | 0.064091606 | 2.844098207 | 11.94296126 | 2.573629848 | 0.042972348 | 0.594255995 | -3.674162623 |
| **A_51_P472323** | Fbh1 | 0.369335622 | 0.016207327 | 0.722463917 | 11.89778979 | 2.573106976 | 0.04300222 | 0.594255995 | -3.674504191 |
| **A_55_P2158281** | NA | 0.254325895 | 0.011106974 | 0.497544815 | 16.99279726 | 2.572541488 | 0.043034551 | 0.594255995 | -3.674873647 |
| **A_51_P254234** | Chchd4 | -0.131781944 | -0.257811245 | -0.005752643 | 9.724174172 | -2.572489106 | 0.043037548 | 0.594255995 | -3.674907873 |
| **A_55_P2004427** | Fbxl15 | 0.150232506 | 0.006552773 | 0.293912239 | 9.466704548 | 2.572394434 | 0.043042963 | 0.594255995 | -3.674969731 |
| **A_55_P1959624** | NA | 0.464162123 | 0.020033101 | 0.908291145 | 9.684476633 | 2.57116358 | 0.043113441 | 0.594255995 | -3.675774104 |
| **A_66_P137654** | Smpd2 | -0.137145254 | -0.268388163 | -0.005902344 | 8.255472061 | -2.570834361 | 0.043132312 | 0.594255995 | -3.675989292 |
| **A_55_P2009543** | NA | -0.195514304 | -0.382700209 | -0.0083284 | 16.61281349 | -2.569653451 | 0.043200077 | 0.594255995 | -3.676761315 |
| **A_52_P246753** | Chmp5 | -0.29440788 | -0.576307841 | -0.01250792 | 13.1079133 | -2.569351837 | 0.043217402 | 0.594255995 | -3.676958532 |
| **A_55_P2039125** | Fam219a | -0.199874225 | -0.391280358 | -0.008468092 | 9.373592525 | -2.569035503 | 0.043235582 | 0.594255995 | -3.67716539 |
| **A_52_P543430** | NA | -0.300282146 | -0.587844873 | -0.012719419 | 11.88709635 | -2.569011694 | 0.04323695 | 0.594255995 | -3.67718096 |
| **A_66_P125722** | Cyb5a | -0.309574577 | -0.606119023 | -0.013030131 | 14.80095472 | -2.568293521 | 0.043278254 | 0.594255995 | -3.67765065 |
| **A_51_P196972** | Slc4a1 | 0.490075727 | 0.020623307 | 0.959528146 | 10.45212272 | 2.568270561 | 0.043279576 | 0.594255995 | -3.677665667 |

| | | | | | | | | | |
|---|---|---|---|---|---|---|---|---|---|
| **A_51_P333159** | Rpa1 | 0.168300871 | 0.007005149 | 0.329596592 | 9.777671679 | 2.567040252 | 0.043350434 | 0.594255995 | -3.678470495 |
| **A_51_P493987** | Moxd1 | -0.126219995 | -0.247195008 | -0.005244982 | 8.063631023 | -2.566856828 | 0.043361009 | 0.594255995 | -3.678590506 |
| **A_55_P2088131** | Zfp975 | -0.148069743 | -0.290026466 | -0.006113021 | 8.985619561 | -2.566135132 | 0.043402642 | 0.594255995 | -3.679062749 |
| **A_55_P2166123** | Atp9a | 0.230276114 | 0.009452825 | 0.451099403 | 9.024778087 | 2.565506864 | 0.043438921 | 0.594255995 | -3.679473925 |
| **A_55_P2133427** | Vkorc1 | 0.210774477 | 0.008572454 | 0.412976499 | 11.40340173 | 2.564494004 | 0.043497475 | 0.594255995 | -3.680136935 |
| **A_52_P264902** | Dnajc3 | -0.263366575 | -0.516045085 | -0.010688064 | 10.95798098 | -2.56425677 | 0.043511202 | 0.594255995 | -3.680292249 |
| **A_55_P1999408** | NA | 0.722138996 | 0.028957006 | 1.415320985 | 13.17786807 | 2.562965089 | 0.043586022 | 0.594255995 | -3.681138058 |
| **A_52_P52263** | Ilrun | -0.201129618 | -0.394217583 | -0.008041653 | 8.372468766 | -2.562654077 | 0.043604058 | 0.594255995 | -3.681341752 |
| **A_52_P603654** | Stx8 | -0.13844657 | -0.271374549 | -0.005518591 | 9.436768695 | -2.562329403 | 0.043622894 | 0.594255995 | -3.681554411 |
| **A_51_P483473** | St3gal5 | -0.160036436 | -0.313722914 | -0.006349957 | 11.13317008 | -2.561842184 | 0.043651177 | 0.594255995 | -3.681873566 |
| **A_51_P466875** | Sbds | -0.158042677 | -0.309842585 | -0.00624277 | 12.55256306 | -2.561368323 | 0.043678704 | 0.594255995 | -3.682184008 |
| **A_55_P1973779** | Pde4c | 0.330632611 | 0.013032286 | 0.648232937 | 16.18843354 | 2.561143493 | 0.04369177 | 0.594255995 | -3.682331314 |
| **A_51_P345902** | Dohh | 0.269689006 | 0.010600897 | 0.528777115 | 12.08330685 | 2.560854619 | 0.043708565 | 0.594255995 | -3.682520592 |
| **A_55_P2107402** | NA | -0.168884621 | -0.331146841 | -0.0066224 | 8.220653988 | -2.560600648 | 0.043723336 | 0.594255995 | -3.682687013 |
| **A_55_P2056300** | Sf3b3 | 0.1668865 | 0.006499313 | 0.327273686 | 11.49252206 | 2.559886442 | 0.043764904 | 0.594255995 | -3.683155066 |
| **A_55_P1962411** | Gm6813 | -0.148184299 | -0.290611281 | -0.005757316 | 16.04833696 | -2.559641117 | 0.043779192 | 0.594255995 | -3.683315859 |
| **A_52_P547234** | Tomm70a | -0.212815069 | -0.417365007 | -0.00826513 | 10.12464915 | -2.559600494 | 0.043781559 | 0.594255995 | -3.683342485 |
| **A_55_P2027212** | NA | 0.244626626 | 0.009416551 | 0.479836701 | 9.438417564 | 2.558685853 | 0.043834875 | 0.594255995 | -3.683942055 |
| **A_55_P2122180** | Snta1 | 0.718353808 | 0.027575192 | 1.409132423 | 11.47727011 | 2.558401357 | 0.043851473 | 0.594255995 | -3.684128575 |
| **A_55_P1955242** | NA | 0.277719224 | 0.010632267 | 0.544806182 | 16.82104305 | 2.558128913 | 0.043867374 | 0.594255995 | -3.684307207 |
| **A_55_P2161640** | Slc38a1 | -0.190022444 | -0.372807286 | -0.007237601 | 10.60677778 | -2.557607414 | 0.043897828 | 0.594255995 | -3.684649168 |
| **A_55_P2161818** | NA | 0.247576177 | 0.009406332 | 0.485746022 | 16.70533089 | 2.557356314 | 0.0439125 | 0.594255995 | -3.684813836 |
| **A_55_P2157927** | Tmem241 | 0.181550807 | 0.006871973 | 0.356229641 | 9.030109428 | 2.556978423 | 0.04393459 | 0.594255995 | -3.685061672 |
| **A_55_P2106079** | NA | 0.502562657 | 0.019005787 | 0.986119528 | 8.957855597 | 2.556888679 | 0.043939837 | 0.594255995 | -3.685120533 |
| **A_51_P123920** | Mydgf | 0.529745987 | 0.01983218 | 1.039659795 | 12.35613044 | 2.555877691 | 0.043999001 | 0.594255995 | -3.718789721 |
| **A_66_P110798** | NA | -0.193134412 | -0.379252267 | -0.007016557 | 12.25068433 | -2.552941044 | 0.044171335 | 0.594255995 | -3.687710955 |
| **A_55_P2033420** | Sppl2a | -0.228181918 | -0.448076608 | -0.008287229 | 9.953193421 | -2.552910853 | 0.044173111 | 0.594255995 | -3.687730776 |
| **A_55_P2078238** | NA | 0.144060272 | 0.005194939 | 0.282925605 | 8.440208991 | 2.552228496 | 0.044213259 | 0.594255995 | -3.68817879 |
| **A_52_P652104** | Brd3 | -0.28761429 | -0.564878236 | -0.010350344 | 10.3427928 | -2.552032628 | 0.04422479 | 0.594255995 | -3.688307404 |
| **A_55_P2015337** | NA | -0.20107035 | -0.394964424 | -0.007176277 | 14.16744614 | -2.551247912 | 0.044271021 | 0.594255995 | -3.688822739 |
| **A_52_P206861** | Klf8 | -0.181266353 | -0.356068824 | -0.006463882 | 10.9973798 | -2.551166443 | 0.044275824 | 0.594255995 | -3.688876247 |
| **A_55_P2085979** | Igf1 | 0.252172092 | 0.008907714 | 0.49543647 | 9.510610982 | 2.550278827 | 0.044328185 | 0.594255995 | -3.689459287 |
| **A_55_P2135601** | Polr2i | 0.412599532 | 0.014487867 | 0.810711196 | 10.48304514 | 2.54972294 | 0.044361011 | 0.594255995 | -3.689824491 |
| **A_55_P2125947** | Agpat2 | -0.159166592 | -0.312746074 | -0.005587109 | 9.431821542 | -2.549692932 | 0.044362784 | 0.594255995 | -3.689844207 |
| **A_52_P407697** | NA | 0.41702388 | 0.01458267 | 0.81946509 | 8.439881748 | 2.549339302 | 0.044383681 | 0.594255995 | -3.69007656 |
| **A_55_P2128158** | Trappc13 | -0.188646101 | -0.370740367 | -0.006551834 | 11.23940388 | -2.548711767 | 0.044420789 | 0.594255995 | -3.690488932 |
| **A_51_P143901** | Tm9sf2 | -0.267156452 | -0.525053074 | -0.009259829 | 13.6485353 | -2.548526625 | 0.044431743 | 0.594255995 | -3.690610606 |
| **A_51_P314763** | Rabggtb | -0.141571654 | -0.278245676 | -0.004897632 | 9.605810807 | -2.548352454 | 0.044442051 | 0.594255995 | -3.690725075 |
| **A_55_P2080902** | NA | 0.313062595 | 0.010824988 | 0.615300202 | 8.39877669 | 2.548307559 | 0.044444708 | 0.594255995 | -3.690754582 |
| **A_55_P2070940** | Krtap20-2 | 0.24987556 | 0.008606739 | 0.491144382 | 9.280399308 | 2.547954927 | 0.044465587 | 0.594255995 | -3.690986356 |
| **A_55_P2126890** | NA | 0.821522254 | 0.028295744 | 1.614748763 | 9.187973549 | 2.547952195 | 0.044465749 | 0.594255995 | -3.690988152 |
| **A_66_P137265** | Gm9933 | 0.153464134 | 0.005277384 | 0.301650884 | 8.615422327 | 2.547807914 | 0.044474294 | 0.594255995 | -3.69108299 |
| **A_55_P1976789** | Dgcr6 | 0.36179568 | 0.012411927 | 0.711179433 | 10.68954222 | 2.547591755 | 0.044487101 | 0.594255995 | -3.691225079 |
| **A_55_P1991016** | Basp1 | -0.203108879 | -0.399266357 | -0.006951401 | 10.73534155 | -2.547376888 | 0.044499834 | 0.594255995 | -3.691366328 |
| **A_55_P1990810** | 4932412D23Rik | 0.280492573 | 0.009587173 | 0.551397974 | 8.348835378 | 2.547257604 | 0.044506905 | 0.594255995 | -3.691444745 |
| **A_52_P167382** | Tmem8 | -0.133339864 | -0.262128131 | -0.004551597 | 8.567968537 | -2.547140333 | 0.044513857 | 0.594255995 | -3.691521841 |
| **A_55_P1955502** | Tasor2 | -0.178928514 | -0.351804891 | -0.006052138 | 9.44166218 | -2.546320506 | 0.044562494 | 0.594255995 | -3.692060871 |
| **A_55_P2360661** | Cbx5 | -0.164577069 | -0.323617751 | -0.005536388 | 9.007623301 | -2.545835046 | 0.044591321 | 0.594255995 | -3.692380107 |
| **A_66_P104422** | Ckap4 | 0.229948748 | 0.00769489 | 0.452202606 | 11.75360704 | 2.545369897 | 0.04461896 | 0.594255995 | -3.692686022 |
| **A_55_P2116853** | Brpf1 | 0.146424946 | 0.004808045 | 0.288041848 | 11.64971574 | 2.543719086 | 0.044717199 | 0.594255995 | -3.693771989 |

| | | | | | | | | | |
|---|---|---|---|---|---|---|---|---|---|
| **A_55_P2028511** | NA | -0.200008137 | -0.393465322 | -0.006550952 | 11.28880209 | -2.543501297 | 0.044730177 | 0.594255995 | -3.69391529 |
| **A_52_P444253** | Gm6900 | -0.157932095 | -0.310694801 | -0.00516939 | 15.45670008 | -2.543444224 | 0.044733578 | 0.594255995 | -3.693952845 |
| **A_55_P2024933** | Zfp706 | -0.281610823 | -0.554118575 | -0.009103072 | 14.2759557 | -2.542375211 | 0.044797342 | 0.594255995 | -3.694656358 |
| **A_51_P404825** | Ccdc58 | 0.149931345 | 0.004816434 | 0.295046255 | 11.5816697 | 2.54184791 | 0.04482883 | 0.594255995 | -3.695003439 |
| **A_51_P258409** | Hey1 | -0.262856216 | -0.517294754 | -0.008417679 | 9.257200583 | -2.541584331 | 0.044844578 | 0.594255995 | -3.695176949 |
| **A_52_P364140** | Itga5 | 0.173022003 | 0.005539597 | 0.340504408 | 9.371860375 | 2.541565498 | 0.044845704 | 0.594255995 | -3.695189347 |
| **A_52_P105765** | Cdc42bpg | -0.189154341 | -0.372252912 | -0.00605577 | 8.514008887 | -2.541560902 | 0.044845978 | 0.594255995 | -3.695192372 |
| **A_55_P1996008** | Gm14399 | 0.347586786 | 0.011073728 | 0.684099844 | 9.515263405 | 2.541151161 | 0.044870472 | 0.594255995 | -3.695462121 |
| **A_51_P146044** | Ccdc92 | 0.141292564 | 0.004489453 | 0.278095676 | 9.154770406 | 2.540928795 | 0.044883771 | 0.594255995 | -3.695608526 |
| **A_52_P199299** | Urgcp | -0.191187856 | -0.376349346 | -0.006026365 | 10.35505778 | -2.540263669 | 0.044923574 | 0.594255995 | -3.696046484 |
| **A_55_P1966169** | Ppdpf | 0.223570048 | 0.007005817 | 0.440134279 | 12.69058546 | 2.539779749 | 0.044952557 | 0.594255995 | -3.69636517 |
| **A_55_P2135967** | NA | -0.195874982 | -0.385692213 | -0.006057751 | 13.99604282 | -2.538706525 | 0.045016905 | 0.594255995 | -3.697072075 |
| **A_55_P2095926** | NA | -0.163426951 | -0.321829055 | -0.005024847 | 8.549669698 | -2.538235396 | 0.045045184 | 0.594255995 | -3.697382453 |
| **A_52_P267106** | Ksr2 | 0.395760045 | 0.012150931 | 0.77936916 | 8.52288393 | 2.538120244 | 0.045052099 | 0.594255995 | -3.69745832 |
| **A_55_P1994047** | NA | -0.182619879 | -0.359705273 | -0.005534485 | 10.68113836 | -2.537081809 | 0.045114506 | 0.594255995 | -3.698142578 |
| **A_55_P2104617** | Kctd5 | -0.182107329 | -0.358729308 | -0.005485351 | 8.862515768 | -2.53659915 | 0.045143544 | 0.594255995 | -3.698460675 |
| **A_55_P2016024** | Kif3c | 0.987897266 | 0.029342179 | 1.946452354 | 10.0789843 | 2.53550149 | 0.045209656 | 0.594255995 | -3.699184226 |
| **A_55_P2082276** | NA | 0.234538449 | 0.006914214 | 0.462162684 | 10.26807031 | 2.53492265 | 0.04524456 | 0.594255995 | -3.69956586 |
| **A_55_P2068391** | Gm9222 | -0.136611215 | -0.269209898 | -0.004012532 | 9.060603212 | -2.534640123 | 0.045261607 | 0.594255995 | -3.699752152 |
| **A_55_P2143953** | Pank2 | -0.162188298 | -0.319625581 | -0.004751014 | 9.909568201 | -2.534434617 | 0.045274012 | 0.594255995 | -3.699887665 |
| **A_55_P1995737** | Gm14288 | 0.216213984 | 0.00620838 | 0.426219587 | 11.51694065 | 2.532923414 | 0.045365337 | 0.594255995 | -3.700884377 |
| **A_55_P1983036** | Tdrd3 | -0.121137021 | -0.238796212 | -0.003477831 | 9.770849031 | -2.532912561 | 0.045365993 | 0.594255995 | -3.700891536 |
| **A_55_P2178006** | NA | 0.292831148 | 0.008352968 | 0.577309328 | 9.02185269 | 2.532430131 | 0.045395189 | 0.594255995 | -3.701209798 |
| **A_55_P2126489** | Atxn7l3b | -0.155382017 | -0.306383071 | -0.004380963 | 12.14753778 | -2.531569987 | 0.045447294 | 0.594255995 | -3.701777332 |
| **A_55_P2106844** | NA | 0.172355559 | 0.004843139 | 0.33986798 | 8.462928665 | 2.5313223 | 0.045462309 | 0.594255995 | -3.701940781 |
| **A_65_P12359** | Pin1rt1 | 0.227695021 | 0.006394479 | 0.448995564 | 10.60686428 | 2.531280186 | 0.045464863 | 0.594255995 | -3.701968572 |
| **A_55_P2013038** | Serpinb6c | 0.320130008 | 0.00898106 | 0.631278955 | 13.17708288 | 2.53120437 | 0.04546946 | 0.594255995 | -3.702018605 |
| **A_55_P2168549** | Ap5b1 | 0.191963195 | 0.00538157 | 0.37854482 | 9.646363546 | 2.531152201 | 0.045472624 | 0.594255995 | -3.702053034 |
| **A_52_P681016** | NA | 1.330576449 | 0.037227048 | 2.623925851 | 10.36705807 | 2.531005734 | 0.045481508 | 0.594255995 | -3.702149696 |
| **A_55_P2162935** | Ntn1 | 0.187195973 | 0.005222795 | 0.369169151 | 9.144704388 | 2.530802679 | 0.045493828 | 0.594255995 | -3.702283709 |
| **A_55_P2154797** | 2810006K23Rik | 0.21946779 | 0.006107054 | 0.432828526 | 8.353100773 | 2.530611358 | 0.045505438 | 0.594255995 | -3.702409983 |
| **A_55_P2115968** | NA | -0.23163113 | -0.456819179 | -0.006443082 | 9.166827594 | -2.530583967 | 0.045507101 | 0.594255995 | -3.702428063 |
| **A_55_P2092859** | Anxa4 | 0.202419538 | 0.005605542 | 0.399233535 | 13.91932278 | 2.530262696 | 0.045526606 | 0.594255995 | -3.70264012 |
| **A_51_P389664** | Scrt2 | 0.490667214 | 0.013571897 | 0.967762532 | 8.654665036 | 2.530177847 | 0.045531759 | 0.594255995 | -3.702696128 |
| **A_55_P2176430** | 0610010K14Rik | -0.16712366 | -0.329654935 | -0.004592386 | 10.52733432 | -2.529706641 | 0.045560386 | 0.594255995 | -3.703007188 |
| **A_55_P2009007** | NA | -0.195322639 | -0.385295367 | -0.005349911 | 14.04959213 | -2.529475564 | 0.045574431 | 0.594255995 | -3.703159742 |
| **A_55_P2066409** | Gabpb1 | 0.301644366 | 0.008237171 | 0.595051562 | 10.21552189 | 2.529260851 | 0.045587486 | 0.594255995 | -3.703301501 |
| **A_55_P2066792** | Sp110 | -0.167135615 | -0.329794113 | -0.004477117 | 16.16416114 | -2.527908847 | 0.045669783 | 0.594255995 | -3.704194293 |
| **A_55_P2066304** | Limk2 | 0.171639195 | 0.004587554 | 0.338690837 | 8.452591279 | 2.527754457 | 0.04567919 | 0.594255995 | -3.704296262 |
| **A_55_P1988951** | NA | 0.453862249 | 0.012030761 | 0.895693737 | 15.06547217 | 2.52718224 | 0.045714076 | 0.594255995 | -3.704674224 |
| **A_55_P2017112** | Mpnd | 0.138421881 | 0.003654986 | 0.273188776 | 9.123534353 | 2.526915316 | 0.045730359 | 0.594255995 | -3.704850551 |
| **A_55_P2088360** | NA | 0.266572577 | 0.007036043 | 0.526109112 | 13.66024054 | 2.52688881 | 0.045731977 | 0.594255995 | -3.704868061 |
| **A_55_P2068104** | NA | 0.544154995 | 0.0142862 | 1.074023789 | 10.42909735 | 2.526524068 | 0.045754238 | 0.594255995 | -3.705109025 |
| **A_51_P315890** | Kcnk6 | -0.376147985 | -0.74243184 | -0.00986413 | 10.55979986 | -2.526446619 | 0.045758966 | 0.594255995 | -3.705160193 |
| **A_52_P123693** | E130304I02Rik | 0.371937004 | 0.009709345 | 0.734164663 | 12.39959304 | 2.526137244 | 0.045777859 | 0.594255995 | -3.705364599 |
| **A_51_P289697** | Nek4 | -0.153290418 | -0.30261128 | -0.003969555 | 9.108273346 | -2.525594838 | 0.045811002 | 0.594255995 | -3.705723006 |
| **A_55_P2024215** | Ctnnbl1 | -0.331531989 | -0.654506955 | -0.008557023 | 13.20775729 | -2.525374208 | 0.045824491 | 0.594255995 | -3.705868805 |
| **A_55_P2050207** | Ankrd27 | 0.126211942 | 0.003192746 | 0.249231138 | 11.2241945 | 2.524042869 | 0.045905976 | 0.594255995 | -3.706748758 |
| **A_66_P133642** | NA | 0.762001093 | 0.019215301 | 1.504786884 | 11.47014187 | 2.523836225 | 0.045918637 | 0.594255995 | -3.706885365 |
| **A_55_P2123234** | NA | 0.571962873 | 0.014319021 | 1.129606724 | 10.13808892 | 2.523365051 | 0.045947521 | 0.594255995 | -3.70719687 |

| | | | | | | | | | |
|---|---|---|---|---|---|---|---|---|---|
| **A_55_P2034320** | NA | 0.144398722 | 0.003609058 | 0.285188385 | 12.23876608 | 2.523258481 | 0.045954056 | 0.594255995 | -3.707267331 |
| **A_55_P2085664** | Meaf6 | 0.227319818 | 0.005678082 | 0.448961555 | 10.68703555 | 2.523218842 | 0.045956487 | 0.594255995 | -3.70729354 |
| **A_66_P137219** | Elane | 1.027237573 | 0.025520931 | 2.028954215 | 10.6371964 | 2.522871729 | 0.045977782 | 0.594255995 | -3.707523055 |
| **A_51_P113906** | Gkn2 | 0.529112479 | 0.013128495 | 1.045096463 | 13.02097505 | 2.522789103 | 0.045982853 | 0.594255995 | -3.733794045 |
| **A_55_P1987483** | Pgm2 | 0.255403957 | 0.006285107 | 0.504522807 | 13.21615701 | 2.522261982 | 0.046015215 | 0.594255995 | -3.707926273 |
| **A_66_P136228** | Rai14 | -0.299644609 | -0.591919558 | -0.00736966 | 11.88479374 | -2.522226233 | 0.046017411 | 0.594255995 | -3.707949916 |
| **A_55_P2100770** | Usp48 | -0.256597585 | -0.506946206 | -0.006248964 | 10.16440517 | -2.521601909 | 0.046055774 | 0.594255995 | -3.708362837 |
| **A_55_P2002723** | Inpp5a | -0.211788414 | -0.418463436 | -0.005113392 | 10.74959841 | -2.521061084 | 0.046089034 | 0.594255995 | -3.708720582 |
| **A_55_P2301941** | A530030E21Rik | 0.518397174 | 0.012442445 | 1.024351902 | 13.82748069 | 2.520694008 | 0.046111623 | 0.594255995 | -3.708963421 |
| **A_66_P106509** | NA | 0.656868828 | 0.015654663 | 1.298082993 | 14.06940801 | 2.520256295 | 0.046138575 | 0.594255995 | -3.709253018 |
| **A_51_P320304** | Plod1 | -0.214730203 | -0.424353077 | -0.00510733 | 8.958653917 | -2.520133967 | 0.04614611 | 0.594255995 | -3.709333957 |
| **A_51_P181365** | Atp5md | 0.505520767 | 0.01201608 | 0.999025455 | 14.8547041 | 2.520094826 | 0.046148521 | 0.594255995 | -3.709359856 |
| **A_55_P2083028** | NA | -0.166084545 | -0.328258162 | -0.003910929 | 9.987845641 | -2.519522162 | 0.046183815 | 0.594255995 | -3.709738798 |
| **A_52_P644674** | Trappc6b | -0.164539704 | -0.325222787 | -0.003856621 | 11.38297594 | -2.51924102 | 0.046201153 | 0.594255995 | -3.709924853 |
| **A_55_P2147616** | NA | 1.035660838 | 0.023936819 | 2.047384856 | 13.11734704 | 2.518399689 | 0.046253078 | 0.594255995 | -3.710481706 |
| **A_55_P2007966** | Tmem215 | 0.771209274 | 0.017726639 | 1.524691908 | 12.07525781 | 2.518072085 | 0.046273313 | 0.594255995 | -3.710698567 |
| **A_55_P1964348** | Gdpd2 | 0.362327408 | 0.008306151 | 0.716348665 | 9.305800584 | 2.517914681 | 0.046283039 | 0.594255995 | -3.710802769 |
| **A_52_P654108** | Dync1li2 | -0.155895228 | -0.30827317 | -0.003517287 | 8.288980329 | -2.516980692 | 0.046340795 | 0.594255995 | -3.711421149 |
| **A_51_P161021** | Ifit2 | -0.215039254 | -0.425257124 | -0.004821383 | 8.161761771 | -2.516617862 | 0.046363252 | 0.594255995 | -3.71166141 |
| **A_55_P2045765** | Rnf224 | 1.075069047 | 0.023874599 | 2.126263495 | 9.712089728 | 2.516068511 | 0.046397276 | 0.594255995 | -3.712025222 |
| **A_55_P2037732** | Foxl2 | -0.302349506 | -0.59799405 | -0.006704963 | 13.26183597 | -2.515987961 | 0.046402267 | 0.594255995 | -3.71207857 |
| **A_55_P2018211** | NA | -0.389899292 | -0.771267792 | -0.008530792 | 14.37307863 | -2.515224709 | 0.046449588 | 0.594255995 | -3.712584126 |
| **A_55_P1960049** | 2810408A11Rik | 0.550038426 | 0.011887757 | 1.088189095 | 10.64321416 | 2.514538614 | 0.046492169 | 0.594255995 | -3.713038653 |
| **A_66_P102260** | Irs2 | -0.29045413 | -0.574652602 | -0.006255658 | 11.69737194 | -2.51434565 | 0.046504153 | 0.594255995 | -3.713166502 |
| **A_51_P368855** | Pfdn6 | -0.218201414 | -0.431706738 | -0.00469609 | 11.24264587 | -2.514305324 | 0.046506657 | 0.594255995 | -3.71319322 |
| **A_66_P126293** | Itgb3 | -0.344402388 | -0.681421871 | -0.007382904 | 9.779298731 | -2.514087035 | 0.046520218 | 0.594255995 | -3.713337857 |
| **A_52_P665817** | Abhd13 | -0.137608482 | -0.272272692 | -0.002944271 | 8.268324162 | -2.513982081 | 0.04652674 | 0.594255995 | -3.713407401 |
| **A_55_P1985890** | Tiparp | -0.200454327 | -0.39663127 | -0.004277384 | 10.27563184 | -2.513834231 | 0.046535928 | 0.594255995 | -3.713505372 |
| **A_51_P359237** | Tmem9 | 0.293829795 | 0.006246976 | 0.581412615 | 11.58499255 | 2.513634098 | 0.046548369 | 0.594255995 | -3.713637992 |
| **A_51_P226932** | Tubgcp2 | -0.113105516 | -0.223813871 | -0.002397161 | 8.72911697 | -2.513463305 | 0.046558989 | 0.594255995 | -3.713751175 |
| **A_52_P218537** | Hnrnpc | -0.160295919 | -0.31720619 | -0.003385648 | 10.99091481 | -2.51327641 | 0.046570614 | 0.594255995 | -3.713875033 |
| **A_55_P1981099** | NA | -0.214633503 | -0.424809094 | -0.004457913 | 14.13546686 | -2.512374638 | 0.046626743 | 0.594255995 | -3.714472731 |
| **A_55_P2089362** | Wfdc13 | 1.238872689 | 0.025672618 | 2.452072761 | 10.24424304 | 2.51225324 | 0.046634305 | 0.594255995 | -3.714553204 |
| **A_52_P437084** | NA | 0.239691131 | 0.004947149 | 0.474435112 | 12.27069184 | 2.512040638 | 0.046647551 | 0.594255995 | -3.714694139 |
| **A_55_P2131865** | Krtap10-10 | 0.59882136 | 0.012354661 | 1.18528806 | 10.127526 | 2.512019979 | 0.046648839 | 0.594255995 | -3.714707835 |
| **A_51_P463497** | Rnh1 | -0.248916066 | -0.492746382 | -0.00508575 | 10.0004318 | -2.511506984 | 0.046680817 | 0.594255995 | -3.715047933 |
| **A_55_P1958919** | Olfr1347 | 0.157146335 | 0.003208671 | 0.311084 | 8.558136323 | 2.511473076 | 0.046682932 | 0.594255995 | -3.715070415 |
| **A_52_P575771** | Mtfr1 | -0.202353635 | -0.400576512 | -0.004130759 | 10.76317603 | -2.511460775 | 0.046683699 | 0.594255995 | -3.715078571 |
| **A_66_P108770** | Oxct1 | 0.159821461 | 0.003252957 | 0.316389966 | 11.44754752 | 2.511307293 | 0.046693272 | 0.594255995 | -3.715180333 |
| **A_51_P155747** | Scyl3 | -0.137930218 | -0.273091391 | -0.002769046 | 8.850961036 | -2.510594859 | 0.046737735 | 0.594255995 | -3.715652743 |
| **A_55_P2074035** | Haao | 0.450385825 | 0.008823603 | 0.891948046 | 8.53594682 | 2.509354199 | 0.046815271 | 0.594255995 | -3.716475605 |
| **A_52_P462203** | Dclre1c | -0.169129297 | -0.33495892 | -0.003299675 | 8.325249834 | -2.509145783 | 0.046828309 | 0.594255995 | -3.71661386 |
| **A_55_P2021585** | Tff1 | 0.651850025 | 0.012531486 | 1.291168564 | 9.547999038 | 2.508415937 | 0.046873999 | 0.594255995 | -3.717098061 |
| **A_55_P2155546** | Nlrc3 | 0.235733978 | 0.004371453 | 0.467096502 | 9.212341333 | 2.506676752 | 0.046983066 | 0.594255995 | -3.71825222 |
| **A_55_P2023692** | Neurl1a | 0.190947749 | 0.003531571 | 0.378363927 | 9.115029086 | 2.506551487 | 0.046990932 | 0.594255995 | -3.718335367 |
| **A_55_P2049186** | Plvap | -0.199257461 | -0.394833312 | -0.003681611 | 8.900370293 | -2.506504726 | 0.046993868 | 0.594255995 | -3.718366405 |
| **A_55_P2039436** | Ilkap | -0.195292917 | -0.386983779 | -0.003602056 | 11.89597309 | -2.506422309 | 0.046999045 | 0.594255995 | -3.718421113 |
| **A_55_P2060793** | NA | -0.181749516 | -0.360204113 | -0.003294919 | 12.51380699 | -2.505616991 | 0.047049657 | 0.594255995 | -3.718955729 |
| **A_55_P2074811** | Zfp609 | 0.59912063 | 0.010691441 | 1.187549818 | 9.651508442 | 2.504893292 | 0.047095188 | 0.594255995 | -3.719436247 |
| **A_51_P107362** | Socs2 | -0.155520024 | -0.308274687 | -0.002765361 | 9.152979456 | -2.504730482 | 0.047105438 | 0.594255995 | -3.719544359 |

| | | | | | | | | | |
|---|---|---|---|---|---|---|---|---|---|
| **A_55_P2137937** | Fxyd1 | 0.843807039 | 0.014783872 | 1.672830205 | 13.15471187 | 2.504065242 | 0.047147342 | 0.594255995 | -3.71998615 |
| **A_55_P2159289** | NA | -0.111758458 | -0.221608872 | -0.001908044 | 8.072832182 | -2.502925163 | 0.04721925 | 0.594255995 | -3.720743444 |
| **A_52_P497388** | NA | 0.295403892 | 0.005027383 | 0.585780401 | 15.28949564 | 2.502787033 | 0.04722797 | 0.594255995 | -3.72083521 |
| **A_52_P186033** | Spn | 0.246846228 | 0.004195944 | 0.489496512 | 10.18182884 | 2.502734928 | 0.04723126 | 0.594255995 | -3.720869826 |
| **A_55_P1971897** | Mdk | 0.45881332 | 0.007768575 | 0.909858065 | 10.26619556 | 2.502566075 | 0.047241922 | 0.594255995 | -3.720982008 |
| **A_55_P2106911** | Wnt1 | 0.685574836 | 0.011548298 | 1.359601375 | 9.42044655 | 2.502344131 | 0.047255942 | 0.594255995 | -3.721129469 |
| **A_55_P2077213** | Cenpb | 0.331006318 | 0.005524799 | 0.656487837 | 10.96393859 | 2.501952797 | 0.047280671 | 0.594255995 | -3.721389493 |
| **A_51_P418317** | Mis12 | -0.1210695 | -0.240124807 | -0.002014192 | 8.10265575 | -2.501814754 | 0.047289398 | 0.594255995 | -3.721481222 |
| **A_55_P2025974** | Ruvbl2 | 0.233725376 | 0.003874766 | 0.463575987 | 10.92306117 | 2.501666245 | 0.047298788 | 0.594255995 | -3.721579909 |
| **A_51_P145772** | Rnf113a2 | -0.134268567 | -0.266322829 | -0.002214306 | 9.541833358 | -2.501445801 | 0.04731273 | 0.594255995 | -3.721726403 |
| **A_55_P2086329** | Krt85 | 0.265192956 | 0.004367857 | 0.526018056 | 8.668835986 | 2.501392053 | 0.04731613 | 0.594255995 | -3.721762123 |
| **A_55_P1976017** | Cnih4 | 0.397025993 | 0.006406049 | 0.787645937 | 13.31208013 | 2.500539335 | 0.047370106 | 0.594255995 | -3.72232887 |
| **A_55_P2153141** | NA | 0.177382445 | 0.002848345 | 0.351916545 | 12.188028 | 2.50034254 | 0.047382572 | 0.594255995 | -3.722459683 |
| **A_52_P400224** | Ccs | 0.153188775 | 0.0024387 | 0.30393885 | 11.77771536 | 2.499991709 | 0.047404804 | 0.594255995 | -3.722692901 |
| **A_55_P2069714** | Sqstm1 | -0.168068617 | -0.333470365 | -0.002666869 | 10.10331706 | -2.499860044 | 0.047413151 | 0.594255995 | -3.722780432 |
| **A_51_P221362** | Tnks2 | -0.226270523 | -0.448720906 | -0.003820139 | 12.81959608 | -2.635996953 | 0.047424428 | 0.594255995 | -3.785743827 |
| **A_55_P2084303** | Nid1 | -0.193171309 | -0.383296529 | -0.003046088 | 8.742507053 | -2.499608849 | 0.047429079 | 0.594255995 | -3.722947433 |
| **A_66_P111773** | Gatad2a | -0.199796513 | -0.396445868 | -0.003147157 | 9.466088196 | -2.499565597 | 0.047431822 | 0.594255995 | -3.722976189 |
| **A_52_P153848** | Prpf4b | -0.121175403 | -0.240460631 | -0.001890175 | 8.329265439 | -2.499176753 | 0.047456491 | 0.594255995 | -3.723234724 |
| **A_55_P2126470** | Rpl7 | -0.494685345 | -0.98174026 | -0.00763043 | 14.44115752 | -2.498735446 | 0.047484505 | 0.594255995 | -3.723528169 |
| **A_55_P2049867** | Ccrl2 | 0.241797935 | 0.003716972 | 0.479878897 | 9.159738108 | 2.498601986 | 0.047492981 | 0.594255995 | -3.723616919 |
| **A_55_P2208682** | B230334C09Rik | 0.19420281 | 0.002973064 | 0.385432557 | 8.51228268 | 2.498441733 | 0.04750316 | 0.594255995 | -3.723723489 |
| **A_51_P375146** | Cd36 | -0.272590301 | -0.541066993 | -0.00411361 | 10.15021965 | -2.497888083 | 0.047538344 | 0.594255995 | -3.724091704 |
| **A_52_P27103** | Necap2 | -0.203004053 | -0.402949107 | -0.003058999 | 9.281474646 | -2.497831889 | 0.047541917 | 0.594255995 | -3.724129079 |
| **A_51_P398260** | Tppp | -0.131302977 | -0.260637234 | -0.001968721 | 8.150307381 | -2.497641867 | 0.047554 | 0.594255995 | -3.724255469 |
| **A_55_P2135261** | Ccdc174 | -0.197568489 | -0.392180226 | -0.002956752 | 10.40485237 | -2.497570819 | 0.047558519 | 0.594255995 | -3.724302727 |
| **A_55_P1976908** | NA | 0.668116237 | 0.009864039 | 1.326368436 | 13.47419517 | 2.497059389 | 0.04759106 | 0.594255995 | -3.72464293 |
| **A_55_P1979743** | NA | 0.232802154 | 0.003381653 | 0.462222656 | 13.8054393 | 2.496456096 | 0.047629476 | 0.594255995 | -3.725044291 |
| **A_55_P1979833** | Cited1 | -0.153942556 | -0.305662115 | -0.002222996 | 16.85225515 | -2.496239675 | 0.047643265 | 0.594255995 | -3.725188286 |
| **A_55_P2160322** | Anapc15 | -0.119633649 | -0.237542544 | -0.001724754 | 8.772678005 | -2.496180252 | 0.047647052 | 0.594255995 | -3.725227824 |
| **A_52_P139740** | Mdp1 | -0.160920152 | -0.319527229 | -0.002313075 | 9.272075446 | -2.496071574 | 0.047653978 | 0.594255995 | -3.725300136 |
| **A_55_P2137586** | Nphp4 | 0.719867362 | 0.010334284 | 1.42940044 | 8.886585494 | 2.496025398 | 0.047656922 | 0.594255995 | -3.725330861 |
| **A_55_P2044359** | Rxrg | 0.766868848 | 0.010942108 | 1.522795587 | 11.18192688 | 2.495804429 | 0.047671009 | 0.594255995 | -3.725477897 |
| **A_51_P118885** | Amacr | 0.186883991 | 0.002653913 | 0.371114069 | 10.00613274 | 2.49563305 | 0.047681938 | 0.594255995 | -3.735286276 |
| **A_55_P2021555** | Tbrg1 | -0.207546609 | -0.412207583 | -0.002885635 | 16.18439665 | -2.494880611 | 0.047729954 | 0.594255995 | -3.726092699 |
| **A_55_P2004168** | NA | 0.540370416 | 0.007455215 | 1.073285617 | 10.433623 | 2.494609772 | 0.047747249 | 0.594255995 | -3.726272967 |
| **A_51_P504065** | Spa17 | -0.129393892 | -0.257007759 | -0.001780026 | 9.053818393 | -2.494508983 | 0.047753687 | 0.594255995 | -3.726340055 |
| **A_51_P214306** | Haus4 | -0.375810205 | -0.746115085 | -0.005505325 | 11.05767488 | -2.630021001 | 0.047761606 | 0.594255995 | -3.788685274 |
| **A_55_P2172196** | Thnsl1 | 0.490848822 | 0.006649672 | 0.975047972 | 8.372643217 | 2.493979579 | 0.047787519 | 0.594255995 | -3.726692463 |
| **A_55_P2082478** | Kidins220 | 0.289172568 | 0.003899291 | 0.574445845 | 9.60651551 | 2.493820347 | 0.047797699 | 0.594255995 | -3.726798467 |
| **A_52_P22617** | NA | 0.381878469 | 0.005028175 | 0.758728763 | 8.788890658 | 2.493018365 | 0.047849009 | 0.594255995 | -3.727332424 |
| **A_55_P2093286** | Apod | -0.25012322 | -0.49704561 | -0.00320083 | 9.304803655 | -2.492084144 | 0.047908854 | 0.594255995 | -3.727954549 |
| **A_55_P2189201** | 5830426C09Rik | 0.262373846 | 0.003290375 | 0.521457318 | 8.732797063 | 2.491437503 | 0.047950323 | 0.594255995 | -3.728385244 |
| **A_55_P1965571** | Rpl8 | -0.276607363 | -0.549834499 | -0.003380227 | 14.24493454 | -2.490629166 | 0.048002215 | 0.594255995 | -3.728923727 |
| **A_51_P360004** | Ocstamp | 0.56272329 | 0.006822872 | 1.118623708 | 9.084825996 | 2.490388217 | 0.048017694 | 0.594255995 | -3.729084258 |
| **A_51_P411297** | Nup50 | -0.121757031 | -0.242038908 | -0.001475154 | 10.08325124 | -2.490365071 | 0.048019182 | 0.594255995 | -3.733930279 |
| **A_65_P05152** | Flrt2 | -0.153707026 | -0.3055521 | -0.001861952 | 8.294770705 | -2.490360241 | 0.048019492 | 0.594255995 | -3.729102897 |
| **A_51_P325374** | Tent2 | -0.121499659 | -0.241537016 | -0.001462302 | 12.35152428 | -2.490163126 | 0.04803216 | 0.594255995 | -3.729234231 |
| **A_51_P483373** | Txndc5 | -0.199627506 | -0.397026101 | -0.00222891 | 9.497486566 | -2.487971983 | 0.048173215 | 0.594255995 | -3.730694546 |
| **A_55_P1955975** | Afg3l2 | -0.118208601 | -0.235122941 | -0.001294261 | 10.63556343 | -2.48742766 | 0.048208324 | 0.594255995 | -3.731057431 |

| | | | | | | | | | |
|---|---|---|---|---|---|---|---|---|---|
| **A_55_P2077751** | NA | 0.933846293 | 0.010004041 | 1.857688545 | 9.846010174 | 2.486833685 | 0.048246666 | 0.594255995 | -3.731453468 |
| **A_55_P2155041** | Ssbp2 | 0.589700333 | 0.006252073 | 1.173148593 | 11.545052 | 2.486555672 | 0.048264623 | 0.594255995 | -3.731638854 |
| **A_55_P2112701** | NA | -0.224592991 | -0.446812698 | -0.002373283 | 10.72141796 | -2.486467516 | 0.048270319 | 0.594255995 | -3.731697641 |
| **A_55_P2081571** | Raver1 | 0.802842851 | 0.008352395 | 1.597333306 | 14.803357 | 2.486056665 | 0.048296873 | 0.594255995 | -3.731971634 |
| **A_51_P334308** | Tspan14 | -0.300244216 | -0.597373524 | -0.003114908 | 13.75079749 | -2.485983951 | 0.048301574 | 0.594255995 | -3.732020129 |
| **A_65_P05358** | NA | -0.180835096 | -0.359667221 | -0.002002971 | 12.49553649 | -2.620518716 | 0.048303213 | 0.594255995 | -3.793374912 |
| **A_55_P2107035** | Mcmbp | -0.212290976 | -0.42252058 | -0.002061371 | 10.71874596 | -2.484315917 | 0.048409552 | 0.594255995 | -3.73313281 |
| **A_55_P1997145** | Nlrx1 | 0.2432918 | 0.002347368 | 0.484236232 | 8.768212015 | 2.484161006 | 0.048419593 | 0.594255995 | -3.733236166 |
| **A_55_P1986049** | Fip1l1 | -0.127806466 | -0.254401501 | -0.001211431 | 8.308440362 | -2.483735334 | 0.048447195 | 0.594255995 | -3.733520193 |
| **A_52_P157274** | Abi1 | -0.128781466 | -0.256343134 | -0.001219797 | 8.805909647 | -2.483718284 | 0.048448301 | 0.594255995 | -3.73353157 |
| **A_55_P2101133** | Tomm7 | -0.24618633 | -0.490094454 | -0.002278205 | 14.5422663 | -2.48317216 | 0.048483739 | 0.594255995 | -3.733896011 |
| **A_55_P2048209** | Ganab | 0.182125583 | 0.001641854 | 0.362609312 | 12.61456161 | 2.482573205 | 0.048522637 | 0.594255995 | -3.734295758 |
| **A_55_P2130975** | Ncoa1 | -0.161379776 | -0.321323357 | -0.001436196 | 9.25398343 | -2.482283951 | 0.048541434 | 0.594255995 | -3.734488828 |
| **A_52_P675395** | Ackr3 | -0.150690998 | -0.300071731 | -0.001310264 | 10.16518781 | -2.481772023 | 0.048574719 | 0.594255995 | -3.734830558 |
| **A_51_P340947** | Selenoo | 0.135726797 | 0.001167778 | 0.270285815 | 9.294590771 | 2.481543841 | 0.048589564 | 0.594255995 | -3.734982891 |
| **A_55_P2004333** | Rpl24 | 0.205227969 | 0.001741825 | 0.408714113 | 15.60781164 | 2.481251968 | 0.048608558 | 0.594255995 | -3.735177755 |
| **A_55_P2021854** | NA | 0.131976364 | 0.001120057 | 0.262832671 | 8.438332534 | 2.481250797 | 0.048608635 | 0.594255995 | -3.735178537 |
| **A_55_P2022263** | NA | -0.233699972 | -0.465434261 | -0.001965682 | 16.52905163 | -2.481061453 | 0.048620961 | 0.594255995 | -3.735304956 |
| **A_52_P546513** | NA | 0.448636696 | 0.003759272 | 0.893514119 | 8.796554306 | 2.480981863 | 0.048626143 | 0.594255995 | -3.735358097 |
| **A_55_P2121142** | Tbc1d22b | 0.248281933 | 0.002072698 | 0.494491168 | 10.42606883 | 2.480903905 | 0.04863122 | 0.594255995 | -3.73541015 |
| **A_66_P129048** | Faap20 | 0.161143695 | 0.001264386 | 0.321023003 | 12.06386144 | 2.479649047 | 0.048713015 | 0.594255995 | -3.736248149 |
| **A_55_P2340206** | E230006M18Rik | 0.175379068 | 0.001347254 | 0.349410881 | 8.983478158 | 2.47923831 | 0.048739819 | 0.594255995 | -3.736522491 |
| **A_52_P496202** | Pcsk1n | 0.285321492 | 0.002084298 | 0.568558686 | 8.911415094 | 2.478297082 | 0.048801302 | 0.594255995 | -3.737151263 |
| **A_52_P65108** | Smad1 | -0.171267209 | -0.3413013 | -0.001233118 | 8.828761082 | -2.47803467 | 0.048818457 | 0.594255995 | -3.737326586 |
| **A_51_P508115** | Trank1 | -0.16774534 | -0.334283314 | -0.001207366 | 8.855445068 | -2.478028805 | 0.048818841 | 0.594255995 | -3.737330505 |
| **A_55_P2157872** | Srsf5 | -0.171957515 | -0.342719572 | -0.001195458 | 11.96200919 | -2.477416042 | 0.048858927 | 0.594255995 | -3.737739948 |
| **A_55_P2024080** | Rpl7a | 0.403087337 | 0.002785456 | 0.803389217 | 15.30415106 | 2.477311893 | 0.048865744 | 0.594255995 | -3.737809545 |
| **A_55_P1960208** | Vcan | 0.199922246 | 0.00136393 | 0.398480562 | 8.693537184 | 2.477092387 | 0.048880114 | 0.594255995 | -3.737956234 |
| **A_55_P2032099** | Zfp715 | 0.193306211 | 0.001277273 | 0.38533515 | 8.334170903 | 2.476556787 | 0.048915197 | 0.594255995 | -3.73831419 |
| **A_55_P2244142** | Klf13 | 0.259380002 | 0.001605053 | 0.51715495 | 10.94961799 | 2.475511469 | 0.048983744 | 0.594255995 | -3.73901293 |
| **A_55_P2127683** | 4933434E20Rik | -0.116699337 | -0.232681254 | -0.00071742 | 9.132942039 | -2.475410731 | 0.048990355 | 0.594255995 | -3.739080277 |
| **A_55_P1991896** | Ceacam5 | 0.274215812 | 0.001677317 | 0.546754307 | 16.69446821 | 2.475333975 | 0.048995393 | 0.594255995 | -3.739131592 |
| **A_66_P111798** | Znrf2 | -0.203128301 | -0.405024936 | -0.001231665 | 8.997230788 | -2.475201257 | 0.049004106 | 0.594255995 | -3.739220322 |
| **A_52_P295067** | Snn | -0.164170691 | -0.327354514 | -0.000986867 | 8.90830124 | -2.475071128 | 0.04901265 | 0.594255995 | -3.739307325 |
| **A_55_P2116385** | Gpr183 | 0.592435071 | 0.003433692 | 1.18143645 | 8.484631901 | 2.474535059 | 0.049047864 | 0.594255995 | -3.739665759 |
| **A_55_P2065751** | Krtap19-5 | 0.533820714 | 0.00286067 | 1.064780758 | 9.92912366 | 2.473447769 | 0.04911937 | 0.594255995 | -3.740392891 |
| **A_55_P2057503** | Arl4c | 0.869822307 | 0.004228674 | 1.73541594 | 11.16502213 | 2.472211663 | 0.049200797 | 0.594255995 | -3.741219763 |
| **A_55_P1971854** | Vcpkmt | 0.252981587 | 0.001200664 | 0.50476251 | 10.36172363 | 2.471924801 | 0.049219714 | 0.594255995 | -3.741411687 |
| **A_52_P21550** | Gcnt1 | -0.324901622 | -0.648319663 | -0.00148358 | 11.76113602 | -2.471478286 | 0.049249175 | 0.594255995 | -3.741710452 |
| **A_51_P467410** | Npm1 | -0.242286671 | -0.483485918 | -0.001087425 | 15.27459865 | -2.47128447 | 0.049261969 | 0.594255995 | -3.741840145 |
| **A_55_P2184113** | Aurkaip1 | 0.367220531 | 0.001592184 | 0.732848879 | 15.60846803 | 2.470906194 | 0.049286948 | 0.594255995 | -3.742093285 |
| **A_55_P2016462** | Cxcl10 | -0.8371754 | -1.670752627 | -0.003598172 | 9.021699597 | -2.470812441 | 0.049293141 | 0.594255995 | -3.742156027 |
| **A_55_P2145879** | Tpt1 | -0.168326967 | -0.335940764 | -0.000713169 | 17.16987602 | -2.47066063 | 0.049303172 | 0.594255995 | -3.742257627 |
| **A_55_P2201676** | Cyp2r1 | 0.58241293 | 0.002450029 | 1.162375831 | 10.09871555 | 2.470585895 | 0.04930811 | 0.594255995 | -3.742307644 |
| **A_55_P1972182** | Siah1b | 0.241367531 | 0.000970876 | 0.481764186 | 9.474401101 | 2.470128749 | 0.04933833 | 0.594255995 | -3.742613615 |
| **A_55_P1965101** | Xlr3b | -0.119336021 | -0.238194245 | -0.000477796 | 12.11002498 | -2.470082608 | 0.049341381 | 0.594255995 | -3.7426445 |
| **A_51_P123297** | Ppp1r7 | -0.145488155 | -0.290397315 | -0.000578995 | 8.387713328 | -2.470022785 | 0.049345337 | 0.594255995 | -3.742684542 |
| **A_55_P1953028** | Tnfsf12 | 0.324282825 | 0.001281693 | 0.647283957 | 9.518828069 | 2.469955144 | 0.049349811 | 0.594255995 | -3.742729819 |
| **A_55_P2034740** | Exosc2 | -0.170537098 | -0.340446573 | -0.000627623 | 17.4020489 | -2.469280545 | 0.049394452 | 0.594255995 | -3.743181408 |
| **A_55_P2043833** | Srgap2 | 0.77292059 | 0.002610672 | 1.543230508 | 9.973799953 | 2.468530799 | 0.049444116 | 0.594255995 | -3.743683382 |

| | | | | | | | | |
|---|---|---|---|---|---|---|---|---|
| **A_55_P2011943** | Gramd1a | 0.191554392 | 0.000640679 | 0.382468105 | 11.81072171 | 2.468448969 | 0.04944954 | 0.594255995 | -3.743738174 |
| **A_55_P2009582** | Nkg7 | 0.129461934 | 0.000432523 | 0.258491346 | 9.61741964 | 2.468439797 | 0.049450148 | 0.594255995 | -3.743744316 |
| **A_52_P257812** | Lpl | -0.171703165 | -0.342848709 | -0.000557621 | 8.314920856 | -2.468208642 | 0.049465473 | 0.594255995 | -3.7438991 |
| **A_51_P427401** | Fam90a1b | -0.23835656 | -0.475948147 | -0.000764973 | 10.33688985 | -2.46811399 | 0.049471749 | 0.594255995 | -3.743962483 |
| **A_66_P108178** | Lin52 | 0.25787518 | 0.000764848 | 0.514985512 | 9.494833657 | 2.467511456 | 0.049511724 | 0.594255995 | -3.744365994 |
| **A_55_P2122429** | Cym | 0.426562177 | 0.001251522 | 0.851872831 | 8.738982839 | 2.467432291 | 0.049516978 | 0.594255995 | -3.744419014 |
| **A_51_P214343** | Sephs1 | -0.192017405 | -0.38347193 | -0.000562881 | 8.310460514 | -2.467425936 | 0.0495174 | 0.594255995 | -3.74442327 |
| **A_52_P8903** | Tmem251 | -0.229407262 | -0.458162915 | -0.00065161 | 10.40803367 | -2.467200765 | 0.049532349 | 0.594255995 | -3.744574082 |
| **A_55_P2065829** | Cfap300 | 0.463776033 | 0.001107899 | 0.926444167 | 14.53482953 | 2.466084057 | 0.049606559 | 0.594255995 | -3.745322128 |
| **A_55_P2140390** | NA | -0.377411945 | -0.754182164 | -0.000641726 | 12.43672489 | -2.464383182 | 0.049719815 | 0.594255995 | -3.746461849 |
| **A_66_P134728** | Psmg1 | 0.157395354 | 0.000256226 | 0.314534482 | 11.7584326 | 2.464204418 | 0.049731734 | 0.594255995 | -3.746581661 |
| **A_55_P1972097** | NA | -0.107226981 | -0.214284678 | -0.000169284 | 16.00299167 | -2.464083064 | 0.049739827 | 0.594255995 | -3.746662997 |
| **A_55_P2055217** | NA | -0.243365829 | -0.48634819 | -0.000383468 | 10.32770214 | -2.464075524 | 0.04974033 | 0.594255995 | -3.746668051 |
| **A_55_P1952182** | Vmn2r111 | 0.444213205 | 0.000624708 | 0.887801702 | 8.729053025 | 2.463657616 | 0.049768211 | 0.594255995 | -3.746948168 |
| **A_55_P2036615** | Lrif1 | 0.195951398 | 0.000180982 | 0.391721814 | 9.584595189 | 2.462467264 | 0.049847718 | 0.594255995 | -3.747746187 |
| **A_51_P490795** | Mxd1 | -0.148901268 | -0.297673384 | -0.000129153 | 9.241263918 | -2.462328669 | 0.049856983 | 0.594255995 | -3.747839116 |
| **A_51_P409988** | Arl13b | -0.138975911 | -0.277875031 | -7.68E-05 | 9.742427628 | -2.46155305 | 0.049908871 | 0.594255995 | -3.748359226 |
| **A_55_P2002782** | Vps26a | -0.222382158 | -0.444698386 | -6.59E-05 | 11.86851555 | -2.460922512 | 0.049951095 | 0.594255995 | -3.748782116 |
| **A_66_P112853** | Eif1a | 0.295739131 | 4.18E-05 | 0.591436458 | 12.89863803 | 2.460540719 | 0.04997668 | 0.594255995 | -3.749038206 |
| **A_51_P103209** | Ttc7 | 0.217037724 | 5.25E-06 | 0.4340702 | 9.8598755 | 2.460252408 | 0.04999601 | 0.594255995 | -3.749231607 |
| **A_55_P2275249** | Th | 0.122015645 | -7.17E-06 | 0.244038456 | 8.81437829 | 2.460048428 | 0.050009691 | 0.594255995 | -3.749368445 |
| **A_55_P2066548** | Baiap2l2 | 1.248403657 | -0.000594005 | 2.497401319 | 11.46529096 | 2.459022881 | 0.050078533 | 0.594255995 | -3.750056521 |
| **A_51_P189208** | 4933417G07Rik | -0.127557443 | -0.25517627 | 6.14E-05 | 8.844182205 | -2.459009575 | 0.050079426 | 0.594255995 | -3.750065449 |
| **A_55_P2050298** | NA | -0.179628144 | -0.359359864 | 0.000103576 | 14.09859149 | -2.458775149 | 0.050095177 | 0.594255995 | -3.750222757 |
| **A_66_P102581** | Ube2z | -0.27282726 | -0.545840506 | 0.000185985 | 12.3145195 | -2.458516953 | 0.050112531 | 0.594255995 | -3.750396025 |
| **A_55_P1952788** | NA | 0.18169878 | -0.000150036 | 0.363547596 | 9.041580616 | 2.458163112 | 0.050136323 | 0.594255995 | -3.750633492 |
| **A_51_P505868** | Lhfp | -0.127746856 | -0.255636092 | 0.000142379 | 8.775646237 | -2.457453982 | 0.050184042 | 0.594255995 | -3.760347133 |
| **A_55_P2082156** | Rab11fip3 | 0.194763037 | -0.000267958 | 0.389794032 | 10.5802632 | 2.456812792 | 0.05022723 | 0.594255995 | -3.751539884 |
| **A_52_P589169** | Aopep | -0.255146579 | -0.510697959 | 0.000404801 | 10.686815 | -2.456295897 | 0.050262074 | 0.594255995 | -3.751886918 |
| **A_51_P249537** | Wdr24 | -0.177572188 | -0.355428059 | 0.000283683 | 9.306026281 | -2.456268862 | 0.050263897 | 0.594255995 | -3.75190507 |
| **A_52_P214752** | NA | -0.200151959 | -0.400684953 | 0.000381035 | 10.20440991 | -2.455518275 | 0.050314543 | 0.594255995 | -3.752409075 |
| **A_52_P284495** | Ankrd33b | -0.246011829 | -0.492539604 | 0.000515946 | 9.61238662 | -2.455044092 | 0.050346566 | 0.594255995 | -3.752727523 |
| **A_55_P2048454** | Rps2 | -0.394022443 | -0.788873627 | 0.000828741 | 13.9118036 | -2.45502929 | 0.050347566 | 0.594255995 | -3.752737465 |
| **A_55_P2052062** | Cd200 | -0.113163512 | -0.226575203 | 0.000248179 | 9.535617888 | -2.454809272 | 0.050362432 | 0.594255995 | -3.752885235 |
| **A_66_P107734** | NA | 0.632717852 | -0.001391265 | 1.26682697 | 14.32950355 | 2.454795137 | 0.050363387 | 0.594255995 | -3.752894728 |
| **A_52_P288614** | Rars | -0.110161634 | -0.220594964 | 0.000271696 | 8.192467014 | -2.454140165 | 0.05040767 | 0.594255995 | -3.753334669 |
| **A_55_P1952882** | Cyp4f18 | 0.57915644 | -0.001443111 | 1.159755992 | 10.20273155 | 2.454077972 | 0.050411877 | 0.594255995 | -3.753376447 |
| **A_52_P173442** | Wscd2 | -0.320411688 | -0.641701328 | 0.000877952 | 10.64726795 | -2.45347022 | 0.050453008 | 0.594255995 | -3.753784733 |
| **A_52_P139588** | Edem3 | -0.172583897 | -0.345658588 | 0.000490795 | 10.67394685 | -2.45321645 | 0.050470193 | 0.594255995 | -3.753955232 |
| **A_55_P2197338** | Tnfsf13os | 0.241444545 | -0.000716037 | 0.483605128 | 9.7091213 | 2.452918445 | 0.050490382 | 0.594255995 | -3.754155463 |
| **A_51_P497563** | Vps18 | -0.289915827 | -0.580695751 | 0.000864097 | 10.30265177 | -2.452882069 | 0.050492847 | 0.594255995 | -3.754179904 |
| **A_55_P2011672** | NA | 0.764927927 | -0.002319205 | 1.53217506 | 14.43885475 | 2.452756336 | 0.050501367 | 0.594255995 | -3.75426439 |
| **A_55_P2069217** | Prr13 | -0.304136025 | -0.609195014 | 0.000922963 | 9.133863677 | -2.45274954 | 0.050501828 | 0.594255995 | -3.754268956 |
| **A_55_P1963579** | NA | 0.375584354 | -0.001145975 | 0.752314683 | 10.28859587 | 2.45270926 | 0.050504558 | 0.594255995 | -3.754296022 |
| **A_55_P1995537** | Mpo | -0.178973052 | -0.358508578 | 0.000562475 | 7.832171564 | -2.452485267 | 0.050519743 | 0.594255995 | -3.75444654 |
| **A_55_P1957173** | NA | 0.490506114 | -0.001572779 | 0.982585007 | 9.178161444 | 2.45232966 | 0.050530294 | 0.594255995 | -3.754551108 |
| **A_55_P2147250** | Caprin2 | -0.115515826 | -0.231430905 | 0.000399253 | 9.049111886 | -2.45171912 | 0.050571717 | 0.594255995 | -3.754961427 |
| **A_55_P1992045** | Msi2 | 0.179396783 | -0.000627637 | 0.359421203 | 10.89585714 | 2.451615704 | 0.050578736 | 0.594255995 | -3.755030934 |
| **A_51_P133422** | Nck2 | 0.146021612 | -0.000513013 | 0.292556236 | 8.304073119 | 2.451579854 | 0.05058117 | 0.594255995 | -3.755055029 |
| **A_55_P2020572** | Psme3 | -0.305720141 | -0.612585482 | 0.001145201 | 11.16487428 | -2.451011637 | 0.050619761 | 0.594255995 | -3.755436967 |

| | | | | | | | | |
|---|---|---|---|---|---|---|---|---|
| **A_52_P650855** | Myo1d | -0.350503769 | -0.702478439 | 0.001470902 | 13.98424293 | -2.449911769 | 0.050694548 | 0.594255995 | -3.7561764 |
| **A_55_P2013655** | Cd4 | 1.173748728 | -0.005046112 | 2.352543569 | 11.91440254 | 2.449661471 | 0.050711583 | 0.594255995 | -3.756344698 |
| **A_55_P2065303** | NA | 0.300155268 | -0.001296121 | 0.601606656 | 15.43217426 | 2.449615064 | 0.050714742 | 0.594255995 | -3.756375902 |
| **A_51_P231738** | Dnajc4 | 0.221538233 | -0.001000547 | 0.444077013 | 10.92837768 | 2.449131742 | 0.050747657 | 0.594255995 | -3.756700914 |
| **A_51_P110814** | Nudt9 | -0.142839824 | -0.286358114 | 0.000678465 | 13.27272707 | -2.448562649 | 0.050786442 | 0.594255995 | -3.766539103 |
| **A_51_P380986** | Epc1 | -0.173929434 | -0.348699499 | 0.000840631 | 9.71392378 | -2.448359567 | 0.05080029 | 0.594255995 | -3.757220239 |
| **A_55_P1999102** | Pi16 | 0.350963344 | -0.001716985 | 0.703643673 | 9.231873987 | 2.448215738 | 0.0508101 | 0.594255995 | -3.757316981 |
| **A_55_P2080592** | Spata31d1b | 0.980813604 | -0.004862625 | 1.966489833 | 10.28351641 | 2.448056072 | 0.050820992 | 0.594255995 | -3.757424378 |
| **A_55_P2029252** | NA | -0.190548976 | -0.382062336 | 0.000964384 | 9.189023345 | -2.44780437 | 0.050838168 | 0.594255995 | -3.75759369 |
| **A_55_P2092329** | Adrm1 | -0.355573821 | -0.713035248 | 0.001887607 | 12.52901924 | -2.447201645 | 0.050879323 | 0.594255995 | -3.757999163 |
| **A_55_P2011169** | NA | 0.497185779 | -0.002969592 | 0.997341151 | 8.817185403 | 2.445585912 | 0.050989822 | 0.594255995 | -3.759086385 |
| **A_55_P2138070** | NA | 0.441725852 | -0.002695206 | 0.88614691 | 13.35875024 | 2.445272991 | 0.051011252 | 0.594255995 | -3.759296994 |
| **A_55_P1955237** | NA | -0.186416773 | -0.373976424 | 0.001142877 | 16.35550476 | -2.445201957 | 0.051016118 | 0.594255995 | -3.759344804 |
| **A_52_P272945** | Ccnh | 0.163623182 | -0.001007914 | 0.328254278 | 8.235064554 | 2.445130975 | 0.051020981 | 0.594255995 | -3.759392581 |
| **A_55_P2382070** | 4930593A02Rik | 0.431630134 | -0.002700112 | 0.86596038 | 8.968378176 | 2.444898566 | 0.051036906 | 0.594255995 | -3.759549016 |
| **A_55_P2015632** | Gm5766 | -0.265321631 | -0.53234431 | 0.001701048 | 14.92291097 | -2.444520437 | 0.051062828 | 0.594255995 | -3.759803553 |
| **A_55_P2089677** | Synrg | 0.919234612 | -0.006127987 | 1.844597212 | 11.27047662 | 2.443900889 | 0.051105329 | 0.594255995 | -3.760220645 |
| **A_51_P374782** | Epn3 | 0.326092778 | -0.002254673 | 0.654440228 | 12.2035753 | 2.443299436 | 0.051146625 | 0.594255995 | -3.760625611 |
| **A_55_P1991985** | Ptprs | 0.279292979 | -0.001944721 | 0.56053068 | 11.13089564 | 2.443181007 | 0.051154761 | 0.594255995 | -3.760705357 |
| **A_55_P2041320** | NA | -0.235518092 | -0.472678753 | 0.00164257 | 15.55599767 | -2.443153666 | 0.051156639 | 0.594255995 | -3.760723768 |
| **A_55_P2000039** | Prf1 | 0.371114111 | -0.002689005 | 0.744917226 | 8.578043146 | 2.442495171 | 0.051201902 | 0.594255995 | -3.761167216 |
| **A_66_P122158** | Pisd-ps3 | -0.224432817 | -0.450503197 | 0.001637564 | 8.828697928 | -2.442372256 | 0.051210355 | 0.594255995 | -3.761249998 |
| **A_55_P2012669** | NA | -0.148082724 | -0.297253383 | 0.001087936 | 16.71978159 | -2.442250164 | 0.051218753 | 0.594255995 | -3.761332227 |
| **A_55_P2183498** | Spcs2 | 0.589146397 | -0.004745871 | 1.183038665 | 10.68490558 | 2.440533189 | 0.051337012 | 0.594255995 | -3.762488849 |
| **A_55_P1963576** | Gm11595 | 0.159890433 | -0.001313527 | 0.321094392 | 8.556022421 | 2.4401467 | 0.051363671 | 0.594255995 | -3.762749263 |
| **A_55_P2068451** | NA | 0.708080933 | -0.005904832 | 1.422066699 | 15.45499973 | 2.439846533 | 0.051384386 | 0.594255995 | -3.76295153 |
| **A_55_P2183777** | NA | 0.436597943 | -0.003703619 | 0.876899506 | 10.26705294 | 2.439498877 | 0.05140839 | 0.594255995 | -3.763185813 |
| **A_66_P115053** | Anp32b | -0.255532764 | -0.51329632 | 0.002230792 | 12.48412326 | -2.43890139 | 0.05144967 | 0.594255995 | -3.763588497 |
| **A_52_P740562** | Rps21 | 0.295757418 | -0.002631539 | 0.594146375 | 15.54067287 | 2.438496084 | 0.051477693 | 0.594255995 | -3.763861689 |
| **A_51_P492125** | Ciapin1 | -0.171967894 | -0.345540355 | 0.001604566 | 10.57733976 | -2.437450002 | 0.051550093 | 0.594255995 | -3.764566899 |
| **A_55_P1968523** | Btbd1 | -0.252905481 | -0.5082058 | 0.002394838 | 10.56963652 | -2.437115135 | 0.051573292 | 0.594255995 | -3.764792683 |
| **A_55_P2158448** | Gdf3 | 0.360473992 | -0.003423406 | 0.72437139 | 8.708106906 | 2.437048368 | 0.051577919 | 0.594255995 | -3.764837702 |
| **A_55_P2145109** | Rpn2 | -0.150124725 | -0.301678345 | 0.001428895 | 16.31541173 | -2.436997446 | 0.051581448 | 0.594255995 | -3.764872038 |
| **A_55_P2158384** | Aanat | 0.138887941 | -0.001328982 | 0.279104863 | 9.059185219 | 2.436875104 | 0.051589927 | 0.594255995 | -3.764954533 |
| **A_55_P2079485** | Ptges3 | -0.15779943 | -0.31711951 | 0.00152065 | 12.04989386 | -2.436711302 | 0.051601283 | 0.594255995 | -3.765064987 |
| **A_51_P335306** | Ppm1a | -0.124618396 | -0.250459713 | 0.001222921 | 9.14589741 | -2.43628486 | 0.051630859 | 0.594255995 | -3.765352564 |
| **A_66_P132062** | Gm5608 | 1.017958882 | -0.010006621 | 2.045924386 | 9.090530694 | 2.436244425 | 0.051633665 | 0.594255995 | -3.765379833 |
| **A_55_P2388197** | 1010001B22Rik | 0.349293135 | -0.003768345 | 0.702354614 | 10.2804202 | 2.433934441 | 0.051794195 | 0.594255995 | -3.76693808 |
| **A_66_P104390** | Arl3 | -0.243347091 | -0.489319561 | 0.00262538 | 9.120829295 | -2.433934115 | 0.051794217 | 0.594255995 | -3.766938299 |
| **A_52_P318438** | Nab1 | -0.192503341 | -0.387112671 | 0.00210599 | 9.740988534 | -2.433569618 | 0.051819596 | 0.594255995 | -3.76718425 |
| **A_55_P1980292** | Purb | -0.254741523 | -0.512287861 | 0.002804815 | 10.41779147 | -2.433400115 | 0.051831402 | 0.594255995 | -3.767298632 |
| **A_55_P2080308** | Gm12538 | -0.122624526 | -0.24661451 | 0.001365459 | 8.362017806 | -2.433099652 | 0.051852337 | 0.594255995 | -3.767501397 |
| **A_52_P368306** | Tmem100 | -0.311614465 | -0.626711686 | 0.003482755 | 9.476527239 | -2.433000512 | 0.051859247 | 0.594255995 | -3.767568304 |
| **A_55_P2008347** | Prelid1 | 0.169289754 | -0.00191626 | 0.340495767 | 15.04757626 | 2.432656678 | 0.051883218 | 0.594255995 | -3.767800358 |
| **A_51_P200517** | Rasgef1a | -0.19481343 | -0.391859458 | 0.002232597 | 8.951540893 | -2.432318105 | 0.051906833 | 0.594255995 | -3.768028879 |
| **A_55_P2020286** | Tmem179 | -0.453744078 | -0.912852471 | 0.005364314 | 8.54836768 | -2.43144753 | 0.051967609 | 0.594255995 | -3.768616554 |
| **A_52_P227445** | Parp2 | -0.126792642 | -0.255104609 | 0.001519324 | 8.372071409 | -2.431062113 | 0.051994539 | 0.594255995 | -3.768876762 |
| **A_52_P167189** | Isca1 | -0.204423512 | -0.411351018 | 0.002503995 | 11.38896226 | -2.430422532 | 0.052039261 | 0.594255995 | -3.769308614 |
| **A_55_P1955821** | Ccl21b | 0.917446396 | -0.01172334 | 1.846616131 | 12.04198024 | 2.429152644 | 0.052128176 | 0.594255995 | -3.770166232 |
| **A_66_P132493** | Siglece | 0.707101037 | -0.009036044 | 1.423238119 | 10.85541382 | 2.429150796 | 0.052128306 | 0.594255995 | -3.770167481 |

| | | | | | | | | |
|---|---|---|---|---|---|---|---|---|
| **A_55_P1960231** | Txn2 | 0.241905796 | -0.003113811 | 0.486925404 | 13.03143625 | 2.428927755 | 0.052143939 | 0.594255995 | -3.770318136 |
| **A_55_P2183110** | Fmr1 | -0.139455662 | -0.280711275 | 0.001799951 | 8.577037866 | -2.428843877 | 0.05214982 | 0.594255995 | -3.770374794 |
| **A_55_P2403453** | BC037032 | 0.132673895 | -0.00173148 | 0.26707927 | 10.08411828 | 2.428499421 | 0.052173976 | 0.594255995 | -3.770607479 |
| **A_51_P160482** | Vsig2 | -0.123684507 | -0.248994519 | 0.001625504 | 7.882805928 | -2.428279633 | 0.052189396 | 0.594255995 | -3.770755958 |
| **A_55_P2021413** | Celf4 | 0.630496735 | -0.008392521 | 1.269385991 | 9.969459562 | 2.427875541 | 0.052217758 | 0.594255995 | -3.771028962 |
| **A_51_P127681** | Clic4 | -0.350982743 | -0.706640041 | 0.004674554 | 14.889533 | -2.427857558 | 0.052219021 | 0.594255995 | -3.789334268 |
| **A_51_P321734** | Bhmt2 | -0.142223639 | -0.286341706 | 0.001894429 | 8.413885926 | -2.427853728 | 0.05221929 | 0.594255995 | -3.7710437 |
| **A_55_P2423586** | Eya1 | -0.296330987 | -0.596613696 | 0.003951722 | 14.75815514 | -2.427816764 | 0.052221885 | 0.594255995 | -3.771068674 |
| **A_55_P2177513** | Chrna2 | 0.360777822 | -0.004834849 | 0.726390492 | 8.642712968 | 2.427659409 | 0.052232935 | 0.594255995 | -3.771174991 |
| **A_55_P2038247** | Tyro3 | -0.248813187 | -0.500993538 | 0.003367165 | 12.14056795 | -2.427343899 | 0.052255098 | 0.594255995 | -3.771388177 |
| **A_51_P288876** | Tmem45a | 0.166816338 | -0.002331683 | 0.335964359 | 8.294485209 | 2.426279481 | 0.052329942 | 0.594255995 | -3.772107496 |
| **A_55_P2099479** | H2afz | 0.222878693 | -0.003174989 | 0.448932375 | 12.0307642 | 2.42563879 | 0.052375046 | 0.594255995 | -3.772540546 |
| **A_55_P2070331** | NA | -0.109480031 | -0.220520025 | 0.001559963 | 10.38238666 | -2.4256305 | 0.05237563 | 0.594255995 | -3.77254615 |
| **A_55_P2138222** | NA | 0.163783379 | -0.002358253 | 0.329925011 | 9.415816059 | 2.425272365 | 0.052400861 | 0.594255995 | -3.772788244 |
| **A_55_P2067221** | Iqsec3 | 2.010325397 | -0.02918621 | 4.049837005 | 13.29190332 | 2.424986588 | 0.052421003 | 0.594255995 | -3.772981438 |
| **A_55_P2155593** | NA | -0.139515587 | -0.28112584 | 0.002094667 | 9.084667946 | -2.423802296 | 0.052504562 | 0.594255995 | -3.773782186 |
| **A_52_P343641** | Siah1a | -0.113681105 | -0.229070762 | 0.001708552 | 8.072721267 | -2.423765328 | 0.052507172 | 0.594255995 | -3.773807185 |
| **A_55_P2062891** | Atf2 | -0.182182952 | -0.367113565 | 0.002747661 | 11.16211016 | -2.423639872 | 0.052516033 | 0.594255995 | -3.773892024 |
| **A_66_P115453** | Coro1c | -0.153764317 | -0.309919953 | 0.002391319 | 12.28405177 | -2.422518288 | 0.052595314 | 0.594255995 | -3.774650586 |
| **A_55_P2129523** | NA | 0.549945484 | -0.008688838 | 1.108579806 | 9.196308325 | 2.42192778 | 0.052637107 | 0.594255995 | -3.77505004 |
| **A_52_P503208** | NA | -0.276677646 | -0.557733471 | 0.004378179 | 10.91519898 | -2.421868976 | 0.05264127 | 0.594255995 | -3.775089821 |
| **A_55_P1999785** | NA | -0.131133631 | -0.26437205 | 0.002104789 | 10.6144045 | -2.421328853 | 0.05267953 | 0.594255995 | -3.775455241 |
| **A_55_P1993544** | NA | 0.128840951 | -0.002073386 | 0.259755288 | 8.810151283 | 2.421229049 | 0.052686603 | 0.594255995 | -3.775522768 |
| **A_52_P26953** | Ergic2 | -0.178857742 | -0.360599214 | 0.00288373 | 10.00959537 | -2.421156513 | 0.052691744 | 0.594255995 | -3.775571846 |
| **A_51_P108935** | Iars2 | -0.166286121 | -0.335254852 | 0.00268261 | 9.417164804 | -2.421133978 | 0.052693342 | 0.594255995 | -3.775587093 |
| **A_55_P2082979** | Irak2 | 0.404364701 | -0.006533239 | 0.815262641 | 13.11983371 | 2.421076076 | 0.052697446 | 0.594255995 | -3.775626271 |
| **A_55_P2121760** | NA | 0.36170251 | -0.005874481 | 0.729279502 | 8.618561721 | 2.420875011 | 0.052711701 | 0.594255995 | -3.775762319 |
| **A_52_P636830** | G3bp2 | -0.263925588 | -0.532156402 | 0.004305226 | 10.00112128 | -2.4207057 | 0.052723708 | 0.594255995 | -3.775876887 |
| **A_55_P2147402** | NA | 0.29162921 | -0.004801344 | 0.588059765 | 8.446584439 | 2.420344683 | 0.05274932 | 0.594255995 | -3.776121189 |
| **A_55_P1973486** | Atp6v1f | 0.342862377 | -0.005676801 | 0.691401554 | 14.85610026 | 2.420122738 | 0.052765072 | 0.594255995 | -3.776271389 |
| **A_52_P86928** | Cog2 | -0.125694361 | -0.253479878 | 0.002091155 | 9.026162118 | -2.419932916 | 0.052778549 | 0.594255995 | -3.776399856 |
| **A_55_P2181251** | Ccdc9b | -0.29045677 | -0.585747756 | 0.004834217 | 11.72527538 | -2.419917026 | 0.052779677 | 0.594255995 | -3.776410611 |
| **A_55_P2027107** | Rgs3 | 0.414332792 | -0.006942522 | 0.835608105 | 9.433411704 | 2.419649488 | 0.052798677 | 0.594255995 | -3.776591684 |
| **A_51_P494006** | Scaf8 | -0.224941778 | -0.453675389 | 0.003791833 | 11.35857463 | -2.419409053 | 0.052815759 | 0.594255995 | -3.776754422 |
| **A_65_P17700** | Apobec3 | 0.746694278 | -0.012614797 | 1.506003354 | 10.76147818 | 2.419320447 | 0.052822055 | 0.594255995 | -3.776814397 |
| **A_66_P107174** | NA | 0.297293172 | -0.005082872 | 0.599669216 | 8.706628735 | 2.418837636 | 0.052856378 | 0.594255995 | -3.777141219 |
| **A_55_P1990121** | Aqp5 | 0.105064529 | -0.00180597 | 0.211935027 | 7.715669513 | 2.418618916 | 0.052871934 | 0.594255995 | -3.777289286 |
| **A_51_P307220** | Ankrd54 | -0.127676332 | -0.257572866 | 0.002220203 | 8.380494396 | -2.418143083 | 0.052905794 | 0.594255995 | -3.777611433 |
| **A_55_P2084278** | Ighmbp2 | 0.230330714 | -0.004007918 | 0.464669345 | 8.449336577 | 2.418115983 | 0.052907723 | 0.594255995 | -3.777629781 |
| **A_55_P2152074** | Ctps2 | -0.17525457 | -0.353576809 | 0.003067669 | 10.62606383 | -2.417870337 | 0.052925213 | 0.594255995 | -3.777796101 |
| **A_55_P2182106** | Rpl39 | 0.21154278 | -0.003813067 | 0.426898627 | 15.20614071 | 2.416633005 | 0.053013403 | 0.594255995 | -3.778634 |
| **A_52_P144263** | Rbms1 | -0.152069202 | -0.306891466 | 0.002753063 | 10.13141863 | -2.416445551 | 0.053026777 | 0.594255995 | -3.778760959 |
| **A_55_P2102454** | NA | 0.2802328 | -0.005106273 | 0.565571872 | 10.0287389 | 2.416166638 | 0.053046682 | 0.594255995 | -3.778949872 |
| **A_55_P2011440** | Ralgapb | -0.205393624 | -0.414546826 | 0.003759577 | 11.35341455 | -2.415970373 | 0.053060695 | 0.594255995 | -3.779082813 |
| **A_55_P2034081** | Hrh1 | 0.298778638 | -0.005603454 | 0.603160731 | 9.776691227 | 2.414902542 | 0.053136999 | 0.594255995 | -3.77980621 |
| **A_55_P2024322** | Rps27 | 0.240389205 | -0.004533003 | 0.485311413 | 16.38187835 | 2.414659833 | 0.053154358 | 0.594255995 | -3.779970655 |
| **A_55_P2089655** | Eif2s2 | -0.278753636 | -0.562767184 | 0.005259912 | 13.86531329 | -2.414630307 | 0.053156471 | 0.594255995 | -3.779990661 |
| **A_55_P1954768** | NA | 0.514295247 | -0.009717804 | 1.038308299 | 9.456497556 | 2.41456872 | 0.053160877 | 0.594255995 | -3.78003239 |
| **A_55_P2076278** | NA | -0.138888532 | -0.280408494 | 0.002631431 | 16.60921596 | -2.414447936 | 0.053169519 | 0.594255995 | -3.780114231 |
| **A_55_P2030501** | Ptpa | 0.614485353 | -0.011700429 | 1.240671136 | 14.22735933 | 2.414223627 | 0.053185573 | 0.594255995 | -3.780266223 |

| | | | | | | | | | |
|---|---|---|---|---|---|---|---|---|---|
| **A_55_P2014450** | B430105G09Rik | -0.269205764 | -0.543567739 | 0.005156211 | 8.541559642 | -2.413957378 | 0.053204634 | 0.594255995 | -3.780446645 |
| **A_55_P2158112** | Rps27 | 0.173115921 | -0.003340959 | 0.349572801 | 16.63436953 | 2.413612676 | 0.053229324 | 0.594255995 | -3.780680244 |
| **A_52_P570700** | Dpp8 | -0.201787498 | -0.407526424 | 0.003951428 | 8.801619847 | -2.41294237 | 0.053277369 | 0.594255995 | -3.781134549 |
| **A_55_P2129029** | Smarcb1 | 0.31491413 | -0.006243817 | 0.636072077 | 11.8591614 | 2.412362884 | 0.053318941 | 0.594255995 | -3.781527353 |
| **A_52_P167511** | NA | -0.151605183 | -0.306217222 | 0.003006856 | 9.079008346 | -2.412347698 | 0.053320031 | 0.594255995 | -3.781537647 |
| **A_66_P124755** | Fut8 | -0.295401624 | -0.596719559 | 0.005916311 | 11.23997347 | -2.41188757 | 0.053353066 | 0.594255995 | -3.781849581 |
| **A_55_P1956063** | Pbld1 | 0.182368038 | -0.003666876 | 0.368402951 | 17.08812806 | 2.41170082 | 0.05336648 | 0.594255995 | -3.781976192 |
| **A_55_P2001593** | Rnf180 | 0.27182153 | -0.005538239 | 0.549181299 | 9.082274727 | 2.4110685 | 0.053411924 | 0.594255995 | -3.782404926 |
| **A_51_P327295** | Akr1a1 | -0.178959793 | -0.361658017 | 0.003738432 | 13.59278679 | -2.409851631 | 0.053499494 | 0.594255995 | -3.783230165 |
| **A_66_P117682** | NA | -0.14250738 | -0.288012325 | 0.002997564 | 9.137582297 | -2.4095102 | 0.053524092 | 0.594255995 | -3.78346175 |
| **A_55_P2158404** | Cmpk2 | -0.189930547 | -0.383881727 | 0.004020634 | 8.637966184 | -2.409192784 | 0.05354697 | 0.594255995 | -3.783677062 |
| **A_55_P2017714** | Kat5 | 0.270175348 | -0.005785888 | 0.546136584 | 11.94250756 | 2.408611757 | 0.053588875 | 0.594255995 | -3.784071225 |
| **A_52_P748958** | Cfl1 | 0.136148509 | -0.002937095 | 0.275234113 | 13.1600028 | 2.408240588 | 0.053615663 | 0.594255995 | -3.784323049 |
| **A_55_P2139625** | Gtf2a2 | -0.134857109 | -0.272638357 | 0.002924139 | 10.50583491 | -2.4079801 | 0.053634471 | 0.594255995 | -3.784499791 |
| **A_55_P2033160** | Pomgnt1 | 0.316858336 | -0.006894686 | 0.640611357 | 13.10581916 | 2.407800329 | 0.053647455 | 0.594255995 | -3.784621772 |
| **A_55_P2100120** | Nme1 | 0.30136748 | -0.00656734 | 0.6093023 | 11.33980207 | 2.407724267 | 0.05365295 | 0.594255995 | -3.784673385 |
| **A_55_P2033864** | Plekha8 | 0.202626159 | -0.004415685 | 0.409668004 | 10.05772627 | 2.407723143 | 0.053653031 | 0.594255995 | -3.784674147 |
| **A_51_P185869** | Hbq1a | 0.162826966 | -0.003561979 | 0.329215911 | 8.000641797 | 2.407526222 | 0.053667259 | 0.594255995 | -3.784807773 |
| **A_55_P2094100** | BC085271 | -0.12931208 | -0.261462194 | 0.002838035 | 9.131394503 | -2.407358201 | 0.053679402 | 0.594255995 | -3.784921793 |
| **A_55_P2152049** | Xrcc3 | 0.275133184 | -0.00604698 | 0.556313348 | 8.612916393 | 2.407284712 | 0.053684714 | 0.594255995 | -3.784971664 |
| **A_55_P2010843** | Fahd1 | -0.151813206 | -0.307019634 | 0.003393222 | 8.422389305 | -2.406406607 | 0.05374823 | 0.594255995 | -3.785567623 |
| **A_55_P2025293** | Ss18 | -0.259541119 | -0.52488653 | 0.005804291 | 12.28780386 | -2.406377485 | 0.053750338 | 0.594255995 | -3.785587389 |
| **A_51_P199567** | Med26 | -0.181718828 | -0.367504562 | 0.004066906 | 10.36189732 | -2.406338547 | 0.053753157 | 0.594255995 | -3.785613819 |
| **A_55_P2116604** | NA | 0.198051041 | -0.004452728 | 0.400554809 | 12.78466922 | 2.406097279 | 0.053770624 | 0.594255995 | -3.785777586 |
| **A_55_P2035717** | Pgap2 | 0.189681273 | -0.004273564 | 0.383636111 | 10.83029554 | 2.405985487 | 0.053778719 | 0.594255995 | -3.785853472 |
| **A_55_P2047800** | NA | 0.248634171 | -0.005628383 | 0.502896724 | 11.84515677 | 2.405733821 | 0.053796949 | 0.594255995 | -3.78602431 |
| **A_55_P1998159** | Deaf1 | -0.110936372 | -0.224392861 | 0.002520118 | 8.339642724 | -2.40554662 | 0.053810513 | 0.594255995 | -3.786151393 |
| **A_51_P297694** | Prpf6 | -0.23865485 | -0.482790247 | 0.005480546 | 11.96072263 | -2.404964541 | 0.053852711 | 0.594255995 | -3.786546577 |
| **A_51_P513541** | Rfk | -0.214361853 | -0.433655392 | 0.004931686 | 11.3064801 | -2.404865708 | 0.05385988 | 0.594255995 | -3.786613681 |
| **A_55_P2004137** | Gnai2 | 0.559645097 | -0.012924393 | 1.132214588 | 14.71584047 | 2.404659916 | 0.053874809 | 0.594255995 | -3.786753412 |
| **A_51_P181451** | C1qa | -0.176059218 | -0.356191293 | 0.004072857 | 8.279018711 | -2.404566978 | 0.053881553 | 0.594255995 | -3.786816518 |
| **A_55_P2102579** | NA | -0.232306972 | -0.469988885 | 0.005374942 | 13.99979026 | -2.40455808 | 0.053882199 | 0.594255995 | -3.786822559 |
| **A_55_P2073915** | Nos1 | 0.222093714 | -0.005150991 | 0.44933842 | 8.633684844 | 2.404427337 | 0.053891687 | 0.594255995 | -3.786911337 |
| **A_55_P2055742** | Adtrp | -0.152558991 | -0.308658863 | 0.003540881 | 8.908633026 | -2.404387285 | 0.053894595 | 0.594255995 | -3.786938534 |
| **A_55_P2387284** | NA | 0.234797802 | -0.005475443 | 0.475071048 | 14.55018522 | 2.404129049 | 0.053913342 | 0.594255995 | -3.787113892 |
| **A_55_P2006768** | NA | 0.220400893 | -0.005163661 | 0.445965447 | 13.36858235 | 2.403873775 | 0.053931882 | 0.594255995 | -3.787287248 |
| **A_55_P2105694** | Prr29 | 0.25277557 | -0.00592845 | 0.51147959 | 9.011547663 | 2.403815238 | 0.053936134 | 0.594255995 | -3.787327002 |
| **A_55_P2024591** | Atxn7l1 | 0.132286081 | -0.003105127 | 0.267677289 | 9.765514757 | 2.403769669 | 0.053939444 | 0.594255995 | -3.787357949 |
| **A_55_P2016034** | Nlrc5 | 0.196493941 | -0.004686475 | 0.397674357 | 8.829646454 | 2.402883001 | 0.0540039 | 0.594255995 | -3.787960169 |
| **A_55_P2401403** | NA | -0.190234399 | -0.385046782 | 0.004577985 | 9.379779421 | -2.402379716 | 0.054040523 | 0.594255995 | -3.788302048 |
| **A_51_P150763** | Ropn1l | 0.158471199 | -0.003821867 | 0.320764264 | 9.921303168 | 2.402257406 | 0.054049427 | 0.594255995 | -3.788385138 |
| **A_55_P2142863** | Parp9 | -0.232120583 | -0.469892609 | 0.005651443 | 10.65909997 | -2.401718243 | 0.054088696 | 0.594255995 | -3.788751438 |
| **A_52_P684378** | Gpx1 | -0.121136534 | -0.245231007 | 0.002957939 | 14.56227128 | -2.401551293 | 0.054100861 | 0.594255995 | -3.78886487 |
| **A_55_P2143522** | Fstl3 | -0.199637861 | -0.404162954 | 0.004887232 | 12.76396615 | -2.401405341 | 0.054111499 | 0.594255995 | -3.788964039 |
| **A_51_P411019** | Tmem127 | -0.167521227 | -0.339154858 | 0.004112404 | 12.81310827 | -2.401245794 | 0.05412313 | 0.594255995 | -3.789072448 |
| **A_55_P2070799** | Tyw1 | 0.176534219 | -0.004340129 | 0.357408567 | 9.880903978 | 2.401159912 | 0.054129392 | 0.594255995 | -3.789130805 |
| **A_55_P2017729** | Gm2138 | 0.21116331 | -0.005230444 | 0.427557064 | 8.219293914 | 2.400727693 | 0.054160919 | 0.594255995 | -3.789424514 |
| **A_55_P2167447** | Cnnm3 | 0.443554265 | -0.011139981 | 0.898248512 | 13.71955032 | 2.399918336 | 0.054220006 | 0.594255995 | -3.789974574 |
| **A_52_P365342** | Lrrn2 | 0.241783793 | -0.006078945 | 0.48964653 | 8.582311568 | 2.399855576 | 0.05422459 | 0.594255995 | -3.790017231 |
| **A_55_P2155438** | Rps27 | 0.126150669 | -0.003174561 | 0.2554759 | 16.84223587 | 2.399802271 | 0.054228485 | 0.594255995 | -3.790053462 |

| | | | | | | | | |
|---|---|---|---|---|---|---|---|---|
| **A_55_P2176688** | NA | 0.472780043 | -0.011912403 | 0.95747249 | 14.67402926 | 2.399728159 | 0.054233899 | 0.594255995 | -3.790103836 |
| **A_51_P159896** | Dnajc1 | -0.201146271 | -0.407409028 | 0.005116486 | 11.85538283 | -2.399166183 | 0.054274977 | 0.594255995 | -3.790485841 |
| **A_55_P2059164** | H3f3b | -0.405608209 | -0.82157263 | 0.010356211 | 14.03901176 | -2.398941816 | 0.054291387 | 0.594255995 | -3.790638367 |
| **A_55_P2109122** | H2bc4 | -0.136674589 | -0.276927348 | 0.00357817 | 16.02292294 | -2.397427737 | 0.054402258 | 0.594255995 | -3.791667838 |
| **A_51_P428505** | Krtcap2 | -0.180882107 | -0.366507593 | 0.004743379 | 14.55253819 | -2.397326399 | 0.054409687 | 0.594255995 | -3.791736753 |
| **A_51_P195875** | Notch4 | -0.146363038 | -0.296564257 | 0.003838182 | 9.285571884 | -2.397326131 | 0.054409707 | 0.594255995 | -3.791736935 |
| **A_55_P2069730** | NA | 0.270852756 | -0.007190298 | 0.54889581 | 8.582183728 | 2.396571395 | 0.05446507 | 0.594255995 | -3.792250238 |
| **A_55_P2101441** | Gpbar1 | 0.194131593 | -0.005209859 | 0.393473044 | 8.381919213 | 2.395894908 | 0.054514745 | 0.594255995 | -3.792710393 |
| **A_52_P229709** | Ube2d3 | -0.174549635 | -0.353787067 | 0.004687797 | 11.49075393 | -2.395848737 | 0.054518137 | 0.594255995 | -3.792741801 |
| **A_55_P2038187** | BC051019 | 0.338799222 | -0.009118858 | 0.686717302 | 11.3404886 | 2.395711788 | 0.054528199 | 0.594255995 | -3.792834964 |
| **A_51_P234462** | Gpn1 | -0.151055673 | -0.306194071 | 0.004082724 | 9.421371071 | -2.395448852 | 0.054547524 | 0.594255995 | -3.793013841 |
| **A_55_P2051124** | Scaf1 | 0.947833774 | -0.026171856 | 1.921839404 | 14.56401999 | 2.39408671 | 0.054647754 | 0.594255995 | -3.79394067 |
| **A_52_P162957** | Frat2 | -0.250188876 | -0.50729967 | 0.006921917 | 8.687488148 | -2.393959785 | 0.054657103 | 0.594255995 | -3.794027046 |
| **A_55_P1956938** | Rpl24 | 0.156075173 | -0.004319304 | 0.31646965 | 16.5220674 | 2.393941754 | 0.054658431 | 0.594255995 | -3.794039317 |
| **A_52_P93933** | Mcl1 | -0.174864077 | -0.354585365 | 0.004857211 | 11.82739634 | -2.393702865 | 0.054676033 | 0.594255995 | -3.794201895 |
| **A_66_P119418** | Rb1cc1 | -0.14736855 | -0.298839692 | 0.004102591 | 9.502092193 | -2.393558659 | 0.054686661 | 0.594255995 | -3.794300038 |
| **A_55_P2057291** | Kansl3 | -0.383628434 | -0.778126574 | 0.010869707 | 12.05949516 | -2.392406595 | 0.054771647 | 0.594255995 | -3.79508422 |
| **A_55_P2150299** | NA | 0.463313241 | -0.013466628 | 0.940093109 | 10.42678587 | 2.390704866 | 0.054897436 | 0.594255995 | -3.796242891 |
| **A_55_P2061134** | NA | 0.723218618 | -0.021038085 | 1.467475322 | 16.25220911 | 2.390650042 | 0.054901493 | 0.594255995 | -3.796280227 |
| **A_55_P2142222** | Serpina3h | 0.570561673 | -0.016886823 | 1.158010169 | 8.449546091 | 2.389472087 | 0.05498875 | 0.594255995 | -3.797082522 |
| **A_51_P262721** | 0610009L18Rik | -0.137377856 | -0.278822145 | 0.004066433 | 10.21372262 | -2.389463941 | 0.054989354 | 0.594255995 | -3.797088071 |
| **A_51_P417854** | Rab20 | -0.156886104 | -0.318427372 | 0.004655163 | 8.343109547 | -2.389297097 | 0.055001725 | 0.594255995 | -3.797201723 |
| **A_51_P450924** | Pole4 | 0.181209689 | -0.005412439 | 0.367831817 | 10.39808656 | 2.388842079 | 0.055035478 | 0.594255995 | -3.797511696 |
| **A_55_P1980677** | Fam24b | 0.130740798 | -0.003919878 | 0.265401474 | 9.794994455 | 2.388578414 | 0.055055046 | 0.594255995 | -3.797691327 |
| **A_55_P2142724** | NA | 0.194359365 | -0.005901618 | 0.394620348 | 8.648848977 | 2.387691922 | 0.055120892 | 0.594255995 | -3.798295351 |
| **A_55_P2057792** | NA | -0.225789479 | -0.458509361 | 0.006930403 | 11.54642763 | -2.386928316 | 0.055177678 | 0.594255995 | -3.798815735 |
| **A_55_P1975922** | Olfr629 | 0.676577612 | -0.020793157 | 1.373948381 | 10.65368109 | 2.386838566 | 0.055184356 | 0.594255995 | -3.798876903 |
| **A_52_P400857** | Zfp595 | 0.119066058 | -0.003662741 | 0.241794858 | 8.588071039 | 2.386770451 | 0.055189425 | 0.594255995 | -3.798923327 |
| **A_55_P1960456** | Larp1 | -0.229332036 | -0.465723222 | 0.00705915 | 12.45680899 | -2.386726255 | 0.055192714 | 0.594255995 | -3.79895345 |
| **A_51_P250551** | Dhx38 | -0.125130563 | -0.254113798 | 0.003852673 | 8.528190762 | -2.386708029 | 0.05519407 | 0.594255995 | -3.798965872 |
| **A_55_P2143075** | Ubc | -0.314731891 | -0.639210211 | 0.009746429 | 16.88402589 | -2.386295538 | 0.05522478 | 0.594255995 | -3.799247025 |
| **A_52_P423247** | Pde4b | -0.133215278 | -0.270569016 | 0.00413846 | 7.871879693 | -2.386067453 | 0.055241769 | 0.594255995 | -3.799402497 |
| **A_55_P2100899** | Arnt | 0.157688107 | -0.00491026 | 0.320286473 | 8.737297196 | 2.385898278 | 0.055254373 | 0.594255995 | -3.799517818 |
| **A_55_P2114402** | Notum | 0.862948326 | -0.02695018 | 1.752846832 | 12.12849774 | 2.385687065 | 0.055270113 | 0.594255995 | -3.799661801 |
| **A_55_P2057519** | Rpl17 | 0.754892258 | -0.023775262 | 1.533559778 | 14.91968425 | 2.385075189 | 0.05531574 | 0.594255995 | -3.800078951 |
| **A_51_P229911** | Adcy4 | -0.199654233 | -0.405615825 | 0.006307358 | 9.394312317 | -2.384852078 | 0.055332387 | 0.594255995 | -3.800231072 |
| **A_66_P101703** | Krtap5-5 | 0.455930091 | -0.014422828 | 0.92628301 | 9.017081162 | 2.384753943 | 0.05533971 | 0.594255995 | -3.800297984 |
| **A_51_P406466** | Ankrd35 | -0.214405613 | -0.435600739 | 0.006789512 | 9.460381498 | -2.384678092 | 0.055345372 | 0.594255995 | -3.800349703 |
| **A_55_P2069392** | NA | -0.206049619 | -0.418703719 | 0.006604481 | 12.49243026 | -2.383785746 | 0.055412021 | 0.594255995 | -3.80095821 |
| **A_51_P247157** | Jpt1 | -0.115855246 | -0.235448005 | 0.003737513 | 9.273182586 | -2.383306956 | 0.055447816 | 0.594255995 | -3.801284752 |
| **A_51_P453351** | Lpgat1 | -0.22446556 | -0.456193179 | 0.007262058 | 11.87167134 | -2.383093498 | 0.055463782 | 0.594255995 | -3.801430344 |
| **A_55_P2098771** | Taf13 | -0.171831306 | -0.349226145 | 0.005563533 | 10.51665634 | -2.38303529 | 0.055468137 | 0.594255995 | -3.801470047 |
| **A_55_P2026785** | Dcaf1 | -0.296072632 | -0.601826633 | 0.009681369 | 9.518749793 | -2.38229357 | 0.055523659 | 0.594255995 | -3.801976004 |
| **A_55_P1973046** | Cttnbp2nl | -0.147457041 | -0.299738347 | 0.004824264 | 11.36627537 | -2.382254119 | 0.055526614 | 0.594255995 | -3.802002917 |
| **A_55_P2002466** | Zfp747 | 0.124608545 | -0.004101523 | 0.253318613 | 9.87800403 | 2.381795497 | 0.055560975 | 0.594255995 | -3.802315803 |
| **A_55_P2120254** | Avpi1 | 0.365968952 | -0.012067049 | 0.744004954 | 12.40822712 | 2.381662644 | 0.055570933 | 0.594255995 | -3.802406444 |
| **A_51_P114910** | Cstb | -0.21161828 | -0.430245489 | 0.00700893 | 15.18805861 | -2.381322038 | 0.055596472 | 0.594255995 | -3.802638842 |
| **A_66_P130813** | Samd4 | -0.184208154 | -0.374519475 | 0.006103166 | 9.25389846 | -2.381296049 | 0.055598421 | 0.594255995 | -3.802656575 |
| **A_55_P2164998** | NA | 0.365096576 | -0.012101956 | 0.742295108 | 12.52442025 | 2.381260615 | 0.055601079 | 0.594255995 | -3.802680752 |
| **A_51_P467346** | Muc1 | -0.158403258 | -0.322086067 | 0.005279551 | 8.24500531 | -2.380839961 | 0.055632639 | 0.594255995 | -3.802967793 |

| | | | | | | | | | |
|---|---|---|---|---|---|---|---|---|---|
| **A_55_P2134852** | Cnot2 | -0.131887912 | -0.268206373 | 0.004430549 | 10.66359568 | -2.380233052 | 0.055678207 | 0.594255995 | -3.803381972 |
| **A_55_P1996998** | Polr2a | 0.198474066 | -0.006706599 | 0.403654731 | 9.271312474 | 2.379778279 | 0.055712378 | 0.594255995 | -3.803692362 |
| **A_55_P2109505** | NA | -0.212226517 | -0.431684913 | 0.007231878 | 15.27932201 | -2.379121438 | 0.055761771 | 0.594255995 | -3.804140717 |
| **A_55_P2147842** | Zglp1 | 0.543335832 | -0.018656532 | 1.105328196 | 10.50631154 | 2.378521577 | 0.055806919 | 0.594255995 | -3.804550232 |
| **A_55_P1952901** | Gm833 | 0.19191479 | -0.006601107 | 0.390430688 | 8.408806027 | 2.378385875 | 0.055817138 | 0.594255995 | -3.80464288 |
| **A_55_P2118309** | NA | 0.130401472 | -0.004511748 | 0.265314692 | 8.19884912 | 2.37791951 | 0.055852272 | 0.594255995 | -3.804961302 |
| **A_66_P126603** | NA | 0.449601987 | -0.015600474 | 0.914804449 | 13.59333184 | 2.377690821 | 0.055869509 | 0.594255995 | -3.805117457 |
| **A_55_P2000548** | Cfl1 | 0.549102906 | -0.019192246 | 1.117398057 | 14.78748895 | 2.377108223 | 0.055913446 | 0.594255995 | -3.805515303 |
| **A_51_P301994** | Trmt10c | -0.18844478 | -0.383477136 | 0.006587575 | 8.709113703 | -2.377095387 | 0.055914414 | 0.594255995 | -3.805524069 |
| **A_55_P1954860** | NA | 0.508838197 | -0.017793272 | 1.035469665 | 13.3833182 | 2.377070495 | 0.055916293 | 0.594255995 | -3.805541069 |
| **A_55_P1968841** | NA | 0.387250295 | -0.013631162 | 0.788131751 | 12.5112805 | 2.376539037 | 0.055956407 | 0.594255995 | -3.805904036 |
| **A_52_P358349** | Mtdh | -0.195007134 | -0.396932072 | 0.006917803 | 11.75184739 | -2.375908472 | 0.056004041 | 0.594255995 | -3.806334741 |
| **A_52_P654965** | Eif3j1 | -0.21045482 | -0.42845455 | 0.007544909 | 12.08649485 | -2.375046328 | 0.056069238 | 0.594255995 | -3.806923717 |
| **A_55_P2133115** | NA | 0.311665405 | -0.011235305 | 0.634566115 | 14.03694501 | 2.374590691 | 0.056103727 | 0.594255995 | -3.807235028 |
| **A_55_P2087785** | NA | 0.287685429 | -0.010386427 | 0.585757285 | 8.581583286 | 2.374466559 | 0.056113127 | 0.594255995 | -3.807319846 |
| **A_55_P1961014** | Selenbp1 | 0.93756616 | -0.033915757 | 1.909048076 | 10.92894412 | 2.374304227 | 0.056125421 | 0.594255995 | -3.807430769 |
| **A_55_P2085060** | Vgf | 0.127983085 | -0.004639884 | 0.260606053 | 10.14303611 | 2.374121784 | 0.056139243 | 0.594255995 | -3.807555437 |
| **A_55_P2003338** | Clint1 | -0.212036732 | -0.431828746 | 0.007755282 | 11.05992597 | -2.37338589 | 0.056195029 | 0.594255995 | -3.808058342 |
| **A_55_P2075013** | Tpt1 | -0.16449506 | -0.335021225 | 0.006031105 | 16.62198558 | -2.373181744 | 0.056210515 | 0.594255995 | -3.808197868 |
| **A_51_P253697** | Rpain | 0.214722135 | -0.007936458 | 0.437380729 | 9.397218915 | 2.372501634 | 0.056262139 | 0.594255995 | -3.808662738 |
| **A_51_P434670** | Qk | -0.142979578 | -0.291246587 | 0.00528743 | 11.70301686 | -2.372458643 | 0.056265404 | 0.594255995 | -3.808692125 |
| **A_66_P127262** | NA | 0.950633382 | -0.035195448 | 1.936462212 | 10.11467209 | 2.372360636 | 0.056272847 | 0.594255995 | -3.808759121 |
| **A_55_P2104653** | Vma21 | -0.122724341 | -0.250036838 | 0.004588155 | 8.372632629 | -2.371531179 | 0.056335888 | 0.594255995 | -3.809326177 |
| **A_55_P2041988** | NA | -0.371374394 | -0.756665454 | 0.013916667 | 8.546540527 | -2.371331041 | 0.056351109 | 0.594255995 | -3.809463015 |
| **A_55_P2052470** | Uqcrh | 0.187888927 | -0.007078468 | 0.382856322 | 14.48462927 | 2.37087338 | 0.056385934 | 0.594255995 | -3.809775948 |
| **A_55_P2112474** | NA | 0.210935369 | -0.007982142 | 0.429852879 | 16.63665479 | 2.370489682 | 0.056415148 | 0.594255995 | -3.81003833 |
| **A_55_P1959828** | Slc35g1 | 0.160879042 | -0.006159827 | 0.327917911 | 8.341395069 | 2.369469347 | 0.056492911 | 0.594255995 | -3.810736159 |
| **A_55_P2009604** | NA | 0.144212512 | -0.005527923 | 0.293952948 | 8.519237744 | 2.369370702 | 0.056500436 | 0.594255995 | -3.810803632 |
| **A_55_P1962876** | Adam11 | 0.330636464 | -0.01288601 | 0.674158939 | 9.201489957 | 2.367907627 | 0.056612154 | 0.594255995 | -3.811804536 |
| **A_55_P2056896** | Kif2a | 0.163568951 | -0.006375489 | 0.333513391 | 10.13014798 | 2.367898441 | 0.056612856 | 0.594255995 | -3.811810821 |
| **A_55_P2022074** | Klf10 | 0.112761116 | -0.004427065 | 0.229949297 | 11.23242206 | 2.367253224 | 0.056662198 | 0.594255995 | -3.812252315 |
| **A_52_P141583** | Macroh2a1 | -0.293349811 | -0.598337198 | 0.011637576 | 9.573037664 | -2.36631794 | 0.056733803 | 0.594255995 | -3.812892394 |
| **A_51_P167763** | Kcnk13 | 0.128027023 | -0.005090036 | 0.261144082 | 8.520646427 | 2.366121792 | 0.056748832 | 0.594255995 | -3.813026646 |
| **A_55_P2073526** | H2-Ke6 | 0.184880317 | -0.007462718 | 0.377223351 | 11.02102947 | 2.364739885 | 0.056854834 | 0.594255995 | -3.813972639 |
| **A_51_P416825** | Tspan15 | 0.163529617 | -0.00660832 | 0.333667554 | 8.065973123 | 2.364636677 | 0.056862759 | 0.594255995 | -3.814043302 |
| **A_51_P183630** | Bcor | 0.178671269 | -0.007279754 | 0.364622292 | 10.34016876 | 2.363879387 | 0.056920945 | 0.594255995 | -3.814561833 |
| **A_55_P1993813** | NA | -0.174470684 | -0.35605384 | 0.007112472 | 15.80444168 | -2.36382905 | 0.056924814 | 0.594255995 | -3.814596302 |
| **A_55_P2115846** | Gm10836 | -0.191681071 | -0.391222172 | 0.007860031 | 9.730814131 | -2.363284599 | 0.056966688 | 0.594255995 | -3.814969152 |
| **A_55_P2122220** | Kcnk4 | 0.124914815 | -0.005143345 | 0.254972976 | 8.778344184 | 2.362900899 | 0.056996218 | 0.594255995 | -3.81523194 |
| **A_55_P1996374** | Prmt9 | -0.142143161 | -0.290152837 | 0.005866516 | 11.19688693 | -2.362680628 | 0.057013178 | 0.594255995 | -3.815382809 |
| **A_55_P2138549** | Cc2d1a | 0.222259264 | -0.009254175 | 0.453772702 | 10.61405131 | 2.361852812 | 0.057076962 | 0.594255995 | -3.81594986 |
| **A_51_P306066** | Eif4g2 | -0.153884383 | -0.314235804 | 0.006467038 | 11.80519625 | -2.360972339 | 0.057144886 | 0.594255995 | -3.816553084 |
| **A_55_P1972923** | Polr1d | -0.179313223 | -0.366165962 | 0.007539516 | 12.88492062 | -2.360924028 | 0.057148616 | 0.594255995 | -3.816586186 |
| **A_55_P2008512** | B4galt7 | 0.171095042 | -0.007215831 | 0.349405915 | 9.322667665 | 2.360634568 | 0.057170966 | 0.594255995 | -3.816784525 |
| **A_51_P223720** | Rtcb | -0.11965336 | -0.244384741 | 0.005078021 | 10.73239257 | -2.360034392 | 0.057217337 | 0.594255995 | -3.817195803 |
| **A_55_P2092611** | Slc45a3 | 0.523251174 | -0.02245735 | 1.068959699 | 10.27788016 | 2.358949464 | 0.057301262 | 0.594255995 | -3.817939391 |
| **A_55_P2159480** | Fam160a1 | -0.121594316 | -0.248425914 | 0.005237282 | 9.859340936 | -2.358603679 | 0.057328037 | 0.594255995 | -3.81817642 |
| **A_55_P1997320** | Zim1 | -0.361674441 | -0.738980771 | 0.015631888 | 11.15009958 | -2.358266557 | 0.057354154 | 0.594255995 | -3.818407526 |
| **A_55_P2016193** | Prxl2c | -0.17308606 | -0.353654451 | 0.007482332 | 9.60505522 | -2.35824827 | 0.057355571 | 0.594255995 | -3.818420062 |
| **A_55_P2055213** | Abcf1 | -0.144158555 | -0.294551642 | 0.006234532 | 8.283048003 | -2.358205838 | 0.05735886 | 0.594255995 | -3.818449152 |

| | | | | | | | | | |
|---|---|---|---|---|---|---|---|---|---|
| **A_55_P2167299** | NA | -0.127655935 | -0.260849579 | 0.00553771 | 15.20200758 | -2.357906994 | 0.057382024 | 0.594255995 | -3.818654033 |
| **A_52_P43848** | Sco1 | -0.170079974 | -0.347579732 | 0.007419784 | 8.433927311 | -2.357352768 | 0.057425009 | 0.594255995 | -3.819034032 |
| **A_51_P219918** | Tmem125 | -0.246617539 | -0.504020653 | 0.010785574 | 12.62374714 | -2.357107165 | 0.057444069 | 0.594255995 | -3.81920244 |
| **A_51_P152747** | Nrip1 | -0.218897596 | -0.447393594 | 0.009598402 | 10.11325471 | -2.356847911 | 0.057464195 | 0.594255995 | -3.819380218 |
| **A_65_P13527** | Osbpl8 | -0.109726445 | -0.224286151 | 0.004833261 | 10.79156011 | -2.356397656 | 0.057499166 | 0.594255995 | -3.819688991 |
| **A_55_P1957900** | Dlg1 | -0.199988093 | -0.40879275 | 0.008816563 | 9.242161995 | -2.356313783 | 0.057505683 | 0.594255995 | -3.819746513 |
| **A_55_P2051778** | Sult2b1 | 0.142170638 | -0.006300027 | 0.290641303 | 8.561138127 | 2.355800027 | 0.057545619 | 0.594255995 | -3.820098875 |
| **A_55_P2163857** | Trim12c | -0.280942343 | -0.574359057 | 0.01247437 | 8.718600759 | -2.355599833 | 0.057561188 | 0.594255995 | -3.820236189 |
| **A_51_P184197** | Smarcd2 | -0.12755971 | -0.260789734 | 0.005670314 | 9.315266641 | -2.355486288 | 0.057570021 | 0.594255995 | -3.820314072 |
| **A_55_P2095909** | Lamtor3 | -0.201622458 | -0.41226282 | 0.009017904 | 8.393824484 | -2.354867503 | 0.05761818 | 0.594255995 | -3.820738544 |
| **A_55_P1983544** | Tmed5 | -0.170671064 | -0.348977314 | 0.007635187 | 10.53160116 | -2.354845884 | 0.057619864 | 0.594255995 | -3.820753375 |
| **A_55_P2002079** | NA | -0.293120057 | -0.599487577 | 0.013247462 | 14.60312582 | -2.353813123 | 0.05770034 | 0.594255995 | -3.821461945 |
| **A_55_P2091055** | NA | 0.505303028 | -0.023114967 | 1.033721022 | 15.01561097 | 2.352574933 | 0.057796979 | 0.594255995 | -3.822311652 |
| **A_55_P1992299** | Cep295 | 0.361506088 | -0.016649367 | 0.739661544 | 8.562684291 | 2.351875936 | 0.057851609 | 0.594255995 | -3.822791431 |
| **A_51_P430383** | Dnajc30 | 0.133810712 | -0.006185683 | 0.273807108 | 10.480851 | 2.351490299 | 0.057881772 | 0.594255995 | -3.823056154 |
| **A_55_P1976978** | Zfp707 | 0.297172731 | -0.013783641 | 0.608129103 | 8.998042682 | 2.351140906 | 0.057909114 | 0.594255995 | -3.823296015 |
| **A_51_P157524** | E2f4 | 0.354251981 | -0.016475036 | 0.724978998 | 9.346603505 | 2.350862419 | 0.057930917 | 0.594255995 | -3.82348721 |
| **A_55_P2043009** | Psmg2 | 0.286636091 | -0.013364983 | 0.586637166 | 11.29446724 | 2.350591848 | 0.057952109 | 0.594255995 | -3.823672981 |
| **A_55_P2072801** | Lypd5 | 0.114233786 | -0.005330905 | 0.233798478 | 8.634877766 | 2.350502885 | 0.057959079 | 0.594255995 | -3.823734064 |
| **A_55_P2096535** | Icmt | 0.348435223 | -0.016393818 | 0.713264265 | 9.73763807 | 2.349642624 | 0.058026518 | 0.594255995 | -3.824324786 |
| **A_52_P473953** | Ctdspl | -0.223247342 | -0.457006312 | 0.010511628 | 11.00872771 | -2.349563439 | 0.05803273 | 0.594255995 | -3.824379166 |
| **A_55_P2110887** | NA | -0.241885003 | -0.495175321 | 0.011405314 | 10.22957026 | -2.349413813 | 0.05804447 | 0.594255995 | -3.824481923 |
| **A_55_P2015767** | Yipf6 | 0.131734328 | -0.006263273 | 0.269731928 | 9.111694269 | 2.348532568 | 0.058113662 | 0.594255995 | -3.825087187 |
| **A_52_P447196** | Col4a6 | 0.172504053 | -0.008216561 | 0.353224667 | 8.219145967 | 2.348338905 | 0.05812888 | 0.594255995 | -3.825220215 |
| **A_55_P2106394** | Knstrn | 1.174402342 | -0.05609582 | 2.404900503 | 14.36135498 | 2.348037899 | 0.05815254 | 0.594255995 | -3.825426987 |
| **A_51_P329356** | Mcts2 | 0.170635323 | -0.008158733 | 0.349429378 | 10.22868111 | 2.347929353 | 0.058161074 | 0.594255995 | -3.825501554 |
| **A_52_P193029** | Hadha | -0.181784271 | -0.372265739 | 0.008697196 | 9.068609999 | -2.347862924 | 0.058166298 | 0.594255995 | -3.825547189 |
| **A_55_P2115136** | Vsir | -0.118264787 | -0.242209249 | 0.005679675 | 9.358349653 | -2.347456169 | 0.058198295 | 0.594255995 | -3.825826633 |
| **A_52_P82765** | NA | 0.624654558 | -0.030193739 | 1.279502856 | 9.572078554 | 2.34675836 | 0.058253229 | 0.594255995 | -3.826306086 |
| **A_55_P2052924** | Selenow | 0.269138097 | -0.013065586 | 0.55134178 | 12.40588506 | 2.346289853 | 0.058290143 | 0.594255995 | -3.826628027 |
| **A_55_P2107528** | NA | 0.195121447 | -0.009507155 | 0.399750048 | 8.03950451 | 2.345891026 | 0.058321586 | 0.594255995 | -3.82690211 |
| **A_55_P2037797** | Tex44 | 0.277393339 | -0.01355377 | 0.568340449 | 8.947313908 | 2.345584833 | 0.058345737 | 0.594255995 | -3.827112547 |
| **A_55_P1974976** | Sprr2a2 | 0.232449979 | -0.011458637 | 0.476358594 | 16.25420424 | 2.344614965 | 0.058422307 | 0.594255995 | -3.827779195 |
| **A_55_P2019214** | NA | 0.121356793 | -0.006019102 | 0.248732688 | 8.255872654 | 2.343937375 | 0.058475865 | 0.594255995 | -3.828245018 |
| **A_55_P2158107** | Ddx31 | -0.198667558 | -0.407189051 | 0.009853936 | 9.429759151 | -2.343933517 | 0.05847617 | 0.594255995 | -3.82824767 |
| **A_55_P2103668** | Zfp316 | 0.206764759 | -0.010293814 | 0.423823332 | 9.967478384 | 2.343520403 | 0.058508848 | 0.594255995 | -3.828531705 |
| **A_52_P22331** | Akap10 | 0.212072103 | -0.010561369 | 0.434705576 | 8.725590244 | 2.343485367 | 0.058511621 | 0.594255995 | -3.828555795 |
| **A_55_P2086128** | Fa2h | -0.278296674 | -0.570524963 | 0.013931615 | 10.87371295 | -2.342906322 | 0.058557459 | 0.594255995 | -3.828953956 |
| **A_55_P2000454** | Tspan18 | 0.147026706 | -0.007399997 | 0.301453408 | 9.141393207 | 2.342302544 | 0.058605295 | 0.594255995 | -3.829369173 |
| **A_66_P131530** | Rpp30 | -0.111552214 | -0.228765768 | 0.005661341 | 8.166049347 | -2.341367145 | 0.058679487 | 0.594255995 | -3.830012544 |
| **A_55_P2131303** | NA | -0.205185927 | -0.420833831 | 0.010461977 | 16.28267747 | -2.340838721 | 0.058721442 | 0.594255995 | -3.830376048 |
| **A_55_P1955244** | NA | 0.60585857 | -0.031023763 | 1.242740902 | 13.3632929 | 2.340352185 | 0.058760099 | 0.594255995 | -3.830710771 |
| **A_55_P2142172** | Olfr1229 | 0.349048487 | -0.017902683 | 0.715999657 | 9.454734075 | 2.340165896 | 0.058774908 | 0.594255995 | -3.830838941 |
| **A_51_P224517** | Jade1 | -0.305627206 | -0.626940558 | 0.015686145 | 12.08031509 | -2.340089148 | 0.05878101 | 0.594255995 | -3.830891746 |
| **A_55_P2170094** | Lrrc27 | 0.110982358 | -0.005697638 | 0.227662355 | 9.253996342 | 2.340058446 | 0.058783451 | 0.594255995 | -3.83091287 |
| **A_52_P624149** | Hsp90b1 | -0.173590532 | -0.356096063 | 0.008914999 | 16.44368719 | -2.340017827 | 0.058786681 | 0.594255995 | -3.830940818 |
| **A_55_P1960735** | Gdf15 | 0.162544199 | -0.008394948 | 0.333483347 | 9.041592169 | 2.339371013 | 0.058838139 | 0.594255995 | -3.831385887 |
| **A_55_P2105152** | Trim59 | -0.243965213 | -0.500563887 | 0.012633461 | 9.579070069 | -2.339066991 | 0.058862342 | 0.594255995 | -3.831595102 |
| **A_55_P2065040** | NA | 0.338302934 | -0.017606312 | 0.69421218 | 14.46618988 | 2.33849075 | 0.058908245 | 0.594255995 | -3.831991682 |
| **A_52_P255804** | NA | -0.140669652 | -0.288706254 | 0.00736695 | 10.92963468 | -2.337762941 | 0.058966275 | 0.594255995 | -3.832492638 |

| | | | | | | | | | |
|---|---|---|---|---|---|---|---|---|---|
| **A_51_P453856** | Csnk1a1 | -0.276104182 | -0.566698468 | 0.014490104 | 12.60721637 | -2.337518609 | 0.05898577 | 0.594255995 | -3.832660829 |
| **A_55_P2108042** | Dgcr2 | 0.177212481 | -0.00932379 | 0.363748752 | 10.9716614 | 2.33722315 | 0.059009353 | 0.594255995 | -3.832864226 |
| **A_55_P2144566** | NA | 0.259231261 | -0.013668269 | 0.532130792 | 15.29117839 | 2.336973283 | 0.059029304 | 0.594255995 | -3.833036246 |
| **A_51_P468055** | Smyd1 | 0.325554649 | -0.017327955 | 0.668437252 | 9.993896945 | 2.335864322 | 0.059117939 | 0.594255995 | -3.833799808 |
| **A_51_P439403** | Padi1 | 0.18326199 | -0.009755873 | 0.376279853 | 8.496873464 | 2.335845196 | 0.059119469 | 0.594255995 | -3.833812978 |
| **A_51_P283876** | Bphl | 0.283402945 | -0.015100084 | 0.581905973 | 10.07816514 | 2.335741513 | 0.059127763 | 0.594255995 | -3.833884376 |
| **A_55_P1998209** | NA | 0.413921346 | -0.022082789 | 0.849925481 | 16.46577409 | 2.335588771 | 0.059139984 | 0.594255995 | -3.833989561 |
| **A_55_P2185667** | NA | 0.338264626 | -0.01818399 | 0.694713241 | 8.348697774 | 2.334687804 | 0.059212125 | 0.594255995 | -3.834610067 |
| **A_55_P1973448** | Ybx2 | 0.14607877 | -0.007892798 | 0.300050338 | 8.108860903 | 2.334079989 | 0.059260846 | 0.594255995 | -3.835028738 |
| **A_51_P290139** | Dlst | -0.266786073 | -0.54821149 | 0.014639344 | 11.25822657 | -2.33221723 | 0.05941042 | 0.594255995 | -3.83631214 |
| **A_55_P2080603** | Hspa8 | 0.388156799 | -0.021360447 | 0.797674044 | 16.686392 | 2.331869085 | 0.059438419 | 0.594255995 | -3.836552058 |
| **A_51_P279712** | Rell1 | -0.149029618 | -0.30627914 | 0.008219904 | 12.13234263 | -2.331591251 | 0.059460773 | 0.594255995 | -3.836743532 |
| **A_55_P2128499** | Mgl2 | 0.329776929 | -0.018214392 | 0.677768251 | 13.40379802 | 2.331422694 | 0.059474339 | 0.594255995 | -3.836859702 |
| **A_55_P2336943** | Dph6 | -0.175197429 | -0.360093002 | 0.009698144 | 8.91550318 | -2.331150851 | 0.059496225 | 0.594255995 | -3.837047064 |
| **A_55_P2149906** | NA | 0.2797018 | -0.015521651 | 0.574925251 | 14.96152647 | 2.330845952 | 0.059520782 | 0.594255995 | -3.837257222 |
| **A_55_P1973334** | Mapk1 | 0.253797131 | -0.014108282 | 0.521702543 | 12.95694929 | 2.330635633 | 0.059537728 | 0.594255995 | -3.837402196 |
| **A_52_P606774** | Stk4 | 0.186426986 | -0.010369683 | 0.383223654 | 10.81875352 | 2.330559522 | 0.059543861 | 0.594255995 | -3.837454661 |
| **A_55_P2168098** | Plrg1 | 0.193053725 | -0.010739239 | 0.396846688 | 11.06318642 | 2.330548598 | 0.059544742 | 0.594255995 | -3.837462191 |
| **A_55_P2081530** | Sh3bgrl3 | 0.19322273 | -0.010771101 | 0.397216561 | 13.96797633 | 2.33029199 | 0.059565426 | 0.594255995 | -3.837639083 |
| **A_55_P2125912** | Cct6a | -0.137122732 | -0.281911067 | 0.007665602 | 9.871496162 | -2.329941673 | 0.059593677 | 0.594255995 | -3.837880587 |
| **A_51_P452779** | Pygl | 0.222410092 | -0.012626276 | 0.457446461 | 12.9793335 | 2.328030065 | 0.059748081 | 0.594255995 | -3.839198718 |
| **A_55_P2142151** | Mapk1ip1 | -0.271321361 | -0.558097973 | 0.015455251 | 10.16915482 | -2.327605745 | 0.059782411 | 0.594255995 | -3.83949137 |
| **A_55_P1998461** | Rpl17 | 0.338297688 | -0.01935138 | 0.695946756 | 9.945401688 | 2.327078831 | 0.05982507 | 0.594255995 | -3.839854814 |
| **A_55_P1996509** | Rpl29 | -0.245533788 | -0.505171176 | 0.014103601 | 16.18820962 | -2.326554309 | 0.059867567 | 0.594255995 | -3.840216645 |
| **A_55_P1973279** | NA | 0.14097366 | -0.00811798 | 0.290065301 | 16.97418406 | 2.32623639 | 0.059893341 | 0.594255995 | -3.840435974 |
| **A_55_P2063490** | Csprs | 0.357697604 | -0.020693404 | 0.736088612 | 9.223124948 | 2.325650164 | 0.059940896 | 0.594255995 | -3.84084044 |
| **A_66_P102076** | Cep68 | -0.211110085 | -0.43443486 | 0.01221469 | 9.385607677 | -2.325633302 | 0.059942264 | 0.594255995 | -3.840852074 |
| **A_55_P2049582** | Apbb3 | 0.169579597 | -0.009882474 | 0.349041668 | 9.527967182 | 2.324716981 | 0.060016678 | 0.594255995 | -3.841484382 |
| **A_55_P2065113** | Gm11559 | 0.134621985 | -0.007869578 | 0.277113547 | 9.033334062 | 2.324320443 | 0.060048911 | 0.594255995 | -3.841758048 |
| **A_55_P1970642** | NA | 0.172609553 | -0.010137644 | 0.355356749 | 9.386645371 | 2.323717165 | 0.060097983 | 0.594255995 | -3.842174434 |
| **A_66_P124674** | Ubxn2a | -0.1356535 | -0.27928086 | 0.007973861 | 9.469240598 | -2.323608659 | 0.060106813 | 0.594255995 | -3.842249331 |
| **A_51_P125679** | Rec114 | -0.131859283 | -0.27149226 | 0.007773695 | 8.391047881 | -2.323228213 | 0.060137786 | 0.594255995 | -3.849022157 |
| **A_55_P2087281** | NA | 0.442415383 | -0.026084238 | 0.910915004 | 9.672207828 | 2.323218932 | 0.060138542 | 0.594255995 | -3.842518353 |
| **A_55_P2124831** | Brdt | 0.423698533 | -0.024988433 | 0.872385499 | 8.810788122 | 2.323178978 | 0.060141795 | 0.594255995 | -3.842545934 |
| **A_55_P1953136** | NA | -0.39154842 | -0.80626064 | 0.023163799 | 13.9312555 | -2.32277855 | 0.060174415 | 0.594255995 | -3.842822367 |
| **A_52_P520408** | Serp1 | 0.181538687 | -0.010881521 | 0.373958896 | 11.74955246 | 2.321066975 | 0.060314053 | 0.594255995 | -3.844004185 |
| **A_55_P1987993** | NA | -0.225621559 | -0.46479409 | 0.013550972 | 15.58915662 | -2.320803976 | 0.060335539 | 0.594255995 | -3.844185816 |
| **A_55_P1993493** | Efr3a | -0.207397951 | -0.427282679 | 0.012486777 | 10.6676268 | -2.32048389 | 0.0603617 | 0.594255995 | -3.844406885 |
| **A_66_P100440** | Vezf1 | -0.222291891 | -0.457981403 | 0.013397621 | 9.931425914 | -2.3203448 | 0.060373072 | 0.594255995 | -3.844502952 |
| **A_55_P1964682** | Rn28s1 | -0.176297121 | -0.363245234 | 0.010650991 | 16.72126629 | -2.320028388 | 0.06039895 | 0.594255995 | -3.844721504 |
| **A_51_P333099** | Mrps15 | -0.188957762 | -0.389358764 | 0.01144324 | 9.892744862 | -2.319711685 | 0.060424863 | 0.594255995 | -3.844940269 |
| **A_55_P1980883** | Ephb6 | 1.129667883 | -0.06855118 | 2.327886945 | 12.64914339 | 2.319443084 | 0.06044685 | 0.594255995 | -3.845125817 |
| **A_55_P2100715** | Auh | 0.142855767 | -0.008731277 | 0.294442811 | 8.515150677 | 2.318488028 | 0.060525094 | 0.594255995 | -3.845785646 |
| **A_55_P2048202** | Ss18 | -0.227957156 | -0.469889737 | 0.013975425 | 12.53295025 | -2.31807794 | 0.060558724 | 0.594255995 | -3.846069004 |
| **A_51_P476711** | Mtrex | -0.167510615 | -0.345299117 | 0.010277886 | 11.00840748 | -2.317970092 | 0.060567571 | 0.594255995 | -3.846143527 |
| **A_55_P2103756** | Hic2 | 0.276988008 | -0.017033821 | 0.571009837 | 9.657226334 | 2.317664425 | 0.060592654 | 0.594255995 | -3.846354753 |
| **A_55_P1977096** | Elk4 | 0.431484818 | -0.026581805 | 0.889551441 | 11.38188083 | 2.31742685 | 0.060612157 | 0.594255995 | -3.846518933 |
| **A_55_P2126448** | Snhg16 | 0.229527584 | -0.014144651 | 0.473199819 | 9.985969894 | 2.317383988 | 0.060615676 | 0.594255995 | -3.846548554 |
| **A_52_P278354** | Bmp7 | 0.252966191 | -0.015610526 | 0.521542907 | 8.742823726 | 2.317198742 | 0.060630889 | 0.594255995 | -3.846676578 |
| **A_55_P2082625** | NA | 0.3137525 | -0.019395932 | 0.646900932 | 9.65629586 | 2.316960257 | 0.06065048 | 0.594255995 | -3.846841402 |

| | | | | | | | | | |
|---|---|---|---|---|---|---|---|---|---|
| **A_55_P2161126** | NA | -0.19493403 | -0.401963311 | 0.012095251 | 8.768956633 | -2.316461313 | 0.060691487 | 0.594255995 | -3.847186261 |
| **A_55_P2010487** | Unk | -0.278161592 | -0.57364183 | 0.017318646 | 11.04380342 | -2.315996434 | 0.060729721 | 0.594255995 | -3.847507605 |
| **A_55_P2098354** | Shf | 0.206708089 | -0.012898705 | 0.426314882 | 9.157122866 | 2.315692364 | 0.060754743 | 0.594255995 | -3.847717806 |
| **A_55_P1974317** | Surf6 | -0.161532236 | -0.333173443 | 0.010108971 | 9.224681432 | -2.315297526 | 0.06078725 | 0.594255995 | -3.847990773 |
| **A_51_P465232** | Mcfd2 | -0.191596605 | -0.395218164 | 0.012024954 | 14.39473327 | -2.314905214 | 0.060819567 | 0.594255995 | -3.848262014 |
| **A_52_P539310** | Serp2 | 0.206157104 | -0.012939676 | 0.425253884 | 9.814068126 | 2.314895943 | 0.060820331 | 0.594255995 | -3.848268424 |
| **A_52_P522977** | Ssbp2 | 0.232708095 | -0.014613157 | 0.480029348 | 8.114636391 | 2.314830614 | 0.060825714 | 0.594255995 | -3.848313595 |
| **A_55_P2007663** | Alkbh8 | -0.140391362 | -0.28959921 | 0.008816486 | 8.982855385 | -2.314823504 | 0.0608263 | 0.594255995 | -3.848318511 |
| **A_51_P367162** | Psmb1 | -0.19815007 | -0.408748941 | 0.012448801 | 11.95397895 | -2.314767383 | 0.060830925 | 0.594255995 | -3.848357315 |
| **A_52_P561377** | Fam160b1 | -0.303778704 | -0.626756289 | 0.01919888 | 11.97500784 | -2.313950722 | 0.060898269 | 0.594255995 | -3.848922027 |
| **A_55_P2424988** | C78228 | 0.299947897 | -0.018994975 | 0.618890768 | 8.416032649 | 2.313673564 | 0.060921142 | 0.594255995 | -3.849113699 |
| **A_55_P2063426** | NA | 0.225159368 | -0.014303407 | 0.464622144 | 8.696815302 | 2.313242552 | 0.06095673 | 0.594255995 | -3.849411791 |
| **A_55_P2106459** | Zfhx3 | 0.264669728 | -0.016938755 | 0.546278212 | 10.23259409 | 2.31221227 | 0.061041887 | 0.594255995 | -3.850124442 |
| **A_51_P292877** | Snrnp200 | -0.119431158 | -0.246515171 | 0.007652855 | 9.747196068 | -2.312042891 | 0.061055898 | 0.594255995 | -3.850241616 |
| **A_55_P2182159** | Ywhaq | 0.118722932 | -0.007607783 | 0.245053647 | 12.95054562 | 2.312037233 | 0.061056367 | 0.594255995 | -3.85024553 |
| **A_55_P2032079** | Dbp | 0.125182328 | -0.008024468 | 0.258389124 | 8.963237261 | 2.311989226 | 0.061060339 | 0.594255995 | -3.850278741 |
| **A_51_P183102** | Rnf139 | -0.247776092 | -0.511453638 | 0.015901455 | 11.18217292 | -2.311827432 | 0.061073727 | 0.594255995 | -3.850390672 |
| **A_55_P2167050** | Paip2 | -0.22164785 | -0.457579425 | 0.014283725 | 12.03637578 | -2.311248372 | 0.061121669 | 0.594255995 | -3.850791302 |
| **A_55_P2038101** | Dock11 | -0.209303796 | -0.432226573 | 0.013618982 | 9.979684291 | -2.309892779 | 0.061234056 | 0.594255995 | -3.851729358 |
| **A_55_P1983588** | Pmepa1 | -0.199166736 | -0.411345338 | 0.013011866 | 11.04245825 | -2.309321434 | 0.061281489 | 0.594255995 | -3.852124795 |
| **A_55_P2094731** | Sox13 | 0.977783265 | -0.064102788 | 2.019669317 | 11.07944203 | 2.308827776 | 0.061322504 | 0.594255995 | -3.852466499 |
| **A_55_P2074080** | NA | -0.194234334 | -0.401206915 | 0.012738247 | 13.62410691 | -2.308778925 | 0.061326564 | 0.594255995 | -3.852500315 |
| **A_55_P2022434** | Gpi1 | 0.208990792 | -0.013753417 | 0.431735002 | 13.73543354 | 2.30828746 | 0.061367427 | 0.594255995 | -3.852840535 |
| **A_55_P2006662** | Zfp967 | -0.14823038 | -0.306250792 | 0.009790032 | 8.909210536 | -2.307773565 | 0.061410186 | 0.594255995 | -3.853196317 |
| **A_66_P121288** | Tceal6 | 0.113114744 | -0.007501929 | 0.233731416 | 8.128564513 | 2.307177652 | 0.061459808 | 0.594255995 | -3.853608925 |
| **A_51_P267740** | Zc3h15 | -0.120754341 | -0.249528232 | 0.00801955 | 10.37254596 | -2.306981415 | 0.061476158 | 0.594255995 | -3.853744808 |
| **A_52_P336768** | NA | -0.222082201 | -0.458960309 | 0.014795907 | 9.742127425 | -2.306524065 | 0.061514281 | 0.594255995 | -3.85406152 |
| **A_55_P1964228** | Elac1 | 0.175688223 | -0.011714615 | 0.363091061 | 9.367896385 | 2.306405416 | 0.061524175 | 0.594255995 | -3.854143688 |
| **A_51_P254895** | Cyp4a10 | 0.169845607 | -0.011346816 | 0.35103803 | 8.063949284 | 2.306128209 | 0.061547297 | 0.594255995 | -3.854335668 |
| **A_55_P2152976** | NA | -0.176045754 | -0.363884612 | 0.011793104 | 14.87013387 | -2.305734397 | 0.061580162 | 0.594255995 | -3.854608422 |
| **A_55_P1984314** | Trim43c | -0.290905121 | -0.601348236 | 0.019537994 | 9.385276767 | -2.30535864 | 0.061611537 | 0.594255995 | -3.85486869 |
| **A_66_P105791** | Prkca | -0.158162249 | -0.326966123 | 0.010641624 | 9.074697457 | -2.305099027 | 0.061633224 | 0.594255995 | -3.855048522 |
| **A_55_P1996941** | Ube2c | 0.266499991 | -0.018014361 | 0.551014344 | 8.942622386 | 2.304422905 | 0.061689742 | 0.594255995 | -3.855516907 |
| **A_55_P2269459** | Prl8a6 | -0.179293485 | -0.370740347 | 0.012153378 | 17.47995873 | -2.304015613 | 0.061723814 | 0.594255995 | -3.855799088 |
| **A_52_P427759** | NA | 0.297866984 | -0.020275998 | 0.616009966 | 13.00308164 | 2.303399055 | 0.06177543 | 0.594255995 | -3.856226295 |
| **A_51_P286737** | Ccl2 | -0.576672731 | -1.192620066 | 0.039274604 | 9.198785251 | -2.303323817 | 0.061781732 | 0.594255995 | -3.863931961 |
| **A_52_P31490** | Vamp3 | -0.180645249 | -0.373648177 | 0.01235768 | 8.916436617 | -2.302670558 | 0.061836475 | 0.594255995 | -3.856731128 |
| **A_51_P425080** | Hdgf | -0.173272027 | -0.358401655 | 0.011857601 | 12.23518592 | -2.30261692 | 0.061840972 | 0.594255995 | -3.8567683 |
| **A_55_P2029021** | NA | 0.409914685 | -0.02812531 | 0.847954679 | 15.95090636 | 2.30223088 | 0.061873349 | 0.594255995 | -3.857035848 |
| **A_55_P2027337** | NA | 0.570554633 | -0.039234195 | 1.180343462 | 12.20145513 | 2.301902561 | 0.061900899 | 0.594255995 | -3.857263408 |
| **A_55_P1978905** | NA | 0.475395728 | -0.03272066 | 0.983512116 | 8.701841429 | 2.301766343 | 0.061912332 | 0.594255995 | -3.857357825 |
| **A_55_P2173333** | Plcb4 | 0.243301126 | -0.016806002 | 0.503408254 | 9.798301171 | 2.301235301 | 0.061956928 | 0.594255995 | -3.857725932 |
| **A_55_P2031471** | Rufy4 | 0.36893257 | -0.025510111 | 0.763375251 | 10.41076313 | 2.301082864 | 0.061969736 | 0.594255995 | -3.857831605 |
| **A_52_P437850** | Fam13c | -0.267941933 | -0.554435375 | 0.018551508 | 8.465583873 | -2.300886338 | 0.061986252 | 0.594255995 | -3.857967846 |
| **A_55_P1991960** | Nwd1 | 0.156764738 | -0.01086865 | 0.324398126 | 8.156348069 | 2.300684255 | 0.06200324 | 0.594255995 | -3.858107944 |
| **A_55_P2010672** | Kif17 | 0.663359618 | -0.046016494 | 1.37273573 | 10.02207795 | 2.300602747 | 0.062010093 | 0.594255995 | -3.858164453 |
| **A_55_P2175275** | Hmbox1 | 0.568480656 | -0.039452078 | 1.17641339 | 10.89934355 | 2.300537547 | 0.062015575 | 0.594255995 | -3.858209656 |
| **A_55_P2021055** | Elp5 | -0.197464555 | -0.408640368 | 0.013711258 | 11.39694305 | -2.300457106 | 0.06202234 | 0.594255995 | -3.858265426 |
| **A_51_P459368** | Appbp2 | -0.282221862 | -0.584044172 | 0.019600448 | 12.63808881 | -2.300427114 | 0.062024863 | 0.594255995 | -3.858286221 |
| **A_55_P2060404** | Pus3 | -0.227771058 | -0.47141133 | 0.015869215 | 9.371705252 | -2.299951219 | 0.062064902 | 0.594255995 | -3.858616182 |

| | | | | | | | | | |
|---|---|---|---|---|---|---|---|---|---|
| **A_55_P2013534** | Slc25a19 | 0.101561283 | -0.007076401 | 0.210198967 | 10.1700677 | 2.299941778 | 0.062065696 | 0.594255995 | -3.858622728 |
| **A_55_P1982096** | Drap1 | 0.250547486 | -0.017488862 | 0.518583834 | 14.09317051 | 2.299670002 | 0.062088574 | 0.594255995 | -3.858811178 |
| **A_55_P2027323** | Golga7 | -0.112942146 | -0.233774443 | 0.007890151 | 12.9415723 | -2.299546352 | 0.062098986 | 0.594255995 | -3.858896919 |
| **A_55_P2050932** | Tlr7 | 1.003012091 | -0.070254364 | 2.076278546 | 9.564999245 | 2.299152486 | 0.062132163 | 0.594255995 | -3.859170049 |
| **A_66_P117022** | Atp6v1e1 | -0.287945384 | -0.596206136 | 0.020315368 | 12.34099774 | -2.298058341 | 0.062224425 | 0.594255995 | -3.859928899 |
| **A_55_P2135031** | NA | 0.464770392 | -0.032859214 | 0.962399997 | 8.939814074 | 2.297742762 | 0.062251062 | 0.594255995 | -3.860147799 |
| **A_51_P181565** | Hbegf | 0.157596472 | -0.011151359 | 0.326344303 | 10.69469021 | 2.29761604 | 0.062261761 | 0.594255995 | -3.860235703 |
| **A_55_P1983358** | Gm4593 | 0.252513142 | -0.017933163 | 0.522959446 | 8.388710119 | 2.297058719 | 0.062308841 | 0.594255995 | -3.860622328 |
| **A_55_P2113459** | Washc1 | 0.294933089 | -0.020974803 | 0.610840981 | 11.49656153 | 2.296847635 | 0.062326682 | 0.594255995 | -3.860768772 |
| **A_52_P679152** | Clk1 | -0.386621623 | -0.800811341 | 0.027568094 | 9.922487773 | -2.296444692 | 0.062360754 | 0.594255995 | -3.861048337 |
| **A_51_P422223** | 1700006J14Rik | 0.636240219 | -0.045390704 | 1.317871143 | 9.052617974 | 2.296365415 | 0.062367459 | 0.594255995 | -3.861103344 |
| **A_66_P128694** | NA | 0.199046119 | -0.014206651 | 0.41229889 | 8.578273002 | 2.296297724 | 0.062373186 | 0.594255995 | -3.86115031 |
| **A_51_P505795** | Tapbpl | -0.182284388 | -0.377598289 | 0.013029513 | 8.40270906 | -2.296071899 | 0.062392293 | 0.594255995 | -3.861307004 |
| **A_51_P261107** | Ogt | -0.289580707 | -0.599927984 | 0.020766569 | 9.459267025 | -2.295571633 | 0.062434644 | 0.594255995 | -3.861654147 |
| **A_55_P1955034** | Myo1c | -0.221361949 | -0.458641452 | 0.015917554 | 11.94241205 | -2.295154413 | 0.062469987 | 0.594255995 | -3.861943687 |
| **A_55_P2004201** | NA | -0.214942082 | -0.445373185 | 0.01548902 | 11.51807713 | -2.294824708 | 0.062497931 | 0.594255995 | -3.862172511 |
| **A_52_P358388** | Ppp4r2 | -0.158078646 | -0.327558124 | 0.011400831 | 8.930321445 | -2.294696514 | 0.0625088 | 0.594255995 | -3.862261484 |
| **A_55_P2103698** | Isg15 | -0.37363661 | -0.774271443 | 0.026998224 | 13.52762935 | -2.294403938 | 0.062533613 | 0.594255995 | -3.862464556 |
| **A_51_P270286** | Dync1li1 | -0.157797587 | -0.327004538 | 0.011409363 | 8.70277639 | -2.294305905 | 0.062541929 | 0.594255995 | -3.862532601 |
| **A_65_P16059** | Tgfbr3 | -0.271387344 | -0.562412081 | 0.019637394 | 13.31942817 | -2.294187169 | 0.062552003 | 0.594255995 | -3.862615018 |
| **A_55_P2008891** | Tmsb15b1 | 0.115482667 | -0.008363374 | 0.239328708 | 8.50206657 | 2.29405507 | 0.062563213 | 0.594255995 | -3.862706713 |
| **A_55_P1997751** | Prpf8 | -0.373686154 | -0.774474444 | 0.027102137 | 14.72512638 | -2.293829557 | 0.062582355 | 0.594255995 | -3.862863255 |
| **A_55_P2029726** | NA | -0.406466662 | -0.842616591 | 0.029683266 | 14.49529569 | -2.292758376 | 0.062673362 | 0.594255995 | -3.863606916 |
| **A_52_P127024** | Tnks | -0.123311956 | -0.25563831 | 0.009014397 | 8.967998191 | -2.292598512 | 0.062686956 | 0.594255995 | -3.863717913 |
| **A_55_P2021099** | Tmem181a | 0.193412595 | -0.016712067 | 0.403537257 | 10.81550363 | 2.590062442 | 0.062719095 | 0.594255995 | -4.041941494 |
| **A_55_P1980651** | Ubr5 | -0.276725771 | -0.573747166 | 0.020295623 | 13.11742802 | -2.292086679 | 0.062730499 | 0.594255995 | -3.864073312 |
| **A_55_P2078780** | Etv4 | 0.112121254 | -0.008255194 | 0.232497701 | 8.327977678 | 2.291477442 | 0.062782371 | 0.594255995 | -3.864496391 |
| **A_66_P122729** | Zmat2 | -0.277888672 | -0.576258699 | 0.020481355 | 10.93214774 | -2.29131508 | 0.062796202 | 0.594255995 | -3.864609149 |
| **A_51_P298666** | Scaf4 | -0.244348142 | -0.506750853 | 0.018054569 | 9.499012029 | -2.2909198 | 0.062829889 | 0.594255995 | -3.864883681 |
| **A_55_P2011425** | Atp10b | -0.168106894 | -0.348638323 | 0.012424535 | 8.394267005 | -2.290877497 | 0.062833495 | 0.594255995 | -3.864913062 |
| **A_55_P2068148** | Pias1 | -0.106986815 | -0.221895897 | 0.007922268 | 8.437192449 | -2.290577876 | 0.062859043 | 0.594255995 | -3.865121171 |
| **A_55_P1957413** | Lsp1 | 0.201959228 | -0.014973859 | 0.418892315 | 11.46403121 | 2.290377495 | 0.062876135 | 0.594255995 | -3.865260358 |
| **A_55_P2031786** | Fcsk | 0.176300423 | -0.013099442 | 0.365700287 | 11.19530947 | 2.290038865 | 0.062905031 | 0.594255995 | -3.865495584 |
| **A_65_P10180** | Rad23b | -0.216749433 | -0.449654963 | 0.016156096 | 11.43224427 | -2.289535251 | 0.062948031 | 0.594255995 | -3.865845443 |
| **A_55_P1996816** | NA | -0.242912127 | -0.503964578 | 0.018140325 | 11.98818288 | -2.289236086 | 0.062973589 | 0.594255995 | -3.866053287 |
| **A_55_P2112997** | Ddx18 | 0.146443469 | -0.010952739 | 0.303839677 | 10.7832933 | 2.288995328 | 0.062994165 | 0.594255995 | -3.866220561 |
| **A_52_P217474** | Ndufa6 | -0.213586278 | -0.44319405 | 0.016021494 | 12.19096036 | -2.288526398 | 0.063034261 | 0.594255995 | -3.866546387 |
| **A_55_P1954925** | Endov | 0.178775062 | -0.013414484 | 0.370964609 | 16.78353454 | 2.288475875 | 0.063038583 | 0.594255995 | -3.866581493 |
| **A_55_P1956687** | Rab37 | 0.125159993 | -0.009457956 | 0.259777942 | 8.065493738 | 2.28734526 | 0.063135374 | 0.594255995 | -3.8673672 |
| **A_52_P220485** | Uhrf2 | -0.156770864 | -0.325392495 | 0.011850767 | 9.478309971 | -2.287289988 | 0.06314011 | 0.594255995 | -3.867405614 |
| **A_51_P292016** | Uts2r | 0.15542209 | -0.011763323 | 0.322607503 | 8.7901146 | 2.287091418 | 0.063157127 | 0.594255995 | -3.867543627 |
| **A_55_P2145543** | Eef2k | -0.183120812 | -0.380114731 | 0.013873107 | 9.02716667 | -2.286936199 | 0.063170432 | 0.594255995 | -3.867651512 |
| **A_51_P256066** | Tiam2 | 0.262397188 | -0.019992337 | 0.544786713 | 9.14855937 | 2.286018583 | 0.063249148 | 0.594255995 | -3.868289366 |
| **A_55_P2024580** | Dtnbp1 | 0.226026583 | -0.017266963 | 0.469320128 | 12.4133683 | 2.285588778 | 0.063286053 | 0.594255995 | -3.868588169 |
| **A_55_P2004811** | NA | 0.131851821 | -0.010079884 | 0.273783527 | 13.63728808 | 2.285471848 | 0.063296097 | 0.594255995 | -3.868669464 |
| **A_51_P453948** | Drg1 | -0.10740585 | -0.223031079 | 0.008219379 | 10.68227039 | -2.285306695 | 0.063310286 | 0.594255995 | -3.868784288 |
| **A_55_P2169514** | Fgf23 | 0.465793675 | -0.035884888 | 0.967472238 | 12.2544125 | 2.284216194 | 0.06340406 | 0.594255995 | -3.869542558 |
| **A_55_P2286493** | 4933405E24Rik | 0.349879366 | -0.026968955 | 0.726727686 | 9.560931495 | 2.284130484 | 0.063411437 | 0.594255995 | -3.869602163 |
| **A_55_P2172299** | Timm17b | 0.299507405 | -0.023117081 | 0.622131892 | 12.52431376 | 2.283912186 | 0.063430228 | 0.594255995 | -3.869753975 |
| **A_51_P416509** | H1f1 | 0.146464125 | -0.011397017 | 0.304325268 | 8.171600273 | 2.282575672 | 0.063545404 | 0.594255995 | -3.870683566 |

| | | | | | | | | | |
|---|---|---|---|---|---|---|---|---|---|
| **A_55_P2145005** | NA | 0.285165699 | -0.022204148 | 0.592535547 | 9.389775994 | 2.282470574 | 0.063554471 | 0.594255995 | -3.870756676 |
| **A_51_P165475** | C1galt1 | -0.178946945 | -0.371843585 | 0.013949696 | 10.24641228 | -2.282279277 | 0.063570976 | 0.594255995 | -3.87088975 |
| **A_55_P1957747** | Inip | 0.227687089 | -0.017848763 | 0.473222941 | 9.557945295 | 2.281353859 | 0.063650887 | 0.594255995 | -3.871533579 |
| **A_55_P2145307** | Ints11 | 0.115963339 | -0.009120068 | 0.241046746 | 9.652556028 | 2.280815591 | 0.063697415 | 0.594255995 | -3.871908112 |
| **A_55_P2183549** | NA | 0.231440091 | -0.01822373 | 0.481103911 | 10.37579541 | 2.280615869 | 0.063714688 | 0.594255995 | -3.87204709 |
| **A_51_P483511** | Dock8 | -0.122181382 | -0.254013277 | 0.009650514 | 9.097750382 | -2.280098973 | 0.063759415 | 0.594255995 | -3.872406799 |
| **A_55_P2117472** | Acap2 | -0.141241733 | -0.293647866 | 0.0111644 | 10.48937797 | -2.279973274 | 0.063770296 | 0.594255995 | -3.872494278 |
| **A_55_P2095732** | Fbrsl1 | -0.218915466 | -0.455151487 | 0.017320555 | 9.488082061 | -2.279814383 | 0.063784054 | 0.594255995 | -3.87260486 |
| **A_55_P2157360** | Tagap1 | -0.125280399 | -0.260474247 | 0.00991345 | 8.219093522 | -2.279792698 | 0.063785932 | 0.594255995 | -3.872619952 |
| **A_55_P2108171** | Glud1 | -0.30037094 | -0.624619245 | 0.023877366 | 12.12625918 | -2.279026429 | 0.063852326 | 0.594255995 | -3.873153294 |
| **A_51_P124345** | Cpne5 | 0.376564274 | -0.030019242 | 0.783147791 | 9.035169512 | 2.278549724 | 0.063893666 | 0.594255995 | -3.873485129 |
| **A_51_P450812** | Cog4 | 0.136677702 | -0.010919888 | 0.284275291 | 11.00053166 | 2.27817754 | 0.063925962 | 0.594255995 | -3.873744227 |
| **A_51_P141860** | Fbxw11 | -0.351976515 | -0.737206769 | 0.033253739 | 13.41977869 | -2.570962897 | 0.063968774 | 0.594255995 | -3.95851216 |
| **A_55_P2128626** | Gm4349 | -0.153099046 | -0.318485856 | 0.012287764 | 8.229170335 | -2.277407654 | 0.063992822 | 0.594255995 | -3.874280244 |
| **A_52_P262219** | Fos | -0.225043293 | -0.46815471 | 0.018068124 | 10.23253853 | -2.27735053 | 0.063997786 | 0.594255995 | -3.874320018 |
| **A_55_P2028310** | Trappc1 | -0.11999928 | -0.249682109 | 0.009683549 | 8.224091506 | -2.276487801 | 0.064072801 | 0.594255995 | -3.874920771 |
| **A_51_P457492** | Tmem223 | 0.123625987 | -0.010106307 | 0.257358282 | 11.99387784 | 2.274273231 | 0.064265781 | 0.594255995 | -3.876463294 |
| **A_51_P391926** | Ubxn6 | 0.144867011 | -0.011846207 | 0.301580229 | 11.86041428 | 2.274222929 | 0.064270171 | 0.594255995 | -3.876498339 |
| **A_51_P270666** | Uspl1 | -0.315284017 | -0.656351969 | 0.025783935 | 11.37404441 | -2.274208115 | 0.064271465 | 0.594255995 | -3.87650866 |
| **A_51_P128065** | Prcc | 0.217501085 | -0.017796842 | 0.452799011 | 10.26696762 | 2.274115351 | 0.064279562 | 0.594255995 | -3.876573287 |
| **A_55_P2060707** | Il15ra | -0.213630205 | -0.444767487 | 0.017507077 | 11.85989162 | -2.27385003 | 0.064302728 | 0.594255995 | -3.87675814 |
| **A_55_P2136413** | NA | 0.138051762 | -0.011376477 | 0.287480002 | 11.49871798 | 2.272890106 | 0.064386615 | 0.594255995 | -3.877427003 |
| **A_55_P2123982** | Rhoh | 0.640456221 | -0.05279317 | 1.333705612 | 9.057399126 | 2.272841312 | 0.064390882 | 0.594255995 | -3.877461005 |
| **A_55_P1999246** | Gm10318 | 0.309358516 | -0.025528182 | 0.644245215 | 12.21733569 | 2.272654103 | 0.064407257 | 0.594255995 | -3.877591465 |
| **A_55_P2121437** | NA | -0.180628781 | -0.376173497 | 0.014915935 | 14.05234958 | -2.272532109 | 0.064417929 | 0.594255995 | -3.877676481 |
| **A_52_P198019** | Fuz | -0.123385113 | -0.256968821 | 0.010198596 | 8.26661131 | -2.272366761 | 0.064432398 | 0.594255995 | -3.877791713 |
| **A_55_P1996329** | Kat5 | -0.103914028 | -0.216427783 | 0.008599727 | 9.213992197 | -2.272153803 | 0.064451037 | 0.594255995 | -3.877940129 |
| **A_66_P114769** | NA | 0.197458626 | -0.01637232 | 0.411289572 | 10.68911509 | 2.271824174 | 0.064479899 | 0.594255995 | -3.878169868 |
| **A_52_P630632** | Rcbtb2 | -0.155737081 | -0.324399233 | 0.012925071 | 10.5772951 | -2.27166118 | 0.064494176 | 0.594255995 | -3.878283474 |
| **A_55_P2092310** | Phgdh | 0.647773669 | -0.053781616 | 1.349328954 | 11.58295527 | 2.271593163 | 0.064500135 | 0.594255995 | -3.878330883 |
| **A_52_P590392** | Rab10 | -0.156209442 | -0.325391792 | 0.012972909 | 12.04712245 | -2.271545227 | 0.064504335 | 0.594255995 | -3.878364295 |
| **A_55_P2027230** | NA | 0.440397715 | -0.036714245 | 0.917509675 | 8.939070128 | 2.270878596 | 0.064562769 | 0.594255995 | -3.878828977 |
| **A_55_P2142808** | NA | 0.455365727 | -0.038034741 | 0.948766195 | 16.28917138 | 2.270544127 | 0.064592108 | 0.594255995 | -3.879062143 |
| **A_52_P304323** | Isca1 | 0.227560474 | -0.019018765 | 0.474139713 | 10.11480812 | 2.27043715 | 0.064601495 | 0.594255995 | -3.879136722 |
| **A_55_P2031966** | Fbxo39 | 0.183028353 | -0.015300255 | 0.38135696 | 8.580862182 | 2.270398922 | 0.06460485 | 0.594255995 | -3.879163373 |
| **A_55_P1977699** | NA | 0.266387202 | -0.022437739 | 0.555212143 | 13.1936741 | 2.269069644 | 0.064721613 | 0.594255995 | -3.880090207 |
| **A_55_P2082361** | Ppt1 | -0.194084435 | -0.40455977 | 0.016390901 | 8.797664803 | -2.268603824 | 0.064762583 | 0.594255995 | -3.880415051 |
| **A_55_P2058494** | NA | 0.225797369 | -0.019153856 | 0.470748593 | 13.17070909 | 2.267819184 | 0.064831654 | 0.594255995 | -3.880962288 |
| **A_51_P449935** | Ftsj3 | -0.19176498 | -0.399803123 | 0.016273162 | 8.794096193 | -2.267751671 | 0.064837601 | 0.594255995 | -3.881009377 |
| **A_51_P454103** | Manba | 0.185826321 | -0.01584975 | 0.387502392 | 11.3081639 | 2.266846016 | 0.064917428 | 0.594255995 | -3.881641119 |
| **A_55_P2061135** | Rpl17 | 0.432345064 | -0.036967559 | 0.901657688 | 14.2544582 | 2.266404545 | 0.064956377 | 0.594255995 | -3.881949105 |
| **A_55_P1978151** | NA | 0.188929569 | -0.016157964 | 0.394017101 | 16.59646182 | 2.266364907 | 0.064959876 | 0.594255995 | -3.881976759 |
| **A_55_P1972842** | Klf3 | -0.172896489 | -0.360585149 | 0.014792172 | 11.29125088 | -2.266299496 | 0.064965649 | 0.594255995 | -3.882022394 |
| **A_51_P208031** | Cnot6l | -0.132000726 | -0.275302041 | 0.01130059 | 8.834126442 | -2.266184702 | 0.064975783 | 0.594255995 | -3.882102484 |
| **A_52_P652193** | Sar1a | -0.266142863 | -0.555170851 | 0.022885125 | 13.94656594 | -2.265395781 | 0.065045469 | 0.594255995 | -3.882652947 |
| **A_55_P2028570** | Sgsh | -0.184317388 | -0.384515781 | 0.015881005 | 15.08275197 | -2.265034823 | 0.065077379 | 0.594255995 | -3.882904827 |
| **A_55_P2163093** | Nxf7 | -0.101421447 | -0.211586713 | 0.008743818 | 8.466797635 | -2.264927373 | 0.065086881 | 0.594255995 | -3.882979811 |
| **A_55_P1971709** | Dnajc8 | -0.20882102 | -0.43564694 | 0.018004901 | 10.07741354 | -2.264908667 | 0.065088535 | 0.594255995 | -3.882992865 |
| **A_55_P2026669** | Atp5l | 0.305659773 | -0.026393977 | 0.637713523 | 15.01553318 | 2.264639404 | 0.065112354 | 0.594255995 | -3.883180775 |
| **A_55_P2183208** | Prl2c1 | -0.304028809 | -0.634386666 | 0.026329048 | 16.28142966 | -2.264119061 | 0.065158408 | 0.594255995 | -3.883543931 |

| | | | | | | | | | |
|---|---|---|---|---|---|---|---|---|---|
| **A_55_P1999847** | Nutf2 | 0.121621328 | -0.010543694 | 0.25378635 | 13.33599977 | 2.26392676 | 0.065175437 | 0.594255995 | -3.88367815 |
| **A_55_P1970920** | Timm17b | 0.1765147 | -0.015304035 | 0.368333434 | 12.55860873 | 2.263909282 | 0.065176985 | 0.594255995 | -3.883690348 |
| **A_55_P2140745** | NA | -0.133049711 | -0.277636065 | 0.011536643 | 8.010277907 | -2.26389246 | 0.065178475 | 0.594255995 | -3.88370209 |
| **A_55_P1953133** | NA | -0.370861414 | -0.773985209 | 0.03226238 | 13.44937475 | -2.263301336 | 0.065230852 | 0.594255995 | -3.884114703 |
| **A_52_P604629** | Csrnp1 | -0.172141957 | -0.359272708 | 0.014988795 | 12.5500538 | -2.263136432 | 0.065245471 | 0.594255995 | -3.884229816 |
| **A_55_P2131766** | Rsrp1 | -0.536901602 | -1.120827059 | 0.047023855 | 13.3431447 | -2.262072155 | 0.065339904 | 0.594255995 | -3.884972828 |
| **A_55_P1957168** | Ube2s | 0.627989256 | -0.055151653 | 1.311130165 | 12.47928372 | 2.261575464 | 0.065384025 | 0.594255995 | -3.885319636 |
| **A_51_P283754** | Cpsf7 | -0.325500679 | -0.679647824 | 0.028646465 | 10.98075924 | -2.261191362 | 0.065418165 | 0.594255995 | -3.88558785 |
| **A_51_P439876** | Map4k4 | -0.153011739 | -0.319496008 | 0.01347253 | 13.0158252 | -2.261104896 | 0.065425853 | 0.594255995 | -3.885648231 |
| **A_55_P2012615** | NA | 0.327856869 | -0.028920131 | 0.684633869 | 15.26325612 | 2.26077114 | 0.065455537 | 0.594255995 | -3.885881308 |
| **A_55_P2062538** | 1700027A15Rik | 0.291347901 | -0.025759759 | 0.608455562 | 8.346786655 | 2.260342878 | 0.065493647 | 0.594255995 | -3.886180403 |
| **A_51_P514961** | Tiparp | -0.177683533 | -0.371083481 | 0.015716415 | 10.54274886 | -2.260268289 | 0.065500287 | 0.594255995 | -3.886232498 |
| **A_52_P259335** | Fkbp15 | 0.172543177 | -0.015291704 | 0.360378059 | 11.28906477 | 2.259907738 | 0.065532393 | 0.594255995 | -3.886484325 |
| **A_55_P2004930** | Hddc2 | -0.157497607 | -0.328967798 | 0.013972584 | 9.901061617 | -2.259719276 | 0.065549181 | 0.594255995 | -3.886615963 |
| **A_55_P1971574** | Acox3 | 0.81555697 | -0.072367095 | 1.703481034 | 11.57252353 | 2.259683634 | 0.065552356 | 0.594255995 | -3.886640859 |
| **A_51_P487048** | Rhox6 | -0.264311409 | -0.552374793 | 0.023751976 | 16.4236328 | -2.257340188 | 0.065761498 | 0.594255995 | -3.888278099 |
| **A_55_P2005972** | NA | -0.150827227 | -0.31522514 | 0.013570687 | 8.847499605 | -2.257109388 | 0.065782133 | 0.594255995 | -3.888439383 |
| **A_52_P650387** | Ccnjl | -0.157663212 | -0.329526443 | 0.014200018 | 10.61744204 | -2.256922067 | 0.065798886 | 0.594255995 | -3.888570289 |
| **A_65_P20371** | Bclaf1 | -0.223706016 | -0.467637096 | 0.020225064 | 10.419917 | -2.25621087 | 0.065862532 | 0.594255995 | -3.889067336 |
| **A_55_P1961135** | Atg4d | 0.207553069 | -0.018834283 | 0.433940422 | 12.2629026 | 2.255517298 | 0.065924662 | 0.594255995 | -3.889552125 |
| **A_55_P1990944** | Casp1 | -0.188313725 | -0.393744599 | 0.017117148 | 8.753439672 | -2.255201881 | 0.065952938 | 0.594255995 | -3.889772612 |
| **A_51_P195240** | Gpatch11 | -0.226996558 | -0.474673472 | 0.020680356 | 10.4370254 | -2.254773425 | 0.065991366 | 0.594255995 | -3.890072138 |
| **A_55_P2102936** | Pnrc1 | -0.186653402 | -0.390349206 | 0.017042402 | 10.17079582 | -2.254358548 | 0.066028599 | 0.594255995 | -3.890362193 |
| **A_55_P2112892** | Dab1 | 0.378770058 | -0.03466347 | 0.792203587 | 8.655078542 | 2.253923178 | 0.066067695 | 0.594255995 | -3.890666598 |
| **A_55_P2129311** | 2210016L21Rik | 0.451387541 | -0.041383337 | 0.944158418 | 9.909585999 | 2.253583722 | 0.066098194 | 0.594255995 | -3.890903957 |
| **A_55_P2090324** | Zfp956 | 0.492518404 | -0.045255239 | 1.030292048 | 10.27905357 | 2.253160419 | 0.066136248 | 0.594255995 | -3.891199965 |
| **A_55_P2060444** | Apopt1 | 0.238904957 | -0.02196696 | 0.499776874 | 16.88494845 | 2.253030105 | 0.066147967 | 0.594255995 | -3.891291096 |
| **A_55_P2094986** | Atp1a3 | 0.143583865 | -0.013225145 | 0.300392875 | 8.503157172 | 2.252702241 | 0.066177462 | 0.594255995 | -3.891520386 |
| **A_51_P438952** | Bag4 | -0.10900039 | -0.228060699 | 0.010059918 | 8.702256623 | -2.252320619 | 0.06621181 | 0.594255995 | -3.891787286 |
| **A_51_P363914** | Cct2 | -0.151710787 | -0.317445964 | 0.01402439 | 13.16390002 | -2.252013176 | 0.066239495 | 0.594255995 | -3.892002322 |
| **A_65_P20683** | NA | 0.826252489 | -0.076541202 | 1.729046181 | 10.14842717 | 2.251611345 | 0.066275698 | 0.594255995 | -3.892283391 |
| **A_51_P245525** | NA | -0.270874587 | -0.5669669 | 0.025217725 | 15.84450163 | -2.250662079 | 0.066361304 | 0.594255995 | -3.892947456 |
| **A_55_P2053798** | NA | 0.199729428 | -0.018614512 | 0.418073368 | 14.45254489 | 2.250453676 | 0.066380113 | 0.594255995 | -3.89309326 |
| **A_55_P2020602** | Pacsin1 | 0.711392137 | -0.066317514 | 1.489101788 | 13.16187284 | 2.250405265 | 0.066384483 | 0.594255995 | -3.893127131 |
| **A_55_P2033845** | NA | -0.269358797 | -0.563853708 | 0.025136114 | 14.90284296 | -2.25020732 | 0.066402355 | 0.594255995 | -3.893265624 |
| **A_51_P304859** | Zfand6 | -0.144322024 | -0.302111993 | 0.013467944 | 9.665180622 | -2.250206551 | 0.066402424 | 0.594255995 | -3.893266162 |
| **A_55_P2462377** | Nsf | -0.255974894 | -0.535845051 | 0.023895263 | 11.91530356 | -2.250142092 | 0.066408245 | 0.594255995 | -3.893311263 |
| **A_66_P121976** | Slc4a11 | -0.13634842 | -0.285426411 | 0.01272957 | 9.094996904 | -2.250120327 | 0.066410211 | 0.594255995 | -3.893326491 |
| **A_51_P395293** | Ccz1 | -0.174033787 | -0.36433633 | 0.016268756 | 12.05153416 | -2.249873711 | 0.066432487 | 0.594255995 | -3.893499047 |
| **A_55_P2092296** | Fbxo2 | 0.16628592 | -0.01555758 | 0.34812942 | 9.128661714 | 2.249711655 | 0.066447129 | 0.594255995 | -3.893612441 |
| **A_51_P245796** | Ddit4 | -0.221460156 | -0.463673174 | 0.020752862 | 12.02583254 | -2.249403072 | 0.066475019 | 0.594255995 | -3.893828372 |
| **A_55_P2078680** | Tpm1 | -0.242210286 | -0.507128264 | 0.022707691 | 10.01997142 | -2.249315185 | 0.066482965 | 0.594255995 | -3.893889873 |
| **A_51_P248865** | Foxf2 | -0.107521728 | -0.225141618 | 0.010098162 | 8.116769794 | -2.248975007 | 0.066513729 | 0.594255995 | -3.89412793 |
| **A_55_P2027361** | Irak1bp1 | 0.308538469 | -0.029104027 | 0.646180965 | 13.05329186 | 2.248129793 | 0.06659023 | 0.594255995 | -3.894719472 |
| **A_51_P459661** | Lipa | -0.242320989 | -0.507571604 | 0.022929627 | 10.28271968 | -2.24752119 | 0.066645372 | 0.594255995 | -3.89514547 |
| **A_55_P2086866** | NA | 0.512795649 | -0.048537175 | 1.074128473 | 13.74081364 | 2.247465616 | 0.06665041 | 0.594255995 | -3.895184372 |
| **A_51_P478952** | N4bp2l1 | -0.134797794 | -0.282354484 | 0.012758897 | 8.22735999 | -2.247465538 | 0.066650417 | 0.594255995 | -3.895184427 |
| **A_55_P2130725** | Rhox8 | 0.15140159 | -0.014353328 | 0.317156508 | 8.489424024 | 2.247155757 | 0.066678504 | 0.594255995 | -3.895401281 |
| **A_66_P109800** | Gcsh | 0.35002583 | -0.033211603 | 0.733263263 | 11.14804897 | 2.246991006 | 0.066693447 | 0.594255995 | -3.895516615 |
| **A_55_P2139089** | Slc22a28 | 0.104674282 | -0.009932237 | 0.219280802 | 8.67928703 | 2.246983227 | 0.066694153 | 0.594255995 | -3.895522061 |

| | | | | | | | | | |
|---|---|---|---|---|---|---|---|---|---|
| **A_55_P2112030** | Tmem145 | -0.337635128 | -0.707311933 | 0.032041678 | 8.727994194 | -2.246956088 | 0.066696615 | 0.594255995 | -3.89554106 |
| **A_55_P2006375** | Dmrtc1c2 | 0.366100686 | -0.034784036 | 0.766985407 | 10.72653611 | 2.246726462 | 0.066717449 | 0.594255995 | -3.895701818 |
| **A_55_P2024515** | Crtc2 | -0.224587083 | -0.470513606 | 0.02133944 | 10.47705596 | -2.24671802 | 0.066718215 | 0.594255995 | -3.895707727 |
| **A_55_P2043451** | NA | -0.133653128 | -0.280042576 | 0.01273632 | 11.32927381 | -2.246148768 | 0.066769893 | 0.594255995 | -3.896106279 |
| **A_55_P2028444** | Ube2j2 | -0.158347262 | -0.331811194 | 0.015116669 | 13.34437345 | -2.245797204 | 0.06680183 | 0.594255995 | -3.896352439 |
| **A_55_P2101340** | Ramp3 | -0.609860809 | -1.277964694 | 0.058243077 | 13.18590979 | -2.24572147 | 0.066808712 | 0.594255995 | -3.896405469 |
| **A_52_P79639** | Vat1 | -0.350454482 | -0.734409205 | 0.033500241 | 10.39794394 | -2.245539854 | 0.066825218 | 0.594255995 | -3.896532642 |
| **A_55_P2074771** | Exosc7 | -0.159400773 | -0.334060691 | 0.015259146 | 10.75700445 | -2.245258415 | 0.066850805 | 0.594255995 | -3.896729722 |
| **A_66_P125560** | Macroh2a2 | 0.279569635 | -0.0267683 | 0.58590757 | 10.54707646 | 2.245217313 | 0.066854543 | 0.594255995 | -3.896758505 |
| **A_55_P2064595** | NA | -0.189385694 | -0.396920799 | 0.01814941 | 8.646477657 | -2.24504353 | 0.066870348 | 0.594255995 | -3.896880203 |
| **A_52_P179238** | Macroh2a1 | -0.167583059 | -0.35123405 | 0.016067931 | 8.313472108 | -2.244946536 | 0.066879172 | 0.594255995 | -3.896948129 |
| **A_55_P2069530** | NA | 0.251925853 | -0.024161108 | 0.528012813 | 10.04705146 | 2.244894858 | 0.066883873 | 0.594255995 | -3.89698432 |
| **A_55_P2006554** | Eif4e2 | -0.255665881 | -0.535948852 | 0.02461709 | 11.64657945 | -2.244115604 | 0.066954809 | 0.594255995 | -3.897530083 |
| **A_51_P378558** | Rpap2 | -0.177690416 | -0.372537572 | 0.017156741 | 9.363290081 | -2.243567257 | 0.067004772 | 0.594255995 | -3.897914171 |
| **A_52_P404363** | Cdc42bpb | -0.129198803 | -0.270923605 | 0.012525999 | 8.749727647 | -2.242754799 | 0.067078871 | 0.594255995 | -3.898483321 |
| **A_55_P2181084** | Rpl17 | 0.42627298 | -0.041333263 | 0.893879224 | 15.8176957 | 2.242728319 | 0.067081287 | 0.594255995 | -3.898501873 |
| **A_55_P2108738** | Nck1 | -0.158293904 | -0.331942894 | 0.015355086 | 10.71095507 | -2.242647882 | 0.067088628 | 0.594255995 | -3.898558226 |
| **A_51_P357735** | Inhbe | 0.198455897 | -0.019340478 | 0.416252272 | 8.084990951 | 2.241725977 | 0.067172825 | 0.594255995 | -3.899204157 |
| **A_55_P2090441** | NA | 0.411160107 | -0.040145599 | 0.862465814 | 8.670010679 | 2.241348088 | 0.067207369 | 0.594255995 | -3.899468953 |
| **A_55_P2058143** | Pclo | 0.554876376 | -0.054264311 | 1.164017064 | 8.616174615 | 2.241030613 | 0.067236405 | 0.594255995 | -3.89969143 |
| **A_51_P418116** | Tmem119 | -0.225399646 | -0.47284355 | 0.022044258 | 9.702095606 | -2.241019493 | 0.067237422 | 0.594255995 | -3.899699223 |
| **A_55_P2057913** | Hmgb1-ps7 | -0.235654696 | -0.494468156 | 0.023158765 | 12.10403217 | -2.240053557 | 0.067325846 | 0.594255995 | -3.900376197 |
| **A_55_P2108122** | Arcn1 | -0.126418124 | -0.265286552 | 0.012450303 | 12.78638598 | -2.239623354 | 0.067365267 | 0.594255995 | -3.90067774 |
| **A_55_P2028821** | Hnrnpf | -0.312935093 | -0.656752923 | 0.030882738 | 13.58286481 | -2.239211086 | 0.067403066 | 0.594255995 | -3.900966733 |
| **A_66_P109220** | Etfbkmt | -0.173899429 | -0.364974959 | 0.017176101 | 8.832456302 | -2.239042028 | 0.067418573 | 0.594255995 | -3.901085246 |
| **A_55_P2151209** | Krtap19-4 | 0.501206912 | -0.049560708 | 1.051974532 | 12.30577933 | 2.238812973 | 0.067439589 | 0.594255995 | -3.901245823 |
| **A_55_P2105808** | Wdr62 | 1.374751594 | -0.136275223 | 2.885778411 | 12.86303336 | 2.238315092 | 0.067485294 | 0.594255995 | -3.90159488 |
| **A_52_P535484** | Gvin1 | -0.177789513 | -0.37320449 | 0.017625465 | 9.098164522 | -2.238295679 | 0.067487077 | 0.594255995 | -3.901608491 |
| **A_55_P1964078** | NA | -0.252950782 | -0.531346234 | 0.025444671 | 12.65848705 | -2.235337235 | 0.067759327 | 0.594255995 | -3.903683238 |
| **A_51_P465350** | Ly86 | -0.210213774 | -0.441595128 | 0.021167581 | 9.069075867 | -2.235125803 | 0.067778828 | 0.594255995 | -3.903831555 |
| **A_55_P2020331** | Camk1g | 0.342301478 | -0.034797014 | 0.719399971 | 8.829071237 | 2.233176974 | 0.067958846 | 0.594255995 | -3.905198886 |
| **A_55_P1993473** | Gm5935 | 0.127873469 | -0.013007968 | 0.268754906 | 8.569148063 | 2.233036582 | 0.067971833 | 0.594255995 | -3.905297405 |
| **A_55_P1971328** | Lage3 | 0.348432606 | -0.035488383 | 0.732353595 | 10.99343605 | 2.232780837 | 0.067995498 | 0.594255995 | -3.905476878 |
| **A_51_P170807** | Map3k6 | 0.227882713 | -0.023213528 | 0.478978955 | 10.05846182 | 2.232751207 | 0.067998241 | 0.594255995 | -3.905497672 |
| **A_51_P489313** | Sri | -0.313239221 | -0.658495434 | 0.032016992 | 14.55497009 | -2.232049364 | 0.068063232 | 0.594255995 | -3.905990244 |
| **A_55_P2121408** | Tpm2 | -0.261435629 | -0.549609428 | 0.02673817 | 12.51535688 | -2.231924223 | 0.068074827 | 0.594255995 | -3.906078078 |
| **A_52_P92557** | Kdm2a | -0.282024007 | -0.592937649 | 0.028889635 | 9.041099599 | -2.231595431 | 0.068105301 | 0.594255995 | -3.906308858 |
| **A_55_P2016604** | Glrx3 | 0.28488984 | -0.029194923 | 0.598974602 | 12.68390878 | 2.231512151 | 0.068113022 | 0.594255995 | -3.906367315 |
| **A_55_P2114993** | Actg1 | 0.490216211 | -0.050293133 | 1.030725554 | 16.7104783 | 2.231277705 | 0.068134763 | 0.594255995 | -3.906531883 |
| **A_55_P1960491** | Snta1 | 0.368478278 | -0.037834059 | 0.774790614 | 12.2048631 | 2.23111032 | 0.068150289 | 0.594255995 | -3.906649382 |
| **A_52_P200458** | Fut4 | -0.111021396 | -0.23345616 | 0.011413368 | 8.208347534 | -2.230853738 | 0.068174097 | 0.594255995 | -3.906829502 |
| **A_55_P2061525** | Cdc27 | -0.152762015 | -0.321244071 | 0.015720041 | 9.326899891 | -2.230647197 | 0.068193267 | 0.594255995 | -3.906974499 |
| **A_55_P2075829** | NA | -0.176860583 | -0.371966058 | 0.018244892 | 8.685863827 | -2.230132973 | 0.06824102 | 0.594255995 | -3.907335519 |
| **A_55_P1969356** | Ccm2l | -0.095392516 | -0.200631424 | 0.009846393 | 8.285940819 | -2.230011645 | 0.068252293 | 0.594255995 | -3.907420704 |
| **A_55_P2049025** | NA | 0.225458292 | -0.023349048 | 0.474265631 | 14.9824455 | 2.229318847 | 0.068316695 | 0.594255995 | -3.907907154 |
| **A_52_P632742** | Shisa5 | 0.119383673 | -0.012386327 | 0.251153674 | 8.733451615 | 2.228935762 | 0.068352333 | 0.594255995 | -3.908176163 |
| **A_55_P2132671** | NA | 0.348160069 | -0.03615688 | 0.732477018 | 8.86760219 | 2.228735781 | 0.068370945 | 0.594255995 | -3.9083166 |
| **A_55_P2057906** | Cetn2 | 0.142221573 | -0.01478211 | 0.299225257 | 10.83121638 | 2.228562405 | 0.068387085 | 0.594255995 | -3.908438358 |
| **A_55_P1998416** | Ifi47 | -0.328134106 | -0.69038542 | 0.034117207 | 10.46031205 | -2.228489374 | 0.068393885 | 0.594255995 | -3.908489647 |
| **A_66_P130914** | NA | -0.131066489 | -0.275766517 | 0.01363354 | 14.79840413 | -2.228395188 | 0.068402656 | 0.594255995 | -3.908555793 |

| | | | | | | | | | |
|---|---|---|---|---|---|---|---|---|---|
| **A_55_P2176292** | NA | 0.142588753 | -0.014848601 | 0.300026108 | 13.56949552 | 2.228161422 | 0.068424429 | 0.594255995 | -3.908719971 |
| **A_55_P2135980** | Ms4a4c | -0.13622871 | -0.286652777 | 0.014195357 | 15.50396157 | -2.228027155 | 0.068436938 | 0.594255995 | -3.908814273 |
| **A_52_P273929** | Mrpl17 | 0.230505174 | -0.024036083 | 0.48504643 | 11.04989067 | 2.227879295 | 0.068450717 | 0.594255995 | -3.908918123 |
| **A_55_P2045127** | NA | 0.497112686 | -0.051939684 | 1.046165057 | 8.891019327 | 2.227461666 | 0.068489649 | 0.594255995 | -3.909211462 |
| **A_51_P209327** | Apln | -0.140690314 | -0.296081492 | 0.014700865 | 8.680095603 | -2.227445063 | 0.068491197 | 0.594255995 | -3.909223125 |
| **A_55_P2106241** | 4930432K21Rik | 0.181732221 | -0.019034691 | 0.382499132 | 8.819736849 | 2.22694227 | 0.0685381 | 0.594255995 | -3.90957631 |
| **A_52_P113916** | NA | 0.277493984 | -0.029076127 | 0.584064096 | 9.157732711 | 2.22686005 | 0.068545773 | 0.594255995 | -3.909634068 |
| **A_52_P72497** | Acbd3 | -0.103876321 | -0.218638744 | 0.010886102 | 11.52787778 | -2.226824613 | 0.06854908 | 0.594255995 | -3.909658962 |
| **A_55_P2094469** | Kdr | 0.409594426 | -0.042927036 | 0.862115888 | 12.52507499 | 2.226814393 | 0.068550034 | 0.594255995 | -3.909666142 |
| **A_51_P209319** | Slc35b1 | 0.199018568 | -0.020940163 | 0.418977298 | 13.48300378 | 2.225981518 | 0.068627816 | 0.594255995 | -3.910251271 |
| **A_66_P132295** | Tfpi | -0.261994686 | -0.551562293 | 0.027572921 | 15.29724684 | -2.225930853 | 0.068632551 | 0.594255995 | -3.910286868 |
| **A_55_P1958314** | Mospd1 | -0.142593295 | -0.300202816 | 0.015016226 | 12.49555387 | -2.225798364 | 0.068644933 | 0.594255995 | -3.910379956 |
| **A_55_P2001064** | NA | -0.208804256 | -0.439604683 | 0.02199617 | 11.95365468 | -2.225727046 | 0.068651599 | 0.594255995 | -3.910430066 |
| **A_55_P2155833** | NA | 0.24514206 | -0.025846254 | 0.516130374 | 16.69419242 | 2.225545261 | 0.068668594 | 0.594255995 | -3.910557793 |
| **A_51_P193336** | Nucb2 | -0.266959564 | -0.562077543 | 0.028158414 | 11.80761888 | -2.225455838 | 0.068676956 | 0.594255995 | -3.910620626 |
| **A_52_P570543** | Sdk2 | 0.24234426 | -0.025595984 | 0.510284503 | 8.336254011 | 2.225173874 | 0.068703329 | 0.594255995 | -3.910818754 |
| **A_51_P148494** | Rnft1 | -0.166651675 | -0.350937369 | 0.017634019 | 10.01858334 | -2.224780781 | 0.068740114 | 0.594255995 | -3.911094984 |
| **A_55_P2144436** | Rrp1 | -0.193872102 | -0.408282828 | 0.020538625 | 10.91332202 | -2.224528489 | 0.068763733 | 0.594255995 | -3.911272282 |
| **A_51_P157976** | Mmab | 0.123263553 | -0.01307139 | 0.259598496 | 8.500203919 | 2.224316912 | 0.068783547 | 0.594255995 | -3.911420973 |
| **A_51_P314907** | Dbf4 | -0.296923283 | -0.625422944 | 0.031576378 | 8.73290641 | -2.22371175 | 0.068840254 | 0.594255995 | -3.911846296 |
| **A_52_P150114** | NA | 0.579597203 | -0.061670754 | 1.22086516 | 9.056263449 | 2.223596105 | 0.068851096 | 0.594255995 | -3.91192758 |
| **A_51_P174081** | Zdhhc14 | -0.217080521 | -0.457316958 | 0.023155916 | 9.443157593 | -2.223059769 | 0.068901402 | 0.594255995 | -3.912304573 |
| **A_52_P648573** | Gorab | 0.149188082 | -0.015916055 | 0.314292218 | 8.824196505 | 2.223030077 | 0.068904188 | 0.594255995 | -3.912325445 |
| **A_55_P1990067** | NA | -0.124119675 | -0.26151969 | 0.013280339 | 12.32013054 | -2.222404025 | 0.068962959 | 0.594255995 | -3.912765545 |
| **A_55_P1963878** | NA | -0.273849943 | -0.577004885 | 0.029305 | 12.01627012 | -2.222374088 | 0.068965771 | 0.594255995 | -3.912786592 |
| **A_55_P2043062** | Hs3st6 | 0.160711352 | -0.017207012 | 0.338629716 | 8.10914019 | 2.222260365 | 0.068976453 | 0.594255995 | -3.912866541 |
| **A_55_P1986983** | Gne | -0.185654635 | -0.391234238 | 0.019924968 | 11.67530747 | -2.221748704 | 0.069024534 | 0.594255995 | -3.913226271 |
| **A_55_P2088625** | Fbln7 | -0.321179662 | -0.676930951 | 0.034571626 | 16.19296915 | -2.22111333 | 0.06908429 | 0.594255995 | -3.913673021 |
| **A_66_P120507** | Plcd3 | 0.166290133 | -0.017900116 | 0.350480383 | 10.35236944 | 2.221104584 | 0.069085112 | 0.594255995 | -3.913679171 |
| **A_55_P2216996** | Tmem87a | 0.142566116 | -0.01537147 | 0.300503702 | 8.95077313 | 2.220751611 | 0.069118332 | 0.594255995 | -3.913927377 |
| **A_51_P439626** | Map2k2 | -0.229879467 | -0.484559755 | 0.02480082 | 10.56950415 | -2.220618808 | 0.069130835 | 0.594255995 | -3.914020767 |
| **A_55_P2167189** | 4921504E06Rik | 0.691434495 | -0.074631641 | 1.45750063 | 9.943865691 | 2.220516172 | 0.069140499 | 0.594255995 | -3.914092943 |
| **A_51_P358872** | Disc1 | 0.176070178 | -0.019028894 | 0.37116925 | 7.973123146 | 2.220239188 | 0.069166588 | 0.594255995 | -3.914287733 |
| **A_51_P271425** | Lhfpl4 | 0.187152857 | -0.020311318 | 0.394617032 | 9.896261421 | 2.219333208 | 0.069251991 | 0.594255995 | -3.91492493 |
| **A_55_P1999364** | Krt85 | -0.142378645 | -0.300237887 | 0.015480597 | 12.03471338 | -2.218932061 | 0.06928984 | 0.594255995 | -3.915207097 |
| **A_52_P587648** | Phf3 | -0.207058299 | -0.436697098 | 0.0225805 | 11.50811237 | -2.218280889 | 0.069351326 | 0.594255995 | -3.915665171 |
| **A_51_P333869** | Nsmce3 | -0.165390688 | -0.348878809 | 0.018097433 | 8.921454504 | -2.217544095 | 0.069420964 | 0.594255995 | -3.916183537 |
| **A_55_P2041060** | Rbp3 | 0.683112159 | -0.074834848 | 1.441059166 | 11.6553548 | 2.217289172 | 0.069445075 | 0.594255995 | -3.916362901 |
| **A_66_P101108** | Tnrc18 | 0.159351471 | -0.017492171 | 0.336195114 | 11.3455114 | 2.216847349 | 0.069486884 | 0.594255995 | -3.916673786 |
| **A_55_P2041265** | Utp6 | 0.208520137 | -0.022890126 | 0.4399304 | 16.17004967 | 2.216841016 | 0.069487484 | 0.594255995 | -3.916678243 |
| **A_55_P2006930** | 2610005L07Rik | -0.124583895 | -0.262845273 | 0.013677482 | 12.21416294 | -2.216818765 | 0.06948959 | 0.594255995 | -3.9166939 |
| **A_51_P518592** | Atxn7 | -0.20485504 | -0.432210236 | 0.022500156 | 9.411954519 | -2.216720472 | 0.069498895 | 0.594255995 | -3.916763066 |
| **A_55_P2017640** | Phf7 | 0.128478391 | -0.014200757 | 0.27115754 | 9.331703716 | 2.215331621 | 0.069630514 | 0.594255995 | -3.917740489 |
| **A_55_P1999269** | Grk2 | 0.231307967 | -0.025619991 | 0.488235925 | 12.2770586 | 2.214870758 | 0.069674246 | 0.594255995 | -3.918064878 |
| **A_51_P161929** | Tspo2 | 0.310134385 | -0.034357643 | 0.654626414 | 10.02132492 | 2.214827496 | 0.069678353 | 0.594255995 | -3.91809533 |
| **A_55_P2016252** | Ctnnal1 | -0.187632434 | -0.396064442 | 0.020799575 | 8.736825154 | -2.214688556 | 0.069691543 | 0.594255995 | -3.918193132 |
| **A_55_P1959496** | Sbno2 | -0.138121837 | -0.291569846 | 0.015326171 | 8.467939557 | -2.214472303 | 0.069712079 | 0.594255995 | -3.918345359 |
| **A_55_P2011220** | Armcx1 | 0.190699106 | -0.021199178 | 0.402597391 | 8.894981137 | 2.214065068 | 0.069750768 | 0.594255995 | -3.91863204 |
| **A_55_P2146535** | Cnr2 | 0.220503312 | -0.024544318 | 0.465550942 | 11.58891009 | 2.213776508 | 0.069778195 | 0.594255995 | -3.918835189 |
| **A_52_P201232** | Xpo5 | 0.239584571 | -0.026676186 | 0.505845327 | 9.073508265 | 2.213710616 | 0.06978446 | 0.594255995 | -3.918881578 |

| | | | | | | | | |
|---|---|---|---|---|---|---|---|---|
| A_55_P2180056 | NA | 0.193539145 | -0.021553681 | 0.408631972 | 9.808087301 | 2.213665799 | 0.069788721 | 0.594255995 | -3.918913131 |
| A_55_P2000813 | Cops2 | 0.181152686 | -0.02022031 | 0.382525682 | 9.587630182 | 2.213159474 | 0.069836882 | 0.594255995 | -3.919269619 |
| A_51_P505322 | Mpnd | 0.176618963 | -0.019730837 | 0.372968763 | 10.17716418 | 2.212972571 | 0.069854668 | 0.594255995 | -3.919401219 |
| A_51_P281700 | Khsrp | 1.048659204 | -0.117198081 | 2.214516488 | 13.40697177 | 2.212881436 | 0.069863343 | 0.594255995 | -3.91946539 |
| A_55_P2048794 | Cd82 | -0.135284204 | -0.285712655 | 0.015144247 | 16.05418749 | -2.212515233 | 0.06989821 | 0.594255995 | -3.919723252 |
| A_51_P382152 | Procr | -0.356959019 | -0.753893636 | 0.039975599 | 14.95726317 | -2.212424943 | 0.06990681 | 0.594255995 | -3.919786832 |
| A_51_P165087 | Snai1 | 0.107089462 | -0.012039665 | 0.226218588 | 10.90244594 | 2.211556007 | 0.069989628 | 0.594255995 | -3.920398766 |
| A_55_P2141884 | Tagln2 | -0.319795154 | -0.675620122 | 0.036029814 | 11.28479944 | -2.211080844 | 0.070034958 | 0.594255995 | -3.92073343 |
| A_66_P137383 | Ddx55 | -0.098915408 | -0.208995938 | 0.011165122 | 8.40268859 | -2.210663287 | 0.070074818 | 0.594255995 | -3.921027542 |
| A_55_P2068701 | NA | 0.157466398 | -0.017786466 | 0.332719261 | 8.351777636 | 2.210507196 | 0.070089725 | 0.594255995 | -3.921137492 |
| A_55_P2076994 | Defa-rs10 | 0.33475995 | -0.037844244 | 0.707364143 | 8.831838064 | 2.210318806 | 0.07010772 | 0.594255995 | -3.921270197 |
| A_55_P2219903 | AW061147 | 0.181512781 | -0.020536241 | 0.383561804 | 11.13407056 | 2.21013917 | 0.070124884 | 0.594255995 | -3.921396739 |
| A_66_P100021 | Ncaph2 | -0.171048784 | -0.361526359 | 0.019428791 | 9.817958224 | -2.209252219 | 0.070209693 | 0.594255995 | -3.922021597 |
| A_51_P117477 | Slc27a1 | 0.335296959 | -0.038164455 | 0.708758373 | 9.365362432 | 2.208782948 | 0.070254607 | 0.594255995 | -3.922352235 |
| A_66_P140711 | Rps10 | 0.173143705 | -0.019723528 | 0.366010938 | 16.4750309 | 2.208601784 | 0.070271954 | 0.594255995 | -3.922479886 |
| A_51_P246727 | Mlxip | -0.157714868 | -0.333435198 | 0.018005462 | 8.803853276 | -2.208105348 | 0.070319512 | 0.594255995 | -3.922829702 |
| A_55_P1962665 | Exoc3l2 | 0.184462429 | -0.02109002 | 0.390014879 | 8.464392845 | 2.207773066 | 0.070351364 | 0.594255995 | -3.923063863 |
| A_55_P2117487 | NA | -0.290966021 | -0.615208639 | 0.033276598 | 13.97360574 | -2.207706516 | 0.070357745 | 0.594255995 | -3.923110763 |
| A_51_P155458 | Dok7 | -0.163189215 | -0.345063449 | 0.018685019 | 8.00827739 | -2.207442699 | 0.070383046 | 0.594255995 | -3.923296686 |
| A_55_P2092904 | Ddb2 | -0.144846694 | -0.306282721 | 0.016589333 | 8.503031053 | -2.207380943 | 0.07038897 | 0.594255995 | -3.92334021 |
| A_55_P1983754 | Pcp4l1 | -0.434483931 | -0.918730464 | 0.049762601 | 9.689010643 | -2.207376236 | 0.070389422 | 0.594255995 | -3.923343527 |
| A_55_P2046411 | NA | 0.486255259 | -0.055757361 | 1.02826788 | 12.09884623 | 2.207110495 | 0.07041492 | 0.594255995 | -3.923530817 |
| A_55_P2132865 | Sumo2 | -0.185804214 | -0.392928966 | 0.021320539 | 11.88340739 | -2.206951143 | 0.070430214 | 0.594255995 | -3.92364313 |
| A_55_P2111018 | NA | -0.151670975 | -0.320829739 | 0.01748779 | 8.679814202 | -2.205855891 | 0.070535428 | 0.594255995 | -3.924415152 |
| A_55_P2180869 | Ocel1 | 0.326653623 | -0.037676726 | 0.690983972 | 10.42353442 | 2.205775419 | 0.070543165 | 0.594255995 | -3.92447188 |
| A_52_P286928 | NA | -0.154310693 | -0.326447234 | 0.017825849 | 9.174623159 | -2.205424074 | 0.070576955 | 0.594255995 | -3.924719569 |
| A_55_P2026624 | NA | 0.209104038 | -0.02417917 | 0.442387246 | 14.05509169 | 2.205200612 | 0.070598455 | 0.594255995 | -3.924877111 |
| A_55_P2125557 | Irx2 | -0.113577396 | -0.240299019 | 0.013144228 | 8.3125968 | -2.205008868 | 0.070616908 | 0.594255995 | -3.925012297 |
| A_66_P127160 | Eif2b3 | -0.121137874 | -0.256321463 | 0.014045716 | 8.591256524 | -2.204576311 | 0.070658556 | 0.594255995 | -3.925317278 |
| A_65_P18858 | Btf3 | -0.206868404 | -0.437763387 | 0.024026579 | 11.49142316 | -2.204188998 | 0.070695869 | 0.594255995 | -3.925590378 |
| A_55_P1967617 | Med16 | 0.373716591 | -0.043430676 | 0.790863858 | 17.05366961 | 2.204053534 | 0.070708924 | 0.594255995 | -3.925685899 |
| A_55_P2120993 | Zfp282 | 0.44976488 | -0.052281108 | 0.951810868 | 10.66286582 | 2.203998032 | 0.070714274 | 0.594255995 | -3.925725036 |
| A_55_P1972927 | Elk3 | -0.251011551 | -0.531214856 | 0.029191755 | 10.65887275 | -2.203888484 | 0.070724834 | 0.594255995 | -3.925802285 |
| A_51_P225179 | Smndc1 | 0.246331025 | -0.028650189 | 0.52131224 | 9.512265265 | 2.20386634 | 0.070726969 | 0.594255995 | -3.925817901 |
| A_52_P58181 | Suclg2 | -0.16001245 | -0.338676813 | 0.018651912 | 9.162985242 | -2.203357685 | 0.070776026 | 0.594255995 | -3.926176605 |
| A_55_P2094901 | Phyhd1 | 0.191711435 | -0.022414202 | 0.405837073 | 16.9426023 | 2.202665316 | 0.070842857 | 0.594255995 | -3.926664913 |
| A_55_P2017491 | Rbm43 | -0.246565522 | -0.521962581 | 0.028831537 | 8.997195344 | -2.202633362 | 0.070845942 | 0.594255995 | -3.926687451 |
| A_52_P428496 | Rhoa | -0.129569899 | -0.274316564 | 0.015176766 | 10.72920037 | -2.202240352 | 0.070883908 | 0.594255995 | -3.926964654 |
| A_51_P501312 | Natd1 | -0.249182555 | -0.5275524 | 0.02918729 | 14.3455633 | -2.202239814 | 0.07088396 | 0.594255995 | -3.926965034 |
| A_55_P2046488 | Nbeal2 | -0.112146093 | -0.237428921 | 0.013136736 | 8.03406544 | -2.202225366 | 0.070885357 | 0.594255995 | -3.926975225 |
| A_55_P1967835 | NA | 0.327667494 | -0.038389678 | 0.693724665 | 9.853908257 | 2.202183999 | 0.070889354 | 0.594255995 | -3.927004404 |
| A_55_P2335903 | NA | 0.149491273 | -0.017527079 | 0.316509625 | 10.88921571 | 2.202017717 | 0.070905425 | 0.594255995 | -3.927121695 |
| A_52_P305685 | NA | -0.13171393 | -0.278880363 | 0.015452503 | 9.071694113 | -2.201872197 | 0.070919493 | 0.594255995 | -3.927224344 |
| A_55_P1953931 | Fam193a | 0.245581861 | -0.028850708 | 0.520014429 | 10.44575918 | 2.201556314 | 0.070950039 | 0.594255995 | -3.927447174 |
| A_55_P1955382 | Rgma | 0.144222232 | -0.016976051 | 0.305420516 | 9.01565669 | 2.201106027 | 0.070993606 | 0.594255995 | -3.927764834 |
| A_65_P05252 | Cdc25a | -0.151093958 | -0.319975083 | 0.017787167 | 9.310471615 | -2.20107656 | 0.070996458 | 0.594255995 | -3.927785623 |
| A_55_P2148896 | NA | -0.117103876 | -0.247999857 | 0.013792106 | 10.63330525 | -2.200969967 | 0.071006776 | 0.594255995 | -3.927860824 |
| A_55_P2037717 | Slc22a18 | 0.534677583 | -0.063026716 | 1.132381881 | 11.53692975 | 2.20077052 | 0.071026086 | 0.594255995 | -3.928001537 |
| A_52_P483280 | Lamp2 | -0.216972657 | -0.459543245 | 0.02559793 | 12.67290609 | -2.200574271 | 0.071045091 | 0.594255995 | -3.928139998 |
| A_51_P109541 | E130308A19Rik | -0.103104022 | -0.218380936 | 0.012172892 | 9.522901751 | -2.200404021 | 0.071061583 | 0.594255995 | -3.92826012 |

| | | | | | | | | | |
|---|---|---|---|---|---|---|---|---|---|
| **A_55_P1969431** | Etfrf1 | -0.154255048 | -0.326732684 | 0.018222589 | 9.105227037 | -2.200268877 | 0.071074678 | 0.594255995 | -3.928355475 |
| **A_55_P1986123** | Sccpdh | 0.17543926 | -0.020765287 | 0.371643808 | 11.14612776 | 2.199818662 | 0.071118317 | 0.594255995 | -3.92867315 |
| **A_55_P2060333** | S100a5 | 0.606677873 | -0.071846976 | 1.285202721 | 9.736752186 | 2.19969041 | 0.071130754 | 0.594255995 | -3.928763651 |
| **A_51_P400555** | Cts3 | -0.282954819 | -0.599488731 | 0.033579093 | 17.29418587 | -2.199206513 | 0.071177697 | 0.594255995 | -3.929105126 |
| **A_51_P407165** | Abhd5 | -0.172607822 | -0.365747691 | 0.020532047 | 9.659759612 | -2.198658117 | 0.071230937 | 0.594255995 | -3.929492149 |
| **A_55_P2082869** | Ezh1 | -0.266562035 | -0.564939298 | 0.031815227 | 12.82141733 | -2.19786865 | 0.071307652 | 0.594255995 | -3.930049366 |
| **A_55_P1967138** | Cenpx | 0.29052862 | -0.03467799 | 0.615735229 | 10.67233348 | 2.197853396 | 0.071309136 | 0.594255995 | -3.930060133 |
| **A_55_P2101666** | Vamp5 | 0.348721752 | -0.04174997 | 0.739193474 | 10.31568282 | 2.197144463 | 0.0713781 | 0.594255995 | -3.930560569 |
| **A_51_P520927** | Ccdc97 | -0.167935246 | -0.355985529 | 0.020115037 | 10.50172609 | -2.197035254 | 0.07138873 | 0.594255995 | -3.930637664 |
| **A_55_P2155654** | Pvt1 | 0.209034024 | -0.025046694 | 0.443114741 | 8.255838063 | 2.196951674 | 0.071396866 | 0.594255995 | -3.930696668 |
| **A_55_P2019019** | Rpl36a | 0.108181263 | -0.012972836 | 0.229335363 | 13.86408751 | 2.196762455 | 0.07141529 | 0.594255995 | -3.930830252 |
| **A_51_P220260** | Phf1 | -0.246449202 | -0.522504402 | 0.029605998 | 9.262891856 | -2.196345441 | 0.07145591 | 0.594255995 | -3.931124667 |
| **A_55_P1986386** | Senp3 | -0.107129515 | -0.227135178 | 0.012876148 | 9.120283714 | -2.196223634 | 0.07146778 | 0.594255995 | -3.931210667 |
| **A_55_P2200628** | 4733401D01Rik | 0.753082371 | -0.090534847 | 1.59669959 | 10.53021003 | 2.196171288 | 0.071472881 | 0.594255995 | -3.931247625 |
| **A_55_P2162404** | NA | 0.135247059 | -0.016275301 | 0.286769419 | 8.422510723 | 2.195938982 | 0.071495526 | 0.594255995 | -3.931411648 |
| **A_55_P2088385** | Tnnt1 | 0.296405047 | -0.035727337 | 0.628537431 | 9.190285018 | 2.195551026 | 0.071533359 | 0.594255995 | -3.931685584 |
| **A_55_P1985288** | Bhlha9 | 0.561567508 | -0.06771208 | 1.190847096 | 8.445061919 | 2.195469914 | 0.071541272 | 0.594255995 | -3.931742859 |
| **A_55_P1960416** | Nrros | -0.229307278 | -0.486302258 | 0.027687701 | 10.3626415 | -2.19514071 | 0.071573396 | 0.594255995 | -3.931975326 |
| **A_51_P104933** | Trub2 | 0.135771571 | -0.016444943 | 0.287988086 | 12.02763151 | 2.194402218 | 0.071645512 | 0.594255995 | -3.971869534 |
| **A_55_P2112125** | Tnk2 | -0.238742569 | -0.5064179 | 0.028932762 | 12.95229463 | -2.194273096 | 0.071658129 | 0.594255995 | -3.932588049 |
| **A_52_P316474** | St3gal4 | -0.202352311 | -0.429262559 | 0.024557938 | 9.097538881 | -2.19393229 | 0.071691442 | 0.594255995 | -3.932828756 |
| **A_55_P2130575** | NA | 0.283676935 | -0.034448608 | 0.601802479 | 14.26333452 | 2.193787954 | 0.071705555 | 0.594255995 | -3.932930702 |
| **A_55_P2125963** | NA | 0.219108847 | -0.026620073 | 0.464837766 | 9.567445966 | 2.193677618 | 0.071716346 | 0.594255995 | -3.933008635 |
| **A_55_P2070174** | NA | 0.403215803 | -0.049046378 | 0.855477984 | 9.662574535 | 2.19339291 | 0.071744197 | 0.594255995 | -3.933209738 |
| **A_55_P2031032** | NA | 0.137175567 | -0.016769546 | 0.29112068 | 11.24349532 | 2.192199224 | 0.07186109 | 0.594255995 | -3.934052997 |
| **A_55_P2027831** | Cryba4 | 0.641607135 | -0.078473963 | 1.361688232 | 8.856163453 | 2.192082713 | 0.07187251 | 0.594255995 | -3.934135313 |
| **A_52_P24428** | Ankrd11 | -0.266994069 | -0.568859337 | 0.034871198 | 10.71982326 | -2.292126337 | 0.071873484 | 0.594255995 | -3.964689266 |
| **A_52_P277139** | Ankrd17 | -0.104006005 | -0.22077853 | 0.012766521 | 12.31632705 | -2.191224634 | 0.071956675 | 0.594255995 | -3.934741599 |
| **A_55_P2081981** | NA | 0.155831891 | -0.019160541 | 0.330824324 | 8.274417328 | 2.190817679 | 0.071996626 | 0.594255995 | -3.935029166 |
| **A_52_P378084** | NA | 0.124739894 | -0.015384454 | 0.264864242 | 8.792367899 | 2.190084797 | 0.072068632 | 0.594255995 | -3.935547091 |
| **A_55_P1969197** | NA | -0.110406414 | -0.234451001 | 0.013638173 | 16.68006733 | -2.189705204 | 0.072105957 | 0.594255995 | -3.935815372 |
| **A_55_P2061341** | Ift74 | -0.129499744 | -0.275026739 | 0.01602725 | 10.25421237 | -2.18924574 | 0.072151162 | 0.594255995 | -3.936140124 |
| **A_55_P2178510** | Mc3r | 1.081592537 | -0.133872386 | 2.29705746 | 13.86315452 | 2.189225081 | 0.072153195 | 0.594255995 | -3.936154726 |
| **A_52_P310530** | Rbfox2 | 0.154083397 | -0.019075446 | 0.32724224 | 10.43899581 | 2.189174253 | 0.072158198 | 0.594255995 | -3.936190654 |
| **A_51_P350073** | Ndel1 | -0.109862766 | -0.233355104 | 0.013629571 | 13.33626869 | -2.188666971 | 0.072208146 | 0.594255995 | -3.936549235 |
| **A_55_P2097072** | Gm7219 | 0.318355228 | -0.039499642 | 0.676210098 | 8.614552098 | 2.18863942 | 0.07221086 | 0.594255995 | -3.936568711 |
| **A_55_P2039196** | Gfra4 | 0.619900081 | -0.07695124 | 1.316751402 | 14.92722503 | 2.188521052 | 0.072222521 | 0.594255995 | -3.936652387 |
| **A_55_P2138960** | Tpk1 | 0.106809061 | -0.013268478 | 0.2268866 | 8.62711217 | 2.18834344 | 0.072240021 | 0.594255995 | -3.936777944 |
| **A_52_P214630** | Sox9 | -0.177658132 | -0.377420652 | 0.022104389 | 8.415103415 | -2.187964365 | 0.072277386 | 0.594255995 | -3.940049394 |
| **A_55_P2157048** | Mllt3 | 0.156848409 | -0.019540836 | 0.333237655 | 8.239757239 | 2.187646665 | 0.072308717 | 0.594255995 | -3.937270545 |
| **A_55_P2062787** | Mfap3 | -0.225885823 | -0.4799334 | 0.028161755 | 12.06880473 | -2.187474899 | 0.072325662 | 0.594255995 | -3.937391987 |
| **A_55_P2184314** | NA | -0.249306711 | -0.529731074 | 0.031117653 | 11.9733553 | -2.187194421 | 0.072353341 | 0.594255995 | -3.937590298 |
| **A_55_P1992388** | NA | -0.192727961 | -0.409529371 | 0.024073449 | 14.59322505 | -2.187015132 | 0.07237104 | 0.594255995 | -3.937717068 |
| **A_55_P2084448** | Cybc1 | 0.170258699 | -0.021341376 | 0.361858774 | 12.14782916 | 2.186164305 | 0.072455091 | 0.594255995 | -3.938318713 |
| **A_55_P2057994** | Mif4gd | 0.192550296 | -0.024220299 | 0.409320891 | 9.492298231 | 2.185309657 | 0.072539621 | 0.594255995 | -3.938923141 |
| **A_55_P2185595** | Samd4b | -0.154341982 | -0.328103956 | 0.019419992 | 13.11232831 | -2.185236744 | 0.072546837 | 0.594255995 | -3.938974711 |
| **A_51_P146813** | Mocs3 | 0.225924566 | -0.028462909 | 0.480312042 | 10.3871915 | 2.184926813 | 0.072577519 | 0.594255995 | -3.939193924 |
| **A_55_P1985298** | Krtap5-2 | 0.138720969 | -0.017482586 | 0.294924523 | 8.52620353 | 2.184843652 | 0.072585755 | 0.594255995 | -3.939252746 |
| **A_55_P2048110** | Tmem97 | 0.193566265 | -0.024398097 | 0.411530628 | 10.12493599 | 2.184808323 | 0.072589253 | 0.594255995 | -3.939277735 |
| **A_55_P2048848** | Smn1 | 0.179968545 | -0.022707746 | 0.382644836 | 10.5846974 | 2.184554182 | 0.072614427 | 0.594255995 | -3.9394575 |

| | | | | | | | | | |
|---|---|---|---|---|---|---|---|---|---|
| **A_55_P1989901** | Ddb1 | -0.267711679 | -0.569332101 | 0.033908743 | 16.32548041 | -2.183613336 | 0.0727077 | 0.594255995 | -3.940123063 |
| **A_55_P2058718** | Ephb4 | 0.177345227 | -0.022490014 | 0.377180467 | 8.937298264 | 2.18331596 | 0.072737206 | 0.594255995 | -3.94033345 |
| **A_55_P2037398** | NA | 0.492284376 | -0.062444959 | 1.04701371 | 16.3547002 | 2.183253087 | 0.072743446 | 0.594255995 | -3.940377932 |
| **A_52_P89683** | NA | -0.16573773 | -0.352512893 | 0.021037434 | 9.559843171 | -2.183088906 | 0.072759744 | 0.594255995 | -3.940494091 |
| **A_51_P414396** | Mmrn2 | -0.214121441 | -0.455430507 | 0.027187625 | 10.17424872 | -2.18300978 | 0.0727676 | 0.594255995 | -3.940550075 |
| **A_51_P214796** | Suz12 | -0.155436214 | -0.330636999 | 0.01976457 | 10.50600658 | -2.182656164 | 0.072802718 | 0.594255995 | -3.940800276 |
| **A_55_P2009567** | Ptbp3 | -0.189852818 | -0.403888708 | 0.024183071 | 12.1231854 | -2.182225419 | 0.07284552 | 0.594255995 | -3.941105067 |
| **A_52_P445828** | Abcf2 | -0.150514963 | -0.320213519 | 0.019183593 | 8.845292184 | -2.182080123 | 0.072859963 | 0.594255995 | -3.941207882 |
| **A_55_P1995449** | NA | 0.497332787 | -0.06339925 | 1.058064823 | 11.92265918 | 2.182030839 | 0.072864863 | 0.594255995 | -3.941242757 |
| **A_55_P2069306** | Ptpn18 | -0.211919097 | -0.450907736 | 0.027069543 | 9.986233554 | -2.181534072 | 0.072914271 | 0.594255995 | -3.941594302 |
| **A_55_P2140195** | Gm15517 | 1.396765804 | -0.178891772 | 2.972423381 | 12.46037006 | 2.180875708 | 0.072979805 | 0.594255995 | -3.942060245 |
| **A_55_P2109589** | Slc1a4 | -0.195731991 | -0.416586478 | 0.025122497 | 11.27093748 | -2.180342642 | 0.073032911 | 0.594255995 | -3.942437545 |
| **A_52_P101333** | Sap18 | -0.130600393 | -0.277982643 | 0.016781857 | 8.317994287 | -2.180060095 | 0.073061076 | 0.594255995 | -3.942637543 |
| **A_55_P2182433** | Atg10 | -0.103565776 | -0.220454771 | 0.01332322 | 8.405210382 | -2.179775657 | 0.073089441 | 0.594255995 | -3.942838889 |
| **A_55_P2016739** | Rbm12b2 | 0.191420293 | -0.024670949 | 0.407511535 | 8.919762238 | 2.179314825 | 0.07313542 | 0.594255995 | -3.943165118 |
| **A_55_P2230506** | 4931402H11Rik | -0.228420686 | -0.48628492 | 0.029443548 | 15.35369685 | -2.179282266 | 0.073138669 | 0.594255995 | -3.943188167 |
| **A_51_P206251** | Ahcyl1 | -0.164049086 | -0.34924452 | 0.021146349 | 9.388823655 | -2.179278328 | 0.073139062 | 0.594255995 | -3.943190955 |
| **A_55_P2010962** | Surf6 | 0.116286101 | -0.015019437 | 0.24759164 | 10.28239963 | 2.178782755 | 0.073188543 | 0.594255995 | -3.943541806 |
| **A_55_P2098499** | Gnpda2 | -0.117854434 | -0.250956828 | 0.01524796 | 9.266258936 | -2.178357838 | 0.073230997 | 0.594255995 | -3.943842655 |
| **A_52_P482983** | Cd164 | -0.218453863 | -0.465219911 | 0.028312185 | 15.76083697 | -2.17792784 | 0.073273984 | 0.594255995 | -3.944147122 |
| **A_55_P2013078** | Thoc5 | -0.141995442 | -0.302421598 | 0.018430714 | 9.395999824 | -2.177551276 | 0.073311651 | 0.594255995 | -3.944413771 |
| **A_55_P2133187** | NA | -0.187272253 | -0.398872441 | 0.024327936 | 8.236088201 | -2.177341481 | 0.073332645 | 0.594255995 | -3.944562336 |
| **A_55_P2029061** | Tnfrsf9 | -0.221500311 | -0.471780999 | 0.028780377 | 11.98608065 | -2.177289426 | 0.073337855 | 0.594255995 | -3.9445992 |
| **A_55_P2017774** | NA | -0.165649834 | -0.352844763 | 0.021545095 | 17.16202555 | -2.177038398 | 0.073362986 | 0.594255995 | -3.944776972 |
| **A_55_P2085672** | NA | 0.117823054 | -0.015341558 | 0.250987666 | 14.19240904 | 2.176760317 | 0.073390834 | 0.594255995 | -3.944973909 |
| **A_55_P2040110** | Cdv3 | -0.204200068 | -0.435051744 | 0.026651608 | 8.938750546 | -2.176165969 | 0.073450393 | 0.594255995 | -3.945394858 |
| **A_66_P119420** | Foxk1 | -0.186361369 | -0.397072374 | 0.024349637 | 10.35716201 | -2.175894498 | 0.073477613 | 0.594255995 | -3.94558714 |
| **A_51_P284577** | Hyal2 | -0.19737843 | -0.420555776 | 0.025798917 | 12.15114481 | -2.175798853 | 0.073487206 | 0.594255995 | -3.945654887 |
| **A_65_P16796** | Psd4 | -0.131936722 | -0.281129965 | 0.017256522 | 8.419427536 | -2.175633289 | 0.073503814 | 0.594255995 | -3.945772162 |
| **A_55_P2031999** | Dglucy | 0.256743155 | -0.033604601 | 0.547090911 | 8.792548203 | 2.17545229 | 0.073521975 | 0.594255995 | -3.945900373 |
| **A_55_P2010966** | Zfp688 | -0.151598195 | -0.323045387 | 0.019848997 | 9.29189236 | -2.175368414 | 0.073530393 | 0.594255995 | -3.945959788 |
| **A_66_P100268** | NA | 0.251800338 | -0.032983335 | 0.536584011 | 9.909927708 | 2.175256051 | 0.073541671 | 0.594255995 | -3.946039383 |
| **A_52_P40241** | Zfp746 | -0.109923753 | -0.234269926 | 0.014422419 | 9.248693925 | -2.174844894 | 0.073582954 | 0.594255995 | -3.94633065 |
| **A_55_P2023200** | Miox | 0.125889697 | -0.016539051 | 0.268318445 | 7.904806797 | 2.17451143 | 0.073616454 | 0.594255995 | -3.946566892 |
| **A_55_P1954436** | NA | -0.290765184 | -0.619751925 | 0.038221557 | 9.031342595 | -2.174368614 | 0.073630806 | 0.594255995 | -3.946668073 |
| **A_52_P555235** | Rgs19 | -0.132101157 | -0.281601784 | 0.017399471 | 8.187994489 | -2.173865992 | 0.07368134 | 0.594255995 | -3.947024185 |
| **A_52_P488612** | Btbd9 | -0.142592609 | -0.303973144 | 0.018787926 | 9.313644967 | -2.173777184 | 0.073690272 | 0.594255995 | -3.947087109 |
| **A_52_P634122** | Rexo1 | -0.167992245 | -0.358119133 | 0.022134643 | 8.45841643 | -2.173776338 | 0.073690357 | 0.594255995 | -3.947087709 |
| **A_55_P2012960** | Gpx7 | 0.262986028 | -0.034665362 | 0.560637419 | 8.538501108 | 2.173671562 | 0.073700897 | 0.594255995 | -3.947161948 |
| **A_52_P432919** | Rab3d | 0.224729326 | -0.029639749 | 0.479098402 | 8.897223271 | 2.17352481 | 0.073715662 | 0.594255995 | -3.947265931 |
| **A_66_P124445** | Rmc1 | -0.164306336 | -0.350302293 | 0.02168962 | 8.681875511 | -2.173301461 | 0.07373814 | 0.594255995 | -3.947424193 |
| **A_55_P2042993** | Arl6ip6 | -0.258945353 | -0.552084249 | 0.034193544 | 15.16518131 | -2.173220712 | 0.073746269 | 0.594255995 | -3.947481412 |
| **A_55_P1964658** | NA | -0.132275682 | -0.282057122 | 0.017505757 | 11.32879164 | -2.172657024 | 0.073803036 | 0.594255995 | -3.947880863 |
| **A_55_P2107447** | Rab37 | 0.128042363 | -0.016986436 | 0.273071162 | 8.04833615 | 2.172043876 | 0.073864835 | 0.594255995 | -3.948315402 |
| **A_52_P322421** | Mpzl2 | 0.278107895 | -0.036901297 | 0.593117088 | 10.78791696 | 2.171997165 | 0.073869546 | 0.594255995 | -3.948348508 |
| **A_55_P2170536** | Zfp949 | -0.108391427 | -0.231199196 | 0.014416342 | 8.44918474 | -2.171392104 | 0.073930587 | 0.594255995 | -3.948777359 |
| **A_55_P1998801** | NA | -0.152402606 | -0.325133851 | 0.020328639 | 9.774266243 | -2.170654249 | 0.074005095 | 0.594255995 | -3.949300385 |
| **A_55_P2075769** | Ube2n | -0.204153566 | -0.435551986 | 0.027244854 | 9.260698269 | -2.170529755 | 0.074017674 | 0.594255995 | -3.949388639 |
| **A_52_P180335** | Mrpl28 | -0.171380736 | -0.365701903 | 0.022940431 | 8.966962692 | -2.169756787 | 0.074095824 | 0.594255995 | -3.949936628 |
| **A_55_P2152009** | Tesc | 0.226899439 | -0.030440986 | 0.484239864 | 9.032557321 | 2.169174901 | 0.074154712 | 0.594255995 | -3.950349195 |

| | | | | | | | | |
|---|---|---|---|---|---|---|---|---|
| A_55_P2077872 | Deaf1 | -0.16285289 | -0.347621396 | 0.021915617 | 10.522113 | -2.168386442 | 0.074234582 | 0.594255995 | -3.950908283 |
| A_55_P2020786 | Prkar1a | 0.182490747 | -0.024573891 | 0.389555385 | 12.84253915 | 2.168223639 | 0.074251085 | 0.594255995 | -3.951023734 |
| A_55_P2030433 | Gpi1 | 0.227666068 | -0.030670536 | 0.486002672 | 13.95831777 | 2.168111057 | 0.074262499 | 0.594255995 | -3.951103572 |
| A_55_P2144305 | Gm3211 | 0.116279054 | -0.015667234 | 0.248225341 | 8.83193464 | 2.168070884 | 0.074266572 | 0.594255995 | -3.951132061 |
| A_55_P1996817 | NA | -0.128564694 | -0.274498746 | 0.017369357 | 10.64344913 | -2.167375927 | 0.074337076 | 0.594255995 | -3.951624929 |
| A_55_P2169839 | Rufy3 | -0.266883248 | -0.569841792 | 0.036075296 | 11.98914362 | -2.167241324 | 0.074350739 | 0.594255995 | -3.951720396 |
| A_55_P2203141 | 2900011F02Rik | 0.137671667 | -0.018628664 | 0.293971999 | 9.42189604 | 2.166974675 | 0.074377814 | 0.594255995 | -3.951909523 |
| A_66_P128631 | Cinp | 0.395630248 | -0.053535663 | 0.844796158 | 9.362248571 | 2.166964832 | 0.074378814 | 0.594255995 | -3.951916504 |
| A_55_P1962289 | NA | -0.353528 | -0.754916692 | 0.047860692 | 8.711803801 | -2.166845001 | 0.074390984 | 0.594255995 | -3.952001499 |
| A_55_P2164615 | NA | -0.166696579 | -0.355967828 | 0.02257467 | 15.9962645 | -2.166761954 | 0.074399421 | 0.594255995 | -3.952060405 |
| A_51_P129803 | Cacybp | -0.123724279 | -0.264250984 | 0.016802427 | 11.42055272 | -2.1660338 | 0.07447343 | 0.594255995 | -3.95257692 |
| A_55_P2166743 | Sp3os | 0.165461056 | -0.022473345 | 0.353395457 | 11.19034187 | 2.166001093 | 0.074476756 | 0.594255995 | -3.952600121 |
| A_55_P2014645 | Knop1 | -0.203704744 | -0.435167307 | 0.027757818 | 10.83925571 | -2.165157784 | 0.074562568 | 0.594255995 | -3.953198396 |
| A_55_P2077909 | Tpgs2 | 0.207407557 | -0.028265199 | 0.443080313 | 8.461331748 | 2.165131897 | 0.074565204 | 0.594255995 | -3.953216762 |
| A_55_P2091348 | NA | -0.139271126 | -0.297579315 | 0.019037063 | 9.715379737 | -2.1643469 | 0.074645178 | 0.594255995 | -3.953773739 |
| A_55_P2036733 | Stk36 | 0.186958832 | -0.025590961 | 0.399508626 | 8.313559949 | 2.163986078 | 0.074681967 | 0.594255995 | -3.954029774 |
| A_51_P347862 | Actn1 | 0.091425756 | -0.012519862 | 0.195371374 | 8.850696612 | 2.163871845 | 0.074693618 | 0.594255995 | -3.954110836 |
| A_51_P193446 | Prpf40a | -0.113941426 | -0.243517602 | 0.01563475 | 8.869774475 | -2.16334435 | 0.074747443 | 0.594255995 | -3.954485172 |
| A_55_P1988259 | Gng5 | -0.111862755 | -0.239092145 | 0.015366635 | 15.76756876 | -2.163053347 | 0.074777154 | 0.594255995 | -3.954691695 |
| A_51_P500550 | Mfsd1 | -0.225300605 | -0.481604493 | 0.031003284 | 9.076474924 | -2.162600633 | 0.0748234 | 0.594255995 | -3.955013001 |
| A_66_P118368 | Catip | 0.284348373 | -0.039195376 | 0.607892122 | 9.053326183 | 2.162155362 | 0.074868914 | 0.594255995 | -3.955329046 |
| A_55_P1990525 | Ube2v1 | -0.353979367 | -0.756756372 | 0.048797638 | 14.19841942 | -2.162133191 | 0.074871181 | 0.594255995 | -3.955344783 |
| A_51_P184910 | Phpt1 | 0.205175619 | -0.028330004 | 0.438681242 | 11.62170994 | 2.161710705 | 0.074914394 | 0.594255995 | -3.955644676 |
| A_55_P2023042 | Ift43 | -0.191896915 | -0.410303916 | 0.026510087 | 11.98332673 | -2.161576441 | 0.074928132 | 0.594255995 | -3.955739985 |
| A_55_P2019751 | Paox | 0.134955253 | -0.018663667 | 0.288574174 | 9.908794697 | 2.161296002 | 0.074956836 | 0.594255995 | -3.955939063 |
| A_55_P2174576 | Nova1 | 0.413031415 | -0.057248236 | 0.883311066 | 8.350416013 | 2.160707905 | 0.075017066 | 0.594255995 | -3.95635657 |
| A_55_P1958140 | B9d1 | 0.184909821 | -0.025666829 | 0.395486471 | 10.18116717 | 2.160324193 | 0.075056391 | 0.594255995 | -3.956628998 |
| A_55_P2139814 | Fgf13 | 0.479787649 | -0.066606027 | 1.026181324 | 8.418697827 | 2.160292526 | 0.075059637 | 0.594255995 | -3.956651482 |
| A_55_P2025153 | Hmgb1 | -0.481613534 | -1.030113344 | 0.066886277 | 13.98259034 | -2.160187078 | 0.075070448 | 0.594255995 | -3.956726351 |
| A_55_P2109744 | NA | -0.111811443 | -0.239160467 | 0.015537581 | 16.03628912 | -2.160030066 | 0.075086549 | 0.594255995 | -3.956837834 |
| A_55_P2024327 | Kmt5c | 0.159322461 | -0.022152563 | 0.340797484 | 11.61385123 | 2.159878424 | 0.075102102 | 0.594255995 | -3.956945505 |
| A_55_P2155901 | Snhg10 | 0.110187179 | -0.015329274 | 0.235703631 | 8.735374596 | 2.15973054 | 0.075117273 | 0.594255995 | -3.957050511 |
| A_55_P2040774 | NA | -0.45090505 | -0.964586356 | 0.062776255 | 13.80391485 | -2.159536268 | 0.075137208 | 0.594255995 | -3.957188459 |
| A_55_P2078994 | Lrrc31 | 0.215328828 | -0.030003639 | 0.460661295 | 16.76318318 | 2.15931655 | 0.075159761 | 0.594255995 | -3.95734448 |
| A_52_P587738 | P2ry2 | 0.132792012 | -0.018514061 | 0.284098085 | 9.009001457 | 2.159159639 | 0.075175871 | 0.594255995 | -3.957455906 |
| A_55_P1988766 | NA | 0.469844259 | -0.065569443 | 1.005257961 | 8.759038212 | 2.15890537 | 0.075201984 | 0.594255995 | -3.957636472 |
| A_55_P2120961 | NA | 0.197514829 | -0.027620059 | 0.422649717 | 10.72772184 | 2.15837086 | 0.075256908 | 0.594255995 | -3.95801607 |
| A_52_P46419 | Grk6 | -0.132308976 | -0.283183983 | 0.01856603 | 8.75834148 | -2.157452143 | 0.075351409 | 0.594255995 | -3.958668597 |
| A_52_P51548 | Pard3 | -0.128744222 | -0.275599879 | 0.018111436 | 10.48238626 | -2.156781889 | 0.07542043 | 0.594255995 | -3.959144707 |
| A_52_P555672 | Vrk1 | -0.112783838 | -0.241435622 | 0.015867946 | 8.346489147 | -2.156752048 | 0.075423504 | 0.594255995 | -3.959165905 |
| A_55_P2152304 | Atp5h | 0.467759649 | -0.065813713 | 1.001333011 | 15.41599945 | 2.156739925 | 0.075424753 | 0.594255995 | -3.959174517 |
| A_52_P40367 | Mrpl42 | -0.206644622 | -0.442371706 | 0.029082462 | 9.748450258 | -2.156670441 | 0.075431913 | 0.594255995 | -3.959223878 |
| A_52_P331212 | Cspp1 | -0.140080137 | -0.299880326 | 0.019720052 | 8.828401053 | -2.156594193 | 0.07543977 | 0.594255995 | -3.959278044 |
| A_55_P2077033 | Ube2e1 | -0.14845779 | -0.317829765 | 0.020914186 | 13.05271572 | -2.156406345 | 0.075459131 | 0.594255995 | -3.959411493 |
| A_55_P1968606 | Synrg | 0.193372361 | -0.02726952 | 0.414014241 | 17.17393001 | 2.156133327 | 0.075487279 | 0.594255995 | -3.959605454 |
| A_55_P2167323 | Cort | 0.311177655 | -0.043929461 | 0.666284771 | 8.354865531 | 2.155848272 | 0.07551668 | 0.594255995 | -3.959807975 |
| A_51_P430950 | Rdx | -0.225677948 | -0.483236668 | 0.031880771 | 11.26348768 | -2.155668771 | 0.075535199 | 0.594255995 | -3.959935508 |
| A_52_P275069 | Epp13 | 0.742427334 | -0.105216309 | 1.590070977 | 9.146985796 | 2.154814091 | 0.075623445 | 0.594255995 | -3.960542794 |
| A_51_P477941 | Fbxw4 | -0.178753809 | -0.382879466 | 0.025371848 | 9.776466241 | -2.154402637 | 0.075665965 | 0.594255995 | -3.960835177 |
| A_55_P2137496 | NA | 0.200708836 | -0.028500454 | 0.429918127 | 8.094433622 | 2.154286402 | 0.075677981 | 0.594255995 | -3.960917778 |

| | | | | | | | | | |
|---|---|---|---|---|---|---|---|---|---|
| **A_51_P475613** | Fahd2a | -0.099778016 | -0.213746701 | 0.01419067 | 9.502620498 | -2.15386504 | 0.075721558 | 0.594255995 | -3.961217224 |
| **A_55_P2112225** | NA | 0.499457076 | -0.071055548 | 1.0699697 | 9.847469112 | 2.153783645 | 0.075729979 | 0.594255995 | -3.961275072 |
| **A_52_P45834** | Pafah1b2 | -0.153972724 | -0.329851364 | 0.021905917 | 9.11539665 | -2.153772638 | 0.075731118 | 0.594255995 | -3.961282894 |
| **A_55_P2152468** | NA | -0.131761602 | -0.282327028 | 0.018803825 | 9.181669755 | -2.15294418 | 0.075816882 | 0.594255995 | -3.961871713 |
| **A_55_P2167025** | Nectin2 | 0.269423711 | -0.038458345 | 0.577305768 | 14.03278645 | 2.152883843 | 0.075823133 | 0.594255995 | -3.9619146 |
| **A_55_P2066799** | Sctr | -0.300063515 | -0.642989209 | 0.04286218 | 9.366174512 | -2.152694139 | 0.075842787 | 0.594255995 | -3.962049442 |
| **A_55_P2008021** | Srpx2 | 0.63629043 | -0.090904713 | 1.363485572 | 10.04125933 | 2.1526508 | 0.075847278 | 0.594255995 | -3.962080248 |
| **A_55_P2182358** | Arid3a | 0.199627461 | -0.028521313 | 0.427776236 | 12.41997577 | 2.152639506 | 0.075848448 | 0.594255995 | -3.962088275 |
| **A_52_P327615** | Rab18 | -0.181553554 | -0.389095525 | 0.025988417 | 11.80486529 | -2.152127416 | 0.075901535 | 0.594255995 | -3.962452292 |
| **A_55_P2207335** | Nron | 0.394844199 | -0.056564272 | 0.846252669 | 8.82148959 | 2.151915533 | 0.075923511 | 0.594255995 | -3.962602916 |
| **A_51_P393598** | Atp6v1d | 0.198414201 | -0.028455926 | 0.425284328 | 13.27568784 | 2.151615188 | 0.075954673 | 0.594255995 | -3.962816433 |
| **A_55_P2142156** | Gm16441 | 0.123781785 | -0.017758274 | 0.265321845 | 8.697075309 | 2.151525664 | 0.075963964 | 0.594255995 | -3.962880078 |
| **A_55_P1965539** | Ndufb4 | 0.128865919 | -0.018511262 | 0.276243099 | 14.5022887 | 2.151181197 | 0.075999726 | 0.594255995 | -3.963124978 |
| **A_52_P645159** | Cpsf6 | -0.168036756 | -0.360219608 | 0.024146097 | 8.549926164 | -2.151091164 | 0.076009075 | 0.594255995 | -3.963188989 |
| **A_51_P479583** | Obsl1 | -0.141458063 | -0.303250784 | 0.020334658 | 10.37791074 | -2.150987519 | 0.07601984 | 0.594255995 | -3.96326268 |
| **A_55_P1968863** | Synpo2l | -0.231782605 | -0.496897174 | 0.033331964 | 13.31811611 | -2.150881127 | 0.076030892 | 0.594255995 | -3.963338323 |
| **A_66_P119801** | Optn | -0.160279543 | -0.343661233 | 0.023102147 | 8.657999938 | -2.150261547 | 0.076095285 | 0.594255995 | -3.963778866 |
| **A_52_P598912** | Erc1 | -0.18674872 | -0.400429808 | 0.026932368 | 8.347246948 | -2.15011016 | 0.076111027 | 0.594255995 | -3.963886514 |
| **A_55_P1979178** | NA | -0.191240759 | -0.410103179 | 0.027621661 | 12.63888986 | -2.149702818 | 0.076153402 | 0.594255995 | -3.964176176 |
| **A_55_P2425437** | D030036P13Rik | 0.173172246 | -0.025015002 | 0.371359494 | 9.43772017 | 2.149669755 | 0.076156843 | 0.594255995 | -3.964199689 |
| **A_55_P2077623** | NA | 0.39323001 | -0.056827001 | 0.84328702 | 9.759937663 | 2.149553636 | 0.076168927 | 0.594255995 | -3.964282265 |
| **A_55_P2049976** | NA | 0.464484913 | -0.067140966 | 0.996110792 | 8.728548488 | 2.149486202 | 0.076175946 | 0.594255995 | -3.96433022 |
| **A_51_P214747** | Parp12 | -0.16603917 | -0.356083685 | 0.024005346 | 8.839474848 | -2.14943529 | 0.076181246 | 0.594255995 | -3.964366427 |
| **A_55_P2061991** | Pnldc1 | 0.538930236 | -0.077924651 | 1.155785123 | 9.203214782 | 2.149407218 | 0.076184168 | 0.594255995 | -3.96438639 |
| **A_55_P2182931** | Sim2 | 0.190734949 | -0.027593479 | 0.409063377 | 8.677737989 | 2.14926097 | 0.076199394 | 0.594255995 | -3.964490396 |
| **A_55_P2065731** | Medag | 0.132702966 | -0.019217464 | 0.284623396 | 17.13657271 | 2.148986121 | 0.076228018 | 0.594255995 | -3.964685865 |
| **A_55_P2017418** | Cfh | 0.168223122 | -0.024374333 | 0.360820577 | 8.881691425 | 2.148841123 | 0.076243122 | 0.594255995 | -3.964788988 |
| **A_52_P152952** | Leng1 | -0.142080185 | -0.304748315 | 0.020587946 | 8.847645648 | -2.1488208 | 0.07624524 | 0.594255995 | -3.964803443 |
| **A_55_P2092884** | NA | 0.418983489 | -0.060934698 | 0.898901676 | 9.154592114 | 2.147824854 | 0.076349077 | 0.594255995 | -3.965511829 |
| **A_55_P1997282** | Fam135a | -0.222538909 | -0.477444982 | 0.032367164 | 9.471879361 | -2.147805429 | 0.076351104 | 0.594255995 | -3.965525647 |
| **A_55_P2083149** | NA | 0.273069367 | -0.039737478 | 0.585876213 | 10.06536056 | 2.147661831 | 0.076366088 | 0.594255995 | -3.965627792 |
| **A_55_P2118906** | Atp5j2 | 0.252792218 | -0.036860414 | 0.54244485 | 14.94305846 | 2.147115387 | 0.076423135 | 0.594255995 | -3.966016515 |
| **A_55_P2015541** | Hif3a | 0.371215012 | -0.054165021 | 0.796595045 | 10.19418813 | 2.146928559 | 0.076442649 | 0.594255995 | -3.966149426 |
| **A_55_P2052475** | NA | 0.285141871 | -0.041705486 | 0.611989228 | 15.87764532 | 2.14627408 | 0.076511051 | 0.594255995 | -3.966615055 |
| **A_55_P2175615** | Csde1 | -0.222144538 | -0.476782528 | 0.032493451 | 13.07273886 | -2.146256418 | 0.076512897 | 0.594255995 | -3.966627621 |
| **A_51_P214127** | Cpa3 | 0.249815009 | -0.036571601 | 0.53620162 | 7.877710001 | 2.146026013 | 0.076536993 | 0.594255995 | -3.966791554 |
| **A_51_P392244** | Tbcd | 0.210582689 | -0.030859724 | 0.452025102 | 10.64354261 | 2.145745786 | 0.07656631 | 0.594255995 | -3.966990942 |
| **A_65_P10351** | Atp2b1 | -0.146822516 | -0.315162706 | 0.021517674 | 9.271974755 | -2.14572475 | 0.076568511 | 0.594255995 | -3.96700591 |
| **A_51_P315673** | Usp11 | 0.207116188 | -0.030368641 | 0.444601017 | 9.050778104 | 2.14559296 | 0.076582303 | 0.594255995 | -3.967099685 |
| **A_55_P2121231** | Nipbl | -0.162475168 | -0.348778545 | 0.02382821 | 8.403376584 | -2.145534137 | 0.07658846 | 0.594255995 | -3.967141541 |
| **A_66_P121361** | C330021F23Rik | 0.117016648 | -0.017179708 | 0.251213004 | 9.186087229 | 2.14524103 | 0.076619146 | 0.594255995 | -3.967350109 |
| **A_52_P146457** | NA | 0.509712857 | -0.074838454 | 1.094264168 | 15.87707472 | 2.145221358 | 0.076621206 | 0.594255995 | -3.967364107 |
| **A_55_P2142393** | Cacna1a | -0.183416821 | -0.393892327 | 0.027058686 | 8.406933591 | -2.143911042 | 0.076758543 | 0.594255995 | -3.968296609 |
| **A_55_P2055291** | Gpn2 | -0.154109946 | -0.330975232 | 0.022755341 | 8.933537841 | -2.143666533 | 0.076784199 | 0.594255995 | -3.968470636 |
| **A_55_P1957028** | Kat7 | -0.244094025 | -0.524235255 | 0.036047205 | 12.65191903 | -2.14362731 | 0.076788315 | 0.594255995 | -3.968498553 |
| **A_55_P2066533** | NA | 0.224007078 | -0.033162302 | 0.481176458 | 15.68001541 | 2.14294807 | 0.076859636 | 0.594255995 | -3.968982033 |
| **A_55_P2006280** | Ogdh | 0.200621562 | -0.029823695 | 0.431066819 | 12.74810817 | 2.141800407 | 0.076980298 | 0.594255995 | -3.969799041 |
| **A_55_P2025403** | Gphn | -0.12152141 | -0.261184224 | 0.018141405 | 9.123407228 | -2.140627858 | 0.077103776 | 0.594255995 | -3.970633906 |
| **A_55_P2179604** | Vash1 | -0.489860891 | -1.052857615 | 0.073135834 | 13.90104685 | -2.140602669 | 0.077106431 | 0.594255995 | -3.970651842 |
| **A_52_P561671** | Msx1 | 0.201153752 | -0.030033925 | 0.432341429 | 8.44818109 | 2.140585697 | 0.07710822 | 0.594255995 | -3.970663928 |

| | | | | | | | | |
|---|---|---|---|---|---|---|---|---|
| A_55_P2109222 | Rps3 | 0.274548046 | -0.041022197 | 0.590118289 | 15.49014294 | 2.140382916 | 0.077129597 | 0.594255995 | -3.970808325 |
| A_51_P450296 | Thap3 | -0.182543318 | -0.392405361 | 0.027318726 | 8.970461843 | -2.13993807 | 0.077176512 | 0.594255995 | -3.971125107 |
| A_51_P109369 | Fbxo32 | -0.294611497 | -0.633319608 | 0.044096614 | 11.01241229 | -2.139898911 | 0.077180643 | 0.594255995 | -3.995740534 |
| A_55_P1963764 | 1-Sep | 0.354265843 | -0.05303656 | 0.761568246 | 8.58067618 | 2.139840842 | 0.07718677 | 0.594255995 | -3.971194348 |
| A_55_P2114838 | NA | -0.281316692 | -0.604775437 | 0.042142054 | 14.48094104 | -2.139664918 | 0.077205334 | 0.594255995 | -3.971319634 |
| A_55_P2166421 | Rpusd3 | 0.215074801 | -0.032253982 | 0.462403584 | 9.517762524 | 2.139360796 | 0.077237438 | 0.594255995 | -3.971536225 |
| A_55_P2187918 | Cep55 | 0.11390343 | -0.017085388 | 0.244892247 | 8.072074593 | 2.139300245 | 0.077243831 | 0.594255995 | -3.971579349 |
| A_55_P1980721 | NA | -0.185965619 | -0.399856778 | 0.027925539 | 12.7519369 | -2.138991172 | 0.077276474 | 0.594255995 | -3.971799478 |
| A_51_P352303 | Homer2 | -0.142893466 | -0.307290955 | 0.021504024 | 8.967976075 | -2.138387228 | 0.0773403 | 0.594255995 | -3.972229648 |
| A_52_P238044 | Psmc3ip | -0.116991365 | -0.251596702 | 0.017613972 | 8.004373787 | -2.138260885 | 0.077353659 | 0.594255995 | -3.972319643 |
| A_55_P2180445 | NA | 0.550320809 | -0.082930407 | 1.183572025 | 11.15629366 | 2.138006718 | 0.077380541 | 0.594255995 | -3.972500693 |
| A_66_P102607 | Btrc | -0.115402871 | -0.248212156 | 0.017406415 | 8.420268451 | -2.137752062 | 0.077407485 | 0.594255995 | -3.972682098 |
| A_55_P1976983 | Rpl36 | 0.158675493 | -0.023952887 | 0.341303873 | 16.13153466 | 2.137522788 | 0.077431751 | 0.594255995 | -3.972845427 |
| A_55_P2051592 | Gfm2 | 0.100672823 | -0.015203553 | 0.216549199 | 9.461380565 | 2.137403447 | 0.077444385 | 0.594255995 | -3.972930444 |
| A_51_P397375 | Gatb | -0.126751959 | -0.272661828 | 0.01915791 | 8.919583972 | -2.137170528 | 0.077469049 | 0.594255995 | -3.973096379 |
| A_55_P1974827 | Pde1b | 0.250054228 | -0.037825614 | 0.537934071 | 9.189588073 | 2.136938922 | 0.077493582 | 0.594255995 | -3.973261384 |
| A_55_P1967983 | Use1 | 0.283288557 | -0.042924474 | 0.609501588 | 12.37264335 | 2.136470445 | 0.077543231 | 0.594255995 | -3.97359516 |
| A_55_P2184414 | Zfp330 | -0.164722128 | -0.354441211 | 0.024996955 | 11.0615818 | -2.136043484 | 0.077588508 | 0.594255995 | -3.973899378 |
| A_51_P113072 | Cdc123 | -0.171320744 | -0.368648936 | 0.026007449 | 11.29465067 | -2.135944562 | 0.077599002 | 0.594255995 | -3.976660201 |
| A_55_P2143015 | Sema6c | 0.240523413 | -0.036554931 | 0.517601757 | 8.930994905 | 2.135619796 | 0.077633465 | 0.594255995 | -3.974201282 |
| A_55_P1965945 | NA | -0.165679433 | -0.35656384 | 0.025204975 | 16.31850674 | -2.135341336 | 0.077663026 | 0.594255995 | -3.974399711 |
| A_55_P1954850 | Pars2 | 0.222492424 | -0.03386914 | 0.478853988 | 10.00610125 | 2.135165175 | 0.077681734 | 0.594255995 | -3.974525247 |
| A_55_P2002819 | Rabgap1l | -0.131865378 | -0.283824022 | 0.020093265 | 8.437914344 | -2.134885263 | 0.077711468 | 0.594255995 | -3.974724725 |
| A_55_P1963150 | Creg1 | -0.218760355 | -0.470858196 | 0.033337485 | 17.18400959 | -2.134856348 | 0.077714541 | 0.594255995 | -3.974745331 |
| A_55_P1954755 | NA | 0.21181997 | -0.032300362 | 0.455940301 | 16.45033292 | 2.134676721 | 0.077733629 | 0.594255995 | -3.974873347 |
| A_55_P2047345 | Sat2 | 0.139663948 | -0.021300327 | 0.300628223 | 9.297148902 | 2.134636746 | 0.077737878 | 0.594255995 | -3.974901836 |
| A_55_P2064568 | Anks1 | 0.123597266 | -0.018913329 | 0.266107861 | 10.85142788 | 2.133687796 | 0.077838806 | 0.594255995 | -3.975578182 |
| A_55_P2213418 | 4933417E11Rik | -0.221267647 | -0.47640475 | 0.033869456 | 8.869420411 | -2.133602251 | 0.077847911 | 0.594255995 | -3.975639157 |
| A_55_P1992019 | Ptpru | -0.298483755 | -0.642702297 | 0.045734788 | 12.64774025 | -2.133318015 | 0.077878172 | 0.594255995 | -3.975841762 |
| A_51_P321651 | Fam192a | -0.170179333 | -0.366443421 | 0.026084755 | 10.60705826 | -2.13321751 | 0.077888875 | 0.594255995 | -3.975913404 |
| A_51_P498257 | NA | -0.201795104 | -0.434545552 | 0.030955344 | 9.672082192 | -2.132992178 | 0.077912876 | 0.594255995 | -3.97607403 |
| A_55_P1984333 | Emd | -0.104636731 | -0.225362915 | 0.016089453 | 8.441122566 | -2.132317414 | 0.077984795 | 0.594255995 | -3.976555059 |
| A_51_P473219 | Siae | -0.219340333 | -0.4724599 | 0.033779234 | 10.20261276 | -2.131876008 | 0.078031879 | 0.594255995 | -3.976869755 |
| A_52_P6828 | Xk | -0.146357749 | -0.315274433 | 0.022558936 | 8.295280761 | -2.131632512 | 0.078057864 | 0.594255995 | -3.977043363 |
| A_52_P630493 | Dnajb6 | -0.164637676 | -0.354666013 | 0.025390661 | 10.05739339 | -2.131473919 | 0.078074794 | 0.594255995 | -3.977156439 |
| A_55_P2026889 | NA | -0.19413041 | -0.418243542 | 0.029982721 | 12.97829186 | -2.131058794 | 0.078119126 | 0.594255995 | -3.977452434 |
| A_55_P2083904 | Tmem185a | -0.123196544 | -0.265468577 | 0.019075488 | 8.605740989 | -2.130336234 | 0.078196351 | 0.594255995 | -3.977967679 |
| A_55_P2068199 | NA | 0.121678793 | -0.018841481 | 0.262199067 | 8.684074044 | 2.130321089 | 0.078197971 | 0.594255995 | -3.977978479 |
| A_65_P02321 | Usp36 | -0.162735983 | -0.350695265 | 0.0252233 | 8.521043428 | -2.130045968 | 0.078227396 | 0.594255995 | -3.978174678 |
| A_55_P1999710 | Smad2 | -0.147056981 | -0.316916022 | 0.022802061 | 10.36245057 | -2.129933966 | 0.078239379 | 0.594255995 | -3.978254552 |
| A_52_P371718 | Tgs1 | -0.136137657 | -0.293386971 | 0.021111658 | 9.36198985 | -2.129897348 | 0.078243297 | 0.594255995 | -3.978280667 |
| A_55_P2108933 | Dao | 0.349778062 | -0.054259465 | 0.753815589 | 8.784886411 | 2.129805901 | 0.078253082 | 0.594255995 | -3.978345884 |
| A_55_P1964812 | Tfap2a | 0.161503781 | -0.025067481 | 0.348075042 | 10.28374178 | 2.129644472 | 0.078270359 | 0.594255995 | -3.978461012 |
| A_51_P246543 | Ebf3 | -0.12844402 | -0.276865709 | 0.01997767 | 8.333940623 | -2.129049112 | 0.07833411 | 0.594255995 | -3.978885635 |
| A_66_P103092 | NA | 0.216053243 | -0.033609259 | 0.465715746 | 12.02758304 | 2.129004768 | 0.078338861 | 0.594255995 | -3.978917263 |
| A_55_P1989469 | NA | 0.215904938 | -0.033619755 | 0.465429631 | 15.5795771 | 2.128718374 | 0.078369549 | 0.594255995 | -3.979121539 |
| A_66_P121196 | Katnb1 | 0.195931914 | -0.030556256 | 0.422420085 | 10.46383525 | 2.128280281 | 0.078416516 | 0.594255995 | -3.979434033 |
| A_55_P2000533 | Polk | -0.109019471 | -0.235071402 | 0.017032459 | 8.463046189 | -2.127765354 | 0.078471757 | 0.594255995 | -3.979801358 |
| A_51_P282268 | Snapc1 | -0.128350611 | -0.276759921 | 0.020058699 | 9.547013312 | -2.127678272 | 0.078481103 | 0.594255995 | -3.979863481 |
| A_55_P2127538 | Ppp1r21 | -0.138894599 | -0.299524443 | 0.021735246 | 8.428800622 | -2.127297756 | 0.078521956 | 0.594255995 | -3.980134943 |

| | | | | | | | | | |
|---|---|---|---|---|---|---|---|---|---|
| **A_55_P1966383** | NA | 0.415751266 | -0.065080901 | 0.896583434 | 11.07973975 | 2.127204433 | 0.078531978 | 0.594255995 | -3.980201523 |
| **A_55_P1979728** | Atf4 | -0.310210686 | -0.672268041 | 0.051846668 | 15.59783776 | -2.220391024 | 0.078535787 | 0.594255995 | -4.00440981 |
| **A_55_P1982034** | NA | 0.200639096 | -0.031488448 | 0.432766639 | 8.564052567 | 2.126464069 | 0.078611537 | 0.594255995 | -3.980729752 |
| **A_55_P1971513** | NA | 0.249642631 | -0.03918543 | 0.538470693 | 16.40886445 | 2.126417458 | 0.078616549 | 0.594255995 | -3.98076301 |
| **A_51_P487105** | Bud31 | -0.208863042 | -0.45051148 | 0.032785396 | 10.16100691 | -2.126408844 | 0.078617475 | 0.594255995 | -3.980769156 |
| **A_51_P131653** | Tsacc | 0.377554739 | -0.059297451 | 0.814406929 | 9.457878931 | 2.126251201 | 0.078634428 | 0.594255995 | -3.980881638 |
| **A_55_P1985005** | Ghrh | 0.323665356 | -0.050865154 | 0.698195866 | 16.0750343 | 2.126073028 | 0.078653592 | 0.594255995 | -3.981008772 |
| **A_55_P1982449** | NA | 0.35536429 | -0.05587115 | 0.76659973 | 8.33694254 | 2.125946898 | 0.078667162 | 0.594255995 | -3.981098773 |
| **A_55_P2038597** | NA | 0.256184478 | -0.040294145 | 0.552663101 | 12.08272355 | 2.125830289 | 0.078679709 | 0.594255995 | -3.98118198 |
| **A_55_P2054744** | Sh3bgrl2 | 0.10511224 | -0.016547696 | 0.226772176 | 9.255340571 | 2.125567342 | 0.07870801 | 0.594255995 | -3.981369616 |
| **A_55_P2170414** | Zfp180 | 0.180355066 | -0.028422441 | 0.389132573 | 9.189574 | 2.125268477 | 0.07874019 | 0.594255995 | -3.98158289 |
| **A_66_P122927** | NA | 0.891377457 | -0.140654256 | 1.923409171 | 10.29055116 | 2.124896429 | 0.078780269 | 0.594255995 | -3.981848402 |
| **A_51_P395736** | Srp72 | 0.198140831 | -0.031283504 | 0.427565165 | 11.4929722 | 2.12472957 | 0.07879825 | 0.594255995 | -3.981967485 |
| **A_66_P106478** | Cct3 | -0.142871297 | -0.308314603 | 0.02257201 | 10.57023438 | -2.124540171 | 0.078818666 | 0.594255995 | -3.982102657 |
| **A_51_P275454** | Trim30a | -0.114999467 | -0.24816815 | 0.018169216 | 8.551241192 | -2.124530089 | 0.078819753 | 0.594255995 | -3.982109853 |
| **A_55_P2060878** | NA | 0.289109925 | -0.045708576 | 0.623928426 | 11.10310686 | 2.124333593 | 0.07884094 | 0.594255995 | -3.982250095 |
| **A_65_P13209** | Ctsd | -0.116290113 | -0.250984464 | 0.018404238 | 9.602842927 | -2.124039431 | 0.078872668 | 0.594255995 | -3.982460049 |
| **A_55_P2022404** | Gtf2h5 | 0.602264306 | -0.095437328 | 1.299965939 | 11.90553412 | 2.123667634 | 0.078912789 | 0.594255995 | -3.982725428 |
| **A_51_P327874** | Pth1r | -0.169718989 | -0.366339011 | 0.026901032 | 11.62680835 | -2.123595819 | 0.078920541 | 0.594255995 | -3.982776688 |
| **A_55_P2138090** | U2af1l4 | 0.307652385 | -0.048872109 | 0.66417688 | 12.30002073 | 2.12295152 | 0.078990124 | 0.59435455 | -3.983236607 |
| **A_55_P2136334** | NA | 0.313937742 | -0.049918174 | 0.677793657 | 10.66140766 | 2.122673769 | 0.079020141 | 0.59435455 | -3.983434887 |
| **A_52_P46085** | Mvp | -0.232342228 | -0.501635138 | 0.036950682 | 9.528761545 | -2.122620693 | 0.079025878 | 0.59435455 | -3.983472777 |
| **A_55_P1970399** | NA | -0.253996991 | -0.548485484 | 0.040491502 | 14.23574087 | -2.121921952 | 0.079101447 | 0.594691511 | -3.983971629 |
| **A_55_P1970740** | Mrps10 | 0.3611866 | -0.057724511 | 0.78009771 | 11.56263249 | 2.121186789 | 0.079181036 | 0.594785348 | -3.984496535 |
| **A_55_P2029750** | Ndufs5 | 0.238380079 | -0.038121206 | 0.514881364 | 14.52627592 | 2.121006349 | 0.079200583 | 0.594785348 | -3.984625377 |
| **A_55_P1963206** | NA | 0.127055726 | -0.020332147 | 0.274443599 | 8.448525662 | 2.120809463 | 0.079221917 | 0.594785348 | -3.984765966 |
| **A_51_P249360** | Suox | -0.246569451 | -0.532638828 | 0.039499926 | 11.84721477 | -2.12049406 | 0.079256106 | 0.594785348 | -3.984991192 |
| **A_55_P1953231** | Map2k5 | -0.181316757 | -0.391729799 | 0.029096285 | 12.22683342 | -2.119993121 | 0.079310438 | 0.594785348 | -3.985348928 |
| **A_55_P1994564** | Sap30bp | 0.16466475 | -0.026430941 | 0.355760442 | 8.668144454 | 2.119917249 | 0.07931867 | 0.594785348 | -3.985403112 |
| **A_55_P2021923** | Fkbp1a | 0.27311866 | -0.043853963 | 0.590091283 | 12.70927614 | 2.119819012 | 0.07932933 | 0.594785348 | -3.98547327 |
| **A_55_P1954393** | Susd4 | -0.27936226 | -0.603674815 | 0.044950295 | 10.3066817 | -2.119205811 | 0.079395906 | 0.594867252 | -3.985911219 |
| **A_52_P280899** | Wdr33 | -0.137600639 | -0.29735981 | 0.022158531 | 9.265988752 | -2.118965166 | 0.079422049 | 0.594867252 | -3.986083098 |
| **A_51_P456187** | Pdcd2 | -0.206414866 | -0.446116438 | 0.033286706 | 10.72307457 | -2.118552609 | 0.079466888 | 0.594867252 | -3.986377778 |
| **A_51_P283083** | Armc1 | 0.216420501 | -0.034917569 | 0.467758571 | 11.73733129 | 2.118406427 | 0.079482782 | 0.594867252 | -3.986482197 |
| **A_55_P2077497** | NA | -0.476683179 | -1.035497095 | 0.082130738 | 11.26353241 | -2.210612878 | 0.079494134 | 0.594867252 | -4.009884 |
| **A_51_P150014** | Rrp7a | 0.246830748 | -0.039955814 | 0.53361731 | 8.520893086 | 2.117432744 | 0.079588733 | 0.595344666 | -3.987177756 |
| **A_52_P493091** | Fmr1 | -0.185339395 | -0.400748439 | 0.03006965 | 8.668503625 | -2.116766576 | 0.079661305 | 0.595510674 | -3.987653693 |
| **A_51_P470989** | Paip1 | -0.111470259 | -0.241031877 | 0.018091358 | 8.731258252 | -2.116663464 | 0.079672544 | 0.595510674 | -3.987727364 |
| **A_55_P2096834** | Fbxl19 | -0.322336181 | -0.69704952 | 0.052377158 | 13.48900712 | -2.116308939 | 0.079711199 | 0.595569296 | -3.987980673 |
| **A_51_P492346** | Tmem174 | 0.270415366 | -0.044046515 | 0.584877246 | 9.988078637 | 2.115594947 | 0.079789107 | 0.595609243 | -3.988490858 |
| **A_52_P464003** | Atg3 | -0.137602292 | -0.297652945 | 0.02244836 | 10.1657099 | -2.115131545 | 0.079839713 | 0.595609243 | -3.988822009 |
| **A_55_P2047009** | NA | 0.497411411 | -0.081159618 | 1.07598244 | 11.15862719 | 2.11508694 | 0.079844586 | 0.595609243 | -3.988853886 |
| **A_51_P330369** | Ssx2ip | 0.097594983 | -0.015940884 | 0.211130851 | 9.113584617 | 2.114772105 | 0.079878988 | 0.595609243 | -3.989078884 |
| **A_55_P2038303** | NA | -0.169987832 | -0.367773165 | 0.027797501 | 9.206837575 | -2.114428064 | 0.0799166 | 0.595609243 | -3.989324765 |
| **A_55_P1973548** | NA | 0.568906644 | -0.093093642 | 1.23090693 | 16.31601123 | 2.114228835 | 0.079938389 | 0.595609243 | -3.989467156 |
| **A_55_P2056419** | Coq6 | 0.159927608 | -0.026205505 | 0.34606072 | 10.96673448 | 2.113824684 | 0.079982607 | 0.595609243 | -3.98975602 |
| **A_55_P2019719** | Oas2 | -0.243347148 | -0.526654998 | 0.039960703 | 8.512557597 | -2.113181571 | 0.080053023 | 0.595609243 | -3.990215712 |
| **A_55_P2061021** | Smg6 | -0.208420276 | -0.451079645 | 0.034239092 | 11.85516838 | -2.113061165 | 0.080066213 | 0.595609243 | -3.990301782 |
| **A_55_P2115028** | Fastkd1 | 0.100184456 | -0.016467277 | 0.216836189 | 9.235613586 | 2.112896929 | 0.080084209 | 0.595609243 | -3.990419185 |
| **A_55_P2157992** | NA | -0.263416959 | -0.570303186 | 0.043469268 | 14.32724992 | -2.111715935 | 0.080213735 | 0.595609243 | -3.991263489 |

| | | | | | | | | | |
|---|---|---|---|---|---|---|---|---|---|
| **A_52_P486482** | R3hcc1l | -0.12023895 | -0.260320134 | 0.019842235 | 8.225090421 | -2.111711236 | 0.080214251 | 0.595609243 | -3.991266849 |
| **A_51_P420655** | Reep4 | -0.131193835 | -0.284053331 | 0.021665662 | 9.438549327 | -2.11149552 | 0.080237933 | 0.595609243 | -3.991421081 |
| **A_51_P366024** | Ercc3 | -0.124503853 | -0.269579716 | 0.020572011 | 9.051613826 | -2.111333263 | 0.080255751 | 0.595609243 | -3.991537094 |
| **A_55_P2118090** | NA | -0.237451337 | -0.514199293 | 0.03929662 | 10.03157895 | -2.110859654 | 0.080307783 | 0.595609243 | -3.991875737 |
| **A_55_P2079568** | NA | 0.197519175 | -0.032691962 | 0.427730312 | 8.836476843 | 2.110824352 | 0.080311662 | 0.595609243 | -3.99190098 |
| **A_55_P2170134** | Eif2b4 | 0.222059961 | -0.036857202 | 0.480977124 | 11.92403073 | 2.109981181 | 0.080404385 | 0.595609243 | -3.992503927 |
| **A_55_P2158376** | NA | 0.287503728 | -0.047872669 | 0.622880125 | 12.89338256 | 2.10901733 | 0.080510514 | 0.595609243 | -3.993193256 |
| **A_55_P2175579** | Olfr1466 | 0.756569039 | -0.125986965 | 1.639125042 | 15.15174872 | 2.108994532 | 0.080513026 | 0.595609243 | -3.993209562 |
| **A_55_P2155572** | NA | 0.581631065 | -0.096987384 | 1.260249514 | 13.8655557 | 2.108584913 | 0.080558173 | 0.595609243 | -3.993502542 |
| **A_66_P127567** | Nxf1 | -0.320318116 | -0.694201535 | 0.053565303 | 12.61950335 | -2.107727489 | 0.080652762 | 0.595609243 | -3.994115868 |
| **A_55_P2213828** | 4933412E12Rik | 0.134944293 | -0.022579099 | 0.292467686 | 8.567469358 | 2.107553603 | 0.080671958 | 0.595609243 | -3.994240259 |
| **A_55_P2067629** | Ush1g | 0.215810072 | -0.036135239 | 0.467755382 | 8.535915302 | 2.107339913 | 0.080695555 | 0.595609243 | -3.994393129 |
| **A_55_P2049448** | Phtf1 | 0.152274605 | -0.025513497 | 0.330062708 | 11.09620586 | 2.107142718 | 0.080717337 | 0.595609243 | -3.994534202 |
| **A_51_P240723** | 1700022A21Rik | 0.345080937 | -0.057857673 | 0.748019547 | 8.848439669 | 2.106935534 | 0.080740229 | 0.595609243 | -3.994682426 |
| **A_55_P2131279** | Atp6v1a | 0.173898448 | -0.029208765 | 0.377005661 | 11.39315239 | 2.106393577 | 0.080800141 | 0.595609243 | -3.995070172 |
| **A_66_P120572** | Gm14488 | 0.206072691 | -0.034627978 | 0.446773361 | 8.707824034 | 2.10626159 | 0.080814739 | 0.595609243 | -3.995164607 |
| **A_55_P2130965** | NA | -0.231699231 | -0.502343215 | 0.038944752 | 14.45285685 | -2.106179489 | 0.080823821 | 0.595609243 | -3.99522335 |
| **A_51_P311038** | Adamts15 | -0.415726456 | -0.901407614 | 0.069954702 | 11.31962796 | -2.105840972 | 0.080861277 | 0.595609243 | -3.995465566 |
| **A_55_P2105002** | Pcna | -0.196389888 | -0.425828501 | 0.033048726 | 12.03286262 | -2.105822569 | 0.080863314 | 0.595609243 | -3.995478734 |
| **A_55_P1971425** | Serpinb9f | -0.370009133 | -0.802315151 | 0.062296884 | 14.54892819 | -2.105670082 | 0.080880193 | 0.595609243 | -3.995587845 |
| **A_55_P2217309** | NA | 0.503148456 | -0.084719002 | 1.091015915 | 8.569739234 | 2.105648557 | 0.080882576 | 0.595609243 | -3.995603247 |
| **A_52_P638794** | Stk31 | -0.1144953 | -0.248280753 | 0.019290154 | 9.041673519 | -2.105464519 | 0.080902953 | 0.595609243 | -3.995734938 |
| **A_51_P473888** | Il6st | -0.320106627 | -0.694189502 | 0.053976248 | 11.66368768 | -2.105212799 | 0.080930832 | 0.595609243 | -3.995915065 |
| **A_66_P112305** | Myo1f | -0.192712142 | -0.417922283 | 0.032498 | 8.310741804 | -2.105185146 | 0.080933895 | 0.595609243 | -3.995934853 |
| **A_51_P116813** | Cyp17a1 | 0.665310637 | -0.112204959 | 1.442826233 | 13.15992097 | 2.105157147 | 0.080936997 | 0.595609243 | -3.995954889 |
| **A_55_P2027225** | Fau | -0.105927529 | -0.229724683 | 0.017869625 | 17.32618901 | -2.105073885 | 0.080946221 | 0.595609243 | -3.996014472 |
| **A_55_P2094378** | NA | 0.134088093 | -0.022631259 | 0.290807444 | 9.19409207 | 2.104925601 | 0.080962652 | 0.595609243 | -3.996120586 |
| **A_55_P1992475** | NA | 0.479059841 | -0.080920659 | 1.039040341 | 11.47130296 | 2.104679764 | 0.0809899 | 0.595609243 | -3.996296515 |
| **A_51_P251357** | Ctps | -0.170016471 | -0.368796687 | 0.028763745 | 12.00737404 | -2.104199927 | 0.081043111 | 0.595609243 | -3.99663992 |
| **A_55_P2019118** | NA | 0.262618556 | -0.044501708 | 0.56973882 | 9.957668253 | 2.103711109 | 0.081097355 | 0.595609243 | -3.996989775 |
| **A_55_P2114963** | Exosc10 | -0.131915332 | -0.286186982 | 0.022356319 | 8.54692204 | -2.103673376 | 0.081101543 | 0.595609243 | -3.997016782 |
| **A_66_P124239** | Gm6557 | 0.149562564 | -0.025348252 | 0.32447338 | 9.006173689 | 2.103659267 | 0.08110311 | 0.595609243 | -3.997026881 |
| **A_55_P2148519** | Teddm2 | 1.230812268 | -0.208664989 | 2.670289526 | 10.17460938 | 2.103566141 | 0.081113449 | 0.595609243 | -3.997093535 |
| **A_52_P267543** | Ginm1 | -0.106704662 | -0.231513088 | 0.018103763 | 8.432566758 | -2.103336001 | 0.081139005 | 0.595609243 | -3.997258262 |
| **A_51_P206268** | Seh1l | -0.145683579 | -0.316085574 | 0.024718417 | 10.34484479 | -2.103318725 | 0.081140924 | 0.595609243 | -3.997270627 |
| **A_55_P2141585** | NA | -0.12902925 | -0.279966579 | 0.021908079 | 10.75961772 | -2.10310364 | 0.081164817 | 0.595609243 | -3.997424583 |
| **A_51_P242356** | Fam114a2 | -0.193170726 | -0.419203665 | 0.032862213 | 12.09319173 | -2.102513256 | 0.081230437 | 0.595820166 | -3.997847194 |
| **A_55_P1980401** | Cbx7 | 0.396343581 | -0.067488958 | 0.860176121 | 10.00591236 | 2.102227824 | 0.081262182 | 0.595820166 | -3.998051526 |
| **A_51_P213153** | Tmem208 | 0.267651379 | -0.045607338 | 0.580910096 | 13.8701064 | 2.102013418 | 0.081286035 | 0.595820166 | -3.998205017 |
| **A_55_P1999172** | Nat10 | 0.112310691 | -0.019164512 | 0.243785894 | 10.33739369 | 2.101582343 | 0.081334017 | 0.59586416 | -3.998513633 |
| **A_55_P2071858** | Mgmt | 0.288724586 | -0.049306395 | 0.626755567 | 8.758034896 | 2.101340467 | 0.081360951 | 0.59586416 | -3.998686806 |
| **A_55_P2072030** | NA | -0.161976832 | -0.351659982 | 0.027706319 | 15.59684877 | -2.100841603 | 0.081416533 | 0.59586416 | -3.999043987 |
| **A_51_P389386** | Nsmce2 | -0.137351 | -0.298200121 | 0.023498121 | 9.732415189 | -2.100788329 | 0.081422471 | 0.59586416 | -3.999082131 |
| **A_55_P2008599** | Pcx | -0.145422847 | -0.315751674 | 0.02490598 | 11.56580004 | -2.100456303 | 0.081459488 | 0.59586416 | -3.999319873 |
| **A_55_P2040371** | Rnaset2a | -0.208731793 | -0.453259196 | 0.03579561 | 14.64368675 | -2.100052885 | 0.081504488 | 0.59586416 | -3.999608748 |
| **A_51_P290904** | Raver2 | 0.439083778 | -0.075331268 | 0.953498824 | 8.461404374 | 2.099920693 | 0.08151924 | 0.59586416 | -3.99970341 |
| **A_52_P12877** | Hspa8 | -0.136789711 | -0.297100802 | 0.023521381 | 10.36864903 | -2.099225161 | 0.081596899 | 0.59586416 | -4.000201504 |
| **A_55_P2177233** | Abhd5 | 0.132393759 | -0.022789541 | 0.28757706 | 11.51847482 | 2.098899734 | 0.08163326 | 0.59586416 | -4.000434569 |
| **A_51_P155345** | Chmp2a | -0.130697791 | -0.283894659 | 0.022499077 | 11.5386997 | -2.098879587 | 0.081635511 | 0.59586416 | -4.000448998 |
| **A_51_P148959** | Dhx36 | -0.19826524 | -0.430748489 | 0.034218009 | 9.899731688 | -2.098089824 | 0.081723826 | 0.59586416 | -4.001014655 |

| | | | | | | | | | |
|---|---|---|---|---|---|---|---|---|---|
| **A_51_P182471** | Celf2 | 0.18464135 | -0.031892116 | 0.401174816 | 9.76573382 | 2.097843574 | 0.081751383 | 0.59586416 | -4.001191041 |
| **A_52_P189957** | Dbnl | -0.115760025 | -0.251539478 | 0.020019428 | 9.244833009 | -2.097460168 | 0.081794307 | 0.59586416 | -4.00146568 |
| **A_52_P451378** | Twsg1 | -0.33341181 | -0.724518174 | 0.057694554 | 12.804979 | -2.097274418 | 0.081815111 | 0.59586416 | -4.00159874 |
| **A_51_P291417** | Thbd | 0.249334928 | -0.043170417 | 0.541840273 | 12.68947951 | 2.0970968 | 0.08183501 | 0.59586416 | -4.001725978 |
| **A_51_P105589** | Eci1 | -0.132873707 | -0.288780461 | 0.023033048 | 10.05024785 | -2.096733737 | 0.081875698 | 0.59586416 | -4.001986071 |
| **A_55_P2059323** | Gm13315 | -0.24684902 | -0.536533421 | 0.042835382 | 10.73787933 | -2.096406315 | 0.081912411 | 0.59586416 | -4.002220642 |
| **A_55_P1967613** | NA | 0.197979201 | -0.034377935 | 0.430336337 | 8.25222932 | 2.096199997 | 0.081935553 | 0.59586416 | -4.002368457 |
| **A_66_P134383** | NA | 0.344187921 | -0.059773599 | 0.748149441 | 11.9314203 | 2.096161741 | 0.081939845 | 0.59586416 | -4.002395866 |
| **A_66_P125071** | Ssb | -0.122246511 | -0.265727166 | 0.021234144 | 10.09919766 | -2.096101392 | 0.081946616 | 0.59586416 | -4.002439103 |
| **A_55_P1962110** | NA | -0.229573853 | -0.499099652 | 0.039951945 | 14.83118122 | -2.095517276 | 0.08201218 | 0.59586416 | -4.002857615 |
| **A_55_P1983135** | NA | 0.171835026 | -0.029919652 | 0.373589705 | 8.538415353 | 2.095353215 | 0.082030604 | 0.59586416 | -4.002975169 |
| **A_51_P170911** | Ptpn9 | -0.210164896 | -0.456966246 | 0.036636455 | 8.933744146 | -2.094989289 | 0.08207149 | 0.59586416 | -4.00323594 |
| **A_55_P1989816** | Zbtb7b | 0.544696345 | -0.094982404 | 1.184375094 | 14.09658286 | 2.094892301 | 0.08208239 | 0.59586416 | -4.003305438 |
| **A_55_P2032222** | Tm9sf2 | -0.281907469 | -0.61301415 | 0.049199213 | 11.78010057 | -2.094632322 | 0.082111614 | 0.59586416 | -4.003491736 |
| **A_51_P103718** | Ip6k1 | -0.108064203 | -0.234991383 | 0.018862978 | 10.8342249 | -2.094577258 | 0.082117805 | 0.59586416 | -4.01371303 |
| **A_55_P2089710** | Ednrb | -0.160672963 | -0.349397212 | 0.028051285 | 8.295394163 | -2.094518793 | 0.082124379 | 0.59586416 | -4.003573091 |
| **A_55_P1994020** | Casc4 | -0.13657008 | -0.297028794 | 0.023888634 | 8.992661485 | -2.09392643 | 0.082191017 | 0.596095387 | -4.003997601 |
| **A_55_P2039190** | Gfra4 | 0.186361156 | -0.032623017 | 0.405345328 | 8.934821242 | 2.093687364 | 0.082217926 | 0.596095387 | -4.004168934 |
| **A_55_P2006861** | Trio | 0.353546014 | -0.062006076 | 0.769098104 | 8.589099188 | 2.093098358 | 0.082284264 | 0.596273223 | -4.004591084 |
| **A_55_P2020090** | Pwwp3b | -0.240220478 | -0.522599366 | 0.04215841 | 11.52774008 | -2.092892715 | 0.082307438 | 0.596273223 | -4.004738479 |
| **A_66_P109368** | Prex2 | -0.130139261 | -0.28314215 | 0.022863627 | 8.090125706 | -2.092559765 | 0.082344972 | 0.596273223 | -4.00497713 |
| **A_66_P135106** | Slco3a1 | -0.355631618 | -0.773820823 | 0.062557586 | 9.498063841 | -2.092168753 | 0.082389074 | 0.596273223 | -4.005257411 |
| **A_55_P1972719** | Pmm1 | -0.148009665 | -0.32207155 | 0.026052219 | 9.021957028 | -2.091970512 | 0.082411442 | 0.596273223 | -4.005399519 |
| **A_55_P2160594** | NA | -0.122458817 | -0.266500559 | 0.021582925 | 15.93786725 | -2.091562559 | 0.082457493 | 0.596273223 | -4.005691967 |
| **A_66_P108822** | NA | -0.12984544 | -0.282614016 | 0.022923137 | 15.35601989 | -2.091037546 | 0.082516797 | 0.596273223 | -4.006068354 |
| **A_55_P2071683** | NA | -0.206354641 | -0.449164408 | 0.036455125 | 12.88593678 | -2.090822927 | 0.082541053 | 0.596273223 | -4.006222224 |
| **A_52_P189132** | Thap1 | 0.172699134 | -0.03055066 | 0.375948929 | 9.998627621 | 2.090399093 | 0.082588974 | 0.596273223 | -4.006526102 |
| **A_55_P1970596** | NA | 0.312142218 | -0.055255311 | 0.679539747 | 16.84014903 | 2.090188453 | 0.082612801 | 0.596273223 | -4.006677132 |
| **A_55_P2118810** | NA | -0.415394253 | -0.904345309 | 0.073556802 | 10.58194239 | -2.090086495 | 0.082624337 | 0.596273223 | -4.006750238 |
| **A_52_P671769** | Arhgap18 | -0.194997767 | -0.424542202 | 0.034546667 | 9.475758652 | -2.089931418 | 0.082641886 | 0.596273223 | -4.006861432 |
| **A_51_P284486** | Gstm2 | 0.274445309 | -0.048624133 | 0.597514751 | 10.89253611 | 2.089917266 | 0.082643487 | 0.596273223 | -4.00687158 |
| **A_55_P2155826** | Rplp1 | -0.134801049 | -0.293544005 | 0.023941907 | 15.91058007 | -2.089142049 | 0.082731271 | 0.596436992 | -4.007427469 |
| **A_51_P115326** | Surf4 | -0.205710387 | -0.447958461 | 0.036537686 | 14.76749743 | -2.089128018 | 0.082732861 | 0.596436992 | -4.00743753 |
| **A_52_P268279** | Fbxl17 | -0.139473958 | -0.303738847 | 0.02479093 | 9.842081229 | -2.088899499 | 0.082758757 | 0.596436992 | -4.007601406 |
| **A_55_P2066484** | Crtc3 | -0.182064948 | -0.396590313 | 0.032460416 | 9.39677315 | -2.087934433 | 0.08286821 | 0.597003215 | -4.00829353 |
| **A_55_P2018479** | Carnmt1 | -0.154031997 | -0.335557971 | 0.027493976 | 11.0603709 | -2.08757139 | 0.082909423 | 0.597077582 | -4.008553919 |
| **A_55_P2033461** | NA | -0.170814 | -0.372218118 | 0.030590117 | 13.90787335 | -2.086528308 | 0.083027951 | 0.597139173 | -4.009302126 |
| **A_51_P246705** | Nop14 | -0.125257091 | -0.273078655 | 0.022564473 | 8.336008956 | -2.084652595 | 0.083241529 | 0.597139173 | -4.010647833 |
| **A_52_P680710** | Kpna3 | -0.142755 | -0.311229877 | 0.025719877 | 10.38513473 | -2.084612531 | 0.083246097 | 0.597139173 | -4.010676581 |
| **A_55_P2151751** | Matr3 | -0.100013968 | -0.218048718 | 0.018020782 | 9.12619261 | -2.084586585 | 0.083249056 | 0.597139173 | -4.010695198 |
| **A_51_P480881** | Thap11 | -0.121534016 | -0.264971184 | 0.021903152 | 11.82592694 | -2.084516369 | 0.083257063 | 0.597139173 | -4.01074558 |
| **A_55_P1982122** | Homer1 | -0.094039428 | -0.205054423 | 0.016975567 | 7.978099938 | -2.083998967 | 0.083316087 | 0.597139173 | -4.011116847 |
| **A_55_P2015753** | Enho | 0.220000854 | -0.039722556 | 0.479724263 | 12.44199029 | 2.083926677 | 0.083324337 | 0.597139173 | -4.011168722 |
| **A_55_P2110507** | NA | 0.104323398 | -0.01884514 | 0.227491936 | 8.185437823 | 2.08377633 | 0.083341498 | 0.597139173 | -4.01127661 |
| **A_55_P2187053** | Tpm1 | 0.179349899 | -0.032421485 | 0.391121282 | 9.815703714 | 2.083545674 | 0.083367833 | 0.597139173 | -4.011442132 |
| **A_51_P208680** | Chtf18 | 0.140604122 | -0.025427342 | 0.306635586 | 8.489653247 | 2.083419957 | 0.08338219 | 0.597139173 | -4.011532351 |
| **A_55_P2054743** | Sh3bgrl2 | -0.098683079 | -0.215223204 | 0.017857045 | 8.535176798 | -2.083225955 | 0.083404351 | 0.597139173 | -4.011671576 |
| **A_51_P324838** | Evpl | 0.159170229 | -0.028818692 | 0.34715915 | 10.79327933 | 2.083045469 | 0.083424973 | 0.597139173 | -4.011801104 |
| **A_55_P1958295** | NA | 0.288989242 | -0.052338689 | 0.630317172 | 11.9983386 | 2.082950795 | 0.083435792 | 0.597139173 | -4.011869049 |
| **A_55_P1970229** | Csnk1d | -0.28340699 | -0.618187415 | 0.051373435 | 13.90284762 | -2.082666176 | 0.083468327 | 0.597139173 | -4.012073318 |

| | | | | | | | | | |
|---|---|---|---|---|---|---|---|---|---|
| **A_66_P133043** | Mbd2 | -0.185425596 | -0.404464646 | 0.033613455 | 11.87835867 | -2.08265483 | 0.083469624 | 0.597139173 | -4.012081461 |
| **A_51_P492298** | Aqp2 | 0.456968896 | -0.082839738 | 0.996777531 | 9.460222486 | 2.082648496 | 0.083470348 | 0.597139173 | -4.012086007 |
| **A_51_P244061** | Snapc5 | -0.159429693 | -0.347764104 | 0.028904717 | 10.74885844 | -2.082613584 | 0.08347434 | 0.597139173 | -4.012111064 |
| **A_55_P2037513** | Pld3 | 0.149708911 | -0.027159953 | 0.326577775 | 11.74895373 | 2.082406098 | 0.083498068 | 0.597139173 | -4.012259981 |
| **A_51_P407904** | Trappc4 | 0.167885054 | -0.030545094 | 0.366315202 | 13.47779313 | 2.081486227 | 0.083603346 | 0.597139173 | -4.012920238 |
| **A_55_P2185347** | NA | 0.228459024 | -0.041604204 | 0.498522251 | 8.436820624 | 2.081191416 | 0.083637115 | 0.597139173 | -4.013131861 |
| **A_55_P2125868** | NA | -0.248623592 | -0.542577833 | 0.045330649 | 11.02438496 | -2.08080685 | 0.083681186 | 0.597139173 | -4.013407924 |
| **A_55_P2057912** | NA | -0.214510367 | -0.46818541 | 0.039164677 | 11.48877806 | -2.080365797 | 0.083731761 | 0.597139173 | -4.013724554 |
| **A_51_P192130** | Stk10 | -0.317368586 | -0.692765813 | 0.058028641 | 9.889913778 | -2.079898009 | 0.083785434 | 0.597139173 | -4.014060396 |
| **A_55_P2006792** | Ntrk2 | 0.315094188 | -0.057626486 | 0.687814863 | 9.491287774 | 2.079821545 | 0.083794211 | 0.597139173 | -4.014115294 |
| **A_55_P2002577** | Ephx1 | 0.134203083 | -0.024555782 | 0.292961948 | 8.679569233 | 2.079666385 | 0.083812024 | 0.597139173 | -4.014226695 |
| **A_55_P2025090** | Gins3 | 0.579323349 | -0.106037172 | 1.264683871 | 12.7279353 | 2.079558354 | 0.083824428 | 0.597139173 | -4.01430426 |
| **A_51_P296528** | Cers4 | 0.142534417 | -0.026107348 | 0.311176182 | 8.993944685 | 2.07933167 | 0.083850463 | 0.597139173 | -4.014467018 |
| **A_52_P293120** | Prrg3 | -0.14782099 | -0.322731995 | 0.027090014 | 8.608430037 | -2.079161076 | 0.083870061 | 0.597139173 | -4.014589508 |
| **A_51_P370350** | Galnt2 | 0.103665051 | -0.018999321 | 0.226329422 | 11.86825603 | 2.07913692 | 0.083872837 | 0.597139173 | -4.014606852 |
| **A_65_P05238** | Knop1 | -0.186414969 | -0.407015835 | 0.034185898 | 9.893623954 | -2.078943715 | 0.083895039 | 0.597139173 | -4.01474558 |
| **A_55_P2094686** | Smim10l1 | 0.109523659 | -0.020161969 | 0.239209287 | 8.609792681 | 2.077711571 | 0.084036773 | 0.597139173 | -4.015630382 |
| **A_51_P215922** | Casp6 | -0.13294996 | -0.290395045 | 0.024495124 | 10.13765744 | -2.077438941 | 0.084068167 | 0.597139173 | -4.015826176 |
| **A_55_P1999202** | Vmn1r217 | 0.148346099 | -0.027348058 | 0.324040256 | 8.912371225 | 2.077246205 | 0.084090368 | 0.597139173 | -4.015964597 |
| **A_51_P378667** | Wrap53 | 0.159406616 | -0.029418425 | 0.348231657 | 10.10782732 | 2.076901585 | 0.084130079 | 0.597139173 | -4.016212107 |
| **A_51_P387608** | Hif1a | -0.191516397 | -0.418389911 | 0.035357118 | 10.29504486 | -2.076783988 | 0.084143635 | 0.597139173 | -4.016296569 |
| **A_55_P2016618** | Neurl1a | 0.185892902 | -0.034319372 | 0.406105176 | 9.250939938 | 2.076779788 | 0.084144119 | 0.597139173 | -4.016299586 |
| **A_55_P2098471** | Rps19 | 0.193717226 | -0.035766571 | 0.423201022 | 15.40771026 | 2.076755539 | 0.084146914 | 0.597139173 | -4.016317003 |
| **A_55_P1961235** | Rpl23 | -0.111848344 | -0.244351978 | 0.02065529 | 16.18046087 | -2.076686464 | 0.084154878 | 0.597139173 | -4.016366615 |
| **A_51_P427812** | Tmem108 | 0.236969048 | -0.043785721 | 0.517723817 | 8.557809386 | 2.076508175 | 0.084175437 | 0.597139173 | -4.016494672 |
| **A_52_P66313** | Psmf1 | -0.104100643 | -0.227439468 | 0.019238181 | 8.387543322 | -2.076456183 | 0.084181433 | 0.597139173 | -4.016532016 |
| **A_55_P2115141** | Lcor | 0.126123961 | -0.023310151 | 0.275558073 | 8.804994638 | 2.076428669 | 0.084184606 | 0.597139173 | -4.016551779 |
| **A_55_P1971093** | NA | -0.2398273 | -0.524035576 | 0.044380975 | 10.31241027 | -2.076017748 | 0.084232014 | 0.597256271 | -4.016846938 |
| **A_52_P658122** | Ets2 | -0.259042413 | -0.566081397 | 0.04799657 | 13.03137469 | -2.075613662 | 0.08427866 | 0.597328661 | -4.017137204 |
| **A_51_P328300** | Pdia4 | -0.272333482 | -0.595160331 | 0.050493368 | 11.766971 | -2.075393985 | 0.08430403 | 0.597328661 | -4.01729501 |
| **A_51_P352283** | Cyb561d1 | 0.125541406 | -0.023300588 | 0.2743834 | 9.603639018 | 2.075060057 | 0.084342609 | 0.597383027 | -4.017534896 |
| **A_55_P2076360** | NA | 0.109706161 | -0.020403712 | 0.239816035 | 8.158919079 | 2.074387698 | 0.084420342 | 0.597500449 | -4.018017935 |
| **A_51_P249848** | Dsg2 | -0.235516253 | -0.514951928 | 0.043919422 | 9.91641445 | -2.073519842 | 0.084520785 | 0.597500449 | -4.018641482 |
| **A_51_P310576** | Smim8 | -0.105153237 | -0.229918616 | 0.019612143 | 8.106481038 | -2.073469809 | 0.08452658 | 0.597500449 | -4.018677433 |
| **A_55_P2112737** | Actb | -0.282265592 | -0.617178765 | 0.052647581 | 17.19988404 | -2.073456239 | 0.084528151 | 0.597500449 | -4.018687183 |
| **A_51_P427663** | Cnn2 | -0.265524858 | -0.58059076 | 0.049541044 | 11.90823513 | -2.073351543 | 0.084540278 | 0.597500449 | -4.018762412 |
| **A_52_P410636** | Trp73 | 0.218652725 | -0.040800478 | 0.478105928 | 8.682777905 | 2.073313715 | 0.08454466 | 0.597500449 | -4.018789593 |
| **A_55_P2063871** | NA | 0.249146479 | -0.046668615 | 0.544961573 | 11.5020892 | 2.07206601 | 0.084689322 | 0.597609264 | -4.0196862 |
| **A_55_P1971759** | Zfp688 | 0.100627856 | -0.018855606 | 0.220111317 | 9.762282663 | 2.071951502 | 0.084702611 | 0.597609264 | -4.019768493 |
| **A_55_P2036967** | Cox17 | 0.391316546 | -0.073586025 | 0.856219116 | 9.09541831 | 2.070786983 | 0.084837878 | 0.597609264 | -4.020605459 |
| **A_52_P370311** | Plbd2 | 0.144199242 | -0.027142849 | 0.315541334 | 12.38245379 | 2.070465881 | 0.084875215 | 0.597609264 | -4.020836263 |
| **A_51_P351860** | C1qb | -0.119712947 | -0.261973116 | 0.022547222 | 8.323298358 | -2.070269888 | 0.084898013 | 0.597609264 | -4.020977145 |
| **A_55_P2076524** | 1700028I16Rik | 0.424390746 | -0.079985857 | 0.92876735 | 10.56503072 | 2.070046666 | 0.084923986 | 0.597609264 | -4.021137604 |
| **A_51_P255441** | Mrpl43 | -0.160953042 | -0.352271825 | 0.030365742 | 13.06631946 | -2.069715925 | 0.084962484 | 0.597609264 | -4.021375358 |
| **A_55_P2274782** | Prl7a1 | 0.409042611 | -0.082502586 | 0.900587808 | 12.08391399 | 2.156528375 | 0.085025997 | 0.597609264 | -4.040414466 |
| **A_55_P2165160** | Arhgef1 | -0.173757362 | -0.380365073 | 0.032850349 | 8.51286097 | -2.069025538 | 0.085042902 | 0.597609264 | -4.021871677 |
| **A_52_P340073** | Efnb2 | -0.111600342 | -0.244304644 | 0.021103959 | 10.25559542 | -2.068948544 | 0.085051875 | 0.597609264 | -4.021927031 |
| **A_55_P2091928** | Raet1b | 0.285095713 | -0.053927689 | 0.624119116 | 9.234137275 | 2.068855567 | 0.085062712 | 0.597609264 | -4.021993876 |
| **A_55_P2151156** | Farsa | 0.135069019 | -0.025555748 | 0.295693786 | 12.03152245 | 2.06877089 | 0.085072583 | 0.597609264 | -4.022054754 |
| **A_55_P2182392** | Adat3 | 0.160222912 | -0.030317516 | 0.35076334 | 9.780008447 | 2.068743505 | 0.085075776 | 0.597609264 | -4.022074442 |

| | | | | | | | | |
|---|---|---|---|---|---|---|---|---|
| **A_55_P2090924** | Tbc1d8b | -0.171472169 | -0.375475175 | 0.032530838 | 8.6625077 | -2.0678843 | 0.085176007 | 0.597609264 | -4.022692204 |
| **A_55_P2023727** | Limch1 | -0.133306478 | -0.291945944 | 0.025332987 | 9.616168397 | -2.067327024 | 0.08524108 | 0.597609264 | -4.023092917 |
| **A_55_P1955733** | NA | 0.280483469 | -0.053388899 | 0.614355837 | 11.91357918 | 2.066788118 | 0.085304057 | 0.597609264 | -4.023480445 |
| **A_55_P1957815** | Tmub1 | 0.214494884 | -0.040846433 | 0.4698362 | 12.0246338 | 2.066640842 | 0.085321276 | 0.597609264 | -4.023586356 |
| **A_52_P559919** | Eif2ak2 | -0.154207609 | -0.337792445 | 0.029377226 | 9.440149434 | -2.066513098 | 0.085336214 | 0.597609264 | -4.023678223 |
| **A_55_P2124120** | Ywhae | -0.176360311 | -0.386343595 | 0.033622973 | 14.4890923 | -2.066261555 | 0.085365637 | 0.597609264 | -4.023859123 |
| **A_55_P2116059** | Crybg1 | 0.145723751 | -0.027807621 | 0.319255122 | 10.97955511 | 2.06595808 | 0.085401149 | 0.597609264 | -4.024077378 |
| **A_55_P2151672** | NA | 0.598500352 | -0.114420844 | 1.311421549 | 9.861159694 | 2.065342329 | 0.085473247 | 0.597609264 | -4.024520242 |
| **A_55_P2064442** | Pabpn1 | -0.180532223 | -0.395580303 | 0.034515856 | 13.05354395 | -2.065324637 | 0.08547532 | 0.597609264 | -4.024532967 |
| **A_55_P2029842** | Hnrnpa1 | 0.097598553 | -0.018667271 | 0.213864377 | 11.34164146 | 2.06519216 | 0.08549084 | 0.597609264 | -4.024628253 |
| **A_55_P1980051** | Il17d | 0.601790068 | -0.115125341 | 1.318705477 | 10.89905664 | 2.065124619 | 0.085498754 | 0.597609264 | -4.024676833 |
| **A_51_P161946** | Epop | 0.410994103 | -0.078686588 | 0.900674794 | 12.61602856 | 2.064865528 | 0.085529119 | 0.597609264 | -4.024863193 |
| **A_55_P2007210** | Abca15 | 0.391721152 | -0.075005611 | 0.858447916 | 10.08754432 | 2.064826099 | 0.085533741 | 0.597609264 | -4.024891554 |
| **A_55_P2051825** | Ephx3 | 0.808724317 | -0.15492157 | 1.772370204 | 9.666354135 | 2.064677346 | 0.085551181 | 0.597609264 | -4.024998552 |
| **A_52_P621418** | NA | -0.317751769 | -0.696441216 | 0.060937679 | 12.22554596 | -2.064305343 | 0.08559481 | 0.597609264 | -4.025266145 |
| **A_55_P2029642** | NA | 0.173842267 | -0.033349908 | 0.381034442 | 13.12405302 | 2.064197231 | 0.085607494 | 0.597609264 | -4.025343915 |
| **A_55_P1972128** | Ccndbp1 | -0.142262456 | -0.311874854 | 0.027349943 | 9.304589524 | -2.063487628 | 0.085690792 | 0.597609264 | -4.025854392 |
| **A_55_P1966843** | Gm12070 | 0.153995625 | -0.029609145 | 0.337600395 | 16.58491985 | 2.063448272 | 0.085695414 | 0.597609264 | -4.025882705 |
| **A_55_P2022971** | Tspan9 | -0.158380298 | -0.347270059 | 0.030509464 | 8.326978207 | -2.062822683 | 0.085768924 | 0.597609264 | -4.026332783 |
| **A_51_P245533** | Tex19.1 | -0.239055884 | -0.524230122 | 0.046118353 | 12.9302978 | -2.062330727 | 0.085826775 | 0.597609264 | -4.026686742 |
| **A_51_P222092** | Igbp1 | -0.126745154 | -0.277955621 | 0.024465314 | 9.250772508 | -2.06214248 | 0.085848923 | 0.597609264 | -4.02682219 |
| **A_55_P1993109** | Taf6 | 0.174395099 | -0.033667541 | 0.382457739 | 11.77754691 | 2.062098157 | 0.085854139 | 0.597609264 | -4.026854082 |
| **A_55_P2128324** | Ubap2l | 0.192696342 | -0.037201245 | 0.422593929 | 9.231308832 | 2.062092866 | 0.085854761 | 0.597609264 | -4.026857889 |
| **A_55_P2000553** | Slc6a8 | -0.225403355 | -0.494342668 | 0.043535959 | 13.7636587 | -2.061936311 | 0.085873186 | 0.597609264 | -4.026970538 |
| **A_55_P2060487** | Zfp593 | 0.125945978 | -0.024327265 | 0.276219222 | 9.986557849 | 2.061919977 | 0.085875109 | 0.597609264 | -4.026982291 |
| **A_52_P676461** | Lpgat1 | -0.185026987 | -0.405926058 | 0.035872084 | 8.817600779 | -2.060679024 | 0.0860213 | 0.597609264 | -4.027875287 |
| **A_52_P594049** | Actr5 | 0.109422206 | -0.021221774 | 0.240066187 | 9.292813637 | 2.060559815 | 0.086035357 | 0.597609264 | -4.027961077 |
| **A_55_P2147276** | Orai1 | -0.189227567 | -0.415263393 | 0.036808259 | 9.327786299 | -2.059568733 | 0.086152315 | 0.597609264 | -4.02867437 |
| **A_55_P2093563** | Ufm1 | 0.211609025 | -0.041222819 | 0.464440869 | 12.30573395 | 2.059072213 | 0.08621097 | 0.597609264 | -4.029031753 |
| **A_55_P2010469** | NA | -0.205250885 | -0.450512349 | 0.040010579 | 10.90060515 | -2.05885085 | 0.086237133 | 0.597609264 | -4.029191091 |
| **A_55_P1988102** | Hmbs | -0.093876445 | -0.206144167 | 0.018391277 | 8.258637057 | -2.057173335 | 0.086435666 | 0.597609264 | -4.030398715 |
| **A_51_P461429** | Cyp7b1 | -0.111953714 | -0.245848981 | 0.021941554 | 7.989351761 | -2.057038596 | 0.086451633 | 0.597609264 | -4.030495722 |
| **A_55_P2258366** | C030009H01Rik | 0.29996562 | -0.0588196 | 0.65875084 | 12.41283275 | 2.056866479 | 0.086472033 | 0.597609264 | -4.030619643 |
| **A_51_P189814** | Cldn5 | -0.321550532 | -0.706326965 | 0.063225902 | 10.02767612 | -2.055937603 | 0.086582213 | 0.597609264 | -4.031288458 |
| **A_55_P1957277** | NA | 0.175170715 | -0.034446382 | 0.384787812 | 9.435269337 | 2.055909358 | 0.086585565 | 0.597609264 | -4.031308796 |
| **A_55_P2043762** | N6amt1 | 0.190199709 | -0.037412137 | 0.417811555 | 8.775367941 | 2.055815562 | 0.086596699 | 0.597609264 | -4.031376336 |
| **A_55_P2106414** | Haus7 | -0.10078815 | -0.221404994 | 0.019828695 | 8.062703588 | -2.055751768 | 0.086604273 | 0.597609264 | -4.031422273 |
| **A_52_P430247** | Nacc1 | -0.146200351 | -0.321164246 | 0.028763544 | 9.339367962 | -2.055744518 | 0.086605134 | 0.597609264 | -4.031427493 |
| **A_51_P487813** | Lxn | 0.208499026 | -0.041029511 | 0.458027564 | 11.11536969 | 2.055667991 | 0.08661422 | 0.597609264 | -4.031482599 |
| **A_55_P2139913** | Ssb | -0.216236784 | -0.475071076 | 0.042597509 | 11.42958986 | -2.055308041 | 0.08665697 | 0.597609264 | -4.031741801 |
| **A_55_P2437253** | Nae1 | -0.174801135 | -0.38405815 | 0.034455879 | 9.056621679 | -2.055102028 | 0.086681448 | 0.597609264 | -4.031890156 |
| **A_55_P1966444** | NA | -0.14763457 | -0.324375299 | 0.029106159 | 15.7266995 | -2.055041439 | 0.086688648 | 0.597609264 | -4.031933789 |
| **A_52_P59579** | Tmbim6 | -0.281172471 | -0.617796476 | 0.055451534 | 12.04441103 | -2.054929268 | 0.08670198 | 0.597609264 | -4.032014568 |
| **A_51_P286488** | Ier3 | -0.787637634 | -1.730667058 | 0.155391789 | 10.74374929 | -2.054803887 | 0.086716884 | 0.597609264 | -4.032104862 |
| **A_51_P519555** | Gnb1l | 0.099110556 | -0.0195541 | 0.217775213 | 9.738166717 | 2.054791173 | 0.086718395 | 0.597609264 | -4.032114019 |
| **A_55_P2068356** | 1700016K19Rik | 0.102507123 | -0.020226424 | 0.22524067 | 8.3856207 | 2.054754409 | 0.086722766 | 0.597609264 | -4.032140495 |
| **A_55_P2415372** | Mta1 | -0.114849151 | -0.252368612 | 0.022670311 | 8.113565887 | -2.054626044 | 0.086738029 | 0.597609264 | -4.03223294 |
| **A_55_P2076757** | Znfx1 | -0.266487819 | -0.585592487 | 0.05261685 | 11.65899686 | -2.054534159 | 0.086748955 | 0.597609264 | -4.032299113 |
| **A_55_P2084238** | NA | 0.205045054 | -0.040490555 | 0.450580664 | 11.95816261 | 2.054489733 | 0.086754239 | 0.597609264 | -4.032331108 |
| **A_55_P1972699** | Rars2 | -0.239970658 | -0.527342347 | 0.047401032 | 10.74546414 | -2.054392039 | 0.086765859 | 0.597609264 | -4.032401467 |

| | | | | | | | | | |
|---|---|---|---|---|---|---|---|---|---|
| **A_55_P2173892** | Isg20 | -0.314483508 | -0.691095838 | 0.062128822 | 13.08244505 | -2.05434086 | 0.086771947 | 0.597609264 | -4.032438326 |
| **A_55_P2210301** | Gm12992 | 0.23463248 | -0.046359528 | 0.515624488 | 9.337643462 | 2.054297448 | 0.086777111 | 0.597609264 | -4.032469592 |
| **A_52_P660032** | Cpsf2 | -0.112044061 | -0.246236401 | 0.022148279 | 9.370947303 | -2.054141137 | 0.086795709 | 0.597609264 | -4.032582169 |
| **A_51_P509140** | U2af1l4 | -0.154563888 | -0.339685941 | 0.030558166 | 9.330530667 | -2.054087957 | 0.086802037 | 0.597609264 | -4.03262047 |
| **A_55_P2087315** | Shroom1 | -0.128103606 | -0.28154588 | 0.025338669 | 10.02322737 | -2.053929295 | 0.08682092 | 0.597609264 | -4.032734743 |
| **A_55_P2059680** | Gm11938 | 0.259047245 | -0.051264423 | 0.569358912 | 8.349872095 | 2.05376163 | 0.086840879 | 0.597609264 | -4.032855503 |
| **A_55_P1977422** | NA | 0.212261383 | -0.042032102 | 0.466554867 | 17.00468645 | 2.053548288 | 0.086866283 | 0.597609264 | -4.033009165 |
| **A_55_P2078516** | NA | -0.215238109 | -0.473151475 | 0.042675257 | 8.590042107 | -2.053120697 | 0.08691722 | 0.597609264 | -4.033317152 |
| **A_52_P28792** | Isg20l2 | -0.186008855 | -0.408920893 | 0.036903184 | 9.823335078 | -2.052906921 | 0.086942698 | 0.597609264 | -4.033471139 |
| **A_51_P244923** | AI837181 | -0.099800025 | -0.219412398 | 0.019812347 | 9.151771081 | -2.052691632 | 0.086968363 | 0.597609264 | -4.033626218 |
| **A_51_P234714** | Ceacam14 | -0.147108263 | -0.323426825 | 0.029210298 | 15.71178243 | -2.052618311 | 0.086977106 | 0.597609264 | -4.033679034 |
| **A_52_P508089** | Psap | -0.383555459 | -0.843346704 | 0.076235785 | 14.03503007 | -2.052280102 | 0.087017445 | 0.597609264 | -4.033922667 |
| **A_55_P2185548** | Cdk8 | -0.16969577 | -0.373138962 | 0.033747422 | 10.51639508 | -2.052092906 | 0.087039781 | 0.597609264 | -4.03405752 |
| **A_55_P1961458** | Speer1 | 0.279960933 | -0.055681221 | 0.615603086 | 8.610044278 | 2.052060196 | 0.087043685 | 0.597609264 | -4.034081085 |
| **A_55_P1996643** | Polr2j | -0.133812167 | -0.294281452 | 0.026657117 | 12.48957505 | -2.051506285 | 0.087109813 | 0.597609264 | -4.034480132 |
| **A_55_P2018747** | Cox8a | -0.25410713 | -0.558845857 | 0.050631598 | 15.39637 | -2.05143785 | 0.087117987 | 0.597609264 | -4.034529436 |
| **A_55_P2000027** | Spink2 | 0.176729194 | -0.035273334 | 0.388731723 | 9.047859193 | 2.050861912 | 0.087186806 | 0.597609264 | -4.034944383 |
| **A_52_P551707** | Kdm6a | -0.11145053 | -0.245163702 | 0.022262643 | 8.582742038 | -2.050581832 | 0.087220293 | 0.597609264 | -4.035146183 |
| **A_55_P2076560** | Nr5a1 | 0.105354978 | -0.021065737 | 0.231775693 | 8.223446585 | 2.050246036 | 0.087260459 | 0.597609264 | -4.035388135 |
| **A_55_P1970910** | Stard13 | -0.138287994 | -0.304242372 | 0.027666384 | 9.547638533 | -2.050052229 | 0.08728365 | 0.597609264 | -4.035527784 |
| **A_51_P473229** | Zbtb7b | 0.187006562 | -0.037425415 | 0.411438539 | 10.20016254 | 2.049940584 | 0.087297012 | 0.597609264 | -4.035608232 |
| **A_52_P685938** | Efcab14 | -0.217613668 | -0.478805906 | 0.043578569 | 9.996082658 | -2.049722492 | 0.08732312 | 0.597609264 | -4.035765384 |
| **A_55_P2090484** | Pde10a | -0.15514541 | -0.341362302 | 0.031071483 | 8.687757081 | -2.049693948 | 0.087326537 | 0.597609264 | -4.035785953 |
| **A_55_P2141395** | Ly6g6c | -0.906644802 | -1.995043433 | 0.181753829 | 12.0576835 | -2.049360457 | 0.087366477 | 0.597609264 | -4.03602627 |
| **A_55_P2127804** | Map3k7 | 0.098788249 | -0.019879399 | 0.217455898 | 8.504918505 | 2.04805737 | 0.087522716 | 0.597609264 | -4.036965374 |
| **A_55_P2056192** | NA | 0.231855438 | -0.04666056 | 0.510371436 | 12.25875554 | 2.048029951 | 0.087526007 | 0.597609264 | -4.036985136 |
| **A_55_P2113066** | Zkscan5 | -0.171279798 | -0.377032816 | 0.03447322 | 10.8533278 | -2.047995933 | 0.087530089 | 0.597609264 | -4.037009653 |
| **A_51_P349825** | 1700021F05Rik | 0.169355385 | -0.034098634 | 0.372809404 | 12.63638324 | 2.047867721 | 0.087545479 | 0.597609264 | -4.037102062 |
| **A_55_P2279690** | 2900076A07Rik | 0.211827894 | -0.042721785 | 0.466377572 | 8.009398443 | 2.047291854 | 0.087614633 | 0.597609264 | -4.037517129 |
| **A_55_P1977787** | NA | 0.136446918 | -0.027611098 | 0.300504934 | 8.068517547 | 2.046140441 | 0.087753072 | 0.597609264 | -4.038347117 |
| **A_52_P258218** | Proser1 | -0.14834898 | -0.326719862 | 0.030021902 | 8.443218797 | -2.046113723 | 0.087756287 | 0.597609264 | -4.038366378 |
| **A_52_P389230** | Smarcc2 | -0.117259953 | -0.258256812 | 0.023736905 | 8.212891263 | -2.046017975 | 0.08776781 | 0.597609264 | -4.038435403 |
| **A_55_P1971159** | NA | -0.164571358 | -0.36248346 | 0.033340744 | 8.961587582 | -2.045742957 | 0.087800915 | 0.597609264 | -4.038633666 |
| **A_55_P1999666** | Cntrl | 0.181786084 | -0.036832993 | 0.400405161 | 9.573689173 | 2.045699038 | 0.087806203 | 0.597609264 | -4.038665329 |
| **A_66_P117289** | Cdk13 | -0.128879024 | -0.283892726 | 0.026134679 | 8.879629035 | -2.045414412 | 0.08784048 | 0.597609264 | -4.038870527 |
| **A_66_P126836** | Zbtb44 | -0.207994354 | -0.458198255 | 0.042209546 | 10.83555526 | -2.04515691 | 0.087871502 | 0.597609264 | -4.039056176 |
| **A_51_P381218** | Gulp1 | 0.199635851 | -0.040587361 | 0.439859063 | 9.035732008 | 2.044526425 | 0.087947507 | 0.597609264 | -4.039510755 |
| **A_55_P2128493** | Rap1a | -0.127265834 | -0.28041487 | 0.025883202 | 9.184280753 | -2.044404006 | 0.087962272 | 0.597609264 | -4.039599023 |
| **A_55_P2071300** | Rnf126 | 0.251265251 | -0.05111096 | 0.553641462 | 10.95966601 | 2.04434399 | 0.087969511 | 0.597609264 | -4.039642296 |
| **A_55_P2054332** | Ephb3 | -0.245461218 | -0.540871051 | 0.049948615 | 14.56801873 | -2.044217493 | 0.087984772 | 0.597609264 | -4.039733506 |
| **A_55_P2042481** | Gm10653 | 0.109534405 | -0.022290681 | 0.241359491 | 15.34961679 | 2.044191859 | 0.087987865 | 0.597609264 | -4.03975199 |
| **A_52_P612803** | Ccng1 | -0.225149167 | -0.49614821 | 0.045849877 | 10.3265788 | -2.043956973 | 0.088016211 | 0.597609264 | -4.039921357 |
| **A_55_P2185397** | Copz2 | -0.099725261 | -0.219777626 | 0.020327104 | 8.418884436 | -2.043636375 | 0.088054915 | 0.597609264 | -4.040152535 |
| **A_55_P1961761** | Dcc | -0.762027268 | -1.679390171 | 0.155335634 | 11.08897994 | -2.043612272 | 0.088057826 | 0.597609264 | -4.040169916 |
| **A_55_P2134427** | NA | -0.200962253 | -0.442908737 | 0.04098423 | 9.989247964 | -2.04345153 | 0.088077239 | 0.597609264 | -4.040285828 |
| **A_55_P2151907** | NA | 0.39774876 | -0.081135084 | 0.876632604 | 14.57452415 | 2.043373758 | 0.088086633 | 0.597609264 | -4.040341911 |
| **A_55_P2001023** | Lrrc19 | 0.705719968 | -0.144014634 | 1.555454569 | 13.61589955 | 2.043234746 | 0.088103427 | 0.597609264 | -4.040442156 |
| **A_52_P527625** | Colec12 | -0.257471046 | -0.567597861 | 0.05265577 | 10.56454973 | -2.042482008 | 0.08819442 | 0.597609264 | -4.040985003 |
| **A_52_P394539** | Tex2 | -0.163230047 | -0.359881819 | 0.033421726 | 12.50120359 | -2.042073657 | 0.088243823 | 0.597609264 | -4.041279509 |
| **A_55_P1958306** | Thrap3 | -0.186595109 | -0.411452261 | 0.038262044 | 12.01476039 | -2.041562648 | 0.088305686 | 0.597609264 | -4.041648074 |

| | | | | | | | | | |
|---|---|---|---|---|---|---|---|---|---|
| **A_55_P1961461** | Bnip3l | -0.160747957 | -0.354461004 | 0.032965089 | 13.61403837 | -2.041529947 | 0.088309646 | 0.597609264 | -4.04167166 |
| **A_55_P1974347** | Fam126b | 0.284529562 | -0.058356389 | 0.627415514 | 9.174413227 | 2.041488167 | 0.088314706 | 0.597609264 | -4.041701795 |
| **A_51_P417701** | Apaf1 | -0.17264146 | -0.380726281 | 0.035443361 | 8.399445927 | -2.041145023 | 0.088356276 | 0.597609264 | -4.041949301 |
| **A_52_P592101** | Washc2 | -0.13681498 | -0.301768384 | 0.028138423 | 9.562134865 | -2.040523195 | 0.088431658 | 0.597609264 | -4.042397843 |
| **A_55_P2091506** | Srp54c | -0.14989065 | -0.330628154 | 0.030846854 | 12.21085912 | -2.040306554 | 0.088457936 | 0.597609264 | -4.042554119 |
| **A_55_P2024595** | Atxn7l1 | 0.112983866 | -0.023252748 | 0.24922048 | 8.401778778 | 2.040289303 | 0.088460028 | 0.597609264 | -4.042566564 |
| **A_55_P1970424** | Fam221a | -0.197412946 | -0.435461532 | 0.040635639 | 8.858839437 | -2.040230276 | 0.08846719 | 0.597609264 | -4.042609145 |
| **A_55_P2055299** | NA | 0.302663085 | -0.062305036 | 0.667631206 | 9.024797297 | 2.040204427 | 0.088470326 | 0.597609264 | -4.042627792 |
| **A_55_P2025490** | Tnfrsf18 | 0.200596511 | -0.041378429 | 0.442571452 | 8.37917355 | 2.039492659 | 0.08855673 | 0.597609264 | -4.043141267 |
| **A_55_P2114938** | Irf9 | -0.109348404 | -0.241277975 | 0.022581167 | 11.83820012 | -2.03910439 | 0.088603899 | 0.597609264 | -4.043421386 |
| **A_55_P2186210** | NA | 0.25882085 | -0.053646568 | 0.571288269 | 13.59888522 | 2.037809974 | 0.088761338 | 0.597609264 | -4.044355336 |
| **A_51_P181002** | Vps50 | -0.103102097 | -0.227583033 | 0.021378839 | 8.989845887 | -2.037669835 | 0.0887784 | 0.597609264 | -4.044456458 |
| **A_55_P2033312** | Lgi4 | 0.15810007 | -0.032800785 | 0.349000926 | 8.883023074 | 2.037479984 | 0.08880152 | 0.597609264 | -4.044593454 |
| **A_52_P559545** | Cercam | 0.331341962 | -0.068764945 | 0.731448869 | 8.164295036 | 2.037368346 | 0.088815118 | 0.597609264 | -4.044674012 |
| **A_55_P2064598** | Cnot6 | -0.419967349 | -0.960977302 | 0.121042603 | 12.5727869 | -2.535381109 | 0.088836309 | 0.597609264 | -4.175248962 |
| **A_51_P461779** | Ppp2r2c | 0.320880872 | -0.066648291 | 0.708410036 | 10.37177686 | 2.037082426 | 0.088849954 | 0.597609264 | -4.044880339 |
| **A_52_P465886** | Gramd3 | 0.159446505 | -0.033226592 | 0.352119603 | 9.57439179 | 2.035931156 | 0.088990364 | 0.597609264 | -4.045711188 |
| **A_55_P1961870** | NA | 0.321955624 | -0.067244529 | 0.711155777 | 13.63933796 | 2.035130095 | 0.089088196 | 0.597609264 | -4.046289362 |
| **A_66_P115406** | Pabpc1 | -0.237594367 | -0.524873889 | 0.049685155 | 14.81163016 | -2.034701164 | 0.089140625 | 0.597609264 | -4.046598968 |
| **A_51_P108901** | Ccdc86 | -0.13271016 | -0.293194824 | 0.027774503 | 10.37478651 | -2.034416183 | 0.089175476 | 0.597609264 | -4.046804678 |
| **A_55_P1997300** | Mtfr2 | -0.250830343 | -0.554219189 | 0.052558503 | 14.32022467 | -2.0339938 | 0.089227156 | 0.597609264 | -4.047109582 |
| **A_51_P359983** | Arhgap17 | -0.18800579 | -0.415413653 | 0.039402074 | 10.54954202 | -2.033924875 | 0.089235592 | 0.597609264 | -4.047159338 |
| **A_52_P522222** | NA | -0.097295972 | -0.214992384 | 0.02040044 | 8.444348948 | -2.033765139 | 0.089255146 | 0.597609264 | -4.047274651 |
| **A_51_P377376** | Gnl3 | -0.123770476 | -0.273497335 | 0.025956384 | 9.451524181 | -2.033698221 | 0.08926334 | 0.597609264 | -4.04732296 |
| **A_55_P2286503** | A930001C03Rik | 0.233217632 | -0.048913653 | 0.515348916 | 9.332192415 | 2.033664454 | 0.089267474 | 0.597609264 | -4.047347336 |
| **A_55_P2066230** | Hck | -0.262431696 | -0.579920774 | 0.055057382 | 8.89373408 | -2.033558452 | 0.089280455 | 0.597609264 | -4.047423861 |
| **A_55_P2171008** | Clk4 | -0.268208019 | -0.592704082 | 0.056288043 | 9.925664335 | -2.033440601 | 0.089294888 | 0.597609264 | -4.04750894 |
| **A_55_P1995633** | NA | -0.239896376 | -0.530190311 | 0.050397559 | 11.7925509 | -2.033081964 | 0.089338827 | 0.597609264 | -4.047767855 |
| **A_55_P2114776** | Greb1 | 0.308477614 | -0.0648827 | 0.681837927 | 9.019092406 | 2.032659636 | 0.089390596 | 0.597609264 | -4.048072764 |
| **A_55_P2007771** | Catsper2 | 0.104351732 | -0.021958512 | 0.230661976 | 8.421129878 | 2.032498576 | 0.089410347 | 0.597609264 | -4.048189049 |
| **A_55_P1964225** | Elac2 | 0.211509672 | -0.044517356 | 0.4675367 | 8.967229879 | 2.032420562 | 0.089419916 | 0.597609264 | -4.048245376 |
| **A_55_P2047571** | Rps9 | -0.21034435 | -0.464970196 | 0.044281497 | 15.17955159 | -2.032345439 | 0.089429131 | 0.597609264 | -4.048299615 |
| **A_55_P2064876** | Fam89b | -0.216159145 | -0.477852936 | 0.045534645 | 11.89484179 | -2.032120048 | 0.089456784 | 0.597609264 | -4.048462353 |
| **A_55_P2014927** | Zfp341 | 0.096192372 | -0.020279728 | 0.212664473 | 9.128082372 | 2.03183245 | 0.089492083 | 0.597609264 | -4.048670012 |
| **A_52_P78257** | Plekha2 | -0.101586568 | -0.224597148 | 0.021424012 | 9.284167541 | -2.031715921 | 0.089506389 | 0.597609264 | -4.048754153 |
| **A_55_P2050988** | NA | 0.315550532 | -0.066550684 | 0.697651749 | 9.308419661 | 2.03170037 | 0.089508298 | 0.597609264 | -4.048765382 |
| **A_55_P1981670** | 8-Sep | -0.109585482 | -0.242343357 | 0.023172394 | 8.412818771 | -2.030775385 | 0.089621944 | 0.597609264 | -4.049433317 |
| **A_55_P2032362** | NA | -0.125944531 | -0.278545481 | 0.02665642 | 8.439727418 | -2.030445026 | 0.089662568 | 0.597609264 | -4.049671888 |
| **A_55_P1981884** | Ercc6l2 | 0.143238483 | -0.030325492 | 0.316802458 | 9.387178861 | 2.030342424 | 0.089675188 | 0.597609264 | -4.049745984 |
| **A_66_P104118** | NA | -0.093578848 | -0.20698101 | 0.019823314 | 8.084409932 | -2.030137821 | 0.089700361 | 0.597609264 | -4.049893744 |
| **A_55_P2114562** | NA | 0.221050273 | -0.046837631 | 0.488938178 | 8.508457887 | 2.030051774 | 0.08971095 | 0.597609264 | -4.049955887 |
| **A_55_P1982733** | Dnajc5 | 0.208689929 | -0.044239471 | 0.46161933 | 9.74971939 | 2.029884561 | 0.08973153 | 0.597609264 | -4.050076649 |
| **A_55_P2040936** | Eng | -0.177830223 | -0.393366311 | 0.037705865 | 9.508309355 | -2.029806974 | 0.089741081 | 0.597609264 | -4.050132684 |
| **A_52_P456158** | Fchsd1 | 0.539985669 | -0.114599696 | 1.194571035 | 11.29427632 | 2.029481543 | 0.089781153 | 0.597609264 | -4.05036772 |
| **A_66_P120260** | Taf10 | 0.099898951 | -0.021214738 | 0.221012641 | 13.44146116 | 2.029256097 | 0.089808924 | 0.597609264 | -4.050530549 |
| **A_51_P167374** | Gpatch1 | -0.131836044 | -0.291675626 | 0.028003538 | 9.844411375 | -2.029172605 | 0.089819211 | 0.597609264 | -4.050590852 |
| **A_55_P2062568** | Caprin2 | -0.182818294 | -0.404527521 | 0.038890933 | 9.19140894 | -2.028640291 | 0.089884825 | 0.597609264 | -4.050975337 |
| **A_55_P1954161** | Plagl2 | 0.094640762 | -0.020136978 | 0.209418502 | 8.046359494 | 2.028568708 | 0.089893653 | 0.597609264 | -4.051027042 |
| **A_51_P157297** | Efcab14 | -0.24183855 | -0.535149387 | 0.051472287 | 11.88780452 | -2.028460634 | 0.089906981 | 0.597609264 | -4.051105106 |
| **A_51_P362881** | Fbxo38 | -0.129324832 | -0.286180212 | 0.027530547 | 11.71164872 | -2.028390976 | 0.089915573 | 0.597609264 | -4.051155422 |

| | | | | | | | | | |
|---|---|---|---|---|---|---|---|---|---|
| **A_52_P678895** | Dynlt3 | -0.129210109 | -0.285928119 | 0.027507901 | 10.22675007 | -2.028367988 | 0.089918409 | 0.597609264 | -4.051172027 |
| **A_55_P2060922** | Unc5a | 0.234140402 | -0.049862256 | 0.51814306 | 9.477125839 | 2.028257625 | 0.089932024 | 0.597609264 | -4.051251746 |
| **A_52_P37431** | NA | -0.180905942 | -0.40034505 | 0.038533166 | 12.43583559 | -2.028186866 | 0.089940754 | 0.597609264 | -4.051302858 |
| **A_52_P436628** | Vegfb | 0.138936812 | -0.029606161 | 0.307479785 | 8.678623293 | 2.028036851 | 0.089959266 | 0.597609264 | -4.051411222 |
| **A_66_P112573** | Isg20 | -0.235109141 | -0.520324857 | 0.050106575 | 15.00431125 | -2.027987271 | 0.089965385 | 0.597609264 | -4.051447036 |
| **A_51_P385635** | Mrpl53 | 0.251227264 | -0.053642827 | 0.556097355 | 12.47223551 | 2.02731443 | 0.090048466 | 0.597609264 | -4.051933089 |
| **A_55_P2013501** | Hmgb1 | -0.146881185 | -0.32513304 | 0.03137067 | 13.02537872 | -2.027221824 | 0.090059907 | 0.597609264 | -4.051999989 |
| **A_55_P1993955** | Pltp | 0.147300415 | -0.031463636 | 0.326064466 | 8.996252341 | 2.027182952 | 0.09006471 | 0.597609264 | -4.052028071 |
| **A_55_P2044091** | Tdg | -0.146134299 | -0.323492904 | 0.031224307 | 8.343562985 | -2.027071452 | 0.090078487 | 0.597609264 | -4.052108622 |
| **A_55_P2443935** | Xpo7 | 0.095283782 | -0.020366568 | 0.210934132 | 8.510768273 | 2.026941418 | 0.090094558 | 0.597609264 | -4.052202564 |
| **A_55_P2139665** | NA | -0.177532012 | -0.393021748 | 0.037957723 | 15.40546249 | -2.026838995 | 0.090107219 | 0.597609264 | -4.052276559 |
| **A_55_P2025225** | Gnaz | 0.184763798 | -0.039516344 | 0.409043941 | 8.427488811 | 2.026726854 | 0.090121083 | 0.597609264 | -4.052357576 |
| **A_55_P2292737** | 2310002F09Rik | -0.140343014 | -0.310715009 | 0.030028981 | 13.28801683 | -2.026570676 | 0.090140394 | 0.597609264 | -4.05247041 |
| **A_55_P1963645** | NA | -0.173683949 | -0.384538656 | 0.037170757 | 10.53063007 | -2.026495059 | 0.090149746 | 0.597609264 | -4.052525041 |
| **A_55_P1961255** | Ppp1cc | 0.115698009 | -0.024798798 | 0.256194815 | 13.22448048 | 2.025949406 | 0.090217256 | 0.597609264 | -4.052919276 |
| **A_55_P2035972** | Mark3 | -0.154362044 | -0.34181114 | 0.033087052 | 11.84357824 | -2.025938858 | 0.090218562 | 0.597609264 | -4.052926897 |
| **A_55_P2111563** | NA | 0.1795083 | -0.038483307 | 0.397499907 | 8.869153233 | 2.02588097 | 0.090225727 | 0.597609264 | -4.052968722 |
| **A_55_P2073294** | Hadhb | -0.217937627 | -0.482645053 | 0.046769798 | 12.39208503 | -2.025514038 | 0.090271159 | 0.597609264 | -4.053233846 |
| **A_51_P376857** | Det1 | -0.107207122 | -0.237446709 | 0.023032465 | 9.153951095 | -2.025115467 | 0.090320535 | 0.597609264 | -4.053521842 |
| **A_55_P1979986** | Crcp | 0.227935016 | -0.04897353 | 0.504843562 | 11.12428545 | 2.025087772 | 0.090323967 | 0.597609264 | -4.053541854 |
| **A_55_P2062793** | Mfap2 | 0.462275416 | -0.099356071 | 1.023906903 | 9.869950627 | 2.024969624 | 0.090338609 | 0.597609264 | -4.053627227 |
| **A_55_P2172430** | Zxdb | 0.200191805 | -0.043032781 | 0.443416392 | 10.88142414 | 2.024920537 | 0.090344693 | 0.597609264 | -4.053662697 |
| **A_55_P1989048** | Tm9sf1 | 0.218773877 | -0.047062465 | 0.48461022 | 10.99136326 | 2.024651475 | 0.090378049 | 0.597609264 | -4.053857125 |
| **A_55_P2180899** | Aldh3a2 | 0.156549689 | -0.033681889 | 0.346781268 | 9.098134361 | 2.024597803 | 0.090384704 | 0.597609264 | -4.05389591 |
| **A_55_P2093794** | Usp3 | 0.261192535 | -0.056234643 | 0.578619714 | 10.05660103 | 2.024350993 | 0.090415316 | 0.597609264 | -4.054074264 |
| **A_51_P332799** | Clptm1 | -0.195958874 | -0.434115581 | 0.042197833 | 12.45955249 | -2.024283253 | 0.090423719 | 0.597609264 | -4.054123217 |
| **A_55_P1961463** | Bnip3l | -0.181130162 | -0.401274327 | 0.039014003 | 13.87590058 | -2.024196921 | 0.09043443 | 0.597609264 | -4.054185605 |
| **A_66_P118592** | Atrx | -0.215223276 | -0.476919806 | 0.046473253 | 8.293744518 | -2.023300732 | 0.090545694 | 0.598140026 | -4.054833275 |
| **A_52_P279184** | Srp54a | -0.166081312 | -0.368197667 | 0.036035044 | 9.239940294 | -2.021568537 | 0.090761145 | 0.598816776 | -4.056085299 |
| **A_51_P241769** | Rhd | 0.362528694 | -0.078696695 | 0.803754083 | 8.60146801 | 2.021394384 | 0.090782835 | 0.598816776 | -4.056211189 |
| **A_51_P516728** | Hap1 | -0.331527273 | -0.73503704 | 0.071982494 | 11.29368652 | -2.021316745 | 0.090792506 | 0.598816776 | -4.056267313 |
| **A_55_P2124115** | Invs | -0.138566701 | -0.307270008 | 0.030136606 | 9.960786428 | -2.020712116 | 0.090867859 | 0.598816776 | -4.056704402 |
| **A_55_P2072315** | NA | 0.153255036 | -0.033379633 | 0.339889705 | 12.09589147 | 2.020187114 | 0.09093334 | 0.598816776 | -4.057083953 |
| **A_51_P467448** | Pif1 | 0.124093995 | -0.027030067 | 0.275218056 | 9.202048115 | 2.020162531 | 0.090936407 | 0.598816776 | -4.057101726 |
| **A_55_P2041581** | NA | -0.123638345 | -0.274227027 | 0.026950337 | 8.50187251 | -2.019900679 | 0.090969086 | 0.598816776 | -4.05729104 |
| **A_51_P475076** | Vbp1 | -0.165063031 | -0.366107094 | 0.035981031 | 8.934739884 | -2.019890039 | 0.090970414 | 0.598816776 | -4.057298732 |
| **A_52_P80944** | Zfp36 | 0.176085923 | -0.038394526 | 0.390566372 | 8.872594156 | 2.019789407 | 0.090982976 | 0.598816776 | -4.057371489 |
| **A_51_P357573** | Cald1 | -0.157344544 | -0.349039805 | 0.034350717 | 10.86640113 | -2.019340129 | 0.091039081 | 0.598816776 | -4.057696325 |
| **A_55_P2038237** | Rps3 | 0.264985391 | -0.057854124 | 0.587824906 | 9.785962609 | 2.019316567 | 0.091042024 | 0.598816776 | -4.057713361 |
| **A_52_P256426** | NA | -0.102436718 | -0.227258137 | 0.022384701 | 8.326417821 | -2.018997144 | 0.091081936 | 0.598816776 | -4.05794432 |
| **A_55_P2107182** | Angptl8 | 0.368702521 | -0.080579804 | 0.817984847 | 8.665427534 | 2.018951736 | 0.091087612 | 0.598816776 | -4.057977153 |
| **A_52_P383653** | Cpne8 | -0.191077643 | -0.423928629 | 0.041773343 | 8.448121158 | -2.018835615 | 0.091102126 | 0.598816776 | -4.058061116 |
| **A_66_P139333** | Atf6 | -0.187485759 | -0.415981704 | 0.041010187 | 8.890503508 | -2.018640346 | 0.091126539 | 0.598816776 | -4.058202311 |
| **A_52_P136782** | Rgs5 | -0.265713815 | -0.589642082 | 0.058214452 | 9.1262467 | -2.01806175 | 0.091198916 | 0.598816776 | -4.058620701 |
| **A_55_P2124381** | NA | 0.208341294 | -0.045662632 | 0.462345219 | 12.39875842 | 2.017920681 | 0.091216572 | 0.598816776 | -4.058722713 |
| **A_55_P2162324** | Bmx | 0.105877692 | -0.023247887 | 0.235003271 | 8.256140996 | 2.017257538 | 0.091299613 | 0.598816776 | -4.059202278 |
| **A_55_P2066559** | Endou | -0.447720669 | -0.993757208 | 0.098315869 | 9.993818586 | -2.01722621 | 0.091303538 | 0.598816776 | -4.059224935 |
| **A_55_P2128646** | Gmnn | -0.225101473 | -0.499640903 | 0.049437957 | 9.360490772 | -2.017171263 | 0.091310422 | 0.598816776 | -4.059264672 |
| **A_55_P2053541** | NA | -0.17014634 | -0.377688493 | 0.037395814 | 15.41469562 | -2.016905061 | 0.091343782 | 0.598816776 | -4.059457193 |
| **A_55_P2150048** | Sypl | -0.147435266 | -0.327309429 | 0.032438897 | 11.07244401 | -2.016516385 | 0.091392513 | 0.598816776 | -4.059738297 |

| | | | | | | | | | |
|---|---|---|---|---|---|---|---|---|---|
| **A_55_P1990633** | Iigp1 | -0.091742337 | -0.203697051 | 0.020212376 | 7.822780944 | -2.016028105 | 0.09145377 | 0.598816776 | -4.060091455 |
| **A_55_P2022805** | Rubcn | -0.207927227 | -0.461688582 | 0.045834128 | 13.693006 | -2.01583527 | 0.091477973 | 0.598816776 | -4.060230932 |
| **A_55_P2143000** | NA | 0.221117113 | -0.048766151 | 0.491000377 | 8.44866619 | 2.015652046 | 0.091500976 | 0.598816776 | -4.06036346 |
| **A_51_P181722** | Rbks | -0.145317874 | -0.322717688 | 0.03208194 | 9.646727143 | -2.015278352 | 0.09154791 | 0.598816776 | -4.060633765 |
| **A_66_P117058** | Aasdh | -0.236854623 | -0.526016125 | 0.05230688 | 9.325065493 | -2.015164742 | 0.091562184 | 0.598816776 | -4.060715944 |
| **A_51_P146505** | Atp11b | -0.149914924 | -0.332945325 | 0.033115477 | 8.894736001 | -2.015073082 | 0.091573702 | 0.598816776 | -4.060782247 |
| **A_55_P2069337** | NA | -0.117033374 | -0.25993871 | 0.025871963 | 8.9937894 | -2.014793036 | 0.0916089 | 0.598816776 | -4.060984825 |
| **A_55_P2122555** | Csnk1e | -0.134788253 | -0.299377907 | 0.029801401 | 9.919718215 | -2.014738453 | 0.091615762 | 0.598816776 | -4.061024309 |
| **A_55_P2006118** | Rbp4 | 0.419267138 | -0.092701688 | 0.931235964 | 15.60851327 | 2.014728219 | 0.091617049 | 0.598816776 | -4.061031712 |
| **A_55_P2005828** | Hbb-bh1 | 0.40583067 | -0.089871584 | 0.901532924 | 8.107956355 | 2.01415614 | 0.091689001 | 0.598816776 | -4.06144556 |
| **A_55_P2024175** | Map4k3 | -0.219061676 | -0.486647421 | 0.04852407 | 11.49344808 | -2.01406088 | 0.091700988 | 0.598816776 | -4.061514474 |
| **A_66_P136857** | Phrf1 | -0.287138114 | -0.637913558 | 0.063637331 | 11.12225805 | -2.013867173 | 0.091725368 | 0.598816776 | -4.06165461 |
| **A_55_P2027259** | Zcchc2 | -0.208980583 | -0.46428387 | 0.046322703 | 10.28534626 | -2.013810933 | 0.091732447 | 0.598816776 | -4.061695297 |
| **A_55_P2072164** | NA | 0.29627971 | -0.06578678 | 0.6583462 | 8.683916212 | 2.013180626 | 0.091811828 | 0.599101192 | -4.062151312 |
| **A_55_P1962214** | Kpna3 | -0.11328972 | -0.251764365 | 0.025184925 | 9.710334552 | -2.012748009 | 0.091866352 | 0.599101192 | -4.062464319 |
| **A_51_P108020** | Zmynd15 | -0.096832284 | -0.215211279 | 0.021546712 | 7.938806538 | -2.012401749 | 0.091910016 | 0.599101192 | -4.062714856 |
| **A_55_P2074154** | Dhx57 | 0.157474284 | -0.035056059 | 0.350004628 | 9.408204371 | 2.012239269 | 0.091930513 | 0.599101192 | -4.062832421 |
| **A_51_P340816** | Ipp | -0.111800408 | -0.24849336 | 0.024892545 | 9.279388226 | -2.012178139 | 0.091938225 | 0.599101192 | -4.062876653 |
| **A_55_P2111302** | Cp | -0.254343473 | -0.565415528 | 0.056728581 | 9.072344984 | -2.011540421 | 0.092018723 | 0.599101192 | -4.063338108 |
| **A_55_P2083814** | Zzz3 | -0.273122885 | -0.60724056 | 0.06099479 | 11.76449901 | -2.011072851 | 0.092077789 | 0.599101192 | -4.063676463 |
| **A_51_P122845** | Naa50 | 0.108765414 | -0.024304537 | 0.241835365 | 9.848848311 | 2.010851419 | 0.092105774 | 0.599101192 | -4.067881237 |
| **A_55_P2127934** | Palm3 | 0.290823704 | -0.065091084 | 0.646738493 | 10.51640309 | 2.010263243 | 0.092180153 | 0.599101192 | -4.064262371 |
| **A_66_P101862** | NA | 0.187070545 | -0.041874396 | 0.416015485 | 8.258142522 | 2.010219695 | 0.092185663 | 0.599101192 | -4.064293887 |
| **A_51_P142091** | Asb8 | -0.167758845 | -0.373077947 | 0.037560258 | 11.65172711 | -2.010135035 | 0.092196374 | 0.599101192 | -4.064355158 |
| **A_55_P2409306** | Vps35l | 0.543995398 | -0.121842926 | 1.209833722 | 9.291130002 | 2.009997886 | 0.09221373 | 0.599101192 | -4.064454418 |
| **A_55_P2016681** | Alkbh7 | 0.134448207 | -0.030136984 | 0.299033398 | 9.607528806 | 2.009710132 | 0.092250154 | 0.599101192 | -4.064662681 |
| **A_51_P401987** | Tmem37 | -0.147107784 | -0.327199619 | 0.032984051 | 16.63761235 | -2.009605415 | 0.092263413 | 0.599101192 | -4.064738472 |
| **A_51_P386526** | Rcc1l | -0.125795527 | -0.279806919 | 0.028215865 | 8.864252166 | -2.009469948 | 0.092280568 | 0.599101192 | -4.06483652 |
| **A_55_P2065991** | S100a11 | -0.306362662 | -0.681479751 | 0.068754427 | 13.10544948 | -2.009269294 | 0.092305984 | 0.599101192 | -4.064981751 |
| **A_51_P322273** | Ptp4a3 | -0.209061204 | -0.4650592 | 0.046936792 | 9.691625062 | -2.009120775 | 0.092324801 | 0.599101192 | -4.06508925 |
| **A_55_P1982872** | Naa10 | -0.097393181 | -0.216656595 | 0.021870233 | 10.01136611 | -2.009048764 | 0.092333926 | 0.599101192 | -4.065141372 |
| **A_55_P2058953** | Rpl13a | 0.144943361 | -0.032580362 | 0.322467084 | 15.95989586 | 2.008681566 | 0.09238047 | 0.599202049 | -4.06540716 |
| **A_51_P319879** | Acadm | -0.202469529 | -0.450621574 | 0.045682516 | 13.19890841 | -2.00729396 | 0.092556572 | 0.599948941 | -4.066411638 |
| **A_66_P126726** | Cwc27 | -0.104629185 | -0.23290415 | 0.023645781 | 8.6920525 | -2.006689127 | 0.092633438 | 0.599948941 | -4.066849516 |
| **A_55_P2347529** | 3010003L21Rik | 0.116460642 | -0.026346609 | 0.259267893 | 8.586744915 | 2.006310219 | 0.092681625 | 0.599948941 | -4.067123847 |
| **A_55_P2075294** | NA | 0.120615519 | -0.02731332 | 0.268544358 | 7.973735079 | 2.005947234 | 0.092727811 | 0.599948941 | -4.067386658 |
| **A_66_P119643** | Atrx | -0.165550524 | -0.368594411 | 0.037493364 | 8.000964543 | -2.00590242 | 0.092733515 | 0.599948941 | -4.067419106 |
| **A_55_P2129477** | NA | -0.126921907 | -0.28258913 | 0.028745316 | 8.452143353 | -2.005896746 | 0.092734237 | 0.599948941 | -4.067423215 |
| **A_55_P2030647** | Dusp13 | 0.149963095 | -0.033968723 | 0.333894913 | 8.616744027 | 2.005841878 | 0.092741221 | 0.599948941 | -4.067462942 |
| **A_55_P1981274** | NA | -0.174645258 | -0.388852366 | 0.039561849 | 16.84387363 | -2.005820591 | 0.092743931 | 0.599948941 | -4.067478354 |
| **A_55_P2075894** | NA | 0.307588077 | -0.069832175 | 0.685008328 | 10.89175814 | 2.004995763 | 0.092848987 | 0.600159378 | -4.068075598 |
| **A_66_P104340** | Atxn7l2 | 0.122933913 | -0.027911477 | 0.273779303 | 9.169548927 | 2.004974372 | 0.092851713 | 0.600159378 | -4.068091088 |
| **A_55_P2149122** | Farp2 | -0.126342125 | -0.281386207 | 0.028701956 | 9.949835832 | -2.0047589 | 0.092879178 | 0.600159378 | -4.068247116 |
| **A_55_P2045258** | Ehd1 | -0.262387905 | -0.584419474 | 0.059643664 | 10.63208113 | -2.004539075 | 0.092907206 | 0.600159378 | -4.0684063 |
| **A_51_P125446** | Lzic | -0.094166394 | -0.209748927 | 0.021416138 | 9.779398666 | -2.004346945 | 0.09293171 | 0.600159378 | -4.070448802 |
| **A_55_P2015605** | Foxp4 | 0.21802289 | -0.049695575 | 0.485741355 | 11.68607671 | 2.003516523 | 0.093037696 | 0.600517617 | -4.069146816 |
| **A_52_P494115** | Gatad2b | -0.088952523 | -0.198185617 | 0.02028057 | 8.855377593 | -2.003425524 | 0.093049318 | 0.600517617 | -4.06921272 |
| **A_55_P2165046** | Klhl22 | -0.28755727 | -0.640728983 | 0.065614443 | 11.61236884 | -2.003122936 | 0.093087972 | 0.60056656 | -4.069431865 |
| **A_52_P297457** | 0610030E20Rik | -0.176826429 | -0.394048579 | 0.040395722 | 9.949886721 | -2.002683085 | 0.093144191 | 0.6005999 | -4.069750435 |
| **A_55_P1957003** | Timm10 | -0.097470631 | -0.217213464 | 0.022272202 | 11.50645555 | -2.002596316 | 0.093155285 | 0.6005999 | -4.06981328 |

| | | | | | | | | | |
|---|---|---|---|---|---|---|---|---|---|
| **A_52_P210338** | Ndufb6 | -0.136828133 | -0.304943518 | 0.031287252 | 12.2622846 | -2.002336693 | 0.093188488 | 0.600613633 | -4.070001325 |
| **A_55_P2113733** | NA | 0.178652429 | -0.0410049 | 0.398309758 | 10.62148861 | 2.000932274 | 0.093368307 | 0.601572005 | -4.071018628 |
| **A_55_P2049080** | Fam160a2 | -0.201117433 | -0.448500782 | 0.046265916 | 11.61870477 | -2.000084833 | 0.093476983 | 0.601806964 | -4.071632549 |
| **A_51_P433026** | Plpp6 | -0.1381249 | -0.308049796 | 0.031799997 | 9.258316307 | -1.999788769 | 0.09351498 | 0.601806964 | -4.071847042 |
| **A_52_P390114** | Eif4b | -0.111556392 | -0.248806096 | 0.025693312 | 8.319983821 | -1.999641803 | 0.093533848 | 0.601806964 | -4.071953518 |
| **A_55_P2146136** | Gla | 0.150803512 | -0.034737628 | 0.336344651 | 16.52911972 | 1.999587429 | 0.093540829 | 0.601806964 | -4.071992913 |
| **A_55_P2005691** | NA | 0.248475399 | -0.057333993 | 0.554284791 | 15.25810577 | 1.998949121 | 0.093622827 | 0.601806964 | -4.072455385 |
| **A_52_P560628** | 1300002E11Rik | -0.13827518 | -0.308471329 | 0.031920968 | 8.80217301 | -1.998773895 | 0.093645349 | 0.601806964 | -4.072582347 |
| **A_51_P390538** | Mpeg1 | -0.199826177 | -0.445810726 | 0.046158372 | 9.230030345 | -1.998544001 | 0.093674907 | 0.601806964 | -4.072748922 |
| **A_51_P193185** | Mb | -0.089633777 | -0.199983096 | 0.020715542 | 8.039737637 | -1.998348375 | 0.093700066 | 0.601806964 | -4.072890671 |
| **A_52_P54856** | Ubald1 | -0.093670557 | -0.208992403 | 0.021651289 | 9.792271854 | -1.998299961 | 0.093706293 | 0.601806964 | -4.072925751 |
| **A_55_P2004096** | Asb8 | -0.087221214 | -0.194634688 | 0.02019226 | 8.219026136 | -1.997710381 | 0.093782164 | 0.601806964 | -4.073352973 |
| **A_52_P110257** | Wdr83 | -0.124773212 | -0.278449001 | 0.028902578 | 8.432759833 | -1.997492075 | 0.093810273 | 0.601806964 | -4.073511169 |
| **A_52_P373973** | Mex3c | -0.166144989 | -0.370827915 | 0.038537936 | 11.49442597 | -1.996984967 | 0.093875601 | 0.601806964 | -4.073878657 |
| **A_52_P469956** | Zc3h6 | -0.141802031 | -0.316513214 | 0.032909151 | 9.148893591 | -1.996783189 | 0.093901608 | 0.601806964 | -4.074024886 |
| **A_52_P376214** | Cstf3 | -0.093517563 | -0.208768956 | 0.021733831 | 9.663782225 | -1.996255648 | 0.093969636 | 0.601806964 | -4.074407209 |
| **A_66_P106072** | Orai1 | -0.187366375 | -0.418306206 | 0.043573456 | 9.023042913 | -1.996006602 | 0.094001768 | 0.601806964 | -4.074587706 |
| **A_51_P175699** | Mtfp1 | -0.181184308 | -0.404504577 | 0.042135961 | 9.311333716 | -1.9960049 | 0.094001988 | 0.601806964 | -4.07458894 |
| **A_55_P1971347** | Mta3 | -0.137837283 | -0.307738556 | 0.03206399 | 8.470430176 | -1.995902096 | 0.094015255 | 0.601806964 | -4.074663449 |
| **A_55_P2340101** | NA | 0.187284775 | -0.043573119 | 0.418142668 | 9.957574891 | 1.995845447 | 0.094022567 | 0.601806964 | -4.074704506 |
| **A_55_P1962065** | NA | 0.347550827 | -0.08091569 | 0.776017345 | 14.08094529 | 1.995586697 | 0.094055971 | 0.601806964 | -4.074892045 |
| **A_55_P2068121** | Sun5 | 0.267290458 | -0.062257729 | 0.596838645 | 8.418048927 | 1.995417106 | 0.094077871 | 0.601806964 | -4.075014964 |
| **A_55_P1958354** | Rps29 | -0.148680053 | -0.331992408 | 0.034632302 | 16.95667692 | -1.995400761 | 0.094079982 | 0.601806964 | -4.075026811 |
| **A_55_P1996716** | Cachd1 | 0.153940831 | -0.035864893 | 0.343746555 | 8.548480901 | 1.99532519 | 0.094089743 | 0.601806964 | -4.075081586 |
| **A_55_P2025855** | Hnrnpul1 | -0.163322272 | -0.364730776 | 0.038086231 | 8.867451206 | -1.994971858 | 0.094135393 | 0.601899774 | -4.075337689 |
| **A_51_P135268** | Corin | -0.185152579 | -0.413525116 | 0.043219958 | 8.362334673 | -1.994596494 | 0.094183915 | 0.602010875 | -4.075609771 |
| **A_55_P1952202** | NA | 0.093739377 | -0.021919126 | 0.20939788 | 8.774026495 | 1.993947218 | 0.094267903 | 0.602147998 | -4.076080424 |
| **A_55_P1998213** | NA | -0.154738153 | -0.345681056 | 0.036204749 | 15.85640273 | -1.993714889 | 0.094297975 | 0.602147998 | -4.076248843 |
| **A_55_P2169149** | Washc3 | -0.156332293 | -0.34925088 | 0.036586294 | 11.37797593 | -1.99362646 | 0.094309424 | 0.602147998 | -4.076312947 |
| **A_52_P119187** | NA | -0.196228972 | -0.438429443 | 0.0459715 | 11.2136062 | -1.9932295 | 0.094360833 | 0.602147998 | -4.076600722 |
| **A_55_P2162472** | NA | 0.195809509 | -0.045903122 | 0.437522139 | 10.14116512 | 1.992983005 | 0.094392771 | 0.602147998 | -4.076779423 |
| **A_55_P1984932** | Zc3h18 | -0.139106912 | -0.310862842 | 0.032649018 | 11.94408802 | -1.992535792 | 0.094450743 | 0.602147998 | -4.07710365 |
| **A_55_P1993629** | NA | -0.349467424 | -0.780985704 | 0.082050855 | 15.16929289 | -1.992400605 | 0.094468274 | 0.602147998 | -4.077201662 |
| **A_55_P1989071** | Oas1d | 0.370334975 | -0.087004024 | 0.827673974 | 8.592381715 | 1.992166607 | 0.094498628 | 0.602147998 | -4.077371317 |
| **A_55_P2133195** | Gm4951 | -0.235478488 | -0.526291243 | 0.055334267 | 8.607165916 | -1.992080804 | 0.09450976 | 0.602147998 | -4.077433528 |
| **A_55_P1993012** | NA | 0.576941745 | -0.135593188 | 1.289476678 | 12.19422808 | 1.992025832 | 0.094516893 | 0.602147998 | -4.077473385 |
| **A_55_P2081353** | NA | -0.228016321 | -0.50975219 | 0.053719549 | 13.37242989 | -1.991099452 | 0.094637179 | 0.602522342 | -4.078145082 |
| **A_55_P1953377** | NA | -0.140340007 | -0.313753262 | 0.033073248 | 14.35382181 | -1.990986735 | 0.094651826 | 0.602522342 | -4.078226815 |
| **A_51_P159503** | Rnf213 | -0.355548478 | -0.794916787 | 0.083819832 | 10.5292831 | -1.990853291 | 0.094669168 | 0.602522342 | -4.078323578 |
| **A_55_P2138257** | Ube2d2a | 0.186054056 | -0.043908626 | 0.416016738 | 11.67849642 | 1.990448479 | 0.094721798 | 0.602658866 | -4.078617124 |
| **A_55_P2054280** | Fam178b | 0.277697836 | -0.065587675 | 0.620983346 | 9.2012355 | 1.990151714 | 0.094760399 | 0.602662043 | -4.078832328 |
| **A_55_P2154132** | Tuba1b | -0.107231359 | -0.23980914 | 0.025346423 | 15.88942256 | -1.989849473 | 0.094799729 | 0.602662043 | -4.079051509 |
| **A_52_P237159** | Spty2d1 | -0.131819817 | -0.2948082 | 0.031168565 | 10.27350112 | -1.989725743 | 0.094815835 | 0.602662043 | -4.079141238 |
| **A_55_P1960764** | Rps4x | -0.252755285 | -0.565396239 | 0.059885669 | 14.61865862 | -1.988948516 | 0.094917067 | 0.602848789 | -4.079704909 |
| **A_55_P2016249** | Ppid | -0.092469673 | -0.206854053 | 0.021914708 | 8.585474875 | -1.988848762 | 0.094930068 | 0.602848789 | -4.079777257 |
| **A_51_P389751** | Relb | -0.151993485 | -0.340022376 | 0.036035407 | 9.334274135 | -1.988701265 | 0.094949294 | 0.602848789 | -4.079884232 |
| **A_55_P2076592** | NA | 0.193703553 | -0.045943357 | 0.433350464 | 8.254573934 | 1.988542676 | 0.094969971 | 0.602848789 | -4.079999254 |
| **A_51_P499169** | Tram1 | -0.170707548 | -0.381957564 | 0.040542468 | 10.97367511 | -1.988040088 | 0.095035527 | 0.603066875 | -4.080363784 |
| **A_55_P2067772** | Cplane1 | -0.228545471 | -0.511431038 | 0.054340097 | 8.4943912 | -1.987609167 | 0.095091772 | 0.603163245 | -4.080676347 |
| **A_55_P2127020** | Nup54 | -0.103881851 | -0.232473743 | 0.02471004 | 10.07336777 | -1.987445645 | 0.095113124 | 0.603163245 | -4.080794959 |

| | | | | | | | | |
|---|---|---|---|---|---|---|---|---|
| **A_55_P2008898** | NA | -0.339075076 | -0.75890678 | 0.080756629 | 11.63827246 | -1.986963083 | 0.095176163 | 0.603365058 | -4.081145001 |
| **A_51_P369803** | Psmb9 | -0.155840913 | -0.348853454 | 0.037171628 | 9.121213411 | -1.986392736 | 0.095250725 | 0.603639759 | -4.08155874 |
| **A_55_P2010768** | Dpm3 | 0.139854319 | -0.033396651 | 0.31310529 | 12.82482807 | 1.98595485 | 0.09530801 | 0.603804828 | -4.081876405 |
| **A_55_P1953809** | Plaat3 | 0.138104382 | -0.033044866 | 0.309253629 | 11.09583882 | 1.985187934 | 0.095408424 | 0.604025992 | -4.082432798 |
| **A_51_P229816** | Eif2b2 | -0.16949887 | -0.379558473 | 0.040560733 | 12.11149279 | -1.985150462 | 0.095413333 | 0.604025992 | -4.082459984 |
| **A_51_P342535** | Rnf166 | -0.231998203 | -0.519538901 | 0.055542494 | 11.70020079 | -1.984972356 | 0.095436669 | 0.604025992 | -4.082589205 |
| **A_52_P585652** | Fndc3b | -0.190216332 | -0.426007394 | 0.04557473 | 11.90335434 | -1.984676042 | 0.095475507 | 0.604073998 | -4.082804194 |
| **A_55_P1974582** | Eif4h | -0.185271034 | -0.41509664 | 0.044554573 | 14.60532096 | -1.983253698 | 0.095662153 | 0.604406463 | -4.083836252 |
| **A_55_P2135351** | Fam89b | -0.165145842 | -0.370031968 | 0.039740285 | 9.616436556 | -1.98300703 | 0.095694559 | 0.604406463 | -4.084015249 |
| **A_66_P140658** | Olfr787 | 0.199292036 | -0.047966813 | 0.446550885 | 8.277352681 | 1.982929452 | 0.095704754 | 0.604406463 | -4.084071545 |
| **A_51_P473242** | Zfp426 | -0.103340367 | -0.231561025 | 0.024880291 | 8.545878489 | -1.982810273 | 0.095720417 | 0.604406463 | -4.08415803 |
| **A_51_P360963** | Mbtd1 | -0.102910506 | -0.230626646 | 0.024805633 | 9.8645026 | -1.982362596 | 0.095779276 | 0.604406463 | -4.084482909 |
| **A_55_P1952828** | NA | -0.115590115 | -0.259076553 | 0.027896323 | 8.376909414 | -1.981887527 | 0.095841776 | 0.604406463 | -4.084827679 |
| **A_51_P214269** | Epha1 | -0.280549637 | -0.628823256 | 0.067723981 | 9.642946752 | -1.981793029 | 0.095854213 | 0.604406463 | -4.084896261 |
| **A_55_P2099540** | H2afj | -0.115656402 | -0.259251029 | 0.027938226 | 8.353568881 | -1.981529983 | 0.095888842 | 0.604406463 | -4.08508717 |
| **A_55_P2183473** | Cldn3 | 0.367908247 | -0.088873459 | 0.824689954 | 11.10653773 | 1.981526951 | 0.095889241 | 0.604406463 | -4.08508937 |
| **A_55_P2153067** | 2610028H24Rik | -0.258233996 | -0.578926798 | 0.062458806 | 17.26048498 | -1.981040559 | 0.095953306 | 0.604406463 | -4.085442386 |
| **A_55_P2001494** | Kitl | -0.197095879 | -0.441868568 | 0.047676809 | 9.856725007 | -1.980996687 | 0.095959086 | 0.604406463 | -4.085474229 |
| **A_55_P2103603** | NA | 0.513783218 | -0.124309695 | 1.151876132 | 8.707997245 | 1.980911881 | 0.095970262 | 0.604406463 | -4.085535781 |
| **A_55_P1997837** | Osbp2 | -0.171433195 | -0.384378186 | 0.041511796 | 9.121830522 | -1.980599446 | 0.096011444 | 0.604406463 | -4.085762553 |
| **A_51_P341130** | Iqgap1 | -0.387357821 | -0.868581459 | 0.093865817 | 12.84176434 | -1.980316202 | 0.096048794 | 0.604406463 | -4.085968143 |
| **A_55_P2093241** | Zc3h4 | -0.140825039 | -0.315775265 | 0.034125187 | 11.58676404 | -1.980316173 | 0.096048798 | 0.604406463 | -4.085968164 |
| **A_55_P2138422** | Obox1 | 0.18802771 | -0.045632599 | 0.42168802 | 8.021249394 | 1.979730501 | 0.096126075 | 0.604406463 | -4.086393286 |
| **A_52_P428745** | Camk2d | -0.127648701 | -0.286277583 | 0.030980182 | 11.6530844 | -1.979717838 | 0.096127746 | 0.604406463 | -4.086402478 |
| **A_55_P2020080** | NA | -0.512090695 | -1.148466756 | 0.124285366 | 12.12084889 | -1.979712902 | 0.096128398 | 0.604406463 | -4.086406061 |
| **A_52_P18596** | NA | -0.271067514 | -0.607934779 | 0.06579975 | 11.23619481 | -1.979647321 | 0.096137055 | 0.604406463 | -4.086453666 |
| **A_51_P211506** | Muc20 | 0.094073999 | -0.022847517 | 0.210995516 | 8.106628907 | 1.979449063 | 0.096163231 | 0.604406463 | -4.086597582 |
| **A_51_P136521** | Lypd2 | 0.114047767 | -0.027716834 | 0.255812369 | 8.066424866 | 1.979193016 | 0.096197049 | 0.604406463 | -4.08678345 |
| **A_51_P446796** | Camta1 | -0.254254959 | -0.570323951 | 0.061814033 | 8.649330187 | -1.979049712 | 0.096215981 | 0.604406463 | -4.086887479 |
| **A_51_P193863** | Gtf3c4 | -0.101436643 | -0.227554084 | 0.024680797 | 8.476881829 | -1.978740689 | 0.096256819 | 0.604466552 | -4.087111814 |
| **A_51_P517870** | Nfatc1 | -0.196059372 | -0.439905298 | 0.047786555 | 9.318048513 | -1.978068217 | 0.096345748 | 0.604505586 | -4.087600014 |
| **A_51_P197259** | Cct7 | -0.113142025 | -0.253908296 | 0.027624247 | 11.95387006 | -1.977399865 | 0.096434215 | 0.604505586 | -4.088085253 |
| **A_55_P2016478** | Ddx39b | -0.140372517 | -0.315027182 | 0.034282147 | 8.460518862 | -1.977293155 | 0.096448347 | 0.604505586 | -4.08816273 |
| **A_51_P217236** | Derl1 | 0.276204523 | -0.067511615 | 0.619920662 | 11.70005234 | 1.976969756 | 0.09649119 | 0.604505586 | -4.088397538 |
| **A_51_P130994** | NA | -0.309819483 | -0.695452458 | 0.075813491 | 10.42436489 | -1.976531438 | 0.096549287 | 0.604505586 | -4.088715796 |
| **A_52_P633481** | Akap11 | -0.15543792 | -0.348937453 | 0.038061612 | 8.535510011 | -1.976269738 | 0.096583991 | 0.604505586 | -4.088905819 |
| **A_55_P2077618** | Csgalnact1 | -0.150663143 | -0.338224125 | 0.036897839 | 9.942813432 | -1.976212705 | 0.096591556 | 0.604505586 | -4.088947232 |
| **A_55_P2032708** | Hmgb1 | -0.151565323 | -0.340260052 | 0.037129406 | 12.88129608 | -1.976101486 | 0.09660631 | 0.604505586 | -4.089027992 |
| **A_55_P2149951** | Prx | -0.299868612 | -0.673239619 | 0.073502394 | 10.52830112 | -1.975875528 | 0.096636292 | 0.604505586 | -4.089192068 |
| **A_55_P2136832** | Gm6981 | 0.236961918 | -0.058090155 | 0.532013992 | 14.05795196 | 1.975827604 | 0.096642652 | 0.604505586 | -4.089226868 |
| **A_55_P2054728** | Ldlrad3 | -0.286556143 | -0.643418689 | 0.070306404 | 14.37950056 | -1.975504007 | 0.096685608 | 0.604505586 | -4.089461852 |
| **A_55_P2009121** | Agrn | -0.372141943 | -0.835629484 | 0.091345598 | 12.71161976 | -1.975330275 | 0.096708678 | 0.604505586 | -4.089588012 |
| **A_55_P2046550** | Adamts8 | 0.206645588 | -0.050729044 | 0.46402022 | 8.195494051 | 1.975284068 | 0.096714815 | 0.604505586 | -4.089621567 |
| **A_55_P2097518** | Zdhhc22 | 0.568719808 | -0.139648976 | 1.277088591 | 14.87309403 | 1.975186475 | 0.096727778 | 0.604505586 | -4.089692438 |
| **A_55_P1996751** | NA | -0.094635016 | -0.212509597 | 0.023239565 | 9.597759044 | -1.975153543 | 0.096732153 | 0.604505586 | -4.089716353 |
| **A_51_P181746** | Zfyve1 | -0.171542085 | -0.385264548 | 0.042180379 | 9.338226666 | -1.974647935 | 0.096799342 | 0.604729953 | -4.090083531 |
| **A_55_P1954511** | Btbd10 | -0.137419032 | -0.308669231 | 0.033831167 | 10.73080404 | -1.974171893 | 0.096862645 | 0.604787878 | -4.090429254 |
| **A_55_P1952076** | Npb | 0.599727106 | -0.147671032 | 1.347125243 | 10.1395969 | 1.974107641 | 0.096871192 | 0.604787878 | -4.090475917 |
| **A_55_P2124183** | NA | -0.085802981 | -0.192779686 | 0.021173724 | 8.359023463 | -1.973250959 | 0.096985227 | 0.605247859 | -4.091098118 |
| **A_55_P2083624** | Wbp2 | 0.204451147 | -0.050474297 | 0.459376591 | 13.41684301 | 1.973083797 | 0.097007495 | 0.605247859 | -4.091219533 |

| | | | | | | | | | |
|---|---|---|---|---|---|---|---|---|---|
| **A_55_P2004752** | Prl2c5 | -0.166720295 | -0.37468656 | 0.04124597 | 17.36475218 | -1.972262611 | 0.097116958 | 0.605735296 | -4.091816008 |
| **A_55_P2186787** | NA | 0.109748551 | -0.027230989 | 0.246728092 | 16.83669098 | 1.971116323 | 0.097269966 | 0.606276352 | -4.092648696 |
| **A_55_P2121185** | Omp | 0.088483212 | -0.021954826 | 0.198921249 | 7.81427539 | 1.971112266 | 0.097270508 | 0.606276352 | -4.092651643 |
| **A_55_P2028193** | Mgat1 | -0.30275007 | -0.680729291 | 0.07522915 | 14.82333176 | -1.970541069 | 0.097346843 | 0.606276352 | -4.093066605 |
| **A_55_P1980863** | NA | 0.121709294 | -0.03025641 | 0.273674999 | 8.223575552 | 1.970367886 | 0.09737 | 0.606276352 | -4.093192423 |
| **A_55_P2002426** | Rps6kb2 | -0.097666889 | -0.219633808 | 0.024300031 | 8.368770274 | -1.970037355 | 0.09741421 | 0.606276352 | -4.09343256 |
| **A_51_P427674** | Cpt1a | -0.093501218 | -0.210272726 | 0.02327029 | 9.969720027 | -1.969924316 | 0.097429335 | 0.606276352 | -4.096936582 |
| **A_55_P2138150** | Pias2 | -0.095424454 | -0.214606033 | 0.023757124 | 9.664607547 | -1.969789028 | 0.097447439 | 0.606276352 | -4.093612978 |
| **A_51_P407683** | Ints3 | 0.162026657 | -0.040363462 | 0.364416776 | 10.85264571 | 1.969546906 | 0.097479849 | 0.606276352 | -4.093788893 |
| **A_51_P433430** | Phc1 | -0.137885333 | -0.310147396 | 0.034376729 | 8.736302392 | -1.969235215 | 0.097521587 | 0.606276352 | -4.094015359 |
| **A_51_P210603** | Cyp4f13 | -0.095477813 | -0.214763922 | 0.023808296 | 9.320820546 | -1.969163396 | 0.097531206 | 0.606276352 | -4.094067541 |
| **A_55_P1988168** | Proser3 | 0.285557913 | -0.071230271 | 0.642346096 | 13.84847429 | 1.969032565 | 0.097548733 | 0.606276352 | -4.094162602 |
| **A_55_P2154240** | Rps29 | -0.154409445 | -0.347416192 | 0.038597302 | 16.58345581 | -1.968205925 | 0.097659545 | 0.606529243 | -4.094763255 |
| **A_55_P2183854** | Sctr | 0.330101635 | -0.082637457 | 0.742840726 | 8.441017698 | 1.967620026 | 0.097738162 | 0.606529243 | -4.095189008 |
| **A_51_P151433** | Ncbp2 | -0.14931604 | -0.336016375 | 0.037384296 | 8.785145461 | -1.967571512 | 0.097744675 | 0.606529243 | -4.095224262 |
| **A_55_P2024307** | Spin1 | -0.187523551 | -0.422002849 | 0.046955747 | 13.78468544 | -1.967525983 | 0.097750787 | 0.606529243 | -4.095257348 |
| **A_66_P104365** | Ccdc167 | 0.174657574 | -0.043754726 | 0.393069875 | 9.920463421 | 1.967340329 | 0.097775715 | 0.606529243 | -4.095392262 |
| **A_55_P2034953** | NA | -0.169659372 | -0.381822926 | 0.042504182 | 10.57654839 | -1.967325566 | 0.097777698 | 0.606529243 | -4.09540299 |
| **A_52_P200359** | Aco2 | 0.175349299 | -0.044001745 | 0.394700343 | 13.46955585 | 1.966679049 | 0.097864559 | 0.606873294 | -4.09587283 |
| **A_55_P2061645** | NA | -0.267013741 | -0.601158396 | 0.067130915 | 9.853013279 | -1.965930925 | 0.097965168 | 0.607045749 | -4.096416543 |
| **A_55_P2074005** | Tctn1 | 0.092016769 | -0.023143879 | 0.207177417 | 8.700810871 | 1.965767005 | 0.097987227 | 0.607045749 | -4.09653568 |
| **A_55_P2320085** | 4833415N18Rik | 0.312880204 | -0.078764344 | 0.704524752 | 8.730756169 | 1.965419065 | 0.098034065 | 0.607045749 | -4.096788568 |
| **A_55_P2081148** | Foxk2 | 0.112919766 | -0.028426831 | 0.254266363 | 9.585384533 | 1.965412775 | 0.098034912 | 0.607045749 | -4.096793139 |
| **A_55_P1988070** | Eef1g | -0.138213817 | -0.311231553 | 0.034803918 | 10.3864809 | -1.965305191 | 0.098049399 | 0.607045749 | -4.096871335 |
| **A_55_P2416494** | 8430426J06Rik | -0.120990734 | -0.272540825 | 0.030559357 | 8.501570988 | -1.964106676 | 0.098210936 | 0.607682493 | -4.0977425 |
| **A_55_P1996518** | Arhgap42 | -0.119423526 | -0.269012935 | 0.030165882 | 8.275239703 | -1.964075638 | 0.098215123 | 0.607682493 | -4.097765062 |
| **A_55_P2019083** | Rps6kb1 | -0.166798531 | -0.375818768 | 0.042221705 | 12.55700712 | -1.96323845 | 0.098328124 | 0.607684306 | -4.098373644 |
| **A_66_P117495** | Capza1 | -0.166866311 | -0.375990882 | 0.042258259 | 9.484940224 | -1.963056351 | 0.098352721 | 0.607684306 | -4.098506024 |
| **A_55_P2151863** | NA | 0.165490767 | -0.041987052 | 0.372968585 | 11.80247644 | 1.96232645 | 0.098451373 | 0.607684306 | -4.09903666 |
| **A_51_P269078** | Habp4 | -0.184424195 | -0.415643541 | 0.04679515 | 8.399723977 | -1.962288657 | 0.098456483 | 0.607684306 | -4.099064137 |
| **A_51_P425632** | Mad2l1bp | -0.159242431 | -0.358893458 | 0.040408595 | 12.04316685 | -1.96225939 | 0.098460441 | 0.607684306 | -4.099085415 |
| **A_55_P2279685** | 2900076A07Rik | 0.507081031 | -0.128726831 | 1.142888894 | 9.402244894 | 1.962097723 | 0.098482308 | 0.607684306 | -4.099202952 |
| **A_55_P2113165** | Camkk1 | 0.121436956 | -0.030828683 | 0.273702595 | 8.282770053 | 1.962086388 | 0.098483841 | 0.607684306 | -4.099211192 |
| **A_51_P158552** | Zscan12 | 0.223484198 | -0.056758595 | 0.503726991 | 9.806458066 | 1.961921072 | 0.098506206 | 0.607684306 | -4.099331384 |
| **A_55_P2011967** | Akirin2 | -0.137587216 | -0.310128946 | 0.034954514 | 12.35641696 | -1.961792628 | 0.098523586 | 0.607684306 | -4.099424769 |
| **A_55_P2121543** | Rps20 | 0.131231618 | -0.033360361 | 0.295823598 | 17.18878591 | 1.961548174 | 0.098556673 | 0.607684306 | -4.099602502 |
| **A_55_P2153862** | Irak1 | -0.165824986 | -0.373807886 | 0.042157915 | 9.205594162 | -1.961514403 | 0.098561245 | 0.607684306 | -4.099627055 |
| **A_55_P1952676** | Gtf2a2 | -0.196264093 | -0.442533627 | 0.050005441 | 11.1973352 | -1.960646628 | 0.098678794 | 0.607825674 | -4.100258014 |
| **A_55_P2073397** | NA | -0.245824338 | -0.55430051 | 0.062651833 | 9.785454566 | -1.960525158 | 0.09869526 | 0.607825674 | -4.100346338 |
| **A_55_P1972841** | Rps28 | -0.103317071 | -0.232981502 | 0.026347359 | 16.88640748 | -1.960290307 | 0.098727102 | 0.607825674 | -4.100517108 |
| **A_55_P2155616** | Gm9457 | -0.236072385 | -0.532360952 | 0.060216181 | 14.16379306 | -1.96019582 | 0.098739916 | 0.607825674 | -4.100585814 |
| **A_55_P2021054** | Elp5 | -0.285124157 | -0.643012847 | 0.072764533 | 11.77859414 | -1.959996084 | 0.09876701 | 0.607825674 | -4.100731054 |
| **A_55_P2091494** | Tango2 | 0.218493394 | -0.055794648 | 0.492781436 | 10.71596574 | 1.959749669 | 0.098800445 | 0.607825674 | -4.10091024 |
| **A_51_P513072** | Rdh13 | -0.095246989 | -0.21485042 | 0.024356442 | 8.502283801 | -1.959191006 | 0.098876291 | 0.607825674 | -4.101316499 |
| **A_52_P220573** | Serpini1 | 0.10892984 | -0.027863185 | 0.245722865 | 8.082496809 | 1.959079568 | 0.098891427 | 0.607825674 | -4.101397539 |
| **A_51_P207412** | Slf1 | 0.162903343 | -0.041681804 | 0.36748849 | 9.4219984 | 1.958957705 | 0.098907982 | 0.607825674 | -4.101486161 |
| **A_51_P205740** | Ube3b | -0.22439764 | -0.506270275 | 0.057474994 | 11.81956991 | -1.958549417 | 0.098963468 | 0.607825674 | -4.101783086 |
| **A_55_P1964347** | NA | -0.130657903 | -0.294827771 | 0.033511965 | 8.522708526 | -1.957994178 | 0.099038974 | 0.607825674 | -4.102186895 |
| **A_52_P220229** | Plekhb2 | 0.237026878 | -0.060851171 | 0.534904926 | 9.535917363 | 1.957619392 | 0.099089974 | 0.607825674 | -4.102459478 |
| **A_55_P1966194** | Plek | -0.216579843 | -0.488773185 | 0.0556135 | 8.634996107 | -1.957535733 | 0.099101361 | 0.607825674 | -4.102520325 |

| | | | | | | | | |
|---|---|---|---|---|---|---|---|---|
| **A_55_P2009066** | Cfap74 | -0.235237001 | -0.530920425 | 0.060446424 | 17.32313875 | -1.957256831 | 0.099139335 | 0.607825674 | -4.102723177 |
| **A_52_P299311** | NA | 0.264000999 | -0.067839299 | 0.595841298 | 14.73068385 | 1.957246876 | 0.099140691 | 0.607825674 | -4.102730418 |
| **A_55_P2143499** | Pgbd5 | -0.099391244 | -0.224340345 | 0.025557857 | 7.96666409 | -1.956969931 | 0.099178413 | 0.607825674 | -4.102931852 |
| **A_55_P2059931** | Prom1 | 0.346329445 | -0.089069354 | 0.781728244 | 12.07295892 | 1.956912254 | 0.099186271 | 0.607825674 | -4.102973804 |
| **A_55_P2353462** | 9530036O11Rik | 0.128237816 | -0.032996647 | 0.289472279 | 7.941804923 | 1.956714212 | 0.099213257 | 0.607825674 | -4.103117852 |
| **A_52_P473975** | Cpsf4 | -0.16085689 | -0.363145801 | 0.04143202 | 9.935158229 | -1.956305863 | 0.099268923 | 0.607825674 | -4.103414877 |
| **A_55_P2170509** | Yipf4 | 0.23727376 | -0.084825563 | 0.559373083 | 11.39481305 | 2.405984791 | 0.099285424 | 0.607825674 | -4.203640544 |
| **A_55_P2128225** | Rpl18 | -0.270766285 | -0.611305032 | 0.069772462 | 14.85738086 | -1.956127754 | 0.099293213 | 0.607825674 | -4.103544433 |
| **A_51_P287198** | Krt23 | 0.293342622 | -0.075599751 | 0.662284995 | 9.679522182 | 1.956076318 | 0.099300229 | 0.607825674 | -4.103581848 |
| **A_51_P517381** | Cers2 | -0.096422972 | -0.217701865 | 0.024855922 | 11.21054167 | -1.955980182 | 0.099313343 | 0.607825674 | -4.103651779 |
| **A_51_P470851** | Cplx3 | 0.270760878 | -0.06983165 | 0.611353406 | 8.359990035 | 1.955779823 | 0.09934068 | 0.607825674 | -4.103797524 |
| **A_52_P342131** | Zfp532 | -0.137237186 | -0.309908102 | 0.03543373 | 8.233316099 | -1.955337709 | 0.099401029 | 0.607825674 | -4.104119135 |
| **A_55_P2172619** | NA | 0.191364959 | -0.049422268 | 0.432152187 | 16.36139916 | 1.955231268 | 0.099415564 | 0.607825674 | -4.104196566 |
| **A_55_P2076134** | NA | 0.20085528 | -0.051889971 | 0.453600531 | 12.28700454 | 1.955101967 | 0.099433224 | 0.607825674 | -4.104290628 |
| **A_51_P267239** | Litaf | -0.361555232 | -0.816612101 | 0.093501638 | 14.00504192 | -1.954691114 | 0.099489357 | 0.607854522 | -4.104589515 |
| **A_52_P683211** | NA | -0.086190577 | -0.194675523 | 0.022294369 | 8.275543322 | -1.954607107 | 0.099500838 | 0.607854522 | -4.104650629 |
| **A_66_P118934** | Plaa | -0.156450288 | -0.353393062 | 0.040492487 | 11.30155231 | -1.954364103 | 0.099534058 | 0.607865344 | -4.104827416 |
| **A_55_P2277014** | AA522020 | 0.267775743 | -0.069429739 | 0.604981224 | 12.29067545 | 1.953645537 | 0.099632355 | 0.60792129 | -4.105350196 |
| **A_52_P17098** | Mpp7 | -0.23838358 | -0.53861283 | 0.06184567 | 9.688421282 | -1.953405922 | 0.099665155 | 0.60792129 | -4.105524531 |
| **A_55_P1964525** | NA | -0.146738362 | -0.33156218 | 0.038085456 | 16.63992319 | -1.95323677 | 0.099688316 | 0.60792129 | -4.105647601 |
| **A_55_P2121364** | Tdg | 0.1254264 | -0.032581797 | 0.283434597 | 8.480231647 | 1.952893237 | 0.099735371 | 0.60792129 | -4.105897553 |
| **A_55_P2155002** | Svbp | -0.154898583 | -0.35004831 | 0.040251144 | 11.78859598 | -1.95275905 | 0.099753757 | 0.60792129 | -4.105995188 |
| **A_51_P346871** | Ift20 | -0.209977813 | -0.474586137 | 0.054630512 | 12.27715274 | -1.952266344 | 0.099821297 | 0.60792129 | -4.106353692 |
| **A_65_P19319** | Soga1 | 0.177066244 | -0.046076224 | 0.400208713 | 10.52020977 | 1.952192801 | 0.099831382 | 0.60792129 | -4.106407205 |
| **A_55_P2079167** | Dnm1 | 0.383640777 | -0.099840324 | 0.867121877 | 9.750753992 | 1.952155563 | 0.099836489 | 0.60792129 | -4.106434301 |
| **A_55_P1968668** | Gpatch2l | -0.111272734 | -0.25152445 | 0.028978983 | 9.548407831 | -1.951864819 | 0.099876371 | 0.60792129 | -4.106645862 |
| **A_55_P2254779** | NA | 0.154713758 | -0.040317763 | 0.349745278 | 9.018023489 | 1.951611144 | 0.099911182 | 0.60792129 | -4.106830454 |
| **A_55_P2068228** | NA | 0.205202472 | -0.053500603 | 0.463905546 | 10.70006974 | 1.951417344 | 0.099937784 | 0.60792129 | -4.106971479 |
| **A_55_P1997400** | Slc7a4 | 0.134804709 | -0.035171562 | 0.304780981 | 8.05116736 | 1.95112875 | 0.099977412 | 0.60792129 | -4.107181488 |
| **A_55_P2169276** | 4732491K20Rik | 0.224670908 | -0.058629167 | 0.507970982 | 8.67950481 | 1.951054111 | 0.099987664 | 0.60792129 | -4.107235803 |
| **A_55_P2006708** | Bmp1 | 0.351358183 | -0.091761347 | 0.794477713 | 10.0198334 | 1.950735304 | 0.100031464 | 0.60792129 | -4.107467806 |
| **A_55_P2023747** | Mfsd10 | 0.302868161 | -0.079229248 | 0.684965569 | 11.27335577 | 1.950063219 | 0.100123862 | 0.60792129 | -4.107956916 |
| **A_51_P362738** | Dusp18 | -0.160976101 | -0.36406445 | 0.042112247 | 8.159539919 | -1.950049162 | 0.100125796 | 0.60792129 | -4.107967146 |
| **A_51_P407866** | 1700020D05Rik | -0.117198433 | -0.265060543 | 0.030663677 | 8.444288827 | -1.949997563 | 0.100132893 | 0.60792129 | -4.108004699 |
| **A_51_P370050** | Nagpa | 0.188185095 | -0.049286709 | 0.425656899 | 9.218221157 | 1.949585715 | 0.100189562 | 0.60792129 | -4.108304436 |
| **A_52_P78123** | Dcaf12 | 0.152918146 | -0.040064038 | 0.34590033 | 10.93842799 | 1.949444919 | 0.100208942 | 0.60792129 | -4.108406907 |
| **A_55_P2256686** | NA | -0.229813299 | -0.519867921 | 0.060241322 | 8.412035818 | -1.949236481 | 0.100237641 | 0.60792129 | -4.108558611 |
| **A_52_P319438** | Ankrd37 | -0.521749629 | -1.180360344 | 0.136861086 | 10.21902509 | -1.948958179 | 0.100275971 | 0.60792129 | -4.108761167 |
| **A_52_P106259** | Egfr | -0.284700179 | -0.644094675 | 0.074694317 | 10.86390654 | -1.948881719 | 0.100286504 | 0.60792129 | -4.1121071 |
| **A_51_P344947** | Rab1b | 0.124013129 | -0.032560287 | 0.280586544 | 12.41701891 | 1.948582515 | 0.100327734 | 0.60792129 | -4.109034593 |
| **A_55_P2116988** | 1700024P12Rik | 0.329938555 | -0.086658539 | 0.746535648 | 9.897847777 | 1.948435326 | 0.100348022 | 0.60792129 | -4.109141726 |
| **A_55_P2091741** | Lamp3 | -0.283541726 | -0.641650896 | 0.074567444 | 11.07039268 | -1.947918131 | 0.100419346 | 0.60792129 | -4.109518183 |
| **A_55_P2150389** | Dnajc21 | -0.179028073 | -0.405164862 | 0.047108717 | 8.888449129 | -1.947686604 | 0.100451291 | 0.60792129 | -4.109686712 |
| **A_66_P112005** | Olfr1252 | 0.220802574 | -0.058146443 | 0.499751591 | 9.403189183 | 1.947369926 | 0.100495001 | 0.60792129 | -4.109917229 |
| **A_55_P1987054** | Krtap5-1 | 0.092191592 | -0.02428293 | 0.208666114 | 8.535732848 | 1.947285109 | 0.100506712 | 0.60792129 | -4.10997897 |
| **A_55_P2048607** | Hp | -0.370183182 | -0.837887248 | 0.097520885 | 8.766006522 | -1.947218561 | 0.100515901 | 0.60792129 | -4.110027413 |
| **A_66_P108247** | Ucp3 | -0.224296612 | -0.507700946 | 0.059107722 | 9.252239394 | -1.947087144 | 0.100534049 | 0.60792129 | -4.110123076 |
| **A_51_P446417** | Ufl1 | -0.10254357 | -0.232126872 | 0.027039732 | 9.661847451 | -1.946832347 | 0.100569246 | 0.60792129 | -4.110308557 |
| **A_55_P2099890** | Atxn7 | -0.191569835 | -0.43366499 | 0.05052532 | 13.03741772 | -1.946750028 | 0.10058062 | 0.60792129 | -4.110368482 |
| **A_51_P298538** | Prmt7 | 0.138472553 | -0.03656218 | 0.313507287 | 11.33214577 | 1.946294814 | 0.10064354 | 0.60792129 | -4.110699868 |

| | | | | | | | | | |
|---|---|---|---|---|---|---|---|---|---|
| A_55_P2112005 | Tff1 | 0.524734859 | -0.138635735 | 1.188105453 | 9.376326547 | 1.946044929 | 0.100678096 | 0.60792129 | -4.110881784 |
| A_55_P2020453 | NA | 0.151609219 | -0.040071357 | 0.343289795 | 8.882376815 | 1.945882746 | 0.100700531 | 0.60792129 | -4.110999855 |
| A_51_P386304 | Ccnl2 | -0.344816111 | -0.780775285 | 0.091143063 | 10.91409712 | -1.945856868 | 0.100704111 | 0.60792129 | -4.111018695 |
| A_66_P114543 | Jmjd6 | 0.149535655 | -0.03954166 | 0.33861297 | 8.500127831 | 1.945693794 | 0.100726674 | 0.60792129 | -4.111137417 |
| A_51_P151484 | Atp1b1 | -0.247687081 | -0.560890815 | 0.065516653 | 12.7098417 | -1.945564294 | 0.100744596 | 0.60792129 | -4.111231697 |
| A_55_P2121877 | Gpr137b | 0.213085766 | -0.056389365 | 0.482560898 | 9.299704657 | 1.945382081 | 0.100769818 | 0.60792129 | -4.111364355 |
| A_51_P201971 | Kmt5a | -0.295130527 | -0.668584352 | 0.078323298 | 11.11662894 | -1.944224381 | 0.100930217 | 0.608517946 | -4.112207252 |
| A_55_P2050592 | NA | 0.123815947 | -0.032869455 | 0.280501348 | 8.312076653 | 1.944093783 | 0.100948328 | 0.608517946 | -4.112302343 |
| A_55_P2260052 | Gm17753 | 0.234524803 | -0.0622757 | 0.531325306 | 9.114856706 | 1.943986792 | 0.100963167 | 0.608517946 | -4.112380246 |
| A_55_P2138906 | Lrfn1 | 0.111018597 | -0.029498127 | 0.25153532 | 8.608332866 | 1.943734219 | 0.100998207 | 0.608539381 | -4.112564153 |
| A_55_P2162920 | NA | 0.163770554 | -0.043608189 | 0.371149298 | 16.54738989 | 1.942856581 | 0.101120057 | 0.609083697 | -4.11320322 |
| A_55_P2031407 | NA | -0.114231839 | -0.258913087 | 0.03044941 | 17.07142437 | -1.942424207 | 0.101180142 | 0.609255753 | -4.113518077 |
| A_51_P236469 | Metap1d | -0.129198851 | -0.292908241 | 0.03451054 | 7.986132209 | -1.941575228 | 0.101298225 | 0.60966884 | -4.114136338 |
| A_55_P2132800 | Zfp367 | 0.119359053 | -0.031889827 | 0.270607932 | 8.835729893 | 1.941477496 | 0.101311827 | 0.60966884 | -4.114207513 |
| A_51_P467751 | Hgsnat | -0.188686498 | -0.427903846 | 0.050530851 | 12.77072787 | -1.940516386 | 0.101445692 | 0.610063621 | -4.114907489 |
| A_55_P2046109 | NA | 0.393688737 | -0.105551426 | 0.8929289 | 8.328595223 | 1.940048723 | 0.101510894 | 0.610063621 | -4.115248106 |
| A_52_P335508 | Tiprl | 0.104404502 | -0.028000704 | 0.236809709 | 9.603929884 | 1.939917796 | 0.101529155 | 0.610063621 | -4.115343467 |
| A_55_P2162815 | NA | 0.263904034 | -0.070807572 | 0.59861564 | 11.90463526 | 1.939744011 | 0.101553399 | 0.610063621 | -4.115470046 |
| A_55_P2085551 | Dennd4b | 0.109930258 | -0.029507875 | 0.249368391 | 9.050816419 | 1.939567286 | 0.101578059 | 0.610063621 | -4.115598768 |
| A_52_P569348 | Dbt | -0.153403259 | -0.347990566 | 0.041184048 | 8.452190702 | -1.93949758 | 0.101587788 | 0.610063621 | -4.115649541 |
| A_55_P2162678 | Dnajc10 | 0.103581962 | -0.027819075 | 0.234982998 | 10.71779611 | 1.939342451 | 0.101609441 | 0.610063621 | -4.115762535 |
| A_52_P254795 | Ndufc1 | 0.225023864 | -0.060542573 | 0.5105903 | 13.57323499 | 1.938610579 | 0.101711663 | 0.610063621 | -4.116295641 |
| A_52_P185044 | Adipor2 | -0.28866907 | -0.65502525 | 0.07768711 | 14.19247167 | -1.938500396 | 0.101727061 | 0.610063621 | -4.116375902 |
| A_55_P2043207 | Magef1 | 0.621734601 | -0.167352025 | 1.410821228 | 10.0820194 | 1.938427303 | 0.101737278 | 0.610063621 | -4.116429147 |
| A_55_P2032258 | Pdlim1 | -0.158461464 | -0.359592183 | 0.042669255 | 15.79191869 | -1.938270659 | 0.101759175 | 0.610063621 | -4.116543253 |
| A_55_P2185526 | 1700016P03Rik | 0.192952828 | -0.051970788 | 0.437876444 | 9.029075678 | 1.9381601 | 0.101774634 | 0.610063621 | -4.116623791 |
| A_55_P2014665 | Racgap1 | 0.139194672 | -0.037512175 | 0.315901519 | 8.144763332 | 1.937931393 | 0.101806618 | 0.610063621 | -4.116790396 |
| A_55_P2142296 | Ywhaq | -0.189749849 | -0.430647474 | 0.051147777 | 12.79575221 | -1.937840737 | 0.1018193 | 0.610063621 | -4.116856437 |
| A_55_P2013756 | NA | 0.281077631 | -0.075890826 | 0.638046087 | 8.944969151 | 1.937160503 | 0.101914503 | 0.610414312 | -4.117351985 |
| A_55_P1966760 | NA | -0.134535782 | -0.305437345 | 0.036365782 | 17.1884527 | -1.936693676 | 0.101979891 | 0.610414312 | -4.117692082 |
| A_66_P139493 | Tmem250-ps | -0.126853148 | -0.287997512 | 0.034291215 | 10.21130208 | -1.936668522 | 0.101983415 | 0.610414312 | -4.117710408 |
| A_55_P2015599 | NA | 0.106173407 | -0.028725145 | 0.24107196 | 12.49919417 | 1.936322213 | 0.102031951 | 0.610414312 | -4.117962712 |
| A_55_P1966124 | Erh | 0.26181862 | -0.07084448 | 0.59448172 | 13.93926458 | 1.936266189 | 0.102039805 | 0.610414312 | -4.118003529 |
| A_55_P1959903 | Map7 | -0.104190771 | -0.23658753 | 0.028205988 | 8.789071758 | -1.936070016 | 0.102067311 | 0.610414312 | -4.118146455 |
| A_55_P2008061 | Itpr2 | -0.364945062 | -0.828791626 | 0.098901502 | 11.57679936 | -1.93562985 | 0.102129056 | 0.610421983 | -4.118467156 |
| A_55_P2225082 | Mettl4 | 0.117673393 | -0.031891424 | 0.267238209 | 8.480438286 | 1.935610616 | 0.102131755 | 0.610421983 | -4.11848117 |
| A_55_P1960655 | NA | 0.633423075 | -0.171959272 | 1.438805422 | 12.45517231 | 1.93491075 | 0.10223001 | 0.610633223 | -4.118991107 |
| A_52_P87843 | Aldh1a3 | 0.570121251 | -0.154803127 | 1.295045629 | 12.02765352 | 1.934833899 | 0.102240805 | 0.610633223 | -4.119047104 |
| A_51_P272553 | Bhlhe40 | -0.227059705 | -0.515803249 | 0.06168384 | 10.31460805 | -1.93462568 | 0.102270058 | 0.610633223 | -4.119198823 |
| A_66_P105422 | Lonrf3 | 0.303920318 | -0.082667792 | 0.690508427 | 11.10881777 | 1.934106594 | 0.102343024 | 0.610633223 | -4.119577067 |
| A_51_P430033 | Ccdc13 | 0.108776792 | -0.029602713 | 0.247156298 | 9.51224658 | 1.933898321 | 0.102372314 | 0.610633223 | -4.119728834 |
| A_55_P2064063 | Casp14 | 0.227005814 | -0.061830725 | 0.515842352 | 8.467731396 | 1.933543786 | 0.102422194 | 0.610633223 | -4.119987186 |
| A_52_P467726 | Nsg1 | -0.137717769 | -0.312947114 | 0.037511576 | 8.354946119 | -1.933536185 | 0.102423263 | 0.610633223 | -4.119992725 |
| A_55_P2054663 | Cox4i2 | 0.157222525 | -0.042838434 | 0.357283484 | 9.066944192 | 1.933399422 | 0.102442511 | 0.610633223 | -4.120092387 |
| A_55_P2062058 | Dbndd2 | -0.108211987 | -0.245913011 | 0.029489037 | 8.413909374 | -1.933336115 | 0.102451422 | 0.610633223 | -4.12013852 |
| A_51_P424641 | Sirt4 | -0.10490483 | -0.238433518 | 0.028623859 | 8.874997029 | -1.932813994 | 0.102524945 | 0.61082455 | -4.120519012 |
| A_51_P411355 | Taf1c | -0.107883917 | -0.245215538 | 0.029447704 | 8.797857806 | -1.932659392 | 0.102546726 | 0.61082455 | -4.12063168 |
| A_51_P398191 | Auts2 | -0.159064778 | -0.361582138 | 0.043452582 | 9.062780479 | -1.932328372 | 0.102593376 | 0.610914161 | -4.120872919 |
| A_55_P2185203 | Snrnp40 | -0.11691594 | -0.265788624 | 0.031956744 | 8.37045555 | -1.932092298 | 0.102626659 | 0.610924142 | -4.121044968 |
| A_55_P2079967 | Map3k5 | 0.50215774 | -0.137463206 | 1.141778685 | 10.79393549 | 1.931464129 | 0.102715274 | 0.6112634 | -4.121502786 |

| | | | | | | | | | |
|---|---|---|---|---|---|---|---|---|---|
| **A_51_P416419** | Calr | -0.373892294 | -0.850335996 | 0.102551409 | 14.75656034 | -1.930652387 | 0.102829899 | 0.611357384 | -4.122094427 |
| **A_55_P2016623** | S100a11-ps | -0.259915853 | -0.591145341 | 0.071313634 | 11.84115507 | -1.930513931 | 0.102849463 | 0.611357384 | -4.122195344 |
| **A_51_P519385** | Sacm1l | -0.098990762 | -0.225148804 | 0.02716728 | 10.14334002 | -1.930407034 | 0.10286457 | 0.611357384 | -4.12227326 |
| **A_55_P1970087** | Rpl35a | 0.215290682 | -0.059119857 | 0.489701221 | 15.86080774 | 1.930161324 | 0.102899304 | 0.611357384 | -4.122452357 |
| **A_55_P2055975** | Ppip5k1 | 0.192452708 | -0.052932637 | 0.437838052 | 9.636217692 | 1.929499046 | 0.102992982 | 0.611357384 | -4.122935104 |
| **A_52_P387564** | Tmpo | -0.162607016 | -0.369943656 | 0.044729623 | 10.49211776 | -1.929444939 | 0.103000639 | 0.611357384 | -4.122974545 |
| **A_51_P206971** | Mnat1 | -0.092677046 | -0.21086042 | 0.025506328 | 9.548180614 | -1.929234241 | 0.103030463 | 0.611357384 | -4.123128133 |
| **A_55_P2024679** | NA | 0.230304784 | -0.063404001 | 0.52401357 | 11.01413177 | 1.929101977 | 0.103049188 | 0.611357384 | -4.123224547 |
| **A_51_P175555** | Pelo | -0.2813677 | -0.645376794 | 0.082641394 | 12.8813459 | -2.003143626 | 0.103060049 | 0.611357384 | -4.129178046 |
| **A_51_P386358** | Wwp2 | -0.248776687 | -0.566116963 | 0.068563589 | 11.32636218 | -1.928651004 | 0.103113062 | 0.611357384 | -4.123553293 |
| **A_55_P2034345** | Terf2 | -0.186519483 | -0.424451715 | 0.05141275 | 10.46774612 | -1.928590776 | 0.103121595 | 0.611357384 | -4.123597199 |
| **A_51_P176071** | Cmtr2 | 0.131038717 | -0.036175133 | 0.298252567 | 8.978009838 | 1.927953471 | 0.103211936 | 0.611357384 | -4.124061797 |
| **A_66_P128537** | Isg15 | -0.183121505 | -0.416808768 | 0.050565757 | 9.783643108 | -1.927851028 | 0.103226465 | 0.611357384 | -4.12413648 |
| **A_55_P2074499** | Kcp | 0.16400078 | -0.045286846 | 0.373288406 | 8.361379875 | 1.927842382 | 0.103227691 | 0.611357384 | -4.124142783 |
| **A_55_P2153621** | Ahnak | 0.241399547 | -0.06667548 | 0.549474574 | 10.88247155 | 1.927742929 | 0.103241798 | 0.611357384 | -4.124215288 |
| **A_66_P135176** | Trak2 | -0.134753796 | -0.306731 | 0.037223408 | 8.747772232 | -1.927699293 | 0.103247989 | 0.611357384 | -4.1242471 |
| **A_55_P1962946** | Syk | -0.163006514 | -0.371056561 | 0.045043533 | 8.99512741 | -1.927552892 | 0.10326876 | 0.611357384 | -4.124353831 |
| **A_52_P75465** | Gdpd1 | -0.169967972 | -0.38698793 | 0.047051985 | 8.601962041 | -1.926799754 | 0.103375682 | 0.611540792 | -4.124902916 |
| **A_55_P1971419** | Serpinb9g | -0.290868136 | -0.662387562 | 0.08065129 | 14.2035863 | -1.926121966 | 0.103472001 | 0.611540792 | -4.125397092 |
| **A_52_P49406** | Pde12 | -0.152334568 | -0.346940334 | 0.042271197 | 9.177014824 | -1.925803297 | 0.103517318 | 0.611540792 | -4.125629442 |
| **A_55_P2076174** | NA | -0.141257388 | -0.321727654 | 0.039212878 | 12.99197512 | -1.925638122 | 0.103540815 | 0.611540792 | -4.125749878 |
| **A_55_P2180944** | Tmco4 | -0.159356819 | -0.362963133 | 0.044249495 | 11.30559633 | -1.925522397 | 0.103557281 | 0.611540792 | -4.125834259 |
| **A_55_P2071930** | NA | -0.089292516 | -0.203384024 | 0.024798992 | 8.645830555 | -1.92544405 | 0.103568429 | 0.611540792 | -4.125891385 |
| **A_51_P122238** | Uchl5 | -0.089929633 | -0.204844535 | 0.024985269 | 10.63735465 | -1.925287682 | 0.103590684 | 0.611540792 | -4.129010014 |
| **A_55_P1976033** | Cnih2 | 0.088990386 | -0.024768045 | 0.202748816 | 8.272291776 | 1.924547618 | 0.103696078 | 0.611540792 | -4.126545044 |
| **A_51_P510939** | Chchd7 | -0.143220106 | -0.326335077 | 0.039894865 | 9.049305171 | -1.924195976 | 0.103746194 | 0.611540792 | -4.126801465 |
| **A_55_P2125457** | NA | 0.189312462 | -0.052734904 | 0.431359827 | 13.56985178 | 1.924190235 | 0.103747012 | 0.611540792 | -4.126805652 |
| **A_52_P417620** | Pacs1 | -0.117493178 | -0.267726355 | 0.032739999 | 8.434721294 | -1.924048268 | 0.103767252 | 0.611540792 | -4.126909177 |
| **A_55_P2014124** | Gjb3 | -0.190212538 | -0.433446037 | 0.05302096 | 13.71724264 | -1.923910733 | 0.103786864 | 0.611540792 | -4.127009473 |
| **A_55_P2062727** | Nasp | -0.160293746 | -0.365280815 | 0.044693324 | 9.054774653 | -1.923797134 | 0.103803066 | 0.611540792 | -4.127092313 |
| **A_55_P2029846** | Fam214a | -0.231690037 | -0.528006969 | 0.064626896 | 11.79517064 | -1.923623403 | 0.103827849 | 0.611540792 | -4.127219006 |
| **A_52_P315976** | Tpm2 | -0.285902226 | -0.651650261 | 0.079845809 | 12.63412087 | -1.923112531 | 0.103900759 | 0.611540792 | -4.127591566 |
| **A_52_P657317** | Tmem160 | -0.232261733 | -0.529434663 | 0.064911198 | 10.5838502 | -1.92281534 | 0.103943197 | 0.611540792 | -4.127808303 |
| **A_51_P123047** | Smc1a | 0.138238641 | -0.03864687 | 0.315124151 | 11.65996756 | 1.922677115 | 0.103962941 | 0.611540792 | -4.12790911 |
| **A_55_P1985094** | Wrap73 | 0.196875154 | -0.055135209 | 0.448885517 | 10.22549395 | 1.92194818 | 0.104067124 | 0.611540792 | -4.128440737 |
| **A_55_P2028655** | Slc31a1 | 0.133369726 | -0.037358835 | 0.304098287 | 11.3084286 | 1.921853343 | 0.104080686 | 0.611540792 | -4.128509905 |
| **A_51_P276039** | Txndc15 | -0.177980858 | -0.405817339 | 0.049855624 | 13.05416232 | -1.921848694 | 0.104081351 | 0.611540792 | -4.128513295 |
| **A_51_P194628** | Tango6 | 0.129309954 | -0.036237924 | 0.294857832 | 8.911906982 | 1.921664208 | 0.104107739 | 0.611540792 | -4.12864785 |
| **A_55_P1978424** | Bcl2a1d | -0.274055013 | -0.624922949 | 0.076812923 | 8.831711792 | -1.921601067 | 0.104116772 | 0.611540792 | -4.128693902 |
| **A_55_P1971889** | F3 | -0.139346273 | -0.317758765 | 0.039066219 | 12.29086689 | -1.92149501 | 0.104131946 | 0.611540792 | -4.128771256 |
| **A_55_P2046378** | Eif4g1 | -0.210915939 | -0.481104089 | 0.05927221 | 13.29980869 | -1.920490962 | 0.104275711 | 0.611540792 | -4.129503595 |
| **A_55_P2011322** | NA | 0.219558182 | -0.061719596 | 0.500835959 | 10.70473879 | 1.920363161 | 0.104294025 | 0.611540792 | -4.129596816 |
| **A_51_P304397** | Cpm | 0.262397641 | -0.073779472 | 0.598574754 | 10.57013068 | 1.920264029 | 0.104308233 | 0.611540792 | -4.129669125 |
| **A_55_P2132237** | Fastkd3 | -0.126345631 | -0.288229182 | 0.03553792 | 9.343510785 | -1.920112473 | 0.104329957 | 0.611540792 | -4.129779675 |
| **A_52_P601980** | Kdm5b | -0.141410405 | -0.322612172 | 0.039791362 | 8.689159167 | -1.919941967 | 0.104354404 | 0.611540792 | -4.129904048 |
| **A_55_P2087539** | Clpsl2 | 0.192717354 | -0.054233415 | 0.439668123 | 8.278910814 | 1.919904403 | 0.104359791 | 0.611540792 | -4.129931448 |
| **A_55_P2279997** | 4930549C15Rik | 0.127297956 | -0.035839049 | 0.29043496 | 8.041715749 | 1.919720972 | 0.104386098 | 0.611540792 | -4.130065252 |
| **A_55_P2109337** | Gba2 | -0.24879933 | -0.567654181 | 0.070055521 | 11.30079642 | -1.919664532 | 0.104394194 | 0.611540792 | -4.130106422 |
| **A_55_P2145908** | Phb | 0.2564649 | -0.072222926 | 0.585152727 | 12.26855675 | 1.919612103 | 0.104401715 | 0.611540792 | -4.130144667 |
| **A_52_P412529** | Fbxo3 | -0.097468551 | -0.222385194 | 0.027448092 | 8.527930907 | -1.919611608 | 0.104401786 | 0.611540792 | -4.130145028 |

| | | | | | | | | | |
|---|---|---|---|---|---|---|---|---|---|
| **A_55_P1963080** | Rps12 | 0.16413219 | -0.046223315 | 0.374487694 | 16.88098627 | 1.919592506 | 0.104404527 | 0.611540792 | -4.130158962 |
| **A_55_P2084533** | Zbtb40 | 0.157634904 | -0.044426749 | 0.359696558 | 10.28446243 | 1.919276951 | 0.104449807 | 0.611540792 | -4.130389149 |
| **A_52_P526852** | Eri1 | -0.103186921 | -0.235476611 | 0.029102769 | 8.428072102 | -1.918968384 | 0.104494103 | 0.611540792 | -4.130614244 |
| **A_55_P2173183** | Cdk5r1 | -0.12144215 | -0.277138988 | 0.034254688 | 9.042159671 | -1.918928607 | 0.104499815 | 0.611540792 | -4.130643261 |
| **A_55_P2080542** | Rbbp7 | -0.213771287 | -0.4878729 | 0.060330326 | 12.51523784 | -1.918699415 | 0.104532731 | 0.611540792 | -4.130810457 |
| **A_55_P2080907** | Rpl26 | -0.184099397 | -0.420161058 | 0.051962264 | 15.33262758 | -1.918651382 | 0.104539631 | 0.611540792 | -4.130845497 |
| **A_52_P132165** | Hsd17b11 | -0.290899697 | -0.678421823 | 0.096622429 | 12.19234546 | -2.112269236 | 0.104565281 | 0.611540792 | -4.200937719 |
| **A_55_P2002389** | Nelfb | -0.194706827 | -0.44449765 | 0.055083996 | 13.4587592 | -1.917669954 | 0.104680708 | 0.611713176 | -4.131561481 |
| **A_55_P1986541** | NA | 0.260060201 | -0.073590484 | 0.593710886 | 15.45124533 | 1.917569157 | 0.104695207 | 0.611713176 | -4.131635018 |
| **A_51_P447258** | Alox12b | 0.298721143 | -0.084651295 | 0.682093581 | 8.442724064 | 1.916965244 | 0.104782125 | 0.611713176 | -4.13207562 |
| **A_55_P1952547** | Sh3rf1 | -0.124611784 | -0.284538158 | 0.035314589 | 8.172364158 | -1.916938536 | 0.104785971 | 0.611713176 | -4.132095106 |
| **A_55_P2178803** | Zbtb5 | -0.090536236 | -0.206731213 | 0.025658741 | 10.26996147 | -1.916921133 | 0.104788477 | 0.611713176 | -4.132107803 |
| **A_55_P2058157** | Ppp4r3b | -0.12059067 | -0.275377583 | 0.034196243 | 9.681120238 | -1.916675685 | 0.104823825 | 0.611713176 | -4.132286882 |
| **A_55_P1993858** | Rmrp | 0.123420838 | -0.034999244 | 0.28184092 | 9.95606644 | 1.916670329 | 0.104824597 | 0.611713176 | -4.13229079 |
| **A_55_P2074864** | NA | 0.223000109 | -0.063287623 | 0.509287841 | 12.82771264 | 1.916335306 | 0.104872865 | 0.611713176 | -4.132535228 |
| **A_55_P2149942** | Ninj2 | 0.140901399 | -0.040044388 | 0.321847186 | 8.441659036 | 1.915737461 | 0.104959056 | 0.611713176 | -4.13297144 |
| **A_55_P2149821** | Srsf11 | -0.242624968 | -0.554207114 | 0.068957178 | 11.50218439 | -1.915720252 | 0.104961538 | 0.611713176 | -4.132983997 |
| **A_55_P2026562** | Cdkn1b | 0.140511967 | -0.039952769 | 0.320976703 | 11.91172663 | 1.915535155 | 0.104988238 | 0.611713176 | -4.133119055 |
| **A_51_P156934** | Dnajc15 | 0.282069946 | -0.080227481 | 0.644367372 | 11.76089668 | 1.915405493 | 0.105006946 | 0.611713176 | -4.133213665 |
| **A_55_P2051039** | Gm11937 | 0.533320672 | -0.151761118 | 1.218402461 | 9.857460888 | 1.915204516 | 0.105035949 | 0.611713176 | -4.133360313 |
| **A_55_P1966337** | Polr2h | -0.183488694 | -0.419217442 | 0.052240055 | 10.62375739 | -1.914987402 | 0.105067291 | 0.611713176 | -4.133518739 |
| **A_52_P227267** | Atp1a2 | -0.217931595 | -0.497911498 | 0.062048308 | 8.730758201 | -1.914972326 | 0.105069468 | 0.611713176 | -4.13352974 |
| **A_52_P421897** | NA | -0.126208547 | -0.288394788 | 0.035977693 | 8.326686747 | -1.914449546 | 0.105144974 | 0.611916926 | -4.133911216 |
| **A_55_P2026710** | 4930445E18Rik | 0.15408425 | -0.043940417 | 0.352108916 | 8.521054055 | 1.914291717 | 0.105167781 | 0.611916926 | -4.134026388 |
| **A_52_P305890** | Yipf5 | -0.268202084 | -0.626422732 | 0.090018565 | 12.19708981 | -2.106755041 | 0.10520537 | 0.611951427 | -4.148174574 |
| **A_51_P425696** | Nt5c2 | -0.119295773 | -0.272676249 | 0.034084704 | 8.873531426 | -1.913480906 | 0.105285021 | 0.612082158 | -4.134618075 |
| **A_55_P2006133** | Tmc6 | 0.285731416 | -0.081679278 | 0.65314211 | 9.406445995 | 1.913266042 | 0.105316112 | 0.612082158 | -4.134774877 |
| **A_55_P1978122** | Mutyh | 0.125559017 | -0.035924178 | 0.287042211 | 8.730709125 | 1.912888852 | 0.105370713 | 0.612082158 | -4.135050146 |
| **A_51_P197175** | Pigk | -0.119969525 | -0.274297333 | 0.034358283 | 9.118633533 | -1.912475654 | 0.105430559 | 0.612082158 | -4.135351701 |
| **A_55_P1975903** | Dnmt3b | 0.252604264 | -0.072379692 | 0.57758822 | 8.876011228 | 1.91226431 | 0.105461183 | 0.612082158 | -4.135505945 |
| **A_55_P2171047** | Oaz1 | 0.121933413 | -0.034938369 | 0.278805195 | 14.59063301 | 1.912260546 | 0.105461728 | 0.612082158 | -4.135508692 |
| **A_52_P283802** | Tapt1 | -0.128509316 | -0.2938626 | 0.036843968 | 9.425808704 | -1.912013487 | 0.105497539 | 0.612082158 | -4.135689004 |
| **A_55_P2123802** | NA | -0.087372208 | -0.199800286 | 0.025055869 | 8.616939944 | -1.911911087 | 0.105512385 | 0.612082158 | -4.13576374 |
| **A_52_P642488** | Kcnk1 | -0.359261098 | -0.821671722 | 0.103149525 | 10.33428483 | -1.911399876 | 0.105586532 | 0.612082158 | -4.136136851 |
| **A_55_P2158697** | H4c9 | -0.115862704 | -0.265005757 | 0.033280348 | 8.856246843 | -1.911216114 | 0.105613198 | 0.612082158 | -4.136270975 |
| **A_52_P520564** | Wasl | -0.122362865 | -0.279884789 | 0.035159059 | 8.606832171 | -1.91107527 | 0.10563364 | 0.612082158 | -4.136373774 |
| **A_55_P2120682** | Golga7 | -0.158807091 | -0.363249055 | 0.045634873 | 10.47137595 | -1.911036624 | 0.10563925 | 0.612082158 | -4.136401981 |
| **A_52_P324754** | Nupr1l | -0.167060238 | -0.382127176 | 0.048006701 | 9.029662596 | -1.91103484 | 0.105639509 | 0.612082158 | -4.136403284 |
| **A_55_P2014555** | Lrp8 | 0.262885292 | -0.075637837 | 0.60140842 | 9.315579357 | 1.910500281 | 0.105717138 | 0.612348385 | -4.136793459 |
| **A_55_P2002562** | Sec11a | -0.119364904 | -0.273167651 | 0.034437843 | 11.55096941 | -1.909333205 | 0.10588682 | 0.612549761 | -4.137645358 |
| **A_55_P2083184** | Gm10272 | 0.194362696 | -0.056092352 | 0.444817745 | 8.355680014 | 1.909203791 | 0.105905652 | 0.612549761 | -4.137739827 |
| **A_55_P2094242** | NA | 0.145832091 | -0.04208853 | 0.333752712 | 8.407441419 | 1.909184185 | 0.105908505 | 0.612549761 | -4.137754139 |
| **A_51_P141521** | Sap130 | -0.28254568 | -0.646664027 | 0.081572668 | 11.89178318 | -1.909041067 | 0.105929336 | 0.612549761 | -4.137858612 |
| **A_52_P648304** | Pigs | -0.205016886 | -0.469227054 | 0.059193281 | 12.45243489 | -1.909014689 | 0.105933176 | 0.612549761 | -4.137877868 |
| **A_55_P1998761** | Coq8b | 0.109050164 | -0.031489879 | 0.249590208 | 9.873912196 | 1.908953745 | 0.105942048 | 0.612549761 | -4.137922357 |
| **A_51_P400659** | Bcap29 | -0.121467741 | -0.278048291 | 0.035112809 | 10.11800722 | -1.908500607 | 0.106008038 | 0.612628786 | -4.138253148 |
| **A_55_P1995447** | NA | 0.202561057 | -0.058593957 | 0.463716071 | 12.90283769 | 1.908212557 | 0.106050008 | 0.612628786 | -4.138463429 |
| **A_52_P487686** | Cstdc5 | -0.544337479 | -1.246235016 | 0.157560057 | 10.30606719 | -1.907935473 | 0.106090396 | 0.612628786 | -4.13866571 |
| **A_55_P1970474** | Tmem67 | -0.15655993 | -0.35844869 | 0.045328829 | 8.744811057 | -1.907821078 | 0.106107075 | 0.612628786 | -4.138749223 |
| **A_55_P1977431** | Cck | -0.341156722 | -0.781118236 | 0.098804792 | 9.711749149 | -1.907692657 | 0.106125801 | 0.612628786 | -4.138842977 |

| | | | | | | | | | |
|---|---|---|---|---|---|---|---|---|---|
| **A_51_P329490** | Slc25a40 | -0.181329258 | -0.415215587 | 0.052557071 | 9.556607558 | -1.90735798 | 0.106174621 | 0.612628786 | -4.139087311 |
| **A_55_P1959064** | Oas1b | -0.128131923 | -0.293406651 | 0.037142804 | 8.741644889 | -1.907304608 | 0.106182408 | 0.612628786 | -4.139126276 |
| **A_55_P2036567** | Vav1 | -0.187056284 | -0.428407374 | 0.054294807 | 8.68923424 | -1.906743168 | 0.106264362 | 0.612628786 | -4.139536173 |
| **A_55_P2183894** | Gm5617 | 0.159868494 | -0.04640719 | 0.366144177 | 10.76482221 | 1.906707223 | 0.106269611 | 0.612628786 | -4.139562416 |
| **A_51_P184806** | Elmod2 | 0.131122713 | -0.038064618 | 0.300310044 | 9.757574056 | 1.906686322 | 0.106272664 | 0.612628786 | -4.139577677 |
| **A_66_P100249** | Snhg12 | 0.151406414 | -0.044004802 | 0.346817631 | 10.45062749 | 1.906180181 | 0.106346606 | 0.612860888 | -4.139947216 |
| **A_51_P286563** | Gna13 | -0.303508181 | -0.695278286 | 0.088261924 | 11.7554509 | -1.90593582 | 0.106382323 | 0.612860888 | -4.14012563 |
| **A_55_P2002602** | NA | 0.172611364 | -0.050216946 | 0.395439674 | 15.49851602 | 1.905759881 | 0.106408047 | 0.612860888 | -4.140254091 |
| **A_51_P334281** | Gucy1b2 | 0.272121279 | -0.079291574 | 0.623534132 | 12.56734394 | 1.905083542 | 0.106506991 | 0.612861899 | -4.140747925 |
| **A_55_P2137773** | NA | 0.178650253 | -0.052073005 | 0.409373511 | 16.70639301 | 1.90494053 | 0.106527925 | 0.612861899 | -4.14085235 |
| **A_55_P2058107** | Pigyl | 0.250289558 | -0.07296826 | 0.573547376 | 12.10150577 | 1.904859104 | 0.106539845 | 0.612861899 | -4.140911805 |
| **A_55_P2169415** | Cgn | -0.295486851 | -0.677181418 | 0.086207717 | 11.29152972 | -1.904545461 | 0.106585775 | 0.612861899 | -4.141140826 |
| **A_55_P2083401** | Acp1 | -0.186112203 | -0.426555527 | 0.054331122 | 11.03902935 | -1.904282113 | 0.106624354 | 0.612861899 | -4.141333124 |
| **A_55_P2047932** | Cyb5r4 | 0.361078706 | -0.105428295 | 0.827585708 | 10.86017458 | 1.904201375 | 0.106636185 | 0.612861899 | -4.14139208 |
| **A_52_P161495** | Bcl6 | 0.143892692 | -0.042028593 | 0.329813976 | 11.08470833 | 1.904051929 | 0.106658087 | 0.612861899 | -4.141501208 |
| **A_55_P1962982** | NA | -0.26433485 | -0.605919866 | 0.077250165 | 9.772187531 | -1.903815144 | 0.106692799 | 0.612861899 | -4.141674116 |
| **A_55_P2049979** | Dbi | 0.270937623 | -0.079180769 | 0.621056015 | 15.21211161 | 1.903809785 | 0.106693584 | 0.612861899 | -4.141678029 |
| **A_55_P2043734** | Cdkl5 | 0.199554277 | -0.0583672 | 0.457475754 | 9.294264797 | 1.903455363 | 0.106745562 | 0.612978303 | -4.141936843 |
| **A_51_P424854** | Ahsg | -1.009462179 | -2.314512009 | 0.295587652 | 10.20059547 | -1.902970783 | 0.106816669 | 0.613088227 | -4.142290714 |
| **A_55_P2114908** | Nek3 | 0.119290318 | -0.034936563 | 0.273517199 | 8.834803563 | 1.90289263 | 0.106828142 | 0.613088227 | -4.142347787 |
| **A_51_P154585** | Vmn1r195 | 0.251982397 | -0.073858252 | 0.577823046 | 8.583302471 | 1.902541345 | 0.106879724 | 0.613129906 | -4.142604326 |
| **A_55_P2011922** | 9-Mar | 0.498488368 | -0.146155424 | 1.143132161 | 10.58947095 | 1.902411171 | 0.106898845 | 0.613129906 | -4.142699392 |
| **A_51_P202964** | Csnk2a2 | -0.17379138 | -0.398622458 | 0.051039699 | 10.08398399 | -1.901695813 | 0.107003985 | 0.613136388 | -4.14322183 |
| **A_55_P2034928** | BC147527 | 0.25283244 | -0.074269324 | 0.579934204 | 9.551543779 | 1.901599582 | 0.107018136 | 0.613136388 | -4.143292111 |
| **A_55_P2003891** | Txlna | 0.188947473 | -0.055506383 | 0.433401329 | 11.19148017 | 1.901574558 | 0.107021816 | 0.613136388 | -4.143310387 |
| **A_51_P247184** | Npr3 | -0.183425865 | -0.420740332 | 0.053888601 | 8.649117027 | -1.901540264 | 0.10702686 | 0.613136388 | -4.143335433 |
| **A_55_P2144597** | 9030025P20Rik | -0.205452505 | -0.471309658 | 0.060404649 | 8.341362793 | -1.901219468 | 0.107074052 | 0.613224995 | -4.143569726 |
| **A_55_P2002113** | Rpl36 | 0.088631211 | -0.026092296 | 0.203354718 | 15.72210634 | 1.900655616 | 0.107157052 | 0.61339401 | -4.143981546 |
| **A_55_P2008422** | Mpc1 | 0.177862636 | -0.05239134 | 0.408116612 | 14.0491825 | 1.900407555 | 0.107193587 | 0.61339401 | -4.144162726 |
| **A_51_P380309** | Ncam1 | 0.337055623 | -0.099291411 | 0.773402656 | 11.15560862 | 1.900372388 | 0.107198767 | 0.61339401 | -4.144188412 |
| **A_52_P300416** | Dcp1a | -0.120527451 | -0.276659013 | 0.035604112 | 9.025352766 | -1.899172568 | 0.107375664 | 0.614224391 | -4.145064787 |
| **A_55_P2007756** | Krt16 | 0.17424848 | -0.051557586 | 0.400054546 | 9.520594043 | 1.898464832 | 0.107480147 | 0.614544872 | -4.145581763 |
| **A_52_P178998** | Fam168b | -0.153183425 | -0.351716302 | 0.045349452 | 13.43947115 | -1.898228558 | 0.107515051 | 0.614544872 | -4.145754359 |
| **A_51_P338397** | Kctd10 | -0.259193047 | -0.595160318 | 0.076774223 | 9.687010073 | -1.897997078 | 0.107549258 | 0.614544872 | -4.145923455 |
| **A_55_P2022444** | Pdss2 | -0.112420764 | -0.258162348 | 0.033320819 | 8.316977681 | -1.897720345 | 0.107590166 | 0.614544872 | -4.14612561 |
| **A_66_P133202** | Nfyc | 0.203267781 | -0.060269631 | 0.466805194 | 11.79289646 | 1.897559629 | 0.107613931 | 0.614544872 | -4.146243016 |
| **A_55_P1964852** | Kbtbd2 | -0.165053074 | -0.379051479 | 0.048945331 | 12.62357251 | -1.897502008 | 0.107622453 | 0.614544872 | -4.14628511 |
| **A_55_P2045085** | NA | 0.138016369 | -0.040998618 | 0.317031356 | 14.78269606 | 1.896751203 | 0.107733553 | 0.614997591 | -4.146833605 |
| **A_55_P2005213** | Ces2c | 0.214467068 | -0.063828673 | 0.492762809 | 8.302783329 | 1.895934011 | 0.107854607 | 0.615345431 | -4.147430627 |
| **A_55_P2083426** | NA | -0.269067192 | -0.618269669 | 0.080135285 | 12.7737194 | -1.895625727 | 0.10790031 | 0.615345431 | -4.14765586 |
| **A_55_P2063594** | Gm5801 | 0.343811244 | -0.102404614 | 0.790027102 | 12.29199601 | 1.895589254 | 0.107905718 | 0.615345431 | -4.147682508 |
| **A_65_P11092** | Tspan15 | -0.093502093 | -0.2148988 | 0.027894614 | 8.036389207 | -1.894888187 | 0.108009728 | 0.615345431 | -4.148194726 |
| **A_55_P2129925** | Mybl2 | -0.16115536 | -0.370459723 | 0.048149004 | 8.679291786 | -1.894242752 | 0.108105573 | 0.615345431 | -4.148666317 |
| **A_55_P2428968** | Lrrc2 | 0.391341095 | -0.116931264 | 0.899613454 | 9.510274076 | 1.894210006 | 0.108110438 | 0.615345431 | -4.148690244 |
| **A_55_P2032643** | Ajap1 | 0.096053336 | -0.028701226 | 0.220807897 | 8.120337405 | 1.89419716 | 0.108112347 | 0.615345431 | -4.14869963 |
| **A_51_P174854** | Maged1 | -0.255138024 | -0.586534841 | 0.076258793 | 15.5806296 | -1.894069967 | 0.108131246 | 0.615345431 | -4.148792567 |
| **A_55_P2126207** | Fthl17a | -0.155460282 | -0.357425627 | 0.046505063 | 16.90801423 | -1.893702531 | 0.10818586 | 0.615345431 | -4.149061047 |
| **A_51_P201721** | Crtap | -0.128264942 | -0.294906749 | 0.038376865 | 12.36161012 | -1.893621453 | 0.108197915 | 0.615345431 | -4.149120291 |
| **A_66_P114784** | Pla2g7 | -0.399181465 | -0.917836495 | 0.119473564 | 9.810225949 | -1.893480938 | 0.10821881 | 0.615345431 | -4.149222965 |
| **A_55_P2179398** | Alg1 | 0.180693458 | -0.054083863 | 0.415470779 | 10.73249246 | 1.893457015 | 0.108222368 | 0.615345431 | -4.149240446 |

| | | | | | | | | | |
|---|---|---|---|---|---|---|---|---|---|
| **A_55_P2008106** | Otud6b | 0.11648593 | -0.034884359 | 0.267856219 | 8.805665158 | 1.893223968 | 0.108257033 | 0.615345431 | -4.149410736 |
| **A_51_P464300** | Gdf1 | 0.090294417 | -0.027046903 | 0.207635736 | 7.938734091 | 1.893124132 | 0.108271887 | 0.615345431 | -4.149483688 |
| **A_55_P2017709** | Igsf9 | -0.246152417 | -0.566071836 | 0.073767003 | 12.94878295 | -1.892921761 | 0.108302002 | 0.615345431 | -4.149631566 |
| **A_55_P1953132** | Camkv | 0.318029778 | -0.095376378 | 0.731435933 | 15.66375826 | 1.892605116 | 0.108349139 | 0.615345431 | -4.14986295 |
| **A_52_P451644** | Ndn | 0.103916055 | -0.031171102 | 0.239003211 | 8.200167992 | 1.892508131 | 0.108363581 | 0.615345431 | -4.149933821 |
| **A_52_P355084** | Metrnl | -0.119443918 | -0.274718912 | 0.035831077 | 12.17021356 | -1.892481664 | 0.108367523 | 0.615345431 | -4.149953162 |
| **A_51_P228295** | Mpzl1 | -0.27827597 | -0.640166053 | 0.083614113 | 14.28933143 | -1.891769358 | 0.108473654 | 0.615767181 | -4.150473689 |
| **A_51_P128336** | Serinc5 | -0.29342934 | -0.675144093 | 0.088285413 | 12.41083209 | -1.891183864 | 0.108560968 | 0.615806062 | -4.154791668 |
| **A_55_P2027386** | Gpr156 | -0.265171054 | -0.610149891 | 0.079807784 | 10.48194486 | -1.891049179 | 0.108581063 | 0.615806062 | -4.150999993 |
| **A_55_P2016682** | Alkbh7 | 0.219166909 | -0.065995141 | 0.504328959 | 11.08746954 | 1.890829711 | 0.108613817 | 0.615806062 | -4.151160383 |
| **A_55_P2182736** | Limd2 | -0.152578841 | -0.351120122 | 0.04596244 | 8.769155273 | -1.890656605 | 0.108639658 | 0.615806062 | -4.151286893 |
| **A_51_P249414** | Ess2 | 0.167922189 | -0.05060451 | 0.386448889 | 10.04219412 | 1.890482862 | 0.1086656 | 0.615806062 | -4.15141387 |
| **A_52_P322141** | Ccdc88b | -0.107488361 | -0.247377228 | 0.032400506 | 8.446337608 | -1.890372766 | 0.108682043 | 0.615806062 | -4.151494332 |
| **A_55_P2032608** | Dipk2a | -0.209987822 | -0.483293624 | 0.063317979 | 9.826844072 | -1.890228999 | 0.108703517 | 0.615806062 | -4.151599403 |
| **A_55_P2019014** | Tmem120b | 0.173893601 | -0.052513625 | 0.400300828 | 10.49841833 | 1.889567806 | 0.108802335 | 0.615820009 | -4.152082643 |
| **A_52_P271377** | Efr3a | -0.154939588 | -0.356674158 | 0.046794982 | 8.570184414 | -1.889518872 | 0.108809651 | 0.615820009 | -4.152118408 |
| **A_51_P176377** | Usp25 | -0.129221664 | -0.297485023 | 0.039041695 | 10.00411512 | -1.889360962 | 0.108833267 | 0.615820009 | -4.152233821 |
| **A_52_P268287** | Zfp507 | -0.190789587 | -0.439222498 | 0.057643323 | 9.348950341 | -1.889359944 | 0.108833419 | 0.615820009 | -4.152234565 |
| **A_51_P400585** | Api5 | -0.088553471 | -0.203891881 | 0.026784939 | 10.420082 | -1.888864446 | 0.108907553 | 0.615877091 | -4.152596722 |
| **A_52_P515036** | Htatip2 | -0.167053308 | -0.384646022 | 0.050539406 | 9.41881382 | -1.888773561 | 0.108921156 | 0.615877091 | -4.15266315 |
| **A_52_P175679** | Gm13023 | 0.175078563 | -0.053002862 | 0.403159989 | 8.709806161 | 1.888479256 | 0.108965218 | 0.615877091 | -4.152878262 |
| **A_52_P476560** | Tpcn1 | -0.183659848 | -0.422925292 | 0.055605596 | 13.28591442 | -1.888440923 | 0.108970958 | 0.615877091 | -4.152906281 |
| **A_52_P359621** | Luzp1 | -0.177080839 | -0.407836496 | 0.053674819 | 12.01929371 | -1.887940818 | 0.109045876 | 0.616120357 | -4.153271824 |
| **A_55_P2131520** | Vash1 | -0.392663639 | -0.90449487 | 0.119167593 | 13.86065039 | -1.887396158 | 0.109127527 | 0.616351064 | -4.153669946 |
| **A_55_P2080649** | Zfp691 | 0.134633841 | -0.040888013 | 0.310155694 | 8.965687026 | 1.887088212 | 0.109173719 | 0.616351064 | -4.153895046 |
| **A_55_P1985633** | Lrp1 | 0.089581052 | -0.027209145 | 0.206371248 | 8.423307588 | 1.88703055 | 0.109182371 | 0.616351064 | -4.153937196 |
| **A_52_P451574** | Xpr1 | 0.125342009 | -0.038120968 | 0.288804985 | 11.30502194 | 1.886454833 | 0.109268788 | 0.616641118 | -4.154358042 |
| **A_52_P503334** | Erlin2 | -0.125646199 | -0.289534979 | 0.038242581 | 11.47088053 | -1.886119887 | 0.109319096 | 0.616641118 | -4.154602893 |
| **A_52_P180842** | Exoc3 | -0.14091361 | -0.32473041 | 0.04290319 | 9.651516783 | -1.885979214 | 0.109340231 | 0.616641118 | -4.154705728 |
| **A_55_P2007510** | Pef1 | 0.262511087 | -0.079950837 | 0.604973011 | 12.74950426 | 1.885838604 | 0.109361361 | 0.616641118 | -4.154808518 |
| **A_55_P1974681** | Nsun4 | -0.443283712 | -1.021774158 | 0.135206734 | 9.807014536 | -1.885188345 | 0.109459132 | 0.616982569 | -4.155283887 |
| **A_52_P272534** | NA | -0.322983485 | -0.744539237 | 0.098572268 | 9.934048189 | -1.884926667 | 0.109498502 | 0.616982569 | -4.155475191 |
| **A_55_P1970380** | NA | -0.158356836 | -0.365059706 | 0.048346034 | 8.509880489 | -1.884774821 | 0.109521354 | 0.616982569 | -4.155586202 |
| **A_55_P1967648** | Flcn | 0.1496351 | -0.045702751 | 0.344972951 | 10.90331391 | 1.884587189 | 0.109549598 | 0.616982569 | -4.155723376 |
| **A_55_P2199717** | 9330102E08Rik | 0.112093931 | -0.034281662 | 0.258469523 | 8.455634602 | 1.884007362 | 0.109636925 | 0.617294529 | -4.156147285 |
| **A_55_P2062652** | Zfp941 | 0.176559755 | -0.054051934 | 0.407171445 | 8.369521213 | 1.883560455 | 0.10970428 | 0.617402649 | -4.156474027 |
| **A_55_P2063558** | NA | 0.171394316 | -0.05248299 | 0.395271623 | 15.56464242 | 1.883456117 | 0.109720011 | 0.617402649 | -4.156550311 |
| **A_51_P327121** | Was | 0.114403101 | -0.035053836 | 0.263860038 | 9.337087849 | 1.883175869 | 0.109762276 | 0.617404938 | -4.156755211 |
| **A_52_P572702** | Gxylt1 | -0.281032332 | -0.648203223 | 0.086138558 | 11.70018323 | -1.883029864 | 0.109784302 | 0.617404938 | -4.156861962 |
| **A_55_P1998389** | NA | -0.262885409 | -0.606420888 | 0.080650069 | 14.49158221 | -1.882625992 | 0.109845251 | 0.617568023 | -4.157157256 |
| **A_66_P136788** | Ubald2 | -0.175087396 | -0.403964066 | 0.053789273 | 13.84472968 | -1.882012579 | 0.109937888 | 0.617595246 | -4.15760577 |
| **A_52_P399054** | Tmed2 | 0.161602918 | -0.049669884 | 0.37287572 | 9.830987516 | 1.881805653 | 0.109969155 | 0.617595246 | -4.157757073 |
| **A_51_P350048** | Gstt2 | 0.225730083 | -0.069390141 | 0.520850307 | 8.89359601 | 1.881740067 | 0.109979067 | 0.617595246 | -4.15780503 |
| **A_51_P296448** | Casp2 | -0.202060028 | -0.466342766 | 0.06222271 | 11.25362286 | -1.880965262 | 0.110096232 | 0.617595246 | -4.15837158 |
| **A_55_P2070825** | Nudt5 | 0.358675057 | -0.110474057 | 0.827824171 | 10.8192875 | 1.880872854 | 0.110110214 | 0.617595246 | -4.158439152 |
| **A_55_P2018847** | Crlf2 | 0.256933876 | -0.079189825 | 0.593057577 | 14.93717037 | 1.880578189 | 0.110154812 | 0.617595246 | -4.158654624 |
| **A_55_P2006869** | Fyb | -0.096788881 | -0.22343095 | 0.029853189 | 7.901190254 | -1.880254479 | 0.110203826 | 0.617595246 | -4.158891339 |
| **A_66_P135694** | NA | -0.159655088 | -0.368563887 | 0.04925371 | 15.76886378 | -1.880161678 | 0.110217881 | 0.617595246 | -4.158959201 |
| **A_66_P139310** | Cog1 | -0.128376336 | -0.296361075 | 0.039608402 | 11.04662593 | -1.880114564 | 0.110225018 | 0.617595246 | -4.158993653 |
| **A_55_P2108404** | NA | -0.17585182 | -0.405961727 | 0.054258088 | 10.2140318 | -1.88009897 | 0.11022738 | 0.617595246 | -4.159005056 |

| | | | | | | | | | |
|---|---|---|---|---|---|---|---|---|---|
| **A_51_P355040** | NA | 0.090083573 | -0.027796609 | 0.207963755 | 8.576054691 | 1.880069782 | 0.110231801 | 0.617595246 | -4.159026401 |
| **A_55_P2076489** | Rac1 | -0.185936534 | -0.429286373 | 0.057413305 | 14.28146916 | -1.879761848 | 0.110278459 | 0.617595246 | -4.159251586 |
| **A_51_P300434** | Dst | -0.358751711 | -0.82834547 | 0.110842048 | 9.890295107 | -1.8794935 | 0.110319134 | 0.617595246 | -4.159447825 |
| **A_55_P2021921** | Fkbp1a | 0.339125176 | -0.104796901 | 0.783047253 | 14.69354268 | 1.879413972 | 0.110331192 | 0.617595246 | -4.159505983 |
| **A_52_P549827** | Mgst1 | -0.149335561 | -0.344836833 | 0.04616571 | 10.83297088 | -1.879242458 | 0.1103572 | 0.617595246 | -4.159631411 |
| **A_55_P2124097** | Gm6981 | -0.268288499 | -0.619520988 | 0.082943991 | 14.1061145 | -1.879215284 | 0.110361321 | 0.617595246 | -4.159651283 |
| **A_52_P16563** | Lsm14b | -0.180483664 | -0.416828211 | 0.055860882 | 10.13353741 | -1.878717484 | 0.110436845 | 0.617649715 | -4.160015331 |
| **A_55_P2404157** | Rbm27 | -0.118617964 | -0.273971582 | 0.036735654 | 8.898624128 | -1.878444016 | 0.110478356 | 0.617649715 | -4.160215326 |
| **A_55_P2005433** | NA | 0.60042458 | -0.185990728 | 1.386839888 | 9.869077284 | 1.878346315 | 0.110493191 | 0.617649715 | -4.160286778 |
| **A_55_P2167769** | Arid1a | -0.171277835 | -0.395616078 | 0.053060407 | 11.02639301 | -1.878308896 | 0.110498873 | 0.617649715 | -4.160314144 |
| **A_55_P2111039** | NA | -0.117834831 | -0.272206015 | 0.036536354 | 10.76605764 | -1.877917924 | 0.110558259 | 0.617725015 | -4.16060008 |
| **A_55_P1968231** | Rbpms | -0.090039701 | -0.208005635 | 0.027926233 | 8.386562226 | -1.877788166 | 0.110577975 | 0.617725015 | -4.16069498 |
| **A_55_P1980326** | Pcbp1 | -0.248417709 | -0.573934433 | 0.077099015 | 14.85478758 | -1.877493359 | 0.110622783 | 0.617725015 | -4.160910591 |
| **A_55_P2088490** | Acad10 | 0.426801027 | -0.132496212 | 0.986098266 | 12.20991045 | 1.877378946 | 0.110640178 | 0.617725015 | -4.16099427 |
| **A_52_P622391** | Usp9x | -0.113068008 | -0.261351402 | 0.035215385 | 9.556231265 | -1.875928967 | 0.110860862 | 0.617822199 | -4.162054791 |
| **A_55_P2061381** | Ghitm | 0.170048523 | -0.05296906 | 0.393066107 | 13.04273562 | 1.875870801 | 0.110869723 | 0.617822199 | -4.162097336 |
| **A_55_P2158522** | Pbx1 | 0.298408751 | -0.093000513 | 0.689818014 | 14.31765782 | 1.875640567 | 0.110904808 | 0.617822199 | -4.162265738 |
| **A_55_P2127692** | Timm50 | 0.726474406 | -0.226413236 | 1.679362049 | 9.721867361 | 1.875632662 | 0.110906013 | 0.617822199 | -4.16227152 |
| **A_52_P468041** | Alkbh5 | -0.166024034 | -0.383796112 | 0.051748044 | 10.25522672 | -1.875590094 | 0.110912501 | 0.617822199 | -4.162302656 |
| **A_55_P2111950** | Dnajc7 | -0.127671882 | -0.295139907 | 0.039796144 | 11.74414189 | -1.875566741 | 0.11091606 | 0.617822199 | -4.162319737 |
| **A_52_P565940** | Nsd1 | -0.146674205 | -0.339079158 | 0.045730748 | 9.970840532 | -1.875455042 | 0.110933087 | 0.617822199 | -4.162401439 |
| **A_66_P112855** | AI413582 | 0.178533445 | -0.055683939 | 0.412750829 | 12.97036221 | 1.875295112 | 0.110957471 | 0.617822199 | -4.162518421 |
| **A_55_P1997097** | Bod1 | -0.09212743 | -0.212990071 | 0.028735212 | 9.004133477 | -1.875279633 | 0.110959831 | 0.617822199 | -4.162529743 |
| **A_52_P639567** | Ate1 | -0.128636312 | -0.297448377 | 0.040175753 | 10.90452822 | -1.874689131 | 0.11104991 | 0.617822199 | -4.162961677 |
| **A_51_P122425** | Ctf1 | -0.126922267 | -0.293492255 | 0.039647721 | 9.022567487 | -1.874606976 | 0.111062449 | 0.617822199 | -4.165415932 |
| **A_55_P2071477** | Prl6a1 | 0.239107986 | -0.074707177 | 0.552923148 | 9.360276972 | 1.874516728 | 0.111076224 | 0.617822199 | -4.163087787 |
| **A_55_P2005552** | Arhgef10l | 0.256387651 | -0.080139115 | 0.592914417 | 11.19166877 | 1.874332579 | 0.111104337 | 0.617822199 | -4.163222491 |
| **A_55_P1957344** | Hnrnpa3 | 0.20039909 | -0.062673492 | 0.463471672 | 12.19865806 | 1.874085153 | 0.111142121 | 0.617822199 | -4.163403482 |
| **A_51_P323610** | Selenos | -0.203895303 | -0.471636108 | 0.063845503 | 12.75714825 | -1.87353503 | 0.111226176 | 0.617822199 | -4.163805906 |
| **A_55_P2035023** | Zfp653 | 0.151215893 | -0.047352913 | 0.349784698 | 9.368217484 | 1.873508104 | 0.111230292 | 0.617822199 | -4.163825602 |
| **A_51_P393078** | Tigd2 | -0.115847404 | -0.267973554 | 0.036278746 | 9.268583055 | -1.873490937 | 0.111232916 | 0.617822199 | -4.163838161 |
| **A_66_P140976** | Rpl15 | 0.141094065 | -0.044195503 | 0.326383632 | 9.887408771 | 1.873384578 | 0.111249175 | 0.617822199 | -4.163915965 |
| **A_51_P336385** | Dido1 | 0.10996898 | -0.034467961 | 0.254405921 | 10.50912793 | 1.873100491 | 0.111292615 | 0.617822199 | -4.164123785 |
| **A_51_P462102** | Adgra3 | 0.272497108 | -0.085415015 | 0.630409231 | 11.97228033 | 1.873072774 | 0.111296854 | 0.617822199 | -4.164144061 |
| **A_55_P1970841** | Nr2c2 | -0.106694581 | -0.246857242 | 0.033468081 | 9.136103527 | -1.872747341 | 0.11134664 | 0.617921102 | -4.164382131 |
| **A_55_P2394308** | Fst | 0.160039833 | -0.050306415 | 0.370386081 | 10.70492234 | 1.8718131 | 0.111489685 | 0.618515343 | -4.165065597 |
| **A_52_P1128182** | Tmem185a | -0.097663702 | -0.226058112 | 0.030730709 | 9.383815213 | -1.871355197 | 0.111559863 | 0.618515343 | -4.165400599 |
| **A_52_P329054** | Cep70 | -0.244588957 | -0.566146587 | 0.076968674 | 9.923798454 | -1.871316245 | 0.111565835 | 0.618515343 | -4.165429096 |
| **A_52_P526396** | Atl1 | -0.101503279 | -0.234962156 | 0.031955599 | 8.707098187 | -1.871120543 | 0.111595843 | 0.618515343 | -4.165572274 |
| **A_55_P2001438** | Coq2 | 0.218439105 | -0.068798099 | 0.50567631 | 11.1288942 | 1.870935698 | 0.111624195 | 0.618515343 | -4.16570751 |
| **A_55_P1957918** | Asap2 | -0.198921106 | -0.460512668 | 0.062670456 | 9.805505993 | -1.870795416 | 0.111645715 | 0.618515343 | -4.165810144 |
| **A_51_P318640** | Rps3 | 0.107314759 | -0.033878514 | 0.248508032 | 15.39649227 | 1.86988377 | 0.111785673 | 0.618994993 | -4.166477146 |
| **A_55_P1997866** | Mtr | 0.172669571 | -0.054531983 | 0.399871124 | 10.11357745 | 1.869707526 | 0.111812751 | 0.618994993 | -4.166606098 |
| **A_51_P133097** | Gm4013 | -0.132364873 | -0.306542288 | 0.041812542 | 8.439052409 | -1.869605898 | 0.111828368 | 0.618994993 | -4.166680456 |
| **A_52_P301495** | Vamp2 | -0.161147538 | -0.373292817 | 0.050997742 | 9.81691148 | -1.86878554 | 0.111954508 | 0.619335058 | -4.167280702 |
| **A_55_P1986551** | Mcam | -0.080331404 | -0.186088451 | 0.025425642 | 7.81281042 | -1.8687242 | 0.111963946 | 0.619335058 | -4.167325585 |
| **A_55_P1983006** | Hdac5 | 0.320811871 | -0.101627675 | 0.743251417 | 12.78076588 | 1.868336191 | 0.112023662 | 0.619335058 | -4.167609496 |
| **A_51_P248705** | Cse1l | -0.11234547 | -0.260285998 | 0.035595058 | 9.562860003 | -1.868261067 | 0.112035227 | 0.619335058 | -4.167664466 |
| **A_55_P2046661** | Tnfrsf14 | -0.087758697 | -0.203328424 | 0.02781103 | 7.987789346 | -1.868165039 | 0.112050013 | 0.619335058 | -4.167734732 |
| **A_52_P539161** | Rdh11 | 0.239710817 | -0.076000731 | 0.555422364 | 10.51314657 | 1.867954646 | 0.112082414 | 0.619337043 | -4.167888683 |

| | | | | | | | | | |
|---|---|---|---|---|---|---|---|---|---|
| **A_51_P495171** | Pbdc1 | -0.104870144 | -0.243005824 | 0.033265537 | 10.58516992 | -1.867734561 | 0.112116318 | 0.619347329 | -4.168049727 |
| **A_52_P556290** | Arih1 | -0.109599764 | -0.253999236 | 0.034799708 | 9.652052505 | -1.86729604 | 0.112183902 | 0.619543609 | -4.168370615 |
| **A_51_P438666** | Keap1 | -0.249940498 | -0.579311713 | 0.079430717 | 10.76911293 | -1.866896117 | 0.112245572 | 0.619701244 | -4.168663264 |
| **A_55_P2067016** | Phykpl | -0.167751545 | -0.388838073 | 0.053334983 | 8.715924157 | -1.866695207 | 0.112276567 | 0.619701244 | -4.168810286 |
| **A_55_P1952696** | NA | 0.175424752 | -0.055905133 | 0.406754637 | 12.01676027 | 1.86564192 | 0.112439197 | 0.620421706 | -4.16958108 |
| **A_55_P2151463** | NA | 0.236590036 | -0.075443429 | 0.5486235 | 9.99515813 | 1.865367643 | 0.112481585 | 0.620478467 | -4.169781802 |
| **A_55_P1962707** | NA | -0.191354159 | -0.443774395 | 0.061066076 | 10.89092344 | -1.86501746 | 0.112535726 | 0.620600014 | -4.170038078 |
| **A_51_P201785** | Morn5 | 0.101627776 | -0.032448762 | 0.235704314 | 8.075453336 | 1.864785123 | 0.112571662 | 0.620621122 | -4.170208113 |
| **A_51_P494929** | Diablo | 0.096582988 | -0.030895917 | 0.224061892 | 9.98857287 | 1.863938055 | 0.112702776 | 0.620697651 | -4.17082805 |
| **A_55_P2026291** | Smarce1 | -0.17027417 | -0.395022907 | 0.054474567 | 10.94111571 | -1.863891706 | 0.112709954 | 0.620697651 | -4.170861972 |
| **A_55_P2279957** | 5730414N17Rik | 0.110537102 | -0.035384491 | 0.256458695 | 8.170906239 | 1.863621335 | 0.112751838 | 0.620697651 | -4.171059853 |
| **A_51_P455241** | Rrp36 | -0.110868805 | -0.259618108 | 0.037880499 | 10.07586134 | -1.931543561 | 0.112818165 | 0.620697651 | -4.171605739 |
| **A_52_P119350** | 4732419C18Rik | -0.194454999 | -0.45125605 | 0.062346053 | 8.972367037 | -1.862908298 | 0.112862372 | 0.620697651 | -4.171581726 |
| **A_55_P2126950** | Zfp467 | 0.098825273 | -0.031687224 | 0.22933777 | 8.404402814 | 1.862880884 | 0.112866624 | 0.620697651 | -4.171601791 |
| **A_55_P1957159** | NA | -0.148907784 | -0.345577499 | 0.047761931 | 15.03092469 | -1.862726419 | 0.112890584 | 0.620697651 | -4.171714846 |
| **A_52_P222350** | Flnb | -0.185805643 | -0.431289437 | 0.05967815 | 13.03664027 | -1.862109594 | 0.112986314 | 0.620697651 | -4.17216632 |
| **A_52_P676063** | Tmem126b | -0.127184276 | -0.295219678 | 0.040851125 | 8.588883388 | -1.862094849 | 0.112988604 | 0.620697651 | -4.172177113 |
| **A_51_P144868** | Mboat7 | -0.105103253 | -0.243972846 | 0.033766341 | 11.12675781 | -1.861993481 | 0.113004344 | 0.620697651 | -4.172251309 |
| **A_55_P2176908** | Tdpoz8 | 0.134845901 | -0.043324064 | 0.313015866 | 10.71348204 | 1.861968879 | 0.113008164 | 0.620697651 | -4.172269316 |
| **A_55_P2132099** | NA | 0.17890946 | -0.057516197 | 0.415335116 | 9.168452007 | 1.861692135 | 0.113051149 | 0.620697651 | -4.172471879 |
| **A_55_P2085156** | NA | -0.166869977 | -0.387389336 | 0.053649382 | 12.57024541 | -1.861661198 | 0.113055955 | 0.620697651 | -4.172494524 |
| **A_55_P2070277** | Mfn1 | -0.13685287 | -0.317738669 | 0.04403293 | 11.06066299 | -1.86130952 | 0.113110605 | 0.620697651 | -4.17275194 |
| **A_55_P2304507** | Noxa1 | 0.102550512 | -0.033022035 | 0.238123059 | 7.896537422 | 1.860952303 | 0.113166142 | 0.620697651 | -4.173013415 |
| **A_55_P2118740** | NA | 0.389172351 | -0.125344868 | 0.90368957 | 13.25338938 | 1.860849403 | 0.113182145 | 0.620697651 | -4.173088736 |
| **A_55_P1968381** | Cast | -0.244358506 | -0.567444787 | 0.078727775 | 10.59929355 | -1.860707499 | 0.113204218 | 0.620697651 | -4.173192608 |
| **A_55_P2047753** | Tec | -0.115655803 | -0.26858468 | 0.037273074 | 8.616116251 | -1.860574612 | 0.113224892 | 0.620697651 | -4.17328988 |
| **A_55_P2174022** | Brd4 | -0.230413369 | -0.535114026 | 0.074287288 | 13.1079968 | -1.860387643 | 0.113253987 | 0.620697651 | -4.173426741 |
| **A_55_P2159865** | Zfp59 | 0.102589047 | -0.033082707 | 0.238260802 | 8.46110226 | 1.860290288 | 0.113269139 | 0.620697651 | -4.173498005 |
| **A_55_P2069515** | Dennd4a | -0.249014725 | -0.578340802 | 0.080311352 | 10.38971524 | -1.860236113 | 0.113277572 | 0.620697651 | -4.173537662 |
| **A_52_P629451** | Gon4l | -0.221638967 | -0.514774772 | 0.071496837 | 8.927157925 | -1.860143348 | 0.113292013 | 0.620697651 | -4.173605567 |
| **A_55_P1972748** | Czib | 0.55153132 | -0.178019514 | 1.281082154 | 13.99705691 | 1.859875121 | 0.113333779 | 0.620750527 | -4.173801913 |
| **A_55_P2085932** | Lamtor2 | -0.178019045 | -0.41357001 | 0.057531919 | 12.06820444 | -1.859305455 | 0.113422533 | 0.621060662 | -4.174218925 |
| **A_55_P1993268** | Ap1s1 | -0.173065867 | -0.402099426 | 0.055967692 | 13.97081259 | -1.859008877 | 0.113468767 | 0.621127791 | -4.174436034 |
| **A_51_P251736** | Reep5 | -0.145320124 | -0.337655355 | 0.047015107 | 9.241810485 | -1.858814617 | 0.113499061 | 0.621127791 | -4.174578242 |
| **A_55_P1961127** | H19 | -0.213475794 | -0.496055157 | 0.069103569 | 17.45401016 | -1.858563305 | 0.113538264 | 0.621166462 | -4.174762217 |
| **A_55_P1984416** | NA | 0.414159102 | -0.134127537 | 0.96244574 | 11.03121818 | 1.85835513 | 0.113570748 | 0.621168363 | -4.174914616 |
| **A_51_P300207** | Rsl24d1 | -0.111357386 | -0.258820566 | 0.036105794 | 9.126613317 | -1.857824115 | 0.113653651 | 0.621445946 | -4.175303361 |
| **A_52_P258338** | Mob3b | 0.174668677 | -0.056707481 | 0.406044835 | 8.541314999 | 1.857229563 | 0.113746545 | 0.621777987 | -4.175738633 |
| **A_55_P2066958** | Sh3glb1 | -0.257612846 | -0.604527902 | 0.08930221 | 11.76071804 | -1.924396019 | 0.113843866 | 0.622094067 | -4.175872065 |
| **A_51_P272563** | Naa25 | -0.116646264 | -0.271227635 | 0.037935107 | 9.865486809 | -1.856448228 | 0.113868737 | 0.622094067 | -4.176310666 |
| **A_52_P352563** | 1700073E17Rik | -0.260918179 | -0.606733152 | 0.084896795 | 14.05136687 | -1.85622111 | 0.11390428 | 0.62209928 | -4.176476948 |
| **A_51_P141104** | Med18 | 0.11806172 | -0.038431034 | 0.274554473 | 8.302231206 | 1.856025912 | 0.113934836 | 0.62209928 | -4.176619861 |
| **A_55_P2097729** | Gtpbp4 | -0.144867259 | -0.336911952 | 0.047177433 | 9.05103636 | -1.855825326 | 0.113966245 | 0.62209928 | -4.17676672 |
| **A_55_P2141709** | Gabarapl2 | -0.134062231 | -0.311891267 | 0.043766804 | 13.34704961 | -1.854696844 | 0.114143108 | 0.622630776 | -4.177592966 |
| **A_55_P2037930** | NA | 0.192919333 | -0.06298511 | 0.448823776 | 8.400403986 | 1.854671886 | 0.114147023 | 0.622630776 | -4.17761124 |
| **A_55_P1971458** | Jak2 | 0.163287754 | -0.053362694 | 0.379938202 | 10.84241417 | 1.854228225 | 0.114216634 | 0.622630776 | -4.177936089 |
| **A_55_P2074591** | Ppm1h | 0.122089687 | -0.03990133 | 0.284080704 | 10.21966348 | 1.854202706 | 0.114220639 | 0.622630776 | -4.177954774 |
| **A_51_P446012** | Phf13 | -0.243579923 | -0.566771076 | 0.07961123 | 13.83060263 | -1.854176996 | 0.114224674 | 0.622630776 | -4.177973599 |
| **A_51_P156438** | Slc25a33 | 0.133415943 | -0.043665702 | 0.310497589 | 10.68890745 | 1.8535459 | 0.114323774 | 0.622751683 | -4.178435701 |
| **A_55_P2121856** | Ier5l | 0.167308102 | -0.054761535 | 0.389377739 | 9.613212018 | 1.853518616 | 0.11432806 | 0.622751683 | -4.178455679 |

| | | | | | | | | | |
|---|---|---|---|---|---|---|---|---|---|
| **A_51_P378550** | Oplah | -0.175468975 | -0.408383807 | 0.057445858 | 8.857129353 | -1.853413643 | 0.114344553 | 0.622751683 | -4.178532544 |
| **A_55_P2064074** | Vps37d | 0.104414244 | -0.034198464 | 0.243026952 | 8.307386012 | 1.853215246 | 0.11437573 | 0.622751683 | -4.178677817 |
| **A_55_P2103626** | Slc27a4 | 0.147648547 | -0.048414568 | 0.343711661 | 8.417874094 | 1.85268866 | 0.114458521 | 0.622938223 | -4.179063409 |
| **A_55_P2093689** | Pold3 | -0.090144595 | -0.209854517 | 0.029565327 | 11.09369396 | -1.852587413 | 0.114474446 | 0.622938223 | -4.179137549 |
| **A_55_P1982578** | Kdm3a | -0.095918569 | -0.223377965 | 0.031540827 | 10.688996 | -1.851398892 | 0.114661554 | 0.62356175 | -4.180007878 |
| **A_55_P2025174** | Rps27a | 0.23136011 | -0.076094526 | 0.538814746 | 16.19160611 | 1.851299142 | 0.114677271 | 0.62356175 | -4.180080926 |
| **A_51_P473576** | Apex1 | -0.085997203 | -0.200282299 | 0.028287894 | 9.574100046 | -1.851244958 | 0.11468581 | 0.62356175 | -4.180120605 |
| **A_51_P393426** | Pros1 | -0.454589618 | -1.058838906 | 0.14965967 | 11.01519005 | -1.850855563 | 0.114747191 | 0.623680108 | -4.180405763 |
| **A_66_P109315** | Snapc2 | 0.13307383 | -0.043825576 | 0.309973236 | 11.35138913 | 1.85069753 | 0.114772112 | 0.623680108 | -4.180521493 |
| **A_55_P2043287** | Tle5 | 0.198704415 | -0.065538523 | 0.462947352 | 8.773716659 | 1.850006656 | 0.11488112 | 0.624033308 | -4.181027442 |
| **A_55_P2001188** | Phf6 | -0.117370714 | -0.273474275 | 0.038732848 | 8.984017398 | -1.84976303 | 0.114919584 | 0.624033308 | -4.181205861 |
| **A_55_P2044953** | Lst1 | 0.410156362 | -0.135380245 | 0.955692969 | 11.70454211 | 1.849671978 | 0.114933963 | 0.624033308 | -4.181272543 |
| **A_55_P1975535** | Nutf2 | -0.168120402 | -0.391757718 | 0.055516913 | 12.3414395 | -1.84946158 | 0.114967196 | 0.624038454 | -4.181426629 |
| **A_55_P1993019** | NA | 0.170901425 | -0.056468223 | 0.398271073 | 10.51257694 | 1.849193497 | 0.115009554 | 0.624093115 | -4.181622963 |
| **A_55_P2092156** | NA | -0.490909789 | -1.144099285 | 0.162279707 | 13.99343921 | -1.848977657 | 0.115043669 | 0.624103027 | -4.181781039 |
| **A_55_P2110215** | Ckap5 | 0.143172322 | -0.047354201 | 0.333698844 | 11.41025366 | 1.848727025 | 0.115083296 | 0.624142825 | -4.181964596 |
| **A_55_P2014978** | Dvl3 | 0.23410434 | -0.077527312 | 0.545735993 | 13.39261419 | 1.84814936 | 0.115174681 | 0.624356729 | -4.182387673 |
| **A_51_P437938** | Coa5 | -0.12441626 | -0.290052988 | 0.041220468 | 10.98029604 | -1.847947638 | 0.115206609 | 0.624356729 | -4.182535415 |
| **A_55_P2065449** | Kdm5b | -0.283930955 | -0.661955814 | 0.094093904 | 13.00598783 | -1.84782801 | 0.115225548 | 0.624356729 | -4.182623031 |
| **A_52_P587831** | Ddx56 | 0.101303238 | -0.033602105 | 0.236208582 | 9.866718611 | 1.847410207 | 0.115291717 | 0.624356729 | -4.182929037 |
| **A_66_P123186** | Mapk1ip1l | -0.136906703 | -0.319242558 | 0.045429152 | 10.17698494 | -1.8472335 | 0.115319714 | 0.624356729 | -4.183058462 |
| **A_66_P124858** | Diras2 | 0.210813466 | -0.069958372 | 0.491585304 | 11.20467316 | 1.847200197 | 0.115324992 | 0.624356729 | -4.183082854 |
| **A_55_P1994204** | Ythdc1 | -0.155604046 | -0.362866395 | 0.051658302 | 11.74079476 | -1.847011648 | 0.115354874 | 0.624356729 | -4.183220953 |
| **A_55_P1958103** | NA | 0.31478369 | -0.104612009 | 0.73417939 | 14.64596169 | 1.846534442 | 0.115430538 | 0.624356729 | -4.183570479 |
| **A_55_P2038972** | Erich1 | 0.142207343 | -0.047278102 | 0.331692789 | 8.778199227 | 1.846355523 | 0.11545892 | 0.624356729 | -4.183701528 |
| **A_55_P2125456** | NA | 0.326569823 | -0.108612645 | 0.761752291 | 14.00419353 | 1.846179068 | 0.115486917 | 0.624356729 | -4.183830774 |
| **A_55_P2125823** | NA | -0.289702528 | -0.675772197 | 0.096367141 | 12.01989429 | -1.846102305 | 0.115499099 | 0.624356729 | -4.183887 |
| **A_51_P148069** | Lpgat1 | -0.215846345 | -0.503526555 | 0.071833865 | 10.93283408 | -1.845881743 | 0.115534108 | 0.624356729 | -4.184048554 |
| **A_55_P2141654** | NA | -0.130177812 | -0.303695962 | 0.043340338 | 15.48953114 | -1.84570047 | 0.115562889 | 0.624356729 | -4.184181332 |
| **A_51_P275697** | Rflnb | -0.236720463 | -0.552265697 | 0.078824772 | 9.911538535 | -1.845624462 | 0.115574959 | 0.624356729 | -4.184237005 |
| **A_55_P2104790** | NA | -0.237774935 | -0.554804253 | 0.079254382 | 11.66074379 | -1.845167556 | 0.115647541 | 0.624523309 | -4.184571682 |
| **A_55_P1985825** | NA | 0.246107639 | -0.082072406 | 0.574287684 | 8.360264781 | 1.844939323 | 0.115683814 | 0.624523309 | -4.184738861 |
| **A_55_P2370394** | Ppp2r5e | -0.13015399 | -0.303723153 | 0.043415172 | 8.706356578 | -1.844820363 | 0.115702725 | 0.624523309 | -4.184825999 |
| **A_55_P1956267** | Syt11 | -0.332057624 | -0.774949701 | 0.110834452 | 9.429018232 | -1.844525693 | 0.115749581 | 0.624601802 | -4.185041846 |
| **A_55_P1965416** | Atp9a | -0.217091722 | -0.506710632 | 0.072527188 | 15.04608969 | -1.844104435 | 0.115816599 | 0.624789017 | -4.185350424 |
| **A_55_P2001510** | NA | -0.144522593 | -0.337451479 | 0.048406294 | 16.73880198 | -1.842924947 | 0.11600445 | 0.624927037 | -4.186214445 |
| **A_52_P108089** | Mosmo | -0.23059877 | -0.538474025 | 0.077276484 | 10.48275116 | -1.842686127 | 0.116042523 | 0.624927037 | -4.186389395 |
| **A_52_P334670** | Clock | -0.114117592 | -0.266483791 | 0.038248607 | 9.180102902 | -1.842608742 | 0.116054862 | 0.624927037 | -4.186446085 |
| **A_52_P223571** | Mlec | -0.223710528 | -0.522416221 | 0.074995164 | 9.291407707 | -1.842519476 | 0.116069097 | 0.624927037 | -4.186511478 |
| **A_55_P2151601** | Samd9l | -0.138791787 | -0.324112353 | 0.046528779 | 8.993329688 | -1.842507709 | 0.116070974 | 0.624927037 | -4.186520098 |
| **A_66_P132232** | Ercc1 | -0.114202092 | -0.266703956 | 0.038299772 | 9.517903859 | -1.842332746 | 0.116098881 | 0.624927037 | -4.186648271 |
| **A_55_P2041843** | Nde1 | -0.081195839 | -0.189623088 | 0.027231411 | 8.094738881 | -1.842317573 | 0.116101302 | 0.624927037 | -4.186659386 |
| **A_55_P2156776** | NA | 0.186386954 | -0.062510369 | 0.435284277 | 8.327466081 | 1.842317377 | 0.116101333 | 0.624927037 | -4.18665953 |
| **A_52_P476400** | NA | -0.197315387 | -0.460834854 | 0.066204079 | 9.918753342 | -1.842117871 | 0.116133164 | 0.624927037 | -4.186805683 |
| **A_52_P527842** | Reps1 | -0.106784892 | -0.249477759 | 0.035907975 | 9.170503828 | -1.841097172 | 0.116296149 | 0.625629909 | -4.187553441 |
| **A_55_P2052240** | Crlf1 | 0.335474726 | -0.112875052 | 0.783824504 | 10.38590889 | 1.840822913 | 0.116339981 | 0.625666463 | -4.187754367 |
| **A_55_P2081378** | NA | -0.139349953 | -0.325603632 | 0.046903726 | 15.68826624 | -1.840649641 | 0.116367683 | 0.625666463 | -4.187881309 |
| **A_55_P1987912** | Chpf2 | -0.245953067 | -0.574773514 | 0.082867379 | 9.960626284 | -1.840189684 | 0.116441248 | 0.625707901 | -4.188218287 |
| **A_55_P1961140** | Mettl3 | -0.193139344 | -0.451365493 | 0.065086806 | 10.48866066 | -1.840092671 | 0.11645677 | 0.625707901 | -4.188289362 |
| **A_51_P115005** | Edn1 | -0.152111059 | -0.355507877 | 0.051285759 | 8.650324464 | -1.839864323 | 0.116493314 | 0.625707901 | -4.190435098 |

| | | | | | | | | | |
|---|---|---|---|---|---|---|---|---|---|
| **A_66_P122758** | Tsr1 | -0.15370081 | -0.359241746 | 0.051840126 | 9.606277998 | -1.839699917 | 0.116519632 | 0.625707901 | -4.188577111 |
| **A_55_P2122491** | Gm6222 | -0.136002409 | -0.317899961 | 0.045895142 | 15.99810957 | -1.839453919 | 0.116559022 | 0.625707901 | -4.188757342 |
| **A_55_P2172334** | Nup155 | 0.107919917 | -0.036423664 | 0.252263499 | 9.191710645 | 1.839387757 | 0.116569618 | 0.625707901 | -4.188805816 |
| **A_66_P110909** | Ppp1r12a | -0.210474548 | -0.492090439 | 0.071141343 | 12.2046771 | -1.838703026 | 0.11667934 | 0.62612045 | -4.189307496 |
| **A_55_P2064230** | Tmx3 | -0.113857738 | -0.266216401 | 0.038500925 | 8.61082105 | -1.83850393 | 0.116711262 | 0.62612045 | -4.189453369 |
| **A_51_P256304** | Mrpl37 | -0.086285111 | -0.201781537 | 0.029211314 | 8.131647922 | -1.837961809 | 0.116798228 | 0.626273927 | -4.189850575 |
| **A_55_P2039694** | NA | 0.191822972 | -0.064970285 | 0.448616228 | 16.26967339 | 1.837748866 | 0.116832405 | 0.626273927 | -4.190006598 |
| **A_55_P1995074** | Ubald2 | -0.276168974 | -0.645882004 | 0.093544056 | 14.45373953 | -1.83771979 | 0.116837073 | 0.626273927 | -4.190027902 |
| **A_55_P1989328** | Rps12 | -0.15965791 | -0.373460556 | 0.054144736 | 16.69067925 | -1.837158081 | 0.116927279 | 0.62658369 | -4.190439473 |
| **A_51_P336391** | Tmem18 | 0.146731929 | -0.04979526 | 0.343259117 | 11.08942591 | 1.836839232 | 0.116978514 | 0.626684506 | -4.190673101 |
| **A_51_P377760** | Rnasel | -0.117858429 | -0.275783658 | 0.0400668 | 8.254769138 | -1.836023751 | 0.117109654 | 0.627213217 | -4.191270635 |
| **A_55_P2048660** | NA | 0.164317109 | -0.05592673 | 0.384560947 | 11.46527048 | 1.835473757 | 0.117198182 | 0.627266047 | -4.191673647 |
| **A_55_P1959838** | NA | 0.171085586 | -0.058239858 | 0.40041103 | 14.62351733 | 1.835398367 | 0.117210323 | 0.627266047 | -4.19172889 |
| **A_55_P2047325** | Mbtps1 | 0.214279534 | -0.077777453 | 0.506336522 | 13.01620043 | 1.901354841 | 0.117216874 | 0.627266047 | -4.189661137 |
| **A_55_P1967533** | Gm5069 | -0.150935352 | -0.353306108 | 0.051435404 | 11.36802485 | -1.834899921 | 0.117290619 | 0.627486959 | -4.192094138 |
| **A_52_P134381** | Psmd12 | -0.150469869 | -0.352260502 | 0.051320763 | 9.856655831 | -1.834499952 | 0.117355092 | 0.627658154 | -4.192387229 |
| **A_51_P202033** | Wls | -0.289482214 | -0.677770529 | 0.098806101 | 9.925311121 | -1.834157928 | 0.117410252 | 0.627769476 | -4.192637863 |
| **A_55_P2119420** | Ndufb2 | 0.26615282 | -0.090880243 | 0.623185884 | 13.46168847 | 1.833968194 | 0.117440862 | 0.627769476 | -4.192776901 |
| **A_51_P492087** | Pole3 | -0.12943251 | -0.303154725 | 0.044289705 | 9.937603655 | -1.83297769 | 0.117600792 | 0.628109777 | -4.19350276 |
| **A_66_P120901** | Atxn10 | -0.235516453 | -0.551674518 | 0.080641612 | 14.17559093 | -1.832677935 | 0.117649235 | 0.628109777 | -4.19372243 |
| **A_55_P2140423** | Arfip2 | -0.227265109 | -0.532362726 | 0.077832509 | 9.537374857 | -1.832580714 | 0.11766495 | 0.628109777 | -4.193793677 |
| **A_51_P490296** | Slx4 | 0.140921026 | -0.048267658 | 0.33010971 | 10.35991148 | 1.832524562 | 0.117674028 | 0.628109777 | -4.193834828 |
| **A_51_P419117** | Rab15 | -0.233697488 | -0.547536948 | 0.080141972 | 15.30369693 | -1.831958621 | 0.117765561 | 0.628109777 | -4.194249577 |
| **A_55_P1957633** | Gpsm3 | 0.258735568 | -0.088827167 | 0.606298302 | 11.56115661 | 1.831437453 | 0.117849914 | 0.628109777 | -4.194631522 |
| **A_66_P134294** | Uba1y | 0.19509739 | -0.066989141 | 0.45718392 | 8.622942072 | 1.831369262 | 0.117860956 | 0.628109777 | -4.194681497 |
| **A_55_P2084686** | Pdik1l | 0.112677722 | -0.038701024 | 0.264056467 | 8.797156162 | 1.831227571 | 0.117883901 | 0.628109777 | -4.194785339 |
| **A_55_P2004511** | Cd300lf | -0.123730456 | -0.290010153 | 0.042549241 | 7.954684246 | -1.83065519 | 0.11797664 | 0.628109777 | -4.195204828 |
| **A_55_P2064478** | Cdc27 | 0.189032785 | -0.065018026 | 0.443083596 | 9.889375972 | 1.830567338 | 0.11799088 | 0.628109777 | -4.195269213 |
| **A_51_P114693** | Parm1 | -0.261798817 | -0.613651325 | 0.090053691 | 14.50752568 | -1.830527222 | 0.117997383 | 0.628109777 | -4.202892842 |
| **A_66_P124146** | Ttc16 | 0.164525153 | -0.056607221 | 0.385657526 | 8.730428731 | 1.830413197 | 0.11801587 | 0.628109777 | -4.195382183 |
| **A_51_P486512** | Letmd1 | 0.152600526 | -0.052510074 | 0.357711126 | 11.05751067 | 1.830362414 | 0.118024104 | 0.628109777 | -4.195419401 |
| **A_66_P128384** | Hdac1 | 0.097388163 | -0.03354089 | 0.228317216 | 11.35765072 | 1.829950372 | 0.118090935 | 0.628109777 | -4.195721388 |
| **A_55_P2070054** | Senp3 | 0.179694811 | -0.061891723 | 0.421281345 | 12.12214626 | 1.829919462 | 0.11809595 | 0.628109777 | -4.195744042 |
| **A_65_P15340** | Zfp251 | -0.109575501 | -0.256908867 | 0.037757865 | 8.728701791 | -1.829706868 | 0.118130448 | 0.628109777 | -4.195899855 |
| **A_52_P453374** | Lrrn4cl | 0.191170423 | -0.065874628 | 0.448215474 | 8.236672024 | 1.829703077 | 0.118131064 | 0.628109777 | -4.195902633 |
| **A_52_P570652** | Flywch1 | -0.19569967 | -0.458843448 | 0.067444109 | 8.547380895 | -1.829642117 | 0.118140957 | 0.628109777 | -4.195947311 |
| **A_52_P126055** | Pon2 | -0.10036608 | -0.23532909 | 0.034596929 | 8.506148026 | -1.829537742 | 0.1181579 | 0.628109777 | -4.196023809 |
| **A_55_P2009013** | Dpf2 | -0.217245826 | -0.509408473 | 0.074916821 | 10.81280918 | -1.829346247 | 0.11818899 | 0.628109777 | -4.19616416 |
| **A_55_P1958133** | NA | 0.594655926 | -0.205179234 | 1.394491087 | 11.96391296 | 1.829087252 | 0.118231051 | 0.628109777 | -4.196353985 |
| **A_55_P2156785** | Ppfia3 | 0.452167105 | -0.156096347 | 1.060430557 | 10.46162862 | 1.828842919 | 0.118270745 | 0.628109777 | -4.196533064 |
| **A_66_P111089** | Ube2f | -0.117725306 | -0.276114358 | 0.040663747 | 9.223253797 | -1.828579423 | 0.118313568 | 0.628109777 | -4.196726191 |
| **A_55_P2004727** | NA | 0.202700795 | -0.070020883 | 0.475422472 | 8.263827273 | 1.82854206 | 0.118319641 | 0.628109777 | -4.196753575 |
| **A_66_P132249** | Akr1c13 | 0.237090483 | -0.081910936 | 0.556091901 | 8.70799949 | 1.828481916 | 0.118329418 | 0.628109777 | -4.196797657 |
| **A_55_P2164139** | NA | 0.372737356 | -0.12902331 | 0.874498021 | 12.63908013 | 1.827576101 | 0.118476764 | 0.628109777 | -4.197461579 |
| **A_51_P438692** | Vti1b | -0.219332447 | -0.514615927 | 0.075951032 | 13.52540517 | -1.827396957 | 0.118505926 | 0.628109777 | -4.197592886 |
| **A_55_P1960703** | NA | 0.122172756 | -0.042323218 | 0.286668729 | 9.550837412 | 1.827209149 | 0.118536507 | 0.628109777 | -4.197730545 |
| **A_55_P2016387** | Eif3k | 0.096443286 | -0.03341658 | 0.226303152 | 15.08023035 | 1.827116385 | 0.118551614 | 0.628109777 | -4.197798539 |
| **A_55_P1971903** | Pla2r1 | 0.179198686 | -0.062136303 | 0.420533674 | 8.472041537 | 1.826769254 | 0.118608164 | 0.628109777 | -4.198052979 |
| **A_51_P514647** | Paip2 | -0.104361312 | -0.244913401 | 0.036190776 | 12.79082118 | -1.826717507 | 0.118616597 | 0.628109777 | -4.198090909 |
| **A_51_P292196** | Prl7d1 | -0.176870694 | -0.41510137 | 0.061359982 | 17.01965361 | -1.826532315 | 0.118646779 | 0.628109777 | -4.198226653 |

| | | | | | | | | | |
|---|---|---|---|---|---|---|---|---|---|
| **A_66_P129873** | Npm1 | -0.133426378 | -0.313152807 | 0.046300051 | 14.12598496 | -1.826412681 | 0.118666281 | 0.628109777 | -4.198314343 |
| **A_66_P110791** | Ppp1r11 | 0.221037601 | -0.076750325 | 0.518825527 | 11.99894624 | 1.826115475 | 0.118714743 | 0.628109777 | -4.198532195 |
| **A_55_P2174203** | Lims2 | -0.16281481 | -0.382199753 | 0.056570133 | 13.24800246 | -1.825812817 | 0.118764114 | 0.628109777 | -4.198754045 |
| **A_55_P2067757** | Psg27 | -0.288905935 | -0.678201862 | 0.100389992 | 16.51877733 | -1.825768738 | 0.118771306 | 0.628109777 | -4.198786355 |
| **A_55_P2004806** | Hexim2 | 0.217996617 | -0.075777183 | 0.511770417 | 12.95806928 | 1.825600961 | 0.118798685 | 0.628109777 | -4.198909338 |
| **A_55_P2105181** | Bhmt | -0.109523455 | -0.257122365 | 0.038075456 | 7.810517586 | -1.825547533 | 0.118807405 | 0.628109777 | -4.198948502 |
| **A_55_P2161690** | Tomm40 | -0.09427969 | -0.221339282 | 0.032779902 | 8.303814391 | -1.825491653 | 0.118816526 | 0.628109777 | -4.198989463 |
| **A_51_P268529** | Csad | -0.242974994 | -0.570466086 | 0.084516098 | 9.765151166 | -1.82528738 | 0.118849874 | 0.628109777 | -4.199139198 |
| **A_55_P2013223** | S100z | 0.20904532 | -0.072728007 | 0.490818647 | 8.497550526 | 1.825196944 | 0.118864641 | 0.628109777 | -4.19920549 |
| **A_51_P490305** | Ifi30 | 0.372221438 | -0.129515505 | 0.873958381 | 11.0656422 | 1.825132783 | 0.118875119 | 0.628109777 | -4.199252521 |
| **A_52_P528963** | Eps15l1 | 0.111730711 | -0.03889052 | 0.262351941 | 10.26799598 | 1.824969176 | 0.11890184 | 0.628109777 | -4.19937245 |
| **A_55_P2023118** | Prss28 | 0.213919876 | -0.074493609 | 0.502333361 | 8.614080161 | 1.824755746 | 0.118936709 | 0.628122307 | -4.199528901 |
| **A_55_P1970631** | Vps54 | -0.177935006 | -0.417911562 | 0.062041551 | 10.20909858 | -1.824155017 | 0.119034904 | 0.628383927 | -4.19996926 |
| **A_55_P2032755** | Gm10586 | -0.124680943 | -0.292844282 | 0.043482396 | 15.88511516 | -1.824054963 | 0.119051267 | 0.628383927 | -4.200042604 |
| **A_55_P2414479** | NA | 0.153085257 | -0.053439858 | 0.359610372 | 8.360410228 | 1.823600315 | 0.119125648 | 0.628559756 | -4.200375887 |
| **A_55_P2047200** | NA | -0.523858051 | -1.230644161 | 0.182928059 | 8.692115134 | -1.823453866 | 0.119149617 | 0.628559756 | -4.200483243 |
| **A_55_P1978226** | Prkn | 0.405040621 | -0.141716656 | 0.951797897 | 9.270848417 | 1.822523644 | 0.119301976 | 0.629191785 | -4.201165165 |
| **A_55_P2026440** | Ceacam-ps1 | -0.547738427 | -1.28738692 | 0.191910065 | 8.817638417 | -1.821868375 | 0.119409417 | 0.62958664 | -4.201645538 |
| **A_51_P196629** | Fam102b | -0.331430522 | -0.779105086 | 0.116244042 | 12.81365998 | -1.821374467 | 0.119490464 | 0.629758697 | -4.202007625 |
| **A_55_P2076533** | Pmvk | 0.233373618 | -0.081869784 | 0.54861702 | 11.70279251 | 1.821272444 | 0.119507212 | 0.629758697 | -4.202082419 |
| **A_51_P371882** | Ccdc71 | -0.193940228 | -0.455994503 | 0.068114046 | 12.24704385 | -1.82073113 | 0.119596113 | 0.630055401 | -4.202479267 |
| **A_51_P233101** | Ltbp3 | -0.421392384 | -0.990857354 | 0.148072586 | 11.71391151 | -1.820492235 | 0.119635368 | 0.630084709 | -4.202654408 |
| **A_55_P2140171** | NA | 0.146965114 | -0.051662815 | 0.345593042 | 10.10806059 | 1.820300566 | 0.119666872 | 0.630084709 | -4.202794927 |
| **A_66_P126984** | Gm10523 | 0.085044037 | -0.029911035 | 0.199999109 | 8.442175023 | 1.820056599 | 0.119706984 | 0.630124264 | -4.202973789 |
| **A_52_P456077** | Commd2 | 0.271759598 | -0.095719588 | 0.639238784 | 11.17605586 | 1.81937117 | 0.119819751 | 0.630404455 | -4.203476309 |
| **A_55_P2112120** | Tnk2 | -0.108716451 | -0.255735579 | 0.038302678 | 8.373813361 | -1.819242462 | 0.119840938 | 0.630404455 | -4.203570673 |
| **A_52_P428446** | Tsn | -0.137969289 | -0.324578432 | 0.048639854 | 11.5204271 | -1.818941246 | 0.119890536 | 0.630404455 | -4.203791513 |
| **A_51_P398723** | Flt1 | -0.372060667 | -0.875299949 | 0.131178614 | 14.13106516 | -1.818898186 | 0.119897628 | 0.630404455 | -4.203823083 |
| **A_51_P514675** | Zcchc10 | 0.123151675 | -0.04344181 | 0.289745161 | 9.483486257 | 1.818659821 | 0.119936893 | 0.630404455 | -4.203997845 |
| **A_55_P2010020** | NA | 0.138985514 | -0.04903913 | 0.327010157 | 15.5876157 | 1.81854447 | 0.1199559 | 0.630404455 | -4.204082417 |
| **A_55_P2106569** | NA | 0.256396398 | -0.090536411 | 0.603329206 | 14.1021512 | 1.818175119 | 0.120016777 | 0.630483326 | -4.204353217 |
| **A_55_P2153904** | Gm6132 | -0.26196796 | -0.616471394 | 0.092535474 | 11.33710329 | -1.818012628 | 0.12004357 | 0.630483326 | -4.204472353 |
| **A_55_P1998957** | Oas1c | 0.205196543 | -0.072520469 | 0.482913555 | 9.766642969 | 1.817760736 | 0.120085114 | 0.630483326 | -4.204657037 |
| **A_55_P2024841** | Gstt1 | 0.535662801 | -0.189356091 | 1.260681693 | 9.056631021 | 1.817654466 | 0.120102645 | 0.630483326 | -4.204734953 |
| **A_55_P2008277** | Tcp10c | 0.124011428 | -0.043855425 | 0.291878281 | 8.369800659 | 1.817464441 | 0.120134 | 0.630483326 | -4.204874278 |
| **A_66_P105307** | Adnp | -0.231471249 | -0.544913796 | 0.081971299 | 11.75292107 | -1.816804804 | 0.120242905 | 0.630883582 | -4.205357926 |
| **A_55_P1978726** | NA | 0.412723939 | -0.146239423 | 0.9716873 | 11.07385561 | 1.816542155 | 0.120286295 | 0.630939974 | -4.205550504 |
| **A_55_P2175550** | Atf2 | -0.120249843 | -0.283165765 | 0.042666078 | 9.162051944 | -1.815892574 | 0.120393674 | 0.631200067 | -4.206026792 |
| **A_55_P2085142** | Spp1 | 0.267793378 | -0.095025366 | 0.630612122 | 8.967326935 | 1.815847116 | 0.120401192 | 0.631200067 | -4.206060123 |
| **A_55_P2081630** | Gpa33 | 0.141721214 | -0.050343089 | 0.333785517 | 9.548919999 | 1.815337466 | 0.120485512 | 0.63147084 | -4.206433817 |
| **A_55_P2002567** | NA | -0.144655082 | -0.340758771 | 0.051448608 | 15.41524862 | -1.814751204 | 0.120582578 | 0.63169705 | -4.206863691 |
| **A_55_P2164218** | NA | -0.140105073 | -0.330055278 | 0.049845133 | 13.17581633 | -1.814609813 | 0.120606 | 0.63169705 | -4.206967367 |
| **A_66_P109183** | Apold1 | -0.187232127 | -0.441173821 | 0.066709566 | 8.452341777 | -1.813909115 | 0.120722138 | 0.63169705 | -4.207481163 |
| **A_66_P119735** | BC049265 | 0.121957032 | -0.043458493 | 0.287372556 | 8.132780872 | 1.813843204 | 0.120733068 | 0.63169705 | -4.207529494 |
| **A_55_P2017066** | NA | 0.638383411 | -0.227509248 | 1.50427607 | 13.96475675 | 1.813788727 | 0.120742103 | 0.63169705 | -4.20756944 |
| **A_55_P2058201** | Scgb2b20 | 0.383844494 | -0.136802703 | 0.904491692 | 9.504147372 | 1.813764694 | 0.120746089 | 0.63169705 | -4.207587062 |
| **A_55_P2131433** | NA | 0.363476334 | -0.12956207 | 0.856514738 | 9.342627626 | 1.813696245 | 0.120757442 | 0.63169705 | -4.207637255 |
| **A_55_P1956989** | BC004004 | -0.165678165 | -0.390544826 | 0.059188495 | 13.43756048 | -1.812630861 | 0.120934289 | 0.632450991 | -4.208418485 |
| **A_51_P158018** | Bend5 | -0.341071208 | -0.804069155 | 0.121926739 | 12.30863052 | -1.812321143 | 0.120985748 | 0.632472045 | -4.2086456 |
| **A_55_P1998396** | Eif2ak4 | -0.16754192 | -0.394990622 | 0.059906782 | 10.21139413 | -1.812212781 | 0.121003757 | 0.632472045 | -4.208725063 |

| | | | | | | | | | |
|---|---|---|---|---|---|---|---|---|---|
| **A_55_P2112947** | Srp9 | -0.101854097 | -0.240183911 | 0.036475717 | 9.677157165 | -1.811473038 | 0.12112677 | 0.632762153 | -4.209267524 |
| **A_55_P2140941** | Tinagl1 | -0.167125549 | -0.394176507 | 0.059925409 | 9.956419204 | -1.810875829 | 0.121226171 | 0.632762153 | -4.209705471 |
| **A_55_P1981739** | Donson | -0.087932732 | -0.207396203 | 0.03153074 | 10.01215537 | -1.810858832 | 0.121229001 | 0.632762153 | -4.209717936 |
| **A_51_P423290** | Mmrn1 | -0.084244347 | -0.198699818 | 0.030211125 | 7.849866477 | -1.810812031 | 0.121236795 | 0.632762153 | -4.209752256 |
| **A_55_P2102715** | NA | -0.222966115 | -0.526016129 | 0.080083899 | 15.05879467 | -1.810063125 | 0.12136157 | 0.632762153 | -4.210301457 |
| **A_55_P2136572** | Cdc42se2 | -0.134904519 | -0.318282604 | 0.048473566 | 12.73840101 | -1.809873534 | 0.121393178 | 0.632762153 | -4.210440494 |
| **A_55_P2144781** | Erich5 | 0.635537351 | -0.22844695 | 1.499521651 | 10.96129428 | 1.809690854 | 0.121423641 | 0.632762153 | -4.210574462 |
| **A_55_P2140076** | NA | 0.163793455 | -0.058890281 | 0.38647719 | 16.43444628 | 1.809577584 | 0.121442533 | 0.632762153 | -4.210657528 |
| **A_52_P106739** | Ddx46 | -0.156835029 | -0.370088239 | 0.056418181 | 10.26023285 | -1.809325296 | 0.121484623 | 0.632762153 | -4.210842546 |
| **A_55_P2173837** | Mpp6 | -0.134871733 | -0.318274482 | 0.048531016 | 11.79441427 | -1.809190336 | 0.121507145 | 0.632762153 | -4.21094152 |
| **A_55_P2187043** | Tpm1 | -0.134551928 | -0.317535861 | 0.048432005 | 10.26871831 | -1.809031501 | 0.121533656 | 0.632762153 | -4.211058004 |
| **A_51_P453995** | Gm11961 | 0.156673124 | -0.056437812 | 0.369784061 | 10.15542588 | 1.808664144 | 0.121594994 | 0.632762153 | -4.211327411 |
| **A_55_P2129034** | Rheb | -0.140628542 | -0.331921526 | 0.050664441 | 11.43284396 | -1.808604459 | 0.121604962 | 0.632762153 | -4.211371183 |
| **A_51_P147714** | Sugp2 | -0.094822555 | -0.223811514 | 0.034166405 | 8.130597699 | -1.808540654 | 0.12161562 | 0.632762153 | -4.211417976 |
| **A_55_P1972892** | NA | 0.129945753 | -0.046833749 | 0.306725256 | 10.46192492 | 1.808420188 | 0.121635744 | 0.632762153 | -4.211506322 |
| **A_55_P2031989** | Rin2 | -0.293333788 | -0.699421418 | 0.112753841 | 10.56477119 | -1.871942107 | 0.121673819 | 0.632762153 | -4.20733977 |
| **A_55_P2102693** | Acvr1 | -0.150581111 | -0.355461028 | 0.054298805 | 9.223549805 | -1.808174218 | 0.121676845 | 0.632762153 | -4.211686711 |
| **A_55_P1971244** | Ndufa4l2 | 0.446475727 | -0.160998515 | 1.05394997 | 11.61076143 | 1.808169539 | 0.121677627 | 0.632762153 | -4.211690143 |
| **A_55_P2119927** | NA | 0.190919027 | -0.068848247 | 0.450686302 | 8.395016376 | 1.808147845 | 0.121681252 | 0.632762153 | -4.211706053 |
| **A_55_P1998001** | DXBay18 | -0.224924986 | -0.531003244 | 0.081153272 | 11.8781468 | -1.807899916 | 0.121722695 | 0.632807417 | -4.211887879 |
| **A_51_P260513** | Polr1d | -0.136003978 | -0.32112898 | 0.049121025 | 11.68346525 | -1.807405897 | 0.121805316 | 0.632987229 | -4.212250187 |
| **A_55_P1985984** | Actg1 | -0.291925139 | -0.68933678 | 0.105486502 | 12.72363434 | -1.807174436 | 0.121844045 | 0.632987229 | -4.21241994 |
| **A_52_P31641** | Tmem145 | -0.286659455 | -0.676927322 | 0.103608412 | 8.658851722 | -1.807060275 | 0.121863151 | 0.632987229 | -4.212503666 |
| **A_51_P406020** | Abhd4 | -0.204053479 | -0.481881855 | 0.073774897 | 14.11864587 | -1.806910186 | 0.121888275 | 0.632987229 | -4.212613741 |
| **A_52_P550147** | NA | -0.23509973 | -0.555288676 | 0.085089215 | 8.8452877 | -1.806404307 | 0.121972994 | 0.633157104 | -4.212984757 |
| **A_55_P2143204** | Pbxip1 | -0.297491798 | -0.702828743 | 0.107845148 | 10.3744292 | -1.805626704 | 0.122103331 | 0.633157104 | -4.213555065 |
| **A_52_P98614** | Slc6a2 | -0.306082735 | -0.723167 | 0.11100153 | 11.20572164 | -1.805444698 | 0.122133857 | 0.633157104 | -4.213688553 |
| **A_55_P2087766** | Dnaja3 | -0.169768499 | -0.401119948 | 0.06158295 | 11.46793693 | -1.805319399 | 0.122154877 | 0.633157104 | -4.213780451 |
| **A_51_P108108** | Lrrc25 | -0.103161487 | -0.243752331 | 0.037429357 | 7.989448159 | -1.805218258 | 0.122171847 | 0.633157104 | -4.213854631 |
| **A_51_P520718** | Ptger3 | -0.194461952 | -0.459482443 | 0.07055854 | 9.528651731 | -1.805195942 | 0.122175592 | 0.633157104 | -4.213870998 |
| **A_55_P2044104** | Fgfr1op | 0.15082841 | -0.054726868 | 0.356383688 | 10.64720331 | 1.805193172 | 0.122176056 | 0.633157104 | -4.21387303 |
| **A_55_P1957018** | Caskin2 | -0.132367549 | -0.312777899 | 0.048042802 | 10.26141531 | -1.805050014 | 0.122200081 | 0.633157104 | -4.213978027 |
| **A_55_P1975660** | Hemgn | -0.290991709 | -0.687629551 | 0.105646133 | 14.88696176 | -1.804910334 | 0.122223526 | 0.633157104 | -4.214080473 |
| **A_55_P2083419** | Pkp4 | -0.123234316 | -0.291223402 | 0.044754769 | 9.346465488 | -1.804761252 | 0.122248555 | 0.633157104 | -4.214189816 |
| **A_55_P2115507** | Egfl8 | 0.322222841 | -0.117141036 | 0.761586718 | 9.690308675 | 1.80426838 | 0.122331335 | 0.633278501 | -4.21455131 |
| **A_51_P468762** | Alkbh6 | 0.1153924 | -0.041961646 | 0.272746446 | 10.29822323 | 1.804132599 | 0.12235415 | 0.633278501 | -4.214650898 |
| **A_55_P2117319** | Ntan1 | 0.114979689 | -0.04181991 | 0.271779288 | 11.80576871 | 1.804036602 | 0.122370283 | 0.633278501 | -4.214721307 |
| **A_55_P2148534** | Nr1d2 | -0.122514407 | -0.28962694 | 0.044598126 | 9.55496611 | -1.803629394 | 0.12243874 | 0.633319774 | -4.215019977 |
| **A_55_P2064901** | NA | -0.084856246 | -0.200612488 | 0.030899996 | 15.34148231 | -1.803468491 | 0.1224658 | 0.633319774 | -4.215137993 |
| **A_55_P2039699** | Camp | -0.326110101 | -0.771038189 | 0.118817987 | 8.301941494 | -1.80319872 | 0.122511182 | 0.633319774 | -4.21533586 |
| **A_66_P102510** | Fam114a2 | -0.10609217 | -0.25083955 | 0.038655211 | 8.63689086 | -1.803191206 | 0.122512446 | 0.633319774 | -4.215341372 |
| **A_51_P149714** | Ms4a6d | -0.145474794 | -0.344003754 | 0.053054166 | 8.169623826 | -1.802739799 | 0.122588422 | 0.633319774 | -4.215672465 |
| **A_52_P193236** | Tedc2 | -0.209611617 | -0.495673479 | 0.076450245 | 9.876516772 | -1.802704532 | 0.12259436 | 0.633319774 | -4.215698333 |
| **A_51_P248444** | Poglut1 | -0.144411925 | -0.341560298 | 0.052736447 | 8.598040366 | -1.802100577 | 0.12269609 | 0.633319774 | -4.216141321 |
| **A_52_P120925** | Metap1 | 0.118233406 | -0.043190005 | 0.279656817 | 8.913909978 | 1.801950444 | 0.122721391 | 0.633319774 | -4.216251441 |
| **A_51_P210928** | Tsta3 | 0.151585322 | -0.055408975 | 0.358579618 | 9.910692298 | 1.801639664 | 0.122773781 | 0.633319774 | -4.216479396 |
| **A_51_P485899** | Srsf1 | -0.115004822 | -0.272056916 | 0.042047271 | 9.914096354 | -1.801529948 | 0.122792282 | 0.633319774 | -4.216559871 |
| **A_55_P1965076** | NA | 0.137460017 | -0.050270678 | 0.325190712 | 11.65665323 | 1.801400458 | 0.122814121 | 0.633319774 | -4.216654852 |
| **A_55_P2175405** | Zfp467 | 0.086628795 | -0.031701346 | 0.204958936 | 8.215409428 | 1.801092651 | 0.122866049 | 0.633319774 | -4.216880628 |
| **A_55_P2166011** | Tmem128 | 0.13303098 | -0.048715769 | 0.314777729 | 9.524312461 | 1.800757795 | 0.122922565 | 0.633319774 | -4.217126246 |

| | | | | | | | | | |
|---|---|---|---|---|---|---|---|---|---|
| **A_55_P2042612** | C1s1 | -0.33851 | -0.800985667 | 0.123965668 | 9.211822934 | -1.800743176 | 0.122925033 | 0.633319774 | -4.217136969 |
| **A_55_P2069917** | Uba52 | -0.100461515 | -0.23772027 | 0.03679724 | 17.04001709 | -1.80064803 | 0.122941096 | 0.633319774 | -4.21720676 |
| **A_52_P311031** | NA | 0.146684847 | -0.05372968 | 0.347099373 | 16.74151864 | 1.800633052 | 0.122943625 | 0.633319774 | -4.217217746 |
| **A_55_P2123962** | Micos10 | -0.1698815 | -0.401993206 | 0.062230206 | 14.98823371 | -1.800603987 | 0.122948533 | 0.633319774 | -4.217239065 |
| **A_55_P2108983** | NA | -0.151227349 | -0.357864855 | 0.055410156 | 17.89663045 | -1.800488508 | 0.122968033 | 0.633319774 | -4.217323771 |
| **A_51_P467233** | Polr3gl | 0.233470627 | -0.085659232 | 0.552600486 | 10.73048418 | 1.7998403 | 0.123077549 | 0.633447246 | -4.217799241 |
| **A_52_P24986** | Gpat4 | 0.090372374 | -0.033169139 | 0.213913888 | 12.54503789 | 1.799666114 | 0.123106994 | 0.633447246 | -4.21792701 |
| **A_55_P2095365** | Meaf6 | 0.113377948 | -0.041630856 | 0.268386751 | 8.351020999 | 1.79945666 | 0.12314241 | 0.633447246 | -4.21808065 |
| **A_55_P1994519** | Tpd52l2 | -0.088509152 | -0.209532428 | 0.032514124 | 8.388462153 | -1.799237275 | 0.123179517 | 0.633447246 | -4.218241575 |
| **A_52_P486279** | Popdc3 | -0.27417356 | -0.649071137 | 0.100724018 | 8.492180465 | -1.79921101 | 0.12318396 | 0.633447246 | -4.218260841 |
| **A_55_P2033666** | NA | 0.09147236 | -0.033632958 | 0.216577679 | 16.32117656 | 1.798801646 | 0.123253231 | 0.633447246 | -4.218561123 |
| **A_55_P1980543** | Ube2i | -0.211808681 | -0.501501482 | 0.077884121 | 11.14400429 | -1.798768252 | 0.123258884 | 0.633447246 | -4.218585619 |
| **A_55_P2042685** | NA | -0.183991729 | -0.435675824 | 0.067692365 | 10.9145755 | -1.798505184 | 0.123303422 | 0.633447246 | -4.21877859 |
| **A_55_P2115582** | Slc20a1 | -0.302605396 | -0.716572038 | 0.111361246 | 14.68749519 | -1.798375944 | 0.123325308 | 0.633447246 | -4.218873393 |
| **A_55_P2052775** | Morc2a | -0.143240394 | -0.339195835 | 0.052715048 | 10.94044634 | -1.798362929 | 0.123327512 | 0.633447246 | -4.21888294 |
| **A_55_P2079599** | NA | -0.145752616 | -0.345176285 | 0.053671052 | 10.68008356 | -1.798079222 | 0.123375572 | 0.633447246 | -4.219091052 |
| **A_52_P58524** | Neu1 | -0.124797048 | -0.295591232 | 0.045997136 | 13.75378987 | -1.797630375 | 0.123451643 | 0.633447246 | -4.219420304 |
| **A_52_P636151** | Golph3 | -0.163009811 | -0.386108572 | 0.06008895 | 11.56889936 | -1.797569745 | 0.123461922 | 0.633447246 | -4.21946478 |
| **A_55_P2112510** | NA | 0.237820168 | -0.087670084 | 0.563310419 | 14.50592436 | 1.797545359 | 0.123466057 | 0.633447246 | -4.219482668 |
| **A_51_P224564** | Ppm1f | -0.148970482 | -0.352869724 | 0.054928759 | 11.52348889 | -1.797437408 | 0.123484362 | 0.633447246 | -4.219561856 |
| **A_55_P2026380** | Pdcd2l | 0.21430983 | -0.079116602 | 0.507736262 | 10.55011335 | 1.796850817 | 0.123583874 | 0.633783376 | -4.219992157 |
| **A_51_P272123** | Ndufa10 | -0.10552091 | -0.250023538 | 0.038981719 | 10.38946561 | -1.796519532 | 0.12364011 | 0.633783376 | -4.220235177 |
| **A_55_P2022094** | Mta3 | -0.137466514 | -0.325747989 | 0.050814961 | 12.04131738 | -1.796215715 | 0.123691705 | 0.633783376 | -4.220458049 |
| **A_51_P421094** | NA | -0.165884085 | -0.393095056 | 0.061326886 | 10.65791937 | -1.796158207 | 0.123701474 | 0.633783376 | -4.220500235 |
| **A_66_P140307** | NA | 0.134844266 | -0.049859032 | 0.319547564 | 13.23567229 | 1.796085454 | 0.123713833 | 0.633783376 | -4.220553605 |
| **A_55_P2125714** | Atp2b4 | 0.0904487 | -0.033560748 | 0.214458149 | 8.120858584 | 1.794389491 | 0.12400229 | 0.634692591 | -4.221797745 |
| **A_55_P2115166** | AI661453 | -0.192721386 | -0.456999332 | 0.07155656 | 13.3102432 | -1.794064908 | 0.124057572 | 0.634692591 | -4.22203586 |
| **A_55_P2075274** | Rab11fip4 | -0.194696226 | -0.461709408 | 0.072316956 | 9.867795102 | -1.793882502 | 0.12408865 | 0.634692591 | -4.222169674 |
| **A_55_P2170345** | Gm6729 | -0.094844858 | -0.224925961 | 0.035236245 | 8.04093583 | -1.793778203 | 0.124106423 | 0.634692591 | -4.222246189 |
| **A_55_P2176350** | 0610010F05Rik | -0.104810138 | -0.248580905 | 0.038960628 | 10.30328389 | -1.793502012 | 0.124153501 | 0.634692591 | -4.222448805 |
| **A_51_P447668** | Ptpn12 | -0.104324307 | -0.247433613 | 0.038784999 | 9.260270905 | -1.793439764 | 0.124164114 | 0.634692591 | -4.222494471 |
| **A_51_P226417** | Rraga | -0.145117392 | -0.344189823 | 0.053955039 | 11.01951336 | -1.793401416 | 0.124170652 | 0.634692591 | -4.222522604 |
| **A_52_P99888** | Cxcl16 | -0.100647573 | -0.238756578 | 0.037461432 | 8.387190969 | -1.792876903 | 0.124260118 | 0.634692591 | -4.222907396 |
| **A_52_P657123** | Gm10270 | -0.114526574 | -0.271694402 | 0.042641253 | 10.06205911 | -1.792717194 | 0.124287372 | 0.634692591 | -4.223024562 |
| **A_55_P2277825** | E430010N07Rik | 0.125039696 | -0.046558095 | 0.296637487 | 9.460895756 | 1.792690762 | 0.124291883 | 0.634692591 | -4.223043953 |
| **A_52_P582150** | Gosr2 | -0.119115354 | -0.28259599 | 0.044365281 | 12.27456181 | -1.792547176 | 0.124316392 | 0.634692591 | -4.223149291 |
| **A_51_P163444** | Carhsp1 | -0.18119375 | -0.429901094 | 0.067513593 | 11.77777947 | -1.792353913 | 0.124349387 | 0.634692591 | -4.224705053 |
| **A_55_P1999958** | NA | -0.256928644 | -0.609678107 | 0.095820819 | 9.182340754 | -1.791906428 | 0.124425819 | 0.634692591 | -4.223619364 |
| **A_55_P2071970** | Nav2 | 0.166922463 | -0.062257725 | 0.39610265 | 8.803409551 | 1.79187156 | 0.124431776 | 0.634692591 | -4.223644944 |
| **A_55_P2104327** | Pxdn | -0.216239196 | -0.513130958 | 0.080652565 | 11.74806805 | -1.791865613 | 0.124432793 | 0.634692591 | -4.223649307 |
| **A_51_P392867** | Pex12 | -0.152816512 | -0.362640753 | 0.057007728 | 9.423052031 | -1.791776298 | 0.124448054 | 0.634692591 | -4.223714831 |
| **A_55_P2088525** | Myh10 | -0.12161896 | -0.288608882 | 0.045370963 | 9.762568693 | -1.791761431 | 0.124450595 | 0.634692591 | -4.223725738 |
| **A_55_P2272007** | Lrrfip2 | -0.104252525 | -0.247412172 | 0.038907122 | 8.683782521 | -1.791575548 | 0.124482364 | 0.634692591 | -4.223862109 |
| **A_51_P249594** | Rev1 | -0.111592022 | -0.264882631 | 0.041698587 | 8.516363287 | -1.790963599 | 0.124587008 | 0.634996604 | -4.22431106 |
| **A_55_P2169069** | Gm10405 | -0.190112764 | -0.451300872 | 0.071075344 | 12.818261 | -1.790717339 | 0.124629144 | 0.634996604 | -4.224491729 |
| **A_55_P2045476** | Hmgxb4 | 0.086994159 | -0.032559973 | 0.206548291 | 9.505883423 | 1.790171617 | 0.124722568 | 0.634996604 | -4.224892099 |
| **A_52_P639101** | Lrrc14 | -0.081029676 | -0.192403236 | 0.030343884 | 8.273960966 | -1.789909876 | 0.124767401 | 0.634996604 | -4.225084127 |
| **A_55_P2102664** | NA | 0.198879875 | -0.074509782 | 0.472269532 | 8.654672228 | 1.789690449 | 0.124804998 | 0.634996604 | -4.225245112 |
| **A_51_P511734** | Snip1 | -0.093834266 | -0.222832532 | 0.035163999 | 8.526076384 | -1.789562025 | 0.124827008 | 0.634996604 | -4.225339332 |
| **A_51_P127464** | Ppp1r37 | -0.184196579 | -0.437425235 | 0.069032078 | 9.390527619 | -1.78952542 | 0.124833282 | 0.634996604 | -4.225366188 |

| | | | | | | | | |
|---|---|---|---|---|---|---|---|---|
| **A_55_P2137461** | NA | -0.128136621 | -0.304302344 | 0.048029103 | 15.92548467 | -1.789455975 | 0.124845186 | 0.634996604 | -4.225417137 |
| **A_55_P2093822** | Ube2d2a | -0.110631862 | -0.26273999 | 0.041476266 | 13.23486723 | -1.789356859 | 0.124862178 | 0.634996604 | -4.225489855 |
| **A_55_P2150962** | Cpne7 | 0.17891462 | -0.067103087 | 0.424932327 | 8.59390686 | 1.789157719 | 0.124896324 | 0.634996604 | -4.225635957 |
| **A_51_P100573** | Map3k3 | -0.090021117 | -0.21381495 | 0.033772715 | 8.4085846 | -1.789017353 | 0.124920398 | 0.634996604 | -4.227835951 |
| **A_51_P490388** | Nudt8 | -0.178838853 | -0.424788706 | 0.067111 | 10.09010611 | -1.788893444 | 0.124941653 | 0.634996604 | -4.225829848 |
| **A_51_P115085** | Nup153 | 0.250801937 | -0.09415254 | 0.595756414 | 10.69763765 | 1.788703117 | 0.124974308 | 0.634996604 | -4.225969486 |
| **A_52_P159256** | Lpp | -0.13516425 | -0.321090913 | 0.050762412 | 8.992526424 | -1.788501585 | 0.125008896 | 0.634996604 | -4.226117345 |
| **A_55_P1966585** | NA | -0.253453998 | -0.602124802 | 0.095216806 | 16.47267028 | -1.788350856 | 0.12503477 | 0.634996604 | -4.226227932 |
| **A_55_P2015238** | Dctn4 | 0.100495958 | -0.037775737 | 0.238767654 | 11.89893372 | 1.788069814 | 0.125083029 | 0.635074825 | -4.226434126 |
| **A_55_P2170474** | Yipf2 | -0.138755889 | -0.329830634 | 0.052318856 | 11.19294295 | -1.786558736 | 0.125342814 | 0.63573278 | -4.227542789 |
| **A_51_P265348** | Ldb3 | 0.139258165 | -0.052509834 | 0.331026165 | 8.123867347 | 1.786543908 | 0.125345366 | 0.63573278 | -4.227553668 |
| **A_55_P1967133** | Ppp1r3b | 0.449959504 | -0.169709033 | 1.069628042 | 13.05849422 | 1.786418249 | 0.125366994 | 0.63573278 | -4.227645865 |
| **A_52_P218955** | D3Ertd751e | -0.14586071 | -0.346762881 | 0.055041461 | 8.354621734 | -1.786170266 | 0.125409687 | 0.63573278 | -4.227827811 |
| **A_55_P2019520** | Cma2 | 0.083979787 | -0.031693447 | 0.199653022 | 8.435851053 | 1.786121725 | 0.125418045 | 0.63573278 | -4.227863425 |
| **A_55_P2103284** | Twf2 | -0.117047509 | -0.278279179 | 0.044184161 | 8.876721061 | -1.785998076 | 0.125439339 | 0.63573278 | -4.227954147 |
| **A_51_P474701** | Fbp1 | -0.267103754 | -0.635040466 | 0.100832958 | 7.904522158 | -1.785977698 | 0.125442849 | 0.63573278 | -4.227969098 |
| **A_55_P2083582** | NA | 0.32202138 | -0.121760525 | 0.765803284 | 14.7454719 | 1.78518932 | 0.125578705 | 0.636129051 | -4.228547539 |
| **A_52_P650473** | NA | 0.191909635 | -0.072570569 | 0.45638984 | 8.705986907 | 1.785142013 | 0.125586862 | 0.636129051 | -4.228582249 |
| **A_51_P159610** | Ccdc120 | -0.166127578 | -0.395144607 | 0.062889451 | 8.76689929 | -1.784609169 | 0.125678772 | 0.636208027 | -4.228973205 |
| **A_55_P2080151** | Hspa2 | 0.177067552 | -0.067054231 | 0.421189335 | 10.16414239 | 1.784438617 | 0.125708205 | 0.636208027 | -4.229098343 |
| **A_51_P162176** | Trappc3 | -0.373731759 | -0.889004577 | 0.141541059 | 10.72633544 | -1.784398853 | 0.125715068 | 0.636208027 | -4.229127518 |
| **A_55_P2099338** | Akt1s1 | -0.104204911 | -0.24788357 | 0.039473749 | 8.160884685 | -1.784288522 | 0.125734113 | 0.636208027 | -4.229208471 |
| **A_55_P1966159** | Dlg5 | -0.260521709 | -0.619855196 | 0.098811779 | 13.94329344 | -1.783673614 | 0.125840306 | 0.636229789 | -4.229659646 |
| **A_55_P2074762** | Lect2 | -0.083803602 | -0.199412131 | 0.031804928 | 7.889048545 | -1.783372102 | 0.125892409 | 0.636229789 | -4.229880874 |
| **A_52_P287683** | Pik3c3 | -0.118731719 | -0.282555833 | 0.045092395 | 8.3385662 | -1.783027709 | 0.125951949 | 0.636229789 | -4.230133568 |
| **A_66_P139295** | Tomm40 | 0.318421779 | -0.120932376 | 0.757775934 | 11.70096805 | 1.783024004 | 0.125952589 | 0.636229789 | -4.230136286 |
| **A_52_P366932** | Gna11 | -0.173527551 | -0.412970088 | 0.065914987 | 10.95038944 | -1.782938213 | 0.125967425 | 0.636229789 | -4.230199234 |
| **A_55_P2001238** | Luc7l3 | -0.29722794 | -0.70745355 | 0.11299767 | 12.10255116 | -1.782526626 | 0.126038626 | 0.636229789 | -4.230501231 |
| **A_55_P2037101** | Exoc2 | -0.086440129 | -0.205742443 | 0.032862186 | 10.62879362 | -1.782525285 | 0.126038858 | 0.636229789 | -4.230502215 |
| **A_55_P2072095** | Mucl3 | 0.310628445 | -0.118180478 | 0.739437369 | 10.58056662 | 1.782159509 | 0.126102167 | 0.636229789 | -4.230770602 |
| **A_55_P2137910** | NA | -0.211368384 | -0.50316512 | 0.080428351 | 14.32642752 | -1.782086426 | 0.12611482 | 0.636229789 | -4.230824225 |
| **A_66_P100105** | Patj | -0.156993798 | -0.37373529 | 0.059747693 | 10.05036307 | -1.782007811 | 0.126128433 | 0.636229789 | -4.230881909 |
| **A_55_P2028532** | Ate1 | 0.172380818 | -0.065638972 | 0.410400608 | 9.876918906 | 1.78174288 | 0.126174317 | 0.636229789 | -4.231076301 |
| **A_55_P1991723** | Alkbh3 | -0.129666716 | -0.308730971 | 0.04939754 | 11.45410372 | -1.781512085 | 0.126214302 | 0.636229789 | -4.231245648 |
| **A_51_P258281** | Gal | 0.212946247 | -0.081146557 | 0.507039051 | 9.125105677 | 1.781372549 | 0.126238482 | 0.636229789 | -4.231348032 |
| **A_55_P1956843** | NA | -0.246167478 | -0.586149037 | 0.093814081 | 13.30755212 | -1.781330392 | 0.126245789 | 0.636229789 | -4.231378965 |
| **A_55_P1980119** | Cptp | -0.17745119 | -0.426993703 | 0.072091323 | 10.08084732 | -1.842826521 | 0.126258429 | 0.636229789 | -4.224919226 |
| **A_55_P2121936** | Wiz | 0.202201443 | -0.077076124 | 0.481479009 | 10.69367425 | 1.781219174 | 0.126265066 | 0.636229789 | -4.231460571 |
| **A_52_P98232** | Hira | -0.120607682 | -0.287265939 | 0.046050575 | 8.268138554 | -1.780398819 | 0.12640735 | 0.636583683 | -4.232062514 |
| **A_51_P159814** | Smpd4 | -0.126928943 | -0.302333708 | 0.048475823 | 8.948370657 | -1.780280509 | 0.126427883 | 0.636583683 | -4.232149325 |
| **A_55_P2061465** | Ppp1r21 | -0.139861966 | -0.333142936 | 0.053419004 | 9.648721292 | -1.780244679 | 0.126434102 | 0.636583683 | -4.232175616 |
| **A_51_P272817** | Map2k3 | -0.136326794 | -0.324770366 | 0.052116778 | 9.416825853 | -1.779791201 | 0.126512838 | 0.636814233 | -4.232508363 |
| **A_55_P1981949** | Klhl5 | -0.099457524 | -0.236968599 | 0.03805355 | 9.601364973 | -1.779381753 | 0.126583971 | 0.6370064 | -4.232808803 |
| **A_55_P2157966** | Map1a | 0.090579875 | -0.034732335 | 0.215892086 | 7.826633244 | 1.778310077 | 0.126770339 | 0.637070187 | -4.233595173 |
| **A_55_P2021119** | Sun2 | -0.284748412 | -0.678686144 | 0.109189319 | 12.41103817 | -1.778291262 | 0.126773613 | 0.637070187 | -4.233608979 |
| **A_55_P2075479** | Rpl37a | -0.116162019 | -0.27692171 | 0.044597672 | 16.86189832 | -1.777690507 | 0.126878207 | 0.637070187 | -4.234049802 |
| **A_55_P1965592** | NA | 0.08918709 | -0.034245577 | 0.212619758 | 9.093640812 | 1.777628659 | 0.12688898 | 0.637070187 | -4.234095186 |
| **A_55_P1989511** | Tbca | 0.259502107 | -0.09971637 | 0.618720583 | 11.82175817 | 1.77726171 | 0.126952915 | 0.637070187 | -4.234364448 |
| **A_55_P2120277** | Gm11619 | -0.105568777 | -0.251725494 | 0.04058794 | 15.47130865 | -1.776993647 | 0.12699964 | 0.637070187 | -4.23456115 |
| **A_51_P501773** | Slc35f3 | -0.280181502 | -0.668090969 | 0.107727965 | 8.658769266 | -1.77696242 | 0.127005084 | 0.637070187 | -4.234584064 |

| | | | | | | | | | |
|---|---|---|---|---|---|---|---|---|---|
| **A_55_P2060863** | NA | 0.117174714 | -0.045059301 | 0.279408728 | 11.18343412 | 1.776892475 | 0.12701728 | 0.637070187 | -4.234635389 |
| **A_52_P36928** | Morn2 | -0.093820637 | -0.22373031 | 0.036089036 | 8.369930566 | -1.776748879 | 0.12704232 | 0.637070187 | -4.234740759 |
| **A_55_P1999637** | Gm15455 | 0.076753086 | -0.029539327 | 0.183045499 | 10.30318502 | 1.776489891 | 0.127087494 | 0.637070187 | -4.234930803 |
| **A_55_P1976108** | Rps27a | -0.104876488 | -0.250116563 | 0.040363587 | 16.60423927 | -1.776482085 | 0.127088856 | 0.637070187 | -4.234936531 |
| **A_55_P1954212** | Snrpd2 | 0.164795199 | -0.063454665 | 0.393045063 | 12.26302859 | 1.776246317 | 0.127129995 | 0.637070187 | -4.235109537 |
| **A_51_P320089** | Gclm | -0.085343808 | -0.203560003 | 0.032872388 | 8.717151972 | -1.776086854 | 0.127157827 | 0.637070187 | -4.235226551 |
| **A_55_P2077526** | NA | -0.366812822 | -0.874922418 | 0.141296775 | 14.38117218 | -1.776054438 | 0.127163485 | 0.637070187 | -4.235250337 |
| **A_55_P2214625** | 4930519A11Rik | 0.320995691 | -0.123649733 | 0.765641114 | 8.719677963 | 1.776047345 | 0.127164723 | 0.637070187 | -4.235255542 |
| **A_55_P2081885** | NA | 0.172570257 | -0.066490533 | 0.411631046 | 16.58473952 | 1.775933738 | 0.127184556 | 0.637070187 | -4.235338907 |
| **A_55_P2109857** | Rgs2 | 0.132852842 | -0.051189305 | 0.316894989 | 9.930289726 | 1.775917231 | 0.127187438 | 0.637070187 | -4.23535102 |
| **A_55_P2003793** | Tspan3 | -0.380289315 | -0.907111473 | 0.146532843 | 13.89981647 | -1.77590305 | 0.127189914 | 0.637070187 | -4.235361426 |
| **A_52_P524895** | Eif3a | -0.160345012 | -0.382507692 | 0.061817668 | 10.8977102 | -1.77563424 | 0.127236856 | 0.637140204 | -4.235558679 |
| **A_55_P2098076** | Trim25 | -0.196780712 | -0.469475185 | 0.07591376 | 13.98497521 | -1.775314733 | 0.127292673 | 0.637141922 | -4.235793134 |
| **A_52_P500244** | Atrn | -0.108472944 | -0.258799236 | 0.041853348 | 10.61491545 | -1.775234163 | 0.127306752 | 0.637141922 | -4.235852256 |
| **A_55_P1960765** | NA | 0.395137987 | -0.152602744 | 0.942878718 | 12.54615498 | 1.774773394 | 0.127387299 | 0.637141922 | -4.236190372 |
| **A_55_P1958280** | Arrdc1 | 0.124207235 | -0.047980307 | 0.296394778 | 9.611089063 | 1.774656605 | 0.127407722 | 0.637141922 | -4.236276073 |
| **A_52_P474636** | Stk4 | -0.077266098 | -0.184399041 | 0.029866846 | 8.449954063 | -1.774332899 | 0.127464348 | 0.637141922 | -4.236513612 |
| **A_55_P2168243** | NA | -0.104695122 | -0.249862335 | 0.040472091 | 9.28870351 | -1.774300075 | 0.127470091 | 0.637141922 | -4.236537698 |
| **A_55_P2038046** | NA | -0.163491267 | -0.390222215 | 0.06323968 | 9.114494269 | -1.773997164 | 0.127523104 | 0.637141922 | -4.236759978 |
| **A_55_P2046374** | NA | -0.147737373 | -0.352636464 | 0.057161719 | 8.284148936 | -1.773860656 | 0.127547001 | 0.637141922 | -4.23686015 |
| **A_55_P2185359** | NA | -0.327600264 | -0.781957153 | 0.126756625 | 13.06301167 | -1.773847537 | 0.127549298 | 0.637141922 | -4.236869777 |
| **A_55_P2108693** | NA | -0.178008647 | -0.424917856 | 0.068900563 | 12.24438305 | -1.773670623 | 0.127580276 | 0.637141922 | -4.236999599 |
| **A_52_P616392** | Sbno2 | -0.174173578 | -0.415778485 | 0.067431328 | 8.692230906 | -1.773559193 | 0.127599792 | 0.637141922 | -4.237081368 |
| **A_55_P1983708** | Insrr | 0.209119416 | -0.081004906 | 0.499243737 | 8.162927716 | 1.773288437 | 0.127647224 | 0.63721415 | -4.237280054 |
| **A_52_P397204** | Csk | -0.123170328 | -0.294125905 | 0.04778525 | 9.326967685 | -1.772523437 | 0.127781332 | 0.637489979 | -4.237841428 |
| **A_52_P600318** | Scamp2 | -0.109439756 | -0.261349514 | 0.042470002 | 10.99252105 | -1.772387217 | 0.127805227 | 0.637489979 | -4.237941389 |
| **A_55_P2172832** | Kcne1l | 0.082249819 | -0.031925591 | 0.196425229 | 7.991822372 | 1.772276733 | 0.12782461 | 0.637489979 | -4.238022465 |
| **A_55_P2067960** | Cplx2 | -0.373693818 | -0.892454695 | 0.145067059 | 14.07259818 | -1.772220926 | 0.127834402 | 0.637489979 | -4.238063417 |
| **A_55_P2054942** | Iffo2 | -0.180618302 | -0.431391809 | 0.070155206 | 9.195501652 | -1.771941023 | 0.127883525 | 0.637570458 | -4.238268818 |
| **A_55_P2173623** | Samd1 | -0.233111065 | -0.556847603 | 0.090625474 | 12.31993157 | -1.771496637 | 0.127961554 | 0.63762448 | -4.238594921 |
| **A_55_P2067031** | Agxt2 | 0.137148278 | -0.053329685 | 0.327626242 | 7.975363034 | 1.771392431 | 0.127979858 | 0.63762448 | -4.238671391 |
| **A_51_P397920** | Ilrun | -0.166841182 | -0.398632296 | 0.064949932 | 11.15271275 | -1.770824971 | 0.128079578 | 0.63762448 | -4.239087812 |
| **A_55_P2178349** | Ywhaq | -0.130604175 | -0.312074002 | 0.050865651 | 11.24763963 | -1.770605465 | 0.128118173 | 0.63762448 | -4.239248894 |
| **A_55_P2168504** | Mcmbp | -0.134259894 | -0.320848348 | 0.052328559 | 9.450532643 | -1.770234084 | 0.128183497 | 0.63762448 | -4.239521427 |
| **A_51_P482410** | Ceacam15 | -0.638853591 | -1.52676151 | 0.249054327 | 10.65143406 | -1.770119451 | 0.128203667 | 0.63762448 | -4.239605549 |
| **A_55_P2183015** | P2rx4 | 0.295897773 | -0.115392664 | 0.70718821 | 13.3559213 | 1.769955096 | 0.128232591 | 0.63762448 | -4.239726159 |
| **A_55_P1957967** | Dhx32 | -0.283902889 | -0.678528076 | 0.110722298 | 11.71297333 | -1.769922188 | 0.128238383 | 0.63762448 | -4.239750308 |
| **A_55_P2037294** | NA | -0.081869404 | -0.195711655 | 0.031972848 | 8.065919178 | -1.769242291 | 0.128358109 | 0.63762448 | -4.240249245 |
| **A_52_P163515** | NA | -0.080891789 | -0.193376246 | 0.031592668 | 8.195029505 | -1.769216928 | 0.128362577 | 0.63762448 | -4.240267858 |
| **A_55_P2110435** | Zscan20 | 0.185703538 | -0.072617367 | 0.444024443 | 8.38589207 | 1.76860068 | 0.128471193 | 0.63762448 | -4.240720089 |
| **A_66_P130646** | H4c8 | 0.177141299 | -0.069292516 | 0.423575114 | 8.520275009 | 1.768433317 | 0.128500708 | 0.63762448 | -4.240842908 |
| **A_55_P1973854** | Sema6b | -0.076308089 | -0.182474605 | 0.029858427 | 8.034179702 | -1.768284652 | 0.12852693 | 0.63762448 | -4.240952006 |
| **A_51_P486810** | Gpx2 | 0.169607279 | -0.066403953 | 0.405618511 | 8.817887554 | 1.767994775 | 0.128578075 | 0.63762448 | -4.241164732 |
| **A_55_P2143885** | Etl4 | -0.171758867 | -0.410796739 | 0.067279005 | 11.1412425 | -1.767753138 | 0.128620724 | 0.63762448 | -4.241342058 |
| **A_55_P2005982** | NA | -0.209554096 | -0.50119649 | 0.082088298 | 12.7246022 | -1.767724831 | 0.128625721 | 0.63762448 | -4.241362831 |
| **A_51_P366161** | Mpg | 0.188548471 | -0.073909712 | 0.451006654 | 10.95693981 | 1.767388644 | 0.128685084 | 0.63762448 | -4.241609542 |
| **A_52_P462125** | Smarca5-ps | -0.211562122 | -0.50607524 | 0.082950997 | 10.57252987 | -1.767268077 | 0.12870638 | 0.63762448 | -4.241698021 |
| **A_55_P1961049** | Cep295 | -0.143560652 | -0.343417927 | 0.056296622 | 8.869876564 | -1.76719562 | 0.12871918 | 0.63762448 | -4.241751194 |
| **A_55_P2117711** | Rnf185 | -0.080101802 | -0.191642727 | 0.031439123 | 11.99907569 | -1.766758579 | 0.128796411 | 0.63762448 | -4.242071919 |
| **A_65_P19458** | Rock2 | -0.129267509 | -0.309303672 | 0.050768654 | 9.669429999 | -1.766439606 | 0.128852807 | 0.63762448 | -4.242305999 |

| | | | | | | | | | |
|---|---|---|---|---|---|---|---|---|---|
| **A_51_P120295** | Cyp2d26 | -0.282476365 | -0.675915624 | 0.110962894 | 8.009252846 | -1.766337079 | 0.128870939 | 0.63762448 | -4.24238124 |
| **A_51_P324814** | Krt18 | -0.138942497 | -0.332483731 | 0.054598736 | 17.08499735 | -1.766162901 | 0.128901749 | 0.63762448 | -4.242509062 |
| **A_55_P2027640** | NA | -0.100726132 | -0.241066529 | 0.039614266 | 16.92379294 | -1.76574757 | 0.128975246 | 0.63762448 | -4.242813857 |
| **A_55_P2015363** | C1ra | -0.16276902 | -0.38960012 | 0.064062079 | 8.671759074 | -1.765380458 | 0.129040244 | 0.63762448 | -4.243083266 |
| **A_55_P2033255** | NA | 0.13912086 | -0.054755495 | 0.332997214 | 15.50149025 | 1.76537337 | 0.129041499 | 0.63762448 | -4.243088468 |
| **A_52_P99848** | Pik3cd | 0.152143404 | -0.059910344 | 0.364197151 | 8.627562396 | 1.765128547 | 0.129084864 | 0.63762448 | -4.243268135 |
| **A_55_P2073841** | NA | -0.176628853 | -0.422824707 | 0.069567 | 13.86928813 | -1.765021822 | 0.129103772 | 0.63762448 | -4.243346456 |
| **A_55_P2125351** | Sema6d | -0.14673052 | -0.351281764 | 0.057820724 | 11.11581169 | -1.764767495 | 0.129148842 | 0.63762448 | -4.243533098 |
| **A_55_P2109599** | NA | 0.171533064 | -0.0675949 | 0.410661028 | 11.66875341 | 1.764764029 | 0.129149456 | 0.63762448 | -4.243535642 |
| **A_51_P138760** | Abcd4 | 0.27293974 | -0.107588869 | 0.653468349 | 11.85271326 | 1.764609543 | 0.12917684 | 0.63762448 | -4.243649013 |
| **A_55_P2184264** | Timm23 | 0.391569805 | -0.154351431 | 0.937491041 | 11.87017477 | 1.764608511 | 0.129177023 | 0.63762448 | -4.243649771 |
| **A_55_P2037982** | Agap1 | 0.193299653 | -0.076207959 | 0.462807266 | 10.10256891 | 1.76453063 | 0.129190831 | 0.63762448 | -4.243706925 |
| **A_51_P291460** | Gm6498 | -0.434265439 | -1.039750951 | 0.171220073 | 14.03723115 | -1.764495985 | 0.129196974 | 0.63762448 | -4.24373235 |
| **A_51_P109050** | Paqr4 | 0.117152558 | -0.046193003 | 0.280498119 | 9.213374158 | 1.764467251 | 0.129202068 | 0.63762448 | -4.243753437 |
| **A_55_P2266880** | Kirrel | -0.14212291 | -0.340305264 | 0.056059444 | 10.14966725 | -1.764283088 | 0.129234727 | 0.63762448 | -4.243888588 |
| **A_55_P2090633** | Ptp4a1 | -0.095825042 | -0.229457664 | 0.037807579 | 9.341041852 | -1.76415075 | 0.129258201 | 0.63762448 | -4.243985706 |
| **A_51_P452203** | Vash2 | -0.077002275 | -0.184391041 | 0.030386491 | 8.052849517 | -1.76406209 | 0.129273929 | 0.63762448 | -4.244050771 |
| **A_51_P107234** | Abcf1 | -0.200383966 | -0.479850964 | 0.079083032 | 12.7680101 | -1.764012272 | 0.129282767 | 0.63762448 | -4.244087331 |
| **A_55_P2032267** | Gm10767 | -0.112218832 | -0.268733014 | 0.044295351 | 11.89673412 | -1.763929442 | 0.129297464 | 0.63762448 | -4.244148117 |
| **A_55_P2181262** | NA | -0.161797021 | -0.38746175 | 0.063867707 | 11.89660978 | -1.763908289 | 0.129301218 | 0.63762448 | -4.244163641 |
| **A_51_P226364** | Coa3 | 0.199335866 | -0.078703382 | 0.477375113 | 13.24485462 | 1.763796618 | 0.129321035 | 0.63762448 | -4.244245592 |
| **A_51_P418809** | Airn | -0.113828661 | -0.272609352 | 0.044952031 | 8.270145823 | -1.763693436 | 0.129339349 | 0.63762448 | -4.244321315 |
| **A_55_P2097329** | Sgsm3 | 0.334565921 | -0.132211079 | 0.801342921 | 13.07752283 | 1.76336175 | 0.129398236 | 0.63762448 | -4.244564729 |
| **A_51_P433091** | Purb | -0.175635329 | -0.420683725 | 0.069413066 | 11.7702361 | -1.763312064 | 0.129407059 | 0.63762448 | -4.244601193 |
| **A_51_P143232** | Wdr59 | 0.173811966 | -0.068706757 | 0.416330688 | 8.984143894 | 1.763208058 | 0.129425531 | 0.63762448 | -4.244677519 |
| **A_66_P115034** | Bend5 | 0.120025133 | -0.047463301 | 0.287513567 | 10.26219981 | 1.763017145 | 0.129459444 | 0.63762448 | -4.244817625 |
| **A_52_P122843** | Bicral | -0.136358749 | -0.326667749 | 0.053950251 | 8.826278346 | -1.762758613 | 0.129505384 | 0.63762448 | -4.245007355 |
| **A_55_P2024575** | Sec24c | -0.335922753 | -0.80476119 | 0.132915683 | 14.16972806 | -1.762728296 | 0.129510772 | 0.63762448 | -4.245029603 |
| **A_51_P322994** | Mrpl22 | -0.158848806 | -0.380579405 | 0.062881794 | 9.676634759 | -1.762493342 | 0.129552537 | 0.637667681 | -4.245202029 |
| **A_52_P368262** | Bcl7c | -0.126050985 | -0.302072562 | 0.049970592 | 8.896955877 | -1.761771171 | 0.129680992 | 0.638077915 | -4.245732013 |
| **A_52_P540219** | Timp2 | -0.828446335 | -2.007199123 | 0.350306453 | 14.29929291 | -1.821342955 | 0.129755083 | 0.638077915 | -4.237937955 |
| **A_55_P2133910** | Ttll10 | 0.112730815 | -0.044732518 | 0.270194148 | 8.733916928 | 1.761296082 | 0.129765566 | 0.638077915 | -4.246080669 |
| **A_52_P661722** | 2-Mar | 0.150705973 | -0.05980296 | 0.361214907 | 14.64260202 | 1.761282817 | 0.129767928 | 0.638077915 | -4.246090404 |
| **A_55_P1980831** | Arpc2 | -0.100217206 | -0.24023194 | 0.039797528 | 16.32147577 | -1.760912241 | 0.129833936 | 0.63811396 | -4.246362362 |
| **A_52_P582394** | Mrps11 | -0.086606244 | -0.207663254 | 0.034450767 | 8.691648862 | -1.760063845 | 0.129985178 | 0.63811396 | -4.246984982 |
| **A_51_P414115** | Naprt | 0.245745294 | -0.097754626 | 0.589245215 | 8.717383956 | 1.760061053 | 0.129985676 | 0.63811396 | -4.246987031 |
| **A_51_P203501** | Vars | -0.084451834 | -0.202500539 | 0.033596871 | 8.383287703 | -1.760017643 | 0.12999342 | 0.63811396 | -4.247018889 |
| **A_52_P494982** | Dhx40 | -0.095176063 | -0.228218402 | 0.037866276 | 8.69577854 | -1.759977142 | 0.130000644 | 0.63811396 | -4.247048612 |
| **A_55_P2132894** | Sdcbp | -0.114891099 | -0.275509045 | 0.045726848 | 8.822063943 | -1.759792559 | 0.130033576 | 0.63811396 | -4.247184074 |
| **A_52_P648715** | Triml1 | -0.228176038 | -0.547183241 | 0.090831166 | 9.214524013 | -1.759700299 | 0.13005004 | 0.63811396 | -4.247251782 |
| **A_55_P2051791** | Cbr4 | -0.103216923 | -0.247545179 | 0.041111334 | 8.574203142 | -1.759416679 | 0.130100664 | 0.63811396 | -4.247459925 |
| **A_55_P1968628** | NA | 0.179051559 | -0.071320546 | 0.429423664 | 8.703655099 | 1.759386804 | 0.130105998 | 0.63811396 | -4.24748185 |
| **A_55_P2070009** | Krt10 | 0.124736149 | -0.04968814 | 0.299160438 | 8.692484939 | 1.759359271 | 0.130110913 | 0.63811396 | -4.247502056 |
| **A_55_P1997926** | NA | 0.170411362 | -0.067934714 | 0.408757437 | 11.29953963 | 1.758975154 | 0.130179511 | 0.63811396 | -4.247783952 |
| **A_55_P2080708** | Cstf2 | 0.170358314 | -0.067993435 | 0.408710064 | 8.751395653 | 1.75838574 | 0.130284841 | 0.63811396 | -4.248216514 |
| **A_55_P2082186** | NA | 0.131505676 | -0.052495472 | 0.315506824 | 12.09679442 | 1.758300615 | 0.130300061 | 0.63811396 | -4.248278986 |
| **A_55_P2108248** | Art4 | 0.20370756 | -0.081357297 | 0.488772417 | 9.436259228 | 1.758055692 | 0.130343859 | 0.63811396 | -4.24845873 |
| **A_55_P2150044** | Cdkl2 | 0.099861852 | -0.039884382 | 0.239608086 | 9.59664195 | 1.758039645 | 0.130346729 | 0.63811396 | -4.248470507 |
| **A_55_P1984070** | NA | -0.2555193 | -0.613137357 | 0.102098756 | 12.29828494 | -1.757816084 | 0.13038672 | 0.63811396 | -4.248634575 |
| **A_66_P134739** | Clpp | 0.219176479 | -0.087626061 | 0.525979019 | 9.768344809 | 1.757535714 | 0.130436891 | 0.63811396 | -4.248840335 |

| | | | | | | | | | |
|---|---|---|---|---|---|---|---|---|---|
| **A_55_P2016089** | Uggt1 | -0.223958245 | -0.53748662 | 0.08957013 | 13.68797487 | -1.757354457 | 0.130469337 | 0.63811396 | -4.248973357 |
| **A_55_P2136813** | Rab10os | -0.132443759 | -0.317860683 | 0.052973165 | 10.43432769 | -1.757321775 | 0.130475188 | 0.63811396 | -4.248997342 |
| **A_55_P2180096** | Gm10136 | 0.277906306 | -0.111234633 | 0.667047244 | 16.26724224 | 1.756955015 | 0.130540865 | 0.63811396 | -4.249266502 |
| **A_55_P1969396** | NA | 0.330851533 | -0.132490411 | 0.794193477 | 11.00782501 | 1.756712525 | 0.130584307 | 0.63811396 | -4.249444461 |
| **A_55_P2060445** | Apopt1 | -0.33594935 | -0.806442241 | 0.13454354 | 14.49918855 | -1.756668861 | 0.130592131 | 0.63811396 | -4.249476506 |
| **A_52_P336825** | Matr3 | -0.127719309 | -0.306611228 | 0.051172609 | 9.667676942 | -1.756446814 | 0.130631925 | 0.63811396 | -4.249639464 |
| **A_55_P2005025** | Mbnl1 | 0.127953807 | -0.051292788 | 0.307200401 | 10.21011433 | 1.756189841 | 0.130677993 | 0.63811396 | -4.249828052 |
| **A_55_P2197998** | D4Ertd58e | 0.142482778 | -0.057136355 | 0.342101912 | 8.306240986 | 1.756019651 | 0.130708513 | 0.63811396 | -4.249952953 |
| **A_52_P632691** | Zfp395 | -0.100380639 | -0.241018522 | 0.040257243 | 8.916420272 | -1.755968825 | 0.130717628 | 0.63811396 | -4.249990254 |
| **A_52_P317923** | Zfp948 | 0.204544833 | -0.082038647 | 0.491128314 | 10.13897822 | 1.755927276 | 0.130725081 | 0.63811396 | -4.250020746 |
| **A_51_P259029** | Dusp26 | 0.104570073 | -0.041991578 | 0.251131725 | 7.757700267 | 1.755319699 | 0.130834105 | 0.63811396 | -4.250466639 |
| **A_55_P2135039** | Hyi | 0.209425208 | -0.084106255 | 0.50295667 | 11.04776231 | 1.755268098 | 0.130843369 | 0.63811396 | -4.250504509 |
| **A_55_P1956998** | Dis3l2 | -0.100260494 | -0.240804346 | 0.040283359 | 8.031964402 | -1.755040521 | 0.130884231 | 0.63811396 | -4.250671525 |
| **A_55_P1957468** | She | -0.128806684 | -0.309374548 | 0.051761179 | 9.371613649 | -1.754959529 | 0.130898777 | 0.63811396 | -4.250730964 |
| **A_55_P2049717** | Agrp | 0.297470227 | -0.119541824 | 0.714482278 | 10.33731271 | 1.754947232 | 0.130900985 | 0.63811396 | -4.250739989 |
| **A_51_P415247** | NA | 0.074690652 | -0.030015917 | 0.17939722 | 8.098201931 | 1.754936814 | 0.130902856 | 0.63811396 | -4.250747634 |
| **A_55_P1976152** | Vrk2 | -0.171865525 | -0.412819482 | 0.069088433 | 9.32002923 | -1.754784816 | 0.130930159 | 0.63811396 | -4.250859184 |
| **A_55_P2114680** | Car4 | 0.694174853 | -0.279100417 | 1.667450123 | 14.13792529 | 1.75469788 | 0.130945777 | 0.63811396 | -4.250922985 |
| **A_51_P379069** | Dkk1 | -0.240468886 | -0.577640065 | 0.096702292 | 8.749092812 | -1.754597923 | 0.130963737 | 0.63811396 | -4.250996342 |
| **A_55_P2131340** | Churc1 | 0.248697534 | -0.100116459 | 0.597511527 | 11.4107071 | 1.754069284 | 0.131058762 | 0.638416031 | -4.251384305 |
| **A_55_P2002379** | Golga2 | 0.195227002 | -0.07864046 | 0.469094464 | 12.96717095 | 1.753753745 | 0.131115513 | 0.638440144 | -4.251615875 |
| **A_55_P2179453** | Lig3 | -0.129919501 | -0.312194493 | 0.052355492 | 8.341715058 | -1.753542984 | 0.131153433 | 0.638440144 | -4.25177055 |
| **A_52_P48777** | Copg2 | -0.228272951 | -0.548559369 | 0.092013468 | 9.835272231 | -1.753416513 | 0.131176193 | 0.638440144 | -4.251863366 |
| **A_55_P1962299** | Hba-a2 | 0.117052882 | -0.047217878 | 0.281323642 | 17.92506888 | 1.753036692 | 0.131244569 | 0.638440144 | -4.252142113 |
| **A_52_P361993** | Zc2hc1a | -0.115734645 | -0.278158695 | 0.046689405 | 9.308956303 | -1.753001193 | 0.131250961 | 0.638440144 | -4.252168165 |
| **A_51_P359137** | Doc2g | -0.096218536 | -0.231268237 | 0.038831166 | 8.20613775 | -1.75280772 | 0.131285806 | 0.638440144 | -4.252310152 |
| **A_55_P2152292** | Map2k2 | -0.205119349 | -0.493027563 | 0.082788866 | 9.461585609 | -1.752757103 | 0.131294923 | 0.638440144 | -4.2523473 |
| **A_55_P1983558** | Tmed1 | 0.14947661 | -0.060371197 | 0.359324416 | 11.58034124 | 1.752419061 | 0.13135583 | 0.638442373 | -4.252595385 |
| **A_66_P133498** | Pdcd6ip | -0.143319257 | -0.344546182 | 0.057907668 | 12.33902533 | -1.752215912 | 0.131392446 | 0.638442373 | -4.252744474 |
| **A_55_P2175469** | Chd2 | -0.228550014 | -0.549446959 | 0.09234693 | 9.464221339 | -1.752204672 | 0.131394473 | 0.638442373 | -4.252752723 |
| **A_55_P2408588** | Arntl | -0.092922386 | -0.223432546 | 0.037587775 | 9.942743972 | -1.751641354 | 0.13149606 | 0.638571534 | -4.253166135 |
| **A_55_P2028894** | Gata4 | 0.127791001 | -0.051699515 | 0.307281516 | 9.223911723 | 1.751571735 | 0.13150862 | 0.638571534 | -4.253217228 |
| **A_55_P1967478** | Hbb-y | 1.038660277 | -0.42025685 | 2.497577405 | 9.656666309 | 1.751507748 | 0.131520165 | 0.638571534 | -4.253264187 |
| **A_55_P2136862** | Dpy30 | -0.114382197 | -0.275071426 | 0.046307032 | 9.171387911 | -1.751220493 | 0.131572007 | 0.638639366 | -4.253475 |
| **A_52_P491544** | Nedd9 | -0.141743818 | -0.340931466 | 0.05744383 | 8.96772015 | -1.750696593 | 0.131666609 | 0.638639366 | -4.253859485 |
| **A_55_P2409570** | Luc7l2 | 0.163212622 | -0.06616839 | 0.392593633 | 15.81372817 | 1.750513406 | 0.131699703 | 0.638639366 | -4.253993924 |
| **A_55_P2115119** | Rnaseh2a | 0.113385963 | -0.045988486 | 0.272760411 | 9.590376499 | 1.750288986 | 0.131740258 | 0.638639366 | -4.254158623 |
| **A_55_P1998943** | Oas1a | -0.296362019 | -0.712949283 | 0.120225244 | 12.77235915 | -1.750192104 | 0.131757769 | 0.638639366 | -4.254229723 |
| **A_55_P2065716** | Glud1 | -0.15352992 | -0.369371163 | 0.062311324 | 11.78417589 | -1.749958508 | 0.1318 | 0.638639366 | -4.254401156 |
| **A_51_P340704** | Prpf31 | 0.130730248 | -0.053079342 | 0.314539838 | 11.68182723 | 1.749754349 | 0.13183692 | 0.638639366 | -4.254550986 |
| **A_55_P2035892** | Prkcsh | -0.176456258 | -0.424614046 | 0.07170153 | 16.04573822 | -1.74935648 | 0.1319089 | 0.638639366 | -4.254842977 |
| **A_66_P136099** | Coasy | 0.152315372 | -0.061902132 | 0.366532875 | 11.83824515 | 1.749274417 | 0.131923751 | 0.638639366 | -4.254903202 |
| **A_52_P621567** | Tcf20 | -0.265205789 | -0.638194057 | 0.107782479 | 11.8646368 | -1.749270577 | 0.131924446 | 0.638639366 | -4.25490602 |
| **A_55_P2034663** | Csf2rb2 | 0.209538956 | -0.085182952 | 0.504260864 | 8.776717992 | 1.749127707 | 0.131950305 | 0.638639366 | -4.25501087 |
| **A_51_P200610** | Zbtb12 | -0.13083445 | -0.314867899 | 0.053198998 | 8.55639496 | -1.749018941 | 0.131969995 | 0.638639366 | -4.255090692 |
| **A_51_P384754** | Ldb1 | -0.125882363 | -0.302954614 | 0.051189888 | 9.227337215 | -1.748974754 | 0.131977995 | 0.638639366 | -4.255123121 |
| **A_55_P2159994** | Atf1 | -0.155046392 | -0.373236032 | 0.063143247 | 10.99953723 | -1.748222496 | 0.132114264 | 0.638639366 | -4.255675192 |
| **A_51_P381873** | Bub3 | 0.128322016 | -0.052263037 | 0.308907069 | 11.00164435 | 1.748189611 | 0.132120224 | 0.638639366 | -4.255699325 |
| **A_52_P344376** | Eif4a2 | -0.166373675 | -0.400510509 | 0.067763158 | 9.401803622 | -1.748171489 | 0.132123509 | 0.638639366 | -4.255712625 |
| **A_55_P1983868** | NA | -0.128781337 | -0.310025441 | 0.052462766 | 9.06419448 | -1.748067535 | 0.132142352 | 0.638639366 | -4.255788916 |

| | | | | | | | | | |
|---|---|---|---|---|---|---|---|---|---|
| **A_55_P1959338** | Spindoc | -0.130544864 | -0.314284368 | 0.05319464 | 11.34227102 | -1.747939565 | 0.132165552 | 0.638639366 | -4.255882831 |
| **A_55_P2171912** | Gja3 | 0.228605409 | -0.093176888 | 0.550387706 | 8.40621315 | 1.747807175 | 0.132189557 | 0.638639366 | -4.25597999 |
| **A_51_P354744** | Cox15 | 0.203285384 | -0.082861607 | 0.489432375 | 9.932873048 | 1.74777746 | 0.132194946 | 0.638639366 | -4.256001797 |
| **A_55_P2048164** | Msantd3 | 0.112120041 | -0.045753985 | 0.269994068 | 8.893365562 | 1.747196399 | 0.13230036 | 0.638988921 | -4.256428229 |
| **A_55_P2073329** | NA | 0.132298776 | -0.054028931 | 0.318626483 | 10.13758225 | 1.746817558 | 0.132369133 | 0.639133369 | -4.256706254 |
| **A_55_P2169721** | Rps17 | -0.233879392 | -0.563348344 | 0.09558956 | 15.14971042 | -1.746411668 | 0.132442855 | 0.639133369 | -4.257004129 |
| **A_55_P1968355** | Tle1 | -0.111474673 | -0.268515134 | 0.045565787 | 10.84388252 | -1.746360141 | 0.132452217 | 0.639133369 | -4.257041944 |
| **A_51_P238734** | Mfsd11 | -0.10121375 | -0.243803724 | 0.041376223 | 9.954413801 | -1.746303372 | 0.132462532 | 0.639133369 | -4.257083606 |
| **A_55_P2044557** | Rpl29 | 0.112367974 | -0.045969741 | 0.270705688 | 16.71693955 | 1.745932064 | 0.132530017 | 0.639293525 | -4.257356101 |
| **A_55_P1987944** | Rai1 | -0.31739049 | -0.764670321 | 0.129889342 | 10.58093673 | -1.745756858 | 0.132561873 | 0.639293525 | -4.257484681 |
| **A_55_P2050628** | Cyp4a31 | 0.092876473 | -0.038101745 | 0.22385469 | 7.982250553 | 1.744519389 | 0.132787082 | 0.639975645 | -4.258392833 |
| **A_55_P2168914** | NA | 0.493837223 | -0.202604111 | 1.190278558 | 8.81739908 | 1.744489846 | 0.132792463 | 0.639975645 | -4.258414514 |
| **A_52_P579876** | Suds3 | -0.129429354 | -0.311978741 | 0.053120033 | 8.767182965 | -1.744301551 | 0.132826766 | 0.639975645 | -4.2585527 |
| **A_55_P2145617** | Krt80 | 0.083105471 | -0.034122941 | 0.200333884 | 8.091443402 | 1.744077974 | 0.132867507 | 0.639975645 | -4.258716778 |
| **A_51_P199866** | Pin1 | 0.157071974 | -0.064515969 | 0.378659916 | 10.97895204 | 1.743900649 | 0.132899828 | 0.639975645 | -4.258846912 |
| **A_55_P1989534** | Cklf | 0.135412268 | -0.055673031 | 0.326497567 | 10.96251494 | 1.743411471 | 0.132989032 | 0.639975645 | -4.259205907 |
| **A_52_P10017** | Tmem64 | -0.124554564 | -0.300373297 | 0.051264169 | 9.400102809 | -1.742864659 | 0.133088815 | 0.639975645 | -4.259607197 |
| **A_55_P2114288** | Mrpl24 | 0.12411817 | -0.051089018 | 0.299325357 | 13.05658672 | 1.742820292 | 0.133096915 | 0.639975645 | -4.259639757 |
| **A_55_P2058152** | Ppp4r3b | -0.147170021 | -0.354933243 | 0.060593201 | 9.605680461 | -1.742688813 | 0.133120919 | 0.639975645 | -4.259736246 |
| **A_55_P1998081** | Trmt1 | 0.216555129 | -0.08923111 | 0.522341367 | 12.62645495 | 1.742287016 | 0.133194304 | 0.639975645 | -4.260031113 |
| **A_52_P641477** | Wars2 | -0.093092877 | -0.224551704 | 0.03836595 | 8.180671402 | -1.742191385 | 0.133211776 | 0.639975645 | -4.260101293 |
| **A_51_P198775** | BC055324 | -0.183617483 | -0.442954371 | 0.075719406 | 9.892220879 | -1.741882662 | 0.133268196 | 0.639975645 | -4.260327855 |
| **A_55_P2144771** | NA | 0.096573079 | -0.039827919 | 0.232974076 | 8.505172353 | 1.741837731 | 0.133276409 | 0.639975645 | -4.260360828 |
| **A_55_P2091858** | Hipk2 | -0.239937751 | -0.57887467 | 0.098999169 | 9.051076199 | -1.741601815 | 0.133319541 | 0.639975645 | -4.26053396 |
| **A_55_P2108451** | Galns | 0.332790435 | -0.137318036 | 0.802898906 | 13.28372753 | 1.741573956 | 0.133324636 | 0.639975645 | -4.260554405 |
| **A_55_P1958678** | Brd2 | 0.180127223 | -0.074328835 | 0.434583281 | 13.0387155 | 1.741549094 | 0.133329182 | 0.639975645 | -4.26057265 |
| **A_52_P384901** | Wdr43 | -0.080639179 | -0.194554999 | 0.033276641 | 8.958497948 | -1.741531046 | 0.133332483 | 0.639975645 | -4.260585895 |
| **A_51_P185135** | Gpr108 | 0.121468594 | -0.050145602 | 0.293082791 | 11.81773648 | 1.741325489 | 0.133370079 | 0.639975645 | -4.260736746 |
| **A_55_P2025730** | Hif3a | 0.122543897 | -0.050595092 | 0.295682885 | 7.942056555 | 1.741269422 | 0.133380336 | 0.639975645 | -4.260777892 |
| **A_51_P153004** | Phax | -0.135718819 | -0.327482126 | 0.056044488 | 9.992351483 | -1.741180222 | 0.133396655 | 0.639975645 | -4.260843352 |
| **A_55_P2024709** | 4933427D14Rik | -0.161325262 | -0.389281336 | 0.066630811 | 10.00347078 | -1.741086611 | 0.133413784 | 0.639975645 | -4.260912051 |
| **A_66_P120305** | Zmynd11 | -0.147868381 | -0.356821358 | 0.061084596 | 10.14755938 | -1.740988558 | 0.133431727 | 0.639975645 | -4.260984008 |
| **A_55_P2024868** | Psg18 | -0.227216397 | -0.548409333 | 0.093976538 | 17.23563731 | -1.740375049 | 0.133544051 | 0.640355486 | -4.26143424 |
| **A_55_P2121252** | Amd2 | 0.093342155 | -0.038620883 | 0.225305193 | 9.861621136 | 1.740182032 | 0.133579409 | 0.640366169 | -4.261575887 |
| **A_55_P2041457** | NA | 0.318770656 | -0.132100337 | 0.769641649 | 10.81394895 | 1.739382928 | 0.13372589 | 0.640909428 | -4.262162317 |
| **A_55_P2033075** | NA | 0.179512738 | -0.074430487 | 0.433455962 | 16.3313288 | 1.739113007 | 0.133775404 | 0.640987797 | -4.262360401 |
| **A_55_P1972118** | NA | -0.246622311 | -0.595545358 | 0.102300736 | 14.73060926 | -1.73888904 | 0.133816501 | 0.640989661 | -4.26252476 |
| **A_55_P2104761** | NA | 0.218382551 | -0.090659866 | 0.527424969 | 15.66612242 | 1.738477227 | 0.133892101 | 0.640989661 | -4.262826971 |
| **A_55_P2113439** | Caln1 | 0.115461262 | -0.047935519 | 0.278858042 | 8.127703391 | 1.738449053 | 0.133897274 | 0.640989661 | -4.262847647 |
| **A_55_P2051727** | Rnf8 | 0.12965223 | -0.053833535 | 0.313137996 | 11.41144182 | 1.738388247 | 0.133908441 | 0.640989661 | -4.262892269 |
| **A_51_P175758** | Atg101 | -0.093240269 | -0.225221165 | 0.038740626 | 11.51114772 | -1.738047379 | 0.133971055 | 0.641045848 | -4.263142416 |
| **A_51_P285413** | Rbbp6 | -0.112804057 | -0.272485073 | 0.046876959 | 8.053921611 | -1.737963277 | 0.133986509 | 0.641045848 | -4.263204134 |
| **A_51_P120717** | Lmnb1 | 0.186842495 | -0.077712376 | 0.451397366 | 9.696536304 | 1.737516986 | 0.134068541 | 0.641158991 | -4.264270887 |
| **A_52_P413947** | Mthfr | -0.157895035 | -0.381483305 | 0.065693235 | 10.72110277 | -1.737355208 | 0.13409829 | 0.641158991 | -4.263650365 |
| **A_55_P2029581** | NA | 0.222301256 | -0.092521278 | 0.537123789 | 9.374846276 | 1.737181794 | 0.134130185 | 0.641158991 | -4.263777624 |
| **A_66_P100349** | Esyt2 | -0.286358288 | -0.691984319 | 0.119267743 | 11.6589103 | -1.736813168 | 0.13419801 | 0.641158991 | -4.264048139 |
| **A_55_P1959056** | Pom121 | -0.252343598 | -0.609803272 | 0.105116076 | 12.31189717 | -1.736738371 | 0.134211776 | 0.641158991 | -4.264103027 |
| **A_52_P239707** | Nfyb | -0.094008807 | -0.227189755 | 0.03917214 | 8.503232435 | -1.736583246 | 0.134240331 | 0.641158991 | -4.264216865 |
| **A_51_P504815** | Stfa3 | -0.390937363 | -0.944785979 | 0.162911252 | 8.760619333 | -1.736541906 | 0.134247942 | 0.641158991 | -4.264247202 |
| **A_55_P2044045** | Ybx1 | 0.178845256 | -0.074569857 | 0.432260368 | 12.64896981 | 1.736257267 | 0.134300355 | 0.641158991 | -4.264456082 |

| | | | | | | | | | |
|---|---|---|---|---|---|---|---|---|---|
| **A_55_P2051899** | Atp2c1 | -0.158876644 | -0.38404468 | 0.066291392 | 10.17749704 | -1.73589112 | 0.134367808 | 0.641158991 | -4.264724776 |
| **A_55_P2174501** | Stx17 | -0.082935578 | -0.200476106 | 0.03460495 | 8.370846033 | -1.735890803 | 0.134367866 | 0.641158991 | -4.264725008 |
| **A_55_P2158319** | Ntan1 | -0.083027399 | -0.200724711 | 0.034669912 | 10.6290157 | -1.735497751 | 0.134440312 | 0.641158991 | -4.265013445 |
| **A_55_P2174401** | NA | 0.247230007 | -0.103244959 | 0.597704972 | 8.625749935 | 1.73545494 | 0.134448205 | 0.641158991 | -4.265044861 |
| **A_55_P2166414** | Uqcr10 | 0.258378002 | -0.107950561 | 0.624706566 | 14.43198647 | 1.735217487 | 0.134491992 | 0.641158991 | -4.265219112 |
| **A_52_P220783** | Tmem229b | 0.135895166 | -0.056798582 | 0.328588914 | 11.48331931 | 1.735024239 | 0.134527638 | 0.641158991 | -4.265360925 |
| **A_52_P460836** | Lrrn4 | -0.441446308 | -1.067411966 | 0.18451935 | 9.061913112 | -1.734988276 | 0.134534273 | 0.641158991 | -4.265387315 |
| **A_55_P2029471** | NA | 0.158047542 | -0.066080521 | 0.382175606 | 12.06472721 | 1.734844967 | 0.134560714 | 0.641158991 | -4.265492481 |
| **A_51_P113784** | Prpf19 | -0.180619048 | -0.436766564 | 0.075528468 | 8.536292605 | -1.734772639 | 0.134574061 | 0.641158991 | -4.265877103 |
| **A_52_P423174** | Acaa1b | -0.26321708 | -0.636558697 | 0.110124536 | 10.17094968 | -1.734510074 | 0.134622524 | 0.641170299 | -4.265738235 |
| **A_51_P398878** | Uck2 | 0.227819025 | -0.095380406 | 0.551018456 | 10.86019096 | 1.734157605 | 0.134687609 | 0.641170299 | -4.265996888 |
| **A_55_P2280821** | Dlc1 | -0.180424105 | -0.436413588 | 0.075565378 | 9.813694892 | -1.733970082 | 0.134722248 | 0.641170299 | -4.266134497 |
| **A_51_P461884** | Abl2 | 0.144695593 | -0.060625849 | 0.350017034 | 10.66038849 | 1.733764723 | 0.134760191 | 0.641170299 | -4.266285195 |
| **A_55_P1959228** | Slc4a4 | -0.124333679 | -0.300775015 | 0.052107656 | 8.78371246 | -1.733634782 | 0.134784206 | 0.641170299 | -4.266380549 |
| **A_55_P2404434** | NA | -0.113227988 | -0.273913771 | 0.047457795 | 8.095445285 | -1.733586435 | 0.134793142 | 0.641170299 | -4.266416027 |
| **A_55_P1974407** | Cdk14 | -0.211900984 | -0.512635225 | 0.088833257 | 9.202097938 | -1.733481688 | 0.134812504 | 0.641170299 | -4.266492893 |
| **A_55_P2060608** | NA | 0.11243622 | -0.047160485 | 0.272032925 | 13.32276848 | 1.733211165 | 0.134862523 | 0.641170299 | -4.266691409 |
| **A_55_P2168587** | Cdk9 | -0.131693561 | -0.318632211 | 0.055245088 | 8.691891862 | -1.733143824 | 0.134874977 | 0.641170299 | -4.266740825 |
| **A_52_P525070** | Vamp4 | -0.1063187 | -0.257256919 | 0.044619519 | 8.641659509 | -1.732924334 | 0.134915577 | 0.641205605 | -4.266901891 |
| **A_51_P189733** | Sult2a8 | 0.082668927 | -0.034711953 | 0.200049808 | 9.266578719 | 1.732663 | 0.134963933 | 0.641277744 | -4.267093663 |
| **A_66_P139460** | Mia3 | -0.103879504 | -0.251440075 | 0.043681067 | 8.847202471 | -1.731923493 | 0.135100859 | 0.641736833 | -4.267636325 |
| **A_55_P2320283** | Hmga2-ps1 | 0.087678475 | -0.036881105 | 0.212238054 | 8.60552977 | 1.731749284 | 0.135133135 | 0.641736833 | -4.267764162 |
| **A_51_P314019** | Banf1 | -0.117275962 | -0.283909791 | 0.049357866 | 10.21525815 | -1.731470099 | 0.135184876 | 0.641736833 | -4.267969031 |
| **A_66_P139799** | Cenpb | -0.148872911 | -0.360418924 | 0.062673101 | 8.48641323 | -1.731330584 | 0.135210739 | 0.641736833 | -4.268071408 |
| **A_66_P134181** | Aym1 | 0.092163606 | -0.038805849 | 0.223133061 | 8.509812021 | 1.731245272 | 0.135226557 | 0.641736833 | -4.268134011 |
| **A_55_P2030225** | NA | 0.119899218 | -0.050558516 | 0.290356953 | 15.3515933 | 1.730488843 | 0.135366885 | 0.642090628 | -4.268689082 |
| **A_51_P190886** | Ankrd13a | -0.136952646 | -0.331668462 | 0.05776317 | 11.14848675 | -1.730367553 | 0.135389399 | 0.642090628 | -4.268778085 |
| **A_55_P2144285** | Nnt | 0.275419633 | -0.116220877 | 0.667060143 | 9.510926962 | 1.730120892 | 0.135435196 | 0.642090628 | -4.268959085 |
| **A_52_P269016** | Nek9 | -0.230558769 | -0.558430534 | 0.097312997 | 11.94833446 | -1.730002729 | 0.13545714 | 0.642090628 | -4.269045793 |
| **A_52_P14453** | Atxn1 | -0.081592713 | -0.197631976 | 0.03444655 | 8.181193379 | -1.729878405 | 0.135480233 | 0.642090628 | -4.269137021 |
| **A_51_P445662** | Hsd17b4 | -0.247644087 | -0.599935405 | 0.104647231 | 12.92618583 | -1.729398926 | 0.135569329 | 0.642090628 | -4.269488861 |
| **A_51_P323880** | COX2 | 0.156762918 | -0.066279366 | 0.379805202 | 17.23896835 | 1.729120657 | 0.135621063 | 0.642090628 | -4.269693053 |
| **A_66_P116311** | Kif5b | -0.187269564 | -0.453744086 | 0.079204958 | 9.291710296 | -1.728942983 | 0.135654105 | 0.642090628 | -4.269823428 |
| **A_55_P2091247** | NA | -0.199547386 | -0.483508185 | 0.084413412 | 12.55048889 | -1.728848026 | 0.135671767 | 0.642090628 | -4.269893106 |
| **A_55_P2027653** | Arhgap25 | -0.08895506 | -0.215549412 | 0.037639293 | 8.503082067 | -1.728723305 | 0.135694969 | 0.642090628 | -4.269984626 |
| **A_55_P2023707** | Camta1 | -0.160131066 | -0.388036592 | 0.067774459 | 8.452172502 | -1.728581672 | 0.135721322 | 0.642090628 | -4.270088554 |
| **A_55_P2067700** | Tas2r113 | 0.87006813 | -0.368379122 | 2.108515381 | 13.33256227 | 1.728402597 | 0.135754649 | 0.642090628 | -4.270219956 |
| **A_55_P1978451** | Ttc1 | 0.123303042 | -0.05220808 | 0.298814163 | 10.54899963 | 1.72837634 | 0.135759536 | 0.642090628 | -4.270239223 |
| **A_55_P2180477** | Rpl31 | 0.19306451 | -0.081751645 | 0.467880666 | 16.15816755 | 1.728340675 | 0.135766175 | 0.642090628 | -4.270265394 |
| **A_52_P283524** | Rcan3 | -0.089859304 | -0.217786079 | 0.038067471 | 8.662097269 | -1.728107522 | 0.135809581 | 0.642138796 | -4.270436478 |
| **A_55_P2186439** | Chtop | -0.157255622 | -0.38117073 | 0.066659486 | 10.36556709 | -1.727793942 | 0.135867982 | 0.642222279 | -4.270666577 |
| **A_52_P571746** | Sh2d5 | -0.120242574 | -0.291500748 | 0.0510156 | 8.376546031 | -1.727333197 | 0.135953836 | 0.642222279 | -4.271004661 |
| **A_55_P2040048** | Swi5 | -0.121349263 | -0.294190687 | 0.051492161 | 13.55831673 | -1.727263001 | 0.135966921 | 0.642222279 | -4.271056169 |
| **A_52_P42380** | Tmem106c | -0.081938344 | -0.198652611 | 0.034775923 | 8.418734383 | -1.727159322 | 0.135986249 | 0.642222279 | -4.271132246 |
| **A_51_P284966** | Dolk | 0.158437974 | -0.067248558 | 0.384124506 | 11.30717664 | 1.727121145 | 0.135993367 | 0.642222279 | -4.27116026 |
| **A_55_P2067086** | Togaram1 | 0.103663882 | -0.044084836 | 0.251412599 | 11.1607031 | 1.726127656 | 0.136178723 | 0.642693855 | -4.271889251 |
| **A_51_P424878** | Prxl2b | 0.17608985 | -0.074886201 | 0.427065901 | 9.662534222 | 1.726120873 | 0.13617999 | 0.642693855 | -4.271894229 |
| **A_51_P520936** | Bcar3 | -0.209058158 | -0.507035045 | 0.088918729 | 12.7618825 | -1.72605132 | 0.136192976 | 0.642693855 | -4.271945264 |
| **A_55_P2165299** | Mybpc1 | 0.131482628 | -0.055954584 | 0.31891984 | 7.89915843 | 1.725765266 | 0.136246396 | 0.642789016 | -4.27215516 |
| **A_51_P110035** | Pik3r4 | -0.116644963 | -0.282996166 | 0.04970624 | 9.620779228 | -1.725079866 | 0.136374479 | 0.642945466 | -4.272934531 |

| | | | | | | | | |
|---|---|---|---|---|---|---|---|---|
| **A_51_P427516** | Thsd1 | -0.098904174 | -0.239958065 | 0.042149717 | 8.665441151 | -1.725038178 | 0.136382273 | 0.642945466 | -4.272688667 |
| **A_52_P505501** | Nsmce2 | -0.091190465 | -0.221253971 | 0.03887304 | 8.465267635 | -1.724896895 | 0.136408691 | 0.642945466 | -4.272792334 |
| **A_52_P652316** | Mapre2 | -0.099637896 | -0.241765816 | 0.042490025 | 10.8729744 | -1.724702957 | 0.136444962 | 0.642945466 | -4.272934636 |
| **A_55_P2042071** | Slc25a3 | -0.179864045 | -0.436433437 | 0.076705347 | 13.64435916 | -1.724680584 | 0.136449147 | 0.642945466 | -4.272951052 |
| **A_51_P348456** | Rest | -0.167329563 | -0.40604117 | 0.071382043 | 9.242381291 | -1.724520277 | 0.136479137 | 0.642945466 | -4.273068678 |
| **A_55_P2036803** | Dhx30 | -0.161647121 | -0.392419043 | 0.069124801 | 9.695918361 | -1.723273346 | 0.136712631 | 0.643309624 | -4.273983604 |
| **A_55_P2060255** | NA | -0.198425914 | -0.514352106 | 0.117500278 | 13.2548693 | -2.051378192 | 0.136732956 | 0.643309624 | -4.293068847 |
| **A_55_P2165544** | Aebp2 | -0.23040817 | -0.559394244 | 0.098577904 | 9.53918322 | -1.723016846 | 0.13676071 | 0.643309624 | -4.274171807 |
| **A_55_P1998384** | Wsb2 | -0.268836814 | -0.65275656 | 0.115082931 | 11.89577711 | -1.722730943 | 0.13681432 | 0.643309624 | -4.274381584 |
| **A_52_P510877** | Bcl2l1 | -0.086118019 | -0.209122238 | 0.0368862 | 8.293807766 | -1.722436359 | 0.136869579 | 0.643309624 | -4.274597729 |
| **A_55_P2037777** | NA | -0.183649932 | -0.445968062 | 0.078668198 | 9.571341854 | -1.722390526 | 0.136878178 | 0.643309624 | -4.274631358 |
| **A_55_P2129234** | Calm2 | -0.15935638 | -0.386976603 | 0.068263843 | 15.40531573 | -1.722375239 | 0.136881047 | 0.643309624 | -4.274642574 |
| **A_55_P2103412** | Golt1a | 0.310234626 | -0.13293477 | 0.753404022 | 10.08977168 | 1.72222413 | 0.136909403 | 0.643309624 | -4.274753447 |
| **A_51_P161691** | Coq7 | -0.255547131 | -0.6206458 | 0.109551537 | 10.39346995 | -1.721987221 | 0.136953872 | 0.643309624 | -4.274927273 |
| **A_55_P2151091** | Phactr4 | 0.159592572 | -0.068461803 | 0.387646946 | 12.31459752 | 1.721644295 | 0.137018265 | 0.643309624 | -4.275178885 |
| **A_55_P2066007** | Crocc | 0.197505425 | -0.084777707 | 0.479788557 | 8.99944496 | 1.721326538 | 0.137077959 | 0.643309624 | -4.275412029 |
| **A_52_P474765** | Denr | -0.12573684 | -0.305451094 | 0.053977414 | 9.880769045 | -1.721270713 | 0.137088449 | 0.643309624 | -4.275452989 |
| **A_55_P2126087** | NA | 0.22551611 | -0.096859395 | 0.547891614 | 14.23994987 | 1.721015172 | 0.137136477 | 0.643309624 | -4.275640483 |
| **A_51_P137611** | Card14 | 0.084990704 | -0.036506763 | 0.206488171 | 8.492458071 | 1.720970259 | 0.13714492 | 0.643309624 | -4.275673437 |
| **A_55_P2159026** | Rpl22l1 | 0.227239117 | -0.097611053 | 0.552089287 | 14.83185429 | 1.720953588 | 0.137148054 | 0.643309624 | -4.275685668 |
| **A_55_P2236291** | Plpp1 | -0.215192774 | -0.522843492 | 0.092457945 | 9.138081615 | -1.720833742 | 0.137170586 | 0.643309624 | -4.2757736 |
| **A_51_P507280** | Patl1 | -0.123953305 | -0.301182283 | 0.053275672 | 9.392526414 | -1.720650019 | 0.137205135 | 0.643309624 | -4.275908399 |
| **A_51_P485594** | Tab3 | -0.129331396 | -0.314272113 | 0.055609322 | 11.11585694 | -1.720444188 | 0.137243851 | 0.643309624 | -4.276059419 |
| **A_55_P2115192** | Sike1 | -0.174048251 | -0.42294508 | 0.074848577 | 9.764553822 | -1.72036051 | 0.137259593 | 0.643309624 | -4.276120814 |
| **A_55_P1966998** | Arhgdib | -0.192829058 | -0.468600772 | 0.082942656 | 10.43258346 | -1.720251415 | 0.13728012 | 0.643309624 | -4.276200858 |
| **A_51_P290685** | Ppp6r3 | -0.115025384 | -0.279540382 | 0.049489615 | 8.659425503 | -1.720114497 | 0.137305886 | 0.643309624 | -4.276301315 |
| **A_55_P2121344** | Nudcd3 | -0.16922385 | -0.411272349 | 0.072824648 | 12.33379873 | -1.719999587 | 0.137327515 | 0.643309624 | -4.276385625 |
| **A_55_P1972997** | A530016L24Rik | 0.101205673 | -0.043553396 | 0.245964741 | 7.840598222 | 1.719999173 | 0.137327593 | 0.643309624 | -4.276385928 |
| **A_51_P260169** | Gstm5 | 0.194583479 | -0.083767156 | 0.472934114 | 9.89957585 | 1.719819662 | 0.137361387 | 0.643309624 | -4.276517636 |
| **A_52_P559498** | Clk3 | -0.168831445 | -0.410463148 | 0.072800258 | 9.577589893 | -1.718971141 | 0.137521237 | 0.643309624 | -4.277140191 |
| **A_51_P275591** | Zfp292 | -0.148185308 | -0.36030887 | 0.063938254 | 9.00889026 | -1.718641914 | 0.137583307 | 0.643309624 | -4.27738174 |
| **A_55_P2452259** | Gls | -0.130619935 | -0.317611104 | 0.056371235 | 9.42573184 | -1.718531622 | 0.137604107 | 0.643309624 | -4.27746266 |
| **A_55_P1984981** | NA | -0.074744823 | -0.181747524 | 0.032257878 | 7.812145566 | -1.718523752 | 0.137605592 | 0.643309624 | -4.277468434 |
| **A_55_P1975185** | Sqle | 0.292685095 | -0.126330879 | 0.71170107 | 12.52918092 | 1.718459058 | 0.137617794 | 0.643309624 | -4.277515899 |
| **A_51_P219325** | Ddx28 | -0.100403304 | -0.244169274 | 0.043362667 | 8.779348201 | -1.718149958 | 0.137676109 | 0.643309624 | -4.27774268 |
| **A_55_P2103561** | Prdx6 | -0.13938358 | -0.33896815 | 0.06020099 | 13.37010312 | -1.718121273 | 0.137681522 | 0.643309624 | -4.277763725 |
| **A_51_P132459** | Rpl27a | 0.127287097 | -0.054985309 | 0.309559503 | 10.92745422 | 1.718037423 | 0.137697346 | 0.643309624 | -4.277825244 |
| **A_51_P306658** | Lsm8 | -0.103815014 | -0.252482982 | 0.044852955 | 10.66179623 | -1.717955544 | 0.1377128 | 0.643309624 | -4.277885317 |
| **A_51_P279854** | Ndufa11 | -0.098921757 | -0.240585252 | 0.042741738 | 10.66458504 | -1.717920377 | 0.137719438 | 0.643309624 | -4.277911118 |
| **A_55_P2078269** | NA | -0.18237522 | -0.443574217 | 0.078823777 | 11.29767014 | -1.717763962 | 0.137748966 | 0.643309624 | -4.278025877 |
| **A_51_P333949** | Ints7 | -0.090836167 | -0.220948387 | 0.039276053 | 8.454826019 | -1.717551921 | 0.137789004 | 0.643309624 | -4.278181445 |
| **A_55_P2084496** | Ssbp3 | -0.123237557 | -0.299780814 | 0.053305701 | 10.20418551 | -1.717359065 | 0.13782543 | 0.643309624 | -4.278322938 |
| **A_55_P2069755** | Tymp | 0.12039338 | -0.052078134 | 0.292864894 | 10.08575535 | 1.717332528 | 0.137830443 | 0.643309624 | -4.278342407 |
| **A_55_P2095933** | Zranb1 | -0.152241167 | -0.370352882 | 0.065870549 | 9.609718621 | -1.717205509 | 0.13785444 | 0.643309624 | -4.278435597 |
| **A_55_P2186480** | NA | -0.141408909 | -0.344052135 | 0.061234316 | 9.46379606 | -1.716776841 | 0.137935455 | 0.643532322 | -4.278750096 |
| **A_55_P2011727** | Neurl1b | -0.143453643 | -0.349060768 | 0.062153481 | 13.89344896 | -1.716495175 | 0.137988713 | 0.643536776 | -4.278956743 |
| **A_55_P1993589** | NA | 0.313491533 | -0.135924127 | 0.762907194 | 15.55135822 | 1.716116541 | 0.138060338 | 0.643536776 | -4.279234531 |
| **A_55_P2035315** | Rasgef1b | -0.182669615 | -0.44455026 | 0.07921103 | 11.76921899 | -1.716058444 | 0.138071331 | 0.643536776 | -4.279277153 |
| **A_51_P249313** | Herc3 | -0.176535472 | -0.429677227 | 0.076606283 | 10.2069105 | -1.715684234 | 0.138142161 | 0.643536776 | -4.279551693 |
| **A_51_P325856** | Mcemp1 | -0.115503264 | -0.281155944 | 0.050149416 | 8.235819238 | -1.71539821 | 0.138196322 | 0.643536776 | -4.279761533 |

| | | | | | | | | | |
|---|---|---|---|---|---|---|---|---|---|
| **A_52_P559817** | Fbln7 | -0.293527591 | -0.714521649 | 0.127466466 | 16.13095795 | -1.715308059 | 0.138213397 | 0.643536776 | -4.279827672 |
| **A_55_P2089310** | 2900097C17Rik | -0.079113336 | -0.192599733 | 0.03437306 | 8.93628959 | -1.715043177 | 0.13826358 | 0.643536776 | -4.280022 |
| **A_52_P177293** | Snx24 | 0.125974546 | -0.054734337 | 0.306683429 | 9.336028379 | 1.715032928 | 0.138265522 | 0.643536776 | -4.28002952 |
| **A_52_P528707** | Mrpl57 | 0.18642778 | -0.081010617 | 0.453866177 | 13.64783739 | 1.714968038 | 0.138277819 | 0.643536776 | -4.280077126 |
| **A_51_P220723** | B230118H07Rik | -0.120259446 | -0.292781149 | 0.052262257 | 11.92389568 | -1.714923005 | 0.138286353 | 0.643536776 | -4.280110164 |
| **A_55_P2053374** | Sorbs1 | -0.229678412 | -0.559216129 | 0.099859304 | 12.64930667 | -1.714684462 | 0.138331568 | 0.643536776 | -4.280285168 |
| **A_52_P523330** | Tmem147 | -0.144969603 | -0.352985874 | 0.063046669 | 11.06283953 | -1.714544671 | 0.138358072 | 0.643536776 | -4.280387723 |
| **A_51_P202633** | Ebi3 | -0.149008363 | -0.362851883 | 0.064835157 | 8.444032988 | -1.714287708 | 0.138406803 | 0.643536776 | -4.280576239 |
| **A_55_P2187225** | NA | 0.160717553 | -0.069975249 | 0.391410355 | 9.710617281 | 1.71395111 | 0.138470663 | 0.643536776 | -4.280823176 |
| **A_55_P2024245** | Mthfr | 0.162361426 | -0.070695866 | 0.395418717 | 9.997624667 | 1.713915179 | 0.138477481 | 0.643536776 | -4.280849537 |
| **A_55_P2046241** | Arl6ip6 | 0.099565103 | -0.04338546 | 0.242515665 | 8.02077355 | 1.713524984 | 0.138551549 | 0.643536776 | -4.281135792 |
| **A_51_P345316** | Cep76 | -0.085470277 | -0.208202615 | 0.037262061 | 8.10975543 | -1.71326785 | 0.13860038 | 0.643536776 | -4.28132443 |
| **A_52_P374292** | Tefm | -0.078614286 | -0.191519794 | 0.034291222 | 8.070777313 | -1.712992689 | 0.138652652 | 0.643536776 | -4.281526292 |
| **A_55_P2074197** | Zbtb24 | -0.102626738 | -0.250020769 | 0.044767293 | 9.054262269 | -1.71297014 | 0.138656937 | 0.643536776 | -4.281542834 |
| **A_55_P2178084** | Dnmt3a | -0.16354091 | -0.398440005 | 0.071358185 | 11.5525436 | -1.71282988 | 0.13868359 | 0.643536776 | -4.281645731 |
| **A_51_P349073** | Endog | -0.119022004 | -0.28998297 | 0.051938962 | 9.361641014 | -1.712771615 | 0.138694664 | 0.643536776 | -4.281688475 |
| **A_51_P130808** | Tpbpb | 0.215763321 | -0.094187504 | 0.525714145 | 17.65776101 | 1.71259229 | 0.138728751 | 0.643536776 | -4.281820029 |
| **A_55_P2102471** | Gm16378 | 0.475289327 | -0.207526003 | 1.158104657 | 14.70953778 | 1.712473905 | 0.138751259 | 0.643536776 | -4.281906877 |
| **A_52_P134083** | Cyhr1 | -0.136238718 | -0.332000139 | 0.059522704 | 11.27196095 | -1.712153116 | 0.138812266 | 0.643536776 | -4.282142208 |
| **A_65_P13402** | Rnf14 | -0.111672029 | -0.272140003 | 0.048795944 | 9.155743831 | -1.71208453 | 0.138825313 | 0.643536776 | -4.282192523 |
| **A_66_P105764** | Tor1b | -0.094352928 | -0.229938064 | 0.041232208 | 9.99452579 | -1.712034309 | 0.138834867 | 0.643536776 | -4.282229365 |
| **A_52_P58257** | Mrps6 | -0.25585527 | -0.623519772 | 0.111809233 | 13.67077904 | -1.712031805 | 0.138835344 | 0.643536776 | -4.282231202 |
| **A_55_P2184526** | NA | 0.249793768 | -0.109341678 | 0.608929215 | 8.391621489 | 1.711167372 | 0.138999898 | 0.643884076 | -4.282865343 |
| **A_52_P727235** | Trabd2b | -0.116791786 | -0.284706826 | 0.051123254 | 8.578006884 | -1.711164904 | 0.139000368 | 0.643884076 | -4.282867153 |
| **A_55_P2006236** | Ugdh | 0.194264924 | -0.085113138 | 0.473642986 | 11.52766458 | 1.710689761 | 0.139090897 | 0.643884076 | -4.28321571 |
| **A_55_P2127439** | Kri1 | -0.14941327 | -0.364314873 | 0.065488332 | 11.33600193 | -1.710482678 | 0.139130371 | 0.643884076 | -4.283367622 |
| **A_55_P2089565** | Tbx22 | -0.118109584 | -0.287987683 | 0.051768515 | 7.83652526 | -1.71047571 | 0.1391317 | 0.643884076 | -4.283372733 |
| **A_55_P2176145** | Rgs17 | 0.227037461 | -0.099570614 | 0.553645536 | 12.27200457 | 1.710171902 | 0.139189631 | 0.643884076 | -4.283595598 |
| **A_55_P2168185** | NA | -0.18381017 | -0.448235043 | 0.080614702 | 16.55690427 | -1.710158631 | 0.139192162 | 0.643884076 | -4.283605333 |
| **A_55_P2043682** | Ubac1 | -0.218749831 | -0.533455934 | 0.095956272 | 11.11544864 | -1.710061477 | 0.139210693 | 0.643884076 | -4.283676603 |
| **A_51_P426754** | Anxa5 | -0.130280739 | -0.31771699 | 0.057155512 | 14.48855447 | -1.709998729 | 0.139222663 | 0.643884076 | -4.283722633 |
| **A_52_P244723** | Tbcel | -0.098424179 | -0.240045884 | 0.043197527 | 9.456451652 | -1.709783579 | 0.139263713 | 0.643884076 | -4.28388046 |
| **A_55_P2135049** | NA | 0.132717917 | -0.058295193 | 0.323731028 | 12.84029121 | 1.709367905 | 0.139343055 | 0.643884076 | -4.284185382 |
| **A_52_P1029978** | Lasp1 | -0.277578585 | -0.677108337 | 0.121951167 | 15.23490555 | -1.709251599 | 0.139365263 | 0.643884076 | -4.2842707 |
| **A_55_P2166990** | Ppcs | 0.305181749 | -0.134084775 | 0.744448273 | 10.93346702 | 1.709226482 | 0.139370059 | 0.643884076 | -4.284289125 |
| **A_66_P111534** | 5430431A17Rik | 0.124440167 | -0.05467769 | 0.303558024 | 8.27920812 | 1.70919205 | 0.139376635 | 0.643884076 | -4.284314383 |
| **A_55_P2178023** | NA | 0.131522129 | -0.057838379 | 0.320882637 | 12.30077021 | 1.708750218 | 0.139461038 | 0.64394421 | -4.284638489 |
| **A_52_P123354** | Tusc2 | 0.126631181 | -0.055712218 | 0.308974581 | 10.12599338 | 1.708518847 | 0.139505257 | 0.64394421 | -4.284808211 |
| **A_55_P2023652** | Vps35l | -0.283792009 | -0.692490294 | 0.124906277 | 12.74448776 | -1.708309315 | 0.139545313 | 0.64394421 | -4.284961912 |
| **A_55_P2147591** | Med27 | -0.082483907 | -0.201277485 | 0.03630967 | 8.107733296 | -1.708226389 | 0.13956117 | 0.64394421 | -4.285022742 |
| **A_55_P1992319** | Col4a3bp | 0.125505121 | -0.055284153 | 0.306294396 | 8.668994278 | 1.707882341 | 0.139626974 | 0.64394421 | -4.285275114 |
| **A_51_P469285** | Nrp1 | -0.07931658 | -0.193583101 | 0.03494994 | 8.070525075 | -1.70771008 | 0.139659932 | 0.64394421 | -4.285401473 |
| **A_55_P2118843** | Hk1 | -0.1051628 | -0.256674268 | 0.046348668 | 8.398466542 | -1.707598628 | 0.13968126 | 0.64394421 | -4.285483227 |
| **A_55_P2082444** | Npepps | -0.176553339 | -0.430986306 | 0.077879627 | 11.71104107 | -1.707150141 | 0.139767118 | 0.64394421 | -4.285812205 |
| **A_51_P206225** | Uroc1 | -0.090463509 | -0.220842861 | 0.039915844 | 8.220300793 | -1.707000988 | 0.139795683 | 0.64394421 | -4.285921612 |
| **A_55_P2084338** | 2310009B15Rik | 0.153700515 | -0.067859319 | 0.375260349 | 9.279050776 | 1.706685329 | 0.139856155 | 0.64394421 | -4.286153154 |
| **A_51_P341010** | Ercc8 | 0.123245795 | -0.05441911 | 0.300910699 | 9.290952441 | 1.706630986 | 0.139866568 | 0.64394421 | -4.286193016 |
| **A_51_P333034** | Ier3ip1 | -0.23297352 | -0.5688483 | 0.102901261 | 12.13466634 | -1.706468704 | 0.139897669 | 0.64394421 | -4.286312052 |
| **A_55_P1998406** | NA | 0.091912461 | -0.040626842 | 0.224451764 | 8.570977815 | 1.706077965 | 0.139972581 | 0.64394421 | -4.286598663 |
| **A_55_P1988048** | Emcn | -0.083435431 | -0.203753262 | 0.036882399 | 8.574054734 | -1.706041868 | 0.139979504 | 0.64394421 | -4.28662514 |

| | | | | | | | | | |
|---|---|---|---|---|---|---|---|---|---|
| **A_55_P1965635** | Setdb1 | -0.084859053 | -0.207235911 | 0.037517806 | 9.222752213 | -1.705956851 | 0.139995809 | 0.64394421 | -4.286687501 |
| **A_55_P1992208** | Gpi1 | 0.214855446 | -0.094996938 | 0.52470783 | 13.6113153 | 1.705927962 | 0.14000135 | 0.64394421 | -4.286708691 |
| **A_66_P106661** | Slc7a1 | -0.1926526 | -0.470500261 | 0.085195062 | 15.82058028 | -1.705836061 | 0.140018979 | 0.64394421 | -4.286776101 |
| **A_52_P585124** | Cxcr4 | -0.11591037 | -0.283080542 | 0.051259803 | 8.038866407 | -1.705817883 | 0.140022466 | 0.64394421 | -4.286789434 |
| **A_55_P2081829** | Rps4x | 0.396562303 | -0.17541713 | 0.968541736 | 14.89930906 | 1.705690294 | 0.140046944 | 0.64394421 | -4.286883021 |
| **A_51_P370561** | Dcakd | -0.086001636 | -0.210064088 | 0.038060816 | 9.884775707 | -1.705436354 | 0.140095675 | 0.64394421 | -4.287069286 |
| **A_55_P2058928** | Ralgds | -0.24862955 | -0.60729875 | 0.110039649 | 13.14328354 | -1.705406149 | 0.140101473 | 0.64394421 | -4.287091441 |
| **A_55_P2047788** | Katnal1 | -0.078581555 | -0.191949417 | 0.034786308 | 8.347439504 | -1.705296191 | 0.14012258 | 0.64394421 | -4.287172094 |
| **A_55_P2058122** | Kyat3 | 0.098709101 | -0.043754292 | 0.241172494 | 9.804889261 | 1.704602324 | 0.140255843 | 0.643956899 | -4.287681037 |
| **A_52_P14666** | NA | -0.078044856 | -0.190691432 | 0.03460172 | 7.88634504 | -1.704493901 | 0.140276678 | 0.643956899 | -4.287760564 |
| **A_55_P1983968** | Slc8b1 | -0.109081595 | -0.266535944 | 0.048372754 | 9.756867848 | -1.704378247 | 0.140298905 | 0.643956899 | -4.287845394 |
| **A_55_P1979973** | Nuak2 | 0.180875108 | -0.08025344 | 0.442003656 | 8.780823943 | 1.704094254 | 0.1403535 | 0.643956899 | -4.288053696 |
| **A_55_P2004711** | Cep95 | 0.093433764 | -0.041460022 | 0.22832755 | 9.803386334 | 1.704045021 | 0.140362967 | 0.643956899 | -4.288089806 |
| **A_55_P2029902** | Gab3 | -0.781612261 | -1.910068234 | 0.346843712 | 10.81702719 | -1.704024784 | 0.140366859 | 0.643956899 | -4.28810465 |
| **A_55_P2087726** | Cops9 | 0.161519491 | -0.071679836 | 0.394718818 | 15.33680908 | 1.703989085 | 0.140373724 | 0.643956899 | -4.288130834 |
| **A_51_P435339** | Epor | 0.101650739 | -0.045163221 | 0.248464698 | 9.27798144 | 1.703383165 | 0.140490292 | 0.643956899 | -4.288575256 |
| **A_55_P2126557** | NA | 0.186290171 | -0.082776006 | 0.455356347 | 10.82068475 | 1.703334708 | 0.140499618 | 0.643956899 | -4.288610798 |
| **A_52_P426870** | Slf2 | -0.27622937 | -0.683237316 | 0.130778576 | 10.32036222 | -1.758802388 | 0.140509891 | 0.643956899 | -4.276041328 |
| **A_55_P2164502** | NA | 0.480128831 | -0.213362929 | 1.173620591 | 10.81075531 | 1.703278419 | 0.140510453 | 0.643956899 | -4.288652083 |
| **A_55_P1982747** | Slc8a3 | -0.689860641 | -1.686404441 | 0.306683159 | 11.15570399 | -1.703076433 | 0.140549338 | 0.643956899 | -4.288800231 |
| **A_55_P2269497** | Pdp1 | 0.079115906 | -0.035177807 | 0.193409618 | 8.244547111 | 1.702984233 | 0.140567091 | 0.643956899 | -4.288867856 |
| **A_55_P1992959** | Pfkfb4 | -0.170665034 | -0.417261909 | 0.07593184 | 9.89171958 | -1.702653 | 0.140630888 | 0.643956899 | -4.289110799 |
| **A_55_P2067747** | Psg21 | -0.200699492 | -0.49074398 | 0.089344995 | 16.99644958 | -1.702357709 | 0.140687786 | 0.643956899 | -4.289327379 |
| **A_55_P2037424** | NA | -0.232799085 | -0.569234867 | 0.103636697 | 13.98294227 | -1.702347641 | 0.140689727 | 0.643956899 | -4.289334764 |
| **A_55_P1973995** | NA | 0.540151924 | -0.240467686 | 1.320771533 | 11.3919252 | 1.702337372 | 0.140691706 | 0.643956899 | -4.289342295 |
| **A_55_P2115732** | Kif1c | 0.111746091 | -0.049804246 | 0.273296427 | 8.130607017 | 1.701741672 | 0.14080656 | 0.644330019 | -4.289779204 |
| **A_52_P500488** | Lars | -0.226687399 | -0.55446365 | 0.101088852 | 12.68016044 | -1.70144948 | 0.14086293 | 0.644345654 | -4.289993506 |
| **A_55_P1985385** | Acp2 | 0.125921162 | -0.056160821 | 0.308003145 | 8.630875458 | 1.701378388 | 0.140876648 | 0.644345654 | -4.290045647 |
| **A_55_P2062678** | Hmox2 | -0.088410163 | -0.216264564 | 0.039444237 | 9.759239746 | -1.701201186 | 0.140910848 | 0.644349604 | -4.290175611 |
| **A_55_P1990780** | Ptp4a1 | -0.173824874 | -0.425232134 | 0.077582386 | 13.30983116 | -1.70099592 | 0.140950474 | 0.644378361 | -4.290326157 |
| **A_55_P2094060** | Gzma | -0.118862431 | -0.290844809 | 0.053119946 | 8.072240063 | -1.700316712 | 0.141081671 | 0.644825636 | -4.290824296 |
| **A_55_P2015292** | Ltc4s | 0.360403853 | -0.16116664 | 0.881974346 | 10.15305864 | 1.699986898 | 0.141145421 | 0.644964502 | -4.291066183 |
| **A_51_P137800** | Acp7 | -0.165654314 | -0.40546086 | 0.074152232 | 9.34467973 | -1.699459733 | 0.141247376 | 0.64520791 | -4.291452804 |
| **A_55_P2118575** | Tgs1 | 0.119499388 | -0.053501303 | 0.292500079 | 9.406327074 | 1.69936632 | 0.14126545 | 0.64520791 | -4.291521312 |
| **A_55_P2121846** | NA | -0.118806129 | -0.290841657 | 0.053229399 | 15.92557512 | -1.698986249 | 0.14133901 | 0.645271091 | -4.29180005 |
| **A_51_P247359** | Ptprcap | 0.149517788 | -0.067130184 | 0.36616576 | 8.851354338 | 1.697881586 | 0.141553021 | 0.645271091 | -4.292610177 |
| **A_55_P1957762** | Tsr1 | -0.118521802 | -0.290266108 | 0.053222503 | 9.544775417 | -1.697794277 | 0.141569949 | 0.645271091 | -4.292674206 |
| **A_55_P2172989** | Ptpn23 | 0.162867478 | -0.073146085 | 0.398881041 | 11.77072599 | 1.69772199 | 0.141583966 | 0.645271091 | -4.292727218 |
| **A_55_P1977245** | Slc22a13 | 0.10801995 | -0.048557957 | 0.264597858 | 8.510065406 | 1.697237634 | 0.141677922 | 0.645271091 | -4.293082421 |
| **A_52_P309718** | Tasor | -0.090765979 | -0.222335141 | 0.040803184 | 8.608156897 | -1.697220027 | 0.141681339 | 0.645271091 | -4.293095333 |
| **A_52_P37681** | Fkbp9 | -0.180814303 | -0.442917776 | 0.081289171 | 12.80294308 | -1.697184933 | 0.141688149 | 0.645271091 | -4.293121069 |
| **A_55_P2078138** | Otop1 | 0.354371102 | -0.159335226 | 0.86807743 | 9.594469063 | 1.697119983 | 0.141700753 | 0.645271091 | -4.2931687 |
| **A_55_P2186619** | Rbmxl1 | -0.116576901 | -0.285578623 | 0.052424821 | 11.41196053 | -1.697033987 | 0.141717444 | 0.645271091 | -4.293231764 |
| **A_55_P2012789** | Impa1 | 0.153314318 | -0.068948156 | 0.375576793 | 9.913647066 | 1.697015207 | 0.141721089 | 0.645271091 | -4.293245537 |
| **A_55_P1986902** | Med23 | 0.14529087 | -0.06535177 | 0.355933511 | 9.192614421 | 1.696919333 | 0.1417397 | 0.645271091 | -4.293315845 |
| **A_55_P1992487** | NA | 0.179387202 | -0.080709224 | 0.439483628 | 10.20258486 | 1.696782724 | 0.141766221 | 0.645271091 | -4.293416026 |
| **A_55_P2152946** | NA | 0.102428165 | -0.046084387 | 0.250940717 | 9.064247151 | 1.696779443 | 0.141766858 | 0.645271091 | -4.293418432 |
| **A_55_P2000943** | Nrip3 | -0.172227322 | -0.421948689 | 0.077494045 | 9.064002879 | -1.696740822 | 0.141774357 | 0.645271091 | -4.293446755 |
| **A_52_P39247** | Hbp1 | -0.19016694 | -0.465904646 | 0.085570766 | 12.62433791 | -1.696711575 | 0.141780037 | 0.645271091 | -4.293468203 |
| **A_52_P558699** | Ppp4r3a | -0.124184165 | -0.304280879 | 0.055912549 | 9.22445996 | -1.6964052 | 0.14183954 | 0.645389942 | -4.293692877 |

| | | | | | | | | | |
|---|---|---|---|---|---|---|---|---|---|
| **A_55_P2127782** | NA | 0.25851955 | -0.116589489 | 0.633628589 | 14.35952594 | 1.695528228 | 0.14201 | 0.645698875 | -4.29433598 |
| **A_55_P1957708** | NA | 0.108064937 | -0.048742949 | 0.264872823 | 8.267462582 | 1.695454216 | 0.142024395 | 0.645698875 | -4.294390254 |
| **A_51_P346165** | Agpat4 | -0.253400648 | -0.621120366 | 0.114319069 | 13.19136552 | -1.695352328 | 0.142044214 | 0.645698875 | -4.29446497 |
| **A_66_P126313** | Rhof | -0.109938058 | -0.269478299 | 0.049602182 | 9.115475376 | -1.695301639 | 0.142054075 | 0.645698875 | -4.29450214 |
| **A_55_P2031501** | Hnrnpa3 | -0.226359558 | -0.561593888 | 0.108874772 | 14.43506375 | -1.74984843 | 0.142122424 | 0.645698875 | -4.281518985 |
| **A_52_P152283** | Gabpa | -0.074649039 | -0.183006393 | 0.033708315 | 9.147434653 | -1.694864543 | 0.142139134 | 0.645698875 | -4.294822665 |
| **A_55_P1954987** | Slc10a1 | -0.278875633 | -0.683682206 | 0.125930941 | 17.18317693 | -1.694853543 | 0.142141275 | 0.645698875 | -4.294830731 |
| **A_51_P440743** | Celsr1 | 0.180188799 | -0.081426916 | 0.441804513 | 8.787768585 | 1.694467041 | 0.142216532 | 0.645888943 | -4.295114151 |
| **A_55_P1993909** | Taz | 0.166493834 | -0.075387911 | 0.408375579 | 11.28424748 | 1.693418208 | 0.142420946 | 0.646220231 | -4.29588324 |
| **A_55_P2039479** | Twistnb | -0.08906942 | -0.21847607 | 0.040337229 | 8.163481918 | -1.693328418 | 0.142438459 | 0.646220231 | -4.29594908 |
| **A_52_P479961** | NA | -0.316533586 | -0.776462518 | 0.143395345 | 14.8315136 | -1.693160904 | 0.142471138 | 0.646220231 | -4.296071912 |
| **A_55_P2095360** | Tfpi | -0.140874606 | -0.345606743 | 0.06385753 | 13.22572056 | -1.692839794 | 0.1425338 | 0.646220231 | -4.296307369 |
| **A_55_P1975843** | NA | -0.180673965 | -0.443268183 | 0.081920253 | 9.580021645 | -1.692698381 | 0.142561404 | 0.646220231 | -4.296411061 |
| **A_52_P339054** | Nap1l4 | -0.101252795 | -0.248489502 | 0.045983913 | 9.232185205 | -1.691843095 | 0.142728468 | 0.646220231 | -4.297038195 |
| **A_52_P615401** | Tpra1 | 0.292518908 | -0.132864117 | 0.717901934 | 9.574697336 | 1.691776356 | 0.142741512 | 0.646220231 | -4.297087131 |
| **A_55_P2130256** | Polr2l | 0.245468538 | -0.111514705 | 0.602451781 | 9.297652852 | 1.691675921 | 0.142761145 | 0.646220231 | -4.297160773 |
| **A_55_P2182412** | Adat1 | 0.117009422 | -0.053180807 | 0.287199651 | 8.231096253 | 1.691435238 | 0.142808202 | 0.646220231 | -4.297337249 |
| **A_51_P487818** | Fabp1 | -0.209917298 | -0.515259299 | 0.095424704 | 7.815323953 | -1.691339696 | 0.142826887 | 0.646220231 | -4.297407303 |
| **A_55_P2058864** | Mvk | -0.101412803 | -0.248953523 | 0.046127916 | 8.872504844 | -1.691025105 | 0.142888425 | 0.646220231 | -4.297637968 |
| **A_66_P133112** | Gm14635 | 0.146190921 | -0.066532148 | 0.35891399 | 8.454593431 | 1.690732788 | 0.14294563 | 0.646220231 | -4.297852299 |
| **A_55_P2147220** | Ak2 | -0.262101324 | -0.643502595 | 0.119299947 | 11.00201382 | -1.690659859 | 0.142959905 | 0.646220231 | -4.297905771 |
| **A_55_P2043668** | Zfp617 | -0.095395594 | -0.234224593 | 0.043433406 | 9.376720711 | -1.690508211 | 0.142989593 | 0.646220231 | -4.298016961 |
| **A_52_P123485** | Tcf25 | -0.176102001 | -0.432418159 | 0.080214157 | 13.29443628 | -1.690275392 | 0.143035184 | 0.646220231 | -4.298187665 |
| **A_66_P123582** | Mog | 0.101877686 | -0.04641949 | 0.250174863 | 8.500862454 | 1.690111487 | 0.143067289 | 0.646220231 | -4.29830784 |
| **A_55_P2329680** | Dyrk2 | -0.120283537 | -0.295378397 | 0.054811323 | 9.759798257 | -1.690059352 | 0.143077502 | 0.646220231 | -4.298346065 |
| **A_55_P2142668** | NA | 0.810632257 | -0.369525783 | 1.990790296 | 12.61656955 | 1.689868362 | 0.143114923 | 0.646220231 | -4.298486098 |
| **A_66_P133450** | Dnajb9 | -0.149436435 | -0.367029032 | 0.068156162 | 8.742145203 | -1.689590838 | 0.143169316 | 0.646220231 | -4.298689575 |
| **A_51_P208410** | Ppp1r12c | -0.160856647 | -0.395098129 | 0.073384835 | 11.14347504 | -1.689446206 | 0.143197671 | 0.646220231 | -4.298795617 |
| **A_55_P2414924** | Luc7l | 0.130220323 | -0.05942119 | 0.319861836 | 9.018057107 | 1.689330097 | 0.143220437 | 0.646220231 | -4.298880746 |
| **A_55_P2002215** | Arfip1 | 0.223533313 | -0.102004455 | 0.549071081 | 10.40384031 | 1.689312657 | 0.143223857 | 0.646220231 | -4.298893533 |
| **A_55_P2069176** | NA | 0.168796457 | -0.077030053 | 0.414622967 | 12.03427077 | 1.6892883 | 0.143228634 | 0.646220231 | -4.298911391 |
| **A_55_P2122718** | Gpr19 | 0.109062981 | -0.049797047 | 0.267923008 | 9.798038974 | 1.689008725 | 0.143283471 | 0.646220231 | -4.299116369 |
| **A_55_P2068152** | Anapc15 | 0.098857559 | -0.045153805 | 0.242868924 | 11.41558567 | 1.688815796 | 0.143321325 | 0.646220231 | -4.299257819 |
| **A_55_P1953382** | Prl8a8 | -0.112660735 | -0.276784032 | 0.051462562 | 17.44703122 | -1.688773907 | 0.143329545 | 0.646220231 | -4.29928853 |
| **A_55_P2099061** | NA | -0.178242734 | -0.437963077 | 0.08147761 | 12.09258241 | -1.688398774 | 0.143403181 | 0.646220231 | -4.299563564 |
| **A_55_P1965079** | Gm10499 | 0.087208772 | -0.039872337 | 0.214289881 | 7.797435949 | 1.688295011 | 0.143423555 | 0.646220231 | -4.299639639 |
| **A_55_P2094706** | Sox17 | -0.106770091 | -0.262360791 | 0.048820609 | 9.855515788 | -1.688243714 | 0.143433628 | 0.646220231 | -4.299677247 |
| **A_55_P1978061** | Hnrnpa3 | -0.105997383 | -0.260466858 | 0.048472092 | 8.717272482 | -1.688191219 | 0.143443938 | 0.646220231 | -4.299715735 |
| **A_51_P189905** | Dad1 | -0.205529545 | -0.505048753 | 0.093989664 | 13.36387334 | -1.688179974 | 0.143446146 | 0.646220231 | -4.299723979 |
| **A_51_P416439** | Zdhhc6 | -0.122222901 | -0.300353382 | 0.05590758 | 10.35254128 | -1.68804302 | 0.143473046 | 0.646220231 | -4.299824387 |
| **A_52_P311104** | Brd7 | -0.108732377 | -0.267217546 | 0.049752793 | 12.67095893 | -1.687871632 | 0.143506716 | 0.646220231 | -4.29995004 |
| **A_51_P176265** | Trip11 | -0.124069685 | -0.304941212 | 0.056801842 | 8.507132139 | -1.68758104 | 0.143563822 | 0.646220231 | -4.300163085 |
| **A_52_P156485** | Ankrd39 | 0.159135984 | -0.072910729 | 0.391182697 | 10.61349689 | 1.687182788 | 0.143642121 | 0.646220231 | -4.300455058 |
| **A_55_P2045224** | NA | 0.194561998 | -0.08914238 | 0.478266376 | 12.23917759 | 1.68717893 | 0.14364288 | 0.646220231 | -4.300457886 |
| **A_55_P1964509** | Lcn12 | 0.126345737 | -0.057898136 | 0.31058961 | 9.681864057 | 1.687083981 | 0.143661553 | 0.646220231 | -4.300527497 |
| **A_55_P2011525** | Rnf214 | -0.090194669 | -0.221721516 | 0.041332178 | 8.650550878 | -1.687079784 | 0.143662379 | 0.646220231 | -4.300530574 |
| **A_55_P2108845** | NA | 0.079619461 | -0.036489935 | 0.195728858 | 8.440677924 | 1.687023101 | 0.143673528 | 0.646220231 | -4.30057213 |
| **A_51_P316949** | Ccdc6 | -0.242207151 | -0.595423774 | 0.111009472 | 10.47877693 | -1.686999642 | 0.143678142 | 0.646220231 | -4.300589328 |
| **A_55_P2131579** | NA | 0.101270021 | -0.046417893 | 0.248957935 | 13.76863866 | 1.686961244 | 0.143685696 | 0.646220231 | -4.300617479 |
| **A_55_P2094611** | NA | 0.136750575 | -0.062724867 | 0.336226018 | 8.787924446 | 1.68658754 | 0.143759228 | 0.646220231 | -4.300891449 |

| | | | | | | | | | |
|---|---|---|---|---|---|---|---|---|---|
| **A_52_P4846** | Zfp451 | -0.139518734 | -0.343045167 | 0.064007699 | 9.659828264 | -1.686478729 | 0.143780645 | 0.646220231 | -4.30097122 |
| **A_66_P136372** | Mrpl9 | -0.110030269 | -0.270548229 | 0.050487692 | 9.588414641 | -1.686388779 | 0.143798352 | 0.646220231 | -4.301037163 |
| **A_55_P1955147** | Camk1d | -0.127994675 | -0.314780169 | 0.05879082 | 8.45777566 | -1.685846068 | 0.143905232 | 0.646220231 | -4.301435028 |
| **A_52_P597860** | Wasf2 | -0.122559418 | -0.301417812 | 0.056298977 | 9.550992025 | -1.685801843 | 0.143913945 | 0.646220231 | -4.30146745 |
| **A_66_P121656** | Pcgf1 | -0.087547961 | -0.215318617 | 0.040222694 | 8.495141843 | -1.685714711 | 0.143931112 | 0.646220231 | -4.301531326 |
| **A_55_P1989405** | Elf1 | -0.21823809 | -0.536776839 | 0.100300658 | 11.31487167 | -1.685533725 | 0.143966778 | 0.646220231 | -4.301664005 |
| **A_55_P1958102** | NA | 0.12914352 | -0.059413314 | 0.317700354 | 15.20970465 | 1.684998449 | 0.144072314 | 0.646220231 | -4.302056409 |
| **A_55_P2176838** | Mitf | -0.287056287 | -0.706204021 | 0.132091446 | 13.94387946 | -1.684880503 | 0.144095578 | 0.646220231 | -4.302142872 |
| **A_55_P2149020** | Rpl35 | 0.087707597 | -0.040363625 | 0.215778818 | 17.03478988 | 1.684825099 | 0.144106507 | 0.646220231 | -4.302183487 |
| **A_51_P399175** | Ppm1j | -0.077330172 | -0.190254583 | 0.035594238 | 7.859953767 | -1.684729992 | 0.144125271 | 0.646220231 | -4.302253208 |
| **A_55_P2041470** | NA | 0.197378807 | -0.090869171 | 0.485626786 | 8.515752029 | 1.684625666 | 0.144145856 | 0.646220231 | -4.302329687 |
| **A_55_P2008619** | Lancl2 | -0.18349361 | -0.45146725 | 0.08448003 | 10.14876198 | -1.684604798 | 0.144149974 | 0.646220231 | -4.302344984 |
| **A_66_P106375** | Strap | -0.115481389 | -0.284142245 | 0.053179467 | 9.926945601 | -1.684483886 | 0.144173835 | 0.646220231 | -4.30243362 |
| **A_66_P114501** | Zcrb1 | 0.13486351 | -0.062119231 | 0.331846251 | 13.08757874 | 1.684362041 | 0.144197885 | 0.646220231 | -4.302522941 |
| **A_51_P243304** | Galk1 | 0.141271574 | -0.065087236 | 0.347630384 | 11.78183777 | 1.684228195 | 0.144224309 | 0.646220231 | -4.302621059 |
| **A_55_P2030859** | NA | -0.171847853 | -0.422873393 | 0.079177687 | 14.99730053 | -1.684206599 | 0.144228572 | 0.646220231 | -4.302636889 |
| **A_51_P323620** | Thyn1 | 0.151777351 | -0.069954298 | 0.373508999 | 13.22285903 | 1.684024655 | 0.144264499 | 0.646231404 | -4.302770265 |
| **A_52_P227065** | Bud31 | -0.143713355 | -0.353686338 | 0.066259629 | 10.83798667 | -1.683847941 | 0.144299402 | 0.646237985 | -4.302899806 |
| **A_51_P234174** | Pms2 | 0.131982466 | -0.060892095 | 0.324857026 | 8.995936327 | 1.68348965 | 0.144370193 | 0.646405247 | -4.30316245 |
| **A_55_P1961760** | NA | -0.575778246 | -1.435183819 | 0.283627327 | 11.36291582 | -1.736229651 | 0.14461117 | 0.647235044 | -4.289859959 |
| **A_51_P339540** | Cdkn1c | -0.123173372 | -0.303316324 | 0.05696958 | 16.21922131 | -1.682165484 | 0.144632112 | 0.647235044 | -4.304133097 |
| **A_55_P2174506** | Stx17 | -0.084124201 | -0.207165955 | 0.038917554 | 8.303434505 | -1.682044948 | 0.144655978 | 0.647235044 | -4.304221452 |
| **A_51_P397768** | Csnk2a1 | -0.090498579 | -0.222935363 | 0.041938206 | 8.527567811 | -1.681133857 | 0.144836488 | 0.64767305 | -4.304889277 |
| **A_55_P2009091** | Mad1l1 | 0.106978719 | -0.049580028 | 0.263537466 | 9.600551043 | 1.681083243 | 0.144846522 | 0.64767305 | -4.304926376 |
| **A_52_P478745** | Paxbp1 | -0.105699065 | -0.260388748 | 0.048990618 | 8.205600666 | -1.681043535 | 0.144854395 | 0.64767305 | -4.304955481 |
| **A_55_P2154977** | Rpap3 | -0.114964406 | -0.283244377 | 0.053315564 | 11.30117876 | -1.680738459 | 0.144914894 | 0.647692555 | -4.305179094 |
| **A_55_P1993293** | Ndrg3 | -0.134288321 | -0.330879852 | 0.062303209 | 9.712544345 | -1.680515814 | 0.144959062 | 0.647692555 | -4.305342287 |
| **A_55_P2073010** | Gm4832 | 0.204014756 | -0.094676771 | 0.502706284 | 14.05779721 | 1.680381302 | 0.144985753 | 0.647692555 | -4.305440879 |
| **A_55_P2005050** | NA | -0.141447005 | -0.348546366 | 0.065652357 | 11.49110342 | -1.680289672 | 0.145003937 | 0.647692555 | -4.30550804 |
| **A_55_P2001143** | Ehmt2 | -0.212286595 | -0.523126451 | 0.098553261 | 14.11479675 | -1.680176998 | 0.145026301 | 0.647692555 | -4.305590626 |
| **A_55_P1972490** | Gm4055 | -0.214622866 | -0.528920192 | 0.099674459 | 12.9911521 | -1.67998138 | 0.145065136 | 0.647716335 | -4.305734005 |
| **A_55_P2131068** | NA | -0.185075141 | -0.456140556 | 0.085990274 | 13.67773077 | -1.679744171 | 0.145112241 | 0.647762179 | -4.305907868 |
| **A_65_P09032** | Rab43 | 0.165104338 | -0.076733248 | 0.406941924 | 12.26755898 | 1.679592195 | 0.145142428 | 0.647762179 | -4.306019258 |
| **A_55_P1976984** | NA | 0.124984311 | -0.058133854 | 0.308102476 | 14.85100858 | 1.679164469 | 0.145227421 | 0.647991878 | -4.306332754 |
| **A_51_P302149** | Cfl2 | -0.118522171 | -0.292231118 | 0.055186777 | 8.858055466 | -1.678597497 | 0.145340157 | 0.648047195 | -4.306748302 |
| **A_55_P2154595** | Mrpl39 | 0.160549432 | -0.074769712 | 0.395868576 | 10.55455796 | 1.678497413 | 0.145360067 | 0.648047195 | -4.306821655 |
| **A_55_P2146105** | NA | 0.211314639 | -0.098423179 | 0.521052458 | 16.28974629 | 1.678434942 | 0.145372495 | 0.648047195 | -4.306867441 |
| **A_55_P2044947** | NA | 0.076392278 | -0.035602769 | 0.188387326 | 8.438890393 | 1.678107615 | 0.145437633 | 0.648047195 | -4.307107341 |
| **A_55_P2009993** | Tomm20 | 0.308829174 | -0.143963258 | 0.761621606 | 12.6395181 | 1.677985963 | 0.145461849 | 0.648047195 | -4.307196501 |
| **A_55_P1981734** | Tut7 | -0.255191735 | -0.629344182 | 0.118960712 | 12.54803339 | -1.677981538 | 0.14546273 | 0.648047195 | -4.307199744 |
| **A_55_P2083806** | Sp8 | 0.146157639 | -0.068140706 | 0.360455983 | 8.406744197 | 1.677922374 | 0.145474509 | 0.648047195 | -4.307243105 |
| **A_55_P2077185** | NA | -0.110073619 | -0.271516909 | 0.051369671 | 8.887271333 | -1.677383662 | 0.145581801 | 0.648347144 | -4.307637923 |
| **A_52_P1055590** | Bnip1 | -0.085184228 | -0.21013277 | 0.039764315 | 8.994395755 | -1.677247522 | 0.145608927 | 0.648347144 | -4.307737697 |
| **A_66_P119968** | Clec4a1 | -0.156738895 | -0.386748285 | 0.073270494 | 8.67616439 | -1.676487729 | 0.145760408 | 0.648774559 | -4.308294527 |
| **A_55_P1987494** | Gabrr2 | 0.162002611 | -0.075747088 | 0.399752311 | 10.46002296 | 1.676375103 | 0.145782875 | 0.648774559 | -4.308377065 |
| **A_55_P2023762** | Epha4 | -0.078094604 | -0.192711467 | 0.036522258 | 7.966309385 | -1.676261141 | 0.145805613 | 0.648774559 | -4.308460583 |
| **A_51_P118765** | Eef1a1 | 0.161347587 | -0.075573853 | 0.398269027 | 16.44595537 | 1.675433802 | 0.145970784 | 0.648965998 | -4.309066893 |
| **A_55_P2021908** | NA | -0.139361189 | -0.344022988 | 0.065300611 | 9.636659028 | -1.675229132 | 0.146011672 | 0.648965998 | -4.309216881 |
| **A_55_P2067463** | 1700110I01Rik | -0.228708746 | -0.56460186 | 0.107184368 | 12.21132226 | -1.675138941 | 0.146029694 | 0.648965998 | -4.309282974 |
| **A_55_P1998688** | Gm2012 | 0.131051612 | -0.061420339 | 0.323523564 | 8.752241265 | 1.675112899 | 0.146034898 | 0.648965998 | -4.309302059 |

| | | | | | | | | | |
|---|---|---|---|---|---|---|---|---|---|
| **A_55_P2000958** | Sdr39u1 | 0.142562588 | -0.066819657 | 0.351944833 | 10.6434968 | 1.675077408 | 0.14604199 | 0.648965998 | -4.309328067 |
| **A_55_P2119100** | Yipf1 | 0.191555751 | -0.089789752 | 0.472901255 | 12.77036609 | 1.675036905 | 0.146050085 | 0.648965998 | -4.309357749 |
| **A_55_P1996973** | Gvin1 | -0.111548612 | -0.275417066 | 0.052319842 | 9.45753303 | -1.674703688 | 0.146116695 | 0.649112756 | -4.309601935 |
| **A_55_P2108424** | NA | -0.171932724 | -0.424561368 | 0.08069592 | 10.16716737 | -1.674345639 | 0.146188302 | 0.649281638 | -4.309864316 |
| **A_51_P191611** | Cat | 0.125067269 | -0.058722765 | 0.308857303 | 8.196673924 | 1.674136523 | 0.146230139 | 0.649318255 | -4.310017556 |
| **A_55_P2122314** | Emc1 | 0.103282227 | -0.04851912 | 0.255083573 | 9.588255084 | 1.673860002 | 0.14628548 | 0.64937757 | -4.310220188 |
| **A_55_P2032559** | NA | 0.315990012 | -0.148581938 | 0.780561962 | 14.06958063 | 1.673360579 | 0.146385482 | 0.64937757 | -4.310586155 |
| **A_51_P230507** | Ell2 | -0.103488743 | -0.255639269 | 0.048661783 | 8.583176991 | -1.673357816 | 0.146386035 | 0.64937757 | -4.31058818 |
| **A_51_P353252** | Mal2 | 0.243145741 | -0.114374404 | 0.600665887 | 8.606827554 | 1.673151674 | 0.146427331 | 0.64937757 | -4.310739235 |
| **A_52_P351882** | Impad1 | -0.153487883 | -0.379184946 | 0.07220918 | 12.4388746 | -1.67308248 | 0.146441195 | 0.64937757 | -4.310789937 |
| **A_52_P599238** | Rspry1 | -0.142185551 | -0.351265459 | 0.066894357 | 8.973069101 | -1.673063126 | 0.146445074 | 0.64937757 | -4.310804119 |
| **A_52_P613808** | Zbtb7a | -0.102517527 | -0.253284927 | 0.048249874 | 7.942986836 | -1.672860922 | 0.146485597 | 0.649408279 | -4.310952287 |
| **A_55_P2037787** | NA | -0.238400852 | -0.589041163 | 0.112239458 | 10.02709955 | -1.672688705 | 0.146520119 | 0.649412376 | -4.311078479 |
| **A_55_P2137840** | Etfbkmt | 0.139264345 | -0.065636087 | 0.344164777 | 9.407334122 | 1.672115331 | 0.146635112 | 0.64955343 | -4.311498616 |
| **A_55_P2034400** | Cdk8 | -0.155867316 | -0.385211209 | 0.073476578 | 10.585738 | -1.672002947 | 0.146657661 | 0.64955343 | -4.311580964 |
| **A_51_P221337** | Ranbp10 | -0.213266325 | -0.527089645 | 0.100556994 | 10.43133634 | -1.671884367 | 0.146681458 | 0.64955343 | -4.311667852 |
| **A_51_P159453** | Serpina3n | -0.135893414 | -0.335864721 | 0.064077893 | 7.800006119 | -1.671859923 | 0.146686364 | 0.64955343 | -4.311685763 |
| **A_55_P2064004** | Rapgef2 | -0.124090484 | -0.306732666 | 0.058551698 | 11.66719152 | -1.67150067 | 0.146758483 | 0.649636989 | -4.311948996 |
| **A_55_P2024337** | Tmem53 | 0.133211678 | -0.062863665 | 0.329287021 | 9.488303493 | 1.671431102 | 0.146772452 | 0.649636989 | -4.311999969 |
| **A_55_P1968493** | Telo2 | -0.198647789 | -0.491075178 | 0.0937796 | 9.023414774 | -1.671224728 | 0.146813901 | 0.649671678 | -4.312151183 |
| **A_55_P2067226** | Arpc5l | -0.152101837 | -0.376040334 | 0.07183666 | 10.79789251 | -1.670993898 | 0.146860274 | 0.649708919 | -4.312320314 |
| **A_55_P2105180** | Bhmt | -0.120107822 | -0.296957165 | 0.05674152 | 7.787844027 | -1.670848246 | 0.146889543 | 0.649708919 | -4.312427033 |
| **A_55_P2046918** | NA | 0.102296737 | -0.048361736 | 0.25295521 | 15.07282249 | 1.670465015 | 0.14696658 | 0.64986022 | -4.312707825 |
| **A_52_P439159** | Thap12 | -0.088512776 | -0.218880267 | 0.041854714 | 9.162982193 | -1.670343617 | 0.146990992 | 0.64986022 | -4.312796772 |
| **A_55_P1984886** | Hcst | 0.103934022 | -0.049174453 | 0.257042498 | 8.060934978 | 1.670043045 | 0.14705145 | 0.649978842 | -4.313016995 |
| **A_55_P2109554** | Serinc3 | -0.192951344 | -0.477232847 | 0.091330159 | 11.31169794 | -1.669815039 | 0.147097328 | 0.650032979 | -4.313184049 |
| **A_55_P1982201** | Sgsm2 | 0.196134138 | -0.09287824 | 0.485146516 | 9.907477202 | 1.669574913 | 0.14714566 | 0.650097932 | -4.313359981 |
| **A_55_P1966372** | Srek1 | -0.16297581 | -0.403219665 | 0.077268044 | 14.15232818 | -1.668937318 | 0.147274067 | 0.650227966 | -4.313827116 |
| **A_55_P2096368** | NA | 0.327190794 | -0.155130197 | 0.809511785 | 11.23180129 | 1.668914451 | 0.147278674 | 0.650227966 | -4.31384387 |
| **A_55_P1964664** | Fus | -0.144416878 | -0.357308133 | 0.068474376 | 12.75521336 | -1.668896082 | 0.147282375 | 0.650227966 | -4.313857328 |
| **A_51_P110341** | Scgb3a1 | 0.15516329 | -0.073588275 | 0.383914856 | 8.654988837 | 1.66876072 | 0.147309652 | 0.650227966 | -4.313915283 |
| **A_55_P2156741** | Stx4a | -0.107719244 | -0.26657609 | 0.051137602 | 13.49867076 | -1.668232294 | 0.147416181 | 0.65024434 | -4.31434364 |
| **A_55_P2082241** | NA | -0.152836022 | -0.37823861 | 0.072566565 | 13.57997595 | -1.668153427 | 0.147432086 | 0.65024434 | -4.31440142 |
| **A_55_P2115981** | Xlr5c | -0.132863786 | -0.32882684 | 0.063099268 | 10.9596286 | -1.668021287 | 0.147458739 | 0.65024434 | -4.314498227 |
| **A_66_P127175** | Ankrd16 | -0.235679471 | -0.583288877 | 0.111929935 | 10.9164659 | -1.668012876 | 0.147460436 | 0.65024434 | -4.31450439 |
| **A_55_P2065369** | H3f3a | -0.193870651 | -0.479833162 | 0.092091861 | 12.99548607 | -1.667908139 | 0.147481566 | 0.65024434 | -4.314581122 |
| **A_55_P1993976** | Aimp2 | 0.155975845 | -0.07413689 | 0.38608858 | 12.59776465 | 1.667576845 | 0.14754842 | 0.650282953 | -4.31482383 |
| **A_52_P615375** | H2aw | -0.225123444 | -0.564232037 | 0.11398515 | 10.81227145 | -1.720410207 | 0.147557609 | 0.650282953 | -4.299562421 |
| **A_55_P2007377** | Elf2 | -0.13238377 | -0.327743021 | 0.062975481 | 10.86192962 | -1.667131769 | 0.147638282 | 0.650490163 | -4.31514989 |
| **A_51_P125183** | Coq5 | -0.101440998 | -0.251193561 | 0.048311565 | 8.659636343 | -1.66651187 | 0.147763528 | 0.650893627 | -4.315739365 |
| **A_55_P2048928** | Cbfb | -0.148538005 | -0.367860033 | 0.070784024 | 11.24458681 | -1.666189889 | 0.147828623 | 0.650957382 | -4.315839887 |
| **A_55_P2325663** | Cers3 | 0.121786939 | -0.058045052 | 0.301618929 | 8.727709992 | 1.66610714 | 0.147845357 | 0.650957382 | -4.315900505 |
| **A_52_P683306** | Fam13b | -0.140502525 | -0.348010098 | 0.067005049 | 9.23739212 | -1.6657865 | 0.147910215 | 0.650968288 | -4.31613539 |
| **A_52_P84275** | Derl2 | 0.167049343 | -0.079697486 | 0.413796171 | 9.818989099 | 1.665567949 | 0.147954439 | 0.650968288 | -4.316295488 |
| **A_52_P672313** | Exoc5 | -0.170768076 | -0.423025796 | 0.081489644 | 11.1383832 | -1.66544917 | 0.14797848 | 0.650968288 | -4.316382498 |
| **A_55_P2005285** | Klhl22 | -0.175946301 | -0.435856404 | 0.083963802 | 13.56573166 | -1.665429073 | 0.147982547 | 0.650968288 | -4.31639722 |
| **A_55_P1998545** | Ncdn | 0.219548127 | -0.104871054 | 0.543967309 | 10.52824062 | 1.664916188 | 0.148086399 | 0.651163372 | -4.316772922 |
| **A_52_P30877** | Gm5176 | -0.183707586 | -0.455172694 | 0.087757522 | 12.57132758 | -1.664877321 | 0.148094272 | 0.651163372 | -4.316801393 |
| **A_51_P324082** | Med1 | -0.214164113 | -0.530728409 | 0.102400183 | 9.602895682 | -1.664385529 | 0.148193924 | 0.651453345 | -4.317161636 |
| **A_55_P2102335** | Zfp423 | 0.118482516 | -0.056742042 | 0.293707074 | 8.42042688 | 1.663521654 | 0.148369129 | 0.65183527 | -4.317794414 |

| | | | | | | | | | |
|---|---|---|---|---|---|---|---|---|---|
| **A_55_P2002460** | Gm2399 | -0.098766472 | -0.244835672 | 0.047302728 | 8.688863215 | -1.663489462 | 0.148375661 | 0.65183527 | -4.317817994 |
| **A_51_P189899** | Olfr1134 | 0.150826706 | -0.072247682 | 0.373901094 | 8.537246933 | 1.66340384 | 0.148393038 | 0.65183527 | -4.31788071 |
| **A_52_P544043** | Pcsk5 | -0.073264475 | -0.181638286 | 0.035109335 | 7.87297694 | -1.663176209 | 0.148439245 | 0.65183527 | -4.318047442 |
| **A_55_P2148912** | S100a16 | 0.273247542 | -0.130998074 | 0.677493159 | 12.00612051 | 1.662953509 | 0.148484464 | 0.65183527 | -4.31821056 |
| **A_55_P2177910** | Lepr | -0.172304325 | -0.427229345 | 0.082620696 | 16.01187254 | -1.662849248 | 0.148505639 | 0.65183527 | -4.318286927 |
| **A_51_P516452** | Eif3f | -0.20137462 | -0.499354075 | 0.096604834 | 11.25716868 | -1.662599237 | 0.148556427 | 0.65183527 | -4.318470047 |
| **A_52_P523445** | Cdhr5 | -0.158382043 | -0.392760552 | 0.075996467 | 8.850148591 | -1.662483388 | 0.148579967 | 0.65183527 | -4.3185549 |
| **A_55_P2027906** | Nsa2 | 0.124401296 | -0.059699567 | 0.30850216 | 12.34698519 | 1.662410387 | 0.148594802 | 0.65183527 | -4.318608369 |
| **A_66_P122362** | Aldh16a1 | 0.163179745 | -0.078325761 | 0.40468525 | 11.18594502 | 1.662296064 | 0.148618037 | 0.65183527 | -4.318692103 |
| **A_55_P2078204** | Prr5 | 0.192486712 | -0.092501004 | 0.477474427 | 11.15230492 | 1.661666163 | 0.148746122 | 0.652249048 | -4.319153458 |
| **A_55_P2179413** | Lgals3bp | -0.264864188 | -0.657127652 | 0.127399275 | 12.59091403 | -1.661171788 | 0.148846725 | 0.65254215 | -4.319515541 |
| **A_55_P1970935** | Usp7 | -0.147587614 | -0.366200528 | 0.0710253 | 11.80972188 | -1.660899145 | 0.148902234 | 0.652589261 | -4.319715223 |
| **A_55_P2138961** | Tpk1 | 0.146143548 | -0.070344924 | 0.362632021 | 9.344684457 | 1.660787377 | 0.148924995 | 0.652589261 | -4.319797081 |
| **A_55_P1987023** | NA | -0.127046946 | -0.315280797 | 0.061186905 | 9.36679703 | -1.660487705 | 0.14898604 | 0.652708785 | -4.320016555 |
| **A_55_P1978023** | NA | -0.109630125 | -0.272122505 | 0.052862254 | 9.117901936 | -1.659839426 | 0.149118179 | 0.653105789 | -4.320491331 |
| **A_52_P153189** | Arl2bp | 0.114976066 | -0.055453118 | 0.285405249 | 9.181189848 | 1.659711653 | 0.149144237 | 0.653105789 | -4.320584906 |
| **A_55_P2012296** | Gopc | 0.131614924 | -0.063529767 | 0.326759614 | 9.713404316 | 1.659271901 | 0.149233952 | 0.653278795 | -4.320906954 |
| **A_55_P2048912** | Themis2 | -0.073045834 | -0.181356034 | 0.035264366 | 7.804203261 | -1.659186697 | 0.14925134 | 0.653278795 | -4.320969352 |
| **A_51_P235123** | Nfkbie | -0.107288739 | -0.266441289 | 0.051863811 | 8.0983343 | -1.658477957 | 0.149396058 | 0.653764187 | -4.321488376 |
| **A_52_P534583** | Ahsp | 0.474680976 | -0.229658163 | 1.179020115 | 11.56198538 | 1.65801772 | 0.149490107 | 0.654027678 | -4.321825408 |
| **A_51_P268343** | Aagab | -0.171908183 | -0.427112549 | 0.083296184 | 11.64013787 | -1.657210252 | 0.14965525 | 0.654469174 | -4.322416698 |
| **A_55_P2108988** | Kcnip3 | 0.095002887 | -0.046040579 | 0.236046353 | 8.155356394 | 1.657116323 | 0.149674472 | 0.654469174 | -4.322485479 |
| **A_55_P2042870** | Aoc2 | 0.20466566 | -0.099210032 | 0.508541351 | 15.94566909 | 1.656983494 | 0.149701658 | 0.654469174 | -4.322582744 |
| **A_55_P1968433** | Gpat3 | 0.116813803 | -0.056637232 | 0.290264837 | 12.15900884 | 1.656862351 | 0.149726457 | 0.654469174 | -4.322671451 |
| **A_51_P320606** | Prl5a1 | -0.422111061 | -1.049001374 | 0.204779251 | 10.81950699 | -1.656549193 | 0.149790581 | 0.654543804 | -4.32290076 |
| **A_55_P2016852** | Zyx | 0.21915651 | -0.10637084 | 0.54468386 | 13.5332203 | 1.656288769 | 0.149843927 | 0.654543804 | -4.323091452 |
| **A_55_P2112379** | NA | -0.234944288 | -0.583923473 | 0.114034897 | 14.26843974 | -1.656282944 | 0.14984512 | 0.654543804 | -4.323095718 |
| **A_55_P2006615** | Rbbp9 | -0.108863395 | -0.270587828 | 0.052861038 | 10.45957121 | -1.656057455 | 0.149891325 | 0.654597701 | -4.323260827 |
| **A_66_P126842** | Frg2f1 | -0.107400727 | -0.267027681 | 0.052226226 | 8.960233366 | -1.655275013 | 0.150051762 | 0.65515033 | -4.323833738 |
| **A_55_P2042184** | NA | -0.295959824 | -0.735882021 | 0.143962373 | 9.721992563 | -1.655106895 | 0.150086256 | 0.655152946 | -4.323956834 |
| **A_55_P1981002** | NA | 0.118281209 | -0.057559975 | 0.294122392 | 9.224380033 | 1.654871662 | 0.150134533 | 0.655215711 | -4.324129068 |
| **A_55_P2184766** | Tmc6 | -0.15088561 | -0.375261887 | 0.073490668 | 8.804442492 | -1.654398191 | 0.150231749 | 0.65549198 | -4.324475731 |
| **A_55_P2097759** | NA | -0.132441213 | -0.329429307 | 0.064546881 | 13.05112325 | -1.654064092 | 0.150300385 | 0.655544474 | -4.324720344 |
| **A_55_P1985569** | Tomm6 | 0.180637979 | -0.088058725 | 0.449334684 | 13.20048339 | 1.653925301 | 0.150328906 | 0.655544474 | -4.324821961 |
| **A_55_P2034705** | Nmi | -0.141491466 | -0.352008931 | 0.069025998 | 10.02166413 | -1.653526958 | 0.150410794 | 0.655544474 | -4.325113603 |
| **A_66_P116807** | Selenot | -0.765649672 | -1.904878685 | 0.373579341 | 10.98795082 | -1.653439191 | 0.150428843 | 0.655544474 | -4.325177861 |
| **A_55_P2001347** | Rpl7a | 0.169460009 | -0.082699893 | 0.421619912 | 15.99842446 | 1.653333105 | 0.150450661 | 0.655544474 | -4.325255529 |
| **A_55_P2022201** | NA | -0.237327216 | -0.59048071 | 0.115826277 | 8.27237325 | -1.653305847 | 0.150456267 | 0.655544474 | -4.325275485 |
| **A_55_P2103429** | Psmc3 | -0.086772766 | -0.215912029 | 0.042366497 | 14.80421369 | -1.653081636 | 0.150502391 | 0.655544474 | -4.325439635 |
| **A_52_P189618** | Armc8 | -0.077476737 | -0.192785525 | 0.037832052 | 9.200820268 | -1.653019869 | 0.1505151 | 0.655544474 | -4.325484856 |
| **A_55_P2077501** | NA | -0.604472671 | -1.504240146 | 0.295294805 | 10.93732039 | -1.65278188 | 0.150564078 | 0.65561006 | -4.32565909 |
| **A_66_P139546** | Igfbp6 | -0.942717728 | -2.346144024 | 0.460708568 | 12.6675599 | -1.652575187 | 0.150606627 | 0.655647633 | -4.325810411 |
| **A_55_P2055477** | Scaf11 | 0.088754892 | -0.04340717 | 0.220916954 | 10.2284593 | 1.652169706 | 0.150690132 | 0.655863444 | -4.32610726 |
| **A_51_P423880** | Smarcd3 | 0.103907973 | -0.050840283 | 0.258656228 | 8.536085441 | 1.651932402 | 0.150739023 | 0.65592854 | -4.326280986 |
| **A_55_P2003043** | P4ha1 | -0.24666484 | -0.614078045 | 0.120748365 | 8.878472014 | -1.651663803 | 0.15079438 | 0.656021737 | -4.32647762 |
| **A_52_P511821** | Wdfy1 | 0.129705158 | -0.063582904 | 0.32299322 | 8.978066624 | 1.650902326 | 0.150951425 | 0.656557176 | -4.327035061 |
| **A_51_P491504** | Aldh5a1 | -0.171293148 | -0.426598186 | 0.08401189 | 9.248764702 | -1.65063013 | 0.1510076 | 0.656611096 | -4.327234317 |
| **A_55_P2097656** | Rps11 | 0.250460894 | -0.122866724 | 0.623788513 | 15.12955244 | 1.650513081 | 0.151031762 | 0.656611096 | -4.32732 |
| **A_55_P2012146** | NA | 0.082273887 | -0.040392873 | 0.204940647 | 7.730746467 | 1.650077272 | 0.151121759 | 0.65673036 | -4.327639018 |
| **A_55_P2247235** | Rab26os | 0.097167931 | -0.047710938 | 0.2420468 | 8.342711759 | 1.650011881 | 0.151135267 | 0.65673036 | -4.327686885 |

| | | | | | | | | | |
|---|---|---|---|---|---|---|---|---|---|
| **A_55_P2006668** | Pi4kb | 0.090856885 | -0.044642709 | 0.226356478 | 9.261578086 | 1.649639373 | 0.15121224 | 0.65673036 | -4.327959561 |
| **A_51_P267508** | Borcs7 | 0.134059904 | -0.065897605 | 0.334017413 | 9.728589632 | 1.649416553 | 0.1512583 | 0.65673036 | -4.328122662 |
| **A_55_P1967216** | Adam10 | -0.137406414 | -0.342361396 | 0.067548568 | 12.89182531 | -1.649368478 | 0.15126824 | 0.65673036 | -4.328157852 |
| **A_55_P2094237** | NA | 0.155879468 | -0.076640143 | 0.388399079 | 15.88270528 | 1.649295559 | 0.151283317 | 0.65673036 | -4.328211227 |
| **A_52_P381665** | Aff1 | -0.285404822 | -0.711180572 | 0.140370929 | 10.13011714 | -1.649109699 | 0.151321753 | 0.65673036 | -4.328347272 |
| **A_55_P2152387** | Wwox | 0.135636188 | -0.066740796 | 0.338013172 | 8.220498306 | 1.648859381 | 0.151373535 | 0.65673036 | -4.328530497 |
| **A_51_P173384** | Tada1 | -0.119030447 | -0.296650465 | 0.058589572 | 10.02784953 | -1.648675993 | 0.151411482 | 0.65673036 | -4.328664729 |
| **A_55_P2073607** | Commd5 | 0.226977797 | -0.111741769 | 0.565697364 | 11.89079826 | 1.648588459 | 0.151429598 | 0.65673036 | -4.328728799 |
| **A_51_P512591** | Cdc42se1 | -0.146119127 | -0.364283481 | 0.072045228 | 10.72140781 | -1.647754238 | 0.151602352 | 0.65673036 | -4.329339396 |
| **A_52_P655948** | Mosmo | -0.136072622 | -0.339241591 | 0.067096348 | 9.501631366 | -1.647716676 | 0.151610135 | 0.65673036 | -4.329366888 |
| **A_55_P2090390** | NA | 0.119519901 | -0.058936567 | 0.297976369 | 9.292764409 | 1.647696022 | 0.151614415 | 0.65673036 | -4.329382005 |
| **A_55_P2016712** | Msl1 | -0.156809736 | -0.404347757 | 0.090728284 | 10.92505054 | -1.782516754 | 0.15166481 | 0.65673036 | -4.32955227 |
| **A_55_P2154690** | Smarca4 | -0.152416329 | -0.380043035 | 0.075210378 | 8.406257695 | -1.647318005 | 0.151692765 | 0.65673036 | -4.329658679 |
| **A_51_P101858** | Dlg1 | -0.137523477 | -0.342931647 | 0.067884693 | 8.524623487 | -1.647131578 | 0.15173142 | 0.65673036 | -4.329795126 |
| **A_52_P183181** | Adar | -0.14595355 | -0.363974709 | 0.07206761 | 9.748274807 | -1.646968073 | 0.151765329 | 0.65673036 | -4.329914794 |
| **A_55_P2171273** | Mier3 | -0.11214179 | -0.279659427 | 0.055375846 | 9.055409127 | -1.646933682 | 0.151772463 | 0.65673036 | -4.329939964 |
| **A_55_P2135750** | NA | 0.76609568 | -0.378558123 | 1.910749483 | 9.534877915 | 1.646561744 | 0.15184963 | 0.65673036 | -4.330212178 |
| **A_55_P2134804** | Cinp | 0.116105555 | -0.057402758 | 0.289613867 | 8.590417637 | 1.646273074 | 0.151909548 | 0.65673036 | -4.330423447 |
| **A_55_P2167005** | NA | -0.216527757 | -0.540108816 | 0.107053301 | 14.50516892 | -1.646264637 | 0.151911299 | 0.65673036 | -4.330429621 |
| **A_55_P2019397** | Mtch1 | 0.155432979 | -0.076892196 | 0.387758153 | 15.20340006 | 1.645947812 | 0.151977088 | 0.65673036 | -4.330661492 |
| **A_55_P1975210** | Fubp3 | 0.122231727 | -0.060479721 | 0.304943176 | 9.304708269 | 1.64583901 | 0.151999687 | 0.65673036 | -4.330741119 |
| **A_55_P2076805** | Ddx5 | -0.280860396 | -0.700696085 | 0.138975293 | 9.101984325 | -1.645812335 | 0.152005228 | 0.65673036 | -4.330760641 |
| **A_55_P2386592** | Cep63 | -0.122604102 | -0.305884084 | 0.06067588 | 10.89905262 | -1.645732059 | 0.152021904 | 0.65673036 | -4.33081939 |
| **A_52_P509185** | Zfp318 | -0.120759085 | -0.301282401 | 0.059764231 | 8.922543341 | -1.645718967 | 0.152024624 | 0.65673036 | -4.330828971 |
| **A_52_P122673** | Cep41 | -0.072068744 | -0.179820242 | 0.035682753 | 8.310129623 | -1.645480747 | 0.152074124 | 0.65673036 | -4.331003309 |
| **A_55_P1999294** | Gpbp1 | -0.229718967 | -0.573184967 | 0.113747032 | 13.26990981 | -1.645440818 | 0.152082422 | 0.65673036 | -4.331032531 |
| **A_51_P455906** | Pold4 | 0.230714831 | -0.114339176 | 0.575768839 | 9.776389935 | 1.644968557 | 0.152180603 | 0.65673036 | -4.331378141 |
| **A_55_P2047614** | Zfp809 | -0.087665598 | -0.21878492 | 0.043453723 | 8.526110486 | -1.644870348 | 0.152201028 | 0.65673036 | -4.331450011 |
| **A_55_P2176212** | NA | 0.309726034 | -0.153582324 | 0.773034392 | 12.28793486 | 1.644662309 | 0.152244304 | 0.65673036 | -4.331602255 |
| **A_55_P2162452** | Mpdu1 | 0.155923453 | -0.077333921 | 0.389180827 | 13.5409381 | 1.644542971 | 0.152269133 | 0.65673036 | -4.331689586 |
| **A_51_P435333** | Dctn2 | -0.115398054 | -0.288051237 | 0.05725513 | 10.95352371 | -1.644345432 | 0.152310242 | 0.65673036 | -4.331834143 |
| **A_55_P2043627** | Fam89a | -0.284887823 | -0.711128876 | 0.14135323 | 12.21184218 | -1.64432543 | 0.152314405 | 0.65673036 | -4.331848781 |
| **A_55_P1981040** | Ddx6 | -0.208584906 | -0.520699559 | 0.103529746 | 10.80676344 | -1.64413655 | 0.152353723 | 0.65673036 | -4.331987 |
| **A_51_P205385** | Uox | -0.132434756 | -0.330662144 | 0.065792631 | 7.910727897 | -1.643642958 | 0.152456517 | 0.65673036 | -4.332348195 |
| **A_51_P388847** | Stap2 | -0.114279148 | -0.285332498 | 0.056774202 | 9.503198455 | -1.643631949 | 0.15245881 | 0.65673036 | -4.332356252 |
| **A_55_P2115320** | Metap1d | -0.081546178 | -0.203606788 | 0.040514431 | 7.883205308 | -1.643604197 | 0.152464592 | 0.65673036 | -4.332376559 |
| **A_52_P337220** | Mgat4b | -0.140311234 | -0.350341401 | 0.069718934 | 9.658247081 | -1.643538671 | 0.152478244 | 0.65673036 | -4.332424508 |
| **A_65_P20249** | Prkcb | -0.104671965 | -0.261357434 | 0.052013503 | 7.9355554 | -1.643504213 | 0.152485424 | 0.65673036 | -4.332449723 |
| **A_52_P237669** | Nup54 | -0.076212027 | -0.190298995 | 0.037874942 | 8.145431511 | -1.643450516 | 0.152496612 | 0.65673036 | -4.332489016 |
| **A_55_P1959403** | Ccdc77 | -0.135653517 | -0.338736507 | 0.067429473 | 8.798867445 | -1.643337143 | 0.152520238 | 0.65673036 | -4.332571977 |
| **A_55_P2045886** | Stk3 | -0.169136132 | -0.422359848 | 0.084087584 | 9.88133156 | -1.643240688 | 0.152540342 | 0.65673036 | -4.332642557 |
| **A_55_P2109454** | Gtf3c5 | 0.129518421 | -0.064404714 | 0.323441556 | 10.61341193 | 1.643126808 | 0.15256408 | 0.65673036 | -4.332725888 |
| **A_55_P2081437** | NA | 0.123877433 | -0.061612684 | 0.30936755 | 8.931690768 | 1.643011431 | 0.152588134 | 0.65673036 | -4.332810313 |
| **A_51_P179258** | Kif26b | -0.094684585 | -0.236486143 | 0.047116973 | 7.972101763 | -1.642734734 | 0.152645835 | 0.65673843 | -4.333012781 |
| **A_55_P1967330** | Gm5069 | -0.239451294 | -0.598071109 | 0.119168522 | 14.08870499 | -1.642676591 | 0.152657963 | 0.65673843 | -4.333055325 |
| **A_55_P2037657** | NA | 0.120485112 | -0.059985323 | 0.300955546 | 8.787104125 | 1.64246636 | 0.152701821 | 0.65678093 | -4.333209154 |
| **A_55_P2092622** | NA | 0.096559271 | -0.048131634 | 0.241250175 | 8.350259976 | 1.641806266 | 0.152839608 | 0.657227315 | -4.333692144 |
| **A_51_P306527** | Ywhah | 0.149959043 | -0.074782983 | 0.374701068 | 14.38974717 | 1.641562915 | 0.152890435 | 0.657299647 | -4.3338702 |
| **A_55_P1985143** | NA | -0.168947022 | -0.422195221 | 0.084301178 | 8.952495886 | -1.641244697 | 0.152956923 | 0.657439264 | -4.33410303 |
| **A_52_P509710** | Zwint | -0.091700095 | -0.2291982 | 0.045798009 | 12.05177938 | -1.640749347 | 0.153060477 | 0.657738097 | -4.334465455 |

| | | | | | | | | | |
|---|---|---|---|---|---|---|---|---|---|
| **A_55_P1971360** | Ciz1 | -0.112044287 | -0.280071197 | 0.055982623 | 9.129757005 | -1.640514372 | 0.153109623 | 0.65773999 | -4.334637371 |
| **A_51_P196509** | Plekhj1 | -0.123192799 | -0.307948963 | 0.061563365 | 8.639907978 | -1.640421864 | 0.153128975 | 0.65773999 | -4.334705053 |
| **A_55_P1979403** | Tigar | 0.177959596 | -0.088964523 | 0.444883714 | 9.042745184 | 1.640222466 | 0.153170697 | 0.657773027 | -4.334850938 |
| **A_55_P1984546** | Mmp1a | 0.447675621 | -0.223993375 | 1.119344616 | 8.764844668 | 1.639748741 | 0.153269862 | 0.657971429 | -4.335197521 |
| **A_55_P2158072** | NA | 0.319878344 | -0.160111164 | 0.799867853 | 9.258196153 | 1.639540911 | 0.153313386 | 0.657971429 | -4.335349568 |
| **A_52_P182331** | NA | -0.095807693 | -0.239595884 | 0.047980499 | 14.75687452 | -1.63925426 | 0.153373437 | 0.657971429 | -4.335559278 |
| **A_55_P2071846** | NA | -0.150258314 | -0.375766306 | 0.075249677 | 10.54632485 | -1.639252059 | 0.153373898 | 0.657971429 | -4.335560888 |
| **A_55_P2147566** | Tac2 | 0.812905233 | -0.407189337 | 2.032999804 | 8.586692222 | 1.639138262 | 0.153397744 | 0.657971429 | -4.33564414 |
| **A_55_P1969543** | Pin4 | -0.234773897 | -0.587237902 | 0.117690109 | 11.46185129 | -1.638717904 | 0.153485861 | 0.657971429 | -4.33595166 |
| **A_51_P153486** | Dnajb1 | -0.258351599 | -0.646220447 | 0.129517249 | 14.83958622 | -1.638684766 | 0.153492809 | 0.657971429 | -4.335975902 |
| **A_55_P1971184** | Ube2l3 | 0.15439558 | -0.077419269 | 0.386210428 | 13.50140897 | 1.63856161 | 0.153518636 | 0.657971429 | -4.336065998 |
| **A_55_P2305294** | Ptpn21 | -0.131159085 | -0.328108926 | 0.065790756 | 8.643139457 | -1.638369702 | 0.153558888 | 0.657971429 | -4.336206388 |
| **A_52_P105015** | Clcn7 | -0.120110362 | -0.300500402 | 0.060279678 | 9.472082131 | -1.63808745 | 0.153618109 | 0.657971429 | -4.336412866 |
| **A_51_P519364** | Fam83f | 0.233064562 | -0.116974264 | 0.583103388 | 9.757433454 | 1.638057669 | 0.153624359 | 0.657971429 | -4.336434652 |
| **A_66_P107973** | Prpf38b | -0.084577513 | -0.211604555 | 0.042449528 | 8.180830247 | -1.63805278 | 0.153625385 | 0.657971429 | -4.336438229 |
| **A_51_P176352** | Ndrg2 | 0.194473489 | -0.097665838 | 0.486612817 | 8.397905582 | 1.637719595 | 0.153695323 | 0.658125098 | -4.336681961 |
| **A_66_P105801** | Igf1r | 0.158873648 | -0.079810993 | 0.397558288 | 10.43354939 | 1.637557494 | 0.153729361 | 0.658125098 | -4.33680054 |
| **A_51_P346826** | Rragc | -0.094678064 | -0.236949058 | 0.04759293 | 11.50092508 | -1.63720162 | 0.153804112 | 0.658299309 | -4.337060863 |
| **A_51_P163233** | Khdc4 | -0.138963472 | -0.347808292 | 0.069881348 | 8.826215546 | -1.636990324 | 0.153848511 | 0.65832649 | -4.337215424 |
| **A_51_P396983** | Ssu2 | 0.074413798 | -0.037430662 | 0.186258258 | 7.933440727 | 1.636847257 | 0.15387858 | 0.65832649 | -4.337320075 |
| **A_51_P468294** | Polg | -0.196750709 | -0.492553491 | 0.099052073 | 10.58717563 | -1.636376431 | 0.153977577 | 0.658533716 | -4.33766447 |
| **A_55_P2101392** | Tmem237 | 0.096586707 | -0.048632978 | 0.241806391 | 8.657883217 | 1.636292849 | 0.153995157 | 0.658533716 | -4.337725607 |
| **A_55_P2131481** | NA | -0.240760403 | -0.602783459 | 0.121262654 | 13.56430425 | -1.636130699 | 0.154029269 | 0.658533896 | -4.337844212 |
| **A_51_P428715** | Mrpl13 | -0.112715181 | -0.282222577 | 0.056792215 | 11.18799274 | -1.63592325 | 0.154072921 | 0.658574855 | -4.33799595 |
| **A_55_P2163493** | Tgm3 | 0.282435009 | -0.14235195 | 0.707221969 | 10.32792942 | 1.635748445 | 0.154109714 | 0.658586482 | -4.338123809 |
| **A_55_P2065834** | Dhx8 | 0.223739303 | -0.112890513 | 0.560369119 | 10.82510639 | 1.635154765 | 0.154234732 | 0.658975053 | -4.338558038 |
| **A_51_P482633** | Rps6ka4 | -0.182586719 | -0.457352825 | 0.092179387 | 9.634408619 | -1.634839753 | 0.154301108 | 0.659047818 | -4.338788438 |
| **A_51_P504314** | Prpf38b | -0.162845915 | -0.40794742 | 0.082255589 | 8.961282809 | -1.634556948 | 0.154360721 | 0.659047818 | -4.338995279 |
| **A_55_P2130219** | Tpr | -0.205773118 | -0.515502751 | 0.103956515 | 8.944661147 | -1.634462814 | 0.154380568 | 0.659047818 | -4.339064127 |
| **A_52_P533792** | Sucla2 | -0.112568569 | -0.282018058 | 0.056880919 | 12.19780662 | -1.634353684 | 0.154403581 | 0.659047818 | -4.339143942 |
| **A_52_P232314** | NA | -0.097631205 | -0.244603435 | 0.049341025 | 9.009505349 | -1.634265186 | 0.154422245 | 0.659047818 | -4.339208666 |
| **A_52_P631568** | Thoc6 | -0.08050715 | -0.201721301 | 0.040707 | 8.865488164 | -1.633993397 | 0.154479578 | 0.659146969 | -4.339407443 |
| **A_52_P141488** | Grk5 | -0.076971274 | -0.192875521 | 0.038932972 | 8.466491171 | -1.633798494 | 0.154520706 | 0.659176941 | -4.339549986 |
| **A_55_P2035554** | Strip1 | -0.177390069 | -0.444772227 | 0.089992089 | 12.89499878 | -1.632172442 | 0.154864236 | 0.660321026 | -4.340739139 |
| **A_55_P2067483** | NA | -0.198846642 | -0.498632526 | 0.100939241 | 10.919435 | -1.63183501 | 0.154935616 | 0.660321026 | -4.340985894 |
| **A_55_P2129691** | AB124611 | 0.51240734 | -0.260144606 | 1.284959286 | 12.78435429 | 1.631761996 | 0.154951065 | 0.660321026 | -4.341039286 |
| **A_55_P2314496** | Secisbp2l | -0.114756581 | -0.287781052 | 0.058267889 | 9.505793672 | -1.63169595 | 0.154965041 | 0.660321026 | -4.341087582 |
| **A_55_P2042615** | NA | -0.430422512 | -1.079432597 | 0.218587573 | 9.221141079 | -1.631596239 | 0.154986144 | 0.660321026 | -4.341160497 |
| **A_55_P2089288** | Rnf121 | -0.141097471 | -0.35389048 | 0.071695538 | 11.47883491 | -1.63128949 | 0.155051081 | 0.660321026 | -4.341384805 |
| **A_55_P2142116** | NA | -0.1690882 | -0.424095419 | 0.08591902 | 15.3306718 | -1.631285545 | 0.155051917 | 0.660321026 | -4.34138769 |
| **A_55_P2288117** | NA | -0.173144168 | -0.434310674 | 0.088022339 | 8.570021408 | -1.63102099 | 0.155107943 | 0.660321026 | -4.341581141 |
| **A_55_P2152836** | NA | 0.111324132 | -0.056603779 | 0.279252042 | 10.02903509 | 1.63093103 | 0.155126999 | 0.660321026 | -4.341646922 |
| **A_51_P442284** | Dnajc12 | 0.17944228 | -0.091241835 | 0.450126395 | 8.992964919 | 1.630914418 | 0.155130518 | 0.660321026 | -4.341659069 |
| **A_51_P405668** | 5-Mar | -0.111113383 | -0.278764705 | 0.05653794 | 10.96296481 | -1.630529077 | 0.15521217 | 0.66042431 | -4.341940836 |
| **A_55_P2000628** | NA | -0.187460566 | -0.470340156 | 0.095419025 | 9.134773498 | -1.630337324 | 0.155252817 | 0.66042431 | -4.342081046 |
| **A_55_P2061170** | Tbc1d2 | -0.118124359 | -0.296378263 | 0.060129545 | 11.3588735 | -1.630307697 | 0.155259098 | 0.66042431 | -4.342102709 |
| **A_52_P338180** | Timm13 | -0.191594274 | -0.48080751 | 0.097618962 | 12.99905595 | -1.629797038 | 0.155367399 | 0.66042431 | -4.342476093 |
| **A_55_P2053398** | NA | -0.096422354 | -0.241975132 | 0.049130424 | 17.29310634 | -1.629770286 | 0.155373075 | 0.66042431 | -4.342495654 |
| **A_55_P1955309** | Zfp800 | 0.119394128 | -0.06085078 | 0.299639036 | 8.763675068 | 1.629630429 | 0.15540275 | 0.66042431 | -4.342597913 |
| **A_51_P467770** | Snx19 | -0.098547301 | -0.24733316 | 0.050238558 | 9.15598152 | -1.629492025 | 0.155432122 | 0.66042431 | -4.342699109 |

| | | | | | | | | | |
|---|---|---|---|---|---|---|---|---|---|
| **A_55_P2117984** | Vangl2 | 0.259644475 | -0.132364794 | 0.651653745 | 8.999736064 | 1.6294908 | 0.155432382 | 0.66042431 | -4.342700004 |
| **A_51_P133562** | Serpina6 | -0.195631895 | -0.491020996 | 0.099757207 | 7.892762503 | -1.629349894 | 0.15546229 | 0.66042431 | -4.342803028 |
| **A_51_P404084** | Psmb2 | -0.128093774 | -0.321526559 | 0.065339011 | 9.962904225 | -1.629172608 | 0.155499929 | 0.660439051 | -4.34293265 |
| **A_55_P1996946** | Cdc20 | 0.188182261 | -0.096036415 | 0.472400937 | 9.353784737 | 1.628903038 | 0.155557176 | 0.660537049 | -4.343129743 |
| **A_51_P484764** | Fam3a | -0.136178447 | -0.34189832 | 0.069541426 | 10.73985329 | -1.628550737 | 0.155632023 | 0.660670766 | -4.343387318 |
| **A_52_P131372** | NA | -0.114316751 | -0.287023424 | 0.058389923 | 16.56271902 | -1.62843308 | 0.155657027 | 0.660670766 | -4.343473339 |
| **A_52_P12001** | Erp44 | 0.139859756 | -0.071465349 | 0.35118486 | 10.28196852 | 1.628211564 | 0.155704113 | 0.660725533 | -4.34363529 |
| **A_55_P1995045** | Zc3hav1 | -0.131757172 | -0.330860679 | 0.067346334 | 9.060696163 | -1.628037933 | 0.15574103 | 0.660737132 | -4.34376223 |
| **A_55_P2013357** | Renbp | 0.113179984 | -0.057872717 | 0.284232685 | 9.905260155 | 1.627829278 | 0.155785406 | 0.660758557 | -4.343914775 |
| **A_51_P311694** | Ypel5 | -0.078719328 | -0.197705935 | 0.040267279 | 9.831744772 | -1.627617919 | 0.155830368 | 0.660758557 | -4.344069295 |
| **A_55_P2140751** | NA | -0.19036112 | -0.478112794 | 0.097390555 | 12.54447724 | -1.627532069 | 0.155848635 | 0.660758557 | -4.344132058 |
| **A_51_P185906** | Abi3 | -0.204347265 | -0.513292336 | 0.104597805 | 8.89136279 | -1.627259155 | 0.155906717 | 0.660859854 | -4.344331574 |
| **A_55_P2184106** | Gpn2 | -0.093587583 | -0.23512017 | 0.047945003 | 8.338519369 | -1.62678797 | 0.156007045 | 0.661140139 | -4.344676032 |
| **A_52_P467140** | Snd1 | -0.108706877 | -0.273125744 | 0.05571199 | 10.14829499 | -1.626576648 | 0.156052061 | 0.661185948 | -4.344830514 |
| **A_51_P430900** | Dusp1 | -0.149689443 | -0.376153753 | 0.076774867 | 10.25231109 | -1.626149868 | 0.156143013 | 0.66142632 | -4.345142496 |
| **A_51_P211573** | Wdr47 | 0.117886211 | -0.06050943 | 0.296281852 | 9.841967291 | 1.62572818 | 0.15623293 | 0.661537416 | -4.345450748 |
| **A_55_P2132711** | Tmem248 | 0.14402871 | -0.073943269 | 0.36200069 | 11.20895182 | 1.625614509 | 0.156257176 | 0.661537416 | -4.345533839 |
| **A_51_P430327** | Gmpr2 | -0.086347115 | -0.217049629 | 0.044355399 | 8.628085661 | -1.625298201 | 0.156324665 | 0.661537416 | -4.345765051 |
| **A_55_P1974421** | Itga3 | -0.119345049 | -0.3000619 | 0.061371802 | 9.611811335 | -1.624706502 | 0.156450989 | 0.661537416 | -4.346197554 |
| **A_55_P2058791** | 4833420G17Rik | -0.198641697 | -0.499470076 | 0.102186681 | 8.734758772 | -1.624503972 | 0.15649425 | 0.661537416 | -4.34634559 |
| **A_66_P124817** | Ing1 | -0.172040031 | -0.43259866 | 0.088518598 | 8.875603555 | -1.624400875 | 0.156516276 | 0.661537416 | -4.346420946 |
| **A_52_P724940** | Ndufa12 | -0.088494713 | -0.222534599 | 0.045545173 | 13.60803452 | -1.624248365 | 0.156548865 | 0.661537416 | -4.346532418 |
| **A_55_P1969078** | Adar | 0.161644256 | -0.083205136 | 0.406493647 | 10.81872176 | 1.624165982 | 0.156566471 | 0.661537416 | -4.346592633 |
| **A_55_P2080562** | Arhgap26 | 0.558804678 | -0.28774109 | 1.405350445 | 10.45573033 | 1.623972808 | 0.156607763 | 0.661537416 | -4.346733825 |
| **A_55_P2170834** | NA | 0.200981911 | -0.103500679 | 0.505464501 | 10.80940751 | 1.623916406 | 0.156619821 | 0.661537416 | -4.346775049 |
| **A_51_P491470** | Ddx47 | -0.146881293 | -0.369421465 | 0.075658878 | 9.471167453 | -1.623780169 | 0.15664895 | 0.661537416 | -4.346874624 |
| **A_55_P2129373** | Capn11 | 0.263015083 | -0.135512796 | 0.661542963 | 9.371002377 | 1.623645112 | 0.156677833 | 0.661537416 | -4.346973336 |
| **A_55_P2024463** | Tbkbp1 | -0.147413188 | -0.370782341 | 0.075955966 | 8.262268423 | -1.623612187 | 0.156684875 | 0.661537416 | -4.3469974 |
| **A_55_P2010861** | Mrps23 | 0.253487607 | -0.130623278 | 0.637598491 | 12.67214835 | 1.623563505 | 0.156695287 | 0.661537416 | -4.347032982 |
| **A_51_P187842** | Eif4e3 | 0.127855107 | -0.065891513 | 0.321601727 | 8.693889735 | 1.623503043 | 0.15670822 | 0.661537416 | -4.347077172 |
| **A_55_P2049282** | Ly6h | 0.074864338 | -0.038592694 | 0.188321369 | 7.81967471 | 1.623352126 | 0.156740507 | 0.661537416 | -4.347187473 |
| **A_52_P217875** | H3f3b | -0.094248546 | -0.237086721 | 0.048589629 | 8.207342389 | -1.623302771 | 0.156751067 | 0.661537416 | -4.347223545 |
| **A_52_P626756** | Trnt1 | -0.101033934 | -0.254183865 | 0.052115997 | 9.879346809 | -1.623004121 | 0.156814981 | 0.661617097 | -4.347441818 |
| **A_55_P1973699** | Htra2 | 0.289576389 | -0.149400818 | 0.728553596 | 11.78281194 | 1.622894695 | 0.156838405 | 0.661617097 | -4.347521792 |
| **A_55_P1962771** | Cyfip2 | 0.217620219 | -0.112339846 | 0.547580283 | 9.690248656 | 1.622583389 | 0.156905065 | 0.661753871 | -4.347749308 |
| **A_51_P333518** | Sh3rf2 | -0.11328948 | -0.285095884 | 0.058516924 | 8.51401839 | -1.622256035 | 0.156975189 | 0.66181517 | -4.347988547 |
| **A_52_P418644** | Plp2 | -0.221136325 | -0.556526473 | 0.114253824 | 11.5074409 | -1.622104947 | 0.157007565 | 0.66181517 | -4.348098965 |
| **A_55_P2078474** | Fam50a | 0.134156208 | -0.069322709 | 0.337635125 | 12.99359284 | 1.622036112 | 0.157022317 | 0.66181517 | -4.34814927 |
| **A_55_P2186833** | NA | 0.129751995 | -0.067150777 | 0.326654768 | 15.50340095 | 1.621180519 | 0.157205794 | 0.662319925 | -4.348774527 |
| **A_55_P2068330** | Cryzl1 | -0.08072471 | -0.203250347 | 0.041800927 | 9.246141484 | -1.620871876 | 0.157272032 | 0.662319925 | -4.349000071 |
| **A_51_P193146** | Ms4a6c | -0.090008916 | -0.226650994 | 0.046633161 | 7.91944554 | -1.620579123 | 0.157334884 | 0.662319925 | -4.349213999 |
| **A_52_P652212** | Psmd14 | -0.071877949 | -0.181005325 | 0.037249427 | 9.575971709 | -1.620433182 | 0.157366225 | 0.662319925 | -4.349320643 |
| **A_55_P2104583** | C87436 | -0.104739118 | -0.263758941 | 0.054280706 | 9.215178148 | -1.620417063 | 0.157369687 | 0.662319925 | -4.349332422 |
| **A_55_P2095345** | Rara | 0.104488506 | -0.05415218 | 0.263129191 | 10.6237312 | 1.620403242 | 0.157372656 | 0.662319925 | -4.349342521 |
| **A_55_P2156126** | Sema6c | -0.277330937 | -0.698402543 | 0.143740669 | 9.549697997 | -1.620360041 | 0.157381935 | 0.662319925 | -4.349374089 |
| **A_55_P2185543** | Tle2 | 0.274976316 | -0.142584267 | 0.692536899 | 11.6157603 | 1.620111697 | 0.157435287 | 0.66240023 | -4.34955556 |
| **A_51_P220806** | Gdf9 | 0.164284783 | -0.085214198 | 0.413783764 | 9.413860446 | 1.619935508 | 0.157473148 | 0.662415337 | -4.349684303 |
| **A_55_P2119219** | NA | -0.088165121 | -0.2221012 | 0.045770958 | 12.85145816 | -1.619453156 | 0.157576845 | 0.662669504 | -4.350036756 |
| **A_55_P2007678** | Xntrpc | 0.441667889 | -0.229345093 | 1.11268087 | 9.959223602 | 1.619325169 | 0.157604372 | 0.662669504 | -4.350130273 |
| **A_55_P2002968** | Coro2a | 0.144833403 | -0.075227957 | 0.364894764 | 8.481017405 | 1.619176177 | 0.157636421 | 0.662669504 | -4.350239137 |

| | | | | | | | | | |
|---|---|---|---|---|---|---|---|---|---|
| **A_51_P289881** | Txndc12 | -0.10721573 | -0.270227773 | 0.055796313 | 11.0202126 | -1.618109772 | 0.157865996 | 0.663479731 | -4.351018299 |
| **A_55_P1988433** | Stard7 | -0.121957808 | -0.3074008 | 0.063485184 | 12.49441179 | -1.61796211 | 0.15789781 | 0.663479731 | -4.351126183 |
| **A_55_P1987156** | Crhr2 | 0.097783093 | -0.050918959 | 0.246485145 | 7.982288555 | 1.617766996 | 0.157939856 | 0.663512168 | -4.351268734 |
| **A_55_P2069022** | Vstm4 | 0.144671632 | -0.075442962 | 0.364786225 | 8.770674167 | 1.61697649 | 0.15811032 | 0.663584694 | -4.351846261 |
| **A_51_P129720** | Hectd3 | -0.132610445 | -0.334384575 | 0.069163685 | 9.865488724 | -1.616893483 | 0.158128229 | 0.663584694 | -4.351906903 |
| **A_52_P464437** | Trp53bp1 | -0.100067017 | -0.252325001 | 0.052190967 | 8.827763883 | -1.616888384 | 0.15812933 | 0.663584694 | -4.351910628 |
| **A_55_P2187266** | Pop1 | -0.16568586 | -0.417823269 | 0.086451548 | 9.396420136 | -1.616654911 | 0.158179715 | 0.663584694 | -4.352081191 |
| **A_52_P512817** | Shisa5 | -0.164411619 | -0.414646945 | 0.085823707 | 15.5825321 | -1.616415658 | 0.158231363 | 0.663584694 | -4.352255974 |
| **A_55_P1968774** | Ptk2b | 0.122044886 | -0.063718429 | 0.307808202 | 8.50206422 | 1.616325397 | 0.158250852 | 0.663584694 | -4.352321913 |
| **A_55_P1995126** | NA | 0.146258411 | -0.076362403 | 0.368879225 | 16.91501853 | 1.616308465 | 0.158254509 | 0.663584694 | -4.352334282 |
| **A_51_P103757** | Kat8 | 0.132577231 | -0.069237349 | 0.33439181 | 11.43928171 | 1.616164522 | 0.158285594 | 0.663584694 | -4.352105599 |
| **A_55_P1963084** | Rps13 | 0.141132611 | -0.073733321 | 0.355998543 | 16.18370899 | 1.615953935 | 0.158331083 | 0.663584694 | -4.352593272 |
| **A_55_P2001233** | Ifi208 | -0.139904494 | -0.352902725 | 0.073093737 | 10.60954223 | -1.61593851 | 0.158334416 | 0.663584694 | -4.352604541 |
| **A_55_P2040988** | NA | 0.173536464 | -0.090665175 | 0.437738104 | 9.772221551 | 1.615936908 | 0.158334762 | 0.663584694 | -4.352605711 |
| **A_55_P2177633** | NA | -0.13634318 | -0.343980241 | 0.071293881 | 8.59162307 | -1.615465579 | 0.158436621 | 0.663867643 | -4.352950015 |
| **A_55_P1963578** | NA | 0.186114894 | -0.09736181 | 0.469591599 | 15.5510363 | 1.615224593 | 0.158488724 | 0.663905007 | -4.353126049 |
| **A_55_P1988498** | Naa20 | 0.158776611 | -0.083095047 | 0.400648269 | 11.26789665 | 1.614993244 | 0.15853876 | 0.663905007 | -4.353295042 |
| **A_51_P312748** | Oxsm | -0.114191881 | -0.288165192 | 0.05978143 | 9.099303426 | -1.61481123 | 0.158578136 | 0.663905007 | -4.353427995 |
| **A_51_P141772** | Uhrf2 | -0.176024505 | -0.444204568 | 0.092155559 | 10.74296345 | -1.614789082 | 0.158582928 | 0.663905007 | -4.353444173 |
| **A_55_P1972817** | Trmt11 | -0.083033561 | -0.209577789 | 0.043510666 | 9.355798914 | -1.61428603 | 0.158691808 | 0.663970899 | -4.35381162 |
| **A_55_P2082609** | NA | -0.132194593 | -0.333707435 | 0.069318248 | 10.82182119 | -1.613913036 | 0.158772585 | 0.663970899 | -4.35408406 |
| **A_55_P1978216** | Cd300c2 | -0.219231597 | -0.553439915 | 0.114976721 | 8.497273375 | -1.613819861 | 0.15879277 | 0.663970899 | -4.354152115 |
| **A_66_P123055** | Gm10845 | -0.330062273 | -0.833311283 | 0.173186737 | 14.12654436 | -1.613548857 | 0.158851491 | 0.663970899 | -4.354350055 |
| **A_51_P363525** | Fbrsl1 | -0.219486019 | -0.554175231 | 0.115203193 | 11.11217683 | -1.613371237 | 0.158889989 | 0.663970899 | -4.354479785 |
| **A_52_P218181** | Zdhhc5 | -0.205605514 | -0.51913208 | 0.107921053 | 11.95805785 | -1.613353641 | 0.158893804 | 0.663970899 | -4.354492637 |
| **A_55_P1994074** | Mrtfa | -0.086768389 | -0.219086858 | 0.04555008 | 8.927966809 | -1.613281779 | 0.158909382 | 0.663970899 | -4.354545123 |
| **A_55_P2127825** | Bcas3 | 0.086640721 | -0.045506249 | 0.218787691 | 9.058626799 | 1.612998677 | 0.158970769 | 0.663970899 | -4.35475189 |
| **A_55_P2053444** | Map4 | -0.214094471 | -0.540651575 | 0.112462632 | 12.04115824 | -1.612929855 | 0.158985695 | 0.663970899 | -4.354802155 |
| **A_55_P2353004** | Pitpnm2os1 | 0.355334991 | -0.186771232 | 0.897441213 | 8.949623589 | 1.612585484 | 0.159060405 | 0.663970899 | -4.355053664 |
| **A_55_P1973213** | Trmt61a | 0.103636937 | -0.054496587 | 0.261770461 | 9.827218623 | 1.612351716 | 0.159111138 | 0.663970899 | -4.355224392 |
| **A_52_P243054** | Nufip2 | -0.145700058 | -0.368035225 | 0.076635109 | 8.77990987 | -1.612206722 | 0.159142614 | 0.663970899 | -4.355330284 |
| **A_55_P1962602** | NA | -0.36648039 | -0.925722776 | 0.192761996 | 11.20744151 | -1.612203367 | 0.159143342 | 0.663970899 | -4.355332734 |
| **A_52_P145767** | Vcpip1 | -0.186191593 | -0.470323139 | 0.097939953 | 10.0079655 | -1.612166068 | 0.15915144 | 0.663970899 | -4.355359974 |
| **A_51_P388478** | Efnb1 | -0.129075823 | -0.32606336 | 0.067911715 | 12.97571304 | -1.612038138 | 0.159179217 | 0.663970899 | -4.355453403 |
| **A_55_P2011674** | Rpl17 | -0.129906066 | -0.328173447 | 0.068361314 | 15.97910791 | -1.611934263 | 0.159201775 | 0.663970899 | -4.355529263 |
| **A_51_P417251** | 6330403K07Rik | 0.125024757 | -0.065829799 | 0.315879314 | 8.289386801 | 1.61162001 | 0.159270038 | 0.663970899 | -4.35575876 |
| **A_66_P130777** | Ephb4 | -0.095639158 | -0.241666269 | 0.050387953 | 8.002262357 | -1.611281475 | 0.159343607 | 0.663970899 | -4.356005984 |
| **A_55_P2020143** | Slc39a3 | 0.157013114 | -0.082737598 | 0.396763827 | 9.509187946 | 1.611184162 | 0.15936476 | 0.663970899 | -4.356077048 |
| **A_55_P2068812** | Pkm | 0.276400683 | -0.145650923 | 0.698452288 | 15.0574295 | 1.611175013 | 0.159366749 | 0.663970899 | -4.35608373 |
| **A_52_P373798** | Zdhhc3 | -0.100000644 | -0.252697569 | 0.052696281 | 8.289398829 | -1.611171123 | 0.159367595 | 0.663970899 | -4.35608657 |
| **A_51_P176883** | Edrf1 | -0.084109235 | -0.212554377 | 0.044335907 | 8.315916525 | -1.6109986 | 0.159405104 | 0.663970899 | -4.356212556 |
| **A_52_P311297** | Als2 | -0.083066584 | -0.209933935 | 0.043800768 | 8.382789461 | -1.610814901 | 0.159445054 | 0.663970899 | -4.356346702 |
| **A_55_P1958350** | Nipsnap3b | 0.156273442 | -0.082428095 | 0.394974979 | 11.06915207 | 1.610642396 | 0.159482577 | 0.663970899 | -4.356472672 |
| **A_55_P1971474** | NA | -0.228277175 | -0.576985448 | 0.120431098 | 10.82597857 | -1.610532158 | 0.15950656 | 0.663970899 | -4.356553171 |
| **A_55_P2014326** | NA | 0.128738889 | -0.06796513 | 0.325442907 | 8.961333618 | 1.610147589 | 0.159590254 | 0.663970899 | -4.35683399 |
| **A_51_P299934** | BC048403 | -0.111417149 | -0.281657038 | 0.05882274 | 9.511540076 | -1.610126052 | 0.159594943 | 0.663970899 | -4.356849717 |
| **A_55_P2088690** | NA | -0.472501026 | -1.194507486 | 0.249505434 | 11.74829241 | -1.610018386 | 0.159618382 | 0.663970899 | -4.356928334 |
| **A_55_P2159807** | Rfc4 | 0.220068554 | -0.11623384 | 0.556370948 | 9.74951017 | 1.609893676 | 0.159645537 | 0.663970899 | -4.357019398 |
| **A_51_P302358** | Ltb | -0.083699539 | -0.211607147 | 0.044208068 | 8.12723767 | -1.60988871 | 0.159646618 | 0.663970899 | -4.357023024 |
| **A_52_P551011** | Smim1 | 0.235951884 | -0.124686984 | 0.596590752 | 9.728677958 | 1.60960785 | 0.159707789 | 0.663970899 | -4.357228103 |

| | | | | | | | | | |
|---|---|---|---|---|---|---|---|---|---|
| **A_51_P319379** | Tmx4 | -0.138226743 | -0.34950812 | 0.073054634 | 9.186750942 | -1.609533492 | 0.159723988 | 0.663970899 | -4.357282398 |
| **A_51_P343309** | Cmas | -0.13716079 | -0.346873445 | 0.072551864 | 10.56121706 | -1.6090684 | 0.159825344 | 0.663970899 | -4.357621991 |
| **A_55_P2185330** | NA | -0.146347862 | -0.370117171 | 0.077421446 | 9.413842517 | -1.608996231 | 0.159841077 | 0.663970899 | -4.357674685 |
| **A_55_P2025837** | NA | -0.130618662 | -0.330340641 | 0.069103317 | 13.78393331 | -1.608972168 | 0.159846323 | 0.663970899 | -4.357692254 |
| **A_52_P297642** | Pdia5 | -0.193662911 | -0.489832662 | 0.102506841 | 12.45654573 | -1.608699465 | 0.159905789 | 0.663970899 | -4.357891365 |
| **A_55_P2052330** | NA | -0.171361404 | -0.433449218 | 0.090726411 | 15.50337151 | -1.608552886 | 0.15993776 | 0.663970899 | -4.357998387 |
| **A_55_P2062733** | NA | -0.224663547 | -0.568388202 | 0.119061109 | 13.84873271 | -1.608018675 | 0.160054334 | 0.663970899 | -4.358388419 |
| **A_55_P2050044** | NA | 0.118371144 | -0.06274866 | 0.299490947 | 10.72706285 | 1.607863097 | 0.160088299 | 0.663970899 | -4.358502005 |
| **A_51_P493105** | Mettl15 | 0.121798148 | -0.064565727 | 0.308162022 | 8.416593847 | 1.607859573 | 0.160089068 | 0.663970899 | -4.358504577 |
| **A_55_P1993423** | NA | -0.284058455 | -0.718743818 | 0.150626909 | 15.2858648 | -1.607688355 | 0.160126455 | 0.663970899 | -4.35862958 |
| **A_51_P425737** | Tbc1d7 | -0.140915488 | -0.356565889 | 0.074734914 | 8.895248463 | -1.607598605 | 0.160146057 | 0.663970899 | -4.358695105 |
| **A_55_P2055632** | Rtf2 | -0.104535651 | -0.26451248 | 0.055441178 | 8.616096002 | -1.60759448 | 0.160146958 | 0.663970899 | -4.358698116 |
| **A_55_P2043862** | Stmn1 | -0.207950528 | -0.526192113 | 0.110291057 | 11.64852693 | -1.607578767 | 0.16015039 | 0.663970899 | -4.358709587 |
| **A_55_P2026738** | Rgs4 | -0.205388759 | -0.519710676 | 0.108933157 | 8.498718599 | -1.607574729 | 0.160151272 | 0.663970899 | -4.358712536 |
| **A_51_P301930** | Lrrc17 | 0.392416913 | -0.208203153 | 0.993036978 | 8.522256474 | 1.607374383 | 0.160195037 | 0.663970899 | -4.358858801 |
| **A_66_P132510** | Scamp3 | 0.131887207 | -0.069985316 | 0.333759729 | 12.46831714 | 1.60729141 | 0.160213165 | 0.663970899 | -4.358919376 |
| **A_52_P69998** | Kdelr2 | -0.169591539 | -0.429215536 | 0.090032458 | 11.79875042 | -1.607046758 | 0.16026663 | 0.663972466 | -4.359097984 |
| **A_55_P2054352** | Rbm6 | -0.18075966 | -0.457526436 | 0.096007116 | 10.59572425 | -1.606781133 | 0.160324698 | 0.663972466 | -4.359291899 |
| **A_55_P2024041** | Psg19 | -0.215920498 | -0.546581053 | 0.114740058 | 14.92976901 | -1.606499687 | 0.160386246 | 0.663972466 | -4.35949736 |
| **A_51_P260639** | Elp3 | -0.090238858 | -0.228446549 | 0.047968833 | 10.43028824 | -1.606314367 | 0.160426785 | 0.663972466 | -4.359632645 |
| **A_55_P2136378** | Dazap2 | -0.275769728 | -0.698149004 | 0.146609548 | 14.29443713 | -1.606250043 | 0.160440858 | 0.663972466 | -4.359679601 |
| **A_55_P2138130** | NA | 0.295862994 | -0.157320898 | 0.749046886 | 13.32449044 | 1.606147205 | 0.16046336 | 0.663972466 | -4.359754673 |
| **A_55_P2062469** | Col12a1 | 0.441843849 | -0.235098773 | 1.118786472 | 10.39332562 | 1.605780269 | 0.160543675 | 0.663972466 | -4.36002253 |
| **A_55_P2026635** | Asnsd1 | 0.124716714 | -0.06638076 | 0.315814187 | 11.69919736 | 1.605605611 | 0.160581917 | 0.663972466 | -4.360150024 |
| **A_51_P309184** | Pxmp2 | 0.19611411 | -0.104418801 | 0.49664702 | 9.922535791 | 1.605410011 | 0.160624755 | 0.663972466 | -4.360292804 |
| **A_51_P200083** | Grina | -0.217146211 | -0.549913344 | 0.115620921 | 10.52564468 | -1.605391634 | 0.160628781 | 0.663972466 | -4.360306218 |
| **A_55_P2002702** | Ddx18 | 0.088081658 | -0.046900548 | 0.223063863 | 10.31040845 | 1.605381012 | 0.160631107 | 0.663972466 | -4.360313972 |
| **A_51_P434240** | Mrpl11 | 0.123161075 | -0.065624507 | 0.311946656 | 13.04928158 | 1.604995466 | 0.160715581 | 0.663972466 | -4.360595397 |
| **A_55_P2148688** | NA | 0.102777261 | -0.054782712 | 0.260337235 | 9.748086117 | 1.604797742 | 0.160758919 | 0.663972466 | -4.36073972 |
| **A_51_P392291** | Pdk3 | -0.130108309 | -0.329613188 | 0.069396571 | 9.10861659 | -1.60442962 | 0.160839635 | 0.663972466 | -4.361008416 |
| **A_55_P2095563** | Slc8a2 | 0.107929783 | -0.057570225 | 0.273429792 | 8.076635889 | 1.604399243 | 0.160846297 | 0.663972466 | -4.361030588 |
| **A_66_P114381** | Ypel2 | -0.160787759 | -0.40735394 | 0.085778422 | 9.16278498 | -1.604311281 | 0.160865591 | 0.663972466 | -4.361094791 |
| **A_55_P2092671** | Nppc | 0.179253892 | -0.09564861 | 0.454156395 | 9.66190549 | 1.604202041 | 0.160889554 | 0.663972466 | -4.361174524 |
| **A_51_P401501** | Tmem213 | 0.073263552 | -0.039103116 | 0.185630221 | 7.766856042 | 1.604056395 | 0.16092151 | 0.663972466 | -4.361280828 |
| **A_51_P111544** | Srp14 | 0.119806867 | -0.063947159 | 0.303560892 | 12.79761496 | 1.604035627 | 0.160926067 | 0.663972466 | -4.361295986 |
| **A_66_P131205** | NA | -0.250906208 | -0.635752282 | 0.133939866 | 14.43082675 | -1.603959912 | 0.160942682 | 0.663972466 | -4.361351249 |
| **A_52_P287052** | Znrd2 | -0.104173482 | -0.26397278 | 0.055625816 | 9.198379457 | -1.603804678 | 0.160976751 | 0.663972466 | -4.361464549 |
| **A_51_P154973** | Stard3 | -0.104850597 | -0.26572221 | 0.056021015 | 11.90413387 | -1.603469326 | 0.161050376 | 0.663972466 | -4.361709307 |
| **A_55_P2167123** | Cyp4b1-ps2 | 0.086384362 | -0.046188344 | 0.218957067 | 7.881748641 | 1.603061456 | 0.161139965 | 0.663972466 | -4.362006984 |
| **A_52_P592951** | Ostc | -0.157383368 | -0.39892932 | 0.084162585 | 12.63424009 | -1.602980475 | 0.161157759 | 0.663972466 | -4.362066086 |
| **A_55_P2004208** | Defa-rs2 | 0.544406718 | -0.291143009 | 1.379956445 | 9.774865217 | 1.602951334 | 0.161164162 | 0.663972466 | -4.362087353 |
| **A_55_P2002918** | Klc2 | 0.080611764 | -0.043146015 | 0.204369542 | 10.81472763 | 1.602489091 | 0.161265766 | 0.663972466 | -4.3624247 |
| **A_55_P2157019** | Phb2 | -0.097055547 | -0.246065064 | 0.051953971 | 9.774580508 | -1.602416896 | 0.161281641 | 0.663972466 | -4.362477387 |
| **A_55_P2037862** | NA | 0.157105218 | -0.084105923 | 0.398316358 | 8.199110895 | 1.602368537 | 0.161292275 | 0.663972466 | -4.362512679 |
| **A_55_P2128672** | Ikbip | -0.113716699 | -0.288313519 | 0.06088012 | 10.72434964 | -1.602348877 | 0.161296598 | 0.663972466 | -4.362527027 |
| **A_55_P1999541** | Ccdc42 | 0.285663035 | -0.15294997 | 0.72427604 | 9.1879117 | 1.602292146 | 0.161309075 | 0.663972466 | -4.362568427 |
| **A_52_P155554** | Cdc42ep2 | 0.165906226 | -0.088860576 | 0.420673027 | 10.60109028 | 1.602097755 | 0.161351833 | 0.663972466 | -4.362710289 |
| **A_51_P382369** | Zfp608 | -0.150568907 | -0.381791003 | 0.080653189 | 8.345502309 | -1.602046538 | 0.1613631 | 0.663972466 | -4.362747665 |
| **A_52_P602091** | Csf1r | -0.196877709 | -0.499260563 | 0.105505145 | 10.401953 | -1.601800956 | 0.161417137 | 0.663972466 | -4.362926881 |
| **A_55_P2121827** | Dgcr2 | 0.32132023 | -0.172200274 | 0.814840735 | 8.987087853 | 1.601776921 | 0.161422426 | 0.663972466 | -4.362944421 |

| | | | | | | | | | |
|---|---|---|---|---|---|---|---|---|---|
| **A_55_P2030899** | NA | 0.101473253 | -0.054385678 | 0.257332184 | 11.13811592 | 1.601729048 | 0.161432962 | 0.663972466 | -4.362979356 |
| **A_52_P598309** | Zfas1 | 0.237692688 | -0.127509317 | 0.602894692 | 11.15510081 | 1.601223044 | 0.161544367 | 0.663972466 | -4.363348604 |
| **A_52_P584188** | Frat1 | -0.134225496 | -0.340459105 | 0.072008112 | 8.598833027 | -1.601196902 | 0.161550124 | 0.663972466 | -4.36336768 |
| **A_52_P337716** | NA | -0.154457633 | -0.391787667 | 0.0828724 | 11.43889456 | -1.60112721 | 0.161565474 | 0.663972466 | -4.363418535 |
| **A_55_P2072556** | Ncoa6 | -0.090013348 | -0.228347045 | 0.048320349 | 12.19739348 | -1.600840622 | 0.161628611 | 0.663972466 | -4.363627661 |
| **A_55_P2302577** | AI426330 | -0.139666573 | -0.354316843 | 0.074983698 | 9.652479454 | -1.60077465 | 0.161643148 | 0.663972466 | -4.3636758 |
| **A_51_P446558** | Timm29 | 0.163147485 | -0.087594223 | 0.413889194 | 8.368399834 | 1.600747997 | 0.161649022 | 0.663972466 | -4.363695248 |
| **A_55_P2144386** | Adgre1 | -0.150329323 | -0.381375532 | 0.080716885 | 8.354373385 | -1.600715018 | 0.16165629 | 0.663972466 | -4.363719313 |
| **A_55_P1965378** | Uqcrq | 0.138554774 | -0.074430911 | 0.351540459 | 15.87711825 | 1.600443119 | 0.161716222 | 0.664077514 | -4.363917712 |
| **A_55_P2100149** | Mapt | 0.133297336 | -0.071654524 | 0.338249196 | 9.15172055 | 1.60006921 | 0.161798675 | 0.664237414 | -4.364190541 |
| **A_55_P2011151** | Tmprss3 | -0.230133924 | -0.584002556 | 0.123734707 | 11.33645674 | -1.599954897 | 0.161823891 | 0.664237414 | -4.364273949 |
| **A_55_P2119962** | Shld1 | 0.286425747 | -0.154175101 | 0.727026596 | 11.29572293 | 1.599321918 | 0.161963584 | 0.664543522 | -4.364735789 |
| **A_55_P2110136** | Arhgap29 | -0.202645566 | -0.514372882 | 0.10908175 | 9.94561261 | -1.599305419 | 0.161967227 | 0.664543522 | -4.364747827 |
| **A_55_P2030775** | Fnbp1 | -0.090598602 | -0.229980356 | 0.048783151 | 10.67231204 | -1.599133555 | 0.162005177 | 0.664558164 | -4.36487322 |
| **A_55_P2090414** | NA | -0.1347806 | -0.342223905 | 0.072662705 | 9.555921331 | -1.598442891 | 0.16215777 | 0.664703854 | -4.365377114 |
| **A_55_P1964108** | B230118H07Rik | -0.097823511 | -0.248389196 | 0.052742173 | 9.86873884 | -1.598403449 | 0.162166489 | 0.664703854 | -4.36540589 |
| **A_55_P2120956** | NA | 0.163820745 | -0.088394441 | 0.416035932 | 9.937152269 | 1.597963399 | 0.162263789 | 0.664703854 | -4.365726926 |
| **A_51_P370615** | Neurog2 | -0.206614277 | -0.52474954 | 0.111520986 | 9.08426727 | -1.597782575 | 0.162303787 | 0.664703854 | -4.365858842 |
| **A_55_P2058137** | Pde4dip | 0.188841699 | -0.101945479 | 0.479628877 | 11.78507736 | 1.597687395 | 0.162324845 | 0.664703854 | -4.365928278 |
| **A_55_P1984307** | Adgrf5 | -0.292960995 | -0.744079435 | 0.158157445 | 11.0762875 | -1.597674801 | 0.162327632 | 0.664703854 | -4.365937465 |
| **A_55_P2005081** | Rpl10 | -0.134952536 | -0.342790704 | 0.072885632 | 17.21412175 | -1.5974413 | 0.162379304 | 0.664703854 | -4.366107807 |
| **A_52_P470401** | Tmem186 | -0.123716016 | -0.314255142 | 0.066823109 | 8.602781733 | -1.597389861 | 0.162390689 | 0.664703854 | -4.366145331 |
| **A_55_P1963105** | NA | -0.168635458 | -0.428358686 | 0.09108777 | 15.54332894 | -1.597376414 | 0.162393665 | 0.664703854 | -4.366155141 |
| **A_51_P164203** | Nme4 | 0.366619744 | -0.198073657 | 0.931313146 | 9.465347957 | 1.597247807 | 0.162422134 | 0.664703854 | -4.366248958 |
| **A_55_P1959753** | NA | 0.211959824 | -0.114540067 | 0.538459716 | 9.176123261 | 1.597127812 | 0.162448701 | 0.664703854 | -4.366336494 |
| **A_51_P293938** | Rasl11b | -0.071224123 | -0.180938192 | 0.038489946 | 10.66814477 | -1.597106766 | 0.162453361 | 0.664703854 | -4.366351846 |
| **A_52_P348250** | Klhl29 | -0.156448672 | -0.397542837 | 0.084645493 | 8.855345813 | -1.596446413 | 0.16259964 | 0.66471533 | -4.36683355 |
| **A_55_P1985793** | Dock6 | 0.200954092 | -0.108728791 | 0.510636975 | 11.33671814 | 1.596426089 | 0.162604144 | 0.66471533 | -4.366848375 |
| **A_55_P2074226** | Fbxo8 | -0.101583313 | -0.258139729 | 0.054973103 | 8.559676241 | -1.596322612 | 0.162627078 | 0.66471533 | -4.366923855 |
| **A_51_P145511** | Prl2a1 | -0.309286029 | -0.785951088 | 0.167379031 | 16.64243578 | -1.596306004 | 0.16263076 | 0.66471533 | -4.36693597 |
| **A_51_P227392** | Rhou | 0.161652494 | -0.08748648 | 0.410791467 | 14.13450225 | 1.596283044 | 0.162635849 | 0.66471533 | -4.366952717 |
| **A_65_P15812** | Pum1 | -0.106838942 | -0.271515136 | 0.057837252 | 10.46931884 | -1.59612875 | 0.162670053 | 0.66471533 | -4.367065264 |
| **A_55_P1968703** | Gfra2 | 0.138019327 | -0.074753319 | 0.350791974 | 8.515367446 | 1.595854433 | 0.162730881 | 0.66471533 | -4.367265356 |
| **A_52_P328497** | Cdk10 | -0.139067416 | -0.353456055 | 0.075321222 | 9.290209965 | -1.595852627 | 0.162731282 | 0.66471533 | -4.367266673 |
| **A_51_P247873** | Ndufb8 | 0.36674054 | -0.198711001 | 0.932192081 | 14.22662003 | 1.59563183 | 0.162780258 | 0.664774901 | -4.367427723 |
| **A_51_P273203** | Gpaa1 | -0.080760404 | -0.205296712 | 0.043775904 | 8.07996949 | -1.595407612 | 0.162830007 | 0.664837603 | -4.367591265 |
| **A_55_P2046338** | Mapk14 | -0.179949218 | -0.457467885 | 0.09756945 | 12.01677679 | -1.595243282 | 0.162866478 | 0.664846074 | -4.367711124 |
| **A_55_P1955227** | Rabac1 | -0.191846343 | -0.487743719 | 0.104051034 | 15.28121589 | -1.595076703 | 0.162903456 | 0.66485661 | -4.367832621 |
| **A_55_P1956707** | Rpl36a | 0.173251376 | -0.094016079 | 0.440518832 | 13.65264232 | 1.594776315 | 0.162970158 | 0.664988428 | -4.368051712 |
| **A_51_P104527** | Atrip | -0.09731715 | -0.247512221 | 0.052877922 | 9.798473673 | -1.594053386 | 0.163130793 | 0.665503396 | -4.368578961 |
| **A_51_P438805** | Txnip | -0.318194213 | -0.809338661 | 0.172950235 | 12.6382036 | -1.593867446 | 0.163172133 | 0.66553158 | -4.368714567 |
| **A_55_P2047778** | Krtap4-8 | 0.091051435 | -0.049526674 | 0.231629544 | 10.33100606 | 1.593449346 | 0.163265126 | 0.665713352 | -4.369019479 |
| **A_55_P2004099** | NA | 0.126992622 | -0.069087898 | 0.323073142 | 8.436071595 | 1.593357409 | 0.163285581 | 0.665713352 | -4.369086525 |
| **A_55_P2115127** | Mphosph10 | -0.1093677 | -0.278265714 | 0.059530315 | 8.697004256 | -1.593065732 | 0.163350494 | 0.665837556 | -4.369299231 |
| **A_51_P254656** | Hdc | 0.116850358 | -0.063680865 | 0.297381581 | 8.163525594 | 1.592380631 | 0.163503059 | 0.666191551 | -4.369798823 |
| **A_55_P2022724** | Inafm2 | -0.161164006 | -0.410194595 | 0.087866584 | 9.696962861 | -1.592151973 | 0.163554008 | 0.666191551 | -4.369965559 |
| **A_55_P2156068** | NA | 0.115372704 | -0.06290739 | 0.293652798 | 9.46356907 | 1.592096469 | 0.163566378 | 0.666191551 | -4.370006032 |
| **A_52_P684138** | Dpf3 | -0.158729449 | -0.404030506 | 0.086571608 | 8.876203409 | -1.591942039 | 0.1636008 | 0.666191551 | -4.37011864 |
| **A_51_P256093** | Map2k6 | 0.101706435 | -0.055474981 | 0.25888785 | 9.592450044 | 1.591902253 | 0.163609669 | 0.666191551 | -4.370147651 |
| **A_55_P1973011** | Ppm1l | -0.123620873 | -0.3147115 | 0.067469754 | 8.843655092 | -1.59155475 | 0.163687155 | 0.666366683 | -4.370401037 |

| | | | | | | | | | |
|---|---|---|---|---|---|---|---|---|---|
| **A_52_P251183** | Tmem177 | 0.144575544 | -0.078935909 | 0.368086997 | 9.166143355 | 1.59134453 | 0.163734046 | 0.666417219 | -4.370554319 |
| **A_55_P1995698** | Thumpd1 | -0.131392626 | -0.334554416 | 0.071769165 | 10.65937564 | -1.591102371 | 0.163788078 | 0.66649679 | -4.370730885 |
| **A_51_P427768** | Diaph1 | 0.116709512 | -0.063824841 | 0.297243865 | 8.658208449 | 1.590433676 | 0.163937368 | 0.666867716 | -4.371218434 |
| **A_55_P2018249** | Lmo4 | -0.082489318 | -0.210093408 | 0.045114772 | 8.497871827 | -1.590385037 | 0.163948233 | 0.666867716 | -4.371253896 |
| **A_51_P426195** | Nppb | -0.828510295 | -2.110459347 | 0.453438756 | 10.1964544 | -1.589997009 | 0.164034928 | 0.667017322 | -4.371536796 |
| **A_55_P2048418** | Mdfic | -0.157856743 | -0.402126391 | 0.086412904 | 11.54413452 | -1.589874327 | 0.164062347 | 0.667017322 | -4.371626238 |
| **A_55_P2070766** | NA | -0.637053232 | -1.622910627 | 0.348804163 | 11.47307404 | -1.589757155 | 0.164088539 | 0.667017322 | -4.371711662 |
| **A_52_P660945** | Ctsf | -0.092915043 | -0.23675563 | 0.050925545 | 9.190741741 | -1.589182399 | 0.164217074 | 0.667399458 | -4.372130674 |
| **A_66_P138046** | Gm12298 | 0.251061713 | -0.137959959 | 0.640083386 | 8.515392559 | 1.587727088 | 0.164542961 | 0.668449579 | -4.373191543 |
| **A_52_P213181** | Cul4b | -0.119676748 | -0.305117476 | 0.065763979 | 11.81018301 | -1.587719655 | 0.164544627 | 0.668449579 | -4.373196961 |
| **A_66_P107790** | Myl12a | -0.103131988 | -0.263034587 | 0.05677061 | 15.73158723 | -1.586744616 | 0.164763315 | 0.669061878 | -4.373907656 |
| **A_51_P266546** | Zc2hc1a | -0.080026364 | -0.204104928 | 0.044052199 | 9.529045589 | -1.586738983 | 0.164764579 | 0.669061878 | -4.373911761 |
| **A_66_P123787** | Lars2 | -0.111316882 | -0.283933655 | 0.061299891 | 9.75155369 | -1.586526029 | 0.164812378 | 0.669063004 | -4.374066973 |
| **A_51_P335419** | Csl | -0.107684628 | -0.274738034 | 0.059368778 | 8.556957271 | -1.585869841 | 0.16495975 | 0.669063004 | -4.374545218 |
| **A_55_P2038121** | Rbx1 | 0.172696182 | -0.095282854 | 0.440675218 | 13.9713303 | 1.585444631 | 0.165055314 | 0.669063004 | -4.374855106 |
| **A_55_P2021892** | Ccdc114 | 0.08727761 | -0.048161228 | 0.222716449 | 9.203195574 | 1.585363259 | 0.165073608 | 0.669063004 | -4.374914409 |
| **A_55_P1976112** | C130074G19Rik | 0.135428 | -0.07474651 | 0.345602509 | 10.28518555 | 1.58524935 | 0.16509922 | 0.669063004 | -4.374997422 |
| **A_55_P2121082** | Cul2 | -0.159368623 | -0.406753102 | 0.088015855 | 10.45225372 | -1.584891501 | 0.165179706 | 0.669063004 | -4.375258205 |
| **A_52_P486322** | Fzr1 | -0.225428573 | -0.575359266 | 0.124502119 | 10.1637823 | -1.58487892 | 0.165182537 | 0.669063004 | -4.375267373 |
| **A_55_P2007768** | Gm8709 | -0.179387162 | -0.457849484 | 0.09907516 | 14.71651081 | -1.584871598 | 0.165184184 | 0.669063004 | -4.375272709 |
| **A_55_P1972688** | Wnk1 | -0.270730348 | -0.690989755 | 0.149529059 | 12.39581836 | -1.584851814 | 0.165188635 | 0.669063004 | -4.375287127 |
| **A_52_P375638** | Gtpbp1 | -0.097413171 | -0.248642097 | 0.053815754 | 8.590184702 | -1.584717958 | 0.165218753 | 0.669063004 | -4.375384672 |
| **A_55_P1963309** | G3bp1 | 0.193953308 | -0.107209477 | 0.495116092 | 12.03187434 | 1.584400785 | 0.165290137 | 0.669063004 | -4.375615802 |
| **A_55_P2092073** | Taf11 | -0.125512281 | -0.320404718 | 0.069380157 | 9.623410934 | -1.584383814 | 0.165293958 | 0.669063004 | -4.375628169 |
| **A_52_P515826** | Med13 | -0.088624934 | -0.226249535 | 0.048999666 | 8.212525103 | -1.584269339 | 0.16531973 | 0.669063004 | -4.375711588 |
| **A_52_P486702** | G3bp1 | -0.089878673 | -0.229461301 | 0.049703954 | 9.018788872 | -1.584143241 | 0.165348123 | 0.669063004 | -4.375803475 |
| **A_55_P2107745** | Yy1 | 0.178081932 | -0.098491225 | 0.454655088 | 11.69605895 | 1.584086872 | 0.165360818 | 0.669063004 | -4.375844551 |
| **A_55_P1967144** | NA | -0.134683447 | -0.34385885 | 0.074491957 | 16.56421309 | -1.584064169 | 0.16536593 | 0.669063004 | -4.375861094 |
| **A_55_P2033287** | NA | -0.089321259 | -0.228045578 | 0.04940306 | 15.43274746 | -1.584059163 | 0.165367058 | 0.669063004 | -4.375864742 |
| **A_52_P255841** | Dbr1 | 0.223569986 | -0.123696384 | 0.570836355 | 9.481985885 | 1.583871469 | 0.165409334 | 0.669063004 | -4.376001512 |
| **A_51_P441426** | Pf4 | 0.33074524 | -0.18301311 | 0.84450359 | 11.28107093 | 1.583812888 | 0.165422531 | 0.669063004 | -4.376044198 |
| **A_51_P436483** | Kin | -0.194585253 | -0.496895927 | 0.10772542 | 10.35244206 | -1.583527489 | 0.165486839 | 0.669106416 | -4.376252158 |
| **A_66_P126300** | Mrpl47 | -0.130193866 | -0.332480495 | 0.072092764 | 10.23537536 | -1.583406807 | 0.165514039 | 0.669106416 | -4.376340093 |
| **A_51_P245405** | Ppp3cc | -0.092827631 | -0.237088498 | 0.051433236 | 9.580450147 | -1.583061891 | 0.165591801 | 0.669106416 | -4.376591411 |
| **A_66_P126523** | Phb2 | -0.199441598 | -0.509397417 | 0.110514221 | 11.3465705 | -1.583015309 | 0.165602306 | 0.669106416 | -4.376625352 |
| **A_51_P438821** | Pycard | 0.260503303 | -0.144354206 | 0.665360812 | 9.09164591 | 1.582997388 | 0.165606348 | 0.669106416 | -4.37663841 |
| **A_51_P339713** | Mrps26 | -0.096977488 | -0.24771613 | 0.053761154 | 10.62945855 | -1.582761564 | 0.165659539 | 0.669181449 | -4.376810234 |
| **A_55_P1959425** | Slc16a3 | 0.145664154 | -0.080804686 | 0.372132993 | 13.66251506 | 1.582389524 | 0.165743489 | 0.669192574 | -4.377081301 |
| **A_55_P1956488** | Dmtn | 0.085167529 | -0.047252116 | 0.217587174 | 7.918489213 | 1.582307146 | 0.165762083 | 0.669192574 | -4.377141319 |
| **A_55_P2028600** | Myh9 | -0.300827851 | -0.768564466 | 0.166908764 | 15.42395223 | -1.582289097 | 0.165766157 | 0.669192574 | -4.377154469 |
| **A_55_P1986199** | Tut1 | -0.111442161 | -0.284751595 | 0.061867274 | 8.337498598 | -1.581963586 | 0.16583965 | 0.669260659 | -4.377391625 |
| **A_52_P684130** | 2510002D24Rik | -0.081766188 | -0.208929483 | 0.045397107 | 8.34893293 | -1.581907699 | 0.165852271 | 0.669260659 | -4.377432342 |
| **A_55_P2020128** | Dhrs3 | 0.136887009 | -0.076043681 | 0.349817699 | 13.57886343 | 1.58158718 | 0.165924673 | 0.669304203 | -4.377665853 |
| **A_55_P2088750** | Mtif3 | 0.081066452 | -0.045078878 | 0.207211782 | 8.14037711 | 1.581026516 | 0.166051394 | 0.669304203 | -4.378074304 |
| **A_55_P1965892** | Gnpda1 | 0.243885632 | -0.13563269 | 0.623403953 | 13.21167943 | 1.58096637 | 0.166064993 | 0.669304203 | -4.37811812 |
| **A_51_P424781** | Sf3b6 | 0.143508275 | -0.079840365 | 0.366856916 | 12.08088962 | 1.580748563 | 0.16611425 | 0.669304203 | -4.378276788 |
| **A_55_P1988774** | Slc17a5 | 0.162476793 | -0.090401892 | 0.415355478 | 12.37829594 | 1.58069572 | 0.166126203 | 0.669304203 | -4.378315283 |
| **A_55_P2410106** | 9130230N09Rik | 0.117925997 | -0.065646763 | 0.301498756 | 8.524575292 | 1.580412596 | 0.166190257 | 0.669304203 | -4.378521528 |
| **A_55_P1976207** | Arf5 | 0.275330023 | -0.153293499 | 0.703953544 | 12.81754294 | 1.580326176 | 0.166209813 | 0.669304203 | -4.378584481 |
| **A_55_P2015430** | Gpn2 | 0.144805379 | -0.080627251 | 0.370238009 | 9.865445608 | 1.580291045 | 0.166217764 | 0.669304203 | -4.378610072 |

| | | | | | | | | |
|---|---|---|---|---|---|---|---|---|
| **A_55_P2028415** | Hacd2 | -0.158943174 | -0.406402817 | 0.088516469 | 11.73873644 | -1.58018037 | 0.166242813 | 0.669304203 | -4.378690693 |
| **A_55_P2120737** | NA | 0.237034322 | -0.132020267 | 0.606088911 | 8.790329452 | 1.580118977 | 0.16625671 | 0.669304203 | -4.378735413 |
| **A_55_P2075268** | Tmem216 | -0.108942565 | -0.278564908 | 0.060679779 | 11.42856653 | -1.580096822 | 0.166261725 | 0.669304203 | -4.378751552 |
| **A_51_P319460** | Osmr | -0.190977658 | -0.488341731 | 0.106386416 | 10.99689952 | -1.580022343 | 0.166278586 | 0.669304203 | -4.378805805 |
| **A_55_P2207593** | 8430437O03Rik | -0.173226314 | -0.443000772 | 0.096548143 | 8.233969104 | -1.57972758 | 0.166345332 | 0.669433461 | -4.379020516 |
| **A_55_P1957937** | NA | 0.091694751 | -0.051206078 | 0.234595581 | 7.89480295 | 1.57862469 | 0.166595295 | 0.670141221 | -4.379823832 |
| **A_55_P2067308** | NA | -0.126612661 | -0.323950784 | 0.070725462 | 12.11630714 | -1.578466271 | 0.166631229 | 0.670141221 | -4.379939213 |
| **A_55_P2006255** | Txnip | -0.174175802 | -0.445650934 | 0.09729933 | 8.796153411 | -1.578435827 | 0.166638136 | 0.670141221 | -4.379961386 |
| **A_55_P2003378** | Atrnl1 | -0.2009332 | -0.514148948 | 0.112282548 | 8.5627295 | -1.578255366 | 0.16667908 | 0.670141221 | -4.38009282 |
| **A_55_P1976351** | Gpcpd1 | -0.222659155 | -0.569831908 | 0.124513598 | 9.701105418 | -1.577844088 | 0.16677243 | 0.670141221 | -4.380392353 |
| **A_55_P1970755** | Hdac9 | -0.151935099 | -0.388884919 | 0.085014721 | 8.08180664 | -1.577505539 | 0.16684931 | 0.670141221 | -4.380638909 |
| **A_55_P2160204** | NA | -0.150995058 | -0.386489467 | 0.084499351 | 10.82486354 | -1.577434354 | 0.166865479 | 0.670141221 | -4.380690751 |
| **A_55_P2116290** | Szrd1 | -0.118416495 | -0.303111319 | 0.06627833 | 10.45800673 | -1.577344795 | 0.166885824 | 0.670141221 | -4.380755973 |
| **A_55_P1986103** | Suco | 0.108220178 | -0.060575943 | 0.2770163 | 9.064373995 | 1.577302331 | 0.166895472 | 0.670141221 | -4.380786897 |
| **A_51_P270364** | Mmaa | 0.206475983 | -0.115625274 | 0.52857724 | 10.68735889 | 1.577052987 | 0.166952131 | 0.670141221 | -4.38096848 |
| **A_55_P1990538** | Ndfip2 | 0.133210525 | -0.07460172 | 0.341022771 | 8.520046891 | 1.577017701 | 0.16696015 | 0.670141221 | -4.380994176 |
| **A_55_P2008538** | Chmp4b | 0.536685051 | -0.300573095 | 1.373943198 | 12.97421103 | 1.576991237 | 0.166966165 | 0.670141221 | -4.381013448 |
| **A_52_P660242** | Isy1 | 0.201877465 | -0.113067515 | 0.516822445 | 10.04056105 | 1.576965946 | 0.166971914 | 0.670141221 | -4.381031865 |
| **A_55_P2072945** | NA | 0.091486156 | -0.051345908 | 0.23431822 | 15.70943715 | 1.575791777 | 0.167238999 | 0.670678148 | -4.381886882 |
| **A_51_P408471** | Cdipt | -0.195772502 | -0.501452037 | 0.109907033 | 12.38886275 | -1.575630902 | 0.167275625 | 0.670678148 | -4.382004022 |
| **A_66_P134878** | D11Wsu47e | -0.149754506 | -0.383586502 | 0.084077491 | 11.18508192 | -1.575596922 | 0.167283362 | 0.670678148 | -4.382028764 |
| **A_55_P1981205** | Xrcc4 | 0.072066049 | -0.040473447 | 0.184605545 | 8.798751283 | 1.575414753 | 0.167324846 | 0.670678148 | -4.382161407 |
| **A_55_P2056344** | Unc93a2 | 0.884207738 | -0.496904878 | 2.265320354 | 14.06997325 | 1.575050134 | 0.167407909 | 0.670678148 | -4.38242689 |
| **A_55_P2108754** | Ubfd1 | 0.133999426 | -0.075357752 | 0.343356604 | 12.26791885 | 1.574650748 | 0.167498936 | 0.670678148 | -4.382717677 |
| **A_51_P138790** | Plekhf2 | -0.16804464 | -0.430602304 | 0.094513023 | 10.44959726 | -1.574595948 | 0.16751143 | 0.670678148 | -4.382757576 |
| **A_55_P2092953** | Traf5 | -0.094728279 | -0.242735187 | 0.053278629 | 8.565779384 | -1.574587585 | 0.167513337 | 0.670678148 | -4.382763665 |
| **A_55_P2087865** | Tinf2 | -0.086947481 | -0.222800136 | 0.048905173 | 9.012062307 | -1.574555741 | 0.167520597 | 0.670678148 | -4.382786849 |
| **A_55_P2158102** | Camp | -0.164901239 | -0.422590094 | 0.092787616 | 8.121125783 | -1.574336069 | 0.167570691 | 0.670678148 | -4.382946783 |
| **A_55_P2021689** | Chrdl2 | 0.265425701 | -0.149368197 | 0.680219599 | 9.222792736 | 1.574272021 | 0.167585299 | 0.670678148 | -4.382993414 |
| **A_55_P2091196** | Hrct1 | 0.077771242 | -0.04376892 | 0.199311404 | 11.43867104 | 1.574230731 | 0.167594717 | 0.670678148 | -4.383023474 |
| **A_66_P111217** | NA | -0.210564425 | -0.539639975 | 0.118511125 | 13.68446594 | -1.574195062 | 0.167602853 | 0.670678148 | -4.383049443 |
| **A_52_P489778** | Ablim1 | -0.083628587 | -0.214350736 | 0.047093561 | 8.2632852 | -1.573891346 | 0.167672148 | 0.670678148 | -4.383270558 |
| **A_55_P2038587** | NA | 0.15964975 | -0.089914727 | 0.409214227 | 8.227566385 | 1.573818468 | 0.16768878 | 0.670678148 | -4.383323614 |
| **A_51_P343889** | Slc35f6 | -0.125773916 | -0.322386727 | 0.070838896 | 8.946766258 | -1.573794169 | 0.167694325 | 0.670678148 | -4.383341304 |
| **A_55_P1988158** | Esd | -0.221827499 | -0.568594775 | 0.124939777 | 13.70389747 | -1.573788758 | 0.16769556 | 0.670678148 | -4.383345244 |
| **A_51_P168792** | Xylb | -0.153735742 | -0.394105807 | 0.086634324 | 12.34687474 | -1.573488699 | 0.167764057 | 0.670813293 | -4.383563688 |
| **A_55_P2076635** | Vps25 | -0.106534421 | -0.273171066 | 0.060102223 | 9.095243467 | -1.572854694 | 0.167908873 | 0.671160659 | -4.384025226 |
| **A_55_P1983958** | NA | -0.275102275 | -0.705419412 | 0.155214862 | 12.36173605 | -1.572804357 | 0.167920376 | 0.671160659 | -4.384061869 |
| **A_55_P2056190** | Rps3a1 | -0.106455013 | -0.27303242 | 0.060122394 | 15.90684259 | -1.572241243 | 0.168049108 | 0.671509967 | -4.384471775 |
| **A_55_P1967481** | Hbb-y | 0.877922117 | -0.496249427 | 2.25209366 | 9.148649136 | 1.571752653 | 0.168160879 | 0.671509967 | -4.384827416 |
| **A_55_P2091230** | NA | -0.119868658 | -0.307511297 | 0.067773982 | 10.66199396 | -1.571604529 | 0.168194778 | 0.671509967 | -4.384935232 |
| **A_51_P257550** | Marcksl1 | -0.332897615 | -0.85404301 | 0.18824778 | 12.07457927 | -1.571523728 | 0.168213273 | 0.671509967 | -4.384994043 |
| **A_52_P674414** | Mkln1 | -0.148269871 | -0.380422234 | 0.083882492 | 9.614704643 | -1.57126329 | 0.168272898 | 0.671509967 | -4.385183604 |
| **A_52_P228031** | Tmem179b | -0.121779395 | -0.312459022 | 0.068900233 | 11.06403218 | -1.57122608 | 0.168281418 | 0.671509967 | -4.385210686 |
| **A_55_P2001250** | Cpz | 0.441639205 | -0.249923885 | 1.133202295 | 9.031647784 | 1.571104151 | 0.168309341 | 0.671509967 | -4.38529943 |
| **A_51_P483557** | Ube2b | -0.140609941 | -0.360912319 | 0.079692437 | 10.83210988 | -1.570239881 | 0.168507393 | 0.671509967 | -4.385928447 |
| **A_52_P55717** | Adnp | -0.077015536 | -0.197686439 | 0.043655368 | 8.278335097 | -1.570163715 | 0.168524858 | 0.671509967 | -4.385983878 |
| **A_55_P2048759** | Mgarp | 0.113876856 | -0.064557238 | 0.29231095 | 8.392761962 | 1.570098115 | 0.168539901 | 0.671509967 | -4.386031619 |
| **A_55_P1963718** | Lias | -0.176969407 | -0.45426552 | 0.100326706 | 11.07206127 | -1.570086488 | 0.168542567 | 0.671509967 | -4.386040081 |
| **A_55_P1954986** | Elk1 | -0.165266379 | -0.424239998 | 0.09370724 | 17.45427607 | -1.569994568 | 0.168563648 | 0.671509967 | -4.386106976 |

| | | | | | | | | | |
|---|---|---|---|---|---|---|---|---|---|
| **A_55_P2092968** | Traf3 | -0.128871041 | -0.330824291 | 0.073082209 | 11.06081533 | -1.569906018 | 0.168583959 | 0.671509967 | -4.386171419 |
| **A_51_P420236** | Srrt | -0.079980837 | -0.205337747 | 0.045376074 | 7.954279269 | -1.569664464 | 0.168639376 | 0.671509967 | -4.386347207 |
| **A_55_P2015118** | NA | 0.165076978 | -0.09365673 | 0.423810686 | 10.32495149 | 1.569649403 | 0.168642832 | 0.671509967 | -4.386358167 |
| **A_51_P494775** | Rcor1 | -0.253627091 | -0.651151368 | 0.143897187 | 12.80063655 | -1.569643936 | 0.168644086 | 0.671509967 | -4.386362146 |
| **A_65_P03031** | Ncapd3 | -0.067519662 | -0.173354855 | 0.038315531 | 7.931512541 | -1.569528907 | 0.168670483 | 0.671509967 | -4.386445856 |
| **A_52_P579252** | NA | -0.183613229 | -0.47145175 | 0.104225292 | 12.65780969 | -1.569365916 | 0.168707892 | 0.671509967 | -4.386564467 |
| **A_55_P2067682** | Atp5l | 0.295285135 | -0.167647487 | 0.758217757 | 13.82592468 | 1.56925298 | 0.168733818 | 0.671509967 | -4.386646651 |
| **A_51_P455807** | Ehd4 | -0.095650858 | -0.245618324 | 0.054316608 | 13.40864258 | -1.569137419 | 0.16876035 | 0.671509967 | -4.386730744 |
| **A_51_P216219** | Ptk2 | 0.150507099 | -0.08546968 | 0.386483877 | 10.6491428 | 1.569122602 | 0.168763752 | 0.671509967 | -4.386741527 |
| **A_55_P1952638** | Fgd6 | 0.182197361 | -0.103472483 | 0.467867205 | 10.36666796 | 1.56908636 | 0.168772074 | 0.671509967 | -4.3867679 |
| **A_52_P273821** | Abhd5 | -0.096779377 | -0.248536532 | 0.054977778 | 9.681640655 | -1.568927263 | 0.16880861 | 0.671517109 | -4.386883673 |
| **A_51_P174434** | Cenpj | -0.15151295 | -0.389148081 | 0.086122181 | 9.553368734 | -1.568585774 | 0.168887059 | 0.671628327 | -4.387132163 |
| **A_55_P2175291** | NA | -0.112471399 | -0.288884399 | 0.063941601 | 15.17776895 | -1.568486106 | 0.168909961 | 0.671628327 | -4.387204687 |
| **A_51_P170946** | Hnrnpm | -0.163715324 | -0.420527131 | 0.093096483 | 12.76444073 | -1.568351876 | 0.168940811 | 0.671628327 | -4.387302358 |
| **A_55_P2005743** | Ubxn11 | -0.083511388 | -0.21452882 | 0.047506044 | 7.97883402 | -1.568143423 | 0.168988729 | 0.671680678 | -4.387454035 |
| **A_55_P2117741** | Nck2 | -0.146006637 | -0.375111423 | 0.08309815 | 12.13662018 | -1.567861143 | 0.169053638 | 0.67180053 | -4.387659427 |
| **A_55_P1983149** | Rps29 | -0.085591293 | -0.21992137 | 0.048738784 | 17.08180967 | -1.567564738 | 0.169121822 | 0.671856968 | -4.38787509 |
| **A_55_P2061061** | Set | -0.093330336 | -0.23981397 | 0.053153299 | 11.18680958 | -1.567483162 | 0.169140592 | 0.671856968 | -4.387934443 |
| **A_55_P2136743** | Slc36a4 | -0.145263205 | -0.373286792 | 0.082760383 | 8.733556952 | -1.567274293 | 0.169188659 | 0.671856968 | -4.38808641 |
| **A_51_P515496** | Rbm17 | -0.180111663 | -0.462852098 | 0.102628772 | 11.37142135 | -1.567195148 | 0.169206877 | 0.671856968 | -4.388143993 |
| **A_66_P122087** | Zfp622 | -0.187828291 | -0.482801015 | 0.107144434 | 14.47088528 | -1.566564603 | 0.16935208 | 0.672034479 | -4.388602737 |
| **A_55_P2149763** | Tapbp | -0.169133354 | -0.434747062 | 0.096480355 | 12.47950174 | -1.566563266 | 0.169352388 | 0.672034479 | -4.38860371 |
| **A_51_P248044** | Lamb2 | -0.237050401 | -0.609327337 | 0.135226534 | 10.71949344 | -1.566548077 | 0.169355887 | 0.672034479 | -4.38861476 |
| **A_55_P1989452** | Nenf | 0.121729251 | -0.069488039 | 0.312946541 | 13.63628674 | 1.566162975 | 0.16944463 | 0.672202907 | -4.388894921 |
| **A_51_P148060** | Ptgfrn | -0.114963518 | -0.295572869 | 0.065645832 | 8.183301167 | -1.565989978 | 0.169484509 | 0.672202907 | -4.389020773 |
| **A_55_P1966987** | Limk1 | 0.240800408 | -0.137592805 | 0.619193622 | 12.1358113 | 1.565608044 | 0.169572585 | 0.672202907 | -4.389298614 |
| **A_55_P1988708** | Gstz1 | 0.182698711 | -0.10440219 | 0.469799611 | 9.957827507 | 1.565561351 | 0.169583355 | 0.672202907 | -4.389332581 |
| **A_66_P131036** | NA | 0.308884021 | -0.176539579 | 0.794307621 | 9.517819782 | 1.565466285 | 0.169605286 | 0.672202907 | -4.389401735 |
| **A_55_P2189571** | NA | -0.175171873 | -0.450501367 | 0.100157621 | 8.932999794 | -1.5652395 | 0.169657614 | 0.672202907 | -4.389566705 |
| **A_52_P456640** | Fgr | -0.070687444 | -0.181798909 | 0.04042402 | 7.765677848 | -1.565137768 | 0.169681092 | 0.672202907 | -4.389640707 |
| **A_55_P2017082** | Hnrnpa2b1 | 0.080980987 | -0.046326749 | 0.208288723 | 12.49557291 | 1.564939078 | 0.169726956 | 0.672202907 | -4.389785235 |
| **A_51_P517012** | Tysnd1 | -0.100105053 | -0.257480019 | 0.057269912 | 10.46104085 | -1.56491054 | 0.169733544 | 0.672202907 | -4.389805993 |
| **A_55_P2100973** | Irf6 | 0.183347554 | -0.104902808 | 0.471597915 | 9.612691089 | 1.564856153 | 0.169746101 | 0.672202907 | -4.389845554 |
| **A_55_P2218223** | Pacs2 | -0.185150363 | -0.476349594 | 0.106048868 | 9.701176576 | -1.564240434 | 0.169888318 | 0.672242594 | -4.39029341 |
| **A_51_P456914** | Bmt2 | -0.094754586 | -0.243796217 | 0.054287045 | 8.765927988 | -1.564090244 | 0.169923026 | 0.672242594 | -4.39040265 |
| **A_55_P1952502** | Mapk7 | 0.140976485 | -0.080772607 | 0.362725578 | 10.96026273 | 1.564062099 | 0.169929531 | 0.672242594 | -4.39042312 |
| **A_52_P303928** | Rab2a | -0.180319933 | -0.463969592 | 0.103329726 | 11.27150217 | -1.563977985 | 0.169948972 | 0.672242594 | -4.390484299 |
| **A_51_P314264** | Ssr2 | -0.108544677 | -0.279333563 | 0.06224421 | 11.98383189 | -1.563572726 | 0.170042671 | 0.672242594 | -4.39077905 |
| **A_51_P300378** | Wwc2 | -0.203710785 | -0.524294065 | 0.116872495 | 13.23131629 | -1.563299962 | 0.170105764 | 0.672242594 | -4.390977429 |
| **A_55_P2162369** | Cacfd1 | -0.075731499 | -0.194912201 | 0.043449203 | 8.507120167 | -1.563290824 | 0.170107879 | 0.672242594 | -4.390984075 |
| **A_52_P103977** | Ints8 | -0.075258597 | -0.193697938 | 0.043180745 | 8.296163817 | -1.563253082 | 0.17011661 | 0.672242594 | -4.391011524 |
| **A_55_P2168781** | NA | -0.222587469 | -0.572942939 | 0.127768 | 11.90593521 | -1.563007179 | 0.170173512 | 0.672242594 | -4.391190361 |
| **A_55_P2051521** | NA | 0.139173997 | -0.07990507 | 0.358253063 | 8.460793869 | 1.562882687 | 0.170202327 | 0.672242594 | -4.391280898 |
| **A_55_P2058493** | NA | 0.108958193 | -0.062564757 | 0.280481143 | 15.47478127 | 1.562812293 | 0.170218622 | 0.672242594 | -4.391332092 |
| **A_51_P506417** | Krt14 | -0.257548738 | -0.662986902 | 0.147889427 | 9.785292358 | -1.562802012 | 0.170221002 | 0.672242594 | -4.391339569 |
| **A_55_P1959606** | Tom1l1 | 0.269684911 | -0.154888582 | 0.694258403 | 10.87277644 | 1.562690363 | 0.170246851 | 0.672242594 | -4.391420764 |
| **A_51_P102860** | Herpud2 | -0.085221558 | -0.219433928 | 0.048990812 | 10.12297759 | -1.562162061 | 0.170369211 | 0.672242594 | -4.38956957 |
| **A_55_P2022840** | Csnk2a1 | -0.12612183 | -0.324754973 | 0.072511313 | 11.43361279 | -1.562095969 | 0.170384525 | 0.672242594 | -4.391853017 |
| **A_51_P448391** | Nkiras1 | -0.07976863 | -0.205419039 | 0.04588178 | 8.683738986 | -1.561843042 | 0.17044314 | 0.672242594 | -4.392036941 |
| **A_55_P2160716** | NA | -0.104049299 | -0.26794652 | 0.059847922 | 15.74875654 | -1.561840688 | 0.170443686 | 0.672242594 | -4.392038653 |

| | | | | | | | | | |
|---|---|---|---|---|---|---|---|---|---|
| **A_52_P401484** | Inha | -0.09049471 | -0.233068851 | 0.052079431 | 8.186962839 | -1.561534536 | 0.170514662 | 0.672242594 | -4.392261276 |
| **A_55_P1984496** | Usp48 | 0.14614636 | -0.084112548 | 0.376405267 | 10.73792555 | 1.561495458 | 0.170523723 | 0.672242594 | -4.392289691 |
| **A_66_P109296** | NA | 0.174905318 | -0.100704432 | 0.450515067 | 10.47432962 | 1.56126851 | 0.170576358 | 0.672242594 | -4.392454714 |
| **A_55_P1981011** | Slc35c2 | 0.073487419 | -0.042314487 | 0.189289325 | 9.478992564 | 1.561228423 | 0.170585657 | 0.672242594 | -4.392483863 |
| **A_55_P2031112** | Dhps | 0.098798815 | -0.056923661 | 0.254521291 | 10.96214904 | 1.560880301 | 0.170666429 | 0.672242594 | -4.392736988 |
| **A_55_P2012734** | Fndc10 | 0.156869837 | -0.090385959 | 0.404125634 | 9.315141364 | 1.560853446 | 0.170672661 | 0.672242594 | -4.392756514 |
| **A_55_P2091364** | Cep164 | 0.121328162 | -0.069932793 | 0.312589117 | 11.04154037 | 1.56064621 | 0.170720763 | 0.672242594 | -4.392907195 |
| **A_55_P2045278** | Gm10845 | -0.189544512 | -0.488362457 | 0.109273433 | 17.32693444 | -1.560535683 | 0.170746423 | 0.672242594 | -4.392987557 |
| **A_55_P2069554** | Brix1 | -0.14047046 | -0.361928388 | 0.080987467 | 11.62386156 | -1.560496999 | 0.170755405 | 0.672242594 | -4.393015684 |
| **A_55_P2013730** | Islr2 | -0.232593048 | -0.599293839 | 0.134107744 | 10.27561157 | -1.560465046 | 0.170762824 | 0.672242594 | -4.393038916 |
| **A_55_P2083954** | Scfd1 | -0.139330155 | -0.359001009 | 0.080340699 | 9.652241721 | -1.560421207 | 0.170773004 | 0.672242594 | -4.39307079 |
| **A_66_P106191** | Eef2 | -0.215451095 | -0.555160207 | 0.124258017 | 14.35039512 | -1.560309213 | 0.170799012 | 0.672242594 | -4.393152217 |
| **A_55_P2000833** | E2f8 | 0.210595337 | -0.121483922 | 0.542674595 | 11.04797738 | 1.560185229 | 0.170827809 | 0.672242594 | -4.39324236 |
| **A_55_P2131844** | NA | 0.369296657 | -0.213089668 | 0.951682982 | 10.49021977 | 1.560031511 | 0.170863519 | 0.672242594 | -4.393354121 |
| **A_55_P1963175** | Lctl | 0.156784659 | -0.090470855 | 0.404040173 | 8.387197904 | 1.560007705 | 0.17086905 | 0.672242594 | -4.393371429 |
| **A_51_P454217** | Nfkb2 | -0.112597615 | -0.29018779 | 0.064992559 | 8.424564081 | -1.559837731 | 0.170908545 | 0.672261145 | -4.393495006 |
| **A_55_P2179974** | Spsb1 | 0.174658558 | -0.100862253 | 0.450179369 | 9.433218515 | 1.559569111 | 0.17097098 | 0.672276562 | -4.393690298 |
| **A_55_P2172375** | NA | 0.125752922 | -0.07263773 | 0.324143574 | 8.537079233 | 1.559430571 | 0.171003189 | 0.672276562 | -4.393791017 |
| **A_55_P2143832** | Col4a4 | 0.076636259 | -0.044271376 | 0.197543895 | 7.809543998 | 1.559372002 | 0.171016807 | 0.672276562 | -4.393833596 |
| **A_55_P1955072** | Alad | -0.116225178 | -0.299659613 | 0.067209257 | 10.33249258 | -1.558793255 | 0.171151431 | 0.672643665 | -4.394254329 |
| **A_55_P2009427** | Nrsn2 | 0.476348516 | -0.275515701 | 1.228212733 | 10.63150364 | 1.558671389 | 0.171179791 | 0.672643665 | -4.394342919 |
| **A_51_P437786** | Rbm28 | -0.167149009 | -0.431155787 | 0.096857769 | 11.46550564 | -1.557607004 | 0.171427683 | 0.67348083 | -4.395116624 |
| **A_51_P388374** | Ppil4 | -0.093703539 | -0.241729642 | 0.054322564 | 10.33530152 | -1.557352231 | 0.171487069 | 0.673577232 | -4.395301806 |
| **A_51_P273609** | Itpka | -0.237690559 | -0.613264783 | 0.137883665 | 12.69818903 | -1.556988183 | 0.171571961 | 0.673773757 | -4.395566407 |
| **A_51_P232474** | Zswim1 | 0.221585124 | -0.128609907 | 0.571780155 | 9.076759452 | 1.556681575 | 0.171643489 | 0.673818227 | -4.395789251 |
| **A_51_P346104** | Sgta | -0.140617261 | -0.362855809 | 0.081621287 | 11.71306413 | -1.556640789 | 0.171653006 | 0.673818227 | -4.395818894 |
| **A_51_P377526** | Sts | 0.279239985 | -0.16219329 | 0.72067326 | 10.48770577 | 1.556258378 | 0.171742262 | 0.674031713 | -4.396096821 |
| **A_55_P2141415** | Ppp6r3 | -0.106557845 | -0.275053522 | 0.061937832 | 8.133387748 | -1.555843211 | 0.171839214 | 0.674181061 | -4.396398542 |
| **A_52_P121491** | Msrb3 | -0.102156549 | -0.263697458 | 0.059384361 | 8.916771486 | -1.555796719 | 0.171850074 | 0.674181061 | -4.39643233 |
| **A_52_P551526** | NA | 0.084757737 | -0.04932112 | 0.218836593 | 15.63920768 | 1.555207045 | 0.171987875 | 0.674435391 | -4.396860849 |
| **A_55_P1996087** | Gm14326 | -0.107464764 | -0.27747229 | 0.062542761 | 11.31932926 | -1.555131461 | 0.172005546 | 0.674435391 | -4.396915774 |
| **A_52_P15212** | Ctbp2 | -0.163084911 | -0.421092571 | 0.094922749 | 10.92118297 | -1.555071434 | 0.172019581 | 0.674435391 | -4.396959394 |
| **A_55_P2012369** | Eea1 | -0.106734072 | -0.27561191 | 0.062143766 | 10.49667519 | -1.554889683 | 0.172062083 | 0.674465219 | -4.397091466 |
| **A_55_P1990092** | Snrpa | 0.085630441 | -0.049881228 | 0.22114211 | 10.69566387 | 1.554607107 | 0.172128182 | 0.674587517 | -4.397296799 |
| **A_55_P2160323** | NA | -0.084688638 | -0.218758615 | 0.04938134 | 8.302635182 | -1.554042079 | 0.172260424 | 0.674968931 | -4.397707358 |
| **A_52_P221024** | NA | -0.188275699 | -0.486411007 | 0.109859609 | 11.91819181 | -1.553638663 | 0.1723549 | 0.675070143 | -4.398000471 |
| **A_55_P1985788** | Usp2 | 0.14014674 | -0.081806139 | 0.362099618 | 10.81827423 | 1.5534289 | 0.172404044 | 0.675070143 | -4.398152876 |
| **A_55_P2048259** | Golph3l | 0.116233476 | -0.067862034 | 0.300328986 | 11.45821122 | 1.553306614 | 0.1724327 | 0.675070143 | -4.398241722 |
| **A_55_P2102624** | Eaf2 | 0.142550177 | -0.083227987 | 0.36832834 | 8.821446763 | 1.553298728 | 0.172434548 | 0.675070143 | -4.398247452 |
| **A_55_P1984690** | Ptprr | 0.185873394 | -0.108543564 | 0.480290352 | 8.587986466 | 1.553186373 | 0.172460881 | 0.675070143 | -4.398329081 |
| **A_52_P254149** | Irgq | -0.096532205 | -0.24947525 | 0.05641084 | 9.476926368 | -1.552786176 | 0.172554707 | 0.675140379 | -4.39861983 |
| **A_55_P2093920** | Pzp | -0.347312741 | -0.897588189 | 0.202962707 | 8.071147748 | -1.552779334 | 0.172556311 | 0.675140379 | -4.398624801 |
| **A_55_P1996504** | Abcg2 | -0.242029891 | -0.625526019 | 0.141466237 | 12.13041555 | -1.552662932 | 0.172583611 | 0.675140379 | -4.398709366 |
| **A_55_P2014611** | Ccdc12 | -0.118246421 | -0.305658187 | 0.069165345 | 10.54823196 | -1.552245163 | 0.172681623 | 0.67538711 | -4.399012864 |
| **A_55_P1993121** | NA | -0.202783085 | -0.524211883 | 0.118645713 | 15.24821528 | -1.552087156 | 0.172718707 | 0.675395486 | -4.399127648 |
| **A_55_P1978716** | Zim1 | -0.213791157 | -0.552723211 | 0.125140898 | 9.451730205 | -1.551837546 | 0.172777305 | 0.67548797 | -4.399308973 |
| **A_55_P2052831** | 0610009B22Rik | -0.202259285 | -0.523013408 | 0.118494838 | 9.791388563 | -1.551334256 | 0.172895514 | 0.675739043 | -4.399674567 |
| **A_55_P1989061** | Tsc22d3 | -0.195143253 | -0.5046259 | 0.114339394 | 14.12438159 | -1.551266454 | 0.172911444 | 0.675739043 | -4.399723817 |
| **A_55_P2041758** | Scand1 | -0.207435332 | -0.536450242 | 0.121579578 | 13.79298838 | -1.551087562 | 0.172953483 | 0.675766702 | -4.39985376 |
| **A_55_P2067563** | NA | -0.275144144 | -0.711678425 | 0.161390137 | 9.425377059 | -1.550640356 | 0.173058617 | 0.676040826 | -4.400178588 |

| | | | | | | | | | |
|---|---|---|---|---|---|---|---|---|---|
| **A_51_P346923** | P3h1 | -0.119006251 | -0.307856604 | 0.069844103 | 8.390893435 | -1.550319225 | 0.173134149 | 0.676193874 | -4.400411832 |
| **A_55_P2023912** | NA | 0.271641796 | -0.15946478 | 0.702748373 | 11.30886254 | 1.550176358 | 0.173167762 | 0.676193874 | -4.400515596 |
| **A_51_P466490** | Armc5 | -0.121431573 | -0.314177192 | 0.071314046 | 9.797013856 | -1.549944929 | 0.173222225 | 0.676199109 | -4.400683681 |
| **A_51_P419439** | Gnmt | 0.144670895 | -0.084972583 | 0.374314373 | 8.097707849 | 1.549873365 | 0.17323907 | 0.676199109 | -4.400735656 |
| **A_52_P86158** | Mtx2 | -0.152693678 | -0.395172421 | 0.089785065 | 10.94549898 | -1.549232312 | 0.17339003 | 0.676492628 | -4.40120122 |
| **A_51_P285779** | Asphd2 | 0.076827768 | -0.045183481 | 0.198839016 | 7.937744738 | 1.54912872 | 0.173414437 | 0.676492628 | -4.401276451 |
| **A_55_P1997872** | Apip | 0.268809282 | -0.158096133 | 0.695714696 | 10.38816296 | 1.549108228 | 0.173419265 | 0.676492628 | -4.401291333 |
| **A_52_P485766** | Zdhhc17 | -0.082860911 | -0.214492472 | 0.048770651 | 8.679462027 | -1.54866981 | 0.173522595 | 0.676636844 | -4.40160971 |
| **A_55_P2055087** | Prkar2b | -0.140866266 | -0.364657007 | 0.082924476 | 9.935423663 | -1.548581441 | 0.17354343 | 0.676636844 | -4.401673882 |
| **A_55_P2100864** | Tma7 | 0.16688713 | -0.098255584 | 0.432029844 | 14.95886001 | 1.54850393 | 0.173561706 | 0.676636844 | -4.401730168 |
| **A_51_P169693** | Bst2 | -0.280131891 | -0.725263569 | 0.164999787 | 13.7230884 | -1.548257574 | 0.173619807 | 0.676636844 | -4.401909061 |
| **A_52_P625431** | Hnrnpa0 | -0.075354936 | -0.195098253 | 0.04438838 | 8.09276776 | -1.548208997 | 0.173631266 | 0.676636844 | -4.401944335 |
| **A_55_P1953884** | Dpm1 | -0.139112244 | -0.360246105 | 0.082021618 | 11.71590622 | -1.547673226 | 0.173757696 | 0.676732071 | -4.40233337 |
| **A_51_P396297** | Hsdl1 | -0.077527966 | -0.20077188 | 0.045715948 | 8.227334501 | -1.547611933 | 0.173772165 | 0.676732071 | -4.402377875 |
| **A_66_P121369** | Gars | -0.19717653 | -0.510625718 | 0.116272657 | 14.4724621 | -1.547594702 | 0.173776233 | 0.676732071 | -4.402390386 |
| **A_52_P504743** | Cdyl | -0.081392544 | -0.210788216 | 0.048003127 | 8.651226263 | -1.547512046 | 0.173795747 | 0.676732071 | -4.402450402 |
| **A_55_P2096827** | Art3 | -0.085729219 | -0.222040384 | 0.050581945 | 8.103050204 | -1.547271782 | 0.173852484 | 0.676733228 | -4.402624853 |
| **A_51_P124798** | Tmem192 | 0.10848602 | -0.064050018 | 0.281022058 | 9.518508901 | 1.546903129 | 0.173939573 | 0.676733228 | -4.399773533 |
| **A_55_P2137701** | Rex2 | 0.236633741 | -0.139772101 | 0.613039582 | 9.201968162 | 1.546640855 | 0.174001557 | 0.676733228 | -4.403082934 |
| **A_55_P2099019** | NA | 0.108576406 | -0.064150663 | 0.281303475 | 11.55838768 | 1.546479688 | 0.174039656 | 0.676733228 | -4.403199943 |
| **A_55_P2117202** | Mtx1 | -0.087989277 | -0.227965742 | 0.051987188 | 12.01768741 | -1.546478522 | 0.174039931 | 0.676733228 | -4.40320079 |
| **A_55_P1985498** | Gm12005 | -0.147133573 | -0.381201078 | 0.086933931 | 13.48694484 | -1.546464023 | 0.174043359 | 0.676733228 | -4.403211316 |
| **A_55_P2088203** | Hcn2 | 0.136132144 | -0.080447486 | 0.352711775 | 9.459912978 | 1.546365816 | 0.174066579 | 0.676733228 | -4.403282615 |
| **A_55_P1983426** | Abhd14a | 0.182722427 | -0.107987677 | 0.473432532 | 9.495847804 | 1.546325406 | 0.174076135 | 0.676733228 | -4.403311953 |
| **A_55_P2030284** | Trem1 | 0.145197667 | -0.085842175 | 0.376237509 | 8.838039991 | 1.546115455 | 0.174125788 | 0.676790138 | -4.403464374 |
| **A_52_P585751** | Gns | -0.422171856 | -1.094103734 | 0.249760023 | 14.32706952 | -1.54572843 | 0.174217354 | 0.6770099 | -4.40374534 |
| **A_52_P594927** | Atl3 | -0.168798357 | -0.437545171 | 0.099948458 | 8.933766688 | -1.545233274 | 0.17433457 | 0.677259804 | -4.404104787 |
| **A_55_P2037221** | Klhdc10 | -0.194143636 | -0.510650638 | 0.122363367 | 11.26822785 | -1.589607553 | 0.17435174 | 0.677259804 | -4.380195893 |
| **A_55_P2035454** | Oaz1 | 0.126666544 | -0.075075768 | 0.328408855 | 14.60606447 | 1.544664235 | 0.174469367 | 0.67758055 | -4.404517844 |
| **A_55_P2058628** | Hnrnpa3 | -0.199445772 | -0.517180066 | 0.118288522 | 9.546845001 | -1.544293722 | 0.17455719 | 0.677612527 | -4.40478678 |
| **A_55_P2116794** | NA | 0.49396489 | -0.293064252 | 1.280994032 | 12.97154813 | 1.544096471 | 0.174603961 | 0.677612527 | -4.40492995 |
| **A_51_P303180** | Fam114a1 | -0.100525325 | -0.260691902 | 0.059641253 | 10.61575053 | -1.544090506 | 0.174605376 | 0.677612527 | -4.40493428 |
| **A_55_P2144110** | Mink1 | 0.213880758 | -0.126906143 | 0.554667659 | 10.17400658 | 1.544037998 | 0.174617828 | 0.677612527 | -4.40497239 |
| **A_55_P2063166** | Ogfod1 | 0.512407753 | -0.304358021 | 1.329173526 | 14.48943245 | 1.543431377 | 0.174761753 | 0.677877271 | -4.405412669 |
| **A_55_P1978089** | Dcaf17 | -0.094265728 | -0.244554052 | 0.056022595 | 9.397967425 | -1.543113069 | 0.174837318 | 0.677877271 | -4.405643681 |
| **A_51_P359293** | Ubxn2b | -0.066885572 | -0.173538022 | 0.039766878 | 8.371445643 | -1.542875108 | 0.174893829 | 0.677877271 | -4.405816376 |
| **A_51_P116906** | Rapgef3 | -0.153596605 | -0.39852273 | 0.091329521 | 10.99026398 | -1.542821445 | 0.174906576 | 0.677877271 | -4.40585532 |
| **A_66_P127463** | NA | 0.383673082 | -0.22816354 | 0.995509704 | 12.51916303 | 1.542748119 | 0.174923994 | 0.677877271 | -4.405908533 |
| **A_55_P2020463** | Rufy2 | -0.187542949 | -0.486631057 | 0.111545159 | 10.72481085 | -1.542661915 | 0.174944473 | 0.677877271 | -4.405971092 |
| **A_55_P2072468** | Sirt6 | 0.105380144 | -0.062705429 | 0.273465716 | 9.974261714 | 1.542401759 | 0.175006293 | 0.677877271 | -4.406159885 |
| **A_51_P448946** | Sertad2 | -0.162306295 | -0.421194477 | 0.096581888 | 11.20626247 | -1.542383248 | 0.175010692 | 0.677877271 | -4.406173318 |
| **A_52_P412321** | Pcgf6 | -0.090605989 | -0.235133567 | 0.05392159 | 9.01045156 | -1.542323021 | 0.175025007 | 0.677877271 | -4.406217024 |
| **A_52_P655084** | Tbc1d17 | -0.106726361 | -0.276982363 | 0.063529641 | 10.41372816 | -1.542191957 | 0.175056161 | 0.677877271 | -4.406312132 |
| **A_51_P233027** | Rmdn3 | -0.152100045 | -0.394749113 | 0.090549023 | 9.788829902 | -1.542126063 | 0.175071827 | 0.677877271 | -4.406359949 |
| **A_55_P1989673** | Slco2a1 | -0.27322525 | -0.709214905 | 0.162764405 | 15.48664596 | -1.541749479 | 0.17516138 | 0.678088187 | -4.406633214 |
| **A_55_P2043377** | NA | -0.073499404 | -0.19082117 | 0.043822362 | 8.493616668 | -1.541254611 | 0.175279128 | 0.678340768 | -4.406992294 |
| **A_55_P2018557** | NA | -0.123626699 | -0.320989947 | 0.07373655 | 11.5623096 | -1.541044395 | 0.175329169 | 0.678340768 | -4.407144823 |
| **A_52_P339996** | Slc6a6 | -0.243989805 | -0.633509065 | 0.145529455 | 12.1829167 | -1.541032883 | 0.175331909 | 0.678340768 | -4.407153175 |
| **A_55_P2068233** | Phactr1 | -0.212611957 | -0.552114503 | 0.12689059 | 11.97632371 | -1.540684848 | 0.175414788 | 0.678525604 | -4.407405694 |
| **A_51_P143915** | Rnf181 | -0.130485588 | -0.338879113 | 0.077907937 | 10.99255618 | -1.540449582 | 0.175470834 | 0.678539531 | -4.407576386 |

| | | | | | | | | | |
|---|---|---|---|---|---|---|---|---|---|
| **A_55_P2152437** | Mitf | -0.162147985 | -0.421172769 | 0.096876798 | 12.14348206 | -1.540066243 | 0.175562191 | 0.678539531 | -4.407854501 |
| **A_52_P633128** | Aasdhppt | 0.117923105 | -0.070466214 | 0.306312424 | 10.43662829 | 1.539968345 | 0.175585529 | 0.678539531 | -4.407925525 |
| **A_55_P1985495** | Gm12005 | -0.196864482 | -0.511379985 | 0.117651022 | 13.89621606 | -1.539906927 | 0.175600172 | 0.678539531 | -4.407970082 |
| **A_51_P413545** | Txndc11 | -0.129988284 | -0.337678631 | 0.077702063 | 11.9656376 | -1.539774286 | 0.1756318 | 0.678539531 | -4.408066309 |
| **A_55_P2008740** | Fcgr1 | -0.113682603 | -0.295330329 | 0.067965122 | 8.009351657 | -1.53968972 | 0.175651967 | 0.678539531 | -4.408127659 |
| **A_55_P2010758** | Npm3 | 0.119327821 | -0.071352407 | 0.31000805 | 8.303682299 | 1.53959046 | 0.175675641 | 0.678539531 | -4.408199667 |
| **A_55_P2020200** | Snx6 | -0.09563915 | -0.248475815 | 0.057197515 | 8.740905602 | -1.539491583 | 0.175699227 | 0.678539531 | -4.408271397 |
| **A_55_P2054883** | Trappc1 | -0.101917679 | -0.264830917 | 0.060995558 | 8.933835121 | -1.539083968 | 0.17579649 | 0.678565056 | -4.408567092 |
| **A_55_P2024993** | Hoxb4 | 0.126397985 | -0.075651873 | 0.328447843 | 9.312312759 | 1.539043034 | 0.17580626 | 0.678565056 | -4.408596786 |
| **A_52_P402897** | Cdh4 | -0.104106012 | -0.270538752 | 0.062326728 | 8.217723756 | -1.538885152 | 0.175843949 | 0.678565056 | -4.408711313 |
| **A_55_P1963201** | Chchd7 | 0.163339038 | -0.097792817 | 0.424470893 | 11.76795441 | 1.53886068 | 0.175849791 | 0.678565056 | -4.408729065 |
| **A_51_P175232** | Fbxl4 | -0.066893564 | -0.173846206 | 0.040059078 | 9.736066111 | -1.538728438 | 0.175881366 | 0.678565056 | -4.408824992 |
| **A_51_P158772** | Elof1 | 0.25189352 | -0.150905099 | 0.654692138 | 12.77202615 | 1.538502424 | 0.175935343 | 0.678637846 | -4.408988936 |
| **A_52_P24843** | Pten | -0.132855637 | -0.345387253 | 0.079675979 | 8.775233995 | -1.537891173 | 0.176081401 | 0.678809945 | -4.409432298 |
| **A_55_P2063396** | Zfand2a | -0.337466436 | -0.877355809 | 0.202422937 | 14.06243302 | -1.537782692 | 0.176107334 | 0.678809945 | -4.40951098 |
| **A_55_P1963031** | Ampd2 | 0.117121072 | -0.070253022 | 0.304495166 | 9.258422121 | 1.537781581 | 0.1761076 | 0.678809945 | -4.409511786 |
| **A_55_P2398534** | Gid8 | 0.190550109 | -0.114308785 | 0.495409003 | 10.75040853 | 1.5377279 | 0.176120434 | 0.678809945 | -4.40955072 |
| **A_51_P188772** | Arhgef28 | 0.084914307 | -0.050977238 | 0.220805852 | 8.748027336 | 1.537296352 | 0.176223643 | 0.679042788 | -4.409863714 |
| **A_52_P137765** | Lmna | -0.203629708 | -0.529530291 | 0.122270875 | 12.63773099 | -1.537181552 | 0.176251108 | 0.679042788 | -4.409946973 |
| **A_51_P375533** | Polr3b | -0.116817023 | -0.303832305 | 0.07019826 | 8.929969229 | -1.536732227 | 0.176358645 | 0.679321691 | -4.410272838 |
| **A_66_P116804** | Smchd1 | -0.190723439 | -0.496129448 | 0.11468257 | 11.23721939 | -1.536369422 | 0.17644552 | 0.679437825 | -4.410535944 |
| **A_55_P2138816** | Mplkip | -0.101730756 | -0.264667589 | 0.061206078 | 9.382975142 | -1.536038712 | 0.176524745 | 0.679437825 | -4.410775765 |
| **A_52_P155124** | Zfp229 | -0.191769774 | -0.498930891 | 0.115391342 | 8.589027092 | -1.535971236 | 0.176540914 | 0.679437825 | -4.410824695 |
| **A_55_P2321453** | D11Ertd726e | -0.220595135 | -0.573963667 | 0.132773398 | 8.212355938 | -1.535809042 | 0.176579785 | 0.679437825 | -4.410942309 |
| **A_51_P116932** | Lad1 | -0.108056058 | -0.281151377 | 0.065039262 | 8.873416574 | -1.535793968 | 0.176583398 | 0.679437825 | -4.41095324 |
| **A_52_P51078** | Ctsh | 0.574031299 | -0.345705547 | 1.493768146 | 13.36233562 | 1.535469347 | 0.176661221 | 0.679437825 | -4.41118863 |
| **A_55_P2039110** | NA | 0.251372526 | -0.151422538 | 0.65416759 | 15.34653707 | 1.535333877 | 0.176693708 | 0.679437825 | -4.41128686 |
| **A_51_P493649** | Sult1e1 | -1.131356145 | -2.9442383 | 0.68152601 | 12.52480504 | -1.535320077 | 0.176697018 | 0.679437825 | -4.411296866 |
| **A_51_P464234** | Fkbp2 | -0.18057109 | -0.469934226 | 0.108792046 | 12.79381464 | -1.535232587 | 0.176718002 | 0.679437825 | -4.411360304 |
| **A_55_P1962926** | Snapc3 | -0.074615945 | -0.194194397 | 0.044962508 | 8.13737229 | -1.535139601 | 0.176740307 | 0.679437825 | -4.411427726 |
| **A_66_P125401** | NA | 0.116686544 | -0.070366511 | 0.3037396 | 9.287259483 | 1.534705801 | 0.176844399 | 0.679662582 | -4.411742259 |
| **A_55_P2072110** | 3-Sep | -0.176375301 | -0.459130704 | 0.106380103 | 14.85761649 | -1.534602906 | 0.176869098 | 0.679662582 | -4.411816862 |
| **A_51_P262101** | Ptcd2 | 0.130243652 | -0.078598552 | 0.339085855 | 8.828038739 | 1.534290021 | 0.176944222 | 0.679791291 | -4.41204371 |
| **A_55_P2179221** | Man1a2 | 0.106623225 | -0.064361524 | 0.277607975 | 11.51742104 | 1.534135086 | 0.176981433 | 0.679791291 | -4.412156039 |
| **A_55_P2156895** | Tmem39b | 0.176059424 | -0.106309765 | 0.458428613 | 9.876388012 | 1.533949755 | 0.177025954 | 0.679791291 | -4.412290402 |
| **A_55_P2172599** | Fam174a | 0.245815522 | -0.148478627 | 0.64010967 | 10.93787905 | 1.533762565 | 0.177070933 | 0.679791291 | -4.412426109 |
| **A_51_P234728** | Lpar6 | -0.136965995 | -0.35666934 | 0.08273735 | 10.90508811 | -1.533717073 | 0.177081865 | 0.679791291 | -4.412459089 |
| **A_55_P2069939** | Ubxn7 | 0.156048991 | -0.094286414 | 0.406384395 | 8.737986071 | 1.533584999 | 0.177113609 | 0.679791291 | -4.412554837 |
| **A_51_P167232** | Cask | -0.071181159 | -0.185443453 | 0.043081135 | 7.948716461 | -1.532608676 | 0.177348431 | 0.680082182 | -4.413262581 |
| **A_51_P406638** | Rfc1 | 0.106170036 | -0.064261391 | 0.276601462 | 10.47339439 | 1.53257398 | 0.177356781 | 0.680082182 | -4.413287731 |
| **A_51_P370252** | Capn10 | -0.097086637 | -0.252938256 | 0.058764982 | 10.1936137 | -1.532559351 | 0.177360302 | 0.680082182 | -4.413298335 |
| **A_55_P1992004** | NA | 0.131910067 | -0.07989783 | 0.343717964 | 10.87991445 | 1.532162945 | 0.177455734 | 0.680082182 | -4.413585668 |
| **A_65_P03719** | Fuca2 | -0.191305884 | -0.498490976 | 0.115879208 | 9.359735638 | -1.532136139 | 0.177462189 | 0.680082182 | -4.413605098 |
| **A_51_P310254** | Fam118a | -0.124124557 | -0.323450661 | 0.075201546 | 8.813984785 | -1.532013874 | 0.177491635 | 0.680082182 | -4.413693718 |
| **A_55_P2041090** | Tyw5 | -0.117793576 | -0.306956725 | 0.071369572 | 10.69490741 | -1.53198403 | 0.177498823 | 0.680082182 | -4.413715349 |
| **A_51_P162671** | Folr2 | 0.261389659 | -0.158375747 | 0.681155064 | 9.452462689 | 1.531972329 | 0.177501641 | 0.680082182 | -4.413723831 |
| **A_55_P2019054** | Acacb | -0.195916096 | -0.510550572 | 0.118718379 | 10.10732759 | -1.531909022 | 0.17751689 | 0.680082182 | -4.413769716 |
| **A_55_P1990970** | Casp9 | -0.183545792 | -0.47835422 | 0.111262637 | 11.17725932 | -1.531699951 | 0.177567258 | 0.680082182 | -4.41392125 |
| **A_55_P2087561** | Eno1b | -0.150721344 | -0.392813665 | 0.091370976 | 9.447665048 | -1.531661896 | 0.177576428 | 0.680082182 | -4.413948832 |
| **A_55_P2117903** | Stard5 | 0.250106627 | -0.151692624 | 0.651905877 | 9.510026391 | 1.531388 | 0.177642437 | 0.680200211 | -4.414147344 |

| | | | | | | | | | |
|---|---|---|---|---|---|---|---|---|---|
| **A_55_P1957583** | Gps2 | 0.191507228 | -0.116226366 | 0.499240822 | 9.831445551 | 1.531014926 | 0.177732386 | 0.680218991 | -4.414417728 |
| **A_55_P2130890** | Parva | 0.1830033 | -0.111113339 | 0.477119939 | 12.46945876 | 1.53076488 | 0.177792696 | 0.680218991 | -4.414598941 |
| **A_55_P1976839** | Spata5 | -0.119821975 | -0.312401053 | 0.072757103 | 9.343427923 | -1.530722738 | 0.177802863 | 0.680218991 | -4.414629482 |
| **A_52_P638459** | Ccl5 | -0.280443223 | -0.731231669 | 0.170345223 | 8.353193108 | -1.530528203 | 0.1778498 | 0.680218991 | -4.414770461 |
| **A_55_P2030802** | Rpl3 | -0.205958917 | -0.537045955 | 0.125128121 | 14.44307637 | -1.530409259 | 0.177878504 | 0.680218991 | -4.414856658 |
| **A_66_P118233** | Hnrnpa0 | -0.139890821 | -0.364828962 | 0.08504732 | 12.27696531 | -1.530013565 | 0.177974028 | 0.680218991 | -4.415143402 |
| **A_55_P2028064** | Kdm4b | 0.090496872 | -0.055018025 | 0.236011768 | 10.07512654 | 1.530013539 | 0.177974034 | 0.680218991 | -4.415143421 |
| **A_52_P147666** | Slc30a7 | 0.121389882 | -0.073803217 | 0.316582981 | 10.01962104 | 1.52998507 | 0.177980909 | 0.680218991 | -4.415164051 |
| **A_55_P2070992** | Aldoc | 0.264598483 | -0.160874347 | 0.690071313 | 9.965187326 | 1.529976223 | 0.177983045 | 0.680218991 | -4.415170461 |
| **A_55_P2117689** | Rmnd1 | -0.148519644 | -0.387383005 | 0.090343718 | 10.74869502 | -1.529690331 | 0.178052095 | 0.680218991 | -4.415377626 |
| **A_51_P220135** | Nfatc1 | -0.194763336 | -0.508020872 | 0.1184942 | 8.516338037 | -1.529589309 | 0.1780765 | 0.680218991 | -4.415450828 |
| **A_52_P585831** | Bap1 | -0.125876654 | -0.328367192 | 0.076613884 | 10.43947311 | -1.529359619 | 0.178132001 | 0.680218991 | -4.415617261 |
| **A_55_P2036250** | Cr1l | 0.164264758 | -0.099985127 | 0.428514642 | 13.65969059 | 1.52932136 | 0.178141248 | 0.680218991 | -4.415644983 |
| **A_51_P481569** | Ddx42 | 0.121221331 | -0.07381491 | 0.316257573 | 11.61613241 | 1.529089454 | 0.178197303 | 0.680218991 | -4.415813016 |
| **A_55_P2066897** | A230045G11Rik | -0.094217227 | -0.245810455 | 0.057376001 | 8.59113832 | -1.529042937 | 0.178208549 | 0.680218991 | -4.415846721 |
| **A_55_P1968143** | Rbm25 | -0.19793975 | -0.516421597 | 0.120542097 | 10.06267551 | -1.52903525 | 0.178210408 | 0.680218991 | -4.41585229 |
| **A_55_P2431118** | Itsn1 | -0.089556435 | -0.233683871 | 0.054571 | 10.6476353 | -1.528689572 | 0.178294001 | 0.68029527 | -4.416102749 |
| **A_55_P2043434** | Atmin | -0.129722458 | -0.343572623 | 0.084127706 | 11.55135687 | -1.572011305 | 0.178307962 | 0.68029527 | -4.39107582 |
| **A_55_P2000683** | Eci2 | 0.19931209 | -0.12153403 | 0.520158211 | 12.37628952 | 1.528290856 | 0.178390466 | 0.68029527 | -4.416391624 |
| **A_51_P507622** | Itpkc | -0.071338087 | -0.186181936 | 0.043505761 | 9.074172365 | -1.528209468 | 0.178410163 | 0.68029527 | -4.416450588 |
| **A_51_P428364** | Psma6 | -0.1580138 | -0.412411316 | 0.096383716 | 11.68676009 | -1.528098379 | 0.178437052 | 0.68029527 | -4.416531071 |
| **A_66_P106968** | NA | -0.120504954 | -0.31451671 | 0.073506803 | 10.53346133 | -1.528079733 | 0.178441565 | 0.68029527 | -4.416544579 |
| **A_55_P2127303** | Spsb2 | 0.119976357 | -0.073263875 | 0.313216588 | 12.44494177 | 1.527450989 | 0.178593825 | 0.680642647 | -4.417000072 |
| **A_51_P367285** | Rnf135 | -0.14282426 | -0.372870299 | 0.087221778 | 8.799737113 | -1.527412663 | 0.17860311 | 0.680642647 | -4.417027836 |
| **A_55_P1985882** | Fastkd1 | -0.084915854 | -0.22173516 | 0.051903451 | 8.541971267 | -1.526899891 | 0.178727381 | 0.680915809 | -4.417399286 |
| **A_52_P605556** | Nxnl2 | -0.117674852 | -0.307285715 | 0.071936011 | 8.616036869 | -1.526826215 | 0.178745244 | 0.680915809 | -4.417452654 |
| **A_55_P2431862** | Srpk2 | 0.180292918 | -0.110248109 | 0.470833946 | 10.27706546 | 1.526653098 | 0.178787221 | 0.680941518 | -4.417578053 |
| **A_55_P2123421** | Dnal4 | -0.086410003 | -0.225675491 | 0.052855486 | 8.377102477 | -1.52647493 | 0.178830434 | 0.680971918 | -4.417707108 |
| **A_55_P2181114** | NA | -0.092450845 | -0.241485834 | 0.056584143 | 12.99313252 | -1.526130988 | 0.17891388 | 0.681051427 | -4.417956233 |
| **A_55_P2125972** | Gorasp1 | 0.153614552 | -0.094024412 | 0.401253515 | 9.929183461 | 1.526098421 | 0.178921783 | 0.681051427 | -4.417979821 |
| **A_55_P2106358** | NA | -0.153614325 | -0.401278719 | 0.09405007 | 7.867325689 | -1.525939461 | 0.178960364 | 0.68106416 | -4.418094955 |
| **A_51_P418420** | Ddx3x | -0.119275641 | -0.311610457 | 0.073059175 | 10.81123887 | -1.525678465 | 0.179023726 | 0.681142196 | -4.418283988 |
| **A_55_P1992680** | Ubtf | -0.118388595 | -0.309331238 | 0.072554048 | 15.93809754 | -1.525373152 | 0.179097874 | 0.681142196 | -4.418505111 |
| **A_55_P1962906** | Als2cl | -0.172253553 | -0.450078187 | 0.105571081 | 9.680865175 | -1.525339794 | 0.179105977 | 0.681142196 | -4.418529271 |
| **A_55_P2163729** | Tvp23a | 0.089636992 | -0.054943072 | 0.234217056 | 8.438954156 | 1.52527455 | 0.179121826 | 0.681142196 | -4.418576522 |
| **A_55_P2146546** | NA | 0.161901487 | -0.099269742 | 0.423072715 | 8.384190584 | 1.525087168 | 0.179167354 | 0.68118131 | -4.418712228 |
| **A_55_P2040367** | Bola3 | 0.128455518 | -0.078847674 | 0.335758709 | 10.61739633 | 1.524459665 | 0.179319895 | 0.681230362 | -4.419166657 |
| **A_51_P445841** | Deptor | -0.081896379 | -0.21406279 | 0.050270032 | 7.913572538 | -1.524448531 | 0.179322603 | 0.681230362 | -4.41917472 |
| **A_55_P2089761** | NA | 0.197817047 | -0.121446371 | 0.517080464 | 8.85214546 | 1.524346574 | 0.179347399 | 0.681230362 | -4.419248552 |
| **A_66_P128313** | Mtmr2 | -0.085202236 | -0.222733158 | 0.052328685 | 10.6400681 | -1.524122254 | 0.179401967 | 0.681230362 | -4.419410991 |
| **A_55_P2417224** | NA | 0.0721898 | -0.044346189 | 0.188725789 | 8.08060534 | 1.523999892 | 0.17943174 | 0.681230362 | -4.419499596 |
| **A_55_P2090996** | Ebna1bp2 | -0.11301721 | -0.295493585 | 0.069459165 | 12.33375888 | -1.523726783 | 0.179498208 | 0.681230362 | -4.419697355 |
| **A_55_P1981296** | Nup85 | -0.098598498 | -0.257803859 | 0.060606862 | 11.36608276 | -1.523637936 | 0.179519836 | 0.681230362 | -4.419761688 |
| **A_55_P2002376** | Srm | 0.080924947 | -0.049797533 | 0.211647426 | 9.777954707 | 1.523004925 | 0.179674002 | 0.681230362 | -4.420220024 |
| **A_55_P2079619** | Rnf43 | 0.094602528 | -0.058218977 | 0.247424032 | 8.463136408 | 1.52295627 | 0.179685857 | 0.681230362 | -4.420255252 |
| **A_55_P2022921** | Nsa2 | 0.257883244 | -0.158748326 | 0.674514814 | 12.32507846 | 1.522790337 | 0.179726292 | 0.681230362 | -4.42037539 |
| **A_52_P109010** | Hipk3 | -0.105283945 | -0.275388142 | 0.064820253 | 12.12068871 | -1.522706781 | 0.179746656 | 0.681230362 | -4.420435885 |
| **A_55_P1955517** | NA | -0.130777301 | -0.342096037 | 0.080541436 | 10.80866538 | -1.522521822 | 0.179791743 | 0.681230362 | -4.420569794 |
| **A_55_P2095118** | Dancr | -0.222962063 | -0.583299142 | 0.137375016 | 9.245979354 | -1.522268227 | 0.179853577 | 0.681230362 | -4.42075339 |
| **A_55_P1975447** | NA | 0.179162446 | -0.110398038 | 0.468722931 | 10.00600437 | 1.522217996 | 0.179865828 | 0.681230362 | -4.420789755 |

| | | | | | | | | | |
|---|---|---|---|---|---|---|---|---|---|
| **A_55_P2144391** | NA | -0.134327507 | -0.351444421 | 0.082789408 | 12.23978818 | -1.52209044 | 0.179896939 | 0.681230362 | -4.420882099 |
| **A_66_P126425** | Eif1 | -0.136261848 | -0.356507435 | 0.083983739 | 15.16762228 | -1.522075596 | 0.17990056 | 0.681230362 | -4.420892846 |
| **A_55_P2040227** | Ptchd4 | 0.12364823 | -0.076210336 | 0.323506797 | 9.652160565 | 1.522068858 | 0.179902204 | 0.681230362 | -4.420897724 |
| **A_52_P467690** | Sptbn1 | -0.169742503 | -0.444132153 | 0.104647147 | 9.991394071 | -1.521920757 | 0.179938334 | 0.681230362 | -4.42100494 |
| **A_52_P5213** | Chd1l | -0.110339251 | -0.288732894 | 0.068054392 | 8.701892126 | -1.521667695 | 0.180000086 | 0.681230362 | -4.421188136 |
| **A_55_P2005300** | Ranbp3 | 0.157915726 | -0.097410381 | 0.413241833 | 12.62936975 | 1.521595868 | 0.180017617 | 0.681230362 | -4.421240132 |
| **A_55_P2060962** | Tm2d3 | 0.179093988 | -0.11049465 | 0.468682625 | 11.4759652 | 1.521488422 | 0.180043845 | 0.681230362 | -4.421317912 |
| **A_66_P114451** | Adgrg6 | -0.267753873 | -0.7007179 | 0.165210155 | 11.4092449 | -1.521433971 | 0.180057137 | 0.681230362 | -4.421357328 |
| **A_55_P2109114** | NA | 0.203615292 | -0.125660265 | 0.532890849 | 13.28625581 | 1.5213182 | 0.180085403 | 0.681230362 | -4.421441133 |
| **A_55_P2154914** | Acap1 | 0.081516513 | -0.050329671 | 0.213362697 | 7.894071047 | 1.521062958 | 0.180147735 | 0.681230362 | -4.421625893 |
| **A_55_P2000369** | Aqp4 | -0.068660516 | -0.179730311 | 0.04240928 | 7.882631555 | -1.520828533 | 0.180205001 | 0.681230362 | -4.42179558 |
| **A_51_P311904** | Ptk2b | 0.07457088 | -0.046081504 | 0.195223265 | 8.278857642 | 1.520556369 | 0.180271508 | 0.681230362 | -4.421992579 |
| **A_55_P2060663** | NA | 0.256823596 | -0.158788183 | 0.672435375 | 15.87719878 | 1.520254293 | 0.180345351 | 0.681230362 | -4.42221122 |
| **A_55_P2018492** | Pgap3 | 0.075629478 | -0.046768111 | 0.198027068 | 9.797400044 | 1.52015336 | 0.180370031 | 0.681230362 | -4.422284272 |
| **A_55_P1967534** | NA | -0.270054428 | -0.707186917 | 0.167078061 | 14.52051739 | -1.519873278 | 0.180438532 | 0.681230362 | -4.422486984 |
| **A_55_P1973036** | NA | -0.118952593 | -0.311516521 | 0.073611335 | 8.980956671 | -1.519735956 | 0.180472126 | 0.681230362 | -4.422586369 |
| **A_55_P2348582** | NA | -0.105533042 | -0.276416534 | 0.06535045 | 8.09345725 | -1.519348879 | 0.180566852 | 0.681230362 | -4.422866503 |
| **A_55_P2040518** | NA | -0.09655001 | -0.252897723 | 0.059797702 | 9.14630551 | -1.519252487 | 0.180590449 | 0.681230362 | -4.422936261 |
| **A_51_P265576** | Rbm18 | -0.077185013 | -0.202174657 | 0.047804631 | 8.271207501 | -1.519246038 | 0.180592028 | 0.681230362 | -4.422940928 |
| **A_55_P2151461** | NA | 0.212814212 | -0.131809404 | 0.557437828 | 12.58635362 | 1.519234297 | 0.180594902 | 0.681230362 | -4.422949426 |
| **A_52_P577662** | Ednrb | -0.150405193 | -0.393967124 | 0.093156737 | 9.238387542 | -1.519226709 | 0.18059676 | 0.681230362 | -4.422954916 |
| **A_55_P1968235** | Rab33b | 0.094693914 | -0.058669548 | 0.248057375 | 10.42841936 | 1.519040409 | 0.180642375 | 0.681230362 | -4.423089738 |
| **A_51_P260504** | Arhgef4 | 0.08773625 | -0.054365521 | 0.229838022 | 8.044075309 | 1.518968405 | 0.180660008 | 0.681230362 | -4.423141845 |
| **A_55_P1963863** | Arhgef17 | -0.13028015 | -0.341309727 | 0.080749428 | 9.014647636 | -1.51881222 | 0.180698262 | 0.681230362 | -4.423254869 |
| **A_55_P2004636** | Ppih | 0.155044135 | -0.096152405 | 0.406240676 | 9.784073194 | 1.518486209 | 0.180778135 | 0.681230362 | -4.423490782 |
| **A_51_P348119** | Ube2q1 | 0.111137128 | -0.068929812 | 0.291204068 | 10.15704619 | 1.5184285 | 0.180792277 | 0.681230362 | -4.423532541 |
| **A_51_P414837** | Faf1 | -0.103633764 | -0.271553653 | 0.064286124 | 10.29811831 | -1.518337433 | 0.180814596 | 0.681230362 | -4.423598438 |
| **A_52_P620748** | Ncl | 0.111600837 | -0.069229784 | 0.292431458 | 8.742357394 | 1.518324646 | 0.180817731 | 0.681230362 | -4.423607691 |
| **A_55_P2027822** | Sgf29 | 0.103812384 | -0.064404338 | 0.272029106 | 10.71849351 | 1.518270529 | 0.180830995 | 0.681230362 | -4.42364685 |
| **A_65_P10357** | Srsf6 | 0.123230617 | -0.076455975 | 0.322917209 | 10.80734432 | 1.518234588 | 0.180839805 | 0.681230362 | -4.423672857 |
| **A_55_P2118734** | Rnmt | -0.134813562 | -0.353328674 | 0.083701551 | 9.872403822 | -1.517823485 | 0.180940606 | 0.681230362 | -4.423970322 |
| **A_55_P2105820** | Tbc1d31 | 0.08323592 | -0.051685145 | 0.218156985 | 8.819579457 | 1.517749812 | 0.180958676 | 0.681230362 | -4.424023629 |
| **A_52_P299358** | Lclat1 | -0.073830263 | -0.193512574 | 0.045852048 | 9.322633801 | -1.517656943 | 0.180981456 | 0.681230362 | -4.424090824 |
| **A_55_P2045549** | Dpy19l1 | -0.125902499 | -0.330000313 | 0.078195315 | 9.37529067 | -1.517627406 | 0.180988702 | 0.681230362 | -4.424112195 |
| **A_55_P2034390** | Ankhd1 | -0.083047974 | -0.217681931 | 0.051585983 | 8.324142243 | -1.517552047 | 0.18100719 | 0.681230362 | -4.42416672 |
| **A_51_P202498** | Sar1a | -0.085153867 | -0.223204835 | 0.052897101 | 11.79485591 | -1.517518801 | 0.181015347 | 0.681230362 | -4.424190775 |
| **A_55_P2117081** | Trgv4 | 0.12195245 | -0.075764937 | 0.319669837 | 7.753921314 | 1.517451542 | 0.18103185 | 0.681230362 | -4.424239438 |
| **A_51_P140211** | Ndufv3 | 0.084047804 | -0.052224793 | 0.2203204 | 8.92077135 | 1.517354307 | 0.18105571 | 0.681230362 | -4.42430979 |
| **A_52_P302544** | Col8a2 | 0.132254757 | -0.082186554 | 0.346696067 | 8.218350028 | 1.517301933 | 0.181068564 | 0.681230362 | -4.424347683 |
| **A_51_P255387** | Med31 | 0.163771037 | -0.101805555 | 0.42934763 | 11.40724702 | 1.517107897 | 0.181116191 | 0.681230362 | -4.424488069 |
| **A_66_P102090** | Pkmyt1 | 0.088919099 | -0.055291487 | 0.233129686 | 10.34812798 | 1.516935358 | 0.181158551 | 0.681230362 | -4.424612899 |
| **A_66_P120125** | Nrep | -0.129733435 | -0.340157388 | 0.080690519 | 9.091335759 | -1.516791563 | 0.181193861 | 0.681230362 | -4.42471693 |
| **A_55_P1968108** | NA | 0.125363366 | -0.077977598 | 0.32870433 | 13.65635368 | 1.516753229 | 0.181203275 | 0.681230362 | -4.424744663 |
| **A_55_P2156462** | Nt5c3b | -0.073974962 | -0.193970092 | 0.046020168 | 8.106216531 | -1.5166672 | 0.181224405 | 0.681230362 | -4.424806902 |
| **A_55_P1977548** | Mapk1 | -0.121954059 | -0.319819683 | 0.075911564 | 10.8436531 | -1.516334706 | 0.18130609 | 0.681307088 | -4.42504744 |
| **A_55_P2022364** | Nxn | 0.168962424 | -0.105253507 | 0.443178355 | 10.0421263 | 1.515886246 | 0.181416319 | 0.681307088 | -4.425371857 |
| **A_55_P2168296** | Rnu7 | 0.077736882 | -0.048425644 | 0.203899408 | 8.125276948 | 1.515883782 | 0.181416925 | 0.681307088 | -4.425373639 |
| **A_52_P230167** | Setbp1 | -0.114816472 | -0.301180553 | 0.07154761 | 8.608126087 | -1.515692657 | 0.181463922 | 0.681307088 | -4.425511893 |
| **A_55_P2033530** | Gm6981 | -0.207641286 | -0.544684885 | 0.129402313 | 10.14774952 | -1.515642553 | 0.181476244 | 0.681307088 | -4.425548137 |
| **A_55_P1966833** | Xaf1 | -0.230723854 | -0.605352704 | 0.143904996 | 10.4812502 | -1.515166784 | 0.181593292 | 0.681307088 | -4.425892277 |

| | | | | | | | | | |
|---|---|---|---|---|---|---|---|---|---|
| **A_51_P370163** | Ctr9 | -0.121268281 | -0.318174961 | 0.0756384 | 10.19304975 | -1.515151056 | 0.181597162 | 0.681307088 | -4.425903654 |
| **A_52_P174775** | Ripk1 | -0.066353615 | -0.174099945 | 0.041392714 | 8.537273617 | -1.515065017 | 0.181618337 | 0.681307088 | -4.425965887 |
| **A_55_P2002752** | Zswim6 | -0.080684631 | -0.211708962 | 0.050339701 | 8.104726397 | -1.514983927 | 0.181638297 | 0.681307088 | -4.426024539 |
| **A_55_P2068570** | NA | 0.156811297 | -0.09786057 | 0.411483163 | 8.312120412 | 1.514835723 | 0.18167478 | 0.681307088 | -4.426131734 |
| **A_55_P2443568** | Ube4a | -0.15659997 | -0.410930904 | 0.097730964 | 10.35053414 | -1.514822162 | 0.181678119 | 0.681307088 | -4.426141542 |
| **A_55_P2136757** | Tomm34 | 0.140290739 | -0.087572297 | 0.368153775 | 11.41170711 | 1.514691843 | 0.181710206 | 0.681307088 | -4.426235798 |
| **A_55_P2255449** | NA | -0.197518993 | -0.518341319 | 0.123303332 | 8.603071066 | -1.514654028 | 0.181719518 | 0.681307088 | -4.426263149 |
| **A_51_P205481** | Mtmr6 | -0.120362983 | -0.315913577 | 0.075187612 | 11.69220074 | -1.514268765 | 0.181814413 | 0.681307088 | -4.42654179 |
| **A_55_P2029574** | Por | 0.104164554 | -0.065070666 | 0.273399774 | 13.57235231 | 1.514252747 | 0.18181836 | 0.681307088 | -4.426553374 |
| **A_55_P2188464** | Uhrf1bp1l | -0.239091858 | -0.627545365 | 0.149361649 | 11.10446187 | -1.514240658 | 0.181821338 | 0.681307088 | -4.426562117 |
| **A_55_P1967504** | Abhd11 | 0.10651659 | -0.066567742 | 0.279600923 | 10.16228061 | 1.514009712 | 0.181878247 | 0.681307088 | -4.426729142 |
| **A_55_P2139763** | Trim56 | -0.238749309 | -0.626749852 | 0.149251235 | 10.34790041 | -1.51383643 | 0.181920958 | 0.681307088 | -4.426854459 |
| **A_55_P1991440** | Zscan21 | 0.181430117 | -0.113454815 | 0.476315049 | 12.4908962 | 1.513651734 | 0.181966493 | 0.681307088 | -4.426988028 |
| **A_55_P2048656** | Atp5l | 0.249668203 | -0.156133722 | 0.655470129 | 14.74979193 | 1.513625014 | 0.181973081 | 0.681307088 | -4.427007351 |
| **A_66_P137297** | Otof | 0.743132114 | -0.464767357 | 1.951031584 | 11.81744879 | 1.513576588 | 0.181985022 | 0.681307088 | -4.427042371 |
| **A_55_P2018934** | Adgre1 | -0.066630792 | -0.174960002 | 0.041698418 | 7.991943205 | -1.513207766 | 0.182075992 | 0.681515656 | -4.427309084 |
| **A_51_P473272** | Tspan6 | 0.146219866 | -0.091584845 | 0.384024577 | 9.759224966 | 1.512707951 | 0.182199339 | 0.681845309 | -4.427670505 |
| **A_51_P365656** | Tmem101 | 0.09291548 | -0.058283137 | 0.244114097 | 9.052533 | 1.51185249 | 0.182410636 | 0.682503909 | -4.428289041 |
| **A_66_P130887** | Pcdh18 | -0.201538036 | -0.52961247 | 0.126536399 | 9.37857495 | -1.511310833 | 0.182544544 | 0.682872748 | -4.428680647 |
| **A_51_P436447** | Zfp871 | -0.111424654 | -0.292848459 | 0.069999152 | 8.589376737 | -1.51097119 | 0.182628557 | 0.682895695 | -4.428926187 |
| **A_55_P2250246** | Taok3 | -0.068285165 | -0.179468591 | 0.042898261 | 7.943284229 | -1.510968726 | 0.182629167 | 0.682895695 | -4.428927968 |
| **A_55_P2087186** | Ier3ip1 | -0.14922329 | -0.392220662 | 0.093774083 | 15.20559292 | -1.510790328 | 0.182673309 | 0.682895695 | -4.429056935 |
| **A_55_P1969042** | Cops6 | 0.115101863 | -0.072344788 | 0.302548513 | 12.47105772 | 1.510684698 | 0.182699451 | 0.682895695 | -4.429133295 |
| **A_51_P224175** | 2210016F16Rik | 0.142757565 | -0.089796599 | 0.375311729 | 8.640959123 | 1.510233762 | 0.18281109 | 0.682895695 | -4.429459264 |
| **A_55_P1979221** | Txndc9 | -0.132662502 | -0.348774393 | 0.08344939 | 10.56084784 | -1.510214659 | 0.182815821 | 0.682895695 | -4.429473072 |
| **A_55_P2169396** | NA | -0.163532751 | -0.429941729 | 0.102876228 | 15.19238543 | -1.51016725 | 0.182827562 | 0.682895695 | -4.429507342 |
| **A_52_P282838** | Spin2c | 0.157441172 | -0.099052137 | 0.413934481 | 8.777582788 | 1.51011992 | 0.182839284 | 0.682895695 | -4.429541554 |
| **A_55_P2085716** | Atg12 | -0.208274323 | -0.54761561 | 0.131066965 | 11.02862574 | -1.509969556 | 0.18287653 | 0.682895695 | -4.429650242 |
| **A_55_P2155834** | NA | -0.140470783 | -0.36935658 | 0.088415014 | 16.59093841 | -1.509858759 | 0.182903979 | 0.682895695 | -4.429730329 |
| **A_51_P301804** | St3gal1 | -0.148634825 | -0.390861715 | 0.093592065 | 12.38602178 | -1.509619115 | 0.182963363 | 0.682985484 | -4.429903544 |
| **A_52_P407871** | Lsm12 | -0.116674842 | -0.306877868 | 0.073528183 | 10.64100356 | -1.509138033 | 0.183082629 | 0.683298732 | -4.430251255 |
| **A_51_P144264** | Klf2 | -0.134356193 | -0.353420117 | 0.084707732 | 12.04952654 | -1.508884467 | 0.183145521 | 0.683401499 | -4.430434516 |
| **A_52_P322535** | Mplkip | -0.075068332 | -0.197483625 | 0.04734696 | 8.097422507 | -1.508656112 | 0.183202177 | 0.683437837 | -4.430599551 |
| **A_52_P420308** | NA | -0.10990188 | -0.289134257 | 0.069330496 | 11.01777295 | -1.508543447 | 0.183230135 | 0.683437837 | -4.430680974 |
| **A_55_P1992640** | Hdac3 | 0.080856239 | -0.051042269 | 0.212754747 | 9.653300726 | 1.508144021 | 0.183329288 | 0.683437837 | -4.430969627 |
| **A_52_P39083** | Ccbe1 | -0.526826836 | -1.386457965 | 0.332804293 | 9.592001395 | -1.507734659 | 0.18343096 | 0.683437837 | -4.431265445 |
| **A_55_P2017369** | NA | -0.193939634 | -0.510480335 | 0.122601067 | 11.52515709 | -1.507322478 | 0.183533386 | 0.683437837 | -4.431563284 |
| **A_51_P423578** | Slfn2 | -0.118819908 | -0.312759848 | 0.075120033 | 8.313275026 | -1.50727021 | 0.183546378 | 0.683437837 | -4.431601051 |
| **A_55_P2075366** | Tead3 | 0.176495159 | -0.111593056 | 0.464583374 | 8.829872963 | 1.507219374 | 0.183559015 | 0.683437837 | -4.431637783 |
| **A_52_P90124** | Prep | 0.14747179 | -0.093246848 | 0.388190428 | 11.62227498 | 1.507191368 | 0.183565978 | 0.683437837 | -4.43165802 |
| **A_55_P2014282** | NA | -0.132665807 | -0.34921824 | 0.083886625 | 16.7430712 | -1.507179924 | 0.183568823 | 0.683437837 | -4.431666288 |
| **A_55_P2156425** | Upk1a | 0.123521874 | -0.078114535 | 0.325158284 | 8.304173805 | 1.50710698 | 0.183586958 | 0.683437837 | -4.431718995 |
| **A_52_P544555** | Hsbp1 | -0.181659429 | -0.478212484 | 0.114893626 | 12.13835988 | -1.507039742 | 0.183603676 | 0.683437837 | -4.431767577 |
| **A_52_P80593** | NA | 0.141038789 | -0.089212755 | 0.371290334 | 11.93455734 | 1.506971996 | 0.183620522 | 0.683437837 | -4.431816527 |
| **A_51_P288618** | Dusp11 | 0.074758939 | -0.047288818 | 0.196806696 | 8.349533063 | 1.506962648 | 0.183622846 | 0.683437837 | -4.431823281 |
| **A_52_P253179** | Igfbp3 | -0.287000461 | -0.755620517 | 0.181619594 | 9.670002158 | -1.506714218 | 0.183684635 | 0.683437837 | -4.432002778 |
| **A_51_P265106** | Dlat | -0.073974925 | -0.194762974 | 0.046813124 | 8.816715779 | -1.506710213 | 0.183685631 | 0.683437837 | -4.432005671 |
| **A_55_P1954544** | Zfp446 | -0.121144628 | -0.319004831 | 0.076715574 | 8.955740426 | -1.506311789 | 0.183784766 | 0.683675087 | -4.432293531 |
| **A_52_P481423** | Cttnbp2nl | -0.151355955 | -0.398648253 | 0.095936344 | 11.70230877 | -1.50576807 | 0.183920135 | 0.684047005 | -4.43268634 |
| **A_55_P2147191** | NA | -0.092100393 | -0.242639417 | 0.05843863 | 16.24190694 | -1.505156139 | 0.184072598 | 0.684410197 | -4.433128394 |

| | | | | | | | | | |
|---|---|---|---|---|---|---|---|---|---|
| **A_55_P2010191** | Serpina11 | 0.124304296 | -0.078881 | 0.327489591 | 8.283237807 | 1.505091926 | 0.184088603 | 0.684410197 | -4.433174779 |
| **A_51_P174016** | Slc39a7 | -0.2322752 | -0.612008309 | 0.147457908 | 14.90865846 | -1.504851142 | 0.184148632 | 0.684501713 | -4.433348708 |
| **A_55_P1993453** | Ftsj1 | 0.149897449 | -0.095219912 | 0.39501481 | 10.5458688 | 1.504490095 | 0.184238678 | 0.684609667 | -4.433609498 |
| **A_55_P2170419** | NA | -0.131588019 | -0.346784838 | 0.083608799 | 15.57793685 | -1.504352688 | 0.184272958 | 0.684609667 | -4.433708746 |
| **A_52_P161707** | Cand1 | 0.120366274 | -0.076507622 | 0.317240169 | 9.351686171 | 1.504131629 | 0.18432812 | 0.684609667 | -4.433868411 |
| **A_55_P1969181** | Paox | 0.109369515 | -0.069525383 | 0.288264413 | 9.921243771 | 1.504068082 | 0.184343981 | 0.684609667 | -4.433914308 |
| **A_55_P2101944** | NA | -0.568339342 | -1.49800977 | 0.361331086 | 11.54835766 | -1.504000106 | 0.184360948 | 0.684609667 | -4.433963405 |
| **A_55_P2114839** | NA | 0.232262234 | -0.147694126 | 0.612218594 | 14.20781578 | 1.503882977 | 0.184390187 | 0.684609667 | -4.434048001 |
| **A_55_P1954555** | NA | -0.165608927 | -0.436572414 | 0.10535456 | 8.306396952 | -1.50363399 | 0.184452357 | 0.684708969 | -4.434227825 |
| **A_51_P340402** | Eefsec | 0.167966031 | -0.10695465 | 0.442886713 | 9.226248343 | 1.503083861 | 0.184589788 | 0.684956332 | -4.434625122 |
| **A_55_P2015530** | NA | 0.071296868 | -0.045403794 | 0.187997529 | 7.797798195 | 1.503025317 | 0.184604419 | 0.684956332 | -4.4346674 |
| **A_51_P352594** | St5 | -0.158696806 | -0.418485343 | 0.101091732 | 9.016907016 | -1.502855981 | 0.184646745 | 0.684956332 | -4.434789686 |
| **A_66_P113760** | Acvr2a | -0.084865992 | -0.223797794 | 0.05406581 | 8.806624705 | -1.502799997 | 0.18466074 | 0.684956332 | -4.434830113 |
| **A_51_P153557** | Hspa12a | -0.086972744 | -0.22940351 | 0.055458023 | 7.862746332 | -1.502271823 | 0.184792825 | 0.685314758 | -4.435211511 |
| **A_55_P2136765** | Uqcr10 | -0.128473723 | -0.338920113 | 0.081972668 | 11.77429395 | -1.501903367 | 0.184885021 | 0.685525141 | -4.435477559 |
| **A_51_P254986** | Serf1 | -0.147834342 | -0.39008694 | 0.094418256 | 12.81484499 | -1.501329621 | 0.18502867 | 0.685884443 | -4.435891813 |
| **A_55_P2057102** | Sft2d3 | -0.086821309 | -0.229116488 | 0.055473869 | 8.310499607 | -1.501085083 | 0.185089927 | 0.685884443 | -4.436068364 |
| **A_55_P2056739** | Col16a1 | 0.416814764 | -0.266407311 | 1.100036839 | 10.22612762 | 1.500895193 | 0.185137508 | 0.685884443 | -4.436205456 |
| **A_55_P1985219** | Pnpla6 | 0.156841165 | -0.100254716 | 0.413937047 | 10.74928791 | 1.500838993 | 0.185151592 | 0.685884443 | -4.436246028 |
| **A_51_P207153** | 1110008P14Rik | 0.134582665 | -0.086043698 | 0.355209029 | 10.53196731 | 1.500724185 | 0.185180367 | 0.685884443 | -4.436328913 |
| **A_55_P2074927** | Gm7932 | -0.307870434 | -0.812626214 | 0.196885347 | 9.266433105 | -1.50056857 | 0.185219376 | 0.685884443 | -4.436441254 |
| **A_55_P2125743** | NA | -0.193029118 | -0.509510942 | 0.123452706 | 15.36848812 | -1.500524932 | 0.185230317 | 0.685884443 | -4.436472757 |
| **A_51_P418339** | Nudcd2 | -0.168682741 | -0.445296416 | 0.107930934 | 9.45786224 | -1.500258743 | 0.185297066 | 0.685970855 | -4.436664917 |
| **A_55_P2025103** | Snx3 | -0.098585607 | -0.260322711 | 0.063151497 | 11.23742225 | -1.499591661 | 0.185464443 | 0.685970855 | -4.437146449 |
| **A_55_P2089820** | Adk | -0.069745018 | -0.184171632 | 0.044681597 | 8.247212878 | -1.499530493 | 0.185479798 | 0.685970855 | -4.437190601 |
| **A_51_P455647** | Car2 | 0.213358051 | -0.136746596 | 0.563462698 | 12.87980942 | 1.499271629 | 0.185544792 | 0.685970855 | -4.437377448 |
| **A_55_P2134251** | Hmgb1 | -0.145145896 | -0.383330211 | 0.093038418 | 11.85259705 | -1.499204118 | 0.185561746 | 0.685970855 | -4.437426176 |
| **A_55_P2048348** | Aig1 | -0.082409544 | -0.217645634 | 0.052826545 | 9.364487922 | -1.499181004 | 0.185567551 | 0.685970855 | -4.43744286 |
| **A_55_P1964573** | Arhgef6 | -0.207329016 | -0.547611813 | 0.132953781 | 15.10081248 | -1.49895728 | 0.185623747 | 0.685970855 | -4.437604335 |
| **A_55_P1991069** | Milr1 | 0.187185185 | -0.120058818 | 0.494429189 | 10.75394701 | 1.498846716 | 0.185651524 | 0.685970855 | -4.437684134 |
| **A_51_P405423** | Smcr8 | 0.097499645 | -0.062553803 | 0.257553093 | 9.95006951 | 1.498673972 | 0.185694931 | 0.685970855 | -4.437808809 |
| **A_55_P2430472** | Dido1 | -0.153156586 | -0.404594187 | 0.098281015 | 8.642919057 | -1.49856166 | 0.185723158 | 0.685970855 | -4.437889867 |
| **A_55_P1998848** | NA | 0.083671629 | -0.05373119 | 0.221074449 | 13.68279718 | 1.498137743 | 0.185829735 | 0.685970855 | -4.438195804 |
| **A_66_P106716** | Akap8 | 0.102513116 | -0.06585305 | 0.270879283 | 9.442618641 | 1.497937783 | 0.185880027 | 0.685970855 | -4.438340106 |
| **A_55_P1959985** | Alas1 | 0.112369027 | -0.072185482 | 0.296923535 | 9.216144375 | 1.497928635 | 0.185882328 | 0.685970855 | -4.438346709 |
| **A_55_P1996573** | Ttc13 | -0.177404521 | -0.468774442 | 0.113965399 | 13.02784057 | -1.497921765 | 0.185884056 | 0.685970855 | -4.438351666 |
| **A_55_P1999548** | Tma7 | 0.189793164 | -0.121945463 | 0.501531791 | 11.36842413 | 1.497818222 | 0.185910104 | 0.685970855 | -4.438426387 |
| **A_55_P1971430** | Serpinb9e | -0.106058608 | -0.280269078 | 0.068151863 | 14.06168142 | -1.49775518 | 0.185925965 | 0.685970855 | -4.438471881 |
| **A_51_P353095** | Zfyve27 | -0.13454218 | -0.355556631 | 0.086472271 | 11.51424513 | -1.497638351 | 0.185955361 | 0.685970855 | -4.438556187 |
| **A_51_P509384** | Aldh8a1 | 0.124699126 | -0.080147707 | 0.329545959 | 8.240762097 | 1.497625817 | 0.185958515 | 0.685970855 | -4.438565232 |
| **A_55_P1975650** | Fbxo33 | -0.174643861 | -0.461541352 | 0.11225363 | 9.422907774 | -1.497599673 | 0.185965095 | 0.685970855 | -4.438584098 |
| **A_55_P2061690** | Atp5d | -0.086905467 | -0.229679185 | 0.055868251 | 12.57217474 | -1.497503996 | 0.185989173 | 0.685970855 | -4.43865314 |
| **A_51_P475995** | Thra | 0.181872885 | -0.11692696 | 0.48067273 | 10.04040824 | 1.497465243 | 0.185998927 | 0.685970855 | -4.438681104 |
| **A_55_P2182586** | Esrp1 | 0.118242069 | -0.076058453 | 0.312542591 | 10.24503951 | 1.497156551 | 0.186076637 | 0.685980474 | -4.438903852 |
| **A_55_P2320263** | NA | -0.447541481 | -1.183009291 | 0.28792633 | 11.35226099 | -1.497058557 | 0.186101313 | 0.685980474 | -4.438974561 |
| **A_55_P2094881** | Fgfr2 | 0.20924502 | -0.134666829 | 0.55315687 | 8.91352604 | 1.496846115 | 0.186154818 | 0.685980474 | -4.439127849 |
| **A_55_P2048813** | NA | -0.246060026 | -0.6504899 | 0.158369847 | 11.12779519 | -1.496811124 | 0.186163632 | 0.685980474 | -4.439153096 |
| **A_51_P282227** | As3mt | -0.186148426 | -0.492143015 | 0.119846162 | 10.50953554 | -1.496631172 | 0.186208968 | 0.685980474 | -4.439282936 |
| **A_52_P345247** | Tfb1m | 0.155166551 | -0.099903121 | 0.410236223 | 9.585217782 | 1.496609322 | 0.186214473 | 0.685980474 | -4.439298701 |
| **A_51_P141818** | Mrpl32 | 0.149316031 | -0.096229365 | 0.394861427 | 10.75156883 | 1.496042065 | 0.186357454 | 0.686376377 | -4.43970797 |

| | | | | | | | | | |
|---|---|---|---|---|---|---|---|---|---|
| **A_52_P612382** | Cdc25b | -0.157853585 | -0.417468041 | 0.101760871 | 9.752832453 | -1.495873058 | 0.186400073 | 0.686402556 | -4.4398299 |
| **A_65_P08971** | F3 | -0.126336266 | -0.334141841 | 0.081469309 | 8.434211644 | -1.495684543 | 0.186447623 | 0.686446877 | -4.439965901 |
| **A_55_P2054913** | Olfr406 | 0.075520516 | -0.048718771 | 0.199759803 | 7.859057862 | 1.495461241 | 0.186503961 | 0.686523532 | -4.440126993 |
| **A_51_P227777** | Top2b | -0.11408544 | -0.301808551 | 0.073637671 | 12.14319126 | -1.495139243 | 0.186585229 | 0.686691905 | -4.440359277 |
| **A_55_P2089158** | NA | -0.119127036 | -0.315194431 | 0.076940358 | 8.464720915 | -1.49476914 | 0.186678678 | 0.686905039 | -4.44062625 |
| **A_52_P514352** | Kcnk5 | -0.337114225 | -0.892086138 | 0.217857688 | 10.29620318 | -1.49442883 | 0.186764644 | 0.687090559 | -4.44087172 |
| **A_55_P1964093** | Rangrf | -0.070622629 | -0.186912959 | 0.0456677 | 8.699961912 | -1.494064831 | 0.186856634 | 0.687298169 | -4.441134264 |
| **A_55_P2002859** | Aifm2 | -0.092367203 | -0.244515533 | 0.059781128 | 8.135512781 | -1.493549991 | 0.186986818 | 0.687468368 | -4.441505582 |
| **A_51_P152990** | Grem2 | 0.141131164 | -0.091365654 | 0.373627982 | 9.163954585 | 1.493396307 | 0.187025695 | 0.687468368 | -4.441616419 |
| **A_55_P2034227** | NA | -0.206348517 | -0.546298639 | 0.133601605 | 14.77454893 | -1.493328361 | 0.187042886 | 0.687468368 | -4.441665421 |
| **A_55_P1996431** | NA | 0.435423693 | -0.281980069 | 1.152827455 | 9.245237553 | 1.493198589 | 0.187075723 | 0.687468368 | -4.441759009 |
| **A_55_P1965655** | Mtmr14 | 0.108488675 | -0.07025956 | 0.28723691 | 8.730998963 | 1.493178765 | 0.18708074 | 0.687468368 | -4.441773305 |
| **A_55_P2055881** | NA | -0.075554305 | -0.200073441 | 0.048964831 | 16.08360851 | -1.492767866 | 0.18718475 | 0.687588164 | -4.442069623 |
| **A_55_P2146483** | Zmynd8 | 0.134599656 | -0.087237625 | 0.356436937 | 10.27625019 | 1.492720787 | 0.187196671 | 0.687588164 | -4.442103573 |
| **A_55_P2039071** | Dhdh | -0.08461929 | -0.224099463 | 0.054860884 | 8.673709423 | -1.492540278 | 0.187242383 | 0.687588164 | -4.44223374 |
| **A_55_P2005753** | Manbal | 0.106145144 | -0.068822786 | 0.281113073 | 11.37392913 | 1.492487971 | 0.187255631 | 0.687588164 | -4.442271458 |
| **A_55_P2073840** | Rpl7 | -0.128663208 | -0.340876765 | 0.083550349 | 16.01384201 | -1.491593264 | 0.187482376 | 0.688289999 | -4.442916587 |
| **A_55_P2010341** | Rpl11 | 0.077776538 | -0.050522964 | 0.20607604 | 15.85493568 | 1.491395409 | 0.187532553 | 0.68834347 | -4.443059238 |
| **A_55_P2128181** | NA | -0.147709267 | -0.391402683 | 0.095984149 | 8.793801845 | -1.491190439 | 0.187584547 | 0.688403592 | -4.443207017 |
| **A_55_P1963124** | NA | 0.14222226 | -0.092521575 | 0.376966095 | 15.52233752 | 1.490536251 | 0.187750585 | 0.688666499 | -4.443678638 |
| **A_55_P2081105** | NA | -0.206609993 | -0.547642964 | 0.134422977 | 8.790007939 | -1.490473019 | 0.187766641 | 0.688666499 | -4.443724222 |
| **A_52_P175086** | Usp48 | -0.112840188 | -0.299110112 | 0.073429735 | 10.56055779 | -1.490356715 | 0.187796176 | 0.688666499 | -4.443808063 |
| **A_55_P2010611** | Slc30a5 | 0.092759347 | -0.060363331 | 0.245882025 | 10.83732068 | 1.490346769 | 0.187798702 | 0.688666499 | -4.443815233 |
| **A_52_P349298** | Rprd1b | -0.100602657 | -0.266691869 | 0.065486556 | 8.578236817 | -1.490174706 | 0.187842407 | 0.688696107 | -4.443939268 |
| **A_55_P2005833** | Adamts8 | -0.118660453 | -0.314588537 | 0.07726763 | 8.128831183 | -1.489973262 | 0.187893585 | 0.688753103 | -4.444084478 |
| **A_55_P2018631** | NA | -0.147102901 | -0.390160797 | 0.095954995 | 12.66157488 | -1.488951894 | 0.188153276 | 0.689478326 | -4.444820662 |
| **A_55_P1993599** | Oaz2 | -0.084347789 | -0.223719004 | 0.055023425 | 13.91709846 | -1.48891458 | 0.188162769 | 0.689478326 | -4.444847555 |
| **A_51_P173564** | Ahsa1 | -0.105942473 | -0.281030931 | 0.069145986 | 12.87514623 | -1.488612804 | 0.188239566 | 0.689482469 | -4.445065048 |
| **A_55_P2040748** | Scrib | -0.177364639 | -0.470532029 | 0.115802751 | 11.53396797 | -1.488403018 | 0.18829297 | 0.689482469 | -4.445216237 |
| **A_55_P2052472** | NA | -0.111982106 | -0.297078487 | 0.073114276 | 13.65307253 | -1.488400694 | 0.188293562 | 0.689482469 | -4.445217912 |
| **A_55_P2149288** | Cmya5 | -0.169731027 | -0.450290839 | 0.110828785 | 7.896679354 | -1.488349549 | 0.188306584 | 0.689482469 | -4.445254771 |
| **A_55_P2129319** | Mre11a | -0.119351516 | -0.316675356 | 0.077972325 | 9.477172482 | -1.488050062 | 0.188382852 | 0.689631088 | -4.445470596 |
| **A_55_P1981479** | Irgm1 | -0.115173099 | -0.305608306 | 0.075262108 | 8.178687472 | -1.48789736 | 0.188421751 | 0.689642874 | -4.445580637 |
| **A_55_P1963980** | Carmil1 | 0.079833082 | -0.052191233 | 0.211857396 | 8.476042408 | 1.487640989 | 0.188487075 | 0.689751357 | -4.44576538 |
| **A_51_P430620** | Slc25a12 | -0.145098731 | -0.385108758 | 0.094911296 | 10.00107478 | -1.487316482 | 0.18856979 | 0.689923429 | -4.445999211 |
| **A_51_P172131** | Lurap1l | -0.172938203 | -0.459052142 | 0.113175735 | 8.860773512 | -1.487034656 | 0.188641654 | 0.689952595 | -4.446202279 |
| **A_55_P2163837** | Med14 | 0.089178832 | -0.058364757 | 0.236722421 | 8.709031674 | 1.486998735 | 0.188650815 | 0.689952595 | -4.446228161 |
| **A_55_P2090349** | Mettl5 | 0.08506073 | -0.055710156 | 0.225831615 | 8.415114411 | 1.486570204 | 0.188760142 | 0.689952595 | -4.446536918 |
| **A_55_P2114789** | Gtf2a2 | 0.064254237 | -0.042092373 | 0.170600847 | 7.911950704 | 1.486439657 | 0.188793459 | 0.689952595 | -4.446630974 |
| **A_51_P303725** | Tmem38a | 0.144017201 | -0.094359217 | 0.382393619 | 9.324132138 | 1.4863471 | 0.188817084 | 0.689952595 | -4.446697658 |
| **A_55_P2074206** | Dlgap4 | -0.2235415 | -0.593552301 | 0.146469301 | 13.37649596 | -1.486322057 | 0.188823477 | 0.689952595 | -4.4467157 |
| **A_55_P2153021** | Ttn | -0.078089704 | -0.207350971 | 0.051171562 | 7.844764312 | -1.486259126 | 0.188839541 | 0.689952595 | -4.446761039 |
| **A_55_P2022885** | Zfp866 | 0.10380451 | -0.06803304 | 0.27564206 | 8.440067143 | 1.486165974 | 0.188863323 | 0.689952595 | -4.446828149 |
| **A_55_P2413458** | Tfcp2l1 | -0.09773622 | -0.259577712 | 0.064105272 | 8.049263358 | -1.485712679 | 0.188979092 | 0.690245061 | -4.447154708 |
| **A_51_P440790** | Dpp7 | 0.20192008 | -0.132551209 | 0.536391369 | 11.67152213 | 1.485216715 | 0.189105833 | 0.690313543 | -4.447511982 |
| **A_55_P2101177** | Ncaph2 | 0.152388438 | -0.100039417 | 0.404816292 | 11.23657124 | 1.485196456 | 0.189111012 | 0.690313543 | -4.447526575 |
| **A_55_P2092616** | Tbc1d25 | 0.140849421 | -0.092474918 | 0.37417376 | 9.966899924 | 1.48512902 | 0.189128251 | 0.690313543 | -4.447575152 |
| **A_51_P286496** | Il2rb | -0.183651423 | -0.487889461 | 0.120586615 | 10.92399102 | -1.485080341 | 0.189140697 | 0.690313543 | -4.447610216 |
| **A_51_P486681** | Ap3b2 | 0.155930287 | -0.102505732 | 0.414366305 | 8.907869449 | 1.484385142 | 0.189318516 | 0.690774913 | -4.448110957 |
| **A_55_P2157914** | NA | 0.120701312 | -0.079390876 | 0.320793501 | 8.815011092 | 1.484058498 | 0.18940212 | 0.690774913 | -4.448346216 |

| | | | | | | | | | |
|---|---|---|---|---|---|---|---|---|---|
| **A_51_P516362** | Vps11 | -0.162929311 | -0.433026502 | 0.107167881 | 10.21338818 | -1.484049251 | 0.189404487 | 0.690774913 | -4.448352876 |
| **A_55_P1978236** | Mrps7 | 0.212645538 | -0.139874245 | 0.565165321 | 12.86014436 | 1.484027484 | 0.189410059 | 0.690774913 | -4.448368553 |
| **A_55_P2184741** | Katnal1 | -0.07543433 | -0.200531233 | 0.049662573 | 10.13181687 | -1.483513983 | 0.189541563 | 0.69093805 | -4.448738367 |
| **A_55_P2058117** | Slc35d1 | 0.140532125 | -0.092535079 | 0.373599329 | 9.376123014 | 1.483418226 | 0.189566095 | 0.69093805 | -4.448807326 |
| **A_52_P320971** | Mt3 | -0.170024016 | -0.452024865 | 0.111976833 | 9.609503157 | -1.483300074 | 0.189596369 | 0.69093805 | -4.448892412 |
| **A_55_P2117425** | Syt5 | 0.070187929 | -0.046225795 | 0.186601653 | 7.864672421 | 1.483294583 | 0.189597776 | 0.69093805 | -4.448896367 |
| **A_51_P150394** | Vps28 | 0.129770606 | -0.085552376 | 0.345093588 | 14.1122566 | 1.48270622 | 0.189748599 | 0.691293036 | -4.449320047 |
| **A_52_P244064** | Ube2g2 | -0.16355646 | -0.434978757 | 0.107865838 | 11.32831414 | -1.482488532 | 0.18980443 | 0.691293036 | -4.449476794 |
| **A_55_P2015934** | NA | 0.102879813 | -0.067864039 | 0.273623666 | 15.99046665 | 1.482361935 | 0.189836905 | 0.691293036 | -4.449567949 |
| **A_52_P460957** | Crem | -0.082144213 | -0.218485997 | 0.054197571 | 9.138926266 | -1.48223534 | 0.189869386 | 0.691293036 | -4.4496591 |
| **A_55_P2057065** | Gtf3c5 | 0.194224114 | -0.128150198 | 0.516598426 | 10.80697757 | 1.48221732 | 0.18987401 | 0.691293036 | -4.449672075 |
| **A_55_P1975087** | NA | -0.144545316 | -0.38455395 | 0.095463318 | 12.2564868 | -1.481652371 | 0.190019025 | 0.691690721 | -4.45007883 |
| **A_55_P2069012** | Pidd1 | 0.069193585 | -0.045724391 | 0.184111561 | 8.288222233 | 1.481313652 | 0.190106019 | 0.691877093 | -4.450322686 |
| **A_55_P2027788** | Baiap2 | 0.221489725 | -0.146466617 | 0.589446067 | 9.291227652 | 1.480902458 | 0.190211677 | 0.692131309 | -4.450618703 |
| **A_55_P2037338** | Plcd3 | 0.110684764 | -0.07324332 | 0.294612847 | 10.4066083 | 1.480501867 | 0.190314664 | 0.69237571 | -4.450907069 |
| **A_51_P228658** | Isyna1 | -0.155845749 | -0.414847535 | 0.103156036 | 11.48577041 | -1.480339632 | 0.190356387 | 0.69239718 | -4.451023849 |
| **A_55_P2167530** | NA | 0.258904279 | -0.171434309 | 0.689242866 | 8.76686799 | 1.480123998 | 0.190411857 | 0.692468633 | -4.451179063 |
| **A_65_P19784** | Asb7 | -0.121225499 | -0.32277208 | 0.080321083 | 9.822324378 | -1.479747809 | 0.190508664 | 0.69268791 | -4.451449832 |
| **A_55_P2120526** | NA | 0.367494443 | -0.24358681 | 0.978575696 | 9.175201687 | 1.479520471 | 0.190567188 | 0.69268791 | -4.451613456 |
| **A_55_P2113359** | Zfp523 | 0.147065823 | -0.097510063 | 0.39164171 | 10.34000752 | 1.479337567 | 0.190614286 | 0.69268791 | -4.451745094 |
| **A_66_P111840** | NA | -0.163572663 | -0.435600899 | 0.108455573 | 14.00518968 | -1.479332854 | 0.1906155 | 0.69268791 | -4.451748486 |
| **A_52_P334796** | Taar4 | 0.146741619 | -0.097333826 | 0.390817065 | 8.069131572 | 1.479102869 | 0.190674737 | 0.692768428 | -4.451914005 |
| **A_55_P1968450** | Gm5766 | 0.202950111 | -0.13466395 | 0.540564172 | 14.70629351 | 1.478897009 | 0.190727774 | 0.692768428 | -4.452062155 |
| **A_55_P1990336** | Add1 | -0.208378373 | -0.555069175 | 0.138312429 | 13.61794309 | -1.478698003 | 0.190779059 | 0.692768428 | -4.452205369 |
| **A_51_P318830** | Syt10 | 0.901087232 | -0.598109916 | 2.40028438 | 8.704012476 | 1.478690394 | 0.19078102 | 0.692768428 | -4.452210845 |
| **A_55_P2000289** | Mab21l2 | 0.130172867 | -0.086430589 | 0.346776324 | 8.35711799 | 1.478509951 | 0.190827533 | 0.692807173 | -4.452340696 |
| **A_55_P2051864** | Golgb1 | -0.200776527 | -0.534924663 | 0.133371609 | 13.7217876 | -1.478233557 | 0.190898799 | 0.692866806 | -4.452539588 |
| **A_55_P2134487** | NA | 0.179995431 | -0.119585985 | 0.479576848 | 11.43951316 | 1.478140701 | 0.190922747 | 0.692866806 | -4.452606406 |
| **A_51_P371119** | Cops4 | -0.150944101 | -0.402198337 | 0.100310135 | 11.89769135 | -1.477991432 | 0.19096125 | 0.692866806 | -4.452713815 |
| **A_51_P318551** | Nle1 | 0.083348166 | -0.055398647 | 0.222094979 | 8.135612047 | 1.477890293 | 0.190987343 | 0.692866806 | -4.45278659 |
| **A_55_P1988113** | Cyth2 | 0.128839451 | -0.085722332 | 0.343401233 | 10.42877299 | 1.47728966 | 0.191142364 | 0.693299074 | -4.453218753 |
| **A_55_P2066478** | Crtc3 | -0.1480901 | -0.394772263 | 0.098592062 | 11.32266161 | -1.476921606 | 0.191237417 | 0.693411942 | -4.453483553 |
| **A_52_P599162** | Tug1 | -0.158526804 | -0.42261708 | 0.105563473 | 9.59453904 | -1.476792422 | 0.19127079 | 0.693411942 | -4.453576493 |
| **A_55_P1993518** | Tbc1d14 | -0.147019013 | -0.39199916 | 0.097961133 | 11.24490142 | -1.476426317 | 0.191365397 | 0.693411942 | -4.453839871 |
| **A_55_P2136385** | Rps3a1 | -0.181321572 | -0.483493368 | 0.120850224 | 14.75373497 | -1.47626632 | 0.191406757 | 0.693411942 | -4.453954969 |
| **A_55_P1966734** | Cyp2u1 | -0.150990219 | -0.402637522 | 0.100657083 | 15.73201743 | -1.476133712 | 0.191441043 | 0.693411942 | -4.454050362 |
| **A_55_P2004152** | NA | 0.151978137 | -0.101337305 | 0.405293578 | 12.65842121 | 1.476007667 | 0.191473637 | 0.693411942 | -4.454141031 |
| **A_52_P543824** | Srsf6 | -0.159165846 | -0.424481802 | 0.10615011 | 8.949667867 | -1.475895726 | 0.191502588 | 0.693411942 | -4.454221554 |
| **A_55_P1960576** | Kdm5b | -0.10476582 | -0.279408089 | 0.069876449 | 8.028499417 | -1.475840469 | 0.191516881 | 0.693411942 | -4.454261303 |
| **A_55_P2119957** | Mlxipl | 0.082977717 | -0.055347309 | 0.221302743 | 8.159229435 | 1.475808084 | 0.191525258 | 0.693411942 | -4.454284597 |
| **A_52_P281033** | Socs5 | -0.091546887 | -0.244159625 | 0.061065851 | 8.46643339 | -1.475781155 | 0.191532224 | 0.693411942 | -4.454303968 |
| **A_55_P2032740** | Iars2 | 0.07007095 | -0.046769255 | 0.186911155 | 8.55740436 | 1.47541726 | 0.191626381 | 0.69350651 | -4.454565716 |
| **A_51_P224575** | Gnpat | 0.095337443 | -0.063639058 | 0.254313945 | 11.64806787 | 1.475365859 | 0.191639684 | 0.69350651 | -4.454602687 |
| **A_51_P225592** | Tpm4 | -0.141366668 | -0.377148303 | 0.094414967 | 15.60948067 | -1.475048193 | 0.191721919 | 0.69350651 | -4.454831168 |
| **A_55_P2003789** | Taf4 | -0.126021168 | -0.341576242 | 0.089533906 | 10.1837387 | -1.515079188 | 0.191722786 | 0.69350651 | -4.426280046 |
| **A_55_P1964068** | Slc25a38 | 0.192440256 | -0.128538929 | 0.513419441 | 11.05913397 | 1.474987092 | 0.191737741 | 0.69350651 | -4.454875114 |
| **A_55_P2408848** | Lgr4 | -0.086432266 | -0.230623383 | 0.057758852 | 8.127679397 | -1.474709756 | 0.191809568 | 0.693517349 | -4.455074577 |
| **A_51_P187665** | Rmi1 | 0.114637856 | -0.076608864 | 0.305884577 | 9.960833834 | 1.474698451 | 0.191812497 | 0.693517349 | -4.455082708 |
| **A_52_P276794** | Tox4 | -0.156161189 | -0.416715004 | 0.104392625 | 9.388066651 | -1.474500198 | 0.191863858 | 0.69354694 | -4.455225288 |
| **A_51_P353592** | Commd4 | 0.125725745 | -0.084062446 | 0.335513936 | 10.89982992 | 1.474389883 | 0.191892443 | 0.69354694 | -4.455304623 |

| | | | | | | | | | |
|---|---|---|---|---|---|---|---|---|---|
| **A_52_P438036** | Lpar4 | -0.154328569 | -0.411935775 | 0.103278637 | 8.818806291 | -1.473864252 | 0.192028701 | 0.693909658 | -4.455682621 |
| **A_55_P2018701** | Kxd1 | 0.087915304 | -0.058871591 | 0.2347022 | 9.579215013 | 1.473487174 | 0.192126505 | 0.693969552 | -4.45595377 |
| **A_55_P1958532** | Hr | -0.096915205 | -0.258735424 | 0.064905015 | 9.131407963 | -1.473425884 | 0.192142406 | 0.693969552 | -4.455997842 |
| **A_55_P1976237** | Dnajb3 | 0.092338821 | -0.061844434 | 0.246522076 | 9.019049636 | 1.473385113 | 0.192152985 | 0.693969552 | -4.456027157 |
| **A_55_P2001228** | Dctn6 | 0.260367748 | -0.174464701 | 0.695200196 | 13.24421919 | 1.473107378 | 0.192225061 | 0.694100169 | -4.456226859 |
| **A_55_P1981090** | Thap7 | 0.182857338 | -0.122603722 | 0.488318397 | 10.34312634 | 1.472738709 | 0.192320775 | 0.694315147 | -4.45649193 |
| **A_55_P2068673** | Stmn2 | -0.120345089 | -0.321398911 | 0.080708733 | 8.564086809 | -1.47260138 | 0.192356439 | 0.694315147 | -4.456590665 |
| **A_51_P192162** | NA | -0.10839624 | -0.289506722 | 0.072714241 | 9.690996565 | -1.472447424 | 0.192396429 | 0.694329832 | -4.456701353 |
| **A_55_P2178203** | Derl2 | 0.14790119 | -0.099261488 | 0.395063869 | 11.20279789 | 1.47216992 | 0.19246853 | 0.694342517 | -4.456900858 |
| **A_55_P2060330** | Cyren | 0.111458322 | -0.074805034 | 0.297721678 | 9.635361988 | 1.472157381 | 0.192471789 | 0.694342517 | -4.456909873 |
| **A_55_P2233271** | Zfp626 | -0.091675233 | -0.244970755 | 0.061620288 | 8.761628256 | -1.471267765 | 0.192703103 | 0.694997375 | -4.457549384 |
| **A_52_P188678** | Nectin2 | 0.135054609 | -0.090808239 | 0.360917457 | 10.64917382 | 1.471071471 | 0.192754177 | 0.694997375 | -4.45769048 |
| **A_52_P131836** | Bysl | 0.126066556 | -0.084817748 | 0.336950861 | 10.19420807 | 1.470702384 | 0.192850245 | 0.694997375 | -4.457955768 |
| **A_55_P2176963** | Hsph1 | -0.255330712 | -0.682482945 | 0.171821522 | 11.58767473 | -1.470582985 | 0.192881332 | 0.694997375 | -4.458041585 |
| **A_55_P2038737** | Slc29a1 | 0.176612366 | -0.11886179 | 0.472086522 | 11.78995867 | 1.47051944 | 0.192897879 | 0.694997375 | -4.458087256 |
| **A_55_P1973347** | A130040M12Rik | -0.509589116 | -1.362295564 | 0.343117331 | 9.702560576 | -1.470245166 | 0.192969314 | 0.694997375 | -4.458284381 |
| **A_55_P2100305** | Lrba | 0.113615562 | -0.07650273 | 0.303733854 | 11.07951125 | 1.470222556 | 0.192975204 | 0.694997375 | -4.458300631 |
| **A_55_P2072443** | Kansl1 | 0.170121257 | -0.114551576 | 0.454794089 | 9.277246464 | 1.470217954 | 0.192976403 | 0.694997375 | -4.458303938 |
| **A_51_P513530** | Spag5 | 0.17894957 | -0.120496549 | 0.478395688 | 8.418914733 | 1.470215963 | 0.192976921 | 0.694997375 | -4.458305369 |
| **A_55_P2170076** | Taf9-ps | 0.089791793 | -0.060511707 | 0.240095293 | 11.54232629 | 1.469727132 | 0.193104303 | 0.69532658 | -4.458656675 |
| **A_55_P1961089** | Npb | 0.519429736 | -0.350153213 | 1.389012686 | 9.719090093 | 1.469551993 | 0.193149961 | 0.695361445 | -4.458782535 |
| **A_55_P2347976** | Mir17hg | -0.245038776 | -0.655351685 | 0.165274133 | 9.563006424 | -1.469226646 | 0.193234804 | 0.695523891 | -4.459016328 |
| **A_52_P598714** | Afg3l1 | -0.113394812 | -0.303305582 | 0.076515958 | 10.45275914 | -1.468969418 | 0.193301907 | 0.695523891 | -4.459201164 |
| **A_52_P320261** | Cpa2 | -0.084579628 | -0.22624685 | 0.057087594 | 8.075087529 | -1.468809785 | 0.193343562 | 0.695523891 | -4.459315867 |
| **A_51_P452798** | Zfp524 | 0.11603699 | -0.078342414 | 0.310416394 | 10.06462842 | 1.468640062 | 0.193387859 | 0.695523891 | -4.459437817 |
| **A_52_P72839** | Sec23a | 0.079566275 | -0.053734901 | 0.21286745 | 9.697264893 | 1.468467055 | 0.193433022 | 0.695523891 | -4.459562124 |
| **A_66_P122110** | Mitd1 | -0.161295103 | -0.431522254 | 0.108932049 | 9.473010415 | -1.468457433 | 0.193435534 | 0.695523891 | -4.459569037 |
| **A_51_P262701** | Cpb2 | -0.065623062 | -0.175572877 | 0.044326752 | 7.797161382 | -1.468355297 | 0.193462202 | 0.695523891 | -4.45964242 |
| **A_51_P452875** | Pigyl | 0.184391955 | -0.124590918 | 0.493374829 | 10.26375497 | 1.46817128 | 0.193510257 | 0.695523891 | -4.459774631 |
| **A_55_P1991124** | Tom1l2 | 0.219433667 | -0.148279187 | 0.587146521 | 11.55709658 | 1.468126951 | 0.193521835 | 0.695523891 | -4.45980648 |
| **A_51_P211786** | Chst13 | -0.176131328 | -0.471306367 | 0.119043711 | 8.615563579 | -1.468000299 | 0.193554918 | 0.695523891 | -4.459897473 |
| **A_51_P413710** | Ap3d1 | -0.164528452 | -0.440311318 | 0.111254414 | 11.06599773 | -1.467718925 | 0.193628435 | 0.695658739 | -4.46009962 |
| **A_51_P504442** | Sf3a1 | -0.10539484 | -0.282100371 | 0.07131069 | 10.85885624 | -1.467365729 | 0.193720754 | 0.695861076 | -4.460353352 |
| **A_55_P2110925** | Flrt3 | 0.164731308 | -0.111517344 | 0.440979959 | 8.957414885 | 1.467050762 | 0.193803115 | 0.695886498 | -4.460579609 |
| **A_51_P112308** | Tcim | -0.078727235 | -0.210752113 | 0.053297642 | 9.683377781 | -1.467027956 | 0.19380908 | 0.695886498 | -4.460595991 |
| **A_55_P2006148** | Aldh3b2 | 0.199581333 | -0.135156473 | 0.534319139 | 9.602543974 | 1.466845311 | 0.193856856 | 0.695886498 | -4.460727188 |
| **A_55_P2160563** | Mrps17 | 0.163094705 | -0.11045841 | 0.436647819 | 11.16909074 | 1.46678804 | 0.19387184 | 0.695886498 | -4.460768327 |
| **A_51_P118132** | Skil | -0.190926485 | -0.511191161 | 0.129338191 | 11.12637541 | -1.466649372 | 0.193908123 | 0.695887505 | -4.456490647 |
| **A_51_P455671** | Psmd5 | 0.08317528 | -0.056389357 | 0.222739917 | 9.486272316 | 1.466182544 | 0.194030316 | 0.696114954 | -4.461203235 |
| **A_55_P1953341** | Wfdc2 | 0.923232963 | -0.625966609 | 2.472432535 | 10.70066879 | 1.466132081 | 0.194043529 | 0.696114954 | -4.46123948 |
| **A_52_P545810** | Lrrfip1 | 0.162131157 | -0.110026963 | 0.434289276 | 10.6983141 | 1.465596261 | 0.194183878 | 0.696365421 | -4.461624304 |
| **A_55_P2162344** | Lrsam1 | 0.184659816 | -0.125370163 | 0.494689795 | 10.645414 | 1.465338198 | 0.194251508 | 0.696365421 | -4.461809633 |
| **A_51_P400543** | Aif1 | -0.256508061 | -0.687178341 | 0.174162218 | 8.891066085 | -1.465295716 | 0.194262643 | 0.696365421 | -4.46184014 |
| **A_51_P431554** | Ten1 | -0.088486879 | -0.237063371 | 0.060089613 | 8.919081317 | -1.465203471 | 0.194286823 | 0.696365421 | -4.461906384 |
| **A_55_P2015405** | 9930111J21Rik1 | -0.075140573 | -0.201318424 | 0.051037278 | 7.890523852 | -1.465077293 | 0.194319904 | 0.696365421 | -4.461996994 |
| **A_52_P354744** | Slc2a3 | 0.279982533 | -0.190182605 | 0.75014767 | 10.47931022 | 1.465040659 | 0.194329509 | 0.696365421 | -4.462023301 |
| **A_55_P2090519** | Ppdpf | 0.16534915 | -0.112361202 | 0.443059502 | 12.64512045 | 1.46480246 | 0.194391975 | 0.696460147 | -4.462194349 |
| **A_55_P2084646** | Csrnp3 | 0.131423743 | -0.089330442 | 0.352177927 | 8.417908572 | 1.464650653 | 0.194431795 | 0.696473716 | -4.462303356 |
| **A_55_P2103115** | Tor1aip2 | -0.096659274 | -0.259064141 | 0.065745593 | 12.56702117 | -1.464244666 | 0.194538326 | 0.696588143 | -4.462594869 |
| **A_51_P464918** | Mefv | -0.079616919 | -0.213402621 | 0.054168782 | 7.783784674 | -1.464080083 | 0.194581529 | 0.696588143 | -4.46271304 |

| | | | | | | | | | |
|---|---|---|---|---|---|---|---|---|---|
| **A_55_P2006754** | Mfsd12 | 0.096774175 | -0.065880677 | 0.259429027 | 12.48527703 | 1.46373217 | 0.194672883 | 0.696588143 | -4.462962831 |
| **A_55_P2128169** | Trappc13 | -0.077399338 | -0.207494613 | 0.052695936 | 9.241614897 | -1.463675788 | 0.194687691 | 0.696588143 | -4.463003311 |
| **A_52_P89477** | Bcl9 | -0.120343901 | -0.322642312 | 0.08195451 | 8.326214126 | -1.463527128 | 0.194726741 | 0.696588143 | -4.463110039 |
| **A_51_P420400** | Lef1 | -0.063769579 | -0.170981934 | 0.043442777 | 7.799268143 | -1.463315167 | 0.194782432 | 0.696588143 | -4.463262209 |
| **A_51_P424221** | Irf2bp2 | -0.138835338 | -0.372256243 | 0.094585566 | 9.545890548 | -1.463286744 | 0.1947899 | 0.696588143 | -4.463282614 |
| **A_52_P66130** | Myo5b | -0.125194706 | -0.335707579 | 0.085318167 | 8.787404047 | -1.463108289 | 0.194836801 | 0.696588143 | -4.463410725 |
| **A_51_P199608** | Ttll1 | 0.100865284 | -0.068750043 | 0.27048061 | 8.798434575 | 1.463004916 | 0.194863973 | 0.696588143 | -4.463484934 |
| **A_55_P2004213** | Gprasp2 | 0.110173784 | -0.075101215 | 0.295448783 | 8.011841203 | 1.462953791 | 0.194877413 | 0.696588143 | -4.463521635 |
| **A_55_P1963017** | Stfa1 | -0.467479883 | -1.253657276 | 0.31869751 | 9.794921563 | -1.462889552 | 0.194894302 | 0.696588143 | -4.463567749 |
| **A_55_P2005440** | Dennd5a | 0.254046494 | -0.336327313 | 0.844420302 | 11.65209692 | 1.981382032 | 0.194896201 | 0.696588143 | -4.408038596 |
| **A_55_P2028029** | Abcb11 | -0.068717403 | -0.184323226 | 0.046888421 | 7.859723348 | -1.462366351 | 0.195031903 | 0.69694429 | -4.463943315 |
| **A_55_P2062622** | Nono | -0.210919595 | -0.565929937 | 0.144090746 | 13.03057302 | -1.461655713 | 0.195218946 | 0.697184181 | -4.464453376 |
| **A_55_P2097553** | C77080 | -0.196218584 | -0.526537126 | 0.134099959 | 13.28725467 | -1.461424373 | 0.195279871 | 0.697184181 | -4.464619408 |
| **A_55_P2132549** | Cd48 | -0.075537255 | -0.202712901 | 0.051638392 | 7.863406663 | -1.461256333 | 0.195324137 | 0.697184181 | -4.464740006 |
| **A_52_P484118** | Urb2 | -0.118384876 | -0.317734624 | 0.080964872 | 8.746155503 | -1.460998246 | 0.195392142 | 0.697184181 | -4.464925221 |
| **A_55_P1980621** | NA | -0.135044269 | -0.362457493 | 0.092368955 | 13.37310161 | -1.46093067 | 0.195409952 | 0.697184181 | -4.464973715 |
| **A_55_P2045903** | NA | -0.071503502 | -0.191922917 | 0.048915912 | 7.999015117 | -1.460830969 | 0.195436231 | 0.697184181 | -4.465045263 |
| **A_55_P2035590** | NA | 0.141010662 | -0.096473622 | 0.378494946 | 13.12321386 | 1.4607848 | 0.195448401 | 0.697184181 | -4.465078394 |
| **A_51_P462746** | Tex10 | 0.182489906 | -0.124858975 | 0.489838786 | 10.09655202 | 1.460751612 | 0.19545715 | 0.697184181 | -4.46510221 |
| **A_52_P119117** | Tmem258 | 0.195892528 | -0.134072165 | 0.525857222 | 13.07960523 | 1.460560537 | 0.195507527 | 0.697184181 | -4.465239324 |
| **A_55_P2327958** | Mtx3 | -0.109437316 | -0.293808032 | 0.0749334 | 8.444093491 | -1.460301913 | 0.195575733 | 0.697184181 | -4.465424904 |
| **A_55_P2121613** | Sox4 | -0.243720919 | -0.654323801 | 0.166881964 | 8.99637978 | -1.460292909 | 0.195578108 | 0.697184181 | -4.465431365 |
| **A_55_P2045602** | Gpkow | -0.073872694 | -0.19833301 | 0.050587622 | 9.507606686 | -1.460233135 | 0.195593875 | 0.697184181 | -4.465474256 |
| **A_52_P492901** | Stx4a | -0.085937671 | -0.230738406 | 0.058863065 | 11.14223696 | -1.460097893 | 0.195629553 | 0.697184181 | -4.465571296 |
| **A_51_P366121** | Klf14 | -0.131737712 | -0.353717503 | 0.090242078 | 8.78365318 | -1.460043661 | 0.195643862 | 0.697184181 | -4.465610209 |
| **A_55_P2161077** | NA | 0.093608693 | -0.064167194 | 0.25138458 | 9.49419468 | 1.459636501 | 0.19575132 | 0.697184181 | -4.465902345 |
| **A_51_P401504** | Col9a2 | 0.244223742 | -0.167431862 | 0.655879347 | 8.719880938 | 1.45956356 | 0.195770577 | 0.697184181 | -4.465954678 |
| **A_55_P2033520** | Med6 | -0.13450255 | -0.361240207 | 0.092235107 | 10.02735166 | -1.459405657 | 0.195812269 | 0.697184181 | -4.466067966 |
| **A_55_P2002703** | NA | 0.208790388 | -0.143203854 | 0.56078463 | 9.69655929 | 1.459298394 | 0.195840595 | 0.697184181 | -4.466144921 |
| **A_55_P2015666** | NA | 0.107836193 | -0.073980757 | 0.289653143 | 16.08388375 | 1.459147997 | 0.195880319 | 0.697184181 | -4.466252819 |
| **A_55_P2444965** | Tmem245 | -0.068787606 | -0.184767546 | 0.047192335 | 8.577820121 | -1.459138357 | 0.195882866 | 0.697184181 | -4.466259735 |
| **A_51_P150678** | Tnfaip8l2 | -0.078785566 | -0.211624516 | 0.054053384 | 7.84585843 | -1.459117917 | 0.195888265 | 0.697184181 | -4.466274399 |
| **A_52_P276607** | Ptp4a2 | -0.10001734 | -0.268685834 | 0.068651154 | 13.67518839 | -1.45884952 | 0.195959177 | 0.697184181 | -4.466466946 |
| **A_55_P1964724** | NA | -0.090059237 | -0.241942345 | 0.061823872 | 8.418881076 | -1.458773776 | 0.195979193 | 0.697184181 | -4.466521283 |
| **A_51_P507801** | F13a1 | 0.275211368 | -0.188981527 | 0.739404263 | 9.225367387 | 1.458602801 | 0.196024382 | 0.697184181 | -4.466643934 |
| **A_51_P371174** | Bag3 | -0.137049034 | -0.368224745 | 0.094126678 | 12.15847102 | -1.458488261 | 0.19605466 | 0.697184181 | -4.466726098 |
| **A_55_P2187261** | NA | -0.107049198 | -0.287627645 | 0.073529249 | 8.443510374 | -1.458433616 | 0.196069107 | 0.697184181 | -4.466765297 |
| **A_55_P2160081** | Ccdc28b | 0.084814302 | -0.0582717 | 0.227900304 | 7.954578297 | 1.458280632 | 0.196109558 | 0.697184181 | -4.466875036 |
| **A_55_P2061591** | Psmd1 | -0.314562895 | -0.845260999 | 0.216135209 | 14.32196456 | -1.458240382 | 0.196120202 | 0.697184181 | -4.466903907 |
| **A_55_P2028788** | Slc39a11 | 0.252247398 | -0.173345836 | 0.677840632 | 12.17545156 | 1.458146445 | 0.196145045 | 0.697184181 | -4.466971288 |
| **A_55_P2109342** | Rnps1 | -0.108982254 | -0.292969435 | 0.075004927 | 9.459075656 | -1.457261139 | 0.196379322 | 0.697668491 | -4.46760627 |
| **A_55_P2097309** | Nudt4 | -0.170826483 | -0.459230766 | 0.117577801 | 10.63560191 | -1.457211719 | 0.196392408 | 0.697668491 | -4.467641713 |
| **A_55_P2019929** | Ythdf3 | -0.089855648 | -0.241572861 | 0.061861565 | 8.573516548 | -1.457067558 | 0.196430584 | 0.697668491 | -4.467745102 |
| **A_55_P2176724** | Fbxw7 | -0.243222292 | -0.653939757 | 0.167495172 | 10.06897196 | -1.456898748 | 0.196475296 | 0.697668491 | -4.467866166 |
| **A_55_P1989663** | Slco3a1 | -0.249279247 | -0.670280437 | 0.171721943 | 9.862502832 | -1.456706184 | 0.196526312 | 0.697668491 | -4.46800426 |
| **A_55_P2015519** | Mtln | 0.25475817 | -0.175503524 | 0.685019863 | 9.198725925 | 1.456681486 | 0.196532856 | 0.697668491 | -4.468021972 |
| **A_55_P2098598** | Btg1 | -0.181867928 | -0.489025994 | 0.125290139 | 12.9704782 | -1.456677314 | 0.196533961 | 0.697668491 | -4.468024964 |
| **A_55_P2073975** | NA | -0.100691455 | -0.270775516 | 0.069392605 | 16.21569573 | -1.456458672 | 0.196591902 | 0.69774603 | -4.468181753 |
| **A_55_P2045268** | Tmem59 | -0.105441595 | -0.28360195 | 0.07271876 | 14.11557821 | -1.456029122 | 0.196705781 | 0.69802204 | -4.468489771 |
| **A_55_P2120354** | Cib2 | -0.125851233 | -0.338543619 | 0.086841154 | 10.27273292 | -1.455709421 | 0.196790578 | 0.698104382 | -4.468719005 |

| | | | | | | | | | |
|---|---|---|---|---|---|---|---|---|---|
| **A_52_P652913** | Akip1 | -0.117730849 | -0.316705026 | 0.081243327 | 10.5328011 | -1.455669308 | 0.19680122 | 0.698104382 | -4.468747766 |
| **A_52_P670263** | Tfpi | 0.143815455 | -0.099268275 | 0.386899185 | 11.69588098 | 1.455522187 | 0.196840255 | 0.698114732 | -4.468853251 |
| **A_52_P583155** | Ykt6 | -0.095315282 | -0.256480827 | 0.065850263 | 9.402907745 | -1.454988286 | 0.196981974 | 0.698215829 | -4.469236033 |
| **A_66_P106418** | Prkacb | -0.099250649 | -0.26707848 | 0.068577182 | 9.572483571 | -1.454918062 | 0.197000622 | 0.698215829 | -4.469286377 |
| **A_66_P116635** | Igf2os | -0.21892161 | -0.589114466 | 0.151271247 | 11.94415323 | -1.454888668 | 0.197008428 | 0.698215829 | -4.46930745 |
| **A_51_P443293** | Decr2 | -0.094502488 | -0.254306644 | 0.065301667 | 10.76648664 | -1.454870505 | 0.197013251 | 0.698215829 | -4.469320472 |
| **A_51_P294095** | Prkar1a | -0.150582745 | -0.4052644 | 0.10409891 | 9.608982295 | -1.454610467 | 0.19708232 | 0.69833257 | -4.46950689 |
| **A_55_P2029622** | Ppp1r3f | -0.108449013 | -0.291921427 | 0.075023401 | 9.597883789 | -1.454199504 | 0.197191523 | 0.698547938 | -4.469801489 |
| **A_55_P2065819** | Fbxo22 | 0.171041873 | -0.125869117 | 0.467952864 | 11.29980776 | 1.492884654 | 0.197215381 | 0.698547938 | -4.439994545 |
| **A_51_P186476** | Slc11a1 | -0.106495402 | -0.286692745 | 0.073701941 | 8.106004797 | -1.453957249 | 0.197255922 | 0.698563524 | -4.469975141 |
| **A_55_P1996299** | Pofut1 | 0.101170053 | -0.070097761 | 0.272437866 | 11.73197383 | 1.453266914 | 0.197439542 | 0.699005986 | -4.470469941 |
| **A_65_P11504** | Zfp85 | -0.06816829 | -0.183592908 | 0.047256328 | 8.24069569 | -1.452958196 | 0.197521709 | 0.699005986 | -4.470691198 |
| **A_55_P2056140** | Flad1 | 0.159775737 | -0.110781313 | 0.430332788 | 10.27231194 | 1.452851205 | 0.197550192 | 0.699005986 | -4.470767875 |
| **A_55_P2004732** | Ttc39a | -0.106007092 | -0.285524179 | 0.073509995 | 9.632443984 | -1.45277478 | 0.19757054 | 0.699005986 | -4.470822646 |
| **A_55_P1963389** | NA | 0.134452308 | -0.093247144 | 0.362151759 | 11.03726152 | 1.452698334 | 0.197590896 | 0.699005986 | -4.470877431 |
| **A_51_P381321** | Ddx17 | -0.178087338 | -0.47970902 | 0.123534343 | 14.22535745 | -1.452578625 | 0.197622776 | 0.699005986 | -4.470963219 |
| **A_52_P357469** | Frk | -0.104420157 | -0.281288676 | 0.072448362 | 8.466015579 | -1.452455936 | 0.197655455 | 0.699005986 | -4.47105114 |
| **A_55_P1982568** | Blcap | 0.301210264 | -0.209027473 | 0.811448001 | 11.17633602 | 1.452333495 | 0.197688072 | 0.699005986 | -4.471138882 |
| **A_55_P2035951** | Haus8 | 0.092298544 | -0.064077383 | 0.248674471 | 10.57439625 | 1.452091944 | 0.197752434 | 0.699005986 | -4.471311975 |
| **A_55_P2160682** | Set | 0.115945176 | -0.08051858 | 0.312408931 | 10.0164583 | 1.451909024 | 0.197801187 | 0.699005986 | -4.471443048 |
| **A_55_P2091191** | Slc28a2 | 0.14681452 | -0.101998207 | 0.395627247 | 8.458850047 | 1.451662244 | 0.197866977 | 0.699005986 | -4.471619874 |
| **A_51_P216702** | Eogt | -0.122439982 | -0.329967919 | 0.085087955 | 10.39510289 | -1.451496025 | 0.197911302 | 0.699005986 | -4.471738972 |
| **A_55_P2068167** | Nufip2 | -0.0824077 | -0.222091719 | 0.057276318 | 11.32470839 | -1.451410418 | 0.197934134 | 0.699005986 | -4.471800308 |
| **A_51_P466964** | Pgp | -0.133985311 | -0.361189135 | 0.093218513 | 11.62252172 | -1.450810584 | 0.198094182 | 0.699005986 | -4.472230061 |
| **A_55_P1984745** | Pappa2 | -0.237829315 | -0.641170146 | 0.165511516 | 16.11467654 | -1.450649055 | 0.198137301 | 0.699005986 | -4.47234578 |
| **A_51_P412225** | Tbc1d16 | -0.145147926 | -0.391310953 | 0.101015101 | 10.24153707 | -1.450631737 | 0.198141925 | 0.699005986 | -4.472358187 |
| **A_55_P2080333** | NA | -0.102208786 | -0.275555818 | 0.071138246 | 8.668116093 | -1.450577657 | 0.198156364 | 0.699005986 | -4.472396929 |
| **A_51_P341025** | Eif3k | 0.172403815 | -0.120002907 | 0.464810537 | 10.92262032 | 1.450536567 | 0.198167335 | 0.699005986 | -4.472426366 |
| **A_52_P1042732** | Akap5 | 0.079319753 | -0.05521309 | 0.213852596 | 7.897673801 | 1.450514904 | 0.198173119 | 0.699005986 | -4.472441884 |
| **A_51_P127915** | Rnasek | -0.128016563 | -0.345166254 | 0.089133128 | 11.23992442 | -1.450360992 | 0.198214221 | 0.699005986 | -4.46679613 |
| **A_55_P2052769** | Mast2 | -0.310208188 | -0.836422359 | 0.216005982 | 12.09107281 | -1.450306795 | 0.198228696 | 0.699005986 | -4.472590967 |
| **A_55_P2003966** | Rybp | -0.197111397 | -0.531483307 | 0.137260513 | 13.19307884 | -1.450277512 | 0.198236517 | 0.699005986 | -4.472611944 |
| **A_55_P2086094** | Slc35b2 | 0.147740381 | -0.102926961 | 0.398407722 | 11.52201877 | 1.450008744 | 0.198308316 | 0.699005986 | -4.472804471 |
| **A_51_P411770** | Nip7 | 0.189289686 | -0.131882336 | 0.510461708 | 12.08646433 | 1.449967966 | 0.198319212 | 0.699005986 | -4.472833681 |
| **A_55_P2156394** | Btbd16 | -0.167671728 | -0.452175468 | 0.116832013 | 9.211245051 | -1.449909922 | 0.198334722 | 0.699005986 | -4.472875259 |
| **A_51_P367060** | Ifrd1 | -0.12838975 | -0.34624723 | 0.089467729 | 13.32959427 | -1.449863251 | 0.198347194 | 0.699005986 | -4.472908689 |
| **A_55_P2022604** | Exoc3l4 | 0.321934065 | -0.224357886 | 0.868226017 | 13.32288435 | 1.449810681 | 0.198361243 | 0.699005986 | -4.472946345 |
| **A_51_P295085** | Ogn | 0.277803834 | -0.193646151 | 0.749253819 | 8.917246797 | 1.449678747 | 0.198396506 | 0.699005986 | -4.473040847 |
| **A_55_P2071329** | Rhog | 0.141870133 | -0.098912668 | 0.382652934 | 11.41772018 | 1.449554927 | 0.198429606 | 0.699005986 | -4.473129536 |
| **A_51_P177984** | Fam234a | -0.111587734 | -0.301054394 | 0.077878925 | 12.87869838 | -1.448948082 | 0.198591901 | 0.699420547 | -4.473564173 |
| **A_51_P201982** | Angpt2 | -0.415539669 | -1.121144889 | 0.29006555 | 11.01445238 | -1.448838134 | 0.198621318 | 0.699420547 | -4.473642916 |
| **A_55_P2146560** | H2-Ab1 | 0.229708829 | -0.160414583 | 0.61983224 | 10.05045258 | 1.448587846 | 0.198688301 | 0.699420547 | -4.473822162 |
| **A_51_P202430** | Pdap1 | 0.216548443 | -0.151246345 | 0.58434323 | 13.31123554 | 1.448500529 | 0.198711674 | 0.699420547 | -4.473884693 |
| **A_55_P2065971** | Setd3 | -0.093986933 | -0.253625017 | 0.065651151 | 11.30102749 | -1.448438743 | 0.198728214 | 0.699420547 | -4.47392894 |
| **A_55_P1998411** | Chchd3 | -0.154805701 | -0.417859144 | 0.108247742 | 10.80726269 | -1.447811837 | 0.19889611 | 0.699884019 | -4.474377858 |
| **A_55_P2011991** | St7 | -0.142490021 | -0.384748117 | 0.099768075 | 10.51825162 | -1.447022604 | 0.199107668 | 0.700220334 | -4.474942949 |
| **A_55_P2326727** | 9430085M18Rik | 0.16560362 | -0.115953125 | 0.447160366 | 10.52321082 | 1.447015069 | 0.199109688 | 0.700220334 | -4.474948343 |
| **A_51_P459340** | Pacrgl | 0.088678776 | -0.062095183 | 0.239452735 | 9.181482652 | 1.446979953 | 0.199119106 | 0.700220334 | -4.474973484 |
| **A_52_P180826** | Ndufaf7 | -0.081471737 | -0.220019701 | 0.057076228 | 10.76729468 | -1.446691695 | 0.199196431 | 0.700220334 | -4.475179856 |
| **A_55_P1962315** | NA | -0.161853491 | -0.437099145 | 0.113392162 | 15.10667279 | -1.446674297 | 0.199201099 | 0.700220334 | -4.475192312 |

| | | | | | | | | | |
|---|---|---|---|---|---|---|---|---|---|
| **A_52_P128068** | Kazn | -0.159486956 | -0.430713666 | 0.111739754 | 9.876599236 | -1.446644685 | 0.199209044 | 0.700220334 | -4.475213511 |
| **A_51_P107020** | Kif5a | -0.125286549 | -0.338392388 | 0.087819291 | 9.00429457 | -1.446366182 | 0.199283783 | 0.700314137 | -4.470596834 |
| **A_51_P361580** | Rab11b | -0.114477807 | -0.309210787 | 0.080255174 | 11.96620646 | -1.446275243 | 0.199308193 | 0.700314137 | -4.475477988 |
| **A_52_P681488** | Stx17 | -0.123515731 | -0.333644196 | 0.086612734 | 8.431153244 | -1.446127377 | 0.19934789 | 0.70032631 | -4.475583838 |
| **A_55_P1992257** | Ap1s1 | -0.132826028 | -0.358816647 | 0.093164591 | 14.14720655 | -1.445978836 | 0.199387774 | 0.700339141 | -4.475690167 |
| **A_55_P2045812** | Sigmar1 | 0.268828347 | -0.18860294 | 0.726259633 | 12.16665103 | 1.445833753 | 0.199426738 | 0.700348732 | -4.47579402 |
| **A_51_P235878** | Guk1 | 0.176449797 | -0.123865658 | 0.476765251 | 10.72460809 | 1.445481852 | 0.199521274 | 0.700396812 | -4.476045904 |
| **A_51_P256682** | Kpna1 | 0.114487811 | -0.080389778 | 0.309365399 | 10.03677482 | 1.445328338 | 0.199562527 | 0.700396812 | -4.476155781 |
| **A_55_P2151952** | NA | -0.235404992 | -0.636115713 | 0.165305729 | 13.52482468 | -1.445286246 | 0.19957384 | 0.700396812 | -4.476185908 |
| **A_52_P312102** | NA | -0.353167029 | -0.954391604 | 0.248057546 | 11.84351599 | -1.445148881 | 0.199610763 | 0.700396812 | -4.476284224 |
| **A_55_P1957593** | Ino80b | 0.193838778 | -0.136181098 | 0.523858654 | 9.571287726 | 1.445006265 | 0.199649103 | 0.700396812 | -4.476386296 |
| **A_55_P2118784** | Rcor2 | -0.135079017 | -0.365083089 | 0.094925054 | 9.412613778 | -1.444845908 | 0.199692222 | 0.700396812 | -4.476501062 |
| **A_55_P2118362** | Tmem121 | -0.06435176 | -0.173928988 | 0.045225468 | 7.79580539 | -1.444805154 | 0.199703181 | 0.700396812 | -4.476530228 |
| **A_55_P1975181** | Acyp1 | 0.078410384 | -0.055114985 | 0.211935754 | 8.124165591 | 1.444704271 | 0.199730313 | 0.700396812 | -4.476602427 |
| **A_55_P2065465** | Mettl1 | -0.126581627 | -0.342193484 | 0.089030229 | 9.28251512 | -1.444332547 | 0.199830317 | 0.700575027 | -4.476868447 |
| **A_55_P2124712** | Ces2c | 0.215625015 | -0.151691547 | 0.582941577 | 8.37402465 | 1.444201509 | 0.19986558 | 0.700575027 | -4.476962219 |
| **A_51_P374305** | Rchy1 | -0.129006059 | -0.348806668 | 0.09079455 | 10.85138737 | -1.443944096 | 0.199934869 | 0.700575027 | -4.477146419 |
| **A_51_P408363** | Cfp | -0.140314114 | -0.379417527 | 0.098789299 | 8.793924604 | -1.443725892 | 0.199993622 | 0.700575027 | -4.477302557 |
| **A_51_P138060** | Rnf13 | 0.121409239 | -0.085485407 | 0.328303885 | 8.5568173 | 1.443682349 | 0.200005348 | 0.700575027 | -4.477333713 |
| **A_55_P1967443** | NA | 0.255528697 | -0.179934483 | 0.690991877 | 10.13325436 | 1.443635003 | 0.200018099 | 0.700575027 | -4.477367591 |
| **A_55_P2161923** | Rabgap1 | -0.084131279 | -0.227511075 | 0.059248517 | 8.193672417 | -1.443572817 | 0.200034848 | 0.700575027 | -4.477412086 |
| **A_51_P251587** | Psmd9 | -0.101346649 | -0.274110424 | 0.071417126 | 9.180830668 | -1.443197842 | 0.200135868 | 0.700801848 | -4.47768038 |
| **A_55_P2062015** | Acyp1 | 0.084386315 | -0.05950467 | 0.2282773 | 8.119022026 | 1.442804879 | 0.200241786 | 0.701031675 | -4.477961526 |
| **A_55_P1994927** | Rab3il1 | 0.264553885 | -0.186586508 | 0.715694278 | 9.674882657 | 1.442685253 | 0.20027404 | 0.701031675 | -4.478047108 |
| **A_51_P223404** | Plin3 | 0.185429279 | -0.130847278 | 0.501705836 | 14.47911786 | 1.442382586 | 0.200355666 | 0.701159739 | -4.478263633 |
| **A_51_P253527** | Rffl | -0.077678421 | -0.210180831 | 0.054823989 | 8.857447569 | -1.442267363 | 0.200386749 | 0.701159739 | -4.47834606 |
| **A_55_P2370250** | Syn3 | -0.075015009 | -0.203010942 | 0.052980924 | 7.731829018 | -1.441853577 | 0.20049841 | 0.701159739 | -4.478642053 |
| **A_66_P123683** | Nabp1 | -0.109231394 | -0.295624268 | 0.077161479 | 8.943605686 | -1.441741297 | 0.200528719 | 0.701159739 | -4.478722367 |
| **A_52_P229728** | Cdc42 | -0.09324872 | -0.252370367 | 0.065872927 | 11.84609367 | -1.441726159 | 0.200532806 | 0.701159739 | -4.478733195 |
| **A_51_P363642** | 1700088E04Rik | -0.087246964 | -0.236133685 | 0.061639757 | 8.534384545 | -1.441662237 | 0.200550063 | 0.701159739 | -4.478778918 |
| **A_55_P2088014** | Cdc42ep4 | -0.14342682 | -0.38819343 | 0.101339791 | 12.14573224 | -1.441608578 | 0.200564551 | 0.701159739 | -4.478817299 |
| **A_55_P2175284** | Ei24 | -0.079418341 | -0.215033394 | 0.056196713 | 9.390320458 | -1.440728255 | 0.200802371 | 0.701674375 | -4.479446922 |
| **A_51_P171200** | Golm1 | 0.124668064 | -0.088227598 | 0.337563726 | 13.21915696 | 1.440646954 | 0.200824348 | 0.701674375 | -4.479505065 |
| **A_55_P2119458** | Slc25a44 | 0.078484426 | -0.055552201 | 0.212521053 | 8.142486654 | 1.440552721 | 0.200849823 | 0.701674375 | -4.479572456 |
| **A_52_P103550** | Tfdp1 | -0.093047821 | -0.251958902 | 0.065863259 | 9.356016091 | -1.440526297 | 0.200856967 | 0.701674375 | -4.479591353 |
| **A_55_P1961650** | Gm2066 | 0.088748651 | -0.062842473 | 0.240339774 | 8.601988172 | 1.440313897 | 0.200914401 | 0.701700674 | -4.479743245 |
| **A_51_P101196** | Psme1 | -0.080573415 | -0.21820848 | 0.05706165 | 11.52588049 | -1.440229962 | 0.200937101 | 0.701700674 | -4.478523853 |
| **A_51_P275123** | Psd3 | -0.125297071 | -0.339354065 | 0.088759924 | 10.07333889 | -1.440060232 | 0.200983013 | 0.701734222 | -4.47992464 |
| **A_55_P2006281** | Ogdh | 0.157856154 | -0.111930137 | 0.427642445 | 12.57797542 | 1.439497127 | 0.2011354 | 0.702080833 | -4.480327283 |
| **A_52_P434820** | Gnpda2 | -0.084823887 | -0.229811359 | 0.060163586 | 8.506819786 | -1.439318314 | 0.201183813 | 0.702080833 | -4.480455134 |
| **A_55_P1969441** | Eef1akmt2 | 0.157170457 | -0.111482434 | 0.425823347 | 10.44603654 | 1.439290835 | 0.201191253 | 0.702080833 | -4.480474781 |
| **A_51_P103929** | Met | 0.072471516 | -0.051461254 | 0.196404286 | 7.924757137 | 1.438634105 | 0.201369158 | 0.702444164 | -4.477205775 |
| **A_55_P2039379** | Slc27a6 | -0.168138113 | -0.455679331 | 0.119403105 | 9.681058502 | -1.438583993 | 0.201382739 | 0.702444164 | -4.480980128 |
| **A_55_P2054857** | Ube2v2 | 0.181383266 | -0.128839318 | 0.491605851 | 11.54224662 | 1.438444038 | 0.201420673 | 0.702444164 | -4.481080179 |
| **A_51_P305532** | Eif2s3x | -0.191514915 | -0.519082667 | 0.136052837 | 9.710231634 | -1.438370026 | 0.201440737 | 0.702444164 | -4.481133088 |
| **A_55_P2160383** | NA | -0.111565996 | -0.3024659 | 0.079333908 | 11.9098101 | -1.437789479 | 0.201598177 | 0.702707852 | -4.481548078 |
| **A_55_P2169332** | NA | 0.132872575 | -0.094503649 | 0.360248799 | 14.73424124 | 1.437670845 | 0.201630363 | 0.702707852 | -4.481632876 |
| **A_55_P2090060** | Mab21l3 | 0.199802441 | -0.142107221 | 0.541712103 | 11.23095262 | 1.437667908 | 0.20163116 | 0.702707852 | -4.481634975 |
| **A_52_P330694** | Tle1 | -0.146961526 | -0.39846751 | 0.104544458 | 9.382582343 | -1.437555082 | 0.201661775 | 0.702707852 | -4.481715619 |
| **A_55_P2030568** | Ifit1bl2 | -0.157857938 | -0.428073149 | 0.112357273 | 9.322446165 | -1.437228418 | 0.20175044 | 0.702773793 | -4.481949098 |

| | | | | | | | | | |
|---|---|---|---|---|---|---|---|---|---|
| **A_51_P129149** | Gatad2a | -0.112159507 | -0.304151348 | 0.079832333 | 10.02774858 | -1.437217457 | 0.201753416 | 0.702773793 | -4.481956932 |
| **A_55_P2044710** | Asb10 | 0.078706572 | -0.056084845 | 0.21349799 | 8.154311463 | 1.436540659 | 0.201937233 | 0.703154828 | -4.482440621 |
| **A_55_P1990086** | Mks1 | -0.112810959 | -0.306052562 | 0.080430643 | 9.187086174 | -1.436216215 | 0.202025406 | 0.703154828 | -4.482672471 |
| **A_55_P2083252** | Mtus2 | -0.190529409 | -0.5169395 | 0.135880681 | 10.11535063 | -1.436043543 | 0.202072347 | 0.703154828 | -4.482795859 |
| **A_55_P2436715** | Paics | 0.085323302 | -0.060861693 | 0.231508297 | 10.88542222 | 1.435932478 | 0.202102546 | 0.703154828 | -4.482875221 |
| **A_55_P2274233** | Agap3 | 0.247534929 | -0.342838878 | 0.837908737 | 13.2885237 | 1.930596453 | 0.202102584 | 0.703154828 | -4.416367658 |
| **A_55_P2025911** | Sms | -0.14691032 | -0.398658315 | 0.104837676 | 8.843796269 | -1.435672713 | 0.202173192 | 0.703154828 | -4.483060832 |
| **A_55_P2029235** | Btn1a1 | -0.158312332 | -0.429605694 | 0.112981031 | 13.21843892 | -1.435637323 | 0.202182818 | 0.703154828 | -4.483086119 |
| **A_51_P395921** | Uchl4 | -0.088541572 | -0.240274335 | 0.063191191 | 11.59737818 | -1.435611816 | 0.202189757 | 0.703154828 | -4.483104344 |
| **A_51_P136888** | Rb1 | 0.164802102 | -0.117652536 | 0.44725674 | 10.68158308 | 1.435433901 | 0.20223816 | 0.703154828 | -4.481927076 |
| **A_55_P2157423** | NA | -0.103626579 | -0.281251524 | 0.073998365 | 8.072879131 | -1.435279128 | 0.202280276 | 0.703154828 | -4.483342046 |
| **A_66_P107379** | Mdk | 0.215791352 | -0.154099266 | 0.58568197 | 9.608102268 | 1.435257692 | 0.202286109 | 0.703154828 | -4.483357361 |
| **A_52_P199586** | Kpna6 | -0.145703062 | -0.395462939 | 0.104056815 | 12.11837244 | -1.435209066 | 0.202299343 | 0.703154828 | -4.483392102 |
| **A_55_P2077558** | Sod3 | 0.189334494 | -0.13529729 | 0.513966278 | 12.15938641 | 1.434854514 | 0.20239586 | 0.703363822 | -4.483645406 |
| **A_55_P2054714** | NA | 0.073858724 | -0.052801331 | 0.20051878 | 9.255018279 | 1.434601535 | 0.202464753 | 0.703369974 | -4.483826133 |
| **A_55_P2110235** | Trip12 | 0.128517804 | -0.091880097 | 0.348915705 | 10.69882347 | 1.434580768 | 0.202470409 | 0.703369974 | -4.483840969 |
| **A_55_P2101357** | Aftph | 0.100598338 | -0.07210182 | 0.273298496 | 11.95814387 | 1.433069434 | 0.202882446 | 0.704674717 | -4.484920482 |
| **A_51_P386983** | Tnni3 | -0.078047625 | -0.212067494 | 0.055972245 | 7.779192113 | -1.432714521 | 0.202979319 | 0.704884522 | -4.485173945 |
| **A_65_P10399** | Gna13 | -0.103144182 | -0.280323317 | 0.074034953 | 8.595937899 | -1.432192254 | 0.203121948 | 0.704941481 | -4.485546896 |
| **A_55_P1998797** | NA | -0.138246091 | -0.375745122 | 0.09925294 | 12.17585686 | -1.432056597 | 0.203159011 | 0.704941481 | -4.485643763 |
| **A_51_P503542** | Zfp414 | 0.091637131 | -0.065795127 | 0.249069389 | 10.80545664 | 1.432012876 | 0.203170957 | 0.704941481 | -4.485674981 |
| **A_52_P552550** | Agtrap | 0.199948253 | -0.143587905 | 0.543484412 | 9.422185341 | 1.431905387 | 0.20320033 | 0.704941481 | -4.485751733 |
| **A_55_P2175272** | NA | -0.151966711 | -0.413070543 | 0.109137121 | 13.7215756 | -1.431872614 | 0.203209286 | 0.704941481 | -4.485775133 |
| **A_52_P649064** | NA | -0.169033237 | -0.459488216 | 0.121421742 | 12.80514021 | -1.431734356 | 0.203247074 | 0.704941481 | -4.485873852 |
| **A_55_P2036459** | Med24 | -0.075919 | -0.206374251 | 0.054536252 | 8.29409374 | -1.431719939 | 0.203251015 | 0.704941481 | -4.485884146 |
| **A_55_P1991688** | Rad51ap1 | 0.130036087 | -0.093477387 | 0.35354956 | 8.15201222 | 1.431295635 | 0.203367027 | 0.705002025 | -4.486187088 |
| **A_51_P217706** | Ribc1 | 0.074716719 | -0.053726311 | 0.203159748 | 8.324644195 | 1.431121198 | 0.203414739 | 0.705002025 | -4.486311626 |
| **A_55_P2022293** | Zfp579 | 0.11108016 | -0.079895378 | 0.302055697 | 9.973816551 | 1.430961396 | 0.203458457 | 0.705002025 | -4.486425711 |
| **A_55_P1969982** | Styx | 0.133492112 | -0.09607027 | 0.363054494 | 9.207538705 | 1.430619183 | 0.203552107 | 0.705002025 | -4.48667001 |
| **A_55_P2143934** | NA | 0.087829172 | -0.06322768 | 0.238886025 | 8.287306192 | 1.43043301 | 0.203603072 | 0.705002025 | -4.48680291 |
| **A_52_P354390** | Snrpn | 0.103044129 | -0.074182849 | 0.280271108 | 8.58977657 | 1.430416738 | 0.203607527 | 0.705002025 | -4.486814525 |
| **A_55_P1982902** | Tceal3 | 0.10286576 | -0.074069629 | 0.279801149 | 8.243804534 | 1.430293939 | 0.203641151 | 0.705002025 | -4.486902182 |
| **A_55_P2101570** | Zhx3 | -0.198695021 | -0.540464691 | 0.14307465 | 9.862549595 | -1.430285143 | 0.203643559 | 0.705002025 | -4.48690846 |
| **A_55_P2095153** | Pdf | 0.118388699 | -0.085278377 | 0.322055776 | 9.740288841 | 1.430074238 | 0.20370132 | 0.705002025 | -4.487059005 |
| **A_52_P537566** | Cenpt | -0.085217147 | -0.231821165 | 0.061386871 | 9.589895288 | -1.4300469 | 0.203708808 | 0.705002025 | -4.487078518 |
| **A_55_P1969650** | Rasgrp1 | 0.265497965 | -0.191268898 | 0.722264829 | 8.685407129 | 1.429999119 | 0.203721896 | 0.705002025 | -4.487112624 |
| **A_55_P2139385** | Gm12070 | 0.13512023 | -0.097360882 | 0.367601342 | 15.5143014 | 1.429887482 | 0.203752479 | 0.705002025 | -4.487192307 |
| **A_55_P2102365** | Spag7 | -0.118259386 | -0.321746078 | 0.085227306 | 16.12181196 | -1.429778533 | 0.203782329 | 0.705002025 | -4.48727007 |
| **A_55_P1962359** | Trim11 | -0.167657683 | -0.456156963 | 0.120841598 | 10.77662725 | -1.429709781 | 0.203801169 | 0.705002025 | -4.487319142 |
| **A_51_P355151** | Camk2n2 | -0.145781747 | -0.396775996 | 0.105212501 | 9.335304232 | -1.428922069 | 0.204017129 | 0.705002025 | -4.487881322 |
| **A_55_P2000798** | Ccdc144b | 0.098405597 | -0.07104139 | 0.267852583 | 8.770983197 | 1.42874628 | 0.204065353 | 0.705002025 | -4.48800677 |
| **A_55_P2013268** | NA | 0.103357118 | -0.074619041 | 0.281333277 | 9.0767187 | 1.428721972 | 0.204072022 | 0.705002025 | -4.488024116 |
| **A_55_P2412369** | Ubxn7 | -0.098484234 | -0.268071957 | 0.071103489 | 8.755809817 | -1.428701383 | 0.204077671 | 0.705002025 | -4.488038808 |
| **A_55_P1994132** | Tmem184a | -0.135822502 | -0.369706774 | 0.09806177 | 9.356624802 | -1.428696138 | 0.20407911 | 0.705002025 | -4.488042552 |
| **A_66_P135842** | Fip1l1 | -0.11294449 | -0.307436331 | 0.081547351 | 11.08170779 | -1.428672954 | 0.204085471 | 0.705002025 | -4.488059096 |
| **A_55_P2086475** | NA | -0.097262051 | -0.264767077 | 0.070242976 | 11.49407892 | -1.428514789 | 0.204128873 | 0.705002025 | -4.488171961 |
| **A_55_P2108850** | 2-Mar | 0.085283254 | -0.061595767 | 0.232162275 | 9.767350207 | 1.42847669 | 0.204139329 | 0.705002025 | -4.488199148 |
| **A_52_P78023** | Limd2 | -0.241655123 | -0.657951883 | 0.174641636 | 13.33023254 | -1.428111576 | 0.204239557 | 0.705002025 | -4.488459676 |
| **A_55_P2143802** | NA | 0.106272658 | -0.076805653 | 0.289350968 | 8.404022448 | 1.428084182 | 0.204247078 | 0.705002025 | -4.488479222 |
| **A_55_P2182077** | NA | -0.149721478 | -0.407717671 | 0.108274714 | 14.50296042 | -1.427709909 | 0.204349871 | 0.705002025 | -4.488746267 |

| | | | | | | | | | |
|---|---|---|---|---|---|---|---|---|---|
| **A_52_P77332** | Ireb2 | -0.075809832 | -0.206447423 | 0.05482776 | 8.575542662 | -1.42766571 | 0.204362013 | 0.705002025 | -4.488777801 |
| **A_55_P2074536** | Ppm1b | -0.205218263 | -0.558874742 | 0.148438215 | 12.59542068 | -1.427590184 | 0.204382763 | 0.705002025 | -4.488831687 |
| **A_55_P2031949** | Bend6 | -0.084034848 | -0.228894842 | 0.060825145 | 9.207850823 | -1.427184501 | 0.204494252 | 0.705002025 | -4.489121114 |
| **A_51_P185846** | Dnal4 | -0.153375115 | -0.417768958 | 0.111018728 | 10.73547197 | -1.427160206 | 0.20450093 | 0.705002025 | -4.489138446 |
| **A_55_P2175206** | Rbbp6 | -0.098459316 | -0.2681975 | 0.071278869 | 8.743996237 | -1.427073769 | 0.204524693 | 0.705002025 | -4.48920011 |
| **A_55_P2448065** | Sfxn2 | -0.094255016 | -0.256747412 | 0.06823738 | 8.335605928 | -1.427054611 | 0.20452996 | 0.705002025 | -4.489213777 |
| **A_55_P2089304** | 2900097C17Rik | -0.09536672 | -0.259786391 | 0.069052951 | 9.337945524 | -1.426961423 | 0.204555582 | 0.705002025 | -4.489280256 |
| **A_51_P451338** | Jag2 | 0.162297798 | -0.117522491 | 0.442118088 | 9.002059244 | 1.42692974 | 0.204564294 | 0.705002025 | -4.489302858 |
| **A_55_P2162651** | NA | 0.101541834 | -0.073541706 | 0.276625373 | 14.5835312 | 1.426818882 | 0.204594779 | 0.705002025 | -4.48938194 |
| **A_55_P2383248** | 4930422N03Rik | -0.193347866 | -0.526765966 | 0.140070235 | 9.878586271 | -1.426656347 | 0.204639483 | 0.705002025 | -4.489497884 |
| **A_66_P139703** | NA | 0.242657953 | -0.175828937 | 0.661144842 | 10.66837738 | 1.426533042 | 0.204673403 | 0.705002025 | -4.489585841 |
| **A_55_P2154782** | NA | 0.243547995 | -0.187692437 | 0.674788428 | 8.890478679 | 1.463575625 | 0.204702939 | 0.705002025 | -4.45808632 |
| **A_52_P56394** | Psmg4 | 0.139350869 | -0.100999411 | 0.379701148 | 9.538966852 | 1.426376618 | 0.204716441 | 0.705002025 | -4.48969742 |
| **A_66_P113892** | Smim6 | 0.143859543 | -0.104269801 | 0.391988887 | 7.970207101 | 1.426361841 | 0.204720507 | 0.705002025 | -4.48970796 |
| **A_55_P2047986** | Ankrd23 | -0.181861606 | -0.495542543 | 0.131819331 | 9.105358649 | -1.426336706 | 0.204727423 | 0.705002025 | -4.489725889 |
| **A_52_P306357** | NA | -0.061087727 | -0.16646688 | 0.044291426 | 7.704997113 | -1.426160571 | 0.204775897 | 0.705021792 | -4.489851522 |
| **A_55_P2088331** | Ctdsp2 | -0.136390217 | -0.371768205 | 0.098987771 | 11.64094156 | -1.4255634 | 0.204940321 | 0.705021792 | -4.490277442 |
| **A_55_P2011747** | Lsg1 | -0.106286424 | -0.289725678 | 0.07715283 | 10.75062608 | -1.425458848 | 0.204969121 | 0.705021792 | -4.490352007 |
| **A_55_P1954897** | NA | 0.176418266 | -0.128077109 | 0.480913642 | 14.3915606 | 1.425384432 | 0.204989622 | 0.705021792 | -4.490405078 |
| **A_55_P1964038** | Gm10032 | 0.122648328 | -0.089050504 | 0.33434716 | 8.766542066 | 1.425319851 | 0.205007415 | 0.705021792 | -4.490451135 |
| **A_55_P1999078** | NA | -0.11684883 | -0.318548124 | 0.084850465 | 9.639072315 | -1.425243771 | 0.205028378 | 0.705021792 | -4.490505392 |
| **A_55_P2149382** | Gm3448 | 0.086126061 | -0.062542058 | 0.234794181 | 8.041196728 | 1.425233102 | 0.205031318 | 0.705021792 | -4.490513 |
| **A_51_P208121** | Klhl25 | 0.092774085 | -0.067393927 | 0.252942097 | 9.396362788 | 1.425017045 | 0.205090861 | 0.705021792 | -4.490667077 |
| **A_55_P2144796** | Gfer | -0.104911262 | -0.286058792 | 0.076236268 | 8.958728133 | -1.424816249 | 0.205146213 | 0.705021792 | -4.490810266 |
| **A_55_P1967659** | Lactb | 0.17475454 | -0.126993769 | 0.47650285 | 10.6096676 | 1.424796323 | 0.205151706 | 0.705021792 | -4.490824475 |
| **A_51_P378858** | Atg10 | 0.065040043 | -0.047267758 | 0.177347844 | 8.788648916 | 1.424754576 | 0.205163216 | 0.705021792 | -4.490854243 |
| **A_55_P2038594** | Tmem234 | -0.170606417 | -0.465227814 | 0.12401498 | 12.37929674 | -1.424623951 | 0.205199235 | 0.705021792 | -4.490947389 |
| **A_51_P285770** | Fem1a | 0.087958481 | -0.063967359 | 0.23988432 | 11.01839538 | 1.424345135 | 0.205276135 | 0.705021792 | -4.491146197 |
| **A_55_P2016968** | NA | 0.174805169 | -0.127138048 | 0.476748385 | 16.01429427 | 1.424289117 | 0.205291589 | 0.705021792 | -4.49118614 |
| **A_52_P141112** | Csnk1g3 | -0.101372513 | -0.276487316 | 0.07374229 | 8.408721335 | -1.424185356 | 0.205320216 | 0.705021792 | -4.491260122 |
| **A_55_P2038831** | NA | -0.134642229 | -0.367246299 | 0.097961842 | 14.51117607 | -1.424075924 | 0.205350411 | 0.705021792 | -4.491338147 |
| **A_51_P488718** | Zfand2b | 0.164816638 | -0.119917927 | 0.449551202 | 11.06671822 | 1.424065689 | 0.205353236 | 0.705021792 | -4.491345445 |
| **A_51_P149373** | Fbxl20 | -0.112238151 | -0.30621369 | 0.081737388 | 10.80223642 | -1.423517137 | 0.205504661 | 0.705416374 | -4.491736536 |
| **A_51_P367563** | Rsph3a | -0.063853857 | -0.174221276 | 0.046513561 | 8.151218193 | -1.42336216 | 0.205547461 | 0.705438009 | -4.49184702 |
| **A_55_P2166995** | NA | -0.090042658 | -0.245706391 | 0.065621075 | 16.2866823 | -1.423082337 | 0.20562476 | 0.705489843 | -4.492046499 |
| **A_55_P2139673** | Rpl11 | 0.142663208 | -0.104001814 | 0.389328231 | 15.88043017 | 1.42289738 | 0.205675867 | 0.705489843 | -4.492178345 |
| **A_55_P1995243** | Cobl | -0.112386349 | -0.306726092 | 0.081953394 | 9.842081218 | -1.42272545 | 0.205723385 | 0.705489843 | -4.492300899 |
| **A_51_P440443** | Exosc5 | 0.124539419 | -0.090816974 | 0.339895812 | 11.34539149 | 1.422716043 | 0.205725985 | 0.705489843 | -4.492307605 |
| **A_51_P485260** | Rgl1 | 0.177807929 | -0.129688696 | 0.485304554 | 12.42767372 | 1.422590595 | 0.205760664 | 0.705489843 | -4.492397024 |
| **A_55_P1993483** | NA | -0.230244863 | -0.628445777 | 0.16795605 | 10.23932725 | -1.42251502 | 0.205781558 | 0.705489843 | -4.492450893 |
| **A_51_P408561** | Oard1 | 0.088198043 | -0.064414089 | 0.240810175 | 10.21219899 | 1.421801774 | 0.205978845 | 0.705868371 | -4.492959246 |
| **A_51_P478688** | Samd10 | 0.241721818 | -0.176565783 | 0.660009418 | 10.92437331 | 1.421706745 | 0.206005144 | 0.705868371 | -4.493026971 |
| **A_51_P487175** | Acsm3 | 0.102036408 | -0.074564741 | 0.278637557 | 7.918851883 | 1.421447416 | 0.206076927 | 0.705868371 | -4.493211783 |
| **A_51_P382484** | Itgav | -0.195217095 | -0.533170395 | 0.142736204 | 9.748954102 | -1.421118585 | 0.206167983 | 0.705868371 | -4.493446112 |
| **A_55_P2002642** | Deup1 | 0.131202959 | -0.095955685 | 0.358361602 | 8.12821135 | 1.420965471 | 0.206210393 | 0.705868371 | -4.493555218 |
| **A_55_P2056286** | Sf3b5 | 0.127215141 | -0.093044577 | 0.347474858 | 12.48713706 | 1.420930667 | 0.206220034 | 0.705868371 | -4.493580018 |
| **A_55_P2090953** | NA | 0.064422788 | -0.047125597 | 0.175971172 | 16.9153369 | 1.420840705 | 0.206244958 | 0.705868371 | -4.493644122 |
| **A_55_P2360266** | 4833411C07Rik | -0.065815645 | -0.179780028 | 0.048148737 | 7.699526904 | -1.420787627 | 0.206259664 | 0.705868371 | -4.493681942 |
| **A_52_P207614** | NA | -0.179574776 | -0.49052361 | 0.131374058 | 10.53164192 | -1.420775843 | 0.206262929 | 0.705868371 | -4.493690339 |
| **A_55_P1978681** | Tspan8 | 0.431481954 | -0.315672715 | 1.178636623 | 9.468319288 | 1.420761843 | 0.206266808 | 0.705868371 | -4.493700315 |

| | | | | | | | | | |
|---|---|---|---|---|---|---|---|---|---|
| **A_51_P194479** | 1110059G10Rik | 0.112830464 | -0.082576841 | 0.308237768 | 9.755764927 | 1.420544172 | 0.206327129 | 0.705868371 | -4.493855412 |
| **A_55_P2010699** | Map11 | -0.101890036 | -0.278351353 | 0.074571281 | 9.318524036 | -1.420533119 | 0.206330193 | 0.705868371 | -4.493863287 |
| **A_51_P455946** | Rac3 | 0.212747439 | -0.155759972 | 0.58125485 | 8.600178595 | 1.420323517 | 0.206388294 | 0.705942193 | -4.494012627 |
| **A_55_P2091368** | Cep164 | 0.089631357 | -0.065645716 | 0.24490843 | 9.343224881 | 1.420109387 | 0.206447666 | 0.705964119 | -4.494165189 |
| **A_55_P2156409** | Rpl26 | 0.196659661 | -0.144085243 | 0.537404565 | 13.44398278 | 1.41989124 | 0.206508168 | 0.705964119 | -4.494320605 |
| **A_52_P20727** | Nhlrc1 | -0.084193988 | -0.230078542 | 0.061690566 | 8.038097655 | -1.419844988 | 0.206520998 | 0.705964119 | -4.494353556 |
| **A_52_P513347** | Phkb | -0.08066797 | -0.220453621 | 0.059117681 | 8.118541124 | -1.419736337 | 0.206551139 | 0.705964119 | -4.49443096 |
| **A_52_P487024** | Smg7 | -0.065252303 | -0.178337651 | 0.047833044 | 9.290626174 | -1.419576082 | 0.206595604 | 0.705964119 | -4.494545124 |
| **A_51_P375201** | Plk3 | -0.09962976 | -0.272303588 | 0.073044068 | 11.27648396 | -1.419488015 | 0.206620043 | 0.705964119 | -4.494607861 |
| **A_51_P105927** | Rasl12 | 0.184756349 | -0.135504013 | 0.505016712 | 9.034037209 | 1.41927105 | 0.206680263 | 0.705964119 | -4.494762416 |
| **A_55_P2140328** | NA | 0.188546025 | -0.138292994 | 0.515385044 | 11.92386999 | 1.419229554 | 0.206691783 | 0.705964119 | -4.494791976 |
| **A_55_P2045163** | Ehmt1 | 0.31216966 | -0.22904747 | 0.85338679 | 12.85676558 | 1.41901936 | 0.206750142 | 0.705964119 | -4.494941701 |
| **A_52_P57526** | Sin3b | -0.15603648 | -0.426571043 | 0.114498083 | 17.4621766 | -1.418967829 | 0.206764452 | 0.705964119 | -4.494978407 |
| **A_52_P112958** | Ywhab | -0.118618904 | -0.324317148 | 0.087079341 | 10.77515251 | -1.418706257 | 0.206837102 | 0.705964119 | -4.495164719 |
| **A_55_P2030736** | Scmh1 | -0.094192186 | -0.257559748 | 0.069175376 | 8.236055528 | -1.418463652 | 0.206904505 | 0.705964119 | -4.495337514 |
| **A_55_P2091105** | NA | -0.072570755 | -0.198449714 | 0.053308205 | 16.02776227 | -1.418331205 | 0.206941312 | 0.705964119 | -4.495431845 |
| **A_51_P106259** | Camk1 | -0.103341086 | -0.282618174 | 0.075936001 | 8.177256144 | -1.418134416 | 0.206996009 | 0.705964119 | -4.494177915 |
| **A_52_P613241** | Icam1 | -0.07251272 | -0.198311836 | 0.053286396 | 7.834362668 | -1.418096454 | 0.207006563 | 0.705964119 | -4.495599033 |
| **A_55_P2024220** | NA | -0.096257729 | -0.263275515 | 0.070760057 | 8.542126532 | -1.417888405 | 0.207064407 | 0.705964119 | -4.495747199 |
| **A_52_P424272** | Gatc | -0.062968258 | -0.172226602 | 0.046290085 | 8.028816534 | -1.417869412 | 0.207069689 | 0.705964119 | -4.495760724 |
| **A_55_P2034998** | Tjp1 | -0.138917148 | -0.386434674 | 0.108600378 | 11.99787904 | -1.454453986 | 0.207088731 | 0.705964119 | -4.463711059 |
| **A_55_P2072906** | Gm6788 | 0.245467259 | -0.180471946 | 0.671406464 | 14.07905152 | 1.417800481 | 0.207088858 | 0.705964119 | -4.495809813 |
| **A_65_P16603** | Bad | -0.096121887 | -0.262938894 | 0.070695119 | 8.470510301 | -1.417591593 | 0.207146957 | 0.705964119 | -4.495958567 |
| **A_55_P2076683** | Sort1 | 0.118902499 | -0.087457374 | 0.325262372 | 8.456017171 | 1.417538599 | 0.207161699 | 0.705964119 | -4.495996305 |
| **A_55_P2086536** | Safb2 | -0.083922667 | -0.229615602 | 0.061770269 | 8.13287487 | -1.417130823 | 0.207275167 | 0.705993976 | -4.496286671 |
| **A_55_P1963109** | Prh1 | 0.082886792 | -0.061037988 | 0.226811573 | 8.626270829 | 1.416833834 | 0.207357843 | 0.705993976 | -4.496498135 |
| **A_55_P2072842** | NA | 0.184383672 | -0.135802214 | 0.504569557 | 10.87364082 | 1.416737659 | 0.207384623 | 0.705993976 | -4.496566611 |
| **A_51_P480904** | Blmh | -0.120348047 | -0.329348706 | 0.088652612 | 9.93755743 | -1.416643434 | 0.207410863 | 0.705993976 | -4.496633698 |
| **A_51_P265545** | Maco1 | -0.095945164 | -0.262567333 | 0.070677005 | 10.49051011 | -1.416639891 | 0.20741185 | 0.705993976 | -4.49663622 |
| **A_55_P2141311** | AU022252 | 0.076449491 | -0.056318174 | 0.209217155 | 8.89426334 | 1.416613721 | 0.207419138 | 0.705993976 | -4.496654852 |
| **A_51_P294555** | Ifitm6 | -0.108749558 | -0.297635959 | 0.080136843 | 8.037682999 | -1.416432789 | 0.207469535 | 0.705993976 | -4.496783669 |
| **A_52_P488305** | Ndel1 | -0.078754024 | -0.215561477 | 0.058053429 | 8.400064743 | -1.416224685 | 0.207527514 | 0.705993976 | -4.496931826 |
| **A_55_P1996983** | Btg1c | -0.139051388 | -0.380634011 | 0.102531236 | 9.612981426 | -1.416050682 | 0.207576004 | 0.705993976 | -4.4970557 |
| **A_51_P348397** | Hexim1 | -0.108550813 | -0.297164116 | 0.080062491 | 9.19424561 | -1.415891322 | 0.207620422 | 0.705993976 | -4.497169146 |
| **A_55_P1967591** | NA | -0.629055159 | -1.722104414 | 0.463994097 | 10.91574859 | -1.415852978 | 0.207631111 | 0.705993976 | -4.497196443 |
| **A_55_P1983523** | Cd300ld | -0.088611364 | -0.24258366 | 0.065360931 | 8.122301557 | -1.415845947 | 0.207633071 | 0.705993976 | -4.497201448 |
| **A_55_P1981210** | Pex26 | -0.115893439 | -0.317289772 | 0.085502894 | 8.623898602 | -1.415717028 | 0.207669013 | 0.705993976 | -4.49729322 |
| **A_51_P466886** | Scoc | -0.11875059 | -0.325118689 | 0.08761751 | 8.584346507 | -1.415671124 | 0.207681812 | 0.705993976 | -4.497325896 |
| **A_51_P123077** | Nubp1 | 0.098091852 | -0.072410699 | 0.268594403 | 10.6831617 | 1.415374012 | 0.207764672 | 0.706151459 | -4.493248808 |
| **A_55_P2144003** | NA | 0.177014831 | -0.130779129 | 0.484808792 | 13.42284911 | 1.414877121 | 0.207903315 | 0.706498448 | -4.497891059 |
| **A_52_P399677** | Tprkb | -0.096940242 | -0.265537503 | 0.07165702 | 8.701298455 | -1.414564463 | 0.207990597 | 0.706670811 | -4.498113582 |
| **A_66_P106606** | 2900097C17Rik | -0.177490488 | -0.486253204 | 0.131272228 | 10.03248975 | -1.414227882 | 0.208084595 | 0.706829481 | -4.498353116 |
| **A_52_P315345** | Lats2 | -0.178809297 | -0.489953495 | 0.132334901 | 10.71535534 | -1.413831172 | 0.208195435 | 0.706829481 | -4.498635421 |
| **A_55_P2182790** | NA | 0.174945467 | -0.129497364 | 0.479388298 | 13.9501111 | 1.4137288 | 0.208224046 | 0.706829481 | -4.498708267 |
| **A_51_P312485** | Fpr1 | -0.089213668 | -0.244471722 | 0.066044386 | 7.810309769 | -1.413664723 | 0.208241957 | 0.706829481 | -4.498753862 |
| **A_55_P1975685** | Nup37 | 0.184664592 | -0.136715323 | 0.506044506 | 11.10533185 | 1.413624495 | 0.208253201 | 0.706829481 | -4.498782487 |
| **A_51_P313157** | Arhgef5 | -0.112791406 | -0.309088994 | 0.083506182 | 11.67389109 | -1.413611954 | 0.208256707 | 0.706829481 | -4.498791411 |
| **A_51_P117865** | Fam20c | -0.138745968 | -0.380329616 | 0.102837681 | 9.692004987 | -1.412934396 | 0.208446193 | 0.707297634 | -4.499273501 |
| **A_55_P2041708** | Gas2 | 0.101518168 | -0.075258883 | 0.27829522 | 7.941816048 | 1.412820706 | 0.208478003 | 0.707297634 | -4.499354386 |
| **A_55_P2132631** | NA | 0.165923849 | -0.123035956 | 0.454883653 | 11.08070509 | 1.41266941 | 0.208520342 | 0.707297634 | -4.499462023 |

| | | | | | | | | |
|---|---|---|---|---|---|---|---|---|
| **A_55_P2003643** | Tstd2 | -0.127430115 | -0.349363891 | 0.094503661 | 10.63136054 | -1.412595558 | 0.208541012 | 0.707297634 | -4.499514564 |
| **A_55_P2102857** | Krtcap3 | 0.282258781 | -0.20954707 | 0.774064632 | 10.9964632 | 1.411961757 | 0.208718478 | 0.707775343 | -4.499965432 |
| **A_52_P85864** | Eif6 | 0.181266972 | -0.134605562 | 0.497139505 | 11.44593541 | 1.411809105 | 0.208761241 | 0.707796182 | -4.500074015 |
| **A_51_P150521** | Mgat2 | -0.12049471 | -0.330547998 | 0.089558578 | 9.678044383 | -1.411262042 | 0.208914561 | 0.708191784 | -4.500463124 |
| **A_52_P460929** | BC048507 | 0.197245289 | -0.146761847 | 0.541252425 | 12.98234874 | 1.410614525 | 0.209096168 | 0.70868312 | -4.500923628 |
| **A_55_P1957850** | Snx9 | -0.164813694 | -0.452384743 | 0.122757355 | 11.60229518 | -1.409994099 | 0.209270313 | 0.709148996 | -4.501364809 |
| **A_51_P278653** | Rprm | 0.07020168 | -0.052304458 | 0.192707819 | 7.796443392 | 1.409804266 | 0.209323623 | 0.709205311 | -4.501499788 |
| **A_55_P2177173** | Gnl1 | -0.126089344 | -0.346147193 | 0.093968504 | 11.70517061 | -1.409648023 | 0.209367509 | 0.709229685 | -4.501610878 |
| **A_55_P2078765** | Kmt2e | -0.24438587 | -0.671109358 | 0.182337617 | 9.831093087 | -1.408960144 | 0.209560825 | 0.709711457 | -4.502099929 |
| **A_55_P2058613** | NA | -0.113964465 | -0.312972177 | 0.085043247 | 8.191618155 | -1.408862831 | 0.209588186 | 0.709711457 | -4.502169108 |
| **A_55_P2096257** | Myl6 | 0.088222999 | -0.065846623 | 0.242292621 | 14.75278665 | 1.408750116 | 0.209619882 | 0.709711457 | -4.502249235 |
| **A_51_P260871** | Sdhd | -0.156116041 | -0.428911102 | 0.116679019 | 12.96356164 | -1.407927174 | 0.20985143 | 0.709968212 | -4.502834196 |
| **A_51_P464738** | Slc2a1 | 0.180522331 | -0.134944145 | 0.495988808 | 16.7886613 | 1.407819194 | 0.20988183 | 0.709968212 | -4.502910943 |
| **A_52_P387954** | Dctn5 | 0.08730311 | -0.065279825 | 0.239886045 | 11.28215361 | 1.407644253 | 0.209931089 | 0.709968212 | -4.503035279 |
| **A_51_P438479** | Zfp281 | -0.068081092 | -0.187092478 | 0.050930294 | 9.987004572 | -1.407366355 | 0.210009361 | 0.709968212 | -4.50323278 |
| **A_55_P2094852** | Rabep1 | -0.118937868 | -0.326861136 | 0.088985399 | 10.72417453 | -1.407298492 | 0.210028479 | 0.709968212 | -4.503281009 |
| **A_55_P1958730** | NA | 0.312825904 | -0.234065143 | 0.859716951 | 12.8576803 | 1.407249353 | 0.210042323 | 0.709968212 | -4.50331593 |
| **A_55_P2015520** | Plekha1 | -0.2793482 | -0.767716981 | 0.209020581 | 12.87904199 | -1.40723668 | 0.210045894 | 0.709968212 | -4.503324937 |
| **A_55_P2050807** | Mtg2 | 0.11719583 | -0.087702951 | 0.32209461 | 10.20173505 | 1.40715503 | 0.210068899 | 0.709968212 | -4.503382962 |
| **A_55_P2108938** | Hps5 | -0.183984338 | -0.505662565 | 0.137693888 | 10.73227299 | -1.407110983 | 0.210081311 | 0.709968212 | -4.503414263 |
| **A_55_P1956752** | NA | -0.073196685 | -0.201183024 | 0.054789655 | 8.07692562 | -1.407009261 | 0.210109977 | 0.709968212 | -4.50348655 |
| **A_55_P2367117** | Pitpnb | -0.097839105 | -0.268920588 | 0.073242377 | 8.719434981 | -1.406949891 | 0.21012671 | 0.709968212 | -4.50352874 |
| **A_55_P1962384** | Churc1 | 0.216491488 | -0.162074912 | 0.595057888 | 11.57747845 | 1.406915206 | 0.210136486 | 0.709968212 | -4.503553388 |
| **A_51_P468418** | Ice2 | 0.087818012 | -0.065775712 | 0.241411736 | 8.413494622 | 1.40662812 | 0.210217419 | 0.710117526 | -4.503757389 |
| **A_55_P2262136** | Tek | -0.092884421 | -0.255381498 | 0.069612656 | 8.157135088 | -1.406262804 | 0.210320446 | 0.710341413 | -4.504016963 |
| **A_55_P2040031** | NA | 0.167841273 | -0.125855804 | 0.46153835 | 12.40697645 | 1.405944908 | 0.210410138 | 0.710430575 | -4.504242828 |
| **A_52_P577384** | Il18bp | -0.072067234 | -0.198183647 | 0.054049179 | 8.015263664 | -1.405838411 | 0.210440193 | 0.710430575 | -4.504318491 |
| **A_52_P651078** | Fyttd1 | 0.078330388 | -0.058752679 | 0.215413456 | 8.419583567 | 1.405774393 | 0.210458262 | 0.710430575 | -4.504363973 |
| **A_55_P2048563** | Cdk2ap1 | -0.160065647 | -0.440232651 | 0.120101357 | 12.41098367 | -1.405562988 | 0.21051794 | 0.710430575 | -4.504514163 |
| **A_55_P2020306** | Aasdh | -0.154729441 | -0.425565 | 0.116106117 | 9.727001046 | -1.405518081 | 0.210530619 | 0.710430575 | -4.504546065 |
| **A_55_P1968977** | Stk38l | -0.148470366 | -0.408374872 | 0.111434139 | 9.014796184 | -1.40538442 | 0.210568361 | 0.710433905 | -4.504641019 |
| **A_55_P2044282** | Mpp6 | -0.090078331 | -0.247837256 | 0.067680595 | 12.50718143 | -1.404738715 | 0.210750775 | 0.710773053 | -4.505099695 |
| **A_55_P2009988** | Trib3 | -0.254653841 | -0.700649586 | 0.191341904 | 11.74814605 | -1.404716482 | 0.210757058 | 0.710773053 | -4.505115487 |
| **A_55_P2057653** | NA | 0.164265015 | -0.123441557 | 0.451971586 | 15.64091088 | 1.404638145 | 0.210779199 | 0.710773053 | -4.505171129 |
| **A_55_P2103097** | Ccr8 | 0.206201955 | -0.155082007 | 0.567485917 | 8.152219234 | 1.404149205 | 0.21091744 | 0.711115159 | -4.5055184 |
| **A_55_P2182810** | Arl16 | 0.07688431 | -0.057840673 | 0.211609292 | 8.883927708 | 1.403972967 | 0.21096729 | 0.71115918 | -4.505643565 |
| **A_55_P2028345** | Ccser2 | -0.167673835 | -0.461683687 | 0.126336017 | 11.99754553 | -1.403048153 | 0.211229052 | 0.71191741 | -4.5063003 |
| **A_55_P2136786** | Zkscan1 | 0.201959484 | -0.152278126 | 0.556197094 | 14.3118299 | 1.402615862 | 0.211351512 | 0.712174223 | -4.506607239 |
| **A_55_P2192577** | H2bfm | 0.198014641 | -0.149332334 | 0.545361617 | 8.344318145 | 1.402500241 | 0.211384276 | 0.712174223 | -4.506689328 |
| **A_55_P1953630** | E330016L19Rik | -0.076477172 | -0.210692047 | 0.057737704 | 8.642577427 | -1.401846074 | 0.211569738 | 0.712174223 | -4.507153743 |
| **A_55_P2096599** | Tsc22d4 | -0.070728563 | -0.194855689 | 0.053398562 | 8.040914791 | -1.4018363 | 0.21157251 | 0.712174223 | -4.507160681 |
| **A_52_P488427** | Sec14l2 | -0.15057298 | -0.414827885 | 0.113681925 | 8.483727616 | -1.401822907 | 0.211576309 | 0.712174223 | -4.507170189 |
| **A_55_P2090943** | Larp4 | 0.088846509 | -0.067085811 | 0.24477883 | 11.63388686 | 1.401759121 | 0.211594401 | 0.712174223 | -4.507215468 |
| **A_66_P135130** | Tmem161a | 0.262867329 | -0.198571262 | 0.72430592 | 9.984921459 | 1.401496001 | 0.211669048 | 0.712174223 | -4.507402246 |
| **A_52_P546552** | Pelp1 | -0.076081553 | -0.209635826 | 0.057472721 | 9.154727253 | -1.401492375 | 0.211670077 | 0.712174223 | -4.507404819 |
| **A_66_P105843** | Cyth2 | 0.103559873 | -0.07823277 | 0.285352516 | 12.12961856 | 1.401471814 | 0.211675912 | 0.712174223 | -4.507419414 |
| **A_55_P2060369** | Akap9 | 0.120004326 | -0.090701215 | 0.330709867 | 10.41903745 | 1.401167666 | 0.211762231 | 0.712174223 | -4.507635301 |
| **A_55_P2024342** | Tmem33 | 0.109582304 | -0.082824367 | 0.301988975 | 10.41138635 | 1.401165597 | 0.211762818 | 0.712174223 | -4.507636769 |
| **A_55_P2149219** | Aar2 | 0.181351631 | -0.137082424 | 0.499785687 | 10.70601966 | 1.401106417 | 0.211779618 | 0.712174223 | -4.507678775 |
| **A_51_P208555** | Decr1 | -0.09062614 | -0.249783695 | 0.068531414 | 8.842778614 | -1.400862113 | 0.211848981 | 0.712174223 | -4.507852172 |

| | | | | | | | | | |
|---|---|---|---|---|---|---|---|---|---|
| **A_55_P2039637** | Ift88 | -0.107236947 | -0.295569546 | 0.081095653 | 10.19787024 | -1.40083861 | 0.211855656 | 0.712174223 | -4.507868853 |
| **A_55_P2063471** | NA | 0.117961787 | -0.089240132 | 0.325163705 | 16.03230912 | 1.400608421 | 0.211921033 | 0.712174223 | -4.508032223 |
| **A_66_P118579** | 3300002A11Rik | -0.100273041 | -0.276408203 | 0.075862121 | 8.073747073 | -1.400577952 | 0.211929688 | 0.712174223 | -4.508053847 |
| **A_55_P2172022** | B4galt1 | -0.093458963 | -0.257648385 | 0.070730458 | 9.415600531 | -1.400376934 | 0.211986797 | 0.712174223 | -4.508196506 |
| **A_51_P384673** | Lrrc59 | -0.133772553 | -0.368803989 | 0.101258882 | 10.78058902 | -1.400264978 | 0.21201861 | 0.712174223 | -4.508275958 |
| **A_55_P2004588** | Nabp2 | 0.073039901 | -0.055296336 | 0.201376138 | 10.58098646 | 1.40016764 | 0.212046273 | 0.712174223 | -4.508345033 |
| **A_55_P2000148** | Dnaic1 | 0.123656346 | -0.093636545 | 0.340949238 | 8.870742265 | 1.400038743 | 0.212082909 | 0.712174223 | -4.508436502 |
| **A_52_P474089** | Capn6 | 0.220154514 | -0.16670829 | 0.607017317 | 8.665236657 | 1.400037868 | 0.212083158 | 0.712174223 | -4.508437123 |
| **A_51_P343323** | Cox6c | -0.195098951 | -0.537961441 | 0.14776354 | 14.7463868 | -1.399922913 | 0.212115837 | 0.712174223 | -4.508518697 |
| **A_51_P509098** | Asah1 | -0.125007083 | -0.344727639 | 0.094713473 | 10.07865362 | -1.399693976 | 0.212180931 | 0.712221322 | -4.508681148 |
| **A_55_P1991585** | Blvrb | 0.243476952 | -0.184524447 | 0.671478352 | 11.08849652 | 1.399528772 | 0.212227916 | 0.712221322 | -4.508798371 |
| **A_52_P605834** | Tm9sf3 | -0.072030558 | -0.198665005 | 0.054603888 | 11.85523245 | -1.39937492 | 0.21227168 | 0.712221322 | -4.508907534 |
| **A_51_P506683** | Fbxw7 | -0.192317863 | -0.530430015 | 0.14579429 | 10.45412742 | -1.39935533 | 0.212277254 | 0.712221322 | -4.508921434 |
| **A_55_P2051229** | Gpr132 | -0.078331676 | -0.216059919 | 0.059396567 | 7.993908961 | -1.399212175 | 0.212317984 | 0.712234347 | -4.509023004 |
| **A_55_P2042526** | Apoo | -0.168738013 | -0.465577869 | 0.128101844 | 10.39420647 | -1.398491659 | 0.212523092 | 0.71261517 | -4.509534173 |
| **A_55_P2062851** | 7-Mar | -0.192338937 | -0.530731781 | 0.146053906 | 11.94199872 | -1.398347806 | 0.212564064 | 0.71261517 | -4.50963622 |
| **A_55_P2129291** | Dclre1c | -0.109274927 | -0.301542296 | 0.082992441 | 8.827338676 | -1.398247678 | 0.212592587 | 0.71261517 | -4.509707248 |
| **A_51_P387188** | Vrk3 | -0.11335934 | -0.312832652 | 0.086113972 | 10.7306955 | -1.398111066 | 0.212631508 | 0.71261517 | -4.509804154 |
| **A_55_P2180909** | Ccdc166 | -0.086163692 | -0.237784251 | 0.065456867 | 8.153645444 | -1.398090779 | 0.212637288 | 0.71261517 | -4.509818544 |
| **A_55_P2099232** | Bdp1 | -0.117496663 | -0.324262853 | 0.089269528 | 10.01314038 | -1.398025743 | 0.21265582 | 0.71261517 | -4.509864676 |
| **A_66_P114627** | S100a16 | 0.2227381 | -0.169272606 | 0.614748806 | 10.60817239 | 1.397866659 | 0.212701157 | 0.71261517 | -4.509977517 |
| **A_51_P115817** | Ralgps2 | 0.087255448 | -0.066335784 | 0.24084668 | 10.0174722 | 1.397639909 | 0.212765792 | 0.71261517 | -4.507112371 |
| **A_51_P474422** | Ift27 | 0.110473476 | -0.08399454 | 0.304941493 | 10.92580657 | 1.397587476 | 0.212780741 | 0.71261517 | -4.510175537 |
| **A_55_P2024649** | Rpe | 0.091889254 | -0.069890052 | 0.253668561 | 9.645485444 | 1.397368408 | 0.212843208 | 0.71261517 | -4.51033091 |
| **A_51_P363634** | Snrpe | -0.103279562 | -0.285125686 | 0.078566562 | 11.47753392 | -1.397267324 | 0.212872038 | 0.71261517 | -4.510402601 |
| **A_55_P2085880** | Aco1 | 0.09541605 | -0.072585444 | 0.263417545 | 8.335692335 | 1.397260726 | 0.21287392 | 0.71261517 | -4.510407281 |
| **A_51_P178347** | Zfp326 | -0.132791555 | -0.366666941 | 0.101083831 | 9.511129496 | -1.396867146 | 0.212986206 | 0.712867597 | -4.510686403 |
| **A_55_P2003513** | Hsph1 | -0.128447963 | -0.354797951 | 0.097902026 | 10.12609226 | -1.396098003 | 0.213205794 | 0.713349681 | -4.511231802 |
| **A_55_P2107025** | Flad1 | -0.095103025 | -0.262717083 | 0.072511033 | 8.185511863 | -1.395895964 | 0.21326351 | 0.713349681 | -4.511375054 |
| **A_65_P00881** | NA | -0.091240134 | -0.252052216 | 0.069571948 | 8.508009357 | -1.395842451 | 0.2132788 | 0.713349681 | -4.511412995 |
| **A_52_P121** | Osgepl1 | 0.110123714 | -0.083972196 | 0.304219623 | 8.801913246 | 1.395833539 | 0.213281346 | 0.713349681 | -4.511419313 |
| **A_55_P2088401** | H2-T9 | -0.257089408 | -0.710254149 | 0.196075333 | 8.4572416 | -1.395716573 | 0.213314768 | 0.713349681 | -4.511502242 |
| **A_52_P641367** | Desi1 | -0.168842596 | -0.466489599 | 0.128804407 | 9.373754084 | -1.395563718 | 0.213358453 | 0.713372348 | -4.511610612 |
| **A_65_P20104** | Mical2 | -0.072677842 | -0.200822879 | 0.055467195 | 8.02165836 | -1.395305784 | 0.213432188 | 0.71339856 | -4.511793474 |
| **A_52_P424826** | Txnl4b | -0.111374325 | -0.307766734 | 0.085018084 | 10.8257317 | -1.395177775 | 0.213468789 | 0.71339856 | -4.511884222 |
| **A_55_P2165539** | Aebp2 | -0.151444241 | -0.418499622 | 0.11561114 | 9.472463144 | -1.395149001 | 0.213477017 | 0.71339856 | -4.511904619 |
| **A_55_P2080870** | Cmtm7 | 0.097082477 | -0.074156838 | 0.268321791 | 12.63206537 | 1.39478263 | 0.21358181 | 0.713625378 | -4.512164331 |
| **A_55_P2005465** | Zfp422 | -0.083886099 | -0.231897872 | 0.064125674 | 8.762634178 | -1.394321422 | 0.213713796 | 0.713881315 | -4.512491242 |
| **A_55_P2055708** | Vmn1r48 | 0.130808421 | -0.100040003 | 0.361656844 | 8.033651929 | 1.394048722 | 0.213791871 | 0.713881315 | -4.512684521 |
| **A_55_P2179448** | NA | -0.211291787 | -0.584245162 | 0.161661588 | 9.535057031 | -1.393789657 | 0.213866066 | 0.713881315 | -4.512868126 |
| **A_52_P247788** | Ino80c | 0.09887874 | -0.075666277 | 0.273423756 | 8.37346997 | 1.39368502 | 0.213896041 | 0.713881315 | -4.512942281 |
| **A_55_P2170139** | Rheb | -0.141306768 | -0.390748509 | 0.108134972 | 11.96894899 | -1.393679781 | 0.213897541 | 0.713881315 | -4.512945994 |
| **A_55_P2168312** | NA | -0.174596031 | -0.482813198 | 0.133621137 | 13.74566622 | -1.393627486 | 0.213912523 | 0.713881315 | -4.512983054 |
| **A_55_P2246900** | NA | 0.067372985 | -0.05156294 | 0.186308911 | 9.054551526 | 1.393612063 | 0.213916942 | 0.713881315 | -4.512993984 |
| **A_55_P1957613** | Ak2 | 0.153892713 | -0.117868866 | 0.425654293 | 13.34415568 | 1.393154117 | 0.214048182 | 0.71410102 | -4.513318504 |
| **A_55_P2140835** | NA | 0.2632057 | -0.201603288 | 0.728014688 | 9.335027546 | 1.393124521 | 0.214056667 | 0.71410102 | -4.513339475 |
| **A_51_P446477** | Cldnd1 | 0.113396805 | -0.086904475 | 0.313698086 | 11.69704744 | 1.392791975 | 0.214152019 | 0.714113084 | -4.513575109 |
| **A_55_P1979508** | Clic3 | 0.175685247 | -0.134670024 | 0.486040517 | 8.11340656 | 1.392660732 | 0.214189661 | 0.714113084 | -4.513668101 |
| **A_66_P101600** | Lamb1 | -0.229189482 | -0.634074461 | 0.175695498 | 14.12365259 | -1.392618564 | 0.214201757 | 0.714113084 | -4.513697978 |
| **A_51_P253633** | Mrps9 | -0.084167948 | -0.232860966 | 0.06452507 | 9.467871724 | -1.392596579 | 0.214208063 | 0.714113084 | -4.513713555 |

| | | | | | | | | | |
|---|---|---|---|---|---|---|---|---|---|
| **A_55_P1969002** | Tbx3 | -0.185981732 | -0.514664104 | 0.14270064 | 11.44997522 | -1.392076298 | 0.214357359 | 0.714295362 | -4.514082166 |
| **A_51_P294113** | Chd2 | -0.075324146 | -0.208456118 | 0.057807826 | 8.354030371 | -1.391941605 | 0.214396026 | 0.714295362 | -4.514177588 |
| **A_55_P2052685** | Dguok | 0.097878156 | -0.075136544 | 0.270892856 | 11.02271224 | 1.391784317 | 0.214441186 | 0.714295362 | -4.514289012 |
| **A_55_P2087404** | Mapre3 | 0.161847929 | -0.124275908 | 0.447971766 | 10.67929511 | 1.391625153 | 0.214486894 | 0.714295362 | -4.514401763 |
| **A_55_P2076777** | Mdfi | -0.205363533 | -0.568468464 | 0.157741398 | 14.08659849 | -1.391426731 | 0.214543888 | 0.714295362 | -4.514542317 |
| **A_55_P1954266** | Zfp672 | 0.095055694 | -0.073019701 | 0.26313109 | 9.347961375 | 1.391371678 | 0.214559704 | 0.714295362 | -4.514581314 |
| **A_52_P364232** | Pik3c2a | 0.088103954 | -0.067695212 | 0.24390312 | 8.588607784 | 1.391231603 | 0.214599949 | 0.714295362 | -4.514680532 |
| **A_51_P509489** | Kras | -0.096274347 | -0.266543648 | 0.073994954 | 10.13784566 | -1.391052085 | 0.214651538 | 0.714295362 | -4.514807686 |
| **A_55_P2025468** | Tst | 0.169922596 | -0.13061164 | 0.470456833 | 9.4053664 | 1.390997485 | 0.214667231 | 0.714295362 | -4.514846359 |
| **A_51_P314285** | Tmem86a | -0.13604281 | -0.37673795 | 0.104652329 | 10.08342089 | -1.390520631 | 0.214804329 | 0.714295362 | -4.515184089 |
| **A_55_P2004501** | Immp1l | 0.253931001 | -0.195343657 | 0.703205659 | 11.94946004 | 1.390506315 | 0.214808447 | 0.714295362 | -4.515194228 |
| **A_55_P2009732** | Prr9 | 0.206709627 | -0.159062449 | 0.572481703 | 8.618655623 | 1.39033456 | 0.214857848 | 0.714295362 | -4.515315864 |
| **A_51_P461031** | Ostm1 | -0.177872567 | -0.492632318 | 0.136887184 | 11.37538108 | -1.390269331 | 0.214876612 | 0.714295362 | -4.515362058 |
| **A_51_P304527** | Cluap1 | -0.101587239 | -0.281400001 | 0.078225524 | 10.33047805 | -1.389913604 | 0.21497897 | 0.714295362 | -4.515613965 |
| **A_51_P462918** | Ehhadh | -0.101509898 | -0.281197123 | 0.078177328 | 8.810802912 | -1.389825733 | 0.215004261 | 0.714295362 | -4.515676188 |
| **A_51_P217498** | Slc2a4 | 0.080542909 | -0.062034196 | 0.223120013 | 8.025218259 | 1.389781996 | 0.21501685 | 0.714295362 | -4.514167414 |
| **A_51_P464822** | H1f4 | 0.175037934 | -0.134817113 | 0.484892981 | 9.59613224 | 1.389769474 | 0.215020455 | 0.714295362 | -4.515716026 |
| **A_52_P260818** | Card10 | -0.088530031 | -0.245273747 | 0.068213684 | 9.296528441 | -1.389535491 | 0.215087818 | 0.714295362 | -4.515881704 |
| **A_52_P352525** | Qser1 | -0.069802341 | -0.193393645 | 0.053788964 | 8.007359285 | -1.389476579 | 0.215104781 | 0.714295362 | -4.515923417 |
| **A_51_P466824** | Tyk2 | -0.082977689 | -0.229911244 | 0.063955866 | 8.368328258 | -1.389343117 | 0.215143216 | 0.714295362 | -4.516017914 |
| **A_55_P1982019** | Rbfox2 | 0.07510202 | -0.057895332 | 0.208099373 | 10.37948913 | 1.38924163 | 0.215172447 | 0.714295362 | -4.516089769 |
| **A_55_P2024675** | Shld2 | -0.149995028 | -0.415621943 | 0.115631888 | 9.665607209 | -1.389229337 | 0.215175988 | 0.714295362 | -4.516098473 |
| **A_55_P2045036** | NA | 0.101045105 | -0.077911357 | 0.280001567 | 8.480137032 | 1.389111342 | 0.215209978 | 0.714295362 | -4.516182015 |
| **A_55_P1963483** | Vps16 | -0.108304515 | -0.300152463 | 0.083543434 | 11.93490386 | -1.388860302 | 0.215282311 | 0.714295362 | -4.516359746 |
| **A_55_P2001714** | Blvra | -0.079662799 | -0.220790845 | 0.061465246 | 8.811308968 | -1.388709477 | 0.215325779 | 0.714295362 | -4.516466523 |
| **A_55_P2135725** | C8g | 0.08698339 | -0.0671265 | 0.24109328 | 7.944174697 | 1.388593036 | 0.215359343 | 0.714295362 | -4.516548955 |
| **A_55_P2077522** | NA | -0.21241437 | -0.588771136 | 0.163942395 | 10.3018545 | -1.388523806 | 0.215379301 | 0.714295362 | -4.516597964 |
| **A_55_P2180292** | Dmrtc1c2 | 0.195446662 | -0.150870119 | 0.541763442 | 9.76742642 | 1.388429666 | 0.215406442 | 0.714295362 | -4.516664606 |
| **A_55_P2045482** | NA | 0.191860393 | -0.148119789 | 0.531840574 | 14.72520158 | 1.388356158 | 0.215427638 | 0.714295362 | -4.516716641 |
| **A_51_P374464** | Gstp1 | 0.15615334 | -0.120575454 | 0.432882134 | 10.03834925 | 1.388244911 | 0.215459718 | 0.714295362 | -4.51679539 |
| **A_55_P1987600** | Trmt112 | 0.175516337 | -0.135537959 | 0.486570634 | 12.71046497 | 1.388195099 | 0.215474084 | 0.714295362 | -4.51683065 |
| **A_51_P463765** | Timp3 | -0.424421839 | -1.176606155 | 0.327762477 | 12.80091342 | -1.388169865 | 0.215481362 | 0.714295362 | -4.516848513 |
| **A_55_P2302285** | Tarbp2 | 0.092372818 | -0.071366066 | 0.256111703 | 11.50146444 | 1.387910718 | 0.215556117 | 0.714295362 | -4.517031947 |
| **A_51_P132081** | Rsl1 | -0.089472155 | -0.248097059 | 0.069152749 | 8.522165928 | -1.387668373 | 0.215626047 | 0.714295362 | -4.517203478 |
| **A_52_P409731** | Scamp5 | -0.105457738 | -0.29243455 | 0.081519075 | 8.557205142 | -1.38758585 | 0.215649864 | 0.714295362 | -4.517261886 |
| **A_55_P1991783** | NA | -0.529025318 | -1.46701123 | 0.408960594 | 11.52386268 | -1.387552116 | 0.2156596 | 0.714295362 | -4.517285761 |
| **A_55_P2031133** | Tmem220 | -0.164183688 | -0.455298457 | 0.126931081 | 8.978466836 | -1.387506195 | 0.215672855 | 0.714295362 | -4.517318263 |
| **A_51_P158044** | NA | -0.120371779 | -0.333811559 | 0.093068001 | 8.889785577 | -1.387453633 | 0.215688028 | 0.714295362 | -4.517355463 |
| **A_65_P08768** | Fbln1 | -0.225760089 | -0.626119867 | 0.174599689 | 11.34461196 | -1.387285639 | 0.215736528 | 0.714295362 | -4.517474359 |
| **A_51_P288138** | Fpr2 | -0.110456498 | -0.306396064 | 0.085483068 | 7.792772325 | -1.386878105 | 0.215854226 | 0.714295362 | -4.517762767 |
| **A_55_P2054311** | M6pr | 0.171483036 | -0.132736363 | 0.475702436 | 10.94644487 | 1.386766758 | 0.215886393 | 0.714295362 | -4.517841562 |
| **A_55_P1956973** | Tln1 | -0.313956102 | -0.870999506 | 0.243087303 | 14.50147003 | -1.386593162 | 0.215936553 | 0.714295362 | -4.517964404 |
| **A_55_P2043612** | Prdm6 | -0.078369891 | -0.21742135 | 0.060681569 | 8.647290598 | -1.386573356 | 0.215942276 | 0.714295362 | -4.517978418 |
| **A_52_P208727** | Zmym4 | -0.131645331 | -0.365256741 | 0.101966079 | 9.029126123 | -1.386374538 | 0.215999738 | 0.714295362 | -4.518119102 |
| **A_55_P2117043** | Mtmr1 | -0.090389612 | -0.250791685 | 0.070012462 | 9.714810074 | -1.386365384 | 0.216002384 | 0.714295362 | -4.518125579 |
| **A_55_P2169888** | Prag1 | -0.075594457 | -0.209747654 | 0.058558739 | 8.700516243 | -1.386302772 | 0.216020484 | 0.714295362 | -4.518169882 |
| **A_55_P1960375** | NA | -0.148270125 | -0.411409148 | 0.114868898 | 14.2214419 | -1.386237229 | 0.216039431 | 0.714295362 | -4.518216258 |
| **A_55_P1969885** | NA | -0.092015749 | -0.255326467 | 0.071294969 | 8.500769323 | -1.386170465 | 0.216058734 | 0.714295362 | -4.518263498 |
| **A_55_P2068265** | Mif | 0.159988559 | -0.123971637 | 0.443948755 | 13.67356521 | 1.38611934 | 0.216073516 | 0.714295362 | -4.518299671 |
| **A_51_P284608** | Cd74 | -0.166099978 | -0.460980607 | 0.128780651 | 8.568310963 | -1.38577427 | 0.216173312 | 0.714503068 | -4.518543814 |

| | | | | | | | | | |
|---|---|---|---|---|---|---|---|---|---|
| **A_55_P2070913** | Rpl14 | -0.11878403 | -0.329708128 | 0.092140069 | 16.45055553 | -1.385482409 | 0.216257752 | 0.714654996 | -4.518750297 |
| **A_55_P1995768** | Zfp967 | -0.094547744 | -0.262458752 | 0.073363264 | 9.908703112 | -1.385291489 | 0.216313004 | 0.714654996 | -4.51888536 |
| **A_51_P212419** | Lrrc41 | -0.107006512 | -0.297051677 | 0.083038654 | 9.970534747 | -1.385232091 | 0.216330197 | 0.714654996 | -4.518927379 |
| **A_51_P275514** | Mprip | -0.152721316 | -0.424113842 | 0.118671211 | 11.7515947 | -1.384429791 | 0.216562543 | 0.715243993 | -4.519494884 |
| **A_55_P2183672** | Pebp1 | -0.074785761 | -0.207696484 | 0.058124961 | 17.03206526 | -1.38429313 | 0.216602142 | 0.715243993 | -4.519591541 |
| **A_52_P459929** | Itga1 | -0.130373014 | -0.362084518 | 0.101338491 | 9.16484951 | -1.38423323 | 0.216619501 | 0.715243993 | -4.519633906 |
| **A_51_P239113** | Zfp35 | 0.121371869 | -0.094418718 | 0.337162456 | 10.8569861 | 1.383740673 | 0.216762292 | 0.715554188 | -4.519982251 |
| **A_55_P2041521** | Doxl2 | -0.905688367 | -2.516068054 | 0.70469132 | 15.14715651 | -1.383629041 | 0.216794666 | 0.715554188 | -4.520061194 |
| **A_52_P443167** | Naa35 | 0.079648647 | -0.062020898 | 0.221318192 | 11.20790679 | 1.383155692 | 0.216931989 | 0.715554188 | -4.520395912 |
| **A_55_P2154982** | Cd8b1 | 0.153422666 | -0.119477165 | 0.426322497 | 8.173487663 | 1.383105864 | 0.216946449 | 0.715554188 | -4.520431145 |
| **A_51_P209372** | Msmo1 | 0.324167017 | -0.25254824 | 0.900882275 | 12.37274216 | 1.38285469 | 0.217019353 | 0.715554188 | -4.52060874 |
| **A_51_P220682** | Retreg3 | -0.12179265 | -0.33847168 | 0.09488638 | 10.63295352 | -1.382844542 | 0.217022299 | 0.715554188 | -4.520615916 |
| **A_51_P265869** | Hspa9 | -0.10019965 | -0.27848879 | 0.07808949 | 11.37389724 | -1.382644327 | 0.217080429 | 0.715554188 | -4.520757473 |
| **A_55_P1962811** | Anp32b | -0.104327182 | -0.289975674 | 0.08132131 | 13.32478313 | -1.382532067 | 0.217113028 | 0.715554188 | -4.52083684 |
| **A_55_P2033120** | Srxn1 | -0.142510619 | -0.396106115 | 0.111084878 | 11.5307564 | -1.382530916 | 0.217113363 | 0.715554188 | -4.520837654 |
| **A_55_P2036514** | Dis3 | 0.079786656 | -0.062199388 | 0.2217727 | 9.647555572 | 1.382463805 | 0.217132853 | 0.715554188 | -4.520885101 |
| **A_55_P2181888** | Tmem54 | 0.146589343 | -0.114380076 | 0.407558762 | 9.605684157 | 1.381916944 | 0.217291735 | 0.715554188 | -4.521271699 |
| **A_66_P137152** | Zbtb8os | -0.128421092 | -0.357082764 | 0.10024058 | 12.5264886 | -1.381694875 | 0.217356284 | 0.715554188 | -4.521428676 |
| **A_51_P496400** | Med9 | -0.119029609 | -0.330973049 | 0.092913831 | 11.24372128 | -1.381669562 | 0.217363643 | 0.715554188 | -4.521446569 |
| **A_52_P392509** | Gin1 | -0.094708352 | -0.263361045 | 0.073944342 | 8.470526506 | -1.381542215 | 0.217400668 | 0.715554188 | -4.521536584 |
| **A_55_P1989125** | Rps17 | -0.099310183 | -0.276169208 | 0.077548843 | 15.10871219 | -1.381451738 | 0.217426977 | 0.715554188 | -4.521600536 |
| **A_66_P118256** | Strn3 | 0.125586768 | -0.098086944 | 0.34926048 | 11.45889702 | 1.381332094 | 0.217461772 | 0.715554188 | -4.521685103 |
| **A_52_P137691** | Trappc5 | -0.083543611 | -0.232343167 | 0.065255945 | 8.183615535 | -1.38127697 | 0.217477804 | 0.715554188 | -4.521724065 |
| **A_55_P2103101** | Snx9 | -0.127990564 | -0.356004928 | 0.100023801 | 9.992754068 | -1.380972105 | 0.217566493 | 0.715554188 | -4.521939536 |
| **A_55_P2088315** | Vmn1r87 | 0.380292267 | -0.297208078 | 1.057792612 | 9.094219747 | 1.380947399 | 0.217573682 | 0.715554188 | -4.521956997 |
| **A_51_P148675** | Dag1 | -0.266397623 | -0.74104144 | 0.208246195 | 11.88419194 | -1.380802864 | 0.217615742 | 0.715554188 | -4.522059145 |
| **A_55_P1962092** | Mpc1 | 0.149197981 | -0.116636868 | 0.415032831 | 12.89871168 | 1.380766356 | 0.217626367 | 0.715554188 | -4.522084946 |
| **A_55_P2067895** | Krba1 | 0.091747109 | -0.071767962 | 0.25526218 | 9.836954317 | 1.380396228 | 0.217734113 | 0.715554188 | -4.522346513 |
| **A_51_P295312** | Gabarap | -0.168711585 | -0.469397213 | 0.131974044 | 12.3470764 | -1.380388702 | 0.217736305 | 0.715554188 | -4.522351831 |
| **A_52_P639229** | Fgd4 | 0.227644141 | -0.178121055 | 0.633409336 | 9.483812646 | 1.380228043 | 0.217783089 | 0.715554188 | -4.522465361 |
| **A_55_P2124523** | Gm13620 | 0.187009616 | -0.146456148 | 0.520475379 | 7.950625367 | 1.379691054 | 0.217939528 | 0.715554188 | -4.522844795 |
| **A_66_P101519** | Abcc9 | -0.087753637 | -0.244250763 | 0.06874349 | 8.057251912 | -1.37951974 | 0.217989458 | 0.715554188 | -4.522965835 |
| **A_55_P2097206** | Camk4 | -0.131663139 | -0.366478864 | 0.103152585 | 7.875205268 | -1.379450728 | 0.218009574 | 0.715554188 | -4.523014593 |
| **A_51_P292566** | Xpot | -0.218262439 | -0.60756601 | 0.171041132 | 10.17699945 | -1.379303316 | 0.21805255 | 0.715554188 | -4.523118741 |
| **A_55_P2064038** | NA | 0.199190684 | -0.156161325 | 0.554542692 | 8.949680867 | 1.379048088 | 0.218126975 | 0.715554188 | -4.523299052 |
| **A_55_P2156093** | NA | 0.150965817 | -0.118358662 | 0.420290297 | 13.83417196 | 1.379024418 | 0.218133879 | 0.715554188 | -4.523315773 |
| **A_55_P1969882** | Morf4l2 | -0.195111152 | -0.543204776 | 0.152982471 | 16.00898217 | -1.378971177 | 0.218149408 | 0.715554188 | -4.523353386 |
| **A_55_P2424459** | Atg9a | -0.159070104 | -0.44287385 | 0.124733642 | 8.157321656 | -1.378921698 | 0.21816384 | 0.715554188 | -4.523388339 |
| **A_55_P2088530** | Htt | -0.168386245 | -0.468851467 | 0.132078976 | 11.14402802 | -1.378737431 | 0.218217596 | 0.715554188 | -4.523518509 |
| **A_55_P1998107** | NA | -0.109610295 | -0.305208739 | 0.085988149 | 11.55171109 | -1.378653461 | 0.218242096 | 0.715554188 | -4.523577826 |
| **A_55_P2026025** | NA | -0.091508386 | -0.254806712 | 0.071789941 | 15.88584555 | -1.378631901 | 0.218248387 | 0.715554188 | -4.523593055 |
| **A_66_P111385** | Klhl28 | 0.087251342 | -0.068451302 | 0.242953985 | 10.73970766 | 1.378622268 | 0.218251198 | 0.715554188 | -4.523599859 |
| **A_55_P2015877** | Clps | 0.115292766 | -0.090460846 | 0.321046379 | 9.171682244 | 1.37855391 | 0.218271146 | 0.715554188 | -4.523648147 |
| **A_51_P232858** | Rpl14 | 0.117491622 | -0.092198751 | 0.327181995 | 9.179679038 | 1.378470797 | 0.218295402 | 0.715554188 | -4.523706855 |
| **A_51_P486091** | Ift52 | 0.094514514 | -0.074172801 | 0.263201829 | 10.11796438 | 1.37843167 | 0.218306821 | 0.715554188 | -4.523734492 |
| **A_51_P361830** | Lgi2 | -0.087446233 | -0.243520253 | 0.068627787 | 8.127172511 | -1.37841393 | 0.218311999 | 0.715554188 | -4.523747023 |
| **A_66_P125701** | NA | -0.135575952 | -0.377582341 | 0.106430437 | 8.555637621 | -1.378240455 | 0.218362637 | 0.715554188 | -4.523869556 |
| **A_51_P146633** | NA | 0.126143281 | -0.099029671 | 0.351316233 | 9.249804942 | 1.378215293 | 0.218369983 | 0.715554188 | -4.523887329 |
| **A_51_P184062** | Hbs1l | 0.13335847 | -0.104716851 | 0.37143379 | 10.98177146 | 1.378083044 | 0.218408596 | 0.715554188 | -4.523980737 |
| **A_52_P647607** | Ubr4 | 0.088762807 | -0.06972383 | 0.247249443 | 8.597336459 | 1.377867766 | 0.218471464 | 0.715554188 | -4.524132785 |

| | | | | | | | | | |
|---|---|---|---|---|---|---|---|---|---|
| **A_66_P100789** | Kansl2 | -0.105508136 | -0.293912216 | 0.082895945 | 9.258057297 | -1.377732197 | 0.218511063 | 0.715554188 | -4.524228532 |
| **A_55_P1968043** | Jmjd1c | -0.297742847 | -0.844199121 | 0.248713426 | 11.52597722 | -1.412004248 | 0.218547164 | 0.715554188 | -4.489838327 |
| **A_55_P1982451** | Hmgn5 | -0.189366203 | -0.52755623 | 0.148823824 | 11.21266325 | -1.377560998 | 0.218561078 | 0.715554188 | -4.524349438 |
| **A_66_P134481** | Skp2 | -0.102810559 | -0.286431437 | 0.080810318 | 8.134600863 | -1.377478489 | 0.218585187 | 0.715554188 | -4.524407707 |
| **A_51_P165683** | Hirip3 | -0.093816726 | -0.261374675 | 0.073741222 | 8.700154954 | -1.377477148 | 0.218585579 | 0.715554188 | -4.524408654 |
| **A_55_P2108143** | NA | 0.236148258 | -0.185639917 | 0.657936433 | 11.85084265 | 1.377398194 | 0.218608651 | 0.715554188 | -4.524464412 |
| **A_55_P2066727** | Serpinb6a | 0.155456091 | -0.122253369 | 0.433165552 | 14.12530469 | 1.377165807 | 0.218676573 | 0.715554188 | -4.524628518 |
| **A_55_P2152956** | Plekhm3 | -0.111126272 | -0.30965324 | 0.087400696 | 8.819297681 | -1.377102918 | 0.218694957 | 0.715554188 | -4.524672927 |
| **A_52_P391081** | Hinfp | 0.096498807 | -0.075950754 | 0.268948367 | 8.664487881 | 1.376667357 | 0.218822324 | 0.715554188 | -4.524980483 |
| **A_51_P329441** | Bsg | -0.181467378 | -0.505777356 | 0.142842599 | 12.36053871 | -1.376598899 | 0.218842348 | 0.715554188 | -4.525028819 |
| **A_51_P184100** | Capn7 | -0.088101128 | -0.245565552 | 0.069363296 | 9.193800652 | -1.376474543 | 0.218878728 | 0.715554188 | -4.525116622 |
| **A_55_P2059556** | NA | 0.212387337 | -0.167231043 | 0.592005717 | 15.84473458 | 1.37641866 | 0.218895078 | 0.715554188 | -4.525156078 |
| **A_55_P2020458** | NA | -0.129261897 | -0.360316049 | 0.101792255 | 11.87958681 | -1.376340569 | 0.218917927 | 0.715554188 | -4.525211213 |
| **A_55_P1987904** | Chpf2 | -0.184331315 | -0.513857293 | 0.145194663 | 8.674099928 | -1.376190726 | 0.218961777 | 0.715554188 | -4.525317005 |
| **A_51_P177171** | Tie1 | -0.081765841 | -0.227938501 | 0.064406818 | 8.716175117 | -1.376178997 | 0.21896521 | 0.715554188 | -4.525325285 |
| **A_52_P627925** | Usp36 | -0.123924371 | -0.345518724 | 0.097669982 | 8.562680226 | -1.375837672 | 0.219065127 | 0.715554188 | -4.525566253 |
| **A_55_P1969914** | Tcf25 | -0.189899709 | -0.529514917 | 0.149715499 | 14.57084156 | -1.37564487 | 0.219121585 | 0.715554188 | -4.525702358 |
| **A_55_P2019058** | Acaca | 0.114087372 | -0.089957128 | 0.318131872 | 9.592393514 | 1.375567313 | 0.219144299 | 0.715554188 | -4.525757107 |
| **A_55_P2182483** | Cd47 | -0.125531679 | -0.350077426 | 0.099014068 | 12.12244568 | -1.375364042 | 0.219203842 | 0.715554188 | -4.525900594 |
| **A_51_P193563** | Cdk7 | -0.102170459 | -0.284937702 | 0.080596785 | 9.637603091 | -1.375295891 | 0.219223809 | 0.715554188 | -4.525948699 |
| **A_51_P259296** | Lpl | -0.181893392 | -0.507348424 | 0.14356164 | 9.570071681 | -1.374975927 | 0.219317572 | 0.715554188 | -4.526174543 |
| **A_55_P1976694** | 11-Sep | 0.092252397 | -0.072840539 | 0.257345333 | 8.522677178 | 1.374732912 | 0.21938881 | 0.715554188 | -4.526346061 |
| **A_55_P2084344** | NA | -0.080531737 | -0.224670286 | 0.063606812 | 11.85427506 | -1.37453589 | 0.219446582 | 0.715554188 | -4.526485111 |
| **A_55_P2150294** | Synj2bp | -0.15791428 | -0.440572939 | 0.12474438 | 10.45650297 | -1.374447869 | 0.219472396 | 0.715554188 | -4.526547231 |
| **A_55_P1974412** | Kat2a | 0.086643518 | -0.068448116 | 0.241735153 | 9.592110928 | 1.37441178 | 0.21948298 | 0.715554188 | -4.526572699 |
| **A_55_P2083484** | Mettl21a | 0.094342383 | -0.074531148 | 0.263215914 | 8.803284011 | 1.374404029 | 0.219485254 | 0.715554188 | -4.52657817 |
| **A_55_P2236818** | A430103D13Rik | -0.081762299 | -0.228120637 | 0.064596039 | 8.122493177 | -1.374373549 | 0.219494194 | 0.715554188 | -4.52659968 |
| **A_55_P2057946** | Hsp90aa1 | -0.162877143 | -0.454460002 | 0.128705716 | 14.99906978 | -1.374254968 | 0.219528978 | 0.715554188 | -4.526683363 |
| **A_55_P1963414** | NA | -0.140264027 | -0.391433692 | 0.110905638 | 13.71222897 | -1.373878349 | 0.219639487 | 0.715554188 | -4.526949132 |
| **A_55_P2153580** | Zfp655 | -0.133778848 | -0.373340926 | 0.105783231 | 8.548240336 | -1.373847543 | 0.219648529 | 0.715554188 | -4.52697087 |
| **A_55_P2145696** | P3h4 | 0.123082972 | -0.097331844 | 0.343497788 | 11.62703819 | 1.373808986 | 0.219659846 | 0.715554188 | -4.526998077 |
| **A_66_P119884** | Wdr91 | -0.100248342 | -0.279774699 | 0.079278014 | 9.322827957 | -1.37378303 | 0.219667465 | 0.715554188 | -4.527016392 |
| **A_55_P1980521** | Ccdc85a | -0.063962346 | -0.178517322 | 0.05059263 | 7.85274935 | -1.373661057 | 0.219703269 | 0.715554188 | -4.527102458 |
| **A_55_P2126855** | E230025N22Rik | 0.182175796 | -0.144120282 | 0.508471874 | 8.430833248 | 1.373561107 | 0.219732613 | 0.715554188 | -4.527172983 |
| **A_55_P2040838** | Gm14548 | -0.071051207 | -0.198313138 | 0.056210724 | 7.916719025 | -1.373542537 | 0.219738066 | 0.715554188 | -4.527186086 |
| **A_55_P2004084** | Iqce | -0.08553212 | -0.238732916 | 0.067668676 | 8.836890639 | -1.373527557 | 0.219742464 | 0.715554188 | -4.527196655 |
| **A_55_P2016216** | NA | -0.173166885 | -0.483381164 | 0.137047394 | 11.28894128 | -1.373321516 | 0.219802969 | 0.715554188 | -4.527342031 |
| **A_55_P2173968** | Ctdspl | -0.194257316 | -0.542266904 | 0.153752271 | 13.88307177 | -1.373268123 | 0.21981865 | 0.715554188 | -4.527379702 |
| **A_51_P356467** | Cldn4 | -0.123605449 | -0.345050352 | 0.097839454 | 9.597367688 | -1.373223066 | 0.219831884 | 0.715554188 | -4.527411492 |
| **A_55_P1987770** | Psenen | 0.231257055 | -0.183071092 | 0.645585202 | 13.53417482 | 1.373155486 | 0.219851735 | 0.715554188 | -4.527459171 |
| **A_55_P1972054** | NA | 0.117201255 | -0.092784938 | 0.327187448 | 8.449977232 | 1.373126932 | 0.219860123 | 0.715554188 | -4.527479317 |
| **A_55_P2129614** | Itprip | -0.197387762 | -0.551163648 | 0.156388123 | 13.21929123 | -1.372654254 | 0.219999017 | 0.715885691 | -4.527812783 |
| **A_52_P354123** | Rhebl1 | -0.109595572 | -0.30604552 | 0.086854375 | 9.020376616 | -1.372493372 | 0.22004631 | 0.715919057 | -4.527926274 |
| **A_52_P200207** | Rmnd5a | -0.118683484 | -0.331466895 | 0.094099927 | 8.813945637 | -1.372213485 | 0.220128607 | 0.716060437 | -4.528123705 |
| **A_55_P2106424** | NA | 0.077224817 | -0.061240914 | 0.215690549 | 16.72069352 | 1.372093633 | 0.220163856 | 0.716060437 | -4.528208245 |
| **A_51_P179241** | Tssc4 | -0.066143929 | -0.184753363 | 0.052465504 | 10.23530466 | -1.371955175 | 0.220204584 | 0.716072411 | -4.528305905 |
| **A_55_P1960698** | Pih1d2 | -0.067369029 | -0.188202032 | 0.053463973 | 7.986626333 | -1.371651821 | 0.220293841 | 0.716242163 | -4.528519864 |
| **A_51_P142187** | Abhd11 | 0.104481589 | -0.082978939 | 0.291942118 | 10.15340707 | 1.371194604 | 0.220428432 | 0.716455079 | -4.528842315 |
| **A_55_P2005470** | Mfap4 | 0.456266294 | -0.362375407 | 1.274907995 | 9.672189292 | 1.371177527 | 0.22043346 | 0.716455079 | -4.528854358 |
| **A_55_P2143946** | Slc18a2 | -0.107265873 | -0.299764982 | 0.085233235 | 8.754138528 | -1.370888121 | 0.220518693 | 0.716574906 | -4.529058443 |

| | | | | | | | | | |
|---|---|---|---|---|---|---|---|---|---|
| **A_52_P515057** | Slc25a24 | 0.087157504 | -0.069296921 | 0.243611928 | 11.46125559 | 1.37052227 | 0.220626482 | 0.716574906 | -4.529316417 |
| **A_52_P376135** | Mknk2 | -0.139196761 | -0.38908566 | 0.110692137 | 12.88935259 | -1.370412561 | 0.220658815 | 0.716574906 | -4.529393772 |
| **A_51_P350403** | Exoc4 | -0.065812546 | -0.183963601 | 0.052338509 | 8.201315771 | -1.370377601 | 0.220669119 | 0.716574906 | -4.529418422 |
| **A_55_P2087374** | NA | 0.121390531 | -0.096545533 | 0.339326595 | 15.46138333 | 1.370329071 | 0.220683423 | 0.716574906 | -4.529452639 |
| **A_55_P2009507** | Ocrl | -0.119570362 | -0.334256171 | 0.095115446 | 10.35646264 | -1.370217062 | 0.220716442 | 0.716574906 | -4.529531613 |
| **A_51_P513392** | Azi2 | 0.075644819 | -0.060178201 | 0.211467839 | 9.197566088 | 1.37017163 | 0.220729836 | 0.716574906 | -4.529563645 |
| **A_55_P2026823** | Uspl1 | -0.129097349 | -0.360918242 | 0.102723544 | 11.07168877 | -1.370042104 | 0.220768025 | 0.716578531 | -4.529654966 |
| **A_55_P1960326** | NA | 0.084827255 | -0.06753729 | 0.2371918 | 9.721206048 | 1.369684864 | 0.220873385 | 0.716616759 | -4.529906819 |
| **A_51_P136699** | Ttc7b | 0.16117448 | -0.128348539 | 0.4506975 | 11.61863639 | 1.369564035 | 0.220909031 | 0.716616759 | -4.529991999 |
| **A_55_P2077826** | Ppp1cc | 0.115102517 | -0.091741068 | 0.321946102 | 11.40239866 | 1.369026728 | 0.221067608 | 0.716616759 | -4.53037075 |
| **A_55_P1985464** | Gm12271 | -0.153255397 | -0.428685032 | 0.122174238 | 12.41482331 | -1.368908036 | 0.221102651 | 0.716616759 | -4.53045441 |
| **A_51_P291682** | Tmed4 | -0.082240034 | -0.230044559 | 0.065564491 | 9.687770993 | -1.368877901 | 0.221111549 | 0.716616759 | -4.530475651 |
| **A_55_P1968178** | Smagp | -0.161558398 | -0.45197415 | 0.128857353 | 12.54657575 | -1.368606296 | 0.221191762 | 0.716616759 | -4.530667083 |
| **A_55_P2119877** | Pnisr | 0.075852774 | -0.060506634 | 0.212212183 | 8.954091508 | 1.368533793 | 0.221213179 | 0.716616759 | -4.530718183 |
| **A_55_P2065327** | Rpl27a | -0.079573476 | -0.222625415 | 0.063478464 | 15.06771986 | -1.368496655 | 0.22122415 | 0.716616759 | -4.530744357 |
| **A_51_P278550** | Tecr | -0.069763858 | -0.195182801 | 0.055655085 | 10.18207339 | -1.368473895 | 0.221230874 | 0.716616759 | -4.530760398 |
| **A_51_P334942** | Aldh1a1 | 0.388768235 | -0.31018825 | 1.08772472 | 8.507878472 | 1.368389702 | 0.221255748 | 0.716616759 | -4.530819734 |
| **A_52_P244132** | Isoc1 | -0.210759484 | -0.589683083 | 0.168164116 | 11.44388711 | -1.368373438 | 0.221260554 | 0.716616759 | -4.530831196 |
| **A_55_P2027117** | Rgs3 | 0.154688181 | -0.123459511 | 0.432835874 | 9.510509387 | 1.368203935 | 0.221310641 | 0.716616759 | -4.530950653 |
| **A_51_P435321** | Ap1s2 | 0.117874291 | -0.094086803 | 0.329835385 | 8.574722686 | 1.368144926 | 0.22132808 | 0.716616759 | -4.530992238 |
| **A_66_P121480** | Sardh | 0.066541283 | -0.05312406 | 0.186206627 | 7.926179608 | 1.368018417 | 0.221365472 | 0.716616759 | -4.53108139 |
| **A_55_P2060284** | Itpripl2 | -0.095734668 | -0.267926194 | 0.076456858 | 8.333720568 | -1.367812676 | 0.221426294 | 0.716616759 | -4.531226373 |
| **A_55_P2052892** | Xiap | -0.130465887 | -0.365130346 | 0.104198571 | 12.17376968 | -1.367788089 | 0.221433564 | 0.716616759 | -4.531243698 |
| **A_55_P2152245** | Rpl19 | -0.114340964 | -0.320023195 | 0.091341267 | 15.51076118 | -1.367647692 | 0.221475079 | 0.716616759 | -4.531342629 |
| **A_52_P484405** | Twf1 | -0.150504017 | -0.421244142 | 0.120236109 | 11.47056603 | -1.367617434 | 0.221484027 | 0.716616759 | -4.531363951 |
| **A_52_P198727** | Cfap410 | -0.060767426 | -0.170082595 | 0.048547743 | 8.570792794 | -1.367601511 | 0.221488736 | 0.716616759 | -4.53137517 |
| **A_55_P2117764** | Vangl2 | 0.091017664 | -0.072744629 | 0.254779957 | 8.500552983 | 1.36735391 | 0.221561972 | 0.716616759 | -4.531549633 |
| **A_52_P466701** | Fuom | -0.089315675 | -0.250035497 | 0.071404146 | 8.259079029 | -1.367185389 | 0.22161183 | 0.716616759 | -4.53166837 |
| **A_52_P398925** | Stfa2l1 | -0.31097747 | -0.870627311 | 0.248672371 | 9.045293496 | -1.367041517 | 0.221654404 | 0.716616759 | -4.531769736 |
| **A_66_P118772** | Tmem136 | -0.192350163 | -0.538519591 | 0.153819265 | 11.26786116 | -1.367014154 | 0.221662502 | 0.716616759 | -4.531789014 |
| **A_55_P1963144** | Mtmr14 | 0.118990228 | -0.095163042 | 0.333143498 | 10.79157407 | 1.366959819 | 0.221678583 | 0.716616759 | -4.531827295 |
| **A_55_P2083889** | Pea15a | 0.256611362 | -0.205258309 | 0.718481033 | 12.61809117 | 1.366864928 | 0.22170667 | 0.716616759 | -4.531894148 |
| **A_55_P2153721** | Ccdc127 | -0.083250712 | -0.23310834 | 0.066606916 | 8.439783367 | -1.366715965 | 0.221750767 | 0.71663514 | -4.531999093 |
| **A_55_P2185573** | Rbms1 | -0.301073015 | -0.843075262 | 0.240929232 | 14.67520279 | -1.366595252 | 0.221786508 | 0.71663514 | -4.532084132 |
| **A_55_P2073795** | Mecr | -0.110105505 | -0.308340102 | 0.088129092 | 8.398673141 | -1.366465732 | 0.221824862 | 0.716639271 | -4.532175374 |
| **A_55_P1971025** | Paqr6 | 0.074501973 | -0.059644374 | 0.20864832 | 7.906242316 | 1.366337817 | 0.221862746 | 0.716641882 | -4.532265482 |
| **A_55_P2093974** | Mphosph10 | 0.172933212 | -0.13850014 | 0.484366564 | 9.273601112 | 1.366099875 | 0.221933233 | 0.716732536 | -4.532433091 |
| **A_52_P578436** | NA | 0.179228013 | -0.14358122 | 0.502037246 | 9.049271657 | 1.365932079 | 0.221982953 | 0.716732536 | -4.532551282 |
| **A_55_P2090600** | Mageb16-ps1 | 0.084059186 | -0.067347736 | 0.235466108 | 8.393383783 | 1.365867626 | 0.222002053 | 0.716732536 | -4.532596681 |
| **A_52_P402319** | Med20 | 0.093210484 | -0.07472327 | 0.261144237 | 9.269872115 | 1.365513286 | 0.222107089 | 0.716951891 | -4.532846251 |
| **A_51_P108334** | Slc25a4 | -0.12914307 | -0.361856131 | 0.10356999 | 14.40516862 | -1.365273036 | 0.222178331 | 0.717049407 | -4.526145319 |
| **A_55_P2114915** | Ube2i | -0.127899953 | -0.358399243 | 0.102599338 | 14.46357524 | -1.365117244 | 0.222224539 | 0.717049407 | -4.533125168 |
| **A_52_P155708** | Creg1 | -0.093594211 | -0.262298454 | 0.075110033 | 10.8675768 | -1.364872685 | 0.222297094 | 0.717049407 | -4.533297389 |
| **A_52_P398989** | Cytip | -0.061989116 | -0.173734074 | 0.049755842 | 7.844649978 | -1.364761211 | 0.222330173 | 0.717049407 | -4.533375888 |
| **A_55_P2063495** | Dph3 | 0.115630446 | -0.092813304 | 0.324074196 | 8.650810072 | 1.364748062 | 0.222334075 | 0.717049407 | -4.533385147 |
| **A_55_P1972019** | 4632415L05Rik | -0.063278411 | -0.177374071 | 0.05081725 | 7.869531367 | -1.364443627 | 0.222424438 | 0.717049407 | -4.533599513 |
| **A_52_P242251** | Utp25 | 0.077736452 | -0.062429925 | 0.21790283 | 8.133145436 | 1.364426137 | 0.22242963 | 0.717049407 | -4.533611828 |
| **A_55_P1958902** | NA | 0.347391119 | -0.279040959 | 0.973823198 | 11.24718683 | 1.364312589 | 0.222463342 | 0.717049407 | -4.533691778 |
| **A_51_P225083** | Snx17 | 0.090873849 | -0.073002668 | 0.254750366 | 12.65262713 | 1.364241825 | 0.222484355 | 0.717049407 | -4.533741603 |
| **A_51_P293688** | Rab32 | -0.191120525 | -0.535796308 | 0.153555258 | 9.79742137 | -1.364161293 | 0.22250827 | 0.717049407 | -4.533798304 |

| | | | | | | | | |
|---|---|---|---|---|---|---|---|---|
| **A_55_P2005754** | Manbal | 0.158043801 | -0.127081114 | 0.443168716 | 11.52540134 | 1.363676823 | 0.222652188 | 0.717091563 | -4.534139387 |
| **A_65_P08227** | Snx5 | -0.079070397 | -0.221724075 | 0.063583281 | 8.526483999 | -1.363641187 | 0.222662777 | 0.717091563 | -4.534164475 |
| **A_55_P1952414** | Odf2 | -0.086479581 | -0.24251188 | 0.069552717 | 9.934199015 | -1.363541109 | 0.222692519 | 0.717091563 | -4.534234928 |
| **A_55_P2142298** | Ywhaq | -0.188745499 | -0.529309914 | 0.151818917 | 13.45923067 | -1.36347286 | 0.222712803 | 0.717091563 | -4.534282973 |
| **A_55_P2074317** | Ap2b1 | -0.16819539 | -0.471700428 | 0.135309649 | 9.720841654 | -1.363381338 | 0.222740007 | 0.717091563 | -4.534347401 |
| **A_55_P1971594** | Pqbp1 | 0.155691999 | -0.125261092 | 0.43664509 | 9.561443177 | 1.363332044 | 0.22275466 | 0.717091563 | -4.534382101 |
| **A_55_P1990608** | Ube2j1 | -0.126413128 | -0.354546055 | 0.101719799 | 9.650816686 | -1.363243291 | 0.222781046 | 0.717091563 | -4.534444576 |
| **A_55_P2148116** | Tsr2 | 0.07328133 | -0.058990379 | 0.205553039 | 8.357478373 | 1.362999017 | 0.222853681 | 0.717175108 | -4.534616522 |
| **A_52_P337427** | Izumo4 | -0.083651776 | -0.234653568 | 0.067350016 | 8.514770582 | -1.362894466 | 0.222884776 | 0.717175108 | -4.534690113 |
| **A_51_P505617** | Il18r1 | -0.059189229 | -0.166041941 | 0.047663482 | 7.711538718 | -1.362781722 | 0.222918312 | 0.717175108 | -4.534769469 |
| **A_55_P1978860** | Trp53bp2 | 0.152085614 | -0.122535456 | 0.426706683 | 9.917098439 | 1.362459004 | 0.22301433 | 0.717191771 | -4.534996605 |
| **A_51_P337897** | Prrc1 | -0.111211884 | -0.312033448 | 0.089609681 | 8.160968023 | -1.362416873 | 0.223026868 | 0.717191771 | -4.535026256 |
| **A_55_P2072213** | NA | 0.105921216 | -0.085363026 | 0.297205459 | 10.76966705 | 1.362300532 | 0.223061495 | 0.717191771 | -4.535108135 |
| **A_52_P100199** | Abhd10 | -0.062619254 | -0.17570693 | 0.050468422 | 8.04476632 | -1.362265543 | 0.223071909 | 0.717191771 | -4.535132759 |
| **A_55_P1971249** | Eya3 | -0.138023317 | -0.387323242 | 0.111276608 | 11.54506304 | -1.362070154 | 0.223130075 | 0.717227961 | -4.535270263 |
| **A_52_P594917** | Pip4k2c | -0.123063101 | -0.345359856 | 0.099233654 | 12.13765365 | -1.361958564 | 0.223163301 | 0.717227961 | -4.535348792 |
| **A_55_P1994302** | NA | 0.071613883 | -0.057756804 | 0.20098457 | 8.022037829 | 1.361853848 | 0.223194484 | 0.717227961 | -4.535422481 |
| **A_55_P2023953** | Arpc3 | -0.125615907 | -0.35258781 | 0.101355997 | 14.47804259 | -1.361575412 | 0.223277419 | 0.717350692 | -4.535618409 |
| **A_55_P2064547** | Tuba1c | 0.078726848 | -0.06354191 | 0.220995606 | 15.96738842 | 1.361389786 | 0.223332724 | 0.717350692 | -4.535749022 |
| **A_55_P2007022** | Adgre5 | -0.199197978 | -0.559198947 | 0.16080299 | 11.33274427 | -1.361289266 | 0.223362678 | 0.717350692 | -4.535819748 |
| **A_52_P431859** | Aopep | -0.176952859 | -0.496765831 | 0.142860114 | 10.07858016 | -1.361227361 | 0.223381127 | 0.717350692 | -4.535863304 |
| **A_55_P1955876** | Ifitm7 | -0.118337869 | -0.332323648 | 0.095647911 | 9.161530204 | -1.360529596 | 0.223589173 | 0.717526296 | -4.536354207 |
| **A_51_P341571** | Klhdc2 | -0.128975495 | -0.362201497 | 0.104250506 | 10.78702941 | -1.360502678 | 0.223597202 | 0.717526296 | -4.536373143 |
| **A_52_P616591** | Eri3 | 0.07213292 | -0.058307558 | 0.202573397 | 8.259182661 | 1.360474151 | 0.223605712 | 0.717526296 | -4.536393211 |
| **A_55_P1953311** | Stx2 | 0.118583464 | -0.095861187 | 0.333028115 | 10.43748708 | 1.360435877 | 0.223617129 | 0.717526296 | -4.536420135 |
| **A_55_P2035286** | Uhrf1 | 0.255432236 | -0.206511446 | 0.717375918 | 9.50207734 | 1.360366212 | 0.223637913 | 0.717526296 | -4.536469142 |
| **A_55_P2041723** | Mid1ip1 | 0.222723768 | -0.180087829 | 0.625535366 | 9.918914304 | 1.360297073 | 0.223658541 | 0.717526296 | -4.536517777 |
| **A_51_P190805** | Ap1m1 | 0.067330464 | -0.054460472 | 0.189121401 | 10.94594896 | 1.360084222 | 0.223722057 | 0.717610959 | -4.536667501 |
| **A_55_P1956812** | Fam83g | 0.240841591 | -0.194965398 | 0.67664858 | 11.93844347 | 1.359585298 | 0.223871004 | 0.717969577 | -4.537018426 |
| **A_51_P123676** | Synpo | -0.101653137 | -0.28563761 | 0.082331335 | 8.749754776 | -1.359279532 | 0.223962331 | 0.71800408 | -4.535538934 |
| **A_55_P2165314** | Trio | -0.195287589 | -0.548774622 | 0.158199444 | 10.02889191 | -1.359159171 | 0.22399829 | 0.71800408 | -4.537318114 |
| **A_51_P327232** | Lenep | -0.084688752 | -0.237987794 | 0.068610291 | 8.382234377 | -1.359112642 | 0.224012192 | 0.71800408 | -4.537350836 |
| **A_55_P2051991** | Brsk1 | 0.064108828 | -0.051942727 | 0.180160383 | 8.005812663 | 1.359051879 | 0.224030349 | 0.71800408 | -4.537393567 |
| **A_55_P2121618** | Tox4 | 1.121404904 | -0.908950303 | 3.151760112 | 11.85178542 | 1.358812679 | 0.224101837 | 0.718114125 | -4.537561776 |
| **A_55_P2084723** | NA | -0.119097342 | -0.334816423 | 0.096621738 | 8.432639072 | -1.358259255 | 0.224267314 | 0.718525262 | -4.537950914 |
| **A_52_P249733** | Tcap | -0.059523667 | -0.167357022 | 0.048309688 | 7.744826659 | -1.358018618 | 0.2243393 | 0.718529402 | -4.538120101 |
| **A_55_P2115955** | Raet1e | 0.218522082 | -0.177387305 | 0.614431469 | 8.653523927 | 1.357902828 | 0.224373946 | 0.718529402 | -4.538201507 |
| **A_55_P2172999** | Ptpn13 | -0.127531646 | -0.358593857 | 0.103530565 | 9.896125298 | -1.357870032 | 0.22438376 | 0.718529402 | -4.538224565 |
| **A_55_P2065231** | Gstm3 | 0.114959067 | -0.093374568 | 0.323292701 | 8.513097404 | 1.357541146 | 0.224482197 | 0.718529402 | -4.538455774 |
| **A_55_P1991618** | Ppp1r16a | -0.165031944 | -0.464115545 | 0.134051656 | 11.3088486 | -1.357514821 | 0.224490079 | 0.718529402 | -4.53847428 |
| **A_55_P2020373** | Chd4 | 0.149502332 | -0.121438355 | 0.420443018 | 12.70571328 | 1.357509579 | 0.224491648 | 0.718529402 | -4.538477965 |
| **A_51_P437240** | Emp2 | 0.162050565 | -0.131683803 | 0.455784933 | 10.0549323 | 1.357265936 | 0.2245646 | 0.71863806 | -4.538649236 |
| **A_55_P2451183** | Pdpk1 | -0.171676699 | -0.482886556 | 0.139533158 | 10.27294691 | -1.357147884 | 0.224599955 | 0.71863806 | -4.538732217 |
| **A_55_P2139009** | NA | 0.088205512 | -0.071711145 | 0.248122168 | 15.88138195 | 1.356972934 | 0.224652359 | 0.718686768 | -4.53885519 |
| **A_51_P419017** | Pde10a | -0.122481746 | -0.344587482 | 0.09962399 | 10.73794715 | -1.356690415 | 0.224737008 | 0.718838596 | -4.539053762 |
| **A_52_P646034** | Nfatc2ip | 0.072091044 | -0.058670786 | 0.202852874 | 8.607010434 | 1.356342862 | 0.224841183 | 0.719052816 | -4.539298026 |
| **A_52_P69707** | Ergic1 | 0.107669411 | -0.087657213 | 0.302996035 | 8.96884099 | 1.356126047 | 0.224906192 | 0.719141735 | -4.539450395 |
| **A_66_P128293** | Pheta2 | 0.070140615 | -0.057143675 | 0.197424904 | 8.129128105 | 1.355701037 | 0.225033676 | 0.719430355 | -4.539749054 |
| **A_55_P1998942** | Oas1a | -0.216058208 | -0.608175631 | 0.176059216 | 10.26436275 | -1.355575753 | 0.225071269 | 0.719431545 | -4.539837087 |
| **A_55_P2054157** | Hspa4 | -0.129619331 | -0.364886201 | 0.105647539 | 13.3070101 | -1.35543334 | 0.225114007 | 0.71944918 | -4.539937151 |

| | | | | | | | | | |
|---|---|---|---|---|---|---|---|---|---|
| **A_55_P2002472** | Zfp777 | -0.155045291 | -0.457722296 | 0.147631714 | 10.80537535 | -1.441390525 | 0.225225761 | 0.719631406 | -4.474813788 |
| **A_51_P285446** | Lig1 | 0.11596348 | -0.094585231 | 0.326512191 | 10.3235798 | 1.354995381 | 0.225245486 | 0.719631406 | -4.540244858 |
| **A_55_P2081560** | Phf20 | 0.167606405 | -0.136804243 | 0.472017054 | 9.916300568 | 1.354565265 | 0.225374679 | 0.719925163 | -4.540547023 |
| **A_55_P1965507** | Trnau1ap | 0.134803565 | -0.11010044 | 0.37970757 | 12.44233708 | 1.354174571 | 0.225492088 | 0.719942071 | -4.540821466 |
| **A_55_P2186621** | Rbmxl1 | -0.121303307 | -0.341683105 | 0.099076491 | 12.0276518 | -1.35416013 | 0.225496429 | 0.719942071 | -4.54083161 |
| **A_51_P170619** | Ddx54 | -0.123040356 | -0.346587002 | 0.10050629 | 11.7276458 | -1.354093285 | 0.225516523 | 0.719942071 | -4.540878562 |
| **A_51_P462972** | Ap5m1 | -0.105458507 | -0.297067314 | 0.086150299 | 9.725614754 | -1.354051918 | 0.225528959 | 0.719942071 | -4.540907618 |
| **A_66_P108792** | Rad1 | 0.092379455 | -0.075503463 | 0.260262373 | 10.36141317 | 1.353748696 | 0.225620133 | 0.720084298 | -4.541120593 |
| **A_52_P56751** | Lcp1 | -0.114412105 | -0.322349613 | 0.093525403 | 10.86485527 | -1.353655972 | 0.225648021 | 0.720084298 | -4.541185716 |
| **A_55_P2163961** | Kcnj5 | -0.071222365 | -0.20067943 | 0.058234701 | 8.082252165 | -1.353504783 | 0.225693499 | 0.720093232 | -4.541291899 |
| **A_55_P2105366** | NA | 0.129375461 | -0.105815115 | 0.364566036 | 9.128690637 | 1.35332205 | 0.225748476 | 0.720093232 | -4.54142023 |
| **A_52_P114620** | Elmo3 | 0.211905431 | -0.173358512 | 0.597169374 | 9.968516503 | 1.353171633 | 0.22579374 | 0.720093232 | -4.541525862 |
| **A_55_P2160291** | Olfm1 | 0.11532713 | -0.094351546 | 0.325005805 | 7.956936867 | 1.353151369 | 0.225799839 | 0.720093232 | -4.541540093 |
| **A_55_P1969367** | Col4a5 | 0.37282516 | -0.305216471 | 1.05086679 | 8.749574728 | 1.352751475 | 0.225920219 | 0.720280532 | -4.541820903 |
| **A_55_P1961750** | Dtd2 | 0.095120013 | -0.077876274 | 0.2681163 | 9.051044454 | 1.3527087 | 0.225933099 | 0.720280532 | -4.541850938 |
| **A_55_P2094721** | Sox11 | -0.292670702 | -0.825042752 | 0.239701348 | 8.91612833 | -1.352487205 | 0.225999804 | 0.720343443 | -4.542006461 |
| **A_52_P83623** | Rapgef6 | -0.068236973 | -0.192369368 | 0.055895422 | 8.55742941 | -1.352395698 | 0.226027367 | 0.720343443 | -4.54207071 |
| **A_55_P2010111** | Zxdc | -0.126583062 | -0.356895326 | 0.103729202 | 9.937744287 | -1.352158785 | 0.226098743 | 0.720409238 | -4.542237046 |
| **A_55_P2011172** | NA | -0.150045664 | -0.423111722 | 0.123020394 | 13.80153077 | -1.351838753 | 0.226195194 | 0.720409238 | -4.542461724 |
| **A_55_P1971278** | Kdm2a | -0.080434936 | -0.226826709 | 0.065956838 | 9.184280556 | -1.351752592 | 0.226221167 | 0.720409238 | -4.542522211 |
| **A_55_P2115068** | Psmb5 | 0.194569848 | -0.159636644 | 0.548776339 | 12.73951448 | 1.35141329 | 0.226323476 | 0.720409238 | -4.542760393 |
| **A_55_P2007601** | Sftpd | -0.168308644 | -0.474708206 | 0.138090918 | 9.56941823 | -1.351410984 | 0.226324171 | 0.720409238 | -4.542762012 |
| **A_55_P1988728** | Fbxl19 | -0.164943375 | -0.46523406 | 0.135347311 | 11.5609288 | -1.351332362 | 0.226347884 | 0.720409238 | -4.542817201 |
| **A_52_P105840** | Fnbp1 | -0.067406414 | -0.190139026 | 0.055326199 | 8.560855587 | -1.351171284 | 0.226396473 | 0.720409238 | -4.542930265 |
| **A_55_P2279575** | 5430400D12Rik | -0.194730512 | -0.549312058 | 0.159851034 | 8.593875062 | -1.351098586 | 0.226418405 | 0.720409238 | -4.542981292 |
| **A_52_P520607** | Ankrd22 | 0.084526147 | -0.069395064 | 0.238447358 | 7.99554474 | 1.351019957 | 0.226442129 | 0.720409238 | -4.543036481 |
| **A_55_P2039038** | Zfp664 | -0.096127491 | -0.271183 | 0.078928018 | 9.766534053 | -1.350955326 | 0.226461631 | 0.720409238 | -4.543081844 |
| **A_51_P442206** | Ankrd42 | -0.069637024 | -0.196457516 | 0.057183467 | 8.215239472 | -1.350889847 | 0.226481391 | 0.720409238 | -4.543127801 |
| **A_52_P8043** | Srsf2 | -0.088895443 | -0.250800196 | 0.073009311 | 9.444085402 | -1.350793807 | 0.226510375 | 0.720409238 | -4.543195207 |
| **A_55_P1995996** | Zfp984 | 0.091422558 | -0.0751083 | 0.257953415 | 8.97798911 | 1.350603314 | 0.226567876 | 0.720409238 | -4.543328901 |
| **A_51_P184849** | Pfkfb2 | 0.080877518 | -0.066456259 | 0.228211294 | 8.648460888 | 1.350500208 | 0.226599004 | 0.720409238 | -4.543401261 |
| **A_52_P706060** | Mex3a | -0.093971329 | -0.265161362 | 0.077218704 | 8.086796457 | -1.350473472 | 0.226607076 | 0.720409238 | -4.543420024 |
| **A_55_P2167107** | Cdk14 | -0.10260795 | -0.289558337 | 0.084342438 | 10.56351088 | -1.350279897 | 0.226665529 | 0.72047265 | -4.54355587 |
| **A_51_P233160** | Lysmd2 | 0.158138956 | -0.130031106 | 0.446309018 | 9.787769505 | 1.350078964 | 0.226726218 | 0.72047265 | -4.543696873 |
| **A_52_P561927** | Ncor1 | -0.142316131 | -0.401660945 | 0.117028682 | 8.984349944 | -1.350037167 | 0.226738845 | 0.72047265 | -4.543726203 |
| **A_55_P2162890** | Zfp780b | -0.121827083 | -0.343901639 | 0.100247474 | 8.455972674 | -1.349628386 | 0.226862364 | 0.720625549 | -4.544013037 |
| **A_55_P2001948** | Cab39l | 0.138650497 | -0.114113131 | 0.391414125 | 10.48572466 | 1.349509709 | 0.226898236 | 0.720625549 | -4.544096305 |
| **A_55_P2132517** | Gmppa | -0.082436747 | -0.23273036 | 0.067856866 | 9.819540236 | -1.349427272 | 0.226923156 | 0.720625549 | -4.544154144 |
| **A_51_P401958** | Taldo1 | -0.132941472 | -0.375355077 | 0.109472133 | 15.45733002 | -1.349188578 | 0.226995327 | 0.720625549 | -4.54432161 |
| **A_55_P2021358** | Klhl15 | -0.074574551 | -0.210559034 | 0.061409932 | 8.447326247 | -1.349181747 | 0.226997392 | 0.720625549 | -4.544326402 |
| **A_51_P495269** | Lor | -0.571364514 | -1.613460758 | 0.47073173 | 8.296874622 | -1.348883979 | 0.227087454 | 0.720625549 | -4.5445353 |
| **A_65_P09657** | Zfp213 | 0.105294516 | -0.086792916 | 0.297381948 | 9.063775147 | 1.34857767 | 0.227180133 | 0.720625549 | -4.544750174 |
| **A_55_P2068988** | NA | 0.132732937 | -0.10941007 | 0.374875944 | 9.866476425 | 1.348577584 | 0.227180159 | 0.720625549 | -4.544750234 |
| **A_55_P2128969** | Mettl27 | -0.140994902 | -0.398276117 | 0.116286312 | 9.815037621 | -1.348231584 | 0.227284888 | 0.720625549 | -4.544992931 |
| **A_55_P2099961** | H2ac11 | 0.389153144 | -0.320985974 | 1.099292261 | 14.84449616 | 1.348175002 | 0.227302019 | 0.720625549 | -4.545032618 |
| **A_51_P245090** | Aqp3 | -0.309490556 | -0.874272757 | 0.255291645 | 12.41995464 | -1.348141764 | 0.227312083 | 0.720625549 | -4.545055931 |
| **A_52_P246749** | Ghitm | -0.10760435 | -0.303973494 | 0.088764794 | 12.10455528 | -1.348111281 | 0.227321312 | 0.720625549 | -4.545077311 |
| **A_51_P218518** | Fiz1 | -0.059133233 | -0.167066461 | 0.048799995 | 8.170518644 | -1.34786259 | 0.227396626 | 0.720625549 | -4.545251735 |
| **A_52_P515282** | 1110051M20Rik | -0.0927329 | -0.262009781 | 0.076543982 | 8.171272334 | -1.347737625 | 0.227434478 | 0.720625549 | -4.545339377 |
| **A_51_P496715** | Pomk | -0.074790033 | -0.211349422 | 0.061769356 | 8.648936868 | -1.347383807 | 0.227541683 | 0.720625549 | -4.545587506 |

| | | | | | | | | | |
|---|---|---|---|---|---|---|---|---|---|
| **A_55_P2182006** | Cacng6 | 0.121187542 | -0.100096932 | 0.342472017 | 8.441940177 | 1.347336875 | 0.227555906 | 0.720625549 | -4.545620418 |
| **A_51_P466759** | Mau2 | 0.067564982 | -0.055812197 | 0.190942162 | 9.47167166 | 1.347274199 | 0.227574902 | 0.720625549 | -4.54566437 |
| **A_55_P1962380** | NA | 0.173069359 | -0.14299478 | 0.489133497 | 11.77122353 | 1.347144323 | 0.227614271 | 0.720625549 | -4.545755443 |
| **A_51_P173709** | Gprc5b | -0.271294377 | -0.766769823 | 0.224181069 | 8.656693341 | -1.347062724 | 0.227639008 | 0.720625549 | -4.545812661 |
| **A_52_P506529** | Rabgap1l | -0.088546649 | -0.25028525 | 0.073191951 | 9.920589098 | -1.346876004 | 0.227695624 | 0.720625549 | -4.545943588 |
| **A_55_P2053320** | Zfr2 | 0.161826183 | -0.133786429 | 0.457438795 | 9.223829183 | 1.346774838 | 0.227726303 | 0.720625549 | -4.546014521 |
| **A_52_P279599** | Ube2k | -0.111730482 | -0.315842365 | 0.092381401 | 10.39085599 | -1.346705234 | 0.227747414 | 0.720625549 | -4.546063324 |
| **A_55_P1976212** | Selplg | 0.063725787 | -0.052702523 | 0.180154097 | 7.946475839 | 1.346560208 | 0.227791405 | 0.720625549 | -4.546165007 |
| **A_55_P2085485** | Pidd1 | -0.062018169 | -0.175328988 | 0.051292649 | 7.768219476 | -1.346532153 | 0.227799916 | 0.720625549 | -4.546184677 |
| **A_55_P2175060** | Plpp5 | -0.138464072 | -0.391496423 | 0.11456828 | 9.524560959 | -1.346263929 | 0.2278813 | 0.720625549 | -4.546372727 |
| **A_52_P55190** | BC005537 | -0.196704343 | -0.55617041 | 0.162761723 | 8.766044913 | -1.34624844 | 0.227886 | 0.720625549 | -4.546383586 |
| **A_55_P1959541** | NA | -0.097659968 | -0.276148521 | 0.080828584 | 9.981939115 | -1.34609396 | 0.227932885 | 0.720625549 | -4.546491885 |
| **A_55_P2022895** | Slc12a8 | -0.104523386 | -0.295559754 | 0.086512983 | 8.58027013 | -1.346066692 | 0.227941162 | 0.720625549 | -4.546511 |
| **A_55_P2014066** | NA | 0.098952378 | -0.08190678 | 0.279811537 | 8.490826291 | 1.346030481 | 0.227952153 | 0.720625549 | -4.546536385 |
| **A_55_P2129786** | Slc38a10 | 0.118414365 | -0.098019781 | 0.33484851 | 8.835635541 | 1.346008413 | 0.227958852 | 0.720625549 | -4.546551855 |
| **A_51_P208020** | Ulk3 | 0.116240205 | -0.096226698 | 0.328707109 | 9.145236729 | 1.345966476 | 0.227971583 | 0.720625549 | -4.546581254 |
| **A_66_P116632** | Mafg | -0.204475894 | -0.578239405 | 0.169287617 | 13.01413063 | -1.345904912 | 0.227990273 | 0.720625549 | -4.546624412 |
| **A_55_P2072035** | Thy1 | 0.219044448 | -0.181423505 | 0.619512401 | 8.732998764 | 1.345654741 | 0.228066236 | 0.720625549 | -4.546799778 |
| **A_55_P2282969** | Gm2a | -0.116740504 | -0.330175995 | 0.096694988 | 9.814442978 | -1.345625126 | 0.22807523 | 0.720625549 | -4.546820536 |
| **A_55_P2317341** | NA | -0.174914816 | -0.494731681 | 0.144902049 | 8.252826903 | -1.345533139 | 0.228103168 | 0.720625549 | -4.546885015 |
| **A_51_P276939** | Gmip | 0.076762868 | -0.06360764 | 0.217133377 | 8.66182957 | 1.345378499 | 0.228150141 | 0.720625549 | -4.546993407 |
| **A_55_P1990964** | Osbpl11 | -0.116940346 | -0.330788908 | 0.096908215 | 11.00978383 | -1.345324979 | 0.228166401 | 0.720625549 | -4.54703092 |
| **A_55_P2030126** | Tfpt | 0.151227402 | -0.125408155 | 0.427862959 | 9.651905054 | 1.344905138 | 0.228293985 | 0.720769419 | -4.547325174 |
| **A_55_P2199040** | NA | 0.264152361 | -0.219077013 | 0.747381736 | 9.21582589 | 1.344839122 | 0.228314052 | 0.720769419 | -4.54737144 |
| **A_52_P186672** | NA | -0.110660602 | -0.313103356 | 0.091782152 | 9.218427497 | -1.344806985 | 0.228323822 | 0.720769419 | -4.547393963 |
| **A_52_P441070** | Apob | 0.307784793 | -0.255449327 | 0.871018913 | 12.94262661 | 1.344396477 | 0.228448647 | 0.720899679 | -4.547681639 |
| **A_55_P2090415** | Nat9 | -0.087233222 | -0.24687918 | 0.072412736 | 8.670749914 | -1.344290563 | 0.228480862 | 0.720899679 | -4.547755857 |
| **A_55_P2071271** | Abracl | 0.173515023 | -0.144038781 | 0.491068827 | 11.83700578 | 1.344277488 | 0.22848484 | 0.720899679 | -4.547765019 |
| **A_55_P1971433** | Serpinb9d | 0.213595397 | -0.177338549 | 0.604529342 | 11.84448117 | 1.344180744 | 0.22851427 | 0.720899679 | -4.547832809 |
| **A_55_P1954896** | Rpl17 | 0.096909733 | -0.080494964 | 0.274314431 | 15.23387854 | 1.3439139 | 0.228595465 | 0.721038143 | -4.548019783 |
| **A_55_P2110177** | NA | 0.116680578 | -0.096997849 | 0.330359005 | 12.94504269 | 1.343405297 | 0.228750294 | 0.721098354 | -4.54837612 |
| **A_55_P2010773** | Tbx2 | 0.12742798 | -0.105949285 | 0.360805245 | 9.546200478 | 1.343307428 | 0.228780098 | 0.721098354 | -4.548444683 |
| **A_52_P527463** | Fam91a1 | -0.10626634 | -0.300890706 | 0.088358027 | 12.82389683 | -1.343283478 | 0.228787392 | 0.721098354 | -4.548461461 |
| **A_51_P308961** | Hmgcl | -0.175796056 | -0.497823809 | 0.146231696 | 12.47035996 | -1.343027763 | 0.228865284 | 0.721098354 | -4.548640598 |
| **A_55_P2120189** | Gas2l3 | 0.065608989 | -0.054582556 | 0.185800535 | 8.148787754 | 1.342946128 | 0.228890155 | 0.721098354 | -4.548697783 |
| **A_55_P2312783** | D030029J20Rik | 0.143489009 | -0.119401505 | 0.406379522 | 10.4766916 | 1.342804797 | 0.228933219 | 0.721098354 | -4.548796783 |
| **A_55_P2112967** | Diaph1 | -0.109638104 | -0.3105228 | 0.091246592 | 8.769363086 | -1.342714957 | 0.228960598 | 0.721098354 | -4.548859712 |
| **A_51_P280532** | Supt16 | -0.062822574 | -0.177929739 | 0.052284592 | 7.997130379 | -1.342710938 | 0.228961823 | 0.721098354 | -4.548862528 |
| **A_55_P2001155** | Rpl31 | 0.101359588 | -0.084381384 | 0.287100559 | 14.71564312 | 1.34253707 | 0.229014817 | 0.721098354 | -4.548984311 |
| **A_51_P106144** | Zfp64 | 0.353921484 | -0.294776751 | 1.00261972 | 11.5707238 | 1.342249878 | 0.229102378 | 0.721098354 | -4.545582842 |
| **A_55_P1992989** | Mrps26 | 0.202457196 | -0.168625862 | 0.573540253 | 13.12948686 | 1.342243329 | 0.229104375 | 0.721098354 | -4.549190048 |
| **A_55_P2179069** | NA | 0.250523461 | -0.208685199 | 0.709732121 | 9.647585316 | 1.342169905 | 0.229126766 | 0.721098354 | -4.549241471 |
| **A_52_P665386** | Ube2m | -0.120163601 | -0.340429478 | 0.100102276 | 9.98767525 | -1.342130898 | 0.229138662 | 0.721098354 | -4.54926879 |
| **A_55_P2043932** | Tmem8b | -0.061161501 | -0.173281345 | 0.050958343 | 8.206806428 | -1.342038001 | 0.229166995 | 0.721098354 | -4.54933385 |
| **A_51_P244856** | Atp2a1 | -0.056925105 | -0.161280711 | 0.047430501 | 7.742982103 | -1.342014536 | 0.229174152 | 0.721098354 | -4.549350283 |
| **A_51_P195506** | Csf1 | -0.26110583 | -0.739816378 | 0.217604719 | 10.46244225 | -1.341877076 | 0.229216084 | 0.721112904 | -4.545959092 |
| **A_55_P2016166** | Wfdc2 | 0.580030133 | -0.483869148 | 1.643929414 | 9.043441183 | 1.341279243 | 0.229398532 | 0.721333462 | -4.549865185 |
| **A_55_P2454896** | Amn1 | 0.10299665 | -0.085934325 | 0.291927626 | 10.53784943 | 1.341186265 | 0.229426919 | 0.721333462 | -4.549930288 |
| **A_51_P172155** | Hal | 0.228014425 | -0.190253193 | 0.646282043 | 8.579272501 | 1.341149657 | 0.229438096 | 0.721333462 | -4.54995592 |
| **A_55_P1963631** | Yae1d1 | 0.141526058 | -0.118101821 | 0.401153936 | 10.10579826 | 1.341078648 | 0.229459779 | 0.721333462 | -4.550005639 |

| | | | | | | | | | |
|---|---|---|---|---|---|---|---|---|---|
| **A_51_P359272** | Alcam | 0.066951101 | -0.055873803 | 0.189776006 | 7.876698505 | 1.341036059 | 0.229472785 | 0.721333462 | -4.550035459 |
| **A_55_P1973462** | Pex11b | 0.169171547 | -0.141308439 | 0.479651533 | 10.35535539 | 1.340487825 | 0.22964026 | 0.721668758 | -4.550419284 |
| **A_55_P2177026** | Mafg | -0.174482033 | -0.494719172 | 0.145755107 | 9.463221722 | -1.340442462 | 0.229654122 | 0.721668758 | -4.550451041 |
| **A_55_P2050773** | LOC100048452 | -0.072169322 | -0.204645753 | 0.060307108 | 8.096830837 | -1.340241843 | 0.229715439 | 0.721729292 | -4.550591483 |
| **A_51_P158120** | B3galnt2 | -0.114393367 | -0.324401271 | 0.095614537 | 10.54642372 | -1.340091228 | 0.229761482 | 0.721729292 | -4.550696914 |
| **A_52_P543040** | Utp14a | -0.075382599 | -0.213781061 | 0.063015863 | 8.140109149 | -1.340012986 | 0.229785403 | 0.721729292 | -4.550751682 |
| **A_52_P115579** | Fbxo45 | -0.073013279 | -0.207094454 | 0.061067896 | 8.22962757 | -1.33968658 | 0.229885224 | 0.721920919 | -4.550980152 |
| **A_55_P2078830** | NA | -0.126792948 | -0.359657461 | 0.106071566 | 9.049151264 | -1.339556235 | 0.229925097 | 0.721920919 | -4.551071382 |
| **A_66_P112006** | H2-T10 | -0.217963375 | -0.618340178 | 0.182413429 | 8.42863134 | -1.339318226 | 0.229997919 | 0.721920919 | -4.55123796 |
| **A_55_P2010347** | Zdhhc6 | 0.10930009 | -0.091480649 | 0.310080829 | 8.839717885 | 1.339268439 | 0.230013155 | 0.721920919 | -4.551272803 |
| **A_55_P2066041** | Hnrnpa1 | -0.067449065 | -0.19135687 | 0.056458739 | 8.234378845 | -1.339203075 | 0.23003316 | 0.721920919 | -4.551318549 |
| **A_52_P453785** | Cthrc1 | 0.235523774 | -0.197262058 | 0.668309606 | 8.916837337 | 1.338846785 | 0.230142227 | 0.722036509 | -4.551567883 |
| **A_55_P1978927** | Srgap1 | -0.075854728 | -0.21524381 | 0.063534354 | 8.472713773 | -1.338822677 | 0.230149608 | 0.722036509 | -4.551584753 |
| **A_55_P1999790** | Pkdcc | 0.17205495 | -0.1441347 | 0.488244601 | 10.84182863 | 1.338716712 | 0.230182056 | 0.722036509 | -4.551658904 |
| **A_55_P2055875** | Josd2 | -0.088677152 | -0.251692984 | 0.074338679 | 8.536739004 | -1.338292726 | 0.230311928 | 0.72220531 | -4.551955574 |
| **A_55_P2092526** | Tgif1 | -0.259161146 | -0.735618223 | 0.217295931 | 11.67982193 | -1.338182272 | 0.230345772 | 0.72220531 | -4.552032855 |
| **A_65_P12392** | Timp2 | -0.439947142 | -1.248863225 | 0.368968942 | 11.9433021 | -1.338031055 | 0.230392113 | 0.72220531 | -4.552138653 |
| **A_51_P203710** | Taf8 | -0.131008552 | -0.371908791 | 0.109891688 | 10.34942928 | -1.337924406 | 0.230424802 | 0.72220531 | -4.552213267 |
| **A_55_P2050902** | Fetub | -0.221720869 | -0.629430836 | 0.185989097 | 11.92940038 | -1.337902323 | 0.230431571 | 0.72220531 | -4.552228717 |
| **A_55_P2163622** | Rps27 | 0.104063549 | -0.08732638 | 0.295453479 | 16.596052 | 1.337669162 | 0.230503052 | 0.72220531 | -4.552391834 |
| **A_51_P371152** | Sos2 | 0.108740413 | -0.091264262 | 0.308745088 | 10.86876731 | 1.337580701 | 0.230530177 | 0.72220531 | -4.552453718 |
| **A_55_P2023937** | Impg1 | 0.349679038 | -0.293503384 | 0.99286146 | 11.01312565 | 1.337533274 | 0.230544721 | 0.72220531 | -4.552486896 |
| **A_55_P2159850** | NA | 0.15333927 | -0.128737322 | 0.435415862 | 13.34018833 | 1.337382102 | 0.230591085 | 0.72220531 | -4.552592645 |
| **A_51_P430082** | Tst | 0.239867671 | -0.201491083 | 0.681226425 | 10.03872625 | 1.33705458 | 0.230691563 | 0.72220531 | -4.552821743 |
| **A_52_P72237** | Actg1 | -0.223319524 | -0.634373983 | 0.187734934 | 12.13197826 | -1.336584727 | 0.230835775 | 0.72220531 | -4.553150367 |
| **A_52_P234910** | Gm15032 | 0.146049019 | -0.122795964 | 0.414894003 | 12.53165323 | 1.336490486 | 0.230864711 | 0.72220531 | -4.553216277 |
| **A_51_P361201** | Atp6v0d1 | -0.157811992 | -0.448318035 | 0.13269405 | 11.0998128 | -1.336453938 | 0.230875933 | 0.72220531 | -4.553241836 |
| **A_55_P2157676** | Wdr34 | -0.194410119 | -0.552297893 | 0.163477656 | 15.61023306 | -1.336414455 | 0.230888057 | 0.72220531 | -4.553269449 |
| **A_52_P354373** | Lockd | 0.096962701 | -0.081535055 | 0.275460457 | 8.17713547 | 1.336414275 | 0.230888112 | 0.72220531 | -4.553269575 |
| **A_66_P125734** | Tnip2 | -0.098057591 | -0.278584915 | 0.082469733 | 9.192541457 | -1.336310677 | 0.230919927 | 0.72220531 | -4.553342025 |
| **A_52_P641849** | Khnyn | -0.093989332 | -0.267034167 | 0.079055503 | 9.12794052 | -1.336254205 | 0.230937271 | 0.72220531 | -4.553381517 |
| **A_55_P2102065** | Gm10639 | 0.126493662 | -0.106465034 | 0.359452359 | 8.113309796 | 1.335854022 | 0.231060212 | 0.72220531 | -4.553661358 |
| **A_55_P2176953** | Usp51 | 0.265006371 | -0.223137263 | 0.753150006 | 8.895714498 | 1.335604422 | 0.231136923 | 0.72220531 | -4.553835885 |
| **A_52_P602669** | Serpinb6d | 0.124439428 | -0.104780288 | 0.353659144 | 9.585965715 | 1.335596273 | 0.231139428 | 0.72220531 | -4.553841583 |
| **A_52_P547187** | Tab2 | -0.061011771 | -0.173402479 | 0.051378937 | 11.51717095 | -1.33552612 | 0.231160992 | 0.72220531 | -4.553890633 |
| **A_55_P2107750** | Hectd1 | -0.086295569 | -0.245289692 | 0.072698554 | 8.861224758 | -1.335293052 | 0.231232649 | 0.72220531 | -4.554053587 |
| **A_55_P2376423** | Igkv4-72 | 0.065124974 | -0.054900235 | 0.185150183 | 8.452396954 | 1.334886236 | 0.231357772 | 0.72220531 | -4.554337997 |
| **A_55_P1957954** | Rsu1 | -0.086509932 | -0.24595123 | 0.072931366 | 11.06255058 | -1.334855679 | 0.231367173 | 0.72220531 | -4.554359359 |
| **A_55_P2113466** | NA | 0.080289655 | -0.067710026 | 0.228289335 | 9.255071032 | 1.334651798 | 0.231429906 | 0.72220531 | -4.554501882 |
| **A_66_P132890** | NA | -0.115426481 | -0.328200098 | 0.097347135 | 7.979998115 | -1.334617591 | 0.231440432 | 0.72220531 | -4.554525794 |
| **A_66_P119680** | Polr2e | 0.228071588 | -0.192401046 | 0.648544221 | 13.13768519 | 1.334450945 | 0.231491721 | 0.72220531 | -4.554642282 |
| **A_52_P24783** | Pds5b | -0.062144851 | -0.176726023 | 0.052436322 | 8.454898437 | -1.334323244 | 0.231531031 | 0.72220531 | -4.554731543 |
| **A_51_P422208** | Dpagt1 | 0.105890847 | -0.089356494 | 0.301138189 | 11.79545942 | 1.334266116 | 0.231548619 | 0.72220531 | -4.554771474 |
| **A_55_P2019833** | NA | -0.31022447 | -0.882240099 | 0.261791159 | 11.25922273 | -1.334250332 | 0.231553478 | 0.72220531 | -4.554782506 |
| **A_55_P2065519** | NA | -0.108861157 | -0.309593559 | 0.091871246 | 10.27141269 | -1.334211333 | 0.231565485 | 0.72220531 | -4.554809765 |
| **A_55_P2066523** | Akt2 | 0.06554037 | -0.055314883 | 0.186395623 | 8.430137649 | 1.334174146 | 0.231576935 | 0.72220531 | -4.554835757 |
| **A_51_P278887** | Hint2 | 0.094224167 | -0.079523605 | 0.267971939 | 11.63620739 | 1.334173235 | 0.231577215 | 0.72220531 | -4.554836393 |
| **A_65_P10673** | Tmem44 | -0.075647441 | -0.215151309 | 0.063856426 | 8.15179896 | -1.334065516 | 0.231610385 | 0.72220531 | -4.554911682 |
| **A_55_P2032845** | Rbm10 | -0.079895499 | -0.227237933 | 0.067446935 | 10.26390063 | -1.334023979 | 0.231623176 | 0.72220531 | -4.554940713 |
| **A_51_P295610** | Cyc1 | 0.107295632 | -0.090581245 | 0.305172508 | 10.9831728 | 1.334001008 | 0.231630251 | 0.72220531 | -4.554956769 |

| | | | | | | | | | |
|---|---|---|---|---|---|---|---|---|---|
| **A_55_P2017413** | Gm4788 | 0.056516356 | -0.047726764 | 0.160759476 | 7.823885814 | 1.333815977 | 0.231687241 | 0.72220531 | -4.555086087 |
| **A_55_P2078955** | Aqp8 | -0.182327509 | -0.518656483 | 0.154001466 | 9.368043042 | -1.333696703 | 0.231723984 | 0.72220531 | -4.555169444 |
| **A_55_P1996419** | NA | 0.182908019 | -0.154493576 | 0.520309615 | 15.75022949 | 1.333689642 | 0.23172616 | 0.72220531 | -4.555174379 |
| **A_55_P1954921** | Fbxw5 | -0.069845904 | -0.198701845 | 0.059010037 | 11.10600751 | -1.33353881 | 0.231772633 | 0.72220531 | -4.555279788 |
| **A_55_P2081313** | Gnas | -0.133168707 | -0.37886944 | 0.112532025 | 15.4419632 | -1.333413648 | 0.231811203 | 0.72220531 | -4.555367254 |
| **A_55_P2089070** | 4933431K14Rik | 0.095758148 | -0.080944276 | 0.272460573 | 8.43352662 | 1.333221765 | 0.231870346 | 0.72220531 | -4.555501341 |
| **A_66_P102062** | NA | -0.07732727 | -0.220024289 | 0.065369749 | 9.571884399 | -1.333174318 | 0.231884972 | 0.72220531 | -4.555534495 |
| **A_52_P140789** | Hspa13 | -0.089262697 | -0.254004685 | 0.075479291 | 11.78793098 | -1.333014478 | 0.231934251 | 0.72220531 | -4.555646185 |
| **A_55_P1978047** | Aven | -0.06298867 | -0.179268048 | 0.053290708 | 7.885041037 | -1.332689278 | 0.232034541 | 0.72220531 | -4.555873406 |
| **A_51_P483379** | Rnaseh1 | -0.077095112 | -0.219428977 | 0.065238753 | 9.774455034 | -1.332563041 | 0.232073482 | 0.72220531 | -4.555961604 |
| **A_55_P1962344** | Trim21 | -0.078925901 | -0.224641766 | 0.066789963 | 9.179065355 | -1.332544973 | 0.232079056 | 0.72220531 | -4.555974227 |
| **A_55_P2137941** | Fxyd2 | 0.518033446 | -0.438584603 | 1.474651495 | 11.94218694 | 1.332258171 | 0.232167551 | 0.72220531 | -4.556174596 |
| **A_51_P384441** | Tmem201 | -0.076542173 | -0.217914454 | 0.064830107 | 8.000009085 | -1.332004496 | 0.232245851 | 0.72220531 | -4.556351809 |
| **A_55_P2007699** | Nanog | 0.062524472 | -0.05297208 | 0.178021023 | 8.831484677 | 1.331834244 | 0.232298414 | 0.72220531 | -4.556470738 |
| **A_55_P2145804** | Aen | -0.170456534 | -0.485345757 | 0.14443269 | 11.76781107 | -1.331757089 | 0.232322238 | 0.72220531 | -4.556524632 |
| **A_66_P118600** | Lama1 | 0.173836427 | -0.147298962 | 0.494971816 | 14.40504608 | 1.33174717 | 0.232325301 | 0.72220531 | -4.556531561 |
| **A_52_P140497** | NA | -0.184987367 | -0.526725581 | 0.156750847 | 9.252564585 | -1.331734617 | 0.232329178 | 0.72220531 | -4.556540329 |
| **A_52_P289685** | Ssbp1 | -0.16111961 | -0.458786049 | 0.136546829 | 11.36023138 | -1.331642636 | 0.232357583 | 0.72220531 | -4.556604578 |
| **A_66_P125212** | Mapk1ip1l | -0.115740091 | -0.329570211 | 0.09809003 | 11.60237221 | -1.331631623 | 0.232360985 | 0.72220531 | -4.556612271 |
| **A_55_P1962603** | Cplx2 | -0.283715011 | -0.807905724 | 0.240475703 | 12.22585197 | -1.331564336 | 0.232381767 | 0.72220531 | -4.55665927 |
| **A_55_P1980986** | Lca5l | 0.091259976 | -0.077357782 | 0.259877733 | 8.117559144 | 1.331515425 | 0.232396874 | 0.72220531 | -4.556693433 |
| **A_51_P139678** | Sprr1a | 0.344168351 | -0.291861893 | 0.980198594 | 8.794397771 | 1.331258295 | 0.232476311 | 0.72220531 | -4.556873025 |
| **A_55_P1985565** | NA | -0.125709641 | -0.358045119 | 0.106625837 | 15.53325289 | -1.331135351 | 0.232514301 | 0.72220531 | -4.556958891 |
| **A_55_P2048867** | NA | 0.524986128 | -0.445309579 | 1.495281835 | 8.894935597 | 1.331106736 | 0.232523144 | 0.72220531 | -4.556978876 |
| **A_55_P1998601** | Slc17a9 | 0.071566013 | -0.060707836 | 0.203839862 | 8.091050625 | 1.331073371 | 0.232533455 | 0.72220531 | -4.557002178 |
| **A_55_P2081223** | Sptbn1 | 0.151741171 | -0.128731549 | 0.432213891 | 14.88049425 | 1.331011988 | 0.232552427 | 0.72220531 | -4.557045047 |
| **A_55_P2044463** | NA | 0.091356724 | -0.077533782 | 0.26024723 | 8.356579559 | 1.330774422 | 0.232625862 | 0.722303897 | -4.557210954 |
| **A_55_P2181597** | Mfap2 | 0.075916895 | -0.064441386 | 0.216275176 | 8.097141398 | 1.330667536 | 0.232658909 | 0.722303897 | -4.557285596 |
| **A_55_P2007566** | Trmt61b | 0.067484257 | -0.057325635 | 0.192294149 | 8.202543141 | 1.330217399 | 0.232798129 | 0.722514357 | -4.557599916 |
| **A_51_P380750** | Cbfa2t3 | -0.116450227 | -0.331837985 | 0.098937532 | 9.049457275 | -1.330112834 | 0.23283048 | 0.722514357 | -4.557672927 |
| **A_52_P36218** | Upf2 | -0.077947448 | -0.222122928 | 0.066228031 | 9.023820824 | -1.330085818 | 0.23283884 | 0.722514357 | -4.55769179 |
| **A_51_P297480** | Dph1 | -0.077542836 | -0.221024071 | 0.065938399 | 9.19228685 | -1.329583864 | 0.232994199 | 0.722880397 | -4.558042239 |
| **A_55_P1978471** | NA | -0.11357502 | -0.3237532 | 0.096603159 | 10.08161218 | -1.329426589 | 0.233042896 | 0.722883862 | -4.558152034 |
| **A_52_P198473** | NA | -0.092803665 | -0.264554415 | 0.078947085 | 9.608668215 | -1.329338698 | 0.233070114 | 0.722883862 | -4.558213389 |
| **A_52_P68714** | NA | -0.093669163 | -0.267090239 | 0.079751913 | 8.377130383 | -1.328813178 | 0.233232915 | 0.723272743 | -4.558580219 |
| **A_55_P2068882** | Ccng2 | -0.097421988 | -0.277808155 | 0.082964178 | 8.101869151 | -1.328687724 | 0.233271794 | 0.723277272 | -4.558667783 |
| **A_55_P2172227** | Ubn2 | -0.07703726 | -0.219718873 | 0.065644352 | 8.167706087 | -1.328317772 | 0.23338648 | 0.72343212 | -4.558925983 |
| **A_52_P1114640** | 2410003L11Rik | 0.175059156 | -0.149205167 | 0.499323478 | 8.647270338 | 1.328173543 | 0.233431205 | 0.72343212 | -4.559026637 |
| **A_55_P2296088** | Sp3 | -0.108562355 | -0.309674738 | 0.092550028 | 9.436579381 | -1.328035264 | 0.233474093 | 0.72343212 | -4.559123136 |
| **A_65_P07062** | Serf2 | -0.165595953 | -0.472388056 | 0.141196149 | 12.4103358 | -1.327928546 | 0.233507196 | 0.72343212 | -4.559197607 |
| **A_55_P2091985** | Gpr157 | 0.284479955 | -0.242600534 | 0.811560445 | 15.72625556 | 1.327834332 | 0.233536424 | 0.72343212 | -4.559263352 |
| **A_55_P2039225** | Rap1gap | 0.11187056 | -0.095406621 | 0.319147741 | 10.98122853 | 1.327802501 | 0.2335463 | 0.72343212 | -4.559285563 |
| **A_52_P163849** | Ivns1abp | -0.060825672 | -0.173555604 | 0.05190426 | 7.877546413 | -1.327445915 | 0.233656959 | 0.723522796 | -4.559534378 |
| **A_55_P2091974** | Gpr137 | 0.115874013 | -0.098906057 | 0.330654082 | 10.46108492 | 1.327275967 | 0.233709715 | 0.723522796 | -4.559652954 |
| **A_55_P1953485** | Copz1 | -0.15774818 | -0.450169636 | 0.134673277 | 13.08088993 | -1.327163055 | 0.233744771 | 0.723522796 | -4.559731732 |
| **A_55_P1995127** | NA | 1.785825927 | -1.524613693 | 5.096265547 | 12.39794059 | 1.327157959 | 0.233746354 | 0.723522796 | -4.559735288 |
| **A_55_P1996583** | Ppdpf | 0.120787039 | -0.10313719 | 0.344711268 | 10.75013728 | 1.327053437 | 0.23377881 | 0.723522796 | -4.55980821 |
| **A_55_P2113071** | Ago4 | -0.094412153 | -0.269449676 | 0.08062537 | 9.677132886 | -1.326984673 | 0.233800165 | 0.723522796 | -4.559856184 |
| **A_55_P2055134** | NA | -0.138359735 | -0.394920951 | 0.11820148 | 11.21965066 | -1.326746289 | 0.233874209 | 0.723619232 | -4.560022488 |
| **A_55_P2027819** | Rpl41 | -0.141957381 | -0.405248983 | 0.121334222 | 17.08414804 | -1.326447703 | 0.233966983 | 0.723619232 | -4.560230776 |

| | | | | | | | | | |
|---|---|---|---|---|---|---|---|---|---|
| **A_51_P488092** | Pde6d | -0.081560171 | -0.232833227 | 0.069712885 | 9.279132477 | -1.326434199 | 0.233971179 | 0.723619232 | -4.560240196 |
| **A_55_P2051914** | Zfp568 | -0.164276721 | -0.469034689 | 0.140481248 | 11.8626922 | -1.326142269 | 0.234061918 | 0.723619232 | -4.560443824 |
| **A_51_P280158** | Sf3b2 | 0.232653512 | -0.198965813 | 0.664272836 | 11.91496874 | 1.326104945 | 0.234073521 | 0.723619232 | -4.560469858 |
| **A_55_P2357860** | Atxn7l1 | 0.065276 | -0.055848238 | 0.186400238 | 8.108172569 | 1.325841595 | 0.234155407 | 0.723619232 | -4.560653535 |
| **A_52_P604195** | Mbp | -0.161057135 | -0.459913935 | 0.137799666 | 11.60490934 | -1.325824343 | 0.234160773 | 0.723619232 | -4.560665568 |
| **A_55_P2052563** | Id1 | 0.157679336 | -0.143347293 | 0.458705965 | 10.99304583 | 1.357437832 | 0.23416463 | 0.723619232 | -4.523264442 |
| **A_66_P135804** | NA | 0.358516677 | -0.306817374 | 1.023850728 | 12.96441682 | 1.325680216 | 0.2342056 | 0.723619232 | -4.560766086 |
| **A_55_P2072373** | Msn | -0.331247335 | -0.945975178 | 0.283480507 | 13.8075128 | -1.325679903 | 0.234205697 | 0.723619232 | -4.560766304 |
| **A_55_P1986113** | Ppil1 | 0.20526841 | -0.175781049 | 0.586317868 | 9.593055695 | 1.325286982 | 0.234327943 | 0.723772714 | -4.561040319 |
| **A_55_P2174967** | NA | -0.079982404 | -0.228458358 | 0.06849355 | 8.905082855 | -1.32527953 | 0.234330263 | 0.723772714 | -4.561045516 |
| **A_55_P1956048** | H2-T3 | -0.076179013 | -0.217624827 | 0.065266801 | 7.703964584 | -1.324995509 | 0.234418664 | 0.723875869 | -4.561243568 |
| **A_55_P2038002** | BC005624 | 0.144440357 | -0.123798807 | 0.412679521 | 10.84464566 | 1.324754886 | 0.234493582 | 0.723875869 | -4.561411346 |
| **A_55_P2044197** | Vps29 | 0.176890848 | -0.151615856 | 0.505397551 | 11.5602315 | 1.324738903 | 0.234498559 | 0.723875869 | -4.56142249 |
| **A_55_P2111399** | Pcp2 | 0.190180443 | -0.16301933 | 0.543380216 | 8.671994785 | 1.324691049 | 0.234513461 | 0.723875869 | -4.561455856 |
| **A_55_P1976504** | NA | 0.134544997 | -0.115373903 | 0.384463897 | 11.65783839 | 1.324456245 | 0.234586593 | 0.723901571 | -4.561619563 |
| **A_51_P246903** | Rps27l | -0.208417805 | -0.595586385 | 0.178750775 | 14.29071691 | -1.324353353 | 0.234618647 | 0.723901571 | -4.561691298 |
| **A_55_P1953157** | Mettl18 | 0.135164158 | -0.115977227 | 0.386305543 | 9.461442441 | 1.3240745 | 0.234705535 | 0.723901571 | -4.561885699 |
| **A_55_P2060607** | NA | 0.112203363 | -0.09630078 | 0.320707505 | 14.54378732 | 1.323915746 | 0.234755015 | 0.723901571 | -4.561996367 |
| **A_55_P1959152** | Gm10324 | 0.190620028 | -0.16361023 | 0.544850286 | 9.97161939 | 1.323890413 | 0.234762912 | 0.723901571 | -4.562014027 |
| **A_55_P2073383** | NA | -0.107135713 | -0.306236248 | 0.091964822 | 14.48660129 | -1.323826285 | 0.234782902 | 0.723901571 | -4.562058729 |
| **A_52_P649224** | Sash1 | -0.277728079 | -0.808868024 | 0.253411866 | 11.79359819 | -1.355067432 | 0.23486639 | 0.723901571 | -4.524711441 |
| **A_55_P2014867** | Iws1 | -0.082934636 | -0.237098067 | 0.071228795 | 8.359561752 | -1.323499364 | 0.234884836 | 0.723901571 | -4.562286607 |
| **A_55_P2030638** | Dusp23 | 0.086283506 | -0.074118656 | 0.246685668 | 9.865260456 | 1.323386586 | 0.234920009 | 0.723901571 | -4.562365214 |
| **A_52_P429534** | Elavl1 | -0.118920189 | -0.339996313 | 0.102155935 | 8.759683122 | -1.323374958 | 0.234923636 | 0.723901571 | -4.562373319 |
| **A_55_P2145720** | Thoc2 | -0.111890529 | -0.319907977 | 0.096126918 | 8.604618439 | -1.323313454 | 0.23494282 | 0.723901571 | -4.562416186 |
| **A_55_P2034097** | Elavl1 | 0.085240711 | -0.077829894 | 0.248311317 | 12.74359869 | 1.35463242 | 0.23499539 | 0.723901571 | -4.524976941 |
| **A_55_P1974567** | Plch2 | 0.166823719 | -0.143370358 | 0.477017796 | 8.463264423 | 1.323102414 | 0.235008658 | 0.723901571 | -4.562563273 |
| **A_55_P2162234** | Tm9sf3 | -0.085507009 | -0.244548069 | 0.07353405 | 9.071567767 | -1.322700807 | 0.235133993 | 0.723914674 | -4.562843154 |
| **A_55_P2253980** | Armt1 | 0.082943448 | -0.071332514 | 0.23721941 | 8.679684625 | 1.322674518 | 0.235142199 | 0.723914674 | -4.562861475 |
| **A_52_P160518** | Sfmbt1 | -0.116745668 | -0.333911009 | 0.100419672 | 8.685210596 | -1.322572307 | 0.235174108 | 0.723914674 | -4.562932701 |
| **A_55_P2157586** | NA | -0.074187431 | -0.212195542 | 0.06382068 | 8.179522297 | -1.322497571 | 0.235197442 | 0.723914674 | -4.562984779 |
| **A_55_P2133661** | NA | 0.177060996 | -0.15232115 | 0.506443142 | 14.75675492 | 1.322488823 | 0.235200173 | 0.723914674 | -4.562990875 |
| **A_52_P663742** | Ganc | -0.115121291 | -0.329309959 | 0.099067377 | 8.507038476 | -1.322294908 | 0.235260728 | 0.723973522 | -4.563125997 |
| **A_55_P2034864** | Tubb2b | 0.122745784 | -0.10564713 | 0.351138697 | 9.48223162 | 1.322187726 | 0.235294204 | 0.723973522 | -4.56320068 |
| **A_51_P510849** | Tmem50b | 0.144659098 | -0.124603337 | 0.413921532 | 11.46883525 | 1.321719043 | 0.235440637 | 0.724308781 | -4.563527225 |
| **A_52_P635338** | Fes | -0.083738145 | -0.239631467 | 0.072155178 | 9.381142378 | -1.321493355 | 0.235511179 | 0.7244105 | -4.563684454 |
| **A_51_P253883** | Fam49a | 0.072273775 | -0.062342701 | 0.206890251 | 8.322884239 | 1.320844487 | 0.235714098 | 0.724849203 | -4.564136445 |
| **A_66_P139267** | Rbpj | -0.080077972 | -0.229237308 | 0.069081365 | 10.00825729 | -1.320783958 | 0.235733035 | 0.724849203 | -4.564178605 |
| **A_55_P2022231** | Gm826 | 0.190346097 | -0.164245142 | 0.544937337 | 8.048476892 | 1.3206421 | 0.235777423 | 0.724849203 | -4.564277409 |
| **A_66_P117659** | Nfatc3 | -0.107380154 | -0.307442734 | 0.092682427 | 9.446927113 | -1.320466288 | 0.235832444 | 0.724849203 | -4.564399857 |
| **A_55_P2167451** | Cnnm2 | -0.249717585 | -0.715017886 | 0.215582717 | 11.56421259 | -1.320337491 | 0.235872759 | 0.724849203 | -4.564489557 |
| **A_51_P459091** | Ybey | 0.079675014 | -0.068786099 | 0.228136126 | 8.261225806 | 1.320318167 | 0.235878808 | 0.724849203 | -4.564503014 |
| **A_55_P1973427** | Tex14 | -0.208998031 | -0.598475684 | 0.180479622 | 9.585383327 | -1.320166822 | 0.235926191 | 0.724879564 | -4.564608413 |
| **A_55_P1988202** | Ifi203 | -0.12971001 | -0.371471085 | 0.112051064 | 8.690625761 | -1.319946352 | 0.235995229 | 0.724976444 | -4.564761943 |
| **A_51_P110371** | Wwox | -0.063967903 | -0.18325614 | 0.055320334 | 7.89520955 | -1.319269918 | 0.236207163 | 0.725348997 | -4.565232941 |
| **A_55_P1968649** | Atp1b2 | 0.100484474 | -0.086957958 | 0.287926905 | 8.228350789 | 1.318864613 | 0.236334231 | 0.725348997 | -4.565515112 |
| **A_52_P965502** | Ssr1 | 0.10225614 | -0.088500433 | 0.293012713 | 8.754245382 | 1.318800329 | 0.23635439 | 0.725348997 | -4.565559863 |
| **A_55_P1967025** | NA | 0.082258645 | -0.071197134 | 0.235714424 | 8.502983874 | 1.318765163 | 0.236365419 | 0.725348997 | -4.565584344 |
| **A_52_P8922** | Sntb2 | -0.145277423 | -0.416328382 | 0.125773537 | 9.222610304 | -1.318609926 | 0.23641411 | 0.725348997 | -4.565692408 |
| **A_55_P2157033** | Bace2 | -0.305099677 | -0.874339929 | 0.264140575 | 10.5462967 | -1.318606794 | 0.236415092 | 0.725348997 | -4.565694589 |

| | | | | | | | | | |
|---|---|---|---|---|---|---|---|---|---|
| **A_55_P1990066** | NA | 0.143528346 | -0.124267179 | 0.411323871 | 9.340613052 | 1.318571026 | 0.236426312 | 0.725348997 | -4.565719486 |
| **A_55_P2003548** | NA | -0.093839784 | -0.268936702 | 0.081257135 | 16.64280295 | -1.318492483 | 0.236450952 | 0.725348997 | -4.56577416 |
| **A_51_P517051** | Castor1 | -0.138691281 | -0.397479272 | 0.120096709 | 8.78231374 | -1.318481998 | 0.236454241 | 0.725348997 | -4.565781458 |
| **A_55_P2040529** | Mtfr1 | -0.118391216 | -0.339390123 | 0.102607691 | 8.929330589 | -1.317948741 | 0.236621593 | 0.725620029 | -4.566152624 |
| **A_55_P1964375** | Gm9731 | 0.067155196 | -0.058218128 | 0.19252852 | 10.78552363 | 1.317782219 | 0.236673875 | 0.725620029 | -4.566268519 |
| **A_55_P2073945** | Eif2b3 | -0.057940736 | -0.166112783 | 0.050231312 | 8.137416327 | -1.317765458 | 0.236679137 | 0.725620029 | -4.566280183 |
| **A_52_P523712** | Tbx5 | -0.135722741 | -0.389205507 | 0.117760025 | 8.342906453 | -1.31726559 | 0.236836142 | 0.725620029 | -4.566628044 |
| **A_55_P2114333** | NA | 0.223590533 | -0.19400327 | 0.641184336 | 15.04819328 | 1.317250977 | 0.236840733 | 0.725620029 | -4.566638213 |
| **A_55_P2061520** | NA | -0.080496744 | -0.230844881 | 0.069851392 | 8.428225656 | -1.317193046 | 0.236858935 | 0.725620029 | -4.566678524 |
| **A_55_P2186961** | NA | 0.07290407 | -0.063264196 | 0.209072336 | 8.935498478 | 1.317179707 | 0.236863126 | 0.725620029 | -4.566687805 |
| **A_55_P2026420** | Pou6f1 | -0.095399086 | -0.273597689 | 0.082799516 | 7.972933814 | -1.317070686 | 0.236897385 | 0.725620029 | -4.566763666 |
| **A_51_P316173** | Vmn1r192 | 0.064872303 | -0.05633489 | 0.186079496 | 7.854841714 | 1.316740177 | 0.23700127 | 0.725620029 | -4.566993631 |
| **A_55_P2046253** | Tmem239 | -0.20363412 | -0.584125059 | 0.176856818 | 14.47938666 | -1.316665309 | 0.237024808 | 0.725620029 | -4.56704572 |
| **A_51_P288549** | Jmjd7 | 0.07707293 | -0.066938143 | 0.221084004 | 8.507444036 | 1.31666456 | 0.237025044 | 0.725620029 | -4.567046242 |
| **A_55_P2069765** | Kiss1 | 0.242417455 | -0.210540638 | 0.695375549 | 8.742251368 | 1.316664199 | 0.237025158 | 0.725620029 | -4.567046493 |
| **A_55_P1956097** | Cyb5d2 | 0.095975405 | -0.083357433 | 0.275308243 | 10.36094163 | 1.316646824 | 0.23703062 | 0.725620029 | -4.567058581 |
| **A_55_P2076797** | Ddx5 | -0.181339336 | -0.520227304 | 0.157548632 | 9.90977314 | -1.316452018 | 0.237091878 | 0.725692622 | -4.567194113 |
| **A_55_P2018307** | NA | -0.214212377 | -0.614638978 | 0.186214223 | 11.83161598 | -1.316105803 | 0.237200781 | 0.725856847 | -4.567434965 |
| **A_55_P1994887** | Zfpm1 | -0.166336488 | -0.477287425 | 0.144614448 | 13.1335057 | -1.316027071 | 0.237225553 | 0.725856847 | -4.567489734 |
| **A_55_P2044109** | NA | 0.081664061 | -0.071016606 | 0.234344729 | 15.79400904 | 1.315879394 | 0.237272023 | 0.725856847 | -4.56759246 |
| **A_51_P426458** | Ube2d2a | -0.108391852 | -0.311068319 | 0.094284614 | 10.88711153 | -1.315716973 | 0.237323143 | 0.725856847 | -4.567705437 |
| **A_51_P507778** | Sdr42e1 | -0.210262453 | -0.60347503 | 0.182950124 | 12.16019919 | -1.31553828 | 0.237379395 | 0.725856847 | -4.567829727 |
| **A_55_P1962693** | Pla2g5 | 0.111758338 | -0.097287031 | 0.320803707 | 8.191074774 | 1.315250711 | 0.237469946 | 0.725856847 | -4.568029733 |
| **A_55_P1994128** | Tmem184a | 0.114091047 | -0.099319887 | 0.327501981 | 10.23750128 | 1.315237136 | 0.237474221 | 0.725856847 | -4.568039174 |
| **A_52_P129697** | Stradb | -0.061131257 | -0.175479477 | 0.053216963 | 8.355685395 | -1.315234158 | 0.237475159 | 0.725856847 | -4.568041246 |
| **A_51_P511899** | Mthfsd | -0.078178573 | -0.224429 | 0.068071854 | 8.58097161 | -1.315102971 | 0.237516479 | 0.725856847 | -4.568132482 |
| **A_55_P2048655** | NA | 0.1286541 | -0.112032181 | 0.369340381 | 15.3421078 | 1.315047556 | 0.237533935 | 0.725856847 | -4.568171019 |
| **A_52_P659258** | Ppat | -0.064065845 | -0.183927462 | 0.055795772 | 8.288267259 | -1.314969223 | 0.237558612 | 0.725856847 | -4.568225495 |
| **A_51_P419226** | S100a14 | 0.366919299 | -0.319773639 | 1.053612236 | 10.62375741 | 1.314550083 | 0.237690693 | 0.726145629 | -4.56851696 |
| **A_52_P72434** | Khk | 0.219327586 | -0.191214245 | 0.629869417 | 9.55761876 | 1.314331772 | 0.237759513 | 0.726241093 | -4.568668757 |
| **A_52_P580582** | Nppa | 0.092875571 | -0.081043915 | 0.266795058 | 8.057116775 | 1.313779309 | 0.237933753 | 0.72649429 | -4.563396079 |
| **A_55_P2018283** | Hmgb1 | -0.066080652 | -0.189833349 | 0.057672044 | 8.542815435 | -1.313677663 | 0.237965823 | 0.72649429 | -4.569123524 |
| **A_55_P2119011** | Osbpl9 | -0.081131427 | -0.233079572 | 0.070816717 | 8.801295671 | -1.31359921 | 0.237990579 | 0.72649429 | -4.569178062 |
| **A_66_P102047** | Tbc1d13 | -0.119640176 | -0.343728746 | 0.104448395 | 9.206163901 | -1.313489181 | 0.238025301 | 0.72649429 | -4.56925455 |
| **A_51_P199168** | Cidea | -0.071457232 | -0.205311707 | 0.062397244 | 8.188669498 | -1.313355975 | 0.238067344 | 0.72649429 | -4.569347146 |
| **A_55_P2018091** | NA | 0.235699349 | -0.205815887 | 0.677214585 | 14.03787491 | 1.313354149 | 0.238067921 | 0.72649429 | -4.569348415 |
| **A_55_P1967291** | Ncaph | 0.156928413 | -0.137095902 | 0.450952728 | 8.893370185 | 1.313068851 | 0.23815799 | 0.726654426 | -4.569546725 |
| **A_55_P1968295** | Pacsin3 | 0.155917111 | -0.136272813 | 0.448107036 | 11.72741857 | 1.3127974 | 0.238243716 | 0.726696685 | -4.569735395 |
| **A_51_P151271** | Eif4b | -0.058018161 | -0.166745531 | 0.05070921 | 8.715985902 | -1.312786901 | 0.238247033 | 0.726696685 | -4.569742692 |
| **A_55_P2153292** | Tubb4b | -0.124239697 | -0.357092081 | 0.108612686 | 14.21701536 | -1.312649753 | 0.238290356 | 0.726714151 | -4.56983801 |
| **A_51_P308469** | Rngtt | 0.058958619 | -0.051565045 | 0.169482283 | 8.952484679 | 1.312384797 | 0.238374072 | 0.726854779 | -4.570022145 |
| **A_55_P1973357** | NA | 0.072518656 | -0.063504361 | 0.208541673 | 15.53171915 | 1.311615396 | 0.238617324 | 0.72708873 | -4.570556775 |
| **A_55_P2152912** | NA | -0.116157971 | -0.334044315 | 0.101728374 | 9.199227495 | -1.311560011 | 0.238634843 | 0.72708873 | -4.570595256 |
| **A_55_P2133337** | NA | 0.361969173 | -0.317014709 | 1.040953055 | 8.847011328 | 1.311539224 | 0.238641419 | 0.72708873 | -4.570609698 |
| **A_51_P117226** | NA | -0.082611728 | -0.237575703 | 0.072352247 | 8.420807241 | -1.311535719 | 0.238642527 | 0.72708873 | -4.566695823 |
| **A_55_P1973770** | Unc5b | -0.168232832 | -0.48381864 | 0.147352976 | 13.35162043 | -1.311482365 | 0.238659405 | 0.72708873 | -4.570649202 |
| **A_55_P1958394** | Slit2 | 0.069671882 | -0.06103009 | 0.200373855 | 8.124400954 | 1.311428343 | 0.238676496 | 0.72708873 | -4.570686735 |
| **A_55_P1985945** | NA | 0.227066031 | -0.198998628 | 0.65313069 | 14.35605834 | 1.311130195 | 0.238770838 | 0.72714424 | -4.570893867 |
| **A_51_P105380** | Csta2 | -0.256149467 | -0.736813456 | 0.224514522 | 9.692480008 | -1.311055366 | 0.238794522 | 0.72714424 | -4.57094585 |
| **A_55_P2088601** | Fbln1 | -0.188389951 | -0.541914458 | 0.165134557 | 9.837880166 | -1.311014123 | 0.238807576 | 0.72714424 | -4.570974501 |

| | | | | | | | | | |
|---|---|---|---|---|---|---|---|---|---|
| **A_55_P2073617** | NA | 0.104652686 | -0.091789403 | 0.301094775 | 9.06365171 | 1.310644771 | 0.238924512 | 0.727357966 | -4.571231069 |
| **A_55_P1967295** | Pemt | 0.07419974 | -0.065089121 | 0.2134886 | 8.387470568 | 1.310554723 | 0.238953028 | 0.727357966 | -4.571293616 |
| **A_51_P289889** | Npnt | -0.06052815 | -0.174185802 | 0.053129502 | 7.783811252 | -1.310170694 | 0.239074678 | 0.727573002 | -4.571560345 |
| **A_51_P141926** | Fxyd4 | 0.183522131 | -0.161115285 | 0.528159547 | 9.762559799 | 1.310072052 | 0.239105934 | 0.727573002 | -4.571628853 |
| **A_55_P2110290** | Tmem254a | 0.144000722 | -0.126451056 | 0.4144525 | 10.78659259 | 1.309917645 | 0.239154868 | 0.727573002 | -4.571736086 |
| **A_52_P347591** | Ndufa7 | 0.137493547 | -0.12077046 | 0.395757555 | 14.83643165 | 1.309747547 | 0.239208784 | 0.727573002 | -4.571854211 |
| **A_55_P1981756** | Vmn2r123 | -0.272324185 | -0.783854081 | 0.239205711 | 8.851849226 | -1.30973778 | 0.23921188 | 0.727573002 | -4.571860994 |
| **A_55_P2107155** | Rasd2 | 0.21998798 | -0.193335327 | 0.633311286 | 8.336392562 | 1.309417736 | 0.239313356 | 0.727767126 | -4.572083234 |
| **A_51_P222522** | Mesd | 0.140934461 | -0.123887576 | 0.405756499 | 13.11939548 | 1.309279118 | 0.239357319 | 0.727786316 | -4.572179485 |
| **A_51_P415809** | Tusc1 | 0.10785016 | -0.094860123 | 0.310560443 | 8.67081521 | 1.308923233 | 0.239470222 | 0.728015087 | -4.57242658 |
| **A_55_P2151897** | NA | 0.139566592 | -0.122850883 | 0.401984068 | 15.30569572 | 1.308452269 | 0.239619707 | 0.728347182 | -4.57275354 |
| **A_55_P1989703** | Galnt11 | 0.140472465 | -0.123689134 | 0.404634064 | 10.02623422 | 1.308249817 | 0.239683992 | 0.728347182 | -4.572894076 |
| **A_55_P1972426** | Pdzd11 | 0.126039088 | -0.110985081 | 0.363063258 | 9.6181845 | 1.308223011 | 0.239692505 | 0.728347182 | -4.572912684 |
| **A_51_P494481** | Snx30 | -0.075325391 | -0.216993165 | 0.066342384 | 9.14239496 | -1.3080956 | 0.23973297 | 0.728355641 | -4.573001123 |
| **A_55_P1968365** | NA | 0.111116544 | -0.097922957 | 0.320156045 | 8.357897404 | 1.307734342 | 0.23984774 | 0.728375608 | -4.573251868 |
| **A_55_P2028883** | Rnft2 | 0.060549223 | -0.053384807 | 0.174483252 | 8.656481862 | 1.307447557 | 0.239938884 | 0.728375608 | -4.573450903 |
| **A_55_P2047509** | Ufd1 | 0.115753998 | -0.102079309 | 0.333587304 | 12.6345678 | 1.307316907 | 0.239980417 | 0.728375608 | -4.573541573 |
| **A_55_P2020395** | 2-Mar | -0.100596332 | -0.289913487 | 0.088720824 | 8.149609847 | -1.307258085 | 0.239999119 | 0.728375608 | -4.573582393 |
| **A_51_P179907** | Appl2 | -0.110578399 | -0.318698211 | 0.097541413 | 9.392018476 | -1.307151833 | 0.240032903 | 0.728375608 | -4.573656127 |
| **A_55_P2038299** | Rpl31 | 0.098031251 | -0.086473815 | 0.282536317 | 14.94990821 | 1.30714996 | 0.240033498 | 0.728375608 | -4.573657426 |
| **A_55_P1986660** | Anapc2 | 0.173542158 | -0.153108586 | 0.500192902 | 13.92025592 | 1.307044895 | 0.240066909 | 0.728375608 | -4.573730334 |
| **A_51_P483617** | 0610040J01Rik | 0.240881324 | -0.212530204 | 0.694292852 | 8.651568908 | 1.307012479 | 0.240077218 | 0.728375608 | -4.573752828 |
| **A_51_P362089** | Lrrc3b | 0.060841773 | -0.053681256 | 0.175364802 | 7.902454537 | 1.307007851 | 0.24007869 | 0.728375608 | -4.573756039 |
| **A_55_P2046793** | Eogt | 0.077398269 | -0.068333404 | 0.223129942 | 9.01010728 | 1.30661146 | 0.240204786 | 0.728643802 | -4.574031085 |
| **A_51_P181341** | Necab2 | 0.067342378 | -0.059480714 | 0.194165469 | 7.866940676 | 1.306349172 | 0.240288255 | 0.728782626 | -4.574213063 |
| **A_51_P369971** | Fra10ac1 | -0.095742272 | -0.276095707 | 0.084611162 | 10.62733569 | -1.306015934 | 0.24039434 | 0.728826975 | -4.574444248 |
| **A_51_P117881** | Leap2 | -0.064882 | -0.187103652 | 0.057339652 | 7.682373236 | -1.306006212 | 0.240397435 | 0.728826975 | -4.574450992 |
| **A_55_P2006504** | Eif4a2 | 0.112947252 | -0.099838141 | 0.325732645 | 13.47277006 | 1.305879247 | 0.240437866 | 0.728826975 | -4.574539069 |
| **A_55_P1988907** | Eml4 | 0.061240742 | -0.05413729 | 0.176618775 | 10.98606549 | 1.305829519 | 0.240453703 | 0.728826975 | -4.574573565 |
| **A_55_P2095854** | Rab3c | 0.229362848 | -0.202809436 | 0.661535131 | 12.97155404 | 1.305675708 | 0.240502693 | 0.728861171 | -4.574680259 |
| **A_52_P178904** | Seh1l | -0.07469858 | -0.215484576 | 0.066087415 | 8.508324498 | -1.30533522 | 0.240611173 | 0.728941512 | -4.574916429 |
| **A_55_P2172747** | Mfn2 | 0.074365049 | -0.065842607 | 0.214572704 | 9.43300959 | 1.304867166 | 0.240760367 | 0.728941512 | -4.575241045 |
| **A_51_P239766** | Plcd1 | 0.116412316 | -0.103090059 | 0.33591469 | 11.4427648 | 1.304754699 | 0.240796229 | 0.728941512 | -4.57531904 |
| **A_55_P1993609** | Slx4ip | -0.076659886 | -0.221208548 | 0.067888777 | 9.149928725 | -1.304737828 | 0.240801609 | 0.728941512 | -4.575330739 |
| **A_51_P389864** | B3gnt9 | 0.085423008 | -0.075651741 | 0.246497757 | 8.400034757 | 1.304717714 | 0.240808023 | 0.728941512 | -4.575344688 |
| **A_52_P651248** | Zdhhc2 | -0.074060761 | -0.213715738 | 0.065594217 | 8.214862655 | -1.304670711 | 0.240823012 | 0.728941512 | -4.575377283 |
| **A_52_P94830** | Galnt16 | 0.194152401 | -0.171964805 | 0.560269606 | 9.694975113 | 1.304643302 | 0.240831754 | 0.728941512 | -4.57539629 |
| **A_55_P2084032** | AW146154 | -0.084734261 | -0.244527086 | 0.075058564 | 9.320702157 | -1.304580653 | 0.240851735 | 0.728941512 | -4.575439734 |
| **A_51_P159415** | Sin3a | -0.108188635 | -0.31222026 | 0.09584299 | 9.637416604 | -1.304527735 | 0.240868614 | 0.728941512 | -4.575476429 |
| **A_55_P2096127** | NA | -0.276694516 | -0.798729338 | 0.245340305 | 10.80553062 | -1.303977934 | 0.241044041 | 0.729278486 | -4.57585765 |
| **A_51_P325485** | Utp3 | 0.121487001 | -0.107737509 | 0.35071151 | 11.75157197 | 1.303880893 | 0.241075016 | 0.729278486 | -4.57592493 |
| **A_52_P238242** | Sertad2 | -0.073844595 | -0.213193866 | 0.065504677 | 9.602200267 | -1.303716522 | 0.241127491 | 0.729278486 | -4.576038887 |
| **A_55_P2138174** | Psd3 | -0.060782832 | -0.175503751 | 0.053938087 | 8.070966277 | -1.303489313 | 0.241200043 | 0.729278486 | -4.576196401 |
| **A_51_P283499** | Drd4 | -0.141907512 | -0.409767488 | 0.125952463 | 8.838122812 | -1.303367012 | 0.241239104 | 0.729278486 | -4.576281182 |
| **A_52_P202029** | Trak2 | -0.17786899 | -0.513613094 | 0.157875113 | 9.759352307 | -1.303349857 | 0.241244584 | 0.729278486 | -4.576293075 |
| **A_55_P2095251** | Ell3 | 0.198408338 | -0.17615634 | 0.572973016 | 8.290332877 | 1.303173567 | 0.2413009 | 0.729278486 | -4.576415277 |
| **A_52_P485450** | Vta1 | -0.109450636 | -0.316098602 | 0.097197329 | 10.61919492 | -1.303035716 | 0.241344944 | 0.729278486 | -4.576510829 |
| **A_51_P249051** | Pycr2 | 0.091134213 | -0.08095057 | 0.263218995 | 11.50832637 | 1.302891168 | 0.241391136 | 0.729278486 | -4.57661102 |
| **A_66_P113505** | Fgd4 | -0.313728879 | -0.906201537 | 0.27874378 | 10.13050324 | -1.302732798 | 0.241441754 | 0.729278486 | -4.576720786 |
| **A_55_P2074081** | NA | -0.172103684 | -0.497120184 | 0.152912816 | 13.58550916 | -1.302728517 | 0.241443123 | 0.729278486 | -4.576723753 |

| | | | | | | | | | |
|---|---|---|---|---|---|---|---|---|---|
| **A_66_P133240** | Hnrnpll | -0.074473616 | -0.215121242 | 0.066174011 | 10.70136039 | -1.302684343 | 0.241457243 | 0.729278486 | -4.576754369 |
| **A_55_P2023952** | Arpc3 | -0.076790278 | -0.221817651 | 0.068237096 | 9.668713195 | -1.302643032 | 0.241470449 | 0.729278486 | -4.576783001 |
| **A_55_P2168267** | NA | 0.323307213 | -0.287438296 | 0.934052722 | 12.18882502 | 1.302339686 | 0.241567441 | 0.729457439 | -4.576993231 |
| **A_55_P2121886** | Map3k9 | 0.091171671 | -0.081124882 | 0.263468224 | 8.461183101 | 1.301824642 | 0.241732201 | 0.729760323 | -4.577350135 |
| **A_51_P179498** | Fam92a | -0.112192764 | -0.32422725 | 0.099841722 | 9.597827876 | -1.301749766 | 0.241756162 | 0.729760323 | -4.577402016 |
| **A_66_P103301** | Dicer1 | 0.08369777 | -0.07450337 | 0.24189891 | 9.124263263 | 1.301587716 | 0.241808026 | 0.729760323 | -4.577514297 |
| **A_51_P111164** | Rnd1 | -0.170620964 | -0.493154916 | 0.151912988 | 8.888240674 | -1.301445891 | 0.241853425 | 0.729760323 | -4.57761256 |
| **A_52_P453884** | Foxo1 | -0.066067525 | -0.19096583 | 0.058830779 | 10.16826282 | -1.301369608 | 0.241877847 | 0.729760323 | -4.577665412 |
| **A_55_P2007196** | Aldoa | -0.123534776 | -0.357082077 | 0.110012526 | 16.07576998 | -1.301318308 | 0.241894272 | 0.729760323 | -4.577700953 |
| **A_55_P2157627** | NA | -0.17987838 | -0.519978344 | 0.160221584 | 10.34145396 | -1.30119248 | 0.241934563 | 0.729767973 | -4.577788126 |
| **A_55_P1965278** | Dmrtc1a | -0.265705877 | -0.768258724 | 0.23684697 | 11.03742239 | -1.300734282 | 0.242081332 | 0.730096751 | -4.578105538 |
| **A_55_P2302383** | 3300002A11Rik | -0.167171865 | -0.483402823 | 0.149059093 | 8.339153936 | -1.300552737 | 0.242139506 | 0.730158271 | -4.57823129 |
| **A_55_P2038947** | Lig3 | 0.080079619 | -0.071482302 | 0.231641541 | 9.226012556 | 1.299873413 | 0.242357299 | 0.730636595 | -4.578701786 |
| **A_51_P297679** | Hcls1 | -0.090849166 | -0.262800744 | 0.081102411 | 9.585595669 | -1.299822182 | 0.242373731 | 0.730636595 | -4.578737264 |
| **A_55_P2073308** | Bin1 | 0.098803676 | -0.088287146 | 0.285894499 | 10.30505552 | 1.299241199 | 0.242560146 | 0.731066132 | -4.579139571 |
| **A_51_P168524** | NA | -0.107994111 | -0.31250314 | 0.096514918 | 8.912239554 | -1.299142379 | 0.242591866 | 0.731066132 | -4.579207993 |
| **A_52_P445360** | Krt20 | 0.358415807 | -0.32057016 | 1.037401774 | 9.039236249 | 1.298660164 | 0.242746704 | 0.731418714 | -4.579541848 |
| **A_55_P2174601** | Plekha7 | 0.224339464 | -0.213219441 | 0.661898369 | 11.56840637 | 1.328676438 | 0.242813836 | 0.731506957 | -4.54078839 |
| **A_52_P536735** | Hpcal1 | -0.10760768 | -0.311565916 | 0.096350556 | 9.139716006 | -1.297989509 | 0.242962198 | 0.731559878 | -4.58000609 |
| **A_55_P2158478** | Nkiras2 | 0.119121331 | -0.106670363 | 0.344913026 | 12.18052162 | 1.297928409 | 0.242981839 | 0.731559878 | -4.580048381 |
| **A_52_P243516** | Rmdn2 | -0.091049386 | -0.263658728 | 0.081559956 | 8.555684249 | -1.29772266 | 0.243047989 | 0.731559878 | -4.580190785 |
| **A_55_P2002854** | Aifm2 | -0.095365796 | -0.276169839 | 0.085438247 | 8.482850066 | -1.297638324 | 0.243075108 | 0.731559878 | -4.580249153 |
| **A_55_P2018611** | Igfbp7 | 0.177799486 | -0.159294478 | 0.514893451 | 10.22846789 | 1.297623461 | 0.243079888 | 0.731559878 | -4.58025944 |
| **A_51_P230439** | Ppfibp2 | -0.122588413 | -0.355009371 | 0.109832545 | 13.37792437 | -1.297607353 | 0.243085068 | 0.731559878 | -4.580270588 |
| **A_55_P2107347** | Ro60 | -0.087853602 | -0.254423748 | 0.078716544 | 8.937031735 | -1.297572311 | 0.243096337 | 0.731559878 | -4.580294839 |
| **A_66_P118849** | NA | 0.06038206 | -0.054128639 | 0.174892758 | 7.957773728 | 1.29727194 | 0.243192955 | 0.731641108 | -4.58050271 |
| **A_55_P2059890** | Fdx1 | -0.19099233 | -0.553228023 | 0.171243363 | 13.97329151 | -1.297160897 | 0.243228681 | 0.731641108 | -4.580579552 |
| **A_51_P464981** | Map1lc3a | -0.219297294 | -0.63524423 | 0.196649642 | 10.43026358 | -1.29707326 | 0.243256881 | 0.731641108 | -4.580640196 |
| **A_55_P2075636** | Epc2 | -0.087621803 | -0.253823487 | 0.078579882 | 9.687345754 | -1.297017772 | 0.243274737 | 0.731641108 | -4.580678592 |
| **A_55_P1982030** | H3f3b | 0.193643663 | -0.173722934 | 0.56101026 | 12.39270313 | 1.296799359 | 0.243345035 | 0.731738671 | -4.580829721 |
| **A_52_P238816** | Ppt2 | -0.111276047 | -0.32245689 | 0.099904797 | 9.604458573 | -1.296332265 | 0.243495432 | 0.732077027 | -4.581152892 |
| **A_51_P142896** | Cd59a | -0.210901356 | -0.611218687 | 0.189415975 | 10.18717736 | -1.296116809 | 0.243564834 | 0.732171799 | -4.581301945 |
| **A_51_P343429** | Slc25a37 | 0.203016177 | -0.182393443 | 0.588425798 | 12.48552373 | 1.295917213 | 0.243629142 | 0.732218501 | -4.581440019 |
| **A_55_P1955243** | Rps24 | -0.133333527 | -0.386483457 | 0.119816403 | 15.51947814 | -1.295778351 | 0.243673892 | 0.732218501 | -4.581536075 |
| **A_55_P1983758** | Sdhc | -0.078475099 | -0.227476806 | 0.070526608 | 10.79694772 | -1.295715914 | 0.243694015 | 0.732218501 | -4.581579264 |
| **A_55_P2069350** | Psph | 0.276543339 | -0.248600053 | 0.801686731 | 12.10870052 | 1.29555084 | 0.243747225 | 0.73226455 | -4.581693444 |
| **A_55_P1971639** | Abitram | 0.080779572 | -0.072689897 | 0.234249041 | 9.650656776 | 1.294937244 | 0.243945102 | 0.732745126 | -4.582117815 |
| **A_52_P153327** | 2-Sep | -0.067031571 | -0.194447845 | 0.060384702 | 8.478608183 | -1.294266359 | 0.244161618 | 0.733281529 | -4.582581723 |
| **A_55_P1986185** | NA | 0.159188007 | -0.143465313 | 0.461841327 | 14.27634185 | 1.293999372 | 0.244247831 | 0.733411823 | -4.582766315 |
| **A_55_P2004833** | NA | -0.199801772 | -0.579808345 | 0.180204802 | 11.75549435 | -1.29353263 | 0.244398612 | 0.733411823 | -4.583088983 |
| **A_51_P213592** | Nrde2 | -0.131703045 | -0.382238563 | 0.118832474 | 8.779478419 | -1.293289269 | 0.244477263 | 0.733411823 | -4.583257206 |
| **A_51_P206405** | Ptprz1 | -0.195614127 | -0.567728863 | 0.176500608 | 7.815359263 | -1.29327985 | 0.244480307 | 0.733411823 | -4.583263717 |
| **A_51_P505229** | Gtpbp8 | 0.139865329 | -0.126200985 | 0.405931642 | 9.319114014 | 1.293270409 | 0.244483359 | 0.733411823 | -4.583270243 |
| **A_51_P275101** | Chst11 | -0.2984228 | -0.866134919 | 0.269289318 | 9.916404993 | -1.293221748 | 0.244499089 | 0.733411823 | -4.583303878 |
| **A_55_P2017526** | Dand5 | 0.110470595 | -0.099686986 | 0.320628176 | 8.38597257 | 1.293215189 | 0.244501209 | 0.733411823 | -4.583308412 |
| **A_55_P2074836** | Mapk8ip1 | 0.155195357 | -0.140055624 | 0.450446338 | 10.51225317 | 1.293172732 | 0.244514934 | 0.733411823 | -4.583337758 |
| **A_55_P1993758** | Nsa2 | 0.109872417 | -0.099225682 | 0.318970517 | 11.20723149 | 1.292729789 | 0.244658164 | 0.733411823 | -4.583643901 |
| **A_55_P2003266** | Fam107a | -0.260778376 | -0.75709034 | 0.235533587 | 9.926738346 | -1.292665017 | 0.244679115 | 0.733411823 | -4.583688665 |
| **A_55_P1992470** | NA | -0.110630107 | -0.32118984 | 0.099929625 | 13.55895746 | -1.292609004 | 0.244697234 | 0.733411823 | -4.583727376 |
| **A_55_P1981259** | Fkbp14 | -0.119598591 | -0.347240324 | 0.108043143 | 9.904647765 | -1.292538062 | 0.244720184 | 0.733411823 | -4.583776403 |

| | | | | | | | | | |
|---|---|---|---|---|---|---|---|---|---|
| A_55_P2144120 | Bcl2l2 | -0.081536884 | -0.236746293 | 0.073672526 | 8.737844188 | -1.292424628 | 0.244756885 | 0.733411823 | -4.583854793 |
| A_55_P1982414 | NA | 0.057183246 | -0.051669212 | 0.166035705 | 8.609912151 | 1.292408269 | 0.244762178 | 0.733411823 | -4.583866098 |
| A_55_P1956302 | H2-Q4 | -0.502670753 | -1.459676319 | 0.454334813 | 13.13044622 | -1.292225529 | 0.244821313 | 0.733411823 | -4.583992376 |
| A_55_P2108837 | Tuba1c | -0.12099756 | -0.351400987 | 0.109405867 | 16.08173486 | -1.291983129 | 0.244899774 | 0.733411823 | -4.584159872 |
| A_55_P2034250 | Zscan26 | 0.086149593 | -0.077911535 | 0.250210722 | 10.51529667 | 1.291863711 | 0.244938435 | 0.733411823 | -4.584242384 |
| A_52_P330984 | Lin9 | -0.153799172 | -0.446691413 | 0.13909307 | 9.225359804 | -1.291859527 | 0.24493979 | 0.733411823 | -4.584245275 |
| A_55_P2149149 | NA | -0.114242306 | -0.331819392 | 0.103334781 | 8.435277476 | -1.291763369 | 0.244970925 | 0.733411823 | -4.584311713 |
| A_52_P491861 | Stag1 | -0.126239471 | -0.366705561 | 0.11422662 | 10.38584094 | -1.291547805 | 0.245040737 | 0.733411823 | -4.584460646 |
| A_55_P2083919 | Robo2 | 0.064957307 | -0.05877655 | 0.188691164 | 7.935217812 | 1.291542267 | 0.24504253 | 0.733411823 | -4.584464472 |
| A_52_P572197 | Pank3 | -0.120742199 | -0.350756998 | 0.109272601 | 8.597833913 | -1.291434735 | 0.245077362 | 0.733411823 | -4.584538762 |
| A_55_P2099105 | Rhobtb1 | 0.18873372 | -0.170868288 | 0.548335729 | 10.15608515 | 1.291209031 | 0.245150486 | 0.733411823 | -4.584694686 |
| A_55_P2176871 | NA | -0.074324923 | -0.215970721 | 0.067320875 | 15.59345274 | -1.290921799 | 0.245243572 | 0.733411823 | -4.5848931 |
| A_55_P2086811 | Sp140 | -0.090139832 | -0.261931072 | 0.081651407 | 8.97777068 | -1.290877099 | 0.245258061 | 0.733411823 | -4.584923976 |
| A_51_P120729 | Polr2f | -0.148368588 | -0.43113867 | 0.134401493 | 11.25327045 | -1.290855624 | 0.245265022 | 0.733411823 | -4.58493881 |
| A_51_P192800 | Grn | -0.215758945 | -0.626997544 | 0.195479654 | 12.68034809 | -1.29075585 | 0.245297366 | 0.733411823 | -4.585007726 |
| A_55_P2105436 | Foxp1 | -0.103798101 | -0.301662132 | 0.094065929 | 8.059245383 | -1.290600179 | 0.245347839 | 0.733411823 | -4.585115249 |
| A_55_P2143337 | Rpp21 | -0.122933865 | -0.357287169 | 0.111419439 | 12.35972945 | -1.290534495 | 0.245369139 | 0.733411823 | -4.585160616 |
| A_55_P2025611 | Psme2 | -0.106852363 | -0.31055228 | 0.096847555 | 11.44579409 | -1.290513167 | 0.245376055 | 0.733411823 | -4.585175346 |
| A_51_P347452 | Htatsf1 | -0.106454048 | -0.309397152 | 0.096489056 | 13.67956366 | -1.290497135 | 0.245381254 | 0.733411823 | -4.585186419 |
| A_52_P425706 | Rnf168 | 0.077066373 | -0.069874168 | 0.224006913 | 9.90953476 | 1.2903052 | 0.245443505 | 0.73346966 | -4.585318978 |
| A_51_P294515 | Gde1 | -0.146723109 | -0.434774361 | 0.141328144 | 9.820452699 | -1.320014901 | 0.245476498 | 0.73346966 | -4.54605063 |
| A_55_P2091206 | NA | -0.250861642 | -0.729355604 | 0.227632321 | 9.235122715 | -1.289813627 | 0.245603002 | 0.733491924 | -4.585658447 |
| A_55_P2101591 | Vta1 | -0.087828767 | -0.255356697 | 0.079699163 | 9.285238204 | -1.289789168 | 0.24561094 | 0.733491924 | -4.585675337 |
| A_51_P447189 | 1700030C10Rik | -0.365091658 | -1.061534746 | 0.33135143 | 9.576023485 | -1.28969032 | 0.245643025 | 0.733491924 | -4.585743593 |
| A_55_P2052578 | Bclaf3 | -0.197984246 | -0.575669981 | 0.179701489 | 12.41598278 | -1.289642139 | 0.245658665 | 0.733491924 | -4.585776862 |
| A_51_P158210 | Mcm2 | 0.11173163 | -0.101430251 | 0.32489351 | 8.925868819 | 1.289542779 | 0.245690921 | 0.733491924 | -4.585845468 |
| A_51_P488768 | Slc25a44 | 0.142824061 | -0.129669519 | 0.415317642 | 8.892458891 | 1.289478978 | 0.245711636 | 0.733491924 | -4.58588952 |
| A_55_P2260648 | 6330418K02Rik | 0.079540452 | -0.072231402 | 0.231312307 | 8.908614382 | 1.289335612 | 0.245758188 | 0.733517607 | -4.585988507 |
| A_51_P430563 | Mtpn | 0.103132126 | -0.09368927 | 0.299953521 | 9.461740883 | 1.289112518 | 0.245830645 | 0.73357011 | -4.586142533 |
| A_55_P2073814 | Gpatch8 | -0.063254511 | -0.183981631 | 0.05747261 | 8.360116241 | -1.289008614 | 0.245864398 | 0.73357011 | -4.586214266 |
| A_55_P1986252 | NA | -0.080351853 | -0.233720086 | 0.07301638 | 16.13990267 | -1.288930934 | 0.245889634 | 0.73357011 | -4.586267893 |
| A_55_P2181162 | NA | -0.180469271 | -0.525015427 | 0.164076886 | 14.07125036 | -1.288620442 | 0.24599053 | 0.733715537 | -4.586482232 |
| A_55_P1964233 | Tnks | -0.096037624 | -0.279399988 | 0.087324741 | 8.552702892 | -1.28854731 | 0.246014299 | 0.733715537 | -4.586532713 |
| A_55_P2183378 | Snx11 | 0.075192358 | -0.068409082 | 0.218793799 | 11.28211957 | 1.288202309 | 0.246126462 | 0.733936806 | -4.586770846 |
| A_55_P2125561 | Mrps36 | 0.11379927 | -0.103699936 | 0.331298475 | 11.91920465 | 1.287214616 | 0.246447818 | 0.734648318 | -4.587452456 |
| A_66_P103490 | Srsf1 | -0.118820874 | -0.345927131 | 0.108285382 | 9.38536829 | -1.287160813 | 0.246465334 | 0.734648318 | -4.587489581 |
| A_51_P432460 | Ppp1r14a | 0.217586691 | -0.198311434 | 0.633484817 | 9.479351353 | 1.287106633 | 0.246482974 | 0.734648318 | -4.587526964 |
| A_51_P151126 | Cd52 | -0.234406943 | -0.682554721 | 0.213740836 | 10.17543607 | -1.286821729 | 0.246575752 | 0.734648318 | -4.587723534 |
| A_52_P209311 | Snx12 | 0.268076378 | -0.285031893 | 0.821184648 | 11.33232614 | 1.363800683 | 0.246578164 | 0.734648318 | -4.51533975 |
| A_55_P2130855 | Nap1l1 | -0.131685098 | -0.383474633 | 0.120104438 | 10.39661185 | -1.286672792 | 0.246624265 | 0.734648318 | -4.587826288 |
| A_52_P171791 | Smg1 | -0.180644865 | -0.526077162 | 0.164787433 | 10.07373227 | -1.286565324 | 0.246659276 | 0.734648318 | -4.587900428 |
| A_51_P249215 | Ptger2 | -0.07925423 | -0.230822073 | 0.072313613 | 7.731860487 | -1.286425217 | 0.246704926 | 0.734648318 | -4.587997081 |
| A_51_P215438 | Prodh | -0.353576074 | -1.029830694 | 0.322678546 | 10.99021203 | -1.286298572 | 0.246746197 | 0.734648318 | -4.588084445 |
| A_51_P518488 | Atpif1 | 0.109490301 | -0.099922407 | 0.318903009 | 15.65872335 | 1.286298551 | 0.246746204 | 0.734648318 | -4.58808446 |
| A_55_P2114482 | Rpgr | 0.074511686 | -0.068022534 | 0.217045905 | 8.846135297 | 1.28609903 | 0.246811235 | 0.734648318 | -4.588222088 |
| A_55_P1979970 | NA | 0.110243017 | -0.100647276 | 0.32113331 | 9.453717866 | 1.286067203 | 0.24682161 | 0.734648318 | -4.588244042 |
| A_55_P1973360 | NA | -0.113081567 | -0.329421265 | 0.103258132 | 14.23175399 | -1.285952003 | 0.246859166 | 0.734648318 | -4.588323502 |
| A_55_P2170963 | Gspt1 | 0.178906891 | -0.163445751 | 0.521259533 | 12.36673916 | 1.285649393 | 0.246957845 | 0.734734776 | -4.588532217 |
| A_55_P2173098 | Commd1 | 0.127981956 | -0.116940885 | 0.372904798 | 10.56126226 | 1.28554895 | 0.246990606 | 0.734734776 | -4.588601491 |
| A_55_P2091201 | Thoc2 | -0.14921701 | -0.434804745 | 0.136370726 | 8.676607585 | -1.285428554 | 0.24702988 | 0.734734776 | -4.588684522 |

| | | | | | | | | | |
|---|---|---|---|---|---|---|---|---|---|
| **A_52_P231737** | Actl6a | -0.086705842 | -0.252658328 | 0.079246643 | 9.245738935 | -1.28538659 | 0.247043571 | 0.734734776 | -4.588713462 |
| **A_65_P09031** | Cdc42se2 | -0.119718018 | -0.348873821 | 0.109437785 | 9.243503834 | -1.285280213 | 0.247078278 | 0.734734776 | -4.588786822 |
| **A_52_P126146** | Riok1 | -0.123730902 | -0.360638127 | 0.113176323 | 9.40824515 | -1.284899137 | 0.247202648 | 0.734872932 | -4.589049601 |
| **A_55_P1962139** | Mff | 0.163844307 | -0.149889083 | 0.477577698 | 11.36569679 | 1.284812569 | 0.247230908 | 0.734872932 | -4.589109292 |
| **A_51_P259571** | Angptl6 | 0.126800681 | -0.116005165 | 0.369606526 | 9.227687059 | 1.284788409 | 0.247238796 | 0.734872932 | -4.58912595 |
| **A_55_P2123101** | Cbarp | 0.089388746 | -0.081821433 | 0.260598924 | 13.37908048 | 1.284465448 | 0.247344256 | 0.735073357 | -4.589348625 |
| **A_55_P2130535** | Dnmt3b | 0.198103449 | -0.181383204 | 0.577590102 | 9.902077064 | 1.284294712 | 0.247400024 | 0.735083092 | -4.589466335 |
| **A_55_P2066299** | Gpr137b | 0.182230104 | -0.166875222 | 0.53133543 | 11.40061456 | 1.284200432 | 0.247430824 | 0.735083092 | -4.589531331 |
| **A_55_P1981681** | Syt8 | -0.116843607 | -0.340701899 | 0.107014685 | 8.773807158 | -1.284106169 | 0.247461621 | 0.735083092 | -4.589596314 |
| **A_51_P145433** | Lin7c | 0.061970165 | -0.056786228 | 0.180726557 | 8.523550602 | 1.283792455 | 0.247564143 | 0.735274635 | -4.58981257 |
| **A_55_P2056293** | Sf3b4 | -0.078810943 | -0.229873904 | 0.072252017 | 8.362550665 | -1.28350539 | 0.247657989 | 0.735440356 | -4.590010437 |
| **A_55_P2171206** | Matn4 | 0.074413084 | -0.068269021 | 0.217095188 | 7.973016483 | 1.283065892 | 0.247801729 | 0.735652583 | -4.590313341 |
| **A_55_P2105271** | Pdlim7 | -0.210451407 | -0.613981581 | 0.193078768 | 12.08196935 | -1.283054134 | 0.247805576 | 0.735652583 | -4.590321444 |
| **A_51_P465582** | Hdhd3 | 0.194706668 | -0.178764452 | 0.568177788 | 8.885242293 | 1.282605105 | 0.247952511 | 0.735793135 | -4.590630875 |
| **A_55_P2143025** | Sema3c | 0.246528756 | -0.226479421 | 0.719536934 | 8.64571219 | 1.282236393 | 0.248073222 | 0.735793135 | -4.590884926 |
| **A_52_P623751** | Fem1b | -0.15234589 | -0.444654473 | 0.139962694 | 13.10167579 | -1.282207571 | 0.248082661 | 0.735793135 | -4.590904785 |
| **A_55_P1964053** | Golga4 | 0.118052584 | -0.108476821 | 0.344581989 | 12.71717732 | 1.282094615 | 0.248119653 | 0.735793135 | -4.590982608 |
| **A_55_P2040170** | Pmp22 | 0.167267698 | -0.153763241 | 0.488298638 | 12.74425513 | 1.281841577 | 0.248202538 | 0.735793135 | -4.591156934 |
| **A_55_P1960167** | Bcat2 | 0.10106557 | -0.092949032 | 0.295080173 | 8.766788674 | 1.281557142 | 0.248295737 | 0.735793135 | -4.591352875 |
| **A_52_P229997** | Cbx8 | -0.094353365 | -0.275483719 | 0.086776989 | 9.522843487 | -1.281549303 | 0.248298306 | 0.735793135 | -4.591358275 |
| **A_55_P2026400** | Capns1 | 0.08421347 | -0.077464719 | 0.245891659 | 12.38781662 | 1.281442984 | 0.248333151 | 0.735793135 | -4.591431511 |
| **A_65_P15295** | NA | -0.130104995 | -0.379929093 | 0.119719104 | 8.736714344 | -1.281235028 | 0.248401319 | 0.735793135 | -4.591574752 |
| **A_55_P2029051** | NA | 0.106954565 | -0.098417424 | 0.312326554 | 11.2934437 | 1.281230532 | 0.248402793 | 0.735793135 | -4.591577849 |
| **A_51_P290074** | Fabp7 | 0.085158026 | -0.078365363 | 0.248681415 | 7.872792263 | 1.281193927 | 0.248414794 | 0.735793135 | -4.591603062 |
| **A_55_P2007535** | Edf1 | -0.096884586 | -0.282929994 | 0.089160822 | 15.18862919 | -1.281164495 | 0.248424444 | 0.735793135 | -4.591623334 |
| **A_55_P2157838** | NA | -0.415734506 | -1.214067421 | 0.38259841 | 11.43711747 | -1.281153595 | 0.248428018 | 0.735793135 | -4.591630841 |
| **A_55_P2148935** | NA | -0.406880007 | -1.188303926 | 0.374543913 | 10.82190026 | -1.280999064 | 0.248478689 | 0.735793135 | -4.591737275 |
| **A_55_P2044364** | Mms22l | -0.20730238 | -0.605479182 | 0.190874422 | 14.25897161 | -1.280847713 | 0.248528326 | 0.735793135 | -4.591841513 |
| **A_55_P1978191** | C2cd4c | 0.11496157 | -0.105860834 | 0.335783973 | 8.213949911 | 1.280792325 | 0.248546493 | 0.735793135 | -4.591879658 |
| **A_55_P2186929** | NA | 0.252402435 | -0.232479651 | 0.737284521 | 10.55189927 | 1.280638529 | 0.248596945 | 0.735793135 | -4.591985574 |
| **A_52_P445387** | Clk4 | -0.195498658 | -0.571109886 | 0.18011257 | 10.45631384 | -1.280484656 | 0.248647431 | 0.735793135 | -4.592091538 |
| **A_55_P2003996** | H2-T24 | -0.0651392 | -0.190291156 | 0.060012755 | 7.925197997 | -1.280483383 | 0.248647848 | 0.735793135 | -4.592092414 |
| **A_51_P248122** | Bbc3 | -0.096177278 | -0.280969854 | 0.088615298 | 10.38515644 | -1.280433785 | 0.248664124 | 0.735793135 | -4.592126569 |
| **A_55_P1999082** | Clec14a | -0.075489895 | -0.220537373 | 0.069557583 | 8.239346356 | -1.280406297 | 0.248673144 | 0.735793135 | -4.592145497 |
| **A_55_P2153296** | Pycrl | 0.083158789 | -0.076630382 | 0.242947959 | 8.010019225 | 1.280353745 | 0.24869039 | 0.735793135 | -4.592181685 |
| **A_55_P2179030** | Ostf1 | -0.14112581 | -0.412343001 | 0.130091381 | 13.50799038 | -1.280142738 | 0.248759647 | 0.735885402 | -4.59232698 |
| **A_51_P459350** | Dstn | -0.162697817 | -0.475471787 | 0.150076153 | 12.34949957 | -1.279735701 | 0.248893293 | 0.735890426 | -4.592607231 |
| **A_55_P1971840** | Muc13 | 0.295521804 | -0.272609805 | 0.863653414 | 12.38724719 | 1.279704625 | 0.248903499 | 0.735890426 | -4.592628626 |
| **A_55_P2113774** | Yif1b | 0.105070814 | -0.096955242 | 0.307096871 | 10.07845664 | 1.279510558 | 0.248967244 | 0.735890426 | -4.592762231 |
| **A_66_P101631** | Zfp27 | 0.074165787 | -0.068437962 | 0.216769537 | 7.965341571 | 1.279504536 | 0.248969222 | 0.735890426 | -4.592766376 |
| **A_52_P207635** | Zfc3h1 | -0.156843574 | -0.458431836 | 0.144744689 | 9.462700247 | -1.279444517 | 0.24898894 | 0.735890426 | -4.592807695 |
| **A_52_P106463** | Jak1 | -0.092832109 | -0.271373285 | 0.085709068 | 12.49667331 | -1.279172121 | 0.249078444 | 0.735890426 | -4.592995208 |
| **A_52_P161237** | Cpa1 | 0.066554297 | -0.061455765 | 0.19456436 | 8.063871618 | 1.279090153 | 0.249105383 | 0.735890426 | -4.593051631 |
| **A_51_P143805** | Tmem42 | 0.138691732 | -0.128080996 | 0.40546446 | 10.75616594 | 1.279022852 | 0.249127504 | 0.735890426 | -4.593097956 |
| **A_55_P2073248** | Slc25a34 | 0.232604768 | -0.214815777 | 0.680025312 | 8.287709706 | 1.279003855 | 0.249133748 | 0.735890426 | -4.593111032 |
| **A_55_P2004876** | Srpx2 | 0.064508302 | -0.059597619 | 0.188614223 | 7.84507319 | 1.278769514 | 0.249210786 | 0.735890426 | -4.593272329 |
| **A_52_P574993** | Pappa2 | -0.161619761 | -0.472569858 | 0.149330335 | 8.987483005 | -1.278712553 | 0.249229515 | 0.735890426 | -4.593311533 |
| **A_52_P289256** | Arid1b | -0.181273939 | -0.530045016 | 0.167497139 | 11.07825895 | -1.278686471 | 0.249238091 | 0.735890426 | -4.593329484 |
| **A_55_P2240478** | NA | 0.085156223 | -0.078701589 | 0.249014034 | 8.309573075 | 1.278552017 | 0.249282306 | 0.735890426 | -4.593422022 |
| **A_55_P2058871** | Otud5 | -0.067283494 | -0.196761715 | 0.062194727 | 13.24517584 | -1.27844184 | 0.249318543 | 0.735890426 | -4.593497847 |

| | | | | | | | | | |
|---|---|---|---|---|---|---|---|---|---|
| **A_55_P2140646** | Creg1 | 0.170453284 | -0.157572435 | 0.498479002 | 17.62756487 | 1.278399641 | 0.249332423 | 0.735890426 | -4.593526888 |
| **A_55_P2129407** | NA | 0.076664147 | -0.070995044 | 0.224323337 | 8.178453649 | 1.277323746 | 0.249686542 | 0.736653684 | -4.594267196 |
| **A_51_P414653** | Plvap | -0.189507113 | -0.554519746 | 0.175505521 | 12.23710416 | -1.277281971 | 0.249700301 | 0.736653684 | -4.594295937 |
| **A_51_P228276** | Nipsnap2 | -0.104643249 | -0.306219536 | 0.096933039 | 9.201879231 | -1.277147136 | 0.249744714 | 0.736653684 | -4.594388696 |
| **A_52_P411331** | Vac14 | -0.079159836 | -0.231649567 | 0.073329894 | 8.128892379 | -1.27712514 | 0.24975196 | 0.736653684 | -4.594403828 |
| **A_51_P221625** | Stk24 | -0.119810526 | -0.350632597 | 0.111011546 | 11.10040417 | -1.276987962 | 0.249797153 | 0.736653684 | -4.594498195 |
| **A_66_P118165** | NA | 0.161344166 | -0.149539392 | 0.472227724 | 10.19892088 | 1.276805295 | 0.249857344 | 0.736653684 | -4.594623849 |
| **A_51_P155313** | Gsto1 | 0.166079083 | -0.153928231 | 0.486086397 | 12.99363088 | 1.27680389 | 0.249857807 | 0.736653684 | -4.594624816 |
| **A_66_P106098** | Kif3a | -0.071549772 | -0.209447212 | 0.066347668 | 9.260020811 | -1.276501167 | 0.249957586 | 0.736835471 | -4.594833039 |
| **A_51_P283292** | Gm14326 | -0.073536471 | -0.215299044 | 0.068226102 | 8.849110185 | -1.276175377 | 0.250065008 | 0.736842516 | -4.595057107 |
| **A_55_P2061025** | Hspa4 | -0.092235893 | -0.270052218 | 0.085580432 | 9.498892199 | -1.276137555 | 0.250077481 | 0.736842516 | -4.595083119 |
| **A_55_P2024858** | Fars2 | -0.154087151 | -0.451286474 | 0.143112173 | 11.37113945 | -1.275521465 | 0.250280743 | 0.736842516 | -4.59550678 |
| **A_55_P1959639** | NA | -0.19990622 | -0.585494008 | 0.185681568 | 13.52317271 | -1.27547573 | 0.250295838 | 0.736842516 | -4.595538228 |
| **A_55_P2052623** | Pgap2 | 0.111216512 | -0.103305084 | 0.325738107 | 10.72929268 | 1.275461674 | 0.250300478 | 0.736842516 | -4.595547893 |
| **A_66_P108567** | Xrcc6 | -0.133928239 | -0.392263464 | 0.124406985 | 10.27125185 | -1.275433133 | 0.250309898 | 0.736842516 | -4.595567517 |
| **A_55_P1955791** | Fopnl | 0.168732218 | -0.156751338 | 0.494215775 | 11.88653169 | 1.275375666 | 0.250328867 | 0.736842516 | -4.59560703 |
| **A_66_P114102** | Zfp36l2 | -0.110176531 | -0.322716418 | 0.102363356 | 12.31469275 | -1.27531601 | 0.25034856 | 0.736842516 | -4.595648047 |
| **A_52_P289213** | Notch2 | -0.12788451 | -0.374607425 | 0.118838405 | 11.02269698 | -1.275197991 | 0.250387523 | 0.736842516 | -4.59572919 |
| **A_52_P138806** | Dlgap3 | 0.055178841 | -0.051282764 | 0.161640446 | 7.860736686 | 1.275113155 | 0.250415535 | 0.736842516 | -4.595787516 |
| **A_55_P2071799** | Slbp | -0.057911093 | -0.169646988 | 0.053824802 | 8.218199662 | -1.275082284 | 0.250425729 | 0.736842516 | -4.595808741 |
| **A_52_P509906** | Ift172 | -0.082146424 | -0.240650603 | 0.076357754 | 8.535640462 | -1.275020337 | 0.250446185 | 0.736842516 | -4.595851329 |
| **A_51_P293069** | Flvcr1 | -0.106528539 | -0.312083368 | 0.099026289 | 10.20618361 | -1.274991977 | 0.25045555 | 0.736842516 | -4.595870827 |
| **A_55_P2114995** | NA | -0.173853522 | -0.509558448 | 0.161851403 | 15.46909823 | -1.274074854 | 0.250758588 | 0.737370527 | -4.596501249 |
| **A_52_P515094** | Tmem176a | -0.102960875 | -0.301782636 | 0.095860886 | 8.833727071 | -1.274023595 | 0.250775535 | 0.737370527 | -4.596536479 |
| **A_51_P158388** | Cdk19 | 0.11455769 | -0.106660792 | 0.335776172 | 9.917630898 | 1.274007553 | 0.250780839 | 0.737370527 | -4.596547505 |
| **A_51_P297963** | Recql | -0.113564992 | -0.332870037 | 0.105740053 | 8.578082117 | -1.273987054 | 0.250787617 | 0.737370527 | -4.596561594 |
| **A_52_P141338** | Acot10 | -0.155071694 | -0.454620784 | 0.144477395 | 8.819701174 | -1.273601881 | 0.250914996 | 0.737384651 | -4.596826299 |
| **A_52_P522379** | Map3k20 | -0.063782369 | -0.187012462 | 0.059447723 | 7.740711302 | -1.273365378 | 0.250993238 | 0.737384651 | -4.596988817 |
| **A_55_P1990134** | Cox18 | -0.173414487 | -0.508477832 | 0.161648858 | 9.630525942 | -1.273290849 | 0.251017899 | 0.737384651 | -4.597040029 |
| **A_55_P2018549** | Gcsh | 0.192729798 | -0.179770357 | 0.565229953 | 11.29794086 | 1.272892045 | 0.251149895 | 0.737384651 | -4.597314043 |
| **A_55_P2039603** | Drosha | -0.227913355 | -0.668431596 | 0.212604887 | 12.43660312 | -1.272843588 | 0.251165937 | 0.737384651 | -4.597347335 |
| **A_51_P265008** | Ppig | -0.08996066 | -0.263847045 | 0.083925726 | 10.99716088 | -1.272788419 | 0.251184203 | 0.737384651 | -4.597385238 |
| **A_55_P2061437** | Lsm5 | 0.137984053 | -0.128741794 | 0.4047099 | 10.38557172 | 1.272720257 | 0.251206772 | 0.737384651 | -4.597432067 |
| **A_55_P1969645** | Sdcbp | -0.114680989 | -0.336392952 | 0.107030974 | 11.31392573 | -1.272540071 | 0.251266442 | 0.737384651 | -4.597555853 |
| **A_55_P2413924** | Gm10404 | -0.067937357 | -0.199317296 | 0.063442581 | 9.280303927 | -1.272180568 | 0.251385532 | 0.737384651 | -4.597802809 |
| **A_55_P2041509** | 1110004F10Rik | -0.072733699 | -0.21339811 | 0.067930711 | 11.32467053 | -1.272098114 | 0.251412853 | 0.737384651 | -4.597859446 |
| **A_55_P1954643** | 2310081J21Rik | 0.074128245 | -0.069243258 | 0.217499748 | 9.250986408 | 1.272008592 | 0.251442519 | 0.737384651 | -4.597920935 |
| **A_51_P343252** | Cd151 | -0.08072083 | -0.236847925 | 0.075406265 | 15.35997484 | -1.271968929 | 0.251455664 | 0.737384651 | -4.597948179 |
| **A_66_P126492** | Brk1 | -0.140599456 | -0.412550867 | 0.131351954 | 12.29843124 | -1.271924956 | 0.251470237 | 0.737384651 | -4.597978381 |
| **A_51_P246924** | Tppp3 | 0.182061397 | -0.170107068 | 0.534229862 | 8.118078734 | 1.271851975 | 0.251494426 | 0.737384651 | -4.598028507 |
| **A_55_P2059372** | Ppip5k2 | 0.058093354 | -0.054282413 | 0.170469121 | 7.935265698 | 1.271812086 | 0.251507648 | 0.737384651 | -4.598055904 |
| **A_55_P1990127** | Dolpp1 | 0.164536368 | -0.15376393 | 0.482836666 | 11.49520374 | 1.271727388 | 0.251535725 | 0.737384651 | -4.598114076 |
| **A_51_P246224** | Tram2 | -0.243435148 | -0.714369288 | 0.227498991 | 11.22650424 | -1.271722256 | 0.251537426 | 0.737384651 | -4.5981176 |
| **A_55_P2025829** | NA | -0.184376988 | -0.541084016 | 0.17233004 | 15.70615404 | -1.271640095 | 0.251564664 | 0.737384651 | -4.598174028 |
| **A_52_P313856** | Slc25a26 | -0.055356755 | -0.162456992 | 0.051743482 | 8.234601042 | -1.271596594 | 0.251579087 | 0.737384651 | -4.598203903 |
| **A_55_P2166528** | Mrps36 | 0.117064434 | -0.109445376 | 0.343574245 | 11.57452696 | 1.271472927 | 0.251620093 | 0.737384651 | -4.598288833 |
| **A_55_P2082708** | Pwp2 | 0.089178506 | -0.08338714 | 0.261744153 | 10.35027223 | 1.271378944 | 0.25165126 | 0.737384651 | -4.598353375 |
| **A_55_P2046842** | Tmem253 | 0.239181572 | -0.223654049 | 0.702017193 | 8.557192196 | 1.271364567 | 0.251656028 | 0.737384651 | -4.598363248 |
| **A_52_P502141** | Hectd2 | -0.061408336 | -0.180252214 | 0.057435542 | 8.196166979 | -1.27121695 | 0.251704989 | 0.737384651 | -4.598464618 |
| **A_55_P1995152** | NA | -0.141460497 | -0.415231704 | 0.132310709 | 14.69566474 | -1.271207873 | 0.251708 | 0.737384651 | -4.598470852 |

| | | | | | | | | | |
|---|---|---|---|---|---|---|---|---|---|
| **A_52_P83557** | Arfgef2 | -0.075923263 | -0.222927907 | 0.071081382 | 8.856436367 | -1.27061205 | 0.251905708 | 0.737852012 | -4.598879962 |
| **A_55_P2008907** | Actg1 | -0.211959422 | -0.622496952 | 0.198578107 | 14.44136155 | -1.270190985 | 0.25204551 | 0.73791414 | -4.599169032 |
| **A_55_P2152467** | NA | -0.057457769 | -0.168750151 | 0.053834612 | 8.417164061 | -1.270142627 | 0.25206157 | 0.73791414 | -4.599202229 |
| **A_55_P1959818** | Tmem40 | -0.160723618 | -0.472105715 | 0.150658479 | 11.55627656 | -1.269858187 | 0.252156054 | 0.73791414 | -4.59939748 |
| **A_55_P2184385** | NA | 0.068433962 | -0.064155097 | 0.201023021 | 7.946642834 | 1.269793669 | 0.25217749 | 0.73791414 | -4.599441765 |
| **A_55_P2125564** | Lrrc58 | -0.074678831 | -0.219370776 | 0.070013114 | 11.72704702 | -1.269761984 | 0.252188017 | 0.73791414 | -4.599463513 |
| **A_51_P156631** | Glrx5 | -0.14478449 | -0.425342254 | 0.135773274 | 13.19718869 | -1.269605844 | 0.252239902 | 0.73791414 | -4.599570683 |
| **A_55_P2059640** | Abhd14b | 0.240740227 | -0.225772856 | 0.70725331 | 9.436186921 | 1.269562253 | 0.252254389 | 0.73791414 | -4.599600601 |
| **A_55_P1957154** | Mageb16 | -0.100993626 | -0.296719713 | 0.094732461 | 10.26686515 | -1.269446534 | 0.25229285 | 0.73791414 | -4.599680023 |
| **A_55_P2131438** | H2ac19 | -0.103701658 | -0.304692539 | 0.097289223 | 14.71394878 | -1.269341582 | 0.252327737 | 0.73791414 | -4.599752053 |
| **A_55_P2114118** | Cadm1 | -0.070214884 | -0.206306025 | 0.065876257 | 8.646569736 | -1.2693123 | 0.252337472 | 0.73791414 | -4.599772149 |
| **A_55_P2161865** | Pdlim7 | -0.056810866 | -0.166941117 | 0.053319385 | 7.976389988 | -1.269094449 | 0.252409904 | 0.73791414 | -4.599921654 |
| **A_52_P495553** | Zbtb10 | -0.121245799 | -0.356299889 | 0.113808292 | 10.17690144 | -1.269018778 | 0.252435068 | 0.73791414 | -4.599973582 |
| **A_55_P2041828** | Tubb3 | 0.216357022 | -0.203155767 | 0.635869811 | 12.39910479 | 1.268805207 | 0.252506101 | 0.73791414 | -4.600120136 |
| **A_55_P1993933** | NA | -0.217561257 | -0.639514282 | 0.204391768 | 11.03741606 | -1.268488743 | 0.252611388 | 0.73791414 | -4.600337278 |
| **A_51_P408989** | L3hypdh | -0.082067404 | -0.241241273 | 0.077106465 | 8.533436545 | -1.268434617 | 0.2526294 | 0.73791414 | -4.600374414 |
| **A_51_P206346** | Reep3 | -0.22690479 | -0.667028462 | 0.213218881 | 11.76290963 | -1.268347042 | 0.252658545 | 0.73791414 | -4.600434499 |
| **A_51_P292550** | Chdh | -0.190481954 | -0.56006171 | 0.179097803 | 9.095162571 | -1.267987069 | 0.252778375 | 0.73791414 | -4.600681459 |
| **A_55_P2011390** | Tead1 | -0.079163427 | -0.232761654 | 0.074434799 | 8.355281454 | -1.267965826 | 0.252785448 | 0.73791414 | -4.600696031 |
| **A_55_P2120141** | NA | -0.497174026 | -1.461830077 | 0.467482026 | 11.17522388 | -1.267958681 | 0.252787827 | 0.73791414 | -4.600700933 |
| **A_55_P2113155** | Scnm1 | -0.116215175 | -0.341708754 | 0.109278403 | 9.460127641 | -1.267937439 | 0.2527949 | 0.73791414 | -4.600715506 |
| **A_51_P350214** | Amz2 | 0.107599814 | -0.101187286 | 0.316386914 | 10.22287468 | 1.267876702 | 0.252815124 | 0.73791414 | -4.60075717 |
| **A_55_P2045414** | Hipk4 | 0.098192879 | -0.092349249 | 0.288735007 | 9.281701819 | 1.267821596 | 0.252833475 | 0.73791414 | -4.600794972 |
| **A_55_P2060072** | Clk2 | -0.131109835 | -0.385566534 | 0.123346864 | 9.952321045 | -1.267624265 | 0.252899198 | 0.73791414 | -4.600930332 |
| **A_55_P1993594** | Dcp1b | 0.073986852 | -0.06961118 | 0.217584885 | 9.206795901 | 1.267579558 | 0.25291409 | 0.73791414 | -4.600960998 |
| **A_65_P06872** | Fam210a | -0.06695732 | -0.196915535 | 0.063000896 | 8.211829629 | -1.267545288 | 0.252925506 | 0.73791414 | -4.600984504 |
| **A_52_P18922** | Chid1 | 0.084303962 | -0.079324768 | 0.247932691 | 8.959857949 | 1.267528079 | 0.252931239 | 0.73791414 | -4.600996308 |
| **A_55_P1989765** | NA | -0.243812739 | -0.71706769 | 0.229442213 | 8.301640032 | -1.267448697 | 0.252957685 | 0.73791414 | -4.601050757 |
| **A_51_P375783** | Prap1 | 0.674684786 | -0.63528694 | 1.984656511 | 12.12186024 | 1.267092026 | 0.25307654 | 0.738048667 | -4.601295383 |
| **A_51_P103509** | Cyb561d2 | 0.131600923 | -0.123968196 | 0.387170042 | 11.07536494 | 1.266834036 | 0.253162542 | 0.738048667 | -4.60147231 |
| **A_51_P317673** | Zfp764 | 0.070501114 | -0.066422618 | 0.207424846 | 9.264736156 | 1.26673688 | 0.253194936 | 0.738048667 | -4.601538936 |
| **A_51_P403049** | Heatr5b | -0.064064024 | -0.188499705 | 0.060371658 | 8.370948238 | -1.266596969 | 0.253241591 | 0.738048667 | -4.601634877 |
| **A_55_P2015989** | Rpl22 | 0.069593163 | -0.065587897 | 0.204774223 | 15.26210574 | 1.266542867 | 0.253259635 | 0.738048667 | -4.601671975 |
| **A_55_P1998299** | Itgb4 | 0.213567301 | -0.20128148 | 0.628416083 | 9.952101163 | 1.266525985 | 0.253265265 | 0.738048667 | -4.601683551 |
| **A_52_P260105** | Man1a | 0.092381863 | -0.087069912 | 0.271833637 | 8.569648341 | 1.266508535 | 0.253271085 | 0.738048667 | -4.601695517 |
| **A_52_P485376** | Ube2k | -0.10947019 | -0.322157724 | 0.103217345 | 12.67807298 | -1.266260315 | 0.253353886 | 0.738178665 | -4.601865714 |
| **A_55_P2075048** | Atic | -0.095392041 | -0.280759702 | 0.089975619 | 9.739270796 | -1.266039739 | 0.253427485 | 0.73818461 | -4.602016944 |
| **A_55_P2046802** | Mlkl | 0.148291967 | -0.139883179 | 0.436467112 | 8.732905929 | 1.265989974 | 0.253444093 | 0.73818461 | -4.602051063 |
| **A_55_P1968568** | Rapgef5 | -0.09785949 | -0.288049519 | 0.092330539 | 8.446637866 | -1.26585618 | 0.253488747 | 0.73818461 | -4.602142788 |
| **A_51_P337856** | Ubl7 | 0.100648164 | -0.094970888 | 0.296267217 | 9.650981446 | 1.265796439 | 0.253508689 | 0.73818461 | -4.602183743 |
| **A_55_P2129348** | Pilra | 0.074072423 | -0.069932345 | 0.21807719 | 7.906570677 | 1.265461225 | 0.253620607 | 0.738293434 | -4.602413535 |
| **A_52_P458870** | Slc25a16 | -0.09550641 | -0.281192518 | 0.090179698 | 8.668062428 | -1.265383798 | 0.253646464 | 0.738293434 | -4.602466609 |
| **A_52_P325443** | Arhgap29 | -0.118876967 | -0.350025453 | 0.112271518 | 10.91058441 | -1.26524849 | 0.253691656 | 0.738293434 | -4.602559355 |
| **A_55_P2000304** | Terf1 | -0.101337426 | -0.298400588 | 0.095725735 | 9.111185796 | -1.265125434 | 0.253732762 | 0.738293434 | -4.602643699 |
| **A_55_P2166399** | Stk11 | -0.122057683 | -0.359416351 | 0.115300985 | 9.751982995 | -1.265112623 | 0.253737042 | 0.738293434 | -4.60265248 |
| **A_55_P2019585** | Klhl17 | -0.13320296 | -0.392291327 | 0.125885407 | 8.855177684 | -1.26483864 | 0.253828586 | 0.738391191 | -4.602840259 |
| **A_55_P2021530** | Clic4 | 0.183520289 | -0.173544185 | 0.540584762 | 8.944714145 | 1.264464396 | 0.253953678 | 0.738391191 | -4.603096728 |
| **A_55_P2134790** | Qsox1 | 0.128698372 | -0.121703631 | 0.379100374 | 13.2380633 | 1.264458028 | 0.253955807 | 0.738391191 | -4.603101092 |
| **A_55_P2174895** | Mrpl43 | -0.097730419 | -0.287882261 | 0.092421424 | 12.25989746 | -1.264440464 | 0.253961679 | 0.738391191 | -4.603113127 |
| **A_55_P2050997** | Serinc4 | 0.06568444 | -0.062127096 | 0.193495976 | 8.863729545 | 1.264333398 | 0.253997477 | 0.738391191 | -4.603186493 |

| | | | | | | | | | |
|---|---|---|---|---|---|---|---|---|---|
| **A_55_P1964213** | Tprkb | -0.148657114 | -0.437945067 | 0.140630838 | 9.544594121 | -1.264225408 | 0.254033588 | 0.738391191 | -4.60326049 |
| **A_51_P418908** | Larp1 | -0.084572515 | -0.249153053 | 0.080008022 | 9.263196878 | -1.264212074 | 0.254038047 | 0.738391191 | -4.603269627 |
| **A_66_P106212** | Secisbp2 | -0.081068188 | -0.238865858 | 0.076729481 | 9.167781437 | -1.26391843 | 0.254136265 | 0.738442905 | -4.603470824 |
| **A_55_P2176565** | B3galt5 | -0.075639663 | -0.222881982 | 0.071602656 | 8.753318365 | -1.263822548 | 0.254168343 | 0.738442905 | -4.603536515 |
| **A_55_P2030428** | Sirt1 | -0.058777797 | -0.173196897 | 0.055641304 | 10.69061251 | -1.263816247 | 0.254170451 | 0.738442905 | -4.603540832 |
| **A_55_P2142008** | 4933434E20Rik | -0.202506219 | -0.596785785 | 0.191773347 | 8.300300943 | -1.263581498 | 0.254249003 | 0.738490009 | -4.603701657 |
| **A_55_P2037538** | NA | -0.122847915 | -0.362040739 | 0.11634491 | 9.32056441 | -1.263539446 | 0.254263077 | 0.738490009 | -4.603730465 |
| **A_55_P1974134** | Mrpl58 | 0.079852117 | -0.075655513 | 0.235359747 | 11.71202019 | 1.26329243 | 0.254345761 | 0.738619172 | -4.60389968 |
| **A_55_P1984118** | Gfra4 | 0.143557732 | -0.136081997 | 0.42319746 | 8.631334288 | 1.262981179 | 0.25444998 | 0.738810826 | -4.604112878 |
| **A_66_P109175** | Farsb | -0.07316319 | -0.215701442 | 0.069375062 | 10.8911612 | -1.262787772 | 0.25451476 | 0.738887923 | -4.604245347 |
| **A_55_P2063146** | NA | 0.112985781 | -0.107162543 | 0.333134105 | 8.690030471 | 1.262634267 | 0.254566185 | 0.738926233 | -4.604350479 |
| **A_51_P267346** | Topors | 0.07834251 | -0.074319433 | 0.231004452 | 10.84809606 | 1.262512999 | 0.254606817 | 0.738933207 | -4.60443353 |
| **A_55_P2020552** | Iscu | -0.161183142 | -0.475328868 | 0.152962584 | 11.59101501 | -1.262285594 | 0.254683026 | 0.738948172 | -4.60458926 |
| **A_55_P2005655** | Ntsr1 | -0.215857051 | -0.636567528 | 0.204853425 | 8.549528463 | -1.262269463 | 0.254688433 | 0.738948172 | -4.604600306 |
| **A_55_P1958165** | Ms4a7 | -0.228851052 | -0.675049208 | 0.217347104 | 9.652948502 | -1.26181099 | 0.254842144 | 0.739283176 | -4.604914239 |
| **A_55_P2177712** | NA | 0.056191765 | -0.053381794 | 0.165765324 | 7.993645777 | 1.261641796 | 0.25489889 | 0.739336831 | -4.605030081 |
| **A_55_P2060036** | Ulk2 | -0.163061794 | -0.481063757 | 0.154940169 | 10.07698164 | -1.261512557 | 0.254942243 | 0.73935163 | -4.605118564 |
| **A_51_P196526** | Bscl2 | -0.097986416 | -0.28913933 | 0.093166499 | 8.165937917 | -1.261113315 | 0.255076209 | 0.739510386 | -4.605391876 |
| **A_52_P13815** | Laptm5 | -0.076248815 | -0.225007568 | 0.072509937 | 7.949648834 | -1.261013498 | 0.255109712 | 0.739510386 | -4.605460204 |
| **A_51_P302651** | Topbp1 | -0.083079452 | -0.245165321 | 0.079006417 | 9.021667676 | -1.26100739 | 0.255111762 | 0.739510386 | -4.605464384 |
| **A_66_P128079** | Krt8 | 0.054192035 | -0.051556716 | 0.159940786 | 16.70036959 | 1.26075116 | 0.255197784 | 0.739541209 | -4.605639771 |
| **A_51_P515605** | Col3a1 | 0.527716278 | -0.502430181 | 1.557862736 | 13.21843833 | 1.260290549 | 0.255352484 | 0.739541209 | -4.605955016 |
| **A_55_P2142873** | Mink1 | 0.084001297 | -0.079976423 | 0.247979017 | 10.08460877 | 1.260289474 | 0.255352845 | 0.739541209 | -4.605955752 |
| **A_55_P2081360** | Btbd1 | 0.096626274 | -0.092001834 | 0.285254382 | 10.9349557 | 1.260253718 | 0.255364858 | 0.739541209 | -4.605980222 |
| **A_55_P1958230** | Dhodh | 0.071576691 | -0.068172493 | 0.211325874 | 9.127588727 | 1.260060794 | 0.255429681 | 0.739541209 | -4.606112245 |
| **A_52_P279152** | Vamp8 | -0.148088602 | -0.437265852 | 0.141088649 | 12.60345043 | -1.259872719 | 0.255492888 | 0.739541209 | -4.606240943 |
| **A_66_P114054** | Sco2 | -0.099449678 | -0.293654161 | 0.094754806 | 10.43114059 | -1.259833901 | 0.255505936 | 0.739541209 | -4.606267504 |
| **A_51_P300572** | Slc30a1 | -0.126086692 | -0.37230862 | 0.120135236 | 10.92134505 | -1.259829249 | 0.255507499 | 0.739541209 | -4.606270688 |
| **A_55_P2148478** | Klhl25 | -0.11611038 | -0.342851351 | 0.11063059 | 11.42528754 | -1.25982496 | 0.255508941 | 0.739541209 | -4.606273622 |
| **A_52_P141161** | Hnrnpf | -0.114873881 | -0.339224884 | 0.109477122 | 9.115341368 | -1.259686402 | 0.255555518 | 0.739541209 | -4.60636843 |
| **A_55_P2149873** | Mapk1ip1l | 0.091212204 | -0.086929924 | 0.269354332 | 9.113733405 | 1.259666204 | 0.255562309 | 0.739541209 | -4.60638225 |
| **A_55_P1953708** | Cry2 | 0.058931812 | -0.056174278 | 0.174037902 | 8.311173458 | 1.259565204 | 0.255596266 | 0.739541209 | -4.606451355 |
| **A_55_P2072841** | NA | 0.136832378 | -0.130444569 | 0.404109325 | 9.144307849 | 1.259495259 | 0.255619785 | 0.739541209 | -4.606499211 |
| **A_66_P101423** | Ogfod3 | 0.115518489 | -0.110206921 | 0.3412439 | 10.574387 | 1.259041984 | 0.255772243 | 0.739619508 | -4.606809313 |
| **A_55_P2185489** | Zfp865 | 0.094256063 | -0.089935867 | 0.278447992 | 8.865299097 | 1.258948197 | 0.255803798 | 0.739619508 | -4.606873471 |
| **A_55_P2042729** | NA | -0.084173789 | -0.2486688 | 0.080321223 | 12.00584273 | -1.258906009 | 0.255817993 | 0.739619508 | -4.60690233 |
| **A_55_P2177386** | Cradd | -0.090614018 | -0.267735489 | 0.086507453 | 9.381565864 | -1.258616265 | 0.255915506 | 0.739619508 | -4.607100522 |
| **A_52_P225570** | Parva | -0.110915452 | -0.327730333 | 0.105899429 | 10.3148225 | -1.258554794 | 0.255936198 | 0.739619508 | -4.607142568 |
| **A_52_P272811** | Cpsf1 | -0.072527593 | -0.214303263 | 0.069248076 | 8.846358692 | -1.258550724 | 0.255937568 | 0.739619508 | -4.607145352 |
| **A_55_P2023732** | Limch1 | -0.162427672 | -0.479998816 | 0.155143472 | 10.21379323 | -1.258311453 | 0.256018124 | 0.739619508 | -4.607309002 |
| **A_52_P308465** | Plxnb1 | 0.067944601 | -0.064917125 | 0.200806328 | 8.401737678 | 1.258126254 | 0.256080491 | 0.739619508 | -4.607435661 |
| **A_52_P382914** | Trip6 | 0.089579906 | -0.085603791 | 0.264763602 | 10.85746456 | 1.258015749 | 0.256117711 | 0.739619508 | -4.607511233 |
| **A_51_P366227** | Zfp799 | -0.067224551 | -0.198710814 | 0.064261712 | 7.849605512 | -1.257814779 | 0.256185413 | 0.739619508 | -4.607648664 |
| **A_55_P1977498** | Htr2b | -0.416212675 | -1.230317997 | 0.397892648 | 12.40873664 | -1.257777642 | 0.256197926 | 0.739619508 | -4.60767406 |
| **A_55_P2018002** | Abcg4 | 0.062665601 | -0.059921507 | 0.185252709 | 8.001435054 | 1.257631981 | 0.256247007 | 0.739619508 | -4.607773662 |
| **A_51_P502764** | Hdac10 | -0.115999409 | -0.342919426 | 0.110920608 | 8.816874006 | -1.257627812 | 0.256248412 | 0.739619508 | -4.607776512 |
| **A_55_P2104689** | NA | -0.137981249 | -0.407910941 | 0.131948443 | 8.156529844 | -1.257588555 | 0.256261642 | 0.739619508 | -4.607803356 |
| **A_51_P204845** | Celf1 | -0.159454274 | -0.471392399 | 0.152483851 | 11.36040421 | -1.257583614 | 0.256263307 | 0.739619508 | -4.607806734 |
| **A_55_P2045928** | NA | -0.275272771 | -0.813843813 | 0.263298271 | 11.96186717 | -1.257446219 | 0.256309613 | 0.739619508 | -4.607900678 |
| **A_51_P413461** | Parg | -0.07344243 | -0.217134294 | 0.070249433 | 10.79092017 | -1.257430605 | 0.256314876 | 0.739619508 | -4.607911355 |

| | | | | | | | | | |
|---|---|---|---|---|---|---|---|---|---|
| **A_55_P2113598** | Ikzf3 | 0.060292915 | -0.057678327 | 0.178264158 | 7.807036043 | 1.257358997 | 0.256339014 | 0.739619508 | -4.607960315 |
| **A_55_P1953683** | Nsmaf | -0.111048529 | -0.328383122 | 0.106286065 | 11.30782896 | -1.257051621 | 0.256442647 | 0.739619508 | -4.608170464 |
| **A_55_P2006525** | Adamtsl4 | 0.18035765 | -0.172655237 | 0.533370537 | 9.11964731 | 1.256936018 | 0.256481632 | 0.739619508 | -4.608249494 |
| **A_52_P288251** | Tmem204 | -0.076955265 | -0.227585743 | 0.073675213 | 8.545623016 | -1.256882408 | 0.256499713 | 0.739619508 | -4.608286144 |
| **A_55_P2036907** | Mycbpap | 0.085295237 | -0.081669953 | 0.252260427 | 8.708625456 | 1.256805314 | 0.256525717 | 0.739619508 | -4.608338846 |
| **A_55_P2131190** | NA | -0.13069023 | -0.386516729 | 0.125136269 | 8.028465856 | -1.256801694 | 0.256526938 | 0.739619508 | -4.608341321 |
| **A_52_P195018** | Arap3 | 0.099603134 | -0.095390508 | 0.294596776 | 10.63466721 | 1.256671359 | 0.256570905 | 0.739635945 | -4.608430415 |
| **A_52_P694988** | Zfp933 | -0.09168372 | -0.271235689 | 0.087868249 | 8.550436199 | -1.256235951 | 0.256717832 | 0.739851664 | -4.608728027 |
| **A_52_P466090** | Samhd1 | -0.109242333 | -0.323183065 | 0.104698399 | 9.417591082 | -1.256222748 | 0.256722288 | 0.739851664 | -4.60873705 |
| **A_55_P2010931** | Cnn2 | 0.078978651 | -0.075736529 | 0.23369383 | 10.17397379 | 1.255873645 | 0.256840148 | 0.739955425 | -4.608975639 |
| **A_51_P136460** | Zmat2 | -0.086774601 | -0.256813697 | 0.083264495 | 12.29911614 | -1.255489257 | 0.256969975 | 0.739955425 | -4.609238312 |
| **A_51_P254583** | Foxred1 | -0.096552234 | -0.285763429 | 0.09265896 | 9.577617793 | -1.25540734 | 0.25699765 | 0.739955425 | -4.609294287 |
| **A_52_P517224** | Trim63 | 0.096965095 | -0.093055239 | 0.286985429 | 8.118043448 | 1.255406902 | 0.256997798 | 0.739955425 | -4.609294586 |
| **A_52_P657240** | Cdc40 | -0.061753241 | -0.18278171 | 0.059275229 | 8.525563998 | -1.255282206 | 0.25703993 | 0.739955425 | -4.609379788 |
| **A_55_P2007496** | NA | -0.093529237 | -0.276851853 | 0.089793379 | 8.48911841 | -1.255164096 | 0.257079843 | 0.739955425 | -4.609460487 |
| **A_52_P87210** | Dcun1d5 | -0.074403939 | -0.220242617 | 0.071434739 | 11.62196303 | -1.25514058 | 0.25708779 | 0.739955425 | -4.609476554 |
| **A_55_P2006911** | Pik3c3 | -0.078721728 | -0.233053052 | 0.075609597 | 8.36417037 | -1.254901669 | 0.257168544 | 0.739955425 | -4.60963978 |
| **A_51_P396818** | Rex1bd | -0.123160709 | -0.364616624 | 0.118295205 | 9.729169757 | -1.254883753 | 0.257174601 | 0.739955425 | -4.60965202 |
| **A_51_P160664** | Cox7b | 0.245702964 | -0.236030712 | 0.72743664 | 13.85509833 | 1.25479434 | 0.257204829 | 0.739955425 | -4.609713104 |
| **A_51_P477676** | Mcph1 | -0.070679094 | -0.209269025 | 0.067910836 | 8.260100295 | -1.254666958 | 0.2572479 | 0.739955425 | -4.609800124 |
| **A_55_P2136902** | Cisd3 | -0.107305811 | -0.317717906 | 0.103106284 | 11.78386546 | -1.254647438 | 0.257254501 | 0.739955425 | -4.609813459 |
| **A_51_P430929** | Fam20a | -0.205550168 | -0.608758874 | 0.197658537 | 8.99224572 | -1.254171998 | 0.257415317 | 0.739955425 | -4.61013822 |
| **A_51_P281593** | Plpp5 | -0.148153721 | -0.438780006 | 0.142472564 | 11.51082374 | -1.254142357 | 0.257425346 | 0.739955425 | -4.610158465 |
| **A_55_P2117333** | NA | -0.109805157 | -0.325207654 | 0.105597341 | 10.53347122 | -1.254125981 | 0.257430887 | 0.739955425 | -4.610169651 |
| **A_55_P2073099** | Trex1 | -0.116492436 | -0.345034812 | 0.112049939 | 10.92436881 | -1.254007559 | 0.257470959 | 0.739955425 | -4.610250533 |
| **A_55_P1982249** | Phactr2 | 0.081770409 | -0.078663846 | 0.242204664 | 9.349875041 | 1.253915381 | 0.257502154 | 0.739955425 | -4.610313488 |
| **A_55_P1982593** | Rad51d | -0.053443866 | -0.158311599 | 0.051423867 | 8.13729008 | -1.253791001 | 0.257544252 | 0.739955425 | -4.610398434 |
| **A_55_P1956898** | NA | 0.109232462 | -0.105106611 | 0.323571535 | 11.65686102 | 1.253774807 | 0.257549734 | 0.739955425 | -4.610409493 |
| **A_55_P2090880** | Stk40 | -0.12107218 | -0.35867963 | 0.116535271 | 12.24427525 | -1.253584084 | 0.2576143 | 0.739955425 | -4.610539741 |
| **A_55_P1953508** | Mob3c | 0.097488339 | -0.093848431 | 0.28882511 | 9.488054708 | 1.253497285 | 0.257643689 | 0.739955425 | -4.610599014 |
| **A_52_P475040** | Etaa1 | -0.065048464 | -0.192731344 | 0.062634416 | 8.458509233 | -1.253353383 | 0.257692419 | 0.739955425 | -4.610697279 |
| **A_55_P2030030** | Adssl1 | -0.223706099 | -0.662827401 | 0.215415203 | 11.85342417 | -1.253321477 | 0.257703224 | 0.739955425 | -4.610719065 |
| **A_55_P2075213** | Kbtbd11 | 0.093801371 | -0.09033111 | 0.277933853 | 8.362052564 | 1.253279525 | 0.257717433 | 0.739955425 | -4.610747711 |
| **A_55_P1997741** | Slc9a8 | 0.08319871 | -0.080126213 | 0.246523633 | 11.66502153 | 1.253237249 | 0.257731752 | 0.739955425 | -4.610776578 |
| **A_55_P1959595** | Fmo5 | 0.240770181 | -0.231975163 | 0.713515525 | 9.566829805 | 1.252981337 | 0.257818444 | 0.739955425 | -4.610951312 |
| **A_55_P2015299** | NA | -0.101858258 | -0.301859721 | 0.098143204 | 15.68363432 | -1.252945666 | 0.25783053 | 0.739955425 | -4.610975667 |
| **A_55_P2093696** | NA | 0.070128869 | -0.06759676 | 0.207854499 | 8.081520809 | 1.252711992 | 0.257909714 | 0.739955425 | -4.611135202 |
| **A_51_P462814** | Sars2 | -0.082740601 | -0.24524185 | 0.079760648 | 8.645792764 | -1.252654008 | 0.257929366 | 0.739955425 | -4.611174787 |
| **A_66_P134412** | Mettl25 | 0.119452316 | -0.115152951 | 0.354057582 | 10.09967182 | 1.252639146 | 0.257934403 | 0.739955425 | -4.611184933 |
| **A_55_P2070576** | Ccl27a | 0.108410288 | -0.10451983 | 0.321340406 | 8.574530177 | 1.252571614 | 0.257957293 | 0.739955425 | -4.611231035 |
| **A_55_P2430221** | Shld2 | -0.078364635 | -0.232291115 | 0.075561844 | 8.542551236 | -1.252494833 | 0.257983321 | 0.739955425 | -4.611283451 |
| **A_51_P375406** | Rprd1a | -0.146898599 | -0.435566897 | 0.141769698 | 9.791482524 | -1.251952141 | 0.25816735 | 0.740357795 | -4.611653889 |
| **A_55_P1988754** | H4f16 | -0.333116412 | -0.987769274 | 0.321536451 | 10.61147145 | -1.251855269 | 0.258200212 | 0.740357795 | -4.611720006 |
| **A_55_P2003494** | NA | -0.139624613 | -0.414061064 | 0.134811838 | 13.52847249 | -1.251668577 | 0.258263554 | 0.740393346 | -4.611847421 |
| **A_55_P2078266** | H3f3a | -0.132393214 | -0.392631855 | 0.127845427 | 11.84094861 | -1.251592941 | 0.25828922 | 0.740393346 | -4.61189904 |
| **A_51_P421804** | Timm9 | -0.118805293 | -0.352363959 | 0.114753373 | 9.629124948 | -1.251436928 | 0.258342168 | 0.740435314 | -4.612005509 |
| **A_55_P2019814** | Rnf20 | -0.120852468 | -0.35845771 | 0.116752774 | 11.42150253 | -1.251320816 | 0.25838158 | 0.740438483 | -4.612084744 |
| **A_55_P2060303** | Mras | 0.109147786 | -0.105487337 | 0.323782909 | 10.57622013 | 1.251074872 | 0.25846508 | 0.740567971 | -4.612252568 |
| **A_55_P2299524** | Scube1 | -0.198144773 | -0.587900806 | 0.191611261 | 8.023014298 | -1.250716661 | 0.258586738 | 0.740590742 | -4.612496975 |
| **A_55_P1996223** | Sf3b4 | 0.152815363 | -0.147792309 | 0.453423035 | 11.51331939 | 1.25065096 | 0.258609057 | 0.740590742 | -4.6125418 |

| | | | | | | | | | |
|---|---|---|---|---|---|---|---|---|---|
| **A_55_P2024260** | Vps33b | 0.101049483 | -0.097743393 | 0.299842359 | 9.307715888 | 1.25055398 | 0.258642005 | 0.740590742 | -4.612607962 |
| **A_51_P268843** | Rasip1 | -0.075028456 | -0.222636167 | 0.072579256 | 9.357775269 | -1.250506991 | 0.25865797 | 0.740590742 | -4.612640019 |
| **A_55_P2112687** | Sipa1l1 | 0.09906038 | -0.095829733 | 0.293950494 | 10.1266243 | 1.250487472 | 0.258664602 | 0.740590742 | -4.612653335 |
| **A_55_P2023136** | Rnf214 | 0.116740485 | -0.113028501 | 0.346509472 | 8.727259544 | 1.249969021 | 0.258840817 | 0.740866307 | -4.613006997 |
| **A_55_P1987839** | Erbin | -0.101954655 | -0.302625487 | 0.098716178 | 10.99707209 | -1.249948065 | 0.258847942 | 0.740866307 | -4.613021291 |
| **A_55_P2012399** | NA | 0.194922581 | -0.188888056 | 0.578733217 | 12.49929668 | 1.249436847 | 0.259021806 | 0.740866307 | -4.613369957 |
| **A_51_P210877** | Pcnp | -0.080281497 | -0.238360317 | 0.077797324 | 11.51749882 | -1.249427142 | 0.259025108 | 0.740866307 | -4.613376576 |
| **A_55_P2266295** | Fzd3 | 0.074378668 | -0.07208056 | 0.220837895 | 8.43963532 | 1.249398036 | 0.25903501 | 0.740866307 | -4.613396425 |
| **A_55_P2429432** | 2610301B20Rik | -0.077454325 | -0.229989602 | 0.075080951 | 8.733244006 | -1.249236153 | 0.25909009 | 0.740866307 | -4.613506821 |
| **A_55_P2137431** | Rtel1 | -0.113582686 | -0.337294722 | 0.11012935 | 9.265835779 | -1.249084867 | 0.259141574 | 0.740866307 | -4.613609985 |
| **A_51_P359262** | Sec11c | -0.129523278 | -0.384670843 | 0.125624286 | 11.34145643 | -1.248893955 | 0.259206555 | 0.740866307 | -4.613740163 |
| **A_52_P614472** | Rpusd1 | -0.07236984 | -0.214942748 | 0.070203067 | 8.241217302 | -1.248791038 | 0.259241591 | 0.740866307 | -4.613810336 |
| **A_51_P428134** | Lrig3 | -0.100307921 | -0.297922433 | 0.097306592 | 9.400329381 | -1.248778909 | 0.259245721 | 0.740866307 | -4.613818606 |
| **A_51_P336161** | Man2b1 | -0.217777476 | -0.646836594 | 0.211281642 | 11.17402255 | -1.24871977 | 0.259265855 | 0.740866307 | -4.613858928 |
| **A_55_P2084741** | Zkscan6 | 0.083480572 | -0.081002417 | 0.247963561 | 9.464417999 | 1.248629499 | 0.259296593 | 0.740866307 | -4.613920476 |
| **A_55_P2037408** | Lrrc57 | 0.168005095 | -0.163017937 | 0.499028128 | 9.817158101 | 1.248628961 | 0.259296776 | 0.740866307 | -4.613920842 |
| **A_52_P229959** | 6-Sep | 0.058838276 | -0.057091873 | 0.174768424 | 8.536507905 | 1.248626957 | 0.259297458 | 0.740866307 | -4.613922208 |
| **A_55_P2068429** | Zfp958 | -0.118421571 | -0.351800596 | 0.114957454 | 10.86817873 | -1.248355157 | 0.259390025 | 0.74097336 | -4.614107512 |
| **A_55_P1972234** | NA | -0.100958476 | -0.299932243 | 0.098015291 | 15.47524323 | -1.248291833 | 0.259411596 | 0.74097336 | -4.614150682 |
| **A_51_P471458** | Sult5a1 | 0.067763912 | -0.065816576 | 0.2013444 | 7.794336142 | 1.248028799 | 0.259501211 | 0.741033739 | -4.614329989 |
| **A_51_P352264** | Sirt7 | -0.14728661 | -0.437699679 | 0.143126459 | 11.44232642 | -1.24771752 | 0.259607299 | 0.741033739 | -4.614542165 |
| **A_55_P2021149** | NA | 0.07784133 | -0.075661524 | 0.231344185 | 8.712417557 | 1.247564349 | 0.259659516 | 0.741033739 | -4.614646562 |
| **A_55_P2047882** | Zc3h7b | 0.104790937 | -0.101874683 | 0.311456558 | 9.820778142 | 1.24745432 | 0.259697031 | 0.741033739 | -4.614721552 |
| **A_55_P2148182** | NA | -0.074149445 | -0.22038563 | 0.07208674 | 9.485944297 | -1.247447338 | 0.259699411 | 0.741033739 | -4.61472631 |
| **A_51_P124089** | Gtpbp2 | -0.137557225 | -0.408857399 | 0.13374295 | 9.059010234 | -1.247390682 | 0.259718731 | 0.741033739 | -4.614764923 |
| **A_52_P661071** | Snhg3 | -0.095271367 | -0.283179905 | 0.092637172 | 10.4175052 | -1.247340553 | 0.259735825 | 0.741033739 | -4.614799087 |
| **A_51_P251892** | Magoh | 0.152612862 | -0.148395767 | 0.45362149 | 12.74956116 | 1.247329962 | 0.259739437 | 0.741033739 | -4.614806304 |
| **A_51_P349192** | NA | 0.072683227 | -0.070694869 | 0.216061324 | 12.10868951 | 1.247155353 | 0.25979899 | 0.741078671 | -4.614925298 |
| **A_52_P236398** | NA | -0.177075473 | -0.526409253 | 0.172258307 | 8.178872214 | -1.247058971 | 0.259831867 | 0.741078671 | -4.614990978 |
| **A_52_P127362** | Vmp1 | 0.132584642 | -0.129058618 | 0.394227902 | 13.28673421 | 1.246673798 | 0.259963291 | 0.741212035 | -4.615253434 |
| **A_55_P2094362** | Erp29 | 0.084266426 | -0.082043506 | 0.250576359 | 9.683714946 | 1.246538088 | 0.260009611 | 0.741212035 | -4.615345899 |
| **A_55_P1969746** | Pitpnm1 | -0.205111309 | -0.610061743 | 0.199839125 | 9.866519482 | -1.24611149 | 0.260155261 | 0.741212035 | -4.615636532 |
| **A_55_P2088535** | Snrpd2 | 0.063052859 | -0.061443326 | 0.187549044 | 12.97432636 | 1.245999605 | 0.260193473 | 0.741212035 | -4.61571275 |
| **A_55_P2319035** | AW011956 | 0.151055509 | -0.14728539 | 0.449396408 | 10.768084 | 1.245641125 | 0.260315937 | 0.741212035 | -4.615956935 |
| **A_66_P105736** | Net1 | -0.124636139 | -0.370800327 | 0.12152805 | 9.761735187 | -1.245627754 | 0.260320506 | 0.741212035 | -4.615966042 |
| **A_55_P2015692** | Elob | -0.103456065 | -0.307844551 | 0.100932421 | 13.73976976 | -1.24528481 | 0.260437712 | 0.741212035 | -4.616199616 |
| **A_55_P1955479** | NA | -0.128618322 | -0.382719156 | 0.125482512 | 10.24138233 | -1.24527684 | 0.260440436 | 0.741212035 | -4.616205044 |
| **A_55_P2146050** | Dennd5b | 0.124377357 | -0.121378745 | 0.370133459 | 12.1951329 | 1.245105572 | 0.260498988 | 0.741212035 | -4.616321681 |
| **A_55_P2010003** | Zfp113 | 0.054189326 | -0.052892506 | 0.161271157 | 8.512342947 | 1.244993601 | 0.260537273 | 0.741212035 | -4.616397933 |
| **A_55_P2132472** | Prpf4b | -0.064119306 | -0.190833281 | 0.062594669 | 8.326932548 | -1.244897117 | 0.260570267 | 0.741212035 | -4.616463636 |
| **A_55_P2057916** | Pex5 | 0.142924519 | -0.139530543 | 0.425379581 | 11.72311729 | 1.244877282 | 0.260577051 | 0.741212035 | -4.616477142 |
| **A_55_P2268221** | Arhgap10 | -0.071256696 | -0.212095339 | 0.069581948 | 8.455039823 | -1.244723848 | 0.260629529 | 0.741212035 | -4.616581621 |
| **A_52_P652513** | Lman1 | 0.088140055 | -0.086071385 | 0.262351495 | 13.63789673 | 1.244703208 | 0.260636589 | 0.741212035 | -4.616595675 |
| **A_55_P2139992** | Mrps18b | -0.140151244 | -0.417179518 | 0.136877031 | 13.68871937 | -1.244635036 | 0.260659909 | 0.741212035 | -4.616642094 |
| **A_55_P1961690** | Tctex1d2 | -0.074838666 | -0.222774715 | 0.073097384 | 10.09688642 | -1.24457531 | 0.260680341 | 0.741212035 | -4.616682761 |
| **A_55_P1967867** | Xpa | -0.067110704 | -0.199773767 | 0.065552359 | 8.76500285 | -1.244545949 | 0.260690386 | 0.741212035 | -4.616702753 |
| **A_55_P2158227** | Angpt1 | -0.07033446 | -0.209370789 | 0.068701868 | 8.332484649 | -1.244540492 | 0.260692253 | 0.741212035 | -4.616706468 |
| **A_55_P2000439** | Ptprz1 | -0.063848668 | -0.190074757 | 0.062377421 | 7.792022593 | -1.244434023 | 0.260728681 | 0.741212035 | -4.616778959 |
| **A_55_P1977776** | Rs5-8s1 | -0.257817754 | -0.767549741 | 0.251914232 | 9.653720422 | -1.244342966 | 0.260759839 | 0.741212035 | -4.616840955 |
| **A_55_P2068248** | NA | -0.393999093 | -1.17323334 | 0.385235155 | 10.78448176 | -1.243931178 | 0.260900788 | 0.741212035 | -4.617121295 |

| | | | | | | | | | |
|---|---|---|---|---|---|---|---|---|---|
| **A_51_P137094** | Sall2 | 0.054972001 | -0.053762448 | 0.16370645 | 8.082790693 | 1.243779954 | 0.260952566 | 0.741212035 | -4.617224237 |
| **A_55_P2055123** | Frrs1 | -0.133239196 | -0.396790341 | 0.130311948 | 8.75694647 | -1.243759071 | 0.260959717 | 0.741212035 | -4.617238452 |
| **A_66_P137157** | Arl6ip4 | 0.075750436 | -0.074109809 | 0.225610681 | 12.30829386 | 1.243563206 | 0.261026796 | 0.741212035 | -4.617371774 |
| **A_55_P2012689** | Stk16 | -0.070387354 | -0.209639018 | 0.06886431 | 9.439856335 | -1.24355045 | 0.261031165 | 0.741212035 | -4.617380456 |
| **A_51_P239036** | Ubn2 | -0.175589421 | -0.522984788 | 0.171805946 | 10.90828739 | -1.243493411 | 0.261050702 | 0.741212035 | -4.617419279 |
| **A_55_P2115297** | Ap3s1 | 0.145808061 | -0.142674127 | 0.434290248 | 12.81859514 | 1.243459642 | 0.261062269 | 0.741212035 | -4.617442264 |
| **A_55_P2169417** | BC021767 | -0.160098385 | -0.476897573 | 0.156700803 | 10.52543181 | -1.243288896 | 0.261120764 | 0.741212035 | -4.617558476 |
| **A_52_P319123** | Vps13b | -0.14566574 | -0.433928122 | 0.142596641 | 10.02470961 | -1.243193165 | 0.261153566 | 0.741212035 | -4.617623629 |
| **A_52_P377723** | Peg10 | -0.161191 | -0.480191457 | 0.157809457 | 11.18804745 | -1.243136009 | 0.261173151 | 0.741212035 | -4.617662528 |
| **A_51_P328489** | 1700025G04Rik | -0.098480728 | -0.293378642 | 0.096417186 | 12.44598959 | -1.243120483 | 0.261178472 | 0.741212035 | -4.617673094 |
| **A_55_P1973254** | Cmtm7 | 0.247824447 | -0.242763332 | 0.738412226 | 12.57191479 | 1.242786661 | 0.26129289 | 0.741212035 | -4.617900267 |
| **A_51_P280492** | Ndufa9 | 0.078159482 | -0.076577004 | 0.232895968 | 11.44429976 | 1.242676561 | 0.261330636 | 0.741212035 | -4.617975187 |
| **A_55_P2086087** | NA | 0.132992388 | -0.130372607 | 0.396357382 | 14.00336253 | 1.242332643 | 0.261448576 | 0.741212035 | -4.618209195 |
| **A_55_P1954604** | NA | -0.116091582 | -0.345988098 | 0.113804935 | 9.786680351 | -1.242331511 | 0.261448964 | 0.741212035 | -4.618209965 |
| **A_52_P228079** | Atf1 | 0.067964193 | -0.066633678 | 0.202562064 | 9.012309349 | 1.242256101 | 0.261474831 | 0.741212035 | -4.618261272 |
| **A_51_P445487** | Mturn | 0.093893692 | -0.092063309 | 0.279850693 | 8.564995145 | 1.242204355 | 0.261492581 | 0.741212035 | -4.618296478 |
| **A_55_P2071691** | Kank3 | 0.121753863 | -0.119404641 | 0.362912368 | 12.71189745 | 1.242079324 | 0.261535476 | 0.741212035 | -4.618381541 |
| **A_52_P83913** | Ssx2ip | -0.156653931 | -0.466954478 | 0.153646616 | 7.827993088 | -1.242018085 | 0.261556488 | 0.741212035 | -4.618423203 |
| **A_51_P503261** | Lrrc8c | -0.088073032 | -0.262534168 | 0.086388104 | 8.368364391 | -1.241976601 | 0.261570722 | 0.741212035 | -4.618451425 |
| **A_55_P1992814** | Shq1 | -0.083805661 | -0.249816943 | 0.08220562 | 8.472201817 | -1.241952306 | 0.261579058 | 0.741212035 | -4.618467952 |
| **A_55_P1975667** | Rhox4g | -0.137127022 | -0.408786358 | 0.134532315 | 11.3082712 | -1.241845508 | 0.261615708 | 0.741212035 | -4.618540605 |
| **A_55_P2091671** | Psmg1 | -0.105423266 | -0.314275696 | 0.103429165 | 9.931459522 | -1.241841288 | 0.261617156 | 0.741212035 | -4.618543476 |
| **A_55_P2098578** | Tubd1 | 0.088669787 | -0.087019662 | 0.264359236 | 8.732942853 | 1.241649866 | 0.261682858 | 0.741212035 | -4.61867369 |
| **A_55_P1980583** | Leprot | -0.171262225 | -0.510625514 | 0.168101064 | 13.47738635 | -1.241554776 | 0.261715501 | 0.741212035 | -4.618738372 |
| **A_51_P462516** | Senp7 | -0.078602344 | -0.234356522 | 0.077151834 | 8.13516056 | -1.241552122 | 0.261716412 | 0.741212035 | -4.618740177 |
| **A_55_P2066553** | Oacyl | -0.403236713 | -1.202272875 | 0.39579945 | 11.01309423 | -1.241545938 | 0.261718535 | 0.741212035 | -4.618744383 |
| **A_55_P2118037** | Sirpa | -0.20516446 | -0.611738938 | 0.201410019 | 9.12634468 | -1.241455567 | 0.261749562 | 0.741212035 | -4.618805853 |
| **A_55_P2050513** | Pgbd1 | -0.062821685 | -0.187317797 | 0.061674427 | 7.98264943 | -1.24143205 | 0.261757636 | 0.741212035 | -4.618821849 |
| **A_55_P2078459** | Kpna2 | -0.10265508 | -0.306126504 | 0.100816343 | 10.98641104 | -1.24121263 | 0.261832984 | 0.741316793 | -4.618971087 |
| **A_55_P2112922** | Impdh2 | 0.109905617 | -0.107970707 | 0.327781941 | 11.64430791 | 1.241020663 | 0.26189892 | 0.741327045 | -4.619101645 |
| **A_55_P2098226** | C1rl | 0.095863271 | -0.094185192 | 0.285911735 | 8.542308826 | 1.240957897 | 0.261920482 | 0.741327045 | -4.619144331 |
| **A_66_P120249** | Nr1h3 | 0.136902018 | -0.134546161 | 0.408350196 | 9.709284829 | 1.240772274 | 0.261984258 | 0.741327045 | -4.619270564 |
| **A_66_P106238** | Gm5106 | -0.225255872 | -0.671960916 | 0.221449172 | 10.07872142 | -1.240579007 | 0.262050674 | 0.741327045 | -4.619401986 |
| **A_55_P2102175** | Lztr1 | 0.075086758 | -0.073819869 | 0.223993386 | 10.84284757 | 1.240562053 | 0.262056502 | 0.741327045 | -4.619413514 |
| **A_51_P310030** | Naca | 0.179410381 | -0.176391837 | 0.535212598 | 14.57441893 | 1.240532312 | 0.262066724 | 0.741327045 | -4.619433738 |
| **A_55_P2175692** | Reep3 | 0.1753229 | -0.172404573 | 0.523050373 | 11.70043852 | 1.240420125 | 0.262105285 | 0.741327636 | -4.61951002 |
| **A_51_P288257** | Cdk5rap3 | 0.064801798 | -0.063755859 | 0.193359454 | 11.10311713 | 1.240104463 | 0.262213815 | 0.741504771 | -4.619724642 |
| **A_52_P478729** | Atg2a | -0.076245758 | -0.227517555 | 0.07502604 | 8.625086994 | -1.240014833 | 0.262244638 | 0.741504771 | -4.619785578 |
| **A_55_P2069460** | NA | -0.092293213 | -0.275483517 | 0.09089709 | 15.60130488 | -1.239471227 | 0.262431649 | 0.741595204 | -4.620155117 |
| **A_55_P1965427** | Uqcrh | 0.080665208 | -0.079448238 | 0.240778653 | 15.19068165 | 1.239446017 | 0.262440325 | 0.741595204 | -4.620172253 |
| **A_51_P403536** | Ltbp4 | 0.303019181 | -0.298453856 | 0.904492219 | 10.38574183 | 1.239433185 | 0.262444741 | 0.741595204 | -4.620180975 |
| **A_55_P2077555** | C330021F23Rik | 0.065765549 | -0.064802301 | 0.196333399 | 8.80807795 | 1.239171344 | 0.262534865 | 0.741595204 | -4.620358946 |
| **A_55_P1991151** | Rpl13 | 0.107272168 | -0.105707802 | 0.320252139 | 15.0410112 | 1.239131677 | 0.262548521 | 0.741595204 | -4.620385906 |
| **A_51_P417053** | Tuft1 | 0.154099723 | -0.151854125 | 0.460053572 | 10.7426534 | 1.239124949 | 0.262550837 | 0.741595204 | -4.620390479 |
| **A_55_P1992216** | Chchd2 | -0.090937248 | -0.271497885 | 0.089623389 | 16.98926303 | -1.239047323 | 0.262577562 | 0.741595204 | -4.620443237 |
| **A_55_P2070131** | NA | 0.19295377 | -0.190182029 | 0.576089568 | 14.47026807 | 1.23899541 | 0.262595436 | 0.741595204 | -4.620478518 |
| **A_55_P1999715** | Tcrb-J | 0.119993215 | -0.118300092 | 0.358286521 | 8.208930764 | 1.238836542 | 0.262650142 | 0.741595204 | -4.620586485 |
| **A_66_P108019** | Heatr1 | 0.103179735 | -0.101728728 | 0.308088199 | 11.20499618 | 1.238807071 | 0.262660291 | 0.741595204 | -4.620606514 |
| **A_55_P2059105** | Gm9920 | 0.16040725 | -0.158227291 | 0.47904179 | 9.845407623 | 1.238512239 | 0.262761846 | 0.741773582 | -4.620806865 |
| **A_55_P2029121** | Pex11g | -0.106073534 | -0.316828278 | 0.104681211 | 8.642840493 | -1.238222926 | 0.262861533 | 0.741946636 | -4.621003447 |

| | | | | | | | | | |
|---|---|---|---|---|---|---|---|---|---|
| **A_52_P24160** | Mrpl51 | -0.115724732 | -0.345722711 | 0.114273247 | 13.46326362 | -1.23785942 | 0.262986831 | 0.742191919 | -4.621250415 |
| **A_66_P140067** | Wdyhv1 | -0.104800407 | -0.313126069 | 0.103525255 | 9.910023556 | -1.237625826 | 0.263067377 | 0.742310851 | -4.621409104 |
| **A_52_P253567** | Hsd3b6 | -0.057403986 | -0.171524685 | 0.056716713 | 7.797516287 | -1.237504503 | 0.263109219 | 0.742320551 | -4.621491517 |
| **A_55_P2123529** | NA | -0.079106495 | -0.236399212 | 0.078186222 | 10.96053651 | -1.237293384 | 0.263182044 | 0.74241765 | -4.621634922 |
| **A_55_P2043382** | Llgl2 | 0.193893732 | -0.191710484 | 0.579497948 | 12.47439501 | 1.237061125 | 0.263262182 | 0.742476672 | -4.621792673 |
| **A_55_P1998186** | NA | 0.078286418 | -0.07741191 | 0.233984747 | 12.37535046 | 1.237005516 | 0.263281372 | 0.742476672 | -4.621830441 |
| **A_55_P2040490** | NA | -0.186666199 | -0.557945452 | 0.184613054 | 8.737062307 | -1.236898793 | 0.263318205 | 0.742476672 | -4.621902923 |
| **A_55_P2156274** | Plekha2 | 0.068107451 | -0.067411876 | 0.203626777 | 8.337683239 | 1.236410126 | 0.263486914 | 0.742844014 | -4.62223477 |
| **A_55_P2113768** | NA | -0.093667225 | -0.280068051 | 0.092733601 | 11.04700347 | -1.236257627 | 0.263539583 | 0.742884147 | -4.622338319 |
| **A_52_P448006** | Nit2 | -0.067006815 | -0.200374112 | 0.066360481 | 8.930634611 | -1.236057838 | 0.263608598 | 0.742892877 | -4.62247397 |
| **A_55_P1974189** | Ptges | -0.162078913 | -0.484681632 | 0.160523805 | 9.009406996 | -1.236026142 | 0.263619548 | 0.742892877 | -4.622495489 |
| **A_55_P2131820** | A530006G24Rik | -0.132398805 | -0.395992677 | 0.131195067 | 12.2218435 | -1.235713862 | 0.263727458 | 0.743022559 | -4.622707498 |
| **A_51_P188795** | Akr7a5 | -0.08287248 | -0.247871238 | 0.082126279 | 9.00143382 | -1.235659523 | 0.263746239 | 0.743022559 | -4.622744387 |
| **A_55_P2008081** | Rhoa | -0.123927667 | -0.370687148 | 0.122831815 | 14.0971272 | -1.23555928 | 0.263780889 | 0.743022559 | -4.622812437 |
| **A_55_P2076832** | Nrd1 | 0.08488054 | -0.084190525 | 0.253951605 | 8.441089742 | 1.23511674 | 0.263933904 | 0.74324614 | -4.623112826 |
| **A_55_P2045114** | Tmem242 | 0.138932853 | -0.137822674 | 0.41568838 | 10.1534203 | 1.235030874 | 0.263963602 | 0.74324614 | -4.623171105 |
| **A_51_P297165** | Sf3a3 | 0.096980652 | -0.096225147 | 0.290186451 | 12.35690633 | 1.23490658 | 0.264006597 | 0.74324614 | -4.623255463 |
| **A_55_P2112569** | Ern1 | 0.055449594 | -0.055022922 | 0.16592211 | 8.631358648 | 1.234847386 | 0.264027075 | 0.74324614 | -4.623295637 |
| **A_52_P475450** | Rbm48 | -0.108341228 | -0.324202884 | 0.107520427 | 8.34784552 | -1.234773828 | 0.264052524 | 0.74324614 | -4.623345558 |
| **A_55_P1994175** | Ppil3 | 0.134859756 | -0.133900661 | 0.403620173 | 9.663449517 | 1.234486163 | 0.26415207 | 0.743418077 | -4.623540774 |
| **A_55_P2180086** | Lrrc28 | -0.147391383 | -0.441177697 | 0.146394931 | 10.51900481 | -1.234268647 | 0.264227362 | 0.743521718 | -4.623688372 |
| **A_55_P2129437** | Plpp2 | 0.157555821 | -0.156559996 | 0.471671639 | 13.56498989 | 1.233996168 | 0.264321706 | 0.743600501 | -4.62387325 |
| **A_51_P350453** | Pdk4 | -0.178948217 | -0.535731861 | 0.177835427 | 10.78078064 | -1.233933066 | 0.264343559 | 0.743600501 | -4.623916063 |
| **A_51_P237754** | H2-T23 | -0.485728991 | -1.454228135 | 0.482770153 | 13.64392619 | -1.233854494 | 0.264370771 | 0.743600501 | -4.62396937 |
| **A_51_P498720** | Cacna2d1 | -0.079450719 | -0.23788998 | 0.078988542 | 7.870906815 | -1.233684722 | 0.264429578 | 0.743657692 | -4.624084547 |
| **A_55_P1986743** | NA | -0.168510811 | -0.50458446 | 0.167562838 | 13.38975523 | -1.233566228 | 0.26447063 | 0.743664942 | -4.624164933 |
| **A_55_P2002963** | Coro1a | -0.136148504 | -0.407743 | 0.135445992 | 9.378855974 | -1.23327825 | 0.264570422 | 0.743813187 | -4.62436028 |
| **A_55_P2159485** | NA | -0.123370065 | -0.369513153 | 0.122773022 | 8.565583295 | -1.23308017 | 0.264639081 | 0.743813187 | -4.624494635 |
| **A_55_P1982227** | NA | 0.236985276 | -0.235865544 | 0.709836097 | 12.04538363 | 1.233009381 | 0.264663622 | 0.743813187 | -4.624542648 |
| **A_55_P2147886** | Fam118a | -0.251694797 | -0.753959932 | 0.250570339 | 11.62312066 | -1.232850364 | 0.264718756 | 0.743813187 | -4.624650498 |
| **A_51_P277006** | Chst8 | -0.200075625 | -0.599381427 | 0.199230178 | 11.00039199 | -1.232700929 | 0.264770578 | 0.743813187 | -4.624751843 |
| **A_55_P2054777** | Maml1 | 0.115804369 | -0.115343733 | 0.34695247 | 10.85493189 | 1.232547812 | 0.264823685 | 0.743813187 | -4.624855681 |
| **A_66_P139250** | Kctd3 | -0.118796852 | -0.355925041 | 0.118331337 | 11.34903316 | -1.232511301 | 0.26483635 | 0.743813187 | -4.624880441 |
| **A_51_P396917** | Zfyve21 | 0.216084476 | -0.21527392 | 0.647442871 | 11.6901877 | 1.232407904 | 0.26487222 | 0.743813187 | -4.624950556 |
| **A_55_P2001683** | Atp5g1 | 0.144668627 | -0.144130521 | 0.433467775 | 14.43834361 | 1.232388437 | 0.264878973 | 0.743813187 | -4.624963756 |
| **A_55_P2072611** | 1-Mar | 0.219931324 | -0.219165455 | 0.659028103 | 8.03850547 | 1.232241982 | 0.264929788 | 0.743813187 | -4.625063066 |
| **A_55_P2084626** | Rab5b | -0.117979508 | -0.353536972 | 0.117577956 | 12.58156267 | -1.23219339 | 0.26494665 | 0.743813187 | -4.625096014 |
| **A_51_P234308** | Rpa2 | 0.07322 | -0.072986707 | 0.219426708 | 8.52136693 | 1.232059247 | 0.264993203 | 0.743835844 | -4.625186969 |
| **A_55_P2177721** | Adra1b | 0.433308444 | -0.432132171 | 1.298749059 | 11.79773185 | 1.231768356 | 0.265094179 | 0.743913386 | -4.625384191 |
| **A_51_P241074** | Map2k1 | 0.107068144 | -0.106791422 | 0.32092771 | 10.84714458 | 1.23168813 | 0.265122033 | 0.743913386 | -4.62543858 |
| **A_55_P2141851** | Pls3 | -0.222881227 | -0.668084843 | 0.222322388 | 11.89206968 | -1.231640528 | 0.265138562 | 0.743913386 | -4.625470851 |
| **A_55_P2151556** | C8g | 0.389198963 | -0.38828894 | 1.166686867 | 9.083107662 | 1.231536243 | 0.265174775 | 0.743913386 | -4.625541548 |
| **A_55_P2069415** | Fxr1 | -0.073794496 | -0.221244174 | 0.073655183 | 10.96840636 | -1.231258668 | 0.265271185 | 0.744020637 | -4.625729708 |
| **A_55_P2042277** | Pbrm1 | -0.077447115 | -0.232211787 | 0.077317557 | 8.729748101 | -1.231126203 | 0.265317205 | 0.744020637 | -4.625819497 |
| **A_51_P450682** | Tepp | 0.095441001 | -0.095293694 | 0.286175696 | 8.719937887 | 1.231046475 | 0.265344907 | 0.744020637 | -4.625873536 |
| **A_55_P2068733** | Gm21953 | 0.078874749 | -0.078761144 | 0.236510642 | 8.318645337 | 1.230982966 | 0.265366975 | 0.744020637 | -4.625916582 |
| **A_55_P1992650** | Bloc1s1 | 0.112312168 | -0.112321697 | 0.336946033 | 12.84544102 | 1.230044276 | 0.265693341 | 0.744827642 | -4.626552701 |
| **A_52_P282035** | Rnf5 | -0.067041132 | -0.201227614 | 0.06714535 | 9.97742847 | -1.229141085 | 0.266007696 | 0.745191532 | -4.627164571 |
| **A_52_P67643** | Mtor | 0.093409473 | -0.093556619 | 0.280375564 | 8.108753771 | 1.229128344 | 0.266012133 | 0.745191532 | -4.627173201 |
| **A_51_P242733** | Tex261 | -0.128315034 | -0.385149 | 0.128518931 | 11.89115604 | -1.229119904 | 0.266015072 | 0.745191532 | -4.627178918 |

| | | | | | | | | | |
|---|---|---|---|---|---|---|---|---|---|
| **A_55_P1976734** | Cacna2d2 | 0.078339834 | -0.078507885 | 0.235187553 | 8.233033878 | 1.228778497 | 0.266133986 | 0.745191532 | -4.627410154 |
| **A_55_P2179309** | P2ry10b | 0.10976258 | -0.110028239 | 0.3295534 | 8.204476257 | 1.22860965 | 0.266192814 | 0.745191532 | -4.627524504 |
| **A_51_P461761** | Zcchc8 | 0.057718394 | -0.057860683 | 0.173297471 | 8.813288743 | 1.228582094 | 0.266202415 | 0.745191532 | -4.627543166 |
| **A_51_P506787** | Nap1l1 | 0.090156555 | -0.09039407 | 0.270707181 | 9.463527429 | 1.228478265 | 0.266238597 | 0.745191532 | -4.627613479 |
| **A_55_P2100050** | NA | -0.142297055 | -0.427286993 | 0.142692884 | 10.17897677 | -1.228387946 | 0.266270074 | 0.745191532 | -4.627674642 |
| **A_52_P88793** | Zfp933 | 0.092455517 | -0.092733601 | 0.277644636 | 8.897240108 | 1.228249317 | 0.266318394 | 0.745191532 | -4.627768515 |
| **A_51_P461724** | Atp6ap2 | -0.090172004 | -0.270792983 | 0.090448976 | 10.90858477 | -1.228210175 | 0.266332039 | 0.745191532 | -4.627795019 |
| **A_51_P404193** | Sp5 | 0.063246763 | -0.063446204 | 0.189939731 | 7.806390764 | 1.228160036 | 0.266349518 | 0.745191532 | -4.62782897 |
| **A_52_P429450** | Ngp | -0.057967442 | -0.174085301 | 0.058150417 | 7.705211819 | -1.228158114 | 0.266350188 | 0.745191532 | -4.627830271 |
| **A_55_P2161465** | Gm10516 | 0.078643975 | -0.078899887 | 0.236187837 | 8.750821334 | 1.228098312 | 0.266371037 | 0.745191532 | -4.627870764 |
| **A_51_P313581** | Fabp2 | 0.279448966 | -0.280412051 | 0.839309984 | 8.189409667 | 1.227980419 | 0.266412142 | 0.745191532 | -4.627950588 |
| **A_55_P2027521** | NA | 0.071202254 | -0.071464858 | 0.213869366 | 7.872876151 | 1.227832245 | 0.266463814 | 0.745191532 | -4.628050911 |
| **A_51_P371091** | Rcsd1 | -0.15204871 | -0.456717398 | 0.152619979 | 11.08478861 | -1.227789962 | 0.26647856 | 0.745191532 | -4.628079538 |
| **A_51_P345046** | Metap2 | -0.077313828 | -0.232244088 | 0.077616431 | 10.91858452 | -1.227693885 | 0.266512071 | 0.745191532 | -4.628144584 |
| **A_55_P2000390** | Arpc5 | -0.180400545 | -0.541911964 | 0.181110873 | 14.63367075 | -1.227679461 | 0.266517102 | 0.745191532 | -4.628154349 |
| **A_55_P1975430** | St3gal3 | -0.095257174 | -0.286191441 | 0.095677093 | 9.845226855 | -1.227391123 | 0.266617694 | 0.745364971 | -4.628349546 |
| **A_52_P266320** | Akap17b | 0.056210921 | -0.056502061 | 0.168923904 | 8.216920397 | 1.226919094 | 0.266782442 | 0.74571769 | -4.628669055 |
| **A_52_P495372** | Twnk | -0.062831613 | -0.188861741 | 0.063198515 | 8.144996182 | -1.226515361 | 0.266923424 | 0.746003883 | -4.628942293 |
| **A_51_P149349** | Nfxl1 | -0.0684944 | -0.205970962 | 0.068982162 | 8.710491393 | -1.225732121 | 0.267197113 | 0.746446223 | -4.629472266 |
| **A_51_P234692** | Neat1 | -0.418502919 | -1.258526735 | 0.421520898 | 11.93528669 | -1.225677051 | 0.267216366 | 0.746446223 | -4.629509523 |
| **A_55_P2228327** | 4921529L05Rik | 0.281671592 | -0.283777361 | 0.847120546 | 9.412562788 | 1.225515496 | 0.267272852 | 0.746446223 | -4.629618817 |
| **A_55_P2079579** | Pira7 | -0.071187434 | -0.214094645 | 0.071719776 | 7.992302028 | -1.225514236 | 0.267273293 | 0.746446223 | -4.62961967 |
| **A_55_P2120433** | Wdr12 | -0.072110701 | -0.21688473 | 0.072663328 | 8.367567792 | -1.225400977 | 0.267312899 | 0.746446223 | -4.629696288 |
| **A_51_P181319** | Dcxr | 0.236871508 | -0.23868735 | 0.712430365 | 8.472868257 | 1.225399538 | 0.267313403 | 0.746446223 | -4.629697262 |
| **A_55_P2040046** | Swi5 | 0.066419907 | -0.066956469 | 0.199796283 | 15.53280553 | 1.225147876 | 0.267401427 | 0.746584165 | -4.629867496 |
| **A_51_P169047** | Tdp2 | -0.125775982 | -0.378426856 | 0.126874892 | 9.943752276 | -1.224746124 | 0.267542001 | 0.746724972 | -4.630139226 |
| **A_51_P446085** | Map10 | 0.065885029 | -0.066464249 | 0.198234307 | 8.640859283 | 1.224713 | 0.267553594 | 0.746724972 | -4.630161629 |
| **A_55_P2063505** | NA | -0.378731558 | -1.139549382 | 0.382086266 | 11.2784843 | -1.224672536 | 0.267567756 | 0.746724972 | -4.630188995 |
| **A_55_P2034420** | NA | 0.110529418 | -0.111531633 | 0.332590469 | 10.37418136 | 1.224544732 | 0.267612492 | 0.746732246 | -4.630275426 |
| **A_55_P2039210** | Cpt1b | 0.075967039 | -0.076668373 | 0.22860245 | 8.033895151 | 1.224444369 | 0.267647628 | 0.746732246 | -4.630343298 |
| **A_55_P2133170** | Luc7l2 | -0.122438803 | -0.368506184 | 0.123628579 | 8.792017347 | -1.224148739 | 0.267751146 | 0.746913249 | -4.630543206 |
| **A_55_P2419711** | 4932412D23Rik | 0.182940349 | -0.184867209 | 0.550747908 | 8.943407361 | 1.223652262 | 0.267925072 | 0.747078863 | -4.630878882 |
| **A_51_P448032** | Cox6b1 | -0.108937206 | -0.327959382 | 0.11008497 | 11.82888564 | -1.223650259 | 0.267925774 | 0.747078863 | -4.630880236 |
| **A_55_P2042146** | Fech | 0.156557029 | -0.158208674 | 0.471322733 | 12.93366658 | 1.223641869 | 0.267928714 | 0.747078863 | -4.630885909 |
| **A_51_P318544** | Fdx2 | 0.098682814 | -0.099746709 | 0.297112337 | 11.37417398 | 1.223501203 | 0.267978011 | 0.747078863 | -4.630981004 |
| **A_51_P401343** | Cldn14 | 0.057728538 | -0.058357879 | 0.173814955 | 7.912834199 | 1.223427715 | 0.268003768 | 0.747078863 | -4.631030684 |
| **A_51_P238183** | Etnk2 | 0.127133182 | -0.128613437 | 0.382879801 | 8.73078816 | 1.222976689 | 0.268161898 | 0.74741187 | -4.631335556 |
| **A_55_P2041584** | NA | 0.065837716 | -0.066645047 | 0.19832048 | 8.112071222 | 1.222600423 | 0.268293878 | 0.747580178 | -4.631589856 |
| **A_51_P273679** | Dmgdh | -0.105474947 | -0.317740084 | 0.10679019 | 9.154114636 | -1.222474501 | 0.26833806 | 0.747580178 | -4.631674954 |
| **A_55_P2023181** | Lsm2 | 0.159620315 | -0.161620689 | 0.480861318 | 11.91516802 | 1.222436623 | 0.268351352 | 0.747580178 | -4.631700551 |
| **A_55_P1955548** | Ezr | 0.117292731 | -0.118776762 | 0.353362224 | 13.08792684 | 1.22236356 | 0.268376991 | 0.747580178 | -4.631749924 |
| **A_55_P2066269** | Aak1 | -0.090772739 | -0.273552937 | 0.092007458 | 8.263470351 | -1.221786896 | 0.268579429 | 0.74785868 | -4.632139568 |
| **A_55_P2127973** | Gm9112 | 0.266342191 | -0.270006732 | 0.802691114 | 15.32173694 | 1.221691966 | 0.268612767 | 0.74785868 | -4.632203703 |
| **A_55_P2037812** | Palmd | -0.076498487 | -0.230562614 | 0.077565639 | 8.801983129 | -1.221575985 | 0.268653503 | 0.74785868 | -4.632282057 |
| **A_55_P1985079** | Mcrip1 | 0.078434278 | -0.079557706 | 0.236426263 | 13.61344166 | 1.221349655 | 0.268733012 | 0.74785868 | -4.632434952 |
| **A_52_P379337** | Rtn4 | -0.095354259 | -0.287436945 | 0.096728427 | 10.03675296 | -1.221296293 | 0.26875176 | 0.74785868 | -4.632470998 |
| **A_55_P2017759** | Evi2a | -0.135337431 | -0.407966777 | 0.137291915 | 8.320172576 | -1.221277875 | 0.268758232 | 0.74785868 | -4.63248344 |
| **A_51_P244504** | LTO1 | -0.09564804 | -0.288339518 | 0.097043437 | 11.62307161 | -1.221188575 | 0.268789611 | 0.74785868 | -4.632543761 |
| **A_51_P191469** | Rnf31 | -0.097492943 | -0.293954201 | 0.098968315 | 10.64805023 | -1.22085876 | 0.268905532 | 0.74785868 | -4.63276653 |
| **A_55_P2042273** | Pbrm1 | -0.089843589 | -0.270910996 | 0.091223817 | 8.493643955 | -1.220719766 | 0.268954397 | 0.74785868 | -4.632860403 |

| | | | | | | | | | |
|---|---|---|---|---|---|---|---|---|---|
| **A_51_P253537** | Atf6b | -0.068701669 | -0.207166774 | 0.069763435 | 8.476006403 | -1.220663939 | 0.268974026 | 0.74785868 | -4.632898106 |
| **A_51_P502054** | Gtf2b | 0.150802235 | -0.153183855 | 0.454788326 | 12.71907657 | 1.220459099 | 0.269046059 | 0.74785868 | -4.63303644 |
| **A_55_P2097269** | NA | -0.082153894 | -0.247766163 | 0.083458376 | 10.76229836 | -1.220407325 | 0.269064269 | 0.74785868 | -4.633071403 |
| **A_55_P2013598** | Ccnq | 0.090698515 | -0.092146981 | 0.273544011 | 10.32139272 | 1.220351872 | 0.269083773 | 0.74785868 | -4.633108849 |
| **A_52_P80567** | Nbas | -0.101713287 | -0.306769156 | 0.103342582 | 8.398991381 | -1.220322584 | 0.269094075 | 0.74785868 | -4.633128627 |
| **A_55_P2098603** | NA | -0.220482592 | -0.664979858 | 0.224014673 | 8.333020005 | -1.220321817 | 0.269094345 | 0.74785868 | -4.633129144 |
| **A_55_P2064811** | Nxt1 | 0.068140208 | -0.069257895 | 0.20553831 | 11.50274084 | 1.220090034 | 0.269175886 | 0.74785868 | -4.633285654 |
| **A_51_P179131** | Vasp | -0.136543888 | -0.41189647 | 0.138808694 | 10.13755129 | -1.219978771 | 0.269215036 | 0.74785868 | -4.633360779 |
| **A_55_P2116833** | 9430038I01Rik | 0.237027192 | -0.240979457 | 0.715033841 | 9.860245954 | 1.219925747 | 0.269233695 | 0.74785868 | -4.633396579 |
| **A_66_P107703** | Smox | -0.118812844 | -0.358426539 | 0.12080085 | 8.647888541 | -1.219890703 | 0.269246028 | 0.74785868 | -4.63342024 |
| **A_55_P1959485** | NA | 0.0748684 | -0.076122792 | 0.225859593 | 10.95850798 | 1.219877166 | 0.269250792 | 0.74785868 | -4.63342938 |
| **A_55_P2039289** | Hspb6 | 0.112279914 | -0.114298272 | 0.3388581 | 8.960531988 | 1.219138759 | 0.269510769 | 0.748473227 | -4.633927859 |
| **A_55_P1967266** | Plxnb2 | 0.119208647 | -0.121437872 | 0.359855166 | 12.65486456 | 1.218701478 | 0.26966483 | 0.748530326 | -4.634222995 |
| **A_55_P2176325** | Prex1 | -0.189283557 | -0.571404359 | 0.192837245 | 10.74295524 | -1.218656674 | 0.269680619 | 0.748530326 | -4.634253232 |
| **A_51_P159402** | Gcat | 0.308471023 | -0.314314851 | 0.931256896 | 10.28644755 | 1.218554009 | 0.269716803 | 0.748530326 | -4.634322516 |
| **A_51_P443403** | Smoc1 | -0.169211965 | -0.510860575 | 0.172436644 | 9.169206107 | -1.218486089 | 0.269740743 | 0.748530326 | -4.634368352 |
| **A_51_P409429** | Aars | 0.159166662 | -0.162210069 | 0.480543394 | 11.44412378 | 1.218447561 | 0.269754324 | 0.748530326 | -4.634394352 |
| **A_52_P263518** | Gng2 | 0.13385036 | -0.136415586 | 0.404116306 | 9.311481361 | 1.218421007 | 0.269763684 | 0.748530326 | -4.63441227 |
| **A_66_P110343** | Snx10 | -0.078917464 | -0.238292534 | 0.080457606 | 8.32005345 | -1.218209255 | 0.269838339 | 0.748630007 | -4.634555158 |
| **A_55_P1952312** | NA | -0.136222947 | -0.411358704 | 0.13891281 | 12.58563929 | -1.218070427 | 0.269887294 | 0.748632324 | -4.634648832 |
| **A_52_P38639** | Fermt3 | -0.05332189 | -0.161027373 | 0.054383593 | 7.763049251 | -1.217970824 | 0.269922421 | 0.748632324 | -4.634716036 |
| **A_55_P2034230** | Polr2c | 0.190130704 | -0.193950833 | 0.57421224 | 10.41901288 | 1.217861743 | 0.269960896 | 0.748632324 | -4.634789632 |
| **A_51_P432563** | Tlr9 | 0.060215448 | -0.061434706 | 0.181865603 | 11.22131776 | 1.217767621 | 0.269994098 | 0.748632324 | -4.634853134 |
| **A_51_P117739** | Vegfd | 0.384901768 | -0.392829748 | 1.162633285 | 9.096804933 | 1.217557194 | 0.270068341 | 0.74865067 | -4.627933829 |
| **A_51_P240614** | Tm4sf1 | -0.102599108 | -0.309915348 | 0.104717131 | 11.36022926 | -1.217529314 | 0.270078179 | 0.74865067 | -4.635013903 |
| **A_55_P2039771** | Itgav | -0.136661399 | -0.412842849 | 0.139520052 | 8.94051935 | -1.217364178 | 0.270136456 | 0.748684202 | -4.6351253 |
| **A_55_P2055587** | Enpp4 | -0.209799746 | -0.633818657 | 0.214219166 | 8.791909779 | -1.217275536 | 0.270167743 | 0.748684202 | -4.635185094 |
| **A_51_P336827** | Cyb5b | -0.144204303 | -0.435777946 | 0.147369339 | 9.538341491 | -1.216743742 | 0.27035551 | 0.748881146 | -4.635543777 |
| **A_55_P1953583** | Ace2 | 0.13052384 | -0.133390702 | 0.394438382 | 11.24980841 | 1.216734113 | 0.270358911 | 0.748881146 | -4.63555027 |
| **A_55_P2013058** | Shb | 0.151956595 | -0.155298677 | 0.459211867 | 12.63313016 | 1.216716428 | 0.270365157 | 0.748881146 | -4.635562198 |
| **A_52_P403527** | NA | 0.085350216 | -0.087238878 | 0.257939309 | 9.939780055 | 1.216635372 | 0.270393787 | 0.748881146 | -4.635616861 |
| **A_55_P1953063** | Hspe1 | 0.206421444 | -0.211068607 | 0.623911495 | 11.18596285 | 1.216404015 | 0.270475521 | 0.749000192 | -4.635772879 |
| **A_51_P485458** | Txlna | -0.084498847 | -0.255474292 | 0.086476598 | 8.308279049 | -1.215867369 | 0.270665189 | 0.749333824 | -4.636134721 |
| **A_51_P485391** | Parn | -0.076274644 | -0.2306378 | 0.078088511 | 9.753149761 | -1.21564203 | 0.270744866 | 0.749333824 | -4.63628664 |
| **A_55_P1992450** | Vdac2 | -0.15013948 | -0.454037787 | 0.153758827 | 14.17457128 | -1.215446342 | 0.270814075 | 0.749333824 | -4.636418558 |
| **A_55_P2026156** | NA | 0.110848776 | -0.113544227 | 0.335241778 | 13.42954377 | 1.215320306 | 0.270858658 | 0.749333824 | -4.636503516 |
| **A_52_P144925** | Taf1d | 0.074564728 | -0.076378101 | 0.225507557 | 9.038319425 | 1.21531852 | 0.27085929 | 0.749333824 | -4.63650472 |
| **A_51_P508029** | Pcyox1 | -0.255811305 | -0.773699264 | 0.262076654 | 10.51975746 | -1.215214891 | 0.270895952 | 0.749333824 | -4.636574572 |
| **A_52_P235102** | Zfp975 | -0.051393389 | -0.15544474 | 0.052657963 | 8.672983097 | -1.215146642 | 0.270920099 | 0.749333824 | -4.636620575 |
| **A_51_P226053** | Sra1 | 0.079934882 | -0.081910852 | 0.241780616 | 12.80647025 | 1.215078248 | 0.2709443 | 0.749333824 | -4.636666673 |
| **A_55_P2029417** | 2610318N02Rik | 0.055050354 | -0.056427162 | 0.16652787 | 7.858149925 | 1.214904104 | 0.271005928 | 0.749333824 | -4.636784045 |
| **A_55_P2152592** | Agpat3 | 0.167472948 | -0.182237222 | 0.517183117 | 12.28735318 | 1.241042016 | 0.271030139 | 0.749333824 | -4.593641443 |
| **A_55_P1960646** | Ube2w | 0.066007633 | -0.067666236 | 0.199681502 | 8.596832156 | 1.214833618 | 0.271030876 | 0.749333824 | -4.63683155 |
| **A_51_P431785** | Myom2 | -0.093831716 | -0.283866825 | 0.096203392 | 8.147512804 | -1.214744607 | 0.271062384 | 0.749333824 | -4.636891538 |
| **A_55_P1967708** | NA | -0.099703704 | -0.301652246 | 0.102244837 | 9.261682493 | -1.214618069 | 0.271107181 | 0.749333824 | -4.636976815 |
| **A_55_P2066468** | Noa1 | 0.11098029 | -0.113832389 | 0.335792969 | 9.750876703 | 1.214490768 | 0.271152254 | 0.749333824 | -4.637062601 |
| **A_66_P140886** | Inafm1 | -0.195436329 | -0.591361682 | 0.200489023 | 10.98188735 | -1.214398292 | 0.271185001 | 0.749333824 | -4.637124916 |
| **A_55_P2148400** | NA | -0.30139138 | -0.912102011 | 0.30931925 | 12.32502566 | -1.214128102 | 0.271280698 | 0.749333824 | -4.637306975 |
| **A_51_P292030** | Nus1 | 0.070798512 | -0.072662092 | 0.214259117 | 11.65429735 | 1.214117275 | 0.271284533 | 0.749333824 | -4.63731427 |
| **A_55_P2110915** | Flrt1 | -0.088831613 | -0.268836975 | 0.091173749 | 8.257813055 | -1.214091082 | 0.271293812 | 0.749333824 | -4.637331918 |

| | | | | | | | | | |
|---|---|---|---|---|---|---|---|---|---|
| **A_55_P1998651** | Entpd5 | 0.189618205 | -0.194812815 | 0.574049225 | 10.11597888 | 1.213474824 | 0.271512201 | 0.749820044 | -4.637747091 |
| **A_55_P1987938** | NA | -0.057447817 | -0.173948856 | 0.059053223 | 11.97367436 | -1.213145497 | 0.27162897 | 0.749820044 | -4.63796892 |
| **A_55_P2133845** | Eid2b | -0.058535228 | -0.177251109 | 0.060180652 | 8.192067362 | -1.213047093 | 0.271663869 | 0.749820044 | -4.638035199 |
| **A_51_P521161** | Caml | -0.085592268 | -0.259199628 | 0.088015092 | 11.16552468 | -1.212929518 | 0.271705573 | 0.749820044 | -4.638114386 |
| **A_55_P2002133** | Ago2 | 0.071572421 | -0.073637965 | 0.216782807 | 9.374322273 | 1.212598955 | 0.271822853 | 0.749820044 | -4.638337005 |
| **A_55_P2005055** | Pepd | -0.205604651 | -0.622762449 | 0.211553148 | 13.28862157 | -1.212555792 | 0.27183817 | 0.749820044 | -4.638366071 |
| **A_66_P103999** | Rsrc1 | -0.077009362 | -0.233258243 | 0.079239518 | 8.363088253 | -1.212539166 | 0.27184407 | 0.749820044 | -4.638377268 |
| **A_52_P130787** | Kcna6 | -0.07593325 | -0.230003041 | 0.078136541 | 7.860322446 | -1.212505338 | 0.271856075 | 0.749820044 | -4.638400047 |
| **A_52_P532227** | S1pr1 | -0.142174935 | -0.430695801 | 0.14634593 | 9.493905694 | -1.212313597 | 0.27192413 | 0.749820044 | -4.638529159 |
| **A_52_P399934** | Dusp2 | -0.171554826 | -0.519742603 | 0.176632951 | 9.27399563 | -1.212156186 | 0.271980012 | 0.749820044 | -4.638635148 |
| **A_55_P1970062** | Igsf9b | 0.135955188 | -0.139988135 | 0.411898512 | 8.763710143 | 1.212118442 | 0.271993412 | 0.749820044 | -4.638660561 |
| **A_52_P429265** | Ppp6r2 | 0.064447706 | -0.066362667 | 0.195258079 | 8.230466133 | 1.212088814 | 0.272003932 | 0.749820044 | -4.638680509 |
| **A_52_P301223** | Brwd1 | -0.0918673 | -0.278362613 | 0.094628013 | 10.14237318 | -1.211887188 | 0.272075529 | 0.749820044 | -4.638816257 |
| **A_55_P2013765** | Raly | 0.155025193 | -0.159718537 | 0.469768922 | 12.44819066 | 1.211753706 | 0.272122938 | 0.749820044 | -4.63890612 |
| **A_65_P20641** | NA | -0.086650569 | -0.262582356 | 0.089281218 | 8.420376477 | -1.211703239 | 0.272140864 | 0.749820044 | -4.638940095 |
| **A_55_P2118520** | Col1a1 | 0.876500966 | -0.903266033 | 2.656267965 | 13.74779109 | 1.211597622 | 0.272178383 | 0.749820044 | -4.639011194 |
| **A_55_P1965674** | Alg2 | -0.077425072 | -0.234649643 | 0.079799498 | 10.71710691 | -1.21151938 | 0.272206181 | 0.749820044 | -4.639063863 |
| **A_55_P2146374** | Nmnat1 | -0.070042479 | -0.212282667 | 0.072197709 | 9.002549347 | -1.21145798 | 0.272227996 | 0.749820044 | -4.639105194 |
| **A_51_P184321** | Aimp1 | 0.132889482 | -0.136986722 | 0.402765686 | 12.33373319 | 1.211421222 | 0.272241057 | 0.749820044 | -4.639129937 |
| **A_55_P1993404** | NA | -0.146767744 | -0.44485692 | 0.151321432 | 12.61389694 | -1.211305179 | 0.272282294 | 0.749820044 | -4.639208048 |
| **A_55_P2041121** | Rn4.5s | -0.207224805 | -0.628106021 | 0.213656411 | 16.85675315 | -1.211299001 | 0.272284489 | 0.749820044 | -4.639212206 |
| **A_51_P496432** | Acsl1 | 0.226818251 | -0.233968794 | 0.687605295 | 10.92080588 | 1.211007687 | 0.272388033 | 0.749998332 | -4.639408278 |
| **A_55_P2046593** | Gm15226 | 0.199773883 | -0.206182208 | 0.605729974 | 10.24752746 | 1.21067845 | 0.272505097 | 0.750061257 | -4.639629851 |
| **A_55_P1957459** | Lilrb4a | -0.220234129 | -0.667813501 | 0.227345242 | 9.156854077 | -1.210552761 | 0.272549799 | 0.750061257 | -4.639714431 |
| **A_52_P676198** | Fam210b | 0.171883425 | -0.177481038 | 0.521247888 | 8.807076008 | 1.210387515 | 0.272608579 | 0.750061257 | -4.639825624 |
| **A_55_P2190299** | Fggy | 0.098401032 | -0.10160826 | 0.298410324 | 8.732256834 | 1.210371369 | 0.272614323 | 0.750061257 | -4.639836488 |
| **A_55_P2056808** | Tanc1 | 0.085605256 | -0.088395689 | 0.259606201 | 8.584815328 | 1.210369542 | 0.272614973 | 0.750061257 | -4.639837717 |
| **A_55_P2057786** | Rft1 | -0.079164046 | -0.240087418 | 0.081759327 | 9.451146662 | -1.21025816 | 0.2726546 | 0.750061257 | -4.639912661 |
| **A_55_P2123220** | Slc35e1 | 0.25742504 | -0.26591712 | 0.780767199 | 10.64936782 | 1.210136135 | 0.27269802 | 0.750061257 | -4.639994764 |
| **A_55_P2177112** | Prrg2 | 0.12390713 | -0.128026137 | 0.375840397 | 8.472936119 | 1.209984873 | 0.272751851 | 0.750061257 | -4.640096532 |
| **A_55_P2021266** | Hpse | 0.354999364 | -0.366814382 | 1.07681311 | 9.639598085 | 1.209961607 | 0.272760131 | 0.750061257 | -4.640112185 |
| **A_51_P343833** | Traf1 | -0.05664871 | -0.171855891 | 0.058558471 | 8.050947304 | -1.209705449 | 0.272851316 | 0.750205275 | -4.640284513 |
| **A_51_P353386** | Micos13 | 0.150338875 | -0.155552167 | 0.456229917 | 11.99317063 | 1.209131956 | 0.273055558 | 0.75066006 | -4.640670266 |
| **A_55_P1993805** | Zfp947 | 0.079516607 | -0.082307994 | 0.241341207 | 10.69958602 | 1.208877953 | 0.27314606 | 0.750802076 | -4.640841093 |
| **A_52_P227307** | Mllt1 | 0.091269447 | -0.094503379 | 0.277042274 | 10.4500685 | 1.208682947 | 0.27321556 | 0.750834066 | -4.640972232 |
| **A_55_P2134246** | Serpinb9 | 0.088255027 | -0.091404498 | 0.267914551 | 9.024912684 | 1.208532594 | 0.273269155 | 0.750834066 | -4.641073336 |
| **A_55_P2052939** | Dyrk2 | -0.059018848 | -0.179164108 | 0.061126412 | 8.875059327 | -1.20851835 | 0.273274233 | 0.750834066 | -4.641082913 |
| **A_55_P1964403** | Agfg2 | 0.078481161 | -0.081305911 | 0.238268233 | 8.158314306 | 1.208350552 | 0.273334059 | 0.750891703 | -4.641195741 |
| **A_52_P409055** | Dcaf8 | -0.148169322 | -0.449929912 | 0.153591269 | 11.06109044 | -1.20799444 | 0.273461063 | 0.751048083 | -4.641435167 |
| **A_51_P172532** | Nit1 | 0.089043553 | -0.092305128 | 0.270392235 | 11.51550588 | 1.207973043 | 0.273468695 | 0.751048083 | -4.641449552 |
| **A_55_P2064906** | NA | -0.120047716 | -0.364563411 | 0.124467978 | 15.79796499 | -1.207859242 | 0.273509293 | 0.751052866 | -4.641526057 |
| **A_55_P2093013** | NA | 0.108042429 | -0.112071866 | 0.328156723 | 13.88268154 | 1.20757817 | 0.273609586 | 0.751077749 | -4.641715 |
| **A_55_P2093665** | NA | 0.077276712 | -0.080163753 | 0.234717177 | 9.659269314 | 1.207539746 | 0.273623299 | 0.751077749 | -4.641740828 |
| **A_55_P2032558** | NA | 0.087950725 | -0.091256164 | 0.267157614 | 13.51314149 | 1.207407548 | 0.273670484 | 0.751077749 | -4.641829687 |
| **A_52_P82660** | Mcm9 | -0.057985394 | -0.176141272 | 0.060170483 | 8.049410001 | -1.20734795 | 0.273691758 | 0.751077749 | -4.641869746 |
| **A_55_P2127497** | Trim35 | 0.162702105 | -0.168870631 | 0.494274841 | 12.14653737 | 1.20721194 | 0.273740314 | 0.751077749 | -4.64196116 |
| **A_55_P2000823** | Rbm39 | -0.098461111 | -0.299129157 | 0.102206935 | 12.94274219 | -1.20713453 | 0.273767952 | 0.751077749 | -4.642013186 |
| **A_55_P2009008** | NA | -0.099900178 | -0.303511688 | 0.103711333 | 12.89361498 | -1.207071784 | 0.273790357 | 0.751077749 | -4.642055356 |
| **A_51_P468780** | Hmg20a | -0.0784063 | -0.238231574 | 0.081418974 | 10.52734677 | -1.206909393 | 0.27384835 | 0.751130234 | -4.64216449 |
| **A_51_P238722** | Cd93 | -0.195085841 | -0.592817712 | 0.20264603 | 12.28197095 | -1.206714469 | 0.273917975 | 0.751214606 | -4.64229548 |

| | | | | | | | | | |
|---|---|---|---|---|---|---|---|---|---|
| **A_65_P06446** | Tfrc | 0.209894514 | -0.2180808 | 0.637869828 | 12.6689861 | 1.206567245 | 0.273970572 | 0.751252261 | -4.642394408 |
| **A_52_P6451** | Sec24a | 0.122716453 | -0.127569251 | 0.373002158 | 9.930920924 | 1.206246079 | 0.274085342 | 0.751420165 | -4.6426102 |
| **A_55_P2063465** | Thbs3 | 0.150063488 | -0.156014907 | 0.456141882 | 8.774289715 | 1.206178339 | 0.274109555 | 0.751420165 | -4.642655712 |
| **A_51_P104392** | Rpp25 | -0.081107547 | -0.246573868 | 0.084358774 | 7.999412005 | -1.205926447 | 0.274199605 | 0.751560433 | -4.642824936 |
| **A_52_P241508** | Zbtb46 | 0.101705838 | -0.105845705 | 0.30925738 | 8.066635829 | 1.20556069 | 0.274330408 | 0.751587509 | -4.643070629 |
| **A_51_P174720** | Ccdc43 | -0.065042942 | -0.197780219 | 0.067694336 | 8.112067676 | -1.205525589 | 0.274342964 | 0.751587509 | -4.643094206 |
| **A_55_P2112111** | Ube2b | 0.140345852 | -0.146099838 | 0.426791542 | 13.65385186 | 1.20538686 | 0.274392592 | 0.751587509 | -4.643187386 |
| **A_51_P415220** | Zmat3 | 0.094842996 | -0.098734947 | 0.288420939 | 9.969783324 | 1.205364941 | 0.274400434 | 0.751587509 | -4.643202108 |
| **A_55_P2419514** | Ccny | -0.105811898 | -0.321779501 | 0.110155704 | 9.073344073 | -1.205355245 | 0.274403904 | 0.751587509 | -4.64320862 |
| **A_55_P2065325** | Tmem94 | -0.063451961 | -0.192979781 | 0.066075858 | 7.895158332 | -1.2051779 | 0.27446736 | 0.751654804 | -4.643327729 |
| **A_52_P394395** | Abcd3 | 0.16602083 | -0.173005818 | 0.505047477 | 12.11469108 | 1.204752701 | 0.274619555 | 0.751681228 | -4.64361327 |
| **A_55_P1983095** | Adam19 | -1.967899251 | -5.986773741 | 2.050975239 | 12.68362151 | -1.204668572 | 0.274649676 | 0.751681228 | -4.643669762 |
| **A_55_P2090817** | Macrod2 | 0.060088604 | -0.062625804 | 0.182803012 | 8.248461212 | 1.204663416 | 0.274651522 | 0.751681228 | -4.643673224 |
| **A_55_P2048730** | NA | -0.074988911 | -0.22818116 | 0.078203338 | 14.84966717 | -1.20428539 | 0.274786907 | 0.751681228 | -4.64392704 |
| **A_55_P1961715** | Nme5 | -0.070871962 | -0.215719667 | 0.073975743 | 7.973561261 | -1.203738081 | 0.27498302 | 0.751681228 | -4.644294455 |
| **A_55_P2211341** | Map3k4 | -0.143375689 | -0.436409836 | 0.149658458 | 12.2492119 | -1.203722694 | 0.274988536 | 0.751681228 | -4.644304783 |
| **A_55_P1969845** | Lrpprc | 0.123372055 | -0.128784042 | 0.375528152 | 11.49041723 | 1.203695088 | 0.274998431 | 0.751681228 | -4.644323313 |
| **A_51_P363338** | Prkd1 | -0.065178503 | -0.198417778 | 0.068060773 | 7.88265026 | -1.203486658 | 0.275073152 | 0.751681228 | -4.644463213 |
| **A_55_P1958653** | Arf2 | -0.302291442 | -0.920283861 | 0.315700977 | 14.22881541 | -1.203405156 | 0.275102374 | 0.751681228 | -4.644517915 |
| **A_55_P2230913** | Etv6 | -0.071974592 | -0.219182513 | 0.075233328 | 9.360714763 | -1.202865857 | 0.275295809 | 0.751681228 | -4.644879836 |
| **A_55_P2332731** | D130062J21Rik | -0.178192506 | -0.542690933 | 0.18630592 | 8.315039346 | -1.202715595 | 0.275349726 | 0.751681228 | -4.644980663 |
| **A_55_P2069525** | Zmym6 | -0.16238357 | -0.494598569 | 0.169831429 | 9.998818838 | -1.20251918 | 0.275420216 | 0.751681228 | -4.645112451 |
| **A_55_P2000613** | Slc6a1 | 0.074160826 | -0.077571717 | 0.22589337 | 8.399856934 | 1.202444352 | 0.275447075 | 0.751681228 | -4.645162656 |
| **A_55_P2014807** | Rpl4 | 0.125867276 | -0.131669935 | 0.383404487 | 15.92510768 | 1.202380733 | 0.275469913 | 0.751681228 | -4.645205339 |
| **A_51_P148037** | Sh3pxd2b | -0.104355285 | -0.317905709 | 0.10919514 | 8.15643143 | -1.202217848 | 0.275528391 | 0.751681228 | -4.645314616 |
| **A_55_P2040775** | NA | 0.174225519 | -0.182313531 | 0.530764569 | 13.26884311 | 1.202191981 | 0.275537679 | 0.751681228 | -4.645331969 |
| **A_52_P342896** | Mrps33 | 0.097085197 | -0.101601303 | 0.295771696 | 9.235060141 | 1.202136595 | 0.275557566 | 0.751681228 | -4.645369126 |
| **A_55_P1954758** | Prickle1 | 0.110355701 | -0.115505486 | 0.336216889 | 11.06745928 | 1.202049441 | 0.275588864 | 0.751681228 | -4.645427592 |
| **A_55_P2160473** | LOC634869 | 0.102982167 | -0.107816749 | 0.313781082 | 15.40130899 | 1.201884726 | 0.275648021 | 0.751681228 | -4.645538083 |
| **A_51_P342652** | Cd79b | -0.08653323 | -0.263687409 | 0.090620949 | 8.348136139 | -1.20171277 | 0.275709791 | 0.751681228 | -4.645653425 |
| **A_51_P108862** | Cops7a | -0.106080846 | -0.323275681 | 0.11111399 | 12.81224729 | -1.201590931 | 0.275753565 | 0.751681228 | -4.640734126 |
| **A_51_P275496** | BC026762 | -0.147532093 | -0.449618698 | 0.154554513 | 9.609860862 | -1.201501167 | 0.275785819 | 0.751681228 | -4.645795351 |
| **A_55_P2006422** | Gm6900 | 0.116581863 | -0.122131381 | 0.355295106 | 16.5786004 | 1.201499623 | 0.275786374 | 0.751681228 | -4.645796386 |
| **A_55_P2136622** | Camta2 | -0.086314629 | -0.263060513 | 0.090431254 | 8.332043787 | -1.201446026 | 0.275805634 | 0.751681228 | -4.645832333 |
| **A_51_P159284** | Bud13 | -0.085157956 | -0.259562499 | 0.089246587 | 8.546973033 | -1.201258841 | 0.275872909 | 0.751681228 | -4.645957869 |
| **A_55_P2028365** | Lrrc36 | 0.066163213 | -0.069340536 | 0.201666962 | 7.848979571 | 1.201252872 | 0.275875054 | 0.751681228 | -4.645961872 |
| **A_55_P2037608** | Acp5 | -0.323169861 | -0.985038454 | 0.338698732 | 13.50389116 | -1.201235728 | 0.275881217 | 0.751681228 | -4.645973369 |
| **A_55_P2073647** | Elp2 | 0.095642195 | -0.100239746 | 0.291524137 | 11.17738365 | 1.201224827 | 0.275885135 | 0.751681228 | -4.64598068 |
| **A_55_P1972575** | Tmeff1 | -0.134292505 | -0.409345445 | 0.140760434 | 12.58722903 | -1.201170476 | 0.275904672 | 0.751681228 | -4.646017128 |
| **A_52_P194316** | Atp13a3 | -0.131852705 | -0.401921162 | 0.138215752 | 9.444784199 | -1.201114315 | 0.275924861 | 0.751681228 | -4.64605479 |
| **A_55_P1999439** | Ino80e | 0.082227064 | -0.086218172 | 0.250672299 | 9.851287168 | 1.200950798 | 0.275983651 | 0.751681228 | -4.64616444 |
| **A_55_P2103944** | Mrpl30 | 0.074920573 | -0.078561247 | 0.228402393 | 11.84409507 | 1.200917878 | 0.275995488 | 0.751681228 | -4.646186514 |
| **A_55_P2081555** | Phf23 | 0.1050495 | -0.110155237 | 0.320254237 | 10.21477222 | 1.200912393 | 0.27599746 | 0.751681228 | -4.646190192 |
| **A_51_P496751** | Nop58 | -0.124428696 | -0.379341207 | 0.130483815 | 11.51159571 | -1.200877095 | 0.276010153 | 0.751681228 | -4.646213861 |
| **A_51_P307082** | Hey2 | -0.073739524 | -0.224810475 | 0.077331427 | 7.692403121 | -1.200849355 | 0.276020128 | 0.751681228 | -4.646232461 |
| **A_52_P597461** | Skil | -0.098036937 | -0.2988932 | 0.102819327 | 8.666935307 | -1.200807844 | 0.276035056 | 0.751681228 | -4.646260296 |
| **A_66_P116126** | Sdhb | 0.077077367 | -0.080839614 | 0.234994348 | 13.35732857 | 1.200790383 | 0.276041335 | 0.751681228 | -4.646272003 |
| **A_55_P2143095** | NA | -0.058403201 | -0.17806283 | 0.061256428 | 8.234456117 | -1.200765392 | 0.276050323 | 0.751681228 | -4.64628876 |
| **A_55_P2425938** | 2810468N07Rik | 0.060130132 | -0.063068585 | 0.183328849 | 8.045816777 | 1.200757021 | 0.276053334 | 0.751681228 | -4.646294373 |
| **A_51_P200477** | Hspa4 | -0.717043493 | -2.186322618 | 0.752235631 | 10.67907295 | -1.200633216 | 0.276097862 | 0.751681228 | -4.646377383 |

| | | | | | | | | | |
|---|---|---|---|---|---|---|---|---|---|
| **A_51_P427624** | Pgc | -0.081100409 | -0.247283404 | 0.085082585 | 8.874183104 | -1.200620155 | 0.27610256 | 0.751681228 | -4.64638614 |
| **A_55_P2067445** | Arf5 | 0.160638839 | -0.178872143 | 0.50014982 | 12.70633079 | 1.226159063 | 0.276110338 | 0.751681228 | -4.602519822 |
| **A_51_P428578** | Retreg1 | -0.158239886 | -0.48253403 | 0.166054258 | 8.88024681 | -1.200455366 | 0.27616184 | 0.751715562 | -4.646496623 |
| **A_55_P2049122** | Tm6sf2 | 0.158249332 | -0.166122296 | 0.48262096 | 8.626135603 | 1.200240256 | 0.276239238 | 0.751770475 | -4.646640834 |
| **A_51_P295215** | Alkbh5 | 0.084299364 | -0.088508473 | 0.257107201 | 11.50857398 | 1.200134794 | 0.276277191 | 0.751770475 | -4.646711531 |
| **A_55_P2017267** | Nanp | 0.085155448 | -0.089423607 | 0.259734503 | 8.688934367 | 1.200022708 | 0.276317533 | 0.751770475 | -4.646786666 |
| **A_66_P135185** | Etv1 | -0.063316936 | -0.193139259 | 0.066505386 | 7.976882576 | -1.19988516 | 0.276367046 | 0.751770475 | -4.646878865 |
| **A_55_P2183172** | Rdx | -0.165491778 | -0.504816411 | 0.173832856 | 12.66104736 | -1.199858952 | 0.276376481 | 0.751770475 | -4.646896433 |
| **A_52_P430169** | Dip2b | -0.07908188 | -0.241277935 | 0.083114174 | 9.434516861 | -1.1995155 | 0.27650015 | 0.751874285 | -4.647126627 |
| **A_55_P2057821** | Golga1 | -0.064420893 | -0.196554196 | 0.06771241 | 9.977353404 | -1.199454044 | 0.276522284 | 0.751874285 | -4.647167814 |
| **A_51_P125648** | Vwce | 0.059232666 | -0.062266297 | 0.18073163 | 8.071722089 | 1.199382951 | 0.27654789 | 0.751874285 | -4.639896802 |
| **A_51_P306710** | Cldn15 | 0.055215684 | -0.058049442 | 0.16848081 | 7.88788035 | 1.199320905 | 0.27657024 | 0.751874285 | -4.647257039 |
| **A_55_P2040044** | NA | 0.085134922 | -0.089571289 | 0.259841134 | 8.965540144 | 1.198860247 | 0.276736224 | 0.75218591 | -4.647565721 |
| **A_55_P1973399** | NA | -0.137643417 | -0.42016483 | 0.144877997 | 13.51445927 | -1.198597137 | 0.276831066 | 0.75218591 | -4.647742004 |
| **A_55_P2074381** | NA | -0.279462315 | -0.85310725 | 0.294182621 | 12.08963761 | -1.198530942 | 0.276854931 | 0.75218591 | -4.647786352 |
| **A_55_P2044724** | Zfp869 | 0.073154334 | -0.077008892 | 0.22331756 | 9.770048114 | 1.198520966 | 0.276858528 | 0.75218591 | -4.647793036 |
| **A_51_P210474** | Mrpl18 | -0.063225542 | -0.193019343 | 0.066568259 | 11.26462402 | -1.19841649 | 0.276896199 | 0.75218591 | -4.647863028 |
| **A_55_P2433438** | Tcf12 | -0.101116109 | -0.308715415 | 0.106483198 | 8.901081119 | -1.19829463 | 0.276940144 | 0.75218591 | -4.647944663 |
| **A_55_P2115477** | Parva | -0.091010758 | -0.277870391 | 0.095848876 | 9.24748296 | -1.198247136 | 0.276957272 | 0.75218591 | -4.647976478 |
| **A_51_P368543** | Abhd8 | -0.1134798 | -0.346518371 | 0.119558772 | 8.597754585 | -1.198008542 | 0.277043336 | 0.752313942 | -4.6481363 |
| **A_55_P2151466** | Casc3 | 0.104906339 | -0.110600203 | 0.320412882 | 11.89205246 | 1.197596276 | 0.277192099 | 0.752612175 | -4.648412422 |
| **A_51_P466591** | 1600012P17Rik | -0.165079958 | -0.504232496 | 0.17407258 | 17.53691378 | -1.197480475 | 0.277233897 | 0.752619944 | -4.648489974 |
| **A_55_P2082826** | Fam160b1 | 0.109424005 | -0.115409498 | 0.334257507 | 11.54088281 | 1.197348962 | 0.277281373 | 0.752643121 | -4.648578044 |
| **A_51_P209807** | Ccnyl1 | 0.082800451 | -0.087378029 | 0.25297893 | 8.656171421 | 1.197008472 | 0.277404323 | 0.752871125 | -4.648806041 |
| **A_55_P2002632** | Rap1b | 0.229562024 | -0.242393033 | 0.701517082 | 14.26242134 | 1.196653912 | 0.277532403 | 0.753006164 | -4.649043427 |
| **A_55_P2161170** | Srp68 | -0.107565611 | -0.328723074 | 0.113591852 | 12.08144477 | -1.196577996 | 0.277559833 | 0.753006164 | -4.649094251 |
| **A_51_P408649** | Shisa2 | -0.092341659 | -0.282203195 | 0.097519877 | 8.338172587 | -1.196547228 | 0.277570951 | 0.753006164 | -4.649114848 |
| **A_55_P2313861** | Zbed6 | -0.077310244 | -0.23630165 | 0.081681162 | 8.686248135 | -1.196279209 | 0.277667815 | 0.753099853 | -4.649294265 |
| **A_55_P1966445** | Krt81 | 0.123050485 | -0.130016832 | 0.376117802 | 9.947519859 | 1.1962348 | 0.277683868 | 0.753099853 | -4.649323992 |
| **A_51_P145415** | Lpcat3 | -0.096144999 | -0.293925981 | 0.101635983 | 11.03410036 | -1.19594535 | 0.277788515 | 0.753099853 | -4.649517731 |
| **A_55_P2049577** | Aggf1 | 0.116568091 | -0.123227265 | 0.356363448 | 12.46163862 | 1.195936386 | 0.277791756 | 0.753099853 | -4.649523731 |
| **A_55_P2008884** | Cltc | -0.193346461 | -0.591141276 | 0.204448354 | 14.28814755 | -1.195766197 | 0.277853303 | 0.753099853 | -4.649637635 |
| **A_52_P491569** | 1700017B05Rik | -0.098014891 | -0.299696137 | 0.103666354 | 9.631195125 | -1.19562699 | 0.277903654 | 0.753099853 | -4.649730797 |
| **A_52_P634447** | Hs1bp3 | 0.119213973 | -0.126108952 | 0.364536898 | 9.901433672 | 1.195523706 | 0.277941017 | 0.753099853 | -4.649799916 |
| **A_55_P1978608** | NA | -0.084548793 | -0.258540648 | 0.089443062 | 9.088516051 | -1.195494707 | 0.277951508 | 0.753099853 | -4.649819322 |
| **A_55_P2041893** | NA | 0.152197315 | -0.161032007 | 0.465426638 | 15.23251502 | 1.195401355 | 0.277985282 | 0.753099853 | -4.64988179 |
| **A_55_P2127702** | Racgap1 | 0.08091347 | -0.085614087 | 0.247441027 | 8.517275286 | 1.195374198 | 0.277995108 | 0.753099853 | -4.649899963 |
| **A_51_P341918** | Tsc22d1 | -0.195018512 | -0.596451294 | 0.206414269 | 14.45137209 | -1.195176836 | 0.278066528 | 0.753187769 | -4.650032025 |
| **A_51_P147373** | Dpp3 | -0.125675798 | -0.384401078 | 0.133049482 | 11.4415226 | -1.19503864 | 0.278116546 | 0.75320288 | -4.65012449 |
| **A_55_P2078285** | Uck1 | -0.163910782 | -0.501375495 | 0.17355393 | 13.08063673 | -1.194946107 | 0.278150042 | 0.75320288 | -4.6501864 |
| **A_51_P351363** | Tyms-ps | 0.125304063 | -0.132713667 | 0.383321794 | 8.589261458 | 1.194771258 | 0.278213344 | 0.753268767 | -4.650303379 |
| **A_51_P223036** | Fbxo30 | 0.094259739 | -0.099886761 | 0.28840624 | 9.500666584 | 1.194444103 | 0.278331821 | 0.753460884 | -4.650522235 |
| **A_55_P2171378** | Pdzk1ip1 | 0.327770248 | -0.347448749 | 1.002989245 | 9.095054226 | 1.194246675 | 0.278403339 | 0.753460884 | -4.650654294 |
| **A_51_P456366** | Pard6a | 0.132346163 | -0.140323436 | 0.405015762 | 8.520915012 | 1.194108524 | 0.278453393 | 0.753460884 | -4.650746697 |
| **A_55_P1974954** | NA | -0.053098008 | -0.162504546 | 0.05630853 | 15.39317915 | -1.193999419 | 0.278492929 | 0.753460884 | -4.650819669 |
| **A_51_P373890** | Guca1b | 0.058869855 | -0.062444645 | 0.180184355 | 8.977291111 | 1.19384905 | 0.278547426 | 0.753460884 | -4.650920234 |
| **A_55_P2046478** | Zfp81 | 0.059671163 | -0.063295196 | 0.182637522 | 8.140621404 | 1.193843369 | 0.278549485 | 0.753460884 | -4.650924033 |
| **A_66_P110817** | Mterf3 | -0.064088149 | -0.196159084 | 0.067982785 | 9.036775457 | -1.193822176 | 0.278557167 | 0.753460884 | -4.650938207 |
| **A_65_P09177** | Strn3 | -0.176621927 | -0.540747817 | 0.187503964 | 9.986732071 | -1.193334565 | 0.278733956 | 0.753678311 | -4.651264274 |
| **A_51_P294643** | Cdr2 | 0.169114174 | -0.179537954 | 0.517766303 | 9.586684631 | 1.193319814 | 0.278739306 | 0.753678311 | -4.651274138 |

| | | | | | | | | | |
|---|---|---|---|---|---|---|---|---|---|
| **A_55_P2080396** | Mrpl4 | 0.153216983 | -0.162675462 | 0.469109428 | 12.92597821 | 1.193264802 | 0.278759257 | 0.753678311 | -4.651310921 |
| **A_51_P282508** | Rhoc | -0.23059333 | -0.706052747 | 0.244866088 | 16.22751043 | -1.193170341 | 0.278793519 | 0.753678311 | -4.651374078 |
| **A_51_P228254** | Zdhhc21 | -0.10925547 | -0.334571811 | 0.116060872 | 11.29073319 | -1.192942913 | 0.278876024 | 0.753726337 | -4.651526131 |
| **A_55_P2128326** | Ubap2l | -0.067215537 | -0.205837635 | 0.071406561 | 9.0418057 | -1.192906396 | 0.278889273 | 0.753726337 | -4.651550543 |
| **A_55_P2172239** | NA | 0.111225391 | -0.118221543 | 0.340672324 | 16.04581303 | 1.192589127 | 0.27900441 | 0.753830055 | -4.651762636 |
| **A_55_P2087528** | Tmem131l | -0.144139844 | -0.441486876 | 0.153207189 | 10.61822648 | -1.19258571 | 0.27900565 | 0.753830055 | -4.65176492 |
| **A_52_P408657** | Mycbp | 0.069822623 | -0.074232989 | 0.213878234 | 8.438356107 | 1.192436166 | 0.279059935 | 0.753871345 | -4.651864881 |
| **A_55_P2058942** | Aldh3b1 | 0.108141955 | -0.115080721 | 0.331364632 | 8.748007277 | 1.19185952 | 0.27926934 | 0.75421932 | -4.652250278 |
| **A_55_P2003720** | NA | 0.170360145 | -0.181316477 | 0.522036767 | 13.96637987 | 1.191773336 | 0.279300649 | 0.75421932 | -4.652307871 |
| **A_55_P1957291** | Rpl39 | 0.084219419 | -0.089637873 | 0.25807671 | 15.59952002 | 1.191759146 | 0.279305805 | 0.75421932 | -4.652317354 |
| **A_51_P168395** | Ttc30a1 | 0.073871801 | -0.078642154 | 0.226385757 | 8.634780477 | 1.191621325 | 0.279355879 | 0.754249167 | -4.652409449 |
| **A_55_P2035205** | Nifk | 0.086810896 | -0.092507231 | 0.266129023 | 11.52134963 | 1.191020423 | 0.279574296 | 0.75473346 | -4.652810931 |
| **A_55_P2106834** | Ndufs3 | 0.178173237 | -0.189931419 | 0.546277894 | 12.99665203 | 1.190804104 | 0.27965296 | 0.754840395 | -4.652955439 |
| **A_51_P256282** | Ppp6c | 0.10065797 | -0.107418631 | 0.308734571 | 11.32350216 | 1.190129127 | 0.279898538 | 0.75538082 | -4.653406266 |
| **A_55_P2010008** | Zfp113 | 0.096033272 | -0.10249835 | 0.294564895 | 8.54839352 | 1.190039016 | 0.279931337 | 0.75538082 | -4.653466444 |
| **A_55_P2108206** | Rrp7a | 0.160918318 | -0.171847793 | 0.493684429 | 11.84784266 | 1.18969478 | 0.280056665 | 0.755613523 | -4.653696312 |
| **A_55_P2033462** | Rps23-ps1 | -0.085648372 | -0.262861887 | 0.091565142 | 16.30488945 | -1.189026237 | 0.280300204 | 0.756095592 | -4.654142654 |
| **A_55_P1990440** | R3hdm1 | -0.13528134 | -0.415200794 | 0.144638113 | 10.73849285 | -1.188978439 | 0.280317623 | 0.756095592 | -4.654174561 |
| **A_55_P2080163** | Carhsp1 | 0.12703636 | -0.135844139 | 0.38991686 | 12.71410707 | 1.188882227 | 0.280352688 | 0.756095592 | -4.654238785 |
| **A_51_P208801** | Ndufab1 | 0.104682978 | -0.111965222 | 0.321331179 | 9.350749716 | 1.188748952 | 0.280401267 | 0.756121107 | -4.654327745 |
| **A_66_P137495** | Btaf1 | -0.087052425 | -0.267247501 | 0.093142651 | 9.80412478 | -1.188521706 | 0.280484116 | 0.756239013 | -4.65447942 |
| **A_52_P64687** | Camk2n1 | -0.147767488 | -0.453683552 | 0.158148576 | 8.960215953 | -1.18835383 | 0.280545334 | 0.756298572 | -4.654591461 |
| **A_55_P2078940** | Nsfl1c | -0.084118329 | -0.258321477 | 0.090084819 | 13.17463253 | -1.187965429 | 0.280687012 | 0.756432578 | -4.654850651 |
| **A_55_P2056260** | NA | -0.098312556 | -0.301928056 | 0.105302944 | 15.37093441 | -1.187865625 | 0.280723428 | 0.756432578 | -4.654917247 |
| **A_55_P2134712** | NA | -0.26397982 | -0.810720163 | 0.282760522 | 11.27306011 | -1.187842258 | 0.280731955 | 0.756432578 | -4.654932838 |
| **A_51_P452629** | Tlr2 | -0.109125007 | -0.335169061 | 0.116919047 | 8.34083012 | -1.187682502 | 0.280790255 | 0.756432578 | -4.647556452 |
| **A_55_P2085157** | Eef1g | -0.115772942 | -0.355606477 | 0.124060593 | 12.25861789 | -1.187589429 | 0.280824226 | 0.756432578 | -4.655101529 |
| **A_55_P1961099** | Atg4c | 0.063292114 | -0.067838752 | 0.194422979 | 9.990837377 | 1.187445911 | 0.280876615 | 0.756432578 | -4.655197279 |
| **A_51_P314521** | Smim11 | 0.129816462 | -0.139155256 | 0.39878818 | 12.04931036 | 1.187387065 | 0.280898098 | 0.756432578 | -4.655236537 |
| **A_51_P272283** | Cmbl | 0.287279247 | -0.307981739 | 0.882540233 | 9.792074916 | 1.187315117 | 0.280924367 | 0.756432578 | -4.655284535 |
| **A_51_P508959** | Prr15 | 0.202186501 | -0.216804569 | 0.621177571 | 9.365188898 | 1.187179947 | 0.280973724 | 0.756432578 | -4.655374706 |
| **A_55_P2012406** | NA | -0.137939677 | -0.423800411 | 0.147921056 | 14.68787385 | -1.187145265 | 0.280986389 | 0.756432578 | -4.655397842 |
| **A_51_P280446** | Sdf2l1 | 0.12563289 | -0.134802925 | 0.386068705 | 11.89927877 | 1.186784338 | 0.281118222 | 0.756573763 | -4.655638588 |
| **A_55_P2150179** | C1s2 | -0.162549125 | -0.499537994 | 0.174439744 | 9.714004413 | -1.186692622 | 0.281151731 | 0.756573763 | -4.655699759 |
| **A_52_P163795** | Tubb5 | -0.157536016 | -0.484135366 | 0.169063333 | 13.86621011 | -1.186680228 | 0.28115626 | 0.756573763 | -4.655708025 |
| **A_51_P251205** | Psd | 0.303018658 | -0.325333909 | 0.931371225 | 9.018709514 | 1.186410934 | 0.281254669 | 0.756576052 | -4.655887621 |
| **A_55_P2113291** | Sik3 | -0.12167131 | -0.395379078 | 0.152036458 | 11.93821829 | -1.250845216 | 0.281290998 | 0.756576052 | -4.574303618 |
| **A_55_P2007693** | NA | -0.064831958 | -0.199294114 | 0.069630197 | 9.796129764 | -1.186200864 | 0.281331456 | 0.756576052 | -4.656027707 |
| **A_55_P2044297** | Ciao2a | 0.164910664 | -0.177129092 | 0.50695042 | 12.84753778 | 1.186154648 | 0.281348352 | 0.756576052 | -4.656058524 |
| **A_55_P2045512** | NA | 0.157052224 | -0.168707221 | 0.482811669 | 15.43470242 | 1.186086156 | 0.281373393 | 0.756576052 | -4.656104195 |
| **A_52_P367565** | Mrps2 | 0.08452734 | -0.090807703 | 0.259862382 | 9.52415454 | 1.186035369 | 0.281391962 | 0.756576052 | -4.65613806 |
| **A_55_P1988083** | Prc1 | 0.199234563 | -0.214080234 | 0.61254936 | 9.210987938 | 1.185913166 | 0.281436648 | 0.756590956 | -4.656219541 |
| **A_55_P1960431** | Ubn1 | -0.120144673 | -0.369459585 | 0.129170238 | 12.37691919 | -1.185565166 | 0.281563934 | 0.756827879 | -4.656451554 |
| **A_51_P154842** | Oas1f | -0.168836459 | -0.519256345 | 0.181583427 | 11.9770918 | -1.185350134 | 0.28164261 | 0.756934094 | -4.656594902 |
| **A_51_P494863** | Vmac | -0.104605498 | -0.321783051 | 0.112572056 | 9.294102976 | -1.184973768 | 0.28178036 | 0.757129442 | -4.656845772 |
| **A_52_P376106** | Slc30a9 | -0.086122475 | -0.264939678 | 0.092694728 | 9.029744696 | -1.184885452 | 0.281812692 | 0.757129442 | -4.656904634 |
| **A_55_P2027594** | Taf3 | 0.139838003 | -0.15052643 | 0.430202436 | 9.083240691 | 1.184816129 | 0.281838073 | 0.757129442 | -4.656950837 |
| **A_55_P1967643** | Plb1 | 0.388494612 | -0.418286301 | 1.195275526 | 11.04844044 | 1.184673157 | 0.281890425 | 0.757129442 | -4.65704612 |
| **A_55_P2015862** | Erg | -0.077226755 | -0.237631285 | 0.083177775 | 8.12707872 | -1.184459782 | 0.281968572 | 0.757129442 | -4.657188314 |
| **A_52_P205797** | Vgll4 | -0.069439478 | -0.213689474 | 0.074810518 | 7.867539241 | -1.184294734 | 0.282029033 | 0.757129442 | -4.657298295 |

| | | | | | | | | | |
|---|---|---|---|---|---|---|---|---|---|
| **A_52_P504787** | Epha7 | 0.05419172 | -0.058383457 | 0.166766897 | 7.744652436 | 1.184293808 | 0.282029373 | 0.757129442 | -4.657298912 |
| **A_51_P237695** | Gnai3 | -0.165401782 | -0.509044025 | 0.17824046 | 13.09963595 | -1.184139324 | 0.282085974 | 0.757129442 | -4.657401847 |
| **A_55_P2171303** | Bin3 | -0.105408452 | -0.324439296 | 0.113622392 | 8.240398449 | -1.183966253 | 0.282149396 | 0.757129442 | -4.657517159 |
| **A_51_P480241** | Elf3 | -0.144384139 | -0.444447527 | 0.155679249 | 9.968253858 | -1.183792659 | 0.282213023 | 0.757129442 | -4.657632813 |
| **A_51_P387868** | Tekt4 | 0.057420634 | -0.061924618 | 0.176765886 | 8.148983346 | 1.183673709 | 0.282256628 | 0.757129442 | -4.657712056 |
| **A_55_P2088028** | Mtss1 | -0.064636309 | -0.198985292 | 0.069712673 | 8.815153597 | -1.183617375 | 0.282277281 | 0.757129442 | -4.657749584 |
| **A_52_P282987** | Yeats2 | -0.118857867 | -0.365929698 | 0.128213964 | 9.301865225 | -1.183515257 | 0.282314723 | 0.757129442 | -4.657817609 |
| **A_52_P454994** | Pi4ka | -0.155135373 | -0.477633761 | 0.167363016 | 12.26937981 | -1.183456903 | 0.282336121 | 0.757129442 | -4.65785648 |
| **A_51_P219385** | Six5 | -0.093305842 | -0.287277466 | 0.100665782 | 8.645245665 | -1.183422433 | 0.282348762 | 0.757129442 | -4.657879441 |
| **A_52_P200265** | Mocs1 | 0.058681397 | -0.063320287 | 0.180683081 | 9.132165412 | 1.1833243 | 0.28238475 | 0.757129442 | -4.657944808 |
| **A_52_P582309** | Nub1 | 0.076249932 | -0.082278158 | 0.234778021 | 11.63728939 | 1.183320524 | 0.282386135 | 0.757129442 | -4.657947323 |
| **A_55_P1989797** | NA | -0.167265471 | -0.515047646 | 0.180516704 | 14.67741555 | -1.183227192 | 0.282420367 | 0.757129442 | -4.658009489 |
| **A_52_P671543** | Pomc | -0.079832149 | -0.245934257 | 0.086269959 | 8.959376801 | -1.182420203 | 0.282716499 | 0.757815321 | -4.658546907 |
| **A_55_P1994694** | Serinc2 | -0.166821083 | -0.5139476 | 0.180305433 | 8.72734561 | -1.18231258 | 0.282756012 | 0.757815321 | -4.658618566 |
| **A_55_P2009187** | Iffo1 | 0.075578155 | -0.081707318 | 0.232863627 | 8.4465171 | 1.182161567 | 0.282811464 | 0.757815321 | -4.658719111 |
| **A_55_P1978311** | NA | 0.121265434 | -0.131139626 | 0.373670493 | 13.60825232 | 1.181974568 | 0.282880143 | 0.757815321 | -4.658843607 |
| **A_55_P1984009** | Rsph3b | -0.080137146 | -0.246945117 | 0.086670825 | 11.97421654 | -1.181914978 | 0.282902031 | 0.757815321 | -4.658883278 |
| **A_55_P2085181** | Chaf1b | 0.063719528 | -0.068919374 | 0.196358431 | 9.353780993 | 1.181872955 | 0.282917468 | 0.757815321 | -4.658911253 |
| **A_55_P2372228** | NA | -0.051448349 | -0.158551686 | 0.055654989 | 7.848490253 | -1.181782623 | 0.282950653 | 0.757815321 | -4.658971387 |
| **A_55_P2127688** | 4933434E20Rik | 0.145964258 | -0.157968935 | 0.44989745 | 12.94332035 | 1.181510413 | 0.283050675 | 0.757978178 | -4.659152584 |
| **A_55_P2361437** | Pitpnc1 | -0.151108151 | -0.465862641 | 0.163646338 | 8.373471026 | -1.181095792 | 0.283203082 | 0.758156701 | -4.659428539 |
| **A_55_P2184567** | NA | 0.061121325 | -0.066197706 | 0.188440357 | 8.507177822 | 1.181050861 | 0.283219602 | 0.758156701 | -4.659458441 |
| **A_55_P2167160** | Hdhd2 | -0.079183434 | -0.244132669 | 0.085765801 | 8.309334991 | -1.181008954 | 0.283235011 | 0.758156701 | -4.65948633 |
| **A_51_P475298** | Zfp110 | -0.070430204 | -0.217217054 | 0.076356645 | 9.027884545 | -1.180431968 | 0.28344724 | 0.758280575 | -4.659870261 |
| **A_55_P2012011** | Tceal7 | 0.276055129 | -0.299313496 | 0.851423754 | 9.072269591 | 1.180371751 | 0.283469396 | 0.758280575 | -4.659910325 |
| **A_55_P2083841** | Atp13a1 | -0.205611226 | -0.634159268 | 0.222936816 | 12.40690967 | -1.180365401 | 0.283471733 | 0.758280575 | -4.65991455 |
| **A_55_P2087825** | Ncoa2 | -0.107050011 | -0.330184987 | 0.116084965 | 11.25586292 | -1.180288645 | 0.283499978 | 0.758280575 | -4.659965616 |
| **A_66_P106289** | Vps54 | -0.066515341 | -0.205202396 | 0.072171713 | 9.718598754 | -1.179926789 | 0.283633168 | 0.758280575 | -4.660206341 |
| **A_51_P295442** | Nek7 | -0.146659426 | -0.45245138 | 0.159132529 | 11.03969121 | -1.179921426 | 0.283635142 | 0.758280575 | -4.660209909 |
| **A_51_P155152** | Ank | 0.135159701 | -0.146665831 | 0.416985234 | 11.54228947 | 1.179875139 | 0.283652183 | 0.758280575 | -4.660240699 |
| **A_55_P2084763** | NA | -0.168465254 | -0.519752065 | 0.182821558 | 10.5850833 | -1.179825174 | 0.283670579 | 0.758280575 | -4.660273935 |
| **A_55_P2174942** | NA | 0.180468768 | -0.195880242 | 0.556817778 | 13.91265943 | 1.179724068 | 0.283707808 | 0.758280575 | -4.660341187 |
| **A_55_P2014269** | Zfp771 | 0.165812258 | -0.180047458 | 0.511671973 | 12.83372943 | 1.179467056 | 0.283802461 | 0.758280575 | -4.660512129 |
| **A_55_P1986406** | Ddx54 | -0.091690789 | -0.282945503 | 0.099563924 | 10.89132398 | -1.179458674 | 0.283805548 | 0.758280575 | -4.660517704 |
| **A_55_P1961620** | Vax2 | 0.207428118 | -0.225269863 | 0.640126098 | 11.4169172 | 1.17937501 | 0.283836367 | 0.758280575 | -4.660573347 |
| **A_52_P68702** | Frmd4b | 0.069740308 | -0.075760053 | 0.21524067 | 8.266833454 | 1.179204031 | 0.283899357 | 0.758280575 | -4.660687054 |
| **A_55_P2014337** | Bach1 | -0.136657984 | -0.421779507 | 0.148463538 | 11.06735138 | -1.179163893 | 0.283914146 | 0.758280575 | -4.660713747 |
| **A_51_P473919** | Mycbp2 | -0.111498589 | -0.344132417 | 0.121135238 | 9.967579351 | -1.17914081 | 0.283922652 | 0.758280575 | -4.660729097 |
| **A_55_P2120055** | Fgfr4 | 0.162864361 | -0.176995126 | 0.502723849 | 9.198392322 | 1.178951189 | 0.283992529 | 0.758280575 | -4.660855189 |
| **A_51_P412835** | Daxx | 0.153463247 | -0.166783657 | 0.473710151 | 12.77095029 | 1.17893159 | 0.283999752 | 0.758280575 | -4.660868222 |
| **A_55_P1990196** | NA | -0.097055241 | -0.299618987 | 0.105508505 | 11.26284097 | -1.178762838 | 0.284061953 | 0.758280575 | -4.660980429 |
| **A_51_P332676** | Top1mt | 0.103866072 | -0.11291856 | 0.320650703 | 9.423795228 | 1.178730115 | 0.284074016 | 0.758280575 | -4.661002187 |
| **A_55_P1980035** | Tbc1d9b | -0.184927209 | -0.570899439 | 0.201045021 | 11.11746997 | -1.178728866 | 0.284074476 | 0.758280575 | -4.661003017 |
| **A_51_P472481** | Dync2li1 | -0.077162229 | -0.238230272 | 0.083905813 | 8.330841276 | -1.178594879 | 0.284123873 | 0.758280575 | -4.661092101 |
| **A_55_P2046398** | Eif4a1 | -0.177358586 | -0.547593439 | 0.192876267 | 14.09824271 | -1.178539332 | 0.284144353 | 0.758280575 | -4.661129031 |
| **A_51_P249118** | Abcb4 | 0.173016029 | -0.188250158 | 0.534282216 | 10.27280453 | 1.178224874 | 0.28426032 | 0.758368297 | -4.661338083 |
| **A_55_P2041894** | NA | -0.088324815 | -0.27275911 | 0.09610948 | 16.59245051 | -1.178176128 | 0.284278301 | 0.758368297 | -4.661370488 |
| **A_52_P91235** | Ccni | -0.210869691 | -0.651277784 | 0.229538403 | 12.37049217 | -1.177953189 | 0.284360546 | 0.758368297 | -4.661518679 |
| **A_51_P502152** | Slc19a1 | 0.091664079 | -0.099788951 | 0.283117109 | 9.321452133 | 1.177893695 | 0.284382497 | 0.758368297 | -4.661558223 |
| **A_55_P1972648** | Pheta2 | 0.05732174 | -0.062410888 | 0.177054368 | 7.960155145 | 1.177812107 | 0.284412603 | 0.758368297 | -4.661612452 |

| | | | | | | | | | |
|---|---|---|---|---|---|---|---|---|---|
| **A_51_P517096** | Casp8ap2 | -0.084415331 | -0.260740579 | 0.091909918 | 9.493094266 | -1.177812027 | 0.284412633 | 0.758368297 | -4.661612505 |
| **A_55_P1962234** | Bad | 0.169907487 | -0.185052641 | 0.524867615 | 12.92320071 | 1.177611685 | 0.284486571 | 0.758377459 | -4.661745657 |
| **A_55_P1955857** | Ap5z1 | -0.344117918 | -1.063130549 | 0.374894713 | 11.24905077 | -1.177443101 | 0.284548801 | 0.758377459 | -4.661857694 |
| **A_52_P583973** | Lrpap1 | -0.101468513 | -0.313487569 | 0.110550544 | 13.43543297 | -1.177404146 | 0.284563182 | 0.758377459 | -4.661883582 |
| **A_55_P1995141** | Casz1 | 0.091131837 | -0.099309218 | 0.281572892 | 8.254256894 | 1.177277134 | 0.284610077 | 0.758377459 | -4.661967985 |
| **A_55_P1963767** | Npr2 | 0.197936156 | -0.215699294 | 0.611571606 | 12.02316361 | 1.177271262 | 0.284612245 | 0.758377459 | -4.661971887 |
| **A_52_P203560** | Fzd10 | -0.062657871 | -0.193625298 | 0.068309556 | 7.774065313 | -1.177013654 | 0.284707378 | 0.758424441 | -4.662143062 |
| **A_55_P2132512** | Ngef | -0.138579555 | -0.42823907 | 0.15107996 | 8.821639145 | -1.177011015 | 0.284708353 | 0.758424441 | -4.662144815 |
| **A_52_P206492** | Pop4 | -0.088021441 | -0.27206571 | 0.096022827 | 10.92869765 | -1.176617601 | 0.284853691 | 0.75870704 | -4.662406195 |
| **A_51_P403814** | Slc4a5 | -0.135928328 | -0.420224963 | 0.148368308 | 9.518979159 | -1.176271074 | 0.284981761 | 0.758820257 | -4.662636392 |
| **A_55_P2091284** | Ltbp3 | -0.257920888 | -0.797426006 | 0.281584231 | 10.78755443 | -1.176142947 | 0.285029126 | 0.758820257 | -4.662721497 |
| **A_51_P230987** | Pi4k2b | -0.181698802 | -0.561783906 | 0.198386301 | 10.07952065 | -1.176089516 | 0.285048881 | 0.758820257 | -4.662756987 |
| **A_55_P2032081** | Dbp | 0.069814309 | -0.076235067 | 0.215863685 | 9.272924062 | 1.176017821 | 0.285075389 | 0.758820257 | -4.662804606 |
| **A_51_P155196** | Abtb2 | 0.075678685 | -0.082645009 | 0.23400238 | 10.13976876 | 1.175971578 | 0.285092488 | 0.758820257 | -4.662835319 |
| **A_52_P486045** | Coq4 | 0.065732666 | -0.071798106 | 0.203263439 | 8.755932303 | 1.175846229 | 0.285138843 | 0.758839142 | -4.66291857 |
| **A_55_P2138306** | NA | -0.104289664 | -0.322560763 | 0.113981436 | 11.40658582 | -1.175477159 | 0.285275363 | 0.759097947 | -4.663163665 |
| **A_55_P2077746** | Mphosph8 | -0.103708301 | -0.320797165 | 0.113380562 | 10.6403 | -1.175290265 | 0.285344517 | 0.759177449 | -4.663287766 |
| **A_55_P2102390** | Spag9 | -0.185442743 | -0.573661444 | 0.202775957 | 12.49275204 | -1.17517503 | 0.285387163 | 0.759186413 | -4.663364279 |
| **A_55_P2027812** | Arxes2 | 0.234672772 | -0.256678716 | 0.726024259 | 8.89162662 | 1.175004665 | 0.285450222 | 0.759249668 | -4.663477391 |
| **A_51_P223946** | Pds5b | -0.059610202 | -0.184449309 | 0.065228905 | 8.262903959 | -1.174732818 | 0.285550868 | 0.759340501 | -4.663657865 |
| **A_52_P286098** | Epb41l4b | 0.11056337 | -0.120991215 | 0.342117955 | 12.16333528 | 1.174700211 | 0.285562942 | 0.759340501 | -4.66367951 |
| **A_51_P277336** | Cavin2 | 0.133283558 | -0.145955902 | 0.412523018 | 11.93190748 | 1.174272664 | 0.2857213 | 0.759567482 | -4.663963307 |
| **A_55_P2015715** | Rab4a | 0.158760768 | -0.173899122 | 0.491420658 | 10.19983333 | 1.174118457 | 0.285778435 | 0.759567482 | -4.664065655 |
| **A_55_P2048779** | Lratd2 | 0.064119268 | -0.070234682 | 0.198473218 | 7.773593651 | 1.17410593 | 0.285783076 | 0.759567482 | -4.664073969 |
| **A_55_P1971076** | Atp11a | 0.081781657 | -0.089629298 | 0.253192611 | 13.05354877 | 1.173779429 | 0.285904081 | 0.759567482 | -4.664290647 |
| **A_55_P2032659** | Tox | 0.074726653 | -0.081901997 | 0.231355302 | 7.801209935 | 1.173744281 | 0.28591711 | 0.759567482 | -4.66431397 |
| **A_55_P2144248** | Zfp790 | -0.117496783 | -0.363796206 | 0.12880264 | 8.985078355 | -1.173631466 | 0.285958933 | 0.759567482 | -4.664388831 |
| **A_55_P2005680** | Ipo11 | 0.109531368 | -0.12008205 | 0.339144786 | 10.78391751 | 1.173573821 | 0.285980305 | 0.759567482 | -4.664427081 |
| **A_55_P2150058** | Zfp593 | -0.056699307 | -0.175571898 | 0.062173284 | 9.931384376 | -1.173451612 | 0.286025618 | 0.759567482 | -4.66450817 |
| **A_55_P1955239** | Rps6 | -0.086861375 | -0.268976997 | 0.095254246 | 13.96236191 | -1.173406972 | 0.286042172 | 0.759567482 | -4.664537788 |
| **A_55_P1967633** | Lsm5 | 0.132353301 | -0.145176953 | 0.409883556 | 9.516137849 | 1.173258225 | 0.286097337 | 0.759567482 | -4.664636478 |
| **A_55_P1996314** | Amy2a5 | -0.060129791 | -0.18622561 | 0.065966028 | 7.691161267 | -1.1731625 | 0.286132842 | 0.759567482 | -4.664699986 |
| **A_55_P2001558** | Zfp128 | -0.060479351 | -0.187312625 | 0.066353923 | 8.272700927 | -1.173121732 | 0.286147965 | 0.759567482 | -4.664727033 |
| **A_55_P1979217** | Abtb1 | 0.147028231 | -0.161317533 | 0.455373994 | 11.81469733 | 1.173091553 | 0.28615916 | 0.759567482 | -4.664747054 |
| **A_55_P2099379** | Tfe3 | -0.08039547 | -0.249044739 | 0.0882538 | 9.207256877 | -1.172779252 | 0.286275032 | 0.759718848 | -4.664954225 |
| **A_55_P2028798** | Nqo2 | 0.159665647 | -0.175287525 | 0.49461882 | 9.278204479 | 1.172725999 | 0.286294795 | 0.759718848 | -4.664989549 |
| **A_51_P239286** | Bcl2l12 | 0.094527634 | -0.103802747 | 0.292858015 | 9.155143437 | 1.172569799 | 0.286352767 | 0.759768379 | -4.665093156 |
| **A_55_P2112637** | NA | 0.053491686 | -0.05876539 | 0.165748763 | 7.682923719 | 1.172307989 | 0.28644996 | 0.759826535 | -4.665266798 |
| **A_51_P234359** | Sct | -0.118029699 | -0.365745007 | 0.12968561 | 17.81705883 | -1.172215922 | 0.286484145 | 0.759826535 | -4.665327856 |
| **A_55_P2124619** | Smurf2 | -0.065905815 | -0.204228604 | 0.072416974 | 10.19186931 | -1.172193107 | 0.286492617 | 0.759826535 | -4.665342987 |
| **A_55_P2062911** | Epha5 | -0.154748216 | -0.479586108 | 0.170089676 | 13.2936027 | -1.172001399 | 0.286563812 | 0.759911087 | -4.665470118 |
| **A_51_P183275** | Sympk | 0.169533352 | -0.18640436 | 0.525471065 | 10.52406281 | 1.171791404 | 0.286641816 | 0.76001367 | -4.665609365 |
| **A_66_P130582** | Ncoa1 | -0.06834036 | -0.211844208 | 0.075163488 | 8.981236713 | -1.171609484 | 0.286709406 | 0.760068425 | -4.665729986 |
| **A_66_P123643** | Ptbp2 | -0.057088299 | -0.176973347 | 0.062796748 | 7.989508353 | -1.171524158 | 0.286741112 | 0.760068425 | -4.665786557 |
| **A_51_P144512** | Rbm19 | -0.054295727 | -0.168330162 | 0.059738709 | 9.520288015 | -1.171382676 | 0.286793692 | 0.760103562 | -4.665880357 |
| **A_51_P326030** | Zfp959 | -0.07891405 | -0.244704935 | 0.086876835 | 8.070475614 | -1.171016045 | 0.286929985 | 0.760319599 | -4.666123401 |
| **A_55_P1970901** | Gramd2 | 0.124507564 | -0.137116507 | 0.386131635 | 8.709397576 | 1.170812093 | 0.287005827 | 0.760319599 | -4.666258587 |
| **A_55_P1968618** | NA | 0.056082681 | -0.061765172 | 0.173930534 | 7.734417467 | 1.170782589 | 0.2870168 | 0.760319599 | -4.666278143 |
| **A_51_P488673** | Rack1 | 0.09937554 | -0.109466833 | 0.308217914 | 17.0466815 | 1.170658023 | 0.287063131 | 0.760319599 | -4.666360703 |
| **A_55_P2122020** | Klf4 | -0.135999311 | -0.421814005 | 0.149815384 | 11.41927209 | -1.170634494 | 0.287071883 | 0.760319599 | -4.666376297 |

| | | | | | | | | | |
|---|---|---|---|---|---|---|---|---|---|
| **A_52_P469381** | Comtd1 | 0.123043959 | -0.135575194 | 0.381663112 | 10.84320175 | 1.170492875 | 0.287124566 | 0.760354945 | -4.666470154 |
| **A_51_P111952** | Nudt19 | 0.088120343 | -0.097161882 | 0.273402568 | 10.27916854 | 1.170069304 | 0.287282187 | 0.760580627 | -4.664145035 |
| **A_55_P1975095** | Timmdc1 | 0.057363896 | -0.063260274 | 0.177988066 | 9.776162395 | 1.169966602 | 0.287320416 | 0.760580627 | -4.666818891 |
| **A_66_P111090** | Sctr | -0.181927167 | -0.564488133 | 0.200633798 | 10.74677227 | -1.16994667 | 0.287327836 | 0.760580627 | -4.666832098 |
| **A_51_P205215** | Klhl30 | 0.251772118 | -0.277795189 | 0.781339425 | 8.155762757 | 1.169649208 | 0.287438588 | 0.760680749 | -4.667029178 |
| **A_55_P2013336** | Melk | 0.22643853 | -0.249849451 | 0.702726511 | 8.368288362 | 1.169633685 | 0.287444368 | 0.760680749 | -4.667039461 |
| **A_55_P2008691** | NA | 0.101488062 | -0.112052221 | 0.315028344 | 15.75308827 | 1.169241733 | 0.287590359 | 0.760788483 | -4.667299107 |
| **A_55_P2172146** | Pds5a | -0.122424464 | -0.380018893 | 0.135169965 | 9.081657787 | -1.169232579 | 0.28759377 | 0.760788483 | -4.667305171 |
| **A_55_P1965308** | Ccdc66 | -0.081728371 | -0.253697487 | 0.090240745 | 8.694666327 | -1.16920738 | 0.287603158 | 0.760788483 | -4.667321863 |
| **A_55_P1998962** | Oas1b | 0.097309866 | -0.107493261 | 0.302112993 | 13.44736809 | 1.168932556 | 0.287705564 | 0.760955233 | -4.667503891 |
| **A_51_P407323** | F5 | -0.051251878 | -0.159140411 | 0.056636656 | 7.674970671 | -1.168701647 | 0.28779163 | 0.761015465 | -4.667656817 |
| **A_51_P391764** | Cbx6 | 0.215723739 | -0.238404855 | 0.669852334 | 10.47626999 | 1.168660203 | 0.28780708 | 0.761015465 | -4.667684263 |
| **A_55_P2117033** | Mtmr3 | 0.070301778 | -0.077718829 | 0.218322385 | 9.959742589 | 1.168458493 | 0.287882285 | 0.761110201 | -4.667817837 |
| **A_55_P2131088** | Vwa5a | -0.12255913 | -0.380671273 | 0.135553012 | 9.877933143 | -1.168170937 | 0.287989525 | 0.761222152 | -4.668008241 |
| **A_55_P2095529** | NA | 0.076074382 | -0.084144989 | 0.236293752 | 14.12762756 | 1.168133754 | 0.288003394 | 0.761222152 | -4.668032859 |
| **A_51_P220062** | Mmp15 | -0.136089182 | -0.422805657 | 0.150627293 | 9.738702774 | -1.167723764 | 0.28815636 | 0.761303185 | -4.66830429 |
| **A_55_P1955287** | NA | 0.099431773 | -0.110066775 | 0.308930322 | 16.27703804 | 1.167651739 | 0.288183239 | 0.761303185 | -4.668351968 |
| **A_51_P198645** | Parvb | -0.05968675 | -0.185445112 | 0.066071613 | 7.978889593 | -1.167643374 | 0.288186361 | 0.761303185 | -4.668357506 |
| **A_55_P2090995** | Ebna1bp2 | -0.084307019 | -0.261941755 | 0.093327717 | 9.12836786 | -1.167629341 | 0.288191598 | 0.761303185 | -4.668366795 |
| **A_55_P1984284** | Gapdh | -0.154985708 | -0.481598686 | 0.17162727 | 14.61391098 | -1.167420666 | 0.288269489 | 0.761404886 | -4.668504924 |
| **A_51_P265219** | Ppm1k | -0.05469544 | -0.169985723 | 0.060594843 | 8.230843983 | -1.167152434 | 0.288369637 | 0.761407021 | -4.668682459 |
| **A_55_P1959119** | NA | -0.07632223 | -0.237227343 | 0.084582882 | 13.94620181 | -1.166944961 | 0.288447119 | 0.761407021 | -4.668819765 |
| **A_51_P271370** | Wsb1 | -0.087847487 | -0.273066966 | 0.097371991 | 8.252855081 | -1.16684146 | 0.28848578 | 0.761407021 | -4.668888259 |
| **A_52_P658034** | Tspyl1 | -0.120003443 | -0.373036316 | 0.13302943 | 11.37167165 | -1.166771796 | 0.288511803 | 0.761407021 | -4.668934359 |
| **A_55_P2070352** | Csf3r | -0.431270319 | -1.340688452 | 0.478147813 | 13.9851472 | -1.166689057 | 0.288542714 | 0.761407021 | -4.668989109 |
| **A_55_P2124528** | Capns1 | 0.10716267 | -0.118823517 | 0.333148856 | 13.8930969 | 1.166623697 | 0.288567134 | 0.761407021 | -4.669032357 |
| **A_55_P1968448** | NA | -0.103133547 | -0.320631135 | 0.114364041 | 14.20721817 | -1.166580387 | 0.288583317 | 0.761407021 | -4.669061015 |
| **A_52_P672803** | Ctsa | 0.105498844 | -0.116987983 | 0.32798567 | 9.272112556 | 1.166574723 | 0.288585433 | 0.761407021 | -4.669064763 |
| **A_51_P279308** | Acbd5 | -0.136696553 | -0.425051876 | 0.151658769 | 10.01607939 | -1.166269061 | 0.288699665 | 0.761604452 | -4.669267 |
| **A_55_P1991416** | NA | 0.260513146 | -0.289094605 | 0.810120897 | 8.454380108 | 1.166127287 | 0.288752662 | 0.7616321 | -4.669360795 |
| **A_55_P1994773** | Gm16381 | 0.204118901 | -0.226598779 | 0.634836582 | 9.893883133 | 1.165895662 | 0.288839264 | 0.7616321 | -4.669514022 |
| **A_55_P1975537** | Nutf2 | 0.064941025 | -0.072098797 | 0.201980847 | 10.9707331 | 1.16584688 | 0.288857506 | 0.7616321 | -4.66954629 |
| **A_52_P358093** | Fbxo10 | -0.137770938 | -0.428504705 | 0.15296283 | 10.22631149 | -1.165819461 | 0.288867759 | 0.7616321 | -4.669564428 |
| **A_55_P2096630** | Ssxb9 | 0.071481927 | -0.079391676 | 0.22235553 | 7.978383407 | 1.165607018 | 0.288947216 | 0.761737689 | -4.669704948 |
| **A_51_P368591** | Tle6 | 0.095276187 | -0.105861273 | 0.296413648 | 9.558239255 | 1.165361245 | 0.289039162 | 0.76187617 | -4.669867499 |
| **A_55_P2002757** | Blnk | 0.396337431 | -0.440512983 | 1.233187844 | 10.48581984 | 1.165162283 | 0.289113613 | 0.761968508 | -4.669999078 |
| **A_55_P2127243** | Tpsb2 | 0.060309429 | -0.067067813 | 0.187686672 | 8.341863187 | 1.164829979 | 0.289237998 | 0.762048016 | -4.670218816 |
| **A_55_P2106956** | Gm5523 | -0.119709821 | -0.372552269 | 0.133132627 | 16.72828657 | -1.164793556 | 0.289251634 | 0.762048016 | -4.670242899 |
| **A_51_P352381** | Dhrs4 | -0.088179582 | -0.274442382 | 0.098083218 | 8.182198887 | -1.164691943 | 0.28928968 | 0.762048016 | -4.670310084 |
| **A_55_P1980966** | Mettl23 | 0.135086701 | -0.150266297 | 0.4204397 | 10.44283261 | 1.164660425 | 0.289301482 | 0.762048016 | -4.670330923 |
| **A_51_P121275** | Cenpo | 0.064837804 | -0.072157591 | 0.201833198 | 8.598830274 | 1.164371297 | 0.289409763 | 0.762181242 | -4.667890242 |
| **A_55_P2000758** | Gkap1 | 0.10787946 | -0.120072002 | 0.335830923 | 10.50105657 | 1.164301737 | 0.289435819 | 0.762181242 | -4.670568058 |
| **A_55_P1957875** | NA | 0.140537906 | -0.156445022 | 0.437520834 | 10.3123398 | 1.164209546 | 0.289470355 | 0.762181242 | -4.670629002 |
| **A_52_P530280** | Nsa2 | 0.121375008 | -0.135154124 | 0.37790414 | 10.60510113 | 1.164023486 | 0.289540068 | 0.76226096 | -4.670751991 |
| **A_51_P399625** | Cnih1 | -0.095707421 | -0.298045606 | 0.106630764 | 13.86192724 | -1.163688994 | 0.289665429 | 0.762487142 | -4.670973073 |
| **A_51_P470311** | Cby1 | -0.057414078 | -0.17881734 | 0.063989185 | 8.16368758 | -1.163475374 | 0.289745515 | 0.76257308 | -4.671114249 |
| **A_55_P2036633** | Ssrp1 | -0.119047456 | -0.370803812 | 0.132708899 | 12.61551964 | -1.163345838 | 0.289794087 | 0.76257308 | -4.671199851 |
| **A_52_P562612** | Srd5a1 | 0.181946347 | -0.2028763 | 0.566768993 | 8.523506255 | 1.16319327 | 0.289851304 | 0.76257308 | -4.671300666 |
| **A_55_P2070721** | Rapgef1 | 0.146709773 | -0.163665253 | 0.457084799 | 11.98217313 | 1.162897504 | 0.289962251 | 0.76257308 | -4.671496088 |
| **A_55_P2154943** | NA | 0.236742783 | -0.264106712 | 0.737592278 | 9.347353229 | 1.162890096 | 0.289965031 | 0.76257308 | -4.671500983 |

| | | | | | | | | |
|---|---|---|---|---|---|---|---|---|
| **A_51_P325592** | 2410004B18Rik | 0.14959539 | -0.166908236 | 0.466099016 | 11.16176053 | 1.162809799 | 0.289995158 | 0.76257308 | -4.671554033 |
| **A_51_P474459** | Socs3 | -0.111419396 | -0.347164729 | 0.124325937 | 10.78999116 | -1.162751368 | 0.290017083 | 0.76257308 | -4.671592636 |
| **A_55_P1960298** | Tmem80 | 0.124688205 | -0.139136066 | 0.388512476 | 8.676110141 | 1.162732442 | 0.290024185 | 0.76257308 | -4.67160514 |
| **A_55_P1978351** | Apobec2 | -0.118143799 | -0.36813754 | 0.131849941 | 7.688258364 | -1.162655265 | 0.290053147 | 0.76257308 | -4.671656126 |
| **A_66_P118752** | Rabif | 0.406537778 | -0.453924704 | 1.26700026 | 9.055491841 | 1.162353248 | 0.290166507 | 0.762609079 | -4.671855634 |
| **A_52_P496655** | Leprotl1 | -0.11408537 | -0.35558607 | 0.127415329 | 8.390533258 | -1.162199614 | 0.290224188 | 0.762609079 | -4.671957113 |
| **A_52_P113672** | Chml | 0.243457885 | -0.271963904 | 0.758879674 | 9.276557609 | 1.162064498 | 0.290274923 | 0.762609079 | -4.672046354 |
| **A_55_P2088202** | Hcn2 | 0.108693696 | -0.121448986 | 0.338836379 | 14.24618567 | 1.161920329 | 0.290329067 | 0.762609079 | -4.672141571 |
| **A_65_P10450** | Tmpo | -0.066102561 | -0.206078708 | 0.073873585 | 7.924622353 | -1.161805473 | 0.290372208 | 0.762609079 | -4.672217423 |
| **A_55_P2096189** | Rfx8 | -0.197702266 | -0.616401952 | 0.220997421 | 8.531766715 | -1.161657694 | 0.290427723 | 0.762609079 | -4.672315013 |
| **A_55_P2126805** | Wdr81 | 0.096417655 | -0.107786359 | 0.300621668 | 11.51677731 | 1.161612969 | 0.290444527 | 0.762609079 | -4.672344547 |
| **A_55_P1960479** | NA | -0.413868605 | -1.290505061 | 0.462767852 | 11.05642098 | -1.161481023 | 0.290494105 | 0.762609079 | -4.672431675 |
| **A_51_P383218** | Tmem161a | 0.05968349 | -0.066739058 | 0.186106037 | 8.722184155 | 1.16144549 | 0.290507457 | 0.762609079 | -4.672455137 |
| **A_51_P436719** | Eftud2 | -0.064067086 | -0.199776063 | 0.07164189 | 9.011089691 | -1.161436743 | 0.290510744 | 0.762609079 | -4.672460913 |
| **A_55_P2089293** | Rnf121 | 0.222802375 | -0.249175326 | 0.694780077 | 10.86743323 | 1.161361697 | 0.290538947 | 0.762609079 | -4.672510465 |
| **A_55_P2020926** | Trip11 | -0.192340065 | -0.599788598 | 0.215108469 | 8.945855883 | -1.161358123 | 0.29054029 | 0.762609079 | -4.672512824 |
| **A_52_P194250** | Ets1 | -0.092982042 | -0.290079697 | 0.104115613 | 8.759883867 | -1.160611274 | 0.290821087 | 0.763242469 | -4.673005875 |
| **A_55_P1979356** | Morf4l1 | -0.112483165 | -0.371526556 | 0.146560225 | 13.87089747 | -1.221848958 | 0.290940425 | 0.763452005 | -4.589367597 |
| **A_55_P2055854** | Ppara | 0.08418442 | -0.094346978 | 0.262715819 | 8.004284988 | 1.160075571 | 0.29102264 | 0.763518953 | -4.67335944 |
| **A_55_P1974452** | Lime1 | -0.10649992 | -0.332372054 | 0.119372214 | 9.07880818 | -1.159994125 | 0.291053294 | 0.763518953 | -4.673413187 |
| **A_55_P2078675** | NA | -0.07221074 | -0.22538828 | 0.0809668 | 8.258761702 | -1.159780674 | 0.291133644 | 0.763518953 | -4.673554038 |
| **A_55_P2072041** | NA | -0.126667353 | -0.39537229 | 0.142037583 | 8.93536481 | -1.159733533 | 0.291151392 | 0.763518953 | -4.673585145 |
| **A_55_P2172233** | NA | 0.085404768 | -0.095773062 | 0.266582598 | 15.76680914 | 1.159701523 | 0.291163444 | 0.763518953 | -4.673606266 |
| **A_55_P2030938** | Trim59 | -0.15287785 | -0.477242703 | 0.171487003 | 9.686212828 | -1.159524529 | 0.29123009 | 0.763570648 | -4.673723046 |
| **A_55_P2146171** | Cox20 | 0.071468735 | -0.085065742 | 0.228003212 | 11.48826362 | 1.183194575 | 0.291262165 | 0.763570648 | -4.62795984 |
| **A_51_P266774** | Mfn2 | -0.074449027 | -0.232436771 | 0.083538716 | 11.18495567 | -1.159323913 | 0.291305647 | 0.763581074 | -4.673855403 |
| **A_52_P646755** | Tex30 | -0.245015402 | -0.765053715 | 0.275022911 | 12.49476896 | -1.159116821 | 0.29138366 | 0.763617675 | -4.673992021 |
| **A_52_P495104** | NA | -0.162595808 | -0.507725861 | 0.182534245 | 11.77551148 | -1.15903281 | 0.291415313 | 0.763617675 | -4.67404744 |
| **A_51_P301636** | Kazn | -0.099476341 | -0.310658467 | 0.111705785 | 12.04159874 | -1.15886223 | 0.291479591 | 0.763617675 | -4.674159958 |
| **A_55_P2093221** | Rnf111 | 0.069373923 | -0.077907291 | 0.216655136 | 11.55534878 | 1.158825551 | 0.291493414 | 0.763617675 | -4.674184151 |
| **A_55_P2027870** | Abhd17a | 0.276379992 | -0.310425761 | 0.863185744 | 10.7469342 | 1.158727728 | 0.291530283 | 0.763617675 | -4.674248672 |
| **A_52_P535012** | NA | -0.086771965 | -0.271030884 | 0.097486955 | 15.16616812 | -1.158564114 | 0.291591957 | 0.763617675 | -4.674356582 |
| **A_55_P2018825** | NA | -0.082378256 | -0.257308883 | 0.092552372 | 8.527019814 | -1.15855298 | 0.291596154 | 0.763617675 | -4.674363925 |
| **A_55_P2102350** | Stambpl1 | 0.094244291 | -0.10591272 | 0.294401301 | 8.912353039 | 1.158386285 | 0.291659001 | 0.763635629 | -4.674473859 |
| **A_51_P167426** | Herc1 | -0.070619869 | -0.220630985 | 0.079391246 | 7.9747424 | -1.158170855 | 0.29174024 | 0.763635629 | -4.674615923 |
| **A_55_P2049072** | NA | 0.119061618 | -0.133857524 | 0.371980761 | 13.0089708 | 1.158135153 | 0.291753705 | 0.763635629 | -4.674639465 |
| **A_51_P271336** | Wipf2 | -0.114610325 | -0.35809485 | 0.128874199 | 9.731387769 | -1.158034624 | 0.291791623 | 0.763635629 | -4.674705752 |
| **A_52_P208613** | Fancb | -0.04945101 | -0.15450962 | 0.055607601 | 7.931430723 | -1.158010969 | 0.291800546 | 0.763635629 | -4.67472135 |
| **A_55_P2049429** | Phtf2 | -0.08025973 | -0.250807437 | 0.090287978 | 9.699924885 | -1.157766472 | 0.291892785 | 0.763665002 | -4.674882555 |
| **A_55_P2156345** | Zfp51 | -0.059576289 | -0.186177007 | 0.06702443 | 9.247024506 | -1.157727732 | 0.291907403 | 0.763665002 | -4.674908096 |
| **A_51_P165914** | Zfp770 | 0.051443026 | -0.057880088 | 0.160766139 | 8.72475795 | 1.157667062 | 0.291930296 | 0.763665002 | -4.674948095 |
| **A_51_P413111** | Ppm1l | -0.104615395 | -0.327034141 | 0.117803351 | 9.135197773 | -1.157159901 | 0.292121729 | 0.763690699 | -4.675282417 |
| **A_51_P313337** | Smim10l1 | -0.124510987 | -0.389280596 | 0.140258622 | 9.232172602 | -1.156934322 | 0.292206911 | 0.763690699 | -4.675431098 |
| **A_55_P2012439** | Tnfrsf19 | -0.068498065 | -0.214163627 | 0.077167497 | 8.300469442 | -1.156886029 | 0.29222515 | 0.763690699 | -4.675462926 |
| **A_66_P115539** | Gm11240 | 0.068931871 | -0.07766045 | 0.215524191 | 10.95780966 | 1.156852554 | 0.292237793 | 0.763690699 | -4.675484988 |
| **A_55_P2135556** | Rpgrip1 | 0.064213547 | -0.072356959 | 0.200784054 | 10.92541791 | 1.156748394 | 0.292277136 | 0.763690699 | -4.675553633 |
| **A_51_P323521** | Snx21 | 0.112051073 | -0.126264095 | 0.350366241 | 12.05661359 | 1.156733995 | 0.292282575 | 0.763690699 | -4.675563122 |
| **A_55_P2062598** | E2f7 | 0.123670167 | -0.139372822 | 0.386713155 | 10.00963997 | 1.156664428 | 0.292308854 | 0.763690699 | -4.675608968 |
| **A_51_P480928** | Zfyve16 | -0.064334412 | -0.201193482 | 0.072524659 | 9.363623099 | -1.156482088 | 0.292377744 | 0.763690699 | -4.675729125 |
| **A_55_P2095181** | NA | 0.143364542 | -0.161624348 | 0.448353432 | 10.13723825 | 1.156450093 | 0.292389834 | 0.763690699 | -4.675750208 |

| | | | | | | | | | |
|---|---|---|---|---|---|---|---|---|---|
| **A_55_P2038414** | Acot8 | 0.183902329 | -0.207325735 | 0.575130394 | 11.01175769 | 1.156448752 | 0.292390341 | 0.763690699 | -4.675751092 |
| **A_55_P1977761** | Ttc14 | -0.147711505 | -0.461963606 | 0.166540596 | 8.147913736 | -1.156392581 | 0.292411566 | 0.763690699 | -4.675788105 |
| **A_55_P2005853** | Nacc2 | -0.101209568 | -0.316531413 | 0.114112278 | 10.0780228 | -1.156385503 | 0.292414241 | 0.763690699 | -4.675792769 |
| **A_51_P419389** | Bmpr2 | -0.141400739 | -0.4422725 | 0.159471023 | 10.33051826 | -1.156217166 | 0.292477859 | 0.763753653 | -4.675903686 |
| **A_55_P2070347** | Rps2 | 0.129663882 | -0.146333467 | 0.405661232 | 15.01660574 | 1.15580155 | 0.292634981 | 0.763948019 | -4.676177502 |
| **A_66_P119155** | NA | -0.093563349 | -0.292724267 | 0.105597569 | 11.73772381 | -1.155768365 | 0.292647529 | 0.763948019 | -4.676199363 |
| **A_55_P1972406** | Tor1b | -0.076554522 | -0.239518791 | 0.086409747 | 9.697929364 | -1.155706665 | 0.292670862 | 0.763948019 | -4.676240008 |
| **A_55_P1981664** | 8-Sep | -0.170115745 | -0.532307005 | 0.192075516 | 13.65752971 | -1.155515317 | 0.292743232 | 0.764033747 | -4.676366051 |
| **A_55_P1957098** | Mtfr1l | -0.119605171 | -0.374286292 | 0.13507595 | 12.8777489 | -1.155373404 | 0.292796915 | 0.764070685 | -4.676459524 |
| **A_55_P2005778** | Uap1 | -0.143996915 | -0.450698008 | 0.162704177 | 12.22735776 | -1.155066608 | 0.292912998 | 0.76427043 | -4.676661582 |
| **A_55_P2048334** | Ubr3 | -0.076134552 | -0.238317755 | 0.086048651 | 9.406290904 | -1.154901873 | 0.292975346 | 0.764305096 | -4.676770066 |
| **A_55_P2037425** | Gapdh | -0.179417096 | -0.561640898 | 0.202806706 | 14.76096041 | -1.154822559 | 0.293005368 | 0.764305096 | -4.676822295 |
| **A_55_P2118674** | Tnip1 | -0.157639946 | -0.493605887 | 0.178325995 | 12.93885808 | -1.154357128 | 0.293181598 | 0.764490606 | -4.67712875 |
| **A_51_P346668** | Irf5 | -0.049171223 | -0.153967814 | 0.055625368 | 7.803651211 | -1.154338065 | 0.293188817 | 0.764490606 | -4.6771413 |
| **A_66_P137102** | Spryd7 | -0.069803313 | -0.218574375 | 0.078967749 | 8.730134517 | -1.154321371 | 0.29319514 | 0.764490606 | -4.677152291 |
| **A_55_P2020472** | Rufy2 | -0.067212004 | -0.210474796 | 0.076050789 | 9.684189875 | -1.15420405 | 0.293239578 | 0.764503345 | -4.677229528 |
| **A_51_P266143** | Adpgk | -0.08197323 | -0.256759046 | 0.092812587 | 9.631253495 | -1.153811919 | 0.293388148 | 0.764787528 | -4.677487657 |
| **A_51_P158876** | Dctpp1 | 0.212416862 | -0.240604711 | 0.665438435 | 12.0731675 | 1.153557557 | 0.293484555 | 0.764870508 | -4.677655074 |
| **A_52_P522372** | Aard | -0.097752635 | -0.306251949 | 0.110746678 | 8.361637345 | -1.153434689 | 0.293531134 | 0.764870508 | -4.677735937 |
| **A_55_P1988010** | Gm10406 | -0.179158254 | -0.561299923 | 0.202983416 | 9.178472132 | -1.153404354 | 0.293542634 | 0.764870508 | -4.677755901 |
| **A_55_P1998912** | Txnrd2 | 0.117357428 | -0.132984473 | 0.367699329 | 11.90418982 | 1.153310378 | 0.293578266 | 0.764870508 | -4.677817746 |
| **A_55_P2062108** | Apold1 | -0.117572527 | -0.368402151 | 0.133257098 | 7.97150758 | -1.153177568 | 0.293628627 | 0.764898616 | -4.677905143 |
| **A_51_P311096** | Malsu1 | -0.086007635 | -0.269513986 | 0.097498717 | 13.09437259 | -1.153068386 | 0.293670034 | 0.764903394 | -4.677976988 |
| **A_55_P2043187** | Exosc9 | -0.097773101 | -0.306414255 | 0.110868052 | 13.0659964 | -1.152891879 | 0.293736985 | 0.764974694 | -4.678093128 |
| **A_55_P1982883** | Gys1 | 0.106281514 | -0.120553112 | 0.333116139 | 10.85643422 | 1.152703321 | 0.293808521 | 0.765057914 | -4.678217188 |
| **A_51_P498023** | Scyl1 | 0.079462653 | -0.090169378 | 0.249094684 | 11.91484953 | 1.152456023 | 0.293902364 | 0.765199191 | -4.67837988 |
| **A_55_P2171897** | Notch1 | -0.089023652 | -0.279111471 | 0.101064167 | 7.933605986 | -1.152179867 | 0.294007189 | 0.765273705 | -4.678561537 |
| **A_66_P119999** | Atg7 | 0.107671123 | -0.122235287 | 0.337577533 | 9.817484421 | 1.15217202 | 0.294010168 | 0.765273705 | -4.678566699 |
| **A_55_P2126537** | Fam53c | -0.089378988 | -0.280277501 | 0.101519526 | 12.976805 | -1.151866233 | 0.294126278 | 0.765408202 | -4.678767822 |
| **A_55_P1982171** | H2afz | 0.149805106 | -0.170164248 | 0.46977446 | 14.00836325 | 1.151827369 | 0.294141038 | 0.765408202 | -4.678793383 |
| **A_52_P317653** | Car1 | 0.056687039 | -0.064405062 | 0.177779141 | 8.181417446 | 1.151694049 | 0.294191675 | 0.765436921 | -4.678881061 |
| **A_51_P199899** | Fto | -0.075279963 | -0.236132704 | 0.085572778 | 9.222477074 | -1.151383747 | 0.294309562 | 0.765548245 | -4.679085113 |
| **A_51_P401683** | Tm6sf1 | -0.061308986 | -0.192310793 | 0.069692821 | 8.647377361 | -1.151372925 | 0.294313674 | 0.765548245 | -4.679092229 |
| **A_52_P494380** | Dgkq | 0.097774924 | -0.111196897 | 0.306746745 | 9.723704339 | 1.151089054 | 0.294421556 | 0.765680435 | -4.679278877 |
| **A_51_P257684** | Stau1 | -0.088345756 | -0.277174425 | 0.100482913 | 9.611901337 | -1.151030743 | 0.29444372 | 0.765680435 | -4.679317213 |
| **A_55_P1974352** | Fam126b | -0.110423373 | -0.346529687 | 0.125682942 | 10.16052111 | -1.150595228 | 0.294609309 | 0.766007982 | -4.679603518 |
| **A_52_P238858** | Shisa9 | 0.076572616 | -0.087195005 | 0.240340237 | 7.913574991 | 1.15030924 | 0.294718089 | 0.766042119 | -4.679791495 |
| **A_55_P2185605** | Cd48 | 0.12963256 | -0.147686656 | 0.406951777 | 8.549239738 | 1.150014449 | 0.294830252 | 0.766042119 | -4.679985236 |
| **A_52_P460703** | Rassf3 | -0.080720796 | -0.253414316 | 0.091972724 | 9.591574633 | -1.149948945 | 0.294855181 | 0.766042119 | -4.680028283 |
| **A_65_P01834** | Lima1 | -0.097742337 | -0.306896464 | 0.111411791 | 9.833928297 | -1.149702408 | 0.294949019 | 0.766042119 | -4.680190286 |
| **A_51_P144543** | NA | 0.061885104 | -0.070544272 | 0.19431448 | 7.799430792 | 1.149664067 | 0.294963615 | 0.766042119 | -4.680215479 |
| **A_55_P1970700** | Rpsa | 0.086899049 | -0.099059523 | 0.272857621 | 16.38627171 | 1.149656196 | 0.294966611 | 0.766042119 | -4.680220651 |
| **A_55_P1966102** | Nme5 | -0.134177358 | -0.421314152 | 0.152959436 | 9.423007412 | -1.149633874 | 0.294975109 | 0.766042119 | -4.680235318 |
| **A_52_P385606** | Ckb | -0.19519444 | -0.612908443 | 0.222519563 | 9.724284361 | -1.149628632 | 0.294977105 | 0.766042119 | -4.680238763 |
| **A_51_P501107** | Cul5 | -0.130071447 | -0.408446204 | 0.14830331 | 9.200344487 | -1.149532582 | 0.295013674 | 0.766042119 | -4.680301872 |
| **A_51_P187082** | G6pdx | 0.069150786 | -0.078845235 | 0.217146806 | 9.932900996 | 1.149519235 | 0.295018756 | 0.766042119 | -4.680310642 |
| **A_52_P606679** | Eloa | -0.10116358 | -0.317727016 | 0.115399855 | 8.967921627 | -1.149233353 | 0.295127623 | 0.766086503 | -4.680498465 |
| **A_55_P2070084** | Zfp988 | -0.186907471 | -0.587029423 | 0.21321448 | 10.67108999 | -1.149220719 | 0.295132436 | 0.766086503 | -4.680506765 |
| **A_55_P2004347** | NA | -0.370831521 | -1.164848902 | 0.423185861 | 11.89159891 | -1.148988801 | 0.295220779 | 0.766086503 | -4.680659116 |
| **A_55_P2146780** | Sdhc | -0.084246194 | -0.264636445 | 0.096144057 | 11.99897164 | -1.148963921 | 0.295230258 | 0.766086503 | -4.680675459 |

| | | | | | | | | | |
|---|---|---|---|---|---|---|---|---|---|
| **A_51_P283344** | ccdc198 | 0.189804533 | -0.216642715 | 0.596251781 | 8.042757006 | 1.148871765 | 0.29526537 | 0.766086503 | -4.680735994 |
| **A_66_P111216** | 4632427E13Rik | -0.114363173 | -0.359264999 | 0.130538652 | 8.048632282 | -1.148850025 | 0.295273654 | 0.766086503 | -4.680750273 |
| **A_55_P2109943** | Rps15a | 0.059319185 | -0.067725448 | 0.186363818 | 15.67459095 | 1.148703685 | 0.295329419 | 0.76612835 | -4.680846394 |
| **A_55_P1961320** | Tes | -0.076341848 | -0.239868318 | 0.087184622 | 12.06173435 | -1.148533766 | 0.29539418 | 0.766193519 | -4.680957994 |
| **A_55_P2086983** | Atxn2 | -0.146925418 | -0.461801985 | 0.16795115 | 12.21302555 | -1.147957356 | 0.295613958 | 0.766660698 | -4.681336511 |
| **A_55_P2144666** | Prpsap1 | -0.136079577 | -0.427986678 | 0.155827524 | 12.06143348 | -1.146878612 | 0.296025642 | 0.767625387 | -4.682044654 |
| **A_55_P2059412** | Ghr | 0.27483789 | -0.314981428 | 0.864657209 | 9.991519835 | 1.146375184 | 0.296217933 | 0.768020984 | -4.682375018 |
| **A_55_P2277019** | Loxl3 | 0.055189203 | -0.063261186 | 0.173639592 | 8.005908723 | 1.146269643 | 0.29625826 | 0.76802252 | -4.682444268 |
| **A_66_P127432** | Tial1 | 0.050714117 | -0.058143107 | 0.159571341 | 10.86738901 | 1.146148198 | 0.296304669 | 0.76803982 | -4.682523949 |
| **A_55_P1982186** | Sgsm1 | 0.057788542 | -0.066283326 | 0.181860409 | 8.993411386 | 1.145875883 | 0.296408753 | 0.768194527 | -4.682702604 |
| **A_55_P2085425** | Ophn1 | -0.097472093 | -0.306771608 | 0.111827422 | 8.334219362 | -1.145727221 | 0.296465589 | 0.768194527 | -4.682800127 |
| **A_66_P122219** | Kcnd3 | 0.055446886 | -0.063617789 | 0.174511561 | 8.071673259 | 1.145680159 | 0.296483583 | 0.768194527 | -4.682830998 |
| **A_55_P1953419** | Rab13 | -0.048570031 | -0.152895031 | 0.055754969 | 8.081351163 | -1.14537883 | 0.296598818 | 0.768280415 | -4.683028647 |
| **A_55_P2076190** | NA | -0.441441873 | -1.389703833 | 0.506820087 | 14.82953048 | -1.145287076 | 0.296633915 | 0.768280415 | -4.683088825 |
| **A_55_P2017804** | Cox5b | 0.116502589 | -0.133757976 | 0.366763154 | 14.23647089 | 1.145281693 | 0.296635974 | 0.768280415 | -4.683092356 |
| **A_55_P2094412** | Ssbp1 | -0.094560766 | -0.297728963 | 0.10860743 | 9.030874407 | -1.145049918 | 0.296724645 | 0.768345614 | -4.683244359 |
| **A_51_P198137** | Ubxn8 | -0.112996932 | -0.355806782 | 0.129812918 | 9.260577758 | -1.144905163 | 0.296780037 | 0.768345614 | -4.683339285 |
| **A_55_P2026125** | Srrm2 | -0.109402113 | -0.344508665 | 0.125704438 | 8.251827031 | -1.144801375 | 0.296819757 | 0.768345614 | -4.683407342 |
| **A_52_P440729** | Atf7ip | 0.095887938 | -0.110176663 | 0.30195254 | 9.766518978 | 1.144800345 | 0.296820151 | 0.768345614 | -4.683408018 |
| **A_55_P2177953** | Aqp11 | 0.097007481 | -0.111495478 | 0.30551044 | 8.127480456 | 1.144622211 | 0.296888335 | 0.768419206 | -4.68352482 |
| **A_51_P407690** | Sppl2b | -0.08039224 | -0.253233742 | 0.092449262 | 11.66345794 | -1.144287785 | 0.297016378 | 0.768536168 | -4.683744077 |
| **A_55_P2018480** | Carnmt1 | -0.093720431 | -0.295228121 | 0.10778726 | 10.96422267 | -1.144226 | 0.297040039 | 0.768536168 | -4.683784581 |
| **A_55_P1973526** | Fam216b | 0.051887965 | -0.059682832 | 0.163458763 | 7.843922392 | 1.144156064 | 0.297066824 | 0.768536168 | -4.683830428 |
| **A_52_P108845** | Clip3 | 0.119465977 | -0.137429939 | 0.376361892 | 8.471795422 | 1.144079492 | 0.297096152 | 0.768536168 | -4.683880623 |
| **A_55_P2144931** | NA | 0.080014007 | -0.092069368 | 0.252097381 | 13.02187566 | 1.143921619 | 0.297156628 | 0.768536168 | -4.683984108 |
| **A_55_P1958200** | P2rx1 | 0.11967925 | -0.13774778 | 0.377106281 | 9.07453761 | 1.14375729 | 0.297219587 | 0.768536168 | -4.684091817 |
| **A_55_P1971522** | Kif16b | 0.127376407 | -0.146615955 | 0.401368768 | 9.949845962 | 1.143719962 | 0.29723389 | 0.768536168 | -4.684116283 |
| **A_52_P97670** | Atad2b | -0.113568254 | -0.357868538 | 0.13073203 | 9.266418474 | -1.143673718 | 0.297251611 | 0.768536168 | -4.684146592 |
| **A_51_P173086** | Eif3l | -0.15127975 | -0.476756648 | 0.174197148 | 15.02476136 | -1.143483214 | 0.29732462 | 0.768622119 | -4.684271444 |
| **A_55_P1999301** | Cyp2e1 | 0.060291821 | -0.069446124 | 0.190029766 | 7.758024267 | 1.143300907 | 0.297394502 | 0.768638459 | -4.684390914 |
| **A_55_P1985591** | Rtl8a | -0.071011221 | -0.223821095 | 0.081798654 | 8.700891391 | -1.143259247 | 0.297410473 | 0.768638459 | -4.684418214 |
| **A_52_P229783** | Ptgr2 | 0.060636297 | -0.069869503 | 0.191142098 | 9.207127439 | 1.143067878 | 0.297483847 | 0.768725305 | -4.684543611 |
| **A_51_P180001** | Ago2 | -0.12825847 | -0.40435842 | 0.147841479 | 11.19496288 | -1.142849103 | 0.297567748 | 0.76878069 | -4.684686953 |
| **A_55_P2100375** | Col11a1 | 0.057644994 | -0.066451294 | 0.181741281 | 7.788980124 | 1.142804574 | 0.297584827 | 0.76878069 | -4.684716127 |
| **A_55_P1955722** | Med15 | -0.106348891 | -0.335420838 | 0.122723057 | 11.18244112 | -1.14216861 | 0.297828847 | 0.769190577 | -4.685132729 |
| **A_51_P245546** | Synrg | 0.096372172 | -0.11121173 | 0.303956074 | 9.047143218 | 1.14216051 | 0.297831956 | 0.769190577 | -4.685138034 |
| **A_52_P585233** | Ccpg1 | -0.15007323 | -0.473351028 | 0.173204568 | 10.94495473 | -1.142079966 | 0.297862873 | 0.769190577 | -4.685190788 |
| **A_52_P468033** | Setd1b | -0.111600009 | -0.352025718 | 0.1288257 | 10.24010524 | -1.141964194 | 0.297907317 | 0.769202583 | -4.685266612 |
| **A_66_P100840** | NA | -0.092564234 | -0.292074143 | 0.106945676 | 8.10256912 | -1.141426368 | 0.29811386 | 0.769441226 | -4.685618807 |
| **A_52_P413502** | Rtf2 | -0.060030097 | -0.189425004 | 0.06936481 | 8.4226211 | -1.141355734 | 0.298140995 | 0.769441226 | -4.685665055 |
| **A_52_P182520** | Timm9 | -0.153114357 | -0.483232396 | 0.177003683 | 8.225705601 | -1.14107928 | 0.298247217 | 0.769441226 | -4.685846053 |
| **A_55_P1990678** | NA | 0.101365121 | -0.117191155 | 0.319921398 | 8.716476206 | 1.141023065 | 0.298268821 | 0.769441226 | -4.685882856 |
| **A_51_P323878** | Coro7 | 0.151488546 | -0.175181436 | 0.478158528 | 10.51034552 | 1.140879383 | 0.298324044 | 0.769441226 | -4.685976915 |
| **A_52_P24631** | Azin1 | 0.179966054 | -0.208129549 | 0.568061657 | 10.12497095 | 1.140830268 | 0.298342923 | 0.769441226 | -4.686009067 |
| **A_55_P2081133** | Gapdh | -0.166689311 | -0.526200152 | 0.192821529 | 15.91633043 | -1.140682884 | 0.298399582 | 0.769441226 | -4.686105542 |
| **A_51_P423825** | Morn1 | -0.074247014 | -0.234383869 | 0.085889842 | 8.119693493 | -1.14066169 | 0.29840773 | 0.769441226 | -4.686119414 |
| **A_55_P1953650** | Zdhhc18 | -0.230748033 | -0.728460038 | 0.266963971 | 12.58738625 | -1.140588676 | 0.298435802 | 0.769441226 | -4.686167206 |
| **A_51_P165098** | Gga2 | -0.17432462 | -0.550334378 | 0.201685139 | 12.07406823 | -1.140587934 | 0.298436088 | 0.769441226 | -4.686167691 |
| **A_52_P357484** | Nfrkb | -0.049745262 | -0.157043757 | 0.057553234 | 8.1371015 | -1.140583933 | 0.298437626 | 0.769441226 | -4.68617031 |
| **A_55_P2148906** | Cltb | 0.06814292 | -0.083642306 | 0.219928146 | 11.04671413 | 1.163432842 | 0.298478032 | 0.769442769 | -4.639557592 |

| | | | | | | | | | |
|---|---|---|---|---|---|---|---|---|---|
| **A_55_P1969530** | Zfp146 | -0.069051132 | -0.218072183 | 0.079969919 | 8.538305447 | -1.139967169 | 0.298674851 | 0.769847473 | -4.686573951 |
| **A_55_P2152305** | Atp5h | 0.072732484 | -0.08428115 | 0.229746117 | 14.76858636 | 1.139620407 | 0.298808296 | 0.769902131 | -4.686800841 |
| **A_51_P201661** | Xrcc5 | 0.057100757 | -0.06616978 | 0.180371294 | 9.464700731 | 1.139598156 | 0.298816861 | 0.769902131 | -4.686815399 |
| **A_51_P353392** | Cript | -0.099783568 | -0.315203465 | 0.115636329 | 9.523622074 | -1.139573598 | 0.298826313 | 0.769902131 | -4.686831466 |
| **A_55_P1970597** | Actg1 | -0.052415335 | -0.165581296 | 0.060750627 | 17.35055899 | -1.139493121 | 0.298857292 | 0.769902131 | -4.686884117 |
| **A_51_P285271** | Ccdc61 | -0.122362809 | -0.386586547 | 0.14186093 | 9.563338325 | -1.139322745 | 0.298922886 | 0.769902131 | -4.686995578 |
| **A_52_P360112** | Clip2 | -0.087280216 | -0.275757967 | 0.101197535 | 9.333553313 | -1.139265339 | 0.29894499 | 0.769902131 | -4.687033131 |
| **A_55_P2112230** | Pcyt2 | 0.229292318 | -0.265887997 | 0.724472632 | 11.43687407 | 1.139187724 | 0.298974877 | 0.769902131 | -4.687083903 |
| **A_51_P360492** | Mcm6 | 0.100395899 | -0.116489415 | 0.317281213 | 11.74559401 | 1.138819754 | 0.299116605 | 0.769945483 | -4.687324588 |
| **A_55_P2144766** | Pigc | -0.135001634 | -0.426704142 | 0.156700875 | 8.515829756 | -1.138591726 | 0.299204462 | 0.769945483 | -4.68747372 |
| **A_55_P2048176** | NA | -0.071127055 | -0.224819589 | 0.082565478 | 8.447721607 | -1.138547678 | 0.299221436 | 0.769945483 | -4.687502526 |
| **A_52_P198009** | Mrpl44 | 0.099232068 | -0.115199165 | 0.3136633 | 10.95044074 | 1.138500335 | 0.29923968 | 0.769945483 | -4.687533486 |
| **A_55_P2131238** | Ttc39a | 0.165648151 | -0.192394372 | 0.523690673 | 10.91154856 | 1.138206721 | 0.29935285 | 0.769945483 | -4.687725481 |
| **A_55_P1955656** | Ctla2a | -0.420063593 | -1.328048365 | 0.487921178 | 17.14203967 | -1.138166088 | 0.299368514 | 0.769945483 | -4.687752049 |
| **A_55_P2032723** | Dnajc10 | -0.071653782 | -0.226555438 | 0.083247874 | 8.47275334 | -1.138026096 | 0.299422487 | 0.769945483 | -4.687843581 |
| **A_55_P2027392** | Gpr146 | -0.167797475 | -0.530555424 | 0.194960474 | 10.63032921 | -1.137987907 | 0.299437212 | 0.769945483 | -4.687868549 |
| **A_51_P232708** | Npff | 0.054925475 | -0.063837307 | 0.173688257 | 8.919411413 | 1.137791337 | 0.299513016 | 0.769945483 | -4.687997061 |
| **A_55_P2003813** | Scn3b | 0.059292346 | -0.06891911 | 0.187503802 | 7.994698841 | 1.137734591 | 0.299534902 | 0.769945483 | -4.688034158 |
| **A_55_P2142191** | Rpl10 | 0.059684945 | -0.069377839 | 0.188747728 | 16.96645746 | 1.137713552 | 0.299543017 | 0.769945483 | -4.688047912 |
| **A_55_P2080905** | NA | 0.070893998 | -0.082414824 | 0.224202819 | 16.43851813 | 1.13765737 | 0.299564687 | 0.769945483 | -4.688084639 |
| **A_51_P103397** | Vwf | -0.158232929 | -0.500425706 | 0.183959847 | 10.3384856 | -1.137614695 | 0.299581149 | 0.769945483 | -4.688112536 |
| **A_66_P108152** | Cbr1 | 0.069619397 | -0.080962747 | 0.220201542 | 10.86814862 | 1.137433311 | 0.299651124 | 0.769945483 | -4.688231101 |
| **A_52_P587948** | Zfp11 | 0.064101983 | -0.074546479 | 0.202750444 | 8.063190171 | 1.13743234 | 0.299651499 | 0.769945483 | -4.688231736 |
| **A_55_P2137653** | NA | -0.129849115 | -0.410746803 | 0.151048572 | 11.95722769 | -1.1372606 | 0.299717766 | 0.769945483 | -4.688343988 |
| **A_55_P1954774** | NA | 0.098538663 | -0.114656717 | 0.311734043 | 13.04558234 | 1.137098378 | 0.299780373 | 0.769945483 | -4.688450012 |
| **A_51_P225771** | NA | 0.13035724 | -0.151700858 | 0.412415338 | 12.27640227 | 1.137013829 | 0.299813007 | 0.769945483 | -4.688505268 |
| **A_55_P2168124** | NA | -0.066597744 | -0.210698301 | 0.077502812 | 12.97686186 | -1.1370067 | 0.299815759 | 0.769945483 | -4.688509927 |
| **A_51_P176474** | Kndc1 | -0.073733309 | -0.233280555 | 0.085813937 | 8.112974398 | -1.136955786 | 0.299835413 | 0.769945483 | -4.6885432 |
| **A_55_P2107499** | Galnt18 | 0.169318463 | -0.197073978 | 0.535710904 | 9.050165897 | 1.136912326 | 0.29985219 | 0.769945483 | -4.688571601 |
| **A_51_P177518** | Emc6 | -0.081443278 | -0.257686787 | 0.094800231 | 11.61668133 | -1.136871235 | 0.299868053 | 0.769945483 | -4.688598454 |
| **A_51_P296429** | Tmem170 | 0.101965175 | -0.118762442 | 0.322692793 | 8.618458337 | 1.136486701 | 0.300016537 | 0.770211037 | -4.688849719 |
| **A_55_P1997486** | Snu13 | 0.080530537 | -0.093819391 | 0.254880466 | 11.47391031 | 1.136339195 | 0.300073512 | 0.770211037 | -4.688946093 |
| **A_55_P2155906** | Chmp3 | 0.074513982 | -0.086816441 | 0.235844405 | 11.96678071 | 1.136293873 | 0.30009102 | 0.770211037 | -4.688975703 |
| **A_55_P2115990** | 2010001K21Rik | 0.075880147 | -0.088430067 | 0.240190361 | 8.48296775 | 1.136142393 | 0.300149542 | 0.770249502 | -4.689074665 |
| **A_51_P357422** | Coprs | 0.115049699 | -0.134098448 | 0.364197846 | 8.912116584 | 1.136048799 | 0.300185705 | 0.770249502 | -4.689135806 |
| **A_55_P2253885** | AI503301 | -0.078384763 | -0.248158602 | 0.091389075 | 8.130709531 | -1.135873708 | 0.300253368 | 0.770268402 | -4.68925018 |
| **A_55_P2289800** | Fryl | 0.076140631 | -0.088779922 | 0.241061183 | 11.33525178 | 1.135823503 | 0.300272772 | 0.770268402 | -4.689282974 |
| **A_55_P2083559** | Hrk | -0.151654596 | -0.480259127 | 0.176949934 | 9.600234002 | -1.135406024 | 0.300434165 | 0.770391589 | -4.689555641 |
| **A_52_P659312** | Spsb4 | 0.23947819 | -0.27953625 | 0.75849263 | 8.452385294 | 1.135156367 | 0.300530715 | 0.770391589 | -4.689718675 |
| **A_55_P2161055** | Trp63 | -0.148940644 | -0.471827722 | 0.173946433 | 10.52861316 | -1.1348324 | 0.300656042 | 0.770391589 | -4.689930209 |
| **A_55_P1997628** | NA | -0.177016045 | -0.560822832 | 0.206790741 | 14.69565584 | -1.134668889 | 0.300719313 | 0.770391589 | -4.690036962 |
| **A_55_P2071893** | NA | -0.08819814 | -0.279463742 | 0.103067462 | 10.69959992 | -1.134466606 | 0.300797603 | 0.770391589 | -4.690169018 |
| **A_55_P2070406** | NA | -0.057336583 | -0.181690805 | 0.067017638 | 8.186939656 | -1.134332667 | 0.300849451 | 0.770391589 | -4.690256451 |
| **A_55_P1973926** | Ipo13 | -0.099443623 | -0.31513818 | 0.116250935 | 10.5129623 | -1.134245106 | 0.30088335 | 0.770391589 | -4.690313606 |
| **A_66_P129800** | Rab8b | 0.125076884 | -0.146220465 | 0.396374234 | 10.59229144 | 1.134228793 | 0.300889666 | 0.770391589 | -4.690324254 |
| **A_55_P2082841** | NA | -0.295448982 | -0.936400957 | 0.345502992 | 11.23670537 | -1.134034251 | 0.300964995 | 0.770391589 | -4.690451232 |
| **A_55_P1963489** | NA | -0.067183785 | -0.212945231 | 0.078577661 | 9.230294503 | -1.133942315 | 0.3010006 | 0.770391589 | -4.690511236 |
| **A_51_P308590** | Ppif | -0.059308652 | -0.18798833 | 0.069371027 | 8.056972978 | -1.133906506 | 0.301014469 | 0.770391589 | -4.690534606 |
| **A_65_P05040** | Ston2 | 0.174100969 | -0.203650506 | 0.551852443 | 9.725634731 | 1.133872395 | 0.30102768 | 0.770391589 | -4.690556868 |
| **A_51_P239203** | Mapk13 | 0.079512631 | -0.093010097 | 0.252035359 | 10.20644073 | 1.133858783 | 0.301032953 | 0.770391589 | -4.690565752 |

| | | | | | | | | | |
|---|---|---|---|---|---|---|---|---|---|
| **A_66_P125496** | NA | 0.169432561 | -0.198243397 | 0.537108518 | 12.54514907 | 1.13370694 | 0.301091771 | 0.770391589 | -4.690664845 |
| **A_51_P499864** | Tubg1 | -0.085802934 | -0.272009474 | 0.100403606 | 10.04273425 | -1.133643156 | 0.301116481 | 0.770391589 | -4.690706468 |
| **A_55_P2183055** | E4f1 | -0.117499482 | -0.372497549 | 0.137498585 | 10.33922284 | -1.133621899 | 0.301124717 | 0.770391589 | -4.69072034 |
| **A_55_P2062508** | Ube2e1 | -0.150635829 | -0.477560498 | 0.176288839 | 12.85893679 | -1.13357368 | 0.301143399 | 0.770391589 | -4.690751805 |
| **A_51_P396811** | Fam122a | -0.060835246 | -0.192877863 | 0.071207371 | 8.828084714 | -1.133470719 | 0.301183293 | 0.770391589 | -4.690818991 |
| **A_55_P1965620** | Bud31 | -0.088014446 | -0.279061355 | 0.103032463 | 12.86956254 | -1.133399739 | 0.301210799 | 0.770391589 | -4.690865305 |
| **A_55_P2068125** | Myct1 | 0.079404012 | -0.092958387 | 0.251766411 | 8.05303744 | 1.133363126 | 0.301224987 | 0.770391589 | -4.690889195 |
| **A_51_P504522** | Derl1 | 0.167941939 | -0.196616426 | 0.532500303 | 12.83608624 | 1.133342716 | 0.301232897 | 0.770391589 | -4.690902512 |
| **A_52_P467266** | Mtf1 | 0.063844297 | -0.074772901 | 0.202461496 | 10.4201101 | 1.133115438 | 0.301320989 | 0.770391589 | -4.691050799 |
| **A_55_P1993517** | Tbc1d14 | 0.105540021 | -0.123609181 | 0.334689222 | 8.700352192 | 1.133099348 | 0.301327226 | 0.770391589 | -4.691061296 |
| **A_52_P159885** | Ddx49 | -0.076800876 | -0.24356177 | 0.089960018 | 8.739762226 | -1.133029258 | 0.301354398 | 0.770391589 | -4.691107023 |
| **A_55_P2151026** | Ccdc50 | -0.112193208 | -0.355810722 | 0.131424305 | 8.38631504 | -1.132992995 | 0.301368456 | 0.770391589 | -4.69113068 |
| **A_52_P533146** | Ddit3 | -0.141193425 | -0.447799436 | 0.165412586 | 10.56985321 | -1.132929724 | 0.301392987 | 0.770391589 | -4.691171956 |
| **A_55_P2145480** | NA | 0.1404346 | -0.164526334 | 0.445395535 | 10.98883689 | 1.132919563 | 0.301396927 | 0.770391589 | -4.691178584 |
| **A_51_P352738** | Mpv17l | 0.084048671 | -0.098487899 | 0.266585242 | 8.680083187 | 1.132791884 | 0.301446435 | 0.770416255 | -4.691261875 |
| **A_55_P1975380** | Nol11 | 0.075789249 | -0.088880177 | 0.240458675 | 9.688462489 | 1.132305967 | 0.301634913 | 0.770796039 | -4.691578814 |
| **A_55_P2011235** | Rpl13 | -0.111992678 | -0.355378223 | 0.131392867 | 15.58433709 | -1.132045838 | 0.301735854 | 0.770952057 | -4.691748456 |
| **A_55_P1981381** | Lamc1 | -0.275686182 | -0.874965186 | 0.323592822 | 14.12220484 | -1.131761978 | 0.301846035 | 0.771131643 | -4.691933551 |
| **A_52_P429749** | Tmem115 | -0.082434773 | -0.261652739 | 0.096783193 | 9.559825146 | -1.131613361 | 0.301903735 | 0.771177123 | -4.69203045 |
| **A_55_P2116105** | Ap2a1 | 0.122373872 | -0.143768413 | 0.388516156 | 11.12268965 | 1.131211946 | 0.302059629 | 0.771456345 | -4.692292142 |
| **A_55_P2056393** | Stk11ip | -0.110972848 | -0.35233811 | 0.130392414 | 10.59998423 | -1.131126378 | 0.302092869 | 0.771456345 | -4.69234792 |
| **A_65_P15675** | Ipo9 | -0.062586032 | -0.198730931 | 0.073558866 | 8.456305165 | -1.130954701 | 0.302159569 | 0.771524744 | -4.692459822 |
| **A_55_P1980262** | Rtl8b | -0.211320946 | -0.671123667 | 0.248481776 | 13.14141428 | -1.130681199 | 0.302265856 | 0.771537801 | -4.692638078 |
| **A_55_P2077635** | NA | -0.07943838 | -0.252288594 | 0.093411835 | 10.85638703 | -1.130653721 | 0.302276535 | 0.771537801 | -4.692655985 |
| **A_52_P257502** | Igfbp4 | -0.076845219 | -0.24405597 | 0.090365533 | 8.029414171 | -1.130633408 | 0.302284431 | 0.771537801 | -4.692669224 |
| **A_65_P19112** | Zfp397 | -0.054019366 | -0.171629492 | 0.063590761 | 8.018952596 | -1.129988245 | 0.302535288 | 0.772076126 | -4.693089617 |
| **A_66_P119421** | Prdx1 | -0.092945815 | -0.295385643 | 0.109494014 | 8.958638267 | -1.129543706 | 0.302708239 | 0.772350887 | -4.693379212 |
| **A_66_P118093** | NA | 0.132112875 | -0.155676064 | 0.419901814 | 8.436117418 | 1.1293803 | 0.302771834 | 0.772350887 | -4.693485648 |
| **A_51_P130332** | Hspa12b | -0.098101649 | -0.311805995 | 0.115602697 | 9.595569297 | -1.129359256 | 0.302780025 | 0.772350887 | -4.693499356 |
| **A_51_P259902** | Arl6ip5 | 0.121344857 | -0.143006113 | 0.385695827 | 11.63414507 | 1.129300784 | 0.302802785 | 0.772350887 | -4.69353744 |
| **A_51_P409039** | Echs1 | 0.148298879 | -0.174848683 | 0.471446441 | 11.60890711 | 1.129031733 | 0.302907529 | 0.772516115 | -4.693712667 |
| **A_51_P483013** | Arel1 | -0.093022995 | -0.295750528 | 0.109704538 | 10.8246952 | -1.128877316 | 0.302967659 | 0.772539063 | -4.693813226 |
| **A_55_P1953053** | NA | -0.148117814 | -0.470949069 | 0.174713441 | 11.17543915 | -1.128758106 | 0.303014087 | 0.772539063 | -4.693890853 |
| **A_55_P2109382** | Adora2a | 0.187812726 | -0.221564051 | 0.597189502 | 8.735881272 | 1.128680383 | 0.303044359 | 0.772539063 | -4.693941461 |
| **A_55_P2184364** | B4galt3 | -0.153974563 | -0.489622021 | 0.181672894 | 12.66577056 | -1.128586321 | 0.303081 | 0.772539063 | -4.694002708 |
| **A_55_P2170514** | Sfn | -0.066720164 | -0.2121935 | 0.078753172 | 9.082415225 | -1.128347492 | 0.303174049 | 0.772539063 | -4.694158204 |
| **A_55_P1987775** | Pla2g4d | 0.320227161 | -0.377982567 | 1.018436888 | 10.77247924 | 1.128343763 | 0.303175502 | 0.772539063 | -4.694160631 |
| **A_52_P625171** | Asph | -0.052652964 | -0.167480545 | 0.062174616 | 8.220855819 | -1.12809527 | 0.303272343 | 0.772539063 | -4.694322401 |
| **A_51_P309733** | Cyp11a1 | 0.171901156 | -0.203064485 | 0.546866797 | 10.33169633 | 1.127863354 | 0.303362747 | 0.772539063 | -4.694473364 |
| **A_55_P2067181** | Usp21 | 0.102523668 | -0.121146519 | 0.326193856 | 9.372461093 | 1.127678233 | 0.303434926 | 0.772539063 | -4.694593854 |
| **A_55_P2114863** | Mgll | -0.134559593 | -0.428136277 | 0.15901709 | 10.64255273 | -1.127618698 | 0.303458141 | 0.772539063 | -4.694632602 |
| **A_51_P226853** | Gga1 | 0.0589197 | -0.069639427 | 0.187478827 | 8.458811824 | 1.127526544 | 0.30349408 | 0.772539063 | -4.694692577 |
| **A_51_P184949** | F12 | -0.048117096 | -0.153126226 | 0.056892035 | 7.686206905 | -1.127305194 | 0.303580418 | 0.772539063 | -4.694836625 |
| **A_51_P164207** | Mta2 | -0.093588402 | -0.29785019 | 0.110673386 | 10.90321136 | -1.127208004 | 0.303618333 | 0.772539063 | -4.69489987 |
| **A_55_P2259500** | D130012P04Rik | 0.063198832 | -0.074763812 | 0.201161476 | 8.148377145 | 1.126981289 | 0.303706795 | 0.772539063 | -4.695047388 |
| **A_55_P2082057** | NA | 0.061973634 | -0.073335479 | 0.197282748 | 16.15140407 | 1.126805821 | 0.303775275 | 0.772539063 | -4.695161551 |
| **A_52_P425092** | NA | -0.195408657 | -0.622112009 | 0.231294695 | 8.518830376 | -1.126644521 | 0.303838237 | 0.772539063 | -4.695266489 |
| **A_55_P2084696** | H2ac1 | -0.049413053 | -0.157314158 | 0.058488051 | 8.105125154 | -1.126639483 | 0.303840204 | 0.772539063 | -4.695269766 |
| **A_55_P2109559** | Trim6 | -0.125653202 | -0.400045664 | 0.14873926 | 9.047886175 | -1.126602073 | 0.303854808 | 0.772539063 | -4.695294103 |
| **A_55_P2180491** | Rrbp1 | -0.1176517 | -0.374578415 | 0.139275016 | 10.63230558 | -1.126569797 | 0.303867409 | 0.772539063 | -4.695315099 |

| | | | | | | | | | |
|---|---|---|---|---|---|---|---|---|---|
| **A_55_P2175451** | NA | 0.155378353 | -0.183995441 | 0.494752146 | 12.52968958 | 1.126370772 | 0.303945118 | 0.772539063 | -4.695444565 |
| **A_51_P418526** | Sfxn1 | 0.093194873 | -0.110417283 | 0.296807029 | 9.49335501 | 1.126049495 | 0.304070597 | 0.772539063 | -4.695653531 |
| **A_55_P2037186** | Alkbh2 | 0.090854608 | -0.107645124 | 0.28935434 | 10.53317486 | 1.126046165 | 0.304071897 | 0.772539063 | -4.695655696 |
| **A_55_P1977747** | Picalm | -0.163286161 | -0.520047612 | 0.193475289 | 13.29416053 | -1.126005784 | 0.304087672 | 0.772539063 | -4.695681959 |
| **A_51_P292447** | Klhl36 | -0.054153357 | -0.172478319 | 0.064171606 | 8.284860799 | -1.125947574 | 0.304110412 | 0.772539063 | -4.695719816 |
| **A_52_P650108** | Uri1 | -0.082745069 | -0.263551381 | 0.098061242 | 8.76015345 | -1.125894509 | 0.304131144 | 0.772539063 | -4.695754326 |
| **A_55_P1983733** | Aldh1l1 | 0.230208658 | -0.272973718 | 0.733391034 | 8.755652276 | 1.125551561 | 0.304265157 | 0.772539063 | -4.695977339 |
| **A_65_P01415** | Kremen1 | -0.316500978 | -1.008302861 | 0.375300904 | 12.18726904 | -1.125544009 | 0.304268108 | 0.772539063 | -4.695982249 |
| **A_55_P1972263** | Nop56 | 0.109612401 | -0.130009997 | 0.3492348 | 10.84909442 | 1.125385833 | 0.304329936 | 0.772539063 | -4.696085096 |
| **A_55_P1981241** | Grasp | 0.161441558 | -0.191502264 | 0.514385379 | 11.10592884 | 1.125327465 | 0.304352753 | 0.772539063 | -4.696123046 |
| **A_65_P07361** | Trim14 | -0.060135625 | -0.191608793 | 0.071337544 | 8.259692882 | -1.125288448 | 0.304368006 | 0.772539063 | -4.696148413 |
| **A_55_P2076248** | NA | 0.090587095 | -0.107470382 | 0.288644572 | 9.137062468 | 1.125237649 | 0.304387867 | 0.772539063 | -4.69618144 |
| **A_55_P1986863** | Ccdc126 | 0.140637537 | -0.166875747 | 0.448150821 | 10.14487773 | 1.125139919 | 0.304426079 | 0.772539063 | -4.696244977 |
| **A_55_P2004920** | NA | -0.054152832 | -0.172564847 | 0.064259182 | 9.19254296 | -1.125108924 | 0.304438198 | 0.772539063 | -4.696265127 |
| **A_55_P2092984** | NA | 0.176520605 | -0.209473038 | 0.562514249 | 14.00769413 | 1.125082626 | 0.304448482 | 0.772539063 | -4.696282223 |
| **A_51_P192751** | Mrpl14 | 0.196811234 | -0.233627306 | 0.627249774 | 13.47219932 | 1.12488441 | 0.304526 | 0.772539063 | -4.696411077 |
| **A_51_P270899** | Zfp61 | -0.107198368 | -0.341650194 | 0.127253458 | 8.834257615 | -1.124873584 | 0.304530235 | 0.772539063 | -4.696418114 |
| **A_55_P2172440** | Scarf2 | 0.162731531 | -0.193205991 | 0.518669052 | 11.51383889 | 1.124778745 | 0.304567331 | 0.772539063 | -4.696479761 |
| **A_55_P1965391** | Klhl18 | 0.102737346 | -0.12200244 | 0.327477132 | 10.83688203 | 1.124650399 | 0.304617539 | 0.772539063 | -4.696563185 |
| **A_55_P2097017** | Eid1 | -0.175086636 | -0.558097625 | 0.207924352 | 11.32121791 | -1.124633274 | 0.304624239 | 0.772539063 | -4.696574315 |
| **A_51_P176783** | Atxn7l3b | -0.091918241 | -0.293013224 | 0.109176741 | 11.32869628 | -1.124526346 | 0.304666074 | 0.772539063 | -4.696643813 |
| **A_55_P2157551** | NA | -0.077141309 | -0.24593393 | 0.091651311 | 9.878068005 | -1.124353078 | 0.304733876 | 0.772539063 | -4.696756421 |
| **A_52_P59457** | Nprl2 | 0.071488057 | -0.084935735 | 0.22791185 | 9.540854505 | 1.124345657 | 0.30473678 | 0.772539063 | -4.696761244 |
| **A_55_P2124273** | Myo1f | -0.053668819 | -0.171113644 | 0.063776006 | 7.89407599 | -1.124235548 | 0.304779874 | 0.772539063 | -4.6968328 |
| **A_51_P520857** | Selenok-ps1 | 0.202784342 | -0.240986779 | 0.646555463 | 14.02925604 | 1.124202494 | 0.304792811 | 0.772539063 | -4.69685428 |
| **A_51_P112445** | Wdr3 | -0.094593489 | -0.301628248 | 0.112441269 | 9.699393424 | -1.124053921 | 0.304850969 | 0.772539063 | -4.680315611 |
| **A_51_P310649** | Arsa | -0.21108075 | -0.688293215 | 0.266131715 | 11.9237782 | -1.146270016 | 0.304873201 | 0.772539063 | -4.649572578 |
| **A_55_P2146185** | Rs1 | -0.349125745 | -1.113363793 | 0.415112302 | 11.33163215 | -1.123886316 | 0.304916588 | 0.772539063 | -4.69705973 |
| **A_66_P100365** | NA | -0.049853835 | -0.159005041 | 0.05929737 | 8.149215844 | -1.123671072 | 0.305000876 | 0.772539063 | -4.697199577 |
| **A_55_P1968600** | NA | 0.094136956 | -0.111988707 | 0.30026262 | 9.108352126 | 1.123562532 | 0.305043387 | 0.772539063 | -4.697270092 |
| **A_66_P100232** | Paxbp1 | -0.103881206 | -0.331352293 | 0.12358988 | 9.216529528 | -1.123517767 | 0.305060921 | 0.772539063 | -4.697299174 |
| **A_55_P2115235** | Bckdhb | -0.085775775 | -0.273602535 | 0.102050986 | 9.744079047 | -1.123508452 | 0.305064569 | 0.772539063 | -4.697305225 |
| **A_55_P2361647** | 4831440E17Rik | -0.050858876 | -0.162230111 | 0.060512359 | 8.234351402 | -1.123473644 | 0.305078204 | 0.772539063 | -4.697327837 |
| **A_51_P419300** | Pdcd2l | -0.055903129 | -0.17833273 | 0.066526471 | 11.2625136 | -1.123359723 | 0.305122832 | 0.772539063 | -4.697401841 |
| **A_55_P2084433** | NA | 0.116715349 | -0.138955636 | 0.372386333 | 12.40234729 | 1.123092924 | 0.305227371 | 0.772539063 | -4.697575142 |
| **A_55_P2109377** | Camk2b | -0.142425878 | -0.45442577 | 0.169574015 | 8.135908979 | -1.123061717 | 0.3052396 | 0.772539063 | -4.697595411 |
| **A_51_P109421** | Eif2d | 0.049078784 | -0.058438361 | 0.15659593 | 9.220194204 | 1.123014161 | 0.305258238 | 0.772539063 | -4.697626299 |
| **A_51_P237040** | Nog | -0.064483489 | -0.205750997 | 0.076784018 | 8.30567407 | -1.122988763 | 0.305268192 | 0.772539063 | -4.697642795 |
| **A_52_P10781** | Lman1 | -0.157039779 | -0.501124075 | 0.187044516 | 11.09690298 | -1.122829949 | 0.30533044 | 0.772539063 | -4.697745938 |
| **A_55_P2143693** | Sytl1 | 0.219032725 | -0.260892495 | 0.698957946 | 8.822103422 | 1.122805669 | 0.305339958 | 0.772539063 | -4.697761706 |
| **A_55_P1990210** | Scpep1 | 0.263841272 | -0.314284768 | 0.841967311 | 14.1394154 | 1.122766288 | 0.305355396 | 0.772539063 | -4.697787281 |
| **A_55_P2123822** | Rbbp7 | -0.200443192 | -0.639659313 | 0.238772928 | 15.77974373 | -1.122747774 | 0.305362654 | 0.772539063 | -4.697799305 |
| **A_51_P282667** | Hexa | -0.298485788 | -0.952582666 | 0.355611089 | 12.88572159 | -1.122666452 | 0.305394536 | 0.772539063 | -4.697852115 |
| **A_52_P339742** | Cyb5r3 | -0.078898948 | -0.251832728 | 0.094034831 | 9.009165988 | -1.122433303 | 0.305485957 | 0.772610761 | -4.698003512 |
| **A_55_P2108690** | Gm8709 | -0.082800807 | -0.264295535 | 0.09869392 | 10.96918351 | -1.122379485 | 0.305507063 | 0.772610761 | -4.698038457 |
| **A_52_P135707** | Creb3 | 0.105546895 | -0.125834056 | 0.336927846 | 13.57662068 | 1.122243304 | 0.305560474 | 0.772610761 | -4.698126877 |
| **A_51_P492595** | Fbxo42 | -0.072093944 | -0.230158515 | 0.085970628 | 10.55231465 | -1.12210477 | 0.305614817 | 0.772610761 | -4.69821682 |
| **A_66_P105288** | Rplp0 | 0.062162587 | -0.074159701 | 0.198484876 | 15.87014879 | 1.121841177 | 0.30571824 | 0.772610761 | -4.698387942 |
| **A_52_P679101** | Tjp2 | -0.13230829 | -0.422477018 | 0.157860438 | 12.70068801 | -1.121774628 | 0.305744356 | 0.772610761 | -4.698431141 |
| **A_55_P2016388** | Eif3k | -0.085872919 | -0.27421134 | 0.102465502 | 14.38176558 | -1.121725162 | 0.305763768 | 0.772610761 | -4.698463251 |

| | | | | | | | | | |
|---|---|---|---|---|---|---|---|---|---|
| **A_55_P2166745** | Sp3os | -0.095992178 | -0.306526536 | 0.114542181 | 10.99079633 | -1.121713704 | 0.305768265 | 0.772610761 | -4.698470688 |
| **A_55_P2136732** | Cd99 | 0.134855247 | -0.160924989 | 0.430635483 | 12.58138253 | 1.121677121 | 0.305782623 | 0.772610761 | -4.698494435 |
| **A_55_P2020661** | Orai3 | -0.07539443 | -0.240795274 | 0.090006414 | 8.27142051 | -1.121426221 | 0.30588111 | 0.77271027 | -4.698657284 |
| **A_55_P2099760** | NA | 0.10563093 | -0.126160864 | 0.337422725 | 9.736876214 | 1.121146099 | 0.3059911 | 0.77271027 | -4.69883908 |
| **A_51_P160439** | Crybg3 | -0.132403222 | -0.422958059 | 0.158151615 | 11.77076848 | -1.121087747 | 0.306014016 | 0.77271027 | -4.698876947 |
| **A_52_P145349** | Stradb | 0.057700991 | -0.068923837 | 0.18432582 | 10.3632157 | 1.121072155 | 0.306020139 | 0.77271027 | -4.698887065 |
| **A_55_P1997831** | Dbndd1 | 0.099980417 | -0.119440548 | 0.319401382 | 8.204670736 | 1.121000965 | 0.3060481 | 0.77271027 | -4.698933261 |
| **A_52_P42834** | Usp25 | -0.117515983 | -0.375446492 | 0.140414526 | 9.020874516 | -1.120891006 | 0.306091291 | 0.77271027 | -4.699004612 |
| **A_52_P644354** | Zfp106 | -0.06853435 | -0.218960896 | 0.081892197 | 10.0378691 | -1.120864141 | 0.306101844 | 0.77271027 | -4.699022044 |
| **A_52_P312371** | Zfp345 | 0.140964289 | -0.168474324 | 0.450402901 | 9.21718608 | 1.120737136 | 0.306151738 | 0.772735302 | -4.69910445 |
| **A_55_P2009792** | Hdac6 | 0.144860072 | -0.173190578 | 0.462910722 | 11.44236616 | 1.120525055 | 0.30623507 | 0.772834082 | -4.699242047 |
| **A_51_P189959** | Oxsr1 | -0.108044647 | -0.345302125 | 0.129212831 | 9.418929365 | -1.120346883 | 0.306305094 | 0.772834082 | -4.699357635 |
| **A_55_P1962314** | NA | 0.078428649 | -0.093796686 | 0.250653985 | 15.15779479 | 1.120332305 | 0.306310823 | 0.772834082 | -4.699367091 |
| **A_55_P1986706** | Pih1d1 | 0.099472147 | -0.1189926 | 0.317936894 | 10.51056509 | 1.120183806 | 0.306369196 | 0.772856387 | -4.699463421 |
| **A_55_P1963443** | Vps18 | -0.06590132 | -0.210646464 | 0.078843824 | 8.491250223 | -1.120106386 | 0.306399632 | 0.772856387 | -4.699513639 |
| **A_55_P1980539** | Cetn4 | -0.097554866 | -0.311844582 | 0.116734851 | 8.580749228 | -1.119996764 | 0.306442733 | 0.772864245 | -4.699584742 |
| **A_52_P484956** | Nnat | 0.294712095 | -0.35292672 | 0.94235091 | 8.566632511 | 1.119526177 | 0.306627812 | 0.773218555 | -4.699889937 |
| **A_52_P253748** | Zbtb45 | 0.210331437 | -0.251915474 | 0.672578348 | 8.940040911 | 1.119436168 | 0.306663223 | 0.773218555 | -4.699948303 |
| **A_55_P2051622** | Gm8479 | -0.099970505 | -0.319703845 | 0.119762835 | 8.328090371 | -1.119296362 | 0.306718232 | 0.773256384 | -4.700038957 |
| **A_55_P2166543** | Pam16 | 0.081609222 | -0.097835223 | 0.261053667 | 12.24012658 | 1.118866787 | 0.306887306 | 0.773581734 | -4.700317468 |
| **A_52_P355075** | Rab40c | -0.087321154 | -0.279388476 | 0.104746169 | 12.33848583 | -1.118497831 | 0.307032583 | 0.77362649 | -4.700556633 |
| **A_55_P2169849** | Thoc6 | -0.05107235 | -0.163422129 | 0.06127743 | 8.693460438 | -1.118362961 | 0.307085704 | 0.77362649 | -4.700644049 |
| **A_51_P502798** | Nxpe4 | -0.056956274 | -0.182269554 | 0.068357006 | 7.815393299 | -1.11818493 | 0.307155835 | 0.77362649 | -4.700759431 |
| **A_52_P375312** | Jaml | -0.122715883 | -0.392827689 | 0.147395922 | 10.23704312 | -1.117702891 | 0.307345791 | 0.77362649 | -4.701071794 |
| **A_55_P2076772** | Hspa5 | 0.057597492 | -0.069186886 | 0.184381869 | 17.00070004 | 1.117653006 | 0.307365455 | 0.77362649 | -4.701104116 |
| **A_52_P549166** | Arsj | 0.057818018 | -0.069453329 | 0.185089365 | 7.841031417 | 1.117639446 | 0.3073708 | 0.77362649 | -4.701112902 |
| **A_55_P2154430** | Cd320 | 0.192966031 | -0.231817339 | 0.617749401 | 16.14436798 | 1.117590037 | 0.307390277 | 0.77362649 | -4.701144914 |
| **A_66_P121770** | Ilf2 | 0.109454821 | -0.131520156 | 0.350429799 | 9.80228869 | 1.117460323 | 0.307441417 | 0.77362649 | -4.701228953 |
| **A_55_P2043767** | NA | -0.104025804 | -0.333072444 | 0.125020836 | 15.52320916 | -1.117342506 | 0.307487872 | 0.77362649 | -4.701305281 |
| **A_55_P2175767** | Rangrf | 0.068572811 | -0.082415378 | 0.219561 | 8.490019269 | 1.117321459 | 0.307496171 | 0.77362649 | -4.701318915 |
| **A_51_P453043** | Aacs | 0.074021032 | -0.089001748 | 0.237043812 | 11.0846949 | 1.117058722 | 0.307599791 | 0.77362649 | -4.701489112 |
| **A_51_P444335** | Acp6 | -0.090380611 | -0.28945824 | 0.108697018 | 10.52042158 | -1.116919767 | 0.307654605 | 0.77362649 | -4.701579116 |
| **A_51_P154997** | Brms1 | -0.090831362 | -0.290903866 | 0.109241141 | 9.513463929 | -1.116908473 | 0.307659061 | 0.77362649 | -4.701586431 |
| **A_51_P443702** | Mrps34 | 0.077622241 | -0.093356064 | 0.248600545 | 12.66489262 | 1.116900101 | 0.307662364 | 0.77362649 | -4.701591853 |
| **A_51_P196590** | Hadh | -0.144506269 | -0.462847992 | 0.173835454 | 12.4539478 | -1.116766268 | 0.307715166 | 0.77362649 | -4.701678534 |
| **A_66_P100561** | Bnip2 | -0.084539925 | -0.270802785 | 0.101722935 | 13.21594315 | -1.116618334 | 0.307773541 | 0.77362649 | -4.701774341 |
| **A_55_P2117664** | Rnf165 | -0.115117452 | -0.368755204 | 0.1385203 | 8.592371541 | -1.116596948 | 0.307781981 | 0.77362649 | -4.701788191 |
| **A_55_P1999923** | Nyap1 | -0.17353067 | -0.555874562 | 0.208813221 | 9.527097708 | -1.116583618 | 0.307787241 | 0.77362649 | -4.701796824 |
| **A_55_P2073940** | Abcc5 | 0.084519036 | -0.10170834 | 0.270746411 | 8.468728043 | 1.116555135 | 0.307798482 | 0.77362649 | -4.701815269 |
| **A_55_P2005719** | NA | -0.16317668 | -0.522730717 | 0.196377356 | 11.9473913 | -1.11651121 | 0.307815818 | 0.77362649 | -4.701843714 |
| **A_52_P296913** | Cnnm1 | -0.147919953 | -0.473869894 | 0.178029987 | 9.291267814 | -1.116464756 | 0.307834153 | 0.77362649 | -4.701873797 |
| **A_55_P2082733** | Cybb | -0.050658591 | -0.162289908 | 0.060972726 | 7.779190964 | -1.116442137 | 0.30784308 | 0.77362649 | -4.701888444 |
| **A_52_P568805** | Ubxn1 | 0.179264935 | -0.215766669 | 0.574296538 | 14.50169238 | 1.116433008 | 0.307846684 | 0.77362649 | -4.701894355 |
| **A_55_P1960813** | NA | -0.186732128 | -0.598282438 | 0.224818182 | 13.72047661 | -1.116259777 | 0.307915066 | 0.77362649 | -4.702006529 |
| **A_51_P510106** | B3gntl1 | 0.096657257 | -0.116401553 | 0.309716068 | 8.518657178 | 1.116102635 | 0.307977109 | 0.77362649 | -4.702108276 |
| **A_55_P2086500** | Dennd5a | 0.161833831 | -0.194921515 | 0.518589178 | 12.08785029 | 1.116009751 | 0.308013786 | 0.77362649 | -4.702168413 |
| **A_55_P2185461** | Rpl9 | 0.074776958 | -0.090094591 | 0.239648506 | 16.51063254 | 1.11581254 | 0.308091671 | 0.77362649 | -4.702296088 |
| **A_55_P1953252** | Ctnnd1 | -0.14911388 | -0.477904972 | 0.179677212 | 13.37983159 | -1.115750757 | 0.308116074 | 0.77362649 | -4.702336084 |
| **A_55_P2036883** | H2bc18 | 0.107058953 | -0.129003239 | 0.343121145 | 8.577733099 | 1.115746977 | 0.308117567 | 0.77362649 | -4.702338531 |
| **A_55_P2131672** | Mical2 | -0.15151434 | -0.485613296 | 0.182584616 | 8.082646938 | -1.115700898 | 0.308135769 | 0.77362649 | -4.70236836 |

| | | | | | | | | | |
|---|---|---|---|---|---|---|---|---|---|
| **A_55_P2017636** | Thbs1 | 0.143326396 | -0.172724812 | 0.459377605 | 10.63525446 | 1.115675482 | 0.308145809 | 0.77362649 | -4.702384813 |
| **A_55_P1993840** | Tmem132c | 0.06538865 | -0.078825565 | 0.209602866 | 7.840568471 | 1.115484305 | 0.308221338 | 0.773715616 | -4.702508563 |
| **A_55_P2128263** | Phf21a | -0.157791407 | -0.505967475 | 0.19038466 | 10.6040079 | -1.114945391 | 0.308434332 | 0.77398822 | -4.702857347 |
| **A_51_P119544** | Aga | -0.075864822 | -0.243270851 | 0.091541206 | 9.906247379 | -1.114906673 | 0.308449639 | 0.77398822 | -4.67182855 |
| **A_51_P165504** | Twist2 | 0.147477938 | -0.177964035 | 0.472919912 | 8.400253674 | 1.114865961 | 0.308465736 | 0.77398822 | -4.702908746 |
| **A_55_P2116650** | A130040M12Rik | -0.530986713 | -1.702852503 | 0.640879077 | 10.89593702 | -1.11474348 | 0.308514165 | 0.77398822 | -4.702988001 |
| **A_55_P2027278** | Gas2l1 | 0.110599119 | -0.133511472 | 0.354709711 | 12.70083248 | 1.114638934 | 0.308555508 | 0.77398822 | -4.703055647 |
| **A_55_P2077783** | Tubb2a-ps2 | -0.053469323 | -0.171499444 | 0.064560798 | 8.297893427 | -1.114502372 | 0.308609519 | 0.77398822 | -4.703144003 |
| **A_55_P1969750** | Stim1 | -0.222292384 | -0.713024426 | 0.268439658 | 13.40854089 | -1.114421113 | 0.308641661 | 0.77398822 | -4.703196576 |
| **A_55_P2025328** | Tacr2 | -0.270833245 | -0.868788025 | 0.327121536 | 11.78173788 | -1.114301701 | 0.308688899 | 0.77398822 | -4.703273829 |
| **A_52_P355014** | Rbm8a | -0.134555558 | -0.431632819 | 0.162521703 | 11.26994069 | -1.11429811 | 0.30869032 | 0.77398822 | -4.703276151 |
| **A_65_P16483** | Pdzk1 | 0.299229306 | -0.361516556 | 0.959975169 | 10.28882507 | 1.114137615 | 0.30875382 | 0.774047028 | -4.703379976 |
| **A_52_P551903** | Slc33a1 | -0.074013282 | -0.237476793 | 0.089450229 | 8.345381977 | -1.113930259 | 0.308835877 | 0.774106405 | -4.703514104 |
| **A_51_P502964** | Tmem63a | -0.098753963 | -0.316869749 | 0.119361822 | 9.845121649 | -1.113875368 | 0.308857602 | 0.774106405 | -4.703549608 |
| **A_51_P264053** | Pigf | 0.11042827 | -0.133509711 | 0.35436625 | 10.01533868 | 1.113704579 | 0.308925207 | 0.77417546 | -4.70366007 |
| **A_55_P2012349** | Fermt2 | 0.078111495 | -0.094484434 | 0.250707424 | 11.89980314 | 1.113406017 | 0.309043418 | 0.774371302 | -4.703853151 |
| **A_55_P1978481** | Nr1h2 | -0.11255538 | -0.361413821 | 0.136303061 | 12.09191488 | -1.112712705 | 0.309318071 | 0.774782508 | -4.704301418 |
| **A_55_P2089488** | Coq10b | -0.08846524 | -0.284064397 | 0.107133917 | 9.757798798 | -1.112691687 | 0.3093264 | 0.774782508 | -4.704315004 |
| **A_51_P412508** | Irx4 | 0.166532465 | -0.201676693 | 0.534741622 | 9.438313328 | 1.112688212 | 0.309327778 | 0.774782508 | -4.704317251 |
| **A_55_P1955276** | Specc1 | -0.057727027 | -0.185383377 | 0.069929323 | 8.357553989 | -1.112515144 | 0.309396372 | 0.774830948 | -4.704429125 |
| **A_55_P1992403** | Simc1 | -0.375974277 | -1.207454291 | 0.455505736 | 9.445409562 | -1.112437146 | 0.30942729 | 0.774830948 | -4.704479542 |
| **A_52_P175157** | NA | -0.116240555 | -0.373402494 | 0.140921383 | 9.307309962 | -1.112039332 | 0.309585022 | 0.775119015 | -4.704736655 |
| **A_51_P343517** | Ly6d | -0.080217588 | -0.257700093 | 0.097264918 | 8.335440125 | -1.111944753 | 0.309622532 | 0.775119015 | -4.704797775 |
| **A_51_P204053** | Hba-x | 0.232597901 | -0.282178577 | 0.747374378 | 8.227470424 | 1.111619764 | 0.309751452 | 0.775295975 | -4.705007775 |
| **A_51_P430973** | Paqr7 | 0.192200786 | -0.233191602 | 0.617593175 | 9.718092324 | 1.111564348 | 0.30977344 | 0.775295975 | -4.705043581 |
| **A_52_P204618** | Rnf222 | 0.205227005 | -0.249113211 | 0.65956722 | 8.856743375 | 1.111277419 | 0.309887306 | 0.775400866 | -4.705228957 |
| **A_55_P2138095** | NA | -0.075401508 | -0.242337134 | 0.091534118 | 15.56161959 | -1.111220293 | 0.30990998 | 0.775400866 | -4.705265861 |
| **A_55_P2013465** | L2hgdh | -0.051358414 | -0.165077834 | 0.062361006 | 8.245087423 | -1.111082042 | 0.309964861 | 0.775400866 | -4.70535517 |
| **A_55_P2151116** | Micu1 | 0.122605206 | -0.148877851 | 0.394088264 | 12.39733556 | 1.111054454 | 0.309975813 | 0.775400866 | -4.705372991 |
| **A_55_P1954271** | Col26a1 | 0.078932158 | -0.0959709 | 0.253835215 | 8.293728447 | 1.110262663 | 0.310290289 | 0.776087085 | -4.705884362 |
| **A_55_P1987231** | Fuom | -0.049086444 | -0.157904064 | 0.059731177 | 8.227531395 | -1.109766237 | 0.310487591 | 0.776467295 | -4.706204879 |
| **A_51_P394735** | Arhgef18 | -0.159540066 | -0.513315142 | 0.194235011 | 12.12751515 | -1.10946012 | 0.310609308 | 0.776467295 | -4.706402486 |
| **A_66_P113884** | NA | 0.072864695 | -0.08871697 | 0.234446359 | 8.429398954 | 1.109415512 | 0.310627048 | 0.776467295 | -4.706431279 |
| **A_51_P296878** | Triqk | 0.142843651 | -0.173927651 | 0.459614952 | 8.795385214 | 1.109390074 | 0.310637165 | 0.776467295 | -4.706447698 |
| **A_55_P2057523** | Ppm1b | -0.085670686 | -0.275666862 | 0.104325489 | 9.446524232 | -1.109319253 | 0.310665332 | 0.776467295 | -4.70649341 |
| **A_52_P31814** | 1110038B12Rik | 0.133017411 | -0.16199558 | 0.428030403 | 10.99076536 | 1.109268074 | 0.310685689 | 0.776467295 | -4.706526443 |
| **A_55_P1953894** | Mccc2 | 0.113058933 | -0.137710587 | 0.363828452 | 8.759446958 | 1.10917302 | 0.3107235 | 0.776467295 | -4.706587792 |
| **A_55_P2120866** | NA | -0.217061181 | -0.698759006 | 0.264636644 | 9.748690392 | -1.108604505 | 0.310949727 | 0.77659336 | -4.706954661 |
| **A_55_P2068247** | NA | -0.214859492 | -0.69167333 | 0.261954346 | 10.74847737 | -1.108599957 | 0.310951537 | 0.77659336 | -4.706957595 |
| **A_55_P1973829** | NA | 0.127753775 | -0.155767708 | 0.411275258 | 8.379836454 | 1.108554201 | 0.31096975 | 0.77659336 | -4.706987118 |
| **A_52_P262676** | NA | -0.118486779 | -0.381445682 | 0.144472125 | 10.47096574 | -1.108539508 | 0.310975599 | 0.77659336 | -4.706996598 |
| **A_55_P2044932** | Gpr84 | -0.078833331 | -0.253797761 | 0.096131099 | 7.733696308 | -1.108483608 | 0.310997852 | 0.77659336 | -4.707032665 |
| **A_52_P494514** | Insr | -0.080380743 | -0.25878647 | 0.098024985 | 8.602742204 | -1.108440501 | 0.311015014 | 0.77659336 | -4.707060477 |
| **A_55_P2061338** | Slc16a11 | 0.246917578 | -0.301169254 | 0.79500441 | 8.316418013 | 1.108336928 | 0.31105625 | 0.776596003 | -4.707127298 |
| **A_66_P111011** | Gata3 | -0.178364142 | -0.574607891 | 0.217879606 | 11.88095901 | -1.107424914 | 0.311419556 | 0.777372866 | -4.707715554 |
| **A_55_P2059318** | Dgkz | -0.088256369 | -0.284340244 | 0.107827507 | 10.11169923 | -1.107320488 | 0.311461178 | 0.777372866 | -4.707782895 |
| **A_55_P2087911** | Mthfsd | 0.116444189 | -0.142281831 | 0.37517021 | 8.805945746 | 1.10725303 | 0.311488067 | 0.777372866 | -4.707826393 |
| **A_55_P2150025** | Otub1 | 0.074415767 | -0.090952315 | 0.239783849 | 9.230750377 | 1.107088748 | 0.311553559 | 0.777406635 | -4.707932321 |
| **A_66_P102954** | Gm2155 | 0.414857539 | -0.507223012 | 1.336938091 | 11.81322663 | 1.106876808 | 0.311638068 | 0.777406635 | -4.708068967 |
| **A_55_P2137314** | Kctd21 | 0.100627086 | -0.123037794 | 0.324291966 | 9.105429968 | 1.106843613 | 0.311651306 | 0.777406635 | -4.708090368 |

| | | | | | | | | | |
|---|---|---|---|---|---|---|---|---|---|
| **A_55_P1962781** | NA | 0.126919675 | -0.155235934 | 0.409075284 | 8.93501752 | 1.1066478 | 0.311729405 | 0.777406635 | -4.708216602 |
| **A_55_P2060722** | Uso1 | -0.057755433 | -0.186153595 | 0.07064273 | 12.10562963 | -1.106631928 | 0.311735736 | 0.777406635 | -4.708226834 |
| **A_55_P2046728** | C920009B18Rik | -0.327131068 | -1.054399645 | 0.400137509 | 11.31465948 | -1.106613927 | 0.311742916 | 0.777406635 | -4.708238438 |
| **A_55_P2016842** | Me1 | -0.134442928 | -0.433379508 | 0.164493652 | 10.60501761 | -1.106440496 | 0.311812104 | 0.777453548 | -4.708350232 |
| **A_51_P419959** | Eaf1 | 0.145342839 | -0.177851929 | 0.468537606 | 9.991263654 | 1.106365133 | 0.311842173 | 0.777453548 | -4.708398808 |
| **A_55_P1959218** | Usp51 | 0.233682451 | -0.28603722 | 0.753402122 | 9.274641975 | 1.106180777 | 0.311915739 | 0.777536667 | -4.70851763 |
| **A_52_P263251** | 3110040N11Rik | -0.095263319 | -0.307195977 | 0.116669338 | 8.914079592 | -1.105851949 | 0.312046991 | 0.777763545 | -4.708729542 |
| **A_55_P2106150** | Cenpk | 0.099390905 | -0.12174792 | 0.32052973 | 8.284034321 | 1.105734374 | 0.312093933 | 0.777780252 | -4.708805306 |
| **A_55_P2212161** | 2310034O05Rik | 0.132795153 | -0.162764358 | 0.428354665 | 8.876783885 | 1.105366882 | 0.312240692 | 0.777929164 | -4.709042083 |
| **A_55_P2232754** | Smarcal1 | 0.108434873 | -0.132916766 | 0.349786511 | 8.473318636 | 1.105319638 | 0.312259563 | 0.777929164 | -4.70907252 |
| **A_51_P214487** | 2-Sep | 0.108894166 | -0.133487915 | 0.351276247 | 14.25067589 | 1.105282429 | 0.312274426 | 0.777929164 | -4.709096491 |
| **A_55_P2053202** | Itsn2 | -0.089626015 | -0.289165026 | 0.109912996 | 10.93499027 | -1.105033478 | 0.312373887 | 0.77803403 | -4.709256862 |
| **A_52_P108447** | Glis2 | -0.103564874 | -0.334152565 | 0.127022816 | 9.978464511 | -1.104957375 | 0.312404297 | 0.77803403 | -4.709305883 |
| **A_55_P2025675** | Dnah8 | 0.05709111 | -0.07004845 | 0.18423067 | 8.187147229 | 1.104732029 | 0.312494358 | 0.77803403 | -4.709451027 |
| **A_51_P340829** | AA986860 | -0.077577456 | -0.250366506 | 0.095211593 | 10.00633026 | -1.104557895 | 0.312563966 | 0.77803403 | -4.709563175 |
| **A_51_P509211** | Rnf34 | -0.097150156 | -0.31355616 | 0.119255849 | 11.3153683 | -1.104443129 | 0.31260985 | 0.77803403 | -4.709637083 |
| **A_55_P2110037** | Akap7 | -0.088550487 | -0.285811735 | 0.10871076 | 9.749789445 | -1.104379521 | 0.312635283 | 0.77803403 | -4.709678044 |
| **A_55_P2014304** | Kank1 | -0.073521779 | -0.237326117 | 0.090282559 | 13.8831276 | -1.104230587 | 0.312694839 | 0.77803403 | -4.709773947 |
| **A_51_P470328** | Selenop | 0.253600698 | -0.311420427 | 0.818621823 | 16.13652317 | 1.104218255 | 0.312699771 | 0.77803403 | -4.709781888 |
| **A_55_P1986630** | Gm3627 | 0.059521184 | -0.07311664 | 0.192159009 | 7.906042227 | 1.104010837 | 0.312782731 | 0.77803403 | -4.709915439 |
| **A_55_P2093595** | Mmd | 0.097582158 | -0.119901074 | 0.315065389 | 10.43858462 | 1.103859506 | 0.31284327 | 0.77803403 | -4.710012868 |
| **A_55_P2060567** | Ddx39 | 0.121357207 | -0.149114308 | 0.391828722 | 11.50200204 | 1.10385798 | 0.31284388 | 0.77803403 | -4.710013851 |
| **A_55_P2032147** | Wnt9a | -0.105772536 | -0.341517939 | 0.129972867 | 9.883408625 | -1.103821498 | 0.312858476 | 0.77803403 | -4.710037337 |
| **A_65_P17218** | Mndal | -0.081282969 | -0.262451901 | 0.099885964 | 8.493814145 | -1.103786288 | 0.312872564 | 0.77803403 | -4.710060005 |
| **A_51_P401907** | Cstdc4 | -0.277718775 | -0.896727505 | 0.341289954 | 9.056862414 | -1.103767574 | 0.312880052 | 0.77803403 | -4.710072053 |
| **A_51_P311990** | P4hb | -0.16982252 | -0.548398736 | 0.208753697 | 14.74338436 | -1.103598537 | 0.312947692 | 0.778102127 | -4.710180869 |
| **A_55_P2153555** | Zkscan3 | -0.07403114 | -0.239083554 | 0.091021274 | 9.284882852 | -1.103473019 | 0.312997926 | 0.778126934 | -4.710261664 |
| **A_55_P1983848** | Tc2n | -0.082087036 | -0.265122889 | 0.100948817 | 7.891115842 | -1.103335444 | 0.313052993 | 0.778163748 | -4.710350216 |
| **A_55_P1952779** | Dcp1b | -0.087877912 | -0.283844852 | 0.108089028 | 9.013892132 | -1.103230044 | 0.313095187 | 0.778168557 | -4.710418053 |
| **A_51_P324640** | C330018D20Rik | 0.120589387 | -0.148383139 | 0.389561913 | 9.871458343 | 1.102986833 | 0.313192568 | 0.77831051 | -4.710574576 |
| **A_55_P2000047** | 9930104L06Rik | -0.06339519 | -0.204814043 | 0.078023663 | 8.179601693 | -1.102854344 | 0.313245627 | 0.778342296 | -4.710659834 |
| **A_55_P2026761** | NA | 0.278290151 | -0.342793392 | 0.899373693 | 11.83263597 | 1.102343581 | 0.313450246 | 0.778750618 | -4.710988465 |
| **A_51_P208580** | Nop53 | -0.097520833 | -0.315207887 | 0.120166221 | 8.820666378 | -1.102132887 | 0.313534686 | 0.778860294 | -4.711124006 |
| **A_55_P2370096** | NA | 0.058940895 | -0.072641166 | 0.190522955 | 8.327099752 | 1.102019309 | 0.313580213 | 0.778873288 | -4.711197065 |
| **A_55_P1956807** | Gm14391 | -0.088117413 | -0.284873175 | 0.108638349 | 9.816114405 | -1.101801706 | 0.313667452 | 0.778894815 | -4.711337029 |
| **A_52_P302041** | NA | 0.158963851 | -0.195989639 | 0.513917341 | 10.11423582 | 1.101783051 | 0.313674932 | 0.778894815 | -4.711349027 |
| **A_55_P1972104** | Acsm3 | 0.074233909 | -0.091540288 | 0.240008106 | 7.824305217 | 1.101677709 | 0.313717173 | 0.778894815 | -4.711416778 |
| **A_55_P2084631** | Hist1h2an | 0.284621256 | -0.351104099 | 0.920346611 | 14.02258519 | 1.101455513 | 0.313806286 | 0.778894815 | -4.711559672 |
| **A_55_P1984123** | Kmt2c | -0.155388856 | -0.502491703 | 0.191713991 | 10.87665767 | -1.101363951 | 0.313843014 | 0.778894815 | -4.711618551 |
| **A_55_P2015178** | Nap1l1 | 0.111101509 | -0.137100193 | 0.35930321 | 12.7395225 | 1.101246053 | 0.313890311 | 0.778894815 | -4.711694362 |
| **A_51_P140803** | Slco1b2 | 0.472053176 | -0.582519363 | 1.526625716 | 8.779986548 | 1.101244188 | 0.313891059 | 0.778894815 | -4.711695561 |
| **A_52_P69043** | Nkap | 0.101927141 | -0.125789739 | 0.329644021 | 9.432014093 | 1.101193855 | 0.313911253 | 0.778894815 | -4.711727924 |
| **A_55_P2043317** | Bag1 | -0.209652914 | -0.678112428 | 0.258806601 | 14.77729735 | -1.101027082 | 0.313978171 | 0.778953488 | -4.711835153 |
| **A_55_P2046245** | Hes1 | 0.129142358 | -0.159444509 | 0.417729225 | 8.649547814 | 1.100934067 | 0.314015499 | 0.778953488 | -4.711894954 |
| **A_55_P2093705** | Meig1 | -0.049621785 | -0.160526285 | 0.061282716 | 7.832443623 | -1.100759327 | 0.314085634 | 0.778964167 | -4.712007292 |
| **A_55_P2103352** | Ndufb5 | 0.166935671 | -0.206259107 | 0.540130449 | 12.79529406 | 1.100481515 | 0.314197165 | 0.778964167 | -4.712185872 |
| **A_55_P1975772** | Ppp2r5a | -0.074213888 | -0.240124451 | 0.091696674 | 11.95496214 | -1.100475338 | 0.314199645 | 0.778964167 | -4.712189843 |
| **A_55_P2116149** | NA | -0.316244349 | -1.023329949 | 0.39084125 | 11.11602739 | -1.100322377 | 0.314261068 | 0.778964167 | -4.712288157 |
| **A_55_P2072391** | Acp1 | 0.057403925 | -0.070946061 | 0.185753912 | 8.452832637 | 1.100309666 | 0.314266172 | 0.778964167 | -4.712296327 |
| **A_55_P2444515** | NA | -0.146228467 | -0.473268093 | 0.180811158 | 11.32606815 | -1.100020336 | 0.314382384 | 0.778964167 | -4.712482271 |

| | | | | | | | | | |
|---|---|---|---|---|---|---|---|---|---|
| **A_55_P2171715** | Mrps33 | -0.081476534 | -0.26371899 | 0.100765921 | 12.1197654 | -1.09989734 | 0.314431797 | 0.778964167 | -4.71256131 |
| **A_51_P117604** | Rab5a | 0.050669944 | -0.062670016 | 0.164009904 | 11.9204181 | 1.099857777 | 0.314447693 | 0.778964167 | -4.712586733 |
| **A_52_P225260** | Ywhaz | -0.062277687 | -0.201599691 | 0.077044317 | 8.706941243 | -1.099719494 | 0.314503258 | 0.778964167 | -4.712675587 |
| **A_55_P1989928** | Lin52 | 0.083145841 | -0.102880736 | 0.269172419 | 8.84140961 | 1.099599919 | 0.314551312 | 0.778964167 | -4.712752416 |
| **A_51_P499698** | Asprv1 | -0.178159016 | -0.576773478 | 0.220455446 | 8.137052143 | -1.099572621 | 0.314562283 | 0.778964167 | -4.712769955 |
| **A_55_P1991500** | Stn1 | 0.141220506 | -0.174751556 | 0.457192568 | 10.82010831 | 1.09955825 | 0.314568059 | 0.778964167 | -4.712779188 |
| **A_55_P2029420** | 2610318N02Rik | 0.057267836 | -0.070870186 | 0.185405859 | 7.998996379 | 1.099516935 | 0.314584665 | 0.778964167 | -4.712805732 |
| **A_52_P629333** | Map3k21 | 0.094668626 | -0.117161095 | 0.306498347 | 8.618861808 | 1.099482558 | 0.314598482 | 0.778964167 | -4.712827819 |
| **A_55_P1970027** | Itch | -0.085525446 | -0.276922259 | 0.105871367 | 10.81737561 | -1.099334373 | 0.31465805 | 0.778964167 | -4.712923019 |
| **A_55_P2085536** | Smarcal1 | -0.076589664 | -0.248009665 | 0.094830338 | 10.72818556 | -1.099202808 | 0.314710945 | 0.778964167 | -4.713007536 |
| **A_51_P145631** | Svip | 0.065441078 | -0.081043072 | 0.211925229 | 7.844002239 | 1.099079157 | 0.314760665 | 0.778964167 | -4.713086965 |
| **A_55_P1960999** | Pigr | 0.175654621 | -0.21754104 | 0.568850283 | 12.29619943 | 1.099056517 | 0.314769769 | 0.778964167 | -4.713101507 |
| **A_55_P2024021** | Atp5l | -0.12537997 | -0.406047885 | 0.155287944 | 14.99955943 | -1.099017376 | 0.314785509 | 0.778964167 | -4.713126649 |
| **A_55_P2167803** | Abi3bp | 0.059888401 | -0.0741948 | 0.193971601 | 7.724540401 | 1.098847721 | 0.314853742 | 0.77903328 | -4.713235617 |
| **A_52_P114494** | Stat6 | -0.105004787 | -0.340140092 | 0.130130518 | 9.315403657 | -1.098652679 | 0.314932201 | 0.779127673 | -4.713360882 |
| **A_55_P1995622** | Pomp | -0.069032919 | -0.223655437 | 0.085589598 | 9.3261241 | -1.098380117 | 0.31504187 | 0.779299246 | -4.713535912 |
| **A_55_P2014706** | Naip5 | -0.147191811 | -0.476925256 | 0.182541633 | 14.57911322 | -1.098221177 | 0.315105837 | 0.779335583 | -4.713637968 |
| **A_55_P1991199** | Nos1ap | 0.143597426 | -0.178110754 | 0.465305607 | 8.890903446 | 1.098129895 | 0.315142579 | 0.779335583 | -4.713696577 |
| **A_55_P2026794** | Unc50 | -0.111750804 | -0.362135696 | 0.138634089 | 12.99850966 | -1.098023659 | 0.315185345 | 0.779335583 | -4.713764784 |
| **A_66_P129029** | Taf10 | 0.078077521 | -0.096873066 | 0.253028108 | 11.85702811 | 1.09794295 | 0.315217838 | 0.779335583 | -4.7138166 |
| **A_51_P209782** | Cyp2c23 | 0.168343588 | -0.208945857 | 0.545633032 | 7.972763289 | 1.09771876 | 0.31530811 | 0.779459068 | -4.713960521 |
| **A_51_P246471** | Pbx2 | 0.082238522 | -0.10209423 | 0.266571273 | 10.84918665 | 1.097594576 | 0.315358123 | 0.779483013 | -4.714040235 |
| **A_52_P381806** | Terf2ip | -0.059401139 | -0.19257571 | 0.073773432 | 8.643766136 | -1.097343589 | 0.315459225 | 0.779633212 | -4.714201331 |
| **A_55_P2167645** | NA | 0.143916216 | -0.178789829 | 0.46662226 | 8.855500022 | 1.097164618 | 0.315531333 | 0.779711728 | -4.714316191 |
| **A_55_P2057212** | Tpr | 0.096642779 | -0.120092739 | 0.313378297 | 11.13873201 | 1.097004691 | 0.315595781 | 0.779738652 | -4.714418822 |
| **A_51_P333071** | Anapc16 | 0.067692464 | -0.084127068 | 0.219511996 | 10.92331548 | 1.096937379 | 0.315622909 | 0.779738652 | -4.714462015 |
| **A_66_P135391** | Igfbp2 | -0.225002381 | -0.729684545 | 0.279679783 | 12.79012116 | -1.096827474 | 0.315667208 | 0.779748431 | -4.714532538 |
| **A_52_P15490** | Nectin3 | -0.0612291 | -0.198607504 | 0.076149304 | 8.004722737 | -1.096499836 | 0.315799299 | 0.779940157 | -4.714742751 |
| **A_52_P310981** | Eif4e | -0.080747501 | -0.26193245 | 0.100437448 | 11.091987 | -1.096417943 | 0.315832322 | 0.779940157 | -4.714795289 |
| **A_55_P1992811** | Nudc | -0.061910327 | -0.20088707 | 0.077066417 | 9.166470704 | -1.095948454 | 0.316021699 | 0.779940157 | -4.715096444 |
| **A_52_P681705** | Zfp955b | -0.128320299 | -0.416376873 | 0.159736274 | 9.661527818 | -1.095939896 | 0.316025152 | 0.779940157 | -4.715101933 |
| **A_55_P2031547** | Vmn2r121 | -0.150226018 | -0.487462804 | 0.187010769 | 8.403435154 | -1.095921319 | 0.316032647 | 0.779940157 | -4.715113848 |
| **A_55_P2007423** | Ppp1r12a | 0.092892453 | -0.115667021 | 0.301451927 | 9.339971229 | 1.095770666 | 0.316093437 | 0.779940157 | -4.71521047 |
| **A_55_P2141301** | Synj2bp | 0.104515352 | -0.130155904 | 0.339186608 | 10.55817206 | 1.095694176 | 0.316124306 | 0.779940157 | -4.715259524 |
| **A_55_P1990314** | Elavl4 | 0.110283703 | -0.137378586 | 0.357945992 | 9.367220782 | 1.095520785 | 0.316194289 | 0.779940157 | -4.715370715 |
| **A_51_P356967** | Lgi3 | 0.066152025 | -0.082409334 | 0.214713385 | 7.689426918 | 1.095485025 | 0.316208723 | 0.779940157 | -4.715393646 |
| **A_55_P2114779** | Greb1 | 0.264281968 | -0.329280663 | 0.857844598 | 9.209703206 | 1.095393459 | 0.316245687 | 0.779940157 | -4.71545236 |
| **A_66_P110088** | NA | -0.282521821 | -0.917251772 | 0.352208131 | 11.53823937 | -1.095045506 | 0.316386184 | 0.779940157 | -4.715675454 |
| **A_55_P1979341** | Cyba | 0.29702479 | -0.370386545 | 0.964436126 | 12.41294151 | 1.094884437 | 0.316451239 | 0.779940157 | -4.715778711 |
| **A_52_P75384** | B230219D22Rik | -0.153271881 | -0.497687787 | 0.191144025 | 10.76805029 | -1.09483444 | 0.316471434 | 0.779940157 | -4.715810762 |
| **A_55_P2017724** | NA | 0.181951325 | -0.226915994 | 0.590818644 | 9.939558542 | 1.094818149 | 0.316478015 | 0.779940157 | -4.715821205 |
| **A_51_P451106** | Brca2 | 0.064988514 | -0.081051387 | 0.211028415 | 8.328953987 | 1.094798613 | 0.316485907 | 0.779940157 | -4.715833728 |
| **A_51_P328133** | Gle1 | -0.063829329 | -0.207279876 | 0.079621219 | 8.40516435 | -1.094680121 | 0.316533776 | 0.779940157 | -4.715909683 |
| **A_52_P424462** | Ero1lb | -0.087310015 | -0.283531337 | 0.108911307 | 8.309371894 | -1.094679608 | 0.316533983 | 0.779940157 | -4.715910012 |
| **A_55_P2152926** | Mybbp1a | -0.076848705 | -0.249592578 | 0.095895168 | 12.18111464 | -1.094467986 | 0.316619491 | 0.779940157 | -4.716045652 |
| **A_51_P217382** | Kif1c | -0.064254544 | -0.20869316 | 0.080184072 | 11.20605929 | -1.094434288 | 0.316633109 | 0.779940157 | -4.71606725 |
| **A_55_P1955346** | NA | 0.137031488 | -0.171011525 | 0.445074501 | 13.80111958 | 1.094405267 | 0.316644837 | 0.779940157 | -4.716085851 |
| **A_55_P2148609** | NA | 0.055051132 | -0.068723101 | 0.178825364 | 8.56048097 | 1.09422132 | 0.316719183 | 0.779940157 | -4.716203738 |
| **A_66_P115949** | Myo9a | -0.073882487 | -0.240001034 | 0.092236061 | 8.636839657 | -1.0941895 | 0.316732045 | 0.779940157 | -4.71622413 |
| **A_55_P2171493** | BC030867 | 0.066274182 | -0.082743631 | 0.215291995 | 7.906078296 | 1.094146193 | 0.316749551 | 0.779940157 | -4.716251882 |

| | | | | | | | | | |
|---|---|---|---|---|---|---|---|---|---|
| **A_52_P434052** | Wdr37 | -0.047383651 | -0.153939691 | 0.05917239 | 8.97345923 | -1.094005752 | 0.316806327 | 0.779940157 | -4.716341877 |
| **A_55_P2240116** | 2310005E17Rik | 0.120285577 | -0.150246775 | 0.390817929 | 8.198958566 | 1.093864455 | 0.316863458 | 0.779940157 | -4.716432415 |
| **A_55_P2171406** | Tm7sf2 | 0.196249424 | -0.245200226 | 0.637699075 | 8.124976231 | 1.09369538 | 0.316931832 | 0.779940157 | -4.716540744 |
| **A_51_P396854** | Timm10b | 0.094338107 | -0.117893595 | 0.306569809 | 12.80068773 | 1.093568682 | 0.316983077 | 0.779940157 | -4.716621916 |
| **A_55_P1968878** | Rhpn2 | 0.234221755 | -0.292736052 | 0.761179562 | 11.58248805 | 1.093504441 | 0.317009063 | 0.779940157 | -4.716663071 |
| **A_55_P1984601** | Vapb | 0.086637944 | -0.108289221 | 0.281565108 | 8.763155554 | 1.093465121 | 0.317024969 | 0.779940157 | -4.71668826 |
| **A_55_P2185900** | Nrg4 | 0.063377513 | -0.079225984 | 0.20598101 | 8.433605463 | 1.093387686 | 0.317056295 | 0.779940157 | -4.716737865 |
| **A_55_P2020461** | Hmgn2 | -0.091352688 | -0.296903088 | 0.114197712 | 10.27299988 | -1.093382624 | 0.317058343 | 0.779940157 | -4.716741108 |
| **A_55_P2193419** | NA | -0.196870658 | -0.639853394 | 0.246112077 | 8.219755252 | -1.093360439 | 0.317067319 | 0.779940157 | -4.716755319 |
| **A_52_P250578** | Tcaf1 | -0.049494088 | -0.16087115 | 0.061882975 | 7.86172377 | -1.093268228 | 0.317104628 | 0.779940157 | -4.716814387 |
| **A_55_P2042818** | Zfp639 | 0.114893617 | -0.143678217 | 0.373465452 | 11.03986969 | 1.093160296 | 0.317148301 | 0.779940157 | -4.716883522 |
| **A_55_P2049647** | H2-K1 | -0.511664785 | -1.663205989 | 0.639876419 | 13.26610441 | -1.093138547 | 0.317157103 | 0.779940157 | -4.716897452 |
| **A_52_P346458** | Tollip | -0.119552921 | -0.388695321 | 0.14958948 | 11.45467915 | -1.092816471 | 0.317287461 | 0.780161473 | -4.717103731 |
| **A_55_P1974984** | NA | 0.074554996 | -0.093319818 | 0.24242981 | 8.282210204 | 1.092597919 | 0.317375944 | 0.780207746 | -4.717243688 |
| **A_55_P2128451** | Sfpq | -0.139877676 | -0.454846859 | 0.175091507 | 11.1024793 | -1.092570592 | 0.31738701 | 0.780207746 | -4.717261187 |
| **A_55_P1968998** | Tmub2 | 0.162041792 | -0.202988535 | 0.527072118 | 11.07592462 | 1.092112187 | 0.317572672 | 0.780564876 | -4.71755469 |
| **A_51_P149422** | Gng13 | 0.238987734 | -0.299439559 | 0.777415028 | 8.009083562 | 1.091987605 | 0.317623146 | 0.780589675 | -4.717634445 |
| **A_55_P2156401** | NA | -0.115298286 | -0.375123389 | 0.144526817 | 13.55744772 | -1.091719091 | 0.317731956 | 0.780619524 | -4.717806325 |
| **A_55_P2065308** | Myo9b | -0.069847519 | -0.227271089 | 0.087576052 | 8.470055448 | -1.091566977 | 0.31779361 | 0.780619524 | -4.717903686 |
| **A_55_P2079089** | Fuca1 | -0.119967845 | -0.390361268 | 0.150425578 | 11.48846613 | -1.09153558 | 0.317806337 | 0.780619524 | -4.717923781 |
| **A_55_P2031432** | Usp47 | -0.055463773 | -0.180486709 | 0.069559163 | 8.325154034 | -1.091412397 | 0.317856275 | 0.780619524 | -4.718002619 |
| **A_55_P1999641** | Frem1 | 0.214658517 | -0.269216643 | 0.698533677 | 8.432240804 | 1.091400026 | 0.31786129 | 0.780619524 | -4.718010536 |
| **A_55_P2162160** | Anp32a | -0.114540234 | -0.372756553 | 0.143676084 | 11.03258827 | -1.091298468 | 0.317902466 | 0.780619524 | -4.718075529 |
| **A_55_P2164957** | Slc9a3r2 | 0.059906748 | -0.075150097 | 0.194963592 | 8.267982565 | 1.091260175 | 0.317917993 | 0.780619524 | -4.718100034 |
| **A_55_P1997335** | Vps9d1 | 0.120253975 | -0.150921237 | 0.391429188 | 10.67469277 | 1.090984589 | 0.318029755 | 0.780741256 | -4.718276379 |
| **A_51_P268094** | Serpine2 | -0.270983456 | -0.882142934 | 0.340176022 | 9.799062492 | -1.090830793 | 0.318092141 | 0.780741256 | -4.718374783 |
| **A_51_P101635** | Stub1 | -0.091871867 | -0.299084457 | 0.115340722 | 12.23528503 | -1.090775988 | 0.318114374 | 0.780741256 | -4.718409846 |
| **A_51_P391380** | Mindy2 | -0.095433159 | -0.310687134 | 0.119820817 | 9.088708491 | -1.090730055 | 0.318133009 | 0.780741256 | -4.718439233 |
| **A_55_P2000158** | Pik3r3 | -0.125525582 | -0.408694954 | 0.157643789 | 11.30642471 | -1.090573976 | 0.318196338 | 0.780741256 | -4.718539084 |
| **A_55_P2032916** | Slc6a20a | 0.359370491 | -0.451397548 | 1.17013853 | 11.80281255 | 1.090473098 | 0.318237274 | 0.780741256 | -4.718603617 |
| **A_65_P01641** | Rnf6 | -0.149927548 | -0.488200322 | 0.188345225 | 12.62496374 | -1.090394264 | 0.318269268 | 0.780741256 | -4.718654045 |
| **A_51_P297336** | Chmp7 | -0.105413079 | -0.34326862 | 0.132442462 | 10.99311805 | -1.090310984 | 0.318303069 | 0.780741256 | -4.718707315 |
| **A_55_P2159432** | Fndc4 | 0.153244036 | -0.192590024 | 0.499078096 | 10.80490954 | 1.090146793 | 0.318369719 | 0.780741256 | -4.718812334 |
| **A_52_P109403** | Cebpz | 0.13826109 | -0.1737614 | 0.450283581 | 10.38310541 | 1.090142425 | 0.318371493 | 0.780741256 | -4.718815128 |
| **A_51_P108226** | Wfdc21 | -0.069541839 | -0.226527079 | 0.0874434 | 7.83568583 | -1.089824372 | 0.318500633 | 0.780777337 | -4.716054536 |
| **A_55_P1955715** | Gm14326 | -0.154140078 | -0.502099997 | 0.193819841 | 10.04551886 | -1.089821866 | 0.31850165 | 0.780777337 | -4.719020137 |
| **A_55_P2030269** | Dnah6 | 0.052336298 | -0.065812874 | 0.17048547 | 7.876815923 | 1.089786639 | 0.318515956 | 0.780777337 | -4.719042664 |
| **A_51_P357433** | Ap1g2 | 0.088721853 | -0.111582099 | 0.289025806 | 8.414520965 | 1.089708277 | 0.318547782 | 0.780777337 | -4.719092774 |
| **A_51_P382121** | Ormdl1 | 0.179303543 | -0.22565519 | 0.584262275 | 10.85573227 | 1.089299402 | 0.318713884 | 0.781066447 | -4.719354205 |
| **A_55_P2053858** | Scamp3 | 0.138042487 | -0.173768909 | 0.449853882 | 12.95339442 | 1.089155664 | 0.318772293 | 0.781066447 | -4.719446097 |
| **A_55_P2035424** | Hpgd | -0.512945429 | -1.67166867 | 0.645777813 | 9.09335411 | -1.089082071 | 0.318802202 | 0.781066447 | -4.719493142 |
| **A_51_P357744** | Jund | 0.380527286 | -0.556335233 | 1.317389804 | 13.25769282 | 1.142911095 | 0.318827372 | 0.781066447 | -4.606927814 |
| **A_55_P1993719** | Tmem216 | 0.148146046 | -0.18660505 | 0.482897143 | 10.10325012 | 1.088772693 | 0.318927961 | 0.781131002 | -4.7196909 |
| **A_55_P1974617** | Lysmd1 | -0.075287942 | -0.245420757 | 0.094844873 | 10.86240568 | -1.088695684 | 0.31895927 | 0.781131002 | -4.71974012 |
| **A_52_P247943** | Parl | -0.054871277 | -0.178871689 | 0.069129134 | 8.322037795 | -1.088657097 | 0.31897496 | 0.781131002 | -4.719764783 |
| **A_55_P2070869** | Lcn2 | 0.143426406 | -0.180731704 | 0.467584516 | 8.834461166 | 1.08853247 | 0.319025637 | 0.781156136 | -4.719844433 |
| **A_55_P2157616** | NA | 0.072005923 | -0.090756118 | 0.234767964 | 16.53077098 | 1.088389284 | 0.31908387 | 0.7811789 | -4.719935938 |
| **A_55_P2028661** | Dusp12 | 0.095084632 | -0.119878294 | 0.310047558 | 10.93141991 | 1.088218057 | 0.319153517 | 0.7811789 | -4.720045355 |
| **A_55_P2042698** | Hsd17b10 | 0.141501138 | -0.178469216 | 0.461471492 | 11.39645097 | 1.087976099 | 0.319251957 | 0.7811789 | -4.720199954 |
| **A_55_P1982628** | Tma16 | 0.146695833 | -0.185049182 | 0.478440848 | 9.135510777 | 1.087883865 | 0.319289489 | 0.7811789 | -4.720258882 |

| | | | | | | | | | |
|---|---|---|---|---|---|---|---|---|---|
| **A_55_P2014928** | Zfp341 | 0.068561926 | -0.086509984 | 0.223633837 | 8.701410382 | 1.087724818 | 0.319354217 | 0.7811789 | -4.720360491 |
| **A_51_P512820** | Dera | 0.128853951 | -0.162593688 | 0.420301591 | 9.664485715 | 1.08769307 | 0.319367139 | 0.7811789 | -4.720380772 |
| **A_55_P2150908** | Atp5c1 | 0.185888103 | -0.234563701 | 0.606339908 | 14.31607394 | 1.087688504 | 0.319368997 | 0.7811789 | -4.720383689 |
| **A_55_P2112599** | Lgals8 | -0.135545482 | -0.442132924 | 0.171041961 | 9.847791298 | -1.087676752 | 0.31937378 | 0.7811789 | -4.720391197 |
| **A_66_P120484** | Mbtd1 | -0.098489721 | -0.321274051 | 0.124294608 | 10.39670867 | -1.08761561 | 0.319398668 | 0.7811789 | -4.720430255 |
| **A_55_P1977563** | Galk2 | -0.081025905 | -0.264356192 | 0.102304382 | 8.690616957 | -1.087323653 | 0.319517529 | 0.781336317 | -4.720616745 |
| **A_55_P2129706** | Srsf4 | -0.112285046 | -0.366357798 | 0.141787707 | 10.87351971 | -1.087258947 | 0.319543877 | 0.781336317 | -4.720658072 |
| **A_52_P472302** | Fxyd6 | -0.145741075 | -0.475584513 | 0.184102363 | 16.31802718 | -1.087034387 | 0.31963533 | 0.781461079 | -4.720801488 |
| **A_55_P1989321** | Sat1 | -0.131488415 | -0.429123161 | 0.166146332 | 12.39532541 | -1.086858541 | 0.31970696 | 0.781537349 | -4.720913782 |
| **A_51_P293781** | Fkbp1b | 0.132370006 | -0.167326135 | 0.432066146 | 9.660758909 | 1.086619764 | 0.319804245 | 0.78167631 | -4.721066248 |
| **A_55_P2000219** | Ubap2 | 0.076087658 | -0.096212137 | 0.248387454 | 12.44876946 | 1.086422172 | 0.319884769 | 0.78172756 | -4.721192402 |
| **A_52_P272854** | NA | 0.147476923 | -0.186501876 | 0.481455723 | 8.688706977 | 1.086361418 | 0.319909532 | 0.78172756 | -4.721231189 |
| **A_66_P120208** | Cnnm3 | -0.052244556 | -0.170568374 | 0.066079262 | 9.299457598 | -1.086270619 | 0.319946543 | 0.78172756 | -4.721289155 |
| **A_51_P505696** | Inpp5k | -0.099472827 | -0.324825437 | 0.125879783 | 9.996556107 | -1.085953005 | 0.320076035 | 0.781728379 | -4.721491898 |
| **A_52_P76196** | NA | 0.122282831 | -0.154762491 | 0.399328154 | 11.75432056 | 1.08588498 | 0.320103775 | 0.781728379 | -4.721535316 |
| **A_55_P1989102** | Hmgn1 | -0.071532297 | -0.233597253 | 0.090532659 | 11.62075097 | -1.085880961 | 0.320105413 | 0.781728379 | -4.721537881 |
| **A_55_P2139211** | NA | 0.088617262 | -0.11217141 | 0.289405934 | 13.40011111 | 1.085796112 | 0.320140017 | 0.781728379 | -4.721592036 |
| **A_55_P2037525** | NA | -0.089744199 | -0.293090448 | 0.113602051 | 13.99795649 | -1.085773854 | 0.320149095 | 0.781728379 | -4.721606241 |
| **A_55_P1983177** | Acsl4 | 0.138301987 | -0.175132226 | 0.4517362 | 12.32245162 | 1.085553375 | 0.320239027 | 0.781849205 | -4.721746949 |
| **A_55_P2068877** | Ibtk | 0.101923614 | -0.129109159 | 0.332956386 | 10.41787164 | 1.085351439 | 0.320321414 | 0.781868099 | -4.721875809 |
| **A_55_P2091861** | Hipk2 | -0.084312398 | -0.275450916 | 0.106826119 | 8.222369623 | -1.085206516 | 0.320380552 | 0.781868099 | -4.72196828 |
| **A_55_P2278604** | A330009N23Rik | 0.124036664 | -0.157158294 | 0.405231622 | 8.624715959 | 1.08520481 | 0.320381248 | 0.781868099 | -4.721969369 |
| **A_55_P1960097** | Epb41l3 | 0.09566659 | -0.121301423 | 0.312634604 | 11.09076471 | 1.084760209 | 0.320562731 | 0.781868099 | -4.722253015 |
| **A_55_P1969523** | Sumo2 | -0.095931258 | -0.313502597 | 0.121640082 | 9.915492214 | -1.084744898 | 0.320568982 | 0.781868099 | -4.722262782 |
| **A_51_P296608** | Gadd45a | -0.13051068 | -0.426542605 | 0.165521245 | 10.18175785 | -1.084617648 | 0.320620941 | 0.781868099 | -4.722343952 |
| **A_55_P1963364** | NA | -0.087845405 | -0.287125187 | 0.111434377 | 8.894482263 | -1.084488553 | 0.32067366 | 0.781868099 | -4.722426294 |
| **A_55_P2178808** | Zbtb5 | 0.084736267 | -0.107491382 | 0.276963915 | 8.864782614 | 1.084482721 | 0.320676042 | 0.781868099 | -4.722430014 |
| **A_55_P2144341** | Smim40 | 0.115263714 | -0.146234752 | 0.37676218 | 9.541263795 | 1.084407783 | 0.320706648 | 0.781868099 | -4.72247781 |
| **A_55_P1995992** | NA | -0.100065493 | -0.327118673 | 0.126987686 | 8.931915417 | -1.084241226 | 0.320774683 | 0.781868099 | -4.722584036 |
| **A_51_P389531** | Prmt1 | -0.171003558 | -0.559020019 | 0.217012902 | 8.861809682 | -1.084236844 | 0.320776473 | 0.781868099 | -4.72258683 |
| **A_55_P2117830** | NA | 0.075585043 | -0.095969508 | 0.247139593 | 8.240061879 | 1.08393386 | 0.320900265 | 0.781868099 | -4.722780041 |
| **A_51_P515242** | Strip2 | -0.099069701 | -0.323928413 | 0.125789012 | 8.559529406 | -1.083927648 | 0.320902804 | 0.781868099 | -4.722784003 |
| **A_51_P126626** | Zfp503 | 0.076358851 | -0.096958004 | 0.249675706 | 8.453529462 | 1.083896342 | 0.320915597 | 0.781868099 | -4.70159457 |
| **A_55_P2056070** | Hfe | 0.118390531 | -0.150334175 | 0.387115237 | 9.219260847 | 1.083873342 | 0.320924996 | 0.781868099 | -4.722818631 |
| **A_52_P442676** | Cux1 | -0.092929708 | -0.303876321 | 0.118016904 | 11.22755117 | -1.083805079 | 0.320952894 | 0.781868099 | -4.722862156 |
| **A_55_P1958912** | NA | -0.127914431 | -0.477113177 | 0.221284316 | 9.960611449 | -1.196409472 | 0.320979467 | 0.781868099 | -4.579812533 |
| **A_55_P2048323** | Cdk10 | 0.126716044 | -0.160946236 | 0.414378324 | 10.39602382 | 1.083721905 | 0.320986889 | 0.781868099 | -4.722915188 |
| **A_55_P2029310** | Arf4 | -0.080882933 | -0.264518868 | 0.102753001 | 11.52183437 | -1.083598478 | 0.321037341 | 0.781868099 | -4.72299388 |
| **A_55_P2012037** | Lrrc8e | -0.216939461 | -0.709497494 | 0.275618573 | 8.931727371 | -1.083553382 | 0.321055776 | 0.781868099 | -4.72302263 |
| **A_55_P2031100** | Ahcy | -0.074737798 | -0.244459951 | 0.094984356 | 7.792694815 | -1.083355335 | 0.321136748 | 0.781918845 | -4.723148884 |
| **A_55_P1994917** | Wdr5 | 0.116727107 | -0.148361048 | 0.381815262 | 11.76640483 | 1.083304535 | 0.32115752 | 0.781918845 | -4.723181267 |
| **A_65_P19250** | Ap2m1 | -0.071407181 | -0.233613488 | 0.090799125 | 8.780091067 | -1.08303706 | 0.321266911 | 0.782086664 | -4.723351758 |
| **A_55_P2106901** | Nkain4 | 0.072757651 | -0.092571257 | 0.238086558 | 8.307008038 | 1.082677307 | 0.321414088 | 0.782283596 | -4.723581032 |
| **A_55_P1957245** | NA | 0.226298641 | -0.287967518 | 0.740564801 | 11.13332996 | 1.082587885 | 0.321450681 | 0.782283596 | -4.723638016 |
| **A_51_P149562** | Apbb2 | -0.096834211 | -0.31691234 | 0.123243919 | 12.00431839 | -1.082483023 | 0.321493595 | 0.782283596 | -4.723704835 |
| **A_51_P234627** | Nubpl | 0.07374198 | -0.093859807 | 0.241343767 | 9.332906275 | 1.082443687 | 0.321509694 | 0.782283596 | -4.723729899 |
| **A_55_P2023942** | Ccdc77 | -0.094139316 | -0.308121236 | 0.119842604 | 8.744167533 | -1.082338535 | 0.321552734 | 0.782289843 | -4.723796898 |
| **A_52_P533779** | Ankrd40 | -0.068593523 | -0.2245454 | 0.087358355 | 9.11881637 | -1.082085709 | 0.321656238 | 0.782292963 | -4.723957975 |
| **A_55_P2014500** | Phf24 | 0.047792312 | -0.060890327 | 0.156474951 | 7.841154719 | 1.081849939 | 0.321752784 | 0.782292963 | -4.724108168 |
| **A_51_P397548** | Mtrf1l | -0.109747014 | -0.359335229 | 0.139841201 | 9.41538985 | -1.08177714 | 0.3217826 | 0.782292963 | -4.724154539 |

| | | | | | | | | | |
|---|---|---|---|---|---|---|---|---|---|
| **A_52_P83959** | Taf7 | -0.081973903 | -0.268440025 | 0.104492219 | 9.977697563 | -1.081545611 | 0.32187744 | 0.782292963 | -4.724302007 |
| **A_51_P236267** | St8sia4 | 0.050415994 | -0.064272659 | 0.165104646 | 8.818000418 | 1.081476397 | 0.321905796 | 0.782292963 | -4.724346089 |
| **A_55_P2045693** | NA | 0.057320348 | -0.073089015 | 0.187729711 | 7.941492028 | 1.081357278 | 0.321954603 | 0.782292963 | -4.724421951 |
| **A_55_P2149500** | Kifc2 | -0.176771998 | -0.578977827 | 0.225433832 | 8.976118219 | -1.081270294 | 0.321990247 | 0.782292963 | -4.724477344 |
| **A_55_P2023384** | NA | 0.062109526 | -0.079217274 | 0.203436326 | 8.613487346 | 1.081192067 | 0.322022306 | 0.782292963 | -4.724527159 |
| **A_55_P2064776** | 4930412O13Rik | -0.13733871 | -0.44987612 | 0.175198701 | 8.510333699 | -1.081085684 | 0.322065907 | 0.782292963 | -4.7245949 |
| **A_55_P2077068** | Malt1 | -0.076693219 | -0.251225369 | 0.09783893 | 9.209373296 | -1.08106223 | 0.32207552 | 0.782292963 | -4.724609835 |
| **A_55_P1972913** | NA | -0.253386419 | -0.830121406 | 0.323348567 | 10.47188667 | -1.080876811 | 0.322151528 | 0.782292963 | -4.724727894 |
| **A_55_P2154870** | Sumo2 | -0.09022064 | -0.295575106 | 0.115133827 | 12.56702666 | -1.080863652 | 0.322156923 | 0.782292963 | -4.724736272 |
| **A_55_P2051429** | NA | 0.068046483 | -0.086852835 | 0.222945802 | 8.438850692 | 1.080750246 | 0.322203419 | 0.782292963 | -4.724808473 |
| **A_55_P1974273** | Ndufs7 | -0.108355167 | -0.355014029 | 0.138303695 | 13.18585143 | -1.080742089 | 0.322206763 | 0.782292963 | -4.724813666 |
| **A_55_P2104894** | Col5a2 | 0.16186781 | -0.206643888 | 0.530379508 | 9.018487632 | 1.080633373 | 0.322251342 | 0.782292963 | -4.724882878 |
| **A_66_P120603** | Trps1 | -0.067029678 | -0.219635404 | 0.085576049 | 7.925946068 | -1.080601242 | 0.322264518 | 0.782292963 | -4.724903332 |
| **A_51_P502119** | F11 | -0.342921848 | -1.123670771 | 0.437827076 | 12.41829215 | -1.080570045 | 0.322277312 | 0.782292963 | -4.724923192 |
| **A_52_P52156** | Isoc2a | 0.259358892 | -0.331188897 | 0.849906681 | 8.264140726 | 1.080476331 | 0.322315745 | 0.782292963 | -4.724982848 |
| **A_51_P337269** | Aldob | 0.641070741 | -0.818662007 | 2.10080349 | 9.680394768 | 1.08044277 | 0.32232951 | 0.782292963 | -4.725004211 |
| **A_55_P2051384** | Ccdc88c | 0.052730503 | -0.067354781 | 0.172815787 | 8.502100957 | 1.080292319 | 0.322391223 | 0.782292963 | -4.725099977 |
| **A_55_P1994539** | Cldn12 | 0.135357017 | -0.172905778 | 0.443619811 | 10.83829868 | 1.08026132 | 0.322403939 | 0.782292963 | -4.725119707 |
| **A_55_P2122841** | NA | 0.216897836 | -0.277225276 | 0.711020948 | 14.14384879 | 1.079914105 | 0.322546404 | 0.782500573 | -4.725340687 |
| **A_55_P2011551** | Anapc5 | -0.105429888 | -0.34562672 | 0.134766944 | 9.289523655 | -1.07985547 | 0.322570468 | 0.782500573 | -4.725378001 |
| **A_55_P2133266** | Rtl8c | 0.237693157 | -0.30394794 | 0.779334254 | 13.67649631 | 1.079628234 | 0.322663738 | 0.78262861 | -4.725522598 |
| **A_55_P1983450** | S100a4 | -0.287674953 | -0.943350542 | 0.368000637 | 9.419669783 | -1.079399465 | 0.32275766 | 0.782758195 | -4.725668153 |
| **A_52_P214612** | NA | -0.07030496 | -0.23057987 | 0.08996995 | 8.839899917 | -1.079169313 | 0.322852173 | 0.782863421 | -4.725814571 |
| **A_55_P2076713** | Immp1l | 0.18507709 | -0.236903302 | 0.607057481 | 10.45730942 | 1.079020148 | 0.322913441 | 0.782863421 | -4.725909459 |
| **A_52_P418489** | Tinagl1 | -0.134648385 | -0.441656383 | 0.172359613 | 11.90720649 | -1.078997961 | 0.322922554 | 0.782863421 | -4.725923572 |
| **A_55_P2156288** | NA | 0.087226104 | -0.111674625 | 0.286126834 | 8.439179383 | 1.078895205 | 0.322964766 | 0.782867566 | -4.725988932 |
| **A_55_P2067583** | Cd68 | 0.223214894 | -0.285931604 | 0.732361391 | 9.274636876 | 1.078573068 | 0.32309713 | 0.78309021 | -4.726193813 |
| **A_52_P563375** | Lgals2 | 0.295333685 | -0.378898064 | 0.969565435 | 10.86836047 | 1.077638127 | 0.323481544 | 0.783828628 | -4.726788256 |
| **A_52_P556082** | Trmt2b | 0.079948096 | -0.10256989 | 0.262466081 | 8.506237686 | 1.077634833 | 0.3234829 | 0.783828628 | -4.72679035 |
| **A_55_P2347756** | AU040972 | 0.138716567 | -0.17799971 | 0.455432843 | 8.658896132 | 1.077524396 | 0.323528332 | 0.783835853 | -4.726860548 |
| **A_55_P1966928** | Lnx1 | -0.121099542 | -0.397616913 | 0.155417829 | 10.40741727 | -1.077430448 | 0.323566986 | 0.783835853 | -4.726920262 |
| **A_55_P1959076** | Zfp930 | 0.057159971 | -0.073416919 | 0.18773686 | 8.309476294 | 1.076948263 | 0.323765434 | 0.784161189 | -4.727226698 |
| **A_55_P2027587** | Supt4b | -0.103965774 | -0.341475746 | 0.133544198 | 10.47311427 | -1.076905774 | 0.323782926 | 0.784161189 | -4.727253698 |
| **A_55_P2003246** | Asxl2 | -0.123317798 | -0.40506296 | 0.158427364 | 9.640658472 | -1.076808458 | 0.323822992 | 0.784161189 | -4.727315533 |
| **A_51_P174275** | Colec11 | 0.170093981 | -0.21856268 | 0.558750641 | 8.555744928 | 1.076693257 | 0.323870426 | 0.78417781 | -4.727388729 |
| **A_51_P412348** | Scnn1b | -0.265331373 | -0.871740481 | 0.341077734 | 9.048184449 | -1.076445515 | 0.323972453 | 0.784326597 | -4.727546124 |
| **A_51_P519570** | 1600002K03Rik | 0.113369382 | -0.145794703 | 0.372533467 | 10.95309308 | 1.076192908 | 0.324076512 | 0.78442701 | -4.727706591 |
| **A_55_P1983162** | Pstpip2 | 0.073280484 | -0.094246808 | 0.240807775 | 8.334717865 | 1.076147802 | 0.324095096 | 0.78442701 | -4.727735242 |
| **A_51_P507290** | Klf5 | 0.099173496 | -0.127576492 | 0.325923485 | 10.045682 | 1.076012989 | 0.324150644 | 0.784463228 | -4.72782087 |
| **A_52_P594756** | Asb4 | 0.165719882 | -0.213283235 | 0.544722998 | 8.992839941 | 1.075724343 | 0.324269605 | 0.784652878 | -4.728004188 |
| **A_66_P109673** | NA | -0.080367973 | -0.264196509 | 0.103460564 | 9.892370325 | -1.075571403 | 0.324332651 | 0.784707199 | -4.728101308 |
| **A_55_P2426549** | NA | -0.147773716 | -0.485858979 | 0.190311546 | 8.214305762 | -1.075325932 | 0.324433862 | 0.78474242 | -4.728257173 |
| **A_55_P1978628** | Hnrnpa2b1 | -0.060185076 | -0.197895426 | 0.077525274 | 8.191779916 | -1.075205287 | 0.324483616 | 0.78474242 | -4.728333771 |
| **A_52_P134666** | Srpr | -0.0842756 | -0.277118527 | 0.108567328 | 10.39702569 | -1.07514564 | 0.324508216 | 0.78474242 | -4.728371639 |
| **A_55_P1972157** | Espnl | 0.109291197 | -0.140823045 | 0.359405439 | 8.776042979 | 1.075018467 | 0.324560672 | 0.78474242 | -4.728452375 |
| **A_55_P1989296** | Tnxb | 0.114744541 | -0.147873957 | 0.37736304 | 8.023327262 | 1.074919354 | 0.324601558 | 0.78474242 | -4.728515292 |
| **A_55_P2411907** | B230213L16Rik | -0.181111507 | -0.595644349 | 0.233421335 | 11.07408243 | -1.074870795 | 0.324621592 | 0.78474242 | -4.728546117 |
| **A_55_P1974193** | Ptges | -0.180411617 | -0.593351669 | 0.232528435 | 13.38161601 | -1.074847014 | 0.324631403 | 0.78474242 | -4.728561213 |
| **A_55_P2069974** | Kctd1 | -0.08935779 | -0.293916716 | 0.115201136 | 8.316922809 | -1.074689856 | 0.324696248 | 0.784786281 | -4.728660968 |
| **A_55_P2038540** | Hbb-b2 | 0.113468157 | -0.146320443 | 0.373256758 | 16.80055398 | 1.074541207 | 0.324757592 | 0.784786281 | -4.728755315 |

| | | | | | | | | |
|---|---|---|---|---|---|---|---|---|
| **A_55_P2066707** | Trim27 | 0.140917145 | -0.181730024 | 0.463564314 | 12.72629549 | 1.074496836 | 0.324775905 | 0.784786281 | -4.728783475 |
| **A_51_P246317** | Mt2 | -0.366779964 | -1.206684499 | 0.473124572 | 16.69350716 | -1.074347654 | 0.324837481 | 0.784786281 | -4.728878152 |
| **A_51_P327451** | Alas2 | -0.158157298 | -0.520349111 | 0.204034514 | 12.65628597 | -1.074285644 | 0.324863079 | 0.784786281 | -4.728917503 |
| **A_52_P481182** | Stard5 | 0.104752635 | -0.135185504 | 0.344690773 | 8.396818651 | 1.074075555 | 0.324949818 | 0.784786281 | -4.729050816 |
| **A_55_P1960960** | Zdhhc9 | -0.24236777 | -0.79754177 | 0.31280623 | 12.99167939 | -1.074026286 | 0.324970162 | 0.784786281 | -4.729082078 |
| **A_55_P2069875** | Trim44 | 0.085597179 | -0.110475831 | 0.281670189 | 13.39604911 | 1.074016122 | 0.324974359 | 0.784786281 | -4.729088528 |
| **A_55_P1967693** | Wipi2 | 0.104002665 | -0.13427734 | 0.342282671 | 8.837703775 | 1.073806506 | 0.325060927 | 0.784836865 | -4.729221522 |
| **A_55_P2003688** | Tmem143 | 0.104336354 | -0.134716565 | 0.343389272 | 10.19197564 | 1.073768768 | 0.325076514 | 0.784836865 | -4.729245465 |
| **A_51_P400463** | Agmat | 0.085463757 | -0.110398023 | 0.281325536 | 8.728960263 | 1.073498512 | 0.325188158 | 0.7849787 | -4.72941691 |
| **A_52_P342159** | Nfatc4 | -0.059125103 | -0.19463489 | 0.076384684 | 8.786516544 | -1.073421794 | 0.325219856 | 0.7849787 | -4.729465575 |
| **A_51_P383270** | Fras1 | 0.152562615 | -0.197127489 | 0.502252719 | 7.973643833 | 1.073331671 | 0.325257096 | 0.7849787 | -4.72952274 |
| **A_55_P2013601** | Ldb2 | 0.067445612 | -0.087186307 | 0.222077531 | 9.228662805 | 1.073059283 | 0.325369672 | 0.785072872 | -4.7296955 |
| **A_55_P2106351** | NA | 0.096867589 | -0.12523499 | 0.318970169 | 10.5430053 | 1.072985993 | 0.325399968 | 0.785072872 | -4.729741979 |
| **A_51_P384629** | Ctsd | -0.152827864 | -0.503315488 | 0.19765976 | 15.14425156 | -1.072751227 | 0.325497028 | 0.785072872 | -4.729890853 |
| **A_55_P1971966** | Uba1 | 0.113270464 | -0.155315835 | 0.381856762 | 12.67228144 | 1.092906244 | 0.32553286 | 0.785072872 | -4.680338596 |
| **A_55_P2164724** | Gnaq | 0.09454877 | -0.12231757 | 0.31141511 | 10.91880902 | 1.072587908 | 0.325564564 | 0.785072872 | -4.729994409 |
| **A_55_P2005903** | Lhx6 | -0.349596606 | -1.151536739 | 0.452343526 | 11.49214034 | -1.072492894 | 0.325603859 | 0.785072872 | -4.730054651 |
| **A_55_P2032192** | Tpm1 | -0.094917945 | -0.312651357 | 0.122815468 | 11.08076926 | -1.07248792 | 0.325605917 | 0.785072872 | -4.730057804 |
| **A_55_P2122898** | Rnf25 | -0.186788139 | -0.615308316 | 0.241732038 | 11.08501675 | -1.072376237 | 0.325652112 | 0.785072872 | -4.730128611 |
| **A_52_P88983** | Dock5 | -0.167758879 | -0.552637559 | 0.217119801 | 11.46679607 | -1.072335848 | 0.325668819 | 0.785072872 | -4.730154217 |
| **A_55_P2111419** | Eny2 | -0.066727955 | -0.219829267 | 0.086373357 | 12.56395941 | -1.072254963 | 0.325702279 | 0.785072872 | -4.730205494 |
| **A_55_P2151082** | NA | -0.088399048 | -0.291303286 | 0.11450519 | 11.37144477 | -1.071829321 | 0.325878407 | 0.785324314 | -4.730475296 |
| **A_55_P2017870** | Pclaf | -0.276512753 | -0.911211943 | 0.358186437 | 12.13122805 | -1.071806497 | 0.325887854 | 0.785324314 | -4.730489762 |
| **A_52_P78373** | Sorbs3 | -0.076091959 | -0.250787592 | 0.098603675 | 8.538813904 | -1.071583152 | 0.325980307 | 0.785437697 | -4.73063131 |
| **A_55_P2006579** | Rbbp6 | -0.164542929 | -0.542346111 | 0.213260253 | 9.612294911 | -1.071476811 | 0.326024333 | 0.785437697 | -4.730698699 |
| **A_52_P483301** | Mios | -0.059844204 | -0.197261145 | 0.077572737 | 8.621211499 | -1.071398375 | 0.32605681 | 0.785437697 | -4.730748402 |
| **A_55_P2087985** | NA | -0.14119979 | -0.465682696 | 0.183283116 | 13.30188852 | -1.070560932 | 0.326403725 | 0.785983674 | -4.731278948 |
| **A_55_P1997291** | Fam53a | -0.08328569 | -0.274682071 | 0.10811069 | 12.16559915 | -1.070547234 | 0.326409402 | 0.785983674 | -4.731287624 |
| **A_52_P199776** | Snhg7 | -0.073420117 | -0.242153701 | 0.095313467 | 8.318542263 | -1.070490214 | 0.326433034 | 0.785983674 | -4.731323739 |
| **A_55_P2099785** | Usp15 | -0.15271669 | -0.503719074 | 0.198285695 | 11.67369468 | -1.07039876 | 0.32647094 | 0.785983674 | -4.731381662 |
| **A_52_P504478** | Adprh | -0.100736874 | -0.33227768 | 0.130803931 | 12.2482907 | -1.070360555 | 0.326486777 | 0.785983674 | -4.731405859 |
| **A_51_P449048** | Cndp2 | -0.076990429 | -0.253999495 | 0.100018636 | 10.16865527 | -1.070065579 | 0.326609071 | 0.786180165 | -4.731592663 |
| **A_55_P2079560** | Lilra6 | -0.05815939 | -0.191891476 | 0.075572696 | 8.033286168 | -1.069925123 | 0.326667315 | 0.786222455 | -4.731681601 |
| **A_51_P353008** | Minpp1 | 0.05621476 | -0.073087088 | 0.185516609 | 10.09698411 | 1.069583741 | 0.326808916 | 0.786354964 | -4.731897743 |
| **A_52_P482203** | Tsen2 | 0.065177695 | -0.084747726 | 0.215103116 | 9.586859172 | 1.06952979 | 0.326831299 | 0.786354964 | -4.731931898 |
| **A_55_P2044369** | Hs2st1 | 0.159512455 | -0.207430897 | 0.526455807 | 10.0248204 | 1.069460473 | 0.326860059 | 0.786354964 | -4.731975779 |
| **A_51_P488175** | Phf20l1 | 0.066981088 | -0.087124083 | 0.221086259 | 8.441433011 | 1.069311288 | 0.326921963 | 0.786354964 | -4.732070216 |
| **A_51_P172853** | Cd14 | -0.213424389 | -0.704541025 | 0.277692246 | 8.53777261 | -1.069125198 | 0.326999194 | 0.786354964 | -4.732188004 |
| **A_55_P2054027** | Chst5 | -0.190910521 | -0.630243696 | 0.248422653 | 8.433642142 | -1.069067257 | 0.327023243 | 0.786354964 | -4.732224676 |
| **A_55_P1988358** | Lrrc47 | -0.191489569 | -0.632169209 | 0.249190071 | 11.58190336 | -1.069033459 | 0.327037273 | 0.786354964 | -4.732246067 |
| **A_66_P129730** | Pdia3 | -0.191617062 | -0.632613293 | 0.24937917 | 14.29494547 | -1.068977245 | 0.327060608 | 0.786354964 | -4.732281644 |
| **A_51_P113722** | Fbxl14 | -0.077180487 | -0.254849931 | 0.100488956 | 10.59634633 | -1.068720006 | 0.327167409 | 0.786354964 | -4.7293893 |
| **A_52_P520397** | Timm8b | 0.093875775 | -0.122234876 | 0.309986426 | 12.80063442 | 1.068677163 | 0.327185199 | 0.786354964 | -4.732471548 |
| **A_55_P2184905** | Rpap2 | -0.046994369 | -0.155185071 | 0.061196332 | 8.695487051 | -1.068624316 | 0.327207145 | 0.786354964 | -4.732504988 |
| **A_51_P100289** | Srf | 0.11233506 | -0.146290591 | 0.37096071 | 10.31972126 | 1.068594384 | 0.327219575 | 0.786354964 | -4.732523928 |
| **A_55_P2135331** | Evl | 0.091179377 | -0.118803359 | 0.301162113 | 11.20917216 | 1.068272852 | 0.327353127 | 0.786354964 | -4.732727365 |
| **A_51_P438701** | Atpaf2 | -0.059466238 | -0.196418573 | 0.077486097 | 8.712808009 | -1.068243323 | 0.327365394 | 0.786354964 | -4.732746046 |
| **A_52_P404005** | NA | 0.08383664 | -0.109265402 | 0.276938682 | 10.75035882 | 1.068110439 | 0.327420604 | 0.786354964 | -4.732830112 |
| **A_52_P393314** | P2rx7 | -0.055606699 | -0.183699931 | 0.072486533 | 7.937043155 | -1.067997153 | 0.327467677 | 0.786354964 | -4.732901775 |
| **A_52_P69789** | Exosc1 | -0.051982619 | -0.171728487 | 0.067763248 | 8.602070399 | -1.06798902 | 0.327471057 | 0.786354964 | -4.73290692 |

| | | | | | | | | | |
|---|---|---|---|---|---|---|---|---|---|
| **A_66_P109562** | Gm12271 | -0.058607184 | -0.193620251 | 0.076405882 | 9.563819748 | -1.067933522 | 0.32749412 | 0.786354964 | -4.732942026 |
| **A_51_P464238** | Atf4 | -0.101370166 | -0.334896735 | 0.132156403 | 8.807780805 | -1.067930583 | 0.327495342 | 0.786354964 | -4.732943885 |
| **A_55_P2002210** | Babam2 | 0.124864818 | -0.162837351 | 0.412566988 | 11.85192431 | 1.06774148 | 0.327573937 | 0.786377222 | -4.733063495 |
| **A_55_P1992084** | Ptprg | 0.062868693 | -0.081991568 | 0.207728953 | 8.226756128 | 1.067712509 | 0.327585979 | 0.786377222 | -4.733081819 |
| **A_52_P632093** | Gtf3c3 | -0.084661342 | -0.279767893 | 0.110445208 | 9.330368432 | -1.067535836 | 0.327659425 | 0.786439866 | -4.733193556 |
| **A_55_P1984152** | 1600029O15Rik | 0.106455232 | -0.138929675 | 0.351840138 | 15.24803871 | 1.067304466 | 0.327755629 | 0.786439866 | -4.73333987 |
| **A_55_P2262753** | 9530053J19Rik | -0.171449555 | -0.566689221 | 0.223790111 | 8.233492186 | -1.067197996 | 0.327799907 | 0.786439866 | -4.733407194 |
| **A_51_P118742** | Drg2 | 0.066662328 | -0.087017455 | 0.22034211 | 8.801198874 | 1.067168263 | 0.327812273 | 0.786439866 | -4.733425994 |
| **A_55_P2226660** | 5033418A18Rik | -0.082116028 | -0.27142333 | 0.107191274 | 7.889521365 | -1.067160479 | 0.327815511 | 0.786439866 | -4.733430916 |
| **A_51_P259318** | Fbrs | -0.053606107 | -0.17721861 | 0.070006396 | 8.413624175 | -1.066893404 | 0.327926606 | 0.786602382 | -4.733599775 |
| **A_55_P2083464** | Lpin3 | 0.087981947 | -0.114916592 | 0.290880487 | 10.28953865 | 1.06680198 | 0.327964643 | 0.786602382 | -4.733657573 |
| **A_55_P2175245** | Tusc3 | 0.208277405 | -0.272129866 | 0.688684676 | 10.13945473 | 1.066600418 | 0.328048516 | 0.786705927 | -4.733784991 |
| **A_51_P383001** | Meox2 | 0.188203684 | -0.24597207 | 0.622379438 | 8.889082251 | 1.066428436 | 0.328120095 | 0.78675741 | -4.733893699 |
| **A_55_P2045952** | Mysm1 | -0.067738058 | -0.224017124 | 0.088541008 | 9.022655917 | -1.06635325 | 0.328151391 | 0.78675741 | -4.73394122 |
| **A_51_P422629** | Tbpl1 | -0.082100392 | -0.271554664 | 0.107353879 | 11.21859702 | -1.06612958 | 0.328244508 | 0.786883058 | -4.73408258 |
| **A_51_P250058** | Epas1 | -0.107442291 | -0.355437557 | 0.140552976 | 16.06916212 | -1.065862127 | 0.328355882 | 0.787029462 | -4.734251589 |
| **A_55_P2127557** | Abracl | -0.061708846 | -0.204153455 | 0.080735763 | 11.65001975 | -1.065787376 | 0.328387015 | 0.787029462 | -4.734298821 |
| **A_51_P183853** | Polr2m | 0.104141871 | -0.136333497 | 0.34461724 | 13.71535992 | 1.065427596 | 0.328536896 | 0.787215925 | -4.734526128 |
| **A_55_P2051322** | Efhd2 | -0.11946196 | -0.395368236 | 0.156444317 | 11.15715238 | -1.065214864 | 0.328625545 | 0.787215925 | -4.734660512 |
| **A_55_P2044468** | Marf1 | 0.152673648 | -0.199987258 | 0.505334555 | 9.799549085 | 1.065064543 | 0.328688198 | 0.787215925 | -4.73475546 |
| **A_55_P1989504** | Tmem164 | -0.104761636 | -0.346773887 | 0.137250615 | 10.67917566 | -1.064961931 | 0.328730972 | 0.787215925 | -4.734820271 |
| **A_55_P2183587** | Gli1 | 0.103691227 | -0.135865669 | 0.343248124 | 7.990217432 | 1.064884489 | 0.328763256 | 0.787215925 | -4.734869181 |
| **A_55_P2215880** | Ephb2 | 0.056361641 | -0.073852339 | 0.186575621 | 8.200937776 | 1.064866538 | 0.32877074 | 0.787215925 | -4.734880518 |
| **A_55_P2009752** | Hlf | 0.055267265 | -0.072425585 | 0.182960115 | 7.897639604 | 1.064806164 | 0.328795912 | 0.787215925 | -4.734918647 |
| **A_55_P2006226** | NA | -0.069436087 | -0.229872456 | 0.091000283 | 16.26640556 | -1.064759627 | 0.328815315 | 0.787215925 | -4.734948036 |
| **A_55_P2173073** | Zfp931 | -0.059211276 | -0.196027532 | 0.077604981 | 8.407480435 | -1.064721145 | 0.328831361 | 0.787215925 | -4.734972338 |
| **A_55_P2018944** | Snx27 | -0.088594352 | -0.29332952 | 0.116140817 | 9.836585961 | -1.064590895 | 0.328885676 | 0.78724845 | -4.735054589 |
| **A_52_P267256** | LOC100861738 | -0.164017729 | -0.543367833 | 0.215332374 | 12.22181101 | -1.063701449 | 0.329256777 | 0.787748871 | -4.735616114 |
| **A_55_P2250296** | 6330566A10Rik | 0.10135138 | -0.133076773 | 0.335779533 | 9.257223673 | 1.063626291 | 0.329288151 | 0.787748871 | -4.735663551 |
| **A_51_P176517** | Ntmt1 | -0.105196224 | -0.348525382 | 0.138132934 | 10.52324038 | -1.06359224 | 0.329302366 | 0.787748871 | -4.735685042 |
| **A_55_P2132467** | Nfx1 | 0.095394196 | -0.125291404 | 0.316079795 | 9.36184389 | 1.063450107 | 0.329361706 | 0.787748871 | -4.735774744 |
| **A_55_P2083489** | Fam129a | 0.074766315 | -0.098215604 | 0.247748234 | 9.176706868 | 1.063345579 | 0.329405352 | 0.787748871 | -4.735840709 |
| **A_55_P2011678** | Pdzk1ip1 | 0.464724385 | -0.610576965 | 1.540025735 | 12.51572632 | 1.063247654 | 0.329446245 | 0.787748871 | -4.735902504 |
| **A_51_P312497** | Atp7a | -0.199517296 | -0.661230709 | 0.262196117 | 10.99915954 | -1.0631076 | 0.329504739 | 0.787748871 | -4.735990879 |
| **A_66_P120414** | Adcy5 | 0.054574988 | -0.071736773 | 0.180886749 | 7.760945949 | 1.062965143 | 0.329564245 | 0.787748871 | -4.736080763 |
| **A_55_P2042778** | Pde5a | 0.072569114 | -0.095396258 | 0.240534486 | 8.006575246 | 1.062921588 | 0.32958244 | 0.787748871 | -4.736108243 |
| **A_55_P2014224** | Sema5a | 0.12188455 | -0.160300239 | 0.40406934 | 9.097474485 | 1.062635259 | 0.329702075 | 0.787748871 | -4.736288881 |
| **A_55_P1978502** | H2-Q1 | 0.183565499 | -0.241464696 | 0.608595693 | 8.742868701 | 1.062528133 | 0.329746844 | 0.787748871 | -4.736356457 |
| **A_51_P128147** | Chmp1a | -0.059740426 | -0.198069271 | 0.07858842 | 11.69190937 | -1.062489685 | 0.329762913 | 0.787748871 | -4.727210905 |
| **A_55_P2048210** | Ganab | -0.104373263 | -0.346067485 | 0.137320959 | 12.13334344 | -1.062410012 | 0.329796214 | 0.787748871 | -4.736430965 |
| **A_66_P115161** | Cep85l | -0.068304763 | -0.22648737 | 0.089877844 | 8.366618124 | -1.062334839 | 0.329827636 | 0.787748871 | -4.736478379 |
| **A_55_P2130877** | Parl | -0.069009439 | -0.228834203 | 0.090815325 | 9.867370234 | -1.062266746 | 0.329856101 | 0.787748871 | -4.736521326 |
| **A_55_P1967227** | Tmem158 | -0.081479259 | -0.270196971 | 0.107238453 | 11.10087721 | -1.062193337 | 0.32988679 | 0.787748871 | -4.736567625 |
| **A_55_P2038422** | Adamts17 | 0.068872687 | -0.090646972 | 0.228392345 | 8.133735656 | 1.062189433 | 0.329888422 | 0.787748871 | -4.736570087 |
| **A_52_P241484** | Rcl1 | -0.054845965 | -0.181890563 | 0.072198632 | 8.996019196 | -1.062081022 | 0.329933749 | 0.787748871 | -4.736638458 |
| **A_55_P2086825** | Rgl3 | 0.161415338 | -0.212497154 | 0.535327829 | 9.104713242 | 1.062047615 | 0.329947718 | 0.787748871 | -4.736659527 |
| **A_55_P2121980** | Plcb4 | -0.085301854 | -0.282918069 | 0.112314361 | 8.500833359 | -1.061952417 | 0.329987526 | 0.787748871 | -4.73671956 |
| **A_55_P1956457** | 6030458C11Rik | -0.093189394 | -0.309093155 | 0.122714368 | 9.639884564 | -1.061879997 | 0.330017812 | 0.787748871 | -4.736765227 |
| **A_55_P1989916** | Lin54 | 0.064040129 | -0.084334542 | 0.2124148 | 9.39100887 | 1.061846137 | 0.330031973 | 0.787748871 | -4.736786579 |
| **A_55_P2033876** | NA | 0.110161492 | -0.145126693 | 0.365449677 | 11.58167826 | 1.061617957 | 0.330127416 | 0.787748871 | -4.736930453 |

| | | | | | | | | | |
|---|---|---|---|---|---|---|---|---|---|
| **A_55_P2045755** | Plcxd2 | 0.098086911 | -0.129225811 | 0.325399634 | 10.08525514 | 1.061589168 | 0.33013946 | 0.787748871 | -4.736948605 |
| **A_55_P1980165** | NA | 0.176170724 | -0.232100364 | 0.584441812 | 14.33751448 | 1.061583783 | 0.330141713 | 0.787748871 | -4.736952 |
| **A_52_P660385** | Trmt6 | -0.074800249 | -0.248163959 | 0.098563461 | 8.81582746 | -1.061485375 | 0.330182883 | 0.787748871 | -4.737014043 |
| **A_55_P2171116** | Lgals3 | -0.12232323 | -0.405861481 | 0.161215021 | 16.6622582 | -1.061369117 | 0.330231527 | 0.787748871 | -4.737087336 |
| **A_55_P2117155** | Apoc3 | 0.498763604 | -0.657564983 | 1.655092191 | 8.601554486 | 1.061164359 | 0.330317214 | 0.787748871 | -4.737216411 |
| **A_51_P513254** | Rac1 | -0.095398129 | -0.316570984 | 0.125774726 | 13.09937702 | -1.061151022 | 0.330322796 | 0.787748871 | -4.737224818 |
| **A_55_P2131105** | NA | 0.070281069 | -0.092682277 | 0.233244415 | 11.00430633 | 1.061005386 | 0.330383754 | 0.787748871 | -4.737316615 |
| **A_55_P2032128** | Dnase1l1 | 0.067902779 | -0.089549549 | 0.225355107 | 9.314738862 | 1.06098104 | 0.330393946 | 0.787748871 | -4.73733196 |
| **A_55_P2049365** | NA | 0.135716716 | -0.178985437 | 0.450418868 | 11.94294176 | 1.060969234 | 0.330398888 | 0.787748871 | -4.737339401 |
| **A_55_P2031481** | Fam131a | -0.111793061 | -0.371070175 | 0.147484053 | 10.97806227 | -1.060766575 | 0.330483732 | 0.787853979 | -4.737467127 |
| **A_55_P2134122** | NA | 0.090547237 | -0.119509001 | 0.300603475 | 8.440902469 | 1.060495387 | 0.330597295 | 0.787988056 | -4.737638022 |
| **A_55_P2077608** | Hdac1 | -0.060567868 | -0.201097303 | 0.079961567 | 9.134744825 | -1.060337567 | 0.330663398 | 0.787988056 | -4.737737465 |
| **A_66_P126080** | Rpl39 | 0.093528531 | -0.123477701 | 0.310534763 | 8.655868514 | 1.060330047 | 0.330666548 | 0.787988056 | -4.737742203 |
| **A_55_P2138097** | NA | -0.093828973 | -0.31156355 | 0.123905604 | 13.67451958 | -1.060177847 | 0.330730309 | 0.787988056 | -4.737838097 |
| **A_51_P309158** | Snx20 | 0.246247379 | -0.325198698 | 0.817693456 | 9.33404542 | 1.060145622 | 0.33074381 | 0.787988056 | -4.737858398 |
| **A_52_P346033** | Trmt1l | -0.058206312 | -0.19336022 | 0.076947596 | 8.418607103 | -1.059523612 | 0.331004498 | 0.788498712 | -4.738250209 |
| **A_52_P87503** | Hbq1b | -0.386785578 | -1.285013064 | 0.511441907 | 8.122818851 | -1.059383234 | 0.331063355 | 0.788498712 | -4.738338618 |
| **A_66_P116373** | Pwwp2b | 0.076774875 | -0.101533198 | 0.255082948 | 9.933877897 | 1.059295856 | 0.331099995 | 0.788498712 | -4.738393644 |
| **A_55_P1999992** | Galnt12 | 0.097412572 | -0.128836992 | 0.323662135 | 8.743547865 | 1.059244998 | 0.331121322 | 0.788498712 | -4.738425671 |
| **A_55_P2056674** | Lrrk1 | -0.140078485 | -0.465465168 | 0.185308199 | 9.597217218 | -1.059109399 | 0.331178192 | 0.788512824 | -4.738511057 |
| **A_55_P2033780** | Fabp3 | 0.110064274 | -0.145682692 | 0.36581124 | 17.02003727 | 1.058778336 | 0.331317071 | 0.788512824 | -4.7387195 |
| **A_52_P563617** | Ssbp4 | -0.073571972 | -0.244541598 | 0.097397654 | 10.52864368 | -1.058674855 | 0.331360491 | 0.788512824 | -4.738784646 |
| **A_51_P346893** | Extl1 | -0.093573169 | -0.311030717 | 0.123884379 | 8.945700381 | -1.05863443 | 0.331377454 | 0.788512824 | -4.738810094 |
| **A_52_P452667** | Prom2 | -0.097918756 | -0.325494278 | 0.129656766 | 8.77640151 | -1.058545431 | 0.331414803 | 0.788512824 | -4.73886612 |
| **A_55_P2016004** | Cd22 | 0.110882111 | -0.146833709 | 0.368597932 | 8.639241007 | 1.058496852 | 0.33143519 | 0.788512824 | -4.738896699 |
| **A_55_P2064492** | Pcmtd2 | -0.075522999 | -0.251059659 | 0.100013661 | 9.47819291 | -1.058474888 | 0.331444408 | 0.788512824 | -4.738910525 |
| **A_51_P138895** | Ccdc102a | 0.10657186 | -0.141144258 | 0.354287977 | 9.107988098 | 1.058418551 | 0.331468054 | 0.788512824 | -4.738945987 |
| **A_51_P282092** | Urod | 0.093432463 | -0.123767175 | 0.310632101 | 12.15101586 | 1.05829773 | 0.331518769 | 0.788512824 | -4.739022034 |
| **A_55_P1979818** | Srp54b | -0.069752616 | -0.23191042 | 0.092405188 | 8.295342519 | -1.058258608 | 0.331535192 | 0.788512824 | -4.739046658 |
| **A_51_P370510** | Clba1 | -0.142425417 | -0.473579214 | 0.18872838 | 8.580016031 | -1.05810051 | 0.331601566 | 0.788543645 | -4.739146159 |
| **A_55_P1973387** | Srrm1 | -0.224992875 | -0.748157715 | 0.298171965 | 12.63736348 | -1.058033403 | 0.331629742 | 0.788543645 | -4.739188392 |
| **A_52_P352362** | NA | -0.215675843 | -0.717342268 | 0.285990581 | 8.577119162 | -1.057683264 | 0.33177679 | 0.788700233 | -4.739408721 |
| **A_52_P144285** | Fam72a | -0.199770996 | -0.6645608 | 0.265018808 | 8.339078924 | -1.057413876 | 0.331889961 | 0.788700233 | -4.739578208 |
| **A_51_P257541** | Plppr1 | -0.097981409 | -0.325953393 | 0.129990576 | 7.681144782 | -1.057380659 | 0.331903918 | 0.788700233 | -4.739599106 |
| **A_66_P132037** | Jmjd4 | 0.052050312 | -0.069065349 | 0.173165973 | 8.348227114 | 1.057285304 | 0.331943986 | 0.788700233 | -4.739659093 |
| **A_52_P625808** | Sf3b1 | -0.048172948 | -0.160282731 | 0.063936834 | 8.51390915 | -1.057131176 | 0.332008759 | 0.788700233 | -4.739756046 |
| **A_55_P2183493** | Spcs3 | 0.172390778 | -0.228811102 | 0.573592658 | 11.24797885 | 1.057110126 | 0.332017607 | 0.788700233 | -4.739769288 |
| **A_51_P487918** | Rinl | -0.068031223 | -0.226363624 | 0.090301179 | 10.69168121 | -1.057079475 | 0.332030489 | 0.788700233 | -4.739788567 |
| **A_51_P314111** | Psip1 | -0.127684004 | -0.424895323 | 0.169527316 | 10.40169585 | -1.056915601 | 0.332099373 | 0.788700233 | -4.73989164 |
| **A_55_P2094118** | Fmnl3 | -0.090752609 | -0.30204799 | 0.120542773 | 9.799184099 | -1.05666732 | 0.332203759 | 0.788700233 | -4.740047787 |
| **A_55_P2048378** | Zfp975 | -0.056931228 | -0.189495821 | 0.075633365 | 8.978655442 | -1.056555154 | 0.332250926 | 0.788700233 | -4.740118322 |
| **A_55_P2012181** | Eps15l1 | -0.094099073 | -0.313212246 | 0.125014099 | 10.44133715 | -1.056540193 | 0.332257218 | 0.788700233 | -4.74012773 |
| **A_51_P516133** | H1f2 | 0.139484138 | -0.185323732 | 0.464292008 | 12.45869864 | 1.056494989 | 0.332276229 | 0.788700233 | -4.740156155 |
| **A_52_P612137** | Runx1t1 | 0.080635134 | -0.107137257 | 0.268407525 | 8.539368858 | 1.05648112 | 0.332282062 | 0.788700233 | -4.740164877 |
| **A_55_P1976928** | Unc13d | 0.17374392 | -0.230885124 | 0.578372964 | 9.334091248 | 1.056383784 | 0.332323001 | 0.788700233 | -4.740226081 |
| **A_55_P2061724** | NA | -0.05332757 | -0.177522535 | 0.070867395 | 8.184890136 | -1.056372207 | 0.33232787 | 0.788700233 | -4.74023336 |
| **A_55_P1993864** | Sh2b3 | 0.049222134 | -0.065416965 | 0.163861233 | 8.594078576 | 1.056323255 | 0.332348461 | 0.788700233 | -4.740264139 |
| **A_51_P154933** | Zdhhc7 | -0.06307209 | -0.209986648 | 0.083842469 | 10.34843888 | -1.056188779 | 0.332405031 | 0.788737643 | -4.740348689 |
| **A_55_P2060111** | NA | -0.097686983 | -0.325263334 | 0.129889369 | 13.63604343 | -1.05603601 | 0.332469306 | 0.788793324 | -4.740444732 |
| **A_55_P2048036** | Zfx | -0.074666127 | -0.248638637 | 0.099306383 | 11.37849472 | -1.055874159 | 0.332537413 | 0.788827593 | -4.740546478 |

| | | | | | | | | | |
|---|---|---|---|---|---|---|---|---|---|
| **A_51_P400581** | Anp32e | -0.106057527 | -0.353203173 | 0.141088119 | 8.424498019 | -1.055741751 | 0.332593138 | 0.788827593 | -4.740629707 |
| **A_55_P2129271** | Tead2 | 0.150340101 | -0.200040014 | 0.500720217 | 11.33655064 | 1.055612564 | 0.332647516 | 0.788827593 | -4.740710907 |
| **A_52_P664506** | Abhd2 | -0.064749086 | -0.215667413 | 0.086169242 | 8.977085956 | -1.055506278 | 0.33269226 | 0.788827593 | -4.740777708 |
| **A_55_P2031207** | Ggnbp2 | -0.113510945 | -0.378118777 | 0.151096887 | 12.07626078 | -1.055368694 | 0.332750187 | 0.788827593 | -4.740864174 |
| **A_55_P1973259** | Gamt | 0.152503889 | -0.203012307 | 0.508020084 | 9.350279577 | 1.055335848 | 0.332764017 | 0.788827593 | -4.740884816 |
| **A_51_P259975** | Aspa | 0.087959523 | -0.117110966 | 0.293030012 | 8.038236277 | 1.055234206 | 0.332806818 | 0.788827593 | -4.740948689 |
| **A_51_P481788** | Mapk1 | 0.062926323 | -0.08378242 | 0.209635065 | 10.57615041 | 1.055226092 | 0.332810235 | 0.788827593 | -4.727773588 |
| **A_55_P2075731** | NA | -0.48281808 | -1.608863917 | 0.643227758 | 11.07634446 | -1.054864357 | 0.332962597 | 0.788841823 | -4.741181078 |
| **A_66_P124788** | Tbcb | -0.058614853 | -0.195322572 | 0.078092866 | 11.254809 | -1.054833235 | 0.332975708 | 0.788841823 | -4.741200631 |
| **A_51_P405397** | Ecm1 | -0.41423823 | -1.38040076 | 0.5519243 | 12.73554457 | -1.054797642 | 0.332990704 | 0.788841823 | -4.741222992 |
| **A_55_P2034555** | Pdcd5 | -0.129221734 | -0.43062746 | 0.172183993 | 13.5271789 | -1.054758971 | 0.333006996 | 0.788841823 | -4.741247287 |
| **A_55_P2060349** | Lzts2 | -0.122685286 | -0.40885351 | 0.163482939 | 13.05863671 | -1.054727409 | 0.333020295 | 0.788841823 | -4.741267116 |
| **A_55_P2131153** | Smarcc1 | 0.34363992 | -0.458072043 | 1.145351884 | 14.71305408 | 1.054518998 | 0.333108115 | 0.78885497 | -4.741398038 |
| **A_51_P277444** | Rnf19a | -0.06713405 | -0.223760315 | 0.089492215 | 10.4543273 | -1.054502027 | 0.333115268 | 0.78885497 | -4.741408699 |
| **A_55_P1995892** | NA | 0.045710671 | -0.060950722 | 0.152372064 | 9.185870037 | 1.054337153 | 0.333184757 | 0.78885497 | -4.741512261 |
| **A_51_P269203** | Trpv4 | 0.066420878 | -0.088567275 | 0.221409032 | 8.160060552 | 1.054326867 | 0.333189093 | 0.78885497 | -4.741518721 |
| **A_55_P2115310** | Spata21 | 0.216599668 | -0.289125783 | 0.72232512 | 9.797558702 | 1.053688255 | 0.333458365 | 0.789313545 | -4.741919763 |
| **A_55_P2048937** | Kif5c | 0.132885659 | -0.177385271 | 0.44315659 | 10.49801018 | 1.053673822 | 0.333464453 | 0.789313545 | -4.741928825 |
| **A_55_P2091473** | Bnip3 | 0.165422041 | -0.220909019 | 0.5517531 | 14.34600418 | 1.053423282 | 0.333570144 | 0.78938004 | -4.742086124 |
| **A_65_P10433** | Rpp14 | -0.096971183 | -0.323456729 | 0.129514363 | 9.445937623 | -1.053346761 | 0.333602431 | 0.78938004 | -4.742134163 |
| **A_55_P1966573** | Gemin4 | 0.048058057 | -0.064189358 | 0.160305472 | 9.760287882 | 1.053316823 | 0.333615063 | 0.78938004 | -4.742152957 |
| **A_55_P2077353** | Ceacam12 | -0.085104728 | -0.283957154 | 0.113747697 | 10.34270714 | -1.05291172 | 0.333786034 | 0.789422795 | -4.742407237 |
| **A_52_P161806** | Ovca2 | 0.073906799 | -0.098790904 | 0.246604503 | 9.304181288 | 1.052851197 | 0.333811583 | 0.789422795 | -4.742445222 |
| **A_51_P237575** | Akirin1 | 0.147264207 | -0.196880352 | 0.491408767 | 11.32696606 | 1.052750504 | 0.333854094 | 0.789422795 | -4.742508416 |
| **A_55_P2060415** | NA | -0.065981418 | -0.220188753 | 0.088225916 | 16.07207679 | -1.052654324 | 0.333894703 | 0.789422795 | -4.742568775 |
| **A_51_P513449** | Pogz | -0.06946781 | -0.231836348 | 0.092900728 | 9.564642716 | -1.052569766 | 0.333930409 | 0.789422795 | -4.742621837 |
| **A_51_P129012** | B2m | -0.357466666 | -1.19299335 | 0.478060018 | 15.45846567 | -1.052554005 | 0.333937065 | 0.789422795 | -4.742631728 |
| **A_55_P2043996** | Cops7a | -0.059038174 | -0.197041142 | 0.078964795 | 8.097947063 | -1.052479511 | 0.333968524 | 0.789422795 | -4.742678472 |
| **A_52_P428125** | Atr | -0.088821708 | -0.296449612 | 0.118806196 | 9.960719948 | -1.052452644 | 0.33397987 | 0.789422795 | -4.74269533 |
| **A_51_P214423** | Tead1 | -0.155271647 | -0.518248653 | 0.20770536 | 11.24744869 | -1.052403314 | 0.334000705 | 0.789422795 | -4.742726283 |
| **A_55_P2036337** | Ifnar1 | 0.074423048 | -0.099593459 | 0.248439556 | 10.17580194 | 1.05217062 | 0.334098996 | 0.789504043 | -4.74287228 |
| **A_51_P506312** | Cdkn2aipnl | -0.055684386 | -0.185891225 | 0.074522453 | 11.04288173 | -1.052128545 | 0.334116771 | 0.789504043 | -4.742898677 |
| **A_55_P1977649** | Hoxd8 | -0.071418473 | -0.238465846 | 0.095628901 | 8.669665681 | -1.051816722 | 0.33424853 | 0.78971884 | -4.743094288 |
| **A_55_P2000668** | Stxbp2 | -0.1491279 | -0.498054521 | 0.199798722 | 10.45550872 | -1.051462913 | 0.334398081 | 0.789952764 | -4.743316198 |
| **A_55_P2133680** | NA | -0.095892845 | -0.320297884 | 0.128512195 | 10.58745774 | -1.051290548 | 0.334470958 | 0.789952764 | -4.743424291 |
| **A_55_P1985514** | Dlg3 | -0.170077794 | -0.568146353 | 0.227990765 | 13.66006955 | -1.051135976 | 0.334536322 | 0.789952764 | -4.743521217 |
| **A_52_P614731** | Gng12 | -0.107357256 | -0.358673182 | 0.14395867 | 10.20572285 | -1.050946369 | 0.334616517 | 0.789952764 | -4.743640102 |
| **A_55_P2144055** | NA | 0.126006867 | -0.168975907 | 0.42098964 | 15.28557895 | 1.050912894 | 0.334630676 | 0.789952764 | -4.74366109 |
| **A_55_P2107686** | Ttll1 | -0.050701646 | -0.169402712 | 0.06799942 | 7.921789181 | -1.050840022 | 0.334661503 | 0.789952764 | -4.743706777 |
| **A_55_P2101387** | Nubpl | 0.16410435 | -0.22012816 | 0.548336859 | 8.666998171 | 1.050739715 | 0.334703939 | 0.789952764 | -4.743769661 |
| **A_52_P222230** | Gm40812 | -0.109884796 | -0.367184229 | 0.147414637 | 8.580688026 | -1.050673891 | 0.334731789 | 0.789952764 | -4.743810926 |
| **A_52_P148678** | Plcg1 | -0.123946376 | -0.414178357 | 0.166285605 | 10.327132 | -1.05064919 | 0.33474224 | 0.789952764 | -4.74382641 |
| **A_55_P2062029** | Tsen34 | 0.16432752 | -0.220472775 | 0.549127814 | 15.17968066 | 1.050616138 | 0.334756226 | 0.789952764 | -4.74384713 |
| **A_52_P142191** | Aph1b | -0.077357685 | -0.258579422 | 0.103864051 | 9.294672233 | -1.050176614 | 0.334942247 | 0.79029525 | -4.744122619 |
| **A_66_P136364** | NA | -0.167218317 | -0.559148297 | 0.224711664 | 12.53410982 | -1.049650033 | 0.335165225 | 0.790483002 | -4.74445259 |
| **A_55_P1966377** | 4930502E18Rik | -0.178198856 | -0.595876488 | 0.239478776 | 9.836491537 | -1.049621833 | 0.335177169 | 0.790483002 | -4.744470258 |
| **A_55_P1954156** | Wdr43 | 0.052618599 | -0.070714772 | 0.17595197 | 10.01118289 | 1.049609716 | 0.335182302 | 0.790483002 | -4.744477849 |
| **A_52_P185485** | Arpc4 | -0.060484656 | -0.202257472 | 0.08128816 | 10.65266393 | -1.049594177 | 0.335188884 | 0.790483002 | -4.744487585 |
| **A_55_P2080021** | Papss2 | -0.310295937 | -1.037674321 | 0.417082446 | 11.90503359 | -1.049505847 | 0.335226301 | 0.790483002 | -4.744542925 |
| **A_55_P2059090** | Tagap | -0.137542139 | -0.460015595 | 0.184931318 | 9.374547877 | -1.049327281 | 0.335301952 | 0.790564947 | -4.74465479 |

| | | | | | | | | | |
|---|---|---|---|---|---|---|---|---|---|
| **A_55_P2227154** | A530041M06Rik | -0.181965223 | -0.608693372 | 0.244762926 | 9.145948318 | -1.049074341 | 0.335409138 | 0.790721213 | -4.744813229 |
| **A_51_P499891** | Krit1 | -0.045531863 | -0.152331429 | 0.061267703 | 8.841114673 | -1.048854137 | 0.335502473 | 0.790844794 | -4.744951145 |
| **A_55_P1957227** | Las1l | -0.119794538 | -0.400811993 | 0.161222916 | 11.48764767 | -1.048752202 | 0.335545687 | 0.790846137 | -4.745014983 |
| **A_55_P2079430** | NA | 0.052988241 | -0.071324046 | 0.177300527 | 8.780725137 | 1.048659775 | 0.335584873 | 0.790846137 | -4.745072863 |
| **A_55_P2074796** | Cdkn2c | 0.058889852 | -0.079290634 | 0.197070338 | 8.187735127 | 1.048486661 | 0.335658279 | 0.790922696 | -4.745181264 |
| **A_55_P2018076** | Arap1 | 0.13250951 | -0.178469232 | 0.443488251 | 12.0272712 | 1.048299813 | 0.335737523 | 0.790938942 | -4.745298253 |
| **A_52_P430934** | Dcun1d1 | -0.079681549 | -0.266701887 | 0.107338788 | 8.754760956 | -1.048185376 | 0.335786065 | 0.790938942 | -4.745369899 |
| **A_55_P1985559** | Gatad1 | 0.098695543 | -0.132953458 | 0.330344544 | 8.795087001 | 1.048180972 | 0.335787933 | 0.790938942 | -4.745372656 |
| **A_55_P2084652** | H2ac15 | 0.210827372 | -0.284196849 | 0.705851593 | 14.9569219 | 1.047779048 | 0.335958466 | 0.791070789 | -4.745624253 |
| **A_55_P1981301** | Nup88 | -0.092634193 | -0.310178784 | 0.124910397 | 11.17761077 | -1.047592062 | 0.336037827 | 0.791070789 | -4.745741285 |
| **A_55_P2103452** | Npy1r | 0.06638839 | -0.089527344 | 0.222304123 | 7.752072552 | 1.047541785 | 0.336059168 | 0.791070789 | -4.74577275 |
| **A_51_P493700** | Cep350 | -0.103376652 | -0.346161772 | 0.139408467 | 10.66671898 | -1.047537461 | 0.336061004 | 0.791070789 | -4.745775457 |
| **A_51_P405565** | Msh6 | -0.065063954 | -0.217871068 | 0.08774316 | 11.10127518 | -1.047528976 | 0.336064606 | 0.791070789 | -4.745780767 |
| **A_55_P2181288** | Ppp1r2 | -0.099650094 | -0.333706108 | 0.134405921 | 9.781933644 | -1.047434969 | 0.336104512 | 0.791070789 | -4.745839598 |
| **A_51_P172344** | Foxn3 | -0.10519403 | -0.352312143 | 0.141924083 | 10.68681459 | -1.047262797 | 0.336177611 | 0.791070789 | -4.745947338 |
| **A_55_P2301063** | 9930024M15Rik | -0.137289219 | -0.459849276 | 0.185270838 | 8.578421415 | -1.047116517 | 0.336239727 | 0.791070789 | -4.746038868 |
| **A_51_P459070** | Aplf | -0.119882986 | -0.401586304 | 0.161820331 | 9.558368188 | -1.046971247 | 0.336301424 | 0.791070789 | -4.746129759 |
| **A_55_P2114498** | Gm8801 | -0.073644281 | -0.246700657 | 0.099412094 | 9.35395806 | -1.046937095 | 0.33631593 | 0.791070789 | -4.746151126 |
| **A_55_P2143968** | Gps1 | -0.064281782 | -0.215339261 | 0.086775697 | 12.92646812 | -1.04692324 | 0.336321815 | 0.791070789 | -4.746159794 |
| **A_55_P2029643** | NA | 0.081354568 | -0.109828387 | 0.272537524 | 11.35002252 | 1.046892134 | 0.336335027 | 0.791070789 | -4.746179254 |
| **A_51_P424959** | Bcl6b | -0.052552513 | -0.176074739 | 0.070969712 | 7.89710552 | -1.046688723 | 0.336421439 | 0.79112318 | -4.746306505 |
| **A_55_P2120615** | Ssh3 | -0.147158501 | -0.493096685 | 0.198779682 | 10.74292766 | -1.04654045 | 0.336484439 | 0.79112318 | -4.746399253 |
| **A_51_P331661** | Ubiad1 | -0.097340615 | -0.326178318 | 0.131497089 | 10.2565687 | -1.046491408 | 0.336505278 | 0.79112318 | -4.746429929 |
| **A_55_P1987261** | Fam214b | -0.162830015 | -0.545664373 | 0.220004343 | 11.20622436 | -1.046387922 | 0.336549257 | 0.79112318 | -4.746494656 |
| **A_65_P19149** | Nln | -0.107762553 | -0.361144414 | 0.145619307 | 9.915429821 | -1.046312745 | 0.336581207 | 0.79112318 | -4.746541674 |
| **A_55_P2051949** | Elmod3 | -0.11202273 | -0.375434371 | 0.151388911 | 10.32942752 | -1.046261759 | 0.336602878 | 0.79112318 | -4.746573562 |
| **A_52_P675617** | Agbl3 | 0.093790264 | -0.126779696 | 0.314360225 | 9.381970893 | 1.046117717 | 0.336664107 | 0.791170885 | -4.746663643 |
| **A_51_P142153** | Filip1l | 0.129312169 | -0.174846841 | 0.433471178 | 10.82001013 | 1.045942654 | 0.336738535 | 0.791249591 | -4.746773115 |
| **A_52_P385615** | Rnpc3 | -0.071091657 | -0.238338701 | 0.096155387 | 8.924991805 | -1.045753553 | 0.336818945 | 0.791342336 | -4.746891354 |
| **A_52_P680495** | Pdcd10 | 0.220436507 | -0.298693112 | 0.739566125 | 12.00043865 | 1.044664593 | 0.337282306 | 0.792296186 | -4.74757201 |
| **A_55_P2016652** | Cdk1 | 0.099301515 | -0.134581101 | 0.333184131 | 7.864328334 | 1.04454486 | 0.337333285 | 0.792296186 | -4.747646826 |
| **A_55_P2164841** | NA | -0.174389706 | -0.585139339 | 0.236359928 | 11.16561921 | -1.044510531 | 0.337347902 | 0.792296186 | -4.747668276 |
| **A_51_P139184** | Fank1 | -0.045840137 | -0.153849345 | 0.062169071 | 7.843399469 | -1.04412932 | 0.337510259 | 0.792411131 | -4.747906438 |
| **A_51_P398467** | Bsdc1 | -0.065767188 | -0.22075933 | 0.089224953 | 11.43867862 | -1.043923705 | 0.337597857 | 0.792411131 | -4.748034876 |
| **A_51_P157982** | Tcaf1 | -0.068943007 | -0.231443069 | 0.093557054 | 8.06005545 | -1.043772514 | 0.33766228 | 0.792411131 | -4.74812931 |
| **A_55_P2177539** | Sod1 | -0.103542469 | -0.34762554 | 0.140540602 | 15.36885736 | -1.043638329 | 0.337719465 | 0.792411131 | -4.748213114 |
| **A_51_P517495** | Cox8a | -0.055944808 | -0.187829912 | 0.075940297 | 16.15714168 | -1.043597909 | 0.337736692 | 0.792411131 | -4.748238357 |
| **A_51_P416585** | Hdlbp | -0.162053785 | -0.544091983 | 0.219984414 | 12.52028494 | -1.04356992 | 0.337748622 | 0.792411131 | -4.748255836 |
| **A_55_P2025712** | Immt | -0.058233248 | -0.1955226 | 0.079056105 | 9.610997517 | -1.0435261 | 0.337767299 | 0.792411131 | -4.748283202 |
| **A_55_P2115260** | NA | -0.080260519 | -0.269483696 | 0.108962659 | 8.315233829 | -1.043510431 | 0.337773978 | 0.792411131 | -4.748292987 |
| **A_55_P2131393** | Eef1akmt2 | 0.086470425 | -0.117424264 | 0.290365113 | 10.22061936 | 1.043351974 | 0.337841527 | 0.792411131 | -4.748391936 |
| **A_55_P2138436** | NA | 0.066702213 | -0.090595694 | 0.22400012 | 15.39253753 | 1.043245357 | 0.337886982 | 0.792411131 | -4.748458508 |
| **A_55_P1952850** | NA | 0.063041544 | -0.0856497 | 0.211732788 | 13.4852243 | 1.043063166 | 0.33796467 | 0.792411131 | -4.748572261 |
| **A_55_P2068554** | NA | 0.045402599 | -0.061686914 | 0.152492113 | 16.39544213 | 1.043044735 | 0.33797253 | 0.792411131 | -4.748583768 |
| **A_55_P1990225** | Fnta | 0.101976315 | -0.138571685 | 0.342524314 | 12.52418104 | 1.042957775 | 0.338009617 | 0.792411131 | -4.748638058 |
| **A_51_P348652** | Spast | 0.078412541 | -0.106552562 | 0.263377644 | 11.28124629 | 1.042953369 | 0.338011496 | 0.792411131 | -4.748640808 |
| **A_55_P2033510** | Ppm1m | -0.072549193 | -0.243683523 | 0.098585137 | 10.36643054 | -1.042952693 | 0.338011784 | 0.792411131 | -4.748641231 |
| **A_55_P2087840** | Gpr50 | -0.080209103 | -0.269438537 | 0.10902033 | 8.499881757 | -1.042807473 | 0.338073725 | 0.792452982 | -4.748731887 |
| **A_52_P199614** | Wdfy4 | -0.068043298 | -0.228588645 | 0.092502049 | 8.219911507 | -1.042693807 | 0.338122213 | 0.792452982 | -4.74880284 |
| **A_51_P289414** | Spg11 | -0.098769062 | -0.331859252 | 0.134321127 | 10.43535924 | -1.042476079 | 0.338215109 | 0.792452982 | -4.748938738 |

| | | | | | | | | | |
|---|---|---|---|---|---|---|---|---|---|
| **A_55_P2001073** | Arid2 | 0.07466741 | -0.10161109 | 0.250945911 | 9.048902228 | 1.042079626 | 0.338384313 | 0.792452982 | -4.749186151 |
| **A_51_P462299** | Aida | -0.076868777 | -0.25835665 | 0.104619097 | 9.727099729 | -1.042009121 | 0.338414411 | 0.792452982 | -4.749230145 |
| **A_51_P175205** | Rps6kb1 | -0.076796445 | -0.25812591 | 0.104533019 | 8.294500084 | -1.041938061 | 0.338444748 | 0.792452982 | -4.749274483 |
| **A_51_P118603** | Wdr82 | -0.073411228 | -0.24675389 | 0.099931434 | 10.40227412 | -1.041900366 | 0.338460843 | 0.792452982 | -4.74612896 |
| **A_55_P1965333** | Naa60 | 0.090890435 | -0.123760179 | 0.305541049 | 11.45077821 | 1.041730095 | 0.338533548 | 0.792452982 | -4.749404236 |
| **A_51_P104418** | Dusp10 | -0.066191003 | -0.222519717 | 0.090137711 | 8.516160203 | -1.04166811 | 0.338560018 | 0.792452982 | -4.749442907 |
| **A_55_P2294037** | NA | -0.127012482 | -0.426995712 | 0.172970747 | 8.637948323 | -1.041642259 | 0.338571059 | 0.792452982 | -4.749459034 |
| **A_55_P2026109** | Rpap1 | -0.118269051 | -0.397635969 | 0.161097867 | 10.93961384 | -1.041514447 | 0.338625647 | 0.792452982 | -4.749538767 |
| **A_51_P503131** | Wnt11 | 0.047176589 | -0.06426404 | 0.158617217 | 7.901478427 | 1.041482904 | 0.338639121 | 0.792452982 | -4.749558444 |
| **A_51_P514412** | Gsk3b | -0.097793213 | -0.3288281 | 0.133241674 | 8.667992688 | -1.041358614 | 0.338692213 | 0.792452982 | -4.749635972 |
| **A_55_P2020592** | Trim3 | -0.048721068 | -0.163842216 | 0.06640008 | 8.113198796 | -1.041192067 | 0.338763368 | 0.792452982 | -4.749739852 |
| **A_66_P102719** | Spop | -0.184818091 | -0.621554174 | 0.251917992 | 11.41313334 | -1.041105087 | 0.338800534 | 0.792452982 | -4.7497941 |
| **A_55_P1980180** | Tardbp | -0.185374006 | -0.638549662 | 0.267801649 | 13.38243963 | -1.060064636 | 0.338841633 | 0.792452982 | -4.698964381 |
| **A_51_P440327** | Efhc1 | -0.101620192 | -0.341784853 | 0.138544469 | 9.002559311 | -1.040974454 | 0.338856358 | 0.792452982 | -4.749875568 |
| **A_55_P2159830** | Yaf2 | -0.080662303 | -0.271298181 | 0.109973575 | 8.967016909 | -1.040962638 | 0.338861408 | 0.792452982 | -4.749882937 |
| **A_55_P2116391** | Tesk2 | -0.18650966 | -0.627310933 | 0.254291612 | 10.40510685 | -1.040944691 | 0.338869078 | 0.792452982 | -4.749894129 |
| **A_55_P2045837** | NA | -0.085406012 | -0.287283858 | 0.116471834 | 8.171290872 | -1.040803977 | 0.338929221 | 0.792452982 | -4.749981877 |
| **A_55_P2098880** | Galnt10 | 0.078831997 | -0.107520838 | 0.265184831 | 11.2848983 | 1.040724279 | 0.338963289 | 0.792452982 | -4.750031572 |
| **A_55_P1991600** | Aplp2 | -0.118087033 | -0.397250913 | 0.161076847 | 14.86080941 | -1.040667875 | 0.338987401 | 0.792452982 | -4.750066742 |
| **A_55_P1970507** | Pef1 | 0.08620609 | -0.117608228 | 0.290020408 | 12.01530999 | 1.040572683 | 0.339028098 | 0.792452982 | -4.750126095 |
| **A_55_P1964867** | Ndufs6 | 0.079164256 | -0.108015309 | 0.26634382 | 14.91411317 | 1.04049468 | 0.339061449 | 0.792452982 | -4.750174727 |
| **A_52_P668984** | Tmem243 | -0.067141723 | -0.225901508 | 0.091618061 | 8.773282733 | -1.040449834 | 0.339080624 | 0.792452982 | -4.750202687 |
| **A_55_P2000399** | Rhox4a | -0.097694721 | -0.328706285 | 0.133316844 | 11.28671948 | -1.04041484 | 0.339095588 | 0.792452982 | -4.750224503 |
| **A_51_P497061** | Ppp2r2d | -0.117546645 | -0.395690615 | 0.160597324 | 12.49545251 | -1.039704094 | 0.339399623 | 0.792944863 | -4.75066752 |
| **A_55_P2146384** | Filip1 | -0.063900615 | -0.215109185 | 0.087307955 | 7.87520824 | -1.039675469 | 0.339411873 | 0.792944863 | -4.750685359 |
| **A_55_P2092085** | Tmsb10 | -0.124227825 | -0.418200593 | 0.169744943 | 16.35112842 | -1.039635126 | 0.339429137 | 0.792944863 | -4.750710499 |
| **A_51_P439612** | Dnajb2 | -0.096666664 | -0.325633339 | 0.132300011 | 10.4444532 | -1.038660502 | 0.339846442 | 0.793707894 | -4.751317694 |
| **A_55_P2148243** | Wdr12 | 0.14042461 | -0.192205678 | 0.473054899 | 10.0167562 | 1.038605451 | 0.339870026 | 0.793707894 | -4.751351981 |
| **A_55_P2073218** | Hps4 | 0.087453657 | -0.119708537 | 0.29461585 | 11.04672753 | 1.038572063 | 0.33988433 | 0.793707894 | -4.751372775 |
| **A_55_P2106690** | Tal1 | -0.061798995 | -0.20821446 | 0.08461647 | 8.09607455 | -1.038397475 | 0.339959134 | 0.793707894 | -4.751481505 |
| **A_52_P513123** | Dna2 | -0.108021471 | -0.363961028 | 0.147918085 | 8.925442433 | -1.038345388 | 0.339981454 | 0.793707894 | -4.751513941 |
| **A_55_P2124461** | NA | -0.3198253 | -1.077635297 | 0.437984697 | 11.65031115 | -1.038297117 | 0.34000214 | 0.793707894 | -4.751544 |
| **A_66_P133244** | Vcl | -0.063944952 | -0.215483409 | 0.087593504 | 8.16884974 | -1.038131985 | 0.340072912 | 0.79377724 | -4.751646826 |
| **A_52_P981697** | Zfp874a | -0.054722544 | -0.184447265 | 0.075002176 | 8.244636259 | -1.037797698 | 0.340216218 | 0.794015851 | -4.751854953 |
| **A_51_P476209** | Fscn3 | -0.071919689 | -0.242479575 | 0.098640196 | 8.173204786 | -1.037385252 | 0.340393097 | 0.794332751 | -4.75211169 |
| **A_66_P128593** | Lims1 | -0.077900837 | -0.262749074 | 0.106947401 | 10.16843779 | -1.036802343 | 0.340643209 | 0.794795096 | -4.752474436 |
| **A_51_P208603** | Dio2 | -0.232988251 | -0.785901464 | 0.319924962 | 8.347870583 | -1.03668357 | 0.340694189 | 0.794795096 | -4.752548335 |
| **A_55_P2096166** | NA | -0.094454884 | -0.318640332 | 0.129730565 | 9.814143288 | -1.036540228 | 0.340755724 | 0.794795096 | -4.752637514 |
| **A_55_P2063795** | Bag6 | -0.072046307 | -0.243053229 | 0.098960614 | 12.29516935 | -1.036494974 | 0.340775153 | 0.794795096 | -4.752665667 |
| **A_52_P496260** | Exosc8 | -0.08628421 | -0.291111107 | 0.118542686 | 9.854775906 | -1.036366836 | 0.340830171 | 0.794795096 | -4.752745378 |
| **A_55_P2088630** | Ap3s2 | 0.087616193 | -0.120376279 | 0.295608664 | 9.488242912 | 1.036348741 | 0.340837941 | 0.794795096 | -4.752756634 |
| **A_55_P2157620** | NA | 0.091293103 | -0.125508649 | 0.308094855 | 9.733826342 | 1.035963241 | 0.341003508 | 0.795037057 | -4.752996409 |
| **A_65_P06061** | Rpl37 | -0.070422256 | -0.237667863 | 0.09682335 | 17.07490411 | -1.035915618 | 0.341023966 | 0.795037057 | -4.753026026 |
| **A_52_P642453** | Obscn | -0.126099651 | -0.425624702 | 0.1734254 | 7.69993364 | -1.035737968 | 0.34110029 | 0.795055104 | -4.753136501 |
| **A_51_P336721** | Tipin | 0.110308417 | -0.151727313 | 0.372344147 | 10.41932178 | 1.035660233 | 0.341133692 | 0.795055104 | -4.753184839 |
| **A_55_P2096967** | Gjb1 | 0.348680954 | -0.479733857 | 1.177095766 | 10.40987641 | 1.035498642 | 0.341203134 | 0.795055104 | -4.753285314 |
| **A_55_P2123471** | Kpna2 | 0.19542234 | -0.284927379 | 0.675772058 | 12.17956691 | 1.054306174 | 0.341222565 | 0.795055104 | -4.702204043 |
| **A_55_P2154242** | Gfpt1 | -0.159546881 | -0.538670523 | 0.219576761 | 12.89918004 | -1.035324792 | 0.341277857 | 0.795055104 | -4.753393402 |
| **A_55_P1996907** | BC067074 | 0.070424685 | -0.096941591 | 0.237790961 | 7.807729096 | 1.035204438 | 0.341329595 | 0.795055104 | -4.753468223 |
| **A_55_P2184449** | NA | -0.234081196 | -0.790406954 | 0.322244562 | 8.754410939 | -1.035157713 | 0.341349683 | 0.795055104 | -4.75349727 |

| | | | | | | | | | |
|---|---|---|---|---|---|---|---|---|---|
| **A_55_P2026535** | Erc2 | -0.112108077 | -0.378554786 | 0.154338633 | 8.49243901 | -1.035131926 | 0.341360769 | 0.795055104 | -4.7535133 |
| **A_55_P2014882** | Dhrs9 | -0.182750095 | -0.617134086 | 0.251633896 | 8.742723609 | -1.03503006 | 0.341404568 | 0.795061312 | -4.753576622 |
| **A_55_P2050697** | Nfkbil1 | 0.128985975 | -0.177675043 | 0.435646993 | 11.36077722 | 1.034792044 | 0.341506923 | 0.795135959 | -4.753724562 |
| **A_52_P672740** | Rala | -0.076739137 | -0.259189475 | 0.105711202 | 8.43625437 | -1.034764207 | 0.341518896 | 0.795135959 | -4.753741864 |
| **A_55_P2178486** | Spin1 | -0.067624603 | -0.228463103 | 0.093213898 | 8.819934234 | -1.034388956 | 0.341680322 | 0.795415987 | -4.753975061 |
| **A_55_P2028054** | Incenp | 0.081221157 | -0.112007765 | 0.27445008 | 10.25517301 | 1.034108733 | 0.341800909 | 0.795600888 | -4.754149172 |
| **A_55_P2143561** | Gm6813 | -0.048376372 | -0.163502164 | 0.06674942 | 15.26299256 | -1.033784044 | 0.341940675 | 0.79566434 | -4.754350878 |
| **A_55_P1954021** | Scarb1 | 0.161403285 | -0.222744144 | 0.545550714 | 11.05624442 | 1.033674023 | 0.341988045 | 0.79566434 | -4.754419218 |
| **A_52_P30989** | Cdkn3 | 0.062004717 | -0.08560151 | 0.209610944 | 7.848591707 | 1.03344939 | 0.342084779 | 0.79566434 | -4.754558736 |
| **A_52_P584335** | Folr1 | 0.163698496 | -0.226050436 | 0.553447428 | 9.40556389 | 1.033305924 | 0.342146571 | 0.79566434 | -4.754647833 |
| **A_55_P2022337** | Dhcr7 | 0.110025712 | -0.151990011 | 0.372041436 | 8.337332018 | 1.033084863 | 0.342241802 | 0.79566434 | -4.754785105 |
| **A_51_P134262** | 1700052K11Rik | 0.070062299 | -0.096788636 | 0.236913234 | 10.48216713 | 1.03305847 | 0.342253174 | 0.79566434 | -4.754801494 |
| **A_55_P2037689** | NA | -0.069663054 | -0.235592107 | 0.096265999 | 9.907394374 | -1.032878498 | 0.342330722 | 0.79566434 | -4.754913237 |
| **A_55_P2151021** | NA | 0.091943295 | -0.127056443 | 0.310943034 | 11.20549754 | 1.032869927 | 0.342334415 | 0.79566434 | -4.754918558 |
| **A_51_P389506** | NA | -0.094962781 | -0.321182665 | 0.131257104 | 12.99719885 | -1.03274193 | 0.342389577 | 0.79566434 | -4.754998023 |
| **A_55_P2124976** | Grhl1 | -0.095402179 | -0.322680876 | 0.131876518 | 9.383776722 | -1.032687037 | 0.342413236 | 0.79566434 | -4.755032101 |
| **A_52_P418665** | Ttpal | 0.061681612 | -0.085267403 | 0.208630627 | 11.54782181 | 1.032662012 | 0.342424023 | 0.79566434 | -4.755047636 |
| **A_55_P2012955** | NA | -0.053359097 | -0.180484708 | 0.073766513 | 15.44206885 | -1.032629634 | 0.342437979 | 0.79566434 | -4.755067736 |
| **A_55_P2136020** | Uba2 | -0.075780239 | -0.256347484 | 0.104787006 | 13.03084897 | -1.032490733 | 0.342497855 | 0.79566434 | -4.755153959 |
| **A_55_P2012498** | Cdca7 | -0.173962224 | -0.588484776 | 0.240560328 | 9.612192032 | -1.032466455 | 0.342508321 | 0.79566434 | -4.755169029 |
| **A_55_P2084571** | NA | -0.048805626 | -0.165143332 | 0.06753208 | 15.61078618 | -1.032092349 | 0.342669634 | 0.79566434 | -4.75540122 |
| **A_51_P152826** | Golt1b | 0.124743917 | -0.172611368 | 0.422099202 | 11.01411973 | 1.032078847 | 0.342675457 | 0.79566434 | -4.755409599 |
| **A_55_P2033407** | Meg3 | -0.194367119 | -0.657715327 | 0.268981089 | 8.712239256 | -1.032011348 | 0.342704569 | 0.79566434 | -4.755451487 |
| **A_51_P472249** | Slc7a7 | 0.275375854 | -0.381134172 | 0.931885879 | 11.52213144 | 1.031938124 | 0.342736153 | 0.79566434 | -4.755496926 |
| **A_52_P553890** | Itgb3 | -0.414197933 | -1.401745864 | 0.573349998 | 13.87489021 | -1.031855556 | 0.342771769 | 0.79566434 | -4.755548161 |
| **A_55_P2013833** | Ddx3x | 0.083360738 | -0.11539179 | 0.282113267 | 12.87007055 | 1.031853529 | 0.342772644 | 0.79566434 | -4.755549419 |
| **A_55_P2131954** | Gm2590 | -0.161849308 | -0.547756063 | 0.224057446 | 8.338335033 | -1.031805008 | 0.342793575 | 0.79566434 | -4.755579526 |
| **A_55_P2330435** | NA | -0.046263136 | -0.156583354 | 0.064057082 | 7.670667843 | -1.031689757 | 0.342843298 | 0.79566434 | -4.755651037 |
| **A_55_P2126572** | Tmem25 | 0.090034112 | -0.124665634 | 0.304733859 | 8.288133015 | 1.031679304 | 0.342847808 | 0.79566434 | -4.755657522 |
| **A_55_P1962731** | Clcn4 | 0.134521661 | -0.186284255 | 0.455327577 | 12.4625261 | 1.031618248 | 0.342874152 | 0.79566434 | -4.755695403 |
| **A_55_P2036854** | Pnkd | -0.0999841 | -0.338426326 | 0.138458125 | 11.40092628 | -1.031613314 | 0.342876281 | 0.79566434 | -4.755698465 |
| **A_55_P1973006** | Bag1 | -0.147951531 | -0.50080412 | 0.204901058 | 14.07059225 | -1.03156196 | 0.34289844 | 0.79566434 | -4.755730325 |
| **A_66_P107680** | Med8 | -0.052369679 | -0.177310783 | 0.072571424 | 8.382250074 | -1.031201988 | 0.343053801 | 0.795908818 | -4.755953633 |
| **A_55_P2177968** | Snrpd2 | 0.090413843 | -0.125312144 | 0.30613983 | 12.46992115 | 1.031102001 | 0.343096965 | 0.795908818 | -4.756015652 |
| **A_55_P1997126** | Ctse | 0.14051942 | -0.194835587 | 0.475874426 | 8.22329161 | 1.030862441 | 0.343200399 | 0.795908818 | -4.756164229 |
| **A_52_P339722** | Csnk1a1 | -0.113251421 | -0.383534187 | 0.157031345 | 11.54737886 | -1.030847612 | 0.343206802 | 0.795908818 | -4.756173425 |
| **A_55_P1967905** | Dcbld1 | -0.069086154 | -0.233966359 | 0.095794052 | 8.528498651 | -1.030840937 | 0.343209685 | 0.795908818 | -4.756177565 |
| **A_55_P2024376** | NA | 0.098458218 | -0.136554982 | 0.333471418 | 16.16890448 | 1.030691936 | 0.343274033 | 0.795915741 | -4.756269965 |
| **A_55_P2000938** | Ahcy | 0.071719477 | -0.099478204 | 0.242917157 | 8.936318223 | 1.030643332 | 0.343295025 | 0.795915741 | -4.756300105 |
| **A_55_P2052719** | Nadk2 | 0.120648749 | -0.167399174 | 0.408696672 | 9.443837767 | 1.030450748 | 0.343378213 | 0.795962511 | -4.756419518 |
| **A_55_P2139809** | Fgf13 | 0.047329621 | -0.065674381 | 0.160333624 | 7.759873806 | 1.030405972 | 0.343397557 | 0.795962511 | -4.75644728 |
| **A_55_P1979082** | Shc1 | 0.065065071 | -0.090301793 | 0.220431934 | 11.70012273 | 1.030288069 | 0.343448496 | 0.79598513 | -4.756520378 |
| **A_55_P1975385** | Dctn1 | 0.08895586 | -0.123498184 | 0.301409904 | 13.16657823 | 1.030098426 | 0.343530444 | 0.796079601 | -4.756637945 |
| **A_51_P286373** | Ttll12 | -0.081606346 | -0.276580274 | 0.113367581 | 10.59125706 | -1.029713857 | 0.343696671 | 0.79636933 | -4.756876315 |
| **A_51_P398235** | Dnajc11 | 0.059244446 | -0.08232695 | 0.200815842 | 8.841358841 | 1.029535419 | 0.343773821 | 0.796452618 | -4.750641473 |
| **A_55_P2008747** | Snrpd3 | -0.044312241 | -0.150228872 | 0.061604389 | 11.29537894 | -1.029268601 | 0.34388921 | 0.796518484 | -4.757152237 |
| **A_55_P1977418** | Ugt2b34 | 0.33544171 | -0.466379761 | 1.13726318 | 8.429961879 | 1.029220777 | 0.343909896 | 0.796518484 | -4.75718187 |
| **A_51_P455897** | Pimreg | 0.046799746 | -0.065083908 | 0.158683399 | 7.711894996 | 1.02907261 | 0.343973989 | 0.796518484 | -4.75727367 |
| **A_55_P2100222** | NA | -0.301239164 | -1.021501786 | 0.419023457 | 10.91006651 | -1.028939216 | 0.344031701 | 0.796518484 | -4.757356312 |
| **A_52_P381953** | Irf2bp1 | -0.051201482 | -0.173627539 | 0.071224574 | 8.093227727 | -1.028911069 | 0.34404388 | 0.796518484 | -4.757373749 |

| | | | | | | | | | |
|---|---|---|---|---|---|---|---|---|---|
| **A_55_P1979694** | NA | -0.090842731 | -0.308056318 | 0.126370857 | 13.74638643 | -1.028898075 | 0.344049502 | 0.796518484 | -4.757381798 |
| **A_55_P2379599** | Asph | -0.055376134 | -0.187816406 | 0.077064139 | 8.866552378 | -1.028659708 | 0.344152653 | 0.796659215 | -4.757529454 |
| **A_55_P2171143** | Hnrnpc | 0.065348574 | -0.090956341 | 0.221653489 | 13.77240214 | 1.028567131 | 0.344192721 | 0.796659215 | -4.757586795 |
| **A_51_P507172** | 2310022A10Rik | 0.078375124 | -0.109131091 | 0.265881338 | 10.5452831 | 1.02832818 | 0.344296159 | 0.796669588 | -4.757734786 |
| **A_55_P2041823** | Hyal1 | -0.161976425 | -0.549494817 | 0.225541968 | 9.323008815 | -1.028320875 | 0.344299322 | 0.796669588 | -4.75773931 |
| **A_55_P2279035** | 9030419F21Rik | -0.230133094 | -0.780738649 | 0.320472461 | 8.663044961 | -1.028271148 | 0.344320851 | 0.796669588 | -4.757770104 |
| **A_55_P2021792** | Dyrk1b | -0.073816509 | -0.250491251 | 0.102858234 | 8.50368814 | -1.027893681 | 0.344484312 | 0.796952397 | -4.758003835 |
| **A_55_P2004126** | NA | 0.098705036 | -0.137579437 | 0.334989508 | 10.6088211 | 1.02771641 | 0.344561101 | 0.797034647 | -4.758113585 |
| **A_51_P451075** | Atp2a2 | 0.119040418 | -0.165989454 | 0.404070289 | 9.059902915 | 1.027479647 | 0.344663681 | 0.797082163 | -4.75826015 |
| **A_55_P2043466** | Eif5 | -0.066708747 | -0.226447525 | 0.093030032 | 13.146118 | -1.027404788 | 0.344696119 | 0.797082163 | -4.758306487 |
| **A_55_P2124751** | Col16a1 | 0.234588844 | -0.327162269 | 0.796339956 | 8.966984006 | 1.027383476 | 0.344705355 | 0.797082163 | -4.758319678 |
| **A_51_P124254** | Col4a1 | -0.221540213 | -0.752136923 | 0.309056496 | 13.799742 | -1.02720513 | 0.344782649 | 0.797165529 | -4.758430063 |
| **A_55_P2018989** | Elk3 | 0.062636638 | -0.087417283 | 0.212690559 | 8.027840401 | 1.026952255 | 0.344892268 | 0.797310465 | -4.758586557 |
| **A_52_P281702** | Igfbp5 | -0.09647589 | -0.327614445 | 0.134662666 | 8.447161178 | -1.026870222 | 0.344927834 | 0.797310465 | -4.758637319 |
| **A_51_P365008** | Tns1 | 0.130176814 | -0.18176698 | 0.442120608 | 12.84298341 | 1.026659549 | 0.345019188 | 0.7973497 | -4.758767673 |
| **A_55_P2034275** | Arfgap2 | 0.066432228 | -0.092762782 | 0.225627238 | 11.8863009 | 1.026640819 | 0.345027311 | 0.7973497 | -4.758779261 |
| **A_51_P464911** | Msrb2 | 0.091379374 | -0.127621592 | 0.31038034 | 9.408332293 | 1.026529204 | 0.345075719 | 0.797366237 | -4.758848316 |
| **A_66_P101107** | NA | -0.174516094 | -0.5928219 | 0.243789712 | 8.584325951 | -1.026386081 | 0.345137801 | 0.797389474 | -4.758936858 |
| **A_52_P431894** | Pknox1 | -0.084364985 | -0.286597215 | 0.117867246 | 9.905359603 | -1.026315816 | 0.345168282 | 0.797389474 | -4.758980324 |
| **A_51_P103222** | Slc39a4 | -0.179020843 | -0.608257111 | 0.250215425 | 14.49706671 | -1.026068489 | 0.345275593 | 0.79751364 | -4.759133309 |
| **A_51_P199098** | Tmem107 | -0.11333213 | -0.385084982 | 0.158420721 | 11.10655421 | -1.02600176 | 0.34530455 | 0.79751364 | -4.75917458 |
| **A_51_P267063** | Ugt3a2 | 0.10955534 | -0.153181835 | 0.372292515 | 7.894330651 | 1.025843679 | 0.345373157 | 0.797576793 | -4.759272347 |
| **A_55_P2114044** | Commd8 | 0.113274557 | -0.158423116 | 0.384972229 | 10.5683325 | 1.025688806 | 0.345440383 | 0.797636742 | -4.759368121 |
| **A_55_P2060996** | NA | -0.091810578 | -0.312051886 | 0.128430729 | 8.514618014 | -1.025564805 | 0.345494216 | 0.797665754 | -4.759444797 |
| **A_55_P1955672** | NA | 0.117826166 | -0.164850847 | 0.400503179 | 15.06762723 | 1.025463994 | 0.345537986 | 0.797671532 | -4.75950713 |
| **A_65_P11773** | Adi1 | 0.064728868 | -0.090586692 | 0.220044428 | 9.812089192 | 1.025302956 | 0.345607915 | 0.797737687 | -4.759606695 |
| **A_55_P2311208** | C130045F17Rik | -0.163266791 | -0.555083955 | 0.228550372 | 8.435248638 | -1.025140908 | 0.345678294 | 0.797804865 | -4.759706875 |
| **A_55_P2379480** | NA | -0.099197803 | -0.337280874 | 0.138885268 | 8.068364749 | -1.025044453 | 0.345720191 | 0.7978063 | -4.759766501 |
| **A_55_P2052495** | Tmx2 | 0.140768294 | -0.197133293 | 0.478669882 | 11.8845429 | 1.024905394 | 0.345780601 | 0.797850452 | -4.759852458 |
| **A_52_P670812** | Snord123 | 0.084966978 | -0.119074502 | 0.289008458 | 9.896292279 | 1.024473836 | 0.345968133 | 0.797973657 | -4.760119172 |
| **A_55_P2131483** | Hmgb1-ps7 | -0.076197267 | -0.259194946 | 0.106800411 | 11.34577671 | -1.024384457 | 0.346006982 | 0.797973657 | -4.760174403 |
| **A_52_P641758** | Ggct | 0.084404624 | -0.118320588 | 0.287129835 | 8.860153404 | 1.024301099 | 0.346043218 | 0.797973657 | -4.76022591 |
| **A_55_P2153941** | Zfp386 | 0.069849845 | -0.097923765 | 0.237623454 | 9.948631763 | 1.024261759 | 0.34606032 | 0.797973657 | -4.760250219 |
| **A_51_P220993** | Pfn1 | 0.048039379 | -0.0673612 | 0.163439957 | 10.14717682 | 1.024138186 | 0.346114045 | 0.797973657 | -4.760326569 |
| **A_55_P2117710** | Snhg1 | -0.067407841 | -0.229344459 | 0.094528777 | 8.619162467 | -1.024081489 | 0.346138697 | 0.797973657 | -4.760361598 |
| **A_51_P223443** | Cd81 | -0.250240893 | -0.872321544 | 0.371839758 | 13.43763078 | -1.042465449 | 0.346163055 | 0.797973657 | -4.708839952 |
| **A_51_P212754** | Tgfbi | 0.308982087 | -0.433340864 | 1.051305038 | 11.5561017 | 1.024022684 | 0.346164267 | 0.797973657 | -4.760397928 |
| **A_51_P177552** | Ndufb9 | 0.143939703 | -0.202027184 | 0.48990659 | 14.35927997 | 1.023564539 | 0.346363533 | 0.798145501 | -4.760680932 |
| **A_51_P459108** | Insl6 | 0.16243624 | -0.228008464 | 0.552880944 | 9.115480695 | 1.023511096 | 0.346386784 | 0.798145501 | -4.760713939 |
| **A_51_P182257** | Catsperz | 0.05277797 | -0.074083793 | 0.179639733 | 7.834462581 | 1.023507664 | 0.346388277 | 0.798145501 | -4.760716059 |
| **A_55_P1987979** | Plrg1 | -0.061231434 | -0.208417877 | 0.08595501 | 11.31443123 | -1.023471562 | 0.346403984 | 0.798145501 | -4.760738356 |
| **A_51_P155073** | Nudt14 | 0.091435954 | -0.128438688 | 0.311310596 | 8.508924444 | 1.023083354 | 0.346572921 | 0.798437167 | -4.760978089 |
| **A_52_P94845** | Nicn1 | -0.053795351 | -0.183167909 | 0.075577206 | 8.302048759 | -1.022990849 | 0.346613186 | 0.798437167 | -4.761035206 |
| **A_55_P2152566** | Sp110 | 0.086463125 | -0.121505877 | 0.294432128 | 9.178490057 | 1.022825349 | 0.346685233 | 0.798507969 | -4.761137387 |
| **A_55_P2119985** | Baiap2 | 0.069491967 | -0.097677127 | 0.236661061 | 10.32252946 | 1.022698877 | 0.346740299 | 0.798529579 | -4.761215465 |
| **A_55_P2159852** | NA | -0.113426958 | -0.386315439 | 0.159461524 | 9.134859273 | -1.022586939 | 0.346789042 | 0.798529579 | -4.761284566 |
| **A_55_P1956572** | Dcaf8 | 0.078552847 | -0.110477695 | 0.267583389 | 12.49166663 | 1.022348852 | 0.346892736 | 0.798529579 | -4.761431526 |
| **A_55_P1990457** | Rpl15 | 0.092502946 | -0.130098423 | 0.315104314 | 15.45338722 | 1.02234363 | 0.346895011 | 0.798529579 | -4.761434749 |
| **A_55_P1952016** | R3hdm2 | -0.077794345 | -0.265003169 | 0.109414478 | 10.87471228 | -1.02232947 | 0.346901178 | 0.798529579 | -4.761443489 |
| **A_55_P2037918** | NA | 0.063404718 | -0.08920418 | 0.216013616 | 8.322188641 | 1.022141169 | 0.346983208 | 0.798623295 | -4.761559703 |

| | | | | | | | | | |
|---|---|---|---|---|---|---|---|---|---|
| **A_51_P214470** | Tdrp | -0.159532831 | -0.543557981 | 0.22449232 | 10.89883162 | -1.022020405 | 0.347035825 | 0.7986493 | -4.761634229 |
| **A_55_P2022509** | Mgat5b | 0.277142448 | -0.390112966 | 0.944397861 | 10.16126602 | 1.021833427 | 0.347117304 | 0.798741711 | -4.761749606 |
| **A_51_P445912** | NA | -0.061132443 | -0.208334842 | 0.086069956 | 8.477458001 | -1.021706197 | 0.347172756 | 0.79876361 | -4.761828108 |
| **A_55_P1984289** | Cdan1 | 0.074385094 | -0.104743475 | 0.253513662 | 8.151376356 | 1.021621967 | 0.34720947 | 0.79876361 | -4.761880076 |
| **A_55_P2179582** | Nmur2 | 0.089194656 | -0.125639241 | 0.304028553 | 7.839278511 | 1.021421966 | 0.34729666 | 0.798869111 | -4.762003461 |
| **A_51_P293607** | Mrps14 | 0.055913492 | -0.078801113 | 0.190628096 | 8.349832535 | 1.021106634 | 0.347434164 | 0.799090308 | -4.762197968 |
| **A_52_P359965** | Cpd | -0.174667398 | -0.595670436 | 0.246335641 | 10.24467394 | -1.02069452 | 0.347613937 | 0.799337748 | -4.762452121 |
| **A_55_P2029498** | Osbpl1a | -0.146720386 | -0.500374542 | 0.20693377 | 10.55781144 | -1.02065944 | 0.347629243 | 0.799337748 | -4.762473752 |
| **A_55_P2029612** | Dhx35 | 0.085179965 | -0.120154298 | 0.290514227 | 9.681771957 | 1.020575637 | 0.34766581 | 0.799337748 | -4.762525425 |
| **A_66_P138915** | Fyn | 0.048058661 | -0.06782229 | 0.163939611 | 8.453332174 | 1.020302095 | 0.347785191 | 0.799414693 | -4.762694075 |
| **A_52_P5652** | Sae1 | -0.086633374 | -0.295544006 | 0.122277258 | 8.758074265 | -1.02022004 | 0.347821008 | 0.799414693 | -4.762744661 |
| **A_55_P2009064** | Psmc1 | 0.174478806 | -0.246267468 | 0.595225079 | 11.64128329 | 1.020214672 | 0.347823352 | 0.799414693 | -4.76274797 |
| **A_55_P1954006** | Ldhb | -0.165305777 | -0.564046043 | 0.23343449 | 14.85496204 | -1.019922326 | 0.347950987 | 0.799529426 | -4.762928176 |
| **A_51_P431894** | Radil | -0.051847001 | -0.176926355 | 0.073232353 | 8.064261881 | -1.019781602 | 0.348012439 | 0.799529426 | -4.76301491 |
| **A_51_P138859** | Tmem68 | -0.09006555 | -0.307372655 | 0.127241554 | 10.35791084 | -1.019656624 | 0.348067023 | 0.799529426 | -4.763091932 |
| **A_55_P2008722** | Zfp984 | 0.101457912 | -0.143338592 | 0.346254417 | 9.362579544 | 1.019647062 | 0.348071199 | 0.799529426 | -4.763097825 |
| **A_55_P2132681** | Fzr1 | -0.057942775 | -0.197759939 | 0.081874389 | 9.358410604 | -1.019548678 | 0.348114173 | 0.799529426 | -4.763158454 |
| **A_55_P2091836** | Fnip1 | 0.075350791 | -0.106475264 | 0.257176845 | 11.07509653 | 1.019532002 | 0.348121457 | 0.799529426 | -4.76316873 |
| **A_55_P2127425** | Sys1 | -0.077968295 | -0.26615225 | 0.110215659 | 11.11179308 | -1.019306074 | 0.34822016 | 0.7996611 | -4.763307944 |
| **A_51_P341177** | Ddah2 | -0.152384731 | -0.520316977 | 0.215547515 | 13.39452053 | -1.018926281 | 0.348386134 | 0.799890332 | -4.763541926 |
| **A_51_P221428** | Cbx6 | -0.105863297 | -0.361479287 | 0.149752692 | 9.178571973 | -1.018888273 | 0.348402748 | 0.799890332 | -4.763565339 |
| **A_55_P1979997** | Spaca9 | -0.107700812 | -0.36778478 | 0.152383155 | 9.168751339 | -1.018766276 | 0.348456077 | 0.799902553 | -4.763640486 |
| **A_66_P117189** | Rad54l2 | -0.043729174 | -0.149337891 | 0.061879544 | 8.989558267 | -1.018686767 | 0.348490837 | 0.799902553 | -4.763689459 |
| **A_51_P481210** | Ofd1 | -0.075129996 | -0.256593677 | 0.106333684 | 10.09736479 | -1.018574539 | 0.348539907 | 0.799920192 | -4.763758582 |
| **A_55_P2039523** | Phf10 | -0.094356093 | -0.322397091 | 0.133684904 | 11.14678024 | -1.017949379 | 0.348813345 | 0.800357047 | -4.764143541 |
| **A_55_P2078073** | Pqlc2 | 0.102237637 | -0.144860311 | 0.349335585 | 10.64624996 | 1.017913391 | 0.348829091 | 0.800357047 | -4.764165697 |
| **A_55_P2115255** | Fer | 0.098859314 | -0.140102111 | 0.33782074 | 9.585111245 | 1.01779182 | 0.348882287 | 0.800357047 | -4.764240541 |
| **A_52_P206526** | Gtf2ird2 | 0.055392263 | -0.078507516 | 0.189292042 | 9.146233369 | 1.01774367 | 0.348903358 | 0.800357047 | -4.764270182 |
| **A_55_P2056871** | Akap13 | -0.162441985 | -0.555184306 | 0.230300336 | 11.63550928 | -1.017559348 | 0.348984029 | 0.800357047 | -4.764383644 |
| **A_52_P445253** | Dst | -0.163152043 | -0.557645454 | 0.231341367 | 8.814271711 | -1.017470736 | 0.349022816 | 0.800357047 | -4.764438186 |
| **A_51_P169880** | Zbtb3 | 0.050863594 | -0.072129756 | 0.173856944 | 8.400477542 | 1.017406657 | 0.349050867 | 0.800357047 | -4.764477626 |
| **A_51_P498831** | Derl2 | 0.089174084 | -0.126473679 | 0.304821846 | 13.43493693 | 1.017332368 | 0.34908339 | 0.800357047 | -4.764523348 |
| **A_55_P2323913** | NA | 0.055200349 | -0.078295317 | 0.188696016 | 9.315878557 | 1.017287768 | 0.349102916 | 0.800357047 | -4.764550797 |
| **A_55_P2060642** | Spidr | -0.124121866 | -0.424517735 | 0.176274003 | 9.276961837 | -1.016537728 | 0.349431424 | 0.800823746 | -4.765012299 |
| **A_52_P162298** | Ydjc | 0.144168921 | -0.204750188 | 0.49308803 | 9.199094254 | 1.016520302 | 0.34943906 | 0.800823746 | -4.765023019 |
| **A_51_P349691** | Ano10 | -0.074139355 | -0.253573807 | 0.105295097 | 9.756476623 | -1.016511121 | 0.349443082 | 0.800823746 | -4.765028666 |
| **A_55_P2099378** | Tfe3 | -0.10740525 | -0.367367889 | 0.152557389 | 12.20159938 | -1.016444653 | 0.349472208 | 0.800823746 | -4.765069554 |
| **A_55_P2054708** | NA | -0.043581384 | -0.149090007 | 0.06192724 | 7.927499958 | -1.016207089 | 0.349576319 | 0.800899963 | -4.765215679 |
| **A_51_P414706** | Ddt | 0.091518033 | -0.1300491 | 0.313085166 | 11.87998184 | 1.016179668 | 0.349588338 | 0.800899963 | -4.765232544 |
| **A_51_P509229** | Tnrc6a | -0.171518967 | -0.58682684 | 0.243788906 | 9.766321158 | -1.01604081 | 0.349649206 | 0.800943745 | -4.765317944 |
| **A_55_P2027240** | Engase | 0.129036009 | -0.183437083 | 0.4415091 | 10.7961304 | 1.015938596 | 0.349694016 | 0.800943745 | -4.765380804 |
| **A_55_P2168431** | Amer1 | 0.109304798 | -0.155409703 | 0.374019298 | 9.056894395 | 1.015852506 | 0.349731761 | 0.800943745 | -4.765433744 |
| **A_51_P394471** | Tcf3 | -0.060582614 | -0.207317791 | 0.086152564 | 9.122001562 | -1.015740867 | 0.349780713 | 0.800960953 | -4.765502392 |
| **A_55_P2433218** | Bloc1s6 | 0.074642152 | -0.106172515 | 0.255456819 | 11.23405271 | 1.015592906 | 0.3498456 | 0.80101464 | -4.765593367 |
| **A_55_P1963639** | Sycn | 0.327076153 | -0.465565291 | 1.119717598 | 9.365534857 | 1.015175827 | 0.350028558 | 0.801110147 | -4.765849772 |
| **A_55_P2073789** | NA | -0.269215316 | -0.921636496 | 0.383205864 | 9.241954895 | -1.015174911 | 0.35002896 | 0.801110147 | -4.765850335 |
| **A_51_P161086** | Ppic | -0.151767305 | -0.519571255 | 0.216036645 | 12.88622769 | -1.015151818 | 0.350039092 | 0.801110147 | -4.765864531 |
| **A_55_P1987988** | Atg2b | 0.061548196 | -0.087621179 | 0.21071757 | 8.495264137 | 1.015090636 | 0.350065938 | 0.801110147 | -4.765902137 |
| **A_66_P116360** | NA | 0.061423376 | -0.087471547 | 0.210318299 | 8.050048859 | 1.014899312 | 0.350149898 | 0.801110147 | -4.766019729 |
| **A_52_P136709** | Pctp | -0.043489102 | -0.148916753 | 0.061938548 | 7.847898277 | -1.014834158 | 0.350178494 | 0.801110147 | -4.766059772 |

| | | | | | | | | | |
|---|---|---|---|---|---|---|---|---|---|
| A_55_P2136662 | 8030462N17Rik | -0.04364539 | -0.149456403 | 0.062165624 | 8.093249017 | -1.014791137 | 0.350197376 | 0.801110147 | -4.766086211 |
| A_55_P2010223 | Gabarap | -0.088391186 | -0.302700766 | 0.125918395 | 14.28714149 | -1.014697372 | 0.350238534 | 0.801110147 | -4.766143833 |
| A_52_P182592 | Ube2w | -0.063495238 | -0.217450929 | 0.090460453 | 9.046186992 | -1.014646049 | 0.350261064 | 0.801110147 | -4.766175371 |
| A_55_P2041626 | Ceacam12 | -0.123090829 | -0.421590357 | 0.175408698 | 17.69770363 | -1.014498042 | 0.350326043 | 0.801110147 | -4.766266318 |
| A_55_P2060183 | Lamtor3 | -0.08973586 | -0.307361816 | 0.127890096 | 13.09812802 | -1.014435643 | 0.350353441 | 0.801110147 | -4.766304658 |
| A_52_P113700 | Sfxn1 | 0.094966907 | -0.135372315 | 0.325306129 | 10.63082275 | 1.014316665 | 0.350405686 | 0.801110147 | -4.766377759 |
| A_51_P479914 | Pik3cb | -0.128115951 | -0.438871378 | 0.182639476 | 11.58100426 | -1.014270154 | 0.350426111 | 0.801110147 | -4.766406334 |
| A_55_P2220283 | Stk35os1 | 0.045523474 | -0.064931205 | 0.155978153 | 8.257672507 | 1.013959113 | 0.350562729 | 0.801283643 | -4.766597411 |
| A_52_P99411 | Itm2b | -0.067682039 | -0.231908745 | 0.096544667 | 14.16173741 | -1.013908617 | 0.350584912 | 0.801283643 | -4.766628428 |
| A_55_P2119892 | NA | -0.100866158 | -0.345665492 | 0.143933176 | 7.708443064 | -1.013688247 | 0.350681736 | 0.801410177 | -4.76676378 |
| A_55_P1987121 | Rtraf | -0.065992342 | -0.226182923 | 0.094198238 | 13.74136933 | -1.01350462 | 0.350762433 | 0.801420741 | -4.766876551 |
| A_55_P2124701 | NA | 0.074037868 | -0.105692899 | 0.253768635 | 12.44373586 | 1.013446066 | 0.350788168 | 0.801420741 | -4.766912508 |
| A_52_P172563 | Dffa | -0.060320185 | -0.206759358 | 0.086118987 | 8.431066802 | -1.013385214 | 0.350814914 | 0.801420741 | -4.766949875 |
| A_55_P2162910 | Rtn1 | -0.206557905 | -0.725651132 | 0.312535322 | 11.0072046 | -1.031208633 | 0.35091597 | 0.801420741 | -4.715115964 |
| A_51_P162437 | Bcr | 0.133339096 | -0.190451401 | 0.457129594 | 12.82762479 | 1.013123925 | 0.35092978 | 0.801420741 | -4.767110309 |
| A_66_P126138 | Ncoa1 | -0.081280924 | -0.278673152 | 0.116111304 | 8.359671878 | -1.013042686 | 0.350965499 | 0.801420741 | -4.767160185 |
| A_55_P2048767 | Efemp1 | 0.079636532 | -0.113767077 | 0.273040141 | 7.917151481 | 1.013017456 | 0.350976593 | 0.801420741 | -4.767175675 |
| A_55_P2099358 | Cars | -0.157413298 | -0.539828123 | 0.225001527 | 12.22868646 | -1.012688461 | 0.351121281 | 0.80165642 | -4.767377634 |
| A_55_P2140212 | NA | -0.52145237 | -1.788789515 | 0.745884774 | 12.14687989 | -1.012258996 | 0.351310227 | 0.801833873 | -4.76764121 |
| A_55_P1996046 | Zfp931 | -0.095106524 | -0.326253364 | 0.136040316 | 9.138614341 | -1.012258685 | 0.351310364 | 0.801833873 | -4.767641401 |
| A_55_P2014511 | Amd2 | -0.094877405 | -0.325474172 | 0.135719361 | 10.28755149 | -1.012228936 | 0.351323455 | 0.801833873 | -4.767659656 |
| A_55_P2052936 | NA | -0.068600148 | -0.23545584 | 0.098255543 | 15.37488711 | -1.011470431 | 0.351657375 | 0.8024098 | -4.768125006 |
| A_51_P363791 | Ap3m1 | -0.067443368 | -0.231518949 | 0.096632214 | 10.86640466 | -1.011263797 | 0.351748386 | 0.8024098 | -4.768251742 |
| A_55_P2186395 | NA | -0.062611845 | -0.214933493 | 0.089709803 | 8.056007585 | -1.011262811 | 0.351748821 | 0.8024098 | -4.768252347 |
| A_52_P604613 | S1pr2 | 0.054356233 | -0.077889156 | 0.186601622 | 10.44926679 | 1.011202138 | 0.351775547 | 0.8024098 | -4.768289557 |
| A_51_P431734 | Fam185a | -0.045265463 | -0.155395497 | 0.064864571 | 8.007011386 | -1.011184394 | 0.351783364 | 0.8024098 | -4.768300439 |
| A_55_P2008889 | Tmsb15b2 | -0.109901082 | -0.377375927 | 0.157573762 | 9.448013967 | -1.01085343 | 0.351929188 | 0.802646035 | -4.768503391 |
| A_52_P71544 | Slc25a26 | -0.071159926 | -0.244363254 | 0.102043402 | 9.39401845 | -1.010760861 | 0.351969982 | 0.802646035 | -4.768560148 |
| A_55_P2073977 | NA | -0.082919672 | -0.284811608 | 0.118972264 | 15.4484302 | -1.010433565 | 0.352114251 | 0.802852067 | -4.768760801 |
| A_51_P382805 | Zpbp2 | 0.086982316 | -0.124815162 | 0.298779794 | 8.58000615 | 1.010367444 | 0.352143402 | 0.802852067 | -4.768801333 |
| A_66_P127411 | Acat2 | 0.093443231 | -0.1341312 | 0.321017662 | 8.185646233 | 1.010167857 | 0.352231407 | 0.802860226 | -4.768923669 |
| A_55_P2152547 | Myl3 | 0.203272467 | -0.291877881 | 0.698422816 | 9.337520681 | 1.009975021 | 0.352316453 | 0.802860226 | -4.769041854 |
| A_51_P368285 | Rflna | 0.302919551 | -0.434981256 | 1.040820358 | 10.55826256 | 1.009946764 | 0.352328916 | 0.802860226 | -4.769059171 |
| A_51_P224468 | Maip1 | 0.110489244 | -0.158671477 | 0.379649964 | 10.66942109 | 1.00989793 | 0.352350456 | 0.802860226 | -4.769089098 |
| A_51_P130666 | Adk | 0.132160863 | -0.189796713 | 0.45411844 | 10.85805274 | 1.009888391 | 0.352354664 | 0.802860226 | -4.769094943 |
| A_52_P389347 | Snap29 | -0.071218742 | -0.244737051 | 0.102299567 | 11.69727909 | -1.009759987 | 0.352411307 | 0.802894644 | -4.769173628 |
| A_55_P2076448 | Tmem255a | 0.088547766 | -0.127257596 | 0.304353128 | 7.95522331 | 1.009449367 | 0.352548362 | 0.803011239 | -4.769363948 |
| A_55_P2080378 | Ttc30a2 | -0.05367129 | -0.184505279 | 0.077162699 | 8.004435516 | -1.009231071 | 0.352644707 | 0.803011239 | -4.769497679 |
| A_51_P244213 | Hapln3 | 0.092644069 | -0.133205539 | 0.318493677 | 9.941103206 | 1.009177231 | 0.352668472 | 0.803011239 | -4.769530659 |
| A_55_P2099418 | Neil2 | -0.233449113 | -0.802576604 | 0.335678378 | 10.73881017 | -1.009140943 | 0.352684491 | 0.803011239 | -4.769552887 |
| A_55_P2001193 | Stx18 | -0.083602784 | -0.287418909 | 0.120213342 | 11.0631014 | -1.009139854 | 0.352684971 | 0.803011239 | -4.769553554 |
| A_51_P278489 | Itfg2 | -0.058357906 | -0.200637827 | 0.083922014 | 8.406059001 | -1.009079193 | 0.35271175 | 0.803011239 | -4.769590711 |
| A_55_P2100550 | Mtif2 | -0.333878252 | -1.147984675 | 0.480228172 | 13.35361935 | -1.008965025 | 0.352762154 | 0.803031407 | -4.769660639 |
| A_55_P2050508 | Spta1 | 0.066069252 | -0.095067733 | 0.227206236 | 7.882084264 | 1.008726242 | 0.352867593 | 0.803101083 | -4.769806879 |
| A_51_P194249 | Stmn4 | -0.421133819 | -1.448260528 | 0.60599289 | 9.653673596 | -1.008707522 | 0.35287586 | 0.803101083 | -4.769818343 |
| A_52_P663526 | Nmrk1 | -0.071427009 | -0.24570339 | 0.102849372 | 8.74468192 | -1.008307722 | 0.353052459 | 0.803342245 | -4.770063148 |
| A_51_P185757 | Casz1 | -0.049174727 | -0.169184768 | 0.070835315 | 8.413924721 | -1.008076597 | 0.353154584 | 0.803342245 | -4.770204644 |
| A_51_P182572 | Phactr1 | -0.1348377 | -0.463924577 | 0.194249177 | 12.99389355 | -1.008021821 | 0.35317879 | 0.803342245 | -4.770238175 |
| A_51_P372895 | Zfp433 | 0.078070836 | -0.112474537 | 0.268616209 | 9.326816213 | 1.007997801 | 0.353189406 | 0.803342245 | -4.770252878 |
| A_55_P2093994 | Maged2 | -0.152638153 | -0.525214681 | 0.219938375 | 14.45265868 | -1.007898444 | 0.353233318 | 0.803342245 | -4.770313697 |

| | | | | | | | | | |
|---|---|---|---|---|---|---|---|---|---|
| **A_51_P394676** | Mef2d | -0.075178946 | -0.258686791 | 0.108328899 | 12.04996212 | -1.007884491 | 0.353239485 | 0.803342245 | -4.770322237 |
| **A_51_P348022** | Nudt21 | -0.104009499 | -0.357910165 | 0.149891167 | 10.2280062 | -1.007809222 | 0.353272755 | 0.803342245 | -4.770368308 |
| **A_55_P1991134** | Fkbp1a | 0.089884996 | -0.129607106 | 0.309377098 | 14.46146747 | 1.007482404 | 0.353417242 | 0.803443718 | -4.770568324 |
| **A_55_P1999197** | Clasp1 | -0.093543167 | -0.321990917 | 0.134904582 | 10.70526263 | -1.007382378 | 0.353461473 | 0.803443718 | -4.770629533 |
| **A_66_P124555** | Rps15 | 0.109564786 | -0.158030349 | 0.377159921 | 13.23579948 | 1.007307212 | 0.353494714 | 0.803443718 | -4.770675527 |
| **A_55_P2087280** | NA | 0.165229407 | -0.238380911 | 0.568839724 | 14.66126724 | 1.007150207 | 0.353564155 | 0.803443718 | -4.770771592 |
| **A_55_P2098450** | Snrnp200 | -0.078004758 | -0.268578794 | 0.112569278 | 11.55078022 | -1.006993174 | 0.353633619 | 0.803443718 | -4.770867665 |
| **A_51_P117995** | Pfkm | 0.065217741 | -0.094117776 | 0.224553259 | 9.029851172 | 1.006983424 | 0.353637933 | 0.803443718 | -4.767559574 |
| **A_52_P101238** | B230354K17Rik | 0.146402744 | -0.211297237 | 0.504102725 | 8.834989135 | 1.006930425 | 0.35366138 | 0.803443718 | -4.770906053 |
| **A_55_P2053929** | NA | 0.234323862 | -0.338309656 | 0.80695738 | 13.07684158 | 1.006720504 | 0.353754262 | 0.803443718 | -4.771034464 |
| **A_51_P156434** | Slc25a33 | -0.05768674 | -0.198659946 | 0.083286466 | 8.366605216 | -1.006719739 | 0.353754601 | 0.803443718 | -4.771034933 |
| **A_51_P215995** | Zswim6 | -0.079023415 | -0.272159435 | 0.114112604 | 8.938389178 | -1.006611022 | 0.353802713 | 0.803443718 | -4.77110143 |
| **A_55_P2014075** | Ercc2 | 0.065784072 | -0.095027733 | 0.226595878 | 10.11060748 | 1.006403159 | 0.353894714 | 0.803443718 | -4.77122856 |
| **A_55_P2121541** | Gm6607 | -0.062578413 | -0.215559371 | 0.090402545 | 15.04863444 | -1.006366869 | 0.353910778 | 0.803443718 | -4.771250753 |
| **A_55_P2068134** | NA | 0.050061488 | -0.072329695 | 0.172452671 | 7.914365703 | 1.006289143 | 0.353945187 | 0.803443718 | -4.771298285 |
| **A_55_P2181222** | Ifi208 | -0.069657212 | -0.239959557 | 0.100645132 | 8.796081504 | -1.006270235 | 0.353953557 | 0.803443718 | -4.771309847 |
| **A_55_P2155996** | NA | -0.077053883 | -0.2654573 | 0.111349533 | 14.61521968 | -1.006178234 | 0.353994289 | 0.803443718 | -4.771366106 |
| **A_52_P570240** | Kbtbd11 | 0.111467911 | -0.161138896 | 0.384074718 | 9.180310091 | 1.005963747 | 0.354089264 | 0.803443718 | -4.771497253 |
| **A_55_P2094574** | Rapgef6 | 0.071468053 | -0.103315908 | 0.246252013 | 10.40047538 | 1.00595727 | 0.354092132 | 0.803443718 | -4.771501213 |
| **A_52_P489729** | Sec61a1 | 0.099677168 | -0.144099508 | 0.343453845 | 14.40397248 | 1.005941452 | 0.354099138 | 0.803443718 | -4.771510884 |
| **A_52_P236448** | Ngfr | 0.048484129 | -0.070114141 | 0.167082399 | 7.75972488 | 1.005750847 | 0.354183557 | 0.803443718 | -4.771627413 |
| **A_55_P2056285** | Sf3b5 | 0.062793411 | -0.090807886 | 0.216394707 | 12.32990075 | 1.00574609 | 0.354185664 | 0.803443718 | -4.771630321 |
| **A_55_P2076176** | NA | -0.08471968 | -0.291957787 | 0.122518427 | 14.10905656 | -1.005735669 | 0.35419028 | 0.803443718 | -4.771636692 |
| **A_51_P407774** | Slc22a21 | -0.068976988 | -0.237748851 | 0.099794874 | 8.66587841 | -1.005479795 | 0.354303634 | 0.803550891 | -4.771793102 |
| **A_65_P12530** | NA | -0.068855464 | -0.237359964 | 0.099649036 | 8.032287292 | -1.005300891 | 0.354382907 | 0.803550891 | -4.771902448 |
| **A_51_P201480** | Stat3 | -0.130841317 | -0.451059936 | 0.189377301 | 11.78986575 | -1.005234746 | 0.35441222 | 0.803550891 | -4.765171362 |
| **A_51_P335081** | Gnb1 | 0.140928214 | -0.203998714 | 0.485855142 | 12.51338502 | 1.005171138 | 0.35444041 | 0.803550891 | -4.771981746 |
| **A_55_P2091245** | NA | -0.087481823 | -0.301660123 | 0.126696477 | 16.30393472 | -1.004873794 | 0.354572213 | 0.803550891 | -4.772163443 |
| **A_52_P595642** | Smim7 | -0.086633469 | -0.298738797 | 0.12547186 | 10.38853647 | -1.004854745 | 0.354580658 | 0.803550891 | -4.772175083 |
| **A_52_P184149** | Mthfd2 | -0.067093702 | -0.231370333 | 0.097182928 | 8.664161016 | -1.004789609 | 0.354609536 | 0.803550891 | -4.77221488 |
| **A_51_P140819** | Wdr36 | -0.083269362 | -0.287151981 | 0.120613256 | 9.431497909 | -1.004787443 | 0.354610497 | 0.803550891 | -4.772216203 |
| **A_55_P2163319** | Peg3 | -0.126681259 | -0.436857469 | 0.183494951 | 8.491971497 | -1.004784781 | 0.354611677 | 0.803550891 | -4.77221783 |
| **A_52_P501816** | Slc6a14 | 0.051071875 | -0.074004688 | 0.176148438 | 7.867542842 | 1.00455803 | 0.354712224 | 0.803684512 | -4.772356361 |
| **A_55_P2064538** | Zfp985 | -0.074156235 | -0.255832823 | 0.107520354 | 8.520796674 | -1.004194568 | 0.354873439 | 0.803955545 | -4.772578377 |
| **A_55_P2001218** | Cd80 | -0.083529785 | -0.288213504 | 0.121153933 | 8.400271638 | -1.003985016 | 0.354966413 | 0.803993826 | -4.772706357 |
| **A_51_P480119** | Prelid2 | 0.081385609 | -0.118047143 | 0.280818361 | 9.292706741 | 1.003968984 | 0.354973527 | 0.803993826 | -4.772716147 |
| **A_55_P1965931** | Nat8f1 | 0.055523926 | -0.080574784 | 0.191622636 | 9.078864017 | 1.003680115 | 0.355101728 | 0.804034768 | -4.772892542 |
| **A_55_P2096395** | Cdc42bpa | -0.139987441 | -0.483135843 | 0.20316096 | 10.06367491 | -1.003636064 | 0.355121281 | 0.804034768 | -4.772919438 |
| **A_55_P2129354** | Pilra | -0.051636874 | -0.178213753 | 0.074940005 | 7.879285266 | -1.003632515 | 0.355122856 | 0.804034768 | -4.772921605 |
| **A_55_P2152901** | 2900040C04Rik | 0.296343379 | -0.430175086 | 1.022861843 | 11.47539956 | 1.003500827 | 0.355181315 | 0.804034768 | -4.773002006 |
| **A_55_P2184897** | Lbr | -0.043461818 | -0.150017627 | 0.063093991 | 7.859062356 | -1.003459661 | 0.35519959 | 0.804034768 | -4.773027139 |
| **A_55_P2029740** | Gm10240 | 0.106287724 | -0.154339535 | 0.366914983 | 14.87780809 | 1.00330375 | 0.355268815 | 0.804097297 | -4.773122319 |
| **A_51_P267494** | Cdc42ep3 | -0.09387976 | -0.324206402 | 0.136446883 | 10.25799936 | -1.002759893 | 0.35551037 | 0.804223035 | -4.773454262 |
| **A_52_P588483** | Fbln1 | -0.199923627 | -0.690438754 | 0.2905915 | 12.71508575 | -1.002722775 | 0.355526861 | 0.804223035 | -4.773476913 |
| **A_55_P1984386** | Depdc5 | 0.050471846 | -0.073372086 | 0.174315778 | 8.641865371 | 1.002636744 | 0.355565085 | 0.804223035 | -4.773529411 |
| **A_51_P497692** | Mageh1 | -0.058969544 | -0.203691176 | 0.085752088 | 9.161008029 | -1.002451756 | 0.355647289 | 0.804223035 | -4.773642286 |
| **A_55_P2291776** | 4833447P13Rik | -0.071858903 | -0.248231287 | 0.104513481 | 8.427879435 | -1.002349462 | 0.355692751 | 0.804223035 | -4.773704698 |
| **A_55_P2136204** | Rnf38 | 0.095539084 | -0.138986486 | 0.330064653 | 12.14891159 | 1.002213008 | 0.355753404 | 0.804223035 | -4.773787946 |
| **A_55_P2109485** | Milr1 | 0.149220647 | -0.217099656 | 0.51554095 | 16.40075812 | 1.002160064 | 0.355776939 | 0.804223035 | -4.773820244 |
| **A_55_P1971330** | NA | 0.054744356 | -0.079677921 | 0.189166634 | 8.46840925 | 1.001929735 | 0.355879341 | 0.804223035 | -4.773960744 |

| | | | | | | | | | |
|---|---|---|---|---|---|---|---|---|---|
| **A_51_P225948** | Snx33 | -0.091465069 | -0.316064046 | 0.133133908 | 10.03529405 | -1.001882186 | 0.355900484 | 0.804223035 | -4.773989746 |
| **A_55_P1975370** | Apoc1 | 0.704651969 | -1.025765974 | 2.435069912 | 13.53188364 | 1.00182721 | 0.355924931 | 0.804223035 | -4.774023277 |
| **A_52_P349467** | Tmlhe | -0.06001072 | -0.207381185 | 0.087359746 | 7.938611365 | -1.001815032 | 0.355930347 | 0.804223035 | -4.774030705 |
| **A_52_P507382** | Unc93b1 | -0.075560535 | -0.261121417 | 0.110000348 | 8.910519841 | -1.001792455 | 0.355940386 | 0.804223035 | -4.774044474 |
| **A_55_P2090406** | H13 | -0.12606551 | -0.435674732 | 0.183543711 | 10.55105575 | -1.001732033 | 0.355967257 | 0.804223035 | -4.774081326 |
| **A_55_P2008651** | Qdpr | 0.150985499 | -0.219855344 | 0.521826342 | 11.50866438 | 1.001651955 | 0.356002872 | 0.804223035 | -4.774130162 |
| **A_55_P2135025** | NA | 0.057881153 | -0.084304669 | 0.200066976 | 10.57782809 | 1.001497903 | 0.356071394 | 0.804223035 | -4.774224107 |
| **A_51_P334979** | Apoc2 | 0.652273594 | -0.950222396 | 2.254769585 | 12.96210486 | 1.001387138 | 0.356120669 | 0.804223035 | -4.774291649 |
| **A_51_P469661** | Stk38 | -0.112425139 | -0.388654856 | 0.163804577 | 10.68407918 | -1.001295352 | 0.356161506 | 0.804223035 | -4.774347615 |
| **A_55_P2046408** | NA | 0.146915098 | -0.214114106 | 0.507944303 | 8.607300951 | 1.001136415 | 0.356232226 | 0.804223035 | -4.774444517 |
| **A_55_P2383877** | 2810040C05Rik | -0.112873855 | -0.390270204 | 0.164522495 | 8.256320311 | -1.001063849 | 0.356264519 | 0.804223035 | -4.774488758 |
| **A_51_P283016** | Slc1a6 | 0.093520298 | -0.136326257 | 0.323366854 | 8.540925968 | 1.001006844 | 0.356289889 | 0.804223035 | -4.774523509 |
| **A_55_P2023441** | Elp4 | -0.085256078 | -0.294794405 | 0.124282249 | 8.589120193 | -1.000992998 | 0.356296051 | 0.804223035 | -4.77453195 |
| **A_55_P2001048** | Rom1 | 0.127629117 | -0.186112691 | 0.441370925 | 10.91654424 | 1.000798237 | 0.35638274 | 0.804223035 | -4.774650672 |
| **A_51_P150087** | Abhd10 | 0.075031142 | -0.109417909 | 0.259480193 | 8.522260896 | 1.000770035 | 0.356395294 | 0.804223035 | -4.774667862 |
| **A_55_P2003123** | Jkamp | 0.094135329 | -0.137287557 | 0.325558214 | 13.56978033 | 1.000726733 | 0.356414571 | 0.804223035 | -4.774694256 |
| **A_55_P2043122** | Arsg | 0.179237142 | -0.261423807 | 0.619898091 | 9.180680547 | 1.00067398 | 0.356438056 | 0.804223035 | -4.77472641 |
| **A_55_P2010936** | Fbxo17 | 0.135572719 | -0.197743184 | 0.468888622 | 9.722756285 | 1.000657451 | 0.356445415 | 0.804223035 | -4.774736484 |
| **A_55_P2015675** | AU015836 | -0.163446554 | -0.565311068 | 0.238417959 | 12.97194726 | -1.000611004 | 0.356466094 | 0.804223035 | -4.774764792 |
| **A_55_P2097244** | Bptf | -0.164225874 | -0.568042577 | 0.239590829 | 12.52895526 | -1.000521597 | 0.356505903 | 0.804223035 | -4.774819283 |
| **A_51_P384640** | Mrpl10 | -0.062330706 | -0.215604948 | 0.090943537 | 9.690890163 | -1.000465295 | 0.356530973 | 0.804223035 | -4.774853595 |
| **A_51_P416919** | Ube2j2 | -0.090482669 | -0.313112593 | 0.132147255 | 10.0182516 | -0.999887241 | 0.356788453 | 0.804605034 | -4.775205814 |
| **A_55_P2031631** | Igf1 | 0.150631744 | -0.220013476 | 0.521276964 | 8.450273415 | 0.999832531 | 0.35681283 | 0.804605034 | -4.775239143 |
| **A_52_P330540** | Mis18a | 0.126601722 | -0.184923758 | 0.438127202 | 9.395411673 | 0.999804761 | 0.356825203 | 0.804605034 | -4.77525606 |
| **A_51_P266640** | Nob1 | -0.045731679 | -0.158280228 | 0.06681687 | 7.91877699 | -0.999646413 | 0.356895768 | 0.804614135 | -4.775352519 |
| **A_66_P111437** | Ube2j2 | 0.098014543 | -0.143214491 | 0.339243577 | 15.08941329 | 0.999608882 | 0.356912494 | 0.804614135 | -4.775375379 |
| **A_52_P627269** | Ces2b | 0.190160113 | -0.278108016 | 0.658428241 | 8.229296344 | 0.999065563 | 0.357154705 | 0.804929038 | -4.775706268 |
| **A_66_P140613** | NA | -0.099667178 | -0.345097312 | 0.145762957 | 8.622074269 | -0.999064295 | 0.35715527 | 0.804929038 | -4.77570704 |
| **A_52_P627816** | Tgm1 | -0.088643921 | -0.306960304 | 0.129672463 | 10.23981796 | -0.99892249 | 0.357218508 | 0.804929038 | -4.775793383 |
| **A_55_P2145389** | Flrt3 | 0.10686981 | -0.156334289 | 0.370073909 | 8.682766715 | 0.99892194 | 0.357218754 | 0.804929038 | -4.775793719 |
| **A_55_P2024799** | NA | -0.05717386 | -0.198000242 | 0.083652522 | 11.94861969 | -0.998809475 | 0.357268914 | 0.804948226 | -4.775862191 |
| **A_55_P2071536** | NA | 0.057597782 | -0.084326865 | 0.199522429 | 7.947726448 | 0.998428796 | 0.357438742 | 0.805027059 | -4.776093931 |
| **A_55_P1972230** | NA | -0.185961678 | -0.644262363 | 0.272339007 | 10.39213253 | -0.998256423 | 0.357515662 | 0.805027059 | -4.776198846 |
| **A_55_P2000623** | Stag1 | -0.054937972 | -0.190334776 | 0.080458833 | 11.07823997 | -0.998236326 | 0.357524631 | 0.805027059 | -4.776211078 |
| **A_52_P24951** | Ado | -0.061734122 | -0.213884211 | 0.090415967 | 9.082438351 | -0.99821072 | 0.357536059 | 0.805027059 | -4.776226662 |
| **A_55_P1968438** | Arxes1 | 0.115822102 | -0.169641906 | 0.40128611 | 8.143756917 | 0.998180882 | 0.357549376 | 0.805027059 | -4.776244821 |
| **A_55_P2068154** | Anapc15 | 0.100845666 | -0.147708738 | 0.349400071 | 10.26868796 | 0.998170981 | 0.357553795 | 0.805027059 | -4.776250847 |
| **A_52_P589119** | Rhno1 | -0.046700506 | -0.161848782 | 0.06844777 | 7.819353578 | -0.997776586 | 0.357729855 | 0.80532965 | -4.776490844 |
| **A_52_P75146** | Lyrm2 | -0.069694752 | -0.241629744 | 0.10224024 | 10.15352514 | -0.997252121 | 0.357964087 | 0.805679938 | -4.776809902 |
| **A_51_P346641** | NA | -0.044394053 | -0.153915626 | 0.06512752 | 7.827136612 | -0.997227597 | 0.357975043 | 0.805679938 | -4.776824819 |
| **A_52_P537852** | Fam120b | -0.05300581 | -0.183783277 | 0.077771656 | 8.022603075 | -0.997148227 | 0.358010502 | 0.805679938 | -4.776873094 |
| **A_51_P260658** | Amdhd2 | 0.180964199 | -0.265563651 | 0.627492048 | 12.10161558 | 0.997041595 | 0.358058145 | 0.80569335 | -4.776937948 |
| **A_55_P1981415** | Gsk3b | -0.119317755 | -0.413767619 | 0.175132109 | 12.34018569 | -0.996925898 | 0.358109844 | 0.805708084 | -4.77700831 |
| **A_55_P2173768** | Pkhd1 | 0.296775301 | -0.435663418 | 1.029214019 | 8.416863502 | 0.996840381 | 0.358148061 | 0.805708084 | -4.777060314 |
| **A_55_P2021345** | Gm14717 | 0.103945085 | -0.152667787 | 0.360557956 | 9.110757867 | 0.996539882 | 0.358282377 | 0.805916451 | -4.777243033 |
| **A_55_P2057405** | Cpt1a | -0.046728385 | -0.162119175 | 0.068662405 | 7.942161488 | -0.996273981 | 0.358401263 | 0.806002086 | -4.777404687 |
| **A_55_P2062034** | Tsen34 | 0.135140201 | -0.198583424 | 0.468863827 | 14.94741238 | 0.996246416 | 0.358413589 | 0.806002086 | -4.777421444 |
| **A_52_P417859** | Gid4 | -0.086071039 | -0.298652756 | 0.126510678 | 9.30041262 | -0.996093943 | 0.358481777 | 0.806002086 | -4.777514126 |
| **A_55_P2040425** | Guf1 | -0.053911901 | -0.187067319 | 0.079243516 | 7.982564301 | -0.99608172 | 0.358487244 | 0.806002086 | -4.777521556 |
| **A_55_P2156293** | Pcyox1 | -0.250789438 | -0.8706215 | 0.369042624 | 12.44459709 | -0.995415428 | 0.358785342 | 0.806578491 | -4.777926466 |

| | | | | | | | | | |
|---|---|---|---|---|---|---|---|---|---|
| **A_51_P426276** | Pdk2 | 0.065490557 | -0.096406998 | 0.227388112 | 8.213338508 | 0.995193564 | 0.358884647 | 0.806707913 | -4.778061259 |
| **A_51_P460774** | Smyd4 | 0.140314545 | -0.206843849 | 0.487472938 | 8.944820846 | 0.994361233 | 0.359257391 | 0.807451874 | -4.778566777 |
| **A_55_P2117169** | Zmym1 | -0.112562164 | -0.400904919 | 0.17578059 | 10.83178794 | -1.011657427 | 0.359301601 | 0.807457347 | -4.725938149 |
| **A_51_P500424** | Dnttip1 | -0.050385716 | -0.175098192 | 0.07432676 | 11.3084655 | -0.993954933 | 0.359439457 | 0.807673247 | -4.778813452 |
| **A_55_P1975011** | Ufc1 | 0.077283084 | -0.114029695 | 0.268595864 | 12.76716242 | 0.993824337 | 0.359497995 | 0.807684376 | -4.778892728 |
| **A_51_P328400** | Cops2 | -0.064119289 | -0.222856034 | 0.094617455 | 10.84118297 | -0.993757439 | 0.359527983 | 0.807684376 | -4.778933334 |
| **A_55_P2422650** | 5031425E22Rik | 0.576701839 | -0.851275019 | 2.004678697 | 12.96856784 | 0.993571968 | 0.359611135 | 0.807777295 | -4.779045905 |
| **A_55_P2052824** | Lrrc14b | 0.205262785 | -0.303178818 | 0.713704388 | 7.999627067 | 0.993203636 | 0.359776314 | 0.808054423 | -4.779269425 |
| **A_55_P2046178** | NA | 0.121365716 | -0.179329032 | 0.422060465 | 16.09106334 | 0.992977352 | 0.359877821 | 0.808137736 | -4.77940672 |
| **A_55_P1967341** | Gm9104 | -0.079248146 | -0.275601199 | 0.117104906 | 12.37854407 | -0.992934544 | 0.359897027 | 0.808137736 | -4.77943269 |
| **A_55_P2121608** | NA | -0.234762661 | -0.816665835 | 0.347140514 | 10.8001661 | -0.992538723 | 0.360074649 | 0.8084314 | -4.779672798 |
| **A_55_P2102095** | 1810010H24Rik | 0.073130657 | -0.108161852 | 0.254423166 | 7.872891898 | 0.992404624 | 0.36013484 | 0.8084314 | -4.77975413 |
| **A_51_P296100** | Rhot2 | 0.10717616 | -0.158538943 | 0.372891262 | 10.29105844 | 0.992318561 | 0.360173475 | 0.8084314 | -4.779806324 |
| **A_55_P2063970** | Ube2l3 | 0.127733359 | -0.188963291 | 0.44443001 | 13.44777338 | 0.992270377 | 0.360195107 | 0.8084314 | -4.779835545 |
| **A_52_P406828** | Dkc1 | -0.045569063 | -0.158580454 | 0.067442329 | 7.907099965 | -0.992012251 | 0.360311007 | 0.808597638 | -4.779992069 |
| **A_55_P2087352** | NA | -0.102463224 | -0.356605887 | 0.15167944 | 12.33311621 | -0.99188107 | 0.36036992 | 0.808615078 | -4.780071607 |
| **A_55_P2009910** | Stxbp3 | -0.068269164 | -0.237625015 | 0.101086687 | 11.32749695 | -0.991730212 | 0.360437679 | 0.808615078 | -4.780163066 |
| **A_51_P489522** | Ctla2b | -0.605024797 | -2.105941324 | 0.895891731 | 16.79668898 | -0.991712524 | 0.360445625 | 0.808615078 | -4.780173789 |
| **A_51_P497724** | Apol7a | 0.225170727 | -0.333472798 | 0.783814252 | 8.29330323 | 0.991622388 | 0.360486116 | 0.808615078 | -4.780228431 |
| **A_51_P104820** | Alkbh5 | -0.073027319 | -0.25424457 | 0.108189932 | 8.359572087 | -0.991413849 | 0.360579809 | 0.808731392 | -4.780354839 |
| **A_55_P2097828** | Gmfg | 0.139881491 | -0.207398266 | 0.487161247 | 10.9126211 | 0.990945903 | 0.360790122 | 0.809109208 | -4.780638429 |
| **A_55_P2125006** | Gpatch1 | -0.134738071 | -0.469285249 | 0.199809106 | 10.33738782 | -0.990836783 | 0.360839179 | 0.809125346 | -4.780704548 |
| **A_52_P380301** | Unc5c | 0.050740174 | -0.075269671 | 0.176750019 | 7.752715748 | 0.990641771 | 0.360926863 | 0.809215017 | -4.780822701 |
| **A_55_P2020577** | Pcolce | 0.318474375 | -0.4725372 | 1.10948595 | 11.62434572 | 0.990514457 | 0.360984117 | 0.809215017 | -4.780899829 |
| **A_51_P155843** | Igsf10 | 0.053979348 | -0.080098216 | 0.188056912 | 7.835301654 | 0.990468547 | 0.361004764 | 0.809215017 | -4.78092764 |
| **A_55_P2114569** | Zfp24 | -0.090729707 | -0.316153896 | 0.134694482 | 11.00782711 | -0.990189133 | 0.36113045 | 0.809402883 | -4.781096887 |
| **A_66_P102467** | Casz1 | -0.043451037 | -0.151437937 | 0.064535864 | 7.694832484 | -0.989915737 | 0.361253462 | 0.809438811 | -4.781262461 |
| **A_55_P1981050** | Gemin8 | -0.044801943 | -0.156150331 | 0.066546445 | 7.868483846 | -0.989878926 | 0.361270027 | 0.809438811 | -4.781284752 |
| **A_52_P249920** | Zscan29 | -0.060091433 | -0.209441195 | 0.089258329 | 8.471751132 | -0.989867784 | 0.361275042 | 0.809438811 | -4.781291499 |
| **A_55_P2036942** | Gm4535 | 0.082243954 | -0.122181007 | 0.286668914 | 8.727552432 | 0.989781245 | 0.361313988 | 0.809438811 | -4.781343901 |
| **A_55_P2102540** | Olfml3 | 0.04284481 | -0.063660959 | 0.149350579 | 7.706486732 | 0.989678765 | 0.361360113 | 0.809448327 | -4.781405953 |
| **A_55_P2150053** | Yipf5 | -0.061202499 | -0.213377431 | 0.090972434 | 10.39996781 | -0.989453065 | 0.361461713 | 0.809564472 | -4.781542601 |
| **A_55_P2072985** | Sptan1 | 0.106000636 | -0.157581273 | 0.369582545 | 13.05021459 | 0.989377511 | 0.36149573 | 0.809564472 | -4.78158834 |
| **A_55_P1960157** | Bcat1 | -0.106366575 | -0.370900259 | 0.15816711 | 9.204986136 | -0.989221065 | 0.361566174 | 0.809583596 | -4.781683044 |
| **A_55_P2153937** | NA | -0.078176583 | -0.272611298 | 0.116258132 | 8.468107686 | -0.989172513 | 0.361588038 | 0.809583596 | -4.781712432 |
| **A_55_P2048279** | Tlr13 | -0.046760442 | -0.163086291 | 0.069565408 | 9.198125707 | -0.98894362 | 0.361691128 | 0.809636986 | -4.78185097 |
| **A_52_P222593** | Cebpzos | 0.168803187 | -0.251148511 | 0.588754884 | 11.23393709 | 0.988895643 | 0.361712739 | 0.809636986 | -4.781880006 |
| **A_66_P127956** | Zfp947 | 0.138493248 | -0.20607866 | 0.483065156 | 11.74403545 | 0.98882149 | 0.361746143 | 0.809636986 | -4.781924881 |
| **A_52_P77227** | Rnf212 | 0.04240723 | -0.063130326 | 0.147944785 | 7.769697229 | 0.988557728 | 0.361864981 | 0.809636986 | -4.782084488 |
| **A_55_P1958887** | DXBay18 | -0.135466913 | -0.47260521 | 0.201671384 | 12.43073903 | -0.988540139 | 0.361872907 | 0.809636986 | -4.78209513 |
| **A_55_P1961645** | Vcp | 0.093928013 | -0.139866304 | 0.32772233 | 14.01130676 | 0.988394561 | 0.361938512 | 0.809636986 | -4.782183211 |
| **A_51_P419286** | Batf3 | 0.2261854 | -0.336823744 | 0.789194544 | 9.961093303 | 0.988367105 | 0.361950886 | 0.809636986 | -4.782199822 |
| **A_55_P2180406** | Cdkn2aipnl | 0.08691799 | -0.129445314 | 0.303281294 | 8.677460371 | 0.98831465 | 0.361974528 | 0.809636986 | -4.782231556 |
| **A_55_P2040815** | Lrrc45 | -0.125173011 | -0.436811719 | 0.186465697 | 9.72026871 | -0.988162724 | 0.362043008 | 0.809636986 | -4.782323463 |
| **A_51_P499838** | Bst1 | 0.353494696 | -0.526713186 | 1.233702579 | 11.57154156 | 0.988022448 | 0.362106247 | 0.809636986 | -4.782408316 |
| **A_52_P218379** | Kif26a | -0.124765633 | -0.435434505 | 0.185903239 | 9.841981012 | -0.988021504 | 0.362106673 | 0.809636986 | -4.782408887 |
| **A_66_P120558** | Pam16 | -0.092282926 | -0.322117871 | 0.137552019 | 12.13990483 | -0.987812366 | 0.362200972 | 0.809636986 | -4.78253538 |
| **A_55_P2083153** | 1700001J03Rik | -0.117160578 | -0.408961551 | 0.174640394 | 8.158721566 | -0.987788429 | 0.362211766 | 0.809636986 | -4.782549856 |
| **A_55_P2133840** | Sipa1 | 0.071027434 | -0.105879082 | 0.247933949 | 11.16172937 | 0.987760053 | 0.362224563 | 0.809636986 | -4.782567018 |
| **A_55_P2005720** | NA | 0.21698027 | -0.323524919 | 0.75748546 | 11.43799995 | 0.987619239 | 0.36228807 | 0.809636986 | -4.782652175 |

| | | | | | | | | | |
|---|---|---|---|---|---|---|---|---|---|
| **A_52_P339791** | Hdgf | 0.070321928 | -0.104869109 | 0.245512966 | 8.301664938 | 0.987524888 | 0.362330627 | 0.809636986 | -4.782709229 |
| **A_52_P214927** | Zmynd8 | 0.082875248 | -0.123589788 | 0.289340285 | 10.00219301 | 0.987523615 | 0.362331201 | 0.809636986 | -4.782709999 |
| **A_51_P315085** | Mia3 | -0.144928902 | -0.506014745 | 0.21615694 | 12.64611766 | -0.987446797 | 0.362365853 | 0.809636986 | -4.782756448 |
| **A_55_P2153923** | Slc17a5 | -0.066261613 | -0.231428841 | 0.098905616 | 8.513796701 | -0.986977572 | 0.362577572 | 0.809898459 | -4.783040128 |
| **A_55_P2102738** | NA | -0.363291494 | -1.268854556 | 0.542271568 | 14.39651639 | -0.986973958 | 0.362579203 | 0.809898459 | -4.783042313 |
| **A_55_P2124016** | Nipsnap1 | 0.146644219 | -0.218998 | 0.512286438 | 8.123999901 | 0.986683293 | 0.362710404 | 0.809898459 | -4.783217999 |
| **A_55_P2047698** | Pop7 | 0.15006843 | -0.224161506 | 0.524298365 | 12.42176083 | 0.986551989 | 0.362769685 | 0.809898459 | -4.783297353 |
| **A_55_P2143041** | Phgdh | 0.142781867 | -0.213282255 | 0.498845988 | 9.970853495 | 0.986538421 | 0.362775811 | 0.809898459 | -4.783305552 |
| **A_55_P2110567** | Proca1 | 0.115177777 | -0.172048363 | 0.402403917 | 8.804137333 | 0.98653817 | 0.362775924 | 0.809898459 | -4.783305703 |
| **A_66_P105711** | Ap4s1 | -0.052536542 | -0.183555677 | 0.078482593 | 8.3351093 | -0.986497336 | 0.362794361 | 0.809898459 | -4.78333038 |
| **A_55_P2099141** | NA | -0.112693564 | -0.393763081 | 0.168375954 | 14.39008496 | -0.986403325 | 0.362836812 | 0.809898459 | -4.783387189 |
| **A_51_P409724** | Secisbp2 | -0.073369499 | -0.256370211 | 0.109631214 | 8.628958019 | -0.98635201 | 0.362859985 | 0.809898459 | -4.783418197 |
| **A_55_P1984523** | NA | -0.256200726 | -0.895399602 | 0.38299815 | 11.55957237 | -0.986083103 | 0.362981439 | 0.810040206 | -4.783580671 |
| **A_55_P2072776** | Fpgs | -0.068518218 | -0.239475233 | 0.102438798 | 10.11504693 | -0.986025835 | 0.363007308 | 0.810040206 | -4.783615269 |
| **A_51_P109449** | Ltb4r1 | -0.051297072 | -0.17931712 | 0.076722976 | 7.949989797 | -0.98578852 | 0.363114526 | 0.810185925 | -4.783758629 |
| **A_51_P289588** | Ube2e3 | -0.082928949 | -0.290004959 | 0.124147061 | 12.66238107 | -0.985247944 | 0.363358849 | 0.810596451 | -4.784085106 |
| **A_51_P504588** | Tm2d2 | -0.062395913 | -0.218214048 | 0.093422223 | 10.33600807 | -0.9851612 | 0.363398067 | 0.810596451 | -4.784137483 |
| **A_55_P2000133** | Cdh2 | 0.083678728 | -0.1253042 | 0.292661656 | 8.547426075 | 0.985084358 | 0.36343281 | 0.810596451 | -4.784183881 |
| **A_51_P411345** | Mogat2 | 0.444760951 | -0.666156875 | 1.555678778 | 10.00309411 | 0.984949304 | 0.363493881 | 0.810596451 | -4.78426542 |
| **A_52_P240542** | Id2 | -0.09999065 | -0.349764038 | 0.149782738 | 12.04153785 | -0.984877893 | 0.363526176 | 0.810596451 | -4.784308533 |
| **A_51_P501815** | Smdt1 | 0.103985848 | -0.155782553 | 0.363754249 | 14.36680052 | 0.984820501 | 0.363552132 | 0.810596451 | -4.784343179 |
| **A_55_P2346639** | Sec14l3 | -0.048939301 | -0.171206171 | 0.073327569 | 7.731784948 | -0.984732189 | 0.363592075 | 0.810596451 | -4.78439649 |
| **A_55_P2134597** | NA | 0.051514991 | -0.077234813 | 0.180264796 | 8.200749584 | 0.984365118 | 0.363758139 | 0.810851559 | -4.784618048 |
| **A_55_P2048769** | Zfp454 | 0.041910909 | -0.062864316 | 0.146686134 | 7.840801939 | 0.984096398 | 0.363879747 | 0.810851559 | -4.784780211 |
| **A_55_P2079561** | Lilra6 | -0.056394644 | -0.197417376 | 0.084628087 | 8.11951878 | -0.98382511 | 0.36400255 | 0.810851559 | -4.784943897 |
| **A_51_P443754** | Hivep2 | 0.064565502 | -0.096890506 | 0.226021511 | 8.807302603 | 0.983819632 | 0.36400503 | 0.810851559 | -4.784947202 |
| **A_52_P381484** | Spon2 | 0.131397613 | -0.19719016 | 0.459985386 | 9.624547555 | 0.983796423 | 0.364015537 | 0.810851559 | -4.784961204 |
| **A_52_P437230** | Rbx1 | 0.118373739 | -0.177664413 | 0.414411891 | 14.60793472 | 0.983732103 | 0.364044658 | 0.810851559 | -4.785000008 |
| **A_55_P2150633** | Gadd45gip1 | 0.062492829 | -0.093797127 | 0.218782785 | 10.88616207 | 0.983712707 | 0.36405344 | 0.810851559 | -4.785011709 |
| **A_51_P320888** | Cln5 | 0.155399424 | -0.233265861 | 0.544064709 | 12.80239811 | 0.983655026 | 0.364079558 | 0.810851559 | -4.785046506 |
| **A_55_P2111770** | Klhl13 | -0.111066726 | -0.388855507 | 0.166722055 | 15.26069637 | -0.983645096 | 0.364084054 | 0.810851559 | -4.785052496 |
| **A_55_P1990648** | Ndufaf4 | 0.076992362 | -0.115613162 | 0.269597886 | 8.527566792 | 0.98344045 | 0.364176728 | 0.810928996 | -4.78517594 |
| **A_55_P2110671** | NA | -0.0840079 | -0.294175894 | 0.126160094 | 11.95621673 | -0.983383034 | 0.364202733 | 0.810928996 | -4.785210571 |
| **A_51_P238523** | Shisa4 | -0.078478507 | -0.274855112 | 0.117898098 | 8.769174387 | -0.983173464 | 0.364297661 | 0.811046936 | -4.785336965 |
| **A_51_P349727** | Slc25a45 | -0.0428783 | -0.150200129 | 0.064443529 | 7.987611404 | -0.982921099 | 0.364412001 | 0.811159509 | -4.785489147 |
| **A_55_P2003133** | Ngrn | -0.140909921 | -0.493615013 | 0.211795171 | 12.41253006 | -0.982876618 | 0.364432157 | 0.811159509 | -4.785515968 |
| **A_55_P2068607** | Rnf19b | -0.096904294 | -0.339501592 | 0.145693004 | 11.37785335 | -0.982711923 | 0.364506795 | 0.811232222 | -4.785615267 |
| **A_52_P116628** | Fam234b | -0.041680179 | -0.146037491 | 0.062677133 | 7.832616673 | -0.982597947 | 0.364558455 | 0.811253784 | -4.785683981 |
| **A_52_P150525** | Ndufaf6 | 0.101816572 | -0.153194327 | 0.35682747 | 9.437445057 | 0.982265501 | 0.364709169 | 0.81146951 | -4.785884377 |
| **A_55_P2062148** | Arl1 | 0.060215066 | -0.090623545 | 0.211053677 | 10.88328705 | 0.982113777 | 0.364777969 | 0.81146951 | -4.785975823 |
| **A_55_P2174258** | Pin4 | -0.172576759 | -0.604884578 | 0.25973106 | 11.60679131 | -0.982106037 | 0.364781479 | 0.81146951 | -4.785980487 |
| **A_52_P250590** | Dnajc13 | -0.117498101 | -0.411922121 | 0.176925919 | 9.563710197 | -0.981808467 | 0.364916446 | 0.81146951 | -4.786159808 |
| **A_55_P1975090** | NA | 0.111552023 | -0.167977342 | 0.391081388 | 11.18579538 | 0.98179129 | 0.364924238 | 0.81146951 | -4.786170158 |
| **A_52_P541270** | Crebl2 | -0.065333554 | -0.229058053 | 0.098390944 | 10.11378978 | -0.981729362 | 0.364952331 | 0.81146951 | -4.786207473 |
| **A_52_P504330** | Ankib1 | -0.063996695 | -0.22437684 | 0.09638345 | 9.639132571 | -0.98169393 | 0.364968406 | 0.81146951 | -4.786228821 |
| **A_55_P1967191** | Gm4632 | 0.163349175 | -0.24603616 | 0.572734511 | 8.591730805 | 0.981643573 | 0.364991252 | 0.81146951 | -4.786259162 |
| **A_55_P1984640** | Ube2f | -0.064632168 | -0.226634746 | 0.097370409 | 11.92612983 | -0.981512794 | 0.365050591 | 0.811477229 | -4.786337953 |
| **A_55_P2122546** | Csnk1e | 0.079186946 | -0.119311528 | 0.277685421 | 11.54739313 | 0.981444138 | 0.365081745 | 0.811477229 | -4.786379314 |
| **A_55_P2181356** | Cdh11 | -0.04880882 | -0.17118386 | 0.07356622 | 8.103668485 | -0.981238606 | 0.365175023 | 0.811477229 | -4.786503124 |
| **A_51_P254912** | Rfc2 | -0.061465446 | -0.215596458 | 0.092665566 | 9.626614419 | -0.981092982 | 0.365241125 | 0.811477229 | -4.786590837 |

| | | | | | | | | | |
|---|---|---|---|---|---|---|---|---|---|
| **A_52_P159429** | AU021092 | -0.051935763 | -0.182207827 | 0.0783363 | 7.962437868 | -0.98080888 | 0.36537011 | 0.811477229 | -4.786761935 |
| **A_52_P558939** | Rab3gap1 | -0.065323785 | -0.229186864 | 0.098539294 | 8.741285158 | -0.980752428 | 0.365395744 | 0.811477229 | -4.786795929 |
| **A_55_P2164258** | Prl7b1 | -0.773643718 | -2.714371404 | 1.167083968 | 14.1510547 | -0.980721204 | 0.365409924 | 0.811477229 | -4.786814731 |
| **A_55_P2156625** | NA | 0.104696168 | -0.157963642 | 0.367355979 | 12.79703119 | 0.980632596 | 0.365450164 | 0.811477229 | -4.786868086 |
| **A_51_P211305** | Alpk3 | 0.089611294 | -0.135225939 | 0.314448526 | 7.860871978 | 0.980536308 | 0.365493895 | 0.811477229 | -4.786926061 |
| **A_52_P650028** | Dock10 | -0.059674774 | -0.209420603 | 0.090071054 | 8.022234408 | -0.980404315 | 0.36555385 | 0.811477229 | -4.787005529 |
| **A_52_P160955** | Sox11 | -0.155286737 | -0.54497319 | 0.234399716 | 8.489081468 | -0.98036595 | 0.365571278 | 0.811477229 | -4.787028626 |
| **A_66_P129622** | Phldb3 | -0.144088648 | -0.505690019 | 0.217512723 | 12.51436411 | -0.980322253 | 0.365591129 | 0.811477229 | -4.787054932 |
| **A_51_P101573** | Klc4 | 0.081127093 | -0.122473456 | 0.284727642 | 10.77465761 | 0.980293523 | 0.365604181 | 0.811477229 | -4.787072228 |
| **A_51_P221651** | Coq8a | -0.04833029 | -0.16962995 | 0.07296937 | 7.971201841 | -0.980232233 | 0.365632026 | 0.811477229 | -4.787109124 |
| **A_51_P224593** | Arl8a | 0.067146629 | -0.101401286 | 0.235694545 | 10.7997243 | 0.980099109 | 0.365692513 | 0.811477229 | -4.787189257 |
| **A_52_P517001** | Rpf1 | -0.073159775 | -0.256814002 | 0.110494452 | 9.801099256 | -0.980032764 | 0.365722661 | 0.811477229 | -4.787229192 |
| **A_52_P456750** | Aph1b | -0.066760676 | -0.234362134 | 0.100840783 | 9.477051133 | -0.979968449 | 0.365751888 | 0.811477229 | -4.787267901 |
| **A_55_P2079855** | Prl6a1 | 0.056399505 | -0.08519213 | 0.197991141 | 8.928654171 | 0.979956638 | 0.365757255 | 0.811477229 | -4.78727501 |
| **A_66_P117786** | Crxos | 0.05361519 | -0.080997008 | 0.188227389 | 7.981513213 | 0.979879332 | 0.365792389 | 0.811477229 | -4.787321538 |
| **A_55_P2160198** | NA | 0.075775963 | -0.114563569 | 0.266115496 | 11.30251317 | 0.979426004 | 0.365998469 | 0.811841223 | -4.787594331 |
| **A_55_P2029775** | Gm38396 | 0.104945442 | -0.158782786 | 0.368673671 | 8.945371598 | 0.978985202 | 0.366198944 | 0.81195654 | -4.787859513 |
| **A_52_P131559** | Srprb | -0.048922832 | -0.171873198 | 0.074027535 | 8.713125308 | -0.978928388 | 0.366224789 | 0.81195654 | -4.787893686 |
| **A_55_P2270137** | B930036G03Rik | 0.122698225 | -0.185675024 | 0.431071473 | 9.84249265 | 0.97888291 | 0.366245478 | 0.81195654 | -4.78792104 |
| **A_52_P142208** | Akirin1 | -0.051085053 | -0.179475738 | 0.077305633 | 10.21258403 | -0.978880085 | 0.366246763 | 0.81195654 | -4.787922739 |
| **A_55_P1986441** | Gcdh | 0.11783309 | -0.178329006 | 0.413995186 | 8.587633815 | 0.978829284 | 0.366269875 | 0.81195654 | -4.787953294 |
| **A_55_P2106039** | Map3k6 | 0.110189584 | -0.166781588 | 0.387160755 | 9.85794618 | 0.978757576 | 0.3663025 | 0.81195654 | -4.787996421 |
| **A_55_P1985758** | Psmd9 | -0.049685336 | -0.174610465 | 0.075239793 | 9.030517402 | -0.978470161 | 0.366433291 | 0.812153319 | -4.788169263 |
| **A_65_P02543** | Srp72 | 0.074212531 | -0.112402893 | 0.260827955 | 9.413261013 | 0.978360409 | 0.366483245 | 0.812170906 | -4.788235257 |
| **A_55_P2156002** | Sumo2 | -0.141895107 | -0.498822379 | 0.215032165 | 14.04412114 | -0.978040526 | 0.36662887 | 0.812279306 | -4.788427574 |
| **A_51_P190111** | Mcm5 | 0.087489415 | -0.13260029 | 0.30757912 | 8.461831412 | 0.977968678 | 0.366661584 | 0.812279306 | -4.788470764 |
| **A_52_P243249** | Mapk3 | 0.085309195 | -0.129303072 | 0.299921463 | 10.20312278 | 0.977936071 | 0.366676432 | 0.812279306 | -4.788490366 |
| **A_55_P2062797** | 4921524J17Rik | -0.087505845 | -0.307688448 | 0.132676759 | 9.102012136 | -0.977739639 | 0.366765889 | 0.812279306 | -4.788608436 |
| **A_55_P1978136** | Tnip2 | 0.059555361 | -0.090299918 | 0.20941064 | 9.387185696 | 0.977727838 | 0.366771264 | 0.812279306 | -4.788615529 |
| **A_55_P1982334** | Llph | 0.104591786 | -0.158593422 | 0.367776993 | 11.40017506 | 0.977699211 | 0.366784302 | 0.812279306 | -4.788632735 |
| **A_55_P2176248** | NA | 0.065464812 | -0.099316668 | 0.230246292 | 15.69366184 | 0.977391793 | 0.366924343 | 0.812392099 | -4.788817482 |
| **A_51_P365980** | 2310022B05Rik | -0.152158801 | -0.535185851 | 0.230868249 | 10.5158632 | -0.977320021 | 0.366957044 | 0.812392099 | -4.78886061 |
| **A_55_P2045741** | Fyb | -0.042118235 | -0.148142849 | 0.06390638 | 7.730759092 | -0.977310629 | 0.366961323 | 0.812392099 | -4.788866254 |
| **A_55_P2206605** | 5830444B04Rik | 0.090908224 | -0.138043185 | 0.319859632 | 9.669787434 | 0.97685255 | 0.367170091 | 0.812761187 | -4.789141464 |
| **A_51_P500344** | Mat2a | -0.133162672 | -0.468589808 | 0.202264463 | 12.85041485 | -0.976682646 | 0.367247548 | 0.812839557 | -4.789243521 |
| **A_51_P112627** | St6galnac2 | 0.128584868 | -0.195400835 | 0.452570572 | 11.69414488 | 0.976412165 | 0.367370883 | 0.812852958 | -4.782394493 |
| **A_51_P345699** | Tnnt3 | -0.043989097 | -0.154829125 | 0.06685093 | 8.118087604 | -0.976377106 | 0.367386872 | 0.812852958 | -4.789427023 |
| **A_55_P2022342** | Stim1 | -0.177434239 | -0.62454947 | 0.269680993 | 12.78239444 | -0.976308625 | 0.367418104 | 0.812852958 | -4.789468147 |
| **A_55_P2067112** | Gm4924 | -0.215245058 | -0.757644017 | 0.327153901 | 9.844018546 | -0.976300486 | 0.367421817 | 0.812852958 | -4.789473035 |
| **A_51_P165984** | Aup1 | -0.075557573 | -0.266022393 | 0.114907247 | 10.1017094 | -0.975960838 | 0.367576754 | 0.81301694 | -4.789676971 |
| **A_55_P2083654** | Prmt2 | -0.080775023 | -0.284393455 | 0.122843409 | 13.30335363 | -0.975953587 | 0.367580063 | 0.81301694 | -4.789681324 |
| **A_55_P2039881** | 5-Sep | 0.368244457 | -0.560315889 | 1.296804804 | 9.310875397 | 0.975652695 | 0.367717366 | 0.813089301 | -4.789861953 |
| **A_51_P128667** | Lynx1 | 0.123349288 | -0.187692997 | 0.434391573 | 8.74146829 | 0.975632761 | 0.367726463 | 0.813089301 | -4.789873919 |
| **A_55_P2111907** | Noxo1 | -0.074234516 | -0.261447382 | 0.112978351 | 8.49916883 | -0.975527125 | 0.367774678 | 0.813089301 | -4.789937324 |
| **A_55_P2058646** | Nhlh1 | 0.070503242 | -0.10731183 | 0.248318315 | 8.725872793 | 0.975460484 | 0.367805097 | 0.813089301 | -4.789977322 |
| **A_52_P541833** | Vps37b | -0.071347686 | -0.251299796 | 0.108604424 | 10.3385158 | -0.975421028 | 0.367823108 | 0.813089301 | -4.790001003 |
| **A_51_P158584** | Poll | -0.042953487 | -0.151314425 | 0.065407451 | 9.138691313 | -0.975202568 | 0.367922843 | 0.813134603 | -4.790132106 |
| **A_55_P2042156** | Aatk | 0.052997936 | -0.080730583 | 0.186726456 | 8.421828899 | 0.974998806 | 0.368015888 | 0.813134603 | -4.790254372 |
| **A_52_P338110** | Chmp1b | -0.081440948 | -0.286945765 | 0.12406387 | 9.845991774 | -0.974967131 | 0.368030353 | 0.813134603 | -4.790273378 |
| **A_52_P52128** | Leo1 | -0.078004292 | -0.274845266 | 0.118836682 | 11.16723647 | -0.974927133 | 0.368048621 | 0.813134603 | -4.790297376 |

| | | | | | | | | | |
|---|---|---|---|---|---|---|---|---|---|
| **A_55_P2038126** | Tpmt | -0.07304722 | -0.257420323 | 0.111325884 | 8.113966862 | -0.974709698 | 0.368147936 | 0.813134603 | -4.790427824 |
| **A_55_P2012478** | Zfp618 | 0.104425517 | -0.159154232 | 0.368005265 | 8.14169551 | 0.97468382 | 0.368159758 | 0.813134603 | -4.790443348 |
| **A_52_P476699** | Gemin4 | 0.045048471 | -0.06866021 | 0.158757152 | 8.044060546 | 0.974665507 | 0.368168123 | 0.813134603 | -4.790454334 |
| **A_52_P708507** | Myadml2 | -0.044607755 | -0.157207047 | 0.067991537 | 7.650929123 | -0.97463919 | 0.368180146 | 0.813134603 | -4.790470121 |
| **A_52_P502267** | Epc2 | -0.136137086 | -0.479812605 | 0.207538433 | 10.45966388 | -0.974534045 | 0.368228183 | 0.813147784 | -4.790533192 |
| **A_52_P512427** | Mkrn2os | 0.213628691 | -0.325818962 | 0.753076344 | 8.59793255 | 0.974270234 | 0.368348731 | 0.813321067 | -4.790691421 |
| **A_55_P2038687** | Tmco1 | 0.120719205 | -0.18417237 | 0.425610781 | 13.32596474 | 0.974092291 | 0.368430059 | 0.813407723 | -4.790798134 |
| **A_55_P2049313** | NA | 0.083034925 | -0.12678279 | 0.292852639 | 8.462115598 | 0.973616235 | 0.368647709 | 0.813795291 | -4.791083566 |
| **A_51_P130544** | Gas5 | -0.052114005 | -0.183887809 | 0.079659799 | 7.930743687 | -0.972958974 | 0.368948371 | 0.814366 | -4.791477503 |
| **A_55_P1982291** | Clca3a1 | -0.077037114 | -0.271849203 | 0.117774975 | 8.68591909 | -0.972866535 | 0.368990672 | 0.814366375 | -4.791532894 |
| **A_55_P1962334** | Trim71 | -0.13813417 | -0.487507894 | 0.211239555 | 9.240379991 | -0.972702528 | 0.369065734 | 0.814367236 | -4.791631163 |
| **A_55_P2107477** | Rab5c | 0.20256409 | -0.309779059 | 0.714907239 | 12.88833312 | 0.972681571 | 0.369075327 | 0.814367236 | -4.791643719 |
| **A_55_P2106608** | NA | -0.063716201 | -0.224910814 | 0.097478413 | 8.073275406 | -0.97245275 | 0.369180074 | 0.814505382 | -4.791780803 |
| **A_52_P763938** | NA | 0.059490616 | -0.091035628 | 0.210016859 | 8.507039182 | 0.972311456 | 0.369244766 | 0.814520703 | -4.791865441 |
| **A_55_P2059055** | Zfp943 | -0.097161844 | -0.343046048 | 0.148722359 | 9.367971309 | -0.972152246 | 0.369317672 | 0.814520703 | -4.791960802 |
| **A_51_P520084** | Lypla2 | 0.066462963 | -0.101739046 | 0.234664971 | 13.45902027 | 0.972115086 | 0.36933469 | 0.814520703 | -4.791983058 |
| **A_55_P2070239** | Preb | -0.058842771 | -0.207800488 | 0.090114947 | 9.268204299 | -0.971850069 | 0.369456076 | 0.814520703 | -4.792141768 |
| **A_52_P97572** | Aplnr | 0.078836619 | -0.120742177 | 0.278415415 | 7.965025518 | 0.971813113 | 0.369473006 | 0.814520703 | -4.792163898 |
| **A_55_P2019776** | Gxylt1 | 0.05105947 | -0.078225622 | 0.180344561 | 9.323912392 | 0.971621279 | 0.369560895 | 0.814520703 | -4.792278761 |
| **A_55_P2046907** | Wdr83os | 0.129859481 | -0.198963828 | 0.458682789 | 10.96243825 | 0.971583721 | 0.369578104 | 0.814520703 | -4.792301248 |
| **A_55_P1974695** | NA | -0.132286393 | -0.467296793 | 0.202724006 | 8.416070379 | -0.971462521 | 0.369633643 | 0.814520703 | -4.79237381 |
| **A_52_P182659** | Cs | -0.0639781 | -0.226086839 | 0.098130638 | 13.71803896 | -0.970943762 | 0.369871433 | 0.814520703 | -4.792684325 |
| **A_51_P173100** | Pttg1ip | 0.081957176 | -0.125713113 | 0.289627464 | 13.03777302 | 0.970916274 | 0.369884036 | 0.814520703 | -4.792700775 |
| **A_51_P297865** | Kctd6 | -0.063341152 | -0.22384099 | 0.097158685 | 9.733257176 | -0.970913472 | 0.369885321 | 0.814520703 | -4.792702452 |
| **A_55_P1974362** | Hp1bp3 | 0.048749145 | -0.074782082 | 0.172280371 | 13.24328825 | 0.970866264 | 0.369906967 | 0.814520703 | -4.792730704 |
| **A_55_P2015258** | Trim13 | 0.085237333 | -0.138506922 | 0.308981587 | 9.198003643 | 0.987252042 | 0.370003456 | 0.814520703 | -4.739302093 |
| **A_51_P401527** | Mrm3 | 0.077899151 | -0.119551061 | 0.275349363 | 8.521582568 | 0.970608932 | 0.370024978 | 0.814520703 | -4.792884691 |
| **A_55_P2104876** | NA | -0.056148763 | -0.19846918 | 0.086171654 | 8.922488828 | -0.970604159 | 0.370027167 | 0.814520703 | -4.792887547 |
| **A_55_P2068734** | Ccl27a | -0.218381467 | -0.791667189 | 0.354904255 | 14.34149862 | -0.987176333 | 0.370037062 | 0.814520703 | -4.739343291 |
| **A_55_P1955457** | Grb10 | -0.074478574 | -0.263273118 | 0.11431597 | 10.90849559 | -0.970534723 | 0.370059015 | 0.814520703 | -4.792929092 |
| **A_51_P155323** | Hc | -0.172340089 | -0.60925117 | 0.264570991 | 8.415342941 | -0.970425986 | 0.370108893 | 0.814520703 | -4.79299415 |
| **A_55_P2124085** | NA | -0.061929398 | -0.218935959 | 0.095077162 | 10.67090184 | -0.970394271 | 0.370123442 | 0.814520703 | -4.793013124 |
| **A_55_P2224830** | D230044B12Rik | -0.107872131 | -0.381357387 | 0.165613125 | 8.039912148 | -0.970385957 | 0.370127256 | 0.814520703 | -4.793018098 |
| **A_55_P1969311** | Gramd1b | 0.361120813 | -0.554431393 | 1.27667302 | 10.9874704 | 0.970372699 | 0.370133338 | 0.814520703 | -4.79302603 |
| **A_55_P2036377** | NA | -0.055593739 | -0.196552508 | 0.08536503 | 16.53210971 | -0.970293112 | 0.370169851 | 0.814520703 | -4.793073642 |
| **A_51_P481821** | Spcs3 | -0.102864356 | -0.363736271 | 0.158007559 | 10.8656174 | -0.970078205 | 0.370268458 | 0.814520703 | -4.793202196 |
| **A_55_P2139181** | Cald1 | -0.082949748 | -0.293354838 | 0.127455343 | 10.51526724 | -0.969902304 | 0.370349184 | 0.814520703 | -4.793307405 |
| **A_55_P1999182** | Cdk12 | 0.057810618 | -0.08882921 | 0.204450445 | 8.700238496 | 0.96989525 | 0.370352422 | 0.814520703 | -4.793311623 |
| **A_55_P2183551** | NA | 0.097298524 | -0.149509433 | 0.344106482 | 14.12756003 | 0.969876102 | 0.37036121 | 0.814520703 | -4.793323075 |
| **A_55_P2335648** | Eprn | 0.115877501 | -0.178066981 | 0.409821983 | 8.229953491 | 0.969846428 | 0.37037483 | 0.814520703 | -4.793340822 |
| **A_51_P479408** | Hyal1 | -0.179102732 | -0.633480494 | 0.27527503 | 8.510501916 | -0.969737757 | 0.370424712 | 0.814520703 | -4.793405812 |
| **A_51_P108190** | Rpl23 | 0.079077264 | -0.12154072 | 0.279695247 | 10.35456643 | 0.969730233 | 0.370428166 | 0.814520703 | -4.776151557 |
| **A_55_P2121349** | NA | 0.100298076 | -0.154170005 | 0.354766158 | 9.429548507 | 0.969680027 | 0.370451213 | 0.814520703 | -4.793440335 |
| **A_52_P235569** | Ipo5 | -0.089514541 | -0.316674202 | 0.13764512 | 8.944322264 | -0.969463675 | 0.370550544 | 0.814595807 | -4.793569703 |
| **A_55_P1998380** | Traf6 | -0.141027584 | -0.498949677 | 0.216894509 | 8.580537451 | -0.969359168 | 0.370598532 | 0.814595807 | -4.793632187 |
| **A_55_P2062513** | Pdzrn3 | -0.252442505 | -0.893150094 | 0.388265085 | 11.06913151 | -0.96933027 | 0.370611802 | 0.814595807 | -4.793649464 |
| **A_55_P2133943** | NA | -0.153307882 | -0.542523871 | 0.235908107 | 9.119239487 | -0.969042833 | 0.370743819 | 0.814793323 | -4.793821296 |
| **A_51_P184728** | Cnksr3 | -0.09317539 | -0.329765521 | 0.143414742 | 9.230709254 | -0.968888395 | 0.370814766 | 0.814856595 | -4.793913607 |
| **A_55_P2044167** | NA | 0.050103282 | -0.077149881 | 0.177356445 | 8.216632944 | 0.968649702 | 0.370924439 | 0.814970818 | -4.794056261 |
| **A_55_P2086153** | Bcdin3d | -0.095997109 | -0.33982678 | 0.147832562 | 10.16910188 | -0.968591749 | 0.370951071 | 0.814970818 | -4.794090893 |

| | | | | | | | | |
|---|---|---|---|---|---|---|---|---|
| **A_66_P123432** | NA | 0.088597567 | -0.136496162 | 0.313691296 | 8.00275302 | 0.968339313 | 0.371067094 | 0.815076654 | -4.794241732 |
| **A_55_P2134387** | Olfr544 | 0.043891323 | -0.067633128 | 0.155415774 | 11.1475517 | 0.968228231 | 0.371118158 | 0.815076654 | -4.794308099 |
| **A_55_P1952235** | Spry1 | -0.092620241 | -0.327965111 | 0.142724629 | 10.48211518 | -0.968211715 | 0.37112575 | 0.815076654 | -4.794317967 |
| **A_55_P2049071** | NA | -0.12384942 | -0.438798838 | 0.191099998 | 13.63110956 | -0.967436191 | 0.371482414 | 0.815691077 | -4.794781177 |
| **A_55_P2069856** | Tnpo3 | 0.071891434 | -0.110942041 | 0.254724909 | 11.47587124 | 0.967365507 | 0.371514935 | 0.815691077 | -4.794823385 |
| **A_51_P240864** | Sppl3 | -0.082466274 | -0.292246506 | 0.127313957 | 13.81408215 | -0.967121363 | 0.371627281 | 0.815691077 | -4.794969155 |
| **A_55_P2107830** | Gemin2 | -0.05436264 | -0.192659157 | 0.083933877 | 8.095685425 | -0.96707122 | 0.371650358 | 0.815691077 | -4.79499909 |
| **A_55_P1990170** | NA | 0.065378704 | -0.100954248 | 0.231711655 | 14.99096862 | 0.967001558 | 0.37168242 | 0.815691077 | -4.795040678 |
| **A_55_P1982420** | Fam8a1 | -0.051457133 | -0.182379555 | 0.07946529 | 10.56347784 | -0.966942638 | 0.37170954 | 0.815691077 | -4.795075851 |
| **A_52_P590235** | Vars2 | -0.059536493 | -0.211025183 | 0.091952197 | 8.357039809 | -0.966879169 | 0.371738755 | 0.815691077 | -4.795113738 |
| **A_51_P286357** | Adgre4 | -0.041530934 | -0.147206094 | 0.064144226 | 7.706143944 | -0.966869695 | 0.371743117 | 0.815691077 | -4.795119393 |
| **A_55_P2002898** | Ctsj | -0.062374113 | -0.221137395 | 0.096389169 | 17.36464456 | -0.966548111 | 0.371891174 | 0.815923327 | -4.795311335 |
| **A_51_P423976** | Crem | -0.065153149 | -0.231044523 | 0.100738225 | 8.53908675 | -0.966230559 | 0.372037421 | 0.816116471 | -4.79550083 |
| **A_51_P132718** | 2310010J17Rik | -0.073031042 | -0.258991877 | 0.112929793 | 11.53654874 | -0.966173612 | 0.372063653 | 0.816116471 | -4.795534809 |
| **A_52_P340669** | Bhlha15 | -0.123067264 | -0.436528747 | 0.190394218 | 11.39142091 | -0.965889683 | 0.372194461 | 0.816152299 | -4.795704202 |
| **A_55_P2095266** | NA | -0.081562907 | -0.289320836 | 0.126195021 | 14.17240388 | -0.965837924 | 0.37221831 | 0.816152299 | -4.795735079 |
| **A_55_P1991906** | Acadvl | -0.051963752 | -0.18441206 | 0.080484556 | 12.99202039 | -0.965213189 | 0.372506271 | 0.816152299 | -4.796107676 |
| **A_55_P2155574** | NA | 0.099889962 | -0.154750312 | 0.354530237 | 13.70282984 | 0.965081343 | 0.372567066 | 0.816152299 | -4.796186291 |
| **A_55_P2033947** | Fnd3c2 | -0.195133113 | -0.692604649 | 0.302338423 | 14.1448824 | -0.965010193 | 0.372599877 | 0.816152299 | -4.796228713 |
| **A_55_P2087589** | Rps11 | -0.231509543 | -0.821729045 | 0.358709959 | 14.54132139 | -0.964993762 | 0.372607454 | 0.816152299 | -4.796238509 |
| **A_52_P138598** | Pum2 | -0.065310841 | -0.231830134 | 0.101208452 | 8.651911634 | -0.964916824 | 0.372642937 | 0.816152299 | -4.796284379 |
| **A_55_P2091691** | Nudt3 | -0.082597312 | -0.293196241 | 0.128001616 | 12.17596895 | -0.964892482 | 0.372654164 | 0.816152299 | -4.796298891 |
| **A_55_P2043798** | Mrto4 | -0.087411256 | -0.310306552 | 0.135484039 | 10.34186941 | -0.964796284 | 0.372698534 | 0.816152299 | -4.796356239 |
| **A_55_P2106827** | NA | 0.089360274 | -0.138523258 | 0.317243807 | 10.12305996 | 0.964718736 | 0.372734305 | 0.816152299 | -4.796402467 |
| **A_52_P274555** | Snx14 | -0.116983266 | -0.415312631 | 0.181346099 | 10.48139338 | -0.964710273 | 0.372738209 | 0.816152299 | -4.796407512 |
| **A_55_P2118931** | Foxj3 | 0.121944087 | -0.189041888 | 0.432930063 | 12.0840844 | 0.964692956 | 0.372746198 | 0.816152299 | -4.796417834 |
| **A_51_P386880** | Stip1 | -0.065473575 | -0.232470981 | 0.101523831 | 10.74672912 | -0.964551654 | 0.372811386 | 0.816152299 | -4.796502059 |
| **A_55_P2100628** | Gm12216 | 0.287541603 | -0.445877896 | 1.020961103 | 10.90682352 | 0.964533688 | 0.372819675 | 0.816152299 | -4.796512768 |
| **A_52_P15197** | Dcun1d2 | 0.0726801 | -0.112727755 | 0.258087955 | 10.94453007 | 0.964398556 | 0.372882026 | 0.816152299 | -4.796593307 |
| **A_66_P123989** | Ikzf5 | 0.047698843 | -0.073998793 | 0.169396479 | 8.41417522 | 0.96426159 | 0.372945232 | 0.816152299 | -4.796674932 |
| **A_55_P2247852** | Ap2s1 | 0.110997278 | -0.172253962 | 0.394248518 | 11.59873017 | 0.96407245 | 0.373032528 | 0.816152299 | -4.796787638 |
| **A_55_P2123502** | Jam2 | -0.488059546 | -1.733553198 | 0.757434107 | 9.469851169 | -0.964051991 | 0.373041972 | 0.816152299 | -4.796799829 |
| **A_55_P2093418** | Zmpste24 | -0.063543681 | -0.225711793 | 0.098624432 | 11.83843805 | -0.963997854 | 0.373066962 | 0.816152299 | -4.796832086 |
| **A_52_P675887** | NA | -0.131562015 | -0.467367313 | 0.204243283 | 14.2025931 | -0.96385596 | 0.373132468 | 0.816152299 | -4.796916626 |
| **A_55_P2005898** | NA | 0.044488746 | -0.069068512 | 0.158046004 | 7.861395877 | 0.963838858 | 0.373140364 | 0.816152299 | -4.796926815 |
| **A_55_P2028961** | Idi1 | 0.30516403 | -0.47391147 | 1.084239531 | 9.898278497 | 0.963658058 | 0.373223845 | 0.816152299 | -4.797034522 |
| **A_51_P397673** | Pcsk9 | 0.133611705 | -0.207510786 | 0.474734197 | 7.918513691 | 0.963614474 | 0.373243971 | 0.816152299 | -4.797060485 |
| **A_55_P1973858** | NA | 0.103059613 | -0.160062173 | 0.366181399 | 9.670165405 | 0.963609032 | 0.373246484 | 0.816152299 | -4.797063727 |
| **A_51_P498772** | 1700019D03Rik | 0.099192977 | -0.154084404 | 0.352470358 | 8.392888031 | 0.963504351 | 0.373294828 | 0.816152299 | -4.79712608 |
| **A_55_P2113984** | Mospd2 | -0.053877644 | -0.191451237 | 0.08369595 | 8.366995875 | -0.963479935 | 0.373306104 | 0.816152299 | -4.797140623 |
| **A_55_P2119817** | Plekhj1 | 0.067925585 | -0.105529388 | 0.241380559 | 8.56696706 | 0.963420306 | 0.373333645 | 0.816152299 | -4.797176139 |
| **A_55_P2069768** | Kiss1 | 0.071025436 | -0.11036588 | 0.252416752 | 7.955985473 | 0.963311134 | 0.373384072 | 0.816152299 | -4.79724116 |
| **A_55_P2095535** | NA | 0.138495419 | -0.215223362 | 0.4922142 | 8.994296414 | 0.963266489 | 0.373404695 | 0.816152299 | -4.797267748 |
| **A_55_P2102060** | Gstm3 | 0.044632727 | -0.069360913 | 0.158626368 | 7.691897371 | 0.963256539 | 0.373409292 | 0.816152299 | -4.797273674 |
| **A_55_P2068566** | NA | 0.085614562 | -0.133054855 | 0.304283978 | 13.75224101 | 0.963227238 | 0.373422827 | 0.816152299 | -4.797291124 |
| **A_55_P2228918** | Rslcan18 | -0.086771996 | -0.308416797 | 0.134872805 | 8.667468186 | -0.963143955 | 0.373461303 | 0.816152299 | -4.79734072 |
| **A_55_P2138739** | NA | -0.187413536 | -0.666143606 | 0.291316534 | 8.846841139 | -0.963117801 | 0.373473386 | 0.816152299 | -4.797356295 |
| **A_66_P114862** | Ankrd11 | -0.106836064 | -0.379776272 | 0.166104144 | 10.3894992 | -0.962985006 | 0.373534744 | 0.816194106 | -4.797435369 |
| **A_55_P2104810** | Mrpl27 | 0.09141941 | -0.14219293 | 0.32503175 | 11.56861988 | 0.96274617 | 0.373645117 | 0.816279622 | -4.797577569 |
| **A_66_P112065** | Sertad2 | -0.221424803 | -0.787303338 | 0.344453733 | 9.520454108 | -0.962658409 | 0.37368568 | 0.816279622 | -4.797629816 |

| | | | | | | | | | |
|---|---|---|---|---|---|---|---|---|---|
| **A_55_P2043982** | Mtm1 | -0.056899142 | -0.202316819 | 0.088518536 | 8.583738758 | -0.962626189 | 0.373700573 | 0.816279622 | -4.797648996 |
| **A_55_P2362635** | Slc9a3 | -0.105262215 | -0.374362271 | 0.163837841 | 8.427535403 | -0.962338539 | 0.373833554 | 0.816477826 | -4.797820218 |
| **A_55_P2231747** | NA | -0.236175559 | -0.84009428 | 0.367743161 | 9.201983385 | -0.962111982 | 0.373938318 | 0.816614365 | -4.797955052 |
| **A_55_P2033906** | Mrpl22 | 0.155338081 | -0.241912411 | 0.552588573 | 12.55798713 | 0.962016796 | 0.37398234 | 0.81661824 | -4.798011695 |
| **A_55_P2093727** | Gemin6 | -0.093451145 | -0.332480015 | 0.145577725 | 9.366685728 | -0.961841322 | 0.374063506 | 0.816703208 | -4.798116108 |
| **A_52_P80444** | Dpm2 | 0.089881248 | -0.140090904 | 0.319853399 | 12.28190418 | 0.961530375 | 0.374207369 | 0.816797235 | -4.798301101 |
| **A_51_P183894** | Fbxo15 | -0.128629539 | -0.45783931 | 0.200580232 | 9.25578829 | -0.961251786 | 0.374336297 | 0.816797235 | -4.798466812 |
| **A_55_P2149931** | Arap2 | -0.076065055 | -0.27074722 | 0.118617111 | 8.990184175 | -0.961231903 | 0.374345501 | 0.816797235 | -4.798478638 |
| **A_55_P2143219** | Rasgrp2 | 0.255676899 | -0.398796536 | 0.910150335 | 9.189499235 | 0.961100117 | 0.374406503 | 0.816797235 | -4.798557016 |
| **A_55_P2058861** | Mvk | 0.067609028 | -0.105463718 | 0.240681773 | 11.21073563 | 0.961048198 | 0.374430538 | 0.816797235 | -4.798587892 |
| **A_55_P2035300** | Rhot1 | 0.080982472 | -0.126331952 | 0.288296897 | 11.4085624 | 0.961016126 | 0.374445386 | 0.816797235 | -4.798606965 |
| **A_55_P1963712** | Cyb5b | 0.058769634 | -0.091683322 | 0.209222591 | 11.38898613 | 0.960995659 | 0.374454862 | 0.816797235 | -4.798619136 |
| **A_55_P2361547** | 2610201A13Rik | 0.069085825 | -0.10780451 | 0.245976159 | 8.379973962 | 0.960846485 | 0.37452393 | 0.816797235 | -4.798707842 |
| **A_51_P345792** | Mfsd13a | 0.220935595 | -0.344776302 | 0.786647492 | 9.770868521 | 0.960814484 | 0.374538748 | 0.816797235 | -4.798726869 |
| **A_52_P573552** | Trib1 | -0.085458618 | -0.304279877 | 0.133362642 | 9.924232719 | -0.960805573 | 0.374542874 | 0.816797235 | -4.798732168 |
| **A_55_P2033958** | Sltm | -0.104767662 | -0.373047942 | 0.163512618 | 10.35407862 | -0.96074396 | 0.374571405 | 0.816797235 | -4.798768802 |
| **A_55_P1988424** | Tpi1 | -0.134008134 | -0.477289324 | 0.209273057 | 10.25737581 | -0.960395937 | 0.374732596 | 0.816829986 | -4.798975704 |
| **A_55_P1974961** | Plekhg3 | -0.08356916 | -0.297646112 | 0.130507792 | 13.4714428 | -0.960384823 | 0.374737745 | 0.816829986 | -4.798982311 |
| **A_55_P2094966** | H3c14 | -0.057122256 | -0.203451799 | 0.089207288 | 8.972971076 | -0.960378646 | 0.374740606 | 0.816829986 | -4.798985982 |
| **A_55_P1998442** | Sema3f | -0.281955057 | -1.004260783 | 0.440350669 | 14.00111102 | -0.960346579 | 0.374755462 | 0.816829986 | -4.799005044 |
| **A_52_P82247** | Cycs | 0.085152685 | -0.133012218 | 0.303317587 | 11.01639127 | 0.960246257 | 0.374801939 | 0.816839179 | -4.799064675 |
| **A_51_P505493** | Elovl5 | -0.109516059 | -0.400260259 | 0.181228142 | 11.42352735 | -0.976150605 | 0.374958091 | 0.816879226 | -4.745325517 |
| **A_55_P2014531** | Hes7 | 0.0957127 | -0.149599591 | 0.341024991 | 9.479021172 | 0.959885483 | 0.374969118 | 0.816879226 | -4.799279088 |
| **A_55_P2110978** | Atp6ap1 | -0.079017509 | -0.28156557 | 0.123530552 | 8.694884625 | -0.959763888 | 0.375025476 | 0.816879226 | -4.799351342 |
| **A_66_P101407** | Acat3 | 0.057458312 | -0.08984806 | 0.204764684 | 7.813465528 | 0.95962266 | 0.375090944 | 0.816879226 | -4.799435255 |
| **A_52_P522255** | Pitx1 | 0.134789526 | -0.210779319 | 0.48035837 | 11.24053743 | 0.959601078 | 0.375100949 | 0.816879226 | -4.799448078 |
| **A_55_P1982817** | NA | 0.091807429 | -0.143577728 | 0.327192586 | 11.03644064 | 0.959550672 | 0.375124318 | 0.816879226 | -4.799478025 |
| **A_51_P499061** | Ube2o | -0.144729052 | -0.515818329 | 0.226360226 | 13.61176662 | -0.959503301 | 0.375146281 | 0.816879226 | -4.799506169 |
| **A_65_P01783** | Med28 | -0.081198504 | -0.289399381 | 0.127002374 | 9.387470731 | -0.959477142 | 0.375158409 | 0.816879226 | -4.799521709 |
| **A_55_P1979893** | Tef | -0.076559884 | -0.272893448 | 0.11977368 | 12.66755811 | -0.95934735 | 0.375218591 | 0.816918242 | -4.799598813 |
| **A_52_P633379** | Plpbp | 0.078113465 | -0.122292255 | 0.278519186 | 12.22603067 | 0.95892569 | 0.37541416 | 0.817251977 | -4.79984926 |
| **A_55_P2183955** | Olfr97 | 0.044843376 | -0.070224185 | 0.159910938 | 7.902004577 | 0.958770269 | 0.375486265 | 0.817316895 | -4.799941555 |
| **A_55_P2052170** | NA | 0.094295938 | -0.147853157 | 0.336445033 | 11.39907736 | 0.958030417 | 0.375829659 | 0.817904756 | -4.800380782 |
| **A_51_P484526** | Wif1 | -0.045264421 | -0.16150503 | 0.070976188 | 7.659408081 | -0.958006064 | 0.375840967 | 0.817904756 | -4.800395236 |
| **A_52_P610987** | Slc28a3 | 0.145902688 | -0.228970432 | 0.520775808 | 9.164245061 | 0.957520664 | 0.376066399 | 0.818134986 | -4.800683284 |
| **A_55_P2017769** | B020031M17Rik | -0.10351055 | -0.369477616 | 0.162456517 | 8.347059802 | -0.957471632 | 0.376089176 | 0.818134986 | -4.800712375 |
| **A_55_P2111355** | Egfl7 | -0.121806376 | -0.434802642 | 0.191189891 | 13.74311625 | -0.957414558 | 0.376115691 | 0.818134986 | -4.800746237 |
| **A_55_P2348676** | B130052P14Rik | -0.084938097 | -0.303201177 | 0.133324984 | 8.411237465 | -0.957395558 | 0.376124518 | 0.818134986 | -4.800757509 |
| **A_51_P456957** | Pex11a | -0.079048096 | -0.282191274 | 0.124095083 | 11.15912316 | -0.957322642 | 0.376158395 | 0.818134986 | -4.797287061 |
| **A_55_P2185840** | Nnmt | 0.055354969 | -0.086950271 | 0.197660209 | 7.977473428 | 0.95698446 | 0.376315549 | 0.818296101 | -4.801001373 |
| **A_55_P2105220** | Hscb | -0.071284336 | -0.25454107 | 0.111972398 | 11.41194383 | -0.956981031 | 0.376317142 | 0.818296101 | -4.801003407 |
| **A_55_P2022663** | NA | 0.088809679 | -0.139593851 | 0.31721321 | 16.21258048 | 0.956591799 | 0.376498082 | 0.818597462 | -4.801234238 |
| **A_52_P582858** | Dgkd | 0.104547742 | -0.164366341 | 0.373461825 | 11.21976014 | 0.956467625 | 0.376555821 | 0.818630915 | -4.801307867 |
| **A_55_P1971269** | NA | -0.071154358 | -0.254195911 | 0.111887194 | 11.21184205 | -0.956359065 | 0.376606305 | 0.818648591 | -4.801372232 |
| **A_55_P2033725** | Ascl2 | 0.243602571 | -0.383486486 | 0.870691628 | 10.35585837 | 0.955700489 | 0.376912679 | 0.819222441 | -4.801762603 |
| **A_51_P348548** | Ecsit | 0.10647486 | -0.167714393 | 0.380664112 | 11.02491611 | 0.955357267 | 0.377072425 | 0.819350306 | -4.801965981 |
| **A_51_P280437** | Slc12a6 | 0.057607738 | -0.090741156 | 0.205956631 | 9.013969974 | 0.955356959 | 0.377072568 | 0.819350306 | -4.801966164 |
| **A_55_P2171788** | NA | -0.496764354 | -1.776188971 | 0.782660264 | 10.96530804 | -0.955223251 | 0.377134814 | 0.819350306 | -4.80204538 |
| **A_55_P2020962** | NA | 0.085113268 | -0.134100411 | 0.304326948 | 16.18739423 | 0.955209822 | 0.377141066 | 0.819350306 | -4.802053336 |
| **A_55_P2069579** | NA | 0.131518988 | -0.207365237 | 0.470403213 | 8.498907512 | 0.954786497 | 0.377338196 | 0.819585575 | -4.802304093 |

| | | | | | | | | | |
|---|---|---|---|---|---|---|---|---|---|
| **A_55_P2176391** | NA | -0.102968852 | -0.368312193 | 0.162374489 | 11.94668326 | -0.954699822 | 0.377378568 | 0.819585575 | -4.802355426 |
| **A_55_P2279807** | NA | -0.178072578 | -0.636970412 | 0.280825256 | 9.524096335 | -0.954663243 | 0.377395607 | 0.819585575 | -4.802377089 |
| **A_55_P2064333** | Lama3 | 0.168248663 | -0.265357256 | 0.601854582 | 8.135645789 | 0.95460913 | 0.377420815 | 0.819585575 | -4.802409135 |
| **A_55_P2025687** | Muc4 | -0.187048405 | -0.669156389 | 0.295059578 | 9.300900788 | -0.954506411 | 0.377468669 | 0.819585575 | -4.802469963 |
| **A_55_P2258567** | D930043N17Rik | -0.148033592 | -0.52961295 | 0.233545765 | 8.353372572 | -0.954431071 | 0.37750377 | 0.819585575 | -4.802514575 |
| **A_55_P2132812** | NA | 0.176943397 | -0.279227127 | 0.633113921 | 10.86618269 | 0.954281059 | 0.37757367 | 0.819630151 | -4.802603398 |
| **A_51_P378051** | Aoc3 | 0.142451203 | -0.224825679 | 0.509728085 | 8.842749455 | 0.954205008 | 0.37760911 | 0.819630151 | -4.802648424 |
| **A_55_P2473316** | Mapre1 | -0.095792273 | -0.342804794 | 0.151220249 | 11.6759432 | -0.95407095 | 0.377671589 | 0.81967372 | -4.802727788 |
| **A_51_P438083** | Slc6a13 | 0.300322274 | -0.474192856 | 1.074837403 | 8.779856531 | 0.95395261 | 0.377726749 | 0.819701395 | -4.802797841 |
| **A_55_P2183750** | Ncoa3 | 0.096186185 | -0.151896894 | 0.344269263 | 11.70497387 | 0.953860182 | 0.377769835 | 0.819702868 | -4.802852552 |
| **A_52_P435561** | Prr15l | 0.057429121 | -0.090759328 | 0.20561757 | 7.791323564 | 0.953425979 | 0.377972296 | 0.819927854 | -4.803109523 |
| **A_55_P1973668** | Slc39a13 | -0.152606119 | -0.546391964 | 0.241179726 | 12.01563342 | -0.953412869 | 0.37797841 | 0.819927854 | -4.80311728 |
| **A_52_P555723** | Ywhag | -0.062207514 | -0.222736275 | 0.098321246 | 11.53606601 | -0.953364902 | 0.378000782 | 0.819927854 | -4.803145663 |
| **A_55_P2040888** | Dhx15 | -0.055513332 | -0.198834781 | 0.087808117 | 8.848458734 | -0.952917429 | 0.378209531 | 0.820288603 | -4.803410401 |
| **A_55_P1979595** | Selenon | 0.127766486 | -0.202163091 | 0.457696063 | 11.8087469 | 0.952719082 | 0.37830209 | 0.820397297 | -4.803527723 |
| **A_52_P240453** | Polr3k | -0.079676443 | -0.285493759 | 0.126140872 | 9.786884873 | -0.95239519 | 0.378453273 | 0.820450617 | -4.803719272 |
| **A_55_P2114953** | Usp18 | -0.079218342 | -0.283852458 | 0.125415773 | 10.87258129 | -0.952394491 | 0.3784536 | 0.820450617 | -4.803719685 |
| **A_55_P2064557** | Furin | 0.10775528 | -0.170629038 | 0.386139599 | 12.88154926 | 0.952276257 | 0.3785088 | 0.820450617 | -4.803789599 |
| **A_52_P263870** | Cdc23 | -0.048602533 | -0.174170949 | 0.076965884 | 7.968010488 | -0.952242689 | 0.378524473 | 0.820450617 | -4.803809447 |
| **A_55_P2422243** | 2900072G11Rik | -0.120974513 | -0.433547215 | 0.191598189 | 10.27754816 | -0.952164532 | 0.378560967 | 0.820450617 | -4.803855658 |
| **A_51_P314161** | Qtrt1 | -0.046758813 | -0.167595582 | 0.074077956 | 8.060077789 | -0.951992524 | 0.378641292 | 0.820450617 | -4.803957351 |
| **A_55_P2035981** | Mindy1 | 0.142990343 | -0.226543714 | 0.512524399 | 12.27841585 | 0.95196592 | 0.378653717 | 0.820450617 | -4.803973079 |
| **A_52_P628117** | Abcc5 | 0.167977951 | -0.266144538 | 0.602100439 | 8.619280193 | 0.951939084 | 0.37866625 | 0.820450617 | -4.803988944 |
| **A_55_P2021953** | Olfr700 | 0.23414098 | -0.371104291 | 0.839386252 | 9.013093777 | 0.951733136 | 0.378762448 | 0.820558791 | -4.804110683 |
| **A_55_P2009677** | Zmiz1 | -0.141044418 | -0.505670401 | 0.223581566 | 13.38991262 | -0.951650438 | 0.378801081 | 0.820558791 | -4.804159563 |
| **A_52_P37498** | Dpcd | -0.049414944 | -0.177203709 | 0.07837382 | 9.200859748 | -0.951337905 | 0.378947112 | 0.820783138 | -4.804344265 |
| **A_55_P1966089** | NA | 0.048878624 | -0.077552049 | 0.175309298 | 8.230271073 | 0.951120817 | 0.379048572 | 0.820871335 | -4.804472538 |
| **A_51_P298802** | Bfsp2 | -0.040904192 | -0.146718533 | 0.064910149 | 7.74136058 | -0.951026127 | 0.379092834 | 0.820871335 | -4.804528482 |
| **A_51_P416059** | Slc49a4 | 0.052232626 | -0.082894681 | 0.187359933 | 8.099062581 | 0.950972378 | 0.37911796 | 0.820871335 | -4.804560236 |
| **A_55_P2389528** | D130061D10Rik | -0.153872008 | -0.551978911 | 0.244234895 | 8.333987712 | -0.950887363 | 0.379157705 | 0.820871335 | -4.80461046 |
| **A_55_P2072193** | Firre | -0.055498858 | -0.199104271 | 0.088106556 | 8.292110797 | -0.950785158 | 0.37920549 | 0.820882845 | -4.804670836 |
| **A_55_P2057686** | Cdc42ep5 | 0.10741746 | -0.170564163 | 0.385399083 | 8.769958066 | 0.950665986 | 0.379261215 | 0.820911536 | -4.804741228 |
| **A_66_P124659** | Mark1 | -0.112013966 | -0.401967946 | 0.177940015 | 9.673220348 | -0.950412765 | 0.379379641 | 0.821075925 | -4.804890784 |
| **A_55_P1967196** | Lamb3 | 0.328280055 | -0.521857319 | 1.17841743 | 9.947670616 | 0.95000207 | 0.379571776 | 0.821399783 | -4.805133291 |
| **A_55_P2410304** | Iigp1 | 0.10174663 | -0.161878462 | 0.365371722 | 8.803643579 | 0.94951636 | 0.379799104 | 0.821623369 | -4.805420008 |
| **A_52_P324767** | Tmed8 | -0.10176532 | -0.365439035 | 0.161908396 | 8.858269491 | -0.949515648 | 0.379799438 | 0.821623369 | -4.805420428 |
| **A_51_P339962** | Cwf19l2 | -0.06491785 | -0.233121057 | 0.103285357 | 8.458431334 | -0.949508854 | 0.379802618 | 0.821623369 | -4.805424438 |
| **A_55_P1976109** | NA | 0.137370512 | -0.218603073 | 0.493344097 | 14.71365263 | 0.949390557 | 0.379858001 | 0.82165122 | -4.805494255 |
| **A_55_P2105321** | Acin1 | 0.154280493 | -0.245624974 | 0.554185959 | 11.64142165 | 0.949123746 | 0.379982938 | 0.821736688 | -4.805651702 |
| **A_55_P2121985** | Plcb4 | -0.047833011 | -0.171827886 | 0.076161865 | 8.124741225 | -0.949058849 | 0.380013331 | 0.821736688 | -4.805689995 |
| **A_55_P2151437** | NA | -0.057858955 | -0.207863002 | 0.092145093 | 8.460488964 | -0.948935664 | 0.380071028 | 0.821736688 | -4.805762674 |
| **A_55_P1999842** | NA | 0.072363463 | -0.115259471 | 0.259986396 | 9.712407656 | 0.948860969 | 0.380106016 | 0.821736688 | -4.805806742 |
| **A_51_P198076** | Mien1 | 0.120503881 | -0.192030831 | 0.433038592 | 11.9303074 | 0.948575576 | 0.380239724 | 0.821736688 | -4.805975095 |
| **A_55_P2021471** | Pwwp3a | 0.115763533 | -0.184508015 | 0.416035082 | 7.938897358 | 0.94847689 | 0.380285967 | 0.821736688 | -4.806033302 |
| **A_55_P2000224** | Cd63 | 0.043681633 | -0.069621922 | 0.156985189 | 17.03047494 | 0.948471956 | 0.380288279 | 0.821736688 | -4.806036212 |
| **A_51_P423091** | Flot1 | -0.054140289 | -0.194582984 | 0.086302407 | 11.3634736 | -0.948397881 | 0.380322992 | 0.821736688 | -4.8060799 |
| **A_55_P2046699** | Tvp23b | -0.093152378 | -0.334829823 | 0.148525066 | 10.59957671 | -0.948259038 | 0.380388065 | 0.821736688 | -4.806161782 |
| **A_55_P2151685** | Pira11 | -0.055030854 | -0.197810286 | 0.087748579 | 7.948538153 | -0.948221423 | 0.380405695 | 0.821736688 | -4.806183964 |
| **A_55_P1975670** | Prps1l3 | 0.098230883 | -0.156641322 | 0.353103089 | 9.946960036 | 0.948188614 | 0.380421074 | 0.821736688 | -4.806203311 |
| **A_51_P182303** | Col1a2 | 0.590748314 | -0.942147706 | 2.123644334 | 14.50986647 | 0.948110502 | 0.38045769 | 0.821736688 | -4.806249372 |

| | | | | | | | | | |
|---|---|---|---|---|---|---|---|---|---|
| **A_51_P163261** | Gm6985 | 0.111106813 | -0.177226663 | 0.399440288 | 9.526274824 | 0.948014075 | 0.380502894 | 0.821736688 | -4.806306229 |
| **A_55_P1965629** | Hcfc1 | -0.077256901 | -0.277781543 | 0.123267741 | 11.09722183 | -0.947847997 | 0.380580761 | 0.821736688 | -4.806404147 |
| **A_55_P1956064** | NA | -0.127986013 | -0.460226816 | 0.20425479 | 12.38759393 | -0.947717075 | 0.380642154 | 0.821736688 | -4.806481329 |
| **A_66_P131389** | Ugt2b38 | 0.048543995 | -0.077484224 | 0.174572214 | 7.732738938 | 0.947625801 | 0.380684958 | 0.821736688 | -4.806535133 |
| **A_51_P311785** | Man2b2 | 0.129442009 | -0.20668912 | 0.465573139 | 8.913874608 | 0.947405005 | 0.380788522 | 0.821736688 | -4.806665276 |
| **A_55_P1999972** | NA | -0.123573781 | -0.444474746 | 0.197327184 | 8.244490323 | -0.947380571 | 0.380799983 | 0.821736688 | -4.806679677 |
| **A_51_P278266** | Gltpd2 | 0.345138485 | -0.551199776 | 1.241476745 | 8.867694219 | 0.947306716 | 0.38083463 | 0.821736688 | -4.806723204 |
| **A_52_P413646** | Bmp6 | -0.05795876 | -0.208480653 | 0.092563133 | 8.331599143 | -0.947302267 | 0.380836718 | 0.821736688 | -4.806725826 |
| **A_55_P2082271** | NA | -0.052765237 | -0.189809753 | 0.084279279 | 16.14660381 | -0.947229893 | 0.380870672 | 0.821736688 | -4.806768478 |
| **A_55_P1991802** | Kat14 | 0.083086864 | -0.132735147 | 0.298908875 | 11.39096669 | 0.947121716 | 0.380921429 | 0.821736688 | -4.806832225 |
| **A_51_P505823** | Endod1 | -0.081428069 | -0.292954597 | 0.130098458 | 11.13123105 | -0.947062111 | 0.380949397 | 0.821736688 | -4.806867348 |
| **A_51_P163797** | Bdp1 | -0.078953959 | -0.284064143 | 0.126156224 | 8.907696962 | -0.947012809 | 0.380972533 | 0.821736688 | -4.806896399 |
| **A_55_P2173927** | Insr | -0.055050805 | -0.198076682 | 0.087975071 | 8.088829655 | -0.946930752 | 0.381011041 | 0.821736688 | -4.806944747 |
| **A_55_P1984770** | Fam220a | -0.100315196 | -0.360955831 | 0.16032544 | 12.39512268 | -0.946877423 | 0.38103607 | 0.821736688 | -4.806976168 |
| **A_55_P2052485** | Ushbp1 | 0.055833596 | -0.089237232 | 0.200904424 | 9.222224203 | 0.946857609 | 0.381045369 | 0.821736688 | -4.806987842 |
| **A_55_P2039215** | Acyp2 | 0.145938263 | -0.23332699 | 0.525203516 | 8.44902617 | 0.946662732 | 0.381136841 | 0.821806523 | -4.807102649 |
| **A_51_P362661** | Spin4 | 0.05839001 | -0.093445036 | 0.210225057 | 8.458875776 | 0.946097052 | 0.38140246 | 0.821806523 | -4.807435822 |
| **A_66_P123527** | Ndufb4 | 0.090715495 | -0.145179347 | 0.326610336 | 13.48655389 | 0.946089434 | 0.381406038 | 0.821806523 | -4.807440308 |
| **A_66_P116154** | Cstf2 | -0.065361283 | -0.235341866 | 0.104619299 | 8.668592715 | -0.94599844 | 0.381448778 | 0.821806523 | -4.807493889 |
| **A_51_P270891** | 1700056E22Rik | 0.044467298 | -0.071190519 | 0.160125115 | 9.012019695 | 0.945877544 | 0.38150557 | 0.821806523 | -4.807565074 |
| **A_55_P2154049** | Myo18a | -0.101498686 | -0.365501723 | 0.162504351 | 8.598485357 | -0.945846496 | 0.381520156 | 0.821806523 | -4.807583353 |
| **A_52_P284958** | Ubtd2 | 0.075895544 | -0.121521466 | 0.273312554 | 9.633138054 | 0.945803401 | 0.381540402 | 0.821806523 | -4.807608726 |
| **A_55_P2128153** | Col6a1 | 0.497646015 | -0.796842893 | 1.792134924 | 10.55288309 | 0.945782688 | 0.381550134 | 0.821806523 | -4.807620921 |
| **A_55_P2228715** | Mm2pr | -0.183731376 | -0.661671972 | 0.294209219 | 9.098764988 | -0.945754837 | 0.381563219 | 0.821806523 | -4.807637318 |
| **A_51_P211934** | Larp7 | -0.062287458 | -0.224321372 | 0.099746456 | 9.621858613 | -0.945722777 | 0.381578282 | 0.821806523 | -4.807656192 |
| **A_51_P202801** | Abcb9 | -0.097562937 | -0.351369745 | 0.156243872 | 10.9383762 | -0.945694273 | 0.381591675 | 0.821806523 | -4.807672973 |
| **A_52_P584415** | Cars2 | -0.06187574 | -0.222865393 | 0.099113913 | 8.031818942 | -0.945565467 | 0.381652201 | 0.821806523 | -4.807748799 |
| **A_51_P479786** | Smtn | -0.05802257 | -0.209017267 | 0.092972126 | 9.965278247 | -0.9453757 | 0.381741384 | 0.821806523 | -4.807860499 |
| **A_52_P77764** | Slc7a2 | 0.071607275 | -0.114741888 | 0.257956438 | 8.130377546 | 0.945363578 | 0.381747082 | 0.821806523 | -4.807867634 |
| **A_51_P437478** | Zfp566 | -0.077250903 | -0.278302583 | 0.123800777 | 9.200715524 | -0.945289907 | 0.38178171 | 0.821806523 | -4.807910994 |
| **A_55_P2035043** | Gm2436 | 0.074487689 | -0.119387911 | 0.268363289 | 10.07293195 | 0.945214793 | 0.381817018 | 0.821806523 | -4.807955201 |
| **A_55_P2045818** | NA | -0.072333142 | -0.260623799 | 0.115957516 | 14.0082404 | -0.945099904 | 0.381871028 | 0.821806523 | -4.808022813 |
| **A_52_P303891** | Nr1d2 | -0.066044828 | -0.237984086 | 0.105894431 | 8.615486618 | -0.945002429 | 0.381916857 | 0.821806523 | -4.808080173 |
| **A_66_P125110** | NA | -0.235539452 | -0.848748165 | 0.377669261 | 10.38786948 | -0.944984113 | 0.381925469 | 0.821806523 | -4.808090951 |
| **A_55_P1985768** | Ttyh2 | 0.12044587 | -0.193127467 | 0.434019206 | 8.759749853 | 0.944978544 | 0.381928087 | 0.821806523 | -4.808094227 |
| **A_55_P1978618** | Grcc10 | 0.084711099 | -0.135900779 | 0.305322976 | 12.65453051 | 0.944671006 | 0.382072709 | 0.822022856 | -4.808275171 |
| **A_52_P477819** | Hmox2 | -0.069984477 | -0.252266286 | 0.112297332 | 11.69034353 | -0.944555658 | 0.382126964 | 0.822022856 | -4.808343029 |
| **A_52_P267391** | Trim12a | -0.080013384 | -0.288430237 | 0.128403469 | 8.795594603 | -0.944493484 | 0.38215621 | 0.822022856 | -4.808379603 |
| **A_55_P2056606** | NA | -0.064084074 | -0.23103566 | 0.102867512 | 8.140615342 | -0.94434074 | 0.382228067 | 0.822039703 | -4.808469447 |
| **A_52_P446120** | Commd6 | 0.161678484 | -0.259545629 | 0.582902597 | 12.16579053 | 0.944296035 | 0.3822491 | 0.822039703 | -4.808495741 |
| **A_52_P279687** | Lrp8 | -0.080918165 | -0.291771905 | 0.129935576 | 9.326663125 | -0.944134519 | 0.382325099 | 0.822107354 | -4.808590732 |
| **A_51_P239750** | Inhba | -1.297240771 | -4.67785402 | 2.083372477 | 13.61532783 | -0.944048407 | 0.382365622 | 0.822107354 | -4.808641373 |
| **A_55_P2074942** | Slc6a4 | -0.104277968 | -0.376138948 | 0.167583012 | 8.51698995 | -0.943658473 | 0.382549163 | 0.822230343 | -4.808870646 |
| **A_55_P2120652** | Snrpa1 | 0.093811449 | -0.150789023 | 0.33841192 | 11.19663814 | 0.943556075 | 0.382597372 | 0.822230343 | -4.808930844 |
| **A_52_P235861** | Peg3 | -0.181058686 | -0.653209184 | 0.291091812 | 11.34485701 | -0.943426509 | 0.38265838 | 0.822230343 | -4.809007008 |
| **A_55_P2053783** | Ppp2r3a | -0.052130618 | -0.188073386 | 0.083812151 | 7.995048635 | -0.943421835 | 0.382660581 | 0.822230343 | -4.809009756 |
| **A_55_P2041397** | Ezh2 | 0.078356321 | -0.12597841 | 0.282691053 | 9.498772752 | 0.94341116 | 0.382665607 | 0.822230343 | -4.80901603 |
| **A_55_P2016014** | Cpsf4l | -0.076833472 | -0.2772301 | 0.123563155 | 8.7494412 | -0.943255217 | 0.382739046 | 0.822230343 | -4.80910769 |
| **A_55_P2158946** | Lama2 | -0.088276134 | -0.318536213 | 0.141983945 | 8.681335458 | -0.943178345 | 0.382775251 | 0.822230343 | -4.80915287 |
| **A_51_P228865** | Zfp72 | -0.049740193 | -0.179485852 | 0.080005467 | 8.776333217 | -0.943156557 | 0.382785513 | 0.822230343 | -4.809165675 |

| | | | | | | | | | |
|---|---|---|---|---|---|---|---|---|---|
| **A_55_P1977149** | Susd6 | 0.125873572 | -0.202492222 | 0.454239366 | 12.5454524 | 0.943074082 | 0.382824362 | 0.822230343 | -4.809214145 |
| **A_55_P2068414** | 6030426L16Rik | -0.100538645 | -0.362836708 | 0.161759418 | 8.764862764 | -0.942990041 | 0.382863951 | 0.822230343 | -4.809263532 |
| **A_51_P406365** | Maf1 | 0.066965189 | -0.107757135 | 0.241687514 | 12.08874888 | 0.942909182 | 0.382902044 | 0.822230343 | -4.809311047 |
| **A_51_P377856** | Gstt3 | 0.142608113 | -0.229504232 | 0.514720457 | 9.474814298 | 0.942842863 | 0.382933289 | 0.822230343 | -4.809350015 |
| **A_55_P2011341** | NA | -0.055624521 | -0.200829437 | 0.089580395 | 7.986999543 | -0.942440906 | 0.383122709 | 0.822545689 | -4.809586166 |
| **A_55_P1979998** | Spaca9 | 0.173095515 | -0.278854685 | 0.625045715 | 10.52066062 | 0.942246201 | 0.383214489 | 0.822551765 | -4.809700532 |
| **A_55_P1966620** | Rab5a | 0.063980717 | -0.10307548 | 0.231036914 | 10.36566116 | 0.942227288 | 0.383223405 | 0.822551765 | -4.809711641 |
| **A_52_P362111** | Fam3c | 0.133170531 | -0.214625143 | 0.480966205 | 11.58999062 | 0.942004808 | 0.3833283 | 0.822551765 | -4.809842302 |
| **A_55_P2092989** | NA | -0.093157212 | -0.336453108 | 0.150138683 | 9.874503895 | -0.941999916 | 0.383330606 | 0.822551765 | -4.809845175 |
| **A_55_P2134032** | NA | 0.096863058 | -0.156155136 | 0.349881253 | 8.704916575 | 0.941836654 | 0.383407596 | 0.822551765 | -4.809941044 |
| **A_65_P19138** | 7-Mar | -0.064622565 | -0.233425744 | 0.104180614 | 11.49953187 | -0.94183047 | 0.383410512 | 0.822551765 | -4.809944675 |
| **A_51_P494361** | Mbnl2 | -0.181901364 | -0.657066983 | 0.293264255 | 12.71853992 | -0.941803087 | 0.383423426 | 0.822551765 | -4.809960754 |
| **A_52_P596541** | Kdelr1 | -0.075361068 | -0.27226346 | 0.121541324 | 9.999366023 | -0.941597323 | 0.383520479 | 0.822552705 | -4.810081564 |
| **A_55_P2003586** | Unkl | -0.094848429 | -0.342677538 | 0.15298068 | 8.814041663 | -0.941557807 | 0.383539119 | 0.822552705 | -4.810104763 |
| **A_51_P211341** | Eif3j1 | -0.093074504 | -0.336275328 | 0.150126319 | 11.10841816 | -0.941531498 | 0.38355153 | 0.822552705 | -4.810120208 |
| **A_55_P2106078** | F10 | 0.123555445 | -0.199332696 | 0.446443585 | 17.44585961 | 0.941410323 | 0.383608697 | 0.822553831 | -4.810191342 |
| **A_52_P158211** | NA | 0.04707902 | -0.075960736 | 0.170118776 | 9.678932481 | 0.941349984 | 0.383637166 | 0.822553831 | -4.810226761 |
| **A_66_P115443** | Snx4 | 0.052260392 | -0.084337525 | 0.188858309 | 8.708213921 | 0.94123431 | 0.383691747 | 0.822579612 | -4.810294658 |
| **A_66_P121249** | Ddi2 | -0.073856577 | -0.266975907 | 0.119262752 | 9.659928502 | -0.940876443 | 0.383860646 | 0.822776885 | -4.810504681 |
| **A_55_P2050799** | Nup35 | 0.092269543 | -0.149069918 | 0.333609004 | 10.21693563 | 0.940587478 | 0.383997068 | 0.822776885 | -4.810674229 |
| **A_51_P343900** | Xrcc1 | 0.060524893 | -0.097790415 | 0.218840202 | 10.34423924 | 0.940546529 | 0.384016403 | 0.822776885 | -4.810698253 |
| **A_52_P533083** | Dnajc16 | -0.085926964 | -0.310703337 | 0.138849408 | 9.566918976 | -0.940476557 | 0.384049445 | 0.822776885 | -4.810739303 |
| **A_55_P2136957** | Wbp1l | -0.166139069 | -0.600771818 | 0.26849368 | 14.79653307 | -0.940412709 | 0.384079596 | 0.822776885 | -4.810776758 |
| **A_51_P139400** | Taf1 | -0.113952538 | -0.412090037 | 0.184184962 | 8.858834614 | -0.940321917 | 0.384122474 | 0.822776885 | -4.810830016 |
| **A_55_P2096043** | Acot11 | 0.057990673 | -0.093736808 | 0.209718153 | 8.468410239 | 0.940292694 | 0.384136276 | 0.822776885 | -4.810847158 |
| **A_52_P426513** | Sh3bgrl | -0.161473828 | -0.583970885 | 0.261023228 | 9.53437139 | -0.940259257 | 0.384152069 | 0.822776885 | -4.810866771 |
| **A_51_P343356** | Isoc2b | 0.060351764 | -0.097564181 | 0.218267709 | 9.776770016 | 0.940227926 | 0.384166868 | 0.822776885 | -4.810885148 |
| **A_55_P2045481** | NA | 0.122255867 | -0.197726934 | 0.442238667 | 13.98028563 | 0.939966199 | 0.384290505 | 0.822828612 | -4.811038649 |
| **A_51_P173735** | Dock11 | -0.097102515 | -0.351305455 | 0.157100424 | 10.31151796 | -0.939764584 | 0.384385768 | 0.822828612 | -4.811156876 |
| **A_51_P245989** | Ccr2 | -0.057331375 | -0.207425985 | 0.092763234 | 7.913811536 | -0.939715583 | 0.384408923 | 0.822828612 | -4.811185608 |
| **A_55_P2079390** | 4930579G24Rik | -0.040443122 | -0.146339937 | 0.065453694 | 8.14430514 | -0.939573874 | 0.384475895 | 0.822828612 | -4.811268694 |
| **A_55_P2031021** | Bex1 | -0.127817736 | -0.462509591 | 0.206874119 | 15.52687499 | -0.939539707 | 0.384492043 | 0.822828612 | -4.811288725 |
| **A_52_P73475** | Fam78a | 0.240665951 | -0.389540179 | 0.870872081 | 8.559295222 | 0.939509533 | 0.384506305 | 0.822828612 | -4.811306415 |
| **A_51_P464576** | Psen1 | -0.054819133 | -0.198395886 | 0.088757621 | 10.56620611 | -0.93932784 | 0.384592191 | 0.822828612 | -4.804100756 |
| **A_51_P450573** | Tgfbr2 | -0.138870583 | -0.502645933 | 0.224904768 | 12.26378267 | -0.939174199 | 0.384664829 | 0.822828612 | -4.811502985 |
| **A_51_P186798** | NA | -0.17447899 | -0.631550245 | 0.282592265 | 8.865709198 | -0.939135791 | 0.384682989 | 0.822828612 | -4.811525496 |
| **A_55_P2052116** | NA | -0.077083968 | -0.279019198 | 0.124851261 | 9.469812053 | -0.939120097 | 0.38469041 | 0.822828612 | -4.811534695 |
| **A_51_P190437** | Ccdc181 | -0.056198488 | -0.20342299 | 0.091026014 | 10.27012083 | -0.939104019 | 0.384698012 | 0.822828612 | -4.811544118 |
| **A_51_P440460** | Hip1r | -0.121601968 | -0.440211514 | 0.197007578 | 8.80404004 | -0.938968416 | 0.384762135 | 0.822828612 | -4.811623591 |
| **A_51_P450527** | Tagln | 0.418581786 | -0.678171907 | 1.515335478 | 13.37639084 | 0.938945499 | 0.384772972 | 0.822828612 | -4.811637021 |
| **A_51_P509999** | Prl8a9 | 0.13245848 | -0.214615663 | 0.479532622 | 17.38426952 | 0.938915847 | 0.384786995 | 0.822828612 | -4.811654398 |
| **A_55_P1968053** | NA | 0.061278141 | -0.09933627 | 0.221892552 | 15.89524015 | 0.938620933 | 0.384926488 | 0.82303585 | -4.811827207 |
| **A_55_P2127265** | NA | -0.078757315 | -0.285260948 | 0.127746317 | 14.31241605 | -0.938279808 | 0.385087886 | 0.823289874 | -4.812027051 |
| **A_51_P365903** | Ankzf1 | -0.07535072 | -0.272945115 | 0.122243675 | 9.651410988 | -0.938170879 | 0.385139436 | 0.823309019 | -4.812090856 |
| **A_55_P2005675** | NA | 0.05733882 | -0.093064308 | 0.207741948 | 15.20188018 | 0.937909735 | 0.385263041 | 0.823335976 | -4.812243801 |
| **A_55_P1998578** | Ifitm2 | -0.06003 | -0.217493691 | 0.097433692 | 10.12019852 | -0.937901164 | 0.385267098 | 0.823335976 | -4.81224882 |
| **A_55_P1962951** | Stx5a | 0.049052418 | -0.079623398 | 0.177728234 | 13.2915773 | 0.937848419 | 0.385292067 | 0.823335976 | -4.812279708 |
| **A_55_P2327177** | Exoc8 | -0.125872084 | -0.456106149 | 0.204361982 | 8.876870404 | -0.937727632 | 0.385349253 | 0.823335976 | -4.812350438 |
| **A_55_P2132132** | Gm5532 | 0.084103918 | -0.136556333 | 0.304764169 | 8.000256907 | 0.937694318 | 0.385365026 | 0.823335976 | -4.812369944 |
| **A_55_P2046092** | NA | 0.088387508 | -0.143576179 | 0.320351195 | 9.663692431 | 0.937432593 | 0.385488962 | 0.823364065 | -4.81252318 |

| | | | | | | | | |
|---|---|---|---|---|---|---|---|---|
| **A_51_P482425** | Lins1 | -0.086566078 | -0.313763336 | 0.140631179 | 9.161442699 | -0.937375983 | 0.385515773 | 0.823364065 | -4.812556321 |
| **A_55_P2106489** | Ankrd52 | 0.060137955 | -0.097699928 | 0.217975839 | 8.776155944 | 0.937360335 | 0.385523185 | 0.823364065 | -4.812565481 |
| **A_52_P343690** | Kpna4 | 0.180052958 | -0.292554848 | 0.652660764 | 11.69269986 | 0.937278237 | 0.38556207 | 0.823364065 | -4.81261354 |
| **A_52_P275408** | Ppp4c | -0.041657561 | -0.151008601 | 0.06769348 | 10.53124544 | -0.937216826 | 0.385591159 | 0.823364065 | -4.812649487 |
| **A_52_P644972** | Mzt1 | 0.044945401 | -0.073075531 | 0.162966333 | 9.33993903 | 0.936904621 | 0.385739071 | 0.823501442 | -4.812832216 |
| **A_55_P2065513** | Ebf1 | 0.060289741 | -0.098024068 | 0.21860355 | 8.100508469 | 0.936901176 | 0.385740704 | 0.823501442 | -4.812834232 |
| **A_55_P2060747** | Esco1 | 0.188954652 | -0.30732357 | 0.685232875 | 10.77030775 | 0.936702188 | 0.385835 | 0.823548993 | -4.812950676 |
| **A_51_P201945** | Cfap97 | -0.043308939 | -0.157060686 | 0.070442808 | 8.736226983 | -0.936674356 | 0.385848191 | 0.823548993 | -4.812966961 |
| **A_51_P113195** | Upk1b | 0.182973169 | -0.297845149 | 0.663791488 | 8.160040714 | 0.936214943 | 0.386065972 | 0.823922841 | -4.813235734 |
| **A_55_P2099650** | Slc24a3 | 0.090773547 | -0.147825801 | 0.329372895 | 9.552594687 | 0.935964157 | 0.386184896 | 0.824085654 | -4.813382418 |
| **A_51_P422751** | Clca1 | -0.244979222 | -0.88915161 | 0.399193166 | 9.662924555 | -0.935613133 | 0.3863514 | 0.824349952 | -4.813587688 |
| **A_55_P2184816** | Eif2s1 | 0.119635461 | -0.194985503 | 0.434256425 | 12.08122025 | 0.935494921 | 0.386407485 | 0.824378619 | -4.813656804 |
| **A_51_P519251** | Nupr1 | -0.226943404 | -0.8239859 | 0.370099092 | 11.18043404 | -0.935150442 | 0.386570958 | 0.824575906 | -4.813858182 |
| **A_52_P563450** | Brix1 | -0.040988088 | -0.148822992 | 0.066846816 | 8.788945312 | -0.93512027 | 0.386585279 | 0.824575906 | -4.813875818 |
| **A_55_P1969072** | Cbx3 | 0.059294208 | -0.096718602 | 0.215307018 | 12.80875244 | 0.935020593 | 0.386632592 | 0.824585829 | -4.813934078 |
| **A_55_P2224582** | Fam219b | 0.051934147 | -0.084758433 | 0.188626728 | 8.526233438 | 0.934710729 | 0.386779702 | 0.824661054 | -4.814115165 |
| **A_55_P2062811** | Tomm5 | 0.164931963 | -0.26919101 | 0.599054937 | 13.38078234 | 0.934676283 | 0.386796058 | 0.824661054 | -4.814135293 |
| **A_66_P110161** | Eppk1 | 0.118280046 | -0.19307412 | 0.429634213 | 9.73277917 | 0.934600409 | 0.386832088 | 0.824661054 | -4.814179627 |
| **A_51_P126198** | Dapk2 | 0.055972828 | -0.091394542 | 0.203340198 | 11.14218455 | 0.93442636 | 0.386914747 | 0.824661054 | -4.814281317 |
| **A_55_P2134690** | Ankrd26 | 0.040604229 | -0.066324348 | 0.147532807 | 8.222619841 | 0.93421459 | 0.387015338 | 0.824661054 | -4.81440503 |
| **A_55_P2049757** | NA | -0.061840289 | -0.224722871 | 0.101042294 | 13.65789078 | -0.934041181 | 0.387097723 | 0.824661054 | -4.81450632 |
| **A_51_P317225** | Cmtm2a | 0.06533876 | -0.106791723 | 0.237469243 | 9.518536237 | 0.933861054 | 0.387183314 | 0.824661054 | -4.814611521 |
| **A_55_P2052172** | NA | 0.112027091 | -0.183189668 | 0.407243849 | 10.44789107 | 0.933579298 | 0.387317226 | 0.824661054 | -4.814776052 |
| **A_55_P2087399** | Tex22 | 0.213800892 | -0.349705518 | 0.777307302 | 8.084653621 | 0.93342583 | 0.38739018 | 0.824661054 | -4.814865656 |
| **A_55_P1987514** | NA | 0.052762743 | -0.086302805 | 0.19182829 | 16.56368788 | 0.933419729 | 0.38739308 | 0.824661054 | -4.814869217 |
| **A_55_P1961414** | NA | -0.131677891 | -0.478751171 | 0.215395389 | 12.82001445 | -0.933385061 | 0.387409562 | 0.824661054 | -4.814889457 |
| **A_55_P2143030** | Timd2 | 0.114102143 | -0.186662514 | 0.4148668 | 8.085439049 | 0.933332016 | 0.387434782 | 0.824661054 | -4.814920425 |
| **A_51_P519189** | Eif3i | 0.095304052 | -0.155941158 | 0.346549262 | 9.515064526 | 0.933217207 | 0.387489371 | 0.824661054 | -4.814987446 |
| **A_51_P198694** | Gckr | -0.053385745 | -0.194127438 | 0.087355948 | 8.027198353 | -0.933193481 | 0.387500653 | 0.824661054 | -4.815001296 |
| **A_51_P223458** | Polr3d | -0.068609052 | -0.249485013 | 0.112266909 | 9.166109871 | -0.933189257 | 0.387502662 | 0.824661054 | -4.815003762 |
| **A_55_P2019934** | Ythdf3 | 0.054128677 | -0.088575609 | 0.196832963 | 9.904962844 | 0.933167403 | 0.387513054 | 0.824661054 | -4.815016519 |
| **A_52_P574653** | Bid | -0.074481054 | -0.270845266 | 0.121883158 | 8.965007574 | -0.933152531 | 0.387520126 | 0.824661054 | -4.8150252 |
| **A_55_P2120577** | Cited2 | -0.23137362 | -0.841384433 | 0.378637193 | 14.42086932 | -0.93313713 | 0.387527449 | 0.824661054 | -4.81503419 |
| **A_55_P2077628** | NA | 0.129629486 | -0.212139395 | 0.471398367 | 8.256316903 | 0.933126333 | 0.387532584 | 0.824661054 | -4.815040492 |
| **A_52_P222096** | 4931428F04Rik | -0.069645522 | -0.253272743 | 0.113981698 | 8.181412076 | -0.933093798 | 0.387548056 | 0.824661054 | -4.815059483 |
| **A_66_P100853** | NA | 0.287670319 | -0.470838835 | 1.046179472 | 14.41275554 | 0.933046722 | 0.387570444 | 0.824661054 | -4.81508696 |
| **A_51_P267194** | Faim | -0.103787977 | -0.377471145 | 0.169895191 | 11.22032289 | -0.932970949 | 0.387606481 | 0.824661054 | -4.815131186 |
| **A_51_P380401** | Cars2 | -0.055232972 | -0.20090061 | 0.090434667 | 8.829372131 | -0.932834271 | 0.387671491 | 0.824708591 | -4.815210954 |
| **A_55_P1996952** | NA | -0.136449016 | -0.496388355 | 0.223490323 | 11.51371088 | -0.932631882 | 0.387767772 | 0.824798381 | -4.815329058 |
| **A_52_P558859** | Rras2 | -0.192839422 | -0.701577218 | 0.315898374 | 10.8533637 | -0.932547537 | 0.387807902 | 0.824798381 | -4.815378272 |
| **A_55_P2102951** | NA | -0.058793682 | -0.213911589 | 0.096324225 | 11.31153994 | -0.932476478 | 0.387841714 | 0.824798381 | -4.815419733 |
| **A_52_P414420** | Dcun1d1 | 0.073446777 | -0.120358615 | 0.267252169 | 10.18487282 | 0.93234372 | 0.387904889 | 0.824821317 | -4.815497186 |
| **A_55_P2214487** | E330013P08Rik | -0.105005741 | -0.382119656 | 0.172108173 | 8.200176694 | -0.932231719 | 0.387958193 | 0.824821317 | -4.815562525 |
| **A_55_P1970650** | Dgkh | -0.057480055 | -0.209179614 | 0.094219503 | 8.518834113 | -0.932184815 | 0.387980517 | 0.824821317 | -4.815589885 |
| **A_55_P2115995** | Sms | -0.084506397 | -0.307564716 | 0.138551921 | 8.997265316 | -0.932052393 | 0.38804355 | 0.824864596 | -4.815667127 |
| **A_55_P1998656** | Entpd5 | 0.09123331 | -0.14961639 | 0.33208301 | 8.605059481 | 0.931915393 | 0.38810877 | 0.824912515 | -4.815747032 |
| **A_52_P574697** | Khdrbs1 | -0.053956024 | -0.196414292 | 0.088502244 | 11.59144976 | -0.931797293 | 0.388164999 | 0.824922008 | -4.815815908 |
| **A_55_P2107422** | Kif21a | 0.077140549 | -0.126546458 | 0.280827556 | 9.376882359 | 0.931726744 | 0.388198592 | 0.824922008 | -4.815857049 |
| **A_51_P418560** | Lnx2 | -0.062442869 | -0.227347854 | 0.102462116 | 10.98116723 | -0.931575863 | 0.388270443 | 0.824983995 | -4.815945031 |
| **A_51_P275435** | Mab21l4 | 0.050488945 | -0.082928658 | 0.183906549 | 7.725143672 | 0.931005674 | 0.388542066 | 0.825428654 | -4.816277433 |

| | | | | | | | | | |
|---|---|---|---|---|---|---|---|---|---|
| **A_51_P502456** | H2-K2 | -0.402604032 | -1.466544993 | 0.66133693 | 12.85157462 | -0.930957282 | 0.388565126 | 0.825428654 | -4.816305639 |
| **A_66_P120736** | Bnip3 | 0.104918699 | -0.172392209 | 0.382229608 | 8.692906042 | 0.930797283 | 0.388641375 | 0.825446059 | -4.816398886 |
| **A_55_P2047361** | Ppih | -0.068336051 | -0.248971864 | 0.112299762 | 8.706235866 | -0.93071172 | 0.388682156 | 0.825446059 | -4.816448749 |
| **A_55_P2014041** | Rab14 | -0.096511893 | -0.351637286 | 0.158613501 | 13.57642776 | -0.930671273 | 0.388701435 | 0.825446059 | -4.816472318 |
| **A_55_P2061319** | Mthfs | -0.051985572 | -0.189425576 | 0.085454433 | 8.371801942 | -0.9305481 | 0.388760149 | 0.825480053 | -4.816544091 |
| **A_51_P508349** | Cdc16 | 0.05501295 | -0.09046537 | 0.20049127 | 12.82336219 | 0.930327409 | 0.388865366 | 0.825576574 | -4.816672671 |
| **A_55_P2137286** | Gm10845 | 0.085964515 | -0.14143067 | 0.313359699 | 14.37636059 | 0.930051755 | 0.388996817 | 0.825576574 | -4.816833246 |
| **A_55_P2028259** | Rhbdf2 | 0.058405008 | -0.096089257 | 0.212899274 | 9.40982606 | 0.930051269 | 0.388997049 | 0.825576574 | -4.81683353 |
| **A_55_P1993365** | NA | 0.092595622 | -0.152351552 | 0.337542797 | 9.342943576 | 0.930009067 | 0.389017177 | 0.825576574 | -4.81685811 |
| **A_55_P2098100** | Arhgef9 | -0.050918119 | -0.18561461 | 0.083778372 | 7.699817377 | -0.930004899 | 0.389019165 | 0.825576574 | -4.816860538 |
| **A_51_P227004** | Cks1b | 0.100478141 | -0.165379073 | 0.366335355 | 11.09137882 | 0.929805915 | 0.38911408 | 0.825687348 | -4.816976429 |
| **A_55_P2071137** | Asap1 | -0.057645302 | -0.210206099 | 0.094915494 | 11.0410237 | -0.929587206 | 0.389218426 | 0.825710592 | -4.817103789 |
| **A_66_P125831** | Pnn | -0.1033933 | -0.377053724 | 0.170267124 | 10.26612527 | -0.929500368 | 0.389259861 | 0.825710592 | -4.817154352 |
| **A_51_P168632** | Prim1 | 0.095596681 | -0.157456391 | 0.348649752 | 9.046674828 | 0.929395066 | 0.389310112 | 0.825710592 | -4.817215661 |
| **A_51_P142164** | Hspa13 | -0.056445475 | -0.205878764 | 0.092987815 | 8.40533573 | -0.929289303 | 0.389360588 | 0.825710592 | -4.817277235 |
| **A_51_P137388** | Zadh2 | -0.086973234 | -0.317231588 | 0.143285119 | 9.818968587 | -0.929264591 | 0.389372383 | 0.825710592 | -4.817291621 |
| **A_52_P444601** | Zfp277 | -0.091498693 | -0.333748569 | 0.150751184 | 9.111292027 | -0.929224148 | 0.389391686 | 0.825710592 | -4.817315165 |
| **A_55_P2052834** | Lst1 | -0.09578056 | -0.349418912 | 0.157857792 | 10.01637227 | -0.929034008 | 0.38948245 | 0.825710592 | -4.817425845 |
| **A_55_P2168664** | Atp5k | 0.158277315 | -0.260914861 | 0.577469492 | 13.43115888 | 0.928912208 | 0.3895406 | 0.825710592 | -4.817496737 |
| **A_51_P181538** | Brwd1 | -0.122084229 | -0.445424707 | 0.201256249 | 9.285292136 | -0.928899334 | 0.389546747 | 0.825710592 | -4.81750423 |
| **A_52_P38157** | Lypla1 | -0.098705768 | -0.360131541 | 0.162720006 | 8.35109043 | -0.928887868 | 0.389552221 | 0.825710592 | -4.817510903 |
| **A_65_P16753** | Bex1 | -0.171015353 | -0.624078002 | 0.282047296 | 13.20148189 | -0.928637044 | 0.389671994 | 0.825800323 | -4.81765687 |
| **A_51_P167048** | Hs6st1 | -0.081346417 | -0.29685741 | 0.134164576 | 9.057999883 | -0.928620277 | 0.389680002 | 0.825800323 | -4.817666627 |
| **A_52_P299363** | NA | -0.054549524 | -0.199093535 | 0.089994487 | 8.630050652 | -0.928453221 | 0.389759791 | 0.825878864 | -4.81776383 |
| **A_51_P272106** | Cirbp | -0.113239865 | -0.413383605 | 0.186903876 | 11.44380407 | -0.928194976 | 0.389883159 | 0.826049718 | -4.81791407 |
| **A_55_P2058746** | Mterf1a | 0.045559439 | -0.075233418 | 0.166352296 | 8.829635633 | 0.927910901 | 0.390018901 | 0.826051824 | -4.818079306 |
| **A_51_P170641** | Dcaf4 | -0.148809029 | -0.54336415 | 0.245746092 | 10.95363284 | -0.92787775 | 0.390034744 | 0.826051824 | -4.818098586 |
| **A_51_P415475** | Dok4 | 0.098294679 | -0.162350865 | 0.358940223 | 8.987330859 | 0.927788247 | 0.390077521 | 0.826051824 | -4.818150639 |
| **A_55_P2135632** | Rps18 | -0.073251644 | -0.267508049 | 0.121004761 | 14.84023966 | -0.927707767 | 0.390115989 | 0.826051824 | -4.818197442 |
| **A_55_P2008874** | Dnmt3b | 0.057401945 | -0.094822978 | 0.209626869 | 8.18439214 | 0.927705239 | 0.390117197 | 0.826051824 | -4.818198911 |
| **A_55_P1953866** | Sema4c | -0.081394571 | -0.297291855 | 0.134502713 | 15.20666837 | -0.927507487 | 0.390211731 | 0.826051824 | -4.818313901 |
| **A_52_P215539** | C030006K11Rik | -0.048483919 | -0.177129493 | 0.080161656 | 8.2053803 | -0.927197015 | 0.390360184 | 0.826051824 | -4.818494403 |
| **A_55_P2049262** | NA | -0.286827231 | -1.047976852 | 0.47432239 | 8.882249331 | -0.927084901 | 0.390413803 | 0.826051824 | -4.818559574 |
| **A_52_P630016** | Bet1l | -0.06674779 | -0.243905749 | 0.110410169 | 10.60707633 | -0.926926689 | 0.390489478 | 0.826051824 | -4.818651533 |
| **A_52_P370022** | NA | -0.104697394 | -0.382594889 | 0.173200101 | 13.91032162 | -0.926873366 | 0.390514986 | 0.826051824 | -4.818682524 |
| **A_55_P1954116** | C8b | -0.042620261 | -0.15576998 | 0.070529458 | 7.668176777 | -0.926684265 | 0.390605455 | 0.826051824 | -4.81879242 |
| **A_52_P467046** | Wdr48 | -0.069605002 | -0.254394871 | 0.115184866 | 9.562710888 | -0.926683562 | 0.390605791 | 0.826051824 | -4.818792828 |
| **A_66_P139647** | Vwa3a | -0.040276077 | -0.147204593 | 0.066652438 | 7.697628521 | -0.926665067 | 0.39061464 | 0.826051824 | -4.818803576 |
| **A_51_P485405** | Gtsf1 | 0.059591394 | -0.098618277 | 0.217801065 | 8.64398332 | 0.92665843 | 0.390617816 | 0.826051824 | -4.818807433 |
| **A_55_P2079142** | Fam124b | 0.065238457 | -0.107964644 | 0.238441558 | 8.51006018 | 0.926653095 | 0.390620368 | 0.826051824 | -4.818810533 |
| **A_55_P2093634** | Ddi2 | -0.076628082 | -0.280076821 | 0.126820657 | 10.07409865 | -0.926620952 | 0.390635748 | 0.826051824 | -4.818829211 |
| **A_66_P106133** | Magi3 | -0.107804866 | -0.394048688 | 0.178438956 | 9.227914849 | -0.926555426 | 0.390667102 | 0.826051824 | -4.818867286 |
| **A_51_P449824** | NA | -0.103261039 | -0.377444429 | 0.170922352 | 8.733891356 | -0.92654072 | 0.39067414 | 0.826051824 | -4.818875831 |
| **A_51_P257156** | Ddrgk1 | -0.123012671 | -0.449657701 | 0.203632359 | 11.72067373 | -0.926494736 | 0.390696144 | 0.826051824 | -4.81890255 |
| **A_51_P138141** | Angel2 | -0.081967808 | -0.299698896 | 0.135763279 | 10.37486062 | -0.926172849 | 0.390850205 | 0.826107874 | -4.819089555 |
| **A_55_P2080804** | Slirp | 0.086250036 | -0.142856778 | 0.315356849 | 12.28372195 | 0.926169427 | 0.390851843 | 0.826107874 | -4.819091544 |
| **A_52_P107234** | Prpf39 | -0.053169559 | -0.194416872 | 0.088077754 | 9.117240403 | -0.926087507 | 0.390891059 | 0.826107874 | -4.819139129 |
| **A_51_P160413** | Yod1 | -0.043742558 | -0.159947291 | 0.072462175 | 8.284760743 | -0.926082175 | 0.390893612 | 0.826107874 | -4.819142227 |
| **A_55_P2108599** | Tlcd1 | -0.049345789 | -0.180466256 | 0.081774677 | 8.446041379 | -0.925867369 | 0.390996456 | 0.826234886 | -4.819266991 |
| **A_55_P2020636** | NA | 0.039190337 | -0.064957376 | 0.143338051 | 7.712587398 | 0.92576003 | 0.391047856 | 0.826253171 | -4.819329329 |

| | | | | | | | | |
|---|---|---|---|---|---|---|---|---|
| **A_52_P379277** | Enpp3 | 0.204989844 | -0.340175499 | 0.750155187 | 8.747062685 | 0.92506717 | 0.391379759 | 0.826864068 | -4.819731599 |
| **A_52_P27332** | Ppp2r5c | -0.051573702 | -0.188754164 | 0.08560676 | 10.42707353 | -0.924922211 | 0.391449227 | 0.826914256 | -4.819815736 |
| **A_55_P2463930** | Zmym5 | -0.079392141 | -0.290588966 | 0.131804683 | 9.923752445 | -0.924824433 | 0.39149609 | 0.826914256 | -4.819872484 |
| **A_55_P2140031** | Fam129b | 0.103923533 | -0.172594131 | 0.380441196 | 15.63931314 | 0.924613405 | 0.391597245 | 0.826914256 | -4.819994946 |
| **A_55_P2048022** | Gm6578 | -0.105430632 | -0.385967192 | 0.175105929 | 10.82611464 | -0.924584278 | 0.391611209 | 0.826914256 | -4.820011848 |
| **A_55_P1973208** | Tax1bp3 | -0.096168685 | -0.352085108 | 0.159747737 | 9.137972437 | -0.924495252 | 0.391653891 | 0.826914256 | -4.820063504 |
| **A_55_P1997375** | NA | 0.061243935 | -0.101751623 | 0.224239493 | 8.433370653 | 0.924392649 | 0.391703086 | 0.826914256 | -4.820123034 |
| **A_52_P412452** | Cntn6 | 0.048180386 | -0.080058971 | 0.176419743 | 7.880920607 | 0.924310975 | 0.39174225 | 0.826914256 | -4.820170418 |
| **A_52_P73369** | Hnrnpu | -0.067604729 | -0.247546328 | 0.112336869 | 11.5539648 | -0.92430365 | 0.391745763 | 0.826914256 | -4.820174668 |
| **A_51_P203192** | Mbd4 | -0.045585012 | -0.166949929 | 0.075779906 | 7.981308259 | -0.924055524 | 0.391864762 | 0.827075123 | -4.820318604 |
| **A_55_P2071097** | Adrm1 | 0.087974313 | -0.146274124 | 0.32222275 | 14.44286599 | 0.923949732 | 0.391915507 | 0.827091914 | -4.820379965 |
| **A_55_P2152246** | Rpl19 | 0.06302108 | -0.104824936 | 0.230867096 | 16.97452712 | 0.923727701 | 0.392022026 | 0.827226391 | -4.820508733 |
| **A_55_P2165869** | Cebpb | -0.062520222 | -0.229089783 | 0.10404934 | 14.66442159 | -0.923408843 | 0.392175036 | 0.827431944 | -4.820693621 |
| **A_55_P1959994** | Unc5cl | 0.194264092 | -0.323348101 | 0.711876286 | 8.228176173 | 0.923330535 | 0.392212621 | 0.827431944 | -4.820739021 |
| **A_55_P1989276** | Romo1 | -0.05897663 | -0.216136811 | 0.09818355 | 13.74427664 | -0.923222965 | 0.392264254 | 0.827431944 | -4.820801382 |
| **A_51_P310548** | Osgep | -0.055740704 | -0.204286682 | 0.092805274 | 10.24918797 | -0.92316794 | 0.392290669 | 0.827431944 | -4.820833279 |
| **A_51_P437289** | Smco4 | 0.087637818 | -0.145947987 | 0.321223624 | 9.166852552 | 0.923026716 | 0.392358468 | 0.827481804 | -4.82091514 |
| **A_55_P1987302** | Eny2 | -0.074265822 | -0.272229045 | 0.123697401 | 11.83723922 | -0.922940364 | 0.392399928 | 0.827481804 | -4.82096519 |
| **A_52_P239023** | Zfp955b | -0.047087714 | -0.172644577 | 0.078469148 | 7.989570679 | -0.922648581 | 0.392540048 | 0.827630638 | -4.821134287 |
| **A_51_P468073** | Ggt1 | -0.234815337 | -0.860960571 | 0.391329896 | 9.625863969 | -0.92261507 | 0.392556143 | 0.827630638 | -4.821153705 |
| **A_55_P2086978** | Col23a1 | -0.140517082 | -0.515272107 | 0.234237942 | 8.536017481 | -0.922466966 | 0.392627283 | 0.827690342 | -4.82123952 |
| **A_55_P1970537** | Bcl7b | 0.077478272 | -0.129322635 | 0.284279178 | 10.58889861 | 0.921714989 | 0.392988637 | 0.828228711 | -4.821675097 |
| **A_55_P1963062** | Rps16 | -0.059580126 | -0.218629407 | 0.099469155 | 17.02543032 | -0.92159237 | 0.393047584 | 0.828228711 | -4.821746101 |
| **A_66_P125067** | Snrpd1 | 0.118585288 | -0.197980264 | 0.435150841 | 10.39691817 | 0.921586963 | 0.393050184 | 0.828228711 | -4.821749232 |
| **A_51_P158400** | Rc3h2 | -0.087366933 | -0.32059657 | 0.145862703 | 10.6489312 | -0.921578896 | 0.393054062 | 0.828228711 | -4.821753903 |
| **A_51_P359549** | Fem1c | -0.078041052 | -0.286424583 | 0.130342478 | 9.924429024 | -0.92135901 | 0.393159788 | 0.828238057 | -4.821881214 |
| **A_51_P129435** | Arid2 | -0.073094319 | -0.268304016 | 0.122115378 | 8.300338604 | -0.921194632 | 0.393238838 | 0.828238057 | -4.821976374 |
| **A_52_P380329** | Tent4a | -0.052840356 | -0.193961649 | 0.088280937 | 10.14476097 | -0.921175447 | 0.393248065 | 0.828238057 | -4.82198748 |
| **A_55_P2008136** | Ctr9 | -0.091628509 | -0.336344261 | 0.153087242 | 9.633028162 | -0.921165914 | 0.39325265 | 0.828238057 | -4.821992998 |
| **A_52_P451834** | Stk35 | 0.070412832 | -0.117654365 | 0.258480028 | 9.785827012 | 0.92110242 | 0.393283189 | 0.828238057 | -4.822029752 |
| **A_51_P400773** | Hdgfl2 | -0.056721491 | -0.208231262 | 0.094788281 | 10.5849675 | -0.921035048 | 0.393315595 | 0.828238057 | -4.822068748 |
| **A_51_P346884** | Snrnp27 | 0.073798622 | -0.12339215 | 0.270989395 | 11.85011922 | 0.920726895 | 0.393463843 | 0.828424005 | -4.822247091 |
| **A_51_P394802** | Fam111a | -0.071403599 | -0.262205942 | 0.119398744 | 8.279172955 | -0.920673331 | 0.393489616 | 0.828424005 | -4.822278087 |
| **A_51_P198835** | Dnajb9 | 0.150958604 | -0.252473655 | 0.554390863 | 12.30958654 | 0.92056914 | 0.393539754 | 0.828434171 | -4.822338377 |
| **A_51_P469008** | Ehd1 | -0.082366482 | -0.302519419 | 0.137786455 | 15.22748985 | -0.920439395 | 0.393602194 | 0.828434171 | -4.822413446 |
| **A_51_P501844** | Cyp26b1 | -0.569797121 | -2.092857783 | 0.953263541 | 11.63867884 | -0.92039068 | 0.39362564 | 0.828434171 | -4.82244163 |
| **A_51_P421958** | Sgms1 | 0.050411714 | -0.084361071 | 0.185184499 | 12.32949571 | 0.92023431 | 0.393700908 | 0.828434171 | -4.822532093 |
| **A_55_P2028710** | Rnf170 | -0.082543126 | -0.303221274 | 0.138135023 | 10.82892923 | -0.920218035 | 0.393708743 | 0.828434171 | -4.822541508 |
| **A_55_P1961965** | Plagl1 | 0.093777022 | -0.157068665 | 0.344622709 | 9.068989965 | 0.919727055 | 0.393945149 | 0.828841383 | -4.822825479 |
| **A_51_P330615** | Zfp119b | -0.157279221 | -0.578049254 | 0.263490812 | 10.89849245 | -0.919593115 | 0.39400966 | 0.828886886 | -4.822902929 |
| **A_55_P1991406** | Fbxo34 | 0.092740011 | -0.155417623 | 0.340897646 | 9.877887326 | 0.919408822 | 0.394098436 | 0.828907407 | -4.823009484 |
| **A_51_P307644** | Gtf3a | -0.063558516 | -0.233633667 | 0.106516634 | 10.20597333 | -0.919394817 | 0.394105183 | 0.828907407 | -4.823017581 |
| **A_52_P686091** | Phf2 | -0.091816925 | -0.337544855 | 0.153911005 | 9.307651918 | -0.919257931 | 0.394171133 | 0.828955917 | -4.823096717 |
| **A_55_P2058467** | Gm4415 | -0.303436255 | -1.115632201 | 0.508759692 | 12.17009853 | -0.919127615 | 0.394233926 | 0.828997776 | -4.823172047 |
| **A_52_P561736** | NA | 0.056624802 | -0.094959727 | 0.208209331 | 11.03780823 | 0.919011572 | 0.394289848 | 0.829025179 | -4.82323912 |
| **A_55_P2013680** | NA | -0.065918919 | -0.242415029 | 0.110577191 | 16.33252641 | -0.9188489 | 0.394368251 | 0.829071369 | -4.823333136 |
| **A_52_P283055** | Slc5a3 | 0.062678781 | -0.105162243 | 0.230519804 | 9.730234489 | 0.918737798 | 0.394421806 | 0.829071369 | -4.823397342 |
| **A_51_P510663** | Supt3 | -0.045624648 | -0.167807677 | 0.076558381 | 8.163564411 | -0.918666338 | 0.394456255 | 0.829071369 | -4.823438635 |
| **A_55_P2069949** | Dynlrb1 | 0.061447827 | -0.103119834 | 0.226015488 | 13.59457394 | 0.91861006 | 0.394483387 | 0.829071369 | -4.823471154 |
| **A_55_P2101739** | NA | -0.094708708 | -0.348424882 | 0.159007466 | 11.04845239 | -0.918355691 | 0.394606036 | 0.829195303 | -4.82361812 |

| | | | | | | | | | |
|---|---|---|---|---|---|---|---|---|---|
| **A_52_P502771** | Rad54b | -0.04303211 | -0.158349795 | 0.072285575 | 7.850359885 | -0.918049049 | 0.39475393 | 0.829195303 | -4.823795252 |
| **A_55_P2192492** | Slmap | -0.050318244 | -0.185164439 | 0.084527951 | 8.265200857 | -0.91802803 | 0.394764069 | 0.829195303 | -4.823807392 |
| **A_55_P2018403** | Sycp1 | -0.090772457 | -0.334067658 | 0.152522743 | 8.863270299 | -0.917888047 | 0.394831599 | 0.829195303 | -4.82388824 |
| **A_55_P2022082** | Klf13 | -0.092403175 | -0.340071854 | 0.155265504 | 11.09384382 | -0.917878019 | 0.394836436 | 0.829195303 | -4.823894031 |
| **A_55_P2108067** | Dgcr8 | 0.076314836 | -0.128241435 | 0.280871108 | 10.09208445 | 0.917836536 | 0.39485645 | 0.829195303 | -4.823917988 |
| **A_55_P2274378** | AW549542 | 0.041783046 | -0.070223673 | 0.153789765 | 7.828442122 | 0.917751664 | 0.3948974 | 0.829195303 | -4.823967 |
| **A_51_P377452** | Ncf4 | 0.127259053 | -0.213894862 | 0.468412967 | 8.975303869 | 0.91771428 | 0.394915438 | 0.829195303 | -4.823988588 |
| **A_51_P156263** | Zfp511 | -0.057132726 | -0.21029767 | 0.096032218 | 10.05702955 | -0.917687318 | 0.394928448 | 0.829195303 | -4.824004157 |
| **A_52_P124173** | Zdhhc8 | 0.044126466 | -0.074208409 | 0.162461341 | 8.03858958 | 0.917393277 | 0.395070352 | 0.829403154 | -4.82417393 |
| **A_55_P2125820** | Gm14288 | 0.065833435 | -0.110801606 | 0.242468476 | 10.55082761 | 0.916935554 | 0.395291327 | 0.829715921 | -4.824438139 |
| **A_55_P2046393** | Rcn3 | 0.119314012 | -0.200822635 | 0.439450659 | 13.15497426 | 0.916906857 | 0.395305185 | 0.829715921 | -4.824454701 |
| **A_55_P2162698** | Prpf40b | -0.056036325 | -0.206415877 | 0.094343226 | 8.36983903 | -0.916748114 | 0.395381845 | 0.82976162 | -4.82454631 |
| **A_52_P622230** | Gm4890 | 0.04730955 | -0.079667376 | 0.174286476 | 8.618443757 | 0.916628104 | 0.395439808 | 0.82976162 | -4.824615558 |
| **A_55_P2125496** | Gm10731 | 0.07375705 | -0.124256456 | 0.271770557 | 9.650973702 | 0.916384825 | 0.395557329 | 0.82976162 | -4.82475592 |
| **A_55_P1954400** | NA | -0.07104566 | -0.261784718 | 0.119693398 | 10.1849617 | -0.916362022 | 0.395568346 | 0.82976162 | -4.824769075 |
| **A_55_P2162765** | Dnajc24 | 0.114312654 | -0.192649597 | 0.421274905 | 10.18652485 | 0.916175134 | 0.395658646 | 0.82976162 | -4.824876882 |
| **A_55_P1976574** | Tmod4 | -0.111235194 | -0.409943233 | 0.187472846 | 8.36128009 | -0.91614553 | 0.395672951 | 0.82976162 | -4.824893958 |
| **A_55_P2036285** | Elp4 | -0.05362934 | -0.197677931 | 0.090419251 | 8.299046055 | -0.915930664 | 0.395776792 | 0.82976162 | -4.825017885 |
| **A_55_P2032695** | Tut4 | -0.097092769 | -0.357900929 | 0.163715391 | 11.88482191 | -0.915872199 | 0.395805051 | 0.82976162 | -4.825051602 |
| **A_55_P1972481** | NA | -0.355204585 | -1.309367413 | 0.598958243 | 13.30697139 | -0.915851863 | 0.39581488 | 0.82976162 | -4.825063329 |
| **A_51_P391805** | Dkkl1 | 0.215701475 | -0.363737833 | 0.795140782 | 12.10740887 | 0.915828858 | 0.395826 | 0.82976162 | -4.825076596 |
| **A_51_P431737** | Cth | 0.127926457 | -0.215751139 | 0.471604054 | 8.632319203 | 0.915752924 | 0.395862706 | 0.82976162 | -4.825120384 |
| **A_52_P462350** | Dr1 | -0.066314846 | -0.251005344 | 0.118375652 | 9.804361777 | -0.930499825 | 0.395906122 | 0.82976162 | -4.769708546 |
| **A_55_P2137023** | NA | 0.083841928 | -0.141446446 | 0.309130302 | 8.649885467 | 0.915570183 | 0.395951053 | 0.82976162 | -4.825225754 |
| **A_55_P2038298** | Rpl31 | -0.039418785 | -0.145343763 | 0.066506194 | 15.89513421 | -0.91553301 | 0.395969026 | 0.82976162 | -4.825247186 |
| **A_51_P267370** | Itm2c | -0.10202373 | -0.376189313 | 0.172141854 | 10.77522439 | -0.915498048 | 0.39598593 | 0.82976162 | -4.825267343 |
| **A_52_P558087** | Poc5 | 0.059117211 | -0.099756862 | 0.217991283 | 9.983159258 | 0.915440389 | 0.396013811 | 0.82976162 | -4.825300585 |
| **A_55_P2139889** | Zfp783 | -0.058400861 | -0.215391861 | 0.098590138 | 9.259716073 | -0.915195047 | 0.39613246 | 0.829803044 | -4.825442017 |
| **A_55_P2012047** | Gigyf1 | -0.103849161 | -0.383025044 | 0.175326721 | 9.741904499 | -0.915154162 | 0.396152235 | 0.829803044 | -4.825465583 |
| **A_51_P482503** | Tpd52l1 | -0.05040569 | -0.185923246 | 0.085111866 | 8.2303371 | -0.915067575 | 0.396194117 | 0.829803044 | -4.82551549 |
| **A_55_P2065567** | Chac2 | 0.096189386 | -0.162462107 | 0.35484088 | 8.368013224 | 0.914916219 | 0.396267336 | 0.829803044 | -4.825602721 |
| **A_55_P2140953** | NA | 0.106940479 | -0.180649282 | 0.39453024 | 16.10411733 | 0.914824671 | 0.396311629 | 0.829803044 | -4.825655479 |
| **A_55_P2019190** | NA | 0.052764364 | -0.089146852 | 0.19467558 | 15.22153839 | 0.914730479 | 0.396357203 | 0.829803044 | -4.825709756 |
| **A_55_P2018559** | Morf4l1 | -0.086959439 | -0.320851774 | 0.146932896 | 11.32012233 | -0.914681515 | 0.396380897 | 0.829803044 | -4.82573797 |
| **A_55_P1969522** | NA | -0.059860339 | -0.220878309 | 0.101157632 | 9.018020195 | -0.914605872 | 0.396417501 | 0.829803044 | -4.825781555 |
| **A_55_P1967539** | Hunk | 0.098622819 | -0.166681536 | 0.363927173 | 9.802202353 | 0.914538928 | 0.396449898 | 0.829803044 | -4.825820124 |
| **A_51_P209302** | Polr1e | 0.047994431 | -0.08112738 | 0.177116243 | 8.814676639 | 0.914450926 | 0.39649249 | 0.829803044 | -4.825870825 |
| **A_55_P2452914** | N4bp2 | -0.106708173 | -0.39381827 | 0.180401924 | 10.9208183 | -0.91436245 | 0.396535314 | 0.829803044 | -4.825921795 |
| **A_55_P2003129** | Nemp1 | -0.050629695 | -0.186891952 | 0.085632562 | 8.162932842 | -0.914110922 | 0.396657078 | 0.829803044 | -4.82606668 |
| **A_55_P2015994** | Fgf9 | 0.06829402 | -0.115510214 | 0.252098254 | 8.004446872 | 0.914105517 | 0.396659695 | 0.829803044 | -4.826069793 |
| **A_55_P2097840** | NA | 0.058173269 | -0.098422609 | 0.214769148 | 8.48324432 | 0.913928681 | 0.396745319 | 0.829803044 | -4.826171638 |
| **A_66_P119074** | Tbc1d22b | 0.042081017 | -0.07120751 | 0.155369544 | 8.639244093 | 0.913838521 | 0.396788979 | 0.829803044 | -4.826223559 |
| **A_52_P609200** | Serinc3 | -0.106286912 | -0.392444917 | 0.179871092 | 15.1740848 | -0.91378296 | 0.396815887 | 0.829803044 | -4.826255553 |
| **A_51_P324410** | Vti1a | -0.064820144 | -0.239342674 | 0.109702385 | 9.904094059 | -0.913750566 | 0.396831576 | 0.829803044 | -4.826274207 |
| **A_55_P2038347** | Acot3 | -0.049200562 | -0.181674002 | 0.083272879 | 8.081080066 | -0.913714274 | 0.396849153 | 0.829803044 | -4.826295104 |
| **A_51_P494686** | Sumf1 | -0.047431924 | -0.175147694 | 0.080283846 | 11.22138399 | -0.913682645 | 0.396864472 | 0.829803044 | -4.826313316 |
| **A_55_P1994817** | Taf15 | -0.068317807 | -0.252286853 | 0.115651238 | 9.691665713 | -0.913604708 | 0.396902222 | 0.829803044 | -4.82635819 |
| **A_55_P2037712** | Smc4 | 0.106177047 | -0.179778234 | 0.392132327 | 9.903048351 | 0.913485556 | 0.396959941 | 0.829803044 | -4.82642679 |
| **A_52_P629895** | Adh1 | 0.196241691 | -0.33229669 | 0.724780072 | 9.127792897 | 0.913448173 | 0.396978051 | 0.829803044 | -4.826448312 |
| **A_51_P373461** | Brcc3 | -0.084489826 | -0.312099938 | 0.143120285 | 10.06795449 | -0.913233907 | 0.397081864 | 0.829930292 | -4.826571654 |

| | | | | | | | | | |
|---|---|---|---|---|---|---|---|---|---|
| **A_52_P88091** | Dsg2 | -0.082414113 | -0.304461603 | 0.139633377 | 8.715230906 | -0.913113755 | 0.397140087 | 0.829949343 | -4.826640811 |
| **A_55_P1994913** | Ppp2cb | 0.057246259 | -0.09700456 | 0.211497079 | 12.85003643 | 0.913037887 | 0.397176854 | 0.829949343 | -4.826684476 |
| **A_55_P1973451** | Stra6 | 0.150081048 | -0.254383177 | 0.554545273 | 11.98226542 | 0.912882545 | 0.397252145 | 0.829970607 | -4.826773874 |
| **A_52_P336958** | Jcad | -0.240889612 | -0.890250131 | 0.408470907 | 12.22076149 | -0.9126439 | 0.397367832 | 0.829970607 | -4.826911193 |
| **A_55_P2067787** | Psg29 | -0.297959427 | -1.101212565 | 0.505293712 | 13.9487911 | -0.912586126 | 0.397395843 | 0.829970607 | -4.826944434 |
| **A_52_P413855** | Slc35a3 | 0.052404138 | -0.088870094 | 0.19367837 | 8.877018647 | 0.912581773 | 0.397397953 | 0.829970607 | -4.826946938 |
| **A_55_P2053733** | NA | 0.093959514 | -0.159344363 | 0.347263391 | 15.94004599 | 0.912573991 | 0.397401726 | 0.829970607 | -4.826951415 |
| **A_55_P1983016** | NA | -0.128850684 | -0.476262577 | 0.218561209 | 8.238446086 | -0.912454487 | 0.397459671 | 0.830001943 | -4.827020167 |
| **A_55_P2020706** | NA | 0.040764769 | -0.06918527 | 0.150714809 | 7.741752509 | 0.912134254 | 0.397614977 | 0.830207047 | -4.827204371 |
| **A_66_P115378** | Orc2 | 0.042780416 | -0.072613705 | 0.158174537 | 8.185078302 | 0.912074853 | 0.397643791 | 0.830207047 | -4.827238535 |
| **A_55_P2181655** | Tns3 | 0.062339886 | -0.10585257 | 0.230532343 | 8.399085876 | 0.91186103 | 0.397747522 | 0.83033393 | -4.8273615 |
| **A_55_P2163353** | Sox3 | 0.097639081 | -0.165940678 | 0.36121884 | 9.038278069 | 0.911340752 | 0.39800001 | 0.830680149 | -4.827660624 |
| **A_55_P1973159** | Serpinf1 | 0.263192667 | -0.447368355 | 0.973753688 | 8.892110583 | 0.911258447 | 0.398039963 | 0.830680149 | -4.827707934 |
| **A_52_P480360** | Dut | 0.116194781 | -0.197531748 | 0.42992131 | 10.04602818 | 0.911180757 | 0.398077678 | 0.830680149 | -4.827752587 |
| **A_51_P481693** | Ero1l | -0.105806957 | -0.391491731 | 0.179877818 | 9.186850704 | -0.91116345 | 0.398086081 | 0.830680149 | -4.827762535 |
| **A_52_P385206** | Harbi1 | 0.056968816 | -0.096864877 | 0.21080251 | 8.650505071 | 0.9110766 | 0.398128248 | 0.830680149 | -4.82781245 |
| **A_51_P203182** | Apobr | -0.105020859 | -0.388742165 | 0.178700447 | 10.84714771 | -0.910652696 | 0.398334107 | 0.831019962 | -4.828056036 |
| **A_55_P1973178** | Haus5 | -0.045329405 | -0.167822317 | 0.077163508 | 8.661591621 | -0.910412514 | 0.398450782 | 0.831173664 | -4.828194018 |
| **A_55_P1986053** | Pcnx | 0.079934515 | -0.136128899 | 0.295997929 | 8.947746803 | 0.910169487 | 0.398568866 | 0.831175256 | -4.828333609 |
| **A_55_P2026547** | Gal3st2 | 0.094230114 | -0.160479717 | 0.348939945 | 8.508574858 | 0.910150416 | 0.398578133 | 0.831175256 | -4.828344563 |
| **A_55_P2210575** | A830073O21Rik | 0.114708942 | -0.195369829 | 0.424787713 | 14.18661754 | 0.910111083 | 0.398597248 | 0.831175256 | -4.828367153 |
| **A_55_P2099520** | H2afv | 0.070395528 | -0.119907483 | 0.260698539 | 8.628330866 | 0.910056958 | 0.398623551 | 0.831175256 | -4.828398237 |
| **A_55_P2147141** | Spata2 | -0.095308088 | -0.353006112 | 0.162389937 | 11.59721246 | -0.909887774 | 0.39870578 | 0.83120569 | -4.828495391 |
| **A_55_P1984825** | Bloc1s2 | 0.122981565 | -0.209597228 | 0.455560359 | 10.94165476 | 0.909734419 | 0.398780326 | 0.83120569 | -4.828583446 |
| **A_55_P1982245** | Phactr2 | -0.041185509 | -0.152566505 | 0.070195486 | 7.886550498 | -0.909709042 | 0.398792663 | 0.83120569 | -4.828598016 |
| **A_55_P2083121** | Vill | 0.261778991 | -0.446293166 | 0.969851147 | 10.98097793 | 0.9095497 | 0.398870134 | 0.83120569 | -4.828689497 |
| **A_55_P2028159** | C9orf72 | -0.065965014 | -0.244417361 | 0.112487333 | 11.31718579 | -0.90941174 | 0.398937217 | 0.83120569 | -4.828768693 |
| **A_55_P2040863** | Cnn3 | -0.105181395 | -0.389741059 | 0.17937827 | 12.76347609 | -0.9093577 | 0.398963497 | 0.83120569 | -4.828799712 |
| **A_52_P480939** | Tnpo1 | -0.063471799 | -0.235193362 | 0.108249763 | 10.98050364 | -0.909337588 | 0.398973278 | 0.83120569 | -4.828811257 |
| **A_55_P2027147** | Mrps18c | -0.099611237 | -0.369167491 | 0.169945018 | 11.863765 | -0.909134379 | 0.399072112 | 0.83120569 | -4.828927888 |
| **A_55_P2169039** | Olfr558 | 0.045509282 | -0.077644943 | 0.168663506 | 7.755587727 | 0.909117106 | 0.399080514 | 0.83120569 | -4.828937802 |
| **A_55_P2181444** | NA | -0.100255587 | -0.371570218 | 0.171059044 | 13.01713003 | -0.909085086 | 0.399096089 | 0.83120569 | -4.828956177 |
| **A_55_P2070657** | Emc10 | -0.050483942 | -0.187122078 | 0.086154194 | 8.189043449 | -0.908971974 | 0.399151113 | 0.83120569 | -4.829021088 |
| **A_51_P304109** | Cyp2c39 | 0.045736197 | -0.078056865 | 0.169529259 | 7.661898889 | 0.908935164 | 0.399169021 | 0.83120569 | -4.829042211 |
| **A_55_P1958517** | Parp6 | 0.087657608 | -0.14961824 | 0.324933456 | 11.34327121 | 0.908877272 | 0.399197187 | 0.83120569 | -4.82907543 |
| **A_55_P2034372** | Eif4g3 | -0.132541032 | -0.491354518 | 0.226272455 | 10.44984513 | -0.908763242 | 0.399252668 | 0.83123167 | -4.829140858 |
| **A_51_P329928** | Phlda3 | -0.146703876 | -0.543924405 | 0.250516652 | 12.76139134 | -0.908613253 | 0.399325655 | 0.831294086 | -4.82922691 |
| **A_55_P1978962** | Slc9a3r2 | 0.11342556 | -0.19378374 | 0.420634861 | 9.81782885 | 0.908334349 | 0.399461401 | 0.831487123 | -4.829386899 |
| **A_55_P2015304** | NA | -0.085542586 | -0.317305078 | 0.146219905 | 12.54662858 | -0.908047128 | 0.399601232 | 0.831638665 | -4.829551624 |
| **A_55_P1984098** | Mst1 | 0.213310737 | -0.364702587 | 0.791324061 | 8.513201589 | 0.90791257 | 0.399666753 | 0.831638665 | -4.829628783 |
| **A_55_P2000487** | Srebf2 | 0.27079021 | -0.463000474 | 1.004580893 | 9.674952887 | 0.907883092 | 0.399681108 | 0.831638665 | -4.829645686 |
| **A_55_P2055763** | Gm5878 | -0.263540082 | -0.977776182 | 0.450696019 | 11.01698627 | -0.90776627 | 0.399738001 | 0.831638665 | -4.829712667 |
| **A_51_P144926** | Cops8 | -0.063808226 | -0.23675145 | 0.109134998 | 9.611816208 | -0.907699893 | 0.39977033 | 0.831638665 | -4.829750723 |
| **A_55_P1963339** | Washc5 | 0.065162438 | -0.111459986 | 0.241784862 | 12.03944344 | 0.907654669 | 0.399792357 | 0.831638665 | -4.82977665 |
| **A_51_P342707** | Pold2 | 0.04275861 | -0.073159236 | 0.158676455 | 8.084005078 | 0.907491234 | 0.39987197 | 0.831714764 | -4.82987034 |
| **A_51_P246677** | Rec8 | 0.185167124 | -0.316914835 | 0.687249083 | 10.68624032 | 0.907315706 | 0.399957487 | 0.831803127 | -4.829970951 |
| **A_66_P135941** | Fam221a | -0.048729502 | -0.180905576 | 0.083446573 | 8.588623115 | -0.90700208 | 0.400110319 | 0.832031453 | -4.830150685 |
| **A_55_P1971962** | Tmem176b | -0.047341395 | -0.175783813 | 0.081101024 | 8.164902426 | -0.906779591 | 0.400218767 | 0.832132952 | -4.830278166 |
| **A_55_P1979708** | Ubap1 | -0.101699431 | -0.377637707 | 0.174238846 | 10.82616726 | -0.906725307 | 0.40024523 | 0.832132952 | -4.830309266 |
| **A_52_P484081** | Ube2v1 | 0.043713932 | -0.074907841 | 0.162335705 | 9.321127794 | 0.906618595 | 0.400297255 | 0.832151607 | -4.830370399 |

| | | | | | | | | | |
|---|---|---|---|---|---|---|---|---|---|
| **A_55_P2052709** | NA | -0.049978403 | -0.185614236 | 0.085657431 | 8.077658383 | -0.906519382 | 0.400345629 | 0.832162669 | -4.830427232 |
| **A_51_P373369** | Med4 | 0.124689277 | -0.213737772 | 0.463116326 | 10.76651416 | 0.906427771 | 0.4003903 | 0.832166033 | -4.830479707 |
| **A_52_P637282** | Wipf1 | -0.104552226 | -0.388353781 | 0.17924933 | 10.13170942 | -0.906332751 | 0.400436638 | 0.83217286 | -4.830534131 |
| **A_52_P77204** | Lrrc20 | 0.14600814 | -0.250389362 | 0.542405642 | 10.28822892 | 0.906181771 | 0.400510273 | 0.832236409 | -4.830620599 |
| **A_52_P245365** | Rtf1 | -0.058191059 | -0.216201772 | 0.099819655 | 8.468733608 | -0.9060223 | 0.400588062 | 0.832303896 | -4.830711919 |
| **A_55_P2057720** | Zfp937 | 0.066719933 | -0.114466626 | 0.247906492 | 9.05128362 | 0.905938647 | 0.400628871 | 0.832303896 | -4.830759818 |
| **A_51_P411917** | Gata6 | 0.094601603 | -0.162365698 | 0.351568905 | 9.165440353 | 0.905711323 | 0.400739785 | 0.832340266 | -4.830889968 |
| **A_55_P2410626** | A430061O12Rik | -0.138131626 | -0.513358125 | 0.237094873 | 8.188003709 | -0.905667505 | 0.400761168 | 0.832340266 | -4.830915053 |
| **A_51_P330630** | Usp3 | 0.075622198 | -0.12980778 | 0.281052177 | 8.300191711 | 0.905638006 | 0.400775563 | 0.832340266 | -4.830931939 |
| **A_51_P446469** | Dok2 | -0.199300791 | -0.740961807 | 0.342360226 | 10.04716769 | -0.905212629 | 0.400983187 | 0.832638103 | -4.831175409 |
| **A_51_P397564** | Snrnp40 | 0.055954574 | -0.096135017 | 0.208044166 | 10.08640668 | 0.905118134 | 0.401029321 | 0.832638103 | -4.831229484 |
| **A_55_P2042823** | Sh3bp2 | -0.066775185 | -0.248283929 | 0.11473356 | 9.115822356 | -0.905079458 | 0.401048204 | 0.832638103 | -4.831251616 |
| **A_55_P1997429** | Gm13698 | -0.076959841 | -0.286190175 | 0.132270493 | 9.390683333 | -0.904916857 | 0.401127601 | 0.832664063 | -4.831344654 |
| **A_55_P2035265** | Trpm7 | 0.05058053 | -0.086953934 | 0.188114993 | 10.85102123 | 0.904775847 | 0.401196464 | 0.832664063 | -4.831425328 |
| **A_52_P104824** | Diaph3 | -0.052668207 | -0.195879707 | 0.090543293 | 8.127458797 | -0.904773355 | 0.401197681 | 0.832664063 | -4.831426754 |
| **A_51_P407606** | Ndc1 | -0.128379671 | -0.477588293 | 0.220828951 | 10.20805337 | -0.904441461 | 0.4013598 | 0.832664063 | -4.831616604 |
| **A_51_P340668** | Bcl9l | -0.048102937 | -0.178951021 | 0.082745146 | 8.429720089 | -0.904426742 | 0.401366991 | 0.832664063 | -4.831625023 |
| **A_52_P351925** | Ank3 | -0.052540543 | -0.195508673 | 0.090427586 | 8.050548108 | -0.904116693 | 0.401518486 | 0.832664063 | -4.831802333 |
| **A_52_P167411** | Tnk1 | 0.09319275 | -0.160397248 | 0.346782747 | 8.330836256 | 0.904105621 | 0.401523897 | 0.832664063 | -4.831808664 |
| **A_51_P296057** | Upf1 | 0.055999489 | -0.096407751 | 0.20840673 | 9.332642782 | 0.903956701 | 0.401596679 | 0.832664063 | -4.831893814 |
| **A_51_P109835** | Uqcrc1 | -0.090366216 | -0.336323831 | 0.1555914 | 10.97070024 | -0.903888758 | 0.401629888 | 0.832664063 | -4.83193266 |
| **A_55_P1961084** | Map3k1 | 0.121976522 | -0.2100421 | 0.453995144 | 11.77453755 | 0.90382212 | 0.401662461 | 0.832664063 | -4.831970757 |
| **A_51_P224227** | Rlf | -0.047737051 | -0.17768919 | 0.082215089 | 10.32372907 | -0.903735437 | 0.401704835 | 0.832664063 | -4.832020311 |
| **A_55_P2039354** | NA | -0.049844 | -0.185532061 | 0.085844062 | 7.852663297 | -0.90373356 | 0.401705753 | 0.832664063 | -4.832021384 |
| **A_55_P1955528** | NA | 0.057540951 | -0.099105197 | 0.214187098 | 10.69304309 | 0.90370457 | 0.401719925 | 0.832664063 | -4.832037957 |
| **A_55_P1964302** | Timm8a1 | -0.04302868 | -0.160179701 | 0.074122342 | 8.324783553 | -0.903610159 | 0.401766082 | 0.832664063 | -4.832091924 |
| **A_51_P257065** | Prpf40b | -0.073446207 | -0.273413059 | 0.126520644 | 8.366876353 | -0.903608963 | 0.401766667 | 0.832664063 | -4.832092607 |
| **A_55_P1989333** | NA | 0.078219142 | -0.134773871 | 0.291212155 | 10.39472076 | 0.903476486 | 0.401831441 | 0.832664063 | -4.832168328 |
| **A_66_P133928** | Fundc2 | 0.120304351 | -0.20732305 | 0.447931752 | 10.44976034 | 0.903379604 | 0.401878817 | 0.832664063 | -4.832223699 |
| **A_55_P2159895** | Zfp52 | -0.074216061 | -0.276330318 | 0.127898197 | 8.150819593 | -0.903379249 | 0.40187899 | 0.832664063 | -4.832223901 |
| **A_55_P2424767** | Hecw1 | -0.11805985 | -0.439597707 | 0.203478007 | 12.87180391 | -0.903315115 | 0.401910355 | 0.832664063 | -4.832260554 |
| **A_51_P332822** | Abce1 | 0.091155004 | -0.157113783 | 0.339423791 | 10.73055761 | 0.903290736 | 0.401922278 | 0.832664063 | -4.832274486 |
| **A_52_P63709** | Fam49b | -0.041600559 | -0.154922818 | 0.071721701 | 7.825614475 | -0.903135893 | 0.401998011 | 0.832731707 | -4.832362969 |
| **A_51_P270213** | Ceacam11 | -0.109924845 | -0.409412661 | 0.189562971 | 17.56598874 | -0.902996084 | 0.402066401 | 0.832760927 | -4.832442852 |
| **A_51_P370825** | Ccdc124 | 0.085898612 | -0.148147153 | 0.319944376 | 11.46290637 | 0.902930911 | 0.402098284 | 0.832760927 | -4.832480088 |
| **A_51_P411271** | Nfe2l2 | -0.045375111 | -0.169037137 | 0.078286915 | 11.39951643 | -0.902714688 | 0.402204077 | 0.832836883 | -4.83260361 |
| **A_51_P418935** | Neurl2 | 0.077164177 | -0.133166652 | 0.287495007 | 9.037271019 | 0.902572211 | 0.402273799 | 0.832836883 | -4.832684993 |
| **A_51_P465809** | Slc30a9 | 0.050066976 | -0.086404352 | 0.186538305 | 8.360217768 | 0.902566284 | 0.4022767 | 0.832836883 | -4.832688378 |
| **A_55_P2095727** | Pml | 0.109314675 | -0.188760194 | 0.407389544 | 10.2951041 | 0.902240401 | 0.402436209 | 0.832836883 | -4.832874489 |
| **A_55_P2111005** | Pafah1b1 | -0.056061599 | -0.208956085 | 0.096832887 | 11.15920487 | -0.902075367 | 0.402517006 | 0.832836883 | -4.832968723 |
| **A_55_P2136817** | Coro2b | 0.083025524 | -0.14341045 | 0.309461498 | 9.408849479 | 0.90205987 | 0.402524593 | 0.832836883 | -4.832977571 |
| **A_55_P1994117** | Ifnl3 | 0.047590998 | -0.082208004 | 0.177389999 | 8.171846035 | 0.902033405 | 0.402537552 | 0.832836883 | -4.832992681 |
| **A_55_P2035613** | Tmbim1 | -0.101588664 | -0.37867666 | 0.175499332 | 13.81438139 | -0.90197957 | 0.402563911 | 0.832836883 | -4.833023417 |
| **A_55_P1954910** | Fbxw2 | -0.066737119 | -0.248768823 | 0.115294585 | 8.698033742 | -0.901964789 | 0.402571149 | 0.832836883 | -4.833031856 |
| **A_55_P2279140** | F830014O18Rik | 0.049138219 | -0.084895733 | 0.183172172 | 7.936744027 | 0.90193191 | 0.402587249 | 0.832836883 | -4.833050627 |
| **A_52_P175952** | Map3k7 | -0.044556322 | -0.166104704 | 0.076992059 | 8.582854154 | -0.901839637 | 0.402632436 | 0.832836883 | -4.833103304 |
| **A_55_P2036220** | Chadl | -0.134316831 | -0.500775023 | 0.23214136 | 10.20416929 | -0.901727194 | 0.402687504 | 0.832836883 | -4.83316749 |
| **A_66_P116299** | Pias1 | -0.089916673 | -0.33530125 | 0.155467904 | 10.95447522 | -0.901492527 | 0.40280245 | 0.832836883 | -4.83330143 |
| **A_51_P154684** | Alyref | 0.064629376 | -0.129198238 | 0.258456989 | 11.46513366 | 0.938246217 | 0.40290761 | 0.832836883 | -4.731911413 |
| **A_52_P93837** | Mme | -0.071382969 | -0.266240907 | 0.12347497 | 10.20541591 | -0.901250804 | 0.402920879 | 0.832836883 | -4.833439373 |

| | | | | | | | | | |
|---|---|---|---|---|---|---|---|---|---|
| **A_55_P2005080** | NA | -0.071556446 | -0.266890455 | 0.123777562 | 16.79555784 | -0.901239181 | 0.402926574 | 0.832836883 | -4.833446005 |
| **A_52_P217675** | Slf1 | -0.046031591 | -0.171688181 | 0.079624998 | 8.063903564 | -0.901238809 | 0.402926757 | 0.832836883 | -4.833446217 |
| **A_55_P1974467** | Ndufb6 | -0.086862621 | -0.32399897 | 0.150273728 | 12.41351875 | -0.90116427 | 0.402963282 | 0.832836883 | -4.833488749 |
| **A_51_P351481** | Ccnl1 | -0.065296349 | -0.243569735 | 0.112977036 | 8.761424593 | -0.901097015 | 0.40299624 | 0.832836883 | -4.833527122 |
| **A_55_P2040168** | Pmp22 | 0.108093831 | -0.187036882 | 0.403224544 | 10.08514892 | 0.901064053 | 0.403012393 | 0.832836883 | -4.833545928 |
| **A_51_P216922** | Ddx59 | -0.059793473 | -0.223058898 | 0.103471953 | 8.51061724 | -0.90100814 | 0.403039795 | 0.832836883 | -4.833577828 |
| **A_52_P135822** | Myo18a | -0.145169176 | -0.541591943 | 0.25125359 | 12.5418932 | -0.900917427 | 0.403084256 | 0.83283972 | -4.833629579 |
| **A_51_P499441** | Prkd2 | -0.086590025 | -0.323082175 | 0.149902125 | 8.618203618 | -0.900783245 | 0.403150028 | 0.832886585 | -4.833706122 |
| **A_55_P2178247** | Fam20b | 0.049967305 | -0.086544486 | 0.186479095 | 9.080756881 | 0.900502505 | 0.403287665 | 0.833012407 | -4.833866245 |
| **A_55_P2013396** | Gm14305 | -0.041133739 | -0.153514438 | 0.07124696 | 8.579399432 | -0.900483215 | 0.403297124 | 0.833012407 | -4.833877246 |
| **A_52_P472660** | Smurf2 | -0.073976892 | -0.276131456 | 0.128177672 | 9.64182197 | -0.900288483 | 0.403392617 | 0.833103212 | -4.833988292 |
| **A_55_P2033445** | Tnfrsf1b | -0.199902413 | -0.746212993 | 0.346408166 | 9.078755268 | -0.900217788 | 0.403427289 | 0.833103212 | -4.834028602 |
| **A_51_P101006** | Ak8 | -0.116844529 | -0.436290381 | 0.202601323 | 8.762489271 | -0.899871073 | 0.403597365 | 0.833181066 | -4.811582439 |
| **A_66_P108815** | Pgls | 0.083464807 | -0.144740752 | 0.311670366 | 9.51338572 | 0.899800722 | 0.403631882 | 0.833181066 | -4.834266368 |
| **A_66_P119191** | Inpp5e | 0.066547664 | -0.115406445 | 0.248501772 | 8.616858938 | 0.899787815 | 0.403638215 | 0.833181066 | -4.834273725 |
| **A_55_P2198568** | 4930506A18Rik | 0.106778322 | -0.185181804 | 0.398738448 | 8.830860706 | 0.899764207 | 0.403649798 | 0.833181066 | -4.834287181 |
| **A_55_P2080756** | Olfr704 | 0.059962569 | -0.104002334 | 0.223927471 | 8.065309764 | 0.899701605 | 0.403680516 | 0.833181066 | -4.834322863 |
| **A_55_P2160387** | Cxcl11 | 0.058661861 | -0.101794809 | 0.219118532 | 16.89513526 | 0.8994297 | 0.403813955 | 0.833225839 | -4.834477823 |
| **A_55_P1992849** | Adrb3 | -0.047238221 | -0.176485969 | 0.082009527 | 7.9122224 | -0.899165652 | 0.403943569 | 0.833225839 | -4.834628275 |
| **A_55_P1952537** | Rrp7a | 0.116360671 | -0.202018906 | 0.434740248 | 12.63666393 | 0.89914592 | 0.403953257 | 0.833225839 | -4.834639518 |
| **A_51_P515452** | Fkbpl | 0.04914655 | -0.085331031 | 0.183624132 | 10.96186128 | 0.899108935 | 0.403971415 | 0.833225839 | -4.834660589 |
| **A_55_P2034175** | Usp28 | 0.057000952 | -0.099040507 | 0.21304241 | 9.312944977 | 0.898692813 | 0.404175755 | 0.833225839 | -4.834897623 |
| **A_51_P497317** | Tcp11l2 | 0.09428845 | -0.163924035 | 0.352500936 | 9.860272331 | 0.898360034 | 0.404339224 | 0.833225839 | -4.835087129 |
| **A_55_P2075474** | Gid8 | -0.049383377 | -0.1846276 | 0.085860846 | 8.665925181 | -0.89832032 | 0.404358737 | 0.833225839 | -4.835109742 |
| **A_55_P2099700** | Klc4 | 0.14209222 | -0.247070109 | 0.531254548 | 11.66284764 | 0.898273666 | 0.404381659 | 0.833225839 | -4.835136305 |
| **A_55_P2081483** | Atp5g2 | -0.058912792 | -0.220267457 | 0.102441874 | 15.57224092 | -0.898250026 | 0.404393274 | 0.833225839 | -4.835149765 |
| **A_52_P580533** | Spata33 | -0.040033164 | -0.149682454 | 0.069616126 | 7.764874055 | -0.898221109 | 0.404407483 | 0.833225839 | -4.835166229 |
| **A_51_P446978** | Cfdp1 | 0.079973985 | -0.139111108 | 0.299059078 | 12.7126775 | 0.898059416 | 0.40448694 | 0.833225839 | -4.835258281 |
| **A_55_P2316612** | 6720407P12Rik | -0.097120574 | -0.363193071 | 0.168951924 | 8.089664488 | -0.898008434 | 0.404511995 | 0.833225839 | -4.835287304 |
| **A_51_P443508** | Ppa1 | -0.056074936 | -0.209721514 | 0.097571643 | 10.91339716 | -0.89787329 | 0.404578417 | 0.833225839 | -4.83536423 |
| **A_55_P2115318** | Fmr1nb | -0.139852538 | -0.523071283 | 0.243366206 | 12.19922187 | -0.897827229 | 0.404601057 | 0.833225839 | -4.835390447 |
| **A_55_P1999648** | Plin2 | -0.090775905 | -0.339541164 | 0.157989354 | 13.01419478 | -0.897738851 | 0.404644501 | 0.833225839 | -4.835440748 |
| **A_55_P1984035** | Litaf | -0.082146235 | -0.307276805 | 0.142984335 | 8.983059438 | -0.897681665 | 0.404672613 | 0.833225839 | -4.835473294 |
| **A_51_P165165** | Hibadh | -0.07894627 | -0.295316928 | 0.137424388 | 11.7534634 | -0.897640446 | 0.404692877 | 0.833225839 | -4.835496751 |
| **A_55_P1992054** | Armc10 | -0.074624458 | -0.279178006 | 0.12992909 | 11.45120143 | -0.897518352 | 0.404752906 | 0.833225839 | -4.835566231 |
| **A_52_P591153** | Pld4 | -0.039247781 | -0.146884204 | 0.068388641 | 7.792456043 | -0.897067286 | 0.404974734 | 0.833225839 | -4.835822864 |
| **A_51_P437978** | Agtr2 | 0.041843093 | -0.072925484 | 0.15661167 | 7.672383171 | 0.896953542 | 0.405030687 | 0.833225839 | -4.835887564 |
| **A_55_P2178036** | Serpina6 | -0.038001071 | -0.142235015 | 0.066232872 | 7.679861562 | -0.896924386 | 0.40504503 | 0.833225839 | -4.835904148 |
| **A_55_P2086770** | Uevld | 0.091980369 | -0.160327118 | 0.344287856 | 9.025319979 | 0.896879659 | 0.405067034 | 0.833225839 | -4.835929588 |
| **A_51_P356283** | Fbxo31 | 0.087657268 | -0.152805579 | 0.328120116 | 10.90683385 | 0.896827898 | 0.4050925 | 0.833225839 | -4.835959028 |
| **A_66_P105244** | Tor1aip1 | 0.061959727 | -0.108036138 | 0.231955593 | 9.224399505 | 0.89668582 | 0.405162407 | 0.833225839 | -4.836039831 |
| **A_51_P331870** | Rnf145 | 0.084836872 | -0.147929157 | 0.317602902 | 10.05134745 | 0.896673252 | 0.405168591 | 0.833225839 | -4.836046978 |
| **A_51_P315682** | Igf2bp2 | 0.070598932 | -0.123114333 | 0.264312197 | 10.76379395 | 0.896618992 | 0.405195291 | 0.833225839 | -4.836077834 |
| **A_51_P111462** | Arl15 | 0.223180229 | -0.389317636 | 0.835678093 | 11.45282693 | 0.896438092 | 0.405284318 | 0.833225839 | -4.794337926 |
| **A_55_P2003473** | Zc4h2 | 0.041238569 | -0.071938581 | 0.15441572 | 7.934947861 | 0.89642508 | 0.405290723 | 0.833225839 | -4.836188097 |
| **A_65_P17866** | Vma21 | -0.045914798 | -0.171934212 | 0.080104615 | 8.723111018 | -0.896363963 | 0.405320804 | 0.833225839 | -4.836222845 |
| **A_55_P2086148** | NA | -0.099679591 | -0.373274307 | 0.173915125 | 11.83518533 | -0.896329532 | 0.405337751 | 0.833225839 | -4.836242421 |
| **A_52_P303077** | Rogdi | -0.069715631 | -0.26108698 | 0.121655718 | 9.114600088 | -0.896236045 | 0.40538377 | 0.833225839 | -4.836295571 |
| **A_55_P1984976** | Wnt5b | 0.048646957 | -0.084893966 | 0.182187879 | 8.925372446 | 0.896211405 | 0.4053959 | 0.833225839 | -4.836309578 |
| **A_55_P1960496** | Ppp1r2 | 0.065289166 | -0.113946452 | 0.244524785 | 8.831743974 | 0.89616085 | 0.405420788 | 0.833225839 | -4.836338318 |

| | | | | | | | | | |
|---|---|---|---|---|---|---|---|---|---|
| **A_55_P2008918** | Cbll1 | -0.071481272 | -0.267775204 | 0.124812659 | 9.065241477 | -0.895889742 | 0.405554271 | 0.833225839 | -4.836492418 |
| **A_51_P265806** | Clca3a2 | -0.085177736 | -0.319092632 | 0.148737159 | 8.353795241 | -0.895854293 | 0.405571727 | 0.833225839 | -4.836512566 |
| **A_55_P1992555** | Gys2 | 0.193270118 | -0.337585495 | 0.724125732 | 8.000445368 | 0.895689456 | 0.405652906 | 0.833225839 | -4.836606243 |
| **A_51_P358906** | U2af2 | -0.04835966 | -0.18120517 | 0.084485851 | 8.232810793 | -0.895582325 | 0.405705672 | 0.833225839 | -4.83666712 |
| **A_55_P2084122** | Dus4l | -0.05567028 | -0.208601308 | 0.097260748 | 8.665254358 | -0.89556469 | 0.405714359 | 0.833225839 | -4.836677141 |
| **A_55_P2072284** | Dnaaf3 | 0.045269008 | -0.079094036 | 0.169632051 | 7.979567645 | 0.895527228 | 0.405732812 | 0.833225839 | -4.836698427 |
| **A_55_P1969625** | Hnrnpr | -0.111350062 | -0.417315927 | 0.194615804 | 11.77369872 | -0.895337238 | 0.405826407 | 0.833225839 | -4.836806372 |
| **A_55_P2057112** | Prl8a2 | -0.954977875 | -3.579057186 | 1.669101436 | 16.25280763 | -0.895334905 | 0.405827556 | 0.833225839 | -4.836807697 |
| **A_55_P1994477** | Prrc2c | -0.088788749 | -0.332782576 | 0.155205079 | 9.522110476 | -0.895258097 | 0.405865399 | 0.833225839 | -4.836851332 |
| **A_66_P130882** | Eif3e | 0.086275503 | -0.150812185 | 0.32336319 | 9.992401741 | 0.895256866 | 0.405866005 | 0.833225839 | -4.836852031 |
| **A_51_P518926** | Borcs6 | -0.074984979 | -0.281061687 | 0.131091728 | 9.215933779 | -0.895188581 | 0.405899652 | 0.833225839 | -4.836890822 |
| **A_55_P2113356** | Miip | -0.040728668 | -0.152687614 | 0.071230278 | 7.89813285 | -0.894974306 | 0.406005244 | 0.833225839 | -4.837012532 |
| **A_51_P464029** | Fads3 | -0.274773074 | -1.030166055 | 0.480619908 | 12.55867508 | -0.894891513 | 0.406046049 | 0.833225839 | -4.837059555 |
| **A_55_P2096165** | NA | -0.075913028 | -0.284617879 | 0.132791823 | 10.92482737 | -0.894855548 | 0.406063776 | 0.833225839 | -4.83707998 |
| **A_55_P2003483** | Gldc | -0.131456819 | -0.492939393 | 0.230025754 | 9.201166816 | -0.894674208 | 0.406153165 | 0.833225839 | -4.837182959 |
| **A_51_P198411** | Grb2 | -0.133172103 | -0.49944552 | 0.233101314 | 11.95885991 | -0.894493153 | 0.406242429 | 0.833225839 | -4.837285762 |
| **A_51_P165435** | Cox4i1 | -0.108208597 | -0.405864143 | 0.189446949 | 15.10763638 | -0.894369437 | 0.406303431 | 0.833225839 | -4.837356 |
| **A_55_P1998309** | Sms | 0.147806871 | -0.258780821 | 0.554394563 | 10.98370761 | 0.894354214 | 0.406310938 | 0.833225839 | -4.837364642 |
| **A_55_P2019690** | Slc43a1 | 0.054054488 | -0.094642699 | 0.202751675 | 7.913198658 | 0.894330755 | 0.406322506 | 0.833225839 | -4.83737796 |
| **A_52_P264905** | Cab39l | 0.080762462 | -0.141424567 | 0.302949491 | 10.24200377 | 0.894252185 | 0.406361253 | 0.833225839 | -4.837422562 |
| **A_55_P2082653** | Zfp711 | 0.045394343 | -0.079505498 | 0.170294184 | 7.914329942 | 0.89414718 | 0.406413041 | 0.833225839 | -4.837482167 |
| **A_51_P280085** | Zfp655 | -0.052613321 | -0.197413639 | 0.092186998 | 14.05529887 | -0.893913218 | 0.406528447 | 0.833225839 | -4.837614956 |
| **A_51_P434758** | Fam83e | 0.36794435 | -0.644770108 | 1.380658808 | 8.729445162 | 0.893849272 | 0.406559994 | 0.833225839 | -4.837651245 |
| **A_55_P1955568** | Extl2 | -0.078755167 | -0.29551831 | 0.138007976 | 10.41411776 | -0.893846161 | 0.406561529 | 0.833225839 | -4.837653011 |
| **A_52_P192307** | Rbak | -0.049079714 | -0.184197433 | 0.086038005 | 8.922574298 | -0.893632347 | 0.406667025 | 0.833225839 | -4.837774338 |
| **A_55_P1966420** | Scrn2 | 0.091856208 | -0.161027696 | 0.344740112 | 9.266032037 | 0.893627423 | 0.406669455 | 0.833225839 | -4.837777132 |
| **A_55_P2159934** | Rnf186 | 0.232767393 | -0.408071585 | 0.873606371 | 8.238054806 | 0.893598408 | 0.406683773 | 0.833225839 | -4.837793595 |
| **A_55_P2064328** | Lama2 | -0.073835931 | -0.277161908 | 0.129490045 | 8.680524295 | -0.893396105 | 0.406783611 | 0.833225839 | -4.837908369 |
| **A_51_P132170** | Ccdc141 | 0.037729796 | -0.066184128 | 0.14164372 | 7.684462698 | 0.893264083 | 0.406848776 | 0.833225839 | -4.837983261 |
| **A_55_P2111478** | Polq | 0.038384851 | -0.067335726 | 0.144105427 | 7.857157511 | 0.893242746 | 0.406859308 | 0.833225839 | -4.837995364 |
| **A_55_P2181542** | Sult1d1 | 0.049059191 | -0.086068085 | 0.184186467 | 7.726413188 | 0.893195497 | 0.406882632 | 0.833225839 | -4.838022165 |
| **A_55_P1970771** | Gm13141 | 0.057022962 | -0.100051115 | 0.214097039 | 12.38633293 | 0.893129469 | 0.406915227 | 0.833225839 | -4.838059616 |
| **A_51_P112762** | Slc5a3 | -0.073754444 | -0.276928891 | 0.129420003 | 9.263001946 | -0.893075699 | 0.406941773 | 0.833225839 | -4.822896863 |
| **A_66_P114585** | NA | 0.056866683 | -0.099791798 | 0.213525164 | 9.131964302 | 0.893044597 | 0.406957129 | 0.833225839 | -4.838107752 |
| **A_55_P2125229** | Arhgef2 | -0.141000623 | -0.529464792 | 0.247463547 | 11.93377055 | -0.892974848 | 0.406991566 | 0.833225839 | -4.838147309 |
| **A_51_P324923** | Man2a1 | -0.211950195 | -0.795898169 | 0.37199778 | 13.39183763 | -0.892953465 | 0.407002124 | 0.833225839 | -4.838159436 |
| **A_55_P1999841** | Cox7c | -0.092013144 | -0.345526763 | 0.161500476 | 12.76030785 | -0.892930662 | 0.407013383 | 0.833225839 | -4.838172367 |
| **A_51_P161354** | Sesn2 | 0.09502355 | -0.166829756 | 0.356876857 | 11.06873873 | 0.892775686 | 0.407089911 | 0.833225839 | -4.838260249 |
| **A_55_P1991778** | Fam219a | 0.17590278 | -0.308920997 | 0.660726558 | 14.50190024 | 0.892602207 | 0.407175588 | 0.833225839 | -4.838358611 |
| **A_55_P2042923** | Sgk2 | 0.292427101 | -0.513574235 | 1.098428437 | 12.74302717 | 0.892587951 | 0.40718263 | 0.833225839 | -4.838366694 |
| **A_55_P2113650** | Clybl | 0.116634627 | -0.204898021 | 0.438167275 | 8.686960202 | 0.892424716 | 0.407263262 | 0.833225839 | -4.838459235 |
| **A_55_P2032600** | Rps24 | -0.070097965 | -0.263343751 | 0.123147822 | 16.90039569 | -0.892410226 | 0.40727042 | 0.833225839 | -4.838467449 |
| **A_51_P183995** | Paf1 | -0.067891684 | -0.255077031 | 0.119293663 | 12.67402829 | -0.892306168 | 0.407321827 | 0.833225839 | -4.838526434 |
| **A_55_P2056729** | Igfbp2 | -0.156790341 | -0.589102622 | 0.275521939 | 11.66069667 | -0.892258915 | 0.407345173 | 0.833225839 | -4.838553219 |
| **A_51_P475580** | Ugp2 | 0.057333514 | -0.100758317 | 0.215425345 | 10.28084039 | 0.892212486 | 0.407368113 | 0.833225839 | -4.838579535 |
| **A_55_P2100679** | Dnm2 | -0.051932533 | -0.195140295 | 0.091275228 | 8.459307459 | -0.892158698 | 0.40739469 | 0.833225839 | -4.838610021 |
| **A_55_P2102961** | NA | -0.067727425 | -0.254506438 | 0.119051588 | 9.140480079 | -0.892083797 | 0.407431702 | 0.833225839 | -4.838652472 |
| **A_55_P2457154** | G2e3 | -0.04960621 | -0.186433747 | 0.087221327 | 8.510870528 | -0.891931913 | 0.407506761 | 0.833225839 | -4.838738545 |
| **A_51_P406429** | Pdk1 | 0.161059305 | -0.283251637 | 0.605370247 | 9.734392502 | 0.89180104 | 0.407571445 | 0.833225839 | -4.838812704 |
| **A_52_P344290** | F2r | -0.111282477 | -0.418294978 | 0.195730024 | 15.16394963 | -0.891743365 | 0.407599953 | 0.833225839 | -4.838845382 |

| | | | | | | | | | |
|---|---|---|---|---|---|---|---|---|---|
| **A_52_P355398** | Dars | -0.098037142 | -0.368513104 | 0.17243882 | 8.854919338 | -0.89172539 | 0.407608839 | 0.833225839 | -4.838855567 |
| **A_51_P223929** | Ash1l | -0.065024031 | -0.244441301 | 0.114393239 | 9.13026634 | -0.891617959 | 0.407661946 | 0.833225839 | -4.838916434 |
| **A_55_P2169539** | NA | 0.064892477 | -0.11417328 | 0.243958234 | 8.013789319 | 0.891560817 | 0.407690196 | 0.833225839 | -4.838948806 |
| **A_52_P418734** | Rnf220 | -0.07663709 | -0.288146819 | 0.134872638 | 13.12045745 | -0.891410658 | 0.407764438 | 0.833225839 | -4.839033869 |
| **A_55_P2150976** | NA | 0.160037614 | -0.281675307 | 0.601750535 | 15.49222174 | 0.891355867 | 0.407791531 | 0.833225839 | -4.839064905 |
| **A_55_P2076288** | Zfp661 | -0.046595139 | -0.17520422 | 0.082013942 | 8.31676256 | -0.891329205 | 0.407804715 | 0.833225839 | -4.839080007 |
| **A_52_P95759** | Sh3rf3 | -0.041700425 | -0.156802467 | 0.073401617 | 7.865974628 | -0.891305559 | 0.407816408 | 0.833225839 | -4.8390934 |
| **A_51_P491227** | Suclg1 | -0.072889088 | -0.274109907 | 0.128331731 | 11.77420263 | -0.891166323 | 0.407885265 | 0.833225839 | -4.83917226 |
| **A_51_P417321** | Zfp236 | -0.108843447 | -0.409332491 | 0.191645597 | 9.520741 | -0.891133579 | 0.40790146 | 0.833225839 | -4.839190805 |
| **A_55_P2136678** | NA | 0.142508275 | -0.250948978 | 0.535965528 | 9.295950129 | 0.891069727 | 0.407933041 | 0.833225839 | -4.839226965 |
| **A_55_P2105200** | Abraxas1 | 0.075496144 | -0.13296259 | 0.283954878 | 8.584239557 | 0.890992071 | 0.407971452 | 0.833225839 | -4.839270941 |
| **A_55_P2174935** | Mn1 | -0.104366668 | -0.392545878 | 0.183812542 | 9.406393086 | -0.890980781 | 0.407977037 | 0.833225839 | -4.839277334 |
| **A_55_P2023488** | NA | -0.194080104 | -0.730003423 | 0.341843215 | 10.71437066 | -0.890938087 | 0.407998156 | 0.833225839 | -4.83930151 |
| **A_55_P2002122** | Rgs12 | 0.181000703 | -0.318859428 | 0.680860834 | 11.37605394 | 0.890842495 | 0.408045445 | 0.833225839 | -4.839355638 |
| **A_55_P2273929** | Gm20558 | -0.047076332 | -0.177091767 | 0.082939103 | 7.756476897 | -0.890793144 | 0.408069861 | 0.833225839 | -4.83938358 |
| **A_51_P117581** | Cables1 | -0.087685216 | -0.329871643 | 0.15450121 | 10.86582977 | -0.890729307 | 0.408101444 | 0.833225839 | -4.835650919 |
| **A_55_P1968833** | Rpgr | 0.042622452 | -0.075113866 | 0.16035877 | 8.971871294 | 0.890629638 | 0.408150759 | 0.833225839 | -4.839476149 |
| **A_55_P2090205** | Tpm3 | -0.054788393 | -0.20616672 | 0.096589935 | 8.148686801 | -0.890418186 | 0.408255399 | 0.833225839 | -4.839595847 |
| **A_51_P267314** | Tbc1d10c | 0.044228737 | -0.077978674 | 0.166436148 | 7.907888529 | 0.890381565 | 0.408273524 | 0.833225839 | -4.839616576 |
| **A_55_P2088095** | NA | -0.055561197 | -0.209088308 | 0.097965914 | 11.00872908 | -0.890339577 | 0.408294305 | 0.833225839 | -4.839640341 |
| **A_55_P1953972** | Pdhb | -0.110357923 | -0.415317728 | 0.194601882 | 13.41105362 | -0.890287098 | 0.40832028 | 0.833225839 | -4.839670043 |
| **A_55_P2087162** | Specc1l | -0.042360736 | -0.15942561 | 0.074704138 | 8.27538883 | -0.890237857 | 0.408344653 | 0.833225839 | -4.839697912 |
| **A_55_P2087384** | Gps1 | 0.077216743 | -0.136181775 | 0.290615261 | 13.33071275 | 0.890203392 | 0.408361713 | 0.833225839 | -4.839717417 |
| **A_55_P1991792** | Slc25a46 | 0.058216788 | -0.102680982 | 0.219114558 | 12.00667328 | 0.890158572 | 0.4083839 | 0.833225839 | -4.839742782 |
| **A_55_P2004046** | NA | -0.042704271 | -0.160741241 | 0.075332699 | 7.865287807 | -0.890066426 | 0.408429517 | 0.833225839 | -4.839794927 |
| **A_52_P85983** | Mrpl45 | 0.065821308 | -0.116116404 | 0.24775902 | 11.24453337 | 0.890047009 | 0.40843913 | 0.833225839 | -4.839805915 |
| **A_55_P1999154** | Gm4987 | 0.117229343 | -0.206888527 | 0.441347213 | 13.05334278 | 0.889820729 | 0.408551168 | 0.833225839 | -4.83993395 |
| **A_51_P394788** | Actn4 | 0.134268097 | -0.236968199 | 0.505504392 | 14.66260694 | 0.889798287 | 0.408562281 | 0.833225839 | -4.839946646 |
| **A_55_P2091551** | Arhgap9 | 0.11712095 | -0.20672346 | 0.440965359 | 11.01893238 | 0.889748664 | 0.408586854 | 0.833225839 | -4.839974721 |
| **A_51_P206655** | Lsm1 | -0.090523357 | -0.340837223 | 0.15979051 | 10.464431 | -0.889702692 | 0.408609621 | 0.833225839 | -4.840000729 |
| **A_55_P2089118** | Zfp976 | -0.085203476 | -0.320810548 | 0.150403596 | 8.892170922 | -0.889688864 | 0.408616469 | 0.833225839 | -4.840008552 |
| **A_51_P417074** | Arhgap8 | -0.093589437 | -0.35250979 | 0.165330916 | 13.28259809 | -0.889262153 | 0.408827835 | 0.833568906 | -4.840249911 |
| **A_55_P2151797** | Dis3l2 | -0.083638636 | -0.315143064 | 0.147865791 | 10.7879182 | -0.888826113 | 0.409043908 | 0.833921496 | -4.840496467 |
| **A_51_P299062** | Kel | 0.056046383 | -0.0991391 | 0.211231867 | 8.151728492 | 0.888516838 | 0.409197217 | 0.834146067 | -4.840671295 |
| **A_51_P488510** | Appl1 | -0.063756116 | -0.240325901 | 0.112813669 | 8.928939261 | -0.88833061 | 0.409289552 | 0.834246309 | -4.840776547 |
| **A_51_P384091** | Rad21 | 0.081106942 | -0.143572689 | 0.305786573 | 13.75868995 | 0.888103308 | 0.409402273 | 0.834388079 | -4.840904993 |
| **A_55_P1998066** | Tmem121 | -0.040014331 | -0.150876773 | 0.070848111 | 7.809158518 | -0.88797407 | 0.409466374 | 0.834430737 | -4.840978014 |
| **A_51_P117236** | Zfp354a | 0.077792126 | -0.137785957 | 0.29337021 | 9.273899746 | 0.887769454 | 0.409567878 | 0.83451926 | -4.822212188 |
| **A_55_P2078335** | Mapk4 | 0.197141368 | -0.349213254 | 0.743495989 | 9.638900484 | 0.887712443 | 0.409596162 | 0.83451926 | -4.841125815 |
| **A_55_P2011410** | Pdzk1 | 0.242284826 | -0.429310614 | 0.913880265 | 12.07902378 | 0.887539397 | 0.409682024 | 0.834606223 | -4.841223558 |
| **A_55_P2133652** | Cep162 | -0.058952905 | -0.222384898 | 0.104479089 | 9.17320968 | -0.887436514 | 0.409733079 | 0.834622266 | -4.841281664 |
| **A_55_P1953087** | Mcm3 | -0.087367105 | -0.329625624 | 0.154891414 | 9.531475384 | -0.887233741 | 0.409833717 | 0.834700471 | -4.841396172 |
| **A_66_P106814** | NA | 0.091711006 | -0.162606561 | 0.346028574 | 13.4661361 | 0.887185145 | 0.409857839 | 0.834700471 | -4.841423612 |
| **A_55_P2146555** | NA | -0.069955421 | -0.263973232 | 0.124062391 | 9.544017639 | -0.887051705 | 0.40992408 | 0.8347032 | -4.841498955 |
| **A_55_P1984283** | NA | 0.096456111 | -0.171073062 | 0.363985283 | 14.76019617 | 0.887008464 | 0.409945547 | 0.8347032 | -4.841523369 |
| **A_52_P11833** | NA | 0.148136732 | -0.262789045 | 0.55906251 | 10.32321255 | 0.886887509 | 0.4100056 | 0.834737543 | -4.841591653 |
| **A_55_P2076906** | NA | -0.143347245 | -0.541056708 | 0.254362217 | 10.14545951 | -0.886732426 | 0.410082606 | 0.834744663 | -4.841679196 |
| **A_55_P2131173** | Sphk2 | 0.062590644 | -0.111103965 | 0.236285254 | 11.47063077 | 0.886527566 | 0.410184346 | 0.834744663 | -4.841794821 |
| **A_55_P1955587** | Hspd1 | -0.072344795 | -0.273114055 | 0.128424465 | 13.50068068 | -0.886501008 | 0.410197537 | 0.834744663 | -4.841809809 |
| **A_51_P261517** | Tyrobp | -0.12580198 | -0.474941427 | 0.223337468 | 9.729473824 | -0.886457092 | 0.41021935 | 0.834744663 | -4.841834593 |

| | | | | | | | | | |
|---|---|---|---|---|---|---|---|---|---|
| **A_52_P25812** | Prdm2 | -0.049227803 | -0.185851961 | 0.087396355 | 8.589077102 | -0.886445662 | 0.410225027 | 0.834744663 | -4.841841043 |
| **A_55_P1984841** | NA | 0.065351806 | -0.116076758 | 0.246780371 | 10.15752683 | 0.886178265 | 0.410357864 | 0.834840586 | -4.841991928 |
| **A_66_P103926** | Eif3m | 0.076527467 | -0.135945017 | 0.288999952 | 12.50742916 | 0.886102185 | 0.410395665 | 0.834840586 | -4.842034853 |
| **A_55_P2162204** | Kctd15 | 0.066996732 | -0.119016984 | 0.253010448 | 12.36812654 | 0.886089956 | 0.410401741 | 0.834840586 | -4.842041752 |
| **A_66_P139618** | Stfa2 | -0.230070266 | -0.869002448 | 0.408861916 | 9.080423795 | -0.885879995 | 0.410506076 | 0.834964952 | -4.842160198 |
| **A_51_P388737** | Slc11a2 | -0.090445684 | -0.341749803 | 0.160858436 | 9.15412027 | -0.885436459 | 0.410726546 | 0.835144941 | -4.842410349 |
| **A_55_P2182700** | Prkab1 | 0.038616404 | -0.068679645 | 0.145912454 | 8.030156807 | 0.885436178 | 0.410726685 | 0.835144941 | -4.842410508 |
| **A_55_P2058923** | Bud23 | 0.091035798 | -0.161913327 | 0.343984924 | 11.15041174 | 0.885417673 | 0.410735886 | 0.835144941 | -4.842420943 |
| **A_55_P1971579** | Acox3 | -0.267645242 | -1.011368886 | 0.476078402 | 9.21297574 | -0.885354301 | 0.410767394 | 0.835144941 | -4.842456677 |
| **A_51_P287069** | Serpinh1 | 0.074209343 | -0.132088687 | 0.280507372 | 15.41385271 | 0.884978394 | 0.410954331 | 0.835319734 | -4.842668607 |
| **A_55_P2027136** | H3c7 | -0.039114706 | -0.147854184 | 0.069624773 | 7.998701847 | -0.884956627 | 0.410965158 | 0.835319734 | -4.842680876 |
| **A_55_P1985077** | Mcrip1 | -0.055256009 | -0.208924971 | 0.098412952 | 8.400178029 | -0.884631751 | 0.411126773 | 0.835319734 | -4.842863984 |
| **A_51_P296249** | Foxs1 | 0.05263595 | -0.093778269 | 0.199050168 | 8.729159254 | 0.884439995 | 0.411222187 | 0.835319734 | -4.842972041 |
| **A_55_P2113539** | NA | -0.037436422 | -0.141593249 | 0.066720406 | 7.938177425 | -0.884251395 | 0.411316048 | 0.835319734 | -4.843078303 |
| **A_52_P550049** | Nfya | -0.04359281 | -0.164887836 | 0.077702215 | 8.71961665 | -0.884180725 | 0.411351222 | 0.835319734 | -4.843118117 |
| **A_51_P326229** | Ddx25 | -0.066215248 | -0.250461492 | 0.118030996 | 8.132723945 | -0.884155252 | 0.411363901 | 0.835319734 | -4.843132467 |
| **A_52_P438082** | Cebpg | -0.046335988 | -0.175286944 | 0.082614969 | 9.369330728 | -0.884021895 | 0.411430285 | 0.835319734 | -4.84320759 |
| **A_52_P464629** | Nme7 | -0.059010802 | -0.223245538 | 0.105223934 | 8.610855816 | -0.883966207 | 0.411458008 | 0.835319734 | -4.843238958 |
| **A_55_P2101906** | Rpl10a | -0.074212159 | -0.280768922 | 0.132344604 | 15.78306897 | -0.883903414 | 0.41148927 | 0.835319734 | -4.843274326 |
| **A_55_P2161675** | Zfp133-ps | 0.397210836 | -0.708500211 | 1.502921882 | 9.640123856 | 0.883789021 | 0.411546226 | 0.835319734 | -4.843338754 |
| **A_55_P1979252** | Glod4 | 0.06769924 | -0.12075529 | 0.256153769 | 11.60619375 | 0.883784486 | 0.411548484 | 0.835319734 | -4.843341308 |
| **A_55_P2056100** | Pus1 | 0.095975002 | -0.171228611 | 0.363178615 | 9.265591738 | 0.883659534 | 0.411610704 | 0.835319734 | -4.843411676 |
| **A_55_P2093874** | Pbxip1 | -0.213312293 | -0.807207395 | 0.380582808 | 12.66385457 | -0.883639874 | 0.411620495 | 0.835319734 | -4.843422747 |
| **A_55_P1958457** | Nbeal2 | -0.057079177 | -0.216008189 | 0.101849835 | 8.20243825 | -0.883575535 | 0.411652536 | 0.835319734 | -4.843458977 |
| **A_55_P2031043** | Stat5b | 0.155246038 | -0.277060316 | 0.587552392 | 10.28683929 | 0.883482739 | 0.411698752 | 0.835319734 | -4.843511229 |
| **A_52_P385489** | Ppp1r3a | 0.037165644 | -0.066332152 | 0.140663439 | 7.691481856 | 0.883445416 | 0.411717341 | 0.835319734 | -4.843532244 |
| **A_55_P1953796** | Fam98c | 0.079845629 | -0.142515802 | 0.302207059 | 10.44427257 | 0.883407026 | 0.411736463 | 0.835319734 | -4.843553859 |
| **A_55_P2053850** | NA | 0.106151711 | -0.189471825 | 0.401775248 | 10.35246458 | 0.883399512 | 0.411740206 | 0.835319734 | -4.843558089 |
| **A_66_P127036** | NA | -0.058643473 | -0.221967196 | 0.10468025 | 15.06965285 | -0.883363755 | 0.411758017 | 0.835319734 | -4.843578221 |
| **A_52_P677891** | Fam174b | -0.076091251 | -0.288098372 | 0.13591587 | 8.379795706 | -0.882985234 | 0.411946597 | 0.835319734 | -4.8437913 |
| **A_55_P1962354** | Lexm | -0.074647483 | -0.282647552 | 0.133352586 | 8.546385443 | -0.882918981 | 0.411979611 | 0.835319734 | -4.843828589 |
| **A_55_P2118675** | Tnip1 | -0.109007863 | -0.412780752 | 0.194765026 | 12.17902259 | -0.882831818 | 0.412023048 | 0.835319734 | -4.843877645 |
| **A_55_P1955437** | Cmtm5 | -0.080303597 | -0.304096823 | 0.143489629 | 9.417450192 | -0.882789638 | 0.412044069 | 0.835319734 | -4.843901382 |
| **A_52_P511381** | Tesk1 | -0.093721675 | -0.35491519 | 0.167471841 | 11.22755194 | -0.882768469 | 0.412054619 | 0.835319734 | -4.843913295 |
| **A_51_P118763** | Ahctf1 | 0.06173518 | -0.110324465 | 0.233794825 | 10.5446948 | 0.88271978 | 0.412078885 | 0.835319734 | -4.824404185 |
| **A_51_P394394** | Tspan2 | -0.068418734 | -0.259110583 | 0.122273116 | 9.585015412 | -0.882697839 | 0.412089821 | 0.835319734 | -4.843953041 |
| **A_55_P2169218** | NA | 0.059275775 | -0.105937313 | 0.224488862 | 9.029286881 | 0.882677292 | 0.412100062 | 0.835319734 | -4.843964603 |
| **A_55_P2009875** | NA | 0.087900606 | -0.157099098 | 0.33290031 | 9.738267275 | 0.882664122 | 0.412106626 | 0.835319734 | -4.843972014 |
| **A_51_P386389** | Actr6 | 0.137164469 | -0.245343604 | 0.519672543 | 10.6087405 | 0.882206362 | 0.412334838 | 0.835553448 | -4.844229553 |
| **A_51_P134812** | Chac1 | -0.20922964 | -0.792826217 | 0.374366938 | 8.559657186 | -0.882022439 | 0.412426557 | 0.835553448 | -4.844333004 |
| **A_55_P2088416** | Lamtor5 | 0.10003174 | -0.179000533 | 0.379064013 | 12.92480531 | 0.881967434 | 0.41245399 | 0.835553448 | -4.84436394 |
| **A_55_P1987746** | NA | 0.058716067 | -0.105070938 | 0.222503072 | 15.26920673 | 0.881955517 | 0.412459934 | 0.835553448 | -4.844370642 |
| **A_52_P231170** | Megf9 | -0.05926417 | -0.224595772 | 0.106067431 | 8.249923346 | -0.881871889 | 0.412501645 | 0.835553448 | -4.844417673 |
| **A_55_P1990663** | Ddx5 | -0.085253819 | -0.323102855 | 0.152595217 | 12.44934121 | -0.881823381 | 0.412525841 | 0.835553448 | -4.844444952 |
| **A_55_P2043182** | Gimap6 | 0.11643957 | -0.208458625 | 0.441337764 | 9.277380934 | 0.881703281 | 0.412585752 | 0.835553448 | -4.844512486 |
| **A_55_P2153570** | Zfp655 | -0.074061603 | -0.280729258 | 0.132606052 | 9.146883385 | -0.8816369 | 0.412618869 | 0.835553448 | -4.84454981 |
| **A_55_P2065621** | Hook2 | 0.189245461 | -0.338898443 | 0.717389365 | 10.78814915 | 0.881540691 | 0.41266687 | 0.835553448 | -4.844603903 |
| **A_52_P48569** | Slc38a4 | -0.104161973 | -0.394904539 | 0.186580593 | 13.16050002 | -0.881393293 | 0.412740418 | 0.835553448 | -4.844686769 |
| **A_52_P90363** | Ifi27l2a | 0.206820513 | -0.370484953 | 0.784125979 | 11.05759908 | 0.881367648 | 0.412753216 | 0.835553448 | -4.844701186 |
| **A_52_P69756** | Mterf4 | 0.113687557 | -0.203729597 | 0.431104711 | 10.28442697 | 0.881153769 | 0.412859957 | 0.835553448 | -4.844821407 |

| | | | | | | | | | |
|---|---|---|---|---|---|---|---|---|---|
| **A_55_P1967015** | Fgfr1 | 0.08877651 | -0.159102163 | 0.336655182 | 9.889390417 | 0.881105817 | 0.412883891 | 0.835553448 | -4.844848358 |
| **A_55_P1969993** | NA | -0.043639488 | -0.165550868 | 0.078271892 | 16.40003476 | -0.880652483 | 0.413110216 | 0.835553448 | -4.845103103 |
| **A_55_P1967855** | Kctd20 | -0.093316343 | -0.354023691 | 0.167391005 | 11.43664393 | -0.8805897 | 0.413141567 | 0.835553448 | -4.845138376 |
| **A_55_P2017684** | Macf1 | 0.09022786 | -0.161889023 | 0.342344742 | 11.84737811 | 0.88045647 | 0.413208104 | 0.835553448 | -4.845213222 |
| **A_51_P308844** | Nrn1 | 0.068933608 | -0.123708824 | 0.261576039 | 7.993890097 | 0.880335509 | 0.413268519 | 0.835553448 | -4.845281169 |
| **A_55_P2108483** | NA | -0.293042449 | -1.112000249 | 0.525915351 | 11.47465404 | -0.880315147 | 0.41327869 | 0.835553448 | -4.845292607 |
| **A_55_P2134785** | Zscan22 | 0.0913224 | -0.1638959 | 0.3465407 | 9.464369803 | 0.880308038 | 0.413282241 | 0.835553448 | -4.845296599 |
| **A_51_P330090** | Trappc9 | 0.057892925 | -0.103931172 | 0.219717022 | 10.72090476 | 0.880139399 | 0.413366484 | 0.835553448 | -4.845391316 |
| **A_51_P162098** | Ndufaf4 | -0.044407198 | -0.168536809 | 0.079722413 | 9.595019477 | -0.880130644 | 0.413370857 | 0.835553448 | -4.845396233 |
| **A_52_P367520** | Nexn | 0.109884855 | -0.197290615 | 0.417060324 | 10.49625065 | 0.880076595 | 0.41339786 | 0.835553448 | -4.845426587 |
| **A_55_P2067453** | Tspan17 | -0.182037131 | -0.691007063 | 0.326932801 | 10.39481376 | -0.879907499 | 0.413482349 | 0.835553448 | -4.845521544 |
| **A_55_P2064896** | Gm8580 | 0.083221249 | -0.149467539 | 0.315910037 | 12.9941475 | 0.879889093 | 0.413491546 | 0.835553448 | -4.845531878 |
| **A_55_P2114809** | Gtf2e2 | 0.06840642 | -0.122877633 | 0.259690472 | 11.15671435 | 0.879806692 | 0.413532723 | 0.835553448 | -4.845578146 |
| **A_55_P2024654** | Trrap | -0.092679031 | -0.351838792 | 0.166480729 | 10.95069645 | -0.879798219 | 0.413536958 | 0.835553448 | -4.845582903 |
| **A_55_P1999962** | Podn | -0.155046325 | -0.588605229 | 0.27851258 | 8.596825472 | -0.879797105 | 0.413537514 | 0.835553448 | -4.845583529 |
| **A_51_P395111** | Nme3 | 0.071073263 | -0.127686835 | 0.26983336 | 10.16395896 | 0.879723542 | 0.413574278 | 0.835553448 | -4.845624831 |
| **A_55_P2122605** | Cbr2 | 0.074402778 | -0.133679336 | 0.282484892 | 7.946709526 | 0.879677663 | 0.413597207 | 0.835553448 | -4.845650588 |
| **A_51_P348433** | Rasal1 | -0.07612827 | -0.289042771 | 0.136786231 | 9.021866118 | -0.879649947 | 0.41361106 | 0.835553448 | -4.845666149 |
| **A_66_P106611** | Gna14 | 0.176513122 | -0.317217987 | 0.67024423 | 9.24308926 | 0.879540147 | 0.413665941 | 0.835553448 | -4.845727788 |
| **A_55_P2024555** | Plpp1 | -0.150099424 | -0.570026986 | 0.269828138 | 13.27000505 | -0.879374381 | 0.413748806 | 0.835553448 | -4.845820836 |
| **A_55_P1968285** | Tle5 | 0.108416844 | -0.194917574 | 0.411751263 | 13.74166217 | 0.8793145 | 0.413778743 | 0.835553448 | -4.845854446 |
| **A_55_P2002712** | Mllt10 | -0.086667417 | -0.32917072 | 0.155835886 | 9.89397814 | -0.879239839 | 0.413816072 | 0.835553448 | -4.845896349 |
| **A_55_P1988970** | Mkks | 0.061574641 | -0.110731389 | 0.23388067 | 9.811056984 | 0.879165372 | 0.413853306 | 0.835553448 | -4.845938141 |
| **A_55_P2045367** | G630025P09Rik | 0.109004772 | -0.196031366 | 0.41404091 | 9.616547839 | 0.879150807 | 0.413860588 | 0.835553448 | -4.845946314 |
| **A_52_P154833** | Ankrd49 | 0.037906599 | -0.068176285 | 0.143989484 | 9.514232746 | 0.879100787 | 0.413885601 | 0.835553448 | -4.845974384 |
| **A_55_P2128597** | Nr1h3 | 0.085917763 | -0.154543823 | 0.326379349 | 9.380872248 | 0.879035502 | 0.413918248 | 0.835553448 | -4.846011019 |
| **A_55_P2022149** | NA | -0.040957185 | -0.155586126 | 0.073671756 | 8.077922247 | -0.8790326 | 0.413919699 | 0.835553448 | -4.846012647 |
| **A_55_P2181246** | NA | -0.083795736 | -0.318356523 | 0.15076505 | 9.829151912 | -0.878892335 | 0.413989848 | 0.835553448 | -4.846091351 |
| **A_52_P2732** | Mfn1 | -0.084119657 | -0.319589534 | 0.151350219 | 11.23417791 | -0.87888348 | 0.413994277 | 0.835553448 | -4.846096319 |
| **A_55_P2058165** | Morn4 | 0.118954437 | -0.214080534 | 0.451989408 | 8.473356218 | 0.878739136 | 0.414066475 | 0.835611913 | -4.846177302 |
| **A_51_P451151** | Ube2c | 0.106337328 | -0.191416491 | 0.404091147 | 9.078834871 | 0.878612883 | 0.414129633 | 0.835652122 | -4.846248128 |
| **A_51_P232705** | Commd9 | 0.072165087 | -0.129987163 | 0.274317337 | 8.836642505 | 0.878249119 | 0.414311646 | 0.83593213 | -4.846452156 |
| **A_52_P581390** | Kif1c | -0.038875801 | -0.147812951 | 0.070061349 | 8.137906849 | -0.877955507 | 0.414458601 | 0.836122873 | -4.846616794 |
| **A_55_P2008042** | Gm15348 | -0.077703023 | -0.29547211 | 0.140066065 | 10.25819961 | -0.87783086 | 0.414521 | 0.836122873 | -4.846686676 |
| **A_55_P1996578** | Fgfr3 | 0.082882935 | -0.149411188 | 0.315177059 | 9.041957391 | 0.877800986 | 0.414535956 | 0.836122873 | -4.846703424 |
| **A_55_P2143657** | Gm5784 | 0.1567907 | -0.282691812 | 0.596273213 | 9.530806514 | 0.877703569 | 0.414584729 | 0.836133997 | -4.846758034 |
| **A_66_P135700** | Cyp2d11 | 0.051705332 | -0.093260703 | 0.196671366 | 9.682895274 | 0.877482034 | 0.414695661 | 0.836270467 | -4.846882208 |
| **A_52_P635182** | Rras | -0.099584953 | -0.378864628 | 0.179694723 | 11.99551533 | -0.877250356 | 0.414811695 | 0.836286891 | -4.847012043 |
| **A_51_P302566** | Maob | 0.477591668 | -0.861890095 | 1.817073432 | 10.76887254 | 0.877180765 | 0.414846554 | 0.836286891 | -4.847051039 |
| **A_55_P1997766** | Acp5 | -0.145862687 | -0.554977195 | 0.263251822 | 13.2609052 | -0.877139139 | 0.414867406 | 0.836286891 | -4.847074363 |
| **A_55_P2147971** | Ranbp1 | -0.057779271 | -0.219847506 | 0.104288964 | 11.16863287 | -0.877088305 | 0.414892871 | 0.836286891 | -4.847102845 |
| **A_55_P2248330** | 2700068H02Rik | 0.051860361 | -0.093625572 | 0.197346294 | 8.135211592 | 0.876967899 | 0.414953195 | 0.836286891 | -4.847170305 |
| **A_52_P381430** | Tbc1d4 | -0.073070757 | -0.278113019 | 0.131971506 | 8.604574843 | -0.876737095 | 0.415068845 | 0.836286891 | -4.847299599 |
| **A_51_P150430** | Mkrn2 | -0.050209526 | -0.191107187 | 0.090688135 | 9.208216964 | -0.876701001 | 0.415086933 | 0.836286891 | -4.847319817 |
| **A_55_P1993698** | Celf1 | 0.123573688 | -0.223221588 | 0.470368964 | 10.61626569 | 0.876641446 | 0.415116779 | 0.836286891 | -4.847353175 |
| **A_55_P2103857** | Hgs | 0.041240421 | -0.074500373 | 0.156981216 | 12.18017291 | 0.87660874 | 0.415133171 | 0.836286891 | -4.847371493 |
| **A_52_P303176** | 1810037I17Rik | 0.123207542 | -0.222575679 | 0.468990763 | 9.582115856 | 0.876602169 | 0.415136464 | 0.836286891 | -4.847375174 |
| **A_55_P2098210** | 4931406P16Rik | -0.095840064 | -0.364933363 | 0.173253236 | 12.28379386 | -0.876220424 | 0.415327824 | 0.836585194 | -4.847588951 |
| **A_66_P118307** | NA | 0.110573293 | -0.200065822 | 0.421212408 | 12.04270194 | 0.87571596 | 0.415580802 | 0.836837078 | -4.847871354 |
| **A_51_P199624** | Sfswap | -0.058115678 | -0.221395112 | 0.105163755 | 8.450057653 | -0.875650881 | 0.415613446 | 0.836837078 | -4.847907777 |

| | | | | | | | | | |
|---|---|---|---|---|---|---|---|---|---|
| **A_55_P2162537** | Acrbp | 0.066224629 | -0.119878299 | 0.252327557 | 8.431821711 | 0.875458349 | 0.415710033 | 0.836837078 | -4.848015524 |
| **A_55_P1988373** | Cenpu | 0.053967632 | -0.097693123 | 0.205628386 | 7.857969279 | 0.875445897 | 0.41571628 | 0.836837078 | -4.848022492 |
| **A_55_P2010014** | Map7d1 | -0.148092627 | -0.564335413 | 0.268150159 | 13.98206878 | -0.875297886 | 0.415790545 | 0.836837078 | -4.848105311 |
| **A_55_P2103351** | Ndufa5 | -0.063064137 | -0.240323317 | 0.114195043 | 12.001307 | -0.8752717 | 0.415803685 | 0.836837078 | -4.848119963 |
| **A_52_P62617** | Gm9895 | -0.043458886 | -0.165619958 | 0.078702185 | 7.908799704 | -0.87521534 | 0.415831967 | 0.836837078 | -4.848151496 |
| **A_51_P279639** | Col6a2 | 0.24178858 | -0.437879487 | 0.921456647 | 8.633875999 | 0.875201557 | 0.415838883 | 0.836837078 | -4.848159207 |
| **A_51_P329370** | Snrnp25 | 0.204256501 | -0.369952719 | 0.778465721 | 12.43146314 | 0.875134672 | 0.415872449 | 0.836837078 | -4.848196627 |
| **A_51_P123795** | Zfp637 | -0.071385976 | -0.272094198 | 0.129322246 | 11.62611343 | -0.875017828 | 0.415931091 | 0.836837078 | -4.82503727 |
| **A_55_P1983268** | 4930444P10Rik | 0.038125858 | -0.069077601 | 0.145329318 | 7.855796384 | 0.874943462 | 0.415968418 | 0.836837078 | -4.84830359 |
| **A_55_P2035495** | Zbtb20 | -0.10964516 | -0.41795159 | 0.19866127 | 9.386783982 | -0.874935515 | 0.415972407 | 0.836837078 | -4.848308035 |
| **A_51_P394814** | Svep1 | 0.091639876 | -0.166101344 | 0.349381095 | 9.852578559 | 0.87472145 | 0.416079866 | 0.836877851 | -4.848427764 |
| **A_51_P418820** | Tfap2c | -0.084415683 | -0.32186909 | 0.153037724 | 13.66037174 | -0.874608906 | 0.416136372 | 0.836877851 | -4.848490704 |
| **A_55_P1971298** | Lonp1 | -0.102300323 | -0.39012411 | 0.185523463 | 12.84204416 | -0.874418802 | 0.41623183 | 0.836877851 | -4.848597005 |
| **A_55_P2054842** | Clpb | -0.076904533 | -0.293281483 | 0.139472417 | 10.45624805 | -0.874399914 | 0.416241316 | 0.836877851 | -4.848607566 |
| **A_55_P2075230** | BC051226 | 0.101988941 | -0.184974617 | 0.388952499 | 10.0593616 | 0.874370503 | 0.416256086 | 0.836877851 | -4.84862401 |
| **A_51_P329413** | Pomgnt1 | 0.125623672 | -0.227897925 | 0.479145268 | 11.03291322 | 0.874227968 | 0.416327672 | 0.836877851 | -4.848703699 |
| **A_52_P261322** | Tanc1 | -0.044748006 | -0.170677163 | 0.081181151 | 8.111416784 | -0.874211572 | 0.416335907 | 0.836877851 | -4.848712865 |
| **A_55_P2036240** | NA | -0.238500669 | -0.909753891 | 0.432752552 | 9.824516691 | -0.874122667 | 0.416380564 | 0.836877851 | -4.848762565 |
| **A_55_P2287269** | AI415730 | 0.04656014 | -0.084489417 | 0.177609697 | 8.363529659 | 0.874073361 | 0.416405332 | 0.836877851 | -4.848790126 |
| **A_55_P2016962** | NA | -0.176772949 | -0.674346368 | 0.320800469 | 9.856242221 | -0.874032938 | 0.416425639 | 0.836877851 | -4.848812722 |
| **A_51_P428742** | Pus7l | 0.060885504 | -0.110533165 | 0.232304173 | 8.335490129 | 0.873825971 | 0.416529619 | 0.836956765 | -4.8489284 |
| **A_55_P2197847** | Lhx1os | 0.090582899 | -0.164459321 | 0.345625119 | 8.599658616 | 0.873782411 | 0.416551507 | 0.836956765 | -4.848952744 |
| **A_55_P2142451** | Echdc2 | -0.102782111 | -0.392242405 | 0.186678182 | 11.82473521 | -0.873569979 | 0.416658259 | 0.83702403 | -4.849071454 |
| **A_51_P235548** | Mtmr9 | 0.067156951 | -0.122015666 | 0.256329569 | 8.852295876 | 0.873377224 | 0.41675514 | 0.83702403 | -4.84917915 |
| **A_55_P2032997** | Srp19 | 0.07127908 | -0.129526831 | 0.27208499 | 11.92300993 | 0.873282496 | 0.416802757 | 0.83702403 | -4.84923207 |
| **A_51_P376057** | Bcorl1 | 0.1226418 | -0.222880613 | 0.468164213 | 8.737241523 | 0.873235645 | 0.41682631 | 0.83702403 | -4.849258243 |
| **A_55_P1979620** | Eif3h | 0.130104219 | -0.23647798 | 0.496686417 | 12.55346946 | 0.873150627 | 0.416869052 | 0.83702403 | -4.849305734 |
| **A_55_P2092103** | NA | -0.081941862 | -0.31283773 | 0.148954006 | 8.843858755 | -0.873089632 | 0.416899718 | 0.83702403 | -4.849339804 |
| **A_55_P1985433** | Nrg1 | -0.081374443 | -0.310683868 | 0.147934982 | 8.051349711 | -0.873042301 | 0.416923516 | 0.83702403 | -4.84936624 |
| **A_51_P188126** | Gca | -0.119558044 | -0.456473157 | 0.21735707 | 8.536295959 | -0.873026587 | 0.416931417 | 0.83702403 | -4.849375017 |
| **A_55_P2031436** | Ly6e | -0.088754748 | -0.338981267 | 0.161471772 | 12.46082845 | -0.872624537 | 0.417133611 | 0.837034474 | -4.849599537 |
| **A_55_P2141889** | NA | -0.111673149 | -0.426527556 | 0.203181259 | 9.003498581 | -0.872585811 | 0.417153091 | 0.837034474 | -4.849621159 |
| **A_52_P8324** | Tmem178 | -0.116874624 | -0.446400717 | 0.21265147 | 8.289679589 | -0.872568596 | 0.41716175 | 0.837034474 | -4.849630771 |
| **A_52_P58471** | Tmem199 | -0.055268117 | -0.211105139 | 0.100568905 | 8.828508761 | -0.872515582 | 0.417188418 | 0.837034474 | -4.84966037 |
| **A_51_P221632** | 1700037H04Rik | 0.148577432 | -0.270426269 | 0.567581132 | 12.13418136 | 0.872376888 | 0.417258192 | 0.837034474 | -4.849737799 |
| **A_55_P1995507** | Aqp11 | 0.173694287 | -0.316239349 | 0.663627924 | 9.074393 | 0.872202729 | 0.417345821 | 0.837034474 | -4.849835015 |
| **A_52_P479262** | NA | 0.410450344 | -0.747328582 | 1.56822927 | 10.26263358 | 0.87217603 | 0.417359256 | 0.837034474 | -4.849849918 |
| **A_51_P169327** | Etd | 0.218883067 | -0.39854446 | 0.836310594 | 11.82944125 | 0.872158345 | 0.417368155 | 0.837034474 | -4.849859788 |
| **A_52_P649332** | Sdf4 | -0.077040212 | -0.294356088 | 0.140275664 | 11.88610026 | -0.872158019 | 0.417368319 | 0.837034474 | -4.84985997 |
| **A_55_P2360427** | BC005561 | -0.066243241 | -0.253108944 | 0.120622462 | 8.290173074 | -0.872129825 | 0.417382507 | 0.837034474 | -4.849875707 |
| **A_55_P1990046** | NA | -0.145506189 | -0.555993403 | 0.264981026 | 8.339279732 | -0.87206929 | 0.41741297 | 0.837034474 | -4.849909492 |
| **A_52_P346581** | Colgalt1 | 0.072760422 | -0.132569098 | 0.278089942 | 11.02192137 | 0.871792197 | 0.417552435 | 0.837082284 | -4.850064124 |
| **A_55_P2017850** | Rpsa | 0.102134998 | -0.186101711 | 0.390371707 | 11.3842162 | 0.87175502 | 0.417571149 | 0.837082284 | -4.850084868 |
| **A_66_P103398** | Ogfrl1 | -0.040212673 | -0.153704265 | 0.073278918 | 8.317645714 | -0.871702759 | 0.417597458 | 0.837082284 | -4.850114027 |
| **A_55_P2068862** | Pigv | 0.049303818 | -0.089849558 | 0.188457195 | 10.45352216 | 0.871677765 | 0.41761004 | 0.837082284 | -4.850127972 |
| **A_51_P404236** | Atp6v0b | -0.09575907 | -0.366073753 | 0.174555614 | 11.25600105 | -0.871524188 | 0.417687361 | 0.837150456 | -4.850213654 |
| **A_55_P2025601** | Lsm6 | 0.084623472 | -0.154292292 | 0.323539235 | 13.75858814 | 0.871395264 | 0.417752278 | 0.837172489 | -4.850285573 |
| **A_55_P2417569** | Prrx2 | 0.127379559 | -0.232275417 | 0.487034536 | 8.086660073 | 0.871330329 | 0.417784977 | 0.837172489 | -4.850321794 |
| **A_55_P2103105** | Hacd1 | 0.158373252 | -0.289040955 | 0.605787459 | 11.68739707 | 0.870845732 | 0.418029067 | 0.837532571 | -4.850592043 |
| **A_55_P2066414** | Dffa | -0.056816483 | -0.217334699 | 0.103701733 | 8.625255146 | -0.870801535 | 0.418051335 | 0.837532571 | -4.850616686 |

| | | | | | | | | |
|---|---|---|---|---|---|---|---|---|
| **A_51_P514319** | Slc13a4 | -0.097093857 | -0.371439936 | 0.177252223 | 15.53542074 | -0.870687192 | 0.418108947 | 0.837539553 | -4.850680435 |
| **A_55_P2070039** | Zfyve9 | -0.063476736 | -0.242848385 | 0.115894913 | 8.008391944 | -0.870622626 | 0.418141481 | 0.837539553 | -4.85071643 |
| **A_51_P124719** | Ccdc93 | -0.040415219 | -0.154651532 | 0.073821094 | 9.108499506 | -0.870382034 | 0.418262731 | 0.837653107 | -4.850850542 |
| **A_55_P2167512** | NA | -0.074388276 | -0.284678734 | 0.135902182 | 13.09722058 | -0.870270158 | 0.418319121 | 0.837653107 | -4.850912895 |
| **A_55_P2003211** | Igsf5 | 0.21138607 | -0.386294908 | 0.809067048 | 8.784819202 | 0.870113875 | 0.418397905 | 0.837653107 | -4.85099999 |
| **A_55_P2046867** | Gata5 | 0.151447957 | -0.276774134 | 0.579670048 | 10.52775608 | 0.870088672 | 0.418410611 | 0.837653107 | -4.851014034 |
| **A_55_P1967231** | Stbd1 | 0.157420715 | -0.287694141 | 0.602535571 | 9.821185203 | 0.870079534 | 0.418415217 | 0.837653107 | -4.851019126 |
| **A_55_P2027157** | NA | 0.058622302 | -0.107151396 | 0.224396001 | 8.103491519 | 0.869994296 | 0.418458192 | 0.837653107 | -4.851066623 |
| **A_51_P273508** | Ube2g1 | 0.047395772 | -0.08666239 | 0.181453933 | 8.641238873 | 0.86979219 | 0.418560102 | 0.837770345 | -4.851179227 |
| **A_55_P1980601** | Oscp1 | 0.041716289 | -0.076320017 | 0.159752595 | 8.157953139 | 0.869479247 | 0.418717938 | 0.837999485 | -4.85135355 |
| **A_66_P132446** | Usp15 | 0.08966112 | -0.164171131 | 0.34349337 | 8.919068351 | 0.869013494 | 0.418952926 | 0.838015328 | -4.851612915 |
| **A_55_P1974288** | Rela | -0.089438422 | -0.342673812 | 0.163796968 | 13.90937947 | -0.868898188 | 0.419011118 | 0.838015328 | -4.851677111 |
| **A_55_P2015941** | NA | -0.047994025 | -0.18388849 | 0.08790044 | 8.243599833 | -0.868869537 | 0.419025578 | 0.838015328 | -4.851693062 |
| **A_51_P155873** | Ppp1r3g | 0.225904394 | -0.41379047 | 0.865599258 | 10.85777456 | 0.868802332 | 0.419059497 | 0.838015328 | -4.851730474 |
| **A_55_P2157610** | Rpusd3 | 0.094428188 | -0.172971847 | 0.361828222 | 10.32603618 | 0.868779088 | 0.41907123 | 0.838015328 | -4.851743413 |
| **A_55_P2115467** | Tmem63b | 0.052098502 | -0.095444808 | 0.199641812 | 9.231268949 | 0.868710113 | 0.419106046 | 0.838015328 | -4.851781808 |
| **A_55_P2121913** | Adi1 | 0.07481984 | -0.137071902 | 0.286711583 | 9.488932963 | 0.868704176 | 0.419109043 | 0.838015328 | -4.851785113 |
| **A_55_P2119257** | Serpine1 | -0.122512769 | -0.469528848 | 0.22450331 | 16.34748204 | -0.868562192 | 0.419180718 | 0.838015328 | -4.851864142 |
| **A_55_P2069083** | Galnt12 | 0.066231795 | -0.121385006 | 0.253848596 | 8.95701442 | 0.868488277 | 0.419218035 | 0.838015328 | -4.85190528 |
| **A_55_P2160286** | Olfm1 | 0.053227212 | -0.097551641 | 0.204006066 | 7.912162615 | 0.868485253 | 0.419219562 | 0.838015328 | -4.851906963 |
| **A_66_P112546** | Yeats4 | -0.049484189 | -0.189665961 | 0.090697583 | 8.484898398 | -0.868448514 | 0.419238111 | 0.838015328 | -4.851927409 |
| **A_55_P2040026** | Itga4 | -0.13532861 | -0.518702594 | 0.248045375 | 8.651312859 | -0.868432655 | 0.419246118 | 0.838015328 | -4.851936235 |
| **A_51_P479769** | Ampd2 | 0.09278952 | -0.170220961 | 0.355800002 | 11.63354759 | 0.867950658 | 0.419489534 | 0.838331593 | -4.852204423 |
| **A_55_P2005229** | NA | -0.115709674 | -0.443688142 | 0.212268794 | 12.5183066 | -0.867947588 | 0.419491084 | 0.838331593 | -4.85220613 |
| **A_66_P130602** | Pdzd8 | -0.054204215 | -0.207919681 | 0.099511251 | 8.370991814 | -0.867530311 | 0.419701901 | 0.838495996 | -4.852438224 |
| **A_51_P338728** | Sdhaf1 | 0.072365686 | -0.132854106 | 0.277585478 | 10.20403976 | 0.867526212 | 0.419703973 | 0.838495996 | -4.852440504 |
| **A_52_P222624** | Hspb2 | 0.20487492 | -0.376125774 | 0.785875615 | 10.24411809 | 0.867523623 | 0.419705281 | 0.838495996 | -4.852441944 |
| **A_52_P526265** | Wdr73 | 0.075329197 | -0.138361301 | 0.289019695 | 11.40777309 | 0.867255951 | 0.419840557 | 0.838495996 | -4.852590785 |
| **A_55_P1988260** | NA | 0.090397869 | -0.166103979 | 0.346899718 | 14.09891156 | 0.867035456 | 0.419952015 | 0.838495996 | -4.852713369 |
| **A_55_P1985453** | NA | 0.194063134 | -0.356680028 | 0.744806296 | 8.63116496 | 0.866888197 | 0.420026465 | 0.838495996 | -4.852795226 |
| **A_51_P316553** | Kdr | -0.328270117 | -1.259956341 | 0.603416107 | 11.78485114 | -0.866823824 | 0.420059013 | 0.838495996 | -4.825693247 |
| **A_55_P2103055** | Gsr | 0.058779183 | -0.108057148 | 0.225615514 | 9.798834889 | 0.86676642 | 0.42008804 | 0.838495996 | -4.85286291 |
| **A_66_P105830** | 1700028B04Rik | 0.080471667 | -0.147938401 | 0.308881734 | 14.44406884 | 0.866756118 | 0.420093249 | 0.838495996 | -4.852868636 |
| **A_55_P2262287** | 5430434G16Rik | -0.117589932 | -0.451378497 | 0.216198633 | 8.518582021 | -0.866698107 | 0.420122584 | 0.838495996 | -4.852900877 |
| **A_52_P565847** | AU018091 | 0.0865961 | -0.159262437 | 0.332454638 | 8.449588153 | 0.86652721 | 0.420209012 | 0.838495996 | -4.852995846 |
| **A_51_P372156** | 4930563E22Rik | -0.137041293 | -0.526121221 | 0.252038635 | 8.849305334 | -0.86652637 | 0.420209437 | 0.838495996 | -4.852996313 |
| **A_55_P1979605** | Eif3m | -0.080896708 | -0.310632374 | 0.148838957 | 12.61310975 | -0.866306534 | 0.420320635 | 0.838495996 | -4.853118461 |
| **A_55_P2172852** | Hacd4 | 0.04862264 | -0.089461673 | 0.186706954 | 8.161746864 | 0.866290109 | 0.420328944 | 0.838495996 | -4.853127586 |
| **A_55_P2038514** | NA | 0.066125938 | -0.121668339 | 0.253920214 | 7.919912582 | 0.866280731 | 0.420333688 | 0.838495996 | -4.853132796 |
| **A_55_P2010196** | Serpina10 | 0.053366269 | -0.098191711 | 0.20492425 | 7.763809419 | 0.866277821 | 0.42033516 | 0.838495996 | -4.853134413 |
| **A_55_P2054310** | M6pr | 0.16403726 | -0.301832148 | 0.629906668 | 11.72906758 | 0.866258436 | 0.420344967 | 0.838495996 | -4.853145182 |
| **A_66_P139167** | Safb | 0.095636625 | -0.175990158 | 0.367263408 | 10.80549358 | 0.866205257 | 0.420371871 | 0.838495996 | -4.853174726 |
| **A_51_P415945** | Psma2 | -0.095094236 | -0.365207967 | 0.175019495 | 15.17257927 | -0.866117266 | 0.420416389 | 0.838495996 | -4.853223606 |
| **A_51_P345467** | Pi4k2a | 0.069552928 | -0.128034931 | 0.267140788 | 9.882989556 | 0.866012831 | 0.420469231 | 0.838495996 | -4.853281617 |
| **A_65_P15503** | Map3k2 | 0.081002567 | -0.149171014 | 0.311176147 | 10.81563406 | 0.86578981 | 0.420582092 | 0.838495996 | -4.853405484 |
| **A_55_P2090070** | Myh14 | 0.271460056 | -0.499944216 | 1.042864328 | 10.8852416 | 0.865751111 | 0.420601678 | 0.838495996 | -4.853426975 |
| **A_55_P2020341** | Camk2g | -0.377774666 | -1.491458508 | 0.735909175 | 11.59494503 | -0.87906501 | 0.420623465 | 0.838495996 | -4.796381392 |
| **A_55_P2104791** | NA | -0.069969005 | -0.268814805 | 0.128876795 | 12.55873401 | -0.8656821 | 0.420636608 | 0.838495996 | -4.853465298 |
| **A_55_P1984861** | NA | 0.096869286 | -0.178438113 | 0.372176685 | 14.61482621 | 0.865640123 | 0.420657855 | 0.838495996 | -4.853488608 |
| **A_55_P2073377** | Mki67 | 0.08785126 | -0.161953124 | 0.337655643 | 8.695738599 | 0.865201174 | 0.420880083 | 0.838676269 | -4.853732308 |

| | | | | | | | | | |
|---|---|---|---|---|---|---|---|---|---|
| **A_51_P521171** | Hif1an | -0.047783762 | -0.183671975 | 0.088104451 | 8.307536573 | -0.86510279 | 0.420929905 | 0.838676269 | -4.853786918 |
| **A_55_P2058601** | Gm7097 | 0.088509749 | -0.163200864 | 0.340220363 | 10.92163683 | 0.865084929 | 0.42093895 | 0.838676269 | -4.853796832 |
| **A_55_P1972799** | Eif4enif1 | -0.06601355 | -0.253750364 | 0.121723264 | 10.53738191 | -0.865073104 | 0.420944939 | 0.838676269 | -4.853803395 |
| **A_52_P495031** | 9130401M01Rik | -0.081192101 | -0.312106099 | 0.149721897 | 11.98083722 | -0.865033014 | 0.420965242 | 0.838676269 | -4.853825646 |
| **A_55_P2002319** | Ap5s1 | -0.072513689 | -0.278893573 | 0.133866195 | 10.66740196 | -0.86441401 | 0.421278824 | 0.839179054 | -4.854169121 |
| **A_55_P1969276** | Hhip | 0.1356814 | -0.250501537 | 0.521864337 | 8.415901505 | 0.864363458 | 0.421304441 | 0.839179054 | -4.854197164 |
| **A_55_P1977792** | Folr1 | 0.2252474 | -0.416020735 | 0.866515535 | 14.80091906 | 0.86415031 | 0.421412465 | 0.839307732 | -4.854315392 |
| **A_55_P2140151** | Flcn | 0.070959627 | -0.131123174 | 0.273042428 | 11.40055171 | 0.863875454 | 0.421551794 | 0.839498725 | -4.854467818 |
| **A_51_P224771** | Tmem184b | -0.14018647 | -0.539522348 | 0.259149408 | 12.08603927 | -0.86364832 | 0.421666958 | 0.83964156 | -4.854593755 |
| **A_55_P2119463** | Ggh | 0.107455123 | -0.198755286 | 0.413665532 | 12.13788947 | 0.863329017 | 0.421828893 | 0.83987749 | -4.854770756 |
| **A_52_P638513** | Zbtb11os1 | -0.040987717 | -0.157808324 | 0.075832889 | 9.07501172 | -0.863184118 | 0.421902394 | 0.839937314 | -4.854851064 |
| **A_51_P137778** | NA | 0.086878653 | -0.160834625 | 0.334591932 | 8.646353883 | 0.862845335 | 0.422074282 | 0.840192976 | -4.855038793 |
| **A_55_P2085974** | Igf1 | 0.130766312 | -0.242197953 | 0.503730577 | 8.210614818 | 0.862576887 | 0.422210522 | 0.840377631 | -4.855187512 |
| **A_55_P2150283** | E030030I06Rik | 0.134500899 | -0.249183483 | 0.51818528 | 12.35540092 | 0.862422797 | 0.422288738 | 0.840385707 | -4.855272862 |
| **A_55_P2054409** | Pira2 | -0.040629626 | -0.156535234 | 0.075275982 | 7.771396019 | -0.862397589 | 0.422301535 | 0.840385707 | -4.855286824 |
| **A_52_P110070** | 5730416F02Rik | -0.120858965 | -0.465874819 | 0.224156889 | 9.077915187 | -0.861804944 | 0.42260247 | 0.840897998 | -4.855614985 |
| **A_55_P2156464** | Nt5c3b | -0.057006277 | -0.21977021 | 0.105757656 | 8.395256347 | -0.8616555 | 0.422678381 | 0.840937033 | -4.855697711 |
| **A_55_P2164189** | Sfpq | -0.088876358 | -0.342722632 | 0.164969916 | 9.64007186 | -0.861359844 | 0.42282859 | 0.840937033 | -4.855861345 |
| **A_52_P406371** | Med14 | 0.075015237 | -0.139263956 | 0.289294431 | 8.557345439 | 0.861268667 | 0.422874921 | 0.840937033 | -4.8559118 |
| **A_51_P361426** | Arl8b | -0.047056833 | -0.181483124 | 0.087369457 | 11.36128166 | -0.861207189 | 0.422906163 | 0.840937033 | -4.855945819 |
| **A_52_P232637** | Dhh | -0.051554976 | -0.198895317 | 0.095785366 | 8.15731506 | -0.860831356 | 0.42309719 | 0.840937033 | -4.856153746 |
| **A_55_P2121426** | NA | -0.078886466 | -0.304370239 | 0.146597306 | 13.37646648 | -0.860709057 | 0.423159365 | 0.840937033 | -4.856221394 |
| **A_55_P2133220** | Arhgef39 | 0.060403139 | -0.112254481 | 0.233060759 | 8.398857315 | 0.860682395 | 0.423172921 | 0.840937033 | -4.85623614 |
| **A_51_P382253** | Rnf2 | -0.07877671 | -0.3040369 | 0.146483479 | 10.94554475 | -0.86036465 | 0.423334495 | 0.840937033 | -4.856411861 |
| **A_55_P2118183** | Tnks2 | -0.056831183 | -0.21938443 | 0.105722064 | 9.508730787 | -0.860122308 | 0.423457757 | 0.840937033 | -4.856545852 |
| **A_55_P2037702** | Gm10638 | 0.101505576 | -0.188836214 | 0.391847366 | 8.821445186 | 0.860101121 | 0.423468535 | 0.840937033 | -4.856557565 |
| **A_55_P2012206** | Ndufb11 | 0.174272227 | -0.324211034 | 0.672755489 | 14.05708774 | 0.860095679 | 0.423471303 | 0.840937033 | -4.856560574 |
| **A_55_P2155942** | Ezh1 | -0.076228911 | -0.294276305 | 0.141818483 | 10.23201443 | -0.860078272 | 0.423480158 | 0.840937033 | -4.856570197 |
| **A_52_P463346** | Serinc3 | -0.086428796 | -0.333667915 | 0.160810322 | 10.97360677 | -0.860023746 | 0.423507896 | 0.840937033 | -4.85660034 |
| **A_55_P2143790** | Gm14410 | -0.07964274 | -0.30751087 | 0.148225389 | 9.749211968 | -0.859867966 | 0.423587151 | 0.840937033 | -4.85668645 |
| **A_51_P244558** | Rab3gap2 | 0.114350637 | -0.212844071 | 0.441545345 | 10.18987211 | 0.859807995 | 0.423617666 | 0.840937033 | -4.856719597 |
| **A_55_P2005343** | NA | -0.088643512 | -0.342311525 | 0.165024502 | 15.47270232 | -0.859706891 | 0.423669113 | 0.840937033 | -4.856775476 |
| **A_51_P165330** | Mrpl33 | 0.112072512 | -0.208674716 | 0.432819741 | 14.01300255 | 0.859617719 | 0.423714491 | 0.840937033 | -4.856824756 |
| **A_52_P23225** | Gpc3 | 0.176754329 | -0.32911521 | 0.682623867 | 11.50949264 | 0.859608484 | 0.423719191 | 0.840937033 | -4.856829859 |
| **A_51_P466378** | Itgal | 0.043824777 | -0.081611702 | 0.169261256 | 7.928406847 | 0.859537882 | 0.423755123 | 0.840937033 | -4.856868874 |
| **A_55_P2084812** | Sec31a | -0.086039089 | -0.33230418 | 0.160226002 | 9.425385747 | -0.859532126 | 0.423758052 | 0.840937033 | -4.856872054 |
| **A_55_P2134566** | NA | 0.039019785 | -0.072667418 | 0.150706987 | 9.050836485 | 0.859509377 | 0.423769631 | 0.840937033 | -4.856884625 |
| **A_55_P2329923** | Camkk2 | -0.155111616 | -0.599219437 | 0.288996205 | 10.64604 | -0.859260928 | 0.423896096 | 0.840937033 | -4.857021898 |
| **A_52_P114722** | Ptpn6 | 0.057998951 | -0.108061709 | 0.224059612 | 9.880462917 | 0.859255936 | 0.423898637 | 0.840937033 | -4.857024656 |
| **A_55_P2092760** | Rhbdd3 | 0.051083897 | -0.095189575 | 0.197357369 | 11.29904411 | 0.859186835 | 0.423933817 | 0.840937033 | -4.857062831 |
| **A_55_P1993940** | Sh3bgrl3 | 0.055858832 | -0.104090324 | 0.215807988 | 11.38196231 | 0.859169908 | 0.423942434 | 0.840937033 | -4.857072181 |
| **A_55_P1993836** | Dusp7 | 0.087494898 | -0.163059173 | 0.338048968 | 9.637288138 | 0.85911327 | 0.423971271 | 0.840937033 | -4.857103469 |
| **A_65_P19780** | Klf9 | -0.161208316 | -0.622987854 | 0.300571221 | 10.99779124 | -0.858859098 | 0.424100695 | 0.840937033 | -4.857243858 |
| **A_55_P2081373** | Glis1 | 0.16901979 | -0.315167158 | 0.653206738 | 11.99478297 | 0.858803179 | 0.424129173 | 0.840937033 | -4.857274741 |
| **A_55_P2082050** | Ankrd12 | -0.098376209 | -0.380205933 | 0.183453515 | 8.66667153 | -0.858761274 | 0.424150515 | 0.840937033 | -4.857297883 |
| **A_51_P453401** | Cipc | 0.057990976 | -0.108169274 | 0.224151225 | 10.55598232 | 0.858622852 | 0.424221018 | 0.840937033 | -4.857374321 |
| **A_55_P2018292** | Map2k4 | 0.092747796 | -0.173018841 | 0.358514432 | 9.823083935 | 0.85856326 | 0.424251373 | 0.840937033 | -4.857407226 |
| **A_55_P2154817** | Ccdc28a | -0.081911538 | -0.316656592 | 0.152833515 | 9.737938044 | -0.858455518 | 0.424306259 | 0.840937033 | -4.857466713 |
| **A_55_P1958840** | Itga6 | 0.067420901 | -0.125823819 | 0.26066562 | 10.12374796 | 0.858333528 | 0.424368409 | 0.840937033 | -4.857534062 |
| **A_51_P264527** | Dipk1b | -0.120316782 | -0.465180379 | 0.224546816 | 8.037946886 | -0.858317596 | 0.424376526 | 0.840937033 | -4.857542856 |

| | | | | | | | | | |
|---|---|---|---|---|---|---|---|---|---|
| **A_55_P1986351** | Tbc1d32 | -0.040640144 | -0.157143372 | 0.075863084 | 8.191470482 | -0.858195922 | 0.424438523 | 0.840937033 | -4.857610023 |
| **A_51_P186547** | Pah | -0.039421894 | -0.152436341 | 0.073592553 | 7.64145586 | -0.858168732 | 0.424452378 | 0.840937033 | -4.857625031 |
| **A_51_P362054** | Atp5e | 0.156690668 | -0.292514579 | 0.605895916 | 14.44866503 | 0.858158434 | 0.424457626 | 0.840937033 | -4.857630715 |
| **A_55_P2057681** | Cdyl2 | -0.048504968 | -0.187564559 | 0.090554623 | 8.372239875 | -0.858132668 | 0.424470756 | 0.840937033 | -4.857644937 |
| **A_51_P413059** | Mcpt8 | 0.038749585 | -0.072344704 | 0.149843874 | 7.744617179 | 0.858113003 | 0.424480777 | 0.840937033 | -4.857655791 |
| **A_55_P2091486** | Tmem219 | 0.068610093 | -0.128101311 | 0.265321497 | 10.7030261 | 0.858079707 | 0.424497745 | 0.840937033 | -4.857674169 |
| **A_55_P2137984** | NA | 0.080507657 | -0.15033645 | 0.311351765 | 9.716825841 | 0.858000534 | 0.424538094 | 0.840937033 | -4.857717867 |
| **A_51_P268734** | Kptn | 0.110611716 | -0.206590298 | 0.427813729 | 9.599225056 | 0.857895432 | 0.424591661 | 0.840937033 | -4.85777587 |
| **A_51_P115715** | Asb2 | -0.039231398 | -0.151739702 | 0.073276906 | 7.730596041 | -0.857863855 | 0.424607756 | 0.840937033 | -4.857793296 |
| **A_65_P07196** | COX3 | 0.068040072 | -0.127102616 | 0.263182759 | 16.68976244 | 0.857791314 | 0.424644732 | 0.840937033 | -4.857833325 |
| **A_55_P2052210** | Gdi2 | 0.088303652 | -0.164959649 | 0.341566953 | 14.09773108 | 0.857779306 | 0.424650853 | 0.840937033 | -4.857839952 |
| **A_51_P340699** | Rasl11a | -0.15035177 | -0.581598225 | 0.280894685 | 9.477873763 | -0.857733099 | 0.424674408 | 0.840937033 | -4.857865449 |
| **A_55_P2018181** | A730008H23Rik | 0.047289161 | -0.088364953 | 0.182943275 | 8.060972749 | 0.857625723 | 0.424729148 | 0.840937033 | -4.857924694 |
| **A_55_P2168048** | Gramd4 | -0.049816536 | -0.192736325 | 0.093103254 | 8.358475439 | -0.857531967 | 0.424776948 | 0.840937033 | -4.857976421 |
| **A_55_P2115567** | Slc26a1 | 0.169155296 | -0.316159305 | 0.654469897 | 8.397324684 | 0.857494622 | 0.42479599 | 0.840937033 | -4.857997024 |
| **A_52_P459276** | Ift81 | -0.050299058 | -0.194609923 | 0.094011806 | 8.015829514 | -0.85749183 | 0.424797413 | 0.840937033 | -4.857998564 |
| **A_55_P2022773** | Glcci1 | 0.06403261 | -0.119798599 | 0.247863819 | 8.605724531 | 0.856941395 | 0.425078138 | 0.841402032 | -4.858302158 |
| **A_55_P2157378** | Xlr5a | -0.069792487 | -0.270178626 | 0.130593652 | 9.10352772 | -0.856860578 | 0.425119367 | 0.841402032 | -4.858346722 |
| **A_66_P132888** | Dhfr | -0.040378922 | -0.156330531 | 0.075572687 | 8.139615049 | -0.856736177 | 0.425182836 | 0.84144149 | -4.858415313 |
| **A_51_P148311** | Chic2 | -0.064132846 | -0.248339668 | 0.120073976 | 11.00753259 | -0.856532739 | 0.425286644 | 0.841448286 | -4.858527467 |
| **A_55_P2133096** | Tgds | -0.053636606 | -0.207696975 | 0.100423764 | 8.379032237 | -0.856523953 | 0.425291127 | 0.841448286 | -4.85853231 |
| **A_55_P2048368** | Fggy | 0.042215089 | -0.079050364 | 0.163480542 | 7.930448173 | 0.85644559 | 0.425331119 | 0.841448286 | -4.858575506 |
| **A_51_P337125** | Inpp5d | -0.0673421 | -0.260805981 | 0.126121781 | 8.986709132 | -0.856359113 | 0.425375255 | 0.841448286 | -4.858623171 |
| **A_55_P1970670** | Scp2 | -0.113657877 | -0.440201552 | 0.212885798 | 12.79917326 | -0.856302924 | 0.425403934 | 0.841448286 | -4.85865414 |
| **A_51_P512061** | Doxl1 | -0.492512749 | -1.908066258 | 0.923040761 | 11.65903576 | -0.855973558 | 0.425572074 | 0.841554114 | -4.858835645 |
| **A_55_P1965000** | Ddx58 | 0.060338723 | -0.113092202 | 0.233769649 | 11.23834647 | 0.855930967 | 0.42559382 | 0.841554114 | -4.858859112 |
| **A_55_P2021350** | NA | -0.046676265 | -0.180844717 | 0.087492187 | 15.85103361 | -0.855883884 | 0.425617861 | 0.841554114 | -4.858885053 |
| **A_55_P2143195** | Dyrk1a | 0.08904724 | -0.166932947 | 0.345027427 | 9.650529267 | 0.85582166 | 0.425649634 | 0.841554114 | -4.858919335 |
| **A_55_P1974640** | NA | -0.06984713 | -0.270645321 | 0.13095106 | 8.652897978 | -0.855771733 | 0.425675129 | 0.841554114 | -4.858946841 |
| **A_55_P2047967** | Bfar | -0.243515699 | -0.943668629 | 0.456637231 | 14.29426363 | -0.855663914 | 0.425730191 | 0.841576893 | -4.859006237 |
| **A_55_P2026818** | Slc4a7 | -0.093951495 | -0.364173686 | 0.176270696 | 8.513786133 | -0.855365731 | 0.425882497 | 0.841756038 | -4.859170474 |
| **A_51_P369200** | Tpx2 | 0.07540704 | -0.14151092 | 0.292325 | 8.237953704 | 0.855235161 | 0.425949202 | 0.841756038 | -4.859242379 |
| **A_66_P126094** | Pycrl | -0.050274864 | -0.194897662 | 0.094347935 | 8.649945631 | -0.855230738 | 0.425951462 | 0.841756038 | -4.859244815 |
| **A_52_P134556** | Cd2bp2 | -0.052700899 | -0.204322535 | 0.098920737 | 9.609351425 | -0.855117924 | 0.426009102 | 0.841783883 | -4.859306935 |
| **A_51_P232371** | Stab1 | -0.06982992 | -0.270800493 | 0.131140653 | 9.608843963 | -0.854827014 | 0.426157765 | 0.841968105 | -4.859467096 |
| **A_55_P2098120** | NA | 0.073567294 | -0.138174801 | 0.285309389 | 8.337802407 | 0.854765015 | 0.426189453 | 0.841968105 | -4.859501225 |
| **A_55_P2143233** | Rasgrp2 | 0.18174075 | -0.341460133 | 0.704941632 | 8.621893313 | 0.854580562 | 0.426283738 | 0.842068306 | -4.859602751 |
| **A_55_P1967618** | Med16 | 0.092027933 | -0.172979296 | 0.357035161 | 10.67219992 | 0.854340726 | 0.426406357 | 0.842074585 | -4.859734739 |
| **A_52_P637172** | Pyroxd2 | -0.087790863 | -0.340613928 | 0.165032203 | 8.812297061 | -0.854283042 | 0.426435852 | 0.842074585 | -4.85976648 |
| **A_55_P2177639** | NA | -0.053147761 | -0.20621025 | 0.099914727 | 16.20395219 | -0.854250754 | 0.426452363 | 0.842074585 | -4.859784246 |
| **A_55_P1984344** | Rhox2h | 0.209530735 | -0.394074371 | 0.813135841 | 9.257350382 | 0.854012044 | 0.426574441 | 0.842074585 | -4.859915578 |
| **A_55_P1970785** | NA | 0.128997181 | -0.242642312 | 0.500636674 | 11.4956412 | 0.853940326 | 0.426611123 | 0.842074585 | -4.85995503 |
| **A_55_P2070194** | Caprin1 | 0.051588072 | -0.097049587 | 0.20022573 | 11.59411661 | 0.853865768 | 0.42664926 | 0.842074585 | -4.859996043 |
| **A_55_P2010152** | Sell | -0.049670816 | -0.192786296 | 0.093444664 | 7.886560053 | -0.853854452 | 0.426655049 | 0.842074585 | -4.860002268 |
| **A_55_P2049289** | NA | -0.042429116 | -0.164694944 | 0.079836711 | 16.3705902 | -0.853744774 | 0.426711156 | 0.842074585 | -4.860062593 |
| **A_55_P2275279** | Mid2 | -0.079207539 | -0.307472715 | 0.149057637 | 8.485774292 | -0.853681794 | 0.426743376 | 0.842074585 | -4.860097232 |
| **A_66_P124958** | Ccdc47 | -0.05700887 | -0.221330878 | 0.107313137 | 8.765683233 | -0.853524252 | 0.426823982 | 0.842074585 | -4.86018387 |
| **A_55_P2090364** | NA | 0.058921672 | -0.110952786 | 0.22879613 | 15.71423141 | 0.853328281 | 0.426924267 | 0.842074585 | -4.860291627 |
| **A_55_P1981494** | H2-Bl | -0.273004381 | -1.060230706 | 0.514221944 | 9.687024944 | -0.853177061 | 0.427001662 | 0.842074585 | -4.860374765 |
| **A_52_P326664** | Unc93a | -0.29736022 | -1.154993543 | 0.560273104 | 9.379107072 | -0.853002659 | 0.427090935 | 0.842074585 | -4.860470635 |

| | | | | | | | | | |
|---|---|---|---|---|---|---|---|---|---|
| **A_55_P2130373** | Dcaf11 | -0.049459524 | -0.192125803 | 0.093206756 | 11.9204807 | -0.852899298 | 0.427143851 | 0.842074585 | -4.860527448 |
| **A_55_P2054240** | Med8 | -0.070633608 | -0.274378913 | 0.133111697 | 13.36454286 | -0.852889847 | 0.427148689 | 0.842074585 | -4.860532642 |
| **A_55_P2419854** | NA | -0.049758545 | -0.193298506 | 0.093781416 | 7.905672912 | -0.852833037 | 0.427177775 | 0.842074585 | -4.860563865 |
| **A_55_P2159555** | NA | -0.422357729 | -1.640749225 | 0.796033768 | 11.80861063 | -0.852830551 | 0.427179048 | 0.842074585 | -4.860565232 |
| **A_55_P1957378** | Unc119 | 0.190206946 | -0.358566073 | 0.738979966 | 12.63777693 | 0.852712806 | 0.427239337 | 0.842074585 | -4.860629941 |
| **A_51_P416556** | Ssr3 | 0.084066019 | -0.15849081 | 0.326622848 | 13.85172857 | 0.852660489 | 0.427266126 | 0.842074585 | -4.86065869 |
| **A_55_P1995623** | Pomp | -0.086919371 | -0.337731416 | 0.163892674 | 13.56714618 | -0.852584338 | 0.427305123 | 0.842074585 | -4.860700536 |
| **A_55_P2010906** | Rbm45 | -0.049462883 | -0.192238333 | 0.093312567 | 9.502012926 | -0.852305032 | 0.427448177 | 0.842074585 | -4.860853992 |
| **A_55_P2140107** | Cyp21a1 | 0.484731464 | -0.914457678 | 1.883920605 | 9.648489437 | 0.852302863 | 0.427449288 | 0.842074585 | -4.860855183 |
| **A_55_P2068486** | Arfgap1 | 0.044605203 | -0.084151804 | 0.173362209 | 10.8978947 | 0.852282967 | 0.427459479 | 0.842074585 | -4.860866113 |
| **A_52_P429364** | Ccdc69 | 0.124327115 | -0.234595498 | 0.483249728 | 8.516537818 | 0.852185612 | 0.427509351 | 0.842074585 | -4.860919593 |
| **A_55_P2058962** | Mcm10 | 0.084720947 | -0.159863851 | 0.329305745 | 8.22065101 | 0.852178362 | 0.427513066 | 0.842074585 | -4.860923576 |
| **A_55_P1959515** | Cep135 | -0.04472751 | -0.173868884 | 0.084413864 | 8.115021884 | -0.852076292 | 0.427565358 | 0.842074585 | -4.86097964 |
| **A_55_P2088692** | NA | -0.279044406 | -1.084747348 | 0.526658536 | 12.70102884 | -0.852054813 | 0.427576363 | 0.842074585 | -4.860991438 |
| **A_51_P477121** | Pmaip1 | 0.161895702 | -0.30558892 | 0.629380324 | 10.65441835 | 0.851995212 | 0.4276069 | 0.842074585 | -4.861024172 |
| **A_55_P2056521** | Rpl21 | 0.038499607 | -0.072680623 | 0.149679837 | 7.919617274 | 0.851918187 | 0.427646368 | 0.842074585 | -4.861066475 |
| **A_55_P2106525** | Nmnat3 | -0.047264779 | -0.183759148 | 0.089229589 | 10.18111499 | -0.851906759 | 0.427652223 | 0.842074585 | -4.861072751 |
| **A_51_P367780** | Adamtsl2 | -0.039990311 | -0.155484465 | 0.075503844 | 7.709277869 | -0.851851588 | 0.427680495 | 0.842074585 | -4.861103049 |
| **A_55_P2083098** | Tatdn3 | 0.045765541 | -0.086407942 | 0.177939023 | 9.12152794 | 0.851850586 | 0.427681008 | 0.842074585 | -4.861103599 |
| **A_52_P525837** | Trmt6 | -0.073768187 | -0.286849004 | 0.13931263 | 8.809134088 | -0.851714262 | 0.427750871 | 0.84212135 | -4.861178457 |
| **A_55_P1985040** | Gpbp1l1 | -0.047567615 | -0.184980491 | 0.089845262 | 8.787566316 | -0.85163422 | 0.427791895 | 0.84212135 | -4.861222407 |
| **A_55_P2068772** | Ncbp2 | 0.118276899 | -0.22355555 | 0.460109349 | 9.910710897 | 0.851247416 | 0.427990183 | 0.84238738 | -4.861434751 |
| **A_55_P2079757** | NA | 0.081188468 | -0.153474446 | 0.315851382 | 8.652252923 | 0.851175378 | 0.428027119 | 0.84238738 | -4.86147429 |
| **A_55_P2153431** | Ralgds | -0.088507959 | -0.344440743 | 0.167424826 | 11.62009932 | -0.850796248 | 0.428221552 | 0.84238738 | -4.861682344 |
| **A_52_P557293** | Tmprss2 | 0.127379125 | -0.240982998 | 0.495741248 | 10.8781726 | 0.850731388 | 0.428254821 | 0.84238738 | -4.86171793 |
| **A_55_P1987196** | Gm3448 | -0.04384925 | -0.170662325 | 0.082963825 | 8.169666714 | -0.85068211 | 0.428280099 | 0.84238738 | -4.861744966 |
| **A_52_P366572** | Slain2 | -0.048632685 | -0.189282818 | 0.092017448 | 9.032919953 | -0.850662459 | 0.428290179 | 0.84238738 | -4.861755747 |
| **A_51_P476851** | Fance | -0.047083556 | -0.183265919 | 0.089098808 | 12.07104166 | -0.850584668 | 0.428330087 | 0.84238738 | -4.861798423 |
| **A_51_P365912** | Ciao3 | 0.067473204 | -0.127710622 | 0.262657031 | 11.00397141 | 0.850465443 | 0.428391255 | 0.84238738 | -4.861863825 |
| **A_55_P2154107** | Gcm1 | -0.233094953 | -0.90741363 | 0.441223723 | 10.31900428 | -0.85042662 | 0.428411174 | 0.84238738 | -4.861885121 |
| **A_51_P494125** | Alpl | -0.195745171 | -0.762063614 | 0.370573272 | 12.58270542 | -0.850353521 | 0.428448682 | 0.84238738 | -4.861925215 |
| **A_55_P1962831** | Ssr1 | -0.128573435 | -0.514550598 | 0.257403728 | 13.40292892 | -0.863256227 | 0.42845933 | 0.84238738 | -4.804396895 |
| **A_52_P49042** | Arfgef3 | -0.074406024 | -0.289686274 | 0.140874226 | 8.607885616 | -0.850301748 | 0.428475249 | 0.84238738 | -4.861953611 |
| **A_55_P1965303** | Dsc2 | 0.089697832 | -0.169854802 | 0.349250466 | 11.28130516 | 0.850208941 | 0.428522874 | 0.84238738 | -4.86200451 |
| **A_55_P2124228** | Uaca | -0.129206354 | -0.503094418 | 0.244681711 | 14.26413138 | -0.850181071 | 0.428537178 | 0.84238738 | -4.862019794 |
| **A_51_P223709** | Pask | -0.074624385 | -0.290596695 | 0.141347924 | 8.620192951 | -0.850064457 | 0.428597028 | 0.842407528 | -4.862083743 |
| **A_55_P2059650** | Atpaf2 | 0.060601857 | -0.11482132 | 0.236025034 | 10.27448649 | 0.849900578 | 0.428681146 | 0.842407528 | -4.8621736 |
| **A_55_P2440032** | Usp24 | -0.110043009 | -0.428588911 | 0.208502893 | 8.218691723 | -0.849883893 | 0.42868971 | 0.842407528 | -4.862182748 |
| **A_51_P290207** | Insig1 | 0.073322686 | -0.138975282 | 0.285620654 | 8.011084814 | 0.84969232 | 0.428788061 | 0.842407528 | -4.862287774 |
| **A_55_P2022678** | C1qtnf1 | -0.19767606 | -0.770041696 | 0.374689576 | 11.50908078 | -0.849668833 | 0.42880012 | 0.842407528 | -4.862300649 |
| **A_52_P196568** | Meg3 | -0.105329716 | -0.410315186 | 0.199655755 | 8.215391132 | -0.849651689 | 0.428808923 | 0.842407528 | -4.862310047 |
| **A_55_P1990285** | Rplp2 | -0.057694724 | -0.224787627 | 0.109398178 | 16.95362624 | -0.849468464 | 0.428903006 | 0.842487405 | -4.862410477 |
| **A_55_P1981829** | Rhox8 | -0.435088015 | -1.695510745 | 0.825334714 | 10.9753528 | -0.849239249 | 0.429020727 | 0.842487405 | -4.862536094 |
| **A_55_P2087796** | Eva1b | -0.084967903 | -0.331123468 | 0.161187663 | 11.73893638 | -0.849208633 | 0.429036452 | 0.842487405 | -4.86255287 |
| **A_51_P383991** | 4-Sep | 0.13953763 | -0.264746945 | 0.543822204 | 13.61512754 | 0.849128334 | 0.429077699 | 0.842487405 | -4.86259687 |
| **A_51_P497882** | Creb3l4 | -0.040452537 | -0.157685788 | 0.076780715 | 8.122429906 | -0.848914814 | 0.429187391 | 0.842487405 | -4.862713854 |
| **A_55_P2010066** | Capn3 | -0.052071022 | -0.202977986 | 0.098835942 | 7.980736693 | -0.848898922 | 0.429195556 | 0.842487405 | -4.86272256 |
| **A_55_P2004781** | Ass1 | 0.177710475 | -0.337491449 | 0.692912399 | 12.97037319 | 0.848603297 | 0.429347464 | 0.842487405 | -4.862884492 |
| **A_51_P503192** | Rbm15b | 0.072634459 | -0.137947347 | 0.283216266 | 8.709579765 | 0.848576543 | 0.429361214 | 0.842487405 | -4.862899145 |
| **A_51_P321531** | Purg | -0.08554939 | -0.333580269 | 0.16248149 | 10.97235891 | -0.84855564 | 0.429371957 | 0.842487405 | -4.862910593 |

| | | | | | | | | | |
|---|---|---|---|---|---|---|---|---|---|
| **A_52_P262967** | Rab10 | -0.067356615 | -0.262643828 | 0.127930598 | 10.32918825 | -0.848546423 | 0.429376694 | 0.842487405 | -4.862915641 |
| **A_55_P2024431** | Gaa | -0.071134922 | -0.277383913 | 0.135114069 | 15.08597635 | -0.848516298 | 0.429392177 | 0.842487405 | -4.862932139 |
| **A_55_P2096144** | Gnao1 | 0.046846556 | -0.088987254 | 0.182680366 | 7.889968127 | 0.848474806 | 0.429413502 | 0.842487405 | -4.862954862 |
| **A_55_P2178317** | Gm9199 | 0.072081197 | -0.136951928 | 0.281114322 | 8.592050532 | 0.848351908 | 0.429476673 | 0.842487405 | -4.863022162 |
| **A_52_P373893** | NA | 0.157637446 | -0.299516115 | 0.614791007 | 11.56675647 | 0.848333168 | 0.429486306 | 0.842487405 | -4.863032424 |
| **A_55_P2030578** | Tns1 | 0.158092253 | -0.300398215 | 0.616582721 | 12.9909255 | 0.848299948 | 0.429503383 | 0.842487405 | -4.863050614 |
| **A_55_P2185821** | Cyp3a13 | 0.128045622 | -0.243375416 | 0.49946666 | 7.889124109 | 0.84813971 | 0.429585761 | 0.842517565 | -4.863138348 |
| **A_55_P2112617** | Smad4 | -0.057371198 | -0.22383443 | 0.109092033 | 10.31828278 | -0.847900253 | 0.429708886 | 0.842517565 | -4.863269433 |
| **A_55_P2108389** | D830030K20Rik | 0.130550986 | -0.248303248 | 0.509405219 | 9.856131588 | 0.8477683 | 0.429776746 | 0.842517565 | -4.863341658 |
| **A_55_P2032272** | Habp2 | 0.347799911 | -0.661532682 | 1.357132504 | 9.497179076 | 0.847743233 | 0.429789639 | 0.842517565 | -4.863355377 |
| **A_55_P2120714** | NA | 0.038201889 | -0.072676625 | 0.149080404 | 8.230711152 | 0.847630559 | 0.429847591 | 0.842517565 | -4.863417041 |
| **A_52_P576100** | Sh2b1 | -0.061609496 | -0.240443397 | 0.117224404 | 10.78454853 | -0.84755321 | 0.429887378 | 0.842517565 | -4.86345937 |
| **A_51_P208769** | Metrn | 0.041917326 | -0.079778372 | 0.163613024 | 7.872096756 | 0.84739814 | 0.429967151 | 0.842517565 | -4.863544221 |
| **A_55_P2122334** | Nfic | -0.107418428 | -0.419320055 | 0.204483198 | 9.920005859 | -0.847286561 | 0.430024558 | 0.842517565 | -4.863605269 |
| **A_55_P1981674** | NA | 0.051989297 | -0.098974111 | 0.202952705 | 8.52539554 | 0.847249689 | 0.43004353 | 0.842517565 | -4.863625442 |
| **A_51_P400040** | Ubxn2a | -0.055730479 | -0.217590598 | 0.106129641 | 9.840880571 | -0.847075418 | 0.430133206 | 0.842517565 | -4.863720776 |
| **A_51_P207622** | Fmod | 0.060959436 | -0.116104275 | 0.238023147 | 8.376908237 | 0.846994402 | 0.430174899 | 0.842517565 | -4.86376509 |
| **A_55_P2077548** | Git1 | 0.079585309 | -0.151584672 | 0.31075529 | 11.98823097 | 0.846975076 | 0.430184846 | 0.842517565 | -4.863775661 |
| **A_55_P2017160** | Cdc14a | -0.060697893 | -0.237022074 | 0.115626289 | 8.257045525 | -0.846897598 | 0.430224722 | 0.842517565 | -4.863818037 |
| **A_55_P2008609** | Rxrb | -0.045892135 | -0.179216606 | 0.087432336 | 11.90747139 | -0.846832574 | 0.430258191 | 0.842517565 | -4.863853599 |
| **A_51_P495077** | Ift22 | -0.067542978 | -0.263793918 | 0.128707963 | 8.294759815 | -0.846715713 | 0.430318345 | 0.842517565 | -4.863917507 |
| **A_55_P2130565** | Prdm11 | -0.065936728 | -0.257523827 | 0.12565037 | 8.244774292 | -0.846701441 | 0.430325692 | 0.842517565 | -4.863925311 |
| **A_51_P197883** | Exo5 | 0.068896478 | -0.13134285 | 0.269135806 | 10.15048148 | 0.846480205 | 0.430439593 | 0.842517565 | -4.86404628 |
| **A_55_P2032498** | Arpin | 0.131963367 | -0.251671707 | 0.515598441 | 9.821092701 | 0.84626084 | 0.430552552 | 0.842517565 | -4.864166204 |
| **A_55_P2061159** | Ing1 | -0.057272249 | -0.223804242 | 0.109259743 | 12.57906286 | -0.846088366 | 0.430641381 | 0.842517565 | -4.864260478 |
| **A_55_P1978050** | Aven | -0.054917343 | -0.21462664 | 0.104791954 | 9.847768356 | -0.845957375 | 0.430708853 | 0.842517565 | -4.864332068 |
| **A_55_P1966538** | Ctu2 | -0.066850925 | -0.261278448 | 0.127576598 | 10.91631532 | -0.845899639 | 0.430738595 | 0.842517565 | -4.86436362 |
| **A_55_P2125588** | Pdgfa | -0.118151475 | -0.461797206 | 0.225494255 | 13.79593143 | -0.8458578 | 0.430760149 | 0.842517565 | -4.864386484 |
| **A_55_P1986691** | Card6 | -0.132242223 | -0.517016669 | 0.252532223 | 8.401816367 | -0.845537907 | 0.430924972 | 0.842517565 | -4.864561267 |
| **A_55_P1991964** | Iqsec2 | 0.073398582 | -0.140166491 | 0.286963654 | 8.766222734 | 0.845525293 | 0.430931472 | 0.842517565 | -4.864568159 |
| **A_51_P102507** | Vps33a | 0.048906053 | -0.093415629 | 0.191227735 | 8.659128505 | 0.845397016 | 0.43099758 | 0.842517565 | -4.864638233 |
| **A_52_P179068** | Gucy1b1 | -0.138390726 | -0.541165255 | 0.264383804 | 9.110409492 | -0.845306389 | 0.431044289 | 0.842517565 | -4.864687736 |
| **A_51_P109939** | Naxe | 0.066816834 | -0.127651255 | 0.261284924 | 12.9522076 | 0.845291905 | 0.431051754 | 0.842517565 | -4.864695647 |
| **A_51_P477456** | Cnot10 | -0.052879381 | -0.206801837 | 0.101043074 | 9.049639933 | -0.845188431 | 0.43110509 | 0.842517565 | -4.864752162 |
| **A_52_P546660** | 1700029I15Rik | -0.052945421 | -0.207080982 | 0.101190139 | 8.581363865 | -0.84507397 | 0.431164094 | 0.842517565 | -4.864814672 |
| **A_55_P1978052** | Gatb | -0.037974082 | -0.148551148 | 0.072602983 | 7.963950621 | -0.844872922 | 0.43126775 | 0.842517565 | -4.864924456 |
| **A_55_P2004179** | Col2a1 | 0.069366842 | -0.13262547 | 0.271359154 | 7.763765503 | 0.844862915 | 0.431272909 | 0.842517565 | -4.864929919 |
| **A_55_P1978201** | Incenp | 0.111736951 | -0.213655828 | 0.437129731 | 9.956802347 | 0.8448081 | 0.431301174 | 0.842517565 | -4.864959848 |
| **A_55_P1976376** | Gimap4 | -0.036552002 | -0.143016841 | 0.069912836 | 7.886788783 | -0.844644847 | 0.431385361 | 0.842517565 | -4.865048974 |
| **A_52_P550843** | Desi2 | -0.060190923 | -0.235523233 | 0.115141386 | 10.79268039 | -0.844574985 | 0.431421391 | 0.842517565 | -4.865087111 |
| **A_55_P2113081** | Tmem132a | -0.143883221 | -0.563006639 | 0.275240198 | 11.60706925 | -0.844573374 | 0.431422222 | 0.842517565 | -4.865087991 |
| **A_55_P1967335** | NA | -0.096852823 | -0.379104048 | 0.185398402 | 11.21004015 | -0.844200512 | 0.431614559 | 0.842517565 | -4.865291495 |
| **A_51_P401964** | Aph1a | 0.083905785 | -0.160637047 | 0.328448617 | 11.29521942 | 0.844123772 | 0.431654152 | 0.842517565 | -4.865333371 |
| **A_52_P302433** | Plau | -0.113795298 | -0.445462661 | 0.217872066 | 8.903927407 | -0.844093861 | 0.431669585 | 0.842517565 | -4.865349692 |
| **A_55_P2126014** | NA | 0.073043423 | -0.139872381 | 0.285959228 | 16.15669486 | 0.843999874 | 0.431718082 | 0.842517565 | -4.865400975 |
| **A_55_P1978866** | NA | -0.305767233 | -1.197060901 | 0.585526435 | 11.80356123 | -0.843993856 | 0.431721187 | 0.842517565 | -4.865404258 |
| **A_55_P2364315** | D030002E05Rik | -0.087674398 | -0.343248577 | 0.16789978 | 8.408735149 | -0.843966067 | 0.431735527 | 0.842517565 | -4.86541942 |
| **A_51_P498403** | Cnpy3 | 0.038597741 | -0.073961068 | 0.151156551 | 9.75539382 | 0.843629129 | 0.431909423 | 0.842517565 | -4.865603227 |
| **A_55_P2110355** | Zfp637 | -0.078184239 | -0.306194662 | 0.149826183 | 9.211946297 | -0.843594384 | 0.431927358 | 0.842517565 | -4.865622178 |
| **A_52_P547662** | P2ry1 | -0.07934129 | -0.310730719 | 0.152048139 | 9.071089998 | -0.843577337 | 0.431936158 | 0.842517565 | -4.865631476 |

| | | | | | | | | | |
|---|---|---|---|---|---|---|---|---|---|
| **A_55_P1963533** | Pcdhac2 | 0.060283019 | -0.11555788 | 0.236123919 | 7.784767214 | 0.843420713 | 0.432017014 | 0.842517565 | -4.865716897 |
| **A_55_P2166384** | Cnot1 | -0.102611711 | -0.401976893 | 0.19675347 | 11.60083988 | -0.843266419 | 0.432096678 | 0.842517565 | -4.865801036 |
| **A_51_P101660** | Ncbp3 | -0.036264705 | -0.142068497 | 0.069539086 | 8.256548705 | -0.843241722 | 0.43210943 | 0.842517565 | -4.857767112 |
| **A_66_P129419** | Cox7a2l | 0.087487909 | -0.167762398 | 0.342738217 | 15.11875281 | 0.843239474 | 0.432110591 | 0.842517565 | -4.865815728 |
| **A_51_P446825** | Trmt9b | -0.050588258 | -0.198192815 | 0.097016299 | 8.567513769 | -0.843177717 | 0.43214248 | 0.842517565 | -4.865849401 |
| **A_55_P2109486** | Milr1 | 0.046401362 | -0.089001743 | 0.181804468 | 17.25156767 | 0.843084817 | 0.432190455 | 0.842517565 | -4.865900052 |
| **A_55_P1975475** | Mdm2 | -0.068345658 | -0.267797744 | 0.131106428 | 12.77741039 | -0.843027046 | 0.432220291 | 0.842517565 | -4.865931548 |
| **A_55_P1965725** | Gapdh | -0.07084376 | -0.277621075 | 0.135933555 | 11.38053894 | -0.842884124 | 0.432294108 | 0.842517565 | -4.866009461 |
| **A_52_P177490** | Cdx2 | 0.176068221 | -0.337849103 | 0.689985544 | 10.8750303 | 0.842862789 | 0.432305128 | 0.842517565 | -4.86602109 |
| **A_55_P1958255** | Rnf114 | -0.037073117 | -0.145285538 | 0.071139305 | 8.549681608 | -0.842851658 | 0.432310878 | 0.842517565 | -4.866027158 |
| **A_55_P2341110** | Chn1os3 | -0.144825076 | -0.567561909 | 0.277911756 | 8.301310199 | -0.842835539 | 0.432319204 | 0.842517565 | -4.866035944 |
| **A_51_P172251** | Ifrd2 | 0.061646522 | -0.1183215 | 0.241614543 | 10.07216871 | 0.842718248 | 0.432379793 | 0.842517565 | -4.866099875 |
| **A_55_P2100108** | Zfp148 | 0.065085503 | -0.124928811 | 0.255099817 | 8.496333797 | 0.842688589 | 0.432395115 | 0.842517565 | -4.866116039 |
| **A_55_P2134042** | NA | -0.057882806 | -0.226873819 | 0.111108207 | 16.92315352 | -0.842665335 | 0.432407128 | 0.842517565 | -4.866128713 |
| **A_52_P328078** | Atp5b | -0.043345944 | -0.169896545 | 0.083204657 | 15.87134856 | -0.842662016 | 0.432408843 | 0.842517565 | -4.866130522 |
| **A_55_P2000117** | Ceacam13 | 0.059784316 | -0.114761488 | 0.23433012 | 16.83251431 | 0.842649589 | 0.432415263 | 0.842517565 | -4.866137295 |
| **A_55_P1981994** | Krt17 | 0.157955185 | -0.303333986 | 0.619244356 | 8.07916493 | 0.842422176 | 0.432532763 | 0.842517565 | -4.866261223 |
| **A_55_P2122380** | Zfp14 | -0.042327424 | -0.165943653 | 0.081288806 | 8.29037771 | -0.842394468 | 0.432547081 | 0.842517565 | -4.866276321 |
| **A_55_P2117400** | Cacng7 | -0.06951901 | -0.272551282 | 0.133513263 | 8.464284792 | -0.84237926 | 0.432554939 | 0.842517565 | -4.866284608 |
| **A_55_P2062627** | Tmem238 | 0.038095534 | -0.073165097 | 0.149356165 | 7.762097073 | 0.842367706 | 0.43256091 | 0.842517565 | -4.866290903 |
| **A_55_P2087375** | Gm4987 | 0.082584263 | -0.158609788 | 0.323778314 | 15.73364899 | 0.842364136 | 0.432562755 | 0.842517565 | -4.866292849 |
| **A_51_P482051** | Cyp3a16 | -0.050613874 | -0.198478329 | 0.097250581 | 7.680729184 | -0.842121883 | 0.432687954 | 0.842517565 | -4.866424832 |
| **A_55_P2186558** | Tmem242 | 0.09505965 | -0.182712806 | 0.372832105 | 9.666627761 | 0.841930407 | 0.43278693 | 0.842517565 | -4.866529132 |
| **A_52_P175066** | Cd2ap | 0.08154391 | -0.156750845 | 0.319838666 | 12.2408072 | 0.841872282 | 0.432816979 | 0.842517565 | -4.866560791 |
| **A_55_P1960506** | Wdr83os | 0.06953665 | -0.133670504 | 0.272743804 | 11.33615067 | 0.841867867 | 0.432819262 | 0.842517565 | -4.866563195 |
| **A_55_P2078088** | Pqlc1 | 0.197734028 | -0.380154399 | 0.775622455 | 10.22726908 | 0.841795459 | 0.432856697 | 0.842517565 | -4.866602631 |
| **A_55_P2052897** | Arhgap30 | -0.042482125 | -0.166647224 | 0.081682973 | 7.861767601 | -0.841735927 | 0.432887476 | 0.842517565 | -4.866635052 |
| **A_55_P2197638** | 1110046J04Rik | 0.110000075 | -0.211527939 | 0.431528088 | 9.514161442 | 0.841672865 | 0.432920083 | 0.842517565 | -4.866669394 |
| **A_55_P2020361** | Lzts3 | 0.163519647 | -0.314448916 | 0.641488211 | 9.879827082 | 0.841665976 | 0.432923645 | 0.842517565 | -4.866673145 |
| **A_55_P2337138** | Ago2 | -0.143482296 | -0.562914736 | 0.275950143 | 11.19567389 | -0.841599493 | 0.432958023 | 0.842517565 | -4.866709348 |
| **A_55_P2154709** | Pter | -0.054579766 | -0.214230744 | 0.105071212 | 8.310317629 | -0.841064399 | 0.433234788 | 0.842517565 | -4.867000654 |
| **A_55_P2185652** | Cln3 | 0.038232767 | -0.073623335 | 0.150088869 | 9.870268624 | 0.840901674 | 0.43331898 | 0.842517565 | -4.867089216 |
| **A_55_P1981830** | NA | -0.191842288 | -0.753121088 | 0.369436511 | 9.618376962 | -0.840881642 | 0.433329345 | 0.842517565 | -4.867100118 |
| **A_55_P1957957** | Dhx34 | 0.058505788 | -0.112718915 | 0.229730491 | 9.449665878 | 0.840623597 | 0.433462881 | 0.842517565 | -4.86724053 |
| **A_51_P496804** | Adat2 | -0.063195095 | -0.248172579 | 0.121782388 | 9.004284839 | -0.840492166 | 0.433530907 | 0.842517565 | -4.867312035 |
| **A_55_P2203478** | Scn5a | -0.050290068 | -0.197496013 | 0.096915876 | 8.439130617 | -0.840477403 | 0.433538549 | 0.842517565 | -4.867320066 |
| **A_55_P1996963** | Mtcl1 | 0.118261026 | -0.227934687 | 0.46445674 | 10.2359056 | 0.840405955 | 0.433575532 | 0.842517565 | -4.867358934 |
| **A_51_P377508** | Srebf2 | -0.079227351 | -0.311198819 | 0.152744118 | 8.326281671 | -0.840252328 | 0.433655062 | 0.842517565 | -4.867442498 |
| **A_51_P331328** | Gpihbp1 | -0.047261515 | -0.185673008 | 0.091149978 | 10.13058495 | -0.840049059 | 0.433760308 | 0.842517565 | -4.867553049 |
| **A_55_P2066976** | Acsm1 | 0.064154679 | -0.123782291 | 0.252091648 | 8.255460598 | 0.83981819 | 0.433879866 | 0.842517565 | -4.867678588 |
| **A_51_P458778** | Hpgd | -0.55843438 | -2.194660489 | 1.077791729 | 10.47431191 | -0.839649421 | 0.433967281 | 0.842517565 | -4.867770343 |
| **A_55_P2028300** | Ppm1d | 0.061637592 | -0.118964033 | 0.242239217 | 10.04036832 | 0.839640099 | 0.433972109 | 0.842517565 | -4.867775411 |
| **A_51_P286321** | Trir | -0.05307266 | -0.208598738 | 0.102453418 | 12.6312246 | -0.839531118 | 0.434028564 | 0.842517565 | -4.867834654 |
| **A_51_P288479** | Slc25a29 | -0.288346628 | -1.133342193 | 0.556648937 | 9.206355978 | -0.839517224 | 0.434035762 | 0.842517565 | -4.867842206 |
| **A_52_P639551** | Tinf2 | -0.06518853 | -0.256237318 | 0.125860259 | 9.211975358 | -0.839452374 | 0.434069358 | 0.842517565 | -4.867877456 |
| **A_51_P350332** | Rbpms | -0.040297523 | -0.15840315 | 0.077808103 | 8.932427215 | -0.839415401 | 0.434088514 | 0.842517565 | -4.867897553 |
| **A_55_P2072228** | Cdk9 | 0.078959798 | -0.152482502 | 0.310402097 | 12.49855727 | 0.839329433 | 0.434133055 | 0.842517565 | -4.867944277 |
| **A_51_P475501** | Ndufa8 | -0.102457037 | -0.40277788 | 0.197863805 | 11.97462301 | -0.839315964 | 0.434140034 | 0.842517565 | -4.867951597 |
| **A_55_P2007733** | Epn1 | -0.065531174 | -0.257631846 | 0.126569499 | 16.18019421 | -0.839243962 | 0.434177343 | 0.842517565 | -4.867990728 |
| **A_51_P141554** | Med22 | 0.092934464 | -0.179506929 | 0.365375857 | 10.82563137 | 0.839214288 | 0.434192719 | 0.842517565 | -4.868006854 |

| | | | | | | | | | |
|---|---|---|---|---|---|---|---|---|---|
| **A_55_P2161695** | Poglut2 | -0.127329469 | -0.500607785 | 0.245948847 | 9.661675226 | -0.83919972 | 0.434200268 | 0.842517565 | -4.868014771 |
| **A_55_P1955039** | Myo1b | -0.092212477 | -0.362606437 | 0.178181482 | 9.208363319 | -0.838999818 | 0.434303866 | 0.842517565 | -4.868123395 |
| **A_51_P377403** | Mettl14 | 0.036786809 | -0.071085565 | 0.144659184 | 8.389180554 | 0.83897891 | 0.434314702 | 0.842517565 | -4.868134755 |
| **A_55_P1994224** | Ythdf1 | 0.068590061 | -0.132571745 | 0.269751868 | 12.29449026 | 0.838851 | 0.434381001 | 0.842517565 | -4.868204249 |
| **A_55_P1956557** | Rab3a | 0.063553128 | -0.122840353 | 0.249946608 | 9.105906699 | 0.838832743 | 0.434390465 | 0.842517565 | -4.868214167 |
| **A_52_P378914** | AU040320 | -0.056463282 | -0.22206747 | 0.109140905 | 8.354527077 | -0.838810711 | 0.434401886 | 0.842517565 | -4.868226136 |
| **A_51_P356525** | Trappc11 | -0.061038595 | -0.240065464 | 0.117988274 | 9.355955399 | -0.83879431 | 0.434410387 | 0.842517565 | -4.868235046 |
| **A_51_P167360** | Ptpn7 | 0.036900799 | -0.071331144 | 0.145132742 | 7.715434764 | 0.838782721 | 0.434416395 | 0.842517565 | -4.868241342 |
| **A_55_P1982454** | Eps8 | 0.06283734 | -0.121519558 | 0.247194238 | 10.52785116 | 0.838547292 | 0.434538451 | 0.842517565 | -4.868369224 |
| **A_66_P128997** | Pml | -0.046212454 | -0.181794853 | 0.089369946 | 8.526741855 | -0.838542105 | 0.43454114 | 0.842517565 | -4.868372041 |
| **A_51_P244531** | NA | 0.087455032 | -0.169144628 | 0.344054693 | 9.446744481 | 0.838490005 | 0.434568154 | 0.842517565 | -4.868400337 |
| **A_55_P2067767** | Psg25 | -0.162240785 | -0.638334936 | 0.313853367 | 15.22596299 | -0.838371208 | 0.434629756 | 0.842517565 | -4.868464854 |
| **A_65_P06453** | Nup35 | 0.080660533 | -0.15609803 | 0.317419095 | 10.28651202 | 0.838155416 | 0.43474167 | 0.842517565 | -4.868582029 |
| **A_52_P817257** | Smim22 | 0.464801735 | -0.899578241 | 1.829181711 | 10.49030503 | 0.83811105 | 0.434764682 | 0.842517565 | -4.868606118 |
| **A_51_P246339** | Rfc5 | 0.093945936 | -0.181847182 | 0.369739053 | 9.976563241 | 0.838038044 | 0.434802551 | 0.842517565 | -4.868645754 |
| **A_52_P152219** | Rpgrip1l | -0.043448796 | -0.171004621 | 0.084107029 | 8.526345031 | -0.838005004 | 0.43481969 | 0.842517565 | -4.868663691 |
| **A_51_P520639** | Mslnl | -0.064308283 | -0.253105681 | 0.124489116 | 7.700476229 | -0.83799238 | 0.434826239 | 0.842517565 | -4.868670545 |
| **A_55_P1964638** | Cxadr | 0.225581477 | -0.436723259 | 0.887886214 | 10.20655637 | 0.837943505 | 0.434851593 | 0.842517565 | -4.868697077 |
| **A_55_P2062437** | Begain | 0.045549667 | -0.088187549 | 0.179286883 | 7.760472688 | 0.837919105 | 0.434864251 | 0.842517565 | -4.868710322 |
| **A_51_P413916** | Pgrmc2 | -0.060820099 | -0.239397899 | 0.117757701 | 9.412835059 | -0.837893493 | 0.434877538 | 0.842517565 | -4.868724226 |
| **A_66_P138765** | Trappc10 | -0.061211594 | -0.240984773 | 0.118561585 | 10.36814653 | -0.837679631 | 0.434988498 | 0.842517565 | -4.868840307 |
| **A_52_P313071** | Gosr1 | -0.083892249 | -0.330297646 | 0.162513147 | 10.54438715 | -0.837607945 | 0.435025696 | 0.842517565 | -4.868879213 |
| **A_52_P271** | Dnajc18 | -0.083339031 | -0.328143777 | 0.161465716 | 9.466154576 | -0.83752499 | 0.435068744 | 0.842517565 | -4.868924231 |
| **A_52_P86073** | Mrpl1 | 0.094184143 | -0.18250938 | 0.370877666 | 9.098079028 | 0.837428931 | 0.435118597 | 0.842517565 | -4.868976357 |
| **A_55_P1952915** | Sh3bgr | 0.086422114 | -0.167468496 | 0.340312724 | 8.538215184 | 0.837427869 | 0.435119148 | 0.842517565 | -4.868976933 |
| **A_55_P2137203** | B4galt3 | -0.120801066 | -0.475720921 | 0.23411879 | 12.07137708 | -0.837355027 | 0.435156954 | 0.842517565 | -4.869016458 |
| **A_55_P2063626** | Matk | -0.083417941 | -0.328518134 | 0.161682251 | 10.0478239 | -0.837307497 | 0.435181625 | 0.842517565 | -4.869042247 |
| **A_55_P2432388** | Prr14l | -0.09977234 | -0.392949164 | 0.193404484 | 10.03578572 | -0.837239452 | 0.435216945 | 0.842517565 | -4.869079164 |
| **A_51_P461452** | Ube2r2 | -0.060029782 | -0.236428747 | 0.116369184 | 13.64914407 | -0.837220572 | 0.435226745 | 0.842517565 | -4.869089407 |
| **A_55_P2158483** | Nkiras2 | 0.039582247 | -0.076733411 | 0.155897905 | 8.265681608 | 0.837204254 | 0.435235216 | 0.842517565 | -4.869098261 |
| **A_51_P366061** | Fscn1 | -0.051702961 | -0.203637688 | 0.100231766 | 9.412519126 | -0.837196742 | 0.435239115 | 0.842517565 | -4.869102336 |
| **A_51_P242414** | Micall1 | 0.160902879 | -0.312029315 | 0.633835072 | 11.79321423 | 0.837016653 | 0.435332608 | 0.842517565 | -4.86920003 |
| **A_55_P2183808** | NA | 0.128664682 | -0.249559949 | 0.506889314 | 14.4469503 | 0.836909904 | 0.435388034 | 0.842517565 | -4.869257932 |
| **A_55_P1959137** | Usp32 | -0.058564601 | -0.230723566 | 0.113594363 | 11.92806684 | -0.836902208 | 0.43539203 | 0.842517565 | -4.869262105 |
| **A_55_P2110512** | Lmo7 | -0.042939232 | -0.169216629 | 0.083338166 | 9.072139975 | -0.83656137 | 0.435569033 | 0.842517565 | -4.869446943 |
| **A_55_P2072980** | Sptan1 | -0.06551997 | -0.258232741 | 0.127192801 | 8.268704752 | -0.836435307 | 0.435634513 | 0.842517565 | -4.869515294 |
| **A_52_P479500** | Npepps | -0.072333586 | -0.285090795 | 0.140423623 | 11.01612824 | -0.836420902 | 0.435641996 | 0.842517565 | -4.869523104 |
| **A_66_P116252** | Dusp16 | -0.092908933 | -0.366220513 | 0.180402646 | 11.84098048 | -0.836312531 | 0.435698293 | 0.842517565 | -4.869581856 |
| **A_55_P2069935** | Fxyd2 | 0.353953855 | -0.687288303 | 1.395196013 | 11.97745928 | 0.836303792 | 0.435702833 | 0.842517565 | -4.869586593 |
| **A_51_P311362** | 4930449E01Rik | 0.037477574 | -0.072774085 | 0.147729232 | 7.764540948 | 0.836287292 | 0.435711405 | 0.842517565 | -4.869595538 |
| **A_55_P2012759** | Bicdl1 | -0.102524707 | -0.404186659 | 0.199137244 | 12.62872282 | -0.836136468 | 0.435789766 | 0.842517565 | -4.869677295 |
| **A_51_P310164** | 2810459M11Rik | 0.068586155 | -0.13322438 | 0.270396691 | 7.851746188 | 0.836106862 | 0.435805149 | 0.842517565 | -4.869693342 |
| **A_52_P681** | Cnot4 | 0.064667304 | -0.125642356 | 0.254976964 | 8.946046528 | 0.835974608 | 0.435873873 | 0.842517565 | -4.869765022 |
| **A_51_P456465** | Cldn10 | 0.088553311 | -0.1720555 | 0.349162122 | 7.925463827 | 0.835958799 | 0.435882088 | 0.842517565 | -4.86977359 |
| **A_55_P2087622** | Il4i1 | 0.169385133 | -0.329139066 | 0.667909333 | 8.860322027 | 0.835907474 | 0.435908761 | 0.842517565 | -4.869801405 |
| **A_51_P342271** | Setd6 | -0.110772538 | -0.436833038 | 0.215287962 | 10.32150897 | -0.835801372 | 0.435963903 | 0.842517565 | -4.869858902 |
| **A_55_P2061110** | Snapin | -0.063924754 | -0.252088718 | 0.124239209 | 9.031029059 | -0.835798866 | 0.435965206 | 0.842517565 | -4.86986026 |
| **A_55_P2037897** | Brd3 | -0.080818162 | -0.318711501 | 0.157075178 | 11.71597338 | -0.835787454 | 0.435971137 | 0.842517565 | -4.869866443 |
| **A_51_P451346** | Klf6 | -0.160864707 | -0.634395812 | 0.312666398 | 14.4783173 | -0.835759694 | 0.435985566 | 0.842517565 | -4.869881486 |
| **A_52_P114282** | Armcx5 | 0.052704608 | -0.102466802 | 0.207876019 | 8.800419481 | 0.83561465 | 0.436060958 | 0.842517565 | -4.869960074 |

| | | | | | | | | | |
|---|---|---|---|---|---|---|---|---|---|
| **A_55_P2150343** | Piezo1 | 0.087681117 | -0.170478664 | 0.345840897 | 12.38640685 | 0.835577337 | 0.436080355 | 0.842517565 | -4.86998029 |
| **A_55_P1997948** | Ppp4r3b | -0.084755611 | -0.334306139 | 0.164794918 | 10.81524041 | -0.835562858 | 0.436087882 | 0.842517565 | -4.869988134 |
| **A_55_P2103145** | Ppp1r2 | 0.088956068 | -0.172978485 | 0.35089062 | 9.075315665 | 0.835510569 | 0.436115065 | 0.842517565 | -4.870016462 |
| **A_51_P425373** | Smg9 | 0.045641673 | -0.088767983 | 0.180051329 | 9.02445092 | 0.835411114 | 0.436166771 | 0.842517565 | -4.870070339 |
| **A_55_P2175341** | NA | 0.094193132 | -0.183219031 | 0.371605295 | 9.74752686 | 0.83533928 | 0.43620412 | 0.842517565 | -4.87010925 |
| **A_55_P1994882** | Ccdc174 | -0.061688266 | -0.243373184 | 0.119996651 | 10.78840923 | -0.835319951 | 0.43621417 | 0.842517565 | -4.87011972 |
| **A_52_P445474** | Lmtk2 | -0.087514789 | -0.345267636 | 0.170238058 | 9.735195115 | -0.835308965 | 0.436219883 | 0.842517565 | -4.870125671 |
| **A_55_P2179796** | NA | 0.173447757 | -0.337407439 | 0.684302952 | 13.66744761 | 0.835295295 | 0.436226991 | 0.842517565 | -4.870133075 |
| **A_52_P96847** | Smlr1 | 0.393786333 | -0.766078483 | 1.553651149 | 9.666383484 | 0.835261431 | 0.436244599 | 0.842517565 | -4.870151417 |
| **A_51_P230287** | Sorcs2 | 0.0401191 | -0.078069687 | 0.158307887 | 8.050045638 | 0.835110744 | 0.436322959 | 0.842517565 | -4.870233029 |
| **A_52_P584078** | Pop1 | 0.074453621 | -0.144890025 | 0.293797267 | 9.665327153 | 0.835083552 | 0.436337101 | 0.842517565 | -4.870247755 |
| **A_65_P12993** | Kif1b | -0.04261251 | -0.168170785 | 0.082945765 | 8.366067369 | -0.834950906 | 0.436406089 | 0.842517565 | -4.870319585 |
| **A_55_P2245422** | NA | -0.15361902 | -0.606266666 | 0.299028625 | 8.613842502 | -0.834937349 | 0.436413141 | 0.842517565 | -4.870326926 |
| **A_52_P18807** | Eif3c | -0.128597445 | -0.507546958 | 0.250352067 | 13.52857419 | -0.834872492 | 0.436446876 | 0.842517565 | -4.870362044 |
| **A_55_P2112693** | Sipa1l1 | 0.047135634 | -0.091763897 | 0.186035165 | 8.807960771 | 0.834867846 | 0.436449293 | 0.842517565 | -4.870364559 |
| **A_52_P260555** | Vil1 | 0.310305559 | -0.604342121 | 1.224953238 | 9.052870181 | 0.83465093 | 0.436562135 | 0.84265124 | -4.870481998 |
| **A_55_P2077866** | Nktr | 0.113036144 | -0.220220328 | 0.446292616 | 11.29216973 | 0.834464574 | 0.436659097 | 0.842754238 | -4.870582873 |
| **A_66_P118004** | Rab3ip | 0.092177096 | -0.179612804 | 0.363966996 | 12.03488049 | 0.834370363 | 0.436708122 | 0.842764705 | -4.870633865 |
| **A_51_P174645** | Asl | 0.090706015 | -0.176788139 | 0.358200168 | 9.313444857 | 0.834239893 | 0.436776022 | 0.842791303 | -4.870704474 |
| **A_55_P1953003** | Atp6v0a4 | 0.315587865 | -0.615159043 | 1.246334774 | 9.41585879 | 0.834176319 | 0.436809109 | 0.842791303 | -4.870738877 |
| **A_51_P392687** | Vim | 0.263118399 | -0.513009507 | 1.039246304 | 13.48280489 | 0.834040388 | 0.436879863 | 0.842825355 | -4.87081243 |
| **A_55_P1962305** | Plac8 | 0.131135678 | -0.255745681 | 0.518017037 | 15.51062129 | 0.833896641 | 0.436954694 | 0.842825355 | -4.870890202 |
| **A_65_P05316** | Zfp182 | -0.080057688 | -0.316248424 | 0.156133048 | 8.922715169 | -0.833891115 | 0.436957571 | 0.842825355 | -4.870893192 |
| **A_51_P483329** | Sdad1 | 0.050003678 | -0.097534833 | 0.197542188 | 10.86919181 | 0.833807345 | 0.437001184 | 0.842825372 | -4.87093851 |
| **A_55_P2060680** | Acsf2 | -0.145052149 | -0.573192983 | 0.283088685 | 11.52362412 | -0.833502064 | 0.437160148 | 0.842950902 | -4.871103634 |
| **A_55_P1970636** | Unc45a | -0.090617282 | -0.358096482 | 0.176861918 | 12.40117176 | -0.833470397 | 0.43717664 | 0.842950902 | -4.871120761 |
| **A_51_P291749** | Pecr | 0.218313897 | -0.42620035 | 0.862828143 | 9.94395144 | 0.833331931 | 0.437248757 | 0.842950902 | -4.87119564 |
| **A_55_P1963547** | Prdm16 | 0.038790371 | -0.075751837 | 0.15333258 | 7.938008826 | 0.833158338 | 0.437339182 | 0.842950902 | -4.871289502 |
| **A_55_P2130576** | NA | -0.069941827 | -0.276478535 | 0.136594882 | 13.81941656 | -0.833122536 | 0.437357833 | 0.842950902 | -4.871308859 |
| **A_55_P2001290** | Ube2e3 | -0.071140961 | -0.281266178 | 0.138984256 | 12.02742306 | -0.832934241 | 0.437455935 | 0.842950902 | -4.871410652 |
| **A_55_P1985764** | NA | -0.075768629 | -0.299573888 | 0.14803663 | 13.38380301 | -0.832891259 | 0.437478331 | 0.842950902 | -4.871433886 |
| **A_51_P157270** | Plekhd1 | 0.302784118 | -0.591668115 | 1.19723635 | 9.467094112 | 0.832808409 | 0.437521503 | 0.842950902 | -4.871478669 |
| **A_51_P149623** | Gabrd | 0.088484065 | -0.172918893 | 0.349887022 | 8.987605186 | 0.832767429 | 0.437542858 | 0.842950902 | -4.871500818 |
| **A_66_P117730** | Hapln1 | 0.039300219 | -0.07680326 | 0.155403697 | 7.721204578 | 0.83275816 | 0.437547688 | 0.842950902 | -4.871505828 |
| **A_51_P377948** | Gimap9 | -0.039510058 | -0.156250095 | 0.077229979 | 7.87565807 | -0.832639485 | 0.437609536 | 0.842950902 | -4.871569967 |
| **A_51_P392705** | Xpc | -0.068876116 | -0.272404086 | 0.134651854 | 9.668423272 | -0.832556496 | 0.437652789 | 0.842950902 | -4.871614814 |
| **A_52_P615952** | Qk | -0.069962772 | -0.276703071 | 0.136777527 | 9.725374105 | -0.832551358 | 0.437655467 | 0.842950902 | -4.87161759 |
| **A_55_P2157307** | NA | -0.097684632 | -0.386364188 | 0.190994924 | 10.10663349 | -0.832490679 | 0.437687095 | 0.842950902 | -4.87165038 |
| **A_51_P431903** | Pdcd6 | 0.082813457 | -0.161937311 | 0.327564225 | 13.53549354 | 0.832426726 | 0.437720431 | 0.842950902 | -4.871684936 |
| **A_52_P309337** | Metap2 | 0.049333821 | -0.096533851 | 0.195201492 | 8.099122564 | 0.832060416 | 0.437911409 | 0.843047856 | -4.871882834 |
| **A_65_P15518** | NA | -0.170779305 | -0.67574716 | 0.334188549 | 8.418123637 | -0.832033234 | 0.437925583 | 0.843047856 | -4.871897516 |
| **A_51_P348325** | Poc1b | -0.055587415 | -0.219955812 | 0.108780981 | 8.960925355 | -0.832007659 | 0.437938919 | 0.843047856 | -4.87191133 |
| **A_55_P2151061** | NA | -0.071523364 | -0.283031569 | 0.139984841 | 8.071330996 | -0.831935915 | 0.437976332 | 0.843047856 | -4.87195008 |
| **A_52_P566605** | Hsd17b7 | 0.159234268 | -0.311665374 | 0.63013391 | 9.298298838 | 0.831911903 | 0.437988855 | 0.843047856 | -4.87196305 |
| **A_51_P351194** | Cnfn | -0.197002621 | -0.779804458 | 0.385799217 | 8.209228541 | -0.831611055 | 0.43814577 | 0.843257099 | -4.872125514 |
| **A_51_P170758** | Grwd1 | -0.049433779 | -0.195712061 | 0.096844502 | 9.156518108 | -0.831405951 | 0.438252771 | 0.843257099 | -4.872236252 |
| **A_52_P244964** | Akap11 | -0.060125625 | -0.238067456 | 0.117816207 | 9.272103049 | -0.831286462 | 0.438315116 | 0.843257099 | -4.872300756 |
| **A_52_P413584** | NA | -0.050489538 | -0.199925496 | 0.09894642 | 9.068713137 | -0.831218973 | 0.438350332 | 0.843257099 | -4.872337185 |
| **A_55_P1997696** | Plbd1 | -0.167639512 | -0.682649738 | 0.347370715 | 14.53642152 | -0.843549585 | 0.438384884 | 0.843257099 | -4.81426225 |
| **A_52_P678816** | Ctsm | -0.048819268 | -0.193331223 | 0.095692687 | 16.38960726 | -0.831106447 | 0.438409053 | 0.843257099 | -4.872397921 |

| | | | | | | | | |
|---|---|---|---|---|---|---|---|---|
| **A_55_P2117681** | Rnf115 | -0.099674073 | -0.394731877 | 0.19538373 | 10.55902748 | -0.831082743 | 0.438421424 | 0.843257099 | -4.872410714 |
| **A_55_P2079272** | Kars | -0.074893217 | -0.296621201 | 0.146834767 | 11.75303459 | -0.830981091 | 0.438474476 | 0.843257099 | -4.872465574 |
| **A_51_P425798** | Mtus2 | -0.054604365 | -0.216271382 | 0.107062653 | 8.233709864 | -0.830950389 | 0.438490501 | 0.843257099 | -4.872482143 |
| **A_55_P1972915** | NA | 0.115477897 | -0.226544838 | 0.457500632 | 15.4173831 | 0.830640404 | 0.438652318 | 0.843257099 | -4.872649404 |
| **A_51_P433733** | Nucb1 | -0.086291519 | -0.341881398 | 0.16929836 | 13.00229694 | -0.830603249 | 0.438671716 | 0.843257099 | -4.872669449 |
| **A_55_P2008258** | Larp1b | -0.089709849 | -0.355431795 | 0.176012098 | 10.15418893 | -0.830580751 | 0.438683463 | 0.843257099 | -4.872681587 |
| **A_55_P2153382** | Ermap | -0.03524868 | -0.139663298 | 0.069165939 | 7.694697445 | -0.830521177 | 0.438714568 | 0.843257099 | -4.872713725 |
| **A_55_P1997991** | Ecpas | -0.067098864 | -0.265877717 | 0.131679989 | 8.257586853 | -0.830451268 | 0.438751071 | 0.843257099 | -4.872751436 |
| **A_51_P146753** | Csf2rb2 | -0.360868927 | -1.429937072 | 0.708199219 | 9.373593224 | -0.830449564 | 0.438751961 | 0.843257099 | -4.872752355 |
| **A_51_P380514** | Tjap1 | -0.053643368 | -0.212625403 | 0.105338666 | 9.069458621 | -0.830112877 | 0.438927795 | 0.843269378 | -4.872933945 |
| **A_66_P102163** | NA | 0.069872393 | -0.137227952 | 0.276972738 | 9.333403357 | 0.830030321 | 0.438970918 | 0.843269378 | -4.872978462 |
| **A_55_P2023523** | Aloxe3 | 0.098777569 | -0.194012414 | 0.391567552 | 9.295425885 | 0.829986981 | 0.438993557 | 0.843269378 | -4.873001832 |
| **A_52_P650379** | Strap | 0.042060747 | -0.082614754 | 0.166736248 | 10.14709517 | 0.829975021 | 0.438999805 | 0.843269378 | -4.873008281 |
| **A_55_P2087984** | Dnaja1 | -0.100686759 | -0.399174592 | 0.197801073 | 15.11022614 | -0.829879228 | 0.439049848 | 0.843269378 | -4.87305993 |
| **A_55_P2074085** | Gimap8 | 0.039791582 | -0.078183297 | 0.157766461 | 8.435560435 | 0.829795037 | 0.439093834 | 0.843269378 | -4.873105321 |
| **A_51_P427232** | Zbed5 | -0.093197028 | -0.369512168 | 0.183118111 | 9.304336405 | -0.829786847 | 0.439098112 | 0.843269378 | -4.873109736 |
| **A_55_P2132403** | Szt2 | 0.061336996 | -0.120521833 | 0.243195825 | 9.643766346 | 0.829769135 | 0.439107367 | 0.843269378 | -4.873119284 |
| **A_55_P2154307** | Peak1 | 0.181794558 | -0.357454862 | 0.721043977 | 11.36024389 | 0.829392978 | 0.439303933 | 0.843410706 | -4.87332204 |
| **A_55_P2061391** | Rhobtb3 | 0.041924678 | -0.082450663 | 0.166300019 | 7.792710631 | 0.829286537 | 0.439359568 | 0.843410706 | -4.873379402 |
| **A_52_P682382** | Scd1 | -0.048930894 | -0.194091904 | 0.096230115 | 8.671942838 | -0.829282187 | 0.439361842 | 0.843410706 | -4.873381746 |
| **A_55_P2018949** | Fbxo7 | 0.104250618 | -0.205028367 | 0.413529603 | 12.90471418 | 0.829272741 | 0.439366779 | 0.843410706 | -4.873386836 |
| **A_52_P674165** | Rsbn1l | -0.035569916 | -0.141108449 | 0.069968617 | 7.913467242 | -0.829164984 | 0.439423107 | 0.843410706 | -4.873444901 |
| **A_55_P1973447** | NA | 0.035430588 | -0.069706418 | 0.140567593 | 7.80244419 | 0.829071362 | 0.43947205 | 0.843410706 | -4.873495345 |
| **A_55_P1989733** | Nfkbib | 0.064726072 | -0.127348945 | 0.256801088 | 11.4605702 | 0.829043914 | 0.4394864 | 0.843410706 | -4.873510133 |
| **A_55_P1985360** | Man1c1 | -0.139715924 | -0.554440291 | 0.275008444 | 9.955358825 | -0.828811018 | 0.439608172 | 0.843560644 | -4.873635598 |
| **A_51_P247799** | Casp8 | -0.052050316 | -0.206615299 | 0.102514666 | 8.67933772 | -0.828478852 | 0.439781893 | 0.843793941 | -4.873814498 |
| **A_55_P2183597** | Tbc1d2 | -0.09465246 | -0.375774729 | 0.186469809 | 11.77912505 | -0.828334633 | 0.439857333 | 0.843793941 | -4.873892156 |
| **A_55_P2412319** | A830052D11Rik | 0.060070647 | -0.118343399 | 0.238484693 | 8.457236785 | 0.82832817 | 0.439860714 | 0.843793941 | -4.873895636 |
| **A_51_P277795** | Resf1 | -0.041492497 | -0.164752129 | 0.081767135 | 10.67977718 | -0.828166894 | 0.439945089 | 0.843872048 | -4.873982467 |
| **A_51_P139320** | Pcbd1 | 0.34430264 | -0.678743737 | 1.367349017 | 11.56874212 | 0.827969225 | 0.44004852 | 0.843986689 | -4.874088877 |
| **A_55_P2140711** | Tufm | -0.085571352 | -0.339870775 | 0.168728072 | 10.4816489 | -0.827851003 | 0.440110389 | 0.8440216 | -4.874152509 |
| **A_55_P2031676** | Gstm6 | -0.035087631 | -0.139403143 | 0.069227881 | 7.643804181 | -0.827512031 | 0.440287816 | 0.844155997 | -4.874334923 |
| **A_55_P2022297** | Ptrh2 | -0.038176603 | -0.151681363 | 0.075328157 | 8.256732903 | -0.827470215 | 0.440309708 | 0.844155997 | -4.874357423 |
| **A_55_P2050483** | Abcc12 | -0.085683745 | -0.340453744 | 0.169086254 | 8.979629815 | -0.827407245 | 0.440342675 | 0.844155997 | -4.874391302 |
| **A_55_P1963737** | Aktip | 0.074238019 | -0.146505894 | 0.294981932 | 8.965359099 | 0.827383396 | 0.440355161 | 0.844155997 | -4.874404133 |
| **A_51_P383194** | Pde9a | -0.094136093 | -0.374103843 | 0.185831657 | 9.119933724 | -0.82721295 | 0.440444407 | 0.844171765 | -4.874495827 |
| **A_66_P112164** | Rab29 | 0.142053403 | -0.280437046 | 0.564543852 | 10.87377767 | 0.827187397 | 0.440457788 | 0.844171765 | -4.874509572 |
| **A_55_P2137887** | Mpdz | -0.041580025 | -0.165260974 | 0.082100925 | 7.894960911 | -0.827086815 | 0.44051046 | 0.844171765 | -4.874563674 |
| **A_51_P248644** | Tdp1 | -0.052804879 | -0.209884489 | 0.104274731 | 10.10244966 | -0.827034072 | 0.440538082 | 0.844171765 | -4.874592042 |
| **A_51_P171616** | Wnt10a | 0.288672295 | -0.570210761 | 1.14755535 | 8.659647852 | 0.826875707 | 0.440621028 | 0.844205845 | -4.874677212 |
| **A_51_P454666** | Arl6ip4 | 0.046989746 | -0.092826178 | 0.18680567 | 9.580578702 | 0.826828847 | 0.440645573 | 0.844205845 | -4.87470241 |
| **A_66_P124420** | NA | -0.068563312 | -0.272589902 | 0.135463277 | 8.450623457 | -0.826749965 | 0.440686894 | 0.844205845 | -4.874744828 |
| **A_51_P513785** | Ctcf | -0.061431173 | -0.244253236 | 0.12139089 | 10.78830214 | -0.82666465 | 0.440731588 | 0.844207795 | -4.8747907 |
| **A_51_P443872** | Slu7 | -0.080473234 | -0.32004991 | 0.159103442 | 9.458801563 | -0.826372935 | 0.440884434 | 0.844416887 | -4.874947527 |
| **A_51_P497402** | Tonsl | -0.071638544 | -0.284980868 | 0.141703779 | 8.527895441 | -0.826111934 | 0.44102122 | 0.844595181 | -4.875087808 |
| **A_55_P2112934** | C030034I22Rik | -0.074758908 | -0.29750096 | 0.147983143 | 8.194850154 | -0.825714479 | 0.441229579 | 0.844860718 | -4.87530137 |
| **A_55_P2014997** | Amz2 | 0.064239462 | -0.127205443 | 0.255684367 | 10.47949669 | 0.825519326 | 0.44133191 | 0.844860718 | -4.875406202 |
| **A_51_P404377** | Rnd2 | -0.086504329 | -0.344320016 | 0.171311357 | 9.200986317 | -0.82546311 | 0.441361391 | 0.844860718 | -4.875436397 |
| **A_55_P1975832** | 1810009N02Rik | 0.081519658 | -0.161441072 | 0.324480389 | 10.04309338 | 0.825458854 | 0.441363623 | 0.844860718 | -4.875438683 |
| **A_51_P163252** | Schip1 | -0.073875583 | -0.294063157 | 0.146311991 | 8.035193724 | -0.825424348 | 0.441381719 | 0.844860718 | -4.875457216 |

| | | | | | | | | | |
|---|---|---|---|---|---|---|---|---|---|
| **A_51_P232399** | Acy3 | 0.142881637 | -0.283019548 | 0.568782821 | 8.450220651 | 0.825347294 | 0.441422132 | 0.844860718 | -4.875498601 |
| **A_55_P1967105** | Adgrb2 | -0.048811954 | -0.194326311 | 0.096702402 | 8.020936456 | -0.82525757 | 0.441469193 | 0.844867132 | -4.875546786 |
| **A_52_P215106** | Pik3r6 | 0.116587643 | -0.231051708 | 0.464226994 | 9.487673655 | 0.82507372 | 0.441565635 | 0.844922971 | -4.875645508 |
| **A_55_P2237432** | Smarca1 | 0.132583814 | -0.262804269 | 0.527971897 | 8.456864035 | 0.824966087 | 0.441622103 | 0.844922971 | -4.875703297 |
| **A_55_P2056995** | Gcgr | -0.107199943 | -0.426894362 | 0.212494475 | 12.70350369 | -0.824951973 | 0.441629509 | 0.844922971 | -4.875710875 |
| **A_55_P2161958** | Gm10324 | 0.122107947 | -0.242095111 | 0.486311006 | 11.03345276 | 0.824839604 | 0.441688468 | 0.844952138 | -4.8757712 |
| **A_55_P2125551** | Mmadhc | -0.082440296 | -0.328397414 | 0.163516822 | 12.83612742 | -0.824611356 | 0.441808246 | 0.845061246 | -4.875893717 |
| **A_55_P2039566** | Arhgap23 | 0.068908851 | -0.136689509 | 0.274507212 | 12.90705131 | 0.824564298 | 0.441832943 | 0.845061246 | -4.875918973 |
| **A_55_P2174339** | NA | 0.036726161 | -0.072883466 | 0.146335789 | 8.009443553 | 0.82432031 | 0.441961013 | 0.845213058 | -4.876049907 |
| **A_51_P292073** | Haghl | -0.107838686 | -0.429737259 | 0.214059888 | 9.639444532 | -0.824184982 | 0.442032058 | 0.845213058 | -4.876122517 |
| **A_55_P1972192** | Phka1 | -0.040804007 | -0.162635035 | 0.081027022 | 9.368315062 | -0.823975049 | 0.442142286 | 0.845213058 | -4.87623514 |
| **A_52_P290783** | BC016579 | 0.10414594 | -0.206845542 | 0.415137421 | 9.027717528 | 0.823878202 | 0.442193143 | 0.845213058 | -4.876287088 |
| **A_51_P476509** | Zbtb38 | -0.069634438 | -0.277626072 | 0.138357195 | 8.361972603 | -0.823658861 | 0.442308342 | 0.845213058 | -4.876404726 |
| **A_55_P1975510** | H6pd | 0.091088083 | -0.180989917 | 0.363166082 | 12.32270576 | 0.823639749 | 0.442318381 | 0.845213058 | -4.876414975 |
| **A_52_P27271** | Dhdds | -0.053978458 | -0.215231652 | 0.107274735 | 9.503142298 | -0.82353359 | 0.442374145 | 0.845213058 | -4.876471902 |
| **A_51_P409010** | Comp | 0.189165804 | -0.376047915 | 0.754379523 | 8.202280036 | 0.823377697 | 0.442456043 | 0.845213058 | -4.876555489 |
| **A_51_P189269** | Mbnl2 | -0.097230563 | -0.387790842 | 0.193329717 | 10.20889124 | -0.823257542 | 0.442519174 | 0.845213058 | -4.876619906 |
| **A_55_P2026748** | Zmat1 | 0.06012804 | -0.119568626 | 0.239824705 | 8.749570004 | 0.823201567 | 0.442548586 | 0.845213058 | -4.876649913 |
| **A_51_P286865** | Wdr33 | -0.059233895 | -0.236292954 | 0.117825165 | 10.07305643 | -0.823040678 | 0.442633132 | 0.845213058 | -4.876736153 |
| **A_55_P2117843** | Coasy | 0.043170249 | -0.085872347 | 0.172212844 | 11.69029692 | 0.82303939 | 0.442633809 | 0.845213058 | -4.876736843 |
| **A_55_P2098995** | Cfhr2 | 0.080122889 | -0.159408717 | 0.319654495 | 9.913871173 | 0.822930084 | 0.442691256 | 0.845213058 | -4.876795427 |
| **A_55_P2124096** | NA | -0.063562525 | -0.253590406 | 0.126465357 | 13.33497673 | -0.822911205 | 0.442701179 | 0.845213058 | -4.876805545 |
| **A_52_P359257** | NA | -0.060752515 | -0.242397274 | 0.120892243 | 8.235687097 | -0.822830833 | 0.442743423 | 0.845213058 | -4.876848617 |
| **A_55_P2147310** | Mrtfa | 0.052839539 | -0.105157424 | 0.210836502 | 11.758362 | 0.82277189 | 0.442774407 | 0.845213058 | -4.876880203 |
| **A_55_P1955861** | Cux2 | -0.036683378 | -0.146388331 | 0.073021576 | 7.697707477 | -0.82264458 | 0.442841332 | 0.845213058 | -4.87694842 |
| **A_55_P2192729** | 5830469G19Rik | -0.275721059 | -1.100407149 | 0.548965031 | 9.654194483 | -0.822527509 | 0.442902881 | 0.845213058 | -4.877011144 |
| **A_55_P2248651** | Aak1 | -0.062535631 | -0.249581621 | 0.12451036 | 10.32036226 | -0.822523457 | 0.442905011 | 0.845213058 | -4.877013314 |
| **A_51_P143142** | Mrpl12 | 0.108405056 | -0.215862795 | 0.432672907 | 11.95805014 | 0.822460042 | 0.442938354 | 0.845213058 | -4.877047288 |
| **A_51_P375390** | Gemin7 | 0.059838298 | -0.119169792 | 0.238846387 | 11.67150195 | 0.822386054 | 0.442977258 | 0.845213058 | -4.877086923 |
| **A_52_P125485** | Slc25a48 | -0.091098299 | -0.3636287 | 0.181432101 | 8.426422464 | -0.822364735 | 0.442988468 | 0.845213058 | -4.877098343 |
| **A_51_P401184** | Rarres1 | 0.219035567 | -0.436262869 | 0.874334004 | 9.953405446 | 0.822327234 | 0.443008188 | 0.845213058 | -4.877118431 |
| **A_55_P1977766** | Ttc14 | -0.131441668 | -0.524705069 | 0.261821734 | 8.317226371 | -0.822278042 | 0.443034057 | 0.845213058 | -4.877144781 |
| **A_55_P1980932** | Elob | -0.057317258 | -0.228838848 | 0.114204332 | 15.10071796 | -0.822121058 | 0.443116617 | 0.845213058 | -4.877228861 |
| **A_55_P2236607** | Ccnc | 0.04029493 | -0.080304131 | 0.160893991 | 8.477886091 | 0.822007247 | 0.443176479 | 0.845213058 | -4.87728981 |
| **A_52_P637812** | Prrg4 | -0.073129891 | -0.292003583 | 0.145743801 | 9.177988785 | -0.82199755 | 0.44318158 | 0.845213058 | -4.877295003 |
| **A_51_P268697** | Slc1a3 | -0.047766194 | -0.190734669 | 0.09520228 | 7.916793699 | -0.821957797 | 0.443202491 | 0.845213058 | -4.87731629 |
| **A_55_P2102621** | Eaf2 | -0.124370354 | -0.496636371 | 0.247895663 | 10.59094821 | -0.821925852 | 0.443219295 | 0.845213058 | -4.877333396 |
| **A_51_P508191** | Vps26c | -0.067993428 | -0.27161652 | 0.135629664 | 12.02182818 | -0.821502849 | 0.443441853 | 0.845213058 | -4.877559859 |
| **A_55_P1966660** | LOC547349 | -0.202090219 | -0.807321695 | 0.403141256 | 10.96581315 | -0.821472354 | 0.4434579 | 0.845213058 | -4.877576182 |
| **A_55_P2088715** | Ss18l2 | 0.067867352 | -0.135388492 | 0.271123195 | 9.186896294 | 0.821461144 | 0.443463799 | 0.845213058 | -4.877582182 |
| **A_55_P2046084** | Dcun1d4 | 0.096409706 | -0.192349047 | 0.385168459 | 10.68819462 | 0.821400126 | 0.443495911 | 0.845213058 | -4.87761484 |
| **A_55_P1952379** | Fkbp5 | 0.100659076 | -0.200831267 | 0.402149419 | 10.12550398 | 0.821388652 | 0.44350195 | 0.845213058 | -4.877620982 |
| **A_52_P655285** | Zfp462 | -0.059949483 | -0.23951428 | 0.119615314 | 8.367237228 | -0.821359726 | 0.443517174 | 0.845213058 | -4.877636464 |
| **A_51_P237514** | Eif1ax | -0.136719805 | -0.546270107 | 0.272830497 | 10.6251949 | -0.821283963 | 0.443557048 | 0.845213058 | -4.87767701 |
| **A_52_P512749** | Ears2 | -0.050063993 | -0.200040182 | 0.099912197 | 8.858011302 | -0.821244227 | 0.443577963 | 0.845213058 | -4.877698276 |
| **A_52_P647919** | Usf2 | -0.048550865 | -0.194007556 | 0.096905826 | 8.700771183 | -0.82116878 | 0.443617676 | 0.845213058 | -4.87773865 |
| **A_55_P2146650** | NA | 0.080336617 | -0.160349087 | 0.321022322 | 9.012842498 | 0.82116874 | 0.443617697 | 0.845213058 | -4.877738671 |
| **A_55_P2039601** | Drosha | -0.092364011 | -0.379648665 | 0.194920644 | 12.1186371 | -0.833183068 | 0.443676399 | 0.845241584 | -4.819394024 |
| **A_51_P490817** | Me2 | 0.094877255 | -0.189450597 | 0.379205108 | 8.958245836 | 0.820940857 | 0.443737663 | 0.845274986 | -4.877860602 |
| **A_55_P2066116** | Bcl3 | -0.129150168 | -0.516226624 | 0.257926287 | 12.09012803 | -0.82085677 | 0.443781936 | 0.845276018 | -4.877905588 |

| | | | | | | | | |
|---|---|---|---|---|---|---|---|---|
| **A_55_P2083764** | NA | 0.049382669 | -0.098703563 | 0.197468901 | 8.026944761 | 0.820406404 | 0.444019111 | 0.845644437 | -4.878146473 |
| **A_55_P2160331** | Bicd2 | -0.074197994 | -0.29675861 | 0.148362622 | 11.04781581 | -0.820187247 | 0.444134559 | 0.845780974 | -4.878263658 |
| **A_52_P374669** | Kdm5a | -0.0500526 | -0.200239554 | 0.100134353 | 11.28508613 | -0.819905122 | 0.444283209 | 0.845880362 | -4.878414479 |
| **A_51_P379373** | Lactb2 | 0.095807379 | -0.191691933 | 0.38330669 | 9.329870795 | 0.819844167 | 0.44431533 | 0.845880362 | -4.87844706 |
| **A_66_P101835** | Cd38 | -0.060727994 | -0.242962034 | 0.121506047 | 8.472605422 | -0.819839033 | 0.444318036 | 0.845880362 | -4.878449804 |
| **A_52_P248604** | Cdh5 | -0.09212815 | -0.368654511 | 0.184398211 | 13.40788331 | -0.819643454 | 0.444421112 | 0.845993271 | -4.878554331 |
| **A_51_P325165** | Il27 | 0.050950327 | -0.102024061 | 0.203924716 | 8.560060305 | 0.819402747 | 0.444547997 | 0.846112878 | -4.878682953 |
| **A_52_P276343** | Pum3 | 0.050138608 | -0.100406844 | 0.200684059 | 8.104454246 | 0.819358178 | 0.444571493 | 0.846112878 | -4.878706764 |
| **A_55_P2143516** | NA | -0.285379443 | -1.142417916 | 0.571659031 | 9.60738786 | -0.819202993 | 0.444653313 | 0.84618528 | -4.87878967 |
| **A_55_P2160128** | Pls1 | 0.15669008 | -0.313990443 | 0.627370603 | 8.632602796 | 0.819001012 | 0.444759822 | 0.846248639 | -4.878897558 |
| **A_55_P2022499** | Pnisr | -0.103823454 | -0.415708552 | 0.208061644 | 8.891826233 | -0.818973807 | 0.44477417 | 0.846248639 | -4.878912088 |
| **A_55_P2067146** | NA | -0.035978556 | -0.144073851 | 0.072116738 | 7.719365767 | -0.818853307 | 0.444837722 | 0.846286253 | -4.878976442 |
| **A_52_P537571** | Trap1a | -0.115106844 | -0.461042905 | 0.230829217 | 13.30961869 | -0.818605151 | 0.444968622 | 0.846366686 | -4.879108951 |
| **A_66_P106113** | Rhoj | -0.220127007 | -0.881726113 | 0.441472099 | 10.53887837 | -0.818554466 | 0.444995362 | 0.846366686 | -4.879136013 |
| **A_55_P1956443** | Ttc5 | 0.045880941 | -0.092020877 | 0.18378276 | 9.495083727 | 0.818524137 | 0.445011363 | 0.846366686 | -4.879152205 |
| **A_55_P2136340** | NA | 0.225377374 | -0.452215349 | 0.902970096 | 14.71277486 | 0.818296595 | 0.445131422 | 0.846479005 | -4.879273672 |
| **A_55_P1967315** | Ppp1r9b | -0.106649365 | -0.427340504 | 0.214041773 | 11.14432691 | -0.818164213 | 0.445201283 | 0.846479005 | -4.87934433 |
| **A_55_P2393734** | 4933421H12Rik | -0.163254896 | -0.654178722 | 0.327668931 | 9.847316559 | -0.818128018 | 0.445220385 | 0.846479005 | -4.879363648 |
| **A_51_P497821** | Zfp790 | -0.037420269 | -0.149976075 | 0.075135536 | 8.339754474 | -0.817914996 | 0.44533282 | 0.846479005 | -4.879477326 |
| **A_55_P2120080** | NA | -0.06288074 | -0.252033343 | 0.126271863 | 7.999369609 | -0.817851552 | 0.445366311 | 0.846479005 | -4.879511178 |
| **A_52_P308507** | Chd6 | -0.055151293 | -0.221055933 | 0.110753347 | 8.216309264 | -0.817836202 | 0.445374414 | 0.846479005 | -4.879519368 |
| **A_51_P104897** | Itpr3 | 0.179304173 | -0.360082018 | 0.718690364 | 10.86232259 | 0.817823785 | 0.445380969 | 0.846479005 | -4.862341895 |
| **A_55_P2128113** | Tbc1d12 | -0.058462114 | -0.234349788 | 0.117425561 | 7.951937119 | -0.817726872 | 0.445432131 | 0.846479005 | -4.879577699 |
| **A_51_P273755** | Exoc7 | 0.078627801 | -0.157947632 | 0.315203234 | 9.348984397 | 0.817665454 | 0.445464558 | 0.846479005 | -4.879610465 |
| **A_55_P2073219** | Hps4 | -0.099592747 | -0.399304767 | 0.200119273 | 11.19734758 | -0.81750932 | 0.445546998 | 0.846552435 | -4.879693753 |
| **A_51_P382849** | Emb | 0.167247265 | -0.336199324 | 0.670693855 | 10.72816587 | 0.817287366 | 0.445664211 | 0.84655604 | -4.879812133 |
| **A_55_P2082439** | Spout1 | 0.06769871 | -0.13609022 | 0.271487639 | 11.03631321 | 0.817276415 | 0.445669995 | 0.84655604 | -4.879817973 |
| **A_55_P2016326** | NA | -0.072685378 | -0.291490563 | 0.146119806 | 10.04927255 | -0.817256928 | 0.445680287 | 0.84655604 | -4.879828365 |
| **A_55_P2160381** | NA | -0.056284719 | -0.225772968 | 0.11320353 | 12.55389442 | -0.816996267 | 0.445817973 | 0.846734361 | -4.879967356 |
| **A_51_P164835** | Ppl | 0.082296462 | -0.165551178 | 0.330144102 | 9.616384692 | 0.816893688 | 0.445872165 | 0.846754085 | -4.880022045 |
| **A_66_P130469** | Rce1 | -0.039429904 | -0.158194693 | 0.079334885 | 9.106096267 | -0.816783929 | 0.445930156 | 0.846768039 | -4.880080556 |
| **A_52_P660922** | Dock1 | 0.054072915 | -0.108819231 | 0.216965061 | 10.55482065 | 0.816674137 | 0.445988171 | 0.846768039 | -4.88013908 |
| **A_51_P383032** | Clec4d | -0.155660682 | -0.624739043 | 0.31341768 | 8.511933097 | -0.816399428 | 0.44613335 | 0.846768039 | -4.880285486 |
| **A_51_P328932** | Tmem165 | -0.03921015 | -0.157370611 | 0.07895031 | 12.76545154 | -0.816385904 | 0.446140499 | 0.846768039 | -4.880292692 |
| **A_52_P149577** | Rab25 | 0.047496037 | -0.095637671 | 0.190629745 | 7.846839316 | 0.816365444 | 0.446151313 | 0.846768039 | -4.880303595 |
| **A_52_P146164** | Ube2n | 0.057501801 | -0.115794316 | 0.230797918 | 9.193016278 | 0.81632252 | 0.446174001 | 0.846768039 | -4.880326467 |
| **A_66_P110633** | Apol9b | -0.199308457 | -0.799991485 | 0.40137457 | 9.507096726 | -0.816299498 | 0.44618617 | 0.846768039 | -4.880338734 |
| **A_55_P1975793** | Srd5a3 | 0.058790531 | -0.118496883 | 0.236077944 | 10.5978684 | 0.815828062 | 0.446435419 | 0.846875658 | -4.880589881 |
| **A_55_P2009111** | NA | 0.041127215 | -0.082896285 | 0.165150716 | 7.774228451 | 0.815820259 | 0.446439545 | 0.846875658 | -4.880594036 |
| **A_55_P2065799** | Naa50 | 0.052708995 | -0.106250254 | 0.211668244 | 10.19361767 | 0.815770683 | 0.446465762 | 0.846875658 | -4.880620441 |
| **A_55_P2059352** | Col18a1 | 0.174262384 | -0.351289686 | 0.699814455 | 11.18354903 | 0.815749964 | 0.446476719 | 0.846875658 | -4.880631475 |
| **A_52_P676956** | Tirap | -0.058716345 | -0.235798451 | 0.11836576 | 8.793279838 | -0.815743271 | 0.446480259 | 0.846875658 | -4.88063504 |
| **A_55_P1965313** | Mctp2 | -0.047762784 | -0.191835766 | 0.096310197 | 8.338863052 | -0.815598192 | 0.446556988 | 0.846875658 | -4.8807123 |
| **A_51_P305437** | Rcn1 | -0.131919596 | -0.529925117 | 0.266085926 | 12.15175842 | -0.815435056 | 0.446643278 | 0.846875658 | -4.880799166 |
| **A_55_P2069246** | Ddx39b | 0.090194343 | -0.192305613 | 0.3726943 | 12.75842446 | 0.827391391 | 0.44665381 | 0.846875658 | -4.822243396 |
| **A_55_P2064025** | A430073D23Rik | -0.165430449 | -0.664638181 | 0.333777283 | 13.22505161 | -0.815273468 | 0.446728762 | 0.846875658 | -4.880885194 |
| **A_51_P279465** | Farsa | -0.068152639 | -0.273817359 | 0.13751208 | 9.243199313 | -0.815252321 | 0.44673995 | 0.846875658 | -4.880896451 |
| **A_55_P1980551** | Dennd1a | -0.108030219 | -0.434041348 | 0.21798091 | 10.3470514 | -0.815233457 | 0.446749931 | 0.846875658 | -4.880906494 |
| **A_55_P2423646** | Nf2 | -0.050479976 | -0.202836454 | 0.101876503 | 9.648686978 | -0.815130932 | 0.446804176 | 0.846875658 | -4.880961069 |
| **A_55_P2078460** | NA | 0.090111599 | -0.181878114 | 0.362101312 | 8.944846396 | 0.815074642 | 0.446833962 | 0.846875658 | -4.88099103 |

| | | | | | | | | | |
|---|---|---|---|---|---|---|---|---|---|
| **A_51_P440865** | Fam110b | -0.040902019 | -0.16439171 | 0.082587673 | 9.233526042 | -0.814860378 | 0.44694735 | 0.846875658 | -4.881105065 |
| **A_52_P273169** | Sh3d19 | 0.046356255 | -0.093607745 | 0.186320255 | 8.253964062 | 0.814819031 | 0.446969233 | 0.846875658 | -4.881127067 |
| **A_55_P2064766** | NA | -0.065125717 | -0.261762823 | 0.131511389 | 10.84603462 | -0.814809726 | 0.446974157 | 0.846875658 | -4.881132019 |
| **A_51_P366138** | Mertk | -0.086859023 | -0.349124753 | 0.175406708 | 8.925055256 | -0.814784119 | 0.446987711 | 0.846875658 | -4.881145645 |
| **A_66_P103792** | AI606181 | -0.073263936 | -0.294510687 | 0.147982816 | 10.64600567 | -0.814671466 | 0.447047339 | 0.846905617 | -4.881205588 |
| **A_55_P2157922** | Ndufaf1 | 0.100834304 | -0.20375887 | 0.405427478 | 10.30365218 | 0.814436636 | 0.447171654 | 0.846995617 | -4.881330521 |
| **A_55_P2091147** | H4c12 | -0.122240362 | -0.491504725 | 0.247024002 | 9.967498184 | -0.814416177 | 0.447182486 | 0.846995617 | -4.881341405 |
| **A_52_P256279** | Zbtb11 | -0.10128808 | -0.407337829 | 0.20476167 | 9.215830499 | -0.814208199 | 0.447292609 | 0.847077155 | -4.881452029 |
| **A_52_P208521** | Khdc1a | -0.094060242 | -0.378284084 | 0.190163601 | 9.415163644 | -0.814169347 | 0.447313184 | 0.847077155 | -4.881472692 |
| **A_55_P2176240** | Pdxk | 0.082050891 | -0.165955298 | 0.330057079 | 8.30689056 | 0.813935415 | 0.447437077 | 0.847192392 | -4.881597092 |
| **A_55_P2066384** | Akt1 | 0.069461246 | -0.140503601 | 0.279426093 | 13.79269897 | 0.813888935 | 0.447461696 | 0.847192392 | -4.881621807 |
| **A_51_P251352** | Slc25a13 | 0.093233257 | -0.188627004 | 0.375093519 | 10.15643759 | 0.813778423 | 0.447520236 | 0.847220238 | -4.881680563 |
| **A_55_P2335529** | 4921531C22Rik | -0.07881238 | -0.317166884 | 0.159542124 | 8.501720024 | -0.813467567 | 0.447684929 | 0.84737826 | -4.881845808 |
| **A_55_P1988531** | Myh11 | -0.172949964 | -0.696015253 | 0.350115325 | 8.217405346 | -0.813455384 | 0.447691385 | 0.84737826 | -4.881852284 |
| **A_51_P258848** | Mfsd13b | -0.066014306 | -0.265701721 | 0.133673108 | 9.059484157 | -0.813310792 | 0.447768007 | 0.847440302 | -4.881929129 |
| **A_55_P2121081** | Cul2 | 0.085333533 | -0.172834418 | 0.343501484 | 9.260021132 | 0.813179759 | 0.447837451 | 0.8474663 | -4.88199876 |
| **A_51_P245789** | Pcolce2 | 0.06563069 | -0.132943043 | 0.264204423 | 7.812832237 | 0.813119421 | 0.447869432 | 0.8474663 | -4.882030821 |
| **A_55_P2108486** | NA | -0.201687024 | -0.812090554 | 0.408716507 | 10.48614819 | -0.812886822 | 0.44799273 | 0.847551154 | -4.882154398 |
| **A_55_P2175880** | Mgst1 | 0.173921094 | -0.352460449 | 0.700302637 | 10.60761999 | 0.812869386 | 0.448001973 | 0.847551154 | -4.88216366 |
| **A_55_P2031403** | NA | 0.343434606 | -0.6961143 | 1.382983512 | 9.602021838 | 0.812771172 | 0.448054044 | 0.847566707 | -4.882215832 |
| **A_55_P2184811** | Eif2s1 | 0.043847751 | -0.088897207 | 0.176592708 | 8.395144513 | 0.81264048 | 0.44812334 | 0.847614839 | -4.882285249 |
| **A_55_P2128229** | NA | 0.154449417 | -0.313218314 | 0.622117148 | 8.913151505 | 0.812490012 | 0.448203131 | 0.847662027 | -4.882365159 |
| **A_55_P2249220** | A430102J17Rik | -0.158937044 | -0.640241222 | 0.322367135 | 8.633530487 | -0.812408879 | 0.448246159 | 0.847662027 | -4.882408243 |
| **A_55_P2067777** | Cplane1 | 0.079005946 | -0.160264049 | 0.318275942 | 9.649347212 | 0.812345353 | 0.448279851 | 0.847662027 | -4.882441975 |
| **A_55_P2130178** | Fn1 | 0.040021792 | -0.081218745 | 0.161262329 | 17.24092817 | 0.812115582 | 0.448401731 | 0.847729853 | -4.882563966 |
| **A_55_P2013645** | Tubg2 | -0.080707185 | -0.325205459 | 0.16379109 | 9.128952671 | -0.812092615 | 0.448413915 | 0.847729853 | -4.882576158 |
| **A_51_P447866** | Sash3 | -0.048524161 | -0.19554469 | 0.098496368 | 7.99676726 | -0.811987258 | 0.448469809 | 0.847729853 | -4.882632085 |
| **A_51_P162676** | Armc7 | -0.045892394 | -0.184945975 | 0.093161187 | 8.962848151 | -0.811947031 | 0.448491152 | 0.847729853 | -4.882653437 |
| **A_55_P2093301** | NA | 0.062301671 | -0.126542782 | 0.251146123 | 8.591086577 | 0.811642212 | 0.4486529 | 0.847841917 | -4.88281521 |
| **A_55_P2208260** | D230019N24Rik | -0.19428124 | -0.783195719 | 0.394633239 | 9.523050644 | -0.811610764 | 0.44866959 | 0.847841917 | -4.882831898 |
| **A_55_P2044533** | NA | -0.07639981 | -0.307993887 | 0.155194268 | 8.223369817 | -0.811584958 | 0.448683285 | 0.847841917 | -4.882845591 |
| **A_55_P1973748** | Nrbp1 | 0.05145614 | -0.104548757 | 0.207461036 | 10.549815 | 0.811461901 | 0.448748599 | 0.847841917 | -4.882910884 |
| **A_55_P2187034** | NA | -0.087336182 | -0.35213532 | 0.177462956 | 10.47915525 | -0.811422038 | 0.448769758 | 0.847841917 | -4.882932033 |
| **A_55_P2115003** | Prpf4 | -0.049321823 | -0.198887405 | 0.100243759 | 8.592715924 | -0.811290928 | 0.448839356 | 0.847890531 | -4.883001588 |
| **A_55_P2281624** | 2610034E01Rik | -0.138009926 | -0.556827843 | 0.280807991 | 8.312474328 | -0.810688912 | 0.449159027 | 0.848246355 | -4.88332086 |
| **A_55_P2012592** | Ildr1 | -0.068929802 | -0.278119857 | 0.140260253 | 9.782491955 | -0.810653314 | 0.449177935 | 0.848246355 | -4.883339733 |
| **A_52_P161630** | St6gal1 | -0.08289905 | -0.33449335 | 0.16869525 | 11.45427791 | -0.810621125 | 0.449195033 | 0.848246355 | -4.883356799 |
| **A_55_P2040971** | Tom1 | 0.117700339 | -0.23955286 | 0.474953539 | 12.80980874 | 0.810533092 | 0.449241794 | 0.848246355 | -4.883403469 |
| **A_55_P2124586** | Xlr4a | -0.078395431 | -0.316372101 | 0.159581238 | 12.04911759 | -0.810448711 | 0.44928662 | 0.848246355 | -4.8834482 |
| **A_55_P2046213** | NA | -0.048308252 | -0.194953972 | 0.098337468 | 16.06396901 | -0.810440423 | 0.449291023 | 0.848246355 | -4.883452593 |
| **A_55_P2010871** | Daam2 | 0.149922035 | -0.305298702 | 0.605142773 | 9.177228279 | 0.810237976 | 0.449398582 | 0.848366559 | -4.883559896 |
| **A_55_P2002667** | Trim26 | -0.058788707 | -0.237318257 | 0.119740844 | 12.31476331 | -0.810126722 | 0.449457699 | 0.848395299 | -4.883618855 |
| **A_55_P2040773** | NA | 0.054817008 | -0.111713941 | 0.221347956 | 9.268641607 | 0.80982193 | 0.449619684 | 0.848618188 | -4.883780352 |
| **A_51_P477364** | Rhob | 0.06035629 | -0.123050402 | 0.243762982 | 8.371931933 | 0.809611228 | 0.449731689 | 0.848725566 | -4.883891969 |
| **A_55_P2086511** | Ppid | -0.055974653 | -0.226082528 | 0.114133222 | 10.87820006 | -0.809535972 | 0.449771698 | 0.848725566 | -4.883931829 |
| **A_51_P219789** | H2-Q2 | -0.316878117 | -1.280075181 | 0.646318947 | 13.28214281 | -0.809368432 | 0.449860779 | 0.848725566 | -4.88402056 |
| **A_55_P1990331** | Add1 | -0.037255428 | -0.150512656 | 0.0760018 | 8.232357794 | -0.809268796 | 0.449913761 | 0.848725566 | -4.884073322 |
| **A_55_P2059686** | Rcc1 | 0.115135336 | -0.23489854 | 0.465169213 | 10.55005528 | 0.809222072 | 0.449938608 | 0.848725566 | -4.884098063 |
| **A_51_P498212** | Ube2j1 | -0.086084195 | -0.347797804 | 0.175629413 | 8.786542389 | -0.809219395 | 0.449940032 | 0.848725566 | -4.884099481 |
| **A_55_P1996852** | NA | -0.061847546 | -0.24993245 | 0.126237358 | 9.345968737 | -0.80897983 | 0.450067446 | 0.848743773 | -4.884226317 |

| | | | | | | | | | |
|---|---|---|---|---|---|---|---|---|---|
| **A_55_P1980636** | Aurka | 0.093895464 | -0.191682649 | 0.379473577 | 8.876438347 | 0.808888862 | 0.450115835 | 0.848743773 | -4.884274472 |
| **A_51_P213359** | Has2 | -0.087134655 | -0.35216472 | 0.177895411 | 7.86854506 | -0.808844307 | 0.450139537 | 0.848743773 | -4.884298057 |
| **A_51_P265495** | Ly6a | -0.85691085 | -3.46429387 | 1.75047217 | 13.05009507 | -0.808537136 | 0.450302965 | 0.848743773 | -4.884460627 |
| **A_52_P445622** | 2610002M06Rik | -0.052366617 | -0.211707565 | 0.106974332 | 8.289716682 | -0.808530266 | 0.45030662 | 0.848743773 | -4.884464262 |
| **A_52_P538470** | Spats2l | 0.052871012 | -0.108050559 | 0.213792582 | 8.299257625 | 0.808299896 | 0.450429216 | 0.848743773 | -4.884586156 |
| **A_51_P140731** | Cops5 | 0.054935539 | -0.112275218 | 0.222146296 | 13.16189119 | 0.808273499 | 0.450443265 | 0.848743773 | -4.884600121 |
| **A_51_P481679** | Angptl3 | -0.099636335 | -0.40297094 | 0.20369827 | 8.106521749 | -0.808099707 | 0.45053577 | 0.848743773 | -4.884692059 |
| **A_51_P314186** | Syne1 | -0.063897094 | -0.258516163 | 0.130721975 | 8.019536648 | -0.807727514 | 0.450733923 | 0.848743773 | -4.884888905 |
| **A_55_P2165839** | Cebpe | -0.050149876 | -0.202906173 | 0.102606422 | 7.825882398 | -0.80768106 | 0.45075866 | 0.848743773 | -4.88491347 |
| **A_55_P2126951** | Zfp467 | 0.044045257 | -0.090117535 | 0.178208048 | 7.95025726 | 0.80767422 | 0.450762302 | 0.848743773 | -4.884917086 |
| **A_55_P2183688** | NA | 0.071538135 | -0.146375422 | 0.289451693 | 15.09807462 | 0.807648756 | 0.450775862 | 0.848743773 | -4.884930551 |
| **A_55_P1955427** | Cux1 | -0.100755602 | -0.407677476 | 0.206166272 | 8.886852827 | -0.80762643 | 0.450787751 | 0.848743773 | -4.884942355 |
| **A_51_P503625** | Gsta3 | 0.168061417 | -0.343891536 | 0.680014369 | 9.512865952 | 0.807620122 | 0.45079111 | 0.848743773 | -4.884945691 |
| **A_55_P2032760** | Ctif | -0.122648567 | -0.496296974 | 0.250999841 | 10.71437222 | -0.807548295 | 0.450829361 | 0.848743773 | -4.884983668 |
| **A_55_P2170978** | Ankrd13d | 0.098812587 | -0.202284915 | 0.399910088 | 9.340881566 | 0.807373107 | 0.450922666 | 0.848743773 | -4.885076284 |
| **A_55_P2300421** | NA | 0.206696989 | -0.423168415 | 0.836562392 | 9.251120372 | 0.807338303 | 0.450941204 | 0.848743773 | -4.885094682 |
| **A_51_P242201** | Naaa | -0.108457009 | -0.438969258 | 0.222055239 | 10.91422046 | -0.80730795 | 0.450957372 | 0.848743773 | -4.885110727 |
| **A_55_P2053838** | Tnfaip3 | -0.063757937 | -0.258063709 | 0.130547836 | 9.262295849 | -0.807267959 | 0.450978675 | 0.848743773 | -4.885131865 |
| **A_55_P1973809** | Hbb-bt | 0.12593349 | -0.257947974 | 0.509814955 | 17.80708021 | 0.807073821 | 0.451082098 | 0.848743773 | -4.885234473 |
| **A_51_P357341** | Clic1 | -0.084594505 | -0.342471611 | 0.173282601 | 15.27232047 | -0.807046442 | 0.451096685 | 0.848743773 | -4.885248942 |
| **A_52_P482124** | Fam32a | -0.056127005 | -0.227224454 | 0.114970445 | 10.52104207 | -0.807044519 | 0.451097709 | 0.848743773 | -4.885249958 |
| **A_55_P2262593** | 4930429F24Rik | 0.051071343 | -0.104621088 | 0.206763775 | 8.023584091 | 0.80701005 | 0.451116074 | 0.848743773 | -4.885268174 |
| **A_55_P1992234** | Slx | 0.058090323 | -0.119002588 | 0.235183233 | 8.70487207 | 0.806996735 | 0.451123169 | 0.848743773 | -4.88527521 |
| **A_51_P291078** | Sel1l3 | 0.154358928 | -0.316249377 | 0.624967234 | 9.131465498 | 0.80694016 | 0.451153313 | 0.848743773 | -4.885305106 |
| **A_55_P1983235** | Tmed10 | -0.059699088 | -0.241709773 | 0.122311597 | 16.10142991 | -0.806937644 | 0.451154654 | 0.848743773 | -4.885306436 |
| **A_51_P214825** | Orc5 | -0.052201835 | -0.211362983 | 0.106959313 | 8.528899037 | -0.806896568 | 0.451176541 | 0.848743773 | -4.885328141 |
| **A_55_P2029517** | Wnk2 | 0.051006976 | -0.104512051 | 0.206526002 | 8.0844651 | 0.806891626 | 0.451179175 | 0.848743773 | -4.885330752 |
| **A_52_P653966** | Atf7ip | -0.065257701 | -0.264287262 | 0.133771859 | 10.39398509 | -0.806646682 | 0.451309708 | 0.84883266 | -4.885460166 |
| **A_55_P2178343** | Rap1gds1 | -0.096366305 | -0.390277152 | 0.197544541 | 11.64387381 | -0.80663815 | 0.451314256 | 0.84883266 | -4.885464673 |
| **A_55_P2025073** | Csf1r | 0.081068973 | -0.166254877 | 0.328392823 | 12.01487015 | 0.806413588 | 0.451433953 | 0.848975177 | -4.885583292 |
| **A_65_P06572** | Smad5 | 0.096659191 | -0.198373022 | 0.391691405 | 8.483548209 | 0.806014554 | 0.451646703 | 0.849276757 | -4.885794012 |
| **A_51_P216605** | Hbp1 | -0.071618444 | -0.290236999 | 0.147000111 | 9.052737332 | -0.805948008 | 0.45168219 | 0.849276757 | -4.885829146 |
| **A_51_P328883** | Zfp810 | -0.042196654 | -0.171025096 | 0.086631789 | 8.330708996 | -0.805815129 | 0.451753057 | 0.849314129 | -4.885899295 |
| **A_55_P2071466** | Ncf1 | -0.037202501 | -0.15079999 | 0.076394988 | 7.845730873 | -0.805698531 | 0.451815247 | 0.849314129 | -4.885960842 |
| **A_52_P518997** | Epha2 | 0.06673486 | -0.137065434 | 0.270535154 | 9.326145809 | 0.805595647 | 0.451870128 | 0.849314129 | -4.886015146 |
| **A_66_P127024** | Mbd3 | 0.055250581 | -0.11348112 | 0.223982282 | 12.39168738 | 0.805581215 | 0.451877826 | 0.849314129 | -4.886022762 |
| **A_55_P2166639** | Cdk4 | 0.103963189 | -0.21358313 | 0.421509508 | 10.50181948 | 0.805455724 | 0.451944773 | 0.849357367 | -4.88608899 |
| **A_52_P33382** | NA | 0.190578351 | -0.391642242 | 0.772798944 | 12.35692829 | 0.805295301 | 0.452030366 | 0.849396461 | -4.886173641 |
| **A_52_P545643** | Tmem263 | -0.037831444 | -0.153413459 | 0.077750571 | 12.31489724 | -0.805252014 | 0.452053464 | 0.849396461 | -4.886196481 |
| **A_55_P1994052** | Asxl1 | -0.075564364 | -0.306455948 | 0.15532722 | 10.95926916 | -0.805152396 | 0.452106622 | 0.849413773 | -4.886249039 |
| **A_51_P496054** | Zcchc14 | -0.068103863 | -0.276248177 | 0.140040451 | 8.79352604 | -0.804963813 | 0.452207266 | 0.849520289 | -4.886348521 |
| **A_52_P156190** | Ednra | 0.227959397 | -0.468964763 | 0.924883557 | 9.530139806 | 0.804713231 | 0.452341024 | 0.849530261 | -4.886480684 |
| **A_52_P669922** | Hamp | 0.101133543 | -0.208055678 | 0.410322764 | 7.904911824 | 0.804711191 | 0.452342113 | 0.849530261 | -4.886481759 |
| **A_55_P2322709** | 4633401B06Rik | -0.064027953 | -0.259777642 | 0.131721736 | 8.005098286 | -0.804706854 | 0.452344428 | 0.849530261 | -4.886484046 |
| **A_52_P59228** | Zswim3 | -0.043299401 | -0.175705956 | 0.089107154 | 8.575241828 | -0.804528751 | 0.452439515 | 0.849591384 | -4.886577963 |
| **A_51_P402458** | Anxa7 | -0.042173742 | -0.171145724 | 0.086798241 | 8.884231514 | -0.804481242 | 0.452464882 | 0.849591384 | -4.886603013 |
| **A_55_P2142251** | H3c15 | -0.121771515 | -0.494289251 | 0.250746221 | 10.00431504 | -0.804207128 | 0.452611262 | 0.849754317 | -4.886747522 |
| **A_51_P146655** | Ptgr2 | -0.077013344 | -0.312624506 | 0.158597818 | 10.08511604 | -0.804154106 | 0.45263958 | 0.849754317 | -4.88677547 |
| **A_52_P496870** | 2310011J03Rik | -0.036717123 | -0.14906497 | 0.075630725 | 8.338930027 | -0.804031471 | 0.452705083 | 0.849790354 | -4.886840107 |
| **A_51_P163694** | Rufy1 | -0.041380315 | -0.168008972 | 0.085248341 | 9.182057237 | -0.803953551 | 0.452746705 | 0.849790354 | -4.886881172 |

| | | | | | | | | |
|---|---|---|---|---|---|---|---|---|
| **A_51_P122723** | Grpel1 | 0.134841017 | -0.278077805 | 0.547759839 | 12.86714036 | 0.80339015 | 0.453047739 | 0.850164868 | -4.887178009 |
| **A_52_P491073** | Nek8 | 0.040787264 | -0.084171178 | 0.165745706 | 9.469572132 | 0.803023277 | 0.453243843 | 0.850164868 | -4.887371221 |
| **A_66_P134428** | Sfrp1 | 0.037770319 | -0.077955331 | 0.153495968 | 7.814301437 | 0.802953109 | 0.453281356 | 0.850164868 | -4.887408167 |
| **A_55_P1993744** | Myo10 | -0.062423754 | -0.253688058 | 0.128840549 | 8.643355557 | -0.802943757 | 0.453286357 | 0.850164868 | -4.887413092 |
| **A_55_P2289765** | Fam160a2 | 0.049787576 | -0.102761089 | 0.202336241 | 8.721504663 | 0.802937487 | 0.453289709 | 0.850164868 | -4.887416393 |
| **A_55_P2042272** | Pbrm1 | -0.034570488 | -0.140496164 | 0.071355188 | 7.736431133 | -0.802922138 | 0.453297915 | 0.850164868 | -4.887424474 |
| **A_55_P2144075** | Pofut2 | -0.040935386 | -0.166366666 | 0.084495895 | 8.31872506 | -0.802901364 | 0.453309022 | 0.850164868 | -4.887435412 |
| **A_55_P2028571** | Sgsh | 0.080682292 | -0.166578098 | 0.327942682 | 9.704071834 | 0.802773156 | 0.453377574 | 0.850164868 | -4.887502909 |
| **A_55_P2045234** | NA | -0.064380062 | -0.261700699 | 0.132940575 | 9.017279477 | -0.802690359 | 0.453421849 | 0.850164868 | -4.887546495 |
| **A_51_P457331** | Hmgn5 | -0.059310084 | -0.24112407 | 0.122503902 | 10.66460769 | -0.802546884 | 0.453498579 | 0.850164868 | -4.887622015 |
| **A_55_P2060343** | Zfp352 | 0.07940618 | -0.164028988 | 0.322841348 | 9.890185994 | 0.802490961 | 0.453528489 | 0.850164868 | -4.887651449 |
| **A_52_P53906** | Ccnd2 | -0.053372 | -0.216997624 | 0.110253623 | 8.259796522 | -0.802474666 | 0.453537204 | 0.850164868 | -4.887660024 |
| **A_55_P2175545** | Gys2 | 0.059387489 | -0.122688478 | 0.241463456 | 7.710064119 | 0.802438023 | 0.453556803 | 0.850164868 | -4.887679309 |
| **A_52_P321627** | NA | 0.054594648 | -0.112797594 | 0.221986889 | 13.70840059 | 0.802387038 | 0.453584074 | 0.850164868 | -4.887706141 |
| **A_52_P92514** | Supt20 | -0.044382903 | -0.180471943 | 0.091706136 | 8.961604684 | -0.802346057 | 0.453605995 | 0.850164868 | -4.887727707 |
| **A_55_P1985351** | Slc35f2 | 0.220141922 | -0.4550418 | 0.895325643 | 11.36129663 | 0.802139593 | 0.453716446 | 0.850289429 | -4.887836345 |
| **A_55_P2149215** | Camk2a | 0.043013013 | -0.088927755 | 0.174953781 | 7.759857176 | 0.802028908 | 0.453775666 | 0.850317969 | -4.887894578 |
| **A_55_P2083566** | NA | 0.050787867 | -0.105042545 | 0.20661828 | 15.7194047 | 0.801820066 | 0.453887419 | 0.850359396 | -4.888004435 |
| **A_55_P2084666** | H2ac10 | 0.165163217 | -0.341672374 | 0.671998808 | 14.98451808 | 0.801706477 | 0.453948209 | 0.850359396 | -4.888064178 |
| **A_55_P1959465** | Lrrfip1 | -0.081409174 | -0.331254878 | 0.16843653 | 10.01390329 | -0.80162384 | 0.453992439 | 0.850359396 | -4.888107638 |
| **A_66_P119518** | Tuba8 | -0.075546458 | -0.307403346 | 0.15631043 | 9.193340692 | -0.801610265 | 0.453999705 | 0.850359396 | -4.888114777 |
| **A_55_P2110713** | Anxa2 | -0.046408979 | -0.18884909 | 0.096031133 | 16.79388111 | -0.801565226 | 0.454023812 | 0.850359396 | -4.888138461 |
| **A_55_P2285599** | 5730437C12Rik | 0.172021991 | -0.356000783 | 0.700044764 | 8.69171302 | 0.801494375 | 0.454061738 | 0.850359396 | -4.888175718 |
| **A_52_P144297** | Tspyl3 | -0.035289719 | -0.143670877 | 0.073091439 | 8.12472517 | -0.801057298 | 0.454295749 | 0.850558732 | -4.888405499 |
| **A_51_P443723** | Slc35c1 | -0.056714607 | -0.230908753 | 0.117479539 | 11.31880834 | -0.800996343 | 0.454328391 | 0.850558732 | -4.888437538 |
| **A_51_P219266** | Tmprss6 | -0.060834037 | -0.247693553 | 0.12602548 | 7.997670932 | -0.800941096 | 0.454357978 | 0.850558732 | -4.888466574 |
| **A_55_P2018393** | NA | -0.057002642 | -0.232139804 | 0.11813452 | 14.98411862 | -0.800729525 | 0.454471296 | 0.850558732 | -4.888577758 |
| **A_51_P175424** | Car14 | 0.038092476 | -0.078958571 | 0.155143524 | 7.895882067 | 0.800632218 | 0.45452342 | 0.850558732 | -4.888628887 |
| **A_55_P1963344** | Upf3b | -0.05403843 | -0.220090924 | 0.112014064 | 8.531288148 | -0.800620093 | 0.454529915 | 0.850558732 | -4.888635257 |
| **A_55_P1966114** | Epn2 | -0.086725294 | -0.353231928 | 0.179781339 | 13.24664177 | -0.800584028 | 0.454549235 | 0.850558732 | -4.888654206 |
| **A_51_P324651** | Adgrl1 | -0.071104723 | -0.289613144 | 0.147403697 | 8.156962741 | -0.800570231 | 0.454556627 | 0.850558732 | -4.888661455 |
| **A_55_P2097116** | Cnep1r1 | 0.049792319 | -0.103232855 | 0.202817494 | 8.980151732 | 0.800513456 | 0.454587043 | 0.850558732 | -4.888691283 |
| **A_51_P270519** | Cenps | -0.037076751 | -0.151039664 | 0.076886163 | 7.875088693 | -0.800400378 | 0.454647627 | 0.850558732 | -4.888750686 |
| **A_55_P2027213** | Krtap20-2 | -0.128092947 | -0.521821439 | 0.265635545 | 8.805939759 | -0.800382414 | 0.454657252 | 0.850558732 | -4.888760122 |
| **A_51_P286748** | Frzb | -0.035340095 | -0.144010558 | 0.073330368 | 7.732259487 | -0.800065162 | 0.45482726 | 0.850558732 | -4.88892675 |
| **A_55_P2085127** | Ramac | 0.127572489 | -0.264733501 | 0.519878478 | 11.18977666 | 0.800020753 | 0.454851061 | 0.850558732 | -4.88895007 |
| **A_55_P2111459** | Trpv2 | 0.188707231 | -0.391604219 | 0.769018682 | 11.94858413 | 0.800012119 | 0.454855689 | 0.850558732 | -4.888954604 |
| **A_66_P107584** | Fam133b | -0.038078923 | -0.155181763 | 0.079023917 | 8.300188094 | -0.79999338 | 0.454865733 | 0.850558732 | -4.888964444 |
| **A_55_P2150906** | Atp5c1 | 0.102783558 | -0.213377707 | 0.418944823 | 12.79058314 | 0.799805065 | 0.454966674 | 0.850558732 | -4.889063322 |
| **A_55_P1968729** | Rpl10 | 0.044892601 | -0.093233265 | 0.183018467 | 17.03607657 | 0.799592873 | 0.455080433 | 0.850558732 | -4.889174716 |
| **A_51_P128648** | Uqcrc2 | 0.075030785 | -0.155825834 | 0.305887404 | 12.83655919 | 0.799588104 | 0.45508299 | 0.850558732 | -4.889177219 |
| **A_55_P1960616** | Dnajc17 | -0.034321944 | -0.139929702 | 0.071285815 | 8.305959706 | -0.79954923 | 0.455103833 | 0.850558732 | -4.889197624 |
| **A_55_P2131875** | Poldip3 | -0.111456523 | -0.454410847 | 0.231497802 | 15.00543956 | -0.799536636 | 0.455110586 | 0.850558732 | -4.889204235 |
| **A_55_P2064771** | Ly6c1 | -0.957488633 | -3.903798317 | 1.988821051 | 15.12747001 | -0.799510914 | 0.455124378 | 0.850558732 | -4.889217736 |
| **A_51_P151835** | Tmtc3 | 0.073011586 | -0.15166023 | 0.297683402 | 9.150635714 | 0.799488735 | 0.45513627 | 0.850558732 | -4.889229377 |
| **A_52_P229972** | Slc22a1 | 0.044213819 | -0.091871565 | 0.180299203 | 7.924160413 | 0.799310856 | 0.455231658 | 0.850564256 | -4.889322733 |
| **A_52_P456059** | Pfdn1 | 0.196899974 | -0.409161894 | 0.802961842 | 11.87735237 | 0.799278004 | 0.455249276 | 0.850564256 | -4.889339973 |
| **A_51_P281582** | Lrrc42 | -0.062755304 | -0.255999932 | 0.130489324 | 8.336502216 | -0.798936328 | 0.455432545 | 0.850564256 | -4.889519245 |
| **A_55_P1960936** | NA | -0.359732372 | -1.467600316 | 0.748135572 | 13.28563527 | -0.798841628 | 0.45548335 | 0.850564256 | -4.889568923 |
| **A_51_P410451** | Tube1 | 0.036568807 | -0.076090601 | 0.149228216 | 8.001883145 | 0.798569084 | 0.455629587 | 0.850564256 | -4.88971187 |

| | | | | | | | | |
|---|---|---|---|---|---|---|---|---|
| **A_51_P407248** | Antxr1 | -0.406170427 | -1.657521065 | 0.845180211 | 10.6652615 | -0.798543251 | 0.45564345 | 0.850564256 | -4.889725418 |
| **A_55_P2112991** | Slc4a3 | -0.099854029 | -0.407541923 | 0.207833864 | 8.971981062 | -0.798407024 | 0.455716558 | 0.850564256 | -4.889796853 |
| **A_55_P2076787** | Synpo | -0.035565819 | -0.145164072 | 0.074032433 | 8.256931005 | -0.798359228 | 0.455742211 | 0.850564256 | -4.889821914 |
| **A_51_P110672** | Mst1r | 0.121769711 | -0.253487747 | 0.497027169 | 8.530276634 | 0.798323853 | 0.455761198 | 0.850564256 | -4.889840462 |
| **A_52_P671661** | Mak16 | -0.04789895 | -0.195524141 | 0.099726241 | 8.561436523 | -0.798242206 | 0.455805022 | 0.850564256 | -4.889883269 |
| **A_55_P2204491** | AI131754 | 0.104799546 | -0.218207774 | 0.427806866 | 9.134332543 | 0.798208229 | 0.455823261 | 0.850564256 | -4.889901082 |
| **A_55_P2148370** | NA | -0.062116402 | -0.253585654 | 0.12935285 | 7.916877198 | -0.798135108 | 0.455862512 | 0.850564256 | -4.889939415 |
| **A_51_P408071** | Kntc1 | 0.045037657 | -0.093798293 | 0.183873607 | 7.879937379 | 0.798073731 | 0.455895461 | 0.850564256 | -4.889971588 |
| **A_55_P2190152** | 4921509J17Rik | -0.044558101 | -0.181920992 | 0.09280479 | 7.856589369 | -0.798043221 | 0.455911841 | 0.850564256 | -4.889987581 |
| **A_55_P2020338** | Scml4 | 0.068333593 | -0.142354636 | 0.279021822 | 8.097371386 | 0.797926973 | 0.455974253 | 0.850564256 | -4.890048512 |
| **A_55_P1966593** | NA | 0.038237534 | -0.079660497 | 0.156135566 | 8.315926143 | 0.797907396 | 0.455984765 | 0.850564256 | -4.890058773 |
| **A_52_P618187** | Mrs2 | -0.065960253 | -0.269349224 | 0.137428718 | 9.839773039 | -0.797855192 | 0.456012795 | 0.850564256 | -4.890086133 |
| **A_51_P327796** | Itgb5 | -0.208946918 | -0.853312847 | 0.435419012 | 14.23879475 | -0.797760563 | 0.456063608 | 0.850564256 | -4.890135725 |
| **A_55_P2045203** | Chp1 | -0.082207364 | -0.335735278 | 0.171320549 | 13.12409542 | -0.797726661 | 0.456081813 | 0.850564256 | -4.890153491 |
| **A_51_P276943** | Gpc1 | 0.159759629 | -0.332957177 | 0.652476435 | 14.73128866 | 0.7976986 | 0.456096882 | 0.850564256 | -4.890168195 |
| **A_55_P2174816** | Zfp266 | -0.051180168 | -0.209027902 | 0.106667565 | 8.378023663 | -0.797687014 | 0.456103104 | 0.850564256 | -4.890174266 |
| **A_51_P268167** | Tasor | -0.053524095 | -0.218618461 | 0.111570271 | 9.100311783 | -0.79760202 | 0.45614875 | 0.850564256 | -4.890218802 |
| **A_52_P487615** | Otulinl | -0.079238644 | -0.32365067 | 0.165173382 | 8.489804843 | -0.797597213 | 0.456151331 | 0.850564256 | -4.89022132 |
| **A_55_P2038757** | Fblim1 | 0.135186586 | -0.281918885 | 0.552292058 | 12.603247 | 0.797364465 | 0.456276344 | 0.850715294 | -4.890343258 |
| **A_55_P2084611** | B4galt6 | -0.041285151 | -0.168699164 | 0.086128863 | 8.040601822 | -0.797160632 | 0.456385847 | 0.850837387 | -4.890450026 |
| **A_55_P2201822** | Naf1 | 0.081652336 | -0.170368858 | 0.333673531 | 10.04077011 | 0.797077801 | 0.45643035 | 0.850838291 | -4.890493407 |
| **A_55_P2115225** | Fap | -0.076101811 | -0.311030556 | 0.158826935 | 8.125999806 | -0.796944345 | 0.45650206 | 0.850889155 | -4.890563296 |
| **A_55_P2020497** | Dcaf12l1 | 0.215117503 | -0.449051045 | 0.87928605 | 11.47597145 | 0.796831704 | 0.456562592 | 0.850889155 | -4.890622277 |
| **A_55_P2260094** | Tcf7l2 | 0.048369095 | -0.100978424 | 0.197716613 | 8.17108781 | 0.796781262 | 0.4565897 | 0.850889155 | -4.890648687 |
| **A_55_P1981964** | Tnfrsf13b | -0.041775051 | -0.170789982 | 0.087239879 | 7.885556295 | -0.796610788 | 0.456681326 | 0.850973705 | -4.890737936 |
| **A_55_P2125208** | Arid5a | -0.055433839 | -0.22665818 | 0.115790502 | 9.909381833 | -0.796486863 | 0.45674794 | 0.850973705 | -4.890802805 |
| **A_55_P2009915** | Stxbp3 | -0.063856313 | -0.261112879 | 0.133400253 | 8.599076782 | -0.796418855 | 0.4567845 | 0.850973705 | -4.890838402 |
| **A_51_P342773** | Vipr2 | -0.039329045 | -0.160826753 | 0.082168663 | 7.764417058 | -0.79636924 | 0.456811173 | 0.850973705 | -4.890864369 |
| **A_55_P2093679** | Zfp599 | -0.098255273 | -0.401896549 | 0.205386002 | 8.598728232 | -0.796093768 | 0.45695929 | 0.851031944 | -4.891008525 |
| **A_55_P2130905** | NA | -0.064739172 | -0.26482115 | 0.135342806 | 13.65263238 | -0.796027977 | 0.45699467 | 0.851031944 | -4.891042949 |
| **A_51_P419925** | Arhgef40 | -0.060226048 | -0.246363671 | 0.125911575 | 8.012109213 | -0.796011542 | 0.457003508 | 0.851031944 | -4.891051548 |
| **A_55_P2024061** | Gtpbp10 | 0.038946835 | -0.081435455 | 0.159329125 | 9.272502381 | 0.79593708 | 0.457043554 | 0.851031944 | -4.891090505 |
| **A_55_P2052097** | Ip6k2 | -0.049824721 | -0.203860139 | 0.104210697 | 8.373379719 | -0.795780782 | 0.457127619 | 0.851031944 | -4.891172268 |
| **A_55_P2011146** | Reep1 | -0.069470752 | -0.284246039 | 0.145304534 | 9.641120619 | -0.79576871 | 0.457134112 | 0.851031944 | -4.891178583 |
| **A_55_P2183237** | Ccdc191 | 0.052930033 | -0.110727745 | 0.21658781 | 9.395523282 | 0.795673104 | 0.45718554 | 0.851031944 | -4.891228591 |
| **A_55_P2059432** | Crabp2 | -0.780176316 | -3.1927329 | 1.632380268 | 12.78166302 | -0.795581026 | 0.457235074 | 0.851031944 | -4.891276749 |
| **A_55_P2091751** | NA | 0.042257685 | -0.088417801 | 0.172933172 | 7.819648211 | 0.795574295 | 0.457238695 | 0.851031944 | -4.891280269 |
| **A_55_P1955931** | Slc25a42 | -0.137444711 | -0.562602192 | 0.28771277 | 9.235932743 | -0.79533001 | 0.457370129 | 0.851094348 | -4.891408013 |
| **A_55_P2130610** | Smc3 | -0.121710244 | -0.498256898 | 0.25483641 | 11.7159594 | -0.795202075 | 0.457438973 | 0.851094348 | -4.891474903 |
| **A_51_P175771** | Flii | 0.063663524 | -0.133302422 | 0.26062947 | 11.49702086 | 0.795185942 | 0.457447655 | 0.851094348 | -4.891483338 |
| **A_55_P2134571** | NA | 0.054044547 | -0.113193019 | 0.221282113 | 8.681529047 | 0.79503676 | 0.457527943 | 0.851094348 | -4.891561325 |
| **A_51_P367310** | Chaf1b | -0.085924132 | -0.351829743 | 0.179981479 | 8.930176127 | -0.794981124 | 0.457557888 | 0.851094348 | -4.891590407 |
| **A_55_P1978875** | Kmt2b | 0.09852944 | -0.206388596 | 0.403447477 | 10.7616946 | 0.794972423 | 0.457562572 | 0.851094348 | -4.891594955 |
| **A_55_P1980501** | Znhit1 | 0.048424192 | -0.101459788 | 0.198308171 | 9.857009139 | 0.794833803 | 0.457637189 | 0.851094348 | -4.891667407 |
| **A_51_P458218** | Wdr13 | -0.069171196 | -0.283282803 | 0.144940411 | 9.661611025 | -0.794793377 | 0.457658951 | 0.851094348 | -4.891688534 |
| **A_55_P1988760** | Sestd1 | 0.091151553 | -0.191018974 | 0.373322079 | 9.489595298 | 0.794733619 | 0.457691121 | 0.851094348 | -4.891719764 |
| **A_55_P2000643** | NA | 0.061047223 | -0.127946323 | 0.250040769 | 8.358937644 | 0.794672348 | 0.457724108 | 0.851094348 | -4.891751782 |
| **A_66_P137556** | Tle1 | 0.040331625 | -0.0845472 | 0.16521045 | 8.363483343 | 0.794558863 | 0.45778521 | 0.851094348 | -4.891811082 |
| **A_55_P2095325** | AI182371 | -0.03600628 | -0.14749655 | 0.075483991 | 7.652194118 | -0.794530267 | 0.457800607 | 0.851094348 | -4.891826023 |
| **A_55_P2068137** | NA | -0.058931624 | -0.241465355 | 0.123602107 | 11.83798927 | -0.794281487 | 0.457934577 | 0.851150199 | -4.891955992 |

| | | | | | | | | | |
|---|---|---|---|---|---|---|---|---|---|
| **A_52_P4698** | Rfwd3 | -0.053073414 | -0.217476658 | 0.11132983 | 11.65026672 | -0.794210827 | 0.457972634 | 0.851150199 | -4.891992902 |
| **A_55_P2051094** | Rorc | 0.041648089 | -0.087369669 | 0.170665848 | 7.736427333 | 0.794172335 | 0.457993366 | 0.851150199 | -4.892013007 |
| **A_51_P503303** | C1galt1c1 | -0.063332055 | -0.25952872 | 0.132864609 | 10.36508922 | -0.794147414 | 0.458006789 | 0.851150199 | -4.892026024 |
| **A_55_P2121195** | Lcorl | -0.039963538 | -0.163814666 | 0.083887589 | 8.161370216 | -0.793840278 | 0.458172241 | 0.851362614 | -4.892186418 |
| **A_55_P2253746** | Nip7 | 0.063397035 | -0.133096955 | 0.259891026 | 8.954423771 | 0.793759324 | 0.458215857 | 0.851362614 | -4.892228687 |
| **A_55_P2118630** | Lasp1 | -0.066048668 | -0.270780534 | 0.138683199 | 8.395721587 | -0.793684277 | 0.458256294 | 0.851362614 | -4.892267869 |
| **A_65_P03364** | Dennd4a | -0.059212763 | -0.24277288 | 0.124347355 | 7.806164468 | -0.793608227 | 0.458297273 | 0.851362614 | -4.892307572 |
| **A_55_P2044380** | NA | -0.061546083 | -0.252368933 | 0.129276766 | 8.900724373 | -0.793485884 | 0.458363204 | 0.851402991 | -4.892371437 |
| **A_55_P2002155** | Sumo1 | -0.081721607 | -0.335143919 | 0.171700704 | 13.47895405 | -0.793343404 | 0.458439995 | 0.851402991 | -4.892445804 |
| **A_52_P669005** | Lrat | -0.034584831 | -0.141847162 | 0.072677501 | 7.693677345 | -0.793245441 | 0.458492797 | 0.851402991 | -4.89249693 |
| **A_55_P2016675** | Ogfr | -0.063735251 | -0.26142107 | 0.133950568 | 10.42253625 | -0.793182913 | 0.458526503 | 0.851402991 | -4.892529561 |
| **A_55_P2093226** | Rnf121 | -0.048782408 | -0.200093928 | 0.102529112 | 8.923631944 | -0.793159269 | 0.458539248 | 0.851402991 | -4.892541899 |
| **A_55_P2144364** | NA | -0.074792495 | -0.306873562 | 0.157288573 | 9.442371741 | -0.792843495 | 0.458709496 | 0.851625575 | -4.892706654 |
| **A_51_P116581** | Ccdc22 | 0.054569363 | -0.114788728 | 0.223927453 | 10.17979792 | 0.792705914 | 0.458783686 | 0.851625575 | -4.88437086 |
| **A_55_P1962011** | Clec1b | 0.20631466 | -0.434002135 | 0.846631455 | 10.59266579 | 0.792691788 | 0.458791304 | 0.851625575 | -4.89278579 |
| **A_55_P2174953** | Akr1b3 | -0.075262206 | -0.308895166 | 0.158370753 | 8.270008612 | -0.792523222 | 0.458882215 | 0.851691014 | -4.892873707 |
| **A_51_P431870** | Map1s | -0.080889291 | -0.332009223 | 0.170230641 | 10.08138001 | -0.792463026 | 0.458914683 | 0.851691014 | -4.8929051 |
| **A_55_P2281095** | 2610002M06Rik | -0.038990632 | -0.160082027 | 0.082100763 | 8.920296478 | -0.792165918 | 0.459074959 | 0.851906671 | -4.893060018 |
| **A_51_P210956** | Vcam1 | -0.073649393 | -0.302424023 | 0.155125237 | 9.250130349 | -0.792009647 | 0.459159276 | 0.851981342 | -4.893141483 |
| **A_52_P56658** | Emc3 | 0.071824247 | -0.151359785 | 0.295008279 | 10.20461797 | 0.791730043 | 0.459310165 | 0.8521678 | -4.893287213 |
| **A_55_P2067153** | NA | -0.076458456 | -0.314063661 | 0.16114675 | 12.88469741 | -0.791660057 | 0.459347939 | 0.8521678 | -4.893323685 |
| **A_55_P2230968** | NA | -0.185371351 | -0.761527144 | 0.390784443 | 8.557134439 | -0.791538136 | 0.459413749 | 0.852208096 | -4.893387214 |
| **A_51_P203878** | Dynll2 | 0.077062792 | -0.162503116 | 0.316628701 | 11.15397242 | 0.791386956 | 0.459495362 | 0.852277695 | -4.89346598 |
| **A_55_P1956147** | B4galnt2 | 0.101780833 | -0.214703609 | 0.418265275 | 9.86575969 | 0.791193662 | 0.459599724 | 0.852389471 | -4.893566671 |
| **A_55_P2035539** | D3Ertd751e | -0.0509764 | -0.209506332 | 0.107553532 | 8.454621058 | -0.791092109 | 0.459654561 | 0.852409385 | -4.893619566 |
| **A_55_P1978676** | Mospd3 | 0.046430633 | -0.097997137 | 0.190858403 | 10.1069375 | 0.790902711 | 0.459756845 | 0.852436288 | -4.893718201 |
| **A_55_P2043892** | Srd5a3 | -0.069298488 | -0.284875695 | 0.146278719 | 9.154279838 | -0.790842645 | 0.459789287 | 0.852436288 | -4.893749478 |
| **A_52_P202725** | Rragc | 0.058764291 | -0.12404778 | 0.241576362 | 9.09092414 | 0.790820269 | 0.459801373 | 0.852436288 | -4.89376113 |
| **A_55_P2081616** | Timeless | -0.079594066 | -0.327237341 | 0.168049209 | 9.328633777 | -0.790721076 | 0.459854952 | 0.852453858 | -4.893812777 |
| **A_52_P191468** | Ubfd1 | 0.099694788 | -0.210620084 | 0.410009659 | 11.15853204 | 0.790385614 | 0.460036185 | 0.852708039 | -4.893987407 |
| **A_55_P2063598** | Tomm5 | 0.130736298 | -0.276287671 | 0.537760267 | 15.08964779 | 0.790215169 | 0.460128287 | 0.852780872 | -4.894076114 |
| **A_55_P1963491** | Ptma | -0.059156374 | -0.243344394 | 0.125031646 | 12.79295705 | -0.790149609 | 0.460163717 | 0.852780872 | -4.894110231 |
| **A_55_P2038484** | 4933427D06Rik | -0.081325235 | -0.334706653 | 0.172056183 | 13.45176175 | -0.789622884 | 0.460448439 | 0.853183008 | -4.894384257 |
| **A_55_P2045859** | NA | 0.105491246 | -0.223198962 | 0.434181453 | 11.915105 | 0.789584871 | 0.460468991 | 0.853183008 | -4.894404028 |
| **A_55_P1989748** | NA | -0.039984997 | -0.164587531 | 0.084617537 | 8.473528847 | -0.789476767 | 0.460527444 | 0.853209525 | -4.894460249 |
| **A_52_P677147** | Prmt5 | -0.055489413 | -0.228438566 | 0.11745974 | 9.882376711 | -0.789334084 | 0.460604603 | 0.853221658 | -4.894534446 |
| **A_55_P2112085** | Vwa8 | 0.043976566 | -0.093108724 | 0.181061857 | 8.574723541 | 0.7892228 | 0.460664788 | 0.853221658 | -4.894592309 |
| **A_55_P1971329** | Lage3 | -0.055191381 | -0.227236541 | 0.116853779 | 9.314864737 | -0.789219784 | 0.460666419 | 0.853221658 | -4.894593877 |
| **A_55_P2049318** | NA | -0.061875046 | -0.254802103 | 0.131052011 | 8.325579434 | -0.789026441 | 0.460770997 | 0.853333583 | -4.894694391 |
| **A_52_P573255** | Cdc42ep1 | -0.076594683 | -0.315449576 | 0.162260209 | 11.6533669 | -0.788921235 | 0.46082791 | 0.853357221 | -4.894749077 |
| **A_52_P574190** | Psmd12 | -0.090449739 | -0.372565578 | 0.1916661 | 13.12983317 | -0.788767507 | 0.46091108 | 0.853429473 | -4.894828976 |
| **A_55_P1955706** | Spg7 | -0.058261995 | -0.240054598 | 0.123530607 | 12.83881881 | -0.788457541 | 0.461078811 | 0.853481264 | -4.894990042 |
| **A_52_P517762** | Slc44a4 | -0.100162584 | -0.41274639 | 0.212421222 | 9.264182143 | -0.788330283 | 0.461147686 | 0.853481264 | -4.895056156 |
| **A_55_P2111237** | Selenof | 0.11904044 | -0.25246837 | 0.490549251 | 14.29239222 | 0.788305525 | 0.461161087 | 0.853481264 | -4.895069017 |
| **A_52_P360330** | Map1b | -0.07315626 | -0.30147184 | 0.15515932 | 8.21200202 | -0.788288353 | 0.461170381 | 0.853481264 | -4.895077937 |
| **A_55_P2099695** | Msantd4 | 0.067712686 | -0.143631455 | 0.279056827 | 10.76572186 | 0.788222795 | 0.461205867 | 0.853481264 | -4.895111991 |
| **A_66_P129469** | NA | -0.035575513 | -0.146636701 | 0.075485676 | 8.747202577 | -0.788057696 | 0.461295241 | 0.853481264 | -4.895197744 |
| **A_51_P163106** | Bdh1 | -0.139777514 | -0.576149327 | 0.296594298 | 10.22805906 | -0.788042766 | 0.461303324 | 0.853481264 | -4.895205498 |
| **A_52_P618745** | Ermp1 | -0.056909447 | -0.234578057 | 0.120759162 | 9.23406483 | -0.788030138 | 0.461310161 | 0.853481264 | -4.895212056 |
| **A_55_P2236382** | D8Ertd158e | -0.198920618 | -0.820016215 | 0.42217498 | 9.658163071 | -0.787935216 | 0.461361552 | 0.853481264 | -4.895261351 |

| | | | | | | | | | |
|---|---|---|---|---|---|---|---|---|---|
| **A_55_P1985673** | Zbtb37 | -0.046084293 | -0.189981042 | 0.097812455 | 8.329236537 | -0.787900026 | 0.461380606 | 0.853481264 | -4.895279625 |
| **A_51_P475291** | Moap1 | -0.058554295 | -0.241427617 | 0.124319027 | 8.640684546 | -0.787730323 | 0.461472497 | 0.853523974 | -4.895367741 |
| **A_55_P2144641** | Sdk1 | 0.064971779 | -0.137966535 | 0.267910093 | 9.700597034 | 0.787643832 | 0.461519336 | 0.853523974 | -4.895412645 |
| **A_55_P2111855** | Gale | -0.112380275 | -0.46342694 | 0.23866639 | 13.11017397 | -0.787579499 | 0.461554177 | 0.853523974 | -4.895446044 |
| **A_51_P114854** | Clns1a | -0.052838935 | -0.21791292 | 0.112235049 | 9.391020412 | -0.787489164 | 0.461603104 | 0.853523974 | -4.874786687 |
| **A_52_P592111** | Brf1 | -0.049984592 | -0.206153918 | 0.106184733 | 8.943945258 | -0.787425699 | 0.46163748 | 0.853523974 | -4.895525879 |
| **A_51_P507787** | Sergef | -0.049733373 | -0.205129155 | 0.105662408 | 8.125810516 | -0.787368173 | 0.46166864 | 0.853523974 | -4.895555737 |
| **A_52_P494930** | Fnip2 | 0.05277864 | -0.112186256 | 0.217743536 | 10.43686779 | 0.787110704 | 0.461808123 | 0.853677614 | -4.895689353 |
| **A_51_P129866** | Atxn1l | -0.047003239 | -0.193938363 | 0.099931884 | 11.05824823 | -0.786993832 | 0.461871448 | 0.853677614 | -4.895749995 |
| **A_55_P2017645** | Tap2 | -0.166807776 | -0.688286798 | 0.354671245 | 11.13244576 | -0.786952671 | 0.461893751 | 0.853677614 | -4.895771351 |
| **A_55_P1989798** | NA | 0.117427756 | -0.249707934 | 0.484563445 | 14.40761449 | 0.786888719 | 0.461928406 | 0.853677614 | -4.895804529 |
| **A_52_P69558** | Gm8221 | 0.049070471 | -0.104365328 | 0.20250627 | 7.784583066 | 0.786796989 | 0.461978117 | 0.853679602 | -4.895852116 |
| **A_55_P2031939** | Aqp4 | 0.036012033 | -0.076602527 | 0.148626593 | 7.78952596 | 0.786723743 | 0.462017813 | 0.853679602 | -4.895890111 |
| **A_55_P1982762** | Hsd3b7 | -0.053340543 | -0.220192351 | 0.113511265 | 11.65121107 | -0.786494476 | 0.462142083 | 0.853764465 | -4.896009021 |
| **A_55_P2049856** | Araf | 0.092854866 | -0.19761817 | 0.383327902 | 11.18963383 | 0.78644437 | 0.462169245 | 0.853764465 | -4.896035005 |
| **A_66_P130654** | Abcg4 | 0.040023611 | -0.085194051 | 0.165241273 | 7.822078307 | 0.786357148 | 0.462216531 | 0.853764465 | -4.896080235 |
| **A_51_P512561** | Zfp68 | -0.050107318 | -0.206881599 | 0.106666963 | 9.839443451 | -0.786313079 | 0.462240423 | 0.853764465 | -4.896103085 |
| **A_55_P2167035** | Zfand1 | -0.039221753 | -0.161964536 | 0.08352103 | 8.269729599 | -0.786140556 | 0.462333965 | 0.853855648 | -4.896192533 |
| **A_51_P401730** | Ints14 | 0.058070049 | -0.123719774 | 0.239859872 | 10.42246301 | 0.785871957 | 0.462479627 | 0.85394207 | -4.896331764 |
| **A_55_P1967022** | Fgfr1 | -0.278956095 | -1.152246098 | 0.594333908 | 9.664945056 | -0.785862435 | 0.462484791 | 0.85394207 | -4.896336699 |
| **A_51_P431996** | Zfp367 | -0.044241917 | -0.182753346 | 0.094269513 | 8.099542661 | -0.785809877 | 0.462513298 | 0.85394207 | -4.896363939 |
| **A_52_P154880** | Ing3 | 0.074594541 | -0.158988191 | 0.308177274 | 8.637754069 | 0.78566151 | 0.462593776 | 0.854009082 | -4.896440827 |
| **A_55_P2148641** | Rab4a | 0.056841525 | -0.121200067 | 0.234883116 | 8.106478418 | 0.785440718 | 0.462713557 | 0.854148633 | -4.896555228 |
| **A_55_P2234704** | NA | 0.037984801 | -0.0810084 | 0.156978001 | 7.712002136 | 0.785338461 | 0.46276904 | 0.854169478 | -4.896608204 |
| **A_51_P222283** | Mlycd | -0.069881914 | -0.288846077 | 0.14908225 | 11.29988774 | -0.78516496 | 0.462863189 | 0.854193832 | -4.896698076 |
| **A_52_P498985** | Zfp512 | -0.043491331 | -0.179767064 | 0.092784402 | 8.227526172 | -0.785151269 | 0.462870619 | 0.854193832 | -4.896705167 |
| **A_51_P152612** | Nol7 | -0.03560981 | -0.147221606 | 0.076001985 | 8.25916122 | -0.784926021 | 0.46299287 | 0.85423093 | -4.896821821 |
| **A_55_P1965772** | Atp2a3 | 0.069518797 | -0.148406676 | 0.287444269 | 8.27901779 | 0.784807986 | 0.463056941 | 0.85423093 | -4.89688294 |
| **A_55_P2077546** | NA | -0.072929915 | -0.301572134 | 0.155712303 | 8.67465051 | -0.78472673 | 0.463101051 | 0.85423093 | -4.896925011 |
| **A_51_P329251** | Chchd1 | 0.085723601 | -0.183032931 | 0.354480133 | 13.47024251 | 0.784712444 | 0.463108807 | 0.85423093 | -4.896932407 |
| **A_52_P297075** | Heatr5a | -0.055864134 | -0.231012479 | 0.119284212 | 10.60789523 | -0.784686522 | 0.46312288 | 0.85423093 | -4.896945828 |
| **A_65_P12977** | Dnaja2 | -0.071396061 | -0.295277125 | 0.152485004 | 12.08159315 | -0.784559797 | 0.463191683 | 0.85423093 | -4.897011432 |
| **A_51_P272586** | Dym | -0.048984716 | -0.202603992 | 0.10463456 | 9.401586409 | -0.784483913 | 0.463232885 | 0.85423093 | -4.897050712 |
| **A_51_P231687** | Ik | 0.085945875 | -0.183593223 | 0.355484972 | 13.62535992 | 0.784462936 | 0.463244276 | 0.85423093 | -4.89706157 |
| **A_52_P73559** | NA | 0.061496975 | -0.131400278 | 0.254394229 | 8.566785026 | 0.784326476 | 0.463318377 | 0.854286074 | -4.897132198 |
| **A_51_P406008** | Hgh1 | -0.046156816 | -0.190960204 | 0.098646573 | 8.122600337 | -0.784198986 | 0.463387616 | 0.854332243 | -4.897198176 |
| **A_65_P08521** | Ssrp1 | 0.103146981 | -0.220497107 | 0.426791069 | 9.305523421 | 0.784075721 | 0.463454567 | 0.854374184 | -4.89726196 |
| **A_51_P144770** | 2410002F23Rik | 0.136704685 | -0.292380938 | 0.565790307 | 11.36297885 | 0.783806025 | 0.463601075 | 0.854496961 | -4.897401489 |
| **A_55_P1962035** | Coa5 | -0.076803585 | -0.317882061 | 0.164274891 | 9.713988458 | -0.783776466 | 0.463617135 | 0.854496961 | -4.897416779 |
| **A_51_P144160** | Colec10 | 0.314966135 | -0.673860384 | 1.303792653 | 8.624368921 | 0.783633365 | 0.463694887 | 0.854496961 | -4.897490796 |
| **A_55_P1958250** | Rnf114 | 0.035571224 | -0.076116628 | 0.147259077 | 13.40139902 | 0.783541562 | 0.463744773 | 0.854496961 | -4.897538275 |
| **A_55_P2249897** | NA | -0.15112971 | -0.625787036 | 0.323527616 | 8.726440439 | -0.783319293 | 0.463865568 | 0.854496961 | -4.897653212 |
| **A_55_P1959263** | Dnpep | -0.096069746 | -0.397847704 | 0.205708213 | 13.78967875 | -0.783192081 | 0.463934713 | 0.854496961 | -4.897718983 |
| **A_55_P2093483** | NA | 0.079974409 | -0.171255437 | 0.331204256 | 15.56828682 | 0.783157248 | 0.463953647 | 0.854496961 | -4.897736991 |
| **A_52_P216672** | Klk8 | 0.426185407 | -0.912669717 | 1.765040531 | 9.173591649 | 0.783130526 | 0.463968173 | 0.854496961 | -4.897750805 |
| **A_55_P2090798** | Mtus1 | -0.091775544 | -0.380089399 | 0.19653831 | 11.20560501 | -0.783124155 | 0.463971637 | 0.854496961 | -4.897754099 |
| **A_55_P1964960** | Il33 | 0.043534107 | -0.09324188 | 0.180310095 | 7.781445016 | 0.783049015 | 0.464012485 | 0.854496961 | -4.897792942 |
| **A_55_P2072453** | 1-Mar | 0.298265462 | -0.638885642 | 1.235416566 | 9.856193417 | 0.783001347 | 0.464038399 | 0.854496961 | -4.897817582 |
| **A_51_P240453** | Nusap1 | 0.096872222 | -0.20760127 | 0.401345714 | 8.288897966 | 0.782742541 | 0.464179116 | 0.854496961 | -4.897951342 |
| **A_51_P122035** | Nipa1 | 0.064858428 | -0.138997224 | 0.26871408 | 9.439250268 | 0.782731523 | 0.464185107 | 0.854496961 | -4.897957035 |

| | | | | | | | | | |
|---|---|---|---|---|---|---|---|---|---|
| A_52_P605812 | Ptrh1 | 0.111037988 | -0.237991637 | 0.460067612 | 11.03581642 | 0.782669582 | 0.464218791 | 0.854496961 | -4.897989043 |
| A_55_P2083649 | Alas1 | -0.114989316 | -0.514058233 | 0.284079602 | 13.06062473 | -0.810796932 | 0.464402236 | 0.854496961 | -4.744486597 |
| A_55_P2001078 | Mmel1 | -0.038836047 | -0.160982369 | 0.083310274 | 7.721579795 | -0.78221077 | 0.464468343 | 0.854496961 | -4.898226076 |
| A_51_P345649 | Pdgfra | -0.173918603 | -0.720927545 | 0.373090338 | 12.80634671 | -0.78220534 | 0.464471298 | 0.854496961 | -4.889748851 |
| A_55_P2254968 | BB212172 | 0.03582393 | -0.076856103 | 0.148503963 | 8.5663031 | 0.782159688 | 0.464496134 | 0.854496961 | -4.89825246 |
| A_55_P2164070 | Grip1 | -0.036108182 | -0.149683638 | 0.077467274 | 7.803522407 | -0.782150444 | 0.464501163 | 0.854496961 | -4.898257234 |
| A_55_P2053958 | Rabep2 | -0.075928494 | -0.314785401 | 0.162928414 | 10.23241108 | -0.782052919 | 0.464554223 | 0.854496961 | -4.898307602 |
| A_55_P2025765 | Adam8 | 0.1225525 | -0.262996947 | 0.508101947 | 10.07150367 | 0.782008102 | 0.464578607 | 0.854496961 | -4.898330746 |
| A_66_P104309 | Myl2 | -0.143237604 | -0.593864096 | 0.307388887 | 7.912904363 | -0.782004932 | 0.464580332 | 0.854496961 | -4.898332383 |
| A_51_P484289 | Slc7a6 | 0.115164012 | -0.247352537 | 0.477680561 | 9.356278353 | 0.781552418 | 0.464826595 | 0.854496961 | -4.898566015 |
| A_55_P2001976 | Rhox12 | -0.156919371 | -0.650966576 | 0.337127834 | 13.99068019 | -0.781406961 | 0.464905774 | 0.854496961 | -4.898641094 |
| A_52_P1037027 | Apela | 0.250564137 | -0.538388531 | 1.039516805 | 9.613285262 | 0.781334722 | 0.4649451 | 0.854496961 | -4.898678376 |
| A_52_P322096 | Patl2 | 0.132472138 | -0.284699329 | 0.549643605 | 9.321585942 | 0.781230361 | 0.465001917 | 0.854496961 | -4.898732232 |
| A_51_P102503 | Metap1 | 0.069429959 | -0.149215231 | 0.288075149 | 11.57491096 | 0.781225018 | 0.465004827 | 0.854496961 | -4.89873499 |
| A_55_P1994339 | Dhrs7c | -0.034914473 | -0.144871954 | 0.075043008 | 7.674831202 | -0.781177765 | 0.465030554 | 0.854496961 | -4.898759373 |
| A_55_P2097736 | Patz1 | -0.035465385 | -0.147174163 | 0.076243393 | 7.848906506 | -0.781063856 | 0.465092579 | 0.854496961 | -4.898818148 |
| A_55_P2033282 | Cept1 | -0.067202999 | -0.278886252 | 0.144480254 | 12.14950705 | -0.781036479 | 0.465107487 | 0.854496961 | -4.898832272 |
| A_51_P221677 | Trim62 | -0.075358655 | -0.312806674 | 0.162089364 | 8.892258095 | -0.780789119 | 0.465242199 | 0.854496961 | -4.898959879 |
| A_55_P2117928 | Mxd4 | 0.058747726 | -0.126397309 | 0.243892761 | 10.31570525 | 0.78063524 | 0.465326015 | 0.854496961 | -4.899039247 |
| A_55_P2177426 | Abcb8 | -0.040443941 | -0.167923965 | 0.087036083 | 8.03149364 | -0.780513631 | 0.465392262 | 0.854496961 | -4.899101962 |
| A_55_P2050652 | Ikbkg | -0.069649738 | -0.289234317 | 0.14993484 | 8.651267349 | -0.780345295 | 0.465483975 | 0.854496961 | -4.899188762 |
| A_51_P303160 | Arg1 | 0.10774347 | -0.231966351 | 0.447453291 | 7.972052206 | 0.780282777 | 0.465518039 | 0.854496961 | -4.899220996 |
| A_66_P116888 | Arid1a | -0.046080553 | -0.191377064 | 0.099215958 | 8.579405636 | -0.780246193 | 0.465537973 | 0.854496961 | -4.899239857 |
| A_51_P499530 | Ankrd10 | -0.11813351 | -0.490620139 | 0.254353119 | 8.868707452 | -0.780246056 | 0.465538048 | 0.854496961 | -4.899239927 |
| A_55_P1960556 | Acox1 | -0.076839211 | -0.319125865 | 0.165447444 | 13.49927987 | -0.780229854 | 0.465546876 | 0.854496961 | -4.89924828 |
| A_66_P129444 | Pnkp | 0.035200553 | -0.075807804 | 0.146208911 | 8.491166067 | 0.780122805 | 0.465605211 | 0.854496961 | -4.899303465 |
| A_55_P2046744 | Cspg5 | 0.066153967 | -0.142471626 | 0.274779561 | 8.669992714 | 0.780112921 | 0.465610597 | 0.854496961 | -4.899308561 |
| A_55_P2167999 | Ldlr | -0.164825262 | -0.684634363 | 0.35498384 | 11.63110231 | -0.780097808 | 0.465618833 | 0.854496961 | -4.899316351 |
| A_55_P2025443 | Oprm1 | -0.160201673 | -0.665445641 | 0.345042296 | 12.71382108 | -0.780072687 | 0.465632524 | 0.854496961 | -4.8993293 |
| A_55_P2083224 | Pum2 | -0.065287094 | -0.271194042 | 0.140619855 | 10.87674723 | -0.780055486 | 0.465641898 | 0.854496961 | -4.899338167 |
| A_51_P513032 | Trps1 | -0.070768494 | -0.293969574 | 0.152432586 | 8.123229756 | -0.780032728 | 0.465654301 | 0.854496961 | -4.899349897 |
| A_52_P39157 | Bmi1 | -0.053131043 | -0.220704564 | 0.114442479 | 9.134536598 | -0.780031438 | 0.465655004 | 0.854496961 | -4.899350562 |
| A_55_P2016959 | Nme2 | -0.068572399 | -0.284894309 | 0.147749511 | 15.825966 | -0.779862431 | 0.465747119 | 0.854496961 | -4.899437669 |
| A_55_P1956497 | Serpinf2 | 0.386393734 | -0.832775157 | 1.605562624 | 8.895248785 | 0.77971406 | 0.465827997 | 0.854496961 | -4.899514128 |
| A_51_P341379 | Acad9 | -0.044255429 | -0.183905314 | 0.095394456 | 9.707628482 | -0.779641836 | 0.46586737 | 0.854496961 | -4.899551343 |
| A_55_P2000182 | Slc5a6 | -0.129949809 | -0.540060821 | 0.280161202 | 14.11511484 | -0.779548929 | 0.465918023 | 0.854496961 | -4.899599212 |
| A_55_P2110032 | Akap7 | 0.071477637 | -0.154102605 | 0.297057879 | 9.960044038 | 0.779539799 | 0.465923 | 0.854496961 | -4.899603916 |
| A_51_P278103 | Stambp | -0.04512774 | -0.187556611 | 0.097301131 | 10.37723253 | -0.779497479 | 0.465946075 | 0.854496961 | -4.899625719 |
| A_51_P160083 | 1500035N22Rik | -0.033408993 | -0.138852912 | 0.072034926 | 7.769251469 | -0.779490828 | 0.465949701 | 0.854496961 | -4.899629145 |
| A_51_P477902 | Kdm5c | 0.117371513 | -0.253094714 | 0.48783774 | 11.78557213 | 0.779440995 | 0.465976873 | 0.854496961 | -4.899654817 |
| A_55_P2035122 | NA | -0.074014762 | -0.307645374 | 0.15961585 | 9.484061792 | -0.779395266 | 0.466001807 | 0.854496961 | -4.899678374 |
| A_55_P1983468 | Cflar | 0.035450349 | -0.076453895 | 0.147354592 | 8.38928019 | 0.779368983 | 0.466016139 | 0.854496961 | -4.899691913 |
| A_51_P400016 | Kalrn | 0.088454511 | -0.19077723 | 0.367686251 | 9.78653913 | 0.779335326 | 0.466034492 | 0.854496961 | -4.89970925 |
| A_52_P22781 | Zfp866 | -0.056152003 | -0.233414447 | 0.12111044 | 9.172479903 | -0.779323347 | 0.466041024 | 0.854496961 | -4.899715421 |
| A_55_P2014332 | Cntln | -0.052575641 | -0.218591824 | 0.113440543 | 8.305617208 | -0.779118135 | 0.46615294 | 0.854561985 | -4.899821115 |
| A_55_P2089530 | Fkbp10 | 0.099461719 | -0.214614085 | 0.413537523 | 11.43910269 | 0.779095408 | 0.466165336 | 0.854561985 | -4.899832819 |
| A_55_P2068177 | NA | -0.034791656 | -0.14467574 | 0.075092428 | 11.39216897 | -0.778949803 | 0.466244756 | 0.854561985 | -4.899907798 |
| A_66_P106836 | Kmo | 0.037963557 | -0.081940908 | 0.157868022 | 7.821632398 | 0.778934077 | 0.466253334 | 0.854561985 | -4.899915895 |
| A_51_P231630 | Zfp239 | 0.041836705 | -0.09032991 | 0.174003319 | 8.605590072 | 0.778762204 | 0.466347096 | 0.854652795 | -4.900004387 |
| A_52_P532456 | Plagl1 | 0.162173806 | -0.35032642 | 0.674674033 | 15.65721845 | 0.77849497 | 0.466492907 | 0.854838965 | -4.900141947 |

| | | | | | | | | | |
|---|---|---|---|---|---|---|---|---|---|
| **A_52_P568781** | Sec63 | -0.090110793 | -0.375018076 | 0.194796489 | 8.509887354 | -0.778112562 | 0.466701617 | 0.855004311 | -4.900338733 |
| **A_51_P438527** | Cyb5r1 | 0.050655452 | -0.109506681 | 0.210817585 | 10.74041398 | 0.77810017 | 0.466708381 | 0.855004311 | -4.900345109 |
| **A_55_P2119478** | Ndufv2 | 0.084649252 | -0.183000028 | 0.352298532 | 13.1554813 | 0.778083508 | 0.466717476 | 0.855004311 | -4.900353681 |
| **A_55_P2028491** | Hmgn2 | 0.122562939 | -0.265003068 | 0.510128946 | 10.21380513 | 0.778005472 | 0.466760075 | 0.855004311 | -4.900393828 |
| **A_55_P2133624** | NA | 0.085508407 | -0.184982346 | 0.35599916 | 12.34430535 | 0.777724096 | 0.466913699 | 0.85514429 | -4.900538563 |
| **A_52_P631499** | NA | 0.158959652 | -0.343894503 | 0.661813808 | 9.497992591 | 0.777703447 | 0.466924975 | 0.85514429 | -4.900549183 |
| **A_55_P2168663** | Atp5k | 0.072214587 | -0.156279914 | 0.300709087 | 14.04122793 | 0.777532125 | 0.467018531 | 0.855234599 | -4.900637287 |
| **A_55_P2066309** | Slc39a8 | -0.129332483 | -0.538622087 | 0.279957122 | 10.36388557 | -0.777402733 | 0.467089199 | 0.855241433 | -4.900703819 |
| **A_52_P131672** | Rbm7 | 0.077404271 | -0.167564134 | 0.322372677 | 10.18398664 | 0.777363264 | 0.467110756 | 0.855241433 | -4.900724111 |
| **A_55_P2117243** | Npm1 | -0.052691462 | -0.219474341 | 0.114091417 | 11.31261289 | -0.777245012 | 0.467175349 | 0.855278682 | -4.900784906 |
| **A_52_P316933** | Sh3bgrl2 | -0.073344492 | -0.305548349 | 0.158859365 | 11.50045552 | -0.777082693 | 0.467264022 | 0.855289239 | -4.900868344 |
| **A_55_P2081050** | Kcnj13 | 0.20897135 | -0.452626972 | 0.870569672 | 10.25443357 | 0.777072458 | 0.467269614 | 0.855289239 | -4.900873604 |
| **A_55_P1995438** | NA | -0.087167522 | -0.363238925 | 0.18890388 | 14.78617002 | -0.776787884 | 0.467425104 | 0.855492835 | -4.901019853 |
| **A_55_P2039180** | Zswim8 | -0.096630352 | -0.402840106 | 0.209579401 | 12.26381941 | -0.776360991 | 0.467658424 | 0.855567603 | -4.901239168 |
| **A_55_P2065671** | Ccnb1 | 0.12367973 | -0.268255385 | 0.515614846 | 8.476953787 | 0.776342778 | 0.46766838 | 0.855567603 | -4.901248523 |
| **A_52_P322181** | Adrb1 | 0.111653063 | -0.242196202 | 0.465502328 | 10.32553514 | 0.776285561 | 0.46769966 | 0.855567603 | -4.901277911 |
| **A_55_P2145800** | NA | -0.093684305 | -0.390596992 | 0.203228381 | 8.605737598 | -0.776260073 | 0.467713594 | 0.855567603 | -4.901291001 |
| **A_55_P2066399** | Fcho2 | -0.057608446 | -0.240205714 | 0.124988822 | 8.465954058 | -0.776177502 | 0.467758737 | 0.855567603 | -4.901333408 |
| **A_51_P382688** | Zmynd19 | 0.040879958 | -0.088702429 | 0.170462345 | 9.614759897 | 0.776128498 | 0.467785529 | 0.855567603 | -4.901358573 |
| **A_55_P2115257** | Fer | 0.110419789 | -0.239608197 | 0.460447776 | 10.55141981 | 0.776092179 | 0.467805387 | 0.855567603 | -4.901377224 |
| **A_52_P89425** | Pcnt | -0.058366156 | -0.243391819 | 0.126659507 | 10.04897597 | -0.776065339 | 0.467820063 | 0.855567603 | -4.901391006 |
| **A_55_P2083894** | NA | -0.124052725 | -0.517448894 | 0.269343444 | 10.20958635 | -0.775792088 | 0.467969489 | 0.855680521 | -4.901531301 |
| **A_66_P130960** | Psmd11 | -0.065372397 | -0.272692977 | 0.141948182 | 9.809386904 | -0.775748884 | 0.467993118 | 0.855680521 | -4.90155348 |
| **A_55_P1952877** | Snrpf | 0.06818999 | -0.148086729 | 0.284466709 | 11.8615948 | 0.7756754 | 0.46803331 | 0.855680521 | -4.901591201 |
| **A_51_P454993** | Tmcc2 | -0.036871364 | -0.153830673 | 0.080087945 | 7.802423437 | -0.775574597 | 0.468088447 | 0.855680521 | -4.901642941 |
| **A_55_P1954092** | NA | -0.280492247 | -1.170269918 | 0.609285423 | 13.21576673 | -0.775547714 | 0.468103152 | 0.855680521 | -4.901656739 |
| **A_55_P2082185** | NA | 0.103528868 | -0.224932034 | 0.43198977 | 12.48789901 | 0.775437764 | 0.4681633 | 0.855709544 | -4.901713167 |
| **A_55_P2114088** | Zfp385c | 0.101541459 | -0.220809165 | 0.423892083 | 9.432635368 | 0.774968494 | 0.468420072 | 0.856097916 | -4.901953936 |
| **A_55_P2009762** | Mapkapk3 | -0.067563737 | -0.28209876 | 0.146971287 | 8.420588183 | -0.774791097 | 0.468517164 | 0.85619441 | -4.902044926 |
| **A_55_P1975445** | NA | 0.086231855 | -0.18770278 | 0.360166491 | 12.66346019 | 0.774443871 | 0.468707249 | 0.856426411 | -4.902222977 |
| **A_55_P2058794** | Cmip | -0.185048956 | -0.772989788 | 0.402891876 | 13.44290037 | -0.774323037 | 0.468773411 | 0.856426411 | -4.902284926 |
| **A_55_P2026945** | Zranb2 | -0.041121707 | -0.171788215 | 0.089544801 | 10.79023486 | -0.774240731 | 0.46881848 | 0.856426411 | -4.902327117 |
| **A_51_P388298** | Mmachc | -0.07586608 | -0.316974955 | 0.165242794 | 8.999650295 | -0.774111667 | 0.46888916 | 0.856426411 | -4.902393271 |
| **A_55_P2171977** | NA | -0.111061169 | -0.4640349 | 0.241912563 | 13.38423089 | -0.774085649 | 0.46890341 | 0.856426411 | -4.902406606 |
| **A_55_P2171196** | Gm3411 | -0.146896488 | -0.61380338 | 0.320010403 | 8.313261277 | -0.774016632 | 0.468941209 | 0.856426411 | -4.902441977 |
| **A_52_P625624** | Stau2 | -0.041808412 | -0.174700047 | 0.091083223 | 8.327862865 | -0.773989722 | 0.468955949 | 0.856426411 | -4.902455768 |
| **A_55_P2136657** | Thbs3 | 0.113892775 | -0.248175323 | 0.475960873 | 8.838910486 | 0.773882592 | 0.469014629 | 0.856426411 | -4.902510667 |
| **A_65_P09443** | Psma4 | -0.072221829 | -0.301842594 | 0.157398936 | 12.18140138 | -0.77379601 | 0.469062057 | 0.856426411 | -4.902555031 |
| **A_51_P394154** | Ddx51 | -0.070978718 | -0.296660541 | 0.154703106 | 9.384767098 | -0.77375012 | 0.469087196 | 0.856426411 | -4.902578544 |
| **A_55_P2144512** | NA | 0.035783167 | -0.07804477 | 0.149611104 | 8.344587606 | 0.773390928 | 0.469284001 | 0.856704803 | -4.902762546 |
| **A_55_P2036026** | Robo4 | -0.080613776 | -0.337104736 | 0.175877185 | 11.48180241 | -0.773225847 | 0.469374469 | 0.85676238 | -4.90284709 |
| **A_51_P249286** | Rgs16 | -0.194966073 | -0.815338241 | 0.425406094 | 9.439075543 | -0.773171618 | 0.469404191 | 0.85676238 | -4.90287486 |
| **A_55_P1973723** | Etfdh | -0.097239971 | -0.406761367 | 0.212281425 | 11.7242063 | -0.772900002 | 0.469553077 | 0.85679634 | -4.903013928 |
| **A_52_P576778** | Tmem30a | 0.067135875 | -0.168812918 | 0.303084668 | 11.92739041 | 0.800643905 | 0.469611373 | 0.85679634 | -4.795367983 |
| **A_55_P1999384** | NA | -0.077007799 | -0.322163903 | 0.168148306 | 11.19894344 | -0.772789408 | 0.469613709 | 0.85679634 | -4.903070543 |
| **A_51_P194853** | Sec14l4 | -0.034444323 | -0.14411015 | 0.075221505 | 7.728922408 | -0.772708146 | 0.469658262 | 0.85679634 | -4.903112137 |
| **A_55_P1985554** | B4galt4 | -0.08310664 | -0.347762058 | 0.181548777 | 9.51994725 | -0.772545559 | 0.469747414 | 0.85679634 | -4.90319535 |
| **A_55_P2119302** | Cspp1 | -0.106823372 | -0.447005834 | 0.233359091 | 8.589517493 | -0.772544534 | 0.469747976 | 0.85679634 | -4.903195874 |
| **A_51_P166244** | Osbpl1a | -0.119386045 | -0.499632699 | 0.260860609 | 11.10618581 | -0.772426787 | 0.469812548 | 0.85679634 | -4.903256129 |
| **A_52_P479269** | Sdc1 | 0.116583663 | -0.254763208 | 0.487930533 | 10.15513117 | 0.772373011 | 0.469842041 | 0.85679634 | -4.903283646 |

| | | | | | | | | | |
|---|---|---|---|---|---|---|---|---|---|
| **A_55_P1955562** | Extl3 | -0.104282997 | -0.436456673 | 0.227890679 | 11.81930003 | -0.772355872 | 0.469851441 | 0.85679634 | -4.903292415 |
| **A_55_P2021049** | Ugp2 | -0.06034465 | -0.252567704 | 0.131878405 | 12.32366054 | -0.772329205 | 0.469866067 | 0.85679634 | -4.90330606 |
| **A_55_P2059134** | Smarca5 | -0.047936105 | -0.200655943 | 0.104783733 | 10.38288538 | -0.77221183 | 0.469930446 | 0.856832902 | -4.903366112 |
| **A_51_P264695** | Crym | -0.077102177 | -0.322856643 | 0.16865229 | 8.739008459 | -0.771852618 | 0.47012751 | 0.85699928 | -4.903549851 |
| **A_55_P2180744** | Clstn3 | -0.04127394 | -0.172841775 | 0.090293895 | 9.427412531 | -0.771783275 | 0.470165559 | 0.85699928 | -4.903585314 |
| **A_51_P160113** | Copb1 | -0.048705884 | -0.203969759 | 0.106557992 | 12.22446721 | -0.771756272 | 0.470180376 | 0.85699928 | -4.903599122 |
| **A_55_P2052744** | NA | 0.037516628 | -0.082083582 | 0.157116839 | 10.59413198 | 0.771722246 | 0.470199046 | 0.85699928 | -4.903616522 |
| **A_55_P2134581** | Papola | 0.100726834 | -0.22045326 | 0.421906927 | 9.908780103 | 0.771552929 | 0.470291964 | 0.857087816 | -4.903703096 |
| **A_55_P2095488** | Syncrip | -0.08497474 | -0.355979879 | 0.186030399 | 9.161801523 | -0.771403282 | 0.470374098 | 0.857156683 | -4.9037796 |
| **A_55_P2107302** | Aurkaip1 | -0.148448686 | -0.621992273 | 0.3250949 | 14.88369042 | -0.771232926 | 0.47046761 | 0.85724627 | -4.903866678 |
| **A_51_P258620** | Rab4b | -0.038791276 | -0.162555196 | 0.084972644 | 8.221370331 | -0.771097284 | 0.470542076 | 0.857283347 | -4.903936002 |
| **A_55_P2313163** | D10Bwg1070e | -0.224480379 | -0.940790985 | 0.491830226 | 10.07326214 | -0.770985428 | 0.47060349 | 0.857283347 | -4.903993163 |
| **A_66_P134775** | Pter | -0.038530731 | -0.161486305 | 0.084424843 | 7.843412824 | -0.770953512 | 0.470621015 | 0.857283347 | -4.904009471 |
| **A_55_P1953391** | Rbfox1 | 0.033612273 | -0.073665059 | 0.140889606 | 7.731079016 | 0.77083084 | 0.470688375 | 0.857325255 | -4.904072149 |
| **A_52_P576442** | Dap | 0.115132432 | -0.252435591 | 0.482700455 | 12.28344731 | 0.770600205 | 0.470815038 | 0.857447663 | -4.904189971 |
| **A_55_P2048744** | Eif2ak3 | 0.084751811 | -0.185842746 | 0.355346367 | 11.28482525 | 0.770546928 | 0.470844301 | 0.857447663 | -4.904217184 |
| **A_51_P417858** | Zfp825 | -0.036598362 | -0.153472449 | 0.080275724 | 8.207404041 | -0.770393457 | 0.470928603 | 0.857517952 | -4.904295567 |
| **A_51_P469578** | Psma1 | -0.080294869 | -0.336736497 | 0.176146758 | 13.0332842 | -0.770315141 | 0.470971627 | 0.857517952 | -4.904335561 |
| **A_51_P341789** | Sugp1 | -0.114857021 | -0.481780355 | 0.252066313 | 11.49229484 | -0.770107547 | 0.471085683 | 0.857644831 | -4.90444156 |
| **A_52_P229320** | Rptor | -0.052790481 | -0.221506667 | 0.115925705 | 10.72954849 | -0.769782497 | 0.471264311 | 0.857889233 | -4.90460749 |
| **A_55_P2117164** | Tmem106b | 0.047587873 | -0.104527537 | 0.199703282 | 9.753332713 | 0.769648176 | 0.47133814 | 0.857936762 | -4.904676042 |
| **A_55_P2051840** | NA | 0.045428868 | -0.09979934 | 0.190657075 | 8.903591919 | 0.769573492 | 0.471379192 | 0.857936762 | -4.904714153 |
| **A_55_P2090330** | Kcnmb4 | -0.033575539 | -0.14094449 | 0.073793411 | 7.71962935 | -0.76933139 | 0.471512291 | 0.858098208 | -4.904837681 |
| **A_52_P316437** | 1810013L24Rik | -0.054991738 | -0.2308995 | 0.120916024 | 9.204974399 | -0.769097861 | 0.471640701 | 0.858251093 | -4.904956807 |
| **A_52_P112651** | Lman2 | -0.097113146 | -0.407795972 | 0.21356968 | 9.678435759 | -0.769006373 | 0.471691014 | 0.858261848 | -4.905003469 |
| **A_55_P1989772** | Sqor | 0.228225786 | -0.502125879 | 0.95857745 | 10.06426336 | 0.768779599 | 0.471815743 | 0.85840799 | -4.905119113 |
| **A_55_P2096010** | NA | -0.054237544 | -0.22784132 | 0.119366232 | 13.38783113 | -0.768617047 | 0.471905162 | 0.858489872 | -4.905201991 |
| **A_55_P2017967** | Usf2 | -0.095968613 | -0.403219556 | 0.21128233 | 13.98805374 | -0.768431497 | 0.472007248 | 0.858594776 | -4.905296579 |
| **A_55_P2088425** | Tcn2 | -0.092209356 | -0.387474255 | 0.203055543 | 14.96229565 | -0.768302649 | 0.472078146 | 0.858596153 | -4.905362251 |
| **A_55_P2167416** | NA | -0.041972843 | -0.176380631 | 0.092434944 | 7.876710791 | -0.76826867 | 0.472096845 | 0.858596153 | -4.905379569 |
| **A_55_P1980336** | NA | 0.044583157 | -0.098202118 | 0.187368432 | 7.845121543 | 0.768168616 | 0.472151906 | 0.858606112 | -4.905430559 |
| **A_55_P2145557** | NA | 0.115533038 | -0.25451587 | 0.485581947 | 13.13924707 | 0.768097287 | 0.472191162 | 0.858606112 | -4.905466906 |
| **A_52_P88818** | Clybl | -0.040110615 | -0.168626838 | 0.088405607 | 8.212087678 | -0.767839652 | 0.472332972 | 0.858783183 | -4.905598169 |
| **A_55_P2413359** | NA | -0.09442644 | -0.397168632 | 0.208315752 | 8.262887118 | -0.767343515 | 0.472606144 | 0.859062819 | -4.905850855 |
| **A_52_P282700** | Zbtb33 | 0.038705518 | -0.085400905 | 0.162811941 | 8.592179539 | 0.767269245 | 0.472647046 | 0.859062819 | -4.905888671 |
| **A_52_P229316** | Galnt4 | -0.090290972 | -0.379853396 | 0.199271453 | 8.671158559 | -0.767134097 | 0.472721482 | 0.859062819 | -4.905957476 |
| **A_55_P2125774** | Ola1 | -0.067442292 | -0.283750336 | 0.148865753 | 11.03872891 | -0.767059071 | 0.472762808 | 0.859062819 | -4.905995669 |
| **A_52_P89064** | Dusp18 | -0.078039506 | -0.328385983 | 0.172306971 | 8.201116894 | -0.766906095 | 0.472847078 | 0.859062819 | -4.906073535 |
| **A_55_P2098321** | NA | -0.065588138 | -0.276001724 | 0.144825449 | 11.87243145 | -0.766868121 | 0.472867999 | 0.859062819 | -4.906092862 |
| **A_55_P2040763** | Sec62 | -0.058995645 | -0.248287796 | 0.130296507 | 13.41757028 | -0.766754808 | 0.472930428 | 0.859062819 | -4.906150529 |
| **A_55_P2244112** | Amotl1 | -0.062441544 | -0.262800551 | 0.137917464 | 8.746093063 | -0.766714933 | 0.472952399 | 0.859062819 | -4.90617082 |
| **A_65_P11603** | Ldb3 | -0.039830474 | -0.167639546 | 0.087978597 | 7.946068446 | -0.766695586 | 0.472963059 | 0.859062819 | -4.906180666 |
| **A_51_P497113** | Igf2bp1 | -0.07956136 | -0.334887072 | 0.175764351 | 9.661544763 | -0.766614115 | 0.473007951 | 0.859062819 | -4.906222121 |
| **A_55_P1958379** | Slit1 | -0.07511201 | -0.316194324 | 0.165970303 | 13.2352784 | -0.766501833 | 0.473069825 | 0.859062819 | -4.906279249 |
| **A_55_P2124233** | Vsig8 | 0.179413973 | -0.396543811 | 0.755371758 | 8.3434588 | 0.766363434 | 0.473146099 | 0.859062819 | -4.906349657 |
| **A_51_P489153** | Crot | 0.079677668 | -0.176106696 | 0.335462032 | 9.966545385 | 0.766358158 | 0.473149007 | 0.859062819 | -4.906352341 |
| **A_51_P434101** | Herpud1 | 0.085566238 | -0.189163789 | 0.360296266 | 13.0605237 | 0.766241152 | 0.473213498 | 0.859062819 | -4.906411857 |
| **A_51_P142923** | Chka | -0.047847711 | -0.20147768 | 0.105782258 | 8.403990728 | -0.766221595 | 0.473224278 | 0.859062819 | -4.906421805 |
| **A_55_P2045086** | NA | 0.071445728 | -0.157975485 | 0.300866941 | 11.91550524 | 0.766146562 | 0.473265639 | 0.859062819 | -4.906459967 |
| **A_55_P1965233** | Sh3tc1 | 0.125560425 | -0.277678613 | 0.528799462 | 12.29678328 | 0.766053974 | 0.47331668 | 0.859062819 | -4.906507054 |

| | | | | | | | | | |
|---|---|---|---|---|---|---|---|---|---|
| **A_55_P2149719** | NA | 0.076263483 | -0.168669632 | 0.321196599 | 13.99284927 | 0.766016801 | 0.473337173 | 0.859062819 | -4.906525958 |
| **A_55_P2054927** | Fam136a | 0.100954844 | -0.223333998 | 0.425243686 | 12.14651446 | 0.765886331 | 0.473409106 | 0.859062819 | -4.906592301 |
| **A_55_P2092983** | NA | 0.063093237 | -0.139609389 | 0.265795863 | 14.46774233 | 0.765759861 | 0.47347884 | 0.859062819 | -4.906656601 |
| **A_52_P463235** | Ankrd33b | -0.092528802 | -0.38981766 | 0.204760056 | 9.443146664 | -0.765715553 | 0.473503273 | 0.859062819 | -4.906679127 |
| **A_55_P2084706** | Acaca | 0.080469744 | -0.17809267 | 0.339032158 | 9.762364898 | 0.765660757 | 0.473533491 | 0.859062819 | -4.906706983 |
| **A_66_P102049** | Zfp1 | -0.084826421 | -0.357456733 | 0.187803891 | 10.59268282 | -0.765466462 | 0.473640646 | 0.859062819 | -4.906805743 |
| **A_52_P149835** | Irak1 | -0.04424086 | -0.186432402 | 0.097950681 | 8.45021094 | -0.765453766 | 0.473647648 | 0.859062819 | -4.906812195 |
| **A_55_P2359436** | A930026B05Rik | -0.08043846 | -0.338985117 | 0.178108197 | 8.236160562 | -0.765409737 | 0.473671934 | 0.859062819 | -4.906834572 |
| **A_55_P2096422** | Inhbb | -0.171008439 | -0.720694766 | 0.378677888 | 14.33882107 | -0.765370594 | 0.473693525 | 0.859062819 | -4.906854465 |
| **A_51_P229599** | Etnk1 | -0.077670438 | -0.327350779 | 0.172009904 | 8.808979311 | -0.7653156 | 0.473723859 | 0.859062819 | -4.906882412 |
| **A_55_P2064686** | Lrrc61 | -0.057124902 | -0.240799637 | 0.126549833 | 9.371138033 | -0.76514758 | 0.473816549 | 0.859062819 | -4.906967789 |
| **A_51_P396570** | Plod2 | -0.154826182 | -0.652645386 | 0.342993022 | 9.367941448 | -0.765141788 | 0.473819745 | 0.859062819 | -4.906970732 |
| **A_52_P284889** | Prkcz | 0.095448486 | -0.211451346 | 0.402348317 | 9.801631934 | 0.765141144 | 0.4738201 | 0.859062819 | -4.906971059 |
| **A_51_P176387** | Hook3 | -0.044450802 | -0.187471935 | 0.098570331 | 8.629122822 | -0.764625101 | 0.47410486 | 0.85938216 | -4.907233187 |
| **A_52_P312398** | Ssr3 | 0.090807776 | -0.201370579 | 0.38298613 | 10.5760822 | 0.764617375 | 0.474109124 | 0.85938216 | -4.907237111 |
| **A_51_P119031** | Naa30 | -0.04211224 | -0.177620597 | 0.093396118 | 9.921176143 | -0.764559735 | 0.474140939 | 0.85938216 | -4.907266381 |
| **A_51_P148355** | Zfp710 | 0.082310107 | -0.182584516 | 0.347204729 | 11.56774868 | 0.764450178 | 0.474201412 | 0.85938216 | -4.907322011 |
| **A_55_P2004119** | NA | -0.074067625 | -0.312480824 | 0.164345575 | 13.68877721 | -0.764306028 | 0.474280989 | 0.85938216 | -4.907395197 |
| **A_55_P1982107** | NA | 0.090232332 | -0.200269276 | 0.38073394 | 16.30636469 | 0.764157365 | 0.474363067 | 0.85938216 | -4.907470663 |
| **A_55_P2106975** | Clcn5 | 0.061816558 | -0.13720504 | 0.260838156 | 9.687039862 | 0.764141478 | 0.474371839 | 0.85938216 | -4.907478727 |
| **A_51_P501897** | Pitx2 | 0.120337958 | -0.26710839 | 0.507784306 | 8.639979378 | 0.764117649 | 0.474384997 | 0.85938216 | -4.907490823 |
| **A_55_P2132942** | Adal | -0.079742708 | -0.336553899 | 0.177068484 | 9.021913254 | -0.763917038 | 0.474495775 | 0.85938216 | -4.907592638 |
| **A_55_P2002117** | Kbtbd4 | 0.084700428 | -0.188106314 | 0.357507171 | 10.26810911 | 0.763835203 | 0.474540969 | 0.85938216 | -4.907634165 |
| **A_55_P1976470** | 2410018L13Rik | -0.061925621 | -0.261394564 | 0.137543322 | 8.897215892 | -0.763772907 | 0.474575375 | 0.85938216 | -4.907665776 |
| **A_55_P2129972** | NA | 0.112675868 | -0.250306558 | 0.475658293 | 11.42703195 | 0.763685379 | 0.47462372 | 0.85938216 | -4.907710186 |
| **A_55_P1955998** | Nlrp1a | -0.213652332 | -0.902029787 | 0.474725123 | 11.60911387 | -0.763572296 | 0.474686185 | 0.85938216 | -4.907767557 |
| **A_55_P1994388** | Lsm14a | -0.068511172 | -0.289264544 | 0.152242199 | 11.54155437 | -0.763524919 | 0.474712356 | 0.85938216 | -4.907791591 |
| **A_55_P2238079** | A830054O07Rik | -0.055298454 | -0.233497989 | 0.12290108 | 7.914220356 | -0.763441195 | 0.474758609 | 0.85938216 | -4.90783406 |
| **A_66_P140533** | NA | -0.090990796 | -0.384210104 | 0.202228511 | 8.389280027 | -0.763438514 | 0.47476009 | 0.85938216 | -4.907835421 |
| **A_55_P2031442** | Pithd1 | -0.040687286 | -0.171810697 | 0.090436125 | 12.32657887 | -0.763392079 | 0.474785744 | 0.85938216 | -4.907858974 |
| **A_51_P474169** | Proser2 | 0.06585425 | -0.146392845 | 0.278101346 | 11.97759334 | 0.763328042 | 0.474821124 | 0.85938216 | -4.907891453 |
| **A_51_P453475** | Slc1a5 | 0.117917295 | -0.262283886 | 0.498118476 | 9.978681496 | 0.763015235 | 0.474993977 | 0.85938216 | -4.908050079 |
| **A_55_P2404484** | 1700001L05Rik | -0.059380525 | -0.25090907 | 0.13214802 | 8.752412745 | -0.762745559 | 0.475143031 | 0.85938216 | -4.908186794 |
| **A_51_P240986** | Plekhg6 | 0.169217153 | -0.376635851 | 0.715070157 | 9.432311526 | 0.76267207 | 0.475183654 | 0.85938216 | -4.908224043 |
| **A_52_P796840** | Cfhr2 | 0.035100793 | -0.078134153 | 0.14833574 | 7.94185296 | 0.762615478 | 0.47521494 | 0.85938216 | -4.908252727 |
| **A_55_P1976127** | Ect2 | 0.120490706 | -0.268260683 | 0.509242095 | 8.640181637 | 0.762519159 | 0.47526819 | 0.85938216 | -4.908301541 |
| **A_51_P260548** | Glg1 | -0.180764341 | -0.763989845 | 0.402461163 | 13.38952291 | -0.762509778 | 0.475273376 | 0.85938216 | -4.908306295 |
| **A_55_P2105833** | Chchd5 | -0.048092824 | -0.203265368 | 0.107079721 | 10.22650657 | -0.76249071 | 0.475283919 | 0.85938216 | -4.908315958 |
| **A_55_P2070079** | Lgals9 | 0.046948477 | -0.104533552 | 0.198430505 | 10.94670099 | 0.762481932 | 0.475288772 | 0.85938216 | -4.908320407 |
| **A_52_P622442** | Pdcl3 | -0.055644932 | -0.235192723 | 0.123902859 | 11.08814993 | -0.762455873 | 0.47530318 | 0.85938216 | -4.908333612 |
| **A_52_P98778** | Ang4 | 0.082878489 | -0.184615236 | 0.350372214 | 8.443803396 | 0.762249923 | 0.47541706 | 0.85938216 | -4.908437966 |
| **A_51_P100174** | Mns1 | 0.051739044 | -0.115263377 | 0.218741466 | 7.986964247 | 0.762192724 | 0.475448692 | 0.85938216 | -4.891380515 |
| **A_55_P1975110** | Pnpt1 | -0.095659238 | -0.404437263 | 0.213118787 | 10.5147166 | -0.762166215 | 0.475463352 | 0.85938216 | -4.908480374 |
| **A_51_P499776** | Denr | -0.040905935 | -0.172949413 | 0.091137544 | 8.432699043 | -0.762146616 | 0.475474192 | 0.85938216 | -4.908490303 |
| **A_55_P2002578** | Ephx1 | 0.107937964 | -0.240573159 | 0.456449086 | 10.61381834 | 0.76195047 | 0.475582677 | 0.85938216 | -4.90858966 |
| **A_55_P2076303** | 8-Mar | -0.092372659 | -0.390633226 | 0.205887907 | 12.58994234 | -0.761932977 | 0.475592353 | 0.85938216 | -4.90859852 |
| **A_55_P1974650** | Npdc1 | 0.136427204 | -0.304093125 | 0.576947532 | 11.56495027 | 0.761910899 | 0.475604566 | 0.85938216 | -4.908609702 |
| **A_55_P2017075** | Zfp54 | 0.082452134 | -0.183803169 | 0.348707437 | 8.402690874 | 0.761855834 | 0.475635026 | 0.85938216 | -4.90863759 |
| **A_55_P2056774** | Hsd17b14 | 0.060707655 | -0.135345548 | 0.256760858 | 8.545419477 | 0.761795986 | 0.475668133 | 0.85938216 | -4.908667899 |
| **A_51_P375862** | Srpk3 | 0.033544269 | -0.074787151 | 0.14187569 | 7.659558218 | 0.761786132 | 0.475673584 | 0.85938216 | -4.908672889 |

| | | | | | | | | | |
|---|---|---|---|---|---|---|---|---|---|
| **A_55_P2030506** | Rp9 | -0.048198462 | -0.203860176 | 0.107463253 | 11.32004005 | -0.761764153 | 0.475685743 | 0.85938216 | -4.90868402 |
| **A_55_P2071389** | Erh | 0.125629752 | -0.280152667 | 0.531412172 | 13.36451202 | 0.761672788 | 0.475736289 | 0.859387233 | -4.908730285 |
| **A_55_P2036999** | Zic3 | 0.122669538 | -0.273590154 | 0.51892923 | 8.962161937 | 0.761598351 | 0.475777473 | 0.859387233 | -4.908767975 |
| **A_55_P2169529** | NA | -0.053215029 | -0.22520066 | 0.118770602 | 8.828864646 | -0.761221942 | 0.475985768 | 0.859549407 | -4.908958522 |
| **A_51_P284686** | Aak1 | 0.032400328 | -0.072316782 | 0.137117439 | 7.935065267 | 0.76120376 | 0.475995832 | 0.859549407 | -4.908967725 |
| **A_55_P2035504** | Nt5c3 | 0.076021395 | -0.169680811 | 0.321723601 | 10.80078528 | 0.761195028 | 0.476000665 | 0.859549407 | -4.908972144 |
| **A_55_P2132358** | NA | 0.108595115 | -0.242442245 | 0.459632474 | 10.24189268 | 0.761072644 | 0.476068404 | 0.859591422 | -4.909034081 |
| **A_55_P2079103** | Pan2 | -0.088617578 | -0.375167856 | 0.197932701 | 10.74619396 | -0.760831006 | 0.47620217 | 0.859752637 | -4.909156348 |
| **A_52_P173368** | NA | -0.034782407 | -0.147294773 | 0.07772996 | 8.016928634 | -0.760551336 | 0.476357022 | 0.859910876 | -4.909297824 |
| **A_51_P208361** | Ak3 | 0.065039705 | -0.145363741 | 0.275443152 | 11.09630947 | 0.760492397 | 0.476389661 | 0.859910876 | -4.909327634 |
| **A_55_P1957393** | NA | 0.07071448 | -0.158082629 | 0.299511589 | 13.4528506 | 0.760373514 | 0.4764555 | 0.859910876 | -4.909387757 |
| **A_55_P2003648** | Bag5 | -0.081996004 | -0.347311283 | 0.183319275 | 11.7927783 | -0.760325557 | 0.476482061 | 0.859910876 | -4.909412008 |
| **A_52_P37757** | Eps8l1 | -0.044977637 | -0.190537778 | 0.100582504 | 8.154792501 | -0.760192061 | 0.476556003 | 0.859910876 | -4.90947951 |
| **A_51_P329332** | Slc19a2 | -0.094240694 | -0.399230323 | 0.210748936 | 11.53943252 | -0.760190723 | 0.476556744 | 0.859910876 | -4.909480187 |
| **A_52_P123864** | Rab21 | -0.047068316 | -0.199418263 | 0.10528163 | 8.864992084 | -0.760073378 | 0.476621747 | 0.859947891 | -4.909539515 |
| **A_51_P224064** | Klhl38 | -0.032795259 | -0.138964636 | 0.073374118 | 7.830321826 | -0.759942892 | 0.476694037 | 0.859998043 | -4.909605478 |
| **A_55_P1960641** | Trpt1 | 0.071672089 | -0.160426469 | 0.303770648 | 10.67875613 | 0.759708148 | 0.476824105 | 0.860152415 | -4.909724125 |
| **A_52_P175242** | Irs1 | 0.057638389 | -0.12908362 | 0.244360397 | 8.615290047 | 0.759426038 | 0.476980451 | 0.860274973 | -4.909866676 |
| **A_55_P2014169** | Ppa2 | 0.102794848 | -0.230261757 | 0.435851453 | 11.20855125 | 0.759315843 | 0.477041531 | 0.860274973 | -4.909922346 |
| **A_55_P1996553** | Ttc17 | -0.109032815 | -0.462361331 | 0.244295701 | 10.86138877 | -0.759185141 | 0.477113985 | 0.860274973 | -4.909988369 |
| **A_55_P2111293** | 3110009E18Rik | 0.091124593 | -0.204231347 | 0.386480532 | 8.736533169 | 0.759030197 | 0.477199887 | 0.860274973 | -4.910066627 |
| **A_51_P324228** | Satb1 | -0.036129071 | -0.153240597 | 0.080982455 | 8.072961077 | -0.75897298 | 0.477231611 | 0.860274973 | -4.910095522 |
| **A_55_P2027052** | Prdm9 | -0.069422657 | -0.294468082 | 0.155622767 | 8.742765934 | -0.758927356 | 0.477256908 | 0.860274973 | -4.910118562 |
| **A_55_P1984366** | Cdadc1 | -0.070993494 | -0.30113909 | 0.159152101 | 9.279463865 | -0.758900866 | 0.477271597 | 0.860274973 | -4.910131939 |
| **A_51_P233788** | Dmap1 | 0.064891428 | -0.14548546 | 0.275268315 | 10.60878359 | 0.75885442 | 0.477297352 | 0.860274973 | -4.910155392 |
| **A_55_P2009940** | Xlr | 0.044977566 | -0.10092331 | 0.190878441 | 8.627723199 | 0.75841552 | 0.477540774 | 0.860274973 | -4.910376963 |
| **A_51_P152155** | Glt8d1 | -0.044529008 | -0.188977366 | 0.09991935 | 11.83337118 | -0.75840218 | 0.477548174 | 0.860274973 | -4.910383697 |
| **A_51_P163173** | Rbm12b1 | -0.038749228 | -0.164460545 | 0.08696209 | 8.356090039 | -0.7583293 | 0.477588603 | 0.860274973 | -4.910420479 |
| **A_51_P160344** | Cenpv | 0.101409256 | -0.227620938 | 0.430439451 | 10.53335765 | 0.758247535 | 0.477633965 | 0.860274973 | -4.910461742 |
| **A_55_P1986998** | Enoph1 | 0.100730807 | -0.226106211 | 0.427567825 | 11.5481923 | 0.758228734 | 0.477644396 | 0.860274973 | -4.91047123 |
| **A_55_P2135759** | NA | -0.038051625 | -0.16152132 | 0.08541807 | 16.15695756 | -0.758196883 | 0.477662067 | 0.860274973 | -4.910487303 |
| **A_51_P153423** | Fndc1 | 0.050410079 | -0.113167952 | 0.21398811 | 7.862907511 | 0.758161219 | 0.477681854 | 0.860274973 | -4.910505299 |
| **A_55_P1977633** | Irf2bpl | -0.067202413 | -0.285275628 | 0.150870802 | 10.95978215 | -0.758144004 | 0.477691405 | 0.860274973 | -4.910513985 |
| **A_55_P2130109** | Nmt2 | -0.04606728 | -0.195560263 | 0.103425703 | 8.405656101 | -0.758125191 | 0.477701844 | 0.860274973 | -4.910523478 |
| **A_55_P2144095** | NA | 0.101737245 | -0.228427622 | 0.431902113 | 9.24139485 | 0.758085656 | 0.47772378 | 0.860274973 | -4.910543426 |
| **A_51_P319592** | Usp42 | -0.048211964 | -0.204677991 | 0.108254063 | 10.02470745 | -0.75806061 | 0.477737677 | 0.860274973 | -4.910556063 |
| **A_55_P2019699** | Samhd1 | -0.088757076 | -0.376856234 | 0.199342082 | 10.75756874 | -0.757931855 | 0.477809124 | 0.86032348 | -4.910621022 |
| **A_51_P306047** | Sec13 | -0.048937032 | -0.20781962 | 0.109945556 | 9.790114415 | -0.757757913 | 0.477905657 | 0.860417143 | -4.910708764 |
| **A_55_P2066274** | NA | -0.049158653 | -0.208786464 | 0.110469158 | 12.19720981 | -0.757635954 | 0.477973349 | 0.860438857 | -4.910770276 |
| **A_55_P2066301** | Gpr137b-ps | -0.050366903 | -0.213966308 | 0.113232502 | 10.0475255 | -0.757412894 | 0.478097174 | 0.860438857 | -4.910882759 |
| **A_55_P2213348** | 5330421C15Rik | -0.103199547 | -0.438411647 | 0.232012553 | 8.10895382 | -0.757403429 | 0.478102428 | 0.860438857 | -4.910887532 |
| **A_55_P1968763** | Shc4 | 0.177605142 | -0.399388689 | 0.754598973 | 9.236427401 | 0.757274841 | 0.478173821 | 0.860438857 | -4.910952364 |
| **A_55_P2137476** | Mpv17 | -0.07962279 | -0.338309381 | 0.179063801 | 11.48947057 | -0.757238413 | 0.478194047 | 0.860438857 | -4.910970729 |
| **A_51_P372550** | Cgref1 | 0.18241384 | -0.410289066 | 0.775116747 | 12.08620912 | 0.757163888 | 0.478235428 | 0.860438857 | -4.911008298 |
| **A_55_P2143517** | Fstl1 | -0.299070648 | -1.270904004 | 0.672762709 | 13.65624817 | -0.757096351 | 0.478272931 | 0.860438857 | -4.911042342 |
| **A_52_P457411** | Ubl5 | -0.05267125 | -0.223834934 | 0.118492434 | 13.12994085 | -0.7570615 | 0.478292284 | 0.860438857 | -4.911059908 |
| **A_55_P1977533** | Eno3 | -0.051018906 | -0.216837698 | 0.114799886 | 8.963928838 | -0.756948892 | 0.478354821 | 0.860438857 | -4.911116664 |
| **A_55_P2108012** | Fam78b | 0.071019554 | -0.159846668 | 0.301885777 | 9.921507917 | 0.756809733 | 0.478432111 | 0.860438857 | -4.911186794 |
| **A_52_P1077588** | Heatr6 | 0.032479109 | -0.073132099 | 0.138090317 | 8.016815727 | 0.756594645 | 0.478551589 | 0.860438857 | -4.911295168 |
| **A_66_P135742** | Wdr1 | -0.083311966 | -0.354227827 | 0.187603895 | 13.1843705 | -0.756557804 | 0.478572056 | 0.860438857 | -4.911313729 |

| | | | | | | | | | |
|---|---|---|---|---|---|---|---|---|---|
| **A_55_P2145034** | Swt1 | -0.083711009 | -0.35593105 | 0.188509032 | 11.29169719 | -0.756539565 | 0.478582188 | 0.860438857 | -4.911322917 |
| **A_52_P230688** | Rep15 | 0.055334786 | -0.124610945 | 0.235280517 | 8.432239137 | 0.756529471 | 0.478587796 | 0.860438857 | -4.911328002 |
| **A_55_P2104119** | NA | -0.052226861 | -0.222067695 | 0.117613973 | 8.576232003 | -0.756520981 | 0.478592513 | 0.860438857 | -4.911332279 |
| **A_55_P2101503** | NA | -0.047426396 | -0.201699375 | 0.106846584 | 8.681641254 | -0.756309259 | 0.478710149 | 0.860438857 | -4.911438928 |
| **A_52_P485971** | Scap | -0.085608933 | -0.364107293 | 0.192889428 | 12.724874 | -0.756250376 | 0.478742869 | 0.860438857 | -4.911468584 |
| **A_55_P1960748** | Arl13b | -0.05427719 | -0.230859655 | 0.122305274 | 9.843313768 | -0.756203963 | 0.478768661 | 0.860438857 | -4.911491959 |
| **A_51_P433870** | Vegfc | 0.10548845 | -0.237753889 | 0.448730789 | 8.376881507 | 0.756089524 | 0.478832258 | 0.860438857 | -4.911549588 |
| **A_55_P1972275** | Wars | -0.155897241 | -0.663162349 | 0.351367868 | 12.69800774 | -0.756088444 | 0.478832858 | 0.860438857 | -4.911550132 |
| **A_52_P654618** | Znhit3 | 0.105781684 | -0.238447766 | 0.450011134 | 8.929908589 | 0.756017095 | 0.478872513 | 0.860438857 | -4.911586059 |
| **A_55_P2380132** | NA | -0.093869225 | -0.399389843 | 0.211651393 | 8.928882087 | -0.755878291 | 0.478949663 | 0.860438857 | -4.911655944 |
| **A_55_P1960238** | Slc2a6 | 0.044855597 | -0.101148991 | 0.190860185 | 8.445845484 | 0.755821602 | 0.478981175 | 0.860438857 | -4.911684483 |
| **A_55_P2295786** | Mindy2 | 0.033151219 | -0.074784989 | 0.141087428 | 7.891686284 | 0.755616634 | 0.479095122 | 0.860438857 | -4.911787658 |
| **A_55_P2150361** | NA | 0.044183194 | -0.099698951 | 0.188065338 | 12.81538555 | 0.755473727 | 0.479174579 | 0.860438857 | -4.91185958 |
| **A_55_P2316300** | Neo1 | 0.055337678 | -0.124946306 | 0.235621661 | 9.568938395 | 0.755149512 | 0.479354877 | 0.860438857 | -4.912022712 |
| **A_51_P164495** | Dda1 | -0.065556418 | -0.27913944 | 0.148026603 | 11.64791242 | -0.755122926 | 0.479369664 | 0.860438857 | -4.912036087 |
| **A_55_P2093720** | Dnajc5 | 0.049989046 | -0.112887048 | 0.212865141 | 8.631412921 | 0.755069046 | 0.479399633 | 0.860438857 | -4.912063191 |
| **A_51_P345836** | Prkra | 0.053521967 | -0.120911425 | 0.227955359 | 9.180371062 | 0.754869023 | 0.479510898 | 0.860438857 | -4.912163801 |
| **A_66_P136801** | Peg13 | -0.058363433 | -0.24859744 | 0.131870575 | 11.16400072 | -0.754782517 | 0.479559024 | 0.860438857 | -4.912207306 |
| **A_51_P386899** | Flvcr2 | -0.140033799 | -0.59648595 | 0.316418352 | 13.36972857 | -0.754756351 | 0.479573581 | 0.860438857 | -4.912220465 |
| **A_55_P1954633** | NA | -0.087985273 | -0.374796339 | 0.198825793 | 9.518526655 | -0.754715458 | 0.479596333 | 0.860438857 | -4.912241029 |
| **A_55_P1953103** | Nudt7 | 0.112738189 | -0.254792977 | 0.480269354 | 9.113110632 | 0.754650812 | 0.479632301 | 0.860438857 | -4.912273536 |
| **A_51_P120615** | Rabl3 | 0.064067087 | -0.144798959 | 0.272933132 | 8.54934073 | 0.754633871 | 0.479641727 | 0.860438857 | -4.903906826 |
| **A_52_P266530** | Ide | 0.196517768 | -0.444154534 | 0.837190071 | 9.854295096 | 0.754631688 | 0.479642942 | 0.860438857 | -4.912283152 |
| **A_55_P2013986** | Ddx31 | -0.052937095 | -0.225521802 | 0.119647612 | 9.089280635 | -0.75461765 | 0.479650753 | 0.860438857 | -4.91229021 |
| **A_55_P2175955** | Ano9 | 0.097855539 | -0.221177504 | 0.416888583 | 8.057952144 | 0.754603668 | 0.479658533 | 0.860438857 | -4.91229724 |
| **A_55_P2085288** | Acoxl | 0.033712324 | -0.07619968 | 0.143624328 | 7.717356838 | 0.754592919 | 0.479664514 | 0.860438857 | -4.912302645 |
| **A_66_P113245** | Mfng | 0.06832435 | -0.154451231 | 0.291099931 | 9.740152226 | 0.754530998 | 0.47969897 | 0.860438857 | -4.912333778 |
| **A_55_P1982084** | NA | -0.110139509 | -0.469258369 | 0.24897935 | 9.34611384 | -0.754525788 | 0.479701869 | 0.860438857 | -4.912336398 |
| **A_55_P1968738** | Nfkb1 | -0.038589344 | -0.164430817 | 0.08725213 | 8.38740047 | -0.754419251 | 0.479761155 | 0.860438857 | -4.912389958 |
| **A_51_P392967** | Zmynd10 | 0.1328378 | -0.300387777 | 0.566063377 | 8.13088709 | 0.754356696 | 0.479795968 | 0.860438857 | -4.912421404 |
| **A_55_P2001963** | NA | -0.062189959 | -0.265028198 | 0.140648281 | 15.38749255 | -0.754292168 | 0.479831881 | 0.860438857 | -4.91245384 |
| **A_51_P513568** | Stx11 | -0.093216236 | -0.397490503 | 0.211058032 | 8.38713183 | -0.753694764 | 0.480164456 | 0.860528766 | -4.912754034 |
| **A_55_P2088571** | Repin1 | 0.068449354 | -0.154991154 | 0.291889863 | 10.28878555 | 0.753661981 | 0.48018271 | 0.860528766 | -4.912770502 |
| **A_55_P2148873** | Cat | -0.146677935 | -0.625516852 | 0.332160982 | 10.25789827 | -0.753606283 | 0.480213727 | 0.860528766 | -4.91279848 |
| **A_55_P2131506** | NA | 0.052281812 | -0.118400407 | 0.222964032 | 8.814667896 | 0.753583732 | 0.480226285 | 0.860528766 | -4.912809807 |
| **A_55_P2014635** | Ccdc82 | 0.076978882 | -0.174345136 | 0.3283029 | 9.681921123 | 0.753540793 | 0.480250197 | 0.860528766 | -4.912831375 |
| **A_66_P117543** | NA | 0.080764307 | -0.182920125 | 0.34444874 | 8.38761176 | 0.753536244 | 0.480252731 | 0.860528766 | -4.91283366 |
| **A_55_P2047163** | Tmc5 | -0.103899319 | -0.443153078 | 0.235354439 | 10.20586166 | -0.753454789 | 0.480298095 | 0.860528766 | -4.91287457 |
| **A_51_P163378** | Zwint | -0.056558392 | -0.2412361 | 0.128119315 | 14.79458279 | -0.753445328 | 0.480303364 | 0.860528766 | -4.912879321 |
| **A_55_P2071802** | Slbp | -0.039930587 | -0.170334312 | 0.090473137 | 8.11008288 | -0.753329315 | 0.48036798 | 0.860528766 | -4.912937582 |
| **A_55_P2184009** | Rnd2 | -0.05250858 | -0.224043413 | 0.119026252 | 9.968786365 | -0.753090406 | 0.480501064 | 0.860528766 | -4.913057539 |
| **A_55_P2058812** | Fancl | 0.055825335 | -0.126552983 | 0.238203653 | 8.58650637 | 0.753056039 | 0.48052021 | 0.860528766 | -4.913074792 |
| **A_55_P2157200** | NA | -0.09541618 | -0.407136444 | 0.216304084 | 11.14845765 | -0.753054056 | 0.480521315 | 0.860528766 | -4.913075788 |
| **A_55_P2107367** | NA | -0.04992353 | -0.21303208 | 0.11318502 | 9.072028785 | -0.753004757 | 0.480548781 | 0.860528766 | -4.913100536 |
| **A_55_P1973965** | Ppp1r13b | 0.120018206 | -0.272135492 | 0.512171904 | 11.62583599 | 0.752939323 | 0.480585239 | 0.860528766 | -4.913133383 |
| **A_55_P2126109** | Usp16 | 0.066317378 | -0.158835531 | 0.291470288 | 11.46838105 | 0.763308093 | 0.480605463 | 0.860528766 | -4.85288673 |
| **A_55_P2111148** | Gemin5 | 0.047462831 | -0.10765362 | 0.202579281 | 8.843624945 | 0.752774572 | 0.480677039 | 0.860528766 | -4.913216075 |
| **A_55_P2102798** | Prl7a2 | 0.120940719 | -0.274315741 | 0.516197178 | 11.20474413 | 0.752770743 | 0.480679172 | 0.860528766 | -4.913217997 |
| **A_55_P2009449** | Pnma2 | -0.148172881 | -0.632433904 | 0.336088142 | 8.743552522 | -0.75276319 | 0.480683382 | 0.860528766 | -4.913221788 |
| **A_51_P179864** | Prkx | -0.070240741 | -0.299843948 | 0.159362466 | 10.30909899 | -0.75262787 | 0.480758793 | 0.860584063 | -4.913289697 |

| | | | | | | | | |
|---|---|---|---|---|---|---|---|---|
| **A_55_P1999576** | Josd1 | 0.077579112 | -0.176042263 | 0.331200486 | 13.57787547 | 0.752537442 | 0.480809191 | 0.860594579 | -4.913335073 |
| **A_55_P2012819** | Sec61a2 | 0.057166667 | -0.12981038 | 0.244143715 | 9.969312833 | 0.75218339 | 0.481006551 | 0.860716117 | -4.913512691 |
| **A_52_P160970** | Akap10 | -0.059236282 | -0.252987498 | 0.134514934 | 9.577968197 | -0.752163958 | 0.481017384 | 0.860716117 | -4.913522438 |
| **A_55_P2013296** | Prkag2 | 0.06580311 | -0.149431184 | 0.281037403 | 11.38325138 | 0.752149399 | 0.481025501 | 0.860716117 | -4.91352974 |
| **A_55_P2056035** | Traf7 | -0.051875814 | -0.221570443 | 0.117818814 | 13.18410282 | -0.752083384 | 0.481062307 | 0.860716117 | -4.91356285 |
| **A_55_P2049261** | NA | -0.146485776 | -0.625708495 | 0.332736942 | 8.314954582 | -0.752016243 | 0.481099743 | 0.860716117 | -4.913596522 |
| **A_55_P1980548** | NA | -0.258519636 | -1.104393604 | 0.587354333 | 10.87292672 | -0.751894726 | 0.481167503 | 0.860724822 | -4.913657459 |
| **A_52_P217710** | Fzd6 | -0.057142774 | -0.244155811 | 0.129870262 | 8.399390331 | -0.751724323 | 0.481262532 | 0.860724822 | -4.913742898 |
| **A_51_P120830** | Mmp10 | -0.173603001 | -0.741770026 | 0.394564023 | 8.532928157 | -0.751710069 | 0.481270482 | 0.860724822 | -4.913750044 |
| **A_55_P2005783** | Ifih1 | -0.049412259 | -0.211133176 | 0.112308658 | 10.91143485 | -0.751688109 | 0.481282729 | 0.860724822 | -4.913761053 |
| **A_55_P2016346** | Eif3d | 0.036631351 | -0.083274313 | 0.156537015 | 13.92345177 | 0.751592441 | 0.481336089 | 0.860740611 | -4.913809013 |
| **A_52_P125363** | Rbp2 | 0.183697568 | -0.417710841 | 0.785105977 | 8.370417475 | 0.751455166 | 0.481412662 | 0.860797904 | -4.913877822 |
| **A_55_P2081283** | Dlx4 | 0.112625052 | -0.256272362 | 0.481522467 | 9.234241823 | 0.751101375 | 0.481610047 | 0.860995614 | -4.914055115 |
| **A_55_P1990081** | Snrpg | -0.083894797 | -0.358691288 | 0.190901694 | 14.49827574 | -0.75109178 | 0.481615401 | 0.860995614 | -4.914059923 |
| **A_55_P2005930** | NA | -0.078309265 | -0.334835786 | 0.178217256 | 11.47099007 | -0.751017471 | 0.481656867 | 0.860995614 | -4.914097153 |
| **A_52_P655663** | H13 | -0.059052637 | -0.252547559 | 0.134442286 | 8.938403152 | -0.750825273 | 0.481764127 | 0.861107713 | -4.914193433 |
| **A_55_P2146620** | Atf2 | -0.049411343 | -0.211342937 | 0.112520252 | 8.408644185 | -0.750696213 | 0.481836162 | 0.861156835 | -4.914258074 |
| **A_66_P111877** | Snx13 | -0.051980213 | -0.222390406 | 0.118429979 | 9.537958006 | -0.750432531 | 0.481983358 | 0.861340267 | -4.914390117 |
| **A_55_P2042297** | NA | 0.184792659 | -0.421246733 | 0.790832052 | 9.976849207 | 0.750158481 | 0.482136375 | 0.861534066 | -4.914527314 |
| **A_51_P358112** | Fads1 | 0.132072182 | -0.301258865 | 0.565403229 | 8.34488094 | 0.749826372 | 0.482321855 | 0.861753532 | -4.914693526 |
| **A_51_P269687** | Pole2 | -0.038772373 | -0.166007947 | 0.0884632 | 8.218342277 | -0.749692208 | 0.482396798 | 0.861753532 | -4.914760655 |
| **A_55_P1961423** | Gsta3 | 0.146337951 | -0.333890044 | 0.626565946 | 8.737428069 | 0.749684719 | 0.482400982 | 0.861753532 | -4.914764403 |
| **A_55_P1953851** | Stk39 | 0.048226399 | -0.110062449 | 0.206515247 | 10.48491423 | 0.749555298 | 0.482473284 | 0.861753532 | -4.91482915 |
| **A_55_P2053177** | NA | -0.041720897 | -0.178660252 | 0.095218458 | 12.11036426 | -0.749539498 | 0.482482111 | 0.861753532 | -4.914837054 |
| **A_55_P2117574** | Ppih | 0.032440056 | -0.074068905 | 0.138949017 | 7.778905979 | 0.749315315 | 0.482607372 | 0.861897615 | -4.914949188 |
| **A_55_P2080243** | NA | -0.047693262 | -0.204329001 | 0.108942477 | 8.777572047 | -0.749092293 | 0.482732006 | 0.862037106 | -4.915060715 |
| **A_55_P1970929** | Hnf1a | 0.17000197 | -0.388408897 | 0.728412837 | 8.348138433 | 0.748978335 | 0.482795699 | 0.862037106 | -4.915117693 |
| **A_55_P2150757** | Gzmm | 0.088423305 | -0.202039888 | 0.378886499 | 10.88604571 | 0.748936162 | 0.482819272 | 0.862037106 | -4.915138777 |
| **A_52_P820196** | Cxcl15 | -0.031941827 | -0.136880568 | 0.072996914 | 7.64419258 | -0.748846955 | 0.482869137 | 0.862046509 | -4.915183373 |
| **A_51_P394339** | Eif2b1 | -0.055666112 | -0.238619093 | 0.12728687 | 10.49170026 | -0.748549558 | 0.483035403 | 0.862263696 | -4.915332017 |
| **A_55_P2096652** | Kyat1 | 0.039123726 | -0.089498175 | 0.167745627 | 9.291463329 | 0.748332226 | 0.483156931 | 0.862331927 | -4.915440615 |
| **A_55_P2020035** | NA | -0.386796087 | -1.708309827 | 0.934717654 | 11.11223403 | -0.758508533 | 0.483222454 | 0.862331927 | -4.855113955 |
| **A_51_P285487** | Rnf113a1 | -0.0385764 | -0.165421792 | 0.088268992 | 8.450193258 | -0.74819735 | 0.483232363 | 0.862331927 | -4.915507999 |
| **A_55_P2099627** | Lrrc51 | -0.033252282 | -0.14260644 | 0.076101876 | 8.134245123 | -0.74809252 | 0.483290996 | 0.862331927 | -4.915560366 |
| **A_55_P2181191** | Btg1 | 0.056288622 | -0.128844475 | 0.241421719 | 7.925999046 | 0.748007092 | 0.483338781 | 0.862331927 | -4.915603036 |
| **A_51_P154753** | Klc3 | 0.092911962 | -0.212677026 | 0.39850095 | 9.508335235 | 0.748002578 | 0.483341306 | 0.862331927 | -4.915605291 |
| **A_51_P121466** | Tab1 | -0.079342892 | -0.340410489 | 0.181724705 | 10.70711755 | -0.747694555 | 0.483513629 | 0.862559754 | -4.915759113 |
| **A_55_P2177103** | Slc25a5 | -0.094216126 | -0.404274989 | 0.215842737 | 11.69540798 | -0.747567232 | 0.483584872 | 0.862598135 | -4.915822683 |
| **A_52_P436318** | Rps25 | 0.037082527 | -0.084969612 | 0.159134665 | 8.166042044 | 0.747468837 | 0.483639933 | 0.862598135 | -4.915871803 |
| **A_52_P372165** | Rpsa | 0.038838472 | -0.089027474 | 0.166704418 | 7.852916678 | 0.747268026 | 0.48375232 | 0.862598135 | -4.915972036 |
| **A_55_P2120278** | Ube2l3 | -0.05248235 | -0.225315498 | 0.120350799 | 12.38349647 | -0.747059844 | 0.483868851 | 0.862598135 | -4.916075927 |
| **A_52_P127925** | Tfec | 0.086602623 | -0.198655913 | 0.371861159 | 7.878764365 | 0.746898455 | 0.483959203 | 0.862598135 | -4.916156451 |
| **A_55_P2112445** | Tbx4 | 0.075499882 | -0.173188338 | 0.324188102 | 9.055653485 | 0.746896152 | 0.483960492 | 0.862598135 | -4.9161576 |
| **A_55_P2137049** | AA467197 | -0.540844296 | -2.322377795 | 1.240689204 | 12.68250179 | -0.746874141 | 0.483972816 | 0.862598135 | -4.916168581 |
| **A_55_P2173457** | 1700097N02Rik | -0.077309439 | -0.331967578 | 0.177348701 | 9.107942562 | -0.746868463 | 0.483975995 | 0.862598135 | -4.916171414 |
| **A_55_P2021125** | NA | 0.041925795 | -0.096190685 | 0.180042274 | 8.058914896 | 0.746801129 | 0.484013695 | 0.862598135 | -4.916205005 |
| **A_52_P494686** | Kansl1l | -0.033564125 | -0.144147184 | 0.077018934 | 7.941414242 | -0.746716749 | 0.484060943 | 0.862598135 | -4.916247097 |
| **A_52_P508317** | Erlec1 | -0.05926678 | -0.254544519 | 0.136010959 | 8.435782701 | -0.746668379 | 0.484088028 | 0.862598135 | -4.916271224 |
| **A_55_P2102759** | NA | 0.058189777 | -0.133592847 | 0.249972402 | 8.563597583 | 0.746460104 | 0.484204667 | 0.862598135 | -4.916375097 |
| **A_55_P2102529** | Fuca2 | -0.043499777 | -0.186874539 | 0.099874985 | 8.775024081 | -0.746420367 | 0.484226923 | 0.862598135 | -4.916394913 |

| | | | | | | | | | |
|---|---|---|---|---|---|---|---|---|---|
| **A_55_P2092963** | Traf2 | -0.040959231 | -0.175989687 | 0.094071224 | 11.83174672 | -0.746258395 | 0.484317647 | 0.862598135 | -4.916475675 |
| **A_52_P3412** | Med17 | -0.042246726 | -0.181534245 | 0.097040793 | 9.679922075 | -0.746191019 | 0.48435539 | 0.862598135 | -4.916509266 |
| **A_51_P208697** | Ttl | -0.077898315 | -0.334735016 | 0.178938386 | 10.37330478 | -0.746174055 | 0.484364893 | 0.862598135 | -4.916517724 |
| **A_55_P2133806** | Tnrc6c | -0.053883202 | -0.231543677 | 0.123777274 | 9.472254358 | -0.74615961 | 0.484372985 | 0.862598135 | -4.916524925 |
| **A_52_P31543** | Btg2 | -0.05728366 | -0.246160059 | 0.13159274 | 11.36750341 | -0.746143264 | 0.484382142 | 0.862598135 | -4.916533073 |
| **A_51_P314339** | Chtop | 0.075448689 | -0.173322219 | 0.324219596 | 10.3817978 | 0.746141626 | 0.48438306 | 0.862598135 | -4.91653389 |
| **A_55_P1979027** | Tmem65 | 0.049295015 | -0.113270587 | 0.211860617 | 9.285161087 | 0.746008038 | 0.484457901 | 0.862616074 | -4.916600482 |
| **A_52_P306537** | Emc1 | 0.05461712 | -0.125518971 | 0.23475321 | 8.490825077 | 0.74592854 | 0.484502443 | 0.862616074 | -4.916640106 |
| **A_66_P134690** | Spin1 | -0.035033361 | -0.150585842 | 0.08051912 | 8.619645354 | -0.745884683 | 0.484527017 | 0.862616074 | -4.916661965 |
| **A_55_P2130388** | Mical1 | 0.091782135 | -0.211051676 | 0.394615947 | 12.18147722 | 0.745629287 | 0.484670137 | 0.86272739 | -4.916789234 |
| **A_55_P2078633** | C4b | 0.384951334 | -0.885216319 | 1.655118987 | 9.170446094 | 0.745613811 | 0.48467881 | 0.86272739 | -4.916796945 |
| **A_55_P1964359** | Cnot8 | -0.043965905 | -0.189058967 | 0.101127156 | 12.07520567 | -0.745484367 | 0.484751361 | 0.862777077 | -4.916861436 |
| **A_55_P2054261** | C2cd4b | -0.033830923 | -0.145498443 | 0.077836598 | 7.782540338 | -0.745342924 | 0.484830645 | 0.862838738 | -4.916931896 |
| **A_55_P2166688** | Timm10b | 0.069565416 | -0.1600812 | 0.299212032 | 10.56968547 | 0.745250879 | 0.484882244 | 0.862851123 | -4.916977742 |
| **A_55_P1977314** | Ulk1 | -0.058084555 | -0.249855271 | 0.133686161 | 12.70455014 | -0.745156576 | 0.484935114 | 0.862852963 | -4.917024709 |
| **A_51_P462428** | NA | 0.115544673 | -0.265969328 | 0.497058674 | 8.482901146 | 0.745089789 | 0.484972559 | 0.862852963 | -4.917057969 |
| **A_52_P321140** | Defb1 | -0.033869685 | -0.145723906 | 0.077984537 | 7.764567662 | -0.744951393 | 0.485050159 | 0.862873383 | -4.917126883 |
| **A_51_P391159** | Ang | -0.084773986 | -0.364754567 | 0.195206594 | 8.991024916 | -0.744910092 | 0.485073319 | 0.862873383 | -4.917147447 |
| **A_55_P2035038** | NA | -0.229623062 | -0.98827917 | 0.529033046 | 11.41692299 | -0.744628592 | 0.485231191 | 0.863074785 | -4.917287583 |
| **A_55_P2054342** | 2810408I11Rik | 0.062751333 | -0.144601282 | 0.270103947 | 8.626344999 | 0.744530686 | 0.485286108 | 0.863082294 | -4.917336313 |
| **A_55_P2062777** | Mfap5 | -1.089873378 | -4.691533738 | 2.511786982 | 14.37655042 | -0.744461857 | 0.485324717 | 0.863082294 | -4.917370568 |
| **A_51_P509997** | Cox6a2 | 0.063231662 | -0.145769477 | 0.272232802 | 7.94145412 | 0.744312152 | 0.485408701 | 0.863152234 | -4.917445065 |
| **A_66_P136834** | Usp7 | -0.04411774 | -0.189967942 | 0.101732462 | 8.080169621 | -0.744175522 | 0.485485358 | 0.863170791 | -4.917513045 |
| **A_55_P1962039** | Anxa7 | 0.07543875 | -0.1740109 | 0.3248884 | 11.36754714 | 0.74401338 | 0.485576341 | 0.863170791 | -4.917593707 |
| **A_52_P33800** | Fndc3a | -0.131024393 | -0.564289797 | 0.30224101 | 9.768866031 | -0.743990361 | 0.485589258 | 0.863170791 | -4.917605157 |
| **A_51_P399614** | Zkscan14 | -0.077139774 | -0.332227209 | 0.17794766 | 9.145521903 | -0.743975204 | 0.485597764 | 0.863170791 | -4.917612696 |
| **A_55_P2077666** | Porcn | -0.16668139 | -0.717955829 | 0.384593049 | 9.329237564 | -0.743855228 | 0.485665095 | 0.863187884 | -4.91767237 |
| **A_55_P2045642** | Stmn4 | -0.309060388 | -1.331349355 | 0.713228579 | 9.730545422 | -0.743770304 | 0.485712759 | 0.863187884 | -4.917714606 |
| **A_51_P433824** | Tpi1 | 0.134338581 | -0.310047894 | 0.578725056 | 13.71569928 | 0.74371936 | 0.485741353 | 0.863187884 | -4.91773994 |
| **A_51_P519837** | Cherp | -0.084918242 | -0.365856456 | 0.196019972 | 11.11075299 | -0.743634178 | 0.485789167 | 0.863193492 | -4.917782298 |
| **A_55_P2098140** | Tmed2 | -0.064554636 | -0.278152826 | 0.149043555 | 15.10879166 | -0.743530913 | 0.485847135 | 0.863217141 | -4.917833643 |
| **A_55_P2154880** | NA | 0.09140567 | -0.211088963 | 0.393900303 | 10.13878977 | 0.743403543 | 0.485918642 | 0.863264837 | -4.917896966 |
| **A_51_P124606** | Nat2 | 0.039416899 | -0.091044947 | 0.169878745 | 8.279187815 | 0.743306789 | 0.485972965 | 0.863282 | -4.895859125 |
| **A_55_P1962736** | Sec23b | 0.042306678 | -0.097769502 | 0.182382859 | 11.58930596 | 0.743042749 | 0.486121233 | 0.863466028 | -4.918076291 |
| **A_51_P192149** | Slc9a6 | -0.142748231 | -0.615475724 | 0.329979263 | 10.68393351 | -0.742897738 | 0.486202675 | 0.863531335 | -4.918148348 |
| **A_55_P2067752** | Psg26 | 0.060883811 | -0.140817508 | 0.26258513 | 16.98536038 | 0.742612502 | 0.4863629 | 0.863736539 | -4.918290051 |
| **A_55_P2037141** | Jsrp1 | 0.240098706 | -0.555412897 | 1.035610309 | 9.392636668 | 0.742527365 | 0.48641073 | 0.863738205 | -4.918332338 |
| **A_55_P2142371** | Rai1 | -0.160070489 | -0.690481091 | 0.370340113 | 11.08170346 | -0.742451756 | 0.48645321 | 0.863738205 | -4.918369889 |
| **A_51_P267700** | Gkn3 | -0.03562162 | -0.153693538 | 0.082450298 | 8.068342635 | -0.742226082 | 0.486580019 | 0.86381765 | -4.918481954 |
| **A_51_P454196** | Sh2d4a | -0.055450915 | -0.239252515 | 0.128350685 | 8.351695939 | -0.742213065 | 0.486587334 | 0.86381765 | -4.918488417 |
| **A_66_P112482** | Tmem184c | -0.090775033 | -0.39170534 | 0.210155274 | 9.960885399 | -0.742112338 | 0.486643942 | 0.863838805 | -4.918538428 |
| **A_55_P2036280** | Psen2 | -0.061561566 | -0.265674818 | 0.142551686 | 11.35875313 | -0.742006344 | 0.486703514 | 0.863865218 | -4.918591047 |
| **A_55_P2104332** | Pxdn | -0.050913434 | -0.219795053 | 0.117968185 | 8.865741378 | -0.741684445 | 0.486884463 | 0.864107041 | -4.918750814 |
| **A_51_P483544** | Aass | 0.161213006 | -0.373687859 | 0.696113872 | 8.842855734 | 0.741474022 | 0.487002774 | 0.864237662 | -4.918855224 |
| **A_51_P268193** | Slc7a10 | -0.03479711 | -0.150280441 | 0.080686221 | 7.864014959 | -0.741298366 | 0.487101551 | 0.864333598 | -4.918942365 |
| **A_55_P1958833** | Gm9079 | -0.081115018 | -0.350401058 | 0.188171023 | 11.00013609 | -0.74106549 | 0.487232527 | 0.864344921 | -4.919057869 |
| **A_51_P429472** | Mindy3 | -0.054137287 | -0.23389859 | 0.125624015 | 10.2803407 | -0.740916808 | 0.487316162 | 0.864344921 | -4.919131599 |
| **A_51_P103780** | Sft2d2 | -0.059573919 | -0.25739134 | 0.138243501 | 9.875912548 | -0.740902059 | 0.487324459 | 0.864344921 | -4.919138912 |
| **A_55_P1969273** | Zfp758 | 0.035645697 | -0.082717705 | 0.1540091 | 8.426547539 | 0.740898712 | 0.487326342 | 0.864344921 | -4.919140571 |
| **A_52_P87964** | Pla2g12a | -0.06740571 | -0.291232674 | 0.156421254 | 10.06216308 | -0.740889508 | 0.48733152 | 0.864344921 | -4.919145135 |

| | | | | | | | | | |
|---|---|---|---|---|---|---|---|---|---|
| **A_55_P1994294** | NA | 0.28297494 | -0.656791309 | 1.222741189 | 12.73650283 | 0.740793727 | 0.487385404 | 0.864361177 | -4.919192625 |
| **A_55_P2015742** | Cggbp1 | -0.041711821 | -0.180296687 | 0.096873045 | 9.418833682 | -0.7404786 | 0.487562716 | 0.864596306 | -4.919348836 |
| **A_52_P493620** | Fgfr1op2 | 0.059293026 | -0.137832378 | 0.256418429 | 10.15656122 | 0.739997376 | 0.487833572 | 0.864927584 | -4.919587284 |
| **A_55_P2015585** | Zcchc24 | -0.269727416 | -1.16647406 | 0.627019228 | 11.32393702 | -0.739987689 | 0.487839025 | 0.864927584 | -4.919592083 |
| **A_55_P2153391** | NA | -0.079949435 | -0.345904985 | 0.186006114 | 7.842322725 | -0.739563563 | 0.488077831 | 0.865117976 | -4.919802139 |
| **A_52_P516097** | Leng9 | -0.087935806 | -0.380460235 | 0.204588622 | 9.812885292 | -0.739558911 | 0.488080451 | 0.865117976 | -4.919804442 |
| **A_55_P2148002** | 2610005L07Rik | -0.065437699 | -0.283121433 | 0.152246035 | 9.748747884 | -0.739556241 | 0.488081955 | 0.865117976 | -4.919805764 |
| **A_55_P1995503** | Gm5643 | 0.050546859 | -0.117626017 | 0.218719735 | 11.61179576 | 0.739447571 | 0.488143155 | 0.865117976 | -4.919859569 |
| **A_55_P2125491** | Tnfrsf4 | 0.066268213 | -0.154225006 | 0.286761431 | 8.574019837 | 0.739399554 | 0.488170198 | 0.865117976 | -4.919883342 |
| **A_51_P355413** | Nisch | -0.147313741 | -0.637570436 | 0.342942955 | 14.34415735 | -0.739245836 | 0.488256781 | 0.865142767 | -4.919959438 |
| **A_55_P2176731** | Prdx6 | 0.103725349 | -0.241484225 | 0.448934924 | 10.40970333 | 0.739215792 | 0.488273705 | 0.865142767 | -4.919974309 |
| **A_55_P2151071** | Cox7a2l | -0.043426449 | -0.188008609 | 0.101155712 | 15.32736925 | -0.738939302 | 0.48842947 | 0.865339433 | -4.920111147 |
| **A_65_P15215** | Mrpl15 | 0.080630457 | -0.187865902 | 0.349126816 | 10.63003445 | 0.738805099 | 0.488505087 | 0.865375059 | -4.920177551 |
| **A_55_P2044788** | NA | -0.047338375 | -0.204986252 | 0.110309502 | 8.723696175 | -0.738744705 | 0.48853912 | 0.865375059 | -4.920207431 |
| **A_55_P1952220** | Pnpo | 0.126596143 | -0.295090597 | 0.548282883 | 13.27468367 | 0.738583656 | 0.488629878 | 0.865456512 | -4.920287102 |
| **A_51_P511375** | Ranbp2 | 0.060085943 | -0.140138902 | 0.260310787 | 11.10829958 | 0.738285057 | 0.488798184 | 0.865625454 | -4.920434783 |
| **A_55_P1962404** | Ugt1a6b | 0.273229878 | -0.637292538 | 1.183752293 | 9.483599355 | 0.73825553 | 0.488814829 | 0.865625454 | -4.920449383 |
| **A_55_P1985840** | Mettl16 | 0.066634896 | -0.155457743 | 0.288727535 | 9.931863633 | 0.738136571 | 0.488881893 | 0.865652037 | -4.920508204 |
| **A_55_P1966508** | Hoxc6 | 0.139476789 | -0.325444524 | 0.604398102 | 8.724077834 | 0.738059963 | 0.488925085 | 0.865652037 | -4.920546079 |
| **A_55_P2139390** | NA | -0.042719759 | -0.185131937 | 0.099692419 | 15.68006744 | -0.737990598 | 0.488964196 | 0.865652037 | -4.920580371 |
| **A_55_P2004606** | Serpina4-ps1 | -0.037511343 | -0.162592618 | 0.087569932 | 7.643124136 | -0.737801401 | 0.489070882 | 0.865761616 | -4.920673892 |
| **A_51_P375187** | Rpf2 | 0.073368189 | -0.171362312 | 0.318098691 | 10.37269588 | 0.73754558 | 0.489215163 | 0.865814349 | -4.920800316 |
| **A_52_P513842** | Fubp1 | -0.036780638 | -0.159474458 | 0.085913182 | 8.176194791 | -0.737506301 | 0.489237318 | 0.865814349 | -4.920819724 |
| **A_55_P2102050** | Cc2d1b | -0.068048012 | -0.295045455 | 0.158949432 | 12.42374872 | -0.737502736 | 0.489239329 | 0.865814349 | -4.920821485 |
| **A_51_P156243** | Eps15 | -0.050040466 | -0.217028364 | 0.116947432 | 9.956431777 | -0.737234265 | 0.489390781 | 0.865814349 | -4.920954119 |
| **A_55_P2112295** | Hmg20b | -0.075987697 | -0.329578585 | 0.17760319 | 13.04139377 | -0.737188925 | 0.489416361 | 0.865814349 | -4.920976514 |
| **A_55_P1978675** | NA | -0.098596881 | -0.42770516 | 0.230511398 | 9.463218304 | -0.737044196 | 0.489498023 | 0.865814349 | -4.921047996 |
| **A_55_P2002160** | Sumo3 | -0.034482614 | -0.149590972 | 0.080625743 | 9.416412061 | -0.736991517 | 0.489527749 | 0.865814349 | -4.921074012 |
| **A_55_P2110497** | Ddc | 0.177518521 | -0.415133158 | 0.770170201 | 9.876938754 | 0.736908075 | 0.489574837 | 0.865814349 | -4.921115217 |
| **A_55_P1960896** | Fndc3c1 | -0.057137029 | -0.247894699 | 0.133620642 | 9.47657169 | -0.736893636 | 0.489582985 | 0.865814349 | -4.921122347 |
| **A_52_P434974** | Ubqln1 | 0.078732222 | -0.184157328 | 0.341621771 | 11.52767806 | 0.736797845 | 0.489637046 | 0.865814349 | -4.921169645 |
| **A_51_P212068** | Aoc1 | -0.699644697 | -3.035834737 | 1.636545342 | 13.07171548 | -0.736781211 | 0.489646434 | 0.865814349 | -4.921177858 |
| **A_55_P1971448** | Snrpd1 | -0.087502384 | -0.379752182 | 0.204747414 | 13.44576399 | -0.736605286 | 0.489745731 | 0.865814349 | -4.921264709 |
| **A_51_P421846** | Ech1 | 0.079391229 | -0.185777183 | 0.344559642 | 11.41751441 | 0.736579963 | 0.489760025 | 0.865814349 | -4.921277209 |
| **A_51_P142989** | Zfp429 | -0.056702053 | -0.246090306 | 0.132686201 | 9.261500069 | -0.73657149 | 0.489764807 | 0.865814349 | -4.921281392 |
| **A_55_P1993054** | Psip1 | -0.048289154 | -0.209580834 | 0.113002526 | 9.196548299 | -0.736557731 | 0.489772574 | 0.865814349 | -4.921288184 |
| **A_52_P89267** | Castor2 | 0.052882542 | -0.123870135 | 0.229635219 | 8.390334297 | 0.736063844 | 0.490051422 | 0.866217317 | -4.921531911 |
| **A_55_P2038458** | Lancl2 | 0.044512244 | -0.104293514 | 0.193318003 | 8.151959333 | 0.735917143 | 0.49013427 | 0.866217317 | -4.921604282 |
| **A_51_P344008** | Slc25a20 | -0.061207515 | -0.265826403 | 0.143411373 | 8.597585797 | -0.735915907 | 0.490134968 | 0.866217317 | -4.921604892 |
| **A_51_P379976** | Arl4a | 0.188633083 | -0.442092192 | 0.819358359 | 10.59327988 | 0.735777989 | 0.490212865 | 0.866223087 | -4.92167292 |
| **A_51_P472329** | Meak7 | -0.036277921 | -0.157597635 | 0.085041793 | 8.583377976 | -0.735665143 | 0.490276607 | 0.866223087 | -4.921728574 |
| **A_52_P145412** | Rictor | -0.057941153 | -0.25176708 | 0.135884775 | 9.194126489 | -0.735435217 | 0.490406501 | 0.866223087 | -4.92184195 |
| **A_51_P150120** | Mgst2 | 0.220633628 | -0.517455578 | 0.958722834 | 11.22801062 | 0.735414208 | 0.49041837 | 0.866223087 | -4.921852308 |
| **A_55_P1977583** | Anxa11 | 0.134218699 | -0.314891687 | 0.583329084 | 13.55399235 | 0.735239937 | 0.49051684 | 0.866223087 | -4.92193822 |
| **A_51_P138772** | Ddx27 | 0.105782354 | -0.248182034 | 0.459746743 | 10.91084509 | 0.735229325 | 0.490522837 | 0.866223087 | -4.921943451 |
| **A_55_P1984361** | NA | 0.03846722 | -0.090267492 | 0.167201933 | 12.50199533 | 0.73513026 | 0.490578818 | 0.866223087 | -4.921992281 |
| **A_55_P2036380** | NA | 0.04498021 | -0.105567023 | 0.195527443 | 17.06150176 | 0.735051659 | 0.490623239 | 0.866223087 | -4.92203102 |
| **A_55_P2174836** | Vldlr | -0.373257943 | -1.62268314 | 0.876167255 | 9.371503324 | -0.734967205 | 0.49067097 | 0.866223087 | -4.922072641 |
| **A_55_P2167984** | NA | 0.054916953 | -0.128923944 | 0.238757849 | 9.946547854 | 0.734908828 | 0.490703966 | 0.866223087 | -4.922101408 |
| **A_52_P574720** | Oit3 | -0.083243162 | -0.362006146 | 0.195519822 | 9.629494849 | -0.734653629 | 0.490848224 | 0.866223087 | -4.922227144 |

| | | | | | | | | | |
|---|---|---|---|---|---|---|---|---|---|
| **A_51_P237752** | Cavin1 | 0.142390108 | -0.334485014 | 0.61926523 | 14.31849413 | 0.734588824 | 0.490884862 | 0.866223087 | -4.922259069 |
| **A_55_P2097657** | Rps11 | 0.147176932 | -0.345750505 | 0.640104368 | 14.87857035 | 0.7345577 | 0.490902459 | 0.866223087 | -4.9222744 |
| **A_66_P102237** | Nemf | -0.06274747 | -0.272906044 | 0.147411104 | 10.47649028 | -0.734544772 | 0.490909768 | 0.866223087 | -4.922280768 |
| **A_52_P551909** | Arhgap5 | 0.041665202 | -0.097956094 | 0.181286498 | 8.773709281 | 0.734160456 | 0.491127087 | 0.866223087 | -4.922470036 |
| **A_55_P2032598** | NA | -0.055518499 | -0.24158432 | 0.130547322 | 15.63849146 | -0.734074729 | 0.491175572 | 0.866223087 | -4.922512244 |
| **A_55_P1988678** | Znhit6 | -0.037744946 | -0.164248301 | 0.088758409 | 11.25556946 | -0.734050483 | 0.491189285 | 0.866223087 | -4.922524182 |
| **A_52_P318955** | Ap3m1 | 0.055099005 | -0.129569654 | 0.239767663 | 9.434001416 | 0.734039995 | 0.491195217 | 0.866223087 | -4.922529345 |
| **A_55_P1977985** | Pigp | -0.085921283 | -0.373928293 | 0.202085727 | 11.75212668 | -0.733950646 | 0.491245756 | 0.866223087 | -4.922573331 |
| **A_51_P390967** | Slc43a3 | -0.085184512 | -0.370725078 | 0.200356054 | 10.33352524 | -0.733942417 | 0.49125041 | 0.866223087 | -4.922577382 |
| **A_55_P1985159** | Mef2a | -0.060435446 | -0.263017644 | 0.142146751 | 11.06944889 | -0.733938411 | 0.491252677 | 0.866223087 | -4.922579354 |
| **A_55_P1996086** | Gm14326 | -0.053486669 | -0.232800495 | 0.125827156 | 8.556573255 | -0.733839257 | 0.491308766 | 0.866223087 | -4.922628162 |
| **A_51_P349023** | Lsm3 | -0.059924929 | -0.260852716 | 0.141002857 | 10.29777929 | -0.733730706 | 0.491370175 | 0.866223087 | -4.922681589 |
| **A_66_P137516** | NA | 0.046968259 | -0.110516749 | 0.204453267 | 13.44710099 | 0.733726842 | 0.491372362 | 0.866223087 | -4.922683491 |
| **A_52_P542794** | 2610507B11Rik | -0.053696926 | -0.233770044 | 0.126376192 | 11.67046533 | -0.733617534 | 0.491434205 | 0.866223087 | -4.922737284 |
| **A_55_P2067905** | Scfd2 | -0.060135277 | -0.261803447 | 0.141532892 | 10.55161196 | -0.733603046 | 0.491442402 | 0.866223087 | -4.922744414 |
| **A_55_P2041082** | NA | -0.035051516 | -0.152606173 | 0.082503142 | 8.081995571 | -0.733560813 | 0.491466298 | 0.866223087 | -4.922765196 |
| **A_55_P2061456** | Nfyc | 0.068411649 | -0.161031471 | 0.297854769 | 9.46817614 | 0.73354064 | 0.491477713 | 0.866223087 | -4.922775122 |
| **A_52_P185907** | Crabp1 | -0.291988777 | -1.271312007 | 0.687334453 | 8.674863848 | -0.73351545 | 0.491491966 | 0.866223087 | -4.922787517 |
| **A_66_P111449** | 1700018A04Rik | 0.032866732 | -0.077390375 | 0.143123839 | 8.094184627 | 0.733363161 | 0.491578143 | 0.866223087 | -4.922862445 |
| **A_55_P2140626** | Arl5b | 0.0545373 | -0.128422101 | 0.237496701 | 8.754243501 | 0.733344545 | 0.491588678 | 0.866223087 | -4.922871603 |
| **A_52_P134639** | Tgoln2 | 0.172446823 | -0.406165728 | 0.751059374 | 8.832678362 | 0.733223729 | 0.491657053 | 0.866223087 | -4.922931036 |
| **A_55_P2126814** | NA | -0.076862079 | -0.334766258 | 0.1810421 | 8.985832999 | -0.733200769 | 0.491670048 | 0.866223087 | -4.92294233 |
| **A_55_P2050083** | Gak | -0.04958006 | -0.215956782 | 0.116796663 | 12.03663148 | -0.733134476 | 0.49170757 | 0.866223087 | -4.922974938 |
| **A_55_P2128085** | NA | -0.087303754 | -0.380290325 | 0.205682816 | 10.08941665 | -0.733085061 | 0.49173554 | 0.866223087 | -4.922999242 |
| **A_55_P2166069** | 2310079G19Rik | 0.051537474 | -0.121442825 | 0.224517773 | 8.042814325 | 0.732985948 | 0.491791643 | 0.866223087 | -4.923047986 |
| **A_65_P01643** | Chd2 | -0.061874302 | -0.269572703 | 0.145824099 | 9.380651512 | -0.732902702 | 0.491838768 | 0.866223087 | -4.923088922 |
| **A_55_P2427377** | NA | -0.031256948 | -0.136180248 | 0.073666353 | 7.708820369 | -0.732898423 | 0.491841191 | 0.866223087 | -4.923091026 |
| **A_55_P1966204** | Cxcl12 | -0.083594426 | -0.364288143 | 0.197099291 | 9.527092438 | -0.732679079 | 0.491965376 | 0.86636286 | -4.923198871 |
| **A_55_P1979053** | Atp6v0c | -0.036962173 | -0.161088147 | 0.087163802 | 13.24079478 | -0.732595062 | 0.492012949 | 0.866367705 | -4.923240174 |
| **A_55_P1953143** | Wt1 | -0.545828563 | -2.379223633 | 1.287566506 | 10.78752651 | -0.732435461 | 0.492103329 | 0.866447918 | -4.923318623 |
| **A_55_P2064457** | Gng7 | 0.097410217 | -0.229875718 | 0.424696152 | 8.688448918 | 0.732228001 | 0.492220828 | 0.866514479 | -4.923420576 |
| **A_51_P334685** | Esyt1 | -0.126065138 | -0.549720135 | 0.297589859 | 8.973842596 | -0.73206869 | 0.49231107 | 0.866514479 | -4.923498852 |
| **A_51_P440852** | Mtrf1 | -0.048307331 | -0.210650102 | 0.11403544 | 8.77332067 | -0.73206434 | 0.492313534 | 0.866514479 | -4.92350099 |
| **A_51_P485756** | Nts | -0.032271363 | -0.140725108 | 0.076182381 | 7.673387778 | -0.732051989 | 0.492320531 | 0.866514479 | -4.923507057 |
| **A_52_P654437** | NA | -0.052565977 | -0.229276104 | 0.12414415 | 11.03519305 | -0.731833807 | 0.49244414 | 0.866514479 | -4.923614236 |
| **A_55_P2024011** | NA | -0.043849375 | -0.191276303 | 0.103577552 | 13.20845243 | -0.731738256 | 0.49249828 | 0.866514479 | -4.923661167 |
| **A_52_P504342** | Coa7 | -0.076058965 | -0.331781757 | 0.179663827 | 9.043886584 | -0.73172878 | 0.49250365 | 0.866514479 | -4.923665821 |
| **A_51_P254855** | Ptgs2 | -0.427406324 | -1.864987614 | 1.010174967 | 9.675236428 | -0.731438296 | 0.492668268 | 0.866514479 | -4.902163366 |
| **A_52_P604515** | Zfp319 | -0.068326483 | -0.298150513 | 0.161497546 | 8.877389005 | -0.731413207 | 0.492682487 | 0.866514479 | -4.923820781 |
| **A_51_P272147** | Emc9 | -0.076567946 | -0.334136325 | 0.181000432 | 11.60422212 | -0.731347227 | 0.492719885 | 0.866514479 | -4.923853174 |
| **A_55_P1989076** | Slc22a23 | 0.097498997 | -0.230491526 | 0.425489519 | 11.17666505 | 0.73132095 | 0.492734778 | 0.866514479 | -4.923866073 |
| **A_55_P2043559** | NA | -0.07534669 | -0.328855777 | 0.178162397 | 8.794472015 | -0.731206108 | 0.492799876 | 0.866514479 | -4.923922448 |
| **A_51_P147034** | Ica1l | 0.248655053 | -0.588183244 | 1.085493351 | 11.43338775 | 0.731012672 | 0.492909537 | 0.866514479 | -4.924017387 |
| **A_51_P460332** | Apoc4 | 0.102965753 | -0.243578572 | 0.449510077 | 7.891757045 | 0.730976091 | 0.492930278 | 0.866514479 | -4.924035339 |
| **A_51_P381618** | Pla1a | -0.320527529 | -1.441969387 | 0.800914329 | 11.72337955 | -0.740693708 | 0.493025994 | 0.866514479 | -4.863294641 |
| **A_51_P499816** | Pbld1 | 0.157075737 | -0.371751371 | 0.685902844 | 8.250720619 | 0.730742825 | 0.493062545 | 0.866514479 | -4.924149797 |
| **A_51_P392546** | Haus3 | -0.075584914 | -0.330071844 | 0.178902016 | 9.868918244 | -0.730699491 | 0.493087119 | 0.866514479 | -4.924171057 |
| **A_52_P305182** | Ccdc32 | -0.058931408 | -0.257350397 | 0.139487582 | 11.28851427 | -0.730689295 | 0.493092901 | 0.866514479 | -4.924176058 |
| **A_51_P483351** | Lix1l | -0.050588085 | -0.220917148 | 0.119740978 | 8.110731343 | -0.730682396 | 0.493096814 | 0.866514479 | -4.924179443 |
| **A_55_P1961335** | Ctsk | -0.990227346 | -4.324922135 | 2.344467443 | 14.47228741 | -0.730546708 | 0.493173767 | 0.866514479 | -4.924246005 |

| | | | | | | | | |
|---|---|---|---|---|---|---|---|---|
| **A_66_P136813** | Smim41 | 0.032740493 | -0.077520405 | 0.143001391 | 7.859088552 | 0.730521249 | 0.493188206 | 0.866514479 | -4.924258493 |
| **A_55_P2078831** | NA | -0.076827991 | -0.335577993 | 0.18192201 | 10.02766095 | -0.73047992 | 0.493211647 | 0.866514479 | -4.924278764 |
| **A_55_P2053511** | Mrfap1 | 0.083406476 | -0.197500601 | 0.364313553 | 15.02669708 | 0.730476512 | 0.49321358 | 0.866514479 | -4.924280436 |
| **A_51_P321965** | Ssna1 | 0.074066428 | -0.175409833 | 0.323542688 | 10.77880832 | 0.730400959 | 0.493256435 | 0.866514479 | -4.924317491 |
| **A_51_P142046** | Oxld1 | 0.098303956 | -0.232856199 | 0.42946411 | 10.06937906 | 0.730301312 | 0.493312959 | 0.866514479 | -4.924366359 |
| **A_55_P2098734** | Ttc21b | -0.053370221 | -0.233171305 | 0.126430863 | 8.711935782 | -0.730257223 | 0.49333797 | 0.866514479 | -4.924387979 |
| **A_55_P2157902** | Igsf10 | 0.031547212 | -0.074736939 | 0.137831364 | 7.714410192 | 0.730233309 | 0.493351536 | 0.866514479 | -4.924399706 |
| **A_55_P2150442** | NA | 0.155721033 | -0.369292509 | 0.680734574 | 12.38687156 | 0.729702666 | 0.493652634 | 0.866956556 | -4.924659835 |
| **A_51_P233396** | Ppp2r5d | -0.065314172 | -0.285546765 | 0.154918421 | 10.64316229 | -0.729617085 | 0.493701206 | 0.866956556 | -4.924701774 |
| **A_66_P125226** | NA | -0.075035682 | -0.328094677 | 0.178023312 | 12.29159274 | -0.729483079 | 0.493777269 | 0.866956556 | -4.924767437 |
| **A_55_P2021991** | Zmym4 | 0.057536862 | -0.13650961 | 0.251583334 | 9.01040369 | 0.72947361 | 0.493782644 | 0.866956556 | -4.924772076 |
| **A_52_P350950** | Dennd2c | -0.032478742 | -0.142035598 | 0.077078113 | 8.067118754 | -0.729337949 | 0.493859654 | 0.866992874 | -4.924838539 |
| **A_55_P1997936** | Hsd17b7 | 0.109437257 | -0.259744786 | 0.4786193 | 8.816544973 | 0.729279145 | 0.493893038 | 0.866992874 | -4.924867346 |
| **A_51_P113274** | Hps6 | -0.042341775 | -0.185220806 | 0.100537256 | 10.60202005 | -0.729070839 | 0.494011308 | 0.867121738 | -4.924969375 |
| **A_55_P1995652** | Rexo4 | -0.079893396 | -0.349599217 | 0.189812424 | 11.5280563 | -0.728768727 | 0.494182874 | 0.867267714 | -4.925117311 |
| **A_55_P2104259** | Plin5 | 0.050176928 | -0.119211957 | 0.219565814 | 7.848507309 | 0.728766373 | 0.494184211 | 0.867267714 | -4.925118463 |
| **A_55_P2079041** | NA | 0.044196358 | -0.10503483 | 0.193427546 | 15.3044115 | 0.728611544 | 0.494272152 | 0.867318127 | -4.92519426 |
| **A_55_P2077055** | Itih4 | -0.223717677 | -0.979181436 | 0.531746082 | 10.52676747 | -0.728544073 | 0.494310478 | 0.867318127 | -4.925227286 |
| **A_66_P130647** | NA | -0.031133154 | -0.136284517 | 0.07401821 | 7.730493364 | -0.728412464 | 0.494385242 | 0.867318127 | -4.925291701 |
| **A_55_P1984168** | Casp4 | -0.146462959 | -0.641146126 | 0.348220209 | 9.737475221 | -0.728399827 | 0.494392422 | 0.867318127 | -4.925297886 |
| **A_55_P2022539** | NA | -0.090607965 | -0.396782976 | 0.215567046 | 9.171123298 | -0.728057699 | 0.494586816 | 0.867580413 | -4.925465293 |
| **A_55_P2056493** | NA | 0.046238904 | -0.110052281 | 0.202530089 | 8.004653871 | 0.727850545 | 0.494704545 | 0.86770818 | -4.925566627 |
| **A_51_P411130** | Rpp25l | 0.053287976 | -0.126874741 | 0.233450694 | 11.32321998 | 0.72766832 | 0.494808122 | 0.867760201 | -4.925655747 |
| **A_55_P2003096** | Pla2g6 | 0.049407045 | -0.117646984 | 0.216461074 | 10.07555759 | 0.727614072 | 0.494838959 | 0.867760201 | -4.925682275 |
| **A_51_P436189** | Nudt22 | 0.033295772 | -0.079294535 | 0.145886078 | 9.176262588 | 0.727540625 | 0.494880713 | 0.867760201 | -4.925718189 |
| **A_66_P140408** | D8Ertd738e | -0.094240346 | -0.412941406 | 0.224460714 | 10.09946226 | -0.72748246 | 0.49491378 | 0.867760201 | -4.925746628 |
| **A_55_P1963900** | Mib2 | -0.099681851 | -0.436849621 | 0.237485919 | 11.02312638 | -0.727342899 | 0.494993128 | 0.867820606 | -4.925814858 |
| **A_52_P625683** | Pxmp4 | 0.14432694 | -0.344005629 | 0.632659509 | 10.43405685 | 0.72711127 | 0.495124841 | 0.867953948 | -4.925928076 |
| **A_55_P1962454** | NA | -0.056107229 | -0.245962641 | 0.133748182 | 8.508873293 | -0.727051217 | 0.495158993 | 0.867953948 | -4.925957425 |
| **A_55_P2159475** | Sbf1 | 0.103185703 | -0.246105146 | 0.452476552 | 11.68797059 | 0.726777516 | 0.495314668 | 0.868148098 | -4.926091164 |
| **A_55_P1971054** | Paqr9 | -0.032149641 | -0.141038935 | 0.076739653 | 7.644144322 | -0.726373693 | 0.495544414 | 0.868453765 | -4.926288413 |
| **A_55_P2171658** | Stx3 | 0.177717784 | -0.424402475 | 0.779838042 | 12.04080017 | 0.726134066 | 0.495680778 | 0.868453765 | -4.926405419 |
| **A_55_P1953402** | Mlana | 0.037017316 | -0.088401599 | 0.162436231 | 7.816161088 | 0.726124435 | 0.49568626 | 0.868453765 | -4.926410121 |
| **A_55_P2171413** | Me1 | -0.091147578 | -0.399971728 | 0.217676573 | 11.26979324 | -0.726111038 | 0.495693884 | 0.868453765 | -4.926416662 |
| **A_55_P2183389** | NA | 0.039460803 | -0.09425111 | 0.173172715 | 13.6108419 | 0.726047404 | 0.495730101 | 0.868453765 | -4.926447728 |
| **A_55_P2096283** | NA | 0.051358175 | -0.122679698 | 0.225396047 | 12.22289982 | 0.725997255 | 0.495758645 | 0.868453765 | -4.926472209 |
| **A_55_P1957213** | NA | -0.378666003 | -1.662334972 | 0.905002967 | 11.22987849 | -0.725725587 | 0.49591329 | 0.868555083 | -4.926604805 |
| **A_55_P1981724** | Nr2c1 | 0.049793178 | -0.119012367 | 0.218598722 | 9.161668756 | 0.725691939 | 0.495932446 | 0.868555083 | -4.926621226 |
| **A_52_P598300** | Gtf2f2 | -0.058675895 | -0.257619298 | 0.140267508 | 10.05794222 | -0.725603459 | 0.495982821 | 0.868555083 | -4.926664401 |
| **A_51_P493857** | Katna1 | 0.04635063 | -0.110811324 | 0.203512585 | 10.76765511 | 0.725566772 | 0.496003709 | 0.868555083 | -4.926682302 |
| **A_55_P1992179** | Tars2 | -0.081724585 | -0.358854796 | 0.195405626 | 11.78074329 | -0.725501001 | 0.496041159 | 0.868555083 | -4.926714393 |
| **A_51_P108978** | Myadm | 0.08461441 | -0.20251893 | 0.37174775 | 10.15911653 | 0.724986421 | 0.496334221 | 0.86896237 | -4.926965385 |
| **A_51_P211491** | Gusb | -0.106495208 | -0.467905348 | 0.254914933 | 10.1379218 | -0.72493471 | 0.496363678 | 0.86896237 | -4.900549415 |
| **A_55_P1995012** | Plekhm2 | -0.141682067 | -0.737833194 | 0.454469061 | 12.89324527 | -0.776231908 | 0.496538804 | 0.869154385 | -4.735544186 |
| **A_55_P2110081** | Cisd3 | 0.044748449 | -0.107205593 | 0.196702492 | 10.78226542 | 0.72449417 | 0.496614677 | 0.869154385 | -4.927205357 |
| **A_55_P2032388** | Trim24 | -0.048568124 | -0.213507535 | 0.116371288 | 9.507150582 | -0.724429366 | 0.496651606 | 0.869154385 | -4.927236939 |
| **A_55_P2026445** | Tfam | -0.047617581 | -0.209383997 | 0.114148836 | 9.796578662 | -0.724182664 | 0.49679221 | 0.869154385 | -4.92735715 |
| **A_51_P258078** | Nop2 | -0.069233521 | -0.304437784 | 0.165970743 | 11.42365159 | -0.724169769 | 0.49679956 | 0.869154385 | -4.927363433 |
| **A_55_P2094964** | NA | -0.047149866 | -0.207339186 | 0.113039453 | 10.00196581 | -0.724129219 | 0.496822674 | 0.869154385 | -4.927383188 |
| **A_55_P2026270** | Cfi | 0.348809953 | -0.836281473 | 1.533901379 | 11.88978488 | 0.724112719 | 0.496832079 | 0.869154385 | -4.927391227 |

| | | | | | | | | | |
|---|---|---|---|---|---|---|---|---|---|
| **A_51_P202745** | Zc3h13 | -0.074941413 | -0.329557552 | 0.179674726 | 10.01360989 | -0.724110945 | 0.49683309 | 0.869154385 | -4.927392091 |
| **A_55_P2161775** | Prr3 | 0.088955601 | -0.213401058 | 0.39131226 | 10.02614931 | 0.72380724 | 0.497006229 | 0.869231206 | -4.927540023 |
| **A_55_P2016262** | Isl2 | 0.070375744 | -0.168863262 | 0.30961475 | 8.550971914 | 0.723702665 | 0.497065855 | 0.869231206 | -4.92759095 |
| **A_52_P609024** | Cenpw | 0.114459118 | -0.27465077 | 0.503569006 | 8.816731454 | 0.723681202 | 0.497078093 | 0.869231206 | -4.927601401 |
| **A_51_P362066** | Chil1 | -0.048544246 | -0.213574685 | 0.116486193 | 7.921716801 | -0.723673832 | 0.497082296 | 0.869231206 | -4.92760499 |
| **A_51_P255329** | Riok2 | 0.089191189 | -0.214153434 | 0.392535813 | 10.47061766 | 0.723360543 | 0.49726096 | 0.869231206 | -4.927757518 |
| **A_51_P155142** | Cdca8 | 0.103583189 | -0.248744674 | 0.455911051 | 8.693220136 | 0.723288316 | 0.497302155 | 0.869231206 | -4.927792675 |
| **A_55_P2033927** | NA | -0.040187216 | -0.176896839 | 0.096522407 | 16.03370887 | -0.723199299 | 0.497352931 | 0.869231206 | -4.927836002 |
| **A_55_P1976869** | Hsdl2 | -0.061039534 | -0.268743976 | 0.146664908 | 9.723670474 | -0.722993822 | 0.49747015 | 0.869231206 | -4.927935996 |
| **A_55_P2052881** | NA | 0.049140646 | -0.118084213 | 0.216365505 | 9.040896161 | 0.722951541 | 0.497494272 | 0.869231206 | -4.927956569 |
| **A_55_P1983488** | Lsr | -0.111697366 | -0.491930514 | 0.268535782 | 13.50947017 | -0.722706765 | 0.497633938 | 0.869231206 | -4.928075653 |
| **A_55_P1995572** | Meis1 | -0.065166434 | -0.295748816 | 0.165415947 | 9.997869187 | -0.732399301 | 0.497638622 | 0.869231206 | -4.867056444 |
| **A_55_P2149515** | Rsrc1 | 0.040108044 | -0.096434763 | 0.176650851 | 8.840450972 | 0.72265634 | 0.497662714 | 0.869231206 | -4.928100181 |
| **A_52_P348720** | Rhobtb3 | 0.041911715 | -0.100776316 | 0.184599746 | 8.098882759 | 0.722631771 | 0.497676734 | 0.869231206 | -4.928112131 |
| **A_51_P511315** | Pstpip1 | -0.061589281 | -0.271274281 | 0.148095719 | 9.59677542 | -0.722614939 | 0.49768634 | 0.869231206 | -4.928120318 |
| **A_55_P1954306** | Esrra | -0.058624866 | -0.258234789 | 0.140985057 | 12.16789455 | -0.722551654 | 0.497722457 | 0.869231206 | -4.928151098 |
| **A_55_P2162573** | Art3 | -0.034190874 | -0.150607826 | 0.082226078 | 7.82725412 | -0.722542066 | 0.497727929 | 0.869231206 | -4.928155762 |
| **A_55_P2023797** | Cep112 | -0.056113218 | -0.247186788 | 0.134960351 | 8.670218059 | -0.722493133 | 0.497755856 | 0.869231206 | -4.92817956 |
| **A_52_P561936** | Sptssa | 0.147491955 | -0.354950492 | 0.649934401 | 11.24033435 | 0.722189504 | 0.497929169 | 0.869231206 | -4.928327199 |
| **A_51_P415607** | Fkbp7 | 0.14690819 | -0.353569891 | 0.647386272 | 10.30619241 | 0.72215448 | 0.497949163 | 0.869231206 | -4.928344226 |
| **A_55_P1954418** | NA | 0.16699903 | -0.401952307 | 0.735950367 | 11.20220996 | 0.722117698 | 0.497970162 | 0.869231206 | -4.928362107 |
| **A_51_P279437** | Mfsd2a | 0.129155398 | -0.310946561 | 0.569257358 | 11.30247913 | 0.721985416 | 0.498045686 | 0.869231206 | -4.928426409 |
| **A_51_P126302** | Rbmx2 | -0.031240635 | -0.137705157 | 0.075223886 | 7.915542332 | -0.721911754 | 0.498087745 | 0.869231206 | -4.916160166 |
| **A_51_P283300** | Cacnb2 | 0.031692848 | -0.076315982 | 0.139701679 | 7.656508474 | 0.721890244 | 0.498100028 | 0.869231206 | -4.928472666 |
| **A_52_P122696** | Arl2 | -0.03920103 | -0.172800423 | 0.094398364 | 8.472551658 | -0.721875248 | 0.498108591 | 0.869231206 | -4.928479954 |
| **A_55_P2096281** | NA | 0.038141794 | -0.091850569 | 0.168134158 | 15.67210723 | 0.721859113 | 0.498117804 | 0.869231206 | -4.928487796 |
| **A_55_P1971194** | Timm23 | 0.098842546 | -0.2380338 | 0.435718892 | 12.63924453 | 0.721842699 | 0.498127177 | 0.869231206 | -4.928495773 |
| **A_51_P230977** | Cdc42 | 0.098407093 | -0.237006174 | 0.43382036 | 14.84194526 | 0.721797427 | 0.498153028 | 0.869231206 | -4.928517774 |
| **A_51_P390377** | Rcor3 | -0.044775956 | -0.197398777 | 0.107846865 | 9.275877238 | -0.721762903 | 0.498172744 | 0.869231206 | -4.928534551 |
| **A_55_P1974233** | Dtd2 | 0.035970495 | -0.086640629 | 0.15858162 | 8.297948836 | 0.721748196 | 0.498181142 | 0.869231206 | -4.928541698 |
| **A_66_P136186** | Wee1 | 0.12846314 | -0.30972308 | 0.566649361 | 10.13951883 | 0.72125524 | 0.498462707 | 0.869584156 | -4.92878118 |
| **A_55_P1981544** | Lactb | 0.075533904 | -0.182148143 | 0.333215952 | 10.87340504 | 0.721152203 | 0.498521573 | 0.869584156 | -4.928831221 |
| **A_51_P284466** | Tsc22d4 | -0.046285398 | -0.204210094 | 0.111639299 | 9.467279846 | -0.721046233 | 0.49858212 | 0.869584156 | -4.92888268 |
| **A_52_P257523** | 1110065P20Rik | 0.052800388 | -0.127367652 | 0.232968427 | 11.47413564 | 0.720988803 | 0.498614935 | 0.869584156 | -4.928910565 |
| **A_52_P641892** | Hacd3 | -0.093429219 | -0.412244845 | 0.225386407 | 13.14218134 | -0.720961847 | 0.498630338 | 0.869584156 | -4.928923653 |
| **A_55_P2001203** | Dnm2 | 0.071813865 | -0.17325574 | 0.31688347 | 13.8697339 | 0.720921559 | 0.498653359 | 0.869584156 | -4.928943214 |
| **A_55_P1990609** | Ube2j1 | -0.042023968 | -0.185487658 | 0.101439723 | 12.33575668 | -0.720649714 | 0.498808717 | 0.869643294 | -4.929075177 |
| **A_51_P358225** | Serinc1 | -0.102583784 | -0.452854332 | 0.247686765 | 13.97813349 | -0.720517037 | 0.498884553 | 0.869643294 | -4.929139569 |
| **A_55_P2030128** | Tfpt | 0.035062171 | -0.084667468 | 0.154791811 | 8.370170395 | 0.720454066 | 0.498920549 | 0.869643294 | -4.929170127 |
| **A_55_P1953043** | E130311K13Rik | -0.041253879 | -0.182157708 | 0.09964995 | 10.05469922 | -0.720296252 | 0.499010767 | 0.869643294 | -4.929246702 |
| **A_55_P1970105** | Ltbp2 | 0.122837998 | -0.296746976 | 0.542422972 | 9.805868194 | 0.720247843 | 0.499038444 | 0.869643294 | -4.929270188 |
| **A_55_P1979067** | A430073D23Rik | -0.234524205 | -1.035602518 | 0.566554109 | 13.44720549 | -0.720247666 | 0.499038545 | 0.869643294 | -4.929270274 |
| **A_55_P2060184** | Lamtor3 | 0.065274465 | -0.157700534 | 0.288249465 | 12.41195438 | 0.720205303 | 0.499062766 | 0.869643294 | -4.929290826 |
| **A_52_P542172** | Rb1cc1 | -0.058608093 | -0.258821303 | 0.141605117 | 9.865736851 | -0.720168337 | 0.499083901 | 0.869643294 | -4.929308759 |
| **A_66_P107818** | Gdf3 | 0.035639866 | -0.086113212 | 0.157392944 | 7.756217949 | 0.72015383 | 0.499092196 | 0.869643294 | -4.929315797 |
| **A_51_P378009** | Blzf1 | -0.073525158 | -0.324854577 | 0.177804262 | 10.90137434 | -0.719717067 | 0.499341968 | 0.869986707 | -4.929527622 |
| **A_51_P285042** | Rtf1 | 0.066106327 | -0.159891619 | 0.292104274 | 11.26554613 | 0.719627412 | 0.49939325 | 0.869986707 | -4.929571092 |
| **A_55_P1958804** | Hsd3b3 | 0.055856466 | -0.135128429 | 0.246841362 | 7.804935407 | 0.719521208 | 0.499454001 | 0.869986707 | -4.929622579 |
| **A_55_P2083382** | Snf8 | -0.100140928 | -0.442585381 | 0.242303525 | 13.18436864 | -0.71943347 | 0.499504194 | 0.869986707 | -4.92966511 |
| **A_55_P1986276** | Tmem41b | 0.09395654 | -0.227363243 | 0.415276322 | 11.47402156 | 0.719380584 | 0.49953445 | 0.869986707 | -4.929690745 |

| | | | | | | | | | |
|---|---|---|---|---|---|---|---|---|---|
| **A_55_P2121521** | H3c11 | 0.032209664 | -0.077950406 | 0.142369735 | 7.719021571 | 0.719334945 | 0.499560562 | 0.869986707 | -4.929712866 |
| **A_55_P2139112** | Cdc26 | 0.070228665 | -0.169985506 | 0.310442837 | 10.77446644 | 0.719258415 | 0.499604348 | 0.869986707 | -4.929749956 |
| **A_52_P1026830** | Tbc1d8b | -0.037992761 | -0.167965557 | 0.091980035 | 9.300542639 | -0.719146809 | 0.499668209 | 0.870019531 | -4.929804041 |
| **A_55_P2020911** | Rps6ka6 | -0.108590348 | -0.480281331 | 0.263100634 | 12.40102344 | -0.718750839 | 0.499894825 | 0.870260766 | -4.929995878 |
| **A_52_P623775** | Eri2 | -0.032538651 | -0.143926406 | 0.078849103 | 8.088088987 | -0.718672885 | 0.499939447 | 0.870260766 | -4.930033635 |
| **A_52_P298002** | Gch1 | -0.12524173 | -0.553977256 | 0.303493796 | 8.6913417 | -0.71866873 | 0.499941826 | 0.870260766 | -4.930035647 |
| **A_55_P2140516** | Pex13 | 0.100895273 | -0.244566379 | 0.446356925 | 12.65678282 | 0.718522109 | 0.50002576 | 0.870328494 | -4.930106654 |
| **A_55_P2141724** | Gm13275 | 0.033426875 | -0.081045873 | 0.147899624 | 8.116460546 | 0.718394231 | 0.500098974 | 0.870346866 | -4.930168575 |
| **A_55_P2000894** | NA | 0.055827325 | -0.135370116 | 0.247024767 | 14.69071194 | 0.71834638 | 0.500126372 | 0.870346866 | -4.930191743 |
| **A_52_P574945** | 1500026H17Rik | -0.044534179 | -0.197097232 | 0.108028874 | 8.581809704 | -0.71814682 | 0.500240643 | 0.870397853 | -4.930288351 |
| **A_52_P102783** | Dync2h1 | -0.051454802 | -0.227747049 | 0.124837444 | 8.957356404 | -0.718061872 | 0.500289292 | 0.870397853 | -4.930329468 |
| **A_55_P2067712** | Hmcn2 | 0.240075506 | -0.582560201 | 1.062711212 | 10.66008727 | 0.717975226 | 0.500338915 | 0.870397853 | -4.930371404 |
| **A_52_P204689** | Map4k2 | -0.03309045 | -0.146484636 | 0.080303735 | 8.096095935 | -0.717928269 | 0.50036581 | 0.870397853 | -4.930394128 |
| **A_55_P2114898** | Gdap2 | 0.08524322 | -0.206880555 | 0.377366996 | 10.15375599 | 0.717896946 | 0.500383751 | 0.870397853 | -4.930409287 |
| **A_52_P84096** | Pgd | -0.039090273 | -0.173070065 | 0.09488952 | 11.45649416 | -0.717791912 | 0.500443913 | 0.870397853 | -4.930460112 |
| **A_52_P271155** | Mtpap | -0.030686284 | -0.135880464 | 0.074507896 | 10.11724594 | -0.71766497 | 0.500516632 | 0.870397853 | -4.930521531 |
| **A_51_P163578** | Ugt2b35 | 0.032001239 | -0.07774081 | 0.141743288 | 7.643347891 | 0.717402507 | 0.500667006 | 0.870397853 | -4.930648492 |
| **A_51_P511015** | Fzd9 | -0.031586225 | -0.139905906 | 0.076733457 | 7.814503749 | -0.717396916 | 0.500670209 | 0.870397853 | -4.930651196 |
| **A_51_P447988** | Atp6ap1 | -0.080617991 | -0.357086358 | 0.195850376 | 9.857072174 | -0.717390606 | 0.500673825 | 0.870397853 | -4.930654249 |
| **A_66_P129672** | 4930453N24Rik | -0.059093849 | -0.26176238 | 0.143574683 | 10.66827046 | -0.717340116 | 0.500702756 | 0.870397853 | -4.930678668 |
| **A_51_P439299** | Relt | -0.044208488 | -0.195851992 | 0.107435016 | 8.415825445 | -0.717217721 | 0.500772894 | 0.870397853 | -4.930737857 |
| **A_52_P127643** | NA | -0.07791462 | -0.345184137 | 0.189354897 | 9.502526838 | -0.717197374 | 0.500784554 | 0.870397853 | -4.930747696 |
| **A_55_P1955778** | Adamts4 | -0.211286106 | -0.936071122 | 0.51349891 | 10.21693962 | -0.717184501 | 0.500791932 | 0.870397853 | -4.930753921 |
| **A_55_P2143070** | Ass1 | -0.1028205 | -0.455578019 | 0.249937019 | 11.8004149 | -0.71708823 | 0.500847106 | 0.870397853 | -4.93080047 |
| **A_55_P2098683** | St7l | 0.050534126 | -0.122854992 | 0.223923245 | 11.10855365 | 0.717021347 | 0.500885439 | 0.870397853 | -4.930832807 |
| **A_55_P2102155** | Rgs20 | 0.244285732 | -0.593963109 | 1.082534573 | 12.59756346 | 0.71695897 | 0.500921192 | 0.870397853 | -4.930862963 |
| **A_55_P2109997** | Lrig1 | -0.084908254 | -0.376361649 | 0.206545141 | 9.349589402 | -0.716720713 | 0.501057771 | 0.870485213 | -4.930978128 |
| **A_51_P240693** | Tecpr1 | -0.121162303 | -0.537106428 | 0.294781821 | 11.28192551 | -0.716641065 | 0.501103434 | 0.870485213 | -4.931016621 |
| **A_51_P332355** | Kif18a | -0.042831237 | -0.189869855 | 0.104207381 | 7.912530401 | -0.716635588 | 0.501106574 | 0.870485213 | -4.931019268 |
| **A_51_P189916** | Nol12 | -0.044880484 | -0.199005981 | 0.109245014 | 11.03696002 | -0.716394427 | 0.501244851 | 0.870644073 | -4.931135796 |
| **A_55_P2145257** | NA | 0.071150912 | -0.173216432 | 0.315518256 | 8.792258051 | 0.716318991 | 0.501288111 | 0.870644073 | -4.93117224 |
| **A_55_P1967568** | Chtop | -0.055473091 | -0.246046086 | 0.135099903 | 12.23176746 | -0.716127209 | 0.5013981 | 0.870756862 | -4.931264879 |
| **A_52_P375876** | Arf3 | -0.084073031 | -0.442404058 | 0.274257996 | 12.49737479 | -0.766311371 | 0.501585612 | 0.871004248 | -4.738978349 |
| **A_55_P2099840** | NA | 0.032548939 | -0.079367591 | 0.144465469 | 7.798605855 | 0.715503504 | 0.501755915 | 0.871221711 | -4.931566021 |
| **A_51_P153224** | Zdhhc12 | 0.053869027 | -0.131383012 | 0.239121065 | 9.657281418 | 0.715394005 | 0.501818752 | 0.871252551 | -4.931618869 |
| **A_55_P1970866** | Rplp0 | 0.062329596 | -0.152075612 | 0.276734803 | 16.0166185 | 0.71520105 | 0.501929493 | 0.871346618 | -4.931711981 |
| **A_52_P253004** | Ralb | 0.112817645 | -0.27529125 | 0.50092654 | 14.11091701 | 0.715142514 | 0.501963092 | 0.871346618 | -4.931740224 |
| **A_51_P142813** | Aspn | 0.102414827 | -0.249962811 | 0.454792465 | 8.00781551 | 0.715029002 | 0.502028249 | 0.871381468 | -4.931794987 |
| **A_52_P653937** | Rpl21 | -0.032672446 | -0.145103493 | 0.079758602 | 7.83963718 | -0.714931699 | 0.502084107 | 0.871400171 | -4.931841925 |
| **A_66_P124952** | Hspe1 | 0.123297158 | -0.301159471 | 0.547753788 | 13.67902398 | 0.714642613 | 0.502250085 | 0.871574735 | -4.931981348 |
| **A_51_P134301** | Pfdn5 | 0.075478922 | -0.184376733 | 0.335334576 | 11.87172877 | 0.714599449 | 0.50227487 | 0.871574735 | -4.932002161 |
| **A_55_P1961531** | Nemp1 | -0.061271106 | -0.272283744 | 0.149741531 | 8.863806366 | -0.714358836 | 0.502413051 | 0.871676272 | -4.932118167 |
| **A_55_P2152872** | Fgfrl1 | 0.08815437 | -0.215449798 | 0.391758537 | 11.90636654 | 0.714340507 | 0.502423578 | 0.871676272 | -4.932127003 |
| **A_55_P2045622** | Setdb2 | 0.056802831 | -0.138850847 | 0.25245651 | 8.952121498 | 0.714251447 | 0.502474731 | 0.871686777 | -4.932169932 |
| **A_55_P2158866** | Nme6 | 0.047389022 | -0.115905931 | 0.210683975 | 10.68917985 | 0.713960436 | 0.502641902 | 0.87189853 | -4.932310179 |
| **A_55_P2045770** | Gmfb | -0.083338375 | -0.370621601 | 0.20394485 | 13.10345006 | -0.713680653 | 0.502802658 | 0.872022928 | -4.932444972 |
| **A_55_P2134862** | Cmip | 0.093382332 | -0.228612286 | 0.415376949 | 11.06249829 | 0.713485685 | 0.502914702 | 0.872022928 | -4.932538879 |
| **A_55_P1990190** | Arhgap28 | 0.066044541 | -0.161700013 | 0.293789095 | 8.930506685 | 0.71344104 | 0.502940361 | 0.872022928 | -4.93256038 |
| **A_55_P2006519** | Clip1 | -0.051444127 | -0.228889482 | 0.126001228 | 9.851659978 | -0.713247619 | 0.503051536 | 0.872022928 | -4.932653518 |
| **A_52_P647906** | NA | -0.044260965 | -0.1969657 | 0.108443771 | 12.08413363 | -0.713078816 | 0.503148575 | 0.872022928 | -4.932734785 |

| | | | | | | | | |
|---|---|---|---|---|---|---|---|---|
| **A_55_P2040678** | Susd2 | 0.101552592 | -0.248842503 | 0.451947687 | 8.643454209 | 0.713020734 | 0.503181968 | 0.872022928 | -4.932762744 |
| **A_55_P1985895** | Thumpd1 | -0.073419073 | -0.326784081 | 0.179945935 | 11.67317583 | -0.712904615 | 0.503248731 | 0.872022928 | -4.932818635 |
| **A_55_P1977330** | Zbtb1 | 0.04194688 | -0.102811754 | 0.186705515 | 8.431694483 | 0.712893004 | 0.503255407 | 0.872022928 | -4.932824224 |
| **A_55_P2058851** | Fancf | 0.058651627 | -0.143764773 | 0.261068026 | 8.731142052 | 0.712858823 | 0.503275061 | 0.872022928 | -4.932840674 |
| **A_55_P1983368** | Sh3d21 | 0.116885426 | -0.286507978 | 0.520278831 | 11.07817978 | 0.712854237 | 0.503277697 | 0.872022928 | -4.932842881 |
| **A_51_P321331** | Sptlc2 | -0.060955511 | -0.2713331 | 0.149422079 | 8.823819849 | -0.712824573 | 0.503294754 | 0.872022928 | -4.932857157 |
| **A_55_P2028721** | Rnf130 | -0.073191765 | -0.32584355 | 0.17946002 | 11.27839191 | -0.712703699 | 0.503364262 | 0.872022928 | -4.932915324 |
| **A_52_P676271** | Tuba4a | 0.06949552 | -0.170441235 | 0.309432274 | 13.56704829 | 0.712572719 | 0.503439588 | 0.872022928 | -4.932978346 |
| **A_55_P1986158** | Klhdc4 | -0.068629017 | -0.305580439 | 0.168322405 | 11.17852086 | -0.712553741 | 0.503450503 | 0.872022928 | -4.932987477 |
| **A_55_P2157388** | Skint9 | 0.076116573 | -0.186740234 | 0.33897338 | 8.614097178 | 0.712408613 | 0.503533976 | 0.872022928 | -4.933057294 |
| **A_55_P2027412** | NA | -0.043089878 | -0.191896378 | 0.105716622 | 8.783808289 | -0.712397724 | 0.503540239 | 0.872022928 | -4.933062532 |
| **A_51_P470724** | Eif2s2 | 0.048987903 | -0.120197906 | 0.218173711 | 9.905791808 | 0.712351068 | 0.503567076 | 0.872022928 | -4.933084974 |
| **A_52_P516409** | Col4a6 | 0.28712799 | -0.704610315 | 1.278866295 | 8.38098083 | 0.712274843 | 0.503610924 | 0.872022928 | -4.933121637 |
| **A_66_P122155** | Wdr26 | -0.11325353 | -0.504476522 | 0.277969462 | 11.30562621 | -0.712191099 | 0.503659099 | 0.872022928 | -4.933161913 |
| **A_52_P219943** | Epc1 | -0.063053556 | -0.280903313 | 0.154796202 | 8.262393418 | -0.712068321 | 0.503729736 | 0.872022928 | -4.933220955 |
| **A_55_P1993463** | NA | 0.035436366 | -0.087007844 | 0.157880577 | 8.455512001 | 0.712000157 | 0.503768956 | 0.872022928 | -4.933253731 |
| **A_51_P114049** | Tmem109 | -0.090060476 | -0.401272666 | 0.221151714 | 13.65215102 | -0.711945583 | 0.503800357 | 0.872022928 | -4.924336623 |
| **A_51_P279062** | Treml1 | -0.05418862 | -0.241443739 | 0.133066499 | 8.755622579 | -0.711940264 | 0.503803417 | 0.872022928 | -4.933282528 |
| **A_51_P421182** | Sys1 | 0.068120605 | -0.167293515 | 0.303534724 | 8.681429788 | 0.711893702 | 0.50383021 | 0.872022928 | -4.933304914 |
| **A_55_P2101426** | Mrps25 | 0.051046061 | -0.12536597 | 0.227458091 | 10.53644981 | 0.711874108 | 0.503841485 | 0.872022928 | -4.933314334 |
| **A_51_P130660** | Mrps30 | 0.089290949 | -0.219329456 | 0.397911355 | 10.55552535 | 0.711790136 | 0.503889807 | 0.872028479 | -4.933354702 |
| **A_52_P212686** | Lrrk1 | -0.048220545 | -0.214955092 | 0.118514002 | 8.000866392 | -0.711501281 | 0.504056053 | 0.872170509 | -4.933493535 |
| **A_55_P2063785** | Ank1 | 0.034810563 | -0.085557414 | 0.15517854 | 7.8039649 | 0.711490737 | 0.504062122 | 0.872170509 | -4.933498601 |
| **A_55_P1980981** | Apba3 | -0.0790548 | -0.352504706 | 0.194395106 | 10.31514143 | -0.71124566 | 0.504203202 | 0.872336529 | -4.933616357 |
| **A_55_P2019463** | Tnks1bp1 | -0.076782063 | -0.342450743 | 0.188886617 | 10.60979295 | -0.711031074 | 0.504326752 | 0.872348287 | -4.933719436 |
| **A_55_P2383902** | 8030453O22Rik | -0.115254395 | -0.514056145 | 0.283547355 | 8.451244883 | -0.711000003 | 0.504344643 | 0.872348287 | -4.93373436 |
| **A_55_P1958447** | Prepl | 0.042473282 | -0.104503305 | 0.189449868 | 8.179408036 | 0.710946345 | 0.504375541 | 0.872348287 | -4.93376013 |
| **A_55_P2151986** | Gngt2 | 0.105249722 | -0.258974807 | 0.469474251 | 10.52033433 | 0.710920324 | 0.504390525 | 0.872348287 | -4.933772627 |
| **A_55_P2091736** | Rassf2 | -0.049841419 | -0.222348529 | 0.122665691 | 8.420654359 | -0.710808419 | 0.504454969 | 0.872381686 | -4.933826366 |
| **A_52_P655803** | Chpt1 | -0.105309436 | -0.469918295 | 0.259299424 | 10.56504157 | -0.710573868 | 0.50459006 | 0.872396515 | -4.933938981 |
| **A_55_P2056021** | Snap91 | 0.113718584 | -0.280011647 | 0.507448814 | 9.019387864 | 0.71056178 | 0.504597023 | 0.872396515 | -4.933944783 |
| **A_55_P2067645** | Gm5136 | 0.042593233 | -0.104878901 | 0.190065367 | 8.432512141 | 0.710558441 | 0.504598946 | 0.872396515 | -4.933946386 |
| **A_51_P273684** | Gbe1 | -0.07513168 | -0.335362426 | 0.185099065 | 10.02670386 | -0.710286663 | 0.504755511 | 0.87246971 | -4.934076836 |
| **A_66_P118759** | Safb2 | 0.105127973 | -0.259000537 | 0.469256483 | 11.30378645 | 0.710285206 | 0.50475635 | 0.87246971 | -4.934077535 |
| **A_55_P2039463** | Timmdc1 | 0.098539431 | -0.242785521 | 0.439864384 | 12.13221285 | 0.710249891 | 0.504776697 | 0.87246971 | -4.934094483 |
| **A_55_P2021378** | Wapl | -0.044955708 | -0.200733197 | 0.11082178 | 11.60231493 | -0.709985223 | 0.5049292 | 0.872547603 | -4.934221477 |
| **A_51_P288026** | Haus6 | -0.038112928 | -0.170188992 | 0.093963136 | 8.179408555 | -0.70993299 | 0.504959302 | 0.872547603 | -4.934246535 |
| **A_55_P2092286** | NA | -0.034809728 | -0.155455302 | 0.085835845 | 7.822322797 | -0.709836628 | 0.505014836 | 0.872547603 | -4.93429276 |
| **A_51_P407879** | Fam193b | -0.154172567 | -0.688549836 | 0.380204702 | 10.55633213 | -0.709787408 | 0.505043203 | 0.872547603 | -4.934316369 |
| **A_55_P1960989** | Nlgn2 | 0.095985446 | -0.236713014 | 0.428683905 | 9.514653227 | 0.709780002 | 0.505047472 | 0.872547603 | -4.934319921 |
| **A_51_P448618** | Slc16a10 | -0.102747854 | -0.458988059 | 0.253492352 | 10.41947562 | -0.709576115 | 0.505164993 | 0.872644722 | -4.934417704 |
| **A_55_P2066697** | Trim47 | 0.156205091 | -0.38541671 | 0.697826893 | 10.70914113 | 0.709525831 | 0.50519398 | 0.872644722 | -4.934441816 |
| **A_52_P475870** | Tmem29 | 0.076425372 | -0.18861009 | 0.341460834 | 8.790961682 | 0.709418871 | 0.505255642 | 0.872673246 | -4.934493102 |
| **A_52_P572045** | Pfkp | 0.039428985 | -0.09733566 | 0.176193631 | 8.406260267 | 0.709268908 | 0.505342102 | 0.872730388 | -4.934564996 |
| **A_55_P1984721** | Ube2i | -0.047085407 | -0.210424163 | 0.116253348 | 12.16669395 | -0.709195962 | 0.505384163 | 0.872730388 | -4.934599963 |
| **A_52_P306305** | NA | -0.076662693 | -0.370948273 | 0.217622887 | 13.94500692 | -0.733022991 | 0.505474696 | 0.872730388 | -4.824454977 |
| **A_55_P2025228** | Akt2 | 0.046342646 | -0.114455801 | 0.207141093 | 12.30894875 | 0.709035762 | 0.505476542 | 0.872730388 | -4.934676746 |
| **A_51_P196158** | Btd | -0.062117176 | -0.277669656 | 0.153435304 | 10.30219378 | -0.708969975 | 0.505514482 | 0.872730388 | -4.934708273 |
| **A_55_P2030383** | Frmd4a | 0.073638597 | -0.182052625 | 0.329329819 | 11.43059866 | 0.708530987 | 0.505767695 | 0.872925508 | -4.934918593 |
| **A_55_P1962274** | Ccnt1 | -0.073938373 | -0.330701452 | 0.182824707 | 9.04582199 | -0.708445547 | 0.505816988 | 0.872925508 | -4.934959515 |

| | | | | | | | | |
|---|---|---|---|---|---|---|---|---|
| **A_55_P2130600** | Musk | 0.069917221 | -0.172894117 | 0.312728558 | 8.606979871 | 0.70840947 | 0.505837803 | 0.872925508 | -4.934976793 |
| **A_55_P1966189** | Il4ra | -0.04333311 | -0.193831399 | 0.107165178 | 8.009893088 | -0.708365602 | 0.505863113 | 0.872925508 | -4.934997802 |
| **A_55_P2180789** | Huwe1 | 0.100875396 | -0.286753966 | 0.488504758 | 11.86511319 | 0.732269752 | 0.505885601 | 0.872925508 | -4.77365356 |
| **A_55_P2397599** | B930025B16Rik | -0.154992661 | -0.693337317 | 0.383351994 | 9.039373158 | -0.708304323 | 0.505898471 | 0.872925508 | -4.935027147 |
| **A_55_P2146783** | Ropn1l | 0.040864536 | -0.101112949 | 0.18284202 | 8.679075727 | 0.708102709 | 0.506014813 | 0.872993174 | -4.935123682 |
| **A_55_P2026223** | St6galnac4 | -0.041639492 | -0.186331462 | 0.103052478 | 7.872258762 | -0.70799495 | 0.506077003 | 0.872993174 | -4.93517527 |
| **A_51_P259009** | Cacnb3 | -0.107328208 | -0.480321448 | 0.265665031 | 8.769413329 | -0.707916577 | 0.506122237 | 0.872993174 | -4.935212785 |
| **A_66_P107038** | Prcp | -0.102329201 | -0.45812412 | 0.253465718 | 12.46998553 | -0.70756934 | 0.506322681 | 0.872993174 | -4.935378961 |
| **A_55_P1955841** | Ctsa | -0.068584541 | -0.307056061 | 0.169886979 | 16.62254292 | -0.707552842 | 0.506332206 | 0.872993174 | -4.935386855 |
| **A_55_P2156973** | Aar2 | 0.069918505 | -0.173229513 | 0.313066522 | 10.44064418 | 0.707441546 | 0.506396464 | 0.872993174 | -4.935440103 |
| **A_52_P508974** | Igf2bp3 | -0.066205068 | -0.296441807 | 0.164031672 | 9.014531818 | -0.707433742 | 0.50640097 | 0.872993174 | -4.935443837 |
| **A_55_P2112504** | Kcnv2 | -0.091969143 | -0.411823612 | 0.227885325 | 8.617182712 | -0.707389945 | 0.506426259 | 0.872993174 | -4.935464789 |
| **A_55_P2041961** | Bmp2 | 0.040446853 | -0.100229458 | 0.181123164 | 8.593015714 | 0.707347677 | 0.506450665 | 0.872993174 | -4.935485008 |
| **A_55_P2017312** | Eapp | 0.035915995 | -0.089003083 | 0.160835072 | 11.78385388 | 0.707340126 | 0.506455025 | 0.872993174 | -4.93548862 |
| **A_51_P508444** | St13 | -0.076557786 | -0.342866702 | 0.18975113 | 12.29722925 | -0.707249782 | 0.506507195 | 0.872993174 | -4.935531834 |
| **A_51_P507053** | Slc38a1 | -0.078184238 | -0.350164605 | 0.193796128 | 15.22082855 | -0.707213947 | 0.506527889 | 0.872993174 | -4.935548974 |
| **A_66_P119017** | Smarca1 | 0.108675985 | -0.269378639 | 0.48673061 | 8.432117441 | 0.707209676 | 0.506530355 | 0.872993174 | -4.935551017 |
| **A_55_P2059337** | Fastk | 0.072493894 | -0.179717413 | 0.324705202 | 12.11396899 | 0.707141036 | 0.506569995 | 0.872993174 | -4.935583845 |
| **A_51_P169032** | 2810004N23Rik | 0.106810931 | -0.264877963 | 0.478499825 | 11.61551076 | 0.70697699 | 0.506664741 | 0.873025553 | -4.935662293 |
| **A_52_P40954** | Dubr | 0.046718035 | -0.115863727 | 0.209299797 | 10.0130835 | 0.706938938 | 0.50668672 | 0.873025553 | -4.935680487 |
| **A_55_P2166282** | NA | -0.098741228 | -0.442440474 | 0.244958018 | 15.97188985 | -0.706787903 | 0.506773965 | 0.873025553 | -4.935752697 |
| **A_51_P261001** | Mtf2 | 0.060584957 | -0.150318684 | 0.271488599 | 9.84197524 | 0.706724081 | 0.506810834 | 0.873025553 | -4.935783207 |
| **A_52_P130727** | Rtl6 | -0.0402806 | -0.180503592 | 0.099942392 | 8.46368784 | -0.706717532 | 0.506814617 | 0.873025553 | -4.935786337 |
| **A_55_P2020248** | NA | 0.095004404 | -0.235825991 | 0.425834799 | 14.05222656 | 0.706492406 | 0.506944685 | 0.873171789 | -4.935893939 |
| **A_52_P578634** | Cela1 | 0.031154381 | -0.077355139 | 0.139663902 | 7.732261201 | 0.706350814 | 0.507026502 | 0.873234898 | -4.935961601 |
| **A_66_P112862** | Gm17821 | -0.210042198 | -0.94184536 | 0.521760964 | 11.75566665 | -0.706124752 | 0.507157148 | 0.873382084 | -4.936069606 |
| **A_55_P2145521** | Stk38l | -0.104107774 | -0.467145819 | 0.25893027 | 11.03358512 | -0.705505145 | 0.507515345 | 0.87386036 | -4.936365495 |
| **A_51_P321374** | Plpp7 | 0.076440649 | -0.190124846 | 0.343006143 | 8.404060346 | 0.70548794 | 0.507525294 | 0.87386036 | -4.936373708 |
| **A_52_P118310** | Chn1 | 0.032554506 | -0.080982859 | 0.146091871 | 7.83258046 | 0.705409759 | 0.507570504 | 0.87386036 | -4.936411028 |
| **A_52_P264229** | Ing5 | 0.083430133 | -0.20777768 | 0.374637946 | 10.16528766 | 0.704837623 | 0.507901432 | 0.874352224 | -4.936684037 |
| **A_52_P361551** | Hspa14 | 0.111695223 | -0.278262572 | 0.501653017 | 12.73442853 | 0.704670606 | 0.507998063 | 0.874440695 | -4.936763701 |
| **A_52_P79187** | Ankrd12 | -0.08909114 | -0.400212389 | 0.222030109 | 9.217194136 | -0.704488657 | 0.508103348 | 0.874453804 | -4.93685047 |
| **A_51_P204080** | Hk2 | -0.127027062 | -0.570739232 | 0.316685108 | 11.75396111 | -0.704310359 | 0.508206534 | 0.874453804 | -4.936935481 |
| **A_55_P2177941** | Tmem208 | 0.094716061 | -0.236142035 | 0.425574156 | 11.61118234 | 0.704289195 | 0.508218783 | 0.874453804 | -4.936945571 |
| **A_55_P2193512** | NA | -0.032590369 | -0.146434784 | 0.081254046 | 7.794169853 | -0.704282191 | 0.508222836 | 0.874453804 | -4.93694891 |
| **A_55_P2168823** | NA | 0.150452055 | -0.375117521 | 0.676021631 | 11.90735441 | 0.704266564 | 0.508231881 | 0.874453804 | -4.93695636 |
| **A_55_P2000102** | Mettl21a | 0.043270015 | -0.107926519 | 0.194466548 | 10.61238744 | 0.70406762 | 0.508347036 | 0.874538256 | -4.937051189 |
| **A_55_P2133205** | Nmt2 | 0.073298451 | -0.182840489 | 0.32943739 | 10.40408923 | 0.704025437 | 0.508371455 | 0.874538256 | -4.937071294 |
| **A_55_P2161702** | Nrip2 | 0.035542237 | -0.088743264 | 0.159827738 | 7.701943458 | 0.703547553 | 0.508648148 | 0.874936376 | -4.937298989 |
| **A_55_P2363442** | Zfp867 | 0.048186886 | -0.120358722 | 0.216732495 | 8.748471368 | 0.703364727 | 0.50875403 | 0.874968441 | -4.937386067 |
| **A_66_P104221** | NA | -0.112088048 | -0.504246208 | 0.280070111 | 8.747831038 | -0.703181141 | 0.508860368 | 0.874968441 | -4.937473489 |
| **A_52_P24308** | Uchl1 | -0.214679835 | -0.96579684 | 0.536437171 | 8.569253315 | -0.703157836 | 0.508873868 | 0.874968441 | -4.937484585 |
| **A_52_P53470** | NA | -0.03121576 | -0.14047366 | 0.07804214 | 8.006603297 | -0.702894623 | 0.509026355 | 0.874968441 | -4.93760989 |
| **A_55_P2012221** | Armcx2 | 0.086945963 | -0.217377838 | 0.391269765 | 9.689384025 | 0.702882395 | 0.50903344 | 0.874968441 | -4.937615711 |
| **A_55_P2106801** | Bglap3 | 0.202494524 | -0.506282063 | 0.91127111 | 12.02102888 | 0.702866885 | 0.509042426 | 0.874968441 | -4.937623093 |
| **A_51_P385697** | Mgat3 | 0.093970699 | -0.234979706 | 0.422921104 | 9.012101064 | 0.702799097 | 0.509081704 | 0.874968441 | -4.937655357 |
| **A_55_P1977917** | Cln3 | 0.072385594 | -0.181023442 | 0.32579463 | 11.67502678 | 0.702747336 | 0.509111696 | 0.874968441 | -4.937679991 |
| **A_55_P2171158** | Hmgn1 | -0.109299997 | -0.491990395 | 0.273390401 | 15.71677613 | -0.702654363 | 0.509165572 | 0.874968441 | -4.937724236 |
| **A_55_P2153743** | Slc26a6 | 0.10089914 | -0.25242028 | 0.454218559 | 8.934265973 | 0.702569217 | 0.509214915 | 0.874968441 | -4.937764752 |
| **A_51_P278994** | Swap70 | 0.109311741 | -0.273473469 | 0.492096951 | 9.897639123 | 0.702555802 | 0.509222689 | 0.874968441 | -4.937771135 |

| | | | | | | | | |
|---|---|---|---|---|---|---|---|---|
| **A_55_P2298029** | Nsd3 | -0.084729598 | -0.381454098 | 0.211994902 | 10.31268606 | -0.702507397 | 0.509250742 | 0.874968441 | -4.937794166 |
| **A_55_P2044252** | Zfp512b | -0.097679243 | -0.439776097 | 0.244417611 | 10.40948484 | -0.702461242 | 0.509277493 | 0.874968441 | -4.937816125 |
| **A_55_P2065631** | Hook3 | -0.099007485 | -0.445801448 | 0.247786479 | 11.3092491 | -0.702369529 | 0.50933065 | 0.874968441 | -4.937859756 |
| **A_51_P352968** | Marcks | -0.079765271 | -0.359174331 | 0.199643788 | 12.56821891 | -0.702332115 | 0.509352336 | 0.874968441 | -4.937877554 |
| **A_55_P2046852** | Gata1 | 0.142264867 | -0.356120879 | 0.640650613 | 10.30891753 | 0.702265303 | 0.509391064 | 0.874968441 | -4.937909335 |
| **A_51_P349803** | Med11 | 0.098274159 | -0.246171101 | 0.442719418 | 10.40715572 | 0.701921082 | 0.509590623 | 0.875233441 | -4.938073033 |
| **A_55_P2028362** | NA | 0.062727012 | -0.157174519 | 0.282628543 | 8.25250528 | 0.701771148 | 0.509677562 | 0.875304984 | -4.938144316 |
| **A_51_P511386** | Zfp267 | 0.038854078 | -0.097377522 | 0.175085679 | 8.029351392 | 0.701661935 | 0.509740896 | 0.875335978 | -4.938196232 |
| **A_55_P2069995** | NA | 0.048017848 | -0.1204109 | 0.216446596 | 9.084364397 | 0.701383639 | 0.509902305 | 0.875535369 | -4.938328494 |
| **A_52_P217240** | Ppme1 | -0.068448294 | -0.308578271 | 0.171681682 | 9.16601675 | -0.701270249 | 0.50996808 | 0.87557053 | -4.938382372 |
| **A_55_P2153887** | Tatdn3 | -0.062062025 | -0.279841276 | 0.155717226 | 8.932772093 | -0.701097798 | 0.510068126 | 0.875607994 | -4.9384643 |
| **A_55_P1956799** | Smpx | 0.046594201 | -0.11695731 | 0.210145712 | 7.911426123 | 0.700884528 | 0.510191871 | 0.875607994 | -4.938565597 |
| **A_55_P2088103** | Ash2l | 0.083914302 | -0.210676344 | 0.378504949 | 13.22293309 | 0.70078726 | 0.510248315 | 0.875607994 | -4.938611789 |
| **A_52_P497424** | Aanat | -0.036256252 | -0.163616076 | 0.091103571 | 8.038131911 | -0.700357244 | 0.5104979 | 0.875607994 | -4.938815939 |
| **A_55_P2181508** | Gm10653 | 0.059974549 | -0.150730741 | 0.27067984 | 14.85907177 | 0.700262254 | 0.510553044 | 0.875607994 | -4.938861023 |
| **A_55_P2140843** | NA | 0.055269046 | -0.138955236 | 0.249493328 | 10.35460384 | 0.700079895 | 0.510658919 | 0.875607994 | -4.938947559 |
| **A_55_P2059864** | Igsf1 | -0.05496088 | -0.248113661 | 0.138191901 | 11.52228246 | -0.700038414 | 0.510683004 | 0.875607994 | -4.93896724 |
| **A_52_P406507** | Mus81 | -0.033594449 | -0.151676256 | 0.084487358 | 8.286592029 | -0.699928531 | 0.51074681 | 0.875607994 | -4.939019373 |
| **A_55_P2105256** | Trim47 | 0.176930276 | -0.444983904 | 0.798844455 | 10.60242418 | 0.69990784 | 0.510758825 | 0.875607994 | -4.939029189 |
| **A_55_P1970309** | Arhgap44 | 0.099898221 | -0.251275564 | 0.451072006 | 11.29188181 | 0.699849776 | 0.510792544 | 0.875607994 | -4.939056733 |
| **A_52_P335609** | Idnk | 0.055135327 | -0.138684645 | 0.248955299 | 9.274300747 | 0.699842953 | 0.510796506 | 0.875607994 | -4.93905997 |
| **A_51_P225506** | Pnrc2 | 0.047281647 | -0.118942268 | 0.213505562 | 12.44149748 | 0.699790836 | 0.510826773 | 0.875607994 | -4.939084691 |
| **A_55_P1972040** | Nox4 | -0.246568846 | -1.113463557 | 0.620325865 | 9.033265836 | -0.699746946 | 0.510852262 | 0.875607994 | -4.939105509 |
| **A_55_P2366358** | 4930444F02Rik | 0.031419411 | -0.079054739 | 0.141893562 | 7.675335031 | 0.699691398 | 0.510884524 | 0.875607994 | -4.939131855 |
| **A_55_P1956652** | Sfmbt2 | -0.145530318 | -0.657251884 | 0.366191247 | 10.11684005 | -0.699663024 | 0.510901004 | 0.875607994 | -4.939145312 |
| **A_51_P484842** | Asgr2 | 0.129108231 | -0.324879191 | 0.583095652 | 8.266505849 | 0.699647478 | 0.510910033 | 0.875607994 | -4.939152685 |
| **A_51_P482121** | Edn2 | -0.642160707 | -2.900935418 | 1.616614005 | 10.56838426 | -0.699423104 | 0.511040364 | 0.875607994 | -4.939259083 |
| **A_51_P441387** | Kansl1l | -0.041253749 | -0.186382757 | 0.103875258 | 8.006166754 | -0.699323891 | 0.511098001 | 0.875607994 | -4.939306121 |
| **A_55_P2076941** | Lrr1 | 0.0340092 | -0.085640739 | 0.15365914 | 7.865129982 | 0.699283203 | 0.511121638 | 0.875607994 | -4.939325409 |
| **A_55_P2133405** | Dmkn | -0.338007741 | -1.527473909 | 0.851458427 | 15.25553697 | -0.699107105 | 0.511223954 | 0.875607994 | -4.939408882 |
| **A_55_P2030282** | Gm5796 | -0.14304096 | -0.646414554 | 0.360332634 | 9.964657593 | -0.699099754 | 0.511228225 | 0.875607994 | -4.939412366 |
| **A_55_P1999289** | Ovol2 | 0.155191244 | -0.391057643 | 0.70144013 | 12.39184801 | 0.698949521 | 0.511315524 | 0.875607994 | -4.939483565 |
| **A_55_P2107288** | Tgfbr2 | -0.061983825 | -0.280165566 | 0.156197917 | 8.463986247 | -0.698922676 | 0.511331124 | 0.875607994 | -4.939496287 |
| **A_55_P2160862** | Wrn | -0.03210048 | -0.145132244 | 0.080931284 | 8.381702498 | -0.698683012 | 0.511470414 | 0.875607994 | -4.939609841 |
| **A_52_P657338** | Mrps17 | 0.117032959 | -0.295117216 | 0.529183133 | 11.63179217 | 0.698589186 | 0.511524951 | 0.875607994 | -4.939654288 |
| **A_51_P489229** | Eif3g | -0.052298455 | -0.236480316 | 0.131883407 | 14.45640143 | -0.698571981 | 0.511534952 | 0.875607994 | -4.939662437 |
| **A_51_P407004** | Prkrip1 | 0.071090837 | -0.179278675 | 0.321460348 | 10.41082324 | 0.698556192 | 0.511544131 | 0.875607994 | -4.939669916 |
| **A_55_P1955078** | Igflr1 | -0.084424784 | -0.381804148 | 0.21295458 | 8.582431665 | -0.698438696 | 0.511612433 | 0.875607994 | -4.939725567 |
| **A_55_P2022519** | Tmem108 | -0.072341488 | -0.327226463 | 0.182543487 | 11.2354559 | -0.698252284 | 0.51172081 | 0.875607994 | -4.939813844 |
| **A_55_P2019349** | Txnrd2 | 0.030235596 | -0.076296558 | 0.136767751 | 7.719339214 | 0.698243648 | 0.511725832 | 0.875607994 | -4.939817934 |
| **A_55_P2149896** | Lrrc75a | -0.096049275 | -0.434478557 | 0.242380008 | 11.88539743 | -0.698224879 | 0.511736745 | 0.875607994 | -4.939826821 |
| **A_52_P341449** | Pgm3 | -0.111051838 | -0.502387985 | 0.280284308 | 10.13857608 | -0.698143905 | 0.511783828 | 0.875607994 | -4.93986516 |
| **A_66_P137605** | Cgas | 0.038833134 | -0.098058226 | 0.175724494 | 8.313016937 | 0.69790381 | 0.51192345 | 0.875607994 | -4.939978818 |
| **A_55_P2047305** | Adcy5 | 0.036484886 | -0.092157812 | 0.165127584 | 7.768231497 | 0.697745458 | 0.51201555 | 0.875607994 | -4.940053763 |
| **A_55_P1991552** | Atraid | 0.079395729 | -0.200549718 | 0.359341177 | 13.07442988 | 0.697738837 | 0.512019401 | 0.875607994 | -4.940056896 |
| **A_55_P2037922** | Rpl7a | 0.094445108 | -0.238645748 | 0.427535965 | 14.10657834 | 0.69756699 | 0.512119363 | 0.875607994 | -4.940138212 |
| **A_55_P2047598** | Arhgdia | -0.100509372 | -0.454994768 | 0.253976025 | 15.52750777 | -0.697553261 | 0.512127349 | 0.875607994 | -4.940144708 |
| **A_51_P349878** | Txndc17 | -0.053465938 | -0.242036538 | 0.135104662 | 14.59896223 | -0.697545224 | 0.512132025 | 0.875607994 | -4.940148511 |
| **A_55_P2027436** | Bri3 | 0.057777965 | -0.146022877 | 0.261578807 | 15.41561562 | 0.697469842 | 0.512175879 | 0.875607994 | -4.940184175 |
| **A_55_P2105371** | Dlk1 | -0.109533116 | -0.495965214 | 0.276898981 | 11.02151194 | -0.697334921 | 0.512254376 | 0.875607994 | -4.940247999 |

| | | | | | | | | | |
|---|---|---|---|---|---|---|---|---|---|
| **A_55_P2206461** | A930006K02Rik | -0.039506789 | -0.178892509 | 0.099878932 | 7.789124083 | -0.697304724 | 0.512271946 | 0.875607994 | -4.940262283 |
| **A_55_P2052535** | Gm14393 | 0.049841853 | -0.12601996 | 0.225703667 | 8.878408971 | 0.697255256 | 0.512300729 | 0.875607994 | -4.940285681 |
| **A_55_P2295951** | NA | -0.118539913 | -0.536890325 | 0.299810498 | 8.408232472 | -0.697097569 | 0.512392487 | 0.875607994 | -4.940360257 |
| **A_55_P2126662** | Cav1 | 0.05039576 | -0.12746249 | 0.228254011 | 8.081237846 | 0.697090475 | 0.512396615 | 0.875607994 | -4.940363611 |
| **A_52_P91402** | Zcchc17 | 0.054497911 | -0.137851034 | 0.246846856 | 11.95141931 | 0.697042419 | 0.512424581 | 0.875607994 | -4.940386336 |
| **A_55_P1981929** | Scg5 | -0.125283241 | -0.567486801 | 0.316920319 | 9.663741931 | -0.697011442 | 0.512442609 | 0.875607994 | -4.940400983 |
| **A_51_P407074** | Tspan31 | -0.06697622 | -0.303384704 | 0.169432264 | 12.5899313 | -0.696990304 | 0.51245491 | 0.875607994 | -4.940410978 |
| **A_51_P334104** | Dcn | -0.48310335 | -2.18880183 | 1.222595129 | 16.19088024 | -0.696798088 | 0.512566785 | 0.875607994 | -4.940501855 |
| **A_51_P328622** | Tlcd2 | -0.033420234 | -0.151424962 | 0.084584494 | 8.195864232 | -0.696753636 | 0.512592659 | 0.875607994 | -4.940522868 |
| **A_55_P2097259** | Zup1 | 0.03168973 | -0.080240378 | 0.143619838 | 9.792973012 | 0.696531521 | 0.51272196 | 0.875607994 | -4.94062785 |
| **A_55_P2038504** | 6-Mar | -0.048432572 | -0.219514282 | 0.122649139 | 8.841562994 | -0.696471114 | 0.512757129 | 0.875607994 | -4.940656397 |
| **A_51_P114028** | Bcl7c | -0.05570976 | -0.252501209 | 0.141081689 | 9.413269595 | -0.696456868 | 0.512765423 | 0.875607994 | -4.940663129 |
| **A_55_P2187121** | Gpr137b-ps | 0.095498518 | -0.241844595 | 0.432841631 | 12.19150089 | 0.696456422 | 0.512765683 | 0.875607994 | -4.94066334 |
| **A_55_P2071811** | Eml2 | -0.110818413 | -0.502363634 | 0.280726808 | 9.229529081 | -0.696304434 | 0.512854177 | 0.875607994 | -4.940735155 |
| **A_55_P2047188** | Fgf1 | 0.08580755 | -0.217381658 | 0.388996757 | 8.709176193 | 0.696275197 | 0.512871202 | 0.875607994 | -4.940748968 |
| **A_52_P237077** | Esr1 | -0.197223429 | -0.894088612 | 0.499641754 | 8.386240476 | -0.696271953 | 0.51287309 | 0.875607994 | -4.92688737 |
| **A_55_P1989568** | Tasp1 | 0.048057286 | -0.121749124 | 0.217863695 | 8.165448976 | 0.696264613 | 0.512877365 | 0.875607994 | -4.940753968 |
| **A_52_P127206** | Gm9 | -0.057971055 | -0.262822124 | 0.146880015 | 10.58121558 | -0.696212999 | 0.51290742 | 0.875607994 | -4.940778352 |
| **A_51_P416822** | Hhip | 0.029582842 | -0.074953552 | 0.134119237 | 7.674440424 | 0.696212062 | 0.512907966 | 0.875607994 | -4.940778795 |
| **A_51_P400217** | Vpreb1 | -0.084114329 | -0.381389481 | 0.213160823 | 8.55024743 | -0.696114276 | 0.512964911 | 0.875607994 | -4.940824988 |
| **A_55_P2023076** | Arpc1b | 0.12676175 | -0.321261014 | 0.574784514 | 14.28039102 | 0.696077038 | 0.512986597 | 0.875607994 | -4.940842578 |
| **A_55_P1997106** | Large2 | -0.063711173 | -0.288920351 | 0.161498004 | 9.049013897 | -0.69598308 | 0.513041318 | 0.875607994 | -4.940886956 |
| **A_52_P110270** | Nelfe | 0.082858297 | -0.210118584 | 0.375835179 | 10.29607335 | 0.695779801 | 0.513159722 | 0.875607994 | -4.940982952 |
| **A_55_P2069425** | Ttll3 | -0.143540575 | -0.651296445 | 0.364215296 | 8.903125078 | -0.695486798 | 0.513330419 | 0.875607994 | -4.941121281 |
| **A_55_P2111790** | Gem | -0.118322934 | -0.536952637 | 0.300306768 | 8.837444568 | -0.695357359 | 0.513405839 | 0.875607994 | -4.941182375 |
| **A_51_P357207** | Zfp518b | 0.040005582 | -0.101541097 | 0.181552262 | 8.300648428 | 0.695328572 | 0.513422613 | 0.875607994 | -4.941195961 |
| **A_55_P2179793** | NA | 0.097456055 | -0.247362363 | 0.442274473 | 11.27576075 | 0.695324506 | 0.513424982 | 0.875607994 | -4.94119788 |
| **A_51_P332141** | Clcn2 | 0.046749933 | -0.118707513 | 0.212207379 | 8.887755545 | 0.695126487 | 0.513540379 | 0.875607994 | -4.941291323 |
| **A_52_P243599** | Hsd17b12 | -0.138730242 | -0.629775348 | 0.352314864 | 10.36722394 | -0.695054596 | 0.513582278 | 0.875607994 | -4.941325242 |
| **A_55_P2034300** | Tmem40 | 0.054550188 | -0.138566147 | 0.247666524 | 11.43354917 | 0.694938556 | 0.513649912 | 0.875607994 | -4.941379985 |
| **A_51_P410927** | NA | -0.041459377 | -0.188247233 | 0.105328479 | 8.837639484 | -0.694867189 | 0.513691512 | 0.875607994 | -4.94141365 |
| **A_52_P609972** | Eln | -0.102165593 | -0.463898192 | 0.259567005 | 8.364560442 | -0.694842183 | 0.513706089 | 0.875607994 | -4.941425445 |
| **A_55_P2085335** | Mia | 0.03116337 | -0.079185002 | 0.141511742 | 7.908416319 | 0.694780549 | 0.513742018 | 0.875607994 | -4.941454516 |
| **A_55_P2061452** | NA | 0.056269787 | -0.143005454 | 0.255545027 | 15.11788424 | 0.694690067 | 0.513794765 | 0.875607994 | -4.941497189 |
| **A_55_P1961436** | Prkca | 0.039832759 | -0.101252969 | 0.180918487 | 8.6360629 | 0.694586711 | 0.513855024 | 0.875607994 | -4.941545929 |
| **A_51_P118650** | Wdr6 | -0.041855848 | -0.190120992 | 0.106409296 | 8.822239386 | -0.694522378 | 0.513892533 | 0.875607994 | -4.900926091 |
| **A_51_P510891** | Afp | -0.098229591 | -0.446224373 | 0.24976519 | 17.51344081 | -0.694446462 | 0.513936798 | 0.875607994 | -4.941612057 |
| **A_55_P2011490** | NA | 0.047562701 | -0.120942993 | 0.216068395 | 11.97640903 | 0.694418191 | 0.513953283 | 0.875607994 | -4.941625386 |
| **A_55_P2010076** | Miga2 | 0.051262405 | -0.130354813 | 0.232879623 | 10.04947732 | 0.694402259 | 0.513962573 | 0.875607994 | -4.941632897 |
| **A_55_P2064567** | Anks1 | 0.05622514 | -0.14298611 | 0.255436391 | 8.989519527 | 0.694361845 | 0.513986139 | 0.875607994 | -4.94165195 |
| **A_51_P171965** | Sdhaf4 | -0.070724862 | -0.321316594 | 0.179866869 | 10.21128132 | -0.694343758 | 0.513996686 | 0.875607994 | -4.941660476 |
| **A_51_P491595** | AW549877 | 0.048275355 | -0.122773549 | 0.21932426 | 9.775614054 | 0.694343453 | 0.513996864 | 0.875607994 | -4.94166062 |
| **A_51_P332932** | NA | 0.038227207 | -0.097235571 | 0.173689984 | 8.1712452 | 0.694259378 | 0.514045893 | 0.875607994 | -4.941700252 |
| **A_52_P558577** | Zfp263 | 0.041928412 | -0.106656195 | 0.19051302 | 10.18413097 | 0.694230611 | 0.51406267 | 0.875607994 | -4.941713812 |
| **A_55_P2044659** | NA | -0.362870368 | -1.648804392 | 0.923063657 | 9.312888019 | -0.694227767 | 0.514064328 | 0.875607994 | -4.941715152 |
| **A_51_P468140** | Serpind1 | 0.32153132 | -0.817923836 | 1.460986475 | 10.52491423 | 0.694216942 | 0.514070642 | 0.875607994 | -4.941720255 |
| **A_55_P2076273** | NA | -0.058176642 | -0.264349928 | 0.147996644 | 12.58750303 | -0.694201291 | 0.514079769 | 0.875607994 | -4.941727631 |
| **A_51_P178828** | Mbl2 | 0.089109993 | -0.226720091 | 0.404940078 | 7.925550078 | 0.694132019 | 0.51412017 | 0.875607994 | -4.941760281 |
| **A_52_P297332** | Aida | -0.039107784 | -0.177729162 | 0.099513595 | 8.676617941 | -0.69406821 | 0.514157386 | 0.875607994 | -4.941790353 |
| **A_51_P336509** | Zfpm2 | 0.043606318 | -0.110971018 | 0.198183654 | 7.775434273 | 0.694021244 | 0.514184781 | 0.875607994 | -4.941812486 |

| | | | | | | | | | |
|---|---|---|---|---|---|---|---|---|---|
| **A_52_P609334** | Klhdc8b | -0.032968246 | -0.149858454 | 0.083921961 | 7.805674131 | -0.693884013 | 0.514264829 | 0.875607994 | -4.941877149 |
| **A_55_P2136763** | NA | 0.075132141 | -0.191259374 | 0.341523656 | 12.16911847 | 0.693864293 | 0.514276332 | 0.875607994 | -4.94188644 |
| **A_51_P318618** | Anks4b | 0.102466631 | -0.260854171 | 0.465787434 | 7.983529802 | 0.693843232 | 0.514288619 | 0.875607994 | -4.941896364 |
| **A_51_P215077** | Mgst3 | 0.14214385 | -0.361865755 | 0.646153455 | 11.48291737 | 0.693838549 | 0.51429135 | 0.875607994 | -4.94189857 |
| **A_51_P110759** | Slc1a1 | 0.074504093 | -0.189694929 | 0.338703114 | 9.235413834 | 0.693774107 | 0.514328944 | 0.875607994 | -4.937496664 |
| **A_55_P2371984** | 5930436O19Rik | -0.105755099 | -0.480782401 | 0.269272202 | 8.28611252 | -0.693757347 | 0.514338722 | 0.875607994 | -4.941936826 |
| **A_55_P2027930** | Nfx1 | -0.042995856 | -0.195496191 | 0.109504479 | 8.829124234 | -0.693625361 | 0.514415726 | 0.875661961 | -4.941998999 |
| **A_55_P2161858** | Pdlim7 | -0.064830933 | -0.294835413 | 0.165173547 | 8.839013579 | -0.693449979 | 0.51451806 | 0.875749346 | -4.942081601 |
| **A_55_P2097869** | Frem2 | 0.068965386 | -0.175731089 | 0.31366186 | 9.315435495 | 0.693382092 | 0.514557676 | 0.875749346 | -4.94211357 |
| **A_52_P623058** | R3hdm4 | -0.038228011 | -0.173900054 | 0.097444031 | 8.591971338 | -0.693203118 | 0.514662126 | 0.875778323 | -4.94219784 |
| **A_51_P479052** | Tars | -0.059610776 | -0.271180473 | 0.151958921 | 11.38298822 | -0.69317114 | 0.51468079 | 0.875778323 | -4.942212894 |
| **A_55_P1958986** | Spata7 | 0.043477486 | -0.110850243 | 0.197805214 | 8.216055309 | 0.693089976 | 0.514728163 | 0.875778323 | -4.942251103 |
| **A_55_P2101776** | Adora1 | 0.070757795 | -0.180425754 | 0.321941344 | 8.684914439 | 0.69303036 | 0.514762961 | 0.875778323 | -4.942279166 |
| **A_55_P2158030** | Glrx3 | 0.110454424 | -0.281685555 | 0.502594402 | 12.58436333 | 0.69296477 | 0.514801247 | 0.875778323 | -4.942310038 |
| **A_52_P663413** | P4ha2 | 0.099455471 | -0.253763949 | 0.452674891 | 13.34311324 | 0.692712886 | 0.514948298 | 0.875939134 | -4.942428576 |
| **A_52_P365896** | Pa2g4 | -0.031987834 | -0.145614885 | 0.081639217 | 7.896744509 | -0.692583693 | 0.515023731 | 0.875939134 | -4.942489362 |
| **A_55_P1975743** | Dap3 | 0.071110455 | -0.181503755 | 0.323724664 | 13.22405241 | 0.692539967 | 0.515049264 | 0.875939134 | -4.942509932 |
| **A_55_P2177659** | Mast1 | -0.064393602 | -0.293184521 | 0.164397317 | 8.473402144 | -0.692425572 | 0.515116065 | 0.875939134 | -4.942563745 |
| **A_55_P2123331** | Cog2 | 0.042558748 | -0.108655136 | 0.193772631 | 8.19760295 | 0.692414789 | 0.515122362 | 0.875939134 | -4.942568817 |
| **A_52_P602996** | Cdip1 | 0.046605901 | -0.119008026 | 0.212219829 | 8.224821852 | 0.692330105 | 0.515171818 | 0.875946171 | -4.942608649 |
| **A_52_P450934** | Paqr9 | 0.064188009 | -0.163938021 | 0.29231404 | 7.993777845 | 0.692226511 | 0.515232322 | 0.875971989 | -4.942657369 |
| **A_55_P2008879** | Cltc | 0.065129451 | -0.16641203 | 0.296670932 | 9.746696762 | 0.692018614 | 0.515353757 | 0.875999888 | -4.942755126 |
| **A_55_P2134800** | Cinp | -0.051848963 | -0.236185854 | 0.132487928 | 9.226861227 | -0.691985477 | 0.515373114 | 0.875999888 | -4.942770705 |
| **A_51_P160824** | Cspg4 | 0.078374747 | -0.20027636 | 0.357025853 | 8.285444838 | 0.691965657 | 0.515384693 | 0.875999888 | -4.942780024 |
| **A_55_P2135311** | Nol3 | 0.035029116 | -0.089565742 | 0.159623974 | 7.767483149 | 0.691668856 | 0.515558098 | 0.876088916 | -4.942919536 |
| **A_51_P233855** | Adprhl1 | 0.03514597 | -0.089881822 | 0.160173761 | 7.939269093 | 0.691573167 | 0.515614012 | 0.876088916 | -4.942964505 |
| **A_52_P58059** | Usp34 | 0.058591384 | -0.149848109 | 0.267030876 | 9.240446784 | 0.691548924 | 0.515628179 | 0.876088916 | -4.942975897 |
| **A_55_P2380806** | Gm2115 | -0.034727679 | -0.158274479 | 0.088819121 | 7.789672612 | -0.69153381 | 0.515637011 | 0.876088916 | -4.942983 |
| **A_55_P2156731** | H2-Eb1 | -0.174383625 | -0.794866838 | 0.446099587 | 9.410931895 | -0.691424603 | 0.515700831 | 0.876088916 | -4.943034313 |
| **A_51_P134142** | Cyp2c70 | -0.065298719 | -0.297646111 | 0.167048672 | 8.462631417 | -0.691410588 | 0.515709022 | 0.876088916 | -4.943040897 |
| **A_52_P501721** | Ccdc127 | -0.040209353 | -0.183335608 | 0.102916901 | 8.339965919 | -0.691157373 | 0.515857021 | 0.876263323 | -4.943159849 |
| **A_51_P359636** | Lgals3bp | -0.136054273 | -0.620482152 | 0.348373605 | 14.22346419 | -0.690958911 | 0.515973038 | 0.876358722 | -4.943253056 |
| **A_55_P2060737** | Trmt10b | -0.049306591 | -0.224878511 | 0.126265329 | 8.983356801 | -0.69090619 | 0.51600386 | 0.876358722 | -4.943277812 |
| **A_55_P2052016** | Crispld2 | -0.223500483 | -1.019631051 | 0.572630085 | 10.45468596 | -0.690658452 | 0.516148713 | 0.876404454 | -4.943394124 |
| **A_66_P103418** | Tmc8 | 0.074986677 | -0.192173045 | 0.342146398 | 9.174089654 | 0.690529579 | 0.516224075 | 0.876404454 | -4.943454616 |
| **A_66_P135534** | Zfp950 | 0.035631739 | -0.091320788 | 0.162584267 | 8.80476953 | 0.690501831 | 0.516240303 | 0.876404454 | -4.94346764 |
| **A_55_P2148402** | Cplx2 | 0.101894256 | -0.261219018 | 0.46500753 | 12.98355582 | 0.690361782 | 0.516322211 | 0.876404454 | -4.943533366 |
| **A_51_P134452** | Lpar3 | 0.133490107 | -0.342225969 | 0.609206184 | 9.227769084 | 0.69035173 | 0.51632809 | 0.876404454 | -4.943538083 |
| **A_51_P155763** | NA | -0.052618668 | -0.240137573 | 0.134900238 | 8.765861033 | -0.690341453 | 0.516334101 | 0.876404454 | -4.943542906 |
| **A_51_P107433** | Mrpl34 | -0.037626134 | -0.17172807 | 0.096475802 | 11.68686288 | -0.690277492 | 0.516371512 | 0.876404454 | -4.939130173 |
| **A_55_P2010552** | Hsd17b13 | -0.029989783 | -0.136895835 | 0.076916269 | 7.668988388 | -0.690144752 | 0.516449158 | 0.876404454 | -4.943635199 |
| **A_52_P79385** | Magt1 | -0.08189005 | -0.373811064 | 0.210030963 | 11.98601664 | -0.690136414 | 0.516454036 | 0.876404454 | -4.943639111 |
| **A_51_P443359** | Trappc2l | 0.091911833 | -0.235759237 | 0.419582903 | 11.94047676 | 0.690084847 | 0.516484202 | 0.876404454 | -4.943663302 |
| **A_51_P252199** | Mmgt1 | -0.064599825 | -0.294933232 | 0.165733583 | 9.384475013 | -0.68999123 | 0.51653897 | 0.876420449 | -4.943707217 |
| **A_55_P2038412** | Acot8 | 0.059679034 | -0.1531478 | 0.272505868 | 9.523937934 | 0.68986572 | 0.516612403 | 0.87644224 | -4.943766086 |
| **A_55_P2019312** | Car12 | -0.032220459 | -0.147133337 | 0.082692419 | 7.759175567 | -0.689814281 | 0.5166425 | 0.87644224 | -4.94379021 |
| **A_55_P2131288** | Ank2 | -0.044516433 | -0.203320373 | 0.114287506 | 8.671289415 | -0.689649227 | 0.516739083 | 0.876529157 | -4.943867609 |
| **A_55_P2086510** | Ppid | -0.069131848 | -0.315803536 | 0.177539841 | 10.95104874 | -0.689490076 | 0.516832223 | 0.876610217 | -4.943942225 |
| **A_55_P2129316** | NA | -0.545780837 | -2.493535286 | 1.401973612 | 10.12506968 | -0.689371367 | 0.516901702 | 0.876651136 | -4.943997872 |
| **A_55_P1967069** | Lpar6 | 0.068704388 | -0.176568337 | 0.313977113 | 10.37773235 | 0.689135117 | 0.517039995 | 0.876793568 | -4.944108595 |

| | | | | | | | | | |
|---|---|---|---|---|---|---|---|---|---|
| **A_52_P123400** | Dnaaf2 | -0.043112159 | -0.197035241 | 0.110810923 | 8.619917669 | -0.68907292 | 0.517076407 | 0.876793568 | -4.94413774 |
| **A_55_P2159710** | Acvr1 | 0.064963154 | -0.167044504 | 0.296970813 | 12.11415535 | 0.688864727 | 0.517198303 | 0.876825747 | -4.944235282 |
| **A_55_P2420873** | NA | -0.093518995 | -0.427534518 | 0.240496529 | 10.10463341 | -0.688814594 | 0.517227658 | 0.876825747 | -4.944258767 |
| **A_55_P2128283** | Nr2f2 | -0.125387939 | -0.573300505 | 0.322524627 | 9.693765533 | -0.688702532 | 0.51729328 | 0.876825747 | -4.944311257 |
| **A_66_P100601** | NA | 0.042594445 | -0.109564986 | 0.194753876 | 11.63474436 | 0.688689163 | 0.517301109 | 0.876825747 | -4.944317519 |
| **A_51_P151968** | Eif2b5 | -0.044096453 | -0.201629722 | 0.113436816 | 9.314299381 | -0.688653147 | 0.517322201 | 0.876825747 | -4.944334387 |
| **A_55_P2157159** | NA | -0.069938451 | -0.319857503 | 0.179980602 | 10.37486759 | -0.688471242 | 0.517428737 | 0.876869931 | -4.944419572 |
| **A_55_P2059860** | Igsf1 | -0.034216819 | -0.1564908 | 0.088057161 | 7.944432524 | -0.688453719 | 0.517439 | 0.876869931 | -4.944427777 |
| **A_55_P2186808** | NA | -0.088536361 | -0.405092919 | 0.228020197 | 10.0971069 | -0.688080918 | 0.517657386 | 0.877081614 | -4.944602299 |
| **A_55_P2009943** | Spen | -0.065257319 | -0.298605424 | 0.168090786 | 11.7089158 | -0.68800899 | 0.517699529 | 0.877081614 | -4.944635963 |
| **A_55_P1961499** | Ly6c1 | -0.725419795 | -3.319806641 | 1.868967051 | 11.76435035 | -0.687897659 | 0.517764761 | 0.877081614 | -4.944688062 |
| **A_55_P2021455** | NA | -0.074119354 | -0.339204756 | 0.190966047 | 11.51960944 | -0.68788364 | 0.517772976 | 0.877081614 | -4.944694623 |
| **A_55_P2043622** | NA | -0.173591534 | -0.794465821 | 0.447282753 | 13.58181088 | -0.687850456 | 0.517792421 | 0.877081614 | -4.94471015 |
| **A_55_P2018507** | Gorasp2 | -0.045262767 | -0.207176206 | 0.116650672 | 8.890443832 | -0.687744883 | 0.517854287 | 0.877081614 | -4.944759546 |
| **A_51_P332201** | Cst3 | 0.128462072 | -0.331102029 | 0.588026173 | 14.95709572 | 0.687698364 | 0.517881549 | 0.877081614 | -4.944781309 |
| **A_52_P401640** | Bpgm | 0.07705769 | -0.198819714 | 0.352935095 | 10.58598205 | 0.687177641 | 0.518186776 | 0.877487177 | -4.945024845 |
| **A_55_P1970710** | Mllt6 | -0.070844865 | -0.324495242 | 0.182805511 | 10.92647581 | -0.687134937 | 0.518211813 | 0.877487177 | -4.945044811 |
| **A_51_P284244** | Tmed3 | 0.061212005 | -0.158036738 | 0.280460749 | 15.31291147 | 0.686860684 | 0.518372622 | 0.877523509 | -4.945173011 |
| **A_51_P417891** | Trim10 | 0.03751928 | -0.096880851 | 0.171919411 | 7.819220758 | 0.686790008 | 0.518414068 | 0.877523509 | -4.945206042 |
| **A_55_P1966644** | Trpc6 | -0.03876886 | -0.177666792 | 0.100129073 | 7.955727667 | -0.686683177 | 0.518476721 | 0.877523509 | -4.945255965 |
| **A_51_P397925** | Smap2 | 0.059518261 | -0.153753774 | 0.272790297 | 9.105191256 | 0.686571047 | 0.518542487 | 0.877523509 | -4.945308358 |
| **A_51_P493886** | Gpt2 | 0.1227152 | -0.317040675 | 0.562471076 | 9.230337392 | 0.686524234 | 0.518569945 | 0.877523509 | -4.945330229 |
| **A_55_P2180846** | Wdsub1 | 0.032061049 | -0.082847252 | 0.14696935 | 8.179142878 | 0.686428787 | 0.518625932 | 0.877523509 | -4.945374819 |
| **A_55_P2124582** | Xlr4a | -0.106052431 | -0.486167593 | 0.27406273 | 9.521631426 | -0.686395774 | 0.518645298 | 0.877523509 | -4.94539024 |
| **A_55_P2124346** | Ivns1abp | -0.060851338 | -0.278968868 | 0.157266191 | 11.07222844 | -0.686354882 | 0.518669286 | 0.877523509 | -4.945409341 |
| **A_55_P2139615** | Rtl6 | 0.036750715 | -0.094981211 | 0.168482641 | 8.722542003 | 0.686347269 | 0.518673752 | 0.877523509 | -4.945412898 |
| **A_51_P357664** | Hras | -0.061397954 | -0.281484616 | 0.158688708 | 10.6855021 | -0.686324239 | 0.518687262 | 0.877523509 | -4.945423654 |
| **A_51_P444052** | Scp2 | -0.037725216 | -0.173002441 | 0.09755201 | 10.76505378 | -0.686082283 | 0.518829218 | 0.877601056 | -4.945536653 |
| **A_51_P296905** | Ppat | -0.04720681 | -0.216500972 | 0.122087352 | 8.367715792 | -0.686012196 | 0.518870343 | 0.877601056 | -4.945569379 |
| **A_55_P2166262** | Rnf215 | 0.112068686 | -0.289839743 | 0.513977115 | 12.74931813 | 0.686003496 | 0.518875448 | 0.877601056 | -4.945573441 |
| **A_55_P2014570** | Zfp961 | 0.071375108 | -0.184620183 | 0.3273704 | 10.42814576 | 0.685936583 | 0.518914712 | 0.877601056 | -4.945604682 |
| **A_51_P254646** | Jdp2 | 0.068250689 | -0.176617689 | 0.313119067 | 11.79703522 | 0.685714759 | 0.519044893 | 0.877716183 | -4.945708233 |
| **A_55_P2209308** | 9530083O12Rik | -0.058049363 | -0.266332501 | 0.150233774 | 11.34610131 | -0.685665841 | 0.519073604 | 0.877716183 | -4.945731065 |
| **A_55_P2109782** | Psg17 | -0.131487729 | -0.60338382 | 0.340408363 | 15.25586946 | -0.685500863 | 0.519170441 | 0.877803136 | -4.945808058 |
| **A_55_P1956847** | Nolc1 | -0.093567383 | -0.442823397 | 0.255688632 | 10.67042179 | -0.69427363 | 0.519232142 | 0.877804413 | -4.88395175 |
| **A_55_P2035102** | Odc1 | 0.109142104 | -0.282662034 | 0.500946243 | 12.93414455 | 0.685318518 | 0.519277486 | 0.877804413 | -4.945893138 |
| **A_55_P2063711** | Snu13 | -0.059200142 | -0.271735784 | 0.153335501 | 12.435392 | -0.685267501 | 0.519307438 | 0.877804413 | -4.945916939 |
| **A_52_P136914** | Nudt7 | 0.035253055 | -0.091337884 | 0.161843993 | 8.212626314 | 0.685114719 | 0.519397142 | 0.877814774 | -4.945988207 |
| **A_55_P2056342** | Mgst3 | -0.132745376 | -0.609503515 | 0.344012763 | 12.08904377 | -0.684999807 | 0.519464619 | 0.877814774 | -4.946041802 |
| **A_51_P479902** | Mars | -0.073137685 | -0.335813998 | 0.189538629 | 9.658086034 | -0.684998242 | 0.519465537 | 0.877814774 | -4.946042532 |
| **A_51_P347177** | Snx2 | -0.08900828 | -0.408741339 | 0.230724779 | 12.56522748 | -0.68487613 | 0.519537248 | 0.877814774 | -4.946099477 |
| **A_55_P2243566** | Redrum | 0.049391264 | -0.128042621 | 0.22682515 | 7.875867721 | 0.684829947 | 0.519564371 | 0.877814774 | -4.946121012 |
| **A_52_P504302** | Pmpca | -0.044791189 | -0.20575902 | 0.116176642 | 9.645679066 | -0.684577559 | 0.519712611 | 0.877814774 | -4.946238677 |
| **A_51_P216550** | Trp63 | 0.107929912 | -0.279950039 | 0.495809864 | 9.058297681 | 0.684563367 | 0.519720948 | 0.877814774 | -4.946245292 |
| **A_51_P185247** | Gdf10 | 0.035646708 | -0.092478651 | 0.163772066 | 7.75053747 | 0.684468541 | 0.519776652 | 0.877814774 | -4.946289491 |
| **A_52_P651034** | Tmx1 | -0.056273833 | -0.258577616 | 0.146029951 | 10.64916777 | -0.684339568 | 0.519852421 | 0.877814774 | -4.946349598 |
| **A_51_P121031** | 1-Mar | -0.030757046 | -0.141342885 | 0.079828793 | 7.797391075 | -0.684249153 | 0.519905543 | 0.877814774 | -4.934872066 |
| **A_52_P557129** | Slc12a5 | -0.031366485 | -0.14415178 | 0.081418811 | 7.838354231 | -0.684199151 | 0.519934922 | 0.877814774 | -4.946415028 |
| **A_55_P2007991** | Tuba3b | 0.092744325 | -0.240858263 | 0.426346912 | 8.708305935 | 0.683954317 | 0.520078793 | 0.877814774 | -4.946529088 |
| **A_51_P415688** | Gng12 | -0.128973483 | -0.592919185 | 0.33497222 | 12.91871325 | -0.68391548 | 0.520101617 | 0.877814774 | -4.946547178 |

| | | | | | | | | |
|---|---|---|---|---|---|---|---|---|
| **A_55_P2037677** | Cttn | -0.064779595 | -0.297813004 | 0.168253815 | 13.29201246 | -0.683894641 | 0.520113864 | 0.877814774 | -4.946556884 |
| **A_55_P2086912** | NA | 0.051157637 | -0.132888516 | 0.235203789 | 11.65337918 | 0.683837469 | 0.520147465 | 0.877814774 | -4.946583512 |
| **A_66_P102046** | NA | -0.073263311 | -0.336853147 | 0.190326525 | 9.853939234 | -0.683796771 | 0.520171385 | 0.877814774 | -4.946602466 |
| **A_55_P2415197** | 2810013P06Rik | -0.042263033 | -0.194319684 | 0.109793617 | 9.319526222 | -0.683792622 | 0.520173824 | 0.877814774 | -4.946604399 |
| **A_55_P2121076** | Cul2 | 0.036341303 | -0.094413305 | 0.16709591 | 8.972213316 | 0.68377411 | 0.520184705 | 0.877814774 | -4.94661302 |
| **A_55_P1989594** | Mynn | -0.041277832 | -0.189826533 | 0.107270868 | 9.66520523 | -0.683623825 | 0.520273041 | 0.877814774 | -4.946683002 |
| **A_55_P1959800** | NA | 0.039261415 | -0.102041659 | 0.180564489 | 9.324951954 | 0.6835708 | 0.520304211 | 0.877814774 | -4.94670769 |
| **A_51_P154780** | Vav1 | -0.029104074 | -0.133858046 | 0.075649898 | 7.71805094 | -0.683521929 | 0.520332939 | 0.877814774 | -4.946730443 |
| **A_55_P2081318** | Gnas | 0.069689515 | -0.181163333 | 0.320542362 | 8.169156433 | 0.683467026 | 0.520365216 | 0.877814774 | -4.946756004 |
| **A_52_P495869** | Mafb | -0.053995921 | -0.248377838 | 0.140385995 | 8.315667321 | -0.683398876 | 0.520405282 | 0.877814774 | -4.946787729 |
| **A_51_P117130** | Naga | -0.052719748 | -0.242543211 | 0.137103715 | 8.525340281 | -0.683270385 | 0.520480828 | 0.877814774 | -4.931974944 |
| **A_55_P1961567** | Ypel3 | 0.103395142 | -0.268972592 | 0.475762875 | 13.25576632 | 0.683120399 | 0.520569021 | 0.877814774 | -4.946917338 |
| **A_55_P1974671** | Sft2d1 | -0.050669798 | -0.233154946 | 0.131815349 | 9.858203968 | -0.683110272 | 0.520574977 | 0.877814774 | -4.946922051 |
| **A_55_P2100008** | NA | 0.070827583 | -0.184289469 | 0.325944634 | 11.0876593 | 0.683017915 | 0.520629288 | 0.877814774 | -4.946965026 |
| **A_51_P320614** | Txnrd1 | -0.07502772 | -0.345312646 | 0.195257205 | 10.79373007 | -0.682918832 | 0.52068756 | 0.877814774 | -4.947011125 |
| **A_55_P2011061** | Tspan12 | 0.112189619 | -0.291989035 | 0.516368273 | 8.701500855 | 0.682886401 | 0.520706634 | 0.877814774 | -4.947026214 |
| **A_51_P161265** | Ccl25 | 0.055142112 | -0.143516521 | 0.253800745 | 10.54895006 | 0.682881134 | 0.520709732 | 0.877814774 | -4.947028664 |
| **A_55_P1964408** | Cryab | -0.617200658 | -2.840923789 | 1.606522472 | 15.37838084 | -0.682833517 | 0.520737738 | 0.877814774 | -4.947050815 |
| **A_66_P126999** | Fis1 | -0.058241195 | -0.268094624 | 0.151612233 | 13.58033859 | -0.68278406 | 0.520766828 | 0.877814774 | -4.947073822 |
| **A_55_P2049141** | Cops7b | -0.03736813 | -0.172058291 | 0.097322031 | 8.634573579 | -0.682550285 | 0.520904344 | 0.877927734 | -4.947182551 |
| **A_55_P2151138** | Dynlt1c | -0.117302728 | -0.540131586 | 0.30552613 | 12.26332026 | -0.682515713 | 0.520924682 | 0.877927734 | -4.947198628 |
| **A_55_P1967500** | Nell1 | 0.041921891 | -0.109223019 | 0.193066801 | 7.890044048 | 0.682364625 | 0.521013573 | 0.877933458 | -4.94726888 |
| **A_51_P437144** | Dynll2 | -0.085691161 | -0.394645622 | 0.2232633 | 11.78177092 | -0.682355538 | 0.52101892 | 0.877933458 | -4.947273105 |
| **A_55_P2079388** | Snora74a | -0.031187431 | -0.143667772 | 0.081292909 | 7.747443735 | -0.682137851 | 0.521147012 | 0.87807275 | -4.947374301 |
| **A_51_P185175** | Fkbp4 | 0.039497669 | -0.103026091 | 0.182021428 | 8.349050325 | 0.681794283 | 0.521349218 | 0.87833688 | -4.947533964 |
| **A_51_P140742** | Islr | 0.180586228 | -0.471255213 | 0.832427668 | 8.837984165 | 0.681572127 | 0.521479994 | 0.878480634 | -4.94763717 |
| **A_55_P2024530** | Siah2 | -0.080775573 | -0.372401219 | 0.210850073 | 9.101994408 | -0.681433525 | 0.521561595 | 0.878541531 | -4.947701546 |
| **A_51_P429295** | Pus10 | -0.06523819 | -0.300818909 | 0.170342529 | 9.471468377 | -0.681288913 | 0.521646744 | 0.878608392 | -4.947768702 |
| **A_55_P2048398** | Kmt5b | -0.04931167 | -0.227428626 | 0.128805287 | 8.60904008 | -0.681104274 | 0.521755474 | 0.878631111 | -4.94785443 |
| **A_55_P1980817** | Gm6548 | 0.059361395 | -0.155117997 | 0.273840787 | 15.89899071 | 0.680906835 | 0.521871759 | 0.878631111 | -4.947946081 |
| **A_55_P1979330** | Dapp1 | 0.136680846 | -0.357176414 | 0.630538106 | 8.701428563 | 0.680887527 | 0.521883131 | 0.878631111 | -4.947955042 |
| **A_51_P506755** | Tep1 | -0.099379503 | -0.458496304 | 0.259737299 | 9.797216646 | -0.680816789 | 0.521924798 | 0.878631111 | -4.947987872 |
| **A_55_P2130716** | Ube2g2 | 0.046224742 | -0.120818682 | 0.213268166 | 8.835528928 | 0.68079174 | 0.521939553 | 0.878631111 | -4.947999497 |
| **A_51_P161054** | Itgb6 | -0.067746936 | -0.312576536 | 0.177082664 | 8.569331789 | -0.680761361 | 0.521957448 | 0.878631111 | -4.948013595 |
| **A_55_P1979669** | Zfp946 | 0.035904011 | -0.093858433 | 0.165666455 | 8.247788599 | 0.680711546 | 0.521986793 | 0.878631111 | -4.948036712 |
| **A_55_P2178207** | Abcc8 | 0.140922453 | -0.368439396 | 0.650284302 | 9.125587937 | 0.680648583 | 0.522023885 | 0.878631111 | -4.948065928 |
| **A_55_P2089840** | Eif2d | 0.038163312 | -0.099815913 | 0.176142537 | 9.291374006 | 0.680458307 | 0.522135988 | 0.878650376 | -4.948154207 |
| **A_51_P498631** | Gsdme | 0.068244229 | -0.17850015 | 0.314988608 | 8.322729888 | 0.680436849 | 0.522148631 | 0.878650376 | -4.94816416 |
| **A_55_P2133632** | Sipa1l2 | 0.080408417 | -0.21033363 | 0.371150464 | 13.03559899 | 0.680397691 | 0.522171704 | 0.878650376 | -4.948182325 |
| **A_66_P109002** | Kcng2 | -0.028941707 | -0.133666382 | 0.075782968 | 7.676534632 | -0.679898817 | 0.522465709 | 0.879068566 | -4.948413668 |
| **A_52_P65719** | Eme2 | -0.11418647 | -0.527464774 | 0.299091833 | 8.58230775 | -0.679737461 | 0.522560824 | 0.879152074 | -4.948488464 |
| **A_52_P367675** | Acin1 | -0.030919871 | -0.142862116 | 0.081022374 | 8.142359016 | -0.679536549 | 0.522679274 | 0.879221044 | -4.948581576 |
| **A_55_P1965836** | Crebbp | -0.098954244 | -0.457220104 | 0.259311615 | 11.09953573 | -0.679513618 | 0.522692794 | 0.879221044 | -4.948592202 |
| **A_55_P2063256** | Lgals4 | 0.038690825 | -0.101445893 | 0.178827544 | 8.679098549 | 0.679243066 | 0.522852331 | 0.87927284 | -4.948717551 |
| **A_51_P310821** | Hoxa5 | -0.053372142 | -0.246716335 | 0.139972052 | 7.763789803 | -0.679129603 | 0.522919246 | 0.87927284 | -4.948770108 |
| **A_66_P133328** | Tpd52 | -0.087109003 | -0.402761346 | 0.228543341 | 12.09531736 | -0.678927166 | 0.523038648 | 0.87927284 | -4.94886386 |
| **A_55_P1980426** | Nnat | 0.114768159 | -0.301147721 | 0.53068404 | 8.080154322 | 0.678867593 | 0.523073789 | 0.87927284 | -4.948891445 |
| **A_55_P2078008** | Ddx46 | -0.052006479 | -0.240497778 | 0.13648482 | 9.107188278 | -0.678789794 | 0.523119684 | 0.87927284 | -4.948927467 |
| **A_55_P2047703** | Serac1 | 0.037989352 | -0.099721627 | 0.17570033 | 8.358088625 | 0.678675982 | 0.523186827 | 0.87927284 | -4.948980157 |
| **A_55_P2020050** | 6720475J19Rik | -0.035459103 | -0.16400378 | 0.093085574 | 8.060865227 | -0.678645248 | 0.52320496 | 0.87927284 | -4.948994384 |

| | | | | | | | | |
|---|---|---|---|---|---|---|---|---|
| **A_55_P2090394** | NA | 0.058306098 | -0.153073348 | 0.269685544 | 12.59255058 | 0.678610209 | 0.523225632 | 0.87927284 | -4.949010604 |
| **A_51_P371876** | Zmynd8 | -0.085490183 | -0.395428805 | 0.224448439 | 10.03114126 | -0.678593525 | 0.523235476 | 0.87927284 | -4.949018327 |
| **A_55_P2026315** | Gys1 | -0.050717758 | -0.234648867 | 0.133213351 | 8.985955767 | -0.678381539 | 0.52336056 | 0.87927284 | -4.949116438 |
| **A_55_P2083396** | Radil | -0.031011679 | -0.143477732 | 0.081454373 | 7.88744817 | -0.678379934 | 0.523361507 | 0.87927284 | -4.949117181 |
| **A_55_P2048588** | Cdk1 | 0.15447737 | -0.405761224 | 0.714715964 | 8.777198726 | 0.678361213 | 0.523372555 | 0.87927284 | -4.949125845 |
| **A_55_P1976356** | Fgf18 | -0.062202677 | -0.287792212 | 0.163386858 | 8.196005442 | -0.678358525 | 0.523374141 | 0.87927284 | -4.949127088 |
| **A_55_P1992762** | Kdelr1 | -0.046531251 | -0.215304729 | 0.122242227 | 10.05568491 | -0.678281063 | 0.523419854 | 0.87927284 | -4.949162933 |
| **A_51_P109258** | Cys1 | -0.169157574 | -0.782718393 | 0.444403245 | 9.619967856 | -0.6782706 | 0.523426029 | 0.87927284 | -4.949167774 |
| **A_55_P2096807** | Rbms1 | -0.07895911 | -0.36544491 | 0.20752669 | 12.00071156 | -0.678060286 | 0.523550155 | 0.87927284 | -4.949265075 |
| **A_55_P2030060** | Kdm8 | 0.032914545 | -0.086520714 | 0.152349805 | 8.754085554 | 0.677991839 | 0.523590557 | 0.87927284 | -4.949296737 |
| **A_55_P2059010** | Rbp1 | -0.054050274 | -0.250191332 | 0.142090785 | 15.52259406 | -0.677951376 | 0.523614441 | 0.87927284 | -4.949315452 |
| **A_55_P1957004** | Timm10 | -0.050598986 | -0.234219229 | 0.133021257 | 10.99283264 | -0.677938695 | 0.523621927 | 0.87927284 | -4.949321318 |
| **A_51_P520849** | Sfrp2 | 0.163876758 | -0.430839343 | 0.75859286 | 8.183662859 | 0.677917478 | 0.523634451 | 0.87927284 | -4.949331131 |
| **A_55_P1972448** | NA | -0.049161425 | -0.227590245 | 0.129267395 | 8.078149215 | -0.677842229 | 0.523678873 | 0.87927284 | -4.949365933 |
| **A_66_P119376** | Kctd12 | -0.036369569 | -0.168392007 | 0.095652868 | 9.476544839 | -0.677734469 | 0.52374249 | 0.879303273 | -4.949415765 |
| **A_52_P629037** | Gm6987 | -0.032643924 | -0.151201095 | 0.085913248 | 11.00572212 | -0.67739765 | 0.523941368 | 0.879460205 | -4.949571484 |
| **A_52_P10041** | Akr1b3 | 0.133448669 | -0.351243324 | 0.618140662 | 10.65632505 | 0.677356909 | 0.523965427 | 0.879460205 | -4.949590315 |
| **A_55_P2114187** | NA | 0.034787101 | -0.09157141 | 0.161145612 | 7.771959972 | 0.677302847 | 0.523997354 | 0.879460205 | -4.949615302 |
| **A_51_P221031** | Slc16a12 | -0.080104346 | -0.371085402 | 0.21087671 | 9.053048761 | -0.677267952 | 0.524017962 | 0.879460205 | -4.949631429 |
| **A_51_P149621** | Stt3b | -0.047829299 | -0.221642249 | 0.125983651 | 10.23597668 | -0.676988123 | 0.524183243 | 0.879661217 | -4.949760732 |
| **A_55_P2150108** | Pgm2l1 | -0.031727089 | -0.147050065 | 0.083595887 | 9.265211697 | -0.676836152 | 0.524273018 | 0.879735495 | -4.949830937 |
| **A_51_P472217** | Sapcd2 | -0.105078586 | -0.487196787 | 0.277039615 | 8.848748318 | -0.676527819 | 0.524455194 | 0.879964796 | -4.949973336 |
| **A_51_P468599** | Smchd1 | -0.125054362 | -0.628893819 | 0.378785095 | 11.79901859 | -0.698407958 | 0.524615936 | 0.880057612 | -4.838713405 |
| **A_55_P1975041** | Arhgef11 | -0.047597171 | -0.220786452 | 0.12559211 | 10.50430857 | -0.676128581 | 0.524691143 | 0.880057612 | -4.950157642 |
| **A_51_P293753** | Cul7 | -0.111874672 | -0.518967727 | 0.295218383 | 13.85801954 | -0.676094252 | 0.524711434 | 0.880057612 | -4.950173485 |
| **A_55_P2183338** | Marveld3 | 0.108921527 | -0.287477724 | 0.505320779 | 9.672420153 | 0.676005236 | 0.524764052 | 0.880057612 | -4.950214565 |
| **A_55_P2339379** | Sass6 | 0.028994194 | -0.076530267 | 0.134518654 | 7.930420634 | 0.675969433 | 0.524785218 | 0.880057612 | -4.950231087 |
| **A_55_P2149274** | Sh3bp5l | 0.077249642 | -0.203916398 | 0.358415682 | 12.66612202 | 0.675931637 | 0.524807561 | 0.880057612 | -4.950248527 |
| **A_55_P2179335** | Usp33 | -0.041151094 | -0.190945143 | 0.108642955 | 11.28240453 | -0.675858819 | 0.52485061 | 0.880057612 | -4.950282126 |
| **A_55_P2157478** | NA | 0.040404323 | -0.106680296 | 0.187488942 | 8.918624864 | 0.675817977 | 0.524874756 | 0.880057612 | -4.950300969 |
| **A_55_P2166049** | Vmn1r65 | 0.064528482 | -0.170509067 | 0.299566031 | 8.511517146 | 0.67543469 | 0.525101394 | 0.880242788 | -4.950477764 |
| **A_51_P288906** | Clip1 | 0.042103843 | -0.111295289 | 0.195502974 | 10.56790343 | 0.675255293 | 0.525207493 | 0.880242788 | -4.950560484 |
| **A_55_P2134082** | Clip4 | -0.028910871 | -0.134259366 | 0.076437625 | 7.685376905 | -0.675152684 | 0.525268185 | 0.880242788 | -4.95060779 |
| **A_51_P394014** | Qprt | 0.088949394 | -0.235192988 | 0.413091776 | 8.131824799 | 0.675112793 | 0.525291781 | 0.880242788 | -4.950626179 |
| **A_52_P176619** | Atat1 | -0.047407629 | -0.220168124 | 0.125352866 | 9.747975379 | -0.675107542 | 0.525294887 | 0.880242788 | -4.9506286 |
| **A_55_P2053933** | Foxa1 | 0.034515362 | -0.091269987 | 0.160300711 | 8.170179366 | 0.675074238 | 0.525314587 | 0.880242788 | -4.950643951 |
| **A_51_P475228** | Armc6 | -0.167058712 | -0.775907108 | 0.441789685 | 10.14648075 | -0.675039403 | 0.525335194 | 0.880242788 | -4.950660009 |
| **A_66_P108672** | Mtg1 | 0.054345315 | -0.143724239 | 0.252414868 | 9.861050667 | 0.675015192 | 0.525349516 | 0.880242788 | -4.950671168 |
| **A_55_P1981879** | Ercc6l2 | -0.052565567 | -0.244212158 | 0.139081024 | 10.20948445 | -0.674791211 | 0.525482026 | 0.880388496 | -4.950774392 |
| **A_52_P257547** | Dele1 | 0.063030452 | -0.166808732 | 0.292869636 | 11.05732482 | 0.67467639 | 0.525549964 | 0.880426006 | -4.950827298 |
| **A_55_P2081388** | Mdm4 | 0.085980746 | -0.227583896 | 0.399545388 | 8.46594738 | 0.674595261 | 0.525597971 | 0.880430122 | -4.950864676 |
| **A_55_P2176345** | Psma5 | -0.11627085 | -0.540436903 | 0.307895202 | 12.81031666 | -0.674379104 | 0.525725892 | 0.88053199 | -4.950964245 |
| **A_55_P2011031** | Cdk5rap1 | -0.048750691 | -0.226608097 | 0.129106714 | 8.471003051 | -0.67433855 | 0.525749894 | 0.88053199 | -4.950982923 |
| **A_55_P1959303** | Rbm47 | 0.14594372 | -0.386634883 | 0.678522324 | 9.794877764 | 0.674172232 | 0.525848336 | 0.880620558 | -4.951059513 |
| **A_55_P2069961** | NA | 0.038682256 | -0.102522901 | 0.179887414 | 16.3883306 | 0.673954224 | 0.525977392 | 0.880760375 | -4.951159885 |
| **A_51_P327564** | Glb1 | -0.046894641 | -0.218169826 | 0.124380545 | 10.86680349 | -0.67359356 | 0.526190943 | 0.880882964 | -4.951325878 |
| **A_51_P432511** | Afg1l | -0.037850084 | -0.176092286 | 0.100392119 | 8.140280724 | -0.673589587 | 0.526193295 | 0.880882964 | -4.951327706 |
| **A_55_P1953728** | Nes | -0.050447478 | -0.234726857 | 0.133831902 | 9.085814226 | -0.673491126 | 0.526251604 | 0.880882964 | -4.951373009 |
| **A_52_P290725** | Arfgef2 | 0.071408439 | -0.189455226 | 0.332272104 | 9.113715947 | 0.673449621 | 0.526276185 | 0.880882964 | -4.951392105 |
| **A_55_P1959628** | Pfdn4 | 0.060045205 | -0.159308445 | 0.279398856 | 9.26686983 | 0.673445769 | 0.526278466 | 0.880882964 | -4.951393877 |

| | | | | | | | | | |
|---|---|---|---|---|---|---|---|---|---|
| **A_55_P2277171** | Gm10134 | 0.029436656 | -0.078131242 | 0.137004555 | 8.047108347 | 0.673247814 | 0.526395713 | 0.881002921 | -4.951484938 |
| **A_51_P452820** | Rpl31 | -0.061365713 | -0.285652543 | 0.162921116 | 8.861895242 | -0.673117959 | 0.526472634 | 0.881055371 | -4.951544661 |
| **A_55_P1956863** | Egfr | -0.137850633 | -0.642005656 | 0.36630439 | 9.264662769 | -0.672688232 | 0.526727239 | 0.881389679 | -4.951742234 |
| **A_55_P2031979** | Sspn | -0.070576144 | -0.328714696 | 0.187562409 | 9.283032931 | -0.672626877 | 0.526763597 | 0.881389679 | -4.951770435 |
| **A_55_P2126945** | 1700066B19Rik | -0.08200423 | -0.382016991 | 0.21800853 | 8.780430504 | -0.672458818 | 0.526863195 | 0.881480022 | -4.951847669 |
| **A_55_P2142028** | Clip4 | 0.033361949 | -0.088726556 | 0.155450453 | 7.777879457 | 0.672273202 | 0.526973212 | 0.88150972 | -4.951932954 |
| **A_55_P2093535** | Rps8 | -0.04173641 | -0.194481952 | 0.111009132 | 15.86772305 | -0.672226627 | 0.52700082 | 0.88150972 | -4.951954351 |
| **A_55_P1955542** | Olfr522 | 0.030165474 | -0.080237831 | 0.14056878 | 8.214487587 | 0.672198046 | 0.527017762 | 0.88150972 | -4.951967481 |
| **A_55_P2185214** | NA | 0.04168409 | -0.110958862 | 0.194327041 | 16.2070988 | 0.671835164 | 0.527232902 | 0.881613718 | -4.952134144 |
| **A_51_P270949** | H1f5 | 0.141274274 | -0.376110097 | 0.658658646 | 8.837540125 | 0.671767428 | 0.527273067 | 0.881613718 | -4.952165246 |
| **A_55_P1960837** | NA | -0.046707776 | -0.217767452 | 0.124351901 | 12.13432639 | -0.67175468 | 0.527280626 | 0.881613718 | -4.952171099 |
| **A_51_P290819** | Hint3 | 0.061009738 | -0.16246248 | 0.284481956 | 11.84275754 | 0.67165273 | 0.527341083 | 0.881613718 | -4.952217905 |
| **A_52_P506344** | B3gat3 | -0.051203157 | -0.23877438 | 0.136368067 | 8.640141839 | -0.671582988 | 0.527382442 | 0.881613718 | -4.952249921 |
| **A_51_P464588** | Dnajc28 | -0.030331969 | -0.141479174 | 0.080815236 | 7.803036503 | -0.671384357 | 0.527500249 | 0.881613718 | -4.952341089 |
| **A_51_P206235** | Slc39a5 | 0.081558183 | -0.217344862 | 0.380461228 | 7.949925323 | 0.671284107 | 0.527559714 | 0.881613718 | -4.952387094 |
| **A_55_P2106800** | Bglap | 0.086824598 | -0.231438429 | 0.405087625 | 8.539518173 | 0.671159519 | 0.52763362 | 0.881613718 | -4.95244426 |
| **A_65_P09285** | 2610524H06Rik | -0.047728421 | -0.222697658 | 0.127240817 | 9.521604548 | -0.671095807 | 0.527671417 | 0.881613718 | -4.952473491 |
| **A_55_P1958537** | Map6 | -0.296473283 | -1.383347955 | 0.790401388 | 11.79826631 | -0.671081487 | 0.527679913 | 0.881613718 | -4.95248006 |
| **A_55_P2016287** | Tpst1 | 0.03973039 | -0.105933946 | 0.185394727 | 8.330480135 | 0.671025095 | 0.527713369 | 0.881613718 | -4.95250593 |
| **A_55_P2151546** | Adarb1 | -0.068153788 | -0.318027682 | 0.181720105 | 10.64038589 | -0.67102435 | 0.527713811 | 0.881613718 | -4.952506272 |
| **A_52_P664617** | Zfp472 | -0.033367471 | -0.155710442 | 0.0889755 | 8.241767669 | -0.670985959 | 0.527736589 | 0.881613718 | -4.952523883 |
| **A_55_P2034042** | Gga3 | 0.052407901 | -0.139784694 | 0.244600495 | 10.40153113 | 0.670855952 | 0.527813727 | 0.881613718 | -4.952583514 |
| **A_52_P375873** | Srp9 | -0.054443384 | -0.254103739 | 0.145216972 | 12.16497359 | -0.670845378 | 0.527820001 | 0.881613718 | -4.952588363 |
| **A_52_P562467** | Sync | 0.063528154 | -0.169455924 | 0.296512232 | 9.283821015 | 0.670824875 | 0.527832168 | 0.881613718 | -4.952597767 |
| **A_55_P2025660** | Ranbp9 | -0.073619416 | -0.343628042 | 0.19638921 | 9.841534368 | -0.670785851 | 0.527855324 | 0.881613718 | -4.952615663 |
| **A_55_P2109712** | Csnk1g2 | -0.036189408 | -0.16895976 | 0.096580945 | 8.35517846 | -0.67057835 | 0.527978464 | 0.881674474 | -4.952710812 |
| **A_55_P1954081** | Ccdc115 | -0.110258102 | -0.514773464 | 0.294257261 | 11.94901163 | -0.670570826 | 0.527982929 | 0.881674474 | -4.952714261 |
| **A_51_P372312** | Mrpl54 | -0.045868334 | -0.214211959 | 0.122475291 | 12.06956483 | -0.670325057 | 0.528128804 | 0.881800841 | -4.952826926 |
| **A_51_P345110** | Thap4 | -0.053003194 | -0.247585471 | 0.141579082 | 11.79446862 | -0.670143681 | 0.528236475 | 0.881800841 | -4.95291005 |
| **A_55_P1956948** | Rpl27 | -0.054282508 | -0.25356614 | 0.145001124 | 16.18721864 | -0.670127502 | 0.52824608 | 0.881800841 | -4.952917464 |
| **A_55_P1987043** | Tpp1 | -0.052819796 | -0.246807527 | 0.141167935 | 9.829489497 | -0.669871681 | 0.528397971 | 0.881800841 | -4.953034673 |
| **A_51_P124535** | Mest | -0.105782129 | -0.49431729 | 0.282753031 | 13.2656214 | -0.669809251 | 0.528435042 | 0.881800841 | -4.948560119 |
| **A_55_P1968789** | Rasa3 | -0.156705327 | -0.73231899 | 0.418908335 | 11.07436377 | -0.669764046 | 0.528461886 | 0.881800841 | -4.953083977 |
| **A_55_P2062418** | Atp6v0a1 | 0.085188809 | -0.227769952 | 0.39814757 | 12.37415328 | 0.669675782 | 0.528514302 | 0.881800841 | -4.953124403 |
| **A_55_P2144115** | 1700125H20Rik | -0.036822779 | -0.172108814 | 0.098463255 | 8.55097274 | -0.6696267 | 0.528543452 | 0.881800841 | -4.953146882 |
| **A_55_P2003981** | NA | 0.074202295 | -0.198450017 | 0.346854608 | 8.470103821 | 0.669541217 | 0.528594222 | 0.881800841 | -4.953186028 |
| **A_52_P316405** | Lamtor4 | -0.040126706 | -0.187572527 | 0.107319115 | 11.59491804 | -0.669530248 | 0.528600737 | 0.881800841 | -4.953191051 |
| **A_55_P2037902** | Brd3 | -0.056844844 | -0.265785304 | 0.152095615 | 9.173470493 | -0.669326003 | 0.528722055 | 0.881800841 | -4.953284565 |
| **A_55_P1980214** | NA | 0.062692027 | -0.167776846 | 0.2931609 | 9.261887108 | 0.669220442 | 0.528784763 | 0.881800841 | -4.953332887 |
| **A_51_P292757** | Acbd4 | -0.066538302 | -0.311199369 | 0.178122765 | 11.15146078 | -0.669076859 | 0.528870067 | 0.881800841 | -4.953398605 |
| **A_55_P1962483** | NA | 0.044358216 | -0.118748319 | 0.207464752 | 9.541208916 | 0.669070488 | 0.528873852 | 0.881800841 | -4.953401521 |
| **A_52_P72587** | Prkcq | -0.031576886 | -0.147704214 | 0.084550443 | 7.735650462 | -0.668965963 | 0.528935957 | 0.881800841 | -4.953449355 |
| **A_52_P263095** | Ahnak | 0.142121309 | -0.380725362 | 0.664967981 | 14.3687972 | 0.668734941 | 0.529073238 | 0.881800841 | -4.953555056 |
| **A_66_P112356** | Agt | 0.122929364 | -0.329323162 | 0.575181889 | 8.462389033 | 0.668719205 | 0.52908259 | 0.881800841 | -4.953562254 |
| **A_52_P578043** | NA | -0.03840152 | -0.17967979 | 0.10287675 | 7.956191475 | -0.668716756 | 0.529084045 | 0.881800841 | -4.953563374 |
| **A_55_P2014690** | NA | 0.034486328 | -0.092408558 | 0.161381215 | 8.390682178 | 0.668608666 | 0.529148285 | 0.881800841 | -4.953612818 |
| **A_55_P2004016** | Crispld2 | -0.223051741 | -1.043920494 | 0.597817011 | 10.78152494 | -0.668499455 | 0.529213196 | 0.881800841 | -4.953662769 |
| **A_55_P2024953** | Tbc1d4 | 0.050723535 | -0.143480335 | 0.244927405 | 10.58925379 | 0.676864468 | 0.529304424 | 0.881800841 | -4.89144401 |
| **A_51_P352720** | Lmod2 | -0.029503554 | -0.138108247 | 0.079101139 | 7.64819143 | -0.668336075 | 0.529310313 | 0.881800841 | -4.953737482 |
| **A_51_P450100** | Rab5if | -0.057968489 | -0.271359629 | 0.15542265 | 13.27776894 | -0.668320468 | 0.52931959 | 0.881800841 | -4.953744618 |

| | | | | | | | | | |
|---|---|---|---|---|---|---|---|---|---|
| **A_51_P339503** | Cct4 | -0.058431997 | -0.273569401 | 0.156705406 | 11.11687187 | -0.668196155 | 0.529393493 | 0.881800841 | -4.953801455 |
| **A_55_P1987396** | Tbl1xr1 | -0.052146858 | -0.244146495 | 0.139852778 | 8.918732 | -0.668185285 | 0.529399956 | 0.881800841 | -4.953806425 |
| **A_55_P2149269** | Galt | 0.039050038 | -0.1047361 | 0.182836176 | 8.089959451 | 0.66814944 | 0.529421267 | 0.881800841 | -4.953822811 |
| **A_52_P681391** | G0s2 | -0.050153319 | -0.234861203 | 0.134554565 | 8.185167201 | -0.668010684 | 0.529503766 | 0.881800841 | -4.953886238 |
| **A_55_P2004801** | Tacc3 | 0.111189696 | -0.298322755 | 0.520702148 | 9.444406657 | 0.667984825 | 0.529519142 | 0.881800841 | -4.953898057 |
| **A_51_P418016** | Ccnj | 0.040989794 | -0.109977308 | 0.191956895 | 8.491239899 | 0.667978646 | 0.529522816 | 0.881800841 | -4.953900881 |
| **A_55_P2053551** | Mast4 | -0.056784969 | -0.265968185 | 0.152398247 | 8.583997342 | -0.667845059 | 0.529602252 | 0.881800841 | -4.953961933 |
| **A_55_P2412349** | AW111846 | -0.029614705 | -0.138711177 | 0.079481767 | 7.675729526 | -0.667829914 | 0.529611259 | 0.881800841 | -4.953968854 |
| **A_55_P2027022** | Prdm1 | -0.076243954 | -0.357123438 | 0.20463553 | 10.07560037 | -0.667812521 | 0.529621602 | 0.881800841 | -4.953976802 |
| **A_52_P671812** | NA | 0.172622042 | -0.463311397 | 0.80855548 | 13.04417418 | 0.66781128 | 0.52962234 | 0.881800841 | -4.953977369 |
| **A_55_P2331974** | NA | 0.094304261 | -0.253119791 | 0.441728313 | 10.01792656 | 0.667791059 | 0.529634365 | 0.881800841 | -4.953986609 |
| **A_51_P170562** | Cpn2 | -0.029206083 | -0.136809185 | 0.07839702 | 7.646733844 | -0.667755816 | 0.529655324 | 0.881800841 | -4.954002713 |
| **A_55_P2103856** | Hgs | 0.047981255 | -0.128816911 | 0.224779422 | 12.62571274 | 0.667671768 | 0.52970531 | 0.881808107 | -4.954041116 |
| **A_55_P2110351** | Eppk1 | -0.144240825 | -0.675857862 | 0.387376212 | 10.03490978 | -0.667511066 | 0.529800892 | 0.881831671 | -4.954114531 |
| **A_52_P649296** | Nras | -0.045718565 | -0.214226759 | 0.122789629 | 10.3009077 | -0.667483801 | 0.52981711 | 0.881831671 | -4.954126985 |
| **A_55_P2186352** | Ttc12 | 0.083984028 | -0.225592509 | 0.393560566 | 8.460760562 | 0.667417865 | 0.529856331 | 0.881831671 | -4.954157102 |
| **A_55_P2013088** | Arpc4 | -0.060309518 | -0.282647294 | 0.162028257 | 9.569099492 | -0.667331717 | 0.529907578 | 0.881841031 | -4.954196448 |
| **A_51_P430357** | Lrp10 | -0.062068156 | -0.290934722 | 0.16679841 | 11.14014105 | -0.667199411 | 0.529986289 | 0.881879852 | -4.954256867 |
| **A_66_P100093** | Ccdc47 | -0.044436453 | -0.208303691 | 0.119430786 | 11.25355354 | -0.667139128 | 0.530022156 | 0.881879852 | -4.954284393 |
| **A_51_P151775** | Mief1 | -0.045478106 | -0.213253264 | 0.122297052 | 9.494445155 | -0.666874136 | 0.530179834 | 0.882002388 | -4.954405366 |
| **A_55_P1954886** | NA | 0.032257209 | -0.086751385 | 0.151265804 | 15.89116335 | 0.666833837 | 0.530203816 | 0.882002388 | -4.95442376 |
| **A_52_P351021** | NA | 0.056563902 | -0.152136122 | 0.265263925 | 11.40006834 | 0.666785311 | 0.530232694 | 0.882002388 | -4.954445908 |
| **A_55_P2078776** | Kmt2e | -0.076284091 | -0.357803879 | 0.205235698 | 9.585574205 | -0.66664436 | 0.530316582 | 0.882043807 | -4.954510232 |
| **A_55_P2075429** | Ttll4 | 0.039353899 | -0.105890051 | 0.184597849 | 8.443553152 | 0.666590129 | 0.530348861 | 0.882043807 | -4.954534978 |
| **A_52_P252121** | Nfs1 | -0.032051103 | -0.150376398 | 0.086274192 | 7.843996391 | -0.666399323 | 0.530462438 | 0.882110895 | -4.95462203 |
| **A_52_P360650** | Abcb7 | -0.029989959 | -0.140711018 | 0.0807311 | 7.912791115 | -0.666369029 | 0.530480472 | 0.882110895 | -4.954635849 |
| **A_66_P130908** | Cyb5r4 | -0.066215329 | -0.310807439 | 0.178376781 | 9.725117106 | -0.666016914 | 0.530690115 | 0.882383591 | -4.954796438 |
| **A_55_P2016708** | Msl1 | 0.042786659 | -0.115317108 | 0.200890426 | 9.274981158 | 0.665787021 | 0.530827018 | 0.882473132 | -4.954901248 |
| **A_55_P2007640** | Eprs | -0.062361918 | -0.292847606 | 0.16812377 | 10.50388373 | -0.665648053 | 0.530909786 | 0.882473132 | -4.95496459 |
| **A_51_P409893** | Prkar2a | 0.036775598 | -0.099153834 | 0.172705029 | 8.239064723 | 0.665603199 | 0.530936502 | 0.882473132 | -4.954985033 |
| **A_51_P153053** | Smpdl3a | 0.096821464 | -0.261171776 | 0.454814705 | 10.7291603 | 0.665374241 | 0.531072889 | 0.882473132 | -4.955089364 |
| **A_52_P163939** | Cln6 | -0.058224157 | -0.273529547 | 0.157081234 | 9.442161583 | -0.665299912 | 0.531117171 | 0.882473132 | -4.955123228 |
| **A_51_P122660** | Liph | -0.042359493 | -0.199007619 | 0.114288634 | 9.761585677 | -0.665265049 | 0.531137942 | 0.882473132 | -4.955139111 |
| **A_51_P501757** | NA | -0.043933581 | -0.206416877 | 0.118549716 | 8.435125497 | -0.665207354 | 0.531172315 | 0.882473132 | -4.955165393 |
| **A_55_P1982559** | 1700007K13Rik | 0.032822839 | -0.088574337 | 0.154220016 | 7.788694204 | 0.665176233 | 0.531190858 | 0.882473132 | -4.955179569 |
| **A_51_P291839** | Cox14 | -0.097795853 | -0.459503934 | 0.263912228 | 13.88620134 | -0.665168065 | 0.531195725 | 0.882473132 | -4.95518329 |
| **A_52_P421344** | Paqr5 | 0.068746978 | -0.185561867 | 0.323055822 | 8.195684263 | 0.665060736 | 0.531259676 | 0.882473132 | -4.955232175 |
| **A_51_P494675** | Cotl1 | -0.135124935 | -0.634998414 | 0.364748543 | 13.15885473 | -0.665035098 | 0.531274953 | 0.882473132 | -4.955243852 |
| **A_55_P2152962** | Mtr | -0.037098711 | -0.174352261 | 0.100154838 | 8.660487904 | -0.664973598 | 0.531311601 | 0.882473132 | -4.955271859 |
| **A_55_P2109585** | Plekha7 | -0.06008288 | -0.282401107 | 0.162235347 | 9.422574768 | -0.664882397 | 0.531365951 | 0.882473132 | -4.95531339 |
| **A_55_P1956160** | Gm8909 | 0.061395867 | -0.165790524 | 0.288582258 | 9.033596255 | 0.664853547 | 0.531383143 | 0.882473132 | -4.955326526 |
| **A_51_P324287** | Kif23 | -0.042778515 | -0.20110689 | 0.115549859 | 7.840037955 | -0.664715977 | 0.531465134 | 0.882533469 | -4.95538916 |
| **A_55_P2002557** | Srebf1 | -0.038391056 | -0.180501326 | 0.103719214 | 8.568421678 | -0.664620532 | 0.531522023 | 0.882552116 | -4.955432609 |
| **A_55_P2091616** | Krt222 | -0.129857653 | -0.610683774 | 0.350968469 | 11.82443007 | -0.664429119 | 0.531636124 | 0.882665749 | -4.95551973 |
| **A_52_P299846** | Sh3rf1 | -0.029055082 | -0.136688192 | 0.078578028 | 8.272500632 | -0.664118199 | 0.531821498 | 0.88272032 | -4.955661201 |
| **A_55_P2076927** | Ints10 | -0.052876687 | -0.248762383 | 0.143009009 | 9.913288055 | -0.664095713 | 0.531834906 | 0.88272032 | -4.95567143 |
| **A_52_P475886** | Rc3h1 | 0.040619795 | -0.109861177 | 0.191100766 | 8.790381127 | 0.66408749 | 0.531839809 | 0.88272032 | -4.955675171 |
| **A_55_P1973783** | Tex9 | -0.037674834 | -0.177250021 | 0.101900353 | 8.184277838 | -0.664067607 | 0.531851666 | 0.88272032 | -4.955684216 |
| **A_51_P319449** | Acat1 | 0.120285293 | -0.325441823 | 0.566012408 | 10.06816931 | 0.663915239 | 0.531942528 | 0.882743022 | -4.95575352 |
| **A_55_P2014357** | Pdpk1 | -0.057891665 | -0.272421696 | 0.156638365 | 10.12174451 | -0.663891504 | 0.531956683 | 0.882743022 | -4.955764315 |

| | | | | | | | | |
|---|---|---|---|---|---|---|---|---|
| **A_55_P2051313** | Gstk1 | -0.111626821 | -0.525353044 | 0.302099401 | 10.27267935 | -0.663780781 | 0.532022718 | 0.882767014 | -4.955814668 |
| **A_55_P2109033** | Hmgcs2 | 0.085922244 | -0.23258542 | 0.404429908 | 8.21681061 | 0.663674127 | 0.532086331 | 0.882767014 | -4.955863164 |
| **A_55_P2068663** | Stmn1 | -0.116582608 | -0.548769889 | 0.315604674 | 13.9134535 | -0.663637542 | 0.532108153 | 0.882767014 | -4.955879798 |
| **A_55_P1962026** | Pigg | 0.045448137 | -0.123063636 | 0.21395991 | 9.605289114 | 0.663521506 | 0.53217737 | 0.882768855 | -4.955932551 |
| **A_55_P2036559** | Fbxl12os | -0.114938446 | -0.541130133 | 0.311253241 | 10.64486672 | -0.663482558 | 0.532200603 | 0.882768855 | -4.955950255 |
| **A_55_P2147001** | Wdr55 | 0.038345161 | -0.10386257 | 0.180552893 | 9.935301629 | 0.663371059 | 0.532267121 | 0.882780657 | -4.956000936 |
| **A_55_P2043147** | Akr1e1 | 0.061367776 | -0.166240557 | 0.288976109 | 9.212667228 | 0.663317398 | 0.532299136 | 0.882780657 | -4.956025325 |
| **A_55_P2068496** | NA | 0.110048943 | -0.298203398 | 0.518301285 | 12.00036097 | 0.663172266 | 0.53238573 | 0.882780657 | -4.956091278 |
| **A_51_P380178** | Id3 | -0.132669976 | -0.624846152 | 0.3595062 | 10.46624401 | -0.663164432 | 0.532390404 | 0.882780657 | -4.956094838 |
| **A_55_P2151822** | Plxnd1 | 0.234006504 | -0.634219284 | 1.102232292 | 12.28527911 | 0.663077682 | 0.532442168 | 0.882790759 | -4.956134254 |
| **A_55_P2032278** | NA | 0.060259105 | -0.163427672 | 0.283945882 | 8.44453534 | 0.662752737 | 0.532636094 | 0.882940151 | -4.956281862 |
| **A_55_P2129309** | Foxo3 | 0.05310393 | -0.144056105 | 0.250263965 | 8.350411586 | 0.662638917 | 0.532704032 | 0.882940151 | -4.956333551 |
| **A_51_P377210** | Caap1 | -0.035657031 | -0.168047517 | 0.096733455 | 8.36352432 | -0.662609364 | 0.532721673 | 0.882940151 | -4.956346971 |
| **A_55_P2162394** | Pdp1 | -0.031512962 | -0.148522236 | 0.085496312 | 7.829256311 | -0.662579668 | 0.532739399 | 0.882940151 | -4.956360455 |
| **A_51_P376711** | Gm15498 | 0.053413743 | -0.144941092 | 0.251768577 | 8.401031904 | 0.662490087 | 0.532792876 | 0.882940151 | -4.956401129 |
| **A_51_P290626** | Gulo | 0.123782336 | -0.33590575 | 0.583470421 | 10.80472922 | 0.662467517 | 0.532806349 | 0.882940151 | -4.956411376 |
| **A_55_P2123727** | Grb2 | -0.032149333 | -0.151561005 | 0.087262339 | 8.184136339 | -0.662360381 | 0.532870311 | 0.882958612 | -4.956460013 |
| **A_55_P2025038** | Cpe | -0.617725741 | -2.91235666 | 1.676905177 | 14.54508412 | -0.662295831 | 0.53290885 | 0.882958612 | -4.956489314 |
| **A_55_P2297485** | Ago2 | -0.09695043 | -0.457366087 | 0.263465227 | 11.55744788 | -0.661782461 | 0.533215421 | 0.883067653 | -4.956722265 |
| **A_55_P1952905** | Dnm1l | -0.040293913 | -0.190093412 | 0.109505586 | 11.89405009 | -0.661756548 | 0.533230898 | 0.883067653 | -4.95673402 |
| **A_51_P477682** | Prss12 | 0.09101682 | -0.247355569 | 0.429389209 | 10.43786556 | 0.661752976 | 0.533233032 | 0.883067653 | -4.956735641 |
| **A_51_P511949** | Setd7 | -0.160237189 | -0.756201916 | 0.435727537 | 11.97287542 | -0.661472701 | 0.533400456 | 0.883067653 | -4.956862754 |
| **A_55_P1990629** | NA | -0.050310735 | -0.237460736 | 0.136839266 | 10.70736466 | -0.661363146 | 0.533465909 | 0.883067653 | -4.956912429 |
| **A_55_P1993636** | Rab9 | -0.061837031 | -0.291888465 | 0.168214403 | 10.74503279 | -0.66129136 | 0.533508799 | 0.883067653 | -4.956944975 |
| **A_51_P326393** | Gapvd1 | -0.053256451 | -0.251407298 | 0.144894395 | 10.55921035 | -0.6612192 | 0.533551916 | 0.883067653 | -4.956977687 |
| **A_51_P354038** | Tmc6 | -0.054917046 | -0.259248752 | 0.149414659 | 10.01586011 | -0.661211768 | 0.533556357 | 0.883067653 | -4.956981056 |
| **A_55_P2080168** | Dgkk | -0.113934583 | -0.538056991 | 0.310187824 | 10.98984772 | -0.660896592 | 0.533744705 | 0.883067653 | -4.957123901 |
| **A_55_P2052425** | Setd4 | 0.06261368 | -0.170472516 | 0.295699876 | 10.36259607 | 0.660878828 | 0.533755322 | 0.883067653 | -4.95713195 |
| **A_55_P2034858** | Pias4 | -0.035630706 | -0.168300291 | 0.097038879 | 8.356098723 | -0.660727258 | 0.533845916 | 0.883067653 | -4.957200624 |
| **A_51_P392928** | Mthfd1 | 0.080349414 | -0.218855848 | 0.379554675 | 10.19021874 | 0.660667052 | 0.533881904 | 0.883067653 | -4.957227899 |
| **A_55_P1986993** | Brd8 | -0.038805987 | -0.183320901 | 0.105708926 | 7.910203368 | -0.660625351 | 0.533906832 | 0.883067653 | -4.957246789 |
| **A_55_P2086143** | Cyp39a1 | -0.085316131 | -0.403049121 | 0.232416858 | 9.575437646 | -0.660599147 | 0.533922496 | 0.883067653 | -4.957258659 |
| **A_55_P1968982** | Pbx3 | -0.034758688 | -0.164212037 | 0.094694662 | 8.436265594 | -0.660570603 | 0.53393956 | 0.883067653 | -4.957271588 |
| **A_51_P144249** | Ubr1 | -0.034859148 | -0.16469159 | 0.094973294 | 8.187614838 | -0.660545449 | 0.533954597 | 0.883067653 | -4.957282982 |
| **A_52_P138205** | Plekha8 | 0.031268325 | -0.085194487 | 0.147731137 | 8.899450581 | 0.660520813 | 0.533969325 | 0.883067653 | -4.95729414 |
| **A_55_P2096121** | NA | 0.097893516 | -0.266741973 | 0.462529005 | 14.2974233 | 0.660486818 | 0.533989648 | 0.883067653 | -4.957309537 |
| **A_55_P1979699** | Spop | -0.088955998 | -0.420372584 | 0.242460588 | 9.218370305 | -0.660343886 | 0.534075103 | 0.883067653 | -4.957374266 |
| **A_55_P2160712** | NA | -0.061240271 | -0.289424236 | 0.166943694 | 8.458369384 | -0.660269364 | 0.534119661 | 0.883067653 | -4.95740801 |
| **A_55_P1986306** | Ltv1 | 0.133644447 | -0.364474076 | 0.631762971 | 12.00816586 | 0.660066043 | 0.534241243 | 0.883067653 | -4.95750006 |
| **A_51_P265571** | Adm | -0.335706301 | -1.587028474 | 0.915615873 | 11.74125549 | -0.660023677 | 0.534266579 | 0.883067653 | -4.957519237 |
| **A_55_P2097178** | Idua | 0.055621387 | -0.151738305 | 0.262981079 | 10.31059848 | 0.659912928 | 0.534332814 | 0.883067653 | -4.957569364 |
| **A_55_P1986225** | NA | 0.105767446 | -0.288554994 | 0.500089885 | 12.1615591 | 0.659887175 | 0.534348217 | 0.883067653 | -4.95758102 |
| **A_55_P2090617** | Pilra | -0.031064544 | -0.146886359 | 0.08475727 | 7.692741514 | -0.659847821 | 0.534371755 | 0.883067653 | -4.957598829 |
| **A_52_P598634** | 1190007I07Rik | 0.045934825 | -0.125331761 | 0.217201411 | 10.32084781 | 0.659839918 | 0.534376482 | 0.883067653 | -4.957602406 |
| **A_52_P292684** | Exosc4 | 0.036701205 | -0.100139935 | 0.173542345 | 10.08715742 | 0.659831134 | 0.534381736 | 0.883067653 | -4.957606381 |
| **A_55_P2025702** | C330007P06Rik | 0.035516083 | -0.096909654 | 0.16794182 | 8.365951993 | 0.659814458 | 0.53439171 | 0.883067653 | -4.957613928 |
| **A_55_P2047852** | Clcn3 | -0.072780346 | -0.344158656 | 0.198597964 | 11.07548583 | -0.659793673 | 0.534404142 | 0.883067653 | -4.957623333 |
| **A_55_P2099530** | NA | -0.050473176 | -0.238695365 | 0.137749013 | 11.14553097 | -0.659718976 | 0.534448822 | 0.883067653 | -4.957657134 |
| **A_55_P2023449** | NA | -0.12999515 | -0.614801815 | 0.354811515 | 10.00880499 | -0.659671515 | 0.534477213 | 0.883067653 | -4.957678609 |
| **A_51_P386562** | Ebf2 | -0.033184386 | -0.156954363 | 0.090585592 | 7.831546305 | -0.65961061 | 0.534513646 | 0.883067653 | -4.957706164 |

| | | | | | | | | | |
|---|---|---|---|---|---|---|---|---|---|
| **A_55_P1984622** | Smarcc1 | -0.032801024 | -0.155144575 | 0.089542528 | 8.161089232 | -0.659592148 | 0.534524691 | 0.883067653 | -4.957714516 |
| **A_55_P2166555** | NA | -0.131507143 | -0.622015781 | 0.359001496 | 12.10821247 | -0.659586631 | 0.534527991 | 0.883067653 | -4.957717013 |
| **A_66_P132777** | Cep68 | -0.042376546 | -0.200522376 | 0.115769284 | 8.930372509 | -0.65923002 | 0.534741353 | 0.883116564 | -4.95787831 |
| **A_55_P2046037** | Gm7083 | -0.076770435 | -0.363287928 | 0.209747058 | 12.3887206 | -0.659192145 | 0.534764017 | 0.883116564 | -4.957895437 |
| **A_51_P121455** | Slc25a39 | -0.036150326 | -0.171069344 | 0.098768692 | 8.211034243 | -0.659186356 | 0.534767481 | 0.883116564 | -4.957898055 |
| **A_51_P137029** | Plce1 | -0.028439804 | -0.134590246 | 0.077710638 | 7.698748544 | -0.659134366 | 0.534798593 | 0.883116564 | -4.957921563 |
| **A_55_P2072288** | Tmem167b | 0.06008132 | -0.164188173 | 0.284350813 | 10.40280214 | 0.659080445 | 0.534830861 | 0.883116564 | -4.957945943 |
| **A_55_P2025248** | Mxd1 | 0.069433959 | -0.189789926 | 0.328657844 | 9.891893066 | 0.658970657 | 0.534896566 | 0.883116564 | -4.957995577 |
| **A_55_P1961873** | Akt3 | -0.147660408 | -0.6989478 | 0.403626984 | 9.96452216 | -0.658954103 | 0.534906473 | 0.883116564 | -4.958003061 |
| **A_55_P2122648** | Dtymk | 0.113230108 | -0.309530229 | 0.535990445 | 10.20362133 | 0.658926312 | 0.534923107 | 0.883116564 | -4.958015624 |
| **A_55_P2113061** | NA | 0.059174428 | -0.161815826 | 0.280164682 | 8.802957716 | 0.658764382 | 0.535020029 | 0.88320114 | -4.958088815 |
| **A_55_P1972653** | Tada3 | 0.042968428 | -0.11761306 | 0.203549917 | 11.31323331 | 0.658298937 | 0.535298682 | 0.883492981 | -4.958299112 |
| **A_55_P2115392** | Mst1 | 0.146020958 | -0.39968961 | 0.691731525 | 8.345913354 | 0.658297175 | 0.535299737 | 0.883492981 | -4.958299908 |
| **A_52_P489295** | Adamts1 | -0.198764268 | -0.941710783 | 0.544182246 | 10.3373015 | -0.658187952 | 0.535365141 | 0.883492981 | -4.958349239 |
| **A_55_P2064578** | Prkci | 0.062159429 | -0.170190337 | 0.294509195 | 14.19557984 | 0.658163721 | 0.535379651 | 0.883492981 | -4.958360183 |
| **A_51_P477440** | Sec23ip | -0.042542857 | -0.201612029 | 0.116526314 | 10.32717232 | -0.657975618 | 0.535492303 | 0.883545393 | -4.958445123 |
| **A_55_P1983819** | Srsf7 | -0.059551537 | -0.282222683 | 0.163119609 | 11.36278504 | -0.657958034 | 0.535502834 | 0.883545393 | -4.958453062 |
| **A_55_P1977850** | Sall1 | -0.042749516 | -0.202625984 | 0.117126952 | 8.504681076 | -0.657833244 | 0.535577577 | 0.883575227 | -4.9585094 |
| **A_55_P2023697** | Zfp386 | -0.074933303 | -0.355203115 | 0.205336508 | 10.71873494 | -0.657760396 | 0.535621213 | 0.883575227 | -4.958542285 |
| **A_52_P445307** | Cwc15 | -0.075395897 | -0.357438379 | 0.206646584 | 12.99059091 | -0.657661396 | 0.535680518 | 0.883575227 | -4.95858697 |
| **A_52_P243391** | Sema4f | -0.030561808 | -0.144894777 | 0.083771161 | 7.648526489 | -0.657622589 | 0.535703766 | 0.883575227 | -4.958604484 |
| **A_55_P1968778** | Rasa1 | -0.042541544 | -0.201715679 | 0.116632591 | 9.85547709 | -0.65752143 | 0.535764369 | 0.883599786 | -4.958650135 |
| **A_66_P122173** | Mrtfb | -0.071573633 | -0.339459643 | 0.196312377 | 12.18005851 | -0.657312955 | 0.535889279 | 0.883730388 | -4.958744199 |
| **A_55_P2162875** | NA | -0.151741016 | -0.719834868 | 0.416352837 | 12.10543918 | -0.657131158 | 0.53599822 | 0.883834635 | -4.958826206 |
| **A_52_P183152** | Txnl1 | -0.035622276 | -0.169090911 | 0.097846359 | 8.255990557 | -0.656616223 | 0.536306869 | 0.884211588 | -4.959058388 |
| **A_55_P2049949** | NA | -0.045560147 | -0.21633474 | 0.125214446 | 9.062063037 | -0.656343244 | 0.536470536 | 0.884211588 | -4.959181412 |
| **A_55_P2164534** | Dtl | 0.058174953 | -0.159894109 | 0.276244014 | 7.936127134 | 0.656313211 | 0.536488545 | 0.884211588 | -4.959194945 |
| **A_52_P52618** | Csf2rb | -0.389212738 | -1.848356233 | 1.069930757 | 10.14267461 | -0.65623321 | 0.536536518 | 0.884211588 | -4.95923099 |
| **A_51_P161342** | Abcb6 | -0.04326722 | -0.20548255 | 0.11894811 | 8.576093752 | -0.656200049 | 0.536556404 | 0.884211588 | -4.959245931 |
| **A_51_P424256** | Gm6211 | -0.072870771 | -0.346103341 | 0.200361798 | 11.19471226 | -0.656130254 | 0.53659826 | 0.884211588 | -4.959277373 |
| **A_55_P2092869** | Anxa9 | -0.130102421 | -0.617956872 | 0.35775203 | 9.558268645 | -0.656091288 | 0.536621628 | 0.884211588 | -4.959294926 |
| **A_55_P2023777** | Tmx2 | 0.048067462 | -0.132179694 | 0.228314618 | 8.937875074 | 0.656072653 | 0.536632805 | 0.884211588 | -4.95930332 |
| **A_52_P195839** | Ctsc | -0.170024691 | -0.807605946 | 0.467556565 | 9.522982778 | -0.656063106 | 0.53663853 | 0.884211588 | -4.959307621 |
| **A_51_P450123** | Mrpl36 | 0.048205864 | -0.132587187 | 0.228998915 | 11.88480984 | 0.655975022 | 0.536691359 | 0.88422326 | -4.959347295 |
| **A_51_P170959** | NA | 0.186057722 | -0.511867675 | 0.883983119 | 8.520157868 | 0.65585504 | 0.536763325 | 0.884266454 | -4.959401331 |
| **A_55_P2158473** | 1700028P14Rik | -0.080992873 | -0.384933808 | 0.222948062 | 8.527488805 | -0.655581624 | 0.536927345 | 0.88439039 | -4.959524436 |
| **A_51_P272876** | Tent5a | -0.069715705 | -0.331379333 | 0.191947923 | 12.96207079 | -0.655475446 | 0.536991048 | 0.88439039 | -4.959572232 |
| **A_55_P2132507** | Bcl2l2 | -0.057389831 | -0.272838156 | 0.158058494 | 10.19695415 | -0.655331414 | 0.537077471 | 0.88439039 | -4.959637057 |
| **A_52_P510771** | Tbl3 | 0.032521372 | -0.089570031 | 0.154612775 | 7.951428436 | 0.655319269 | 0.537084759 | 0.88439039 | -4.959642523 |
| **A_55_P2010161** | Fastkd5 | -0.052423743 | -0.249237111 | 0.144389625 | 10.10659211 | -0.655303663 | 0.537094123 | 0.88439039 | -4.959649546 |
| **A_55_P2028394** | Atg13 | 0.098016079 | -0.269981392 | 0.466013551 | 13.15868259 | 0.655272067 | 0.537113083 | 0.88439039 | -4.959663764 |
| **A_51_P185713** | Ptdss2 | -0.0375991 | -0.178825871 | 0.103627671 | 8.49900003 | -0.654982332 | 0.537286966 | 0.884601342 | -4.959794122 |
| **A_55_P2049549** | Rab22a | -0.053795617 | -0.255893999 | 0.148302764 | 10.84665097 | -0.654867177 | 0.537356085 | 0.884621955 | -4.95984592 |
| **A_51_P273556** | Fam83d | 0.10285928 | -0.283594907 | 0.489313468 | 9.394490021 | 0.654808981 | 0.537391018 | 0.884621955 | -4.959872094 |
| **A_51_P394115** | Aadac | -0.07414553 | -0.352810879 | 0.204519818 | 7.853664272 | -0.654592718 | 0.537520847 | 0.8846938 | -4.959969344 |
| **A_55_P2126259** | Slc35e2 | -0.095586448 | -0.45483931 | 0.263666414 | 8.773547092 | -0.654583795 | 0.537526203 | 0.8846938 | -4.959973355 |
| **A_55_P2007919** | Akr1c19 | 0.051433014 | -0.141937355 | 0.244803383 | 7.853652999 | 0.654366737 | 0.53765653 | 0.884728203 | -4.960070935 |
| **A_51_P397296** | Marveld3 | -0.055748178 | -0.265350313 | 0.153853956 | 8.471299218 | -0.654341015 | 0.537671976 | 0.884728203 | -4.960082497 |
| **A_55_P2161600** | Diaph2 | -0.07166237 | -0.341107255 | 0.197782515 | 10.1148513 | -0.654320286 | 0.537684423 | 0.884728203 | -4.960091814 |
| **A_55_P2039712** | Gigyf2 | -0.146987426 | -0.699776254 | 0.405801401 | 11.99459012 | -0.6541692 | 0.537775153 | 0.884802174 | -4.960159717 |

| | | | | | | | | | |
|---|---|---|---|---|---|---|---|---|---|
| **A_55_P2310898** | Kantr | -0.052068656 | -0.247990919 | 0.143853607 | 8.716831661 | -0.653825332 | 0.537981689 | 0.8849546 | -4.960314213 |
| **A_51_P119401** | Tug1 | -0.03818988 | -0.181895441 | 0.105515682 | 8.913350508 | -0.653798435 | 0.537997847 | 0.8849546 | -4.960326295 |
| **A_55_P2100968** | Dnah7b | 0.028592792 | -0.07900606 | 0.136191645 | 7.782755968 | 0.653759623 | 0.538021162 | 0.8849546 | -4.960343728 |
| **A_55_P2082434** | Btbd11 | 0.076689834 | -0.211927085 | 0.365306753 | 8.802185479 | 0.653710068 | 0.538050932 | 0.8849546 | -4.960365985 |
| **A_55_P1976690** | Snrnp35 | -0.072370804 | -0.344799458 | 0.20005785 | 10.6907249 | -0.653551436 | 0.538146235 | 0.884978724 | -4.960437224 |
| **A_55_P1958623** | Ncln | 0.064320157 | -0.177809831 | 0.306450145 | 12.05465625 | 0.653533238 | 0.538157169 | 0.884978724 | -4.960445396 |
| **A_55_P2039532** | Pax8 | 0.093781827 | -0.259327735 | 0.446891389 | 9.726360377 | 0.653398862 | 0.53823791 | 0.885036202 | -4.96050573 |
| **A_52_P254174** | Nr3c1 | 0.047722191 | -0.132000985 | 0.227445368 | 8.858636501 | 0.653259077 | 0.538321908 | 0.885093381 | -4.960568481 |
| **A_55_P2071496** | Fcf1 | -0.053422351 | -0.254654749 | 0.147810048 | 12.24146395 | -0.653121912 | 0.538404341 | 0.885093381 | -4.960630046 |
| **A_66_P105046** | Il18 | -0.062432321 | -0.297629807 | 0.172765166 | 8.163594482 | -0.653049297 | 0.538447983 | 0.885093381 | -4.960662634 |
| **A_55_P1995902** | Zfp992 | 0.050891952 | -0.140834442 | 0.242618346 | 9.102891899 | 0.653034868 | 0.538456656 | 0.885093381 | -4.960669109 |
| **A_55_P2160094** | Plscr2 | 0.08130272 | -0.225025947 | 0.387631386 | 8.010368482 | 0.65296003 | 0.538501638 | 0.885093381 | -4.960702692 |
| **A_51_P349888** | Ang2 | 0.074390386 | -0.205989508 | 0.354770281 | 8.463902102 | 0.65273832 | 0.538634912 | 0.885112232 | -4.960802161 |
| **A_52_P637353** | Zbtb43 | -0.047703399 | -0.227517481 | 0.132110682 | 9.599601869 | -0.652671713 | 0.538674954 | 0.885112232 | -4.960832038 |
| **A_55_P2140592** | Dmkn | -0.319985507 | -1.526230112 | 0.886259098 | 14.79005032 | -0.652625573 | 0.538702694 | 0.885112232 | -4.960852734 |
| **A_55_P2168301** | Slc46a1 | 0.06252967 | -0.173204278 | 0.298263618 | 11.2060701 | 0.652579115 | 0.538730626 | 0.885112232 | -4.960873571 |
| **A_55_P2043987** | Luzp1 | -0.059885777 | -0.285676131 | 0.165904577 | 13.12672433 | -0.652510447 | 0.538771912 | 0.885112232 | -4.960904367 |
| **A_55_P2010332** | Ewsr1 | -0.076959366 | -0.367134925 | 0.213216192 | 10.24301376 | -0.652483926 | 0.538787859 | 0.885112232 | -4.96091626 |
| **A_55_P2146175** | NA | 0.052928151 | -0.146669914 | 0.252526216 | 9.426478118 | 0.652378379 | 0.538851324 | 0.885141263 | -4.960963589 |
| **A_55_P2061909** | Tceal9 | -0.080892061 | -0.386010201 | 0.224226078 | 13.31623779 | -0.652239411 | 0.538934893 | 0.885203309 | -4.961025894 |
| **A_51_P299287** | Impa2 | -0.103562703 | -0.494249926 | 0.287124521 | 11.3301877 | -0.652143741 | 0.538992428 | 0.885222589 | -4.961068781 |
| **A_55_P2050857** | D11Wsu47e | -0.032385714 | -0.154604578 | 0.089833151 | 10.10453363 | -0.651905118 | 0.539135953 | 0.885369821 | -4.961175728 |
| **A_51_P296036** | Nrbp2 | 0.098128164 | -0.272228562 | 0.46848489 | 9.938862731 | 0.651842392 | 0.539173685 | 0.885369821 | -4.961203836 |
| **A_51_P202074** | Ncapd2 | -0.05247576 | -0.250673237 | 0.145721718 | 9.449681297 | -0.651373031 | 0.539456077 | 0.885758283 | -4.961414087 |
| **A_52_P436238** | Odc1 | -0.056230439 | -0.268642466 | 0.156181589 | 9.999055442 | -0.651270688 | 0.539517664 | 0.885778898 | -4.961459915 |
| **A_55_P1991075** | Lmf1 | 0.054014372 | -0.150071457 | 0.258100202 | 10.80713875 | 0.651126912 | 0.539604192 | 0.885778898 | -4.961524287 |
| **A_55_P1966721** | NA | -0.044476086 | -0.212523644 | 0.123571472 | 8.783683321 | -0.651123724 | 0.539606111 | 0.885778898 | -4.961525715 |
| **A_51_P493016** | Cmya5 | -0.029692522 | -0.141915242 | 0.082530198 | 7.725285223 | -0.650931761 | 0.539721653 | 0.885893328 | -4.961611642 |
| **A_55_P2171348** | Grm2 | 0.060822254 | -0.169087086 | 0.290731594 | 9.129281455 | 0.650841231 | 0.539776149 | 0.885897416 | -4.961652159 |
| **A_52_P135873** | NA | -0.152687626 | -0.729923295 | 0.424548043 | 10.9295606 | -0.650758494 | 0.539825956 | 0.885897416 | -4.961689184 |
| **A_55_P2057390** | Smap1 | -0.052895123 | -0.25290524 | 0.147114994 | 12.63173927 | -0.650628125 | 0.539904443 | 0.885897416 | -4.961747516 |
| **A_55_P2008176** | Zscan12 | 0.053023481 | -0.14749338 | 0.253540341 | 8.577235631 | 0.650558719 | 0.539946231 | 0.885897416 | -4.961778567 |
| **A_51_P248666** | Cd274 | -0.139512343 | -0.667153429 | 0.388128743 | 9.165505316 | -0.650493844 | 0.539985294 | 0.885897416 | -4.961807588 |
| **A_55_P2140118** | Qpct | -0.456523752 | -2.183241335 | 1.270193831 | 11.46500534 | -0.650445974 | 0.540014118 | 0.885897416 | -4.961829001 |
| **A_52_P284348** | Smim20 | 0.067578338 | -0.188081394 | 0.323238069 | 10.84739528 | 0.650300878 | 0.540101492 | 0.885897416 | -4.961893897 |
| **A_55_P2072403** | Suv39h2 | 0.084903718 | -0.236302857 | 0.406110293 | 13.60026442 | 0.650296542 | 0.540104103 | 0.885897416 | -4.961895836 |
| **A_52_P48546** | Rnf41 | 0.039897301 | -0.111067116 | 0.190861719 | 9.144630063 | 0.65018671 | 0.540170248 | 0.885897416 | -4.961944951 |
| **A_55_P2070005** | NA | 0.046864569 | -0.130514629 | 0.224243767 | 12.00596274 | 0.649996631 | 0.540284732 | 0.885897416 | -4.962029936 |
| **A_52_P674309** | Mocs2 | 0.044113954 | -0.122861256 | 0.211089164 | 11.95459543 | 0.649969764 | 0.540300915 | 0.885897416 | -4.962041947 |
| **A_51_P395164** | Pex16 | -0.075290776 | -0.360295332 | 0.209713779 | 10.25384825 | -0.649918852 | 0.540331582 | 0.885897416 | -4.962064705 |
| **A_51_P214083** | 2310039H08Rik | -0.073637356 | -0.352397346 | 0.205122633 | 10.39921638 | -0.649885599 | 0.540351613 | 0.885897416 | -4.962079569 |
| **A_66_P127224** | Krt10 | -0.071647705 | -0.342955533 | 0.199660123 | 8.390208807 | -0.649694398 | 0.540466798 | 0.885897416 | -4.962165023 |
| **A_55_P2050408** | NA | 0.076311646 | -0.212683079 | 0.36530637 | 9.6552693 | 0.649635976 | 0.540501996 | 0.885897416 | -4.96219113 |
| **A_55_P2073699** | Bnip3l | -0.057805762 | -0.276736211 | 0.161124687 | 12.68029534 | -0.649582217 | 0.540534386 | 0.885897416 | -4.962215151 |
| **A_52_P322927** | NA | -0.029572279 | -0.141574921 | 0.082430362 | 7.699163407 | -0.649569605 | 0.540541985 | 0.885897416 | -4.962220786 |
| **A_55_P2039250** | Mboat2 | -0.163979157 | -0.785048431 | 0.457090118 | 9.81235254 | -0.649557748 | 0.54054913 | 0.885897416 | -4.962226084 |
| **A_55_P2154097** | NA | -0.046748036 | -0.223829574 | 0.130333502 | 14.66423057 | -0.649470227 | 0.540601865 | 0.885908727 | -4.962265186 |
| **A_51_P130079** | Lrmp | -0.027780048 | -0.133038136 | 0.07747804 | 7.731481258 | -0.649301913 | 0.540703291 | 0.885938607 | -4.962340373 |
| **A_55_P2139473** | Ilk | -0.055994045 | -0.268196206 | 0.156208116 | 12.87158132 | -0.649174122 | 0.540780306 | 0.885938607 | -4.962397447 |
| **A_55_P1976395** | Nsdhl | 0.034430405 | -0.096057724 | 0.164918534 | 7.772546072 | 0.649142875 | 0.540799139 | 0.885938607 | -4.962411402 |

| | | | | | | | | | |
|---|---|---|---|---|---|---|---|---|---|
| **A_52_P593268** | Lsm6 | -0.050565636 | -0.242206938 | 0.141075666 | 12.33398828 | -0.649135744 | 0.540803436 | 0.885938607 | -4.962414586 |
| **A_55_P2341890** | NA | -0.117850123 | -0.564563799 | 0.328863552 | 8.40634907 | -0.649037748 | 0.540862501 | 0.885960279 | -4.962458345 |
| **A_55_P2009952** | Me1 | -0.044190517 | -0.211715808 | 0.123334774 | 12.49686933 | -0.648959903 | 0.540909424 | 0.885962058 | -4.962493102 |
| **A_55_P1989813** | Gcm1 | -0.139144881 | -0.666797088 | 0.388507326 | 10.34896778 | -0.64876683 | 0.541025812 | 0.885998531 | -4.962579292 |
| **A_51_P324303** | Mylip | -0.106556546 | -0.510766937 | 0.297653845 | 10.51932476 | -0.648547552 | 0.541158018 | 0.885998531 | -4.962677155 |
| **A_55_P2129856** | Mtfmt | 0.03422454 | -0.095608217 | 0.164057296 | 10.29467202 | 0.648518693 | 0.541175419 | 0.885998531 | -4.962690032 |
| **A_52_P381784** | Nsd2 | -0.087087076 | -0.417457939 | 0.243283787 | 9.427976595 | -0.648516655 | 0.541176648 | 0.885998531 | -4.962690942 |
| **A_55_P2061804** | NA | -0.05147261 | -0.246743296 | 0.143798076 | 9.159940398 | -0.648497489 | 0.541188204 | 0.885998531 | -4.962699494 |
| **A_52_P93066** | Klhl42 | -0.059254931 | -0.28405991 | 0.165550047 | 9.271782116 | -0.648466785 | 0.541206718 | 0.885998531 | -4.962713194 |
| **A_55_P2159506** | Gm5458 | -0.097701856 | -0.468428938 | 0.273025226 | 8.734038946 | -0.648362166 | 0.541269805 | 0.886006313 | -4.962759871 |
| **A_55_P2155062** | Naa40 | 0.05778401 | -0.16149457 | 0.27706259 | 10.05309055 | 0.648306876 | 0.541303148 | 0.886006313 | -4.962784537 |
| **A_55_P2102449** | Nxf7 | 0.053326075 | -0.149143095 | 0.255795246 | 11.26302558 | 0.647962511 | 0.541510847 | 0.886155237 | -4.962938124 |
| **A_55_P1965554** | Orc3 | 0.0659391 | -0.18446649 | 0.316344689 | 10.67736324 | 0.647840593 | 0.541584391 | 0.886155237 | -4.962992484 |
| **A_55_P1996212** | Zbtb2 | -0.062340415 | -0.299091322 | 0.174410491 | 10.09166521 | -0.647809336 | 0.541603248 | 0.886155237 | -4.963006419 |
| **A_55_P2064171** | NA | -0.143957984 | -0.690723365 | 0.402807398 | 11.16213141 | -0.647744761 | 0.541642205 | 0.886155237 | -4.963035207 |
| **A_55_P2037912** | Cyp2d10 | -0.030751787 | -0.147557241 | 0.086053666 | 7.699040109 | -0.647703747 | 0.541666949 | 0.886155237 | -4.96305349 |
| **A_55_P1982111** | Homer1 | -0.031907454 | -0.153109091 | 0.089294184 | 8.969326276 | -0.647668571 | 0.541688172 | 0.886155237 | -4.963069169 |
| **A_55_P2073482** | NA | 0.039060846 | -0.109323424 | 0.187445117 | 11.26835328 | 0.647624016 | 0.541715054 | 0.886155237 | -4.963089029 |
| **A_55_P2057984** | Tbc1d19 | -0.036782784 | -0.176548261 | 0.102982692 | 8.731284348 | -0.647461359 | 0.541813199 | 0.886230188 | -4.96316152 |
| **A_55_P2454099** | Oga | -0.037016862 | -0.177685953 | 0.103652228 | 8.237557002 | -0.647396112 | 0.541852572 | 0.886230188 | -4.963190595 |
| **A_55_P2034517** | NA | -0.060770424 | -0.291943509 | 0.17040266 | 9.077593645 | -0.64673172 | 0.542253593 | 0.886735494 | -4.963486515 |
| **A_51_P155174** | Zfp672 | 0.087334365 | -0.244902846 | 0.419571576 | 9.678115506 | 0.646704762 | 0.542269869 | 0.886735494 | -4.963498517 |
| **A_66_P130601** | Foxp2 | 0.029538509 | -0.082840254 | 0.141917273 | 7.649609034 | 0.646656263 | 0.54229915 | 0.886735494 | -4.963520108 |
| **A_55_P2093143** | Casd1 | 0.068010144 | -0.190926677 | 0.326946966 | 8.815439416 | 0.646173354 | 0.542590764 | 0.886782131 | -4.963735019 |
| **A_51_P135004** | Zfp212 | -0.047414047 | -0.227975827 | 0.133147732 | 10.58611169 | -0.646026549 | 0.542679435 | 0.886782131 | -4.963800326 |
| **A_55_P2089507** | NA | -0.082056566 | -0.394548724 | 0.230435592 | 15.04866096 | -0.646016153 | 0.542685714 | 0.886782131 | -4.96380495 |
| **A_55_P2109257** | Nipal3 | -0.101748206 | -0.489251249 | 0.285754836 | 9.714981397 | -0.645982583 | 0.542705992 | 0.886782131 | -4.963819883 |
| **A_55_P2235931** | Cacna1c | -0.056854923 | -0.273402907 | 0.159693061 | 8.109826372 | -0.645926488 | 0.542739877 | 0.886782131 | -4.963844832 |
| **A_55_P2043554** | Slc12a4 | 0.119510158 | -0.335694845 | 0.574715161 | 11.65453631 | 0.645902489 | 0.542754374 | 0.886782131 | -4.963855506 |
| **A_55_P2076545** | F5 | 0.057552866 | -0.161678417 | 0.276784149 | 7.988630214 | 0.645852865 | 0.542784352 | 0.886782131 | -4.963877575 |
| **A_52_P117999** | Selenot | 0.03024899 | -0.084979439 | 0.145477418 | 8.60367789 | 0.645833243 | 0.542796206 | 0.886782131 | -4.963886301 |
| **A_55_P2148001** | 6820431F20Rik | -0.0822129 | -0.39543643 | 0.23101063 | 10.63375957 | -0.645735634 | 0.542855175 | 0.886782131 | -4.963929706 |
| **A_55_P2368355** | 2610319H10Rik | -0.206484617 | -0.993182014 | 0.58021278 | 9.951869254 | -0.645727307 | 0.542860206 | 0.886782131 | -4.963933409 |
| **A_52_P413492** | Ldlrap1 | -0.040261586 | -0.193732675 | 0.113209503 | 10.34663364 | -0.645406693 | 0.543053931 | 0.886782131 | -4.96407594 |
| **A_55_P2049061** | NA | 0.034541913 | -0.097131953 | 0.16621578 | 9.700055232 | 0.645380688 | 0.543069646 | 0.886782131 | -4.964087498 |
| **A_55_P2009861** | NA | -0.187825798 | -0.903860496 | 0.5282089 | 11.18132128 | -0.645342604 | 0.543092661 | 0.886782131 | -4.964104424 |
| **A_52_P51029** | Ralgapa1 | -0.040980537 | -0.197215716 | 0.115254642 | 9.450087466 | -0.645309374 | 0.543112743 | 0.886782131 | -4.964119193 |
| **A_51_P312846** | Fbxo9 | 0.029248329 | -0.082269971 | 0.140766628 | 10.55295844 | 0.645244153 | 0.543152159 | 0.886782131 | -4.964148176 |
| **A_52_P489202** | 4732465J04Rik | -0.048614982 | -0.233980364 | 0.136750401 | 7.984561206 | -0.645224214 | 0.54316421 | 0.886782131 | -4.964157036 |
| **A_51_P103706** | Cyp2c29 | 0.048869021 | -0.137469411 | 0.235207453 | 7.799416548 | 0.645208921 | 0.543173452 | 0.886782131 | -4.950473402 |
| **A_55_P2075298** | Atp8a1 | 0.052981987 | -0.149046002 | 0.255009975 | 10.11719967 | 0.645187381 | 0.543186471 | 0.886782131 | -4.964173403 |
| **A_51_P508580** | Tcf15 | -0.045886106 | -0.220883016 | 0.129110805 | 8.209789069 | -0.645089514 | 0.543245623 | 0.886782131 | -4.964216887 |
| **A_55_P2042958** | Rad50 | -0.043697336 | -0.210361977 | 0.122967306 | 9.002899347 | -0.645031087 | 0.543280939 | 0.886782131 | -4.964242845 |
| **A_51_P349008** | Ptges2 | -0.043030382 | -0.207155501 | 0.121094737 | 9.735041324 | -0.645014249 | 0.543291117 | 0.886782131 | -4.964250325 |
| **A_55_P2152427** | Zwilch | -0.063473794 | -0.305603111 | 0.178655523 | 8.556327223 | -0.644935444 | 0.543338753 | 0.886785001 | -4.964285332 |
| **A_55_P1954331** | Cavin4 | -0.030397083 | -0.146385393 | 0.085591226 | 8.026899473 | -0.644743333 | 0.543454892 | 0.886899663 | -4.964370658 |
| **A_51_P477419** | Nfic | -0.128202139 | -0.617531571 | 0.361127293 | 10.0060906 | -0.644559622 | 0.543565968 | 0.887006044 | -4.964452233 |
| **A_66_P113268** | Mme | -0.063566553 | -0.306257953 | 0.179124847 | 9.605784726 | -0.644382058 | 0.54367334 | 0.887106364 | -4.96453106 |
| **A_55_P2066878** | Kcnj13 | 0.112839668 | -0.318061821 | 0.543741158 | 9.056124072 | 0.644247836 | 0.543754512 | 0.887163921 | -4.964590635 |
| **A_52_P140881** | Slc26a10 | 0.037911215 | -0.106911873 | 0.182734302 | 8.137658783 | 0.644019568 | 0.543892578 | 0.887314285 | -4.964691929 |

| | | | | | | | | |
|---|---|---|---|---|---|---|---|---|
| **A_51_P370717** | Gsc | 0.10900394 | -0.307554785 | 0.525562665 | 8.977103302 | 0.643776506 | 0.544039617 | 0.887362139 | -4.964799755 |
| **A_52_P221756** | Spock2 | 0.028788976 | -0.081233374 | 0.138811326 | 7.767041325 | 0.643745893 | 0.544058138 | 0.887362139 | -4.964813333 |
| **A_51_P453111** | Hexb | -0.097361862 | -0.469449493 | 0.274725769 | 13.05076631 | -0.643743416 | 0.544059636 | 0.887362139 | -4.964814432 |
| **A_55_P2008936** | Slc2a9 | -0.031588705 | -0.152359007 | 0.089181597 | 7.858786159 | -0.64348856 | 0.54421384 | 0.887455595 | -4.964927449 |
| **A_55_P1956482** | NA | -0.045106526 | -0.217571917 | 0.127358866 | 10.97497635 | -0.643437814 | 0.544244547 | 0.887455595 | -4.964949948 |
| **A_55_P1964832** | Reck | 0.034992741 | -0.098805944 | 0.168791427 | 8.184779266 | 0.643421077 | 0.544254676 | 0.887455595 | -4.964957369 |
| **A_55_P2191002** | 2610027K06Rik | 0.030003061 | -0.084746374 | 0.144752495 | 8.11322404 | 0.643256482 | 0.544354285 | 0.887543145 | -4.965030334 |
| **A_51_P357877** | Fbxo46 | -0.051365416 | -0.247841686 | 0.145110854 | 9.294666464 | -0.643176052 | 0.544402964 | 0.887547645 | -4.965065983 |
| **A_51_P324442** | Wdr35 | 0.080257881 | -0.226889591 | 0.387405352 | 8.927982545 | 0.642850382 | 0.544600098 | 0.887621487 | -4.965210293 |
| **A_52_P278295** | Pear1 | 0.036086251 | -0.10202308 | 0.174195582 | 8.490551766 | 0.642817821 | 0.54461981 | 0.887621487 | -4.965224718 |
| **A_55_P1980556** | NA | 0.040369718 | -0.114158115 | 0.194897551 | 7.740789803 | 0.642714598 | 0.544682304 | 0.887621487 | -4.965270443 |
| **A_51_P139069** | Wdr45 | 0.111987519 | -0.316698619 | 0.540673657 | 11.78067806 | 0.642686747 | 0.544699166 | 0.887621487 | -4.965282779 |
| **A_52_P158710** | Dcbld1 | -0.049427413 | -0.238658204 | 0.139803378 | 8.345828586 | -0.642606679 | 0.544747645 | 0.887621487 | -4.965318242 |
| **A_55_P2134148** | Mzt2 | -0.054224004 | -0.261829351 | 0.153381343 | 10.20871384 | -0.642572614 | 0.544768271 | 0.887621487 | -4.965333329 |
| **A_52_P430262** | Rbm15 | 0.065679461 | -0.18578955 | 0.317148471 | 9.492566256 | 0.642560862 | 0.544775388 | 0.887621487 | -4.965338533 |
| **A_51_P429246** | Fbxl6 | -0.040628621 | -0.1962185 | 0.114961258 | 8.863552251 | -0.642421262 | 0.544859921 | 0.887621487 | -4.965400351 |
| **A_52_P516034** | Ptp4a1 | -0.071648728 | -0.346074435 | 0.20277698 | 9.017153724 | -0.642322081 | 0.544919984 | 0.887621487 | -4.965444264 |
| **A_55_P2032458** | Gm6934 | 0.034163377 | -0.096707028 | 0.165033781 | 8.485814661 | 0.642226925 | 0.544977613 | 0.887621487 | -4.965486389 |
| **A_66_P127458** | Zfp652 | 0.078015931 | -0.220847827 | 0.376879689 | 10.94632979 | 0.642213167 | 0.544985946 | 0.887621487 | -4.96549248 |
| **A_55_P2023572** | NA | 0.035215019 | -0.099701749 | 0.170131787 | 7.994912966 | 0.642142128 | 0.545028973 | 0.887621487 | -4.965523925 |
| **A_51_P196862** | Amdhd1 | 0.028100244 | -0.079562701 | 0.135763188 | 7.714657365 | 0.642115271 | 0.545045239 | 0.887621487 | -4.965535812 |
| **A_55_P2027077** | Shc2 | -0.208098494 | -1.005646973 | 0.589449985 | 8.704594535 | -0.64192015 | 0.545163432 | 0.887697427 | -4.965622163 |
| **A_55_P2124941** | Brinp1 | 0.027980401 | -0.079272842 | 0.135233644 | 7.685633567 | 0.641819144 | 0.545224623 | 0.887697427 | -4.965666856 |
| **A_55_P2133524** | Nbr1 | 0.054021568 | -0.153089354 | 0.26113249 | 8.858637188 | 0.641701935 | 0.545295634 | 0.887697427 | -4.96571871 |
| **A_55_P2133993** | NA | -0.049044397 | -0.237078961 | 0.138990168 | 10.07942392 | -0.641683499 | 0.545306804 | 0.887697427 | -4.965726866 |
| **A_55_P2042520** | Cop1 | -0.042485627 | -0.205396763 | 0.120425509 | 11.352903 | -0.641594195 | 0.545360913 | 0.887697427 | -4.965766368 |
| **A_51_P417839** | Hdac4 | 0.067365451 | -0.190951695 | 0.325682597 | 10.06425625 | 0.641583449 | 0.545367424 | 0.887697427 | -4.965771121 |
| **A_51_P482043** | Epm2aip1 | -0.050887815 | -0.246132006 | 0.144356376 | 9.3454721 | -0.641216734 | 0.545589655 | 0.887923122 | -4.965933284 |
| **A_52_P251615** | Arhgef40 | 0.084185428 | -0.238820518 | 0.407191373 | 9.356610022 | 0.641203035 | 0.545597958 | 0.887923122 | -4.96593934 |
| **A_55_P2055819** | Adam15 | -0.146613613 | -0.709336664 | 0.416109439 | 14.40241186 | -0.640986308 | 0.545729324 | 0.88806214 | -4.966035141 |
| **A_55_P2104422** | Patj | -0.052707645 | -0.255089161 | 0.149673871 | 8.991735267 | -0.640725385 | 0.545887504 | 0.888232534 | -4.966150442 |
| **A_52_P424767** | Rbbp4 | -0.029819623 | -0.144335045 | 0.084695799 | 7.878904975 | -0.6406301 | 0.545945277 | 0.888232534 | -4.966192538 |
| **A_55_P1993512** | NA | -0.059680396 | -0.288902463 | 0.169541671 | 10.10899711 | -0.64053731 | 0.54600154 | 0.888232534 | -4.966233527 |
| **A_52_P234354** | Hic2 | -0.056226696 | -0.272247386 | 0.159793994 | 12.78361545 | -0.64034847 | 0.546116054 | 0.888232534 | -4.966316931 |
| **A_55_P2065824** | Smtn | -0.068195387 | -0.330206293 | 0.193815518 | 12.4262847 | -0.640331395 | 0.546126409 | 0.888232534 | -4.966324471 |
| **A_55_P2470474** | 9530082P21Rik | -0.055126914 | -0.266960301 | 0.156706472 | 8.711070735 | -0.640233567 | 0.546185741 | 0.888232534 | -4.966367669 |
| **A_55_P2018210** | NA | 0.037097402 | -0.105477953 | 0.179672757 | 16.39462422 | 0.640130028 | 0.546248539 | 0.888232534 | -4.966413383 |
| **A_51_P394833** | Tshz1 | -0.078409946 | -0.379771236 | 0.222951344 | 10.08577431 | -0.640107408 | 0.546262259 | 0.888232534 | -4.96642337 |
| **A_55_P2033521** | Med6 | 0.06798801 | -0.193341228 | 0.329317249 | 9.935375114 | 0.640049396 | 0.546297447 | 0.888232534 | -4.966448979 |
| **A_52_P527500** | Helb | 0.03454039 | -0.098250981 | 0.167331762 | 8.530546016 | 0.639921271 | 0.546375168 | 0.888232534 | -4.966505534 |
| **A_55_P2180854** | Mrgprg | -0.628174136 | -3.043797178 | 1.787448906 | 10.62382714 | -0.639764372 | 0.546470354 | 0.888232534 | -4.966574777 |
| **A_51_P360918** | Ehd3 | 0.098950996 | -0.281576013 | 0.479478006 | 9.974233583 | 0.639740501 | 0.546484836 | 0.888232534 | -4.966585311 |
| **A_55_P2009857** | NA | -0.143823265 | -0.697016399 | 0.409369869 | 11.5928609 | -0.63961925 | 0.546558404 | 0.888232534 | -4.966638811 |
| **A_55_P2149363** | 6430548M08Rik | -0.03998458 | -0.19378571 | 0.11381655 | 8.478063469 | -0.639590756 | 0.546575693 | 0.888232534 | -4.966651382 |
| **A_55_P2052634** | Rpl6 | -0.045956728 | -0.22273427 | 0.130820814 | 15.23376627 | -0.63957455 | 0.546585526 | 0.888232534 | -4.966658531 |
| **A_55_P2041075** | Ccnd3 | 0.092003511 | -0.261933829 | 0.445940851 | 12.43799309 | 0.639509768 | 0.546624836 | 0.888232534 | -4.96668711 |
| **A_55_P2053266** | St6galnac6 | 0.16218107 | -0.461777064 | 0.786139204 | 9.884777156 | 0.639460723 | 0.546654597 | 0.888232534 | -4.966708745 |
| **A_55_P2012331** | NA | 0.083462521 | -0.237667829 | 0.404592871 | 11.69581777 | 0.639409831 | 0.54668548 | 0.888232534 | -4.966731193 |
| **A_51_P369516** | Nudt16l1 | 0.085889058 | -0.244596087 | 0.416374203 | 11.70986133 | 0.639374127 | 0.546707148 | 0.888232534 | -4.966746941 |
| **A_55_P2122604** | Cbr2 | 0.029476701 | -0.084006382 | 0.142959785 | 7.712941251 | 0.639023627 | 0.54691988 | 0.888503476 | -4.966901497 |

| | | | | | | | | | |
|---|---|---|---|---|---|---|---|---|---|
| **A_51_P451428** | Marveld2 | 0.061281683 | -0.174685118 | 0.297248485 | 8.755897835 | 0.638923618 | 0.546980588 | 0.888527422 | -4.966945583 |
| **A_55_P2182324** | NA | -0.038095381 | -0.184811007 | 0.108620244 | 8.058275985 | -0.638800314 | 0.547055444 | 0.888551316 | -4.966999932 |
| **A_55_P2005475** | Sult1a1 | -0.053724828 | -0.260650647 | 0.15320099 | 9.444022245 | -0.638747947 | 0.547087237 | 0.888551316 | -4.967023011 |
| **A_51_P465273** | Mettl17 | -0.050711868 | -0.246166784 | 0.144743049 | 8.164316283 | -0.638310764 | 0.547352705 | 0.888883481 | -4.967215624 |
| **A_51_P176086** | Ffar2 | -0.02754511 | -0.133718642 | 0.078628422 | 7.703814762 | -0.638259684 | 0.547383727 | 0.888883481 | -4.967238122 |
| **A_55_P2113683** | Raph1 | 0.048934134 | -0.139754975 | 0.237623243 | 10.69073956 | 0.638019919 | 0.547529358 | 0.889045275 | -4.967343704 |
| **A_66_P134674** | Far1 | 0.058243238 | -0.166392815 | 0.282879291 | 8.992683213 | 0.637874459 | 0.54761772 | 0.889114062 | -4.967407742 |
| **A_55_P2109874** | Rpl23a | -0.06074931 | -0.295170561 | 0.173671941 | 15.72662663 | -0.63754895 | 0.54781549 | 0.889360458 | -4.967551004 |
| **A_55_P2022658** | Cdk11b | -0.058455339 | -0.284225254 | 0.167314575 | 13.09959033 | -0.636982174 | 0.548159953 | 0.889833378 | -4.967800306 |
| **A_55_P2067251** | Tmpo | -0.038214942 | -0.185825945 | 0.109396062 | 8.0566762 | -0.636918163 | 0.548198865 | 0.889833378 | -4.967828451 |
| **A_55_P2016069** | Adamtsl1 | 0.054870498 | -0.15715474 | 0.266895736 | 8.08303811 | 0.636678969 | 0.548344285 | 0.889908504 | -4.967933599 |
| **A_55_P2139611** | Acbd4 | 0.03589575 | -0.102819977 | 0.174611477 | 9.03993903 | 0.636629111 | 0.5483746 | 0.889908504 | -4.967955513 |
| **A_55_P2037613** | Elmsan1 | 0.067403508 | -0.193076799 | 0.327883815 | 9.825357851 | 0.636614855 | 0.548383268 | 0.889908504 | -4.967961778 |
| **A_55_P2079009** | Slco2b1 | 0.04147683 | -0.118896736 | 0.201850395 | 8.204915056 | 0.636270714 | 0.548592543 | 0.890173376 | -4.968112992 |
| **A_55_P2074438** | Gtf2a1 | 0.059116949 | -0.169569693 | 0.287803591 | 11.02444684 | 0.63597549 | 0.548772111 | 0.890331627 | -4.968242657 |
| **A_55_P1999849** | NA | -0.053260094 | -0.259302406 | 0.152782218 | 11.40413382 | -0.635937857 | 0.548795003 | 0.890331627 | -4.968259182 |
| **A_52_P259402** | Dsg3 | 0.027346803 | -0.078456292 | 0.133149898 | 7.65892738 | 0.635883197 | 0.548828255 | 0.890331627 | -4.968283183 |
| **A_55_P2137828** | Gprc5b | -0.188929559 | -0.920259294 | 0.542400175 | 8.85086998 | -0.635558956 | 0.549025527 | 0.890476286 | -4.96842552 |
| **A_55_P1953356** | Triobp | 0.082229621 | -0.236076977 | 0.400536219 | 10.66549669 | 0.635553053 | 0.549029119 | 0.890476286 | -4.96842811 |
| **A_51_P400375** | Erg28 | 0.121444934 | -0.348694344 | 0.591584213 | 14.24625086 | 0.635509476 | 0.549055636 | 0.890476286 | -4.968447235 |
| **A_55_P1954001** | Snrpc | 0.093029083 | -0.267298536 | 0.453356703 | 11.60917105 | 0.635170548 | 0.549261899 | 0.890736073 | -4.968595946 |
| **A_52_P1092823** | Irx1 | 0.071568691 | -0.205784906 | 0.348922288 | 8.129993127 | 0.634831448 | 0.549468316 | 0.890975537 | -4.968744666 |
| **A_52_P226127** | Dido1 | -0.058978988 | -0.287567654 | 0.169609679 | 8.892396726 | -0.634763263 | 0.549509827 | 0.890975537 | -4.968774562 |
| **A_55_P2111163** | S100g | 0.122747773 | -0.353090343 | 0.59858589 | 16.32382819 | 0.634634325 | 0.54958833 | 0.890975537 | -4.968831089 |
| **A_51_P317317** | Gatd1 | 0.046096245 | -0.132615531 | 0.224808021 | 8.748837414 | 0.63457293 | 0.549625713 | 0.890975537 | -4.968858001 |
| **A_51_P153013** | Map2k3os | -0.030750412 | -0.149971932 | 0.088471109 | 8.536389598 | -0.634549403 | 0.549640039 | 0.890975537 | -4.968868313 |
| **A_55_P1984337** | Emd | -0.043087011 | -0.210166932 | 0.123992911 | 11.58462189 | -0.634441032 | 0.549706029 | 0.891007786 | -4.968915811 |
| **A_55_P2050040** | Rpl36al | 0.059514202 | -0.17129659 | 0.290324994 | 14.15335844 | 0.634356895 | 0.549757266 | 0.891016116 | -4.968952682 |
| **A_52_P327402** | Cds1 | 0.052159187 | -0.150229031 | 0.254547404 | 8.997837549 | 0.634037212 | 0.549951973 | 0.891188837 | -4.969092741 |
| **A_52_P168338** | Fundc2 | 0.030434636 | -0.087676172 | 0.148545444 | 7.766296166 | 0.633939241 | 0.550011651 | 0.891188837 | -4.969135652 |
| **A_55_P2034600** | Gm5523 | -0.037961167 | -0.185289619 | 0.109367285 | 8.267122159 | -0.633901957 | 0.550034364 | 0.891188837 | -4.969151981 |
| **A_55_P1958245** | Tcof1 | -0.064782964 | -0.316216761 | 0.186650834 | 9.60664239 | -0.633878936 | 0.550048388 | 0.891188837 | -4.969162062 |
| **A_55_P2243431** | Gdap10 | -0.146015132 | -0.712856189 | 0.420825925 | 8.870550549 | -0.633732133 | 0.550137824 | 0.891188837 | -4.969226346 |
| **A_55_P1999309** | Zfp668 | 0.080098173 | -0.230882513 | 0.391078859 | 10.03287225 | 0.633663009 | 0.550179939 | 0.891188837 | -4.969256611 |
| **A_51_P412966** | Lrrn1 | 0.027831295 | -0.080229219 | 0.135891809 | 7.653136877 | 0.633629736 | 0.550200212 | 0.891188837 | -4.969271178 |
| **A_55_P2134092** | Brsk1 | 0.030416304 | -0.087712214 | 0.148544823 | 7.927697236 | 0.63346241 | 0.550302169 | 0.891188837 | -4.969344424 |
| **A_52_P64707** | Foxa3 | 0.04506989 | -0.129978754 | 0.220118535 | 7.848411879 | 0.633427497 | 0.550323444 | 0.891188837 | -4.969359705 |
| **A_51_P288828** | Slc52a2 | -0.069417953 | -0.339050953 | 0.200215047 | 10.23109396 | -0.633385215 | 0.550349211 | 0.891188837 | -4.96937821 |
| **A_55_P1987245** | Ywhaz | 0.046270493 | -0.133477837 | 0.226018824 | 9.398630388 | 0.633298457 | 0.550402083 | 0.891188837 | -4.969416178 |
| **A_55_P1978506** | H2-Q6 | -0.249955189 | -1.221018783 | 0.721108404 | 12.36318237 | -0.633262322 | 0.550424106 | 0.891188837 | -4.969431991 |
| **A_55_P2168692** | Cep131 | -0.036969778 | -0.1806183 | 0.106678745 | 9.025984011 | -0.633161997 | 0.550485251 | 0.891188837 | -4.969475888 |
| **A_52_P433828** | Tctn3 | 0.044883874 | -0.129526272 | 0.219294019 | 8.736303261 | 0.633122502 | 0.550509323 | 0.891188837 | -4.969493168 |
| **A_55_P1979457** | Prdx4 | -0.069109842 | -0.337710434 | 0.199490751 | 14.17717298 | -0.632997645 | 0.550585429 | 0.891223649 | -4.969547789 |
| **A_52_P148658** | Mettl22 | -0.054782465 | -0.267741888 | 0.158176958 | 8.557526732 | -0.63286907 | 0.550663807 | 0.891223649 | -4.969604027 |
| **A_51_P106859** | Lsm10 | 0.086967816 | -0.251179671 | 0.425115302 | 12.13663359 | 0.632734568 | 0.550745806 | 0.891223649 | -4.965054438 |
| **A_55_P2005190** | Herc4 | -0.067738849 | -0.331177894 | 0.195700196 | 11.11276661 | -0.632596566 | 0.550829947 | 0.891223649 | -4.969723188 |
| **A_51_P516870** | Itm2a | -0.138086531 | -0.675122469 | 0.398949406 | 13.57877048 | -0.632582444 | 0.550838558 | 0.891223649 | -4.969729362 |
| **A_52_P164709** | Poc1a | 0.037320102 | -0.10784927 | 0.182489475 | 8.72832561 | 0.632465716 | 0.550909734 | 0.891223649 | -4.969780391 |
| **A_55_P1979684** | Rhoc | -0.130588381 | -0.638703255 | 0.377526492 | 13.56045308 | -0.63228342 | 0.551020904 | 0.891223649 | -4.969860069 |
| **A_55_P2102469** | Gm6428 | -0.033670418 | -0.164682934 | 0.097342097 | 16.76425652 | -0.63227337 | 0.551027033 | 0.891223649 | -4.969864461 |

| | | | | | | | | | |
|---|---|---|---|---|---|---|---|---|---|
| **A_66_P126189** | Ercc5 | -0.045829579 | -0.22416967 | 0.132510512 | 8.674029045 | -0.632216821 | 0.551061522 | 0.891223649 | -4.969889173 |
| **A_55_P2173438** | Arhgap4 | -0.02819775 | -0.137932001 | 0.081536501 | 7.793180884 | -0.632180969 | 0.551083388 | 0.891223649 | -4.969904839 |
| **A_52_P325116** | Gramd1b | 0.252024922 | -0.728797846 | 1.23284769 | 11.37308724 | 0.632152868 | 0.551100527 | 0.891223649 | -4.969917118 |
| **A_51_P405912** | Lmcd1 | -0.450435273 | -2.203431034 | 1.302560487 | 10.50687944 | -0.632150797 | 0.55110179 | 0.891223649 | -4.969918023 |
| **A_55_P2435504** | Cep295 | -0.032156737 | -0.157312971 | 0.092999497 | 8.299823798 | -0.632104164 | 0.551130233 | 0.891223649 | -4.969938399 |
| **A_55_P2079516** | Mpp1 | 0.050778491 | -0.1469107 | 0.248467682 | 8.277588387 | 0.631925717 | 0.551239083 | 0.891325099 | -4.970016357 |
| **A_55_P2039454** | Gm5088 | -0.026799616 | -0.131164639 | 0.077565407 | 7.928541564 | -0.631746379 | 0.551348489 | 0.891383817 | -4.970094687 |
| **A_55_P2052281** | Rnf208 | 0.063789332 | -0.184654643 | 0.312233308 | 10.47880301 | 0.631667815 | 0.551396422 | 0.891383817 | -4.970128996 |
| **A_51_P292368** | Tmco6 | 0.028888122 | -0.083629164 | 0.141405408 | 8.259595763 | 0.63163942 | 0.551413746 | 0.891383817 | -4.970141394 |
| **A_55_P2159522** | Col14a1 | 0.032923736 | -0.095347401 | 0.161194873 | 7.805997138 | 0.63146507 | 0.551520131 | 0.891481235 | -4.970217517 |
| **A_55_P1979705** | Spop | 0.027104239 | -0.078561518 | 0.132769997 | 7.985712205 | 0.631062126 | 0.551766047 | 0.891686253 | -4.970393379 |
| **A_52_P440709** | Ryr2 | 0.02796636 | -0.0810889 | 0.137021619 | 7.673349358 | 0.63089704 | 0.551866819 | 0.891686253 | -4.970465403 |
| **A_55_P1964615** | Dnttip1 | -0.030946686 | -0.151629913 | 0.08973654 | 8.060705945 | -0.63086496 | 0.551886403 | 0.891686253 | -4.970479397 |
| **A_51_P235117** | Zc3h11a | -0.095640979 | -0.468627038 | 0.277345079 | 13.51467322 | -0.630841969 | 0.551900438 | 0.891686253 | -4.970489426 |
| **A_55_P2079471** | NA | 0.120473618 | -0.349370171 | 0.590317407 | 8.900381549 | 0.630823153 | 0.551911924 | 0.891686253 | -4.970497633 |
| **A_55_P2126063** | Mrps21 | 0.054437817 | -0.157874694 | 0.266750329 | 13.0009083 | 0.630803768 | 0.551923759 | 0.891686253 | -4.970506089 |
| **A_55_P2147487** | Cyth3 | -0.043217754 | -0.211795374 | 0.125359866 | 10.0366442 | -0.630712495 | 0.551979482 | 0.891686306 | -4.970545898 |
| **A_55_P1977353** | Ogfod2 | 0.036263866 | -0.105202463 | 0.177730196 | 11.66126202 | 0.630652592 | 0.552016055 | 0.891686306 | -4.970572022 |
| **A_55_P2024366** | Raver2 | -0.083712036 | -0.410325534 | 0.242901462 | 8.867405851 | -0.630554949 | 0.552075674 | 0.89170809 | -4.970614601 |
| **A_51_P495379** | Fhod1 | -0.086515462 | -0.424160962 | 0.251130038 | 10.61620471 | -0.630379278 | 0.552182945 | 0.89180683 | -4.970691192 |
| **A_55_P2056639** | Bcar1 | -0.043104126 | -0.211377276 | 0.125169024 | 12.84790247 | -0.63019243 | 0.552297055 | 0.8919166 | -4.970772636 |
| **A_52_P495565** | Efnb3 | -0.027384723 | -0.134336831 | 0.079567385 | 7.764774942 | -0.629924018 | 0.552461004 | 0.891937014 | -4.970889598 |
| **A_52_P491244** | Zfp287 | -0.053665754 | -0.263268547 | 0.15593704 | 8.273661564 | -0.629896693 | 0.552477695 | 0.891937014 | -4.970901502 |
| **A_55_P2077188** | Stxbp1 | 0.071372397 | -0.207395529 | 0.350140324 | 9.333469262 | 0.629878294 | 0.552488935 | 0.891937014 | -4.970909518 |
| **A_55_P1964193** | Map1b | 0.043093579 | -0.125249887 | 0.211437046 | 8.439088041 | 0.62977507 | 0.552551995 | 0.891937014 | -4.970954484 |
| **A_52_P601914** | Lemd2 | -0.102137886 | -0.501157886 | 0.296882115 | 13.11212193 | -0.629740118 | 0.552573348 | 0.891937014 | -4.970969709 |
| **A_51_P128987** | Akr1b8 | 0.052791153 | -0.153454056 | 0.259036361 | 10.34009803 | 0.629718482 | 0.552586567 | 0.891937014 | -4.966362398 |
| **A_55_P2098061** | Tchp | -0.048445928 | -0.237816726 | 0.14092487 | 8.802385136 | -0.629380718 | 0.552792948 | 0.892100801 | -4.971126217 |
| **A_55_P1963777** | Gtf3c6 | 0.048667536 | -0.141589889 | 0.238924962 | 11.15034166 | 0.629313296 | 0.55283415 | 0.892100801 | -4.971155569 |
| **A_55_P2016316** | Xrn2 | 0.050689753 | -0.147480673 | 0.248860179 | 10.12852192 | 0.629289518 | 0.552848682 | 0.892100801 | -4.97116592 |
| **A_52_P517063** | Immp2l | -0.040347071 | -0.19810721 | 0.117413068 | 8.125883046 | -0.629193014 | 0.55290766 | 0.892100801 | -4.971207927 |
| **A_55_P2165719** | Pcyt1a | -0.083272708 | -0.408894238 | 0.242348821 | 12.48555483 | -0.629156576 | 0.55292993 | 0.892100801 | -4.971223786 |
| **A_52_P410765** | Sema7a | -0.085635446 | -0.420545334 | 0.249274443 | 11.78549647 | -0.629063887 | 0.552986582 | 0.892100801 | -4.971264126 |
| **A_55_P1971124** | Mbd1 | -0.066518701 | -0.326681905 | 0.193644504 | 10.80408415 | -0.629023755 | 0.553011112 | 0.892100801 | -4.97128159 |
| **A_55_P2079912** | NA | 0.02926533 | -0.085218507 | 0.143749168 | 8.350730247 | 0.628895394 | 0.553089576 | 0.892140861 | -4.971337443 |
| **A_55_P2095301** | NA | 0.035825833 | -0.104336315 | 0.175987981 | 8.01097913 | 0.628832118 | 0.553128257 | 0.892140861 | -4.971364973 |
| **A_55_P1955483** | Grb14 | 0.034115179 | -0.099397 | 0.167627357 | 7.898931608 | 0.628631198 | 0.553251093 | 0.892264529 | -4.971452373 |
| **A_51_P236197** | Polr1a | 0.058985659 | -0.171908742 | 0.289880061 | 9.905087676 | 0.628495537 | 0.553334041 | 0.892274587 | -4.971511371 |
| **A_51_P395842** | Timm21 | 0.07067009 | -0.206027082 | 0.347367262 | 10.0400719 | 0.628347782 | 0.553424393 | 0.892274587 | -4.971575618 |
| **A_55_P2077964** | Vps8 | -0.049085523 | -0.241310024 | 0.143138977 | 8.512203132 | -0.628223019 | 0.553500692 | 0.892274587 | -4.971629857 |
| **A_55_P2373121** | Dcaf17 | 0.042045075 | -0.122654216 | 0.206744367 | 8.728228509 | 0.628047607 | 0.553607978 | 0.892274587 | -4.971706101 |
| **A_55_P2123831** | Thap6 | 0.030456313 | -0.088861434 | 0.149774061 | 9.671521592 | 0.627973693 | 0.553653189 | 0.892274587 | -4.971738223 |
| **A_51_P254045** | Traip | 0.057482891 | -0.167725344 | 0.282691126 | 8.721723031 | 0.627947734 | 0.553669068 | 0.892274587 | -4.971749504 |
| **A_55_P2358002** | 5-Mar | -0.030399139 | -0.149501397 | 0.088703119 | 8.071202077 | -0.627928874 | 0.553680605 | 0.892274587 | -4.971757699 |
| **A_51_P107782** | Kdsr | -0.087152008 | -0.428619602 | 0.254315586 | 11.67616052 | -0.627909517 | 0.553692446 | 0.892274587 | -4.97176611 |
| **A_66_P134040** | Vps25 | -0.037183688 | -0.182905465 | 0.108538088 | 9.361367024 | -0.627765103 | 0.553780789 | 0.892274587 | -4.971828855 |
| **A_55_P1980591** | Hace1 | -0.035445364 | -0.174428145 | 0.103537418 | 8.59956184 | -0.627433351 | 0.55398377 | 0.892274587 | -4.971972951 |
| **A_51_P314153** | Nr2c2ap | 0.04527434 | -0.132256346 | 0.222805025 | 8.549575987 | 0.62740483 | 0.554001223 | 0.892274587 | -4.971985336 |
| **A_52_P476029** | Nol4l | -0.066247791 | -0.32602684 | 0.193531258 | 9.722736189 | -0.627388335 | 0.554011316 | 0.892274587 | -4.971992499 |
| **A_55_P1966749** | Cyp2s1 | -0.080564327 | -0.396502262 | 0.235373607 | 13.61182444 | -0.627350393 | 0.554034535 | 0.892274587 | -4.972008974 |

| | | | | | | | | | |
|---|---|---|---|---|---|---|---|---|---|
| **A_52_P421713** | 9930012K11Rik | -0.083620972 | -0.411602031 | 0.244360087 | 9.280193815 | -0.627242693 | 0.554100444 | 0.892274587 | -4.972055735 |
| **A_51_P439996** | Gp5 | 0.028350852 | -0.082853147 | 0.139554851 | 7.683478449 | 0.627212748 | 0.55411877 | 0.892274587 | -4.972068734 |
| **A_52_P249588** | Cnot11 | 0.037875926 | -0.110699737 | 0.186451588 | 9.071417827 | 0.62716923 | 0.554145404 | 0.892274587 | -4.972087626 |
| **A_55_P2055284** | Chrm2 | -0.027558312 | -0.135667481 | 0.080550858 | 7.691316281 | -0.627132401 | 0.554167944 | 0.892274587 | -4.972103613 |
| **A_55_P2096887** | Steap3 | 0.039158783 | -0.114460438 | 0.192778004 | 8.835045128 | 0.627123092 | 0.554173642 | 0.892274587 | -4.972107654 |
| **A_55_P2145029** | Gm826 | 0.033670565 | -0.098439694 | 0.165780823 | 7.829912307 | 0.627022347 | 0.554235305 | 0.892274587 | -4.972151382 |
| **A_52_P213889** | Tmc7 | -0.142652132 | -0.702365774 | 0.417061511 | 8.99807705 | -0.627020206 | 0.554236616 | 0.892274587 | -4.972152312 |
| **A_55_P1993049** | Gm4076 | -0.058366821 | -0.287484646 | 0.170751005 | 11.13920495 | -0.626723993 | 0.554417942 | 0.892274587 | -4.972280847 |
| **A_55_P2036638** | Ints11 | -0.029123105 | -0.143459783 | 0.085213574 | 9.748465198 | -0.626644541 | 0.554466585 | 0.892274587 | -4.972315315 |
| **A_52_P234958** | Dvl1 | -0.063081718 | -0.310757734 | 0.184594298 | 10.20746898 | -0.626597593 | 0.554495329 | 0.892274587 | -4.972335681 |
| **A_55_P1986556** | Minpp1 | 0.080477974 | -0.235502351 | 0.396458299 | 10.17440286 | 0.626593892 | 0.554497595 | 0.892274587 | -4.972337286 |
| **A_55_P2112240** | Cdk5rap2 | 0.03704557 | -0.108419721 | 0.182510862 | 9.454202553 | 0.626536053 | 0.554533009 | 0.892274587 | -4.972362374 |
| **A_55_P1954724** | A130040M12Rik | -0.327732778 | -1.614963096 | 0.959497539 | 11.31816683 | -0.626372645 | 0.554633068 | 0.892274587 | -4.972433243 |
| **A_52_P335064** | Mustn1 | -0.437645044 | -2.156678459 | 1.281388372 | 9.96412169 | -0.626335256 | 0.554655963 | 0.892274587 | -4.972449456 |
| **A_55_P2138416** | NA | 0.043259308 | -0.126664806 | 0.213183422 | 12.69030835 | 0.6263163 | 0.554667572 | 0.892274587 | -4.972457675 |
| **A_55_P2171373** | Eloc | 0.044144997 | -0.129271652 | 0.217561646 | 13.04657845 | 0.626267482 | 0.554697468 | 0.892274587 | -4.972478843 |
| **A_51_P176505** | Rps19bp1 | -0.034620504 | -0.170621745 | 0.101380736 | 10.96977591 | -0.626267226 | 0.554697625 | 0.892274587 | -4.972478954 |
| **A_52_P217646** | Snapc3 | -0.063764856 | -0.314367837 | 0.186838126 | 9.439754947 | -0.625985553 | 0.554870139 | 0.892274587 | -4.97260106 |
| **A_51_P221256** | Slc34a2 | -0.030119058 | -0.148493574 | 0.088255458 | 7.787746883 | -0.625968279 | 0.55488072 | 0.892274587 | -4.972608547 |
| **A_55_P2106180** | Dnah2 | 0.101093891 | -0.296243602 | 0.498431383 | 8.092680721 | 0.625942625 | 0.554896434 | 0.892274587 | -4.972619665 |
| **A_52_P314129** | Pkia | 0.02687591 | -0.078757833 | 0.132509654 | 7.773033189 | 0.625935633 | 0.554900717 | 0.892274587 | -4.972622695 |
| **A_55_P2325698** | 2010002M09Rik | 0.074046763 | -0.216994165 | 0.365087692 | 8.653745726 | 0.625923382 | 0.554908221 | 0.892274587 | -4.972628005 |
| **A_51_P173622** | Ptpn11 | -0.071497676 | -0.352527444 | 0.209532092 | 14.91413736 | -0.625905493 | 0.554919179 | 0.892274587 | -4.972635758 |
| **A_55_P2028922** | Stoml2 | -0.069691888 | -0.343885559 | 0.204501784 | 12.15106363 | -0.62530797 | 0.555285271 | 0.892758608 | -4.972894607 |
| **A_55_P2035932** | Fgf7 | 0.031161007 | -0.09144666 | 0.153768674 | 7.789856689 | 0.625263417 | 0.555312574 | 0.892758608 | -4.9729139 |
| **A_51_P158922** | Prkaa1 | -0.050467306 | -0.249090539 | 0.148155927 | 9.930597104 | -0.625099622 | 0.555412957 | 0.89284573 | -4.972984817 |
| **A_51_P336952** | Cryl1 | 0.18717996 | -0.549596778 | 0.923956697 | 11.34101967 | 0.625018118 | 0.555462913 | 0.892851779 | -4.973020099 |
| **A_66_P126415** | NA | 0.032776372 | -0.096257759 | 0.161810503 | 7.76809106 | 0.624921472 | 0.555522151 | 0.892872748 | -4.973061931 |
| **A_51_P486068** | Zfp280b | 0.074425631 | -0.218608834 | 0.367460097 | 10.13897245 | 0.624845992 | 0.55556842 | 0.892872868 | -4.973094598 |
| **A_55_P2012641** | NA | -0.028633474 | -0.141463565 | 0.084196618 | 16.45797663 | -0.62433583 | 0.555881205 | 0.893032461 | -4.973315305 |
| **A_55_P2066463** | Enah | -0.085825319 | -0.42409499 | 0.252444351 | 10.36568451 | -0.62419679 | 0.555966471 | 0.893032461 | -4.973375431 |
| **A_55_P2026487** | NA | 0.027163504 | -0.079903711 | 0.134230719 | 7.773594892 | 0.624163611 | 0.555986819 | 0.893032461 | -4.973389777 |
| **A_52_P212418** | Trmt13 | -0.035851831 | -0.17718283 | 0.105479167 | 8.49763628 | -0.624084046 | 0.556035617 | 0.893032461 | -4.973424177 |
| **A_55_P1954820** | D1Ertd622e | -0.033474709 | -0.165447715 | 0.098498297 | 9.335690975 | -0.624023387 | 0.556072821 | 0.893032461 | -4.973450401 |
| **A_51_P479894** | Cpsf3 | -0.032083316 | -0.158571777 | 0.094405145 | 10.33915766 | -0.624018558 | 0.556075783 | 0.893032461 | -4.973452488 |
| **A_55_P2177885** | Dact3 | 0.050357227 | -0.14818485 | 0.248899304 | 8.490370958 | 0.623991121 | 0.556092612 | 0.893032461 | -4.973464349 |
| **A_55_P1993708** | Lrp12 | -0.081115974 | -0.400931993 | 0.238700045 | 10.02302763 | -0.623986705 | 0.55609532 | 0.893032461 | -4.973466258 |
| **A_55_P2113160** | Mal | 0.38958804 | -1.146454667 | 1.925630747 | 11.23901743 | 0.623981176 | 0.556098711 | 0.893032461 | -4.973468648 |
| **A_55_P2287260** | Snap23 | -0.112744531 | -0.557334077 | 0.331845014 | 12.11804261 | -0.623886233 | 0.556156948 | 0.893032461 | -4.973509687 |
| **A_55_P2001891** | Prps1 | 0.078824477 | -0.232022332 | 0.389671286 | 8.935924147 | 0.62385527 | 0.556175941 | 0.893032461 | -4.973523069 |
| **A_55_P2046838** | Amfr | 0.060225753 | -0.177375828 | 0.297827333 | 10.80079849 | 0.62359421 | 0.556336095 | 0.893152544 | -4.973635879 |
| **A_55_P2025695** | C330007P06Rik | -0.058593744 | -0.289782699 | 0.172595211 | 10.87994086 | -0.623524226 | 0.556379033 | 0.893152544 | -4.973666114 |
| **A_55_P2211164** | 5330406M23Rik | -0.136315111 | -0.674195483 | 0.401565261 | 8.541398319 | -0.623487093 | 0.556401816 | 0.893152544 | -4.973682155 |
| **A_66_P121745** | Arhgef28 | -0.029253786 | -0.144695318 | 0.086187746 | 7.98487398 | -0.623432099 | 0.55643556 | 0.893152544 | -4.973705911 |
| **A_55_P2032318** | 4930522L14Rik | 0.046772237 | -0.137835823 | 0.231380297 | 8.277465639 | 0.623313662 | 0.556508235 | 0.893195025 | -4.973757067 |
| **A_55_P2053923** | Rab17 | 0.205393101 | -0.605569862 | 1.016356063 | 8.735288892 | 0.62309461 | 0.556642665 | 0.893336606 | -4.973851659 |
| **A_55_P1984730** | Sgpl1 | -0.136291455 | -0.674584866 | 0.402001955 | 11.54278845 | -0.622900571 | 0.556761763 | 0.893344797 | -4.973935426 |
| **A_55_P1988368** | Upp1 | 0.081082515 | -0.239176438 | 0.401341469 | 10.90107058 | 0.622866675 | 0.556782569 | 0.893344797 | -4.973950057 |
| **A_55_P2001871** | Uros | -0.070599273 | -0.349454428 | 0.208255881 | 10.80031224 | -0.622860397 | 0.556786423 | 0.893344797 | -4.973952767 |
| **A_55_P2001903** | Smim4 | 0.087127558 | -0.257241741 | 0.431496857 | 10.98529571 | 0.622443992 | 0.557042064 | 0.893569726 | -4.974132449 |

| | | | | | | | | | |
|---|---|---|---|---|---|---|---|---|---|
| **A_55_P2080703** | Cstf2 | 0.032452682 | -0.095818002 | 0.160723365 | 8.126972696 | 0.62243262 | 0.557049046 | 0.893569726 | -4.974137354 |
| **A_55_P2022812** | Cyp4f17 | -0.033275166 | -0.16482518 | 0.098274848 | 8.932455802 | -0.622298129 | 0.55713163 | 0.893569726 | -4.974195366 |
| **A_55_P2074972** | Zswim6 | -0.034561651 | -0.171203363 | 0.102080062 | 7.973634972 | -0.622272119 | 0.557147602 | 0.893569726 | -4.974206584 |
| **A_55_P2024358** | Rpl34 | 0.047914065 | -0.141522312 | 0.237350441 | 15.79510123 | 0.622255577 | 0.55715776 | 0.893569726 | -4.974213718 |
| **A_55_P1983959** | Ftl2-ps | -0.093572 | -0.463581979 | 0.27643798 | 14.07378274 | -0.622159356 | 0.55721685 | 0.893590351 | -4.974255214 |
| **A_51_P296796** | Nt5dc2 | -0.028379965 | -0.140634624 | 0.083874694 | 7.876910211 | -0.621980312 | 0.557326814 | 0.893611728 | -4.974332412 |
| **A_51_P271503** | Il1r1 | -0.070543747 | -0.349579431 | 0.208491937 | 9.862275828 | -0.621967856 | 0.557334464 | 0.893611728 | -4.974337782 |
| **A_55_P1990041** | Nxt2 | 0.118085358 | -0.349043175 | 0.585213892 | 9.312635778 | 0.621911832 | 0.557368875 | 0.893611728 | -4.974361934 |
| **A_66_P121474** | Chmp6 | 0.030868864 | -0.091318009 | 0.153055738 | 11.01837573 | 0.621534535 | 0.557600654 | 0.893841411 | -4.974524538 |
| **A_55_P1974845** | Pde1a | 0.031709838 | -0.09381178 | 0.157231457 | 7.745884304 | 0.621505047 | 0.557618771 | 0.893841411 | -4.974537243 |
| **A_55_P2007026** | Gm9112 | 0.20547586 | -0.608042822 | 1.018994542 | 14.16672691 | 0.621387394 | 0.557691061 | 0.893841411 | -4.97458793 |
| **A_51_P171288** | Gli3 | -0.102199865 | -0.506835314 | 0.302435583 | 8.822372687 | -0.621377551 | 0.557697109 | 0.893841411 | -4.97459217 |
| **A_51_P463120** | Hacd4 | -0.048880815 | -0.242486837 | 0.144725206 | 8.368622596 | -0.621138919 | 0.55784375 | 0.893860554 | -4.97469495 |
| **A_55_P2221365** | NA | -0.098325386 | -0.487795375 | 0.291144604 | 8.347483286 | -0.621098989 | 0.55786829 | 0.893860554 | -4.974712144 |
| **A_55_P2151143** | Dynlt1c | 0.157743783 | -0.467115601 | 0.782603167 | 13.5052859 | 0.62106795 | 0.557887366 | 0.893860554 | -4.97472551 |
| **A_55_P2012101** | Bmp8b | 0.059448547 | -0.176044925 | 0.294942019 | 9.650275625 | 0.621057106 | 0.557894031 | 0.893860554 | -4.974730179 |
| **A_52_P456134** | Dgat1 | -0.048455043 | -0.24049088 | 0.143580794 | 9.465564343 | -0.620763056 | 0.55807477 | 0.894076024 | -4.974856769 |
| **A_51_P438967** | Gpnmb | -0.252510383 | -1.253495286 | 0.74847452 | 9.935052331 | -0.62061301 | 0.55816701 | 0.894079337 | -4.974921345 |
| **A_55_P2002572** | Ephx2 | 0.223355477 | -0.662060912 | 1.108771866 | 8.884954155 | 0.620609205 | 0.55816935 | 0.894079337 | -4.974922982 |
| **A_52_P312467** | Hoxb3 | -0.028604004 | -0.142029848 | 0.08482184 | 7.729866623 | -0.620417411 | 0.558287268 | 0.894194118 | -4.975005506 |
| **A_55_P2003369** | Fkbp8 | -0.05088002 | -0.252693528 | 0.150933488 | 11.04328354 | -0.620249189 | 0.558390708 | 0.89428569 | -4.975077871 |
| **A_51_P423127** | Gba | -0.04762273 | -0.236593424 | 0.141347964 | 10.67011907 | -0.619996153 | 0.558546321 | 0.894293688 | -4.975186689 |
| **A_55_P1976224** | Ckb | 0.092879094 | -0.27567734 | 0.461435528 | 11.46324028 | 0.619987789 | 0.558551465 | 0.894293688 | -4.975190285 |
| **A_55_P1968240** | Cbr4 | 0.077201445 | -0.229176639 | 0.38357953 | 10.62194946 | 0.619921784 | 0.558592062 | 0.894293688 | -4.975218664 |
| **A_55_P2010134** | NA | -0.037240838 | -0.185044816 | 0.110563139 | 9.545303633 | -0.619872672 | 0.558622269 | 0.894293688 | -4.975239778 |
| **A_55_P2043277** | Siglecf | 0.027810057 | -0.082565773 | 0.138185887 | 7.684784234 | 0.619864922 | 0.558627036 | 0.894293688 | -4.97524311 |
| **A_55_P1969615** | Pnck | 0.064988746 | -0.192986535 | 0.322964026 | 8.626257603 | 0.619768108 | 0.558686588 | 0.894314953 | -4.975284728 |
| **A_51_P346938** | Lrg1 | -0.08015164 | -0.398453432 | 0.238150152 | 8.380488287 | -0.619501686 | 0.558850487 | 0.894447546 | -4.975399227 |
| **A_55_P2091472** | Bnip3 | 0.103836243 | -0.308535332 | 0.516207819 | 13.97198033 | 0.619483023 | 0.55886197 | 0.894447546 | -4.975407246 |
| **A_55_P2042879** | Mbtd1 | -0.026948081 | -0.133986379 | 0.080090218 | 7.817121089 | -0.619380896 | 0.558924806 | 0.89447405 | -4.975451126 |
| **A_55_P2046509** | Amot | -0.100257209 | -0.498789272 | 0.298274853 | 15.30977854 | -0.61890146 | 0.559219849 | 0.894526915 | -4.975657035 |
| **A_55_P2049479** | NA | -0.072492092 | -0.360684647 | 0.215700463 | 9.362186312 | -0.618838093 | 0.559258851 | 0.894526915 | -4.97568424 |
| **A_51_P437717** | Glmp | -0.063174766 | -0.314332546 | 0.187983013 | 12.22247901 | -0.618822611 | 0.559268382 | 0.894526915 | -4.975690887 |
| **A_55_P1956918** | Adamts5 | -0.192517312 | -0.958048207 | 0.573013583 | 8.507255627 | -0.618694464 | 0.559347263 | 0.894526915 | -4.975745895 |
| **A_55_P1980746** | Dram2 | -0.059668835 | -0.296949575 | 0.177611905 | 10.23766317 | -0.618663128 | 0.559366553 | 0.894526915 | -4.975759345 |
| **A_55_P2060278** | Fam45a | 0.073619978 | -0.219152371 | 0.366392328 | 11.06257569 | 0.618635433 | 0.559383602 | 0.894526915 | -4.975771232 |
| **A_51_P246773** | Sesn3 | 0.083464762 | -0.248469147 | 0.41539867 | 8.914268751 | 0.618615362 | 0.559395958 | 0.894526915 | -4.975779846 |
| **A_55_P2123362** | NA | -0.089540521 | -0.445645041 | 0.266564 | 8.398490563 | -0.61860196 | 0.559404209 | 0.894526915 | -4.975785598 |
| **A_55_P2104978** | Edc4 | 0.037601089 | -0.111959765 | 0.187161943 | 11.89013872 | 0.618517012 | 0.559456506 | 0.894526915 | -4.975822053 |
| **A_52_P430274** | Dnaaf5 | -0.058477413 | -0.291150648 | 0.174195823 | 10.17294458 | -0.618316567 | 0.559579919 | 0.894526915 | -4.975908057 |
| **A_55_P2133901** | NA | -0.057688728 | -0.287240812 | 0.171863355 | 14.00046753 | -0.618271016 | 0.559607967 | 0.894526915 | -4.975927598 |
| **A_55_P2121466** | Ncor1 | -0.03386926 | -0.16866279 | 0.100924269 | 8.177287322 | -0.618167017 | 0.559672008 | 0.894526915 | -4.975972208 |
| **A_55_P1964218** | Tprkb | -0.089616983 | -0.446308924 | 0.267074957 | 14.04683843 | -0.618110592 | 0.559706755 | 0.894526915 | -4.975996409 |
| **A_51_P246816** | Napa | 0.073231274 | -0.218328885 | 0.364791433 | 10.75141565 | 0.617927573 | 0.55981947 | 0.894526915 | -4.976074894 |
| **A_55_P1990573** | Snrnp70 | -0.116916775 | -0.582459769 | 0.348626218 | 11.54448744 | -0.617854475 | 0.559864493 | 0.894526915 | -4.976106236 |
| **A_51_P212560** | Cog7 | -0.113561935 | -0.565907558 | 0.338783688 | 9.496660295 | -0.617634511 | 0.559999986 | 0.894526915 | -4.976200529 |
| **A_52_P359502** | Pdxk | 0.060303905 | -0.179940659 | 0.30054847 | 11.75724601 | 0.617534224 | 0.560061767 | 0.894526915 | -4.97624351 |
| **A_51_P174681** | Bzw2 | 0.054144492 | -0.161580324 | 0.269869308 | 9.901269357 | 0.617480635 | 0.560094783 | 0.894526915 | -4.976266476 |
| **A_51_P227222** | Adamts2 | -0.540403202 | -2.693607353 | 1.612800949 | 12.58793757 | -0.617450104 | 0.560113593 | 0.894526915 | -4.976279559 |
| **A_55_P1963610** | NA | -0.07594296 | -0.378584004 | 0.226698084 | 14.95142129 | -0.61734631 | 0.560177543 | 0.894526915 | -4.976324032 |

| | | | | | | | | | |
|---|---|---|---|---|---|---|---|---|---|
| **A_55_P2037883** | Ino80 | 0.035625091 | -0.106352691 | 0.177602872 | 9.316370995 | 0.617312052 | 0.560198651 | 0.894526915 | -4.976338709 |
| **A_52_P200925** | NA | -0.039909085 | -0.198964705 | 0.119146534 | 13.51619622 | -0.617293809 | 0.560209892 | 0.894526915 | -4.976346525 |
| **A_55_P2032363** | Tomm40l | 0.041508812 | -0.123931419 | 0.206949042 | 8.286064144 | 0.617260287 | 0.560230547 | 0.894526915 | -4.976360886 |
| **A_51_P413740** | Ftcd | 0.05611819 | -0.167573708 | 0.279810088 | 7.760910861 | 0.617195235 | 0.560270632 | 0.894526915 | -4.976388753 |
| **A_55_P2179027** | Gem | -0.074503112 | -0.371499351 | 0.222493127 | 8.140390623 | -0.61715269 | 0.560296849 | 0.894526915 | -4.976406977 |
| **A_55_P1998631** | Capg | 0.103833362 | -0.310096998 | 0.517763721 | 13.55736977 | 0.61713304 | 0.560308958 | 0.894526915 | -4.976415394 |
| **A_55_P1992455** | Vdac3 | -0.058425812 | -0.291387329 | 0.174535705 | 13.62675894 | -0.617006492 | 0.560386945 | 0.894526915 | -4.976469592 |
| **A_55_P2031217** | Vps52 | -0.084065996 | -0.419262795 | 0.251130802 | 11.19756699 | -0.617006396 | 0.560387004 | 0.894526915 | -4.976469633 |
| **A_55_P2003199** | Setd1b | -0.071239714 | -0.355312608 | 0.21283318 | 8.265241818 | -0.616966431 | 0.560411634 | 0.894526915 | -4.976486748 |
| **A_55_P1985025** | Pard3 | 0.045646791 | -0.136373981 | 0.227667563 | 11.92365922 | 0.616962069 | 0.560414323 | 0.894526915 | -4.976488616 |
| **A_51_P178735** | Hibch | -0.084609386 | -0.422018574 | 0.252799802 | 10.52694972 | -0.616922774 | 0.56043854 | 0.894526915 | -4.976505442 |
| **A_55_P2024625** | Ccl21a | 0.040913309 | -0.122242758 | 0.204069377 | 7.8308105 | 0.616922405 | 0.560438768 | 0.894526915 | -4.9765056 |
| **A_55_P1964174** | Nme1 | 0.074337958 | -0.222166812 | 0.370842728 | 10.56657257 | 0.616805316 | 0.560510936 | 0.894531374 | -4.976555734 |
| **A_55_P2114994** | NA | -0.03255435 | -0.162408719 | 0.097300019 | 16.85000309 | -0.616767703 | 0.56053412 | 0.894531374 | -4.976571837 |
| **A_55_P1967414** | Asph | -0.026269984 | -0.131103766 | 0.078563798 | 7.678866529 | -0.616492391 | 0.560703834 | 0.894601482 | -4.976689679 |
| **A_51_P330213** | Asf1b | 0.055509635 | -0.166008851 | 0.277028121 | 8.397613576 | 0.616492165 | 0.560703974 | 0.894601482 | -4.976689776 |
| **A_55_P2007871** | Ptpdc1 | -0.065773082 | -0.328258129 | 0.196711964 | 9.476914837 | -0.616471199 | 0.560716899 | 0.894601482 | -4.976698749 |
| **A_51_P445473** | Fut7 | -0.028430988 | -0.141909052 | 0.085047076 | 7.732169685 | -0.616380939 | 0.560772547 | 0.894616422 | -4.976737372 |
| **A_51_P482029** | Mcoln1 | -0.068895456 | -0.343988747 | 0.206197834 | 13.35569912 | -0.616140486 | 0.560920811 | 0.894761599 | -4.976840242 |
| **A_55_P2136962** | NA | -0.029872252 | -0.149160524 | 0.089416021 | 9.230933074 | -0.616083209 | 0.560956131 | 0.894761599 | -4.976864741 |
| **A_55_P2159070** | 2810025M15Rik | -0.040598423 | -0.202803008 | 0.121606162 | 9.25361685 | -0.615765283 | 0.561152209 | 0.89493593 | -4.977000694 |
| **A_51_P405089** | NA | -0.060142739 | -0.300437229 | 0.180151752 | 8.753095675 | -0.615755854 | 0.561158025 | 0.89493593 | -4.977004725 |
| **A_55_P2038318** | NA | -0.058702522 | -0.293288594 | 0.17588355 | 9.338400402 | -0.61563556 | 0.561232227 | 0.894963586 | -4.97705615 |
| **A_51_P314323** | Naa38 | 0.068189422 | -0.204335424 | 0.340714269 | 13.23351874 | 0.61557372 | 0.561270374 | 0.894963586 | -4.977082582 |
| **A_51_P390078** | NA | -0.038755977 | -0.193665452 | 0.116153499 | 9.309110484 | -0.615502562 | 0.561314271 | 0.894963586 | -4.977112995 |
| **A_66_P137048** | Men1 | 0.075524952 | -0.226447846 | 0.377497751 | 11.85070947 | 0.615306919 | 0.561434975 | 0.895082203 | -4.977196598 |
| **A_55_P2064984** | Cenpi | 0.052988964 | -0.158918921 | 0.264896849 | 8.023840363 | 0.61518746 | 0.561508684 | 0.895112812 | -4.977247635 |
| **A_55_P1990261** | Chchd6 | 0.069449745 | -0.208314247 | 0.347213737 | 10.86511288 | 0.615125701 | 0.561546793 | 0.895112812 | -4.977274017 |
| **A_51_P163953** | Nsg2 | 0.026052273 | -0.078156872 | 0.130261419 | 7.688575968 | 0.61504792 | 0.561594791 | 0.895115504 | -4.97730724 |
| **A_55_P2085596** | Nudt1 | -0.036648604 | -0.183337102 | 0.110039895 | 9.615975185 | -0.614653743 | 0.561838073 | 0.895369867 | -4.977475554 |
| **A_55_P1955921** | NA | -0.03736469 | -0.186924816 | 0.112195435 | 13.86498613 | -0.614631382 | 0.561851875 | 0.895369867 | -4.9774851 |
| **A_55_P2249849** | Sema6a | 0.051616415 | -0.155018409 | 0.258251239 | 9.200220403 | 0.61454471 | 0.561905378 | 0.895369867 | -4.977522096 |
| **A_66_P109049** | Brd9 | 0.050356514 | -0.151253736 | 0.251966765 | 9.395837429 | 0.614486316 | 0.561941426 | 0.895369867 | -4.977547019 |
| **A_55_P2053206** | Kdm4c | 0.04360465 | -0.130993958 | 0.218203258 | 9.62198977 | 0.614414128 | 0.561985991 | 0.895369867 | -4.977577826 |
| **A_55_P2096340** | Rnf170 | 0.055204555 | -0.165877431 | 0.27628654 | 9.094642259 | 0.614314431 | 0.562047544 | 0.89539413 | -4.977620369 |
| **A_55_P1955747** | Gm12070 | -0.049614375 | -0.248357563 | 0.149128813 | 14.81152542 | -0.614164115 | 0.562140356 | 0.895407113 | -4.977684501 |
| **A_55_P1989728** | C2cd2l | -0.119602757 | -0.618239843 | 0.379034329 | 12.44701489 | -0.621593832 | 0.562148343 | 0.895407113 | -4.914261322 |
| **A_55_P2148744** | NA | -0.059074321 | -0.295768035 | 0.177619393 | 11.13478114 | -0.614018108 | 0.562230516 | 0.895464209 | -4.977746782 |
| **A_55_P2086423** | Recql4 | 0.030266549 | -0.09103557 | 0.151568667 | 7.812910263 | 0.613852006 | 0.562333095 | 0.895553794 | -4.977817619 |
| **A_51_P513244** | Zfp949 | -0.03128476 | -0.156685847 | 0.094116327 | 8.402803734 | -0.613762981 | 0.562388079 | 0.895567571 | -4.977855579 |
| **A_55_P2000107** | Atox1 | -0.049574803 | -0.248313762 | 0.149164155 | 15.86636193 | -0.613687327 | 0.562434807 | 0.895568201 | -4.977887834 |
| **A_52_P422755** | Lsm6 | -0.026234395 | -0.131439368 | 0.078970579 | 8.171377201 | -0.613484984 | 0.562559798 | 0.895658944 | -4.977974087 |
| **A_55_P2020203** | Sfrp5 | -0.833753493 | -4.177936657 | 2.510429671 | 13.16474838 | -0.613361869 | 0.562635857 | 0.895658944 | -4.978026555 |
| **A_51_P284823** | Polr1b | -0.043732544 | -0.21914795 | 0.131682861 | 8.50505276 | -0.613346904 | 0.562645102 | 0.895658944 | -4.978032932 |
| **A_55_P2033105** | Cdcp1 | -0.068491941 | -0.343264936 | 0.206281054 | 9.12746975 | -0.61324581 | 0.562707561 | 0.895658944 | -4.978076008 |
| **A_55_P2155727** | Mafa | -0.075957328 | -0.380727704 | 0.228813048 | 10.09310664 | -0.613149096 | 0.562767319 | 0.895658944 | -4.978117213 |
| **A_51_P139030** | Slc38a3 | 0.042962997 | -0.129425169 | 0.215351163 | 7.978135235 | 0.613135243 | 0.562775879 | 0.895658944 | -4.978123115 |
| **A_51_P294807** | Stom | -0.075317478 | -0.377561718 | 0.226926762 | 9.357403895 | -0.613065534 | 0.562818953 | 0.895658944 | -4.978152809 |
| **A_55_P2024853** | Cep57l1 | -0.056274504 | -0.28212646 | 0.169577452 | 8.743883854 | -0.61299507 | 0.562862496 | 0.895658944 | -4.978182823 |
| **A_51_P125135** | Cdca5 | 0.067231497 | -0.202766421 | 0.337229416 | 8.144384014 | 0.612606401 | 0.563102711 | 0.895865824 | -4.973685041 |

| | | | | | | | | | |
|---|---|---|---|---|---|---|---|---|---|
| **A_55_P1966322** | Eif3h | 0.049012686 | -0.147838056 | 0.245863428 | 15.50091469 | 0.612548683 | 0.563138388 | 0.895865824 | -4.978372893 |
| **A_55_P2168058** | Glod4 | -0.033552887 | -0.168359299 | 0.101253524 | 10.29426226 | -0.612334194 | 0.563270984 | 0.895865824 | -4.978464181 |
| **A_55_P2186110** | R3hdm2 | -0.077833649 | -0.403333478 | 0.247666179 | 11.91395278 | -0.619678904 | 0.563309557 | 0.895865824 | -4.915024638 |
| **A_55_P2051962** | Sec61b | 0.048660662 | -0.146870077 | 0.2441914 | 15.0439337 | 0.612254708 | 0.563320126 | 0.895865824 | -4.978498003 |
| **A_55_P1984830** | Fgl1 | 0.056579115 | -0.170819544 | 0.283977775 | 8.546310723 | 0.612121193 | 0.563402678 | 0.895865824 | -4.978554808 |
| **A_55_P1968133** | Ttc4 | 0.087344186 | -0.263703286 | 0.438391659 | 11.27986287 | 0.612121052 | 0.563402765 | 0.895865824 | -4.978554868 |
| **A_55_P1961736** | Rcan2 | -0.03412386 | -0.171284504 | 0.103036784 | 8.025357356 | -0.612065359 | 0.563437203 | 0.895865824 | -4.97857856 |
| **A_51_P349341** | Npc1 | 0.036055822 | -0.10888002 | 0.180991665 | 9.764187607 | 0.612024446 | 0.563462501 | 0.895865824 | -4.978595963 |
| **A_55_P2087225** | Hnrnpa3 | -0.109594289 | -0.550153023 | 0.330964444 | 10.53394844 | -0.612002607 | 0.563476006 | 0.895865824 | -4.978605253 |
| **A_55_P2280868** | 0610005C13Rik | -0.031065996 | -0.155964351 | 0.09383236 | 7.734613468 | -0.611924329 | 0.563524413 | 0.895865824 | -4.978638547 |
| **A_55_P2161400** | Sctr | -0.101231157 | -0.50824905 | 0.305786736 | 10.31788185 | -0.611885078 | 0.563548686 | 0.895865824 | -4.97865524 |
| **A_51_P450518** | Hpf1 | -0.096831998 | -0.486261826 | 0.292597831 | 12.36626314 | -0.611728679 | 0.563645412 | 0.895945902 | -4.978721746 |
| **A_51_P229105** | Cetn3 | 0.063708363 | -0.192583515 | 0.320000242 | 10.85599783 | 0.611548303 | 0.56375698 | 0.895991238 | -4.978798431 |
| **A_51_P121302** | Them4 | 0.03096878 | -0.09365499 | 0.15559255 | 8.894436522 | 0.611353463 | 0.563877509 | 0.895991238 | -4.969342062 |
| **A_55_P2315921** | 5330431K02Rik | -0.135060826 | -0.678589702 | 0.408468051 | 8.651666086 | -0.611330328 | 0.563891821 | 0.895991238 | -4.978891075 |
| **A_51_P234044** | 1190005I06Rik | 0.053738858 | -0.162545572 | 0.270023289 | 9.263925087 | 0.611268957 | 0.563929789 | 0.895991238 | -4.978917154 |
| **A_51_P246387** | Nktr | -0.080397629 | -0.4039843 | 0.243189041 | 9.801484405 | -0.611254094 | 0.563938985 | 0.895991238 | -4.978923469 |
| **A_66_P114782** | Nr2f1 | -0.033602819 | -0.16885308 | 0.101647442 | 7.954314207 | -0.611232958 | 0.563952062 | 0.895991238 | -4.97893245 |
| **A_55_P2122666** | Pnma3 | -0.030203931 | -0.151805287 | 0.091397426 | 8.840238241 | -0.611074564 | 0.564050064 | 0.896047623 | -4.978999745 |
| **A_55_P1953503** | Lrrc4b | -0.02734029 | -0.137445857 | 0.082765276 | 7.671014957 | -0.610889994 | 0.564164276 | 0.896047623 | -4.979078142 |
| **A_55_P2145376** | Fastk | -0.058124319 | -0.29220549 | 0.175956852 | 8.424098038 | -0.610886543 | 0.564166412 | 0.896047623 | -4.979079607 |
| **A_51_P102257** | Tns1 | -0.117627143 | -0.591349303 | 0.356095017 | 8.375237903 | -0.610875927 | 0.564172982 | 0.896047623 | -4.979084116 |
| **A_52_P552547** | Lppos | 0.028012735 | -0.084856794 | 0.140882263 | 8.793120863 | 0.610587576 | 0.564351443 | 0.896194385 | -4.979206553 |
| **A_51_P157462** | Rgn | -0.048515736 | -0.24401671 | 0.146985239 | 7.99347901 | -0.610524165 | 0.564390694 | 0.896194385 | -4.979233472 |
| **A_52_P157880** | Gm1947 | -0.034000037 | -0.171012963 | 0.103012889 | 8.910729738 | -0.610501888 | 0.564404483 | 0.896194385 | -4.979242928 |
| **A_55_P2146249** | NA | -0.083161051 | -0.418422249 | 0.252100146 | 12.31395445 | -0.610247264 | 0.564562108 | 0.896325393 | -4.979350991 |
| **A_51_P479321** | Acss1 | 0.142806601 | -0.432960581 | 0.718573783 | 10.95379898 | 0.610197661 | 0.564592819 | 0.896325393 | -4.979372038 |
| **A_52_P309737** | Cdk12 | -0.027934686 | -0.140578176 | 0.084708805 | 8.121622751 | -0.610108187 | 0.564648216 | 0.896325393 | -4.97941 |
| **A_52_P472324** | Slpi | -0.729719646 | -3.672421358 | 2.212982066 | 12.46826167 | -0.610069004 | 0.564672477 | 0.896325393 | -4.979426623 |
| **A_51_P438200** | Sil1 | -0.115400289 | -0.580853774 | 0.350053195 | 13.91107911 | -0.60995778 | 0.564741348 | 0.896361103 | -4.979473803 |
| **A_51_P153042** | Psg16 | -0.093375864 | -0.470222928 | 0.2834712 | 17.56256888 | -0.609591159 | 0.564968397 | 0.896647848 | -4.97962927 |
| **A_55_P2295671** | NA | -0.033483024 | -0.168643677 | 0.101677629 | 8.738129372 | -0.609457677 | 0.565051076 | 0.896705439 | -4.979685854 |
| **A_51_P427530** | Pgm1 | -0.054130206 | -0.272705848 | 0.164445437 | 9.069568848 | -0.60926619 | 0.565169698 | 0.896753012 | -4.979767009 |
| **A_55_P2454004** | Smim8 | 0.04598766 | -0.139710741 | 0.231686061 | 10.72554725 | 0.6092595 | 0.565173842 | 0.896753012 | -4.979769844 |
| **A_51_P189985** | Ppan | -0.076374665 | -0.384960687 | 0.232211357 | 10.91735799 | -0.608894751 | 0.565399838 | 0.89691709 | -4.979924369 |
| **A_55_P2147816** | NA | -0.029230352 | -0.14733387 | 0.088873166 | 16.74644484 | -0.608892154 | 0.565401447 | 0.89691709 | -4.979925469 |
| **A_51_P323299** | Trim28 | -0.051637936 | -0.260289649 | 0.157013777 | 10.39875671 | -0.608858099 | 0.56542255 | 0.89691709 | -4.979939892 |
| **A_52_P1026777** | Ecscr | -0.036553829 | -0.184274576 | 0.111166919 | 10.24009552 | -0.608780225 | 0.565470809 | 0.89691709 | -4.979972871 |
| **A_66_P114974** | NA | 0.075383004 | -0.229299884 | 0.380065892 | 13.10076157 | 0.608687718 | 0.565528138 | 0.89691709 | -4.980012043 |
| **A_55_P2099448** | Slc12a2 | -0.045702868 | -0.230445169 | 0.139039433 | 10.90269446 | -0.608620067 | 0.565570067 | 0.89691709 | -4.980040687 |
| **A_51_P479434** | Rfc3 | -0.02845874 | -0.143505768 | 0.086588289 | 8.058494534 | -0.608568431 | 0.56560207 | 0.89691709 | -4.980062547 |
| **A_55_P2420724** | Srek1 | -0.032577211 | -0.164425893 | 0.09927147 | 7.888322595 | -0.607865198 | 0.566038039 | 0.897534804 | -4.980360118 |
| **A_55_P2030662** | Eda | -0.070331826 | -0.35508928 | 0.214425629 | 8.382362417 | -0.607639435 | 0.566178044 | 0.897683161 | -4.980455588 |
| **A_52_P28057** | Gm14124 | -0.055597121 | -0.280746546 | 0.169552304 | 8.98017959 | -0.607506078 | 0.566260755 | 0.89774066 | -4.980511967 |
| **A_51_P202014** | Taf12 | -0.0440556 | -0.222557398 | 0.134446199 | 10.32146422 | -0.607194298 | 0.566454154 | 0.897819996 | -4.980643738 |
| **A_51_P262533** | Cxxc1 | -0.030594383 | -0.154589394 | 0.093400628 | 8.836885169 | -0.607025101 | 0.566559126 | 0.897819996 | -4.980715224 |
| **A_52_P478629** | Arcn1 | -0.036336052 | -0.183604614 | 0.11093251 | 8.806506158 | -0.607011411 | 0.56656762 | 0.897819996 | -4.980721007 |
| **A_52_P377576** | NA | 0.054106356 | -0.165230306 | 0.273443019 | 10.39228975 | 0.606884741 | 0.566646215 | 0.897819996 | -4.980774513 |
| **A_51_P365468** | Clpb | 0.074316881 | -0.226966 | 0.375599763 | 10.24432546 | 0.606851155 | 0.566667056 | 0.897819996 | -4.980788699 |
| **A_51_P184969** | Rilpl1 | -0.075957197 | -0.383907147 | 0.231992754 | 10.00537976 | -0.606817298 | 0.566688065 | 0.897819996 | -4.980802998 |

| | | | | | | | | | |
|---|---|---|---|---|---|---|---|---|---|
| **A_55_P1956464** | Stam | -0.054481805 | -0.27539991 | 0.166436301 | 8.836359023 | -0.606721433 | 0.566747554 | 0.897819996 | -4.980843482 |
| **A_55_P2017789** | Il4ra | -0.05348048 | -0.270342159 | 0.1633812 | 8.390862808 | -0.606710679 | 0.566754228 | 0.897819996 | -4.980848023 |
| **A_51_P317191** | Eepd1 | 0.027822625 | -0.085021177 | 0.140666427 | 7.855785701 | 0.606582046 | 0.566834058 | 0.897819996 | -4.980902336 |
| **A_55_P1993348** | NA | -0.033864111 | -0.171237351 | 0.103509129 | 15.89191907 | -0.606466336 | 0.566905873 | 0.897819996 | -4.980951184 |
| **A_52_P642626** | Rbm20 | -0.02674284 | -0.135237325 | 0.081751646 | 7.728421673 | -0.606413724 | 0.566938529 | 0.897819996 | -4.980973392 |
| **A_55_P2075996** | Cpne1 | -0.063745549 | -0.322404977 | 0.19491388 | 12.02814147 | -0.60630439 | 0.567006394 | 0.897819996 | -4.981019538 |
| **A_51_P392005** | Car8 | -0.038390911 | -0.194199963 | 0.117418142 | 7.745934008 | -0.606184585 | 0.567080766 | 0.897819996 | -4.981070095 |
| **A_55_P1979814** | Abcc8 | 0.158662747 | -0.485357843 | 0.802683336 | 9.055776268 | 0.606100133 | 0.567133194 | 0.897819996 | -4.981105728 |
| **A_66_P125161** | 1700057H15Rik | 0.037213133 | -0.113852365 | 0.18827863 | 8.651164542 | 0.606038351 | 0.567171551 | 0.897819996 | -4.981131794 |
| **A_66_P131604** | Otud4 | -0.07953619 | -0.402451101 | 0.243378722 | 13.10988701 | -0.605962634 | 0.567218561 | 0.897819996 | -4.981163735 |
| **A_55_P2157770** | Neu2 | 0.066085513 | -0.202280915 | 0.33445194 | 8.26807084 | 0.605825069 | 0.567303977 | 0.897819996 | -4.981221759 |
| **A_66_P125828** | NA | 0.037766322 | -0.115632383 | 0.191165027 | 9.214968282 | 0.605692446 | 0.567386332 | 0.897819996 | -4.981277688 |
| **A_55_P2002553** | Stx16 | 0.057264268 | -0.175388294 | 0.289916831 | 9.851842369 | 0.605543072 | 0.567479097 | 0.897819996 | -4.981340668 |
| **A_55_P2081055** | 1700056E22Rik | -0.031735557 | -0.160683245 | 0.09721213 | 8.515437834 | -0.605482698 | 0.567516594 | 0.897819996 | -4.98136612 |
| **A_55_P2013011** | NA | -0.10317997 | -0.522436106 | 0.316076165 | 9.756104539 | -0.605459552 | 0.56753097 | 0.897819996 | -4.981375877 |
| **A_52_P40293** | Hgd | 0.054517471 | -0.167027181 | 0.276062123 | 7.831917894 | 0.605401642 | 0.567566938 | 0.897819996 | -4.981400287 |
| **A_55_P2046198** | Slc25a25 | -0.044463724 | -0.225177143 | 0.136249694 | 9.282662085 | -0.605319409 | 0.567618016 | 0.897819996 | -4.981434947 |
| **A_51_P277295** | Apmap | -0.207148071 | -1.049138989 | 0.634842848 | 10.16511907 | -0.605260941 | 0.567654334 | 0.897819996 | -4.981459587 |
| **A_55_P2063376** | Add2 | -0.086372586 | -0.437452468 | 0.264707297 | 8.109796981 | -0.60525605 | 0.567657372 | 0.897819996 | -4.981461648 |
| **A_52_P23379** | Depdc7 | 0.090647261 | -0.277808472 | 0.459102993 | 8.319552717 | 0.605255202 | 0.567657898 | 0.897819996 | -4.981462006 |
| **A_52_P238846** | Bpnt1 | 0.05919351 | -0.181468266 | 0.299855287 | 8.879074923 | 0.605112522 | 0.567746533 | 0.897819996 | -4.981522128 |
| **A_55_P1993807** | Nudt10 | 0.035587478 | -0.109119037 | 0.180293993 | 7.941341277 | 0.605031917 | 0.567796609 | 0.897819996 | -4.981556087 |
| **A_55_P1958857** | Nek6 | -0.067506237 | -0.342005141 | 0.206992668 | 10.23254176 | -0.605023781 | 0.567801663 | 0.897819996 | -4.981559515 |
| **A_52_P88695** | Dgka | -0.044391651 | -0.224942663 | 0.136159361 | 10.53516782 | -0.604881822 | 0.567889863 | 0.897819996 | -4.981619314 |
| **A_55_P2040121** | Cdv3 | 0.053604318 | -0.16442038 | 0.271629016 | 10.50316297 | 0.60487167 | 0.567896171 | 0.897819996 | -4.98162359 |
| **A_55_P2045916** | NA | -0.169142063 | -0.857156661 | 0.518872534 | 8.956727153 | -0.60481581 | 0.567930879 | 0.897819996 | -4.981647117 |
| **A_55_P1976490** | Lman2l | -0.051117461 | -0.259047062 | 0.15681214 | 10.77055633 | -0.604814393 | 0.56793176 | 0.897819996 | -4.981647714 |
| **A_66_P123938** | Smim13 | -0.057037662 | -0.289050487 | 0.174975164 | 11.38897223 | -0.604809886 | 0.56793456 | 0.897819996 | -4.981649612 |
| **A_55_P2082989** | Rab7b | 0.17703439 | -0.543248687 | 0.897317468 | 9.03477889 | 0.604677197 | 0.568017012 | 0.897819996 | -4.98170549 |
| **A_52_P112178** | Srgap2 | -0.043236873 | -0.219227625 | 0.132753878 | 8.321100349 | -0.604412722 | 0.568181375 | 0.897819996 | -4.981816835 |
| **A_51_P512992** | Entpd3 | 0.062744297 | -0.192689249 | 0.318177843 | 8.169841297 | 0.604317943 | 0.568240285 | 0.897819996 | -4.981856727 |
| **A_51_P495780** | Plin4 | 0.061680453 | -0.189433735 | 0.312794642 | 9.586756777 | 0.604290083 | 0.568257602 | 0.897819996 | -4.981868452 |
| **A_55_P2144850** | Psmb5 | -0.0631995 | -0.320502988 | 0.194103988 | 15.99994189 | -0.604278486 | 0.56826481 | 0.897819996 | -4.981873333 |
| **A_51_P115159** | Fam162a | -0.106368016 | -0.539428564 | 0.326692533 | 13.31878222 | -0.604270787 | 0.568269596 | 0.897819996 | -4.981876572 |
| **A_55_P2161585** | Coq10a | -0.034030283 | -0.172592678 | 0.104532112 | 8.673041881 | -0.604211992 | 0.568306142 | 0.897819996 | -4.981901315 |
| **A_51_P429212** | Npl | 0.197719819 | -0.607345015 | 1.002784652 | 9.460470991 | 0.604210837 | 0.568306861 | 0.897819996 | -4.981901801 |
| **A_55_P1957909** | Mrpl49 | 0.072404155 | -0.222478948 | 0.367287257 | 10.00356208 | 0.604063735 | 0.568398305 | 0.897819996 | -4.981963696 |
| **A_52_P391718** | Rpl22 | 0.040898225 | -0.125719475 | 0.207515926 | 10.16454273 | 0.603882563 | 0.568510941 | 0.897819996 | -4.98203991 |
| **A_55_P2063937** | D630033A02Rik | 0.042244209 | -0.129868548 | 0.214356967 | 8.775056599 | 0.603841961 | 0.568536186 | 0.897819996 | -4.982056987 |
| **A_51_P331021** | Ttc32 | -0.05544192 | -0.281341792 | 0.170457952 | 8.68590628 | -0.603797684 | 0.568563716 | 0.897819996 | -4.982075609 |
| **A_55_P2007646** | Cryaa | -0.670090011 | -3.400398459 | 2.060218438 | 11.86789917 | -0.603796504 | 0.56856445 | 0.897819996 | -4.982076105 |
| **A_55_P2108151** | NA | 0.094158377 | -0.289546901 | 0.477863654 | 15.59628556 | 0.603712756 | 0.568616524 | 0.897819996 | -4.982111325 |
| **A_55_P2117375** | NA | -0.036563533 | -0.185568676 | 0.112441609 | 8.550435682 | -0.60369289 | 0.568628877 | 0.897819996 | -4.982119678 |
| **A_55_P2010301** | Serpina1c | 0.422777698 | -1.300243584 | 2.145798981 | 13.33968968 | 0.603657487 | 0.568650892 | 0.897819996 | -4.982134565 |
| **A_51_P136845** | Usp19 | 0.042301188 | -0.13013244 | 0.214734815 | 9.809074574 | 0.603531246 | 0.568729397 | 0.897819996 | -4.982187643 |
| **A_55_P2014319** | Armcx6 | 0.04000567 | -0.123100293 | 0.203111633 | 8.0630347 | 0.60342163 | 0.568797569 | 0.897819996 | -4.982233723 |
| **A_66_P134534** | Atl2 | -0.030989273 | -0.157335301 | 0.095356755 | 8.745906188 | -0.60341897 | 0.568799223 | 0.897819996 | -4.982234841 |
| **A_52_P116264** | Adhfe1 | -0.028221326 | -0.143288272 | 0.086845619 | 7.768520907 | -0.60338707 | 0.568819063 | 0.897819996 | -4.98224825 |
| **A_51_P226429** | Rhobtb2 | -0.036963082 | -0.187738999 | 0.113812835 | 8.217336352 | -0.603122258 | 0.568983777 | 0.897919517 | -4.982359536 |
| **A_51_P215038** | Tmem59l | -0.05084671 | -0.258289059 | 0.15659564 | 11.13049351 | -0.603023997 | 0.569044904 | 0.897919517 | -4.98240082 |

| | | | | | | | | | |
|---|---|---|---|---|---|---|---|---|---|
| **A_55_P1971094** | Esx1 | 0.128154861 | -0.394821887 | 0.651131609 | 11.72577403 | 0.602867493 | 0.56914227 | 0.897919517 | -4.982466562 |
| **A_65_P03022** | Cdc14b | -0.062004644 | -0.31503691 | 0.191027623 | 9.025987192 | -0.602861393 | 0.569146066 | 0.897919517 | -4.982469124 |
| **A_52_P408025** | Mpped2 | 0.075696697 | -0.233211885 | 0.38460528 | 9.731798991 | 0.602859515 | 0.569147234 | 0.897919517 | -4.982469913 |
| **A_55_P2130660** | Slitrk4 | -0.028353106 | -0.14406272 | 0.087356508 | 7.64288258 | -0.602837643 | 0.569160842 | 0.897919517 | -4.9824791 |
| **A_55_P1957419** | Ensa | 0.044352411 | -0.136692806 | 0.225397628 | 8.971579579 | 0.602697433 | 0.569248082 | 0.897955817 | -4.982537983 |
| **A_55_P2081482** | Atp5g2 | -0.062796323 | -0.319148646 | 0.193555999 | 15.43019512 | -0.602651337 | 0.569276764 | 0.897955817 | -4.982557339 |
| **A_55_P2081137** | NA | 0.065381701 | -0.201640429 | 0.332403832 | 10.20361686 | 0.602390514 | 0.569439077 | 0.898138549 | -4.982666837 |
| **A_51_P448489** | Mea1 | 0.034061746 | -0.105065411 | 0.173188904 | 10.98975475 | 0.602315672 | 0.569485657 | 0.898138729 | -4.982698249 |
| **A_55_P2208579** | Onecut2 | 0.027241959 | -0.084048067 | 0.138531985 | 8.049943182 | 0.602214569 | 0.569548585 | 0.89816469 | -4.982740679 |
| **A_51_P413539** | Nek6 | -0.055537321 | -0.282519988 | 0.171445346 | 11.0651626 | -0.601951354 | 0.569712434 | 0.898211001 | -4.982851114 |
| **A_51_P219849** | Alg13 | 0.026826347 | -0.082837415 | 0.13649011 | 7.997728897 | 0.60182131 | 0.569793396 | 0.898211001 | -4.982905661 |
| **A_55_P1993522** | NA | -0.097436485 | -0.495754911 | 0.300881941 | 12.04669133 | -0.601811353 | 0.569799595 | 0.898211001 | -4.982909837 |
| **A_51_P248857** | Inka1 | -0.037662234 | -0.191645741 | 0.116321274 | 8.049577671 | -0.601729119 | 0.569850795 | 0.898211001 | -4.982944324 |
| **A_51_P197378** | Atg12 | -0.054572909 | -0.277745852 | 0.168600034 | 13.47052262 | -0.601595703 | 0.569933868 | 0.898211001 | -4.983000267 |
| **A_66_P106789** | NA | -0.076273822 | -0.388230371 | 0.235682726 | 8.333102028 | -0.601520686 | 0.569980582 | 0.898211001 | -4.983031718 |
| **A_51_P429366** | Hes6 | 0.051146032 | -0.158056263 | 0.260348327 | 9.444544224 | 0.601470962 | 0.570011547 | 0.898211001 | -4.983052564 |
| **A_55_P2026233** | Uba7 | 0.030778704 | -0.09513125 | 0.156688658 | 9.775865537 | 0.601394465 | 0.570059186 | 0.898211001 | -4.983084629 |
| **A_55_P2115281** | Chchd2-ps | -0.04667331 | -0.237636714 | 0.144290094 | 15.9714449 | -0.601295032 | 0.570121112 | 0.898211001 | -4.983126305 |
| **A_55_P2168426** | Pced1b | -0.052723066 | -0.268439655 | 0.162993522 | 10.84232056 | -0.601293185 | 0.570122263 | 0.898211001 | -4.983127079 |
| **A_51_P292008** | Gpx3 | -0.383372284 | -1.951989923 | 1.185245355 | 14.40484911 | -0.601274493 | 0.570133904 | 0.898211001 | -4.983134913 |
| **A_51_P484158** | Steap1 | -0.059671884 | -0.30382827 | 0.184484501 | 8.112775368 | -0.60127179 | 0.570135588 | 0.898211001 | -4.983136045 |
| **A_55_P2180805** | Pex3 | 0.037455943 | -0.115860569 | 0.190772454 | 10.67936942 | 0.601036664 | 0.570282041 | 0.898368507 | -4.983234567 |
| **A_51_P234788** | Cxxc5 | -0.062939714 | -0.320610456 | 0.194731027 | 10.85339845 | -0.60093683 | 0.570344232 | 0.898393258 | -4.98327639 |
| **A_52_P295104** | Smim5 | -0.032241707 | -0.16429116 | 0.099807746 | 7.938410164 | -0.600690246 | 0.570497858 | 0.898522272 | -4.983379663 |
| **A_55_P1972907** | Cab39 | -0.036302295 | -0.184990773 | 0.112386184 | 13.23343913 | -0.60065614 | 0.570519108 | 0.898522272 | -4.983393945 |
| **A_55_P2232023** | NA | -0.105905512 | -0.539795952 | 0.327984929 | 8.348311618 | -0.600492577 | 0.570621026 | 0.898609566 | -4.983462425 |
| **A_51_P275949** | Loxl2 | 0.087141788 | -0.269974161 | 0.444257736 | 9.322443216 | 0.600324936 | 0.570725497 | 0.898700866 | -4.983532596 |
| **A_51_P409452** | Cldn11 | -0.549866027 | -2.804125784 | 1.704393731 | 10.39446974 | -0.60009788 | 0.570867012 | 0.898787769 | -4.983627611 |
| **A_55_P2108609** | Srgap3 | -0.0427462 | -0.217993904 | 0.132501503 | 8.684666644 | -0.600087177 | 0.570873684 | 0.898787769 | -4.983632089 |
| **A_51_P253803** | Mki67 | 0.097115535 | -0.301173361 | 0.495404431 | 9.09908694 | 0.599873491 | 0.571006887 | 0.898924264 | -4.98372148 |
| **A_55_P2205156** | Aamdc | -0.05405876 | -0.275834198 | 0.167716679 | 9.579183201 | -0.599683079 | 0.571125597 | 0.898946305 | -4.983801112 |
| **A_55_P2172420** | Krit1 | 0.041930434 | -0.130119778 | 0.213980645 | 10.19329258 | 0.599574712 | 0.571193165 | 0.898946305 | -4.983846422 |
| **A_51_P440892** | Dph2 | -0.02898915 | -0.147953231 | 0.089974931 | 8.931969045 | -0.599499459 | 0.571240088 | 0.898946305 | -4.983877883 |
| **A_55_P2079054** | Ndufv1 | 0.069011406 | -0.214210792 | 0.352233603 | 13.41144651 | 0.599463507 | 0.571262506 | 0.898946305 | -4.983892913 |
| **A_55_P2038242** | Tchhl1 | -0.625800112 | -3.194299766 | 1.942699542 | 10.71297615 | -0.5994118 | 0.57129475 | 0.898946305 | -4.983914527 |
| **A_51_P232411** | Rag1 | -0.027718602 | -0.141486891 | 0.086049687 | 7.617952816 | -0.599403487 | 0.571299933 | 0.898946305 | -4.983918001 |
| **A_51_P292527** | P4htm | -0.032039768 | -0.163563377 | 0.099483842 | 8.00086077 | -0.599314523 | 0.571355413 | 0.898960421 | -4.983955185 |
| **A_66_P138319** | Acox2 | 0.2676058 | -0.831150593 | 1.366362194 | 9.642666659 | 0.599188225 | 0.57143418 | 0.899011172 | -4.984007965 |
| **A_51_P502082** | Rrm1 | -0.27164527 | -1.387176404 | 0.843885863 | 11.30006985 | -0.599086614 | 0.571497555 | 0.899037702 | -4.984050422 |
| **A_55_P2141429** | Park7 | -0.057325461 | -0.292810759 | 0.178159837 | 12.14801787 | -0.598898079 | 0.571615157 | 0.899096155 | -4.984129182 |
| **A_66_P111433** | NA | 0.062197243 | -0.19330929 | 0.317703776 | 10.78652297 | 0.598877902 | 0.571627743 | 0.899096155 | -4.98413761 |
| **A_55_P1969710** | NA | 0.038263086 | -0.118955758 | 0.19548193 | 13.17226003 | 0.598748666 | 0.571708366 | 0.899122863 | -4.984191584 |
| **A_51_P385351** | Slc44a3 | -0.035510724 | -0.181435924 | 0.110414477 | 8.083931613 | -0.59868501 | 0.571748079 | 0.899122863 | -4.984218166 |
| **A_55_P2145227** | Slmap | -0.06887789 | -0.351947142 | 0.214191362 | 11.61163844 | -0.598626997 | 0.571784274 | 0.899122863 | -4.984242389 |
| **A_55_P2011352** | NA | -0.051818089 | -0.264807472 | 0.161171295 | 11.47233125 | -0.598539196 | 0.571839056 | 0.899135859 | -4.984279047 |
| **A_52_P89099** | Nolc1 | -0.02618352 | -0.13383633 | 0.08146929 | 8.010054429 | -0.598372781 | 0.571942897 | 0.899225984 | -4.984348514 |
| **A_55_P2032543** | Cblc | 0.167594148 | -0.521952974 | 0.85714127 | 9.353140092 | 0.597948889 | 0.572207451 | 0.899502584 | -4.984525387 |
| **A_55_P2125841** | Cog1 | -0.036317169 | -0.185747899 | 0.113113562 | 9.606816977 | -0.597917452 | 0.572227073 | 0.899502584 | -4.9845385 |
| **A_55_P2457153** | G2e3 | -0.032159322 | -0.164505183 | 0.100186539 | 8.647482074 | -0.597813447 | 0.572291996 | 0.899502584 | -4.984581879 |
| **A_52_P109304** | Tbl2 | -0.02919632 | -0.149352662 | 0.090960022 | 9.036385506 | -0.597792664 | 0.572304971 | 0.899502584 | -4.984590547 |

| | | | | | | | | |
|---|---|---|---|---|---|---|---|---|
| **A_55_P2101920** | Eva1c | -0.509252618 | -2.606317404 | 1.587812169 | 12.67520606 | -0.597434895 | 0.572528338 | 0.899682349 | -4.984739714 |
| **A_55_P2159949** | Polr1e | 0.031260908 | -0.097469719 | 0.159991535 | 8.401808376 | 0.597432533 | 0.572529812 | 0.899682349 | -4.984740699 |
| **A_55_P2113886** | NA | -0.059430383 | -0.304227325 | 0.185366559 | 10.50702393 | -0.597271383 | 0.572630441 | 0.899682349 | -4.984807863 |
| **A_55_P2032294** | Zfp932 | 0.041349959 | -0.12901304 | 0.211712957 | 9.152655106 | 0.597130107 | 0.572718668 | 0.899682349 | -4.984866732 |
| **A_51_P478610** | Asb6 | -0.038040057 | -0.194769632 | 0.118689518 | 10.07418932 | -0.597116907 | 0.572726912 | 0.899682349 | -4.984872232 |
| **A_55_P2064321** | Lama5 | -0.126141257 | -0.645885035 | 0.393602521 | 14.10824284 | -0.597086177 | 0.572746104 | 0.899682349 | -4.984885035 |
| **A_55_P1983999** | Desi1 | -0.038404265 | -0.196668633 | 0.119860104 | 9.781623559 | -0.596987819 | 0.572807535 | 0.899682349 | -4.984926011 |
| **A_52_P477431** | Mnt | -0.079990021 | -0.409643133 | 0.249663091 | 12.77119089 | -0.596963525 | 0.57282271 | 0.899682349 | -4.984936131 |
| **A_52_P586141** | Adcy7 | 0.070171084 | -0.219077198 | 0.359419366 | 12.97014026 | 0.596838132 | 0.572901034 | 0.899682349 | -4.98498836 |
| **A_51_P428372** | Ppbp | -0.446939444 | -2.289279092 | 1.395400205 | 9.750348759 | -0.596826569 | 0.572908256 | 0.899682349 | -4.984993175 |
| **A_52_P38964** | Sap25 | -0.043195812 | -0.221265318 | 0.134873694 | 7.996467114 | -0.596789601 | 0.572931349 | 0.899682349 | -4.985008571 |
| **A_55_P2033500** | Ltbp4 | 0.065937545 | -0.206004589 | 0.337879679 | 8.462840302 | 0.596520583 | 0.573099412 | 0.899873154 | -4.985120583 |
| **A_51_P439085** | Hilpda | -0.117954727 | -0.604598518 | 0.368689063 | 11.953516 | -0.596311697 | 0.573229929 | 0.900004979 | -4.985207528 |
| **A_52_P286141** | Cisd2 | 0.070946964 | -0.221832529 | 0.363726457 | 9.857637714 | 0.5961593 | 0.573325162 | 0.900036416 | -4.985270945 |
| **A_51_P115178** | Scara3 | 0.033694263 | -0.10537696 | 0.172765487 | 8.238137111 | 0.596057081 | 0.573389044 | 0.900036416 | -4.985313473 |
| **A_52_P253317** | 4930481A15Rik | -0.041029688 | -0.210390928 | 0.128331551 | 8.98967881 | -0.59600974 | 0.573418631 | 0.900036416 | -4.985333168 |
| **A_55_P2002103** | Arhgap45 | 0.049231017 | -0.153993036 | 0.25245507 | 9.604053342 | 0.595981617 | 0.573436208 | 0.900036416 | -4.985344866 |
| **A_52_P473966** | Kdelr3 | -0.073305389 | -0.375981078 | 0.2293703 | 10.48400972 | -0.59583708 | 0.573526549 | 0.90010512 | -4.985404984 |
| **A_55_P2021886** | Mrpl2 | 0.089109058 | -0.278970915 | 0.457189032 | 13.87414403 | 0.595591962 | 0.573679776 | 0.900191296 | -4.985506909 |
| **A_55_P2138878** | Lrfn4 | 0.053819273 | -0.168500731 | 0.276139277 | 9.500471177 | 0.595564015 | 0.573697247 | 0.900191296 | -4.985518528 |
| **A_55_P1959006** | Imp4 | 0.061965035 | -0.194020441 | 0.317950512 | 11.53328561 | 0.595525743 | 0.573721174 | 0.900191296 | -4.985534438 |
| **A_66_P129893** | 1700047M11Rik | 0.03341331 | -0.104648655 | 0.171475274 | 7.99641734 | 0.595407922 | 0.573794838 | 0.900233801 | -4.985583413 |
| **A_51_P219465** | Tmem11 | 0.043673976 | -0.136818691 | 0.224166643 | 12.95991661 | 0.595295133 | 0.573865361 | 0.900247053 | -4.985630289 |
| **A_52_P305073** | Pias1 | 0.055609051 | -0.174238872 | 0.285456973 | 9.912288974 | 0.595215269 | 0.5739153 | 0.900247053 | -4.985663477 |
| **A_55_P1998097** | Tmem214 | -0.046311966 | -0.237782063 | 0.145158132 | 9.356849347 | -0.595060905 | 0.574011831 | 0.900247053 | -4.985727611 |
| **A_51_P302377** | Fgfbp1 | 0.060066127 | -0.188273792 | 0.308406046 | 7.843280557 | 0.595048355 | 0.57401968 | 0.900247053 | -4.985732825 |
| **A_52_P251699** | Prodh2 | 0.053227751 | -0.166849043 | 0.273304544 | 7.90815319 | 0.595022004 | 0.574036159 | 0.900247053 | -4.985743772 |
| **A_55_P2007802** | Smarca2 | 0.039705761 | -0.12456185 | 0.203973371 | 9.521375895 | 0.594662761 | 0.574260857 | 0.900272824 | -4.985892968 |
| **A_55_P2145656** | NA | 0.145714197 | -0.457167841 | 0.748596235 | 8.401982912 | 0.594618868 | 0.574288314 | 0.900272824 | -4.985911191 |
| **A_51_P391825** | Exosc10 | 0.046286614 | -0.145233052 | 0.23780628 | 9.41529367 | 0.594581235 | 0.574311856 | 0.900272824 | -4.985926815 |
| **A_55_P2147831** | Gimap4 | 0.08979917 | -0.281833159 | 0.461431499 | 8.56123561 | 0.594467338 | 0.57438311 | 0.900272824 | -4.985974096 |
| **A_55_P2100655** | Lsm6 | 0.045578738 | -0.143100129 | 0.234257606 | 13.03315653 | 0.594303385 | 0.574485689 | 0.900272824 | -4.986042142 |
| **A_55_P1957219** | Ptma | -0.0734047 | -0.37727782 | 0.230468421 | 14.88865568 | -0.594293177 | 0.574492076 | 0.900272824 | -4.986046379 |
| **A_51_P425824** | C1qtnf12 | 0.102356582 | -0.321372396 | 0.526085559 | 10.64646179 | 0.594287742 | 0.574495477 | 0.900272824 | -4.986048634 |
| **A_55_P1966765** | NA | 0.055581211 | -0.174547343 | 0.285709765 | 11.37007092 | 0.594191808 | 0.574555504 | 0.900272824 | -4.986088442 |
| **A_52_P343627** | Rbp7 | 0.02639417 | -0.082890534 | 0.135678873 | 7.91970509 | 0.594179675 | 0.574563097 | 0.900272824 | -4.986093476 |
| **A_51_P361220** | Fzd4 | -0.039017611 | -0.200608125 | 0.122572904 | 9.013661728 | -0.594037648 | 0.574651973 | 0.900272824 | -4.986152399 |
| **A_55_P2067513** | Slc10a3 | 0.092355615 | -0.29013392 | 0.474845151 | 10.61338041 | 0.594036201 | 0.574652879 | 0.900272824 | -4.986152999 |
| **A_55_P2091145** | H4c12 | 0.124448283 | -0.390985813 | 0.639882379 | 12.78710649 | 0.593997925 | 0.574676833 | 0.900272824 | -4.986168877 |
| **A_52_P515497** | Atp1a1 | -0.062166621 | -0.319657214 | 0.195323971 | 14.98239372 | -0.593970754 | 0.574693837 | 0.900272824 | -4.986180147 |
| **A_66_P131788** | Ythdf2 | 0.029112379 | -0.091486593 | 0.149711351 | 8.260658358 | 0.593886229 | 0.574746736 | 0.900272824 | -4.986215206 |
| **A_51_P465128** | Chst15 | 0.039858311 | -0.125275805 | 0.204992427 | 8.397220896 | 0.593815119 | 0.574791243 | 0.900272824 | -4.986244696 |
| **A_51_P108459** | Gpr65 | -0.033495651 | -0.172292703 | 0.105301401 | 7.892820816 | -0.593714073 | 0.574854489 | 0.900272824 | -4.972172573 |
| **A_51_P110931** | Znrd1as | -0.036930463 | -0.189962525 | 0.116101599 | 9.352115031 | -0.593706058 | 0.574859506 | 0.900272824 | -4.967495454 |
| **A_55_P2165969** | NA | 0.040822966 | -0.128424289 | 0.210070221 | 8.35684968 | 0.59340621 | 0.575047211 | 0.900272824 | -4.986414222 |
| **A_55_P1965935** | Rwdd1 | -0.052390076 | -0.269607086 | 0.164826935 | 9.630013033 | -0.59336832 | 0.575070933 | 0.900272824 | -4.986429925 |
| **A_51_P171107** | Tmem35a | -0.026074899 | -0.134185605 | 0.082035806 | 7.688686519 | -0.593366604 | 0.575072007 | 0.900272824 | -4.986430636 |
| **A_51_P295582** | Cutc | -0.032739846 | -0.168497678 | 0.103017987 | 8.249652265 | -0.593308947 | 0.575108106 | 0.900272824 | -4.98645453 |
| **A_55_P2178608** | NA | 0.035313661 | -0.111131093 | 0.181758415 | 8.531366947 | 0.59325047 | 0.575144719 | 0.900272824 | -4.986478762 |
| **A_51_P332136** | Zmat5 | -0.040969505 | -0.210886864 | 0.128947855 | 8.383246815 | -0.593187685 | 0.575184031 | 0.900272824 | -4.986504777 |

| | | | | | | | | | |
|---|---|---|---|---|---|---|---|---|---|
| **A_55_P1975874** | Bcl2l15 | 0.030589848 | -0.09628563 | 0.157465326 | 7.781059347 | 0.593155814 | 0.575203987 | 0.900272824 | -4.986517982 |
| **A_52_P509965** | Ager | -0.027992198 | -0.144128003 | 0.088143608 | 7.83380805 | -0.592979971 | 0.5753141 | 0.900272824 | -4.986590826 |
| **A_52_P35064** | Tm7sf2 | 0.278886432 | -0.878194082 | 1.435966945 | 9.75792745 | 0.592970338 | 0.575320133 | 0.900272824 | -4.986594816 |
| **A_55_P2129232** | Calm2 | -0.045902279 | -0.236379697 | 0.14457514 | 14.86216925 | -0.592870597 | 0.575382597 | 0.900272824 | -4.986636126 |
| **A_51_P484054** | F7 | 0.03047301 | -0.09599167 | 0.15693769 | 7.793666144 | 0.592809647 | 0.575420769 | 0.900272824 | -4.986661367 |
| **A_51_P406188** | Asb10 | 0.056613073 | -0.178340577 | 0.291566724 | 8.361954128 | 0.592793862 | 0.575430655 | 0.900272824 | -4.986667904 |
| **A_51_P272250** | Snf8 | -0.030275084 | -0.155928215 | 0.095378047 | 10.78383589 | -0.592763167 | 0.57544988 | 0.900272824 | -4.986680614 |
| **A_55_P1985304** | Krtap5-3 | 0.026275685 | -0.082822763 | 0.135374134 | 8.61759736 | 0.592522218 | 0.575600805 | 0.900402584 | -4.986780369 |
| **A_55_P2181419** | Tcf4 | -0.051389029 | -0.264774282 | 0.161996224 | 7.983985934 | -0.592482017 | 0.575625988 | 0.900402584 | -4.986797009 |
| **A_55_P2425190** | Nhsl2 | 0.030599171 | -0.096526799 | 0.157725141 | 7.91817704 | 0.592167465 | 0.575823057 | 0.900577993 | -4.986927177 |
| **A_55_P2141938** | 1810058I24Rik | -0.057712175 | -0.297485974 | 0.182061624 | 8.995054327 | -0.592154291 | 0.575831311 | 0.900577993 | -4.986932627 |
| **A_51_P184796** | Zfp101 | -0.03448574 | -0.177798161 | 0.108826681 | 7.896761807 | -0.592004328 | 0.575925279 | 0.900652081 | -4.986994663 |
| **A_51_P491987** | Ripk3 | -0.06671685 | -0.34403902 | 0.210605319 | 8.618780198 | -0.591861526 | 0.576014768 | 0.900683986 | -4.987053724 |
| **A_51_P519369** | Agtpbp1 | -0.041345176 | -0.213231271 | 0.130540919 | 8.801001278 | -0.59177043 | 0.576071859 | 0.900683986 | -4.987091394 |
| **A_52_P229210** | NA | 0.059581141 | -0.188127217 | 0.307289499 | 11.27038681 | 0.591748707 | 0.576085474 | 0.900683986 | -4.987100377 |
| **A_55_P2229351** | NA | 0.039181059 | -0.123764353 | 0.202126471 | 8.732283914 | 0.591565989 | 0.576199997 | 0.900729935 | -4.987175917 |
| **A_55_P2022585** | Pira6 | 0.036348724 | -0.114822416 | 0.187519864 | 7.992055042 | 0.591547255 | 0.57621174 | 0.900729935 | -4.987183661 |
| **A_51_P416613** | 1810058I24Rik | -0.052126391 | -0.268940557 | 0.164687774 | 8.705486791 | -0.59147878 | 0.576254663 | 0.900729935 | -4.987211964 |
| **A_66_P105564** | Gjc2 | 0.050037289 | -0.158140821 | 0.258215399 | 8.851053139 | 0.591327227 | 0.576349669 | 0.900805592 | -4.987274597 |
| **A_55_P2092399** | Ccdc160 | -0.064015849 | -0.33041845 | 0.202386753 | 8.874645002 | -0.591177924 | 0.576443274 | 0.900872474 | -4.987336287 |
| **A_55_P2161390** | NA | 0.065864798 | -0.208286775 | 0.340016372 | 10.21399098 | 0.591060296 | 0.576517026 | 0.900872474 | -4.98738488 |
| **A_51_P407193** | Clp1 | -0.035154262 | -0.181484217 | 0.111175693 | 8.243074155 | -0.591035964 | 0.576532283 | 0.900872474 | -4.987394931 |
| **A_55_P2045026** | Mgrn1 | -0.082630587 | -0.426749665 | 0.261488491 | 13.18597557 | -0.590746627 | 0.576713724 | 0.901083144 | -4.987514419 |
| **A_55_P1997895** | Rpl35a | 0.061913453 | -0.195970218 | 0.319797124 | 14.68346038 | 0.590650188 | 0.576774208 | 0.901104807 | -4.987554235 |
| **A_55_P2127174** | Zfp628 | -0.045754072 | -0.236386642 | 0.144878499 | 17.76640259 | -0.590475401 | 0.576883839 | 0.901203243 | -4.987626383 |
| **A_66_P111747** | NA | -0.093978596 | -0.485768092 | 0.2978109 | 12.86219617 | -0.590126785 | 0.577102538 | 0.901472035 | -4.98777023 |
| **A_55_P1997501** | Gm10778 | 0.057504853 | -0.182294877 | 0.297304584 | 9.864784662 | 0.589963269 | 0.577205134 | 0.901547501 | -4.987837676 |
| **A_55_P2048855** | Sprr2a2 | 0.311690317 | -0.988219598 | 1.611600232 | 9.215075684 | 0.589901116 | 0.577244134 | 0.901547501 | -4.987863308 |
| **A_55_P2051596** | NA | -0.157936439 | -0.816974374 | 0.501101496 | 9.29778904 | -0.589577759 | 0.577447061 | 0.901770682 | -4.987996625 |
| **A_55_P2060854** | NA | -0.032450935 | -0.167874533 | 0.102972663 | 10.50424037 | -0.589524736 | 0.577480341 | 0.901770682 | -4.98801848 |
| **A_55_P2111825** | 1700001C02Rik | -0.025446445 | -0.131678359 | 0.080785469 | 7.701721456 | -0.58930656 | 0.577617289 | 0.901911671 | -4.98810839 |
| **A_51_P502872** | 2200002D01Rik | -0.080456819 | -0.416387057 | 0.255473419 | 12.41320747 | -0.589227385 | 0.577666991 | 0.90191642 | -4.988141011 |
| **A_55_P2005345** | Epb41 | -0.027055795 | -0.140068247 | 0.085956658 | 7.761598703 | -0.588983543 | 0.577820081 | 0.901945846 | -4.988241453 |
| **A_51_P104891** | Selenoi | -0.053742305 | -0.278238841 | 0.170754231 | 9.346734627 | -0.58894645 | 0.577843371 | 0.901945846 | -4.973437287 |
| **A_55_P2083128** | Sec16a | 0.060317232 | -0.191648064 | 0.312282528 | 11.05478094 | 0.588938353 | 0.577848455 | 0.901945846 | -4.988260063 |
| **A_55_P2088070** | Spata5 | 0.052354243 | -0.166410082 | 0.271118568 | 9.926278647 | 0.588768475 | 0.577955126 | 0.901945846 | -4.988330013 |
| **A_55_P2029187** | Pde4a | 0.067308705 | -0.213994238 | 0.348611648 | 8.933720491 | 0.588662161 | 0.578021889 | 0.901945846 | -4.988373781 |
| **A_55_P2023703** | Zfp386 | -0.033320048 | -0.172578671 | 0.105938575 | 8.28334717 | -0.588643946 | 0.578033328 | 0.901945846 | -4.988381279 |
| **A_51_P384318** | C1ra | -0.045252428 | -0.234434746 | 0.14392989 | 9.057458167 | -0.588478374 | 0.578137315 | 0.901945846 | -4.988449427 |
| **A_51_P160536** | Cfap161 | -0.025650829 | -0.132887459 | 0.081585801 | 7.682093705 | -0.588474187 | 0.578139945 | 0.901945846 | -4.988451151 |
| **A_52_P161297** | Tcea3 | 0.076879525 | -0.244555936 | 0.398314986 | 8.123832617 | 0.588418159 | 0.578175135 | 0.901945846 | -4.988474207 |
| **A_55_P2139587** | NA | -0.027130808 | -0.140584231 | 0.086322614 | 7.746270445 | -0.588320926 | 0.57823621 | 0.901945846 | -4.988514217 |
| **A_51_P205278** | Tmem161b | -0.039455097 | -0.204504754 | 0.125594559 | 8.87956504 | -0.588108773 | 0.578369482 | 0.901945846 | -4.988601494 |
| **A_51_P489337** | Becn1 | -0.031364084 | -0.162586912 | 0.099858745 | 8.762465541 | -0.588020371 | 0.57842502 | 0.901945846 | -4.988637853 |
| **A_52_P54238** | Lmf2 | 0.042962537 | -0.13679052 | 0.222715594 | 9.631364762 | 0.588007406 | 0.578433166 | 0.901945846 | -4.988643186 |
| **A_55_P2180301** | Clcn5 | 0.072235344 | -0.230035704 | 0.374506393 | 9.646386894 | 0.587925582 | 0.578484574 | 0.901945846 | -4.988676835 |
| **A_51_P491051** | Ints9 | -0.069812507 | -0.361953245 | 0.222328232 | 10.93463477 | -0.587909222 | 0.578494854 | 0.901945846 | -4.988683562 |
| **A_55_P2186085** | Atp11c | 0.038759943 | -0.123462898 | 0.200982784 | 8.557849941 | 0.587814488 | 0.578554378 | 0.901945846 | -4.988722515 |
| **A_55_P2055638** | Gsn | -0.075066926 | -0.389273186 | 0.239139334 | 14.65658946 | -0.587763972 | 0.57858612 | 0.901945846 | -4.988743283 |
| **A_55_P2075385** | Orai2-ps | -0.028967079 | -0.150214743 | 0.092280585 | 8.072734437 | -0.587760626 | 0.578588223 | 0.901945846 | -4.988744659 |

| | | | | | | | | | |
|---|---|---|---|---|---|---|---|---|---|
| **A_52_P375546** | Lrrc46 | -0.037490906 | -0.194420554 | 0.119438743 | 7.870927899 | -0.587746554 | 0.578597065 | 0.901945846 | -4.988750445 |
| **A_55_P1953439** | Lemd3 | 0.056811411 | -0.181004306 | 0.294627127 | 9.972492255 | 0.587711496 | 0.578619095 | 0.901945846 | -4.988764857 |
| **A_55_P2013591** | NA | 0.056563884 | -0.180279046 | 0.293406815 | 8.597179575 | 0.587554237 | 0.57871792 | 0.90202715 | -4.988829497 |
| **A_51_P471779** | Phf5a | -0.040758356 | -0.211459678 | 0.129942966 | 11.57381963 | -0.587420283 | 0.578802108 | 0.902039813 | -4.988884546 |
| **A_55_P2044843** | NA | -0.029272854 | -0.151893755 | 0.093348047 | 15.7519845 | -0.58731315 | 0.578869445 | 0.902039813 | -4.988928565 |
| **A_52_P106620** | Tnfrsf11b | -0.654505138 | -3.39749689 | 2.088486613 | 11.84386792 | -0.58702652 | 0.579049624 | 0.902039813 | -4.989046303 |
| **A_55_P1959718** | Cobll1 | -0.073043714 | -0.379170296 | 0.233082868 | 11.91915001 | -0.58701739 | 0.579055364 | 0.902039813 | -4.989050053 |
| **A_55_P1973339** | Mapk1 | -0.140762847 | -0.730727569 | 0.449201874 | 10.69269806 | -0.586990622 | 0.579072193 | 0.902039813 | -4.989061046 |
| **A_52_P90684** | Klhl31 | 0.025664557 | -0.081921393 | 0.133250508 | 7.693510973 | 0.586877393 | 0.579143381 | 0.902039813 | -4.989107541 |
| **A_55_P2168990** | NA | -0.051541645 | -0.267704292 | 0.164621003 | 15.99771937 | -0.586606383 | 0.579313789 | 0.902039813 | -4.989218794 |
| **A_52_P628162** | Zfp592 | 0.035821341 | -0.114413085 | 0.186055767 | 8.387831457 | 0.586599298 | 0.579318244 | 0.902039813 | -4.989221701 |
| **A_55_P2028837** | Tspan2 | -0.065015267 | -0.337748024 | 0.20771749 | 8.403745685 | -0.586471905 | 0.579398358 | 0.902039813 | -4.989273982 |
| **A_52_P256817** | Zik1 | 0.027472819 | -0.087801 | 0.142746637 | 7.819059078 | 0.586329441 | 0.579487957 | 0.902039813 | -4.989332437 |
| **A_52_P194242** | Sel1l | -0.051172151 | -0.265906688 | 0.163562386 | 11.05949454 | -0.586274402 | 0.579522575 | 0.902039813 | -4.989355017 |
| **A_55_P1956017** | Usp45 | -0.028032001 | -0.145676054 | 0.089612052 | 9.000456815 | -0.586210083 | 0.579563031 | 0.902039813 | -4.989381402 |
| **A_51_P257762** | Kat2b | -0.02771405 | -0.144023777 | 0.088595677 | 8.024012537 | -0.586209868 | 0.579563167 | 0.902039813 | -4.98938149 |
| **A_55_P2168316** | Aldh3b2 | 0.077998163 | -0.249381132 | 0.405377459 | 9.126755172 | 0.586141308 | 0.579606292 | 0.902039813 | -4.989409612 |
| **A_51_P423008** | Klhl7 | -0.034961786 | -0.181708327 | 0.111784756 | 8.219308532 | -0.586131271 | 0.579612605 | 0.902039813 | -4.989413728 |
| **A_55_P2419483** | 4732460I02Rik | -0.0615916 | -0.320116907 | 0.196933707 | 8.146217956 | -0.586121409 | 0.579618809 | 0.902039813 | -4.989417773 |
| **A_55_P2025078** | Csf1r | -0.046359102 | -0.240956473 | 0.148238269 | 16.6477487 | -0.586093914 | 0.579636105 | 0.902039813 | -4.98942905 |
| **A_51_P440210** | Prkg1 | -0.036649962 | -0.190497407 | 0.117197483 | 7.936894211 | -0.586073935 | 0.579648673 | 0.902039813 | -4.989437244 |
| **A_55_P2159189** | Decr1 | -0.07239283 | -0.376407732 | 0.231622071 | 13.17362456 | -0.585827626 | 0.579803628 | 0.902039813 | -4.989538241 |
| **A_55_P2001861** | NA | 0.069220155 | -0.221479698 | 0.359920007 | 9.123354079 | 0.585810182 | 0.579814603 | 0.902039813 | -4.989545392 |
| **A_55_P2076846** | Tead2 | 0.06057546 | -0.193827169 | 0.314978088 | 10.01467566 | 0.585793149 | 0.57982532 | 0.902039813 | -4.989552375 |
| **A_55_P2027999** | Hk1 | -0.104943037 | -0.545679342 | 0.335793267 | 13.23041305 | -0.585792716 | 0.579825592 | 0.902039813 | -4.989552553 |
| **A_55_P2030717** | NA | -0.048578936 | -0.252620127 | 0.155462254 | 10.84595616 | -0.585732493 | 0.579863484 | 0.902039813 | -4.98957724 |
| **A_51_P282616** | Maats1 | 0.034307526 | -0.109801254 | 0.178416306 | 8.454260782 | 0.585690421 | 0.579889956 | 0.902039813 | -4.989594485 |
| **A_51_P443364** | Cbll1 | -0.046884145 | -0.243823596 | 0.150055306 | 10.19779803 | -0.585682762 | 0.579894775 | 0.902039813 | -4.989597625 |
| **A_52_P351816** | Itgb1 | -0.042059447 | -0.21876781 | 0.134648916 | 9.602778927 | -0.585565681 | 0.579968448 | 0.902039813 | -4.98964561 |
| **A_66_P128608** | Fut11 | 0.028580103 | -0.091501951 | 0.148662157 | 8.766864568 | 0.585537673 | 0.579986072 | 0.902039813 | -4.989657088 |
| **A_55_P1998471** | S100a9 | -0.174603734 | -0.908439091 | 0.559231624 | 11.25476065 | -0.585361366 | 0.580097025 | 0.902040156 | -4.989729329 |
| **A_55_P2029678** | Etfa | -0.051336063 | -0.267135742 | 0.164463615 | 12.26147885 | -0.585249339 | 0.580167532 | 0.902040156 | -4.989775222 |
| **A_55_P1971179** | Cd1d1 | 0.025826458 | -0.0827412 | 0.134394116 | 7.828235369 | 0.585239387 | 0.580173796 | 0.902040156 | -4.989779299 |
| **A_52_P147803** | Syne4 | 0.093287773 | -0.298966821 | 0.485542366 | 8.744251962 | 0.585094275 | 0.580265133 | 0.902040156 | -4.989838733 |
| **A_55_P2060752** | Usp12 | -0.048404183 | -0.251963867 | 0.155155501 | 12.83992356 | -0.585005954 | 0.580320729 | 0.902040156 | -4.989874901 |
| **A_51_P297586** | Birc6 | -0.073847733 | -0.384432284 | 0.236736818 | 9.748438016 | -0.584960421 | 0.580349392 | 0.902040156 | -4.989893546 |
| **A_52_P297009** | Itpk1 | -0.090453124 | -0.470916177 | 0.290009928 | 10.20429315 | -0.584898148 | 0.580388595 | 0.902040156 | -4.989919043 |
| **A_55_P2089238** | Fam76a | 0.034465384 | -0.110512113 | 0.17944288 | 11.99991126 | 0.584859681 | 0.580412812 | 0.902040156 | -4.989934791 |
| **A_55_P2108808** | Tubgcp4 | 0.040645563 | -0.130347229 | 0.211638355 | 8.468873377 | 0.584796147 | 0.580452811 | 0.902040156 | -4.989960801 |
| **A_55_P1984497** | Usp48 | 0.045489138 | -0.145880288 | 0.236858564 | 10.51137832 | 0.584795894 | 0.58045297 | 0.902040156 | -4.989960904 |
| **A_55_P2038358** | Acot1 | 0.100511712 | -0.32259466 | 0.523618084 | 9.61550495 | 0.584435069 | 0.580680165 | 0.902257923 | -4.990108572 |
| **A_52_P302463** | Arl5a | -0.039738695 | -0.207022507 | 0.127545118 | 8.401482448 | -0.584425075 | 0.580686459 | 0.902257923 | -4.990112661 |
| **A_51_P487933** | Psmd7 | 0.039047962 | -0.125353089 | 0.203449013 | 12.09548159 | 0.584336412 | 0.580742295 | 0.902272149 | -4.990148934 |
| **A_55_P1990235** | Bin3 | -0.032004238 | -0.16683886 | 0.102830383 | 11.54564078 | -0.583949432 | 0.580986032 | 0.902578285 | -4.990307196 |
| **A_55_P2097820** | NA | -0.065806512 | -0.343144172 | 0.211531147 | 10.54233402 | -0.58375309 | 0.581109721 | 0.902660729 | -4.99038746 |
| **A_51_P394921** | 1700020L24Rik | -0.030838951 | -0.160831103 | 0.099153201 | 8.784608056 | -0.583648844 | 0.581175399 | 0.902660729 | -4.990430066 |
| **A_55_P1956827** | Ogfr | -0.034960019 | -0.182324793 | 0.112404755 | 13.62332767 | -0.58364281 | 0.581179201 | 0.902660729 | -4.990432532 |
| **A_55_P2122830** | Zfp185 | 0.052471889 | -0.168781715 | 0.273725494 | 10.95017722 | 0.583452506 | 0.581299109 | 0.902698079 | -4.990510293 |
| **A_55_P2148708** | NA | -0.05388061 | -0.281098993 | 0.173337773 | 13.03270253 | -0.583388959 | 0.581339153 | 0.902698079 | -4.990536254 |
| **A_66_P104134** | Homer1 | -0.027485658 | -0.143395968 | 0.088424651 | 7.958585576 | -0.583382292 | 0.581343354 | 0.902698079 | -4.990538978 |

| | | | | | | | | | |
|---|---|---|---|---|---|---|---|---|---|
| **A_55_P1972530** | Efl1 | -0.064198337 | -0.33509372 | 0.206697047 | 9.633091195 | -0.583030584 | 0.58156501 | 0.902969723 | -4.990682619 |
| **A_55_P2014189** | NA | 0.087261232 | -0.281001706 | 0.45552417 | 10.12312248 | 0.58295159 | 0.581614801 | 0.902974496 | -4.990714871 |
| **A_66_P115004** | Gabrp | -0.140114205 | -0.73153316 | 0.45130475 | 9.27085635 | -0.582849047 | 0.581679439 | 0.903002319 | -4.990756732 |
| **A_66_P106861** | B3glct | -0.061517071 | -0.321281986 | 0.198247844 | 8.789371423 | -0.582618566 | 0.581824739 | 0.903155347 | -4.990850798 |
| **A_55_P2033435** | Stra6l | -0.058020484 | -0.303079605 | 0.187038637 | 8.146923984 | -0.582478153 | 0.581913269 | 0.903157476 | -4.99090809 |
| **A_55_P2007001** | Ctnnbip1 | -0.060740412 | -0.317291996 | 0.195811171 | 12.08293328 | -0.582468172 | 0.581919562 | 0.903157476 | -4.990912161 |
| **A_55_P1952512** | Sult2a2 | 0.025326322 | -0.081665726 | 0.132318371 | 7.728341009 | 0.582357655 | 0.581989248 | 0.90319311 | -4.990957246 |
| **A_51_P131561** | Plet1 | -0.146231477 | -0.764189763 | 0.47172681 | 14.356147 | -0.582171403 | 0.582106701 | 0.90330286 | -4.991033209 |
| **A_51_P137336** | Cdh1 | -0.089237987 | -0.466558833 | 0.288082858 | 11.54170754 | -0.581846104 | 0.582311872 | 0.903484575 | -4.981467469 |
| **A_55_P2032445** | Slc25a37 | 0.071825914 | -0.23187675 | 0.375528579 | 9.067384228 | 0.58183752 | 0.582317287 | 0.903484575 | -4.991169331 |
| **A_55_P1991931** | Nosip | 0.044253227 | -0.142924794 | 0.231431248 | 10.85559151 | 0.581646684 | 0.58243767 | 0.903555067 | -4.991247104 |
| **A_51_P111259** | Cdk2ap2 | -0.077738621 | -0.406566541 | 0.251089299 | 12.53903585 | -0.581617291 | 0.582456213 | 0.903555067 | -4.991259081 |
| **A_55_P2047599** | Arhgdia | -0.068417942 | -0.357879409 | 0.221043525 | 13.02254652 | -0.581498251 | 0.582531315 | 0.903599052 | -4.991307581 |
| **A_66_P105965** | Drp2 | -0.058241581 | -0.30484277 | 0.188359608 | 8.951114705 | -0.581041498 | 0.582819532 | 0.903930053 | -4.991493597 |
| **A_51_P336622** | Dexi | -0.037079617 | -0.194096028 | 0.119936794 | 11.62531617 | -0.580977557 | 0.582859887 | 0.903930053 | -4.991519628 |
| **A_51_P214985** | Zfp521 | -0.03140154 | -0.164382818 | 0.101579738 | 7.921878789 | -0.580937766 | 0.582885001 | 0.903930053 | -4.991535826 |
| **A_55_P2005570** | Celf2 | -0.044024408 | -0.23050221 | 0.142453395 | 10.77652336 | -0.580811949 | 0.582964413 | 0.903972307 | -4.991587036 |
| **A_55_P2167763** | NA | 0.029327775 | -0.094923954 | 0.153579504 | 7.853175403 | 0.580692001 | 0.583040128 | 0.903972307 | -4.991635849 |
| **A_51_P113773** | Slx1b | 0.03860546 | -0.12495818 | 0.2021691 | 9.032110619 | 0.580672321 | 0.58305255 | 0.903972307 | -4.991643856 |
| **A_55_P2088615** | Fbln5 | -0.07240545 | -0.379279663 | 0.234468762 | 10.60839327 | -0.58047033 | 0.583180067 | 0.903987911 | -4.991726035 |
| **A_55_P2170355** | NA | -0.044353773 | -0.232340317 | 0.143632771 | 8.486095924 | -0.580460896 | 0.583186023 | 0.903987911 | -4.991729872 |
| **A_51_P337543** | Glce | -0.079807327 | -0.418076475 | 0.25846182 | 11.02532714 | -0.580429584 | 0.583205792 | 0.903987911 | -4.991742609 |
| **A_52_P5901** | Saal1 | -0.025682378 | -0.134552043 | 0.083187287 | 8.297138956 | -0.580360057 | 0.583249689 | 0.903987911 | -4.991770888 |
| **A_55_P2181071** | NA | 0.052180651 | -0.169213994 | 0.273575296 | 15.59253634 | 0.579844504 | 0.583575255 | 0.904419989 | -4.991980491 |
| **A_55_P2128869** | Ccdc80 | -0.044672097 | -0.234262686 | 0.144918492 | 7.97535033 | -0.579680542 | 0.583678817 | 0.904507966 | -4.992047119 |
| **A_52_P527800** | Emilin2 | -0.662420486 | -3.475161287 | 2.150320315 | 12.13646267 | -0.579392948 | 0.583860496 | 0.904716973 | -4.992163946 |
| **A_52_P71105** | Sertad3 | -0.048835058 | -0.256252408 | 0.158582291 | 9.483159994 | -0.579236329 | 0.583959449 | 0.904762842 | -4.992227548 |
| **A_55_P2047225** | Uxs1 | 0.039987511 | -0.129862876 | 0.209837898 | 8.842545833 | 0.579197924 | 0.583983714 | 0.904762842 | -4.992243141 |
| **A_52_P573366** | Hnrnpul1 | 0.025788622 | -0.083769272 | 0.135346516 | 7.987566 | 0.579100084 | 0.584045538 | 0.904786102 | -4.992282864 |
| **A_55_P1979402** | Tigar | 0.055056042 | -0.178957852 | 0.289069936 | 8.921152254 | 0.578805307 | 0.584231823 | 0.904985123 | -4.992402506 |
| **A_55_P2071028** | NA | -0.06545461 | -0.343735861 | 0.21282664 | 10.77847928 | -0.578662659 | 0.584321982 | 0.904985123 | -4.992460384 |
| **A_52_P165654** | Star | -0.06479677 | -0.3402825 | 0.210688959 | 9.787729578 | -0.578659937 | 0.584323703 | 0.904985123 | -4.992461489 |
| **A_55_P2101431** | Mrps25 | -0.032484744 | -0.170644099 | 0.10567461 | 8.639714144 | -0.578453322 | 0.584454307 | 0.904985123 | -4.992545299 |
| **A_66_P132855** | Baz1b | 0.025527249 | -0.083047518 | 0.134102015 | 8.64049249 | 0.578421283 | 0.584474561 | 0.904985123 | -4.992558293 |
| **A_55_P2028243** | Dnajc19 | -0.060768744 | -0.319273984 | 0.197736496 | 11.00142825 | -0.578335793 | 0.584528607 | 0.904985123 | -4.992592962 |
| **A_51_P463428** | Pik3ip1 | -0.106634344 | -0.560465092 | 0.347196404 | 10.90188666 | -0.578059239 | 0.584703461 | 0.904985123 | -4.992705083 |
| **A_66_P101538** | Lrrk2 | -0.025961572 | -0.136468321 | 0.084545176 | 7.935559903 | -0.577978061 | 0.584754792 | 0.904985123 | -4.992737986 |
| **A_55_P1980611** | Trmt2b | 0.087806771 | -0.285965369 | 0.461578912 | 9.748107223 | 0.577949968 | 0.584772557 | 0.904985123 | -4.992749372 |
| **A_51_P336691** | Srsf9 | -0.03988627 | -0.209702404 | 0.129929864 | 11.57210255 | -0.577848034 | 0.584837018 | 0.904985123 | -4.99279068 |
| **A_55_P2457912** | Psme4 | -0.036890848 | -0.193970137 | 0.12018844 | 8.430806977 | -0.57778848 | 0.584874681 | 0.904985123 | -4.992814811 |
| **A_52_P65237** | Zbtb7c | 0.088591368 | -0.288664057 | 0.465846793 | 8.4037321 | 0.577730211 | 0.584911532 | 0.904985123 | -4.992838419 |
| **A_55_P2094925** | Srgn | -0.640235884 | -3.366783481 | 2.086311714 | 14.515542 | -0.577691651 | 0.58493592 | 0.904985123 | -4.992854041 |
| **A_55_P2083351** | Hs3st1 | -0.072059534 | -0.378943693 | 0.234824626 | 9.250999902 | -0.577678412 | 0.584944293 | 0.904985123 | -4.992859405 |
| **A_52_P612693** | Polb | -0.035467004 | -0.186517525 | 0.115583516 | 9.32998801 | -0.577658871 | 0.584956652 | 0.904985123 | -4.992867321 |
| **A_66_P122086** | 9030619P08Rik | -0.415213102 | -2.183660942 | 1.353234738 | 10.23367214 | -0.577627628 | 0.584976413 | 0.904985123 | -4.992879977 |
| **A_55_P2151868** | Mest | -0.039275372 | -0.20656365 | 0.128012905 | 8.3652323 | -0.577595721 | 0.584996594 | 0.904985123 | -4.992892902 |
| **A_51_P436201** | Gart | -0.042907093 | -0.225674299 | 0.139860113 | 8.40433158 | -0.577563825 | 0.585016768 | 0.904985123 | -4.992905821 |
| **A_52_P96782** | Wasl | -0.091976137 | -0.484055728 | 0.300103454 | 9.644464811 | -0.577125271 | 0.5852942 | 0.905341836 | -4.993083397 |
| **A_55_P2029294** | Cetn4 | 0.032306035 | -0.105508008 | 0.170120078 | 8.094896737 | 0.576712475 | 0.585555408 | 0.9056734 | -4.993250438 |
| **A_55_P2001998** | Zfp503 | -0.04472625 | -0.235625231 | 0.14617273 | 8.186642166 | -0.576405405 | 0.585749759 | 0.905861813 | -4.99337463 |

| | | | | | | | | | |
|---|---|---|---|---|---|---|---|---|---|
| **A_55_P2091762** | NA | -0.033582607 | -0.176954806 | 0.109789592 | 15.43568935 | -0.576260205 | 0.585841673 | 0.905861813 | -4.993433336 |
| **A_55_P2089913** | Cd6 | -0.18326735 | -0.965698133 | 0.599163432 | 8.642978354 | -0.576246547 | 0.585850319 | 0.905861813 | -4.993438857 |
| **A_55_P1978511** | H2-Q7 | -0.259169476 | -1.365695883 | 0.84735693 | 13.14040269 | -0.576223852 | 0.585864686 | 0.905861813 | -4.993448031 |
| **A_55_P1963061** | Rps16 | 0.037937395 | -0.124062942 | 0.199937733 | 16.86324609 | 0.576130349 | 0.585923881 | 0.905877555 | -4.993485827 |
| **A_55_P1996953** | NA | -0.089650608 | -0.472523746 | 0.29322253 | 10.97984055 | -0.576059715 | 0.5859686 | 0.905877555 | -4.993514374 |
| **A_52_P383862** | Cmc1 | 0.067737224 | -0.221612824 | 0.357087272 | 10.83049034 | 0.575934373 | 0.58604796 | 0.905927785 | -4.993565026 |
| **A_55_P1952744** | Timm8a1 | -0.03148481 | -0.166035719 | 0.103066099 | 9.841601329 | -0.575683259 | 0.586206972 | 0.905994985 | -4.993666474 |
| **A_52_P597775** | Gprc5a | -0.1385234 | -0.730731284 | 0.453684484 | 12.72962846 | -0.575463949 | 0.586345866 | 0.905994985 | -4.993755043 |
| **A_55_P2099526** | H2afv | 0.07329985 | -0.240133509 | 0.386733209 | 12.79615608 | 0.575343263 | 0.586422307 | 0.905994985 | -4.99380377 |
| **A_66_P133520** | Glmn | -0.051330453 | -0.270833375 | 0.16817247 | 8.953327243 | -0.575312686 | 0.586441675 | 0.905994985 | -4.993816114 |
| **A_51_P220896** | Chmp2b | 0.097356145 | -0.319012745 | 0.513725035 | 10.86701576 | 0.575246863 | 0.58648337 | 0.905994985 | -4.993842685 |
| **A_55_P2085295** | Inf2 | 0.101963364 | -0.334164345 | 0.538091073 | 13.23773588 | 0.57517452 | 0.586529196 | 0.905994985 | -4.993871885 |
| **A_55_P1993728** | Tmem116 | 0.075282059 | -0.246732673 | 0.397296792 | 9.844383776 | 0.575155018 | 0.586541551 | 0.905994985 | -4.993879757 |
| **A_55_P2018274** | NA | 0.069928238 | -0.229233004 | 0.369089479 | 11.04534649 | 0.575064317 | 0.586599011 | 0.905994985 | -4.993916361 |
| **A_55_P2008576** | Ubr2 | -0.037365574 | -0.19722166 | 0.122490512 | 8.885279885 | -0.575057992 | 0.586603018 | 0.905994985 | -4.993918914 |
| **A_55_P2092909** | Rassf1 | 0.036815838 | -0.120699817 | 0.194331494 | 12.12346458 | 0.575016264 | 0.586629454 | 0.905994985 | -4.993935752 |
| **A_55_P1958419** | NA | -0.039226944 | -0.207118603 | 0.128664714 | 12.04715579 | -0.574810279 | 0.586759964 | 0.905994985 | -4.994018858 |
| **A_55_P2008427** | Dnd1 | 0.02570351 | -0.084311188 | 0.135718209 | 7.906344027 | 0.574792234 | 0.586771398 | 0.905994985 | -4.994026137 |
| **A_66_P102706** | 2700012I20Rik | -0.039398402 | -0.208045288 | 0.129248485 | 8.210858604 | -0.574737374 | 0.58680616 | 0.905994985 | -4.994048266 |
| **A_51_P342567** | Akap12 | 0.066944605 | -0.219646584 | 0.353535794 | 11.19375346 | 0.574674482 | 0.586846012 | 0.905994985 | -4.994073632 |
| **A_51_P290921** | Sytl2 | -0.038678125 | -0.204286841 | 0.126930591 | 8.060140193 | -0.574581169 | 0.586905145 | 0.905994985 | -4.994111264 |
| **A_55_P2179378** | Zkscan8 | -0.039657025 | -0.209466672 | 0.130152622 | 9.236347848 | -0.574548815 | 0.586925648 | 0.905994985 | -4.994124311 |
| **A_51_P457528** | Ccnb2 | 0.095872566 | -0.314655621 | 0.506400752 | 8.681437695 | 0.574540347 | 0.586931015 | 0.905994985 | -4.994127725 |
| **A_55_P1953387** | Fabp5 | 0.112059684 | -0.36794363 | 0.592062999 | 15.14661225 | 0.574346953 | 0.587053583 | 0.905994985 | -4.994205697 |
| **A_52_P209514** | Cacnb2 | 0.030861981 | -0.101349266 | 0.163073228 | 7.678039272 | 0.574281149 | 0.587095292 | 0.905994985 | -4.994232222 |
| **A_55_P1979575** | Shroom2 | -0.087635132 | -0.463060926 | 0.287790662 | 10.92019167 | -0.574279482 | 0.587096349 | 0.905994985 | -4.994232894 |
| **A_55_P2166538** | Mrps16 | 0.060383095 | -0.198353458 | 0.319119648 | 12.96740673 | 0.574151821 | 0.587177269 | 0.905994985 | -4.994284346 |
| **A_55_P2182016** | Faf2 | -0.081376727 | -0.430079095 | 0.267325642 | 12.54021371 | -0.574135607 | 0.587187547 | 0.905994985 | -4.994290881 |
| **A_55_P2157705** | Uqcr10 | -0.051947328 | -0.274561372 | 0.170666717 | 13.56070458 | -0.574089779 | 0.587216598 | 0.905994985 | -4.994309348 |
| **A_55_P2024155** | Zbtb16 | 0.085545322 | -0.281051757 | 0.4521424 | 9.484311817 | 0.574085302 | 0.587219436 | 0.905994985 | -4.994311153 |
| **A_55_P2125643** | NA | 0.075132132 | -0.246878881 | 0.397143146 | 13.25970118 | 0.574016203 | 0.58726324 | 0.905994985 | -4.994338995 |
| **A_52_P830312** | Iba57 | -0.047657915 | -0.252072547 | 0.156756717 | 9.014966423 | -0.573577656 | 0.587541296 | 0.906351612 | -4.994515636 |
| **A_55_P2151609** | Sorl1 | 0.074790262 | -0.246057179 | 0.395637703 | 8.235316092 | 0.573476518 | 0.587605432 | 0.906378213 | -4.994556357 |
| **A_66_P110490** | Polr3h | -0.049878919 | -0.263908271 | 0.164150432 | 9.13410483 | -0.573340819 | 0.587691491 | 0.906438624 | -4.994610983 |
| **A_55_P2345116** | NA | -0.070822002 | -0.374862532 | 0.233218529 | 8.104483272 | -0.573067632 | 0.587864767 | 0.906615504 | -4.994720922 |
| **A_55_P2155397** | Rps8 | -0.030921024 | -0.163678909 | 0.101836861 | 16.95098409 | -0.57301066 | 0.587900907 | 0.906615504 | -4.994743843 |
| **A_55_P2095578** | NA | -0.068632378 | -0.363339374 | 0.226074618 | 8.79038864 | -0.572938182 | 0.587946885 | 0.906615504 | -4.994773001 |
| **A_52_P529570** | Nsl1 | 0.037833025 | -0.124657063 | 0.200323114 | 7.940712847 | 0.57281365 | 0.588025889 | 0.906630161 | -4.994823092 |
| **A_51_P185693** | Slc2a2 | 0.126817354 | -0.417890386 | 0.671525094 | 8.645292889 | 0.57277533 | 0.588050201 | 0.906630161 | -4.994838504 |
| **A_55_P1974967** | H4c14 | -0.024334036 | -0.128930099 | 0.080262027 | 7.687714711 | -0.572358283 | 0.588314831 | 0.90670846 | -4.995006177 |
| **A_66_P122372** | Ppp4r2 | -0.036075584 | -0.191165005 | 0.119013837 | 9.240463617 | -0.57226918 | 0.588371379 | 0.90670846 | -4.995041987 |
| **A_55_P2143494** | Doc2b | 0.144368976 | -0.476364708 | 0.765102661 | 8.419134335 | 0.572186658 | 0.588423753 | 0.90670846 | -4.995075148 |
| **A_51_P347823** | Ipo4 | 0.033827874 | -0.111620441 | 0.179276189 | 10.59668018 | 0.572183295 | 0.588425888 | 0.90670846 | -4.995076499 |
| **A_55_P2130970** | Parp10 | -0.036035389 | -0.191004136 | 0.118933358 | 12.0507906 | -0.572076699 | 0.588493545 | 0.90670846 | -4.995119328 |
| **A_55_P2057321** | Myh7b | 0.026305646 | -0.08684597 | 0.139457261 | 7.730635278 | 0.571949085 | 0.588574549 | 0.90670846 | -4.995170592 |
| **A_66_P125962** | Hipk1 | -0.03375273 | -0.178943509 | 0.11143805 | 8.043349672 | -0.571924927 | 0.588589884 | 0.90670846 | -4.995180296 |
| **A_65_P10195** | Myl7 | 0.032352769 | -0.10681956 | 0.171525098 | 8.023675673 | 0.571910049 | 0.588599329 | 0.90670846 | -4.995186272 |
| **A_55_P2000127** | Tgfb1i1 | 0.09421899 | -0.311132817 | 0.499570797 | 10.22377253 | 0.57184127 | 0.588642991 | 0.90670846 | -4.995213896 |
| **A_55_P2003824** | Asgr1 | -0.02971333 | -0.157553691 | 0.098127031 | 7.840746941 | -0.571810989 | 0.588662214 | 0.90670846 | -4.995226057 |
| **A_66_P130310** | A4galt | 0.134447118 | -0.444018334 | 0.71291257 | 9.258427993 | 0.571798792 | 0.588669957 | 0.90670846 | -4.995230955 |

| | | | | | | | | | |
|---|---|---|---|---|---|---|---|---|---|
| **A_51_P211325** | Nudt2 | 0.046991474 | -0.155219795 | 0.249202744 | 8.711556659 | 0.571719333 | 0.588720402 | 0.90670846 | -4.995262863 |
| **A_51_P521176** | Rassf7 | 0.087222877 | -0.288170354 | 0.462616109 | 12.11513088 | 0.571627528 | 0.588778688 | 0.90670846 | -4.995299724 |
| **A_55_P2088272** | Golga3 | -0.048669399 | -0.258140817 | 0.160802019 | 12.1289263 | -0.57161073 | 0.588789354 | 0.90670846 | -4.995306468 |
| **A_66_P124497** | Imp3 | -0.074531085 | -0.408562259 | 0.25950009 | 13.16107128 | -0.578229871 | 0.588818268 | 0.90670846 | -4.931082908 |
| **A_55_P1988328** | NA | 0.043699121 | -0.144417656 | 0.231815898 | 15.340939 | 0.571497501 | 0.588861247 | 0.90670846 | -4.995351922 |
| **A_52_P427240** | Ltn1 | -0.028076231 | -0.148955927 | 0.092803465 | 8.233725329 | -0.571418913 | 0.588911149 | 0.90670846 | -4.995383466 |
| **A_55_P2139886** | Sft2d1 | -0.065939849 | -0.349864724 | 0.217985025 | 9.440588666 | -0.57136505 | 0.588945353 | 0.90670846 | -4.995405084 |
| **A_52_P588539** | Snapin | 0.027467007 | -0.090850941 | 0.145784954 | 8.787146181 | 0.571123287 | 0.589098888 | 0.906872603 | -4.995502092 |
| **A_55_P2173066** | NA | 0.050757741 | -0.167939591 | 0.269455074 | 8.689744199 | 0.570989293 | 0.589183993 | 0.906931385 | -4.995555842 |
| **A_55_P2043028** | Rexo5 | -0.052870644 | -0.280745717 | 0.175004429 | 9.004025546 | -0.570803916 | 0.589301746 | 0.90698335 | -4.995630187 |
| **A_52_P619248** | Prl2c5 | -0.039890133 | -0.211855188 | 0.132074923 | 16.84648877 | -0.570682349 | 0.589378973 | 0.90698335 | -4.995678929 |
| **A_55_P2052145** | Ly6g6d | 0.034872188 | -0.11546143 | 0.185205805 | 7.979577536 | 0.57067947 | 0.589380802 | 0.90698335 | -4.995680083 |
| **A_55_P2081168** | Rpl7 | 0.029309463 | -0.0970519 | 0.155670827 | 16.32964311 | 0.57064068 | 0.589405445 | 0.90698335 | -4.995695634 |
| **A_55_P2157632** | 2510039O18Rik | -0.057244598 | -0.304091472 | 0.189602275 | 12.95610135 | -0.570526791 | 0.589477802 | 0.907022484 | -4.995741288 |
| **A_55_P1956223** | Chka | 0.048270763 | -0.159960721 | 0.256502248 | 9.026422384 | 0.570304681 | 0.589618931 | 0.907036034 | -4.995830299 |
| **A_51_P338262** | Tnnt2 | 0.05550612 | -0.183949362 | 0.294961602 | 8.663229335 | 0.570276205 | 0.589637026 | 0.907036034 | -4.995841709 |
| **A_55_P2425661** | Ncbp3 | 0.03558935 | -0.117979384 | 0.189158084 | 10.54663442 | 0.570146442 | 0.589719488 | 0.907036034 | -4.995893696 |
| **A_55_P2199118** | Bend4 | 0.096919014 | -0.321328469 | 0.515166498 | 11.71062288 | 0.570091828 | 0.589754196 | 0.907036034 | -4.995915573 |
| **A_52_P592909** | Dgat2 | -0.122576507 | -0.651590249 | 0.406437235 | 12.7228018 | -0.570045408 | 0.589783698 | 0.907036034 | -4.995934167 |
| **A_52_P355057** | Wdr4 | 0.032670104 | -0.108336615 | 0.173676823 | 10.34695652 | 0.570006588 | 0.58980837 | 0.907036034 | -4.995949715 |
| **A_51_P267454** | Slc25a11 | -0.085645273 | -0.455303815 | 0.284013269 | 12.98536779 | -0.569996012 | 0.589815092 | 0.907036034 | -4.995953951 |
| **A_55_P2158990** | Jun | -0.129015768 | -0.686025298 | 0.427993762 | 12.89413461 | -0.569835274 | 0.589917257 | 0.907120975 | -4.996018319 |
| **A_55_P2115401** | Bst1 | 0.126139099 | -0.418684859 | 0.670963057 | 8.774211701 | 0.569590438 | 0.590072894 | 0.907140142 | -4.996116335 |
| **A_55_P2088720** | NA | -0.308624178 | -1.641826761 | 1.024578404 | 11.55994249 | -0.569512111 | 0.590122691 | 0.907140142 | -4.996147684 |
| **A_55_P2069277** | Mtf1 | -0.038447999 | -0.204541336 | 0.127645338 | 9.244765355 | -0.569496028 | 0.590132916 | 0.907140142 | -4.99615412 |
| **A_52_P447284** | Clic6 | 0.199514623 | -0.662427802 | 1.061457048 | 9.266459662 | 0.569463165 | 0.590153809 | 0.907140142 | -4.996167272 |
| **A_55_P2065050** | NA | -0.033890125 | -0.180330875 | 0.112550626 | 8.415205674 | -0.569351388 | 0.590224877 | 0.907140142 | -4.996211999 |
| **A_52_P648424** | Nedd4 | -0.050281045 | -0.267585353 | 0.167023263 | 12.22385549 | -0.569252731 | 0.590287608 | 0.907140142 | -4.99625147 |
| **A_52_P799815** | Tmem171 | 0.032109387 | -0.106672901 | 0.170891675 | 7.841110065 | 0.569202937 | 0.59031927 | 0.907140142 | -4.99627139 |
| **A_55_P2103963** | Bhmt | -0.025951381 | -0.13815157 | 0.086248808 | 7.664306365 | -0.569031161 | 0.590428507 | 0.907140142 | -4.996340096 |
| **A_55_P2067518** | Slc13a3 | 0.2468122 | -0.820298704 | 1.313923103 | 9.760042522 | 0.569018293 | 0.59043669 | 0.907140142 | -4.996345242 |
| **A_55_P2073705** | Tfdp2 | -0.033718048 | -0.17955548 | 0.112119384 | 9.290423626 | -0.568803925 | 0.590573028 | 0.907140142 | -4.996430957 |
| **A_51_P254471** | Birc2 | -0.060050643 | -0.31978704 | 0.199685754 | 9.555438902 | -0.5687927 | 0.590580168 | 0.907140142 | -4.996435444 |
| **A_52_P679205** | Slc35f6 | 0.034547178 | -0.114897181 | 0.183991537 | 10.27726992 | 0.568724863 | 0.590623317 | 0.907140142 | -4.996462562 |
| **A_55_P1958275** | Bcl11a | 0.026537737 | -0.088264074 | 0.141339547 | 7.880312447 | 0.568701413 | 0.590638233 | 0.907140142 | -4.996471936 |
| **A_51_P518298** | Rnf187 | 0.066286465 | -0.220477213 | 0.353050142 | 12.080838 | 0.568682519 | 0.590650251 | 0.907140142 | -4.996479488 |
| **A_55_P2350022** | Akap6 | -0.024412465 | -0.130024642 | 0.081199713 | 7.721093387 | -0.56867848 | 0.590652821 | 0.907140142 | -4.996481103 |
| **A_55_P2048221** | NA | 0.068848505 | -0.229023898 | 0.366720908 | 15.06015737 | 0.568634767 | 0.590680627 | 0.907140142 | -4.996498575 |
| **A_55_P2383897** | 8030453O22Rik | -0.072940191 | -0.388566212 | 0.24268583 | 8.449498289 | -0.568542926 | 0.59073905 | 0.907140421 | -4.99653528 |
| **A_55_P2036813** | H2bu2 | 0.03377918 | -0.112414075 | 0.179972436 | 7.956926937 | 0.568448252 | 0.590799278 | 0.907140421 | -4.996573111 |
| **A_55_P1980432** | Zfp488 | -0.059770026 | -0.318465292 | 0.19892524 | 9.739905527 | -0.568413161 | 0.590821603 | 0.907140421 | -4.996587132 |
| **A_52_P357402** | Mlst8 | 0.059213534 | -0.197127794 | 0.315554862 | 10.27783561 | 0.568291961 | 0.590898714 | 0.907186754 | -4.996635553 |
| **A_55_P2184128** | Ndufa3 | -0.060540148 | -0.322911565 | 0.201831268 | 14.50303856 | -0.567670233 | 0.591294367 | 0.907516799 | -4.996883802 |
| **A_55_P2010957** | Gm4971 | -0.063028253 | -0.336185076 | 0.21012857 | 13.13142968 | -0.567665339 | 0.591297482 | 0.907516799 | -4.996885755 |
| **A_55_P2131473** | Rmnd5a | -0.03323844 | -0.177303565 | 0.110826685 | 8.439348287 | -0.567611174 | 0.591331959 | 0.907516799 | -4.996907372 |
| **A_55_P1972039** | Nox4 | -0.13642581 | -0.7278391 | 0.45498748 | 8.519658159 | -0.567511445 | 0.591395441 | 0.907516799 | -4.996947167 |
| **A_51_P500718** | Dck | 0.043590818 | -0.145411044 | 0.232592679 | 7.951739288 | 0.56741145 | 0.591459096 | 0.907516799 | -4.996987063 |
| **A_55_P2132014** | Tekt2 | 0.040724757 | -0.135859276 | 0.217308791 | 8.027177699 | 0.567382892 | 0.591477276 | 0.907516799 | -4.996998455 |
| **A_55_P2002689** | Hmgb1 | -0.028969422 | -0.154586944 | 0.0966481 | 11.4598708 | -0.567360081 | 0.591491798 | 0.907516799 | -4.997007555 |
| **A_55_P2081005** | Cox7c | -0.030457673 | -0.162563688 | 0.101648342 | 8.02799255 | -0.567209236 | 0.591587834 | 0.907516799 | -4.997067722 |

| | | | | | | | | | |
|---|---|---|---|---|---|---|---|---|---|
| **A_55_P2031871** | Plekhh3 | 0.072446901 | -0.241794915 | 0.386688717 | 10.6483589 | 0.567185344 | 0.591603046 | 0.907516799 | -4.997077251 |
| **A_52_P585448** | Ptp4a2 | 0.075865831 | -0.253213959 | 0.404945622 | 11.24320564 | 0.567171203 | 0.591612049 | 0.907516799 | -4.99708289 |
| **A_52_P542970** | NA | -0.08054753 | -0.429964113 | 0.268869053 | 10.56863486 | -0.567123805 | 0.591642228 | 0.907516799 | -4.997101792 |
| **A_55_P1997747** | Cox16 | 0.076852861 | -0.256568385 | 0.410274108 | 11.76393273 | 0.567069035 | 0.591677102 | 0.907516799 | -4.997123632 |
| **A_55_P2146127** | Chmp4b | -0.026279025 | -0.140342717 | 0.087784667 | 12.65471076 | -0.566801499 | 0.591847468 | 0.907576784 | -4.997230287 |
| **A_52_P141136** | Fggy | 0.040310535 | -0.134662064 | 0.215283133 | 8.19373436 | 0.566784131 | 0.591858528 | 0.907576784 | -4.99723721 |
| **A_55_P2001198** | NA | 0.046735405 | -0.156153972 | 0.249624782 | 8.832047817 | 0.56670346 | 0.591909906 | 0.907576784 | -4.997269361 |
| **A_51_P338317** | Rhoq | -0.133834069 | -0.714878885 | 0.447210746 | 10.25190008 | -0.566664774 | 0.591934545 | 0.907576784 | -4.997284777 |
| **A_55_P1975714** | Tor1aip1 | -0.041015348 | -0.21912729 | 0.137096594 | 11.47911536 | -0.566529491 | 0.59202071 | 0.907576784 | -4.997338681 |
| **A_51_P253207** | NA | -0.203944712 | -1.090002005 | 0.682112582 | 9.613810393 | -0.566265114 | 0.592189121 | 0.907576784 | -4.997443991 |
| **A_51_P257675** | Tspyl4 | -0.02967761 | -0.158615438 | 0.099260218 | 8.094898749 | -0.566262417 | 0.592190839 | 0.907576784 | -4.997445065 |
| **A_55_P2021476** | NA | 0.069286049 | -0.231739954 | 0.370312052 | 13.58682245 | 0.566253566 | 0.592196479 | 0.907576784 | -4.99744859 |
| **A_55_P2104835** | NA | -0.063843751 | -0.341247148 | 0.213559645 | 8.36566854 | -0.566207718 | 0.592225687 | 0.907576784 | -4.997466848 |
| **A_52_P249856** | Cc2d2a | -0.025173674 | -0.13456365 | 0.084216302 | 7.769439898 | -0.566158771 | 0.592256871 | 0.907576784 | -4.997486339 |
| **A_51_P222657** | Tspo | -0.190196674 | -1.016723916 | 0.636330567 | 10.49614059 | -0.566128359 | 0.592276247 | 0.907576784 | -4.997498449 |
| **A_51_P461005** | Nrk | -0.095526642 | -0.510723285 | 0.319670001 | 13.12493541 | -0.56603051 | 0.592338591 | 0.907576784 | -4.997537407 |
| **A_55_P2000077** | Nrf1 | -0.04503453 | -0.240807047 | 0.150737988 | 9.565137915 | -0.56593046 | 0.59240234 | 0.907576784 | -4.997577235 |
| **A_55_P2017851** | NA | 0.029310668 | -0.098117601 | 0.156738938 | 11.6002342 | 0.565886194 | 0.592430546 | 0.907576784 | -4.997594854 |
| **A_55_P2042723** | Gm2769 | -0.038670495 | -0.206791721 | 0.129450732 | 8.257986366 | -0.5658826 | 0.592432837 | 0.907576784 | -4.997596285 |
| **A_51_P333923** | Tspan1 | 0.141402688 | -0.473409132 | 0.756214507 | 11.88271288 | 0.565828239 | 0.592467477 | 0.907576784 | -4.997617921 |
| **A_52_P97699** | D430019H16Rik | -0.080882771 | -0.432616976 | 0.270851433 | 8.569787261 | -0.565731788 | 0.592528941 | 0.90759901 | -4.997656304 |
| **A_55_P1985422** | NA | -0.333835258 | -1.785963131 | 1.118292615 | 13.85884251 | -0.565583205 | 0.592623634 | 0.907662941 | -4.997715424 |
| **A_55_P2091676** | Fas | 0.102735617 | -0.344207706 | 0.54967894 | 10.08197888 | 0.565506684 | 0.592672405 | 0.907662941 | -4.997745865 |
| **A_52_P60353** | Greb1l | 0.101446277 | -0.339974652 | 0.542867205 | 8.261546678 | 0.56539551 | 0.592743266 | 0.907662941 | -4.997790086 |
| **A_55_P2038519** | NA | -0.03307829 | -0.177024535 | 0.110867954 | 8.659556622 | -0.565342819 | 0.592776852 | 0.907662941 | -4.997811042 |
| **A_55_P2008735** | Cdc123 | 0.040390776 | -0.135393192 | 0.216174743 | 8.833995483 | 0.565291031 | 0.592809865 | 0.907662941 | -4.997831637 |
| **A_55_P2172924** | NA | -0.030233748 | -0.161848902 | 0.101381406 | 8.009083429 | -0.565138967 | 0.592906802 | 0.907662941 | -4.9978921 |
| **A_55_P2009022** | NA | -0.036590821 | -0.19591578 | 0.122734139 | 8.779286605 | -0.565011773 | 0.592987894 | 0.907662941 | -4.997942664 |
| **A_55_P2145833** | Dmpk | 0.104740606 | -0.351350004 | 0.560831215 | 8.893614771 | 0.564980051 | 0.593008119 | 0.907662941 | -4.997955274 |
| **A_51_P118223** | Gm1943 | -0.038642837 | -0.20694862 | 0.129662945 | 8.55640752 | -0.564857803 | 0.593086064 | 0.907662941 | -4.978562774 |
| **A_51_P472726** | Pdlim2 | 0.084545994 | -0.283719698 | 0.452811686 | 13.34025484 | 0.564808123 | 0.593117742 | 0.907662941 | -4.998023602 |
| **A_51_P400752** | NA | -0.197608844 | -1.058372198 | 0.663154511 | 12.23692502 | -0.564796206 | 0.593125341 | 0.907662941 | -4.998028338 |
| **A_55_P1960291** | Tjp3 | 0.139086039 | -0.466773001 | 0.744945079 | 9.887753809 | 0.564782341 | 0.593134182 | 0.907662941 | -4.998033847 |
| **A_51_P245454** | Ufsp2 | 0.038701259 | -0.129944894 | 0.207347413 | 11.44374319 | 0.56457003 | 0.593269573 | 0.907798256 | -4.998118197 |
| **A_51_P247694** | Adgrg3 | -0.087350045 | -0.468072734 | 0.293372645 | 11.42632167 | -0.564447476 | 0.593347734 | 0.907845986 | -4.998166874 |
| **A_55_P2109429** | Rfng | -0.052474536 | -0.281254943 | 0.17630587 | 11.34473188 | -0.564285571 | 0.593451 | 0.90785536 | -4.998231168 |
| **A_55_P2128303** | Ids | 0.024191497 | -0.081282233 | 0.129665227 | 8.716146673 | 0.564270831 | 0.593460402 | 0.90785536 | -4.99823702 |
| **A_65_P02177** | Gstm4 | -0.037001439 | -0.198346883 | 0.124344004 | 7.982980907 | -0.564197396 | 0.593507245 | 0.90785536 | -4.998266176 |
| **A_55_P2129219** | NA | -0.054423119 | -0.291758854 | 0.182912616 | 11.61941559 | -0.564143328 | 0.593541735 | 0.90785536 | -4.99828764 |
| **A_55_P2141878** | Ldha | -0.055741645 | -0.298961359 | 0.187478069 | 11.00803092 | -0.563832585 | 0.593739981 | 0.908051595 | -4.998410967 |
| **A_55_P2018457** | Hdac7 | 0.046713028 | -0.157125368 | 0.250551424 | 10.25666653 | 0.56379496 | 0.593763987 | 0.908051595 | -4.998425896 |
| **A_51_P505521** | NA | -0.045487939 | -0.244039652 | 0.153063775 | 11.08608841 | -0.56362699 | 0.593871166 | 0.908127347 | -4.998492531 |
| **A_51_P507509** | Gimap1 | -0.063524726 | -0.340859264 | 0.213809812 | 8.658047075 | -0.563518274 | 0.593940542 | 0.908127347 | -4.99853565 |
| **A_55_P2009360** | Psg-ps1 | -0.044492551 | -0.238747379 | 0.149762277 | 16.69536863 | -0.563487966 | 0.593959884 | 0.908127347 | -4.99854767 |
| **A_55_P2115245** | Zfyve19 | 0.07324313 | -0.246573923 | 0.393060184 | 11.90187181 | 0.563422833 | 0.594001451 | 0.908127347 | -4.998573499 |
| **A_55_P2171993** | Mrgbp | 0.030269401 | -0.101927082 | 0.162465884 | 8.339900554 | 0.563317298 | 0.594068806 | 0.908140239 | -4.998615344 |
| **A_55_P2247633** | Gpatch2 | -0.057274022 | -0.307433 | 0.192884955 | 8.874634997 | -0.563262391 | 0.59410385 | 0.908140239 | -4.998637112 |
| **A_51_P288103** | Psmd6 | -0.042412004 | -0.227719501 | 0.142895494 | 12.14842106 | -0.563073336 | 0.594224524 | 0.908252873 | -4.99871205 |
| **A_55_P2075359** | Eif4a3l1 | 0.042826049 | -0.144348085 | 0.230000182 | 11.96790185 | 0.562900115 | 0.594335104 | 0.908264742 | -4.998780693 |
| **A_51_P172688** | Stard3nl | -0.064851222 | -0.348358783 | 0.21865634 | 10.37800864 | -0.562759296 | 0.594425008 | 0.908264742 | -4.998836482 |

| | | | | | | | | | |
|---|---|---|---|---|---|---|---|---|---|
| **A_55_P2091631** | Coa4 | 0.04414674 | -0.148875437 | 0.237168917 | 10.26643264 | 0.562678851 | 0.594476371 | 0.908264742 | -4.998868347 |
| **A_55_P2046709** | NA | 0.391708617 | -1.321007745 | 2.104424979 | 13.24357847 | 0.562661036 | 0.594487745 | 0.908264742 | -4.998875403 |
| **A_51_P393897** | Mpi | -0.05398785 | -0.290061586 | 0.182085885 | 8.970227222 | -0.562623058 | 0.594511995 | 0.908264742 | -4.998890445 |
| **A_52_P80965** | Sephs1 | -0.025224242 | -0.135523465 | 0.085074982 | 7.801165975 | -0.562619561 | 0.594514228 | 0.908264742 | -4.99889183 |
| **A_55_P2080758** | Slc8a1 | 0.023810845 | -0.08034362 | 0.127965311 | 7.727775988 | 0.562426901 | 0.594637253 | 0.908316504 | -4.998968121 |
| **A_51_P322542** | Bbx | 0.06880029 | -0.232182874 | 0.369783453 | 11.97340565 | 0.562363632 | 0.594677657 | 0.908316504 | -4.99899317 |
| **A_51_P443569** | Pwwp3a | -0.043037707 | -0.231350593 | 0.145275179 | 9.327838116 | -0.562261371 | 0.594742965 | 0.908316504 | -4.999033652 |
| **A_55_P1985070** | Robo1 | 0.061230302 | -0.206698231 | 0.329158835 | 9.235368447 | 0.56223334 | 0.594760868 | 0.908316504 | -4.999044747 |
| **A_55_P2024763** | Lama3 | 0.03108552 | -0.104949659 | 0.167120698 | 8.48575518 | 0.562180874 | 0.594794377 | 0.908316504 | -4.999065513 |
| **A_52_P58283** | Kcna5 | -0.02381917 | -0.128070257 | 0.080431916 | 7.643140222 | -0.562102089 | 0.594844698 | 0.908316504 | -4.999096693 |
| **A_55_P2105457** | Zfp7 | -0.036361952 | -0.19555241 | 0.122828505 | 8.258960023 | -0.561952134 | 0.594940482 | 0.908316504 | -4.999156028 |
| **A_55_P2117630** | Snrnp25 | 0.124212592 | -0.419583232 | 0.668008416 | 10.67070743 | 0.561951609 | 0.594940818 | 0.908316504 | -4.999156236 |
| **A_55_P2141786** | Pard6g | 0.06184604 | -0.208972675 | 0.332664756 | 13.04993762 | 0.561826719 | 0.595020599 | 0.908316504 | -4.999205644 |
| **A_55_P2102574** | Prorsd1 | 0.044673793 | -0.150976741 | 0.240324327 | 9.785101294 | 0.56174725 | 0.595071368 | 0.908316504 | -4.999237077 |
| **A_51_P227108** | Bri3bp | -0.042658428 | -0.229508532 | 0.144191676 | 12.06272039 | -0.561669278 | 0.595121183 | 0.908316504 | -4.999267915 |
| **A_52_P19532** | Fgd3 | 0.052467628 | -0.177358177 | 0.282293433 | 8.415776859 | 0.561644881 | 0.59513677 | 0.908316504 | -4.999277563 |
| **A_51_P490509** | Bub1b | -0.025940708 | -0.139576752 | 0.087695335 | 7.874114572 | -0.561610071 | 0.595159011 | 0.908316504 | -4.999291329 |
| **A_55_P2369474** | Nr1h4 | 0.142829056 | -0.483000225 | 0.768658337 | 8.581302625 | 0.561474258 | 0.595245789 | 0.908377219 | -4.999345029 |
| **A_52_P234729** | Pkd2 | -0.10730564 | -0.577608513 | 0.362997233 | 10.58650563 | -0.561324607 | 0.595341417 | 0.908378152 | -4.999404187 |
| **A_55_P2067842** | 5-Mar | -0.043585457 | -0.234626267 | 0.147455354 | 10.09190841 | -0.56128652 | 0.595365757 | 0.908378152 | -4.999419241 |
| **A_55_P1953758** | Serinc2 | -0.132132082 | -0.711319366 | 0.447055202 | 10.93148181 | -0.561252672 | 0.595387387 | 0.908378152 | -4.999432619 |
| **A_55_P2301312** | Wdr78 | 0.043490897 | -0.147208598 | 0.234190392 | 8.623407407 | 0.561071212 | 0.595503359 | 0.908441543 | -4.999504326 |
| **A_55_P2169198** | NA | -0.030761198 | -0.165681012 | 0.104158616 | 7.984347558 | -0.560914512 | 0.595603517 | 0.908441543 | -4.999566232 |
| **A_55_P1982384** | NA | -0.045707533 | -0.246219574 | 0.154804509 | 12.40698126 | -0.560810948 | 0.595669716 | 0.908441543 | -4.999607138 |
| **A_51_P491350** | Col4a2 | -0.11566827 | -0.623111177 | 0.391774637 | 13.50931377 | -0.560784777 | 0.595686446 | 0.908441543 | -4.999617475 |
| **A_55_P1958137** | NA | -0.044503625 | -0.239747349 | 0.1507401 | 8.316331421 | -0.560773475 | 0.595693671 | 0.908441543 | -4.999621938 |
| **A_55_P2098802** | Uqcrq | -0.06885441 | -0.370943033 | 0.233234213 | 14.39157867 | -0.560746478 | 0.59571093 | 0.908441543 | -4.9996326 |
| **A_55_P2055682** | Atp5o | 0.072124194 | -0.244390549 | 0.388638937 | 12.40870256 | 0.560603999 | 0.595802016 | 0.908508769 | -4.999688861 |
| **A_55_P2116664** | Hook2 | 0.163515909 | -0.554381098 | 0.881412916 | 11.24947217 | 0.56035988 | 0.595958099 | 0.90861248 | -4.999785228 |
| **A_51_P283708** | Msh2 | -0.046067592 | -0.24832519 | 0.156190007 | 10.93292462 | -0.560350581 | 0.595964045 | 0.90861248 | -4.999788898 |
| **A_51_P486001** | Mob3a | -0.032416896 | -0.1748236 | 0.109989808 | 9.152453791 | -0.560028541 | 0.596169986 | 0.908854772 | -4.999915967 |
| **A_55_P1996837** | Baz2b | -0.035492369 | -0.191455743 | 0.120471005 | 9.844236521 | -0.559862697 | 0.596276057 | 0.908887282 | -4.999981381 |
| **A_55_P2184541** | Cdadc1 | 0.063899539 | -0.216966473 | 0.344765552 | 10.97262452 | 0.559715976 | 0.596369907 | 0.908887282 | -5.000039238 |
| **A_55_P2067727** | Mxra7 | -0.079246151 | -0.42757691 | 0.269084609 | 8.817860523 | -0.559700264 | 0.596379958 | 0.908887282 | -5.000045433 |
| **A_66_P105074** | Enpp5 | -0.069142479 | -0.373162364 | 0.234877406 | 8.90556728 | -0.559515493 | 0.596498159 | 0.908887282 | -5.000118274 |
| **A_51_P503937** | Calm1 | -0.032799268 | -0.177026993 | 0.111428457 | 10.83708966 | -0.559479993 | 0.596520871 | 0.908887282 | -5.000132266 |
| **A_51_P436727** | Arrb1 | 0.053986096 | -0.183454369 | 0.291426562 | 11.13653497 | 0.559366372 | 0.596593565 | 0.908887282 | -5.000177046 |
| **A_55_P2021029** | Tigd3 | 0.02358097 | -0.080142462 | 0.127304403 | 7.777566998 | 0.55931177 | 0.596628501 | 0.908887282 | -5.000198562 |
| **A_51_P336403** | Emc7 | 0.080795619 | -0.274626983 | 0.436218221 | 13.5049473 | 0.559257648 | 0.59666313 | 0.908887282 | -5.000219888 |
| **A_55_P2287910** | Rragd | -0.031105041 | -0.167951936 | 0.105741854 | 8.096260326 | -0.559197203 | 0.596701807 | 0.908887282 | -5.000243702 |
| **A_55_P1971828** | Csnk2a1 | 0.027147738 | -0.092296022 | 0.146591497 | 8.683191755 | 0.559164177 | 0.596722941 | 0.908887282 | -5.000256714 |
| **A_55_P2125786** | Lrrc8e | 0.087225013 | -0.296724434 | 0.471174459 | 8.25146731 | 0.558902637 | 0.596890313 | 0.908887282 | -5.000359728 |
| **A_55_P2330545** | NA | 0.026247488 | -0.089291875 | 0.141786852 | 8.060967668 | 0.558890776 | 0.596897904 | 0.908887282 | -5.000364398 |
| **A_55_P1969987** | Fam204a | -0.044394126 | -0.239825564 | 0.151037312 | 8.515313932 | -0.558856424 | 0.59691989 | 0.908887282 | -5.000377926 |
| **A_52_P37894** | Cox7a2 | 0.055743584 | -0.189666656 | 0.301153823 | 13.95669812 | 0.558819264 | 0.596943673 | 0.908887282 | -5.000392557 |
| **A_55_P2126391** | Srl | -0.024575817 | -0.132781714 | 0.083630081 | 7.781451765 | -0.558761134 | 0.596980879 | 0.908887282 | -5.000415445 |
| **A_55_P2212908** | Pon2 | -0.030558135 | -0.165106364 | 0.103990094 | 7.886799011 | -0.558750626 | 0.596987605 | 0.908887282 | -5.000419582 |
| **A_55_P2049592** | NA | 0.028683714 | -0.097613985 | 0.154981413 | 8.499148068 | 0.55873916 | 0.596994943 | 0.908887282 | -5.000424096 |
| **A_55_P2046703** | Atp5g3 | -0.046339797 | -0.250509885 | 0.157830291 | 15.47369874 | -0.558381697 | 0.597223769 | 0.908887282 | -5.000564789 |
| **A_55_P2094896** | Phyhd1 | -0.082111282 | -0.443947668 | 0.279725104 | 10.68549944 | -0.558289885 | 0.597282549 | 0.908887282 | -5.000600912 |

| | | | | | | | | |
|---|---|---|---|---|---|---|---|---|
| **A_52_P176737** | Lcorl | -0.025481272 | -0.137772248 | 0.086809703 | 7.884733509 | -0.558271445 | 0.597294355 | 0.908887282 | -5.000608167 |
| **A_51_P201773** | Tubgcp3 | -0.053732382 | -0.2905305 | 0.183065736 | 10.25492404 | -0.558247788 | 0.597309502 | 0.908887282 | -5.000617474 |
| **A_51_P216075** | Ddost | -0.057990783 | -0.313557481 | 0.197575915 | 13.68610062 | -0.558243759 | 0.597312081 | 0.908887282 | -5.000619059 |
| **A_55_P2328787** | NA | -0.108179462 | -0.585128461 | 0.368769537 | 8.499739684 | -0.558010071 | 0.597461714 | 0.908887282 | -5.000710974 |
| **A_55_P1968188** | Btf3 | 0.054750255 | -0.186650483 | 0.296150994 | 13.97181379 | 0.557977537 | 0.597482547 | 0.908887282 | -5.000723768 |
| **A_51_P479043** | Nif3l1 | -0.026653459 | -0.14420323 | 0.090896313 | 11.06083398 | -0.557828814 | 0.597577789 | 0.908887282 | -5.000782244 |
| **A_55_P2113026** | Foxo4 | 0.037041093 | -0.126343988 | 0.200426174 | 9.165576606 | 0.557751261 | 0.597627458 | 0.908887282 | -5.000812731 |
| **A_65_P01418** | Wnk2 | 0.105896406 | -0.361284228 | 0.573077041 | 9.721197398 | 0.55765494 | 0.597689149 | 0.908887282 | -5.000850592 |
| **A_55_P2066440** | Myh3 | 0.023970464 | -0.081787093 | 0.129728021 | 7.699658168 | 0.557614673 | 0.59771494 | 0.908887282 | -5.000866418 |
| **A_55_P1989698** | Mmp11 | -0.131811592 | -0.713390482 | 0.449767298 | 10.99482886 | -0.557588918 | 0.597731437 | 0.908887282 | -5.00087654 |
| **A_55_P2011286** | Hopx | -0.121560116 | -0.657911648 | 0.414791417 | 10.24846191 | -0.557584563 | 0.597734226 | 0.908887282 | -5.000878251 |
| **A_52_P294253** | Zfp41 | 0.047204079 | -0.161121935 | 0.255530093 | 8.476241288 | 0.557449058 | 0.597821024 | 0.908887282 | -5.000931498 |
| **A_55_P1999838** | Cox7c | -0.059576215 | -0.322506035 | 0.203353605 | 14.02540191 | -0.557445259 | 0.597823458 | 0.908887282 | -5.000932991 |
| **A_52_P471143** | Pusl1 | 0.025204268 | -0.086033787 | 0.136442323 | 8.696685111 | 0.557429385 | 0.597833626 | 0.908887282 | -5.000939228 |
| **A_51_P234140** | Cdkl1 | -0.031243702 | -0.16917786 | 0.106690456 | 8.164089476 | -0.557262504 | 0.597940533 | 0.908887282 | -5.001004787 |
| **A_55_P2004230** | Emc6 | -0.046463503 | -0.251653036 | 0.158726029 | 11.19647091 | -0.557090707 | 0.598050601 | 0.908887282 | -5.00107226 |
| **A_51_P386058** | Ppp2r5b | -0.085000178 | -0.460377093 | 0.290376737 | 10.5454118 | -0.557084967 | 0.598054279 | 0.908887282 | -5.001074514 |
| **A_52_P50478** | Rnf123 | -0.024928733 | -0.135022696 | 0.085165229 | 8.195074618 | -0.557065002 | 0.598067071 | 0.908887282 | -5.001082354 |
| **A_66_P124011** | Spint2 | -0.059180356 | -0.320542376 | 0.202181664 | 11.98059572 | -0.557062927 | 0.5980684 | 0.908887282 | -5.001083168 |
| **A_55_P1981322** | Grhl2 | -0.039160577 | -0.212116265 | 0.133795111 | 10.98328861 | -0.557036171 | 0.598085544 | 0.908887282 | -5.001093675 |
| **A_52_P139650** | Ska1 | 0.032905542 | -0.112425427 | 0.178236511 | 7.790819455 | 0.55703187 | 0.5980883 | 0.908887282 | -5.001095363 |
| **A_55_P2101473** | NA | 0.050890758 | -0.173893374 | 0.27567489 | 14.83974864 | 0.556983628 | 0.598119211 | 0.908887282 | -5.001114305 |
| **A_55_P2106063** | NA | 0.033151769 | -0.113302839 | 0.179606377 | 11.93461139 | 0.556894372 | 0.598176405 | 0.908902737 | -5.001149347 |
| **A_55_P2056325** | Anxa3 | -0.083475019 | -0.452335261 | 0.285385223 | 10.53654747 | -0.556754635 | 0.598265952 | 0.908967347 | -5.001204199 |
| **A_55_P2239317** | Afap1l1 | -0.058055305 | -0.314666179 | 0.198555569 | 10.93810043 | -0.556590792 | 0.598370956 | 0.909002976 | -5.001268497 |
| **A_51_P450278** | 2010003K11Rik | 0.208798783 | -0.714148344 | 1.131745911 | 9.349081129 | 0.556570655 | 0.598383863 | 0.909002976 | -5.001276399 |
| **A_55_P2048434** | Hsp90ab1 | -0.046206638 | -0.250495758 | 0.158082483 | 16.13765491 | -0.556452747 | 0.598459436 | 0.909002976 | -5.001322659 |
| **A_66_P117695** | Zpr1 | 0.050426365 | -0.172551911 | 0.273404642 | 11.93922111 | 0.556370732 | 0.598512007 | 0.909002976 | -5.001354832 |
| **A_55_P2325378** | NA | -0.065510806 | -0.355211637 | 0.224190025 | 9.300806495 | -0.55632985 | 0.598538213 | 0.909002976 | -5.001370867 |
| **A_55_P1968129** | Eloc | 0.076403825 | -0.261499546 | 0.414307195 | 9.951762301 | 0.556277813 | 0.59857157 | 0.909002976 | -5.001391277 |
| **A_51_P381558** | Rasa4 | -0.049472889 | -0.268441679 | 0.169495902 | 8.331647658 | -0.555845652 | 0.598848641 | 0.90931321 | -5.001560714 |
| **A_55_P2135220** | NA | 0.025000252 | -0.085676323 | 0.135676826 | 7.763995931 | 0.555722314 | 0.598927729 | 0.90931321 | -5.001609049 |
| **A_55_P2112622** | Smad4 | -0.033876623 | -0.183857688 | 0.116104442 | 12.917708 | -0.555690337 | 0.598948236 | 0.90931321 | -5.00162158 |
| **A_51_P441942** | Tial1 | -0.049313587 | -0.267691738 | 0.169064564 | 8.961218585 | -0.555554371 | 0.599035431 | 0.90931321 | -5.001674851 |
| **A_55_P2107357** | Gm15421 | 0.090983101 | -0.311934124 | 0.493900327 | 14.58727222 | 0.555538376 | 0.599045689 | 0.90931321 | -5.001681117 |
| **A_55_P2119969** | Bdh2 | -0.039246611 | -0.213055671 | 0.134562449 | 9.121678806 | -0.555518992 | 0.599058121 | 0.90931321 | -5.001688711 |
| **A_55_P2042229** | NA | 0.105074935 | -0.360421126 | 0.570570996 | 12.15797905 | 0.555331469 | 0.599178394 | 0.909424357 | -5.00176216 |
| **A_55_P2074406** | Ldoc1 | 0.141492412 | -0.485447124 | 0.768431948 | 11.32332329 | 0.555234769 | 0.599240421 | 0.909447087 | -5.001800027 |
| **A_55_P2129920** | Chd7 | -0.092602875 | -0.503064484 | 0.317858734 | 12.31412124 | -0.555035921 | 0.599367981 | 0.909569263 | -5.001877877 |
| **A_55_P1980321** | Pcbp3 | 0.059731197 | -0.205139317 | 0.324601711 | 9.826744758 | 0.554800405 | 0.599519084 | 0.909727145 | -5.001970051 |
| **A_55_P2074736** | Prkar1b | -0.050677741 | -0.275498889 | 0.174143407 | 9.007077707 | -0.554560906 | 0.599672764 | 0.909888913 | -5.00206375 |
| **A_51_P518586** | Emg1 | -0.034606981 | -0.188169741 | 0.118955779 | 9.175345634 | -0.554430314 | 0.599756571 | 0.909944644 | -5.002114826 |
| **A_55_P1957624** | Insig2 | -0.080029872 | -0.435414036 | 0.275354292 | 9.724268771 | -0.554017157 | 0.600021759 | 0.910077165 | -5.002276349 |
| **A_55_P2056654** | Kif22 | 0.047895627 | -0.164801342 | 0.260592596 | 8.443612281 | 0.553992296 | 0.600037719 | 0.910077165 | -5.002286065 |
| **A_52_P202142** | Sv2a | 0.025398068 | -0.087430886 | 0.138227021 | 7.63328271 | 0.553795316 | 0.600164177 | 0.910077165 | -5.002363034 |
| **A_52_P354298** | Ap2a2 | -0.052562504 | -0.286085225 | 0.180960216 | 10.12682814 | -0.553752975 | 0.600191362 | 0.910077165 | -5.002379576 |
| **A_52_P149391** | Usp40 | -0.025632409 | -0.139521792 | 0.088256974 | 8.844106356 | -0.553701049 | 0.600224701 | 0.910077165 | -5.002399861 |
| **A_51_P401668** | Laptm5 | -0.035286627 | -0.192089403 | 0.12151615 | 8.044753207 | -0.553637573 | 0.600265457 | 0.910077165 | -5.002424655 |
| **A_55_P2118491** | Gm10046 | -0.027946208 | -0.152137558 | 0.096245142 | 8.088097099 | -0.553605886 | 0.600285803 | 0.910077165 | -5.002437032 |
| **A_52_P670874** | Ebag9 | 0.062775502 | -0.216304791 | 0.341855795 | 9.827775616 | 0.55338857 | 0.600425351 | 0.910077165 | -5.002521895 |

| | | | | | | | | |
|---|---|---|---|---|---|---|---|---|
| A_51_P125368 | Hars | 0.052795353 | -0.181994357 | 0.287585062 | 9.607748608 | 0.553204623 | 0.600543486 | 0.910077165 | -5.002593704 |
| A_55_P2183735 | Pisd | -0.085500093 | -0.508086985 | 0.337086798 | 12.23474814 | -0.569315591 | 0.600559128 | 0.910077165 | -4.887563368 |
| A_55_P2109752 | Nans | 0.06906068 | -0.238081272 | 0.376202632 | 12.44579799 | 0.553172874 | 0.600563877 | 0.910077165 | -5.002606097 |
| A_51_P294020 | Ticam1 | -0.032460018 | -0.176845812 | 0.111925777 | 9.164101621 | -0.553086994 | 0.600619037 | 0.910077165 | -5.002639614 |
| A_55_P2179726 | Mettl18 | -0.041259862 | -0.224804694 | 0.142284969 | 9.10392206 | -0.55303775 | 0.600650667 | 0.910077165 | -5.002658831 |
| A_55_P2038489 | NA | 0.034275455 | -0.118199496 | 0.186750406 | 8.220273195 | 0.553036623 | 0.600651391 | 0.910077165 | -5.002659271 |
| A_55_P2098335 | Map2k4 | -0.035455434 | -0.193196204 | 0.122285336 | 8.704147744 | -0.552978205 | 0.600688915 | 0.910077165 | -5.002682066 |
| A_55_P2057142 | Cks1brt | -0.029529444 | -0.160908277 | 0.101849388 | 8.184268168 | -0.552966778 | 0.600696255 | 0.910077165 | -5.002686524 |
| A_52_P298088 | Itgb1bp2 | 0.023823355 | -0.082185573 | 0.129832283 | 7.63375459 | 0.552878429 | 0.600753008 | 0.910077165 | -5.002720994 |
| A_51_P388801 | Tbcc | -0.051048922 | -0.278208156 | 0.176110312 | 12.19201442 | -0.552872951 | 0.600756527 | 0.910077165 | -5.002723131 |
| A_55_P2017944 | Etfb | 0.083091929 | -0.286670808 | 0.452854666 | 14.31005998 | 0.552846878 | 0.600773276 | 0.910077165 | -5.002733303 |
| A_51_P384148 | Ryk | -0.053602282 | -0.292172684 | 0.18496812 | 11.06345401 | -0.552759073 | 0.600829684 | 0.910077165 | -5.002767554 |
| A_66_P101755 | Sp1 | 0.062188536 | -0.214638912 | 0.339015984 | 10.70793805 | 0.552675672 | 0.600883265 | 0.910077165 | -5.002800083 |
| A_55_P2125049 | Gm11127 | -0.186983804 | -1.019351063 | 0.645383455 | 13.00833384 | -0.552660168 | 0.600893226 | 0.910077165 | -5.002806129 |
| A_51_P133582 | Cbx4 | 0.084618131 | -0.292205873 | 0.461442135 | 11.37634567 | 0.552451339 | 0.601027402 | 0.910077165 | -5.002887558 |
| A_52_P370203 | Jarid2 | 0.059886353 | -0.206850994 | 0.326623699 | 10.66767094 | 0.552348526 | 0.601093467 | 0.910077165 | -5.002927638 |
| A_55_P2006250 | Myo5a | -0.150243238 | -0.819440067 | 0.518953591 | 9.711169323 | -0.552344742 | 0.601095899 | 0.910077165 | -5.002929113 |
| A_55_P2180201 | Mrpl10 | -0.031255412 | -0.170473989 | 0.107963165 | 8.976553415 | -0.552328177 | 0.601106543 | 0.910077165 | -5.00293557 |
| A_55_P2128606 | Nr1h5 | 0.024309433 | -0.083972838 | 0.132591704 | 7.689287158 | 0.552314749 | 0.601115172 | 0.910077165 | -5.002940804 |
| A_55_P2142580 | NA | 0.060299541 | -0.208331787 | 0.328930868 | 10.66449747 | 0.552238282 | 0.601164312 | 0.910080278 | -5.002970608 |
| A_51_P109144 | Grtp1 | -0.094627476 | -0.516294419 | 0.327039467 | 10.56229761 | -0.552098877 | 0.601253903 | 0.910144623 | -4.993173434 |
| A_55_P2140449 | B3galt4 | -0.033086231 | -0.180613302 | 0.11444084 | 8.947155657 | -0.551753045 | 0.601476191 | 0.910195066 | -5.003159651 |
| A_55_P2261882 | D230040J21Rik | -0.045823937 | -0.250147836 | 0.158499961 | 7.866536554 | -0.551750073 | 0.601478102 | 0.910195066 | -5.003160809 |
| A_55_P1982156 | Nelfcd | -0.050465904 | -0.275559167 | 0.17462736 | 13.759245 | -0.551575186 | 0.601590531 | 0.910195066 | -5.003228907 |
| A_52_P354823 | Irf8 | -0.047149757 | -0.257457958 | 0.163158443 | 9.878877961 | -0.551559562 | 0.601600576 | 0.910195066 | -5.00323499 |
| A_51_P126835 | Arl14ep | -0.036166184 | -0.197483192 | 0.125150825 | 9.409892054 | -0.551558634 | 0.601601173 | 0.910195066 | -5.003235351 |
| A_55_P1999823 | Zmynd19 | -0.033439401 | -0.182631247 | 0.115752445 | 8.133213344 | -0.551420069 | 0.601690261 | 0.910195066 | -5.003289292 |
| A_55_P1960083 | NA | 0.060956146 | -0.211012249 | 0.332924541 | 13.83909512 | 0.55140186 | 0.601701969 | 0.910195066 | -5.003296379 |
| A_55_P2171484 | Lta4h | -0.050002481 | -0.273129709 | 0.173124748 | 11.08327254 | -0.551325579 | 0.601751017 | 0.910195066 | -5.003326068 |
| A_52_P614960 | Skiv2l | -0.039294584 | -0.214669287 | 0.13608012 | 8.737537574 | -0.551232613 | 0.601810796 | 0.910195066 | -5.003362246 |
| A_55_P1991214 | Lair1 | 0.02806026 | -0.097205209 | 0.153325728 | 8.339490547 | 0.551098823 | 0.601896831 | 0.910195066 | -5.003414301 |
| A_55_P2074291 | Fbxo6 | 0.056043367 | -0.194144831 | 0.306231566 | 13.78840617 | 0.55109512 | 0.601899212 | 0.910195066 | -5.003415742 |
| A_52_P33453 | NA | -0.052791316 | -0.288471201 | 0.18288857 | 11.60201832 | -0.551072996 | 0.60191344 | 0.910195066 | -5.003424349 |
| A_55_P1989215 | Entpd8 | 0.047880438 | -0.165894081 | 0.261654956 | 8.178239633 | 0.551025044 | 0.601944279 | 0.910195066 | -5.003443003 |
| A_55_P2097048 | NA | 0.097428877 | -0.337570085 | 0.532427839 | 11.32407134 | 0.55102162 | 0.601946481 | 0.910195066 | -5.003444335 |
| A_52_P117352 | Gja4 | -0.039626252 | -0.216602861 | 0.137350357 | 10.50496677 | -0.550853729 | 0.60205446 | 0.91027458 | -5.003509634 |
| A_55_P2004153 | NA | 0.067814417 | -0.235149564 | 0.370778398 | 10.33125582 | 0.550681133 | 0.602165478 | 0.91027458 | -5.003576746 |
| A_55_P2069485 | Ptpn13 | -0.073205405 | -0.400282751 | 0.253871942 | 10.64938934 | -0.550632501 | 0.602196761 | 0.91027458 | -5.003595653 |
| A_55_P2078433 | Mcoln2 | 0.250416241 | -0.868565927 | 1.369398409 | 9.568779258 | 0.55056486 | 0.602240274 | 0.91027458 | -5.003621948 |
| A_51_P475938 | Mfsd6l | -0.039158255 | -0.214160187 | 0.135843678 | 7.956157232 | -0.550490267 | 0.602288261 | 0.91027458 | -5.003650942 |
| A_55_P1983903 | Thap4 | -0.045772278 | -0.250359237 | 0.158814681 | 8.38170284 | -0.550419413 | 0.602333844 | 0.91027458 | -5.003678479 |
| A_51_P154534 | Bloc1s5 | 0.046785041 | -0.162343655 | 0.255913737 | 10.32363361 | 0.550379881 | 0.602359278 | 0.91027458 | -5.003693841 |
| A_51_P435968 | Tnfaip8 | -0.061835749 | -0.338270575 | 0.214599077 | 10.81773642 | -0.550320935 | 0.602397203 | 0.91027458 | -5.003716747 |
| A_55_P1981217 | Aatf | 0.06577441 | -0.228289428 | 0.359838247 | 9.042597465 | 0.55028098 | 0.60242291 | 0.91027458 | -5.003732272 |
| A_51_P218653 | Jph2 | 0.024507967 | -0.085082977 | 0.13409891 | 7.691230256 | 0.550176175 | 0.602490345 | 0.910305314 | -5.003772989 |
| A_55_P1953301 | Sord | 0.103546964 | -0.359546763 | 0.566640691 | 11.20736257 | 0.55009492 | 0.60254263 | 0.910313154 | -5.003804553 |
| A_55_P2031242 | Sun1 | 0.032701837 | -0.113578775 | 0.178982449 | 8.592466068 | 0.54998968 | 0.602610353 | 0.910344314 | -5.003845427 |
| A_55_P2065370 | NA | -0.051347509 | -0.281136938 | 0.178441919 | 11.28393331 | -0.549741469 | 0.602770096 | 0.910461275 | -5.003941805 |
| A_52_P160936 | Gusb | -0.101517587 | -0.555847374 | 0.352812199 | 9.28664807 | -0.549717091 | 0.602785786 | 0.910461275 | -5.003951268 |
| A_51_P165082 | Tmem191c | -0.071282117 | -0.39033573 | 0.247771495 | 8.75447263 | -0.549649819 | 0.602829085 | 0.910461275 | -5.003977382 |

| | | | | | | | | | |
|---|---|---|---|---|---|---|---|---|---|
| **A_66_P123824** | Sorl1 | 0.025399328 | -0.088352158 | 0.139150814 | 7.767636253 | 0.549331256 | 0.603034152 | 0.91069983 | -5.004101002 |
| **A_51_P335900** | Cox5a | -0.044174561 | -0.242068083 | 0.153718961 | 15.44856302 | -0.549173824 | 0.60313551 | 0.91078174 | -5.004162072 |
| **A_52_P243412** | Ulk2 | -0.045398815 | -0.24883984 | 0.158042211 | 9.351311075 | -0.549003534 | 0.603245157 | 0.910876153 | -5.004228113 |
| **A_52_P356068** | Proz | 0.036615636 | -0.127514393 | 0.200745665 | 7.768248176 | 0.548842459 | 0.60334888 | 0.910961608 | -5.004290563 |
| **A_55_P2061273** | Tbx6 | -0.044595361 | -0.244533657 | 0.155342936 | 8.101398311 | -0.548735245 | 0.603417926 | 0.910994697 | -5.004332122 |
| **A_51_P295237** | Lrp11 | -0.055081705 | -0.302117206 | 0.191953796 | 8.847448378 | -0.548551201 | 0.603536462 | 0.911084042 | -5.004403447 |
| **A_55_P2169714** | Gcn1 | 0.054504256 | -0.189978301 | 0.298986814 | 12.73442948 | 0.548468513 | 0.603589722 | 0.911084042 | -5.004435485 |
| **A_51_P462192** | Olr1 | 0.111348289 | -0.388152761 | 0.61084934 | 11.82160705 | 0.548423815 | 0.603618513 | 0.911084042 | -5.004452802 |
| **A_55_P2003005** | Capza1 | -0.039858845 | -0.218699666 | 0.138981977 | 13.40076164 | -0.548311323 | 0.603690977 | 0.911122268 | -5.004496378 |
| **A_51_P238448** | Ccnd3 | -0.081774668 | -0.448785178 | 0.285235841 | 10.17443584 | -0.548162667 | 0.603786744 | 0.911171433 | -5.004553951 |
| **A_52_P69020** | Slc24a5 | -0.023636108 | -0.129725975 | 0.082453759 | 7.726833843 | -0.548114411 | 0.603817833 | 0.911171433 | -5.004572637 |
| **A_66_P116372** | Soat2 | 0.134428994 | -0.469173449 | 0.738031437 | 8.776674154 | 0.547912391 | 0.603947995 | 0.911261305 | -5.00465085 |
| **A_52_P411601** | Ep300 | -0.043267142 | -0.237602979 | 0.151068696 | 9.067572388 | -0.547740018 | 0.604059069 | 0.911261305 | -5.004717565 |
| **A_55_P2030334** | D0Kist3 | 0.054732807 | -0.191103839 | 0.300569454 | 8.52570706 | 0.547734712 | 0.604062488 | 0.911261305 | -5.004719618 |
| **A_52_P174915** | Gja1 | -0.227696599 | -1.250423604 | 0.795030407 | 9.553140333 | -0.547729312 | 0.604065968 | 0.911261305 | -4.990256737 |
| **A_55_P2067632** | Ush1c | 0.068339278 | -0.238668354 | 0.37534691 | 8.052028387 | 0.547633968 | 0.604127411 | 0.911282872 | -5.004758601 |
| **A_55_P2054628** | Nrxn1 | -0.025022843 | -0.137513263 | 0.087467576 | 7.682276845 | -0.547255687 | 0.604371224 | 0.911439481 | -5.004904922 |
| **A_51_P292754** | Yaf2 | -0.045706657 | -0.251198461 | 0.159785147 | 11.3706196 | -0.547210111 | 0.604400603 | 0.911439481 | -5.004922545 |
| **A_51_P479132** | Zcchc3 | -0.039343521 | -0.216292372 | 0.13760533 | 8.345724958 | -0.547009209 | 0.604530116 | 0.911439481 | -5.005000214 |
| **A_55_P2106961** | 2900097C17Rik | -0.030994813 | -0.170446131 | 0.108456506 | 8.258406125 | -0.546808874 | 0.60465928 | 0.911439481 | -5.005077639 |
| **A_55_P2019577** | 1500011B03Rik | 0.03549613 | -0.124232773 | 0.195225033 | 8.450294577 | 0.546722136 | 0.604715208 | 0.911439481 | -5.005111153 |
| **A_51_P140797** | Dcaf6 | -0.026029651 | -0.143163108 | 0.091103806 | 8.768551981 | -0.546709413 | 0.604723412 | 0.911439481 | -5.005116069 |
| **A_51_P232207** | Hoxb6 | -0.026896653 | -0.147933415 | 0.094140108 | 8.135838376 | -0.546701313 | 0.604728636 | 0.911439481 | -5.005119198 |
| **A_51_P322989** | Ptrhd1 | 0.04754476 | -0.166412787 | 0.261502308 | 11.37517103 | 0.546693885 | 0.604733425 | 0.911439481 | -5.005122068 |
| **A_52_P317393** | Adgrg1 | 0.074417631 | -0.260471864 | 0.409307126 | 13.21274843 | 0.546692956 | 0.604734025 | 0.911439481 | -5.005122427 |
| **A_55_P1975341** | Nasp | 0.044219061 | -0.154783013 | 0.243221135 | 11.0616047 | 0.54666476 | 0.604752206 | 0.911439481 | -5.00513332 |
| **A_55_P2137309** | Kctd11 | -0.216575216 | -1.191473761 | 0.75832333 | 11.18660939 | -0.54653565 | 0.604835464 | 0.911439481 | -5.005183194 |
| **A_55_P1956284** | NA | 0.023417687 | -0.082005558 | 0.128840933 | 7.966668812 | 0.546483163 | 0.604869313 | 0.911439481 | -5.005203466 |
| **A_55_P2061809** | Ndufc2 | 0.048730893 | -0.170665452 | 0.268127239 | 13.62495475 | 0.546442093 | 0.604895799 | 0.911439481 | -5.005219327 |
| **A_51_P262144** | Ddx19b | -0.033601496 | -0.184923458 | 0.117720465 | 8.965719108 | -0.546293229 | 0.604991809 | 0.911439481 | -5.00527681 |
| **A_55_P2118268** | Chpf | -0.043023084 | -0.236787944 | 0.150741777 | 13.39647352 | -0.546255319 | 0.605016261 | 0.911439481 | -5.005291447 |
| **A_55_P1973583** | Hpn | 0.201632858 | -0.706506455 | 1.10977217 | 9.692378782 | 0.546233073 | 0.605030609 | 0.911439481 | -5.005300035 |
| **A_51_P315391** | Txlnb | -0.025725922 | -0.141604941 | 0.090153097 | 7.886940113 | -0.546179387 | 0.605065238 | 0.911439481 | -5.00532076 |
| **A_52_P143477** | Tgoln1 | -0.123776846 | -0.6813567 | 0.433803008 | 12.91654848 | -0.546136877 | 0.605092659 | 0.911439481 | -5.00533717 |
| **A_51_P363719** | Stx7 | -0.058458956 | -0.321828819 | 0.204910908 | 10.9359982 | -0.546077316 | 0.605131078 | 0.911439481 | -5.00536016 |
| **A_55_P1968370** | Gprc5c | -0.134940424 | -0.768226582 | 0.498345734 | 12.54058009 | -0.552194601 | 0.605199909 | 0.911439481 | -4.940704614 |
| **A_52_P187132** | Ybx1 | -0.080826422 | -0.445070976 | 0.283418133 | 11.6464666 | -0.545920555 | 0.605232204 | 0.911439481 | -5.005420657 |
| **A_51_P156274** | Ltbr | 0.028252444 | -0.099080462 | 0.155585351 | 12.59660416 | 0.545864106 | 0.605268621 | 0.911439481 | -5.005442438 |
| **A_55_P1962122** | NA | -0.032634151 | -0.179754526 | 0.114486224 | 15.09771303 | -0.54571848 | 0.605362577 | 0.911509951 | -5.005498619 |
| **A_52_P354271** | 2310057M21Rik | 0.060650247 | -0.212855943 | 0.334156437 | 9.19122822 | 0.545550021 | 0.605471273 | 0.911589919 | -5.005563593 |
| **A_51_P298843** | Pcbd2 | 0.066498863 | -0.233420044 | 0.366417769 | 10.59448498 | 0.545480887 | 0.605515884 | 0.911589919 | -5.005590252 |
| **A_51_P234864** | Sfxn5 | -0.045271636 | -0.249484813 | 0.158941542 | 8.426839333 | -0.545395547 | 0.605570955 | 0.911589919 | -5.005623157 |
| **A_51_P463860** | Fam161b | -0.026762468 | -0.147516354 | 0.093991418 | 8.075894494 | -0.54524816 | 0.605666074 | 0.911589919 | -5.005679975 |
| **A_55_P2124791** | Col18a1 | 0.125817978 | -0.441898846 | 0.693534803 | 11.16119086 | 0.545230448 | 0.605677505 | 0.911589919 | -5.005686802 |
| **A_55_P2008681** | Ktn1 | -0.053990885 | -0.297685905 | 0.189704135 | 11.79945698 | -0.5450583 | 0.605788614 | 0.911589919 | -5.005753147 |
| **A_55_P2088905** | Hpca | 0.07102637 | -0.249585374 | 0.391638114 | 10.91521394 | 0.545016129 | 0.605815835 | 0.911589919 | -5.005769397 |
| **A_55_P2121603** | 2810047C21Rik1 | 0.046688129 | -0.164165586 | 0.257541845 | 9.703547114 | 0.544746411 | 0.605989947 | 0.911589919 | -5.005873301 |
| **A_55_P1964459** | NA | -0.068885161 | -0.380101834 | 0.242331512 | 9.643741502 | -0.544542758 | 0.606121431 | 0.911589919 | -5.005951725 |
| **A_51_P461319** | Gatm | -0.615714588 | -3.397744489 | 2.166315312 | 11.49209977 | -0.54448612 | 0.606158001 | 0.911589919 | -5.005973531 |
| **A_52_P404461** | Tmem60 | -0.02973802 | -0.164109311 | 0.104633271 | 10.59339888 | -0.544470964 | 0.606167787 | 0.911589919 | -5.005979366 |

| | | | | | | | | | |
|---|---|---|---|---|---|---|---|---|---|
| **A_52_P745** | Acox3 | -0.170812733 | -0.94277739 | 0.601151924 | 10.04540008 | -0.544367248 | 0.606234758 | 0.911589919 | -5.006019292 |
| **A_55_P2158498** | Btk | -0.024041681 | -0.132701228 | 0.084617866 | 7.890471182 | -0.544334809 | 0.606255705 | 0.911589919 | -5.006031777 |
| **A_51_P195958** | Phlda1 | 0.045577512 | -0.16042037 | 0.251575393 | 10.29813718 | 0.54432342 | 0.60626306 | 0.911589919 | -5.006036161 |
| **A_55_P2096883** | Steap3 | -0.031274441 | -0.172629395 | 0.110080514 | 8.749166429 | -0.54431171 | 0.606270621 | 0.911589919 | -5.006040668 |
| **A_52_P330395** | Farp1 | -0.05159544 | -0.284809624 | 0.181618744 | 11.5654772 | -0.544283947 | 0.60628855 | 0.911589919 | -5.006051353 |
| **A_55_P1987146** | Adgrf4 | 0.185830784 | -0.654440671 | 1.026102238 | 8.648168701 | 0.544085575 | 0.60641666 | 0.911589919 | -5.006127688 |
| **A_55_P2107207** | Gpr137b-ps | -0.057654133 | -0.318375241 | 0.203066974 | 12.87569843 | -0.544030715 | 0.606452092 | 0.911589919 | -5.006148794 |
| **A_66_P131708** | Znhit1 | -0.045200075 | -0.249610133 | 0.159209982 | 12.50304629 | -0.544008973 | 0.606466134 | 0.911589919 | -5.006157158 |
| **A_51_P342549** | Hand1 | -0.070406127 | -0.388815798 | 0.248003543 | 12.18858066 | -0.543993075 | 0.606476403 | 0.911589919 | -5.006163274 |
| **A_52_P282741** | Sdc3 | -0.029256282 | -0.161569957 | 0.103057392 | 9.521210907 | -0.543980801 | 0.60648433 | 0.911589919 | -5.006167996 |
| **A_55_P2033084** | Zdhhc20 | 0.042325543 | -0.149106202 | 0.233757287 | 8.562494263 | 0.543948444 | 0.606505229 | 0.911589919 | -5.006180442 |
| **A_55_P2069721** | Hspa8 | -0.03343819 | -0.184686827 | 0.117810448 | 15.6347318 | -0.543901742 | 0.606535394 | 0.911589919 | -5.006198406 |
| **A_52_P579531** | Pdlim3 | 0.151896914 | -0.535191639 | 0.838985466 | 8.979722813 | 0.543882895 | 0.606547569 | 0.911589919 | -5.006205655 |
| **A_55_P2050439** | Dlgap5 | -0.038651116 | -0.213533622 | 0.13623139 | 7.73587331 | -0.543731925 | 0.606645088 | 0.911596982 | -5.006263714 |
| **A_51_P234025** | Rbm5 | -0.070390829 | -0.388885575 | 0.248103916 | 12.25975159 | -0.543729597 | 0.606646592 | 0.911596982 | -5.006264609 |
| **A_66_P120277** | 9330182L06Rik | 0.023723987 | -0.083683995 | 0.13113197 | 7.657669434 | 0.543400811 | 0.606859004 | 0.911845281 | -5.006391002 |
| **A_55_P1967592** | Ccnk | -0.029183984 | -0.16133999 | 0.102972022 | 12.4223265 | -0.543283901 | 0.606934544 | 0.911887898 | -5.006435929 |
| **A_55_P2071236** | Aqp5 | 0.027347064 | -0.096541783 | 0.15123591 | 7.741368808 | 0.543059803 | 0.607079357 | 0.912034579 | -5.006522024 |
| **A_55_P1958374** | NA | 0.043199422 | -0.152551686 | 0.23895053 | 11.89549489 | 0.542928783 | 0.607164032 | 0.912090897 | -5.006572345 |
| **A_55_P2102210** | NA | -0.051547192 | -0.285259222 | 0.182164838 | 11.51339359 | -0.542616641 | 0.607365788 | 0.912230414 | -5.006692188 |
| **A_55_P2111277** | NA | 0.037873953 | -0.133898805 | 0.20964671 | 9.660439064 | 0.542444747 | 0.607476909 | 0.912230414 | -5.006758158 |
| **A_55_P2134616** | Med12l | -0.113450034 | -0.62806277 | 0.401162702 | 8.591461997 | -0.542367007 | 0.607527169 | 0.912230414 | -5.006787988 |
| **A_55_P2026753** | Zmat1 | -0.052763213 | -0.302050204 | 0.196523778 | 10.44396369 | -0.548506405 | 0.607542492 | 0.912230414 | -4.942037907 |
| **A_55_P2102578** | NA | -0.054934901 | -0.304167026 | 0.194297223 | 14.19659495 | -0.542267396 | 0.607591571 | 0.912230414 | -5.006826205 |
| **A_55_P2054362** | NA | -0.085121836 | -0.471307949 | 0.301064278 | 9.075876843 | -0.542267391 | 0.607591574 | 0.912230414 | -5.006826207 |
| **A_55_P2162811** | NA | -0.033258602 | -0.184208785 | 0.11769158 | 15.57664177 | -0.542050211 | 0.607732003 | 0.912230414 | -5.006909508 |
| **A_51_P383629** | Vps4a | 0.036127429 | -0.127847364 | 0.200102223 | 8.88232454 | 0.542037243 | 0.607740388 | 0.912230414 | -5.006914481 |
| **A_55_P2097612** | Zfp131 | -0.037394275 | -0.207125329 | 0.132336778 | 9.836767029 | -0.542017088 | 0.607753422 | 0.912230414 | -5.00692221 |
| **A_55_P1972187** | Slc25a14 | -0.0363488 | -0.201340008 | 0.128642407 | 10.39385288 | -0.54199895 | 0.607765151 | 0.912230414 | -5.006929165 |
| **A_55_P2050426** | Ankrd50 | 0.051078396 | -0.180804739 | 0.282961532 | 10.33390525 | 0.541922588 | 0.607814533 | 0.912230414 | -5.006958446 |
| **A_55_P2204804** | Dclk1 | 0.025933735 | -0.091801858 | 0.143669328 | 7.762528769 | 0.541909116 | 0.607823245 | 0.912230414 | -5.006963611 |
| **A_52_P576689** | Zfp944 | 0.038034959 | -0.134709061 | 0.210778978 | 9.130659932 | 0.541687846 | 0.607966349 | 0.912367063 | -5.007048432 |
| **A_66_P119429** | NA | 0.043088339 | -0.152649386 | 0.238826064 | 13.12051257 | 0.541569726 | 0.60804275 | 0.912367063 | -5.007093699 |
| **A_55_P2023314** | Casz1 | -0.026754698 | -0.14829803 | 0.094788634 | 8.289985051 | -0.541549396 | 0.608055901 | 0.912367063 | -5.00710149 |
| **A_51_P442097** | Slc41a3 | -0.040870723 | -0.226572827 | 0.144831381 | 8.934960574 | -0.541457852 | 0.608115116 | 0.912385087 | -5.007136565 |
| **A_55_P2090184** | Fgfr1op2 | -0.0555567 | -0.308052669 | 0.196939269 | 12.20333945 | -0.541316365 | 0.608206644 | 0.912399253 | -5.007190767 |
| **A_51_P293339** | Tob1 | -0.050710195 | -0.281205614 | 0.179785225 | 9.508995849 | -0.541255272 | 0.608246168 | 0.912399253 | -5.007214166 |
| **A_52_P125350** | Anp32e | -0.063235646 | -0.350680002 | 0.224208711 | 8.411697025 | -0.541224357 | 0.608266168 | 0.912399253 | -5.007226007 |
| **A_55_P2021515** | Rpl3 | 0.032829205 | -0.11642303 | 0.182081439 | 15.87056775 | 0.54113881 | 0.608321516 | 0.912411468 | -5.007258768 |
| **A_55_P2024888** | Ctss | -0.109324518 | -0.60688126 | 0.388232223 | 11.28433419 | -0.540560268 | 0.608695897 | 0.912770096 | -5.007480206 |
| **A_51_P141164** | Ddx50 | -0.041081267 | -0.228052451 | 0.145889916 | 9.765341919 | -0.540553046 | 0.608700571 | 0.912770096 | -5.007482969 |
| **A_65_P08291** | Gpalpp1 | -0.039070346 | -0.216915623 | 0.138774932 | 11.13198286 | -0.540473095 | 0.608752319 | 0.912770096 | -5.007513554 |
| **A_51_P369862** | Nceh1 | 0.048873878 | -0.173679598 | 0.271427353 | 10.44855197 | 0.540270903 | 0.608883197 | 0.912770096 | -5.007590884 |
| **A_55_P2020743** | Abcc10 | 0.057371451 | -0.203910666 | 0.318653567 | 9.665344619 | 0.540200907 | 0.608928509 | 0.912770096 | -5.007617649 |
| **A_55_P2012389** | Sfxn3 | 0.125998751 | -0.447925593 | 0.699923095 | 10.12661104 | 0.540108183 | 0.608988537 | 0.912770096 | -5.0076531 |
| **A_55_P2063096** | Txnrd3 | -0.046002559 | -0.25554834 | 0.163543221 | 8.915480048 | -0.540097589 | 0.608995396 | 0.912770096 | -5.00765715 |
| **A_52_P100252** | Fasn | 0.028161889 | -0.100126003 | 0.156449781 | 7.831638218 | 0.540064061 | 0.609017102 | 0.912770096 | -5.007669967 |
| **A_55_P1953920** | Me3 | 0.026018148 | -0.092521312 | 0.144557607 | 7.795687266 | 0.53998612 | 0.609067563 | 0.912770096 | -5.007699759 |
| **A_55_P2121484** | Smo | -0.10138499 | -0.56339335 | 0.360623369 | 9.851926519 | -0.539874723 | 0.60913969 | 0.912770096 | -5.007742334 |
| **A_55_P2018387** | Zkscan17 | -0.033692266 | -0.187238148 | 0.119853616 | 11.84337823 | -0.539835211 | 0.609165274 | 0.912770096 | -5.007757433 |

| | | | | | | | | | |
|---|---|---|---|---|---|---|---|---|---|
| **A_55_P1957088** | Fam53b | 0.05261252 | -0.187212098 | 0.292437138 | 10.98280956 | 0.53971502 | 0.609243101 | 0.912770096 | -5.007803357 |
| **A_55_P2059357** | Myo7a | -0.052473251 | -0.291669119 | 0.186722617 | 9.388098888 | -0.539701298 | 0.609251987 | 0.912770096 | -5.007808599 |
| **A_51_P196844** | Osbpl3 | -0.04369337 | -0.242870066 | 0.155483326 | 9.719350809 | -0.539692248 | 0.609257847 | 0.912770096 | -5.007812057 |
| **A_55_P1970867** | Gm5779 | -0.043084021 | -0.239489239 | 0.153321197 | 12.16134424 | -0.539675083 | 0.609268962 | 0.912770096 | -5.007818614 |
| **A_51_P501018** | Nek2 | 0.024947892 | -0.088815676 | 0.138711461 | 7.734603865 | 0.539510395 | 0.609375615 | 0.912801744 | -5.007881522 |
| **A_55_P2054128** | NA | 0.060982881 | -0.217109364 | 0.339075126 | 12.71493394 | 0.539495992 | 0.609384943 | 0.912801744 | -5.007887022 |
| **A_55_P1988952** | NA | -0.051604031 | -0.286992734 | 0.183784673 | 14.31276198 | -0.539345638 | 0.609482322 | 0.912801744 | -5.007944438 |
| **A_55_P2156186** | BC028528 | 0.050230452 | -0.178910889 | 0.279371794 | 8.633724767 | 0.539302957 | 0.609509967 | 0.912801744 | -5.007960735 |
| **A_55_P2056241** | Pdlim5 | -0.027552263 | -0.153246053 | 0.098141528 | 9.518349024 | -0.53927788 | 0.60952621 | 0.912801744 | -5.007970309 |
| **A_51_P265406** | Ric8a | 0.039734421 | -0.141591012 | 0.221059854 | 12.39416978 | 0.539109932 | 0.609634999 | 0.91284927 | -5.00803442 |
| **A_55_P2149540** | Srf | -0.049401539 | -0.274860376 | 0.176057299 | 10.50886438 | -0.539066539 | 0.609663109 | 0.91284927 | -5.008050982 |
| **A_55_P2164265** | Prl7c1 | -0.266284901 | -1.481753514 | 0.949183711 | 11.5309346 | -0.538979139 | 0.609719729 | 0.91284927 | -5.008084336 |
| **A_51_P217336** | Scamp1 | 0.066530561 | -0.237194473 | 0.370255594 | 12.35119624 | 0.538901953 | 0.609769734 | 0.91284927 | -5.008113788 |
| **A_55_P2007413** | Agps | 0.049625367 | -0.176968362 | 0.276219096 | 12.10160917 | 0.538796802 | 0.60983786 | 0.91284927 | -5.008153905 |
| **A_55_P2176045** | NA | 0.028295301 | -0.100915304 | 0.157505906 | 7.89614703 | 0.538747571 | 0.609869758 | 0.91284927 | -5.008172686 |
| **A_55_P2124090** | Ambra1 | -0.024064512 | -0.133993969 | 0.085864945 | 10.94165887 | -0.538557586 | 0.609992862 | 0.91284927 | -5.008245146 |
| **A_55_P2111735** | Ppp1r2 | 0.055145342 | -0.196774832 | 0.307065515 | 9.971295119 | 0.53853638 | 0.610006604 | 0.91284927 | -5.008253232 |
| **A_55_P2160077** | NA | 0.023567683 | -0.084106665 | 0.131242031 | 16.93059461 | 0.538485232 | 0.610039749 | 0.91284927 | -5.008272736 |
| **A_55_P1993851** | Gm10021 | -0.0971965 | -0.541356097 | 0.346963097 | 8.349680298 | -0.53836986 | 0.610114517 | 0.91284927 | -5.008316722 |
| **A_55_P2039235** | Mat2b | 0.087755388 | -0.313261503 | 0.488772279 | 12.13773281 | 0.538369302 | 0.610114879 | 0.91284927 | -5.008316935 |
| **A_51_P118168** | Zbtb43 | -0.034447334 | -0.191885408 | 0.12299074 | 10.17567212 | -0.538288383 | 0.610167321 | 0.91284927 | -4.993813998 |
| **A_55_P2052479** | NA | 0.06051288 | -0.216058779 | 0.33708454 | 14.49954453 | 0.538281325 | 0.610171895 | 0.91284927 | -5.008350471 |
| **A_55_P2011937** | Fignl1 | 0.08189046 | -0.292645301 | 0.456426221 | 8.845149462 | 0.537909462 | 0.610412931 | 0.913139196 | -5.008492172 |
| **A_55_P1959758** | Top3b | 0.044670658 | -0.159696916 | 0.249038233 | 9.958826956 | 0.537748892 | 0.610517027 | 0.913224239 | -5.008553331 |
| **A_65_P19395** | H2-D1 | -0.268046024 | -1.494699361 | 0.958607314 | 14.32567442 | -0.537596816 | 0.610615626 | 0.9132609 | -5.008611241 |
| **A_55_P1963002** | Ppfia4 | -0.031595403 | -0.176201069 | 0.113010263 | 8.199671216 | -0.537536237 | 0.610654904 | 0.9132609 | -5.008634305 |
| **A_51_P472621** | Zfp719 | -0.03929459 | -0.2191525 | 0.140563319 | 8.06808129 | -0.537492474 | 0.610683281 | 0.9132609 | -5.008650966 |
| **A_55_P2008452** | Wtap | 0.072926788 | -0.260971245 | 0.406824821 | 11.0834499 | 0.537331609 | 0.610787593 | 0.913346232 | -5.008712196 |
| **A_55_P2113523** | Nsd2 | -0.033375099 | -0.186215756 | 0.119465558 | 8.48124433 | -0.537220819 | 0.610859441 | 0.913360304 | -5.008754357 |
| **A_55_P2022574** | Gpank1 | 0.054318717 | -0.194495619 | 0.303133054 | 12.49330755 | 0.537085304 | 0.61094733 | 0.913360304 | -5.008805917 |
| **A_55_P2160253** | 6720475J19Rik | -0.025747078 | -0.143693828 | 0.092199672 | 7.77530449 | -0.537045558 | 0.610973108 | 0.913360304 | -5.008821037 |
| **A_55_P2046453** | Aarsd1 | 0.035606494 | -0.127512051 | 0.198725038 | 10.83535834 | 0.537025654 | 0.610986017 | 0.913360304 | -5.008828608 |
| **A_51_P198434** | H2-K1 | -0.166026807 | -0.927279249 | 0.595225636 | 16.1644692 | -0.536560476 | 0.611287774 | 0.913487476 | -5.009005493 |
| **A_55_P2179296** | Mad2l1 | -0.025562363 | -0.142781321 | 0.091656595 | 7.798265977 | -0.536503182 | 0.611324946 | 0.913487476 | -5.009027269 |
| **A_51_P143190** | Lyl1 | 0.052386863 | -0.187839247 | 0.292612973 | 8.527344006 | 0.536501999 | 0.611325713 | 0.913487476 | -5.009027719 |
| **A_55_P2052261** | Zfp280c | 0.02921457 | -0.104753772 | 0.163182912 | 8.120491154 | 0.536495989 | 0.611329613 | 0.913487476 | -5.009030003 |
| **A_55_P2318584** | Aqp8 | -0.142756671 | -0.79752261 | 0.512009267 | 15.12695349 | -0.536388549 | 0.611399323 | 0.913487476 | -5.009070834 |
| **A_55_P2415472** | 4931408D14Rik | 0.032779656 | -0.117574757 | 0.18313407 | 7.757652628 | 0.536361226 | 0.611417051 | 0.913487476 | -5.009081217 |
| **A_52_P64356** | Sparcl1 | -0.088197095 | -0.492882343 | 0.316488152 | 13.30420274 | -0.536174396 | 0.611538284 | 0.913487476 | -5.009152198 |
| **A_55_P1990653** | Comt | 0.054664866 | -0.196207256 | 0.305536987 | 11.69445135 | 0.53607437 | 0.611603196 | 0.913487476 | -5.009190192 |
| **A_55_P2044582** | Iglon5 | 0.02253334 | -0.080894601 | 0.12596128 | 7.74928558 | 0.535990201 | 0.61165782 | 0.913487476 | -5.009222157 |
| **A_55_P2108032** | Tmem255b | -0.023514032 | -0.131447706 | 0.084419642 | 7.669699035 | -0.535968555 | 0.611671869 | 0.913487476 | -5.009230377 |
| **A_51_P495232** | Gpx8 | -0.163418933 | -0.913582179 | 0.586744313 | 11.58500113 | -0.535939482 | 0.611690738 | 0.913487476 | -5.009241418 |
| **A_52_P293556** | Wdr41 | -0.030241336 | -0.169062235 | 0.108579563 | 8.704224853 | -0.535938904 | 0.611691113 | 0.913487476 | -5.009241637 |
| **A_55_P2046563** | Cym | -0.022396503 | -0.125220661 | 0.080427656 | 7.703825916 | -0.53586354 | 0.611740027 | 0.913487476 | -5.009270253 |
| **A_55_P2104798** | NA | -0.048445359 | -0.270898477 | 0.174007759 | 9.701678252 | -0.535775495 | 0.611797174 | 0.913487476 | -5.009303679 |
| **A_55_P2106255** | Mbnl3 | 0.043558594 | -0.164790884 | 0.251908071 | 13.12075544 | 0.541790665 | 0.611821861 | 0.913487476 | -4.944446483 |
| **A_55_P1954356** | Ttc23 | 0.04333335 | -0.155667982 | 0.242334682 | 9.53195414 | 0.535717018 | 0.611835131 | 0.913487476 | -5.009325877 |
| **A_55_P1962419** | Ccar1 | -0.044938854 | -0.25135132 | 0.161473611 | 9.43295974 | -0.535618093 | 0.611899346 | 0.913487476 | -5.009363424 |
| **A_51_P418168** | Manf | -0.067024598 | -0.374941368 | 0.240892171 | 13.06008361 | -0.535513027 | 0.611967552 | 0.913487476 | -5.009403296 |

| | | | | | | | | | |
|---|---|---|---|---|---|---|---|---|---|
| **A_55_P2186928** | NA | -0.054004068 | -0.302104455 | 0.194096319 | 8.518494746 | -0.535510755 | 0.611969027 | 0.913487476 | -5.009404158 |
| **A_55_P2077263** | Cenpk | 0.043913654 | -0.157919939 | 0.245747246 | 8.023082178 | 0.535272935 | 0.612123428 | 0.913488447 | -5.009494383 |
| **A_55_P2182497** | Maml3 | -0.078501662 | -0.439317399 | 0.282314075 | 11.42480855 | -0.535257232 | 0.612133624 | 0.913488447 | -5.00950034 |
| **A_55_P1952793** | Son | -0.055746019 | -0.311974971 | 0.200482933 | 11.63630771 | -0.535247712 | 0.612139805 | 0.913488447 | -5.00950395 |
| **A_52_P401311** | NA | 0.186805654 | -0.671942576 | 1.045553884 | 9.000512805 | 0.535171928 | 0.612189012 | 0.913488447 | -5.009532693 |
| **A_55_P2066827** | Hexim2 | -0.043035435 | -0.240879597 | 0.154808727 | 9.10826264 | -0.5351458 | 0.612205977 | 0.913488447 | -5.009542602 |
| **A_55_P1958808** | Stoml2 | 0.086859527 | -0.312599324 | 0.486318379 | 12.58912196 | 0.534951705 | 0.612332016 | 0.913605986 | -5.009616197 |
| **A_51_P129317** | Mob4 | 0.073174619 | -0.263524766 | 0.409874003 | 11.52552437 | 0.534671836 | 0.61251378 | 0.9137896 | -4.990330514 |
| **A_55_P2047621** | Cutal | 0.167702512 | -0.60411974 | 0.939524763 | 8.564816473 | 0.534553817 | 0.612590438 | 0.9137896 | -5.009766993 |
| **A_55_P2143279** | Nkd2 | 0.079386251 | -0.286017098 | 0.444789601 | 10.70472084 | 0.534492892 | 0.612630013 | 0.9137896 | -5.009790074 |
| **A_55_P1969477** | Abcb1a | -0.050748212 | -0.284344373 | 0.182847949 | 9.506728347 | -0.534471076 | 0.612644184 | 0.9137896 | -5.009798338 |
| **A_55_P2183438** | Runx1 | -0.25604048 | -1.435087793 | 0.923006833 | 9.18504444 | -0.5342525 | 0.612786178 | 0.913792335 | -5.009881123 |
| **A_55_P1983769** | Birc5 | 0.037190421 | -0.134140569 | 0.208521412 | 7.906103693 | 0.534028378 | 0.612931795 | 0.913792335 | -5.009965978 |
| **A_51_P472888** | Enkd1 | -0.029436652 | -0.165051256 | 0.106177953 | 8.277382499 | -0.53401212 | 0.612942359 | 0.913792335 | -5.009972132 |
| **A_51_P475875** | Lrwd1 | 0.042163853 | -0.152086457 | 0.236414162 | 9.235492541 | 0.534007963 | 0.61294506 | 0.913792335 | -5.009973706 |
| **A_52_P367760** | Calml4 | 0.086696879 | -0.312720903 | 0.486114661 | 9.571091186 | 0.534004886 | 0.612947059 | 0.913792335 | -5.00997487 |
| **A_55_P2049473** | Sh3bp5 | 0.029244562 | -0.105506634 | 0.163995758 | 8.227263514 | 0.533926719 | 0.612997851 | 0.913792335 | -5.010004457 |
| **A_55_P2034720** | Cfap36 | 0.043005902 | -0.155202933 | 0.241214738 | 9.759536873 | 0.53379465 | 0.613083673 | 0.913792335 | -5.010054436 |
| **A_55_P2047842** | Clcn3 | -0.05568119 | -0.312317097 | 0.200954717 | 12.09498821 | -0.533777483 | 0.61309483 | 0.913792335 | -5.010060932 |
| **A_55_P1969131** | Cidec | -0.106852934 | -0.599350037 | 0.385644168 | 8.944720138 | -0.533767266 | 0.613101469 | 0.913792335 | -5.010064798 |
| **A_52_P437795** | Cdh11 | -0.052822593 | -0.296299965 | 0.19065478 | 8.362292445 | -0.533740636 | 0.613118775 | 0.913792335 | -5.010074874 |
| **A_51_P114843** | H1f10 | 0.024262046 | -0.087592965 | 0.136117056 | 7.698872771 | 0.533631104 | 0.613189959 | 0.913808495 | -5.010116313 |
| **A_51_P482711** | Dhcr24 | 0.122381346 | -0.441983184 | 0.686745875 | 13.1265955 | 0.533488028 | 0.613282951 | 0.913808495 | -5.010170432 |
| **A_55_P1956862** | Egfr | -0.127331833 | -0.714626673 | 0.459963006 | 9.520283412 | -0.533396265 | 0.613342595 | 0.913808495 | -5.010205134 |
| **A_55_P2166128** | NA | -0.080875393 | -0.453899576 | 0.292148791 | 8.545241868 | -0.533394554 | 0.613343708 | 0.913808495 | -5.010205781 |
| **A_55_P2116111** | Prag1 | 0.037155945 | -0.134230639 | 0.208542529 | 11.08534974 | 0.533360259 | 0.613366 | 0.913808495 | -5.01021875 |
| **A_55_P2048011** | Larp4b | -0.062886631 | -0.353100751 | 0.227327489 | 13.43213905 | -0.533100336 | 0.613534969 | 0.913989781 | -5.010317013 |
| **A_55_P1984113** | Nup62 | -0.055051475 | -0.30916502 | 0.199062069 | 12.93778295 | -0.53297926 | 0.613613686 | 0.914036601 | -5.010362771 |
| **A_55_P2061076** | Gm3219 | -0.023918589 | -0.134358471 | 0.086521292 | 7.792792943 | -0.532817887 | 0.613718611 | 0.914122449 | -5.010423744 |
| **A_51_P232439** | Ice1 | -0.043078247 | -0.242099926 | 0.155943432 | 9.831673325 | -0.532508815 | 0.613919597 | 0.914351356 | -5.010540479 |
| **A_52_P354682** | Elovl7 | -0.036416471 | -0.204711387 | 0.131878445 | 8.27231668 | -0.532348489 | 0.614023871 | 0.914363557 | -5.01060101 |
| **A_51_P151820** | Urb1 | -0.073745427 | -0.414571013 | 0.267080159 | 8.900289377 | -0.532319122 | 0.614042971 | 0.914363557 | -5.010612095 |
| **A_51_P475249** | Rnf44 | 0.059454201 | -0.215353926 | 0.334262328 | 11.8532193 | 0.532257927 | 0.614082775 | 0.914363557 | -5.010635195 |
| **A_52_P437734** | Fbxo28 | 0.037733604 | -0.136756167 | 0.212223375 | 9.164829821 | 0.532019413 | 0.614237927 | 0.914363557 | -5.010725202 |
| **A_52_P257625** | Esm1 | 0.166662359 | -0.604043031 | 0.93736775 | 8.224597338 | 0.532008159 | 0.614245248 | 0.914363557 | -5.010729448 |
| **A_55_P2001970** | Zswim9 | 0.032011867 | -0.116023674 | 0.180047409 | 9.197811977 | 0.532003116 | 0.614248528 | 0.914363557 | -5.010731351 |
| **A_52_P605517** | Phactr1 | 0.095620887 | -0.346615297 | 0.537857071 | 11.48436536 | 0.531946134 | 0.614285599 | 0.914363557 | -5.010752849 |
| **A_51_P495485** | Pip4p2 | -0.05519647 | -0.310502881 | 0.200109941 | 9.822457706 | -0.531886229 | 0.614324572 | 0.914363557 | -5.010775447 |
| **A_55_P2150555** | Pcgf5 | 0.101773075 | -0.369061633 | 0.572607784 | 13.05697769 | 0.531781948 | 0.614392418 | 0.914363557 | -5.010814779 |
| **A_55_P2035484** | Gucd1 | 0.046580968 | -0.168922696 | 0.262084632 | 11.41565739 | 0.531769 | 0.614400842 | 0.914363557 | -5.010819663 |
| **A_66_P104190** | Gsk3b | 0.075709999 | -0.274614394 | 0.426034392 | 9.622884747 | 0.531682083 | 0.614457396 | 0.914377319 | -5.01085244 |
| **A_66_P138929** | Lox | 0.199859224 | -0.725084016 | 1.124802465 | 8.981330008 | 0.531591806 | 0.614516138 | 0.914394336 | -5.01088648 |
| **A_51_P496720** | Dnmt3l | 0.126169332 | -0.45801197 | 0.710350633 | 10.33858093 | 0.531343428 | 0.61467777 | 0.914526484 | -5.010980107 |
| **A_55_P2045040** | F7 | 0.027011494 | -0.0980752 | 0.152098188 | 7.750916914 | 0.53125943 | 0.614732438 | 0.914526484 | -5.011011762 |
| **A_51_P288505** | Tradd | 0.037046636 | -0.134554362 | 0.208647633 | 11.31200049 | 0.531126698 | 0.614818827 | 0.914526484 | -5.011061773 |
| **A_52_P86517** | Cbwd1 | -0.042658893 | -0.240268956 | 0.15495117 | 10.40755384 | -0.53109191 | 0.614841471 | 0.914526484 | -5.011074878 |
| **A_55_P2006008** | Serpinb1a | -0.053067449 | -0.298893413 | 0.192758514 | 9.710882126 | -0.531091841 | 0.614841516 | 0.914526484 | -5.011074904 |
| **A_55_P1967168** | Lrp4 | -0.038730159 | -0.218213946 | 0.140753628 | 8.037160646 | -0.530876151 | 0.614981917 | 0.914531139 | -5.011156145 |
| **A_55_P1980020** | Rsl1d1 | 0.065009694 | -0.23627008 | 0.366289467 | 12.85339707 | 0.530856705 | 0.614994576 | 0.914531139 | -5.011163468 |
| **A_55_P2003931** | NA | 0.0276422 | -0.100494146 | 0.155778545 | 8.081308378 | 0.530724856 | 0.615080412 | 0.914531139 | -5.011213113 |

| | | | | | | | | | |
|---|---|---|---|---|---|---|---|---|---|
| **A_51_P222590** | Arf6 | -0.047627782 | -0.268413316 | 0.173157752 | 11.08275959 | -0.530711997 | 0.615088783 | 0.914531139 | -5.011217954 |
| **A_66_P104473** | Polg2 | -0.046873283 | -0.264168595 | 0.170422028 | 9.255641055 | -0.530694005 | 0.615100497 | 0.914531139 | -5.011224728 |
| **A_55_P2085170** | 6-Sep | 0.039415194 | -0.143321903 | 0.22215229 | 8.528926914 | 0.53064748 | 0.615130788 | 0.914531139 | -5.011242243 |
| **A_55_P2089506** | NA | 0.064062846 | -0.233026978 | 0.361152671 | 13.07187138 | 0.530502723 | 0.615225039 | 0.914531139 | -5.011296729 |
| **A_55_P2021540** | Myocd | 0.045271442 | -0.164704755 | 0.255247638 | 8.111745109 | 0.530424312 | 0.615276095 | 0.914531139 | -5.011326238 |
| **A_52_P444637** | Rcbtb2 | -0.023986888 | -0.135268189 | 0.087294413 | 8.068879393 | -0.530299085 | 0.615357641 | 0.914531139 | -5.011373357 |
| **A_55_P2081488** | Pglyrp1 | 0.035523158 | -0.129279612 | 0.200325927 | 8.073445228 | 0.530293402 | 0.615361341 | 0.914531139 | -5.011375495 |
| **A_55_P2100700** | Pygo2 | -0.044334972 | -0.258590373 | 0.169920429 | 12.15091311 | -0.536246836 | 0.615367841 | 0.914531139 | -4.946416002 |
| **A_66_P128963** | Cyp2d9 | 0.023612835 | -0.085951383 | 0.133177054 | 7.682453784 | 0.530210784 | 0.615415144 | 0.914531139 | -5.011406576 |
| **A_51_P291501** | Ccno | -0.027358758 | -0.154356506 | 0.09963899 | 8.005167934 | -0.529992254 | 0.615557468 | 0.914531139 | -5.011488766 |
| **A_55_P2080119** | NA | -0.056151355 | -0.316821256 | 0.204518547 | 13.66595892 | -0.529954412 | 0.615582116 | 0.914531139 | -5.011502995 |
| **A_55_P1961943** | Cd68 | -0.256124682 | -1.445316297 | 0.933066932 | 13.79310301 | -0.5298693 | 0.615637554 | 0.914531139 | -5.011534996 |
| **A_55_P2154425** | Cd320 | 0.081956922 | -0.298582584 | 0.462496428 | 8.964048021 | 0.52985258 | 0.615648445 | 0.914531139 | -5.011541282 |
| **A_55_P2029746** | NA | -0.281699028 | -1.589676111 | 1.026278054 | 12.50621845 | -0.529851755 | 0.615648982 | 0.914531139 | -5.011541592 |
| **A_55_P2009813** | NA | 0.03016865 | -0.109935368 | 0.170272668 | 8.018115417 | 0.52975425 | 0.615712497 | 0.914555203 | -5.011578247 |
| **A_55_P2419021** | 9330188P03Rik | -0.023425803 | -0.132243651 | 0.085392045 | 7.643967521 | -0.529618954 | 0.615800634 | 0.914592626 | -5.011629097 |
| **A_51_P456870** | Foxj1 | 0.033857067 | -0.123461093 | 0.191175226 | 7.929868583 | 0.529467898 | 0.615899046 | 0.914592626 | -5.011685858 |
| **A_52_P399095** | Mettl24 | -0.038512661 | -0.217465602 | 0.140440279 | 8.521431016 | -0.529460855 | 0.615903635 | 0.914592626 | -5.011688504 |
| **A_55_P2063381** | Tmem74b | -0.031567806 | -0.17828004 | 0.115144428 | 7.955363561 | -0.529355259 | 0.615972436 | 0.914592626 | -5.011728174 |
| **A_55_P2073709** | Tfdp2 | 0.060080768 | -0.219147721 | 0.339309257 | 9.484611675 | 0.529352433 | 0.615974277 | 0.914592626 | -5.011729235 |
| **A_55_P2119716** | Mapk1ip1 | 0.035973383 | -0.13127065 | 0.203217416 | 9.6964116 | 0.529175606 | 0.616089498 | 0.914603338 | -5.011795649 |
| **A_55_P2381538** | NA | -0.098364368 | -0.555737864 | 0.359009127 | 9.409920515 | -0.529097826 | 0.616140183 | 0.914603338 | -5.011824856 |
| **A_51_P421538** | Slc9b2 | -0.107860528 | -0.609449199 | 0.393728144 | 9.016806501 | -0.529034487 | 0.61618146 | 0.914603338 | -5.011848637 |
| **A_55_P1996746** | NA | 0.207994785 | -0.759289537 | 1.175279108 | 8.61259349 | 0.52901436 | 0.616194577 | 0.914603338 | -5.011856194 |
| **A_55_P2168628** | Sac3d1 | 0.036649229 | -0.133800435 | 0.207098893 | 11.23834912 | 0.528978296 | 0.61621808 | 0.914603338 | -5.011869733 |
| **A_52_P127465** | Ypel1 | -0.061624606 | -0.348312534 | 0.225063322 | 8.365391609 | -0.528827352 | 0.616316457 | 0.914679115 | -5.011926391 |
| **A_51_P406334** | Dnlz | -0.048751302 | -0.275649768 | 0.178147164 | 9.503666248 | -0.528595937 | 0.616467296 | 0.914832735 | -5.012013228 |
| **A_51_P390285** | Larp6 | -0.199034957 | -1.125679766 | 0.727609852 | 8.913692314 | -0.528427274 | 0.616577247 | 0.914921256 | -5.012076496 |
| **A_66_P128440** | NA | 0.048919175 | -0.178864729 | 0.27670308 | 10.36070192 | 0.528354315 | 0.616624812 | 0.914921256 | -5.012103859 |
| **A_66_P111987** | Brwd3 | -0.052536594 | -0.29719804 | 0.192124853 | 9.742491226 | -0.528281661 | 0.61667218 | 0.914921256 | -5.012131104 |
| **A_55_P2051476** | C8a | 0.02280371 | -0.083407137 | 0.129014558 | 7.627654473 | 0.528209008 | 0.616719549 | 0.914921256 | -5.012158345 |
| **A_55_P1982986** | Cfap410 | 0.025844616 | -0.094546358 | 0.146235591 | 8.217479245 | 0.528135455 | 0.616767507 | 0.914921256 | -5.012185921 |
| **A_55_P2028015** | Pawr | 0.121380424 | -0.44450308 | 0.687263927 | 9.895865873 | 0.527704477 | 0.617048557 | 0.914921256 | -5.01234743 |
| **A_51_P206824** | Hjv | 0.02826961 | -0.103542633 | 0.160081853 | 7.623889159 | 0.527634555 | 0.617094162 | 0.914921256 | -5.012373622 |
| **A_55_P2098935** | Sh2d3c | -0.053290396 | -0.301850441 | 0.19526965 | 10.51262243 | -0.527456669 | 0.617210191 | 0.914921256 | -5.012440243 |
| **A_55_P1955970** | Cbfa2t2 | -0.046701091 | -0.26456924 | 0.171167058 | 11.07857022 | -0.527354246 | 0.617277003 | 0.914921256 | -5.012478594 |
| **A_66_P101261** | Gm3367 | -0.02580572 | -0.146208646 | 0.094597205 | 8.550754311 | -0.527288265 | 0.617320045 | 0.914921256 | -5.012503295 |
| **A_51_P470448** | Setx | -0.045912879 | -0.260154537 | 0.168328778 | 11.02564022 | -0.527229589 | 0.617358324 | 0.914921256 | -5.01252526 |
| **A_55_P2343151** | Akap1 | -0.040433182 | -0.229110207 | 0.148243844 | 10.34516685 | -0.527215366 | 0.617367603 | 0.914921256 | -5.012530584 |
| **A_55_P2178632** | Tmem80 | 0.02624877 | -0.096253444 | 0.148750984 | 9.217176573 | 0.52714996 | 0.617410274 | 0.914921256 | -5.012555065 |
| **A_55_P1971769** | NA | 0.044286356 | -0.162406825 | 0.250979537 | 11.77275448 | 0.527124207 | 0.617427076 | 0.914921256 | -5.012564703 |
| **A_51_P114634** | Amz1 | 0.046071888 | -0.168966777 | 0.261110553 | 10.5046566 | 0.527094659 | 0.617446354 | 0.914921256 | -5.007697573 |
| **A_55_P2051176** | NA | 0.063357913 | -0.232370963 | 0.35908679 | 14.76976357 | 0.527079705 | 0.617456111 | 0.914921256 | -5.012581358 |
| **A_55_P2176729** | Prdx6 | 0.071582273 | -0.262630559 | 0.405795105 | 11.47832004 | 0.526928307 | 0.617554894 | 0.914921256 | -5.012638008 |
| **A_55_P2156712** | Gpr1 | -0.059309798 | -0.336234281 | 0.217614685 | 11.66373439 | -0.526907349 | 0.617568569 | 0.914921256 | -5.012645849 |
| **A_55_P1997485** | NA | -0.041016449 | -0.232534614 | 0.150501716 | 10.9279123 | -0.52688672 | 0.61758203 | 0.914921256 | -5.012653566 |
| **A_55_P1973818** | Mmel1 | -0.023094966 | -0.130938477 | 0.084748545 | 7.712413147 | -0.52685665 | 0.617601651 | 0.914921256 | -5.012664815 |
| **A_66_P124139** | Ikzf1 | -0.023070564 | -0.13080115 | 0.084660022 | 7.698641039 | -0.526851653 | 0.617604912 | 0.914921256 | -5.012666684 |
| **A_51_P144581** | Pgrmc1 | -0.075818608 | -0.429866525 | 0.278229309 | 13.51774602 | -0.526845075 | 0.617609204 | 0.914921256 | -5.012669145 |
| **A_55_P2140147** | Ppp4r3b | -0.047275218 | -0.26803904 | 0.173488603 | 10.6404547 | -0.526835224 | 0.617615632 | 0.914921256 | -5.01267283 |

| | | | | | | | | | |
|---|---|---|---|---|---|---|---|---|---|
| **A_55_P1974636** | Gm16430 | -0.035913268 | -0.203651633 | 0.131825096 | 8.575235576 | -0.526734407 | 0.617681421 | 0.914936006 | -5.01271054 |
| **A_51_P160372** | Wbp1 | -0.074863824 | -0.424566177 | 0.274838529 | 11.89584505 | -0.526674894 | 0.617720258 | 0.914936006 | -5.012732797 |
| **A_55_P2259125** | D7Ertd143e | 0.084304737 | -0.309613739 | 0.478223213 | 10.43941165 | 0.526519898 | 0.617821413 | 0.91500876 | -5.012790755 |
| **A_52_P1004509** | BC065397 | -0.091506149 | -0.519151564 | 0.336139265 | 9.163221548 | -0.526423931 | 0.617884049 | 0.91500876 | -5.012826632 |
| **A_65_P05396** | Rad18 | -0.023761603 | -0.134818069 | 0.087294862 | 7.947499271 | -0.526382035 | 0.617911395 | 0.91500876 | -5.012842293 |
| **A_51_P453519** | Zdhhc16 | 0.062927331 | -0.231387175 | 0.357241836 | 12.80602183 | 0.526013399 | 0.618152034 | 0.915294979 | -5.012980044 |
| **A_51_P268903** | Mon1a | -0.035073055 | -0.199150735 | 0.129004625 | 9.774440926 | -0.525887989 | 0.618233911 | 0.915343157 | -5.013026887 |
| **A_51_P477833** | Klhl21 | -0.030775012 | -0.174776713 | 0.113226689 | 10.41388692 | -0.525774806 | 0.618307811 | 0.915343157 | -5.013069156 |
| **A_52_P459564** | Itga2b | -0.022271292 | -0.126500891 | 0.081958307 | 7.700808443 | -0.525682493 | 0.618368088 | 0.915343157 | -5.013103624 |
| **A_51_P208263** | NA | 0.054367537 | -0.200076796 | 0.30881187 | 14.37455804 | 0.525673447 | 0.618373995 | 0.915343157 | -5.013107001 |
| **A_51_P133792** | Prickle3 | -0.031184394 | -0.177185865 | 0.114817078 | 9.054941495 | -0.52547158 | 0.618505818 | 0.915400168 | -5.013182356 |
| **A_55_P2077884** | Kat6b | -0.051270739 | -0.291314981 | 0.188773502 | 8.055995377 | -0.525469423 | 0.618507227 | 0.915400168 | -5.013183161 |
| **A_55_P2041340** | Atp5a1 | -0.061509513 | -0.349602202 | 0.226583175 | 14.08333245 | -0.525265912 | 0.61864014 | 0.915464058 | -5.013259103 |
| **A_55_P2033650** | Cpt2 | 0.041938859 | -0.154504594 | 0.238382312 | 11.78007625 | 0.525228418 | 0.618664629 | 0.915464058 | -5.013273092 |
| **A_55_P2052959** | Ctbp1 | -0.032657648 | -0.185649524 | 0.120334227 | 13.82182404 | -0.525152822 | 0.618714006 | 0.915464058 | -5.013301293 |
| **A_55_P2108784** | Arhgap22 | -0.042763749 | -0.24311495 | 0.157587451 | 11.66795664 | -0.525113265 | 0.618739844 | 0.915464058 | -5.013316048 |
| **A_55_P2084345** | NA | -0.062588233 | -0.355890139 | 0.230713674 | 13.11627731 | -0.524985085 | 0.618823574 | 0.915515163 | -5.013363854 |
| **A_55_P2105638** | Ralgapa2 | 0.03291143 | -0.121339084 | 0.187161944 | 9.012798105 | 0.524915373 | 0.618869115 | 0.915515163 | -5.01338985 |
| **A_52_P547447** | Sirt2 | 0.067518368 | -0.249007143 | 0.38404388 | 10.14875318 | 0.524786172 | 0.618953522 | 0.915525161 | -5.013438021 |
| **A_51_P317695** | Trpc3 | 0.032034849 | -0.118155732 | 0.18222543 | 8.133905013 | 0.524746007 | 0.618979763 | 0.915525161 | -5.013452994 |
| **A_51_P168695** | Mast4 | 0.048209047 | -0.177880349 | 0.274298443 | 8.67524484 | 0.52458699 | 0.61908366 | 0.915525161 | -5.013512263 |
| **A_55_P2179106** | Ppig | 0.033165353 | -0.122390684 | 0.188721391 | 8.743005893 | 0.524525876 | 0.619123592 | 0.915525161 | -5.013535037 |
| **A_52_P449871** | Id4 | -0.046699329 | -0.26573604 | 0.172337381 | 8.122764271 | -0.524521021 | 0.619126765 | 0.915525161 | -5.013536847 |
| **A_55_P2081805** | Slc46a1 | -0.037213593 | -0.211814468 | 0.137387281 | 10.49049704 | -0.52435372 | 0.619236089 | 0.915525161 | -5.01359918 |
| **A_51_P132013** | Cysltr2 | -0.288664371 | -1.643224405 | 1.065895662 | 9.156509266 | -0.524280964 | 0.619283635 | 0.915525161 | -5.013626282 |
| **A_55_P1978176** | Abhd17c | -0.069420464 | -0.395213799 | 0.256372871 | 12.15900738 | -0.524221076 | 0.619322774 | 0.915525161 | -5.013648588 |
| **A_52_P32864** | Prps2 | -0.03513974 | -0.2000951 | 0.129815619 | 7.782342567 | -0.524084459 | 0.619412062 | 0.915525161 | -5.013699465 |
| **A_55_P1987290** | Rnu1b6 | 0.108072377 | -0.399373569 | 0.615518323 | 14.40996831 | 0.523955108 | 0.619496608 | 0.915525161 | -5.013747625 |
| **A_55_P2165414** | Serpina1a | 0.370090495 | -1.36769023 | 2.10787122 | 13.309507 | 0.523940679 | 0.619506039 | 0.915525161 | -5.013752996 |
| **A_51_P134972** | Shkbp1 | -0.043072425 | -0.245327264 | 0.159182414 | 10.09893314 | -0.523925536 | 0.619515937 | 0.915525161 | -5.013758633 |
| **A_55_P2105362** | Tmed5 | 0.033828521 | -0.125035413 | 0.192692455 | 9.454626986 | 0.523874024 | 0.619549609 | 0.915525161 | -5.013777808 |
| **A_55_P2129911** | NA | -0.0396617 | -0.225949341 | 0.146625942 | 15.00305421 | -0.52378908 | 0.619605136 | 0.915525161 | -5.013809425 |
| **A_52_P359819** | Ntf5 | 0.055217899 | -0.204145735 | 0.314581534 | 9.464566268 | 0.523769204 | 0.619618129 | 0.915525161 | -5.013816822 |
| **A_55_P2148385** | NA | -0.029691301 | -0.169160467 | 0.109777865 | 8.594087109 | -0.523745356 | 0.619633719 | 0.915525161 | -5.013825697 |
| **A_55_P2169729** | NA | -0.067917062 | -0.387093217 | 0.251259093 | 13.04670028 | -0.523501118 | 0.619793394 | 0.915691089 | -5.013916571 |
| **A_66_P128446** | Ttpa | -0.022789382 | -0.129986155 | 0.084407392 | 7.643073689 | -0.523022039 | 0.620106668 | 0.91579681 | -5.014094715 |
| **A_55_P2045802** | Nsmf | 0.123459522 | -0.457313836 | 0.704232881 | 10.91723569 | 0.522982395 | 0.620132596 | 0.91579681 | -5.014109451 |
| **A_55_P1965524** | NA | -0.036888414 | -0.210420094 | 0.136643266 | 13.71836714 | -0.522974338 | 0.620137865 | 0.91579681 | -5.014112445 |
| **A_51_P321886** | Cmtm3 | -0.119241383 | -0.680414484 | 0.441931717 | 11.66286462 | -0.522756358 | 0.620280437 | 0.91579681 | -5.014193446 |
| **A_52_P31011** | Cep162 | -0.081145438 | -0.463072252 | 0.300781376 | 8.735777186 | -0.52270075 | 0.620316811 | 0.91579681 | -5.014214106 |
| **A_55_P2006915** | Myo5c | 0.023904192 | -0.088610804 | 0.136419189 | 7.808572098 | 0.522676319 | 0.620332791 | 0.91579681 | -5.014223181 |
| **A_55_P2136289** | NA | 0.094955358 | -0.352068606 | 0.541979322 | 11.51577352 | 0.522586074 | 0.620391825 | 0.91579681 | -5.014256703 |
| **A_55_P2216456** | Rint1 | 0.029885427 | -0.110845469 | 0.170616322 | 9.319913225 | 0.522443306 | 0.620485223 | 0.91579681 | -5.014309725 |
| **A_55_P2164090** | Kcnj15 | -0.049316895 | -0.281570281 | 0.182936491 | 9.104912634 | -0.522399599 | 0.620513817 | 0.91579681 | -5.014325954 |
| **A_55_P1955612** | Nup93 | -0.042898541 | -0.244926263 | 0.159129181 | 11.00057998 | -0.522397054 | 0.620515483 | 0.91579681 | -5.014326899 |
| **A_55_P1964364** | Kctd14 | 0.058835529 | -0.218276314 | 0.335947372 | 17.45151255 | 0.522340546 | 0.620552453 | 0.91579681 | -5.01434788 |
| **A_52_P55772** | Tbxa2r | 0.032712448 | -0.121369832 | 0.186794729 | 8.156965448 | 0.522311414 | 0.620571513 | 0.91579681 | -5.014358695 |
| **A_55_P2132756** | Cacna1g | -0.027312159 | -0.155991288 | 0.10136697 | 7.972505188 | -0.522176208 | 0.620659978 | 0.91579681 | -5.014408885 |
| **A_55_P2093889** | Arhgef19 | 0.025031967 | -0.092934803 | 0.142998737 | 7.976122067 | 0.522040809 | 0.620748576 | 0.91579681 | -5.014459136 |
| **A_66_P122559** | Myct1 | -0.023665306 | -0.13523901 | 0.087908398 | 7.866079818 | -0.521818451 | 0.620894091 | 0.91579681 | -5.014541634 |

| | | | | | | | | |
|---|---|---|---|---|---|---|---|---|
| **A_55_P2068027** | NA | -0.022188071 | -0.126809707 | 0.082433564 | 7.771207467 | -0.521755714 | 0.62093515 | 0.91579681 | -5.014564904 |
| **A_51_P316951** | Wipf3 | 0.050545662 | -0.187790911 | 0.288882235 | 10.42019729 | 0.521749886 | 0.620938965 | 0.91579681 | -5.014567066 |
| **A_55_P2096942** | Nrn1l | -0.379872468 | -2.17164628 | 1.411901344 | 10.12523226 | -0.521583443 | 0.621047905 | 0.91579681 | -5.014628792 |
| **A_55_P2038987** | Gas7 | 0.090525051 | -0.336563273 | 0.517613375 | 8.550568719 | 0.521459089 | 0.621129303 | 0.91579681 | -5.014674897 |
| **A_55_P2032034** | Tbp | 0.043730958 | -0.16262366 | 0.250085576 | 9.607340666 | 0.521367507 | 0.621189254 | 0.91579681 | -5.014708846 |
| **A_55_P2054619** | NA | 0.049758798 | -0.185043884 | 0.28456148 | 14.27903326 | 0.521357935 | 0.62119552 | 0.91579681 | -5.014712394 |
| **A_55_P2106426** | NA | -0.066854431 | -0.382330449 | 0.248621587 | 14.49577108 | -0.521354362 | 0.621197859 | 0.91579681 | -5.014713718 |
| **A_55_P2039611** | Eed | -0.034480331 | -0.197193813 | 0.12823315 | 12.35343338 | -0.521335212 | 0.621210396 | 0.91579681 | -5.014720816 |
| **A_55_P1965070** | Map7d1 | -0.080526332 | -0.460570394 | 0.29951773 | 14.70299636 | -0.521282479 | 0.621244917 | 0.91579681 | -5.01474036 |
| **A_55_P2014640** | Knop1 | 0.060890186 | -0.226488122 | 0.348268493 | 11.40975697 | 0.521269696 | 0.621253286 | 0.91579681 | -5.014745098 |
| **A_51_P516615** | Ndufb10 | 0.047876015 | -0.178090492 | 0.273842521 | 13.82108929 | 0.521246417 | 0.621268526 | 0.91579681 | -5.014753725 |
| **A_55_P2116496** | Tuba1b | 0.079334877 | -0.295160277 | 0.453830031 | 13.51951057 | 0.521179247 | 0.621312502 | 0.91579681 | -5.014778616 |
| **A_55_P1960941** | Wac | -0.039326524 | -0.224966455 | 0.146313407 | 11.95865402 | -0.521174703 | 0.621315477 | 0.91579681 | -5.014780299 |
| **A_55_P1966863** | Mad2l2 | -0.034516296 | -0.197472697 | 0.128440104 | 11.40784381 | -0.521101029 | 0.621363713 | 0.91579681 | -5.014807597 |
| **A_52_P565496** | Prl8a1 | 0.115481007 | -0.429728997 | 0.660691011 | 12.36602352 | 0.521093803 | 0.621368444 | 0.91579681 | -5.014810274 |
| **A_51_P346575** | Rhag | 0.023264119 | -0.086575137 | 0.133103375 | 7.704699799 | 0.521072545 | 0.621382362 | 0.91579681 | -5.01481815 |
| **A_55_P2159575** | Reep6 | 0.050775256 | -0.188968405 | 0.290518916 | 12.158716 | 0.5210437 | 0.621401248 | 0.91579681 | -5.014828836 |
| **A_55_P2120931** | Dleu2 | 0.033252274 | -0.123778773 | 0.190283322 | 8.330393566 | 0.520960736 | 0.621455571 | 0.91579681 | -5.014859569 |
| **A_51_P349015** | Pacrg | -0.027984157 | -0.160144826 | 0.104176511 | 7.890807468 | -0.520929762 | 0.621475853 | 0.91579681 | -5.014871042 |
| **A_52_P613977** | NA | 0.033812672 | -0.125955411 | 0.193580755 | 11.04549506 | 0.520665297 | 0.621649037 | 0.91598218 | -5.014968976 |
| **A_55_P1976764** | Polr3e | -0.038441516 | -0.220252822 | 0.143369791 | 10.30860061 | -0.520174163 | 0.621970727 | 0.916386323 | -5.01515073 |
| **A_51_P109171** | Os9 | -0.126524461 | -0.725047162 | 0.47199824 | 12.55785237 | -0.520071472 | 0.622038 | 0.916415587 | -5.005180685 |
| **A_55_P1976744** | NA | 0.056599415 | -0.211208368 | 0.324407197 | 8.146267319 | 0.519945601 | 0.622120464 | 0.916467225 | -5.015235263 |
| **A_55_P2006940** | Sfmbt1 | -0.044208106 | -0.253435455 | 0.165019243 | 9.074013463 | -0.519819561 | 0.622203046 | 0.916497019 | -5.015281864 |
| **A_51_P494825** | Mfhas1 | -0.063258832 | -0.362703627 | 0.236185962 | 11.19688643 | -0.519724952 | 0.622265038 | 0.916497019 | -5.015316838 |
| **A_51_P173601** | Bcam | -0.029131769 | -0.167038441 | 0.108774903 | 8.222964575 | -0.519697636 | 0.622282936 | 0.916497019 | -5.015326935 |
| **A_51_P402536** | Uvrag | -0.06994554 | -0.401124492 | 0.261233412 | 12.30812169 | -0.519596794 | 0.622349017 | 0.916524507 | -5.015364205 |
| **A_55_P1959683** | Xlr3b | -0.051633832 | -0.296245903 | 0.192978239 | 10.04922458 | -0.519308746 | 0.622537793 | 0.916659019 | -5.015470629 |
| **A_55_P2142830** | Ninj1 | -0.055106448 | -0.316188304 | 0.205975407 | 14.55205573 | -0.519271987 | 0.622561886 | 0.916659019 | -5.015484206 |
| **A_52_P572447** | Agpat5 | 0.092399726 | -0.345418853 | 0.530218305 | 11.16559881 | 0.519213122 | 0.622600468 | 0.916659019 | -5.015505947 |
| **A_55_P1970033** | Per1 | -0.06595159 | -0.378488214 | 0.246585034 | 12.16263934 | -0.519150788 | 0.622641326 | 0.916659019 | -5.015528967 |
| **A_51_P328968** | Zfp322a | 0.033721183 | -0.126096409 | 0.193538775 | 11.00553121 | 0.519095639 | 0.622677476 | 0.916659019 | -5.015549331 |
| **A_55_P2153827** | Fgfr1op | -0.036504961 | -0.209598042 | 0.13658812 | 10.3922635 | -0.518849427 | 0.622838878 | 0.916666088 | -5.015640224 |
| **A_52_P656867** | Fam98b | 0.038936258 | -0.145691209 | 0.223563725 | 9.577859251 | 0.518832373 | 0.622850059 | 0.916666088 | -5.015646518 |
| **A_66_P106654** | Camsap1 | -0.059148683 | -0.339644711 | 0.221347346 | 10.64470752 | -0.518785136 | 0.622881028 | 0.916666088 | -5.015663952 |
| **A_55_P2153631** | NA | -0.038423833 | -0.220650883 | 0.143803218 | 8.329565265 | -0.518748673 | 0.622904933 | 0.916666088 | -5.015677408 |
| **A_51_P314397** | Crip2 | -0.042247334 | -0.242616061 | 0.158121394 | 16.23083015 | -0.51872661 | 0.622919399 | 0.916666088 | -5.015685549 |
| **A_52_P537381** | Dimt1 | -0.048121593 | -0.276460443 | 0.180217257 | 9.925151051 | -0.518476828 | 0.623083177 | 0.916789114 | -5.015777702 |
| **A_55_P2073734** | Bfar | -0.040268828 | -0.231411729 | 0.150874072 | 11.79348715 | -0.518298538 | 0.623200094 | 0.916789114 | -5.015843455 |
| **A_52_P458697** | Higd1a | 0.046323658 | -0.173595024 | 0.26624234 | 11.80146135 | 0.518214889 | 0.623254953 | 0.916789114 | -5.015874298 |
| **A_52_P541175** | Marveld1 | 0.028937175 | -0.108442092 | 0.166316441 | 8.575529715 | 0.51820798 | 0.623259484 | 0.916789114 | -5.015876846 |
| **A_55_P1973349** | H4c14 | 0.053407618 | -0.200168099 | 0.306983336 | 12.26390923 | 0.518160989 | 0.623290303 | 0.916789114 | -5.01589417 |
| **A_55_P2136906** | Vpreb3 | 0.030357571 | -0.113786804 | 0.174501945 | 8.006445344 | 0.518129688 | 0.623310831 | 0.916789114 | -5.015905709 |
| **A_55_P2064240** | Hemk1 | -0.030775299 | -0.176940428 | 0.115389829 | 8.599409812 | -0.517997511 | 0.623397525 | 0.916789114 | -5.015954429 |
| **A_55_P1992571** | NA | 0.076155931 | -0.285582406 | 0.437894268 | 13.06464782 | 0.517938695 | 0.623436105 | 0.916789114 | -5.015976105 |
| **A_66_P132846** | Exosc1 | -0.027325898 | -0.157133529 | 0.102481734 | 9.533281426 | -0.517896975 | 0.62346347 | 0.916789114 | -5.015991479 |
| **A_55_P2124026** | Synj2 | 0.048480447 | -0.181828123 | 0.278789017 | 9.777693897 | 0.517875878 | 0.623477309 | 0.916789114 | -5.015999253 |
| **A_55_P1962084** | H2ac18 | -0.127754826 | -0.734837535 | 0.479327883 | 14.318016 | -0.517724378 | 0.623576692 | 0.916865501 | -5.01605507 |
| **A_51_P367125** | Pold1 | 0.042353615 | -0.159081548 | 0.243788778 | 9.21967854 | 0.517278425 | 0.623869284 | 0.917189886 | -5.016219289 |
| **A_55_P2083063** | Mob1b | -0.033757485 | -0.194320023 | 0.126805053 | 8.27263197 | -0.517243479 | 0.623892215 | 0.917189886 | -5.016232153 |

| | | | | | | | | | |
|---|---|---|---|---|---|---|---|---|---|
| **A_52_P3845** | Rnf217 | 0.051860602 | -0.194874254 | 0.298595459 | 8.44021228 | 0.517101994 | 0.623985061 | 0.917256617 | -5.016284224 |
| **A_55_P2070296** | NA | 0.025735489 | -0.096730851 | 0.148201829 | 7.650982715 | 0.516993221 | 0.624056447 | 0.917267165 | -5.016324248 |
| **A_51_P290576** | Plk2 | 0.036763757 | -0.13824326 | 0.211770774 | 9.493413814 | 0.51681319 | 0.624174607 | 0.917267165 | -5.016390476 |
| **A_51_P112734** | Slc7a8 | -0.063815551 | -0.367637078 | 0.240005976 | 12.41275267 | -0.516746024 | 0.624218693 | 0.917267165 | -5.006391398 |
| **A_51_P437021** | Trak1 | -0.054079253 | -0.311549924 | 0.203391418 | 9.921193969 | -0.516740004 | 0.624222644 | 0.917267165 | -5.016417393 |
| **A_55_P2085225** | NA | -0.038359182 | -0.221005871 | 0.144287508 | 10.99648372 | -0.516685997 | 0.624258095 | 0.917267165 | -5.016437254 |
| **A_55_P2177911** | Lepr | -0.142665679 | -0.822004056 | 0.536672698 | 14.42152202 | -0.516657242 | 0.62427697 | 0.917267165 | -5.016447828 |
| **A_55_P2360501** | NA | -0.074130713 | -0.427243995 | 0.278982568 | 8.153938836 | -0.51647974 | 0.624393493 | 0.917368639 | -5.016513088 |
| **A_55_P1970665** | Kdm3b | -0.026314703 | -0.151700117 | 0.099070711 | 8.231601633 | -0.516321983 | 0.624497064 | 0.917387547 | -5.016571073 |
| **A_51_P286106** | NA | -0.051897213 | -0.299182335 | 0.19538791 | 8.730549936 | -0.516315553 | 0.624501286 | 0.917387547 | -5.016573436 |
| **A_55_P2023081** | Dnaja2 | -0.064444225 | -0.371590059 | 0.242701609 | 9.949683662 | -0.516188754 | 0.624584539 | 0.917438427 | -5.016620029 |
| **A_55_P2036460** | Synb | -0.099237823 | -0.572277097 | 0.37380145 | 11.02595416 | -0.516118224 | 0.62463085 | 0.917438427 | -5.016645942 |
| **A_51_P276063** | Phyh | -0.125267718 | -0.722705516 | 0.47217008 | 11.88889942 | -0.515840735 | 0.624813072 | 0.917581042 | -5.016747861 |
| **A_55_P2096204** | Kdm1b | 0.065047042 | -0.245190014 | 0.375284099 | 11.43967097 | 0.515825783 | 0.624822891 | 0.917581042 | -5.016753351 |
| **A_55_P1983683** | Rtl5 | -0.026338262 | -0.151993567 | 0.099317042 | 7.947066535 | -0.515674259 | 0.624922408 | 0.917629227 | -5.016808982 |
| **A_66_P101942** | Gm9706 | 0.103914665 | -0.391898848 | 0.599728179 | 11.99142819 | 0.515617499 | 0.624959689 | 0.917629227 | -5.016829818 |
| **A_55_P2008895** | Tmsb15b1 | -0.023064345 | -0.133150709 | 0.087022019 | 8.282507239 | -0.515438396 | 0.625077333 | 0.917629227 | -5.01689555 |
| **A_55_P1998947** | Trim16 | 0.06121647 | -0.230988385 | 0.353421326 | 8.42307238 | 0.515406654 | 0.625098184 | 0.917629227 | -5.016907197 |
| **A_51_P376445** | Rhox5 | -0.051717006 | -0.298598177 | 0.195164165 | 16.28407213 | -0.515364584 | 0.62512582 | 0.917629227 | -5.016922633 |
| **A_55_P1956762** | Sgms1 | -0.097807307 | -0.56472979 | 0.369115177 | 8.773941036 | -0.515342163 | 0.625140549 | 0.917629227 | -5.016930859 |
| **A_51_P309920** | Itga8 | 0.05564611 | -0.210047122 | 0.321339341 | 8.584655308 | 0.515256502 | 0.625196823 | 0.917642144 | -5.016962285 |
| **A_55_P2127884** | Btbd1 | -0.024109595 | -0.139300736 | 0.091081546 | 7.945838147 | -0.51492028 | 0.625417727 | 0.917896678 | -5.017085587 |
| **A_51_P125205** | Aqp1 | -0.567948072 | -3.282666996 | 2.146770851 | 12.78282487 | -0.514698524 | 0.625563449 | 0.917994183 | -5.017166873 |
| **A_51_P141390** | Anapc1 | -0.05451621 | -0.315108797 | 0.206076376 | 11.99076234 | -0.514674634 | 0.625579149 | 0.917994183 | -5.017175628 |
| **A_55_P2007738** | Mir22hg | 0.036273188 | -0.137145573 | 0.209691948 | 8.587701536 | 0.514586998 | 0.625636743 | 0.918009003 | -5.017207742 |
| **A_51_P116665** | 2610005L07Rik | -0.074036267 | -0.428081828 | 0.280009295 | 9.137563241 | -0.514463444 | 0.625717947 | 0.918058463 | -5.017253009 |
| **A_55_P1966134** | Prl3c1 | 0.093196254 | -0.352543807 | 0.538936314 | 9.519826452 | 0.514382222 | 0.625771332 | 0.918067102 | -5.017282762 |
| **A_55_P2151308** | G430095P16Rik | 0.024456116 | -0.092537434 | 0.141449665 | 7.857010935 | 0.514274186 | 0.625842345 | 0.9181016 | -5.01732233 |
| **A_52_P366803** | Cyp3a44 | -0.022890504 | -0.132426054 | 0.086645046 | 7.643758415 | -0.514125834 | 0.625939864 | 0.918110031 | -5.017376652 |
| **A_51_P309501** | Praf2 | -0.048181989 | -0.27878289 | 0.182418913 | 8.839220083 | -0.514035226 | 0.62599943 | 0.918110031 | -5.017409824 |
| **A_51_P196925** | Cx3cl1 | -0.051074612 | -0.295532479 | 0.193383255 | 10.19064965 | -0.514008407 | 0.626017062 | 0.918110031 | -5.017419641 |
| **A_66_P125777** | Slc43a2 | 0.104403802 | -0.395334086 | 0.60414169 | 11.44476982 | 0.513976425 | 0.626038088 | 0.918110031 | -5.017431348 |
| **A_55_P1992421** | Hmgb3 | 0.074423368 | -0.28187988 | 0.430726615 | 9.788248128 | 0.513876431 | 0.62610383 | 0.918136783 | -5.017467946 |
| **A_55_P2078395** | Gm1943 | 0.051091995 | -0.19355212 | 0.295736111 | 8.531825889 | 0.513791901 | 0.626159408 | 0.918148627 | -5.017498879 |
| **A_51_P167513** | Leng1 | 0.028938005 | -0.10971026 | 0.167586269 | 10.39738963 | 0.513479737 | 0.626364678 | 0.918359095 | -5.017613073 |
| **A_52_P663585** | 10-Sep | -0.036940869 | -0.213950043 | 0.140068305 | 8.325574283 | -0.513429117 | 0.626397967 | 0.918359095 | -5.017631585 |
| **A_55_P2097474** | Creb5 | 0.074139756 | -0.281216526 | 0.429496037 | 15.81829909 | 0.513282333 | 0.626494504 | 0.918389803 | -5.017685256 |
| **A_55_P1984675** | Abhd6 | 0.071724178 | -0.272073889 | 0.415522244 | 11.88973787 | 0.513252782 | 0.62651394 | 0.918389803 | -5.017696059 |
| **A_55_P2135023** | NA | 0.046112998 | -0.175047416 | 0.267273412 | 11.3356469 | 0.512961923 | 0.626705258 | 0.918561323 | -5.017802364 |
| **A_55_P1966790** | Impdh2 | -0.051900058 | -0.300843717 | 0.1970436 | 11.26995589 | -0.512903831 | 0.626743473 | 0.918561323 | -5.017823589 |
| **A_51_P424338** | Nqo1 | 0.042850954 | -0.162706099 | 0.248408006 | 8.023733971 | 0.512858163 | 0.626773516 | 0.918561323 | -5.017840274 |
| **A_51_P470079** | Il1r2 | -0.271893161 | -1.576908084 | 1.033121762 | 11.14978022 | -0.512568566 | 0.626964047 | 0.918766325 | -5.017946045 |
| **A_55_P1975742** | Dap3 | -0.023459804 | -0.136075463 | 0.089155854 | 10.83193831 | -0.512501062 | 0.627008464 | 0.918766325 | -5.017970692 |
| **A_66_P119174** | Cmtm6 | -0.071749623 | -0.416239146 | 0.2727399 | 12.58697069 | -0.512404302 | 0.627072134 | 0.918789969 | -5.018006017 |
| **A_55_P2138718** | H2al1a | 0.024251639 | -0.092215578 | 0.140718856 | 7.769179583 | 0.512279006 | 0.627154586 | 0.918841128 | -5.018051751 |
| **A_52_P471395** | Ifngr2 | -0.049188754 | -0.285556731 | 0.187179223 | 10.09845429 | -0.511972162 | 0.627356532 | 0.919067335 | -5.018163708 |
| **A_51_P111612** | Arrdc4 | 0.048398234 | -0.184235822 | 0.281032289 | 10.77753691 | 0.511829583 | 0.627450382 | 0.919135159 | -5.008168813 |
| **A_51_P301311** | Ciz1 | 0.036530228 | -0.139102916 | 0.212163373 | 11.25648409 | 0.511699596 | 0.627535949 | 0.919190843 | -5.018263109 |
| **A_52_P556099** | Oip5 | -0.022750221 | -0.132185953 | 0.08668551 | 7.786283653 | -0.511441117 | 0.627706119 | 0.91937043 | -5.018357329 |
| **A_55_P2219765** | Wnt5a | 0.04831002 | -0.184154059 | 0.280774099 | 8.280075372 | 0.511270259 | 0.627818617 | 0.919465529 | -5.018419586 |

| | | | | | | | | | |
|---|---|---|---|---|---|---|---|---|---|
| **A_55_P2023607** | Ikzf1 | -0.024111067 | -0.140221094 | 0.091998959 | 7.710717469 | -0.510876444 | 0.628077959 | 0.919775655 | -5.018563016 |
| **A_55_P2011659** | Anapc5 | -0.062568784 | -0.364191834 | 0.239054266 | 10.0625717 | -0.510343222 | 0.6284292 | 0.920092742 | -5.018757062 |
| **A_52_P244193** | Cd24a | -0.031249915 | -0.181904133 | 0.119404303 | 10.89639479 | -0.51031309 | 0.628449051 | 0.920092742 | -5.018768022 |
| **A_55_P2052106** | BC049762 | 0.040395224 | -0.154406884 | 0.235197333 | 9.844168668 | 0.510158975 | 0.62855059 | 0.920092742 | -5.01882407 |
| **A_55_P2143042** | Gm8096 | 0.083689839 | -0.319919826 | 0.487299505 | 8.945360645 | 0.510129383 | 0.628570088 | 0.920092742 | -5.01883483 |
| **A_66_P137660** | Fam166b | 0.022322527 | -0.085342283 | 0.129987336 | 7.83980127 | 0.510080519 | 0.628602285 | 0.920092742 | -5.018852597 |
| **A_55_P2032946** | Gsta1 | 0.051055947 | -0.195245711 | 0.297357605 | 8.143407746 | 0.509974148 | 0.628672376 | 0.920092742 | -5.018891268 |
| **A_51_P256945** | Opa3 | -0.035703499 | -0.207944074 | 0.136537077 | 8.39251162 | -0.509969813 | 0.628675233 | 0.920092742 | -5.018892844 |
| **A_52_P306351** | Asph | 0.051718289 | -0.197780778 | 0.301217355 | 10.48188726 | 0.509969713 | 0.628675298 | 0.920092742 | -5.01889288 |
| **A_51_P236864** | Parp8 | 0.02881785 | -0.110248531 | 0.167884232 | 8.213885128 | 0.509810283 | 0.62878036 | 0.920121104 | -5.018950826 |
| **A_55_P2140383** | Mis18a | 0.029039799 | -0.111116991 | 0.169196588 | 8.746472516 | 0.509739889 | 0.628826752 | 0.920121104 | -5.018976406 |
| **A_52_P175376** | Tfcp2l1 | -0.04942391 | -0.287969572 | 0.189121752 | 8.403562416 | -0.5097236 | 0.628837487 | 0.920121104 | -5.018982325 |
| **A_66_P108685** | Cideb | 0.121761302 | -0.466280534 | 0.709803138 | 10.33873932 | 0.509413232 | 0.629042052 | 0.920350755 | -5.019095069 |
| **A_55_P2140002** | Tnfaip8 | -0.057961377 | -0.337960324 | 0.22203757 | 10.72699498 | -0.509273951 | 0.629133865 | 0.920415415 | -5.019145644 |
| **A_55_P2175502** | M1ap | 0.045258966 | -0.173496058 | 0.264013991 | 8.227125685 | 0.508997628 | 0.629316036 | 0.920424437 | -5.019245944 |
| **A_55_P2036495** | NA | 0.029576202 | -0.113380435 | 0.172532838 | 13.76295276 | 0.508987644 | 0.629322618 | 0.920424437 | -5.019249567 |
| **A_55_P1963036** | Oxr1 | 0.047400758 | -0.181711169 | 0.276512686 | 10.94335307 | 0.508987076 | 0.629322993 | 0.920424437 | -5.019249774 |
| **A_51_P392303** | Lrrn3 | -0.033880388 | -0.197645166 | 0.129884391 | 8.067056489 | -0.508975679 | 0.629330507 | 0.920424437 | -5.019253909 |
| **A_55_P2003753** | Sh2d5 | 0.041921638 | -0.160823223 | 0.2446665 | 8.857397122 | 0.508695099 | 0.629515516 | 0.920625361 | -5.019355702 |
| **A_51_P111962** | Bean1 | -0.088231809 | -0.515055644 | 0.338592026 | 8.210625657 | -0.508564081 | 0.629601917 | 0.920682058 | -5.019403218 |
| **A_55_P2447879** | Asxl2 | -0.042888108 | -0.250408256 | 0.164632041 | 8.58634811 | -0.508447103 | 0.629679064 | 0.920717612 | -5.019445632 |
| **A_55_P1958562** | Dmrtc1b | -0.091615133 | -0.535026743 | 0.351796476 | 12.17786231 | -0.50831078 | 0.629768976 | 0.920717612 | -5.01949505 |
| **A_55_P2076792** | Anxa6 | -0.055958073 | -0.326791832 | 0.214875686 | 11.82363153 | -0.508310544 | 0.629769132 | 0.920717612 | -5.019495136 |
| **A_51_P324491** | Cmpk1 | 0.050424718 | -0.193668522 | 0.294517958 | 10.43771591 | 0.508226009 | 0.629824891 | 0.920727682 | -5.019525774 |
| **A_55_P2096777** | Tprn | 0.03653692 | -0.140353504 | 0.213427344 | 9.415508448 | 0.508155669 | 0.629871289 | 0.920727682 | -5.019551265 |
| **A_52_P171724** | 2310009B15Rik | -0.040491397 | -0.236574883 | 0.155592088 | 9.811726913 | -0.508031813 | 0.629952991 | 0.920758929 | -5.019596141 |
| **A_55_P1952720** | Kctd2 | -0.069951619 | -0.40875616 | 0.268852922 | 12.17981004 | -0.507946196 | 0.630009473 | 0.920758929 | -5.019627157 |
| **A_55_P1976926** | Tmem104 | -0.079380173 | -0.463881035 | 0.30512069 | 8.851205232 | -0.507906634 | 0.630035573 | 0.920758929 | -5.019641487 |
| **A_51_P256246** | Tspan13 | 0.15610197 | -0.600340175 | 0.912544115 | 11.55362358 | 0.507693766 | 0.630176017 | 0.920829106 | -5.019718575 |
| **A_55_P2133370** | NA | 0.073245999 | -0.281693997 | 0.428185994 | 10.63059303 | 0.507689439 | 0.630178872 | 0.920829106 | -5.019720142 |
| **A_51_P373609** | Yars2 | -0.03249566 | -0.190001243 | 0.125009923 | 9.836418545 | -0.507573075 | 0.630255653 | 0.920871684 | -5.019762269 |
| **A_55_P2081615** | Timeless | 0.030279774 | -0.116514362 | 0.177073911 | 8.004583535 | 0.507473176 | 0.630321573 | 0.920898389 | -5.019798429 |
| **A_55_P1966649** | Glmn | -0.041218966 | -0.241097633 | 0.158659701 | 9.486809787 | -0.50734083 | 0.630408911 | 0.920956378 | -5.019846325 |
| **A_55_P2100866** | NA | 0.052790209 | -0.203287741 | 0.308868159 | 10.20093951 | 0.507166267 | 0.630524118 | 0.921052553 | -5.019909481 |
| **A_55_P1971009** | Gzme | 0.297411497 | -1.145850873 | 1.740673867 | 12.27324593 | 0.506969261 | 0.630654151 | 0.921052553 | -5.019980733 |
| **A_55_P2122987** | Dlx3 | 0.033392169 | -0.128652585 | 0.195436923 | 11.28211633 | 0.506965975 | 0.63065632 | 0.921052553 | -5.019981922 |
| **A_55_P1975396** | Ighmbp2 | -0.05605323 | -0.328095259 | 0.2159888 | 9.114752608 | -0.50691343 | 0.630691004 | 0.921052553 | -5.020000922 |
| **A_55_P2005908** | Lin9 | 0.022264544 | -0.085798629 | 0.130327716 | 7.794142424 | 0.506880108 | 0.630713001 | 0.921052553 | -5.02001297 |
| **A_55_P2175381** | NA | -0.029830661 | -0.174670198 | 0.115008876 | 16.6559833 | -0.506693009 | 0.630836514 | 0.921067166 | -5.020080607 |
| **A_55_P2156504** | NA | -0.052010477 | -0.304545961 | 0.200525007 | 11.52068297 | -0.506684466 | 0.630842154 | 0.921067166 | -5.020083694 |
| **A_52_P405145** | NA | 0.074389279 | -0.286831592 | 0.43561015 | 8.918678492 | 0.506648401 | 0.630865964 | 0.921067166 | -5.020096729 |
| **A_55_P1983046** | Plch1 | 0.023586998 | -0.090966089 | 0.138140085 | 7.725871087 | 0.506564836 | 0.630921135 | 0.921068904 | -5.020126929 |
| **A_52_P367294** | Fsd1l | -0.045661543 | -0.267462374 | 0.176139288 | 8.215084407 | -0.506473327 | 0.630981554 | 0.921068904 | -5.020159994 |
| **A_55_P2006476** | NA | 0.059265883 | -0.228727259 | 0.347259025 | 11.21780693 | 0.506281172 | 0.631108434 | 0.921068904 | -5.020229408 |
| **A_55_P1989986** | Fam149b | 0.023031502 | -0.088887655 | 0.134950659 | 9.234460606 | 0.506275599 | 0.631112115 | 0.921068904 | -5.020231421 |
| **A_55_P2048406** | NA | -0.06596635 | -0.386540655 | 0.254607955 | 11.72620438 | -0.506247519 | 0.631130657 | 0.921068904 | -5.020241562 |
| **A_51_P414548** | Casp7 | -0.034449178 | -0.201871834 | 0.132973479 | 8.956731203 | -0.506213583 | 0.631153067 | 0.921068904 | -5.020253819 |
| **A_52_P432685** | Ctdsp2 | -0.040958915 | -0.240065747 | 0.158147916 | 9.27934005 | -0.506094302 | 0.631231839 | 0.921111239 | -5.020296891 |
| **A_55_P2008591** | Kras | 0.049017552 | -0.189295881 | 0.287330986 | 8.506488165 | 0.506025336 | 0.631277386 | 0.921111239 | -5.020321791 |
| **A_55_P2172001** | Mrgpra2a | 0.021984846 | -0.084922932 | 0.128892624 | 7.713435738 | 0.50592168 | 0.631345846 | 0.921141595 | -5.020359209 |

| | | | | | | | | |
|---|---|---|---|---|---|---|---|---|
| **A_55_P2046812** | Sparc | 0.057493893 | -0.222163227 | 0.337151012 | 17.49531243 | 0.505783896 | 0.631436852 | 0.92120484 | -5.020408937 |
| **A_55_P2128821** | Selenok | 0.03388464 | -0.130973892 | 0.198743172 | 14.36626657 | 0.505662345 | 0.631517142 | 0.921249374 | -5.020452796 |
| **A_51_P455166** | Prl4a1 | -0.104686702 | -0.61408713 | 0.404713725 | 13.48120617 | -0.505593378 | 0.631562701 | 0.921249374 | -5.020477677 |
| **A_55_P1998374** | Ppfibp1 | -0.033465692 | -0.196384582 | 0.129453198 | 12.05785264 | -0.505356123 | 0.631719441 | 0.921298718 | -5.020563248 |
| **A_55_P2159600** | Herc1 | 0.065287571 | -0.252618858 | 0.383194001 | 11.1953553 | 0.505243069 | 0.631794137 | 0.921298718 | -5.020604011 |
| **A_55_P2108883** | NA | 0.031382691 | -0.121470515 | 0.184235897 | 9.092912399 | 0.505108634 | 0.631882966 | 0.921298718 | -5.020652473 |
| **A_52_P460791** | Ppy | 0.038026694 | -0.147237446 | 0.223290835 | 8.421162783 | 0.504970924 | 0.631973965 | 0.921298718 | -5.020702103 |
| **A_51_P342716** | Ppp1cb | -0.05194298 | -0.305023623 | 0.201137663 | 13.07700998 | -0.504936887 | 0.631996458 | 0.921298718 | -5.020714368 |
| **A_65_P12104** | Tmtc1 | -0.050496873 | -0.296536902 | 0.195543156 | 8.552643022 | -0.504926167 | 0.632003543 | 0.921298718 | -5.020718231 |
| **A_55_P2013371** | Ttc29 | -0.021212083 | -0.124587078 | 0.082162912 | 7.662141142 | -0.5048205 | 0.632073375 | 0.921298718 | -5.020756302 |
| **A_52_P432570** | Chdh | -0.049067652 | -0.288206252 | 0.190070948 | 8.644589136 | -0.504794667 | 0.632090447 | 0.921298718 | -5.020765608 |
| **A_55_P1989419** | AI987944 | 0.029377305 | -0.113798055 | 0.172552665 | 8.668813823 | 0.50479243 | 0.632091925 | 0.921298718 | -5.020766414 |
| **A_55_P2068066** | Rapgefl1 | -0.025232558 | -0.148254877 | 0.097789761 | 7.876048187 | -0.504599172 | 0.632219656 | 0.921298718 | -5.020836021 |
| **A_51_P326994** | Calhm2 | -0.04742702 | -0.278689284 | 0.183835244 | 8.551151338 | -0.504533752 | 0.632262897 | 0.921298718 | -5.020859578 |
| **A_51_P260490** | Ralbp1 | -0.057070803 | -0.335369292 | 0.221227685 | 13.07343926 | -0.50451293 | 0.632276661 | 0.921298718 | -5.020867076 |
| **A_55_P2077048** | Itih5 | -0.090087209 | -0.529443176 | 0.349268758 | 9.520110844 | -0.504447259 | 0.63232007 | 0.921298718 | -5.02089072 |
| **A_52_P65494** | Iqgap2 | 0.034828757 | -0.135064174 | 0.204721688 | 8.473604321 | 0.504349778 | 0.63238451 | 0.921298718 | -5.020925813 |
| **A_55_P2101862** | Spryd3 | -0.029966644 | -0.176150912 | 0.116217624 | 10.57963908 | -0.504320511 | 0.632403857 | 0.921298718 | -5.020936347 |
| **A_52_P302395** | Lnpep | 0.085420824 | -0.331283863 | 0.50212551 | 9.831106967 | 0.504318074 | 0.632405468 | 0.921298718 | -5.020937224 |
| **A_52_P131548** | Ajuba | -0.064006489 | -0.376280318 | 0.24826734 | 9.685717958 | -0.504263552 | 0.632441512 | 0.921298718 | -5.020956848 |
| **A_55_P2151666** | NA | -0.044305927 | -0.260473381 | 0.171861528 | 8.458011567 | -0.504243932 | 0.632454482 | 0.921298718 | -5.020963909 |
| **A_55_P2271742** | NA | -0.029329723 | -0.172461207 | 0.11380176 | 8.12648803 | -0.504129322 | 0.632530254 | 0.921303412 | -5.021005153 |
| **A_55_P2073084** | Gtf3c2 | -0.03990839 | -0.234729174 | 0.154912394 | 9.90468977 | -0.503962344 | 0.632640656 | 0.921303412 | -5.021065226 |
| **A_55_P2121096** | Fbn2 | 0.225642915 | -0.875888984 | 1.327174813 | 9.359731147 | 0.503957353 | 0.632643956 | 0.921303412 | -5.021067021 |
| **A_55_P2107957** | Abraxas1 | -0.033271694 | -0.195711182 | 0.129167794 | 8.033953623 | -0.503909404 | 0.632675661 | 0.921303412 | -5.021084268 |
| **A_55_P1956734** | NA | -0.034544669 | -0.203234098 | 0.134144759 | 8.589606921 | -0.503804842 | 0.632744802 | 0.921303412 | -5.021121874 |
| **A_52_P331762** | Lmo1 | -0.025263623 | -0.148681684 | 0.098154438 | 7.778242496 | -0.503600404 | 0.632879998 | 0.921303412 | -5.02119538 |
| **A_51_P236412** | Cdkl2 | -0.038520175 | -0.226738441 | 0.149698091 | 9.384650572 | -0.503495561 | 0.632949338 | 0.921303412 | -5.021233066 |
| **A_55_P2182452** | Tnfsf15 | -0.023103235 | -0.136004361 | 0.089797891 | 7.785744691 | -0.503435321 | 0.632989179 | 0.921303412 | -5.021254716 |
| **A_55_P2040341** | NA | -0.078428088 | -0.461707233 | 0.304851057 | 9.357066449 | -0.503414361 | 0.633003043 | 0.921303412 | -5.021262248 |
| **A_52_P565575** | Arhgap20 | -0.041254883 | -0.242881945 | 0.160372179 | 8.015740944 | -0.503379708 | 0.633025963 | 0.921303412 | -5.021274701 |
| **A_51_P487999** | Sgo1 | 0.047719244 | -0.185517285 | 0.280955773 | 8.069510159 | 0.503345452 | 0.633048621 | 0.921303412 | -5.021287011 |
| **A_51_P464630** | Rad17 | -0.035530242 | -0.209274543 | 0.138214059 | 10.93102113 | -0.50310283 | 0.633209112 | 0.921303412 | -5.021374172 |
| **A_65_P16952** | Ywhaq | 0.041576272 | -0.161853213 | 0.245005758 | 11.56780541 | 0.502806419 | 0.633405212 | 0.921303412 | -5.021480606 |
| **A_55_P2142218** | NA | 0.068577273 | -0.267033491 | 0.404188037 | 14.67662941 | 0.502705333 | 0.633472097 | 0.921303412 | -5.021516891 |
| **A_51_P312437** | Dhrs7 | 0.068012505 | -0.26488434 | 0.40090935 | 12.51637697 | 0.502629826 | 0.633522059 | 0.921303412 | -5.02154399 |
| **A_55_P2083879** | Vsig2 | 0.057159784 | -0.222657571 | 0.336977139 | 10.24163615 | 0.502556731 | 0.633570427 | 0.921303412 | -5.02157022 |
| **A_55_P2132902** | Gprasp1 | 0.047380517 | -0.184569597 | 0.279330632 | 8.974179974 | 0.50254432 | 0.63357864 | 0.921303412 | -5.021574673 |
| **A_52_P573336** | Sbsn | 0.071883077 | -0.280026223 | 0.423792377 | 15.45230927 | 0.502533569 | 0.633585755 | 0.921303412 | -5.021578531 |
| **A_51_P466148** | Bckdha | 0.047312935 | -0.184313631 | 0.2789395 | 10.4956372 | 0.502528485 | 0.633589119 | 0.921303412 | -5.021580355 |
| **A_52_P226929** | Cflar | -0.05156007 | -0.303988714 | 0.200868575 | 8.309296664 | -0.502509209 | 0.633601875 | 0.921303412 | -5.021587271 |
| **A_55_P2084758** | Agfg1 | 0.068007776 | -0.264956048 | 0.4009716 | 10.80126212 | 0.502493774 | 0.633612089 | 0.921303412 | -5.021592809 |
| **A_52_P117420** | Tacc1 | 0.070409926 | -0.274327734 | 0.415147585 | 10.54587384 | 0.502474838 | 0.63362462 | 0.921303412 | -5.021599603 |
| **A_66_P108267** | Foxn4 | -0.022085169 | -0.130217733 | 0.086047395 | 7.659126458 | -0.502473758 | 0.633625335 | 0.921303412 | -5.021599991 |
| **A_51_P376149** | Eef1akmt1 | -0.057449426 | -0.338735219 | 0.223836366 | 11.01108922 | -0.502466443 | 0.633630175 | 0.921303412 | -5.021602615 |
| **A_55_P2038640** | Gcn1 | -0.048573716 | -0.286468115 | 0.189320682 | 9.275427505 | -0.50232672 | 0.633722644 | 0.921303412 | -5.021652737 |
| **A_55_P2019217** | NA | -0.021867741 | -0.128973118 | 0.085237635 | 7.956956757 | -0.502298426 | 0.633741369 | 0.921303412 | -5.021662885 |
| **A_52_P1197913** | Gadd45b | -0.065951432 | -0.388976221 | 0.257073357 | 12.30716923 | -0.502293482 | 0.633744641 | 0.921303412 | -5.021664658 |
| **A_55_P2086949** | Cntf | -0.041785259 | -0.246583458 | 0.16301294 | 8.587161089 | -0.501956552 | 0.633967653 | 0.921531089 | -5.021785465 |
| **A_55_P1959409** | Deup1 | -0.021787589 | -0.128582487 | 0.085007308 | 7.671983092 | -0.501912305 | 0.633996943 | 0.921531089 | -5.021801325 |

| | | | | | | | | | |
|---|---|---|---|---|---|---|---|---|---|
| **A_55_P2003115** | Rnf7 | 0.046609354 | -0.181953794 | 0.275172502 | 12.45085601 | 0.501690684 | 0.634143659 | 0.921531089 | -5.021880741 |
| **A_55_P1954365** | Ttc28 | -0.06109994 | -0.360737307 | 0.238537427 | 9.744311438 | -0.501665198 | 0.634160532 | 0.921531089 | -5.021889872 |
| **A_52_P423810** | Mt1 | -0.075173023 | -0.443835678 | 0.293489633 | 8.323139009 | -0.5016514 | 0.634169668 | 0.921531089 | -5.021894815 |
| **A_55_P2094963** | NA | -0.040278533 | -0.237913479 | 0.157356412 | 9.414152509 | -0.501393931 | 0.634340143 | 0.921531089 | -5.021987033 |
| **A_55_P2034245** | Meox1 | -0.022391735 | -0.132288464 | 0.087504993 | 7.728653946 | -0.501270508 | 0.634421872 | 0.921531089 | -5.022031224 |
| **A_55_P1954718** | Cyb561 | -0.036634425 | -0.216433387 | 0.143164537 | 12.56662375 | -0.5012696 | 0.634422473 | 0.921531089 | -5.022031549 |
| **A_55_P2013138** | NA | 0.052739293 | -0.206114258 | 0.311592844 | 12.21809168 | 0.501244178 | 0.634439308 | 0.921531089 | -5.02204065 |
| **A_55_P2000818** | Cab39l | 0.051760412 | -0.202323954 | 0.305844778 | 8.775966702 | 0.501174474 | 0.634485469 | 0.921531089 | -5.022065602 |
| **A_52_P334562** | Vdr | -0.095665436 | -0.565331917 | 0.374001044 | 8.756107625 | -0.501111827 | 0.634526957 | 0.921531089 | -5.022088026 |
| **A_55_P2182705** | Kpna2 | 0.094971349 | -0.371326556 | 0.561269255 | 11.43141417 | 0.501069891 | 0.634554731 | 0.921531089 | -5.022103034 |
| **A_55_P1974204** | Fam172a | -0.026653714 | -0.157529411 | 0.104221983 | 10.64087275 | -0.501034798 | 0.634577972 | 0.921531089 | -5.022115593 |
| **A_51_P315904** | Gadd45g | -0.171826138 | -1.015605844 | 0.671953569 | 14.77000621 | -0.500990297 | 0.634607446 | 0.921531089 | -5.022131518 |
| **A_55_P2038697** | Zfp235 | -0.032552326 | -0.192443846 | 0.127339194 | 8.466516949 | -0.500870849 | 0.634686561 | 0.921531089 | -5.022174255 |
| **A_55_P2074473** | Gtf2f1 | 0.048605515 | -0.190143993 | 0.287355022 | 13.38189628 | 0.500855243 | 0.634696898 | 0.921531089 | -5.022179838 |
| **A_55_P2048096** | Mzf1 | -0.025967174 | -0.153538123 | 0.101603775 | 8.062008197 | -0.500774336 | 0.634750489 | 0.921531089 | -5.02220878 |
| **A_51_P103650** | Trim8 | -0.049121169 | -0.290468521 | 0.192226183 | 11.62539867 | -0.500720438 | 0.634786192 | 0.921531089 | -5.006732671 |
| **A_55_P1987439** | Pigt | 0.113136823 | -0.442774091 | 0.669047737 | 11.74755911 | 0.500688874 | 0.634807101 | 0.921531089 | -5.022239347 |
| **A_51_P487678** | Dtnbp1 | -0.031862573 | -0.188481785 | 0.124756638 | 9.3612442 | -0.50050103 | 0.634931541 | 0.921594594 | -5.022306517 |
| **A_55_P2015403** | 9930111J21Rik1 | 0.043756866 | -0.171337783 | 0.258851516 | 9.278584246 | 0.500478893 | 0.634946206 | 0.921594594 | -5.022314431 |
| **A_55_P2058359** | Macroh2a1 | -0.02796214 | -0.165530087 | 0.109605807 | 8.328495241 | -0.500060224 | 0.63522361 | 0.921928004 | -5.022464052 |
| **A_55_P1986421** | Tcp11 | 0.021346773 | -0.083693219 | 0.126386766 | 7.72199805 | 0.499973195 | 0.635281282 | 0.921942481 | -5.02249514 |
| **A_55_P1962284** | Klhl24 | -0.022087716 | -0.130851492 | 0.086676059 | 7.808639947 | -0.499615274 | 0.635518499 | 0.922217497 | -5.022622944 |
| **A_51_P483658** | Trim3 | 0.030332427 | -0.119088168 | 0.179753022 | 9.378704125 | 0.499419918 | 0.635647993 | 0.92224126 | -5.022692666 |
| **A_51_P367866** | Egr1 | 0.032424958 | -0.127318695 | 0.192168611 | 8.832601692 | 0.499372907 | 0.635679158 | 0.92224126 | -5.022709441 |
| **A_66_P117389** | Rcn2 | -0.052958826 | -0.313892238 | 0.207974586 | 10.29737299 | -0.499318685 | 0.635715103 | 0.92224126 | -5.022728786 |
| **A_55_P2082570** | NA | -0.024746221 | -0.146692441 | 0.0972 | 8.412374835 | -0.499240375 | 0.635767019 | 0.92224126 | -5.022756723 |
| **A_51_P313483** | Dyrk1a | -0.035218643 | -0.208774993 | 0.138337707 | 10.27070802 | -0.499230692 | 0.635773438 | 0.92224126 | -5.022760177 |
| **A_55_P2063646** | Krcc1 | 0.057944311 | -0.227705298 | 0.34359392 | 12.91080877 | 0.49905261 | 0.635891507 | 0.922343309 | -5.022823691 |
| **A_51_P428086** | Sptbn1 | -0.036094822 | -0.214060794 | 0.14187115 | 8.810319902 | -0.498973058 | 0.635944254 | 0.922350604 | -5.022852058 |
| **A_55_P2172688** | Srbd1 | -0.042200845 | -0.250351839 | 0.165950149 | 8.428463262 | -0.498783207 | 0.636070145 | 0.922463973 | -5.022919738 |
| **A_55_P2162855** | Nipa2 | 0.027628351 | -0.108676881 | 0.163933584 | 10.66130754 | 0.498668121 | 0.636146466 | 0.922505442 | -5.022960754 |
| **A_55_P2097448** | Rbpms | -0.051322713 | -0.304577302 | 0.201931877 | 9.921912235 | -0.498564604 | 0.636215118 | 0.922535785 | -5.022997639 |
| **A_51_P380013** | Bcl2 | -0.028029099 | -0.166368484 | 0.110310286 | 8.181342068 | -0.498462471 | 0.636282856 | 0.922564799 | -5.023034025 |
| **A_55_P2162503** | Tbx20 | 0.021606759 | -0.085095358 | 0.128308876 | 7.688708779 | 0.49817939 | 0.636470627 | 0.922767832 | -5.023134841 |
| **A_55_P1964985** | Gm10240 | 0.040040892 | -0.157756314 | 0.237838099 | 16.10120564 | 0.498026849 | 0.636571821 | 0.922796255 | -5.023189145 |
| **A_55_P2144636** | Ccdc163 | -0.033374839 | -0.198249475 | 0.131499797 | 9.4336218 | -0.498005907 | 0.636585714 | 0.922796255 | -5.023196599 |
| **A_55_P1999800** | Gm14066 | 0.024875194 | -0.098063629 | 0.147814018 | 7.833979505 | 0.497790487 | 0.636728638 | 0.922886691 | -5.02327326 |
| **A_51_P368496** | Tmem98 | 0.049737701 | -0.196097262 | 0.295572663 | 8.987420184 | 0.497749942 | 0.63675554 | 0.922886691 | -5.023287685 |
| **A_55_P2049687** | Efna2 | -0.035456 | -0.210740021 | 0.139828021 | 8.148508544 | -0.497641482 | 0.636827508 | 0.922886691 | -5.023326269 |
| **A_55_P1979090** | Ublcp1 | 0.040923893 | -0.161398876 | 0.243246662 | 12.61530933 | 0.497624034 | 0.636839086 | 0.922886691 | -5.023332475 |
| **A_52_P172014** | Ramp1 | -0.070817533 | -0.421063591 | 0.279428524 | 8.045796295 | -0.497435416 | 0.636964253 | 0.922998879 | -5.023399554 |
| **A_51_P399960** | Mier2 | -0.032582748 | -0.193775811 | 0.128610315 | 8.624747811 | -0.497290921 | 0.637060149 | 0.923068638 | -5.023450926 |
| **A_51_P461040** | Crct1 | -0.363087194 | -2.159677779 | 1.433503392 | 9.858633282 | -0.497199834 | 0.637120605 | 0.923087038 | -5.023483303 |
| **A_51_P509643** | Snca | 0.092360982 | -0.364812533 | 0.549534497 | 11.69276457 | 0.497023179 | 0.637237861 | 0.923187724 | -5.023546081 |
| **A_55_P2052181** | Ccdc82 | 0.039639675 | -0.156662539 | 0.235941889 | 11.39127029 | 0.496791376 | 0.63739174 | 0.923341448 | -5.023628426 |
| **A_55_P2045407** | Ppp4r1 | -0.037436017 | -0.222895437 | 0.148023403 | 12.25779759 | -0.496603646 | 0.637516376 | 0.923385773 | -5.02369509 |
| **A_51_P187716** | Zc3h11a | -0.072367051 | -0.43087775 | 0.286143647 | 12.93278445 | -0.496601377 | 0.637517882 | 0.923385773 | -5.023695895 |
| **A_51_P465772** | Ruvbl1 | 0.029897933 | -0.118250188 | 0.178046054 | 11.48531755 | 0.496494199 | 0.637589044 | 0.923396717 | -5.023733944 |
| **A_52_P276935** | Dnajb6 | 0.070443149 | -0.27872339 | 0.419609687 | 13.48407802 | 0.496335462 | 0.637694448 | 0.923396717 | -5.023790284 |
| **A_55_P2025143** | Tlk1 | -0.034917873 | -0.207998213 | 0.138162468 | 11.00217302 | -0.496328486 | 0.637699081 | 0.923396717 | -5.023792759 |

| | | | | | | | | |
|---|---|---|---|---|---|---|---|---|
| A_52_P406036 | Cdkn2aip | 0.026789501 | -0.106007289 | 0.159586291 | 8.623796955 | 0.496302211 | 0.637716528 | 0.923396717 | -5.023802083 |
| A_55_P1973289 | Mapk9 | 0.053506053 | -0.211801445 | 0.318813552 | 10.93082761 | 0.496160923 | 0.637810356 | 0.923419179 | -5.023852213 |
| A_51_P236013 | Rnf40 | 0.056631308 | -0.224240595 | 0.337503211 | 10.00611785 | 0.496040868 | 0.637890088 | 0.923419179 | -5.0238948 |
| A_51_P240136 | Gng8 | 0.025021979 | -0.099089568 | 0.149133526 | 7.806606707 | 0.49599652 | 0.637919543 | 0.923419179 | -5.023910529 |
| A_55_P2183692 | Rgs19 | -0.023203988 | -0.13831191 | 0.091903934 | 7.99433161 | -0.495937085 | 0.637959018 | 0.923419179 | -5.023931607 |
| A_51_P112677 | Mrpl50 | -0.052441309 | -0.312596071 | 0.207713453 | 10.22031672 | -0.495919181 | 0.63797091 | 0.923419179 | -5.023937956 |
| A_55_P2073489 | NA | -0.035218236 | -0.209979296 | 0.139542825 | 9.006073302 | -0.495783521 | 0.638061019 | 0.923480451 | -5.023986056 |
| A_55_P2128511 | Ociad2 | 0.030251353 | -0.119910507 | 0.180413212 | 7.962357339 | 0.495626277 | 0.638165473 | 0.923562475 | -5.024041795 |
| A_55_P2035087 | Arhgef15 | -0.053349711 | -0.318390179 | 0.211690757 | 11.52363931 | -0.49520959 | 0.638442313 | 0.92383481 | -5.024189423 |
| A_51_P375693 | Tmem135 | 0.043938528 | -0.174400312 | 0.262277369 | 9.675530365 | 0.495089448 | 0.638522145 | 0.92383481 | -5.024231967 |
| A_55_P2002893 | Pfkfb1 | 0.048904557 | -0.194151827 | 0.291960942 | 9.291233945 | 0.495007137 | 0.638576843 | 0.92383481 | -5.02426111 |
| A_51_P338443 | Angptl4 | -0.091722381 | -0.547584004 | 0.364139242 | 11.84271678 | -0.495007127 | 0.63857685 | 0.92383481 | -5.024261113 |
| A_55_P2349198 | NA | -0.063657834 | -0.380053409 | 0.252737741 | 8.412852207 | -0.494983384 | 0.638592628 | 0.92383481 | -5.024269519 |
| A_55_P2032960 | Brap | 0.058636312 | -0.23291955 | 0.350192174 | 12.47855334 | 0.494782159 | 0.638726358 | 0.923959122 | -5.024340742 |
| A_51_P331003 | Sema4g | 0.044326024 | -0.176113982 | 0.264766029 | 8.726209799 | 0.494695002 | 0.638784286 | 0.923973768 | -5.024371583 |
| A_55_P1966432 | Gstm1 | 0.08543682 | -0.33975807 | 0.51063171 | 9.436056326 | 0.49434051 | 0.639019923 | 0.923994322 | -5.024496973 |
| A_55_P1956502 | Hnrnph1 | -0.070056567 | -0.418729984 | 0.27861685 | 13.79604636 | -0.494309751 | 0.639040371 | 0.923994322 | -5.024507849 |
| A_52_P222967 | Adck2 | 0.051783413 | -0.205944445 | 0.309511271 | 8.996644221 | 0.494308942 | 0.639040909 | 0.923994322 | -5.024508135 |
| A_51_P287232 | Qars | 0.039237835 | -0.1560859 | 0.234561571 | 11.98800753 | 0.494218709 | 0.639100897 | 0.923994322 | -5.024540038 |
| A_55_P2031496 | Rufy3 | -0.057620015 | -0.344450065 | 0.229210035 | 8.4914432 | -0.494217232 | 0.639101879 | 0.923994322 | -5.02454056 |
| A_55_P2011887 | Kif13a | -0.057461784 | -0.343529253 | 0.228605685 | 11.00511586 | -0.494173891 | 0.639130693 | 0.923994322 | -5.024555881 |
| A_55_P2171623 | Ptbp1 | 0.033191844 | -0.132051492 | 0.19843518 | 8.671109106 | 0.494170241 | 0.639133121 | 0.923994322 | -5.024557171 |
| A_52_P161087 | 2610008E11Rik | 0.036632967 | -0.145775791 | 0.219041726 | 8.74952469 | 0.494078068 | 0.639194403 | 0.924013807 | -5.024589751 |
| A_55_P2097391 | Tent5a | -0.05907738 | -0.353335872 | 0.235181113 | 9.244783427 | -0.493925425 | 0.639295896 | 0.924036003 | -5.024643693 |
| A_55_P1998506 | Aplf | -0.021522205 | -0.128744461 | 0.085700051 | 8.007172773 | -0.493822626 | 0.639364253 | 0.924036003 | -5.024680013 |
| A_55_P2071286 | Tmc4 | 0.050092046 | -0.19947354 | 0.299657633 | 12.57548653 | 0.493802449 | 0.63937767 | 0.924036003 | -5.024687141 |
| A_55_P1997894 | NA | 0.036584926 | -0.145701251 | 0.218871103 | 14.63778461 | 0.493761936 | 0.639404611 | 0.924036003 | -5.024701452 |
| A_51_P186899 | Egln1 | -0.025277187 | -0.151243175 | 0.100688801 | 7.817719064 | -0.493678948 | 0.639459799 | 0.924036003 | -5.024730764 |
| A_51_P173678 | Slc10a6 | -0.036338386 | -0.217470202 | 0.144793429 | 7.966456536 | -0.49356012 | 0.639538825 | 0.924036003 | -5.024772727 |
| A_55_P1987650 | Ctnnd2 | 0.0390323 | -0.155531022 | 0.233595622 | 8.201581116 | 0.493551337 | 0.639544666 | 0.924036003 | -5.024775828 |
| A_51_P446106 | Lrrc38 | 0.027449145 | -0.109395734 | 0.164294025 | 8.384573633 | 0.493479863 | 0.639592203 | 0.924036003 | -5.024801064 |
| A_66_P122099 | Zc3h8 | -0.025497889 | -0.152662823 | 0.101667045 | 8.068389291 | -0.493294206 | 0.63971569 | 0.924052455 | -5.0248666 |
| A_55_P2016877 | Nsd1 | -0.078847168 | -0.472108525 | 0.314414188 | 11.9267029 | -0.493257834 | 0.639739884 | 0.924052455 | -5.024879437 |
| A_51_P301736 | Ppp2r2a | -0.030824811 | -0.184581231 | 0.122931608 | 11.15823475 | -0.493215066 | 0.639768333 | 0.924052455 | -5.02489453 |
| A_66_P116678 | Rps8 | -0.04672273 | -0.27979795 | 0.186352489 | 8.51047493 | -0.493175253 | 0.639794816 | 0.924052455 | -5.024908579 |
| A_51_P303238 | Tmed10 | -0.106594338 | -0.638531352 | 0.425342677 | 12.87552959 | -0.492995649 | 0.639914297 | 0.924060663 | -5.024971944 |
| A_55_P2155931 | Tnfsfm13 | -0.100444701 | -0.60170923 | 0.400819828 | 9.169816034 | -0.492979907 | 0.63992477 | 0.924060663 | -5.024977496 |
| A_55_P1974554 | Eif4b | -0.034527431 | -0.206866428 | 0.137811567 | 11.75084734 | -0.492889836 | 0.639984695 | 0.924060663 | -5.025009265 |
| A_52_P425839 | Retnlg | -0.040055631 | -0.23999218 | 0.159880918 | 8.131279799 | -0.492879265 | 0.639991727 | 0.924060663 | -5.025012993 |
| A_51_P405227 | Nradd | -0.027336228 | -0.163811106 | 0.10913865 | 8.378662343 | -0.492782225 | 0.640056292 | 0.924074048 | -5.025047213 |
| A_51_P289239 | F8 | -0.021249802 | -0.127355518 | 0.084855914 | 7.686414314 | -0.49270307 | 0.64010896 | 0.924074048 | -5.025075122 |
| A_55_P2039556 | Pak6 | 0.056565734 | -0.225912051 | 0.339043519 | 8.318846258 | 0.492649777 | 0.640144421 | 0.924074048 | -5.02509391 |
| A_55_P1961152 | Pou3f1 | 0.080971266 | -0.32361346 | 0.485555993 | 10.16483836 | 0.492368897 | 0.640331334 | 0.924240882 | -5.025192902 |
| A_51_P146407 | Fam104a | 0.063242502 | -0.252810381 | 0.379295384 | 9.43620587 | 0.492287091 | 0.640385778 | 0.924240882 | -5.025221724 |
| A_52_P660693 | Dsel | -0.072430922 | -0.434495656 | 0.289633811 | 8.94518091 | -0.492160727 | 0.640469882 | 0.924240882 | -5.025266236 |
| A_51_P219444 | Plcl2 | -0.044870761 | -0.269177062 | 0.17943554 | 10.30215178 | -0.492142789 | 0.640481821 | 0.924240882 | -5.025272554 |
| A_52_P432969 | Pcdh19 | -0.049462911 | -0.296738124 | 0.197812302 | 8.568177403 | -0.492116867 | 0.640499075 | 0.924240882 | -5.025281684 |
| A_52_P127682 | Dagla | 0.026856585 | -0.107487126 | 0.161200295 | 7.96811018 | 0.491815949 | 0.640699383 | 0.924460912 | -5.025387633 |
| A_52_P163766 | Atp6v0a2 | -0.049764252 | -0.298742605 | 0.199214102 | 9.969549624 | -0.491728127 | 0.640757849 | 0.92447626 | -5.025418544 |
| A_51_P416689 | Ext1 | -0.040123604 | -0.240931192 | 0.160683983 | 10.05806563 | -0.491574088 | 0.640860404 | 0.924524647 | -5.025472748 |

| | | | | | | | | | |
|---|---|---|---|---|---|---|---|---|---|
| **A_51_P281089** | S100a6 | -0.0894792 | -0.53733441 | 0.358376009 | 16.88967615 | -0.491534071 | 0.640887047 | 0.924524647 | -5.025486827 |
| **A_55_P1987409** | Nbeal1 | -0.039633827 | -0.238106651 | 0.158838997 | 8.389944391 | -0.491285699 | 0.641052429 | 0.924624478 | -5.025574188 |
| **A_66_P119170** | Phf21a | 0.042948966 | -0.17215488 | 0.258052813 | 8.628399198 | 0.491217357 | 0.64109794 | 0.924624478 | -5.025598219 |
| **A_66_P138201** | Rhox10 | -0.096893926 | -0.582176202 | 0.388388349 | 11.44132657 | -0.491214625 | 0.641099759 | 0.924624478 | -5.02559918 |
| **A_55_P2088912** | Gapdh | 0.053620707 | -0.21510405 | 0.322345464 | 14.6154408 | 0.490901117 | 0.641308553 | 0.924856602 | -5.025709381 |
| **A_55_P2094034** | Nrg2 | 0.02530875 | -0.101557574 | 0.152175073 | 7.776500425 | 0.490787505 | 0.641384227 | 0.924896727 | -5.025749301 |
| **A_55_P2015964** | Rif1 | 0.034254402 | -0.137501609 | 0.206010413 | 8.652805744 | 0.490652038 | 0.641474463 | 0.924957844 | -5.025796889 |
| **A_55_P2100895** | Arnt | 0.068158521 | -0.27371323 | 0.410030273 | 10.72711722 | 0.490485424 | 0.641585457 | 0.925029765 | -5.025855403 |
| **A_55_P2175195** | Stxbp4 | 0.02403989 | -0.096552948 | 0.144632727 | 7.875813545 | 0.490433491 | 0.641620056 | 0.925029765 | -5.025873639 |
| **A_55_P1983314** | Crat | 0.022658686 | -0.091079151 | 0.136396524 | 7.702798924 | 0.490116052 | 0.641831559 | 0.92512669 | -5.025985062 |
| **A_65_P09523** | Scrn3 | -0.051287393 | -0.308753821 | 0.206179035 | 9.84036719 | -0.490071196 | 0.641861449 | 0.92512669 | -5.026000802 |
| **A_55_P1993640** | Gm6994 | -0.059257756 | -0.356741386 | 0.238225873 | 13.49741819 | -0.490062301 | 0.641867376 | 0.92512669 | -5.026003923 |
| **A_51_P331207** | Tmem218 | 0.071709423 | -0.288319779 | 0.431738624 | 11.40030996 | 0.490013069 | 0.641900183 | 0.92512669 | -5.026021196 |
| **A_55_P2160732** | Tfcp2 | -0.042719477 | -0.257217135 | 0.171778181 | 9.195307327 | -0.489973432 | 0.641926596 | 0.92512669 | -5.026035102 |
| **A_52_P669964** | Cchcr1 | 0.02181682 | -0.087777162 | 0.131410801 | 7.792667109 | 0.489749386 | 0.642075907 | 0.925272885 | -5.026113685 |
| **A_52_P112110** | Tmem82 | -0.035315481 | -0.212770819 | 0.142139858 | 8.517499869 | -0.489604294 | 0.642172611 | 0.925343254 | -5.026164558 |
| **A_51_P387235** | Nampt | -0.097723518 | -0.588878891 | 0.393431854 | 10.15809455 | -0.489496238 | 0.642244635 | 0.925378051 | -5.026202436 |
| **A_55_P2104612** | NA | 0.05343388 | -0.215191471 | 0.322059231 | 8.707596985 | 0.48937173 | 0.642327631 | 0.92542228 | -5.026246073 |
| **A_55_P2065334** | Gm14326 | -0.046945855 | -0.282985742 | 0.189094031 | 8.648293269 | -0.489306543 | 0.642371086 | 0.92542228 | -5.026268915 |
| **A_55_P2101182** | Pcmt1 | -0.031315879 | -0.188808427 | 0.12617667 | 10.69776244 | -0.489185698 | 0.642451648 | 0.925469363 | -5.026311253 |
| **A_55_P2185870** | Bcl7a | -0.020757221 | -0.125227104 | 0.083712662 | 7.884916566 | -0.488818087 | 0.642696752 | 0.925653335 | -5.026439988 |
| **A_55_P2034350** | Terf2 | 0.047609329 | -0.192034503 | 0.287253161 | 9.636454359 | 0.488759227 | 0.642736001 | 0.925653335 | -5.026460593 |
| **A_55_P2011380** | Acvrl1 | 0.105925359 | -0.427282112 | 0.63913283 | 13.31686864 | 0.488734371 | 0.642752576 | 0.925653335 | -5.026469293 |
| **A_66_P129747** | Dnajc24 | -0.054791521 | -0.330620405 | 0.221037364 | 9.953232467 | -0.488700487 | 0.642775172 | 0.925653335 | -5.026481153 |
| **A_55_P2050268** | Nfkbid | 0.023058207 | -0.093035875 | 0.13915229 | 8.090351054 | 0.488635055 | 0.642818807 | 0.925653335 | -5.026504053 |
| **A_55_P2408355** | BB166591 | -0.135015923 | -0.815083811 | 0.545051965 | 10.22924414 | -0.488429497 | 0.642955897 | 0.925718903 | -5.026575976 |
| **A_51_P325836** | Hpx | 0.163932465 | -0.661797607 | 0.989662537 | 8.478615124 | 0.488422914 | 0.642960288 | 0.925718903 | -5.026578279 |
| **A_52_P561205** | Stard4 | -0.02319981 | -0.140100213 | 0.093700593 | 7.839360259 | -0.488244747 | 0.643079123 | 0.925718903 | -5.026640596 |
| **A_55_P2051337** | Ppfia1 | -0.033786837 | -0.204068157 | 0.136494483 | 10.95334384 | -0.488145951 | 0.643145025 | 0.925718903 | -5.026675143 |
| **A_55_P2078325** | Smco3 | 0.031077487 | -0.125580486 | 0.18773546 | 7.854349253 | 0.488048018 | 0.643210354 | 0.925718903 | -5.026709382 |
| **A_55_P2005813** | Abhd18 | -0.036270371 | -0.219107563 | 0.146566821 | 9.049083664 | -0.488041351 | 0.643214802 | 0.925718903 | -5.026711713 |
| **A_55_P2066180** | Hid1 | 0.044064162 | -0.178087355 | 0.266215679 | 8.689083012 | 0.487983791 | 0.6432532 | 0.925718903 | -5.026731833 |
| **A_52_P384574** | Stard4 | -0.095382659 | -0.576328973 | 0.385563655 | 9.692396005 | -0.487912548 | 0.643300729 | 0.925718903 | -5.026756734 |
| **A_51_P352687** | Rnf216 | 0.072696797 | -0.293863918 | 0.439257512 | 11.8177702 | 0.487908654 | 0.643303327 | 0.925718903 | -5.026758095 |
| **A_55_P2074647** | Snap47 | 0.051856966 | -0.209654682 | 0.313368614 | 9.987804373 | 0.487848786 | 0.643343268 | 0.925718903 | -5.026779017 |
| **A_51_P157042** | Ccn2 | -0.410408962 | -2.48075562 | 1.659937695 | 14.77537102 | -0.487688966 | 0.6434499 | 0.925769119 | -5.026834859 |
| **A_51_P425833** | Lztfl1 | -0.025418984 | -0.153675529 | 0.10283756 | 10.08855553 | -0.487582174 | 0.643521157 | 0.925769119 | -5.026872164 |
| **A_55_P2197914** | C030027H14Rik | -0.062331045 | -0.376854923 | 0.252192833 | 11.8842713 | -0.487550886 | 0.643542034 | 0.925769119 | -5.026883092 |
| **A_55_P2354531** | 2610029K11Rik | -0.025634936 | -0.155029046 | 0.103759173 | 8.030199839 | -0.487401539 | 0.643641694 | 0.925769119 | -5.026935247 |
| **A_52_P222073** | Ackr2 | -0.107104622 | -0.647816872 | 0.433607628 | 9.957452109 | -0.487316556 | 0.643698408 | 0.925769119 | -5.026964918 |
| **A_55_P2053497** | Poli | -0.035596323 | -0.215347843 | 0.144155197 | 9.721333896 | -0.487193773 | 0.643780351 | 0.925769119 | -5.027007778 |
| **A_51_P478311** | Nmd3 | -0.029503785 | -0.178497272 | 0.119489703 | 8.770924935 | -0.487168956 | 0.643796915 | 0.925769119 | -5.02701644 |
| **A_51_P456857** | Ift122 | -0.040226838 | -0.243433021 | 0.162979345 | 9.259583747 | -0.487021506 | 0.64389533 | 0.925769119 | -5.027067897 |
| **A_52_P219372** | Tmem177 | -0.046348605 | -0.280506861 | 0.187809651 | 8.698764853 | -0.486963439 | 0.643934088 | 0.925769119 | -5.027088156 |
| **A_51_P462692** | Ptbp1 | -0.036275199 | -0.219556173 | 0.147005775 | 11.68674169 | -0.486924449 | 0.643960114 | 0.925769119 | -5.027101759 |
| **A_52_P590535** | Fbln2 | -0.315764986 | -1.911195559 | 1.279665588 | 10.74772907 | -0.48691732 | 0.643964873 | 0.925769119 | -5.027104246 |
| **A_55_P1999878** | Sec24b | 0.041280115 | -0.167330705 | 0.249890935 | 12.35885932 | 0.486825403 | 0.64402623 | 0.925769119 | -5.02713631 |
| **A_51_P170008** | Napg | -0.037841179 | -0.229149089 | 0.15346673 | 9.088146797 | -0.486632264 | 0.644155165 | 0.925769119 | -5.027203665 |
| **A_55_P2028171** | Dppa5a | 0.114567753 | -0.464711348 | 0.693846855 | 10.17111969 | 0.486568173 | 0.644197954 | 0.925769119 | -5.027226011 |
| **A_55_P2010197** | Serpina10 | 0.203522136 | -0.825668117 | 1.232712388 | 9.981577014 | 0.486502582 | 0.644241746 | 0.925769119 | -5.027248877 |

| | | | | | | | | | |
|---|---|---|---|---|---|---|---|---|---|
| **A_55_P2157695** | Tsc1 | -0.039140122 | -0.237108006 | 0.158827763 | 10.81200559 | -0.486403392 | 0.644307973 | 0.925769119 | -5.027283451 |
| **A_51_P436878** | Sertad1 | -0.025312655 | -0.153355844 | 0.102730535 | 10.09750258 | -0.486351627 | 0.644342537 | 0.925769119 | -5.027301491 |
| **A_52_P139097** | NA | -0.030080356 | -0.18224355 | 0.122082838 | 10.58591894 | -0.486342835 | 0.644348408 | 0.925769119 | -5.027304555 |
| **A_66_P102299** | Ascc1 | -0.029161828 | -0.17668552 | 0.118361864 | 8.350869581 | -0.486319999 | 0.644363656 | 0.925769119 | -5.027312513 |
| **A_55_P2037081** | Zfp979 | 0.0280307 | -0.11377669 | 0.169838091 | 8.466418438 | 0.486299974 | 0.644377027 | 0.925769119 | -5.027319491 |
| **A_55_P1962154** | Rarb | -0.031363026 | -0.190032053 | 0.127306001 | 8.316584646 | -0.486289579 | 0.644383968 | 0.925769119 | -5.027323114 |
| **A_55_P2000603** | Slc50a1 | -0.082908256 | -0.502458994 | 0.336642483 | 10.26071563 | -0.486163613 | 0.644468084 | 0.925821152 | -5.027367003 |
| **A_55_P2005420** | NA | -0.059907811 | -0.363162571 | 0.243346948 | 8.543104704 | -0.486009761 | 0.644570829 | 0.925899937 | -5.027420593 |
| **A_52_P504421** | Rnf138 | -0.057025206 | -0.345757047 | 0.231706635 | 10.44356165 | -0.485893783 | 0.644648286 | 0.925914041 | -5.027460982 |
| **A_51_P456208** | Tff3 | 0.289289343 | -1.176000057 | 1.754578744 | 10.5627684 | 0.485711281 | 0.644770182 | 0.925914041 | -5.027524519 |
| **A_51_P161037** | Cep170 | -0.026467272 | -0.160554946 | 0.107620402 | 8.597045408 | -0.485612084 | 0.644836443 | 0.925914041 | -5.027559045 |
| **A_55_P2062836** | Ttf2 | -0.027661578 | -0.167817603 | 0.112494447 | 10.00677293 | -0.485550427 | 0.64487763 | 0.925914041 | -5.027580502 |
| **A_55_P2147101** | Sytl4 | 0.023892799 | -0.097184811 | 0.144970409 | 7.870899225 | 0.485481134 | 0.644923919 | 0.925914041 | -5.027604613 |
| **A_55_P2014315** | NA | -0.027607554 | -0.167512821 | 0.112297712 | 8.019488009 | -0.485470711 | 0.644930882 | 0.925914041 | -5.02760824 |
| **A_51_P194609** | Prss34 | -0.020604834 | -0.125030583 | 0.083820915 | 7.756166399 | -0.485434556 | 0.644955035 | 0.925914041 | -5.027620819 |
| **A_55_P2042753** | Cltrn | -0.033184737 | -0.20142619 | 0.135056716 | 7.848022798 | -0.485260041 | 0.645071626 | 0.925914041 | -5.027681524 |
| **A_55_P2425235** | NA | 0.038133359 | -0.155278202 | 0.23154492 | 9.492759755 | 0.485055908 | 0.645208018 | 0.925914041 | -5.027752508 |
| **A_55_P1973698** | Htra2 | 0.062446979 | -0.254380052 | 0.37927401 | 9.848165476 | 0.484906906 | 0.645307584 | 0.925914041 | -5.027804304 |
| **A_55_P2122849** | NA | -0.04990039 | -0.303106755 | 0.203305976 | 10.90937955 | -0.484840043 | 0.645352265 | 0.925914041 | -5.027827542 |
| **A_51_P113205** | F13b | 0.045063643 | -0.183624872 | 0.273752157 | 7.843356412 | 0.484787153 | 0.64538761 | 0.925914041 | -5.007633234 |
| **A_55_P1964309** | Mlf2 | 0.033248318 | -0.135507966 | 0.202004602 | 13.52521116 | 0.48470655 | 0.645441478 | 0.925914041 | -5.027873929 |
| **A_55_P2022104** | Cop1 | 0.024529179 | -0.099984947 | 0.149043306 | 8.294547593 | 0.484655958 | 0.64547529 | 0.925914041 | -5.027891506 |
| **A_51_P161874** | Med30 | 0.03923875 | -0.159978115 | 0.238455615 | 10.04542501 | 0.484571897 | 0.645531472 | 0.925914041 | -5.027920707 |
| **A_52_P502577** | S1pr3 | -0.045463693 | -0.276286883 | 0.185359496 | 8.459681704 | -0.484567674 | 0.645534294 | 0.925914041 | -5.027922174 |
| **A_51_P160907** | Morc3 | -0.038931028 | -0.236612828 | 0.158750772 | 9.182101516 | -0.484505095 | 0.645576121 | 0.925914041 | -5.02794391 |
| **A_55_P1955891** | NA | 0.059839654 | -0.244016395 | 0.363695703 | 13.66377702 | 0.484496172 | 0.645582085 | 0.925914041 | -5.027947009 |
| **A_55_P1957911** | Tom1 | -0.022273935 | -0.135417302 | 0.090869432 | 8.591928236 | -0.484325136 | 0.64569641 | 0.925914041 | -5.028006402 |
| **A_51_P187171** | Smim12 | -0.035723973 | -0.217203692 | 0.145755746 | 12.97650135 | -0.484284771 | 0.645723392 | 0.925914041 | -5.028020416 |
| **A_55_P2059382** | Arl6 | 0.053339464 | -0.217628882 | 0.324307809 | 8.771487818 | 0.484283028 | 0.645724557 | 0.925914041 | -5.028021021 |
| **A_55_P2107647** | NA | 0.050889097 | -0.207666872 | 0.309445065 | 10.28270018 | 0.484216227 | 0.645769213 | 0.925914041 | -5.028044211 |
| **A_51_P273326** | Sec31a | 0.107171747 | -0.499017041 | 0.713360534 | 12.14384279 | 0.497479191 | 0.645789878 | 0.925914041 | -4.848456651 |
| **A_52_P519538** | Uba6 | 0.059065914 | -0.241068239 | 0.359200068 | 9.725750244 | 0.484161973 | 0.645805482 | 0.925914041 | -5.028063043 |
| **A_51_P321743** | NA | -0.02382164 | -0.144892665 | 0.097249385 | 7.799897095 | -0.484061569 | 0.645872606 | 0.925914041 | -5.028097889 |
| **A_55_P2007249** | Spdef | -0.025746195 | -0.15659894 | 0.105106549 | 8.365865172 | -0.484060209 | 0.645873515 | 0.925914041 | -5.028098361 |
| **A_55_P2053810** | NA | -0.031358929 | -0.190738144 | 0.128020286 | 8.979050119 | -0.484059447 | 0.645874024 | 0.925914041 | -5.028098626 |
| **A_66_P140742** | Azin2 | 0.090875401 | -0.371159531 | 0.552910332 | 8.393479158 | 0.483883364 | 0.645991751 | 0.925971651 | -5.028159721 |
| **A_55_P2096226** | Loxl1 | -0.024889177 | -0.151451899 | 0.101673545 | 7.81432012 | -0.483808944 | 0.646041511 | 0.925971651 | -5.028185537 |
| **A_55_P1964896** | NA | -0.027109308 | -0.164968505 | 0.110749889 | 7.750861163 | -0.483784391 | 0.646057927 | 0.925971651 | -5.028194053 |
| **A_51_P120066** | 9330151L19Rik | -0.031113005 | -0.189389878 | 0.127163869 | 8.203074523 | -0.483608198 | 0.646175745 | 0.926071845 | -5.008031933 |
| **A_55_P2091681** | Nudt1 | 0.034697503 | -0.141881954 | 0.21127696 | 9.619860075 | 0.483422888 | 0.646299672 | 0.92618078 | -5.028319399 |
| **A_55_P2181753** | Bdh1 | 0.06627711 | -0.27106575 | 0.40361997 | 9.240661905 | 0.483349422 | 0.646348805 | 0.926182524 | -5.028344862 |
| **A_55_P2170847** | Fbxo27 | 0.036845572 | -0.150765298 | 0.224456443 | 12.28284741 | 0.483166119 | 0.646471407 | 0.92618692 | -5.028408379 |
| **A_51_P373901** | Rsph1 | -0.022645016 | -0.137953067 | 0.092663035 | 7.716013004 | -0.483150197 | 0.646482057 | 0.92618692 | -5.028413895 |
| **A_51_P510764** | Agmo | -0.023452677 | -0.14287834 | 0.095972986 | 7.797408598 | -0.483129913 | 0.646495625 | 0.92618692 | -5.028420922 |
| **A_51_P148122** | Klhdc3 | -0.040538259 | -0.247013531 | 0.165937012 | 11.59709238 | -0.483021223 | 0.646568328 | 0.92621544 | -5.028458572 |
| **A_55_P2107667** | Enah | -0.022824516 | -0.1390931 | 0.093444068 | 7.924921083 | -0.482956882 | 0.646611369 | 0.92621544 | -5.028480855 |
| **A_55_P2133895** | Hnrnph2 | -0.042267033 | -0.257642861 | 0.173108795 | 8.932569611 | -0.482807457 | 0.646711332 | 0.926263768 | -5.028532597 |
| **A_51_P121412** | Slc35a4 | -0.028993089 | -0.176743778 | 0.118757599 | 8.904346564 | -0.482763186 | 0.64674095 | 0.926263768 | -5.013632106 |
| **A_55_P2100241** | Rn18s | 0.042994736 | -0.176317515 | 0.262306987 | 17.40889561 | 0.482304773 | 0.647047678 | 0.926634405 | -5.028706557 |
| **A_52_P15300** | Hoxb2 | -0.021206516 | -0.129407482 | 0.08699445 | 7.867206353 | -0.482177956 | 0.647132546 | 0.926687285 | -5.028750418 |

| | | | | | | | | | |
|---|---|---|---|---|---|---|---|---|---|
| **A_51_P504134** | NA | -0.125759115 | -0.767539304 | 0.516021074 | 13.54006132 | -0.482083568 | 0.647195715 | 0.926709088 | -5.028783056 |
| **A_55_P2034670** | Fbxl12 | -0.053103762 | -0.324222941 | 0.218015417 | 12.80005172 | -0.481874794 | 0.64733545 | 0.926720563 | -5.028855227 |
| **A_55_P2073507** | 2610528J11Rik | -0.035524468 | -0.216899911 | 0.145850976 | 8.471339171 | -0.48185709 | 0.6473473 | 0.926720563 | -5.028861346 |
| **A_51_P487869** | Cyp4f16 | -0.027884122 | -0.170250762 | 0.114482519 | 7.880458999 | -0.481856697 | 0.647347563 | 0.926720563 | -5.028861482 |
| **A_55_P2028773** | Slc22a17 | -0.109585247 | -0.669303405 | 0.450132911 | 10.39661904 | -0.481672508 | 0.647470857 | 0.926828423 | -5.028925129 |
| **A_51_P249930** | Tceal1 | -0.035959217 | -0.219652477 | 0.147734044 | 8.459262004 | -0.481599648 | 0.647519632 | 0.926829604 | -5.0289503 |
| **A_52_P1130990** | Trim67 | -0.020647502 | -0.126150279 | 0.084855274 | 7.627753043 | -0.481473955 | 0.647603779 | 0.92686262 | -5.028993715 |
| **A_52_P536293** | NA | -0.039435732 | -0.240980611 | 0.162109147 | 13.81128534 | -0.481379184 | 0.64766723 | 0.92686262 | -5.029026443 |
| **A_55_P1975640** | Zfp217 | 0.049005105 | -0.201463154 | 0.299473363 | 10.11923709 | 0.481346466 | 0.647689136 | 0.92686262 | -5.02903774 |
| **A_55_P1968553** | Btbd9 | -0.046159791 | -0.282138292 | 0.189818709 | 9.72608392 | -0.48123872 | 0.647761278 | 0.92686262 | -5.02907494 |
| **A_55_P2098999** | NA | 0.058339461 | -0.239923661 | 0.356602583 | 12.87620714 | 0.481207088 | 0.647782458 | 0.92686262 | -5.029085859 |
| **A_51_P504354** | Nrarp | -0.025842358 | -0.157994325 | 0.10630961 | 7.892971368 | -0.481091476 | 0.647859874 | 0.926887075 | -5.029125764 |
| **A_55_P1960775** | Rps15a | -0.02436669 | -0.149079373 | 0.100345992 | 15.5111636 | -0.480678929 | 0.648136159 | 0.926887075 | -5.029268087 |
| **A_51_P503866** | Edem1 | -0.046895795 | -0.286917718 | 0.193126127 | 10.8948295 | -0.480675688 | 0.64813833 | 0.926887075 | -5.029269205 |
| **A_55_P2073855** | Mrpl27 | 0.080408304 | -0.331140397 | 0.491957004 | 12.20648669 | 0.480672005 | 0.648140797 | 0.926887075 | -5.029270475 |
| **A_55_P2104917** | Nphp1 | -0.021981898 | -0.134503296 | 0.0905395 | 7.810903692 | -0.480617112 | 0.648177563 | 0.926887075 | -5.029289404 |
| **A_55_P2039598** | Rpl7a | 0.046677237 | -0.192260025 | 0.285614499 | 11.97092743 | 0.480607366 | 0.648184092 | 0.926887075 | -5.029292764 |
| **A_55_P2096530** | Icmt | -0.049740826 | -0.304375326 | 0.204893673 | 10.05307648 | -0.480579139 | 0.648202998 | 0.926887075 | -5.029302497 |
| **A_55_P2029711** | Cdh2 | 0.047258163 | -0.194705853 | 0.289222179 | 8.760048588 | 0.480502016 | 0.648254657 | 0.926887075 | -5.029329086 |
| **A_52_P282905** | Ces1b | -0.020921867 | -0.128046029 | 0.086202296 | 7.643019115 | -0.480487563 | 0.648264339 | 0.926887075 | -5.029334069 |
| **A_55_P1955827** | NA | 0.040327258 | -0.166188912 | 0.246843429 | 11.5418409 | 0.480411946 | 0.648314992 | 0.926887075 | -5.029360135 |
| **A_51_P183213** | Riiad1 | -0.02411544 | -0.147615395 | 0.099384516 | 7.785542489 | -0.480393969 | 0.648327034 | 0.926887075 | -5.029366331 |
| **A_52_P361081** | Arhgef16 | -0.04580002 | -0.280443997 | 0.188843957 | 11.87541654 | -0.480203613 | 0.648454558 | 0.926966081 | -5.029431929 |
| **A_55_P2179216** | Man1a2 | 0.04460798 | -0.183945854 | 0.273161814 | 10.04005569 | 0.480167996 | 0.64847842 | 0.926966081 | -5.029444201 |
| **A_51_P283649** | Brat1 | 0.075231849 | -0.310283927 | 0.460747626 | 10.54954266 | 0.480096727 | 0.648526169 | 0.926966081 | -5.029468753 |
| **A_51_P140141** | Hspbp1 | 0.046075991 | -0.190162211 | 0.282314192 | 9.22039824 | 0.479836983 | 0.648700207 | 0.927096376 | -5.029558208 |
| **A_52_P595717** | Smug1 | 0.025179135 | -0.103928674 | 0.154286944 | 8.489131487 | 0.479796922 | 0.648727052 | 0.927096376 | -5.029572001 |
| **A_51_P374869** | Abca7 | -0.075202752 | -0.460851206 | 0.310445701 | 11.04717263 | -0.479745937 | 0.648761218 | 0.927096376 | -5.029589554 |
| **A_52_P609120** | Pdxp | 0.024680528 | -0.101951405 | 0.151312461 | 8.411730174 | 0.479490896 | 0.648932137 | 0.927268382 | -5.029677332 |
| **A_55_P1965338** | Anxa8 | -0.115991178 | -0.711207871 | 0.479225515 | 9.654106966 | -0.47942317 | 0.648977529 | 0.927268382 | -5.029700634 |
| **A_52_P48767** | Rassf8 | -0.033573412 | -0.20593908 | 0.138792257 | 10.71227502 | -0.47919676 | 0.649129287 | 0.92741666 | -5.029778513 |
| **A_55_P2046145** | Gm10693 | -0.03636918 | -0.223241691 | 0.15050333 | 8.029072812 | -0.478803433 | 0.649392969 | 0.927713407 | -5.029913729 |
| **A_52_P351669** | Gcc2 | -0.029078714 | -0.178509898 | 0.12035247 | 9.194693765 | -0.478743754 | 0.649432982 | 0.927713407 | -5.029934236 |
| **A_55_P2041131** | NA | 0.05066387 | -0.20974757 | 0.311075311 | 12.98575857 | 0.478638319 | 0.649503676 | 0.927721862 | -5.029970461 |
| **A_55_P1969352** | Get4 | -0.045070655 | -0.276766243 | 0.186624933 | 10.96750865 | -0.478569776 | 0.649549636 | 0.927721862 | -5.029994007 |
| **A_52_P69194** | Rimkla | 0.022705123 | -0.0940287 | 0.139438947 | 7.79180419 | 0.478515838 | 0.649585805 | 0.927721862 | -5.030012534 |
| **A_55_P1970583** | Ttc38 | 0.028708467 | -0.118945531 | 0.176362466 | 9.384204302 | 0.478336978 | 0.649705748 | 0.927721862 | -5.030073955 |
| **A_55_P2130666** | Armc1 | -0.027473121 | -0.168806776 | 0.113860534 | 10.24953028 | -0.478224222 | 0.649781368 | 0.927721862 | -5.030112665 |
| **A_52_P487599** | Thnsl2 | 0.075176154 | -0.311601612 | 0.46195392 | 9.078557517 | 0.478175991 | 0.649813716 | 0.927721862 | -5.030129221 |
| **A_55_P1985623** | Abcc3 | -0.047145798 | -0.289732764 | 0.195441169 | 8.869669228 | -0.478128562 | 0.649845526 | 0.927721862 | -5.0301455 |
| **A_51_P434776** | Jmjd8 | -0.028190313 | -0.173247053 | 0.116866426 | 8.079788102 | -0.478113667 | 0.649855516 | 0.927721862 | -5.030150612 |
| **A_51_P307325** | Rbbp4 | 0.032077676 | -0.132997509 | 0.197152861 | 12.65986745 | 0.478068652 | 0.649885709 | 0.927721862 | -5.030166061 |
| **A_55_P2083839** | Atp13a1 | -0.030152921 | -0.185382582 | 0.125076741 | 10.94156517 | -0.477885483 | 0.650008571 | 0.927721862 | -5.030228909 |
| **A_55_P2348815** | Casd1 | 0.046493182 | -0.192884736 | 0.285871101 | 9.449455146 | 0.477831029 | 0.650045099 | 0.927721862 | -5.030247588 |
| **A_55_P2255399** | F630048H11Rik | -0.045620767 | -0.280538664 | 0.18929713 | 7.978226583 | -0.477766444 | 0.650088424 | 0.927721862 | -5.030269741 |
| **A_52_P118958** | Pex19 | 0.036609164 | -0.151906151 | 0.225124479 | 9.574634765 | 0.477762804 | 0.650090866 | 0.927721862 | -5.03027099 |
| **A_51_P185259** | Gnb5 | -0.021679821 | -0.133324903 | 0.089965261 | 7.763738379 | -0.477733014 | 0.65011085 | 0.927721862 | -5.030281207 |
| **A_55_P2061737** | Tmsb4x | -0.033833931 | -0.208144696 | 0.140476833 | 16.71014267 | -0.477526433 | 0.650249442 | 0.927819393 | -5.030352041 |
| **A_51_P258372** | Igsf6 | -0.033082896 | -0.203548171 | 0.137382379 | 8.301174223 | -0.477459747 | 0.650294184 | 0.927819393 | -5.030374901 |
| **A_55_P2087944** | NA | -0.072049285 | -0.443329166 | 0.299230595 | 9.077369704 | -0.4774165 | 0.650323201 | 0.927819393 | -5.030389725 |

| | | | | | | | | | |
|---|---|---|---|---|---|---|---|---|---|
| **A_51_P507899** | Ttc8 | -0.046670913 | -0.287277491 | 0.193935666 | 9.767384782 | -0.477208268 | 0.650462925 | 0.927876726 | -5.030461083 |
| **A_52_P49601** | Fth1 | -0.058704126 | -0.361405653 | 0.243997402 | 16.5762518 | -0.477115113 | 0.650525436 | 0.927876726 | -5.030492997 |
| **A_55_P1974645** | Entpd2 | 0.213528183 | -0.887579424 | 1.314635791 | 12.49078606 | 0.477083729 | 0.650546498 | 0.927876726 | -5.030503748 |
| **A_55_P2026325** | NA | -0.068889721 | -0.424145431 | 0.286365988 | 14.44903533 | -0.477070458 | 0.650555404 | 0.927876726 | -5.030508294 |
| **A_51_P451458** | Mamdc2 | -0.140410097 | -0.864926314 | 0.584106121 | 8.488890695 | -0.476781495 | 0.650749338 | 0.927995695 | -5.030607245 |
| **A_51_P316523** | Irf2 | -0.029003989 | -0.178666458 | 0.120658479 | 10.0240822 | -0.476775572 | 0.650753313 | 0.927995695 | -5.030609273 |
| **A_55_P2155848** | Gm14207 | -0.023393582 | -0.144117127 | 0.097329962 | 7.8719445 | -0.476731571 | 0.650782847 | 0.927995695 | -5.030624336 |
| **A_65_P18948** | Ppm1h | -0.050080785 | -0.308769968 | 0.208608398 | 9.954084639 | -0.476279646 | 0.65108622 | 0.928322867 | -5.030778969 |
| **A_51_P364485** | Tnfaip2 | 0.09833172 | -0.409629811 | 0.606293252 | 11.33895473 | 0.476246697 | 0.651108341 | 0.928322867 | -5.030790238 |
| **A_51_P351394** | Vps45 | 0.032632175 | -0.136021065 | 0.201285415 | 9.067158179 | 0.476014846 | 0.651264012 | 0.928366691 | -5.030869513 |
| **A_55_P2025851** | Heatr3 | -0.076092055 | -0.469365658 | 0.317181548 | 11.20414951 | -0.476007374 | 0.651269029 | 0.928366691 | -5.030872068 |
| **A_55_P1987449** | Ep400 | 0.045638287 | -0.190248723 | 0.281525296 | 10.69372435 | 0.475986318 | 0.651283167 | 0.928366691 | -5.030879265 |
| **A_51_P371750** | Marco | 0.056971605 | -0.237554815 | 0.351498026 | 8.138459079 | 0.475886475 | 0.651350211 | 0.928393794 | -5.030913392 |
| **A_55_P2026089** | Usp30 | -0.052998229 | -0.327208913 | 0.221212454 | 10.69990742 | -0.475495216 | 0.651612975 | 0.928444589 | -5.031047061 |
| **A_51_P189272** | Tmcc3 | -0.044216894 | -0.273073828 | 0.18464004 | 13.18482767 | -0.475327916 | 0.651725348 | 0.928444589 | -5.031104188 |
| **A_55_P2134735** | Fdxacb1 | -0.042664421 | -0.263509982 | 0.17818114 | 10.03309678 | -0.475276503 | 0.651759883 | 0.928444589 | -5.03112174 |
| **A_55_P2144376** | NA | 0.03133584 | -0.130888439 | 0.193560119 | 8.091997343 | 0.475219936 | 0.651797881 | 0.928444589 | -5.031141049 |
| **A_55_P2085905** | Anpep | -0.315988614 | -1.951860682 | 1.319883455 | 11.96678352 | -0.475216225 | 0.651800374 | 0.928444589 | -5.031142316 |
| **A_55_P1981780** | Cul4b | 0.053866802 | -0.225044702 | 0.332778306 | 12.46514089 | 0.475142554 | 0.651849864 | 0.928444589 | -5.03116746 |
| **A_51_P480982** | Glo1 | 0.041353314 | -0.172800128 | 0.255506756 | 10.24966284 | 0.475066517 | 0.651900945 | 0.928444589 | -5.031193409 |
| **A_55_P2060158** | Ermn | 0.023821014 | -0.099581203 | 0.147223231 | 7.893534148 | 0.47490468 | 0.652009673 | 0.928444589 | -5.031248625 |
| **A_55_P2116621** | Cited2 | 0.075820502 | -0.316995548 | 0.468636552 | 12.65736271 | 0.474861099 | 0.652038954 | 0.928444589 | -5.031263491 |
| **A_55_P2068723** | Ccl27a | 0.114724385 | -0.47966246 | 0.709111231 | 14.19659313 | 0.474849203 | 0.652046947 | 0.928444589 | -5.031267549 |
| **A_51_P466685** | Fbxl22 | -0.02064275 | -0.12761077 | 0.086325269 | 7.712934179 | -0.474769457 | 0.652100528 | 0.928444589 | -5.031294748 |
| **A_66_P124017** | Map7d3 | -0.025312991 | -0.15650194 | 0.105875958 | 7.987221282 | -0.474695779 | 0.652150035 | 0.928444589 | -5.031319874 |
| **A_51_P425962** | Khdrbs3 | 0.109373565 | -0.457522879 | 0.676270008 | 9.612508283 | 0.474654713 | 0.652177629 | 0.928444589 | -5.031333877 |
| **A_51_P116027** | Med27 | -0.030548774 | -0.188897881 | 0.127800334 | 10.9918933 | -0.474621408 | 0.652200009 | 0.928444589 | -5.031345232 |
| **A_51_P127841** | Pdss1 | 0.022349292 | -0.093499438 | 0.138198022 | 8.261733669 | 0.474615213 | 0.652204172 | 0.928444589 | -5.016378756 |
| **A_55_P1955532** | Tfr2 | 0.022059302 | -0.092293438 | 0.136412041 | 7.750867389 | 0.474585376 | 0.652224222 | 0.928444589 | -5.031357517 |
| **A_51_P151902** | Spon1 | 0.057546628 | -0.240798811 | 0.355892067 | 8.021949397 | 0.474536519 | 0.652257053 | 0.928444589 | -5.031374173 |
| **A_51_P135732** | Ccdc136 | -0.060993292 | -0.377273423 | 0.255286839 | 7.99727614 | -0.474437846 | 0.652323363 | 0.928444589 | -5.031407807 |
| **A_55_P2043269** | Cdc42se1 | 0.024526074 | -0.102693324 | 0.151745472 | 9.563525276 | 0.474289886 | 0.652422801 | 0.928444589 | -5.031458229 |
| **A_51_P350666** | Peli2 | -0.038536647 | -0.238458143 | 0.161384849 | 8.26647685 | -0.47422407 | 0.652467036 | 0.928444589 | -5.031480654 |
| **A_55_P2408415** | Kcnq1ot1 | -0.103753813 | -0.64212164 | 0.434614015 | 8.736002903 | -0.47412639 | 0.65253269 | 0.928444589 | -5.03151393 |
| **A_55_P2079324** | Sephs2 | -0.022594089 | -0.139855086 | 0.094666908 | 8.034158164 | -0.474035001 | 0.652594118 | 0.928444589 | -5.031545057 |
| **A_52_P220810** | Trib2 | 0.035454805 | -0.148597179 | 0.219506789 | 8.450253469 | 0.473918607 | 0.652672358 | 0.928444589 | -5.031584693 |
| **A_52_P667913** | Pcdhga9 | 0.048728456 | -0.204256966 | 0.301713878 | 8.286548762 | 0.473866841 | 0.652707157 | 0.928444589 | -5.031602318 |
| **A_51_P399889** | Ap4b1 | 0.048799778 | -0.204590881 | 0.302190437 | 11.37820472 | 0.473801474 | 0.6527511 | 0.928444589 | -5.031624572 |
| **A_55_P2006479** | 3300002I08Rik | -0.038314983 | -0.237286797 | 0.160656832 | 8.233062555 | -0.473746741 | 0.652787895 | 0.928444589 | -5.031643203 |
| **A_51_P438619** | Sobp | -0.069143494 | -0.428303474 | 0.290016486 | 8.546132527 | -0.473622738 | 0.652871263 | 0.928444589 | -5.031685406 |
| **A_55_P2042202** | Pigo | 0.058984751 | -0.247416049 | 0.36538555 | 11.11150597 | 0.473607987 | 0.652881181 | 0.928444589 | -5.031690426 |
| **A_51_P493037** | Dclre1a | -0.023367278 | -0.144756886 | 0.09802233 | 8.022618209 | -0.47358264 | 0.652898222 | 0.928444589 | -5.031699051 |
| **A_55_P2044413** | Cxcl11 | -0.084871706 | -0.525786312 | 0.356042901 | 10.96971474 | -0.473562829 | 0.652911542 | 0.928444589 | -5.031705793 |
| **A_55_P1983858** | Scel | -0.080080767 | -0.496138997 | 0.335977463 | 11.55713388 | -0.473525391 | 0.652936713 | 0.928444589 | -5.031718531 |
| **A_55_P2164111** | Nploc4 | 0.059089884 | -0.24797726 | 0.366157028 | 10.90213186 | 0.473422559 | 0.653005856 | 0.928444589 | -5.031753515 |
| **A_55_P2050448** | Slc25a3 | -0.045703273 | -0.283234783 | 0.191828237 | 10.31722531 | -0.473364013 | 0.653045223 | 0.928444589 | -5.03177343 |
| **A_51_P466221** | Amhr2 | -0.045825244 | -0.283999675 | 0.192349186 | 8.603946566 | -0.473346117 | 0.653057257 | 0.928444589 | -5.031779517 |
| **A_51_P421984** | Timm22 | 0.04316212 | -0.181177743 | 0.267501984 | 11.59344768 | 0.473331582 | 0.653067031 | 0.928444589 | -5.031784461 |
| **A_55_P2095603** | Ccdc65 | 0.021374448 | -0.089738839 | 0.132487735 | 7.744154209 | 0.473258118 | 0.653116431 | 0.928446531 | -5.031809445 |
| **A_52_P141628** | Slc23a1 | -0.033301698 | -0.206462907 | 0.139859512 | 7.872361085 | -0.47313484 | 0.653199333 | 0.928496096 | -5.031851364 |

| | | | | | | | | | |
|---|---|---|---|---|---|---|---|---|---|
| **A_51_P514449** | Nr5a2 | -0.04288051 | -0.26594216 | 0.18018114 | 8.204169941 | -0.472937984 | 0.653331726 | 0.928615996 | -5.03191828 |
| **A_51_P214251** | Rnls | 0.023649303 | -0.099405252 | 0.146703857 | 7.910070887 | 0.472813435 | 0.653415497 | 0.928649374 | -5.031960604 |
| **A_52_P467930** | NA | 0.04765434 | -0.200355215 | 0.295663895 | 9.147494382 | 0.472719164 | 0.653478907 | 0.928649374 | -5.031992633 |
| **A_55_P2091626** | NA | 0.026238027 | -0.110342581 | 0.162818635 | 15.6345744 | 0.472619133 | 0.653546195 | 0.928649374 | -5.032026612 |
| **A_51_P166023** | Hip1 | -0.021110698 | -0.131041969 | 0.088820572 | 8.006346504 | -0.472444198 | 0.653663876 | 0.928649374 | -5.03208602 |
| **A_55_P2043486** | Msx2 | 0.048968704 | -0.206034627 | 0.303972035 | 9.561666154 | 0.472434845 | 0.653670169 | 0.928649374 | -5.032089195 |
| **A_55_P1974788** | Cfl1 | 0.029645208 | -0.124748262 | 0.184038678 | 10.80767584 | 0.472383523 | 0.653704696 | 0.928649374 | -5.03210662 |
| **A_52_P504268** | B3galnt1 | 0.123724284 | -0.520694915 | 0.768143483 | 10.80105598 | 0.472340997 | 0.653733307 | 0.928649374 | -5.032121057 |
| **A_55_P2030524** | Vldlr | -0.316162839 | -1.963021125 | 1.330695446 | 13.43726413 | -0.47230632 | 0.653756638 | 0.928649374 | -5.032132829 |
| **A_55_P2051047** | Rps10 | 0.021349381 | -0.089868092 | 0.132566855 | 17.02371957 | 0.472260288 | 0.653787608 | 0.928649374 | -5.032148454 |
| **A_51_P304125** | Slc29a2 | 0.042498459 | -0.178962433 | 0.263959352 | 8.617251848 | 0.472112286 | 0.65388719 | 0.928722574 | -5.032198682 |
| **A_55_P2454784** | Ifnar1 | -0.021887366 | -0.136006825 | 0.092232093 | 9.097185088 | -0.471848913 | 0.654064418 | 0.928906035 | -5.032288029 |
| **A_51_P161463** | Ccdc103 | -0.025426877 | -0.158052816 | 0.107199063 | 8.189665657 | -0.471665058 | 0.654188152 | 0.928997472 | -5.032350374 |
| **A_55_P1972991** | Pcsk6 | -0.08917391 | -0.55442273 | 0.376074911 | 10.44902615 | -0.471543421 | 0.65427002 | 0.928997472 | -5.032391609 |
| **A_55_P2062444** | Memo1 | 0.040562151 | -0.17106725 | 0.252191552 | 10.92864784 | 0.471535224 | 0.654275537 | 0.928997472 | -5.032394387 |
| **A_55_P1998023** | NA | -0.033030022 | -0.205385926 | 0.139325882 | 8.534217245 | -0.471467605 | 0.65432105 | 0.928997472 | -5.032417305 |
| **A_52_P418934** | Sec22b | -0.064454477 | -0.400894312 | 0.271985358 | 13.63902276 | -0.471318882 | 0.65442116 | 0.929011715 | -5.032467702 |
| **A_51_P268145** | Naa80 | 0.067597222 | -0.285253858 | 0.420448302 | 9.215242407 | 0.471309897 | 0.654427208 | 0.929011715 | -5.032470746 |
| **A_55_P2072666** | Cyp2d37-ps | 0.020130115 | -0.085061729 | 0.125321959 | 7.657171128 | 0.470796641 | 0.654772756 | 0.929351469 | -5.032644556 |
| **A_51_P243808** | Traf3ip3 | -0.021465232 | -0.133637418 | 0.090706953 | 7.724162508 | -0.470781697 | 0.654782819 | 0.929351469 | -5.032649614 |
| **A_55_P2095108** | A430033K04Rik | -0.033542276 | -0.208841678 | 0.141757126 | 8.319793182 | -0.470740166 | 0.654810784 | 0.929351469 | -5.032663671 |
| **A_51_P244492** | Nbl1 | -0.274422553 | -1.709144864 | 1.160299759 | 13.37873456 | -0.470566614 | 0.654927653 | 0.92944909 | -5.032722398 |
| **A_55_P1994027** | Adra2b | 0.062656342 | -0.26504156 | 0.390354243 | 12.66091118 | 0.470392661 | 0.655044802 | 0.92949656 | -5.032781241 |
| **A_52_P17112** | Znrf1 | 0.047225507 | -0.199820407 | 0.294271421 | 10.77004267 | 0.470292572 | 0.655112212 | 0.92949656 | -5.032815089 |
| **A_66_P100565** | Ccdc107 | 0.087907002 | -0.371956793 | 0.547770797 | 13.60390336 | 0.470287476 | 0.655115644 | 0.92949656 | -5.032816813 |
| **A_55_P2111292** | 3110009E18Rik | -0.028258094 | -0.176137553 | 0.119621366 | 9.299289905 | -0.47011507 | 0.655231769 | 0.92949656 | -5.032875101 |
| **A_51_P196395** | Cfap298 | -0.036611448 | -0.228267302 | 0.155044406 | 10.51505529 | -0.469963334 | 0.65533398 | 0.92949656 | -5.032926386 |
| **A_51_P215489** | Slc37a1 | -0.034628931 | -0.215933279 | 0.146675418 | 8.868108084 | -0.469894134 | 0.655380596 | 0.92949656 | -5.032949769 |
| **A_55_P2018941** | Fmr1 | -0.045923309 | -0.28636917 | 0.194522553 | 10.44206835 | -0.469877909 | 0.655391526 | 0.92949656 | -5.032955251 |
| **A_55_P2426272** | Scaper | -0.028325324 | -0.176642681 | 0.119992034 | 8.521205784 | -0.469842246 | 0.655415551 | 0.92949656 | -5.032967301 |
| **A_52_P402705** | Kcnj11 | 0.020086718 | -0.085144582 | 0.125318018 | 7.666397196 | 0.469605536 | 0.655575029 | 0.92949656 | -5.033047257 |
| **A_51_P204913** | Bag2 | 0.069546887 | -0.294809436 | 0.433903209 | 10.0147608 | 0.469591843 | 0.655584254 | 0.92949656 | -5.033051881 |
| **A_51_P338461** | Numa1 | 0.070036089 | -0.312455177 | 0.452527356 | 13.68013913 | 0.474515548 | 0.655648016 | 0.92949656 | -4.967173647 |
| **A_55_P2102424** | Msto1 | 0.070242412 | -0.297896696 | 0.438381521 | 9.665509616 | 0.469414634 | 0.655703658 | 0.92949656 | -5.033111714 |
| **A_51_P415905** | Pola1 | 0.043777696 | -0.185689828 | 0.273245219 | 8.332042479 | 0.469354333 | 0.655744291 | 0.92949656 | -5.033132069 |
| **A_55_P2180071** | Zfand5 | -0.046729819 | -0.291672156 | 0.198212518 | 12.91330672 | -0.46935279 | 0.655745332 | 0.92949656 | -5.03313259 |
| **A_51_P348592** | Slc38a7 | -0.026226013 | -0.163697549 | 0.111245522 | 9.369487523 | -0.469341176 | 0.655753157 | 0.92949656 | -5.03313651 |
| **A_51_P334789** | Nudt16 | -0.026397928 | -0.164782057 | 0.1119862 | 8.493512319 | -0.469302346 | 0.655779324 | 0.92949656 | -5.033149615 |
| **A_55_P2009146** | Gtpbp3 | 0.025553655 | -0.108437385 | 0.159544696 | 7.984629482 | 0.469187504 | 0.655856715 | 0.92949656 | -5.033188372 |
| **A_51_P185292** | 4930581F22Rik | -0.043772373 | -0.273332944 | 0.185788198 | 8.454822539 | -0.469107044 | 0.655910939 | 0.92949656 | -5.033215519 |
| **A_55_P1990755** | Ppp3r1 | -0.042910865 | -0.268050699 | 0.182228969 | 11.73007891 | -0.468904167 | 0.656047674 | 0.92949656 | -5.033283953 |
| **A_55_P2230638** | Ece1 | -0.089145789 | -0.556925316 | 0.378633738 | 8.634514405 | -0.468844455 | 0.656087922 | 0.92949656 | -5.033304089 |
| **A_51_P476579** | NA | -0.073569998 | -0.45973745 | 0.312597454 | 9.704078617 | -0.46869923 | 0.656185812 | 0.92949656 | -5.033353054 |
| **A_52_P17146** | Bmpr1a | -0.050975239 | -0.318558501 | 0.216608024 | 11.92254117 | -0.468672517 | 0.65620382 | 0.92949656 | -5.033362059 |
| **A_51_P430423** | Ada | 0.09304772 | -0.395420066 | 0.581515507 | 15.15201879 | 0.468639588 | 0.656226017 | 0.92949656 | -5.033373159 |
| **A_52_P193925** | Sulf2 | 0.056290097 | -0.239236158 | 0.351816353 | 11.68518774 | 0.468603029 | 0.656250662 | 0.92949656 | -5.033385482 |
| **A_51_P432659** | Pex7 | 0.061999958 | -0.263508145 | 0.387508062 | 10.06919625 | 0.468596193 | 0.65625527 | 0.92949656 | -5.033387786 |
| **A_55_P2442017** | Samd8 | -0.028736213 | -0.179705082 | 0.122232656 | 9.572831686 | -0.468286136 | 0.656464305 | 0.92949656 | -5.033492259 |
| **A_55_P2064233** | Ipo8 | -0.032349155 | -0.202330476 | 0.137632165 | 10.15326083 | -0.468199462 | 0.656522745 | 0.92949656 | -5.033521453 |
| **A_55_P1971649** | Ttc19 | 0.062094315 | -0.264185857 | 0.388374488 | 10.58603609 | 0.468198828 | 0.656523172 | 0.92949656 | -5.033521666 |

| | | | | | | | | | |
|---|---|---|---|---|---|---|---|---|---|
| **A_55_P2046101** | Xlr4b | -0.065558636 | -0.410115571 | 0.2789983 | 11.80653478 | -0.468099389 | 0.656590222 | 0.92949656 | -5.033555153 |
| **A_55_P1958921** | Ankrd29 | 0.029135251 | -0.123991874 | 0.182262375 | 8.150938936 | 0.468096932 | 0.656591879 | 0.92949656 | -5.033555981 |
| **A_51_P379775** | Nrm | -0.048664271 | -0.304459844 | 0.207131303 | 9.073669427 | -0.468043651 | 0.656627807 | 0.92949656 | -5.033573921 |
| **A_55_P2165011** | Fut10 | 0.026599074 | -0.113230347 | 0.166428494 | 7.987196881 | 0.467990586 | 0.656663591 | 0.92949656 | -5.033591787 |
| **A_51_P101075** | Cnppd1 | -0.062868949 | -0.393411289 | 0.267673391 | 14.50156112 | -0.467927171 | 0.656706355 | 0.92949656 | -5.028603909 |
| **A_51_P233815** | Tmem126a | 0.03895602 | -0.165902714 | 0.243814754 | 9.624950801 | 0.467831286 | 0.656771018 | 0.92949656 | -5.033645408 |
| **A_51_P180032** | Sem1 | 0.043691672 | -0.186073008 | 0.273456352 | 15.39659477 | 0.467826219 | 0.656774435 | 0.92949656 | -5.033647113 |
| **A_52_P176160** | Ahcy | 0.022878885 | -0.097457367 | 0.143215137 | 7.792204463 | 0.46774326 | 0.656830383 | 0.92949656 | -5.03367503 |
| **A_52_P1172945** | Ankrd34b | 0.021861311 | -0.09315889 | 0.136881512 | 7.688200755 | 0.467596485 | 0.656929377 | 0.92949656 | -5.033724413 |
| **A_51_P294778** | Zfp954 | 0.041661346 | -0.177561366 | 0.260884059 | 10.08415 | 0.467538003 | 0.656968823 | 0.92949656 | -5.033744085 |
| **A_52_P369123** | Fam189a2 | -0.020873772 | -0.130726733 | 0.088979189 | 7.805649665 | -0.467474939 | 0.65701136 | 0.92949656 | -5.033765296 |
| **A_55_P2183433** | Rab30 | 0.139641305 | -0.595254916 | 0.874537525 | 9.75559299 | 0.467473554 | 0.657012294 | 0.92949656 | -5.033765762 |
| **A_51_P210560** | Pptc7 | -0.034482118 | -0.215958763 | 0.146994526 | 11.86240374 | -0.467457746 | 0.657022957 | 0.92949656 | -5.033771079 |
| **A_55_P2123566** | NA | -0.120422913 | -0.754212689 | 0.513366863 | 8.811314534 | -0.467447739 | 0.657029707 | 0.92949656 | -5.033774444 |
| **A_51_P427934** | Megf8 | -0.058486792 | -0.366343969 | 0.249370386 | 9.334680419 | -0.467388131 | 0.657069916 | 0.92949656 | -5.033794489 |
| **A_55_P2064996** | Capzb | -0.038348774 | -0.240209908 | 0.163512359 | 11.0467036 | -0.467377655 | 0.657076982 | 0.92949656 | -5.033798012 |
| **A_51_P481832** | Rabggta | -0.070585345 | -0.442192644 | 0.301021953 | 10.1045607 | -0.46730397 | 0.657126688 | 0.929498848 | -5.033822787 |
| **A_55_P2004881** | Ldlrad4 | -0.021646103 | -0.135710824 | 0.092418617 | 7.855529414 | -0.466871697 | 0.657418326 | 0.929588824 | -5.033968062 |
| **A_51_P476601** | Ly96 | -0.026113302 | -0.163719177 | 0.111492573 | 9.882553449 | -0.466867869 | 0.657420909 | 0.929588824 | -5.033969348 |
| **A_52_P246165** | Ptprk | -0.028158123 | -0.1765399 | 0.120223655 | 8.865814769 | -0.466866046 | 0.65742214 | 0.929588824 | -5.03396996 |
| **A_55_P1986188** | Def8 | 0.052630838 | -0.224718344 | 0.32998002 | 12.44181861 | 0.466855584 | 0.657429198 | 0.929588824 | -5.033973474 |
| **A_66_P119164** | Zfp354c | -0.027528181 | -0.172594398 | 0.117538035 | 8.106013615 | -0.466853265 | 0.657430763 | 0.929588824 | -5.033974254 |
| **A_55_P1993064** | Mbd6 | 0.054804365 | -0.234124683 | 0.343733414 | 11.67929733 | 0.466651976 | 0.657566589 | 0.929681316 | -5.034041857 |
| **A_52_P667477** | Fyco1 | -0.040944238 | -0.256842872 | 0.174954396 | 8.650527428 | -0.466564899 | 0.657625352 | 0.929681316 | -5.034071094 |
| **A_55_P2009069** | NA | -0.167279265 | -1.049383819 | 0.71482529 | 15.57000277 | -0.466542498 | 0.657640469 | 0.929681316 | -5.034078615 |
| **A_52_P117966** | Atpsckmt | 0.046224262 | -0.197577647 | 0.29002617 | 10.36511035 | 0.46644672 | 0.657705107 | 0.929704695 | -5.034110766 |
| **A_52_P176300** | Med1 | 0.028151787 | -0.120398447 | 0.176702021 | 8.84496534 | 0.466231693 | 0.657850235 | 0.929841842 | -5.034182925 |
| **A_51_P227077** | Mdh1b | 0.028202129 | -0.120664025 | 0.177068283 | 7.934983833 | 0.466074227 | 0.657956523 | 0.929857975 | -5.03423575 |
| **A_51_P156291** | F9 | 0.022822264 | -0.097659353 | 0.14330388 | 8.134233707 | 0.466022723 | 0.65799129 | 0.929857975 | -5.034253024 |
| **A_55_P1990500** | Masp2 | 0.019831961 | -0.084868357 | 0.12453228 | 7.69639747 | 0.466000977 | 0.65800597 | 0.929857975 | -5.034260317 |
| **A_55_P2151006** | C1qtnf7 | -0.022242324 | -0.13971979 | 0.095235141 | 8.053185755 | -0.465794937 | 0.658145064 | 0.929956516 | -5.034329401 |
| **A_51_P192860** | Hint1 | -0.045015336 | -0.282803484 | 0.192772812 | 16.47356438 | -0.465735621 | 0.65818511 | 0.929956516 | -5.034349285 |
| **A_55_P2092661** | Nup210 | -0.075043617 | -0.471517841 | 0.321430606 | 10.12902448 | -0.465658962 | 0.658236867 | 0.929956516 | -5.034374979 |
| **A_55_P2011137** | Armc10 | 0.039718271 | -0.170144181 | 0.249580723 | 9.160849071 | 0.465612631 | 0.658268149 | 0.929956516 | -5.034390505 |
| **A_51_P215559** | Gstp3 | 0.024248242 | -0.103899043 | 0.152395526 | 7.741603215 | 0.465521782 | 0.658329491 | 0.929975204 | -5.034420948 |
| **A_55_P1976012** | Cnih4 | -0.05528976 | -0.347649435 | 0.237069915 | 10.04390997 | -0.465260728 | 0.658505771 | 0.930156244 | -5.034508392 |
| **A_55_P1973683** | Morn4 | -0.105196071 | -0.661534414 | 0.451142272 | 8.888416064 | -0.465189273 | 0.658554026 | 0.930156432 | -5.03453232 |
| **A_52_P613643** | Zfp3 | -0.030241811 | -0.190235983 | 0.12975236 | 8.348030438 | -0.465021254 | 0.6586675 | 0.93024873 | -5.03458857 |
| **A_52_P603421** | Ccdc189 | -0.019744324 | -0.124249182 | 0.084760533 | 7.76849426 | -0.464809464 | 0.65881055 | 0.930315507 | -5.034659447 |
| **A_51_P305320** | B4gat1 | 0.082256978 | -0.353122179 | 0.517636136 | 11.12606465 | 0.464808734 | 0.658811043 | 0.930315507 | -5.034659691 |
| **A_66_P110584** | Tpbpa | -0.040515147 | -0.255009694 | 0.173979401 | 10.52446061 | -0.464697487 | 0.65888619 | 0.930353654 | -5.03469691 |
| **A_55_P1956712** | Med28 | -0.050271794 | -0.316480389 | 0.215936801 | 12.31591733 | -0.464591729 | 0.658957633 | 0.930386565 | -5.034732284 |
| **A_51_P327632** | Igsf3 | -0.046217643 | -0.290999602 | 0.198564316 | 9.79898137 | -0.464512659 | 0.659011049 | 0.930394023 | -5.034758727 |
| **A_52_P578562** | Slc41a1 | 0.055780389 | -0.239694974 | 0.351255751 | 9.131313183 | 0.464439797 | 0.659060274 | 0.930395562 | -5.03478309 |
| **A_51_P339444** | Rab9 | -0.065391052 | -0.412112346 | 0.281330242 | 12.53304741 | -0.463988239 | 0.659365384 | 0.930758307 | -5.034934002 |
| **A_55_P2017929** | Clu | 0.08877776 | -0.382376814 | 0.559932333 | 13.43886813 | 0.463564248 | 0.659651932 | 0.9310948 | -5.03507558 |
| **A_55_P2178658** | NA | -0.047270541 | -0.298268351 | 0.203727269 | 11.36257141 | -0.46332934 | 0.659810719 | 0.931185042 | -5.03515397 |
| **A_66_P106536** | Atp5mpl | 0.043628366 | -0.188031278 | 0.27528801 | 14.20824903 | 0.463327125 | 0.659812217 | 0.931185042 | -5.035154709 |
| **A_51_P136781** | Pnliprp1 | 0.019803972 | -0.085385892 | 0.124993836 | 7.678028928 | 0.463177619 | 0.659913286 | 0.931249234 | -5.035204581 |
| **A_55_P2130930** | Nhp2 | 0.076512031 | -0.329938675 | 0.482962738 | 12.94672784 | 0.463117309 | 0.659954059 | 0.931249234 | -5.035224695 |

| | | | | | | | | | |
|---|---|---|---|---|---|---|---|---|---|
| **A_55_P2069597** | Tspan17 | 0.0592916 | -0.255746797 | 0.374329997 | 8.439326114 | 0.463019032 | 0.660020503 | 0.931275005 | -5.035257466 |
| **A_55_P2101585** | Cd55 | -0.262727246 | -1.659243964 | 1.133789473 | 12.03623655 | -0.462837071 | 0.660143533 | 0.931323193 | -5.035318126 |
| **A_55_P2156435** | NA | -0.100584083 | -0.635272717 | 0.434104551 | 9.995715357 | -0.462804393 | 0.660165629 | 0.931323193 | -5.035329017 |
| **A_51_P441898** | Krt25 | 0.070666447 | -0.305025201 | 0.446358095 | 9.142212067 | 0.462754742 | 0.660199203 | 0.931323193 | -5.035345564 |
| **A_66_P126504** | Trabd2b | -0.044046699 | -0.27835493 | 0.190261533 | 9.252542571 | -0.462482154 | 0.66038354 | 0.931515248 | -5.035436381 |
| **A_55_P2058992** | Vamp4 | 0.030304521 | -0.130967196 | 0.191576238 | 8.757402057 | 0.46229413 | 0.660510705 | 0.931597926 | -5.035498996 |
| **A_55_P2053516** | Cacfd1 | -0.064477256 | -0.407636738 | 0.278682226 | 12.81470019 | -0.462252966 | 0.660538547 | 0.931597926 | -5.035512701 |
| **A_55_P2095814** | Kalrn | -0.021826236 | -0.138081903 | 0.094429431 | 7.74997943 | -0.461885017 | 0.660787443 | 0.931832809 | -5.035635157 |
| **A_52_P420504** | Acta2 | 0.169672531 | -0.734115061 | 1.073460123 | 12.32008406 | 0.461864229 | 0.660801506 | 0.931832809 | -5.025717797 |
| **A_51_P260035** | Sar1b | 0.063038243 | -0.273015614 | 0.3990921 | 11.15719958 | 0.461492216 | 0.661053202 | 0.932119736 | -5.035765786 |
| **A_55_P2280237** | Zfp697 | -0.04430105 | -0.280563895 | 0.191961796 | 8.874976886 | -0.461304563 | 0.661180182 | 0.932131519 | -5.035828157 |
| **A_66_P139393** | Psmd10 | -0.025127057 | -0.159157181 | 0.108903067 | 9.942008374 | -0.461220253 | 0.661237237 | 0.932131519 | -5.035856171 |
| **A_52_P96748** | Ttc30b | 0.033777445 | -0.146404193 | 0.213959083 | 8.536997225 | 0.461195887 | 0.661253727 | 0.932131519 | -5.035864267 |
| **A_52_P337236** | Alyref2 | -0.03380011 | -0.214111591 | 0.14651137 | 8.700333157 | -0.461173033 | 0.661269193 | 0.932131519 | -5.03587186 |
| **A_51_P500541** | Flywch2 | -0.055725634 | -0.353033826 | 0.241582559 | 8.104612634 | -0.461123551 | 0.661302681 | 0.932131519 | -5.035888298 |
| **A_55_P1965298** | Gm5640 | -0.03895069 | -0.247000778 | 0.169099398 | 12.45960563 | -0.460592028 | 0.661662452 | 0.932570624 | -5.036064773 |
| **A_52_P30451** | Ppp1r3c | -0.06572623 | -0.416924305 | 0.285471845 | 9.091233818 | -0.460421673 | 0.66177778 | 0.932583539 | -5.036121295 |
| **A_51_P402435** | Trap1 | -0.023699092 | -0.150363547 | 0.102965364 | 9.430975457 | -0.460305434 | 0.661856479 | 0.932583539 | -5.036159851 |
| **A_55_P2014209** | Scyl2 | 0.045475838 | -0.197586088 | 0.288537763 | 9.041861667 | 0.46029148 | 0.661865927 | 0.932583539 | -5.036164479 |
| **A_52_P32903** | Zzef1 | -0.05172965 | -0.328275557 | 0.224816258 | 10.66232086 | -0.460194543 | 0.661931561 | 0.932583539 | -5.036196625 |
| **A_55_P2119917** | Ikzf4 | -0.061469533 | -0.390088797 | 0.267149731 | 8.700004436 | -0.460188816 | 0.661935439 | 0.932583539 | -5.036198524 |
| **A_51_P302139** | Mastl | -0.027117039 | -0.172120403 | 0.117886325 | 7.843176768 | -0.460079997 | 0.662009123 | 0.932583539 | -5.036234602 |
| **A_55_P2081178** | Msl3l2 | -0.040367416 | -0.256224884 | 0.175490052 | 8.704391055 | -0.46007966 | 0.662009351 | 0.932583539 | -5.036234714 |
| **A_52_P207335** | Tmem246 | 0.037392466 | -0.162650978 | 0.237435909 | 8.668792657 | 0.459863506 | 0.662155726 | 0.932711166 | -5.036306356 |
| **A_55_P2107854** | Hnrnpk | 0.051412991 | -0.223673896 | 0.326499877 | 10.61420021 | 0.459803362 | 0.662196458 | 0.932711166 | -5.036326285 |
| **A_52_P228236** | Tfrc | 0.075149974 | -0.327078137 | 0.477378085 | 13.31567419 | 0.459648215 | 0.662301533 | 0.932791193 | -5.011073339 |
| **A_55_P1962801** | Max | -0.067765909 | -0.430600869 | 0.295069051 | 13.11740772 | -0.459484967 | 0.662412105 | 0.932815852 | -5.036431745 |
| **A_55_P2153016** | Ccdc34 | -0.026294477 | -0.167113754 | 0.1145248 | 7.905453088 | -0.459379478 | 0.66248356 | 0.932815852 | -5.036466671 |
| **A_55_P1977478** | Zfp629 | -0.033363107 | -0.212064507 | 0.145338293 | 9.448956222 | -0.459311897 | 0.662529339 | 0.932815852 | -5.036489043 |
| **A_55_P1963687** | Tsen54 | 0.095513076 | -0.416120259 | 0.60714641 | 11.90926739 | 0.459275375 | 0.662554079 | 0.932815852 | -5.036501132 |
| **A_55_P2030761** | NA | 0.077010501 | -0.335572761 | 0.489593763 | 13.72782443 | 0.45920595 | 0.662601111 | 0.932815852 | -5.036524109 |
| **A_52_P407796** | Mdh1 | 0.052821706 | -0.230215577 | 0.33585899 | 13.82800656 | 0.459132401 | 0.662650937 | 0.932815852 | -5.036548447 |
| **A_55_P1955279** | Specc1 | -0.046119314 | -0.293247525 | 0.201008898 | 12.00082535 | -0.459123657 | 0.662656861 | 0.932815852 | -5.036551341 |
| **A_55_P2100155** | Mapt | 0.024528227 | -0.106961832 | 0.156018287 | 8.434237603 | 0.458925723 | 0.662790964 | 0.932926595 | -5.036616821 |
| **A_55_P2031382** | Crebzf | -0.036305265 | -0.230954982 | 0.158344452 | 8.64943268 | -0.458865067 | 0.662832062 | 0.932926595 | -5.036636883 |
| **A_51_P375969** | Ces1d | 0.021082558 | -0.091976024 | 0.13414114 | 7.694004985 | 0.458763571 | 0.662900834 | 0.932949198 | -5.036670446 |
| **A_51_P473940** | Slc22a15 | 0.046088913 | -0.201132803 | 0.293310629 | 8.574742537 | 0.45864748 | 0.6629795 | 0.932949198 | -5.036708827 |
| **A_52_P532559** | Thrb | 0.027197342 | -0.118695966 | 0.173090651 | 7.859921705 | 0.458627673 | 0.662992922 | 0.932949198 | -5.036715375 |
| **A_51_P236175** | Rit1 | 0.043867429 | -0.191579388 | 0.279314246 | 13.31003929 | 0.458372465 | 0.663165876 | 0.93306807 | -5.036799716 |
| **A_55_P2062039** | Tsen34 | 0.103134432 | -0.450502616 | 0.656771481 | 12.85366313 | 0.458297724 | 0.663216532 | 0.93306807 | -5.036824408 |
| **A_55_P2018322** | Zfp644 | -0.036397215 | -0.231883426 | 0.159088996 | 10.19192796 | -0.458058752 | 0.663378509 | 0.93306807 | -5.036903333 |
| **A_55_P2347384** | NA | 0.02147809 | -0.093880898 | 0.136837078 | 7.83467556 | 0.458050524 | 0.663384087 | 0.93306807 | -5.03690605 |
| **A_55_P2138366** | Xpo4 | 0.038684466 | -0.177923524 | 0.255292455 | 8.751136841 | 0.462820101 | 0.663438666 | 0.93306807 | -4.970857908 |
| **A_55_P2074831** | Fam168a | -0.020364746 | -0.129797239 | 0.089067747 | 10.98293882 | -0.45782749 | 0.66353528 | 0.93306807 | -5.036979676 |
| **A_52_P275627** | Kpnb1 | -0.027758739 | -0.176931621 | 0.121414143 | 9.715320067 | -0.457803405 | 0.663551609 | 0.93306807 | -5.036987625 |
| **A_55_P2114467** | Romo1 | 0.03028894 | -0.13248241 | 0.193060291 | 13.44227585 | 0.45779946 | 0.663554283 | 0.93306807 | -5.036988927 |
| **A_55_P2109128** | H2bc6 | 0.039323103 | -0.172014178 | 0.250660384 | 8.413883299 | 0.457763151 | 0.663578898 | 0.93306807 | -5.03700091 |
| **A_51_P431329** | Car3 | 0.022899656 | -0.100190884 | 0.145990196 | 7.687510132 | 0.457692136 | 0.663627044 | 0.93306807 | -5.037024343 |
| **A_52_P253915** | Cds2 | -0.053984495 | -0.344166614 | 0.236197624 | 10.49137495 | -0.457685928 | 0.663631253 | 0.93306807 | -5.037026391 |
| **A_55_P2084965** | Klk13 | -0.019760095 | -0.125984962 | 0.086464773 | 7.664883338 | -0.457648436 | 0.663656672 | 0.93306807 | -5.037038761 |

| | | | | | | | | | |
|---|---|---|---|---|---|---|---|---|---|
| **A_55_P2131899** | Sox4 | 0.022375245 | -0.097958366 | 0.142708856 | 7.781606933 | 0.457456723 | 0.663786659 | 0.933164793 | -5.037101999 |
| **A_55_P2035937** | Rab6a | 0.032494375 | -0.14229008 | 0.20727883 | 12.67475229 | 0.457377239 | 0.663840555 | 0.933164793 | -5.037128211 |
| **A_55_P2043337** | Colq | 0.079631352 | -0.348797848 | 0.508060552 | 8.378498815 | 0.457271556 | 0.66391222 | 0.933164793 | -5.037163056 |
| **A_55_P2180099** | NA | 0.047868664 | -0.20967756 | 0.305414887 | 13.43531672 | 0.457262178 | 0.66391858 | 0.933164793 | -5.037166148 |
| **A_55_P1969831** | Ptpmt1 | -0.04691866 | -0.299437126 | 0.205599806 | 12.66103482 | -0.457110948 | 0.664021137 | 0.933241079 | -5.037215996 |
| **A_55_P2158873** | Ppid | -0.027031228 | -0.172616041 | 0.118553586 | 9.427850209 | -0.456792387 | 0.664237198 | 0.933330682 | -5.037320952 |
| **A_55_P2015670** | Itga6 | 0.09780127 | -0.451250422 | 0.646852962 | 14.29195116 | 0.461616521 | 0.664243189 | 0.933330682 | -4.971232468 |
| **A_55_P1992174** | Tars2 | -0.032866215 | -0.209888201 | 0.144155771 | 8.689665906 | -0.456763766 | 0.664256612 | 0.933330682 | -5.037330379 |
| **A_52_P152631** | Tmem17 | -0.026739705 | -0.170778276 | 0.117298866 | 8.627186204 | -0.456716778 | 0.664288485 | 0.933330682 | -5.037345853 |
| **A_55_P2003823** | Asgr1 | -0.020315288 | -0.129773653 | 0.089143077 | 7.646853371 | -0.45660766 | 0.664362503 | 0.933330682 | -5.037381783 |
| **A_51_P308814** | Toe1 | -0.025809813 | -0.164906232 | 0.113286606 | 8.966690599 | -0.456497155 | 0.664437468 | 0.933330682 | -5.037418163 |
| **A_52_P273594** | Zc3h14 | -0.047490732 | -0.303432096 | 0.208450633 | 10.00370154 | -0.456496596 | 0.664437847 | 0.933330682 | -5.037418347 |
| **A_55_P2003251** | Magel2 | -0.297765722 | -1.902684554 | 1.307153111 | 9.200498636 | -0.456447456 | 0.664471184 | 0.933330682 | -5.037434521 |
| **A_55_P1989251** | Chfr | -0.057854322 | -0.405377374 | 0.289668729 | 11.41171748 | -0.468440787 | 0.664636739 | 0.933446832 | -4.920308749 |
| **A_66_P139974** | Arl6 | 0.038197021 | -0.167860881 | 0.244254922 | 8.533826513 | 0.456046766 | 0.664743047 | 0.933446832 | -5.037566351 |
| **A_66_P136095** | Ctnna3 | 0.019954598 | -0.087697675 | 0.127606872 | 7.613892884 | 0.456025311 | 0.664757605 | 0.933446832 | -5.037573407 |
| **A_51_P465161** | Hoga1 | 0.077809119 | -0.341987412 | 0.497605649 | 8.24586674 | 0.455995771 | 0.66477765 | 0.933446832 | -5.037583121 |
| **A_55_P1964183** | Dysf | 0.06074041 | -0.267033942 | 0.388514761 | 11.93514348 | 0.455902437 | 0.664840986 | 0.933446832 | -5.037613811 |
| **A_55_P2133568** | Snhg5 | -0.022651615 | -0.144907486 | 0.099604256 | 7.936592103 | -0.455825486 | 0.664893206 | 0.933446832 | -5.037639109 |
| **A_55_P2146525** | Camta2 | -0.062638926 | -0.400836888 | 0.275559036 | 11.50947832 | -0.455661649 | 0.665004396 | 0.933446832 | -5.037692959 |
| **A_55_P2101336** | Ramp2 | 0.109791434 | -0.483043444 | 0.702626311 | 12.15682367 | 0.455621146 | 0.665031885 | 0.933446832 | -5.037706268 |
| **A_52_P422859** | Hspg2 | 0.117671815 | -0.517846378 | 0.753190008 | 13.98570144 | 0.455526481 | 0.665096137 | 0.933446832 | -5.037737372 |
| **A_55_P2102419** | Lime1 | -0.045150616 | -0.289081321 | 0.198780089 | 8.954855166 | -0.455372047 | 0.665200961 | 0.933446832 | -5.037788102 |
| **A_51_P116407** | Pald1 | -0.020291947 | -0.129935465 | 0.089351571 | 7.960227278 | -0.455312863 | 0.665241136 | 0.933446832 | -5.037807539 |
| **A_55_P2076871** | Lef1 | -0.038594021 | -0.247131973 | 0.169943931 | 8.249980664 | -0.455306748 | 0.665245286 | 0.933446832 | -5.037809547 |
| **A_52_P447944** | Epcam | -0.041144074 | -0.263480193 | 0.181192046 | 9.416717265 | -0.455267273 | 0.665272083 | 0.933446832 | -5.037822509 |
| **A_55_P2132845** | NA | 0.045229105 | -0.199190652 | 0.289648863 | 16.14060423 | 0.455250939 | 0.665283171 | 0.933446832 | -5.037827873 |
| **A_55_P2166446** | NA | -0.021221555 | -0.135908526 | 0.093465415 | 8.183436237 | -0.455231488 | 0.665296375 | 0.933446832 | -5.03783426 |
| **A_66_P126153** | 1700011M02Rik | 0.109370289 | -0.48175761 | 0.700498187 | 12.18362266 | 0.45518408 | 0.665328558 | 0.933446832 | -5.037849825 |
| **A_55_P2081164** | Cd300lf | -0.069161761 | -0.443025379 | 0.304701857 | 8.312635911 | -0.455115892 | 0.665374849 | 0.933446832 | -5.03787221 |
| **A_55_P2131522** | Arfrp1 | 0.02445156 | -0.107801283 | 0.156704402 | 10.89710247 | 0.454852633 | 0.665553582 | 0.933567962 | -5.037958607 |
| **A_55_P2013029** | NA | -0.028283081 | -0.181266993 | 0.124700831 | 8.452885914 | -0.454831059 | 0.665568231 | 0.933567962 | -5.037965685 |
| **A_55_P2327027** | Ano6 | 0.029703758 | -0.13099622 | 0.190403737 | 8.356745327 | 0.454741664 | 0.66562893 | 0.933567962 | -5.037995012 |
| **A_51_P140641** | Eva1a | -0.060388676 | -0.387123731 | 0.266346379 | 13.2217806 | -0.454704172 | 0.665654388 | 0.933567962 | -5.038007309 |
| **A_55_P2135271** | Zfp358 | 0.039622908 | -0.174937769 | 0.254183586 | 10.09354221 | 0.454323689 | 0.665912771 | 0.93362086 | -5.03813206 |
| **A_51_P187018** | Magohb | -0.040965401 | -0.262808145 | 0.180877342 | 10.047039 | -0.454298336 | 0.665929989 | 0.93362086 | -5.038140369 |
| **A_51_P144319** | Efna5 | -0.040867647 | -0.262196597 | 0.180461303 | 8.629991702 | -0.454266359 | 0.665951707 | 0.93362086 | -5.038150848 |
| **A_55_P1980267** | Sidt2 | -0.046109438 | -0.295866836 | 0.20364796 | 12.04356418 | -0.4541932 | 0.666001396 | 0.93362086 | -5.038174822 |
| **A_55_P2039274** | Hspb3 | -0.019623368 | -0.125932749 | 0.086686012 | 7.652331822 | -0.454120526 | 0.666050758 | 0.93362086 | -5.038198633 |
| **A_51_P403760** | Braf | 0.049230631 | -0.217477698 | 0.31593896 | 10.60721928 | 0.454117238 | 0.666052991 | 0.93362086 | -5.03819971 |
| **A_55_P2146520** | Carns1 | 0.025552898 | -0.112900915 | 0.164006712 | 8.037374841 | 0.454050763 | 0.666098144 | 0.93362086 | -5.038221487 |
| **A_55_P2055369** | NA | -0.02589737 | -0.16622229 | 0.11442755 | 8.063307854 | -0.454035723 | 0.66610836 | 0.93362086 | -5.038226413 |
| **A_55_P2186005** | Sphk1 | -0.183372129 | -1.17710688 | 0.810362622 | 9.178201208 | -0.453975079 | 0.666149553 | 0.93362086 | -5.038246276 |
| **A_55_P2066081** | Prmt6 | 0.028299534 | -0.12508585 | 0.181684918 | 8.892724682 | 0.453904481 | 0.66619751 | 0.93362086 | -5.038269397 |
| **A_55_P2143139** | Myadml2 | -0.021508561 | -0.138133479 | 0.095116356 | 7.67259647 | -0.453721311 | 0.666321944 | 0.93362086 | -5.03832937 |
| **A_52_P239052** | Zfp148 | -0.048098958 | -0.309003586 | 0.212805671 | 10.93350607 | -0.453547779 | 0.666439841 | 0.93362086 | -5.038386166 |
| **A_55_P2154054** | Myo18a | -0.033449044 | -0.214912693 | 0.148014606 | 11.25375912 | -0.453485312 | 0.666482284 | 0.93362086 | -5.038406607 |
| **A_55_P1982744** | NA | -0.038893881 | -0.249904129 | 0.172116367 | 12.12953871 | -0.453468258 | 0.666493871 | 0.93362086 | -5.038412187 |
| **A_55_P2039105** | Rbck1 | 0.059290415 | -0.262446198 | 0.381027029 | 10.83945749 | 0.453370408 | 0.666560358 | 0.93362086 | -5.038444199 |
| **A_51_P231320** | Mmp8 | -0.02570163 | -0.165173761 | 0.113770501 | 7.816860232 | -0.453359156 | 0.666568003 | 0.93362086 | -5.03844788 |

| | | | | | | | | | |
|---|---|---|---|---|---|---|---|---|---|
| **A_55_P1963434** | Gpc4 | -0.028361757 | -0.182345126 | 0.125621613 | 8.338201117 | -0.453135901 | 0.666719712 | 0.93362086 | -5.038520894 |
| **A_52_P53626** | NA | -0.033156488 | -0.213188882 | 0.146875906 | 8.996927554 | -0.453092666 | 0.666749094 | 0.93362086 | -5.03853503 |
| **A_55_P2089233** | Pou2af1 | 0.021172276 | -0.093797766 | 0.136142318 | 7.816816321 | 0.45305614 | 0.666773917 | 0.93362086 | -5.038546972 |
| **A_55_P1998825** | Rpl9 | 0.031176615 | -0.138124697 | 0.200477927 | 16.91543259 | 0.453041301 | 0.666784002 | 0.93362086 | -5.038551823 |
| **A_52_P558609** | Clec16a | -0.041157536 | -0.264731325 | 0.182416253 | 8.82948863 | -0.45289512 | 0.666883351 | 0.93362086 | -5.038599604 |
| **A_52_P357133** | Selenom | 0.04777411 | -0.21179953 | 0.30734775 | 11.83655771 | 0.452794542 | 0.666951711 | 0.93362086 | -5.038632471 |
| **A_55_P2159746** | Mfsd2b | -0.018984465 | -0.122165154 | 0.084196225 | 7.730243693 | -0.452656846 | 0.667045305 | 0.93362086 | -5.038677456 |
| **A_55_P2185651** | NA | 0.031618643 | -0.140244024 | 0.203481309 | 8.871428724 | 0.45261698 | 0.667072404 | 0.93362086 | -5.038690478 |
| **A_55_P2067608** | Clmp | -0.176024536 | -1.133024177 | 0.780975105 | 9.094393038 | -0.452512517 | 0.667143415 | 0.93362086 | -5.038724596 |
| **A_55_P2015734** | Syvn1 | -0.073147735 | -0.47093403 | 0.324638559 | 14.80366183 | -0.452397537 | 0.667221579 | 0.93362086 | -5.038762139 |
| **A_55_P2095688** | Sh3bp5l | -0.035432051 | -0.228166334 | 0.157302232 | 12.77665418 | -0.452279063 | 0.667302124 | 0.93362086 | -5.038800815 |
| **A_55_P2006698** | Bmp1 | -0.066433249 | -0.427814284 | 0.294947786 | 12.17409111 | -0.452261164 | 0.667314293 | 0.93362086 | -5.038806658 |
| **A_55_P2071191** | Prss8 | -0.070586135 | -0.454573818 | 0.313401549 | 10.46267021 | -0.452242391 | 0.667327056 | 0.93362086 | -5.038812785 |
| **A_55_P2079251** | Ap2m1 | -0.062115983 | -0.400045439 | 0.275813472 | 12.27866105 | -0.452216579 | 0.667344606 | 0.93362086 | -5.038821209 |
| **A_55_P1961210** | Trp53inp2 | -0.080318912 | -0.51729074 | 0.356652916 | 12.84473345 | -0.452203107 | 0.667353765 | 0.93362086 | -5.038825606 |
| **A_66_P106760** | Adam32 | 0.029104385 | -0.129244825 | 0.187453595 | 8.061218245 | 0.452180356 | 0.667369233 | 0.93362086 | -5.038833031 |
| **A_52_P378968** | Rgs2 | -0.019109772 | -0.123095368 | 0.084875824 | 7.687152014 | -0.45211768 | 0.667411848 | 0.93362086 | -5.038853484 |
| **A_51_P283473** | Fibin | -0.026007821 | -0.167531866 | 0.115516224 | 8.693271453 | -0.452108737 | 0.667417928 | 0.93362086 | -5.038856402 |
| **A_51_P186856** | Krt5 | 0.219797875 | -0.97658324 | 1.416178989 | 8.625029106 | 0.451984043 | 0.667502714 | 0.93362086 | -5.038897084 |
| **A_55_P2127131** | Trmt112 | -0.031670348 | -0.204058214 | 0.140717518 | 11.35160956 | -0.451975926 | 0.667508233 | 0.93362086 | -5.038899732 |
| **A_51_P279571** | Syap1 | -0.073326677 | -0.472498259 | 0.325844906 | 11.89306731 | -0.451930395 | 0.667539194 | 0.93362086 | -5.038914584 |
| **A_51_P502348** | Tfip11 | 0.031580647 | -0.14034422 | 0.203505514 | 9.589872064 | 0.45190952 | 0.667553389 | 0.93362086 | -5.038921393 |
| **A_55_P2027327** | Usp1 | 0.059773465 | -0.265656748 | 0.385203677 | 11.41639178 | 0.451876466 | 0.667575866 | 0.93362086 | -5.038932174 |
| **A_55_P2348799** | 1700025G04Rik | -0.063468319 | -0.409122471 | 0.282185832 | 9.315846571 | -0.45173567 | 0.667671612 | 0.933635685 | -5.038978087 |
| **A_55_P1958464** | NA | 0.0806003 | -0.358372526 | 0.519573127 | 10.41652927 | 0.45171882 | 0.667683071 | 0.933635685 | -5.038983581 |
| **A_55_P2084492** | NA | 0.039079628 | -0.17384909 | 0.252008346 | 8.586675984 | 0.451528683 | 0.667812383 | 0.933656337 | -5.039045562 |
| **A_55_P2138177** | Psd3 | -0.028785259 | -0.185656714 | 0.128086195 | 8.581759096 | -0.451435165 | 0.667875989 | 0.933656337 | -5.039076038 |
| **A_52_P610808** | Hexdc | 0.021325664 | -0.094895567 | 0.137546895 | 7.885938848 | 0.451425667 | 0.667882449 | 0.933656337 | -5.039079133 |
| **A_55_P2013113** | Bak1 | 0.060738195 | -0.270283887 | 0.391760277 | 12.09969719 | 0.451413017 | 0.667891053 | 0.933656337 | -5.039083255 |
| **A_51_P214663** | Tpmt | -0.040106655 | -0.258753161 | 0.17853985 | 8.195791205 | -0.451276864 | 0.667983664 | 0.933705747 | -5.039127614 |
| **A_55_P2034481** | Erdr1 | -0.252241219 | -1.628053797 | 1.123571359 | 13.65310179 | -0.451051305 | 0.668137102 | 0.933705747 | -5.039201073 |
| **A_52_P382886** | Gjb2 | -0.06980447 | -0.45056546 | 0.31095652 | 16.0794355 | -0.45102431 | 0.668155467 | 0.933705747 | -5.039209863 |
| **A_55_P2240468** | 1110019D14Rik | -0.026155323 | -0.168827117 | 0.11651647 | 8.513645731 | -0.451015152 | 0.668161697 | 0.933705747 | -5.039212844 |
| **A_65_P16688** | Taok2 | -0.041365835 | -0.267012358 | 0.184280688 | 8.300631166 | -0.451005992 | 0.668167929 | 0.933705747 | -5.039215827 |
| **A_55_P1993168** | Ppargc1b | 0.031836539 | -0.141887567 | 0.205560645 | 8.186538026 | 0.450852961 | 0.668272042 | 0.93374175 | -5.039265642 |
| **A_55_P2060454** | Nsd3 | -0.068801963 | -0.444355238 | 0.306751313 | 11.6357089 | -0.450711289 | 0.668368434 | 0.93374175 | -5.039311746 |
| **A_51_P148196** | Sdf2 | -0.023998336 | -0.154999024 | 0.107002351 | 14.97580537 | -0.450688759 | 0.668383764 | 0.93374175 | -5.039319077 |
| **A_55_P1994997** | Arhgap11a | -0.021327098 | -0.137801708 | 0.095147513 | 7.768350413 | -0.450473925 | 0.66852995 | 0.93374175 | -5.039388961 |
| **A_51_P143031** | Eps8l3 | 0.198360894 | -0.885034658 | 1.281756445 | 10.15788484 | 0.450441267 | 0.668552174 | 0.93374175 | -5.039399582 |
| **A_55_P2069470** | 1110017D15Rik | -0.026364596 | -0.170369229 | 0.117640036 | 8.332952603 | -0.450416017 | 0.668569357 | 0.93374175 | -5.039407794 |
| **A_66_P121583** | Tmem267 | -0.036414009 | -0.235320489 | 0.16249247 | 8.63183467 | -0.450389994 | 0.668587066 | 0.93374175 | -5.039416256 |
| **A_66_P130759** | 4631405J19Rik | -0.02206583 | -0.142604865 | 0.098473206 | 7.675854203 | -0.450361972 | 0.668606136 | 0.93374175 | -5.039425368 |
| **A_55_P2140288** | Phka2 | 0.031199781 | -0.13926864 | 0.201668202 | 8.657168314 | 0.450273892 | 0.668666079 | 0.93374175 | -5.039454004 |
| **A_55_P2107203** | Gpr137b | 0.048292592 | -0.215606169 | 0.312191353 | 10.63181237 | 0.450207088 | 0.668711544 | 0.93374175 | -5.03947572 |
| **A_51_P327778** | Pfdn2 | 0.045541259 | -0.203362448 | 0.294444967 | 9.874560903 | 0.450135052 | 0.668760572 | 0.93374175 | -5.039499134 |
| **A_52_P104761** | Rlim | -0.040393318 | -0.261183103 | 0.180396466 | 9.825267548 | -0.450090367 | 0.668790985 | 0.93374175 | -5.039513656 |
| **A_52_P507310** | Mrps24 | 0.030550919 | -0.136457077 | 0.197558916 | 10.88331528 | 0.45004525 | 0.668821694 | 0.93374175 | -5.039528318 |
| **A_55_P2038207** | Zfp687 | -0.036730278 | -0.237604603 | 0.164144048 | 9.941150157 | -0.449851261 | 0.668953738 | 0.933858646 | -5.039591341 |
| **A_55_P1954481** | Dock9 | -0.046808484 | -0.302890579 | 0.20927361 | 8.648964857 | -0.449691346 | 0.669062599 | 0.933942478 | -5.039643276 |
| **A_55_P2071952** | Wdr92 | -0.100137028 | -0.64805707 | 0.447783013 | 11.55247288 | -0.449621092 | 0.669110426 | 0.933942478 | -5.039666086 |

| | | | | | | | | |
|---|---|---|---|---|---|---|---|---|
| **A_55_P2081865** | Slc2a8 | -0.056145608 | -0.36349512 | 0.251203903 | 11.28245927 | -0.449420031 | 0.669247314 | 0.934028468 | -5.039731351 |
| **A_51_P410949** | Polr3g | 0.093310729 | -0.417521916 | 0.604143374 | 9.990204231 | 0.449388654 | 0.669268678 | 0.934028468 | -5.039741533 |
| **A_55_P2023783** | Leng8 | -0.078585046 | -0.508900143 | 0.351730052 | 9.26152945 | -0.449285589 | 0.669338854 | 0.934044764 | -5.039774976 |
| **A_55_P2036526** | Mbd2 | 0.056217347 | -0.251679714 | 0.364114408 | 9.974592461 | 0.449194018 | 0.669401207 | 0.934044764 | -5.039804683 |
| **A_66_P130241** | Slc25a1 | 0.036464816 | -0.163272516 | 0.236202148 | 9.665299454 | 0.449142286 | 0.669436434 | 0.934044764 | -5.039821463 |
| **A_55_P2081925** | Zfp53 | -0.072669942 | -0.470891235 | 0.32555135 | 9.25021658 | -0.448951577 | 0.669566305 | 0.934044764 | -5.039883308 |
| **A_55_P2042121** | Slc25a22 | 0.045921929 | -0.205724015 | 0.297567872 | 9.651220499 | 0.448951417 | 0.669566414 | 0.934044764 | -5.03988336 |
| **A_55_P2181976** | Akap11 | -0.034530595 | -0.223801916 | 0.154740726 | 8.739738513 | -0.448836751 | 0.669644507 | 0.934044764 | -5.039920533 |
| **A_55_P1999532** | Col9a1 | -0.020153547 | -0.130637539 | 0.090330445 | 7.669739505 | -0.448767396 | 0.669691743 | 0.934044764 | -5.039943013 |
| **A_51_P511482** | Ift57 | -0.036478726 | -0.236461994 | 0.163504542 | 9.635963921 | -0.448761058 | 0.66969606 | 0.934044764 | -5.039945067 |
| **A_55_P2116802** | NA | -0.044648643 | -0.2894722 | 0.200174915 | 10.79690392 | -0.448667096 | 0.669760058 | 0.934044764 | -5.039975518 |
| **A_52_P113830** | 1600025M17Rik | 0.037500222 | -0.168128513 | 0.243128958 | 14.05401192 | 0.448661911 | 0.66976359 | 0.934044764 | -5.039977198 |
| **A_55_P1961094** | Tmem230 | 0.034974114 | -0.156933141 | 0.226881368 | 11.21388705 | 0.448357553 | 0.669970911 | 0.934266485 | -5.04007579 |
| **A_51_P423743** | Cldn6 | 0.063523108 | -0.285100968 | 0.412147184 | 8.188966355 | 0.448273974 | 0.670027849 | 0.934273195 | -5.040102854 |
| **A_55_P1973886** | Socs4 | 0.02529836 | -0.113585447 | 0.164182167 | 8.54704461 | 0.448136088 | 0.670121788 | 0.934273195 | -5.040147492 |
| **A_55_P2121275** | Gm4907 | 0.028992682 | -0.130215545 | 0.188200909 | 10.742817 | 0.448014478 | 0.670204644 | 0.934273195 | -5.040186851 |
| **A_51_P292116** | Ica1 | 0.028012198 | -0.125838711 | 0.181863107 | 9.975396445 | 0.44793633 | 0.670257892 | 0.934273195 | -5.040212139 |
| **A_55_P2040628** | Gid8 | -0.034173788 | -0.221963155 | 0.15361558 | 11.90835488 | -0.447704314 | 0.670415991 | 0.934273195 | -5.040287192 |
| **A_51_P496934** | Hps3 | 0.020526688 | -0.092274067 | 0.133327443 | 8.53357644 | 0.447688599 | 0.6704267 | 0.934273195 | -5.040292275 |
| **A_52_P131490** | Atg4b | -0.032543972 | -0.211386951 | 0.146299007 | 9.561604304 | -0.447680138 | 0.670432466 | 0.934273195 | -5.040295011 |
| **A_55_P2093770** | NA | 0.069160215 | -0.310965845 | 0.449286276 | 11.4582482 | 0.447608014 | 0.670481617 | 0.934273195 | -5.040318333 |
| **A_55_P2024391** | NA | 0.025507203 | -0.114691878 | 0.165706284 | 7.765556098 | 0.447596654 | 0.670489358 | 0.934273195 | -5.040322006 |
| **A_55_P2244677** | 6720422M22Rik | -0.080963649 | -0.526033929 | 0.364106631 | 8.867065439 | -0.447538746 | 0.670528823 | 0.934273195 | -5.040340729 |
| **A_65_P16542** | NA | -0.045375494 | -0.294922389 | 0.204171401 | 12.38981355 | -0.447340643 | 0.67066384 | 0.934273195 | -5.040404762 |
| **A_55_P2012630** | NA | -0.037716694 | -0.245154948 | 0.169721559 | 14.55660647 | -0.447315491 | 0.670680983 | 0.934273195 | -5.04041289 |
| **A_55_P2035804** | Fzd8 | -0.020920908 | -0.135985039 | 0.094143224 | 7.883726074 | -0.447311145 | 0.670683945 | 0.934273195 | -5.040414295 |
| **A_55_P2123381** | Fga | -0.166238538 | -1.080594628 | 0.748117553 | 9.496492591 | -0.447286213 | 0.670700939 | 0.934273195 | -5.040422352 |
| **A_55_P1953489** | Lrrc8b | -0.054321116 | -0.353103657 | 0.244461425 | 9.730515333 | -0.447283245 | 0.670702962 | 0.934273195 | -5.040423311 |
| **A_52_P76034** | Rcc2 | -0.055330186 | -0.359727122 | 0.24906675 | 11.55945308 | -0.447188902 | 0.670767267 | 0.934273195 | -5.040453793 |
| **A_51_P309854** | Kcnn2 | 0.020111511 | -0.090574808 | 0.13079783 | 7.740657002 | 0.447012765 | 0.670887334 | 0.934273195 | -5.030526395 |
| **A_52_P229044** | Slc20a2 | -0.050223684 | -0.326683094 | 0.226235726 | 8.554508182 | -0.446937044 | 0.670938953 | 0.934273195 | -5.040535139 |
| **A_55_P2024461** | Tbkbp1 | 0.037249107 | -0.167860434 | 0.242358648 | 11.20100028 | 0.446785598 | 0.671042201 | 0.934273195 | -5.040584034 |
| **A_55_P2176792** | Sh3gl3 | -0.118793438 | -0.77297825 | 0.535391374 | 8.736092648 | -0.446746499 | 0.671068858 | 0.934273195 | -5.040596654 |
| **A_55_P2112185** | Nhsl1 | 0.026010671 | -0.117238013 | 0.169259356 | 9.657592387 | 0.446714534 | 0.671090651 | 0.934273195 | -5.040606972 |
| **A_52_P372032** | Ddx21 | 0.020718986 | -0.093406939 | 0.134844911 | 9.212632213 | 0.446635618 | 0.671144457 | 0.934273195 | -5.04063244 |
| **A_55_P2092776** | Apoo | -0.043333554 | -0.28204347 | 0.195376362 | 9.970225409 | -0.446604411 | 0.671165735 | 0.934273195 | -5.040642511 |
| **A_55_P2105843** | NA | -0.173367085 | -1.128768298 | 0.782034128 | 11.63765659 | -0.446426556 | 0.671287008 | 0.934273195 | -5.040699891 |
| **A_51_P429682** | Ebp | 0.029814517 | -0.134492641 | 0.194121675 | 12.42607002 | 0.446416721 | 0.671293715 | 0.934273195 | -5.040703063 |
| **A_55_P1972222** | NA | -0.037332415 | -0.243076754 | 0.168411924 | 13.50280993 | -0.446403255 | 0.671302897 | 0.934273195 | -5.040707407 |
| **A_51_P308681** | Mcur1 | 0.024579341 | -0.110885055 | 0.160043737 | 9.24361952 | 0.446389764 | 0.671312097 | 0.934273195 | -5.040711758 |
| **A_51_P414448** | Naa15 | 0.030004755 | -0.135387909 | 0.19539742 | 9.738984773 | 0.446316569 | 0.67136201 | 0.934273195 | -5.040735365 |
| **A_55_P2092864** | Emp2 | -0.030975274 | -0.201733291 | 0.139782743 | 9.183247592 | -0.446275674 | 0.671389898 | 0.934273195 | -5.040748553 |
| **A_52_P274028** | Ssr1 | 0.072683976 | -0.328097166 | 0.473465118 | 9.087917322 | 0.446170201 | 0.671461827 | 0.934273195 | -5.040782561 |
| **A_51_P141610** | Cables2 | 0.027006841 | -0.121932135 | 0.175945817 | 9.310163706 | 0.446102423 | 0.671508051 | 0.934273195 | -5.040804411 |
| **A_55_P1997275** | Xpnpep1 | -0.041475697 | -0.270233664 | 0.187282269 | 13.75723018 | -0.446053172 | 0.671541642 | 0.934273195 | -5.040820286 |
| **A_55_P2052185** | Lamtor1 | 0.030333285 | -0.137007037 | 0.197673607 | 13.37525446 | 0.44595189 | 0.67161072 | 0.934273195 | -5.040852928 |
| **A_52_P460537** | Lpar2 | -0.026792293 | -0.174602024 | 0.121017439 | 8.496179726 | -0.445939569 | 0.671619124 | 0.934273195 | -5.040856898 |
| **A_51_P519276** | Ndufb7 | 0.037731517 | -0.170584654 | 0.246047688 | 13.82306423 | 0.445605402 | 0.671847067 | 0.934366466 | -5.040964546 |
| **A_52_P59264** | Sncaip | 0.022913076 | -0.103659724 | 0.149485876 | 8.000592367 | 0.445360989 | 0.67201381 | 0.934366466 | -5.041043233 |
| **A_55_P1991770** | Pdlim4 | -0.053603674 | -0.349872422 | 0.242665075 | 12.59814787 | -0.445120785 | 0.672177703 | 0.934366466 | -5.041120528 |

| | | | | | | | | | |
|---|---|---|---|---|---|---|---|---|---|
| **A_52_P203948** | Pla2g4e | -0.074074211 | -0.483486094 | 0.335337673 | 8.541944871 | -0.445118609 | 0.672179187 | 0.934366466 | -5.041121228 |
| **A_51_P506748** | Arhgap35 | 0.036367148 | -0.164636131 | 0.237370426 | 11.40364622 | 0.445118106 | 0.67217953 | 0.934366466 | -5.041121389 |
| **A_55_P2085915** | Tnfrsf14 | -0.020620782 | -0.134607914 | 0.09336635 | 7.673223041 | -0.445059909 | 0.672219241 | 0.934366466 | -5.041140111 |
| **A_55_P2013273** | NA | -0.047597872 | -0.310809436 | 0.215613693 | 11.49541554 | -0.444889063 | 0.672335826 | 0.934366466 | -5.041195057 |
| **A_55_P2043662** | Zfp607a | 0.021690752 | -0.098283825 | 0.141665329 | 7.953168496 | 0.444789514 | 0.672403763 | 0.934366466 | -5.041227064 |
| **A_52_P19016** | Clasrp | -0.054202516 | -0.354082302 | 0.245677269 | 11.52490505 | -0.444673675 | 0.67248282 | 0.934366466 | -5.0412643 |
| **A_55_P2091359** | Padi2 | -0.415719599 | -2.715861003 | 1.884421805 | 11.84827061 | -0.444646755 | 0.672501194 | 0.934366466 | -5.041272952 |
| **A_55_P2402134** | NA | -0.109919742 | -0.718214774 | 0.498375291 | 9.143200982 | -0.444560214 | 0.67256026 | 0.934366466 | -5.041300764 |
| **A_51_P435068** | Acadsb | -0.048068157 | -0.314154365 | 0.21801805 | 9.438143315 | -0.444430928 | 0.672648505 | 0.934366466 | -5.041342303 |
| **A_55_P2173982** | Rrm2 | -0.262944519 | -1.718655404 | 1.192766367 | 12.07380046 | -0.444383736 | 0.672680718 | 0.934366466 | -5.041357463 |
| **A_52_P655265** | Bora | 0.021929014 | -0.099533096 | 0.143391123 | 7.819240746 | 0.444168179 | 0.672827865 | 0.934366466 | -5.041426689 |
| **A_51_P408703** | Rnf138rt1 | 0.207313485 | -0.941062268 | 1.355689239 | 9.115615479 | 0.444132651 | 0.67285212 | 0.934366466 | -5.041438096 |
| **A_51_P130905** | Gcm1 | 0.093357289 | -0.423888852 | 0.610603431 | 8.842558804 | 0.444037999 | 0.672916739 | 0.934366466 | -5.041468481 |
| **A_51_P176042** | Pklr | 0.026075774 | -0.118399881 | 0.170551428 | 7.802973023 | 0.444029371 | 0.672922629 | 0.934366466 | -5.041471251 |
| **A_55_P2044242** | Slc13a5 | -0.278584778 | -1.822593404 | 1.265423848 | 9.071061459 | -0.443891495 | 0.673016763 | 0.934366466 | -5.041515501 |
| **A_55_P2058783** | Hilpda | -0.100985863 | -0.660706269 | 0.458734543 | 12.29027131 | -0.443872872 | 0.673029478 | 0.934366466 | -5.041521477 |
| **A_52_P121051** | 2610001J05Rik | -0.029599762 | -0.193757694 | 0.134558169 | 10.36019507 | -0.443604065 | 0.673213025 | 0.934366466 | -5.041607709 |
| **A_55_P1991851** | Speg | -0.033872001 | -0.221748444 | 0.154004442 | 8.586260624 | -0.443545 | 0.67325336 | 0.934366466 | -5.041626651 |
| **A_66_P128434** | Kit | -0.06959695 | -0.455676656 | 0.316482755 | 12.5582443 | -0.443488538 | 0.673291918 | 0.934366466 | -5.041644756 |
| **A_51_P488991** | Oaf | -0.069999729 | -0.458338887 | 0.318339429 | 11.90757405 | -0.443459884 | 0.673311485 | 0.934366466 | -5.041653942 |
| **A_55_P2050192** | NA | -0.031440773 | -0.205908535 | 0.143026988 | 12.31335375 | -0.44335049 | 0.673386195 | 0.934366466 | -5.041689012 |
| **A_51_P374818** | Shd | 0.021394367 | -0.097382835 | 0.14017157 | 7.707362239 | 0.443134457 | 0.673533743 | 0.934366466 | -5.041758244 |
| **A_52_P569038** | Cnot2 | -0.04301815 | -0.28193344 | 0.19589714 | 10.93219674 | -0.442972686 | 0.673644242 | 0.934366466 | -5.041810066 |
| **A_55_P1974442** | Sumf2 | 0.049827698 | -0.226916738 | 0.326572134 | 9.217566855 | 0.442956511 | 0.67365529 | 0.934366466 | -5.041815247 |
| **A_55_P1959989** | D430042O09Rik | -0.041529616 | -0.272225396 | 0.189166164 | 10.37433907 | -0.442881388 | 0.673706608 | 0.934366466 | -5.041839306 |
| **A_52_P216539** | Fbxw8 | -0.03896138 | -0.255396531 | 0.17747377 | 9.175145937 | -0.44286943 | 0.673714776 | 0.934366466 | -5.041843135 |
| **A_51_P217535** | Actr3 | -0.024499092 | -0.160600837 | 0.111602653 | 13.81341515 | -0.442848791 | 0.673728875 | 0.934366466 | -5.041849743 |
| **A_51_P122649** | Degs2 | 0.224325228 | -1.021890647 | 1.470541102 | 10.95620358 | 0.4428473 | 0.673729893 | 0.934366466 | -5.041850221 |
| **A_55_P1972772** | Gsg1 | -0.026526387 | -0.173905737 | 0.120852962 | 8.071594401 | -0.44280308 | 0.673760102 | 0.934366466 | -5.04186438 |
| **A_55_P1958246** | Tcof1 | -0.046447527 | -0.30451396 | 0.211618907 | 10.05950195 | -0.442792481 | 0.673767342 | 0.934366466 | -5.041867774 |
| **A_51_P469053** | Mfsd4a | -0.052284406 | -0.34284322 | 0.238274408 | 9.020118417 | -0.442697722 | 0.673832078 | 0.934366466 | -5.04189811 |
| **A_51_P369252** | Ddias | 0.027474921 | -0.125264509 | 0.180214352 | 7.924949128 | 0.442541955 | 0.673938498 | 0.934366466 | -5.041947964 |
| **A_55_P2090219** | Mrpl48 | 0.056829257 | -0.272549754 | 0.386208268 | 11.64716778 | 0.4471221 | 0.673972147 | 0.934366466 | -4.975675446 |
| **A_51_P432199** | Sap30 | -0.033077859 | -0.216995338 | 0.15083962 | 8.767356828 | -0.442469714 | 0.673987857 | 0.934366466 | -5.04197108 |
| **A_55_P1969904** | Abcc6 | 0.047991858 | -0.21888392 | 0.314867636 | 7.91269026 | 0.442412683 | 0.674026824 | 0.934366466 | -5.041989327 |
| **A_55_P2001583** | Tspan18 | -0.026177221 | -0.171774818 | 0.119420375 | 7.859137736 | -0.442321962 | 0.674088813 | 0.934366466 | -5.042018348 |
| **A_55_P2021114** | Ier5 | -0.035075388 | -0.230186277 | 0.1600355 | 8.304378877 | -0.442272714 | 0.674122465 | 0.934366466 | -5.0420341 |
| **A_55_P1980956** | Osbp | 0.021228078 | -0.096863867 | 0.139320024 | 8.843560487 | 0.442241575 | 0.674143743 | 0.934366466 | -5.042044059 |
| **A_55_P1970154** | Rnf10 | -0.044078353 | -0.28939029 | 0.201233584 | 11.16825816 | -0.442054525 | 0.674271566 | 0.934366466 | -5.042103867 |
| **A_51_P101545** | Hgfac | 0.051075531 | -0.233194786 | 0.335345847 | 8.064684451 | 0.442028772 | 0.674289166 | 0.934366466 | -5.0421121 |
| **A_55_P2084308** | Nid1 | -0.047717759 | -0.313327562 | 0.217892044 | 15.27572906 | -0.44198253 | 0.674320769 | 0.934366466 | -5.042126881 |
| **A_55_P1999811** | NA | 0.022616562 | -0.103289433 | 0.148522557 | 8.973883945 | 0.441925785 | 0.67435955 | 0.934366466 | -5.042145018 |
| **A_51_P196207** | Capsl | -0.022596428 | -0.148432564 | 0.103239708 | 7.727000396 | -0.441777494 | 0.674460902 | 0.934366466 | -5.042192405 |
| **A_55_P2195157** | Dbn1 | -0.042204927 | -0.277245264 | 0.192835411 | 8.632619515 | -0.441763582 | 0.674470411 | 0.934366466 | -5.04219685 |
| **A_55_P2157330** | NA | -0.042452348 | -0.278879837 | 0.193975142 | 15.32974103 | -0.441746283 | 0.674482235 | 0.934366466 | -5.042202377 |
| **A_55_P2058903** | Sgpp1 | -0.148160725 | -0.973310915 | 0.676989464 | 13.31067624 | -0.441742571 | 0.674484772 | 0.934366466 | -5.042203562 |
| **A_55_P2023114** | Hsd3b2 | 0.020444688 | -0.09341992 | 0.134309295 | 7.705408473 | 0.441734062 | 0.674490588 | 0.934366466 | -5.042206281 |
| **A_51_P427476** | Mxra8 | -0.125196376 | -0.822656538 | 0.572263786 | 9.393056199 | -0.44161266 | 0.67457357 | 0.934366466 | -5.042245061 |
| **A_55_P2035662** | Aebp1 | -0.122198729 | -0.803223765 | 0.558826306 | 13.60532774 | -0.44144111 | 0.674690837 | 0.934366466 | -5.042299843 |
| **A_55_P2161347** | Acmsd | 0.018962238 | -0.086718584 | 0.12464306 | 7.672897314 | 0.441430742 | 0.674697925 | 0.934366466 | -5.042303154 |

| | | | | | | | | |
|---|---|---|---|---|---|---|---|---|
| **A_55_P2000515** | Peli1 | -0.036874012 | -0.242388958 | 0.168640933 | 11.62313205 | -0.441414049 | 0.674709337 | 0.934366466 | -5.042308483 |
| **A_55_P1970695** | NA | 0.030254682 | -0.138379039 | 0.198888402 | 8.776009381 | 0.441384758 | 0.67472936 | 0.934366466 | -5.042317834 |
| **A_51_P208377** | Trappc5 | 0.050013409 | -0.228855841 | 0.32888266 | 11.37970003 | 0.441219799 | 0.674842136 | 0.934366466 | -5.042370488 |
| **A_52_P566681** | Gpm6a | 0.02537436 | -0.116133594 | 0.166882315 | 7.750726434 | 0.441147086 | 0.674891849 | 0.934366466 | -5.042393691 |
| **A_55_P2064451** | Pced1a | -0.026666245 | -0.175380973 | 0.122048483 | 9.895111006 | -0.441140613 | 0.674896275 | 0.934366466 | -5.042395757 |
| **A_55_P2430122** | Ilf2 | -0.021541614 | -0.141690841 | 0.098607612 | 9.759147217 | -0.441089206 | 0.674931423 | 0.934366466 | -5.042412159 |
| **A_51_P281380** | Tspan5 | 0.160424929 | -0.734565663 | 1.05541552 | 9.576040241 | 0.440983711 | 0.675003555 | 0.934366466 | -5.042445813 |
| **A_55_P2081303** | Tet2 | 0.057725159 | -0.264322947 | 0.379773265 | 9.504888453 | 0.440974576 | 0.675009801 | 0.934366466 | -5.042448727 |
| **A_55_P2066613** | Lcp1 | -0.035957894 | -0.236600717 | 0.164684928 | 9.905399486 | -0.440899684 | 0.675061011 | 0.934366466 | -5.042472614 |
| **A_55_P2090698** | NA | -0.039061097 | -0.257092155 | 0.178969962 | 13.3555465 | -0.440752954 | 0.675161347 | 0.934366466 | -5.042519402 |
| **A_55_P1968718** | Plekha6 | 0.072584472 | -0.33256763 | 0.477736573 | 9.929497832 | 0.440752503 | 0.675161655 | 0.934366466 | -5.042519546 |
| **A_52_P121960** | NA | -0.044983237 | -0.296071503 | 0.206105029 | 14.2798812 | -0.440751143 | 0.675162585 | 0.934366466 | -5.04251998 |
| **A_55_P1978576** | Dtx2 | 0.032011999 | -0.146751324 | 0.210775322 | 11.43443054 | 0.440558454 | 0.675294361 | 0.934366466 | -5.042581402 |
| **A_55_P2026049** | NA | 0.030457461 | -0.139636764 | 0.200551686 | 8.471137152 | 0.440527771 | 0.675315345 | 0.934366466 | -5.04259118 |
| **A_51_P192847** | Ghdc | 0.031307658 | -0.143595384 | 0.206210699 | 8.92820102 | 0.440374717 | 0.675420026 | 0.934366466 | -5.042639947 |
| **A_51_P405476** | Fcer1g | -0.152975967 | -1.00774218 | 0.701790246 | 10.94176318 | -0.440296287 | 0.675473671 | 0.934366466 | -5.042664931 |
| **A_55_P2049383** | Erp29 | -0.047237512 | -0.31118972 | 0.216714696 | 15.57917634 | -0.440281947 | 0.675483479 | 0.934366466 | -5.042669499 |
| **A_55_P1989981** | Fam129b | 0.040817352 | -0.187288042 | 0.268922747 | 15.5830303 | 0.440228787 | 0.675519842 | 0.934366466 | -5.04268643 |
| **A_55_P2103206** | NA | 0.036668588 | -0.168282943 | 0.241620118 | 8.444538965 | 0.440161629 | 0.67556578 | 0.934366466 | -5.042707817 |
| **A_52_P88773** | Tor2a | 0.062901384 | -0.288696632 | 0.414499401 | 12.2287601 | 0.440132006 | 0.675586043 | 0.934366466 | -5.04271725 |
| **A_51_P434198** | Anapc4 | 0.025947162 | -0.119115369 | 0.171009692 | 13.37313941 | 0.440051772 | 0.675640929 | 0.934366466 | -5.042742796 |
| **A_52_P566348** | Nras | -0.038129188 | -0.251302056 | 0.17504368 | 10.3960774 | -0.440042673 | 0.675647153 | 0.934366466 | -5.042745692 |
| **A_52_P206613** | Adss | -0.035634994 | -0.234886666 | 0.163616678 | 10.34598874 | -0.439991084 | 0.675682445 | 0.934366466 | -5.042762115 |
| **A_55_P2004777** | Micall2 | 0.02503069 | -0.114941133 | 0.165002513 | 8.295271895 | 0.439948019 | 0.675711907 | 0.934366466 | -5.042775823 |
| **A_55_P2263098** | A130078K24Rik | -0.123713042 | -0.815581266 | 0.568155182 | 8.930002667 | -0.439907397 | 0.675739697 | 0.934366466 | -5.042788752 |
| **A_51_P239984** | Exo1 | 0.049912113 | -0.229227466 | 0.329051691 | 8.270563566 | 0.439899733 | 0.67574494 | 0.934366466 | -5.042791192 |
| **A_55_P1996821** | Myo15b | 0.121957606 | -0.560137773 | 0.804052984 | 8.795006208 | 0.439878714 | 0.67575932 | 0.934366466 | -5.042797881 |
| **A_55_P2184478** | NA | 0.03603794 | -0.165565933 | 0.237641812 | 15.26322201 | 0.439774708 | 0.675830476 | 0.934366466 | -5.042830977 |
| **A_55_P2065929** | NA | 0.021847635 | -0.100397711 | 0.144092981 | 8.02444907 | 0.439684611 | 0.67589212 | 0.934366466 | -5.042859642 |
| **A_55_P2171086** | NA | -0.030223978 | -0.199347911 | 0.138899954 | 8.36864698 | -0.439658752 | 0.675909814 | 0.934366466 | -5.042867868 |
| **A_52_P441036** | Col23a1 | -0.024186634 | -0.159563515 | 0.111190247 | 7.78755001 | -0.439541715 | 0.675989894 | 0.934366466 | -5.042905093 |
| **A_55_P2060097** | Prdm5 | 0.027370803 | -0.125832886 | 0.180574492 | 8.150867469 | 0.439528949 | 0.67599863 | 0.934366466 | -5.042909153 |
| **A_52_P429774** | Cnst | -0.02763225 | -0.182299753 | 0.127035252 | 9.4244083 | -0.439527793 | 0.675999421 | 0.934366466 | -5.042909521 |
| **A_55_P2141955** | NA | 0.030414962 | -0.139926256 | 0.20075618 | 15.6266083 | 0.439275204 | 0.676172268 | 0.934366466 | -5.042989828 |
| **A_51_P245631** | Rftn2 | 0.026732957 | -0.123017358 | 0.176483272 | 8.019045367 | 0.439185936 | 0.67623336 | 0.934366466 | -5.043018199 |
| **A_55_P2086692** | Nfib | -0.050475977 | -0.333294422 | 0.232342468 | 8.937819114 | -0.439082537 | 0.676304125 | 0.934366466 | -5.043051055 |
| **A_55_P2131128** | Helq | 0.027465651 | -0.126428259 | 0.181359562 | 8.33891351 | 0.43907391 | 0.67631003 | 0.934366466 | -5.043053796 |
| **A_55_P1954086** | Postn | 0.110280976 | -0.507647212 | 0.728209164 | 10.20306595 | 0.439067972 | 0.676314094 | 0.934366466 | -5.043055682 |
| **A_55_P2117445** | NA | -0.031798738 | -0.209983555 | 0.146386079 | 12.25607156 | -0.43904431 | 0.676330289 | 0.934366466 | -5.0430632 |
| **A_55_P1989312** | Prkd3 | 0.030719867 | -0.141444393 | 0.202884126 | 11.45695661 | 0.438980766 | 0.676373781 | 0.934366466 | -5.043083387 |
| **A_51_P375078** | Lmbrd2 | 0.030312005 | -0.139572262 | 0.200196271 | 8.974185815 | 0.438965776 | 0.67638404 | 0.934366466 | -5.043088148 |
| **A_55_P2060101** | Tprgl | -0.031425898 | -0.207570515 | 0.144718719 | 13.85072829 | -0.438922133 | 0.676413912 | 0.934366466 | -5.04310201 |
| **A_66_P111660** | Mt1 | -0.104781104 | -0.692156589 | 0.482594381 | 17.01860634 | -0.438870427 | 0.676449304 | 0.934366466 | -5.043118432 |
| **A_51_P291388** | Pafah1b1 | -0.046690835 | -0.308468882 | 0.215087212 | 11.50899203 | -0.438800974 | 0.676496844 | 0.934366466 | -5.043140488 |
| **A_51_P157083** | Gas1 | -0.040326147 | -0.266439876 | 0.185787582 | 8.324885683 | -0.438761949 | 0.676523557 | 0.934366466 | -5.043152879 |
| **A_51_P389895** | Slc35f5 | 0.04227677 | -0.194775789 | 0.279329329 | 12.25297756 | 0.438759282 | 0.676525383 | 0.934366466 | -5.043153726 |
| **A_51_P279163** | Plcg2 | 0.091420916 | -0.421230184 | 0.604072015 | 10.13720967 | 0.438725458 | 0.676548536 | 0.934366466 | -5.043164465 |
| **A_55_P2021187** | Malat1 | -0.142745351 | -0.943311819 | 0.657821116 | 8.851754018 | -0.438665765 | 0.676589399 | 0.934366466 | -5.043183416 |
| **A_51_P334318** | Chp2 | 0.018678611 | -0.086136908 | 0.123494131 | 7.709880152 | 0.438417776 | 0.676759171 | 0.934366466 | -5.043262119 |
| **A_52_P270145** | Zfp329 | -0.05099792 | -0.337211418 | 0.235215578 | 9.10713505 | -0.438360603 | 0.676798315 | 0.934366466 | -5.043280258 |

| | | | | | | | | | |
|---|---|---|---|---|---|---|---|---|---|
| **A_51_P368823** | Grb7 | 0.132801061 | -0.612776279 | 0.8783784 | 9.509728901 | 0.438205685 | 0.676904385 | 0.934366466 | -5.043329397 |
| **A_52_P520466** | Kif18b | 0.019393775 | -0.089534071 | 0.12832162 | 7.674301547 | 0.438018639 | 0.677032462 | 0.934366466 | -5.043388706 |
| **A_51_P466829** | Pheta1 | 0.040042313 | -0.184994463 | 0.265079089 | 12.78513672 | 0.437758734 | 0.67721045 | 0.934366466 | -5.043471079 |
| **A_55_P1967332** | NA | 0.03614278 | -0.166997691 | 0.239283252 | 13.23426774 | 0.437717859 | 0.677238444 | 0.934366466 | -5.043484029 |
| **A_55_P2452384** | Mga | -0.026447864 | -0.175100869 | 0.122205141 | 8.78333729 | -0.437709604 | 0.677244098 | 0.934366466 | -5.043486644 |
| **A_55_P2049086** | Map3k7 | -0.031564682 | -0.208981754 | 0.14585239 | 11.03777467 | -0.43769862 | 0.677251621 | 0.934366466 | -5.043490125 |
| **A_55_P2146389** | NA | -0.021442572 | -0.141967014 | 0.099081869 | 8.08920118 | -0.43769433 | 0.677254559 | 0.934366466 | -5.043491484 |
| **A_55_P2168667** | Rbpms2 | -0.050278494 | -0.332910149 | 0.23235316 | 9.900259583 | -0.437653721 | 0.677282371 | 0.934366466 | -5.043504348 |
| **A_55_P2181738** | Lyz1 | -0.087899149 | -0.58202796 | 0.406229663 | 11.00297365 | -0.43763662 | 0.677294084 | 0.934366466 | -5.043509765 |
| **A_55_P2169059** | Strada | 0.035725298 | -0.16510867 | 0.236559266 | 11.27418434 | 0.437630774 | 0.677298088 | 0.934366466 | -5.043511617 |
| **A_55_P2047748** | NA | -0.076800886 | -0.50860149 | 0.354999718 | 11.24698485 | -0.437574645 | 0.677336532 | 0.934366466 | -5.043529396 |
| **A_55_P2118717** | Ninl | -0.020506822 | -0.135809951 | 0.094796306 | 7.906491796 | -0.437548743 | 0.677354273 | 0.934366466 | -5.043537599 |
| **A_55_P2282257** | NA | -0.029879782 | -0.197900644 | 0.13814108 | 9.684788257 | -0.437505363 | 0.677383986 | 0.934366466 | -5.043551338 |
| **A_52_P350616** | Mrm1 | 0.029795595 | -0.137803038 | 0.197394229 | 10.74359102 | 0.437371776 | 0.677475489 | 0.934366466 | -5.043593636 |
| **A_51_P467224** | Fbn1 | 0.091105452 | -0.421367134 | 0.603578038 | 9.484493958 | 0.437363858 | 0.677480913 | 0.934366466 | -5.043596143 |
| **A_55_P2080774** | Qdpr | -0.030963178 | -0.205134778 | 0.143208421 | 9.24818267 | -0.437358283 | 0.677484732 | 0.934366466 | -5.043597908 |
| **A_51_P327828** | Ppib | -0.04249037 | -0.28150561 | 0.19652487 | 14.92523381 | -0.437354987 | 0.67748699 | 0.934366466 | -5.043598951 |
| **A_51_P103865** | Ppp1r35 | 0.03149431 | -0.14575819 | 0.208746809 | 8.606975123 | 0.437128262 | 0.677642305 | 0.934470265 | -5.043670711 |
| **A_51_P519108** | Dzank1 | -0.021030909 | -0.139401131 | 0.097339312 | 8.287700627 | -0.437103975 | 0.677658943 | 0.934470265 | -5.043678396 |
| **A_55_P2179676** | Spp2 | 0.256760539 | -1.188789417 | 1.702310494 | 12.96769055 | 0.436982793 | 0.677741965 | 0.934475167 | -5.043716735 |
| **A_51_P432802** | Rbm4 | 0.037229678 | -0.172383747 | 0.246843104 | 11.17832623 | 0.436957654 | 0.677759189 | 0.934475167 | -5.043724687 |
| **A_52_P18068** | Txlng | -0.034123978 | -0.226356805 | 0.158108849 | 8.343347809 | -0.436718176 | 0.677923274 | 0.934603668 | -5.043800419 |
| **A_52_P270164** | E2f6 | 0.040598401 | -0.188210615 | 0.269407417 | 10.46833282 | 0.436520816 | 0.678058515 | 0.934603668 | -5.043862804 |
| **A_51_P296487** | Lss | 0.066864938 | -0.310045177 | 0.443775053 | 8.370277287 | 0.436445297 | 0.678110267 | 0.934603668 | -5.043886668 |
| **A_51_P490257** | Fbxo4 | 0.04604737 | -0.213518282 | 0.305613022 | 10.48321905 | 0.436442236 | 0.678112365 | 0.934603668 | -5.043887635 |
| **A_55_P2075237** | Ulk3 | 0.061412802 | -0.284861221 | 0.407686826 | 9.630803864 | 0.436323058 | 0.678194041 | 0.934603668 | -5.043925288 |
| **A_55_P1960857** | Sorbs2 | -0.091882214 | -0.61014836 | 0.426383933 | 8.163169924 | -0.436161946 | 0.678304464 | 0.934603668 | -5.043976174 |
| **A_51_P320539** | Otulin | 0.027331297 | -0.126921845 | 0.181584439 | 10.62739201 | 0.435908545 | 0.678478157 | 0.934603668 | -5.044056175 |
| **A_52_P37702** | NA | -0.020942168 | -0.139142452 | 0.097258116 | 7.935156107 | -0.435885355 | 0.678494053 | 0.934603668 | -5.044063494 |
| **A_55_P2324921** | NA | -0.033887583 | -0.225185989 | 0.157410823 | 7.872641598 | -0.435811214 | 0.678544878 | 0.934603668 | -5.044086892 |
| **A_55_P2117420** | NA | -0.033794738 | -0.232711318 | 0.165121842 | 14.17464367 | -0.440279218 | 0.678590796 | 0.934603668 | -4.977729257 |
| **A_55_P2177023** | Mafg | -0.05600641 | -0.37223745 | 0.26022463 | 9.958750628 | -0.435714892 | 0.67861091 | 0.934603668 | -5.044117284 |
| **A_51_P102789** | C1qc | -0.067655549 | -0.449683378 | 0.31437228 | 9.346188133 | -0.435689993 | 0.67862798 | 0.934603668 | -5.008012961 |
| **A_51_P468126** | Trim39 | -0.026854457 | -0.17849915 | 0.124790236 | 8.66920761 | -0.43567067 | 0.678641228 | 0.934603668 | -5.044131235 |
| **A_55_P2178378** | Gas5 | -0.027790648 | -0.184776098 | 0.129194801 | 8.140193546 | -0.435520339 | 0.678744293 | 0.934603668 | -5.044178652 |
| **A_55_P2030373** | Atp1a2 | -0.019541785 | -0.12993297 | 0.0908494 | 7.797085409 | -0.435510872 | 0.678750784 | 0.934603668 | -5.044181637 |
| **A_55_P2131348** | Loxl2 | 0.04873482 | -0.226582599 | 0.324052239 | 9.464547748 | 0.435486643 | 0.678767396 | 0.934603668 | -5.044189278 |
| **A_51_P125260** | Acaa2 | 0.071088878 | -0.330547798 | 0.472725553 | 11.47658739 | 0.435449159 | 0.678793097 | 0.934603668 | -5.044201098 |
| **A_55_P2090505** | NA | 0.051018485 | -0.23733124 | 0.339368209 | 10.70410578 | 0.435288483 | 0.678903268 | 0.934603668 | -5.044251754 |
| **A_55_P1978855** | NA | -0.059007363 | -0.392520347 | 0.27450562 | 15.17664769 | -0.435273901 | 0.678913267 | 0.934603668 | -5.04425635 |
| **A_55_P1953899** | Mccc1 | 0.035261949 | -0.16404551 | 0.234569407 | 9.814807839 | 0.435263171 | 0.678920625 | 0.934603668 | -5.044259732 |
| **A_55_P2073579** | Chchd5 | 0.03131964 | -0.145714213 | 0.208353492 | 9.817958006 | 0.435240801 | 0.678935964 | 0.934603668 | -5.044266783 |
| **A_55_P1957618** | Ak2 | -0.03869564 | -0.25744646 | 0.18005518 | 9.402922414 | -0.435192606 | 0.678969013 | 0.934603668 | -5.044281973 |
| **A_55_P2105873** | Ppp2r5c | -0.031688003 | -0.210855958 | 0.147479952 | 9.386016462 | -0.43511464 | 0.679022478 | 0.934603668 | -5.044306541 |
| **A_51_P478172** | Tcrb-J | 0.032081074 | -0.149336401 | 0.213498549 | 7.915244299 | 0.435049772 | 0.679066962 | 0.934603668 | -5.04432698 |
| **A_55_P2136353** | NA | -0.021636362 | -0.143989774 | 0.100717051 | 9.340129455 | -0.435048129 | 0.679068089 | 0.934603668 | -5.044327497 |
| **A_55_P2088296** | Fam160b2 | 0.068937693 | -0.320973896 | 0.458849283 | 14.64435677 | 0.434970463 | 0.679121352 | 0.934603668 | -5.044351964 |
| **A_55_P2081349** | Focad | 0.035526257 | -0.165434872 | 0.236487385 | 9.591475474 | 0.434917166 | 0.679157904 | 0.934603668 | -5.044368752 |
| **A_51_P462047** | Prpf38a | 0.040360049 | -0.188030954 | 0.268751052 | 10.43904539 | 0.434752269 | 0.679270999 | 0.934692756 | -5.04442068 |
| **A_52_P422494** | Cd300lf | -0.059552155 | -0.396671176 | 0.277566866 | 8.182123988 | -0.434593662 | 0.679379789 | 0.934728368 | -5.04447061 |

| | | | | | | | | | |
|---|---|---|---|---|---|---|---|---|---|
| **A_55_P2058290** | Cluh | -0.020711833 | -0.137976314 | 0.096552648 | 8.032530476 | -0.434531445 | 0.679422466 | 0.934728368 | -5.044490192 |
| **A_66_P118989** | Gm8350 | -0.058996993 | -0.393042954 | 0.275048969 | 8.705329448 | -0.434503033 | 0.679441956 | 0.934728368 | -5.044499133 |
| **A_55_P2037585** | Trf | -0.186882475 | -1.245473379 | 0.871708428 | 15.96813585 | -0.434319754 | 0.679567685 | 0.9348348 | -5.044556799 |
| **A_55_P1975420** | Dhx40 | 0.095028624 | -0.443697151 | 0.633754398 | 12.17980456 | 0.433966143 | 0.679810293 | 0.935099554 | -5.044667993 |
| **A_51_P303620** | Whrn | -0.079051426 | -0.527335726 | 0.369232874 | 8.549494657 | -0.433835755 | 0.679899761 | 0.935099554 | -5.044708973 |
| **A_52_P477369** | Csnk1d | -0.02573109 | -0.171652894 | 0.120190715 | 8.388049299 | -0.433817582 | 0.679912231 | 0.935099554 | -5.044714684 |
| **A_52_P680870** | Ctxn1 | -0.037982451 | -0.25341208 | 0.177447179 | 8.045700409 | -0.433757213 | 0.679953657 | 0.935099554 | -5.044733653 |
| **A_52_P514306** | Spata2 | 0.042652629 | -0.19940115 | 0.284706408 | 9.427257701 | 0.433513976 | 0.68012058 | 0.935255026 | -5.044810058 |
| **A_66_P128431** | Fam122b | -0.032504194 | -0.217016547 | 0.152008159 | 8.750536928 | -0.433394226 | 0.680202768 | 0.935255026 | -5.044847659 |
| **A_51_P185660** | Ccl9 | -0.091352578 | -0.609938329 | 0.427233173 | 8.910413981 | -0.433380525 | 0.680212171 | 0.935255026 | -5.04485196 |
| **A_55_P1987364** | Ap5z1 | 0.038006212 | -0.177780435 | 0.25379286 | 10.12105722 | 0.433310473 | 0.680260252 | 0.935255026 | -5.044873952 |
| **A_52_P891775** | Cdr2l | 0.087772874 | -0.410680734 | 0.586226482 | 10.15402735 | 0.433216249 | 0.680324925 | 0.935277417 | -5.044903525 |
| **A_51_P145520** | Sigirr | 0.093129312 | -0.435842143 | 0.622100767 | 9.814241516 | 0.43313504 | 0.680380668 | 0.935287528 | -5.04492901 |
| **A_55_P2081169** | NA | -0.040202527 | -0.26859908 | 0.188194026 | 15.6791002 | -0.433044942 | 0.680442516 | 0.935305329 | -5.044957279 |
| **A_55_P1973573** | Spn | 0.027818469 | -0.130247864 | 0.185884803 | 7.884065693 | 0.432975194 | 0.680490395 | 0.935305329 | -5.044979159 |
| **A_55_P2063251** | Dusp3 | 0.030214026 | -0.141536906 | 0.201964959 | 10.30352849 | 0.432791441 | 0.680616544 | 0.935389946 | -5.045036787 |
| **A_52_P920129** | Gnai1 | 0.030345716 | -0.142172497 | 0.202863928 | 8.566040237 | 0.432744538 | 0.680648745 | 0.935389946 | -5.045051493 |
| **A_51_P353173** | Fam98a | -0.068744226 | -0.459657401 | 0.32216895 | 10.70649585 | -0.432638415 | 0.680721607 | 0.935408809 | -5.045084762 |
| **A_52_P59206** | Cst6 | -0.050419595 | -0.33716836 | 0.236329171 | 9.452099593 | -0.432580519 | 0.680761359 | 0.935408809 | -5.045102909 |
| **A_55_P1959500** | Ces2e | 0.018522059 | -0.086833915 | 0.123878033 | 7.679007822 | 0.432513096 | 0.680807653 | 0.935408809 | -5.045124038 |
| **A_51_P465740** | Lysmd4 | 0.040827537 | -0.191463388 | 0.273118462 | 9.614644194 | 0.432404399 | 0.68088229 | 0.935444864 | -5.045158097 |
| **A_51_P473259** | Dpyd | 0.021118007 | -0.099155643 | 0.141391656 | 7.72517459 | 0.431968025 | 0.681181969 | 0.93572527 | -5.04529475 |
| **A_52_P449694** | NA | 0.060243422 | -0.282863132 | 0.403349976 | 9.372385901 | 0.431966218 | 0.68118321 | 0.93572527 | -5.045295316 |
| **A_55_P1999361** | Dip2a | -0.035240928 | -0.236005216 | 0.16552336 | 10.26094713 | -0.431847129 | 0.681265006 | 0.935771127 | -5.045332587 |
| **A_51_P520966** | Icosl | -0.047543178 | -0.318488312 | 0.223401956 | 8.982522272 | -0.431694004 | 0.681370185 | 0.935808886 | -5.045380497 |
| **A_55_P2021629** | Abcb7 | 0.018687003 | -0.087815749 | 0.125189754 | 7.832204005 | 0.43166614 | 0.681389325 | 0.935808886 | -5.045389213 |
| **A_55_P2102181** | NA | -0.18414492 | -1.234179928 | 0.865890088 | 14.3969479 | -0.431444688 | 0.681541454 | 0.935884349 | -5.045458469 |
| **A_55_P2135860** | Prss36 | 0.061099518 | -0.287312478 | 0.409511515 | 10.24597204 | 0.431433485 | 0.68154915 | 0.935884349 | -5.045461972 |
| **A_55_P2147846** | Npcd | 0.061797495 | -0.290642634 | 0.414237624 | 9.517126658 | 0.431374713 | 0.681589528 | 0.935884349 | -5.045480347 |
| **A_55_P2120843** | NA | -0.033056901 | -0.221684703 | 0.155570902 | 10.10862784 | -0.431147222 | 0.681745827 | 0.936032469 | -5.045551447 |
| **A_52_P251366** | Neil3 | 0.02036132 | -0.095842662 | 0.136565302 | 7.685387832 | 0.431076232 | 0.681794605 | 0.936032951 | -5.045573627 |
| **A_55_P2078123** | Rora | -0.04261577 | -0.285929973 | 0.200698434 | 9.179872834 | -0.430895579 | 0.681918741 | 0.936136885 | -5.045630056 |
| **A_65_P17492** | Med29 | -0.059494024 | -0.399381037 | 0.280392988 | 10.03472015 | -0.430633628 | 0.68209876 | 0.936317516 | -5.04571184 |
| **A_52_P645862** | Agtr1a | -0.057038638 | -0.382985049 | 0.268907773 | 9.95006103 | -0.430518787 | 0.682177689 | 0.936332435 | -5.04574768 |
| **A_55_P2095663** | Pgr | -0.178030043 | -1.195576674 | 0.839516587 | 8.621562199 | -0.430435557 | 0.682234895 | 0.936332435 | -5.04577365 |
| **A_55_P1986296** | Tagln2 | 0.020539962 | -0.096909994 | 0.137989918 | 16.77544095 | 0.430245112 | 0.682365801 | 0.936332435 | -5.045833055 |
| **A_55_P2081278** | Mrpl55 | -0.054257355 | -0.364552005 | 0.256037295 | 12.93163656 | -0.430183246 | 0.682408328 | 0.936332435 | -5.045852348 |
| **A_55_P1973362** | Kcnj12 | 0.037372101 | -0.176377003 | 0.251121206 | 9.098552502 | 0.43014252 | 0.682436324 | 0.936332435 | -5.045865046 |
| **A_51_P225224** | Htra1 | -0.263076891 | -1.768208953 | 1.242055171 | 14.61623441 | -0.430008715 | 0.68252831 | 0.936332435 | -5.045906761 |
| **A_51_P281806** | Cactin | -0.043922307 | -0.295273601 | 0.207428986 | 12.08743649 | -0.429905682 | 0.682599144 | 0.936332435 | -5.045938874 |
| **A_66_P106231** | Nccrp1 | -0.488848059 | -3.286867285 | 2.309171167 | 11.59997035 | -0.429825685 | 0.682654144 | 0.936332435 | -5.045963802 |
| **A_55_P2040996** | Klf7 | -0.041390503 | -0.278322614 | 0.195541607 | 11.2213046 | -0.429779746 | 0.68268573 | 0.936332435 | -5.045978115 |
| **A_51_P502150** | Slc9a3r1 | 0.029036055 | -0.137187852 | 0.195259962 | 10.6241102 | 0.429747431 | 0.682707948 | 0.936332435 | -5.045988183 |
| **A_51_P116601** | Cfap69 | -0.037814786 | -0.254318071 | 0.1786885 | 8.08732824 | -0.429700953 | 0.682739905 | 0.936332435 | -5.046002662 |
| **A_55_P2015217** | Alg11 | 0.040849743 | -0.193062225 | 0.274761711 | 9.374644386 | 0.429641325 | 0.682780905 | 0.936332435 | -5.046021235 |
| **A_66_P101646** | NA | -0.125024868 | -0.84102793 | 0.590978194 | 15.50693046 | -0.429586563 | 0.68281856 | 0.936332435 | -5.04603829 |
| **A_51_P189361** | Osgin1 | -0.028931428 | -0.194682717 | 0.13681986 | 8.917879867 | -0.429419859 | 0.682933194 | 0.936332435 | -5.046090198 |
| **A_51_P117794** | Bik | -0.027394481 | -0.184345337 | 0.129556375 | 7.956859077 | -0.429406437 | 0.682942424 | 0.936332435 | -5.046094376 |
| **A_55_P2184246** | Rps26 | -0.020516241 | -0.138083751 | 0.097051268 | 15.72645536 | -0.429318541 | 0.683002869 | 0.936332435 | -5.046121736 |
| **A_55_P2128929** | Cc2d2a | 0.027989339 | -0.13241547 | 0.188394148 | 8.292766807 | 0.429283716 | 0.683026819 | 0.936332435 | -5.046132575 |

| | | | | | | | | |
|---|---|---|---|---|---|---|---|---|
| **A_55_P1960621** | NA | 0.018662442 | -0.088306712 | 0.125631596 | 15.76527367 | 0.429219132 | 0.683071235 | 0.936332435 | -5.046152674 |
| **A_55_P2061565** | Zfp229 | -0.063846385 | -0.429820893 | 0.302128122 | 8.647233739 | -0.429194988 | 0.683087841 | 0.936332435 | -5.046160187 |
| **A_52_P251450** | Cldn2 | 0.098142779 | -0.464449373 | 0.66073493 | 8.953466544 | 0.429174435 | 0.683101976 | 0.936332435 | -5.046166582 |
| **A_55_P2000909** | Atp5l | 0.04100396 | -0.194123183 | 0.276131102 | 15.16087874 | 0.429034478 | 0.683198236 | 0.936332435 | -5.046210124 |
| **A_51_P365578** | Ppp2r1a | 0.034790107 | -0.164706143 | 0.234286358 | 9.227229105 | 0.429032499 | 0.683199597 | 0.936332435 | -5.04621074 |
| **A_55_P2096737** | Elmod3 | -0.044232263 | -0.297893283 | 0.209428758 | 10.4603819 | -0.428997323 | 0.683223791 | 0.936332435 | -5.046221681 |
| **A_55_P2120773** | Dctn4 | 0.024045521 | -0.113947553 | 0.162038596 | 11.94206354 | 0.428692681 | 0.683433346 | 0.936445381 | -5.046316405 |
| **A_55_P1979463** | Prdx5 | 0.069942494 | -0.33145295 | 0.471337939 | 13.39663676 | 0.428684557 | 0.683438934 | 0.936445381 | -5.046318931 |
| **A_55_P1984961** | NA | 0.086927769 | -0.412113139 | 0.585968676 | 8.703062643 | 0.428540179 | 0.683538258 | 0.936445381 | -5.0463638 |
| **A_55_P1978597** | Trit1 | 0.027380236 | -0.129806443 | 0.184566915 | 10.43302585 | 0.428539262 | 0.68353889 | 0.936445381 | -5.046364085 |
| **A_55_P2030204** | Supt6 | 0.040550035 | -0.192250436 | 0.273350506 | 13.08273076 | 0.428525373 | 0.683548445 | 0.936445381 | -5.046368401 |
| **A_66_P128918** | Hacl1 | -0.04074608 | -0.274760744 | 0.193268585 | 9.573286409 | -0.428362968 | 0.68366018 | 0.936472401 | -5.046418855 |
| **A_55_P2088725** | Usp14 | 0.052135312 | -0.247311522 | 0.351582146 | 11.72199697 | 0.428332879 | 0.683680882 | 0.936472401 | -5.046428201 |
| **A_55_P2044385** | Fgfbp3 | 0.119394162 | -0.566439716 | 0.805228041 | 9.049665874 | 0.428285451 | 0.683713515 | 0.936472401 | -5.046442931 |
| **A_55_P2124736** | Col14a1 | 0.145301102 | -0.689577231 | 0.980179436 | 9.680824947 | 0.428168666 | 0.683793872 | 0.936516102 | -5.046479196 |
| **A_66_P135165** | Nfib | 0.02164216 | -0.102795597 | 0.146079917 | 7.821492299 | 0.427875663 | 0.6839955 | 0.936521391 | -5.046570141 |
| **A_52_P268231** | NA | 0.040963716 | -0.19458577 | 0.276513202 | 9.34429036 | 0.427844888 | 0.684016679 | 0.936521391 | -5.046579689 |
| **A_55_P2012394** | Sfxn2 | 0.047517905 | -0.22572649 | 0.320762299 | 9.896536248 | 0.42783389 | 0.684024248 | 0.936521391 | -5.046583102 |
| **A_55_P2122618** | Asb11 | -0.104808054 | -0.707544434 | 0.497928326 | 9.333953554 | -0.427795699 | 0.684050532 | 0.936521391 | -5.04659495 |
| **A_51_P417469** | Rnaseh2c | -0.03373473 | -0.227782752 | 0.160313291 | 11.74497997 | -0.427697969 | 0.684117793 | 0.936521391 | -5.046625267 |
| **A_55_P2160032** | Atp5pb | -0.069533081 | -0.469519321 | 0.330453158 | 12.87934473 | -0.427676698 | 0.684132433 | 0.936521391 | -5.046631864 |
| **A_51_P351970** | Hells | -0.019364835 | -0.130779375 | 0.092049705 | 7.77427744 | -0.427603347 | 0.684182918 | 0.936521391 | -5.046654613 |
| **A_55_P1957424** | Sulf1 | -0.01842533 | -0.124435355 | 0.087584694 | 7.69259716 | -0.427599819 | 0.684185347 | 0.936521391 | -5.046655707 |
| **A_55_P1998697** | NA | -0.03331667 | -0.225085626 | 0.158452285 | 12.06366721 | -0.427417649 | 0.684310737 | 0.936626699 | -5.046712188 |
| **A_52_P154101** | Calca | 0.018612748 | -0.088543488 | 0.125768984 | 7.664493322 | 0.427328849 | 0.684371864 | 0.936644039 | -5.046739712 |
| **A_55_P2293013** | Ces2a | -0.021764233 | -0.147094526 | 0.10356606 | 7.714795535 | -0.42722482 | 0.684443477 | 0.936675727 | -5.04677195 |
| **A_55_P2041700** | Hoxa11os | -0.127831706 | -0.864183927 | 0.608520516 | 10.1485114 | -0.427092697 | 0.684534435 | 0.936733882 | -5.046812884 |
| **A_52_P287992** | Tbcel | -0.020187146 | -0.136516512 | 0.09614222 | 8.063488923 | -0.426928079 | 0.684647771 | 0.936822651 | -5.046863868 |
| **A_51_P112223** | Gsta4 | 0.025177837 | -0.119945424 | 0.170301098 | 7.817858114 | 0.426825686 | 0.684718271 | 0.936852797 | -5.041804024 |
| **A_51_P424810** | Ncapg2 | -0.026375042 | -0.178457665 | 0.125707581 | 8.233711312 | -0.426660784 | 0.684831818 | 0.936874152 | -5.046946615 |
| **A_52_P49321** | Adamts9 | 0.037330685 | -0.177931617 | 0.252592987 | 9.658268711 | 0.426645472 | 0.684842362 | 0.936874152 | -5.046951354 |
| **A_51_P517430** | Cd1d1 | 0.041350669 | -0.197172765 | 0.279874102 | 8.655624635 | 0.426501583 | 0.684941448 | 0.936874152 | -5.046995876 |
| **A_51_P331462** | Cgrrf1 | 0.050820413 | -0.242407166 | 0.344047993 | 9.653425878 | 0.426385612 | 0.685021314 | 0.936874152 | -5.047031751 |
| **A_55_P1962466** | NA | -0.020969534 | -0.141989481 | 0.100050413 | 16.49243899 | -0.426285907 | 0.685089981 | 0.936874152 | -5.047062586 |
| **A_55_P2126790** | 1500009L16Rik | -0.0398685 | -0.27008635 | 0.19034935 | 8.282841316 | -0.426049507 | 0.685252805 | 0.936874152 | -5.04713567 |
| **A_51_P489903** | Ogg1 | 0.035503602 | -0.169530627 | 0.240537831 | 10.00494844 | 0.426005505 | 0.685283114 | 0.936874152 | -5.047149269 |
| **A_55_P2133255** | Mad2l1 | 0.057484633 | -0.27450349 | 0.389472757 | 9.053433859 | 0.425988998 | 0.685294485 | 0.936874152 | -5.047154371 |
| **A_51_P386911** | Taco1 | -0.020776976 | -0.140788518 | 0.099234566 | 8.341453943 | -0.425920449 | 0.685341703 | 0.936874152 | -5.047175553 |
| **A_51_P412270** | Rapgef6 | -0.021237361 | -0.143916554 | 0.101441831 | 8.215902321 | -0.425891342 | 0.685361754 | 0.936874152 | -5.047184546 |
| **A_51_P426739** | Gpt | 0.026552342 | -0.126866011 | 0.179970694 | 8.14802236 | 0.425789232 | 0.685432094 | 0.936874152 | -5.047216092 |
| **A_51_P498890** | Degs1 | 0.047066443 | -0.224932274 | 0.319065159 | 13.88649432 | 0.425709835 | 0.685486791 | 0.936874152 | -5.047240616 |
| **A_55_P2096917** | Mreg | -0.102479288 | -0.694832776 | 0.4898742 | 9.829379002 | -0.425622239 | 0.685547139 | 0.936874152 | -5.047267667 |
| **A_52_P540434** | Ppp1cc | 0.055082875 | -0.263329099 | 0.373494849 | 9.855248079 | 0.425594856 | 0.685566004 | 0.936874152 | -5.047276122 |
| **A_55_P1961034** | Gm13272 | 0.025159868 | -0.120306555 | 0.170626291 | 8.334452946 | 0.425514891 | 0.685621097 | 0.936874152 | -5.047300811 |
| **A_51_P282394** | Mrps28 | 0.060027848 | -0.287134327 | 0.407190022 | 11.23722294 | 0.425392211 | 0.685705624 | 0.936874152 | -5.04733868 |
| **A_52_P497188** | Prrg1 | 0.023993541 | -0.114769797 | 0.162756878 | 8.012172571 | 0.425391465 | 0.685706139 | 0.936874152 | -5.04733891 |
| **A_55_P2000638** | Stag2 | -0.037649423 | -0.255403362 | 0.180104515 | 11.28156532 | -0.425364727 | 0.685724561 | 0.936874152 | -5.047347162 |
| **A_51_P175871** | Got2 | -0.022992422 | -0.155976143 | 0.109991299 | 11.88526594 | -0.425358777 | 0.685728662 | 0.936874152 | -5.047348998 |
| **A_52_P539632** | Plxna3 | 0.019960588 | -0.095492875 | 0.135414051 | 7.730333112 | 0.425339318 | 0.685742069 | 0.936874152 | -5.047355004 |
| **A_51_P109335** | Copa | 0.040428198 | -0.193449986 | 0.274306383 | 12.30312442 | 0.425269106 | 0.685790449 | 0.936874152 | -5.047376669 |

| | | | | | | | | |
|---|---|---|---|---|---|---|---|---|
| **A_51_P513224** | Cep164 | 0.044162969 | -0.21133876 | 0.299664698 | 9.226772854 | 0.425239483 | 0.685810861 | 0.936874152 | -5.04738581 |
| **A_55_P2085984** | Igf1 | -0.052190935 | -0.354214995 | 0.249833124 | 8.258763078 | -0.42513093 | 0.685885664 | 0.936874152 | -5.047419298 |
| **A_55_P2062133** | Etv3 | 0.030116431 | -0.144171373 | 0.204404234 | 8.419504043 | 0.425114253 | 0.685897156 | 0.936874152 | -5.047424442 |
| **A_55_P2035167** | Pcdh7 | 0.019547592 | -0.093633868 | 0.132729053 | 7.72068282 | 0.424900402 | 0.686044532 | 0.937009239 | -5.04749039 |
| **A_55_P2098300** | Chmp6 | 0.023059025 | -0.110645102 | 0.156763152 | 8.640332052 | 0.42429244 | 0.686463591 | 0.937025001 | -5.047677706 |
| **A_66_P140531** | Selenok | 0.040379442 | -0.193776398 | 0.274535282 | 12.13966711 | 0.424252568 | 0.686491079 | 0.937025001 | -5.047689983 |
| **A_55_P1992329** | Gmpr | 0.02465783 | -0.118353859 | 0.16766952 | 7.93934162 | 0.424182247 | 0.686539559 | 0.937025001 | -5.047711631 |
| **A_51_P306151** | Sgk3 | -0.032714973 | -0.222473911 | 0.157043964 | 8.496764474 | -0.424144164 | 0.686565815 | 0.937025001 | -5.047723354 |
| **A_51_P465281** | Lgals1 | 0.025242106 | -0.121174577 | 0.171658789 | 15.68927756 | 0.424135066 | 0.686572087 | 0.937025001 | -5.047726154 |
| **A_55_P2005549** | Tnn | -0.018198802 | -0.123777253 | 0.087379649 | 7.660080736 | -0.424069139 | 0.686617541 | 0.937025001 | -5.047746445 |
| **A_52_P177847** | Tril | 0.018272727 | -0.087744435 | 0.12428989 | 7.740110015 | 0.42402978 | 0.686644677 | 0.937025001 | -5.047758557 |
| **A_51_P172054** | Gas6 | 0.037268892 | -0.178992374 | 0.253530157 | 11.53322559 | 0.423971729 | 0.686684702 | 0.937025001 | -5.04777642 |
| **A_55_P2032818** | Trim2 | 0.024831344 | -0.119258766 | 0.168921453 | 9.006844646 | 0.423970081 | 0.686685838 | 0.937025001 | -5.047776927 |
| **A_51_P459944** | Tcf21 | -0.019911603 | -0.135470382 | 0.095647177 | 7.824386468 | -0.423908806 | 0.686728088 | 0.937025001 | -5.04779578 |
| **A_55_P2025463** | Brpf3 | -0.054214864 | -0.368924835 | 0.260495106 | 10.56589634 | -0.423815696 | 0.68679229 | 0.937025001 | -5.047824422 |
| **A_55_P2001553** | Igdcc4 | 0.045471138 | -0.218486556 | 0.309428831 | 8.63600139 | 0.42380947 | 0.686796583 | 0.937025001 | -5.047826337 |
| **A_55_P2033997** | Ddo | 0.018765131 | -0.090211188 | 0.127741451 | 7.731589733 | 0.42363188 | 0.686919045 | 0.937025001 | -5.04788095 |
| **A_55_P2105100** | Lrig1 | -0.081936255 | -0.55780057 | 0.393928061 | 10.14759353 | -0.423606029 | 0.686936873 | 0.937025001 | -5.047888898 |
| **A_55_P2183288** | Cryzl1 | 0.041507311 | -0.199567203 | 0.282581824 | 10.49855202 | 0.42358684 | 0.686950105 | 0.937025001 | -5.047894797 |
| **A_55_P2074924** | Ppp1r13l | 0.042297792 | -0.203396649 | 0.287992234 | 12.45140376 | 0.423537172 | 0.686984358 | 0.937025001 | -5.047910066 |
| **A_51_P304126** | Necap1 | 0.027699231 | -0.133226059 | 0.18862452 | 11.51453807 | 0.423460174 | 0.68703746 | 0.937025001 | -5.047933733 |
| **A_51_P315646** | Nfs1 | -0.02915454 | -0.198560232 | 0.140251153 | 10.32154115 | -0.423396584 | 0.687081316 | 0.937025001 | -5.047953276 |
| **A_55_P1968340** | Plekhh1 | -0.049353376 | -0.336154075 | 0.237447323 | 15.1961082 | -0.423356103 | 0.687109236 | 0.937025001 | -5.047965716 |
| **A_52_P42269** | Fgf10 | -0.018349576 | -0.124982966 | 0.088283814 | 7.655879487 | -0.42335236 | 0.687111817 | 0.937025001 | -5.027608736 |
| **A_55_P2181109** | Il10rb | 0.05288927 | -0.254532351 | 0.360310892 | 9.734681221 | 0.423255225 | 0.687178813 | 0.937025001 | -5.04799671 |
| **A_51_P353882** | Zfp580 | -0.025171887 | -0.171508251 | 0.121164478 | 8.639706846 | -0.423187346 | 0.687225632 | 0.937025001 | -5.048017562 |
| **A_51_P195374** | Baiap2l1 | 0.034525455 | -0.166224479 | 0.235275389 | 8.682856147 | 0.423109876 | 0.687279068 | 0.937025001 | -5.048041356 |
| **A_52_P420394** | Apeh | -0.036127728 | -0.246219113 | 0.173963657 | 11.11397569 | -0.42305961 | 0.68731374 | 0.937025001 | -5.048056792 |
| **A_55_P2010292** | Serpina1e | 0.226001223 | -1.088269532 | 1.540271978 | 10.58310125 | 0.42305332 | 0.68731808 | 0.937025001 | -5.048058724 |
| **A_55_P1955289** | Slc18b1 | -0.034871869 | -0.237694315 | 0.167950577 | 8.21054926 | -0.422988315 | 0.68736292 | 0.937025001 | -5.048078684 |
| **A_51_P455997** | Meg3 | -0.118220107 | -0.80588884 | 0.569448625 | 12.19517126 | -0.422942408 | 0.687394588 | 0.937025001 | -5.048092778 |
| **A_51_P202596** | Sh3tc2 | -0.058667613 | -0.400000926 | 0.2826657 | 10.43257517 | -0.422852501 | 0.687456611 | 0.937025001 | -5.048120378 |
| **A_51_P351413** | Phospho2 | -0.022459349 | -0.153132194 | 0.108213495 | 8.444361243 | -0.422844799 | 0.687461924 | 0.937025001 | -5.048122742 |
| **A_52_P37091** | Magi1 | -0.023302741 | -0.158933841 | 0.11232836 | 7.821616083 | -0.422685041 | 0.68757214 | 0.937059601 | -5.048171769 |
| **A_51_P258698** | Phf8 | -0.057877289 | -0.394770528 | 0.27901595 | 12.80719102 | -0.422654063 | 0.687593513 | 0.937059601 | -5.048181274 |
| **A_55_P1976898** | Pnp2 | -0.087455031 | -0.596637824 | 0.421727763 | 9.411256609 | -0.422552076 | 0.687663878 | 0.937059601 | -5.048212561 |
| **A_52_P82510** | Nmt1 | -0.025586811 | -0.174567829 | 0.123394206 | 11.64133245 | -0.422526932 | 0.687681227 | 0.937059601 | -5.048220273 |
| **A_55_P2170515** | NA | -0.034225134 | -0.233564754 | 0.165114485 | 9.157423966 | -0.422396879 | 0.687770963 | 0.937072753 | -5.048260158 |
| **A_55_P1975322** | NA | 0.034856841 | -0.168173824 | 0.237887506 | 10.59034711 | 0.422372421 | 0.68778784 | 0.937072753 | -5.048267658 |
| **A_52_P639402** | Kcnk3 | 0.021123986 | -0.10198247 | 0.144230443 | 7.826916686 | 0.422147488 | 0.687943057 | 0.937218166 | -5.04833661 |
| **A_55_P2274553** | NA | 0.018007617 | -0.08698432 | 0.122999554 | 7.673750523 | 0.421958238 | 0.688073664 | 0.937269732 | -5.048394598 |
| **A_55_P1963198** | NA | -0.01904006 | -0.130053179 | 0.091973059 | 7.691850557 | -0.421952117 | 0.688077888 | 0.937269732 | -5.048396473 |
| **A_55_P2149993** | NA | -0.132253696 | -0.903663861 | 0.639156469 | 10.0992459 | -0.421785478 | 0.688192901 | 0.93735502 | -5.048447513 |
| **A_55_P2123002** | Tet1 | 0.018111019 | -0.087543752 | 0.12376579 | 7.688605093 | 0.421718774 | 0.688238942 | 0.93735502 | -5.048467938 |
| **A_52_P554703** | Gprin3 | -0.025088522 | -0.17147181 | 0.121294766 | 7.917094243 | -0.421650621 | 0.688285985 | 0.93735502 | -5.048488804 |
| **A_55_P2052290** | Psat1 | 0.154732066 | -0.748387096 | 1.057851228 | 10.62060652 | 0.421506649 | 0.688385367 | 0.937367039 | -5.048532873 |
| **A_55_P1975026** | Gm9104 | 0.034316702 | -0.166008115 | 0.234641518 | 15.23622222 | 0.421444072 | 0.688428565 | 0.937367039 | -5.048552023 |
| **A_55_P1989921** | Eml2 | 0.081708382 | -0.395286079 | 0.558702843 | 12.71676769 | 0.421427078 | 0.688440296 | 0.937367039 | -5.048557223 |
| **A_55_P2042988** | Aoah | 0.030114497 | -0.145759996 | 0.205988989 | 7.81610938 | 0.42125194 | 0.688561206 | 0.93737017 | -5.048610803 |
| **A_51_P462385** | G6pc | 0.018558001 | -0.089829132 | 0.126945135 | 7.755066649 | 0.421233242 | 0.688574115 | 0.93737017 | -5.048616523 |

| | | | | | | | | | |
|---|---|---|---|---|---|---|---|---|---|
| **A_55_P2018017** | Tnfsf10 | -0.074963008 | -0.512829251 | 0.362903234 | 8.952761431 | -0.421186756 | 0.688606209 | 0.93737017 | -5.04863074 |
| **A_55_P2000783** | Axin2 | -0.074207397 | -0.507705313 | 0.359290518 | 8.435013781 | -0.42114277 | 0.688636578 | 0.93737017 | -5.048644192 |
| **A_52_P490272** | Nup98 | -0.026582477 | -0.181947205 | 0.128782251 | 9.285994566 | -0.420932229 | 0.688781949 | 0.937502028 | -5.048708561 |
| **A_55_P1978241** | Artn | 0.026875131 | -0.130307908 | 0.184058171 | 7.812439918 | 0.420643398 | 0.688981401 | 0.937676651 | -5.048796817 |
| **A_55_P2169621** | Prorsd1 | -0.03781207 | -0.259007894 | 0.183383754 | 15.92213625 | -0.420554896 | 0.689042521 | 0.937676651 | -5.048823849 |
| **A_55_P1982718** | Rtf2 | -0.041691595 | -0.28559328 | 0.202210091 | 11.1094798 | -0.420535699 | 0.689055779 | 0.937676651 | -5.048829711 |
| **A_55_P1996344** | Dcst2 | -0.029462371 | -0.201885542 | 0.142960801 | 9.115673674 | -0.420379201 | 0.689163866 | 0.937757716 | -5.048877497 |
| **A_51_P472829** | Aif1l | -0.336330061 | -2.304973282 | 1.63231316 | 13.39790964 | -0.420308171 | 0.689212925 | 0.937758456 | -5.04889918 |
| **A_51_P488554** | Mrnip | 0.068043743 | -0.330351708 | 0.466439194 | 9.829013057 | 0.420187364 | 0.68929637 | 0.937805978 | -5.048936051 |
| **A_51_P315634** | Nedd1 | 0.044616356 | -0.216873049 | 0.306105761 | 8.862204446 | 0.419767839 | 0.689586184 | 0.938134245 | -5.049064015 |
| **A_55_P2040878** | Ung | -0.057228483 | -0.392736107 | 0.278279142 | 14.4805059 | -0.41964205 | 0.689673092 | 0.93814486 | -5.049102361 |
| **A_51_P104747** | Gtf2h4 | -0.057469955 | -0.394532033 | 0.279592124 | 10.37394074 | -0.419469242 | 0.689792494 | 0.93814486 | -5.049155022 |
| **A_51_P292490** | Ttbk2 | -0.042635865 | -0.292697802 | 0.207426073 | 9.031156597 | -0.419465886 | 0.689794814 | 0.93814486 | -5.049156045 |
| **A_51_P329949** | Fam13a | 0.024446466 | -0.118940634 | 0.167833567 | 8.001668175 | 0.419445144 | 0.689809146 | 0.93814486 | -5.049162365 |
| **A_51_P447545** | Igfbp1 | 0.28854493 | -1.40417662 | 1.98126648 | 11.69257928 | 0.419369737 | 0.689861253 | 0.93814486 | -5.049185337 |
| **A_55_P2107432** | Slc18a1 | -0.078286691 | -0.537615135 | 0.381041753 | 9.26786758 | -0.419308591 | 0.689903506 | 0.93814486 | -5.049203961 |
| **A_55_P1965075** | NA | -0.019515619 | -0.134065747 | 0.095034509 | 15.46845873 | -0.419136912 | 0.690022147 | 0.93814486 | -5.049256241 |
| **A_55_P2137973** | Bnip2 | 0.049698636 | -0.242034383 | 0.341431655 | 9.303371494 | 0.419110025 | 0.690040728 | 0.93814486 | -5.049264426 |
| **A_55_P2039284** | Hspb1 | 0.026628391 | -0.129692718 | 0.182949499 | 16.99965072 | 0.419079543 | 0.690061795 | 0.93814486 | -5.049273706 |
| **A_55_P2113524** | Nsd2 | -0.027031823 | -0.185730882 | 0.131667236 | 8.502988608 | -0.419054152 | 0.690079343 | 0.93814486 | -5.049281436 |
| **A_55_P2042396** | Alkbh4 | 0.047311897 | -0.230736375 | 0.325360169 | 9.55017102 | 0.418619375 | 0.690379856 | 0.93845525 | -5.04941372 |
| **A_55_P1979616** | Tmem189 | -0.041000909 | -0.292300507 | 0.21029869 | 11.12059129 | -0.422816345 | 0.690450298 | 0.93845525 | -4.98284208 |
| **A_55_P2173373** | Zfp62 | 0.027626952 | -0.134799388 | 0.190053291 | 8.457618112 | 0.418452024 | 0.690495545 | 0.93845525 | -5.049464605 |
| **A_55_P2042486** | Dpysl3 | -0.024716234 | -0.17003285 | 0.120600382 | 7.867000137 | -0.41844288 | 0.690501866 | 0.93845525 | -5.049467385 |
| **A_51_P191354** | NA | -0.027073379 | -0.186282871 | 0.132136113 | 7.98371006 | -0.418352792 | 0.690564147 | 0.938473908 | -5.049494768 |
| **A_55_P1971963** | Tmem176b | 0.023324314 | -0.113866655 | 0.160515283 | 12.28023466 | 0.418265964 | 0.690624177 | 0.938489505 | -5.049521156 |
| **A_55_P2436438** | Dusp22 | 0.031859259 | -0.155617684 | 0.219336202 | 8.658214249 | 0.418077667 | 0.690754367 | 0.938600433 | -5.049578363 |
| **A_52_P350786** | Phlda2 | 0.115575283 | -0.564651463 | 0.795802029 | 8.608203261 | 0.418003987 | 0.690805313 | 0.938603677 | -5.049600742 |
| **A_55_P1970120** | Vopp1 | -0.184693763 | -1.272164365 | 0.90277684 | 11.27691513 | -0.417834087 | 0.690922798 | 0.938697319 | -5.049652331 |
| **A_55_P2056236** | Pdlim5 | -0.031427162 | -0.216560342 | 0.153706018 | 8.124392283 | -0.417628444 | 0.691065011 | 0.938783947 | -5.049714748 |
| **A_55_P2061045** | Rpl29 | -0.025839519 | -0.178080582 | 0.126401545 | 14.46839822 | -0.417562774 | 0.691110429 | 0.938783947 | -5.049734674 |
| **A_51_P298178** | Dctn3 | 0.030057801 | -0.147049893 | 0.207165494 | 13.67374829 | 0.4175312 | 0.691132266 | 0.938783947 | -5.049744254 |
| **A_52_P212214** | Pomt2 | -0.025831981 | -0.178072049 | 0.126408087 | 8.354241642 | -0.417443702 | 0.691192782 | 0.938800174 | -5.049770797 |
| **A_51_P467110** | Dpp4 | 0.028671314 | -0.140360546 | 0.197703174 | 8.048739959 | 0.417299812 | 0.691292306 | 0.938817546 | -5.049814435 |
| **A_51_P267354** | Lrfn3 | -0.019339316 | -0.133374278 | 0.094695646 | 8.637125005 | -0.417226847 | 0.691342777 | 0.938817546 | -5.049836559 |
| **A_51_P138538** | Eif1b | -0.025124662 | -0.173298758 | 0.123049433 | 11.90116834 | -0.417154674 | 0.691392701 | 0.938817546 | -5.049858439 |
| **A_52_P481686** | Wtip | -0.026384971 | -0.182027462 | 0.129257519 | 12.37429883 | -0.417059117 | 0.691458804 | 0.938817546 | -5.049887402 |
| **A_51_P453088** | Coa6 | -0.0299702 | -0.206774461 | 0.146834061 | 10.78582658 | -0.417028828 | 0.691479757 | 0.938817546 | -5.049896581 |
| **A_66_P100620** | Ensa | 0.078422118 | -0.384243901 | 0.541088138 | 11.57585546 | 0.417003911 | 0.691496994 | 0.938817546 | -5.049904132 |
| **A_55_P2052690** | Synm | 0.018033751 | -0.088377941 | 0.124445443 | 7.739615282 | 0.416932635 | 0.691546303 | 0.938818549 | -5.049925729 |
| **A_55_P2065149** | Kdm2b | -0.072415215 | -0.499802572 | 0.354972142 | 11.6987542 | -0.41684761 | 0.691605126 | 0.938832466 | -5.049951488 |
| **A_55_P2029106** | Bmf | 0.087400638 | -0.450737598 | 0.625538873 | 9.794439415 | 0.420892156 | 0.691763375 | 0.938878839 | -4.983394083 |
| **A_51_P413698** | Mrpl20 | 0.059341293 | -0.291105706 | 0.409788291 | 13.3260074 | 0.416585185 | 0.691786695 | 0.938878839 | -5.050030961 |
| **A_51_P423518** | Amph | -0.019926183 | -0.137610743 | 0.097758377 | 7.836769368 | -0.416556384 | 0.691806623 | 0.938878839 | -5.05003968 |
| **A_65_P15875** | Chpf2 | -0.059825369 | -0.413218854 | 0.293568117 | 11.7067496 | -0.41648178 | 0.691858246 | 0.938878839 | -5.050062264 |
| **A_51_P308298** | Myl9 | 0.108205951 | -0.531126474 | 0.747538376 | 13.05851059 | 0.416383564 | 0.691926209 | 0.938878839 | -5.050091989 |
| **A_55_P2015485** | Cfap45 | -0.057981527 | -0.400597103 | 0.284634049 | 8.247782652 | -0.416343422 | 0.691953988 | 0.938878839 | -5.050104136 |
| **A_55_P2119032** | Rasl2-9 | -0.03782728 | -0.261391343 | 0.185736784 | 10.13593867 | -0.416267284 | 0.692006676 | 0.938878839 | -5.050127172 |
| **A_55_P1987201** | Capn2 | 0.020443393 | -0.100442317 | 0.141329103 | 12.8877926 | 0.416051583 | 0.692155957 | 0.938878839 | -5.050192415 |
| **A_55_P2128734** | Pbxip1 | -0.1303986 | -0.902009374 | 0.641212173 | 12.68533307 | -0.415761059 | 0.692357043 | 0.938878839 | -5.050280239 |

| | | | | | | | | | |
|---|---|---|---|---|---|---|---|---|---|
| **A_55_P2153600** | Mfsd4b3-ps | 0.037377129 | -0.183796747 | 0.258551005 | 8.444988993 | 0.415758629 | 0.692358726 | 0.938878839 | -5.050280974 |
| **A_52_P110291** | Phf12 | -0.026576637 | -0.183862038 | 0.130708764 | 9.051013745 | -0.415700725 | 0.692398807 | 0.938878839 | -5.050298471 |
| **A_55_P2067722** | Camsap3 | 0.087297538 | -0.429361352 | 0.603956429 | 12.67925553 | 0.415687775 | 0.692407771 | 0.938878839 | -5.050302384 |
| **A_55_P2037343** | H2-Q10 | 0.240094733 | -1.180913087 | 1.661102554 | 8.830324582 | 0.415676362 | 0.692415672 | 0.938878839 | -5.050305832 |
| **A_51_P240019** | Gyg | -0.069068052 | -0.47788768 | 0.339751576 | 11.83556512 | -0.415637412 | 0.692442634 | 0.938878839 | -5.0503176 |
| **A_55_P2331804** | AU015791 | 0.071506391 | -0.351752898 | 0.494765681 | 8.499182004 | 0.415630611 | 0.692447342 | 0.938878839 | -5.050319655 |
| **A_51_P204740** | Cd34 | 0.063500644 | -0.312372136 | 0.439373424 | 12.83894788 | 0.415629549 | 0.692448077 | 0.938878839 | -5.050319976 |
| **A_51_P383140** | Slk | -0.040284829 | -0.278819174 | 0.198249516 | 9.818252786 | -0.415489228 | 0.692545217 | 0.938878839 | -5.050362362 |
| **A_51_P191586** | Arpc1a | 0.035449795 | -0.174495837 | 0.245395428 | 11.45504568 | 0.415409143 | 0.69260066 | 0.938878839 | -5.050386547 |
| **A_55_P2054420** | Tcf4 | 0.089356067 | -0.439845774 | 0.618557908 | 11.05905627 | 0.415405134 | 0.692603435 | 0.938878839 | -5.050387757 |
| **A_51_P238383** | B4galt4 | 0.058602031 | -0.288488865 | 0.405692926 | 11.68025182 | 0.415373329 | 0.692625455 | 0.938878839 | -5.050397361 |
| **A_51_P435817** | Zbtb8a | -0.023269822 | -0.161136219 | 0.114596574 | 8.263472823 | -0.415244423 | 0.692714702 | 0.938878839 | -5.050436277 |
| **A_55_P2075080** | Fat2 | 0.032634828 | -0.160729759 | 0.225999416 | 8.275278461 | 0.415215495 | 0.692734731 | 0.938878839 | -5.050445009 |
| **A_51_P258766** | Smo | 0.037161098 | -0.183038743 | 0.257360938 | 9.748056717 | 0.415184084 | 0.69275648 | 0.938878839 | -5.05045449 |
| **A_55_P1974114** | NA | 0.06033291 | -0.297261499 | 0.417927318 | 14.18192688 | 0.415080865 | 0.692827949 | 0.938909866 | -5.050485639 |
| **A_51_P116088** | Rexo2 | 0.024065198 | -0.118615227 | 0.166745623 | 10.40861116 | 0.414948507 | 0.6929196 | 0.938968238 | -5.04541158 |
| **A_51_P155755** | Pld6 | 0.021711686 | -0.107040955 | 0.150464326 | 7.828770211 | 0.41486478 | 0.692977579 | 0.93897747 | -5.050550825 |
| **A_51_P405985** | Fig4 | 0.049936125 | -0.246237899 | 0.34611015 | 9.220465339 | 0.414798366 | 0.69302357 | 0.93897747 | -5.050570854 |
| **A_55_P1981306** | Hdgfl3 | -0.068343428 | -0.473851744 | 0.337164887 | 8.425865469 | -0.414635192 | 0.693136576 | 0.939064756 | -5.050620051 |
| **A_55_P2084512** | NA | -0.019404571 | -0.134574017 | 0.095764875 | 7.810882577 | -0.41451088 | 0.693222672 | 0.939115575 | -5.050657519 |
| **A_51_P141535** | Oat | 0.027208189 | -0.134398791 | 0.188815169 | 10.54640896 | 0.414198654 | 0.693438939 | 0.939138552 | -5.050751579 |
| **A_51_P398406** | Kctd13 | -0.026471567 | -0.183726732 | 0.130783598 | 8.519590091 | -0.414136871 | 0.693481737 | 0.939138552 | -5.050770184 |
| **A_55_P1974243** | Dbh | 0.019187735 | -0.094808583 | 0.133184053 | 7.676167597 | 0.414096973 | 0.693509376 | 0.939138552 | -5.050782197 |
| **A_52_P343856** | Creb3l2 | -0.072717945 | -0.504842751 | 0.359406861 | 10.53150278 | -0.414001164 | 0.693575748 | 0.939138552 | -5.050811041 |
| **A_66_P109577** | Plcb3 | 0.031949407 | -0.157910606 | 0.22180942 | 10.38728381 | 0.413998206 | 0.693577798 | 0.939138552 | -5.050811931 |
| **A_52_P13730** | Cbx2 | 0.048104285 | -0.237777962 | 0.333986532 | 8.808267996 | 0.41396702 | 0.693599402 | 0.939138552 | -5.050821318 |
| **A_55_P2053083** | Gm6098 | -0.035153756 | -0.244095047 | 0.173787535 | 10.56057953 | -0.413920203 | 0.693631837 | 0.939138552 | -5.050835409 |
| **A_66_P134785** | 2610044O15Rik8 | 0.033909486 | -0.167659718 | 0.23547869 | 8.44786524 | 0.413872131 | 0.693665142 | 0.939138552 | -5.050849876 |
| **A_55_P2387379** | NA | 0.028336804 | -0.140113455 | 0.196787063 | 8.614871688 | 0.413855137 | 0.693676916 | 0.939138552 | -5.050854991 |
| **A_55_P2139937** | Lrrc56 | -0.026626107 | -0.184954361 | 0.131702147 | 9.326886507 | -0.413731332 | 0.693762693 | 0.939188899 | -5.050892241 |
| **A_66_P129795** | Ccdc59 | -0.037893277 | -0.263284564 | 0.187498009 | 11.60894582 | -0.413613026 | 0.693844665 | 0.939234087 | -5.050927827 |
| **A_55_P2032238** | Zbtb8os | -0.047388066 | -0.329451402 | 0.23467527 | 11.53486923 | -0.413324844 | 0.69404436 | 0.939296224 | -5.051014473 |
| **A_52_P625244** | Ube3a | -0.018313576 | -0.127322621 | 0.09069547 | 8.623624712 | -0.413313674 | 0.694052101 | 0.939296224 | -5.05101783 |
| **A_51_P406077** | Fmc1 | -0.060982083 | -0.424146578 | 0.302182413 | 13.05818788 | -0.41311221 | 0.694191722 | 0.939296224 | -5.051078368 |
| **A_55_P2027496** | Adck5 | -0.06009083 | -0.417957058 | 0.297775399 | 12.0296105 | -0.413101381 | 0.694199227 | 0.939296224 | -5.051081621 |
| **A_55_P2009588** | Rpl13 | -0.019503112 | -0.135667082 | 0.096660858 | 15.4079688 | -0.413049048 | 0.694235498 | 0.939296224 | -5.051097342 |
| **A_52_P294510** | Fgl1 | -0.017813965 | -0.12393696 | 0.088309031 | 7.729294048 | -0.412971661 | 0.694289135 | 0.939296224 | -5.051120586 |
| **A_51_P460890** | Rgl2 | -0.059671226 | -0.415182286 | 0.295839835 | 10.54652002 | -0.412934344 | 0.694315 | 0.939296224 | -5.051131792 |
| **A_55_P1976278** | Prkcb | -0.034331332 | -0.238931089 | 0.170268426 | 8.185923035 | -0.412814268 | 0.694398229 | 0.939296224 | -5.051167847 |
| **A_55_P1997977** | Rab2b | -0.053011374 | -0.368940524 | 0.262917776 | 10.56395484 | -0.4128084 | 0.694402297 | 0.939296224 | -5.051169609 |
| **A_55_P2186027** | Gapdh | -0.027572407 | -0.191935272 | 0.136790457 | 16.67620452 | -0.412705392 | 0.6944737 | 0.939296224 | -5.05120053 |
| **A_55_P1958434** | Got1 | -0.03760559 | -0.26180776 | 0.186596579 | 12.72395796 | -0.41264992 | 0.694512154 | 0.939296224 | -5.051217179 |
| **A_55_P2179020** | Mrpl15 | -0.033564233 | -0.233695708 | 0.166567241 | 9.911460226 | -0.41260121 | 0.694545921 | 0.939296224 | -5.051231797 |
| **A_66_P139785** | Plpp3 | -0.063555958 | -0.458801051 | 0.331689136 | 12.74806365 | -0.416715698 | 0.694617664 | 0.939296224 | -4.984584399 |
| **A_55_P2133315** | Steap2 | -0.058342174 | -0.40633708 | 0.289652732 | 8.735499562 | -0.412457196 | 0.694645759 | 0.939296224 | -5.051275005 |
| **A_51_P158678** | Fam181b | 0.020262749 | -0.100604151 | 0.141129649 | 7.71741756 | 0.412439398 | 0.694658097 | 0.939296224 | -5.051280344 |
| **A_55_P2429480** | Atad2 | -0.020711559 | -0.144259921 | 0.102836803 | 7.876059101 | -0.412424984 | 0.694668091 | 0.939296224 | -5.051284668 |
| **A_55_P2066234** | Arfip2 | 0.040964571 | -0.203529075 | 0.285458217 | 9.579000531 | 0.412201911 | 0.694822753 | 0.939318311 | -5.051351565 |
| **A_55_P2038183** | Insc | 0.081978519 | -0.407440636 | 0.571397673 | 9.655044826 | 0.412086386 | 0.694902855 | 0.939318311 | -5.051386196 |
| **A_55_P2102515** | Daam1 | -0.044540759 | -0.310490643 | 0.221409126 | 10.21144834 | -0.412028224 | 0.694943185 | 0.939318311 | -5.051403628 |

| | | | | | | | | |
|---|---|---|---|---|---|---|---|---|
| **A_55_P2023818** | Cysltr1 | 0.048224356 | -0.239749639 | 0.336198351 | 8.085495397 | 0.411985878 | 0.694972549 | 0.939318311 | -5.051416318 |
| **A_51_P278427** | 3830406C13Rik | 0.029344681 | -0.145895532 | 0.204584895 | 9.618715425 | 0.411969237 | 0.694984088 | 0.939318311 | -5.051421305 |
| **A_55_P2030209** | Cir1 | -0.050680668 | -0.353375391 | 0.252014054 | 9.706129011 | -0.411914091 | 0.695022329 | 0.939318311 | -5.051437829 |
| **A_51_P342481** | Gpd2 | 0.054570704 | -0.271360187 | 0.380501595 | 10.17503988 | 0.411910817 | 0.695024599 | 0.939318311 | -5.05143881 |
| **A_55_P2064995** | Capzb | -0.017954957 | -0.125268572 | 0.089358658 | 7.719957821 | -0.411622116 | 0.695224814 | 0.939489753 | -5.051525281 |
| **A_52_P450835** | Cdpf1 | -0.035464691 | -0.247473073 | 0.176543691 | 11.35950291 | -0.411540243 | 0.695281598 | 0.939489753 | -5.051549793 |
| **A_55_P2110875** | 2700046G09Rik | -0.039294153 | -0.274225711 | 0.195637405 | 9.594305391 | -0.411486636 | 0.695318779 | 0.939489753 | -5.05156584 |
| **A_65_P18167** | Hnrnpul2 | -0.022309616 | -0.155716705 | 0.111097472 | 9.873243534 | -0.411417121 | 0.695366995 | 0.939489753 | -5.051586647 |
| **A_55_P2132781** | Slc16a2 | -0.027767296 | -0.193826222 | 0.13829163 | 8.066548743 | -0.411377497 | 0.695394479 | 0.939489753 | -5.051598505 |
| **A_55_P1956517** | Polr2l | 0.068335074 | -0.340469154 | 0.477139301 | 14.35740017 | 0.411241989 | 0.695488474 | 0.939492907 | -5.051639051 |
| **A_55_P2401646** | Trdn | 0.019765939 | -0.098495593 | 0.138027472 | 7.66124182 | 0.411190538 | 0.695524165 | 0.939492907 | -5.051654442 |
| **A_55_P2152771** | Lhfpl2 | -0.120295743 | -0.840184775 | 0.59959329 | 9.593680565 | -0.411106045 | 0.695582778 | 0.939492907 | -5.051679714 |
| **A_51_P251717** | Hikeshi | 0.037432183 | -0.186580873 | 0.261445239 | 10.11679848 | 0.411093854 | 0.695591235 | 0.939492907 | -5.051683361 |
| **A_51_P307168** | Ddah1 | -0.022786157 | -0.159177541 | 0.113605227 | 8.56695786 | -0.411010873 | 0.695648802 | 0.939505009 | -5.051708175 |
| **A_55_P2021490** | Gm16386 | 0.062985007 | -0.314127512 | 0.440097525 | 9.394696491 | 0.410899292 | 0.695726213 | 0.939543909 | -5.051741535 |
| **A_55_P1974477** | Msh3 | -0.064451687 | -0.450505274 | 0.3216019 | 9.851065955 | -0.410729465 | 0.69584404 | 0.939574068 | -5.051792293 |
| **A_51_P487668** | H2afy3 | -0.035161329 | -0.245772408 | 0.17544975 | 8.459841357 | -0.410726981 | 0.695845764 | 0.939574068 | -5.051793035 |
| **A_51_P288916** | Tmtc2 | -0.035932279 | -0.251341478 | 0.179476919 | 8.321567848 | -0.410383307 | 0.696084238 | 0.939688 | -5.051895692 |
| **A_55_P2400691** | Arhgap21 | -0.048452857 | -0.338923527 | 0.242017814 | 12.4169081 | -0.410380072 | 0.696086483 | 0.939688 | -5.051896658 |
| **A_51_P365859** | Sowaha | 0.018441214 | -0.092123076 | 0.129005503 | 7.688349312 | 0.410339931 | 0.696114339 | 0.939688 | -5.051908643 |
| **A_55_P2009351** | Prxl2c | -0.038062808 | -0.266276589 | 0.190150972 | 9.538364069 | -0.410325139 | 0.696124604 | 0.939688 | -5.051913059 |
| **A_66_P107129** | 3300005D01Rik | 0.079766378 | -0.398660473 | 0.558193229 | 9.306721161 | 0.410179066 | 0.696225978 | 0.939759213 | -5.051956662 |
| **A_52_P196105** | Ttyh3 | 0.065059088 | -0.325242314 | 0.455360489 | 10.36494253 | 0.410087962 | 0.696289207 | 0.939778931 | -5.05198385 |
| **A_66_P134474** | Ang3 | 0.047463301 | -0.237335231 | 0.332261832 | 8.915752451 | 0.410005188 | 0.696346657 | 0.939787489 | -5.052008546 |
| **A_51_P478003** | Papolg | -0.025070109 | -0.175547347 | 0.125407129 | 9.079372005 | -0.40987797 | 0.696434958 | 0.939787489 | -5.052046495 |
| **A_55_P1956472** | Cybrd1 | 0.022761761 | -0.113929711 | 0.159453233 | 7.764572505 | 0.40966947 | 0.696579688 | 0.939787489 | -5.052108665 |
| **A_52_P134141** | Pyurf | -0.021926931 | -0.153606571 | 0.109752709 | 8.106550404 | -0.409664552 | 0.696583102 | 0.939787489 | -5.052110131 |
| **A_51_P232901** | Cnp | -0.022704293 | -0.159084232 | 0.113675647 | 8.176715054 | -0.40956859 | 0.696649719 | 0.939787489 | -5.052138735 |
| **A_52_P582261** | Ikzf5 | -0.027177573 | -0.1904355 | 0.136080354 | 9.149531435 | -0.409548702 | 0.696663525 | 0.939787489 | -5.052144662 |
| **A_55_P2203996** | 4932433N03Rik | -0.018210302 | -0.127645169 | 0.091224564 | 9.01132893 | -0.409383757 | 0.696778038 | 0.939787489 | -5.052193811 |
| **A_55_P2111444** | Trpv6 | 0.038030377 | -0.190515805 | 0.266576559 | 9.993693658 | 0.409379246 | 0.69678117 | 0.939787489 | -5.052195155 |
| **A_55_P2056473** | Spc24 | 0.021165342 | -0.106032448 | 0.148363132 | 7.856019095 | 0.409368932 | 0.69678833 | 0.939787489 | -5.052198228 |
| **A_51_P335969** | Des | -0.313846537 | -2.200046863 | 1.572353789 | 9.837468832 | -0.40935367 | 0.696798927 | 0.939787489 | -5.041721784 |
| **A_51_P454309** | Zfp772 | -0.033921748 | -0.237838331 | 0.169994835 | 9.442076824 | -0.409255796 | 0.696866881 | 0.939787489 | -5.052231928 |
| **A_51_P388002** | Snrpb2 | 0.049336841 | -0.2599329 | 0.358606583 | 11.34740825 | 0.413412871 | 0.696878995 | 0.939787489 | -4.985518137 |
| **A_55_P1976754** | Ppia | -0.029876329 | -0.209576328 | 0.14982367 | 17.02031362 | -0.409023557 | 0.697028137 | 0.93984023 | -5.052301078 |
| **A_52_P272054** | NA | -0.036175785 | -0.253774503 | 0.181422933 | 9.922668003 | -0.409007049 | 0.6970396 | 0.93984023 | -5.052305992 |
| **A_52_P458000** | AI467606 | -0.019753376 | -0.138586872 | 0.09908012 | 7.878386509 | -0.408951333 | 0.697078289 | 0.93984023 | -5.052322575 |
| **A_51_P417747** | Kap | -0.253127191 | -1.776310986 | 1.270056604 | 8.842418454 | -0.408842139 | 0.697154117 | 0.93984023 | -5.05235507 |
| **A_52_P215170** | Acot4 | 0.026768323 | -0.134321842 | 0.187858489 | 7.837521626 | 0.408809807 | 0.69717657 | 0.93984023 | -5.05236469 |
| **A_51_P463452** | Acsl1 | -0.025926227 | -0.181966998 | 0.130114543 | 7.955827569 | -0.408761896 | 0.697209843 | 0.93984023 | -5.052378945 |
| **A_52_P508920** | Abca6 | -0.017834632 | -0.125227348 | 0.089558085 | 7.639360297 | -0.408562481 | 0.697348338 | 0.939912611 | -5.052438257 |
| **A_51_P111932** | Tor1aip2 | -0.039995314 | -0.280840992 | 0.200850363 | 10.28366872 | -0.408544549 | 0.697360792 | 0.939912611 | -5.052443589 |
| **A_55_P1989013** | Eif6 | 0.044349305 | -0.222893917 | 0.311592527 | 10.7252214 | 0.408271701 | 0.69755031 | 0.940102492 | -5.052524696 |
| **A_51_P385773** | Eif2ak1 | -0.018215865 | -0.128115585 | 0.091683854 | 8.573827172 | -0.407776682 | 0.697894206 | 0.940492494 | -5.052671718 |
| **A_55_P1995874** | Gm14410 | -0.034902583 | -0.24554471 | 0.175739545 | 8.417610855 | -0.407644412 | 0.697986109 | 0.940492494 | -5.052710974 |
| **A_52_P223704** | Faah | 0.019030891 | -0.095840248 | 0.13390203 | 7.709259405 | 0.407584214 | 0.698027937 | 0.940492494 | -5.052728837 |
| **A_65_P15689** | Mmut | -0.033587097 | -0.236335769 | 0.169161575 | 9.431537531 | -0.407552546 | 0.698049943 | 0.940492494 | -5.052738233 |
| **A_66_P109708** | Il1f6 | 0.069913258 | -0.352167693 | 0.491994209 | 8.435361267 | 0.407505009 | 0.698082974 | 0.940492494 | -5.052752335 |
| **A_51_P328001** | Sftpc | 0.032508557 | -0.163808376 | 0.22882549 | 7.748345912 | 0.407388806 | 0.698163724 | 0.940535727 | -5.052786802 |

| | | | | | | | | | |
|---|---|---|---|---|---|---|---|---|---|
| **A_55_P2252184** | NA | 0.072056526 | -0.363293605 | 0.507406658 | 9.435705828 | 0.407196284 | 0.698297517 | 0.940583088 | -5.052843887 |
| **A_55_P2162152** | NA | -0.053918779 | -0.379779804 | 0.271942247 | 9.821812138 | -0.407077211 | 0.698380273 | 0.940583088 | -5.052879181 |
| **A_55_P2113703** | Spib | -0.019223866 | -0.135405694 | 0.096957961 | 7.767003467 | -0.407072435 | 0.698383592 | 0.940583088 | -5.052880596 |
| **A_55_P2295445** | 2600002B07Rik | 0.07191566 | -0.362780002 | 0.506611322 | 10.12143773 | 0.407012105 | 0.698425523 | 0.940583088 | -5.052898475 |
| **A_55_P1972025** | Mycl | 0.023817717 | -0.120186664 | 0.167822099 | 7.98392698 | 0.406905533 | 0.698499597 | 0.940583088 | -5.05293005 |
| **A_55_P2171528** | 3110082I17Rik | -0.019716808 | -0.13905585 | 0.099622233 | 8.109225467 | -0.406465076 | 0.698805778 | 0.940583088 | -5.05306047 |
| **A_51_P351896** | Gask1b | -0.027529772 | -0.194172021 | 0.139112478 | 7.884150298 | -0.406430838 | 0.698829581 | 0.940583088 | -5.053070602 |
| **A_55_P2008790** | Zfp30 | 0.019249918 | -0.097282561 | 0.135782398 | 7.859724736 | 0.406397538 | 0.698852733 | 0.940583088 | -5.053080456 |
| **A_55_P2086885** | Slc45a4 | -0.05577809 | -0.393512858 | 0.281956678 | 12.16987399 | -0.406309549 | 0.698913907 | 0.940583088 | -5.05310649 |
| **A_55_P2078645** | Mettl5 | -0.03178504 | -0.224251501 | 0.160681421 | 9.272416335 | -0.40629069 | 0.698927019 | 0.940583088 | -5.053112069 |
| **A_55_P1994957** | NA | -0.035706716 | -0.252009315 | 0.180595883 | 15.16752309 | -0.406122765 | 0.699043777 | 0.940583088 | -5.053161738 |
| **A_52_P498193** | Aldh1l2 | 0.034641796 | -0.175215162 | 0.244498754 | 8.110016749 | 0.406112346 | 0.699051021 | 0.940583088 | -5.053164819 |
| **A_51_P370678** | Gfi1b | 0.045141543 | -0.22841041 | 0.318693496 | 7.886867163 | 0.405981029 | 0.699142333 | 0.940583088 | -5.053203646 |
| **A_51_P268234** | Il34 | 0.089774167 | -0.454247671 | 0.633796005 | 9.945906506 | 0.405979601 | 0.699143326 | 0.940583088 | -5.053204068 |
| **A_55_P1966804** | Fdps | 0.126980354 | -0.642812782 | 0.896773489 | 12.52904914 | 0.405818333 | 0.699255472 | 0.940583088 | -5.053251734 |
| **A_52_P575668** | NA | -0.020105419 | -0.14201006 | 0.101799222 | 8.812489036 | -0.405753285 | 0.699300708 | 0.940583088 | -5.053270955 |
| **A_55_P1959763** | Dach1 | 0.020583956 | -0.104264574 | 0.145432485 | 7.846329083 | 0.405615526 | 0.699396515 | 0.940583088 | -5.053311653 |
| **A_51_P284426** | Cstad | -0.020226468 | -0.142917428 | 0.102464492 | 7.942191379 | -0.405580114 | 0.699421144 | 0.940583088 | -5.053322112 |
| **A_55_P1967720** | Polr2i | -0.023939551 | -0.169169567 | 0.121290465 | 12.45124528 | -0.40553541 | 0.699452236 | 0.940583088 | -5.053335315 |
| **A_55_P2008896** | NA | 0.045521483 | -0.230636244 | 0.32167921 | 11.73209676 | 0.405535024 | 0.699452504 | 0.940583088 | -5.053335429 |
| **A_55_P2018706** | Cep19 | -0.029616884 | -0.209302574 | 0.150068805 | 11.20932834 | -0.405503907 | 0.699474147 | 0.940583088 | -5.053344619 |
| **A_55_P2094197** | Dnase1l3 | 0.021110049 | -0.106970061 | 0.14919016 | 8.495518729 | 0.405486795 | 0.699486049 | 0.940583088 | -5.053349672 |
| **A_55_P1964559** | Smarca5-ps | -0.04564512 | -0.322620779 | 0.23133054 | 9.525413249 | -0.405435624 | 0.69952164 | 0.940583088 | -5.053364781 |
| **A_55_P2086746** | Fam3a | -0.032636645 | -0.230700262 | 0.165426972 | 8.480944139 | -0.405387141 | 0.699555363 | 0.940583088 | -5.053379095 |
| **A_51_P273538** | Syce2 | 0.084149706 | -0.426556131 | 0.594855543 | 11.34536302 | 0.405369383 | 0.699567714 | 0.940583088 | -5.053384338 |
| **A_51_P255193** | Faf2 | -0.034743401 | -0.245608028 | 0.176121225 | 11.05416926 | -0.405357084 | 0.699576269 | 0.940583088 | -5.053387969 |
| **A_55_P2107362** | Gjc1 | 0.031244837 | -0.158412011 | 0.220901685 | 9.07576503 | 0.405302144 | 0.699614484 | 0.940583088 | -5.053404186 |
| **A_51_P363947** | Cdkn1a | -0.08806693 | -0.622658427 | 0.446524567 | 9.991094258 | -0.405284481 | 0.69962677 | 0.940583088 | -5.027561452 |
| **A_52_P549977** | Fam32a | -0.036951244 | -0.261266429 | 0.18736394 | 10.15104853 | -0.40526543 | 0.699640022 | 0.940583088 | -5.053415022 |
| **A_55_P2005920** | NA | 0.098089761 | -0.497446372 | 0.693625895 | 14.30316951 | 0.405214264 | 0.699675614 | 0.940583088 | -5.053430123 |
| **A_55_P2016540** | Cry2 | -0.059796918 | -0.422922452 | 0.303328615 | 11.15404967 | -0.405126991 | 0.699736324 | 0.940583088 | -5.053455875 |
| **A_55_P1999829** | Thoc7 | -0.039441363 | -0.278971579 | 0.200088854 | 12.17053323 | -0.405098625 | 0.699756056 | 0.940583088 | -5.053464244 |
| **A_55_P1991505** | Gamt | 0.041381812 | -0.210079002 | 0.292842627 | 9.20179675 | 0.404863246 | 0.699919808 | 0.940721508 | -5.05353367 |
| **A_51_P503822** | Slitrk6 | 0.018790091 | -0.09543091 | 0.133011093 | 7.663337123 | 0.404717601 | 0.700021141 | 0.940721508 | -5.053576609 |
| **A_55_P2150831** | Cdh4 | -0.044931743 | -0.31810701 | 0.228243524 | 8.545995325 | -0.404651409 | 0.700067197 | 0.940721508 | -5.05359612 |
| **A_55_P2212498** | NA | -0.059536545 | -0.421627763 | 0.302554672 | 8.432183288 | -0.40451516 | 0.700162001 | 0.940721508 | -5.05363627 |
| **A_52_P393589** | Clmp | -0.145432961 | -1.029990071 | 0.739124149 | 8.994172218 | -0.404488456 | 0.700180583 | 0.940721508 | -5.053644138 |
| **A_51_P148612** | Cox7a1 | 0.051999645 | -0.26428709 | 0.36828638 | 11.81418892 | 0.404472093 | 0.700191969 | 0.940721508 | -5.053648958 |
| **A_51_P243930** | Qrsl1 | -0.021670459 | -0.153532429 | 0.110191511 | 8.252678794 | -0.404313015 | 0.700302669 | 0.940721508 | -5.053695816 |
| **A_51_P271439** | Garem1 | 0.025039854 | -0.127328914 | 0.177408622 | 8.192180761 | 0.404301172 | 0.70031091 | 0.940721508 | -5.053699303 |
| **A_66_P133525** | NA | 0.034842057 | -0.17719446 | 0.246878574 | 8.489338851 | 0.404261411 | 0.700338581 | 0.940721508 | -5.053711012 |
| **A_55_P2073467** | Gtf2i | 0.050062286 | -0.254661377 | 0.35478595 | 11.40149548 | 0.404178923 | 0.700395987 | 0.940721508 | -5.0537353 |
| **A_51_P263220** | Taf5 | -0.021976639 | -0.155765905 | 0.111812627 | 8.360858065 | -0.404118907 | 0.700437756 | 0.940721508 | -5.053752968 |
| **A_55_P2148288** | Tmem79 | -0.036281954 | -0.257163206 | 0.184599298 | 10.23505429 | -0.404111284 | 0.700443062 | 0.940721508 | -5.053755212 |
| **A_65_P10913** | Tgfb2 | -0.190412838 | -1.349988111 | 0.969162436 | 11.87565911 | -0.403986118 | 0.700530175 | 0.940773137 | -5.053792051 |
| **A_55_P2145691** | Hlcs | -0.025594882 | -0.181510527 | 0.130320764 | 9.058585435 | -0.403861627 | 0.700616825 | 0.940824136 | -5.053828681 |
| **A_51_P208240** | Tnfsf14 | 0.017757995 | -0.090462707 | 0.125978696 | 7.699252284 | 0.403694413 | 0.70073322 | 0.940874349 | -5.053877866 |
| **A_55_P1964076** | Dnajb6 | 0.024442012 | -0.124522129 | 0.173406153 | 10.32755858 | 0.403668049 | 0.700751572 | 0.940874349 | -5.053885618 |
| **A_51_P436817** | Cbarp | -0.049449127 | -0.350915899 | 0.252017646 | 9.231715688 | -0.403541625 | 0.700839579 | 0.940927154 | -5.05392279 |
| **A_55_P2086687** | Nfib | 0.018136289 | -0.092480649 | 0.128753228 | 7.787313182 | 0.403362914 | 0.700963995 | 0.940927992 | -5.053975317 |

| | | | | | | | | | |
|---|---|---|---|---|---|---|---|---|---|
| **A_51_P216496** | Tfpi2 | -0.457391973 | -3.247501286 | 2.33271734 | 10.7924189 | -0.403307672 | 0.701002456 | 0.940927992 | -5.053991549 |
| **A_55_P2021659** | Abcb8 | 0.065630494 | -0.334783005 | 0.466043992 | 11.04660334 | 0.403242338 | 0.701047944 | 0.940927992 | -5.054010744 |
| **A_55_P2018994** | Elk3 | -0.025777472 | -0.183056136 | 0.131501191 | 10.34895372 | -0.403217787 | 0.701065037 | 0.940927992 | -5.054017957 |
| **A_52_P70854** | Tob2 | -0.041325861 | -0.29351404 | 0.210862318 | 9.78523871 | -0.40314971 | 0.701112437 | 0.940927992 | -5.054037954 |
| **A_55_P2087385** | Gps1 | 0.031892886 | -0.162768124 | 0.226553896 | 12.96455481 | 0.403073281 | 0.701165654 | 0.940927992 | -5.0540604 |
| **A_65_P20433** | Parpbp | -0.017993184 | -0.127843001 | 0.091856634 | 7.784935959 | -0.402974748 | 0.701234264 | 0.940927992 | -5.054089332 |
| **A_55_P1964752** | Slc23a3 | 0.082538763 | -0.421402257 | 0.586479783 | 11.40797197 | 0.402946517 | 0.701253923 | 0.940927992 | -5.054097621 |
| **A_55_P2062642** | Magt1 | 0.048187242 | -0.246045903 | 0.342420388 | 12.23316494 | 0.402911481 | 0.70127832 | 0.940927992 | -5.054107906 |
| **A_66_P125660** | Nde1 | -0.01935808 | -0.137621487 | 0.098905327 | 7.887287373 | -0.402699472 | 0.701425961 | 0.941060762 | -5.054170127 |
| **A_55_P2098801** | Gm15393 | 0.034003163 | -0.173816253 | 0.24182258 | 15.90255395 | 0.40253381 | 0.701541336 | 0.941150228 | -5.054218725 |
| **A_52_P485007** | Abca2 | -0.039043571 | -0.277748534 | 0.199661393 | 7.915534637 | -0.402399325 | 0.701635004 | 0.941210563 | -5.054258163 |
| **A_51_P338600** | Tmem14c | -0.052045136 | -0.37051015 | 0.266419878 | 12.75230404 | -0.402056958 | 0.701873488 | 0.941396868 | -5.054358509 |
| **A_51_P464182** | Mat1a | -0.017318074 | -0.123291465 | 0.088655318 | 7.64847871 | -0.402042458 | 0.701883589 | 0.941396868 | -5.054362757 |
| **A_51_P308948** | Tamm41 | -0.042835203 | -0.304988023 | 0.219317617 | 10.16185804 | -0.401990193 | 0.701919999 | 0.941396868 | -5.054378068 |
| **A_55_P1960411** | Lrrc23 | 0.018190987 | -0.093163502 | 0.129545476 | 7.763541893 | 0.40189972 | 0.701983028 | 0.941416079 | -5.054404568 |
| **A_55_P2165334** | Nde1 | -0.041219015 | -0.293627881 | 0.211189851 | 11.21156444 | -0.40175581 | 0.70208329 | 0.941465252 | -5.054446708 |
| **A_55_P2077133** | Oxsr1 | -0.022318396 | -0.159069652 | 0.114432861 | 10.2506476 | -0.401514105 | 0.702251701 | 0.941465252 | -5.054517453 |
| **A_52_P172405** | Washc4 | -0.036213427 | -0.258111133 | 0.185684278 | 8.957171518 | -0.4015004 | 0.702261251 | 0.941465252 | -5.054521463 |
| **A_55_P1952359** | Anapc5 | -0.112033865 | -0.827995036 | 0.603927306 | 13.89303487 | -0.405517807 | 0.70229901 | 0.941465252 | -4.98772288 |
| **A_52_P528634** | Ttc27 | -0.018672253 | -0.133104021 | 0.095759515 | 8.33126143 | -0.401438741 | 0.702304216 | 0.941465252 | -5.054539504 |
| **A_55_P2102628** | Usp4 | 0.031663583 | -0.162423368 | 0.225750534 | 11.01518431 | 0.401358886 | 0.702359862 | 0.941465252 | -5.054562864 |
| **A_55_P2082806** | Trib1 | 0.042772852 | -0.219410857 | 0.30495656 | 9.702673712 | 0.40135776 | 0.702360646 | 0.941465252 | -5.054563193 |
| **A_51_P395743** | Spopl | -0.024167192 | -0.172377839 | 0.124043455 | 8.711953998 | -0.401158459 | 0.702499536 | 0.941578196 | -5.054621476 |
| **A_55_P2047461** | Fcho1 | 0.018709183 | -0.096061239 | 0.133479606 | 7.903842763 | 0.401045834 | 0.702578028 | 0.941578196 | -5.0546544 |
| **A_55_P2130792** | Tuft1 | 0.030486703 | -0.15654096 | 0.217514367 | 9.004340484 | 0.401027156 | 0.702591046 | 0.941578196 | -5.05465986 |
| **A_55_P2202539** | BC028789 | 0.018783252 | -0.096512216 | 0.134078719 | 7.837792024 | 0.400799994 | 0.702749376 | 0.941676339 | -5.054726238 |
| **A_51_P194498** | Wfdc1 | 0.049581769 | -0.254809508 | 0.353973045 | 9.956983237 | 0.40073657 | 0.702793585 | 0.941676339 | -5.054744764 |
| **A_55_P2280581** | A130088B03Rik | -0.024853887 | -0.177445511 | 0.127737738 | 7.808497744 | -0.400712398 | 0.702810434 | 0.941676339 | -5.054751824 |
| **A_55_P2074062** | Sdhaf2 | 0.051293917 | -0.263790588 | 0.366378423 | 10.2026878 | 0.400505038 | 0.702954983 | 0.941804731 | -5.054812373 |
| **A_55_P2149440** | Ehmt1 | -0.023038392 | -0.164634206 | 0.118557423 | 10.3819046 | -0.400286465 | 0.703107362 | 0.94189052 | -5.054876165 |
| **A_55_P2033665** | NA | 0.063591505 | -0.327259778 | 0.454442789 | 9.633755882 | 0.400273397 | 0.703116473 | 0.94189052 | -5.054879978 |
| **A_55_P2108476** | Mavs | 0.043795698 | -0.225467272 | 0.313058667 | 10.90497129 | 0.400151068 | 0.703201763 | 0.941931171 | -5.054915665 |
| **A_51_P115471** | Apol8 | -0.017008596 | -0.121596109 | 0.087578917 | 7.653590363 | -0.400090086 | 0.703244282 | 0.941931171 | -5.049792239 |
| **A_52_P571371** | Tmem144 | 0.0571875 | -0.294704079 | 0.409079079 | 9.560927885 | 0.399817133 | 0.703434612 | 0.942102212 | -5.055013033 |
| **A_51_P293269** | Pde7a | 0.041437234 | -0.213570179 | 0.296444647 | 8.945280333 | 0.399767158 | 0.703469461 | 0.942102212 | -5.055027598 |
| **A_52_P365011** | Fes | -0.028056012 | -0.200760423 | 0.144648399 | 8.445150891 | -0.39966091 | 0.703543556 | 0.942103492 | -5.055058558 |
| **A_55_P2042590** | Polm | 0.027854879 | -0.143631892 | 0.199341649 | 10.77222494 | 0.399613186 | 0.703576839 | 0.942103492 | -5.055072462 |
| **A_55_P2116054** | Caprin1 | -0.028982548 | -0.207463585 | 0.149498488 | 8.407437061 | -0.399497117 | 0.703657788 | 0.942103492 | -5.055106271 |
| **A_55_P2288232** | Gucy1a2 | 0.028904229 | -0.149111633 | 0.20692009 | 8.275701476 | 0.399458668 | 0.703684605 | 0.942103492 | -5.055117469 |
| **A_51_P512783** | Rmdn1 | 0.075424417 | -0.389149994 | 0.539998827 | 9.919099161 | 0.399416349 | 0.70371412 | 0.942103492 | -5.055129792 |
| **A_51_P487547** | Ccdc91 | -0.02707436 | -0.193879125 | 0.139730406 | 9.417762405 | -0.399318014 | 0.703782707 | 0.94213006 | -5.055158423 |
| **A_51_P324934** | Mcm3 | -0.040390589 | -0.289327373 | 0.208546195 | 9.247733599 | -0.399172187 | 0.703884424 | 0.942147679 | -5.05520087 |
| **A_51_P335770** | Afap1 | 0.055746464 | -0.287843239 | 0.399336166 | 13.10757195 | 0.399159386 | 0.703893354 | 0.942147679 | -5.055204596 |
| **A_55_P2025626** | NA | 0.030009898 | -0.155002409 | 0.215022206 | 8.27215515 | 0.399055288 | 0.703965969 | 0.94217963 | -5.055234886 |
| **A_51_P242859** | Akr1c12 | 0.0202483 | -0.10460639 | 0.14510299 | 7.774254854 | 0.398981605 | 0.70401737 | 0.942183185 | -5.055256323 |
| **A_52_P134023** | Rtp3 | -0.017353311 | -0.124391214 | 0.089684591 | 7.710169047 | -0.39885398 | 0.704106405 | 0.942237101 | -5.055293443 |
| **A_52_P598835** | Ccdc71l | -0.021181111 | -0.151894528 | 0.109532306 | 9.333249817 | -0.398655475 | 0.704244897 | 0.942312694 | -5.055351157 |
| **A_55_P2149049** | Rpl32 | 0.022298893 | -0.115320277 | 0.159918064 | 16.642219 | 0.398633263 | 0.704260395 | 0.942312694 | -5.055357614 |
| **A_51_P235726** | Icam4 | -0.055307763 | -0.39682294 | 0.286207414 | 8.116706688 | -0.398423775 | 0.704406565 | 0.942411361 | -5.055418489 |
| **A_55_P2010778** | Il11ra1 | 0.139015303 | -0.719455866 | 0.997486472 | 14.4142112 | 0.398387827 | 0.704431649 | 0.942411361 | -5.055428932 |

| | | | | | | | | |
|---|---|---|---|---|---|---|---|---|
| **A_55_P2037919** | NA | 0.027349142 | -0.141664887 | 0.196363171 | 8.981375378 | 0.398098108 | 0.704633827 | 0.94256533 | -5.055513066 |
| **A_55_P2156905** | NA | -0.045210802 | -0.32464287 | 0.234221266 | 13.54482922 | -0.398047707 | 0.704669001 | 0.94256533 | -5.055527697 |
| **A_51_P143296** | Myh8 | 0.037693243 | -0.195326266 | 0.270712752 | 7.859486026 | 0.397960879 | 0.7047296 | 0.94256533 | -5.055552898 |
| **A_55_P2305010** | A730089K16Rik | 0.017102993 | -0.088646668 | 0.122852655 | 7.689109106 | 0.397889341 | 0.704779529 | 0.94256533 | -5.055573657 |
| **A_51_P424561** | Zfp93 | -0.028363066 | -0.203782714 | 0.147056583 | 8.812745822 | -0.397781059 | 0.704855107 | 0.94256533 | -5.055605072 |
| **A_55_P2066096** | Zfand3 | -0.041974422 | -0.301609486 | 0.217660642 | 13.09082847 | -0.397732012 | 0.704889341 | 0.94256533 | -5.055619299 |
| **A_55_P2018482** | Adh5 | -0.02750815 | -0.197694942 | 0.142678643 | 7.905551821 | -0.397653387 | 0.704944222 | 0.94256533 | -5.055642102 |
| **A_55_P2058737** | Mex3d | 0.043696401 | -0.226649439 | 0.314042241 | 10.25891553 | 0.397644649 | 0.704950322 | 0.94256533 | -5.055644636 |
| **A_55_P1973352** | NA | 0.092093921 | -0.477868907 | 0.66205675 | 9.817941478 | 0.397515067 | 0.705040777 | 0.94256533 | -5.055682208 |
| **A_52_P683441** | Capn5 | 0.045289414 | -0.235043546 | 0.325622375 | 8.746810087 | 0.397458422 | 0.70508032 | 0.94256533 | -5.055698629 |
| **A_51_P322640** | Ccl24 | 0.018068147 | -0.093771424 | 0.129907718 | 7.732234676 | 0.397454375 | 0.705083145 | 0.94256533 | -5.055699802 |
| **A_55_P2082614** | Zfp791 | 0.051073988 | -0.265334982 | 0.367482959 | 12.68878787 | 0.397118525 | 0.705317616 | 0.942788735 | -5.055797115 |
| **A_55_P2066657** | Vps13a | -0.019018714 | -0.136858279 | 0.098820851 | 8.037135974 | -0.397062777 | 0.70535654 | 0.942788735 | -5.055813261 |
| **A_55_P1997585** | NA | 0.017504437 | -0.090968369 | 0.125977242 | 7.711115582 | 0.397005417 | 0.70539659 | 0.942788735 | -5.055829871 |
| **A_52_P94521** | 2510009E07Rik | 0.020849477 | -0.108388895 | 0.150087849 | 8.099835081 | 0.396892457 | 0.705475464 | 0.942828959 | -5.055862575 |
| **A_52_P575078** | Babam1 | 0.032384713 | -0.168448715 | 0.233218142 | 11.59995295 | 0.396710066 | 0.705602827 | 0.942933977 | -5.055915363 |
| **A_51_P513941** | Lrpap1 | -0.060039638 | -0.432440922 | 0.312361646 | 12.43852711 | -0.396639588 | 0.705652044 | 0.942934556 | -5.055935755 |
| **A_51_P127320** | Cabs1 | 0.050766776 | -0.264270887 | 0.365804439 | 8.554584825 | 0.396448034 | 0.705785821 | 0.943048122 | -5.033094367 |
| **A_51_P131025** | Ngdn | 0.035455875 | -0.184611519 | 0.255523269 | 13.30884559 | 0.396370811 | 0.705839755 | 0.943054996 | -5.056013491 |
| **A_55_P2126552** | Zdhhc13 | 0.025746516 | -0.134215369 | 0.185708401 | 11.39562153 | 0.395978055 | 0.706114093 | 0.943116439 | -5.056126997 |
| **A_55_P2040161** | Upf3a | -0.025582702 | -0.184541505 | 0.133376102 | 9.132722085 | -0.395941468 | 0.706139651 | 0.943116439 | -5.056137565 |
| **A_55_P2077083** | Tmem132e | 0.104438032 | -0.544559524 | 0.753435588 | 10.491139 | 0.395899344 | 0.706169078 | 0.943116439 | -5.056149731 |
| **A_55_P1962503** | Nup62cl | 0.079852851 | -0.41638964 | 0.576095342 | 11.39467975 | 0.395881895 | 0.706181267 | 0.943116439 | -5.05615477 |
| **A_55_P1999057** | Cps1 | -0.016896253 | -0.121914826 | 0.088122319 | 7.715692709 | -0.395816109 | 0.706227225 | 0.943116439 | -5.056173768 |
| **A_52_P203691** | Arl5c | -0.02484245 | -0.179270949 | 0.129586049 | 7.860992305 | -0.395763859 | 0.706263728 | 0.943116439 | -5.056188855 |
| **A_51_P303089** | Ttc28 | -0.031311304 | -0.225970773 | 0.163348165 | 8.551203776 | -0.395726175 | 0.706290055 | 0.943116439 | -5.056199734 |
| **A_55_P2110101** | Tm2d3 | -0.02824986 | -0.203885802 | 0.147386082 | 11.32267629 | -0.395705484 | 0.70630451 | 0.943116439 | -5.056205707 |
| **A_55_P1989653** | Slco4a1 | -0.088947294 | -0.642130157 | 0.46423557 | 14.11085292 | -0.395578959 | 0.706392908 | 0.943116439 | -5.056242227 |
| **A_55_P2129286** | Zc3hc1 | -0.0318071 | -0.229636483 | 0.166022282 | 9.002538445 | -0.395550965 | 0.706412467 | 0.943116439 | -5.056250306 |
| **A_51_P151038** | Pmpcb | 0.025689055 | -0.134108454 | 0.185486564 | 11.50653692 | 0.395500727 | 0.706447569 | 0.943116439 | -5.056264802 |
| **A_55_P2109658** | NA | 0.032993472 | -0.172301309 | 0.238288253 | 11.06514135 | 0.395384166 | 0.706529013 | 0.943116439 | -5.05629843 |
| **A_51_P441687** | Lrrc10 | 0.016678053 | -0.087103069 | 0.120459176 | 7.720591247 | 0.395363124 | 0.706543716 | 0.943116439 | -5.0563045 |
| **A_55_P2169010** | Ndfip1 | -0.035734575 | -0.258117948 | 0.186648799 | 12.15614399 | -0.395326081 | 0.7065696 | 0.943116439 | -5.056315184 |
| **A_55_P2065732** | Medag | 0.048926811 | -0.255675501 | 0.353529123 | 15.13579388 | 0.39516901 | 0.706679359 | 0.943116439 | -5.056360479 |
| **A_66_P125209** | Kif2c | 0.017690236 | -0.092481657 | 0.127862129 | 7.711259482 | 0.395031732 | 0.706775293 | 0.943116439 | -5.056400052 |
| **A_52_P401504** | Thbs4 | 0.018054794 | -0.094449899 | 0.130559487 | 7.736228107 | 0.394812654 | 0.706928404 | 0.943116439 | -5.056463179 |
| **A_55_P1982533** | Elovl4 | 0.040212415 | -0.210401436 | 0.290826266 | 8.503590393 | 0.394751922 | 0.706970852 | 0.943116439 | -5.056480673 |
| **A_52_P392674** | Acot9 | 0.072562489 | -0.379810282 | 0.52493526 | 10.65568315 | 0.394625258 | 0.707059385 | 0.943116439 | -5.056517151 |
| **A_55_P2289040** | NA | -0.031424653 | -0.227345513 | 0.164496207 | 7.957127068 | -0.394601726 | 0.707075834 | 0.943116439 | -5.056523927 |
| **A_66_P128702** | Nup43 | -0.028575219 | -0.206845035 | 0.149694597 | 9.045398257 | -0.39434915 | 0.707252392 | 0.943116439 | -5.056596629 |
| **A_51_P356642** | Krt19 | -0.032439426 | -0.234818137 | 0.169939285 | 14.2548221 | -0.394346056 | 0.707254555 | 0.943116439 | -5.05659752 |
| **A_55_P2064712** | NA | -0.024690466 | -0.178743488 | 0.129362556 | 15.5748146 | -0.394301321 | 0.707285828 | 0.943116439 | -5.056610392 |
| **A_66_P126332** | Zfp703 | 0.033214259 | -0.17407782 | 0.240506338 | 12.08148256 | 0.394194924 | 0.70736021 | 0.943116439 | -5.056641001 |
| **A_55_P2103422** | Psmc2 | -0.052663199 | -0.381349568 | 0.276023171 | 13.43674351 | -0.394180108 | 0.707370568 | 0.943116439 | -5.056645263 |
| **A_55_P2065726** | Snx29 | -0.034880301 | -0.252583305 | 0.182822704 | 8.302735954 | -0.394171264 | 0.707376751 | 0.943116439 | -5.056647807 |
| **A_52_P556140** | Vapb | 0.030431249 | -0.159516593 | 0.22037909 | 12.2308048 | 0.394143684 | 0.707396033 | 0.943116439 | -5.05665574 |
| **A_51_P512629** | Ttll5 | -0.041097147 | -0.297634029 | 0.215439735 | 9.663088327 | -0.394122314 | 0.707410973 | 0.943116439 | -5.056661886 |
| **A_55_P2044439** | Mfng | 0.032355274 | -0.169659892 | 0.23437044 | 9.03937282 | 0.394030893 | 0.707474891 | 0.943116439 | -5.056688176 |
| **A_55_P2357303** | 5430401H09Rik | -0.02033943 | -0.147340391 | 0.106661531 | 7.92189372 | -0.39400427 | 0.707493505 | 0.943116439 | -5.056695831 |
| **A_51_P386344** | Mtmr10 | -0.038592867 | -0.279621314 | 0.20243558 | 9.695645338 | -0.393919883 | 0.707552508 | 0.943116439 | -5.056720092 |

| | | | | | | | | | |
|---|---|---|---|---|---|---|---|---|---|
| **A_52_P386837** | Trmu | 0.019760451 | -0.103671393 | 0.143192294 | 9.019965749 | 0.393857201 | 0.707596336 | 0.943116439 | -5.05673811 |
| **A_55_P1973643** | Raf1 | 0.027306725 | -0.143308274 | 0.197921723 | 12.26635177 | 0.39375091 | 0.707670659 | 0.943116439 | -5.056768657 |
| **A_55_P2031671** | Gstm6 | -0.017059847 | -0.123671838 | 0.089552145 | 7.776791813 | -0.393675356 | 0.707723492 | 0.943116439 | -5.056790366 |
| **A_55_P2017764** | Cep89 | -0.02844999 | -0.206248579 | 0.149348598 | 9.599915328 | -0.393661532 | 0.707733158 | 0.943116439 | -5.056794337 |
| **A_52_P79947** | Ndst1 | -0.055344751 | -0.401248479 | 0.290558976 | 10.1442098 | -0.393631967 | 0.707753833 | 0.943116439 | -5.056802831 |
| **A_55_P2007656** | Cryab | -0.292229162 | -2.119309163 | 1.534850838 | 11.48882233 | -0.393491316 | 0.707852192 | 0.943116439 | -5.056843229 |
| **A_52_P217437** | Htra3 | -0.168815487 | -1.224680918 | 0.887049943 | 8.92111844 | -0.393344316 | 0.707954998 | 0.943116439 | -5.056885437 |
| **A_55_P2164909** | Gatd3a | -0.046458025 | -0.337056285 | 0.244140236 | 11.89340413 | -0.393311724 | 0.707977793 | 0.943116439 | -5.056894793 |
| **A_52_P174661** | Gatc | 0.019611981 | -0.103068411 | 0.142292373 | 8.018601209 | 0.393292318 | 0.707991365 | 0.943116439 | -5.056900364 |
| **A_55_P2017694** | Gprc5c | -0.074371126 | -0.539684176 | 0.390941923 | 12.70121495 | -0.393213382 | 0.708046574 | 0.943116439 | -5.056923019 |
| **A_51_P386270** | Cyp2d12 | -0.017347366 | -0.125907541 | 0.091212809 | 7.726643681 | -0.393126362 | 0.708107439 | 0.943116439 | -5.056947991 |
| **A_55_P2057296** | Upk3bl | 0.531023801 | -2.792542714 | 3.854590317 | 10.2059877 | 0.393078034 | 0.708141243 | 0.943116439 | -5.056961857 |
| **A_52_P304947** | Cenpn | -0.019777299 | -0.143600492 | 0.104045893 | 7.814637185 | -0.392947159 | 0.708232788 | 0.943116439 | -5.056999398 |
| **A_51_P106952** | Rbm26 | 0.030808889 | -0.16216123 | 0.223779007 | 9.523449999 | 0.392785215 | 0.708346074 | 0.943116439 | -5.046503333 |
| **A_51_P107090** | BC031181 | -0.023510739 | -0.170801022 | 0.123779544 | 10.97934929 | -0.392700408 | 0.708405402 | 0.943116439 | -5.041618643 |
| **A_52_P225898** | Kcnj8 | -0.018747493 | -0.136211038 | 0.098716052 | 7.790977985 | -0.392653314 | 0.708438349 | 0.943116439 | -5.057083646 |
| **A_55_P2048409** | Apool | -0.041376849 | -0.300669415 | 0.217915717 | 10.08108253 | -0.392587541 | 0.708484365 | 0.943116439 | -5.057102496 |
| **A_55_P2083023** | Sac3d1 | 0.030437989 | -0.160323144 | 0.221199122 | 10.67331123 | 0.392550219 | 0.708510476 | 0.943116439 | -5.05711319 |
| **A_55_P2117649** | Slfn9 | -0.031612728 | -0.229740782 | 0.166515327 | 8.263864566 | -0.392541122 | 0.70851684 | 0.943116439 | -5.057115797 |
| **A_52_P58041** | Arpc5 | 0.031098014 | -0.163811402 | 0.22600743 | 10.86559646 | 0.392526515 | 0.70852706 | 0.943116439 | -5.057119982 |
| **A_51_P153683** | Lrrc26 | 0.016656595 | -0.087770944 | 0.121084133 | 7.686778451 | 0.392410246 | 0.708608409 | 0.943116439 | -5.057153292 |
| **A_51_P489996** | Dst | -0.047887716 | -0.348156646 | 0.252381213 | 8.456785914 | -0.392358346 | 0.708644722 | 0.943116439 | -5.057168157 |
| **A_55_P2074281** | Nr2f1 | -0.0261255 | -0.18993971 | 0.13768871 | 8.348022743 | -0.39235772 | 0.708645161 | 0.943116439 | -5.057168336 |
| **A_55_P2010298** | Serpina1d | 0.201487888 | -1.062324776 | 1.465300553 | 10.30085925 | 0.392225121 | 0.708737942 | 0.943116439 | -5.057206308 |
| **A_55_P2003517** | Zfp536 | -0.017215164 | -0.125240666 | 0.090810338 | 7.680331975 | -0.392061351 | 0.708852542 | 0.943116439 | -5.057253189 |
| **A_52_P602898** | Nkrf | 0.020742354 | -0.109442071 | 0.150926779 | 8.009058181 | 0.391983858 | 0.708906771 | 0.943116439 | -5.057275366 |
| **A_55_P2032165** | NA | 0.033742182 | -0.178076476 | 0.245560839 | 14.56647275 | 0.391902569 | 0.708963659 | 0.943116439 | -5.057298625 |
| **A_55_P1966958** | Mef2d | -0.025512453 | -0.185683342 | 0.134658435 | 8.389928836 | -0.391866196 | 0.708989114 | 0.943116439 | -5.05730903 |
| **A_51_P416858** | Myl1 | -0.026815459 | -0.195178268 | 0.141547351 | 7.798557865 | -0.391839512 | 0.709007789 | 0.943116439 | -5.057316664 |
| **A_51_P321150** | Lyz2 | -0.029021977 | -0.211263903 | 0.153219949 | 8.189088702 | -0.391785052 | 0.709045904 | 0.943116439 | -5.057332241 |
| **A_51_P386280** | Olfr187 | -0.018566221 | -0.135169856 | 0.098037414 | 7.937308464 | -0.391724368 | 0.709088376 | 0.943116439 | -5.057349597 |
| **A_55_P2154228** | Ncapd3 | -0.026811043 | -0.195219121 | 0.141597035 | 9.353801974 | -0.391669681 | 0.709126651 | 0.943116439 | -5.057365235 |
| **A_55_P1999334** | Inpp5b | 0.036923857 | -0.195083937 | 0.268931652 | 11.3236178 | 0.39153776 | 0.709218986 | 0.943116439 | -5.057402951 |
| **A_55_P2154235** | 1700008J07Rik | 0.025245692 | -0.133390153 | 0.183881536 | 8.510212632 | 0.391521045 | 0.709230686 | 0.943116439 | -5.057407729 |
| **A_55_P1965459** | Zfp521 | -0.017122381 | -0.124736915 | 0.090492154 | 7.689994604 | -0.391437459 | 0.709289194 | 0.943116439 | -5.057431618 |
| **A_55_P1971174** | Cd1d2 | -0.018225762 | -0.132798419 | 0.096346895 | 7.935792864 | -0.391357687 | 0.709345033 | 0.943116439 | -5.057454414 |
| **A_52_P621064** | NA | -0.0348879 | -0.25422061 | 0.18444481 | 8.991385244 | -0.391327693 | 0.70936603 | 0.943116439 | -5.057462984 |
| **A_55_P1998776** | NA | -0.019377038 | -0.141226608 | 0.102472533 | 15.77318027 | -0.391230356 | 0.709434169 | 0.943116439 | -5.05749079 |
| **A_55_P2106593** | NA | -0.055547361 | -0.40504493 | 0.293950208 | 9.394952858 | -0.391010513 | 0.709588076 | 0.943116439 | -5.05755357 |
| **A_51_P359603** | Itgb7 | 0.120148424 | -0.635864372 | 0.87616122 | 8.604939933 | 0.390983199 | 0.709607199 | 0.943116439 | -5.057561368 |
| **A_51_P195573** | Dnmt1 | 0.029070314 | -0.153859417 | 0.212000046 | 9.268762621 | 0.390962042 | 0.709622012 | 0.943116439 | -5.057567407 |
| **A_51_P517695** | Ly6f | 0.127917287 | -0.677024592 | 0.932859166 | 8.867512635 | 0.390961398 | 0.709622463 | 0.943116439 | -5.057567591 |
| **A_55_P2100088** | Ccdc92b | 0.018884494 | -0.099951402 | 0.13772039 | 7.717433263 | 0.390955086 | 0.709626882 | 0.943116439 | -5.057569393 |
| **A_51_P208152** | 3930402G23Rik | 0.023088515 | -0.122231461 | 0.16840849 | 7.926723523 | 0.390876753 | 0.709681726 | 0.943116439 | -5.057591751 |
| **A_52_P638895** | Vegfa | -0.050285219 | -0.366804738 | 0.266234301 | 10.62197871 | -0.390849004 | 0.709701156 | 0.943116439 | -5.057599671 |
| **A_55_P2068892** | Il6ra | 0.093204059 | -0.493777553 | 0.680185672 | 12.4720881 | 0.390642503 | 0.709845747 | 0.943116439 | -5.057658588 |
| **A_55_P2022861** | Elovl1 | -0.033609314 | -0.24528889 | 0.178070261 | 10.57183774 | -0.390615846 | 0.709864413 | 0.943116439 | -5.057666192 |
| **A_55_P2043367** | Apol9a | -0.079177223 | -0.577872291 | 0.419517845 | 10.13764724 | -0.390601905 | 0.709874175 | 0.943116439 | -5.057670168 |
| **A_52_P35377** | Plagl2 | 0.020202845 | -0.107167858 | 0.147573548 | 8.772883237 | 0.390222353 | 0.710139976 | 0.943116439 | -5.057778373 |
| **A_55_P2343648** | 1700120C14Rik | -0.017593567 | -0.128555705 | 0.09336857 | 7.647989015 | -0.390075125 | 0.710243092 | 0.943116439 | -5.057820319 |

| | | | | | | | | | |
|---|---|---|---|---|---|---|---|---|---|
| **A_55_P2014214** | NA | 0.031091275 | -0.16501794 | 0.22720049 | 13.21664272 | 0.390040495 | 0.710267347 | 0.943116439 | -5.057830183 |
| **A_55_P2250249** | Taok1 | 0.049797004 | -0.264356563 | 0.363950572 | 13.96845025 | 0.389969268 | 0.710317236 | 0.943116439 | -5.057850469 |
| **A_51_P103975** | Mpc2 | 0.046671984 | -0.247805216 | 0.341149184 | 12.38175771 | 0.389918419 | 0.710352854 | 0.943116439 | -5.052705652 |
| **A_51_P146560** | Msln | 0.243339172 | -1.292234187 | 1.77891253 | 12.47453392 | 0.389861743 | 0.710392553 | 0.943116439 | -5.057881086 |
| **A_55_P2057803** | Stum | 0.032367756 | -0.171903571 | 0.236639083 | 7.908479387 | 0.389829178 | 0.710415364 | 0.943116439 | -5.057890358 |
| **A_55_P2174582** | Urah | -0.083997774 | -0.614261995 | 0.446266447 | 8.945590811 | -0.389712751 | 0.710496921 | 0.943116439 | -5.057923499 |
| **A_55_P2257341** | 2410004I01Rik | 0.016671119 | -0.088585006 | 0.121927243 | 7.658520996 | 0.389660638 | 0.710533428 | 0.943116439 | -5.057938329 |
| **A_52_P106789** | Rbm18 | 0.022535767 | -0.119781739 | 0.164853273 | 11.90565184 | 0.389567916 | 0.710598384 | 0.943116439 | -5.057964713 |
| **A_55_P2316335** | Gfm2 | -0.034614788 | -0.253343429 | 0.184113853 | 8.401812594 | -0.389336558 | 0.710760474 | 0.943116439 | -5.058030519 |
| **A_55_P2001068** | NA | -0.033327933 | -0.243947543 | 0.177291678 | 10.74818263 | -0.389294918 | 0.710789648 | 0.943116439 | -5.058042358 |
| **A_51_P254425** | Ahrr | -0.018894909 | -0.138306485 | 0.100516666 | 7.783389638 | -0.389284884 | 0.710796679 | 0.943116439 | -5.058045211 |
| **A_52_P84353** | Homez | -0.033287083 | -0.243654808 | 0.177080642 | 9.981265088 | -0.389283315 | 0.710797778 | 0.943116439 | -5.058045657 |
| **A_52_P155805** | 2010204K13Rik | 0.028027893 | -0.149121537 | 0.205177322 | 8.144572631 | 0.389242138 | 0.71082663 | 0.943116439 | -5.058057364 |
| **A_66_P102395** | Elac2 | -0.020618232 | -0.150947655 | 0.109711191 | 8.159857866 | -0.389204735 | 0.710852837 | 0.943116439 | -5.058067997 |
| **A_52_P591166** | Dpysl2 | -0.200927781 | -1.471069655 | 1.069214093 | 11.85094389 | -0.389185738 | 0.710866147 | 0.943116439 | -5.058073397 |
| **A_55_P2087872** | NA | 0.033594111 | -0.178802277 | 0.245990499 | 13.51017716 | 0.389121463 | 0.710911184 | 0.943116439 | -5.058091666 |
| **A_55_P2023484** | Osbpl5 | -0.214306815 | -1.569407367 | 1.140793737 | 12.41695991 | -0.389075265 | 0.710943556 | 0.943116439 | -5.058104795 |
| **A_55_P2004927** | Dnajc25 | 0.061382713 | -0.326841886 | 0.449607311 | 9.889527947 | 0.388984406 | 0.711007224 | 0.943116439 | -5.058130612 |
| **A_66_P105175** | Bche | 0.01886348 | -0.100465187 | 0.138192148 | 7.685564153 | 0.388907392 | 0.711061192 | 0.943116439 | -5.058152491 |
| **A_55_P2096933** | Cntln | -0.048303466 | -0.353896948 | 0.257290015 | 8.846259797 | -0.388869046 | 0.711088064 | 0.943116439 | -5.058163383 |
| **A_55_P1998164** | Rab28 | -0.026924734 | -0.197300346 | 0.143450879 | 8.250607659 | -0.388788268 | 0.711144673 | 0.943116439 | -5.058186325 |
| **A_55_P1965402** | Uqcrb | 0.042341354 | -0.225628388 | 0.310311097 | 12.94068272 | 0.388730081 | 0.711185451 | 0.943116439 | -5.058202847 |
| **A_55_P1961160** | NA | -0.038391636 | -0.281372008 | 0.204588736 | 13.32250394 | -0.388717947 | 0.711193955 | 0.943116439 | -5.058206293 |
| **A_55_P1982767** | Tmem9b | -0.0558908 | -0.409731982 | 0.297950382 | 12.55801647 | -0.388598494 | 0.711277673 | 0.943116439 | -5.058240205 |
| **A_55_P2037478** | Kifap3 | -0.036376286 | -0.266722478 | 0.193969906 | 10.66693588 | -0.388513824 | 0.711337016 | 0.943116439 | -5.058264236 |
| **A_55_P2054300** | Alg8 | -0.059661666 | -0.437488342 | 0.31816501 | 9.83040787 | -0.388482914 | 0.711358681 | 0.943116439 | -5.058273008 |
| **A_55_P1966908** | Zfp40 | 0.020221196 | -0.107884543 | 0.148326936 | 7.97193254 | 0.388335797 | 0.711461799 | 0.943116439 | -5.058314748 |
| **A_52_P206881** | Prpf18 | -0.022715652 | -0.166681376 | 0.121250072 | 11.14551307 | -0.388181886 | 0.711569687 | 0.943116439 | -5.0583584 |
| **A_51_P428754** | Grik5 | 0.017354581 | -0.092636006 | 0.127345167 | 8.08639816 | 0.388175186 | 0.711574383 | 0.943116439 | -5.0583603 |
| **A_51_P209930** | Rtn2 | 0.046497512 | -0.248209162 | 0.341204187 | 8.987615314 | 0.388158328 | 0.711586201 | 0.943116439 | -5.058365081 |
| **A_51_P156857** | Smim3 | -0.087720799 | -0.643734077 | 0.468292478 | 9.544662245 | -0.388138373 | 0.711600189 | 0.943116439 | -5.058370739 |
| **A_55_P1988033** | Atp6v1g1 | 0.05402197 | -0.28842836 | 0.3964723 | 13.21346179 | 0.388098523 | 0.711628125 | 0.943116439 | -5.058382037 |
| **A_55_P2013823** | Gal3st1 | 0.059260845 | -0.316451786 | 0.434973477 | 8.453820853 | 0.388044212 | 0.711666199 | 0.943116439 | -5.058397434 |
| **A_52_P451888** | Tlk2 | -0.020227724 | -0.148480995 | 0.108025547 | 8.321195638 | -0.388014295 | 0.711687171 | 0.943116439 | -5.058405914 |
| **A_55_P2146749** | Rps13 | 0.041726734 | -0.222869545 | 0.306323013 | 16.21682822 | 0.387971501 | 0.711717173 | 0.943116439 | -5.058418044 |
| **A_55_P2110513** | Lmo7 | 0.038242285 | -0.204282063 | 0.280766632 | 9.948906813 | 0.387933825 | 0.711743586 | 0.943116439 | -5.058428722 |
| **A_55_P2103807** | NA | 0.05287423 | -0.282511634 | 0.388260095 | 14.41076796 | 0.387854171 | 0.71179943 | 0.943116439 | -5.058451293 |
| **A_55_P2036629** | NA | -0.063815264 | -0.468667826 | 0.341037299 | 9.954473654 | -0.387790208 | 0.711844275 | 0.943116439 | -5.058469416 |
| **A_55_P2183404** | Ptn | -0.049528364 | -0.363759274 | 0.264702545 | 8.376990495 | -0.387770035 | 0.711858419 | 0.943116439 | -5.058475131 |
| **A_51_P316801** | Wdr60 | 0.035121198 | -0.187740216 | 0.257982611 | 10.03440557 | 0.387706963 | 0.711902641 | 0.943116439 | -5.058492997 |
| **A_51_P414637** | Col5a1 | 0.141317925 | -0.755417617 | 1.038053467 | 10.20974874 | 0.387705562 | 0.711903623 | 0.943116439 | -5.058493393 |
| **A_51_P373379** | Csnk2b | -0.047442525 | -0.348497725 | 0.253612676 | 13.66606486 | -0.387695556 | 0.711910638 | 0.943116439 | -5.058496228 |
| **A_52_P66371** | Nlrx1 | -0.035005605 | -0.25716083 | 0.18714962 | 9.164908698 | -0.38765931 | 0.711936053 | 0.943116439 | -5.058506493 |
| **A_51_P434426** | Rwdd4a | -0.01761748 | -0.12951171 | 0.09427675 | 8.98087014 | -0.387351517 | 0.712151879 | 0.943289802 | -5.058593632 |
| **A_55_P2003492** | NA | 0.062892855 | -0.336578303 | 0.462364012 | 12.84769638 | 0.387333484 | 0.712164524 | 0.943289802 | -5.058598735 |
| **A_55_P2015410** | 0610010K14Rik | -0.020591753 | -0.151440868 | 0.110257362 | 13.63890705 | -0.387161086 | 0.712285424 | 0.943307941 | -5.058647512 |
| **A_55_P2164047** | NA | 0.044561814 | -0.238700503 | 0.327824132 | 12.30462424 | 0.387028744 | 0.71237824 | 0.943307941 | -5.058684942 |
| **A_55_P1988009** | Gm10406 | -0.062288494 | -0.458275065 | 0.333698077 | 9.644953324 | -0.386987143 | 0.712407417 | 0.943307941 | -5.058696705 |
| **A_55_P2054491** | Slc31a2 | 0.043262085 | -0.231822601 | 0.318346772 | 9.212018617 | 0.386910217 | 0.712461372 | 0.943307941 | -5.058718455 |
| **A_51_P140607** | Ints13 | -0.028502563 | -0.209755984 | 0.152750858 | 9.99288106 | -0.386871611 | 0.71248845 | 0.943307941 | -5.058729368 |

| | | | | | | | | |
|---|---|---|---|---|---|---|---|---|
| A_55_P2006467 | Serf2 | 0.025518545 | -0.136770064 | 0.187807154 | 15.19411765 | 0.386845042 | 0.712507085 | 0.943307941 | -5.058736878 |
| A_51_P349495 | Mboat1 | -0.093513744 | -0.688367062 | 0.501339574 | 9.779532602 | -0.386753916 | 0.712571003 | 0.943307941 | -5.058762633 |
| A_51_P287100 | Cdh16 | 0.01691308 | -0.090709044 | 0.124535204 | 7.718728495 | 0.38662534 | 0.712661194 | 0.943307941 | -5.058798962 |
| A_51_P469789 | Agxt | -0.016804568 | -0.123751099 | 0.090141963 | 7.660368564 | -0.386571492 | 0.712698968 | 0.943307941 | -5.058814174 |
| A_51_P486610 | Afg3l1 | -0.018476761 | -0.136069731 | 0.099116209 | 8.426987599 | -0.386557086 | 0.712709073 | 0.943307941 | -5.058818243 |
| A_55_P2428967 | Lrrc2 | -0.020627672 | -0.151937585 | 0.110682242 | 7.735089261 | -0.386475405 | 0.712766374 | 0.943307941 | -5.058841312 |
| A_55_P2028942 | Ncam1 | -0.018871098 | -0.139023984 | 0.101281788 | 7.848152957 | -0.386395558 | 0.71282239 | 0.943307941 | -5.05886386 |
| A_55_P2125200 | Mterf1b | 0.018841251 | -0.101190848 | 0.138873349 | 8.235217309 | 0.386172628 | 0.712978793 | 0.943307941 | -5.058926787 |
| A_55_P2052759 | Scd3 | -0.044744266 | -0.329840716 | 0.240352183 | 8.392222922 | -0.386113284 | 0.713020431 | 0.943307941 | -5.058943533 |
| A_55_P2000022 | Ccdc151 | -0.016350427 | -0.120573555 | 0.0878727 | 7.764605608 | -0.385952775 | 0.713133054 | 0.943307941 | -5.058988813 |
| A_52_P175028 | NA | 0.039291561 | -0.211169272 | 0.289752394 | 8.683096761 | 0.385947851 | 0.713136509 | 0.943307941 | -5.058990202 |
| A_55_P2110718 | Npm1 | -0.03047021 | -0.224762379 | 0.163821959 | 11.14919868 | -0.38582407 | 0.713223367 | 0.943307941 | -5.059025109 |
| A_55_P2128944 | Bcl2l13 | 0.045398016 | -0.24409943 | 0.334895461 | 10.76757063 | 0.385799176 | 0.713240835 | 0.943307941 | -5.059032128 |
| A_55_P2079973 | NA | -0.039748414 | -0.293228895 | 0.213732067 | 12.70975545 | -0.385784208 | 0.713251339 | 0.943307941 | -5.059036348 |
| A_55_P1997105 | Large2 | 0.028315489 | -0.152305866 | 0.208936844 | 10.03054943 | 0.385677348 | 0.713326329 | 0.943307941 | -5.059066471 |
| A_55_P2139713 | Phf13 | -0.024501532 | -0.180799856 | 0.131796792 | 8.876686563 | -0.385663102 | 0.713336326 | 0.943307941 | -5.059070487 |
| A_51_P263246 | Dusp8 | -0.036064638 | -0.266129335 | 0.194000059 | 10.02868694 | -0.385656589 | 0.713340897 | 0.943307941 | -5.059072322 |
| A_52_P527822 | Slc22a3 | 0.04216904 | -0.226841719 | 0.311179799 | 8.453549749 | 0.385649907 | 0.713345586 | 0.943307941 | -5.059074206 |
| A_51_P278540 | Myg1 | 0.03074757 | -0.165404543 | 0.226899684 | 8.853087658 | 0.385644353 | 0.713349484 | 0.943307941 | -5.059075771 |
| A_55_P2429225 | Psrc1 | -0.021407435 | -0.158020669 | 0.115205799 | 8.253716823 | -0.385514774 | 0.713440423 | 0.943363656 | -5.059112286 |
| A_55_P1967553 | Tmem254a | 0.044322702 | -0.238607954 | 0.327253358 | 11.60913708 | 0.38540326 | 0.713518687 | 0.943371933 | -5.059143701 |
| A_55_P2101001 | Smad3 | 0.093368048 | -0.502696337 | 0.689432433 | 12.22487663 | 0.385366776 | 0.713544294 | 0.943371933 | -5.059153977 |
| A_55_P2178862 | U2surp | 0.029346986 | -0.158057509 | 0.216751481 | 8.605008825 | 0.385258882 | 0.713620023 | 0.943407526 | -5.059184361 |
| A_51_P473953 | Arhgef26 | 0.036099128 | -0.19450332 | 0.266701576 | 8.432331261 | 0.385125222 | 0.713713841 | 0.943440356 | -5.059221991 |
| A_52_P51429 | Dennd1c | 0.017278401 | -0.093122986 | 0.127679787 | 7.856087814 | 0.385033194 | 0.713778441 | 0.943440356 | -5.059247892 |
| A_55_P2029592 | Dus2 | 0.02200634 | -0.118611185 | 0.162623865 | 9.082449015 | 0.385014896 | 0.713791286 | 0.943440356 | -5.059253042 |
| A_55_P2027701 | NA | 0.060842731 | -0.328094224 | 0.449779685 | 12.59829175 | 0.384856345 | 0.713902589 | 0.943508906 | -5.059297651 |
| A_51_P412640 | Ccdc51 | 0.063343068 | -0.341634521 | 0.468320656 | 9.730156011 | 0.38480195 | 0.713940776 | 0.943508906 | -5.059312951 |
| A_55_P2051874 | Psmc2 | 0.02363509 | -0.127502894 | 0.174773075 | 13.10980687 | 0.384727125 | 0.713993307 | 0.943513819 | -5.059333995 |
| A_66_P106090 | Rpa3 | 0.063015268 | -0.340203413 | 0.466233949 | 11.38186182 | 0.384480492 | 0.714166469 | 0.94360886 | -5.059403331 |
| A_51_P146149 | Napsa | 0.019722563 | -0.106548679 | 0.145993804 | 7.853294652 | 0.384262546 | 0.714319504 | 0.94360886 | -5.059464567 |
| A_51_P360880 | Zfp266 | -0.026183275 | -0.193837364 | 0.141470815 | 8.453583765 | -0.384219123 | 0.714349997 | 0.94360886 | -5.059476764 |
| A_51_P327904 | Yipf1 | -0.04042041 | -0.299241584 | 0.218400764 | 8.847524433 | -0.384211244 | 0.71435553 | 0.94360886 | -5.059478977 |
| A_52_P601688 | Abcc1 | -0.041585876 | -0.307928728 | 0.224756975 | 9.503716589 | -0.38412624 | 0.714415223 | 0.94360886 | -5.059502849 |
| A_55_P2040357 | Bola1 | 0.026545334 | -0.14347143 | 0.196562098 | 8.669023948 | 0.384118842 | 0.714420418 | 0.94360886 | -5.059504927 |
| A_55_P2025363 | Ccnl2 | -0.017736687 | -0.131343996 | 0.095870622 | 8.655664835 | -0.384092121 | 0.714439183 | 0.94360886 | -5.05951243 |
| A_55_P2241553 | 9330133O14Rik | -0.019210783 | -0.14226756 | 0.103845995 | 8.187975423 | -0.384068495 | 0.714455775 | 0.94360886 | -5.059519063 |
| A_55_P2098697 | Tnfaip2 | -0.088321991 | -0.654627293 | 0.477983311 | 10.53450399 | -0.383696101 | 0.714717316 | 0.943889792 | -5.059623571 |
| A_51_P386612 | Pir | 0.019124811 | -0.103549943 | 0.141799564 | 7.956690964 | 0.383540396 | 0.714826684 | 0.943969733 | -5.059667239 |
| A_51_P184385 | Pla2g12b | -0.085036279 | -0.630699528 | 0.46062697 | 9.572424731 | -0.383396997 | 0.714927415 | 0.944024814 | -5.059707442 |
| A_55_P1983152 | Gramd4 | -0.028665735 | -0.212652177 | 0.155320706 | 11.1395492 | -0.383306715 | 0.714990837 | 0.944024814 | -5.059732746 |
| A_55_P1985428 | Atg16l2 | -0.056475094 | -0.418983874 | 0.306033686 | 10.07333632 | -0.383272442 | 0.715014914 | 0.944024814 | -5.05974235 |
| A_55_P1991164 | Mlc1 | 0.065803231 | -0.356892592 | 0.488499055 | 8.591661058 | 0.382990877 | 0.71521273 | 0.944055522 | -5.059821224 |
| A_55_P2053616 | NA | -0.038089155 | -0.282793888 | 0.206615579 | 9.061554374 | -0.382937701 | 0.715250091 | 0.944055522 | -5.059836114 |
| A_55_P2162422 | NA | 0.023665202 | -0.128382377 | 0.17571278 | 16.65463436 | 0.38291278 | 0.715267601 | 0.944055522 | -5.059843091 |
| A_55_P2346654 | AU035318 | -0.0692437 | -0.514138282 | 0.375650881 | 8.598045919 | -0.382906126 | 0.715272277 | 0.944055522 | -5.059844954 |
| A_55_P2008933 | Slc17a3 | -0.017194356 | -0.127673191 | 0.093284478 | 7.638529256 | -0.382891746 | 0.71528238 | 0.944055522 | -5.05984898 |
| A_55_P1965846 | Sp2 | 0.039333563 | -0.213475014 | 0.292142139 | 9.791181554 | 0.382772425 | 0.715366221 | 0.944101713 | -5.059882381 |
| A_51_P237668 | Bex2 | 0.244276006 | -1.326280352 | 1.814832363 | 11.53950447 | 0.382645357 | 0.715455509 | 0.944125712 | -5.059917939 |
| A_51_P408946 | Ccne1 | 0.060705877 | -0.329637123 | 0.451048876 | 9.342431397 | 0.382607523 | 0.715482096 | 0.944125712 | -5.059928524 |

| | | | | | | | | | |
|---|---|---|---|---|---|---|---|---|---|
| **A_51_P485421** | NA | -0.01693582 | -0.125903666 | 0.092032025 | 7.733166812 | -0.382364038 | 0.715653206 | 0.944287038 | -5.054235253 |
| **A_51_P107808** | Ttc19 | -0.020745125 | -0.154247326 | 0.112757075 | 8.479704926 | -0.382293402 | 0.715702849 | 0.94428808 | -5.049436181 |
| **A_51_P128775** | Zfp84 | 0.018344787 | -0.099740701 | 0.136430275 | 8.46718003 | 0.38219527 | 0.715771819 | 0.94431462 | -4.999997305 |
| **A_55_P2082604** | Ndor1 | -0.027680416 | -0.205988706 | 0.150627873 | 16.96139194 | -0.381918106 | 0.715966634 | 0.944474197 | -5.060121238 |
| **A_55_P2058127** | Pde4dip | -0.054015927 | -0.401999995 | 0.293968141 | 8.213913693 | -0.381884151 | 0.715990502 | 0.944474197 | -5.060130721 |
| **A_65_P12152** | Rasa2 | -0.036172159 | -0.269261347 | 0.196917028 | 9.235070148 | -0.381787296 | 0.716058587 | 0.944499551 | -5.060157767 |
| **A_55_P2155499** | Tmem9 | 0.030360844 | -0.165336366 | 0.226058055 | 8.883619979 | 0.381679095 | 0.71613465 | 0.944535425 | -5.060187974 |
| **A_52_P650453** | Spink4 | -0.017248499 | -0.128481975 | 0.093984978 | 7.793268923 | -0.381491574 | 0.716266483 | 0.944563394 | -5.060240305 |
| **A_55_P2015177** | NA | -0.042540708 | -0.316975637 | 0.23189422 | 10.44056413 | -0.381359434 | 0.716359388 | 0.944563394 | -5.060277167 |
| **A_51_P487278** | Yipf3 | 0.042224113 | -0.230177507 | 0.314625734 | 11.55174855 | 0.38134672 | 0.716368327 | 0.944563394 | -5.060280713 |
| **A_55_P2209767** | 2610002J23Rik | -0.025954853 | -0.193403796 | 0.141494089 | 10.04421602 | -0.38133383 | 0.71637739 | 0.944563394 | -5.060284308 |
| **A_55_P1959061** | NA | 0.03582 | -0.195294053 | 0.266934053 | 11.84334042 | 0.381301396 | 0.716400195 | 0.944563394 | -5.060293353 |
| **A_55_P2138792** | Fdx1 | -0.046616251 | -0.347466355 | 0.254233852 | 13.57526296 | -0.381203032 | 0.716469358 | 0.944590152 | -5.060320781 |
| **A_55_P2010641** | Slc38a5 | 0.024615334 | -0.134297379 | 0.183528047 | 8.053067396 | 0.381080086 | 0.716555809 | 0.944639697 | -5.060355055 |
| **A_51_P376789** | Pgghg | -0.029688562 | -0.221432204 | 0.162055079 | 8.56493117 | -0.380923145 | 0.716666171 | 0.944643336 | -5.060398789 |
| **A_55_P1972634** | Cmc4 | 0.024976524 | -0.13634014 | 0.186293187 | 8.658570724 | 0.380909602 | 0.716675695 | 0.944643336 | -5.060402562 |
| **A_55_P1990735** | Ift122 | 0.026236941 | -0.14323908 | 0.195712963 | 8.672580865 | 0.380867668 | 0.716705185 | 0.944643336 | -5.060414245 |
| **A_51_P337918** | Aldh4a1 | 0.053737746 | -0.293461542 | 0.400937034 | 10.24521041 | 0.380776188 | 0.716769519 | 0.944663715 | -5.060439726 |
| **A_55_P2176802** | Hipk1 | 0.022413313 | -0.122470965 | 0.167297591 | 8.187790854 | 0.380587007 | 0.716902572 | 0.944712307 | -5.060492404 |
| **A_52_P920158** | Rbm24 | 0.01619222 | -0.08850761 | 0.12089205 | 7.681913972 | 0.380478027 | 0.716979223 | 0.944712307 | -5.060522739 |
| **A_51_P435198** | Cryz | 0.108776032 | -0.594681425 | 0.812233489 | 10.41043469 | 0.380421048 | 0.717019301 | 0.944712307 | -5.060538596 |
| **A_52_P239403** | Ipo7 | -0.024238093 | -0.181027916 | 0.132551729 | 8.424679732 | -0.380320512 | 0.717090019 | 0.944712307 | -5.060566569 |
| **A_55_P2083449** | Spryd7 | -0.02537424 | -0.189514987 | 0.138766507 | 9.803022244 | -0.380317052 | 0.717092452 | 0.944712307 | -5.060567531 |
| **A_55_P2065244** | Rbm26 | 0.020851996 | -0.114052937 | 0.155756929 | 8.809832047 | 0.380267283 | 0.717127461 | 0.944712307 | -5.060581376 |
| **A_55_P2123716** | Tug1 | -0.0302839 | -0.226234616 | 0.165666815 | 8.98833821 | -0.380219263 | 0.71716124 | 0.944712307 | -5.060594733 |
| **A_55_P2055199** | Airn | 0.051475277 | -0.281638384 | 0.384588937 | 10.04478442 | 0.380167871 | 0.717197392 | 0.944712307 | -5.060609026 |
| **A_55_P1971951** | Cd86 | 0.020730922 | -0.11345856 | 0.154920403 | 7.885670005 | 0.380074993 | 0.71726273 | 0.94473399 | -5.060634853 |
| **A_51_P168203** | Aig1 | 0.022764078 | -0.124635466 | 0.170163621 | 8.648262065 | 0.379947054 | 0.717352736 | 0.944776691 | -5.060670419 |
| **A_55_P1986486** | Gtf2ird1 | -0.055346337 | -0.413772942 | 0.303080267 | 10.32219028 | -0.379889957 | 0.717392907 | 0.944776691 | -5.060686288 |
| **A_55_P2001718** | Pex1 | -0.033615373 | -0.251596318 | 0.184365572 | 9.441499684 | -0.37939235 | 0.717743035 | 0.945042573 | -5.060824493 |
| **A_51_P174696** | Casq2 | 0.018391871 | -0.100874857 | 0.137658598 | 7.678167445 | 0.379381162 | 0.717750908 | 0.945042573 | -5.060827599 |
| **A_55_P2115442** | Clec2d | -0.091133766 | -0.68212117 | 0.499853638 | 9.100658531 | -0.379376351 | 0.717754293 | 0.945042573 | -5.060828934 |
| **A_51_P175580** | Trp53inp1 | -0.046913571 | -0.351310002 | 0.257482861 | 11.13245738 | -0.379164873 | 0.717903118 | 0.945042573 | -5.060887616 |
| **A_55_P1967677** | Cped1 | -0.042542915 | -0.318609208 | 0.233523377 | 7.947968363 | -0.379125529 | 0.717930808 | 0.945042573 | -5.060898531 |
| **A_55_P2178064** | NA | -0.023503824 | -0.17605786 | 0.129050212 | 8.369287667 | -0.37903908 | 0.71799165 | 0.945042573 | -5.060922508 |
| **A_55_P2440441** | Wdhd1 | -0.025399018 | -0.190421267 | 0.139623232 | 8.500573102 | -0.378654898 | 0.718262063 | 0.945042573 | -5.061029002 |
| **A_55_P2176361** | Zcchc9 | 0.019713743 | -0.108382146 | 0.147809633 | 8.219067627 | 0.378619578 | 0.718286926 | 0.945042573 | -5.061038788 |
| **A_55_P2000409** | Rab44 | -0.016128742 | -0.120973396 | 0.088715911 | 7.784866469 | -0.378462956 | 0.718397182 | 0.945042573 | -5.06108217 |
| **A_55_P2398264** | Rnf130 | -0.016626312 | -0.124706656 | 0.091454032 | 7.840280434 | -0.378458601 | 0.718400247 | 0.945042573 | -5.061083376 |
| **A_66_P128942** | Mipep | -0.018950247 | -0.142142707 | 0.104242214 | 7.823607616 | -0.378442499 | 0.718411583 | 0.945042573 | -5.061087835 |
| **A_51_P485161** | Cplx2 | 0.038648288 | -0.212625459 | 0.289922035 | 8.794399798 | 0.378401029 | 0.718440778 | 0.945042573 | -5.061099318 |
| **A_55_P1956418** | Efr3b | -0.059159224 | -0.443808473 | 0.325490026 | 11.54040273 | -0.378378751 | 0.718456462 | 0.945042573 | -5.061105487 |
| **A_55_P1995195** | Fosl2 | -0.088130337 | -0.661285414 | 0.485024739 | 11.5493174 | -0.378287902 | 0.718520421 | 0.945042573 | -5.061130638 |
| **A_51_P167105** | B3galt4 | -0.036808076 | -0.276203424 | 0.202587272 | 9.428789808 | -0.378265362 | 0.71853629 | 0.945042573 | -5.061136877 |
| **A_52_P350537** | Mtmr11 | 0.031157513 | -0.171648335 | 0.233963361 | 8.34882536 | 0.377964905 | 0.718747839 | 0.945042573 | -5.061220012 |
| **A_55_P1974283** | Rbmx | 0.047277784 | -0.260522079 | 0.355077646 | 9.884902604 | 0.37788343 | 0.718805209 | 0.945042573 | -5.061242546 |
| **A_51_P157255** | Sdc2 | -0.060168995 | -0.451983423 | 0.331645433 | 9.065986481 | -0.377799602 | 0.718864239 | 0.945042573 | -5.061265725 |
| **A_51_P109840** | Vtn | 0.115243345 | -0.635260702 | 0.865747391 | 8.449949503 | 0.377773925 | 0.71888232 | 0.945042573 | -5.061272824 |
| **A_55_P1970846** | Pemt | -0.019921436 | -0.149666107 | 0.109823234 | 7.902015077 | -0.377746354 | 0.718901736 | 0.945042573 | -5.061280446 |
| **A_55_P2065364** | Tdrkh | -0.019572733 | -0.147054809 | 0.107909343 | 7.926430511 | -0.377721333 | 0.718919356 | 0.945042573 | -5.061287363 |

| | | | | | | | | |
|---|---|---|---|---|---|---|---|---|
| **A_55_P2001589** | Klhl8 | 0.02161699 | -0.119182997 | 0.162416977 | 8.963041748 | 0.377712856 | 0.718925325 | 0.945042573 | -5.061289706 |
| **A_55_P1990870** | Mylk3 | 0.048588849 | -0.267923284 | 0.365100981 | 8.26028091 | 0.377672542 | 0.718953715 | 0.945042573 | -5.061300849 |
| **A_55_P2144511** | NA | -0.019155925 | -0.143942912 | 0.105631061 | 8.287966794 | -0.377661749 | 0.718961316 | 0.945042573 | -5.061303832 |
| **A_55_P2243768** | D930030O05Rik | -0.044885561 | -0.337311508 | 0.247540386 | 8.144127437 | -0.377624286 | 0.718987699 | 0.945042573 | -5.061314186 |
| **A_51_P296416** | Orc6 | 0.047186306 | -0.260274645 | 0.354647257 | 11.14392408 | 0.377567998 | 0.719027339 | 0.945042573 | -5.06132974 |
| **A_51_P483280** | Prnp | -0.122949853 | -0.924286154 | 0.678386448 | 13.30212873 | -0.37746993 | 0.719096406 | 0.945042573 | -5.024459403 |
| **A_52_P432153** | Yipf6 | -0.041591135 | -0.312712132 | 0.229529862 | 9.212084858 | -0.377404246 | 0.719142667 | 0.945042573 | -5.06137498 |
| **A_51_P295708** | Ints1 | 0.035431674 | -0.19563281 | 0.266496158 | 9.324328053 | 0.377248602 | 0.719252291 | 0.945042573 | -5.061417962 |
| **A_52_P337126** | Bves | -0.054885671 | -0.412866366 | 0.303095025 | 8.718145261 | -0.377197261 | 0.719288454 | 0.945042573 | -5.061432136 |
| **A_51_P468329** | Psmb7 | 0.044338477 | -0.245051654 | 0.333728608 | 10.58167318 | 0.376934789 | 0.719473341 | 0.945042573 | -5.061504573 |
| **A_51_P209122** | Agr2 | -0.017389063 | -0.130978669 | 0.096200544 | 7.690436907 | -0.376622919 | 0.719693052 | 0.945042573 | -5.06159058 |
| **A_51_P136337** | Galm | 0.021691088 | -0.120013013 | 0.163395188 | 7.890621026 | 0.376589387 | 0.719716677 | 0.945042573 | -5.061599824 |
| **A_55_P2102182** | NA | -0.073010004 | -0.550026208 | 0.404006201 | 9.056255044 | -0.376546314 | 0.719747025 | 0.945042573 | -5.061611697 |
| **A_55_P1994355** | Ptges3 | 0.027180301 | -0.150424136 | 0.204784738 | 13.67714104 | 0.376504012 | 0.719776829 | 0.945042573 | -5.061623355 |
| **A_55_P2005873** | Zfp280d | 0.025532963 | -0.141333287 | 0.192399214 | 9.304743159 | 0.376445295 | 0.719818201 | 0.945042573 | -5.061639536 |
| **A_55_P1960023** | NA | -0.0378922 | -0.285554469 | 0.209770069 | 8.651024334 | -0.376408253 | 0.719844301 | 0.945042573 | -5.061649743 |
| **A_55_P1989578** | NA | -0.06082812 | -0.458420746 | 0.336764506 | 8.639389077 | -0.376387538 | 0.719858896 | 0.945042573 | -5.06165545 |
| **A_51_P122582** | Daglb | -0.028806842 | -0.217161085 | 0.159547401 | 9.52126815 | -0.376261173 | 0.719947938 | 0.945042573 | -5.040610355 |
| **A_52_P553316** | Snrk | -0.03445171 | -0.259750373 | 0.190846953 | 10.26247366 | -0.376202201 | 0.719989493 | 0.945042573 | -5.061706501 |
| **A_55_P2031337** | Trmt2a | 0.030182266 | -0.16723081 | 0.227595341 | 10.80929572 | 0.376136163 | 0.720036029 | 0.945042573 | -5.061724686 |
| **A_55_P2001481** | Pde6c | 0.040324238 | -0.223439193 | 0.30408767 | 7.888627129 | 0.376115085 | 0.720050882 | 0.945042573 | -5.061730489 |
| **A_55_P1970676** | Sh3yl1 | -0.035136139 | -0.26501337 | 0.194741092 | 8.992942619 | -0.376034112 | 0.720107944 | 0.945042573 | -5.061752781 |
| **A_52_P111031** | Pcdh17 | -0.029043335 | -0.219134042 | 0.161047373 | 8.140687012 | -0.375884792 | 0.720213177 | 0.945042573 | -5.061793877 |
| **A_51_P228171** | Cenpp | 0.03883219 | -0.215329528 | 0.292993909 | 8.129736245 | 0.375881467 | 0.72021552 | 0.945042573 | -5.061794791 |
| **A_55_P1988246** | NA | 0.041581629 | -0.23069704 | 0.313860299 | 8.890350179 | 0.375713712 | 0.720333753 | 0.945042573 | -5.061840943 |
| **A_55_P2109354** | Dot1l | -0.032898872 | -0.24832265 | 0.182524906 | 10.7820942 | -0.375713268 | 0.720334065 | 0.945042573 | -5.061841065 |
| **A_51_P221144** | Vwa8 | 0.027065578 | -0.150184364 | 0.20431552 | 9.562143632 | 0.375664681 | 0.720368311 | 0.945042573 | -5.061854428 |
| **A_51_P189082** | Akr1c6 | 0.016249654 | -0.090176117 | 0.122675425 | 7.717002368 | 0.375635368 | 0.720388972 | 0.945042573 | -5.061862489 |
| **A_51_P237383** | Rnase4 | -0.052093055 | -0.393284915 | 0.289098805 | 9.382598003 | -0.375621403 | 0.720398815 | 0.945042573 | -5.061866329 |
| **A_55_P2049572** | Pld2 | -0.05088833 | -0.384231715 | 0.282455056 | 11.75306019 | -0.375574 | 0.720432227 | 0.945042573 | -5.061879364 |
| **A_51_P362209** | Commd10 | 0.028554244 | -0.158491639 | 0.215600128 | 10.00573155 | 0.375570679 | 0.720434568 | 0.945042573 | -5.061880277 |
| **A_55_P2163928** | Rtl8c | -0.026960068 | -0.203563984 | 0.149643848 | 8.195362564 | -0.375569075 | 0.720435699 | 0.945042573 | -5.061880718 |
| **A_55_P1965368** | Ctc1 | 0.030416664 | -0.168835985 | 0.229669313 | 10.76070198 | 0.375557676 | 0.720443734 | 0.945042573 | -5.061883852 |
| **A_52_P154501** | Ube2q2 | -0.047400206 | -0.357970437 | 0.263170026 | 10.2175692 | -0.375482381 | 0.720496807 | 0.945042573 | -5.061904552 |
| **A_55_P2087013** | Hipk1 | -0.016235319 | -0.122621857 | 0.090151218 | 7.862500543 | -0.375442407 | 0.720524985 | 0.945042573 | -5.06191554 |
| **A_55_P2042525** | Apoo | 0.048519043 | -0.269562439 | 0.366600525 | 10.78691529 | 0.375269274 | 0.720647031 | 0.945042573 | -5.061963118 |
| **A_55_P2103682** | NA | -0.033982767 | -0.256773331 | 0.188807797 | 13.72959729 | -0.37525899 | 0.72065428 | 0.945042573 | -5.061965944 |
| **A_55_P2057132** | Scn8a | -0.04569378 | -0.345276847 | 0.253889286 | 11.91976586 | -0.375239882 | 0.72066775 | 0.945042573 | -5.061971193 |
| **A_55_P2002864** | Aifm1 | -0.045943194 | -0.347162492 | 0.255276105 | 11.53102438 | -0.375238638 | 0.720668627 | 0.945042573 | -5.061971535 |
| **A_52_P45738** | Trim35 | 0.01783716 | -0.099119189 | 0.134793508 | 7.786248642 | 0.375207115 | 0.72069085 | 0.945042573 | -5.061980195 |
| **A_55_P2021510** | NA | 0.033250677 | -0.184771855 | 0.251273209 | 8.310133367 | 0.375204706 | 0.720692549 | 0.945042573 | -5.061980857 |
| **A_55_P2186672** | Rps14 | 0.017266394 | -0.095954182 | 0.130486971 | 17.40141228 | 0.375184994 | 0.720706445 | 0.945042573 | -5.061986271 |
| **A_52_P238468** | Gipr | -0.019158281 | -0.144793258 | 0.106476696 | 7.704807402 | -0.375158801 | 0.720724911 | 0.945042573 | -5.061993466 |
| **A_55_P2273705** | NA | -0.051519793 | -0.389473414 | 0.286433829 | 8.720337206 | -0.375047406 | 0.720803444 | 0.945042573 | -5.062024059 |
| **A_55_P2266178** | Il2rb | 0.041106408 | -0.228569686 | 0.310782502 | 10.62791278 | 0.375004295 | 0.720833839 | 0.945042573 | -5.062035896 |
| **A_55_P2033795** | Fabp6 | 0.089358734 | -0.497222656 | 0.675940123 | 8.923607175 | 0.374781278 | 0.720991081 | 0.945042573 | -5.062097112 |
| **A_52_P216427** | Nkx2-1 | -0.015904342 | -0.120352023 | 0.088543338 | 7.64004936 | -0.37461579 | 0.721107771 | 0.945042573 | -5.056956857 |
| **A_52_P279557** | NA | -0.015848329 | -0.11993713 | 0.088240472 | 7.694621799 | -0.374583503 | 0.721130538 | 0.945042573 | -5.062151371 |
| **A_51_P227445** | NA | -0.017850386 | -0.135093972 | 0.099393201 | 7.772062268 | -0.374565416 | 0.721143292 | 0.945042573 | -5.062156331 |
| **A_55_P1956130** | LOC68395 | -0.171235713 | -1.296494125 | 0.954022698 | 12.90525574 | -0.374378795 | 0.721274895 | 0.945042573 | -5.062207503 |

| | | | | | | | | | |
|---|---|---|---|---|---|---|---|---|---|
| **A_55_P2132207** | 1810037I17Rik | -0.026514299 | -0.200764186 | 0.147735588 | 8.718623786 | -0.374349113 | 0.721295828 | 0.945042573 | -5.062215639 |
| **A_55_P2177614** | Srsf3 | 0.034472169 | -0.192111307 | 0.261055645 | 10.28891264 | 0.374291132 | 0.721336719 | 0.945042573 | -5.062231532 |
| **A_52_P610403** | Sf1 | 0.022275196 | -0.12414481 | 0.168695201 | 8.327283038 | 0.374274532 | 0.721348426 | 0.945042573 | -5.062236081 |
| **A_55_P2107192** | NA | -0.058012012 | -0.43937205 | 0.323348026 | 13.78796266 | -0.374241467 | 0.721371745 | 0.945042573 | -5.062245143 |
| **A_51_P262230** | Ggact | 0.049666691 | -0.276848989 | 0.376182372 | 8.799234794 | 0.374222891 | 0.721384846 | 0.945042573 | -5.062250233 |
| **A_51_P189722** | Cabin1 | 0.026659236 | -0.148728642 | 0.202047114 | 9.33321661 | 0.37395323 | 0.72157504 | 0.945042573 | -5.062324103 |
| **A_55_P2136880** | Ppp1r15a | -0.043349496 | -0.328614331 | 0.241915338 | 13.70125412 | -0.373856537 | 0.721643245 | 0.945042573 | -5.062350579 |
| **A_55_P1952743** | NA | 0.02215301 | -0.123718913 | 0.168024933 | 9.802926591 | 0.373620075 | 0.721810049 | 0.945042573 | -5.062415298 |
| **A_55_P1991351** | S100a13 | -0.02874659 | -0.218047912 | 0.160554732 | 12.18139733 | -0.373595688 | 0.721827252 | 0.945042573 | -5.062421971 |
| **A_51_P480233** | Nmral1 | 0.017170306 | -0.095911769 | 0.130252382 | 7.95467525 | 0.373554037 | 0.721856635 | 0.945042573 | -5.062433366 |
| **A_55_P1979242** | Kcne2 | 0.026889419 | -0.150202365 | 0.203981203 | 7.796932286 | 0.373552948 | 0.721857404 | 0.945042573 | -5.062433664 |
| **A_51_P355943** | Mvd | 0.067268567 | -0.375768869 | 0.510306004 | 8.458608713 | 0.37354327 | 0.721864231 | 0.945042573 | -5.062436311 |
| **A_55_P2041673** | Ttc38 | -0.028556615 | -0.216671021 | 0.159557791 | 17.36047119 | -0.373468377 | 0.721917067 | 0.945042573 | -5.062456797 |
| **A_52_P470956** | NA | 0.019172565 | -0.107163984 | 0.145509114 | 15.79877565 | 0.373353622 | 0.721998029 | 0.945042573 | -5.06248818 |
| **A_51_P131164** | Enkur | -0.020127354 | -0.152769849 | 0.112515141 | 7.943251682 | -0.373313041 | 0.72202666 | 0.945042573 | -5.062499275 |
| **A_51_P116487** | Lsm4 | -0.028314019 | -0.214911616 | 0.158283577 | 12.3725631 | -0.373305719 | 0.722031826 | 0.945042573 | -5.062501277 |
| **A_55_P2170309** | Ppfibp2 | -0.051024457 | -0.387363279 | 0.285314365 | 9.860504414 | -0.373224851 | 0.722088883 | 0.945042573 | -5.062523384 |
| **A_55_P2106763** | Pxdc1 | -0.032535258 | -0.247000182 | 0.181929665 | 9.411468573 | -0.373221926 | 0.722090946 | 0.945042573 | -5.062524184 |
| **A_51_P110841** | Mrps35 | -0.034715323 | -0.263577873 | 0.194147227 | 9.930764922 | -0.373177666 | 0.722122176 | 0.945042573 | -5.046247315 |
| **A_55_P1981714** | Rreb1 | -0.07141524 | -0.542228612 | 0.399398132 | 11.63404871 | -0.37317391 | 0.722124825 | 0.945042573 | -5.062537308 |
| **A_51_P210143** | Syne2 | 0.055777626 | -0.311943751 | 0.423499004 | 9.029205077 | 0.373173087 | 0.722125406 | 0.945042573 | -5.062537533 |
| **A_55_P2044119** | Klc1 | -0.035142107 | -0.266870825 | 0.196586612 | 8.949489634 | -0.373092996 | 0.722181918 | 0.945042573 | -5.06255942 |
| **A_55_P2060966** | Mov10 | 0.051894437 | -0.29037023 | 0.394159104 | 11.73661493 | 0.373016375 | 0.722235983 | 0.945042573 | -5.062580355 |
| **A_52_P10254** | Psmc4 | 0.02370337 | -0.132665417 | 0.180072158 | 11.46036548 | 0.372931611 | 0.722295797 | 0.945042573 | -5.06260351 |
| **A_52_P308669** | Cpn1 | 0.04351889 | -0.243684411 | 0.330722191 | 8.120054754 | 0.37278424 | 0.722399794 | 0.945042573 | -5.062643756 |
| **A_51_P317542** | Slc24a2 | 0.016313413 | -0.091380367 | 0.124007192 | 7.707891369 | 0.372669084 | 0.722481062 | 0.945042573 | -5.062675194 |
| **A_66_P124776** | NA | -0.020095429 | -0.152820296 | 0.112629437 | 8.53155817 | -0.372489603 | 0.722607732 | 0.945042573 | -5.062724175 |
| **A_52_P649285** | NA | 0.020834563 | -0.116781695 | 0.158450821 | 9.623618057 | 0.372463576 | 0.722626102 | 0.945042573 | -5.062731276 |
| **A_51_P314418** | Blm | -0.027200581 | -0.206872787 | 0.152471625 | 8.425781658 | -0.372448684 | 0.722636613 | 0.945042573 | -5.062735339 |
| **A_51_P348183** | Tmem141 | 0.027656088 | -0.15503437 | 0.210346545 | 10.54027788 | 0.37242947 | 0.722650174 | 0.945042573 | -5.06274058 |
| **A_55_P2034067** | Myom3 | -0.053462746 | -0.406667334 | 0.299741841 | 11.10453285 | -0.372386641 | 0.722680404 | 0.945042573 | -5.062752263 |
| **A_55_P2168899** | NA | -0.022579634 | -0.171770999 | 0.126611731 | 8.142837166 | -0.372342294 | 0.722711705 | 0.945042573 | -5.062764359 |
| **A_55_P2179791** | NA | -0.018179623 | -0.13831376 | 0.101954514 | 8.02786475 | -0.372295344 | 0.722744844 | 0.945042573 | -5.062777163 |
| **A_66_P113749** | Il2rg | -0.04131354 | -0.314332606 | 0.231705526 | 9.863507737 | -0.372279051 | 0.722756345 | 0.945042573 | -5.062781606 |
| **A_55_P1987264** | Gnb2 | 0.033802273 | -0.189596749 | 0.257201295 | 15.47040009 | 0.372249222 | 0.7227774 | 0.945042573 | -5.06278974 |
| **A_55_P1955209** | NA | 0.036308804 | -0.203702819 | 0.276320427 | 14.64074049 | 0.3721764 | 0.722828803 | 0.945045856 | -5.062809595 |
| **A_55_P1955632** | Trp53i11 | -0.063651441 | -0.484522511 | 0.35721963 | 13.16520237 | -0.372073146 | 0.72290169 | 0.945077225 | -5.062837741 |
| **A_51_P394515** | Tkt | -0.026969962 | -0.205388235 | 0.15144831 | 11.81009272 | -0.371886296 | 0.723033595 | 0.945185741 | -5.062888656 |
| **A_55_P2174987** | Creb3l3 | 0.100309665 | -0.563765318 | 0.764384648 | 10.16555411 | 0.371616359 | 0.723224174 | 0.94536593 | -5.062962169 |
| **A_51_P142744** | Sulf1 | -0.055444871 | -0.422566809 | 0.311677066 | 8.257201224 | -0.371552516 | 0.723269252 | 0.94536593 | -5.062979548 |
| **A_55_P2073253** | NA | 0.020648813 | -0.11616161 | 0.157459236 | 10.3122541 | 0.371317201 | 0.723435409 | 0.945399346 | -5.063043581 |
| **A_55_P2117411** | NA | 0.018529785 | -0.104243185 | 0.141302755 | 10.93203314 | 0.371310112 | 0.723440415 | 0.945399346 | -5.06304551 |
| **A_55_P2366273** | Amn1 | 0.041294568 | -0.232312423 | 0.314901559 | 8.543706723 | 0.371308505 | 0.72344155 | 0.945399346 | -5.063045947 |
| **A_52_P595824** | Cpeb4 | -0.016856613 | -0.128608807 | 0.094895582 | 7.943810768 | -0.371093554 | 0.723593344 | 0.945533784 | -5.063104404 |
| **A_55_P2037957** | NA | -0.021950869 | -0.167571273 | 0.123669535 | 13.38610273 | -0.370850308 | 0.723765135 | 0.945694335 | -5.063170517 |
| **A_55_P2069842** | D5Ertd579e | -0.03339233 | -0.255075374 | 0.188290714 | 8.362776654 | -0.370581222 | 0.723955196 | 0.945878734 | -5.063243607 |
| **A_55_P1958867** | Gm9790 | 0.063313671 | -0.357207153 | 0.483834495 | 14.13413475 | 0.370406969 | 0.724078287 | 0.945975614 | -5.063290911 |
| **A_51_P328089** | Crb3 | 0.081174303 | -0.458167206 | 0.620515811 | 10.07561918 | 0.370274568 | 0.724171819 | 0.946033867 | -5.06332684 |
| **A_55_P1961009** | Apbb1ip | -0.050227878 | -0.384074536 | 0.28361878 | 9.546051067 | -0.370140803 | 0.724266321 | 0.946036531 | -5.063363127 |
| **A_52_P295313** | Wdr46 | -0.031194059 | -0.238534063 | 0.176145944 | 10.84120102 | -0.370133124 | 0.724271746 | 0.946036531 | -5.06336521 |

| | | | | | | | | | |
|---|---|---|---|---|---|---|---|---|---|
| **A_51_P117109** | Ilvbl | -0.021901249 | -0.167509438 | 0.123706941 | 9.907569487 | -0.370043041 | 0.724335391 | 0.946055732 | -5.063389639 |
| **A_55_P1972605** | Adora3 | 0.01633791 | -0.0923534 | 0.12502922 | 7.70876516 | 0.369803348 | 0.724504748 | 0.946212992 | -5.063454615 |
| **A_66_P105075** | Mmp11 | -0.095312437 | -0.729538375 | 0.5389135 | 9.75033592 | -0.369721528 | 0.724562563 | 0.946224565 | -5.063476785 |
| **A_52_P449417** | Vangl1 | -0.032696166 | -0.25033918 | 0.184946848 | 10.37110771 | -0.369590891 | 0.724654876 | 0.946225631 | -5.063512174 |
| **A_51_P159352** | Sub1 | -0.02432195 | -0.186237299 | 0.1375934 | 8.196156101 | -0.369555377 | 0.724679973 | 0.946225631 | -5.063521792 |
| **A_51_P345747** | E130309D02Rik | 0.02869201 | -0.162348442 | 0.219732461 | 9.492350993 | 0.369491793 | 0.724724906 | 0.946225631 | -5.063539011 |
| **A_52_P454295** | Ttn | -0.015866893 | -0.12152755 | 0.089793763 | 7.678742981 | -0.369443274 | 0.724759195 | 0.946225631 | -5.063552148 |
| **A_51_P111488** | Gfod2 | -0.020673705 | -0.158369896 | 0.117022485 | 8.175994654 | -0.369373354 | 0.724808608 | 0.946226232 | -5.063571077 |
| **A_52_P657435** | Psmc5 | -0.025626982 | -0.196459785 | 0.145205821 | 11.73066912 | -0.369058628 | 0.725031047 | 0.946299166 | -5.063656239 |
| **A_52_P548202** | NA | -0.017732278 | -0.135939095 | 0.100474539 | 7.693483745 | -0.369055068 | 0.725033563 | 0.946299166 | -5.063657202 |
| **A_55_P2068977** | Armcx3 | 0.041826248 | -0.237024172 | 0.320676668 | 8.642165602 | 0.369017337 | 0.725060232 | 0.946299166 | -5.063667407 |
| **A_51_P354433** | Ppie | 0.016859614 | -0.095594013 | 0.129313242 | 8.127172837 | 0.368844511 | 0.725182396 | 0.946299166 | -5.063714138 |
| **A_51_P466288** | Frmd5 | -0.018793678 | -0.144151684 | 0.106564328 | 7.935347258 | -0.368832233 | 0.725191075 | 0.946299166 | -5.063717458 |
| **A_51_P512002** | Rer1 | -0.040191127 | -0.308381909 | 0.227999655 | 13.29265861 | -0.368685029 | 0.725295135 | 0.946299166 | -5.063757243 |
| **A_66_P133383** | NA | -0.016176982 | -0.124182425 | 0.091828461 | 7.650428959 | -0.368486029 | 0.725435819 | 0.946299166 | -5.063811004 |
| **A_51_P519756** | Rusc1 | -0.137261503 | -1.053689222 | 0.779166216 | 11.25878215 | -0.368484901 | 0.725436617 | 0.946299166 | -5.063811309 |
| **A_55_P2077816** | Hmgxb3 | 0.04211556 | -0.239228041 | 0.323459162 | 12.17452214 | 0.368277092 | 0.725583542 | 0.946299166 | -5.063867421 |
| **A_52_P326657** | Fam167b | -0.024688942 | -0.189653598 | 0.140275713 | 8.311567678 | -0.368197423 | 0.725639873 | 0.946299166 | -5.063888925 |
| **A_51_P429335** | Prss16 | -0.02886237 | -0.221713522 | 0.163988782 | 9.098900519 | -0.368195872 | 0.72564097 | 0.946299166 | -5.063889344 |
| **A_52_P14526** | Zyg11b | -0.02606194 | -0.200214978 | 0.148091098 | 8.565636598 | -0.368166993 | 0.725661389 | 0.946299166 | -5.063897138 |
| **A_51_P421664** | Tmem123 | -0.036836245 | -0.28303221 | 0.209359721 | 12.10855386 | -0.368098105 | 0.7257101 | 0.946299166 | -5.063915727 |
| **A_52_P425734** | Afm | -0.087970645 | -0.675968151 | 0.500026861 | 8.863024287 | -0.368070878 | 0.725729353 | 0.946299166 | -5.063923073 |
| **A_65_P13270** | Csnk1g1 | -0.019962013 | -0.15339271 | 0.113468684 | 8.956063894 | -0.36805926 | 0.725737567 | 0.946299166 | -5.063926207 |
| **A_55_P2137186** | Rps25 | -0.031153577 | -0.239396039 | 0.177088884 | 15.88362606 | -0.368050828 | 0.72574353 | 0.946299166 | -5.063928482 |
| **A_51_P202623** | Mterf2 | -0.042670297 | -0.327986367 | 0.242645773 | 9.382899703 | -0.367932876 | 0.725826938 | 0.946299166 | -5.0639603 |
| **A_51_P295967** | Proc | 0.101765127 | -0.578829806 | 0.782360061 | 11.32504781 | 0.367857345 | 0.725880351 | 0.946299166 | -5.063980669 |
| **A_55_P1976864** | Fbxl20 | 0.039424213 | -0.224382902 | 0.303231327 | 10.13792209 | 0.367659412 | 0.72602033 | 0.946299166 | -5.06403403 |
| **A_66_P125173** | Aip | 0.016856235 | -0.095941606 | 0.129654077 | 9.075695246 | 0.367645251 | 0.726030345 | 0.946299166 | -5.064037846 |
| **A_51_P492482** | Cstf2t | -0.022919305 | -0.176362918 | 0.130524308 | 10.2770247 | -0.367469915 | 0.726154354 | 0.946299166 | -5.06408509 |
| **A_55_P1970775** | Hdac1 | 0.033140224 | -0.188835489 | 0.255115936 | 13.18478769 | 0.36729849 | 0.726275606 | 0.946299166 | -5.064131261 |
| **A_52_P162509** | Ahcyl2 | -0.046991586 | -0.361839095 | 0.267855923 | 9.069366473 | -0.367188443 | 0.726353449 | 0.946299166 | -5.064160889 |
| **A_55_P2064928** | Nebl | -0.016051566 | -0.123607321 | 0.091504189 | 7.638692616 | -0.367157937 | 0.726375029 | 0.946299166 | -5.064169101 |
| **A_55_P2143311** | Pdxk | 0.03147047 | -0.179408133 | 0.242349072 | 8.381497143 | 0.367146909 | 0.726382829 | 0.946299166 | -5.064172069 |
| **A_55_P2105858** | Atf5 | 0.117549534 | -0.670159178 | 0.905258246 | 14.01605947 | 0.367133848 | 0.726392068 | 0.946299166 | -5.064175585 |
| **A_52_P356119** | Hus1 | -0.018882695 | -0.145425697 | 0.107660307 | 8.218440002 | -0.367108974 | 0.726409664 | 0.946299166 | -5.064182279 |
| **A_52_P44030** | Exoc3 | 0.03390834 | -0.193400929 | 0.261217609 | 8.617516316 | 0.366993649 | 0.726491247 | 0.946299166 | -5.064213313 |
| **A_51_P473240** | Zfp426 | 0.034144665 | -0.194784785 | 0.263074114 | 10.3917037 | 0.366936024 | 0.726532014 | 0.946299166 | -5.064228817 |
| **A_55_P1994586** | Itsn1 | -0.01859367 | -0.143294753 | 0.106107412 | 8.197926977 | -0.366829345 | 0.726607485 | 0.946299166 | -5.064257512 |
| **A_55_P2154416** | Car15 | 0.022491587 | -0.128368976 | 0.17335215 | 8.246547682 | 0.366786667 | 0.726637679 | 0.946299166 | -5.06426899 |
| **A_52_P218976** | Cyld | -0.026401584 | -0.203498772 | 0.150695604 | 8.340390742 | -0.366764656 | 0.726653252 | 0.946299166 | -5.064274909 |
| **A_55_P2017914** | Csf1 | -0.082554157 | -0.636410722 | 0.471302407 | 9.245762221 | -0.366699911 | 0.72669906 | 0.946299166 | -5.064292317 |
| **A_55_P2006808** | Ntrk3 | 0.017879889 | -0.10209179 | 0.137851567 | 7.709193932 | 0.366652999 | 0.726732251 | 0.946299166 | -5.064304929 |
| **A_55_P1998872** | Slitrk5 | -0.016020492 | -0.12353513 | 0.091494146 | 7.652337348 | -0.366587303 | 0.726778734 | 0.946299166 | -5.064322589 |
| **A_51_P364210** | Ms4a3 | -0.015521774 | -0.119715148 | 0.088671599 | 7.653298137 | -0.366497009 | 0.726842623 | 0.946299166 | -5.064346855 |
| **A_55_P2059342** | Lamp1 | -0.019933004 | -0.153741782 | 0.113875773 | 16.33628915 | -0.366485943 | 0.726850453 | 0.946299166 | -5.064349829 |
| **A_55_P2014229** | Sema4a | 0.138202448 | -0.789752487 | 1.066157382 | 10.42617649 | 0.366402149 | 0.726909745 | 0.946299166 | -5.064372343 |
| **A_55_P2171185** | NA | -0.029494898 | -0.22756391 | 0.168574114 | 10.74064435 | -0.366352812 | 0.726944657 | 0.946299166 | -5.064385597 |
| **A_52_P64364** | Ikbkb | -0.02622898 | -0.20239084 | 0.14993288 | 8.391779275 | -0.366301489 | 0.726980975 | 0.946299166 | -5.064399382 |
| **A_55_P1959953** | Helz2 | -0.049588963 | -0.382665745 | 0.283487819 | 10.0092139 | -0.366277152 | 0.726998196 | 0.946299166 | -5.064405919 |
| **A_55_P2147126** | Zfp712 | 0.020577636 | -0.117638291 | 0.158793563 | 8.267673893 | 0.366274388 | 0.727000153 | 0.946299166 | -5.064406661 |

| | | | | | | | | | |
|---|---|---|---|---|---|---|---|---|---|
| **A_55_P2029046** | Fgd3 | -0.025637536 | -0.197844566 | 0.146569495 | 9.115417304 | -0.366264277 | 0.727007308 | 0.946299166 | -5.064409377 |
| **A_55_P1971074** | Atp11a | 0.041733479 | -0.23866653 | 0.322133489 | 13.55030387 | 0.366164075 | 0.727078217 | 0.946299166 | -5.064436283 |
| **A_55_P2090535** | Thrap3 | 0.018264587 | -0.104452382 | 0.140981555 | 8.089619361 | 0.366162949 | 0.727079014 | 0.946299166 | -5.064436586 |
| **A_55_P2107502** | Fat1 | 0.041950954 | -0.239954078 | 0.323855987 | 11.37887327 | 0.366107124 | 0.72711852 | 0.946299166 | -5.064451573 |
| **A_55_P2179869** | NA | 0.041927848 | -0.239888577 | 0.323744273 | 12.0750518 | 0.366020523 | 0.727179809 | 0.946299166 | -5.064474819 |
| **A_51_P218774** | Rgs10 | -0.035465758 | -0.273954768 | 0.203023251 | 9.000332105 | -0.365855882 | 0.727296333 | 0.946299166 | -5.064518997 |
| **A_51_P162162** | Inmt | 0.190198966 | -1.088817583 | 1.469215514 | 9.013290983 | 0.365848392 | 0.727301634 | 0.946299166 | -5.064521007 |
| **A_55_P2171746** | NA | -0.021668076 | -0.167383774 | 0.124047622 | 10.72727205 | -0.365833252 | 0.72731235 | 0.946299166 | -5.064525068 |
| **A_55_P2076580** | Acaa1a | -0.063143749 | -0.48788227 | 0.361594772 | 14.71884753 | -0.365744562 | 0.727375123 | 0.946317141 | -5.064548858 |
| **A_55_P2074843** | Ccdc122 | -0.027845919 | -0.215204515 | 0.159512677 | 8.49469638 | -0.365642853 | 0.727447115 | 0.946332163 | -5.064576133 |
| **A_52_P88808** | Vps37a | 0.030550363 | -0.175079376 | 0.236180102 | 10.0221659 | 0.365510292 | 0.727540949 | 0.946332163 | -5.064611671 |
| **A_52_P504361** | Naglu | -0.106841166 | -0.826046337 | 0.612364006 | 10.90327097 | -0.365472732 | 0.727567537 | 0.946332163 | -5.064621738 |
| **A_51_P229893** | Cts6 | -0.030134766 | -0.23301144 | 0.172741909 | 17.09153356 | -0.365430559 | 0.727597391 | 0.946332163 | -5.06463304 |
| **A_51_P376347** | Hebp1 | 0.094665145 | -0.542790278 | 0.732120567 | 11.00028197 | 0.365350281 | 0.72765422 | 0.946332163 | -5.064654551 |
| **A_55_P2021011** | Sfi1 | 0.062947539 | -0.360971069 | 0.486866148 | 9.673617528 | 0.365313264 | 0.727680425 | 0.946332163 | -5.064664469 |
| **A_55_P2135098** | Rplp0 | -0.018893003 | -0.146274893 | 0.108488888 | 16.74567755 | -0.364890423 | 0.727979794 | 0.946569335 | -5.064777688 |
| **A_55_P2128526** | Pfn2 | 0.036205393 | -0.207914865 | 0.280325651 | 9.710537208 | 0.364870378 | 0.727993987 | 0.946569335 | -5.064783052 |
| **A_51_P362959** | Fbxo36 | -0.041924907 | -0.324627047 | 0.240777234 | 9.842951066 | -0.364848169 | 0.728009713 | 0.946569335 | -5.064788995 |
| **A_51_P364421** | Vamp7 | -0.047045336 | -0.364430908 | 0.270340236 | 8.871282359 | -0.364668758 | 0.728136753 | 0.946576165 | -5.064836992 |
| **A_52_P537545** | Smpd3 | 0.041010133 | -0.235682897 | 0.317703162 | 8.044516926 | 0.364638162 | 0.728158419 | 0.946576165 | -5.064845175 |
| **A_52_P343343** | Rpl7l1 | 0.019399499 | -0.111489572 | 0.150288571 | 8.473440298 | 0.364633273 | 0.728161881 | 0.946576165 | -5.064846482 |
| **A_55_P2038117** | Tmem104 | -0.111432274 | -0.863440618 | 0.640576069 | 11.06530204 | -0.364550332 | 0.728220615 | 0.946588854 | -5.064868661 |
| **A_51_P200561** | Shtn1 | -0.083130048 | -0.644428974 | 0.478168878 | 10.4266124 | -0.364361919 | 0.728354046 | 0.946609845 | -5.064919026 |
| **A_66_P115693** | Rnf207 | -0.020258088 | -0.157051647 | 0.116535472 | 7.879764437 | -0.364335894 | 0.728372477 | 0.946609845 | -5.064925981 |
| **A_55_P2030721** | Ankle1 | -0.01676724 | -0.129993626 | 0.096459146 | 7.73212623 | -0.36432007 | 0.728383684 | 0.946609845 | -5.06493021 |
| **A_55_P2074656** | Padi2 | -0.343097132 | -2.661454289 | 1.975260025 | 12.03240991 | -0.364087617 | 0.728548321 | 0.946727629 | -5.064992307 |
| **A_51_P224983** | Cln8 | 0.037692942 | -0.217027403 | 0.292413288 | 11.91389779 | 0.3640538 | 0.728572274 | 0.946727629 | -5.065001338 |
| **A_55_P2007470** | Pdgfa | 0.062404077 | -0.359502721 | 0.484310875 | 14.32236516 | 0.363886216 | 0.728690978 | 0.946818225 | -5.065046079 |
| **A_66_P121459** | Cenpa | 0.033425894 | -0.192680451 | 0.25953224 | 8.27759448 | 0.363696773 | 0.728825176 | 0.946928939 | -5.065096633 |
| **A_51_P478895** | Ppp6r1 | 0.029259112 | -0.168703112 | 0.227221336 | 9.13561488 | 0.363620183 | 0.728879434 | 0.946935783 | -5.065117065 |
| **A_51_P447976** | Tent5c | -0.037040658 | -0.287701092 | 0.213619775 | 11.95980268 | -0.363548263 | 0.728930385 | 0.94693833 | -5.065136247 |
| **A_55_P1975947** | Fam234b | -0.040624545 | -0.315790936 | 0.234541846 | 11.66740494 | -0.363213754 | 0.729167385 | 0.947108229 | -5.065225417 |
| **A_51_P371001** | Tm4sf4 | 0.015630103 | -0.090255467 | 0.121515673 | 7.825785806 | 0.363156839 | 0.729207713 | 0.947108229 | -5.065240582 |
| **A_52_P39418** | Ccpg1 | 0.030441522 | -0.175793645 | 0.236676689 | 8.684535479 | 0.363138919 | 0.72922041 | 0.947108229 | -5.065245356 |
| **A_52_P240152** | Snx27 | -0.053325727 | -0.414685587 | 0.308034133 | 10.66216884 | -0.363049663 | 0.729283656 | 0.947108229 | -5.065269131 |
| **A_66_P138999** | NA | -0.021223994 | -0.165104716 | 0.122656727 | 8.034971916 | -0.362905605 | 0.729385739 | 0.947108229 | -5.065307493 |
| **A_51_P336833** | Fabp4 | -0.192012059 | -1.494346667 | 1.110322549 | 13.50599813 | -0.362722993 | 0.72951515 | 0.947108229 | -5.065356101 |
| **A_52_P487436** | Nags | 0.015581872 | -0.090129663 | 0.121293408 | 7.694696277 | 0.362632244 | 0.729579464 | 0.947108229 | -5.065380248 |
| **A_55_P2077153** | Slc27a2 | 0.171603365 | -0.992629399 | 1.335836129 | 10.34798663 | 0.362622833 | 0.729586134 | 0.947108229 | -5.065382752 |
| **A_51_P300297** | Srek1ip1 | -0.043047045 | -0.335120433 | 0.249026343 | 9.438332489 | -0.362593922 | 0.729606624 | 0.947108229 | -5.065390444 |
| **A_55_P1969776** | Pdhb | -0.065141833 | -0.507231104 | 0.376947438 | 12.88146458 | -0.362509308 | 0.729666594 | 0.947108229 | -5.065412951 |
| **A_51_P506937** | Mrps12 | -0.028881594 | -0.224890892 | 0.167127703 | 12.29464796 | -0.362504711 | 0.729669852 | 0.947108229 | -5.065414174 |
| **A_51_P101228** | Gmeb2 | -0.015553378 | -0.12111377 | 0.090007015 | 8.136689951 | -0.362487372 | 0.729682142 | 0.947108229 | -5.065418785 |
| **A_55_P2339650** | Hnrnpl | 0.017167406 | -0.099354626 | 0.133689438 | 8.336640601 | 0.362464758 | 0.729698169 | 0.947108229 | -5.065424799 |
| **A_52_P480351** | Loxl2 | 0.059505901 | -0.344461419 | 0.463473221 | 11.07480462 | 0.362395639 | 0.72974716 | 0.947108229 | -5.065443179 |
| **A_51_P304510** | BC017158 | -0.025255215 | -0.196864316 | 0.146353886 | 10.94788742 | -0.362059473 | 0.729985448 | 0.94721137 | -5.065532524 |
| **A_55_P1969745** | Pitpnm1 | -0.068512316 | -0.534115511 | 0.397090879 | 10.54719017 | -0.362011077 | 0.730019756 | 0.94721137 | -5.06554538 |
| **A_55_P2134012** | Oxa1l | 0.043879184 | -0.254320072 | 0.34207844 | 11.73799889 | 0.362010484 | 0.730020176 | 0.94721137 | -5.065545538 |
| **A_51_P267388** | Dxo | -0.023443573 | -0.182774422 | 0.135887276 | 8.827714827 | -0.361987104 | 0.730036751 | 0.94721137 | -5.065551748 |
| **A_51_P260740** | Pcdh7 | -0.018880845 | -0.147219227 | 0.109457537 | 7.766438683 | -0.36193787 | 0.730071654 | 0.94721137 | -5.065564824 |

| | | | | | | | | | |
|---|---|---|---|---|---|---|---|---|---|
| **A_51_P254299** | Dlk2 | 0.048408766 | -0.280792363 | 0.377609895 | 9.023224011 | 0.361769424 | 0.730191073 | 0.947261411 | -5.06560955 |
| **A_51_P393862** | Spryd4 | -0.021392047 | -0.166877215 | 0.12409312 | 8.884999759 | -0.361745215 | 0.730208237 | 0.947261411 | -5.065615976 |
| **A_55_P2462358** | Fam118b | -0.038977997 | -0.304255279 | 0.226299285 | 11.3007289 | -0.36148362 | 0.730393713 | 0.947438433 | -5.065685391 |
| **A_51_P327491** | Apoa4 | 0.187001895 | -1.086450528 | 1.460454319 | 10.52221912 | 0.361270456 | 0.730544866 | 0.947570911 | -5.06574192 |
| **A_55_P1980041** | Kpna6 | 0.035184053 | -0.204619878 | 0.274987984 | 9.81659282 | 0.360959714 | 0.730765234 | 0.947694161 | -5.065824269 |
| **A_55_P2133165** | Wwc1 | -0.063863887 | -0.499282022 | 0.371554248 | 11.19554727 | -0.360842762 | 0.730848181 | 0.947694161 | -5.065855246 |
| **A_65_P16081** | NA | -0.085470538 | -0.668362322 | 0.497421247 | 9.152687964 | -0.360742794 | 0.730919084 | 0.947694161 | -5.065881716 |
| **A_55_P1978615** | Rpl17 | 0.028924943 | -0.168448059 | 0.226297944 | 15.19478127 | 0.36054039 | 0.731062651 | 0.947694161 | -5.065935288 |
| **A_51_P116755** | Cap2 | 0.016511659 | -0.096163528 | 0.129186846 | 7.735413476 | 0.360521849 | 0.731075803 | 0.947694161 | -5.044751978 |
| **A_66_P103654** | Asxl3 | 0.020055271 | -0.11682333 | 0.156933873 | 7.769612187 | 0.360464207 | 0.731116691 | 0.947694161 | -5.065955445 |
| **A_51_P319453** | Zfp143 | -0.022364352 | -0.175044789 | 0.130316085 | 8.486963593 | -0.360364569 | 0.731187371 | 0.947694161 | -5.065981802 |
| **A_51_P226142** | Dennd6a | 0.053582668 | -0.312253496 | 0.419418831 | 11.41730125 | 0.360335342 | 0.731208105 | 0.947694161 | -5.065989532 |
| **A_66_P109941** | Mamdc4 | 0.024071343 | -0.140298497 | 0.188441183 | 7.902270484 | 0.360285974 | 0.731243127 | 0.947694161 | -5.066002587 |
| **A_55_P2097372** | Tctex1d4 | -0.016774104 | -0.13132335 | 0.097775141 | 7.757200394 | -0.360260189 | 0.731261419 | 0.947694161 | -5.066009406 |
| **A_51_P479818** | Lonrf3 | -0.070467499 | -0.551701191 | 0.410766192 | 11.21339201 | -0.360248348 | 0.731269819 | 0.947694161 | -5.066012537 |
| **A_52_P673499** | Shmt1 | -0.01739513 | -0.136276135 | 0.101485876 | 7.908370159 | -0.359984965 | 0.731456678 | 0.947694161 | -5.066082154 |
| **A_55_P1964842** | Opa1 | 0.02237251 | -0.130558161 | 0.175303181 | 10.37848894 | 0.359906153 | 0.731512597 | 0.947694161 | -5.066102976 |
| **A_55_P1998892** | Smox | -0.052498353 | -0.411445163 | 0.306448458 | 10.74706321 | -0.359819536 | 0.731574054 | 0.947694161 | -5.066125855 |
| **A_55_P2033610** | Fastkd3 | 0.055829063 | -0.325931325 | 0.437589451 | 10.07374321 | 0.359781343 | 0.731601154 | 0.947694161 | -5.066135942 |
| **A_55_P2060604** | Gm5797 | 0.054727057 | -0.319625322 | 0.429079435 | 10.00226763 | 0.359658772 | 0.731688128 | 0.947694161 | -5.066168306 |
| **A_55_P2146254** | Ifitm1 | 0.111232999 | -0.649944601 | 0.8724106 | 9.973653984 | 0.359514831 | 0.73179027 | 0.947694161 | -5.0662063 |
| **A_51_P261059** | Cpne2 | -0.019213474 | -0.150697136 | 0.112270188 | 8.087496091 | -0.359503619 | 0.731798227 | 0.947694161 | -5.066209259 |
| **A_55_P2071626** | C530008M17Rik | -0.015124608 | -0.118632943 | 0.088383727 | 7.699912109 | -0.359482679 | 0.731813086 | 0.947694161 | -5.066214784 |
| **A_52_P11441** | Rab6b | -0.031652731 | -0.248306291 | 0.185000829 | 12.86692739 | -0.35943016 | 0.731850357 | 0.947694161 | -5.066228642 |
| **A_55_P2062190** | Gstm1 | 0.091727798 | -0.53612354 | 0.719579136 | 11.04601913 | 0.35942916 | 0.731851067 | 0.947694161 | -5.066228906 |
| **A_55_P1978441** | Kif3a | -0.040824048 | -0.32037808 | 0.238729983 | 10.72805684 | -0.359268776 | 0.731964889 | 0.947694161 | -5.066271214 |
| **A_51_P135517** | Coch | -0.31637314 | -2.482833684 | 1.850087404 | 10.49790912 | -0.359267543 | 0.731965764 | 0.947694161 | -5.066271539 |
| **A_52_P644690** | Flt3l | 0.017251294 | -0.100902575 | 0.135405163 | 8.057660514 | 0.359205434 | 0.732009843 | 0.947694161 | -5.066287918 |
| **A_55_P2151011** | NA | -0.015322689 | -0.120279325 | 0.089633947 | 7.770971783 | -0.35916519 | 0.732038406 | 0.947694161 | -5.066298529 |
| **A_55_P2082180** | Tnfrsf9 | -0.048176073 | -0.378199807 | 0.281847661 | 13.14492192 | -0.35913306 | 0.732061211 | 0.947694161 | -5.066307 |
| **A_52_P242973** | Sec11a | -0.023702458 | -0.186094885 | 0.138689969 | 13.774279 | -0.359084598 | 0.732095606 | 0.947694161 | -5.066319776 |
| **A_55_P2076462** | Lnx1 | 0.033686639 | -0.197135794 | 0.264509073 | 10.13753573 | 0.359044962 | 0.732123739 | 0.947694161 | -5.066330224 |
| **A_51_P306017** | Dll1 | -0.027260965 | -0.214117729 | 0.159595799 | 8.623382076 | -0.358923234 | 0.73221014 | 0.947694161 | -5.066362303 |
| **A_52_P427818** | Zscan18 | -0.022189113 | -0.174290081 | 0.129911855 | 8.315169602 | -0.358903033 | 0.732224479 | 0.947694161 | -5.066367626 |
| **A_66_P126320** | Mrpl46 | 0.037495948 | -0.219645681 | 0.294637578 | 10.01860107 | 0.358741082 | 0.732339438 | 0.947694161 | -5.066410288 |
| **A_51_P336874** | Atp6v0e | -0.034132104 | -0.268253411 | 0.199989203 | 15.02868956 | -0.358666888 | 0.732392106 | 0.947694161 | -5.066429826 |
| **A_55_P2211319** | Fgf18 | 0.057695097 | -0.338055293 | 0.453445487 | 8.716648918 | 0.358663118 | 0.732394783 | 0.947694161 | -5.066430819 |
| **A_55_P2003309** | NA | 0.036693671 | -0.215049006 | 0.288436348 | 13.99766999 | 0.358594382 | 0.732443578 | 0.947694161 | -5.066448917 |
| **A_51_P348804** | Amotl2 | -0.038982595 | -0.306429757 | 0.228464567 | 10.76248432 | -0.358593089 | 0.732444496 | 0.947694161 | -5.066449257 |
| **A_55_P1991516** | E4f1 | 0.024023615 | -0.140815891 | 0.18886312 | 9.519659905 | 0.358547102 | 0.732477143 | 0.947694161 | -5.066461363 |
| **A_55_P2099952** | Car1 | 0.015921738 | -0.093330467 | 0.125173943 | 7.652027381 | 0.358533243 | 0.732486981 | 0.947694161 | -5.066465011 |
| **A_52_P468343** | Bdh1 | -0.015394095 | -0.12103236 | 0.09024417 | 7.80642045 | -0.358510659 | 0.732503014 | 0.947694161 | -5.066470956 |
| **A_51_P513959** | NA | 0.029490854 | -0.173007836 | 0.231989543 | 9.077244322 | 0.358289676 | 0.732659904 | 0.947724958 | -5.066529104 |
| **A_55_P2022369** | Lbh | 0.016818585 | -0.09867473 | 0.1323119 | 8.529773705 | 0.358262839 | 0.732678958 | 0.947724958 | -5.066536163 |
| **A_66_P106363** | Itgb1bp1 | 0.032469535 | -0.190510957 | 0.255450027 | 9.874721526 | 0.358243535 | 0.732692664 | 0.947724958 | -5.06654124 |
| **A_52_P651948** | Fam229b | -0.01718456 | -0.135219576 | 0.100850456 | 7.772902333 | -0.358176189 | 0.73274048 | 0.947724958 | -5.066558952 |
| **A_55_P1959550** | Carmil2 | 0.017677667 | -0.103759377 | 0.139114711 | 8.516859672 | 0.358131833 | 0.732771975 | 0.947724958 | -5.066570616 |
| **A_66_P100780** | Katnbl1 | 0.037170188 | -0.218248361 | 0.292588736 | 10.49311614 | 0.358023458 | 0.732848927 | 0.947761067 | -5.066599109 |
| **A_55_P1960916** | Egln3 | 0.222450553 | -1.306769265 | 1.751670372 | 9.518350288 | 0.357876133 | 0.732953542 | 0.947831375 | -5.06663783 |
| **A_52_P286360** | Otc | 0.022622708 | -0.13292469 | 0.178170106 | 7.758077528 | 0.357808787 | 0.733001365 | 0.947831375 | -5.066655524 |

| | | | | | | | | | |
|---|---|---|---|---|---|---|---|---|---|
| **A_55_P2006261** | Krt15 | 0.344927178 | -2.027729435 | 2.717583791 | 9.329754724 | 0.35765285 | 0.733112106 | 0.947860283 | -5.066696485 |
| **A_52_P252737** | Rangap1 | 0.020785511 | -0.122209145 | 0.163780167 | 9.587318561 | 0.357610335 | 0.733142298 | 0.947860283 | -5.066707649 |
| **A_55_P2408330** | Rbm33 | -0.028920535 | -0.227902732 | 0.170061662 | 8.408631047 | -0.357570153 | 0.733170836 | 0.947860283 | -5.0667182 |
| **A_55_P2091350** | NA | -0.040350524 | -0.318152785 | 0.237451736 | 14.27043398 | -0.357340771 | 0.733333751 | 0.947977813 | -5.066778408 |
| **A_66_P109595** | Ube4b | -0.034421612 | -0.271429297 | 0.202586073 | 10.59113029 | -0.357304049 | 0.733359834 | 0.947977813 | -5.066788043 |
| **A_55_P2324976** | 5033406O09Rik | 0.02251547 | -0.132545811 | 0.177576751 | 8.621132857 | 0.357229085 | 0.733413081 | 0.947983244 | -5.06680771 |
| **A_55_P1960451** | Thsd7a | -0.098699614 | -0.778794969 | 0.581395742 | 8.320152829 | -0.3570383 | 0.733548601 | 0.947989568 | -5.066857744 |
| **A_51_P143468** | Klhl26 | 0.030156987 | -0.17766021 | 0.237974185 | 10.48925504 | 0.357006097 | 0.733571477 | 0.947989568 | -5.066866187 |
| **A_52_P16419** | Gpd1 | 0.017668142 | -0.104090091 | 0.139426376 | 7.865340582 | 0.356994657 | 0.733579603 | 0.947989568 | -5.066869186 |
| **A_51_P495131** | Prl7c1 | -0.086441295 | -0.682223989 | 0.509341398 | 8.976580168 | -0.356946021 | 0.733614153 | 0.947989568 | -5.066881936 |
| **A_55_P2114799** | NA | 0.024759459 | -0.146006944 | 0.195525862 | 8.713645835 | 0.356703923 | 0.733786144 | 0.948148431 | -5.066945376 |
| **A_55_P2090708** | Fam207a | 0.034607317 | -0.204162744 | 0.273377378 | 12.36539726 | 0.356580202 | 0.733874045 | 0.948158494 | -5.066977781 |
| **A_55_P2172470** | Nrgn | 0.11511768 | -0.679182781 | 0.909418141 | 8.821951503 | 0.356554875 | 0.733892039 | 0.948158494 | -5.066984413 |
| **A_55_P2181371** | NA | -0.015926161 | -0.125854353 | 0.094002032 | 8.29978645 | -0.35642747 | 0.733982562 | 0.948212067 | -5.067017769 |
| **A_55_P1953608** | NA | -0.025710292 | -0.203493825 | 0.152073241 | 9.743125174 | -0.355782544 | 0.734440866 | 0.948384793 | -5.067186446 |
| **A_66_P112436** | Prkar2b | -0.025597857 | -0.202625251 | 0.151429537 | 7.963615527 | -0.355739674 | 0.734471334 | 0.948384793 | -5.067197648 |
| **A_55_P2045066** | Eif5a | -0.039127308 | -0.309781679 | 0.231527062 | 11.76239249 | -0.355659237 | 0.734528505 | 0.948384793 | -5.067218664 |
| **A_55_P2085781** | Smim26 | 0.02345219 | -0.138773914 | 0.185678294 | 11.58649516 | 0.355657386 | 0.73452982 | 0.948384793 | -5.067219147 |
| **A_55_P1967538** | Hunk | 0.049298541 | -0.291725254 | 0.390322337 | 10.19864259 | 0.355646509 | 0.734537551 | 0.948384793 | -5.067221989 |
| **A_52_P432580** | NA | -0.049303634 | -0.390374782 | 0.291767514 | 12.62117441 | -0.355633865 | 0.734546538 | 0.948384793 | -5.067225292 |
| **A_55_P1975690** | Bicc1 | 0.066876296 | -0.395779084 | 0.529531677 | 10.2970266 | 0.355618018 | 0.734557801 | 0.948384793 | -5.067229431 |
| **A_55_P2040907** | Gtf2i | 0.015667812 | -0.092730507 | 0.124066131 | 7.723053345 | 0.355594447 | 0.734574555 | 0.948384793 | -5.067235588 |
| **A_66_P117417** | NA | -0.022136024 | -0.175295781 | 0.131023733 | 8.418427394 | -0.355569182 | 0.734592513 | 0.948384793 | -5.067242187 |
| **A_52_P625270** | 1700024G13Rik | 0.015849158 | -0.093819923 | 0.125518239 | 7.736750838 | 0.355542206 | 0.734611687 | 0.948384793 | -5.067249232 |
| **A_55_P2047290** | Ankrd40 | 0.026821078 | -0.158801314 | 0.212443471 | 8.87410852 | 0.355479887 | 0.734655984 | 0.948384793 | -5.067265505 |
| **A_52_P128964** | Nhej1 | 0.025735986 | -0.152417959 | 0.203889932 | 8.696064147 | 0.355397636 | 0.734714449 | 0.948396927 | -5.06728698 |
| **A_51_P481482** | Dram1 | -0.034515839 | -0.273493808 | 0.20446213 | 8.397815937 | -0.355328246 | 0.734763775 | 0.948397263 | -5.067305093 |
| **A_51_P389957** | Rgs14 | 0.015486555 | -0.091784803 | 0.122757912 | 7.829928713 | 0.355173212 | 0.734873985 | 0.948453915 | -5.067345549 |
| **A_51_P125842** | Tmem51 | -0.064969343 | -0.515348223 | 0.385409538 | 10.08499876 | -0.354894786 | 0.735071929 | 0.948453915 | -5.067418164 |
| **A_55_P2000280** | Mthfd1l | -0.048172944 | -0.382119133 | 0.285773245 | 8.254396254 | -0.354891716 | 0.735074112 | 0.948453915 | -5.067418964 |
| **A_52_P124734** | Cybrd1 | 0.0368151 | -0.218398254 | 0.292028453 | 8.233909749 | 0.354888355 | 0.735076501 | 0.948453915 | -5.06741984 |
| **A_55_P2110245** | Fignl1 | 0.055995427 | -0.332193561 | 0.444184416 | 8.986578798 | 0.35487754 | 0.735084191 | 0.948453915 | -5.06742266 |
| **A_55_P2065501** | Rnf7 | 0.040496325 | -0.240265326 | 0.321257977 | 13.01569122 | 0.354851782 | 0.735102504 | 0.948453915 | -5.067429374 |
| **A_55_P2006340** | Uba52 | -0.021908655 | -0.173830991 | 0.130013682 | 16.89125681 | -0.354783364 | 0.735151149 | 0.948453915 | -5.067447208 |
| **A_55_P2007713** | Tspan4 | 0.094872047 | -0.563419665 | 0.75316376 | 10.41407939 | 0.354559437 | 0.735310371 | 0.948456105 | -5.067505553 |
| **A_51_P503654** | Tmem205 | 0.05360974 | -0.31857731 | 0.42579679 | 8.899055462 | 0.354365643 | 0.735448179 | 0.948456105 | -5.067556018 |
| **A_51_P247249** | Alox5 | -0.085556184 | -0.679542749 | 0.50843038 | 8.283784595 | -0.354359393 | 0.735452623 | 0.948456105 | -5.067557645 |
| **A_55_P2073452** | Glp2r | 0.015991147 | -0.095056973 | 0.127039267 | 7.719366779 | 0.354272602 | 0.735514344 | 0.948456105 | -5.067580237 |
| **A_55_P2178553** | Apoa2 | 0.133976883 | -0.79660065 | 1.064554415 | 15.04683378 | 0.354198297 | 0.735567188 | 0.948456105 | -5.067599575 |
| **A_51_P453246** | Uqcc3 | -0.019375749 | -0.153978116 | 0.115226618 | 12.76568838 | -0.354139986 | 0.735608658 | 0.948456105 | -5.067614748 |
| **A_51_P280401** | Tcta | -0.026754506 | -0.212646582 | 0.15913757 | 8.763389252 | -0.354083122 | 0.7356491 | 0.948456105 | -5.067629542 |
| **A_55_P2026214** | NA | 0.05662039 | -0.336786802 | 0.450027581 | 11.75171392 | 0.354078634 | 0.735652292 | 0.948456105 | -5.067630709 |
| **A_51_P184024** | Tsen15 | 0.027499574 | -0.16357788 | 0.218577029 | 10.67209082 | 0.354067194 | 0.735660428 | 0.948456105 | -5.067633685 |
| **A_55_P2155504** | Lck | 0.026424219 | -0.157201964 | 0.210050403 | 8.113924599 | 0.354027274 | 0.73568882 | 0.948456105 | -5.067644069 |
| **A_55_P1971956** | Ift46 | -0.049197597 | -0.39109958 | 0.292704386 | 12.20655642 | -0.354006661 | 0.735703481 | 0.948456105 | -5.067649431 |
| **A_55_P2079520** | Mpp1 | 0.06230444 | -0.370849036 | 0.495457916 | 10.35356361 | 0.353872126 | 0.735799169 | 0.948456105 | -5.067684416 |
| **A_55_P2136338** | Tomm20 | 0.054655699 | -0.325363183 | 0.434674581 | 13.01620863 | 0.353833901 | 0.735826358 | 0.948456105 | -5.067694354 |
| **A_55_P2186648** | Bard1 | 0.021507049 | -0.128138215 | 0.171152313 | 7.962593908 | 0.353579452 | 0.736007353 | 0.948456105 | -5.067760481 |
| **A_51_P364984** | Gabpa | -0.021436132 | -0.17059665 | 0.127724387 | 8.430426715 | -0.353558834 | 0.73602202 | 0.948456105 | -5.067765837 |
| **A_55_P2041634** | Vps51 | 0.026335998 | -0.156954357 | 0.209626353 | 10.65157705 | 0.353491787 | 0.736069715 | 0.948456105 | -5.067783253 |

| | | | | | | | | | |
|---|---|---|---|---|---|---|---|---|---|
| **A_55_P2075814** | Tnni3k | 0.016754225 | -0.099879936 | 0.133388385 | 7.824152212 | 0.353400969 | 0.736134323 | 0.948456105 | -5.067806839 |
| **A_55_P2055411** | Rps7 | -0.020918355 | -0.166575147 | 0.124738437 | 15.32123528 | -0.353318156 | 0.736193238 | 0.948456105 | -5.067828341 |
| **A_55_P1963300** | Vsig10 | -0.029507888 | -0.235019049 | 0.176003272 | 9.634176741 | -0.353241631 | 0.736247682 | 0.948456105 | -5.067848206 |
| **A_51_P496381** | Ptk7 | -0.030183527 | -0.240413244 | 0.18004619 | 10.15939926 | -0.353219805 | 0.73626321 | 0.948456105 | -5.067853871 |
| **A_55_P2010622** | Sec16b | 0.059871535 | -0.357277217 | 0.477020287 | 8.255122749 | 0.353100725 | 0.736347933 | 0.948456105 | -5.067884773 |
| **A_55_P2186572** | Lypla2 | -0.031026293 | -0.247264464 | 0.185211879 | 13.35296258 | -0.352993483 | 0.736424237 | 0.948456105 | -5.067912594 |
| **A_52_P162099** | Ckap2 | -0.016509963 | -0.13160922 | 0.098589293 | 7.756483872 | -0.352892766 | 0.736495901 | 0.948456105 | -5.067938715 |
| **A_52_P407692** | H2afj | -0.03861857 | -0.307947735 | 0.230710594 | 14.82143115 | -0.352762141 | 0.736588851 | 0.948456105 | -5.067972583 |
| **A_66_P134542** | Anln | 0.047069099 | -0.281195255 | 0.375333453 | 8.794332629 | 0.352761614 | 0.736589226 | 0.948456105 | -5.06797272 |
| **A_51_P404077** | Fzd2 | 0.066420898 | -0.39683943 | 0.529681226 | 8.663504461 | 0.352735196 | 0.736608024 | 0.948456105 | -5.067979568 |
| **A_55_P2114318** | NA | -0.102257979 | -0.815559583 | 0.611043624 | 10.96485472 | -0.352690019 | 0.736640173 | 0.948456105 | -5.067991277 |
| **A_55_P2029687** | Hmox1 | -0.015511467 | -0.123725411 | 0.092702478 | 8.285450697 | -0.352645869 | 0.736671591 | 0.948456105 | -5.068002719 |
| **A_55_P2018417** | Osbpl3 | -0.031543353 | -0.251674393 | 0.188587687 | 9.813299424 | -0.352529721 | 0.736754247 | 0.948456105 | -5.068032814 |
| **A_51_P220162** | Notch3 | -0.048648262 | -0.388383587 | 0.291087062 | 9.335979224 | -0.352286329 | 0.736927468 | 0.948456105 | -5.068095849 |
| **A_55_P1976805** | Isca2 | -0.025978839 | -0.207415355 | 0.155457678 | 9.604419286 | -0.352260701 | 0.736945709 | 0.948456105 | -5.068102484 |
| **A_51_P235705** | Tsfm | -0.02713819 | -0.216700411 | 0.162424031 | 8.801831215 | -0.352207218 | 0.736983775 | 0.948456105 | -5.068116328 |
| **A_55_P2021275** | Iqca | 0.020691324 | -0.123845013 | 0.165227661 | 7.757855351 | 0.352192747 | 0.736994075 | 0.948456105 | -5.068120074 |
| **A_55_P2174632** | Rpl7 | -0.021880357 | -0.174737912 | 0.130977198 | 15.94044138 | -0.352157268 | 0.737019327 | 0.948456105 | -5.068129257 |
| **A_55_P2049597** | Tpp2 | -0.036153586 | -0.288772159 | 0.216464987 | 10.22561436 | -0.352091281 | 0.737066296 | 0.948456105 | -5.068146333 |
| **A_51_P449911** | NA | 0.026043251 | -0.156059305 | 0.208145807 | 11.81572957 | 0.351842517 | 0.737243371 | 0.948456105 | -5.068210683 |
| **A_51_P382970** | Itga9 | -0.027825738 | -0.222434774 | 0.166783298 | 8.622208191 | -0.351765181 | 0.737298424 | 0.948456105 | -5.068230679 |
| **A_52_P589568** | Foxo6 | 0.033250103 | -0.199386428 | 0.265886635 | 8.176764797 | 0.351628648 | 0.737395622 | 0.948456105 | -5.068265972 |
| **A_55_P1992099** | Ptprj | -0.026129333 | -0.208946308 | 0.156687642 | 9.620325167 | -0.35162599 | 0.737397515 | 0.948456105 | -5.068266659 |
| **A_52_P168575** | Fam169b | 0.037987433 | -0.227808795 | 0.303783661 | 8.230132845 | 0.351609257 | 0.737409427 | 0.948456105 | -5.068270983 |
| **A_51_P345340** | Upk3bl | 0.441562612 | -2.648881614 | 3.532006838 | 9.998532974 | 0.351512317 | 0.737478443 | 0.948456105 | -5.068296032 |
| **A_51_P256827** | S100a8 | -0.255939342 | -2.047239114 | 1.535360429 | 12.82448401 | -0.351510209 | 0.737479944 | 0.948456105 | -5.068296576 |
| **A_55_P2061428** | Fads6 | 0.053081697 | -0.318456355 | 0.42461975 | 9.122195956 | 0.351488131 | 0.737495662 | 0.948456105 | -5.06830228 |
| **A_55_P2156490** | Gm3146 | 0.076694217 | -0.460143462 | 0.613531897 | 9.28880428 | 0.35147043 | 0.737508264 | 0.948456105 | -5.068306853 |
| **A_52_P353478** | Scn7a | 0.016035931 | -0.096226638 | 0.128298501 | 7.647008575 | 0.351421536 | 0.737543076 | 0.948456105 | -5.068319483 |
| **A_55_P2082425** | Mtln | 0.075118752 | -0.451002266 | 0.60123977 | 10.50472654 | 0.351262569 | 0.73765626 | 0.948456105 | -5.068360536 |
| **A_55_P1960972** | Lrrc59 | -0.014796786 | -0.118449216 | 0.088855644 | 8.347240245 | -0.351202074 | 0.737699334 | 0.948456105 | -5.068376153 |
| **A_55_P2083939** | Irf3 | 0.024661317 | -0.148106937 | 0.197429572 | 12.36923054 | 0.351173301 | 0.737719822 | 0.948456105 | -5.068383581 |
| **A_55_P2016976** | Zfp729a | -0.024209427 | -0.193811985 | 0.145393131 | 9.044678114 | -0.351173131 | 0.737719943 | 0.948456105 | -5.068383625 |
| **A_51_P374900** | P2ry13 | 0.016016033 | -0.096199171 | 0.128231237 | 7.688476151 | 0.351133622 | 0.737748076 | 0.948456105 | -5.068393822 |
| **A_55_P2067952** | NA | -0.037574893 | -0.300966454 | 0.225816668 | 11.89667775 | -0.350966013 | 0.737867428 | 0.948456105 | -5.068437072 |
| **A_52_P447971** | NA | -0.035500152 | -0.284371836 | 0.213371533 | 11.63034927 | -0.350932736 | 0.737891125 | 0.948456105 | -5.068445657 |
| **A_51_P425680** | Ivd | -0.041234548 | -0.330401711 | 0.247932615 | 9.820838157 | -0.350817644 | 0.737973087 | 0.948456105 | -5.068475342 |
| **A_55_P2053718** | Pink1 | -0.027502241 | -0.220385296 | 0.165380814 | 12.58539646 | -0.350786743 | 0.737995092 | 0.948456105 | -5.06848331 |
| **A_55_P2151273** | Elovl1 | -0.018055879 | -0.144716615 | 0.108604858 | 15.37960632 | -0.350708091 | 0.738051106 | 0.948456105 | -5.068503589 |
| **A_51_P214197** | Stk17b | 0.043155287 | -0.259622495 | 0.345933068 | 9.642684044 | 0.350654299 | 0.738089417 | 0.948456105 | -5.068517456 |
| **A_52_P117576** | Casp3 | -0.015173115 | -0.121648049 | 0.09130182 | 7.753323439 | -0.350587576 | 0.738136937 | 0.948456105 | -5.068534653 |
| **A_52_P590625** | Nlrp10 | 0.016217448 | -0.097612062 | 0.130046958 | 7.655821434 | 0.35050709 | 0.738194261 | 0.948456105 | -5.068555393 |
| **A_55_P2064272** | NA | -0.018691861 | -0.149927984 | 0.112544262 | 8.641101112 | -0.350403402 | 0.738268114 | 0.948456105 | -5.068582107 |
| **A_51_P161554** | Arid3b | 0.043825111 | -0.263880402 | 0.351530624 | 9.547867373 | 0.3503942 | 0.738274668 | 0.948456105 | -5.068584477 |
| **A_51_P151254** | Slc35b3 | -0.049574859 | -0.397764366 | 0.298614647 | 10.76161125 | -0.350279706 | 0.738356221 | 0.948456105 | -5.068613964 |
| **A_55_P2034853** | 2610028H24Rik | 0.039791494 | -0.239691847 | 0.319274835 | 9.009507891 | 0.350270437 | 0.738362824 | 0.948456105 | -5.068616351 |
| **A_55_P2035877** | Faap20 | -0.024458351 | -0.19626487 | 0.147348168 | 10.13973734 | -0.350232705 | 0.738389701 | 0.948456105 | -5.068626067 |
| **A_51_P505967** | Spag6l | -0.014768526 | -0.118510134 | 0.088973082 | 7.664265236 | -0.350229997 | 0.73839163 | 0.948456105 | -5.068626764 |
| **A_52_P75777** | NA | -0.015476321 | -0.12419637 | 0.093243729 | 7.657559607 | -0.350208948 | 0.738406623 | 0.948456105 | -5.068632183 |
| **A_51_P393768** | Osbp | -0.020431121 | -0.163975777 | 0.123113535 | 9.694430439 | -0.350166291 | 0.738437009 | 0.948456105 | -5.068643165 |

| | | | | | | | | |
|---|---|---|---|---|---|---|---|---|
| **A_55_P2106265** | Mbnl2 | -0.029456638 | -0.236427666 | 0.177514389 | 9.620245214 | -0.350140856 | 0.738455128 | 0.948456105 | -5.068649713 |
| **A_52_P547612** | Tmem30b | 0.094007002 | -0.566634203 | 0.754648207 | 9.752495347 | 0.350077104 | 0.738500542 | 0.948456105 | -5.068666121 |
| **A_51_P418737** | Rbm33 | -0.016974953 | -0.136285915 | 0.102336009 | 8.415005607 | -0.350023662 | 0.738538612 | 0.948456105 | -5.068679875 |
| **A_55_P1982076** | Emsy | -0.040021189 | -0.321593573 | 0.241551194 | 11.19883358 | -0.349678634 | 0.73878442 | 0.948708747 | -5.068768619 |
| **A_55_P1967820** | AI661453 | -0.031826696 | -0.255812824 | 0.192159433 | 10.29482941 | -0.349574375 | 0.738858704 | 0.948741108 | -5.068795419 |
| **A_52_P219473** | Cdc6 | 0.016379664 | -0.098919135 | 0.131678463 | 7.737829367 | 0.349501768 | 0.738910437 | 0.94874451 | -5.068814078 |
| **A_55_P2047768** | Serpinb6c | 0.032733312 | -0.197739143 | 0.263205766 | 13.8408303 | 0.349413822 | 0.738973103 | 0.948761947 | -5.068836675 |
| **A_55_P2017183** | NA | 0.041232198 | -0.24916369 | 0.331628087 | 10.93815034 | 0.349313354 | 0.739044693 | 0.948790839 | -5.068862482 |
| **A_52_P15008** | Mapkapk2 | -0.035227511 | -0.283611444 | 0.213156423 | 10.73613701 | -0.348921408 | 0.739324009 | 0.949060209 | -5.068963094 |
| **A_51_P392090** | Tmem19 | 0.022108291 | -0.133812991 | 0.178029573 | 10.84879006 | 0.3488341 | 0.739386234 | 0.949060209 | -5.068985492 |
| **A_55_P1986647** | Podxl2 | 0.022551808 | -0.136523005 | 0.181626621 | 7.802824805 | 0.348778013 | 0.739426208 | 0.949060209 | -5.068999877 |
| **A_55_P2123199** | Akirin1 | -0.016674029 | -0.134300149 | 0.100952092 | 8.678537996 | -0.348743348 | 0.739450915 | 0.949060209 | -5.069008767 |
| **A_55_P2113698** | Nup205 | -0.032594543 | -0.262649783 | 0.197460696 | 10.45182451 | -0.348563524 | 0.739579089 | 0.94916169 | -5.06905487 |
| **A_55_P2123942** | Lmbrd1 | 0.047016764 | -0.284974712 | 0.37900824 | 9.551662716 | 0.34841349 | 0.739686035 | 0.949230543 | -5.069093318 |
| **A_51_P186487** | NA | 0.015079228 | -0.091416601 | 0.121575057 | 7.738990217 | 0.348349889 | 0.739731374 | 0.949230543 | -5.069109612 |
| **A_51_P243207** | Zfp574 | -0.019321262 | -0.155802875 | 0.117160351 | 10.40617714 | -0.348281583 | 0.739780067 | 0.949230543 | -5.069127108 |
| **A_55_P1952618** | Ear2 | -0.42389089 | -3.420669677 | 2.572887896 | 10.94005581 | -0.34799144 | 0.739986915 | 0.949432929 | -5.06920139 |
| **A_55_P1998072** | NA | 0.015095978 | -0.091701865 | 0.121893821 | 7.745443872 | 0.347750645 | 0.7401586 | 0.949590175 | -5.069262993 |
| **A_55_P2035872** | Mical3 | -0.050931382 | -0.453270357 | 0.351407594 | 11.31335429 | -0.356201596 | 0.740259557 | 0.949642657 | -4.950301126 |
| **A_51_P284665** | Plcb1 | -0.015061887 | -0.121851625 | 0.09172785 | 7.715805192 | -0.346991665 | 0.740699855 | 0.949642657 | -5.069456901 |
| **A_51_P424532** | Vnn1 | -0.07613675 | -0.616081742 | 0.463808242 | 12.6795853 | -0.346907731 | 0.740759721 | 0.949642657 | -5.069478321 |
| **A_51_P196973** | Chaf1a | -0.022429158 | -0.181492252 | 0.136633936 | 8.403679963 | -0.346906718 | 0.740760444 | 0.949642657 | -5.069478579 |
| **A_52_P204331** | D630039A03Rik | 0.014625124 | -0.089111969 | 0.118362217 | 7.639646551 | 0.34684437 | 0.740804916 | 0.949642657 | -5.069494487 |
| **A_66_P127484** | Abcf3 | -0.019464061 | -0.157535371 | 0.11860725 | 10.12601926 | -0.346816032 | 0.740825129 | 0.949642657 | -5.069501716 |
| **A_55_P1968723** | Noct | -0.099100789 | -0.802141123 | 0.603939546 | 13.79061998 | -0.346789574 | 0.740844001 | 0.949642657 | -5.069508465 |
| **A_51_P413376** | Zfp42 | 0.026355684 | -0.160644503 | 0.213355871 | 9.750251499 | 0.346737981 | 0.740880803 | 0.949642657 | -5.069521625 |
| **A_55_P1980302** | Psenen | -0.040677003 | -0.329345753 | 0.247991747 | 12.9042865 | -0.34667166 | 0.740928111 | 0.949642657 | -5.069538538 |
| **A_55_P2043117** | Ccdc138 | 0.040895884 | -0.249337984 | 0.331129752 | 15.58795134 | 0.346657558 | 0.740938171 | 0.949642657 | -5.069542134 |
| **A_51_P466162** | C1qbp | 0.042047935 | -0.25643931 | 0.34053518 | 12.21532133 | 0.346567679 | 0.741002286 | 0.949642657 | -5.069565049 |
| **A_52_P485850** | Mgam | -0.016642219 | -0.134798955 | 0.101514516 | 7.728703983 | -0.346514908 | 0.741039931 | 0.949642657 | -5.069578501 |
| **A_55_P2116744** | Xirp1 | 0.016171545 | -0.098646291 | 0.130989381 | 7.763162804 | 0.346506451 | 0.741045964 | 0.949642657 | -5.069580657 |
| **A_66_P106583** | NA | 0.040119176 | -0.24473243 | 0.324970781 | 14.17554934 | 0.346499404 | 0.741050991 | 0.949642657 | -5.069582453 |
| **A_55_P2018919** | Spns1 | -0.030070423 | -0.243611037 | 0.183470192 | 13.02419929 | -0.346440143 | 0.741093268 | 0.949642657 | -5.069597556 |
| **A_52_P36680** | Pggt1b | -0.031134126 | -0.252232819 | 0.189964566 | 8.526039696 | -0.346433333 | 0.741098126 | 0.949642657 | -5.069599292 |
| **A_55_P1992839** | Dsn1 | 0.035651661 | -0.217546711 | 0.288850032 | 8.811686072 | 0.346408084 | 0.741116139 | 0.949642657 | -5.069605726 |
| **A_51_P208931** | N4bp3 | 0.05024966 | -0.306631274 | 0.407130594 | 10.07924556 | 0.346400847 | 0.741121302 | 0.949642657 | -5.06960757 |
| **A_51_P464387** | Hspb8 | -0.138680198 | -1.12373873 | 0.846378334 | 10.68942011 | -0.346355094 | 0.741153943 | 0.949642657 | -5.069619227 |
| **A_66_P115323** | Utp4 | -0.023272686 | -0.188632247 | 0.142086876 | 9.903186336 | -0.346247269 | 0.741230869 | 0.949642657 | -5.069646695 |
| **A_51_P146760** | Pcdhb20 | -0.020014809 | -0.162226379 | 0.12219676 | 7.723284507 | -0.346246739 | 0.741231247 | 0.949642657 | -5.06964683 |
| **A_55_P1967736** | Nkd1 | 0.044282365 | -0.270511742 | 0.359076472 | 9.211400911 | 0.34607751 | 0.741351989 | 0.949659204 | -5.069689923 |
| **A_55_P1971006** | Urah | -0.103856226 | -0.842181338 | 0.634468885 | 9.696629889 | -0.346062118 | 0.741362971 | 0.949659204 | -5.069693842 |
| **A_52_P446457** | Ly6g6f | 0.015211468 | -0.092947344 | 0.123370281 | 7.648585359 | 0.346001826 | 0.74140599 | 0.949659204 | -5.069709189 |
| **A_55_P2038573** | NA | 0.019448648 | -0.118857419 | 0.157754716 | 13.48519757 | 0.345953198 | 0.741440688 | 0.949659204 | -5.069721566 |
| **A_55_P2175314** | NA | 0.030482682 | -0.186390116 | 0.24735548 | 14.8816296 | 0.34579384 | 0.741554399 | 0.949741915 | -5.069762114 |
| **A_55_P2131607** | Trnau1ap | 0.028784773 | -0.176112533 | 0.233682079 | 9.964176855 | 0.345617496 | 0.741680239 | 0.949785827 | -5.069806963 |
| **A_55_P2124425** | Zfp661 | -0.018372404 | -0.149155393 | 0.112410586 | 8.431713979 | -0.345608077 | 0.741686961 | 0.949785827 | -5.069809358 |
| **A_51_P337523** | Adcy2 | 0.017557243 | -0.107480737 | 0.142595224 | 7.7377529 | 0.345448685 | 0.741800712 | 0.949791703 | -5.069849876 |
| **A_52_P63905** | Ddc | 0.015888222 | -0.097286712 | 0.129063156 | 7.709336226 | 0.345377641 | 0.741851416 | 0.949791703 | -5.06986793 |
| **A_65_P13910** | Ndufb2 | 0.061206656 | -0.374798319 | 0.497211632 | 14.34311816 | 0.345363449 | 0.741861545 | 0.949791703 | -5.069871536 |
| **A_55_P1997661** | Smim15 | -0.032881878 | -0.267140834 | 0.201377077 | 11.76775828 | -0.345326238 | 0.741888102 | 0.949791703 | -5.06988099 |

| | | | | | | | | | |
|---|---|---|---|---|---|---|---|---|---|
| **A_52_P469502** | Cda | -0.080406233 | -0.653399962 | 0.492587495 | 10.49428128 | -0.345230385 | 0.741956516 | 0.949816378 | -5.06990534 |
| **A_55_P2091153** | BC051142 | 0.021563426 | -0.13214828 | 0.175275132 | 8.441870284 | 0.345127831 | 0.742029714 | 0.949847175 | -5.069931385 |
| **A_52_P611798** | Ddx5 | -0.036504839 | -0.296801526 | 0.223791848 | 12.2736549 | -0.345025314 | 0.742102889 | 0.949848073 | -5.069957413 |
| **A_55_P2125013** | H4c11 | -0.041821697 | -0.34006971 | 0.256426316 | 8.833587984 | -0.344979474 | 0.74213561 | 0.949848073 | -5.069969049 |
| **A_51_P194306** | Lrrc1 | 0.046892177 | -0.287572904 | 0.381357257 | 11.09997431 | 0.344920315 | 0.742177839 | 0.949848073 | -5.069984063 |
| **A_66_P115061** | Smc6 | 0.035421879 | -0.217367386 | 0.288211144 | 9.566889969 | 0.344732418 | 0.74231197 | 0.94984989 | -5.070031735 |
| **A_55_P2222870** | Mylk | -0.03106422 | -0.252757086 | 0.190628647 | 8.477680827 | -0.344729057 | 0.74231437 | 0.94984989 | -5.070032588 |
| **A_51_P367298** | NA | -0.035825029 | -0.291524182 | 0.219874124 | 8.452184537 | -0.344688206 | 0.742343533 | 0.94984989 | -5.070042949 |
| **A_55_P2031668** | Gstp1 | -0.032605292 | -0.265354294 | 0.20014371 | 9.234210354 | -0.344642976 | 0.742375823 | 0.94984989 | -5.07005442 |
| **A_51_P486239** | Clec3b | 0.018200613 | -0.111795106 | 0.148196332 | 7.873316504 | 0.344449952 | 0.742513629 | 0.949963327 | -5.070103355 |
| **A_51_P264495** | Pgam2 | -0.02058942 | -0.167709877 | 0.126531038 | 8.177646579 | -0.344302519 | 0.742618894 | 0.950032207 | -5.070140715 |
| **A_55_P2104572** | Tmem268 | -0.038911899 | -0.317033128 | 0.239209331 | 9.034927565 | -0.344205215 | 0.742688371 | 0.950032207 | -5.070165364 |
| **A_55_P2145606** | Kmo | -0.016642587 | -0.135607658 | 0.102322484 | 7.65254159 | -0.344168033 | 0.74271492 | 0.950032207 | -5.070174781 |
| **A_55_P2000384** | Pcnx3 | -0.057174591 | -0.46609003 | 0.351740847 | 11.64933858 | -0.343984382 | 0.742846059 | 0.950114705 | -5.07022128 |
| **A_51_P205779** | Cd5l | 0.033322454 | -0.205032074 | 0.271676982 | 7.905148654 | 0.343940039 | 0.742877725 | 0.950114705 | -5.070232504 |
| **A_55_P2170009** | Lactb2 | 0.05565161 | -0.342721634 | 0.454024855 | 10.87680604 | 0.343681958 | 0.743062031 | 0.950259321 | -5.070297801 |
| **A_55_P1984962** | NA | 0.031439448 | -0.19363971 | 0.256518606 | 7.961927618 | 0.343644025 | 0.743089122 | 0.950259321 | -5.070307394 |
| **A_55_P2075884** | Senp8 | 0.017936174 | -0.110499134 | 0.146371481 | 8.056909848 | 0.343569446 | 0.743142386 | 0.950264566 | -5.070326253 |
| **A_51_P323435** | Gtf2h2 | 0.017544331 | -0.108117556 | 0.143206218 | 8.22930554 | 0.343480745 | 0.743205739 | 0.95028271 | -5.070348677 |
| **A_55_P1976958** | NA | -0.036573426 | -0.298744006 | 0.225597153 | 9.738797005 | -0.343202828 | 0.743404248 | 0.950473655 | -5.070418902 |
| **A_51_P426270** | Mgp | 0.147694724 | -0.911400122 | 1.20678957 | 11.19849264 | 0.343083073 | 0.743489793 | 0.950488935 | -5.070449145 |
| **A_52_P68477** | Zfp809 | 0.024322442 | -0.150148114 | 0.198792998 | 8.953554631 | 0.342968472 | 0.743571659 | 0.950488935 | -5.070478077 |
| **A_55_P2163404** | NA | -0.031911001 | -0.26084123 | 0.197019227 | 13.86479149 | -0.34293077 | 0.743598593 | 0.950488935 | -5.070487594 |
| **A_55_P2122688** | Zmym3 | 0.029127986 | -0.179849029 | 0.238105001 | 10.04198246 | 0.342910749 | 0.743612896 | 0.950488935 | -5.070492647 |
| **A_51_P136355** | Gng11 | -0.14796993 | -1.209858973 | 0.913919113 | 13.25231612 | -0.342817902 | 0.743679227 | 0.950495825 | -5.070516077 |
| **A_55_P2427685** | Agl | -0.028357999 | -0.231954029 | 0.175238031 | 11.1571475 | -0.342669492 | 0.743785257 | 0.950495825 | -5.070553515 |
| **A_55_P2015113** | Atg5 | -0.031951858 | -0.261478222 | 0.197574506 | 10.96772936 | -0.342478023 | 0.74392206 | 0.950495825 | -5.070601794 |
| **A_55_P1999314** | Dhx33 | 0.024587518 | -0.152074637 | 0.201249673 | 10.15817796 | 0.342405186 | 0.743974104 | 0.950495825 | -5.070620152 |
| **A_52_P561650** | Vwa1 | 0.019534586 | -0.120839588 | 0.15990876 | 7.981864835 | 0.342362475 | 0.744004622 | 0.950495825 | -5.070630916 |
| **A_51_P213030** | Macrod1 | 0.03364263 | -0.208196156 | 0.275481416 | 8.832897398 | 0.342241876 | 0.744090799 | 0.950495825 | -5.070661302 |
| **A_55_P2158547** | Cyp2c54 | 0.016027673 | -0.099222276 | 0.131277622 | 7.654186501 | 0.342136099 | 0.744166386 | 0.950495825 | -5.070687945 |
| **A_55_P2159585** | Pm20d1 | -0.021880944 | -0.179253757 | 0.135491869 | 7.891002261 | -0.342062534 | 0.744218958 | 0.950495825 | -5.07070647 |
| **A_52_P555680** | Trappc12 | -0.022200503 | -0.181959393 | 0.137558386 | 10.80633524 | -0.341874686 | 0.744353205 | 0.950495825 | -5.070753756 |
| **A_51_P297968** | Pdia6 | -0.022449411 | -0.18403493 | 0.139136108 | 10.93305209 | -0.341799699 | 0.744406798 | 0.950495825 | -5.070772625 |
| **A_51_P457196** | Sfrp4 | -0.376130047 | -3.083723865 | 2.331463771 | 10.85950863 | -0.341761925 | 0.744433796 | 0.950495825 | -5.070782129 |
| **A_51_P419571** | Polr2d | -0.030880805 | -0.25319417 | 0.191432561 | 10.38512305 | -0.341737154 | 0.744451501 | 0.950495825 | -5.070788361 |
| **A_55_P2114596** | Zbtb14 | 0.034465986 | -0.213697794 | 0.282629765 | 10.12087513 | 0.341681507 | 0.744491273 | 0.950495825 | -5.070802358 |
| **A_55_P1994751** | NA | 0.023389581 | -0.14503997 | 0.191819132 | 8.445443188 | 0.341643615 | 0.744518356 | 0.950495825 | -5.070811888 |
| **A_55_P2063151** | NA | -0.052462827 | -0.430292873 | 0.32536722 | 10.51747755 | -0.341605111 | 0.744545878 | 0.950495825 | -5.070821572 |
| **A_55_P2058384** | NA | 0.035512795 | -0.220248921 | 0.291274511 | 9.356345493 | 0.341600485 | 0.744549184 | 0.950495825 | -5.070822735 |
| **A_55_P2049256** | Mapkapk5 | 0.02522262 | -0.156452136 | 0.206897375 | 11.34606573 | 0.341558243 | 0.744579377 | 0.950495825 | -5.070833357 |
| **A_65_P03606** | Rpap2 | 0.019184393 | -0.119073773 | 0.157442558 | 8.31621219 | 0.341370847 | 0.744713328 | 0.950495825 | -5.070880463 |
| **A_55_P1977468** | Dab2ip | -0.059867306 | -0.491344506 | 0.371609894 | 13.28993385 | -0.341350878 | 0.744727602 | 0.950495825 | -5.070885481 |
| **A_55_P2126827** | NA | 0.041984231 | -0.260647683 | 0.344616146 | 14.15152913 | 0.341303423 | 0.744761525 | 0.950495825 | -5.070897406 |
| **A_55_P2164805** | Gm6749 | -0.039958918 | -0.328061105 | 0.248143269 | 12.53752849 | -0.341221451 | 0.744820124 | 0.950495825 | -5.070918 |
| **A_66_P134394** | Pde6c | 0.122806172 | -0.762648766 | 1.008261109 | 8.824670527 | 0.341210897 | 0.744827668 | 0.950495825 | -5.070920651 |
| **A_55_P2014284** | Serpina11 | 0.023838086 | -0.148074725 | 0.195750896 | 8.04676185 | 0.341139729 | 0.744878545 | 0.950495825 | -5.070938526 |
| **A_55_P1969156** | Pam | -0.081811001 | -0.67183186 | 0.508209858 | 8.912377127 | -0.341124966 | 0.744889099 | 0.950495825 | -5.070942234 |
| **A_55_P2083499** | Shisal2a | -0.020162476 | -0.165605983 | 0.125281031 | 8.683999534 | -0.341050495 | 0.744942339 | 0.950495825 | -5.070960935 |
| **A_52_P176999** | Klhl31 | 0.015174838 | -0.094310289 | 0.124659966 | 7.633743449 | 0.340987222 | 0.744987575 | 0.950495825 | -5.07097682 |

| | | | | | | | | | |
|---|---|---|---|---|---|---|---|---|---|
| **A_52_P556337** | Txlng | 0.026333469 | -0.163666223 | 0.21633316 | 8.979022344 | 0.340976411 | 0.744995304 | 0.950495825 | -5.070979534 |
| **A_55_P2062480** | NA | 0.036315971 | -0.22578749 | 0.298419432 | 14.38207348 | 0.340874148 | 0.745068417 | 0.950495825 | -5.071005202 |
| **A_55_P2096297** | NA | -0.019202077 | -0.157819217 | 0.119415062 | 8.41378563 | -0.340800671 | 0.745120951 | 0.950495825 | -5.07102364 |
| **A_51_P474454** | Rnaseh2b | 0.024467354 | -0.152159234 | 0.201093943 | 10.40821832 | 0.3408004 | 0.745121145 | 0.950495825 | -5.071023708 |
| **A_51_P438711** | Tbx18 | 0.0176395 | -0.109732698 | 0.145011698 | 7.867484515 | 0.340706786 | 0.745188079 | 0.950495825 | -5.071047194 |
| **A_51_P250807** | Spata2l | 0.025374344 | -0.157861269 | 0.208609957 | 8.501777669 | 0.340685849 | 0.745203049 | 0.950495825 | -5.071052445 |
| **A_55_P2135035** | NA | 0.022349277 | -0.139091827 | 0.183790382 | 8.114101245 | 0.340579518 | 0.74527908 | 0.950495825 | -5.071079113 |
| **A_52_P269461** | Ddhd2 | 0.034143625 | -0.212505816 | 0.280793066 | 9.179105942 | 0.340563934 | 0.745290223 | 0.950495825 | -5.07108302 |
| **A_55_P2063553** | Gm4234 | -0.058548501 | -0.481772172 | 0.364675171 | 9.544625801 | -0.340341563 | 0.745449237 | 0.950612885 | -5.071138761 |
| **A_51_P509669** | Uap1l1 | -0.060869864 | -0.500935051 | 0.379195322 | 10.31767381 | -0.340294151 | 0.745483143 | 0.950612885 | -5.071150642 |
| **A_66_P103271** | Irf2bp2 | -0.036381551 | -0.310853415 | 0.238090313 | 11.64870129 | -0.343505318 | 0.745551804 | 0.950612885 | -5.00368052 |
| **A_51_P337089** | Gins1 | 0.035454396 | -0.220967668 | 0.29187646 | 8.211138108 | 0.340160485 | 0.745578733 | 0.950612885 | -5.071184126 |
| **A_55_P2064243** | Rhot2 | -0.015148939 | -0.124760301 | 0.094462423 | 7.946593844 | -0.340013232 | 0.745684047 | 0.950627614 | -5.071221 |
| **A_55_P1996202** | NA | 0.046679728 | -0.291081564 | 0.384441019 | 11.50850442 | 0.340006798 | 0.745688649 | 0.950627614 | -5.071222611 |
| **A_55_P2175942** | NA | 0.062253954 | -0.388513376 | 0.513021284 | 12.15852479 | 0.339768937 | 0.745858777 | 0.950736081 | -5.071282142 |
| **A_55_P2083629** | Tle2 | -0.015735495 | -0.129690496 | 0.098219505 | 7.831287187 | -0.339716151 | 0.745896534 | 0.950736081 | -5.071295347 |
| **A_55_P1968908** | Hypk | 0.0160851 | -0.100441058 | 0.132611257 | 11.54744824 | 0.339601415 | 0.745978605 | 0.950736081 | -5.071324044 |
| **A_51_P125691** | Cdca4 | -0.028944819 | -0.238644543 | 0.180754906 | 8.991606693 | -0.339580026 | 0.745993905 | 0.950736081 | -5.071329393 |
| **A_55_P1953236** | Epb42 | 0.015142122 | -0.094591992 | 0.124876236 | 7.663950142 | 0.339480028 | 0.746065438 | 0.950736081 | -5.071354395 |
| **A_55_P1980062** | NA | 0.033863311 | -0.211689932 | 0.279416554 | 8.462081982 | 0.339275819 | 0.746211526 | 0.950736081 | -5.07140543 |
| **A_55_P2435563** | Etl4 | -0.049313292 | -0.406904802 | 0.308278218 | 8.787219362 | -0.339270391 | 0.746215409 | 0.950736081 | -5.071406787 |
| **A_52_P987071** | Gm6803 | -0.032207936 | -0.26576936 | 0.201353489 | 11.75203775 | -0.339258657 | 0.746223804 | 0.950736081 | -5.071409718 |
| **A_55_P2110758** | B430306N03Rik | -0.03813998 | -0.314763788 | 0.238483828 | 8.045012736 | -0.339203301 | 0.746263407 | 0.950736081 | -5.071423547 |
| **A_55_P2017055** | Kctd12b | -0.046035971 | -0.380022308 | 0.287950366 | 9.690396344 | -0.339107792 | 0.746331738 | 0.950736081 | -5.071447402 |
| **A_51_P391716** | Ermap | 0.044427964 | -0.277958793 | 0.366814721 | 8.837259558 | 0.339038001 | 0.746381671 | 0.950736081 | -5.071464829 |
| **A_51_P367843** | Polr2g | 0.039932849 | -0.24990835 | 0.329774047 | 12.77246678 | 0.338952888 | 0.746442569 | 0.950736081 | -5.071486077 |
| **A_51_P249193** | Gsg1l | 0.018349156 | -0.114834382 | 0.151532695 | 8.00258397 | 0.338949281 | 0.74644515 | 0.950736081 | -5.071486978 |
| **A_51_P416546** | Ctsg | 0.082600689 | -0.517082881 | 0.68228426 | 8.415882729 | 0.338868098 | 0.746503238 | 0.950736081 | -5.07150724 |
| **A_55_P2127765** | Crp | 0.014534139 | -0.091050702 | 0.12011898 | 7.648591666 | 0.338654545 | 0.746656047 | 0.950736081 | -5.071560519 |
| **A_51_P333965** | Cisd2 | -0.016215002 | -0.134036163 | 0.10160616 | 7.963194557 | -0.338581216 | 0.746708521 | 0.950736081 | -5.071578806 |
| **A_55_P1996199** | NA | 0.060283827 | -0.377801342 | 0.498368996 | 14.09115775 | 0.338541119 | 0.746737215 | 0.950736081 | -5.071588804 |
| **A_51_P133612** | Cdt1 | -0.023016672 | -0.190307535 | 0.144274192 | 11.0269341 | -0.338485027 | 0.746777356 | 0.950736081 | -5.071602789 |
| **A_55_P1962054** | NA | -0.022400197 | -0.18529144 | 0.140491045 | 15.02872239 | -0.338316572 | 0.746897912 | 0.950736081 | -5.071644774 |
| **A_55_P2070661** | Taf7l | -0.062826614 | -0.519711788 | 0.39405856 | 14.43861696 | -0.338302924 | 0.746907679 | 0.950736081 | -5.071648175 |
| **A_55_P2097665** | Arl4a | 0.054681517 | -0.342988849 | 0.452351883 | 8.756612684 | 0.338287918 | 0.746918419 | 0.950736081 | -5.071651914 |
| **A_51_P324871** | Cyb561a3 | -0.036346201 | -0.300711122 | 0.22801872 | 12.06387352 | -0.338239529 | 0.746953051 | 0.950736081 | -5.071663969 |
| **A_55_P2034027** | Nos3 | -0.029110227 | -0.240853497 | 0.182633042 | 9.390943474 | -0.338224563 | 0.746963761 | 0.950736081 | -5.071667697 |
| **A_55_P1996082** | Zfp936 | 0.036472385 | -0.228860272 | 0.301805042 | 10.88373847 | 0.338175875 | 0.746998608 | 0.950736081 | -5.071679825 |
| **A_55_P2008056** | Cideb | -0.036252693 | -0.300031521 | 0.227526134 | 8.156342699 | -0.338118944 | 0.747039355 | 0.950736081 | -5.071694005 |
| **A_51_P354126** | Reg3g | -0.015160454 | -0.125475726 | 0.095154817 | 7.658910902 | -0.338100447 | 0.747052595 | 0.950736081 | -5.071698611 |
| **A_51_P443443** | 5830417I10Rik | -0.034085653 | -0.282175415 | 0.214004109 | 11.18014244 | -0.338011859 | 0.747116001 | 0.950754176 | -5.071720669 |
| **A_55_P2092364** | Ermard | -0.024046261 | -0.199146388 | 0.151053867 | 9.370020684 | -0.337854923 | 0.747228334 | 0.950811164 | -5.071759732 |
| **A_55_P1960202** | Vcan | -0.071639341 | -0.593378172 | 0.45009949 | 8.923396568 | -0.337806174 | 0.747263229 | 0.950811164 | -5.071771862 |
| **A_55_P2153541** | Sik3 | 0.050125767 | -0.315000995 | 0.415252528 | 12.27822646 | 0.337743133 | 0.747308356 | 0.950811164 | -5.071787547 |
| **A_55_P1987730** | Prxl2a | -0.06478749 | -0.536898199 | 0.407323218 | 11.64749106 | -0.337610908 | 0.74740301 | 0.950840103 | -5.071820435 |
| **A_55_P1982239** | Ctrb1 | -0.015922737 | -0.131965483 | 0.100120008 | 8.083402785 | -0.337573925 | 0.747429486 | 0.950840103 | -5.071829632 |
| **A_55_P1990038** | Mark2 | -0.018858036 | -0.156345687 | 0.118629616 | 8.546012166 | -0.337444164 | 0.747522382 | 0.950895697 | -5.071861892 |
| **A_51_P406583** | Myo1e | -0.025421035 | -0.210945807 | 0.160103736 | 10.90233096 | -0.337101351 | 0.747767826 | 0.951084595 | -5.071947064 |
| **A_51_P269173** | Alg9 | 0.037851116 | -0.23839106 | 0.314093291 | 9.322570182 | 0.337099308 | 0.747769289 | 0.951084595 | -5.071947571 |
| **A_52_P244702** | Tcf7 | -0.015671982 | -0.13011605 | 0.098772087 | 7.798980174 | -0.336899053 | 0.74791268 | 0.951148593 | -5.071997286 |

| | | | | | | | | |
|---|---|---|---|---|---|---|---|---|
| **A_55_P2069787** | NA | -0.049444475 | -0.410518928 | 0.311629977 | 12.50086917 | -0.336891592 | 0.747918023 | 0.951148593 | -5.071999137 |
| **A_51_P221949** | Csnk1g3 | -0.035165433 | -0.292136345 | 0.221805479 | 11.37073324 | -0.33666748 | 0.748078511 | 0.951190446 | -5.07205474 |
| **A_51_P120093** | Snx10 | -0.048929701 | -0.40649881 | 0.308639409 | 9.583096499 | -0.336652412 | 0.748089302 | 0.951190446 | -5.061325839 |
| **A_55_P2052252** | NA | -0.01569021 | -0.130374248 | 0.098993829 | 8.265074798 | -0.336585142 | 0.748137477 | 0.951190446 | -5.07207516 |
| **A_55_P1991239** | NA | 0.08349182 | -0.526926901 | 0.69391054 | 9.323928085 | 0.336500137 | 0.748198355 | 0.951190446 | -5.072096236 |
| **A_55_P1981550** | Park7 | -0.027753112 | -0.23066599 | 0.175159766 | 11.62707675 | -0.336489286 | 0.748206127 | 0.951190446 | -5.072098926 |
| **A_55_P2037544** | NA | 0.029855898 | -0.188467497 | 0.248179293 | 15.64519949 | 0.336433338 | 0.748246197 | 0.951190446 | -5.072112794 |
| **A_51_P127756** | Ddx20 | -0.016655114 | -0.138489602 | 0.105179375 | 9.511134223 | -0.336315216 | 0.748330797 | 0.951235432 | -5.056409248 |
| **A_55_P2009375** | Cdk18 | -0.174210881 | -1.449572906 | 1.101151144 | 10.34492459 | -0.336055463 | 0.748516851 | 0.951367259 | -5.072206405 |
| **A_66_P138053** | Gm9917 | 0.018012631 | -0.113870722 | 0.149895984 | 8.00808295 | 0.336013195 | 0.748547128 | 0.951367259 | -5.07221687 |
| **A_66_P114463** | NA | -0.031900605 | -0.265501751 | 0.201700541 | 11.00716393 | -0.335964285 | 0.748582164 | 0.951367259 | -5.072228978 |
| **A_52_P1016836** | NA | 0.014204677 | -0.089874294 | 0.118283649 | 7.677826376 | 0.335766643 | 0.748723746 | 0.951484634 | -5.072277887 |
| **A_52_P157170** | Rnf157 | 0.031278695 | -0.198037513 | 0.260594903 | 8.503939801 | 0.335569929 | 0.748864675 | 0.951544166 | -5.07232654 |
| **A_51_P399853** | Zfp704 | -0.017834217 | -0.148586136 | 0.112917702 | 8.728109426 | -0.335563823 | 0.748869049 | 0.951544166 | -5.072328049 |
| **A_55_P2030080** | Kdm6b | -0.043871842 | -0.365622285 | 0.277878601 | 10.85000153 | -0.335456246 | 0.748946124 | 0.951545295 | -5.072354644 |
| **A_55_P2026014** | Mc5r | 0.017106204 | -0.108364251 | 0.142576659 | 7.958239748 | 0.335414119 | 0.748976307 | 0.951545295 | -5.072365057 |
| **A_52_P454183** | Olfml2b | -0.022146376 | -0.184613381 | 0.140320628 | 7.981389032 | -0.335356452 | 0.749017625 | 0.951545295 | -5.072379308 |
| **A_55_P2207255** | Srpx | 0.068694605 | -0.435699654 | 0.573088864 | 8.618525144 | 0.335059286 | 0.749230556 | 0.951601526 | -5.072452709 |
| **A_55_P2164348** | Hoxa10 | -0.15616899 | -1.303028812 | 0.990690833 | 9.055797186 | -0.335006802 | 0.749268166 | 0.951601526 | -5.072465667 |
| **A_55_P2160451** | Stk26 | -0.014745108 | -0.123032841 | 0.093542624 | 7.717685702 | -0.334994648 | 0.749276875 | 0.951601526 | -5.072468667 |
| **A_66_P139946** | Sfta3-ps | 0.014473077 | -0.091817032 | 0.120763187 | 7.639595865 | 0.334994121 | 0.749277253 | 0.951601526 | -5.072468797 |
| **A_55_P1968664** | NA | 0.03606708 | -0.228843113 | 0.300977273 | 11.171221 | 0.334951149 | 0.749308047 | 0.951601526 | -5.072479404 |
| **A_55_P1983418** | Amy1 | -0.017467267 | -0.145823777 | 0.110889242 | 7.754521625 | -0.334792894 | 0.749421458 | 0.951679301 | -5.072518457 |
| **A_52_P211223** | Cdca2 | -0.015443936 | -0.128954096 | 0.098066225 | 7.780450161 | -0.334728287 | 0.749467759 | 0.951679301 | -5.072534395 |
| **A_51_P285077** | Hhatl | 0.017628139 | -0.112031398 | 0.147287676 | 7.710758053 | 0.334480781 | 0.749645148 | 0.951721185 | -5.072595426 |
| **A_52_P130912** | Pcdh7 | 0.014125602 | -0.089775312 | 0.118026515 | 7.67787319 | 0.334469682 | 0.749653103 | 0.951721185 | -5.072598161 |
| **A_51_P412338** | Vps41 | 0.021907683 | -0.139249098 | 0.183064464 | 9.927758113 | 0.334439087 | 0.749675033 | 0.951721185 | -5.072605702 |
| **A_55_P2025760** | Adam9 | 0.037148061 | -0.236183964 | 0.310480087 | 11.94597107 | 0.33436037 | 0.749731454 | 0.951721185 | -5.072625101 |
| **A_55_P2011877** | Eif2s3y | -0.289744697 | -2.422034405 | 1.842545012 | 9.188933334 | -0.334301594 | 0.749773583 | 0.951721185 | -5.072639583 |
| **A_52_P574759** | Rbpjl | 0.0145521 | -0.092616112 | 0.121720312 | 7.707792267 | 0.33406336 | 0.749944355 | 0.951721185 | -5.072698257 |
| **A_51_P521052** | Ly6k | 0.045656217 | -0.290639906 | 0.38195234 | 9.958383268 | 0.334000583 | 0.749989357 | 0.951721185 | -5.072713712 |
| **A_55_P2185826** | Cyp3a13 | 0.015768316 | -0.100404 | 0.131940632 | 7.881770196 | 0.333927229 | 0.750041944 | 0.951721185 | -5.072731767 |
| **A_52_P387724** | Tcf7l1 | 0.041956591 | -0.267193148 | 0.35110633 | 9.526102786 | 0.333887741 | 0.750070253 | 0.951721185 | -5.072741485 |
| **A_55_P2006975** | NA | -0.014856856 | -0.124328664 | 0.094614951 | 9.316755952 | -0.333882608 | 0.750073932 | 0.951721185 | -5.072742748 |
| **A_55_P1971815** | Clcn2 | -0.026042621 | -0.217967865 | 0.165882623 | 8.406431454 | -0.333827228 | 0.750113635 | 0.951721185 | -5.072756374 |
| **A_55_P1965902** | Pi4k2a | -0.036556954 | -0.305994221 | 0.232880314 | 10.00970225 | -0.333796281 | 0.750135822 | 0.951721185 | -5.072763988 |
| **A_66_P120770** | Ywhaz | 0.020641116 | -0.131523546 | 0.172805778 | 9.157913121 | 0.333724838 | 0.750187042 | 0.951721185 | -5.072781563 |
| **A_55_P2060459** | Nsd3 | -0.039689318 | -0.332279521 | 0.252900884 | 8.892707656 | -0.333720607 | 0.750190075 | 0.951721185 | -5.072782603 |
| **A_51_P411909** | Sirt5 | 0.031289124 | -0.199458772 | 0.26203702 | 8.952998458 | 0.333599062 | 0.75027722 | 0.951769271 | -5.072812494 |
| **A_51_P212420** | Lama4 | -0.09549149 | -0.79990948 | 0.6089265 | 10.47627977 | -0.333505802 | 0.750344087 | 0.951791629 | -5.072835422 |
| **A_52_P307961** | Abca5 | -0.032350969 | -0.271047646 | 0.206345708 | 8.794028389 | -0.333434154 | 0.750395459 | 0.951794332 | -5.072853032 |
| **A_52_P382876** | Bloc1s3 | 0.018748456 | -0.1196981 | 0.157195012 | 8.354601968 | 0.333159743 | 0.75059223 | 0.951981445 | -5.072920446 |
| **A_51_P288558** | Cfap52 | 0.015973494 | -0.102032653 | 0.13397964 | 7.704839643 | 0.333015498 | 0.750695672 | 0.95205017 | -5.072955861 |
| **A_55_P2176917** | NA | -0.024363001 | -0.204425151 | 0.155699148 | 9.7685511 | -0.332872194 | 0.750798444 | 0.952094045 | -5.072991031 |
| **A_55_P2047155** | Meis2 | -0.056890853 | -0.477556603 | 0.363774897 | 10.21085127 | -0.332716589 | 0.750910045 | 0.952094045 | -5.073029204 |
| **A_55_P2170225** | Sh3bgrl2 | 0.029249441 | -0.187098829 | 0.245597711 | 9.580143357 | 0.33260847 | 0.750987592 | 0.952094045 | -5.073055717 |
| **A_66_P126552** | Prr14l | -0.024419069 | -0.205100181 | 0.156262042 | 8.66322115 | -0.332495302 | 0.751068764 | 0.952094045 | -5.07308346 |
| **A_55_P2085460** | 5530601H04Rik | -0.033961974 | -0.285297218 | 0.217373271 | 8.408651366 | -0.332436491 | 0.751110949 | 0.952094045 | -5.073097874 |
| **A_51_P251209** | Cacna1d | -0.014235115 | -0.119586435 | 0.091116204 | 7.715235203 | -0.332422319 | 0.751121115 | 0.952094045 | -5.073101347 |
| **A_51_P309534** | Pex14 | 0.041726421 | -0.26709181 | 0.350544652 | 11.38102981 | 0.332412515 | 0.751128147 | 0.952094045 | -5.073103749 |

| | | | | | | | | | |
|---|---|---|---|---|---|---|---|---|---|
| **A_55_P2121491** | Bbs9 | -0.020078782 | -0.168693745 | 0.128536181 | 9.392210264 | -0.332386971 | 0.751146471 | 0.952094045 | -5.073110008 |
| **A_55_P2035593** | Hsf1 | -0.027221403 | -0.228752797 | 0.174309991 | 10.84753903 | -0.332305065 | 0.751205224 | 0.952094045 | -5.073130075 |
| **A_66_P102628** | 1700016J18Rik | -0.014533059 | -0.122135308 | 0.093069189 | 7.697201956 | -0.332280507 | 0.75122284 | 0.952094045 | -5.073136091 |
| **A_51_P368483** | 1700123O20Rik | -0.023423401 | -0.196885364 | 0.150038561 | 10.37370553 | -0.332211655 | 0.751272231 | 0.952094215 | -5.073152955 |
| **A_52_P77080** | Gypa | 0.026964046 | -0.172774258 | 0.22670235 | 7.980727382 | 0.33211834 | 0.751339174 | 0.952116625 | -5.073175806 |
| **A_55_P2135880** | Fancl | 0.044712028 | -0.286678457 | 0.376102513 | 9.277091778 | 0.331935344 | 0.751470458 | 0.952160341 | -5.0732206 |
| **A_55_P2177347** | NA | 0.019058387 | -0.122196985 | 0.160313759 | 8.686660067 | 0.331932926 | 0.751472192 | 0.952160341 | -5.073221191 |
| **A_55_P2041270** | Arhgef1 | 0.052791298 | -0.338677651 | 0.444260246 | 13.20936309 | 0.331767761 | 0.751590692 | 0.952198544 | -5.0732616 |
| **A_51_P169516** | Ppp1r3d | 0.041963181 | -0.269224181 | 0.353150543 | 9.032401526 | 0.331753578 | 0.751600868 | 0.952198544 | -5.073265069 |
| **A_52_P362981** | NA | 0.015850404 | -0.101784886 | 0.133485694 | 8.569092724 | 0.331491103 | 0.7517892 | 0.952374718 | -5.073329243 |
| **A_55_P1970810** | NA | 0.036140631 | -0.232278018 | 0.304559281 | 13.13394386 | 0.331247197 | 0.751964226 | 0.952534015 | -5.073388833 |
| **A_55_P1967325** | Tafa5 | -0.034383492 | -0.289935678 | 0.221168693 | 8.439525233 | -0.331008809 | 0.752135308 | 0.952553287 | -5.073447036 |
| **A_55_P2058657** | NA | -0.018691893 | -0.157621636 | 0.120237849 | 16.30365961 | -0.330999415 | 0.75214205 | 0.952553287 | -5.073449328 |
| **A_55_P2124498** | Il17rb | -0.038709957 | -0.326553702 | 0.249133788 | 8.947263569 | -0.330852985 | 0.752247145 | 0.952553287 | -5.073485059 |
| **A_52_P102559** | Dars2 | -0.021668845 | -0.182830985 | 0.139493296 | 8.945876419 | -0.330782019 | 0.75229808 | 0.952553287 | -5.073502369 |
| **A_55_P2019512** | Mocs2 | -0.031595073 | -0.266616674 | 0.203426528 | 11.22661642 | -0.330735449 | 0.752331506 | 0.952553287 | -5.073513727 |
| **A_55_P2051099** | Tnfrsf1a | -0.043331123 | -0.365691647 | 0.279029402 | 11.26740021 | -0.330694712 | 0.752360747 | 0.952553287 | -5.073523662 |
| **A_51_P277745** | Abcc2 | 0.061170378 | -0.394039333 | 0.516380089 | 8.222919878 | 0.330596923 | 0.752430939 | 0.952553287 | -5.073547504 |
| **A_52_P586004** | Hk3 | -0.016217438 | -0.136910985 | 0.10447611 | 7.77062732 | -0.330572977 | 0.752448127 | 0.952553287 | -5.073553341 |
| **A_51_P414243** | Pomgnt2 | -0.015706601 | -0.132602983 | 0.101189782 | 10.85233755 | -0.330559994 | 0.752457447 | 0.952553287 | -5.073556505 |
| **A_55_P2083093** | Haus7 | -0.02067169 | -0.174530349 | 0.13318697 | 8.058005198 | -0.33053937 | 0.752472251 | 0.952553287 | -5.073561533 |
| **A_55_P1991530** | Cep126 | -0.01488621 | -0.125732412 | 0.095959993 | 8.014539378 | -0.330394247 | 0.752576426 | 0.95261809 | -5.073596897 |
| **A_51_P516705** | Ern1 | -0.03065537 | -0.25896631 | 0.19765557 | 9.373504328 | -0.330330747 | 0.752622011 | 0.95261809 | -5.073612366 |
| **A_55_P2009895** | Gm5113 | -0.020603169 | -0.17419609 | 0.132989752 | 8.041517758 | -0.330013713 | 0.752849616 | 0.952843782 | -5.073689558 |
| **A_51_P309988** | Gprc5c | 0.023865739 | -0.154088404 | 0.201819881 | 7.995756253 | 0.32994074 | 0.752902009 | 0.9528477 | -5.073707315 |
| **A_51_P193302** | Mrps7 | -0.024075968 | -0.203682668 | 0.155530732 | 9.61984228 | -0.329784606 | 0.753014114 | 0.952927184 | -5.073745297 |
| **A_55_P2181029** | NA | -0.045599908 | -0.385999307 | 0.294799491 | 8.118258994 | -0.329567477 | 0.753170025 | 0.952931085 | -5.073798088 |
| **A_55_P2133280** | Ap4b1 | -0.033733966 | -0.285581572 | 0.218113641 | 10.46669674 | -0.329532865 | 0.753194879 | 0.952931085 | -5.0738065 |
| **A_55_P2094352** | Zfp523 | -0.021976662 | -0.186055075 | 0.14210175 | 7.908040157 | -0.329518232 | 0.753205387 | 0.952931085 | -5.073810056 |
| **A_51_P341647** | Fancc | -0.018549627 | -0.157046971 | 0.119947717 | 8.524343817 | -0.329505682 | 0.753214399 | 0.952931085 | -5.073813106 |
| **A_55_P2125811** | Esam | 0.044398538 | -0.287228437 | 0.376025514 | 13.24914843 | 0.329372993 | 0.753309686 | 0.952973853 | -5.073845345 |
| **A_52_P518808** | Mmd | 0.020618574 | -0.133412365 | 0.174649513 | 8.512384661 | 0.3293213 | 0.753346809 | 0.952973853 | -5.073857901 |
| **A_55_P2139396** | Unc45b | 0.01457782 | -0.094385191 | 0.12354083 | 7.783407383 | 0.329141502 | 0.753475936 | 0.953074823 | -5.07390156 |
| **A_55_P2091402** | Kpna2 | -0.06211414 | -0.526537278 | 0.402308998 | 11.44697659 | -0.329037799 | 0.753550417 | 0.953099759 | -5.07392673 |
| **A_51_P188271** | Cd248 | 0.06794434 | -0.440261245 | 0.576149926 | 10.61896766 | 0.328914497 | 0.753638978 | 0.953099759 | -5.073956649 |
| **A_55_P1957373** | Pdcl | -0.021328422 | -0.180902932 | 0.138246088 | 9.850138361 | -0.328824651 | 0.753703512 | 0.953099759 | -5.073978442 |
| **A_55_P2125322** | NA | -0.028664857 | -0.243275908 | 0.185946193 | 15.26196477 | -0.328599474 | 0.753865261 | 0.953099759 | -5.074033038 |
| **A_65_P15809** | Gm4759 | 0.055838091 | -0.362260565 | 0.473936747 | 9.159385933 | 0.328564738 | 0.753890213 | 0.953099759 | -5.074041456 |
| **A_51_P160744** | Ndufb3 | 0.059835722 | -0.388220166 | 0.50789161 | 12.87894146 | 0.328547003 | 0.753902954 | 0.953099759 | -5.074045754 |
| **A_55_P2080248** | Cyp2j8 | -0.016851378 | -0.143092637 | 0.109389882 | 7.86423717 | -0.32840008 | 0.7540085 | 0.953099759 | -5.074081352 |
| **A_55_P2021285** | Cpped1 | -0.034919761 | -0.296523463 | 0.226683942 | 10.73029229 | -0.328394998 | 0.754012151 | 0.953099759 | -5.074082583 |
| **A_55_P1956593** | Plekha4 | 0.043995215 | -0.285721493 | 0.373711924 | 10.27641651 | 0.32827186 | 0.754100617 | 0.953099759 | -5.074112406 |
| **A_55_P2088699** | NA | -0.017180629 | -0.14594408 | 0.111582821 | 15.81549256 | -0.328258234 | 0.754110406 | 0.953099759 | -5.074115705 |
| **A_55_P1988183** | Cdkn2b | -0.025802215 | -0.219187681 | 0.167583251 | 9.06347447 | -0.328248174 | 0.754117634 | 0.953099759 | -5.074118141 |
| **A_55_P2073642** | 1600014C10Rik | 0.049356807 | -0.320617274 | 0.419330887 | 8.715715553 | 0.328204791 | 0.754148802 | 0.953099759 | -5.074128645 |
| **A_55_P2013948** | Gsn | -0.054678284 | -0.464633782 | 0.355277215 | 15.27446239 | -0.328131044 | 0.754201787 | 0.953099759 | -5.074146497 |
| **A_55_P1973930** | Krt79 | 0.068853852 | -0.447460056 | 0.58516776 | 9.163466064 | 0.328082888 | 0.754236386 | 0.953099759 | -5.074158152 |
| **A_66_P104312** | Pdcl | 0.026744761 | -0.173843402 | 0.227332924 | 9.716694542 | 0.328021707 | 0.754280344 | 0.953099759 | -5.074172958 |
| **A_55_P1952269** | Xylt2 | 0.034567415 | -0.224696049 | 0.293830879 | 9.900676828 | 0.328015786 | 0.754284599 | 0.953099759 | -5.074174391 |
| **A_55_P2110595** | Slc35a2 | 0.018116416 | -0.117868252 | 0.154101085 | 9.037068369 | 0.327756644 | 0.754470803 | 0.953272725 | -5.074237071 |

| | | | | | | | | | |
|---|---|---|---|---|---|---|---|---|---|
| **A_51_P369897** | Zfp410 | -0.029309954 | -0.249397867 | 0.190777959 | 9.454969084 | -0.327633354 | 0.754559399 | 0.953322349 | -5.074266875 |
| **A_55_P1974533** | Eif4e | -0.02197982 | -0.187190398 | 0.143230759 | 8.745482982 | -0.327307107 | 0.754793857 | 0.953487702 | -5.074345692 |
| **A_55_P2001628** | Rps4l | -0.025447555 | -0.216793271 | 0.165898161 | 14.18264222 | -0.327187332 | 0.754879941 | 0.953487702 | -5.074374608 |
| **A_55_P1977144** | Cyp2c67 | 0.087865677 | -0.57284455 | 0.748575904 | 7.968371999 | 0.327172953 | 0.754890276 | 0.953487702 | -5.074378079 |
| **A_51_P464023** | Chst12 | 0.060791388 | -0.396456319 | 0.518039094 | 9.904169129 | 0.327084289 | 0.754954003 | 0.953487702 | -5.074399478 |
| **A_55_P2113857** | Serpina1e | 0.244915895 | -1.597346117 | 2.087177908 | 13.05554852 | 0.327065502 | 0.754967506 | 0.953487702 | -5.074404011 |
| **A_55_P2102002** | Noc3l | 0.030085511 | -0.196236319 | 0.25640734 | 8.876668975 | 0.32703942 | 0.754986253 | 0.953487702 | -5.074410305 |
| **A_52_P596592** | Rassf9 | -0.014120488 | -0.12038036 | 0.092139384 | 7.722245401 | -0.326926089 | 0.755067714 | 0.953528279 | -5.074437645 |
| **A_55_P1984635** | Parp4 | -0.025422755 | -0.216836612 | 0.165991102 | 11.48567493 | -0.326752111 | 0.755192773 | 0.953536572 | -5.074479598 |
| **A_51_P461108** | Osbpl10 | -0.037026239 | -0.315809916 | 0.241757438 | 9.315577785 | -0.326746858 | 0.75519655 | 0.953536572 | -5.074480865 |
| **A_55_P1988108** | Mrc1 | 0.041211376 | -0.269146514 | 0.351569265 | 8.881327243 | 0.326680706 | 0.755244103 | 0.953536572 | -5.074496811 |
| **A_55_P1992720** | NA | 0.01441845 | -0.094199314 | 0.123036214 | 7.700106072 | 0.326577966 | 0.755317961 | 0.953536572 | -5.074521571 |
| **A_51_P135654** | Eif4ebp2 | 0.034978326 | -0.228525432 | 0.298482084 | 11.72229369 | 0.326573823 | 0.75532094 | 0.953536572 | -5.074522569 |
| **A_55_P2001159** | Oser1 | 0.025507253 | -0.166792908 | 0.217807414 | 12.83231324 | 0.326327144 | 0.755498284 | 0.953651918 | -5.074581986 |
| **A_51_P243514** | Macc1 | 0.016606691 | -0.108598567 | 0.141811948 | 7.789275876 | 0.32630948 | 0.755510984 | 0.953651918 | -5.074586239 |
| **A_55_P1953459** | Yap1 | -0.024354722 | -0.208206836 | 0.159497391 | 13.25480494 | -0.325899516 | 0.755805758 | 0.953926754 | -5.074684887 |
| **A_55_P1994348** | Golim4 | -0.03294659 | -0.28168111 | 0.21578793 | 9.126803681 | -0.325869389 | 0.755827422 | 0.953926754 | -5.074692132 |
| **A_66_P129508** | Hyls1 | -0.014190977 | -0.121409255 | 0.093027301 | 7.800818957 | -0.32562117 | 0.756005921 | 0.95404736 | -5.074751797 |
| **A_52_P239536** | Ppp1r9a | -0.032228034 | -0.275786067 | 0.21133 | 9.355680417 | -0.325537118 | 0.756066368 | 0.95404736 | -5.074771991 |
| **A_52_P144818** | Ppp1r1a | -0.013708958 | -0.117332083 | 0.089914168 | 7.659288938 | -0.325474462 | 0.756111429 | 0.95404736 | -5.074787041 |
| **A_51_P193000** | Bbs4 | -0.016496182 | -0.141192135 | 0.108199771 | 9.337756259 | -0.325461968 | 0.756120415 | 0.95404736 | -5.074790042 |
| **A_55_P2003398** | Rmdn1 | -0.031009337 | -0.26551116 | 0.203492487 | 8.632534543 | -0.325323487 | 0.756220013 | 0.954110746 | -5.074823295 |
| **A_55_P2155764** | NA | 0.022938841 | -0.150621669 | 0.196499352 | 10.04264794 | 0.325154467 | 0.756341581 | 0.954150814 | -5.074863863 |
| **A_55_P1984069** | NA | -0.052303607 | -0.448136393 | 0.343529179 | 12.7331881 | -0.325079091 | 0.756395798 | 0.954150814 | -5.074881948 |
| **A_51_P252784** | Fam216a | -0.042687959 | -0.365850969 | 0.280475052 | 10.32828561 | -0.32497721 | 0.756469083 | 0.954150814 | -5.074906386 |
| **A_55_P2007334** | Mepce | -0.025800659 | -0.221222986 | 0.169621667 | 11.80073383 | -0.324807306 | 0.756591303 | 0.954150814 | -5.074947125 |
| **A_55_P2186446** | NA | -0.028502552 | -0.244429041 | 0.187423937 | 13.91472102 | -0.324748377 | 0.756633696 | 0.954150814 | -5.07496125 |
| **A_51_P114722** | Hao2 | -0.022667338 | -0.194388821 | 0.149054144 | 7.876925077 | -0.324746936 | 0.756634732 | 0.954150814 | -5.069697134 |
| **A_55_P1995745** | Gm14308 | 0.016828563 | -0.110663564 | 0.14432069 | 9.028704049 | 0.324737795 | 0.756641308 | 0.954150814 | -5.074963786 |
| **A_55_P2255628** | 9430013L14Rik | -0.074428988 | -0.638427076 | 0.4895691 | 9.07325575 | -0.324663634 | 0.756694659 | 0.954150814 | -5.074981558 |
| **A_51_P499020** | Fbp2 | 0.062345277 | -0.410112198 | 0.534802752 | 8.419558666 | 0.324645954 | 0.756707379 | 0.954150814 | -5.074985794 |
| **A_51_P166740** | Kcnj4 | -0.013715992 | -0.117673803 | 0.090241818 | 7.73308421 | -0.324593095 | 0.756745407 | 0.954150814 | -5.074998458 |
| **A_55_P1971674** | Golga7b | -0.027589715 | -0.23681892 | 0.181639489 | 9.954564045 | -0.324409883 | 0.756877219 | 0.95421723 | -5.075042337 |
| **A_55_P2315457** | Gripap1 | -0.028325678 | -0.243154166 | 0.186502809 | 9.826492555 | -0.324382645 | 0.756896816 | 0.95421723 | -5.075048859 |
| **A_52_P570266** | Psmb10 | 0.04285776 | -0.282403276 | 0.368118797 | 11.76403649 | 0.324165351 | 0.757053163 | 0.954342153 | -5.075100866 |
| **A_55_P2039842** | Frmpd1 | -0.040676553 | -0.349454933 | 0.268101827 | 12.16379401 | -0.32409059 | 0.757106957 | 0.954342153 | -5.075118752 |
| **A_51_P128463** | Grrp1 | -0.02937168 | -0.252369477 | 0.193626117 | 9.45767217 | -0.324039073 | 0.757144028 | 0.954342153 | -5.042826153 |
| **A_66_P139876** | Psph | -0.053602525 | -0.460870088 | 0.353665038 | 9.962645343 | -0.323798317 | 0.757317279 | 0.954451175 | -5.075188637 |
| **A_55_P2172058** | Trim55 | 0.013725583 | -0.090565636 | 0.118016802 | 7.684950313 | 0.323781639 | 0.757329281 | 0.954451175 | -5.075192623 |
| **A_52_P604526** | Mrpl19 | -0.033270661 | -0.28622842 | 0.219687097 | 9.616915571 | -0.323580686 | 0.757473901 | 0.954571198 | -5.075240635 |
| **A_55_P2022987** | Psmd2 | -0.071917993 | -0.619132399 | 0.475296412 | 13.87397444 | -0.323332382 | 0.757652614 | 0.954649675 | -5.075299921 |
| **A_51_P165293** | Ckmt2 | -0.023018968 | -0.198241059 | 0.152203123 | 7.72269028 | -0.323196135 | 0.757750683 | 0.954649675 | -5.075332434 |
| **A_55_P2129578** | Pkp3 | 0.089648412 | -0.592821534 | 0.772118358 | 13.00130922 | 0.323167912 | 0.757770998 | 0.954649675 | -5.075339167 |
| **A_55_P1999419** | Bag5 | 0.023456374 | -0.155212267 | 0.202125015 | 10.29242875 | 0.322984518 | 0.757903011 | 0.954649675 | -5.075382906 |
| **A_55_P2070179** | Ing4 | 0.030195367 | -0.199825189 | 0.260215923 | 11.57440455 | 0.322955605 | 0.757923825 | 0.954649675 | -5.0753898 |
| **A_52_P504068** | Cdk8 | -0.027516013 | -0.237308138 | 0.182276113 | 8.209805789 | -0.32267512 | 0.758125748 | 0.954649675 | -5.075456643 |
| **A_51_P320980** | Sfxn4 | 0.025510075 | -0.169016004 | 0.220036154 | 8.158339871 | 0.322628755 | 0.758159129 | 0.954649675 | -5.075467687 |
| **A_55_P2126469** | NA | -0.018981956 | -0.16373042 | 0.125766508 | 15.72140675 | -0.322623628 | 0.758162819 | 0.954649675 | -5.075468908 |
| **A_55_P2169391** | Atp6v1a | -0.039435206 | -0.340185277 | 0.261314865 | 12.27438665 | -0.322587502 | 0.758188829 | 0.954649675 | -5.075477512 |
| **A_51_P105709** | Trip13 | 0.037551945 | -0.24884349 | 0.32394738 | 8.81745622 | 0.322578567 | 0.758195262 | 0.954649675 | -5.046554681 |

| | | | | | | | | | |
|---|---|---|---|---|---|---|---|---|---|
| **A_51_P468240** | Dennd4a | 0.056652206 | -0.375420749 | 0.48872516 | 9.240760042 | 0.322573661 | 0.758198794 | 0.954649675 | -5.075480809 |
| **A_55_P1991109** | Gatd3a | 0.034902769 | -0.231441501 | 0.30124704 | 11.79518574 | 0.322393067 | 0.758328821 | 0.954649675 | -5.075523805 |
| **A_55_P2066543** | Orai2 | 0.015982738 | -0.105990133 | 0.13795561 | 8.356132932 | 0.322371842 | 0.758344104 | 0.954649675 | -5.075528857 |
| **A_51_P462271** | Acan | 0.014092027 | -0.093463844 | 0.121647899 | 7.790986629 | 0.322335781 | 0.758370068 | 0.954649675 | -5.075537439 |
| **A_55_P2020153** | Cycs | -0.020414322 | -0.182982821 | 0.142154178 | 12.88924782 | -0.325423293 | 0.758377588 | 0.954649675 | -5.007870495 |
| **A_52_P116372** | Ppp1r15b | -0.025418087 | -0.219468381 | 0.168632208 | 9.453888318 | -0.32225355 | 0.758429278 | 0.954649675 | -5.075557005 |
| **A_55_P2075195** | Cd302 | 0.041020546 | -0.272192901 | 0.354233993 | 9.189649474 | 0.322203461 | 0.758465346 | 0.954649675 | -5.075568922 |
| **A_65_P02323** | Sec22b | -0.035801664 | -0.309481867 | 0.237878538 | 8.436616478 | -0.321831831 | 0.758732963 | 0.954649675 | -5.075657278 |
| **A_52_P12806** | Gm5069 | -0.061771263 | -0.533990679 | 0.410448153 | 14.08079004 | -0.32181909 | 0.758742139 | 0.954649675 | -5.075660305 |
| **A_55_P2198946** | Gm10768 | 0.058018524 | -0.385572551 | 0.5016096 | 8.06333449 | 0.321775549 | 0.758773496 | 0.954649675 | -5.075670651 |
| **A_52_P484838** | Rfxank | 0.018268679 | -0.12142134 | 0.157958698 | 9.417285805 | 0.321744347 | 0.758795968 | 0.954649675 | -5.075678063 |
| **A_55_P1982916** | Tspoap1 | -0.013972138 | -0.120811386 | 0.09286711 | 7.801946951 | -0.321737145 | 0.758801154 | 0.954649675 | -5.075679774 |
| **A_55_P1990217** | Zfp13 | 0.014894863 | -0.099004038 | 0.128793765 | 7.884335182 | 0.321725991 | 0.758809187 | 0.954649675 | -5.075682424 |
| **A_52_P367774** | Zfp821 | -0.028172398 | -0.243657998 | 0.187313201 | 8.955184816 | -0.321643464 | 0.758868624 | 0.954649675 | -5.075702026 |
| **A_55_P2225087** | 5430439M09Rik | 0.015561868 | -0.103470524 | 0.13459426 | 8.501677929 | 0.3216368 | 0.758873423 | 0.954649675 | -5.075703608 |
| **A_55_P2159573** | Reep6 | 0.014481027 | -0.096296984 | 0.125259038 | 7.863227483 | 0.321599202 | 0.758900503 | 0.954649675 | -5.075712537 |
| **A_52_P657360** | Tnni1 | 0.017137681 | -0.113963437 | 0.148238798 | 7.804810233 | 0.321599093 | 0.758900581 | 0.954649675 | -5.075712563 |
| **A_51_P492446** | Ldah | 0.020070066 | -0.133475695 | 0.173615826 | 11.33444582 | 0.321573408 | 0.75891908 | 0.954649675 | -5.075718662 |
| **A_55_P2185504** | Masp2 | -0.013962299 | -0.120826952 | 0.092902353 | 7.667462591 | -0.321434166 | 0.75901937 | 0.954684241 | -5.075751716 |
| **A_65_P11853** | Arhgap28 | -0.034548728 | -0.299081269 | 0.229983814 | 9.603698464 | -0.321308429 | 0.759109937 | 0.954684241 | -5.075781554 |
| **A_52_P444162** | Rab11a | 0.039715094 | -0.264415119 | 0.343845307 | 11.34972517 | 0.321266316 | 0.759140272 | 0.954684241 | -5.075791545 |
| **A_52_P381303** | Gins2 | 0.022675779 | -0.151041337 | 0.196392895 | 8.263335268 | 0.321135831 | 0.759234265 | 0.954684241 | -5.075822493 |
| **A_51_P379597** | Ndufs1 | -0.014741269 | -0.127676062 | 0.098193523 | 12.290577 | -0.321126608 | 0.759240909 | 0.954684241 | -5.07582468 |
| **A_51_P172231** | Gsdmd | -0.086177523 | -0.746400497 | 0.57404545 | 9.218297711 | -0.321123834 | 0.759242907 | 0.954684241 | -5.070555578 |
| **A_55_P2041017** | Gm15365 | -0.054355767 | -0.470990163 | 0.362278629 | 13.62976327 | -0.32096647 | 0.759356268 | 0.954720198 | -5.075862644 |
| **A_55_P2363030** | Cmtm4 | 0.059856103 | -0.399192158 | 0.518904364 | 12.11345327 | 0.320788842 | 0.759484236 | 0.954720198 | -5.075904733 |
| **A_51_P304239** | Loxl1 | 0.05445989 | -0.363284649 | 0.47220443 | 8.572321139 | 0.320726721 | 0.759528991 | 0.954720198 | -5.075919447 |
| **A_51_P302520** | Myom1 | -0.018173262 | -0.157583501 | 0.121236977 | 7.866724961 | -0.320706216 | 0.759543765 | 0.954720198 | -5.075924304 |
| **A_55_P2403325** | NA | -0.022177683 | -0.192309935 | 0.14795457 | 7.793930099 | -0.32069979 | 0.759548394 | 0.954720198 | -5.075925825 |
| **A_52_P42194** | NA | 0.016135476 | -0.107666514 | 0.139937467 | 7.76586502 | 0.320644156 | 0.759588477 | 0.954720198 | -5.075939 |
| **A_55_P2006018** | Slc4a2 | 0.014259133 | -0.095159948 | 0.123678214 | 10.6981322 | 0.320604213 | 0.759617255 | 0.954720198 | -5.075948457 |
| **A_51_P164630** | Fitm1 | 0.013680273 | -0.091406636 | 0.118767181 | 7.637562455 | 0.320269293 | 0.759858579 | 0.954946585 | -5.076027713 |
| **A_55_P1983209** | Rcbtb1 | -0.030931265 | -0.268572786 | 0.206710256 | 10.34238058 | -0.3202171 | 0.759896188 | 0.954946585 | -5.076040057 |
| **A_51_P355301** | Cyp3a11 | 0.018349427 | -0.122676237 | 0.159375092 | 7.780722286 | 0.320105787 | 0.759976402 | 0.954949722 | -5.076066376 |
| **A_55_P2122709** | Arhgef3 | -0.051094911 | -0.443823931 | 0.341634109 | 11.32095556 | -0.320076517 | 0.759997495 | 0.954949722 | -5.076073296 |
| **A_55_P2120545** | Wdtc1 | -0.016812888 | -0.146106324 | 0.112480548 | 8.398443582 | -0.319915297 | 0.760113679 | 0.954991232 | -5.076111397 |
| **A_55_P2055612** | Zfp800 | 0.019346536 | -0.129471435 | 0.168164506 | 8.461381032 | 0.319828376 | 0.760176322 | 0.954991232 | -5.076131932 |
| **A_51_P132978** | Idh1 | 0.039353419 | -0.263365262 | 0.342072101 | 8.519325441 | 0.319825004 | 0.760178752 | 0.954991232 | -5.076132728 |
| **A_51_P512172** | Tet1 | -0.023872431 | -0.207546183 | 0.159801322 | 8.002468583 | -0.319755999 | 0.760228485 | 0.954991641 | -5.076149026 |
| **A_52_P537466** | Zfp330 | -0.033084793 | -0.287799889 | 0.221630304 | 9.429699815 | -0.319552997 | 0.760374798 | 0.955025032 | -5.076196953 |
| **A_55_P2088325** | Ctdsp2 | -0.033151787 | -0.288415555 | 0.22211198 | 10.3580444 | -0.319511825 | 0.760404474 | 0.955025032 | -5.07620667 |
| **A_55_P1967149** | Irx3 | 0.03919827 | -0.262663244 | 0.341059784 | 11.33422338 | 0.319468703 | 0.760435556 | 0.955025032 | -5.076216845 |
| **A_55_P1992676** | Ubtf | 0.020963879 | -0.140496898 | 0.182424656 | 8.402805263 | 0.319428574 | 0.760464481 | 0.955025032 | -5.076226313 |
| **A_55_P1962149** | Dnah10 | -0.018218111 | -0.158554318 | 0.122118095 | 8.613525221 | -0.31937637 | 0.760502111 | 0.955025032 | -5.076238629 |
| **A_51_P518163** | Rrp9 | -0.017134902 | -0.14917792 | 0.114908116 | 8.540970453 | -0.319253267 | 0.760590848 | 0.955028424 | -5.076267663 |
| **A_55_P2053157** | Zc3h3 | 0.03793738 | -0.254427546 | 0.330302306 | 10.57342343 | 0.319235536 | 0.760603629 | 0.955028424 | -5.076271844 |
| **A_55_P2001148** | Pdlim5 | 0.014305852 | -0.09607453 | 0.124686233 | 7.824947276 | 0.318853351 | 0.760879149 | 0.955312314 | -5.07636191 |
| **A_55_P1952517** | Sult2a1 | -0.014006868 | -0.122150449 | 0.094136713 | 7.66083374 | -0.318646715 | 0.761028132 | 0.955338218 | -5.076410563 |
| **A_55_P2102828** | NA | -0.013782246 | -0.120196587 | 0.092632094 | 7.674943338 | -0.318631725 | 0.761038939 | 0.955338218 | -5.076414091 |
| **A_52_P456279** | Cct8 | -0.015654463 | -0.136529202 | 0.105220276 | 12.36058299 | -0.31861908 | 0.761048057 | 0.955338218 | -5.076417067 |

| | | | | | | | | | |
|---|---|---|---|---|---|---|---|---|---|
| A_55_P2078895 | Gin1 | 0.025783706 | -0.173460024 | 0.225027437 | 8.210298593 | 0.318368321 | 0.761228868 | 0.955503136 | -5.076476064 |
| A_51_P440047 | Lgalsl | 0.071487445 | -0.48121447 | 0.624189361 | 11.3212536 | 0.318205711 | 0.761346127 | 0.955587536 | -5.076514298 |
| A_51_P270206 | Cep83 | -0.014601349 | -0.127535433 | 0.098332735 | 9.860502447 | -0.318080552 | 0.761436385 | 0.955587536 | -5.076543713 |
| A_51_P300655 | Tbccd1 | 0.016492107 | -0.111070507 | 0.144054721 | 9.344704709 | 0.318069408 | 0.761444422 | 0.955587536 | -5.076546332 |
| A_55_P2145709 | NA | 0.014197191 | -0.095752489 | 0.12414687 | 7.768177834 | 0.317671031 | 0.761731739 | 0.955664464 | -5.076639884 |
| A_55_P1963873 | NA | -0.036914637 | -0.322835711 | 0.249006436 | 14.15399289 | -0.317630065 | 0.761761286 | 0.955664464 | -5.076649498 |
| A_51_P244690 | Ube2b | -0.021641604 | -0.189307783 | 0.146024574 | 13.25469821 | -0.317550756 | 0.761818492 | 0.955664464 | -5.076668106 |
| A_55_P2061084 | Wfikkn2 | 0.016988037 | -0.114632685 | 0.14860876 | 7.807377461 | 0.31753244 | 0.761831703 | 0.955664464 | -5.076672403 |
| A_55_P2016119 | Sgcb | -0.017086481 | -0.149502921 | 0.115329958 | 9.124583272 | -0.317453334 | 0.761888764 | 0.955664464 | -5.076690959 |
| A_66_P137475 | Dnajb14 | -0.0437478 | -0.382847125 | 0.295351525 | 11.92867193 | -0.317393813 | 0.761931698 | 0.955664464 | -5.076704918 |
| A_55_P2099390 | Ranbp9 | -0.038677055 | -0.338493872 | 0.261139762 | 9.210066295 | -0.317370514 | 0.761948505 | 0.955664464 | -5.076710381 |
| A_51_P245156 | Gdf2 | -0.014928256 | -0.130670195 | 0.100813682 | 7.646191842 | -0.317312732 | 0.761990186 | 0.955664464 | -5.076723929 |
| A_55_P1983414 | NA | -0.030988825 | -0.27127866 | 0.209301009 | 9.021481527 | -0.31727721 | 0.762015811 | 0.955664464 | -5.076732256 |
| A_55_P2008687 | Nlk | -0.028802738 | -0.252149354 | 0.194543878 | 11.02671322 | -0.317266017 | 0.762023885 | 0.955664464 | -5.07673488 |
| A_55_P2004447 | Fgl1 | -0.015919054 | -0.139405418 | 0.10756731 | 7.798867908 | -0.317151972 | 0.762106157 | 0.955664464 | -5.076761608 |
| A_51_P390276 | Cwc25 | -0.020558324 | -0.180052741 | 0.138936093 | 9.385699988 | -0.317111056 | 0.762135674 | 0.955664464 | -5.076771195 |
| A_55_P2057380 | NA | -0.029226154 | -0.255979519 | 0.19752721 | 7.979786898 | -0.317093324 | 0.762148467 | 0.955664464 | -5.07677535 |
| A_66_P132448 | Gnpnat1 | 0.027747975 | -0.187672457 | 0.243168407 | 10.05508886 | 0.316893671 | 0.762292506 | 0.955783073 | -5.076822111 |
| A_52_P252258 | Mapk12 | -0.015409734 | -0.135128784 | 0.104309315 | 7.946325468 | -0.316665723 | 0.762456972 | 0.955927275 | -5.076875466 |
| A_55_P1990548 | Tbc1d22a | 0.031806421 | -0.215435071 | 0.279047912 | 12.19191064 | 0.316491903 | 0.762582393 | 0.956022511 | -5.076916126 |
| A_55_P2010788 | Slc6a9 | -0.015003025 | -0.131668579 | 0.101662528 | 8.281863919 | -0.316377334 | 0.762665066 | 0.956064146 | -5.076942915 |
| A_55_P1953688 | Top1 | -0.029863018 | -0.262169108 | 0.202443072 | 12.10484676 | -0.31625854 | 0.762750791 | 0.956084592 | -5.076970681 |
| A_51_P172502 | Cxcl12 | 0.035601456 | -0.241406344 | 0.312609257 | 12.46064451 | 0.316187669 | 0.762801935 | 0.956084592 | -5.076987242 |
| A_51_P405167 | Maf | -0.056867968 | -0.499652125 | 0.385916188 | 9.545216781 | -0.315969239 | 0.762959574 | 0.956084592 | -5.07703826 |
| A_55_P2005005 | Cdk5rap1 | -0.159481837 | -1.401331897 | 1.082368223 | 11.43439224 | -0.315944813 | 0.762977202 | 0.956084592 | -5.077043964 |
| A_55_P2449750 | Fam177a2 | -0.023498016 | -0.206478502 | 0.159482469 | 8.543921435 | -0.315933433 | 0.762985416 | 0.956084592 | -5.077046621 |
| A_52_P188261 | Camk2d | -0.026582783 | -0.233625968 | 0.180460402 | 9.483842522 | -0.315870207 | 0.763031049 | 0.956084592 | -5.07706138 |
| A_52_P515769 | Pcdh12 | -0.068533183 | -0.602368963 | 0.465302596 | 12.35764587 | -0.315836552 | 0.763055339 | 0.956084592 | -5.077069236 |
| A_55_P1961888 | Twistnb | -0.039014316 | -0.342943364 | 0.264914731 | 9.789392111 | -0.315806421 | 0.763077086 | 0.956084592 | -5.077076268 |
| A_55_P1975837 | NA | -0.026884078 | -0.236480508 | 0.182712352 | 12.53782674 | -0.315558896 | 0.763255749 | 0.956092208 | -5.077134013 |
| A_55_P2008437 | Cdkn2a | 0.057247837 | -0.389149865 | 0.503645539 | 9.337298687 | 0.315505036 | 0.763294627 | 0.956092208 | -5.077146573 |
| A_55_P1979156 | Arap3 | -0.014274083 | -0.125584856 | 0.09703669 | 7.845169756 | -0.315486059 | 0.763308325 | 0.956092208 | -5.077150997 |
| A_52_P7183 | Tmem248 | -0.021601858 | -0.19006821 | 0.146864495 | 11.3662494 | -0.315462026 | 0.763325673 | 0.956092208 | -5.0771566 |
| A_52_P563340 | Smim4 | 0.052365385 | -0.356025043 | 0.460755813 | 11.00535276 | 0.315455359 | 0.763330486 | 0.956092208 | -5.077158155 |
| A_51_P162955 | Serpina7 | -0.013705585 | -0.120691678 | 0.093280508 | 7.638008219 | -0.315166047 | 0.763539338 | 0.956213284 | -5.077225572 |
| A_55_P2111870 | Rogdi | -0.015984425 | -0.140760594 | 0.108791743 | 8.376175819 | -0.315162509 | 0.763541892 | 0.956213284 | -5.077226396 |
| A_55_P2025665 | Mrpl40 | 0.028591203 | -0.194628837 | 0.251811243 | 11.09460555 | 0.315114516 | 0.76357654 | 0.956213284 | -5.077237574 |
| A_55_P1990495 | Stat5a | 0.033110321 | -0.225496238 | 0.291716879 | 9.130382564 | 0.31498728 | 0.763668399 | 0.956213284 | -5.077267201 |
| A_51_P245368 | Abcb1b | -0.021023 | -0.18529714 | 0.14325114 | 10.39616174 | -0.314843438 | 0.763772252 | 0.956213284 | -5.077300679 |
| A_52_P597726 | Ccdc84 | -0.019338645 | -0.170466504 | 0.131789213 | 8.730506594 | -0.314811568 | 0.763795263 | 0.956213284 | -5.077308095 |
| A_55_P2024704 | Cpe | -0.295252586 | -2.602674718 | 2.012169545 | 14.44174345 | -0.314800795 | 0.763803042 | 0.956213284 | -5.077310602 |
| A_51_P223498 | Slc39a10 | -0.023588121 | -0.207947259 | 0.160771016 | 8.75944672 | -0.314773271 | 0.763822914 | 0.956213284 | -5.077317005 |
| A_51_P271865 | Clec4f | -0.013716279 | -0.120961374 | 0.093528817 | 7.71656788 | -0.314650205 | 0.763911774 | 0.956225288 | -5.077345631 |
| A_55_P2185332 | Il17rc | 0.115592003 | -0.788473835 | 1.019657841 | 10.81008467 | 0.314555218 | 0.763980362 | 0.956225288 | -5.077367718 |
| A_51_P436386 | Qrich1 | -0.038192308 | -0.336961715 | 0.260577099 | 10.97535672 | -0.314491521 | 0.764026357 | 0.956225288 | -5.077382526 |
| A_52_P636343 | Wipi2 | -0.02378104 | -0.209834809 | 0.162272729 | 9.221174663 | -0.314457192 | 0.764051147 | 0.956225288 | -5.077390506 |
| A_55_P1987504 | Ccnc | 0.019059825 | -0.130075819 | 0.168195469 | 8.380098413 | 0.314417433 | 0.764079858 | 0.956225288 | -5.077399746 |
| A_51_P456721 | Azgp1 | 0.021745974 | -0.148541594 | 0.192033543 | 7.743175106 | 0.314170275 | 0.764258345 | 0.956386737 | -5.077457163 |
| A_55_P2122255 | Polm | 0.020523691 | -0.14047214 | 0.181519521 | 9.073674198 | 0.313624509 | 0.76465253 | 0.956596073 | -5.077583798 |
| A_55_P2235970 | C130068B02Rik | 0.046788758 | -0.320306656 | 0.413884173 | 8.271069529 | 0.313567992 | 0.764693355 | 0.956596073 | -5.077596899 |

| | | | | | | | | | |
|---|---|---|---|---|---|---|---|---|---|
| **A_51_P464064** | Zc3h13 | -0.017933422 | -0.158656657 | 0.122789813 | 8.304980223 | -0.313520919 | 0.764727358 | 0.956596073 | -5.07760781 |
| **A_51_P119597** | Ap3b1 | 0.030450759 | -0.208507032 | 0.269408549 | 12.45105324 | 0.313506168 | 0.764738014 | 0.956596073 | -5.077611229 |
| **A_55_P2119155** | Set | 0.015959948 | -0.109284492 | 0.141204388 | 9.937396496 | 0.313503352 | 0.764740048 | 0.956596073 | -5.077611881 |
| **A_55_P1964757** | Stam2 | -0.019591603 | -0.173365667 | 0.134182462 | 8.799329228 | -0.313441162 | 0.764784972 | 0.956596073 | -5.077626292 |
| **A_55_P2182806** | NA | -0.031523786 | -0.278990638 | 0.215943066 | 11.45920765 | -0.31339387 | 0.764819135 | 0.956596073 | -5.07763725 |
| **A_55_P2033616** | Dhrs7b | -0.03048761 | -0.269984067 | 0.209008847 | 11.57786334 | -0.313179589 | 0.764973936 | 0.956596073 | -5.077686877 |
| **A_55_P2009206** | Sumo2 | 0.018024333 | -0.123629341 | 0.159678007 | 8.823751933 | 0.313040501 | 0.765074423 | 0.956596073 | -5.077719072 |
| **A_52_P506271** | Cmtm4 | 0.042129797 | -0.289003884 | 0.373263478 | 8.514908758 | 0.313007811 | 0.765098041 | 0.956596073 | -5.077726637 |
| **A_51_P130095** | Fcgr2b | -0.016064031 | -0.142389797 | 0.110261735 | 7.83333892 | -0.312846821 | 0.765214358 | 0.956596073 | -5.071880785 |
| **A_55_P2006277** | Ogdh | 0.085411232 | -0.586302352 | 0.757124816 | 10.64448625 | 0.312823966 | 0.765230872 | 0.956596073 | -5.077769167 |
| **A_51_P237834** | Tmcc1 | -0.04483201 | -0.397544096 | 0.307880075 | 10.01339273 | -0.312706591 | 0.765315682 | 0.956596073 | -5.077796308 |
| **A_51_P113403** | Slc26a11 | 0.020431751 | -0.140314889 | 0.18117839 | 9.25464755 | 0.312703574 | 0.765317862 | 0.956596073 | -5.051175204 |
| **A_55_P2427900** | Unc13c | 0.014061118 | -0.096657374 | 0.12477961 | 7.634069227 | 0.312441596 | 0.765507169 | 0.956596073 | -5.077857547 |
| **A_51_P221495** | Nsun5 | -0.01450355 | -0.128712848 | 0.099705749 | 8.276947138 | -0.312422284 | 0.765521125 | 0.956596073 | -5.077862008 |
| **A_55_P2142590** | Mark3 | -0.02257857 | -0.200409233 | 0.155252092 | 11.93366884 | -0.312362547 | 0.765564294 | 0.956596073 | -5.077875805 |
| **A_55_P2053258** | Poc1a | 0.034205227 | -0.235220102 | 0.303630555 | 9.361939099 | 0.312336844 | 0.765582868 | 0.956596073 | -5.077881741 |
| **A_55_P1975021** | Rad18 | -0.01609384 | -0.142878808 | 0.110691127 | 8.241086511 | -0.312292166 | 0.765615156 | 0.956596073 | -5.077892058 |
| **A_51_P307243** | Ppp5c | 0.033722057 | -0.231993829 | 0.299437942 | 10.72681723 | 0.312223579 | 0.765664723 | 0.956596073 | -5.077907893 |
| **A_55_P2186025** | NA | 0.048910457 | -0.33660406 | 0.434424974 | 14.85273391 | 0.312126143 | 0.765735141 | 0.956596073 | -5.077930382 |
| **A_55_P2055442** | Pafah2 | 0.017183686 | -0.118261961 | 0.152629332 | 9.152682631 | 0.312119165 | 0.765740184 | 0.956596073 | -5.077931993 |
| **A_55_P2040723** | Dnali1 | 0.013252475 | -0.091214834 | 0.117719784 | 7.683873088 | 0.312094234 | 0.765758202 | 0.956596073 | -5.077937746 |
| **A_55_P1978795** | Pja2 | 0.02253114 | -0.155137953 | 0.200200233 | 11.97191349 | 0.311989837 | 0.765833655 | 0.956596073 | -5.077961832 |
| **A_55_P1987186** | Ttll9 | 0.013524047 | -0.093129964 | 0.120178059 | 7.663106703 | 0.311959811 | 0.765855356 | 0.956596073 | -5.077968758 |
| **A_55_P2053459** | Timd2 | 0.083149184 | -0.572994964 | 0.739293332 | 10.35371611 | 0.311765385 | 0.765995885 | 0.956596073 | -5.078013592 |
| **A_51_P140237** | Fhl2 | 0.03826384 | -0.263715949 | 0.340243629 | 13.16980991 | 0.311730889 | 0.76602082 | 0.956596073 | -5.078021543 |
| **A_51_P124126** | Cyp2d22 | 0.015017144 | -0.103501172 | 0.13353546 | 7.722884629 | 0.311724574 | 0.766025385 | 0.956596073 | -5.078022999 |
| **A_66_P140076** | NA | 0.030834205 | -0.21253333 | 0.27420174 | 10.82058953 | 0.311701775 | 0.766041865 | 0.956596073 | -5.078028254 |
| **A_55_P2000521** | Lonrf1 | 0.03018996 | -0.2081171 | 0.268497021 | 10.43529662 | 0.311669853 | 0.766064939 | 0.956596073 | -5.078035611 |
| **A_52_P283910** | Hnrnpdl | 0.017283863 | -0.119168511 | 0.153736237 | 9.870431606 | 0.31162255 | 0.766099132 | 0.956596073 | -5.078046511 |
| **A_55_P2022181** | Arvcf | -0.033070281 | -0.294174798 | 0.228034237 | 9.279223953 | -0.311596564 | 0.766117916 | 0.956596073 | -5.078052499 |
| **A_55_P2102719** | NA | -0.042346031 | -0.376747622 | 0.29205556 | 12.44259802 | -0.311539801 | 0.766158948 | 0.956596073 | -5.078065576 |
| **A_51_P388696** | Ndufs4 | 0.050523129 | -0.348815702 | 0.449861961 | 12.36950264 | 0.311256094 | 0.766364043 | 0.956596073 | -5.078130904 |
| **A_55_P2012729** | Kcnh2 | 0.015485828 | -0.106923006 | 0.137894663 | 7.761054582 | 0.311236727 | 0.766378045 | 0.956596073 | -5.078135361 |
| **A_51_P169745** | Tuba1a | -0.035580538 | -0.316854257 | 0.245693182 | 14.55107689 | -0.311209262 | 0.766397901 | 0.956596073 | -5.078141682 |
| **A_51_P192344** | Ash2l | 0.027128797 | -0.187354481 | 0.241612074 | 11.87619462 | 0.311176118 | 0.766421862 | 0.956596073 | -5.078149309 |
| **A_52_P486253** | Polr2b | -0.019292076 | -0.171833859 | 0.133249707 | 11.26954504 | -0.311142477 | 0.766446184 | 0.956596073 | -5.07815705 |
| **A_52_P140005** | Nipal1 | 0.05663332 | -0.391175723 | 0.504442363 | 8.587723616 | 0.31113461 | 0.766451872 | 0.956596073 | -5.07815886 |
| **A_55_P1987499** | Pttg1 | 0.018761771 | -0.129591112 | 0.167114654 | 12.79137124 | 0.311133659 | 0.766452559 | 0.956596073 | -5.078159079 |
| **A_51_P349546** | Cd109 | -0.207437507 | -1.847701056 | 1.432826041 | 9.835030468 | -0.311130663 | 0.766454725 | 0.956596073 | -5.078159768 |
| **A_51_P347240** | Lrr1 | -0.016866794 | -0.150299428 | 0.11656584 | 7.80595565 | -0.310985147 | 0.766559933 | 0.956665609 | -5.07819324 |
| **A_55_P1959576** | Slitrk4 | 0.013796403 | -0.095435723 | 0.12302853 | 7.669304997 | 0.310731055 | 0.766743655 | 0.956772384 | -5.078251652 |
| **A_55_P1959923** | Cth | 0.042293481 | -0.29256365 | 0.377150613 | 8.325909946 | 0.310729901 | 0.766744489 | 0.956772384 | -5.078251917 |
| **A_52_P313217** | Adgrd1 | -0.154696014 | -1.379784494 | 1.070392465 | 8.867871129 | -0.310656776 | 0.766797365 | 0.956776598 | -5.078268719 |
| **A_51_P401876** | Kif1bp | 0.022514329 | -0.155868844 | 0.200897502 | 11.36309509 | 0.310509068 | 0.766904176 | 0.956848104 | -5.078302645 |
| **A_51_P415395** | C2cd4b | 0.014064887 | -0.097414565 | 0.125544338 | 7.786478977 | 0.310392044 | 0.766988803 | 0.95686068 | -5.078329514 |
| **A_51_P507974** | Brf2 | -0.01616378 | -0.144293194 | 0.111965634 | 8.390477468 | -0.31035822 | 0.767013264 | 0.95686068 | -5.078337278 |
| **A_55_P1973560** | NA | 0.022797127 | -0.158020455 | 0.20361471 | 9.628125054 | 0.310176313 | 0.76714482 | 0.956902731 | -5.078379019 |
| **A_51_P400366** | Rhbg | 0.014653765 | -0.101574566 | 0.130882095 | 7.680423701 | 0.310174703 | 0.767145984 | 0.956902731 | -5.078379388 |
| **A_55_P2042973** | 1010001N08Rik | 0.016349895 | -0.113423114 | 0.146122904 | 7.887507637 | 0.309955798 | 0.767304308 | 0.956980318 | -5.078429588 |
| **A_52_P375493** | Tfdp1 | -0.032677785 | -0.292082865 | 0.226727295 | 9.88400409 | -0.3099155 | 0.767333455 | 0.956980318 | -5.078438825 |

| | | | | | | | | |
|---|---|---|---|---|---|---|---|---|
| **A_51_P196444** | Foxc2 | -0.016765976 | -0.149872633 | 0.116340681 | 7.865272838 | -0.309883342 | 0.767356715 | 0.956980318 | -5.078446196 |
| **A_55_P2069251** | Prr18 | 0.013258388 | -0.092043303 | 0.118560078 | 7.72050612 | 0.309759428 | 0.767446345 | 0.957030348 | -5.078474591 |
| **A_66_P122699** | NA | -0.013232216 | -0.118428116 | 0.091963683 | 7.686319453 | -0.30945888 | 0.767663753 | 0.957055505 | -5.078543417 |
| **A_51_P100828** | Prdx3 | 0.022898349 | -0.159146918 | 0.204943616 | 10.76155875 | 0.309452464 | 0.767668395 | 0.957055505 | -5.073258449 |
| **A_51_P184284** | Dld | -0.027251965 | -0.243921125 | 0.189417196 | 11.47446482 | -0.309435318 | 0.767680798 | 0.957055505 | -5.07854881 |
| **A_51_P346445** | Hspb7 | 0.065759847 | -0.45712087 | 0.588640564 | 8.177593126 | 0.309405002 | 0.767702729 | 0.957055505 | -5.046118123 |
| **A_51_P122740** | Slc37a3 | -0.028077581 | -0.251406323 | 0.195251161 | 10.2953827 | -0.309303071 | 0.76777647 | 0.957055505 | -5.040058305 |
| **A_55_P1971498** | Exoc1 | -0.03158024 | -0.282837702 | 0.219677222 | 11.60090167 | -0.309218609 | 0.767837576 | 0.957055505 | -5.078598393 |
| **A_51_P102911** | Hoxa11 | -0.180159054 | -1.614122228 | 1.253804119 | 10.39250468 | -0.30909164 | 0.767929437 | 0.957055505 | -5.078627428 |
| **A_55_P2006762** | Bzw1 | -0.024394826 | -0.218593142 | 0.16980349 | 12.05524265 | -0.309044788 | 0.767963335 | 0.957055505 | -5.078638139 |
| **A_52_P661587** | Ccdc141 | -0.013098022 | -0.117371264 | 0.09117522 | 7.653087678 | -0.309030968 | 0.767973334 | 0.957055505 | -5.078641298 |
| **A_55_P2038747** | Ano1 | -0.187750292 | -1.682572703 | 1.307072118 | 11.74060095 | -0.309001213 | 0.767994862 | 0.957055505 | -5.078648099 |
| **A_55_P2016356** | Eif3b | 0.034470376 | -0.23999444 | 0.308935191 | 10.33167957 | 0.308978671 | 0.768011172 | 0.957055505 | -5.078653252 |
| **A_55_P1991160** | Mrrf | -0.014717517 | -0.131971106 | 0.102536073 | 7.792702965 | -0.308800183 | 0.768140319 | 0.957146315 | -5.078694034 |
| **A_52_P110052** | Ackr1 | 0.014576241 | -0.101580888 | 0.13073337 | 7.742785458 | 0.308722887 | 0.768196249 | 0.957146315 | -5.078711689 |
| **A_55_P2070045** | Golga7b | -0.177464356 | -1.592371065 | 1.237442352 | 9.109645502 | -0.308569144 | 0.768307501 | 0.957146315 | -5.078746791 |
| **A_55_P2043362** | NA | -0.03167747 | -0.284245931 | 0.22089099 | 8.208582235 | -0.30856065 | 0.768313648 | 0.957146315 | -5.07874873 |
| **A_66_P139387** | Prlr | 0.046216915 | -0.322306057 | 0.414739886 | 9.030150893 | 0.30853579 | 0.768331638 | 0.957146315 | -5.078754404 |
| **A_66_P117578** | Mtmr7 | -0.056823153 | -0.510055243 | 0.396408938 | 10.13924464 | -0.308442232 | 0.768399342 | 0.957168968 | -5.078775755 |
| **A_55_P2151855** | Acp1 | -0.031022101 | -0.278552453 | 0.216508251 | 11.3335228 | -0.308327248 | 0.768482555 | 0.957210936 | -5.078801987 |
| **A_51_P276235** | Pnpla7 | -0.019543651 | -0.175582567 | 0.136495266 | 8.203295312 | -0.308135636 | 0.76862123 | 0.957239575 | -5.078845679 |
| **A_55_P2319370** | Plekha3 | 0.022550634 | -0.157496753 | 0.202598021 | 11.81034586 | 0.308135049 | 0.768621654 | 0.957239575 | -5.078845813 |
| **A_52_P399990** | B4galt2 | 0.017267895 | -0.120637357 | 0.155173147 | 8.053517878 | 0.308054647 | 0.768679847 | 0.957239575 | -5.078864139 |
| **A_52_P298548** | Nbn | -0.018351336 | -0.164959327 | 0.128256655 | 8.19364176 | -0.307949289 | 0.768756103 | 0.957239575 | -5.078888146 |
| **A_51_P207591** | Anxa8 | -0.092752221 | -0.833880712 | 0.64837627 | 12.53149161 | -0.307893111 | 0.768796765 | 0.957239575 | -5.073612325 |
| **A_55_P1997509** | Zbtb32 | 0.050627282 | -0.354004328 | 0.455258892 | 9.229873784 | 0.307817969 | 0.768851155 | 0.957239575 | -5.078918058 |
| **A_51_P354272** | Cluh | -0.017803209 | -0.160093586 | 0.124487169 | 10.50873997 | -0.30781651 | 0.768852211 | 0.957239575 | -5.07891839 |
| **A_55_P1984178** | Pde4dip | -0.042166705 | -0.379333909 | 0.295000499 | 11.80772216 | -0.307675915 | 0.768953981 | 0.957304618 | -5.078950401 |
| **A_55_P2048101** | Gm6762 | -0.024683406 | -0.22214222 | 0.172775408 | 8.601379325 | -0.307537246 | 0.769054362 | 0.957367925 | -5.078981959 |
| **A_55_P2110537** | Slfn9 | 0.042962183 | -0.300800821 | 0.386725188 | 9.485285508 | 0.307465485 | 0.769106311 | 0.957370935 | -5.078998286 |
| **A_55_P1989544** | Cklf | 0.023606995 | -0.165419799 | 0.212633788 | 10.43954104 | 0.307246182 | 0.769265076 | 0.957382075 | -5.079048156 |
| **A_55_P2154208** | NA | -0.02700403 | -0.243274862 | 0.189266801 | 12.51538317 | -0.307184854 | 0.769309477 | 0.957382075 | -5.079062096 |
| **A_51_P117618** | Ethe1 | 0.036596634 | -0.256506091 | 0.329699359 | 9.669040991 | 0.307178244 | 0.769314262 | 0.957382075 | -5.079063599 |
| **A_51_P424054** | Btbd2 | -0.016530971 | -0.148949347 | 0.115887405 | 9.375744003 | -0.307127906 | 0.769350708 | 0.957382075 | -5.079075039 |
| **A_66_P115467** | Sfta2 | 0.016268916 | -0.114057465 | 0.146595297 | 8.478176246 | 0.307111046 | 0.769362914 | 0.957382075 | -5.07907887 |
| **A_55_P2063041** | Prelid3b | 0.017815752 | -0.124986666 | 0.16061817 | 9.070943719 | 0.306928882 | 0.769494809 | 0.957401395 | -5.079120252 |
| **A_66_P105270** | Hao1 | 0.015711254 | -0.11025895 | 0.141681458 | 7.919425013 | 0.306840142 | 0.769559063 | 0.957401395 | -5.079140402 |
| **A_55_P2048518** | Zfpm1 | -0.034971221 | -0.315378662 | 0.24543622 | 13.10075619 | -0.306824778 | 0.769570188 | 0.957401395 | -5.07914389 |
| **A_55_P2103706** | NA | 0.037569343 | -0.26377129 | 0.338909977 | 8.11814398 | 0.306722102 | 0.769644536 | 0.957401395 | -5.079167197 |
| **A_55_P2044684** | Rsph1 | -0.015035056 | -0.135654949 | 0.105584838 | 8.223988161 | -0.306658684 | 0.769690458 | 0.957401395 | -5.079181589 |
| **A_52_P106537** | Chrac1 | -0.023962822 | -0.216246387 | 0.168320743 | 9.469771013 | -0.306594921 | 0.769736631 | 0.957401395 | -5.079196056 |
| **A_51_P279127** | Fam173a | -0.019395509 | -0.175030281 | 0.136239263 | 11.72306112 | -0.306594036 | 0.769737273 | 0.957401395 | -5.079196256 |
| **A_55_P2088995** | Plscr1 | 0.047892826 | -0.336476866 | 0.432262518 | 9.079259509 | 0.306542359 | 0.769774695 | 0.957401395 | -5.079207979 |
| **A_52_P365768** | Nsun2 | 0.036629631 | -0.257601193 | 0.330860456 | 11.62343841 | 0.306276406 | 0.769967296 | 0.957431554 | -5.07926828 |
| **A_55_P2024525** | Prdm8 | 0.013230462 | -0.093048735 | 0.119509658 | 7.680700065 | 0.306263964 | 0.769976307 | 0.957431554 | -5.0792711 |
| **A_55_P1995055** | Prr7 | 0.015174473 | -0.106742403 | 0.137091348 | 8.097696361 | 0.306209703 | 0.770015605 | 0.957431554 | -5.079283396 |
| **A_55_P1956842** | NA | 0.037900101 | -0.266622886 | 0.342423088 | 14.42808077 | 0.306188902 | 0.770030671 | 0.957431554 | -5.079288109 |
| **A_55_P2083963** | Phgr1 | 0.081530638 | -0.573750929 | 0.736812205 | 10.46401806 | 0.3060991 | 0.770095711 | 0.957431554 | -5.079308454 |
| **A_55_P2394490** | NA | 0.013711396 | -0.096490647 | 0.12391344 | 7.773670968 | 0.306098501 | 0.770096144 | 0.957431554 | -5.079308589 |
| **A_55_P2055537** | Slc16a6 | -0.030648807 | -0.277051294 | 0.215753679 | 12.00487153 | -0.306011435 | 0.770159205 | 0.957448371 | -5.079328308 |

| | | | | | | | | | |
|---|---|---|---|---|---|---|---|---|---|
| **A_55_P1972034** | Muc16 | 0.22539435 | -1.587437304 | 2.038226005 | 9.026694951 | 0.305882558 | 0.770252553 | 0.957502836 | -5.079357487 |
| **A_55_P2039541** | Arhgap15 | -0.015421241 | -0.139533474 | 0.108690992 | 7.815337619 | -0.305684838 | 0.770395772 | 0.957603901 | -5.079402229 |
| **A_51_P211822** | Neurl4 | -0.031309563 | -0.283335458 | 0.220716333 | 10.79497332 | -0.305633531 | 0.770432938 | 0.957603901 | -5.079413835 |
| **A_51_P114826** | Cdh13 | -0.203664725 | -1.844914438 | 1.437584989 | 10.12992331 | -0.305288408 | 0.770682958 | 0.957830193 | -5.079491854 |
| **A_55_P2382262** | A830039N20Rik | 0.022610425 | -0.159694198 | 0.204915048 | 8.777435421 | 0.305126697 | 0.770800118 | 0.957830193 | -5.079528382 |
| **A_66_P111301** | Mplkip | -0.024737209 | -0.224195482 | 0.174721064 | 10.33622753 | -0.305117983 | 0.770806431 | 0.957830193 | -5.07953035 |
| **A_55_P2042773** | Ctdnep1 | 0.024562228 | -0.17350478 | 0.222629236 | 11.40121667 | 0.305087756 | 0.770828332 | 0.957830193 | -5.079537176 |
| **A_55_P2001773** | Copg2 | -0.017471449 | -0.158483839 | 0.12354094 | 9.911311 | -0.304818151 | 0.771023679 | 0.957830193 | -5.079598026 |
| **A_51_P396708** | Med21 | -0.02113597 | -0.191736852 | 0.149464911 | 11.17222242 | -0.304796577 | 0.771039312 | 0.957830193 | -5.079602893 |
| **A_52_P127325** | AI182371 | -0.013009934 | -0.118050885 | 0.092031016 | 7.682651617 | -0.304709244 | 0.771102594 | 0.957830193 | -5.079622592 |
| **A_52_P323820** | Ufm1 | 0.020859719 | -0.147593136 | 0.189312574 | 8.400343117 | 0.304648637 | 0.771146512 | 0.957830193 | -5.07963626 |
| **A_55_P2169659** | Vwf | -0.016587762 | -0.150553065 | 0.117377541 | 8.076618004 | -0.304624356 | 0.771164108 | 0.957830193 | -5.079641735 |
| **A_55_P2074488** | Crim1 | 0.091660017 | -0.648676613 | 0.831996647 | 11.9079282 | 0.304592958 | 0.77118686 | 0.957830193 | -5.079648814 |
| **A_55_P2030752** | Nedd4l | -0.016197289 | -0.147040939 | 0.114646362 | 11.97138068 | -0.304550159 | 0.771217876 | 0.957830193 | -5.079658462 |
| **A_55_P2047620** | NA | 0.050708037 | -0.358931787 | 0.460347861 | 13.50997266 | 0.304539613 | 0.771225517 | 0.957830193 | -5.079660839 |
| **A_55_P1970017** | Zer1 | -0.028930012 | -0.262696514 | 0.20483649 | 11.76815711 | -0.304463682 | 0.771280543 | 0.957830193 | -5.079677953 |
| **A_55_P2016981** | Nfu1 | -0.019710814 | -0.179002738 | 0.139581111 | 11.94022697 | -0.304424751 | 0.771308756 | 0.957830193 | -5.079686725 |
| **A_55_P2156101** | Skp1a | 0.051701041 | -0.366247346 | 0.469649427 | 13.53984913 | 0.304330722 | 0.771376901 | 0.957853277 | -5.07970791 |
| **A_55_P1957733** | Cysrt1 | 0.036059647 | -0.255842091 | 0.327961384 | 8.865681798 | 0.303916269 | 0.771677287 | 0.95807067 | -5.079801209 |
| **A_51_P130427** | Psmd3 | -0.020078024 | -0.182707416 | 0.142551368 | 13.98990827 | -0.303732379 | 0.77181058 | 0.95807067 | -5.079842566 |
| **A_51_P121818** | Prkag1 | 0.013943464 | -0.099000669 | 0.126887597 | 10.89226703 | 0.303721937 | 0.771818149 | 0.95807067 | -5.079844914 |
| **A_52_P630867** | Abcc4 | 0.049592442 | -0.352127509 | 0.451312393 | 9.066799264 | 0.303711516 | 0.771825704 | 0.95807067 | -5.079847257 |
| **A_51_P164296** | Adamdec1 | -0.014502847 | -0.131982321 | 0.102976626 | 7.678392004 | -0.30371095 | 0.771826114 | 0.95807067 | -5.079847384 |
| **A_51_P106397** | Prkd3 | -0.01308748 | -0.119113048 | 0.092938089 | 8.426065977 | -0.303678868 | 0.77184937 | 0.95807067 | -5.079854597 |
| **A_55_P2096187** | Hax1 | 0.032834811 | -0.233247849 | 0.298917471 | 12.68442922 | 0.303589753 | 0.771913969 | 0.958089329 | -5.079874627 |
| **A_55_P2375121** | AI225934 | -0.060311219 | -0.549330012 | 0.428707574 | 8.47778902 | -0.303418266 | 0.772038286 | 0.958182101 | -5.079913156 |
| **A_52_P193233** | Trnau1ap | -0.027057235 | -0.246583112 | 0.192468641 | 9.814550948 | -0.303226298 | 0.77217746 | 0.958193292 | -5.079956262 |
| **A_51_P372702** | Il16 | 0.01641983 | -0.116856953 | 0.149696613 | 7.919655946 | 0.303098177 | 0.77227035 | 0.958193292 | -5.079985016 |
| **A_55_P1953093** | Pabpc4 | -0.035464433 | -0.323440484 | 0.252511618 | 10.8433662 | -0.302974315 | 0.772360157 | 0.958193292 | -5.080012804 |
| **A_51_P266958** | Nr1i2 | -0.013510771 | -0.123227562 | 0.09620602 | 7.735221401 | -0.302953649 | 0.772375141 | 0.958193292 | -5.080017439 |
| **A_55_P2062495** | Smyd1 | 0.014307595 | -0.101957057 | 0.130572246 | 7.742547558 | 0.30275275 | 0.772520812 | 0.958193292 | -5.080062483 |
| **A_52_P129428** | Fzd2 | 0.01610874 | -0.114813086 | 0.147030567 | 7.809836615 | 0.302704366 | 0.772555897 | 0.958193292 | -5.080073327 |
| **A_55_P1981804** | Gp1ba | 0.021898863 | -0.156094469 | 0.199892196 | 7.967720109 | 0.302682282 | 0.772571911 | 0.958193292 | -5.080078276 |
| **A_66_P123735** | NA | -0.055714429 | -0.508623095 | 0.397194237 | 9.7421384 | -0.302639921 | 0.772602629 | 0.958193292 | -5.080087769 |
| **A_55_P1966878** | Rnaseh2a | 0.01581797 | -0.112809737 | 0.144445676 | 8.379275161 | 0.30254179 | 0.77267379 | 0.958193292 | -5.080109753 |
| **A_52_P480064** | Tmem134 | -0.031300733 | -0.285850807 | 0.223249341 | 11.79480389 | -0.302517459 | 0.772691435 | 0.958193292 | -5.080115203 |
| **A_55_P2113975** | Psg20 | 0.016674032 | -0.118958824 | 0.152306888 | 8.546653257 | 0.302443944 | 0.772744748 | 0.958193292 | -5.080131666 |
| **A_55_P2132502** | Pkn1 | -0.029575513 | -0.270155433 | 0.211004406 | 11.08361739 | -0.302441986 | 0.772746168 | 0.958193292 | -5.080132105 |
| **A_55_P1962010** | Clec1b | -0.055598818 | -0.507972441 | 0.396774806 | 8.955855126 | -0.302369126 | 0.772799007 | 0.958193292 | -5.080148418 |
| **A_52_P273120** | Myl1 | -0.013101773 | -0.119744928 | 0.093541381 | 7.638420634 | -0.302249965 | 0.772885426 | 0.958193292 | -5.080175089 |
| **A_52_P244803** | D630033O11Rik | 0.033056386 | -0.236094129 | 0.302206902 | 8.129833849 | 0.302154679 | 0.772954534 | 0.958193292 | -5.08019641 |
| **A_55_P2062632** | Smc5 | 0.025200129 | -0.180026989 | 0.230427246 | 11.34598267 | 0.302090577 | 0.773001026 | 0.958193292 | -5.080210749 |
| **A_55_P2059958** | Snx7 | -0.042925079 | -0.392506605 | 0.306656446 | 10.54060653 | -0.302086833 | 0.773003742 | 0.958193292 | -5.080211586 |
| **A_55_P1966982** | Gsr | -0.027413278 | -0.250671099 | 0.195844544 | 11.28876343 | -0.302081024 | 0.773007955 | 0.958193292 | -5.080212886 |
| **A_52_P315030** | NA | 0.030832884 | -0.220298072 | 0.28196384 | 14.01096537 | 0.302052935 | 0.773028328 | 0.958193292 | -5.080219168 |
| **A_55_P2092046** | NA | 0.035687361 | -0.2549967 | 0.326371422 | 9.175085404 | 0.302038552 | 0.77303876 | 0.958193292 | -5.080222385 |
| **A_51_P444447** | Cebpd | -0.078174719 | -0.715123139 | 0.558773701 | 12.74189822 | -0.301947352 | 0.773104909 | 0.958213837 | -5.080242777 |
| **A_55_P2033680** | NA | -0.014360512 | -0.131431338 | 0.102710314 | 9.114165405 | -0.301779961 | 0.773226326 | 0.958264971 | -5.080280191 |
| **A_55_P2008181** | Il3ra | -0.033394329 | -0.305678628 | 0.23888997 | 11.68120783 | -0.301730553 | 0.773262166 | 0.958264971 | -5.080291231 |
| **A_51_P469951** | Srgap3 | -0.033450172 | -0.306230584 | 0.239330239 | 10.65035281 | -0.301685434 | 0.773294894 | 0.958264971 | -5.08030131 |

| | | | | | | | | |
|---|---|---|---|---|---|---|---|---|
| **A_51_P284946** | Rnd3 | 0.022866554 | -0.163671613 | 0.209404722 | 10.23226293 | 0.301579752 | 0.773371557 | 0.958290211 | -5.080324914 |
| **A_55_P2017982** | Pde8b | 0.015717149 | -0.112542127 | 0.143976425 | 7.916229436 | 0.301476967 | 0.773446121 | 0.958290211 | -5.080347863 |
| **A_55_P2015007** | Zfp46 | 0.019059461 | -0.136487361 | 0.174606284 | 11.54534927 | 0.301452326 | 0.773463996 | 0.958290211 | -5.080353364 |
| **A_65_P18981** | Palb2 | 0.015711248 | -0.112579776 | 0.144002271 | 7.896714485 | 0.301289206 | 0.773582336 | 0.958350955 | -5.080389766 |
| **A_66_P103957** | Bbs7 | -0.029242206 | -0.268053598 | 0.209569186 | 8.23115832 | -0.301248061 | 0.773612187 | 0.958350955 | -5.080398945 |
| **A_55_P1975665** | Rhox4e | -0.026681376 | -0.24465904 | 0.191296288 | 9.656799028 | -0.301137879 | 0.773692126 | 0.958377771 | -5.08042352 |
| **A_51_P365516** | Spink1 | 0.195604598 | -1.402716701 | 1.793925897 | 14.661076 | 0.301081545 | 0.773732998 | 0.958377771 | -5.080436081 |
| **A_55_P2013586** | Prss8 | 0.065271036 | -0.468489029 | 0.599031102 | 10.8550353 | 0.300845551 | 0.77390423 | 0.958485859 | -5.080488677 |
| **A_55_P1977681** | Vps13d | -0.029785881 | -0.273379707 | 0.213807945 | 9.218765665 | -0.300824593 | 0.773919437 | 0.958485859 | -5.080493346 |
| **A_66_P108188** | Car5a | -0.019702177 | -0.18087014 | 0.141465785 | 9.042922431 | -0.300749331 | 0.773974049 | 0.958492081 | -5.080510111 |
| **A_55_P2067533** | Foxa2 | 0.061962788 | -0.44503122 | 0.568956796 | 8.739962625 | 0.300674976 | 0.774028004 | 0.958497488 | -5.080526669 |
| **A_55_P2002314** | Zmiz2 | 0.022653183 | -0.162766356 | 0.208072721 | 10.36484922 | 0.300568106 | 0.774105555 | 0.958503424 | -5.080550462 |
| **A_55_P2049147** | NA | 0.020814231 | -0.149609533 | 0.191237995 | 15.12934952 | 0.300468802 | 0.774177619 | 0.958503424 | -5.080572562 |
| **A_55_P2033863** | Psmc1 | 0.018938056 | -0.136126674 | 0.174002785 | 13.82652202 | 0.300463366 | 0.774181564 | 0.958503424 | -5.080573772 |
| **A_52_P197095** | Cdk17 | -0.013681756 | -0.125778756 | 0.098415244 | 8.25167873 | -0.300273509 | 0.774319349 | 0.95861261 | -5.080616006 |
| **A_55_P1985410** | Reps2 | -0.032742111 | -0.301073001 | 0.235588778 | 10.49458243 | -0.300196189 | 0.774375464 | 0.958620683 | -5.080633198 |
| **A_55_P2012166** | Spata6 | -0.02656727 | -0.244475309 | 0.191340768 | 10.09603913 | -0.299945841 | 0.774557167 | 0.958784213 | -5.080688835 |
| **A_55_P2085641** | Lmbr1l | 0.03376233 | -0.243252878 | 0.310777538 | 11.92594977 | 0.299845795 | 0.774629785 | 0.958784436 | -5.080711057 |
| **A_55_P2011300** | Fggy | -0.014093317 | -0.129750666 | 0.101564031 | 7.808125012 | -0.299784487 | 0.774674286 | 0.958784436 | -5.080724671 |
| **A_55_P2060021** | Otub2 | 0.044667715 | -0.321953303 | 0.411288733 | 8.945862312 | 0.29974058 | 0.774706157 | 0.958784436 | -5.080734419 |
| **A_55_P2150901** | Foxo4 | -0.018767888 | -0.172853738 | 0.135317962 | 9.465307891 | -0.299655189 | 0.774768142 | 0.958786179 | -5.080753374 |
| **A_55_P2144476** | NA | -0.034480296 | -0.317655376 | 0.248694784 | 8.768199918 | -0.299560895 | 0.774836592 | 0.958786179 | -5.080774299 |
| **A_51_P267796** | Ddr2 | -0.084776628 | -0.781303483 | 0.611750228 | 9.359388593 | -0.299438358 | 0.774925547 | 0.958786179 | -5.080801481 |
| **A_52_P607507** | Olfm4 | -0.012897472 | -0.118901574 | 0.09310663 | 7.653323344 | -0.299330575 | 0.775003795 | 0.958786179 | -5.080825383 |
| **A_55_P2149575** | Sdr39u1 | -0.016474567 | -0.151896546 | 0.118947412 | 8.198284369 | -0.29929124 | 0.775032351 | 0.958786179 | -5.080834103 |
| **A_55_P2126695** | NA | 0.015657615 | -0.113119083 | 0.144434312 | 8.216639395 | 0.299128287 | 0.775150658 | 0.958786179 | -5.080870218 |
| **A_55_P1992655** | Hdac6 | 0.035970454 | -0.259997004 | 0.331937912 | 11.57296154 | 0.298999956 | 0.775243834 | 0.958786179 | -5.080898646 |
| **A_51_P349961** | Gc | 0.101929316 | -0.736901622 | 0.940760254 | 9.754898407 | 0.298946748 | 0.775282467 | 0.958786179 | -5.08091043 |
| **A_52_P216226** | Masp1 | -0.012677986 | -0.117024048 | 0.091668077 | 7.714269606 | -0.298912001 | 0.775307696 | 0.958786179 | -5.080918124 |
| **A_55_P2455725** | Klhl20 | 0.016223223 | -0.117316312 | 0.149762758 | 9.38328682 | 0.298879714 | 0.775331139 | 0.958786179 | -5.080925272 |
| **A_52_P408757** | Fcgr3 | -0.052741129 | -0.486910192 | 0.381427933 | 8.468249261 | -0.298854441 | 0.77534949 | 0.958786179 | -5.080930867 |
| **A_52_P476731** | Fam110c | 0.016266472 | -0.117668189 | 0.150201133 | 7.905495985 | 0.298792402 | 0.775394537 | 0.958786179 | -5.080944599 |
| **A_52_P313102** | Yipf7 | 0.012865763 | -0.093085306 | 0.118816833 | 7.659041804 | 0.298744124 | 0.775429594 | 0.958786179 | -5.080955284 |
| **A_51_P362176** | Fgg | -0.185179478 | -1.710726545 | 1.340367589 | 13.31788907 | -0.298632044 | 0.77551098 | 0.958786179 | -5.080980082 |
| **A_55_P2167620** | NA | 0.015809171 | -0.114430761 | 0.146049103 | 8.067178607 | 0.298630458 | 0.775512132 | 0.958786179 | -5.080980433 |
| **A_55_P2065966** | Ctdp1 | 0.017679122 | -0.127990873 | 0.163349118 | 9.62094014 | 0.298579346 | 0.775549248 | 0.958786179 | -5.080991739 |
| **A_51_P122246** | Creld2 | -0.046366276 | -0.428423864 | 0.335691313 | 15.45418435 | -0.298567511 | 0.775557842 | 0.958786179 | -5.054271493 |
| **A_55_P1961247** | NA | 0.026828121 | -0.194278725 | 0.247934968 | 14.61361009 | 0.298508866 | 0.775600429 | 0.958786179 | -5.081007326 |
| **A_55_P2035667** | Gk | 0.014101264 | -0.102221627 | 0.130424155 | 7.764321935 | 0.298237341 | 0.775797618 | 0.958916713 | -5.081067342 |
| **A_52_P590396** | Sort1 | -0.022783097 | -0.210730013 | 0.165163818 | 8.920207698 | -0.298226841 | 0.775805244 | 0.958916713 | -5.081069662 |
| **A_51_P206134** | Akna | -0.013293762 | -0.123028461 | 0.096440936 | 7.934027482 | -0.298039002 | 0.775941669 | 0.958940604 | -5.081111148 |
| **A_55_P1959738** | Zfp511 | 0.02011088 | -0.145916586 | 0.186138346 | 9.351259809 | 0.298002771 | 0.775967983 | 0.958940604 | -5.081119147 |
| **A_65_P03728** | St3gal1 | 0.03551881 | -0.257880542 | 0.328918161 | 12.39910421 | 0.297829984 | 0.776093486 | 0.958940604 | -5.081157283 |
| **A_51_P295625** | Arf4 | -0.026181231 | -0.242481794 | 0.190119332 | 11.18940609 | -0.297784144 | 0.776126782 | 0.958940604 | -5.081167397 |
| **A_51_P454691** | 4930430F08Rik | -0.01914378 | -0.17732041 | 0.139032851 | 8.343995081 | -0.297751892 | 0.776150209 | 0.958940604 | -5.081174511 |
| **A_55_P1956647** | Sfmbt2 | 0.038599706 | -0.280489476 | 0.357688888 | 12.63304469 | 0.297605586 | 0.776256486 | 0.958940604 | -5.081206778 |
| **A_55_P2078403** | Rgs19 | 0.012876234 | -0.093581681 | 0.119334149 | 8.962813121 | 0.297563778 | 0.776286857 | 0.958940604 | -5.081215995 |
| **A_51_P426633** | Dnttip2 | 0.040163162 | -0.291991004 | 0.372317328 | 12.77923304 | 0.297479712 | 0.776347925 | 0.958940604 | -5.081234526 |
| **A_66_P103780** | Gm3646 | -0.012585326 | -0.116677726 | 0.091507073 | 7.672566596 | -0.29745045 | 0.776369183 | 0.958940604 | -5.081240975 |
| **A_55_P2060243** | Itgbl1 | -0.092405989 | -0.85683559 | 0.672023611 | 8.436657974 | -0.297393717 | 0.776410397 | 0.958940604 | -5.081253476 |

| | | | | | | | | | |
|---|---|---|---|---|---|---|---|---|---|
| **A_55_P2046943** | Bnip3l | 0.030068576 | -0.218683824 | 0.278820975 | 12.82323678 | 0.297382042 | 0.776418878 | 0.958940604 | -5.081256048 |
| **A_55_P1992079** | Ptprf | -0.052773319 | -0.489543605 | 0.383996966 | 10.46730887 | -0.297255904 | 0.776510517 | 0.958940604 | -5.081283835 |
| **A_55_P2016237** | Hand2 | 0.122227808 | -0.889411475 | 1.133867091 | 9.424718744 | 0.297244277 | 0.776518963 | 0.958940604 | -5.081286395 |
| **A_51_P135389** | Zfp474 | 0.012775377 | -0.092963057 | 0.118513812 | 7.634679306 | 0.297241891 | 0.776520697 | 0.958940604 | -5.081286921 |
| **A_55_P2126925** | Msantd3 | -0.014882108 | -0.138106463 | 0.108342246 | 10.46403654 | -0.29712355 | 0.776606675 | 0.958940604 | -5.081312978 |
| **A_55_P2009086** | Creb3l1 | -0.080186158 | -0.744166153 | 0.583793837 | 12.17614236 | -0.297107471 | 0.776618357 | 0.958940604 | -5.081316518 |
| **A_55_P2070241** | Preb | -0.015803055 | -0.146729429 | 0.115123318 | 12.56999602 | -0.296949835 | 0.776732889 | 0.958963844 | -5.081351209 |
| **A_52_P255224** | NA | -0.041766749 | -0.387806795 | 0.304273296 | 8.456764238 | -0.296943263 | 0.776737664 | 0.958963844 | -5.081352655 |
| **A_52_P30550** | Acp2 | 0.023723322 | -0.172891491 | 0.220338135 | 8.970297853 | 0.296844105 | 0.776809712 | 0.958963844 | -5.081374468 |
| **A_55_P2089358** | Hspe1 | -0.034755711 | -0.322839684 | 0.253328262 | 13.52877103 | -0.296808436 | 0.776835629 | 0.958963844 | -5.081382313 |
| **A_55_P2035963** | Hipk3 | 0.013733683 | -0.100189626 | 0.127656993 | 8.909700405 | 0.296581185 | 0.77700076 | 0.958995555 | -5.081432272 |
| **A_55_P2001168** | NA | -0.03738447 | -0.347558769 | 0.272789828 | 11.70897035 | -0.296520407 | 0.777044926 | 0.958995555 | -5.081445627 |
| **A_51_P108853** | Lyplal1 | 0.013832095 | -0.100944037 | 0.128608226 | 7.943171209 | 0.296486915 | 0.777069264 | 0.958995555 | -5.058708417 |
| **A_51_P255304** | Cav3 | -0.012528241 | -0.116488904 | 0.091432423 | 7.671622929 | -0.29647645 | 0.777076869 | 0.958995555 | -5.081455284 |
| **A_55_P2040873** | NA | 0.026648404 | -0.194625889 | 0.247922697 | 7.900556028 | 0.296284824 | 0.777216128 | 0.958995555 | -5.081497368 |
| **A_51_P391367** | Tfap4 | -0.013337967 | -0.124095405 | 0.09741947 | 7.808641591 | -0.296268796 | 0.777227777 | 0.958995555 | -5.081500887 |
| **A_55_P1990785** | Zranb3 | 0.019984346 | -0.14596539 | 0.185934082 | 8.193959886 | 0.296266493 | 0.77722945 | 0.958995555 | -5.081501393 |
| **A_52_P450188** | Mtmr4 | 0.029488025 | -0.215412868 | 0.274388918 | 11.0101336 | 0.29622689 | 0.777258232 | 0.958995555 | -5.081510087 |
| **A_55_P2107972** | Smad5 | -0.019404338 | -0.180615698 | 0.141807023 | 11.47283505 | -0.296123143 | 0.777333633 | 0.95901439 | -5.081532856 |
| **A_55_P2127258** | Dok7 | -0.014273843 | -0.132882565 | 0.104334879 | 7.683617732 | -0.296069352 | 0.777372728 | 0.95901439 | -5.081544658 |
| **A_55_P1967076** | Rpl11 | -0.018693929 | -0.174123692 | 0.136735834 | 16.53933159 | -0.295893593 | 0.777500473 | 0.959108945 | -5.081583208 |
| **A_55_P2044704** | NA | 0.052199425 | -0.381960022 | 0.486358872 | 11.81610084 | 0.295791459 | 0.77757471 | 0.959108945 | -5.081605599 |
| **A_55_P2052754** | Pcm1 | 0.015747244 | -0.115241993 | 0.146736481 | 8.699491399 | 0.295759095 | 0.777598234 | 0.959108945 | -5.081612693 |
| **A_55_P2073015** | Lin37 | 0.022385335 | -0.16392017 | 0.20869084 | 9.712869499 | 0.295601799 | 0.777712572 | 0.959121235 | -5.081647159 |
| **A_55_P2125241** | NA | -0.048144244 | -0.44886164 | 0.352573151 | 8.299304863 | -0.295580202 | 0.777728272 | 0.959121235 | -5.08165189 |
| **A_55_P1998539** | C1qtnf1 | -0.070676815 | -0.659017638 | 0.517664009 | 12.78894719 | -0.295540597 | 0.777757061 | 0.959121235 | -5.081660565 |
| **A_55_P2019293** | NA | 0.013189425 | -0.096679229 | 0.123058079 | 7.702432118 | 0.295339295 | 0.777903399 | 0.959240497 | -5.081704639 |
| **A_55_P2141093** | Eya2 | -0.019477454 | -0.18187466 | 0.142919751 | 7.794597821 | -0.29506847 | 0.778100293 | 0.959264011 | -5.08176389 |
| **A_55_P2130264** | NA | -0.015470545 | -0.144501862 | 0.113560773 | 16.36501753 | -0.294971212 | 0.778171005 | 0.959264011 | -5.081785155 |
| **A_51_P201338** | Mtss1 | -0.017915292 | -0.167338857 | 0.131508273 | 9.92524164 | -0.294967364 | 0.778173803 | 0.959264011 | -5.081785996 |
| **A_55_P2017038** | Cbx3 | -0.021384391 | -0.199752066 | 0.156983283 | 10.2095721 | -0.294951023 | 0.778185684 | 0.959264011 | -5.081789569 |
| **A_51_P464146** | Sowahb | 0.027930128 | -0.205067969 | 0.260928225 | 8.109754321 | 0.294910147 | 0.778215404 | 0.959264011 | -5.081798503 |
| **A_55_P2391065** | Cep128 | -0.030715659 | -0.28695693 | 0.225525611 | 8.01233773 | -0.2949035 | 0.778220237 | 0.959264011 | -5.081799956 |
| **A_55_P1953414** | Rab12 | 0.025036591 | -0.183924579 | 0.233997761 | 12.62455285 | 0.294766933 | 0.778319536 | 0.959295966 | -5.081829798 |
| **A_52_P420466** | H2ac4 | 0.084869099 | -0.62355356 | 0.793291758 | 12.15006176 | 0.294731334 | 0.778345421 | 0.959295966 | -5.081837575 |
| **A_55_P1973868** | Sema3b | 0.113880811 | -0.837809758 | 1.065571381 | 11.50099845 | 0.294390608 | 0.778593188 | 0.959408275 | -5.081911963 |
| **A_55_P2145449** | Cep68 | -0.01510124 | -0.14130319 | 0.111100709 | 7.972447131 | -0.29438503 | 0.778597244 | 0.959408275 | -5.08191318 |
| **A_55_P2024808** | Abl1 | 0.029768497 | -0.219042114 | 0.278579108 | 11.22356634 | 0.294345345 | 0.778626104 | 0.959408275 | -5.081921839 |
| **A_51_P365952** | Trdmt1 | -0.021563953 | -0.201807036 | 0.158679129 | 9.300743456 | -0.294332991 | 0.778635088 | 0.959408275 | -5.081924534 |
| **A_55_P2032668** | NA | -0.050683509 | -0.474423325 | 0.373056307 | 12.34788415 | -0.294263615 | 0.778685541 | 0.959409282 | -5.081939667 |
| **A_52_P530291** | Pim1 | -0.032985461 | -0.308936005 | 0.242965084 | 8.705868921 | -0.294076596 | 0.778821554 | 0.959462136 | -5.081980445 |
| **A_52_P531175** | Fkbp3 | 0.039562976 | -0.291475259 | 0.370601211 | 11.90393058 | 0.29402209 | 0.778861197 | 0.959462136 | -5.081992325 |
| **A_55_P2145754** | Atp5pb | -0.032244196 | -0.302063845 | 0.237575453 | 12.34446331 | -0.293999875 | 0.778877354 | 0.959462136 | -5.081997166 |
| **A_52_P231635** | Oxsm | 0.027473453 | -0.202609575 | 0.25755648 | 8.496235527 | 0.293763493 | 0.779049284 | 0.959526861 | -5.082048659 |
| **A_51_P394728** | Kti12 | -0.016244629 | -0.152320577 | 0.119831319 | 9.120628201 | -0.293695695 | 0.779098599 | 0.959526861 | -5.082063421 |
| **A_51_P469568** | Cldn13 | 0.013097972 | -0.096628514 | 0.122824458 | 7.655840918 | 0.293671459 | 0.779116227 | 0.959526861 | -5.082068697 |
| **A_51_P169087** | Gls2 | -0.016647051 | -0.156113457 | 0.122819355 | 8.993292856 | -0.293654637 | 0.779128464 | 0.959526861 | -5.082072359 |
| **A_55_P2033917** | Fut10 | 0.017901099 | -0.132163396 | 0.167965595 | 8.573110883 | 0.293474869 | 0.779259231 | 0.959578559 | -5.082111478 |
| **A_52_P628067** | Cdca3 | 0.044443186 | -0.328205831 | 0.417092203 | 8.829892746 | 0.293409636 | 0.779306685 | 0.959578559 | -5.082125668 |
| **A_55_P1958220** | Crebl2 | 0.019694514 | -0.145472004 | 0.184861033 | 8.783697209 | 0.293354276 | 0.779346957 | 0.959578559 | -5.082137708 |

| | | | | | | | | | |
|---|---|---|---|---|---|---|---|---|---|
| **A_55_P2073809** | AU015836 | 0.073495691 | -0.542972842 | 0.689964223 | 13.11254798 | 0.293305445 | 0.77938248 | 0.959578559 | -5.082148326 |
| **A_51_P475378** | Echdc3 | -0.042842978 | -0.402263064 | 0.316577108 | 9.413256617 | -0.293255706 | 0.779418665 | 0.959578559 | -5.08215914 |
| **A_55_P1959633** | Hnf4a | 0.148463955 | -1.097695639 | 1.394623549 | 10.68988543 | 0.293100477 | 0.779531596 | 0.959651509 | -5.082192877 |
| **A_52_P179729** | Txnl4a | -0.030966279 | -0.291015653 | 0.229083094 | 11.24665061 | -0.292955987 | 0.77963672 | 0.959651509 | -5.082224264 |
| **A_55_P2028418** | Hacd2 | 0.03504859 | -0.25929718 | 0.32939436 | 11.05808064 | 0.292942183 | 0.779646763 | 0.959651509 | -5.082227262 |
| **A_51_P279623** | Pagr1a | 0.028018231 | -0.217558836 | 0.273595298 | 12.90218282 | 0.295667092 | 0.779676512 | 0.959651509 | -5.014300362 |
| **A_51_P193173** | Slc25a25 | -0.020650037 | -0.194242842 | 0.152942769 | 10.07055255 | -0.292656563 | 0.779854583 | 0.959756098 | -5.082289261 |
| **A_52_P84027** | Cyp7a1 | 0.013227987 | -0.097979546 | 0.12443552 | 7.653954611 | 0.292636655 | 0.779869068 | 0.959756098 | -5.08229358 |
| **A_51_P233928** | B4galnt1 | 0.055850666 | -0.413776485 | 0.525477817 | 9.484312883 | 0.29257979 | 0.779910447 | 0.959756098 | -5.082305915 |
| **A_55_P1972969** | NA | 0.026095863 | -0.193402374 | 0.245594099 | 12.92957705 | 0.292489166 | 0.779976391 | 0.959758759 | -5.08232557 |
| **A_55_P2072661** | Prex1 | -0.042251065 | -0.397735441 | 0.313233311 | 11.72683236 | -0.292406016 | 0.780036899 | 0.959758759 | -5.082343598 |
| **A_55_P2074246** | Zfp184 | 0.018757339 | -0.139113327 | 0.176628005 | 8.447037241 | 0.292306824 | 0.780109083 | 0.959758759 | -5.082365098 |
| **A_51_P391727** | Slc3a1 | 0.086926786 | -0.644925355 | 0.818778927 | 8.966206089 | 0.292212935 | 0.780177409 | 0.959758759 | -5.082385442 |
| **A_51_P394558** | Dchs1 | -0.016497844 | -0.155412073 | 0.122416384 | 8.217885565 | -0.292179427 | 0.780201795 | 0.959758759 | -5.082392701 |
| **A_51_P290018** | NA | 0.015928098 | -0.118223619 | 0.150079815 | 12.1529989 | 0.29210356 | 0.780257009 | 0.959758759 | -5.082409134 |
| **A_55_P2076631** | Gypc | 0.034727913 | -0.257802215 | 0.327258042 | 9.861953635 | 0.292063475 | 0.780286182 | 0.959758759 | -5.082417814 |
| **A_55_P2028576** | NA | 0.035274519 | -0.261893011 | 0.332442048 | 12.1844012 | 0.292030969 | 0.78030984 | 0.959758759 | -5.082424853 |
| **A_51_P128075** | Tescl | -0.018829834 | -0.177607135 | 0.139947468 | 8.412924816 | -0.291760991 | 0.780506337 | 0.959917119 | -5.082483282 |
| **A_55_P2147876** | Abhd17a | 0.035842259 | -0.26643251 | 0.338117028 | 9.827273665 | 0.291717607 | 0.780537915 | 0.959917119 | -5.082492667 |
| **A_52_P561020** | Slc25a32 | -0.026321462 | -0.24841227 | 0.195769346 | 9.324277393 | -0.29157386 | 0.780642547 | 0.959931494 | -5.082523751 |
| **A_55_P2307621** | 5730526G10Rik | -0.049031859 | -0.462757077 | 0.36469336 | 8.50572456 | -0.291565092 | 0.780648929 | 0.959931494 | -5.082525647 |
| **A_55_P1988866** | NA | 0.02447218 | -0.182236453 | 0.231180813 | 15.0835302 | 0.291261584 | 0.780869867 | 0.96014209 | -5.082591229 |
| **A_51_P506297** | Cst9 | -0.012843589 | -0.121417644 | 0.095730466 | 7.629274628 | -0.291024467 | 0.781042492 | 0.960221894 | -5.082642419 |
| **A_51_P467837** | Ap3m2 | -0.019160413 | -0.181164997 | 0.142844171 | 8.731125016 | -0.290969004 | 0.781082871 | 0.960221894 | -5.082654387 |
| **A_55_P2143336** | Rpp21 | 0.02958017 | -0.220571084 | 0.279731425 | 9.380217372 | 0.290915693 | 0.781121685 | 0.960221894 | -5.082665888 |
| **A_51_P134627** | NA | -0.026555305 | -0.251255614 | 0.198145003 | 9.406400161 | -0.290748039 | 0.781243753 | 0.960221894 | -5.082702045 |
| **A_52_P580707** | Zfp781 | 0.029530689 | -0.220413239 | 0.279474616 | 9.140816545 | 0.290669957 | 0.781300605 | 0.960221894 | -5.082718878 |
| **A_51_P169920** | Noc2l | -0.016872757 | -0.159693258 | 0.125947745 | 9.838304436 | -0.290646201 | 0.781317903 | 0.960221894 | -5.082723998 |
| **A_55_P2006703** | Bmp1 | -0.034146122 | -0.323200481 | 0.254908238 | 12.68346153 | -0.290623696 | 0.78133429 | 0.960221894 | -5.082728849 |
| **A_55_P2047375** | NA | -0.044525235 | -0.421697878 | 0.332647408 | 14.51610247 | -0.290425802 | 0.781478389 | 0.960221894 | -5.082771484 |
| **A_51_P310949** | Ppox | -0.016280093 | -0.154283057 | 0.12172287 | 8.752673844 | -0.290226884 | 0.781623244 | 0.960221894 | -5.082814312 |
| **A_55_P2023772** | Fam107b | 0.037462764 | -0.280266147 | 0.355191676 | 11.36357847 | 0.290076301 | 0.781732907 | 0.960221894 | -5.082846714 |
| **A_66_P118863** | Ncald | 0.012695633 | -0.095008392 | 0.120399658 | 7.812910022 | 0.289995728 | 0.781791587 | 0.960221894 | -5.082864045 |
| **A_65_P13713** | Ank3 | 0.013871889 | -0.103830011 | 0.131573788 | 7.795872083 | 0.289948779 | 0.781825779 | 0.960221894 | -5.082874142 |
| **A_51_P224311** | Popdc2 | -0.020343287 | -0.192981601 | 0.152295027 | 8.171147085 | -0.289903263 | 0.781858929 | 0.960221894 | -5.082883929 |
| **A_55_P2013414** | Dmwd | -0.041373094 | -0.392642033 | 0.309895845 | 12.03079432 | -0.289765992 | 0.781958908 | 0.960221894 | -5.082913435 |
| **A_55_P2110837** | Rnf7 | 0.018023572 | -0.135084914 | 0.171132058 | 11.06898898 | 0.289608143 | 0.782073881 | 0.960221894 | -5.082947349 |
| **A_55_P1993762** | NA | -0.013249203 | -0.125824357 | 0.099325951 | 7.834013169 | -0.289545197 | 0.78211973 | 0.960221894 | -5.082960868 |
| **A_55_P2034784** | Snhg17 | 0.015881722 | -0.119073875 | 0.150837318 | 8.162444465 | 0.289518185 | 0.782139406 | 0.960221894 | -5.082966668 |
| **A_51_P431316** | Ube2w | 0.016929698 | -0.126948488 | 0.160807883 | 8.910792763 | 0.289483233 | 0.782164865 | 0.960221894 | -5.082974173 |
| **A_52_P541161** | Rgs18 | -0.01314729 | -0.124922953 | 0.098628372 | 7.77850346 | -0.289373107 | 0.782245085 | 0.960221894 | -5.082997813 |
| **A_55_P2046153** | NA | 0.031962854 | -0.239882255 | 0.303807964 | 15.88143614 | 0.289263205 | 0.782325144 | 0.960221894 | -5.083021396 |
| **A_55_P2078173** | NA | -0.037464032 | -0.356250902 | 0.281322838 | 14.64177725 | -0.289123407 | 0.782426985 | 0.960221894 | -5.083051383 |
| **A_55_P2114148** | NA | 0.023927792 | -0.179785053 | 0.227640637 | 9.149295352 | 0.288970412 | 0.782538445 | 0.960221894 | -5.083084183 |
| **A_52_P164161** | Cyp51 | 0.037582001 | -0.28239173 | 0.357555733 | 8.371833444 | 0.288958012 | 0.782547479 | 0.960221894 | -5.083086841 |
| **A_52_P195922** | Exosc6 | -0.040032278 | -0.380873408 | 0.300808852 | 12.04552908 | -0.288953175 | 0.782551004 | 0.960221894 | -5.083087878 |
| **A_55_P2159870** | Tead4 | -0.039076444 | -0.371800026 | 0.293647139 | 11.66909592 | -0.288935305 | 0.782564023 | 0.960221894 | -5.083091708 |
| **A_55_P2000853** | Mvb12b | -0.030033695 | -0.29690047 | 0.236833081 | 11.63513427 | -0.291651582 | 0.782568243 | 0.960221894 | -5.015123307 |
| **A_51_P247481** | Adck1 | 0.017841936 | -0.134092853 | 0.169776725 | 9.686667511 | 0.288904243 | 0.782586653 | 0.960221894 | -5.083098364 |
| **A_51_P173212** | Osbpl2 | 0.031824532 | -0.239217173 | 0.302866237 | 12.43148201 | 0.288865095 | 0.782615175 | 0.960221894 | -5.083106753 |

| | | | | | | | | | |
|---|---|---|---|---|---|---|---|---|---|
| **A_55_P2035138** | NA | -0.036983354 | -0.351968915 | 0.278002207 | 13.13426399 | -0.288858272 | 0.782620146 | 0.960221894 | -5.083108214 |
| **A_55_P2253284** | NA | -0.021245426 | -0.202206682 | 0.159715831 | 8.125203078 | -0.288834457 | 0.782637497 | 0.960221894 | -5.083113317 |
| **A_66_P139196** | Baz1a | 0.017990531 | -0.13525344 | 0.171234502 | 8.275480422 | 0.288821646 | 0.782646831 | 0.960221894 | -5.083116061 |
| **A_51_P139108** | Cpxm1 | -0.043743441 | -0.416401888 | 0.328915005 | 11.45459328 | -0.28878268 | 0.782675221 | 0.960221894 | -5.083124409 |
| **A_55_P2170569** | Ggps1 | -0.033554905 | -0.319479138 | 0.252369329 | 10.72400771 | -0.288718229 | 0.782722179 | 0.960221894 | -5.083138213 |
| **A_55_P1961541** | NA | 0.017580687 | -0.132300284 | 0.167461657 | 8.204317689 | 0.288574866 | 0.782826637 | 0.960221894 | -5.083168907 |
| **A_55_P2430367** | Zbtb8b | 0.019567498 | -0.147281112 | 0.186416109 | 10.28269762 | 0.28852395 | 0.782863737 | 0.960221894 | -5.083179806 |
| **A_55_P2154684** | Smarca4 | 0.032940992 | -0.247960968 | 0.313842951 | 12.73974899 | 0.288503484 | 0.782878649 | 0.960221894 | -5.083184185 |
| **A_55_P2094445** | Ms4a10 | 0.055612305 | -0.418696775 | 0.529921386 | 9.995920861 | 0.288455366 | 0.782913711 | 0.960221894 | -5.083194482 |
| **A_55_P2086659** | Rad54l | 0.013556019 | -0.102078212 | 0.12919025 | 7.901114406 | 0.288413061 | 0.782944537 | 0.960221894 | -5.083203533 |
| **A_55_P1956873** | Timm17a | -0.022972239 | -0.218929831 | 0.172985354 | 13.34694349 | -0.288410047 | 0.782946734 | 0.960221894 | -5.083204178 |
| **A_55_P1953578** | Top3a | 0.030058672 | -0.226437567 | 0.286554912 | 9.726618928 | 0.288308839 | 0.783020483 | 0.960221894 | -5.083225827 |
| **A_55_P2060107** | Pkm | 0.093402489 | -0.703843305 | 0.890648283 | 12.72457493 | 0.288227474 | 0.783079775 | 0.960221894 | -5.083243225 |
| **A_55_P2113031** | Foxo4 | -0.025764376 | -0.245704752 | 0.194176 | 8.849768545 | -0.288193266 | 0.783104703 | 0.960221894 | -5.083250539 |
| **A_55_P2270412** | C230096K16Rik | 0.0485795 | -0.366166174 | 0.463325173 | 9.151570023 | 0.288164407 | 0.783125734 | 0.960221894 | -5.083256708 |
| **A_55_P1985808** | Ranbp9 | 0.045513201 | -0.343099413 | 0.434125816 | 11.49037085 | 0.288130779 | 0.78315024 | 0.960221894 | -5.083263896 |
| **A_55_P2007404** | Vbp1 | -0.015809198 | -0.150808268 | 0.119189872 | 8.637961629 | -0.288103292 | 0.783170271 | 0.960221894 | -5.083269771 |
| **A_55_P2008417** | Mnd1 | 0.028667994 | -0.21629897 | 0.273634958 | 8.227918079 | 0.287911458 | 0.783310075 | 0.960332389 | -5.083310756 |
| **A_51_P114005** | Gstm7 | -0.012373546 | -0.118139246 | 0.093392154 | 7.650372635 | -0.287818352 | 0.783377932 | 0.960354668 | -5.078014888 |
| **A_52_P325161** | Zbtb10 | -0.02128037 | -0.203364219 | 0.160803479 | 9.165025078 | -0.287525863 | 0.783591115 | 0.96044667 | -5.083393059 |
| **A_55_P2137376** | Dip2c | 0.033915604 | -0.256345038 | 0.324176245 | 9.718737042 | 0.287462081 | 0.783637606 | 0.96044667 | -5.083406662 |
| **A_52_P193194** | Slc35d2 | -0.034626971 | -0.330976893 | 0.261722952 | 8.805019039 | -0.287460942 | 0.783638435 | 0.96044667 | -5.083406905 |
| **A_55_P2019068** | Flii | 0.019688316 | -0.14882199 | 0.188198623 | 8.449650416 | 0.287442693 | 0.783651737 | 0.96044667 | -5.083410797 |
| **A_55_P2095213** | Dusp22 | 0.040243047 | -0.304392826 | 0.384878921 | 9.537797439 | 0.287276129 | 0.783773152 | 0.960474614 | -5.083446306 |
| **A_51_P370341** | Zbtb48 | -0.015764575 | -0.150770701 | 0.119241551 | 8.578396108 | -0.287275075 | 0.78377392 | 0.960474614 | -5.083446531 |
| **A_55_P1979551** | Pitrm1 | -0.055583388 | -0.531817313 | 0.420650537 | 13.38105215 | -0.2871401 | 0.783872312 | 0.960502397 | -5.083475291 |
| **A_55_P2103880** | Gtf2ird1 | -0.021666421 | -0.207323536 | 0.163990693 | 9.384960096 | -0.28710764 | 0.783895976 | 0.960502397 | -5.083482206 |
| **A_51_P393958** | Fbxo5 | 0.020662536 | -0.15644992 | 0.197774992 | 8.252292227 | 0.287014393 | 0.783963953 | 0.9605248 | -5.083502065 |
| **A_55_P2179299** | Ceacam13 | -0.016062928 | -0.153834897 | 0.12170904 | 16.96576687 | -0.286835582 | 0.784094313 | 0.960613855 | -5.08354013 |
| **A_55_P2046158** | NA | 0.034825949 | -0.264034052 | 0.333685949 | 13.56243442 | 0.286684574 | 0.78420441 | 0.960613855 | -5.083572258 |
| **A_55_P2115713** | Nid1 | 0.0240852 | -0.182677128 | 0.230847528 | 8.828689269 | 0.286581406 | 0.784279631 | 0.960613855 | -5.083594199 |
| **A_52_P151320** | Tnfaip8l1 | 0.073778306 | -0.559589841 | 0.707146452 | 11.00235194 | 0.286577192 | 0.784282704 | 0.960613855 | -5.083595095 |
| **A_51_P165060** | Slc22a5 | -0.03215048 | -0.308171004 | 0.243870044 | 10.05215959 | -0.286559788 | 0.784295393 | 0.960613855 | -5.083598795 |
| **A_51_P346964** | Mrap | 0.06398902 | -0.485606622 | 0.613584662 | 9.233147006 | 0.286438467 | 0.784383853 | 0.960613855 | -5.083624585 |
| **A_55_P2061333** | NA | 0.018131343 | -0.137597551 | 0.173860237 | 9.829979293 | 0.286437546 | 0.784384525 | 0.960613855 | -5.08362478 |
| **A_55_P2117119** | Efhd1 | -0.046595771 | -0.447045585 | 0.353854043 | 9.797789097 | -0.286264551 | 0.784510669 | 0.960707471 | -5.083661536 |
| **A_55_P2083609** | Dkk3 | 0.039505444 | -0.300441922 | 0.379452809 | 10.80039516 | 0.285900179 | 0.784776384 | 0.960971982 | -5.083738883 |
| **A_51_P206585** | Runx1 | -0.10490844 | -1.00802801 | 0.79821113 | 8.73705958 | -0.28578165 | 0.784862827 | 0.9609751 | -5.083764023 |
| **A_51_P432851** | Dhrs11 | -0.016098998 | -0.154699896 | 0.1225019 | 8.055454451 | -0.285760346 | 0.784878364 | 0.9609751 | -5.083768541 |
| **A_55_P1980421** | Wdr81 | 0.021155508 | -0.161091228 | 0.203402245 | 11.43992184 | 0.285583342 | 0.785007461 | 0.961072283 | -5.083806063 |
| **A_55_P2194958** | Nfia | 0.013736873 | -0.104672439 | 0.132146185 | 7.778632502 | 0.285411322 | 0.785132929 | 0.961113577 | -5.083842506 |
| **A_55_P2159869** | Zfp59 | 0.01862349 | -0.14195402 | 0.179201 | 8.651567005 | 0.285328734 | 0.78519317 | 0.961113577 | -5.083859996 |
| **A_66_P123526** | 4930504D19Rik | -0.013935883 | -0.134117153 | 0.106245386 | 7.793141037 | -0.285277079 | 0.785230848 | 0.961113577 | -5.083870932 |
| **A_55_P2010704** | Selenoh | 0.029685417 | -0.226352645 | 0.285723479 | 11.34726615 | 0.285238265 | 0.785259161 | 0.961113577 | -5.083879148 |
| **A_55_P2182423** | NA | 0.041299226 | -0.314960942 | 0.397559395 | 14.32962876 | 0.28519625 | 0.785289809 | 0.961113577 | -5.083888041 |
| **A_55_P2093439** | Mex3d | 0.026018233 | -0.198493815 | 0.250530282 | 10.37129298 | 0.285106629 | 0.785355184 | 0.961132731 | -5.083907006 |
| **A_55_P2182867** | Sgcb | -0.016757246 | -0.161452869 | 0.127938378 | 8.908521607 | -0.284915716 | 0.785494455 | 0.961204184 | -5.083947386 |
| **A_52_P330950** | C2cd2 | -0.039427559 | -0.379911184 | 0.301056065 | 10.66533989 | -0.284887129 | 0.78551531 | 0.961204184 | -5.08395343 |
| **A_55_P2010312** | Serpina1a | 0.165048794 | -1.260879168 | 1.590976755 | 10.60646238 | 0.284763244 | 0.785605689 | 0.961204184 | -5.083979616 |
| **A_51_P333274** | Gzmb | 0.067220822 | -0.513673773 | 0.648115418 | 8.233398704 | 0.284692252 | 0.785657483 | 0.961204184 | -5.083994617 |

| | | | | | | | | | |
|---|---|---|---|---|---|---|---|---|---|
| **A_55_P2064299** | Msrb1 | -0.013102626 | -0.12636187 | 0.100156617 | 8.156244384 | -0.284612425 | 0.785715724 | 0.961204184 | -5.084011481 |
| **A_55_P2096867** | Gap43 | -0.018154043 | -0.175093851 | 0.138785765 | 8.031880247 | -0.284583298 | 0.785736974 | 0.961204184 | -5.084017633 |
| **A_52_P209172** | Mrpl41 | -0.028521175 | -0.275113085 | 0.218070735 | 11.75656659 | -0.284549451 | 0.785761669 | 0.961204184 | -5.084024781 |
| **A_55_P2075794** | Tti2 | -0.020698322 | -0.19973756 | 0.158340917 | 11.36155813 | -0.284417342 | 0.785858059 | 0.9612156 | -5.084052674 |
| **A_51_P359485** | Zfpl1 | -0.013751209 | -0.132755113 | 0.105252695 | 10.21340199 | -0.284281659 | 0.785957062 | 0.9612156 | -5.084081308 |
| **A_55_P1955108** | Ripor1 | 0.032874843 | -0.251731545 | 0.31748123 | 14.84472298 | 0.284176527 | 0.786033775 | 0.9612156 | -5.084103486 |
| **A_51_P106527** | Mcrip2 | 0.024199003 | -0.185417571 | 0.233815577 | 9.877258735 | 0.28401483 | 0.786151768 | 0.9612156 | -5.08413758 |
| **A_52_P277104** | Bank1 | 0.012537927 | -0.096071914 | 0.121147768 | 7.761474975 | 0.284004829 | 0.786159067 | 0.9612156 | -5.084139689 |
| **A_55_P2221647** | AI605517 | -0.017612833 | -0.170185999 | 0.134960334 | 7.814901378 | -0.284001223 | 0.786161698 | 0.9612156 | -5.084140449 |
| **A_55_P2072150** | Vpreb2 | -0.014089487 | -0.136158322 | 0.107979348 | 7.769484335 | -0.283961558 | 0.786190644 | 0.9612156 | -5.084148809 |
| **A_55_P2067976** | Gmeb1 | 0.022392674 | -0.171668264 | 0.216453611 | 9.229074538 | 0.283881433 | 0.786249115 | 0.9612156 | -5.084165694 |
| **A_51_P404413** | Bckdk | 0.016763849 | -0.128616508 | 0.162144207 | 10.20709734 | 0.283685529 | 0.786392084 | 0.9612156 | -5.084206957 |
| **A_51_P105017** | Rad52 | 0.013274549 | -0.101851879 | 0.128400978 | 8.260521027 | 0.283670332 | 0.786403176 | 0.9612156 | -5.073690922 |
| **A_55_P2151951** | NA | -0.032035348 | -0.309893762 | 0.245823067 | 13.55635545 | -0.283644947 | 0.786421703 | 0.9612156 | -5.084215502 |
| **A_51_P150608** | Jagn1 | -0.023374703 | -0.22613273 | 0.179383325 | 11.71991346 | -0.283620229 | 0.786439743 | 0.9612156 | -5.084220706 |
| **A_55_P2020072** | NA | -0.117751243 | -1.139186604 | 0.903684118 | 11.36926037 | -0.28361146 | 0.786446142 | 0.9612156 | -5.084222552 |
| **A_55_P2077613** | Fam102a | -0.041102511 | -0.39775805 | 0.315553029 | 13.0203394 | -0.283523161 | 0.786510587 | 0.9612156 | -5.084241137 |
| **A_66_P138567** | NA | 0.024464835 | -0.187828444 | 0.236758114 | 9.597940607 | 0.283514457 | 0.78651694 | 0.9612156 | -5.084242968 |
| **A_52_P399584** | Ckap2l | -0.012091396 | -0.117058604 | 0.092875812 | 7.69431455 | -0.283394858 | 0.786604231 | 0.961261502 | -5.084268132 |
| **A_55_P2033393** | Slc25a21 | -0.012583785 | -0.121893618 | 0.096726048 | 7.860419025 | -0.283218233 | 0.786733151 | 0.961300156 | -5.084305274 |
| **A_55_P2037156** | Pja1 | -0.023601518 | -0.228619738 | 0.181416702 | 11.37944761 | -0.283215251 | 0.786735328 | 0.961300156 | -5.084305901 |
| **A_51_P407999** | 1500011B03Rik | 0.030704481 | -0.236280604 | 0.297689567 | 9.704478316 | 0.282933211 | 0.786941206 | 0.96148212 | -5.084365165 |
| **A_52_P120803** | Ankrd1 | 0.017304366 | -0.133210256 | 0.167818987 | 7.954716803 | 0.282843469 | 0.787006719 | 0.96148212 | -5.08438401 |
| **A_51_P241995** | Col5a3 | 0.089101143 | -0.686007482 | 0.864209767 | 9.455892974 | 0.282806813 | 0.787033478 | 0.96148212 | -5.084391706 |
| **A_52_P201551** | Nav1 | -0.045303789 | -0.439678379 | 0.349070801 | 9.429931847 | -0.282614711 | 0.787173723 | 0.961514259 | -5.084432021 |
| **A_51_P171772** | Bcl11b | -0.012327842 | -0.119650287 | 0.094994604 | 7.653233361 | -0.282595764 | 0.787187555 | 0.961514259 | -5.084435996 |
| **A_52_P68028** | Ace | 0.025014766 | -0.192778849 | 0.242808381 | 8.457142604 | 0.282566364 | 0.78720902 | 0.961514259 | -5.084442164 |
| **A_55_P2110200** | Ubox5 | -0.021055853 | -0.204587253 | 0.162475548 | 9.147332219 | -0.282248483 | 0.787441112 | 0.961736967 | -5.084508807 |
| **A_55_P2185160** | NA | -0.028252602 | -0.274661094 | 0.218155891 | 12.49086108 | -0.282079768 | 0.787564303 | 0.961814043 | -5.084544148 |
| **A_55_P2052862** | Miga1 | -0.034033694 | -0.330926312 | 0.262858925 | 10.5813515 | -0.282019311 | 0.787608449 | 0.961814043 | -5.084556807 |
| **A_51_P106428** | Nsmce4a | -0.038681818 | -0.376205429 | 0.298841793 | 13.28217397 | -0.281949859 | 0.787659165 | 0.961814043 | -5.067852404 |
| **A_51_P125467** | Igf2r | -0.016714588 | -0.162645105 | 0.129215928 | 12.96758756 | -0.281785558 | 0.787779146 | 0.961814043 | -5.05234554 |
| **A_55_P2106645** | Adcy5 | -0.014905636 | -0.145077811 | 0.115266538 | 7.733004565 | -0.281709519 | 0.787834676 | 0.961814043 | -5.084621634 |
| **A_55_P2159264** | Lifr | 0.018367814 | -0.142080768 | 0.178816395 | 12.01129583 | 0.28163767 | 0.787887147 | 0.961814043 | -5.08463666 |
| **A_55_P2070316** | Ankrd36 | -0.011893302 | -0.115798013 | 0.092011408 | 7.700641422 | -0.281602424 | 0.787912887 | 0.961814043 | -5.084644029 |
| **A_55_P2077313** | Ddr1 | -0.022865742 | -0.222710791 | 0.176979307 | 11.91918952 | -0.281488766 | 0.787995895 | 0.961814043 | -5.084667787 |
| **A_51_P302856** | Ugt2b37 | 0.012387479 | -0.095895802 | 0.12067076 | 7.69793516 | 0.281443147 | 0.788029212 | 0.961814043 | -5.08467732 |
| **A_51_P356931** | Orc1 | -0.011990017 | -0.11683722 | 0.092857185 | 7.660884127 | -0.281340418 | 0.788104241 | 0.961814043 | -5.084698782 |
| **A_52_P582112** | Hcfc1r1 | 0.035817227 | -0.277402144 | 0.349036597 | 12.38815626 | 0.281327705 | 0.788113527 | 0.961814043 | -5.084701438 |
| **A_55_P2111523** | Higd1b | 0.035884802 | -0.277939824 | 0.349709427 | 10.67148332 | 0.281314875 | 0.788122898 | 0.961814043 | -5.084704118 |
| **A_55_P2251082** | Qk | 0.014930705 | -0.115661244 | 0.145522654 | 8.563031311 | 0.281276261 | 0.788151101 | 0.961814043 | -5.084712182 |
| **A_52_P435670** | Npepl1 | -0.021502416 | -0.209764458 | 0.166759626 | 10.99773405 | -0.280991806 | 0.788358874 | 0.961980901 | -5.084771559 |
| **A_55_P2039586** | Grk3 | -0.01343911 | -0.131120103 | 0.104241882 | 7.718773687 | -0.280952796 | 0.788387369 | 0.961980901 | -5.084779697 |
| **A_51_P402160** | Zfp750 | 0.015773214 | -0.122416944 | 0.153963372 | 7.774077665 | 0.28080979 | 0.788491833 | 0.962011388 | -5.084809522 |
| **A_55_P2058240** | Ipo5 | 0.022586708 | -0.175395145 | 0.220568561 | 9.277633555 | 0.280670467 | 0.78859361 | 0.962011388 | -5.084838564 |
| **A_55_P2028496** | NA | 0.030482776 | -0.236721297 | 0.297686849 | 13.61867625 | 0.280660058 | 0.788601215 | 0.962011388 | -5.084840734 |
| **A_52_P182118** | NA | -0.014235524 | -0.13913807 | 0.110667021 | 8.008254716 | -0.280395697 | 0.788794348 | 0.962011388 | -5.084895801 |
| **A_66_P112816** | B4galt6 | -0.02227298 | -0.217707663 | 0.173161704 | 8.806309771 | -0.280379232 | 0.788806377 | 0.962011388 | -5.084899229 |
| **A_52_P64763** | Gen1 | -0.012147621 | -0.118787762 | 0.09449252 | 7.719111589 | -0.280246167 | 0.788903597 | 0.962011388 | -5.084926926 |
| **A_51_P374137** | Bco1 | 0.018364142 | -0.142891089 | 0.179619373 | 7.79095937 | 0.280172817 | 0.78895719 | 0.962011388 | -5.084942189 |

| | | | | | | | | | |
|---|---|---|---|---|---|---|---|---|---|
| **A_55_P1992739** | Dync1i2 | 0.024744198 | -0.192651298 | 0.242139694 | 13.3841179 | 0.280021901 | 0.78906746 | 0.962011388 | -5.084973578 |
| **A_55_P2067652** | Boc | 0.026322044 | -0.204955576 | 0.257599665 | 8.281903818 | 0.279998154 | 0.789084811 | 0.962011388 | -5.084978516 |
| **A_52_P95910** | Ugcg | -0.101162502 | -0.990048845 | 0.787723841 | 10.17965049 | -0.279989981 | 0.789090783 | 0.962011388 | -5.084980215 |
| **A_55_P2037524** | Rps15 | -0.024280104 | -0.237625701 | 0.189065493 | 14.79155603 | -0.279985814 | 0.789093828 | 0.962011388 | -5.084981082 |
| **A_51_P245275** | H2afx | 0.012598784 | -0.098108154 | 0.123305722 | 12.00680856 | 0.279977385 | 0.789099987 | 0.962011388 | -5.084982834 |
| **A_51_P423509** | Faap100 | -0.012757916 | -0.124868222 | 0.099352389 | 7.907736976 | -0.279964768 | 0.789109207 | 0.962011388 | -5.084985457 |
| **A_55_P1984357** | NA | 0.018672739 | -0.145590798 | 0.182936276 | 14.71405171 | 0.279663652 | 0.789329243 | 0.962011388 | -5.085048027 |
| **A_51_P127297** | Hsd11b1 | -0.074632347 | -0.731288731 | 0.582024037 | 14.02113329 | -0.279613472 | 0.789365913 | 0.962011388 | -5.047087049 |
| **A_52_P367791** | Mri1 | 0.06514749 | -0.508250027 | 0.638545007 | 10.34601006 | 0.279518812 | 0.78943509 | 0.962011388 | -5.0850781 |
| **A_55_P2009994** | NA | 0.020763298 | -0.16201308 | 0.203539676 | 11.34261232 | 0.279476594 | 0.789465943 | 0.962011388 | -5.085086863 |
| **A_52_P554650** | Gzmd | 0.150448658 | -1.174000168 | 1.474897483 | 9.233499016 | 0.279461701 | 0.789476828 | 0.962011388 | -5.085089955 |
| **A_55_P2076001** | Rbm12 | 0.023909971 | -0.186594887 | 0.234414829 | 10.34756142 | 0.279438405 | 0.789493853 | 0.962011388 | -5.085094789 |
| **A_55_P2167840** | Gm14151 | -0.013267602 | -0.130097221 | 0.103562017 | 7.863675072 | -0.279388574 | 0.789530271 | 0.962011388 | -5.085105129 |
| **A_55_P1967885** | Nr3c1 | 0.025630244 | -0.209909496 | 0.261169984 | 11.63264821 | 0.281993196 | 0.789539987 | 0.962011388 | -5.017058802 |
| **A_66_P104479** | NA | 0.015656838 | -0.122265997 | 0.153579673 | 16.20068255 | 0.2792782 | 0.789610937 | 0.962011388 | -5.085128026 |
| **A_51_P380005** | Galnt3 | 0.032728439 | -0.255707081 | 0.321163958 | 10.04782612 | 0.279155193 | 0.789700841 | 0.962011388 | -5.085153533 |
| **A_55_P1953593** | Ece2 | -0.016007233 | -0.157142179 | 0.125127713 | 10.28030838 | -0.279029987 | 0.789792355 | 0.962011388 | -5.085179485 |
| **A_55_P2056973** | Treml2 | -0.012390642 | -0.121641313 | 0.096860029 | 7.70574596 | -0.279022265 | 0.789797999 | 0.962011388 | -5.085181086 |
| **A_51_P514256** | Tubb2b | 0.024062585 | -0.18819602 | 0.23632119 | 8.548990286 | 0.278898476 | 0.789888481 | 0.962011388 | -5.085206732 |
| **A_52_P222709** | Pigu | 0.037041599 | -0.289740434 | 0.363823631 | 9.952048581 | 0.278869306 | 0.789909803 | 0.962011388 | -5.085212774 |
| **A_51_P483311** | Mpv17l2 | -0.01762966 | -0.173222457 | 0.137963136 | 9.134355953 | -0.278755615 | 0.789992908 | 0.962011388 | -5.085236317 |
| **A_51_P391445** | Ifngr1 | 0.034673823 | -0.307260984 | 0.376608629 | 11.90540314 | 0.285338859 | 0.790004187 | 0.962011388 | -4.895251028 |
| **A_52_P540470** | Zfp407 | -0.022139809 | -0.217551914 | 0.173272295 | 10.47161385 | -0.278735047 | 0.790007943 | 0.962011388 | -5.085240575 |
| **A_52_P235241** | Fntb | -0.02468287 | -0.242569697 | 0.193203956 | 8.783451887 | -0.278698 | 0.790035024 | 0.962011388 | -5.085248244 |
| **A_52_P411003** | Dlgap5 | -0.012048535 | -0.118420729 | 0.09432366 | 7.679476303 | -0.27866041 | 0.790062503 | 0.962011388 | -5.085256024 |
| **A_55_P1975155** | Galnt6 | 0.084516967 | -0.661701393 | 0.830735327 | 13.22285124 | 0.278642358 | 0.790075699 | 0.962011388 | -5.08525976 |
| **A_52_P366525** | Coq10b | 0.012261616 | -0.096014149 | 0.120537382 | 8.85055137 | 0.278602895 | 0.790104547 | 0.962011388 | -5.085267927 |
| **A_55_P1962651** | NA | -0.013837499 | -0.136220309 | 0.10854531 | 7.771401424 | -0.278167478 | 0.790422869 | 0.962018928 | -5.085357957 |
| **A_55_P2061278** | Sik1 | 0.02620867 | -0.205741529 | 0.258158869 | 9.88756617 | 0.277983742 | 0.790557206 | 0.962018928 | -5.085395907 |
| **A_55_P1970578** | Aamdc | 0.014754305 | -0.115853428 | 0.145362039 | 10.77578842 | 0.27791951 | 0.790604171 | 0.962018928 | -5.085409168 |
| **A_55_P2467801** | Srp54a | 0.026578073 | -0.20871536 | 0.261871506 | 9.729451013 | 0.277896357 | 0.790621101 | 0.962018928 | -5.085413948 |
| **A_51_P487308** | Pstk | 0.024657125 | -0.193657733 | 0.242971982 | 9.392758964 | 0.277861454 | 0.790646621 | 0.962018928 | -5.085421152 |
| **A_51_P393067** | Ap4m1 | -0.014386298 | -0.14176889 | 0.112996294 | 8.826469099 | -0.277848547 | 0.790656058 | 0.962018928 | -5.085423816 |
| **A_55_P1991901** | NA | 0.033306641 | -0.261815067 | 0.328428349 | 16.12134691 | 0.277650746 | 0.790800696 | 0.962018928 | -5.085464625 |
| **A_51_P105408** | Alg14 | 0.015751115 | -0.123835244 | 0.155337474 | 10.79362915 | 0.277611523 | 0.790829377 | 0.962018928 | -5.057807859 |
| **A_55_P2002597** | NA | -0.046674574 | -0.460306195 | 0.366957048 | 10.02272333 | -0.277610437 | 0.790830171 | 0.962018928 | -5.085472938 |
| **A_51_P250123** | Zfp763 | 0.012818501 | -0.100780166 | 0.126417167 | 7.852556237 | 0.277608758 | 0.790831399 | 0.962018928 | -5.085473284 |
| **A_55_P2151056** | Rarg | 0.068138603 | -0.53588847 | 0.672165677 | 10.0248787 | 0.277527476 | 0.790890838 | 0.962018928 | -5.085490043 |
| **A_51_P153995** | Gp9 | 0.027194756 | -0.214007743 | 0.268397255 | 8.355942248 | 0.277378327 | 0.790999909 | 0.962018928 | -5.085520783 |
| **A_52_P411430** | Gm5914 | -0.011674872 | -0.115243768 | 0.091894024 | 7.831765898 | -0.277326865 | 0.791037544 | 0.962018928 | -5.085531385 |
| **A_51_P464394** | Klb | 0.032293591 | -0.254308777 | 0.318895958 | 8.266678575 | 0.277207978 | 0.79112449 | 0.962018928 | -5.085555872 |
| **A_55_P2085616** | Nsa2 | -0.017598345 | -0.173824992 | 0.138628302 | 8.621580562 | -0.277131491 | 0.79118043 | 0.962018928 | -5.085571621 |
| **A_55_P1998354** | Wbp4 | 0.023288172 | -0.18346409 | 0.230040434 | 8.747462657 | 0.277111339 | 0.791195168 | 0.962018928 | -5.085575769 |
| **A_51_P233603** | Psma7 | -0.034823722 | -0.344002153 | 0.274354708 | 13.98677549 | -0.277099131 | 0.791204097 | 0.962018928 | -5.085578282 |
| **A_52_P20357** | Aqr | -0.01903304 | -0.188018012 | 0.149951931 | 9.572636918 | -0.277095357 | 0.791206858 | 0.962018928 | -5.085579059 |
| **A_55_P2013043** | Serpinb6b | 0.022186232 | -0.174872206 | 0.21924467 | 13.45807972 | 0.27698591 | 0.791286905 | 0.962018928 | -5.085601584 |
| **A_51_P124568** | Mpp1 | -0.034318147 | -0.339144173 | 0.270507879 | 11.88538823 | -0.276975243 | 0.791294707 | 0.962018928 | -5.080274118 |
| **A_55_P1952256** | Ccnb1 | -0.011785848 | -0.116489677 | 0.09291798 | 7.687326338 | -0.276928372 | 0.791328989 | 0.962018928 | -5.085613422 |
| **A_55_P2133356** | NA | -0.028616781 | -0.282877361 | 0.225643799 | 15.12807503 | -0.276892319 | 0.791355359 | 0.962018928 | -5.085620838 |
| **A_55_P2131964** | Pdia2 | -0.01291689 | -0.127687879 | 0.101854099 | 7.728018502 | -0.276882177 | 0.791362777 | 0.962018928 | -5.085622925 |

| | | | | | | | | | |
|---|---|---|---|---|---|---|---|---|---|
| **A_51_P288447** | Nup133 | 0.021346332 | -0.168435921 | 0.211128585 | 10.12850744 | 0.276717626 | 0.791483137 | 0.962018928 | -5.085656762 |
| **A_55_P2120906** | NA | 0.0143036 | -0.112867368 | 0.141474568 | 9.375801338 | 0.27671107 | 0.791487933 | 0.962018928 | -5.085658109 |
| **A_52_P283501** | Zc3h7b | -0.044379954 | -0.438987057 | 0.350227149 | 9.421808531 | -0.276688502 | 0.791504441 | 0.962018928 | -5.085662748 |
| **A_55_P2069832** | Prpsap1 | -0.014717396 | -0.145606223 | 0.116171432 | 12.29375544 | -0.2766289 | 0.791548038 | 0.962018928 | -5.085674998 |
| **A_51_P198387** | Uckl1 | -0.019687052 | -0.194844782 | 0.155470678 | 10.43315108 | -0.276516175 | 0.791630496 | 0.962018928 | -5.08569816 |
| **A_52_P109548** | Map7 | 0.029872189 | -0.235934054 | 0.295678431 | 9.237694052 | 0.276484653 | 0.791653555 | 0.962018928 | -5.085704635 |
| **A_52_P453624** | Arl6ip6 | -0.016224983 | -0.160613335 | 0.12816337 | 8.591377361 | -0.27645296 | 0.791676739 | 0.962018928 | -5.085711145 |
| **A_51_P346523** | Zcchc7 | -0.065287263 | -0.64642814 | 0.515853615 | 9.093960053 | -0.276386101 | 0.791725649 | 0.962018928 | -5.085724875 |
| **A_51_P209736** | Atoh8 | -0.012629992 | -0.125060171 | 0.099800187 | 7.822333314 | -0.27636901 | 0.791738152 | 0.962018928 | -5.085728384 |
| **A_55_P1972839** | NA | 0.014220717 | -0.112412181 | 0.140853615 | 16.47842239 | 0.276276603 | 0.791805753 | 0.962018928 | -5.085747354 |
| **A_51_P257885** | Mmd2 | 0.012042975 | -0.095221589 | 0.119307538 | 7.654376595 | 0.276214628 | 0.791851092 | 0.962018928 | -5.085760074 |
| **A_55_P2095133** | Synpo2 | 0.028801423 | -0.227766715 | 0.28536956 | 8.130922971 | 0.276172468 | 0.791881935 | 0.962018928 | -5.085768725 |
| **A_55_P2082529** | Sav1 | -0.042537136 | -0.421540078 | 0.336465806 | 11.36530297 | -0.276118071 | 0.791921732 | 0.962018928 | -5.085779885 |
| **A_55_P2025155** | NA | -0.026336763 | -0.261084568 | 0.208411042 | 12.81761219 | -0.276013306 | 0.79199838 | 0.962018928 | -5.085801373 |
| **A_55_P1969660** | Pex26 | -0.028515914 | -0.282691764 | 0.225659937 | 11.15857258 | -0.27600832 | 0.792002028 | 0.962018928 | -5.085802396 |
| **A_51_P363801** | Pgpep1 | 0.015313675 | -0.121282785 | 0.151910136 | 8.519012945 | 0.275809454 | 0.792147529 | 0.962135201 | -5.085843162 |
| **A_52_P221776** | Kif12 | -0.011651303 | -0.115634109 | 0.092331502 | 7.676214639 | -0.275665328 | 0.792252985 | 0.962202824 | -5.085872689 |
| **A_51_P220407** | Rnpepl1 | 0.016899307 | -0.13397927 | 0.167777883 | 12.66725339 | 0.275556382 | 0.792332703 | 0.962239182 | -5.085894999 |
| **A_52_P168567** | Cebpa | 0.028453657 | -0.225693732 | 0.282601047 | 8.329485319 | 0.275436575 | 0.792420372 | 0.96228519 | -5.085919523 |
| **A_55_P2115871** | Inpp5j | -0.013268966 | -0.131992388 | 0.105454457 | 7.708569947 | -0.274960192 | 0.792768998 | 0.962648068 | -5.086016936 |
| **A_55_P2085412** | Ankle1 | 0.012012074 | -0.095562708 | 0.119586856 | 7.736911937 | 0.274711401 | 0.792951089 | 0.962808693 | -5.086067745 |
| **A_55_P2031167** | Efna1 | -0.025573727 | -0.254968205 | 0.20382075 | 13.4898554 | -0.274271217 | 0.793273296 | 0.962998348 | -5.086157532 |
| **A_55_P2093704** | Meig1 | -0.011763018 | -0.11727757 | 0.093751534 | 7.700383591 | -0.274268273 | 0.79327545 | 0.962998348 | -5.086158132 |
| **A_51_P225793** | Prr5l | 0.067211888 | -0.535693653 | 0.670117428 | 12.29810815 | 0.274262216 | 0.793279885 | 0.962998348 | -5.086159367 |
| **A_51_P167843** | Timm17a | 0.035121252 | -0.279966045 | 0.350208549 | 12.4278025 | 0.274225762 | 0.79330657 | 0.962998348 | -5.086166796 |
| **A_51_P103850** | Fbn2 | 0.05522408 | -0.440839866 | 0.551288026 | 8.189622925 | 0.273879792 | 0.79355985 | 0.963186455 | -5.086237257 |
| **A_55_P2078300** | Fndc7 | -0.016297442 | -0.162756671 | 0.130161787 | 8.01468459 | -0.273761176 | 0.793646693 | 0.963186455 | -5.086261395 |
| **A_55_P1988273** | Rnf103 | 0.017304707 | -0.138220778 | 0.172830192 | 11.57547097 | 0.273735957 | 0.793665158 | 0.963186455 | -5.086266525 |
| **A_52_P5549** | Fam133b | 0.019423498 | -0.155225322 | 0.194072318 | 9.6776969 | 0.273609366 | 0.793757844 | 0.963186455 | -5.086292272 |
| **A_52_P520495** | Vcam1 | -0.01254833 | -0.125378822 | 0.100282161 | 7.73884539 | -0.273607899 | 0.793758918 | 0.963186455 | -5.086292571 |
| **A_55_P2077510** | Zfp583 | -0.021463304 | -0.214478844 | 0.171552236 | 8.744784857 | -0.273573143 | 0.793784366 | 0.963186455 | -5.086299638 |
| **A_55_P2009267** | Gm16060 | -0.038862642 | -0.388392566 | 0.310667282 | 14.10064458 | -0.273537657 | 0.793810349 | 0.963186455 | -5.086306852 |
| **A_52_P180565** | 2310061I04Rik | 0.022643909 | -0.181213543 | 0.226501362 | 10.05921566 | 0.273271273 | 0.794005405 | 0.963261591 | -5.086360981 |
| **A_55_P2185273** | Umps | -0.029715834 | -0.29729341 | 0.237861741 | 11.14900409 | -0.273216784 | 0.794045305 | 0.963261591 | -5.086372046 |
| **A_55_P2089253** | Atg4c | -0.031028131 | -0.310431094 | 0.248374831 | 10.48162015 | -0.273208231 | 0.794051569 | 0.963261591 | -5.086373783 |
| **A_55_P2012051** | Sox1 | -0.028926858 | -0.28944898 | 0.231595263 | 8.354402456 | -0.273165486 | 0.79408287 | 0.963261591 | -5.086382462 |
| **A_51_P487298** | Vasn | -0.119468296 | -1.1956724 | 0.956735809 | 12.03381454 | -0.27310345 | 0.794128298 | 0.963261591 | -5.086395055 |
| **A_52_P94401** | Rtn4rl1 | -0.019957169 | -0.199831564 | 0.159917225 | 8.359097885 | -0.272959842 | 0.794233466 | 0.963261591 | -5.086424198 |
| **A_52_P28806** | Foxm1 | 0.01442802 | -0.115622794 | 0.144478834 | 7.906933448 | 0.27293726 | 0.794250003 | 0.963261591 | -5.086428779 |
| **A_55_P2013143** | Esrp2 | -0.034899088 | -0.349595764 | 0.279797587 | 9.389657609 | -0.272829352 | 0.794329031 | 0.963261591 | -5.086450665 |
| **A_55_P2262560** | 5730471H19Rik | 0.013313715 | -0.106747188 | 0.133374618 | 8.028010678 | 0.272814103 | 0.794340198 | 0.963261591 | -5.086453757 |
| **A_66_P127255** | Ing3 | 0.021778135 | -0.174674604 | 0.218230874 | 9.728431094 | 0.272729273 | 0.794402326 | 0.963261591 | -5.086470956 |
| **A_51_P491976** | Fbxw17 | -0.031503269 | -0.315932366 | 0.252925828 | 10.91757605 | -0.272490125 | 0.794577483 | 0.963261591 | -5.086519413 |
| **A_52_P126266** | Prkab2 | -0.040150531 | -0.402714014 | 0.322412951 | 9.046725541 | -0.272443469 | 0.794611656 | 0.963261591 | -5.086528862 |
| **A_55_P2061219** | Ces3a | -0.012379476 | -0.124180066 | 0.099421113 | 7.673091423 | -0.272412693 | 0.794634198 | 0.963261591 | -5.086535094 |
| **A_51_P424232** | Mecp2 | -0.044938057 | -0.450807027 | 0.360930913 | 9.126054187 | -0.272394042 | 0.794647859 | 0.963261591 | -5.086538871 |
| **A_55_P2089780** | Mtfr1l | -0.015548319 | -0.155986571 | 0.124889932 | 8.747951724 | -0.272374973 | 0.794661827 | 0.963261591 | -5.086542732 |
| **A_55_P2033250** | Fdft1 | -0.021430743 | -0.215008823 | 0.172147337 | 10.21659016 | -0.272364317 | 0.794669632 | 0.963261591 | -5.086544889 |
| **A_55_P2231152** | Gm13372 | -0.025607161 | -0.257063151 | 0.20584883 | 8.656956638 | -0.272183733 | 0.794801909 | 0.963322368 | -5.086581436 |
| **A_52_P190647** | Mxd3 | 0.012011607 | -0.096567515 | 0.120590729 | 7.678289395 | 0.272159789 | 0.794819449 | 0.963322368 | -5.08658628 |

| | | | | | | | | | |
|---|---|---|---|---|---|---|---|---|---|
| **A_52_P447424** | NA | 0.024871297 | -0.200043522 | 0.249786116 | 9.566412764 | 0.272050495 | 0.79489951 | 0.963358996 | -5.086608387 |
| **A_51_P235687** | Alox5ap | -0.023623795 | -0.237408132 | 0.190160541 | 8.9877445 | -0.271858523 | 0.795040143 | 0.963423482 | -5.086647195 |
| **A_55_P2087114** | NA | 0.026308361 | -0.211788834 | 0.264405556 | 10.25814665 | 0.271837066 | 0.795055863 | 0.963423482 | -5.086651531 |
| **A_51_P288341** | Hpd | -0.011522704 | -0.11584346 | 0.092798051 | 7.655349284 | -0.271739554 | 0.795127301 | 0.963423482 | -5.086671232 |
| **A_55_P1976057** | Smurf2 | -0.013688569 | -0.137661369 | 0.110284231 | 8.643644065 | -0.27164443 | 0.795196992 | 0.963423482 | -5.086690444 |
| **A_51_P196127** | Papss1 | -0.03103994 | -0.312165298 | 0.250085418 | 10.22706628 | -0.27163768 | 0.795201937 | 0.963423482 | -5.086691807 |
| **A_66_P102879** | Thra | 0.012495893 | -0.100732417 | 0.125724204 | 7.746434296 | 0.271507263 | 0.795297487 | 0.963478855 | -5.086718135 |
| **A_52_P205255** | Adam11 | 0.011656304 | -0.094040433 | 0.117353042 | 7.758033849 | 0.271311662 | 0.795440804 | 0.963534154 | -5.086757601 |
| **A_55_P2145531** | Mpst | -0.036044798 | -0.362900723 | 0.290811126 | 9.855140416 | -0.271303503 | 0.795446782 | 0.963534154 | -5.086759246 |
| **A_51_P192089** | Pinx1 | -0.02736805 | -0.275600127 | 0.220864027 | 9.175356569 | -0.271240862 | 0.795492681 | 0.963534154 | -5.086771879 |
| **A_51_P242603** | E430018J23Rik | 0.012110262 | -0.097810717 | 0.122031242 | 8.045204141 | 0.271045451 | 0.795635869 | 0.963647203 | -5.086811268 |
| **A_55_P2045658** | Nme6 | -0.01812012 | -0.182746693 | 0.146506453 | 8.941372191 | -0.270788549 | 0.795824129 | 0.963814823 | -5.086863011 |
| **A_52_P413395** | Sln | 0.016104286 | -0.130287265 | 0.162495837 | 7.786597698 | 0.270641646 | 0.795931787 | 0.963884814 | -5.086892577 |
| **A_55_P2042312** | NA | 0.014533437 | -0.11764239 | 0.146709264 | 16.41315905 | 0.270511329 | 0.796027295 | 0.963940081 | -5.086918793 |
| **A_51_P382925** | Fadd | 0.0217445 | -0.176138272 | 0.219627271 | 9.005122322 | 0.270340179 | 0.796152734 | 0.963996396 | -5.086953203 |
| **A_52_P105138** | Frs2 | -0.025459071 | -0.257170113 | 0.206251972 | 12.59372903 | -0.270311784 | 0.796173546 | 0.963996396 | -5.08695891 |
| **A_55_P2005410** | NA | 0.019988368 | -0.162021803 | 0.201998538 | 12.34617927 | 0.270178533 | 0.796271214 | 0.964023374 | -5.086985684 |
| **A_51_P160713** | Alb | 0.204328745 | -1.656951141 | 2.065608632 | 10.72751607 | 0.270076594 | 0.796345935 | 0.964023374 | -5.087006158 |
| **A_55_P2087414** | Sema3e | -0.076774627 | -0.776308674 | 0.62275942 | 8.962816316 | -0.270008865 | 0.79639558 | 0.964023374 | -5.087019757 |
| **A_55_P1987291** | Plppr5 | 0.011950469 | -0.096960547 | 0.120861484 | 7.678835475 | 0.269949351 | 0.796439205 | 0.964023374 | -5.087031703 |
| **A_52_P507214** | Mmp9 | -0.011563014 | -0.117001398 | 0.093875369 | 7.675527422 | -0.269799716 | 0.796548895 | 0.964023374 | -5.076139892 |
| **A_55_P2135701** | BC024978 | 0.023446568 | -0.190375161 | 0.237268296 | 8.341512866 | 0.269771835 | 0.796569333 | 0.964023374 | -5.087067322 |
| **A_51_P468249** | Phex | -0.011644662 | -0.117846139 | 0.094556814 | 7.648986778 | -0.269752517 | 0.796583495 | 0.964023374 | -5.087071197 |
| **A_52_P483129** | Utp23 | 0.022785852 | -0.185037244 | 0.230608948 | 9.744747622 | 0.269737064 | 0.796594823 | 0.964023374 | -5.087074296 |
| **A_55_P1974577** | Nfam1 | -0.016846953 | -0.170620229 | 0.136926323 | 8.414413003 | -0.269531576 | 0.796745467 | 0.964145315 | -5.087115494 |
| **A_55_P2071596** | Ppara | 0.013600956 | -0.110582462 | 0.137784374 | 7.786271437 | 0.269448018 | 0.796806726 | 0.964159083 | -5.087132238 |
| **A_51_P237585** | Btnl9 | -0.012086011 | -0.122477457 | 0.098305435 | 7.64719857 | -0.269349839 | 0.796878706 | 0.964185823 | -5.087151905 |
| **A_66_P104442** | Cog6 | -0.013629009 | -0.138150772 | 0.110892754 | 8.144915143 | -0.269270132 | 0.796937145 | 0.964196174 | -5.087167867 |
| **A_51_P488196** | Bmper | 0.046283011 | -0.376720898 | 0.46928692 | 10.31352132 | 0.26918223 | 0.797001595 | 0.964213797 | -5.087185464 |
| **A_55_P1990490** | Recql | 0.012751129 | -0.103844217 | 0.129346476 | 8.654283951 | 0.269052229 | 0.797096914 | 0.96426876 | -5.08721148 |
| **A_52_P490770** | Zfp26 | -0.016186677 | -0.164279754 | 0.131906399 | 8.158694346 | -0.268900816 | 0.797207938 | 0.964316959 | -5.087241765 |
| **A_55_P2033849** | Sphkap | 0.011405765 | -0.092961538 | 0.115773068 | 7.64762067 | 0.268861816 | 0.797236536 | 0.964316959 | -5.087249562 |
| **A_55_P1999324** | Nudt6 | 0.019424495 | -0.158465589 | 0.19731458 | 8.965544816 | 0.268637825 | 0.797400789 | 0.964455281 | -5.087294328 |
| **A_55_P2019615** | Neil1 | 0.018561721 | -0.151545481 | 0.188668923 | 9.359148256 | 0.2684508 | 0.797537944 | 0.964560593 | -5.087331677 |
| **A_55_P2027969** | Myl6 | 0.017424065 | -0.142341665 | 0.177189796 | 16.3263417 | 0.268308863 | 0.797642039 | 0.964560593 | -5.087360006 |
| **A_55_P2135633** | NA | -0.016995007 | -0.173040307 | 0.139050293 | 15.91515337 | -0.267941396 | 0.797911556 | 0.964560593 | -5.08743328 |
| **A_52_P111866** | Naa50 | -0.013712397 | -0.139629582 | 0.112204788 | 9.321055264 | -0.267915309 | 0.79793069 | 0.964560593 | -5.087438478 |
| **A_55_P1994258** | Gapdh | 0.02334598 | -0.191046899 | 0.237738858 | 10.84714843 | 0.26789889 | 0.797942734 | 0.964560593 | -5.08744175 |
| **A_51_P215887** | Aldh18a1 | -0.025591444 | -0.260672449 | 0.20948956 | 8.856116479 | -0.267822107 | 0.797999055 | 0.964560593 | -5.087457046 |
| **A_51_P249793** | Dusp28 | -0.015521004 | -0.158097634 | 0.127055627 | 8.740468612 | -0.267818531 | 0.798001678 | 0.964560593 | -5.087457758 |
| **A_66_P100696** | Slc26a2 | -0.033479823 | -0.341044265 | 0.274084619 | 9.630538389 | -0.267803464 | 0.79801273 | 0.964560593 | -5.087460759 |
| **A_55_P2163098** | Akr1c18 | 0.079889214 | -0.6541333 | 0.813911729 | 8.635035109 | 0.267761376 | 0.798043602 | 0.964560593 | -5.087469141 |
| **A_51_P193490** | Fbxo21 | 0.041575416 | -0.340459235 | 0.423610067 | 13.13780775 | 0.267733681 | 0.798063918 | 0.964560593 | -5.087474656 |
| **A_55_P2015495** | Abat | 0.06504115 | -0.532978479 | 0.663060778 | 9.512198199 | 0.267572782 | 0.798181945 | 0.964560593 | -5.087506685 |
| **A_55_P2007495** | NA | -0.089929811 | -0.916958847 | 0.737099225 | 10.99270634 | -0.267517432 | 0.798222549 | 0.964560593 | -5.087517699 |
| **A_65_P18433** | Ankrd28 | 0.017313042 | -0.141919925 | 0.176546009 | 10.12174212 | 0.267491236 | 0.798241766 | 0.964560593 | -5.087522911 |
| **A_52_P205789** | Unc119b | -0.033130452 | -0.337911823 | 0.271650919 | 9.260432553 | -0.267428759 | 0.798287599 | 0.964560593 | -5.087535339 |
| **A_51_P173961** | Pdrg1 | -0.015838338 | -0.161551617 | 0.129874942 | 12.69202477 | -0.267411221 | 0.798300464 | 0.964560593 | -5.087538827 |
| **A_55_P2243828** | NA | -0.048873537 | -0.498549411 | 0.400802337 | 8.194429004 | -0.267388882 | 0.798316853 | 0.964560593 | -5.08754327 |
| **A_55_P1996469** | Zfp592 | -0.026025029 | -0.265499857 | 0.213449799 | 11.48845178 | -0.267362512 | 0.798336198 | 0.964560593 | -5.087548514 |

| | | | | | | | | | |
|---|---|---|---|---|---|---|---|---|---|
| **A_55_P2142439** | Echdc2 | 0.03019433 | -0.24781302 | 0.30820168 | 9.634928753 | 0.267201125 | 0.798454598 | 0.964626744 | -5.087580597 |
| **A_51_P317640** | Tgfb2 | -0.109746514 | -1.12042363 | 0.900930602 | 10.7899439 | -0.267145256 | 0.798495587 | 0.964626744 | -5.0875917 |
| **A_55_P2131707** | Lpcat1 | -0.017492567 | -0.178669169 | 0.143684034 | 8.005042696 | -0.267005817 | 0.79859789 | 0.964626744 | -5.087619399 |
| **A_66_P121495** | Psat1 | 0.014162475 | -0.116333822 | 0.144658772 | 7.781959323 | 0.266999307 | 0.798602667 | 0.964626744 | -5.087620692 |
| **A_55_P2011116** | L3mbtl3 | -0.020396611 | -0.208371946 | 0.167578725 | 9.507547103 | -0.266947775 | 0.798640477 | 0.964626744 | -5.087630925 |
| **A_55_P2079636** | Scgb3a2 | 0.012064912 | -0.099167011 | 0.123296835 | 7.63927182 | 0.266847955 | 0.798713717 | 0.9646419 | -5.087650741 |
| **A_55_P2079981** | NA | -0.012798921 | -0.130821588 | 0.105223746 | 7.893299221 | -0.266794638 | 0.798752838 | 0.9646419 | -5.087661323 |
| **A_55_P2115691** | NA | -0.032897389 | -0.336572375 | 0.270777597 | 10.13353529 | -0.266514951 | 0.798958067 | 0.964642647 | -5.087716797 |
| **A_55_P1963817** | Rps6 | -0.014536747 | -0.148811114 | 0.119737619 | 15.55059589 | -0.26634423 | 0.799083347 | 0.964642647 | -5.087750631 |
| **A_51_P148105** | Rad51 | 0.021205128 | -0.174679195 | 0.217089452 | 7.983533343 | 0.266324049 | 0.799098156 | 0.964642647 | -5.087754629 |
| **A_51_P519592** | Srsf10 | -0.017473532 | -0.179014116 | 0.144067051 | 7.993060336 | -0.26611431 | 0.79925208 | 0.964642647 | -5.087796164 |
| **A_55_P1972720** | Pmm1 | -0.029582589 | -0.303127106 | 0.243961928 | 12.41213794 | -0.266058613 | 0.799292956 | 0.964642647 | -5.087807189 |
| **A_52_P370443** | Riok3 | 0.026355212 | -0.217406843 | 0.270117266 | 11.25473905 | 0.26599261 | 0.799341397 | 0.964642647 | -5.08782025 |
| **A_55_P2116525** | NA | -0.012904321 | -0.132408082 | 0.10659944 | 7.671895119 | -0.265657903 | 0.79958706 | 0.964642647 | -5.087886438 |
| **A_55_P2082779** | Gm10240 | -0.027859262 | -0.28589774 | 0.230179217 | 14.78128467 | -0.265616037 | 0.79961779 | 0.964642647 | -5.087894711 |
| **A_55_P2008197** | Esr2 | -0.043215757 | -0.443564256 | 0.357132742 | 10.00687695 | -0.265566375 | 0.799654243 | 0.964642647 | -5.087904523 |
| **A_55_P2341468** | Rbms3 | 0.03890118 | -0.321484503 | 0.399286863 | 9.392629263 | 0.265561069 | 0.799658138 | 0.964642647 | -5.087905572 |
| **A_51_P145785** | Ambp | 0.155017921 | -1.282083274 | 1.592119115 | 9.844082006 | 0.265377268 | 0.799793056 | 0.964642647 | -5.087941871 |
| **A_55_P1987534** | Tnrc6b | -0.011580359 | -0.118939023 | 0.095778306 | 8.191872475 | -0.265371374 | 0.799797383 | 0.964642647 | -5.087943034 |
| **A_55_P2045792** | Pes1 | -0.04448572 | -0.456916648 | 0.367945209 | 11.7258319 | -0.265361894 | 0.799804342 | 0.964642647 | -5.087944906 |
| **A_55_P1966086** | NA | -0.024913256 | -0.255962242 | 0.20613573 | 8.991609181 | -0.265274553 | 0.799868458 | 0.964642647 | -5.087962145 |
| **A_51_P362903** | Mrps22 | 0.028383852 | -0.234864364 | 0.291632069 | 10.72742772 | 0.265262014 | 0.799877663 | 0.964642647 | -5.08796462 |
| **A_51_P460954** | Ccl6 | 0.078576363 | -0.650200928 | 0.807353654 | 9.305443558 | 0.265256634 | 0.799881612 | 0.964642647 | -5.087965682 |
| **A_55_P2023235** | Fen1 | -0.024897317 | -0.25588353 | 0.206088895 | 10.70862448 | -0.265176884 | 0.799940157 | 0.964642647 | -5.087981417 |
| **A_55_P2115704** | Aip | -0.018047769 | -0.185570571 | 0.149475034 | 8.399705076 | -0.265044474 | 0.800037363 | 0.964642647 | -5.088007533 |
| **A_52_P536494** | Mycn | 0.019572749 | -0.162164217 | 0.201309716 | 8.089969237 | 0.264958416 | 0.800100542 | 0.964642647 | -5.0880245 |
| **A_52_P608495** | Aars2 | 0.017620338 | -0.146019802 | 0.181260479 | 9.022117672 | 0.264907081 | 0.800138231 | 0.964642647 | -5.088034618 |
| **A_55_P2048671** | Rpl27a | -0.013561113 | -0.139503906 | 0.112381681 | 16.02258859 | -0.264905614 | 0.800139308 | 0.964642647 | -5.088034907 |
| **A_55_P2008704** | Gstp2 | 0.036708355 | -0.30425031 | 0.377667021 | 10.07934845 | 0.264869747 | 0.80016564 | 0.964642647 | -5.088041976 |
| **A_55_P2171578** | Mkrn1 | 0.024211847 | -0.200764654 | 0.249188348 | 12.19437139 | 0.264764607 | 0.800242834 | 0.964642647 | -5.088062691 |
| **A_55_P2055404** | Slc15a4 | -0.020727055 | -0.213350104 | 0.171895993 | 12.27152914 | -0.264727169 | 0.800270321 | 0.964642647 | -5.088070065 |
| **A_51_P348617** | Borcs8 | 0.035285515 | -0.292634857 | 0.363205886 | 12.00702402 | 0.26472638 | 0.8002709 | 0.964642647 | -5.08807022 |
| **A_51_P370470** | Cyp20a1 | -0.014363362 | -0.147894335 | 0.119167612 | 8.687395 | -0.26463254 | 0.8003398 | 0.964642647 | -5.088088699 |
| **A_55_P2156016** | NA | 0.013489485 | -0.111920231 | 0.138899202 | 7.807090199 | 0.264626515 | 0.800344223 | 0.964642647 | -5.088089885 |
| **A_52_P71686** | Atp6v0d2 | -0.06648795 | -0.684991952 | 0.552016051 | 8.337876199 | -0.264465847 | 0.800462195 | 0.964642647 | -5.088121509 |
| **A_55_P1980311** | Pcbp2 | 0.015633985 | -0.12982192 | 0.161089891 | 9.079633148 | 0.26442804 | 0.800489956 | 0.964642647 | -5.088128948 |
| **A_51_P129464** | Scd2 | -0.05459199 | -0.562509716 | 0.453325735 | 13.74510081 | -0.264426346 | 0.8004912 | 0.964642647 | -5.088129281 |
| **A_55_P2075592** | Clstn1 | 0.015991279 | -0.132823534 | 0.164806092 | 8.16964229 | 0.264366371 | 0.800535239 | 0.964642647 | -5.08814108 |
| **A_51_P137168** | Smarcal1 | -0.02560307 | -0.263868233 | 0.212662093 | 9.559782606 | -0.26436299 | 0.800537721 | 0.964642647 | -5.088141745 |
| **A_55_P2163446** | NA | 0.015163786 | -0.12597955 | 0.156307122 | 9.172896679 | 0.264311733 | 0.800575359 | 0.964642647 | -5.088151826 |
| **A_51_P316935** | Depp1 | -0.074594637 | -0.769136476 | 0.619947203 | 10.17939339 | -0.264227705 | 0.800637063 | 0.964642647 | -5.088168348 |
| **A_55_P2133067** | Ube2v1 | 0.014712563 | -0.122282047 | 0.151707174 | 8.324712268 | 0.264212908 | 0.800647928 | 0.964642647 | -5.088171257 |
| **A_55_P2016670** | Alpk2 | -0.011724744 | -0.120994804 | 0.097545315 | 7.649812195 | -0.263980209 | 0.800818812 | 0.964642647 | -5.088216983 |
| **A_55_P2054098** | Gm9495 | -0.01554041 | -0.160404645 | 0.129323825 | 8.323313048 | -0.263918879 | 0.800863852 | 0.964642647 | -5.088229028 |
| **A_55_P1978466** | Pnma5 | 0.020347734 | -0.169333336 | 0.210028803 | 8.707928763 | 0.263913265 | 0.800867975 | 0.964642647 | -5.088230131 |
| **A_55_P1952577** | Ces1a | 0.017253923 | -0.143589698 | 0.178097543 | 8.712587178 | 0.263908373 | 0.800871568 | 0.964642647 | -5.088231091 |
| **A_55_P1999473** | Ppp1r26 | -0.011394525 | -0.117618926 | 0.094829877 | 7.796097255 | -0.26390103 | 0.80087696 | 0.964642647 | -5.088232533 |
| **A_51_P485479** | Prrc2a | -0.019566625 | -0.201992606 | 0.162859355 | 9.099512587 | -0.263875093 | 0.800896009 | 0.964642647 | -5.088237626 |
| **A_55_P2164469** | Prcd | -0.012807902 | -0.132251984 | 0.10663618 | 7.710044332 | -0.26380469 | 0.800947713 | 0.964642647 | -5.088251447 |
| **A_55_P2152248** | Cpe | -0.115404317 | -1.19206359 | 0.961254955 | 9.108881177 | -0.263701703 | 0.801023351 | 0.964642647 | -5.088271659 |

| | | | | | | | | |
|---|---|---|---|---|---|---|---|---|
| **A_55_P2090718** | Pop5 | 0.01829248 | -0.152440565 | 0.189025524 | 12.18063609 | 0.263587107 | 0.801107516 | 0.964642647 | -5.08829414 |
| **A_55_P2061620** | Chst1 | 0.051112726 | -0.425950816 | 0.528176267 | 9.988190458 | 0.26358578 | 0.801108491 | 0.964642647 | -5.088294401 |
| **A_52_P448466** | BC004004 | -0.012201158 | -0.126115416 | 0.101713101 | 8.126240941 | -0.263506974 | 0.801166372 | 0.964642647 | -5.088309855 |
| **A_55_P2079713** | Gramd1c | 0.027662803 | -0.23068234 | 0.286007947 | 10.42841392 | 0.263429892 | 0.801222989 | 0.964642647 | -5.088324967 |
| **A_55_P2184189** | Ncf2 | 0.027609859 | -0.230269889 | 0.285489606 | 8.850373464 | 0.263400206 | 0.801244793 | 0.964642647 | -5.088330786 |
| **A_55_P2152364** | NA | -0.036205322 | -0.374517095 | 0.302106452 | 8.568462765 | -0.263283997 | 0.801330151 | 0.964642647 | -5.088353558 |
| **A_55_P1997614** | Gm9693 | -0.016179559 | -0.167398723 | 0.135039605 | 7.924484586 | -0.263226136 | 0.801372653 | 0.964642647 | -5.088364893 |
| **A_55_P2062707** | Mthfsl | -0.022537758 | -0.2332206 | 0.188145084 | 9.053629061 | -0.263178681 | 0.801407511 | 0.964642647 | -5.088374187 |
| **A_51_P357085** | Dctn6 | 0.011986517 | -0.100066753 | 0.124039788 | 11.68264506 | 0.263170769 | 0.801413323 | 0.964642647 | -5.088375737 |
| **A_55_P2015936** | NA | 0.026792452 | -0.223720505 | 0.277305409 | 15.41770095 | 0.263118527 | 0.801451699 | 0.964642647 | -5.088385967 |
| **A_55_P2158653** | Cldn7 | 0.077937942 | -0.650885658 | 0.806761543 | 11.83643063 | 0.263084749 | 0.801476511 | 0.964642647 | -5.08839258 |
| **A_55_P2009518** | NA | 0.01818866 | -0.1519702 | 0.188347519 | 15.06931743 | 0.262975501 | 0.801556764 | 0.964642647 | -5.088413964 |
| **A_51_P168630** | Klf1 | 0.011083724 | -0.092624975 | 0.114792422 | 7.682405832 | 0.262929714 | 0.8015904 | 0.964642647 | -5.088422923 |
| **A_52_P937** | Spcs2 | -0.036995818 | -0.383173852 | 0.309182215 | 12.8625833 | -0.262919195 | 0.801598127 | 0.964642647 | -5.088424981 |
| **A_55_P2008407** | Ppp1r14c | 0.05680873 | -0.474998238 | 0.588615699 | 10.30130028 | 0.262802941 | 0.801683532 | 0.964685363 | -5.088447722 |
| **A_51_P123879** | Tecrl | -0.012039604 | -0.124860641 | 0.100781432 | 7.650103802 | -0.26253747 | 0.801878566 | 0.964804946 | -5.055677392 |
| **A_51_P420100** | Emc2 | 0.017270445 | -0.144571391 | 0.179112282 | 9.885094826 | 0.262531791 | 0.801882738 | 0.964804946 | -5.088500725 |
| **A_52_P589699** | Polr3a | -0.017259742 | -0.179201816 | 0.144682332 | 8.505623564 | -0.26220669 | 0.802121604 | 0.964946015 | -5.088564203 |
| **A_51_P276479** | 4930486L24Rik | -0.128096994 | -1.330148977 | 1.073954989 | 13.83211206 | -0.262171122 | 0.802147739 | 0.964946015 | -5.088571144 |
| **A_55_P2170732** | Fndc10 | 0.03631105 | -0.304477598 | 0.377099699 | 9.084261909 | 0.262133698 | 0.802175238 | 0.964946015 | -5.088578445 |
| **A_51_P292357** | Rps3a1 | -0.022375456 | -0.232494176 | 0.187743263 | 14.4737638 | -0.261984934 | 0.802284551 | 0.964946015 | -5.088607459 |
| **A_52_P214891** | Stx6 | 0.039389526 | -0.330545801 | 0.409324852 | 10.47517828 | 0.261953442 | 0.802307692 | 0.964946015 | -5.088613599 |
| **A_55_P1957080** | NA | -0.012384468 | -0.12873732 | 0.103968384 | 7.726207503 | -0.261860199 | 0.802376211 | 0.964946015 | -5.088631775 |
| **A_55_P2419299** | NA | 0.050531312 | -0.424545295 | 0.525607919 | 9.300674241 | 0.261677323 | 0.802510602 | 0.964946015 | -5.088667404 |
| **A_51_P382764** | Akr1c20 | 0.011011005 | -0.092521084 | 0.114543095 | 7.664014323 | 0.26165025 | 0.802530498 | 0.964946015 | -5.088672676 |
| **A_65_P03874** | Nav2 | 0.011288439 | -0.09485298 | 0.117429858 | 7.860623943 | 0.261648453 | 0.802531819 | 0.964946015 | -5.088673026 |
| **A_55_P1959305** | Rbm47 | 0.034357849 | -0.288715152 | 0.35743085 | 11.50426185 | 0.26163417 | 0.802542316 | 0.964946015 | -5.088675808 |
| **A_52_P508991** | Fmo1 | -0.047866788 | -0.498047185 | 0.402313609 | 9.617769922 | -0.261587429 | 0.802576666 | 0.964946015 | -5.088684909 |
| **A_51_P136792** | Calcoco1 | 0.017138244 | -0.144063298 | 0.178339786 | 8.402917998 | 0.261556968 | 0.802599052 | 0.964946015 | -5.088690839 |
| **A_55_P2071326** | Rpl36 | -0.034495351 | -0.359183832 | 0.290193131 | 14.10707978 | -0.261374278 | 0.802733319 | 0.964957077 | -5.088726392 |
| **A_52_P633752** | Apc | 0.013695062 | -0.115250519 | 0.142640643 | 8.035209041 | 0.261292349 | 0.802793535 | 0.964957077 | -5.088742328 |
| **A_55_P1960566** | Ybx3 | 0.033353581 | -0.280843888 | 0.34755105 | 12.46146334 | 0.261161374 | 0.802889801 | 0.964957077 | -5.088767794 |
| **A_52_P507498** | Plxnc1 | 0.012672604 | -0.106735676 | 0.132080883 | 7.725374638 | 0.261096216 | 0.802937693 | 0.964957077 | -5.088780458 |
| **A_51_P411757** | Ankrd13b | -0.047491716 | -0.495072805 | 0.400089374 | 8.811152162 | -0.261044948 | 0.802975377 | 0.964957077 | -5.088790421 |
| **A_55_P2151899** | NA | -0.021021143 | -0.219153524 | 0.177111238 | 11.17085336 | -0.261017738 | 0.802995377 | 0.964957077 | -5.088795708 |
| **A_55_P2054887** | Trappc1 | -0.024466509 | -0.255086902 | 0.206153885 | 12.0216449 | -0.26100177 | 0.803007114 | 0.964957077 | -5.08879881 |
| **A_55_P2023446** | NA | -0.029867975 | -0.311403153 | 0.251667203 | 11.13153845 | -0.261001062 | 0.803007635 | 0.964957077 | -5.088798947 |
| **A_52_P48681** | Cldn1 | -0.036677793 | -0.382614588 | 0.309259002 | 8.469886336 | -0.260840845 | 0.803125404 | 0.964975838 | -5.088830064 |
| **A_52_P11174** | Hip1 | -0.022081119 | -0.230420013 | 0.186257774 | 9.139651619 | -0.260747344 | 0.803194136 | 0.964975838 | -5.088848214 |
| **A_51_P259879** | Fkrp | 0.019240435 | -0.162538167 | 0.201019036 | 8.997384984 | 0.260400181 | 0.80344935 | 0.964975838 | -5.088915552 |
| **A_51_P274504** | Tex264 | 0.039586833 | -0.334632881 | 0.413806547 | 13.61045163 | 0.260251512 | 0.803558651 | 0.964975838 | -5.088944362 |
| **A_52_P664699** | Cul1 | 0.017998422 | -0.152243032 | 0.188239877 | 9.124107105 | 0.260098761 | 0.803670958 | 0.964975838 | -5.088973946 |
| **A_51_P354077** | Svil | -0.031119917 | -0.325502777 | 0.263262942 | 9.742654666 | -0.260072886 | 0.803689982 | 0.964975838 | -5.088978955 |
| **A_51_P327585** | Gstm4 | -0.021241729 | -0.222204961 | 0.179721502 | 8.495650356 | -0.26004136 | 0.803713162 | 0.964975838 | -5.088985059 |
| **A_52_P355095** | Dstyk | 0.021646474 | -0.183202244 | 0.226495192 | 8.69505338 | 0.259969905 | 0.803765701 | 0.964975838 | -5.088998889 |
| **A_51_P371745** | Havcr1 | 0.017558966 | -0.148624661 | 0.183742593 | 8.226103321 | 0.259944046 | 0.803784714 | 0.964975838 | -5.089003893 |
| **A_55_P2084408** | Uba1 | 0.012524953 | -0.106023359 | 0.131073265 | 8.073729588 | 0.259926094 | 0.803797914 | 0.964975838 | -5.089007367 |
| **A_55_P1985012** | Pard3 | 0.01574323 | -0.133283469 | 0.164769928 | 8.259039453 | 0.25989559 | 0.803820343 | 0.964975838 | -5.089013269 |
| **A_55_P2139748** | Phf14 | -0.019729252 | -0.206521931 | 0.167063427 | 8.508538281 | -0.259848327 | 0.803855095 | 0.964975838 | -5.089022412 |
| **A_55_P2019684** | Bspry | 0.031517056 | -0.266901335 | 0.329935448 | 8.595211019 | 0.259829959 | 0.803868602 | 0.964975838 | -5.089025965 |

| | | | | | | | | | |
|---|---|---|---|---|---|---|---|---|---|
| **A_52_P253230** | Ankrd46 | -0.016340288 | -0.171107195 | 0.138426618 | 10.30787605 | -0.259747141 | 0.803929499 | 0.964975838 | -5.089041982 |
| **A_55_P2057469** | Caskin1 | -0.023571732 | -0.246975947 | 0.199832484 | 8.74268871 | -0.259578839 | 0.804053259 | 0.964975838 | -5.089074516 |
| **A_52_P641922** | Nhlrc3 | 0.019182276 | -0.162686147 | 0.201050699 | 9.686795122 | 0.259484847 | 0.804122377 | 0.964975838 | -5.089092677 |
| **A_51_P461475** | Atn1 | -0.019225467 | -0.201527434 | 0.163076501 | 9.677473716 | -0.259450615 | 0.804147552 | 0.964975838 | -5.089099289 |
| **A_51_P345593** | Sptlc1 | 0.028948273 | -0.245610278 | 0.303506824 | 10.72160157 | 0.259392162 | 0.804190537 | 0.964975838 | -5.089110578 |
| **A_55_P2077879** | Kat6b | 0.01676982 | -0.142310253 | 0.175849893 | 8.191086326 | 0.259347335 | 0.804223504 | 0.964975838 | -5.089119234 |
| **A_51_P395473** | Tns2 | -0.027813385 | -0.29165384 | 0.23602707 | 9.95576242 | -0.259347235 | 0.804223577 | 0.964975838 | -5.089119254 |
| **A_55_P2143150** | Qk | -0.024379066 | -0.255669392 | 0.20691126 | 9.569518453 | -0.259315668 | 0.804246792 | 0.964975838 | -5.089125348 |
| **A_52_P402663** | Ninl | -0.011437929 | -0.119996914 | 0.097121056 | 7.855253283 | -0.259209427 | 0.804324925 | 0.964975838 | -5.089145854 |
| **A_55_P2121961** | Psmc3 | 0.022519105 | -0.191258015 | 0.236296226 | 12.07200395 | 0.259154691 | 0.804365181 | 0.964975838 | -5.089156416 |
| **A_52_P609868** | Timd4 | 0.011062568 | -0.094045771 | 0.116170907 | 7.666098768 | 0.258933322 | 0.804527993 | 0.964975838 | -5.08919911 |
| **A_55_P2081585** | Dennd5b | 0.043518112 | -0.370090792 | 0.457127017 | 9.359154598 | 0.258850693 | 0.804588767 | 0.964975838 | -5.089215036 |
| **A_55_P1999760** | Lrrc51 | 0.01761023 | -0.14976733 | 0.18498779 | 10.10713473 | 0.258843318 | 0.804594192 | 0.964975838 | -5.089216458 |
| **A_55_P2090794** | NA | 0.020399942 | -0.173493109 | 0.214292993 | 10.32872764 | 0.258842657 | 0.804594678 | 0.964975838 | -5.089216585 |
| **A_51_P229925** | Chst7 | 0.02557611 | -0.217593408 | 0.268745627 | 8.975441851 | 0.258758433 | 0.804656628 | 0.964975838 | -5.089232814 |
| **A_55_P2031045** | Stat5b | 0.035994506 | -0.306274618 | 0.378263631 | 9.311371688 | 0.258724563 | 0.80468154 | 0.964975838 | -5.089239338 |
| **A_55_P2164629** | Ptn | -0.043069084 | -0.452762975 | 0.366624808 | 9.136815451 | -0.25862786 | 0.804752672 | 0.964975838 | -5.089257963 |
| **A_55_P2108689** | NA | -0.038847741 | -0.408393081 | 0.330697598 | 13.63836741 | -0.258623036 | 0.80475622 | 0.964975838 | -5.089258892 |
| **A_52_P671062** | Uba6 | 0.022899403 | -0.19498615 | 0.240784957 | 9.130919547 | 0.258562116 | 0.804801031 | 0.964975838 | -5.089270621 |
| **A_51_P205390** | Cabcoco1 | 0.011179714 | -0.095203777 | 0.117563205 | 7.698736415 | 0.258538736 | 0.804818229 | 0.964975838 | -5.089275122 |
| **A_52_P97192** | Ergic1 | -0.02358908 | -0.248071112 | 0.200892951 | 10.0162594 | -0.258522644 | 0.804830066 | 0.964975838 | -5.089278219 |
| **A_55_P2075670** | NA | 0.020433633 | -0.17408907 | 0.214956336 | 8.337326212 | 0.258430904 | 0.804897551 | 0.964975838 | -5.089295875 |
| **A_52_P527106** | Arhgap12 | -0.018191943 | -0.191374686 | 0.1549908 | 9.711562885 | -0.258430416 | 0.80489791 | 0.964975838 | -5.089295969 |
| **A_52_P514317** | Slc35a3 | 0.033216058 | -0.283028551 | 0.349460666 | 9.666993819 | 0.258400956 | 0.804919581 | 0.964975838 | -5.089301637 |
| **A_55_P2471798** | NA | 0.041781479 | -0.356077406 | 0.439640364 | 10.20346557 | 0.258359189 | 0.804950306 | 0.964975838 | -5.089309672 |
| **A_51_P386870** | Sprr2f | 0.060567582 | -0.516240208 | 0.637375372 | 8.45158263 | 0.258332046 | 0.804970274 | 0.964975838 | -5.089314893 |
| **A_52_P488437** | Plek2 | -0.012296289 | -0.129480793 | 0.104888215 | 7.786858779 | -0.258150536 | 0.805103802 | 0.965076055 | -5.089349794 |
| **A_52_P576720** | Calm4 | -0.169274785 | -1.783514161 | 1.444964592 | 9.318363974 | -0.25798443 | 0.805226005 | 0.965117472 | -5.089381712 |
| **A_51_P147064** | 1600014C23Rik | 0.064480213 | -0.550555391 | 0.679515817 | 9.823085029 | 0.257926146 | 0.805268886 | 0.965117472 | -5.089392907 |
| **A_55_P2090914** | Tpst2 | -0.015755941 | -0.166091898 | 0.134580016 | 12.19067934 | -0.257840214 | 0.805332109 | 0.965117472 | -5.089409408 |
| **A_55_P2140201** | Dupd1 | -0.060096769 | -0.633554698 | 0.513361159 | 8.72895007 | -0.25782126 | 0.805346054 | 0.965117472 | -5.089413047 |
| **A_51_P199104** | Casq1 | 0.011597194 | -0.099116184 | 0.122310571 | 7.695118032 | 0.257704485 | 0.805431973 | 0.965117472 | -5.08943546 |
| **A_52_P253044** | Syt13 | 0.050390691 | -0.430774581 | 0.531555962 | 10.56602729 | 0.257647066 | 0.805474221 | 0.965117472 | -5.089446477 |
| **A_55_P2137206** | Tgfbr1 | 0.018456328 | -0.157790186 | 0.194702842 | 10.54388031 | 0.257628514 | 0.805487871 | 0.965117472 | -5.089450037 |
| **A_51_P511176** | Triap1 | -0.017981348 | -0.189898607 | 0.153935911 | 11.22218661 | -0.257319047 | 0.805715584 | 0.965226064 | -5.08950937 |
| **A_55_P2057370** | Slc22a19 | -0.011084683 | -0.117069342 | 0.094899977 | 7.644672635 | -0.257305705 | 0.805725402 | 0.965226064 | -5.089511926 |
| **A_51_P487360** | Hpcal1 | -0.016474346 | -0.174026883 | 0.141078191 | 10.07289853 | -0.257247961 | 0.805767894 | 0.965226064 | -5.089522989 |
| **A_55_P2132024** | Tekt1 | 0.044910287 | -0.38477234 | 0.474592914 | 8.28335849 | 0.257138556 | 0.805848404 | 0.965226064 | -5.089543943 |
| **A_51_P430552** | G6pc3 | 0.02338862 | -0.200537095 | 0.247314334 | 11.66863781 | 0.256962525 | 0.805977948 | 0.965226064 | -5.089577639 |
| **A_55_P2095047** | Mat2a | -0.012529513 | -0.132553595 | 0.107494568 | 8.285464095 | -0.256823628 | 0.80608017 | 0.965226064 | -5.089604211 |
| **A_55_P2069221** | Prr11 | -0.012218184 | -0.129263477 | 0.104827108 | 7.721964731 | -0.256815888 | 0.806085866 | 0.965226064 | -5.089605691 |
| **A_55_P2127139** | H3c4 | -0.022646245 | -0.239637872 | 0.194345383 | 13.17593449 | -0.256757053 | 0.806129167 | 0.965226064 | -5.089616942 |
| **A_66_P120567** | Anxa1 | -0.032474394 | -0.343657312 | 0.278708524 | 12.26719481 | -0.256740552 | 0.806141312 | 0.965226064 | -5.089620097 |
| **A_51_P410681** | Irx5 | 0.011445802 | -0.098246639 | 0.121138243 | 7.713998851 | 0.256707579 | 0.806165579 | 0.965226064 | -5.089626401 |
| **A_55_P1992769** | Pafah1b1 | -0.012853876 | -0.136107045 | 0.110399294 | 12.9992608 | -0.256569582 | 0.806267146 | 0.965226064 | -5.089652776 |
| **A_51_P209518** | Galnt1 | 0.018395482 | -0.158088338 | 0.194879303 | 9.205863396 | 0.256433906 | 0.806367009 | 0.965226064 | -5.089678693 |
| **A_52_P258557** | Slc26a9 | -0.010812503 | -0.114554814 | 0.092929808 | 7.670468605 | -0.25641268 | 0.806382632 | 0.965226064 | -5.089682747 |
| **A_51_P348665** | Ramp1 | -0.183174936 | -1.94145828 | 1.575108408 | 11.01315072 | -0.256298668 | 0.806466553 | 0.965226064 | -5.089704514 |
| **A_55_P2107398** | NA | -0.027240287 | -0.288727449 | 0.234246875 | 11.72885109 | -0.256289298 | 0.80647345 | 0.965226064 | -5.089706302 |
| **A_55_P2112479** | Nudt16l1 | 0.023733473 | -0.204179904 | 0.251646849 | 9.409815164 | 0.256189092 | 0.806547211 | 0.965226064 | -5.089725425 |

| | | | | | | | | | |
|---|---|---|---|---|---|---|---|---|---|
| A_51_P209183 | Cxcl14 | -0.219948924 | -2.332431188 | 1.892533341 | 10.05371252 | -0.256152107 | 0.806574436 | 0.965226064 | -5.089732481 |
| A_55_P2080048 | NA | 0.020340647 | -0.175019583 | 0.215700876 | 14.75494546 | 0.256152007 | 0.80657451 | 0.965226064 | -5.0897325 |
| A_52_P86693 | Ifi27 | 0.041876804 | -0.360336003 | 0.444089611 | 13.39732701 | 0.256145541 | 0.806579269 | 0.965226064 | -5.089733734 |
| A_55_P2009329 | Gm8709 | 0.02861268 | -0.246237204 | 0.303462565 | 9.237819827 | 0.256113311 | 0.806602995 | 0.965226064 | -5.089739882 |
| A_51_P369371 | Ubtd1 | 0.040353807 | -0.347394911 | 0.428102525 | 11.11615778 | 0.256037337 | 0.806658921 | 0.965226064 | -5.089754371 |
| A_52_P35057 | Kctd17 | 0.026443005 | -0.227718179 | 0.280604189 | 11.31054104 | 0.2559592 | 0.806716442 | 0.965226064 | -5.089769269 |
| A_51_P174415 | Retreg2 | 0.021916997 | -0.188753596 | 0.232587591 | 10.08020962 | 0.255944795 | 0.806727046 | 0.965226064 | -5.089772015 |
| A_55_P2157814 | Gmds | -0.022840385 | -0.242489683 | 0.196808913 | 11.99702321 | -0.255824871 | 0.806815331 | 0.965230129 | -5.08979487 |
| A_51_P419971 | Tmem106b | -0.023334974 | -0.247815048 | 0.2011451 | 12.28126253 | -0.255740021 | 0.806877797 | 0.965230129 | -5.089811034 |
| A_52_P344424 | Lipg | 0.037918542 | -0.326932785 | 0.402769869 | 9.578151624 | 0.255684773 | 0.806918471 | 0.965230129 | -5.089821557 |
| A_52_P496924 | Mgat4a | 0.014180124 | -0.122290444 | 0.150650693 | 11.26450803 | 0.255629049 | 0.806959496 | 0.965230129 | -5.089832167 |
| A_51_P486618 | Akr1b10 | 0.052849398 | -0.455960311 | 0.561659107 | 9.338804223 | 0.255537016 | 0.807027254 | 0.965230129 | -5.089849687 |
| A_55_P2119030 | Rasl2-9 | -0.022076239 | -0.234619295 | 0.190466817 | 12.28288124 | -0.255533199 | 0.807030064 | 0.965230129 | -5.089850413 |
| A_55_P1979645 | S100a1 | 0.097778709 | -0.844748925 | 1.040306343 | 11.57728298 | 0.255222741 | 0.807258649 | 0.965443783 | -5.089909466 |
| A_66_P136998 | Tmem43 | -0.047410043 | -0.504689465 | 0.40986938 | 9.966179557 | -0.255069101 | 0.80737178 | 0.965477879 | -5.089938665 |
| A_55_P2319468 | NA | -0.018611472 | -0.198199747 | 0.160976802 | 7.886139139 | -0.254959922 | 0.807452175 | 0.965477879 | -5.089959404 |
| A_51_P497100 | Lgals4 | 0.019371379 | -0.16755868 | 0.206301437 | 8.922750904 | 0.25494738 | 0.80746141 | 0.965477879 | -5.089961785 |
| A_52_P546635 | Zeb1 | -0.028561308 | -0.304224264 | 0.247101649 | 9.229755699 | -0.254899415 | 0.80749673 | 0.965477879 | -5.089970893 |
| A_55_P2016351 | Eif3c | 0.021876012 | -0.189315151 | 0.233067175 | 13.87116404 | 0.25483647 | 0.807543083 | 0.965477879 | -5.089982843 |
| A_55_P2032853 | Entpd1 | 0.012724501 | -0.110162833 | 0.135611834 | 8.885559311 | 0.254743316 | 0.807611682 | 0.965477879 | -5.090000522 |
| A_55_P2089477 | Sypl | 0.024447094 | -0.211683238 | 0.260577427 | 12.56202967 | 0.2547092 | 0.807636806 | 0.965477879 | -5.090006995 |
| A_66_P129188 | H2ac20 | 0.02795412 | -0.242335358 | 0.298243599 | 14.85593276 | 0.254440273 | 0.80783486 | 0.965654917 | -5.090057993 |
| A_51_P273433 | Prkg2 | -0.024872713 | -0.26557784 | 0.215832414 | 8.171287854 | -0.2542184 | 0.807998271 | 0.965747296 | -5.090100028 |
| A_52_P546090 | Ssr4 | -0.016559767 | -0.176828351 | 0.143708818 | 15.45804924 | -0.254199667 | 0.808012069 | 0.965747296 | -5.090103575 |
| A_66_P120745 | Armc1 | 0.017379694 | -0.150897325 | 0.185656712 | 10.93291106 | 0.254089357 | 0.808093317 | 0.965773237 | -5.090124459 |
| A_55_P2010116 | Rab27b | -0.068811103 | -0.735510655 | 0.597888449 | 8.818285973 | -0.253920356 | 0.8082178 | 0.965773237 | -5.090156438 |
| A_51_P176365 | Gimap5 | 0.020556771 | -0.17864699 | 0.219760532 | 8.195611523 | 0.253878848 | 0.808248375 | 0.965773237 | -5.090164289 |
| A_55_P2088218 | Uqcrb | -0.024443385 | -0.261403567 | 0.212516797 | 12.92918189 | -0.253778681 | 0.80832216 | 0.965773237 | -5.090183229 |
| A_55_P2121352 | Cdk5 | 0.024886612 | -0.216374417 | 0.26614764 | 10.60495944 | 0.25377437 | 0.808325336 | 0.965773237 | -5.090184044 |
| A_55_P2033010 | Mrrf | -0.012285417 | -0.13141929 | 0.106848455 | 8.008118076 | -0.253701956 | 0.808378678 | 0.965773237 | -5.090197733 |
| A_52_P485891 | Dnah5 | 0.010843891 | -0.094313975 | 0.116001757 | 7.66227424 | 0.253695374 | 0.808383528 | 0.965773237 | -5.090198977 |
| A_55_P2089942 | Xpo1 | -0.023225351 | -0.248547633 | 0.202096931 | 10.85294999 | -0.253587186 | 0.808463225 | 0.965808756 | -5.09021942 |
| A_55_P2110935 | Mlip | 0.011102537 | -0.096680632 | 0.118885706 | 7.651329228 | 0.253419749 | 0.808586574 | 0.965896415 | -5.090251041 |
| A_51_P371255 | Rwdd2b | 0.01183003 | -0.103048072 | 0.126708132 | 8.414478041 | 0.253348162 | 0.808639313 | 0.965899721 | -5.090264555 |
| A_52_P304128 | Mmp14 | -0.060650756 | -0.650276611 | 0.528975099 | 9.055223835 | -0.253063123 | 0.808849317 | 0.966090864 | -5.090318326 |
| A_55_P2029022 | Rpl34 | -0.017802992 | -0.190955706 | 0.155349722 | 16.70510361 | -0.252948931 | 0.808933453 | 0.966105655 | -5.09033985 |
| A_52_P281209 | Kdm4a | -0.020293871 | -0.217830737 | 0.177242996 | 10.3275451 | -0.252746929 | 0.809082294 | 0.966105655 | -5.090377904 |
| A_55_P2122066 | Ppp2r3d | -0.012056733 | -0.129427684 | 0.105314218 | 8.243100425 | -0.252719164 | 0.809102753 | 0.966105655 | -5.090383133 |
| A_51_P341736 | Mmp2 | 0.053169353 | -0.464711319 | 0.571050025 | 9.010830753 | 0.252581091 | 0.809204495 | 0.966105655 | -5.090409124 |
| A_55_P2068289 | Slc17a2 | -0.018453533 | -0.198211413 | 0.161304347 | 7.75930251 | -0.252557783 | 0.809221671 | 0.966105655 | -5.09041351 |
| A_51_P341664 | Zfp87 | -0.012772092 | -0.137213965 | 0.111669781 | 8.8096795 | -0.252501902 | 0.809262849 | 0.966105655 | -5.090424024 |
| A_55_P2245920 | Lrrc8a | -0.030795728 | -0.330926395 | 0.269334939 | 13.35561916 | -0.252434821 | 0.809312282 | 0.966105655 | -5.090436642 |
| A_52_P523480 | Rfx1 | -0.022092108 | -0.237409641 | 0.193225424 | 9.22672916 | -0.252421843 | 0.809321846 | 0.966105655 | -5.090439083 |
| A_55_P2052048 | Pih1d2 | 0.010771168 | -0.094310366 | 0.115852702 | 7.905108127 | 0.25217705 | 0.809502246 | 0.966105655 | -5.090485101 |
| A_55_P2370384 | 1600029I14Rik | -0.041858853 | -0.450378078 | 0.366660372 | 8.682932533 | -0.252083252 | 0.809571374 | 0.966105655 | -5.090502723 |
| A_55_P2037584 | Trf | -0.074216331 | -0.798653064 | 0.650220401 | 16.6523298 | -0.252039252 | 0.809603801 | 0.966105655 | -5.090510986 |
| A_51_P472274 | Sox18 | 0.01973409 | -0.172900559 | 0.212368739 | 8.634988317 | 0.252029781 | 0.809610782 | 0.966105655 | -5.090512765 |
| A_55_P2118103 | Zfp68 | 0.014425668 | -0.126395234 | 0.15524657 | 8.071979937 | 0.252021724 | 0.80961672 | 0.966105655 | -5.090514278 |
| A_51_P163958 | Spr | 0.031232525 | -0.273721277 | 0.336186327 | 10.08501209 | 0.251966153 | 0.809657676 | 0.966105655 | -5.090524712 |
| A_55_P1980190 | NA | -0.025258763 | -0.271900322 | 0.221382796 | 11.76190167 | -0.251950362 | 0.809669315 | 0.966105655 | -5.090527677 |

| | | | | | | | | | |
|---|---|---|---|---|---|---|---|---|---|
| **A_55_P2177488** | H2bc21 | -0.012476808 | -0.1343362 | 0.109382585 | 7.844815705 | -0.251891571 | 0.809712645 | 0.966105655 | -5.090538713 |
| **A_51_P164939** | Tmem150a | -0.012986051 | -0.139870156 | 0.113898053 | 10.67066967 | -0.25179034 | 0.809787257 | 0.966105655 | -5.090557709 |
| **A_52_P98625** | Sowahc | -0.013829889 | -0.148962203 | 0.121302426 | 8.523926304 | -0.251784296 | 0.809791712 | 0.966105655 | -5.090558843 |
| **A_51_P297105** | Ucp2 | -0.017110315 | -0.184346886 | 0.150126255 | 9.523181023 | -0.251707367 | 0.809848414 | 0.966105655 | -5.090573274 |
| **A_52_P434841** | Coa4 | 0.03250727 | -0.285250228 | 0.350264768 | 10.0851998 | 0.251682983 | 0.809866387 | 0.966105655 | -5.090577847 |
| **A_52_P559975** | Cxcr2 | -0.013698862 | -0.147688075 | 0.120290351 | 11.0578248 | -0.251526539 | 0.809981702 | 0.966105655 | -5.090607178 |
| **A_55_P2082778** | NA | 0.032644865 | -0.286770826 | 0.352060556 | 13.38555847 | 0.25143619 | 0.8100483 | 0.966105655 | -5.090624109 |
| **A_66_P123448** | Pigx | 0.022899597 | -0.201246224 | 0.247045418 | 12.19690616 | 0.251342746 | 0.810117182 | 0.966105655 | -5.090641614 |
| **A_55_P2024909** | Dnah17 | 0.014549424 | -0.127868604 | 0.156967453 | 8.029158138 | 0.251333285 | 0.810124156 | 0.966105655 | -5.090643386 |
| **A_55_P2024749** | Hspb11 | 0.022598011 | -0.198635597 | 0.24383162 | 10.57201959 | 0.251297565 | 0.810150488 | 0.966105655 | -5.090650075 |
| **A_55_P1956672** | Rab34 | 0.051999753 | -0.45719568 | 0.561195187 | 10.48191418 | 0.251238357 | 0.810194133 | 0.966105655 | -5.090661161 |
| **A_55_P2052420** | Setd5 | 0.042679814 | -0.375370005 | 0.460729634 | 12.61262037 | 0.251167617 | 0.810246282 | 0.966105655 | -5.090674404 |
| **A_52_P562593** | Tpgs1 | 0.024215519 | -0.212995194 | 0.261426232 | 10.32723821 | 0.251147378 | 0.810261202 | 0.966105655 | -5.090678191 |
| **A_55_P2459897** | A2m | 0.063512585 | -0.559181574 | 0.686206744 | 8.238715479 | 0.250930909 | 0.810420787 | 0.966178216 | -5.090718687 |
| **A_51_P392214** | Pcnx4 | -0.013863488 | -0.149785704 | 0.122058727 | 9.874764577 | -0.250929224 | 0.81042203 | 0.966178216 | -5.090719003 |
| **A_55_P2120466** | Snx11 | 0.014046318 | -0.123778941 | 0.151871577 | 9.030302375 | 0.250728 | 0.810570385 | 0.966233274 | -5.090756616 |
| **A_52_P203981** | Srek1ip1 | -0.034668517 | -0.374936122 | 0.305599088 | 9.98647434 | -0.2506593 | 0.810621037 | 0.966233274 | -5.090769451 |
| **A_51_P470715** | Cish | 0.026241899 | -0.23135861 | 0.283842409 | 9.985983157 | 0.250621147 | 0.810649167 | 0.966233274 | -5.090776577 |
| **A_55_P2467415** | B3galt6 | -0.024133579 | -0.261075512 | 0.212808355 | 8.828437662 | -0.250581477 | 0.810678417 | 0.966233274 | -5.090783986 |
| **A_55_P2062714** | Zfp975 | 0.028664345 | -0.252820902 | 0.310149591 | 9.341089049 | 0.25052758 | 0.810718156 | 0.966233274 | -5.09079405 |
| **A_55_P2131353** | 1110059G10Rik | -0.024242492 | -0.262447854 | 0.21396287 | 9.285311375 | -0.250377266 | 0.810828989 | 0.966277513 | -5.090822106 |
| **A_55_P2041050** | Zmym5 | -0.01975175 | -0.213858952 | 0.174355452 | 9.558711851 | -0.250341643 | 0.810855257 | 0.966277513 | -5.090828753 |
| **A_55_P2073935** | Abcc5 | 0.028486559 | -0.251782557 | 0.308755674 | 8.409419579 | 0.250054057 | 0.811067322 | 0.966415201 | -5.090882378 |
| **A_52_P556281** | G3bp2 | 0.016658469 | -0.147241371 | 0.18055831 | 10.77151043 | 0.250049348 | 0.811070795 | 0.966415201 | -5.090883255 |
| **A_52_P419373** | Lrrc39 | 0.010996144 | -0.09726344 | 0.119255727 | 7.687947332 | 0.249886743 | 0.811190707 | 0.966415461 | -5.090913548 |
| **A_55_P1984422** | Nme2 | 0.033835212 | -0.299363006 | 0.367033431 | 15.24148731 | 0.249824713 | 0.811236453 | 0.966415461 | -5.0909251 |
| **A_51_P439836** | NA | 0.02595224 | -0.229721983 | 0.281626463 | 11.19726528 | 0.249722152 | 0.811312091 | 0.966415461 | -5.090944192 |
| **A_51_P354706** | Lefty1 | -0.011496124 | -0.124810647 | 0.101818399 | 7.808580492 | -0.249594507 | 0.811406231 | 0.966415461 | -5.090967944 |
| **A_51_P193686** | 1700012B09Rik | 0.010726064 | -0.095014479 | 0.116466607 | 7.664466662 | 0.249555993 | 0.811434636 | 0.966415461 | -5.090975108 |
| **A_55_P2139043** | 4833439L19Rik | -0.019011632 | -0.206542114 | 0.168518849 | 11.75871059 | -0.249411633 | 0.811541109 | 0.966415461 | -5.091001952 |
| **A_55_P2165854** | Gemin7 | 0.03268677 | -0.289761115 | 0.355134655 | 10.78881536 | 0.249391494 | 0.811555963 | 0.966415461 | -5.091005695 |
| **A_52_P150236** | Apom | 0.148142378 | -1.314594799 | 1.610879555 | 13.2945394 | 0.249162211 | 0.811725081 | 0.966415461 | -5.091048296 |
| **A_52_P552832** | Ndufa4 | 0.035376285 | -0.314057624 | 0.384810195 | 13.12517704 | 0.249067088 | 0.811795247 | 0.966415461 | -5.091065959 |
| **A_51_P252677** | Card19 | 0.055547636 | -0.493153673 | 0.604248945 | 11.56374168 | 0.249056997 | 0.81180269 | 0.966415461 | -5.091067832 |
| **A_51_P462546** | Gmppa | -0.018349609 | -0.199621969 | 0.162922751 | 10.01515837 | -0.249037297 | 0.811817222 | 0.966415461 | -5.091071489 |
| **A_51_P137419** | Cst7 | 0.01361154 | -0.120874977 | 0.148098056 | 7.775405722 | 0.248999037 | 0.811845445 | 0.966415461 | -5.091078591 |
| **A_55_P2030756** | Nedd4l | 0.012936573 | -0.114922408 | 0.140795555 | 8.112175755 | 0.248918501 | 0.811904853 | 0.966415461 | -5.091093535 |
| **A_51_P305547** | Snai2 | -0.027087333 | -0.294907918 | 0.240733251 | 8.48195026 | -0.248823539 | 0.811974905 | 0.966415461 | -5.091111151 |
| **A_51_P509971** | Plekho1 | -0.107975297 | -1.17642552 | 0.960474925 | 12.02690857 | -0.248621841 | 0.8121237 | 0.966415461 | -5.091148546 |
| **A_51_P191782** | Olfml3 | -0.017467163 | -0.190327281 | 0.155392956 | 8.248721635 | -0.248597481 | 0.812141672 | 0.966415461 | -5.09115306 |
| **A_55_P2004258** | Gm7102 | 0.021077917 | -0.187571535 | 0.229727368 | 9.796216391 | 0.248530444 | 0.812191128 | 0.966415461 | -5.091165481 |
| **A_52_P112888** | Atp6v1h | 0.015860565 | -0.141182103 | 0.172903233 | 9.508409533 | 0.248467822 | 0.812237328 | 0.966415461 | -5.091177081 |
| **A_52_P208222** | Pde1c | -0.011536136 | -0.125766796 | 0.102694524 | 7.629075921 | -0.248454492 | 0.812247162 | 0.966415461 | -5.09117955 |
| **A_51_P344878** | Thop1 | 0.012053756 | -0.107313318 | 0.13142083 | 8.338738683 | 0.248431702 | 0.812263976 | 0.966415461 | -5.09118377 |
| **A_55_P2100209** | Kif2c | -0.011591298 | -0.126484875 | 0.10330228 | 7.682403023 | -0.248202108 | 0.81243337 | 0.966415461 | -5.091226269 |
| **A_55_P2055168** | Gm2016 | 0.017925726 | -0.159761762 | 0.195613213 | 9.66904469 | 0.248192731 | 0.812440289 | 0.966415461 | -5.091228004 |
| **A_55_P1959869** | Ppp1r2 | -0.019725741 | -0.215263615 | 0.175812134 | 11.05652209 | -0.248182751 | 0.812447653 | 0.966415461 | -5.09122985 |
| **A_55_P2007155** | Akr1c14 | 0.021978477 | -0.195955204 | 0.239912157 | 8.18150727 | 0.24810893 | 0.812502121 | 0.966415461 | -5.091243506 |
| **A_51_P302167** | Cobl | -0.012865999 | -0.140495036 | 0.114763038 | 7.909212405 | -0.248006573 | 0.812577645 | 0.966415461 | -5.091262433 |
| **A_52_P662244** | Slc25a47 | 0.014222791 | -0.126867827 | 0.155313409 | 8.338414735 | 0.248002381 | 0.812580739 | 0.966415461 | -5.091263208 |

| | | | | | | | | | |
|---|---|---|---|---|---|---|---|---|---|
| **A_55_P2009498** | Ccdc28a | -0.017292697 | -0.188850615 | 0.154265222 | 10.3858943 | -0.247982546 | 0.812595374 | 0.966415461 | -5.091266875 |
| **A_55_P2158152** | NA | 0.018871742 | -0.168357524 | 0.206101009 | 15.32205937 | 0.247974731 | 0.81260114 | 0.966415461 | -5.09126832 |
| **A_55_P1978103** | Ppip5k2 | -0.012974845 | -0.141703949 | 0.115754259 | 10.18009817 | -0.247967399 | 0.812606551 | 0.966415461 | -5.091269675 |
| **A_55_P1985804** | Thumpd2 | -0.012063169 | -0.131818213 | 0.107691875 | 7.963750482 | -0.247820236 | 0.81271514 | 0.966415461 | -5.09129687 |
| **A_55_P2050226** | Ackr4 | 0.023155491 | -0.20681645 | 0.253127432 | 8.04908925 | 0.24771272 | 0.812794478 | 0.966415461 | -5.091316729 |
| **A_55_P2070823** | Ino80d | -0.02197835 | -0.240294941 | 0.196338241 | 10.33693115 | -0.247672339 | 0.812824277 | 0.966415461 | -5.091324186 |
| **A_55_P2109717** | Kif20b | 0.023470023 | -0.209734671 | 0.256674718 | 8.14127603 | 0.24759701 | 0.812879865 | 0.966415461 | -5.091338092 |
| **A_52_P24439** | Sftpa1 | 0.010410196 | -0.093104944 | 0.113925337 | 7.640425542 | 0.247413963 | 0.813014949 | 0.966415461 | -5.091371867 |
| **A_55_P2028847** | NA | -0.204911823 | -2.483792425 | 2.073968778 | 10.91421411 | -0.253015983 | 0.813127108 | 0.966415461 | -4.90032874 |
| **A_55_P2035320** | Nfil3 | -0.015513478 | -0.169893283 | 0.138866327 | 11.29914477 | -0.24722242 | 0.813156309 | 0.966415461 | -5.091407184 |
| **A_66_P107353** | NA | 0.017625346 | -0.157897799 | 0.193148491 | 15.82459367 | 0.247042928 | 0.813288784 | 0.966415461 | -5.091440255 |
| **A_51_P337944** | Bmp2k | -0.013584951 | -0.148881676 | 0.121711773 | 10.71621649 | -0.247024464 | 0.813302412 | 0.966415461 | -5.091443655 |
| **A_55_P2410007** | NA | 0.044618463 | -0.399771523 | 0.489008448 | 9.780915776 | 0.247012826 | 0.813311002 | 0.966415461 | -5.091445799 |
| **A_52_P14456** | Srrm4 | -0.015943683 | -0.17474832 | 0.142860953 | 7.798245319 | -0.246998686 | 0.813321438 | 0.966415461 | -5.091448403 |
| **A_51_P234847** | 1700001L19Rik | 0.012057873 | -0.108045536 | 0.132161283 | 7.949748879 | 0.246992943 | 0.813325677 | 0.966415461 | -5.09144946 |
| **A_55_P2074908** | Ube4b | -0.011804308 | -0.129411743 | 0.105803128 | 8.597960262 | -0.246930598 | 0.813371693 | 0.966415461 | -5.091460939 |
| **A_55_P2048840** | Ppil2 | -0.012164307 | -0.133370594 | 0.109041981 | 9.193688621 | -0.246905852 | 0.813389958 | 0.966415461 | -5.091465495 |
| **A_55_P1994648** | Eef1d | 0.016980348 | -0.152223848 | 0.186184544 | 14.86283636 | 0.246890638 | 0.813401187 | 0.966415461 | -5.091468295 |
| **A_55_P2099930** | Cers5 | -0.013288267 | -0.145768648 | 0.119192115 | 12.64738031 | -0.246766342 | 0.813492932 | 0.966415461 | -5.091491169 |
| **A_55_P2099080** | Irak1 | -0.022232242 | -0.2439175 | 0.199453016 | 10.27646177 | -0.246726393 | 0.813522419 | 0.966415461 | -5.091498518 |
| **A_51_P372141** | Pnkd | -0.010844919 | -0.118983868 | 0.097294029 | 7.847350258 | -0.246725112 | 0.813523365 | 0.966415461 | -5.091498754 |
| **A_55_P1969857** | Ap2m1 | 0.038323965 | -0.343885874 | 0.420533804 | 14.19545104 | 0.246682158 | 0.813555071 | 0.966415461 | -5.091506655 |
| **A_55_P2019113** | Apol7b | -0.034064953 | -0.373823665 | 0.30569376 | 9.117829886 | -0.246664331 | 0.81356823 | 0.966415461 | -5.091509933 |
| **A_51_P361492** | Pou2f1 | 0.025097449 | -0.22528935 | 0.275484248 | 9.84710497 | 0.246596731 | 0.81361813 | 0.966415461 | -5.091522364 |
| **A_55_P2028660** | Dusp12 | -0.011229852 | -0.123267129 | 0.100807425 | 8.006887862 | -0.246592948 | 0.813620922 | 0.966415461 | -5.091523059 |
| **A_55_P1998811** | NA | -0.089733842 | -0.985834816 | 0.806367132 | 10.04539259 | -0.246359025 | 0.813793601 | 0.966561172 | -5.091566047 |
| **A_51_P166288** | Acy1 | -0.011366915 | -0.124928655 | 0.102194826 | 7.831792726 | -0.246251972 | 0.81387263 | 0.966589636 | -5.091585707 |
| **A_66_P113578** | Washc1 | -0.021364257 | -0.234857742 | 0.192129229 | 11.08643227 | -0.246191084 | 0.813917581 | 0.966589636 | -5.091596885 |
| **A_51_P299632** | Spef1 | 0.016528392 | -0.148705274 | 0.181762059 | 9.367052767 | 0.24609412 | 0.813989166 | 0.96661526 | -5.091614681 |
| **A_52_P198435** | Rasgrp3 | -0.029674614 | -0.326418589 | 0.267069361 | 9.64730455 | -0.246021086 | 0.814043085 | 0.966619903 | -5.09162808 |
| **A_55_P2035677** | Srp19 | -0.01270302 | -0.139781031 | 0.114374991 | 11.4213367 | -0.245926733 | 0.814112745 | 0.966643236 | -5.091645385 |
| **A_51_P185509** | Mapre1 | 0.030225752 | -0.272237877 | 0.332689381 | 9.164894367 | 0.245851644 | 0.814168184 | 0.966649682 | -5.091659152 |
| **A_55_P1958210** | Ephb1 | -0.01140323 | -0.125559393 | 0.102752933 | 7.653163235 | -0.245752361 | 0.814241488 | 0.966665311 | -5.091677349 |
| **A_55_P1953980** | Sdhb | 0.025674184 | -0.231403026 | 0.282751394 | 11.25764985 | 0.245698346 | 0.81428137 | 0.966665311 | -5.091687246 |
| **A_55_P2063126** | Patj | -0.019832879 | -0.218529822 | 0.178864065 | 10.40404805 | -0.245563454 | 0.814380969 | 0.966696929 | -5.091711952 |
| **A_52_P635271** | Btbd6 | -0.011549067 | -0.127271393 | 0.104173258 | 8.055357089 | -0.245526808 | 0.814408029 | 0.966696929 | -5.091718662 |
| **A_55_P1955417** | Atad3a | 0.045339632 | -0.409384597 | 0.500063861 | 12.37005191 | 0.245300854 | 0.814574876 | 0.966763224 | -5.091760012 |
| **A_55_P2119907** | Dnah11 | 0.010376731 | -0.093702092 | 0.114455554 | 7.696519352 | 0.245282948 | 0.814588098 | 0.966763224 | -5.091763287 |
| **A_55_P2071406** | NA | -0.01126942 | -0.124318046 | 0.101779207 | 7.973548917 | -0.245247968 | 0.814613929 | 0.966763224 | -5.091769684 |
| **A_55_P2208071** | Nbeal1 | -0.020068873 | -0.221470809 | 0.181333063 | 9.640160518 | -0.245148087 | 0.814687687 | 0.966791399 | -5.091787947 |
| **A_55_P1975415** | Cpped1 | -0.019221752 | -0.212266375 | 0.173822871 | 10.55456448 | -0.244965218 | 0.814822733 | 0.966825048 | -5.091821365 |
| **A_51_P515623** | Qpctl | -0.024945851 | -0.275516502 | 0.225624799 | 10.75448318 | -0.244927356 | 0.814850695 | 0.966825048 | -5.091828281 |
| **A_51_P343613** | Ehd2 | 0.012079365 | -0.109294454 | 0.133453184 | 8.181481015 | 0.244843309 | 0.814912765 | 0.966825048 | -5.091843629 |
| **A_55_P2077053** | Itih4 | -0.015336773 | -0.169443991 | 0.138770444 | 7.867179113 | -0.244838767 | 0.81491612 | 0.966825048 | -5.091844458 |
| **A_55_P2061495** | Cks1b | 0.038245529 | -0.346328908 | 0.422819967 | 10.83227793 | 0.244663635 | 0.815045464 | 0.966871022 | -5.091876423 |
| **A_51_P479352** | Klhdc9 | -0.010391162 | -0.114895147 | 0.094112823 | 7.87649187 | -0.244624768 | 0.81507417 | 0.966871022 | -5.091883514 |
| **A_55_P2110027** | NA | -0.017649618 | -0.19518217 | 0.159882935 | 12.37003321 | -0.244583113 | 0.815104936 | 0.966871022 | -5.091891113 |
| **A_52_P121342** | Nrxn1 | 0.010861419 | -0.098551543 | 0.120274381 | 7.66259366 | 0.244223221 | 0.815370761 | 0.967111911 | -5.091956709 |
| **A_55_P2048224** | NA | -0.011068342 | -0.122601311 | 0.100464626 | 7.725366761 | -0.244145364 | 0.815428271 | 0.967111911 | -5.091970887 |
| **A_55_P2111153** | NA | -0.013954716 | -0.154596241 | 0.126686808 | 10.22973486 | -0.244104962 | 0.815458116 | 0.967111911 | -5.091978243 |

| | | | | | | | | | |
|---|---|---|---|---|---|---|---|---|---|
| **A_55_P2176300** | Slc33a1 | -0.016834666 | -0.186596953 | 0.152927621 | 8.538774071 | -0.243967766 | 0.815559463 | 0.967113705 | -5.092003213 |
| **A_55_P1966144** | Slfn14 | 0.010521181 | -0.09559452 | 0.116636881 | 7.659069816 | 0.243923699 | 0.815592016 | 0.967113705 | -5.09201123 |
| **A_51_P105124** | P2ry6 | 0.017967024 | -0.163264611 | 0.199198658 | 8.498812265 | 0.243899717 | 0.815609732 | 0.967113705 | -5.081031464 |
| **A_55_P2063266** | Piga | 0.025590654 | -0.232663675 | 0.283844983 | 9.100522609 | 0.243782731 | 0.815696154 | 0.96715685 | -5.092036868 |
| **A_55_P1956627** | Gpr160 | 0.023786207 | -0.216349409 | 0.263921823 | 8.585276519 | 0.24369004 | 0.815764631 | 0.967169806 | -5.092053718 |
| **A_55_P2131920** | Prrx1 | -0.071998013 | -0.799265348 | 0.655269322 | 9.081597154 | -0.243554183 | 0.815865 | 0.967169806 | -5.092078403 |
| **A_52_P18816** | Gosr2 | 0.014444727 | -0.131469327 | 0.16035878 | 9.55714764 | 0.243546205 | 0.815870895 | 0.967169806 | -5.092079852 |
| **A_51_P337662** | Ints6l | -0.020077685 | -0.223085691 | 0.182930321 | 8.532234521 | -0.243315419 | 0.816041406 | 0.967169806 | -5.092121754 |
| **A_55_P2063316** | Mgll | 0.027076101 | -0.246795688 | 0.300947889 | 11.81097358 | 0.243224873 | 0.816108307 | 0.967169806 | -5.092138183 |
| **A_51_P312121** | Xdh | -0.054180299 | -0.602228318 | 0.49386772 | 9.593250256 | -0.243215891 | 0.816114943 | 0.967169806 | -5.092139812 |
| **A_55_P2008870** | NA | -0.023861147 | -0.26525744 | 0.217535146 | 13.29995397 | -0.243181136 | 0.816140622 | 0.967169806 | -5.092146117 |
| **A_55_P2028977** | Tcf20 | -0.018319048 | -0.20386251 | 0.167224414 | 8.470675562 | -0.242899381 | 0.816348814 | 0.967169806 | -5.092197193 |
| **A_55_P2026530** | Erc2 | -0.01572974 | -0.175052955 | 0.143593476 | 8.151939465 | -0.242891118 | 0.81635492 | 0.967169806 | -5.09219869 |
| **A_55_P1959648** | Hspg2 | -0.032729193 | -0.364240638 | 0.298782252 | 8.838998465 | -0.242887931 | 0.816357275 | 0.967169806 | -5.092199267 |
| **A_55_P2127421** | NA | -0.011746717 | -0.13081934 | 0.107325906 | 7.981265828 | -0.242702219 | 0.816494509 | 0.967169806 | -5.0922329 |
| **A_51_P265374** | Vnn3 | 0.01042891 | -0.095295607 | 0.116153428 | 7.668135351 | 0.242679109 | 0.816511586 | 0.967169806 | -5.092237083 |
| **A_52_P250555** | Dynll1 | -0.016238619 | -0.18086599 | 0.148388752 | 8.364705779 | -0.242670067 | 0.816518268 | 0.967169806 | -5.09223872 |
| **A_52_P242724** | NA | 0.017236028 | -0.15756066 | 0.192032717 | 10.02049096 | 0.24259015 | 0.816577327 | 0.967169806 | -5.092253183 |
| **A_52_P285470** | Lrp2 | 0.113722637 | -1.040042343 | 1.267487617 | 11.29964696 | 0.242492735 | 0.816649317 | 0.967169806 | -5.092270807 |
| **A_55_P1986596** | Cacna1h | 0.011739387 | -0.107400345 | 0.130879118 | 7.813602924 | 0.242414141 | 0.816707401 | 0.967169806 | -5.092285021 |
| **A_55_P1992031** | Dydc2 | 0.010573825 | -0.096737832 | 0.117885482 | 7.692681917 | 0.242412146 | 0.816708875 | 0.967169806 | -5.092285382 |
| **A_55_P1959748** | Asns | -0.044254789 | -0.493666479 | 0.405156901 | 12.46684033 | -0.242261872 | 0.816819936 | 0.967169806 | -5.092312546 |
| **A_55_P2018345** | Amz2 | 0.030881907 | -0.283046065 | 0.34480988 | 9.985316334 | 0.242015547 | 0.817001994 | 0.967169806 | -5.092357039 |
| **A_55_P2055597** | Enpp2 | 0.043287203 | -0.396749121 | 0.483323528 | 10.92387983 | 0.242013819 | 0.817003271 | 0.967169806 | -5.092357351 |
| **A_52_P227937** | Cystm1 | 0.048034609 | -0.440362465 | 0.536431683 | 11.69774 | 0.241963786 | 0.817040252 | 0.967169806 | -5.092366382 |
| **A_52_P466853** | Bmyc | -0.022947399 | -0.256294337 | 0.21039954 | 9.103561105 | -0.241936013 | 0.81706078 | 0.967169806 | -5.092371395 |
| **A_51_P195153** | Gtse1 | -0.012913453 | -0.144252939 | 0.118426032 | 7.9195475 | -0.241889072 | 0.817095476 | 0.967169806 | -5.092379866 |
| **A_55_P2004469** | NA | -0.0146924 | -0.164125714 | 0.134740913 | 15.40264271 | -0.241888096 | 0.817096197 | 0.967169806 | -5.092380042 |
| **A_55_P1988789** | Acss2 | -0.044469153 | -0.496861651 | 0.407923344 | 9.720314753 | -0.241831367 | 0.817138129 | 0.967169806 | -5.092390277 |
| **A_55_P2007464** | Rps2 | -0.01579028 | -0.176428037 | 0.144847478 | 15.75923342 | -0.241830653 | 0.817138656 | 0.967169806 | -5.092390406 |
| **A_52_P237652** | 2610035D17Rik | 0.073637885 | -0.675543942 | 0.822819711 | 9.157447384 | 0.241814998 | 0.817150228 | 0.967169806 | -5.09239323 |
| **A_51_P429209** | Smpd1 | -0.025367865 | -0.283618375 | 0.232882645 | 13.28524578 | -0.241663965 | 0.817261869 | 0.967169806 | -5.092420466 |
| **A_55_P1983070** | Uhrf1bp1 | -0.016206458 | -0.181248598 | 0.148835682 | 10.24254408 | -0.241580811 | 0.817323336 | 0.967169806 | -5.092435455 |
| **A_55_P2015984** | Odr4 | 0.014982111 | -0.137652859 | 0.167617082 | 11.77056687 | 0.241483876 | 0.817394993 | 0.967169806 | -5.092452921 |
| **A_52_P200617** | Iars | 0.013057451 | -0.119997274 | 0.146112177 | 10.04233092 | 0.241433358 | 0.817432337 | 0.967169806 | -5.092462021 |
| **A_52_P442756** | Btf3l4 | 0.019887797 | -0.182873792 | 0.222649385 | 10.82541757 | 0.241307128 | 0.817525653 | 0.967169806 | -5.09248475 |
| **A_55_P2144975** | Ccdc90b | 0.046680903 | -0.42926915 | 0.522630956 | 9.139573077 | 0.241294284 | 0.817535149 | 0.967169806 | -5.092487062 |
| **A_51_P263993** | Lipc | 0.010447555 | -0.096087155 | 0.116982264 | 7.651153082 | 0.241264088 | 0.817557472 | 0.967169806 | -5.092492498 |
| **A_51_P389636** | Kcnn4 | -0.045650379 | -0.511235917 | 0.419935159 | 8.144033653 | -0.241220418 | 0.817589756 | 0.967169806 | -5.092500357 |
| **A_52_P60194** | C4bp | 0.011114197 | -0.102268713 | 0.124497108 | 7.662651824 | 0.24115688 | 0.817636729 | 0.967169806 | -5.09251179 |
| **A_51_P133369** | Scly | 0.022252613 | -0.204767341 | 0.249272567 | 10.38533287 | 0.241149376 | 0.817642277 | 0.967169806 | -5.09251314 |
| **A_55_P1984243** | Naip2 | 0.023603566 | -0.217421292 | 0.264628424 | 8.080159827 | 0.240926709 | 0.8178069 | 0.967169806 | -5.092553181 |
| **A_55_P2308835** | E130110O22Rik | -0.017075614 | -0.191452601 | 0.157301373 | 8.404437691 | -0.240910833 | 0.817818638 | 0.967169806 | -5.092556034 |
| **A_51_P501656** | Poglut3 | 0.013530044 | -0.124660732 | 0.151720821 | 8.638476861 | 0.240873665 | 0.817846118 | 0.967169806 | -5.092562714 |
| **A_55_P1992889** | Serping1 | -0.15741978 | -1.765849454 | 1.451009894 | 11.86286811 | -0.240783313 | 0.817912921 | 0.967169806 | -5.092578948 |
| **A_55_P2128704** | NA | -0.013637122 | -0.152977724 | 0.12570348 | 14.01297239 | -0.240776557 | 0.817917917 | 0.967169806 | -5.092580162 |
| **A_51_P381360** | Ndufa13 | -0.022943475 | -0.257422912 | 0.211535963 | 15.81136115 | -0.240726328 | 0.817955056 | 0.967169806 | -5.092589184 |
| **A_55_P1973402** | Tada2a | -0.022787737 | -0.255679809 | 0.210104334 | 9.593016578 | -0.240721933 | 0.817958305 | 0.967169806 | -5.092589973 |
| **A_55_P1962304** | Plac8 | -0.043686025 | -0.49031352 | 0.402941469 | 10.14649337 | -0.240639127 | 0.818019532 | 0.967169806 | -5.092604842 |
| **A_51_P344566** | Plk1 | 0.029445885 | -0.271767943 | 0.330659713 | 8.491889174 | 0.240502095 | 0.818120856 | 0.967169806 | -5.092629437 |

| | | | | | | | | | |
|---|---|---|---|---|---|---|---|---|---|
| **A_55_P2097972** | NA | -0.015451706 | -0.173531552 | 0.142628141 | 8.600871323 | -0.24047453 | 0.818141238 | 0.967169806 | -5.092634383 |
| **A_55_P2087265** | NA | 0.05648567 | -0.521481309 | 0.634452649 | 9.690001712 | 0.240438728 | 0.818167712 | 0.967169806 | -5.092640806 |
| **A_51_P165451** | Pbld2 | 0.024727749 | -0.228346966 | 0.277802465 | 8.027098972 | 0.24038369 | 0.81820841 | 0.967169806 | -5.092650678 |
| **A_52_P207303** | Bmp3 | -0.012109175 | -0.136072889 | 0.111854538 | 7.682832317 | -0.240319582 | 0.818255816 | 0.967169806 | -5.092662174 |
| **A_51_P439824** | Gipc1 | 0.014799884 | -0.136738944 | 0.166338712 | 10.07724 | 0.240272218 | 0.81829084 | 0.967169806 | -5.092670666 |
| **A_55_P2148431** | Slc4a3 | -0.010868908 | -0.122199154 | 0.100461338 | 7.788486922 | -0.24018281 | 0.818356956 | 0.967169806 | -5.092686691 |
| **A_55_P2148348** | NA | -0.022718934 | -0.255472347 | 0.210034479 | 12.84416921 | -0.240138095 | 0.818390023 | 0.967169806 | -5.092694703 |
| **A_52_P394385** | 1600012H06Rik | 0.01683313 | -0.155696212 | 0.189362473 | 10.6756508 | 0.240033069 | 0.818467693 | 0.967169806 | -5.092713517 |
| **A_66_P138584** | Mnd1 | 0.013770103 | -0.127410168 | 0.154950374 | 7.927668953 | 0.239956403 | 0.81852439 | 0.967169806 | -5.092727245 |
| **A_55_P2157205** | Cdx1 | 0.040472757 | -0.374506175 | 0.455451689 | 10.10611651 | 0.239941796 | 0.818535193 | 0.967169806 | -5.092729861 |
| **A_55_P2105794** | Wdr61 | -0.02237964 | -0.251925149 | 0.207165869 | 14.23985567 | -0.239857588 | 0.818597469 | 0.967169806 | -5.092744934 |
| **A_55_P2027836** | Tnfrsf10b | 0.020533583 | -0.190091099 | 0.231158265 | 10.17905818 | 0.23984167 | 0.818609241 | 0.967169806 | -5.092747782 |
| **A_51_P410650** | Dpp9 | -0.012036801 | -0.13554822 | 0.111474617 | 9.38010778 | -0.239758018 | 0.818671109 | 0.967183782 | -5.09276275 |
| **A_51_P272993** | Ntm | 0.010494603 | -0.09729571 | 0.118284916 | 7.684926654 | 0.239527537 | 0.818841575 | 0.967326048 | -5.092803962 |
| **A_52_P101399** | Tmem130 | -0.034338722 | -0.387193306 | 0.318515861 | 10.96267938 | -0.239418404 | 0.818922295 | 0.967362283 | -5.092823463 |
| **A_66_P136899** | Nadk | 0.020931894 | -0.194269211 | 0.236133 | 11.53499213 | 0.239294764 | 0.819013748 | 0.967373533 | -5.092845545 |
| **A_55_P2091116** | Kbtbd12 | 0.010247092 | -0.095132133 | 0.115626317 | 7.652014378 | 0.239229536 | 0.819061996 | 0.967373533 | -5.092857191 |
| **A_52_P632191** | Nfx1 | -0.014659851 | -0.165461563 | 0.136141861 | 8.721831938 | -0.239162148 | 0.819111843 | 0.967373533 | -5.092869219 |
| **A_51_P430259** | Cfap126 | -0.027158729 | -0.306627653 | 0.252310194 | 9.650785663 | -0.239081013 | 0.819171861 | 0.967373533 | -5.092883696 |
| **A_66_P134809** | Gas5 | -0.019504671 | -0.220268108 | 0.181258765 | 11.78390633 | -0.239013912 | 0.819221497 | 0.967373533 | -5.092895665 |
| **A_55_P2071384** | Crh | -0.010411544 | -0.117648583 | 0.096825496 | 7.646305558 | -0.238857824 | 0.819336964 | 0.967373533 | -5.092923495 |
| **A_52_P110088** | Ndufaf3 | 0.011806851 | -0.109802778 | 0.133416479 | 9.459872724 | 0.238855513 | 0.819338674 | 0.967373533 | -5.092923907 |
| **A_55_P2100824** | LOC676689 | -0.104049346 | -1.17576299 | 0.967664298 | 11.93076492 | -0.238852482 | 0.819340916 | 0.967373533 | -5.092924448 |
| **A_55_P2072000** | Pcmtd1 | -0.014165102 | -0.160118473 | 0.131788268 | 8.653980055 | -0.238767247 | 0.819403971 | 0.967373533 | -5.092939637 |
| **A_55_P2094049** | Hectd4 | 0.03828992 | -0.356302297 | 0.432882136 | 11.31129304 | 0.238728959 | 0.819432296 | 0.967373533 | -5.092946458 |
| **A_55_P1981155** | Phldb1 | -0.021227866 | -0.240186151 | 0.197730418 | 12.68734106 | -0.238514139 | 0.819591225 | 0.967402435 | -5.092984711 |
| **A_55_P2106058** | Gm11110 | -0.022892409 | -0.259028685 | 0.213243867 | 8.394817486 | -0.238505255 | 0.819597798 | 0.967402435 | -5.092986292 |
| **A_55_P1954461** | Ptdss1 | -0.016500194 | -0.186793621 | 0.153793233 | 8.406896232 | -0.238374796 | 0.819694319 | 0.967402435 | -5.093009506 |
| **A_55_P2175400** | Zfp467 | -0.012271642 | -0.139002663 | 0.114459379 | 8.320515491 | -0.238225863 | 0.819804512 | 0.967402435 | -5.093035991 |
| **A_55_P1972018** | H4c3 | -0.017584956 | -0.199252457 | 0.164082545 | 8.79054451 | -0.238140474 | 0.819867692 | 0.967402435 | -5.093051169 |
| **A_55_P2013716** | Napa | 0.015705453 | -0.146598269 | 0.178009174 | 12.85826029 | 0.238062583 | 0.819925326 | 0.967402435 | -5.09306501 |
| **A_51_P472932** | Atp8b1 | -0.053132354 | -0.602278365 | 0.496013656 | 8.393201239 | -0.238034765 | 0.81994591 | 0.967402435 | -5.093069952 |
| **A_51_P511707** | Ntpcr | -0.013622867 | -0.154432779 | 0.127187045 | 9.586752238 | -0.238015067 | 0.819960485 | 0.967402435 | -5.093073451 |
| **A_51_P222359** | Emilin1 | 0.019711815 | -0.184106978 | 0.223530607 | 8.485435739 | 0.237931286 | 0.820022479 | 0.967402435 | -5.09308833 |
| **A_51_P307741** | Cckar | 0.010246179 | -0.095702879 | 0.116195237 | 7.653845055 | 0.237921662 | 0.8200296 | 0.967402435 | -5.093090039 |
| **A_51_P275976** | Dok1 | 0.012516799 | -0.117137794 | 0.142171392 | 8.490634304 | 0.237505971 | 0.820337214 | 0.967402435 | -5.093163786 |
| **A_52_P588559** | Ube2d3 | 0.022302882 | -0.208883263 | 0.253489026 | 12.09242323 | 0.23733858 | 0.820461095 | 0.967402435 | -5.093193448 |
| **A_51_P479808** | Mapk6 | 0.013814114 | -0.129400405 | 0.157028633 | 8.890511917 | 0.237304049 | 0.82048665 | 0.967402435 | -5.093199564 |
| **A_66_P111781** | 4930432M17Rik | 0.013480058 | -0.126273122 | 0.153233238 | 9.440430202 | 0.237300808 | 0.82048905 | 0.967402435 | -5.093200138 |
| **A_55_P2145059** | Gbp5 | -0.017678716 | -0.20096505 | 0.165607618 | 8.557772749 | -0.237295662 | 0.820492858 | 0.967402435 | -5.093201049 |
| **A_51_P191463** | H2-M2 | 0.063165871 | -0.591741481 | 0.718073223 | 11.2157395 | 0.237285819 | 0.820500143 | 0.967402435 | -5.065286811 |
| **A_55_P2184877** | NA | -0.016653648 | -0.189330956 | 0.15602366 | 10.59158113 | -0.237270237 | 0.820511675 | 0.967402435 | -5.093205552 |
| **A_55_P2042713** | Pigq | 0.025972493 | -0.243359211 | 0.295304197 | 13.07490645 | 0.237244049 | 0.820531056 | 0.967402435 | -5.093210189 |
| **A_52_P101982** | Tmem129 | 0.022596577 | -0.21185097 | 0.257044125 | 11.04649105 | 0.237118875 | 0.820623699 | 0.967402435 | -5.093232348 |
| **A_55_P2159801** | Rfc4 | 0.014153912 | -0.132699783 | 0.161007607 | 8.09130348 | 0.237115955 | 0.82062586 | 0.967402435 | -5.093232865 |
| **A_51_P378381** | Crnde | -0.011455219 | -0.130309383 | 0.107398946 | 7.765403065 | -0.237114514 | 0.820626927 | 0.967402435 | -5.09323312 |
| **A_55_P1961168** | NA | 0.02040637 | -0.19135628 | 0.232169019 | 10.61512972 | 0.237074884 | 0.820656259 | 0.967402435 | -5.093240133 |
| **A_51_P385952** | Man2a2 | -0.027516813 | -0.313171551 | 0.258137924 | 8.351888546 | -0.236987736 | 0.820720761 | 0.967402435 | -5.09325555 |
| **A_55_P2045931** | Btnl10 | 0.010219456 | -0.095886755 | 0.116325667 | 7.663566333 | 0.236949688 | 0.820748922 | 0.967402435 | -5.09326228 |
| **A_55_P2074776** | Gm8894 | -0.01336959 | -0.152195796 | 0.125456616 | 16.21373362 | -0.236927684 | 0.820765209 | 0.967402435 | -5.093266171 |

| | | | | | | | | |
|---|---|---|---|---|---|---|---|---|
| **A_66_P124179** | Atp6v0d2 | -0.092610142 | -1.054649629 | 0.869429345 | 8.839780822 | -0.236828964 | 0.820838279 | 0.967402435 | -5.093283625 |
| **A_55_P1952127** | Psma3 | 0.025912169 | -0.243298575 | 0.295122912 | 14.28446779 | 0.236799367 | 0.820860186 | 0.967402435 | -5.093288856 |
| **A_51_P190765** | Rtca | 0.018851603 | -0.17714691 | 0.214850116 | 10.2887209 | 0.236627198 | 0.820987628 | 0.967402435 | -5.093319275 |
| **A_51_P466673** | Srsf7 | 0.012012963 | -0.112886883 | 0.136912808 | 8.5245292 | 0.236623234 | 0.820990562 | 0.967402435 | -5.093319975 |
| **A_55_P2044593** | 1810037I17Rik | -0.019395512 | -0.221072425 | 0.1822814 | 13.25499346 | -0.236599726 | 0.821007963 | 0.967402435 | -5.093324127 |
| **A_52_P503730** | Fchsd2 | -0.028813274 | -0.328417807 | 0.27079126 | 9.625175621 | -0.236599263 | 0.821008306 | 0.967402435 | -5.093324209 |
| **A_55_P2138333** | Mad2l1 | -0.010509336 | -0.119845219 | 0.098826548 | 7.75645586 | -0.23647308 | 0.821101712 | 0.96745352 | -5.093346487 |
| **A_55_P2209298** | NA | -0.028177172 | -0.321572235 | 0.265217892 | 8.506440476 | -0.236272818 | 0.821249962 | 0.967488504 | -5.09338182 |
| **A_55_P2186287** | Sub1 | -0.049255825 | -0.562149115 | 0.463637464 | 14.22540796 | -0.236265194 | 0.821255606 | 0.967488504 | -5.093383164 |
| **A_55_P2028046** | Cyp3a59 | 0.010021298 | -0.094344421 | 0.114387018 | 7.655783444 | 0.236230129 | 0.821281565 | 0.967488504 | -5.093389348 |
| **A_52_P441974** | Evc2 | -0.043218279 | -0.493798579 | 0.407362021 | 9.162563481 | -0.235974149 | 0.821471075 | 0.967641902 | -5.093434461 |
| **A_55_P2150737** | NA | -0.032788317 | -0.374786271 | 0.309209637 | 9.419784343 | -0.235865695 | 0.821551371 | 0.967641902 | -5.09345356 |
| **A_55_P2165655** | Zbtb4 | 0.020494209 | -0.193328037 | 0.234316455 | 10.02410231 | 0.235801972 | 0.821598552 | 0.967641902 | -5.093464778 |
| **A_55_P1976494** | Bbof1 | 0.010132431 | -0.095618343 | 0.115883204 | 7.820562169 | 0.235721526 | 0.821658114 | 0.967641902 | -5.093478936 |
| **A_51_P374101** | Pard6b | -0.021699211 | -0.248176033 | 0.204777611 | 8.382094233 | -0.235716156 | 0.82166209 | 0.967641902 | -5.093479881 |
| **A_51_P471520** | Stk25 | -0.013485853 | -0.154412172 | 0.127440465 | 12.73282405 | -0.235426575 | 0.821876507 | 0.967664265 | -5.093530804 |
| **A_52_P589550** | Nr2c1 | -0.017231754 | -0.197304094 | 0.162840585 | 8.423329328 | -0.235424496 | 0.821878047 | 0.967664265 | -5.093531169 |
| **A_52_P171033** | Mcmbp | 0.012065283 | -0.114050538 | 0.138181103 | 8.544022196 | 0.235362405 | 0.821924024 | 0.967664265 | -5.09354208 |
| **A_55_P2044653** | Cyp2b10 | -0.010380563 | -0.118900285 | 0.098139159 | 7.834210938 | -0.235332224 | 0.821946373 | 0.967664265 | -5.093547382 |
| **A_55_P2184180** | Ndufv3 | 0.024865387 | -0.235096637 | 0.284827411 | 14.40328601 | 0.235317637 | 0.821957174 | 0.967664265 | -5.093549945 |
| **A_51_P269792** | Rad51b | -0.018356055 | -0.210291226 | 0.173579116 | 8.050360378 | -0.235284845 | 0.821981457 | 0.967664265 | -5.093555705 |
| **A_51_P217047** | Gng10 | 0.042290762 | -0.400073078 | 0.484654602 | 8.92602492 | 0.235198775 | 0.822045192 | 0.96768036 | -5.09357082 |
| **A_52_P283628** | Rps6ka3 | -0.0159239 | -0.182599782 | 0.150751983 | 8.177694375 | -0.235042192 | 0.822161147 | 0.967753812 | -5.093598304 |
| **A_55_P2154835** | NA | 0.03147298 | -0.298108863 | 0.361054822 | 11.96532831 | 0.234932849 | 0.822242122 | 0.967753812 | -5.093617485 |
| **A_52_P487362** | Ppp4r4 | -0.010444827 | -0.119863079 | 0.098973426 | 7.847001064 | -0.234844631 | 0.822307454 | 0.967753812 | -5.093632955 |
| **A_51_P234253** | Sdcbp2 | -0.026320642 | -0.302051906 | 0.249410622 | 8.724382224 | -0.234844084 | 0.822307859 | 0.967753812 | -5.093633051 |
| **A_55_P2190994** | Cdsn | 0.083632949 | -0.794211699 | 0.961477596 | 8.524879368 | 0.23438451 | 0.822648235 | 0.968045797 | -5.093713547 |
| **A_55_P2055127** | Slc7a14 | -0.07186266 | -0.826375081 | 0.68264976 | 8.28821747 | -0.23431822 | 0.822697336 | 0.968045797 | -5.093725146 |
| **A_55_P2045158** | NA | -0.03046158 | -0.350316829 | 0.289393669 | 10.19059135 | -0.234297744 | 0.822712502 | 0.968045797 | -5.093728727 |
| **A_55_P2050688** | Zbed4 | -0.02525694 | -0.290528927 | 0.240015046 | 11.78957001 | -0.234238626 | 0.822756291 | 0.968045797 | -5.093739067 |
| **A_51_P116651** | Dpt | -0.010215313 | -0.117593892 | 0.097163267 | 7.665547246 | -0.234047049 | 0.822898198 | 0.968100566 | -5.088392987 |
| **A_51_P213691** | Scnn1a | -0.036244897 | -0.417370397 | 0.344880602 | 10.2061781 | -0.233963457 | 0.822960119 | 0.968100566 | -5.093787162 |
| **A_51_P198675** | Ttc36 | 0.010775125 | -0.102601304 | 0.124151553 | 7.664383056 | 0.233813022 | 0.823071558 | 0.968100566 | -5.093813432 |
| **A_55_P2174288** | Snap29 | 0.016034132 | -0.152719114 | 0.184787378 | 10.81763812 | 0.233755843 | 0.823113916 | 0.968100566 | -5.093823412 |
| **A_55_P2061294** | NA | -0.015220056 | -0.175418437 | 0.144978326 | 13.09949952 | -0.233736902 | 0.823127948 | 0.968100566 | -5.093826718 |
| **A_66_P126459** | Gm13429 | -0.00981053 | -0.113117449 | 0.093496388 | 7.671717759 | -0.233631955 | 0.823205695 | 0.968100566 | -5.093845029 |
| **A_55_P2042655** | NA | 0.015728145 | -0.149945296 | 0.181401586 | 11.27890223 | 0.233557472 | 0.823260875 | 0.968100566 | -5.09385802 |
| **A_51_P521374** | Nudt13 | 0.021755525 | -0.207421049 | 0.250932098 | 10.31237466 | 0.233543884 | 0.823270941 | 0.968100566 | -5.093860389 |
| **A_51_P284988** | Nrbf2 | 0.015462597 | -0.147448823 | 0.178374017 | 10.09902179 | 0.233507086 | 0.823298203 | 0.968100566 | -5.093866805 |
| **A_55_P2131027** | NA | -0.024754028 | -0.285625356 | 0.236117301 | 11.79367051 | -0.233447208 | 0.823342565 | 0.968100566 | -5.093877244 |
| **A_55_P2358679** | NA | -0.018245557 | -0.210619124 | 0.17412801 | 10.51235446 | -0.23333554 | 0.823425298 | 0.968100566 | -5.093896703 |
| **A_55_P2017826** | Myb | -0.011813873 | -0.136390789 | 0.112763043 | 7.804730634 | -0.233304917 | 0.823447986 | 0.968100566 | -5.093902038 |
| **A_55_P2009953** | Cd300lg | 0.041066346 | -0.392120818 | 0.47425351 | 9.041161477 | 0.233227441 | 0.823505388 | 0.968100566 | -5.093915533 |
| **A_55_P2032453** | Mrpl23 | 0.037533426 | -0.358621879 | 0.433688731 | 13.72469029 | 0.233089062 | 0.823607917 | 0.968100566 | -5.093939624 |
| **A_66_P113662** | Tmem62 | 0.022484557 | -0.21484922 | 0.259818334 | 10.33554939 | 0.233074065 | 0.823619029 | 0.968100566 | -5.093942234 |
| **A_55_P1979335** | NA | 0.018510884 | -0.177021431 | 0.214043199 | 8.777281812 | 0.232904448 | 0.823744709 | 0.968100566 | -5.093971742 |
| **A_55_P2126627** | Phf20l1 | -0.012423397 | -0.143670771 | 0.118823978 | 7.962043852 | -0.232872872 | 0.823768106 | 0.968100566 | -5.093977233 |
| **A_55_P2349265** | Ptk2 | 0.016728366 | -0.159999576 | 0.193456309 | 9.128141912 | 0.232872108 | 0.823768672 | 0.968100566 | -5.093977366 |
| **A_55_P2009783** | Sfr1 | -0.016401735 | -0.197353398 | 0.164549928 | 13.36277074 | -0.234896905 | 0.82384611 | 0.968100566 | -5.025599273 |
| **A_51_P249127** | Pkd1 | -0.040350668 | -0.466846797 | 0.386145461 | 10.38347176 | -0.232758099 | 0.823853152 | 0.968100566 | -5.093997186 |

| | | | | | | | | | |
|---|---|---|---|---|---|---|---|---|---|
| **A_52_P455089** | NA | -0.010694465 | -0.123794011 | 0.10240508 | 8.898888172 | -0.232630892 | 0.823947415 | 0.968100566 | -5.094019289 |
| **A_55_P2212027** | Utrn | -0.038581227 | -0.446611426 | 0.369448973 | 9.106168937 | -0.232623125 | 0.82395317 | 0.968100566 | -5.094020638 |
| **A_52_P211418** | G2e3 | -0.011698682 | -0.135423641 | 0.112026278 | 7.790888056 | -0.23262092 | 0.823954804 | 0.968100566 | -5.094021021 |
| **A_51_P409349** | Trim29 | -0.035349172 | -0.409433151 | 0.338734807 | 8.303342693 | -0.23247663 | 0.824061731 | 0.968116046 | -5.094046077 |
| **A_51_P108659** | Pon1 | -0.009903724 | -0.114714167 | 0.09490672 | 7.649989205 | -0.232467966 | 0.824068151 | 0.968116046 | -5.088666333 |
| **A_52_P630571** | Epg5 | -0.014075398 | -0.163147118 | 0.134996321 | 9.139855749 | -0.232292186 | 0.824198419 | 0.968167063 | -5.094078084 |
| **A_51_P474053** | Erv3 | -0.32118029 | -3.723045448 | 3.080684868 | 11.74171605 | -0.232274189 | 0.824211756 | 0.968167063 | -5.094081206 |
| **A_55_P2004797** | Tacc2 | -0.04321341 | -0.501233937 | 0.414807116 | 13.03524416 | -0.232114764 | 0.824329909 | 0.968205623 | -5.094108849 |
| **A_66_P129263** | Lekr1 | 0.012369541 | -0.118746997 | 0.143486079 | 8.454091679 | 0.23209472 | 0.824344764 | 0.968205623 | -5.094112323 |
| **A_52_P35048** | Serpinf1 | 0.010266693 | -0.098678652 | 0.119212037 | 7.722280468 | 0.231841437 | 0.824532489 | 0.968242593 | -5.094156199 |
| **A_55_P1995151** | NA | -0.028271918 | -0.328352997 | 0.27180916 | 9.744964577 | -0.231785267 | 0.824574122 | 0.968242593 | -5.094165923 |
| **A_51_P389885** | Spic | -0.010401902 | -0.120837068 | 0.100033265 | 7.754733917 | -0.231725865 | 0.824618151 | 0.968242593 | -5.094176204 |
| **A_55_P1966274** | Pdzd4 | 0.016619883 | -0.159867773 | 0.193107539 | 7.953252305 | 0.231676929 | 0.824654423 | 0.968242593 | -5.094184671 |
| **A_66_P107585** | Taf2 | 0.02126699 | -0.204808776 | 0.247342756 | 10.01810987 | 0.231430812 | 0.824836857 | 0.968242593 | -5.094227231 |
| **A_55_P2114133** | Garnl3 | -0.015188997 | -0.176671213 | 0.146293219 | 8.901943287 | -0.231405433 | 0.82485567 | 0.968242593 | -5.094231618 |
| **A_52_P276955** | Epha3 | 0.012061802 | -0.116215526 | 0.140339131 | 7.741152389 | 0.231329737 | 0.824911782 | 0.968242593 | -5.094244697 |
| **A_52_P633410** | Gabarapl1 | -0.030773687 | -0.358059731 | 0.296512357 | 16.2743534 | -0.231324273 | 0.824915832 | 0.968242593 | -5.094245641 |
| **A_51_P302942** | Rasl10a | -0.02696071 | -0.313775823 | 0.259854403 | 9.385182851 | -0.2312589 | 0.824964293 | 0.968242593 | -5.094256933 |
| **A_52_P883557** | Slc30a10 | -0.010225016 | -0.11900622 | 0.098556187 | 7.652707741 | -0.23124871 | 0.824971847 | 0.968242593 | -5.094258693 |
| **A_55_P2016902** | Entr1 | 0.018996186 | -0.183193487 | 0.221185858 | 9.795889549 | 0.231140792 | 0.825051849 | 0.968242593 | -5.094277327 |
| **A_52_P575178** | Cdon | -0.042821971 | -0.498667403 | 0.41302346 | 9.320277216 | -0.23110972 | 0.825074884 | 0.968242593 | -5.09428269 |
| **A_55_P1962756** | Ttll2 | 0.060235472 | -0.581170133 | 0.701641078 | 8.430195534 | 0.231040827 | 0.825125956 | 0.968242593 | -5.09429458 |
| **A_55_P2373987** | A730009E18Rik | -0.021580545 | -0.251406195 | 0.208245105 | 7.991372185 | -0.231011221 | 0.825147905 | 0.968242593 | -5.094299688 |
| **A_55_P2181271** | Nhsl2 | -0.024907078 | -0.290308689 | 0.240494533 | 8.626126584 | -0.230881102 | 0.825244371 | 0.968242593 | -5.094322131 |
| **A_55_P2031192** | Gtdc1 | -0.037948205 | -0.442659363 | 0.366762953 | 9.884851422 | -0.230682807 | 0.825391387 | 0.968242593 | -5.09435631 |
| **A_55_P2114218** | Zcchc7 | -0.030656051 | -0.357625281 | 0.29631318 | 9.942871537 | -0.230663289 | 0.825405858 | 0.968242593 | -5.094359673 |
| **A_52_P316697** | Ost4 | -0.028738234 | -0.33548205 | 0.278005582 | 12.62422963 | -0.230490707 | 0.825533818 | 0.968242593 | -5.094389394 |
| **A_55_P1956718** | Rab38 | -0.01051325 | -0.122732501 | 0.101706001 | 7.724071599 | -0.230482945 | 0.825539573 | 0.968242593 | -5.09439073 |
| **A_52_P528600** | Hhex | -0.017588451 | -0.205430747 | 0.170253844 | 8.265849832 | -0.230358043 | 0.825632184 | 0.968242593 | -5.094412226 |
| **A_55_P2129975** | Rabgef1 | -0.01559021 | -0.182142208 | 0.150961788 | 11.46374869 | -0.230287981 | 0.825684135 | 0.968242593 | -5.094424279 |
| **A_52_P70027** | Gm20498 | 0.014537291 | -0.140793106 | 0.169867688 | 11.38016209 | 0.230248178 | 0.825713649 | 0.968242593 | -5.094431124 |
| **A_55_P2113210** | Cyth4 | -0.019353288 | -0.226193306 | 0.187486731 | 8.82601128 | -0.230191533 | 0.825755652 | 0.968242593 | -5.094440865 |
| **A_51_P239673** | Hprt | -0.042371023 | -0.495399972 | 0.410657926 | 13.76328661 | -0.230097638 | 0.825825277 | 0.968242593 | -5.089073259 |
| **A_51_P412914** | Efs | -0.03056804 | -0.357461402 | 0.296325322 | 11.9374735 | -0.230054461 | 0.825857295 | 0.968242593 | -5.094464425 |
| **A_55_P2073845** | Mrpl47 | -0.015045864 | -0.175976458 | 0.14588473 | 8.676116243 | -0.230010509 | 0.825889887 | 0.968242593 | -5.094471977 |
| **A_52_P50287** | Bend7 | -0.01434545 | -0.167805699 | 0.139114799 | 8.341094078 | -0.229978606 | 0.825913546 | 0.968242593 | -5.094477458 |
| **A_51_P333349** | Tmem120a | -0.017945137 | -0.209941824 | 0.17405155 | 11.59777538 | -0.229944067 | 0.825939158 | 0.968242593 | -5.09448339 |
| **A_55_P2009201** | NA | -0.011611853 | -0.135968518 | 0.112744811 | 10.7886223 | -0.229721499 | 0.826104212 | 0.968242593 | -5.094521599 |
| **A_55_P2085546** | Dnaja4 | 0.029963052 | -0.291009643 | 0.350935748 | 9.940252464 | 0.229660935 | 0.826149128 | 0.968242593 | -5.09453199 |
| **A_55_P1980125** | Srsf3 | -0.027945502 | -0.327356638 | 0.271465634 | 10.51580503 | -0.229621807 | 0.826178145 | 0.968242593 | -5.094538701 |
| **A_55_P1982683** | Cib1 | 0.044868738 | -0.43597053 | 0.525708006 | 13.74579865 | 0.229568921 | 0.826217368 | 0.968242593 | -5.094547772 |
| **A_55_P2115458** | Pot1a | -0.02238524 | -0.262433776 | 0.217663297 | 11.23753002 | -0.229420304 | 0.82632759 | 0.968242593 | -5.094573248 |
| **A_51_P156927** | Arid4a | -0.01557146 | -0.182564402 | 0.151421482 | 8.444509125 | -0.22940368 | 0.82633992 | 0.968242593 | -5.094576097 |
| **A_51_P422685** | Zmat4 | -0.027412956 | -0.32141666 | 0.266590748 | 8.707292146 | -0.229388813 | 0.826350947 | 0.968242593 | -5.094578645 |
| **A_55_P2187206** | NA | -0.014792412 | -0.173539794 | 0.14395497 | 8.449539979 | -0.229245906 | 0.826456939 | 0.968242593 | -5.094603124 |
| **A_51_P181523** | Tmem214 | -0.016096501 | -0.188880532 | 0.15668753 | 11.00444774 | -0.229190723 | 0.826497869 | 0.968242593 | -5.094612573 |
| **A_55_P1982613** | Hnrnpa3 | -0.019369421 | -0.227323637 | 0.188584796 | 13.42236456 | -0.229149054 | 0.826528776 | 0.968242593 | -5.094619706 |
| **A_55_P2134236** | Foxa2 | 0.073295819 | -0.713795806 | 0.860387444 | 9.958608198 | 0.229098936 | 0.82656595 | 0.968242593 | -5.094628284 |
| **A_52_P215176** | Opn3 | -0.04524805 | -0.531324206 | 0.440828107 | 9.206212601 | -0.229015412 | 0.826627903 | 0.968242593 | -5.094642575 |
| **A_51_P302458** | Plscr3 | -0.02181556 | -0.256194193 | 0.212563074 | 9.629944879 | -0.228990522 | 0.826646365 | 0.968242593 | -5.094646833 |

| | | | | | | | | | |
|---|---|---|---|---|---|---|---|---|---|
| **A_51_P437608** | Tulp3 | -0.020158402 | -0.236767781 | 0.196450977 | 8.306985909 | -0.228953881 | 0.826673544 | 0.968242593 | -5.094653101 |
| **A_51_P181851** | Eml3 | -0.016603497 | -0.195022632 | 0.161815637 | 10.35495134 | -0.228942968 | 0.826681639 | 0.968242593 | -5.094654967 |
| **A_51_P115666** | Fam83h | 0.032700874 | -0.318765614 | 0.384167362 | 11.60860818 | 0.228899372 | 0.826713977 | 0.968242593 | -5.094662422 |
| **A_52_P302496** | Fah | -0.018406927 | -0.216292673 | 0.179478819 | 9.132596307 | -0.228842106 | 0.826756456 | 0.968242593 | -5.094672213 |
| **A_52_P223333** | Mib1 | -0.012198653 | -0.14337118 | 0.118973875 | 9.251299404 | -0.228790584 | 0.826794674 | 0.968242593 | -5.09468102 |
| **A_55_P1952950** | Cic | -0.030121592 | -0.35410795 | 0.293864767 | 13.53434684 | -0.228728538 | 0.8268407 | 0.968242593 | -5.094691623 |
| **A_55_P1989846** | Tm6sf1 | -0.012704002 | -0.149360988 | 0.123952983 | 8.039296809 | -0.228706176 | 0.826857288 | 0.968242593 | -5.094695444 |
| **A_55_P2056014** | NA | 0.016455845 | -0.160588332 | 0.193500021 | 9.146378221 | 0.22866921 | 0.82688471 | 0.968242593 | -5.094701759 |
| **A_52_P22324** | Pln | 0.010091021 | -0.098512167 | 0.11869421 | 7.646893013 | 0.228592359 | 0.82694172 | 0.968242593 | -5.094714885 |
| **A_51_P181891** | Amer2 | -0.01883212 | -0.221578014 | 0.183913774 | 7.94390541 | -0.228515838 | 0.826998486 | 0.968242593 | -5.09472795 |
| **A_55_P2091461** | Casp4 | -0.05235741 | -0.616105153 | 0.511390332 | 9.968547172 | -0.228487532 | 0.827019486 | 0.968242593 | -5.094732782 |
| **A_51_P172842** | Med19 | 0.02814775 | -0.274949829 | 0.331245328 | 10.27562256 | 0.228470629 | 0.827032025 | 0.968242593 | -5.094735667 |
| **A_52_P390944** | Chst3 | -0.01323314 | -0.155826108 | 0.129359828 | 7.817854723 | -0.228314747 | 0.82714767 | 0.968242593 | -5.094762264 |
| **A_55_P2042062** | Mier1 | 0.015715112 | -0.153647783 | 0.185078006 | 8.610172238 | 0.228280263 | 0.827173253 | 0.968242593 | -5.094768145 |
| **A_55_P2035986** | E330020D12Rik | 0.060705951 | -0.59359555 | 0.715007452 | 10.76155058 | 0.228256163 | 0.827191133 | 0.968242593 | -5.094772255 |
| **A_52_P676076** | Rbx1 | -0.024613421 | -0.290101783 | 0.240874941 | 10.89401489 | -0.228084438 | 0.827318538 | 0.968242593 | -5.094801527 |
| **A_51_P277718** | Cenpc1 | -0.021501868 | -0.253459386 | 0.21045565 | 9.5610236 | -0.228053584 | 0.82734143 | 0.968242593 | -5.094806784 |
| **A_51_P322473** | Perm1 | 0.009716747 | -0.095107925 | 0.114541419 | 7.70434404 | 0.228048145 | 0.827345465 | 0.968242593 | -5.094807711 |
| **A_51_P365409** | Rpusd4 | -0.031667277 | -0.373459231 | 0.310124678 | 10.3957534 | -0.227938687 | 0.827426678 | 0.968242593 | -5.094826355 |
| **A_55_P2048478** | Olfml1 | 0.033657272 | -0.329624354 | 0.396938897 | 8.457824108 | 0.227931652 | 0.827431897 | 0.968242593 | -5.094827553 |
| **A_55_P1957249** | Pdgfrb | -0.032092109 | -0.378743026 | 0.314558808 | 11.70283055 | -0.227758749 | 0.827560188 | 0.968286126 | -5.094856985 |
| **A_51_P282523** | Adgra2 | -0.021601505 | -0.254948111 | 0.211745102 | 9.869546344 | -0.227746483 | 0.82756929 | 0.968286126 | -5.094859072 |
| **A_66_P111572** | Fam122b | 0.015105472 | -0.148318764 | 0.178529707 | 10.84112023 | 0.227398185 | 0.827827739 | 0.968412667 | -5.094918291 |
| **A_66_P100186** | Psg28 | 0.041153887 | -0.404113235 | 0.48642101 | 15.25197995 | 0.227383737 | 0.82783846 | 0.968412667 | -5.094920746 |
| **A_55_P2336173** | Mirg | 0.051566715 | -0.506573222 | 0.609706652 | 9.462094038 | 0.227297956 | 0.827902116 | 0.968412667 | -5.094935316 |
| **A_51_P415225** | Zfp105 | -0.013788804 | -0.163036505 | 0.135458898 | 8.782642925 | -0.227294068 | 0.827905001 | 0.968412667 | -5.094935976 |
| **A_55_P2129658** | Smad6 | -0.024804997 | -0.29338727 | 0.243777276 | 9.778771154 | -0.22721186 | 0.827966008 | 0.968412667 | -5.094949935 |
| **A_52_P219966** | Spg21 | -0.015145362 | -0.179217723 | 0.148926998 | 8.5159354 | -0.227098048 | 0.828050469 | 0.968412667 | -5.094969251 |
| **A_55_P1959393** | Hhat | 0.011867902 | -0.116732032 | 0.140467837 | 8.192972911 | 0.227040002 | 0.828093547 | 0.968412667 | -5.094979099 |
| **A_55_P2028248** | Dnajc19 | -0.022369484 | -0.264778299 | 0.22003933 | 11.02076333 | -0.227026592 | 0.828103499 | 0.968412667 | -5.094981373 |
| **A_66_P116195** | Psmd13 | -0.019839643 | -0.234871614 | 0.195192328 | 11.45756575 | -0.226986474 | 0.828133272 | 0.968412667 | -5.094988178 |
| **A_51_P121179** | Erfe | -0.009847231 | -0.116605079 | 0.096910617 | 7.661511811 | -0.226925592 | 0.828178456 | 0.968412667 | -5.078841378 |
| **A_51_P269084** | Chchd10 | -0.031487689 | -0.373248382 | 0.310273004 | 15.63477796 | -0.226666758 | 0.828370559 | 0.968547839 | -5.095042363 |
| **A_51_P490348** | 1110059E24Rik | -0.021721198 | -0.257511706 | 0.21406931 | 10.40272634 | -0.22663481 | 0.828394271 | 0.968547839 | -5.095047774 |
| **A_52_P505827** | Cyb5r4 | -0.01760471 | -0.209020043 | 0.173810623 | 9.004389071 | -0.226267049 | 0.828667243 | 0.968684754 | -5.095110001 |
| **A_52_P158431** | Tshz2 | -0.077368185 | -0.918651466 | 0.763915095 | 9.025888409 | -0.22625038 | 0.828679616 | 0.968684754 | -5.095112819 |
| **A_55_P2153381** | Ermap | 0.020347389 | -0.200989573 | 0.241684351 | 8.558859319 | 0.226164224 | 0.828743569 | 0.968684754 | -5.095127382 |
| **A_55_P2008755** | NA | -0.019327678 | -0.229646745 | 0.19099139 | 12.49562636 | -0.226084187 | 0.828802982 | 0.968684754 | -5.095140906 |
| **A_51_P205099** | Dazap1 | -0.020024649 | -0.238156325 | 0.198107026 | 10.98857073 | -0.225847533 | 0.828978662 | 0.968684754 | -5.095180865 |
| **A_55_P1997898** | Tfeb | 0.017482495 | -0.173035918 | 0.208000908 | 8.791351867 | 0.225754087 | 0.829048034 | 0.968684754 | -5.095196632 |
| **A_55_P1963508** | Slc13a5 | -0.132999325 | -1.583159071 | 1.317160421 | 8.928135466 | -0.225633071 | 0.829137877 | 0.968684754 | -5.095217042 |
| **A_55_P1991475** | Sesn1 | -0.043020117 | -0.51250222 | 0.426461986 | 11.35902631 | -0.225435189 | 0.82928479 | 0.968684754 | -5.095250393 |
| **A_52_P96552** | Vkorc1l1 | 0.027126985 | -0.268927616 | 0.323181585 | 9.84729243 | 0.225423335 | 0.829293591 | 0.968684754 | -5.09525239 |
| **A_55_P2136935** | Rgs7bp | -0.01461948 | -0.174195737 | 0.144956777 | 7.838577198 | -0.225389047 | 0.829319049 | 0.968684754 | -5.095258165 |
| **A_51_P490023** | Tubb2a | 0.022688276 | -0.22499671 | 0.270373262 | 12.12038998 | 0.225356963 | 0.82934287 | 0.968684754 | -5.095263569 |
| **A_51_P509760** | Actr1a | 0.014208791 | -0.140950207 | 0.169367789 | 12.81225851 | 0.22529384 | 0.829389737 | 0.968684754 | -5.095274198 |
| **A_55_P2123841** | Zfhx4 | 0.011067009 | -0.109784613 | 0.131918631 | 7.749820989 | 0.225292603 | 0.829390655 | 0.968684754 | -5.095274406 |
| **A_55_P2077664** | Gm5458 | -0.026807629 | -0.319565474 | 0.265950217 | 9.380323053 | -0.225278122 | 0.829401407 | 0.968684754 | -5.095276844 |
| **A_55_P2049612** | Ube2a | 0.030775619 | -0.305399056 | 0.366950293 | 12.3268029 | 0.225222078 | 0.829443019 | 0.968684754 | -5.095286278 |
| **A_55_P2027609** | Scgb3a2 | 0.009956186 | -0.098839473 | 0.118751844 | 7.669781224 | 0.225138921 | 0.829504763 | 0.968684754 | -5.095300271 |

| | | | | | | | | | |
|---|---|---|---|---|---|---|---|---|---|
| **A_55_P2005859** | Fn3k | 0.009977738 | -0.09907374 | 0.119029216 | 7.699951255 | 0.225096992 | 0.829535896 | 0.968684754 | -5.095307325 |
| **A_55_P2419296** | NA | 0.025217204 | -0.250405158 | 0.300839565 | 8.445993842 | 0.225087634 | 0.829542844 | 0.968684754 | -5.095308899 |
| **A_51_P242166** | Lap3 | -0.030262292 | -0.361059402 | 0.300534818 | 10.99092382 | -0.225065678 | 0.829559147 | 0.968684754 | -5.095312592 |
| **A_52_P49014** | Shh | -0.009583848 | -0.114350065 | 0.095182368 | 7.631561366 | -0.225054571 | 0.829567395 | 0.968684754 | -5.084660361 |
| **A_51_P411264** | Txndc16 | 0.01313092 | -0.13042057 | 0.156682411 | 9.106311924 | 0.225038392 | 0.829579408 | 0.968684754 | -5.095317181 |
| **A_51_P345927** | Ddx41 | -0.011776892 | -0.140584688 | 0.117030904 | 8.641776402 | -0.224935346 | 0.829655923 | 0.968684754 | -5.095334507 |
| **A_55_P2029528** | NA | 0.069791207 | -0.693681961 | 0.833264375 | 10.1964573 | 0.22489308 | 0.829687307 | 0.968684754 | -5.095341611 |
| **A_65_P19832** | Ttr | 0.153028653 | -1.521280437 | 1.827337742 | 14.36236619 | 0.224856933 | 0.829714149 | 0.968684754 | -5.095347686 |
| **A_55_P2098539** | Itgb3bp | -0.024963802 | -0.298229048 | 0.248301444 | 9.432033146 | -0.22474782 | 0.829795174 | 0.968707148 | -5.095366018 |
| **A_55_P1994524** | Tpd52l2 | -0.027328685 | -0.326549859 | 0.271892489 | 12.96427104 | -0.224696121 | 0.829833564 | 0.968707148 | -5.0953747 |
| **A_51_P455266** | Ring1 | 0.0104095 | -0.103631713 | 0.124450713 | 8.285897105 | 0.224562491 | 0.829932799 | 0.968764483 | -5.095397133 |
| **A_55_P2113723** | Pecr | 0.012858478 | -0.12805659 | 0.153773545 | 7.953934549 | 0.224492212 | 0.82998499 | 0.968766901 | -5.095408926 |
| **A_52_P13897** | Hook1 | -0.014527344 | -0.173799978 | 0.144745289 | 8.043993723 | -0.224395547 | 0.830056778 | 0.96879219 | -5.095425141 |
| **A_52_P335259** | Cdkal1 | 0.030738301 | -0.306647655 | 0.368124257 | 9.818277007 | 0.22414137 | 0.830245549 | 0.968941766 | -5.095467744 |
| **A_51_P443475** | Bccip | -0.020790924 | -0.249073434 | 0.207491585 | 10.80074556 | -0.224063092 | 0.830303687 | 0.968941766 | -5.095480855 |
| **A_55_P1970825** | NA | -0.058864703 | -0.705318974 | 0.587589569 | 8.307899284 | -0.22401975 | 0.830335878 | 0.968941766 | -5.095488113 |
| **A_52_P654752** | Trub1 | 0.009969999 | -0.09957588 | 0.119515878 | 7.962607568 | 0.22390729 | 0.830419405 | 0.968941766 | -5.095506937 |
| **A_52_P423364** | Fbxl7 | 0.017886234 | -0.178658803 | 0.214431271 | 8.593313446 | 0.223885516 | 0.830435578 | 0.968941766 | -5.095510581 |
| **A_51_P120398** | Asb17 | -0.009717696 | -0.116659258 | 0.097223867 | 7.687626779 | -0.223555795 | 0.83068049 | 0.969169022 | -5.076602974 |
| **A_55_P1955691** | Alg6 | 0.018274002 | -0.18304932 | 0.219597325 | 9.176344716 | 0.223310299 | 0.830862853 | 0.969323278 | -5.095606713 |
| **A_55_P2054739** | Polr3c | -0.019161096 | -0.230459066 | 0.192136874 | 10.05878044 | -0.223097231 | 0.831021137 | 0.969449427 | -5.095642261 |
| **A_55_P2090782** | NA | -0.031007457 | -0.373394542 | 0.311379627 | 13.28638174 | -0.222801414 | 0.831240909 | 0.969528435 | -5.095691558 |
| **A_52_P330289** | Inpp4b | -0.010241568 | -0.123410937 | 0.102927801 | 7.836402429 | -0.222641812 | 0.831359489 | 0.969528435 | -5.095718129 |
| **A_55_P2016079** | Erich4 | 0.092297944 | -0.928271643 | 1.112867531 | 9.065978381 | 0.222494135 | 0.831469213 | 0.969528435 | -5.095742698 |
| **A_51_P108489** | Cfap20 | 0.011799613 | -0.118699066 | 0.142298293 | 11.44183847 | 0.222449189 | 0.831502609 | 0.969528435 | -5.095750173 |
| **A_55_P2029344** | Rpl23 | -0.010220025 | -0.123269678 | 0.102829627 | 16.8616443 | -0.222408764 | 0.831532646 | 0.969528435 | -5.095756895 |
| **A_51_P222475** | Ehbp1l1 | 0.031164956 | -0.313572115 | 0.375902026 | 8.959696578 | 0.222406611 | 0.831534246 | 0.969528435 | -5.095757253 |
| **A_55_P2056496** | Tk1 | -0.02545272 | -0.307016388 | 0.256110948 | 9.096314922 | -0.222395886 | 0.831542215 | 0.969528435 | -5.095759036 |
| **A_55_P2045225** | NA | 0.021947997 | -0.220848542 | 0.264744536 | 12.16824714 | 0.22239323 | 0.831544189 | 0.969528435 | -5.095759477 |
| **A_55_P2109439** | Lsmem1 | 0.010024441 | -0.100873468 | 0.12092235 | 7.652130403 | 0.222385247 | 0.831550121 | 0.969528435 | -5.095760804 |
| **A_55_P2051334** | H2-K2 | -0.112169917 | -1.354569972 | 1.130230138 | 13.86001391 | -0.222118176 | 0.831748571 | 0.969528435 | -5.095805175 |
| **A_55_P2177755** | Cdk2ap1 | 0.03471644 | -0.349842914 | 0.419275794 | 9.400236932 | 0.22209612 | 0.831764961 | 0.969528435 | -5.095808837 |
| **A_51_P379443** | Bclaf1 | 0.026446773 | -0.266516604 | 0.31941015 | 11.77438154 | 0.222089751 | 0.831769694 | 0.969528435 | -5.095809895 |
| **A_55_P2072908** | NA | -0.03376419 | -0.407791416 | 0.340263037 | 14.04273579 | -0.222086559 | 0.831772066 | 0.969528435 | -5.095810425 |
| **A_55_P1980854** | Ankrd24 | -0.029876236 | -0.361189262 | 0.30143679 | 9.955386769 | -0.221848519 | 0.831948957 | 0.969528435 | -5.095849923 |
| **A_52_P117408** | Tg | 0.009757251 | -0.098474085 | 0.117988587 | 7.710383508 | 0.221790855 | 0.83199181 | 0.969528435 | -5.095859485 |
| **A_51_P329322** | Zfand1 | -0.022940145 | -0.277407366 | 0.231527076 | 10.0211133 | -0.221785667 | 0.831995666 | 0.969528435 | -5.095860346 |
| **A_66_P117251** | Snx6 | -0.03721456 | -0.468282051 | 0.393852932 | 12.1551474 | -0.223726637 | 0.832052706 | 0.969528435 | -5.02740414 |
| **A_51_P256202** | Ctsz | 0.047752053 | -0.48229487 | 0.577798977 | 13.02260614 | 0.221639365 | 0.832104393 | 0.969528435 | -5.095884594 |
| **A_55_P2151956** | NA | 0.03144644 | -0.317817462 | 0.380710341 | 10.05405994 | 0.22150674 | 0.832202959 | 0.969528435 | -5.095906563 |
| **A_52_P407854** | Dda1 | 0.012123768 | -0.122555552 | 0.146803088 | 8.675201902 | 0.221465388 | 0.832233692 | 0.969528435 | -5.09591341 |
| **A_55_P2108773** | Tedc1 | 0.018571545 | -0.18775802 | 0.224901111 | 8.76005933 | 0.221439832 | 0.832252686 | 0.969528435 | -5.095917641 |
| **A_55_P2091846** | Tut4 | -0.025961839 | -0.31444751 | 0.262523833 | 10.38872574 | -0.221401399 | 0.832281251 | 0.969528435 | -5.095924003 |
| **A_51_P290601** | Znhit2 | -0.013804771 | -0.167237968 | 0.139628426 | 10.47334043 | -0.221349757 | 0.832319633 | 0.969528435 | -5.09593255 |
| **A_52_P233441** | Gata2 | -0.018449587 | -0.223607713 | 0.186708539 | 11.35670415 | -0.221241749 | 0.832399909 | 0.969528435 | -5.095950419 |
| **A_55_P1968488** | Cblif | -0.009760858 | -0.118330397 | 0.09880868 | 7.656083117 | -0.221181688 | 0.83244455 | 0.969528435 | -5.095960352 |
| **A_52_P9437** | Psmb3 | -0.020435636 | -0.248139356 | 0.207268084 | 14.28702689 | -0.220793967 | 0.832732746 | 0.969528435 | -5.09602441 |
| **A_51_P341746** | Mpl | -0.013263552 | -0.161089411 | 0.134562308 | 8.106349401 | -0.220738751 | 0.832773791 | 0.969528435 | -5.096033523 |
| **A_66_P140858** | NA | -0.014493945 | -0.176084943 | 0.147097053 | 10.67343175 | -0.220667622 | 0.832826665 | 0.969528435 | -5.09604526 |
| **A_55_P1984481** | Mfn2 | 0.012395452 | -0.125841696 | 0.1506326 | 9.254651853 | 0.220600646 | 0.832876454 | 0.969528435 | -5.096056308 |

| | | | | | | | | |
|---|---|---|---|---|---|---|---|---|
| **A_55_P2058497** | Tmem50a | -0.052329318 | -0.635968571 | 0.531309935 | 11.05519542 | -0.220581835 | 0.832890437 | 0.969528435 | -5.096059411 |
| **A_51_P271725** | Fundc1 | -0.024655989 | -0.299682023 | 0.250370044 | 10.28447854 | -0.22055545 | 0.832910051 | 0.969528435 | -5.096063762 |
| **A_55_P1994275** | Gar1 | -0.012009643 | -0.146003804 | 0.121984517 | 11.44584051 | -0.220502443 | 0.832949456 | 0.969528435 | -5.096072502 |
| **A_51_P342906** | Sh3bp1 | 0.018249171 | -0.185478658 | 0.221977 | 8.568882961 | 0.22037481 | 0.83304434 | 0.969528435 | -5.096093537 |
| **A_55_P2176585** | Donson | -0.011312876 | -0.137636758 | 0.115011005 | 7.995454639 | -0.220321428 | 0.833084025 | 0.969528435 | -5.096102332 |
| **A_55_P2060483** | Rnf141 | 0.016927 | -0.17208995 | 0.20594395 | 8.310967458 | 0.2203172 | 0.833087169 | 0.969528435 | -5.096103028 |
| **A_51_P353856** | Ubac2 | 0.025085644 | -0.255070358 | 0.305241646 | 9.547844233 | 0.220289849 | 0.833107502 | 0.969528435 | -5.096107534 |
| **A_55_P2009973** | Ttc3 | -0.013035857 | -0.15864292 | 0.132571207 | 11.54790159 | -0.220255263 | 0.833133215 | 0.969528435 | -5.09611323 |
| **A_52_P90289** | Oaz2 | -0.015896585 | -0.193539432 | 0.161746261 | 8.981209507 | -0.220153345 | 0.833208987 | 0.969528435 | -5.09613001 |
| **A_55_P1965574** | A2ml1 | 0.009463173 | -0.096340639 | 0.115266985 | 7.702657575 | 0.220041514 | 0.833292131 | 0.969528435 | -5.096148413 |
| **A_52_P550218** | Trmt12 | 0.012220863 | -0.124504363 | 0.14894609 | 8.4587227 | 0.219898562 | 0.833398415 | 0.969528435 | -5.096171925 |
| **A_51_P327983** | 4833427G06Rik | -0.00965898 | -0.117733465 | 0.098415505 | 7.668469041 | -0.21987571 | 0.833415406 | 0.969528435 | -5.096175682 |
| **A_55_P2055042** | Uqcc2 | 0.06888889 | -0.70210847 | 0.839886251 | 12.72217255 | 0.219819118 | 0.833457483 | 0.969528435 | -5.096184985 |
| **A_65_P11306** | NA | -0.012439181 | -0.151766348 | 0.126887986 | 8.982699156 | -0.219646929 | 0.833585514 | 0.969528435 | -5.096213275 |
| **A_52_P552879** | Cracr2b | -0.01371233 | -0.167318698 | 0.139894038 | 8.273005292 | -0.219619656 | 0.833605793 | 0.969528435 | -5.096217754 |
| **A_51_P240594** | Zfp7 | 0.03093801 | -0.315956209 | 0.37783223 | 9.41266017 | 0.219414074 | 0.833758661 | 0.969528435 | -5.096251499 |
| **A_51_P307316** | Syde1 | 0.047010289 | -0.480258461 | 0.574279038 | 10.04303545 | 0.219346167 | 0.833809157 | 0.969528435 | -5.096262638 |
| **A_55_P1995442** | NA | -0.020267849 | -0.247609033 | 0.207073334 | 8.729274057 | -0.219330343 | 0.833820925 | 0.969528435 | -5.096265234 |
| **A_55_P2106175** | Gch1 | -0.039032189 | -0.476970554 | 0.398906176 | 12.23963796 | -0.219269931 | 0.833865849 | 0.969528435 | -5.09627514 |
| **A_55_P2012939** | Arid4b | -0.012978008 | -0.158590611 | 0.132634595 | 9.819025333 | -0.219269505 | 0.833866165 | 0.969528435 | -5.09627521 |
| **A_55_P2102897** | Gm12657 | -0.022539142 | -0.286531013 | 0.241452728 | 12.1407693 | -0.221256948 | 0.83387049 | 0.969528435 | -5.027791664 |
| **A_55_P1956179** | Git2 | 0.020548433 | -0.21002778 | 0.251124645 | 10.39961513 | 0.219246849 | 0.833883013 | 0.969528435 | -5.096278924 |
| **A_55_P1962279** | Plac1 | -0.036669847 | -0.448207377 | 0.374867682 | 12.34121641 | -0.219214268 | 0.833907241 | 0.969528435 | -5.096284265 |
| **A_55_P2086203** | Kcnq1 | 0.013236832 | -0.13535247 | 0.161826134 | 8.306780264 | 0.219162216 | 0.83394595 | 0.969528435 | -5.096292797 |
| **A_52_P33014** | Ppp1r8 | -0.011758518 | -0.143782748 | 0.120265713 | 8.179996721 | -0.219112972 | 0.83398257 | 0.969528435 | -5.096300866 |
| **A_55_P2118391** | Lrp6 | -0.027441435 | -0.335738684 | 0.280855814 | 11.92156793 | -0.218980951 | 0.834080751 | 0.969528435 | -5.096322491 |
| **A_52_P638605** | Ap1m2 | -0.030553509 | -0.37392708 | 0.312820061 | 9.221466331 | -0.21890889 | 0.834134343 | 0.969528435 | -5.096334289 |
| **A_55_P1988689** | Rbpjl | 0.009408103 | -0.096356502 | 0.115172708 | 7.710179348 | 0.218842098 | 0.834184016 | 0.969528435 | -5.096345221 |
| **A_55_P2080451** | NA | 0.038284143 | -0.392145568 | 0.468713855 | 12.60711254 | 0.218819417 | 0.834200885 | 0.969528435 | -5.096348932 |
| **A_51_P327021** | Zbtb25 | -0.023088692 | -0.282697311 | 0.236519927 | 9.867357844 | -0.218801039 | 0.834214553 | 0.969528435 | -5.09635194 |
| **A_52_P684057** | Snupn | 0.019135409 | -0.196060825 | 0.234331644 | 10.83928985 | 0.218762186 | 0.834243449 | 0.969528435 | -5.096358296 |
| **A_55_P2010271** | Samsn1 | -0.010744537 | -0.131599304 | 0.110110229 | 7.781985805 | -0.218722319 | 0.834273099 | 0.969528435 | -5.096364817 |
| **A_55_P2002357** | Tnrc18 | 0.031285267 | -0.320622858 | 0.383193391 | 10.58964254 | 0.218715585 | 0.834278107 | 0.969528435 | -5.096365919 |
| **A_55_P2005330** | Egln2 | 0.012852954 | -0.131735842 | 0.157441749 | 9.641011625 | 0.218694302 | 0.834293937 | 0.969528435 | -5.096369399 |
| **A_55_P2019964** | Lancl1 | -0.016615658 | -0.203538733 | 0.170307416 | 11.11585469 | -0.218687417 | 0.834299057 | 0.969528435 | -5.096370525 |
| **A_52_P304720** | Crlf1 | -0.009476275 | -0.116161926 | 0.097209376 | 7.701372407 | -0.218524835 | 0.834419978 | 0.969599988 | -5.096397103 |
| **A_55_P2021065** | E130308A19Rik | 0.019468308 | -0.199782283 | 0.2387189 | 8.331089451 | 0.218452293 | 0.834473934 | 0.969599988 | -5.096408956 |
| **A_51_P256224** | Zfp768 | -0.02190773 | -0.268726835 | 0.224911374 | 11.03280977 | -0.218367387 | 0.834537086 | 0.969599988 | -5.096422824 |
| **A_55_P2063999** | Gpatch8 | -0.024515424 | -0.300841307 | 0.251810459 | 10.53110065 | -0.218266463 | 0.834612155 | 0.969599988 | -5.096439301 |
| **A_55_P2079684** | NA | 0.031193378 | -0.320519585 | 0.38290634 | 14.5516873 | 0.218194194 | 0.834665911 | 0.969599988 | -5.096451095 |
| **A_52_P67212** | 2900052N01Rik | -0.011418626 | -0.140231883 | 0.11739463 | 7.974925324 | -0.218083326 | 0.834748379 | 0.969599988 | -5.096469181 |
| **A_52_P525317** | Gja5 | 0.017045305 | -0.175281986 | 0.209372597 | 8.384450411 | 0.218038424 | 0.83478178 | 0.969599988 | -5.096476504 |
| **A_52_P608132** | Snx32 | 0.014405194 | -0.148149164 | 0.176959552 | 8.020179292 | 0.218016647 | 0.834797979 | 0.969599988 | -5.096480055 |
| **A_55_P2106803** | Bglap2 | 0.045413396 | -0.46709536 | 0.557922152 | 11.11132368 | 0.21799767 | 0.834812096 | 0.969599988 | -5.096483149 |
| **A_51_P518246** | Fibp | 0.02866284 | -0.294931305 | 0.352256986 | 9.810067719 | 0.217915304 | 0.834873367 | 0.96961144 | -5.096496574 |
| **A_55_P1957922** | Arrdc2 | 0.020123153 | -0.207129275 | 0.247375581 | 9.396543412 | 0.217849546 | 0.834922283 | 0.96961144 | -5.096507289 |
| **A_52_P338956** | Aspg | 0.07848871 | -0.808398145 | 0.965375564 | 9.796966503 | 0.217724917 | 0.835014996 | 0.969622565 | -5.096527588 |
| **A_52_P609109** | Pitpnc1 | -0.010274927 | -0.12639809 | 0.105848236 | 7.927725743 | -0.21768528 | 0.835044484 | 0.969622565 | -5.096534042 |
| **A_51_P210395** | Glrx | 0.038167893 | -0.393305314 | 0.4696411 | 9.298210686 | 0.217627369 | 0.835087566 | 0.969622565 | -5.096543469 |
| **A_51_P100034** | Mif4gd | -0.016896712 | -0.208015691 | 0.174222266 | 10.69826587 | -0.217504154 | 0.835179232 | 0.969622565 | -5.096563517 |

| | | | | | | | | |
|---|---|---|---|---|---|---|---|---|
| **A_51_P472518** | Capn1 | -0.00947236 | -0.116621858 | 0.097677138 | 8.373614815 | -0.217488962 | 0.835190534 | 0.969622565 | -5.096565989 |
| **A_51_P407675** | Zrsr1 | -0.020269642 | -0.249683007 | 0.209143722 | 9.990557579 | -0.217368464 | 0.835280183 | 0.969622565 | -5.096585583 |
| **A_55_P1969058** | Epha7 | 0.010350969 | -0.10683111 | 0.127533047 | 7.737225014 | 0.217314628 | 0.835320236 | 0.969622565 | -5.096594334 |
| **A_52_P481279** | Drc1 | 0.009964293 | -0.102849309 | 0.122777895 | 8.32123265 | 0.217297237 | 0.835333175 | 0.969622565 | -5.096597161 |
| **A_55_P2131677** | NA | -0.012057585 | -0.148641143 | 0.124525973 | 8.012466029 | -0.217185623 | 0.835416217 | 0.969660726 | -5.096615295 |
| **A_51_P445683** | 4931406C07Rik | -0.039516517 | -0.487390441 | 0.408357408 | 11.56609559 | -0.217066118 | 0.835505134 | 0.9697057 | -5.096634702 |
| **A_55_P2036180** | Mcf2l | 0.026740048 | -0.276546825 | 0.330026922 | 11.14015687 | 0.216909081 | 0.835621979 | 0.969783082 | -5.096660188 |
| **A_51_P203675** | Trem3 | -0.009332088 | -0.115223752 | 0.096559577 | 7.677037075 | -0.216813445 | 0.835693141 | 0.969806774 | -5.0966757 |
| **A_51_P342669** | Pgam1 | -0.024133433 | -0.298241195 | 0.249974328 | 14.59698689 | -0.216604233 | 0.835848818 | 0.969806774 | -5.09670961 |
| **A_55_P2006392** | NA | 0.018309613 | -0.18965136 | 0.226270585 | 16.02250098 | 0.216604002 | 0.83584899 | 0.969806774 | -5.096709647 |
| **A_55_P2261772** | C230098O21Rik | -0.039867196 | -0.492758667 | 0.413024276 | 10.04245298 | -0.216566216 | 0.835877109 | 0.969806774 | -5.096715769 |
| **A_55_P2129696** | Smad9 | 0.012651628 | -0.131113048 | 0.156416303 | 7.943441063 | 0.216502727 | 0.835924353 | 0.969806774 | -5.096726051 |
| **A_66_P121965** | 1110032F04Rik | 0.009587265 | -0.099421033 | 0.118595562 | 7.663019058 | 0.216373626 | 0.836020426 | 0.969806774 | -5.096746951 |
| **A_66_P104980** | Tceal5 | 0.078590783 | -0.815071395 | 0.972252961 | 9.358298745 | 0.216355231 | 0.836034115 | 0.969806774 | -5.096749928 |
| **A_55_P1973941** | Slc7a5 | 0.021848862 | -0.226648022 | 0.270345745 | 11.93748267 | 0.216310215 | 0.836067615 | 0.969806774 | -5.096757212 |
| **A_55_P2338535** | Slc35a1 | -0.018390826 | -0.22759216 | 0.190810507 | 9.567990498 | -0.216274819 | 0.836093957 | 0.969806774 | -5.096762939 |
| **A_52_P531651** | Parvg | -0.009183407 | -0.11378116 | 0.095414346 | 7.700181806 | -0.215998456 | 0.836299632 | 0.969987134 | -5.096807618 |
| **A_55_P2013665** | Mybpc2 | -0.009303825 | -0.115326557 | 0.096718906 | 7.69022725 | -0.215889601 | 0.836380648 | 0.970022894 | -5.096825201 |
| **A_55_P2073885** | Bbs5 | -0.013472743 | -0.167057355 | 0.14011187 | 9.475558793 | -0.215812934 | 0.836437709 | 0.970030869 | -5.096837579 |
| **A_55_P2090055** | Phrf1 | -0.024434329 | -0.303170844 | 0.254302186 | 11.07324631 | -0.215663037 | 0.836549278 | 0.970051403 | -5.096861769 |
| **A_51_P138879** | Zgpat | 0.013898587 | -0.144666111 | 0.172463286 | 9.233278671 | 0.215641982 | 0.836564949 | 0.970051403 | -5.096865166 |
| **A_55_P1967928** | Srfbp1 | -0.014906036 | -0.18500789 | 0.155195817 | 9.188623045 | -0.215586863 | 0.836605975 | 0.970051403 | -5.096874056 |
| **A_55_P2043684** | Ubac1 | -0.018912068 | -0.234805075 | 0.19698094 | 11.69874036 | -0.21551108 | 0.836662383 | 0.970058617 | -5.096886275 |
| **A_52_P15388** | Ltf | -0.024650133 | -0.306308089 | 0.257007824 | 8.07258789 | -0.215311091 | 0.836811246 | 0.97016011 | -5.0969185 |
| **A_55_P2418745** | D230034L24Rik | -0.013610112 | -0.169208037 | 0.141987814 | 7.896529277 | -0.215192458 | 0.836899554 | 0.97016011 | -5.096937603 |
| **A_55_P2021996** | Hbs1l | -0.01501684 | -0.186698228 | 0.156664548 | 8.14502005 | -0.215191194 | 0.836900496 | 0.97016011 | -5.096937806 |
| **A_55_P2040280** | Dhps | -0.009948684 | -0.123746076 | 0.103848707 | 8.282617193 | -0.215081227 | 0.836982356 | 0.970181866 | -5.096955504 |
| **A_55_P2069885** | Slc50a1 | -0.027545483 | -0.342711495 | 0.287620528 | 8.802205744 | -0.215020656 | 0.837027446 | 0.970181866 | -5.096965248 |
| **A_51_P241943** | Sap30l | -0.028565289 | -0.355543265 | 0.298412687 | 13.57260308 | -0.214926164 | 0.83709779 | 0.970181866 | -5.096980444 |
| **A_52_P577388** | Epdr1 | -0.11965281 | -1.489725304 | 1.250419685 | 10.09590901 | -0.21485651 | 0.837149644 | 0.970181866 | -5.096991641 |
| **A_55_P2141860** | Aen | 0.012848487 | -0.134342577 | 0.160039551 | 11.85681043 | 0.214753231 | 0.837226532 | 0.970181866 | -5.097008237 |
| **A_55_P2068260** | H3c8 | -0.009340259 | -0.116415232 | 0.097734713 | 7.675189196 | -0.214605144 | 0.837336782 | 0.970181866 | -5.09703202 |
| **A_51_P384113** | Acad12 | -0.017061103 | -0.212660367 | 0.178538161 | 8.995090521 | -0.214589785 | 0.837348217 | 0.970181866 | -5.097034486 |
| **A_55_P2078845** | Qk | 0.015667166 | -0.164045617 | 0.195379949 | 11.14561761 | 0.214476956 | 0.83743222 | 0.970181866 | -5.097052594 |
| **A_51_P188073** | Dnajb11 | -0.031654452 | -0.394791933 | 0.331483029 | 13.96882595 | -0.214453374 | 0.837449778 | 0.970181866 | -5.097056378 |
| **A_55_P2145942** | Atg16l1 | -0.016343723 | -0.203882926 | 0.171195479 | 8.959862257 | -0.214401637 | 0.837488298 | 0.970181866 | -5.097064678 |
| **A_52_P624362** | Aamp | 0.020505864 | -0.214805274 | 0.255817002 | 10.91671993 | 0.214390112 | 0.837496879 | 0.970181866 | -5.097066526 |
| **A_55_P2139901** | Tsr3 | -0.026988841 | -0.336742077 | 0.282764396 | 11.95117669 | -0.214356937 | 0.83752158 | 0.970181866 | -5.097071847 |
| **A_55_P2095311** | NA | -0.128126656 | -1.599355762 | 1.343102451 | 9.539540979 | -0.214253707 | 0.837598441 | 0.970208396 | -5.097088397 |
| **A_51_P336599** | Kcne3 | -0.011395806 | -0.142314438 | 0.119522827 | 8.003388938 | -0.21414737 | 0.837677617 | 0.970208396 | -5.097105438 |
| **A_51_P227962** | Dynlrb2 | -0.011007209 | -0.137502245 | 0.115487826 | 7.742953472 | -0.21407843 | 0.83772895 | 0.970208396 | -5.097116481 |
| **A_51_P297069** | Tmod1 | 0.024593304 | -0.258062282 | 0.30724889 | 8.16076759 | 0.214056526 | 0.83774526 | 0.970208396 | -5.097119989 |
| **A_55_P2414524** | NA | -0.015581925 | -0.194777371 | 0.16361352 | 8.02550837 | -0.213925876 | 0.837842545 | 0.970235966 | -5.097140906 |
| **A_52_P594768** | Aprt | -0.012995658 | -0.162521179 | 0.136529863 | 15.28910062 | -0.213821871 | 0.837919991 | 0.970235966 | -5.097157548 |
| **A_55_P2104941** | Rpl30 | -0.011330873 | -0.141713486 | 0.119051741 | 15.04240411 | -0.213802528 | 0.837934395 | 0.970235966 | -5.097160642 |
| **A_51_P261379** | Spats2 | -0.021352721 | -0.267301356 | 0.224595915 | 9.967423319 | -0.213588551 | 0.838093738 | 0.970235966 | -5.097194853 |
| **A_51_P337195** | Pipox | -0.009218772 | -0.115421576 | 0.096984032 | 7.683429287 | -0.213553282 | 0.838120003 | 0.970235966 | -5.097200488 |
| **A_51_P481238** | Dop1b | -0.035250163 | -0.441504997 | 0.371004672 | 10.25283966 | -0.213467492 | 0.838183891 | 0.970235966 | -5.097214193 |
| **A_51_P491095** | Gzf1 | 0.013975391 | -0.147093956 | 0.175044739 | 9.45637612 | 0.213461835 | 0.838188104 | 0.970235966 | -5.097215096 |
| **A_55_P2026982** | Tdrp | -0.06437985 | -0.806669249 | 0.677909549 | 14.50038114 | -0.213376145 | 0.83825192 | 0.970235966 | -5.097228779 |

| | | | | | | | | |
|---|---|---|---|---|---|---|---|---|
| **A_66_P121787** | Samd9l | -0.02640838 | -0.330935258 | 0.278118497 | 8.980739067 | -0.21334639 | 0.83827408 | 0.970235966 | -5.097233529 |
| **A_55_P2059347** | Ormdl3 | -0.015745469 | -0.197427657 | 0.165936719 | 13.42949112 | -0.21321238 | 0.838373884 | 0.970235966 | -5.097254914 |
| **A_52_P707503** | Tmem212 | -0.009554238 | -0.11980851 | 0.100700034 | 7.676575172 | -0.213191457 | 0.838389467 | 0.970235966 | -5.097258251 |
| **A_55_P2093553** | NA | -0.042828098 | -0.537075964 | 0.451419768 | 8.548951956 | -0.213183285 | 0.838395553 | 0.970235966 | -5.097259555 |
| **A_51_P501803** | Hoxa2 | 0.009172499 | -0.096698346 | 0.115043344 | 7.670779301 | 0.213147612 | 0.838422121 | 0.970235966 | -5.097265244 |
| **A_51_P100787** | Snw1 | 0.017388882 | -0.18347463 | 0.218252394 | 10.58488734 | 0.212980464 | 0.838546612 | 0.970235966 | -5.058768934 |
| **A_55_P2039439** | Ppp1ca | 0.026600768 | -0.280688242 | 0.333889779 | 14.80979439 | 0.212968962 | 0.838555179 | 0.970235966 | -5.097293724 |
| **A_52_P501049** | NA | -0.016772327 | -0.21055024 | 0.177005586 | 8.884603521 | -0.212940471 | 0.838576399 | 0.970235966 | -5.097298263 |
| **A_52_P577729** | Rnf169 | -0.016273991 | -0.204348956 | 0.171800974 | 10.00965692 | -0.212878715 | 0.838622396 | 0.970235966 | -5.097308101 |
| **A_52_P531662** | Ttc9c | -0.009856493 | -0.124018133 | 0.104305148 | 8.969334706 | -0.212408243 | 0.838972836 | 0.97056505 | -5.097382958 |
| **A_55_P2019483** | Ndrg4 | -0.081629222 | -1.027421467 | 0.864163023 | 10.50718727 | -0.212333771 | 0.839028312 | 0.97056505 | -5.097394792 |
| **A_55_P1975682** | Rhox2e | 0.035958512 | -0.380749541 | 0.452666566 | 8.630591169 | 0.212294619 | 0.839057477 | 0.97056505 | -5.097401012 |
| **A_52_P333097** | NA | -0.039076639 | -0.492086185 | 0.413932908 | 8.404284238 | -0.212216431 | 0.839115723 | 0.97057434 | -5.097413431 |
| **A_51_P393518** | Hmcn1 | 0.009030412 | -0.095704543 | 0.113765367 | 7.676864396 | 0.212121692 | 0.839186299 | 0.97059395 | -5.097428472 |
| **A_55_P1953361** | Luc7l2 | 0.023293059 | -0.24694049 | 0.293526609 | 16.08759893 | 0.212058863 | 0.839233105 | 0.97059395 | -5.097438443 |
| **A_55_P2092958** | Traf2 | 0.010461981 | -0.111012001 | 0.131935962 | 7.986210117 | 0.211884807 | 0.839362775 | 0.970685837 | -5.097466051 |
| **A_51_P508485** | Tmtc4 | 0.010826738 | -0.115076212 | 0.136729688 | 9.04473422 | 0.211558698 | 0.839605739 | 0.970907349 | -5.097517718 |
| **A_55_P1988872** | NA | 0.009327971 | -0.099212519 | 0.117868462 | 7.774487328 | 0.211429016 | 0.839702363 | 0.970907349 | -5.097538242 |
| **A_51_P262878** | Tmed9 | -0.021279673 | -0.268894604 | 0.226335258 | 15.59611444 | -0.211425461 | 0.839705011 | 0.970907349 | -5.097538805 |
| **A_55_P2129469** | NA | 0.009355165 | -0.099604645 | 0.118314974 | 7.9318143 | 0.21122935 | 0.839851135 | 0.97097951 | -5.097569819 |
| **A_55_P2131577** | NA | 0.009590946 | -0.102159577 | 0.12134147 | 16.79376213 | 0.211145126 | 0.839913894 | 0.97097951 | -5.09758313 |
| **A_51_P216593** | Wfs1 | 0.028693597 | -0.305710874 | 0.363098069 | 12.03715315 | 0.211097012 | 0.839949746 | 0.97097951 | -5.097590731 |
| **A_51_P330133** | Pspc1 | -0.009401402 | -0.119007784 | 0.100204979 | 8.697360129 | -0.211021136 | 0.840006286 | 0.97097951 | -5.097602716 |
| **A_52_P337259** | Heyl | 0.012637057 | -0.134703786 | 0.1599779 | 8.203922697 | 0.211004621 | 0.840018592 | 0.97097951 | -5.097605324 |
| **A_51_P144349** | Dtx4 | -0.02440516 | -0.309168877 | 0.260358556 | 9.042488467 | -0.210846393 | 0.840136501 | 0.971033244 | -5.0976303 |
| **A_55_P2096657** | NA | -0.018901161 | -0.239484032 | 0.201681709 | 14.94095145 | -0.210807408 | 0.840165553 | 0.971033244 | -5.097636451 |
| **A_55_P2163659** | Rspo3 | 0.01184886 | -0.126517093 | 0.150214812 | 8.479672501 | 0.210676687 | 0.840262969 | 0.971087768 | -5.097657068 |
| **A_52_P89567** | Rhob | 0.014775135 | -0.157862338 | 0.187412607 | 9.125788031 | 0.210554991 | 0.840353662 | 0.971134517 | -5.09767625 |
| **A_55_P2078623** | Stxbp3 | -0.033146447 | -0.420871755 | 0.354578861 | 10.97613909 | -0.210320689 | 0.84052828 | 0.971221545 | -5.097713151 |
| **A_51_P439857** | Dnajc21 | 0.019378151 | -0.207296407 | 0.246052708 | 9.252948219 | 0.210319101 | 0.840529464 | 0.971221545 | -5.097713401 |
| **A_51_P295816** | Cep192 | -0.013937552 | -0.177116542 | 0.149241438 | 9.224989465 | -0.21013163 | 0.840669189 | 0.971304471 | -5.097742898 |
| **A_51_P452153** | Mzb1 | -0.009078388 | -0.115389033 | 0.097232256 | 7.715792362 | -0.210087965 | 0.840701733 | 0.971304471 | -5.092345936 |
| **A_55_P2122588** | Ubr2 | -0.016012249 | -0.203645794 | 0.171621297 | 9.265845775 | -0.209947643 | 0.840806322 | 0.971330536 | -5.09777182 |
| **A_55_P2026492** | NA | 0.010232711 | -0.109706389 | 0.13017181 | 7.830194159 | 0.209893543 | 0.840846646 | 0.971330536 | -5.09778032 |
| **A_51_P492765** | Wdr75 | -0.015069038 | -0.191727503 | 0.161589428 | 10.43044346 | -0.209855436 | 0.840875051 | 0.971330536 | -5.097786306 |
| **A_55_P2029682** | Fry | -0.050813259 | -0.647295508 | 0.545668991 | 9.76888082 | -0.209579445 | 0.841080775 | 0.971510117 | -5.097829627 |
| **A_51_P323812** | Slc6a12 | -0.089873078 | -1.145219587 | 0.96547343 | 12.36474482 | -0.209509492 | 0.841132921 | 0.971512294 | -5.097840598 |
| **A_55_P1974228** | NA | -0.01175582 | -0.149880272 | 0.126368631 | 15.25299642 | -0.209387879 | 0.841223578 | 0.971558947 | -5.097859663 |
| **A_51_P247637** | Rnf144a | 0.015699694 | -0.168914447 | 0.200313835 | 8.292646123 | 0.209216237 | 0.841351533 | 0.971648669 | -5.097886553 |
| **A_51_P150876** | Lypd3 | 0.056038524 | -0.603627377 | 0.715704425 | 8.312270238 | 0.208993036 | 0.841517933 | 0.971782778 | -5.097921487 |
| **A_55_P2386256** | D130062J10Rik | 0.038878685 | -0.419056624 | 0.496813994 | 9.48243621 | 0.208870255 | 0.841609472 | 0.971804264 | -5.097940689 |
| **A_55_P1993503** | Adora2b | 0.025122542 | -0.271235312 | 0.321480397 | 9.087357426 | 0.208552935 | 0.841846061 | 0.971804264 | -5.097990263 |
| **A_55_P2035827** | Vma21 | -0.023387019 | -0.299278848 | 0.252504809 | 10.67366318 | -0.208547601 | 0.841850038 | 0.971804264 | -5.097991095 |
| **A_55_P2172121** | Mxd4 | -0.010471156 | -0.134027333 | 0.11308502 | 8.929427551 | -0.208496779 | 0.841887932 | 0.971804264 | -5.097999028 |
| **A_55_P2003353** | Gemin8 | 0.011132958 | -0.120239465 | 0.142505381 | 8.132816633 | 0.208485343 | 0.841896459 | 0.971804264 | -5.098000813 |
| **A_66_P113868** | Cdh3 | 0.016983195 | -0.183429686 | 0.217396076 | 11.73975097 | 0.208479293 | 0.84190097 | 0.971804264 | -5.098001757 |
| **A_51_P274124** | Mansc1 | 0.013928262 | -0.150434773 | 0.178291297 | 8.839054917 | 0.208478826 | 0.841901318 | 0.971804264 | -5.09800183 |
| **A_51_P422124** | Fam126a | -0.028424155 | -0.36397393 | 0.307125621 | 9.715627012 | -0.208400984 | 0.84195936 | 0.971804264 | -5.098013975 |
| **A_55_P2153031** | Ttn | -0.008780415 | -0.112453821 | 0.094892992 | 7.675469118 | -0.20836119 | 0.841989033 | 0.971804264 | -5.098020183 |
| **A_66_P139116** | Rpl8 | 0.016845742 | -0.182304816 | 0.215996301 | 16.26589294 | 0.208102737 | 0.842181754 | 0.971968661 | -5.098060469 |

| | | | | | | | | | |
|---|---|---|---|---|---|---|---|---|---|
| **A_66_P124549** | H13 | -0.029931467 | -0.383987714 | 0.324124779 | 14.55416238 | -0.207981598 | 0.842272088 | 0.971970795 | -5.098079335 |
| **A_55_P2159299** | Ing1 | -0.022398748 | -0.287582851 | 0.242785355 | 11.67135164 | -0.207799938 | 0.842407559 | 0.971970795 | -5.098107606 |
| **A_55_P2167000** | Dmac2 | -0.010459794 | -0.1343019 | 0.113382311 | 10.95618881 | -0.207789682 | 0.842415207 | 0.971970795 | -5.098109201 |
| **A_55_P2008639** | NA | -0.018810766 | -0.241620261 | 0.203998728 | 11.60013026 | -0.207702613 | 0.84248014 | 0.971970795 | -5.098122742 |
| **A_51_P459128** | Serpinb9e | 0.016517715 | -0.179303512 | 0.212338942 | 11.04229533 | 0.207519717 | 0.842616541 | 0.971970795 | -5.098151167 |
| **A_55_P2065866** | Cygb | 0.038931423 | -0.422633715 | 0.500496561 | 9.111495927 | 0.207508764 | 0.842624709 | 0.971970795 | -5.098152869 |
| **A_55_P2069979** | Large1 | -0.06330109 | -0.814021694 | 0.687419514 | 11.64205352 | -0.207444543 | 0.842672606 | 0.971970795 | -5.098162844 |
| **A_55_P1996872** | Polr2k | 0.028436482 | -0.308976161 | 0.365849126 | 12.34387978 | 0.207340282 | 0.842750366 | 0.971970795 | -5.098179031 |
| **A_55_P2109397** | Scn10a | -0.008912358 | -0.114757181 | 0.096932465 | 7.660090928 | -0.207153453 | 0.842889713 | 0.971970795 | -5.098208018 |
| **A_55_P2095196** | Akr1b10 | 0.02942277 | -0.320203257 | 0.379048798 | 8.773888346 | 0.207037479 | 0.842976215 | 0.971970795 | -5.098225999 |
| **A_55_P1981794** | Sgf29 | -0.011260077 | -0.145085666 | 0.122565512 | 8.164850407 | -0.207000482 | 0.843003811 | 0.971970795 | -5.098231733 |
| **A_55_P2041070** | Tle3 | -0.021456064 | -0.276701886 | 0.233789758 | 12.55646631 | -0.206804785 | 0.843149784 | 0.971970795 | -5.098262046 |
| **A_55_P2119145** | NA | 0.011100881 | -0.120973893 | 0.143175656 | 8.488355901 | 0.206779148 | 0.843168907 | 0.971970795 | -5.098266015 |
| **A_52_P27020** | Sgo2a | -0.010090209 | -0.130154588 | 0.109974171 | 7.725943531 | -0.206754576 | 0.843187237 | 0.971970795 | -5.098269819 |
| **A_55_P2122619** | Asb11 | -0.054640714 | -0.704939275 | 0.595657848 | 9.434805205 | -0.206715353 | 0.843216494 | 0.971970795 | -5.09827589 |
| **A_51_P309589** | 2700099C18Rik | -0.009028514 | -0.116513717 | 0.098456689 | 7.792034117 | -0.206650639 | 0.843264768 | 0.971970795 | -5.098285904 |
| **A_55_P2004037** | NA | 0.009447994 | -0.103053775 | 0.121949763 | 7.644566235 | 0.206609087 | 0.843295764 | 0.971970795 | -5.098292332 |
| **A_55_P2168397** | NA | 0.015331853 | -0.167283395 | 0.197947102 | 16.30555879 | 0.206550752 | 0.84333928 | 0.971970795 | -5.098301354 |
| **A_51_P455866** | Elf5 | -0.008632234 | -0.111468706 | 0.094204238 | 7.717388701 | -0.206511966 | 0.843368214 | 0.971970795 | -5.098307352 |
| **A_51_P227165** | 2310030G06Rik | -0.015696702 | -0.202748356 | 0.171354952 | 11.86843817 | -0.206450539 | 0.843414037 | 0.971970795 | -5.098316848 |
| **A_52_P409833** | Plat | -0.015376756 | -0.198761453 | 0.168007941 | 8.879443438 | -0.206286498 | 0.843536413 | 0.971970795 | -5.098342193 |
| **A_51_P305628** | Atp8a2 | 0.027943307 | -0.30558689 | 0.361473503 | 8.254169078 | 0.206116044 | 0.843663578 | 0.971970795 | -5.098368509 |
| **A_51_P491916** | Rassf6 | 0.030400097 | -0.332527125 | 0.393327319 | 11.05214322 | 0.206074659 | 0.843694454 | 0.971970795 | -5.098374895 |
| **A_55_P2126780** | Pgam5 | -0.028886508 | -0.373792722 | 0.316019707 | 12.55406398 | -0.206045522 | 0.843716192 | 0.971970795 | -5.09837939 |
| **A_55_P2359807** | NA | 0.015293298 | -0.167320224 | 0.19790682 | 10.07423217 | 0.206033279 | 0.843725326 | 0.971970795 | -5.098381279 |
| **A_55_P2001608** | Klhl4 | -0.008952467 | -0.115860256 | 0.097955322 | 7.637632538 | -0.206016759 | 0.843737651 | 0.971970795 | -5.098383827 |
| **A_55_P2006713** | NA | 0.009690301 | -0.106155345 | 0.125535948 | 16.78595645 | 0.205791166 | 0.843905964 | 0.971970795 | -5.098418606 |
| **A_51_P243914** | Skap2 | -0.014513829 | -0.188232409 | 0.159204752 | 10.92646517 | -0.205544036 | 0.844090356 | 0.971970795 | -5.098456662 |
| **A_55_P2178044** | Inppl1 | -0.022893882 | -0.296957139 | 0.251169376 | 12.67394987 | -0.205512283 | 0.844114048 | 0.971970795 | -5.098461549 |
| **A_55_P2068459** | Hspa1a | 0.015304001 | -0.168010036 | 0.198618039 | 8.436986718 | 0.205389595 | 0.844205595 | 0.971970795 | -5.098480422 |
| **A_52_P628590** | Pvr | -0.08898662 | -1.155250538 | 0.977277297 | 10.59344607 | -0.20531901 | 0.844258264 | 0.971970795 | -5.098491276 |
| **A_52_P547491** | Lurap1 | 0.034347222 | -0.377467679 | 0.446162123 | 10.1670526 | 0.205191196 | 0.84435364 | 0.971970795 | -5.098510919 |
| **A_55_P2036863** | Rabl2 | -0.016997617 | -0.220829458 | 0.186834223 | 10.43887285 | -0.205156453 | 0.844379566 | 0.971970795 | -5.098516257 |
| **A_55_P1956195** | Sap18b | -0.029194933 | -0.379308592 | 0.320918725 | 12.63805801 | -0.205148147 | 0.844385764 | 0.971970795 | -5.098517533 |
| **A_52_P123944** | Mospd2 | -0.011817413 | -0.15356487 | 0.129930044 | 8.393999943 | -0.205105022 | 0.844417945 | 0.971970795 | -5.098524157 |
| **A_55_P2145441** | NA | -0.012854881 | -0.167102305 | 0.141392542 | 9.973392963 | -0.205030898 | 0.84447326 | 0.971970795 | -5.098535539 |
| **A_55_P2114074** | Acrbp | -0.031057097 | -0.403729857 | 0.341615663 | 8.844543347 | -0.2050229 | 0.844479228 | 0.971970795 | -5.098536766 |
| **A_52_P505157** | Fsd2 | -0.009786108 | -0.127245542 | 0.107673325 | 7.646508385 | -0.204970461 | 0.844518361 | 0.971970795 | -5.098544816 |
| **A_51_P162471** | Dnajc9 | 0.02644693 | -0.291183685 | 0.344077546 | 10.66456481 | 0.204843449 | 0.844613146 | 0.971970795 | -5.098564304 |
| **A_66_P119350** | 2310040G24Rik | 0.015094372 | -0.166210226 | 0.196398969 | 8.228098221 | 0.204821425 | 0.844629582 | 0.971970795 | -5.098567682 |
| **A_51_P171697** | Pebp1 | -0.019567437 | -0.254655516 | 0.215520642 | 12.24533305 | -0.20477291 | 0.844665788 | 0.971970795 | -5.098575122 |
| **A_51_P443618** | Mettl26 | 0.03184484 | -0.350790997 | 0.414480678 | 11.60565478 | 0.20474938 | 0.844683349 | 0.971970795 | -5.09857873 |
| **A_52_P296109** | NA | -0.009051809 | -0.117851705 | 0.099748087 | 8.027966094 | -0.204680309 | 0.844734896 | 0.971970795 | -5.098589318 |
| **A_55_P1974005** | Tnfrsf23 | -0.053177344 | -0.692400609 | 0.58604592 | 8.67862159 | -0.204664837 | 0.844746443 | 0.971970795 | -5.098591689 |
| **A_51_P410744** | Dgke | -0.02264127 | -0.294821388 | 0.249538847 | 9.032240949 | -0.204650852 | 0.84475688 | 0.971970795 | -5.098593832 |
| **A_55_P2106434** | Zfp114 | 0.010076123 | -0.111103953 | 0.131256198 | 8.035874315 | 0.20456503 | 0.844820931 | 0.971970795 | -5.098606981 |
| **A_55_P1984541** | NA | -0.019167641 | -0.24979274 | 0.211457459 | 8.917662524 | -0.204470777 | 0.844891275 | 0.971970795 | -5.098621416 |
| **A_51_P466910** | Cacna1g | -0.00940742 | -0.122606989 | 0.103792149 | 7.914874135 | -0.204453683 | 0.844904034 | 0.971970795 | -5.098624033 |
| **A_51_P325318** | Serinc2 | -0.038147237 | -0.497568545 | 0.42127407 | 9.998815055 | -0.204277777 | 0.845035324 | 0.971970795 | -5.098650954 |
| **A_55_P2161342** | Pkib | 0.014424537 | -0.159359421 | 0.188208495 | 11.32095661 | 0.204202643 | 0.845091402 | 0.971970795 | -5.098662445 |

| | | | | | | | | | |
|---|---|---|---|---|---|---|---|---|---|
| **A_55_P1999633** | Cit | -0.008736794 | -0.113999868 | 0.096526279 | 7.715339742 | -0.204195061 | 0.845097062 | 0.971970795 | -5.098663604 |
| **A_66_P133404** | Pclaf | -0.008848438 | -0.11547279 | 0.097775913 | 7.670978847 | -0.2041641 | 0.845120171 | 0.971970795 | -5.098668338 |
| **A_55_P1978316** | Adamts14 | -0.025481778 | -0.332544999 | 0.281581442 | 9.932617228 | -0.204160207 | 0.845123077 | 0.971970795 | -5.098668933 |
| **A_52_P463143** | Cdc37 | -0.011874009 | -0.154977341 | 0.131229323 | 14.58177037 | -0.204134676 | 0.845142134 | 0.971970795 | -5.098672837 |
| **A_51_P221510** | Fam81a | -0.009295516 | -0.121377823 | 0.10278679 | 7.71304899 | -0.204035445 | 0.8452162 | 0.971970795 | -5.098688002 |
| **A_55_P2043782** | Trpm1 | -0.021584567 | -0.282018175 | 0.238849041 | 8.149129945 | -0.203899175 | 0.845317917 | 0.971970795 | -5.098708817 |
| **A_66_P129852** | Fxn | -0.01994641 | -0.260617402 | 0.220724582 | 8.886262723 | -0.203896681 | 0.845319779 | 0.971970795 | -5.098709197 |
| **A_51_P151586** | Haspin | -0.009201314 | -0.120292318 | 0.10188969 | 7.720626962 | -0.203769939 | 0.845414387 | 0.971970795 | -5.098728544 |
| **A_55_P2062981** | Abca9 | 0.011940013 | -0.132294273 | 0.156174299 | 7.888670756 | 0.203659861 | 0.845496557 | 0.971970795 | -5.098745337 |
| **A_51_P461894** | Tnnc1 | -0.037518271 | -0.490795448 | 0.415758906 | 9.147600732 | -0.203632984 | 0.845516621 | 0.971970795 | -5.098749436 |
| **A_55_P2386236** | Ceacam2 | -0.030391892 | -0.397580385 | 0.336796601 | 9.465066014 | -0.203628159 | 0.845520223 | 0.971970795 | -5.098750172 |
| **A_55_P1966528** | Pmf1 | 0.013395039 | -0.148457712 | 0.175247789 | 8.777487516 | 0.203607166 | 0.845535894 | 0.971970795 | -5.098753373 |
| **A_55_P2141479** | Rnu2-10 | 0.020242668 | -0.224475625 | 0.264960961 | 10.51275788 | 0.203502846 | 0.84561377 | 0.971970795 | -5.098769276 |
| **A_51_P235088** | Dusp27 | 0.008957328 | -0.09934155 | 0.117256206 | 7.662921183 | 0.203480918 | 0.845630139 | 0.971970795 | -5.098772618 |
| **A_55_P1957038** | Gstp2 | -0.036632541 | -0.47954772 | 0.406282637 | 13.32828999 | -0.203477151 | 0.845632952 | 0.971970795 | -5.098773192 |
| **A_52_P42069** | Prokr1 | 0.008978998 | -0.0996184 | 0.117576396 | 7.666120238 | 0.203412489 | 0.845681224 | 0.971970795 | -5.098783044 |
| **A_52_P682745** | Dock4 | 0.010776545 | -0.119567821 | 0.141120911 | 8.363494354 | 0.203402574 | 0.845688625 | 0.971970795 | -5.098784554 |
| **A_51_P510418** | Aldh1b1 | -0.015229968 | -0.199449651 | 0.168989716 | 8.104641144 | -0.203391178 | 0.845697133 | 0.971970795 | -5.09878629 |
| **A_55_P2104597** | Kctd9 | -0.015801993 | -0.206996511 | 0.175392524 | 12.98789578 | -0.203331938 | 0.845741358 | 0.971970795 | -5.098795312 |
| **A_52_P227391** | Kif15 | 0.008670257 | -0.096248126 | 0.113588641 | 7.695329575 | 0.203305706 | 0.845760942 | 0.971970795 | -5.098799307 |
| **A_55_P2025273** | Trabd | -0.011605372 | -0.152061869 | 0.128851126 | 10.06020376 | -0.203276129 | 0.845783022 | 0.971970795 | -5.09880381 |
| **A_55_P2037116** | Ran | 0.0271564 | -0.301555549 | 0.355868348 | 14.4172647 | 0.203247805 | 0.845804168 | 0.971970795 | -5.098808121 |
| **A_51_P172573** | Sod2 | -0.016794198 | -0.220181205 | 0.18659281 | 12.47400374 | -0.203144568 | 0.845881242 | 0.971995075 | -5.098823832 |
| **A_52_P462472** | Nexmif | 0.008816579 | -0.09798849 | 0.115621648 | 7.677979972 | 0.203084791 | 0.84592587 | 0.971995075 | -5.098832925 |
| **A_55_P2181386** | Adm2 | 0.010625726 | -0.118301845 | 0.139553298 | 7.779812056 | 0.202759863 | 0.846168468 | 0.972132749 | -5.098882306 |
| **A_55_P2011132** | Armc10 | -0.019117806 | -0.251088576 | 0.212852963 | 10.74771469 | -0.202756114 | 0.846171267 | 0.972132749 | -5.098882876 |
| **A_51_P196673** | Khdc1b | -0.019542728 | -0.256785629 | 0.217700173 | 8.449121693 | -0.202656773 | 0.846245441 | 0.972132749 | -5.098897957 |
| **A_66_P127311** | Esco1 | 0.011843344 | -0.132003024 | 0.155689711 | 8.557335862 | 0.202555761 | 0.846320864 | 0.972132749 | -5.098913285 |
| **A_55_P2184925** | Tmem185b | 0.01328244 | -0.148181777 | 0.174746657 | 11.59758109 | 0.202381466 | 0.84645101 | 0.972132749 | -5.098939716 |
| **A_51_P283004** | Tinag | 0.074423518 | -0.830449828 | 0.979296863 | 8.924708253 | 0.202344573 | 0.846478558 | 0.972132749 | -5.098945308 |
| **A_52_P539434** | Lbh | -0.009098617 | -0.119790453 | 0.101593219 | 7.807293936 | -0.202222255 | 0.846569897 | 0.972132749 | -5.09896384 |
| **A_55_P2094520** | Fam171a1 | 0.026758723 | -0.298825248 | 0.352342694 | 12.23480718 | 0.202195522 | 0.84658986 | 0.972132749 | -5.098967889 |
| **A_52_P653825** | Keg1 | 0.008549936 | -0.095513898 | 0.112613769 | 7.653653836 | 0.202130657 | 0.846638298 | 0.972132749 | -5.09897771 |
| **A_52_P432780** | Mrpl50 | -0.012772164 | -0.168387333 | 0.142843004 | 8.382075218 | -0.201921113 | 0.84679478 | 0.972132749 | -5.099009418 |
| **A_55_P2067151** | NA | -0.011457318 | -0.151087898 | 0.128173263 | 8.757623673 | -0.201869902 | 0.846833025 | 0.972132749 | -5.099017162 |
| **A_55_P2057528** | Arl4d | -0.107697904 | -1.420398815 | 1.205003008 | 10.44339516 | -0.201841575 | 0.84685418 | 0.972132749 | -5.099021445 |
| **A_52_P72654** | Slc25a17 | 0.025882161 | -0.289655293 | 0.341419615 | 11.1426365 | 0.201798895 | 0.846886054 | 0.972132749 | -5.099027897 |
| **A_55_P2146177** | Cerk | -0.07162534 | -0.944932743 | 0.801682063 | 11.8457398 | -0.201775633 | 0.846903426 | 0.972132749 | -5.099031413 |
| **A_55_P2258832** | 5230400M03Rik | 0.030765739 | -0.344492477 | 0.406023954 | 8.670855695 | 0.201700186 | 0.846959772 | 0.972132749 | -5.099042813 |
| **A_51_P379478** | Nckap5 | 0.014022997 | -0.15707132 | 0.185117314 | 7.92891784 | 0.201638947 | 0.847005508 | 0.972132749 | -5.099052064 |
| **A_51_P133684** | Csrp3 | -0.008986651 | -0.118649372 | 0.100676069 | 7.744066729 | -0.201608131 | 0.847028523 | 0.972132749 | -5.099056718 |
| **A_52_P219904** | Afmid | 0.009564617 | -0.107196943 | 0.126326177 | 8.275453957 | 0.201528676 | 0.847087864 | 0.972132749 | -5.099068714 |
| **A_51_P114062** | Ncs1 | -0.008789749 | -0.116132154 | 0.098552656 | 7.734218733 | -0.201453269 | 0.847144184 | 0.972132749 | -5.088007288 |
| **A_51_P228817** | NA | -0.011472913 | -0.151646356 | 0.12870053 | 8.847275588 | -0.201361817 | 0.847212488 | 0.972132749 | -5.099093892 |
| **A_55_P1983448** | S100a4 | -0.067248346 | -0.888890195 | 0.754393504 | 10.14580279 | -0.201357688 | 0.847215571 | 0.972132749 | -5.099094514 |
| **A_55_P2049266** | Mettl6 | 0.015260919 | -0.171233708 | 0.201755547 | 9.698006057 | 0.201318429 | 0.847244894 | 0.972132749 | -5.099100435 |
| **A_55_P2142527** | NA | 0.012258917 | -0.137566209 | 0.162084042 | 8.368546531 | 0.201296675 | 0.847261142 | 0.972132749 | -5.099103715 |
| **A_51_P136729** | Idh3g | -0.022835573 | -0.301995191 | 0.256324044 | 12.39791171 | -0.201246571 | 0.847298565 | 0.972132749 | -5.099111269 |
| **A_51_P344399** | Rilpl2 | 0.043296265 | -0.486090226 | 0.572682755 | 10.25559808 | 0.201208692 | 0.847326858 | 0.972132749 | -5.099116979 |
| **A_55_P1985463** | NA | -0.01673201 | -0.221398813 | 0.187934793 | 13.12284027 | -0.201126768 | 0.847388049 | 0.972132749 | -5.099129324 |

| | | | | | | | | | |
|---|---|---|---|---|---|---|---|---|---|
| **A_55_P2041592** | Rdh1 | -0.010038559 | -0.132851066 | 0.112773949 | 8.118518885 | -0.201093448 | 0.847412937 | 0.972132749 | -5.099134343 |
| **A_52_P653902** | Eny2 | -0.018673642 | -0.247222684 | 0.2098754 | 8.65044773 | -0.201010521 | 0.847474879 | 0.972132749 | -5.099146832 |
| **A_55_P2036693** | Cd40 | -0.054845008 | -0.726359319 | 0.616669303 | 8.727523456 | -0.200932873 | 0.847532879 | 0.972132749 | -5.099158521 |
| **A_52_P235278** | Pms1 | 0.015190235 | -0.170936059 | 0.201316529 | 8.439077074 | 0.200782536 | 0.847645178 | 0.972132749 | -5.099181141 |
| **A_55_P2122643** | Dtymk | -0.022063916 | -0.292430839 | 0.248303007 | 10.46902106 | -0.200769715 | 0.847654756 | 0.972132749 | -5.099183069 |
| **A_51_P516850** | Rbbp5 | 0.010688726 | -0.120317937 | 0.14169539 | 8.373623003 | 0.200725121 | 0.847688067 | 0.972132749 | -5.099189775 |
| **A_52_P647740** | Klhl41 | -0.008591566 | -0.113906493 | 0.096723361 | 7.648268073 | -0.20070193 | 0.84770539 | 0.972132749 | -5.099193262 |
| **A_55_P1967683** | Bex4 | 0.089646723 | -1.010490931 | 1.189784377 | 11.91380622 | 0.200473306 | 0.84787618 | 0.972137313 | -5.099227615 |
| **A_51_P351697** | Mrpl35 | -0.008935371 | -0.118627595 | 0.100756854 | 8.161003793 | -0.200403772 | 0.847928125 | 0.972137313 | -5.099238056 |
| **A_55_P2111713** | Spint2 | 0.035090593 | -0.395707458 | 0.465888643 | 10.82093357 | 0.200394656 | 0.847934935 | 0.972137313 | -5.099239424 |
| **A_66_P137943** | Tctn2 | -0.01476652 | -0.196093359 | 0.166560318 | 10.15086856 | -0.200348107 | 0.84796971 | 0.972137313 | -5.099246411 |
| **A_55_P2033116** | Cdcp2 | -0.008459383 | -0.112348101 | 0.095429335 | 7.693320094 | -0.200326982 | 0.847985493 | 0.972137313 | -5.099249582 |
| **A_55_P1960117** | Dipk2a | 0.016216056 | -0.182968694 | 0.215400807 | 8.680496192 | 0.200289563 | 0.848013447 | 0.972137313 | -5.099255197 |
| **A_51_P161362** | Gnl2 | 0.015680977 | -0.176993084 | 0.208355038 | 10.58641575 | 0.200225337 | 0.848061429 | 0.972137313 | -5.099264832 |
| **A_55_P1957474** | Pus7 | 0.016820448 | -0.189968892 | 0.223609788 | 9.816411539 | 0.200114507 | 0.848144231 | 0.972174574 | -5.099281451 |
| **A_55_P2047639** | Cubn | 0.116756498 | -1.319820243 | 1.553333239 | 13.41364217 | 0.199949992 | 0.848267144 | 0.972217833 | -5.099306104 |
| **A_51_P329966** | D6Wsu163e | -0.015824385 | -0.21060661 | 0.178957841 | 9.074844672 | -0.199869565 | 0.848327235 | 0.972217833 | -5.099318149 |
| **A_52_P599317** | Hs6st2 | -0.009998712 | -0.133097349 | 0.113099925 | 8.059384176 | -0.199829674 | 0.848357039 | 0.972217833 | -5.099324121 |
| **A_66_P111285** | Pdp2 | -0.010414977 | -0.138660885 | 0.11783093 | 8.133319744 | -0.199794708 | 0.848383165 | 0.972217833 | -5.099329355 |
| **A_51_P194004** | Glyctk | 0.010594578 | -0.11999636 | 0.141185517 | 7.807559377 | 0.199590463 | 0.848535773 | 0.972270065 | -5.099359911 |
| **A_55_P2169586** | Ccdc80 | 0.033405698 | -0.378506699 | 0.445318095 | 9.179477441 | 0.199519269 | 0.84858897 | 0.972270065 | -5.099370554 |
| **A_55_P2100963** | Hmces | -0.021554667 | -0.287586977 | 0.244477644 | 12.07732922 | -0.199331569 | 0.848729225 | 0.972270065 | -5.099398598 |
| **A_55_P2086835** | NA | 0.044191168 | -0.501286912 | 0.589669247 | 14.20560088 | 0.199309195 | 0.848745944 | 0.972270065 | -5.099401939 |
| **A_51_P232142** | Thumpd3 | -0.016379514 | -0.218627803 | 0.185868775 | 11.14281034 | -0.199244029 | 0.84879464 | 0.972270065 | -5.099411668 |
| **A_55_P2236138** | 4930447F24Rik | -0.048167309 | -0.642983696 | 0.546649077 | 8.342758449 | -0.199222611 | 0.848810645 | 0.972270065 | -5.099414865 |
| **A_55_P2054511** | NA | -0.017181668 | -0.229369918 | 0.195006582 | 9.75720983 | -0.199210929 | 0.848819374 | 0.972270065 | -5.099416609 |
| **A_55_P2011126** | Mfap1a | 0.01033834 | -0.117347041 | 0.138023721 | 8.815255423 | 0.199195167 | 0.848831153 | 0.972270065 | -5.099418961 |
| **A_51_P361448** | Scara5 | -0.009482108 | -0.126681416 | 0.1077172 | 8.141444741 | -0.199043968 | 0.848944141 | 0.972341864 | -5.099441517 |
| **A_55_P1996421** | NA | -0.032153411 | -0.430375545 | 0.366068723 | 12.87631077 | -0.19864188 | 0.849244634 | 0.972549222 | -5.099501419 |
| **A_55_P2152035** | Tcf19 | 0.008916213 | -0.101520419 | 0.119352845 | 8.211101796 | 0.19862616 | 0.849256383 | 0.972549222 | -5.099503759 |
| **A_51_P279100** | Ptgs1 | 0.034830949 | -0.397103075 | 0.466764972 | 11.20566745 | 0.198388754 | 0.849433817 | 0.972549222 | -5.099539068 |
| **A_52_P618132** | Manea | -0.009816149 | -0.131559764 | 0.111927465 | 8.48291054 | -0.198364583 | 0.849451883 | 0.972549222 | -5.099542661 |
| **A_51_P110471** | Ddah1 | -0.012503881 | -0.167622632 | 0.14261487 | 8.444998139 | -0.19831232 | 0.849490945 | 0.972549222 | -5.060061548 |
| **A_51_P314277** | Parp1 | 0.014450507 | -0.164879443 | 0.193780457 | 8.580064514 | 0.198243711 | 0.849542225 | 0.972549222 | -5.09956062 |
| **A_51_P317272** | Bod1 | 0.022517033 | -0.256989536 | 0.302023602 | 12.36981053 | 0.198192997 | 0.849580131 | 0.972549222 | -5.099568152 |
| **A_55_P2157068** | Cdhr2 | 0.070901623 | -0.80926119 | 0.951064436 | 11.10749978 | 0.198181141 | 0.849588993 | 0.972549222 | -5.099569913 |
| **A_51_P460048** | Cnrip1 | 0.026634704 | -0.304021811 | 0.357291218 | 8.867653038 | 0.198170931 | 0.849596624 | 0.972549222 | -5.099571429 |
| **A_55_P1984173** | Pde4dip | -0.028737713 | -0.385676339 | 0.328200913 | 11.78260818 | -0.198074155 | 0.84966896 | 0.972549222 | -5.099585795 |
| **A_52_P282500** | Kif21b | -0.074043959 | -0.993916658 | 0.84582874 | 9.96839053 | -0.198030035 | 0.849701938 | 0.972549222 | -5.099592342 |
| **A_55_P1956308** | NA | 0.014350141 | -0.164051146 | 0.192751427 | 13.15261365 | 0.197891589 | 0.849805424 | 0.972549222 | -5.099612877 |
| **A_55_P1966883** | Rnaseh2a | -0.010074115 | -0.1353264 | 0.11517817 | 9.24089185 | -0.19787477 | 0.849817996 | 0.972549222 | -5.099615371 |
| **A_55_P2120161** | Myh8 | 0.008801528 | -0.100637167 | 0.118240224 | 7.659690231 | 0.197859246 | 0.849829601 | 0.972549222 | -5.099617673 |
| **A_52_P150565** | Usp39 | 0.02287587 | -0.261679683 | 0.307431422 | 9.185451498 | 0.197778789 | 0.849889743 | 0.972560466 | -5.099629598 |
| **A_51_P183051** | Upb1 | -0.019835934 | -0.266805624 | 0.227133757 | 7.966878875 | -0.197596001 | 0.850026382 | 0.972659244 | -5.099656674 |
| **A_55_P2028474** | Adam22 | 0.009331187 | -0.107003345 | 0.125665719 | 7.765583724 | 0.197331956 | 0.850223774 | 0.972672204 | -5.099695743 |
| **A_52_P641216** | Galk2 | -0.018845464 | -0.253801218 | 0.216110291 | 11.09141908 | -0.19732854 | 0.850226328 | 0.972672204 | -5.099696248 |
| **A_55_P2011862** | Mybpc3 | 0.02012701 | -0.23081865 | 0.27107267 | 8.131195013 | 0.197318922 | 0.850233518 | 0.972672204 | -5.09969767 |
| **A_55_P2121744** | Vars2 | 0.009094997 | -0.104523019 | 0.122713013 | 7.914503862 | 0.196935732 | 0.850520001 | 0.972672204 | -5.099754273 |
| **A_55_P1963585** | Mbd5 | 0.014451811 | -0.166137407 | 0.195041029 | 8.410631589 | 0.196879099 | 0.850562343 | 0.972672204 | -5.09976263 |
| **A_55_P1959703** | Zfp646 | -0.035440462 | -0.478453883 | 0.40757296 | 15.43642335 | -0.196812034 | 0.850612486 | 0.972672204 | -5.099772522 |

| | | | | | | | | | |
|---|---|---|---|---|---|---|---|---|---|
| **A_55_P2093649** | NA | -0.023799343 | -0.32130203 | 0.273703345 | 15.22219012 | -0.19680822 | 0.850615337 | 0.972672204 | -5.099773085 |
| **A_51_P390715** | Tgfb1 | -0.022308111 | -0.301186176 | 0.256569953 | 8.489578075 | -0.196796608 | 0.85062402 | 0.972672204 | -5.099774797 |
| **A_51_P278018** | Vps36 | -0.016599339 | -0.224155636 | 0.190956957 | 11.30834233 | -0.196754216 | 0.850655716 | 0.972672204 | -5.099781048 |
| **A_66_P134453** | I830077J02Rik | 0.009025709 | -0.103859645 | 0.121911063 | 7.935288984 | 0.196703866 | 0.850693362 | 0.972672204 | -5.099788471 |
| **A_66_P135194** | Yrdc | -0.013059544 | -0.17640453 | 0.150285442 | 8.692157643 | -0.19669412 | 0.850700649 | 0.972672204 | -5.099789908 |
| **A_55_P2163774** | Crip1 | 0.028463569 | -0.327562759 | 0.384489897 | 16.01630597 | 0.196687337 | 0.850705721 | 0.972672204 | -5.099790907 |
| **A_55_P2004741** | Cyb5rl | 0.009581514 | -0.11028299 | 0.129446017 | 8.378767172 | 0.196658487 | 0.850727292 | 0.972672204 | -5.099795159 |
| **A_55_P1971991** | Kdf1 | -0.017012347 | -0.229857992 | 0.195833299 | 11.27637561 | -0.196638529 | 0.850742215 | 0.972672204 | -5.0997981 |
| **A_65_P06611** | Wtap | 0.016150919 | -0.186112395 | 0.218414234 | 10.19499743 | 0.196448759 | 0.850884111 | 0.972776896 | -5.099826051 |
| **A_55_P1986526** | Cdh26 | 0.011173807 | -0.128893963 | 0.151241577 | 9.021713463 | 0.196260141 | 0.851025151 | 0.972880598 | -5.099853805 |
| **A_55_P2099363** | Stac2 | -0.09395422 | -1.272819796 | 1.084911356 | 8.899709156 | -0.196074523 | 0.851163955 | 0.972951457 | -5.099881092 |
| **A_66_P118513** | 11-Sep | 0.020400952 | -0.235616211 | 0.276418115 | 11.90385528 | 0.196042626 | 0.851187808 | 0.972951457 | -5.099885779 |
| **A_66_P118124** | Fxyd3 | 0.048860662 | -0.565300867 | 0.663022192 | 11.28052226 | 0.19572482 | 0.851425475 | 0.973033327 | -5.099932433 |
| **A_66_P127118** | Trmt112 | 0.012785336 | -0.147932109 | 0.173502781 | 8.4758528 | 0.195712374 | 0.851434783 | 0.973033327 | -5.099934258 |
| **A_55_P2113912** | NA | 0.020321487 | -0.235155271 | 0.275798245 | 13.55722418 | 0.19569208 | 0.85144996 | 0.973033327 | -5.099937235 |
| **A_55_P2064257** | Dnaic2 | -0.008456256 | -0.11478004 | 0.097867527 | 7.674508231 | -0.195666677 | 0.851468958 | 0.973033327 | -5.09994096 |
| **A_52_P631547** | Cytl1 | 0.009327423 | -0.108056192 | 0.126711037 | 7.781395182 | 0.195489456 | 0.851601501 | 0.973033327 | -5.099966936 |
| **A_51_P501069** | Spta1 | 0.008468681 | -0.098121132 | 0.115058494 | 7.724044855 | 0.195465096 | 0.851619719 | 0.973033327 | -5.099970505 |
| **A_55_P2194064** | NA | -0.009079132 | -0.123357091 | 0.105198827 | 7.748190086 | -0.195456905 | 0.851625846 | 0.973033327 | -5.099971705 |
| **A_51_P252410** | Cope | 0.012301626 | -0.14263499 | 0.167238242 | 11.20467129 | 0.195333896 | 0.851717847 | 0.973033327 | -5.099989718 |
| **A_55_P2065239** | NA | -0.011926667 | -0.162207168 | 0.138353834 | 15.66463815 | -0.195247569 | 0.851782414 | 0.973033327 | -5.100002353 |
| **A_52_P76402** | Ago1 | -0.011269948 | -0.15328828 | 0.130748385 | 8.717259139 | -0.195230042 | 0.851795523 | 0.973033327 | -5.100004918 |
| **A_51_P174961** | F10 | 0.056169983 | -0.651978117 | 0.764318083 | 8.878311008 | 0.195141375 | 0.851861842 | 0.973033327 | -5.100017889 |
| **A_55_P2163669** | NA | 0.013327567 | -0.154718488 | 0.181373622 | 13.99272445 | 0.195115475 | 0.851881215 | 0.973033327 | -5.100021677 |
| **A_51_P208870** | Zdhhc24 | 0.019861229 | -0.23073336 | 0.270455818 | 8.846202396 | 0.194986076 | 0.851978002 | 0.973033327 | -5.100040593 |
| **A_55_P2112105** | NA | -0.014228134 | -0.193816789 | 0.165360522 | 10.27143848 | -0.194911829 | 0.852033539 | 0.973033327 | -5.100051441 |
| **A_55_P2097508** | Mcc | -0.061331496 | -0.836008319 | 0.713345326 | 10.52856403 | -0.194774528 | 0.852136243 | 0.973033327 | -5.100071492 |
| **A_55_P2085574** | Nfix | -0.014816076 | -0.202015612 | 0.172383461 | 8.23966609 | -0.194714179 | 0.852181386 | 0.973033327 | -5.1000803 |
| **A_55_P1991693** | Pprc1 | -0.013264703 | -0.180884099 | 0.154354692 | 8.266077915 | -0.194689459 | 0.852199877 | 0.973033327 | -5.100083908 |
| **A_51_P467629** | Efcab10 | -0.008446447 | -0.115181303 | 0.09828841 | 7.651044465 | -0.19468699 | 0.852201724 | 0.973033327 | -5.100084268 |
| **A_55_P2073557** | Slc27a5 | -0.010345745 | -0.141094121 | 0.120402631 | 7.753863741 | -0.194668035 | 0.852215904 | 0.973033327 | -5.100087034 |
| **A_55_P1993128** | Gas8 | -0.01610156 | -0.219838987 | 0.187635868 | 10.22307704 | -0.194431349 | 0.852392959 | 0.973156771 | -5.100121547 |
| **A_55_P2038273** | NA | 0.020572475 | -0.239833941 | 0.28097889 | 8.957283143 | 0.194358715 | 0.852447297 | 0.973156771 | -5.10013213 |
| **A_51_P217395** | Cyp2d34 | 0.008597588 | -0.100269708 | 0.117464884 | 7.636505739 | 0.194289072 | 0.852499397 | 0.973156771 | -5.100142273 |
| **A_66_P115531** | Crebzf | -0.026785884 | -0.366023888 | 0.31245212 | 10.21506452 | -0.194254303 | 0.852525407 | 0.973156771 | -5.100147336 |
| **A_55_P2083988** | Lmo2 | 0.01351305 | -0.157797713 | 0.184823813 | 8.92407121 | 0.194060831 | 0.852670151 | 0.973264518 | -5.100175492 |
| **A_55_P2093949** | Zfp428 | -0.008573308 | -0.11744796 | 0.100301345 | 7.808306515 | -0.193727286 | 0.852919701 | 0.973416733 | -5.100223967 |
| **A_51_P175567** | Dact1 | -0.017715496 | -0.242788501 | 0.207357509 | 8.120151908 | -0.193641785 | 0.852983673 | 0.973416733 | -5.10023638 |
| **A_55_P2093196** | NA | 0.01018191 | -0.119198073 | 0.139561893 | 16.05856105 | 0.193611578 | 0.853006275 | 0.973416733 | -5.100240765 |
| **A_55_P1981366** | Lamc2 | -0.045163371 | -0.619114436 | 0.528787693 | 12.16536814 | -0.193588989 | 0.853023177 | 0.973416733 | -5.100244043 |
| **A_52_P566390** | Zfyve26 | -0.015923327 | -0.218438549 | 0.186591895 | 10.2300258 | -0.193439561 | 0.853134985 | 0.973416733 | -5.100265718 |
| **A_51_P433584** | Bcas3 | -0.012640818 | -0.173455989 | 0.148174353 | 8.56809179 | -0.193382572 | 0.853177627 | 0.973416733 | -5.10027398 |
| **A_66_P116998** | Tro | -0.0262503 | -0.360538306 | 0.308037705 | 10.73904715 | -0.193189113 | 0.853322389 | 0.973416733 | -5.10030201 |
| **A_52_P529486** | Wdr77 | 0.025686301 | -0.301741482 | 0.353114085 | 9.459922128 | 0.192999067 | 0.853464602 | 0.973416733 | -5.100329518 |
| **A_55_P2171091** | Sap30bp | -0.016317759 | -0.224493486 | 0.191857967 | 9.77034685 | -0.192841099 | 0.853582816 | 0.973416733 | -5.100352363 |
| **A_52_P197666** | Med12 | 0.018304217 | -0.215262721 | 0.251871155 | 10.71211377 | 0.192800858 | 0.85361293 | 0.973416733 | -5.100358179 |
| **A_51_P236829** | Zswim7 | -0.011877551 | -0.163486254 | 0.139731153 | 10.75419143 | -0.192740027 | 0.853658454 | 0.973416733 | -5.10036697 |
| **A_55_P2139027** | Plec | 0.023224831 | -0.273225002 | 0.319674665 | 12.45151714 | 0.192739409 | 0.853658916 | 0.973416733 | -5.100367059 |
| **A_66_P128909** | Numbl | -0.042256508 | -0.58163911 | 0.497126094 | 10.1432859 | -0.192737328 | 0.853660474 | 0.973416733 | -5.10036736 |
| **A_51_P105068** | Lypd6b | -0.008680915 | -0.119585876 | 0.102224046 | 7.763252371 | -0.192567809 | 0.853787339 | 0.973416733 | -5.094971915 |

| | | | | | | | | | |
|---|---|---|---|---|---|---|---|---|---|
| **A_55_P2378827** | NA | -0.036078934 | -0.497684186 | 0.425526318 | 8.595448995 | -0.192287973 | 0.853996774 | 0.973416733 | -5.100432209 |
| **A_55_P2006906** | Armcx3 | -0.01371635 | -0.189341658 | 0.161908958 | 8.550429011 | -0.192141256 | 0.854106586 | 0.973416733 | -5.100453351 |
| **A_51_P371675** | Slc7a6os | -0.015918701 | -0.219876484 | 0.188039082 | 9.645297326 | -0.192015594 | 0.854200642 | 0.973416733 | -5.100471446 |
| **A_55_P2076782** | 1700028P14Rik | -0.018761133 | -0.259145854 | 0.221623588 | 8.29402566 | -0.1920089 | 0.854205652 | 0.973416733 | -5.100472409 |
| **A_51_P520106** | Adprhl2 | 0.014542705 | -0.171823082 | 0.200908493 | 9.236211791 | 0.191976548 | 0.854229867 | 0.973416733 | -5.100477066 |
| **A_66_P124732** | Zfp606 | -0.00807728 | -0.111637845 | 0.095483284 | 8.085239297 | -0.191884504 | 0.854298763 | 0.973416733 | -5.10049031 |
| **A_55_P2116924** | Dbi | -0.019602748 | -0.271192314 | 0.231986817 | 15.77001189 | -0.191687372 | 0.854446322 | 0.973416733 | -5.100518653 |
| **A_55_P2017710** | Igsf9 | -0.045550288 | -0.69951248 | 0.608411903 | 11.76495047 | -0.195993198 | 0.85447634 | 0.973416733 | -4.979930846 |
| **A_55_P2077343** | Ahdc1 | -0.01854752 | -0.256772873 | 0.219677834 | 10.0003381 | -0.191543326 | 0.854554149 | 0.973416733 | -5.100539347 |
| **A_55_P2090025** | Mest | -0.030291017 | -0.419454126 | 0.358872093 | 11.25618168 | -0.191492315 | 0.854592334 | 0.973416733 | -5.100546671 |
| **A_51_P195034** | Esco2 | -0.009337209 | -0.129443366 | 0.110768948 | 7.71303721 | -0.191258599 | 0.854767294 | 0.973416733 | -5.100580204 |
| **A_51_P298486** | Cog6 | -0.011729538 | -0.162645984 | 0.139186908 | 9.924120302 | -0.191211277 | 0.85480272 | 0.973416733 | -5.100586989 |
| **A_55_P2171463** | Nptn | -0.010919983 | -0.151428943 | 0.129588978 | 13.27295797 | -0.191199647 | 0.854811427 | 0.973416733 | -5.100588656 |
| **A_51_P404463** | Ecrg4 | -0.025232703 | -0.350029144 | 0.299563738 | 9.10153409 | -0.191126839 | 0.854865933 | 0.973416733 | -5.100599091 |
| **A_66_P135987** | Ocrl | 0.012222202 | -0.145116506 | 0.169560909 | 9.99291213 | 0.191109831 | 0.854878666 | 0.973416733 | -5.100601528 |
| **A_51_P276599** | AU040320 | 0.013921947 | -0.165323438 | 0.193167333 | 11.16688513 | 0.191082612 | 0.854899043 | 0.973416733 | -5.100605428 |
| **A_55_P1977318** | Wbp11 | 0.017357078 | -0.206155931 | 0.240870087 | 11.77302206 | 0.191048211 | 0.854924798 | 0.973416733 | -5.100610356 |
| **A_52_P471502** | Dcaf13 | 0.011045686 | -0.131210358 | 0.153301731 | 9.604768244 | 0.191025413 | 0.854941865 | 0.973416733 | -5.100613621 |
| **A_55_P1952434** | Tm2d1 | -0.019144165 | -0.265714036 | 0.227425706 | 11.45536655 | -0.191014169 | 0.854950283 | 0.973416733 | -5.100615231 |
| **A_55_P1988313** | NA | -0.018397719 | -0.255364166 | 0.218568728 | 8.448507222 | -0.191005682 | 0.854956637 | 0.973416733 | -5.100616447 |
| **A_51_P515965** | Nfe2 | -0.023492729 | -0.326113227 | 0.279127769 | 8.938499796 | -0.190987215 | 0.854970463 | 0.973416733 | -5.100619091 |
| **A_51_P149775** | Spcs1 | 0.012601322 | -0.157100052 | 0.182302697 | 14.23736753 | 0.192433572 | 0.855170016 | 0.973416733 | -5.032004197 |
| **A_55_P1986680** | NA | 0.018344323 | -0.218296831 | 0.254985477 | 8.568636763 | 0.190713125 | 0.855175669 | 0.973416733 | -5.100658311 |
| **A_52_P621617** | Dnajb4 | 0.012885302 | -0.153398667 | 0.179169271 | 8.168097496 | 0.190639716 | 0.855230632 | 0.973416733 | -5.100668806 |
| **A_51_P370380** | Rdh14 | -0.014777206 | -0.205584081 | 0.176029669 | 10.1152722 | -0.190531799 | 0.855311432 | 0.973416733 | -5.100684227 |
| **A_51_P162116** | Mblac1 | -0.011743536 | -0.163391622 | 0.13990455 | 8.365033567 | -0.190515844 | 0.855323379 | 0.973416733 | -5.100686506 |
| **A_55_P2122997** | Tet1 | -0.010502576 | -0.146170148 | 0.125164996 | 7.921069202 | -0.190453494 | 0.855370063 | 0.973416733 | -5.100695411 |
| **A_52_P232580** | Dyrk3 | -0.01302974 | -0.181365578 | 0.155306097 | 8.470732067 | -0.190426916 | 0.855389964 | 0.973416733 | -5.100699206 |
| **A_52_P415155** | Wnt6 | -0.017872513 | -0.248791938 | 0.213046912 | 8.553043222 | -0.190412002 | 0.85540113 | 0.973416733 | -5.100701335 |
| **A_52_P595225** | Vps35 | -0.012125273 | -0.168813444 | 0.144562899 | 9.049818798 | -0.190381377 | 0.855424061 | 0.973416733 | -5.100705707 |
| **A_55_P2029630** | Mpdu1 | 0.015799953 | -0.188468918 | 0.220068824 | 9.39919708 | 0.190292982 | 0.855490249 | 0.973416733 | -5.100718323 |
| **A_55_P2009285** | Fbxo25 | 0.017739804 | -0.211624755 | 0.247104363 | 10.75749654 | 0.190279354 | 0.855500453 | 0.973416733 | -5.100720267 |
| **A_52_P533304** | Eif5 | -0.01442843 | -0.201147478 | 0.172290618 | 13.28337909 | -0.190107656 | 0.85562902 | 0.973416733 | -5.100744752 |
| **A_55_P2003541** | Nrcam | 0.046130922 | -0.57716662 | 0.669428464 | 10.17046561 | 0.191799206 | 0.855640459 | 0.973416733 | -5.032090458 |
| **A_55_P2064043** | Cd44 | 0.029547101 | -0.352893569 | 0.411987771 | 9.284557567 | 0.190072798 | 0.855655122 | 0.973416733 | -5.100749721 |
| **A_55_P2020054** | 2610035D17Rik | 0.009828841 | -0.117442511 | 0.137100194 | 8.403282948 | 0.189994415 | 0.855713817 | 0.973416733 | -5.100760889 |
| **A_55_P2044592** | NA | -0.023615947 | -0.329423892 | 0.282191999 | 13.34461954 | -0.189987819 | 0.855718756 | 0.973416733 | -5.100761829 |
| **A_55_P2045682** | Rassf5 | 0.027170477 | -0.324691189 | 0.379032142 | 11.00461287 | 0.18997413 | 0.855729007 | 0.973416733 | -5.100763779 |
| **A_55_P2322139** | C530042K13Rik | 0.008382951 | -0.100223713 | 0.116989615 | 7.707617449 | 0.189893293 | 0.855789541 | 0.973416733 | -5.100775291 |
| **A_51_P311905** | Elp6 | -0.012452378 | -0.173830681 | 0.148925926 | 8.050514817 | -0.189835004 | 0.85583319 | 0.973416733 | -5.10078359 |
| **A_55_P2011387** | Tifa | 0.013354908 | -0.159782758 | 0.186492575 | 8.135123645 | 0.189766048 | 0.855884829 | 0.973416733 | -5.100793403 |
| **A_55_P2006499** | Esrrg | 0.00893473 | -0.106926055 | 0.124795515 | 7.674609136 | 0.189720445 | 0.85591898 | 0.973416733 | -5.100799892 |
| **A_51_P193813** | Fga | -0.051283437 | -0.716413587 | 0.613846712 | 8.614998333 | -0.189687912 | 0.855943343 | 0.973416733 | -5.100804519 |
| **A_51_P388691** | Bbx | -0.025709952 | -0.359242469 | 0.307822565 | 9.093672238 | -0.189641004 | 0.855978471 | 0.973416733 | -5.10081119 |
| **A_51_P341349** | Mtap | -0.008568232 | -0.119804005 | 0.10266754 | 8.085784676 | -0.189502925 | 0.856081878 | 0.973416733 | -5.100830818 |
| **A_55_P2007228** | Prkaca | -0.00953652 | -0.13334542 | 0.11427238 | 8.104242111 | -0.189499138 | 0.856084714 | 0.973416733 | -5.100831356 |
| **A_51_P145662** | Clec4g | 0.009189807 | -0.110169307 | 0.128548922 | 7.851331759 | 0.189417447 | 0.856145894 | 0.973416733 | -5.100842962 |
| **A_51_P371051** | Glipr1 | 0.027459632 | -0.329284663 | 0.384203928 | 9.460141193 | 0.189368109 | 0.856182844 | 0.973416733 | -5.100849968 |
| **A_55_P2046348** | Itpripl2 | -0.017139829 | -0.240064923 | 0.205785264 | 12.36511099 | -0.18915451 | 0.856342819 | 0.973416733 | -5.100880282 |
| **A_51_P439452** | Insig2 | -0.026235788 | -0.367521042 | 0.315049467 | 11.25798781 | -0.189123611 | 0.856365962 | 0.973416733 | -5.100884665 |

| | | | | | | | | | |
|---|---|---|---|---|---|---|---|---|---|
| **A_55_P1991054** | Tmf1 | -0.013532139 | -0.189615979 | 0.162551701 | 9.818295708 | -0.189067164 | 0.856408239 | 0.973416733 | -5.100892669 |
| **A_51_P308912** | Mybphl | 0.008226046 | -0.098858546 | 0.115310639 | 7.624921329 | 0.188987603 | 0.856467829 | 0.973416733 | -5.100903946 |
| **A_55_P1980411** | Zkscan1 | -0.020248167 | -0.28402529 | 0.243528956 | 10.83439541 | -0.188850328 | 0.856570649 | 0.973416733 | -5.100923394 |
| **A_52_P233335** | Pphln1 | 0.01469047 | -0.176724592 | 0.206105533 | 10.50437899 | 0.188811634 | 0.856599631 | 0.973416733 | -5.100928873 |
| **A_65_P11425** | Bmp2k | 0.016626499 | -0.200195913 | 0.233448911 | 9.088496037 | 0.188653906 | 0.856717775 | 0.973416733 | -5.100951196 |
| **A_55_P2136556** | Pkn2 | -0.024102826 | -0.338463135 | 0.290257484 | 10.03563748 | -0.188629414 | 0.856736121 | 0.973416733 | -5.100954661 |
| **A_55_P2172274** | Smc2 | -0.027582937 | -0.387489353 | 0.332323479 | 9.027483181 | -0.188547197 | 0.856797706 | 0.973416733 | -5.100966288 |
| **A_55_P2037313** | Gm12027 | -0.016536141 | -0.232341164 | 0.199268883 | 12.40622144 | -0.188513204 | 0.85682317 | 0.973416733 | -5.100971094 |
| **A_55_P2049211** | Pisd-ps1 | 0.026156766 | -0.315288405 | 0.367601937 | 12.61822157 | 0.188465665 | 0.85685878 | 0.973416733 | -5.100977814 |
| **A_66_P113487** | Pacsin2 | -0.040435054 | -0.56844717 | 0.487577062 | 13.8688722 | -0.188401043 | 0.856907187 | 0.973416733 | -5.100986945 |
| **A_52_P194680** | NA | -0.014925222 | -0.209876503 | 0.18002606 | 9.411777628 | -0.188349235 | 0.856945996 | 0.973416733 | -5.100994264 |
| **A_52_P66226** | Rab33a | -0.009341772 | -0.131397588 | 0.112714045 | 7.808903976 | -0.188295494 | 0.856986254 | 0.973416733 | -5.101001854 |
| **A_55_P2165324** | Acsl3 | 0.021632059 | -0.261103149 | 0.304367267 | 10.30424815 | 0.188229258 | 0.857035872 | 0.973416733 | -5.101011205 |
| **A_55_P2006093** | Nebl | -0.008401675 | -0.118227745 | 0.101424394 | 7.640814564 | -0.188204336 | 0.857054542 | 0.973416733 | -5.101014723 |
| **A_52_P329917** | Prrg2 | 0.037179654 | -0.44891579 | 0.523275099 | 11.90765004 | 0.188171116 | 0.857079427 | 0.973416733 | -5.101019411 |
| **A_52_P111963** | 1810030O07Rik | -0.01705617 | -0.240174435 | 0.206062095 | 9.677273515 | -0.188068279 | 0.857156466 | 0.973416733 | -5.101033919 |
| **A_66_P121446** | Gm7969 | -0.038928873 | -0.548206681 | 0.470348936 | 9.437742718 | -0.188055585 | 0.857165976 | 0.973416733 | -5.10103571 |
| **A_52_P323658** | Zfp955a | 0.015472189 | -0.187001891 | 0.217946269 | 8.997547638 | 0.187997247 | 0.85720968 | 0.973416733 | -5.101043936 |
| **A_55_P2019650** | NA | 0.015621939 | -0.188978133 | 0.220222012 | 10.70258167 | 0.187844432 | 0.857324164 | 0.973416733 | -5.101065472 |
| **A_55_P2005335** | Kcng2 | -0.00790663 | -0.111536675 | 0.095723415 | 7.685155633 | -0.187704586 | 0.857428936 | 0.973416733 | -5.101085166 |
| **A_66_P101620** | Cfap97 | -0.016745213 | -0.236323753 | 0.202833327 | 8.715439854 | -0.187616032 | 0.857495281 | 0.973416733 | -5.101097629 |
| **A_55_P1952420** | Odf2 | 0.010710983 | -0.129748027 | 0.151169994 | 8.1596261 | 0.187606942 | 0.857502092 | 0.973416733 | -5.101098908 |
| **A_55_P2005555** | Celf2 | 0.010154058 | -0.12304341 | 0.143351527 | 8.0808607 | 0.187548175 | 0.857546121 | 0.973416733 | -5.101107175 |
| **A_52_P466887** | Hnrnpd | 0.013099159 | -0.158735485 | 0.184933804 | 11.6955424 | 0.187543432 | 0.857549675 | 0.973416733 | -5.101107842 |
| **A_55_P2209046** | NA | 0.008106762 | -0.098320852 | 0.114534376 | 7.913489863 | 0.187396833 | 0.857659514 | 0.973416733 | -5.101128454 |
| **A_55_P2170109** | Bivm | -0.01511429 | -0.213591002 | 0.183362422 | 11.00403735 | -0.187347265 | 0.857696653 | 0.973416733 | -5.101135419 |
| **A_51_P102438** | Ugt2b36 | 0.00817223 | -0.099189474 | 0.115533933 | 7.678668217 | 0.187266602 | 0.857757091 | 0.973416733 | -5.10114675 |
| **A_52_P10732** | Map6 | -0.032295503 | -0.4565815 | 0.391990495 | 9.27834769 | -0.187263231 | 0.857759617 | 0.973416733 | -5.101147224 |
| **A_55_P2123933** | D2hgdh | 0.010772307 | -0.130804345 | 0.15234896 | 8.355638694 | 0.18719156 | 0.857813319 | 0.973416733 | -5.101157288 |
| **A_51_P149325** | Psmd8 | 0.033242431 | -0.403879862 | 0.470364724 | 12.09475038 | 0.187093622 | 0.857886703 | 0.973416733 | -5.101171034 |
| **A_51_P413866** | Cfb | -0.063941282 | -0.904871119 | 0.776988555 | 9.66153871 | -0.187064227 | 0.857908729 | 0.973416733 | -5.101175159 |
| **A_55_P2094613** | NA | -0.008471836 | -0.119912627 | 0.102968955 | 8.100929408 | -0.187026226 | 0.857937204 | 0.973416733 | -5.10118049 |
| **A_55_P2177463** | B130024G19Rik | -0.011294806 | -0.160131152 | 0.137541541 | 7.945552942 | -0.186697683 | 0.858183395 | 0.973416733 | -5.101226535 |
| **A_52_P74427** | Mms19 | 0.011089774 | -0.135049287 | 0.157228835 | 8.383614258 | 0.186691925 | 0.85818771 | 0.973416733 | -5.101227341 |
| **A_55_P2011545** | Pyroxd1 | -0.011364035 | -0.161126779 | 0.13839871 | 8.857550154 | -0.186680055 | 0.858196605 | 0.973416733 | -5.101229003 |
| **A_52_P69109** | Slc10a1 | 0.008215292 | -0.100083115 | 0.116513698 | 7.654755843 | 0.18662511 | 0.858237779 | 0.973416733 | -5.101236695 |
| **A_55_P2234744** | Spice1 | -0.011406236 | -0.161788298 | 0.138975826 | 8.768646354 | -0.18660165 | 0.858255359 | 0.973416733 | -5.101239979 |
| **A_55_P2084332** | Pigp | 0.031600224 | -0.385038177 | 0.448238625 | 9.38254233 | 0.186595013 | 0.858260333 | 0.973416733 | -5.101240908 |
| **A_51_P173285** | Nkx2-5 | -0.007890739 | -0.111937943 | 0.096156464 | 7.695712732 | -0.186576282 | 0.85827437 | 0.973416733 | -5.101243529 |
| **A_55_P2134877** | Eml1 | -0.00790339 | -0.112121246 | 0.096314466 | 7.839369625 | -0.186569407 | 0.858279522 | 0.973416733 | -5.101244491 |
| **A_52_P536481** | Mogs | 0.02930821 | -0.357204081 | 0.415820501 | 13.65623852 | 0.186549955 | 0.858294099 | 0.973416733 | -5.101247213 |
| **A_51_P284793** | Ints12 | -0.016784976 | -0.238168911 | 0.20459896 | 8.637756919 | -0.186527888 | 0.858310636 | 0.973416733 | -5.1012503 |
| **A_52_P678056** | Eva1b | -0.02652819 | -0.376469079 | 0.323412699 | 12.188972 | -0.186501396 | 0.858330489 | 0.973416733 | -5.101254006 |
| **A_55_P1997421** | Csta3 | -0.015857148 | -0.225190164 | 0.193475867 | 8.113137392 | -0.186361641 | 0.858435223 | 0.973416733 | -5.101273548 |
| **A_51_P289987** | Psmg3 | 0.020063511 | -0.244904487 | 0.285031509 | 10.31290235 | 0.186287053 | 0.858491121 | 0.973416733 | -5.101283972 |
| **A_55_P2025523** | Arhgap6 | 0.011333064 | -0.138408034 | 0.161074162 | 8.743480591 | 0.186198206 | 0.858557707 | 0.973416733 | -5.101296384 |
| **A_52_P162486** | Bgn | -0.014960189 | -0.212627304 | 0.182706927 | 8.594672534 | -0.186196625 | 0.858558892 | 0.973416733 | -5.101296604 |
| **A_55_P2017600** | Crmp1 | 0.008198815 | -0.100163768 | 0.116561398 | 7.721597033 | 0.186140512 | 0.858600946 | 0.973416733 | -5.10130444 |
| **A_51_P293989** | Senp6 | 0.008273612 | -0.101099891 | 0.117647115 | 7.86786235 | 0.186102491 | 0.858629441 | 0.973416733 | -5.101309748 |
| **A_55_P1986813** | Ibsp | 0.008063526 | -0.09853685 | 0.114663902 | 7.658809332 | 0.186095312 | 0.858634822 | 0.973416733 | -5.10131075 |

| | | | | | | | | | |
|---|---|---|---|---|---|---|---|---|---|
| **A_51_P513661** | Slc4a8 | -0.024728458 | -0.351695126 | 0.30223821 | 9.232919775 | -0.186064157 | 0.858658172 | 0.973416733 | -5.101315098 |
| **A_55_P1990864** | NA | -0.008294524 | -0.117991113 | 0.101402066 | 7.683003603 | -0.186023361 | 0.858688747 | 0.973416733 | -5.101320791 |
| **A_55_P2019909** | Tmem35b | 0.02213619 | -0.270633516 | 0.314905897 | 10.50083637 | 0.186014117 | 0.858695675 | 0.973416733 | -5.101322081 |
| **A_51_P186053** | Rrp15 | -0.011629888 | -0.165552602 | 0.142292825 | 9.518335786 | -0.185883996 | 0.8587932 | 0.973435392 | -5.101340229 |
| **A_55_P2057441** | NA | 0.011787492 | -0.144246503 | 0.167821488 | 15.78486847 | 0.18585376 | 0.858815861 | 0.973435392 | -5.101344445 |
| **A_55_P1997544** | NA | 0.008116896 | -0.09941081 | 0.115644602 | 7.660098432 | 0.185711478 | 0.858922504 | 0.973435392 | -5.101364272 |
| **A_51_P262208** | Itgb2 | -0.010396079 | -0.148277885 | 0.127485726 | 8.165926126 | -0.18549482 | 0.8590849 | 0.973435392 | -5.101394436 |
| **A_55_P1988228** | Aspm | -0.012499236 | -0.178290279 | 0.153291807 | 7.866385298 | -0.185477648 | 0.859097772 | 0.973435392 | -5.101396825 |
| **A_55_P2067142** | NA | 0.008598176 | -0.105484727 | 0.122681079 | 17.17214158 | 0.185419296 | 0.85914151 | 0.973435392 | -5.101404942 |
| **A_55_P2081123** | Srcin1 | 0.009704266 | -0.119071227 | 0.138479759 | 7.845156017 | 0.185395266 | 0.859159523 | 0.973435392 | -5.101408284 |
| **A_55_P2249055** | NA | 0.020055698 | -0.246169568 | 0.286280965 | 10.32526605 | 0.1853351 | 0.859204623 | 0.973435392 | -5.10141665 |
| **A_55_P1961968** | Plagl1 | -0.038995134 | -0.556806776 | 0.478816507 | 15.59095938 | -0.185271139 | 0.859252567 | 0.973435392 | -5.10142554 |
| **A_55_P2097616** | Stxbp1 | 0.010506747 | -0.129046061 | 0.150059554 | 7.940252165 | 0.185224677 | 0.859287395 | 0.973435392 | -5.101431996 |
| **A_55_P2149511** | Mff | -0.014254855 | -0.203609556 | 0.175099847 | 12.11188968 | -0.185206347 | 0.859301135 | 0.973435392 | -5.101434543 |
| **A_51_P330044** | Cyp2j9 | -0.008398947 | -0.120041243 | 0.103243348 | 7.765455336 | -0.185082456 | 0.859394006 | 0.973435392 | -5.101451749 |
| **A_55_P2186668** | Cfap126 | 0.014191125 | -0.174487188 | 0.202869437 | 8.461188303 | 0.185039307 | 0.859426353 | 0.973435392 | -5.101457739 |
| **A_55_P2010386** | Tspan32 | 0.010216986 | -0.125627903 | 0.146061875 | 7.962408078 | 0.18503277 | 0.859431253 | 0.973435392 | -5.101458647 |
| **A_55_P2088965** | Scarb1 | -0.01109444 | -0.158644562 | 0.136455682 | 8.189926621 | -0.184984342 | 0.859467557 | 0.973435392 | -5.101465368 |
| **A_55_P2178127** | 1-Sep | -0.009749745 | -0.139573437 | 0.120073946 | 8.092084819 | -0.184760224 | 0.85963557 | 0.973479161 | -5.101496448 |
| **A_51_P372992** | Vsir | -0.044289469 | -0.634118714 | 0.545539775 | 10.00689185 | -0.184732512 | 0.859656345 | 0.973479161 | -5.101500289 |
| **A_55_P2039572** | NA | 0.008694193 | -0.107150448 | 0.124538835 | 15.65071703 | 0.184638605 | 0.859726747 | 0.973479161 | -5.101513299 |
| **A_51_P483220** | Impact | 0.014822282 | -0.182685574 | 0.212330137 | 11.12051186 | 0.18462897 | 0.85973397 | 0.973479161 | -5.101514634 |
| **A_51_P368903** | Anapc13 | -0.022959907 | -0.328955312 | 0.283035498 | 13.81124475 | -0.184596892 | 0.859758019 | 0.973479161 | -5.101519076 |
| **A_55_P2336558** | Chn2 | 0.017104936 | -0.210997158 | 0.24520703 | 8.364087794 | 0.184485121 | 0.859841816 | 0.973484013 | -5.101534549 |
| **A_51_P196726** | Itih2 | 0.03746218 | -0.46218965 | 0.537114009 | 8.289016368 | 0.184456822 | 0.859863032 | 0.973484013 | -5.101538465 |
| **A_51_P365399** | Smg5 | -0.009266815 | -0.133020934 | 0.114487303 | 11.19063607 | -0.184221374 | 0.860039559 | 0.973539796 | -5.101571025 |
| **A_55_P2139407** | Myl12b | -0.01500388 | -0.215382589 | 0.185374828 | 13.45206445 | -0.184213384 | 0.86004555 | 0.973539796 | -5.101572129 |
| **A_55_P2087647** | Grhpr | 0.015119809 | -0.18700494 | 0.217244557 | 8.241364706 | 0.184033109 | 0.860180718 | 0.973539796 | -5.10159703 |
| **A_66_P139664** | Ppp2r5c | -0.008327222 | -0.119764989 | 0.103110545 | 8.337130486 | -0.183838682 | 0.860326502 | 0.973539796 | -5.101623859 |
| **A_55_P2023864** | Mfap1a | 0.015630841 | -0.193590604 | 0.224852286 | 10.27741896 | 0.183799919 | 0.860355568 | 0.973539796 | -5.101629204 |
| **A_55_P1990324** | Ptprc | -0.00785396 | -0.112993808 | 0.097285888 | 7.761652826 | -0.183776718 | 0.860372965 | 0.973539796 | -5.101632403 |
| **A_55_P1992582** | Hmgcs2 | 0.007924427 | -0.098179543 | 0.114028396 | 7.672176189 | 0.183740706 | 0.860399968 | 0.973539796 | -5.101637368 |
| **A_55_P2064053** | NA | 0.020517029 | -0.254256957 | 0.295291014 | 10.99890384 | 0.183699517 | 0.860430854 | 0.973539796 | -5.101643045 |
| **A_55_P2119721** | NA | 0.009247935 | -0.114671462 | 0.133167332 | 16.71119673 | 0.183600827 | 0.860504858 | 0.973539796 | -5.101656642 |
| **A_55_P2042356** | Rftn1 | 0.03311437 | -0.410858345 | 0.477087084 | 9.847565609 | 0.183497171 | 0.860582587 | 0.973539796 | -5.101670916 |
| **A_51_P212038** | Atp6v0e2 | 0.032787623 | -0.406915855 | 0.4724911 | 9.394152705 | 0.183450623 | 0.860617493 | 0.973539796 | -5.101677323 |
| **A_55_P2068822** | Cdk20 | -0.020030279 | -0.288736231 | 0.248675674 | 8.795436946 | -0.183391359 | 0.860661935 | 0.973539796 | -5.101685479 |
| **A_55_P1955717** | Hccs | 0.018714807 | -0.232554445 | 0.26998406 | 16.41568712 | 0.183237845 | 0.860777057 | 0.973539796 | -5.101706591 |
| **A_51_P336060** | Lcat | -0.008702824 | -0.125597125 | 0.108191476 | 7.663070906 | -0.183162284 | 0.860833722 | 0.973539796 | -5.101716977 |
| **A_52_P658437** | Espl1 | 0.008459871 | -0.105204078 | 0.122123819 | 7.730335039 | 0.183109197 | 0.860873535 | 0.973539796 | -5.101724271 |
| **A_52_P190405** | Ifnar2 | -0.02817692 | -0.40683788 | 0.350484041 | 11.27735632 | -0.183067877 | 0.860904523 | 0.973539796 | -5.101729947 |
| **A_55_P2101231** | Aldh7a1 | 0.043523854 | -0.541394369 | 0.628442078 | 13.92576308 | 0.183063331 | 0.860907932 | 0.973539796 | -5.101730571 |
| **A_51_P200667** | Clmn | 0.021930634 | -0.272970366 | 0.316831635 | 8.432875511 | 0.182954928 | 0.860989231 | 0.973539796 | -5.101745456 |
| **A_52_P336290** | Rap2a | 0.029690521 | -0.369966402 | 0.429347444 | 9.943769371 | 0.182767781 | 0.861129588 | 0.973539796 | -5.101771131 |
| **A_66_P122877** | NA | 0.010725137 | -0.133644095 | 0.155094369 | 16.06567796 | 0.182766828 | 0.861130303 | 0.973539796 | -5.101771262 |
| **A_51_P141836** | Ube3c | 0.012805543 | -0.159636941 | 0.185248027 | 9.614517804 | 0.182693407 | 0.861185369 | 0.973539796 | -5.101781328 |
| **A_55_P2122789** | Homer1 | -0.009492525 | -0.137360601 | 0.118375551 | 8.013243445 | -0.182637013 | 0.861227666 | 0.973539796 | -5.101789057 |
| **A_66_P131645** | Ahnak2 | -0.046607115 | -0.674985432 | 0.581771202 | 8.677039718 | -0.182473664 | 0.861350183 | 0.973539796 | -5.101811431 |
| **A_55_P2021709** | Oxnad1 | 0.015800139 | -0.197237076 | 0.228837353 | 10.30376922 | 0.182462905 | 0.861358253 | 0.973539796 | -5.101812904 |
| **A_55_P2108864** | Fer | 0.023096636 | -0.288443905 | 0.334637177 | 8.792534034 | 0.182390966 | 0.861412212 | 0.973539796 | -5.10182275 |

| | | | | | | | | |
|---|---|---|---|---|---|---|---|---|
| **A_55_P1983768** | Birc5 | 0.012195076 | -0.152343316 | 0.176733469 | 7.870493842 | 0.182341884 | 0.861449026 | 0.973539796 | -5.101829466 |
| **A_51_P433789** | Agr3 | 0.008927061 | -0.111589618 | 0.129443741 | 7.659667198 | 0.182234468 | 0.861529596 | 0.973539796 | -5.101844158 |
| **A_52_P554267** | Ilf3 | -0.011748536 | -0.1705694 | 0.147072327 | 9.153226326 | -0.1819891 | 0.861713649 | 0.973539796 | -5.101877685 |
| **A_51_P313761** | Shmt2 | 0.009897163 | -0.123976073 | 0.143770399 | 9.837796328 | 0.181880496 | 0.861795116 | 0.973539796 | -5.101892511 |
| **A_55_P2064507** | Msra | -0.025805659 | -0.375114427 | 0.323503109 | 9.696219033 | -0.181750089 | 0.861892941 | 0.973539796 | -5.101910301 |
| **A_55_P2335695** | Ndor1 | 0.013178864 | -0.165254696 | 0.191612424 | 8.835682249 | 0.181706556 | 0.861925599 | 0.973539796 | -5.101916237 |
| **A_52_P49378** | Kif1a | 0.010879804 | -0.13644852 | 0.158208128 | 8.215977214 | 0.181678683 | 0.861946508 | 0.973539796 | -5.101920037 |
| **A_51_P319732** | Mrps31 | 0.024283158 | -0.304622884 | 0.353189201 | 10.64215047 | 0.18163623 | 0.861978355 | 0.973539796 | -5.101925824 |
| **A_66_P134149** | Snx30 | -0.012996558 | -0.189052079 | 0.163058962 | 10.53739262 | -0.181613395 | 0.861995486 | 0.973539796 | -5.101928936 |
| **A_51_P181517** | Fcgr4 | -0.013404148 | -0.195020582 | 0.168212286 | 8.938238464 | -0.181573822 | 0.862025173 | 0.973539796 | -5.101934328 |
| **A_51_P158073** | Shfl | 0.017117415 | -0.214856249 | 0.249091078 | 9.596236353 | 0.181538463 | 0.8620517 | 0.973539796 | -5.101939145 |
| **A_51_P333111** | Aox1 | -0.008484389 | -0.12352186 | 0.106553081 | 7.832964851 | -0.181447262 | 0.862120119 | 0.973539796 | -5.101951565 |
| **A_55_P1989539** | Cklf | -0.011800251 | -0.171854611 | 0.148254109 | 7.873599428 | -0.181381459 | 0.862169485 | 0.973539796 | -5.101960523 |
| **A_51_P190845** | Atp6v1b2 | 0.01295057 | -0.162741811 | 0.18864295 | 8.748016718 | 0.1813448 | 0.862196988 | 0.973539796 | -5.101965512 |
| **A_51_P315042** | Avpr1a | -0.008436425 | -0.122906363 | 0.106033514 | 7.663609642 | -0.181316004 | 0.862218591 | 0.973539796 | -5.10196943 |
| **A_55_P2031466** | Lrrc32 | -0.019317827 | -0.281514539 | 0.242878884 | 10.78675246 | -0.181259262 | 0.862261162 | 0.973539796 | -5.101977149 |
| **A_55_P1977473** | Dab2 | -0.051595661 | -0.752301971 | 0.649110649 | 10.78763101 | -0.181153327 | 0.862340639 | 0.973539796 | -5.101991553 |
| **A_55_P2024689** | Rsbn1 | -0.011536829 | -0.168222986 | 0.145149328 | 8.606516917 | -0.181144431 | 0.862347314 | 0.973539796 | -5.101992762 |
| **A_51_P190254** | NA | 0.025901735 | -0.326012513 | 0.377815983 | 9.533683415 | 0.181076113 | 0.86239857 | 0.973539796 | -5.102002047 |
| **A_52_P461343** | Upk3a | 0.008320397 | -0.104747117 | 0.121387912 | 7.767650035 | 0.181040352 | 0.862425401 | 0.973539796 | -5.102006905 |
| **A_55_P2119441** | Vma21 | 0.013700117 | -0.172642 | 0.200042235 | 11.74865266 | 0.180876615 | 0.862548251 | 0.973539796 | -5.102029139 |
| **A_51_P363400** | Prg2 | 0.007688317 | -0.096905603 | 0.112282237 | 7.667601306 | 0.180839805 | 0.862575869 | 0.973539796 | -5.102034135 |
| **A_52_P686136** | Prrc2b | -0.015539692 | -0.226948807 | 0.195869424 | 12.30756263 | -0.180837233 | 0.862577799 | 0.973539796 | -5.102034484 |
| **A_51_P255456** | Cyp1b1 | -0.016882962 | -0.246626302 | 0.212860378 | 7.907963156 | -0.180790194 | 0.862613093 | 0.973539796 | -5.102040866 |
| **A_55_P2132651** | Ccn4 | -0.013768017 | -0.201227962 | 0.173691927 | 7.996850399 | -0.180689152 | 0.862688907 | 0.973539796 | -5.10205457 |
| **A_55_P1976539** | Nup62cl | 0.016801817 | -0.211992789 | 0.245596423 | 10.54522625 | 0.180667336 | 0.862705276 | 0.973539796 | -5.102057528 |
| **A_55_P2079425** | Spred1 | -0.009239003 | -0.135052026 | 0.116574019 | 8.343755701 | -0.180662783 | 0.862708693 | 0.973539796 | -5.102058145 |
| **A_55_P1964477** | Srsf1 | 0.011881917 | -0.149981996 | 0.17374583 | 9.05739218 | 0.180594971 | 0.862759575 | 0.973539796 | -5.102067337 |
| **A_51_P366672** | Slc36a2 | -0.015808562 | -0.231336381 | 0.199719256 | 7.911671251 | -0.180450548 | 0.862867944 | 0.973539796 | -5.102086902 |
| **A_55_P2051859** | Golgb1 | 0.020502671 | -0.259075561 | 0.300080903 | 11.23870396 | 0.180416496 | 0.862893495 | 0.973539796 | -5.102091512 |
| **A_51_P422369** | Odf3b | -0.008787012 | -0.128609327 | 0.111035302 | 7.87157386 | -0.18041502 | 0.862894603 | 0.973539796 | -5.102091712 |
| **A_55_P2122260** | Tmem87b | -0.012341756 | -0.180645085 | 0.155961573 | 9.005249743 | -0.180407008 | 0.862900615 | 0.973539796 | -5.102092797 |
| **A_66_P120472** | Pglyrp2 | -0.008112786 | -0.118761686 | 0.102536115 | 7.650089493 | -0.180381527 | 0.862919736 | 0.973539796 | -5.102096246 |
| **A_52_P128691** | Gpr4 | -0.020908803 | -0.306458998 | 0.264641392 | 8.671948606 | -0.18014237 | 0.863099198 | 0.973539796 | -5.102128598 |
| **A_52_P654043** | Msto1 | 0.017949446 | -0.227229075 | 0.263127967 | 10.78027568 | 0.180109985 | 0.863123501 | 0.973539796 | -5.102132976 |
| **A_55_P2057459** | Mall | 0.120215418 | -1.521942058 | 1.762372893 | 9.859221483 | 0.18010034 | 0.863130739 | 0.973539796 | -5.102134279 |
| **A_55_P2111990** | Tekt1 | 0.032018473 | -0.405367875 | 0.469404822 | 8.229807483 | 0.180096203 | 0.863133843 | 0.973539796 | -5.102134838 |
| **A_66_P113008** | Pla2g2e | -0.127786645 | -1.874527002 | 1.618953712 | 9.600995201 | -0.179980841 | 0.863220415 | 0.973539796 | -5.102150425 |
| **A_52_P32466** | Yif1a | -0.009097758 | -0.133493822 | 0.115298306 | 12.06072165 | -0.179927233 | 0.863260645 | 0.973539796 | -5.102157665 |
| **A_55_P1954277** | Emid1 | 0.016237186 | -0.205858772 | 0.238333144 | 8.994601898 | 0.179861937 | 0.863309646 | 0.973539796 | -5.10216648 |
| **A_55_P2078271** | Rbis | 0.011302635 | -0.14334836 | 0.16595363 | 12.28379003 | 0.179802681 | 0.863354116 | 0.973539796 | -5.102174477 |
| **A_55_P2032936** | Cisd1 | -0.020759874 | -0.304881194 | 0.263361446 | 11.86160569 | -0.179758752 | 0.863387084 | 0.973539796 | -5.102180404 |
| **A_55_P2127357** | NA | 0.026598862 | -0.337460576 | 0.390658301 | 11.3707211 | 0.179746287 | 0.863396439 | 0.973539796 | -5.102182085 |
| **A_55_P2100065** | Fsd1l | -0.015758006 | -0.231468069 | 0.199952057 | 8.430857133 | -0.179721491 | 0.863415048 | 0.973539796 | -5.10218543 |
| **A_55_P2057268** | Magi1 | 0.022595785 | -0.286768686 | 0.331960256 | 9.820723624 | 0.179690933 | 0.863437981 | 0.973539796 | -5.102189551 |
| **A_55_P1961395** | Pdpn | -0.061752833 | -0.907739314 | 0.784233648 | 13.53588361 | -0.179581927 | 0.86351979 | 0.973575246 | -5.102204246 |
| **A_52_P124472** | Kcnd2 | -0.008024573 | -0.118036465 | 0.101987318 | 7.680944816 | -0.179453316 | 0.863616314 | 0.973627281 | -5.102221573 |
| **A_66_P101477** | Bglap3 | -0.028566841 | -0.420454454 | 0.363320772 | 9.309529756 | -0.179336977 | 0.863703631 | 0.97366893 | -5.102237236 |
| **A_66_P132832** | Myzap | 0.009355181 | -0.119304244 | 0.138014606 | 7.815931127 | 0.178887402 | 0.864041073 | 0.973721757 | -5.10229767 |
| **A_55_P2052799** | Cask | -0.02323497 | -0.342785774 | 0.296315834 | 9.020671854 | -0.178883945 | 0.864043667 | 0.973721757 | -5.102298134 |

| | | | | | | | | |
|---|---|---|---|---|---|---|---|---|
| **A_55_P2061094** | Fnta | -0.01850668 | -0.27315994 | 0.236146579 | 8.530270614 | -0.178792148 | 0.864112573 | 0.973721757 | -5.102310456 |
| **A_66_P111666** | Nprl3 | -0.018921383 | -0.279379884 | 0.241537117 | 9.475846888 | -0.17872426 | 0.864163532 | 0.973721757 | -5.102319564 |
| **A_51_P362538** | Syde2 | -0.018048088 | -0.266521003 | 0.230424828 | 9.002504839 | -0.17869866 | 0.864182748 | 0.973721757 | -5.102322997 |
| **A_55_P1998188** | Cdk4 | -0.018222924 | -0.269203981 | 0.232758132 | 11.87774429 | -0.178626667 | 0.86423679 | 0.973721757 | -5.102332651 |
| **A_52_P639774** | Gart | 0.013400246 | -0.171167802 | 0.197968295 | 8.409345095 | 0.178618085 | 0.864243232 | 0.973721757 | -5.102333802 |
| **A_51_P315555** | Nars | -0.026068739 | -0.385408238 | 0.333270761 | 10.51735841 | -0.178477807 | 0.864348534 | 0.973721757 | -5.1023526 |
| **A_51_P161636** | Kng2 | 0.064866482 | -0.830133314 | 0.959866278 | 11.40228168 | 0.178306252 | 0.864477319 | 0.973721757 | -5.10237557 |
| **A_51_P180724** | Mlh1 | -0.011847875 | -0.175331728 | 0.151635979 | 9.543799166 | -0.178293188 | 0.864487126 | 0.973721757 | -5.102377318 |
| **A_55_P2076984** | Farp1 | 0.016889441 | -0.216322848 | 0.250101729 | 8.587623115 | 0.178169353 | 0.864580091 | 0.973721757 | -5.102393884 |
| **A_55_P2029191** | Pde4a | 0.016764151 | -0.214845912 | 0.248374213 | 9.750063645 | 0.178071039 | 0.864653899 | 0.973721757 | -5.102407028 |
| **A_51_P401024** | Lrrc34 | 0.007950286 | -0.101910753 | 0.117811325 | 7.72650262 | 0.178036161 | 0.864680083 | 0.973721757 | -5.102411689 |
| **A_55_P2105242** | Arhgap39 | 0.028678927 | -0.367772163 | 0.425130018 | 11.46205511 | 0.177968218 | 0.864731092 | 0.973721757 | -5.102420766 |
| **A_55_P1995210** | Gm11849 | 0.011564856 | -0.148446117 | 0.17157583 | 16.05089686 | 0.177811417 | 0.864848813 | 0.973721757 | -5.102441702 |
| **A_55_P2127179** | NA | -0.008143339 | -0.12082551 | 0.104538832 | 7.903592296 | -0.177793741 | 0.864862084 | 0.973721757 | -5.102444061 |
| **A_51_P201137** | Lbr | 0.011422159 | -0.146745297 | 0.169589614 | 9.150900005 | 0.177664327 | 0.864959247 | 0.973721757 | -5.102461325 |
| **A_55_P2067262** | Cdv3 | -0.008052372 | -0.119586109 | 0.103481365 | 8.313590298 | -0.177617899 | 0.864994106 | 0.973721757 | -5.102467516 |
| **A_55_P1982494** | H2al1m | 0.007501517 | -0.096417643 | 0.111420677 | 7.765583807 | 0.177591684 | 0.865013788 | 0.973721757 | -5.102471011 |
| **A_55_P2185712** | Copa | -0.028094796 | -0.417457972 | 0.361268379 | 11.9924196 | -0.177517093 | 0.865069793 | 0.973721757 | -5.102480951 |
| **A_52_P821** | Fli1 | -0.010035491 | -0.149131318 | 0.129060335 | 9.034135279 | -0.177498101 | 0.865084053 | 0.973721757 | -5.102483482 |
| **A_51_P509452** | Tmlhe | 0.009552947 | -0.122899366 | 0.142005261 | 7.837523737 | 0.177438151 | 0.865129066 | 0.973721757 | -5.102491468 |
| **A_52_P327467** | Med7 | -0.01461674 | -0.217293223 | 0.188059742 | 11.18141502 | -0.17742562 | 0.865138474 | 0.973721757 | -5.102493137 |
| **A_51_P506309** | Hrg | 0.00773735 | -0.099568672 | 0.115043373 | 7.770569529 | 0.177393344 | 0.865162709 | 0.973721757 | -5.102497435 |
| **A_66_P139568** | Helz | -0.015248991 | -0.226747142 | 0.196249161 | 12.54273598 | -0.177379606 | 0.865173024 | 0.973721757 | -5.102499264 |
| **A_55_P2063163** | Agk | 0.01051071 | -0.135312815 | 0.156334234 | 9.329455823 | 0.177326488 | 0.865212907 | 0.973721757 | -5.102506335 |
| **A_51_P141264** | Zfp24 | 0.01148658 | -0.147909845 | 0.170883004 | 8.212934647 | 0.177288809 | 0.865241199 | 0.973721757 | -5.10251135 |
| **A_51_P112557** | Polr1c | 0.010445132 | -0.134524718 | 0.155414982 | 10.69404003 | 0.177257822 | 0.865264467 | 0.973721757 | -5.102515473 |
| **A_55_P2113673** | Eml1 | 0.025206136 | -0.324845173 | 0.375257445 | 10.79141079 | 0.17715105 | 0.865344639 | 0.973721757 | -5.102529675 |
| **A_66_P116771** | Vps8 | -0.015642621 | -0.233220523 | 0.20193528 | 10.74111242 | -0.176873964 | 0.865552704 | 0.973721757 | -5.102566492 |
| **A_51_P230904** | Copz2 | -0.015165653 | -0.226171523 | 0.195840217 | 9.612965207 | -0.176821772 | 0.865591897 | 0.973721757 | -5.102573421 |
| **A_52_P177341** | Cox19 | -0.019670729 | -0.293386895 | 0.254045437 | 10.54563105 | -0.176802814 | 0.865606133 | 0.973721757 | -5.102575937 |
| **A_55_P2025959** | Chd9 | 0.020060938 | -0.259285472 | 0.299407349 | 9.063160699 | 0.176675899 | 0.86570144 | 0.973721757 | -5.102592775 |
| **A_55_P2275057** | Nrap | -0.007823771 | -0.11678958 | 0.101142037 | 7.664296038 | -0.176642443 | 0.865726564 | 0.973721757 | -5.102597211 |
| **A_55_P1960216** | Cd151 | 0.036050425 | -0.466456695 | 0.538557544 | 10.82653351 | 0.176497001 | 0.865835787 | 0.973721757 | -5.102616489 |
| **A_55_P2205650** | 2610507I01Rik | 0.014153619 | -0.183176583 | 0.211483821 | 8.820210648 | 0.176458711 | 0.865864543 | 0.973721757 | -5.102621561 |
| **A_55_P2053848** | NA | 0.029763625 | -0.385545827 | 0.445073078 | 12.23507525 | 0.17631253 | 0.865974325 | 0.973721757 | -5.102640917 |
| **A_55_P1996818** | NA | 0.017897777 | -0.231929725 | 0.267725279 | 12.32857823 | 0.176249546 | 0.866021627 | 0.973721757 | -5.102649251 |
| **A_51_P141274** | Sec24d | -0.017294378 | -0.258731805 | 0.224143048 | 11.2875985 | -0.176225812 | 0.866039452 | 0.973721757 | -5.102652391 |
| **A_51_P417720** | Itga11 | 0.044309566 | -0.574745789 | 0.663364921 | 9.475078361 | 0.176091007 | 0.866140696 | 0.973721757 | -5.102670218 |
| **A_51_P181400** | Efcab2 | -0.012453488 | -0.186473158 | 0.161566182 | 8.24231468 | -0.17606046 | 0.866163638 | 0.973721757 | -5.102674256 |
| **A_55_P1960311** | Dnase1l1 | -0.013937005 | -0.208752807 | 0.180878798 | 11.88572193 | -0.176000711 | 0.866208513 | 0.973721757 | -5.102682151 |
| **A_55_P2075631** | Slc20a2 | 0.025305606 | -0.328658221 | 0.379269432 | 11.6023158 | 0.175884276 | 0.866295964 | 0.973721757 | -5.10269753 |
| **A_55_P2041539** | Pigx | -0.015507105 | -0.23248639 | 0.20147218 | 12.37075863 | -0.175825401 | 0.866340184 | 0.973721757 | -5.102705302 |
| **A_55_P2036083** | Gm5797 | -0.020253193 | -0.303672863 | 0.263166476 | 9.631154497 | -0.175805592 | 0.866355062 | 0.973721757 | -5.102707917 |
| **A_55_P1955059** | Rpl38 | -0.009609066 | -0.144079619 | 0.124861487 | 16.45470952 | -0.175801732 | 0.866357962 | 0.973721757 | -5.102708426 |
| **A_51_P261999** | Ccdc166 | 0.011223606 | -0.146000473 | 0.168447685 | 8.411913627 | 0.175623454 | 0.866491868 | 0.973721757 | -5.102731943 |
| **A_55_P2027771** | Zfp260 | 0.01212483 | -0.157831946 | 0.182081606 | 11.16426817 | 0.175511806 | 0.866575729 | 0.973721757 | -5.102746658 |
| **A_55_P2102816** | Rpl36a | -0.015956412 | -0.239655102 | 0.207742278 | 13.71294907 | -0.175485389 | 0.866595573 | 0.973721757 | -5.102750139 |
| **A_51_P157214** | Mfsd14a | -0.012568933 | -0.188790221 | 0.163652356 | 10.64085983 | -0.175472554 | 0.866605213 | 0.973721757 | -5.10275183 |
| **A_51_P448814** | Zfp90 | 0.013446047 | -0.175131878 | 0.202023972 | 9.69321516 | 0.175417508 | 0.866646561 | 0.973721757 | -5.10275908 |
| **A_52_P58145** | Aldh1a2 | -0.017796061 | -0.267383685 | 0.231791564 | 8.257844314 | -0.175416318 | 0.866647455 | 0.973721757 | -5.102759236 |

| | | | | | | | | |
|---|---|---|---|---|---|---|---|---|
| **A_51_P303749** | Depdc1b | -0.008000851 | -0.120327464 | 0.104325762 | 7.77359447 | -0.175235736 | 0.866783102 | 0.973721757 | -5.102783005 |
| **A_55_P2022845** | Csnk2a1 | 0.022821986 | -0.298183287 | 0.343827259 | 11.97900245 | 0.174908306 | 0.867029069 | 0.973721757 | -5.102826041 |
| **A_55_P2105512** | Nop56 | -0.016658367 | -0.251128005 | 0.217811272 | 11.2280752 | -0.174789352 | 0.867118432 | 0.973721757 | -5.102841656 |
| **A_51_P378789** | Cxcl13 | 0.007915945 | -0.103538595 | 0.119370485 | 7.7275136 | 0.1747327 | 0.867160992 | 0.973721757 | -5.10284909 |
| **A_55_P2122633** | Airn | -0.058814093 | -0.887013104 | 0.769384919 | 10.83141971 | -0.174709234 | 0.867178621 | 0.973721757 | -5.102852168 |
| **A_55_P2071354** | Togaram2 | 0.007568132 | -0.099051565 | 0.114187829 | 7.646101668 | 0.17463063 | 0.867237673 | 0.973721757 | -5.102862476 |
| **A_55_P2101696** | Gnat2 | -0.011774655 | -0.177681666 | 0.154132356 | 8.659989073 | -0.174603368 | 0.867258154 | 0.973721757 | -5.10286605 |
| **A_55_P2071834** | Tln2 | -0.009039997 | -0.136475396 | 0.118395402 | 8.185716349 | -0.174520873 | 0.867320132 | 0.973721757 | -5.102876861 |
| **A_55_P2042426** | Rpl6 | 0.009016157 | -0.118179831 | 0.136212144 | 16.39000095 | 0.174388243 | 0.867419777 | 0.973721757 | -5.102894233 |
| **A_55_P1988148** | NA | 0.00828458 | -0.108592979 | 0.125162139 | 7.740932771 | 0.174384756 | 0.867422397 | 0.973721757 | -5.10289469 |
| **A_55_P1956667** | Cyfip1 | 0.012178113 | -0.159629714 | 0.18398594 | 10.79718265 | 0.174383832 | 0.867423091 | 0.973721757 | -5.102894811 |
| **A_65_P09090** | Arih2 | -0.015152599 | -0.228974001 | 0.198668803 | 11.3513972 | -0.174343241 | 0.867453588 | 0.973721757 | -5.102900124 |
| **A_55_P2250424** | 9530029O12Rik | -0.055428384 | -0.837814825 | 0.726958057 | 10.11954209 | -0.174293048 | 0.867491298 | 0.973721757 | -5.102906693 |
| **A_55_P2022277** | Nfe2l1 | 0.019144276 | -0.251084565 | 0.289373116 | 8.85428931 | 0.174291581 | 0.867492401 | 0.973721757 | -5.102906885 |
| **A_52_P161400** | Cnpy4 | 0.021909717 | -0.287465102 | 0.331284536 | 9.380233135 | 0.174229209 | 0.867539262 | 0.973721757 | -5.102915046 |
| **A_55_P2106250** | Trappc6a | -0.023831383 | -0.360355394 | 0.312692628 | 12.45230106 | -0.17422174 | 0.867544874 | 0.973721757 | -5.102916023 |
| **A_55_P1984023** | Arrb2 | -0.01059164 | -0.160217067 | 0.139033788 | 8.363418894 | -0.174151395 | 0.867597727 | 0.973721757 | -5.102925223 |
| **A_55_P2133385** | NA | 0.013159307 | -0.172766726 | 0.19908534 | 14.72052453 | 0.174125339 | 0.867617305 | 0.973721757 | -5.102928629 |
| **A_52_P624434** | Clec1a | 0.019067737 | -0.250431276 | 0.28856675 | 8.914319385 | 0.174064874 | 0.867662735 | 0.973721757 | -5.102936533 |
| **A_55_P2062543** | Kifc5b | 0.016551114 | -0.217444104 | 0.250546332 | 8.811468241 | 0.174016091 | 0.867699389 | 0.973721757 | -5.102942907 |
| **A_55_P2086258** | Pan3 | -0.024400679 | -0.614774486 | 0.565973128 | 11.78536185 | -0.190307948 | 0.867712662 | 0.973721757 | -4.822354674 |
| **A_66_P113659** | Ripply1 | -0.042904866 | -0.65003303 | 0.564223297 | 9.486592384 | -0.173858263 | 0.867817977 | 0.973721757 | -5.102963519 |
| **A_55_P2154387** | Bmp4 | -0.031272214 | -0.473814222 | 0.411269794 | 11.11618351 | -0.173849438 | 0.867824608 | 0.973721757 | -5.102964671 |
| **A_52_P576049** | Inhbc | 0.007476314 | -0.098431361 | 0.113383989 | 7.670603758 | 0.173671789 | 0.867958095 | 0.973721757 | -5.102987848 |
| **A_55_P2060991** | Plppr3 | -0.007507462 | -0.113866491 | 0.098851566 | 7.735223274 | -0.173655271 | 0.867970506 | 0.973721757 | -5.102990002 |
| **A_51_P410918** | Tor4a | 0.012450987 | -0.163950924 | 0.188852897 | 9.822025507 | 0.173647946 | 0.867976011 | 0.973721757 | -5.102990957 |
| **A_55_P2005868** | Bbs7 | -0.008867796 | -0.134516067 | 0.116780474 | 7.879486967 | -0.173631435 | 0.867988417 | 0.973721757 | -5.102993109 |
| **A_66_P105689** | Trim34a | -0.011736888 | -0.178052458 | 0.154578682 | 9.745217284 | -0.173615789 | 0.868000174 | 0.973721757 | -5.102995149 |
| **A_55_P1992049** | Gucy1a1 | 0.01494569 | -0.196843541 | 0.226734921 | 10.2803902 | 0.173612611 | 0.868002562 | 0.973721757 | -5.102995563 |
| **A_55_P2097808** | P4ha1 | 0.025150596 | -0.331579563 | 0.381880755 | 10.53646245 | 0.173451326 | 0.868123758 | 0.973721757 | -5.103016578 |
| **A_51_P414305** | Acot12 | -0.007733065 | -0.117465111 | 0.101998981 | 7.652322769 | -0.173375346 | 0.868180854 | 0.973721757 | -5.103026471 |
| **A_52_P316644** | Bet1 | -0.034896454 | -0.530087962 | 0.460295053 | 10.607643 | -0.173371329 | 0.868183872 | 0.973721757 | -5.103026994 |
| **A_51_P211998** | Sgms2 | -0.015722844 | -0.238927452 | 0.207481764 | 8.72832608 | -0.173299421 | 0.868237909 | 0.973721757 | -5.103036353 |
| **A_51_P192964** | Clec12b | 0.027269169 | -0.359871185 | 0.414409523 | 8.401946189 | 0.173289647 | 0.868245254 | 0.973721757 | -5.103037625 |
| **A_55_P2178678** | P2rx5 | -0.057481401 | -0.873682977 | 0.758720176 | 8.790490328 | -0.173260305 | 0.868267304 | 0.973721757 | -5.103041442 |
| **A_51_P518600** | Atp6v1c2 | -0.008368382 | -0.127205747 | 0.110468983 | 7.719382599 | -0.173243781 | 0.868279722 | 0.973721757 | -5.103043592 |
| **A_55_P1996683** | Itih1 | 0.014239222 | -0.188138256 | 0.216616701 | 7.979628042 | 0.173098479 | 0.868388915 | 0.973721757 | -5.103062485 |
| **A_51_P282837** | St14 | -0.016039556 | -0.244081899 | 0.212002786 | 9.411822694 | -0.173039808 | 0.868433007 | 0.973721757 | -5.10307011 |
| **A_52_P46310** | Dcaf7 | -0.009545811 | -0.145328119 | 0.126236497 | 8.517702326 | -0.172957265 | 0.86849504 | 0.973721757 | -5.103080832 |
| **A_55_P2104542** | NA | 0.010929886 | -0.144550322 | 0.166410093 | 8.388027581 | 0.172945662 | 0.86850376 | 0.973721757 | -5.103082339 |
| **A_55_P2053077** | Chuk | 0.0182078 | -0.240849107 | 0.277264707 | 11.51230311 | 0.172914522 | 0.868527162 | 0.973721757 | -5.103086383 |
| **A_52_P137371** | Hmgcr | -0.016489523 | -0.251253181 | 0.218274135 | 8.614192643 | -0.172801051 | 0.86861244 | 0.973721757 | -5.103101111 |
| **A_55_P2035018** | Clec4n | 0.052158841 | -0.690625768 | 0.794943449 | 8.673729613 | 0.17275642 | 0.868645983 | 0.973721757 | -5.103106901 |
| **A_55_P1978696** | NA | 0.008082968 | -0.107046921 | 0.123212856 | 7.699471155 | 0.172723696 | 0.868670577 | 0.973721757 | -5.103111146 |
| **A_55_P1960371** | NA | 0.016399572 | -0.217213703 | 0.250012847 | 15.57124941 | 0.1727047 | 0.868684854 | 0.973721757 | -5.103113609 |
| **A_51_P310896** | Pfkl | -0.016131281 | -0.245933848 | 0.213671285 | 9.124902666 | -0.172696345 | 0.868691133 | 0.973721757 | -5.103114693 |
| **A_55_P2122200** | Prkar2b | -0.008594461 | -0.131095852 | 0.11390693 | 8.061893269 | -0.17260238 | 0.868761755 | 0.973721757 | -5.103126874 |
| **A_55_P2345853** | NA | 0.013983888 | -0.185475364 | 0.21344314 | 11.64281985 | 0.17248166 | 0.868852487 | 0.973721757 | -5.103142515 |
| **A_55_P2055324** | Spint1 | -0.023377547 | -0.35693594 | 0.310180847 | 9.833317901 | -0.172423407 | 0.86889627 | 0.973721757 | -5.103150059 |
| **A_51_P497985** | C2 | -0.042198095 | -0.644685179 | 0.56028899 | 12.00908891 | -0.1723115 | 0.868980381 | 0.973721757 | -5.103164543 |

| | | | | | | | | | |
|---|---|---|---|---|---|---|---|---|---|
| **A_55_P2105286** | Lrrc56 | -0.009640276 | -0.147282441 | 0.128001888 | 9.457088584 | -0.172308682 | 0.868982499 | 0.973721757 | -5.103164908 |
| **A_55_P2093014** | NA | 0.017842061 | -0.236917739 | 0.272601861 | 15.11310223 | 0.172299212 | 0.868989617 | 0.973721757 | -5.103166133 |
| **A_55_P1994492** | Usp31 | -0.017200794 | -0.263192446 | 0.228790857 | 9.805167316 | -0.172027272 | 0.86919402 | 0.973831487 | -5.10320129 |
| **A_52_P578790** | Dok3 | 0.008070946 | -0.107387028 | 0.12352892 | 7.915043953 | 0.171976725 | 0.869232015 | 0.973831487 | -5.103207819 |
| **A_55_P2103796** | Arhgap11a | -0.013164754 | -0.201556044 | 0.175226536 | 7.901768906 | -0.171917896 | 0.869276235 | 0.973831487 | -5.103215415 |
| **A_55_P2128668** | Ccnb1 | 0.007852519 | -0.104530372 | 0.120235409 | 7.74198932 | 0.171900818 | 0.869289072 | 0.973831487 | -5.10321762 |
| **A_51_P292057** | Tbl1xr1 | -0.007949021 | -0.121827342 | 0.1059293 | 9.086031359 | -0.17172825 | 0.869418792 | 0.973912403 | -5.103239885 |
| **A_51_P210340** | 2610306M01Rik | -0.008762538 | -0.13441066 | 0.116885584 | 8.557791055 | -0.171570678 | 0.869537243 | 0.973912403 | -5.103260196 |
| **A_55_P2050390** | Cdh22 | -0.009259483 | -0.142049348 | 0.123530381 | 7.826224185 | -0.171550107 | 0.869552707 | 0.973912403 | -5.103262847 |
| **A_55_P2008066** | Itpr1 | 0.025098953 | -0.334988377 | 0.385186283 | 11.1988029 | 0.171481361 | 0.869604386 | 0.973912403 | -5.103271701 |
| **A_55_P1985850** | Timp1 | -0.036374428 | -0.558263513 | 0.485514658 | 10.01322708 | -0.171469593 | 0.869613232 | 0.973912403 | -5.103273216 |
| **A_51_P467505** | Tec | 0.017105847 | -0.228618862 | 0.262830556 | 9.290151681 | 0.171263542 | 0.869768134 | 0.973993828 | -5.103299734 |
| **A_55_P2218334** | 9430011C21Rik | 0.007693784 | -0.102843023 | 0.118230591 | 7.673322195 | 0.171238824 | 0.869786717 | 0.973993828 | -5.103302912 |
| **A_55_P1962174** | Rab8a | -0.0116589 | -0.179381919 | 0.156064119 | 9.959109477 | -0.171014942 | 0.869955032 | 0.974085094 | -5.103331684 |
| **A_55_P2099560** | Apobec1 | -0.037260618 | -0.573709256 | 0.499188019 | 8.748223325 | -0.170879937 | 0.870056533 | 0.974085094 | -5.103349017 |
| **A_55_P2063992** | Rps5 | -0.008770063 | -0.135067682 | 0.117527556 | 16.74843754 | -0.170834941 | 0.870090363 | 0.974085094 | -5.10335479 |
| **A_55_P1970299** | Mttp | 0.056186951 | -0.752983021 | 0.865356923 | 10.03144905 | 0.170830287 | 0.870093862 | 0.974085094 | -5.103355387 |
| **A_51_P518051** | Dusp4 | 0.036293493 | -0.48751323 | 0.560100216 | 8.856317776 | 0.170461718 | 0.870370979 | 0.974085094 | -5.103402622 |
| **A_55_P2196027** | NA | -0.00780756 | -0.120500438 | 0.104885317 | 7.829838273 | -0.170446484 | 0.870382434 | 0.974085094 | -5.103404572 |
| **A_55_P1990874** | Banp | -0.031967507 | -0.493478094 | 0.42954308 | 10.25793554 | -0.170410468 | 0.870409515 | 0.974085094 | -5.103409182 |
| **A_51_P446315** | Cdc37l1 | 0.01708213 | -0.229719588 | 0.263883848 | 9.782666451 | 0.170279748 | 0.870507805 | 0.974085094 | -5.103425906 |
| **A_52_P376360** | Pdgfc | 0.029274146 | -0.393800949 | 0.452349241 | 8.959863723 | 0.170229939 | 0.870545259 | 0.974085094 | -5.103432275 |
| **A_51_P123805** | Sbno1 | -0.011952643 | -0.184714693 | 0.160809408 | 9.671532836 | -0.170209874 | 0.870560346 | 0.974085094 | -5.10343484 |
| **A_66_P132969** | NA | 0.016136798 | -0.217149471 | 0.249423066 | 14.22990636 | 0.17017562 | 0.870586103 | 0.974085094 | -5.103439219 |
| **A_55_P2033041** | Sirpb1b | 0.012915419 | -0.173822762 | 0.1996536 | 8.075388983 | 0.17015493 | 0.870601661 | 0.974085094 | -5.103441863 |
| **A_51_P145533** | Rnf14 | 0.009543173 | -0.128438224 | 0.147524569 | 13.7392658 | 0.170153703 | 0.870602584 | 0.974085094 | -5.10344202 |
| **A_52_P263201** | Tmem33 | -0.009629161 | -0.148870636 | 0.129612313 | 8.555010786 | -0.170133177 | 0.870618018 | 0.974085094 | -5.103444643 |
| **A_55_P2012979** | Itgb2l | 0.007935315 | -0.106902091 | 0.122772722 | 7.659379654 | 0.170000416 | 0.870717849 | 0.974085094 | -5.1034616 |
| **A_55_P2163009** | Mapkapk5 | -0.015546183 | -0.24059291 | 0.209500544 | 11.79781506 | -0.169949639 | 0.870756032 | 0.974085094 | -5.103468082 |
| **A_51_P116616** | Rbm14 | 0.016129915 | -0.21742137 | 0.2496812 | 11.10016841 | 0.169910016 | 0.870785828 | 0.974085094 | -5.103473139 |
| **A_55_P2093291** | Coil | -0.014802833 | -0.229171282 | 0.199565616 | 9.434633381 | -0.169884258 | 0.870805198 | 0.974085094 | -5.103476426 |
| **A_52_P365005** | Zbtb22 | 0.007972799 | -0.107527831 | 0.123473429 | 8.955457183 | 0.169822664 | 0.870851516 | 0.974085094 | -5.103484284 |
| **A_51_P502225** | Slc22a4 | -0.031310313 | -0.485105991 | 0.422485365 | 11.06212187 | -0.169744697 | 0.870910147 | 0.974085094 | -5.103494226 |
| **A_55_P2102295** | Tmem184c | 0.020169336 | -0.272192207 | 0.31253088 | 9.905703077 | 0.169722933 | 0.870926514 | 0.974085094 | -5.103497 |
| **A_52_P621588** | Ifnlr1 | -0.025954892 | -0.402555384 | 0.350645599 | 10.70108129 | -0.169553794 | 0.871053711 | 0.97416589 | -5.10351855 |
| **A_55_P2050877** | Plekhg2 | -0.021331783 | -0.33098411 | 0.288320545 | 13.27942748 | -0.169481371 | 0.871108176 | 0.97416589 | -5.103527771 |
| **A_55_P2007500** | Tmem151b | 0.007248591 | -0.098031871 | 0.112529052 | 7.670963025 | 0.169385008 | 0.871180647 | 0.97416589 | -5.103540035 |
| **A_51_P484254** | Pcca | -0.017891167 | -0.27782113 | 0.242038795 | 9.051910179 | -0.169336858 | 0.871216858 | 0.97416589 | -5.10354616 |
| **A_51_P338998** | Prim2 | 0.017876993 | -0.24193126 | 0.277685245 | 9.632249794 | 0.169281961 | 0.871258145 | 0.97416589 | -5.103553141 |
| **A_55_P2161219** | Thsd7a | -0.074901135 | -1.164809657 | 1.015007387 | 8.952330819 | -0.169070374 | 0.871417278 | 0.97416589 | -5.103580027 |
| **A_52_P185343** | Gna13 | -0.011720216 | -0.182303303 | 0.158862872 | 9.877705205 | -0.169031945 | 0.87144618 | 0.97416589 | -5.103584906 |
| **A_55_P1985749** | Rad23a | 0.007430789 | -0.100732515 | 0.115594092 | 7.838974962 | 0.169014568 | 0.871459251 | 0.97416589 | -5.103587112 |
| **A_55_P2148171** | Ncmap | -0.024009549 | -0.373608316 | 0.325589218 | 10.11496285 | -0.168959754 | 0.871500477 | 0.97416589 | -5.10359407 |
| **A_51_P342871** | S100a10 | 0.02449175 | -0.332134625 | 0.381118125 | 13.93170273 | 0.168956738 | 0.871502745 | 0.97416589 | -5.103594452 |
| **A_51_P394585** | Snx22 | -0.007155531 | -0.111474782 | 0.09716372 | 7.918823246 | -0.168751089 | 0.871657422 | 0.974282445 | -5.103620535 |
| **A_55_P2003053** | Dct | -0.011513223 | -0.179479299 | 0.156452854 | 7.973767128 | -0.168633744 | 0.871745685 | 0.974283563 | -5.103635404 |
| **A_55_P2144126** | NA | -0.018737584 | -0.292315286 | 0.254840118 | 16.08628909 | -0.168500838 | 0.871845655 | 0.974283563 | -5.103652232 |
| **A_55_P2044207** | Bphl | -0.008959942 | -0.139804569 | 0.121884686 | 7.805247497 | -0.1684684 | 0.871870055 | 0.974283563 | -5.103656337 |
| **A_55_P2063257** | Lgals4 | -0.012430848 | -0.193985801 | 0.169124105 | 8.48401902 | -0.16844643 | 0.87188658 | 0.974283563 | -5.103659117 |
| **A_52_P645632** | Senp8 | 0.011904725 | -0.162002518 | 0.185811969 | 8.126118045 | 0.168411164 | 0.871913108 | 0.974283563 | -5.103663579 |

| | | | | | | | | |
|---|---|---|---|---|---|---|---|---|
| **A_51_P343350** | Amn | 0.086079888 | -1.172655632 | 1.344815409 | 10.33090549 | 0.168242755 | 0.872039788 | 0.974283563 | -5.103684872 |
| **A_51_P286460** | H2ac8 | 0.007716558 | -0.105163024 | 0.120596141 | 7.908707467 | 0.16818119 | 0.872086099 | 0.974283563 | -5.103692651 |
| **A_51_P308254** | Surf2 | -0.007815677 | -0.122164517 | 0.106533163 | 8.373608489 | -0.16815277 | 0.872107478 | 0.974283563 | -5.103696241 |
| **A_51_P403273** | Tax1bp1 | -0.031538284 | -0.492982243 | 0.429905675 | 11.3023145 | -0.168146665 | 0.872112071 | 0.974283563 | -5.103697012 |
| **A_52_P509744** | Nup160 | -0.01413394 | -0.221040092 | 0.192772211 | 8.927123606 | -0.168057931 | 0.872178821 | 0.974285472 | -5.103708217 |
| **A_55_P2349148** | 5830408B19Rik | 0.025069336 | -0.34202347 | 0.392162142 | 8.486963927 | 0.168010383 | 0.87221459 | 0.974285472 | -5.103714219 |
| **A_55_P1961129** | Egfem1 | -0.007375605 | -0.115473485 | 0.100722276 | 7.685850891 | -0.167860926 | 0.872327023 | 0.974312393 | -5.103733072 |
| **A_55_P2089035** | Pole | 0.007087594 | -0.096812278 | 0.110987465 | 7.710945535 | 0.167823573 | 0.872355123 | 0.974312393 | -5.103737782 |
| **A_55_P1989384** | Rnf181 | -0.017162392 | -0.268898552 | 0.234573768 | 11.3541643 | -0.16772638 | 0.872428241 | 0.974312393 | -5.103750031 |
| **A_55_P2121372** | Trp53i13 | -0.019894429 | -0.311806945 | 0.272018087 | 13.02712322 | -0.167667128 | 0.872472817 | 0.974312393 | -5.103757495 |
| **A_55_P2058591** | NA | -0.007109897 | -0.111458168 | 0.097238374 | 7.830960308 | -0.167628253 | 0.872502064 | 0.974312393 | -5.103762391 |
| **A_66_P112623** | Psmc6 | -0.012616217 | -0.197955081 | 0.172722647 | 10.73849747 | -0.167467994 | 0.872622632 | 0.974312393 | -5.103782561 |
| **A_51_P511511** | Stk33 | 0.00858068 | -0.117513602 | 0.134674962 | 7.912039936 | 0.16741542 | 0.872662186 | 0.974312393 | -5.103789174 |
| **A_55_P2016049** | Gp6 | -0.007915421 | -0.124268962 | 0.10843812 | 7.758800028 | -0.167364588 | 0.87270043 | 0.974312393 | -5.103795566 |
| **A_55_P2105035** | NA | 0.016533361 | -0.226572101 | 0.259638824 | 11.80686758 | 0.167315279 | 0.872737528 | 0.974312393 | -5.103801765 |
| **A_55_P2229137** | Tmem116 | -0.019848708 | -0.311714945 | 0.272017528 | 8.690966405 | -0.167308327 | 0.872742758 | 0.974312393 | -5.103802638 |
| **A_55_P2075303** | Zfp609 | -0.008031843 | -0.126229422 | 0.110165736 | 8.471316062 | -0.167176718 | 0.872841778 | 0.974321638 | -5.103819173 |
| **A_51_P195483** | Ube2q2 | -0.027942611 | -0.439399566 | 0.383514345 | 11.23895406 | -0.167075103 | 0.872918233 | 0.974321638 | -5.103831931 |
| **A_55_P1974587** | Vangl1 | -0.009113312 | -0.143381303 | 0.125154679 | 8.791892355 | -0.166983248 | 0.872987345 | 0.974321638 | -5.103843456 |
| **A_65_P15245** | Nrp2 | -0.012538994 | -0.197302212 | 0.172224224 | 8.195836914 | -0.166961496 | 0.873003712 | 0.974321638 | -5.103846185 |
| **A_52_P596755** | Dnph1 | 0.007177522 | -0.098600856 | 0.1129559 | 7.823948268 | 0.166934759 | 0.873023829 | 0.974321638 | -5.103849538 |
| **A_52_P183038** | Impdh1 | -0.031078851 | -0.48921007 | 0.427052367 | 10.46391098 | -0.166895348 | 0.873053483 | 0.974321638 | -5.10385448 |
| **A_52_P197627** | Umps | -0.009890031 | -0.155785703 | 0.136005641 | 8.471080236 | -0.166772488 | 0.873145927 | 0.974368549 | -5.103869878 |
| **A_51_P218975** | Kcns3 | -0.007968864 | -0.125622119 | 0.109684391 | 7.757218575 | -0.166633232 | 0.873250712 | 0.974429223 | -5.103887318 |
| **A_51_P240253** | Rrad | 0.007895922 | -0.108827393 | 0.124619236 | 7.73761637 | 0.166423402 | 0.873408605 | 0.974453178 | -5.103913569 |
| **A_55_P2030486** | Srsf2 | -0.007659844 | -0.120900339 | 0.10558065 | 8.915604165 | -0.16641304 | 0.873416403 | 0.974453178 | -5.103914864 |
| **A_52_P357611** | Neu3 | 0.02515373 | -0.347123679 | 0.397431138 | 8.468890213 | 0.166228264 | 0.873555449 | 0.974453178 | -5.103937952 |
| **A_55_P1984043** | Pdcd4 | 0.031423126 | -0.433937661 | 0.496783913 | 12.65492523 | 0.166122615 | 0.873634953 | 0.974453178 | -5.103951142 |
| **A_55_P2000658** | Gm15645 | -0.01663068 | -0.262979384 | 0.229718023 | 10.74005741 | -0.166084422 | 0.873663695 | 0.974453178 | -5.103955908 |
| **A_52_P139747** | Ndufa1 | -0.014344205 | -0.226853801 | 0.198165392 | 14.82869353 | -0.166060784 | 0.873681484 | 0.974453178 | -5.103958857 |
| **A_55_P2035946** | Penk | -0.027986659 | -0.442692231 | 0.386718913 | 8.211804264 | -0.166027621 | 0.873706441 | 0.974453178 | -5.103962994 |
| **A_55_P2154027** | Ptgis | 0.020608403 | -0.284808456 | 0.326025262 | 9.301290428 | 0.166004742 | 0.873723659 | 0.974453178 | -5.103965848 |
| **A_51_P167527** | Lum | -0.12198776 | -1.931404486 | 1.687428965 | 14.66634353 | -0.165861971 | 0.873831104 | 0.974453178 | -5.103983646 |
| **A_51_P358243** | Tstd3 | 0.007916095 | -0.109526267 | 0.125358458 | 8.578599043 | 0.165827057 | 0.87385738 | 0.974453178 | -5.103987996 |
| **A_55_P2046184** | Ccdc90b | 0.017600839 | -0.243601825 | 0.278803503 | 10.39467996 | 0.16577725 | 0.873894864 | 0.974453178 | -5.103994201 |
| **A_55_P2051716** | Tbx1 | 0.009146385 | -0.126663748 | 0.144956519 | 8.004766463 | 0.16568625 | 0.87396335 | 0.974453178 | -5.104005531 |
| **A_52_P80806** | Syncrip | 0.007682397 | -0.106396732 | 0.121761526 | 9.518118666 | 0.165676039 | 0.873971035 | 0.974453178 | -5.104006803 |
| **A_55_P2070769** | Dus1l | 0.007782588 | -0.107790758 | 0.123355934 | 11.27786462 | 0.165666817 | 0.873977975 | 0.974453178 | -5.10400795 |
| **A_55_P2227355** | Ptpro | 0.011355153 | -0.157452239 | 0.180162545 | 8.577472111 | 0.165489596 | 0.874111356 | 0.974545677 | -5.104029997 |
| **A_55_P2022251** | Cdkn2d | -0.011826571 | -0.187888451 | 0.164235309 | 11.43726243 | -0.165258063 | 0.87428562 | 0.974636641 | -5.104058765 |
| **A_51_P458384** | Slc38a2 | 0.030812079 | -0.427936742 | 0.489560901 | 12.94871142 | 0.165240009 | 0.874299208 | 0.974636641 | -5.104061006 |
| **A_55_P1973848** | Sema4b | 0.03266318 | -0.453821992 | 0.519148352 | 10.70883422 | 0.165180212 | 0.874344216 | 0.974636641 | -5.104068428 |
| **A_55_P2083406** | Myh3 | 0.007290993 | -0.101409192 | 0.115991178 | 7.675239036 | 0.165015815 | 0.874467956 | 0.974718362 | -5.104088821 |
| **A_52_P129443** | Wdpcp | 0.010316555 | -0.143950462 | 0.164583571 | 8.17962745 | 0.164524568 | 0.874837736 | 0.974972607 | -5.104149638 |
| **A_66_P133269** | Hesx1 | -0.007476846 | -0.119338738 | 0.104385047 | 7.784836524 | -0.164439225 | 0.874901981 | 0.974972607 | -5.104160185 |
| **A_52_P554451** | 1300017J02Rik | -0.056774865 | -0.906547108 | 0.792997379 | 10.21308344 | -0.164370066 | 0.874954043 | 0.974972607 | -5.104168728 |
| **A_55_P2059019** | Mapk8ip3 | 0.034779032 | -0.485957819 | 0.555515883 | 11.81924946 | 0.164311644 | 0.874998023 | 0.974972607 | -5.104175942 |
| **A_51_P249608** | Trdn | 0.007099697 | -0.09925061 | 0.113450004 | 7.625909662 | 0.164236707 | 0.875054436 | 0.974972607 | -5.104185192 |
| **A_51_P338485** | Aldh6a1 | -0.019324229 | -0.308795785 | 0.270147328 | 10.32099588 | -0.164234893 | 0.875055802 | 0.974972607 | -5.104185416 |
| **A_55_P2056220** | NA | -0.011926193 | -0.190844808 | 0.166992422 | 10.40803253 | -0.163989282 | 0.875240705 | 0.974972607 | -5.104215703 |

| | | | | | | | | | |
|---|---|---|---|---|---|---|---|---|---|
| **A_66_P134090** | Grpel2 | 0.007354544 | -0.103049034 | 0.117758122 | 8.456827827 | 0.163885974 | 0.875318482 | 0.974972607 | -5.104228429 |
| **A_55_P1962699** | NA | 0.046973973 | -0.658677889 | 0.752625836 | 11.05064345 | 0.163770611 | 0.875405336 | 0.974972607 | -5.10424263 |
| **A_51_P446131** | Gipc2 | -0.04294147 | -0.688037793 | 0.602154853 | 9.43428885 | -0.163765157 | 0.875409442 | 0.974972607 | -5.104243301 |
| **A_55_P2292435** | Slc8a1 | -0.00778863 | -0.124802088 | 0.109224829 | 7.64938106 | -0.163754939 | 0.875417135 | 0.974972607 | -5.104244559 |
| **A_55_P2075358** | NA | 0.010390946 | -0.145817324 | 0.166599217 | 11.98147921 | 0.163651593 | 0.875494943 | 0.974972607 | -5.104257271 |
| **A_52_P578768** | Slc38a9 | 0.008086166 | -0.113581371 | 0.129753702 | 9.453247634 | 0.163507275 | 0.875603602 | 0.974972607 | -5.10427501 |
| **A_55_P1992445** | Vdac1 | 0.016941732 | -0.238051115 | 0.27193458 | 14.88842356 | 0.16345529 | 0.875642742 | 0.974972607 | -5.104281396 |
| **A_66_P106065** | Ttc3 | -0.012854006 | -0.206472617 | 0.180764606 | 8.473913659 | -0.163327965 | 0.87573861 | 0.974972607 | -5.104297029 |
| **A_55_P2046262** | Phgdh | 0.028653045 | -0.403086662 | 0.460392752 | 11.29796125 | 0.163274346 | 0.875778983 | 0.974972607 | -5.104303609 |
| **A_51_P472879** | NA | -0.007306024 | -0.117392554 | 0.102780506 | 7.747646459 | -0.16327364 | 0.875779514 | 0.974972607 | -5.104303696 |
| **A_55_P1970159** | Rnf11 | 0.016091433 | -0.226463674 | 0.25864654 | 11.46197697 | 0.163212512 | 0.875825541 | 0.974972607 | -5.104311194 |
| **A_55_P1964332** | NA | -0.021623444 | -0.347744415 | 0.304497526 | 10.49734077 | -0.163123042 | 0.875892909 | 0.974972607 | -5.104322164 |
| **A_55_P1993549** | Npnt | -0.007147364 | -0.115010327 | 0.100715599 | 7.787206119 | -0.163020681 | 0.875969985 | 0.974972607 | -5.104334707 |
| **A_66_P109395** | Rpia | 0.017178122 | -0.242124478 | 0.276480722 | 11.10637267 | 0.16298137 | 0.875999586 | 0.974972607 | -5.104339522 |
| **A_52_P113537** | Xist | 0.057318428 | -0.807931136 | 0.922567991 | 8.119370464 | 0.162975395 | 0.876004085 | 0.974972607 | -5.104340254 |
| **A_52_P209484** | Tmem88 | 0.012240174 | -0.172586341 | 0.197066689 | 10.19218273 | 0.16292678 | 0.876040692 | 0.974972607 | -5.104346207 |
| **A_52_P632423** | NA | -0.007022143 | -0.113099129 | 0.099054842 | 7.675978932 | -0.16286122 | 0.876090059 | 0.974972607 | -5.104354232 |
| **A_55_P2124341** | Lysmd1 | 0.015917124 | -0.224590346 | 0.256424594 | 11.5777837 | 0.16281904 | 0.876121821 | 0.974972607 | -5.104359394 |
| **A_55_P2010121** | Rab27b | -0.024457555 | -0.394420391 | 0.34550528 | 8.173260659 | -0.162638781 | 0.876257562 | 0.974972607 | -5.104381437 |
| **A_55_P2014427** | Il17re | 0.008907097 | -0.125884283 | 0.143698478 | 8.081679922 | 0.162571059 | 0.87630856 | 0.974972607 | -5.104389712 |
| **A_51_P497768** | Gpat2 | -0.014134398 | -0.228152585 | 0.199883789 | 9.695384402 | -0.162478461 | 0.876378292 | 0.974972607 | -5.104401022 |
| **A_55_P1980080** | Rbms2 | -0.013590461 | -0.21946973 | 0.192288808 | 13.05946001 | -0.16240176 | 0.876436054 | 0.974972607 | -5.104410385 |
| **A_52_P408530** | St3gal2 | 0.016725985 | -0.236812293 | 0.270264263 | 9.889136843 | 0.162299554 | 0.876513023 | 0.974972607 | -5.104422854 |
| **A_55_P2002115** | NA | 0.014374591 | -0.203548741 | 0.232297923 | 14.5444901 | 0.162278476 | 0.876528897 | 0.974972607 | -5.104425425 |
| **A_55_P2059025** | U2af1 | -0.017960852 | -0.290266466 | 0.254344762 | 13.08070372 | -0.16227047 | 0.876534926 | 0.974972607 | -5.104426401 |
| **A_55_P2067200** | NA | -0.010838945 | -0.175178539 | 0.15350065 | 8.586283232 | -0.162260925 | 0.876542114 | 0.974972607 | -5.104427565 |
| **A_55_P1981599** | Ccdc125 | 0.007162894 | -0.101442786 | 0.115768573 | 8.083028042 | 0.162257628 | 0.876544597 | 0.974972607 | -5.104427967 |
| **A_55_P1980341** | Romo1 | -0.008772812 | -0.141832673 | 0.12428705 | 13.51289914 | -0.162203752 | 0.876585172 | 0.974972607 | -5.104434536 |
| **A_55_P2168736** | Relb | -0.011647997 | -0.188386575 | 0.165090581 | 8.841399745 | -0.16213959 | 0.876633492 | 0.974972607 | -5.104442355 |
| **A_55_P1959973** | Dcaf12l2 | 0.044915356 | -0.63664619 | 0.726476902 | 11.27650849 | 0.162128338 | 0.876641967 | 0.974972607 | -5.104443726 |
| **A_55_P2137706** | Tmpo | -0.008596363 | -0.139158903 | 0.121966176 | 9.554319836 | -0.161981471 | 0.876752576 | 0.974972607 | -5.104461613 |
| **A_55_P2014259** | Zfp82 | -0.017277426 | -0.279740463 | 0.245185611 | 9.65920622 | -0.161949666 | 0.87677653 | 0.974972607 | -5.104465484 |
| **A_52_P358076** | Ppp2ca | 0.010367342 | -0.14714465 | 0.167879333 | 12.02062912 | 0.161928373 | 0.876792567 | 0.974972607 | -5.104468075 |
| **A_55_P2089080** | Ssbp2 | 0.009663429 | -0.137195542 | 0.1565224 | 9.608765027 | 0.161882516 | 0.876827104 | 0.974972607 | -5.104473655 |
| **A_55_P2104846** | Rsl24d1 | 0.007811138 | -0.110954533 | 0.126576809 | 8.302599839 | 0.161805233 | 0.87688531 | 0.974972607 | -5.104483055 |
| **A_55_P2344598** | E330037I15Rik | -0.009708014 | -0.157379503 | 0.137963474 | 8.481094981 | -0.16173459 | 0.876938516 | 0.974972607 | -5.104491643 |
| **A_52_P472799** | Ecd | 0.010011316 | -0.142373398 | 0.162396029 | 9.792146839 | 0.161628859 | 0.87701815 | 0.974972607 | -5.10450449 |
| **A_55_P1988699** | Cacna2d1 | -0.011360313 | -0.184374344 | 0.161653719 | 8.020070971 | -0.161539271 | 0.877085627 | 0.974972607 | -5.10451537 |
| **A_55_P2159665** | Zfp534 | 0.015766363 | -0.224615448 | 0.256148175 | 10.4028527 | 0.161361192 | 0.877219759 | 0.974972607 | -5.104536977 |
| **A_55_P1961385** | NA | 0.008893295 | -0.126727571 | 0.144514161 | 8.2968632 | 0.161326364 | 0.877245992 | 0.974972607 | -5.1045412 |
| **A_55_P1970715** | NA | 0.034069344 | -0.486432779 | 0.554571467 | 8.639576922 | 0.161031348 | 0.877468214 | 0.974972607 | -5.104576937 |
| **A_52_P476754** | Ubash3b | -0.022379348 | -0.364344985 | 0.319586289 | 10.21012734 | -0.161003057 | 0.877489525 | 0.974972607 | -5.104580361 |
| **A_52_P588881** | Iqgap3 | 0.013055138 | -0.186474872 | 0.212585148 | 8.278514315 | 0.16096906 | 0.877515134 | 0.974972607 | -5.104584474 |
| **A_55_P2079746** | Pls3 | 0.038879952 | -0.555520165 | 0.633280069 | 12.08985983 | 0.160922214 | 0.877550422 | 0.974972607 | -5.104590141 |
| **A_51_P481920** | Ccna2 | -0.015789916 | -0.257251399 | 0.225671568 | 8.26538768 | -0.160879649 | 0.877582486 | 0.974972607 | -5.104595288 |
| **A_51_P252859** | Ccn1 | -0.050213339 | -0.81859299 | 0.718166313 | 11.53441786 | -0.160772737 | 0.877663023 | 0.974972607 | -5.104608211 |
| **A_51_P215097** | Klf16 | 0.009451153 | -0.135201239 | 0.154103545 | 8.824278287 | 0.160741616 | 0.877686466 | 0.974972607 | -5.104611971 |
| **A_55_P2062070** | Pdzd2 | 0.006845344 | -0.097952483 | 0.111643171 | 7.707083709 | 0.160698628 | 0.87771885 | 0.974972607 | -5.104617164 |
| **A_51_P136303** | Cyp4f15 | -0.007249845 | -0.118252269 | 0.103752578 | 7.653803955 | -0.160681339 | 0.877731874 | 0.974972607 | -5.104619252 |
| **A_55_P2096305** | Rc3h1 | 0.011898247 | -0.170298473 | 0.194094967 | 11.49397548 | 0.160661419 | 0.877746881 | 0.974972607 | -5.104621657 |

| | | | | | | | | | |
|---|---|---|---|---|---|---|---|---|---|
| **A_55_P1963724** | NA | 0.008727639 | -0.124919392 | 0.14237467 | 7.866046067 | 0.160659581 | 0.877748265 | 0.974972607 | -5.104621879 |
| **A_52_P407280** | Adamtsl5 | 0.046473975 | -0.665253784 | 0.758201734 | 8.841609749 | 0.160644211 | 0.877759844 | 0.974972607 | -5.104623735 |
| **A_51_P351238** | Plgrkt | 0.057523526 | -0.823441802 | 0.938488855 | 11.79103349 | 0.160640795 | 0.877762417 | 0.974972607 | -5.104624147 |
| **A_51_P465473** | Btf3l4 | -0.009621384 | -0.15698446 | 0.137741691 | 8.626580427 | -0.160626816 | 0.877772948 | 0.974972607 | -5.104625835 |
| **A_51_P154469** | Ccndbp1 | -0.013980343 | -0.228421183 | 0.200460496 | 11.34939051 | -0.160390815 | 0.877950739 | 0.975052945 | -5.104654306 |
| **A_55_P2050273** | Cct6b | -0.044542528 | -0.727801405 | 0.638716349 | 11.52704542 | -0.160383153 | 0.877956511 | 0.975052945 | -5.104655229 |
| **A_55_P1991911** | NA | -0.02464412 | -0.402923333 | 0.353635092 | 13.27890272 | -0.160276558 | 0.878036816 | 0.975052945 | -5.104668075 |
| **A_55_P2128032** | Hsf2 | 0.010430674 | -0.149772176 | 0.170633525 | 9.418767044 | 0.160181116 | 0.878108721 | 0.975052945 | -5.104679569 |
| **A_55_P2111439** | Gpsm1 | -0.007071844 | -0.115717922 | 0.101574234 | 7.78443541 | -0.160135549 | 0.878143051 | 0.975052945 | -5.104685054 |
| **A_55_P2005808** | Abhd18 | 0.027076895 | -0.388927473 | 0.443081262 | 9.423074537 | 0.16012905 | 0.878147947 | 0.975052945 | -5.104685836 |
| **A_52_P325237** | Rgs6 | 0.007435489 | -0.106951751 | 0.121822728 | 7.697529903 | 0.159919379 | 0.878305916 | 0.975172327 | -5.104711055 |
| **A_51_P142515** | Antxr2 | -0.051684436 | -0.848007133 | 0.744638261 | 9.664507182 | -0.159676075 | 0.878489233 | 0.975238422 | -5.104740278 |
| **A_55_P2147186** | A430005L14Rik | 0.016307346 | -0.234997188 | 0.26761188 | 12.15806934 | 0.159643827 | 0.87851353 | 0.975238422 | -5.104744148 |
| **A_52_P78922** | Ythdc2 | -0.007128358 | -0.11710004 | 0.102843323 | 8.017458927 | -0.159469572 | 0.878644828 | 0.975238422 | -5.104765046 |
| **A_51_P140321** | Mocos | -0.024779369 | -0.407066321 | 0.357507583 | 9.590824736 | -0.159466672 | 0.878647013 | 0.975238422 | -5.104765394 |
| **A_55_P2072032** | NA | 0.008543069 | -0.123260209 | 0.140346347 | 16.12680205 | 0.159461877 | 0.878650626 | 0.975238422 | -5.104765969 |
| **A_52_P367745** | Acads | -0.009874534 | -0.162289505 | 0.142540437 | 8.19230035 | -0.159388928 | 0.878705593 | 0.975238422 | -5.104774711 |
| **A_55_P2089804** | Atf7 | 0.00945917 | -0.136724463 | 0.155642803 | 7.950281454 | 0.1591928 | 0.878853379 | 0.975238422 | -5.104798193 |
| **A_55_P2147291** | NA | 0.007792817 | -0.112710817 | 0.128296452 | 15.81825511 | 0.159097558 | 0.878925147 | 0.975238422 | -5.104809587 |
| **A_55_P2139894** | Cwc15 | 0.012421912 | -0.179738153 | 0.204581977 | 10.25408581 | 0.159035644 | 0.878971803 | 0.975238422 | -5.10481699 |
| **A_52_P529360** | Rab11fip5 | 0.024666273 | -0.35704411 | 0.406376656 | 13.23652418 | 0.158978623 | 0.879014771 | 0.975238422 | -5.104823805 |
| **A_55_P1982951** | Dcaf10 | -0.009442847 | -0.15559094 | 0.136705246 | 8.782965695 | -0.158956741 | 0.879031261 | 0.975238422 | -5.10482642 |
| **A_55_P2008071** | Prkcd | -0.016904804 | -0.27861243 | 0.244802821 | 9.552369987 | -0.158914285 | 0.879063254 | 0.975238422 | -5.104831492 |
| **A_52_P644534** | Dhrsx | -0.009936134 | -0.163785016 | 0.143912748 | 8.519718373 | -0.158888428 | 0.879082739 | 0.975238422 | -5.10483458 |
| **A_52_P625277** | Accs | -0.016293791 | -0.268656939 | 0.236069356 | 8.89708757 | -0.158842012 | 0.879117717 | 0.975238422 | -5.104840123 |
| **A_66_P114333** | Tlr12 | -0.020240233 | -0.333738967 | 0.293258501 | 8.803783269 | -0.158835976 | 0.879122266 | 0.975238422 | -5.104840844 |
| **A_51_P165704** | Mcm7 | -0.012603299 | -0.207985565 | 0.182778967 | 9.962135311 | -0.158696835 | 0.879227121 | 0.975298768 | -5.104857449 |
| **A_51_P275953** | Dcps | -0.01417848 | -0.234141996 | 0.205785037 | 9.809359963 | -0.158579913 | 0.879315235 | 0.975340535 | -5.104871392 |
| **A_55_P2119802** | NA | 0.011268929 | -0.163801811 | 0.186339669 | 15.78116914 | 0.158357353 | 0.879482963 | 0.975362205 | -5.104897904 |
| **A_55_P2083929** | Robo2 | 0.006816552 | -0.099198659 | 0.112831764 | 7.700511019 | 0.158185161 | 0.879612737 | 0.975362205 | -5.104918391 |
| **A_51_P315795** | Tubb4a | 0.013467086 | -0.196027241 | 0.222961412 | 9.34954237 | 0.158150482 | 0.879638874 | 0.975362205 | -5.104922514 |
| **A_55_P2079787** | Prss50 | 0.008536774 | -0.124282283 | 0.141355831 | 7.902428335 | 0.158125736 | 0.879657524 | 0.975362205 | -5.104925456 |
| **A_55_P2058757** | Cap1 | 0.009495617 | -0.138393695 | 0.157384928 | 11.79901447 | 0.157963064 | 0.879780128 | 0.975362205 | -5.104944782 |
| **A_55_P2125466** | Ube2l3 | 0.009800474 | -0.142842398 | 0.162443346 | 11.58837666 | 0.157957302 | 0.879784471 | 0.975362205 | -5.104945466 |
| **A_51_P170696** | Haus1 | -0.015788541 | -0.261916998 | 0.230339917 | 10.86788218 | -0.157815379 | 0.87989144 | 0.975362205 | -5.104962311 |
| **A_55_P2082314** | NA | 0.012037698 | -0.175687053 | 0.199762448 | 8.188415431 | 0.157757878 | 0.879934781 | 0.975362205 | -5.104969132 |
| **A_55_P2005315** | NA | 0.015124107 | -0.220815872 | 0.251064085 | 11.36423615 | 0.157702056 | 0.879976855 | 0.975362205 | -5.104975751 |
| **A_55_P1985259** | C2cd3 | -0.00806995 | -0.134048344 | 0.117908444 | 8.026300354 | -0.157595547 | 0.880057136 | 0.975362205 | -5.104988373 |
| **A_52_P59681** | Rida | 0.023722214 | -0.34661489 | 0.394059318 | 9.432998479 | 0.157589456 | 0.880061727 | 0.975362205 | -5.104989095 |
| **A_55_P1985273** | Lmcd1 | 0.07120434 | -1.041864553 | 1.184273233 | 10.33668501 | 0.157381465 | 0.880218504 | 0.975362205 | -5.105013719 |
| **A_55_P2192662** | Lepr | -0.034131942 | -0.568026036 | 0.499762151 | 10.09258335 | -0.15728056 | 0.880294565 | 0.975362205 | -5.105025654 |
| **A_55_P2108579** | Phkg2 | 0.016365601 | -0.239683166 | 0.272414368 | 10.59121295 | 0.157245576 | 0.880320936 | 0.975362205 | -5.10502979 |
| **A_66_P108468** | 2610528A11Rik | 0.048901735 | -0.716271039 | 0.814074509 | 11.07681281 | 0.157229459 | 0.880333085 | 0.975362205 | -5.105031695 |
| **A_51_P406557** | Myorg | 0.021019031 | -0.308230851 | 0.350268913 | 8.614013937 | 0.157056614 | 0.880463378 | 0.975362205 | -5.105052115 |
| **A_66_P136259** | Rbm4b | -0.007452323 | -0.124190507 | 0.109285862 | 8.072371949 | -0.157053592 | 0.880465657 | 0.975362205 | -5.105052471 |
| **A_55_P2041045** | Csnk1g2 | -0.010469416 | -0.174470035 | 0.153531203 | 12.73166277 | -0.157052961 | 0.880466132 | 0.975362205 | -5.105052546 |
| **A_66_P112024** | NA | 0.029707905 | -0.436314098 | 0.495729907 | 11.72599703 | 0.156832029 | 0.88063268 | 0.975362205 | -5.105078613 |
| **A_52_P89717** | NA | -0.00774407 | -0.129239016 | 0.113750876 | 8.574397276 | -0.156812341 | 0.880647523 | 0.975362205 | -5.105080934 |
| **A_55_P2034260** | Ctdsp1 | -0.015871796 | -0.264985443 | 0.233241851 | 12.00881722 | -0.15674645 | 0.880697195 | 0.975362205 | -5.105088701 |
| **A_55_P2152188** | NA | 0.02043456 | -0.300406223 | 0.341275343 | 16.00453658 | 0.1566913 | 0.880738771 | 0.975362205 | -5.105095199 |

| | | | | | | | | | |
|---|---|---|---|---|---|---|---|---|---|
| **A_51_P299216** | Wdr90 | -0.010350209 | -0.172871865 | 0.152171448 | 8.227388924 | -0.156677643 | 0.880749067 | 0.975362205 | -5.105096807 |
| **A_55_P2112797** | NA | 0.011876473 | -0.174611743 | 0.198364689 | 9.140352566 | 0.156677002 | 0.88074955 | 0.975362205 | -5.105096883 |
| **A_51_P304683** | Clpx | 0.018720944 | -0.275362746 | 0.312804634 | 11.32453415 | 0.156612335 | 0.880798302 | 0.975362205 | -5.105104498 |
| **A_55_P1953082** | NA | -0.073346846 | -1.225690603 | 1.078996912 | 8.683674222 | -0.156591633 | 0.880813909 | 0.975362205 | -5.105106936 |
| **A_51_P363556** | Abi2 | 0.008128605 | -0.119622413 | 0.135879624 | 9.232888587 | 0.156538377 | 0.880854058 | 0.975362205 | -5.105113204 |
| **A_51_P406306** | Hecw2 | -0.011353083 | -0.189823713 | 0.167117547 | 8.042284282 | -0.156500677 | 0.88088248 | 0.975362205 | -5.105117641 |
| **A_51_P229886** | Fpgt | -0.014923322 | -0.249523687 | 0.219677044 | 8.893877493 | -0.156496987 | 0.880885262 | 0.975362205 | -5.105118075 |
| **A_51_P494541** | Abhd17b | 0.012067041 | -0.177655499 | 0.20178958 | 10.13869389 | 0.156477181 | 0.880900194 | 0.975362205 | -5.105120405 |
| **A_55_P2033586** | Mmab | -0.008847334 | -0.148005269 | 0.1303106 | 8.080812122 | -0.156413282 | 0.880948368 | 0.975362205 | -5.105127921 |
| **A_55_P2044202** | Vps29 | -0.017123406 | -0.286531334 | 0.252284522 | 11.27279239 | -0.15636838 | 0.880982221 | 0.975362205 | -5.1051332 |
| **A_51_P243122** | Sec14l3 | 0.006697149 | -0.098687077 | 0.112081374 | 7.699383201 | 0.156344815 | 0.880999987 | 0.975362205 | -5.10513597 |
| **A_55_P2128663** | Uggt1 | 0.012591828 | -0.185929039 | 0.211112695 | 8.864733875 | 0.156045692 | 0.88122551 | 0.975556007 | -5.105171097 |
| **A_52_P255849** | NA | 0.009829447 | -0.145261526 | 0.164920421 | 8.932542594 | 0.155923559 | 0.881317594 | 0.9755623 | -5.10518542 |
| **A_55_P1957267** | Rab9 | -0.018889739 | -0.316972664 | 0.279193186 | 11.94022578 | -0.155904272 | 0.881332137 | 0.9755623 | -5.105187681 |
| **A_55_P2110601** | Slc35a2 | -0.017219248 | -0.289178685 | 0.25474019 | 10.82253079 | -0.155768346 | 0.881434625 | 0.975616469 | -5.105203607 |
| **A_51_P336635** | Xpo1 | 0.01906646 | -0.282274785 | 0.320407706 | 10.11668084 | 0.155661301 | 0.881515338 | 0.975616469 | -5.105216139 |
| **A_55_P2007303** | Snrpg | 0.011880249 | -0.175911951 | 0.199672449 | 13.76688392 | 0.155638544 | 0.881532497 | 0.975616469 | -5.105218803 |
| **A_55_P2129000** | Cyyr1 | -0.012721985 | -0.213960476 | 0.188516506 | 8.243173968 | -0.155529578 | 0.881614661 | 0.975644665 | -5.105231549 |
| **A_55_P2122504** | Zpr1 | 0.008699572 | -0.129010246 | 0.14640939 | 11.32375454 | 0.155418291 | 0.881698576 | 0.975644665 | -5.105244558 |
| **A_52_P440962** | Cdc73 | 0.015882785 | -0.235674859 | 0.26744043 | 9.815005096 | 0.155331063 | 0.881764351 | 0.975644665 | -5.105254748 |
| **A_51_P159612** | Hebp2 | 0.012073324 | -0.179208256 | 0.203354903 | 8.547015178 | 0.155282623 | 0.881800878 | 0.975644665 | -5.105260405 |
| **A_51_P279038** | Ppargc1a | 0.012307559 | -0.182732756 | 0.207347874 | 8.095832736 | 0.155244673 | 0.881829495 | 0.975644665 | -5.105264835 |
| **A_51_P432520** | Wrnip1 | -0.009141917 | -0.154074563 | 0.135790729 | 9.988947814 | -0.155181595 | 0.881877061 | 0.975644665 | -5.105272196 |
| **A_55_P2163873** | Uox | 0.007141254 | -0.106146955 | 0.120429462 | 7.682789983 | 0.155081115 | 0.881952832 | 0.975644665 | -5.105283917 |
| **A_55_P2178496** | Fam53b | 0.025395476 | -0.377811666 | 0.428602617 | 9.18646475 | 0.15495204 | 0.882050169 | 0.975644665 | -5.105298961 |
| **A_55_P1969166** | Fxr2 | 0.010156746 | -0.151149598 | 0.17146309 | 12.21314857 | 0.154907455 | 0.882083791 | 0.975644665 | -5.105304155 |
| **A_51_P112237** | Angel1 | 0.008862593 | -0.131961436 | 0.149686622 | 8.490072962 | 0.154829326 | 0.88214271 | 0.975644665 | -5.094162353 |
| **A_55_P2077228** | Cenpl | 0.013515709 | -0.201386999 | 0.228418417 | 8.247643215 | 0.154726996 | 0.882219881 | 0.975644665 | -5.105325162 |
| **A_52_P276525** | Ankrd9 | -0.017293115 | -0.292327274 | 0.257741043 | 8.935300347 | -0.154687695 | 0.88224952 | 0.975644665 | -5.105329733 |
| **A_51_P499854** | Ghr | -0.029504844 | -0.499295627 | 0.440285939 | 10.92191772 | -0.154510499 | 0.882383155 | 0.975644665 | -5.105350332 |
| **A_55_P2002093** | Dph7 | 0.009131418 | -0.136280163 | 0.154542998 | 8.649816284 | 0.154492847 | 0.882396468 | 0.975644665 | -5.105352382 |
| **A_51_P328669** | Ncbp1 | 0.011870338 | -0.177267736 | 0.201008413 | 8.81274586 | 0.154402135 | 0.882464881 | 0.975644665 | -5.105362917 |
| **A_52_P168097** | Atp2a2 | -0.00688462 | -0.116610875 | 0.102841635 | 7.847643144 | -0.154361359 | 0.882495634 | 0.975644665 | -5.105367651 |
| **A_55_P2175190** | Cep250 | -0.01094398 | -0.185404298 | 0.163516339 | 8.36667215 | -0.154329082 | 0.882519977 | 0.975644665 | -5.105371397 |
| **A_51_P484111** | Matn2 | -0.101334145 | -1.717175292 | 1.514507002 | 10.53111657 | -0.154285924 | 0.882552527 | 0.975644665 | -5.105376404 |
| **A_55_P2033660** | Crls1 | -0.015927615 | -0.269958613 | 0.238103382 | 10.40805415 | -0.154252853 | 0.88257747 | 0.975644665 | -5.10538024 |
| **A_55_P2058270** | Gm6507 | 0.018438251 | -0.275715594 | 0.312592095 | 8.397408588 | 0.154210642 | 0.882609306 | 0.975644665 | -5.105385136 |
| **A_52_P583458** | E2f3 | -0.020382471 | -0.345577557 | 0.304812616 | 9.353054016 | -0.154199163 | 0.882617963 | 0.975644665 | -5.105386467 |
| **A_51_P328014** | Cops3 | -0.01477456 | -0.250760424 | 0.221211303 | 10.19109313 | -0.154027316 | 0.882747576 | 0.97565605 | -5.105406381 |
| **A_55_P1956083** | Gpr68 | -0.01328986 | -0.225631484 | 0.199051765 | 7.902441153 | -0.153976495 | 0.882785907 | 0.97565605 | -5.105412266 |
| **A_55_P2024075** | Prkacb | 0.017760313 | -0.266384052 | 0.301904678 | 12.24050917 | 0.153773229 | 0.882939223 | 0.97565605 | -5.105435786 |
| **A_66_P131398** | Ror2 | 0.022210088 | -0.333143907 | 0.377564083 | 8.420610079 | 0.153765265 | 0.88294523 | 0.97565605 | -5.105436706 |
| **A_51_P295896** | 4930452B06Rik | 0.006554476 | -0.098336608 | 0.111445559 | 7.735942051 | 0.153733515 | 0.882969178 | 0.97565605 | -5.105440377 |
| **A_55_P2123484** | Gm11627 | -0.009311259 | -0.158320424 | 0.139697906 | 7.845743457 | -0.153732116 | 0.882970233 | 0.97565605 | -5.105440539 |
| **A_55_P2083967** | Trmt5 | 0.013585447 | -0.20384534 | 0.231016233 | 9.193850819 | 0.153717052 | 0.882981596 | 0.97565605 | -5.10544228 |
| **A_55_P1998596** | Nupl2 | -0.008616928 | -0.146602375 | 0.12936852 | 9.445552243 | -0.153634349 | 0.883043978 | 0.975667509 | -5.105451837 |
| **A_51_P123705** | Dus3l | 0.008897346 | -0.133698433 | 0.151493125 | 9.837377366 | 0.153505161 | 0.883141425 | 0.975667509 | -5.077973373 |
| **A_55_P2040993** | NA | 0.013820766 | -0.207717235 | 0.235358767 | 8.240106992 | 0.15348044 | 0.883160073 | 0.975667509 | -5.105469609 |
| **A_51_P130757** | Acer3 | -0.020720257 | -0.353122593 | 0.311682078 | 8.984816596 | -0.153355812 | 0.883254082 | 0.975667509 | -5.105483987 |
| **A_51_P462448** | Gpx4 | 0.016552522 | -0.249321421 | 0.282426466 | 15.15486193 | 0.153164305 | 0.883398545 | 0.975667509 | -5.105506059 |

| | | | | | | | | | |
|---|---|---|---|---|---|---|---|---|---|
| **A_52_P190973** | Vcl | -0.015029882 | -0.256513971 | 0.226454206 | 11.17984292 | -0.153121517 | 0.883430822 | 0.975667509 | -5.105510986 |
| **A_52_P558368** | Zdhhc1 | 0.011674775 | -0.175934085 | 0.199283634 | 8.435386516 | 0.153096172 | 0.883449942 | 0.975667509 | -5.105513904 |
| **A_55_P1988481** | NA | -0.010246638 | -0.174992667 | 0.154499392 | 8.40603941 | -0.153015561 | 0.883510753 | 0.975667509 | -5.105523182 |
| **A_55_P2111331** | Klhl32 | 0.007261229 | -0.109493181 | 0.124015638 | 7.853118343 | 0.153005126 | 0.883518624 | 0.975667509 | -5.105524383 |
| **A_65_P02958** | Rnf13 | 0.012970792 | -0.195608018 | 0.221549603 | 11.26848249 | 0.15299086 | 0.883529386 | 0.975667509 | -5.105526024 |
| **A_65_P19089** | Esrrg | -0.006970566 | -0.119110078 | 0.105168945 | 7.701884193 | -0.152925029 | 0.883579048 | 0.975667509 | -5.105533597 |
| **A_52_P417825** | Osgin2 | -0.012724213 | -0.217459024 | 0.192010598 | 9.160791941 | -0.152900319 | 0.883597689 | 0.975667509 | -5.105536438 |
| **A_55_P2027698** | Ndufa5 | -0.010964316 | -0.187489125 | 0.165560493 | 13.66083722 | -0.152807604 | 0.883667633 | 0.97568562 | -5.105547095 |
| **A_55_P1993475** | Gm5168 | 0.021682768 | -0.327736947 | 0.371102483 | 8.575347485 | 0.152663945 | 0.883776012 | 0.97568562 | -5.105563596 |
| **A_55_P2042487** | Dpysl3 | -0.035676415 | -0.610678843 | 0.539326014 | 11.20217832 | -0.152644334 | 0.883790807 | 0.97568562 | -5.105565847 |
| **A_51_P424499** | Txn1 | -0.007011638 | -0.120055809 | 0.106032534 | 15.13577784 | -0.152595051 | 0.883827988 | 0.97568562 | -5.105571504 |
| **A_55_P1972541** | Zfp960 | -0.022169224 | -0.379709071 | 0.335370622 | 10.46540129 | -0.15254403 | 0.88386648 | 0.97568562 | -5.105577358 |
| **A_55_P2049999** | Slc9a1 | -0.011634472 | -0.199615857 | 0.176346914 | 11.89863381 | -0.152265314 | 0.88407676 | 0.975811919 | -5.105609302 |
| **A_55_P2047867** | NA | -0.019556724 | -0.335736109 | 0.296622662 | 10.26145244 | -0.152170935 | 0.884147968 | 0.975811919 | -5.105620106 |
| **A_55_P1962224** | Afap1l2 | 0.023768996 | -0.360721896 | 0.408259888 | 12.65157362 | 0.15208765 | 0.884210806 | 0.975811919 | -5.105629635 |
| **A_55_P2009477** | 9530068E07Rik | -0.011121554 | -0.191058431 | 0.168815324 | 13.23353139 | -0.15205981 | 0.884231812 | 0.975811919 | -5.105632819 |
| **A_55_P1974028** | Stab2 | -0.014022243 | -0.240892679 | 0.212848192 | 10.03198258 | -0.152057816 | 0.884233316 | 0.975811919 | -5.105633047 |
| **A_66_P106385** | Cd55b | -0.070583675 | -1.213427982 | 1.072260632 | 10.46179471 | -0.151944982 | 0.884318451 | 0.975828677 | -5.105645945 |
| **A_52_P512575** | Hopx | -0.027419532 | -0.471498669 | 0.416659604 | 10.30291194 | -0.151903869 | 0.884349472 | 0.975828677 | -5.105650643 |
| **A_55_P2002849** | Aifm3 | -0.010828248 | -0.186352144 | 0.164695647 | 10.3566695 | -0.151771815 | 0.884449111 | 0.975872813 | -5.105665722 |
| **A_52_P248483** | Zfx | -0.00970495 | -0.167142673 | 0.147732773 | 9.401479221 | -0.151653928 | 0.884538063 | 0.975872813 | -5.105679173 |
| **A_52_P424840** | Mrc2 | 0.016057365 | -0.244438397 | 0.276553126 | 8.0692433 | 0.151650125 | 0.884540933 | 0.975872813 | -5.105679607 |
| **A_55_P1986466** | Retreg2 | -0.012196551 | -0.210183905 | 0.185790803 | 8.499297707 | -0.151554471 | 0.88461311 | 0.975896741 | -5.105690513 |
| **A_55_P2175356** | Uros | 0.008393472 | -0.128025151 | 0.144812094 | 8.054544124 | 0.151369065 | 0.884753015 | 0.975958825 | -5.105711633 |
| **A_55_P2076057** | Hmga1 | -0.008221759 | -0.141978661 | 0.125535142 | 7.859678107 | -0.151222957 | 0.884863269 | 0.975958825 | -5.105728258 |
| **A_55_P2062213** | Tmem41a | -0.022592502 | -0.390173451 | 0.344988447 | 11.52045939 | -0.151209993 | 0.884873052 | 0.975958825 | -5.105729733 |
| **A_55_P2000728** | Myef2 | 0.008177193 | -0.124912179 | 0.141266565 | 8.179847778 | 0.151157618 | 0.884912575 | 0.975958825 | -5.105735688 |
| **A_55_P2127971** | Gm9112 | 0.032272484 | -0.493577044 | 0.558122012 | 12.23568375 | 0.150987178 | 0.885041196 | 0.975958825 | -5.105755054 |
| **A_51_P418357** | Mov10l1 | -0.006621699 | -0.114566914 | 0.101323516 | 7.629052687 | -0.150915967 | 0.885094935 | 0.975958825 | -5.105763138 |
| **A_66_P118696** | Nup107 | 0.013918944 | -0.213078188 | 0.240916075 | 11.28118416 | 0.150853389 | 0.885142161 | 0.975958825 | -5.10577024 |
| **A_55_P2002903** | Smoc2 | -0.110275339 | -1.908895584 | 1.688344906 | 11.95823132 | -0.15083707 | 0.885154477 | 0.975958825 | -5.105772091 |
| **A_52_P349844** | Obscn | 0.006415964 | -0.098251201 | 0.111083128 | 7.650177961 | 0.150806693 | 0.885177401 | 0.975958825 | -5.105775537 |
| **A_52_P35961** | NA | 0.009643437 | -0.147683297 | 0.166970172 | 8.410192117 | 0.150799012 | 0.885183198 | 0.975958825 | -5.105776408 |
| **A_51_P338961** | Qser1 | -0.012910631 | -0.22370007 | 0.197878808 | 8.885381827 | -0.150684222 | 0.885269828 | 0.975958825 | -5.105789423 |
| **A_55_P2028288** | Nhlrc4 | 0.013023804 | -0.209006924 | 0.235054531 | 8.26297396 | 0.152010938 | 0.88527529 | 0.975958825 | -5.036941368 |
| **A_52_P251623** | Fam222a | -0.007492293 | -0.129947778 | 0.114963192 | 7.901942035 | -0.150523979 | 0.885390763 | 0.975967346 | -5.105807574 |
| **A_65_P01065** | Phf14 | 0.015586805 | -0.239190925 | 0.270364534 | 10.51477654 | 0.150509804 | 0.885401461 | 0.975967346 | -5.105809179 |
| **A_55_P2052696** | Synm | -0.010960193 | -0.190164685 | 0.168244299 | 7.956312043 | -0.150466032 | 0.885434496 | 0.975967346 | -5.105814133 |
| **A_55_P2025283** | NA | -0.01348748 | -0.234354098 | 0.207379137 | 15.49081364 | -0.150234578 | 0.885609183 | 0.976104231 | -5.105840307 |
| **A_51_P372456** | Wdr86 | 0.012414991 | -0.191151524 | 0.215981506 | 9.364110578 | 0.15004075 | 0.885755477 | 0.976209808 | -5.105862195 |
| **A_55_P2065360** | Siva1 | -0.012749377 | -0.222122629 | 0.196623876 | 12.19328521 | -0.149808658 | 0.885930657 | 0.976328425 | -5.105888368 |
| **A_55_P2067741** | Psg23 | -0.014950267 | -0.260539744 | 0.23063921 | 17.91985017 | -0.149764318 | 0.885964126 | 0.976328425 | -5.105893364 |
| **A_51_P317433** | Mesp2 | 0.006603141 | -0.10200058 | 0.115206862 | 7.862746543 | 0.149580512 | 0.886102867 | 0.976410794 | -5.105914056 |
| **A_52_P569549** | Lig4 | 0.008044481 | -0.124308789 | 0.140397752 | 9.036745801 | 0.149531449 | 0.886139901 | 0.976410794 | -5.105919576 |
| **A_55_P1970234** | Dclk2 | -0.015554351 | -0.271610515 | 0.240501813 | 8.167880033 | -0.149446525 | 0.886204006 | 0.976411598 | -5.105929125 |
| **A_55_P2099064** | Nsd2 | 0.009843557 | -0.152255467 | 0.171942581 | 8.08592937 | 0.14939664 | 0.886241662 | 0.976411598 | -5.105934732 |
| **A_51_P450021** | Higd2a | 0.021914674 | -0.339526764 | 0.383356112 | 12.19820296 | 0.149164758 | 0.886416704 | 0.976478653 | -5.105960769 |
| **A_55_P2093173** | NA | -0.015240978 | -0.266657336 | 0.236175379 | 11.34421538 | -0.149138057 | 0.88643686 | 0.976478653 | -5.105963765 |
| **A_52_P465121** | NA | -0.022624184 | -0.396010318 | 0.35076195 | 11.62628067 | -0.14906782 | 0.886489882 | 0.976478653 | -5.105971642 |
| **A_65_P08239** | Mef2a | -0.007027601 | -0.123025247 | 0.108970045 | 8.433075138 | -0.149048324 | 0.8865046 | 0.976478653 | -5.105973828 |

| | | | | | | | | | |
|---|---|---|---|---|---|---|---|---|---|
| **A_55_P2044381** | NA | -0.006389641 | -0.112134139 | 0.099354857 | 9.192423375 | -0.148657844 | 0.886799385 | 0.976674127 | -5.10601755 |
| **A_55_P2002212** | Babam2 | 0.012611315 | -0.196099258 | 0.221321889 | 9.231116316 | 0.148656908 | 0.886800092 | 0.976674127 | -5.106017655 |
| **A_55_P1992074** | Ptpra | -0.016657035 | -0.292499282 | 0.259185212 | 13.96035311 | -0.148561431 | 0.886872173 | 0.976674127 | -5.106028328 |
| **A_55_P1972277** | Samd11 | -0.006825925 | -0.119876054 | 0.106224205 | 7.671104727 | -0.148545531 | 0.886884178 | 0.976674127 | -5.106030105 |
| **A_55_P1965757** | Eme2 | -0.010363978 | -0.182240378 | 0.161512421 | 9.858402196 | -0.148347219 | 0.887033899 | 0.976783356 | -5.10605225 |
| **A_55_P2079158** | Saxo2 | 0.006717846 | -0.104970696 | 0.118406388 | 7.69350878 | 0.147975758 | 0.887314361 | 0.977014091 | -5.106093651 |
| **A_55_P2003410** | Kng1 | 0.05897239 | -0.922127395 | 1.040072176 | 9.90643543 | 0.14787839 | 0.887387878 | 0.977014091 | -5.106104486 |
| **A_55_P1979281** | Rab1a | -0.013932699 | -0.246030125 | 0.218164727 | 14.01168537 | -0.147684222 | 0.887534489 | 0.977014091 | -5.106126072 |
| **A_55_P2340593** | 1810019D21Rik | -0.009244738 | -0.163280183 | 0.144790707 | 8.486705236 | -0.147653277 | 0.887557856 | 0.977014091 | -5.10612951 |
| **A_55_P2145149** | Cpn2 | -0.006330058 | -0.111815346 | 0.09915523 | 7.656405677 | -0.147633519 | 0.887572774 | 0.977014091 | -5.106131705 |
| **A_55_P2026275** | Ppp1r1b | 0.006516948 | -0.102094141 | 0.115128037 | 7.732678003 | 0.147617977 | 0.88758451 | 0.977014091 | -5.106133431 |
| **A_51_P279232** | Dennd2d | -0.009040051 | -0.159718276 | 0.141638174 | 7.87370961 | -0.147601092 | 0.88759726 | 0.977014091 | -5.106135306 |
| **A_51_P153063** | Fabp12 | 0.00919645 | -0.144295273 | 0.162688174 | 8.0366963 | 0.147402359 | 0.887747324 | 0.977123628 | -5.106157358 |
| **A_51_P278868** | H2-DMb1 | -0.017388034 | -0.307763427 | 0.27298736 | 8.320393996 | -0.147319359 | 0.88781 | 0.977136971 | -5.106166559 |
| **A_55_P2002884** | Zcchc24 | 0.052116582 | -0.819602084 | 0.923835249 | 11.44419735 | 0.147085122 | 0.887986883 | 0.977276004 | -5.106192499 |
| **A_51_P126563** | Otc | 0.006399713 | -0.100733359 | 0.113532786 | 7.659878973 | 0.146962365 | 0.888079586 | 0.977279772 | -5.100750766 |
| **A_55_P1981709** | Rreb1 | -0.021029652 | -0.373283272 | 0.331223968 | 9.362695939 | -0.146874289 | 0.888146099 | 0.977279772 | -5.106215812 |
| **A_51_P210082** | Ercc4 | 0.010906683 | -0.171895354 | 0.19370872 | 10.06538881 | 0.146784713 | 0.888213746 | 0.977279772 | -5.106225706 |
| **A_51_P498388** | Sbk1 | 0.007123827 | -0.112302573 | 0.126550226 | 8.081133384 | 0.146751368 | 0.888238928 | 0.977279772 | -5.106229388 |
| **A_55_P2066518** | Akt2 | -0.015035737 | -0.267246817 | 0.237175343 | 12.52706029 | -0.146666093 | 0.888303329 | 0.977279772 | -5.1062388 |
| **A_55_P2169829** | Armh4 | -0.007216326 | -0.128345059 | 0.113912407 | 8.191744968 | -0.146567653 | 0.888377673 | 0.977279772 | -5.106249659 |
| **A_51_P122085** | Usp20 | -0.018467561 | -0.328483226 | 0.291548104 | 10.96484171 | -0.146553118 | 0.88838865 | 0.977279772 | -5.095088621 |
| **A_55_P2052749** | Pcm1 | -0.011566418 | -0.20584393 | 0.182711094 | 9.645469723 | -0.146468932 | 0.88845223 | 0.977279772 | -5.106260541 |
| **A_55_P2043008** | NA | -0.015526798 | -0.276549281 | 0.245495684 | 8.703904491 | -0.146343406 | 0.888547034 | 0.977279772 | -5.106274367 |
| **A_51_P251129** | Stard8 | -0.026993217 | -0.481301363 | 0.427314929 | 11.354475 | -0.146175063 | 0.888674178 | 0.977279772 | -5.106292891 |
| **A_55_P1962966** | NA | -0.007727806 | -0.137818152 | 0.12236254 | 13.65522155 | -0.146143765 | 0.888697817 | 0.977279772 | -5.106296333 |
| **A_55_P2169963** | Zfp982 | 0.008971049 | -0.142069956 | 0.160012053 | 8.303676841 | 0.14612264 | 0.888713772 | 0.977279772 | -5.106298656 |
| **A_52_P249424** | Vegfa | -0.028420098 | -0.507464257 | 0.450624062 | 9.708666341 | -0.145955067 | 0.888840339 | 0.977279772 | -5.106317068 |
| **A_55_P2133047** | Tollip | -0.008182416 | -0.146113559 | 0.129748726 | 9.163415468 | -0.145944725 | 0.888848151 | 0.977279772 | -5.106318203 |
| **A_51_P244969** | Rmnd5b | 0.010914677 | -0.173135418 | 0.194964773 | 8.995468971 | 0.14589621 | 0.888884794 | 0.977279772 | -5.106323529 |
| **A_51_P480861** | Pga5 | -0.030284165 | -0.541610843 | 0.481042514 | 10.59230188 | -0.145708977 | 0.889026217 | 0.977279772 | -5.106344069 |
| **A_55_P2038767** | Fam49a | -0.013042572 | -0.233319593 | 0.207234449 | 8.664145684 | -0.14566768 | 0.88905741 | 0.977279772 | -5.106348595 |
| **A_55_P2055063** | Mcts1 | 0.007906894 | -0.125694987 | 0.141508776 | 11.06317691 | 0.145600388 | 0.889108239 | 0.977279772 | -5.106355969 |
| **A_52_P102248** | Mex3b | 0.01349125 | -0.214562726 | 0.241545226 | 9.521213053 | 0.145540446 | 0.889153516 | 0.977279772 | -5.106362534 |
| **A_52_P108346** | Myc | -0.018961538 | -0.33953213 | 0.301609054 | 8.533335386 | -0.14551878 | 0.889169882 | 0.977279772 | -5.106364906 |
| **A_51_P416191** | Hectd1 | -0.013384959 | -0.239820717 | 0.213050798 | 10.82313755 | -0.145425715 | 0.88924018 | 0.977279772 | -5.106375092 |
| **A_55_P1975326** | Tmem259 | -0.010736361 | -0.192393868 | 0.170921146 | 13.80730153 | -0.145402852 | 0.889257451 | 0.977279772 | -5.106377594 |
| **A_55_P1970954** | Pknox2 | 0.044198232 | -0.703856657 | 0.79225312 | 8.877650705 | 0.145358553 | 0.889290913 | 0.977279772 | -5.106382439 |
| **A_52_P269158** | Pid1 | 0.011459351 | -0.182551523 | 0.205470226 | 8.001520544 | 0.14531255 | 0.889325663 | 0.977279772 | -5.10638747 |
| **A_55_P1966327** | NA | -0.022717095 | -0.407332377 | 0.361898187 | 9.754217022 | -0.145309973 | 0.88932761 | 0.977279772 | -5.106387751 |
| **A_52_P482897** | Areg | -0.006258043 | -0.112230292 | 0.099714206 | 7.675258161 | -0.145283253 | 0.889347794 | 0.977279772 | -5.106390672 |
| **A_55_P1984158** | Slc44a2 | 0.037695984 | -0.600959663 | 0.67635163 | 12.76632381 | 0.145210321 | 0.889402886 | 0.977279772 | -5.106398643 |
| **A_55_P2163263** | NA | -0.009091745 | -0.163163409 | 0.144979918 | 9.400040169 | -0.145175611 | 0.889429107 | 0.977279772 | -5.106402435 |
| **A_55_P2096081** | Myo5b | 0.019466592 | -0.310503164 | 0.349436348 | 11.52460682 | 0.145139276 | 0.889456555 | 0.977279772 | -5.106406403 |
| **A_55_P2052555** | Dpysl5 | -0.020265669 | -0.364334122 | 0.323802783 | 12.3635209 | -0.14490563 | 0.889633057 | 0.977311739 | -5.106431898 |
| **A_51_P421140** | Tubb6 | -0.037109548 | -0.667862649 | 0.593643552 | 12.0985524 | -0.144742289 | 0.889756453 | 0.977311739 | -5.106449697 |
| **A_55_P2192633** | E330017L17Rik | -0.006534318 | -0.117608097 | 0.104539461 | 7.65554053 | -0.14472977 | 0.889765911 | 0.977311739 | -5.10645106 |
| **A_52_P50417** | 1700084J12Rik | -0.006842413 | -0.123163566 | 0.10947874 | 8.214171689 | -0.144717063 | 0.889775511 | 0.977311739 | -5.106452444 |
| **A_52_P4046** | Baz2a | 0.014146926 | -0.226398834 | 0.254692686 | 10.34031749 | 0.144688339 | 0.88979721 | 0.977311739 | -5.106455571 |
| **A_55_P2051405** | Tet2 | -0.006346028 | -0.114308592 | 0.101616536 | 8.105484822 | -0.144609875 | 0.889856488 | 0.977311739 | -5.106464111 |

| | | | | | | | | | |
|---|---|---|---|---|---|---|---|---|---|
| **A_51_P219028** | Wdr53 | -0.007619242 | -0.137258664 | 0.12202018 | 10.78072245 | -0.144591862 | 0.889870097 | 0.977311739 | -5.106466071 |
| **A_55_P2107247** | Tssk5 | 0.02041541 | -0.327213149 | 0.368043968 | 8.350819343 | 0.144481358 | 0.889953581 | 0.977311739 | -5.106478089 |
| **A_51_P379750** | Myoz1 | -0.006243788 | -0.112585786 | 0.100098211 | 7.701724183 | -0.144448311 | 0.889978548 | 0.977311739 | -5.106481681 |
| **A_55_P1978465** | H2-Q8 | 0.087601992 | -1.404578317 | 1.579782302 | 10.73704249 | 0.144431474 | 0.889991269 | 0.977311739 | -5.106483511 |
| **A_52_P384314** | Tmem198b | 0.014917894 | -0.239329039 | 0.269164828 | 8.730201029 | 0.14435139 | 0.890051772 | 0.977320081 | -5.106492212 |
| **A_51_P322972** | Hkdc1 | 0.041255642 | -0.662178753 | 0.744690037 | 10.4628712 | 0.144287569 | 0.89009999 | 0.977320081 | -5.106499143 |
| **A_55_P2029558** | Adrb2 | 0.00629245 | -0.10121405 | 0.11379895 | 7.733743849 | 0.143997264 | 0.890319327 | 0.977415576 | -5.106530629 |
| **A_55_P2010018** | NA | 0.007734098 | -0.124500694 | 0.139968889 | 16.23474984 | 0.143890815 | 0.890399756 | 0.977415576 | -5.106542159 |
| **A_52_P9703** | Cyp2a4 | 0.006210188 | -0.099982706 | 0.112403082 | 7.676403749 | 0.143872724 | 0.890413425 | 0.977415576 | -5.106544118 |
| **A_52_P514061** | Padi4 | -0.00693004 | -0.125463215 | 0.111603135 | 7.711308437 | -0.143835141 | 0.890441822 | 0.977415576 | -5.106548186 |
| **A_51_P463087** | Cenpm | -0.006083494 | -0.110137879 | 0.09797089 | 7.673772524 | -0.143834109 | 0.890442602 | 0.977415576 | -5.106548298 |
| **A_55_P2181665** | NA | -0.013562428 | -0.24574221 | 0.218617354 | 9.368583785 | -0.143708416 | 0.890537573 | 0.977415576 | -5.106561896 |
| **A_51_P123134** | Ercc6l | -0.006514983 | -0.118072039 | 0.105042072 | 7.765030876 | -0.143676399 | 0.890561765 | 0.977415576 | -5.084348353 |
| **A_51_P239924** | Polr2c | -0.010085693 | -0.182831874 | 0.162660488 | 12.07340906 | -0.143637045 | 0.890591501 | 0.977415576 | -5.106569612 |
| **A_55_P2157023** | Sox17 | 0.013978104 | -0.225641805 | 0.253598012 | 10.36851291 | 0.143514085 | 0.890684411 | 0.977462044 | -5.106582896 |
| **A_51_P307964** | Krt13 | -0.009493466 | -0.172471771 | 0.15348484 | 9.082157524 | -0.143305927 | 0.890841702 | 0.977579156 | -5.10660536 |
| **A_51_P100298** | Stx3 | 0.01471614 | -0.238594184 | 0.268026465 | 8.746422677 | 0.142925656 | 0.891129061 | 0.977677975 | -5.08482571 |
| **A_55_P2128373** | Cops6 | -0.00982407 | -0.178950212 | 0.159302071 | 12.23159294 | -0.14290581 | 0.891144058 | 0.977677975 | -5.106648448 |
| **A_55_P2339601** | NA | -0.007277169 | -0.132586994 | 0.118032656 | 8.307142909 | -0.142871793 | 0.891169764 | 0.977677975 | -5.106652105 |
| **A_52_P260629** | Cpne3 | -0.010803422 | -0.196861819 | 0.175254974 | 9.117076591 | -0.14285033 | 0.891185984 | 0.977677975 | -5.106654413 |
| **A_55_P2092384** | Pik3r2 | -0.009528887 | -0.173705888 | 0.154648115 | 8.804004771 | -0.142790398 | 0.891231275 | 0.977677975 | -5.106660854 |
| **A_55_P2040951** | Actc1 | 0.063746775 | -1.034612266 | 1.162105817 | 9.209482145 | 0.142785154 | 0.891235238 | 0.977677975 | -5.106661417 |
| **A_55_P2223851** | NA | -0.009434197 | -0.172083888 | 0.153215494 | 7.960466056 | -0.142698979 | 0.891300362 | 0.977685142 | -5.106670674 |
| **A_55_P2042953** | Acd | -0.011996952 | -0.21893411 | 0.194940205 | 10.47899542 | -0.142626955 | 0.891354792 | 0.977685142 | -5.106678406 |
| **A_55_P1995205** | Top2a | 0.009118938 | -0.148285077 | 0.166522953 | 7.884204706 | 0.142527153 | 0.891430216 | 0.977685142 | -5.106689115 |
| **A_55_P2068366** | Gm9292 | 0.008645779 | -0.140610173 | 0.157901732 | 8.157819567 | 0.142508787 | 0.891444097 | 0.977685142 | -5.106691084 |
| **A_55_P2031692** | Gstm6 | -0.006938871 | -0.126794742 | 0.112917 | 7.838503343 | -0.142429085 | 0.891504331 | 0.977695729 | -5.10669963 |
| **A_55_P2029549** | Dtd1 | -0.010702334 | -0.195957121 | 0.174552453 | 9.999172957 | -0.142127536 | 0.891732234 | 0.977839261 | -5.106731917 |
| **A_55_P2114009** | Ankra2 | 0.008944355 | -0.145893483 | 0.163782194 | 11.07192725 | 0.142115391 | 0.891741413 | 0.977839261 | -5.106733216 |
| **A_51_P246166** | Wfdc18 | -0.006512665 | -0.119302784 | 0.106277454 | 7.692616082 | -0.142055104 | 0.891786978 | 0.977839261 | -5.106739662 |
| **A_51_P337675** | Cd53 | -0.006141991 | -0.112664111 | 0.100380129 | 7.749394365 | -0.141853003 | 0.891939729 | 0.977917431 | -5.106761253 |
| **A_52_P239726** | Larp4 | 0.016734991 | -0.273557655 | 0.307027638 | 9.760472302 | 0.141826904 | 0.891959455 | 0.977917431 | -5.106764038 |
| **A_55_P2077079** | Rab29 | -0.018352075 | -0.337005226 | 0.300301075 | 9.912922076 | -0.141688999 | 0.892063689 | 0.977976238 | -5.106778751 |
| **A_55_P2017362** | Prr36 | -0.0059948 | -0.11028155 | 0.09829195 | 7.737313123 | -0.141421272 | 0.892266054 | 0.978006615 | -5.106807272 |
| **A_51_P312336** | Slc14a1 | -0.012478043 | -0.229632572 | 0.204676487 | 9.398621302 | -0.141366577 | 0.892307398 | 0.978006615 | -5.106813093 |
| **A_55_P2132039** | NA | -0.022485286 | -0.413822294 | 0.368851721 | 9.656117432 | -0.141356787 | 0.892314798 | 0.978006615 | -5.106814134 |
| **A_55_P2153797** | Slc38a6 | 0.01380378 | -0.226502662 | 0.254110223 | 9.454796506 | 0.141319402 | 0.892343057 | 0.978006615 | -5.106818111 |
| **A_55_P2133716** | Tax1bp3 | -0.02885242 | -0.531142996 | 0.473438156 | 14.29873276 | -0.141317642 | 0.892344387 | 0.978006615 | -5.106818298 |
| **A_51_P206445** | Zfp773 | 0.006940866 | -0.113979534 | 0.127861266 | 8.733065492 | 0.141215782 | 0.892421384 | 0.978029995 | -5.106829127 |
| **A_55_P1964033** | Carm1 | -0.010411663 | -0.191876015 | 0.171052689 | 10.26608214 | -0.141155545 | 0.892466917 | 0.978029995 | -5.106835528 |
| **A_51_P363103** | Blvra | 0.018982807 | -0.312054804 | 0.350020417 | 10.06816921 | 0.141075711 | 0.892527265 | 0.978040245 | -5.106844007 |
| **A_66_P113059** | Itgb1 | -0.022811161 | -0.420855768 | 0.375233446 | 11.8854529 | -0.140988863 | 0.892592916 | 0.978040245 | -5.106853225 |
| **A_51_P514922** | Ccar2 | -0.009212186 | -0.170020358 | 0.151595987 | 9.36343414 | -0.140936578 | 0.89263244 | 0.978040245 | -5.106858772 |
| **A_55_P1999349** | Nudt4 | 0.025546116 | -0.420756227 | 0.47184846 | 13.18225837 | 0.140820176 | 0.892720434 | 0.978040245 | -5.106871114 |
| **A_66_P117477** | Dlk1 | -0.029740557 | -0.549364829 | 0.489883714 | 14.92830223 | -0.140808489 | 0.892729269 | 0.978040245 | -5.106872352 |
| **A_55_P2152225** | Ihh | -0.023531682 | -0.453138017 | 0.406074654 | 8.944628822 | -0.141949011 | 0.892805874 | 0.978059722 | -5.037992743 |
| **A_51_P402994** | Ddx3y | -0.135701712 | -2.509322589 | 2.237919165 | 9.317943956 | -0.140651101 | 0.892848249 | 0.978059722 | -5.106889022 |
| **A_51_P358722** | Lancl3 | -0.006248896 | -0.115668744 | 0.103170951 | 7.722975232 | -0.14050002 | 0.892962465 | 0.978082266 | -5.106905007 |
| **A_52_P342620** | Csrnp2 | -0.011176495 | -0.206893849 | 0.184540859 | 8.901530708 | -0.140490009 | 0.892970032 | 0.978082266 | -5.106906066 |
| **A_52_P408690** | Sft2d2 | -0.019547755 | -0.362311103 | 0.323215592 | 8.842764935 | -0.140304525 | 0.89311026 | 0.978116975 | -5.106925666 |

| | | | | | | | | | |
|---|---|---|---|---|---|---|---|---|---|
| **A_51_P124748** | Tgfb3 | -0.016181666 | -0.300097412 | 0.26773408 | 12.03252412 | -0.140217722 | 0.893175886 | 0.978116975 | -5.106934829 |
| **A_52_P620793** | Eif2a | -0.008196292 | -0.152053189 | 0.135660604 | 8.97178816 | -0.140170271 | 0.89321176 | 0.978116975 | -5.106939836 |
| **A_52_P222354** | Vps13c | 0.00716284 | -0.118604386 | 0.132930066 | 8.335621938 | 0.140115742 | 0.893252986 | 0.978116975 | -5.106945588 |
| **A_65_P19286** | Alg10b | -0.009570343 | -0.177720656 | 0.15857997 | 8.586632228 | -0.140022875 | 0.893323198 | 0.978116975 | -5.106955378 |
| **A_52_P539414** | Gtf2h3 | -0.012048304 | -0.223744456 | 0.199647847 | 9.849006089 | -0.140017437 | 0.893327309 | 0.978116975 | -5.106955951 |
| **A_52_P325527** | Edil3 | -0.006297086 | -0.117051679 | 0.104457508 | 7.683237175 | -0.139877231 | 0.893433314 | 0.978116975 | -5.10697072 |
| **A_52_P551743** | Aff4 | -0.017577693 | -0.326762103 | 0.291606717 | 11.01157574 | -0.139866417 | 0.89344149 | 0.978116975 | -5.106971858 |
| **A_55_P2180425** | Uqcc1 | 0.00988473 | -0.164009364 | 0.183778823 | 9.049063542 | 0.139845702 | 0.893457153 | 0.978116975 | -5.106974039 |
| **A_55_P2017845** | Ehbp1 | 0.007552561 | -0.125517421 | 0.140622543 | 8.159550365 | 0.139631467 | 0.893619133 | 0.978169025 | -5.106996571 |
| **A_55_P1971199** | Slc4a1ap | 0.007354154 | -0.122245619 | 0.136953927 | 10.90506351 | 0.139603931 | 0.893639953 | 0.978169025 | -5.106999465 |
| **A_51_P224445** | Gpn3 | 0.009852381 | -0.163800043 | 0.183504804 | 8.648064234 | 0.139582024 | 0.893656517 | 0.978169025 | -5.107001767 |
| **A_55_P2084980** | Dcaf12l2 | 0.021212643 | -0.352985142 | 0.395410428 | 11.26489944 | 0.1394642 | 0.893745606 | 0.978195131 | -5.10701414 |
| **A_55_P2118302** | Fam110a | 0.013820039 | -0.230053051 | 0.257693128 | 11.05105904 | 0.13941662 | 0.893781583 | 0.978195131 | -5.107019134 |
| **A_55_P2141038** | Gnpnat1 | -0.014197468 | -0.264884881 | 0.236489945 | 8.844032774 | -0.139330927 | 0.893846378 | 0.978200417 | -5.107028124 |
| **A_52_P114889** | Upk3b | -0.022576286 | -0.421366239 | 0.376213667 | 8.022795253 | -0.139276374 | 0.893887628 | 0.978200417 | -5.107033844 |
| **A_55_P2033932** | Zzz3 | -0.009577433 | -0.178858871 | 0.159704005 | 9.485372049 | -0.139190295 | 0.893952717 | 0.978216263 | -5.107042865 |
| **A_55_P2110998** | Gpr153 | -0.028278401 | -0.528467817 | 0.471911016 | 8.743162527 | -0.139087951 | 0.894030106 | 0.978245565 | -5.107053583 |
| **A_51_P369628** | Ptov1 | -0.007867866 | -0.14713429 | 0.131398558 | 10.96761894 | -0.138988764 | 0.894105108 | 0.978249546 | -5.107063964 |
| **A_55_P1960053** | Hvcn1 | 0.008226983 | -0.137437436 | 0.153891403 | 10.42688274 | 0.138949282 | 0.894134964 | 0.978249546 | -5.107068094 |
| **A_55_P2038525** | C3 | -0.016177254 | -0.302776461 | 0.270421952 | 10.15821025 | -0.138866981 | 0.894197199 | 0.978262263 | -5.107076699 |
| **A_51_P348749** | Llgl2 | -0.018987063 | -0.355566456 | 0.31759233 | 11.24849743 | -0.138784008 | 0.894259943 | 0.978275536 | -5.107085369 |
| **A_55_P1979305** | Gle1 | -0.013454081 | -0.252363128 | 0.225454965 | 9.468129335 | -0.138544922 | 0.894440746 | 0.978379431 | -5.107110324 |
| **A_55_P1960145** | Clip4 | -0.005984164 | -0.112353001 | 0.100384673 | 7.685450109 | -0.138407054 | 0.894545007 | 0.978379431 | -5.107124695 |
| **A_55_P2166506** | NA | 0.00768204 | -0.128877162 | 0.144241242 | 15.29002364 | 0.138396389 | 0.894553073 | 0.978379431 | -5.107125806 |
| **A_55_P2045050** | Zfp953 | 0.006677588 | -0.112121838 | 0.125477014 | 8.269275521 | 0.138284806 | 0.894637459 | 0.978379431 | -5.107137426 |
| **A_52_P201206** | Scrn1 | 0.013072972 | -0.219511942 | 0.245657887 | 9.528153498 | 0.138280827 | 0.894640468 | 0.978379431 | -5.10713784 |
| **A_55_P2153620** | Ahnak | 0.021628895 | -0.363371449 | 0.406629238 | 9.695909075 | 0.138210925 | 0.894693333 | 0.978379431 | -5.107145114 |
| **A_55_P2063216** | Dusp14 | 0.019504666 | -0.327778768 | 0.366788099 | 9.098743515 | 0.138173134 | 0.894721913 | 0.978379431 | -5.107149045 |
| **A_51_P229676** | Plscr1 | -0.017303355 | -0.325591997 | 0.290985287 | 9.042966082 | -0.138083553 | 0.894789663 | 0.978379431 | -5.10715836 |
| **A_55_P2021398** | NA | 0.048063761 | -0.80853047 | 0.904657993 | 11.30835249 | 0.138042168 | 0.894820962 | 0.978379431 | -5.107162661 |
| **A_55_P2186693** | NA | 0.011137301 | -0.187428433 | 0.209703036 | 8.629628397 | 0.137989113 | 0.894861087 | 0.978379431 | -5.107168173 |
| **A_55_P1977289** | Zbtb9 | -0.010144613 | -0.191121251 | 0.170832025 | 9.73943886 | -0.137905667 | 0.894924198 | 0.97839309 | -5.107176838 |
| **A_51_P507023** | Nt5dc3 | -0.019799386 | -0.373337683 | 0.333738912 | 10.80560141 | -0.137779438 | 0.895019668 | 0.97839656 | -5.107189936 |
| **A_66_P119771** | Nme8 | -0.006028694 | -0.113771542 | 0.101714153 | 7.633768529 | -0.137658795 | 0.895110915 | 0.97839656 | -5.107202443 |
| **A_55_P2004637** | NA | -0.013340096 | -0.251800733 | 0.225120541 | 9.554032972 | -0.137629466 | 0.895133098 | 0.97839656 | -5.107205482 |
| **A_51_P364786** | U2af1 | -0.022614424 | -0.427408963 | 0.382180115 | 8.802006536 | -0.137442182 | 0.895274752 | 0.97839656 | -5.107224872 |
| **A_55_P2037454** | Etv5 | -0.017451126 | -0.329859021 | 0.294956769 | 10.56173248 | -0.137426541 | 0.895286582 | 0.97839656 | -5.10722649 |
| **A_66_P117913** | NA | 0.008012717 | -0.135454851 | 0.151480284 | 15.57978178 | 0.137402682 | 0.895304628 | 0.97839656 | -5.107228959 |
| **A_55_P2121222** | Clcc1 | -0.00924196 | -0.174771907 | 0.156287987 | 10.86764258 | -0.137358863 | 0.895337772 | 0.97839656 | -5.10723349 |
| **A_55_P2219657** | Tmf1 | -0.006023499 | -0.113911152 | 0.101864155 | 8.601154721 | -0.137355554 | 0.895340275 | 0.97839656 | -5.107233832 |
| **A_51_P281024** | Rad9a | -0.009086265 | -0.17207321 | 0.15390068 | 8.571173033 | -0.13715187 | 0.89549434 | 0.97839656 | -5.107254878 |
| **A_52_P424784** | Clstn2 | 0.009304587 | -0.157615143 | 0.176224318 | 8.037898673 | 0.13713825 | 0.895504642 | 0.97839656 | -5.107256284 |
| **A_55_P2066782** | Fam104a | 0.010136032 | -0.171862399 | 0.192134462 | 13.23964908 | 0.137015429 | 0.895597546 | 0.97839656 | -5.107268958 |
| **A_52_P476431** | Evc | 0.007884728 | -0.133730722 | 0.149500178 | 7.859781239 | 0.136976249 | 0.895627183 | 0.97839656 | -5.107272999 |
| **A_51_P230734** | Slc34a3 | 0.04050213 | -0.687280358 | 0.768284617 | 9.240397462 | 0.136913233 | 0.89567485 | 0.97839656 | -5.107279495 |
| **A_51_P312997** | Zfp346 | 0.007427451 | -0.126052055 | 0.140906956 | 8.343473579 | 0.136897133 | 0.895687029 | 0.97839656 | -5.107281155 |
| **A_55_P2010048** | NA | -0.013874053 | -0.26330971 | 0.235561603 | 8.575460631 | -0.136840291 | 0.895730026 | 0.97839656 | -5.107287011 |
| **A_52_P665202** | NA | -0.017554263 | -0.333176894 | 0.298068368 | 13.68271476 | -0.136830726 | 0.895737261 | 0.97839656 | -5.107287997 |
| **A_51_P205106** | Mlf1 | -0.007284583 | -0.138381522 | 0.123812355 | 8.029136017 | -0.136704034 | 0.895833098 | 0.978445949 | -5.107301041 |
| **A_52_P521507** | Trio | -0.021087752 | -0.401207586 | 0.359032081 | 11.18781209 | -0.136483116 | 0.896000217 | 0.978463317 | -5.107323758 |

| | | | | | | | | | |
|---|---|---|---|---|---|---|---|---|---|
| **A_55_P2005248** | Meis3 | -0.008989139 | -0.171155671 | 0.153177393 | 9.007050472 | -0.136372255 | 0.896084082 | 0.978463317 | -5.107335144 |
| **A_66_P133270** | Mfsd1 | -0.010585984 | -0.201650598 | 0.180478629 | 8.826281545 | -0.136307624 | 0.896132976 | 0.978463317 | -5.107341778 |
| **A_55_P2089020** | Slc35g2 | -0.010970469 | -0.209035936 | 0.187094998 | 8.398045091 | -0.1362654 | 0.896164918 | 0.978463317 | -5.10734611 |
| **A_55_P2045946** | Tepsin | -0.012160107 | -0.231829958 | 0.207509744 | 9.822330129 | -0.136187137 | 0.896224126 | 0.978463317 | -5.107354137 |
| **A_66_P131110** | Zfp473 | -0.012060073 | -0.230047629 | 0.205927484 | 8.073691892 | -0.136109171 | 0.896283109 | 0.978463317 | -5.107362128 |
| **A_55_P2034550** | 1700013F07Rik | 0.006362882 | -0.108651422 | 0.121377186 | 7.715291664 | 0.136104087 | 0.896286955 | 0.978463317 | -5.107362649 |
| **A_55_P1961466** | Dct | 0.016029349 | -0.273782857 | 0.305841554 | 8.229923621 | 0.136071873 | 0.896311326 | 0.978463317 | -5.10736595 |
| **A_55_P2133365** | 1110017D15Rik | -0.005862821 | -0.111869045 | 0.100143404 | 7.709317087 | -0.136064366 | 0.896317006 | 0.978463317 | -5.107366719 |
| **A_55_P1998115** | Klkb1 | 0.006283305 | -0.107429923 | 0.119996534 | 7.759765151 | 0.135939709 | 0.896411315 | 0.978463317 | -5.107379482 |
| **A_55_P1975695** | Snx18 | 0.012307838 | -0.210661736 | 0.235277411 | 11.17084778 | 0.135801734 | 0.896515701 | 0.978463317 | -5.107393596 |
| **A_55_P2066778** | AB124611 | -0.005760356 | -0.110157298 | 0.098636587 | 7.708083919 | -0.135747137 | 0.896557008 | 0.978463317 | -5.107399177 |
| **A_51_P250445** | Zfp276 | 0.014167522 | -0.242657769 | 0.270992814 | 9.363623526 | 0.135714194 | 0.896581931 | 0.978463317 | -5.107402543 |
| **A_55_P2127959** | Zfp36l3 | 0.027598098 | -0.472694833 | 0.52789103 | 12.16507394 | 0.135713783 | 0.896582242 | 0.978463317 | -5.107402585 |
| **A_55_P2168307** | Gm13213 | -0.006079302 | -0.116422049 | 0.104263445 | 15.94958939 | -0.135543625 | 0.896710982 | 0.978463317 | -5.10741996 |
| **A_51_P114297** | Kifc3 | -0.013568111 | -0.259837373 | 0.232701152 | 9.262261518 | -0.135543386 | 0.896711163 | 0.978463317 | -5.107419985 |
| **A_55_P2079928** | Ccdc68 | -0.011263146 | -0.24003975 | 0.217513457 | 11.5281636 | -0.138532167 | 0.896729081 | 0.978463317 | -4.986498204 |
| **A_55_P2106106** | C5ar2 | 0.054560563 | -0.936219675 | 1.0453408 | 10.4971264 | 0.13547859 | 0.896760187 | 0.978463317 | -5.107426595 |
| **A_51_P378856** | Pfkp | -0.024221165 | -0.465010415 | 0.416568084 | 10.12603224 | -0.135186462 | 0.896981218 | 0.978618968 | -5.10745636 |
| **A_55_P1987645** | Unc13b | -0.014776933 | -0.283848785 | 0.254294919 | 9.580519608 | -0.135109288 | 0.897039612 | 0.978618968 | -5.107464213 |
| **A_52_P169901** | Slc25a51 | 0.015461604 | -0.266146944 | 0.297070151 | 10.97572152 | 0.1350759 | 0.897064875 | 0.978618968 | -5.107467609 |
| **A_52_P429876** | Tbx20 | -0.015842084 | -0.304494803 | 0.272810636 | 8.332369362 | -0.135022396 | 0.89710536 | 0.978618968 | -5.107473049 |
| **A_51_P159565** | Arhgef9 | -0.005758463 | -0.110748948 | 0.099232022 | 7.74407755 | -0.134935369 | 0.89717121 | 0.978635571 | -5.107481893 |
| **A_55_P2115176** | Pdhx | -0.016199545 | -0.311950275 | 0.279551185 | 9.402965318 | -0.134755393 | 0.897307394 | 0.978728888 | -5.107500165 |
| **A_55_P2054350** | Fbxo44 | 0.008293557 | -0.143207826 | 0.15979494 | 8.08376199 | 0.134676995 | 0.897366719 | 0.978738365 | -5.107508117 |
| **A_51_P263591** | Pank1 | -0.006003457 | -0.115760268 | 0.103753354 | 7.749739196 | -0.134567161 | 0.897449831 | 0.978773784 | -5.10751925 |
| **A_52_P356204** | Nostrin | 0.016640259 | -0.288149107 | 0.321429625 | 9.348209362 | 0.13431652 | 0.897639498 | 0.978791894 | -5.107544621 |
| **A_51_P256665** | Pon3 | 0.068360592 | -1.183854873 | 1.320576057 | 10.07219351 | 0.134306154 | 0.897647342 | 0.978791894 | -5.107545669 |
| **A_55_P1953241** | Mafk | 0.016228323 | -0.281099239 | 0.313555885 | 10.17962669 | 0.134278857 | 0.897668 | 0.978791894 | -5.107548429 |
| **A_51_P518621** | H3c1 | 0.007514419 | -0.130171401 | 0.145200238 | 7.817677737 | 0.134268869 | 0.897675558 | 0.978791894 | -5.107549439 |
| **A_66_P128586** | Cwc22 | 0.010207394 | -0.176926235 | 0.197341023 | 8.940746032 | 0.134193723 | 0.897732425 | 0.978791894 | -5.107557034 |
| **A_55_P2123491** | B3gnt5 | -0.015553199 | -0.300844086 | 0.269737687 | 8.993893506 | -0.134122303 | 0.897786474 | 0.978791894 | -5.107564248 |
| **A_55_P2094626** | Ralgps1 | 0.01866463 | -0.324016999 | 0.361346258 | 12.37894483 | 0.133997815 | 0.897880683 | 0.978791894 | -5.107576814 |
| **A_51_P495825** | Apof | -0.021004556 | -0.406950634 | 0.364941522 | 7.957042337 | -0.133892436 | 0.897960432 | 0.978791894 | -5.107587442 |
| **A_55_P2008016** | Armc3 | 0.006026669 | -0.104727892 | 0.116781231 | 7.851658586 | 0.133870501 | 0.897977033 | 0.978791894 | -5.107589653 |
| **A_55_P2077114** | NA | 0.019213123 | -0.334796827 | 0.373223072 | 14.07859471 | 0.133521637 | 0.89824106 | 0.978791894 | -5.107624772 |
| **A_52_P679105** | Prss23 | -0.046045828 | -0.894655713 | 0.802564057 | 10.5596152 | -0.133490809 | 0.898264391 | 0.978791894 | -5.107627871 |
| **A_55_P2172314** | Scamp4 | -0.007146207 | -0.139066492 | 0.124774078 | 13.22429282 | -0.13327024 | 0.89843133 | 0.978791894 | -5.107650023 |
| **A_55_P2170210** | Ap1s3 | 0.025450726 | -0.444495161 | 0.495396613 | 12.06248564 | 0.133235967 | 0.898457271 | 0.978791894 | -5.107653462 |
| **A_55_P2066453** | NA | 0.007387604 | -0.129046976 | 0.143822184 | 7.79062818 | 0.133213521 | 0.898474259 | 0.978791894 | -5.107655714 |
| **A_55_P2101246** | Aldh6a1 | -0.005692533 | -0.110870682 | 0.099485616 | 7.76786178 | -0.133152451 | 0.898520482 | 0.978791894 | -5.107661838 |
| **A_55_P2048751** | Prox1 | 0.006339703 | -0.110898436 | 0.123577842 | 7.632484866 | 0.133035995 | 0.898608626 | 0.978791894 | -5.107673509 |
| **A_52_P668715** | Nutf2 | -0.006291849 | -0.122704483 | 0.110120785 | 8.169322425 | -0.132968062 | 0.898660045 | 0.978791894 | -5.107680313 |
| **A_55_P2015232** | NA | 0.010581359 | -0.185339526 | 0.206502244 | 9.94929992 | 0.132870905 | 0.898733585 | 0.978791894 | -5.107690037 |
| **A_51_P173906** | Mto1 | -0.007483012 | -0.146057038 | 0.131091015 | 8.161262814 | -0.132850668 | 0.898748902 | 0.978791894 | -5.107692062 |
| **A_55_P2165074** | Rnf31 | -0.012397223 | -0.242149066 | 0.217354621 | 10.22529781 | -0.132750008 | 0.898825094 | 0.978791894 | -5.107702128 |
| **A_55_P2058348** | Pacc1 | 0.014031403 | -0.246048485 | 0.274111292 | 8.560920184 | 0.13272829 | 0.898841533 | 0.978791894 | -5.107704298 |
| **A_55_P2004452** | Tceal8 | 0.016276024 | -0.285929246 | 0.318481294 | 8.726830713 | 0.13249987 | 0.899014435 | 0.978791894 | -5.107727108 |
| **A_51_P315666** | Nid2 | -0.017245998 | -0.337617744 | 0.303125748 | 10.95378233 | -0.132435156 | 0.899063421 | 0.978791894 | -5.107733564 |
| **A_52_P73848** | Clint1 | 0.010058649 | -0.176822323 | 0.196939621 | 9.364008574 | 0.132416993 | 0.89907717 | 0.978791894 | -5.107735375 |
| **A_55_P1952920** | AI854703 | -0.029912635 | -0.585984503 | 0.526159234 | 8.360143463 | -0.132340541 | 0.899135042 | 0.978791894 | -5.107742996 |

| | | | | | | | | | |
|---|---|---|---|---|---|---|---|---|---|
| **A_55_P2041693** | Gm5834 | 0.005785221 | -0.101770447 | 0.11334089 | 7.688363787 | 0.132329249 | 0.89914359 | 0.978791894 | -5.107744121 |
| **A_51_P481644** | Mbip | -0.013661887 | -0.267825706 | 0.240501931 | 9.621832298 | -0.132241005 | 0.899210389 | 0.978791894 | -5.107752912 |
| **A_51_P509263** | Hoxa7 | 0.011152582 | -0.196422217 | 0.21872738 | 8.57945623 | 0.132181278 | 0.899255602 | 0.978791894 | -5.107758858 |
| **A_55_P2007964** | Cx3cr1 | -0.019917565 | -0.390704266 | 0.350869136 | 9.134426374 | -0.132154287 | 0.899276034 | 0.978791894 | -5.107761544 |
| **A_51_P346453** | Noc4l | 0.008712236 | -0.153533049 | 0.170957522 | 8.718033783 | 0.132107274 | 0.899311623 | 0.978791894 | -5.107766222 |
| **A_55_P2078670** | Cbs | 0.007903844 | -0.139291439 | 0.155099128 | 7.977790694 | 0.132103295 | 0.899314635 | 0.978791894 | -5.107766618 |
| **A_55_P2248320** | C430003N24Rik | 0.014155265 | -0.249514313 | 0.277824844 | 8.291140353 | 0.132076987 | 0.89933455 | 0.978791894 | -5.107769235 |
| **A_52_P181709** | Crkl | -0.007668276 | -0.150547888 | 0.135211336 | 8.554244748 | -0.132037299 | 0.899364595 | 0.978791894 | -5.107773181 |
| **A_55_P1973553** | Bbs12 | 0.007308601 | -0.128909956 | 0.143527158 | 8.085513617 | 0.131997936 | 0.899394393 | 0.978791894 | -5.107777095 |
| **A_52_P615312** | Mrps27 | 0.008811155 | -0.15543275 | 0.173055061 | 8.376612341 | 0.13198141 | 0.899406903 | 0.978791894 | -5.107778737 |
| **A_51_P469968** | H2-M3 | -0.019695853 | -0.38700043 | 0.347608724 | 9.161593098 | -0.131922119 | 0.899451788 | 0.978791894 | -5.107784629 |
| **A_55_P2011385** | NA | 0.008338158 | -0.147257201 | 0.163933516 | 8.078716032 | 0.131838617 | 0.899515002 | 0.978791894 | -5.107792921 |
| **A_51_P399985** | Myo9b | -0.013417415 | -0.263795451 | 0.23696062 | 12.23930244 | -0.131838361 | 0.899515196 | 0.978791894 | -5.107792947 |
| **A_51_P375509** | Spty2d1 | -0.016708151 | -0.328577903 | 0.2951616 | 8.791930727 | -0.131802701 | 0.899542191 | 0.978791894 | -5.107796486 |
| **A_55_P1962661** | Cyp2c67 | -0.006717482 | -0.132159718 | 0.118724753 | 7.759404427 | -0.131744323 | 0.899586386 | 0.978791894 | -5.107802279 |
| **A_55_P2032678** | Lox | 0.026420087 | -0.466962456 | 0.519802631 | 8.22384984 | 0.131740599 | 0.899589206 | 0.978791894 | -5.107802649 |
| **A_51_P379687** | Gm10413 | -0.014913026 | -0.293417978 | 0.263591927 | 8.507434599 | -0.131735252 | 0.899593254 | 0.978791894 | -5.107803179 |
| **A_55_P1960366** | Fam47e | 0.005810631 | -0.102820069 | 0.114441332 | 7.735383788 | 0.131595156 | 0.899699314 | 0.978818479 | -5.10781707 |
| **A_51_P114353** | Ctu1 | 0.00767217 | -0.135852811 | 0.15119715 | 8.567926633 | 0.131510327 | 0.899763536 | 0.978818479 | -5.107825474 |
| **A_55_P1972792** | Stil | -0.005619642 | -0.110784046 | 0.099544763 | 7.86427484 | -0.131464656 | 0.899798112 | 0.978818479 | -5.107829996 |
| **A_55_P2000930** | Ccnyl1 | -0.008477566 | -0.167159653 | 0.150204522 | 10.34739407 | -0.131435419 | 0.899820247 | 0.978818479 | -5.10783289 |
| **A_51_P218953** | Zfp536 | -0.005919324 | -0.116833252 | 0.104994603 | 7.725917033 | -0.131297122 | 0.899924951 | 0.978838716 | -5.107846572 |
| **A_51_P423549** | Shbg | 0.005530584 | -0.09811512 | 0.109176289 | 7.680406297 | 0.131277069 | 0.899940132 | 0.978838716 | -5.107848554 |
| **A_66_P112593** | Sec61b | 0.009612045 | -0.170661412 | 0.189885503 | 11.24060226 | 0.131175639 | 0.900016926 | 0.978867159 | -5.107858578 |
| **A_55_P2102385** | Etl4 | -0.0254056 | -0.502576247 | 0.451765047 | 10.88219276 | -0.130986007 | 0.900160501 | 0.978968229 | -5.107877297 |
| **A_55_P1964787** | Slc35e4 | -0.038661541 | -0.766531972 | 0.689208891 | 10.72178028 | -0.130675521 | 0.900395586 | 0.979164163 | -5.107907889 |
| **A_55_P2175494** | NA | 0.007394988 | -0.131939779 | 0.146729755 | 8.538106237 | 0.130571122 | 0.900474635 | 0.979164163 | -5.107918159 |
| **A_55_P2405784** | BC023202 | -0.020735084 | -0.411519873 | 0.370049705 | 8.453614365 | -0.130538106 | 0.900499635 | 0.979164163 | -5.107921405 |
| **A_55_P2030160** | Tcf7l2 | 0.009503969 | -0.169692204 | 0.188700142 | 8.517946021 | 0.130480446 | 0.900543294 | 0.979164163 | -5.107927072 |
| **A_51_P180140** | H2bc1 | 0.009589437 | -0.171419959 | 0.190598833 | 12.18250714 | 0.130335029 | 0.900653404 | 0.979173191 | -5.107941354 |
| **A_55_P2136752** | Ermn | 0.039855829 | -0.712859426 | 0.792571085 | 10.25240353 | 0.130265766 | 0.900705851 | 0.979173191 | -5.107948151 |
| **A_52_P640386** | Usp53 | -0.038918043 | -0.774045954 | 0.696209868 | 8.717508749 | -0.130243855 | 0.900722443 | 0.979173191 | -5.1079503 |
| **A_51_P435922** | Rsph9 | 0.007214045 | -0.129122173 | 0.143550263 | 8.06320017 | 0.130177749 | 0.9007725 | 0.979173191 | -5.107956783 |
| **A_55_P2091212** | NA | -0.007784054 | -0.155012995 | 0.139444887 | 15.63165818 | -0.1300714 | 0.900853031 | 0.979173191 | -5.107967205 |
| **A_55_P2036317** | NA | 0.006278211 | -0.112471986 | 0.125028408 | 16.63550568 | 0.130068078 | 0.900855546 | 0.979173191 | -5.10796753 |
| **A_51_P286814** | Ncor2 | -0.014579637 | -0.290840942 | 0.261681667 | 12.98175987 | -0.129836207 | 0.901031132 | 0.979182281 | -5.107990224 |
| **A_55_P1986531** | Mpig6b | 0.00562593 | -0.10101865 | 0.112270509 | 7.781146112 | 0.129785055 | 0.901069868 | 0.979182281 | -5.107995225 |
| **A_55_P2069818** | Mapk10 | -0.006717454 | -0.134096062 | 0.120661154 | 7.946665438 | -0.129741037 | 0.901103202 | 0.979182281 | -5.107999526 |
| **A_52_P18081** | Pacc1 | -0.008850385 | -0.17693027 | 0.1592295 | 9.611694907 | -0.129543489 | 0.901252803 | 0.979182281 | -5.108018815 |
| **A_55_P1968464** | Kat6a | 0.013015016 | -0.234185248 | 0.26021528 | 10.08304022 | 0.129528379 | 0.901264245 | 0.979182281 | -5.108020289 |
| **A_55_P1986958** | Brms1l | 0.010285409 | -0.185098966 | 0.205669784 | 11.33331794 | 0.129509288 | 0.901278703 | 0.979182281 | -5.108022151 |
| **A_55_P2139402** | Amy2a5 | -0.005492891 | -0.109884598 | 0.098898816 | 7.663440689 | -0.129450622 | 0.901323131 | 0.979182281 | -5.108027872 |
| **A_51_P286946** | Lhpp | 0.017606408 | -0.317075185 | 0.352288002 | 9.750696491 | 0.129422 | 0.901344807 | 0.979182281 | -5.108030663 |
| **A_52_P455404** | Itpa | 0.018164257 | -0.327203272 | 0.363531785 | 12.31720846 | 0.129391363 | 0.90136801 | 0.979182281 | -5.108033649 |
| **A_51_P486155** | Ocln | -0.028576244 | -0.572081568 | 0.514929079 | 8.505855667 | -0.129351215 | 0.901398414 | 0.979182281 | -5.10803756 |
| **A_52_P606303** | Mrpl3 | 0.015893837 | -0.286468884 | 0.318256559 | 11.84970532 | 0.129321188 | 0.901421155 | 0.979182281 | -5.108040485 |
| **A_55_P2051479** | NA | -0.008434568 | -0.169209618 | 0.152340483 | 9.698306566 | -0.129066442 | 0.901614084 | 0.979335636 | -5.108065273 |
| **A_51_P121447** | 1110038F14Rik | -0.008186652 | -0.164315262 | 0.147941958 | 11.59191896 | -0.129000977 | 0.901663665 | 0.979335636 | -5.108071635 |
| **A_55_P2200618** | NA | -0.033114996 | -0.665421871 | 0.599191878 | 9.816393381 | -0.128844526 | 0.901782157 | 0.979365228 | -5.108086827 |
| **A_55_P2007650** | Nt5dc1 | -0.015847324 | -0.318523816 | 0.286829168 | 9.489134149 | -0.128809059 | 0.901809019 | 0.979365228 | -5.108090268 |

| | | | | | | | | | |
|---|---|---|---|---|---|---|---|---|---|
| **A_55_P2148046** | NA | -0.017656762 | -0.35525844 | 0.319944917 | 8.923932812 | -0.128669502 | 0.901914718 | 0.979365228 | -5.108103801 |
| **A_55_P2084656** | H2ac12 | -0.01669157 | -0.336262485 | 0.302879344 | 15.2323715 | -0.128498813 | 0.902044 | 0.979365228 | -5.108120332 |
| **A_51_P479865** | Glb1l | -0.011243009 | -0.226689804 | 0.204203787 | 8.506582936 | -0.128384229 | 0.902130789 | 0.979365228 | -5.108131417 |
| **A_55_P2181104** | Shisa3 | -0.068549746 | -1.382204966 | 1.245105474 | 8.869031243 | -0.12837889 | 0.902134833 | 0.979365228 | -5.108131933 |
| **A_55_P2173737** | Hspa4l | 0.008844138 | -0.160819129 | 0.178507405 | 9.295291606 | 0.128243945 | 0.902237046 | 0.979365228 | -5.108144975 |
| **A_51_P257743** | Dnajc22 | 0.026994295 | -0.490859451 | 0.544848042 | 8.805205361 | 0.128243108 | 0.90223768 | 0.979365228 | -5.108145055 |
| **A_51_P484919** | Polrmt | -0.009179198 | -0.185468751 | 0.167110354 | 9.740195635 | -0.128099474 | 0.902346477 | 0.979365228 | -5.108158922 |
| **A_51_P330428** | Eif4ebp1 | 0.026834856 | -0.488885481 | 0.542555193 | 14.23187107 | 0.128013029 | 0.902411957 | 0.979365228 | -5.10816726 |
| **A_51_P295420** | Zc3h7a | -0.011943125 | -0.241616758 | 0.217730507 | 11.36908634 | -0.127931062 | 0.902474045 | 0.979365228 | -5.108175161 |
| **A_55_P2015032** | Kazald1 | 0.006079816 | -0.110848445 | 0.123008077 | 8.370239912 | 0.12792049 | 0.902482052 | 0.979365228 | -5.108176179 |
| **A_52_P118560** | Kynu | -0.005545864 | -0.11225065 | 0.101158921 | 7.698186887 | -0.127865833 | 0.902523455 | 0.979365228 | -5.108181445 |
| **A_55_P2007457** | Muc5b | 0.006388184 | -0.11656946 | 0.129345829 | 7.786204244 | 0.12781772 | 0.9025599 | 0.979365228 | -5.108186078 |
| **A_55_P2329313** | C530014P21Rik | 0.012056047 | -0.220071715 | 0.244183809 | 8.054308486 | 0.12777533 | 0.902592011 | 0.979365228 | -5.108190158 |
| **A_55_P2043837** | Cmtr1 | -0.016032076 | -0.324793792 | 0.292729639 | 13.95608407 | -0.127742522 | 0.902616863 | 0.979365228 | -5.108193315 |
| **A_55_P2053988** | Epas1 | -0.018210991 | -0.369083494 | 0.332661511 | 12.27104309 | -0.12768898 | 0.902657421 | 0.979365228 | -5.108198466 |
| **A_55_P2397240** | B430316J06Rik | -0.011556458 | -0.234293037 | 0.21118012 | 8.100003267 | -0.12764458 | 0.902691055 | 0.979365228 | -5.108202736 |
| **A_51_P261351** | Mrpl49 | -0.010540964 | -0.213883255 | 0.192801327 | 10.16431137 | -0.127532764 | 0.902775759 | 0.979365228 | -5.108213482 |
| **A_55_P2071426** | Eef2kmt | -0.008605259 | -0.174620818 | 0.1574103 | 8.531102264 | -0.127521765 | 0.90278409 | 0.979365228 | -5.108214538 |
| **A_55_P1973838** | Slc6a9 | 0.007363814 | -0.134712582 | 0.149440211 | 8.352253734 | 0.127511703 | 0.902791713 | 0.979365228 | -5.108215505 |
| **A_55_P1996927** | 4933409K07Rik | 0.014431525 | -0.264314525 | 0.293177576 | 9.345043924 | 0.127371621 | 0.902897831 | 0.979365228 | -5.108228952 |
| **A_55_P1995007** | Arhgap11a | -0.011796884 | -0.240042393 | 0.216448625 | 8.040790554 | -0.127155233 | 0.903061758 | 0.979365228 | -5.108249695 |
| **A_55_P1984144** | NA | -0.0113561 | -0.231243585 | 0.208531386 | 14.91825205 | -0.127056781 | 0.903136342 | 0.979365228 | -5.108259121 |
| **A_51_P493117** | Slc16a9 | 0.022962636 | -0.422074382 | 0.467999654 | 12.19952583 | 0.126938911 | 0.90322564 | 0.979365228 | -5.108270397 |
| **A_55_P2095880** | NA | 0.006436832 | -0.118398141 | 0.131271806 | 7.881856383 | 0.126854265 | 0.903289768 | 0.979365228 | -5.108278488 |
| **A_55_P2136533** | Kmt2d | -0.010822405 | -0.220757962 | 0.199113152 | 9.20922507 | -0.126825602 | 0.903311483 | 0.979365228 | -5.108281227 |
| **A_52_P183524** | Tmem86b | 0.005581617 | -0.102707953 | 0.113871186 | 7.684875866 | 0.126806799 | 0.903325729 | 0.979365228 | -5.108283023 |
| **A_51_P487913** | NA | 0.005510465 | -0.101399607 | 0.112420538 | 7.698480429 | 0.12680571 | 0.903326554 | 0.979365228 | -5.108283127 |
| **A_55_P2012096** | Bmp8a | -0.017897724 | -0.365152528 | 0.329357079 | 11.14838992 | -0.126799843 | 0.903330998 | 0.979365228 | -5.108283687 |
| **A_55_P2158701** | H4c4 | 0.016011645 | -0.294665947 | 0.326689236 | 11.26082073 | 0.126792972 | 0.903336204 | 0.979365228 | -5.108284344 |
| **A_52_P305851** | 2410002F23Rik | 0.019711951 | -0.362994606 | 0.402418509 | 9.220571076 | 0.126716414 | 0.903394206 | 0.979365228 | -5.108291653 |
| **A_52_P564951** | C2cd3 | 0.006392589 | -0.117729781 | 0.130514959 | 8.511814846 | 0.126705621 | 0.903402383 | 0.979365228 | -5.108292684 |
| **A_52_P137378** | Prkaa2 | 0.00895103 | -0.16493959 | 0.18284165 | 8.510833114 | 0.126638576 | 0.903453178 | 0.979365228 | -5.108299081 |
| **A_52_P633489** | C1qtnf4 | -0.006648989 | -0.135833089 | 0.122535112 | 8.083129409 | -0.1266239 | 0.903464296 | 0.979365228 | -5.108300481 |
| **A_51_P205968** | Snx15 | -0.00744679 | -0.152247316 | 0.137353736 | 9.315430287 | -0.126522605 | 0.903541041 | 0.979393494 | -5.108310139 |
| **A_52_P352735** | Ccdc153 | 0.005350029 | -0.098825986 | 0.109526045 | 7.694157211 | 0.126344857 | 0.903675713 | 0.979448504 | -5.108327068 |
| **A_51_P369762** | Nmrk2 | -0.005350789 | -0.109560562 | 0.098858984 | 7.686606036 | -0.126321862 | 0.903693136 | 0.979448504 | -5.108329256 |
| **A_51_P182296** | Washc3 | 0.017289278 | -0.319950421 | 0.354528976 | 11.84576892 | 0.126126783 | 0.903840943 | 0.979541485 | -5.108347805 |
| **A_55_P2176530** | Ldah | 0.00888982 | -0.164762999 | 0.182542639 | 9.160237986 | 0.125944818 | 0.903978817 | 0.979541485 | -5.108365081 |
| **A_55_P1974783** | Tsnax | 0.006724075 | -0.124667911 | 0.138116062 | 11.95209556 | 0.125902066 | 0.904011211 | 0.979541485 | -5.108369137 |
| **A_55_P1964483** | Cyp2c37 | -0.005878455 | -0.120850104 | 0.109093193 | 7.675291508 | -0.125788701 | 0.90409711 | 0.979541485 | -5.108379884 |
| **A_55_P2273811** | Msh5 | -0.007540378 | -0.155151165 | 0.140070409 | 7.884588465 | -0.125673632 | 0.904184301 | 0.979541485 | -5.108390783 |
| **A_66_P100050** | Ndufaf8 | -0.012527406 | -0.258409008 | 0.233354195 | 10.017717 | -0.125344215 | 0.904433919 | 0.979541485 | -5.108421929 |
| **A_55_P2121662** | Mapk3 | 0.007100438 | -0.132297265 | 0.146498141 | 8.672942401 | 0.125313734 | 0.904457017 | 0.979541485 | -5.108424807 |
| **A_55_P2085400** | NA | 0.007931737 | -0.147788793 | 0.163652268 | 7.939537019 | 0.125311694 | 0.904458563 | 0.979541485 | -5.108424999 |
| **A_55_P2166572** | Strbp | -0.00871548 | -0.179890422 | 0.162459462 | 8.912076961 | -0.125262271 | 0.904496014 | 0.979541485 | -5.108429664 |
| **A_55_P2158011** | Knstrn | 0.006327916 | -0.117984533 | 0.130640365 | 7.821132889 | 0.125231977 | 0.904518971 | 0.979541485 | -5.108432522 |
| **A_51_P434171** | Myo19 | -0.009208322 | -0.190292065 | 0.171875422 | 8.738489077 | -0.125103707 | 0.904616173 | 0.979541485 | -5.108444618 |
| **A_55_P2327518** | 1500032P08Rik | -0.010392 | -0.214801682 | 0.194017682 | 8.151488184 | -0.125073946 | 0.904638726 | 0.979541485 | -5.108447422 |
| **A_66_P121117** | Serpina1d | 0.039031108 | -0.728709459 | 0.806771675 | 8.753311891 | 0.125073571 | 0.90463901 | 0.979541485 | -5.108447457 |
| **A_55_P2100884** | Fjx1 | 0.013842417 | -0.258749969 | 0.286434804 | 8.501233244 | 0.124930185 | 0.90474767 | 0.979541485 | -5.10846096 |

| | | | | | | | | |
|---|---|---|---|---|---|---|---|---|
| **A_55_P2012849** | Dock10 | -0.00631251 | -0.130702514 | 0.118077494 | 7.777153925 | -0.124849204 | 0.904809039 | 0.979541485 | -5.108468579 |
| **A_51_P442152** | Itfg1 | -0.011057696 | -0.228988358 | 0.206872965 | 12.6710267 | -0.124828997 | 0.904824353 | 0.979541485 | -5.10847048 |
| **A_55_P2012523** | Twf1 | 0.01720478 | -0.322196523 | 0.356606084 | 11.21573257 | 0.124711006 | 0.904913771 | 0.979541485 | -5.10848157 |
| **A_55_P2048684** | Rpl27 | 0.015287811 | -0.286367011 | 0.316942633 | 13.60372371 | 0.124682126 | 0.904935658 | 0.979541485 | -5.108484284 |
| **A_55_P1965050** | Cks2 | -0.01787898 | -0.370911408 | 0.335153447 | 8.731537093 | -0.124594053 | 0.905002404 | 0.979541485 | -5.108492553 |
| **A_51_P259861** | Mettl9 | -0.01114314 | -0.231212913 | 0.208926632 | 12.34514303 | -0.124570834 | 0.905020001 | 0.979541485 | -5.108494733 |
| **A_55_P1977008** | Gfi1 | -0.005957205 | -0.123626493 | 0.111712084 | 7.74389926 | -0.124551383 | 0.905034742 | 0.979541485 | -5.108496558 |
| **A_55_P2293351** | Slc2a4rg-ps | -0.013812147 | -0.286663666 | 0.259039372 | 8.28492334 | -0.124538599 | 0.90504443 | 0.979541485 | -5.108497758 |
| **A_55_P1965728** | Abca17 | -0.010055905 | -0.208820984 | 0.188709173 | 8.570686795 | -0.124465863 | 0.905099555 | 0.979541485 | -5.10850458 |
| **A_55_P2024406** | Napepld | -0.008523924 | -0.177121908 | 0.16007406 | 8.796430271 | -0.124381664 | 0.905163367 | 0.979541485 | -5.108512472 |
| **A_55_P2147641** | Cstf1 | 0.007245536 | -0.13633633 | 0.150827401 | 10.95663161 | 0.124148097 | 0.905340386 | 0.979541485 | -5.108534339 |
| **A_52_P472319** | Plxnd1 | -0.017440046 | -0.363235585 | 0.328355492 | 8.398620705 | -0.124078751 | 0.905392944 | 0.979541485 | -5.108540823 |
| **A_51_P289742** | Slc27a5 | -0.005516132 | -0.114896501 | 0.103864237 | 7.679998448 | -0.124069325 | 0.905400088 | 0.979541485 | -5.108541704 |
| **A_55_P2011802** | Pdk2 | 0.010948532 | -0.206273261 | 0.228170324 | 9.161618146 | 0.123999989 | 0.905452639 | 0.979541485 | -5.108548183 |
| **A_55_P2027737** | Il1rl1 | -0.005467262 | -0.114035475 | 0.103100952 | 7.785925917 | -0.123890022 | 0.905535987 | 0.979541485 | -5.108558452 |
| **A_55_P2083794** | Tmem159 | -0.012610535 | -0.263287982 | 0.238066913 | 9.514307206 | -0.123762023 | 0.905633002 | 0.979541485 | -5.108570393 |
| **A_51_P516637** | Bmp5 | 0.005583611 | -0.105558774 | 0.116725995 | 7.660621837 | 0.123596048 | 0.905758804 | 0.979541485 | -5.108585859 |
| **A_51_P120461** | Riox2 | -0.013179658 | -0.275824069 | 0.249464753 | 10.44912655 | -0.123453992 | 0.905866479 | 0.979541485 | -5.081007458 |
| **A_55_P1978642** | Wasf1 | 0.020645538 | -0.390936877 | 0.432227953 | 10.12995197 | 0.123406646 | 0.905902367 | 0.979541485 | -5.108603482 |
| **A_52_P441044** | Rfxap | 0.006966597 | -0.131969678 | 0.145902872 | 8.125328785 | 0.123359953 | 0.90593776 | 0.979541485 | -5.108607823 |
| **A_51_P262670** | Rbm38 | -0.006172205 | -0.129275149 | 0.116930738 | 7.949136794 | -0.123350545 | 0.905944891 | 0.979541485 | -5.108608697 |
| **A_55_P2007273** | Pole | -0.007158973 | -0.14997064 | 0.135652694 | 8.774293536 | -0.123326445 | 0.905963159 | 0.979541485 | -5.108610937 |
| **A_52_P77245** | Snx8 | 0.028426039 | -0.538800967 | 0.595653045 | 11.87736995 | 0.123290216 | 0.90599062 | 0.979541485 | -5.108614302 |
| **A_52_P681310** | Plaur | -0.018663205 | -0.391124578 | 0.353798169 | 11.79214549 | -0.123274753 | 0.906002341 | 0.979541485 | -5.108615739 |
| **A_55_P2169669** | Csf1 | -0.028733351 | -0.602696142 | 0.54522944 | 9.314169522 | -0.123160574 | 0.90608889 | 0.979541485 | -5.108626339 |
| **A_51_P493720** | Tfam | -0.015655767 | -0.328525285 | 0.297213751 | 10.40683976 | -0.123106295 | 0.906130034 | 0.979541485 | -5.108631374 |
| **A_55_P2117614** | Tnfrsf13c | 0.005595961 | -0.106320423 | 0.117512344 | 7.689158542 | 0.123012758 | 0.906200938 | 0.979541485 | -5.108640047 |
| **A_66_P131324** | Platr21 | -0.005224281 | -0.10970967 | 0.099261107 | 7.770789473 | -0.123009924 | 0.906203086 | 0.979541485 | -5.108640309 |
| **A_51_P391616** | Etnppl | 0.005315774 | -0.101021449 | 0.111652997 | 7.669537023 | 0.122984498 | 0.90622236 | 0.979541485 | -5.108642666 |
| **A_51_P205545** | Creld1 | -0.018895574 | -0.397222882 | 0.359431735 | 9.216605555 | -0.122874441 | 0.906305787 | 0.979541485 | -5.108652859 |
| **A_55_P2036007** | Rai2 | -0.014625937 | -0.307552098 | 0.278300224 | 8.206929699 | -0.122838557 | 0.906332989 | 0.979541485 | -5.108656181 |
| **A_55_P2106922** | Arfgap3 | -0.007387717 | -0.155408448 | 0.140633015 | 11.22138152 | -0.12278826 | 0.906371117 | 0.979541485 | -5.108660835 |
| **A_51_P335981** | Narf | 0.009518334 | -0.181324783 | 0.200361452 | 8.411051378 | 0.122702557 | 0.906436085 | 0.979541485 | -5.108668761 |
| **A_55_P2144526** | Ripor2 | -0.01219907 | -0.256827796 | 0.232429655 | 7.944185238 | -0.122684144 | 0.906450043 | 0.979541485 | -5.108670463 |
| **A_51_P320481** | Mcee | 0.014830266 | -0.282630364 | 0.312290897 | 10.70560932 | 0.12265595 | 0.906471416 | 0.979541485 | -5.108673069 |
| **A_51_P320852** | Cd9 | 0.018703594 | -0.356485269 | 0.393892458 | 15.62109839 | 0.122643433 | 0.906480905 | 0.979541485 | -5.108674226 |
| **A_55_P2040011** | Itga7 | 0.029308078 | -0.558832412 | 0.617448568 | 9.922708398 | 0.122595752 | 0.906517051 | 0.979541485 | -5.108678631 |
| **A_51_P286792** | Zfp275 | -0.005510082 | -0.116139092 | 0.105118927 | 8.527989983 | -0.122534452 | 0.906563521 | 0.979541485 | -5.108684292 |
| **A_55_P1961451** | Adamts7 | -0.011053966 | -0.233040244 | 0.210932312 | 7.888613612 | -0.122507071 | 0.906584278 | 0.979541485 | -5.108686819 |
| **A_55_P1971938** | Atp2b2 | 0.005846047 | -0.11158067 | 0.123272764 | 7.989124701 | 0.122479816 | 0.90660494 | 0.979541485 | -5.108689335 |
| **A_55_P2167922** | Dmd | -0.005478756 | -0.115566651 | 0.104609139 | 7.648140268 | -0.122436679 | 0.906637642 | 0.979541485 | -5.108693315 |
| **A_55_P1977538** | Apba3 | -0.013216512 | -0.278859734 | 0.252426711 | 10.31278315 | -0.122401647 | 0.906664199 | 0.979541485 | -5.108696546 |
| **A_51_P269134** | Sftpb | -0.005426358 | -0.114534397 | 0.103681681 | 7.640663672 | -0.122354761 | 0.906699744 | 0.979541485 | -5.108700869 |
| **A_55_P1965150** | Pnkp | -0.008601269 | -0.18158183 | 0.164379291 | 11.28211132 | -0.122330404 | 0.906718209 | 0.979541485 | -5.108703115 |
| **A_51_P212679** | Cwf19l1 | 0.005718524 | -0.109451264 | 0.120888312 | 8.33009862 | 0.12215593 | 0.906850479 | 0.979602037 | -5.108719185 |
| **A_51_P150802** | Mpv17 | 0.00752775 | -0.14412061 | 0.159176111 | 10.88663367 | 0.12212277 | 0.906875619 | 0.979602037 | -5.108722236 |
| **A_51_P171999** | Apoe | 0.042915307 | -0.822210268 | 0.908040882 | 13.15638142 | 0.122040011 | 0.906938361 | 0.979615065 | -5.103259197 |
| **A_55_P2157008** | Ear10 | -0.090255864 | -1.910762672 | 1.730250943 | 9.407794747 | -0.121969793 | 0.906991597 | 0.979617824 | -5.108736304 |
| **A_51_P406835** | Mei1 | 0.011401747 | -0.219368467 | 0.242171961 | 8.672146684 | 0.121551637 | 0.907308629 | 0.979762908 | -5.108774668 |
| **A_66_P103229** | Foxl1 | 0.005263685 | -0.101374929 | 0.111902298 | 7.670627005 | 0.121435186 | 0.907396921 | 0.979762908 | -5.108785328 |

| | | | | | | | | |
|---|---|---|---|---|---|---|---|---|
| **A_55_P2027312** | Cdc34 | 0.025251541 | -0.486502862 | 0.537005943 | 15.322954 | 0.121393506 | 0.907428523 | 0.979762908 | -5.108789142 |
| **A_55_P2136426** | Prss57 | -0.005342606 | -0.113646492 | 0.102961279 | 7.689327796 | -0.121360764 | 0.907453348 | 0.979762908 | -5.108792136 |
| **A_51_P513460** | Cfhr1 | 0.00513436 | -0.098988291 | 0.109257012 | 7.633969521 | 0.121313823 | 0.907488939 | 0.979762908 | -5.108796428 |
| **A_55_P2150786** | Mul1 | 0.006206012 | -0.119672464 | 0.132084489 | 11.35926437 | 0.121291486 | 0.907505875 | 0.979762908 | -5.108798469 |
| **A_52_P661** | Plxna4 | 0.011689549 | -0.225490709 | 0.248869806 | 9.911936558 | 0.121251848 | 0.90753593 | 0.979762908 | -5.108802092 |
| **A_52_P560728** | Serhl | -0.005384451 | -0.114651971 | 0.103883068 | 9.401517286 | -0.121232632 | 0.9075505 | 0.979762908 | -5.108803847 |
| **A_52_P377299** | Zbtb11 | -0.008499814 | -0.181190109 | 0.164190481 | 8.731966923 | -0.121090663 | 0.907658145 | 0.979762908 | -5.108816808 |
| **A_51_P229602** | Plxna2 | 0.020090734 | -0.388344666 | 0.428526134 | 10.99201261 | 0.121015665 | 0.907715012 | 0.979762908 | -5.108823649 |
| **A_51_P321341** | Sult1a1 | 0.014335499 | -0.277124227 | 0.305795224 | 9.73405295 | 0.121005027 | 0.907723078 | 0.979762908 | -5.108824619 |
| **A_55_P1955637** | Trp53i11 | 0.010619205 | -0.205309478 | 0.226547887 | 8.550145165 | 0.120990373 | 0.90773419 | 0.979762908 | -5.108825956 |
| **A_55_P2151442** | C330007P06Rik | -0.006869466 | -0.146672 | 0.132933068 | 10.32824651 | -0.120886307 | 0.907813098 | 0.979793365 | -5.108835439 |
| **A_55_P2012984** | Ldha | -0.010312999 | -0.220947897 | 0.200321899 | 15.94216523 | -0.120454715 | 0.908140367 | 0.979820645 | -5.108874683 |
| **A_55_P2081656** | NA | 0.011020547 | -0.214129196 | 0.236170291 | 7.966648607 | 0.120420621 | 0.908166221 | 0.979820645 | -5.108877778 |
| **A_52_P32081** | Crbn | -0.006520449 | -0.139789278 | 0.126748381 | 8.930565479 | -0.120369944 | 0.90820465 | 0.979820645 | -5.108882375 |
| **A_55_P2014249** | Sema3a | 0.017271128 | -0.335888742 | 0.370430997 | 7.99625484 | 0.12031465 | 0.908246581 | 0.979820645 | -5.10888739 |
| **A_51_P390775** | Ube2e1 | 0.015421932 | -0.300207998 | 0.331051861 | 12.00637599 | 0.120207 | 0.908328215 | 0.979820645 | -5.108897145 |
| **A_55_P2064962** | Pdpn | 0.020148195 | -0.392372477 | 0.432668868 | 8.482934629 | 0.120159912 | 0.908363923 | 0.979820645 | -5.10890141 |
| **A_55_P1975285** | Lrig2 | 0.007656919 | -0.149152015 | 0.164465853 | 9.310619552 | 0.120130257 | 0.908386411 | 0.979820645 | -5.108904094 |
| **A_55_P2016667** | Gstm7 | 0.005168704 | -0.100744268 | 0.111081677 | 7.66300045 | 0.120060924 | 0.90843899 | 0.979820645 | -5.108910369 |
| **A_55_P1974367** | Fabp3 | 0.009005497 | -0.175704392 | 0.193715386 | 17.01220928 | 0.119946258 | 0.908525948 | 0.979820645 | -5.108920738 |
| **A_51_P215066** | Emc4 | -0.020566424 | -0.442645038 | 0.401512189 | 13.02185024 | -0.119876654 | 0.908578733 | 0.979820645 | -5.108927028 |
| **A_52_P54261** | Tmem56 | -0.006631106 | -0.142839937 | 0.129577725 | 7.877671046 | -0.1197705 | 0.908659237 | 0.979820645 | -5.108936613 |
| **A_51_P376238** | Serping1 | -0.076771755 | -1.653854879 | 1.500311369 | 11.04996671 | -0.119761175 | 0.908666309 | 0.979820645 | -5.108937455 |
| **A_55_P2111416** | Rcc2 | -0.013541084 | -0.291814567 | 0.264732399 | 9.765477345 | -0.119715607 | 0.908700867 | 0.979820645 | -5.108941566 |
| **A_66_P131091** | Rabl6 | -0.008373436 | -0.180492872 | 0.163745999 | 11.70332924 | -0.119685897 | 0.908723398 | 0.979820645 | -5.108944246 |
| **A_51_P209280** | Rab31 | -0.043535559 | -0.938454864 | 0.851383746 | 10.46014544 | -0.119682157 | 0.908726235 | 0.979820645 | -5.108944584 |
| **A_55_P2055869** | Dvl2 | 0.005701887 | -0.11172898 | 0.123132755 | 9.087669643 | 0.119455331 | 0.90889826 | 0.979820645 | -5.108965022 |
| **A_51_P414126** | Rab19 | -0.005865965 | -0.126714476 | 0.114982545 | 8.394374335 | -0.119417331 | 0.908927079 | 0.979820645 | -5.108968442 |
| **A_55_P2013236** | S100g | 0.026393856 | -0.517866387 | 0.5706541 | 16.02105171 | 0.119306856 | 0.909010866 | 0.979820645 | -5.108978379 |
| **A_55_P2399688** | Glipr2 | 0.005453399 | -0.107015589 | 0.117922386 | 7.999460163 | 0.119289891 | 0.909023733 | 0.979820645 | -5.108979904 |
| **A_51_P139651** | Nos3 | -0.005218793 | -0.112856045 | 0.10241846 | 7.9298106 | -0.119282461 | 0.909029369 | 0.979820645 | -5.097783791 |
| **A_55_P1963772** | Atp5j | 0.01199073 | -0.235343464 | 0.259324924 | 13.33853945 | 0.119269837 | 0.909038943 | 0.979820645 | -5.108981707 |
| **A_51_P104077** | Usp13 | -0.005138482 | -0.111178069 | 0.100901105 | 7.727164073 | -0.119216392 | 0.909079477 | 0.979820645 | -5.103514298 |
| **A_52_P400999** | Arhgap31 | -0.008753628 | -0.189437803 | 0.171930547 | 11.10402369 | -0.119189261 | 0.909100054 | 0.979820645 | -5.108988947 |
| **A_66_P132663** | Uqcrb | -0.011136174 | -0.241058798 | 0.21878645 | 14.24462203 | -0.119158067 | 0.909123713 | 0.979820645 | -5.108991748 |
| **A_51_P234544** | Azin1 | -0.025254574 | -0.546993141 | 0.496483993 | 12.22267972 | -0.119084783 | 0.909179295 | 0.979820645 | -5.108998327 |
| **A_55_P2162582** | Rab5b | 0.008284046 | -0.162903776 | 0.179471868 | 9.014916489 | 0.119052577 | 0.909203722 | 0.979820645 | -5.109001217 |
| **A_51_P434503** | Dtwd1 | 0.006928094 | -0.136319444 | 0.150175633 | 8.207801151 | 0.118985979 | 0.909254234 | 0.979820645 | -5.10900719 |
| **A_55_P1963563** | Zfp689 | 0.007197435 | -0.141624874 | 0.156019745 | 8.551581902 | 0.118981353 | 0.909257743 | 0.979820645 | -5.109007605 |
| **A_51_P193794** | Lrp1 | -0.045953044 | -0.997164211 | 0.905258123 | 11.5260624 | -0.118852004 | 0.90935585 | 0.979850088 | -5.109019197 |
| **A_55_P2044217** | 9930021J03Rik | -0.012610859 | -0.273739657 | 0.248517939 | 11.26173561 | -0.118811658 | 0.909386452 | 0.979850088 | -5.10902281 |
| **A_55_P1999022** | Zfp760 | -0.006273688 | -0.136353737 | 0.12380636 | 8.466720615 | -0.118653734 | 0.909506236 | 0.979901615 | -5.109036941 |
| **A_52_P670978** | Nkain1 | 0.014031317 | -0.277189461 | 0.305252095 | 8.659429337 | 0.11853463 | 0.909596576 | 0.979901615 | -5.109047586 |
| **A_55_P2090194** | NA | -0.009631216 | -0.209532178 | 0.190269746 | 15.82969233 | -0.118531943 | 0.909598614 | 0.979901615 | -5.109047826 |
| **A_55_P2135486** | NA | -0.019400952 | -0.422250178 | 0.383448274 | 8.811300551 | -0.11848126 | 0.909637057 | 0.979901615 | -5.109052352 |
| **A_55_P1974892** | Zbtb18 | -0.010633578 | -0.231750891 | 0.210483734 | 9.662856961 | -0.118311198 | 0.909766054 | 0.979906234 | -5.109067526 |
| **A_55_P2085445** | Acsm1 | 0.005847923 | -0.115785224 | 0.127481069 | 7.694080486 | 0.118282049 | 0.909788165 | 0.979906234 | -5.109070125 |
| **A_55_P2091323** | Wdr26 | 0.011637389 | -0.230504453 | 0.253779231 | 9.361296866 | 0.118237399 | 0.909822033 | 0.979906234 | -5.109074104 |
| **A_55_P1992617** | Clec1b | -0.033373066 | -0.728075882 | 0.66132975 | 10.09630928 | -0.118186047 | 0.909860986 | 0.979906234 | -5.109078679 |
| **A_66_P108276** | Gsg1l | 0.00682672 | -0.135404455 | 0.149057894 | 7.97285611 | 0.118082744 | 0.909939347 | 0.979906234 | -5.109087876 |

| | | | | | | | | | |
|---|---|---|---|---|---|---|---|---|---|
| **A_51_P126337** | Fgf12 | 0.005184346 | -0.103114988 | 0.11348368 | 7.663689554 | 0.117770721 | 0.910176039 | 0.979906234 | -5.092697796 |
| **A_51_P507942** | Atp13a2 | -0.01104335 | -0.242078663 | 0.219991962 | 10.93996849 | -0.117595757 | 0.910308767 | 0.979906234 | -5.109131126 |
| **A_51_P331752** | Ccl11 | 0.005329744 | -0.106196885 | 0.116856374 | 7.685119174 | 0.117570124 | 0.910328212 | 0.979906234 | -5.109133397 |
| **A_55_P1985670** | Exosc3 | 0.014940742 | -0.297984934 | 0.327866419 | 10.011733 | 0.117462744 | 0.910409672 | 0.979906234 | -5.109142908 |
| **A_55_P2210213** | 4732423E21Rik | 0.035621174 | -0.710632142 | 0.78187449 | 9.303035176 | 0.117433259 | 0.91043204 | 0.979906234 | -5.109145517 |
| **A_51_P138923** | Peg12 | -0.006579987 | -0.144439735 | 0.13127976 | 8.220666829 | -0.117423967 | 0.910439089 | 0.979906234 | -5.10914634 |
| **A_51_P489138** | Sptb | 0.005260305 | -0.104959449 | 0.11548006 | 7.678065064 | 0.117414214 | 0.910446488 | 0.979906234 | -5.109147203 |
| **A_51_P208919** | Acbd6 | -0.019974242 | -0.438699587 | 0.398751103 | 10.7442496 | -0.11735733 | 0.910489642 | 0.979906234 | -5.109152235 |
| **A_55_P2181743** | Elmod1 | -0.01542549 | -0.338879429 | 0.308028448 | 8.911930255 | -0.117326388 | 0.910513116 | 0.979906234 | -5.109154972 |
| **A_52_P517896** | Man2c1 | -0.024966451 | -0.548725989 | 0.498793088 | 11.17935333 | -0.117271917 | 0.910554439 | 0.979906234 | -5.109159787 |
| **A_55_P2047673** | Srr | -0.011069317 | -0.243403405 | 0.221264772 | 10.53193987 | -0.117213338 | 0.91059888 | 0.979906234 | -5.109164964 |
| **A_55_P1975989** | Wdr63 | -0.006589792 | -0.144939401 | 0.131759817 | 7.819797735 | -0.11718255 | 0.910622238 | 0.979906234 | -5.109167683 |
| **A_51_P158638** | Ggnbp1 | 0.00528763 | -0.105749655 | 0.116324915 | 7.806719665 | 0.117155152 | 0.910643023 | 0.979906234 | -5.109170103 |
| **A_55_P2038007** | Csrp1 | 0.005860012 | -0.117256215 | 0.12897624 | 16.78509156 | 0.117098789 | 0.910685783 | 0.979906234 | -5.109175078 |
| **A_55_P1980446** | Sbno1 | 0.010224537 | -0.204661647 | 0.225110721 | 11.0998738 | 0.117058867 | 0.910716071 | 0.979906234 | -5.109178601 |
| **A_51_P256957** | Ankmy2 | -0.006588218 | -0.145132035 | 0.131955599 | 8.560530063 | -0.116990325 | 0.910768071 | 0.979906234 | -5.109184646 |
| **A_51_P106059** | Traf4 | -0.021517958 | -0.474608176 | 0.431572261 | 10.7511343 | -0.116838379 | 0.910883349 | 0.979906234 | -5.109198035 |
| **A_51_P126525** | D230025D16Rik | 0.009963754 | -0.199901891 | 0.2198294 | 10.08822224 | 0.116802147 | 0.910910838 | 0.979906234 | -5.092781837 |
| **A_55_P1964282** | Sbk3 | -0.012007027 | -0.264994455 | 0.2409804 | 8.759692598 | -0.11676313 | 0.91094044 | 0.979906234 | -5.10920466 |
| **A_55_P1981405** | Gsk3a | 0.012763155 | -0.256179914 | 0.281706224 | 12.403894 | 0.116752677 | 0.910948371 | 0.979906234 | -5.109205579 |
| **A_55_P1958349** | Nipsnap3b | 0.01228685 | -0.246690632 | 0.271264331 | 8.800206199 | 0.116720652 | 0.910972668 | 0.979906234 | -5.109208397 |
| **A_55_P1985693** | Fhdc1 | 0.025197445 | -0.506129424 | 0.556524313 | 11.81377217 | 0.11667126 | 0.911010142 | 0.979906234 | -5.109212741 |
| **A_51_P481398** | Kif11 | -0.005124714 | -0.113279556 | 0.103030128 | 7.720147282 | -0.116571617 | 0.911085741 | 0.97993302 | -5.1092215 |
| **A_55_P2077535** | Zfp513 | 0.010101215 | -0.203248476 | 0.223450906 | 11.80851503 | 0.116479835 | 0.911155378 | 0.979953389 | -5.109229561 |
| **A_55_P2141943** | NA | -0.006648001 | -0.147582488 | 0.134286486 | 8.003232666 | -0.116049417 | 0.911481956 | 0.980168657 | -5.109267278 |
| **A_55_P2090239** | Casp8ap2 | -0.009309706 | -0.206909198 | 0.188289785 | 8.414505701 | -0.115909577 | 0.911588063 | 0.980168657 | -5.109279503 |
| **A_51_P128575** | Scgb1a1 | -0.067259676 | -1.495032324 | 1.360512972 | 9.30834436 | -0.115895047 | 0.911599088 | 0.980168657 | -5.109280772 |
| **A_51_P281246** | Cep57 | -0.007753829 | -0.172441298 | 0.15693364 | 8.580291008 | -0.115831 | 0.911647686 | 0.980168657 | -5.109286365 |
| **A_55_P2279977** | NA | 0.006881399 | -0.139400453 | 0.153163251 | 8.616841893 | 0.115732533 | 0.911722403 | 0.980168657 | -5.109294958 |
| **A_51_P465148** | Ctsb | -0.007030598 | -0.156559773 | 0.142498576 | 16.51913113 | -0.115673935 | 0.911766867 | 0.980168657 | -5.109300069 |
| **A_55_P2042918** | Trim37 | 0.007570403 | -0.153446149 | 0.168586955 | 7.956122078 | 0.115669174 | 0.91177048 | 0.980168657 | -5.109300484 |
| **A_55_P2055463** | Snapin | -0.009081316 | -0.202260934 | 0.184098301 | 12.28373628 | -0.115652939 | 0.911782799 | 0.980168657 | -5.109301899 |
| **A_55_P2027067** | Phc2 | 0.012096211 | -0.24534221 | 0.269534633 | 13.53149946 | 0.115596628 | 0.911825529 | 0.980168657 | -5.109306807 |
| **A_55_P1998194** | Snrpn | 0.006433548 | -0.130546805 | 0.143413901 | 8.079548906 | 0.115547733 | 0.911862631 | 0.980168657 | -5.109311066 |
| **A_55_P2045136** | Ly6a2 | 0.013216597 | -0.268758712 | 0.295191906 | 7.914884054 | 0.115312856 | 0.912040863 | 0.980231953 | -5.109331502 |
| **A_51_P225832** | 2700097O09Rik | 0.012201641 | -0.248271425 | 0.272674707 | 10.04207426 | 0.11524566 | 0.912091855 | 0.980231953 | -5.10933734 |
| **A_51_P438149** | Mapre2 | -0.010067797 | -0.225017857 | 0.204882263 | 11.74465068 | -0.115230131 | 0.912103639 | 0.980231953 | -5.109338689 |
| **A_55_P2019890** | Rasef | -0.008166868 | -0.182572936 | 0.1662392 | 9.466505799 | -0.115202814 | 0.912124368 | 0.980231953 | -5.109341062 |
| **A_55_P2172807** | NA | -0.006626465 | -0.148359276 | 0.135106346 | 7.741923673 | -0.11502193 | 0.912261634 | 0.980324964 | -5.109356756 |
| **A_55_P2036673** | Ddx52 | 0.007740179 | -0.15813468 | 0.173615037 | 10.23191227 | 0.114799389 | 0.912430517 | 0.980451939 | -5.109376032 |
| **A_55_P2296118** | 9630010G10Rik | 0.013527605 | -0.276641325 | 0.303696535 | 8.374462364 | 0.114693594 | 0.912510805 | 0.980478277 | -5.109385183 |
| **A_51_P398887** | Ctns | -0.009048395 | -0.203239579 | 0.185142789 | 12.5470778 | -0.114633409 | 0.91255648 | 0.980478277 | -5.109390384 |
| **A_51_P123405** | Bub1 | 0.007287305 | -0.149245186 | 0.163819796 | 7.819838978 | 0.114533263 | 0.912632483 | 0.980505434 | -5.109399034 |
| **A_51_P242076** | Ubr7 | 0.015460937 | -0.31739865 | 0.348320523 | 11.15397956 | 0.11427307 | 0.912829953 | 0.980606235 | -5.109421472 |
| **A_51_P115626** | Shank3 | -0.005256549 | -0.11857267 | 0.108059573 | 7.976915531 | -0.114124309 | 0.912942856 | 0.980606235 | -5.109434278 |
| **A_55_P1955325** | Zfp820 | -0.005850076 | -0.131984964 | 0.120284813 | 8.150021747 | -0.114102565 | 0.912959359 | 0.980606235 | -5.109436148 |
| **A_52_P117313** | Azin1 | 0.006145326 | -0.126418181 | 0.138708834 | 7.948786343 | 0.114048645 | 0.913000283 | 0.980606235 | -5.109440785 |
| **A_51_P153973** | Zfp273 | 0.006622042 | -0.13624647 | 0.149490555 | 8.409946935 | 0.114031436 | 0.913013343 | 0.980606235 | -5.109442264 |
| **A_51_P312310** | Mtbp | 0.005079507 | -0.104531264 | 0.114690278 | 8.843273465 | 0.114008568 | 0.9130307 | 0.980606235 | -5.10944423 |
| **A_52_P213696** | Fndc5 | -0.022726551 | -0.513887215 | 0.468434113 | 12.10170817 | -0.113835866 | 0.913161778 | 0.980657241 | -5.109459061 |

| | | | | | | | | |
|---|---|---|---|---|---|---|---|---|
| **A_55_P1996171** | Pcgf5 | -0.031504746 | -0.712750902 | 0.649741411 | 13.87176104 | -0.113773489 | 0.913209122 | 0.980657241 | -5.109464412 |
| **A_51_P259415** | Tmem70 | -0.01316294 | -0.297863282 | 0.271537403 | 10.73619585 | -0.113745459 | 0.913230397 | 0.980657241 | -5.109466816 |
| **A_51_P342082** | Cnpy2 | -0.008511398 | -0.192775215 | 0.17575242 | 12.35739253 | -0.113639674 | 0.913310689 | 0.980662764 | -5.109475882 |
| **A_55_P2412214** | Mhrt | -0.005441193 | -0.123273906 | 0.112391519 | 7.652656518 | -0.113604995 | 0.91333701 | 0.980662764 | -5.109478852 |
| **A_55_P2145401** | H3c3 | -0.006209514 | -0.140832734 | 0.128413706 | 9.071690183 | -0.113476725 | 0.91343437 | 0.980697788 | -5.109489831 |
| **A_51_P185939** | Gper1 | 0.006590049 | -0.136398953 | 0.149579051 | 7.987770929 | 0.113384894 | 0.913504073 | 0.980697788 | -5.109497683 |
| **A_55_P2045876** | NA | 0.006304196 | -0.13065672 | 0.143265112 | 8.20868706 | 0.113240616 | 0.913613587 | 0.980697788 | -5.109510007 |
| **A_55_P2371801** | A430046D13Rik | 0.005259955 | -0.109116495 | 0.119636406 | 7.920552368 | 0.113139591 | 0.913690271 | 0.980697788 | -5.109518627 |
| **A_52_P323830** | Zfp830 | -0.006599629 | -0.150170939 | 0.13697168 | 9.243645613 | -0.113089178 | 0.913728537 | 0.980697788 | -5.109522926 |
| **A_55_P2163344** | NA | -0.011498373 | -0.261973355 | 0.238976609 | 9.244228765 | -0.112938291 | 0.913843072 | 0.980697788 | -5.109535781 |
| **A_51_P219505** | Slc41a2 | 0.014057152 | -0.292314883 | 0.320429187 | 10.66356346 | 0.112880098 | 0.913887246 | 0.980697788 | -5.109540734 |
| **A_51_P499195** | Nkg7 | -0.005003959 | -0.114161436 | 0.104153518 | 7.692524893 | -0.112779312 | 0.913963752 | 0.980697788 | -5.109549306 |
| **A_51_P392147** | Xab2 | -0.009095811 | -0.207551173 | 0.189359551 | 10.29543792 | -0.112758102 | 0.913979853 | 0.980697788 | -5.109551109 |
| **A_51_P241843** | Tmem202 | 0.036754522 | -0.765573404 | 0.839082449 | 8.458358833 | 0.11270107 | 0.914023146 | 0.980697788 | -5.109555956 |
| **A_55_P1984406** | Lrrk2 | 0.007078137 | -0.147440036 | 0.161596309 | 8.521307971 | 0.11269601 | 0.914026987 | 0.980697788 | -5.109556386 |
| **A_55_P2298319** | NA | 0.004834619 | -0.100737652 | 0.110406891 | 7.65797937 | 0.112663068 | 0.914051993 | 0.980697788 | -5.109559184 |
| **A_51_P318381** | Pgf | -0.031707583 | -0.724913055 | 0.661497889 | 8.873783804 | -0.112530519 | 0.914152614 | 0.980697788 | -5.109570435 |
| **A_55_P1953846** | Abca8b | 0.008634893 | -0.180183584 | 0.19745337 | 7.902552404 | 0.112507541 | 0.914170057 | 0.980697788 | -5.109572384 |
| **A_52_P235479** | Usp38 | -0.007615881 | -0.174194304 | 0.158962542 | 9.887888274 | -0.112478774 | 0.914191895 | 0.980697788 | -5.109574824 |
| **A_55_P1952738** | Ppp3ca | 0.008061713 | -0.168443502 | 0.184566927 | 11.87996769 | 0.112367043 | 0.914276714 | 0.980697788 | -5.109584293 |
| **A_52_P577748** | Lpxn | -0.007574385 | -0.173494993 | 0.158346223 | 8.022995518 | -0.112309432 | 0.914320449 | 0.980697788 | -5.109589172 |
| **A_55_P2158741** | Nos2 | -0.015635975 | -0.358515351 | 0.327243401 | 11.22473323 | -0.112189645 | 0.914411386 | 0.980697788 | -5.109599309 |
| **A_51_P110301** | C3 | 0.014487823 | -0.303343209 | 0.332318855 | 11.03287185 | 0.112143988 | 0.914446046 | 0.980697788 | -5.07950436 |
| **A_51_P200144** | Pak1ip1 | 0.025281699 | -0.52939203 | 0.579955429 | 12.5145952 | 0.112134132 | 0.914453529 | 0.980697788 | -5.109604003 |
| **A_52_P302345** | Cyp4v3 | -0.043058754 | -0.988359682 | 0.902242174 | 9.330556661 | -0.112062558 | 0.914507865 | 0.980697788 | -5.109610052 |
| **A_55_P1957389** | Snx25 | -0.006733718 | -0.154574746 | 0.141107311 | 9.7092915 | -0.112054448 | 0.914514022 | 0.980697788 | -5.109610737 |
| **A_52_P252322** | Pdcd11 | -0.005930343 | -0.136167542 | 0.124306857 | 8.07994961 | -0.112024729 | 0.914536584 | 0.980697788 | -5.109613247 |
| **A_52_P2947** | Ndufaf2 | 0.01195023 | -0.25090872 | 0.27480918 | 10.11637682 | 0.111846566 | 0.914671842 | 0.980788419 | -5.109628282 |
| **A_55_P2078735** | AI662270 | 0.010922374 | -0.230429278 | 0.252274026 | 7.913124815 | 0.111336082 | 0.915059411 | 0.98104838 | -5.109671228 |
| **A_55_P1966583** | Bpifc | -0.011599603 | -0.268014779 | 0.244815572 | 12.74946192 | -0.111293186 | 0.91509198 | 0.98104838 | -5.109674828 |
| **A_66_P140335** | Mbnl3 | 0.015559023 | -0.32852608 | 0.359644126 | 10.54367581 | 0.111246311 | 0.915127569 | 0.98104838 | -5.10967876 |
| **A_55_P1955869** | NA | 0.013070641 | -0.276274557 | 0.302415839 | 11.04090972 | 0.11113472 | 0.915212295 | 0.98104838 | -5.109688115 |
| **A_55_P2005135** | Tardbp | -0.013619704 | -0.315284661 | 0.288045252 | 10.14567147 | -0.111073891 | 0.915258481 | 0.98104838 | -5.10969321 |
| **A_51_P427964** | Lmod3 | -0.005000611 | -0.11582913 | 0.105827907 | 7.91986544 | -0.111004534 | 0.915311142 | 0.98104838 | -5.109699016 |
| **A_55_P2141084** | Tenm4 | 0.042441736 | -0.898270423 | 0.983153894 | 10.35195979 | 0.110995543 | 0.915317968 | 0.98104838 | -5.109699768 |
| **A_65_P12533** | Naa10 | -0.006857048 | -0.158846312 | 0.145132217 | 9.198109942 | -0.110992445 | 0.915320321 | 0.98104838 | -5.109700028 |
| **A_55_P1964965** | Gm1673 | 0.015006153 | -0.318030579 | 0.348042886 | 9.917552592 | 0.110852733 | 0.915426402 | 0.981070419 | -5.109711712 |
| **A_55_P1970414** | Mccc1 | -0.006488834 | -0.150525142 | 0.137547473 | 8.06771661 | -0.110831667 | 0.915442397 | 0.981070419 | -5.109713472 |
| **A_55_P2180551** | Sinhcaf | -0.00549964 | -0.12778132 | 0.116782041 | 8.580927647 | -0.110647601 | 0.915582158 | 0.981134919 | -5.10972884 |
| **A_55_P2067942** | D16Ertd472e | -0.012242458 | -0.284565383 | 0.260080467 | 10.05042042 | -0.110599603 | 0.915618604 | 0.981134919 | -5.109732843 |
| **A_55_P2117028** | Mtmr3 | -0.011673503 | -0.271452688 | 0.248105681 | 11.91446078 | -0.110551854 | 0.915654861 | 0.981134919 | -5.109736824 |
| **A_55_P2003903** | Bfsp1 | 0.004958145 | -0.105467359 | 0.11538365 | 8.098244647 | 0.110463555 | 0.915721908 | 0.98115237 | -5.10974418 |
| **A_51_P165934** | Phb | 0.007444275 | -0.158584246 | 0.173472795 | 10.32527747 | 0.110308466 | 0.915839673 | 0.981224158 | -5.109757088 |
| **A_55_P2049854** | Araf | -0.007316572 | -0.170706336 | 0.156073192 | 8.724077658 | -0.110167112 | 0.915947011 | 0.981284767 | -5.109768836 |
| **A_55_P2088375** | Tnnt3 | 0.006646692 | -0.14212027 | 0.155413653 | 8.374425383 | 0.109917843 | 0.916136298 | 0.981433159 | -5.109789517 |
| **A_55_P2011702** | Ppargc1a | -0.004733811 | -0.110961099 | 0.101493476 | 7.664905607 | -0.109633688 | 0.916352084 | 0.981584103 | -5.109813036 |
| **A_55_P2013118** | NA | -0.008620502 | -0.20226784 | 0.185026837 | 8.360820655 | -0.109519175 | 0.916439047 | 0.981584103 | -5.109822497 |
| **A_66_P121774** | Tmem59 | -0.006664127 | -0.156806242 | 0.143477988 | 16.32608912 | -0.109196797 | 0.916683872 | 0.981584103 | -5.109849079 |
| **A_52_P307893** | Pip4k2b | -0.006564922 | -0.15449457 | 0.141364727 | 9.575373789 | -0.109180099 | 0.916696554 | 0.981584103 | -5.109850453 |
| **A_51_P155522** | Psmb4 | -0.010102312 | -0.237896755 | 0.217692132 | 14.59942645 | -0.10910554 | 0.916753178 | 0.981584103 | -5.109856589 |

| | | | | | | | | | |
|---|---|---|---|---|---|---|---|---|---|
| **A_55_P2300071** | 4833412C15Rik | -0.012169623 | -0.286729886 | 0.26239064 | 8.193036063 | -0.109045713 | 0.916798614 | 0.981584103 | -5.109861509 |
| **A_52_P532033** | Hltf | 0.008727162 | -0.188350127 | 0.205804451 | 9.610960124 | 0.108944581 | 0.916875422 | 0.981584103 | -5.109869821 |
| **A_55_P2177105** | NA | -0.009898043 | -0.233469753 | 0.213673668 | 10.41408817 | -0.108918495 | 0.916895234 | 0.981584103 | -5.109871963 |
| **A_55_P2080794** | Taf9 | -0.007020796 | -0.172617657 | 0.158576064 | 11.07952317 | -0.109871323 | 0.916895502 | 0.981584103 | -5.040868632 |
| **A_55_P2048041** | Slc44a1 | 0.014550032 | -0.314287144 | 0.343387209 | 8.752806582 | 0.108855961 | 0.916942727 | 0.981584103 | -5.109877097 |
| **A_65_P19933** | Zdhhc23 | 0.007901647 | -0.170699197 | 0.18650249 | 8.629518767 | 0.10884369 | 0.916952047 | 0.981584103 | -5.109878104 |
| **A_55_P2024640** | Eral1 | 0.007863842 | -0.169910066 | 0.18563775 | 9.675506629 | 0.108826816 | 0.916964862 | 0.981584103 | -5.109879489 |
| **A_55_P2293967** | Tbc1d5 | 0.006354673 | -0.137501276 | 0.150210623 | 9.222415891 | 0.108676231 | 0.917079231 | 0.981584103 | -5.109891837 |
| **A_51_P230624** | Ccdc71l | 0.008248907 | -0.178634868 | 0.195132682 | 10.03481816 | 0.108591029 | 0.917143942 | 0.981584103 | -5.109898816 |
| **A_55_P1957880** | Jpt2 | 0.0170543 | -0.37003341 | 0.40414201 | 10.00878581 | 0.108391114 | 0.917295782 | 0.981584103 | -5.10991517 |
| **A_55_P2124415** | 6030468B19Rik | -0.004697613 | -0.111424467 | 0.102029242 | 7.664030273 | -0.108286083 | 0.917375556 | 0.981584103 | -5.10992375 |
| **A_52_P516021** | Ptpn1 | -0.010769571 | -0.255471676 | 0.233932534 | 8.738828665 | -0.108275416 | 0.917383659 | 0.981584103 | -5.109924621 |
| **A_52_P150683** | NA | -0.004761557 | -0.113080066 | 0.103556952 | 7.699934238 | -0.10814725 | 0.917481006 | 0.981584103 | -5.109935079 |
| **A_51_P335480** | 1810055G02Rik | 0.010192454 | -0.221961918 | 0.242346827 | 9.619750566 | 0.108011766 | 0.917583914 | 0.981584103 | -5.10994612 |
| **A_55_P2009042** | NA | 0.025728603 | -0.560416546 | 0.611873752 | 11.59384905 | 0.107989167 | 0.91760108 | 0.981584103 | -5.10994796 |
| **A_55_P2052490** | Ushbp1 | -0.006666751 | -0.158576491 | 0.145242988 | 8.024632256 | -0.107968683 | 0.917616639 | 0.981584103 | -5.109949628 |
| **A_55_P2025008** | Hvcn1 | 0.00952714 | -0.20787679 | 0.22693107 | 10.24590681 | 0.107811307 | 0.917736179 | 0.981584103 | -5.109962431 |
| **A_52_P228318** | Smu1 | 0.01488706 | -0.324862097 | 0.354636217 | 13.97756764 | 0.107800235 | 0.917744589 | 0.981584103 | -5.109963331 |
| **A_55_P2144456** | Kmt5b | 0.005862221 | -0.128040499 | 0.13976494 | 8.513859601 | 0.107706502 | 0.917815787 | 0.981584103 | -5.109970947 |
| **A_55_P2159666** | NA | -0.007020108 | -0.167409973 | 0.153369757 | 9.144474158 | -0.107680244 | 0.917835733 | 0.981584103 | -5.109973079 |
| **A_51_P190961** | Gzmf | 0.071822392 | -1.570024928 | 1.713669712 | 10.40897969 | 0.10762081 | 0.917880879 | 0.981584103 | -5.109977904 |
| **A_52_P600535** | Zfp39 | 0.004840628 | -0.10581644 | 0.115497696 | 7.910114399 | 0.107619689 | 0.91788173 | 0.981584103 | -5.109977995 |
| **A_52_P134924** | Wdr20 | -0.011981265 | -0.286026744 | 0.262064214 | 9.028449061 | -0.107559604 | 0.917927371 | 0.981584103 | -5.10998287 |
| **A_55_P2153251** | Plagl1 | -0.005190289 | -0.123941134 | 0.113560556 | 8.318002195 | -0.1075286 | 0.917950922 | 0.981584103 | -5.109985384 |
| **A_51_P317181** | Pcdhb21 | 0.005369322 | -0.11753571 | 0.128274355 | 7.765200412 | 0.107477853 | 0.91798947 | 0.981584103 | -5.109989497 |
| **A_55_P2023424** | Trip10 | 0.013522478 | -0.296238518 | 0.323283475 | 10.35720192 | 0.107398627 | 0.918049652 | 0.981584103 | -5.109995916 |
| **A_55_P2182222** | Rab3c | -0.02717025 | -0.649639699 | 0.5952992 | 12.31536844 | -0.107385279 | 0.918059792 | 0.981584103 | -5.109996997 |
| **A_55_P2029763** | Peg10 | -0.007503352 | -0.179418173 | 0.164411469 | 10.51738047 | -0.10737698 | 0.918066096 | 0.981584103 | -5.109997669 |
| **A_55_P2170256** | Ptgr2 | 0.005783326 | -0.126916133 | 0.138482785 | 8.458582734 | 0.107220469 | 0.918184987 | 0.981584103 | -5.110010332 |
| **A_51_P359046** | Slurp1 | -0.004840068 | -0.115921197 | 0.106241061 | 7.6952201 | -0.107196439 | 0.918203241 | 0.981584103 | -5.110012275 |
| **A_55_P2108201** | Gnptab | -0.006970771 | -0.166987094 | 0.153045551 | 9.084163842 | -0.10717308 | 0.918220986 | 0.981584103 | -5.110014163 |
| **A_65_P07627** | Dach1 | -0.004687721 | -0.112337761 | 0.102962318 | 7.685975589 | -0.107131396 | 0.918252651 | 0.981584103 | -5.110017531 |
| **A_51_P359333** | Fh1 | 0.01821374 | -0.40006907 | 0.43649655 | 11.77356615 | 0.107126837 | 0.918256114 | 0.981584103 | -5.110017899 |
| **A_51_P257951** | Retnla | 0.004747293 | -0.104367618 | 0.113862205 | 7.651983802 | 0.107036312 | 0.918324882 | 0.981584103 | -5.110025209 |
| **A_52_P236705** | Ripply3 | -0.00800843 | -0.192239955 | 0.176223094 | 8.483030198 | -0.106943061 | 0.918395722 | 0.981584103 | -5.110032732 |
| **A_55_P2180884** | Psmd10 | 0.012174384 | -0.268041805 | 0.292390573 | 9.766178027 | 0.10688652 | 0.918438674 | 0.981584103 | -5.110037291 |
| **A_55_P2062936** | Abca3 | 0.014115607 | -0.311127608 | 0.339358822 | 12.7953308 | 0.10677276 | 0.918525095 | 0.981584103 | -5.110046455 |
| **A_51_P443693** | Rasl10b | 0.00483688 | -0.10664374 | 0.116317499 | 7.690784185 | 0.106741939 | 0.918548509 | 0.981584103 | -5.110048936 |
| **A_55_P2153291** | NA | 0.016821307 | -0.37193519 | 0.405577804 | 14.72972498 | 0.106451367 | 0.918769257 | 0.981584103 | -5.110072293 |
| **A_55_P1999561** | Pram1 | 0.005300747 | -0.117347807 | 0.127949301 | 7.936586097 | 0.106327065 | 0.918863692 | 0.981584103 | -5.110082266 |
| **A_55_P2095019** | NA | -0.00576548 | -0.13935617 | 0.127825211 | 8.26472845 | -0.106176504 | 0.918978078 | 0.981584103 | -5.110094329 |
| **A_51_P513085** | Gins4 | -0.007180407 | -0.173731511 | 0.159370696 | 9.664412176 | -0.106064668 | 0.919063045 | 0.981584103 | -5.110103279 |
| **A_55_P2015092** | E430024I08Rik | 0.00596617 | -0.132438491 | 0.14437083 | 8.984262947 | 0.106050822 | 0.919073565 | 0.981584103 | -5.110104387 |
| **A_66_P112399** | BC029722 | -0.014333636 | -0.347052844 | 0.318385572 | 11.20180638 | -0.105985793 | 0.919122971 | 0.981584103 | -5.110109586 |
| **A_55_P2159595** | NA | 0.007549232 | -0.167878867 | 0.182977331 | 7.891407823 | 0.105869965 | 0.919210973 | 0.981584103 | -5.110118838 |
| **A_52_P598605** | Ptch1 | -0.005484505 | -0.132942093 | 0.121973083 | 7.999856021 | -0.105862197 | 0.919216874 | 0.981584103 | -5.110119458 |
| **A_51_P204247** | C8a | 0.00472356 | -0.1051415 | 0.114588621 | 7.64755413 | 0.105774025 | 0.919283866 | 0.981584103 | -5.110126494 |
| **A_55_P2103902** | Mrpl4 | 0.010867895 | -0.241992652 | 0.263728442 | 10.55202558 | 0.105738593 | 0.919310786 | 0.981584103 | -5.11012932 |
| **A_51_P246854** | Acta1 | -0.01408675 | -0.341872995 | 0.313699495 | 8.704343526 | -0.105727809 | 0.91931898 | 0.981584103 | -5.11013018 |
| **A_55_P2140780** | Clip4 | 0.004817154 | -0.107316821 | 0.116951129 | 7.710278668 | 0.105687219 | 0.919349819 | 0.981584103 | -5.110133416 |

| | | | | | | | | | |
|---|---|---|---|---|---|---|---|---|---|
| **A_55_P2099750** | NA | -0.01408022 | -0.341863793 | 0.313703353 | 11.72411537 | -0.105679664 | 0.919355559 | 0.981584103 | -5.110134018 |
| **A_52_P378287** | Lca5 | -0.005669277 | -0.137690255 | 0.126351701 | 8.210927177 | -0.105646196 | 0.919380988 | 0.981584103 | -5.110136685 |
| **A_51_P172085** | Arhgdig | -0.009268826 | -0.225150993 | 0.206613341 | 8.198064984 | -0.10562753 | 0.919395171 | 0.981584103 | -5.110138172 |
| **A_55_P2086075** | Gm2058 | 0.014632879 | -0.326501635 | 0.355767394 | 10.28641612 | 0.105529358 | 0.919469761 | 0.981584103 | -5.110145988 |
| **A_52_P513177** | Sla | -0.004521562 | -0.109955463 | 0.100912339 | 7.77495843 | -0.10550605 | 0.91948747 | 0.981584103 | -5.110147843 |
| **A_66_P103477** | Nkapd1 | -0.006839231 | -0.166489944 | 0.152811482 | 9.702896775 | -0.105391492 | 0.919574512 | 0.981584103 | -5.110156953 |
| **A_51_P387845** | Fam120a | -0.010858809 | -0.26435264 | 0.242635022 | 10.81327923 | -0.105386255 | 0.919578492 | 0.981584103 | -5.110157369 |
| **A_52_P191537** | NA | 0.005715321 | -0.127843034 | 0.139273675 | 10.15212733 | 0.105278262 | 0.919660547 | 0.981584103 | -5.110165947 |
| **A_52_P274496** | Tspan18 | -0.014557396 | -0.355015314 | 0.325900521 | 10.95882705 | -0.105193628 | 0.919724854 | 0.981584103 | -5.110172664 |
| **A_55_P2037146** | Igfbpl1 | -0.004463875 | -0.108911657 | 0.099983908 | 7.644369438 | -0.10514338 | 0.919763035 | 0.981584103 | -5.11017665 |
| **A_52_P282762** | Myd88 | -0.005753039 | -0.140525722 | 0.129019644 | 10.51689215 | -0.105018207 | 0.919858147 | 0.981584103 | -5.071084269 |
| **A_55_P2092668** | Kyat3 | 0.004623396 | -0.103710738 | 0.11295753 | 7.739553891 | 0.104994108 | 0.919876458 | 0.981584103 | -5.110188477 |
| **A_55_P1975045** | Sgcg | -0.004591377 | -0.112247484 | 0.103064731 | 7.819162506 | -0.104923659 | 0.919929989 | 0.981584103 | -5.110194054 |
| **A_55_P2061386** | Chd3 | -0.014679374 | -0.359055997 | 0.329697248 | 11.86032484 | -0.104868014 | 0.919972272 | 0.981584103 | -5.110198456 |
| **A_51_P235801** | Zfp36l1 | -0.010757344 | -0.263172443 | 0.241657755 | 10.97307435 | -0.104847696 | 0.919987711 | 0.981584103 | -5.110200062 |
| **A_65_P15412** | Usp13 | -0.004709149 | -0.115269927 | 0.105851629 | 7.725795625 | -0.104787747 | 0.920033264 | 0.981584103 | -5.110204801 |
| **A_66_P128701** | NA | -0.005507961 | -0.134914034 | 0.123898111 | 14.89447116 | -0.104714156 | 0.920089184 | 0.981584103 | -5.110210615 |
| **A_51_P131408** | Tnfrsf12a | -0.029859597 | -0.731492805 | 0.671773611 | 11.87044527 | -0.104699105 | 0.920100621 | 0.981584103 | -5.110211803 |
| **A_55_P2170670** | NA | -0.005067146 | -0.124144938 | 0.114010645 | 9.059292557 | -0.104689192 | 0.920108153 | 0.981584103 | -5.110212586 |
| **A_52_P593037** | Acsl5 | -0.00993021 | -0.243295857 | 0.223435438 | 9.584381389 | -0.104686495 | 0.920110203 | 0.981584103 | -5.110212799 |
| **A_66_P134501** | Olfm1 | -0.009771255 | -0.239636835 | 0.220094326 | 8.217147437 | -0.104579257 | 0.920191692 | 0.981584103 | -5.110221261 |
| **A_66_P101499** | AW822252 | -0.01337909 | -0.328762649 | 0.302004469 | 8.765048646 | -0.104365436 | 0.920354174 | 0.981584103 | -5.110238108 |
| **A_55_P1987761** | Prkcz | 0.012007381 | -0.271159028 | 0.295173789 | 9.905444738 | 0.104321955 | 0.920387216 | 0.981584103 | -5.11024153 |
| **A_55_P2095039** | Ncmap | 0.021171257 | -0.478419493 | 0.520762007 | 11.86056771 | 0.104256087 | 0.92043727 | 0.981584103 | -5.11024671 |
| **A_51_P433127** | Rdm1 | -0.011393017 | -0.280564697 | 0.257778663 | 11.84281384 | -0.104130643 | 0.920532598 | 0.981584103 | -5.110256568 |
| **A_66_P126563** | Ptges3l | 0.008853961 | -0.20050931 | 0.218217232 | 8.562949241 | 0.10404142 | 0.920600402 | 0.981584103 | -5.110263572 |
| **A_51_P106799** | Pparg | -0.017098185 | -0.421487767 | 0.387291397 | 13.7156498 | -0.104020567 | 0.920616249 | 0.981584103 | -5.086752846 |
| **A_55_P1967880** | Chtf8 | 0.014272366 | -0.323653475 | 0.352198206 | 12.91828428 | 0.103906743 | 0.920702749 | 0.981584103 | -5.110274133 |
| **A_55_P2014987** | Castor2 | 0.022885126 | -0.519405464 | 0.565175715 | 11.10990404 | 0.10382224 | 0.920766968 | 0.981584103 | -5.110280752 |
| **A_55_P2098428** | Bloc1s4 | -0.007340741 | -0.181312442 | 0.166630959 | 9.533237433 | -0.103807914 | 0.920777855 | 0.981584103 | -5.110281874 |
| **A_51_P252363** | Stx12 | -0.00937674 | -0.231727238 | 0.212973759 | 11.94861696 | -0.103748761 | 0.920822809 | 0.981584103 | -5.110286504 |
| **A_55_P1957228** | Sgk3 | -0.007080684 | -0.175012646 | 0.160851277 | 11.27310639 | -0.103731589 | 0.920835859 | 0.981584103 | -5.110287847 |
| **A_51_P436068** | Gpr182 | 0.019577946 | -0.446442224 | 0.485598116 | 8.804789212 | 0.10335502 | 0.921122046 | 0.981584103 | -5.110317254 |
| **A_52_P501875** | Ccdc173 | 0.004525734 | -0.103245601 | 0.11229707 | 7.805487066 | 0.103312996 | 0.921153985 | 0.981584103 | -5.11032053 |
| **A_55_P2071532** | NA | -0.011648865 | -0.289249119 | 0.265951388 | 14.40118936 | -0.103236416 | 0.921212186 | 0.981584103 | -5.110326494 |
| **A_55_P1958160** | Sgce | -0.014549931 | -0.361340029 | 0.332240168 | 11.72613772 | -0.103219891 | 0.921224746 | 0.981584103 | -5.110327781 |
| **A_51_P192589** | Pno1 | 0.012075362 | -0.275757701 | 0.299908425 | 11.44867826 | 0.103211629 | 0.921231025 | 0.981584103 | -5.110328424 |
| **A_55_P1963868** | Plekho2 | -0.054174961 | -1.347110417 | 1.238760495 | 12.63616512 | -0.10308392 | 0.921328087 | 0.981584103 | -5.110338359 |
| **A_55_P1987984** | NA | -0.006991772 | -0.173881432 | 0.159897888 | 8.982154431 | -0.103068747 | 0.921339618 | 0.981584103 | -5.110339539 |
| **A_55_P1966659** | LOC547349 | 0.024090647 | -0.551228978 | 0.599410273 | 8.962960926 | 0.103016893 | 0.92137903 | 0.981584103 | -5.110343569 |
| **A_55_P2048249** | NA | 0.005010329 | -0.114733882 | 0.124754539 | 12.7534313 | 0.102939215 | 0.921438067 | 0.981584103 | -5.110349602 |
| **A_55_P2059765** | NA | 0.00923321 | -0.211528078 | 0.229994499 | 10.60584806 | 0.102896113 | 0.921470827 | 0.981584103 | -5.110352948 |
| **A_55_P2086507** | Bglap3 | 0.02661151 | -0.610046313 | 0.663269333 | 11.25192159 | 0.102833022 | 0.921518779 | 0.981584103 | -5.110357842 |
| **A_51_P279552** | Cav2 | 0.008669715 | -0.198757861 | 0.216097292 | 8.76500213 | 0.102827083 | 0.921523293 | 0.981584103 | -5.110358303 |
| **A_55_P1953226** | H2al1e | -0.00481626 | -0.12005473 | 0.11042221 | 7.842297585 | -0.102820953 | 0.921527952 | 0.981584103 | -5.110358778 |
| **A_55_P2171648** | Ppp1r7 | 0.015819945 | -0.362740523 | 0.394380412 | 10.918911 | 0.10281083 | 0.921535646 | 0.981584103 | -5.110359563 |
| **A_51_P185593** | Synm | 0.004819596 | -0.110543442 | 0.120182635 | 7.725938586 | 0.10278107 | 0.921558265 | 0.981584103 | -5.110361871 |
| **A_51_P502132** | Mmp23 | -0.029031931 | -0.724789576 | 0.666725714 | 9.177320219 | -0.102656652 | 0.92165283 | 0.981584103 | -5.11037151 |
| **A_55_P2029151** | Snap47 | -0.006424705 | -0.16052797 | 0.14767856 | 8.491833887 | -0.102567672 | 0.921720462 | 0.981584103 | -5.110378396 |
| **A_51_P331385** | Paip2b | -0.007459078 | -0.186419171 | 0.171501015 | 8.905290156 | -0.102541129 | 0.921740637 | 0.981584103 | -5.11038045 |

| | | | | | | | | | |
|---|---|---|---|---|---|---|---|---|---|
| A_51_P491667 | Derl3 | -0.017507233 | -0.437694893 | 0.402680428 | 12.99434909 | -0.102504603 | 0.921768399 | 0.981584103 | -5.110383274 |
| A_55_P2128695 | Nckap1 | -0.008045224 | -0.20123767 | 0.185147222 | 10.7668795 | -0.102451226 | 0.92180897 | 0.981584103 | -5.1103874 |
| A_55_P2340448 | B230114P17Rik | 0.00464177 | -0.107008266 | 0.116291806 | 7.77253907 | 0.102280752 | 0.921938546 | 0.981584103 | -5.110400562 |
| A_55_P2016105 | Errfi1 | -0.03046284 | -0.763258332 | 0.702332651 | 13.15988273 | -0.102272004 | 0.921945195 | 0.981584103 | -5.110401237 |
| A_52_P566840 | Adgrf1 | 0.00446934 | -0.103194623 | 0.112133303 | 7.687195316 | 0.102127383 | 0.922055123 | 0.981584103 | -5.110412385 |
| A_55_P2064676 | Gas2l3 | -0.004404831 | -0.110533264 | 0.101723601 | 8.170712818 | -0.102109631 | 0.922068617 | 0.981584103 | -5.110413752 |
| A_55_P1986552 | Mcam | -0.013430554 | -0.337092082 | 0.310230975 | 13.98674857 | -0.102087368 | 0.922085539 | 0.981584103 | -5.110415466 |
| A_55_P2054817 | Rxfp1 | 0.00471983 | -0.109041601 | 0.118481261 | 7.808102932 | 0.102070551 | 0.922098322 | 0.981584103 | -5.110416761 |
| A_55_P2054998 | NA | -0.018979327 | -0.47647508 | 0.438516427 | 10.80304978 | -0.102061723 | 0.922105032 | 0.981584103 | -5.110417441 |
| A_52_P66199 | Fbxo11 | -0.008706639 | -0.218612773 | 0.201199496 | 9.651660961 | -0.102045662 | 0.922117241 | 0.981584103 | -5.110418677 |
| A_51_P151897 | R3hcc1 | 0.006063217 | -0.140215933 | 0.152342367 | 8.648846703 | 0.101974091 | 0.922171644 | 0.981587957 | -5.110424184 |
| A_55_P2164977 | NA | -0.009905464 | -0.249238442 | 0.229427514 | 8.48736607 | -0.101821962 | 0.922287283 | 0.981656987 | -5.110435876 |
| A_55_P2136501 | Midn | 0.013691106 | -0.317450255 | 0.344832466 | 12.57220631 | 0.101717167 | 0.922366942 | 0.981687717 | -5.11044392 |
| A_55_P1996653 | D030056L22Rik | 0.014024351 | -0.325694366 | 0.353743068 | 11.26283401 | 0.101562283 | 0.922484679 | 0.981692136 | -5.110455794 |
| A_66_P125257 | Cep120 | -0.006882357 | -0.173878593 | 0.160113878 | 9.550608852 | -0.101391065 | 0.922614834 | 0.981692136 | -5.110468899 |
| A_51_P114878 | Elp1 | 0.006356568 | -0.147973733 | 0.160686869 | 10.47996036 | 0.101330614 | 0.922660788 | 0.981692136 | -5.070482622 |
| A_51_P424272 | Mt4 | -0.008738418 | -0.221015314 | 0.203538478 | 8.00599885 | -0.101274301 | 0.922703597 | 0.981692136 | -5.110477824 |
| A_51_P442964 | Knl1 | 0.004360466 | -0.101692809 | 0.110413741 | 7.695243757 | 0.101152814 | 0.922795951 | 0.981692136 | -5.110487099 |
| A_55_P2060253 | Gm12657 | 0.006825193 | -0.159263742 | 0.172914127 | 13.32261683 | 0.101098187 | 0.922837478 | 0.981692136 | -5.110491266 |
| A_51_P248638 | Myoz2 | -0.014562746 | -0.369061593 | 0.339936101 | 7.985140868 | -0.101064265 | 0.922863267 | 0.981692136 | -5.110493852 |
| A_55_P2096055 | NA | 0.01670637 | -0.390228408 | 0.423641149 | 13.75952904 | 0.101001182 | 0.922911223 | 0.981692136 | -5.110498659 |
| A_65_P17602 | Pgm5 | -0.014491476 | -0.36781847 | 0.338835517 | 7.953535495 | -0.100903208 | 0.922985705 | 0.981692136 | -5.11050612 |
| A_55_P1972381 | Gjb4 | -0.009412674 | -0.239190648 | 0.2203653 | 8.423514791 | -0.100779867 | 0.923079472 | 0.981692136 | -5.110515502 |
| A_55_P1963529 | Abcb10 | 0.009841531 | -0.230484425 | 0.250167487 | 11.19274557 | 0.100746773 | 0.923104632 | 0.981692136 | -5.110518017 |
| A_52_P18765 | Hsbp1l1 | 0.005258339 | -0.123210101 | 0.133726778 | 7.972547673 | 0.100698095 | 0.923141639 | 0.981692136 | -5.110521715 |
| A_55_P2156062 | NA | -0.011468405 | -0.292105966 | 0.269169156 | 8.829064298 | -0.100537107 | 0.92326403 | 0.981692136 | -5.110533934 |
| A_55_P2117726 | Stk33 | 0.004368342 | -0.102548684 | 0.111285368 | 7.68355044 | 0.100516867 | 0.923279418 | 0.981692136 | -5.110535469 |
| A_51_P453167 | Naxd | 0.006426004 | -0.15090203 | 0.163754038 | 12.57699891 | 0.100485644 | 0.923303156 | 0.981692136 | -5.110537836 |
| A_51_P371867 | 3830417A13Rik | -0.027751987 | -0.707282155 | 0.651778181 | 13.22439497 | -0.100474186 | 0.923311867 | 0.981692136 | -5.110538704 |
| A_55_P2111429 | Btf3l4 | 0.008439776 | -0.198216768 | 0.215096319 | 10.72129569 | 0.100473354 | 0.923312499 | 0.981692136 | -5.110538767 |
| A_55_P1997604 | Pla2g4a | -0.021367943 | -0.544949604 | 0.502213718 | 10.18534317 | -0.100403176 | 0.923365853 | 0.981692136 | -5.110544084 |
| A_55_P2010038 | Tnfsf9 | -0.038646978 | -0.985665642 | 0.908371686 | 9.3891464 | -0.100398256 | 0.923369594 | 0.981692136 | -5.110544457 |
| A_55_P2165028 | NA | 0.012314659 | -0.289516221 | 0.314145539 | 14.58972508 | 0.100375537 | 0.923386866 | 0.981692136 | -5.110546177 |
| A_55_P2020256 | Sik2 | 0.007329918 | -0.172526033 | 0.187185869 | 9.014542954 | 0.100263644 | 0.923471936 | 0.981728581 | -5.110554644 |
| A_55_P2106033 | Myo18b | 0.025065749 | -0.592018938 | 0.642150436 | 8.236912215 | 0.099932116 | 0.923723996 | 0.981942536 | -5.110579677 |
| A_55_P1963807 | Actg2 | 0.077522022 | -1.834392531 | 1.989436576 | 11.13280646 | 0.099752957 | 0.923860214 | 0.981944578 | -5.11059317 |
| A_55_P2079860 | Asap3 | -0.00944114 | -0.242422637 | 0.223540356 | 8.992807548 | -0.099694727 | 0.923904488 | 0.981944578 | -5.11059755 |
| A_66_P120882 | Nfkbib | -0.010009344 | -0.257147981 | 0.237129292 | 9.367192957 | -0.099640099 | 0.923946024 | 0.981944578 | -5.110601657 |
| A_55_P1971010 | Gzme | 0.067235003 | -1.59329993 | 1.727769937 | 12.71985798 | 0.099613128 | 0.923966531 | 0.981944578 | -5.110603684 |
| A_51_P512157 | Slc30a6 | -0.006543986 | -0.168274137 | 0.155186164 | 9.178522508 | -0.099545255 | 0.924018137 | 0.981944578 | -5.110608783 |
| A_52_P596008 | Zeb2 | 0.017338876 | -0.411479344 | 0.446157097 | 9.267295723 | 0.099475672 | 0.924071045 | 0.981944578 | -5.110614006 |
| A_55_P2025820 | NA | -0.010801857 | -0.278141284 | 0.25653757 | 8.217925752 | -0.099404163 | 0.924125418 | 0.981944578 | -5.11061937 |
| A_52_P451073 | Tnfrsf21 | 0.018168518 | -0.431930511 | 0.468267547 | 9.045604239 | 0.099307168 | 0.924199169 | 0.981944578 | -5.11062664 |
| A_55_P2355345 | A130019P10Rik | 0.011820754 | -0.281472724 | 0.305114231 | 8.580893745 | 0.099154386 | 0.92431534 | 0.981944578 | -5.110638076 |
| A_55_P2154252 | Gfpt2 | 0.009904604 | -0.235868325 | 0.255677533 | 8.653853953 | 0.099145326 | 0.924322229 | 0.981944578 | -5.110638754 |
| A_51_P212183 | Dmac1 | -0.007094166 | -0.183162369 | 0.168974038 | 10.80726053 | -0.099126454 | 0.924336579 | 0.981944578 | -5.110640165 |
| A_52_P675530 | Fat4 | -0.005148759 | -0.133003079 | 0.122705562 | 7.864794953 | -0.099073222 | 0.924377056 | 0.981944578 | -5.110644144 |
| A_55_P1994898 | NA | -0.014435474 | -0.373095492 | 0.344224543 | 11.64108979 | -0.099018707 | 0.924418509 | 0.981944578 | -5.110648218 |
| A_55_P2002815 | Rabgap1l | 0.004533727 | -0.108137939 | 0.117205392 | 7.770558843 | 0.098994205 | 0.92443714 | 0.981944578 | -5.110650048 |
| A_55_P2138071 | NA | 0.009746502 | -0.232858242 | 0.252351246 | 11.26571217 | 0.098836796 | 0.924556834 | 0.981995946 | -5.110661793 |

| | | | | | | | | |
|---|---|---|---|---|---|---|---|---|
| **A_51_P204402** | Shcbp1 | -0.008652384 | -0.224109699 | 0.206804931 | 8.024144574 | -0.098796983 | 0.924587108 | 0.981995946 | -5.110664761 |
| **A_55_P1991920** | Dhx32 | 0.00878087 | -0.21047056 | 0.228032301 | 11.44367181 | 0.098529047 | 0.924790853 | 0.982110173 | -5.110684703 |
| **A_55_P2172396** | A530064D06Rik | 0.004730467 | -0.113638747 | 0.123099682 | 7.799931164 | 0.098318318 | 0.924951101 | 0.982110173 | -5.11070035 |
| **A_55_P2093614** | NA | -0.005941169 | -0.1547697 | 0.142887361 | 7.997464719 | -0.098209817 | 0.925033612 | 0.982110173 | -5.110708393 |
| **A_51_P286142** | Znrd1 | -0.005698451 | -0.148505134 | 0.137108231 | 11.14162669 | -0.0981697 | 0.925064119 | 0.982110173 | -5.110711364 |
| **A_52_P57308** | Cand1 | -0.004736046 | -0.123475826 | 0.114003734 | 13.21785534 | -0.098127071 | 0.925096537 | 0.982110173 | -5.110714521 |
| **A_55_P2096465** | NA | -0.005169265 | -0.134860819 | 0.124522289 | 11.39086328 | -0.098058732 | 0.925148507 | 0.982110173 | -5.110719578 |
| **A_55_P2137932** | NA | -0.006946543 | -0.181436602 | 0.167543517 | 8.18582584 | -0.097941596 | 0.925237587 | 0.982110173 | -5.110728238 |
| **A_55_P1970164** | Rnf11 | 0.012797066 | -0.309233803 | 0.334827935 | 10.82230397 | 0.0977647 | 0.925372116 | 0.982110173 | -5.110741296 |
| **A_51_P181922** | Yjefn3 | -0.004541999 | -0.118871463 | 0.109787465 | 7.998122481 | -0.097736783 | 0.925393347 | 0.982110173 | -5.110743355 |
| **A_55_P2129172** | Rab40b | 0.006606356 | -0.159773693 | 0.172986406 | 8.021100967 | 0.097685456 | 0.925432381 | 0.982110173 | -5.110747138 |
| **A_55_P2026218** | Igsf8 | -0.016111578 | -0.421919455 | 0.389696298 | 12.44893684 | -0.097675755 | 0.925439759 | 0.982110173 | -5.110747853 |
| **A_52_P400425** | Pirt | 0.004299366 | -0.104158078 | 0.112756809 | 7.831278057 | 0.097524605 | 0.925554711 | 0.982110173 | -5.110758983 |
| **A_55_P2134356** | Msl3 | -0.006135705 | -0.160971271 | 0.14869986 | 11.31469985 | -0.097490644 | 0.925580539 | 0.982110173 | -5.110761481 |
| **A_55_P1955676** | Kat14 | 0.007194701 | -0.174370645 | 0.188760048 | 10.77880513 | 0.097487505 | 0.925582926 | 0.982110173 | -5.110761712 |
| **A_51_P403693** | Uba5 | -0.006030402 | -0.158397093 | 0.146336289 | 12.07841201 | -0.097370052 | 0.925672253 | 0.982110173 | -5.110770345 |
| **A_52_P541157** | Calu | -0.013191515 | -0.346602517 | 0.320219488 | 9.752850375 | -0.097338331 | 0.925696378 | 0.982110173 | -5.110772674 |
| **A_51_P140871** | Ankhd1 | -0.013419152 | -0.352958495 | 0.326120192 | 10.95471457 | -0.097230859 | 0.925778116 | 0.982110173 | -5.110780562 |
| **A_55_P1956882** | Cpt1b | 0.006184718 | -0.150432491 | 0.162801926 | 7.983986825 | 0.097151513 | 0.925838462 | 0.982110173 | -5.11078638 |
| **A_51_P395555** | Vkorc1 | -0.009203446 | -0.242451757 | 0.224044865 | 13.5513761 | -0.097073595 | 0.925897724 | 0.982110173 | -5.110792089 |
| **A_55_P2108334** | Med10 | -0.010377659 | -0.273400504 | 0.252645186 | 10.63434005 | -0.097067777 | 0.925902148 | 0.982110173 | -5.110792515 |
| **A_52_P278154** | Eif1ad | 0.005537789 | -0.134902573 | 0.145978151 | 10.01304095 | 0.097009359 | 0.925946579 | 0.982110173 | -5.110796791 |
| **A_55_P2127659** | Mtmr2 | 0.004571102 | -0.111388121 | 0.120530325 | 8.398048381 | 0.096980577 | 0.92596847 | 0.982110173 | -5.110798898 |
| **A_55_P2235911** | 0610009E02Rik | 0.005300141 | -0.12927119 | 0.139871473 | 8.084832325 | 0.096895602 | 0.926033099 | 0.982110173 | -5.110805112 |
| **A_55_P2120348** | Pdcd7 | 0.007294108 | -0.177930252 | 0.192518468 | 10.93978856 | 0.096882036 | 0.926043418 | 0.982110173 | -5.110806104 |
| **A_55_P2107292** | Ptch1 | -0.004310966 | -0.113808296 | 0.105186363 | 7.810252457 | -0.096859068 | 0.926060887 | 0.982110173 | -5.110807782 |
| **A_55_P2186562** | Hars2 | 0.014611586 | -0.356613004 | 0.385836176 | 10.88191809 | 0.096834427 | 0.926079628 | 0.982110173 | -5.110809583 |
| **A_51_P488730** | Zfp612 | 0.006661786 | -0.162880904 | 0.176204475 | 8.238582107 | 0.096667557 | 0.926206547 | 0.982110173 | -5.110821763 |
| **A_55_P1955726** | Ptpn5 | -0.00418519 | -0.110712509 | 0.102342128 | 7.65600055 | -0.096654785 | 0.926216262 | 0.982110173 | -5.110822694 |
| **A_55_P1986703** | Aldh3b3 | -0.007435626 | -0.196770738 | 0.181899485 | 9.267368048 | -0.096617448 | 0.92624466 | 0.982110173 | -5.110825416 |
| **A_55_P2105052** | Tfb2m | -0.007266476 | -0.192349895 | 0.177816943 | 10.35507485 | -0.096588514 | 0.926266668 | 0.982110173 | -5.110827525 |
| **A_55_P2115315** | Fmr1nb | -0.008465277 | -0.224092325 | 0.20716177 | 8.777620338 | -0.096584431 | 0.926269773 | 0.982110173 | -5.110827823 |
| **A_52_P138110** | 1110032A03Rik | 0.005434411 | -0.133115296 | 0.143984119 | 8.475591255 | 0.096497498 | 0.926335895 | 0.982126407 | -5.110834154 |
| **A_55_P1991718** | Hmmr | -0.010576684 | -0.280963821 | 0.259810452 | 8.34212813 | -0.096234919 | 0.926535619 | 0.982194686 | -5.110853245 |
| **A_51_P436342** | Hoxb5 | 0.004166252 | -0.102485316 | 0.110817821 | 7.903605923 | 0.096105335 | 0.926634186 | 0.982194686 | -5.110862647 |
| **A_55_P2060938** | NA | 0.007469966 | -0.183860148 | 0.198800081 | 11.4504162 | 0.096051573 | 0.92667508 | 0.982194686 | -5.110866544 |
| **A_51_P501248** | Sphk1 | -0.064256117 | -1.710233933 | 1.5817217 | 11.99335069 | -0.09604166 | 0.92668262 | 0.982194686 | -5.110867262 |
| **A_55_P2107120** | Kansl2 | -0.006707268 | -0.178692679 | 0.165278144 | 8.837475627 | -0.095945186 | 0.926756003 | 0.982194686 | -5.110874249 |
| **A_55_P2001072** | NA | -0.008953174 | -0.238554242 | 0.220647894 | 12.11361411 | -0.095933939 | 0.926764559 | 0.982194686 | -5.110875063 |
| **A_55_P2107045** | Myl4 | 0.020980438 | -0.518066679 | 0.560027555 | 14.19783212 | 0.095754014 | 0.926901421 | 0.982194686 | -5.110888074 |
| **A_51_P324704** | Mrpl16 | 0.009694416 | -0.239556548 | 0.25894538 | 11.18101072 | 0.095687227 | 0.926952225 | 0.982194686 | -5.110892897 |
| **A_52_P105537** | Ccn3 | -0.022071315 | -0.589710655 | 0.545568026 | 8.212726234 | -0.095658788 | 0.926973858 | 0.982194686 | -5.11089495 |
| **A_55_P2012430** | NA | 0.004335599 | -0.107203044 | 0.115874241 | 7.915375445 | 0.095629721 | 0.926995969 | 0.982194686 | -5.110897048 |
| **A_52_P171212** | Irs1 | 0.004017468 | -0.099382016 | 0.107416952 | 7.769679328 | 0.095587975 | 0.927027724 | 0.982194686 | -5.110900059 |
| **A_55_P2139753** | Jade2 | -0.007534766 | -0.201509311 | 0.186439779 | 8.886434589 | -0.09556397 | 0.927045985 | 0.982194686 | -5.11090179 |
| **A_52_P102002** | Srrd | -0.005383174 | -0.144045658 | 0.13327931 | 8.311257543 | -0.095509948 | 0.927087079 | 0.982194686 | -5.110905684 |
| **A_52_P409769** | Mrvi1 | -0.008032672 | -0.21521813 | 0.199152787 | 8.747094878 | -0.095382764 | 0.927183828 | 0.982194686 | -5.110914843 |
| **A_66_P127969** | Bcat1 | -0.021974473 | -0.588807506 | 0.544858559 | 8.924904354 | -0.095374546 | 0.92719008 | 0.982194686 | -5.110915434 |
| **A_51_P343913** | Parp4 | -0.0100171 | -0.268491707 | 0.248457506 | 9.80422131 | -0.095343983 | 0.927213329 | 0.982194686 | -5.110917633 |
| **A_55_P2026475** | NA | 0.005854826 | -0.14576588 | 0.157475532 | 10.65200823 | 0.095000233 | 0.927474829 | 0.982399272 | -5.110942316 |

| | | | | | | | | | |
|---|---|---|---|---|---|---|---|---|---|
| **A_55_P2097022** | Eid1 | 0.004470628 | -0.111380195 | 0.120321451 | 7.940131671 | 0.094937668 | 0.927522424 | 0.982399272 | -5.1109468 |
| **A_51_P303906** | Actr2 | 0.006065128 | -0.151306353 | 0.163436608 | 12.98633906 | 0.094816316 | 0.927614743 | 0.982399272 | -5.110955486 |
| **A_55_P2056315** | Tbca | 0.018610142 | -0.464669101 | 0.501889384 | 13.65155946 | 0.094737234 | 0.927674905 | 0.982399272 | -5.110961142 |
| **A_55_P2016685** | Csnk1g1 | 0.006333185 | -0.158195239 | 0.170861608 | 10.39516935 | 0.094700087 | 0.927703165 | 0.982399272 | -5.110963796 |
| **A_55_P1972084** | Napb | -0.010896547 | -0.294007967 | 0.272214874 | 8.561913953 | -0.094689245 | 0.927711414 | 0.982399272 | -5.110964571 |
| **A_55_P1971439** | Serpinb9c | 0.019726124 | -0.554952624 | 0.594404871 | 11.26966454 | 0.0965871 | 0.927849266 | 0.982478221 | -4.989885975 |
| **A_55_P2065953** | Btbd19 | 0.006054415 | -0.15169208 | 0.16380091 | 8.889580566 | 0.094423834 | 0.927913332 | 0.982478221 | -5.110983506 |
| **A_51_P424518** | Nin | -0.008342116 | -0.225770224 | 0.209085992 | 8.675980444 | -0.094390811 | 0.927938456 | 0.982478221 | -5.110985859 |
| **A_65_P01991** | Prrg4 | -0.010023396 | -0.272125255 | 0.252078463 | 9.105163627 | -0.094083605 | 0.928172179 | 0.98252751 | -5.111007702 |
| **A_66_P124551** | Akap13 | -0.006368287 | -0.173082952 | 0.160346378 | 11.97841279 | -0.093976221 | 0.928253879 | 0.98252751 | -5.111015321 |
| **A_55_P1970655** | Kdm3b | -0.004937449 | -0.134237886 | 0.124362988 | 8.477893193 | -0.09394459 | 0.928277944 | 0.98252751 | -5.111017563 |
| **A_55_P1968035** | 1810009A15Rik | 0.008311293 | -0.209434514 | 0.226057099 | 12.11106748 | 0.09390483 | 0.928308195 | 0.98252751 | -5.111020381 |
| **A_55_P2001342** | Cacul1 | -0.01033423 | -0.2811909 | 0.26052244 | 9.92321383 | -0.093865878 | 0.928337831 | 0.98252751 | -5.111023141 |
| **A_55_P2056926** | Mtss2 | -0.012218743 | -0.332630838 | 0.308193353 | 10.1860147 | -0.093818133 | 0.928374157 | 0.98252751 | -5.111026521 |
| **A_55_P2115607** | Ikbip | 0.014243186 | -0.359383486 | 0.387869858 | 10.91119086 | 0.09378609 | 0.928398537 | 0.98252751 | -5.111028789 |
| **A_55_P2064792** | Arpc3 | 0.014171488 | -0.357817239 | 0.386160214 | 12.74667835 | 0.093724866 | 0.928445119 | 0.98252751 | -5.11103312 |
| **A_55_P2161132** | NA | -0.00810055 | -0.220933479 | 0.20473238 | 8.670331372 | -0.093636426 | 0.928512408 | 0.98252751 | -5.111039372 |
| **A_51_P300337** | Csrp2 | 0.019792927 | -0.500352693 | 0.539938548 | 13.05181274 | 0.093616899 | 0.928527266 | 0.98252751 | -5.111040751 |
| **A_51_P356762** | Mcm4 | 0.009156398 | -0.231525039 | 0.249837836 | 10.15600174 | 0.093594698 | 0.928544157 | 0.98252751 | -5.111042319 |
| **A_51_P133137** | Kif20a | -0.012211626 | -0.334857531 | 0.31043428 | 8.555087011 | -0.093114322 | 0.928909665 | 0.982606315 | -5.111076157 |
| **A_55_P2040777** | Lipo3 | -0.00895269 | -0.245763844 | 0.227858463 | 8.958735425 | -0.093008056 | 0.928990522 | 0.982606315 | -5.111083619 |
| **A_55_P2080264** | NA | -0.00404971 | -0.111199399 | 0.103099979 | 7.790908834 | -0.092982698 | 0.929009817 | 0.982606315 | -5.111085399 |
| **A_55_P2051721** | Psg22 | -0.027847179 | -0.765775855 | 0.710081496 | 11.72550906 | -0.092840183 | 0.929118259 | 0.982606315 | -5.11109539 |
| **A_66_P129976** | Apba1 | -0.008727197 | -0.240040338 | 0.222585944 | 9.913804426 | -0.092820447 | 0.929133277 | 0.982606315 | -5.111096772 |
| **A_55_P1957584** | Gps2 | 0.006583405 | -0.167937815 | 0.181104625 | 9.404147721 | 0.09280503 | 0.929145007 | 0.982606315 | -5.111097852 |
| **A_51_P223575** | Tbc1d23 | -0.007388681 | -0.203288985 | 0.188511622 | 9.484231995 | -0.092789962 | 0.929156473 | 0.982606315 | -5.111098907 |
| **A_55_P2135027** | NA | -0.004209516 | -0.115854524 | 0.107435491 | 9.820744465 | -0.092760276 | 0.929179062 | 0.982606315 | -5.111100985 |
| **A_55_P2077994** | Ube2v2 | -0.008649118 | -0.238096425 | 0.220798188 | 9.109541234 | -0.092738068 | 0.92919596 | 0.982606315 | -5.11110254 |
| **A_55_P2012815** | Acvr2b | -0.016555898 | -0.455891129 | 0.422779333 | 13.71296298 | -0.09270985 | 0.929217433 | 0.982606315 | -5.111104514 |
| **A_55_P2127204** | Amigo1 | -0.004529043 | -0.12475595 | 0.115697863 | 7.917268177 | -0.092677429 | 0.929242103 | 0.982606315 | -5.111106782 |
| **A_51_P418056** | Sc5d | 0.024329172 | -0.621532906 | 0.67019125 | 11.06238244 | 0.092673743 | 0.929244908 | 0.982606315 | -5.111107039 |
| **A_66_P129111** | NA | 0.003976979 | -0.10165098 | 0.109604939 | 7.688313108 | 0.092628284 | 0.929279499 | 0.982606315 | -5.111110218 |
| **A_55_P2091120** | Kbtbd12 | -0.004033566 | -0.111270252 | 0.10320312 | 7.65704109 | -0.092536903 | 0.929349035 | 0.982626087 | -5.111116602 |
| **A_51_P489452** | Cdo1 | -0.042028144 | -1.168390074 | 1.084333786 | 9.713870291 | -0.091797619 | 0.929911611 | 0.983167134 | -5.111168017 |
| **A_51_P308048** | Cmtm8 | 0.014945522 | -0.386944286 | 0.416835329 | 11.59580362 | 0.091489922 | 0.930145774 | 0.983354314 | -5.111189296 |
| **A_52_P390918** | Trip4 | -0.004862006 | -0.135717562 | 0.12599355 | 8.531338101 | -0.091409741 | 0.930206794 | 0.983354314 | -5.111194829 |
| **A_51_P372473** | Auh | -0.006386266 | -0.178422658 | 0.165650126 | 10.92343205 | -0.091326294 | 0.930270301 | 0.983354314 | -5.111200583 |
| **A_52_P398211** | NA | 0.005333714 | -0.138592209 | 0.149259638 | 8.362405888 | 0.091171667 | 0.93038798 | 0.983354314 | -5.11121123 |
| **A_52_P23308** | Fam241a | 0.014917082 | -0.387983888 | 0.417818051 | 8.997662547 | 0.091086648 | 0.930452684 | 0.983354314 | -5.111217077 |
| **A_66_P124806** | Tlr4 | 0.022891861 | -0.595831523 | 0.641615245 | 8.449713188 | 0.091023541 | 0.930500713 | 0.983354314 | -5.111221413 |
| **A_55_P2144746** | Apip | -0.010108325 | -0.283484695 | 0.263268046 | 9.741954147 | -0.090967734 | 0.930543186 | 0.983354314 | -5.111225245 |
| **A_51_P104125** | Fam76b | -0.005450728 | -0.153113577 | 0.14221212 | 9.461557937 | -0.090813927 | 0.930660245 | 0.983354314 | -5.094778923 |
| **A_52_P84901** | Slc44a1 | 0.023926871 | -0.624684221 | 0.672537964 | 9.146932627 | 0.090755031 | 0.93070507 | 0.983354314 | -5.111239829 |
| **A_55_P1953998** | NA | -0.032902699 | -0.925796889 | 0.85999149 | 10.13633205 | -0.090656865 | 0.930779784 | 0.983354314 | -5.111246549 |
| **A_55_P2042521** | Cop1 | -0.005974633 | -0.168220257 | 0.156270991 | 10.88120619 | -0.090595664 | 0.930826364 | 0.983354314 | -5.111250734 |
| **A_51_P483159** | Gchfr | 0.012045082 | -0.315197094 | 0.339287258 | 11.64635149 | 0.090554418 | 0.930857756 | 0.983354314 | -5.111253554 |
| **A_55_P2142758** | Slc9a7 | 0.003986363 | -0.104317568 | 0.112290293 | 7.682029858 | 0.090552769 | 0.930859012 | 0.983354314 | -5.111253666 |
| **A_51_P339822** | Arhgap4 | -0.004242371 | -0.119508962 | 0.11102422 | 7.930474127 | -0.09054706 | 0.930863357 | 0.983354314 | -5.111254056 |
| **A_55_P1983773** | Birc5 | -0.020537139 | -0.579583919 | 0.538509641 | 10.09481437 | -0.09037763 | 0.930992312 | 0.983354314 | -5.111265623 |
| **A_55_P1983739** | NA | -0.006347394 | -0.179184395 | 0.166489607 | 9.996809287 | -0.090349951 | 0.931013379 | 0.983354314 | -5.11126751 |

| | | | | | | | | | |
|---|---|---|---|---|---|---|---|---|---|
| **A_55_P1980566** | Ndufb11 | 0.011568116 | -0.303562212 | 0.326698444 | 14.11016011 | 0.090311195 | 0.931042878 | 0.983354314 | -5.111270152 |
| **A_52_P418515** | Cfap299 | 0.004432201 | -0.116342274 | 0.125206675 | 7.772918778 | 0.09028455 | 0.931063157 | 0.983354314 | -5.111271968 |
| **A_55_P2040743** | Zwilch | -0.00675291 | -0.190956657 | 0.177450838 | 8.042192609 | -0.090190674 | 0.931134609 | 0.983354314 | -5.111278361 |
| **A_51_P173022** | Nsmce1 | 0.008082423 | -0.212489474 | 0.228654321 | 10.92233397 | 0.090148932 | 0.931166381 | 0.983354314 | -5.111281201 |
| **A_52_P403443** | Tns3 | 0.011784522 | -0.310311471 | 0.333880516 | 11.78449993 | 0.090011049 | 0.93127133 | 0.983354314 | -5.111290574 |
| **A_55_P2117130** | Zfp937 | -0.007706976 | -0.218377665 | 0.202963713 | 9.37968617 | -0.090001359 | 0.931278706 | 0.983354314 | -5.111291233 |
| **A_55_P2137591** | Rnf220 | 0.008553729 | -0.22536025 | 0.242467708 | 9.447835491 | 0.089963942 | 0.931307185 | 0.983354314 | -5.111293773 |
| **A_55_P1968399** | Rdh7 | 0.005623353 | -0.14831823 | 0.159564936 | 7.751412209 | 0.089868724 | 0.931379661 | 0.983354314 | -5.111300234 |
| **A_55_P2275437** | Hoxb5os | -0.004014871 | -0.113965522 | 0.10593578 | 7.757183911 | -0.089834457 | 0.931405744 | 0.983354314 | -5.111302558 |
| **A_55_P2049752** | Pamr1 | -0.053596794 | -1.521932584 | 1.414738995 | 10.77631738 | -0.089801294 | 0.931430987 | 0.983354314 | -5.111304806 |
| **A_52_P291971** | Stau2 | 0.004432845 | -0.11707442 | 0.125940111 | 8.911000334 | 0.089753106 | 0.931467666 | 0.983354314 | -5.111308071 |
| **A_52_P28651** | Nectin1 | 0.02735647 | -0.723002196 | 0.777715135 | 10.18837505 | 0.089693363 | 0.93151314 | 0.983354314 | -5.111312116 |
| **A_52_P353417** | Gm2a | 0.016849816 | -0.446156009 | 0.479855641 | 11.86733503 | 0.089531914 | 0.931636032 | 0.983411975 | -5.111323034 |
| **A_55_P1953919** | Me3 | 0.007320701 | -0.194054729 | 0.20869613 | 7.910073048 | 0.08943661 | 0.931708577 | 0.983411975 | -5.11132947 |
| **A_55_P2108903** | Zbtb20 | 0.005207805 | -0.138122431 | 0.148538042 | 7.975291598 | 0.089389414 | 0.931744503 | 0.983411975 | -5.111332655 |
| **A_51_P381260** | Fxyd5 | 0.064563261 | -1.713058756 | 1.842185277 | 12.12291385 | 0.089354247 | 0.931771272 | 0.983411975 | -5.111335027 |
| **A_55_P2382741** | NA | 0.004009619 | -0.106557244 | 0.114576481 | 7.718872408 | 0.089216922 | 0.931875804 | 0.983468601 | -5.11134428 |
| **A_55_P2093543** | Snapc4 | 0.013646296 | -0.363246105 | 0.390538697 | 10.01520431 | 0.089077202 | 0.931982162 | 0.983480818 | -5.11135368 |
| **A_55_P1974855** | Zfp868 | 0.007287379 | -0.194089227 | 0.208663984 | 9.163222521 | 0.089028998 | 0.932018856 | 0.983480818 | -5.11135692 |
| **A_55_P2057777** | Fgfr1 | -0.041490318 | -1.188512151 | 1.105531515 | 14.57840526 | -0.088990622 | 0.932048069 | 0.983480818 | -5.111359498 |
| **A_51_P428483** | Fgb | -0.054164049 | -1.552505333 | 1.444177236 | 12.77092022 | -0.08893435 | 0.932090905 | 0.983480818 | -5.111363276 |
| **A_55_P2095271** | Pkn3 | 0.004935735 | -0.132070781 | 0.141942252 | 8.287438065 | 0.088629809 | 0.932322736 | 0.983616353 | -5.111383681 |
| **A_55_P2153587** | Afg3l1 | -0.006905761 | -0.198597284 | 0.184785762 | 11.4012857 | -0.088629398 | 0.932323049 | 0.983616353 | -5.111383709 |
| **A_55_P2109922** | NA | -0.009047753 | -0.260513218 | 0.242417712 | 9.015256335 | -0.088517991 | 0.932407859 | 0.983616353 | -5.111391156 |
| **A_66_P123155** | Ddo | 0.003840173 | -0.10298265 | 0.110662997 | 7.695315885 | 0.088441468 | 0.932466114 | 0.983616353 | -5.111396266 |
| **A_51_P214319** | Cox11 | 0.011143356 | -0.298868855 | 0.321155567 | 11.65631732 | 0.088431373 | 0.932473799 | 0.983616353 | -5.11139694 |
| **A_51_P250358** | Prpf39 | 0.009067359 | -0.244067758 | 0.262202477 | 8.725871076 | 0.088124687 | 0.932707275 | 0.983671091 | -5.111417372 |
| **A_55_P2092730** | Wdfy1 | 0.024586818 | -0.661915784 | 0.711089419 | 9.808094158 | 0.088110831 | 0.932717824 | 0.983671091 | -5.111418294 |
| **A_55_P2016862** | Ncoa4 | -0.008600084 | -0.24876768 | 0.231567511 | 12.97674177 | -0.088096258 | 0.932728918 | 0.983671091 | -5.111419263 |
| **A_55_P2167294** | NA | -0.003711411 | -0.107437253 | 0.100014431 | 7.667831492 | -0.088028086 | 0.932780818 | 0.983671091 | -5.111423794 |
| **A_55_P2022116** | NA | 0.007996937 | -0.21585516 | 0.231849035 | 15.78870928 | 0.087888427 | 0.932887142 | 0.983671091 | -5.111433065 |
| **A_55_P1982563** | Rhbdf1 | -0.009678188 | -0.28068956 | 0.261333183 | 10.77173386 | -0.087856867 | 0.932911169 | 0.983671091 | -5.111435158 |
| **A_66_P140949** | NA | -0.006226476 | -0.180791596 | 0.168338644 | 12.12464208 | -0.087751389 | 0.932991472 | 0.983671091 | -5.111442148 |
| **A_55_P2163812** | 4-Sep | -0.019127484 | -0.555814078 | 0.517559111 | 10.91905342 | -0.087681153 | 0.933044945 | 0.983671091 | -5.111446797 |
| **A_55_P2021565** | Ntf3 | 0.003911544 | -0.105896434 | 0.113719521 | 7.749923467 | 0.087636179 | 0.933079185 | 0.983671091 | -5.111449773 |
| **A_55_P1960331** | Srbd1 | 0.006322287 | -0.171226837 | 0.183871411 | 9.201498955 | 0.087604182 | 0.933103546 | 0.983671091 | -5.111451889 |
| **A_66_P128055** | Spindoc | -0.006510006 | -0.189769855 | 0.176749843 | 9.021706029 | -0.087394321 | 0.933263323 | 0.983671091 | -5.111465748 |
| **A_51_P230098** | Pbk | 0.006040336 | -0.16418951 | 0.176270181 | 7.898339782 | 0.08729604 | 0.933338151 | 0.983671091 | -5.111472227 |
| **A_55_P1970905** | Lin28b | 0.004147341 | -0.112936452 | 0.121231134 | 7.750521648 | 0.087144937 | 0.933453196 | 0.983671091 | -5.111482174 |
| **A_51_P293982** | Plekho2 | -0.031539453 | -0.92198032 | 0.858901414 | 9.225312262 | -0.087140136 | 0.933456851 | 0.983671091 | -5.111482489 |
| **A_52_P296632** | NA | -0.013683304 | -0.400031215 | 0.372664607 | 11.627235 | -0.087132778 | 0.933462453 | 0.983671091 | -5.111482973 |
| **A_55_P1994939** | Hmgb2 | 0.011720137 | -0.319407753 | 0.342848027 | 11.78958965 | 0.087077527 | 0.93350452 | 0.983671091 | -5.111486606 |
| **A_52_P412585** | Ear1 | 0.070187961 | -1.917320206 | 2.057696128 | 9.530270132 | 0.086880611 | 0.93365445 | 0.983671091 | -5.111499533 |
| **A_51_P369784** | Ces1e | 0.003752172 | -0.102613553 | 0.110117898 | 7.666310111 | 0.086786114 | 0.9337264 | 0.983671091 | -5.111505726 |
| **A_55_P2138343** | Ndufaf5 | -0.006984759 | -0.205029904 | 0.191060386 | 8.671365105 | -0.086767363 | 0.933740677 | 0.983671091 | -5.111506954 |
| **A_52_P476075** | Clspn | 0.004009843 | -0.109776851 | 0.117796537 | 7.691980875 | 0.086697196 | 0.933794102 | 0.983671091 | -5.111511547 |
| **A_55_P2158151** | NA | -0.0056089 | -0.164809353 | 0.153591553 | 15.69528675 | -0.086676739 | 0.933809679 | 0.983671091 | -5.111512886 |
| **A_55_P2079064** | NA | -0.013034682 | -0.38305506 | 0.356985695 | 8.689267793 | -0.086665047 | 0.933818581 | 0.983671091 | -5.11151365 |
| **A_51_P446510** | Emp3 | -0.025499692 | -0.74966897 | 0.698669585 | 9.411420599 | -0.086629141 | 0.93384592 | 0.983671091 | -5.111515999 |
| **A_65_P11718** | Nol8 | 0.00411284 | -0.112725238 | 0.120950918 | 8.540416599 | 0.086601735 | 0.933866788 | 0.983671091 | -5.11151779 |

| | | | | | | | | |
|---|---|---|---|---|---|---|---|---|
| **A_55_P2078710** | Morc4 | 0.018747301 | -0.514058632 | 0.551553233 | 12.73746506 | 0.086564307 | 0.933895286 | 0.983671091 | -5.111520236 |
| **A_52_P239787** | Ccdc25 | 0.005084718 | -0.139470457 | 0.149639893 | 8.586955334 | 0.086537111 | 0.933915994 | 0.983671091 | -5.111522013 |
| **A_51_P354572** | Rab35 | -0.007220593 | -0.212524649 | 0.198083462 | 12.9306687 | -0.086525581 | 0.933924773 | 0.983671091 | -5.111522766 |
| **A_55_P1996674** | Itih3 | 0.035310767 | -0.969230134 | 1.039851668 | 8.928787101 | 0.086478608 | 0.93396054 | 0.983671091 | -5.111525833 |
| **A_55_P2099269** | Ift80 | -0.005644234 | -0.16632696 | 0.155038491 | 8.310419475 | -0.086418161 | 0.934006565 | 0.983671091 | -5.111529777 |
| **A_51_P108266** | Actn2 | -0.003689229 | -0.108835934 | 0.101457477 | 7.677757713 | -0.086319536 | 0.934081663 | 0.983671091 | -5.111536206 |
| **A_55_P2173952** | Myh6 | 0.030648488 | -0.843151784 | 0.904448759 | 8.586660634 | 0.086291107 | 0.93410331 | 0.983671091 | -5.111538058 |
| **A_55_P2087923** | Otud3 | 0.007501406 | -0.206746571 | 0.221749383 | 9.988755116 | 0.086138066 | 0.934219842 | 0.983740211 | -5.111548016 |
| **A_55_P1955015** | Myo1g | 0.00938917 | -0.259247978 | 0.278026318 | 9.88381883 | 0.085986502 | 0.934335252 | 0.983761708 | -5.111557862 |
| **A_51_P354698** | Dync1h1 | -0.011288812 | -0.334310872 | 0.311733249 | 12.85863121 | -0.085977578 | 0.934342047 | 0.983761708 | -5.111558441 |
| **A_55_P2121701** | Mir670hg | -0.003646236 | -0.108123351 | 0.100830878 | 7.628712799 | -0.085860383 | 0.934431287 | 0.983802078 | -5.111566041 |
| **A_55_P1977066** | Klk4 | -0.065295581 | -1.938169652 | 1.80757849 | 10.2355007 | -0.085771771 | 0.934498763 | 0.983819531 | -5.111571781 |
| **A_65_P03442** | Pgk1 | 0.012699938 | -0.352375613 | 0.377775489 | 12.88849087 | 0.085583099 | 0.934642435 | 0.983838887 | -5.111583982 |
| **A_51_P161830** | Enpep | 0.009189583 | -0.255026223 | 0.273405389 | 8.313553476 | 0.085566976 | 0.934654712 | 0.983838887 | -5.111585023 |
| **A_52_P560146** | NA | -0.017493651 | -0.521050417 | 0.486063114 | 8.959629662 | -0.085467539 | 0.934730434 | 0.983838887 | -5.111591442 |
| **A_55_P2401084** | 4930558N01Rik | -0.007129468 | -0.212691978 | 0.198433043 | 8.466941381 | -0.085326191 | 0.934838072 | 0.983838887 | -5.111600553 |
| **A_55_P2174067** | Cerk | 0.010542809 | -0.293447476 | 0.314533094 | 9.248139625 | 0.085322937 | 0.93484055 | 0.983838887 | -5.111600762 |
| **A_55_P2039324** | Ccnf | -0.010843281 | -0.323557712 | 0.30187115 | 10.19059051 | -0.085306467 | 0.934853092 | 0.983838887 | -5.111601823 |
| **A_55_P2141088** | NA | 0.021309306 | -0.594135569 | 0.636754181 | 9.539578578 | 0.085182291 | 0.934947655 | 0.983838887 | -5.111609812 |
| **A_51_P202050** | Dtx1 | -0.019109798 | -0.571332623 | 0.533113027 | 12.41829826 | -0.085135542 | 0.934983256 | 0.983838887 | -5.111612817 |
| **A_55_P2158350** | Gm15427 | 0.004933797 | -0.137901182 | 0.147768777 | 16.26483142 | 0.084979831 | 0.935101835 | 0.983838887 | -5.111622814 |
| **A_51_P282424** | Eapp | -0.008756044 | -0.262399825 | 0.244887737 | 10.35834002 | -0.084928388 | 0.935141011 | 0.983838887 | -5.111626113 |
| **A_55_P2079344** | Zfp362 | 0.010887244 | -0.30450301 | 0.326277498 | 10.20011065 | 0.084925642 | 0.935143102 | 0.983838887 | -5.111626289 |
| **A_51_P326685** | Lrtm1 | -0.003769041 | -0.113008645 | 0.105470562 | 7.653498804 | -0.084882852 | 0.935175688 | 0.983838887 | -5.111629031 |
| **A_55_P2020378** | Ehd4 | 0.009656406 | -0.270438995 | 0.289751807 | 9.76674027 | 0.084816178 | 0.935226464 | 0.983838887 | -5.111633302 |
| **A_55_P2106429** | Herc2 | -0.011288311 | -0.339029685 | 0.316453062 | 10.32228663 | -0.084735787 | 0.935287687 | 0.983838887 | -5.111638446 |
| **A_55_P2112302** | NA | 0.012634308 | -0.354187522 | 0.379456138 | 11.98383025 | 0.084735511 | 0.935287897 | 0.983838887 | -5.111638463 |
| **A_52_P318073** | NA | 0.00385936 | -0.10835524 | 0.116073961 | 7.73064381 | 0.084612614 | 0.935381491 | 0.983838887 | -5.111646318 |
| **A_55_P2092826** | Anxa1 | -0.013934219 | -0.437850371 | 0.409981932 | 14.99354691 | -0.085182966 | 0.935507096 | 0.983838887 | -5.042586274 |
| **A_51_P485862** | Eef1a2 | -0.003663749 | -0.110460166 | 0.103132669 | 7.683171067 | -0.084399164 | 0.935544049 | 0.983838887 | -5.111659934 |
| **A_51_P131358** | Selplg | -0.004955303 | -0.149631927 | 0.13972132 | 8.171871025 | -0.084263798 | 0.935647143 | 0.983838887 | -5.11166855 |
| **A_51_P172752** | Gpatch4 | 0.009228799 | -0.260249402 | 0.278707001 | 10.40795116 | 0.084254038 | 0.935654576 | 0.983838887 | -5.111669171 |
| **A_55_P2130463** | Ptgdr2 | -0.007902429 | -0.238762092 | 0.222957234 | 10.10194866 | -0.084213497 | 0.935685452 | 0.983838887 | -5.111671749 |
| **A_55_P2251181** | 2900076A13Rik | -0.005254216 | -0.158777429 | 0.148268996 | 7.882825848 | -0.084198251 | 0.935697064 | 0.983838887 | -5.111672718 |
| **A_51_P101137** | Samd14 | -0.012548291 | -0.37948478 | 0.354388199 | 8.651888052 | -0.084132314 | 0.935747282 | 0.983838887 | -5.111676907 |
| **A_52_P490863** | Nop10 | 0.00633518 | -0.178919312 | 0.191589672 | 11.54649691 | 0.084131639 | 0.935747795 | 0.983838887 | -5.11167695 |
| **A_51_P108581** | Grk3 | -0.005295203 | -0.160488213 | 0.149897808 | 7.913231269 | -0.083942052 | 0.935892187 | 0.983838887 | -5.089743827 |
| **A_51_P211131** | Traf3ip2 | -0.004867224 | -0.147640335 | 0.137905887 | 8.41499756 | -0.083869503 | 0.935947442 | 0.983838887 | -5.111693571 |
| **A_52_P232830** | Ets1 | 0.004881905 | -0.138522708 | 0.148286518 | 8.502630483 | 0.083752036 | 0.936036908 | 0.983838887 | -5.111701003 |
| **A_51_P487389** | Drc3 | -0.003562215 | -0.108220738 | 0.101096308 | 7.678759303 | -0.083736475 | 0.93604876 | 0.983838887 | -5.111701986 |
| **A_55_P2063336** | Tedc1 | -0.007092997 | -0.215652467 | 0.201466474 | 9.170371105 | -0.083669854 | 0.936099501 | 0.983838887 | -5.111706195 |
| **A_55_P2136318** | Slc1a5 | -0.007294849 | -0.221935706 | 0.207346008 | 8.516251767 | -0.083612857 | 0.936142913 | 0.983838887 | -5.111709794 |
| **A_55_P2427150** | C430002N11Rik | 0.038475246 | -1.094699462 | 1.171649953 | 9.042014328 | 0.083532156 | 0.936204378 | 0.983838887 | -5.111714885 |
| **A_55_P2004571** | Pitpnm2 | 0.008312489 | -0.236698801 | 0.253323779 | 11.05373304 | 0.083466875 | 0.9362541 | 0.983838887 | -5.111719 |
| **A_55_P1976097** | NA | -0.019954223 | -0.608749036 | 0.568840589 | 9.782408555 | -0.083375801 | 0.936323467 | 0.983838887 | -5.111724735 |
| **A_55_P2071581** | NA | -0.00389712 | -0.118911886 | 0.111117645 | 8.150219059 | -0.083360319 | 0.936335259 | 0.983838887 | -5.111725709 |
| **A_55_P2119430** | Usp10 | 0.006094869 | -0.174025891 | 0.186215628 | 10.7143956 | 0.083247219 | 0.936421403 | 0.983838887 | -5.111732821 |
| **A_52_P56634** | Map3k11 | -0.006769747 | -0.206887755 | 0.193348262 | 9.207679705 | -0.083225306 | 0.936438094 | 0.983838887 | -5.111734198 |
| **A_51_P396752** | Arl2bp | -0.006572788 | -0.200928429 | 0.187782852 | 12.37456587 | -0.083199682 | 0.936457611 | 0.983838887 | -5.111735807 |
| **A_51_P446570** | Bbox1 | 0.003902357 | -0.111697131 | 0.119501846 | 7.710443434 | 0.083050126 | 0.936571525 | 0.983838887 | -5.111745192 |

| | | | | | | | | | |
|---|---|---|---|---|---|---|---|---|---|
| **A_51_P103594** | Fam162b | -0.008125263 | -0.248933661 | 0.232683136 | 9.40398641 | -0.083010864 | 0.93660143 | 0.983838887 | -5.111747653 |
| **A_55_P2015426** | Asnsd1 | 0.005217568 | -0.149423676 | 0.159858812 | 11.695697 | 0.083006474 | 0.936604774 | 0.983838887 | -5.111747928 |
| **A_66_P125862** | A930005H10Rik | -0.004448902 | -0.136355705 | 0.127457901 | 8.137479109 | -0.082976441 | 0.93662765 | 0.983838887 | -5.111749809 |
| **A_55_P1952475** | Rpl30 | 0.006155833 | -0.176439096 | 0.188750763 | 15.77477772 | 0.082940623 | 0.936654933 | 0.983838887 | -5.111752052 |
| **A_51_P324351** | Meltf | -0.025872896 | -0.796217833 | 0.74447204 | 10.55535049 | -0.082628331 | 0.936892807 | 0.98400612 | -5.111771568 |
| **A_55_P1968304** | NA | -0.006127792 | -0.189024824 | 0.176769241 | 12.47452533 | -0.08242643 | 0.9370466 | 0.98400612 | -5.111784146 |
| **A_65_P16680** | B4galt5 | 0.015882002 | -0.459046784 | 0.490810788 | 12.66229099 | 0.082270839 | 0.937165119 | 0.98400612 | -5.111793818 |
| **A_51_P391934** | Cpb1 | 0.00357259 | -0.103273393 | 0.110418573 | 7.647595174 | 0.082261034 | 0.937172589 | 0.98400612 | -5.111794427 |
| **A_51_P453909** | Cyp2f2 | 0.003575362 | -0.103551329 | 0.110702053 | 7.669721577 | 0.082109134 | 0.937288298 | 0.98400612 | -5.111803851 |
| **A_52_P120612** | Grap2 | -0.003756461 | -0.116406878 | 0.108893957 | 7.703768251 | -0.082038027 | 0.937342464 | 0.98400612 | -5.111808257 |
| **A_55_P2023617** | Kiss1r | -0.008531854 | -0.264436677 | 0.247372968 | 9.163727415 | -0.082022713 | 0.93735413 | 0.98400612 | -5.111809205 |
| **A_52_P148212** | Mis18bp1 | 0.003595773 | -0.104365332 | 0.111556879 | 7.671503953 | 0.081939658 | 0.937417398 | 0.98400612 | -5.111814345 |
| **A_55_P1978306** | Zfp318 | 0.007609721 | -0.221115582 | 0.236335024 | 9.654970084 | 0.081850942 | 0.93748498 | 0.98400612 | -5.11181983 |
| **A_55_P2092750** | Car9 | -0.003828881 | -0.118953766 | 0.111296005 | 7.985567931 | -0.081822316 | 0.937506786 | 0.98400612 | -5.111821598 |
| **A_55_P2002517** | Fhl1 | 0.003497327 | -0.101763807 | 0.108758461 | 7.89608506 | 0.08174051 | 0.937569104 | 0.98400612 | -5.111826649 |
| **A_51_P149455** | Acadl | -0.006796486 | -0.211489507 | 0.197896534 | 10.58930299 | -0.081686551 | 0.937610209 | 0.98400612 | -5.111829977 |
| **A_51_P380153** | Ubr4 | 0.008454052 | -0.247116095 | 0.264024199 | 11.99389646 | 0.081381173 | 0.937842845 | 0.98400612 | -5.111848773 |
| **A_52_P391983** | Nt5c2 | 0.003632556 | -0.106216982 | 0.113482095 | 8.703698369 | 0.081354822 | 0.93786292 | 0.98400612 | -5.111850392 |
| **A_55_P2091297** | NA | 0.005834207 | -0.170611254 | 0.182279667 | 7.944930659 | 0.081346803 | 0.937869029 | 0.98400612 | -5.111850884 |
| **A_51_P261443** | Mlx | 0.004814786 | -0.14083736 | 0.150466931 | 8.971699373 | 0.081325971 | 0.937884899 | 0.98400612 | -5.111852164 |
| **A_52_P180408** | Bcas2 | 0.013461048 | -0.393800855 | 0.420722951 | 11.93101741 | 0.081315673 | 0.937892743 | 0.98400612 | -5.111852796 |
| **A_55_P2001920** | Flna | 0.030400099 | -0.890399215 | 0.951199414 | 13.42250086 | 0.081223028 | 0.937963322 | 0.98400612 | -5.11185848 |
| **A_51_P370286** | Med24 | 0.005386713 | -0.157812962 | 0.168586388 | 10.44184168 | 0.081203305 | 0.937978348 | 0.98400612 | -5.111859689 |
| **A_51_P453010** | Mphosph6 | 0.007665935 | -0.224594199 | 0.23992607 | 10.49776636 | 0.081200676 | 0.93798035 | 0.98400612 | -5.11185985 |
| **A_55_P2102533** | Fuca2 | -0.007689628 | -0.240736033 | 0.225356778 | 8.626709418 | -0.081176824 | 0.937998521 | 0.98400612 | -5.111861312 |
| **A_55_P2071059** | Nab2 | 0.010382525 | -0.304332472 | 0.325097522 | 10.63563105 | 0.081162371 | 0.938009532 | 0.98400612 | -5.111862198 |
| **A_51_P119429** | Nckap1l | 0.008436608 | -0.247459995 | 0.264333212 | 9.541000095 | 0.081109651 | 0.938049696 | 0.98400612 | -5.089915923 |
| **A_55_P2007630** | Sez6l2 | 0.003987122 | -0.11704919 | 0.125023435 | 7.888857808 | 0.08104254 | 0.938100823 | 0.98400612 | -5.111869534 |
| **A_51_P387220** | Ddx1 | 0.009408528 | -0.2762196 | 0.295036656 | 10.41415003 | 0.081038217 | 0.938104116 | 0.98400612 | -5.111869799 |
| **A_55_P1988795** | Acss2 | 0.004283362 | -0.125823684 | 0.134390408 | 7.909510799 | 0.080994054 | 0.938137761 | 0.98400612 | -5.1118725 |
| **A_55_P2375905** | 1110035E04Rik | -0.010215168 | -0.320985476 | 0.300555139 | 8.577091917 | -0.080867715 | 0.938234011 | 0.984053677 | -5.111880219 |
| **A_55_P2302413** | BC002189 | -0.003593065 | -0.11329819 | 0.106112061 | 7.682160654 | -0.080576291 | 0.938456034 | 0.984142847 | -5.111897978 |
| **A_55_P2118799** | Rmi1 | 0.006618442 | -0.195576985 | 0.208813869 | 8.425405765 | 0.080529243 | 0.938491879 | 0.984142847 | -5.111900839 |
| **A_55_P2064351** | Vipr1 | 0.003572566 | -0.10562074 | 0.112765872 | 7.700964315 | 0.08049213 | 0.938520154 | 0.984142847 | -5.111903095 |
| **A_52_P323852** | Tnfrsf22 | -0.018259044 | -0.576358685 | 0.539840597 | 9.075084427 | -0.0804888 | 0.938522691 | 0.984142847 | -5.111903297 |
| **A_65_P17827** | Cenpl | 0.004129979 | -0.122276077 | 0.130536035 | 8.150398718 | 0.080380206 | 0.938605426 | 0.984176211 | -5.111909891 |
| **A_55_P1994907** | Senp2 | 0.006796118 | -0.201397537 | 0.214989773 | 12.3420528 | 0.080308701 | 0.938659904 | 0.984179945 | -5.111914229 |
| **A_55_P2160416** | Acoxl | 0.003408852 | -0.101126805 | 0.107944509 | 7.66270358 | 0.080225581 | 0.938723232 | 0.984192957 | -5.111919265 |
| **A_55_P2086198** | NA | -0.007381972 | -0.234514243 | 0.219750298 | 10.43159407 | -0.07995815 | 0.938926988 | 0.984245394 | -5.111935436 |
| **A_51_P160544** | Efemp2 | -0.016573118 | -0.526591356 | 0.49344512 | 11.29796012 | -0.079944334 | 0.938937515 | 0.984245394 | -5.11193627 |
| **A_51_P448664** | Tbxas1 | -0.004258559 | -0.135359286 | 0.126842167 | 7.939523195 | -0.079914717 | 0.93896008 | 0.984245394 | -5.111938057 |
| **A_55_P2249536** | Nucks1 | -0.004796569 | -0.152500923 | 0.142907785 | 10.08033872 | -0.079892604 | 0.938976929 | 0.984245394 | -5.111939391 |
| **A_55_P2062262** | Aspscr1 | -0.004397509 | -0.140425011 | 0.131629992 | 9.008217031 | -0.079533338 | 0.939250663 | 0.984478937 | -5.111961012 |
| **A_52_P118461** | Ipmk | 0.007522387 | -0.225519419 | 0.240564193 | 11.04233304 | 0.079412892 | 0.939342436 | 0.984518397 | -5.111968238 |
| **A_52_P599624** | Pfas | -0.004809489 | -0.153997867 | 0.144378889 | 8.101848016 | -0.079310944 | 0.939420116 | 0.984518397 | -5.111974347 |
| **A_55_P2174085** | St8sia5 | -0.007847982 | -0.251373787 | 0.235677823 | 10.86738455 | -0.079283385 | 0.939441115 | 0.984518397 | -5.111975996 |
| **A_55_P2085186** | NA | 0.004626761 | -0.139131396 | 0.148384917 | 9.831470546 | 0.079179673 | 0.939520139 | 0.984547833 | -5.1119822 |
| **A_51_P480499** | Emc8 | -0.008289466 | -0.266632594 | 0.250053662 | 10.43945524 | -0.078940305 | 0.939702532 | 0.984648531 | -5.111996488 |
| **A_51_P286665** | Rbl1 | 0.007925115 | -0.239294208 | 0.255144438 | 9.103812965 | 0.078866459 | 0.939758801 | 0.984648531 | -5.112000887 |
| **A_55_P2088640** | Trip4 | -0.006033534 | -0.194278221 | 0.182211152 | 9.071823909 | -0.078853 | 0.939769057 | 0.984648531 | -5.112001688 |

| | | | | | | | | |
|---|---|---|---|---|---|---|---|---|
| **A_51_P381749** | Prelid3b | 0.009911438 | -0.300137025 | 0.319959901 | 12.10709185 | 0.078645932 | 0.939926842 | 0.984739358 | -5.112013999 |
| **A_51_P335460** | Scin | 0.004357122 | -0.132112358 | 0.140826601 | 7.871206907 | 0.07854767 | 0.940001718 | 0.984739358 | -5.11201983 |
| **A_55_P2050236** | Hikeshi | 0.004617178 | -0.140014118 | 0.149248474 | 11.773806 | 0.078538662 | 0.940008583 | 0.984739358 | -5.112020364 |
| **A_55_P2019203** | Spire1 | -0.005580222 | -0.181106859 | 0.169946415 | 8.616066485 | -0.078212761 | 0.940256926 | 0.984739404 | -5.112039649 |
| **A_55_P2118609** | St6galnac1 | -0.011369542 | -0.36926609 | 0.346527006 | 8.824105184 | -0.078154616 | 0.940301234 | 0.984739404 | -5.112043081 |
| **A_52_P674357** | Thtpa | -0.00584705 | -0.189962308 | 0.178268207 | 9.046021436 | -0.078129712 | 0.940320212 | 0.984739404 | -5.112044551 |
| **A_55_P2423228** | AI464196 | 0.00423029 | -0.129055809 | 0.137516389 | 8.409495881 | 0.078082632 | 0.940356089 | 0.984739404 | -5.112047327 |
| **A_51_P263302** | Rnf24 | -0.003809131 | -0.123829114 | 0.116210852 | 9.037444315 | -0.078080309 | 0.940357859 | 0.984739404 | -5.112047464 |
| **A_55_P1961059** | Usp16 | -0.005416467 | -0.176149753 | 0.165316819 | 11.32658349 | -0.078048954 | 0.940381753 | 0.984739404 | -5.112049312 |
| **A_51_P398164** | Klhl2 | -0.004354475 | -0.141694138 | 0.132985189 | 9.92223333 | -0.078002574 | 0.940417097 | 0.984739404 | -5.112052044 |
| **A_52_P582907** | Entpd4 | -0.003877811 | -0.126335073 | 0.118579452 | 8.195887112 | -0.077906057 | 0.940490648 | 0.984739404 | -5.112057725 |
| **A_51_P340355** | Tarsl2 | 0.00441487 | -0.135041631 | 0.143871371 | 7.866968131 | 0.077884011 | 0.940507448 | 0.984739404 | -5.112059021 |
| **A_55_P1984103** | Gfra3 | 0.009159667 | -0.281120657 | 0.29943999 | 9.569402886 | 0.077630295 | 0.940700796 | 0.984739404 | -5.112073916 |
| **A_55_P2177198** | H2bc3 | -0.00340209 | -0.111240571 | 0.10443639 | 7.760952095 | -0.077614211 | 0.940713053 | 0.984739404 | -5.112074859 |
| **A_51_P380432** | Scx | 0.00448555 | -0.137708607 | 0.146679707 | 7.995134209 | 0.07760739 | 0.940718252 | 0.984739404 | -5.112075259 |
| **A_52_P151853** | Tpd52 | -0.008210586 | -0.268643461 | 0.25222229 | 9.759598514 | -0.07756173 | 0.940753048 | 0.984739404 | -5.112077933 |
| **A_55_P1961913** | Pygm | -0.005239613 | -0.171696354 | 0.161217128 | 8.14377586 | -0.077440294 | 0.940845592 | 0.984739404 | -5.112085039 |
| **A_51_P270733** | Syngr1 | -0.011052727 | -0.362256491 | 0.340151038 | 11.19930802 | -0.077424683 | 0.940857489 | 0.984739404 | -5.112085952 |
| **A_55_P2060672** | Hoxa3 | 0.003852419 | -0.118712749 | 0.126417587 | 8.034648731 | 0.077327791 | 0.94093133 | 0.984739404 | -5.112091612 |
| **A_55_P2126870** | Nsf | 0.011361485 | -0.350296693 | 0.373019663 | 8.595870064 | 0.077286914 | 0.940962482 | 0.984739404 | -5.112093998 |
| **A_55_P2093764** | Eif2s3x | -0.008580457 | -0.281779405 | 0.26461849 | 9.059593577 | -0.077268161 | 0.940976774 | 0.984739404 | -5.112095093 |
| **A_66_P134252** | NA | -0.009922949 | -0.326130079 | 0.306284182 | 12.54898104 | -0.077203724 | 0.941025881 | 0.984739404 | -5.11209885 |
| **A_52_P99807** | Dpy19l3 | -0.004542723 | -0.149625451 | 0.140540005 | 8.282985499 | -0.077031746 | 0.941156947 | 0.984739404 | -5.112108864 |
| **A_55_P1993390** | NA | 0.006116554 | -0.189333249 | 0.201566358 | 14.00946614 | 0.07699114 | 0.941187893 | 0.984739404 | -5.112111225 |
| **A_52_P528463** | Mfsd3 | -0.004060413 | -0.134041329 | 0.125920503 | 9.45204427 | -0.076852811 | 0.941293317 | 0.984739404 | -5.112119259 |
| **A_52_P662013** | Plg | -0.019038537 | -0.62860471 | 0.590527637 | 8.353498878 | -0.07683903 | 0.94130382 | 0.984739404 | -5.112120059 |
| **A_55_P1979674** | Ikzf4 | -0.006573637 | -0.217257997 | 0.204110724 | 8.707750079 | -0.076761346 | 0.941363025 | 0.984739404 | -5.112124563 |
| **A_51_P199135** | Cd83 | 0.013155438 | -0.408507556 | 0.434818432 | 11.39003863 | 0.07675541 | 0.941367549 | 0.984739404 | -5.112124907 |
| **A_55_P2155921** | Hace1 | -0.003428705 | -0.113379168 | 0.106521757 | 8.083338969 | -0.076718884 | 0.941395387 | 0.984739404 | -5.112127024 |
| **A_55_P1976310** | Tbx20 | 0.003441647 | -0.106974982 | 0.113858275 | 7.773893755 | 0.076683329 | 0.941422485 | 0.984739404 | -5.112129083 |
| **A_55_P1964464** | NA | -0.007448119 | -0.246532879 | 0.231636641 | 15.45668323 | -0.076641478 | 0.941454381 | 0.984739404 | -5.112131505 |
| **A_55_P2096486** | Bcs1l | -0.00462872 | -0.153368626 | 0.144111185 | 9.176097396 | -0.076560122 | 0.941516386 | 0.984739404 | -5.112136211 |
| **A_51_P243134** | Adcy6 | 0.00513713 | -0.16027438 | 0.17054864 | 8.237816123 | 0.076405391 | 0.941634315 | 0.984739404 | -5.112145146 |
| **A_55_P2087671** | Abhd12 | -0.005344104 | -0.177498349 | 0.166810142 | 8.874181046 | -0.076370616 | 0.941660819 | 0.984739404 | -5.112147152 |
| **A_55_P2162782** | Ahnak | 0.012333011 | -0.385307568 | 0.40997359 | 8.701104823 | 0.076304049 | 0.941711553 | 0.984739404 | -5.112150988 |
| **A_55_P2032966** | Hmgcs1 | -0.025357999 | -0.843111645 | 0.792395647 | 12.34158399 | -0.076288953 | 0.941723059 | 0.984739404 | -5.112151858 |
| **A_52_P26299** | Slc35f1 | 0.00329145 | -0.102884777 | 0.109467677 | 7.647555122 | 0.076265679 | 0.941740798 | 0.984739404 | -5.112153199 |
| **A_55_P2167112** | Kank2 | -0.013088467 | -0.43656264 | 0.410385706 | 11.1631922 | -0.076038058 | 0.941914285 | 0.984867533 | -5.112166287 |
| **A_51_P379660** | Mllt11 | -0.010043007 | -0.336308668 | 0.316222654 | 10.6736319 | -0.075728887 | 0.942149933 | 0.984962446 | -5.112184001 |
| **A_51_P419246** | Mirt1 | 0.003623323 | -0.114549679 | 0.121796326 | 7.939483916 | 0.075432409 | 0.942375913 | 0.984962446 | -5.112200921 |
| **A_55_P2035400** | Ripk4 | 0.010927573 | -0.345659169 | 0.367514314 | 11.78261169 | 0.07539242 | 0.942406393 | 0.984962446 | -5.112203198 |
| **A_55_P2180764** | Gls | 0.00599094 | -0.189633811 | 0.201615691 | 9.30206652 | 0.075342555 | 0.942444402 | 0.984962446 | -5.112206036 |
| **A_55_P1990866** | Mylk3 | -0.003257227 | -0.109650305 | 0.103135852 | 7.760198008 | -0.075318863 | 0.942462461 | 0.984962446 | -5.112207383 |
| **A_55_P2130104** | Slc52a3 | 0.010623201 | -0.336552255 | 0.357798657 | 11.15491666 | 0.075279296 | 0.94249262 | 0.984962446 | -5.112209633 |
| **A_51_P403942** | A930016O22Rik | -0.00341994 | -0.115203839 | 0.10836396 | 7.629495633 | -0.075267645 | 0.9425015 | 0.984962446 | -5.112210295 |
| **A_52_P286520** | Slc23a2 | -0.011765265 | -0.396365191 | 0.372834661 | 9.19615967 | -0.07525956 | 0.942507663 | 0.984962446 | -5.112210754 |
| **A_51_P260300** | Ikbkg | -0.0050096 | -0.168780318 | 0.158761119 | 8.39394928 | -0.075255098 | 0.942511065 | 0.984962446 | -5.112211008 |
| **A_52_P93910** | Nrp2 | -0.0092105 | -0.310522998 | 0.292101997 | 10.03864844 | -0.075203013 | 0.942550766 | 0.984962446 | -5.112213967 |
| **A_52_P99531** | Ippk | 0.014424505 | -0.457613007 | 0.486462017 | 11.51724582 | 0.075178485 | 0.942569462 | 0.984962446 | -5.112215359 |
| **A_51_P394172** | Ces1c | -0.003222055 | -0.108789379 | 0.102345269 | 7.677796688 | -0.07508836 | 0.942638159 | 0.984962446 | -5.112220472 |

| | | | | | | | | | |
|---|---|---|---|---|---|---|---|---|---|
| **A_52_P617636** | Klf12 | -0.003326455 | -0.112370234 | 0.105717325 | 7.71030048 | -0.075049855 | 0.942667509 | 0.984962446 | -5.112222655 |
| **A_52_P417654** | Tcea1 | 0.005283692 | -0.168263377 | 0.17883076 | 9.088853899 | 0.074901301 | 0.942780745 | 0.985027515 | -5.112231065 |
| **A_51_P387379** | Tshz3 | 0.024725662 | -0.789946935 | 0.839398259 | 9.189896829 | 0.074667908 | 0.942958653 | 0.985160143 | -5.112244245 |
| **A_55_P2018929** | Spns2 | 0.009356048 | -0.299337672 | 0.318049769 | 11.1265035 | 0.074564795 | 0.943037254 | 0.985189011 | -5.112250054 |
| **A_55_P2086664** | Tspan33 | -0.009309708 | -0.317318384 | 0.298698969 | 12.5098499 | -0.074360491 | 0.943192992 | 0.985275065 | -5.112261542 |
| **A_51_P506045** | Cpq | -0.012383307 | -0.422287725 | 0.397521111 | 9.025305053 | -0.074322997 | 0.943221574 | 0.985275065 | -5.112263646 |
| **A_55_P2021665** | Zfp638 | -0.004014567 | -0.137100001 | 0.129070867 | 10.1766354 | -0.07421255 | 0.943305767 | 0.985287959 | -5.11226984 |
| **A_55_P2011912** | Ecm2 | 0.003507923 | -0.11292236 | 0.119938206 | 7.776739913 | 0.074123042 | 0.943374 | 0.985287959 | -5.112274853 |
| **A_52_P289692** | NA | -0.006974594 | -0.238617724 | 0.224668537 | 10.59943964 | -0.074074485 | 0.943411016 | 0.985287959 | -5.11227757 |
| **A_55_P2039416** | Sgk1 | -0.023203331 | -0.794208032 | 0.74780137 | 13.37776551 | -0.074039329 | 0.943437816 | 0.985287959 | -5.112279536 |
| **A_55_P2082203** | Baz1a | 0.005343696 | -0.172509273 | 0.183196665 | 9.297874162 | 0.073917928 | 0.943530362 | 0.985331372 | -5.112286318 |
| **A_55_P2013928** | Dhrs13 | 0.010531324 | -0.340560363 | 0.361623011 | 10.2064467 | 0.073795792 | 0.94362347 | 0.985375367 | -5.112293129 |
| **A_55_P1958961** | 2010001K21Rik | 0.003617129 | -0.117559697 | 0.124793954 | 8.438422104 | 0.073436769 | 0.943897169 | 0.985515019 | -5.112313087 |
| **A_55_P2108868** | Nufip1 | -0.009405729 | -0.324762715 | 0.305951256 | 9.092004784 | -0.07337687 | 0.943942834 | 0.985515019 | -5.112316407 |
| **A_55_P2168383** | Myo18b | -0.04536105 | -1.566707664 | 1.475985563 | 9.157599428 | -0.07335405 | 0.943960231 | 0.985515019 | -5.112317672 |
| **A_66_P113490** | Sh2b2 | -0.009168647 | -0.316677388 | 0.298340093 | 10.3709647 | -0.073352845 | 0.94396115 | 0.985515019 | -5.112317738 |
| **A_55_P2101526** | L1td1 | 0.01514711 | -0.49421984 | 0.52451406 | 10.62889492 | 0.073159072 | 0.944108875 | 0.98557148 | -5.112328458 |
| **A_55_P2106745** | Trim63 | 0.003338766 | -0.108974456 | 0.115651988 | 7.712839861 | 0.073134831 | 0.944127356 | 0.98557148 | -5.112329796 |
| **A_51_P444994** | Tmem203 | 0.003659959 | -0.119548167 | 0.126868085 | 10.79960449 | 0.07308126 | 0.944168198 | 0.98557148 | -5.112332754 |
| **A_51_P161890** | Fcgrt | -0.021132463 | -0.7349333 | 0.692668374 | 10.91705116 | -0.072835353 | 0.944355674 | 0.9856338 | -5.112346302 |
| **A_55_P2040276** | Prcd | 0.003254223 | -0.10671603 | 0.113224477 | 7.724060464 | 0.07280166 | 0.944381361 | 0.9856338 | -5.112348155 |
| **A_55_P2102569** | Cox7a2l | -0.008948473 | -0.311803428 | 0.293906482 | 11.44445423 | -0.072691464 | 0.944465375 | 0.9856338 | -5.112354208 |
| **A_55_P2074453** | Slc16a9 | 0.011026602 | -0.362632709 | 0.384685913 | 9.126077429 | 0.072599738 | 0.944535308 | 0.9856338 | -5.11235924 |
| **A_52_P433864** | Naa16 | -0.004591845 | -0.160255193 | 0.151071504 | 8.455529851 | -0.072572147 | 0.944556343 | 0.9856338 | -5.112360752 |
| **A_55_P2033135** | Sh3bgr | 0.006202367 | -0.20471698 | 0.217121715 | 8.42043183 | 0.072345283 | 0.944729309 | 0.9856338 | -5.112373166 |
| **A_51_P469902** | Ggps1 | -0.008162111 | -0.286079307 | 0.269755086 | 8.882962712 | -0.072253057 | 0.944799625 | 0.9856338 | -5.112378201 |
| **A_55_P1978166** | Zfp945 | 0.003375494 | -0.112007068 | 0.118758056 | 7.814499598 | 0.071972456 | 0.945013566 | 0.9856338 | -5.112393482 |
| **A_55_P1957876** | NA | -0.004660508 | -0.164145337 | 0.154824322 | 10.38436451 | -0.071892406 | 0.945074601 | 0.9856338 | -5.11239783 |
| **A_51_P511680** | Ttc38 | -0.005226718 | -0.184551717 | 0.174098281 | 8.187574377 | -0.071706317 | 0.945216487 | 0.9856338 | -5.11240792 |
| **A_55_P2106096** | Fam171a1 | 0.004413863 | -0.147035422 | 0.155863147 | 10.11856948 | 0.071700262 | 0.945221103 | 0.9856338 | -5.112408248 |
| **A_55_P1996456** | Srrm1 | -0.006518492 | -0.230308097 | 0.217271114 | 9.033990118 | -0.071659927 | 0.945251858 | 0.9856338 | -5.112410431 |
| **A_55_P1999852** | Tchp | -0.00423579 | -0.149732382 | 0.141260802 | 8.799591863 | -0.071622712 | 0.945280233 | 0.9856338 | -5.112412445 |
| **A_51_P123017** | Gmcl1 | 0.007895882 | -0.263399696 | 0.279191459 | 11.28583177 | 0.071602317 | 0.945295783 | 0.9856338 | -5.090450518 |
| **A_51_P235139** | Klhl40 | 0.003182658 | -0.106176066 | 0.112541381 | 7.686185975 | 0.071598792 | 0.945298471 | 0.9856338 | -5.112413738 |
| **A_51_P460734** | Prkcd | -0.009873605 | -0.349156009 | 0.329408799 | 10.47182752 | -0.071595146 | 0.945301251 | 0.9856338 | -5.112413935 |
| **A_55_P2103501** | Ankrd29 | 0.003074272 | -0.102584771 | 0.108733314 | 7.770208366 | 0.071582155 | 0.945311157 | 0.9856338 | -5.112414638 |
| **A_55_P2351193** | Snora47 | 0.003083625 | -0.102910758 | 0.109078008 | 7.726691724 | 0.071572778 | 0.945318306 | 0.9856338 | -5.112415144 |
| **A_52_P181468** | Kmt5a | 0.008345763 | -0.278559443 | 0.295250969 | 10.44737688 | 0.071564358 | 0.945324727 | 0.9856338 | -5.1124156 |
| **A_55_P2305525** | C81189 | -0.007553735 | -0.267267492 | 0.252160021 | 8.267298286 | -0.071554337 | 0.945332367 | 0.9856338 | -5.112416141 |
| **A_55_P2149787** | Fcnb | 0.003273618 | -0.109452223 | 0.115999459 | 7.710316593 | 0.071445307 | 0.945415501 | 0.9856338 | -5.112422028 |
| **A_66_P100159** | Cfl2 | 0.007168263 | -0.239875735 | 0.254212261 | 11.66087544 | 0.071385301 | 0.945461255 | 0.9856338 | -5.112425264 |
| **A_55_P2126368** | Atad5 | 0.004975146 | -0.166604782 | 0.176555073 | 8.344021689 | 0.071335958 | 0.945498878 | 0.9856338 | -5.112427923 |
| **A_52_P179178** | Bend3 | 0.006209854 | -0.208316674 | 0.220736383 | 10.4862225 | 0.071214688 | 0.945591346 | 0.9856338 | -5.112434451 |
| **A_55_P2038162** | Nt5c | 0.010304527 | -0.345816682 | 0.366425736 | 11.20497836 | 0.071186784 | 0.945612623 | 0.9856338 | -5.112435951 |
| **A_55_P2155216** | Ankrd34a | -0.003661133 | -0.130245404 | 0.122923137 | 7.760814964 | -0.071154926 | 0.945636915 | 0.9856338 | -5.112437664 |
| **A_55_P2062603** | 4932438A13Rik | -0.006220154 | -0.221353725 | 0.208913417 | 10.0878142 | -0.071131524 | 0.945654759 | 0.9856338 | -5.112438921 |
| **A_52_P494126** | Usp46 | -0.011700561 | -0.416389363 | 0.392988241 | 9.725800776 | -0.071130304 | 0.945655689 | 0.9856338 | -5.112438986 |
| **A_55_P2129187** | NA | -0.005092471 | -0.181897608 | 0.171712666 | 15.59483503 | -0.070860275 | 0.945861589 | 0.985795247 | -5.112453464 |
| **A_51_P221014** | Cd47 | 0.007062988 | -0.238703922 | 0.252829899 | 12.0964327 | 0.070702414 | 0.945981962 | 0.985867545 | -5.112461902 |
| **A_55_P2137521** | Nmral1 | -0.003083287 | -0.110545353 | 0.10437878 | 7.694165327 | -0.070587513 | 0.946069579 | 0.985905698 | -5.112468032 |

| | | | | | | | | | |
|---|---|---|---|---|---|---|---|---|---|
| **A_51_P195215** | Fam149a | -0.020789052 | -0.746164924 | 0.70458682 | 9.307867781 | -0.070508381 | 0.946129921 | 0.985915426 | -5.112472248 |
| **A_55_P2091308** | Rpl38 | 0.004339199 | -0.147424202 | 0.1561026 | 16.58677444 | 0.070341505 | 0.946257171 | 0.985964634 | -5.112481124 |
| **A_51_P344046** | Nt5m | 0.00584851 | -0.19889313 | 0.21059015 | 9.802256645 | 0.070276195 | 0.946306974 | 0.985964634 | -5.112484591 |
| **A_55_P1965194** | Fbxl5 | 0.005760186 | -0.196042825 | 0.207563197 | 9.368267349 | 0.070222783 | 0.946347704 | 0.985964634 | -5.112487425 |
| **A_51_P363749** | Irf6 | 0.01678825 | -0.57174114 | 0.60531764 | 10.30957993 | 0.070178881 | 0.946381182 | 0.985964634 | -5.112489753 |
| **A_55_P1953713** | Bop1 | -0.006972667 | -0.252021037 | 0.238075703 | 10.18076984 | -0.070002939 | 0.94651535 | 0.98596666 | -5.112499066 |
| **A_55_P1992910** | Gyg | -0.015403614 | -0.557431078 | 0.52662385 | 11.83056455 | -0.069915022 | 0.946582393 | 0.98596666 | -5.112503711 |
| **A_51_P361788** | Vapa | 0.00697153 | -0.238484353 | 0.252427413 | 14.36300986 | 0.069875324 | 0.946612667 | 0.98596666 | -5.112505806 |
| **A_55_P1995924** | Zfp984 | 0.010510767 | -0.35963774 | 0.380659275 | 9.513470851 | 0.06985984 | 0.946624475 | 0.98596666 | -5.112506623 |
| **A_55_P2184339** | Mdga1 | 0.003026479 | -0.103581792 | 0.109634749 | 7.674228079 | 0.069841874 | 0.946638175 | 0.98596666 | -5.112507571 |
| **A_52_P423814** | Cox8b | -0.004095004 | -0.148654214 | 0.140464205 | 7.754406129 | -0.069691171 | 0.946753099 | 0.986033227 | -5.112515512 |
| **A_51_P135340** | Panx1 | -0.005794261 | -0.211279319 | 0.199690798 | 9.308303288 | -0.069372434 | 0.94699617 | 0.986115259 | -5.112532249 |
| **A_51_P334289** | Zfp628 | -0.003724258 | -0.135821327 | 0.128372812 | 8.631144742 | -0.069361054 | 0.947004848 | 0.986115259 | -5.112532845 |
| **A_55_P2040245** | Piezo2 | -0.003119177 | -0.113817428 | 0.107579075 | 8.252713017 | -0.069321571 | 0.947034959 | 0.986115259 | -5.112534913 |
| **A_55_P2202043** | C430045I18Rik | -0.007930401 | -0.289643476 | 0.273782673 | 8.341910636 | -0.069255987 | 0.947084974 | 0.986115259 | -5.112538345 |
| **A_55_P2054854** | Art4 | -0.003086617 | -0.112770604 | 0.106597369 | 7.747413196 | -0.069232296 | 0.947103042 | 0.986115259 | -5.112539584 |
| **A_51_P101460** | Dsp | -0.0078506 | -0.287055873 | 0.271354673 | 12.43340097 | -0.069174877 | 0.947146831 | 0.986115259 | -5.095281713 |
| **A_55_P2117959** | Cyp4b1 | -0.003849699 | -0.140873104 | 0.133173705 | 7.90104299 | -0.069119602 | 0.947188986 | 0.986115259 | -5.112545471 |
| **A_52_P470316** | Laptm4b | -0.008026369 | -0.294939861 | 0.278887122 | 10.51125411 | -0.068823592 | 0.947414734 | 0.986257315 | -5.112560891 |
| **A_55_P2097478** | Gda | -0.043562094 | -1.60164285 | 1.514518663 | 10.26936911 | -0.068784082 | 0.947444867 | 0.986257315 | -5.112562944 |
| **A_51_P194273** | Sgcg | 0.003026012 | -0.105274505 | 0.11132653 | 7.805222727 | 0.068739971 | 0.947478508 | 0.986257315 | -5.112565234 |
| **A_55_P2106666** | Adcy9 | -0.008808858 | -0.324851885 | 0.307234169 | 10.37307367 | -0.06857133 | 0.947607124 | 0.986301391 | -5.112573979 |
| **A_55_P1980931** | NA | -0.006835339 | -0.253120294 | 0.239449616 | 11.8624976 | -0.068279659 | 0.947829573 | 0.986301391 | -5.112589052 |
| **A_55_P2183032** | Zfp787 | -0.00444399 | -0.164619418 | 0.155731439 | 10.08798628 | -0.068256859 | 0.947846962 | 0.986301391 | -5.112590227 |
| **A_55_P2093231** | NA | -0.010499742 | -0.389038676 | 0.368039192 | 8.008601096 | -0.068239718 | 0.947860035 | 0.986301391 | -5.112591111 |
| **A_66_P138780** | Elob | 0.005610626 | -0.196677399 | 0.207898651 | 13.16945518 | 0.068235484 | 0.947863264 | 0.986301391 | -5.112591329 |
| **A_51_P323842** | Ciao2b | -0.005205482 | -0.193216886 | 0.182805922 | 13.09476574 | -0.068115499 | 0.947954776 | 0.986301391 | -5.112597507 |
| **A_51_P519791** | Ska3 | -0.00297309 | -0.110397823 | 0.104451644 | 7.707850413 | -0.068088364 | 0.947975471 | 0.986301391 | -5.112598903 |
| **A_52_P613498** | 4833420G17Rik | -0.008679407 | -0.322306941 | 0.304948128 | 10.89368143 | -0.068083992 | 0.947978806 | 0.986301391 | -5.112599128 |
| **A_55_P2102126** | Lamp2 | -0.004637364 | -0.172211164 | 0.162936436 | 12.34301219 | -0.068082303 | 0.947980094 | 0.986301391 | -5.112599215 |
| **A_55_P2060376** | Gzmg | 0.048168949 | -1.697844935 | 1.794182832 | 10.9622862 | 0.067871686 | 0.948140731 | 0.986415426 | -5.112610028 |
| **A_51_P291819** | 2310033P09Rik | -0.007961185 | -0.297267156 | 0.281344786 | 10.3469161 | -0.067700128 | 0.948271581 | 0.986418686 | -5.112618811 |
| **A_52_P494622** | Nr4a2 | -0.006421849 | -0.240023992 | 0.227180294 | 8.61223377 | -0.06763203 | 0.948323521 | 0.986418686 | -5.112622291 |
| **A_51_P416137** | Slc31a2 | 0.011206035 | -0.396777512 | 0.419189583 | 11.38968286 | 0.067573826 | 0.948367914 | 0.986418686 | -5.112625263 |
| **A_51_P111757** | Asb1 | -0.004489888 | -0.168152683 | 0.159172906 | 9.749674885 | -0.06749238 | 0.948430035 | 0.986418686 | -5.090259862 |
| **A_55_P1982727** | Dnajb5 | -0.003430782 | -0.128565637 | 0.121704073 | 8.084934438 | -0.067450323 | 0.948462113 | 0.986418686 | -5.11263156 |
| **A_55_P2157102** | NA | 0.011672435 | -0.414328188 | 0.437673058 | 14.11631556 | 0.067409389 | 0.948493335 | 0.986418686 | -5.112633645 |
| **A_55_P1982818** | Odc1 | -0.006135268 | -0.230390189 | 0.218119652 | 9.690716073 | -0.067307078 | 0.948571371 | 0.986418686 | -5.11263885 |
| **A_55_P2084251** | Kcna4 | 0.005368315 | -0.190868023 | 0.201604652 | 11.67666463 | 0.067301958 | 0.948575277 | 0.986418686 | -5.11263911 |
| **A_55_P2018904** | Kbtbd6 | 0.004400378 | -0.156558021 | 0.165358776 | 7.971245539 | 0.067258235 | 0.948608626 | 0.986418686 | -5.112641332 |
| **A_55_P2032768** | NA | 0.007938609 | -0.282965494 | 0.298842713 | 13.70723801 | 0.06713728 | 0.948700884 | 0.986418686 | -5.112647471 |
| **A_51_P441970** | Stox2 | 0.005651921 | -0.201678731 | 0.212982574 | 8.621819081 | 0.067065897 | 0.948755332 | 0.986418686 | -5.112651089 |
| **A_52_P367060** | Atxn2l | 0.009976188 | -0.356374527 | 0.376326903 | 12.4284185 | 0.06699413 | 0.948810073 | 0.986418686 | -5.112654722 |
| **A_51_P511236** | Pik3r1 | -0.006195845 | -0.233854283 | 0.221462593 | 8.99743944 | -0.066955454 | 0.948839573 | 0.986418686 | -5.112656679 |
| **A_51_P408082** | Apoa1 | 0.038630237 | -1.381310105 | 1.458570579 | 11.53747075 | 0.066930865 | 0.948858328 | 0.986418686 | -5.107163415 |
| **A_51_P441914** | Hsd17b2 | -0.012430989 | -0.470274747 | 0.445412768 | 12.62600065 | -0.066797092 | 0.948960367 | 0.986471707 | -5.112664678 |
| **A_55_P2130448** | NA | 0.010829769 | -0.389184627 | 0.410844166 | 12.72003601 | 0.066605907 | 0.949106198 | 0.986570245 | -5.112674309 |
| **A_55_P1956978** | Tln1 | 0.00410872 | -0.147953831 | 0.156171272 | 8.161300626 | 0.066474255 | 0.94920662 | 0.986619294 | -5.112680926 |
| **A_55_P2123673** | Trnp1 | 0.007585999 | -0.273440385 | 0.288612383 | 9.762891741 | 0.066410212 | 0.949255472 | 0.986619294 | -5.11268414 |
| **A_55_P2007981** | Fmn2 | -0.009146334 | -0.34932631 | 0.331033643 | 9.174010976 | -0.066146589 | 0.949456563 | 0.986770326 | -5.112697337 |

| | | | | | | | | |
|---|---|---|---|---|---|---|---|---|
| A_52_P442710 | Ndnf | 0.002995598 | -0.108545547 | 0.114536743 | 7.627613176 | 0.066072021 | 0.949513445 | 0.986770326 | -5.11270106 |
| A_55_P2137406 | Bax | -0.007750109 | -0.296557612 | 0.281057393 | 11.72198735 | -0.066018937 | 0.949553938 | 0.986770326 | -5.112703708 |
| A_51_P244950 | Dpys | 0.002889625 | -0.104995256 | 0.110774507 | 7.656131723 | 0.065894643 | 0.949648753 | 0.986803055 | -5.1127099 |
| A_51_P121915 | Rdh16f2 | -0.002805764 | -0.107789278 | 0.102177751 | 7.656477214 | -0.065750515 | 0.949758697 | 0.986803055 | -5.103419086 |
| A_55_P1981499 | Rpl10a | 0.00373689 | -0.136281653 | 0.143755434 | 15.51722062 | 0.065658952 | 0.949828545 | 0.986803055 | -5.11272161 |
| A_55_P1992199 | Psg20 | 0.012718078 | -0.463913778 | 0.489349935 | 12.84646528 | 0.065645897 | 0.949838504 | 0.986803055 | -5.112722257 |
| A_51_P425048 | H2-Q2 | -0.017640846 | -0.679835703 | 0.644554011 | 9.896085463 | -0.065539445 | 0.94991971 | 0.986803055 | -5.112727531 |
| A_55_P2069052 | Sacs | 0.006615141 | -0.242050412 | 0.255280694 | 8.965910185 | 0.065447434 | 0.9499899 | 0.986803055 | -5.112732083 |
| A_51_P234113 | Nod1 | -0.008464258 | -0.327480712 | 0.310552197 | 9.831755567 | -0.065274709 | 0.950121664 | 0.986803055 | -5.11274061 |
| A_55_P2084002 | Aptx | 0.002952281 | -0.108353279 | 0.114257842 | 8.1539209 | 0.065254439 | 0.950137127 | 0.986803055 | -5.112741609 |
| A_55_P1977929 | Itgam | -0.002928239 | -0.113454225 | 0.107597748 | 7.903651941 | -0.065179531 | 0.950194272 | 0.986803055 | -5.112745299 |
| A_55_P2018191 | Ccm2l | 0.00350151 | -0.12894439 | 0.135947411 | 8.334043318 | 0.065040826 | 0.950300087 | 0.986803055 | -5.112752121 |
| A_55_P2012899 | Vsig4 | 0.003047862 | -0.112271857 | 0.118367581 | 7.673534478 | 0.065022083 | 0.950314385 | 0.986803055 | -5.112753041 |
| A_51_P300936 | Rbbp8 | 0.004616747 | -0.170070564 | 0.179304059 | 10.82759297 | 0.065019542 | 0.950316323 | 0.986803055 | -5.112753166 |
| A_55_P2052166 | Akap13 | 0.004695454 | -0.173224752 | 0.182615659 | 10.68305291 | 0.064926419 | 0.950387365 | 0.986803055 | -5.112757736 |
| A_55_P1967761 | Foxc1 | 0.004246628 | -0.156675339 | 0.165168596 | 8.633508337 | 0.064922924 | 0.950390032 | 0.986803055 | -5.112757908 |
| A_55_P1962747 | H2-Ab1 | 0.011590284 | -0.427712467 | 0.450893035 | 10.21709696 | 0.064908162 | 0.950401293 | 0.986803055 | -5.112758632 |
| A_52_P206925 | Parp16 | 0.005025785 | -0.185612878 | 0.195664448 | 9.170816983 | 0.064857777 | 0.950439731 | 0.986803055 | -5.112761101 |
| A_55_P1962339 | Trim41 | -0.005731743 | -0.223301323 | 0.211837838 | 15.21312331 | -0.064812334 | 0.950474398 | 0.986803055 | -5.112763326 |
| A_52_P574214 | Rrp1b | -0.010628357 | -0.414696794 | 0.39344008 | 9.46845819 | -0.064711335 | 0.950551449 | 0.986803055 | -5.112768267 |
| A_55_P2031357 | Zfpm2 | -0.00296143 | -0.115706616 | 0.109783756 | 7.714423098 | -0.06462084 | 0.950620488 | 0.986803055 | -5.112772687 |
| A_55_P2121225 | Cyp3a11 | 0.002936758 | -0.10895273 | 0.114826245 | 7.653534912 | 0.064572555 | 0.950657324 | 0.986803055 | -5.112775043 |
| A_51_P454008 | Lbp | -0.032683345 | -1.280595034 | 1.215228343 | 9.589442992 | -0.064433514 | 0.9507634 | 0.986803055 | -5.112781817 |
| A_55_P1995092 | H2-Q10 | 0.010352532 | -0.385134305 | 0.405839369 | 8.998521931 | 0.064399681 | 0.950789212 | 0.986803055 | -5.112783463 |
| A_55_P1997415 | Exoc6b | -0.006513027 | -0.255608937 | 0.242582884 | 10.47912104 | -0.064325833 | 0.950845551 | 0.986803055 | -5.112787054 |
| A_55_P1956819 | Gm4832 | -0.005768183 | -0.226551488 | 0.215015122 | 15.62733222 | -0.064274982 | 0.950884346 | 0.986803055 | -5.112789523 |
| A_55_P2074718 | Fh1 | -0.012270256 | -0.48231867 | 0.457778157 | 12.31622018 | -0.064221466 | 0.950925175 | 0.986803055 | -5.112792121 |
| A_55_P1971804 | Hdac10 | -0.003165594 | -0.124506374 | 0.118175186 | 8.467207524 | -0.064182649 | 0.950954789 | 0.986803055 | -5.112794003 |
| A_55_P1971872 | NA | -0.003862721 | -0.152076717 | 0.144351276 | 8.883377294 | -0.06411701 | 0.951004866 | 0.986803055 | -5.112797184 |
| A_66_P135882 | Rrp1 | -0.006995436 | -0.276072279 | 0.262081407 | 12.27526356 | -0.06395988 | 0.951124746 | 0.986803055 | -5.112804784 |
| A_51_P167876 | Tfrc | 0.007518618 | -0.281750527 | 0.296787763 | 9.266181593 | 0.063944776 | 0.95113627 | 0.986803055 | -5.112805514 |
| A_55_P2065059 | Wnt2 | -0.003165221 | -0.125018273 | 0.11868783 | 7.730379356 | -0.063905296 | 0.951166391 | 0.986803055 | -5.11280742 |
| A_51_P373619 | Mbl1 | 0.002774172 | -0.104028137 | 0.10957648 | 7.648392157 | 0.063903088 | 0.951168075 | 0.986803055 | -5.112807527 |
| A_52_P73126 | Epb42 | 0.003263134 | -0.122803963 | 0.129330231 | 7.758879773 | 0.063679887 | 0.951338365 | 0.986926753 | -5.112818282 |
| A_55_P2043702 | Tfg | -0.007884076 | -0.31380773 | 0.298039579 | 12.79770621 | -0.063402575 | 0.951549943 | 0.986967714 | -5.112831592 |
| A_55_P2044547 | NA | 0.003386341 | -0.128083145 | 0.134855827 | 8.670397348 | 0.063368713 | 0.951575779 | 0.986967714 | -5.112833213 |
| A_55_P1969566 | Atp9b | 0.003040064 | -0.115062709 | 0.121142837 | 8.404962967 | 0.06332742 | 0.951607284 | 0.986967714 | -5.112835189 |
| A_51_P481159 | Cbr3 | -0.00565618 | -0.22550358 | 0.21419122 | 9.13199969 | -0.063295241 | 0.951631836 | 0.986967714 | -5.112836728 |
| A_55_P2037328 | Ppp1r12b | -0.002658097 | -0.105977247 | 0.100661053 | 7.690280559 | -0.063293511 | 0.951633156 | 0.986967714 | -5.112836811 |
| A_55_P2028898 | Prl2b1 | 0.008630378 | -0.328317987 | 0.345578743 | 17.56925897 | 0.063013791 | 0.951846577 | 0.98705416 | -5.112850155 |
| A_51_P141071 | Cand2 | 0.008429496 | -0.321437483 | 0.338296475 | 8.63061975 | 0.06286833 | 0.951957563 | 0.98705416 | -5.11285707 |
| A_51_P232742 | Mfsd14b | -0.006489729 | -0.260475748 | 0.24749629 | 10.77596371 | -0.06286167 | 0.951962645 | 0.98705416 | -5.112857387 |
| A_52_P550185 | Urm1 | -0.007096317 | -0.284967713 | 0.270775078 | 11.58299944 | -0.06282874 | 0.951987771 | 0.98705416 | -5.11285895 |
| A_55_P2159216 | NA | 0.007243154 | -0.276475463 | 0.29096177 | 10.77919853 | 0.062807141 | 0.95200425 | 0.98705416 | -5.112859975 |
| A_52_P742248 | Fam117b | -0.005014312 | -0.201506999 | 0.191478375 | 11.19301081 | -0.062781848 | 0.952023549 | 0.98705416 | -5.112861175 |
| A_51_P140901 | Pmm2 | -0.003022835 | -0.121871066 | 0.115825396 | 9.130113243 | -0.062573556 | 0.952182478 | 0.98705416 | -5.112871036 |
| A_55_P1978321 | Fbxo8 | 0.007368323 | -0.282433545 | 0.29717019 | 9.916459016 | 0.062551341 | 0.952199428 | 0.98705416 | -5.112872086 |
| A_55_P2066429 | NA | -0.004812163 | -0.194374592 | 0.184750267 | 10.81584346 | -0.062453558 | 0.952274039 | 0.98705416 | -5.112876703 |
| A_55_P2060498 | Zfp553 | 0.005088256 | -0.195400115 | 0.205576628 | 9.736654367 | 0.062437998 | 0.952285911 | 0.98705416 | -5.112877437 |
| A_51_P483324 | Ptpn22 | 0.002782862 | -0.107104573 | 0.112670296 | 7.776701731 | 0.062303541 | 0.952388505 | 0.98705416 | -5.112883772 |

| | | | | | | | | |
|---|---|---|---|---|---|---|---|---|
| **A_51_P155085** | Dennd2a | 0.007006743 | -0.269823979 | 0.283837465 | 12.33167596 | 0.06226888 | 0.952414953 | 0.98705416 | -5.112885403 |
| **A_55_P2011852** | Cdk2 | 0.003387929 | -0.130501804 | 0.137277662 | 8.337899767 | 0.06225241 | 0.95242752 | 0.98705416 | -5.112886177 |
| **A_51_P113182** | Lifr | 0.004369404 | -0.168322092 | 0.1770609 | 9.743995194 | 0.062247284 | 0.952431431 | 0.98705416 | -5.112886418 |
| **A_55_P2015681** | AU015836 | -0.012354758 | -0.503426943 | 0.478717428 | 10.39185492 | -0.061895355 | 0.952699968 | 0.987209617 | -5.11290292 |
| **A_55_P1963454** | Cenpq | 0.002672831 | -0.103728109 | 0.109073771 | 7.780360512 | 0.061800952 | 0.952772003 | 0.987209617 | -5.11290733 |
| **A_55_P2015277** | Cyld | 0.010095207 | -0.391886853 | 0.412077267 | 9.459087568 | 0.061784242 | 0.952784754 | 0.987209617 | -5.11290811 |
| **A_55_P2076471** | Mon1b | -0.005373862 | -0.219360634 | 0.208612911 | 10.00777165 | -0.061782961 | 0.952785731 | 0.987209617 | -5.11290817 |
| **A_55_P2078428** | Tstd1 | -0.003539039 | -0.144982704 | 0.137904626 | 8.015167494 | -0.061556086 | 0.952958852 | 0.987336066 | -5.112918739 |
| **A_55_P2109613** | Ahr | -0.006872769 | -0.28210241 | 0.268356872 | 8.866964767 | -0.061433565 | 0.953052344 | 0.987367233 | -5.11292443 |
| **A_51_P225427** | Pkp2 | -0.009171281 | -0.376751892 | 0.358409331 | 12.63874288 | -0.061382779 | 0.953091098 | 0.987367233 | -5.112926786 |
| **A_55_P2003803** | Scml2 | -0.002565547 | -0.105831811 | 0.100700717 | 7.701398594 | -0.061121029 | 0.953290836 | 0.987521227 | -5.112938898 |
| **A_55_P1979650** | S100a1 | -0.022900951 | -0.945914804 | 0.900112902 | 11.9827641 | -0.061039991 | 0.953352676 | 0.987532362 | -5.112942637 |
| **A_55_P2032538** | Pde3b | -0.004127679 | -0.170920963 | 0.162665605 | 8.043256932 | -0.060883066 | 0.953472426 | 0.987603479 | -5.112949863 |
| **A_55_P1954965** | Npm1 | -0.007190655 | -0.298429247 | 0.284047937 | 13.81005575 | -0.060741946 | 0.953580116 | 0.987662098 | -5.112956346 |
| **A_55_P2157093** | Bcl2l14 | 0.003219255 | -0.127819182 | 0.134257691 | 7.92093354 | 0.060440183 | 0.953810399 | 0.987762893 | -5.112970159 |
| **A_52_P284821** | Slco4c1 | 0.003491955 | -0.138788741 | 0.145772652 | 7.752586762 | 0.060379828 | 0.953856458 | 0.987762893 | -5.112972913 |
| **A_55_P1957927** | Nono | -0.005628976 | -0.235252247 | 0.223994295 | 13.38966584 | -0.060309074 | 0.953910453 | 0.987762893 | -5.112976139 |
| **A_55_P2088440** | Arrdc2 | 0.005739894 | -0.228505534 | 0.239985322 | 9.464869152 | 0.06028398 | 0.953929603 | 0.987762893 | -5.112977282 |
| **A_52_P245415** | AW554918 | -0.002645383 | -0.110611462 | 0.105320696 | 8.272959733 | -0.060279599 | 0.953932946 | 0.987762893 | -5.112977481 |
| **A_55_P2173313** | Plcb1 | 0.002775974 | -0.110912933 | 0.116464881 | 7.750702568 | 0.060071224 | 0.954091967 | 0.987857662 | -5.112986954 |
| **A_55_P2005585** | Trps1 | 0.003438049 | -0.137635499 | 0.144511597 | 7.956754525 | 0.059956406 | 0.954179591 | 0.987857662 | -5.112992159 |
| **A_51_P353232** | Tnnc2 | -0.003944155 | -0.166419003 | 0.158530694 | 8.098684879 | -0.059722357 | 0.954358208 | 0.987857662 | -5.113002739 |
| **A_55_P1966664** | Cdc14b | -0.003083413 | -0.130207131 | 0.124040306 | 9.944609108 | -0.059672499 | 0.954396259 | 0.987857662 | -5.113004988 |
| **A_66_P126485** | Dipk1a | -0.007211264 | -0.304542616 | 0.290120088 | 10.36931898 | -0.059667779 | 0.954399861 | 0.987857662 | -5.113005201 |
| **A_55_P2170787** | Braf | -0.003561378 | -0.150454728 | 0.143331973 | 8.458617308 | -0.059646513 | 0.954416091 | 0.987857662 | -5.113006159 |
| **A_55_P1957835** | Cpeb3 | -0.005638665 | -0.238412592 | 0.227135263 | 9.921414413 | -0.059595176 | 0.95445527 | 0.987857662 | -5.113008471 |
| **A_55_P2146229** | NA | 0.003940114 | -0.158804273 | 0.166684501 | 8.004950554 | 0.059562363 | 0.954480312 | 0.987857662 | -5.113009948 |
| **A_55_P1995135** | Casz1 | 0.003441699 | -0.138811608 | 0.145695006 | 9.031380566 | 0.059522296 | 0.95451089 | 0.987857662 | -5.11301175 |
| **A_55_P2007384** | Sgca | -0.002530664 | -0.107185618 | 0.102124289 | 7.687962966 | -0.05948999 | 0.954535546 | 0.987857662 | -5.113013203 |
| **A_55_P2023391** | Grhl3 | 0.002626454 | -0.106754337 | 0.112007246 | 7.739838797 | 0.059074212 | 0.954852865 | 0.988027623 | -5.113031823 |
| **A_51_P358354** | Jam3 | -0.005931756 | -0.253090772 | 0.241227261 | 9.209470031 | -0.059044025 | 0.954875904 | 0.988027623 | -5.11303317 |
| **A_55_P2142146** | Glul | 0.008353131 | -0.340349112 | 0.357055375 | 12.259107 | 0.058933704 | 0.954960102 | 0.988027623 | -5.113038086 |
| **A_55_P2041355** | Gm4750 | 0.006071329 | -0.247459871 | 0.25960253 | 12.59239547 | 0.058914412 | 0.954974826 | 0.988027623 | -5.113038945 |
| **A_52_P326214** | Cttn | 0.004195097 | -0.171046755 | 0.179436949 | 8.80257765 | 0.058894307 | 0.95499017 | 0.988027623 | -5.11303984 |
| **A_55_P2094641** | NA | 0.003104558 | -0.126712695 | 0.132921812 | 8.205868501 | 0.05883511 | 0.95503535 | 0.988027623 | -5.113042472 |
| **A_51_P350817** | Cnn1 | 0.036089784 | -1.473753966 | 1.545933534 | 10.10887053 | 0.058805974 | 0.955057588 | 0.988027623 | -5.113043767 |
| **A_51_P226269** | Rgcc | -0.013397955 | -0.576191568 | 0.549395658 | 14.72290114 | -0.058567748 | 0.955239407 | 0.988124007 | -5.113054329 |
| **A_51_P175974** | NA | 0.003635601 | -0.149345571 | 0.156616773 | 8.180844381 | 0.058466543 | 0.95531665 | 0.988124007 | -5.113058803 |
| **A_55_P2103536** | Irf2bpl | 0.005584623 | -0.229437727 | 0.240606974 | 9.083693905 | 0.058459338 | 0.95532215 | 0.988124007 | -5.113059121 |
| **A_55_P2001063** | Fcf1 | -0.005817268 | -0.250811892 | 0.239177356 | 12.78286926 | -0.058415982 | 0.955355241 | 0.988124007 | -5.113061035 |
| **A_55_P2032727** | NA | 0.003719265 | -0.153274337 | 0.160712866 | 15.31897477 | 0.058283322 | 0.955456492 | 0.988175854 | -5.113066883 |
| **A_55_P2050602** | Ncam1 | 0.002455898 | -0.101365095 | 0.106276891 | 7.695902965 | 0.058196163 | 0.955523016 | 0.988179949 | -5.113070718 |
| **A_55_P1965154** | Spc25 | 0.011748519 | -0.485354159 | 0.508851197 | 11.04207879 | 0.058144171 | 0.9555627 | 0.988179949 | -5.113073003 |
| **A_52_P700056** | Wfdc17 | 0.00522071 | -0.216541536 | 0.226982956 | 8.19749158 | 0.057917677 | 0.955735573 | 0.988248424 | -5.113082933 |
| **A_55_P1967224** | NA | -0.002942977 | -0.128051555 | 0.122165602 | 8.17127744 | -0.057872054 | 0.955770396 | 0.988248424 | -5.113084928 |
| **A_52_P480301** | Sppl3 | -0.00475054 | -0.206754708 | 0.197253627 | 9.550151908 | -0.05785646 | 0.955782298 | 0.988248424 | -5.11308561 |
| **A_51_P241667** | Prkdc | -0.002645584 | -0.11545516 | 0.110163992 | 8.048523236 | -0.057695869 | 0.955904873 | 0.988271952 | -5.101844451 |
| **A_55_P2070976** | Gm6923 | -0.003325646 | -0.145238734 | 0.138587443 | 7.852976793 | -0.057653102 | 0.955937516 | 0.988271952 | -5.113094483 |
| **A_55_P2092831** | NA | 0.012678907 | -0.52947925 | 0.554837065 | 11.54183556 | 0.057534056 | 0.956028381 | 0.988271952 | -5.113099663 |
| **A_51_P243857** | Copb2 | -0.004654399 | -0.20397365 | 0.194664853 | 12.86292821 | -0.057449137 | 0.956093199 | 0.988271952 | -5.113103351 |

| | | | | | | | | |
|---|---|---|---|---|---|---|---|---|
| **A_55_P2116305** | Tnfrsf25 | 0.004130699 | -0.172772195 | 0.181033594 | 8.038071101 | 0.057445737 | 0.956095794 | 0.988271952 | -5.113103499 |
| **A_55_P2108708** | Kcne3 | 0.003014501 | -0.126314577 | 0.132343578 | 7.948751175 | 0.057344049 | 0.956173412 | 0.988271952 | -5.113107908 |
| **A_55_P1961863** | Pcdhga9 | 0.016132405 | -0.676348355 | 0.708613165 | 11.59226866 | 0.057313981 | 0.956196363 | 0.988271952 | -5.113109211 |
| **A_55_P2170335** | Bmi1 | 0.004163837 | -0.174640613 | 0.182968288 | 10.50795012 | 0.057290763 | 0.956214085 | 0.988271952 | -5.113110216 |
| **A_55_P2111380** | Ctnnd2 | -0.003346322 | -0.147343833 | 0.14065119 | 8.089017738 | -0.0571718 | 0.956304889 | 0.988312956 | -5.113115359 |
| **A_55_P1952166** | Arid3a | -0.011844926 | -0.523533752 | 0.4998439 | 12.89689514 | -0.056950245 | 0.956474005 | 0.988412762 | -5.113124911 |
| **A_51_P116421** | Abcc5 | 0.014793906 | -0.624725254 | 0.654313067 | 10.48334333 | 0.056911295 | 0.956503736 | 0.988412762 | -5.062120758 |
| **A_52_P244463** | D16Ertd472e | 0.005389741 | -0.22835731 | 0.239136793 | 8.822003554 | 0.056727145 | 0.956644301 | 0.988501916 | -5.113134491 |
| **A_52_P16752** | Aox3 | 0.002382339 | -0.101051674 | 0.105816352 | 7.656662867 | 0.056664272 | 0.956692294 | 0.988501916 | -5.113137184 |
| **A_55_P1960178** | NA | -0.003263004 | -0.145233284 | 0.138707275 | 16.26274596 | -0.056544373 | 0.956783817 | 0.988543639 | -5.113142311 |
| **A_51_P114616** | Batf | -0.002688671 | -0.11988959 | 0.114512248 | 7.926577177 | -0.056438542 | 0.956864602 | 0.988544022 | -5.078096094 |
| **A_55_P1959164** | Gm10324 | 0.006733519 | -0.28713844 | 0.300605478 | 9.663960996 | 0.056370657 | 0.956916421 | 0.988544022 | -5.113149721 |
| **A_55_P2101757** | Dscc1 | 0.0029559 | -0.12611244 | 0.13202424 | 7.775277801 | 0.05634289 | 0.956937616 | 0.988544022 | -5.113150903 |
| **A_66_P110742** | Gm9949 | 0.002520029 | -0.10791629 | 0.112956347 | 7.756936689 | 0.056138749 | 0.957093448 | 0.988652162 | -5.113159578 |
| **A_51_P431046** | St8sia3 | 0.002362907 | -0.101413201 | 0.106139014 | 7.665720232 | 0.056016806 | 0.957186534 | 0.98869548 | -5.113164744 |
| **A_51_P358574** | Pdzd3 | 0.007500759 | -0.322384884 | 0.337386402 | 8.708553598 | 0.055938521 | 0.957246295 | 0.988704372 | -5.113168055 |
| **A_52_P674759** | Btbd3 | 0.019334075 | -0.832433456 | 0.871101607 | 11.05950019 | 0.055843353 | 0.957318943 | 0.988726574 | -5.113172074 |
| **A_55_P2108803** | Tubgcp4 | -0.002749139 | -0.124506551 | 0.119008272 | 8.789320682 | -0.055548269 | 0.957544203 | 0.988906385 | -5.113184491 |
| **A_51_P352562** | Mrpl38 | -0.0028673 | -0.130150473 | 0.124415873 | 9.316266621 | -0.055420615 | 0.957641652 | 0.988919192 | -5.113189842 |
| **A_52_P381929** | Ubqln2 | 0.00754235 | -0.327409104 | 0.342493804 | 9.93956558 | 0.055397984 | 0.957658929 | 0.988919192 | -5.113190789 |
| **A_51_P387302** | Zmym2 | -0.003667762 | -0.167524966 | 0.160189441 | 10.10867775 | -0.055068697 | 0.957910307 | 0.989083171 | -5.113204531 |
| **A_66_P134704** | Gm4952 | -0.002452729 | -0.112066313 | 0.107160856 | 7.672953075 | -0.055049617 | 0.957924873 | 0.989083171 | -5.113205324 |
| **A_55_P1962937** | Trem2 | -0.047788833 | -2.185852737 | 2.090275071 | 11.71958218 | -0.054988884 | 0.957971237 | 0.989083171 | -5.113207849 |
| **A_55_P1953030** | Tnfsf12 | -0.004128621 | -0.189326978 | 0.181069735 | 8.061855682 | -0.054845002 | 0.958081079 | 0.989107739 | -5.113213819 |
| **A_52_P387009** | Egln3 | -0.042980315 | -1.971707426 | 1.885746796 | 10.63650153 | -0.054823653 | 0.958097377 | 0.989107739 | -5.113214704 |
| **A_52_P219266** | Frmd8 | 0.004339613 | -0.191239168 | 0.199918393 | 11.36607843 | 0.054588153 | 0.958277164 | 0.989206397 | -5.113224437 |
| **A_52_P239292** | AI597479 | -0.002844382 | -0.131395184 | 0.125706419 | 10.54446207 | -0.054435514 | 0.958393694 | 0.989206397 | -5.113230724 |
| **A_51_P513062** | B630019K06Rik | -0.004854745 | -0.224333153 | 0.214623663 | 8.254055371 | -0.054418153 | 0.958406949 | 0.989206397 | -5.113231438 |
| **A_51_P418259** | Pccb | 0.00351723 | -0.155962602 | 0.162997062 | 8.700857834 | 0.054258045 | 0.958529182 | 0.989206397 | -5.113238011 |
| **A_51_P468464** | Gdpd5 | -0.007258787 | -0.336407543 | 0.32188997 | 10.3547748 | -0.05425515 | 0.958531392 | 0.989206397 | -5.11323813 |
| **A_55_P2095342** | Rara | -0.003373024 | -0.156485658 | 0.149739609 | 9.078645288 | -0.054197292 | 0.958575564 | 0.989206397 | -5.1132405 |
| **A_51_P201904** | Ndufb5 | -0.005676288 | -0.263398622 | 0.252046047 | 9.519619753 | -0.054185302 | 0.958584717 | 0.989206397 | -5.113240991 |
| **A_55_P2022569** | Pak3 | -0.002327152 | -0.108144605 | 0.103490301 | 7.680387362 | -0.054104902 | 0.958646099 | 0.989206397 | -5.11324428 |
| **A_55_P2138714** | Sgcd | -0.002488027 | -0.115786307 | 0.110810253 | 7.677332637 | -0.054025769 | 0.958706514 | 0.989206397 | -5.113247513 |
| **A_55_P2017677** | Cap2 | 0.00237919 | -0.105997255 | 0.110755636 | 7.693607606 | 0.05400866 | 0.958719576 | 0.989206397 | -5.113248211 |
| **A_55_P2161595** | Hpca | -0.008705655 | -0.405645489 | 0.388234179 | 10.60978721 | -0.05395677 | 0.958759192 | 0.989206397 | -5.113250328 |
| **A_55_P2003414** | Kng1 | -0.029382931 | -1.37076534 | 1.311999479 | 14.20646671 | -0.053890433 | 0.958809838 | 0.989206397 | -5.113253031 |
| **A_55_P2135526** | Gzmc | 0.03839193 | -1.716477809 | 1.793261669 | 10.82024652 | 0.053822545 | 0.958861669 | 0.989206397 | -5.113255793 |
| **A_55_P2090557** | Lrrc75a | 0.008154191 | -0.365002162 | 0.381310545 | 12.19550972 | 0.053759994 | 0.958909425 | 0.989206397 | -5.113258335 |
| **A_51_P121228** | Ascc2 | -0.004872119 | -0.228682899 | 0.218938662 | 11.56915847 | -0.053555741 | 0.959065367 | 0.989245664 | -5.113266616 |
| **A_51_P259214** | Slc39a6 | -0.018938395 | -0.890035907 | 0.852159117 | 11.21861909 | -0.05348667 | 0.959118101 | 0.989245664 | -5.113269409 |
| **A_51_P492830** | Cenph | 0.004165039 | -0.187442428 | 0.195772506 | 7.985801516 | 0.053478081 | 0.959124659 | 0.989245664 | -5.113269756 |
| **A_55_P2040893** | Tnni2 | -0.008470639 | -0.398595943 | 0.381654664 | 9.016368891 | -0.053417213 | 0.959171131 | 0.989245664 | -5.113272214 |
| **A_55_P1974877** | Bcl2l1 | -0.005207122 | -0.245476751 | 0.235062508 | 15.61678925 | -0.053317285 | 0.959247425 | 0.989245664 | -5.113276244 |
| **A_55_P1957845** | Tnfaip1 | -0.005722909 | -0.270452097 | 0.259006279 | 13.86195782 | -0.053184387 | 0.959348893 | 0.989245664 | -5.113281591 |
| **A_51_P126437** | Enc1 | -0.005016009 | -0.237079899 | 0.227047882 | 10.24231794 | -0.053176517 | 0.959354901 | 0.989245664 | -5.113281907 |
| **A_55_P2051894** | Zfp598 | 0.004320249 | -0.195846348 | 0.204486846 | 9.768738695 | 0.053099 | 0.959414086 | 0.989245664 | -5.113285019 |
| **A_55_P2036764** | Prr15l | -0.002679199 | -0.126827359 | 0.121468962 | 7.760444783 | -0.053092581 | 0.959418986 | 0.989245664 | -5.113285277 |
| **A_55_P1961265** | NA | 0.006174799 | -0.280338426 | 0.292688023 | 9.010772592 | 0.053020923 | 0.959473698 | 0.989245664 | -5.11328815 |
| **A_51_P237688** | Smim14 | -0.010285942 | -0.487991737 | 0.467419854 | 11.68535723 | -0.052972773 | 0.959510461 | 0.989245664 | -5.113290078 |

| | | | | | | | | | |
|---|---|---|---|---|---|---|---|---|---|
| **A_55_P2026863** | Trim32 | 0.003568005 | -0.162736064 | 0.169872073 | 10.85037878 | 0.052782711 | 0.959655576 | 0.989294918 | -5.113297671 |
| **A_55_P2096266** | Sftpb | 0.002310464 | -0.105393262 | 0.110014189 | 7.704724989 | 0.052776133 | 0.959660598 | 0.989294918 | -5.113297934 |
| **A_52_P510215** | Traf6 | 0.006747469 | -0.309393628 | 0.322888566 | 8.949070448 | 0.052508438 | 0.959864991 | 0.989433633 | -5.113308581 |
| **A_51_P375431** | Phip | 0.002835822 | -0.130139716 | 0.135811361 | 8.187226837 | 0.052465812 | 0.959897536 | 0.989433633 | -5.113310272 |
| **A_52_P362033** | Polr2e | 0.008060229 | -0.370396309 | 0.386516768 | 10.17229706 | 0.052396292 | 0.959950618 | 0.989435583 | -5.113313026 |
| **A_51_P113003** | Cenpq | -0.002483613 | -0.119584806 | 0.114617579 | 7.905487301 | -0.05217853 | 0.960116888 | 0.989505981 | -5.1020706 |
| **A_55_P2079729** | NA | 0.004228935 | -0.195185452 | 0.203643323 | 15.98755977 | 0.052172747 | 0.960121304 | 0.989505981 | -5.113321857 |
| **A_51_P354165** | Apcs | 0.00238316 | -0.110297533 | 0.115063853 | 7.72003649 | 0.052032277 | 0.96022856 | 0.989539969 | -5.113327387 |
| **A_55_P1962901** | Osr1 | 0.003286993 | -0.152238827 | 0.158812812 | 7.790320439 | 0.05199546 | 0.960256672 | 0.989539969 | -5.113328834 |
| **A_55_P2051254** | Pvt1 | -0.003577216 | -0.173072277 | 0.165917845 | 8.099799365 | -0.051922695 | 0.960312232 | 0.989544468 | -5.113331691 |
| **A_55_P1981731** | Nr2c2 | -0.00292389 | -0.142589599 | 0.13674182 | 8.214707291 | -0.051503927 | 0.96063199 | 0.989764244 | -5.113348054 |
| **A_51_P481221** | Bace2 | 0.005427815 | -0.253883344 | 0.264738974 | 8.075312001 | 0.051495942 | 0.960638087 | 0.989764244 | -5.113348365 |
| **A_51_P463440** | Elovl6 | 0.007682553 | -0.35973111 | 0.375096216 | 9.287276596 | 0.051442186 | 0.960679134 | 0.989764244 | -5.113350456 |
| **A_55_P2144210** | Snx5 | 0.008571692 | -0.403248119 | 0.420391503 | 11.86344976 | 0.0512069 | 0.960858796 | 0.989896581 | -5.11335958 |
| **A_55_P1960197** | P2ry14 | -0.005591184 | -0.277114968 | 0.2659326 | 8.019377946 | -0.050659983 | 0.961276424 | 0.990108488 | -5.113380629 |
| **A_55_P2162592** | Lman2 | 0.00267301 | -0.127408352 | 0.132754372 | 15.94681119 | 0.0505539 | 0.961357431 | 0.990108488 | -5.113384686 |
| **A_51_P227770** | Phykpl | 0.004283269 | -0.204189963 | 0.2127565 | 9.356274903 | 0.050546859 | 0.961362808 | 0.990108488 | -5.113384954 |
| **A_55_P1982049** | Icam2 | -0.005105137 | -0.253624718 | 0.243414445 | 11.91941734 | -0.050537752 | 0.961369762 | 0.990108488 | -5.113385302 |
| **A_55_P2054331** | Gm9222 | -0.006624211 | -0.33073031 | 0.317481888 | 8.551576638 | -0.050282413 | 0.961564746 | 0.990108488 | -5.113395028 |
| **A_51_P511060** | Csgalnact2 | 0.004220004 | -0.202520359 | 0.210960366 | 9.986142276 | 0.050217686 | 0.961614174 | 0.990108488 | -5.113397486 |
| **A_55_P1970002** | Sh3kbp1 | 0.003975574 | -0.191252537 | 0.199203684 | 8.219811077 | 0.050098719 | 0.961705022 | 0.990108488 | -5.113401995 |
| **A_66_P109720** | Jpt2 | -0.003660212 | -0.18368605 | 0.176365626 | 8.39096662 | -0.05001964 | 0.96176541 | 0.990108488 | -5.113404986 |
| **A_55_P2116889** | Hax1 | 0.004093405 | -0.197738022 | 0.205924832 | 10.20758712 | 0.049895922 | 0.961859887 | 0.990108488 | -5.113409657 |
| **A_55_P1994000** | Snx16 | -0.002456591 | -0.123671573 | 0.11875839 | 9.970452782 | -0.049859257 | 0.961887887 | 0.990108488 | -5.113411039 |
| **A_55_P2045188** | NA | -0.003323164 | -0.167518464 | 0.160872136 | 9.047658945 | -0.049792076 | 0.96193919 | 0.990108488 | -5.113413568 |
| **A_51_P131800** | Cyba | -0.010310113 | -0.519866168 | 0.499245942 | 9.137422545 | -0.049778365 | 0.96194966 | 0.990108488 | -5.113414084 |
| **A_51_P255565** | Smarcad1 | 0.004344733 | -0.210422782 | 0.219112247 | 9.327185142 | 0.049769542 | 0.961956397 | 0.990108488 | -5.113414416 |
| **A_51_P501550** | NA | -0.005211005 | -0.263085262 | 0.252663251 | 9.014395685 | -0.049714457 | 0.961998464 | 0.990108488 | -5.113416486 |
| **A_55_P2069955** | NA | -0.003545356 | -0.179308546 | 0.172217833 | 16.99604827 | -0.049625067 | 0.962066727 | 0.990108488 | -5.113419841 |
| **A_55_P1997141** | Mybl2 | -0.004729048 | -0.239251844 | 0.229793749 | 9.595917265 | -0.049608692 | 0.962079232 | 0.990108488 | -5.113420455 |
| **A_51_P420547** | Clic5 | 0.01023023 | -0.497128721 | 0.517589181 | 12.42651357 | 0.049606573 | 0.962080851 | 0.990108488 | -5.113420534 |
| **A_51_P219721** | Ola1 | 0.005418612 | -0.263562331 | 0.274399556 | 10.79409331 | 0.049560506 | 0.962116031 | 0.990108488 | -5.11342226 |
| **A_55_P2240823** | Epb41 | 0.002417582 | -0.117637422 | 0.122472586 | 7.776274128 | 0.049541612 | 0.962130459 | 0.990108488 | -5.113422968 |
| **A_55_P2051701** | Pbx1 | 0.015071014 | -0.734252996 | 0.764395024 | 8.997429071 | 0.049481401 | 0.962176441 | 0.990108488 | -5.113425221 |
| **A_55_P1981291** | Spink8 | 0.017490659 | -0.852579881 | 0.887561199 | 9.877751861 | 0.049456214 | 0.962195675 | 0.990108488 | -5.113426162 |
| **A_52_P267824** | Ces2g | 0.002264102 | -0.110388037 | 0.114916241 | 7.717365399 | 0.04944537 | 0.962203956 | 0.990108488 | -5.113426567 |
| **A_51_P469252** | Zfp647 | -0.002747825 | -0.139608183 | 0.134112534 | 7.96550901 | -0.049394718 | 0.962242638 | 0.990108488 | -5.113428459 |
| **A_51_P280893** | Pla2g1b | 0.002201076 | -0.107597864 | 0.112000016 | 7.660938633 | 0.049318068 | 0.962301173 | 0.990116011 | -5.113431318 |
| **A_55_P2040559** | NA | -0.005194761 | -0.276355363 | 0.265965842 | 14.15866793 | -0.049646546 | 0.962375939 | 0.990140231 | -5.044297651 |
| **A_55_P1957855** | NA | 0.009357159 | -0.460055488 | 0.478769805 | 11.86859677 | 0.049040893 | 0.962512848 | 0.990228381 | -5.113441618 |
| **A_52_P163928** | Poldip2 | -0.004494188 | -0.231643625 | 0.222655249 | 10.10886331 | -0.048675312 | 0.962792041 | 0.990462896 | -5.113455115 |
| **A_55_P1965989** | Zfp329 | -0.003803345 | -0.196391356 | 0.188784665 | 8.355530367 | -0.048585387 | 0.962860717 | 0.99048083 | -5.11345842 |
| **A_51_P339200** | Abhd16a | 0.005599188 | -0.278655939 | 0.289854316 | 8.963835619 | 0.048460281 | 0.962956262 | 0.9905264 | -5.113463007 |
| **A_51_P329158** | Zc3h10 | 0.002999177 | -0.149654638 | 0.155652991 | 8.963543099 | 0.048335205 | 0.963051784 | 0.990571942 | -5.113467581 |
| **A_52_P331901** | Trp53rkb | 0.002942198 | -0.147058615 | 0.152943011 | 8.264204808 | 0.048255569 | 0.963112604 | 0.990581787 | -5.113470487 |
| **A_55_P2061268** | Rnpc3 | -0.004037452 | -0.210590887 | 0.202515983 | 8.027604623 | -0.048088819 | 0.963239955 | 0.990660056 | -5.113476557 |
| **A_55_P1991605** | Aplp1 | -0.006083574 | -0.318018636 | 0.305851487 | 9.802223077 | -0.04798039 | 0.963322766 | 0.990692511 | -5.113480493 |
| **A_55_P2018116** | Pabpc4l | -0.00220084 | -0.115347315 | 0.110945634 | 7.725947502 | -0.047853829 | 0.963419426 | 0.990739205 | -5.113485076 |
| **A_52_P91618** | Lysmd3 | -0.00359493 | -0.18886573 | 0.18167587 | 9.387428157 | -0.047736725 | 0.963508863 | 0.990762361 | -5.113489305 |
| **A_55_P2124153** | Rusc2 | -0.011237571 | -0.590950794 | 0.568475652 | 11.4848523 | -0.047690118 | 0.963544459 | 0.990762361 | -5.113490986 |

| | | | | | | | | | |
|---|---|---|---|---|---|---|---|---|---|
| **A_51_P401263** | Eme1 | -0.002354091 | -0.124288458 | 0.119580276 | 7.894606669 | -0.047497005 | 0.96369195 | 0.990861306 | -5.113497931 |
| **A_52_P663757** | Tnik | 0.002344671 | -0.119463527 | 0.124152868 | 8.182850913 | 0.047355942 | 0.963799688 | 0.99091937 | -5.113502987 |
| **A_51_P271665** | Rev3l | 0.005456757 | -0.27906237 | 0.289975884 | 10.12005691 | 0.047183734 | 0.963931214 | 0.99099542 | -5.113509138 |
| **A_52_P428521** | Fgd1 | -0.003568678 | -0.189915721 | 0.182778364 | 8.38810469 | -0.047114444 | 0.963984136 | 0.99099542 | -5.113511607 |
| **A_51_P258435** | Zdhhc3 | 0.00281845 | -0.144940418 | 0.150577319 | 9.121756279 | 0.046927348 | 0.964127036 | 0.99099542 | -5.113518255 |
| **A_52_P506995** | Plet1 | 0.004703123 | -0.242543979 | 0.251950225 | 9.664920213 | 0.046797676 | 0.964226077 | 0.99099542 | -5.113522847 |
| **A_55_P2134496** | Amfr | 0.003593548 | -0.185331139 | 0.192518235 | 12.19943071 | 0.04679548 | 0.964227754 | 0.99099542 | -5.113522925 |
| **A_55_P2163198** | NA | 0.004103612 | -0.211873751 | 0.220080976 | 10.63302 | 0.046744147 | 0.964266961 | 0.99099542 | -5.113524739 |
| **A_51_P385598** | Slc37a4 | 0.011690609 | -0.604643701 | 0.628024919 | 11.1301883 | 0.046664857 | 0.964327522 | 0.99099542 | -5.113527538 |
| **A_51_P463187** | Tbc1d2b | -0.002588745 | -0.139252948 | 0.134075457 | 8.218268858 | -0.046601906 | 0.964375604 | 0.99099542 | -5.113529756 |
| **A_51_P282144** | Adam23 | -0.00405055 | -0.218335853 | 0.210234753 | 8.879475532 | -0.046504047 | 0.964450348 | 0.99099542 | -5.113533199 |
| **A_55_P2098688** | Mcrs1 | -0.003413023 | -0.184690485 | 0.177864439 | 8.722874836 | -0.04631957 | 0.964591252 | 0.99099542 | -5.113539669 |
| **A_55_P2069729** | Ormdl2 | -0.005007154 | -0.271245667 | 0.26123136 | 12.28401075 | -0.046268904 | 0.964629951 | 0.99099542 | -5.113541441 |
| **A_51_P108252** | Gpsm2 | -0.002969348 | -0.161092547 | 0.155153851 | 9.285513641 | -0.046199222 | 0.964683174 | 0.99099542 | -5.102669308 |
| **A_55_P2130090** | Cnn3 | 0.007569549 | -0.395525719 | 0.410664817 | 14.22159911 | 0.046198884 | 0.964683433 | 0.99099542 | -5.113543888 |
| **A_55_P2091446** | Trim33 | 0.005159154 | -0.269682375 | 0.280000684 | 9.464534231 | 0.046181213 | 0.96469693 | 0.99099542 | -5.113544505 |
| **A_51_P114817** | Ccna1 | 0.001987879 | -0.103983506 | 0.107959263 | 7.653324572 | 0.046149865 | 0.964720874 | 0.99099542 | -5.097046199 |
| **A_51_P451588** | Plekhb1 | 0.002071007 | -0.108390208 | 0.112532223 | 7.790659027 | 0.046125492 | 0.964739491 | 0.99099542 | -5.113546448 |
| **A_51_P510722** | Rita1 | -0.002434937 | -0.132328276 | 0.127458401 | 8.121443037 | -0.046117959 | 0.964745244 | 0.99099542 | -5.11354671 |
| **A_55_P2044458** | Pramef8 | 0.003625012 | -0.190640237 | 0.197890261 | 10.80837867 | 0.045907483 | 0.96490601 | 0.991107889 | -5.113554029 |
| **A_55_P2027127** | Triml2 | 0.004514969 | -0.237839065 | 0.246869003 | 9.578226288 | 0.045832515 | 0.964963272 | 0.991114038 | -5.113556628 |
| **A_55_P2023537** | NA | 0.002071993 | -0.109937459 | 0.114081445 | 7.819250424 | 0.04550957 | 0.965209947 | 0.991314722 | -5.113567774 |
| **A_51_P160673** | Kcne1l | 0.002146267 | -0.114339683 | 0.118632218 | 7.699160349 | 0.045329346 | 0.96534761 | 0.991388028 | -5.11357396 |
| **A_55_P2071132** | Tnfrsf23 | -0.032586508 | -1.803033748 | 1.737860732 | 13.45430224 | -0.045281833 | 0.965383903 | 0.991388028 | -5.113575587 |
| **A_51_P142421** | Rspo1 | 0.002160189 | -0.115700834 | 0.120021211 | 7.709518474 | 0.045091081 | 0.965529609 | 0.991484982 | -5.1135821 |
| **A_51_P222386** | Tbc1d10a | -0.009521722 | -0.532214581 | 0.513171137 | 11.63087407 | -0.044816516 | 0.965739338 | 0.991596287 | -5.113591428 |
| **A_55_P1967978** | Nfatc2 | -0.005516078 | -0.308331166 | 0.29729901 | 8.393101549 | -0.044814862 | 0.965740602 | 0.991596287 | -5.113591484 |
| **A_55_P2088495** | NA | 0.012757413 | -0.69222396 | 0.717738785 | 9.405519003 | 0.044519894 | 0.965965919 | 0.991774953 | -5.113601441 |
| **A_52_P245766** | NA | 0.006071919 | -0.331926315 | 0.344070153 | 10.06164801 | 0.044195769 | 0.966213514 | 0.99189021 | -5.113612306 |
| **A_55_P2169445** | Mfsd6 | 0.005309493 | -0.290388749 | 0.301007735 | 9.28515601 | 0.044174689 | 0.966229617 | 0.99189021 | -5.11361301 |
| **A_66_P118195** | Ciao1 | 0.003611192 | -0.197553412 | 0.204775796 | 10.22579273 | 0.044163978 | 0.966237799 | 0.99189021 | -5.113613368 |
| **A_52_P266132** | Fgl2 | 0.00785781 | -0.430858653 | 0.446574274 | 8.228365555 | 0.044064289 | 0.96631395 | 0.99189021 | -5.113616691 |
| **A_55_P2032039** | Bola3 | 0.003731547 | -0.204736646 | 0.212199739 | 10.57308065 | 0.04403705 | 0.966334758 | 0.99189021 | -5.113617598 |
| **A_55_P2003216** | Igsf5 | 0.003127474 | -0.172308117 | 0.178563066 | 7.850106635 | 0.043857633 | 0.966471815 | 0.991978213 | -5.113623557 |
| **A_55_P1964433** | Zmynd12 | -0.002934904 | -0.168511258 | 0.162641449 | 9.011247597 | -0.043607863 | 0.966662617 | 0.992026051 | -5.113631812 |
| **A_52_P595954** | Nelfa | -0.004229104 | -0.242877535 | 0.234419327 | 9.954047116 | -0.043597237 | 0.966670734 | 0.992026051 | -5.113632162 |
| **A_55_P2006677** | Gm7969 | -0.005957416 | -0.342888006 | 0.330973174 | 9.436675647 | -0.043499739 | 0.966745215 | 0.992026051 | -5.113635371 |
| **A_51_P426283** | Dhx30 | -0.004674477 | -0.269123763 | 0.259774809 | 12.33941485 | -0.043487037 | 0.966754918 | 0.992026051 | -5.113635788 |
| **A_52_P409746** | Serpinc1 | -0.014954508 | -0.861488873 | 0.831579857 | 8.71619484 | -0.043460698 | 0.966775039 | 0.992026051 | -5.113636654 |
| **A_55_P2333126** | Oga | -0.005742273 | -0.331642414 | 0.320157868 | 8.684017344 | -0.043347939 | 0.966861177 | 0.992061774 | -5.113640352 |
| **A_66_P119247** | Gatad1 | -0.00826338 | -0.47976685 | 0.46324009 | 10.5727268 | -0.04311635 | 0.967038095 | 0.992190633 | -5.113647919 |
| **A_52_P257728** | Cnnm4 | -0.0024811 | -0.144838948 | 0.139876748 | 9.428503113 | -0.04287775 | 0.967220371 | 0.99220725 | -5.113655672 |
| **A_55_P2028804** | Uchl3 | 0.002904525 | -0.163995793 | 0.169804842 | 9.337808909 | 0.042814123 | 0.967268978 | 0.99220725 | -5.113657733 |
| **A_51_P337771** | Pcdhb17 | -0.002902718 | -0.169916037 | 0.164110601 | 7.797613965 | -0.042758539 | 0.967311441 | 0.99220725 | -5.11365953 |
| **A_55_P2060188** | NA | -0.002004819 | -0.117451145 | 0.113441507 | 8.004855453 | -0.042723246 | 0.967338403 | 0.99220725 | -5.11366067 |
| **A_55_P2062737** | Elovl2 | -0.002612328 | -0.153055324 | 0.147830668 | 7.970121248 | -0.042719378 | 0.967341358 | 0.99220725 | -5.113660795 |
| **A_55_P2075941** | Nf1 | -0.002352588 | -0.137924167 | 0.133218991 | 9.877268798 | -0.042691984 | 0.967362286 | 0.99220725 | -5.113661679 |
| **A_55_P1996988** | Tent4b | -0.003402752 | -0.200078528 | 0.193273024 | 10.40101176 | -0.042564599 | 0.967459602 | 0.992212372 | -5.113665783 |
| **A_55_P1954486** | Mapt | 0.004177699 | -0.23772466 | 0.246080057 | 8.654390204 | 0.042487988 | 0.967518129 | 0.992212372 | -5.113668245 |
| **A_52_P876147** | Crocc2 | 0.001790573 | -0.1021799 | 0.105761046 | 7.672764351 | 0.04236928 | 0.967608818 | 0.992212372 | -5.113672052 |

| | | | | | | | | | |
|---|---|---|---|---|---|---|---|---|---|
| **A_55_P2124871** | Npw | 0.001854174 | -0.105846399 | 0.109554746 | 7.728745884 | 0.042354701 | 0.967619956 | 0.992212372 | -5.113672518 |
| **A_51_P244824** | Dapp1 | -0.002497679 | -0.147594437 | 0.142599079 | 7.878741861 | -0.04234948 | 0.967623944 | 0.992212372 | -5.113672685 |
| **A_51_P380807** | Ckm | 0.002011318 | -0.115568854 | 0.119591489 | 7.67750119 | 0.042083875 | 0.967826857 | 0.992352076 | -5.11368116 |
| **A_55_P2025954** | Acly | -0.003833277 | -0.228175109 | 0.220508556 | 13.25876285 | -0.042036742 | 0.967862866 | 0.992352076 | -5.113682658 |
| **A_55_P2286650** | 6720462K09Rik | -0.004908846 | -0.293679112 | 0.28386142 | 8.1336287 | -0.041821163 | 0.968027563 | 0.992422942 | -5.113689489 |
| **A_51_P240801** | Tmem173 | -0.011789087 | -0.705454197 | 0.681876023 | 9.374794952 | -0.04181186 | 0.968034671 | 0.992422942 | -5.113689783 |
| **A_51_P234833** | Strn4 | 0.002540794 | -0.147944481 | 0.153026069 | 10.6620686 | 0.041537909 | 0.968243966 | 0.992545987 | -5.113698412 |
| **A_55_P1997225** | Agap2 | -0.001810441 | -0.109594219 | 0.105973337 | 7.700905933 | -0.041323792 | 0.968407551 | 0.992545987 | -5.113705116 |
| **A_55_P2086387** | 1700030K09Rik | -0.002266464 | -0.137696468 | 0.133163539 | 8.74277231 | -0.041172111 | 0.968523435 | 0.992545987 | -5.113709845 |
| **A_55_P2058933** | Kif1b | -0.005769559 | -0.350905889 | 0.33936677 | 11.25140935 | -0.041126441 | 0.968558328 | 0.992545987 | -5.113711265 |
| **A_55_P2059986** | Chst14 | -0.004401428 | -0.267745087 | 0.258942232 | 9.742519706 | -0.041118746 | 0.968564207 | 0.992545987 | -5.113711504 |
| **A_55_P2149001** | NA | -0.003559604 | -0.216569009 | 0.209449801 | 13.74976787 | -0.041112326 | 0.968569112 | 0.992545987 | -5.113711704 |
| **A_55_P2070838** | NA | -0.00851507 | -0.518102917 | 0.501072778 | 13.74152806 | -0.041109132 | 0.968571552 | 0.992545987 | -5.113711803 |
| **A_52_P320193** | Clec2h | 0.021900311 | -1.296301342 | 1.340101964 | 10.32250464 | 0.040873102 | 0.968751882 | 0.992545987 | -5.113719115 |
| **A_52_P525183** | Acot2 | -0.002128826 | -0.130669987 | 0.126412334 | 8.146910004 | -0.040744328 | 0.968850268 | 0.992545987 | -5.113723087 |
| **A_51_P140350** | Nadsyn1 | 0.002830483 | -0.168458901 | 0.174119868 | 8.604331266 | 0.040653628 | 0.968919565 | 0.992545987 | -5.113725877 |
| **A_55_P1965659** | Havcr2 | -0.018170105 | -1.119662917 | 1.083322706 | 8.82996298 | -0.040583074 | 0.96897347 | 0.992545987 | -5.113728043 |
| **A_55_P1974900** | Ndufab1 | 0.003099828 | -0.185097758 | 0.191297414 | 13.78524384 | 0.040522176 | 0.969019998 | 0.992545987 | -5.113729909 |
| **A_55_P2082045** | Vps13a | -0.001995085 | -0.123123438 | 0.119133268 | 8.033415805 | -0.040521433 | 0.969020566 | 0.992545987 | -5.113729932 |
| **A_55_P2100495** | Synb | -0.010929631 | -0.676183368 | 0.654324106 | 14.26485658 | -0.040419165 | 0.969098702 | 0.992545987 | -5.11373306 |
| **A_66_P137462** | Nsdhl | 0.003038596 | -0.182014086 | 0.188091279 | 8.220727295 | 0.040396781 | 0.969115804 | 0.992545987 | -5.113733743 |
| **A_55_P2042217** | Fam161a | 0.002044258 | -0.122529029 | 0.126617545 | 7.843243147 | 0.040371964 | 0.969134765 | 0.992545987 | -5.113734501 |
| **A_55_P1957198** | Apob | -0.023109809 | -1.431897326 | 1.385677707 | 13.73136466 | -0.040357107 | 0.969146116 | 0.992545987 | -5.113734954 |
| **A_55_P1978366** | NA | -0.003945438 | -0.24501813 | 0.237127254 | 9.969302615 | -0.040263945 | 0.969217295 | 0.992545987 | -5.113737792 |
| **A_51_P233947** | Tbc1d10b | 0.004250452 | -0.255927629 | 0.264428533 | 13.7898987 | 0.04019144 | 0.969272692 | 0.992545987 | -5.113739997 |
| **A_55_P2092501** | Med1 | 0.003546738 | -0.213924247 | 0.221017723 | 12.7152811 | 0.040123331 | 0.96932473 | 0.992545987 | -5.113742064 |
| **A_55_P1971229** | Slc25a36 | 0.003570271 | -0.215505814 | 0.222646355 | 9.111171237 | 0.040093627 | 0.969347425 | 0.992545987 | -5.113742964 |
| **A_51_P357561** | Fbxw9 | -0.002884374 | -0.179988009 | 0.17421926 | 8.992764462 | -0.040067595 | 0.969367315 | 0.992545987 | -5.113743753 |
| **A_52_P449629** | Ankrd13c | 0.003268737 | -0.197509878 | 0.204047353 | 9.579491289 | 0.040052696 | 0.969378699 | 0.992545987 | -5.113744204 |
| **A_55_P2083725** | NA | -0.003184255 | -0.198827329 | 0.192458818 | 8.84714279 | -0.040041708 | 0.969387094 | 0.992545987 | -5.113744537 |
| **A_51_P239654** | Nr4a1 | -0.005356647 | -0.335246045 | 0.324532751 | 10.42624228 | -0.039947891 | 0.969458775 | 0.992546665 | -5.113747372 |
| **A_55_P1999902** | Pip5k1a | 0.018565956 | -1.126007445 | 1.163139357 | 15.11559776 | 0.039906426 | 0.969490456 | 0.992546665 | -5.113748624 |
| **A_52_P393306** | Snap23 | -0.002964714 | -0.18621661 | 0.180287182 | 9.339774964 | -0.039801871 | 0.969570342 | 0.992575877 | -5.113751773 |
| **A_52_P109319** | E2f5 | -0.001852337 | -0.116592824 | 0.112888149 | 8.084998356 | -0.039716641 | 0.969635463 | 0.992589973 | -5.113754334 |
| **A_51_P275350** | Pten | -0.003648763 | -0.230626709 | 0.223329184 | 10.67745236 | -0.039548602 | 0.969763855 | 0.992668831 | -5.113759368 |
| **A_52_P608322** | Maff | 0.011210592 | -0.689641683 | 0.712062866 | 11.74310185 | 0.039352399 | 0.969913767 | 0.992686522 | -5.113765218 |
| **A_52_P198898** | Samd5 | 0.004869829 | -0.299941857 | 0.309681516 | 8.317522742 | 0.039305316 | 0.969949742 | 0.992686522 | -5.113766617 |
| **A_51_P221132** | L2hgdh | -0.0020084 | -0.127905391 | 0.123888591 | 9.130801956 | -0.039246785 | 0.969994464 | 0.992686522 | -5.113768355 |
| **A_52_P358860** | Gss | -0.003462863 | -0.221076939 | 0.214151212 | 8.891728434 | -0.039148719 | 0.970069394 | 0.992686522 | -5.11377126 |
| **A_55_P2258261** | 1810008I18Rik | 0.001715099 | -0.106300632 | 0.109730831 | 7.706281465 | 0.039063525 | 0.970134489 | 0.992686522 | -5.113773778 |
| **A_52_P275354** | Osbpl7 | -0.003205977 | -0.205404694 | 0.198992739 | 9.974643336 | -0.039007779 | 0.970177084 | 0.992686522 | -5.113775423 |
| **A_52_P579933** | Slc16a6 | -0.003365469 | -0.215879157 | 0.209148218 | 9.298408073 | -0.038960804 | 0.970212977 | 0.992686522 | -5.113776807 |
| **A_55_P2143431** | Nfib | 0.005044966 | -0.313851614 | 0.323941546 | 9.722398063 | 0.038920421 | 0.970243833 | 0.992686522 | -5.113777995 |
| **A_52_P81630** | 2810001G20Rik | 0.004032883 | -0.251025577 | 0.259091342 | 8.92192007 | 0.038899589 | 0.97025975 | 0.992686522 | -5.113778608 |
| **A_51_P481676** | Zfp787 | 0.002254896 | -0.140752569 | 0.145262362 | 9.503539681 | 0.038791541 | 0.970342309 | 0.992686522 | -5.11378178 |
| **A_51_P233153** | Cadps2 | 0.001675486 | -0.104598781 | 0.107949754 | 7.645765916 | 0.038786621 | 0.970346068 | 0.992686522 | -5.113781924 |
| **A_55_P2160043** | A330069E16Rik | -0.001825139 | -0.118187564 | 0.114537287 | 8.017574634 | -0.038588003 | 0.970497831 | 0.992789234 | -5.113787732 |
| **A_52_P498208** | H2ac15 | -0.005548061 | -0.360911514 | 0.349815392 | 10.65462271 | -0.038409409 | 0.970634295 | 0.992876285 | -5.113792929 |
| **A_55_P2102240** | Nprl3 | 0.003742264 | -0.237291408 | 0.244775936 | 8.983126757 | 0.038196703 | 0.970796825 | 0.99298999 | -5.113799087 |
| **A_51_P375987** | Fign | -0.002197692 | -0.144274304 | 0.13987892 | 7.90364451 | -0.038055149 | 0.970904989 | 0.993048076 | -5.113803166 |

| | | | | | | | | | |
|---|---|---|---|---|---|---|---|---|---|
| **A_55_P2011436** | NA | -0.011363219 | -0.748850265 | 0.726123826 | 13.73509266 | -0.037906715 | 0.97101841 | 0.993063839 | -5.113807427 |
| **A_51_P256384** | Atp2b2 | 0.001675735 | -0.107119292 | 0.110470762 | 7.687178036 | 0.037893564 | 0.971028459 | 0.993063839 | -5.113807803 |
| **A_51_P377390** | Ndst2 | 0.00405722 | -0.259772602 | 0.267887043 | 9.470170869 | 0.03783327 | 0.971074531 | 0.993063839 | -5.113809529 |
| **A_55_P2160750** | Col25a1 | -0.00162632 | -0.107656744 | 0.104404103 | 7.681455814 | -0.037735034 | 0.971149596 | 0.993087699 | -5.113812334 |
| **A_55_P2101806** | Esf1 | 0.002946236 | -0.189592697 | 0.195485169 | 10.22601274 | 0.037645938 | 0.971217677 | 0.993087699 | -5.113814872 |
| **A_52_P535907** | Sde2 | 0.002278924 | -0.146837839 | 0.151395688 | 9.422063471 | 0.037598681 | 0.971253788 | 0.993087699 | -5.113816216 |
| **A_55_P2186282** | Agxt2 | 0.001690782 | -0.109566188 | 0.112947752 | 7.71111167 | 0.037387768 | 0.971414954 | 0.993087699 | -5.113822193 |
| **A_55_P2173702** | Ebpl | -0.008348441 | -0.559002912 | 0.542306029 | 12.72300536 | -0.037298846 | 0.971482903 | 0.993087699 | -5.113824703 |
| **A_55_P2031898** | NA | 0.005690571 | -0.370636174 | 0.382017316 | 10.78610127 | 0.037201454 | 0.971557324 | 0.993087699 | -5.113827445 |
| **A_66_P140452** | Map1a | -0.001665702 | -0.111912103 | 0.108580699 | 7.712002789 | -0.037170816 | 0.971580736 | 0.993087699 | -5.113828306 |
| **A_51_P262340** | Rbm3 | -0.00359929 | -0.242754624 | 0.235556044 | 12.68279753 | -0.037025926 | 0.971691453 | 0.993087699 | -5.113832369 |
| **A_51_P114177** | Eci3 | 0.00323594 | -0.212092421 | 0.2185643 | 10.32920869 | 0.036971609 | 0.97173296 | 0.993087699 | -5.097329183 |
| **A_51_P372463** | Fxr1 | 0.003843081 | -0.252082853 | 0.259769016 | 12.07230899 | 0.036943195 | 0.971754673 | 0.993087699 | -5.113834681 |
| **A_66_P114768** | Bach2 | 0.00248196 | -0.163093666 | 0.168057587 | 9.217603271 | 0.036878022 | 0.971804475 | 0.993087699 | -5.1138365 |
| **A_55_P1977589** | Nfu1 | -0.001710507 | -0.11590131 | 0.112480297 | 11.65059895 | -0.036852146 | 0.971824248 | 0.993087699 | -5.113837221 |
| **A_55_P2009572** | Ptbp3 | -0.002245168 | -0.152288202 | 0.147797865 | 8.886447966 | -0.036813087 | 0.971854095 | 0.993087699 | -5.113838308 |
| **A_51_P164420** | Eef1e1 | -0.002982379 | -0.202636263 | 0.196671504 | 9.160650582 | -0.036749741 | 0.971902502 | 0.993087699 | -5.113840069 |
| **A_55_P2037111** | Ran | -0.002379337 | -0.162462558 | 0.157703885 | 9.796470386 | -0.036566152 | 0.972042793 | 0.993087699 | -5.113845156 |
| **A_51_P511250** | Tent5b | 0.010189273 | -0.675416358 | 0.695794903 | 10.15187582 | 0.036562675 | 0.97204545 | 0.993087699 | -5.113845252 |
| **A_51_P254262** | Nxph3 | 0.013121107 | -0.87016062 | 0.896402835 | 8.762778557 | 0.036546047 | 0.972058157 | 0.993087699 | -5.113845711 |
| **A_51_P191909** | Rem2 | -0.001555481 | -0.106306231 | 0.10319527 | 7.678885414 | -0.036532268 | 0.972068687 | 0.993087699 | -5.113846092 |
| **A_55_P1954907** | Rpl12 | -0.002026217 | -0.138504003 | 0.134451569 | 16.1533188 | -0.036525253 | 0.972074048 | 0.993087699 | -5.113846286 |
| **A_51_P352385** | Ism1 | -0.002488199 | -0.170621524 | 0.165645126 | 8.095110744 | -0.036408307 | 0.972163414 | 0.993126507 | -5.113849509 |
| **A_52_P507305** | Iah1 | 0.005663434 | -0.380547244 | 0.391874113 | 9.601959974 | 0.036076528 | 0.972416952 | 0.993307424 | -5.113858597 |
| **A_51_P134062** | Stxbp2 | 0.002480321 | -0.166951012 | 0.171911655 | 11.02374969 | 0.036014999 | 0.972463971 | 0.993307424 | -5.113860274 |
| **A_52_P167278** | Mthfd1l | 0.004899412 | -0.33021325 | 0.340012074 | 9.358681644 | 0.035968495 | 0.972499509 | 0.993307424 | -5.113861539 |
| **A_66_P130911** | Proser2 | 0.00478102 | -0.32278868 | 0.332350719 | 8.997849731 | 0.035907565 | 0.972546071 | 0.993307424 | -5.113863194 |
| **A_51_P441974** | Pip5k1a | -0.003409273 | -0.238493157 | 0.231674611 | 11.50465326 | -0.03567862 | 0.972721029 | 0.993432046 | -5.113869388 |
| **A_55_P2021821** | Prmt1 | 0.00765408 | -0.5210941 | 0.53640226 | 11.74744469 | 0.035613387 | 0.97277088 | 0.993432046 | -5.113871145 |
| **A_51_P246653** | Clec7a | -0.00337114 | -0.237182518 | 0.230440238 | 8.689425039 | -0.035471561 | 0.972879263 | 0.993471282 | -5.113874956 |
| **A_55_P2130695** | Armc4 | 0.001488171 | -0.101851704 | 0.104828045 | 7.669373692 | 0.035428598 | 0.972912096 | 0.993471282 | -5.113876107 |
| **A_55_P2079659** | NA | 0.001544504 | -0.10656449 | 0.109653499 | 7.65628436 | 0.03514766 | 0.97312679 | 0.993534269 | -5.1138836 |
| **A_66_P138829** | Ndufs8 | 0.004811125 | -0.332082808 | 0.341705058 | 10.25812645 | 0.035133599 | 0.973137536 | 0.993534269 | -5.113883974 |
| **A_52_P56397** | Aunip | -0.005354952 | -0.380796866 | 0.370086963 | 8.216249644 | -0.035089887 | 0.973170941 | 0.993534269 | -5.113885134 |
| **A_51_P353502** | Sharpin | -0.002915904 | -0.207744436 | 0.201912627 | 11.83836706 | -0.035022892 | 0.97322214 | 0.993534269 | -5.113886909 |
| **A_55_P1963184** | NA | 0.003157893 | -0.219110389 | 0.225426175 | 8.076489787 | 0.034953371 | 0.973275269 | 0.993534269 | -5.113888747 |
| **A_51_P278034** | Ufsp1 | 0.002923868 | -0.202925906 | 0.208773643 | 9.346260281 | 0.034944319 | 0.973282186 | 0.993534269 | -5.113888986 |
| **A_55_P1954221** | Emp1 | -0.01054486 | -0.76179687 | 0.740707149 | 13.10918036 | -0.034532208 | 0.973597132 | 0.993754252 | -5.113899807 |
| **A_55_P2011237** | NA | 0.003411745 | -0.239727161 | 0.24655065 | 15.666155 | 0.034521623 | 0.973605221 | 0.993754252 | -5.113900084 |
| **A_51_P196895** | Ist1 | -0.003499749 | -0.254505409 | 0.247505911 | 10.14574781 | -0.034302244 | 0.973772878 | 0.993754252 | -5.11390579 |
| **A_55_P1989573** | Tmem175 | 0.00351294 | -0.248459353 | 0.255485233 | 11.14533692 | 0.034299444 | 0.973775018 | 0.993754252 | -5.113905862 |
| **A_51_P283266** | Zfp874b | 0.001667991 | -0.118255119 | 0.121591101 | 8.313835648 | 0.034218426 | 0.973836935 | 0.993754252 | -5.11390796 |
| **A_66_P108345** | Sgcb | 0.002276227 | -0.161481915 | 0.16603437 | 8.886216922 | 0.034196518 | 0.973853679 | 0.993754252 | -5.113908527 |
| **A_55_P2141699** | Rnf149 | 0.005477038 | -0.388615055 | 0.399569131 | 11.30831887 | 0.034191422 | 0.973857573 | 0.993754252 | -5.113908659 |
| **A_51_P137604** | Fcna | 0.001478358 | -0.105267106 | 0.108223823 | 7.783791578 | 0.034072147 | 0.973948729 | 0.993759581 | -5.113911736 |
| **A_51_P365964** | Tor1a | 0.004614184 | -0.329189877 | 0.338418245 | 12.10363434 | 0.034007323 | 0.97399827 | 0.993759581 | -5.113913404 |
| **A_55_P2066505** | Mpo | -0.001426162 | -0.10499891 | 0.102146586 | 7.630274238 | -0.033876035 | 0.974098607 | 0.993759581 | -5.113916772 |
| **A_52_P599055** | Atg14 | -0.004063035 | -0.299178358 | 0.291052288 | 10.47092105 | -0.033870999 | 0.974102455 | 0.993759581 | -5.113916901 |
| **A_55_P2039444** | Cmss1 | -0.00280895 | -0.20697205 | 0.20135415 | 9.150661307 | -0.033848225 | 0.97411986 | 0.993759581 | -5.113917484 |
| **A_51_P263965** | Hmox1 | 0.004248894 | -0.3082554 | 0.316753189 | 11.78293411 | 0.03344946 | 0.974424619 | 0.994018021 | -5.113927626 |

| | | | | | | | | |
|---|---|---|---|---|---|---|---|---|
| **A_55_P1997791** | Fbl | 0.002190486 | -0.159706816 | 0.164087788 | 11.09547947 | 0.033286652 | 0.974549048 | 0.994052359 | -5.113931732 |
| **A_55_P2062549** | Gm6524 | 0.002206055 | -0.160919399 | 0.165331508 | 9.844786267 | 0.033270836 | 0.974561136 | 0.994052359 | -5.11393213 |
| **A_55_P2016732** | Slc30a2 | 0.002920936 | -0.213842935 | 0.219684807 | 9.665055474 | 0.033151583 | 0.974652278 | 0.994092865 | -5.113935124 |
| **A_55_P2102679** | NA | 0.001736837 | -0.12780013 | 0.131273804 | 8.356803774 | 0.032986369 | 0.974778546 | 0.994166098 | -5.113939253 |
| **A_51_P351285** | Pcsk7 | 0.002259174 | -0.166558855 | 0.171077202 | 9.883410835 | 0.032923041 | 0.974826947 | 0.994166098 | -5.113940831 |
| **A_52_P404329** | Saa4 | 0.001465581 | -0.108492169 | 0.111423331 | 7.683827479 | 0.032790886 | 0.974927951 | 0.994188643 | -5.113944113 |
| **A_55_P2128792** | NA | -0.001478618 | -0.112520696 | 0.10956346 | 8.15885765 | -0.03275952 | 0.974951923 | 0.994188643 | -5.11394489 |
| **A_55_P2179314** | Macrod2 | -0.001429831 | -0.109541732 | 0.106682069 | 7.83545635 | -0.032537224 | 0.975121821 | 0.994309437 | -5.113950376 |
| **A_55_P2082658** | NA | 0.00163253 | -0.122130019 | 0.125395078 | 7.922723916 | 0.032451964 | 0.975186984 | 0.994313004 | -5.11395247 |
| **A_52_P381418** | Slc35b4 | -0.001676751 | -0.129003306 | 0.125649805 | 8.646989966 | -0.032398035 | 0.975228203 | 0.994313004 | -5.113953792 |
| **A_55_P2069592** | Pitpnm3 | -0.001507154 | -0.116772002 | 0.113757693 | 7.67568885 | -0.03216844 | 0.975403682 | 0.994393485 | -5.113959395 |
| **A_55_P1994668** | Tti1 | -0.003492719 | -0.27134402 | 0.264358583 | 10.02312124 | -0.032080342 | 0.975471015 | 0.994393485 | -5.113961534 |
| **A_55_P2107785** | Gm13105 | -0.008740563 | -0.682037254 | 0.664556129 | 8.836622072 | -0.031937585 | 0.975580125 | 0.994393485 | -5.113964988 |
| **A_51_P225525** | Gm9079 | 0.002533343 | -0.192808907 | 0.197875593 | 12.91751276 | 0.031905607 | 0.975604566 | 0.994393485 | -5.11396576 |
| **A_55_P2120919** | NA | -0.003321682 | -0.25975572 | 0.253112355 | 10.20540192 | -0.031867763 | 0.975633491 | 0.994393485 | -5.113966672 |
| **A_52_P405206** | Tm7sf3 | 0.003507432 | -0.267529985 | 0.274544849 | 11.01386703 | 0.031836784 | 0.975657168 | 0.994393485 | -5.113967418 |
| **A_55_P2020796** | Man1b1 | -0.002196264 | -0.172265127 | 0.167872598 | 12.51058291 | -0.031770862 | 0.975707553 | 0.994393485 | -5.113969002 |
| **A_55_P2088018** | Fhod3 | -0.006962158 | -0.546328071 | 0.532403756 | 10.75016943 | -0.031756273 | 0.975718704 | 0.994393485 | -5.113969353 |
| **A_55_P2160726** | Tfcp2 | 0.001399928 | -0.107557974 | 0.110357829 | 8.459057075 | 0.031609381 | 0.975830975 | 0.99440258 | -5.113972871 |
| **A_52_P634090** | Jag1 | -0.001990769 | -0.156981637 | 0.153000099 | 8.661454233 | -0.031599764 | 0.975838326 | 0.99440258 | -5.1139731 |
| **A_52_P498355** | Lyrm4 | -0.003197432 | -0.252583473 | 0.246188609 | 10.00186132 | -0.031542666 | 0.975881967 | 0.99440258 | -5.113974463 |
| **A_55_P1993761** | Nsa2 | 0.002862541 | -0.221380942 | 0.227106025 | 10.79027074 | 0.031405167 | 0.975987061 | 0.99443294 | -5.113977734 |
| **A_51_P120254** | Fbxl8 | -0.001713102 | -0.136553461 | 0.133127258 | 8.460471711 | -0.03125593 | 0.976101126 | 0.99443294 | -5.113981268 |
| **A_55_P2126281** | NA | 0.002015516 | -0.156751307 | 0.16078234 | 8.501147786 | 0.03123171 | 0.976119639 | 0.99443294 | -5.11398184 |
| **A_55_P2090179** | Cbx5 | -0.001799047 | -0.144052136 | 0.140454043 | 8.231522261 | -0.031113578 | 0.97620993 | 0.99443294 | -5.113984624 |
| **A_52_P607683** | Ppm1e | 0.006381676 | -0.498615976 | 0.511379329 | 8.107631006 | 0.031089559 | 0.976228288 | 0.99443294 | -5.113985189 |
| **A_52_P468068** | Tchh | -0.015347067 | -1.229890597 | 1.199196463 | 10.97767298 | -0.03108719 | 0.976230099 | 0.99443294 | -5.113985244 |
| **A_51_P144783** | Cct5 | 0.004250524 | -0.333682645 | 0.342183693 | 14.21445117 | 0.030944312 | 0.976339306 | 0.99443294 | -5.113988594 |
| **A_51_P349662** | Taf5l | -0.00247823 | -0.199821684 | 0.194865225 | 10.42947159 | -0.030894984 | 0.976377009 | 0.99443294 | -5.113989747 |
| **A_52_P265877** | Aldh9a1 | -0.004019298 | -0.325861825 | 0.317823229 | 10.92111603 | -0.030723872 | 0.976507797 | 0.99443294 | -5.113993732 |
| **A_55_P2037439** | Depdc1a | -0.001360045 | -0.11036705 | 0.10764696 | 7.64986654 | -0.03069503 | 0.976529842 | 0.99443294 | -5.113994402 |
| **A_55_P1957204** | NA | 0.00158289 | -0.126069318 | 0.129235098 | 16.93039354 | 0.030506442 | 0.976673988 | 0.99443294 | -5.113998764 |
| **A_55_P2065529** | Mok | 0.005711605 | -0.455941862 | 0.467365072 | 9.002058139 | 0.030437657 | 0.976726564 | 0.99443294 | -5.114000348 |
| **A_65_P04472** | Atp6v1c1 | -0.002357647 | -0.193287766 | 0.188572473 | 10.21393948 | -0.030378996 | 0.976771402 | 0.99443294 | -5.114001697 |
| **A_55_P2198364** | Tsen54 | -0.003115442 | -0.255770215 | 0.249539331 | 8.572311287 | -0.03033621 | 0.976804106 | 0.99443294 | -5.114002678 |
| **A_55_P1989865** | Thrap3 | 0.002354004 | -0.188718208 | 0.193426216 | 8.79558265 | 0.030309505 | 0.976824517 | 0.99443294 | -5.114003291 |
| **A_55_P1953400** | Rbfox1 | -0.001329215 | -0.109251108 | 0.106592677 | 7.778392574 | -0.030300856 | 0.976831128 | 0.99443294 | -5.114003489 |
| **A_55_P2047310** | Pdgfb | -0.006883548 | -0.565782952 | 0.552015857 | 13.51500439 | -0.030300363 | 0.976831505 | 0.99443294 | -5.1140035 |
| **A_55_P2227335** | Ptpre | 0.001759326 | -0.141125462 | 0.144644113 | 9.031021408 | 0.030292101 | 0.97683782 | 0.99443294 | -5.114003689 |
| **A_55_P2048284** | Aasdhppt | -0.001885127 | -0.156601448 | 0.152831193 | 8.382436702 | -0.029976002 | 0.977079433 | 0.994500785 | -5.11401089 |
| **A_55_P2065897** | NA | 0.002747337 | -0.223283117 | 0.228777791 | 11.12884693 | 0.029902954 | 0.977135268 | 0.994500785 | -5.114012543 |
| **A_55_P2159259** | Lifr | -0.002923523 | -0.24368632 | 0.237839274 | 13.6918699 | -0.029873512 | 0.977157772 | 0.994500785 | -5.114013208 |
| **A_55_P2096762** | Arhgdig | 0.002540435 | -0.206902151 | 0.211983021 | 7.972431946 | 0.029840926 | 0.97718268 | 0.994500785 | -5.114013943 |
| **A_55_P2142033** | Clip4 | -0.001283122 | -0.107208619 | 0.104642375 | 7.676836841 | -0.029801392 | 0.977212899 | 0.994500785 | -5.114014835 |
| **A_51_P465292** | Hnmt | -0.00131736 | -0.110201275 | 0.107566555 | 7.692341483 | -0.029765277 | 0.977240504 | 0.994500785 | -5.114015648 |
| **A_55_P2006876** | NA | -0.002989439 | -0.250338147 | 0.244359268 | 13.12350543 | -0.029733721 | 0.977264624 | 0.994500785 | -5.114016357 |
| **A_51_P236324** | Gtf2e1 | -0.003464762 | -0.290970583 | 0.284041058 | 10.02575188 | -0.029648039 | 0.977330117 | 0.994515074 | -5.11401828 |
| **A_66_P132070** | Abra | -0.001249267 | -0.105698393 | 0.103199859 | 7.630362685 | -0.029425209 | 0.977500442 | 0.994565276 | -5.114023255 |
| **A_65_P11193** | Flad1 | 0.003726106 | -0.308447328 | 0.31589954 | 9.972785446 | 0.029364893 | 0.977546547 | 0.994565276 | -5.114024596 |
| **A_55_P2003158** | Wwp1 | 0.001947488 | -0.162980554 | 0.166875531 | 8.216112018 | 0.029050226 | 0.977787073 | 0.994565276 | -5.114031543 |

| | | | | | | | | |
|---|---|---|---|---|---|---|---|---|
| **A_55_P2076048** | Nuf2 | 0.001280152 | -0.107258043 | 0.109818348 | 7.759921131 | 0.029016712 | 0.97781269 | 0.994565276 | -5.114032279 |
| **A_55_P2085940** | NA | -0.00255593 | -0.219450444 | 0.214338584 | 11.92555644 | -0.028991421 | 0.977832023 | 0.994565276 | -5.114032833 |
| **A_55_P2136776** | Sult1b1 | -0.001551924 | -0.133417305 | 0.130313456 | 7.650053026 | -0.028954025 | 0.977860607 | 0.994565276 | -5.114033652 |
| **A_55_P2232988** | Scube2 | 0.002954951 | -0.248134605 | 0.254044506 | 9.452285401 | 0.028952813 | 0.977861534 | 0.994565276 | -5.114033679 |
| **A_66_P131979** | Cdc6 | -0.001500821 | -0.129067962 | 0.126066321 | 7.794092236 | -0.02894404 | 0.97786824 | 0.994565276 | -5.114033871 |
| **A_52_P664404** | Zfp286 | -0.001733556 | -0.149156973 | 0.145689861 | 7.989551869 | -0.028929474 | 0.977879374 | 0.994565276 | -5.114034189 |
| **A_52_P376337** | Sdhaf3 | 0.001600463 | -0.134652198 | 0.137853124 | 8.26204132 | 0.028898133 | 0.977903331 | 0.994565276 | -5.114034874 |
| **A_51_P356055** | Grp | -0.001290426 | -0.111590602 | 0.109009751 | 7.635651839 | -0.028782327 | 0.977991852 | 0.994565276 | -5.114037399 |
| **A_55_P2179498** | Ppp1r8 | -0.001418659 | -0.122707556 | 0.119870238 | 9.015783307 | -0.028775716 | 0.977996906 | 0.994565276 | -5.114037543 |
| **A_55_P2087240** | Mipep | -0.001572897 | -0.136727584 | 0.133581789 | 8.404122334 | -0.028631123 | 0.978107432 | 0.99456894 | -5.11404068 |
| **A_51_P341108** | Spint1 | -0.002511609 | -0.219326666 | 0.214303449 | 11.45113449 | -0.028499136 | 0.978208323 | 0.99456894 | -5.11404353 |
| **A_55_P2151398** | Tmem116 | -0.002774665 | -0.243390226 | 0.237840896 | 8.920556964 | -0.02836978 | 0.978307203 | 0.99456894 | -5.11404631 |
| **A_52_P270035** | Tspyl2 | -0.002274885 | -0.20003371 | 0.19548394 | 8.610377304 | -0.028300412 | 0.978360228 | 0.99456894 | -5.114047796 |
| **A_55_P2111985** | Prss35 | -0.006672465 | -0.587206956 | 0.573862025 | 8.511791559 | -0.028276619 | 0.978378416 | 0.99456894 | -5.114048305 |
| **A_55_P2139878** | NA | -0.001194129 | -0.10512588 | 0.102737621 | 7.678990913 | -0.028266518 | 0.978386137 | 0.99456894 | -5.114048521 |
| **A_55_P2368625** | Sp4 | 0.001727633 | -0.149283031 | 0.152738298 | 8.241113382 | 0.028145766 | 0.978478441 | 0.99456894 | -5.114051096 |
| **A_51_P461877** | Thbs2 | -0.009155766 | -0.80980438 | 0.791492849 | 9.318857511 | -0.028133377 | 0.978487911 | 0.99456894 | -5.114051359 |
| **A_55_P1976242** | Arhgef7 | -0.003219865 | -0.284830837 | 0.278391106 | 12.28122703 | -0.028129194 | 0.978491108 | 0.99456894 | -5.114051448 |
| **A_55_P2170681** | Cdca8 | 0.001336077 | -0.115648159 | 0.118320313 | 7.714959056 | 0.028097866 | 0.978515056 | 0.99456894 | -5.114052114 |
| **A_55_P1980826** | Kcnrg | 0.001222007 | -0.106975948 | 0.109419961 | 7.695446341 | 0.027785854 | 0.978753563 | 0.994667886 | -5.114058704 |
| **A_55_P2020612** | Arl4d | 0.013713138 | -1.202447337 | 1.229873614 | 10.24318492 | 0.027740554 | 0.978788191 | 0.994667886 | -5.114059655 |
| **A_55_P1994194** | Heph | 0.007585182 | -0.668595632 | 0.683765996 | 9.816140747 | 0.027597664 | 0.978897419 | 0.994667886 | -5.114062644 |
| **A_55_P2000793** | Rbm34 | -0.002629784 | -0.237104837 | 0.231845268 | 9.736513286 | -0.027592604 | 0.978901288 | 0.994667886 | -5.114062749 |
| **A_66_P115757** | Mthfd2l | 0.001504735 | -0.133273682 | 0.136283151 | 8.254088831 | 0.027466844 | 0.978997421 | 0.994667886 | -5.114065366 |
| **A_51_P275083** | Mrps18a | 0.001992223 | -0.176460365 | 0.180444812 | 11.34675077 | 0.0274653 | 0.978998601 | 0.994667886 | -5.114065398 |
| **A_52_P655687** | Egfl6 | -0.001162366 | -0.105552601 | 0.103227869 | 7.647413587 | -0.027393803 | 0.979053256 | 0.994667886 | -5.114066881 |
| **A_55_P1996698** | Gm16381 | 0.002807526 | -0.250983022 | 0.256598074 | 12.47565558 | 0.027215572 | 0.979189501 | 0.994667886 | -5.11407056 |
| **A_55_P2018666** | Thrsp | 0.004972017 | -0.445276836 | 0.455220871 | 8.678960272 | 0.027167469 | 0.979226273 | 0.994667886 | -5.114071548 |
| **A_55_P2362601** | D4Ertd681e | 0.001742881 | -0.156264843 | 0.159750606 | 7.919392416 | 0.027136799 | 0.979249718 | 0.994667886 | -5.114072178 |
| **A_55_P2404554** | Atcayos | 0.001152184 | -0.103433149 | 0.105737517 | 7.641900688 | 0.027103181 | 0.979275417 | 0.994667886 | -5.114072867 |
| **A_52_P603976** | Jtb | -0.001552003 | -0.142433817 | 0.139329811 | 11.72802923 | -0.027102342 | 0.979276058 | 0.994667886 | -5.114072884 |
| **A_55_P2056542** | Nin | 0.002220156 | -0.19936452 | 0.203804832 | 9.438922225 | 0.027095375 | 0.979281384 | 0.994667886 | -5.114073027 |
| **A_55_P1960683** | F2 | -0.015287008 | -1.414552875 | 1.38397886 | 11.73056672 | -0.026877657 | 0.979447816 | 0.994668632 | -5.114077469 |
| **A_55_P2075140** | Sirt3 | 0.003453789 | -0.312846235 | 0.319753813 | 10.55394006 | 0.026863695 | 0.979458489 | 0.994668632 | -5.114077752 |
| **A_66_P102202** | Gm648 | 0.005005047 | -0.456287196 | 0.466297289 | 12.48530217 | 0.026693232 | 0.979588799 | 0.994668632 | -5.114081203 |
| **A_66_P124052** | Bok | -0.00812458 | -0.757064057 | 0.740814896 | 13.75623623 | -0.026688451 | 0.979592453 | 0.994668632 | -5.114081299 |
| **A_51_P152404** | Ikbip | -0.005047107 | -0.470913954 | 0.460819739 | 9.82417585 | -0.026653234 | 0.979619375 | 0.994668632 | -5.114082009 |
| **A_55_P2283207** | Maz | 0.001217221 | -0.111250072 | 0.113684514 | 7.865809812 | 0.02662639 | 0.979639896 | 0.994668632 | -5.11408255 |
| **A_55_P1973813** | NA | -0.00118003 | -0.110224117 | 0.107864057 | 7.702082407 | -0.026623196 | 0.979642337 | 0.994668632 | -5.114082614 |
| **A_55_P2056615** | NA | -0.002402844 | -0.225337551 | 0.220531864 | 11.80789008 | -0.026516548 | 0.979723865 | 0.994699159 | -5.114084756 |
| **A_55_P2170034** | Rgl1 | -0.004140638 | -0.393138639 | 0.384857363 | 10.63539923 | -0.026187201 | 0.979975635 | 0.994895473 | -5.114091317 |
| **A_66_P111021** | Nlrp6 | -0.00116062 | -0.110661032 | 0.108339791 | 7.731667657 | -0.026076157 | 0.980060524 | 0.994895473 | -5.114093511 |
| **A_55_P2016486** | Tmem131 | 0.001259314 | -0.118084701 | 0.120603328 | 10.82804525 | 0.02595987 | 0.980149421 | 0.994895473 | -5.114095798 |
| **A_55_P2094602** | Lrch1 | -0.001961529 | -0.188407389 | 0.184484332 | 9.9237133 | -0.025882791 | 0.980208345 | 0.994895473 | -5.114097308 |
| **A_55_P1980416** | Zkscan1 | 0.003674435 | -0.345893214 | 0.353242084 | 10.00804272 | 0.025859997 | 0.98022577 | 0.994895473 | -5.114097754 |
| **A_55_P1976664** | Gm16430 | 0.00239874 | -0.225808824 | 0.230606304 | 8.697269567 | 0.025859626 | 0.980226053 | 0.994895473 | -5.114097761 |
| **A_55_P1966035** | Apoa2 | 0.012926331 | -1.221631151 | 1.247483813 | 14.16635493 | 0.025759244 | 0.980302792 | 0.994921117 | -5.11409972 |
| **A_55_P1988623** | Add3 | 0.003169339 | -0.301776142 | 0.30811482 | 10.19799189 | 0.025569113 | 0.980448142 | 0.994970375 | -5.114103408 |
| **A_55_P2114034** | NA | -0.001269245 | -0.124332029 | 0.12179354 | 8.519033259 | -0.02537393 | 0.980597354 | 0.994970375 | -5.114107167 |
| **A_55_P2082688** | Ak1 | -0.009510109 | -0.932587834 | 0.913567616 | 10.31224204 | -0.025346406 | 0.980618396 | 0.994970375 | -5.114107694 |

| | | | | | | | | |
|---|---|---|---|---|---|---|---|---|
| **A_55_P2125982** | Lrrc4b | -0.001081065 | -0.10601618 | 0.103854051 | 7.688687766 | -0.025345451 | 0.980619126 | 0.994970375 | -5.114107713 |
| **A_52_P500002** | Tspyl5 | -0.001089293 | -0.107132521 | 0.104953936 | 7.71333228 | -0.025271492 | 0.980675666 | 0.994970375 | -5.114109128 |
| **A_55_P2036978** | Mrps14 | -0.002502731 | -0.246198047 | 0.241192585 | 12.299665 | -0.025265982 | 0.980679878 | 0.994970375 | -5.114109233 |
| **A_55_P2121456** | Fam43a | -0.002440116 | -0.24043001 | 0.235549779 | 10.20938833 | -0.025224414 | 0.980711656 | 0.994970375 | -5.114110026 |
| **A_52_P179583** | Adnp2 | -0.002606862 | -0.259095964 | 0.25388224 | 8.867268192 | -0.025004506 | 0.980879771 | 0.995039827 | -5.114114201 |
| **A_52_P162184** | Snx12 | 0.002090769 | -0.203754269 | 0.207935807 | 8.910987347 | 0.02498819 | 0.980892245 | 0.995039827 | -5.11411451 |
| **A_55_P2068849** | Nudc | 0.002105512 | -0.205651143 | 0.209862167 | 10.13277862 | 0.024932852 | 0.98093455 | 0.995039827 | -5.114115554 |
| **A_55_P2172624** | Gm1123 | -0.001241932 | -0.124261255 | 0.121777391 | 8.119782738 | -0.024836683 | 0.98100807 | 0.995062184 | -5.114117363 |
| **A_51_P406165** | Nxpe2 | 0.001238022 | -0.122470974 | 0.124947018 | 7.90424494 | 0.024620471 | 0.981173362 | 0.995171808 | -5.114121405 |
| **A_55_P2028214** | NA | -0.001049084 | -0.106133972 | 0.104035803 | 7.895612708 | -0.02456062 | 0.981219117 | 0.995171808 | -5.114122517 |
| **A_55_P1981836** | Ccm2l | -0.001765948 | -0.179276228 | 0.175744333 | 9.456283549 | -0.024475043 | 0.981284541 | 0.995185943 | -5.114124104 |
| **A_55_P2059085** | Zfp457 | -0.001697608 | -0.173701774 | 0.170306558 | 9.338723835 | -0.024281056 | 0.981432844 | 0.995199314 | -5.114127679 |
| **A_52_P656714** | Map3k5 | 0.001924783 | -0.193691457 | 0.197541023 | 8.198174235 | 0.02420728 | 0.981489246 | 0.995199314 | -5.114129031 |
| **A_55_P2072631** | Prr5l | -0.002205135 | -0.226896111 | 0.222485841 | 10.47399932 | -0.024144523 | 0.981537224 | 0.995199314 | -5.114130178 |
| **A_51_P103054** | Adamts19 | 0.001022819 | -0.103199297 | 0.105244936 | 7.634934289 | 0.024143946 | 0.981537665 | 0.995199314 | -5.114130188 |
| **A_55_P2133057** | Pdia2 | -0.001063309 | -0.109619826 | 0.107493208 | 7.745061203 | -0.02409754 | 0.981573142 | 0.995199314 | -5.114131034 |
| **A_51_P155501** | Alg3 | -0.001124992 | -0.116416614 | 0.114166631 | 8.29849999 | -0.024006051 | 0.981643087 | 0.995199314 | -5.114132698 |
| **A_52_P670026** | Rsad2 | -0.002013247 | -0.208504535 | 0.204478041 | 11.51354462 | -0.023986364 | 0.981658137 | 0.995199314 | -5.114133055 |
| **A_52_P417437** | Gab1 | 0.001485402 | -0.151506072 | 0.154476877 | 9.241405753 | 0.023886144 | 0.981734756 | 0.995224791 | -5.114134868 |
| **A_51_P332652** | Pqlc3 | -0.011634311 | -1.226298635 | 1.203030013 | 10.03131555 | -0.023564247 | 0.981980851 | 0.995340211 | -5.114140642 |
| **A_52_P32369** | Hmgn3 | -0.001477396 | -0.155835913 | 0.15288112 | 8.438712419 | -0.023547 | 0.981994036 | 0.995340211 | -5.114140949 |
| **A_66_P112394** | NA | 0.004873979 | -0.506485584 | 0.516233543 | 12.66849198 | 0.023449115 | 0.982068871 | 0.995340211 | -5.114142688 |
| **A_51_P502422** | Acot13 | -0.001448342 | -0.153634878 | 0.150738195 | 12.54718564 | -0.023413372 | 0.982096197 | 0.995340211 | -5.114143321 |
| **A_55_P2056348** | NA | 0.002333017 | -0.243060103 | 0.247726136 | 12.13895289 | 0.023389697 | 0.982114297 | 0.995340211 | -5.11414374 |
| **A_52_P377416** | Disp1 | -0.001526612 | -0.162489497 | 0.159436273 | 8.877077018 | -0.023333083 | 0.982157579 | 0.995340211 | -5.11414474 |
| **A_55_P2148268** | Xxylt1 | -0.001650796 | -0.1766231 | 0.173321509 | 8.988749163 | -0.023210965 | 0.982250942 | 0.995382639 | -5.114146888 |
| **A_51_P135357** | Sike1 | -0.00125337 | -0.134764313 | 0.132257573 | 8.435303692 | -0.02309573 | 0.982339042 | 0.99541973 | -5.114148905 |
| **A_52_P252425** | 2700062C07Rik | 0.001806142 | -0.192761131 | 0.196373415 | 9.215969274 | 0.022837647 | 0.982536354 | 0.995420407 | -5.114153386 |
| **A_65_P02327** | Pogk | 0.001747318 | -0.187529349 | 0.191023985 | 8.634961696 | 0.022711407 | 0.982632868 | 0.995420407 | -5.114155559 |
| **A_55_P1981704** | Ccdc113 | -0.001629533 | -0.178186872 | 0.174927805 | 7.791075154 | -0.022706314 | 0.982636762 | 0.995420407 | -5.114155647 |
| **A_51_P357685** | Dhx16 | 0.002904465 | -0.312508202 | 0.318317132 | 10.99548012 | 0.022654589 | 0.982676307 | 0.995420407 | -5.114156534 |
| **A_55_P1975080** | Ugt2b5 | 0.000992987 | -0.106888073 | 0.108874048 | 7.654395133 | 0.02264476 | 0.982683822 | 0.995420407 | -5.114156702 |
| **A_55_P1987459** | Pigt | 0.001429486 | -0.154227215 | 0.157086187 | 8.571970367 | 0.022593383 | 0.982723102 | 0.995420407 | -5.11415758 |
| **A_55_P2368710** | NA | 0.00320318 | -0.345954769 | 0.352361129 | 8.587279269 | 0.022569844 | 0.982741098 | 0.995420407 | -5.114157982 |
| **A_55_P2128853** | Ear12 | 0.015521451 | -1.677411982 | 1.708454885 | 9.22414815 | 0.022555975 | 0.982751701 | 0.995420407 | -5.114158219 |
| **A_55_P1996623** | Dnajb12 | 0.001984592 | -0.215668899 | 0.219638083 | 10.00025742 | 0.022432349 | 0.982846218 | 0.995463977 | -5.114160321 |
| **A_52_P35304** | Chkb | 0.001624223 | -0.177427766 | 0.180676213 | 9.981147754 | 0.022316996 | 0.98293441 | 0.995501137 | -5.114162272 |
| **A_55_P2144556** | Flrt3 | -0.003658095 | -0.410133752 | 0.402817562 | 9.642243306 | -0.022140612 | 0.983069263 | 0.995585549 | -5.114165236 |
| **A_52_P466147** | Rarres2 | -0.004094606 | -0.464462596 | 0.456273384 | 16.49066469 | -0.021881453 | 0.983267402 | 0.995628014 | -5.114169549 |
| **A_55_P2086682** | Nfib | 0.002585098 | -0.289526635 | 0.294696831 | 10.07369874 | 0.021771944 | 0.983351128 | 0.995628014 | -5.114171356 |
| **A_51_P455338** | Rnase2a | 0.018686598 | -2.096462749 | 2.133835945 | 9.774400078 | 0.021734936 | 0.983379422 | 0.995628014 | -5.114171964 |
| **A_55_P2086954** | NA | -0.007146799 | -0.818030681 | 0.803737082 | 11.35530269 | -0.021683136 | 0.983419027 | 0.995628014 | -5.114172814 |
| **A_55_P2000543** | Cd209f | -0.000955864 | -0.109488147 | 0.107576418 | 7.659808419 | -0.021667387 | 0.983431067 | 0.995628014 | -5.114173073 |
| **A_55_P2093232** | Myh7 | -0.002378264 | -0.275178068 | 0.270421539 | 7.999242096 | -0.021447923 | 0.98359886 | 0.995628014 | -5.114176649 |
| **A_51_P140368** | Cmc2 | 0.001363006 | -0.156330333 | 0.159056346 | 10.38511903 | 0.021264428 | 0.983739153 | 0.995628014 | -5.114179612 |
| **A_51_P491635** | Ly6g6e | 0.001944695 | -0.223190292 | 0.227079681 | 8.136718203 | 0.021250912 | 0.983749487 | 0.995628014 | -5.114179829 |
| **A_52_P364130** | Map3k14 | 0.00121459 | -0.139620464 | 0.142049644 | 8.120479731 | 0.021217197 | 0.983775264 | 0.995628014 | -5.11418037 |
| **A_52_P81783** | Gspt1 | 0.001614955 | -0.185709054 | 0.188938963 | 8.85775534 | 0.021209774 | 0.98378094 | 0.995628014 | -5.114180489 |
| **A_55_P2175301** | NA | 0.001285969 | -0.148583756 | 0.151155693 | 14.71895632 | 0.021109872 | 0.983857321 | 0.995628014 | -5.114182087 |
| **A_55_P2136522** | Tmem128 | 0.001566417 | -0.180992999 | 0.184125832 | 12.36283822 | 0.02110922 | 0.98385782 | 0.995628014 | -5.114182098 |

| | | | | | | | | | |
|---|---|---|---|---|---|---|---|---|---|
| **A_65_P04980** | Ilf3 | 0.002289036 | -0.264799044 | 0.269377117 | 9.485112173 | 0.021084697 | 0.983876569 | 0.995628014 | -5.114182489 |
| **A_52_P260994** | Fgd2 | 0.000947639 | -0.110003425 | 0.111898702 | 7.739176194 | 0.02101263 | 0.983931669 | 0.995628014 | -5.114183636 |
| **A_55_P1993879** | NA | -0.001220944 | -0.144500186 | 0.142058299 | 15.9697061 | -0.020964353 | 0.98396858 | 0.995628014 | -5.114184402 |
| **A_55_P1999818** | NA | -0.002160473 | -0.257690205 | 0.25336926 | 14.08686855 | -0.02080063 | 0.984093758 | 0.995628014 | -5.114186986 |
| **A_55_P2118857** | NA | -0.001088509 | -0.129919958 | 0.12774294 | 8.025708067 | -0.020786403 | 0.984104635 | 0.995628014 | -5.11418721 |
| **A_55_P2148248** | NA | -0.002450227 | -0.292482722 | 0.287582269 | 9.19885524 | -0.020783981 | 0.984106487 | 0.995628014 | -5.114187248 |
| **A_55_P2293381** | Cldn18 | 0.000978759 | -0.114959697 | 0.116917215 | 7.665810283 | 0.02076909 | 0.984117872 | 0.995628014 | -5.114187482 |
| **A_51_P451052** | Pgm5 | 0.004705792 | -0.555791996 | 0.56520358 | 8.398877951 | 0.020655133 | 0.984205001 | 0.995628014 | -5.114189266 |
| **A_55_P1958082** | Prpsap2 | 0.001689055 | -0.200201802 | 0.203579912 | 9.582687729 | 0.020582418 | 0.984260596 | 0.995628014 | -5.1141904 |
| **A_51_P421734** | Slc12a7 | 0.002037335 | -0.241964839 | 0.24603951 | 10.42088315 | 0.020541774 | 0.984291672 | 0.995628014 | -5.114191032 |
| **A_55_P1971849** | Sh3glb2 | -0.001772761 | -0.214145426 | 0.210599905 | 12.90118001 | -0.020536225 | 0.984295914 | 0.995628014 | -5.114191118 |
| **A_51_P274947** | Atad1 | -0.002332711 | -0.284028583 | 0.279363162 | 11.74914964 | -0.020372745 | 0.984420908 | 0.995648263 | -5.114193646 |
| **A_51_P396774** | Sugt1 | -0.001842933 | -0.224989148 | 0.221303283 | 13.98515655 | -0.02031838 | 0.984462474 | 0.995648263 | -5.114194482 |
| **A_55_P2128224** | NA | 0.00144372 | -0.174135052 | 0.177022492 | 15.77343596 | 0.020229265 | 0.984530609 | 0.995648263 | -5.114195848 |
| **A_55_P2373852** | 2310058N22Rik | 0.003108162 | -0.37599224 | 0.382208563 | 9.415484421 | 0.020170586 | 0.984575474 | 0.995648263 | -5.114196744 |
| **A_55_P2040860** | Sun3 | 0.001091706 | -0.132095167 | 0.134278579 | 7.812069682 | 0.020165705 | 0.984579206 | 0.995648263 | -5.114196818 |
| **A_51_P299805** | Slc46a3 | 0.002714454 | -0.333472242 | 0.338901151 | 10.38386513 | 0.019864205 | 0.984809729 | 0.995648263 | -5.11420138 |
| **A_51_P221248** | Tspan32 | 0.000860755 | -0.105817047 | 0.107538557 | 7.696426417 | 0.019850649 | 0.984820094 | 0.995648263 | -5.114201584 |
| **A_55_P2040115** | Cdv3 | 0.001867454 | -0.230032147 | 0.233767056 | 13.82287853 | 0.019811581 | 0.984849965 | 0.995648263 | -5.11420217 |
| **A_51_P382928** | Cstf3 | -0.001565513 | -0.19623097 | 0.193099944 | 9.913742242 | -0.019785042 | 0.984870256 | 0.995648263 | -5.114202567 |
| **A_52_P467449** | Alox12 | 0.001283891 | -0.159083552 | 0.161651335 | 8.069966348 | 0.019696143 | 0.984938228 | 0.995648263 | -5.114203893 |
| **A_51_P441782** | Micu2 | 0.002065956 | -0.255992734 | 0.260124645 | 10.6797591 | 0.019695711 | 0.984938558 | 0.995648263 | -5.1142039 |
| **A_55_P2035812** | Rnf208 | -0.001126394 | -0.142217185 | 0.139964398 | 10.18505161 | -0.019640871 | 0.984980488 | 0.995648263 | -5.114204715 |
| **A_55_P2035657** | Psmb6 | -0.00348402 | -0.441038105 | 0.434070065 | 14.611727 | -0.019589259 | 0.98501995 | 0.995648263 | -5.114205481 |
| **A_55_P1955472** | Bbs1 | 0.001221422 | -0.152351282 | 0.154794127 | 8.079707247 | 0.019566854 | 0.98503708 | 0.995648263 | -5.114205812 |
| **A_51_P180974** | Cavin3 | 0.004966012 | -0.622500894 | 0.632432919 | 10.20994737 | 0.019470905 | 0.985110443 | 0.995653992 | -5.114207228 |
| **A_51_P194377** | Pcbp4 | -0.003299379 | -0.421174955 | 0.414576198 | 9.653047322 | -0.0194247 | 0.985145771 | 0.995653992 | -5.114207908 |
| **A_51_P474468** | Utp11 | -0.001712693 | -0.219738997 | 0.216313611 | 12.4220083 | -0.019325906 | 0.985221309 | 0.995678274 | -5.114209355 |
| **A_55_P2068172** | Nufip2 | 0.002388184 | -0.304206774 | 0.308983143 | 9.248637305 | 0.019163373 | 0.985345581 | 0.995751803 | -5.114211719 |
| **A_55_P2353079** | NA | 0.000800796 | -0.102923681 | 0.104525272 | 7.632458937 | 0.018993699 | 0.985475314 | 0.995830842 | -5.114214167 |
| **A_55_P2105888** | G6pc3 | -0.001357852 | -0.180157426 | 0.177441723 | 10.82190275 | -0.018683364 | 0.985712598 | 0.996018548 | -5.114218586 |
| **A_55_P2145711** | Tox3 | 0.001051704 | -0.137936509 | 0.140039917 | 7.974988657 | 0.018615927 | 0.985764161 | 0.996018581 | -5.114219537 |
| **A_55_P1986711** | NA | 0.00156583 | -0.206173868 | 0.209305527 | 8.82873774 | 0.018543607 | 0.985819457 | 0.996022387 | -5.114220553 |
| **A_55_P2025902** | NA | -0.001972857 | -0.265029795 | 0.261084081 | 9.405487502 | -0.018450793 | 0.985890424 | 0.996042024 | -5.114221851 |
| **A_51_P502614** | Dusp6 | -0.002122873 | -0.28993885 | 0.285693105 | 11.1092535 | -0.01814589 | 0.986123558 | 0.996225487 | -5.114226069 |
| **A_55_P1974830** | Pde1c | -0.000769616 | -0.105679156 | 0.104139924 | 7.698565902 | -0.018047967 | 0.986198431 | 0.996249058 | -5.114227409 |
| **A_55_P2101191** | Arrdc4 | 0.001838174 | -0.253591477 | 0.257267824 | 10.48777347 | 0.01770453 | 0.986461031 | 0.996462255 | -5.11423205 |
| **A_55_P1977653** | Palld | -0.002163931 | -0.30860247 | 0.304274608 | 11.30065621 | -0.017372772 | 0.986714702 | 0.996666412 | -5.114236449 |
| **A_55_P2424042** | Ttn | -0.000744821 | -0.106803621 | 0.105313978 | 7.653654627 | -0.017277247 | 0.986787743 | 0.99666819 | -5.1142377 |
| **A_55_P2126850** | Nudt11 | -0.001878882 | -0.270666104 | 0.266908339 | 8.109244849 | -0.017197293 | 0.986848878 | 0.99666819 | -5.114238742 |
| **A_66_P101764** | NA | 0.004242637 | -0.60372601 | 0.612211283 | 12.82692358 | 0.017168163 | 0.986871152 | 0.99666819 | -5.114239121 |
| **A_55_P2013391** | Gm4402 | -0.002338173 | -0.339596624 | 0.334920279 | 8.857100399 | -0.017056227 | 0.986956742 | 0.996702307 | -5.114240569 |
| **A_55_P2126333** | Cyp2a12 | -0.000876352 | -0.127780848 | 0.126028144 | 7.69765598 | -0.016989109 | 0.987008063 | 0.996702307 | -5.114241432 |
| **A_55_P2085779** | Ifi27l2b | 0.005174493 | -0.747513984 | 0.757862971 | 9.655716262 | 0.016913042 | 0.987066226 | 0.996703812 | -5.114242407 |
| **A_51_P220343** | Ccn4 | -0.004299429 | -0.633955003 | 0.625356146 | 9.447598432 | -0.016798746 | 0.987153621 | 0.996703812 | -5.114243864 |
| **A_66_P136714** | Scpep1 | -0.004144145 | -0.611560803 | 0.603272513 | 14.43727361 | -0.016784847 | 0.987164249 | 0.996703812 | -5.11424404 |
| **A_52_P27122** | Ago4 | -0.001286844 | -0.193388473 | 0.190814786 | 8.536766271 | -0.016480254 | 0.987397154 | 0.996841204 | -5.11424787 |
| **A_55_P2006634** | Map4k5 | 0.0013436 | -0.199331279 | 0.202018479 | 12.04981656 | 0.016471992 | 0.987403471 | 0.996841204 | -5.114247973 |
| **A_52_P8227** | Lypla1 | -0.002395286 | -0.373539073 | 0.368748501 | 9.69356777 | -0.015877581 | 0.987857985 | 0.99724428 | -5.114255242 |
| **A_55_P1954598** | NA | 0.002052407 | -0.317831042 | 0.321935856 | 11.33908341 | 0.015784864 | 0.987928882 | 0.99724428 | -5.114256352 |

| | | | | | | | | | |
|---|---|---|---|---|---|---|---|---|---|
| **A_51_P212630** | Chd8 | 0.002495956 | -0.387594182 | 0.392586093 | 12.66132199 | 0.015741316 | 0.987962181 | 0.99724428 | -5.114256871 |
| **A_51_P345663** | Rrbp1 | 0.001895631 | -0.296908764 | 0.300700026 | 13.13832252 | 0.015607596 | 0.988064431 | 0.99724428 | -5.114258455 |
| **A_52_P425860** | Iqsec1 | 0.001533162 | -0.240627076 | 0.243693401 | 11.58807098 | 0.015575947 | 0.988088632 | 0.99724428 | -5.114258828 |
| **A_66_P122415** | Snhg6 | -0.001759005 | -0.281354504 | 0.277836495 | 13.47293819 | -0.015477684 | 0.988163769 | 0.99724428 | -5.114259982 |
| **A_55_P2008006** | Ap1g1 | 0.001009766 | -0.159495028 | 0.16151456 | 10.23508924 | 0.015477535 | 0.988163883 | 0.99724428 | -5.114259984 |
| **A_55_P1985001** | NA | -0.002725417 | -0.439788879 | 0.434338045 | 8.858207544 | -0.015341139 | 0.988268179 | 0.997297464 | -5.114261573 |
| **A_66_P123014** | Twistnb | 0.001404797 | -0.226459204 | 0.229268798 | 9.788926419 | 0.015167257 | 0.98840114 | 0.997354223 | -5.114263578 |
| **A_52_P195798** | Dmac2l | 0.001291805 | -0.209301561 | 0.21188517 | 8.934372726 | 0.015091114 | 0.988459363 | 0.997354223 | -5.114264449 |
| **A_55_P1964568** | Megf10 | -0.000659583 | -0.108371924 | 0.107052757 | 7.678332985 | -0.015065146 | 0.98847922 | 0.997354223 | -5.114264745 |
| **A_51_P344376** | Gpr162 | 0.000630727 | -0.10674407 | 0.108005523 | 7.851761216 | 0.014451343 | 0.988948575 | 0.997637435 | -5.114271593 |
| **A_52_P2706** | Mypn | 0.000637428 | -0.108042717 | 0.109317573 | 7.68799165 | 0.014429463 | 0.988965306 | 0.997637435 | -5.114271832 |
| **A_55_P2108655** | Mtch2 | -0.001038521 | -0.178179477 | 0.176102435 | 12.90390864 | -0.014423322 | 0.988970002 | 0.997637435 | -5.114271899 |
| **A_52_P508700** | Zfp951 | -0.001095907 | -0.188043104 | 0.18585129 | 8.774088345 | -0.014421948 | 0.988971053 | 0.997637435 | -5.114271914 |
| **A_51_P149699** | P3h3 | 0.00164435 | -0.280058668 | 0.283347369 | 8.47206334 | 0.01436058 | 0.989017979 | 0.997637435 | -5.114272582 |
| **A_51_P255657** | Misp3 | 0.003911955 | -0.67286768 | 0.68069159 | 11.68364869 | 0.014220529 | 0.989125072 | 0.99768888 | -5.114274095 |
| **A_55_P2047559** | Ccp110 | 0.001237181 | -0.215216495 | 0.217690857 | 8.679896588 | 0.014061688 | 0.989246534 | 0.99768888 | -5.114275793 |
| **A_55_P2109983** | Mpg | -0.0007383 | -0.130330265 | 0.128853665 | 8.332021636 | -0.01401599 | 0.989281478 | 0.99768888 | -5.114276278 |
| **A_52_P285024** | Sertad4 | 0.000707331 | -0.123710047 | 0.125124709 | 7.759394894 | 0.01398656 | 0.989303983 | 0.99768888 | -5.11427659 |
| **A_55_P2031011** | S100pbp | -0.002110117 | -0.374885694 | 0.370665461 | 9.323503984 | -0.013926059 | 0.989350246 | 0.99768888 | -5.114277228 |
| **A_55_P1995173** | Odc1 | 0.002175399 | -0.383161781 | 0.387512579 | 13.50193799 | 0.013888879 | 0.989378677 | 0.99768888 | -5.114277619 |
| **A_55_P2007389** | Fam189b | 0.004387107 | -0.782915012 | 0.791689226 | 9.392391612 | 0.013709006 | 0.989516223 | 0.9976911 | -5.114279496 |
| **A_55_P2203051** | Gt(ROSA)26Sor | -0.001362032 | -0.247338841 | 0.244614776 | 9.866932625 | -0.013622677 | 0.989582237 | 0.9976911 | -5.114280388 |
| **A_55_P2087805** | Hsd3b1 | -0.005647956 | -1.026677463 | 1.015381551 | 9.61447228 | -0.013608873 | 0.989592792 | 0.9976911 | -5.11428053 |
| **A_55_P2037368** | Pdxdc1 | -0.001232698 | -0.225782737 | 0.223317341 | 10.04884093 | -0.013505561 | 0.989671793 | 0.9976911 | -5.114281589 |
| **A_55_P2120807** | Dld | -0.00143346 | -0.263080916 | 0.260213997 | 10.37025018 | -0.013478392 | 0.989692568 | 0.9976911 | -5.114281866 |
| **A_51_P495242** | Lat | 0.000571927 | -0.105231358 | 0.106375213 | 7.759151541 | 0.013298754 | 0.989829935 | 0.9976911 | -5.114283684 |
| **A_55_P2030702** | NA | -0.001518845 | -0.283653985 | 0.280616296 | 9.727223279 | -0.013244187 | 0.989871661 | 0.9976911 | -5.114284232 |
| **A_55_P2055137** | NA | 0.000975535 | -0.180929208 | 0.182880279 | 12.50891971 | 0.013193748 | 0.989910232 | 0.9976911 | -5.114284736 |
| **A_65_P14951** | Cblb | -0.001355983 | -0.254777941 | 0.252065975 | 9.608449013 | -0.013163738 | 0.98993318 | 0.9976911 | -5.114285035 |
| **A_51_P327530** | Ciao2a | -0.001814658 | -0.343539032 | 0.339909716 | 11.30623531 | -0.013064355 | 0.990009177 | 0.9976911 | -5.11428602 |
| **A_66_P119123** | A1cf | -0.000583796 | -0.111709018 | 0.110541427 | 7.66370554 | -0.012924606 | 0.990116042 | 0.9976911 | -5.114287393 |
| **A_55_P2062255** | Syne3 | -0.000621695 | -0.119657086 | 0.118413696 | 7.762642318 | -0.01284904 | 0.990173826 | 0.9976911 | -5.114288129 |
| **A_51_P118539** | 2700049A03Rik | 0.000948851 | -0.181161519 | 0.183059222 | 9.277223467 | 0.012818366 | 0.990197282 | 0.9976911 | -5.103025325 |
| **A_55_P2022049** | Klf15 | -0.000969883 | -0.189520385 | 0.18758062 | 8.059354726 | -0.012654955 | 0.990322241 | 0.9976911 | -5.11429 |
| **A_55_P1973033** | Ltbp1 | 0.002614419 | -0.508685473 | 0.513914312 | 9.495187746 | 0.012579655 | 0.990379823 | 0.9976911 | -5.114290719 |
| **A_55_P2135631** | Rps18 | -0.001304467 | -0.256764191 | 0.254155257 | 13.95369488 | -0.01256261 | 0.990392857 | 0.9976911 | -5.114290881 |
| **A_55_P2113633** | Bzw1 | 0.000841303 | -0.165083232 | 0.166765837 | 11.24165848 | 0.012474145 | 0.990460505 | 0.9976911 | -5.114291718 |
| **A_55_P1966332** | Plac9a | -0.005760236 | -1.144860299 | 1.133339828 | 14.206653 | -0.012440778 | 0.990486021 | 0.9976911 | -5.114292032 |
| **A_55_P2174821** | Rab8a | 0.001323489 | -0.260747173 | 0.263394151 | 10.5695435 | 0.012424278 | 0.990498638 | 0.9976911 | -5.114292187 |
| **A_55_P2033215** | Cabp5 | -0.003264769 | -0.656388989 | 0.649859452 | 9.905635075 | -0.012297754 | 0.990595391 | 0.9976911 | -5.11429337 |
| **A_51_P126177** | Map1lc3b | 0.001459516 | -0.291556539 | 0.294475571 | 12.6762385 | 0.012254248 | 0.990628659 | 0.9976911 | -5.085695386 |
| **A_55_P1964628** | NA | -0.001134692 | -0.229597229 | 0.227327846 | 14.71342004 | -0.012218898 | 0.990655692 | 0.9976911 | -5.114294101 |
| **A_66_P105032** | Gm13889 | -0.002421851 | -0.497644455 | 0.492800753 | 10.17359692 | -0.012031399 | 0.990799072 | 0.9976911 | -5.114295819 |
| **A_51_P201884** | Dsp | -0.000758072 | -0.156441399 | 0.154925254 | 7.934299596 | -0.011979474 | 0.990838779 | 0.9976911 | -5.114296291 |
| **A_66_P125392** | Atp8b4 | 0.000514689 | -0.105514111 | 0.106543488 | 7.704650243 | 0.01194235 | 0.990867168 | 0.9976911 | -5.114296626 |
| **A_51_P164014** | Cenpe | -0.00141307 | -0.296619704 | 0.293793565 | 9.169115483 | -0.011776241 | 0.990994192 | 0.9976911 | -5.114298116 |
| **A_55_P2024290** | Fam149a | -0.000858792 | -0.18066191 | 0.178944325 | 7.869973739 | -0.011750604 | 0.991013797 | 0.9976911 | -5.114298344 |
| **A_55_P2162269** | Bri3 | 0.001643905 | -0.346341293 | 0.349629104 | 11.97391703 | 0.011622115 | 0.991112052 | 0.9976911 | -5.114299479 |
| **A_51_P466229** | Pdgfrl | 0.000622613 | -0.131289849 | 0.132535074 | 8.43370036 | 0.011611845 | 0.991119906 | 0.9976911 | -5.114299569 |
| **A_55_P1981105** | Ap1b1 | -0.001407918 | -0.301860572 | 0.299044736 | 13.12721826 | -0.011528439 | 0.991183687 | 0.9976911 | -5.114300299 |

| | | | | | | | | |
|---|---|---|---|---|---|---|---|---|
| **A_55_P2122329** | Nfic | -0.000507838 | -0.109369524 | 0.108353847 | 8.209744215 | -0.011476768 | 0.9912232 | 0.9976911 | -5.114300748 |
| **A_51_P254805** | Kif4 | -0.000508003 | -0.110202825 | 0.109186819 | 7.744327657 | -0.011393291 | 0.991287035 | 0.9976911 | -5.11430147 |
| **A_52_P516091** | Pla2g15 | 0.001325621 | -0.285063281 | 0.287714524 | 9.886295052 | 0.011387606 | 0.991291383 | 0.9976911 | -5.114301519 |
| **A_52_P50329** | Stimate | -0.000785566 | -0.171392493 | 0.169821362 | 8.658450446 | -0.011328045 | 0.99133693 | 0.9976911 | -5.11430203 |
| **A_55_P2102901** | Fbxl4 | 0.00059117 | -0.127964411 | 0.129146752 | 8.499136885 | 0.011313339 | 0.991348175 | 0.9976911 | -5.114302156 |
| **A_51_P476261** | Zfp120 | 0.00059664 | -0.130074568 | 0.131267847 | 8.725616816 | 0.011233146 | 0.9914095 | 0.9976911 | -5.11430284 |
| **A_52_P599264** | Mdfic | -0.000890654 | -0.197225712 | 0.195444405 | 10.68988886 | -0.011160409 | 0.991465122 | 0.9976911 | -5.114303455 |
| **A_55_P2106235** | Syngr2 | 0.000756704 | -0.166807863 | 0.168321272 | 13.69901745 | 0.011109978 | 0.991503687 | 0.9976911 | -5.11430388 |
| **A_52_P601385** | Cd99l2 | 0.002704166 | -0.597630681 | 0.603039013 | 10.82875229 | 0.011081766 | 0.991525261 | 0.9976911 | -5.114304117 |
| **A_55_P1958007** | Ddx24 | -0.000972101 | -0.219156322 | 0.217212121 | 13.0045347 | -0.010961173 | 0.99161748 | 0.9976911 | -5.114305122 |
| **A_52_P493519** | Igll1 | 0.000464713 | -0.104841711 | 0.105771137 | 7.685555516 | 0.010856725 | 0.991697353 | 0.9976911 | -5.114305983 |
| **A_55_P1998751** | Ccdc137 | 0.000656285 | -0.14905649 | 0.15036906 | 8.586508281 | 0.010784573 | 0.991752528 | 0.9976911 | -5.114306573 |
| **A_55_P2227967** | NA | 0.000714097 | -0.162292388 | 0.163720582 | 8.846394517 | 0.01077759 | 0.991757868 | 0.9976911 | -5.11430663 |
| **A_51_P305843** | Chordc1 | 0.000798007 | -0.181671782 | 0.183267797 | 10.3136311 | 0.010759325 | 0.991771836 | 0.9976911 | -5.114306779 |
| **A_55_P1964368** | Kctd14 | -0.000470877 | -0.110660333 | 0.109718578 | 7.73786686 | -0.010513248 | 0.991960014 | 0.9976911 | -5.114308758 |
| **A_55_P2065646** | Zfp189 | 0.000642357 | -0.150084109 | 0.151368823 | 8.626216875 | 0.010484708 | 0.991981839 | 0.9976911 | -5.114308984 |
| **A_55_P2022158** | Gc | 0.00517105 | -1.213298688 | 1.223640787 | 11.28634154 | 0.010440784 | 0.992015428 | 0.9976911 | -5.114309332 |
| **A_55_P2051414** | Axl | -0.001149037 | -0.274097922 | 0.271799848 | 10.25320606 | -0.010356714 | 0.992079718 | 0.9976911 | -5.114309993 |
| **A_51_P451516** | Arhgef25 | -0.001417281 | -0.342755099 | 0.339920536 | 12.62141399 | -0.010215055 | 0.992188046 | 0.9976911 | -5.114311094 |
| **A_55_P2024302** | Spin1 | 0.001074993 | -0.260826527 | 0.262976513 | 9.926775729 | 0.010098031 | 0.992277537 | 0.9976911 | -5.114311993 |
| **A_51_P211968** | Lcmt1 | 0.001637449 | -0.398308416 | 0.401583315 | 10.28685992 | 0.010072467 | 0.992297087 | 0.9976911 | -5.114312188 |
| **A_55_P2066996** | Cav2 | 0.000425494 | -0.103701748 | 0.104552735 | 7.749018019 | 0.010053051 | 0.992311934 | 0.9976911 | -5.114312336 |
| **A_55_P2019557** | Mrgpra2b | -0.000455583 | -0.112005395 | 0.111094229 | 7.739199722 | -0.010047721 | 0.99231601 | 0.9976911 | -5.114312376 |
| **A_55_P1983988** | Enox2 | -0.000686857 | -0.171471659 | 0.170097945 | 8.406375443 | -0.009894327 | 0.992433314 | 0.9976911 | -5.114313532 |
| **A_51_P143470** | Timm44 | -0.000669583 | -0.168145692 | 0.166806526 | 11.04686588 | -0.009836047 | 0.992477882 | 0.9976911 | -5.114313967 |
| **A_55_P2063715** | Cfap53 | 0.00046278 | -0.115449641 | 0.116375201 | 7.874897723 | 0.009822308 | 0.992488388 | 0.9976911 | -5.114314069 |
| **A_51_P492410** | Pmvk | -0.001001622 | -0.254488383 | 0.25248514 | 10.18260987 | -0.00972115 | 0.992565746 | 0.9976911 | -5.114314817 |
| **A_66_P132322** | Mrpl21 | -0.001298065 | -0.330309777 | 0.327713648 | 11.97086804 | -0.009706312 | 0.992577093 | 0.9976911 | -5.114314926 |
| **A_55_P2020326** | Madcam1 | -0.001404758 | -0.358183538 | 0.355374023 | 8.40960945 | -0.009686603 | 0.992592164 | 0.9976911 | -5.11431507 |
| **A_55_P2097578** | NA | -0.000764793 | -0.195372647 | 0.193843061 | 10.34815518 | -0.009668356 | 0.992606118 | 0.9976911 | -5.114315203 |
| **A_55_P1965467** | 2300009A05Rik | -0.001084801 | -0.277382656 | 0.275213053 | 10.29087264 | -0.009659214 | 0.99261311 | 0.9976911 | -5.11431527 |
| **A_55_P2161535** | Ddx23 | -0.000416028 | -0.107950134 | 0.107118077 | 8.06638076 | -0.009518005 | 0.992721096 | 0.9976911 | -5.114316294 |
| **A_55_P2070936** | Tra2b | -0.000963965 | -0.250289115 | 0.248361185 | 8.313426972 | -0.009511837 | 0.992725812 | 0.9976911 | -5.114316338 |
| **A_51_P212023** | Sart3 | -0.000844765 | -0.219987499 | 0.218297968 | 11.24077819 | -0.009483707 | 0.992747324 | 0.9976911 | -5.11431654 |
| **A_51_P279606** | Socs1 | 0.000782311 | -0.20586669 | 0.207431312 | 9.135616492 | 0.009313552 | 0.992877446 | 0.9976911 | -5.114317749 |
| **A_52_P646783** | Fbxl22 | -0.000528143 | -0.142375734 | 0.141319448 | 7.829426253 | -0.009160067 | 0.99299482 | 0.9976911 | -5.114318821 |
| **A_55_P1958480** | NA | -0.001939654 | -0.523615264 | 0.519735956 | 11.06292451 | -0.0091473 | 0.993004583 | 0.9976911 | -5.11431891 |
| **A_55_P2104662** | Ttc33 | 0.000956143 | -0.256817894 | 0.258730179 | 10.80451612 | 0.009125416 | 0.993021319 | 0.9976911 | -5.114319061 |
| **A_55_P1967483** | NA | 0.000583157 | -0.156703611 | 0.157869925 | 10.76080364 | 0.009121415 | 0.993024378 | 0.9976911 | -5.114319088 |
| **A_51_P361951** | Uqcrfs1 | 0.001299871 | -0.350310289 | 0.352910031 | 12.56203518 | 0.00909511 | 0.993044495 | 0.9976911 | -5.11431927 |
| **A_55_P2119737** | NA | -0.000983684 | -0.267110064 | 0.265142696 | 8.511531782 | -0.009093618 | 0.993045635 | 0.9976911 | -5.11431928 |
| **A_52_P552589** | Map4k1 | 0.000601732 | -0.165362208 | 0.166565672 | 8.375051752 | 0.008919869 | 0.993178506 | 0.997702692 | -5.114320463 |
| **A_51_P459320** | Gkap1 | 0.000947091 | -0.261627538 | 0.263521719 | 11.90483218 | 0.008873764 | 0.993213764 | 0.997702692 | -5.114320773 |
| **A_55_P1960472** | Polg | 0.001416471 | -0.394687381 | 0.397520323 | 9.735929671 | 0.008797672 | 0.993271954 | 0.997702692 | -5.114321281 |
| **A_55_P1968276** | Tomm22 | -0.001111551 | -0.312676728 | 0.310453625 | 12.17258929 | -0.008777074 | 0.993287705 | 0.997702692 | -5.114321418 |
| **A_51_P402193** | Map3k1 | -0.001043729 | -0.294804416 | 0.292716957 | 11.59292736 | -0.008741046 | 0.993315257 | 0.997702692 | -5.114321657 |
| **A_55_P2058433** | Cyp2c68 | -0.000572386 | -0.163456139 | 0.162311368 | 7.71245496 | -0.008645303 | 0.993388475 | 0.997724388 | -5.114322286 |
| **A_55_P2186615** | Acp1 | -0.001056644 | -0.306978978 | 0.30486569 | 9.824693652 | -0.008497412 | 0.993501572 | 0.997741856 | -5.114323245 |
| **A_55_P2140453** | Myh6 | -0.00036166 | -0.105191891 | 0.104468571 | 7.668936107 | -0.00848756 | 0.993509106 | 0.997741856 | -5.114323308 |
| **A_55_P1980316** | Pcbp2 | -0.000585161 | -0.176577462 | 0.17540714 | 13.75244855 | -0.008179951 | 0.993744345 | 0.997919283 | -5.114325246 |

| | | | | | | | | | |
|---|---|---|---|---|---|---|---|---|---|
| **A_52_P93902** | Alg5 | -0.001290662 | -0.392261993 | 0.389680669 | 12.57849193 | -0.00812151 | 0.993789036 | 0.997919283 | -5.114325606 |
| **A_55_P2008349** | Prelid1 | 0.000699724 | -0.216233537 | 0.217632984 | 13.80442321 | 0.007935415 | 0.99393135 | 0.998002885 | -5.114326736 |
| **A_51_P333622** | 2010300C02Rik | 0.000465637 | -0.145595266 | 0.14652654 | 9.508584918 | 0.00784301 | 0.994002016 | 0.998002885 | -5.114327287 |
| **A_55_P2182273** | NA | -0.000718939 | -0.227186064 | 0.225748186 | 9.009945364 | -0.007810092 | 0.99402719 | 0.998002885 | -5.114327482 |
| **A_55_P2053032** | Xkr8 | 0.000636578 | -0.204718157 | 0.205991314 | 8.604699202 | 0.007626341 | 0.994167711 | 0.998085901 | -5.114328554 |
| **A_55_P2021216** | Crlf3 | 0.000786703 | -0.2587625 | 0.260335905 | 11.23426381 | 0.00745693 | 0.994297266 | 0.998085901 | -5.11432952 |
| **A_55_P1970788** | C5ar2 | 0.00287344 | -0.94621125 | 0.95195813 | 10.08817433 | 0.007448457 | 0.994303745 | 0.998085901 | -5.114329568 |
| **A_51_P271984** | Tmem45b | -0.001992923 | -0.667091005 | 0.66310516 | 8.395586354 | -0.007371806 | 0.994362364 | 0.998085901 | -5.114329997 |
| **A_55_P2227321** | Ptprd | 0.000316258 | -0.105335136 | 0.105967652 | 7.675893369 | 0.00736436 | 0.994368058 | 0.998085901 | -5.114330039 |
| **A_52_P676359** | Fcnb | -0.00031057 | -0.106473305 | 0.105852166 | 7.640259084 | -0.007197081 | 0.994495983 | 0.998162471 | -5.114330959 |
| **A_55_P1975400** | Flnc | 0.003701205 | -1.278845117 | 1.286247528 | 10.93354136 | 0.007099688 | 0.994570463 | 0.998185393 | -5.114331486 |
| **A_55_P2176385** | St6galnac4 | 0.001019107 | -0.360746658 | 0.362784872 | 9.501651338 | 0.006930452 | 0.994699885 | 0.998222614 | -5.114332383 |
| **A_55_P2326517** | Ube2i | -0.001406476 | -0.501715282 | 0.49890233 | 9.171596532 | -0.006916132 | 0.994710837 | 0.998222614 | -5.114332458 |
| **A_55_P2162136** | Hmgn2 | 0.0010794 | -0.388847651 | 0.39100645 | 12.50576834 | 0.006810328 | 0.99479175 | 0.99822621 | -5.114333007 |
| **A_52_P284778** | Rae1 | 0.000486756 | -0.176231765 | 0.177205276 | 10.60991108 | 0.006776384 | 0.994817708 | 0.99822621 | -5.114333182 |
| **A_51_P134551** | Coq9 | 0.000505505 | -0.188056105 | 0.189067115 | 8.947269069 | 0.0065954 | 0.994956114 | 0.998313265 | -5.114334096 |
| **A_52_P420665** | Uts2b | 0.000326064 | -0.123666554 | 0.124318681 | 7.841593618 | 0.006469577 | 0.995052337 | 0.998357987 | -5.114334718 |
| **A_52_P590474** | Ccnl1 | 0.000517709 | -0.199878973 | 0.200914392 | 8.532225064 | 0.006355719 | 0.99513941 | 0.998393525 | -5.11433527 |
| **A_51_P436020** | Tmem167 | -0.000450128 | -0.180499158 | 0.179598903 | 9.037192388 | -0.006150552 | 0.99529631 | 0.998479467 | -5.11433624 |
| **A_55_P1964313** | Pla2g4b | 0.000543779 | -0.219103272 | 0.220190829 | 11.75549084 | 0.006090683 | 0.995342095 | 0.998479467 | -5.114336517 |
| **A_51_P363630** | Snrpe | -0.00110914 | -0.45280121 | 0.45058293 | 13.52089611 | -0.006041061 | 0.995380043 | 0.998479467 | -5.114336745 |
| **A_52_P597634** | Fzd1 | -0.002261239 | -0.936427961 | 0.931905483 | 10.55357117 | -0.00595513 | 0.995445759 | 0.998492649 | -5.114337134 |
| **A_65_P20799** | Abi2 | 0.000261623 | -0.109038441 | 0.109561688 | 8.05072193 | 0.005888781 | 0.9954965 | 0.998492649 | -5.114337431 |
| **A_55_P2207055** | NA | 0.000271897 | -0.116595657 | 0.117139451 | 7.684315172 | 0.005723732 | 0.99562272 | 0.998525678 | -5.114338156 |
| **A_55_P2160648** | Gm6900 | 0.000466206 | -0.202735081 | 0.203667492 | 16.32335442 | 0.00564443 | 0.995683366 | 0.998525678 | -5.114338497 |
| **A_55_P1982404** | Gpm6b | 0.000528345 | -0.229812905 | 0.230869596 | 9.787458663 | 0.005643069 | 0.995684407 | 0.998525678 | -5.114338502 |
| **A_55_P2234361** | Rnf150 | -0.000520193 | -0.244416045 | 0.24337566 | 8.300680804 | -0.005247215 | 0.995987138 | 0.998777452 | -5.114340132 |
| **A_55_P2156340** | NA | -0.000401603 | -0.195703181 | 0.194899975 | 16.32918601 | -0.005058948 | 0.996131115 | 0.9987819 | -5.114340866 |
| **A_55_P2022211** | Plxdc2 | 0.001342274 | -0.653233842 | 0.65591839 | 11.35520566 | 0.005044872 | 0.99614188 | 0.9987819 | -5.114340919 |
| **A_55_P2087182** | Car4 | 0.000791666 | -0.385889014 | 0.387472346 | 11.40687026 | 0.005036846 | 0.996148018 | 0.9987819 | -5.11434095 |
| **A_55_P2105542** | Pde2a | -0.000826381 | -0.409797996 | 0.408145235 | 11.41382301 | -0.004971143 | 0.996198264 | 0.9987819 | -5.114341199 |
| **A_55_P2007326** | Cyp2a4 | 0.000218143 | -0.109503418 | 0.109939703 | 7.711571769 | 0.004891231 | 0.996259377 | 0.998791364 | -5.114341497 |
| **A_52_P49250** | Erbb2 | 0.000239482 | -0.121918345 | 0.122397309 | 8.278555726 | 0.004823039 | 0.996311528 | 0.998791843 | -5.108837023 |
| **A_55_P2266977** | Ahcyl2 | 0.000760198 | -0.396304364 | 0.39782476 | 11.19373312 | 0.004710151 | 0.996397859 | 0.998799814 | -5.114342154 |
| **A_55_P2139256** | Rps6ka1 | 0.000689987 | -0.362217658 | 0.363597632 | 12.8118221 | 0.004677503 | 0.996422827 | 0.998799814 | -5.11434227 |
| **A_51_P383399** | Aldh1a7 | 0.000199223 | -0.107771807 | 0.108170253 | 7.700695816 | 0.004539433 | 0.996528417 | 0.998853856 | -5.114342751 |
| **A_55_P2133225** | Mmgt2 | 0.000426303 | -0.250901151 | 0.251753756 | 11.55001766 | 0.004172991 | 0.996808655 | 0.999037138 | -5.114343958 |
| **A_51_P301809** | Slit3 | 0.000963236 | -0.576154565 | 0.578081038 | 8.460462829 | 0.004106176 | 0.996859752 | 0.999037138 | -5.114344167 |
| **A_55_P1958088** | Lyn | 0.000221626 | -0.133075097 | 0.13351835 | 8.214006896 | 0.004090447 | 0.996871782 | 0.999037138 | -5.114344215 |
| **A_55_P2095394** | Dhx37 | -0.000224135 | -0.140073488 | 0.139625217 | 8.583748102 | -0.003942927 | 0.996984598 | 0.999037138 | -5.114344663 |
| **A_55_P2032714** | Fhit | -0.000268347 | -0.170530626 | 0.169993932 | 8.329393707 | -0.003877462 | 0.997034663 | 0.999037138 | -5.114344857 |
| **A_55_P2160686** | Tsc22d1 | 0.000894535 | -0.574534827 | 0.576323897 | 12.80762974 | 0.003824499 | 0.997075167 | 0.999037138 | -5.114345011 |
| **A_52_P295533** | Slc25a28 | -0.000289087 | -0.187452383 | 0.186874208 | 9.805640281 | -0.003799948 | 0.997093943 | 0.999037138 | -5.114345082 |
| **A_51_P384718** | Efna4 | -0.000192066 | -0.128067862 | 0.127683729 | 7.975122193 | -0.003695152 | 0.997174086 | 0.999037138 | -5.114345379 |
| **A_55_P2049287** | NA | 0.00031677 | -0.21076167 | 0.21139521 | 13.98931044 | 0.003692068 | 0.997176445 | 0.999037138 | -5.114345388 |
| **A_51_P212801** | Ube2a | -0.000593089 | -0.411039319 | 0.40985314 | 10.68623724 | -0.003554947 | 0.99728131 | 0.999071973 | -5.114345763 |
| **A_55_P1966438** | Gstm2 | -0.000188585 | -0.132500327 | 0.132123156 | 8.03384273 | -0.003506542 | 0.997318328 | 0.999071973 | -5.114345892 |
| **A_52_P542290** | Pdxdc1 | 0.000265333 | -0.189281493 | 0.189812158 | 9.833052544 | 0.003443842 | 0.997366278 | 0.999071973 | -5.114346057 |
| **A_55_P2002482** | Sarnp | -0.00025703 | -0.199993404 | 0.199479344 | 11.41940976 | -0.003165889 | 0.997578845 | 0.999233119 | -5.114346752 |
| **A_55_P2131925** | Kprp | 0.001597628 | -1.312334017 | 1.315529272 | 8.72858352 | 0.002991383 | 0.9977123 | 0.999315009 | -5.114347158 |

| | | | | | | | | | |
|---|---|---|---|---|---|---|---|---|---|
| **A_55_P2054062** | Chst2 | -0.000247761 | -0.222938479 | 0.222442957 | 8.581967963 | -0.002737157 | 0.997906723 | 0.999451424 | -5.114347708 |
| **A_55_P2145136** | NA | 0.000114887 | -0.105425673 | 0.105655447 | 7.710546301 | 0.002678069 | 0.99795191 | 0.999451424 | -5.114347829 |
| **A_55_P1961039** | Pnpla2 | 0.000285383 | -0.276268293 | 0.276839059 | 11.45311506 | 0.002538734 | 0.998058469 | 0.999506354 | -5.114348104 |
| **A_66_P111714** | Utp20 | -0.000151846 | -0.155683698 | 0.155380007 | 8.260574581 | -0.002401887 | 0.998163124 | 0.999542038 | -5.11434836 |
| **A_51_P420415** | Srd5a1 | 0.001094529 | -1.168072854 | 1.170261912 | 12.67546072 | 0.002303137 | 0.998238644 | 0.999542038 | -5.114348535 |
| **A_52_P526372** | Zeb2 | -0.000101788 | -0.109488659 | 0.109285083 | 7.763482304 | -0.002289286 | 0.998249237 | 0.999542038 | -5.114348559 |
| **A_51_P427828** | Mlip | 8.96E-05 | -0.106412958 | 0.106592131 | 7.640027126 | 0.002069442 | 0.998417365 | 0.999638323 | -5.114348922 |
| **A_55_P2147160** | NA | 0.000132782 | -0.173626112 | 0.173891676 | 8.363491544 | 0.001880018 | 0.99856223 | 0.999638323 | -5.114349204 |
| **A_55_P2274592** | Wnt7b | 0.000129438 | -0.170114283 | 0.170373159 | 8.89760384 | 0.001870514 | 0.998569499 | 0.999638323 | -5.114349218 |
| **A_52_P600625** | Rad23b | 0.000128613 | -0.182795616 | 0.183052842 | 9.045932712 | 0.001729747 | 0.998677151 | 0.999638323 | -5.11434941 |
| **A_55_P2033055** | Pdp1 | -0.000112109 | -0.159801949 | 0.159577732 | 8.346019901 | -0.001727155 | 0.998679134 | 0.999638323 | -5.114349413 |
| **A_55_P2136209** | Ublcp1 | -0.000232387 | -0.341642758 | 0.341177984 | 9.837486738 | -0.001674574 | 0.998719346 | 0.999638323 | -5.114349481 |
| **A_55_P2163725** | Slc7a9 | 0.000672752 | -1.02484084 | 1.026186344 | 10.52047852 | 0.001613922 | 0.99876573 | 0.999638323 | -5.114349556 |
| **A_51_P191649** | Ndc80 | 7.43E-05 | -0.119748715 | 0.119897333 | 7.80047759 | 0.001525705 | 0.998833195 | 0.999638323 | -5.114349661 |
| **A_51_P490700** | Ccdc40 | -6.66E-05 | -0.109473122 | 0.109339823 | 7.826245494 | -0.001498726 | 0.998853828 | 0.999638323 | -5.114349691 |
| **A_55_P2159890** | Zbtb7a | -0.000185933 | -0.30774439 | 0.307372525 | 9.290876203 | -0.001487298 | 0.998862568 | 0.999638323 | -5.114349704 |
| **A_51_P209818** | Prtn3 | 6.38E-05 | -0.110598697 | 0.110726332 | 7.696447069 | 0.001418756 | 0.998914986 | 0.999639025 | -5.11434978 |
| **A_55_P2100928** | Ptgds | 0.000507992 | -1.060023357 | 1.06103934 | 8.852039745 | 0.001178425 | 0.999098782 | 0.999701522 | -5.114350016 |
| **A_51_P308446** | 1700094D03Rik | -5.04E-05 | -0.114959746 | 0.114858999 | 8.168044828 | -0.001078482 | 0.999175215 | 0.999701522 | -5.114350101 |
| **A_55_P2081405** | BC052688 | -8.63E-05 | -0.198752979 | 0.198580327 | 13.1911834 | -0.001069019 | 0.999182453 | 0.999701522 | -5.114350109 |
| **A_55_P2003719** | NA | 5.31E-05 | -0.125749192 | 0.125855321 | 15.15021022 | 0.001037732 | 0.999206379 | 0.999701522 | -5.114350134 |
| **A_51_P177432** | NA | -4.53E-05 | -0.111502387 | 0.111411874 | 8.095853906 | -0.000998949 | 0.999236039 | 0.999701522 | -5.114350163 |
| **A_52_P399717** | Ccdc50 | -5.46E-05 | -0.148451733 | 0.148342604 | 8.556914148 | -0.000904597 | 0.999308196 | 0.999708081 | -5.114350231 |
| **A_55_P2162552** | Txndc9 | 5.35E-05 | -0.153943979 | 0.154051033 | 11.60032794 | 0.000855118 | 0.999346036 | 0.999708081 | -5.114350264 |
| **A_51_P490678** | Trim68 | 4.16E-05 | -0.167839178 | 0.167922423 | 8.355020859 | 0.000609951 | 0.999533531 | 0.999779128 | -5.1143504 |
| **A_51_P483261** | Pomt1 | 3.91E-05 | -0.170389872 | 0.170468049 | 8.8228428 | 0.000564258 | 0.999568476 | 0.999779128 | -5.11435042 |
| **A_55_P2031157** | Efna3 | -3.37E-05 | -0.148129065 | 0.148061723 | 8.662796599 | -0.000559348 | 0.999572231 | 0.999779128 | -5.114350422 |
| **A_52_P370935** | Glcci1 | 2.85E-05 | -0.194370015 | 0.194427018 | 8.661769284 | 0.0003607 | 0.999724149 | 0.999879338 | -5.114350491 |
| **A_51_P434697** | Ube2e2 | -2.1E-05 | -0.233902453 | 0.233860421 | 10.93300546 | -0.000221068 | 0.999830935 | 0.999934399 | -5.114350522 |
| **A_66_P121059** | Mirg | -3.61E-05 | -0.64787126 | 0.647799111 | 10.19057444 | -0.000136995 | 0.999895231 | 0.999946964 | -5.114350534 |
| **A_52_P177021** | Pts | 6.95E-06 | -0.324206569 | 0.324220476 | 9.390277609 | 5.28E-05 | 0.999959645 | 0.999959645 | -5.11435054 |

### Rychlik_Arsenic_microarray_Supplemental_TableS5

| #term ID | term description | observed gene count | background gene count | strength | false discovery rate | matching proteins in your network (IDs) | matching proteins in your network (labels) |
|---|---|---|---|---|---|---|---|
| GO:0050789 | Regulation of biological process | 163 | 9973 | 0.18 | 1.95E-08 | 10090.ENSMUSP00000002284,10090.ENSMUSP00000002413,10090.ENSMUSP00000004120,10090.ENSMUSP00000004430,10090.ENSMUSP00000004985,10090.ENSMUSP00000005503,10090.ENSMUSP00000015581,10090.ENSMUSP00000017851,10090.ENSMUSP00000018361,10090.ENSMUSP00000018506,10090.ENSMUSP00000019469,10090.ENSMUSP00000019726,10090.ENSMUSP00000020123,10090.ENSMUSP00000020308,10090.ENSMUSP00000020835,10090.ENSMUSP00000020908,10090.ENSMUSP00000021807,10090.ENSMUSP00000022227,10090.ENSMUSP00000022380,10090.ENSMUSP00000022612,10090.ENSMUSP00000022927,10090.ENSMUSP00000023769,10090.ENSMUSP00000023774,10090.ENSMUSP00000023897,10090.ENSMUSP00000024104,10090.ENSMUSP00000024657,10090.ENSMUSP00000025161,10090.ENSMUSP00000025196,10090.ENSMUSP00000025254,10090.ENSMUSP00000025562,10090.ENSMUSP00000025904,10090.ENSMUSP00000025930,10090.ENSMUSP00000026565,10090.ENSMUSP00000027697,10090.ENSMUSP00000027760,10090.ENSMUSP00000027809,10090.ENSMUSP00000027933,10090.ENSMUSP00000028328,10090.ENSMUSP00000028382,10090.ENSMUSP00000028398,10090.ENSMUSP00000029018,10090.ENSMUSP00000029566,10090.ENSMUSP00000030121,10090.ENSMUSP00000030289,10090.ENSMUSP00000030367,10090.ENSMUSP00000030623,10090.ENSMUSP00000031540,10090.ENSMUSP00000031542,10090.ENSMUSP00000032198,10090.ENSMUSP00000032473,10090.ENSMUSP00000032800,10090.ENSMUSP00000032898,10090.ENSMUSP00000033626,10090.ENSMUSP00000034003, | Plaur,Tmem161a,Fancl,Clcn3,Brpf3,Msh6,Gzmb,Serinc3,Pmp22,Kpna2,G6pc,Plin3,Tmpo,Ddit4,Ppm1d,E2f6,Dek,Cenpk,Psmc6,Pbk,Rad21,Atf1,Slc11a2,Gzma,Gcm1,Phf10,Tapbp,Psmb8,Lims2,Incenp,Prdx5,Smc3,Ifitm3,Cdk18,Rgl1,Opn3,Dtl,Entpd2,Nup35,Ube2e3,Zbp1,Efna1,B4galt1,Usp1,Ssbp3,Sfpq,Oasl1,Oasl2,Usp18,Iigp1,Tyrobp,Ipo5,Serpina7,Fgl1,Map1lc3b,Ddx25,Phip,Tipin,Me1,Csrnp1,Mvk,Hspb8,Zfand1,Slx4,Atg14,Hace1,Serpine1,Cblc,C1qb,Ofd1,Pim3,Card19,Tiparp,Rbm12,Zfp367,Elmod2,Hic1,Atxn2,E2f8,Cenpe,Zfpm1,S100a1,Whsc1,Aspm,Ercc8,Camk2n1,Hilpda,Tob2,Insig1,Adam10,Artn,Nfil3,Azin1,Hmgb2,Ly6c1,Cdc42ep3,Ifnlr1,Frmd4a,Tnfaip8l1,Dchs1,Phf6,Gspt1,Krit1,Birc5,Oas1a,Slc39a8,Rps6ka6,Isg15,Kif20b,Fmr1,Magi1,Rnf213,Hbb-bt,Arrb1,Cox7a1,Mbnl1,Mad2l1,Atp11c,Flcn,Ifit3,Anp32b,Pdpk1,Sirt1,Zbtb8b,Txnip,Crebzf,Trim59,Ect2,Irf1,Map4k5,Hnrnpa3,Bclaf3,Cnot4,Arid5a,Dnm1l,Ube2v2,Rnf186,Arhgap23,Rsl1d1,Bcl3,Srsf3,Irf9,Srsf1,Nmi,Prlr,Ppp1r3g,Xaf1,Cep295,Camsap3,Lsm5,Tgm1,Tfdp2,Odc1,Tcea1,Parp10,Sh3glb1,Zfyve27,Tgif1,Psmb9,Arhgap21,H2-D1,Golph3l,Zfp131 |
| GO:0009987 | Cellular process | 195 | 13330 | 0.13 | 2.93E-08 | 10090.ENSMUSP00000002198,10090.ENSMUSP00000002284,10090.ENSMUSP00000002413,10090.ENSMUSP00000004120,10090.ENSMUSP00000004430,10090.ENSMUSP00000004985,10090.ENSMUSP00000005503,10090.ENSMUSP00000011058,10090.ENSMUSP00000013667,10090.ENSMUSP00000015581,10090.ENSMUSP00000017851,10090.ENSMUSP00000018361,10090.ENSMUSP00000018506,10090.ENSMUSP00000019469,10090.ENSMUSP00000020217,10090.ENSMUSP00000020308,10090.ENSMUSP00000020835,10090.ENSMUSP00000020908,10090.ENSMUSP00000021120,10090.ENSMUSP00000021381,10090.ENSMUSP00000021807,10090.ENSMUSP00000022227,10090.ENSMUSP00000022380,10090.ENSMUSP00000022612,10090.ENSMUSP00000022927,10090.ENSMUSP00000023353,10090.ENSMUSP00000023359,10090.ENSMUSP00000023769,10090.ENSMUSP00000023774,10090.ENSMUSP00000023897,10090.ENSMUSP00000024104,10090.ENSMUSP00000024657,10090.ENSMUSP00000025161,10090.ENSMUSP00000025196,10090.ENSMUSP00000025254,10090.ENSMUSP00000025562,10090.ENSMUSP00000025632,10090.ENSMUSP00000025904,10090.ENSMUSP00000025930,10090.ENSMUSP00000026565,10090.ENSMUSP00000026823,10090.ENSMUSP00000027623,10090.ENSMUSP00000027697,10090.ENSMUSP00000027760,10090.ENSMUSP00000027800,10090.ENSMUSP00000027809,10090.ENSMUSP00000027933,10090.ENSMUSP00000028328,10090.ENSMUSP00000028382,10090.ENSMUSP00000028398,10090.ENSMUSP00000029018,10090.ENSMUSP00000029566,10090.ENSMUSP00000030121,10090.ENSMUSP00000030289, | Sf3a1,Plaur,Tmem161a,Fancl,Clcn3,Brpf3,Msh6,Pdhx,Bcas1,Gzmb,Serinc3,Pmp22,Kpna2,G6pc,Nudt4,Ddit4,Ppm1d,E2f6,Trim47,Pnn,Dek,Cenpk,Psmc6,Pbk,Rad21,Mcm4,Nde1,Atf1,Slc11a2,Gzma,Gcm1,Phf10,Tapbp,Psmb8,Lims2,Incenp,Carnmt1,Prdx5,Smc3,Ifitm3,Pigq,Tsn,Cdk18,Rgl1,Tmem63a,Opn3,Dtl,Entpd2,Nup35,Ube2e3,Zbp1,Efna1,B4galt1,Usp1,Ssbp3,Sfpq,Sh3bgrl3,Oasl2,Fscn3,Usp18,Iigp1,Tyrobp,Ipo5,Polr3e,Dctn6,Fgl1,Heatr3,Dnaja2,Map1lc3b,Ddx25,Phip,Tipin,Me1,Csrnp1,Mvk,Hspb8,Zfand1,Slx4,Wdr55,Atg14,Hace1,Serpine1,Cblc,Ubac1,C1qb,Rbm45,Rtp4,Ofd1,Dis3,Pim3,Smc4,Tiparp,Cdh26,Cbr1,Haus3,Mettl5,Polr2k,Hic1,Atxn2,E2f8,Cenpe,Zfpm1,Whsc1,Aspm,Ercc8,Hilpda,Insig1,Nupl2,Adam10,Artn,Nfil3,Azin1,Hmgb2,Ly6c1,Cdc42ep3,Rabgap1,Srp19,Ifnlr1,Frmd4a,Dchs1,Mrpl18,Prpf38a,Gspt1,Krit1,Birc5,Oas1a,Amd2,Slc39a8,Rps6ka6,Mdc1,Prpf4,Nhlrc4,Isg15,Kif20b,Fmr1,Magi1,Mfap1a,Asgr1,Rnf213,Hbb-bt,Arrb1,Cox7a1,Mbnl1,Mad2l1,Atp11c,Nop56,Flcn,Ifit3,Anp32b,Pdpk1,Mcts2,Sirt1,Txnip,Crebzf,Trim59,Ect2,Irf1,Ninl,Map4k5,Pom121,Hnrnpa3,Ppip5k2,Nmnat3,Cnot4,Arid5a,Dnm1l,Ube2v2,Rnf186,Arhgap23,Rsl1d1,Bcl3,Slc16a11,Srsf3,Srsf1,Nmi,Prlr,Crispld2,Xaf1,Camsap3,Lsm5,Tgm1,Tfdp2,Odc1,Tcea1,Parp10,Sh3glb1,Zfyve27,Tgif1,Ube2d2a,Psmb9,Arhgap21,H2-D1,Golph3l,Tomm20,Metap2 |
| GO:0065007 | Biological regulation | 167 | 10591 | 0.16 | 6.45E-08 | 10090.ENSMUSP00000002284,10090.ENSMUSP00000002413,10090.ENSMUSP00000004120,10090.ENSMUSP00000004430,10090.ENSMUSP00000004985,10090.ENSMUSP00000005503,10090.ENSMUSP00000015581,10090.ENSMUSP00000017851,10090.ENSMUSP00000018361,10090.ENSMUSP00000018506,10090.ENSMUSP00000019469,10090.ENSMUSP00000019726,10090.ENSMUSP00000020123,10090.ENSMUSP00000020308,10090.ENSMUSP00000020835,10090.ENSMUSP00000020908,10090.ENSMUSP00000021807,10090.ENSMUSP00000022227,10090.ENSMUSP00000022380,10090.ENSMUSP00000022612,10090.ENSMUSP00000022927,10090.ENSMUSP00000023359,10090.ENSMUSP00000023769,10090.ENSMUSP00000023774,10090.ENSMUSP00000023897,10090.ENSMUSP00000024104,10090.ENSMUSP00000024657,10090.ENSMUSP00000025161,10090.ENSMUSP00000025196,10090.ENSMUSP00000025254,10090.ENSMUSP00000025562,10090.ENSMUSP00000025904,10090.ENSMUSP00000025930,10090.ENSMUSP00000026565,10090.ENSMUSP00000027697,10090.ENSMUSP00000027760,10090.ENSMUSP00000027809,10090.ENSMUSP00000027933,10090.ENSMUSP00000028328,10090.ENSMUSP00000028382,10090.ENSMUSP00000028398,10090.ENSMUSP00000029018,10090.ENSMUSP00000029566,10090.ENSMUSP00000030121,10090.ENSMUSP00000030289,10090.ENSMUSP00000030367,10090.ENSMUSP00000030623,10090.ENSMUSP00000031540,10090.ENSMUSP00000031542,10090.ENSMUSP00000032198,10090.ENSMUSP00000032473,10090.ENSMUSP00000032800,10090.ENSMUSP00000032898,10090.ENSMUSP00000033626, | Plaur,Tmem161a,Fancl,Clcn3,Brpf3,Msh6,Gzmb,Serinc3,Pmp22,Kpna2,G6pc,Plin3,Tmpo,Ddit4,Ppm1d,E2f6,Dek,Cenpk,Psmc6,Pbk,Rad21,Nde1,Atf1,Slc11a2,Gzma,Gcm1,Phf10,Tapbp,Psmb8,Lims2,Incenp,Prdx5,Smc3,Ifitm3,Cdk18,Rgl1,Opn3,Dtl,Entpd2,Nup35,Ube2e3,Zbp1,Efna1,B4galt1,Usp1,Ssbp3,Sfpq,Oasl1,Oasl2,Usp18,Iigp1,Tyrobp,Ipo5,Serpina7,Fgl1,Dnaja2,Map1lc3b,Ddx25,Phip,Tipin,Me1,Csrnp1,Mvk,Hspb8,Zfand1,Slx4,Atg14,Hace1,Serpine1,Cblc,C1qb,Ofd1,Dis3,Pim3,Card19,Tiparp,Rbm12,Zfp367,Elmod2,Hic1,Atxn2,E2f8,Cenpe,Zfpm1,S100a1,Whsc1,Aspm,Ercc8,Camk2n1,Hilpda,Tob2,Insig1,Adam10,Artn,Nfil3,Azin1,Hmgb2,Ly6c1,Cdc42ep3,Rabgap1,Ifnlr1,Frmd4a,Tnfaip8l1,Dchs1,Phf6,Gspt1,Krit1,Birc5,Oas1a,Slc39a8,Rps6ka6,Isg15,Kif20b,Fmr1,Magi1,Rnf213,Hbb-bt,Arrb1,Cox7a1,Mbnl1,Mad2l1,Atp11c,Flcn,Ifit3,Anp32b,Pdpk1,Sirt1,Zbtb8b,Txnip,Crebzf,Trim59,Ect2,Irf1,Map4k5,Hnrnpa3,Bclaf3,Cnot4,Arid5a,Dnm1l,Ube2v2,Rnf186,Arhgap23,Rsl1d1,Bcl3,Srsf3,Irf9,Srsf1,Nmi,Prlr,Ppp1r3g,Xaf1,Cep295,Camsap3,Lsm5,Tgm1,Tfdp2,Odc1,Tcea1,Parp10,Sh3glb1,Zfyve27,Tgif1,Psmb9,Arhgap21,H2-D1,Golph3l,Zfp131 |
| GO:0019222 | Regulation of metabolic process | 115 | 6085 | 0.24 | 7.81E-08 | 10090.ENSMUSP00000002284,10090.ENSMUSP00000002413,10090.ENSMUSP00000004430,10090.ENSMUSP00000004985,10090.ENSMUSP00000005503,10090.ENSMUSP00000018361,10090.ENSMUSP00000019469,10090.ENSMUSP00000020123,10090.ENSMUSP00000020308,10090.ENSMUSP00000020835,10090.ENSMUSP00000020908,10090.ENSMUSP00000021807,10090.ENSMUSP00000022227,10090.ENSMUSP00000022380,10090.ENSMUSP00000022612,10090.ENSMUSP00000022927,10090.ENSMUSP00000023769,10090.ENSMUSP00000023774,10090.ENSMUSP00000023897,10090.ENSMUSP00000024104,10090.ENSMUSP00000024657,10090.ENSMUSP00000025161,10090.ENSMUSP00000025196,10090.ENSMUSP00000025562,10090.ENSMUSP00000025904,10090.ENSMUSP00000025930,10090.ENSMUSP00000027697,10090.ENSMUSP00000027933,10090.ENSMUSP00000028382,10090.ENSMUSP00000029566,10090.ENSMUSP00000030289,10090.ENSMUSP00000030367,10090.ENSMUSP00000030623,10090.ENSMUSP00000031540,10090.ENSMUSP00000031542,10090.ENSMUSP00000032473,10090.ENSMUSP00000032800,10090.ENSMUSP00000033626,10090.ENSMUSP00000034003,10090.ENSMUSP00000034612,10090.ENSMUSP00000034787,10090.ENSMUSP00000034964,10090.ENSMUSP00000034989,10090.ENSMUSP00000035101,10090.ENSMUSP00000036971,10090.ENSMUSP00000037007,10090.ENSMUSP00000038871,10090.ENSMUSP00000039047,10090.ENSMUSP00000039206,10090.ENSMUSP00000039586,10090.ENSMUSP00000039955,10090.ENSMUSP00000048051,10090.ENSMUSP00000050461,10090.ENSMUSP00000050854, | Plaur,Tmem161a,Clcn3,Brpf3,Msh6,Pmp22,G6pc,Tmpo,Ddit4,Ppm1d,E2f6,Dek,Cenpk,Psmc6,Pbk,Rad21,Atf1,Slc11a2,Gzma,Gcm1,Phf10,Tapbp,Psmb8,Incenp,Prdx5,Smc3,Cdk18,Dtl,Nup35,Efna1,Usp1,Ssbp3,Sfpq,Oasl1,Oasl2,Iigp1,Tyrobp,Serpina7,Fgl1,Ddx25,Phip,Tipin,Me1,Csrnp1,Mvk,Hspb8,Slx4,Atg14,Hace1,Serpine1,Cblc,Tiparp,Rbm12,Zfp367,Hic1,Atxn2,E2f8,Cenpe,Zfpm1,S100a1,Whsc1,Ercc8,Camk2n1,Hilpda,Tob2,Insig1,Nfil3,Azin1,Hmgb2,Phf6,Gspt1,Birc5,Oas1a,Slc39a8,Rps6ka6,Isg15,Fmr1,Hbb-bt,Arrb1,Cox7a1,Mbnl1,Mad2l1,Flcn,Anp32b,Pdpk1,Sirt1,Zbtb8b,Txnip,Crebzf,Ect2,Irf1,Map4k5,Hnrnpa3,Bclaf3,Arid5a,Dnm1l,Ube2v2,Bcl3,Srsf3,Irf9,Srsf1,Nmi,Prlr,Ppp1r3g,Cep295,Camsap3,Lsm5,Tfdp2,Odc1,Tcea1,Parp10,Sh3glb1,Tgif1,Psmb9,Zfp131 |

| | | | | | | | |
|---|---|---|---|---|---|---|---|
| GO:0060255 | Regulation of macromolecule metabolic process | 107 | 5618 | 0.24 | 3.81E-07 | 10090.ENSMUSP00000002284,10090.ENSMUSP00000002413, 10090.ENSMUSP00000004985,10090.ENSMUSP00000005503, 10090.ENSMUSP00000018361,10090.ENSMUSP00000019469, 10090.ENSMUSP00000020123,10090.ENSMUSP00000020308, 10090.ENSMUSP00000020835,10090.ENSMUSP00000020908, 10090.ENSMUSP00000021807,10090.ENSMUSP00000022227, 10090.ENSMUSP00000022380,10090.ENSMUSP00000022612, 10090.ENSMUSP00000022927,10090.ENSMUSP00000023769, 10090.ENSMUSP00000023774,10090.ENSMUSP00000023897, 10090.ENSMUSP00000024104,10090.ENSMUSP00000024657, 10090.ENSMUSP00000025161,10090.ENSMUSP00000025196, 10090.ENSMUSP00000025562,10090.ENSMUSP00000025904, 10090.ENSMUSP00000025930,10090.ENSMUSP00000027697, 10090.ENSMUSP00000027933,10090.ENSMUSP00000028382, 10090.ENSMUSP00000029566,10090.ENSMUSP00000030289, 10090.ENSMUSP00000030367,10090.ENSMUSP00000030623, 10090.ENSMUSP00000031540,10090.ENSMUSP00000031542, 10090.ENSMUSP00000032800,10090.ENSMUSP00000033626, 10090.ENSMUSP00000034612,10090.ENSMUSP00000034787, 10090.ENSMUSP00000034964,10090.ENSMUSP00000035101, 10090.ENSMUSP00000036971,10090.ENSMUSP00000038871, 10090.ENSMUSP00000039047,10090.ENSMUSP00000039206, 10090.ENSMUSP00000039586,10090.ENSMUSP00000039955, 10090.ENSMUSP00000048051,10090.ENSMUSP00000050461, 10090.ENSMUSP00000050854,10090.ENSMUSP00000053483, 10090.ENSMUSP00000056715,10090.ENSMUSP00000056778, 10090.ENSMUSP00000057938,10090.ENSMUSP00000058037, | Plaur,Tmem161a,Brpf3,Msh6,Pmp22,G6pc,Tmpo,Ddit4,Ppm1d,E2f6,Dek,Cenpk,Psmc6,Pbk,Rad21,Atf1,Slc11a2,Gzma,Gcm1,Phf10,Tapbp,Psmb8,Incenp,Prdx5,Smc3,Cdk18,Dtl,Nup35,Efna1,Usp1,Ssbp3,Sfpq,Oasl1,Oasl2,Tyrobp,Serpina7,Ddx25,Phip,Tipin,Csrnp1,Mvk,Slx4,Atg14,Hace1,Serpine1,Cblc,Tiparp,Rbm12,Zfp367,Hic1,Atxn2,E2f8,Cenpe,Zfpm1,S100a1,Whsc1,Ercc8,Camk2n1,Hilpda,Tob2,Insig1,Nfil3,Azin1,Hmgb2,Phf6,Gspt1,Birc5,Oas1a,Slc39a8,Rps6ka6,Isg15,Fmr1,Hbb-bt,Arrb1,Mbnl1,Mad2l1,Flcn,Anp32b,Pdpk1,Sirt1,Zbtb8b,Txnip,Crebzf,Ect2,Irf1,Map4k5,Hnrnpa3,Bclaf3,Arid5a,Dnm1l,Ube2v2,Bcl3,Srsf3,Irf9,Srsf1,Nmi,Prlr,Ppp1r3g,Cep295,Lsm5,Tfdp2,Odc1,Tcea1,Parp10,Tgif1,Psmb9,Zfp131 |
| GO:0031323 | Regulation of cellular metabolic process | 103 | 5479 | 0.24 | 1.94E-06 | 10090.ENSMUSP00000002284,10090.ENSMUSP00000002413, 10090.ENSMUSP00000004430,10090.ENSMUSP00000004985, 10090.ENSMUSP00000005503,10090.ENSMUSP00000018361, 10090.ENSMUSP00000020123,10090.ENSMUSP00000020308, 10090.ENSMUSP00000020835,10090.ENSMUSP00000020908, 10090.ENSMUSP00000021807,10090.ENSMUSP00000022227, 10090.ENSMUSP00000022380,10090.ENSMUSP00000022612, 10090.ENSMUSP00000022927,10090.ENSMUSP00000023769, 10090.ENSMUSP00000023774,10090.ENSMUSP00000023897, 10090.ENSMUSP00000024104,10090.ENSMUSP00000024657, 10090.ENSMUSP00000025562,10090.ENSMUSP00000025904, 10090.ENSMUSP00000025930,10090.ENSMUSP00000027697, 10090.ENSMUSP00000028382,10090.ENSMUSP00000029566, 10090.ENSMUSP00000030289,10090.ENSMUSP00000030367, 10090.ENSMUSP00000030623,10090.ENSMUSP00000031540, 10090.ENSMUSP00000031542,10090.ENSMUSP00000032473, 10090.ENSMUSP00000032800,10090.ENSMUSP00000033626, 10090.ENSMUSP00000034003,10090.ENSMUSP00000034612, 10090.ENSMUSP00000034787,10090.ENSMUSP00000034964, 10090.ENSMUSP00000035101,10090.ENSMUSP00000036971, 10090.ENSMUSP00000037007,10090.ENSMUSP00000038871, 10090.ENSMUSP00000039047,10090.ENSMUSP00000039586, 10090.ENSMUSP00000039955,10090.ENSMUSP00000050461, 10090.ENSMUSP00000050854,10090.ENSMUSP00000053483, 10090.ENSMUSP00000056715,10090.ENSMUSP00000056778, 10090.ENSMUSP00000057938,10090.ENSMUSP00000058037, 10090.ENSMUSP00000058237,10090.ENSMUSP00000058940, | Plaur,Tmem161a,Clcn3,Brpf3,Msh6,Pmp22,Tmpo,Ddit4,Ppm1d,E2f6,Dek,Cenpk,Psmc6,Pbk,Rad21,Atf1,Slc11a2,Gzma,Gcm1,Phf10,Incenp,Prdx5,Smc3,Cdk18,Nup35,Efna1,Usp1,Ssbp3,Sfpq,Oasl1,Oasl2,Iigp1,Tyrobp,Serpina7,Fgl1,Ddx25,Phip,Tipin,Csrnp1,Mvk,Hspb8,Slx4,Atg14,Serpine1,Cblc,Rbm12,Zfp367,Hic1,Atxn2,E2f8,Cenpe,Zfpm1,S100a1,Whsc1,Ercc8,Camk2n1,Tob2,Insig1,Nfil3,Hmgb2,Phf6,Gspt1,Birc5,Oas1a,Slc39a8,Rps6ka6,Isg15,Fmr1,Hbb-bt,Arrb1,Cox7a1,Mbnl1,Mad2l1,Flcn,Anp32b,Pdpk1,Sirt1,Zbtb8b,Txnip,Crebzf,Ect2,Irf1,Map4k5,Hnrnpa3,Bclaf3,Arid5a,Dnm1l,Ube2v2,Bcl3,Srsf3,Irf9,Srsf1,Nmi,Prlr,Ppp1r3g,Cep295,Camsap3,Tfdp2,Tcea1,Parp10,Sh3glb1,Tgif1,Zfp131 |
| GO:0050794 | Regulation of cellular process | 151 | 9541 | 0.16 | 1.94E-06 | 10090.ENSMUSP00000002284,10090.ENSMUSP00000002413, 10090.ENSMUSP00000004120,10090.ENSMUSP00000004430, 10090.ENSMUSP00000004985,10090.ENSMUSP00000005503, 10090.ENSMUSP00000015581,10090.ENSMUSP00000017851, 10090.ENSMUSP00000018361,10090.ENSMUSP00000018506, 10090.ENSMUSP00000020123,10090.ENSMUSP00000020308, 10090.ENSMUSP00000020835,10090.ENSMUSP00000020908, 10090.ENSMUSP00000021807,10090.ENSMUSP00000022227, 10090.ENSMUSP00000022380,10090.ENSMUSP00000022612, 10090.ENSMUSP00000022927,10090.ENSMUSP00000023769, 10090.ENSMUSP00000023774,10090.ENSMUSP00000023897, 10090.ENSMUSP00000024104,10090.ENSMUSP00000024657, 10090.ENSMUSP00000025254,10090.ENSMUSP00000025562, 10090.ENSMUSP00000025904,10090.ENSMUSP00000025930, 10090.ENSMUSP00000026565,10090.ENSMUSP00000027697, 10090.ENSMUSP00000027760,10090.ENSMUSP00000027809, 10090.ENSMUSP00000027933,10090.ENSMUSP00000028328, 10090.ENSMUSP00000028382,10090.ENSMUSP00000029018, 10090.ENSMUSP00000029566,10090.ENSMUSP00000030121, 10090.ENSMUSP00000030289,10090.ENSMUSP00000030367, 10090.ENSMUSP00000030623,10090.ENSMUSP00000031540, 10090.ENSMUSP00000031542,10090.ENSMUSP00000032473, 10090.ENSMUSP00000032800,10090.ENSMUSP00000032898, 10090.ENSMUSP00000033626,10090.ENSMUSP00000034003, 10090.ENSMUSP00000034612,10090.ENSMUSP00000034787, 10090.ENSMUSP00000034964,10090.ENSMUSP00000035101, 10090.ENSMUSP00000036971,10090.ENSMUSP00000037007, | Plaur,Tmem161a,Fancl,Clcn3,Brpf3,Msh6,Gzmb,Serinc3,Pmp22,Kpna2,Tmpo,Ddit4,Ppm1d,E2f6,Dek,Cenpk,Psmc6,Pbk,Rad21,Atf1,Slc11a2,Gzma,Gcm1,Phf10,Lims2,Incenp,Prdx5,Smc3,Ifitm3,Cdk18,Rgl1,Opn3,Dtl,Entpd2,Nup35,Zbp1,Efna1,B4galt1,Usp1,Ssbp3,Sfpq,Oasl1,Oasl2,Iigp1,Tyrobp,Ipo5,Serpina7,Fgl1,Ddx25,Phip,Tipin,Csrnp1,Mvk,Hspb8,Zfand1,Slx4,Atg14,Hace1,Serpine1,Cblc,Ofd1,Pim3,Card19,Tiparp,Rbm12,Zfp367,Hic1,Atxn2,E2f8,Cenpe,Zfpm1,S100a1,Whsc1,Aspm,Ercc8,Camk2n1,Hilpda,Tob2,Insig1,Adam10,Artn,Nfil3,Azin1,Hmgb2,Ly6c1,Cdc42ep3,Ifnlr1,Frmd4a,Tnfaip8l1,Dchs1,Phf6,Gspt1,Krit1,Birc5,Oas1a,Slc39a8,Rps6ka6,Isg15,Kif20b,Fmr1,Magi1,Rnf213,Hbb-bt,Arrb1,Cox7a1,Mbnl1,Mad2l1,Atp11c,Flcn,Ifit3,Anp32b,Pdpk1,Sirt1,Zbtb8b,Txnip,Crebzf,Trim59,Ect2,Irf1,Map4k5,Hnrnpa3,Bclaf3,Cnot4,Arid5a,Dnm1l,Ube2v2,Rnf186,Arhgap23,Rsl1d1,Bcl3,Srsf3,Irf9,Srsf1,Nmi,Prlr,Ppp1r3g,Xaf1,Cep295,Camsap3,Tgm1,Tfdp2,Odc1,Tcea1,Parp10,Sh3glb1,Zfyve27,Tgif1,Arhgap21,H2-D1,Golph3l,Zfp131 |
| GO:0080090 | Regulation of primary metabolic process | 100 | 5290 | 0.24 | 2.46E-06 | 10090.ENSMUSP00000002284,10090.ENSMUSP00000002413, 10090.ENSMUSP00000005503,10090.ENSMUSP00000018361, 10090.ENSMUSP00000020123,10090.ENSMUSP00000020308, 10090.ENSMUSP00000020835,10090.ENSMUSP00000020908, 10090.ENSMUSP00000021807,10090.ENSMUSP00000022227, 10090.ENSMUSP00000022380,10090.ENSMUSP00000022612, 10090.ENSMUSP00000022927,10090.ENSMUSP00000023769, 10090.ENSMUSP00000023774,10090.ENSMUSP00000023897, 10090.ENSMUSP00000024104,10090.ENSMUSP00000024657, 10090.ENSMUSP00000025196,10090.ENSMUSP00000025562, 10090.ENSMUSP00000025904,10090.ENSMUSP00000027697, 10090.ENSMUSP00000027933,10090.ENSMUSP00000028382, 10090.ENSMUSP00000029566,10090.ENSMUSP00000030289, 10090.ENSMUSP00000030367,10090.ENSMUSP00000030623, 10090.ENSMUSP00000031540,10090.ENSMUSP00000031542, 10090.ENSMUSP00000033626,10090.ENSMUSP00000034003, 10090.ENSMUSP00000034612,10090.ENSMUSP00000034787, 10090.ENSMUSP00000035101,10090.ENSMUSP00000036971, 10090.ENSMUSP00000038871,10090.ENSMUSP00000039047, 10090.ENSMUSP00000039206,10090.ENSMUSP00000039586, 10090.ENSMUSP00000039955,10090.ENSMUSP00000048051, 10090.ENSMUSP00000050461,10090.ENSMUSP00000050854, 10090.ENSMUSP00000053483,10090.ENSMUSP00000056778, 10090.ENSMUSP00000057938,10090.ENSMUSP00000058037, 10090.ENSMUSP00000058237,10090.ENSMUSP00000058940, 10090.ENSMUSP00000059211,10090.ENSMUSP00000060349, 10090.ENSMUSP00000061390,10090.ENSMUSP00000061877, | Plaur,Tmem161a,Msh6,Pmp22,Tmpo,Ddit4,Ppm1d,E2f6,Dek,Cenpk,Psmc6,Pbk,Rad21,Atf1,Slc11a2,Gzma,Gcm1,Phf10,Psmb8,Incenp,Prdx5,Cdk18,Dtl,Nup35,Efna1,Usp1,Ssbp3,Sfpq,Oasl1,Oasl2,Serpina7,Fgl1,Ddx25,Phip,Csrnp1,Mvk,Slx4,Atg14,Hace1,Serpine1,Cblc,Tiparp,Rbm12,Zfp367,Hic1,E2f8,Cenpe,Zfpm1,S100a1,Whsc1,Ercc8,Camk2n1,Tob2,Insig1,Nfil3,Azin1,Hmgb2,Phf6,Gspt1,Birc5,Oas1a,Slc39a8,Rps6ka6,Isg15,Fmr1,Hbb-bt,Arrb1,Mbnl1,Mad2l1,Flcn,Anp32b,Pdpk1,Sirt1,Zbtb8b,Txnip,Crebzf,Ect2,Irf1,Map4k5,Hnrnpa3,Bclaf3,Arid5a,Dnm1l,Ube2v2,Bcl3,Srsf3,Irf9,Srsf1,Nmi,Prlr,Ppp1r3g,Cep295,Tfdp2,Odc1,Tcea1,Parp10,Sh3glb1,Tgif1,Psmb9,Zfp131 |

| | | | | | | | |
|---|---|---|---|---|---|---|---|
| GO:0009615 | Response to virus | 17 | 235 | 0.82 | 3.28E-06 | 10090.ENSMUSP00000017851,10090.ENSMUSP00000020308, 10090.ENSMUSP00000026565,10090.ENSMUSP00000029018, 10090.ENSMUSP00000031540,10090.ENSMUSP00000031542, 10090.ENSMUSP00000033173,10090.ENSMUSP00000041091, 10090.ENSMUSP00000074009,10090.ENSMUSP00000079198, 10090.ENSMUSP00000082548,10090.ENSMUSP00000085906, 10090.ENSMUSP00000099889,10090.ENSMUSP00000102824, 10090.ENSMUSP00000104548,10090.ENSMUSP00000113851, 10090.ENSMUSP00000128661 | Serinc3,Ddit4,Ifitm3,Zbp1,Oasl1,Oasl2,Polr3e,Rtp4,Ifnlr1,Oas1a,Isg15,Fmr1,Ifit3,Crebzf,Irf1,Bcl3,Odc1 |
| GO:0051171 | Regulation of nitrogen compound metabolic process | 97 | 5126 | 0.24 | 4.02E-06 | 10090.ENSMUSP00000002284,10090.ENSMUSP00000002413, 10090.ENSMUSP00000005503,10090.ENSMUSP00000018361, 10090.ENSMUSP00000020123,10090.ENSMUSP00000020308, 10090.ENSMUSP00000020835,10090.ENSMUSP00000020908, 10090.ENSMUSP00000021807,10090.ENSMUSP00000022227, 10090.ENSMUSP00000022380,10090.ENSMUSP00000022612, 10090.ENSMUSP00000022927,10090.ENSMUSP00000023769, 10090.ENSMUSP00000023774,10090.ENSMUSP00000023897, 10090.ENSMUSP00000024104,10090.ENSMUSP00000024657, 10090.ENSMUSP00000025196,10090.ENSMUSP00000025562, 10090.ENSMUSP00000025904,10090.ENSMUSP00000027697, 10090.ENSMUSP00000027933,10090.ENSMUSP00000028382, 10090.ENSMUSP00000029566,10090.ENSMUSP00000030289, 10090.ENSMUSP00000030367,10090.ENSMUSP00000030623, 10090.ENSMUSP00000031540,10090.ENSMUSP00000031542, 10090.ENSMUSP00000033626,10090.ENSMUSP00000034612, 10090.ENSMUSP00000034787,10090.ENSMUSP00000035101, 10090.ENSMUSP00000036971,10090.ENSMUSP00000038871, 10090.ENSMUSP00000039047,10090.ENSMUSP00000039206, 10090.ENSMUSP00000039586,10090.ENSMUSP00000039955, 10090.ENSMUSP00000048051,10090.ENSMUSP00000050461, 10090.ENSMUSP00000050854,10090.ENSMUSP00000053483, 10090.ENSMUSP00000056778,10090.ENSMUSP00000057938, 10090.ENSMUSP00000058037,10090.ENSMUSP00000058237, 10090.ENSMUSP00000058940,10090.ENSMUSP00000059211, 10090.ENSMUSP00000060349,10090.ENSMUSP00000061390, 10090.ENSMUSP00000061877,10090.ENSMUSP00000065363, [illegible] | Plaur,Tmem161a,Msh6,Pmp22,Tmpo,Ddit4,Ppm1d,E2f6,Dek,Cenpk,Psmc6,Pbk,Rad21,Atf1,Slc11a2,Gzma,Gcm1,Phf10,Psmb8,Incenp,Prdx5,Cdk18,Dtl,Nup35,Efna1,Usp1,Ssbp3,Sfpq,Oasl1,Oasl2,Serpina7,Ddx25,Phip,Csrnp1,Mvk,Slx4,Atg14,Hace1,Serpine1,Cblc,Tiparp,Rbm12,Zfp367,Hic1,E2f8,Cenpe,Zfpm1,S100a1,Whsc1,Ercc8,Camk2n1,Tob2,Insig1,Nfil3,Azin1,Hmgb2,Phf6,Gspt1,Birc5,Oas1a,Slc39a8,Rps6ka6,Isg15,Fmr1,Hbb-bt,Arrb1,Mbnl1,Mad2l1,Flcn,Anp32b,Pdpk1,Sirt1,Zbtb8b,Txnip,Crebzf,Ect2,Irf1,Map4k5,Hnrnpa3,Bclaf3,Arid5a,Dnm1l,Ube2v2,Bcl3,Srsf3,Irf9,Srsf1,Nmi,Prlr,Cep295,Tfdp2,Odc1,Tcea1,Parp10,Tgif1,Psmb9,Zfp131 |
| GO:0044237 | Cellular metabolic process | 111 | 6445 | 0.2 | 2.70E-05 | 10090.ENSMUSP00000002198,10090.ENSMUSP00000004120, 10090.ENSMUSP00000004985,10090.ENSMUSP00000005503, 10090.ENSMUSP00000011058,10090.ENSMUSP00000017851, 10090.ENSMUSP00000018361,10090.ENSMUSP00000019469, 10090.ENSMUSP00000020217,10090.ENSMUSP00000020308, 10090.ENSMUSP00000020835,10090.ENSMUSP00000021120, 10090.ENSMUSP00000021381,10090.ENSMUSP00000022380, 10090.ENSMUSP00000022612,10090.ENSMUSP00000022927, 10090.ENSMUSP00000023353,10090.ENSMUSP00000023774, 10090.ENSMUSP00000023897,10090.ENSMUSP00000024104, 10090.ENSMUSP00000024657,10090.ENSMUSP00000025161, 10090.ENSMUSP00000025196,10090.ENSMUSP00000025562, 10090.ENSMUSP00000025632,10090.ENSMUSP00000025904, 10090.ENSMUSP00000025930,10090.ENSMUSP00000026823, 10090.ENSMUSP00000027623,10090.ENSMUSP00000027697, 10090.ENSMUSP00000027809,10090.ENSMUSP00000027933, 10090.ENSMUSP00000028328,10090.ENSMUSP00000028398, 10090.ENSMUSP00000030121,10090.ENSMUSP00000030289, 10090.ENSMUSP00000030623,10090.ENSMUSP00000030651, 10090.ENSMUSP00000031542,10090.ENSMUSP00000032198, 10090.ENSMUSP00000033173,10090.ENSMUSP00000034270, 10090.ENSMUSP00000034964,10090.ENSMUSP00000034989, 10090.ENSMUSP00000036971,10090.ENSMUSP00000038871, 10090.ENSMUSP00000039010,10090.ENSMUSP00000039047, 10090.ENSMUSP00000039206,10090.ENSMUSP00000039955, 10090.ENSMUSP00000040220,10090.ENSMUSP00000041906, 10090.ENSMUSP00000044603,10090.ENSMUSP00000047872, [illegible] | Sf3a1,Fancl,Brpf3,Msh6,Pdhx,Serinc3,Pmp22,G6pc,Nudt4,Ddit4,Ppm1d,Trim47,Pnn,Psmc6,Pbk,Rad21,Mcm4,Slc11a2,Gzma,Gcm1,Phf10,Tapbp,Psmb8,Incenp,Carnmt1,Prdx5,Smc3,Pigq,Tsn,Cdk18,Opn3,Dtl,Entpd2,Ube2e3,B4galt1,Usp1,Sfpq,Sh3bgrl3,Oasl2,Usp18,Polr3e,Map1lc3b,Tipin,Me1,Mvk,Slx4,Wdr55,Atg14,Hace1,Cblc,Ubac1,Dis3,Pim3,Smc4,Tiparp,Cbr1,Mettl5,Polr2k,Whsc1,Ercc8,Insig1,Adam10,Nfil3,Azin1,Hmgb2,Mrpl18,Prpf38a,Gspt1,Birc5,Oas1a,Amd2,Slc39a8,Rps6ka6,Mdc1,Prpf4,Nhlrc4,Isg15,Fmr1,Mfap1a,Rnf213,Hbb-bt,Arrb1,Cox7a1,Mbnl1,Pdpk1,Mcts2,Sirt1,Crebzf,Trim59,Map4k5,Hnrnpa3,Ppip5k2,Nmnat3,Cnot4,Ube2v2,Rnf186,Rsl1d1,Bcl3,Srsf3,Srsf1,Nmi,Prlr,Lsm5,Tgm1,Odc1,Tcea1,Parp10,Sh3glb1,Ube2d2a,Psmb9,Metap2 |
| GO:0006950 | Response to stress | 66 | 3045 | 0.3 | 2.77E-05 | 10090.ENSMUSP00000002413,10090.ENSMUSP00000004120, 10090.ENSMUSP00000005503,10090.ENSMUSP00000017851, 10090.ENSMUSP00000018361,10090.ENSMUSP00000020308, 10090.ENSMUSP00000020835,10090.ENSMUSP00000022380, 10090.ENSMUSP00000022927,10090.ENSMUSP00000023353, 10090.ENSMUSP00000023774,10090.ENSMUSP00000025161, 10090.ENSMUSP00000025904,10090.ENSMUSP00000025930, 10090.ENSMUSP00000026565,10090.ENSMUSP00000027933, 10090.ENSMUSP00000028328,10090.ENSMUSP00000029018, 10090.ENSMUSP00000030121,10090.ENSMUSP00000030289, 10090.ENSMUSP00000030623,10090.ENSMUSP00000031540, 10090.ENSMUSP00000031542,10090.ENSMUSP00000032473, 10090.ENSMUSP00000032800,10090.ENSMUSP00000033173, 10090.ENSMUSP00000034138,10090.ENSMUSP00000034270, 10090.ENSMUSP00000034964,10090.ENSMUSP00000037007, 10090.ENSMUSP00000038871,10090.ENSMUSP00000039047, 10090.ENSMUSP00000039586,10090.ENSMUSP00000040246, 10090.ENSMUSP00000041091,10090.ENSMUSP00000047872, 10090.ENSMUSP00000053483,10090.ENSMUSP00000059211, 10090.ENSMUSP00000060791,10090.ENSMUSP00000061877, 10090.ENSMUSP00000065940,10090.ENSMUSP00000074009, 10090.ENSMUSP00000079198,10090.ENSMUSP00000080694, 10090.ENSMUSP00000080949,10090.ENSMUSP00000082548, 10090.ENSMUSP00000085906,10090.ENSMUSP00000099758, 10090.ENSMUSP00000099889,10090.ENSMUSP00000099991, 10090.ENSMUSP00000101082,10090.ENSMUSP00000102710, 10090.ENSMUSP00000103432,10090.ENSMUSP00000103935, | Tmem161a,Fancl,Msh6,Serinc3,Pmp22,Ddit4,Ppm1d,Psmc6,Rad21,Mcm4,Slc11a2,Tapbp,Prdx5,Smc3,Ifitm3,Dtl,Entpd2,Zbp1,B4galt1,Usp1,Sfpq,Oasl1,Oasl2,Iigp1,Tyrobp,Polr3e,Dnaja2,Map1lc3b,Tipin,Hspb8,Slx4,Atg14,Serpine1,C1qb,Rtp4,Smc4,Hic1,Ercc8,Hilpda,Insig1,Hmgb2,Ifnlr1,Oas1a,Rps6ka6,Mdc1,Isg15,Fmr1,Flcn,Ifit3,Pdpk1,Sirt1,Txnip,Trim59,Ect2,Irf1,Map4k5,Fgfr1op2,Nmnat3,Arid5a,Dnm1l,Ube2v2,Rnf186,Bcl3,Nmi,Parp10,Sh3glb1 |

| | | | | | | | |
|---|---|---|---|---|---|---|---|
| GO:0006139 | Nucleobase-containing compound metabolic process | 53 | 2205 | 0.34 | 3.68E-05 | 10090.ENSMUSP00000002198,10090.ENSMUSP00000004120,10090.ENSMUSP00000005503,10090.ENSMUSP00000011058,10090.ENSMUSP00000020217,10090.ENSMUSP00000020835,10090.ENSMUSP00000021381,10090.ENSMUSP00000022927,10090.ENSMUSP00000023353,10090.ENSMUSP00000024104,10090.ENSMUSP00000025930,10090.ENSMUSP00000027623,10090.ENSMUSP00000027933,10090.ENSMUSP00000028328,10090.ENSMUSP00000030289,10090.ENSMUSP00000030623,10090.ENSMUSP00000031542,10090.ENSMUSP00000033173,10090.ENSMUSP00000034964,10090.ENSMUSP00000036971,10090.ENSMUSP00000038871,10090.ENSMUSP00000039010,10090.ENSMUSP00000041906,10090.ENSMUSP00000047872,10090.ENSMUSP00000050907,10090.ENSMUSP00000051968,10090.ENSMUSP00000059211,10090.ENSMUSP00000065363,10090.ENSMUSP00000065940,10090.ENSMUSP00000078207,10090.ENSMUSP00000078940,10090.ENSMUSP00000079198,10090.ENSMUSP00000079703,10090.ENSMUSP00000080640,10090.ENSMUSP00000080949,10090.ENSMUSP00000081572,10090.ENSMUSP00000085906,10090.ENSMUSP00000087372,10090.ENSMUSP00000095866,10090.ENSMUSP00000096686,10090.ENSMUSP00000101082,10090.ENSMUSP00000102824,10090.ENSMUSP00000107595,10090.ENSMUSP00000108557,10090.ENSMUSP00000111443,10090.ENSMUSP00000113431,10090.ENSMUSP00000113851,10090.ENSMUSP00000117045,10090.ENSMUSP00000120595,10090.ENSMUSP00000120647,10090.ENSMUSP00000126565,10090.ENSMUSP00000129157,10090.ENSMUSP00000129765 | Sf3a1,Fancl,Msh6,Pdhx,Nudt4,Ppm1d,Pnn,Rad21,Mcm4,Gcm1,Smc3,Tsn,Dtl,Entpd2,Usp1,Sfpq,Oasl2,Polr3e,Tipin,Mvk,Slx4,Wdr55,Dis3,Smc4,Mettl5,Polr2k,Ercc8,Nfil3,Hmgb2,Prpf38a,Gspt1,Oas1a,Amd2,Slc39a8,Mdc1,Prpf4,Fmr1,Mfap1a,Arrb1,Mbnl1,Sirt1,Crebzf,Hnrnpa3,Nmnat3,Ube2v2,Rsl1d1,Bcl3,Srsf3,Srsf1,Nmi,Lsm5,Tcea1,Parp10 |
| GO:0034641 | Cellular nitrogen compound metabolic process | 62 | 2805 | 0.31 | 3.68E-05 | 10090.ENSMUSP00000002198,10090.ENSMUSP00000004120,10090.ENSMUSP00000005503,10090.ENSMUSP00000011058,10090.ENSMUSP00000019469,10090.ENSMUSP00000020217,10090.ENSMUSP00000020835,10090.ENSMUSP00000021381,10090.ENSMUSP00000022927,10090.ENSMUSP00000023353,10090.ENSMUSP00000023774,10090.ENSMUSP00000024104,10090.ENSMUSP00000025161,10090.ENSMUSP00000025632,10090.ENSMUSP00000025930,10090.ENSMUSP00000027623,10090.ENSMUSP00000027933,10090.ENSMUSP00000028328,10090.ENSMUSP00000030289,10090.ENSMUSP00000030623,10090.ENSMUSP00000031542,10090.ENSMUSP00000033173,10090.ENSMUSP00000034964,10090.ENSMUSP00000036971,10090.ENSMUSP00000038871,10090.ENSMUSP00000039010,10090.ENSMUSP00000041906,10090.ENSMUSP00000047872,10090.ENSMUSP00000050907,10090.ENSMUSP00000051968,10090.ENSMUSP00000059211,10090.ENSMUSP00000063839,10090.ENSMUSP00000065363,10090.ENSMUSP00000065544,10090.ENSMUSP00000065940,10090.ENSMUSP00000078123,10090.ENSMUSP00000078207,10090.ENSMUSP00000078940,10090.ENSMUSP00000079198,10090.ENSMUSP00000079703,10090.ENSMUSP00000080640,10090.ENSMUSP00000080949,10090.ENSMUSP00000081572,10090.ENSMUSP00000085906,10090.ENSMUSP00000087372,10090.ENSMUSP00000095866,10090.ENSMUSP00000096686,10090.ENSMUSP00000100534,10090.ENSMUSP00000101082,10090.ENSMUSP00000102824,10090.ENSMUSP00000107595,10090.ENSMUSP00000108557,10090.ENSMUSP00000111443,10090.ENSMUSP00000113431,[illegible] | Sf3a1,Fancl,Msh6,Pdhx,G6pc,Nudt4,Ppm1d,Pnn,Rad21,Mcm4,Slc11a2,Gcm1,Tapbp,Carnmt1,Smc3,Tsn,Dtl,Entpd2,Usp1,Sfpq,Oasl2,Polr3e,Tipin,Mvk,Slx4,Wdr55,Dis3,Smc4,Mettl5,Polr2k,Ercc8,Adam10,Nfil3,Azin1,Hmgb2,Mrpl18,Prpf38a,Gspt1,Oas1a,Amd2,Slc39a8,Mdc1,Prpf4,Fmr1,Mfap1a,Arrb1,Mbnl1,Mcts2,Sirt1,Crebzf,Hnrnpa3,Nmnat3,Ube2v2,Rsl1d1,Bcl3,Srsf3,Srsf1,Nmi,Lsm5,Odc1,Tcea1,Parp10 |
| GO:0002376 | Immune system process | 47 | 1842 | 0.37 | 3.81E-05 | 10090.ENSMUSP00000003512,10090.ENSMUSP00000005503,10090.ENSMUSP00000015581,10090.ENSMUSP00000017851,10090.ENSMUSP00000018361,10090.ENSMUSP00000020308,10090.ENSMUSP00000023774,10090.ENSMUSP00000025161,10090.ENSMUSP00000025196,10090.ENSMUSP00000026565,10090.ENSMUSP00000029018,10090.ENSMUSP00000029671,10090.ENSMUSP00000030121,10090.ENSMUSP00000030367,10090.ENSMUSP00000030623,10090.ENSMUSP00000031540,10090.ENSMUSP00000031542,10090.ENSMUSP00000032473,10090.ENSMUSP00000032800,10090.ENSMUSP00000033173,10090.ENSMUSP00000034003,10090.ENSMUSP00000040246,10090.ENSMUSP00000041091,10090.ENSMUSP00000048051,10090.ENSMUSP00000048829,10090.ENSMUSP00000058037,10090.ENSMUSP00000063839,10090.ENSMUSP00000064521,10090.ENSMUSP00000065363,10090.ENSMUSP00000065940,10090.ENSMUSP00000074009,10090.ENSMUSP00000079198,10090.ENSMUSP00000082548,10090.ENSMUSP00000095794,10090.ENSMUSP00000099066,10090.ENSMUSP00000099758,10090.ENSMUSP00000099889,10090.ENSMUSP00000101082,10090.ENSMUSP00000103432,10090.ENSMUSP00000104548,10090.ENSMUSP00000110684,10090.ENSMUSP00000113851,10090.ENSMUSP00000120525,10090.ENSMUSP00000120647,10090.ENSMUSP00000129157,10090.ENSMUSP00000133499,10090.ENSMUSP00000134570 | Fcgrt,Msh6,Gzmb,Serinc3,Pmp22,Ddit4,Slc11a2,Tapbp,Psmb8,Ifitm3,Zbp1,Ifi44,B4galt1,Ssbp3,Sfpq,Oasl1,Oasl2,Iigp1,Tyrobp,Polr3e,Fgl1,C1qb,Rtp4,Tiparp,Cdh26,Zfpm1,Adam10,Artn,Nfil3,Hmgb2,Ifnlr1,Oas1a,Isg15,Hbb-bt,Atp11c,Flcn,Ifit3,Sirt1,Trim59,Irf1,Arid5a,Bcl3,Irf9,Nmi,Tcea1,Psmb9,H2-D1 |
| GO:0006807 | Nitrogen compound metabolic process | 103 | 5878 | 0.21 | 3.81E-05 | 10090.ENSMUSP00000002198,10090.ENSMUSP00000004120,10090.ENSMUSP00000004985,10090.ENSMUSP00000005503,10090.ENSMUSP00000011058,10090.ENSMUSP00000015581,10090.ENSMUSP00000017851,10090.ENSMUSP00000018361,10090.ENSMUSP00000019469,10090.ENSMUSP00000020217,10090.ENSMUSP00000020835,10090.ENSMUSP00000021120,10090.ENSMUSP00000021381,10090.ENSMUSP00000022380,10090.ENSMUSP00000022612,10090.ENSMUSP00000022641,10090.ENSMUSP00000022927,10090.ENSMUSP00000023353,10090.ENSMUSP00000023774,10090.ENSMUSP00000023897,10090.ENSMUSP00000024104,10090.ENSMUSP00000024657,10090.ENSMUSP00000025161,10090.ENSMUSP00000025196,10090.ENSMUSP00000025562,10090.ENSMUSP00000025632,10090.ENSMUSP00000025904,10090.ENSMUSP00000025930,10090.ENSMUSP00000026823,10090.ENSMUSP00000027623,10090.ENSMUSP00000027697,10090.ENSMUSP00000027809,10090.ENSMUSP00000027933,10090.ENSMUSP00000028328,10090.ENSMUSP00000028398,10090.ENSMUSP00000030121,10090.ENSMUSP00000030289,10090.ENSMUSP00000030623,10090.ENSMUSP00000031542,10090.ENSMUSP00000032198,10090.ENSMUSP00000033173,10090.ENSMUSP00000034964,10090.ENSMUSP00000036971,10090.ENSMUSP00000038871,10090.ENSMUSP00000039010,10090.ENSMUSP00000039206,10090.ENSMUSP00000039955,10090.ENSMUSP00000040220,10090.ENSMUSP00000041906,10090.ENSMUSP00000044603,10090.ENSMUSP00000047872,10090.ENSMUSP00000048051,10090.ENSMUSP00000050907,10090.ENSMUSP00000051968,[illegible] | Sf3a1,Fancl,Brpf3,Msh6,Pdhx,Gzmb,Serinc3,Pmp22,G6pc,Nudt4,Ppm1d,Trim47,Pnn,Psmc6,Pbk,Adamdec1,Rad21,Mcm4,Slc11a2,Gzma,Gcm1,Phf10,Tapbp,Psmb8,Incenp,Carnmt1,Prdx5,Smc3,Pigq,Tsn,Cdk18,Opn3,Dtl,Entpd2,Ube2e3,B4galt1,Usp1,Sfpq,Oasl2,Usp18,Polr3e,Tipin,Mvk,Slx4,Wdr55,Hace1,Cblc,Ubac1,Dis3,Pim3,Smc4,Tiparp,Mettl5,Polr2k,Whsc1,Ercc8,Adam10,Nfil3,Azin1,Hmgb2,Mrpl18,Prpf38a,Gspt1,Birc5,Oas1a,Amd2,Slc39a8,Rps6ka6,Mdc1,Prpf4,Nhlrc4,Isg15,Fmr1,Mfap1a,Rnf213,Arrb1,Mbnl1,Pdpk1,Mcts2,Sirt1,Crebzf,Cisd3,Trim59,Map4k5,Hnrnpa3,Nmnat3,Cnot4,Ube2v2,Rnf186,Rsl1d1,Bcl3,Srsf3,Srsf1,Nmi,Prlr,Lsm5,Tgm1,Odc1,Tcea1,Parp10,Ube2d2a,Psmb9,Metap2 |

| | | | | | | | |
|---|---|---|---|---|---|---|---|
| GO:0046483 | Heterocycle metabolic process | 55 | 2347 | 0.33 | 3.81E-05 | 10090.ENSMUSP00000002198,10090.ENSMUSP00000004120,10090.ENSMUSP00000005503,10090.ENSMUSP00000011058,10090.ENSMUSP00000019469,10090.ENSMUSP00000020217,10090.ENSMUSP00000020835,10090.ENSMUSP00000021381,10090.ENSMUSP00000022927,10090.ENSMUSP00000023353,10090.ENSMUSP00000023774,10090.ENSMUSP00000024104,10090.ENSMUSP00000025930,10090.ENSMUSP00000027623,10090.ENSMUSP00000027933,10090.ENSMUSP00000028328,10090.ENSMUSP00000030289,10090.ENSMUSP00000030623,10090.ENSMUSP00000031542,10090.ENSMUSP00000033173,10090.ENSMUSP00000034964,10090.ENSMUSP00000036971,10090.ENSMUSP00000038871,10090.ENSMUSP00000039010,10090.ENSMUSP00000041906,10090.ENSMUSP00000047872,10090.ENSMUSP00000050907,10090.ENSMUSP00000051968,10090.ENSMUSP00000059211,10090.ENSMUSP00000065363,10090.ENSMUSP00000065940,10090.ENSMUSP00000078207,10090.ENSMUSP00000078940,10090.ENSMUSP00000079198,10090.ENSMUSP00000079703,10090.ENSMUSP00000080640,10090.ENSMUSP00000080949,10090.ENSMUSP00000081572,10090.ENSMUSP00000085906,10090.ENSMUSP00000087372,10090.ENSMUSP00000095866,10090.ENSMUSP00000096686,10090.ENSMUSP00000101082,10090.ENSMUSP00000102824,10090.ENSMUSP00000107595,10090.ENSMUSP00000108557,10090.ENSMUSP00000111443,10090.ENSMUSP00000113431,10090.ENSMUSP00000113851,10090.ENSMUSP00000117045,10090.ENSMUSP00000120595,10090.ENSMUSP00000120647,10090.ENSMUSP00000126565,10090.ENSMUSP00000129157,[illegible] | Sf3a1,Fancl,Msh6,Pdhx,G6pc,Nudt4,Ppm1d,Pnn,Rad21,Mcm4,Slc11a2,Gcm1,Smc3,Tsn,Dtl,Entpd2,Usp1,Sfpq,Oasl2,Polr3e,Tipin,Mvk,Slx4,Wdr55,Dis3,Smc4,Mettl5,Polr2k,Ercc8,Nfil3,Hmgb2,Prpf38a,Gspt1,Oas1a,Amd2,Slc39a8,Mdc1,Prpf4,Fmr1,Mfap1a,Arrb1,Mbnl1,Sirt1,Crebzf,Hnrnpa3,Nmnat3,Ube2v2,Rsl1d1,Bcl3,Srsf3,Srsf1,Nmi,Lsm5,Tcea1,Parp10 |
| GO:1901360 | Organic cyclic compound metabolic process | 59 | 2614 | 0.32 | 3.81E-05 | 10090.ENSMUSP00000002198,10090.ENSMUSP00000004120,10090.ENSMUSP00000005503,10090.ENSMUSP00000011058,10090.ENSMUSP00000018361,10090.ENSMUSP00000019469,10090.ENSMUSP00000020217,10090.ENSMUSP00000020835,10090.ENSMUSP00000021381,10090.ENSMUSP00000022927,10090.ENSMUSP00000023353,10090.ENSMUSP00000023774,10090.ENSMUSP00000024104,10090.ENSMUSP00000025930,10090.ENSMUSP00000027623,10090.ENSMUSP00000027933,10090.ENSMUSP00000028328,10090.ENSMUSP00000030289,10090.ENSMUSP00000030623,10090.ENSMUSP00000031542,10090.ENSMUSP00000033173,10090.ENSMUSP00000034003,10090.ENSMUSP00000034964,10090.ENSMUSP00000036971,10090.ENSMUSP00000038871,10090.ENSMUSP00000039010,10090.ENSMUSP00000041906,10090.ENSMUSP00000047872,10090.ENSMUSP00000048051,10090.ENSMUSP00000050907,10090.ENSMUSP00000051968,10090.ENSMUSP00000059211,10090.ENSMUSP00000061877,10090.ENSMUSP00000065363,10090.ENSMUSP00000065940,10090.ENSMUSP00000078207,10090.ENSMUSP00000078940,10090.ENSMUSP00000079198,10090.ENSMUSP00000079703,10090.ENSMUSP00000080640,10090.ENSMUSP00000080949,10090.ENSMUSP00000081572,10090.ENSMUSP00000085906,10090.ENSMUSP00000087372,10090.ENSMUSP00000095866,10090.ENSMUSP00000096686,10090.ENSMUSP00000101082,10090.ENSMUSP00000102824,10090.ENSMUSP00000107595,10090.ENSMUSP00000108557,10090.ENSMUSP00000111443,10090.ENSMUSP00000113431,10090.ENSMUSP00000113851,10090.ENSMUSP00000117045,[illegible] | Sf3a1,Fancl,Msh6,Pdhx,Pmp22,G6pc,Nudt4,Ppm1d,Pnn,Rad21,Mcm4,Slc11a2,Gcm1,Smc3,Tsn,Dtl,Entpd2,Usp1,Sfpq,Oasl2,Polr3e,Fgl1,Tipin,Mvk,Slx4,Wdr55,Dis3,Smc4,Tiparp,Mettl5,Polr2k,Ercc8,Insig1,Nfil3,Hmgb2,Prpf38a,Gspt1,Oas1a,Amd2,Slc39a8,Mdc1,Prpf4,Fmr1,Mfap1a,Arrb1,Mbnl1,Sirt1,Crebzf,Hnrnpa3,Nmnat3,Ube2v2,Rsl1d1,Bcl3,Srsf3,Srsf1,Nmi,Lsm5,Tcea1,Parp10 |
| GO:0006725 | Cellular aromatic compound metabolic process | 55 | 2412 | 0.32 | 6.82E-05 | 10090.ENSMUSP00000002198,10090.ENSMUSP00000004120,10090.ENSMUSP00000005503,10090.ENSMUSP00000011058,10090.ENSMUSP00000019469,10090.ENSMUSP00000020217,10090.ENSMUSP00000020835,10090.ENSMUSP00000021381,10090.ENSMUSP00000022927,10090.ENSMUSP00000023353,10090.ENSMUSP00000023774,10090.ENSMUSP00000024104,10090.ENSMUSP00000025930,10090.ENSMUSP00000027623,10090.ENSMUSP00000027933,10090.ENSMUSP00000028328,10090.ENSMUSP00000030289,10090.ENSMUSP00000030623,10090.ENSMUSP00000031542,10090.ENSMUSP00000033173,10090.ENSMUSP00000034964,10090.ENSMUSP00000036971,10090.ENSMUSP00000038871,10090.ENSMUSP00000039010,10090.ENSMUSP00000041906,10090.ENSMUSP00000047872,10090.ENSMUSP00000050907,10090.ENSMUSP00000051968,10090.ENSMUSP00000059211,10090.ENSMUSP00000065363,10090.ENSMUSP00000065940,10090.ENSMUSP00000078207,10090.ENSMUSP00000078940,10090.ENSMUSP00000079198,10090.ENSMUSP00000079703,10090.ENSMUSP00000080640,10090.ENSMUSP00000080949,10090.ENSMUSP00000081572,10090.ENSMUSP00000085906,10090.ENSMUSP00000087372,10090.ENSMUSP00000095866,10090.ENSMUSP00000096686,10090.ENSMUSP00000101082,10090.ENSMUSP00000102824,10090.ENSMUSP00000107595,10090.ENSMUSP00000108557,10090.ENSMUSP00000111443,10090.ENSMUSP00000113431,10090.ENSMUSP00000113851,10090.ENSMUSP00000117045,10090.ENSMUSP00000120595,10090.ENSMUSP00000120647,10090.ENSMUSP00000126565,10090.ENSMUSP00000129157,[illegible] | Sf3a1,Fancl,Msh6,Pdhx,G6pc,Nudt4,Ppm1d,Pnn,Rad21,Mcm4,Slc11a2,Gcm1,Smc3,Tsn,Dtl,Entpd2,Usp1,Sfpq,Oasl2,Polr3e,Tipin,Mvk,Slx4,Wdr55,Dis3,Smc4,Mettl5,Polr2k,Ercc8,Nfil3,Hmgb2,Prpf38a,Gspt1,Oas1a,Amd2,Slc39a8,Mdc1,Prpf4,Fmr1,Mfap1a,Arrb1,Mbnl1,Sirt1,Crebzf,Hnrnpa3,Nmnat3,Ube2v2,Rsl1d1,Bcl3,Srsf3,Srsf1,Nmi,Lsm5,Tcea1,Parp10 |
| GO:0009607 | Response to biotic stimulus | 35 | 1178 | 0.44 | 6.82E-05 | 10090.ENSMUSP00000017851,10090.ENSMUSP00000018361,10090.ENSMUSP00000020308,10090.ENSMUSP00000020835,10090.ENSMUSP00000023897,10090.ENSMUSP00000026565,10090.ENSMUSP00000029018,10090.ENSMUSP00000029671,10090.ENSMUSP00000030623,10090.ENSMUSP00000031540,10090.ENSMUSP00000031542,10090.ENSMUSP00000032198,10090.ENSMUSP00000032473,10090.ENSMUSP00000033173,10090.ENSMUSP00000038871,10090.ENSMUSP00000039586,10090.ENSMUSP00000040246,10090.ENSMUSP00000041091,10090.ENSMUSP00000065940,10090.ENSMUSP00000074009,10090.ENSMUSP00000079198,10090.ENSMUSP00000082548,10090.ENSMUSP00000085906,10090.ENSMUSP00000099889,10090.ENSMUSP00000102710,10090.ENSMUSP00000102824,10090.ENSMUSP00000103432,10090.ENSMUSP00000104548,10090.ENSMUSP00000110684,10090.ENSMUSP00000113851,10090.ENSMUSP00000120647,10090.ENSMUSP00000122219,10090.ENSMUSP00000126565,10090.ENSMUSP00000128661,10090.ENSMUSP00000133499 | Serinc3,Pmp22,Ddit4,Ppm1d,Gzma,Ifitm3,Zbp1,Ifi44,Sfpq,Oasl1,Oasl2,Usp18,Iigp1,Polr3e,Slx4,Serpine1,C1qb,Rtp4,Hmgb2,Ifnlr1,Oas1a,Isg15,Fmr1,Ifit3,Txnip,Crebzf,Trim59,Irf1,Arid5a,Bcl3,Nmi,Prlr,Lsm5,Odc1,Psmb9 |
| GO:0051607 | Defense response to virus | 13 | 170 | 0.85 | 6.82E-05 | 10090.ENSMUSP00000017851,10090.ENSMUSP00000020308,10090.ENSMUSP00000026565,10090.ENSMUSP00000029018,10090.ENSMUSP00000031540,10090.ENSMUSP00000031542,10090.ENSMUSP00000033173,10090.ENSMUSP00000041091,10090.ENSMUSP00000074009,10090.ENSMUSP00000079198,10090.ENSMUSP00000082548,10090.ENSMUSP00000099889,10090.ENSMUSP00000104548 | Serinc3,Ddit4,Ifitm3,Zbp1,Oasl1,Oasl2,Polr3e,Rtp4,Ifnlr1,Oas1a,Isg15,Ifit3,Irf1 |

| | | | | | | | |
|---|---|---|---|---|---|---|---|
| GO:0090304 | Nucleic acid metabolic process | 45 | 1777 | 0.37 | 6.97E-05 | 10090.ENSMUSP00000002198,10090.ENSMUSP00000004120,10090.ENSMUSP00000005503,10090.ENSMUSP00000020835,10090.ENSMUSP00000021381,10090.ENSMUSP00000022927,10090.ENSMUSP00000023353,10090.ENSMUSP00000024104,10090.ENSMUSP00000025930,10090.ENSMUSP00000027623,10090.ENSMUSP00000027933,10090.ENSMUSP00000030289,10090.ENSMUSP00000030623,10090.ENSMUSP00000033173,10090.ENSMUSP00000034964,10090.ENSMUSP00000038871,10090.ENSMUSP00000039010,10090.ENSMUSP00000041906,10090.ENSMUSP00000047872,10090.ENSMUSP00000050907,10090.ENSMUSP00000051968,10090.ENSMUSP00000059211,10090.ENSMUSP00000065363,10090.ENSMUSP00000065940,10090.ENSMUSP00000078207,10090.ENSMUSP00000078940,10090.ENSMUSP00000080640,10090.ENSMUSP00000080949,10090.ENSMUSP00000081572,10090.ENSMUSP00000085906,10090.ENSMUSP00000087372,10090.ENSMUSP00000095866,10090.ENSMUSP00000096686,10090.ENSMUSP00000101082,10090.ENSMUSP00000102824,10090.ENSMUSP00000107595,10090.ENSMUSP00000111443,10090.ENSMUSP00000113431,10090.ENSMUSP00000113851,10090.ENSMUSP00000117045,10090.ENSMUSP00000120595,10090.ENSMUSP00000120647,10090.ENSMUSP00000126565,10090.ENSMUSP00000129157,10090.ENSMUSP00000129765 | Sf3a1,Fancl,Msh6,Ppm1d,Pnn,Rad21,Mcm4,Gcm1,Smc3,Tsn,Dtl,Usp1,Sfpq,Polr3e,Tipin,Slx4,Wdr55,Dis3,Smc4,Mettl5,Polr2k,Ercc8,Nfil3,Hmgb2,Prpf38a,Gspt1,Slc39a8,Mdc1,Prpf4,Fmr1,Mfap1a,Arrb1,Mbnl1,Sirt1,Crebzf,Hnrnpa3,Ube2v2,Rsl1d1,Bcl3,Srsf3,Srsf1,Nmi,Lsm5,Tcea1,Parp10 |
| GO:0008152 | Metabolic process | 119 | 7331 | 0.17 | 8.36E-05 | 10090.ENSMUSP00000002198,10090.ENSMUSP00000004120,10090.ENSMUSP00000004985,10090.ENSMUSP00000005503,10090.ENSMUSP00000011058,10090.ENSMUSP00000015581,10090.ENSMUSP00000017851,10090.ENSMUSP00000018361,10090.ENSMUSP00000019469,10090.ENSMUSP00000020217,10090.ENSMUSP00000020308,10090.ENSMUSP00000020835,10090.ENSMUSP00000021120,10090.ENSMUSP00000021381,10090.ENSMUSP00000022380,10090.ENSMUSP00000022612,10090.ENSMUSP00000022641,10090.ENSMUSP00000022927,10090.ENSMUSP00000023353,10090.ENSMUSP00000023774,10090.ENSMUSP00000023897,10090.ENSMUSP00000024104,10090.ENSMUSP00000024657,10090.ENSMUSP00000025161,10090.ENSMUSP00000025196,10090.ENSMUSP00000025562,10090.ENSMUSP00000025632,10090.ENSMUSP00000025904,10090.ENSMUSP00000025930,10090.ENSMUSP00000026823,10090.ENSMUSP00000027623,10090.ENSMUSP00000027697,10090.ENSMUSP00000027809,10090.ENSMUSP00000027933,10090.ENSMUSP00000028328,10090.ENSMUSP00000028398,10090.ENSMUSP00000030121,10090.ENSMUSP00000030289,10090.ENSMUSP00000030623,10090.ENSMUSP00000030651,10090.ENSMUSP00000031542,10090.ENSMUSP00000032198,10090.ENSMUSP00000033173,10090.ENSMUSP00000033913,10090.ENSMUSP00000034003,10090.ENSMUSP00000034270,10090.ENSMUSP00000034612,10090.ENSMUSP00000034964,10090.ENSMUSP00000034989,10090.ENSMUSP00000036971,10090.ENSMUSP00000038871,10090.ENSMUSP00000039010,10090.ENSMUSP00000039047,10090.ENSMUSP00000039206, [illegible] | Sf3a1,Fancl,Brpf3,Msh6,Pdhx,Gzmb,Serinc3,Pmp22,G6pc,Nudt4,Ddit4,Ppm1d,Trim47,Pnn,Psmc6,Pbk,Adamdec1,Rad21,Mcm4,Slc11a2,Gzma,Gcm1,Phf10,Tapbp,Psmb8,Incenp,Carnmt1,Prdx5,Smc3,Pigq,Tsn,Cdk18,Opn3,Dtl,Entpd2,Ube2e3,B4galt1,Usp1,Sfpq,Sh3bgrl3,Oasl2,Usp18,Polr3e,Dctn6,Fgl1,Map1lc3b,Ddx25,Tipin,Me1,Mvk,Slx4,Wdr55,Atg14,Hace1,Cblc,Ubac1,Dis3,Pim3,Smc4,Tiparp,Cbr1,Mettl5,Polr2k,Whsc1,Ercc8,Insig1,Adam10,Nfil3,Azin1,Hmgb2,Mrpl18,Prpf38a,Gspt1,Birc5,Oas1a,Amd2,Slc39a8,Rps6ka6,Mdc1,Prpf4,Nhlrc4,Isg15,Fmr1,Mfap1a,Rnf213,Hbb-bt,Arrb1,Cox7a1,Mbnl1,Pdpk1,Mcts2,Sirt1,Crebzf,Cisd3,Trim59,Map4k5,Hnrnpa3,Ppip5k2,Nmnat3,Cnot4,Ube2v2,Rnf186,Rsl1d1,Bcl3,Slc16a11,Srsf3,Irf9,Srsf1,Nmi,Prlr,Lsm5,Tgm1,Odc1,Tcea1,Parp10,Sh3glb1,Ube2d2a,Psmb9,Metap2 |
| GO:0032069 | Regulation of nuclease activity | 6 | 22 | 1.4 | 0.00024 | 10090.ENSMUSP00000023897,10090.ENSMUSP00000031540,10090.ENSMUSP00000031542,10090.ENSMUSP00000065940,10090.ENSMUSP00000079198,10090.ENSMUSP00000101082 | Gzma,Oasl1,Oasl2,Hmgb2,Oas1a,Sirt1 |
| GO:0051707 | Response to other organism | 33 | 1145 | 0.42 | 0.00025 | 10090.ENSMUSP00000017851,10090.ENSMUSP00000018361,10090.ENSMUSP00000020308,10090.ENSMUSP00000020835,10090.ENSMUSP00000023897,10090.ENSMUSP00000026565,10090.ENSMUSP00000029018,10090.ENSMUSP00000029671,10090.ENSMUSP00000030623,10090.ENSMUSP00000031540,10090.ENSMUSP00000031542,10090.ENSMUSP00000032198,10090.ENSMUSP00000032473,10090.ENSMUSP00000033173,10090.ENSMUSP00000039586,10090.ENSMUSP00000040246,10090.ENSMUSP00000041091,10090.ENSMUSP00000065940,10090.ENSMUSP00000074009,10090.ENSMUSP00000079198,10090.ENSMUSP00000082548,10090.ENSMUSP00000085906,10090.ENSMUSP00000099889,10090.ENSMUSP00000102824,10090.ENSMUSP00000103432,10090.ENSMUSP00000104548,10090.ENSMUSP00000110684,10090.ENSMUSP00000113851,10090.ENSMUSP00000120647,10090.ENSMUSP00000122219,10090.ENSMUSP00000126565,10090.ENSMUSP00000128661,10090.ENSMUSP00000133499 | Serinc3,Pmp22,Ddit4,Ppm1d,Gzma,Ifitm3,Zbp1,Ifi44,Sfpq,Oasl1,Oasl2,Usp18,Iigp1,Polr3e,Serpine1,C1qb,Rtp4,Hmgb2,Ifnlr1,Oas1a,Isg15,Fmr1,Ifit3,Crebzf,Trim59,Irf1,Arid5a,Bcl3,Nmi,Prlr,Lsm5,Odc1,Psmb9 |
| GO:0050896 | Response to stimulus | 112 | 6908 | 0.17 | 0.00027 | 10090.ENSMUSP00000002284,10090.ENSMUSP00000002413,10090.ENSMUSP00000004120,10090.ENSMUSP00000005503,10090.ENSMUSP00000015581,10090.ENSMUSP00000017851,10090.ENSMUSP00000018361,10090.ENSMUSP00000018506,10090.ENSMUSP00000019469,10090.ENSMUSP00000020308,10090.ENSMUSP00000020835,10090.ENSMUSP00000022380,10090.ENSMUSP00000022612,10090.ENSMUSP00000022927,10090.ENSMUSP00000023353,10090.ENSMUSP00000023769,10090.ENSMUSP00000023774,10090.ENSMUSP00000023897,10090.ENSMUSP00000025161,10090.ENSMUSP00000025904,10090.ENSMUSP00000025930,10090.ENSMUSP00000026565,10090.ENSMUSP00000027760,10090.ENSMUSP00000027809,10090.ENSMUSP00000027933,10090.ENSMUSP00000028328,10090.ENSMUSP00000028382,10090.ENSMUSP00000029018,10090.ENSMUSP00000029566,10090.ENSMUSP00000029671,10090.ENSMUSP00000030121,10090.ENSMUSP00000030289,10090.ENSMUSP00000030623,10090.ENSMUSP00000031540,10090.ENSMUSP00000031542,10090.ENSMUSP00000032198,10090.ENSMUSP00000032473,10090.ENSMUSP00000032800,10090.ENSMUSP00000032898,10090.ENSMUSP00000033173,10090.ENSMUSP00000033626,10090.ENSMUSP00000034003,10090.ENSMUSP00000034138,10090.ENSMUSP00000034270,10090.ENSMUSP00000034787,10090.ENSMUSP00000034964,10090.ENSMUSP00000034989,10090.ENSMUSP00000035101,10090.ENSMUSP00000037007,10090.ENSMUSP00000037459,10090.ENSMUSP00000038871,10090.ENSMUSP00000039047,10090.ENSMUSP00000039586,10090.ENSMUSP00000039955, [illegible] | Plaur,Tmem161a,Fancl,Msh6,Gzmb,Serinc3,Pmp22,Kpna2,G6pc,Ddit4,Ppm1d,Psmc6,Pbk,Rad21,Mcm4,Atf1,Slc11a2,Gzma,Tapbp,Prdx5,Smc3,Ifitm3,Rgl1,Opn3,Dtl,Entpd2,Nup35,Zbp1,Efna1,Ifi44,B4galt1,Usp1,Sfpq,Oasl1,Oasl2,Usp18,Iigp1,Tyrobp,Ipo5,Polr3e,Serpina7,Fgl1,Dnaja2,Map1lc3b,Phip,Tipin,Me1,Csrnp1,Hspb8,Zfand1,Slx4,Atg14,Serpine1,Cblc,C1qb,Rtp4,Smc4,Tiparp,Hic1,Ercc8,Hilpda,Insig1,Adam10,Artn,Nfil3,Hmgb2,Ly6c1,Cdc42ep3,Srp19,Ifnlr1,Dchs1,Oas1a,Rps6ka6,Mdc1,Isg15,Fmr1,Magi1,Asgr1,Hbb-bt,Arrb1,Flcn,Ifit3,Pdpk1,Sirt1,Txnip,Crebzf,Trim59,Ect2,Irf1,Map4k5,Fgfr1op2,Nmnat3,Arid5a,Dnm1l,Ube2v2,Rnf186,Arhgap23,Bcl3,Srsf3,Nmi,Prlr,Xaf1,Lsm5,Odc1,Parp10,Sh3glb1,Zfyve27,Tgif1,Psmb9,Arhgap21,H2-D1,Tomm20 |

| | | | | | | | |
|---|---|---|---|---|---|---|---|
| GO:0071704 | Organic substance metabolic process | 111 | 6824 | 0.17 | 0.00027 | 10090.ENSMUSP00000002198,10090.ENSMUSP00000004120, 10090.ENSMUSP00000004985,10090.ENSMUSP00000005503, 10090.ENSMUSP00000011058,10090.ENSMUSP00000015581, 10090.ENSMUSP00000017851,10090.ENSMUSP00000018361, 10090.ENSMUSP00000019469,10090.ENSMUSP00000020217, 10090.ENSMUSP00000020835,10090.ENSMUSP00000021120, 10090.ENSMUSP00000021381,10090.ENSMUSP00000022380, 10090.ENSMUSP00000022612,10090.ENSMUSP00000022641, 10090.ENSMUSP00000022927,10090.ENSMUSP00000023353, 10090.ENSMUSP00000023774,10090.ENSMUSP00000023897, 10090.ENSMUSP00000024104,10090.ENSMUSP00000024657, 10090.ENSMUSP00000025161,10090.ENSMUSP00000025196, 10090.ENSMUSP00000025562,10090.ENSMUSP00000025632, 10090.ENSMUSP00000025930,10090.ENSMUSP00000026823, 10090.ENSMUSP00000027623,10090.ENSMUSP00000027697, 10090.ENSMUSP00000027809,10090.ENSMUSP00000027933, 10090.ENSMUSP00000028328,10090.ENSMUSP00000028398, 10090.ENSMUSP00000030121,10090.ENSMUSP00000030289, 10090.ENSMUSP00000030623,10090.ENSMUSP00000031542, 10090.ENSMUSP00000032198,10090.ENSMUSP00000033173, 10090.ENSMUSP00000033913,10090.ENSMUSP00000034003, 10090.ENSMUSP00000034612,10090.ENSMUSP00000034964, 10090.ENSMUSP00000034989,10090.ENSMUSP00000036971, 10090.ENSMUSP00000038871,10090.ENSMUSP00000039010, 10090.ENSMUSP00000039206,10090.ENSMUSP00000039955, 10090.ENSMUSP00000040220,10090.ENSMUSP00000041906, 10090.ENSMUSP00000044603,10090.ENSMUSP00000047872, | Sf3a1,Fancl,Brpf3,Msh6,Pdhx,Gzmb,Serinc3,Pmp22,G6pc,Nudt4,Ppm1d,Trim47,Pnn,Psmc6,Pbk,Adamdec1,Rad21,Mcm4,Slc11a2,Gzma,Gcm1,Phf10,Tapbp,Psmb8,Incenp,Carnmt1,Smc3,Pigq,Tsn,Cdk18,Opn3,Dtl,Entpd2,Ube2e3,B4galt1,Usp1,Sfpq,Oasl2,Usp18,Polr3e,Dctn6,Fgl1,Ddx25,Tipin,Me1,Mvk,Slx4,Wdr55,Hace1,Cblc,Ubac1,Dis3,Pim3,Smc4,Tiparp,Mettl5,Polr2k,Whsc1,Ercc8,Insig1,Adam10,Nfil3,Azin1,Hmgb2,Mrpl18,Prpf38a,Gspt1,Birc5,Oas1a,Amd2,Slc39a8,Rps6ka6,Mdc1,Prpf4,Nhlrc4,Isg15,Fmr1,Mfap1a,Rnf213,Arrb1,Mbnl1,Pdpk1,Mcts2,Sirt1,Crebzf,Cisd3,Trim59,Map4k5,Hnrnpa3,Ppip5k2,Nmnat3,Cnot4,Ube2v2,Rnf186,Rsl1d1,Bcl3,Slc16a11,Srsf3,Irf9,Srsf1,Nmi,Prlr,Lsm5,Tgm1,Odc1,Tcea1,Parp10,Sh3glb1,Ube2d2a,Psmb9,Metap2 |
| GO:0019219 | Regulation of nucleobase-containing compound metabolic process | 65 | 3265 | 0.26 | 0.00034 | 10090.ENSMUSP00000002413,10090.ENSMUSP00000005503, 10090.ENSMUSP00000020123,10090.ENSMUSP00000020308, 10090.ENSMUSP00000020835,10090.ENSMUSP00000020908, 10090.ENSMUSP00000021807,10090.ENSMUSP00000022227, 10090.ENSMUSP00000022380,10090.ENSMUSP00000022927, 10090.ENSMUSP00000023769,10090.ENSMUSP00000023897, 10090.ENSMUSP00000024104,10090.ENSMUSP00000024657, 10090.ENSMUSP00000025904,10090.ENSMUSP00000027697, 10090.ENSMUSP00000028382,10090.ENSMUSP00000029566, 10090.ENSMUSP00000030289,10090.ENSMUSP00000030367, 10090.ENSMUSP00000030623,10090.ENSMUSP00000031540, 10090.ENSMUSP00000031542,10090.ENSMUSP00000034787, 10090.ENSMUSP00000035101,10090.ENSMUSP00000038871, 10090.ENSMUSP00000050461,10090.ENSMUSP00000050854, 10090.ENSMUSP00000053483,10090.ENSMUSP00000056778, 10090.ENSMUSP00000058037,10090.ENSMUSP00000058237, 10090.ENSMUSP00000058940,10090.ENSMUSP00000059211, 10090.ENSMUSP00000061390,10090.ENSMUSP00000061877, 10090.ENSMUSP00000065363,10090.ENSMUSP00000065940, 10090.ENSMUSP00000077971,10090.ENSMUSP00000079124, 10090.ENSMUSP00000079198,10090.ENSMUSP00000080640, 10090.ENSMUSP00000085906,10090.ENSMUSP00000095866, 10090.ENSMUSP00000096686,10090.ENSMUSP00000099758, 10090.ENSMUSP00000101082,10090.ENSMUSP00000101661, 10090.ENSMUSP00000102710,10090.ENSMUSP00000102824, 10090.ENSMUSP00000104548,10090.ENSMUSP00000107595, 10090.ENSMUSP00000108083,10090.ENSMUSP00000110684, | Tmem161a,Msh6,Tmpo,Ddit4,Ppm1d,E2f6,Dek,Cenpk,Psmc6,Rad21,Atf1,Gzma,Gcm1,Phf10,Prdx5,Cdk18,Nup35,Efna1,Usp1,Ssbp3,Sfpq,Oasl1,Oasl2,Phip,Csrnp1,Slx4,Rbm12,Zfp367,Hic1,E2f8,Zfpm1,S100a1,Whsc1,Ercc8,Tob2,Insig1,Nfil3,Hmgb2,Phf6,Birc5,Oas1a,Slc39a8,Fmr1,Arrb1,Mbnl1,Flcn,Sirt1,Zbtb8b,Txnip,Crebzf,Irf1,Hnrnpa3,Bclaf3,Arid5a,Ube2v2,Bcl3,Srsf3,Irf9,Srsf1,Nmi,Tfdp2,Tcea1,Parp10,Tgif1,Zfp131 |
| GO:0044238 | Primary metabolic process | 105 | 6369 | 0.18 | 0.00034 | 10090.ENSMUSP00000002198,10090.ENSMUSP00000004120, 10090.ENSMUSP00000004985,10090.ENSMUSP00000005503, 10090.ENSMUSP00000011058,10090.ENSMUSP00000015581, 10090.ENSMUSP00000017851,10090.ENSMUSP00000018361, 10090.ENSMUSP00000019469,10090.ENSMUSP00000020217, 10090.ENSMUSP00000020835,10090.ENSMUSP00000021120, 10090.ENSMUSP00000021381,10090.ENSMUSP00000022380, 10090.ENSMUSP00000022612,10090.ENSMUSP00000022641, 10090.ENSMUSP00000022927,10090.ENSMUSP00000023353, 10090.ENSMUSP00000023897,10090.ENSMUSP00000024104, 10090.ENSMUSP00000024657,10090.ENSMUSP00000025196, 10090.ENSMUSP00000025562,10090.ENSMUSP00000025930, 10090.ENSMUSP00000026823,10090.ENSMUSP00000027623, 10090.ENSMUSP00000027697,10090.ENSMUSP00000027809, 10090.ENSMUSP00000027933,10090.ENSMUSP00000028328, 10090.ENSMUSP00000028398,10090.ENSMUSP00000030121, 10090.ENSMUSP00000030289,10090.ENSMUSP00000030623, 10090.ENSMUSP00000031542,10090.ENSMUSP00000032198, 10090.ENSMUSP00000033173,10090.ENSMUSP00000033913, 10090.ENSMUSP00000034003,10090.ENSMUSP00000034964, 10090.ENSMUSP00000036971,10090.ENSMUSP00000038871, 10090.ENSMUSP00000039010,10090.ENSMUSP00000039206, 10090.ENSMUSP00000039955,10090.ENSMUSP00000040220, 10090.ENSMUSP00000041906,10090.ENSMUSP00000044603, 10090.ENSMUSP00000047872,10090.ENSMUSP00000048051, 10090.ENSMUSP00000050907,10090.ENSMUSP00000051968, 10090.ENSMUSP00000058940,10090.ENSMUSP00000059211, | Sf3a1,Fancl,Brpf3,Msh6,Pdhx,Gzmb,Serinc3,Pmp22,G6pc,Nudt4,Ppm1d,Trim47,Pnn,Psmc6,Pbk,Adamdec1,Rad21,Mcm4,Gzma,Gcm1,Phf10,Psmb8,Incenp,Smc3,Pigq,Tsn,Cdk18,Opn3,Dtl,Entpd2,Ube2e3,B4galt1,Usp1,Sfpq,Oasl2,Usp18,Polr3e,Dctn6,Fgl1,Tipin,Mvk,Slx4,Wdr55,Hace1,Cblc,Ubac1,Dis3,Pim3,Smc4,Tiparp,Mettl5,Polr2k,Whsc1,Ercc8,Insig1,Adam10,Nfil3,Azin1,Hmgb2,Mrpl18,Prpf38a,Gspt1,Birc5,Oas1a,Amd2,Slc39a8,Rps6ka6,Mdc1,Prpf4,Nhlrc4,Isg15,Fmr1,Mfap1a,Rnf213,Arrb1,Mbnl1,Pdpk1,Mcts2,Sirt1,Crebzf,Cisd3,Trim59,Map4k5,Hnrnpa3,Ppip5k2,Nmnat3,Cnot4,Ube2v2,Rnf186,Rsl1d1,Bcl3,Slc16a11,Srsf3,Srsf1,Nmi,Prlr,Lsm5,Tgm1,Odc1,Tcea1,Parp10,Sh3glb1,Ube2d2a,Psmb9,Metap2 |
| GO:0042981 | Regulation of apoptotic process | 38 | 1486 | 0.37 | 0.00044 | 10090.ENSMUSP00000002284,10090.ENSMUSP00000002413, 10090.ENSMUSP00000017851,10090.ENSMUSP00000018361, 10090.ENSMUSP00000023774,10090.ENSMUSP00000023897, 10090.ENSMUSP00000025254,10090.ENSMUSP00000025904, 10090.ENSMUSP00000029018,10090.ENSMUSP00000029566, 10090.ENSMUSP00000030121,10090.ENSMUSP00000030623, 10090.ENSMUSP00000032800,10090.ENSMUSP00000034787, 10090.ENSMUSP00000039586,10090.ENSMUSP00000044603, 10090.ENSMUSP00000047569,10090.ENSMUSP00000063839, 10090.ENSMUSP00000065940,10090.ENSMUSP00000076961, 10090.ENSMUSP00000078985,10090.ENSMUSP00000079124, 10090.ENSMUSP00000085906,10090.ENSMUSP00000095866, 10090.ENSMUSP00000098897,10090.ENSMUSP00000099758, 10090.ENSMUSP00000099889,10090.ENSMUSP00000099990, 10090.ENSMUSP00000099991,10090.ENSMUSP00000101082, 10090.ENSMUSP00000102710,10090.ENSMUSP00000103935, 10090.ENSMUSP00000111415,10090.ENSMUSP00000111443, 10090.ENSMUSP00000113431,10090.ENSMUSP00000113851, 10090.ENSMUSP00000122219,10090.ENSMUSP00000129800 | Plaur,Tmem161a,Serinc3,Pmp22,Slc11a2,Gzma,Lims2,Prdx5,Zbp1,Efna1,B4galt1,Sfpq,Tyrobp,Phip,Serpine1,Pim3,Card19,Adam10,Hmgb2,Tnfaip8l1,Krit1,Birc5,Fmr1,Arrb1,Mad2l1,Flcn,Ifit3,Anp32b,Pdpk1,Sirt1,Txnip,Ect2,Dnm1l,Ube2v2,Rsl1d1,Bcl3,Prlr,Sh3glb1 |

| | | | | | | | |
|---|---|---|---|---|---|---|---|
| GO:0071840 | Cellular component organization or biogenesis | 87 | 4954 | 0.21 | 0.00044 | 10090.ENSMUSP00000002198,10090.ENSMUSP00000004430, 10090.ENSMUSP00000004985,10090.ENSMUSP00000005503, 10090.ENSMUSP00000018361,10090.ENSMUSP00000020308, 10090.ENSMUSP00000020835,10090.ENSMUSP00000021807, 10090.ENSMUSP00000022227,10090.ENSMUSP00000022927, 10090.ENSMUSP00000023353,10090.ENSMUSP00000023359, 10090.ENSMUSP00000023769,10090.ENSMUSP00000023774, 10090.ENSMUSP00000024657,10090.ENSMUSP00000025161, 10090.ENSMUSP00000025254,10090.ENSMUSP00000025562, 10090.ENSMUSP00000025930,10090.ENSMUSP00000028382, 10090.ENSMUSP00000029566,10090.ENSMUSP00000030121, 10090.ENSMUSP00000030367,10090.ENSMUSP00000030623, 10090.ENSMUSP00000031719,10090.ENSMUSP00000032800, 10090.ENSMUSP00000033913,10090.ENSMUSP00000034079, 10090.ENSMUSP00000034270,10090.ENSMUSP00000034787, 10090.ENSMUSP00000037459,10090.ENSMUSP00000038871, 10090.ENSMUSP00000039010,10090.ENSMUSP00000039047, 10090.ENSMUSP00000039206,10090.ENSMUSP00000040246, 10090.ENSMUSP00000041091,10090.ENSMUSP00000041744, 10090.ENSMUSP00000041906,10090.ENSMUSP00000044603, 10090.ENSMUSP00000047872,10090.ENSMUSP00000048829, 10090.ENSMUSP00000049973,10090.ENSMUSP00000050907, 10090.ENSMUSP00000056715,10090.ENSMUSP00000057938, 10090.ENSMUSP00000058037,10090.ENSMUSP00000058940, 10090.ENSMUSP00000059159,10090.ENSMUSP00000063839, 10090.ENSMUSP00000064521,10090.ENSMUSP00000065940, 10090.ENSMUSP00000067217,10090.ENSMUSP00000072386, | Sf3a1,Clcn3,Brpf3,Msh6,Pmp22,Ddit4,Ppm1d,Dek,Cenpk,Rad21,Mcm4,Nde1,Atf1,Slc11a2,Phf10,Tapbp,Lims2,Incenp,Smc3,Nup35,Efna1,B4galt1,Ssbp3,Sfpq,Fscn3,Tyrobp,Dctn6,Heatr3,Map1lc3b,Phip,Zfand1,Slx4,Wdr55,Atg14,Hace1,C1qb,Rtp4,Ofd1,Dis3,Pim3,Smc4,Cdh26,Haus3,Mettl5,Atxn2,Cenpe,Zfpm1,Whsc1,Aspm,Adam10,Artn,Hmgb2,Cdc42ep3,Srp19,Gspt1,Birc5,Slc39a8,Kif20b,Fmr1,Rnf213,Arrb1,Cox7a1,Mbnl1,Mad2l1,Atp11c,Nop56,Flcn,Anp32b,Pdpk1,Mcts2,Sirt1,Ect2,Ninl,Pom121,Dnm1l,Rsl1d1,Bcl3,Srsf1,Prlr,Crispld2,Xaf1,Camsap3,Sh3glb1,Zfyve27,Arhgap21,Golph3l,Tomm20 |
| GO:0007049 | Cell cycle | 32 | 1138 | 0.41 | 0.00048 | 10090.ENSMUSP00000005503,10090.ENSMUSP00000018361, 10090.ENSMUSP00000020835,10090.ENSMUSP00000020908, 10090.ENSMUSP00000022227,10090.ENSMUSP00000022612, 10090.ENSMUSP00000022927,10090.ENSMUSP00000023353, 10090.ENSMUSP00000023359,10090.ENSMUSP00000025562, 10090.ENSMUSP00000025930,10090.ENSMUSP00000027697, 10090.ENSMUSP00000033913,10090.ENSMUSP00000034964, 10090.ENSMUSP00000038871,10090.ENSMUSP00000039206, 10090.ENSMUSP00000044603,10090.ENSMUSP00000047872, 10090.ENSMUSP00000049973,10090.ENSMUSP00000056778, 10090.ENSMUSP00000057938,10090.ENSMUSP00000059159, 10090.ENSMUSP00000068835,10090.ENSMUSP00000079124, 10090.ENSMUSP00000080949,10090.ENSMUSP00000084599, 10090.ENSMUSP00000098897,10090.ENSMUSP00000099990, 10090.ENSMUSP00000102710,10090.ENSMUSP00000103935, 10090.ENSMUSP00000104548,10090.ENSMUSP00000128260 | Msh6,Pmp22,Ppm1d,E2f6,Cenpk,Pbk,Rad21,Mcm4,Nde1,Incenp,Smc3,Cdk18,Dctn6,Tipin,Slx4,Hace1,Pim3,Smc4,Haus3,E2f8,Cenpe,Aspm,Rabgap1,Birc5,Mdc1,Kif20b,Mad2l1,Anp32b,Txnip,Ect2,Irf1,Tfdp2 |
| GO:0050658 | RNA transport | 11 | 148 | 0.83 | 0.00049 | 10090.ENSMUSP00000028382,10090.ENSMUSP00000030623, 10090.ENSMUSP00000034612,10090.ENSMUSP00000062766, 10090.ENSMUSP00000078123,10090.ENSMUSP00000085906, 10090.ENSMUSP00000106801,10090.ENSMUSP00000107595, 10090.ENSMUSP00000117045,10090.ENSMUSP00000120595, 10090.ENSMUSP00000136493 | Nup35,Sfpq,Ddx25,Nupl2,Mrpl18,Fmr1,Pom121,Hnrnpa3,Srsf3,Srsf1,Tomm20 |
| GO:0043170 | Macromolecule metabolic process | 91 | 5313 | 0.2 | 0.00052 | 10090.ENSMUSP00000002198,10090.ENSMUSP00000004120, 10090.ENSMUSP00000004985,10090.ENSMUSP00000005503, 10090.ENSMUSP00000015581,10090.ENSMUSP00000018361, 10090.ENSMUSP00000019469,10090.ENSMUSP00000020835, 10090.ENSMUSP00000021120,10090.ENSMUSP00000021381, 10090.ENSMUSP00000022380,10090.ENSMUSP00000022612, 10090.ENSMUSP00000022641,10090.ENSMUSP00000022927, 10090.ENSMUSP00000023353,10090.ENSMUSP00000023897, 10090.ENSMUSP00000024104,10090.ENSMUSP00000024657, 10090.ENSMUSP00000025196,10090.ENSMUSP00000025562, 10090.ENSMUSP00000025930,10090.ENSMUSP00000026823, 10090.ENSMUSP00000027623,10090.ENSMUSP00000027697, 10090.ENSMUSP00000027809,10090.ENSMUSP00000027933, 10090.ENSMUSP00000028398,10090.ENSMUSP00000030121, 10090.ENSMUSP00000030289,10090.ENSMUSP00000030623, 10090.ENSMUSP00000032198,10090.ENSMUSP00000033173, 10090.ENSMUSP00000034612,10090.ENSMUSP00000034964, 10090.ENSMUSP00000038871,10090.ENSMUSP00000039010, 10090.ENSMUSP00000039206,10090.ENSMUSP00000039955, 10090.ENSMUSP00000040220,10090.ENSMUSP00000041906, 10090.ENSMUSP00000044603,10090.ENSMUSP00000047872, 10090.ENSMUSP00000048051,10090.ENSMUSP00000050907, 10090.ENSMUSP00000051968,10090.ENSMUSP00000058940, 10090.ENSMUSP00000059211,10090.ENSMUSP00000063839, 10090.ENSMUSP00000065363,10090.ENSMUSP00000065940, 10090.ENSMUSP00000078123,10090.ENSMUSP00000078207, 10090.ENSMUSP00000078940,10090.ENSMUSP00000079124, | Sf3a1,Fancl,Brpf3,Msh6,Gzmb,Pmp22,G6pc,Ppm1d,Trim47,Pnn,Psmc6,Pbk,Adamdec1,Rad21,Mcm4,Gzma,Gcm1,Phf10,Psmb8,Incenp,Smc3,Pigq,Tsn,Cdk18,Opn3,Dtl,Ube2e3,B4galt1,Usp1,Sfpq,Usp18,Polr3e,Ddx25,Tipin,Slx4,Wdr55,Hace1,Cblc,Ubac1,Dis3,Pim3,Smc4,Tiparp,Mettl5,Polr2k,Whsc1,Ercc8,Adam10,Nfil3,Hmgb2,Mrpl18,Prpf38a,Gspt1,Birc5,Slc39a8,Rps6ka6,Mdc1,Prpf4,Nhlrc4,Isg15,Fmr1,Mfap1a,Rnf213,Arrb1,Mbnl1,Pdpk1,Mcts2,Sirt1,Crebzf,Cisd3,Trim59,Map4k5,Hnrnpa3,Cnot4,Ube2v2,Rnf186,Rsl1d1,Bcl3,Srsf3,Irf9,Srsf1,Nmi,Prlr,Lsm5,Tgm1,Tcea1,Parp10,Sh3glb1,Ube2d2a,Psmb9,Metap2 |
| GO:0048518 | Positive regulation of biological process | 98 | 5869 | 0.19 | 0.00052 | 10090.ENSMUSP00000002284,10090.ENSMUSP00000002413, 10090.ENSMUSP00000004430,10090.ENSMUSP00000004985, 10090.ENSMUSP00000017851,10090.ENSMUSP00000018361, 10090.ENSMUSP00000018506,10090.ENSMUSP00000019726, 10090.ENSMUSP00000020308,10090.ENSMUSP00000020908, 10090.ENSMUSP00000022227,10090.ENSMUSP00000022380, 10090.ENSMUSP00000022612,10090.ENSMUSP00000022927, 10090.ENSMUSP00000023769,10090.ENSMUSP00000023774, 10090.ENSMUSP00000023897,10090.ENSMUSP00000024104, 10090.ENSMUSP00000024657,10090.ENSMUSP00000025254, 10090.ENSMUSP00000025562,10090.ENSMUSP00000025904, 10090.ENSMUSP00000027933,10090.ENSMUSP00000029018, 10090.ENSMUSP00000029566,10090.ENSMUSP00000030121, 10090.ENSMUSP00000030367,10090.ENSMUSP00000030623, 10090.ENSMUSP00000032800,10090.ENSMUSP00000032898, 10090.ENSMUSP00000034270,10090.ENSMUSP00000034787, 10090.ENSMUSP00000034964,10090.ENSMUSP00000035101, 10090.ENSMUSP00000037007,10090.ENSMUSP00000037459, 10090.ENSMUSP00000038871,10090.ENSMUSP00000039047, 10090.ENSMUSP00000039206,10090.ENSMUSP00000039586, 10090.ENSMUSP00000040246,10090.ENSMUSP00000048051, 10090.ENSMUSP00000050854,10090.ENSMUSP00000053483, 10090.ENSMUSP00000056778,10090.ENSMUSP00000057938, 10090.ENSMUSP00000058037,10090.ENSMUSP00000058237, 10090.ENSMUSP00000058940,10090.ENSMUSP00000059159, 10090.ENSMUSP00000059211,10090.ENSMUSP00000060349, 10090.ENSMUSP00000060791,10090.ENSMUSP00000061390, | Plaur,Tmem161a,Clcn3,Brpf3,Serinc3,Pmp22,Kpna2,Plin3,Ddit4,E2f6,Cenpk,Psmc6,Pbk,Rad21,Atf1,Slc11a2,Gzma,Gcm1,Phf10,Lims2,Incenp,Prdx5,Dtl,Zbp1,Efna1,B4galt1,Ssbp3,Sfpq,Tyrobp,Ipo5,Map1lc3b,Phip,Tipin,Csrnp1,Hspb8,Zfand1,Slx4,Atg14,Hace1,Serpine1,C1qb,Tiparp,Zfp367,Hic1,E2f8,Cenpe,Zfpm1,S100a1,Whsc1,Aspm,Ercc8,Camk2n1,Hilpda,Tob2,Insig1,Adam10,Artn,Nfil3,Azin1,Hmgb2,Cdc42ep3,Frmd4a,Birc5,Isg15,Kif20b,Fmr1,Magi1,Arrb1,Mad2l1,Atp11c,Flcn,Anp32b,Pdpk1,Sirt1,Txnip,Ect2,Irf1,Map4k5,Bclaf3,Arid5a,Dnm1l,Ube2v2,Bcl3,Srsf1,Nmi,Prlr,Ppp1r3g,Cep295,Tgm1,Tfdp2,Odc1,Tcea1,Sh3glb1,Zfyve27,Tgif1,H2-D1,Golph3l,Zfp131 |

| | | | | | | | |
|---|---|---|---|---|---|---|---|
| GO:0002252 | Immune effector process | 18 | 435 | 0.58 | 0.00073 | 10090.ENSMUSP00000005503,10090.ENSMUSP00000015581,10090.ENSMUSP00000017851,10090.ENSMUSP00000020308,10090.ENSMUSP00000026565,10090.ENSMUSP00000029018,10090.ENSMUSP00000031540,10090.ENSMUSP00000031542,10090.ENSMUSP00000032800,10090.ENSMUSP00000033173,10090.ENSMUSP00000040246,10090.ENSMUSP00000041091,10090.ENSMUSP00000074009,10090.ENSMUSP00000079198,10090.ENSMUSP00000082548,10090.ENSMUSP00000099889,10090.ENSMUSP00000104548,10090.ENSMUSP00000113851 | Msh6,Gzmb,Serinc3,Ddit4,Ifitm3,Zbp1,Oasl1,Oasl2,Tyrobp,Polr3e,C1qb,Rtp4,Ifnlr1,Oas1a,Isg15,Ifit3,Irf1,Bcl3 |
| GO:0010941 | Regulation of cell death | 40 | 1661 | 0.34 | 0.00073 | 10090.ENSMUSP00000002284,10090.ENSMUSP00000002413,10090.ENSMUSP00000017851,10090.ENSMUSP00000018361,10090.ENSMUSP00000020308,10090.ENSMUSP00000023774,10090.ENSMUSP00000023897,10090.ENSMUSP00000025254,10090.ENSMUSP00000025904,10090.ENSMUSP00000029018,10090.ENSMUSP00000029566,10090.ENSMUSP00000030121,10090.ENSMUSP00000030623,10090.ENSMUSP00000032800,10090.ENSMUSP00000034787,10090.ENSMUSP00000039586,10090.ENSMUSP00000044603,10090.ENSMUSP00000047569,10090.ENSMUSP00000063839,10090.ENSMUSP00000065940,10090.ENSMUSP00000076961,10090.ENSMUSP00000078985,10090.ENSMUSP00000079124,10090.ENSMUSP00000085906,10090.ENSMUSP00000095866,10090.ENSMUSP00000098897,10090.ENSMUSP00000099758,10090.ENSMUSP00000099889,10090.ENSMUSP00000099990,10090.ENSMUSP00000099991,10090.ENSMUSP00000101082,10090.ENSMUSP00000102710,10090.ENSMUSP00000103935,10090.ENSMUSP00000104548,10090.ENSMUSP00000111415,10090.ENSMUSP00000111443,10090.ENSMUSP00000113431,10090.ENSMUSP00000113851,10090.ENSMUSP00000122219,10090.ENSMUSP00000129800 | Plaur,Tmem161a,Serinc3,Pmp22,Ddit4,Slc11a2,Gzma,Lims2,Prdx5,Zbp1,Efna1,B4galt1,Sfpq,Tyrobp,Phip,Serpine1,Pim3,Card19,Adam10,Hmgb2,Tnfaip8l1,Krit1,Birc5,Fmr1,Arrb1,Mad2l1,Flcn,Ifit3,Anp32b,Pdpk1,Sirt1,Txnip,Ect2,Irf1,Dnm1l,Ube2v2,Rsl1d1,Bcl3,Prlr,Sh3glb1 |
| GO:0009605 | Response to external stimulus | 48 | 2199 | 0.3 | 0.00088 | 10090.ENSMUSP00000017851,10090.ENSMUSP00000018361,10090.ENSMUSP00000019469,10090.ENSMUSP00000020308,10090.ENSMUSP00000020835,10090.ENSMUSP00000023897,10090.ENSMUSP00000026565,10090.ENSMUSP00000027809,10090.ENSMUSP00000029018,10090.ENSMUSP00000029566,10090.ENSMUSP00000029671,10090.ENSMUSP00000030623,10090.ENSMUSP00000031540,10090.ENSMUSP00000031542,10090.ENSMUSP00000032198,10090.ENSMUSP00000032473,10090.ENSMUSP00000033173,10090.ENSMUSP00000033626,10090.ENSMUSP00000034270,10090.ENSMUSP00000039047,10090.ENSMUSP00000039586,10090.ENSMUSP00000040246,10090.ENSMUSP00000041091,10090.ENSMUSP00000059211,10090.ENSMUSP00000064521,10090.ENSMUSP00000065940,10090.ENSMUSP00000074009,10090.ENSMUSP00000079198,10090.ENSMUSP00000082548,10090.ENSMUSP00000085906,10090.ENSMUSP00000090637,10090.ENSMUSP00000095866,10090.ENSMUSP00000099758,10090.ENSMUSP00000099889,10090.ENSMUSP00000101082,10090.ENSMUSP00000102710,10090.ENSMUSP00000102824,10090.ENSMUSP00000103432,10090.ENSMUSP00000104548,10090.ENSMUSP00000110684,10090.ENSMUSP00000111415,10090.ENSMUSP00000113851,10090.ENSMUSP00000120647,10090.ENSMUSP00000122219,10090.ENSMUSP00000126565,10090.ENSMUSP00000128661,10090.ENSMUSP00000129800,10090.ENSMUSP00000133499 | Serinc3,Pmp22,G6pc,Ddit4,Ppm1d,Gzma,Ifitm3,Opn3,Zbp1,Efna1,Ifi44,Sfpq,Oasl1,Oasl2,Usp18,Iigp1,Polr3e,Serpina7,Map1lc3b,Atg14,Serpine1,C1qb,Rtp4,Ercc8,Artn,Hmgb2,Ifnlr1,Oas1a,Isg15,Fmr1,Asgr1,Arrb1,Flcn,Ifit3,Sirt1,Txnip,Crebzf,Trim59,Irf1,Arid5a,Dnm1l,Bcl3,Nmi,Prlr,Lsm5,Odc1,Sh3glb1,Psmb9 |
| GO:0033554 | Cellular response to stress | 37 | 1493 | 0.36 | 0.00092 | 10090.ENSMUSP00000002413,10090.ENSMUSP00000004120,10090.ENSMUSP00000005503,10090.ENSMUSP00000018361,10090.ENSMUSP00000020308,10090.ENSMUSP00000020835,10090.ENSMUSP00000022380,10090.ENSMUSP00000022927,10090.ENSMUSP00000023353,10090.ENSMUSP00000023774,10090.ENSMUSP00000025904,10090.ENSMUSP00000025930,10090.ENSMUSP00000027933,10090.ENSMUSP00000030289,10090.ENSMUSP00000030623,10090.ENSMUSP00000034270,10090.ENSMUSP00000034964,10090.ENSMUSP00000037007,10090.ENSMUSP00000038871,10090.ENSMUSP00000039047,10090.ENSMUSP00000047872,10090.ENSMUSP00000053483,10090.ENSMUSP00000059211,10090.ENSMUSP00000060791,10090.ENSMUSP00000061877,10090.ENSMUSP00000080694,10090.ENSMUSP00000080949,10090.ENSMUSP00000085906,10090.ENSMUSP00000099758,10090.ENSMUSP00000101082,10090.ENSMUSP00000103935,10090.ENSMUSP00000106199,10090.ENSMUSP00000111443,10090.ENSMUSP00000111780,10090.ENSMUSP00000113851,10090.ENSMUSP00000129765,10090.ENSMUSP00000129800 | Tmem161a,Fancl,Msh6,Pmp22,Ddit4,Ppm1d,Psmc6,Rad21,Mcm4,Slc11a2,Prdx5,Smc3,Dtl,Usp1,Sfpq,Map1lc3b,Tipin,Hspb8,Slx4,Atg14,Smc4,Hic1,Ercc8,Hilpda,Insig1,Rps6ka6,Mdc1,Fmr1,Flcn,Sirt1,Ect2,Map4k5,Ube2v2,Rnf186,Bcl3,Parp10,Sh3glb1 |
| GO:0044419 | Interspecies interaction between organisms | 34 | 1309 | 0.38 | 0.00092 | 10090.ENSMUSP00000017851,10090.ENSMUSP00000018361,10090.ENSMUSP00000018506,10090.ENSMUSP00000020308,10090.ENSMUSP00000020835,10090.ENSMUSP00000023897,10090.ENSMUSP00000026565,10090.ENSMUSP00000029018,10090.ENSMUSP00000029671,10090.ENSMUSP00000030623,10090.ENSMUSP00000031540,10090.ENSMUSP00000031542,10090.ENSMUSP00000032198,10090.ENSMUSP00000032473,10090.ENSMUSP00000033173,10090.ENSMUSP00000039586,10090.ENSMUSP00000040246,10090.ENSMUSP00000041091,10090.ENSMUSP00000065940,10090.ENSMUSP00000074009,10090.ENSMUSP00000079198,10090.ENSMUSP00000082548,10090.ENSMUSP00000085906,10090.ENSMUSP00000099889,10090.ENSMUSP00000102824,10090.ENSMUSP00000103432,10090.ENSMUSP00000104548,10090.ENSMUSP00000110684,10090.ENSMUSP00000113851,10090.ENSMUSP00000120647,10090.ENSMUSP00000122219,10090.ENSMUSP00000126565,10090.ENSMUSP00000128661,10090.ENSMUSP00000133499 | Serinc3,Pmp22,Kpna2,Ddit4,Ppm1d,Gzma,Ifitm3,Zbp1,Ifi44,Sfpq,Oasl1,Oasl2,Usp18,Iigp1,Polr3e,Serpine1,C1qb,Rtp4,Hmgb2,Ifnlr1,Oas1a,Isg15,Fmr1,Ifit3,Crebzf,Trim59,Irf1,Arid5a,Bcl3,Nmi,Prlr,Lsm5,Odc1,Psmb9 |

| | | | | | | | |
|---|---|---|---|---|---|---|---|
| GO:0051726 | Regulation of cell cycle | 29 | 1016 | 0.42 | 0.00092 | 10090.ENSMUSP00000018361,10090.ENSMUSP00000020908, 10090.ENSMUSP00000022927,10090.ENSMUSP00000025562, 10090.ENSMUSP00000025930,10090.ENSMUSP00000027697, 10090.ENSMUSP00000027933,10090.ENSMUSP00000030623, 10090.ENSMUSP00000034787,10090.ENSMUSP00000034964, 10090.ENSMUSP00000044603,10090.ENSMUSP00000056778, 10090.ENSMUSP00000057938,10090.ENSMUSP00000059159, 10090.ENSMUSP00000060349,10090.ENSMUSP00000065544, 10090.ENSMUSP00000079124,10090.ENSMUSP00000084599, 10090.ENSMUSP00000098897,10090.ENSMUSP00000099758, 10090.ENSMUSP00000099990,10090.ENSMUSP00000101082, 10090.ENSMUSP00000103935,10090.ENSMUSP00000104548, 10090.ENSMUSP00000106199,10090.ENSMUSP00000123788, 10090.ENSMUSP00000128090,10090.ENSMUSP00000128260, 10090.ENSMUSP00000129800 | Pmp22,E2f6,Rad21,Incenp,Smc3,Cdk18,Dtl,Sfpq,Phip,Tipin,Pim3,E2f8,Cenpe,Aspm,Camk2n1,Azin1,Birc5,Kif20b,Mad2l1,Flcn,Anp32b,Sirt1,Ect2,Irf1,Map4k5,Cep295,Tgm1,Tfdp2,Sh3glb1 |
| GO:0006955 | Immune response | 28 | 979 | 0.42 | 0.0012 | 10090.ENSMUSP00000005503,10090.ENSMUSP00000015581, 10090.ENSMUSP00000017851,10090.ENSMUSP00000018361, 10090.ENSMUSP00000026565,10090.ENSMUSP00000029018, 10090.ENSMUSP00000029671,10090.ENSMUSP00000030623, 10090.ENSMUSP00000031540,10090.ENSMUSP00000031542, 10090.ENSMUSP00000032473,10090.ENSMUSP00000032800, 10090.ENSMUSP00000033173,10090.ENSMUSP00000034003, 10090.ENSMUSP00000040246,10090.ENSMUSP00000065363, 10090.ENSMUSP00000065940,10090.ENSMUSP00000074009, 10090.ENSMUSP00000079198,10090.ENSMUSP00000082548, 10090.ENSMUSP00000099889,10090.ENSMUSP00000101082, 10090.ENSMUSP00000103432,10090.ENSMUSP00000104548, 10090.ENSMUSP00000110684,10090.ENSMUSP00000113851, 10090.ENSMUSP00000120647,10090.ENSMUSP00000134570 | Msh6,Gzmb,Serinc3,Pmp22,Ifitm3,Zbp1,Ifi44,Sfpq,Oasl1,Oasl2,Iigp1,Tyrobp,Polr3e,Fgl1,C1qb,Nfil3,Hmgb2,Ifnlr1,Oas1a,Isg15,Ifit3,Sirt1,Trim59,Irf1,Arid5a,Bcl3,Nmi,H2-D1 |
| GO:0010605 | Negative regulation of macromolecule metabolic process | 54 | 2646 | 0.27 | 0.0012 | 10090.ENSMUSP00000002284,10090.ENSMUSP00000005503, 10090.ENSMUSP00000018361,10090.ENSMUSP00000020308, 10090.ENSMUSP00000020835,10090.ENSMUSP00000020908, 10090.ENSMUSP00000022612,10090.ENSMUSP00000023897, 10090.ENSMUSP00000024657,10090.ENSMUSP00000025904, 10090.ENSMUSP00000029566,10090.ENSMUSP00000030623, 10090.ENSMUSP00000032800,10090.ENSMUSP00000033626, 10090.ENSMUSP00000034964,10090.ENSMUSP00000036971, 10090.ENSMUSP00000038871,10090.ENSMUSP00000039047, 10090.ENSMUSP00000039586,10090.ENSMUSP00000039955, 10090.ENSMUSP00000048051,10090.ENSMUSP00000053483, 10090.ENSMUSP00000056715,10090.ENSMUSP00000056778, 10090.ENSMUSP00000058037,10090.ENSMUSP00000058237, 10090.ENSMUSP00000058940,10090.ENSMUSP00000060349, 10090.ENSMUSP00000061390,10090.ENSMUSP00000065363, 10090.ENSMUSP00000065544,10090.ENSMUSP00000065940, 10090.ENSMUSP00000077971,10090.ENSMUSP00000078940, 10090.ENSMUSP00000079124,10090.ENSMUSP00000080694, 10090.ENSMUSP00000082548,10090.ENSMUSP00000085906, 10090.ENSMUSP00000095866,10090.ENSMUSP00000098897, 10090.ENSMUSP00000099758,10090.ENSMUSP00000099991, 10090.ENSMUSP00000101082,10090.ENSMUSP00000102710, 10090.ENSMUSP00000102824,10090.ENSMUSP00000104548, 10090.ENSMUSP00000110684,10090.ENSMUSP00000113851, 10090.ENSMUSP00000120647,10090.ENSMUSP00000126565, 10090.ENSMUSP00000128260,10090.ENSMUSP00000129765, 10090.ENSMUSP00000130930,10090.ENSMUSP00000136867 | Plaur,Msh6,Pmp22,Ddit4,Ppm1d,E2f6,Pbk,Gzma,Phf10,Prdx5,Efna1,Sfpq,Tyrobp,Serpina7,Tipin,Mvk,Slx4,Atg14,Serpine1,Cblc,Tiparp,Hic1,Atxn2,E2f8,Zfpm1,S100a1,Whsc1,Camk2n1,Tob2,Nfil3,Azin1,Hmgb2,Phf6,Gspt1,Birc5,Rps6ka6,Isg15,Fmr1,Arrb1,Mad2l1,Flcn,Pdpk1,Sirt1,Txnip,Crebzf,Irf1,Arid5a,Bcl3,Nmi,Lsm5, Tfdp2,Parp10,Tgif1,Zfp131 |
| GO:0006974 | Cellular response to dna damage stimulus | 22 | 667 | 0.48 | 0.0016 | 10090.ENSMUSP00000004120,10090.ENSMUSP00000005503, 10090.ENSMUSP00000020308,10090.ENSMUSP00000020835, 10090.ENSMUSP00000022927,10090.ENSMUSP00000023353, 10090.ENSMUSP00000025930,10090.ENSMUSP00000027933, 10090.ENSMUSP00000030289,10090.ENSMUSP00000030623, 10090.ENSMUSP00000034964,10090.ENSMUSP00000038871, 10090.ENSMUSP00000047872,10090.ENSMUSP00000053483, 10090.ENSMUSP00000059211,10090.ENSMUSP00000080694, 10090.ENSMUSP00000080949,10090.ENSMUSP00000085906, 10090.ENSMUSP00000101082,10090.ENSMUSP00000111443, 10090.ENSMUSP00000113851,10090.ENSMUSP00000129765 | Fancl,Msh6,Ddit4,Ppm1d,Rad21,Mcm4,Smc3,Dtl,Usp1,Sfpq,Tipin,Slx4,Smc4,Hic1,Ercc8,Rps6ka6,Mdc1,Fmr1,Sirt1,Ube2v2,Bcl3,Parp10 |
| GO:0009411 | Response to uv | 10 | 143 | 0.81 | 0.0017 | 10090.ENSMUSP00000002413,10090.ENSMUSP00000005503, 10090.ENSMUSP00000022612,10090.ENSMUSP00000027933, 10090.ENSMUSP00000030289,10090.ENSMUSP00000034964, 10090.ENSMUSP00000059211,10090.ENSMUSP00000085906, 10090.ENSMUSP00000101082,10090.ENSMUSP00000113851 | Tmem161a,Msh6,Pbk,Dtl,Usp1,Tipin,Ercc8,Fmr1,Sirt1,Bcl3 |
| GO:0090068 | Positive regulation of cell cycle process | 13 | 252 | 0.68 | 0.0017 | 10090.ENSMUSP00000022927,10090.ENSMUSP00000027933, 10090.ENSMUSP00000030623,10090.ENSMUSP00000034787, 10090.ENSMUSP00000056778,10090.ENSMUSP00000057938, 10090.ENSMUSP00000065544,10090.ENSMUSP00000079124, 10090.ENSMUSP00000084599,10090.ENSMUSP00000098897, 10090.ENSMUSP00000099990,10090.ENSMUSP00000103935, 10090.ENSMUSP00000123788 | Rad21,Dtl,Sfpq,Phip,E2f8,Cenpe,Azin1,Birc5,Kif20b,Mad2l1,Anp32b,Ect2,Cep295 |
| GO:0016043 | Cellular component organization | 81 | 4769 | 0.19 | 0.0029 | 10090.ENSMUSP00000002198,10090.ENSMUSP00000004430, 10090.ENSMUSP00000004985,10090.ENSMUSP00000005503, 10090.ENSMUSP00000018361,10090.ENSMUSP00000020308, 10090.ENSMUSP00000020835,10090.ENSMUSP00000021807, 10090.ENSMUSP00000022227,10090.ENSMUSP00000022927, 10090.ENSMUSP00000023353,10090.ENSMUSP00000023359, 10090.ENSMUSP00000023769,10090.ENSMUSP00000023774, 10090.ENSMUSP00000024657,10090.ENSMUSP00000025161, 10090.ENSMUSP00000025254,10090.ENSMUSP00000025562, 10090.ENSMUSP00000025930,10090.ENSMUSP00000028382, 10090.ENSMUSP00000029566,10090.ENSMUSP00000030121, 10090.ENSMUSP00000030367,10090.ENSMUSP00000030623, 10090.ENSMUSP00000031719,10090.ENSMUSP00000032800, 10090.ENSMUSP00000033913,10090.ENSMUSP00000034270, 10090.ENSMUSP00000034787,10090.ENSMUSP00000037459, 10090.ENSMUSP00000038871,10090.ENSMUSP00000039047, 10090.ENSMUSP00000039206,10090.ENSMUSP00000040246, 10090.ENSMUSP00000041091,10090.ENSMUSP00000041744, 10090.ENSMUSP00000044603,10090.ENSMUSP00000047872, 10090.ENSMUSP00000048829,10090.ENSMUSP00000049973, 10090.ENSMUSP00000056715,10090.ENSMUSP00000057938, 10090.ENSMUSP00000058037,10090.ENSMUSP00000058940, 10090.ENSMUSP00000059159,10090.ENSMUSP00000063839, 10090.ENSMUSP00000064521,10090.ENSMUSP00000065940, 10090.ENSMUSP00000067217,10090.ENSMUSP00000072386, 10090.ENSMUSP00000078940,10090.ENSMUSP00000079124, 10090.ENSMUSP00000080640,10090.ENSMUSP00000084599, [illegible] | Sf3a1,Clcn3,Brpf3,Msh6,Pmp22,Ddit4,Ppm1d,Dek,Cenpk,Rad21,Mcm4,Nde1,Atf1,Slc11a2,Phf10,Tapbp,Lims2,Incenp,Smc3,Nup35,Efna1,B4galt1,Ssbp3,Sfpq,Fscn3,Tyrobp,Dctn6,Map1lc3b,Phip,Zfand1,Slx4,Atg14,Hace1,C1qb,Rtp4,Ofd1,Pim3,Smc4,Cdh26,Haus3,Atxn2,Cenpe,Zfpm1,Whsc1,Aspm,Adam10,Artn,Hmgb2,Cdc42ep3,Srp19,Gspt1,Birc5,Slc39a8,Kif20b,Fmr1,Rnf213,Arrb1,Cox7a1,Mbnl1,Mad2l1,Atp11c,Flcn,Anp32b,Pdpk1,Mcts2,Sirt1,Ect2,Ninl,Pom121,Dnm1l,Bcl3,Srsf1,Prlr,Crispld2,Xaf1,Camsap3,Sh3glb1,Zfyve27,Arhgap21,Golph3l,Tomm20 |

| | | | | | | | |
|---|---|---|---|---|---|---|---|
| GO:0009892 | Negative regulation of metabolic process | 56 | 2895 | 0.25 | 0.0032 | 10090.ENSMUSP00000002284,10090.ENSMUSP00000005503, 10090.ENSMUSP00000018361,10090.ENSMUSP00000020308, 10090.ENSMUSP00000020835,10090.ENSMUSP00000020908, 10090.ENSMUSP00000022612,10090.ENSMUSP00000023897, 10090.ENSMUSP00000024657,10090.ENSMUSP00000025904, 10090.ENSMUSP00000029566,10090.ENSMUSP00000030623, 10090.ENSMUSP00000032800,10090.ENSMUSP00000033626, 10090.ENSMUSP00000034964,10090.ENSMUSP00000036971, 10090.ENSMUSP00000038871,10090.ENSMUSP00000039047, 10090.ENSMUSP00000039586,10090.ENSMUSP00000039955, 10090.ENSMUSP00000048051,10090.ENSMUSP00000053483, 10090.ENSMUSP00000056715,10090.ENSMUSP00000056778, 10090.ENSMUSP00000058037,10090.ENSMUSP00000058237, 10090.ENSMUSP00000058940,10090.ENSMUSP00000060349, 10090.ENSMUSP00000061390,10090.ENSMUSP00000061877, 10090.ENSMUSP00000065363,10090.ENSMUSP00000065544, 10090.ENSMUSP00000065940,10090.ENSMUSP00000077971, 10090.ENSMUSP00000078940,10090.ENSMUSP00000079124, 10090.ENSMUSP00000080694,10090.ENSMUSP00000082548, 10090.ENSMUSP00000085906,10090.ENSMUSP00000095866, 10090.ENSMUSP00000098897,10090.ENSMUSP00000099758, 10090.ENSMUSP00000099991,10090.ENSMUSP00000101082, 10090.ENSMUSP00000102710,10090.ENSMUSP00000102824, 10090.ENSMUSP00000104548,10090.ENSMUSP00000110684, 10090.ENSMUSP00000113851,10090.ENSMUSP00000120647, 10090.ENSMUSP00000125993,10090.ENSMUSP00000126565, 10090.ENSMUSP00000128260,10090.ENSMUSP00000129765, | Plaur,Msh6,Pmp22,Ddit4,Ppm1d,E2f6,Pbk,Gzma,Phf10,Prdx5,Efna1,Sfpq,Tyrobp,Serpina7,Tipin,Mvk,Slx4,Atg14,Serpine1,Cblc,Tiparp,Hic1,Atxn2,E2f8,Zfpm1,S100a1,Whsc1,Camk2n1,Tob2,Insig1,Nfil3,Azin1,Hmgb2,Phf6,Gspt1,Birc5,Rps6ka6,Isg15,Fmr1,Arrb1,Mad2l1,Flcn,Pdpk1,Sirt1,Txnip,Crebzf,Irf1,Arid5a,Bcl3,Nmi,Camsap3,Lsm5,Tfdp2,Parp10,Tgif1,Zfp131 |
| GO:0044260 | Cellular macromolecule metabolic process | 73 | 4161 | 0.21 | 0.0032 | 10090.ENSMUSP00000004120,10090.ENSMUSP00000004985, 10090.ENSMUSP00000005503,10090.ENSMUSP00000018361, 10090.ENSMUSP00000019469,10090.ENSMUSP00000020835, 10090.ENSMUSP00000021120,10090.ENSMUSP00000022380, 10090.ENSMUSP00000022612,10090.ENSMUSP00000022927, 10090.ENSMUSP00000023353,10090.ENSMUSP00000023897, 10090.ENSMUSP00000024104,10090.ENSMUSP00000024657, 10090.ENSMUSP00000025196,10090.ENSMUSP00000025562, 10090.ENSMUSP00000025930,10090.ENSMUSP00000026823, 10090.ENSMUSP00000027697,10090.ENSMUSP00000027809, 10090.ENSMUSP00000027933,10090.ENSMUSP00000028398, 10090.ENSMUSP00000030121,10090.ENSMUSP00000030289, 10090.ENSMUSP00000030623,10090.ENSMUSP00000032198, 10090.ENSMUSP00000033173,10090.ENSMUSP00000034964, 10090.ENSMUSP00000038871,10090.ENSMUSP00000039206, 10090.ENSMUSP00000039955,10090.ENSMUSP00000040220, 10090.ENSMUSP00000041906,10090.ENSMUSP00000044603, 10090.ENSMUSP00000047872,10090.ENSMUSP00000048051, 10090.ENSMUSP00000050907,10090.ENSMUSP00000051968, 10090.ENSMUSP00000058940,10090.ENSMUSP00000059211, 10090.ENSMUSP00000063839,10090.ENSMUSP00000065363, 10090.ENSMUSP00000065940,10090.ENSMUSP00000078123, 10090.ENSMUSP00000078940,10090.ENSMUSP00000079124, 10090.ENSMUSP00000080640,10090.ENSMUSP00000080694, 10090.ENSMUSP00000080949,10090.ENSMUSP00000082102, 10090.ENSMUSP00000082548,10090.ENSMUSP00000091429, 10090.ENSMUSP00000095866,10090.ENSMUSP00000099991, | Fancl,Brpf3,Msh6,Pmp22,G6pc,Ppm1d,Trim47,Psmc6,Pbk,Rad21,Mcm4,Gzma,Gcm1,Phf10,Psmb8,Incenp,Smc3,Pigq,Cdk18,Opn3,Dtl,Ube2e3,B4galt1,Usp1,Sfpq,Usp18,Polr3e,Tipin,Slx4,Hace1,Cblc,Ubac1,Dis3,Pim3,Smc4,Tiparp,Mettl5,Polr2k,Whsc1,Ercc8,Adam10,Nfil3,Hmgb2,Mrpl18,Gspt1,Birc5,Slc39a8,Rps6ka6,Mdc1,Nhlrc4,Isg15,Rnf213,Arrb1,Pdpk1,Mcts2,Sirt1,Crebzf,Trim59,Map4k5,Cnot4,Ube2v2,Rnf186,Bcl3,Nmi,Prlr,Lsm5,Tgm1,Tcea1,Parp10,Sh3glb1,Ube2d2a,Psmb9,Metap2 |
| GO:0048519 | Negative regulation of biological process | 85 | 5108 | 0.18 | 0.0032 | 10090.ENSMUSP00000002284,10090.ENSMUSP00000002413, 10090.ENSMUSP00000005503,10090.ENSMUSP00000018361, 10090.ENSMUSP00000020308,10090.ENSMUSP00000020835, 10090.ENSMUSP00000020908,10090.ENSMUSP00000022612, 10090.ENSMUSP00000022927,10090.ENSMUSP00000023897, 10090.ENSMUSP00000024657,10090.ENSMUSP00000025254, 10090.ENSMUSP00000025904,10090.ENSMUSP00000026565, 10090.ENSMUSP00000027933,10090.ENSMUSP00000029566, 10090.ENSMUSP00000030121,10090.ENSMUSP00000030623, 10090.ENSMUSP00000031540,10090.ENSMUSP00000031542, 10090.ENSMUSP00000032800,10090.ENSMUSP00000033626, 10090.ENSMUSP00000034003,10090.ENSMUSP00000034787, 10090.ENSMUSP00000034964,10090.ENSMUSP00000036971, 10090.ENSMUSP00000038871,10090.ENSMUSP00000039047, 10090.ENSMUSP00000039586,10090.ENSMUSP00000039955, 10090.ENSMUSP00000041744,10090.ENSMUSP00000044603, 10090.ENSMUSP00000048051,10090.ENSMUSP00000053483, 10090.ENSMUSP00000056715,10090.ENSMUSP00000056778, 10090.ENSMUSP00000057938,10090.ENSMUSP00000058037, 10090.ENSMUSP00000058237,10090.ENSMUSP00000058940, 10090.ENSMUSP00000059159,10090.ENSMUSP00000060349, 10090.ENSMUSP00000061390,10090.ENSMUSP00000061877, 10090.ENSMUSP00000063839,10090.ENSMUSP00000065363, 10090.ENSMUSP00000065544,10090.ENSMUSP00000065940, 10090.ENSMUSP00000075172,10090.ENSMUSP00000076961, 10090.ENSMUSP00000077971,10090.ENSMUSP00000078940, 10090.ENSMUSP00000078985,10090.ENSMUSP00000079124, | Plaur,Tmem161a,Msh6,Pmp22,Ddit4,Ppm1d,E2f6,Pbk,Rad21,Gzma,Phf10,Lims2,Prdx5,Ifitm3,Dtl,Efna1,B4galt1,Sfpq,Oasl1,Oasl2,Tyrobp,Serpina7,Fgl1,Phip,Tipin,Mvk,Slx4,Atg14,Serpine1,Cblc,Ofd1,Pim3,Tiparp,Hic1,Atxn2,E2f8,Cenpe,Zfpm1,S100a1,Whsc1,Aspm,Camk2n1,Tob2,Insig1,Adam10,Nfil3,Azin1,Hmgb2,Frmd4a,Tnfaip8l1,Phf6,Gspt1,Krit1,Birc5,Oas1a,Rps6ka6,Isg15,Fmr1,Rnf213,Arrb1,Mad2l1,Flcn,Ifit3,Anp32b,Pdpk1,Sirt1,Txnip,Crebzf,Trim59,Irf1,Arid5a,Dnm1l,Ube2v2,Bcl3,Nmi,Prlr,Xaf1,Camsap3,Lsm5,Tfdp2,Parp10,Sh3glb1,Tgif1,H2-D1,Zfp131 |
| GO:0050790 | Regulation of catalytic activity | 46 | 2202 | 0.28 | 0.0034 | 10090.ENSMUSP00000002284,10090.ENSMUSP00000005503, 10090.ENSMUSP00000018361,10090.ENSMUSP00000022612, 10090.ENSMUSP00000023359,10090.ENSMUSP00000023774, 10090.ENSMUSP00000023897,10090.ENSMUSP00000025196, 10090.ENSMUSP00000025562,10090.ENSMUSP00000025904, 10090.ENSMUSP00000029566,10090.ENSMUSP00000031540, 10090.ENSMUSP00000031542,10090.ENSMUSP00000033626, 10090.ENSMUSP00000034138,10090.ENSMUSP00000038871, 10090.ENSMUSP00000039047,10090.ENSMUSP00000039586, 10090.ENSMUSP00000039955,10090.ENSMUSP00000052287, 10090.ENSMUSP00000057938,10090.ENSMUSP00000058237, 10090.ENSMUSP00000060349,10090.ENSMUSP00000065544, 10090.ENSMUSP00000065940,10090.ENSMUSP00000068835, 10090.ENSMUSP00000078985,10090.ENSMUSP00000079124, 10090.ENSMUSP00000079198,10090.ENSMUSP00000095866, 10090.ENSMUSP00000098897,10090.ENSMUSP00000099758, 10090.ENSMUSP00000099990,10090.ENSMUSP00000099991, 10090.ENSMUSP00000101082,10090.ENSMUSP00000102710, 10090.ENSMUSP00000103935,10090.ENSMUSP00000106199, 10090.ENSMUSP00000111415,10090.ENSMUSP00000112999, 10090.ENSMUSP00000122219,10090.ENSMUSP00000122712, 10090.ENSMUSP00000125993,10090.ENSMUSP00000129800, 10090.ENSMUSP00000133499,10090.ENSMUSP00000133851 | Plaur,Msh6,Pmp22,Pbk,Nde1,Slc11a2,Gzma,Psmb8,Incenp,Prdx5,Efna1,Oasl1,Oasl2,Serpina7,Dnaja2,Slx4,Atg14,Serpine1,Cblc,Elmod2,Cenpe,S100a1,Camk2n1,Azin1,Hmgb2,Rabgap1,Krit1,Birc5,Oas1a,Arrb1,Mad2l1,Flcn,Anp32b,Pdpk1,Sirt1,Txnip,Ect2,Map4k5,Dnm1l,Arhgap23,Prlr,Ppp1r3g,Camsap3,Sh3glb1,Psmb9,Arhgap21 |
| GO:0006913 | Nucleocytoplasmic transport | 11 | 199 | 0.71 | 0.0042 | 10090.ENSMUSP00000018506,10090.ENSMUSP00000028382, 10090.ENSMUSP00000032898,10090.ENSMUSP00000034079, 10090.ENSMUSP00000034612,10090.ENSMUSP00000034787, 10090.ENSMUSP00000062766,10090.ENSMUSP00000063839, 10090.ENSMUSP00000102710,10090.ENSMUSP00000106801, 10090.ENSMUSP00000117045 | Kpna2,Nup35,Ipo5,Heatr3,Ddx25,Phip,Nupl2,Adam10,Txnip,Pom121,Srsf3 |

| | | | | | | | |
|---|---|---|---|---|---|---|---|
| GO:0048522 | Positive regulation of cellular process | 88 | 5407 | 0.17 | 0.0046 | 10090.ENSMUSP00000002284,10090.ENSMUSP00000002413,10090.ENSMUSP00000004430,10090.ENSMUSP00000004985,10090.ENSMUSP00000017851,10090.ENSMUSP00000020308,10090.ENSMUSP00000020908,10090.ENSMUSP00000022227,10090.ENSMUSP00000022380,10090.ENSMUSP00000022612,10090.ENSMUSP00000022927,10090.ENSMUSP00000023769,10090.ENSMUSP00000023774,10090.ENSMUSP00000023897,10090.ENSMUSP00000024104,10090.ENSMUSP00000024657,10090.ENSMUSP00000025254,10090.ENSMUSP00000025562,10090.ENSMUSP00000027933,10090.ENSMUSP00000029018,10090.ENSMUSP00000029566,10090.ENSMUSP00000030121,10090.ENSMUSP00000030367,10090.ENSMUSP00000030623,10090.ENSMUSP00000032800,10090.ENSMUSP00000032898,10090.ENSMUSP00000034787,10090.ENSMUSP00000034964,10090.ENSMUSP00000035101,10090.ENSMUSP00000037007,10090.ENSMUSP00000037459,10090.ENSMUSP00000038871,10090.ENSMUSP00000039047,10090.ENSMUSP00000039586,10090.ENSMUSP00000050854,10090.ENSMUSP00000053483,10090.ENSMUSP00000056778,10090.ENSMUSP00000057938,10090.ENSMUSP00000058037,10090.ENSMUSP00000058237,10090.ENSMUSP00000058940,10090.ENSMUSP00000059159,10090.ENSMUSP00000059211,10090.ENSMUSP00000060349,10090.ENSMUSP00000060791,10090.ENSMUSP00000061877,10090.ENSMUSP00000063839,10090.ENSMUSP00000065363,10090.ENSMUSP00000065544,10090.ENSMUSP00000065940,10090.ENSMUSP00000067217,10090.ENSMUSP00000075172,10090.ENSMUSP00000079124,10090.ENSMUSP00000082548, [illegible] | Plaur,Tmem161a,Clcn3,Brpf3,Serinc3,Ddit4,E2f6,Cenpk,Psmc6,Pbk,Rad21,Atf1,Slc11a2,Gzma,Gcm1,Phf10,Lims2,Incenp,Dtl,Zbp1,Efna1,B4galt1,Ssbp3,Sfpq,Tyrobp,Ipo5,Phip,Tipin,Csrnp1,Hspb8,Zfand1,Slx4,Atg14,Serpine1,Zfp367,Hic1,E2f8,Cenpe,Zfpm1,S100a1,Whsc1,Aspm,Ercc8,Camk2n1,Hilpda,Insig1,Adam10,Nfil3,Azin1,Hmgb2,Cdc42ep3,Frmd4a,Birc5,Isg15,Kif20b,Fmr1,Magi1,Arrb1,Mad2l1,Atp11c,Flcn,Anp32b,Pdpk1,Sirt1,Txnip,Ect2,Irf1,Map4k5,Bclaf3,Arid5a,Dnm1l,Ube2v2,Bcl3,Srsf1,Nmi,Prlr,Ppp1r3g,Cep295,Tgm1,Tfdp2,Odc1,Tcea1,Sh3glb1,Zfyve27,Tgif1,H2-D1,Golph3l,Zfp131 |
| GO:0022402 | Cell cycle process | 23 | 784 | 0.43 | 0.0049 | 10090.ENSMUSP00000005503,10090.ENSMUSP00000018361,10090.ENSMUSP00000020835,10090.ENSMUSP00000020908,10090.ENSMUSP00000022227,10090.ENSMUSP00000022927,10090.ENSMUSP00000023353,10090.ENSMUSP00000023359,10090.ENSMUSP00000025562,10090.ENSMUSP00000025930,10090.ENSMUSP00000027697,10090.ENSMUSP00000033913,10090.ENSMUSP00000034964,10090.ENSMUSP00000038871,10090.ENSMUSP00000047872,10090.ENSMUSP00000049973,10090.ENSMUSP00000057938,10090.ENSMUSP00000059159,10090.ENSMUSP00000079124,10090.ENSMUSP00000098897,10090.ENSMUSP00000099990,10090.ENSMUSP00000103935,10090.ENSMUSP00000104548 | Msh6,Pmp22,Ppm1d,E2f6,Cenpk,Rad21,Mcm4,Nde1,Incenp,Smc3,Cdk18,Dctn6,Tipin,Slx4,Smc4,Haus3,Cenpe,Aspm,Birc5,Mad2l1,Anp32b,Ect2,Irf1 |
| GO:0000278 | Mitotic cell cycle | 18 | 527 | 0.5 | 0.0058 | 10090.ENSMUSP00000020835,10090.ENSMUSP00000020908,10090.ENSMUSP00000022227,10090.ENSMUSP00000022612,10090.ENSMUSP00000023353,10090.ENSMUSP00000023359,10090.ENSMUSP00000025562,10090.ENSMUSP00000027697,10090.ENSMUSP00000033913,10090.ENSMUSP00000034964,10090.ENSMUSP00000047872,10090.ENSMUSP00000056778,10090.ENSMUSP00000057938,10090.ENSMUSP00000079124,10090.ENSMUSP00000098897,10090.ENSMUSP00000099990,10090.ENSMUSP00000103935,10090.ENSMUSP00000128260 | Ppm1d,E2f6,Cenpk,Pbk,Mcm4,Nde1,Incenp,Cdk18,Dctn6,Tipin,Smc4,E2f8,Cenpe,Birc5,Mad2l1,Anp32b,Ect2,Tfdp2 |
| GO:0019941 | Modification-dependent protein catabolic process | 17 | 481 | 0.51 | 0.0063 | 10090.ENSMUSP00000018361,10090.ENSMUSP00000022380,10090.ENSMUSP00000025196,10090.ENSMUSP00000027933,10090.ENSMUSP00000030289,10090.ENSMUSP00000032198,10090.ENSMUSP00000039206,10090.ENSMUSP00000039955,10090.ENSMUSP00000059211,10090.ENSMUSP00000082548,10090.ENSMUSP00000091429,10090.ENSMUSP00000095866,10090.ENSMUSP00000101082,10090.ENSMUSP00000110645,10090.ENSMUSP00000111780,10090.ENSMUSP00000132446,10090.ENSMUSP00000133499 | Pmp22,Psmc6,Psmb8,Dtl,Usp1,Usp18,Hace1,Cblc,Ercc8,Isg15,Rnf213,Arrb1,Sirt1,Cnot4,Rnf186,Ube2d2a,Psmb9 |
| GO:1903900 | Regulation of viral life cycle | 9 | 138 | 0.78 | 0.0064 | 10090.ENSMUSP00000018506,10090.ENSMUSP00000026565,10090.ENSMUSP00000031540,10090.ENSMUSP00000031542,10090.ENSMUSP00000079198,10090.ENSMUSP00000082548,10090.ENSMUSP00000085906,10090.ENSMUSP00000103432,10090.ENSMUSP00000129765 | Kpna2,Ifitm3,Oasl1,Oasl2,Oas1a,Isg15,Fmr1,Trim59,Parp10 |
| GO:1903901 | Negative regulation of viral life cycle | 7 | 77 | 0.92 | 0.0068 | 10090.ENSMUSP00000026565,10090.ENSMUSP00000031540,10090.ENSMUSP00000031542,10090.ENSMUSP00000079198,10090.ENSMUSP00000082548,10090.ENSMUSP00000103432,10090.ENSMUSP00000129765 | Ifitm3,Oasl1,Oasl2,Oas1a,Isg15,Trim59,Parp10 |
| GO:0031324 | Negative regulation of cellular metabolic process | 49 | 2497 | 0.26 | 0.0072 | 10090.ENSMUSP00000002284,10090.ENSMUSP00000005503,10090.ENSMUSP00000020308,10090.ENSMUSP00000020835,10090.ENSMUSP00000020908,10090.ENSMUSP00000022612,10090.ENSMUSP00000023897,10090.ENSMUSP00000024657,10090.ENSMUSP00000025904,10090.ENSMUSP00000029566,10090.ENSMUSP00000030623,10090.ENSMUSP00000033626,10090.ENSMUSP00000034964,10090.ENSMUSP00000036971,10090.ENSMUSP00000038871,10090.ENSMUSP00000039047,10090.ENSMUSP00000039586,10090.ENSMUSP00000039955,10090.ENSMUSP00000053483,10090.ENSMUSP00000056715,10090.ENSMUSP00000056778,10090.ENSMUSP00000058037,10090.ENSMUSP00000058237,10090.ENSMUSP00000058940,10090.ENSMUSP00000060349,10090.ENSMUSP00000061390,10090.ENSMUSP00000061877,10090.ENSMUSP00000065363,10090.ENSMUSP00000065940,10090.ENSMUSP00000077971,10090.ENSMUSP00000079124,10090.ENSMUSP00000080694,10090.ENSMUSP00000082548,10090.ENSMUSP00000085906,10090.ENSMUSP00000095866,10090.ENSMUSP00000098897,10090.ENSMUSP00000099758,10090.ENSMUSP00000099991,10090.ENSMUSP00000101082,10090.ENSMUSP00000102710,10090.ENSMUSP00000102824,10090.ENSMUSP00000104548,10090.ENSMUSP00000110684,10090.ENSMUSP00000113851,10090.ENSMUSP00000125993,10090.ENSMUSP00000128260,10090.ENSMUSP00000129765,10090.ENSMUSP00000130930,10090.ENSMUSP00000136867 | Plaur,Msh6,Ddit4,Ppm1d,E2f6,Pbk,Gzma,Phf10,Prdx5,Efna1,Sfpq,Serpina7,Tipin,Mvk,Slx4,Atg14,Serpine1,Cblc,Hic1,Atxn2,E2f8,Zfpm1,S100a1,Whsc1,Camk2n1,Tob2,Insig1,Nfil3,Hmgb2,Phf6,Birc5,Rps6ka6,Isg15,Fmr1,Arrb1,Mad2l1,Flcn,Pdpk1,Sirt1,Txnip,Crebzf,Irf1,Arid5a,Bcl3,Camsap3,Tfdp2,Parp10,Tgif1,Zfp131 |
| GO:0045071 | Negative regulation of viral genome replication | 6 | 53 | 1.02 | 0.0077 | 10090.ENSMUSP00000026565,10090.ENSMUSP00000031540,10090.ENSMUSP00000031542,10090.ENSMUSP00000079198,10090.ENSMUSP00000082548,10090.ENSMUSP00000129765 | Ifitm3,Oasl1,Oasl2,Oas1a,Isg15,Parp10 |
| GO:0051028 | mRNA transport | 8 | 109 | 0.83 | 0.0077 | 10090.ENSMUSP00000028382,10090.ENSMUSP00000034612,10090.ENSMUSP00000062766,10090.ENSMUSP00000085906,10090.ENSMUSP00000106801,10090.ENSMUSP00000107595,10090.ENSMUSP00000117045,10090.ENSMUSP00000120595 | Nup35,Ddx25,Nupl2,Fmr1,Pom121,Hnrnpa3,Srsf3,Srsf1 |

| | | | | | | | |
|---|---|---|---|---|---|---|---|
| GO:0051603 | Proteolysis involved in cellular protein catabolic process | 18 | 545 | 0.48 | 0.0077 | 10090.ENSMUSP00000018361,10090.ENSMUSP00000022380,10090.ENSMUSP00000023897,10090.ENSMUSP00000025196,10090.ENSMUSP00000027933,10090.ENSMUSP00000030289,10090.ENSMUSP00000032198,10090.ENSMUSP00000039206,10090.ENSMUSP00000039955,10090.ENSMUSP00000059211,10090.ENSMUSP00000082548,10090.ENSMUSP00000091429,10090.ENSMUSP00000095866,10090.ENSMUSP00000101082,10090.ENSMUSP00000110645,10090.ENSMUSP00000111780,10090.ENSMUSP00000132446,10090.ENSMUSP00000133499 | Pmp22,Psmc6,Gzma,Psmb8,Dtl,Usp1,Usp18,Hace1,Cblc,Ercc8,Isg15,Rnf213,Arrb1,Sirt1,Cnot4,Rnf186,Ube2d2a,Psmb9 |
| GO:0065009 | Regulation of molecular function | 74 | 4379 | 0.19 | 0.0077 | 10090.ENSMUSP00000002284,10090.ENSMUSP00000005503,10090.ENSMUSP00000018361,10090.ENSMUSP00000020908,10090.ENSMUSP00000022612,10090.ENSMUSP00000023359,10090.ENSMUSP00000023769,10090.ENSMUSP00000023774,10090.ENSMUSP00000023897,10090.ENSMUSP00000024104,10090.ENSMUSP00000025196,10090.ENSMUSP00000025562,10090.ENSMUSP00000025904,10090.ENSMUSP00000027760,10090.ENSMUSP00000029566,10090.ENSMUSP00000031540,10090.ENSMUSP00000031542,10090.ENSMUSP00000033626,10090.ENSMUSP00000034138,10090.ENSMUSP00000034270,10090.ENSMUSP00000035101,10090.ENSMUSP00000038871,10090.ENSMUSP00000039047,10090.ENSMUSP00000039586,10090.ENSMUSP00000039955,10090.ENSMUSP00000041906,10090.ENSMUSP00000050854,10090.ENSMUSP00000052287,10090.ENSMUSP00000053483,10090.ENSMUSP00000056778,10090.ENSMUSP00000057938,10090.ENSMUSP00000058037,10090.ENSMUSP00000058237,10090.ENSMUSP00000060349,10090.ENSMUSP00000061390,10090.ENSMUSP00000064521,10090.ENSMUSP00000065363,10090.ENSMUSP00000065544,10090.ENSMUSP00000065940,10090.ENSMUSP00000066954,10090.ENSMUSP00000068835,10090.ENSMUSP00000077971,10090.ENSMUSP00000078985,10090.ENSMUSP00000079124,10090.ENSMUSP00000079198,10090.ENSMUSP00000085906,10090.ENSMUSP00000095866,10090.ENSMUSP00000098897,10090.ENSMUSP00000099758,10090.ENSMUSP00000099990,10090.ENSMUSP00000099991,10090.ENSMUSP00000101082,10090.ENSMUSP00000102710,10090.ENSMUSP00000102824,[illegible] | Plaur,Msh6,Pmp22,E2f6,Pbk,Nde1,Atf1,Slc11a2,Gzma,Gcm1,Psmb8,Incenp,Prdx5,Rgl1,Efna1,Oasl1,Oasl2,Serpina7,Dnaja2,Map1lc3b,Csrnp1,Slx4,Atg14,Serpine1,Cblc,Dis3,Zfp367,Elmod2,Hic1,E2f8,Cenpe,Zfpm1,S100a1,Camk2n1,Tob2,Artn,Nfil3,Azin1,Hmgb2,Ly6c1,Rabgap1,Phf6,Krit1,Birc5,Oas1a,Fmr1,Arrb1,Mad2l1,Flcn,Anp32b,Pdpk1,Sirt1,Txnip,Crebzf,Ect2,Irf1,Map4k5,Bclaf3,Arid5a,Dnm1l,Arhgap23,Bcl3,Irf9,Nmi,Prlr,Ppp1r3g,Camsap3,Tfdp2,Parp10,Sh3glb1,Tgif1,Psmb9,Arhgap21,Zfp131 |
| GO:0051252 | Regulation of rna metabolic process | 56 | 3027 | 0.23 | 0.0083 | 10090.ENSMUSP00000020123,10090.ENSMUSP00000020835,10090.ENSMUSP00000020908,10090.ENSMUSP00000022227,10090.ENSMUSP00000022380,10090.ENSMUSP00000022927,10090.ENSMUSP00000023769,10090.ENSMUSP00000024104,10090.ENSMUSP00000024657,10090.ENSMUSP00000025904,10090.ENSMUSP00000027697,10090.ENSMUSP00000028382,10090.ENSMUSP00000029566,10090.ENSMUSP00000030367,10090.ENSMUSP00000030623,10090.ENSMUSP00000031540,10090.ENSMUSP00000031542,10090.ENSMUSP00000034787,10090.ENSMUSP00000035101,10090.ENSMUSP00000050461,10090.ENSMUSP00000050854,10090.ENSMUSP00000053483,10090.ENSMUSP00000056778,10090.ENSMUSP00000058037,10090.ENSMUSP00000058237,10090.ENSMUSP00000058940,10090.ENSMUSP00000061390,10090.ENSMUSP00000061877,10090.ENSMUSP00000065363,10090.ENSMUSP00000065940,10090.ENSMUSP00000077971,10090.ENSMUSP00000079124,10090.ENSMUSP00000079198,10090.ENSMUSP00000080640,10090.ENSMUSP00000085906,10090.ENSMUSP00000095866,10090.ENSMUSP00000096686,10090.ENSMUSP00000099758,10090.ENSMUSP00000101082,10090.ENSMUSP00000101661,10090.ENSMUSP00000102710,10090.ENSMUSP00000102824,10090.ENSMUSP00000104548,10090.ENSMUSP00000107595,10090.ENSMUSP00000108083,10090.ENSMUSP00000110684,10090.ENSMUSP00000113851,10090.ENSMUSP00000117045,10090.ENSMUSP00000120525,10090.ENSMUSP00000120595,10090.ENSMUSP00000120647,10090.ENSMUSP00000128260,10090.ENSMUSP00000129157,10090.ENSMUSP00000129765, | Tmpo,Ppm1d,E2f6,Cenpk,Psmc6,Rad21,Atf1,Gcm1,Phf10,Prdx5,Cdk18,Nup35,Efna1,Ssbp3,Sfpq,Oasl1,Oasl2,Phip,Csrnp1,Rbm12,Zfp367,Hic1,E2f8,Zfpm1,S100a1,Whsc1,Tob2,Insig1,Nfil3,Hmgb2,Phf6,Birc5,Oas1a,Slc39a8,Fmr1,Arrb1,Mbnl1,Flcn,Sirt1,Zbtb8b,Txnip,Crebzf,Irf1,Hnrnpa3,Bclaf3,Arid5a,Bcl3,Srsf3,Irf9,Srsf1,Nmi,Tfdp2,Tcea1,Parp10,Tgif1,Zfp131 |
| GO:0051172 | Negative regulation of nitrogen compound metabolic process | 46 | 2310 | 0.26 | 0.0084 | 10090.ENSMUSP00000002284,10090.ENSMUSP00000005503,10090.ENSMUSP00000020308,10090.ENSMUSP00000020835,10090.ENSMUSP00000020908,10090.ENSMUSP00000022612,10090.ENSMUSP00000023897,10090.ENSMUSP00000024657,10090.ENSMUSP00000025904,10090.ENSMUSP00000029566,10090.ENSMUSP00000030623,10090.ENSMUSP00000033626,10090.ENSMUSP00000036971,10090.ENSMUSP00000038871,10090.ENSMUSP00000039047,10090.ENSMUSP00000039586,10090.ENSMUSP00000039955,10090.ENSMUSP00000053483,10090.ENSMUSP00000056778,10090.ENSMUSP00000058037,10090.ENSMUSP00000058237,10090.ENSMUSP00000058940,10090.ENSMUSP00000060349,10090.ENSMUSP00000061390,10090.ENSMUSP00000065363,10090.ENSMUSP00000065544,10090.ENSMUSP00000065940,10090.ENSMUSP00000077971,10090.ENSMUSP00000079124,10090.ENSMUSP00000080694,10090.ENSMUSP00000082548,10090.ENSMUSP00000085906,10090.ENSMUSP00000095866,10090.ENSMUSP00000098897,10090.ENSMUSP00000099758,10090.ENSMUSP00000099991,10090.ENSMUSP00000101082,10090.ENSMUSP00000102710,10090.ENSMUSP00000102824,10090.ENSMUSP00000104548,10090.ENSMUSP00000110684,10090.ENSMUSP00000113851,10090.ENSMUSP00000128260,10090.ENSMUSP00000129765,10090.ENSMUSP00000130930,10090.ENSMUSP00000136867 | Plaur,Msh6,Ddit4,Ppm1d,E2f6,Pbk,Gzma,Phf10,Prdx5,Efna1,Sfpq,Serpina7,Mvk,Slx4,Atg14,Serpine1,Cblc,Hic1,E2f8,Zfpm1,S100a1,Whsc1,Camk2n1,Tob2,Nfil3,Azin1,Hmgb2,Phf6,Birc5,Rps6ka6,Isg15,Fmr1,Arrb1,Mad2l1,Flcn,Pdpk1,Sirt1,Txnip,Crebzf,Irf1,Arid5a,Bcl3,Tfdp2,Parp10,Tgif1,Zfp131 |
| GO:0045787 | Positive regulation of cell cycle | 14 | 354 | 0.56 | 0.0086 | 10090.ENSMUSP00000022927,10090.ENSMUSP00000027933,10090.ENSMUSP00000030623,10090.ENSMUSP00000034787,10090.ENSMUSP00000056778,10090.ENSMUSP00000057938,10090.ENSMUSP00000065544,10090.ENSMUSP00000079124,10090.ENSMUSP00000084599,10090.ENSMUSP00000098897,10090.ENSMUSP00000099990,10090.ENSMUSP00000103935,10090.ENSMUSP00000123788,10090.ENSMUSP00000128090 | Rad21,Dtl,Sfpq,Phip,E2f8,Cenpe,Azin1,Birc5,Kif20b,Mad2l1,Anp32b,Ect2,Cep295,Tgm1 |
| GO:0009057 | Macromolecule catabolic process | 23 | 827 | 0.41 | 0.0087 | 10090.ENSMUSP00000018361,10090.ENSMUSP00000019469,10090.ENSMUSP00000022380,10090.ENSMUSP00000023897,10090.ENSMUSP00000025196,10090.ENSMUSP00000027933,10090.ENSMUSP00000030289,10090.ENSMUSP00000032198,10090.ENSMUSP00000039206,10090.ENSMUSP00000039955,10090.ENSMUSP00000041906,10090.ENSMUSP00000059211,10090.ENSMUSP00000078940,10090.ENSMUSP00000082548,10090.ENSMUSP00000091429,10090.ENSMUSP00000095866,10090.ENSMUSP00000101082,10090.ENSMUSP00000110645,10090.ENSMUSP00000111780,10090.ENSMUSP00000126565,10090.ENSMUSP00000129800,10090.ENSMUSP00000132446,10090.ENSMUSP00000133499 | Pmp22,G6pc,Psmc6,Gzma,Psmb8,Dtl,Usp1,Usp18,Hace1,Cblc,Dis3,Ercc8,Gspt1,Isg15,Rnf213,Arrb1,Sirt1,Cnot4,Rnf186,Lsm5,Sh3glb1,Ube2d2a,Psmb9 |

| | | | | | | | |
|---|---|---|---|---|---|---|---|
| GO:0098542 | Defense response to other organism | 22 | 775 | 0.42 | 0.0094 | 10090.ENSMUSP00000017851,10090.ENSMUSP00000020308,10090.ENSMUSP00000026565,10090.ENSMUSP00000029018,10090.ENSMUSP00000030623,10090.ENSMUSP00000031540,10090.ENSMUSP00000031542,10090.ENSMUSP00000032473,10090.ENSMUSP00000033173,10090.ENSMUSP00000039586,10090.ENSMUSP00000040246,10090.ENSMUSP00000041091,10090.ENSMUSP00000065940,10090.ENSMUSP00000074009,10090.ENSMUSP00000079198,10090.ENSMUSP00000082548,10090.ENSMUSP00000099889,10090.ENSMUSP00000103432,10090.ENSMUSP00000104548,10090.ENSMUSP00000110684,10090.ENSMUSP00000113851,10090.ENSMUSP00000120647 | Serinc3,Ddit4,Ifitm3,Zbp1,Sfpq,Oasl1,Oasl2,Iigp1,Polr3e,Serpine1,C1qb,Rtp4,Hmgb2,Ifnlr1,Oas1a,Isg15,Ifit3,Trim59,Irf1,Arid5a,Bcl3,Nmi |
| GO:0045087 | Innate immune response | 18 | 558 | 0.47 | 0.0096 | 10090.ENSMUSP00000017851,10090.ENSMUSP00000026565,10090.ENSMUSP00000029018,10090.ENSMUSP00000030623,10090.ENSMUSP00000031540,10090.ENSMUSP00000031542,10090.ENSMUSP00000032473,10090.ENSMUSP00000033173,10090.ENSMUSP00000040246,10090.ENSMUSP00000065940,10090.ENSMUSP00000074009,10090.ENSMUSP00000079198,10090.ENSMUSP00000082548,10090.ENSMUSP00000099889,10090.ENSMUSP00000103432,10090.ENSMUSP00000104548,10090.ENSMUSP00000110684,10090.ENSMUSP00000120647 | Serinc3,Ifitm3,Zbp1,Sfpq,Oasl1,Oasl2,Iigp1,Polr3e,C1qb,Hmgb2,Ifnlr1,Oas1a,Isg15,Ifit3,Trim59,Irf1,Arid5a,Nmi |
| GO:0048523 | Negative regulation of cellular process | 77 | 4672 | 0.18 | 0.0102 | 10090.ENSMUSP00000002284,10090.ENSMUSP00000002413,10090.ENSMUSP00000005503,10090.ENSMUSP00000018361,10090.ENSMUSP00000020308,10090.ENSMUSP00000020835,10090.ENSMUSP00000020908,10090.ENSMUSP00000022612,10090.ENSMUSP00000022927,10090.ENSMUSP00000023897,10090.ENSMUSP00000024657,10090.ENSMUSP00000025254,10090.ENSMUSP00000025904,10090.ENSMUSP00000026565,10090.ENSMUSP00000027933,10090.ENSMUSP00000029566,10090.ENSMUSP00000030121,10090.ENSMUSP00000030623,10090.ENSMUSP00000032800,10090.ENSMUSP00000033626,10090.ENSMUSP00000034003,10090.ENSMUSP00000034787,10090.ENSMUSP00000034964,10090.ENSMUSP00000036971,10090.ENSMUSP00000038871,10090.ENSMUSP00000039047,10090.ENSMUSP00000039586,10090.ENSMUSP00000039955,10090.ENSMUSP00000041744,10090.ENSMUSP00000044603,10090.ENSMUSP00000053483,10090.ENSMUSP00000056715,10090.ENSMUSP00000056778,10090.ENSMUSP00000057938,10090.ENSMUSP00000058037,10090.ENSMUSP00000058237,10090.ENSMUSP00000058940,10090.ENSMUSP00000059159,10090.ENSMUSP00000060349,10090.ENSMUSP00000061390,10090.ENSMUSP00000061877,10090.ENSMUSP00000063839,10090.ENSMUSP00000065363,10090.ENSMUSP00000065940,10090.ENSMUSP00000075172,10090.ENSMUSP00000076961,10090.ENSMUSP00000077971,10090.ENSMUSP00000078985,10090.ENSMUSP00000079124,10090.ENSMUSP00000080694,10090.ENSMUSP00000082548,10090.ENSMUSP00000085906,10090.ENSMUSP00000091429,10090.ENSMUSP00000095866,[illegible] | Plaur,Tmem161a,Msh6,Pmp22,Ddit4,Ppm1d,E2f6,Pbk,Rad21,Gzma,Phf10,Lims2,Prdx5,Ifitm3,Dtl,Efna1,B4galt1,Sfpq,Tyrobp,Serpina7,Fgl1,Phip,Tipin,Mvk,Slx4,Atg14,Serpine1,Cblc,Ofd1,Pim3,Hic1,Atxn2,E2f8,Cenpe,Zfpm1,S100a1,Whsc1,Aspm,Camk2n1,Tob2,Insig1,Adam10,Nfil3,Hmgb2,Frmd4a,Tnfaip8l1,Phf6,Krit1,Birc5,Rps6ka6,Isg15,Fmr1,Rnf213,Arrb1,Mad2l1,Flcn,Ifit3,Anp32b,Pdpk1,Sirt1,Txnip,Crebzf,Trim59,Irf1,Arid5a,Dnm1l,Ube2v2,Bcl3,Prlr,Xaf1,Camsap3,Tfdp2,Parp10,Sh3glb1,Tgif1,H2-D1,Zfp131 |
| GO:0045069 | Regulation of viral genome replication | 7 | 87 | 0.87 | 0.0114 | 10090.ENSMUSP00000026565,10090.ENSMUSP00000031540,10090.ENSMUSP00000031542,10090.ENSMUSP00000079198,10090.ENSMUSP00000082548,10090.ENSMUSP00000085906,10090.ENSMUSP00000129765 | Ifitm3,Oasl1,Oasl2,Oas1a,Isg15,Fmr1,Parp10 |
| GO:0044265 | Cellular macromolecule catabolic process | 21 | 735 | 0.42 | 0.012 | 10090.ENSMUSP00000018361,10090.ENSMUSP00000022380,10090.ENSMUSP00000023897,10090.ENSMUSP00000025196,10090.ENSMUSP00000027933,10090.ENSMUSP00000030289,10090.ENSMUSP00000032198,10090.ENSMUSP00000039206,10090.ENSMUSP00000039955,10090.ENSMUSP00000041906,10090.ENSMUSP00000059211,10090.ENSMUSP00000078940,10090.ENSMUSP00000082548,10090.ENSMUSP00000091429,10090.ENSMUSP00000095866,10090.ENSMUSP00000101082,10090.ENSMUSP00000110645,10090.ENSMUSP00000111780,10090.ENSMUSP00000126565,10090.ENSMUSP00000132446,10090.ENSMUSP00000133499 | Pmp22,Psmc6,Gzma,Psmb8,Dtl,Usp1,Usp18,Hace1,Cblc,Dis3,Ercc8,Gspt1,Isg15,Rnf213,Arrb1,Sirt1,Cnot4,Rnf186,Lsm5,Ube2d2a,Psmb9 |
| GO:0006511 | Ubiquitin-dependent protein catabolic process | 16 | 471 | 0.49 | 0.0131 | 10090.ENSMUSP00000018361,10090.ENSMUSP00000022380,10090.ENSMUSP00000025196,10090.ENSMUSP00000027933,10090.ENSMUSP00000030289,10090.ENSMUSP00000032198,10090.ENSMUSP00000039206,10090.ENSMUSP00000039955,10090.ENSMUSP00000059211,10090.ENSMUSP00000091429,10090.ENSMUSP00000095866,10090.ENSMUSP00000101082,10090.ENSMUSP00000110645,10090.ENSMUSP00000111780,10090.ENSMUSP00000132446,10090.ENSMUSP00000133499 | Pmp22,Psmc6,Psmb8,Dtl,Usp1,Usp18,Hace1,Cblc,Ercc8,Rnf213,Arrb1,Sirt1,Cnot4,Rnf186,Ube2d2a,Psmb9 |
| GO:0045934 | Negative regulation of nucleobase-containing compound metabolic process | 32 | 1415 | 0.32 | 0.0135 | 10090.ENSMUSP00000005503,10090.ENSMUSP00000020308,10090.ENSMUSP00000020835,10090.ENSMUSP00000020908,10090.ENSMUSP00000023897,10090.ENSMUSP00000024657,10090.ENSMUSP00000025904,10090.ENSMUSP00000029566,10090.ENSMUSP00000030623,10090.ENSMUSP00000038871,10090.ENSMUSP00000053483,10090.ENSMUSP00000056778,10090.ENSMUSP00000058037,10090.ENSMUSP00000058237,10090.ENSMUSP00000058940,10090.ENSMUSP00000061390,10090.ENSMUSP00000065363,10090.ENSMUSP00000065940,10090.ENSMUSP00000077971,10090.ENSMUSP00000079124,10090.ENSMUSP00000085906,10090.ENSMUSP00000099758,10090.ENSMUSP00000101082,10090.ENSMUSP00000102710,10090.ENSMUSP00000102824,10090.ENSMUSP00000104548,10090.ENSMUSP00000110684,10090.ENSMUSP00000113851,10090.ENSMUSP00000128260,10090.ENSMUSP00000129765,10090.ENSMUSP00000130930,10090.ENSMUSP00000136867 | Msh6,Ddit4,Ppm1d,E2f6,Gzma,Phf10,Prdx5,Efna1,Sfpq,Slx4,Hic1,E2f8,Zfpm1,S100a1,Whsc1,Tob2,Nfil3,Hmgb2,Phf6,Birc5,Fmr1,Flcn,Sirt1,Txnip,Crebzf,Irf1,Arid5a,Bcl3,Tfdp2,Parp10,Tgif1,Zfp131 |
| GO:0051302 | Regulation of cell division | 10 | 197 | 0.67 | 0.0135 | 10090.ENSMUSP00000025562,10090.ENSMUSP00000034787,10090.ENSMUSP00000056778,10090.ENSMUSP00000057938,10090.ENSMUSP00000059159,10090.ENSMUSP00000084599,10090.ENSMUSP00000099758,10090.ENSMUSP00000102710,10090.ENSMUSP00000103935,10090.ENSMUSP00000129800 | Incenp,Phip,E2f8,Cenpe,Aspm,Kif20b,Flcn,Txnip,Ect2,Sh3glb1 |
| GO:0034654 | Nucleobase-containing compound biosynthetic process | 20 | 689 | 0.43 | 0.0138 | 10090.ENSMUSP00000011058,10090.ENSMUSP00000020217,10090.ENSMUSP00000024104,10090.ENSMUSP00000027933,10090.ENSMUSP00000028328,10090.ENSMUSP00000031542,10090.ENSMUSP00000033173,10090.ENSMUSP00000051968,10090.ENSMUSP00000065363,10090.ENSMUSP00000079198,10090.ENSMUSP00000079703,10090.ENSMUSP00000080640,10090.ENSMUSP00000095866,10090.ENSMUSP00000101082,10090.ENSMUSP00000102824,10090.ENSMUSP00000108557,10090.ENSMUSP00000113851,10090.ENSMUSP00000120647,10090.ENSMUSP00000129157,10090.ENSMUSP00000129765 | Pdhx,Nudt4,Gcm1,Dtl,Entpd2,Oasl2,Polr3e,Polr2k,Nfil3,Oas1a,Amd2,Slc39a8,Arrb1,Sirt1,Crebzf,Nmnat3,Bcl3,Nmi,Tcea1,Parp10 |

| | | | | | | | |
|---|---|---|---|---|---|---|---|
| GO:0010468 | Regulation of gene expression | 68 | 4041 | 0.19 | 0.0162 | 10090.ENSMUSP00000018361,10090.ENSMUSP00000019469,10090.ENSMUSP00000020123,10090.ENSMUSP00000020835,10090.ENSMUSP00000020908,10090.ENSMUSP00000022227,10090.ENSMUSP00000022380,10090.ENSMUSP00000022927,10090.ENSMUSP00000023769,10090.ENSMUSP00000024104,10090.ENSMUSP00000024657,10090.ENSMUSP00000025161,10090.ENSMUSP00000025904,10090.ENSMUSP00000027697,10090.ENSMUSP00000028382,10090.ENSMUSP00000029566,10090.ENSMUSP00000030367,10090.ENSMUSP00000030623,10090.ENSMUSP00000032800,10090.ENSMUSP00000034612,10090.ENSMUSP00000034787,10090.ENSMUSP00000035101,10090.ENSMUSP00000036971,10090.ENSMUSP00000039047,10090.ENSMUSP00000039586,10090.ENSMUSP00000048051,10090.ENSMUSP00000050461,10090.ENSMUSP00000050854,10090.ENSMUSP00000053483,10090.ENSMUSP00000056778,10090.ENSMUSP00000058037,10090.ENSMUSP00000058237,10090.ENSMUSP00000058940,10090.ENSMUSP00000060349,10090.ENSMUSP00000060791,10090.ENSMUSP00000061390,10090.ENSMUSP00000061877,10090.ENSMUSP00000065363,10090.ENSMUSP00000065940,10090.ENSMUSP00000077971,10090.ENSMUSP00000078940,10090.ENSMUSP00000079124,10090.ENSMUSP00000080640,10090.ENSMUSP00000082548,10090.ENSMUSP00000085906,10090.ENSMUSP00000095794,10090.ENSMUSP00000095866,10090.ENSMUSP00000096686,10090.ENSMUSP00000099758,10090.ENSMUSP00000101082,10090.ENSMUSP00000101661,10090.ENSMUSP00000102710,10090.ENSMUSP00000102824,10090.ENSMUSP00000104548,[illegible] | Pmp22,G6pc,Tmpo,Ppm1d,E2f6,Cenpk,Psmc6,Rad21,Atf1,Gcm1,Phf10,Tapbp,Prdx5,Cdk18,Nup35,Efna1,Ssbp3,Sfpq,Tyrobp,Ddx25,Phip,Csrnp1,Mvk,Atg14,Serpine1,Tiparp,Rbm12,Zfp367,Hic1,E2f8,Zfpm1,S100a1,Whsc1,Camk2n1,Hilpda,Tob2,Insig1,Nfil3,Hmgb2,Phf6,Gspt1,Birc5,Slc39a8,Isg15,Fmr1,Hbb-bt,Arrb1,Mbnl1,Flcn,Sirt1,Zbtb8b,Txnip,Crebzf,Irf1,Bclaf3,Arid5a,Dnm1l,Bcl3,Srsf3,Irf9,Srsf1,Nmi,Lsm5,Tfdp2,Tcea1,Parp10,Tgif1,Zfp131 |
| GO:0018130 | Heterocycle biosynthetic process | 21 | 755 | 0.41 | 0.0162 | 10090.ENSMUSP00000011058,10090.ENSMUSP00000020217,10090.ENSMUSP00000023774,10090.ENSMUSP00000024104,10090.ENSMUSP00000027933,10090.ENSMUSP00000028328,10090.ENSMUSP00000031542,10090.ENSMUSP00000033173,10090.ENSMUSP00000051968,10090.ENSMUSP00000065363,10090.ENSMUSP00000079198,10090.ENSMUSP00000079703,10090.ENSMUSP00000080640,10090.ENSMUSP00000095866,10090.ENSMUSP00000101082,10090.ENSMUSP00000102824,10090.ENSMUSP00000108557,10090.ENSMUSP00000113851,10090.ENSMUSP00000120647,10090.ENSMUSP00000129157,10090.ENSMUSP00000129765 | Pdhx,Nudt4,Slc11a2,Gcm1,Dtl,Entpd2,Oasl2,Polr3e,Polr2k,Nfil3,Oas1a,Amd2,Slc39a8,Arrb1,Sirt1,Crebzf,Nmnat3,Bcl3,Nmi,Tcea1,Parp10 |
| GO:0006606 | Protein import into nucleus | 7 | 94 | 0.84 | 0.0164 | 10090.ENSMUSP00000018506,10090.ENSMUSP00000028382,10090.ENSMUSP00000032898,10090.ENSMUSP00000034079,10090.ENSMUSP00000034787,10090.ENSMUSP00000102710,10090.ENSMUSP00000106801 | Kpna2,Nup35,Ipo5,Heatr3,Phip,Txnip,Pom121 |
| GO:0006281 | DNA repair | 15 | 435 | 0.5 | 0.0171 | 10090.ENSMUSP00000004120,10090.ENSMUSP00000005503,10090.ENSMUSP00000022927,10090.ENSMUSP00000023353,10090.ENSMUSP00000025930,10090.ENSMUSP00000027933,10090.ENSMUSP00000030289,10090.ENSMUSP00000030623,10090.ENSMUSP00000038871,10090.ENSMUSP00000047872,10090.ENSMUSP00000059211,10090.ENSMUSP00000080949,10090.ENSMUSP00000101082,10090.ENSMUSP00000111443,10090.ENSMUSP00000129765 | Fancl,Msh6,Rad21,Mcm4,Smc3,Dtl,Usp1,Sfpq,Slx4,Smc4,Ercc8,Mdc1,Sirt1,Ube2v2,Parp10 |
| GO:0051336 | Regulation of hydrolase activity | 28 | 1184 | 0.34 | 0.0173 | 10090.ENSMUSP00000002284,10090.ENSMUSP00000005503,10090.ENSMUSP00000023359,10090.ENSMUSP00000023774,10090.ENSMUSP00000023897,10090.ENSMUSP00000025196,10090.ENSMUSP00000025904,10090.ENSMUSP00000029566,10090.ENSMUSP00000031540,10090.ENSMUSP00000031542,10090.ENSMUSP00000033626,10090.ENSMUSP00000034138,10090.ENSMUSP00000039586,10090.ENSMUSP00000052287,10090.ENSMUSP00000065940,10090.ENSMUSP00000068835,10090.ENSMUSP00000079124,10090.ENSMUSP00000079198,10090.ENSMUSP00000095866,10090.ENSMUSP00000099758,10090.ENSMUSP00000099990,10090.ENSMUSP00000099991,10090.ENSMUSP00000101082,10090.ENSMUSP00000103935,10090.ENSMUSP00000112999,10090.ENSMUSP00000125993,10090.ENSMUSP00000133499,10090.ENSMUSP00000133851 | Plaur,Msh6,Nde1,Slc11a2,Gzma,Psmb8,Prdx5,Efna1,Oasl1,Oasl2,Serpina7,Dnaja2,Serpine1,Elmod2,Hmgb2,Rabgap1,Birc5,Oas1a,Arrb1,Flcn,Anp32b,Pdpk1,Sirt1,Ect2,Arhgap23,Camsap3,Psmb9,Arhgap21 |
| GO:2001244 | Positive regulation of intrinsic apoptotic signaling pathway | 6 | 66 | 0.92 | 0.0179 | 10090.ENSMUSP00000017851,10090.ENSMUSP00000030623,10090.ENSMUSP00000099758,10090.ENSMUSP00000101082,10090.ENSMUSP00000111415,10090.ENSMUSP00000129800 | Serinc3,Sfpq,Flcn,Sirt1,Dnm1l,Sh3glb1 |
| GO:1901362 | Organic cyclic compound biosynthetic process | 23 | 883 | 0.38 | 0.0182 | 10090.ENSMUSP00000011058,10090.ENSMUSP00000020217,10090.ENSMUSP00000023774,10090.ENSMUSP00000024104,10090.ENSMUSP00000027933,10090.ENSMUSP00000028328,10090.ENSMUSP00000031542,10090.ENSMUSP00000033173,10090.ENSMUSP00000036971,10090.ENSMUSP00000051968,10090.ENSMUSP00000061877,10090.ENSMUSP00000065363,10090.ENSMUSP00000079198,10090.ENSMUSP00000079703,10090.ENSMUSP00000080640,10090.ENSMUSP00000095866,10090.ENSMUSP00000101082,10090.ENSMUSP00000102824,10090.ENSMUSP00000108557,10090.ENSMUSP00000113851,10090.ENSMUSP00000120647,10090.ENSMUSP00000129157,10090.ENSMUSP00000129765 | Pdhx,Nudt4,Slc11a2,Gcm1,Dtl,Entpd2,Oasl2,Polr3e,Mvk,Polr2k,Insig1,Nfil3,Oas1a,Amd2,Slc39a8,Arrb1,Sirt1,Crebzf,Nmnat3,Bcl3,Nmi,Tcea1,Parp10 |
| GO:0044085 | Cellular component biogenesis | 43 | 2201 | 0.25 | 0.0185 | 10090.ENSMUSP00000002198,10090.ENSMUSP00000018361,10090.ENSMUSP00000022227,10090.ENSMUSP00000023353,10090.ENSMUSP00000023359,10090.ENSMUSP00000023769,10090.ENSMUSP00000025161,10090.ENSMUSP00000025562,10090.ENSMUSP00000030367,10090.ENSMUSP00000031719,10090.ENSMUSP00000034079,10090.ENSMUSP00000034270,10090.ENSMUSP00000038871,10090.ENSMUSP00000039010,10090.ENSMUSP00000039047,10090.ENSMUSP00000041744,10090.ENSMUSP00000041906,10090.ENSMUSP00000048829,10090.ENSMUSP00000049973,10090.ENSMUSP00000050907,10090.ENSMUSP00000056715,10090.ENSMUSP00000057938,10090.ENSMUSP00000059159,10090.ENSMUSP00000063839,10090.ENSMUSP00000072386,10090.ENSMUSP00000079124,10090.ENSMUSP00000091429,10090.ENSMUSP00000095866,10090.ENSMUSP00000096193,10090.ENSMUSP00000096686,10090.ENSMUSP00000099487,10090.ENSMUSP00000099758,10090.ENSMUSP00000099990,10090.ENSMUSP00000099991,10090.ENSMUSP00000100534,10090.ENSMUSP00000101082,10090.ENSMUSP00000103935,10090.ENSMUSP00000111415,10090.ENSMUSP00000113431,10090.ENSMUSP00000120595,10090.ENSMUSP00000122219,10090.ENSMUSP00000130684,10090.ENSMUSP00000136493 | Sf3a1,Pmp22,Cenpk,Mcm4,Nde1,Atf1,Tapbp,Incenp,Ssbp3,Fscn3,Heatr3,Map1lc3b,Slx4,Wdr55,Atg14,Ofd1,Dis3,Cdh26,Haus3,Mettl5,Atxn2,Cenpe,Aspm,Adam10,Srp19,Birc5,Rnf213,Arrb1,Cox7a1,Mbnl1,Nop56,Flcn,Anp32b,Pdpk1,Mcts2,Sirt1,Ect2,Dnm1l,Rsl1d1,Srsf1,Prlr,Zfyve27,Tomm20 |

| | | | | | | | |
|---|---|---|---|---|---|---|---|
| GO:0019438 | Aromatic compound biosynthetic process | 21 | 769 | 0.4 | 0.0188 | 10090.ENSMUSP00000011058,10090.ENSMUSP00000020217,10090.ENSMUSP00000023774,10090.ENSMUSP00000024104,10090.ENSMUSP00000027933,10090.ENSMUSP00000028328,10090.ENSMUSP00000031542,10090.ENSMUSP00000033173,10090.ENSMUSP00000051968,10090.ENSMUSP00000065363,10090.ENSMUSP00000079198,10090.ENSMUSP00000079703,10090.ENSMUSP00000080640,10090.ENSMUSP00000095866,10090.ENSMUSP00000101082,10090.ENSMUSP00000102824,10090.ENSMUSP00000108557,10090.ENSMUSP00000113851,10090.ENSMUSP00000120647,10090.ENSMUSP00000129157,10090.ENSMUSP00000129765 | Pdhx,Nudt4,Slc11a2,Gcm1,Dtl,Entpd2,Oasl2,Polr3e,Polr2k,Nfil3,Oas1a,Amd2,Slc39a8,Arrb1,Sirt1,Crebzf,Nmnat3,Bcl3,Nmi,Tcea1,Parp10 |
| GO:0035456 | Response to interferon-beta | 5 | 43 | 1.03 | 0.0217 | 10090.ENSMUSP00000026565,10090.ENSMUSP00000032473,10090.ENSMUSP00000099889,10090.ENSMUSP00000104548,10090.ENSMUSP00000123011 | Ifitm3,Iigp1,Ifit3,Irf1,Xaf1 |
| GO:0010467 | Gene expression | 36 | 1743 | 0.28 | 0.0238 | 10090.ENSMUSP00000002198,10090.ENSMUSP00000021381,10090.ENSMUSP00000024104,10090.ENSMUSP00000030623,10090.ENSMUSP00000033173,10090.ENSMUSP00000034612,10090.ENSMUSP00000039010,10090.ENSMUSP00000041906,10090.ENSMUSP00000050907,10090.ENSMUSP00000051968,10090.ENSMUSP00000063839,10090.ENSMUSP00000065363,10090.ENSMUSP00000078123,10090.ENSMUSP00000078207,10090.ENSMUSP00000078940,10090.ENSMUSP00000080640,10090.ENSMUSP00000081572,10090.ENSMUSP00000082548,10090.ENSMUSP00000085906,10090.ENSMUSP00000087372,10090.ENSMUSP00000095866,10090.ENSMUSP00000096686,10090.ENSMUSP00000100534,10090.ENSMUSP00000101082,10090.ENSMUSP00000102824,10090.ENSMUSP00000103209,10090.ENSMUSP00000107595,10090.ENSMUSP00000113431,10090.ENSMUSP00000113851,10090.ENSMUSP00000117045,10090.ENSMUSP00000120525,10090.ENSMUSP00000120595,10090.ENSMUSP00000120647,10090.ENSMUSP00000126565,10090.ENSMUSP00000129157,10090.ENSMUSP00000138006 | Sf3a1,Pnn,Gcm1,Sfpq,Polr3e,Ddx25,Wdr55,Dis3,Mettl5,Polr2k,Adam10,Nfil3,Mrpl18,Prpf38a,Gspt1,Slc39a8,Prpf4,Isg15,Fmr1,Mfap1a,Arrb1,Mbnl1,Mcts2,Sirt1,Crebzf,Cisd3,Hnrnpa3,Rsl1d1,Bcl3,Srsf3,Irf9,Srsf1,Nmi,Lsm5,Tcea1,Metap2 |
| GO:1903047 | Mitotic cell cycle process | 15 | 454 | 0.48 | 0.0241 | 10090.ENSMUSP00000020835,10090.ENSMUSP00000020908,10090.ENSMUSP00000022227,10090.ENSMUSP00000023353,10090.ENSMUSP00000023359,10090.ENSMUSP00000025562,10090.ENSMUSP00000027697,10090.ENSMUSP00000033913,10090.ENSMUSP00000034964,10090.ENSMUSP00000047872,10090.ENSMUSP00000057938,10090.ENSMUSP00000079124,10090.ENSMUSP00000098897,10090.ENSMUSP00000099990,10090.ENSMUSP00000103935 | Ppm1d,E2f6,Cenpk,Mcm4,Nde1,Incenp,Cdk18,Dctn6,Tipin,Smc4,Cenpe,Birc5,Mad2l1,Anp32b,Ect2 |
| GO:0009416 | Response to light stimulus | 12 | 307 | 0.56 | 0.0245 | 10090.ENSMUSP00000002413,10090.ENSMUSP00000005503,10090.ENSMUSP00000022612,10090.ENSMUSP00000027809,10090.ENSMUSP00000027933,10090.ENSMUSP00000030289,10090.ENSMUSP00000034964,10090.ENSMUSP00000059211,10090.ENSMUSP00000085906,10090.ENSMUSP00000095866,10090.ENSMUSP00000101082,10090.ENSMUSP00000113851 | Tmem161a,Msh6,Pbk,Opn3,Dtl,Usp1,Tipin,Ercc8,Fmr1,Arrb1,Sirt1,Bcl3 |
| GO:0008380 | RNA splicing | 12 | 312 | 0.55 | 0.0274 | 10090.ENSMUSP00000002198,10090.ENSMUSP00000021381,10090.ENSMUSP00000030623,10090.ENSMUSP00000078207,10090.ENSMUSP00000081572,10090.ENSMUSP00000085906,10090.ENSMUSP00000087372,10090.ENSMUSP00000096686,10090.ENSMUSP00000107595,10090.ENSMUSP00000117045,10090.ENSMUSP00000120595,10090.ENSMUSP00000126565 | Sf3a1,Pnn,Sfpq,Prpf38a,Prpf4,Fmr1,Mfap1a,Mbnl1,Hnrnpa3,Srsf3,Srsf1,Lsm5 |
| GO:0043933 | Protein-containing complex subunit organization | 27 | 1164 | 0.33 | 0.0274 | 10090.ENSMUSP00000002198,10090.ENSMUSP00000020308,10090.ENSMUSP00000022227,10090.ENSMUSP00000023353,10090.ENSMUSP00000023359,10090.ENSMUSP00000023769,10090.ENSMUSP00000025161,10090.ENSMUSP00000028382,10090.ENSMUSP00000030367,10090.ENSMUSP00000034270,10090.ENSMUSP00000037459,10090.ENSMUSP00000039047,10090.ENSMUSP00000057938,10090.ENSMUSP00000063839,10090.ENSMUSP00000072386,10090.ENSMUSP00000078940,10090.ENSMUSP00000091429,10090.ENSMUSP00000096193,10090.ENSMUSP00000096686,10090.ENSMUSP00000099990,10090.ENSMUSP00000100534,10090.ENSMUSP00000103935,10090.ENSMUSP00000106801,10090.ENSMUSP00000111415,10090.ENSMUSP00000120595,10090.ENSMUSP00000122219,10090.ENSMUSP00000136493 | Sf3a1,Ddit4,Cenpk,Mcm4,Nde1,Atf1,Tapbp,Nup35,Ssbp3,Map1lc3b,Zfand1,Atg14,Cenpe,Adam10,Srp19,Gspt1,Rnf213,Cox7a1,Mbnl1,Anp32b,Mcts2,Ect2,Pom121,Dnm1l,Srsf1,Prlr,Tomm20 |
| GO:0017038 | Protein import | 8 | 142 | 0.71 | 0.0279 | 10090.ENSMUSP00000018506,10090.ENSMUSP00000028382,10090.ENSMUSP00000032898,10090.ENSMUSP00000034079,10090.ENSMUSP00000034787,10090.ENSMUSP00000102710,10090.ENSMUSP00000106801,10090.ENSMUSP00000136493 | Kpna2,Nup35,Ipo5,Heatr3,Phip,Txnip,Pom121,Tomm20 |
| GO:0016070 | RNA metabolic process | 28 | 1240 | 0.32 | 0.0307 | 10090.ENSMUSP00000002198,10090.ENSMUSP00000021381,10090.ENSMUSP00000024104,10090.ENSMUSP00000027623,10090.ENSMUSP00000030623,10090.ENSMUSP00000033173,10090.ENSMUSP00000039010,10090.ENSMUSP00000041906,10090.ENSMUSP00000050907,10090.ENSMUSP00000051968,10090.ENSMUSP00000065363,10090.ENSMUSP00000078207,10090.ENSMUSP00000078940,10090.ENSMUSP00000080640,10090.ENSMUSP00000081572,10090.ENSMUSP00000085906,10090.ENSMUSP00000087372,10090.ENSMUSP00000095866,10090.ENSMUSP00000096686,10090.ENSMUSP00000102824,10090.ENSMUSP00000107595,10090.ENSMUSP00000113431,10090.ENSMUSP00000113851,10090.ENSMUSP00000117045,10090.ENSMUSP00000120595,10090.ENSMUSP00000120647,10090.ENSMUSP00000126565,10090.ENSMUSP00000129157 | Sf3a1,Pnn,Gcm1,Tsn,Sfpq,Polr3e,Wdr55,Dis3,Mettl5,Polr2k,Nfil3,Prpf38a,Gspt1,Slc39a8,Prpf4,Fmr1,Mfap1a,Arrb1,Mbnl1,Crebzf,Hnrnpa3,Rsl1d1,Bcl3,Srsf3,Srsf1,Nmi,Lsm5,Tcea1 |
| GO:0035634 | Response to stilbenoid | 4 | 25 | 1.17 | 0.0307 | 10090.ENSMUSP00000019469,10090.ENSMUSP00000032198,10090.ENSMUSP00000034003,10090.ENSMUSP00000099889 | G6pc,Usp18,Fgl1,Ifit3 |
| GO:0045841 | Negative regulation of mitotic metaphase/anaphase transition | 4 | 25 | 1.17 | 0.0307 | 10090.ENSMUSP00000022927,10090.ENSMUSP00000057938,10090.ENSMUSP00000079124,10090.ENSMUSP00000098897 | Rad21,Cenpe,Birc5,Mad2l1 |
| GO:1904951 | Positive regulation of establishment of protein localization | 13 | 368 | 0.51 | 0.0317 | 10090.ENSMUSP00000032898,10090.ENSMUSP00000037459,10090.ENSMUSP00000060349,10090.ENSMUSP00000075172,10090.ENSMUSP00000084599,10090.ENSMUSP00000095866,10090.ENSMUSP00000099990,10090.ENSMUSP00000101082,10090.ENSMUSP00000103935,10090.ENSMUSP00000111415,10090.ENSMUSP00000123788,10090.ENSMUSP00000129800,10090.ENSMUSP00000134799 | Ipo5,Zfand1,Camk2n1,Frmd4a,Kif20b,Arrb1,Anp32b,Sirt1,Ect2,Dnm1l,Cep295,Sh3glb1,Golph3l |

| | | | | | | |
|---|---|---|---|---|---|---|
| GO:0006259 | DNA metabolic process | 18 | 638 | 0.41 | 0.0335 | 10090.ENSMUSP00000004120,10090.ENSMUSP00000005503,10090.ENSMUSP00000020835,10090.ENSMUSP00000022927,10090.ENSMUSP00000023353,10090.ENSMUSP00000025930,10090.ENSMUSP00000027933,10090.ENSMUSP00000030289,10090.ENSMUSP00000030623,10090.ENSMUSP00000034964,10090.ENSMUSP00000038871,10090.ENSMUSP00000047872,10090.ENSMUSP00000059211,10090.ENSMUSP00000065940,10090.ENSMUSP00000080949,10090.ENSMUSP00000101082,10090.ENSMUSP00000111443,10090.ENSMUSP00000129765 | Fancl,Msh6,Ppm1d,Rad21,Mcm4,Smc3,Dtl,Usp1,Sfpq,Tipin,Slx4,Smc4,Ercc8,Hmgb2,Mdc1,Sirt1,Ube2v2,Parp10 |
| GO:0010564 | Regulation of cell cycle process | 17 | 583 | 0.43 | 0.0335 | 10090.ENSMUSP00000022927,10090.ENSMUSP00000025562,10090.ENSMUSP00000025930,10090.ENSMUSP00000027933,10090.ENSMUSP00000030623,10090.ENSMUSP00000034787,10090.ENSMUSP00000056778,10090.ENSMUSP00000057938,10090.ENSMUSP00000065544,10090.ENSMUSP00000079124,10090.ENSMUSP00000084599,10090.ENSMUSP00000098897,10090.ENSMUSP00000099758,10090.ENSMUSP00000099990,10090.ENSMUSP00000103935,10090.ENSMUSP00000123788,10090.ENSMUSP00000129800 | Rad21,Incenp,Smc3,Dtl,Sfpq,Phip,E2f8,Cenpe,Azin1,Birc5,Kif20b,Mad2l1,Flcn,Anp32b,Ect2,Cep295,Sh3glb1 |
| GO:0060700 | Regulation of ribonuclease activity | 3 | 10 | 1.44 | 0.0368 | 10090.ENSMUSP00000031540,10090.ENSMUSP00000031542,10090.ENSMUSP00000079198 | Oasl1,Oasl2,Oas1a |
| GO:0000398 | mRNA splicing, via spliceosome | 9 | 191 | 0.64 | 0.0369 | 10090.ENSMUSP00000002198,10090.ENSMUSP00000030623,10090.ENSMUSP00000078207,10090.ENSMUSP00000081572,10090.ENSMUSP00000087372,10090.ENSMUSP00000096686,10090.ENSMUSP00000117045,10090.ENSMUSP00000120595,10090.ENSMUSP00000126565 | Sf3a1,Sfpq,Prpf38a,Prpf4,Mfap1a,Mbnl1,Srsf3,Srsf1,Lsm5 |
| GO:0016567 | Protein ubiquitination | 17 | 591 | 0.42 | 0.0369 | 10090.ENSMUSP00000004120,10090.ENSMUSP00000021120,10090.ENSMUSP00000027933,10090.ENSMUSP00000028398,10090.ENSMUSP00000039206,10090.ENSMUSP00000039955,10090.ENSMUSP00000040220,10090.ENSMUSP00000059211,10090.ENSMUSP00000082102,10090.ENSMUSP00000091429,10090.ENSMUSP00000095866,10090.ENSMUSP00000101082,10090.ENSMUSP00000103432,10090.ENSMUSP00000110645,10090.ENSMUSP00000111443,10090.ENSMUSP00000111780,10090.ENSMUSP00000132446 | Fancl,Trim47,Dtl,Ube2e3,Hace1,Cblc,Ubac1,Ercc8,Nhlrc4,Rnf213,Arrb1,Sirt1,Trim59,Cnot4,Ube2v2,Rnf186,Ube2d2a |
| GO:0010942 | Positive regulation of cell death | 19 | 708 | 0.39 | 0.0383 | 10090.ENSMUSP00000002284,10090.ENSMUSP00000017851,10090.ENSMUSP00000020308,10090.ENSMUSP00000023774,10090.ENSMUSP00000023897,10090.ENSMUSP00000029018,10090.ENSMUSP00000030121,10090.ENSMUSP00000030623,10090.ENSMUSP00000032800,10090.ENSMUSP00000063839,10090.ENSMUSP00000095866,10090.ENSMUSP00000099758,10090.ENSMUSP00000099990,10090.ENSMUSP00000101082,10090.ENSMUSP00000102710,10090.ENSMUSP00000103935,10090.ENSMUSP00000104548,10090.ENSMUSP00000111415,10090.ENSMUSP00000129800 | Plaur,Serinc3,Ddit4,Slc11a2,Gzma,Zbp1,B4galt1,Sfpq,Tyrobp,Adam10,Arrb1,Flcn,Anp32b,Sirt1,Txnip,Ect2,Irf1,Dnm1l,Sh3glb1 |
| GO:0010629 | Negative regulation of gene expression | 36 | 1811 | 0.26 | 0.0395 | 10090.ENSMUSP00000018361,10090.ENSMUSP00000020835,10090.ENSMUSP00000020908,10090.ENSMUSP00000024657,10090.ENSMUSP00000025904,10090.ENSMUSP00000029566,10090.ENSMUSP00000030623,10090.ENSMUSP00000032800,10090.ENSMUSP00000036971,10090.ENSMUSP00000039586,10090.ENSMUSP00000048051,10090.ENSMUSP00000053483,10090.ENSMUSP00000056778,10090.ENSMUSP00000058037,10090.ENSMUSP00000058237,10090.ENSMUSP00000058940,10090.ENSMUSP00000065363,10090.ENSMUSP00000065940,10090.ENSMUSP00000077971,10090.ENSMUSP00000078940,10090.ENSMUSP00000079124,10090.ENSMUSP00000085906,10090.ENSMUSP00000095866,10090.ENSMUSP00000099758,10090.ENSMUSP00000101082,10090.ENSMUSP00000102710,10090.ENSMUSP00000102824,10090.ENSMUSP00000104548,10090.ENSMUSP00000110684,10090.ENSMUSP00000113851,10090.ENSMUSP00000120647,10090.ENSMUSP00000126565,10090.ENSMUSP00000128260,10090.ENSMUSP00000129765,10090.ENSMUSP00000130930,10090.ENSMUSP00000136867 | Pmp22,Ppm1d,E2f6,Phf10,Prdx5,Efna1,Sfpq,Tyrobp,Mvk,Serpine1,Tiparp,Hic1,E2f8,Zfpm1,S100a1,Whsc1,Nfil3,Hmgb2,Phf6,Gspt1,Birc5,Fmr1,Arrb1,Flcn,Sirt1,Txnip,Crebzf,Irf1,Arid5a,Bcl3,Nmi,Lsm5,Tfdp2,Parp10,Tgif1,Zfp131 |
| GO:0031327 | Negative regulation of cellular biosynthetic process | 31 | 1475 | 0.29 | 0.0416 | 10090.ENSMUSP00000020835,10090.ENSMUSP00000020908,10090.ENSMUSP00000024657,10090.ENSMUSP00000025904,10090.ENSMUSP00000029566,10090.ENSMUSP00000030623,10090.ENSMUSP00000034964,10090.ENSMUSP00000036971,10090.ENSMUSP00000053483,10090.ENSMUSP00000056778,10090.ENSMUSP00000058037,10090.ENSMUSP00000058237,10090.ENSMUSP00000058940,10090.ENSMUSP00000061390,10090.ENSMUSP00000061877,10090.ENSMUSP00000065363,10090.ENSMUSP00000065940,10090.ENSMUSP00000077971,10090.ENSMUSP00000079124,10090.ENSMUSP00000085906,10090.ENSMUSP00000099758,10090.ENSMUSP00000101082,10090.ENSMUSP00000102710,10090.ENSMUSP00000102824,10090.ENSMUSP00000104548,10090.ENSMUSP00000110684,10090.ENSMUSP00000113851,10090.ENSMUSP00000128260,10090.ENSMUSP00000129765,10090.ENSMUSP00000130930,10090.ENSMUSP00000136867 | Ppm1d,E2f6,Phf10,Prdx5,Efna1,Sfpq,Tipin,Mvk,Hic1,E2f8,Zfpm1,S100a1,Whsc1,Tob2,Insig1,Nfil3,Hmgb2,Phf6,Birc5,Fmr1,Flcn,Sirt1,Txnip,Crebzf,Irf1,Arid5a,Bcl3,Tfdp2,Parp10,Tgif1,Zfp131 |
| GO:0010556 | Regulation of macromolecule biosynthetic process | 56 | 3294 | 0.19 | 0.0436 | 10090.ENSMUSP00000004985,10090.ENSMUSP00000020123,10090.ENSMUSP00000020835,10090.ENSMUSP00000020908,10090.ENSMUSP00000022227,10090.ENSMUSP00000022380,10090.ENSMUSP00000022927,10090.ENSMUSP00000023769,10090.ENSMUSP00000024104,10090.ENSMUSP00000024657,10090.ENSMUSP00000025904,10090.ENSMUSP00000025930,10090.ENSMUSP00000027697,10090.ENSMUSP00000028382,10090.ENSMUSP00000029566,10090.ENSMUSP00000030367,10090.ENSMUSP00000030623,10090.ENSMUSP00000034612,10090.ENSMUSP00000034787,10090.ENSMUSP00000034964,10090.ENSMUSP00000035101,10090.ENSMUSP00000036971,10090.ENSMUSP00000050854,10090.ENSMUSP00000053483,10090.ENSMUSP00000056778,10090.ENSMUSP00000058037,10090.ENSMUSP00000058237,10090.ENSMUSP00000058940,10090.ENSMUSP00000061390,10090.ENSMUSP00000061877,10090.ENSMUSP00000065363,10090.ENSMUSP00000065940,10090.ENSMUSP00000077971,10090.ENSMUSP00000078940,10090.ENSMUSP00000079124,10090.ENSMUSP00000080640,10090.ENSMUSP00000085906,10090.ENSMUSP00000095794,10090.ENSMUSP00000095866,10090.ENSMUSP00000099758,10090.ENSMUSP00000101082,10090.ENSMUSP00000101661,10090.ENSMUSP00000102710,10090.ENSMUSP00000102824,10090.ENSMUSP00000104548,10090.ENSMUSP00000108083,10090.ENSMUSP00000110684,10090.ENSMUSP00000113851,10090.ENSMUSP00000120525,10090.ENSMUSP00000120647,10090.ENSMUSP00000122712,10090.ENSMUSP00000128260,10090.ENSMUSP00000129157,10090.ENSMUSP00000129765,[illegible] | Brpf3,Tmpo,Ppm1d,E2f6,Cenpk,Psmc6,Rad21,Atf1,Gcm1,Phf10,Prdx5,Smc3,Cdk18,Nup35,Efna1,Ssbp3,Sfpq,Ddx25,Phip,Tipin,Csrnp1,Mvk,Zfp367,Hic1,E2f8,Zfpm1,S100a1,Whsc1,Tob2,Insig1,Nfil3,Hmgb2,Phf6,Gspt1,Birc5,Slc39a8,Fmr1,Hbb-bt,Arrb1,Flcn,Sirt1,Zbtb8b,Txnip,Crebzf,Irf1,Bclaf3,Arid5a,Bcl3,Irf9,Nmi,Ppp1r3g,Tfdp2,Tcea1,Parp10,Tgif1,Zfp131 |

| | | | | | | | |
|---|---|---|---|---|---|---|---|
| GO:0032465 | Regulation of cytokinesis | 6 | 84 | 0.82 | 0.0436 | 10090.ENSMUSP00000025562,10090.ENSMUSP00000056778,10090.ENSMUSP00000084599,10090.ENSMUSP00000099758,10090.ENSMUSP00000103935,10090.ENSMUSP00000129800 | Incenp,E2f8,Kif20b,Flcn,Ect2,Sh3glb1 |
| GO:1903507 | Negative regulation of nucleic acid-templated transcription | 27 | 1215 | 0.31 | 0.0436 | 10090.ENSMUSP00000020835,10090.ENSMUSP00000020908,10090.ENSMUSP00000024657,10090.ENSMUSP00000025904,10090.ENSMUSP00000029566,10090.ENSMUSP00000030623,10090.ENSMUSP00000053483,10090.ENSMUSP00000056778,10090.ENSMUSP00000058037,10090.ENSMUSP00000058237,10090.ENSMUSP00000058940,10090.ENSMUSP00000061390,10090.ENSMUSP00000065363,10090.ENSMUSP00000065940,10090.ENSMUSP00000077971,10090.ENSMUSP00000079124,10090.ENSMUSP00000099758,10090.ENSMUSP00000101082,10090.ENSMUSP00000102710,10090.ENSMUSP00000102824,10090.ENSMUSP00000104548,10090.ENSMUSP00000110684,10090.ENSMUSP00000113851,10090.ENSMUSP00000128260,10090.ENSMUSP00000129765,10090.ENSMUSP00000130930,10090.ENSMUSP00000136867 | Ppm1d,E2f6,Phf10,Prdx5,Efna1,Sfpq,Hic1,E2f8,Zfpm1,S100a1,Whsc1,Tob2,Nfil3,Hmgb2,Phf6,Birc5,Flcn,Sirt1,Txnip,Crebzf,Irf1,Arid5a,Bcl3,Tfdp2,Parp10,Tgif1,Zfp131 |
| GO:0009893 | Positive regulation of metabolic process | 62 | 3767 | 0.18 | 0.0449 | 10090.ENSMUSP00000002284,10090.ENSMUSP00000002413,10090.ENSMUSP00000004430,10090.ENSMUSP00000004985,10090.ENSMUSP00000018361,10090.ENSMUSP00000020908,10090.ENSMUSP00000022227,10090.ENSMUSP00000022380,10090.ENSMUSP00000022612,10090.ENSMUSP00000023769,10090.ENSMUSP00000023774,10090.ENSMUSP00000024104,10090.ENSMUSP00000024657,10090.ENSMUSP00000025562,10090.ENSMUSP00000025904,10090.ENSMUSP00000027933,10090.ENSMUSP00000029566,10090.ENSMUSP00000030367,10090.ENSMUSP00000030623,10090.ENSMUSP00000032800,10090.ENSMUSP00000034787,10090.ENSMUSP00000035101,10090.ENSMUSP00000037007,10090.ENSMUSP00000038871,10090.ENSMUSP00000039047,10090.ENSMUSP00000039206,10090.ENSMUSP00000039586,10090.ENSMUSP00000048051,10090.ENSMUSP00000050854,10090.ENSMUSP00000056778,10090.ENSMUSP00000057938,10090.ENSMUSP00000058037,10090.ENSMUSP00000058237,10090.ENSMUSP00000058940,10090.ENSMUSP00000059211,10090.ENSMUSP00000060349,10090.ENSMUSP00000060791,10090.ENSMUSP00000061877,10090.ENSMUSP00000065363,10090.ENSMUSP00000065940,10090.ENSMUSP00000085906,10090.ENSMUSP00000095866,10090.ENSMUSP00000099758,10090.ENSMUSP00000099990,10090.ENSMUSP00000099991,10090.ENSMUSP00000101082,10090.ENSMUSP00000103935,10090.ENSMUSP00000104548,10090.ENSMUSP00000106199,10090.ENSMUSP00000108083,10090.ENSMUSP00000110684,10090.ENSMUSP00000111443,10090.ENSMUSP00000113851,10090.ENSMUSP00000120595,[illegible] | Plaur,Tmem161a,Clcn3,Brpf3,Pmp22,E2f6,Cenpk,Psmc6,Pbk,Atf1,Slc11a2,Gcm1,Phf10,Incenp,Prdx5,Dtl,Efna1,Ssbp3,Sfpq,Tyrobp,Phip,Csrnp1,Hspb8,Slx4,Atg14,Hace1,Serpine1,Tiparp,Zfp367,E2f8,Cenpe,Zfpm1,S100a1,Whsc1,Ercc8,Camk2n1,Hilpda,Insig1,Nfil3,Hmgb2,Fmr1,Arrb1,Flcn,Anp32b,Pdpk1,Sirt1,Ect2,Irf1,Map4k5,Bclaf3,Arid5a,Ube2v2,Bcl3,Srsf1,Nmi,Prlr,Ppp1r3g,Cep295,Tfdp2,Tcea1,Sh3glb1,Zfp131 |
| GO:0010558 | Negative regulation of macromolecule biosynthetic process | 30 | 1419 | 0.29 | 0.0449 | 10090.ENSMUSP00000020835,10090.ENSMUSP00000020908,10090.ENSMUSP00000024657,10090.ENSMUSP00000025904,10090.ENSMUSP00000029566,10090.ENSMUSP00000030623,10090.ENSMUSP00000034964,10090.ENSMUSP00000036971,10090.ENSMUSP00000053483,10090.ENSMUSP00000056778,10090.ENSMUSP00000058037,10090.ENSMUSP00000058237,10090.ENSMUSP00000058940,10090.ENSMUSP00000061390,10090.ENSMUSP00000065363,10090.ENSMUSP00000065940,10090.ENSMUSP00000077971,10090.ENSMUSP00000079124,10090.ENSMUSP00000085906,10090.ENSMUSP00000099758,10090.ENSMUSP00000101082,10090.ENSMUSP00000102710,10090.ENSMUSP00000102824,10090.ENSMUSP00000104548,10090.ENSMUSP00000110684,10090.ENSMUSP00000113851,10090.ENSMUSP00000128260,10090.ENSMUSP00000129765,10090.ENSMUSP00000130930,10090.ENSMUSP00000136867 | Ppm1d,E2f6,Phf10,Prdx5,Efna1,Sfpq,Tipin,Mvk,Hic1,E2f8,Zfpm1,S100a1,Whsc1,Tob2,Nfil3,Hmgb2,Phf6,Birc5,Fmr1,Flcn,Sirt1,Txnip,Crebzf,Irf1,Arid5a,Bcl3,Tfdp2,Parp10,Tgif1,Zfp131 |
| GO:0044271 | Cellular nitrogen compound biosynthetic process | 26 | 1156 | 0.32 | 0.0449 | 10090.ENSMUSP00000011058,10090.ENSMUSP00000020217,10090.ENSMUSP00000023774,10090.ENSMUSP00000024104,10090.ENSMUSP00000027933,10090.ENSMUSP00000028328,10090.ENSMUSP00000031542,10090.ENSMUSP00000033173,10090.ENSMUSP00000051968,10090.ENSMUSP00000065363,10090.ENSMUSP00000065544,10090.ENSMUSP00000078123,10090.ENSMUSP00000078940,10090.ENSMUSP00000079198,10090.ENSMUSP00000079703,10090.ENSMUSP00000080640,10090.ENSMUSP00000095866,10090.ENSMUSP00000100534,10090.ENSMUSP00000101082,10090.ENSMUSP00000102824,10090.ENSMUSP00000108557,10090.ENSMUSP00000113851,10090.ENSMUSP00000120647,10090.ENSMUSP00000128661,10090.ENSMUSP00000129157,10090.ENSMUSP00000129765 | Pdhx,Nudt4,Slc11a2,Gcm1,Dtl,Entpd2,Oasl2,Polr3e,Polr2k,Nfil3,Azin1,Mrpl18,Gspt1,Oas1a,Amd2,Slc39a8,Arrb1,Mcts2,Sirt1,Crebzf,Nmnat3,Bcl3,Nmi,Odc1,Tcea1,Parp10 |
| GO:0051641 | Cellular localization | 40 | 2115 | 0.24 | 0.0449 | 10090.ENSMUSP00000004430,10090.ENSMUSP00000018361,10090.ENSMUSP00000018506,10090.ENSMUSP00000022927,10090.ENSMUSP00000023359,10090.ENSMUSP00000023774,10090.ENSMUSP00000025562,10090.ENSMUSP00000028382,10090.ENSMUSP00000030623,10090.ENSMUSP00000032898,10090.ENSMUSP00000034079,10090.ENSMUSP00000034612,10090.ENSMUSP00000034787,10090.ENSMUSP00000039047,10090.ENSMUSP00000041091,10090.ENSMUSP00000057938,10090.ENSMUSP00000059159,10090.ENSMUSP00000061877,10090.ENSMUSP00000062766,10090.ENSMUSP00000063839,10090.ENSMUSP00000068835,10090.ENSMUSP00000072386,10090.ENSMUSP00000077574,10090.ENSMUSP00000079124,10090.ENSMUSP00000080640,10090.ENSMUSP00000084599,10090.ENSMUSP00000085906,10090.ENSMUSP00000098897,10090.ENSMUSP00000099758,10090.ENSMUSP00000102710,10090.ENSMUSP00000106801,10090.ENSMUSP00000111415,10090.ENSMUSP00000111780,10090.ENSMUSP00000117045,10090.ENSMUSP00000125993,10090.ENSMUSP00000129800,10090.ENSMUSP00000130684,10090.ENSMUSP00000133851,10090.ENSMUSP00000134799,10090.ENSMUSP00000136493 | Clcn3,Pmp22,Kpna2,Rad21,Nde1,Slc11a2,Incenp,Nup35,Sfpq,Ipo5,Heatr3,Ddx25,Phip,Atg14,Rtp4,Cenpe,Aspm,Insig1,Nupl2,Adam10,Rabgap1,Srp19,Dchs1,Birc5,Slc39a8,Kif20b,Fmr1,Mad2l1,Flcn,Txnip,Pom121,Dnm1l,Rnf186,Srsf3,Camsap3,Sh3glb1,Zfyve27,Arhgap21,Golph3l,Tomm20 |
| GO:0008219 | Cell death | 22 | 906 | 0.35 | 0.0456 | 10090.ENSMUSP00000005503,10090.ENSMUSP00000015581,10090.ENSMUSP00000018361,10090.ENSMUSP00000020308,10090.ENSMUSP00000022927,10090.ENSMUSP00000023897,10090.ENSMUSP00000025904,10090.ENSMUSP00000029018,10090.ENSMUSP00000032800,10090.ENSMUSP00000035101,10090.ENSMUSP00000044603,10090.ENSMUSP00000053483,10090.ENSMUSP00000079124,10090.ENSMUSP00000095866,10090.ENSMUSP00000099991,10090.ENSMUSP00000101082,10090.ENSMUSP00000104548,10090.ENSMUSP00000111415,10090.ENSMUSP00000111780,10090.ENSMUSP00000113851,10090.ENSMUSP00000123011,10090.ENSMUSP00000129800 | Msh6,Gzmb,Pmp22,Ddit4,Rad21,Gzma,Prdx5,Zbp1,Tyrobp,Csrnp1,Pim3,Hic1,Birc5,Arrb1,Pdpk1,Sirt1,Irf1,Dnm1l,Rnf186,Bcl3,Xaf1,Sh3glb1 |
| GO:0000012 | Single strand break repair | 3 | 12 | 1.36 | 0.0491 | 10090.ENSMUSP00000047872,10090.ENSMUSP00000059211,10090.ENSMUSP00000101082 | Smc4,Ercc8,Sirt1 |

| | | | | | | | |
|---|---|---|---|---|---|---|---|
| GO:0003183 | Mitral valve morphogenesis | 3 | 12 | 1.36 | 0.0491 | 10090.ENSMUSP00000029566,10090.ENSMUSP00000058037, 10090.ENSMUSP00000077574 | Efna1,Zfpm1,Dchs1 |
| GO:0006596 | Polyamine biosynthetic process | 3 | 12 | 1.36 | 0.0491 | 10090.ENSMUSP00000065544,10090.ENSMUSP00000079703, 10090.ENSMUSP00000128661 | Azin1,Amd2,Odc1 |
| GO:0006607 | NLS-bearing protein import into nucleus | 3 | 12 | 1.36 | 0.0491 | 10090.ENSMUSP00000018506,10090.ENSMUSP00000028382, 10090.ENSMUSP00000032898 | Kpna2,Nup35,Ipo5 |
| GO:0043066 | Negative regulation of apoptotic process | 22 | 913 | 0.35 | 0.0491 | 10090.ENSMUSP00000002284,10090.ENSMUSP00000002413, 10090.ENSMUSP00000025254,10090.ENSMUSP00000025904, 10090.ENSMUSP00000029566,10090.ENSMUSP00000034787, 10090.ENSMUSP00000039586,10090.ENSMUSP00000044603, 10090.ENSMUSP00000065940,10090.ENSMUSP00000078985, 10090.ENSMUSP00000079124,10090.ENSMUSP00000085906, 10090.ENSMUSP00000095866,10090.ENSMUSP00000098897, 10090.ENSMUSP00000099889,10090.ENSMUSP00000099990, 10090.ENSMUSP00000099991,10090.ENSMUSP00000101082, 10090.ENSMUSP00000111443,10090.ENSMUSP00000113851, 10090.ENSMUSP00000122219,10090.ENSMUSP00000129800 | Plaur,Tmem161a,Lims2,Prdx5,Efna1,Phip,Serpine1,Pim3,Hmgb2,Krit1,Birc5,Fmr1,Arrb1,Mad2l1,Ifit3,Anp32b,Pdpk1,Sirt1,Ube2v2,Bcl3,Prlr,Sh3glb1 |
